# The BTB-ZF gene *Bm-mamo* regulates pigmentation in silkworm caterpillars

**DOI:** 10.1101/2023.04.07.536007

**Authors:** Songyuan Wu, Xiaoling Tong, Chenxing Peng, Jiangwen Luo, Chenghao Zhang, Kunpeng Lu, Chunlin Li, Xin Ding, Xiaohui Duan, Yaru Lu, Hai Hu, Duan Tan, Fangyin Dai

**Affiliations:** State Key Laboratory of Resource Insects, Key Laboratory of Sericultural Biology and Genetic Breeding, Ministry of Agriculture and Rural Affairs, College of Sericulture, Textile and Biomass Sciences, Southwest University, Chongqing 400715, China

## Abstract

The color pattern of insects is one of the most diverse adaptive evolutionary phenotypes. However, the molecular regulation of this color pattern is not fully understood. In this study, we found that the transcription factor Bm-mamo is responsible for *black dilute* (*bd*) allele mutations in the silkworm. Bm-mamo belongs to the BTB zinc finger family and is orthologous to mamo in *Drosophila melanogaster*. This gene has a conserved function in gamete production in *Drosophila* and silkworms and has evolved a pleiotropic function in the regulation of color patterns in caterpillars. Using RNAi and clustered regularly interspaced short palindromic repeats (CRISPR) technology, we showed that Bm-mamo is a repressor or has dark melanin patterns in the larval epidermis. Using in vitro binding assays and gene expression profiling in wild-type and mutant larvae, we also showed that *Bm-mamo* likely regulates the expression of related pigment synthesis and cuticular protein genes in a coordinated manner to mediate its role in color pattern formation. This mechanism is consistent with the dual role of this transcription factor in regulating both the structure and shape of the cuticle and the pigments that are embedded within it. This study provides new insight into the regulation of color patterns as well as into the construction of more complex epidermis features in some insects.

## Introduction

Insects often display stunning colors, and the patterns of these colors have been shown to be involved in behavior(*1*), immunity(*2*), thermoregulation(*3*), and UV protection(*4*); in particular, visual antagonism of predators via aposematism(*5*), mimicry(*6*), cryptic color patterns(*7*) or some combination of the above (*8*). In addition, the color patterns are divergent and convergent among populations(*9*). Due to these striking visual features and highly active adaptive evolutionary phenotypes, the genetic basis and evolutionary mechanism of color patterns have long been a topic of interest.

Insect coloration can be pigmentary, structural, or bioluminescent. Pigments are synthesized by insects themselves and form solid particles that are deposited within the cuticle of the body surface and the scales of the wings(*10, 11*). Interestingly, recent studies have shown that bile pigments and carotenoid pigments synthesized through biological synthesis are incorporated into body fluids and fill in the wing membranes of two butterflies (*Siproeta stelenes* and *Philaethria diatonica*) via hemolymph circulation, providing color in the form of liquid pigments(*12*). These pigments form colors by selective absorption and/or scattering of light depending on their physical properties(*13*). However, structural color refers to colors, such as metallic colors and iridescence, generated by optical interference and grating diffraction of the microstructure/nanostructure of the body surface or appendages (such as scales)(*14, 15*). Pigment color and structural color are widely distributed in insects and can only be observed by the naked eye in illuminated environments. However, some insects, such as fireflies, exhibit colors (green to orange) in the dark due to bioluminescence(*16*). Bioluminescence occurs when luciferase catalyzes the oxidation of small molecules of luciferin(*17*). In conclusion, the color patterns of insects have evolved to become highly sophisticated and are closely related to their living environment. For example, cryptic color can deceive animals via high similarity to the surrounding environment. However, the molecular mechanism by which insects form precise color patterns to match their living environment has not been determined.

Recent research has identified the metabolic pathways associated with related pigments, such as melanins, pterins, and ommochromes, in Lepidoptera(*18*). A deficiency of enzymes in the pigment metabolism pathway can lead to changes in color patterns(*19, 20*). In addition to pigment synthesis, the microstructure/nanostructure of the body surface and wing scales are important factors that influence body color patterns. The body surface and wing scales of Lepidoptera are composed mainly of cuticular proteins (CPs)(*21, 22*). There are multiple genes encoding CPs in Lepidopteran genomes. For example, in *Bombyx mori,* more than 220 genes encode CPs(*23*). However, the functions of CPs and the molecular mechanisms underlying their fine-scale localization in the cuticle are still unclear.

In addition, several pleiotropic factors, such as *wnt1*(*24*), *Apontic-like*(*25*), *clawless*(*26*), *abdominal-A*(*27*), *abdominal-B*(*28*), *engrailed*(*29*), *antennapedia*(*30*), *optix*(*31*), *bric* à *brac* (*bab*)(*32*), and *Distal-less* (*Dll*)(*33*), play important roles in the regulation of color patterns. The molecular mechanism by which these factors participate in the regulation of body color patterns needs further study. In addition, there are many undiscovered factors in the gene regulatory network (GRN) involved in the regulation of insect color patterns. The identification of new color pattern regulatory genes and the study of their molecular mechanisms would be helpful for further understanding color patterns.

Silkworms (*B. mori*) have been completely domesticated for more than 5000 years and are famous for their silk fiber production(*34*). Because of this long history of domestication, artificial selection and genetic research, its genetic background has been well studied, and approximately 200 strains of mutants with variable color patterns have been preserved(*35*), which provides a good resource for color pattern research. The *black dilute* (*bd*) mutant, which exhibits recessive Mendelian inheritance, has a dark gray larval body color, and the female is sterile. *Black dilute fertile* (*bd^f^*) is an allele that leads to a lighter body color than *bd* and fertile females (Fig. 1). The two mutations were mapped to a single locus at 22.9 centimorgans (cM) of linkage group 9(*36*). No pigmentation-related genes have been reported at this locus. Thus, this research may reveal a new color pattern-related gene, which stimulated our interest.

**Fig. 1.**
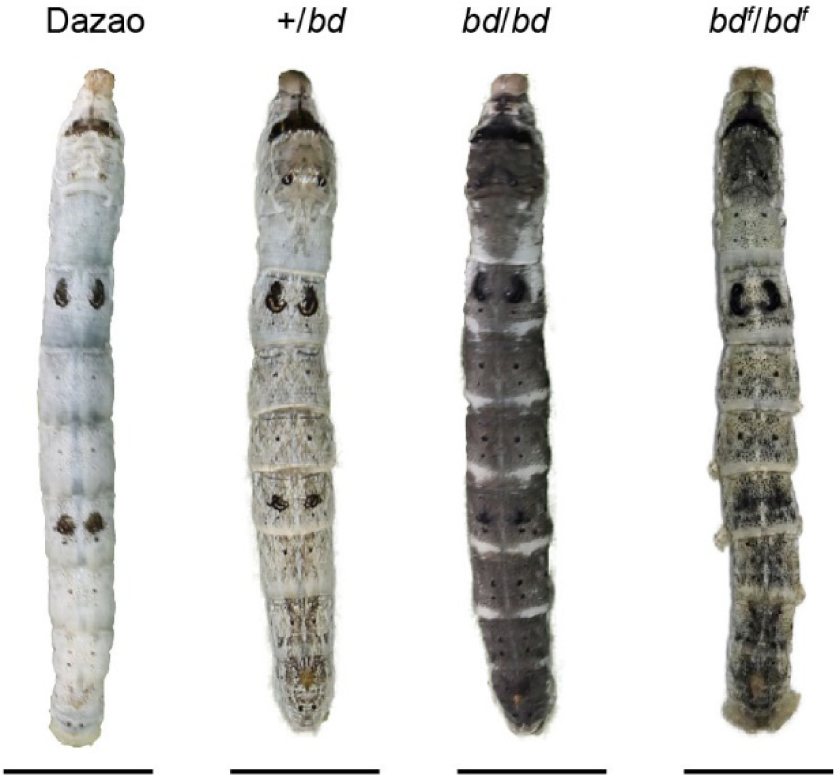
Phenotypes of *bd*, *bd^f^*, and wild-type Dazao larvae. The epidermis of *bd* is dark gray, and the epidermis of *bd^f^* is light gray at the 5th instar (day 3) of silkworm larvae. The bar indicates 1 centimeter.

## Results

### Candidate gene of the *bd* allele

To identify the genomic region responsible for the *bd* alleles, positional cloning of the *bd* locus was performed. Due to the female infertility of the *bd* mutant and the fertility of females with the *bd^f^* allele, we used *bd^f^* and the wild-type Dazao strain as parents for mapping analysis. The 1162 back-crossed filial 1st (BC1M) generation individuals from *bd^f^*and Dazao were subjected to fine mapping with molecular markers (Fig. 2). A genomic region of approximately 390 kilobases (kb) was responsible for the *bd* phenotype (Table S1). According to the SilkDB database(*37*), this region included five predicted genes (*BGIBMGA012516*, *BGIBMGA012517*, *BGIBMGA012518*, *BGIBMGA012519* and *BGIBMGA014089*). In addition, we analyzed the predictive genes for this genomic region from the GenBank(*38*) and SilkBase(*39*) databases (Fig. S1). The number of predicted genes varied among the different databases. We performed sequence alignment analysis of the predicted genes in the three databases to determine their correspondence. Real-time quantitative polymerase chain reaction (qPCR) was subsequently performed, which revealed that *BGIBMGA012517* and *BGIBMGA012518* were significantly downregulated on Day 3 of the fifth instar of the larvae in the *bd* phenotype individuals, while there was no difference in the expression levels of the other genes (Fig. S1). These two genes were predicted to be associated with a single locus (*LOC101738295*) in GenBank. To determine the gene structures of *BGIBMGA012517* and *BGIBMGA012518*, we used forward primers for the *BGIBMGA012517* gene and reverse primers for the *BGIBMGA012518* gene to amplify cDNA from the wild-type Dazao strain. By gene cloning, the predicted genes *BGIBMGA012517* and *BGIBMGA012518* were proven to be one such gene. For convenience, we temporarily called this the *12517/8* gene.

**Fig. 2.**
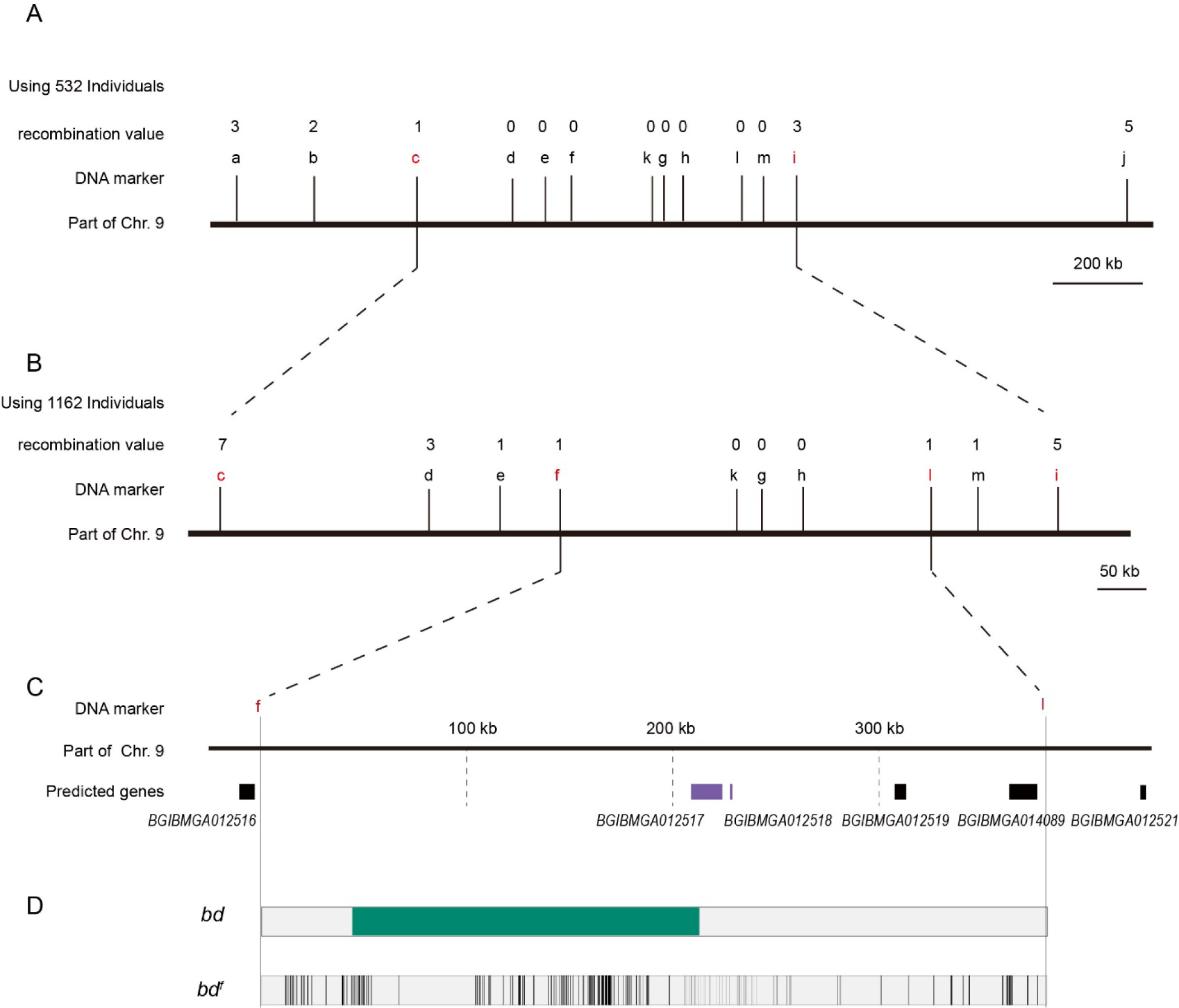
Positional cloning of the *bd* locus. (A) We used 532 BC1 individuals to map the *bd* locus between PCR markers c and i. The numbers above the DNA markers indicate recombination events. (B) A total of 1162 BC1 individuals were used to narrow the *bd* locus to an approximately 400 kb genomic region. (C) Partial enlarged view of the region responsible for *bd*. This region contains 4 predicted genes, *BGIBMGA012517*, *BGIBMGA012518*, *BGIBMGA012519* and *BGIBMGA014089*. (D) Analysis of nucleotide differences in the region responsible for *bd*. The green block indicates the deletion of the genome in *bd* mutants. The black vertical lines indicate the SNPs and indels of *bd^f^* mutants.

The *12517/8* gene produces two transcripts; the open reading frame (ORF) of the long transcript is 2397 bp, and the ORF of the short transcript is 1824 bp, which is the same 5’-terminus as that of the wild-type Dazao strain (Fig. S2). The *12517/8* gene showed significantly lower expression in the *bd^f^* mutant (Fig. S3), and multiple variations were found in the region nearby this gene by comparative genomic analysis between *bd^f^* and Dazao (Table S2). In addition, *12517/8* was completely silenced due to the deletion of DNA fragments (approximately 168 kb) from the first upstream intron and an insertion of 3560 bp in the *bd* mutant (Fig. S4).

To predict the function of the *12517*/8 gene, we performed a BLAST search using its full-length sequence and found a transcription factor, the *maternal gene required for meiosis* (*mamo*) in *D. melanogaster*, that had high sequence similarity to that of the *12517/8* gene (Fig. S5). Therefore, we named the *12517/18* gene *Bm-mamo*; the long transcript was designated *Bm-mamo-L*, and the short transcript was designated *Bm-mamo-S*.

The *mamo* gene belongs to the Broad-complex, Tramtrack and bric à brac/poxvirus zinc finger protein (BTB-ZF) family. In the BTB-ZF family, the zinc finger serves as a recognition motif for DNA-specific sequences, and the BTB domain promotes oligomerization and the recruitment of other regulatory factors(*40*). Most of these factors are transcription repressors, such as nuclear receptor corepressor (N-CoR) and silencing mediator for retinoid and thyroid hormone receptor (SMRT)(*41*), but some are activators, such as p300(*42*). Therefore, these features commonly serve as regulators of gene expression. *mamo* is enriched in embryonic primordial germ cells (PGCs) in *D. melanogaster*. Individuals deficient in *mamo* are able to undergo oogenesis but fail to execute meiosis properly, leading to female infertility in *D. melanogaster*(*43*). *Bm-mamo* was identified as an important candidate gene for further analysis.

### Expression pattern analysis of *Bm-mamo*

To analyze the expression profiles of *Bm-mamo*, we performed quantitative PCR. The expression levels of the *Bm-mamo* gene in the Dazao strain were investigated throughout the body at different developmental stages, from the embryonic stage to the adult stage. The gene was highly expressed in the molting stage of caterpillars, and its expression was upregulated in the later pupal and adult stages (Fig. 3A). This finding suggested that the *Bm-mamo* gene responds to ecdysone and participates in the processes of molting and metamorphosis in silkworms.

**Fig. 3.**
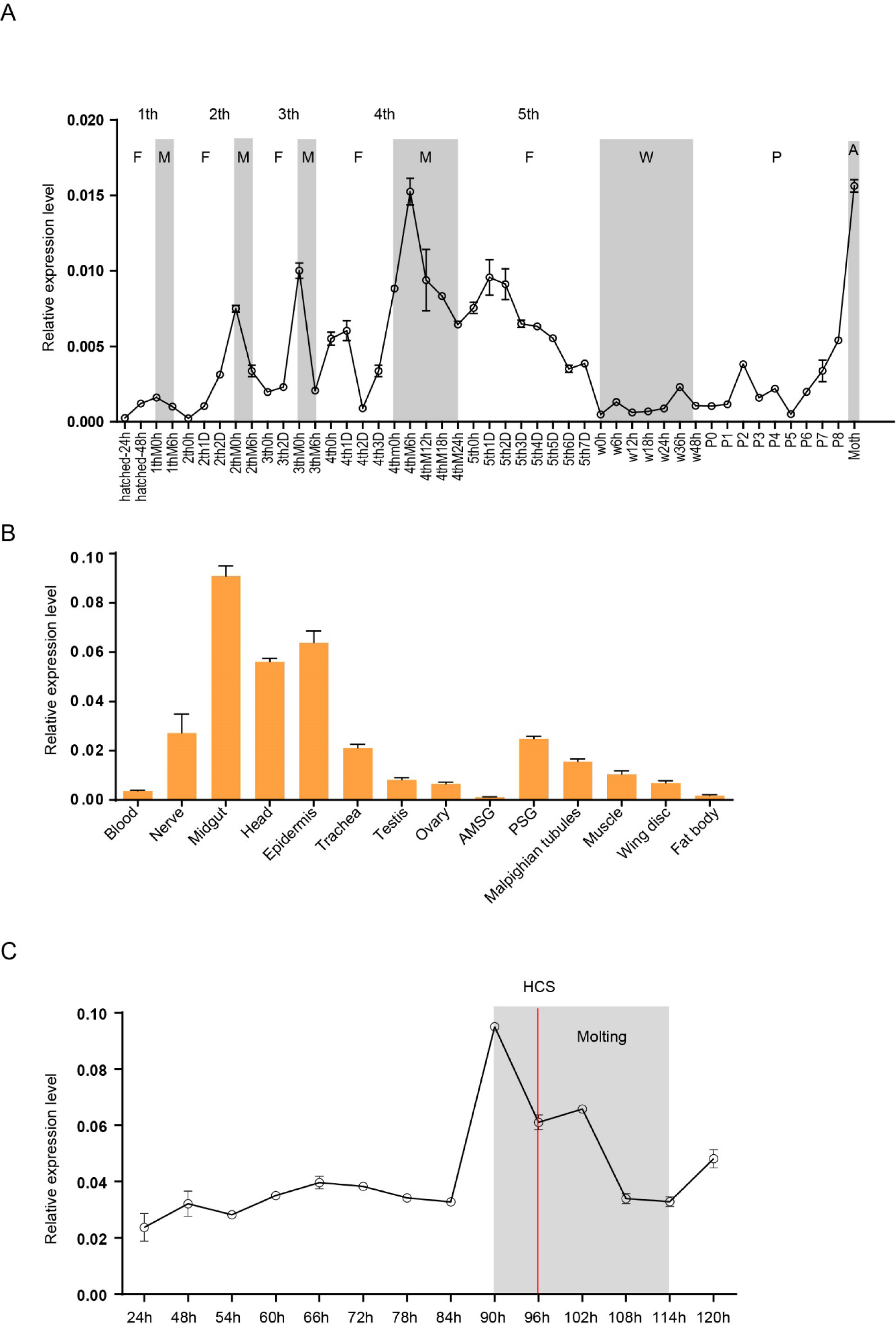
Spatiotemporal expression of *Bm-mamo*. (A) Temporal expression of *Bm-mamo*. In the molting stage and adult stage, this gene was significantly upregulated. M: molting stage, W: wandering stage, P: pupal stage, P1: Day 1 of the pupal stage, A: adult stage, 4th3d indicates the 4th instar on Day 3. 1st to 5th denote the first instar of larvae to fifth instar of larvae, respectively. (B) Tissue-specific expression of 4th-instar molting larvae. *Bm-mamo* had relatively high expression levels in the midgut, head, and epidermis. AMSG: anterior division of silk gland and middle silk gland, PSG: posterior silk gland. (C) Detailed analysis of *Bm-mamo* at the 4th larval stage in the epidermis of the Dazao strain. *Bm-mamo* expression is upregulated during the molting stage. HCS indicates the head capsule stage. The “h” indicates the hour, 90 h: at 90 hours of the 4^th^ instar.

According to the investigation of tissue-specific Dazao strain expression in 5th-instar 3rd-day larvae, the midgut, head, and epidermis exhibited high expression levels; the trachea, nerves, silk glands, testis, ovary, muscle, wing disc and Malpighian tubules exhibited moderate expression levels; and the blood and fat bodies exhibited low expression levels (Fig. 3B). These findings suggested that *Bm-mamo* is involved in the developmental regulation of multiple silkworm tissues. Due to the melanism of the epidermis of the *bd* mutant and the high expression level of the *Bm-mamo* gene in the epidermis, we measured the expression level of this gene in the epidermis of the 4th to 5th instars of the Dazao strain. In the epidermis, the *Bm-mamo* gene was upregulated during the molting period, and the highest expression was observed at the beginning of molting (Fig. 3C).

### Functional analyses of *Bm-mamo*

To study the function of the *Bm-mamo* gene, we carried out an RNA interference (RNAi) experiment. Short interfering RNA (siRNA) was injected into the hemolymph of silkworms, and an electroporation experiment(*44*) was immediately conducted. We found significant melanin pigmentation in the epidermis of the newly molted 5th-instar larvae. These findings indicate that *Bm-mamo* deficiency can cause melanin pigmentation (Fig. 4). The melanistic phenotype of the RNAi individuals was similar to that of the *bd* mutants.

**Fig. 4.**
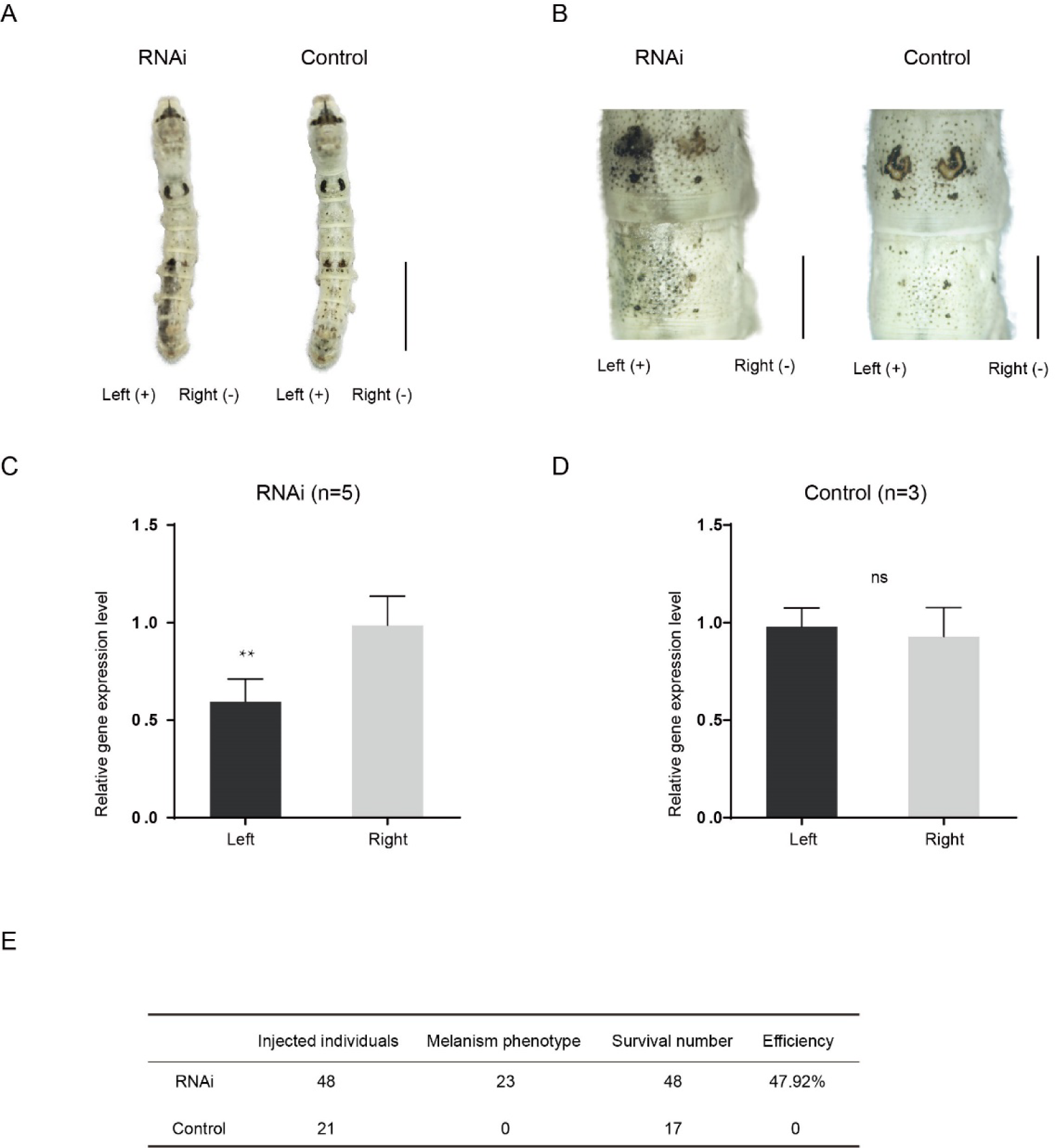
(A) siRNA was introduced by microinjection followed by electroporation. “+” and “-” indicate the positive and negative poles of the electrical current, respectively. Scale bars, 1 cm. (B) Partial magnification of the siRNA experimental group and negative control group. Scale bars, 0.2 cm. (C) and (**D**) Relative expression levels of *Bm-mamo* in the negative control and RNAi groups were determined by qPCR analysis. The means ± s.d.s. Numbers of samples are shown in the upper right in each graph. \*\**P*<0.01, paired Student’s t test (NS, not significant). (E) Statistical analysis of the efficiency of the RNAi.

In addition, gene knockout was performed. Ribonucleoproteins (RNPs) generated from guide RNA (gRNA) and recombinant Cas9 protein were injected into 450 silkworm embryos. In the G0 generation, individuals with a mosaic melanization phenotype were found. These melanistic individuals were raised to moths and subsequently crossed. A homozygous line with a gene knockout was obtained through generations of screening. The gene-edited individuals had a significantly melanistic body color, and the female moths were sterile (Fig. 5).

**Fig. 5.**
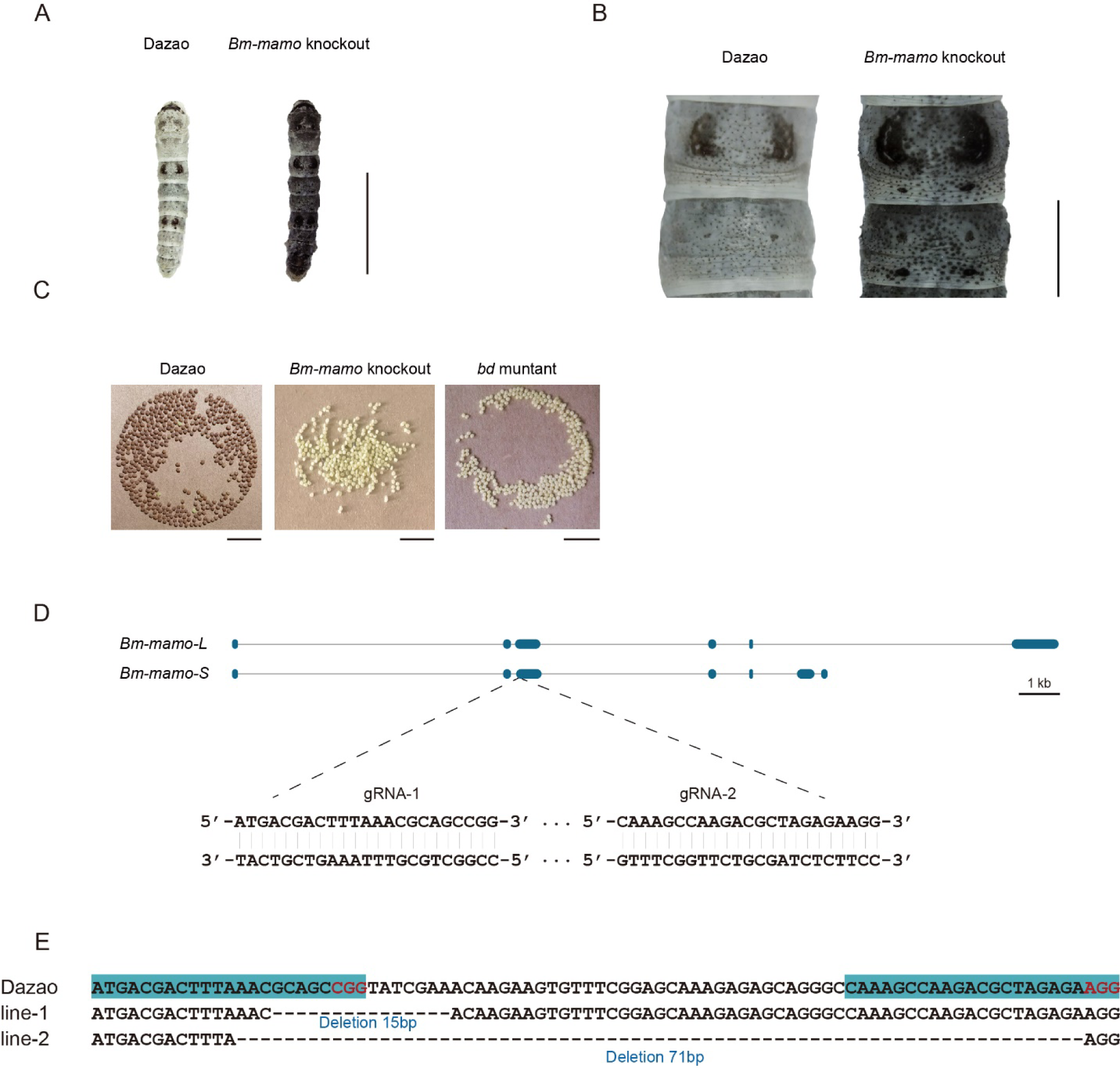
*Bm-mamo* knockout. (A) Larval phenotype of G3 fourth-instar larvae of Dazao targeted for *Bm-mamo*. Scale bars, 1 cm. (B) Partial magnification of the *Bm-mamo* knockout individual and control. Scale bars, 0.2 cm. (C) After 48 hours of egg laying, the pigmentation of the eggs indicates that those produced by knockout homozygous females cannot undergo normal pigmentation and development. (D) Genomic structure of *Bm-mamo*. The open reading frame (blue) and untranslated region (black) are shown. The gRNA 1 and gRNA 2 sequences are shown. (E) Sequences of the *Bm-mamo* knockout individuals. Lines 1 and 2 indicate deletions of 15 and 71 bp, respectively.

These results indicated that the *Bm-mamo* gene negatively regulates melanin pigmentation in caterpillars and participates in the reproductive regulation of female moths.

### Downstream target gene analysis

*Bm-mamo* belongs to the zinc finger protein family, which specifically recognizes downstream DNA sequences according to their zinc fingers. We identified homologous genes of *mamo* in multiple species and conducted a phylogenetic analysis (Fig. S6). Sequence alignment revealed that the amino acid residues of the zinc finger motif of the mamo-S protein were highly conserved among 57 species. The insufficient ability of the different species in GenBank to predict alternative transcripts of mamo may have resulted in the mamo-L protein being found in only 30 species (Fig. S7). As the mamo protein contains a tandem Cys_2_His_2_ zinc finger (C2H2-ZF) motif, it can directly bind to DNA sequences. Previous research has suggested that the ZF-DNA binding interface can be understood as a “canonical binding model”, in which each finger contacts DNA in an antiparallel manner. The binding sequence of the C2H2-ZF motif is determined by the amino acid residue sequence of its α-helical component. The first amino acid residue in the α-helical region of the C2H2-ZF domain is at position 1, and positions −1, 2, 3, and 6 are key amino acids involved in the recognition and binding of DNA. The residues at positions −1, 3, and 6 specifically interact with base 3, base 2, and base 1 of the DNA sense sequence, respectively, while the residue at position 2 interacts with the complementary DNA strand(*45, 46*). To analyze the downstream target genes of mamo, we first predicted the DNA-binding motifs of these genes using online software (http://zf.princeton.edu) based on the canonical binding model(*47*). In addition, the DNA-binding sequence of mamo (<u>TGCGT</u>) in *Drosophila* was confirmed by electrophoretic mobility shift assay (EMSA)(*48*), which has a consensus sequence with the predicted binding site sequence of Bm-mamo-S (G<u>TGCGT</u>GGC), and the predicted sequence was longer. This indicates that the predicted results for the DNA-binding site have good reliability. Furthermore, the predicted DNA-binding sites of Bm-mamo-L and Bm-mamo-S were highly consistent with those of mamo orthologs in different species (Fig. S8). This finding suggested that the protein may regulate similar target genes between species.

C2H2-ZF transcription factors function by recognizing and binding to *cis*-regulatory sequences in the genome, which harbor *cis*-regulatory elements (CREs)(*49*). CREs are broadly classified as promoters, enhancers, silencers or insulators(*50*). CREs are often near their target genes, such as enhancers, which are typically located upstream (5’) or downstream (3’) of the gene they regulate or in introns, but approximately 12% of CREs are located far from their target gene(*51*). Therefore, we first investigated the 2 kb upstream and downstream regions of the predicted genes in silkworms.

The predicted position weight matrices (PWMs) of the recognized sequences of Bm-mamo protein and the Find Individual Motif Occurrences (FIMO) software of MEME were used to perform silkworm whole-genome scanning for possible downstream target genes. The 2 kb upstream and downstream regions of the 14,623 predicted genes in silkworms were investigated. A total of 10,622 genes contained recognition sites within 2 kb of the upstream/downstream region of the Bm-mamo protein in the silkworm genome (Fig. S9, Table S3 & Table S4).

Moreover, we compared the transcriptome data of integument tissue between homozygotes and heterozygotes of the *bd* mutant at the 4^th^ instar/beginning molting stage(*52*). In the integument tissue, 10,072 genes (∼69% of the total predicted genes of silkworm) were expressed in heterozygotes, and 9,853 genes (∼67% of the total predicted genes) were expressed in homozygotes of the *bd* mutant. In addition, there were 191 genes whose expression significantly differed between homozygotes (*bd*/*bd*) and heterozygotes (+/*bd*) according to comparative transcriptome analysis (Table S5)(*52*). Protein functional annotation was performed, and 19 CP genes were significantly differentially expressed between heterozygotes and homozygotes of *bd*. In addition, the orthologs of these CPs were analyzed in *Danaus plexippus*, *Papilio xuthu* and *D. melanogaster* (Table S6). Furthermore, we identified 53 enzyme-encoding genes, 17 antimicrobial peptide genes, 6 transporter genes, 5 transcription factor genes, 5 cytochrome genes, and others. Among the differentially expressed genes (DEGs), CP was significantly enriched, and previous studies have shown that CPs can affect pigmentation(*53*). Therefore, we first investigated the expression of the CP genes. Among them, 18 CP genes had Bm-mamo binding sites in the upstream and downstream 2 kb genomic regions. In addition, we investigated the expression levels of the 18 CP genes in the integument from the 4^th^ instar (Day 1) to the beginning of the 5^th^ instar in the *Bm-mamo* knockout lines (Fig. 7& Fig. S10). The CP genes were significantly upregulated at one or several time points in homozygous (*mamo^-^*/*mamo^-^*) individuals compared with heterozygous (*mamo^-^*/*+*) *Bm-mamo* knockout individuals. Interestingly, the expression of the CP gene *BmorCPH24* was significantly upregulated at the feeding stage in *Bm-mamo* knockout homozygotes. Previous studies have shown that BmorCPH24 deficiency can lead to a marked decrease in pigmentation in silkworm larvae(*53*). Therefore, the expression of some CP genes may be necessary for determining the color patterns of caterpillars.

In addition, the synthesis of pigment substances is an important driver of color patterns(*54*). Eight key genes (*TH*, *DDC*, *aaNAT*, *ebony*, *black*, *tan*, *yellow* and *laccase2*) involved in melanin synthesis(*11*) were investigated in heterozygous and homozygous *Bm-mamo* gene knockout lines. Among them, *yellow*, *tan*, and *DDC* were significantly upregulated during the molting period in the homozygous *Bm-mamo* knockout individuals (Fig. 6). The upstream and/or downstream 2 kb genomic regions of *yellow*, *tan* and *DDC* contain the binding site of the Bm-mamo protein. In addition, the expression of *yellow*, *DDC* and *tan* can promote the generation of melanin(*55*). To explore the interaction between the Bm-mamo protein and its binding sequence, EMSA was conducted. The binding site of Bm-mamo-S (C<u>TGCGT</u>GGT) was located approximately 70 bp upstream of the transcription initiation site of the *Bm*-*yellow* gene. The EMSA results showed that the Bm-mamo-S protein expressed in prokaryotes can bind to the C<u>TGCGT</u>GGT sequence in vitro (Fig. S11). This finding suggested that the Bm-mamo-S protein can bind to the upstream region of the *Bm*-*yellow* gene and regulate its transcription. Therefore, the *Bm-mamo* gene may control the color pattern of caterpillars by regulating key melanin synthesis genes and 18 CP genes.

**Fig. 6.**
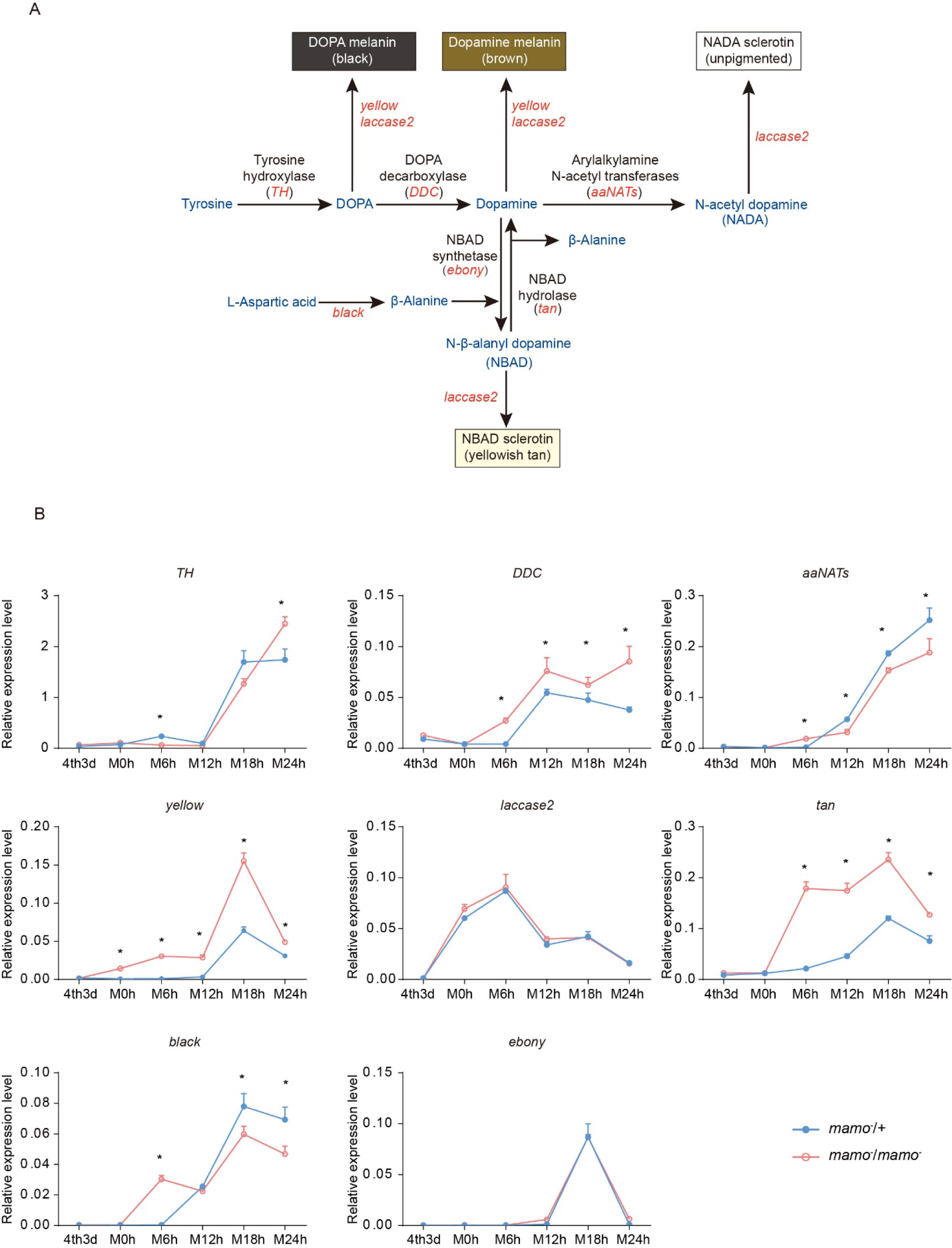
Melanin metabolism pathway and qPCR of related genes. (A) The melanin metabolism pathway. Blue indicates amino acids and catecholamines, red indicates the names of genes, and black indicates the names of enzymes. (B) Relative expression levels of eight genes in the heterozygous *Bm-mamo* knockout group (blue) and homozygous *Bm-mamo* knockout group (red) determined via qPCR analysis. The means ± s.d.s. \**P*<0.05, paired Student’s t test. 4th3d indicates the 4th instar on Day 3, M indicates molting, and h indicates hours.

## Discussion

Insects have evolved many important phenotypes during the process of adapting to the environment. Among these traits, the color pattern is one of the most interesting. The biochemical metabolic pathways of pigments, such as melanin, ommochromes, and pteridines, have been identified in insects(*18*). However, the regulation of pigment metabolism-related genes and the processes involved in the transport and deposition of pigment substances are unclear. In this study, we discovered that the *Bm-mamo* gene negatively regulates melanin pigmentation in caterpillars. When this gene is deficient, the body color of silkworms exhibits substantial melanism, and changes in the expression of some melanin synthesis genes and some CP genes are also significant.

### Structural genes of melanin synthesis

Deficiencies of some genes in the melanin synthesis pathway, such as *TH*, *yellow*, *ebony*, *tan*, and *aaNAT,* can lead to variations in color patterns (*11*). However, they often lead to localized pigmentation changes only in later-instar caterpillars. For example, in *yellow*-mutant silkworms, the eye spot, lunar spot, star spot, spiracle plate, head, and some sclerotized areas appear reddish brown, and the other cuticle is consistent with that in the wild type in later instars of larvae (third instar to fifth instar)(*56*). This situation is highly similar to that of the *tan* mutant in silkworms(*57*). Why do the *yellow* and *tan* phenotypes appear only in a limited cuticle region in later instars of silkworm larvae? One possible reason is that the expression of *yellow* and *tan* is limited to a certain region by transcription regulators. Alternatively, other factors, such as CPs in the cuticle, may limit pigmentation by interacting with pigment substances in the cuticle.

On the one hand, we investigated the expression levels of *yellow* and *tan* in the pigmented region (lunar spot) and nonpigmented region of the epidermis during the 4th molting of the wild-type Dazao strain. The expression level of *yellow* was significantly upregulated in the lunar spot (the epidermis on the dorsum of the fifth body segment) compared with the nonpigmented region (the epidermis on the dorsum of the sixth body segment) at 6 hours and 12 hours after molting.

Moreover, *tan* was significantly upregulated 18 hours after molting in the lunar spot region (Fig. S12). This finding suggested that the upregulated expression of pigment synthesis genes at key time points may be important for pigmentation. However, *yellow* and *tan* were still moderately expressed in the nonpigmented epidermis, although they did not cause significant melanin pigmentation. This finding indicates that pigment synthesis alone cannot determine the predominant color pattern of the cuticle in caterpillars.

### Cuticular proteins participate in pigmentation

On the other hand, synthesized pigment substances need to be transported from epidermal cells and embedded into the cuticle to allow pigmentation. Therefore, the correct cuticle structure and location of cuticular proteins in the cuticle may be important factors affecting pigmentation. Previous studies have shown that a lack of expression of *BmorCPH24*, which encodes an important component of the endocuticle, can lead to dramatic changes in body shape and a significant reduction in the pigmentation of caterpillars(*53*). We crossed *Bo* (*BmorCPH24* null mutation) and *bd* to obtain F_1_ (*Bo*/+*^Bo^*, *bd*/*+*) and then self-crossed F_1_ to observe the phenotype of F_2_. The area of lunar spots and star spots decreased, and light-colored stripes appeared on the body segments; however, the other areas still exhibited significant melanin pigmentation in double-mutation (*Bo*, *bd*) individuals (Fig. S13). However, in previous studies, the introduction of *Bo* into *L* (ectopic expression of *wnt1* results in lunar stripes generated on each body segment)(*24*) and *U* (overexpression of *SoxD* results in excessive melanin pigmentation of the epidermis)(*58*) strains by genetic crosses markedly reduced the pigmentation of *L* and *U*(*53*). Interestingly, there was a more significant decrease in pigmentation in the double mutants (*Bo*, *L*) and (*Bo*, *U*) than in (*Bo*, *bd*). These findings suggested that *Bm-mamo* has a greater ability than does *wnt1* and *SoxD* to regulate pigmentation. On the one hand, mamo may be a stronger regulator of the melanin metabolic pathway, and on the other hand, mamo may regulate other CP genes to reduce the impact of BmorCPH24 deficiency.

How do CPs affect pigmentation? One study showed that some CPs can form “pore canals” to transport macromolecules(*59*). In addition, some CPs can be crosslinked with catecholamines, which are synthesized in the melanin metabolism pathway(*60*). Because there are no live cells in the cuticle, melanin precursor substances may be transported by the pore canals formed by some CPs and fixed to specific positions through cross-linking with CPs. The cuticular protein TcCPR4 is needed for the formation of pore canals in the cuticle of *Tribolium castaneum*(*61*). In contrast, the vertical pore canal is lacking in the less pigmented cuticles of *T. castaneum*(*62*). This finding suggested that the pore canals constructed by TcCPR4 may transport pigments and contribute to cuticle pigmentation in *T. castaneum*. Moreover, the melanin metabolites N-acetyldopamine (NADA) and N-β-alanyldopamine (NBAD) can target and sclerotize the cuticle by cross-linking with specific cuticular proteins (*57*). This finding suggested that pigments interact with specific CPs, thereby affecting pigmentation, hardening properties and the structure of the cuticle. Interestingly, a study showed that in addition to absorbing specific wavelengths, pigments can affect cuticle polymerization, density, and the refractive index, which in turn affects the reflected wavelengths that produce structural color in butterfly wing scales(*63*). This implies that the interaction between pigments and CPs can be very subtle, resulting in the formation of unique nanostructures, such as those on wing scales, that produce brilliant structural colors.

Consequently, to maintain the accuracy of the color pattern, the localization of CPs in the cuticle may be very important. In a previous study employing microarray analysis, different CPs were found in differently colored areas of the epidermis in *Papilio xuthus* larvae(*64*). We investigated whether the CP genes were highly expressed in the black region of *Papilio xuthus* caterpillars. Thirteen orthologous genes were found in silkworms (Table S7). Among them, 11 genes (*BmorCPR67*, *BmorCPR71*, *BmorCPR76*, *BmorCPR79*, *BmorCPR99*, *BmorCPR107*, *BmorCPT4*, *BmorCPH5*, *BmorCPFL4*, *BmorCPG27* and *BmorCPG4*) were significantly upregulated in homozygous (*mamo^-^*/*mamo^-^*) knockout individuals at some time points from the 3rd day of the 4th instar to the 5th instar; *BmorCPG4* was also among the 18 previously detected CP genes, and two genes (*BmorCPG38* and *BmorCPR129*) were not differentially expressed between homozygous (*mamo^-^*/*mamo^-^*) and heterozygous (*mamo^-^*/*+*) individuals (Fig. S14). The expression of the 28 CP genes mentioned above was significantly upregulated in homozygous (*mamo^-^*/*mamo^-^*) gene knockout individuals at several stages. These CPs may be involved in the transportation or cross-linking of melanin in the cuticle. However, there were no differences in the expression levels of these genes during some periods compared with those in the control group, and the expression of some genes was significantly downregulated at some time points in the melanic individuals (Fig. 7). This finding suggested that the regulation of CP genes is complex and may involve other transcription factors and feedback effects. CPs are essential components of the insect cuticle and are involved in cuticular microstructure construction (*65*), body shape development(*66*), wing morphogenesis(*67*), and pigmentation(*53*). CP genes usually account for more than 1% of the total genes in an insect genome and can be categorized into several families, including CPR, CPG, CPH, CPAP1, CPAP3, CPT, CPF and CPFL(*68*). The CPR family is the largest group of CPs and contains a chitin-binding domain called the Rebers and Riddiford motif (R&R)(*69*). The variation in the R&R consensus sequence can be divided into three subfamilies (RR-1, RR-2, and RR-3)(*70*). Among the 28 CPs, 11 RR-1 genes, 6 RR-2 genes, 4 hypothetical cuticular protein (CPH) genes, 3 glycine-rich cuticular protein (CPG) genes, 3 cuticular protein Tweedle motif (CPT) genes, and 1 CPFL (like the CPFs in a conserved C-terminal region) gene were identified. The RR-1 consensus among species is usually more variable than the RR-2 consensus, which suggests that RR-1 may have a species-specific function. RR-2 often clustered into several branches, which may be due to gene duplication events in co-orthologous groups and may result in conserved functions between species (*71*). The classification of CPH was based on the lack of known motifs. In the epidermis of Lepidoptera, CPH genes often exhibit high expression levels. For example, *BmorCPH24* has the highest expression level in the silkworm larval epidermis(*72*). The CPG protein is rich in glycine. The CPH and CPG genes are less commonly found in insects outside the order Lepidoptera (*73*). This finding suggested that these genes may provide species-specific functions for Lepidoptera. CPT contains a Tweedle motif, and the *TweedleD1* mutation has a dramatic effect on body shape in *D. melanogaster*(*74*). The CPFL family members are relatively conserved among species and may be involved in the synthesis of larval cuticles(*75*). CPT and CPFL may have relatively conserved functions among insects. The CP genes are a group of rapidly evolving genes, and their copy numbers may undergo significant changes in different species. In addition, RNAi experiments on 135 CP genes in the brown planthopper (*Nilaparvata lugens*) revealed that deficiency of 32 CP genes leads to significant defects in phenotypes, such as lethality and developmental retardation. These findings suggested that the 32 CP genes are indispensable and that other CP genes may have redundant and complementary functions (*76*). In previous studies, it was found that the construction of the larval cuticle of silkworms requires the precise expression of more than two hundred CP genes(*22*). The production, interaction, and deposition of CPs and pigments are complex and precise processes, and our research showed that *Bm-mamo* plays an important regulatory role in this process in silkworm caterpillars. To further understand the role of CPs, future work should aim to identify the functions of important cuticular protein genes and the deposition mechanism in the cuticle.

**Fig. 7.**
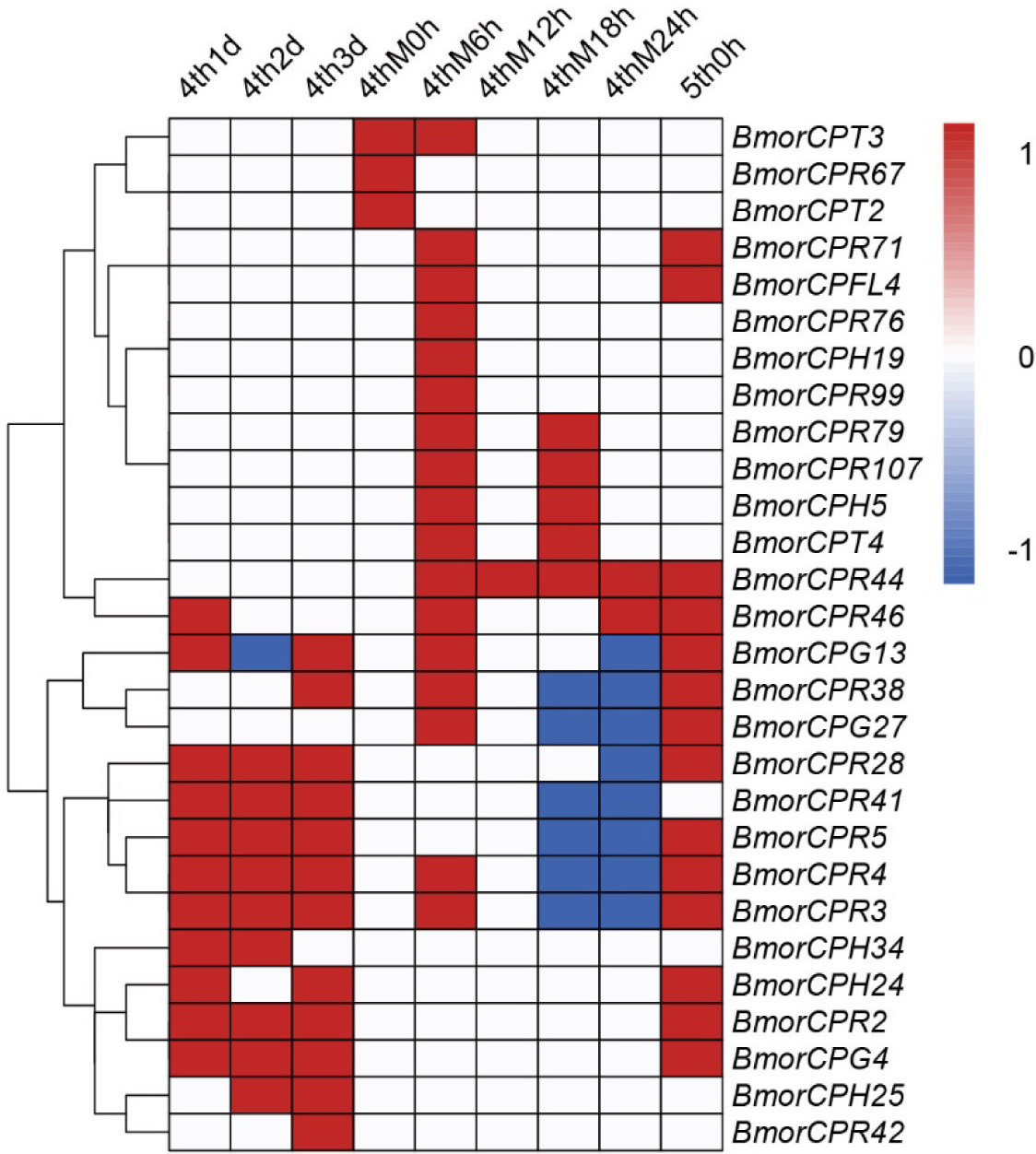
Heatmap of 28 differentially expressed cuticular protein genes. Differentially expressed cuticular protein genes between homozygous (*mamo^-^*/*mamo^-^*) and heterozygous (*mamo^-^*/+) *Bm-mamo* knockout individuals. Red indicates upregulation in homozygous individuals. Blue indicates downregulation in homozygous individuals. White indicates no difference in expression level or no investigation. 4th1d indicates 4-instar Day 1, M indicates molting, and h indicates hours.

### Maturation of pigment granules

In addition, among the 191 DEGs found in the comparative transcriptome data, we also discovered several interesting genes. For example, *BGIBMGA013242*, which encodes a major facilitator superfamily protein (MFS) named *BmMFS* and is responsible for the *cheek and tail spot (cts)* mutant, was significantly upregulated in the *bd* mutant. Deficiency of this gene (*BmMFS*) results in chocolate-colored head and anal plates on the silkworm caterpillar(*77*). In the ommochrome metabolic pathway, *Bm-re*, which encodes an MFS protein, may function in the transportation of several amino acids, such as cysteine or methionine, into pigment granules(*78, 79*). Therefore, the encoded product of *BmMFS* may participate in pigmentation by promoting the maturation of pigment granules. Moreover, *BGIBMGA013576* and *BGIBMGA013656*, which encode MFS domain-containing proteins belonging to the solute carrier family 22 (SLC22) and solute carrier family 2 (SLC2) families, respectively, were significantly upregulated in *bd*. These two genes may participate in pigmentation in a similar manner to that of the *BmMFS* gene. In addition, the *red Malpighian tubule* (*red*) gene encodes a LysM domain-containing protein. *Red* deficiency results in a significant decrease in orange pterin in the wing of *Colias* butterflies. Research suggests that the product of *red* may interact with V-ATPase to modulate vacuolar pH across a variety of endosomal organelles, thereby affecting the maturation of pigment granules in cells(*80*). These findings indicate that the maturation of pigment granules plays an important role in the coloring of Lepidoptera and that *Bm-mamo* may be involved in regulating the maturation of pigment granules in the epidermal cells of silkworms.

*Bm-mamo* may affect the synthesis of melanin in epidermal cells by regulating *yellow*, *DDC*, and *tan*, regulate the maturation of melanin granules in epidermal cells through *BmMFS*, and affect the deposition of melanin granules in the cuticle by regulating CP genes, thereby comprehensively regulating the color pattern of caterpillars.

### The upstream sequence of *Bm-mamo*

Moreover, in *D. melanogaster*, *mamo* is needed for functional gamete production. In the silkworm, *Bm-mamo* has a conserved function in female reproduction. The DNA-binding motifs of the mamo proteins in silkworms and fruit flies are highly conserved, suggesting that they may regulate the same downstream target genes. However, *mamo* has developed new functions in color pattern regulation in silkworm caterpillars. We found that in *D. melanogaster*, the *mamo* gene is expressed mainly during the adult stage; it is not expressed during the 1^st^ instar or 2^nd^ instar larval stage and has very low expression at the 3^rd^ instar (Table S8). In addition, several binding sites of mamo were found near the TSS of *yellow* in *D. melanogaster* (Fig. S15). The *yellow* gene is a key melanin metabolic gene with upstream and intron sequences that have been identified as multiple CREs, and it has been considered a research model for CREs (*81, 82*). This difference may be due to a change in the expression pattern of *mamo,* which led to the development of a new function for this gene in regulating coloration in silkworm caterpillars. Changes in gene expression patterns are generally believed to be the result of the evolution of CREs. Because CREs play important roles in the spatiotemporal expression pattern regulation of genes, many CREs are found in noncoding regions, which are relatively prone to sequence variation compared with the sequences of coding genes(*49*). However, the molecular mechanism underlying the sequence changes in CREs is unclear.

TFs, because they recognize relatively short sequences, generally between 4 and 20 bp in length, can have many binding sites in the genome(*83*). Therefore, one member of the TF family has the potential to regulate many genes in one genome. Single regulation of specific TFs can lead to patterned changes in the expression of multiple downstream genes, enabling organisms to adapt to the environment by altering a given type of trait. For example, the marine form of the three-spined stickleback (*Gasterosteus aculeatus*) has thick armor, whereas the lake population (which was recently derived from the marine form) does not. Research has shown that pelvic loss in different natural populations of three-spined stickleback fishes occurs via regulatory mutations resulting from the deletion of a tissue-specific enhancer (*Pel*) of the *pituitary homeobox transcription factor 1* (*Pitx1*) gene. The researchers genotyped 13 pelvic-reduced populations of three-spined sticklebacks from disparate geographic locations. Nine of the 13 pelvic-reduced stickleback populations had sequence deletions of varying lengths, all of which were located at the *Pel* enhancer (*84*).

We investigated nucleotide diversity in 51 wild silkworms and 171 domesticated silkworms (*34, 85*). The nucleotide diversity of introns and upstream sequences of *Bm-mamo* in wild silkworms was significantly greater than that in domestic silkworms. In addition, the approximately 1 kb genomic region upstream had high fixation indices (*F_ST_*) (Fig. S16). This indicates a significant degree of differentiation between the wild strains in this genomic region. Multiple sequence alignment analysis of this region was performed between 12 wild silkworms and 12 domestic silkworms (Fig. S17 & Table S9). The results showed that the wild silkworm group had a greater degree of nucleotide variation, while the sequences in domestic silkworms were highly conserved. The domestic silkworm group had two different forms: one had a long interspersed nuclear element (LINE) inserted at approximately 4.5 kb, and the other did not have this transposon sequence. It is suggested that this genomic region is prone to variation in wild silkworm and is fixed in domestic silkworms. The larvae of domesticated strains often have a lighter or even pale body color, but the wild-type strains have a darker color. The *Bm-mamo* gene may be involved in the domestication of silkworms.

The function of TFs is to precisely regulate the temporal and spatial expression of target genes. Therefore, the expression of TFs requires strict regulation. There are long intergenic regions upstream of many important TFs, dozens of kilobase pairs (Kb) to hundreds of Kb, which may contain many CREs for better control its expression pattern. It has often been believed that changes in CREs are caused by random mutations. However, recent studies have shown that there is a mutation bias in the genome; compared with that in the intergenic region, the mutation frequency is reduced by half inside gene bodies and by two-thirds in essential genes. In addition, they compared the mutation rates of genes with different functions. The mutation rate in the coding region of essential genes (such as translation) is the lowest, and the mutation rates in the coding region of specialized functional genes (such as environmental response) are the highest. These patterns are mainly affected by the features of the epigenome(*86*). Due to the plasticity of epigenomic features, mutations bias associated with epigenomes may even lead to environmental influences on mutations (*87*). This finding suggested that different functional regions in the genome are subject to distinct controls and that some sequences can undergo procedural changes under environmental changes. Therefore, some variations in the CRE of TFs in response to environmental changes may be mutation bias.

## Materials and Methods

### Silkworm strains

The *bd* and *bd^f^* mutant strains and the wild-type Dazao and N4 strains were obtained from the bank of genetic resources of Southwest University. Silkworms were reared on mulberry leaves or artificial diets at 25°C and 73% relative humidity in the dark for 12 hours and light for 12 hours.

### Positional cloning of the *bd* locus

For mapping of the *bd* locus, F1 heterozygous individuals were obtained from a cross between a *bd^f^* strain and a Dazao strain. Then, an F1 female was crossed with a *bd^f^* male (BC1F), and an F1 male was backcrossed with a *bd^f^* female (BC1M). A total of 1162 BC1M individuals were used for recombination analysis. Genomic DNA was extracted from the parents (Dazao, *bd^f^* and F1) and each BC1 individual using the phenol chloroform extraction method. Available DNA molecular markers were identified through polymorphism screening and linkage analysis. The primers used for mapping are listed in Table S10.

### Phylogenetic analysis

To determine whether *Bm-mamo* orthologs were present in other species, the BlastX program of the National Center for Biotechnology Information (NCBI) (http://www.ncbi.nlm.nih.gov/BLAST/) was used. The *Bm-mamo-L* and *Bm-mamo-S* sequences were subjected to BLAST searches against the nonredundant protein sequence (nr) database.

Sequences with a maximum score and an E-value≤10^-4^ were downloaded. The sequences of multiple species were subjected to multiple sequence alignment of the predicted amino acid sequences by MUSCLE(*88*). A phylogenetic tree was subsequently constructed using the neighborLjoining method with the MEGA7 program (Pearson model). The confidence levels for various phylogenetic lineages were estimated by bootstrap analysis (2,000 replicates).

### Quantitative PCR

Total RNA was isolated from the whole body and integument of the silkworms using TRIzol reagent (Invitrogen, California, USA), purified by phenol chloroform extraction and then reverse transcribed with a PrimeScript™ RT Reagent Kit (TAKARA, Dalian, China) according to the manufacturers’ protocol.

qPCR was performed using a CFX96™ Real-Time PCR Detection System (Bio-Rad, Hercules, CA) with a qPCR system and an iTaq Universal SYBR Green Supermix System (Bio-Rad). The cycling parameters were as follows: 95°C for 3 min, followed by 40 cycles of 95°C for 10 s and annealing for 30 s. The primers used for the target genes are listed in Table S10. The expression levels of the genes in each sample were determined with biological replicates, and each sample was analyzed in triplicate. The gene expression levels were normalized against the expression levels of the ribosomal protein L3 (RpL3). The relative expression levels were analyzed using the classical R=2^−ΔΔCt^ method.

### siRNA for gene knockdown

siRNAs for *Bm-mamo* were designed with the siDirect program (http://sidirect2.rnai.jp). The target siRNAs and negative controls were synthesized by Tsingke Biotechnology Company Limited. The siRNA (5 μl, 1 μl/μg) was injected from the abdominal spiracle into the hemolymph at the fourth instar (Day 3) larval stage. Immediately after injection, phosphate-buffered saline (pH 7.3) droplets were placed nearby, and a 20-voltage pulse for one second and pause for one second were applied 3 times. The phenotype was observed for fifth-instar larvae. The left and right epidermis were separately dissected from the injected larvae, after which RNA was extracted. Then, cDNA was synthesized, and the expression level of the gene was detected via qPCR.

### sgRNA synthesis and RNP complex assembly

CRISPRdirect (http://crispr.dbcls.jp/doc/) online software was used to screen appropriate single guide RNA (sgRNA) target sequences. The gRNAs were synthetized by the Beijing Genomics Institute. The sgRNA templates were transcribed using T7 polymerase with RiboMAX™ Large-Scale RNA Production Systems (Promega, Beijing, China) according to the manufacturer’s instructions. The RNA transcripts were purified using 3 M sodium acetate (pH 5.2) and anhydrous ethanol (1:30) precipitation, washed with 75% ethanol, and eluted in nuclease-free water. All injection mixes contained 300 ng/µL Cas9 nuclease (Invitrogen, California, USA) and 300 ng/µL purified sgRNA. Before injection, mixtures of Cas9 nuclease and gRNA were incubated for 15 min at 37°C to reconstitute active ribonucleoproteins (RNPs)(*89*).

### Microinjection of embryos

For embryo microinjection, microcapillary needles were manufactured using a PC-10 model micropipette puller (Narishige, Tokyo, Japan). Microinjection was performed using an Eppendorf TransferMan NK 2 and a FemtoJet 4i system (Eppendorf, Hamburg, Germany). The eggs used for microinjection were generated by mating female and male wild-type Dazao moths. Within 4 hours of culture, the eggs were allowed to adhere to a clean glass slide. CRISPR/Cas9-messenger RNP mixtures with volumes of approximately 1 nL were injected into the middle of the eggs, and the wound was sealed with glue. All the injected embryos were allowed to develop in a climate chamber at 25°C and 80% humidity(*90*).

### Comparative genomics

The reference genome of Dazao was downloaded from Silkbase (https://silkbase.ab.a.u-tokyo.ac.jp/cgi-bin/index.cgi). The *bd^f^* genome was obtained from the silkworm pangenome project(*34*).

The short reads of the *bd^f^* strains were mapped to the silkworm reference genome by BWA55 v0.7.17 mem with default parameters. The SAMtools56 v1.11 and Picard v2.23.5 (https://broadinstitute.github.io/picard/) programs were used to filter the unmapped and duplicated reads. A GVCF file of the samples was obtained using GATK57 v4.1.8.1 HaplotypeCaller with the parameter-ERC = GVCF. The VCF files of insertions/deletions (indels) and single-nucleotide polymorphisms (SNPs) were used for further analysis via eGPS software.

### Downstream target gene screening

Online software (http://zf.princeton.edu) was used to predict the DNA-binding site for Cys2His2 zinc finger proteins. The confident ZF domains with scores higher than 17.7 were chosen. RF regression on the B1H model was used to predict the DNA-binding sites. Then, the sequence logo and position weight matrices (PWMs) of the DNA-binding sites were obtained. The sequences 2 kb upstream and downstream of the predicted genes were extracted by a Perl script. The FIMO package of the MEME suite was used to search for binding sites in the silkworm genome.

### Analysis of EMSA

Primers with binding sites and flanking sequences were designed according to the FIMO results. A biotin label was added to one end of the upstream primer (probe), and the downstream primer was used as a reference. Primers with the same sequence as the labeled probes were used as competitive probes. The EMSA was conducted with an EMSA reagent kit (Beyotime, Shanghai, China) according to the manufacturers’ protocol.

## Acknowledgments

None

## Author contributions

FY Dai conceived and designed the experiments. SY Wu, CX Peng, JW Luo, KP Lu and CH Zhang performed the study. SY Wu, XL Tong, KP Lu, CL Li, X Ding, YR Lu, XH Duan, D Tan and H Hu analyzed the data. SY Wu wrote the paper. XL Tong and FY Dai edited and revised the manuscript.

## Data and material availability

The authors confirm that the data supporting the findings of this study are available within the article and its supplementary materials.

## Additional information

Supplementary Information is available for this paper.

## References

1. G. A. Sword, S. J. Simpson, O. T. M. El Hadi, H. Wilps, Density-dependent aposematism in the desert locust. P Roy Soc B-Biol Sci 267, 63–68 (2000).

2. A. I. Barnes, M. T. Siva-Jothy, Density-dependent prophylaxis in the mealworm beetle *Tenebrio* molitor L. (Coleoptera: Tenebrionidae): cuticular melanization is an indicator of investment in immunity. P Roy Soc B-Biol Sci 267, 177–182 (2000).

3. N. F. Hadley, A. Savill, T. D. Schultz, Coloration and its thermal consequences in the New Zealand tiger beetle *Neocicindela perhispida*. J Therm Biol 17, 55–61 (1992).

4. Y. G. Hu, Y. H. Shen, Z. Zhang, G. Q. Shi, Melanin and urate act to prevent ultraviolet damage in the integument of the silkworm, Bombyx mori. Arch Insect Biochem 83, 41–55 (2013).

5. M. Stevens, G. D. Ruxton, Linking the evolution and form of warning coloration in nature. P Roy Soc B-Biol Sci 279, 417–426 (2012).

6. K. K. Dasmahapatra et al., Butterfly genome reveals promiscuous exchange of mimicry adaptations among species. Nature 487, 94–98 (2012).

7. N. Gaitonde, J. Joshi, K. Kunte, Evolution of ontogenic change in color defenses of swallowtail butterflies. Ecol Evol 8, 9751–9763 (2018).

8. B. S. Tullberg, S. Merilaita, C. Wiklund, Aposematism and crypsis combined as a result of distance dependence: functional versatility of the colour pattern in the swallowtail butterfly larva. P Roy Soc B-Biol Sci 272, 1315–1321 (2005).

9. P. J. Wittkopp, B. L. Williams, J. E. Selegue, S. B. Carroll, *Drosophila* pigmentation evolution: divergent genotypes underlying convergent phenotypes. P Natl Acad Sci USA 100, 1808–1813 (2003).

10. R. Futahashi, Y. Banno, H. Fujiwara, Caterpillar color patterns are determined by a two-phase melanin gene prepatterning process: new evidence from *tan* and *laccase2*. Evol Dev 12, 157–167 (2010).

11. Y. Matsuoka, A. Monteiro, Melanin pathway genes regulate color and morphology of butterfly wing scales. Cell Rep 24, 56–65 (2018).

12. C. Finet et al., Multi-scale dissection of wing transparency in the clearwing butterfly. J R Soc Interface 20, (2023).

13. A. Gürses, M. Açıkyıldız, K. Güneş, M. S. Gürses, in Dyes and Pigments. (Springer International Publishing, 2016), pp. 13-29.

14. V. J. Lloyd, N. J. Nadeau, The evolution of structural colour in butterflies. Curr Opin Genet Dev 69, 28–34 (2021).

15. A. E. Seago, P. Brady, J. P. Vigneron, T. D. Schultz, Gold bugs and beyond: a review of iridescence and structural colour mechanisms in beetles (Coleoptera). J R Soc Interface 6, S165–S184 (2009).

16. Y. Oba et al., Resurrecting the ancient glow of the fireflies. Sci Adv 6, (2020).

17. A. J. Syed, J. C. Anderson, Applications of bioluminescence in biotechnology and beyond. Chem Soc Rev 50, 5668–5705 (2021).

18. X. L. Tong, L. Qiao, J. W. Luo, X. Ding, S. Y. Wu, The evolution and genetics of lepidopteran egg and caterpillar coloration. Curr Opin Genet Dev 69, 140–146 (2021).

19. F. Y. Dai et al., Mutations of an arylalkylamine-*N*-acetyltransferase, *Bm-iAANAT*, are responsible for silkworm melanism mutant. J Biol Chem 285, 19553–19560 (2010).

20. P. J. Wittkopp, J. R. True, S. B. Carroll, Reciprocal functions of the *Drosophila* yellow and ebony proteins in the development and evolution of pigment patterns. Development 129, 1849–1858 (2002).

21. J. Q. Liu et al., Lepidopteran wing scales contain abundant cross-linked film-forming histidine-rich cuticular proteins. Commun Biol 4, (2021).

22. Z. W. Yan et al., A blueprint of microstructures and stage-specific transcriptome dynamics of cuticle formation in *Bombyx mori*. Int J Mol Sci 23, (2022).

23. R. Futahashi et al., Genome-wide identification of cuticular protein genes in the silkworm, Bombyx mori. Insect Biochem Molec 38, 1138–1146 (2008).

24. J. Yamaguchi et al., Periodic Wnt1 expression in response to ecdysteroid generates twin-spot markings on caterpillars. Nat Commun 4, (2013).

25. S. Yoda et al., The transcription factor Apontic-like controls diverse colouration pattern in caterpillars. Nat Commun 5, (2014).

26. H. Jin, T. Seki, J. Yamaguchi, H. Fujiwara, Prepatterning of *Papilio xuthus* caterpillar camouflage is controlled by three homeobox genes: *clawless*, *abdominal-A*, and *Abdominal-B*. Sci Adv 5, (2019).

27. W. A. Rogers et al., A survey of the trans-regulatory landscape for *Drosophila melanogaster* abdominal pigmentation. Dev Biol 385, 417–432 (2014).

28. S. Jeong, A. Rokas, S. B. Carroll, Regulation of body pigmentation by the Abdominal-B Hox protein and its gain and loss in *Drosophila* evolution. Cell 125, 1387–1399 (2006).

29. H. D. Dufour, S. Koshikawa, C. Finet, Temporal flexibility of gene regulatory network underlies a novel wing pattern in flies. P Natl Acad Sci USA 117, 11589–11596 (2020).

30. A. Prakash, C. Finet, T. Das Banerjee, V. Saranathan, A. Monteiro, *Antennapedia* and *optix* regulate metallic silver wing scale development and cell shape in *Bicyclus anynana* butterflies. Cell Rep 40, (2022).

31. R. D. Reed et al., Drives the repeated convergent evolution of butterfly wing pattern mimicry. Science 333, 1137–1141 (2011).

32. A. Kopp, I. Duncan, S. B. Carroll, Genetic control and evolution of sexually dimorphic characters in *Drosophila*. Nature 408, 553–559 (2000).

33. L. Arnoult et al., Emergence and diversification of fly pigmentation through evolution of a gene regulatory module. Science 339, 1423–1426 (2013).

34. X. L. Tong et al., High-resolution silkworm pan-genome provides genetic insights into artificial selection and ecological adaptation. Nat Commun 13, (2022).

35. L. Cheng., D. Fangyin., X. Zhonghuai., Studies on the mutant systems of the *Bombyx mori* gene bank. Scientia agricultura sinica 36, 968–975 (2003).

36. Y. Kawaguchi, T. Kusakabe, J. M. Lee, K. Koga, Manifestation of the *dilute black* (*bd*) mutation and constitution of the *bd* Locus in *Bombyx mori*. J Fac Agr Kyushu U 52, 355–359 (2007).

37. J. Duan et al., SilkDB v2.0: a platform for silkworm (*Bombyx mori*) genome biology. Nucleic Acids Res 38, D453–D456 (2010).

38. E. W. Sayers et al., GenBank. Nucleic Acids Res 50, D161–d164 (2022).

39. M. Kawamoto, T. Kiuchi, S. Katsuma, SilkBase: an integrated transcriptomic and genomic database for *Bombyx mori* and related species. Database-Oxford 2022, (2022).

40. V. J. Bardwell, R. Treisman, The POZ domain: a conserved protein-protein interaction motif. Gene Dev 8, 1664–1677 (1994).

41. K. D. Huynh, V. J. Bardwell, The BCL-6 POZ domain and other POZ domains interact with the co-repressors N-CoR and SMRT. Oncogene 17, 2473–2484 (1998).

42. P. Staller et al., Repression of p15INK4b expression by Myc through association with Miz-1. Nat Cell Biol 3, 392–399 (2001).

43. M. Mukai et al., MAMO, a maternal BTB/POZ-Zn-finger protein enriched in germline progenitors is required for the production of functional eggs in *Drosophila*. Mech Develop 124, 570–583 (2007).

44. T. Ando, H. Fujiwara, Electroporation-mediated somatic transgenesis for rapid functional analysis in insects. Development 140, 454–458 (2013).

45. C. O. Pabo, E. Peisach, R. A. Grant, Design and selection of novel Cys2His2 zinc finger proteins. Annu Rev Biochem 70, 313–340 (2001).

46. S. A. Wolfe, L. Nekludova, C. O. Pabo, DNA recognition by Cys2His2 zinc finger proteins. Annu Rev Bioph Biom 29, 183–212 (2000).

47. A. V. Persikov, M. Singh, De novo prediction of DNA-binding specificities for Cys2His2 zinc finger proteins. Nucleic Acids Res 42, 97–108 (2014).

48. S. Hira et al., Binding of *Drosophila* maternal Mamo protein to chromatin and specific DNA sequences. Biochem Bioph Res Co 438, 156–160 (2013).

49. P. J. Wittkopp, G. Kalay, *Cis*-regulatory elements: molecular mechanisms and evolutionary processes underlying divergence. Nat Rev Genet 13, 59–69 (2012).

50. S. Preissl, K. J. Gaulton, B. Ren, Characterizing *cis*-regulatory elements using single-cell epigenomics. Nat Rev Genet, (2022).

51. E. Z. Kvon et al., Genome-scale functional characterization of *Drosophila* developmental enhancers *in vivo*. Nature 512, 91-+ (2014).

52. S. Y. Wu et al., Comparative analysis of the integument transcriptomes of the *black dilute* mutant and the wild-type silkworm *Bombyx mori*. Sci Rep-Uk 6, (2016).

53. G. Xiong et al., Body shape and coloration of silkworm larvae are influenced by a novel cuticular protein. Genetics 207, 1053–1066 (2017).

54. P. J. Wittkopp, S. B. Carroll, A. Kopp, Evolution in black and white: genetic control of pigment patterns in *Drosophila*. Trends Genet 19, 495–504 (2003).

55. J. M. Gibert, Phenotypic plasticity in insects. Biologie aujourd’hui 214, 33–44 (2020).

56. R. Futahashi et al., *yellow* and *ebony* are the responsible genes for the larval color mutants of the silkworm *Bombyx mori*. Genetics 180, 1995–2005 (2008).

57. M. Y. Noh, S. Muthukrishnan, K. J. Kramer, Y. Arakane, Cuticle formation and pigmentation in beetles. Curr Opin Insect Sci 17, 1–9 (2016).

58. N. N. Wang et al., Abnormal overexpression of *SoxD* enhances melanin synthesis in the *Ursa* mutant of *Bombyx mori*. Insect Biochem Molec 149, (2022).

59. K. J. Kramer et al., Oxidative conjugation of catechols with proteins in insect skeletal systems. Tetrahedron 57, 385–392 (2001).

60. M. Y. Noh, S. Muthukrishnan, K. J. Kramer, Y. Arakane, *Tribolium castaneum* RR-1 cuticular protein TcCPR4 is required for formation of pore canals in rigid cuticle. Plos Genet 11, (2015).

61. M. Y. Noh et al., Two major cuticular proteins are required for assembly of horizontal laminae and vertical pore canals in rigid cuticle of *Tribolium castaneum*. Insect Biochem Molec 53, 22–29 (2014).

62. S. Mun et al., Cuticular protein with a low complexity sequence becomes cross-linked during insect cuticle sclerotization and is required for the adult molt. Sci Rep-Uk 5, (2015).

63. A. Prakash et al., Nanoscale cuticle density variations correlate with pigmentation and color in butterfly wing scales. arXiv, 2305.16628 (2023).

64. R. Futahashi, H. Shirataki, T. Narita, K. Mita, H. Fujiwara, Comprehensive microarray-based analysis for stage-specific larval camouflage pattern-associated genes in the swallowtail butterfly, *Papilio xuthus*. Bmc Biol 10, (2012).

65. M. Y. Noh, S. Muthukrishnan, K. J. Kramer, Y. Arakane, Development and ultrastructure of the rigid dorsal and flexible ventral cuticles of the elytron of the red flour beetle, Tribolium castaneum. Insect Biochem Molec 91, 21–33 (2017).

66. R. Tajiri, N. Ogawa, H. Fujiwara, T. Kojima, Mechanical control of whole body shape by a single cuticular protein Obstructor-E in *Drosophila melanogaster*. Plos Genet 13, (2017).

67. X. M. Zhao et al., The wing-specific cuticular protein LmACP7 is essential for normal wing morphogenesis in the migratory locust. Insect Biochem Molec 112, (2019).

68. J. H. Willis, Structural cuticular proteins from arthropods: Annotation, nomenclature, and sequence characteristics in the genomics era. Insect Biochem Molec 40, 189–204 (2010).

69. J. E. Rebers, L. M. Riddiford, Structure and expression of a *Manduca sexta* larval cuticle gene homologous to *Drosophila* cuticle genes. J Mol Biol 203, 411–423 (1988).

70. M. V. Karouzou et al., *Drosophila* cuticular proteins with the R&R Consensus: Annotation and classification with a new tool for discriminating RR-1 and RR-2 sequences. Insect Biochem Molec 37, 754–760 (2007).

71. E. H. Chen, Q. L. Hou, Identification and expression analysis of cuticular protein genes in the diamondback moth, Plutella xylostella (Lepidoptera: Plutellidae). Pestic Biochem Phys 178, (2021).

72. X. Fu et al., Genome-Wide Identification and Transcriptome-Based Expression Profile of Cuticular Protein Genes in *Antheraea pernyi*. Int J Mol Sci 24, (2023).

73. C. H. Yang et al., Identification, expression pattern, and feature analysis of cuticular protein genes in the pine moth *Dendrolimus punctatus* (Lepidoptera: Lasiocampidae). Insect Biochem Molec 83, 94–106 (2017).

74. X. Guan, B. W. Middlebrooks, S. Alexander, S. A. Wasserman, Mutation of TweedleD, a member of an unconventional cuticle protein family, alters body shape in *Drosophila*. P Natl Acad Sci USA 103, 16794–16799 (2006).

75. T. Togawa, W. A. Dunn, A. C. Emmons, J. H. Willis, CPF and CPFL, two related gene families encoding cuticular proteins of *Anopheles gambiae* and other insects. Insect Biochem Molec 37, 675–688 (2007).

76. P. L. Pan et al., A comprehensive omics analysis and functional survey of cuticular proteins in the brown planthopper. P Natl Acad Sci USA 115, 5175–5180 (2018).

77. K. Ito et al., Positional cloning of a gene responsible for the *cts* mutation of the silkworm, *Bombyx mori*. Genome 55, 493–504 (2012).

78. M. Osanai-Futahashi et al., Identification of the *Bombyx red egg* gene reveals involvement of a novel transporter family gene in late steps of the insect ommochrome biosynthesis pathway. J Biol Chem 287, 17706–17714 (2012).

79. J. W. Luo et al., Molecular basis of the silkworm mutant *re^l^* causing red egg color and embryonic death. Insect Sci 28, 1290–1299 (2021).

80. J. J. Hanly et al., Genetics of yellow-orange color variation in a pair of sympatric sulphur butterflies. Cell Rep 42, (2023).

81. G. Kalay, J. Lachowiec, U. Rosas, M. R. Dome, P. Wittkopp, Redundant and cryptic enhancer activities of the *Drosophila yellow* gene. Genetics 212, 343–360 (2019).

82. Y. Q. Xin et al., Enhancer evolutionary co-option through shared chromatin accessibility input. P Natl Acad Sci USA 117, 20636–20644 (2020).

83. A. Jolma, J. Taipale, Methods for analysis of transcription factor DNA-binding specificity in vitro. Sub-cellular biochemistry 52, 155–173 (2011).

84. Y. F. Chan et al., Adaptive evolution of pelvic reduction in sticklebacks by recurrent deletion of a *Pitx1* enhancer. Science 327, 302–305 (2010).

85. K. P. Lu et al., SilkMeta: a comprehensive platform for sharing and exploiting pan-genomic and multi-omic silkworm data. Nucleic Acids Res, (2023).

86. J. G. Monroe et al., Mutation bias reflects natural selection in *Arabidopsis thaliana*. Nature 602, 101–105 (2022).

87. E. J. Belfield et al., Thermal stress accelerates *Arabidopsis thaliana* mutation rate. Genome Res 31, 40–50 (2021).

88. R. C. Edgar, MUSCLE: multiple sequence alignment with high accuracy and high throughput. Nucleic Acids Res 32, 1792–1797 (2004).

89. Y. L. Zou et al., Bmmp influences wing morphology by regulating anterior-posterior and proximal-distal axes development. Insect Sci 29, 1569–1582 (2022).

90. D. P. Long et al., Genetic hybridization of highly active exogenous functional proteins into silk-based materials using “light-clothing” strategy. Matter-Us 4, 2039–2058 (2021).

