## Supplemental Figure for "The BTB-ZF gene *Bm-mamo* regulates pigmentation in silkworm caterpillars"

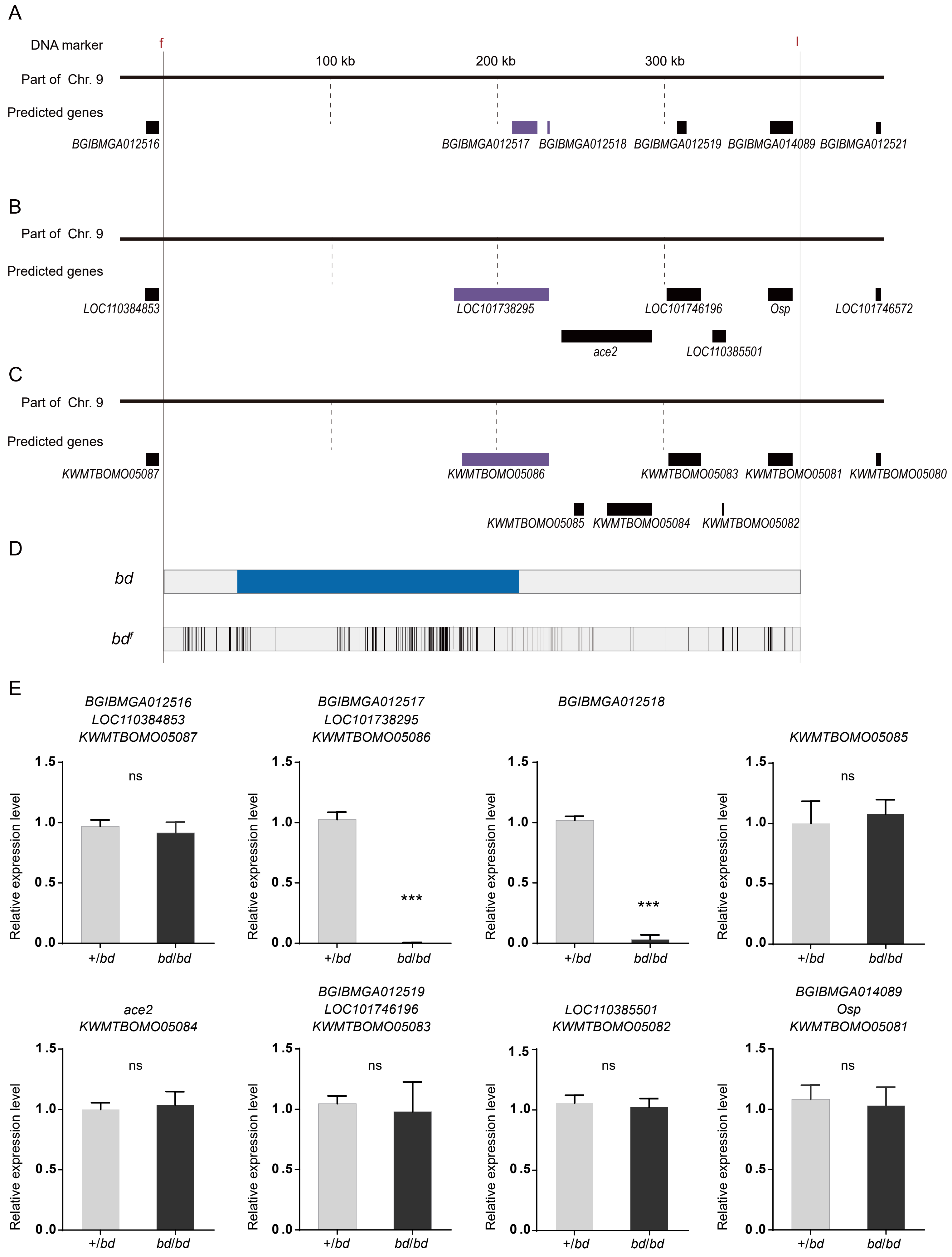


Fig. S1. The predicted genes and qPCR analysis of candidate genes in the responsible genomic region for *bd* mutant.
(A) The predicted genes in SilkDB;(B) the predicted genes in Genbak;(C) the predicted genes in Silkbase;(D) analysis of nucleotide differences in the responsible region of *bd*;(E) investigation of the expression level of candidate genes.

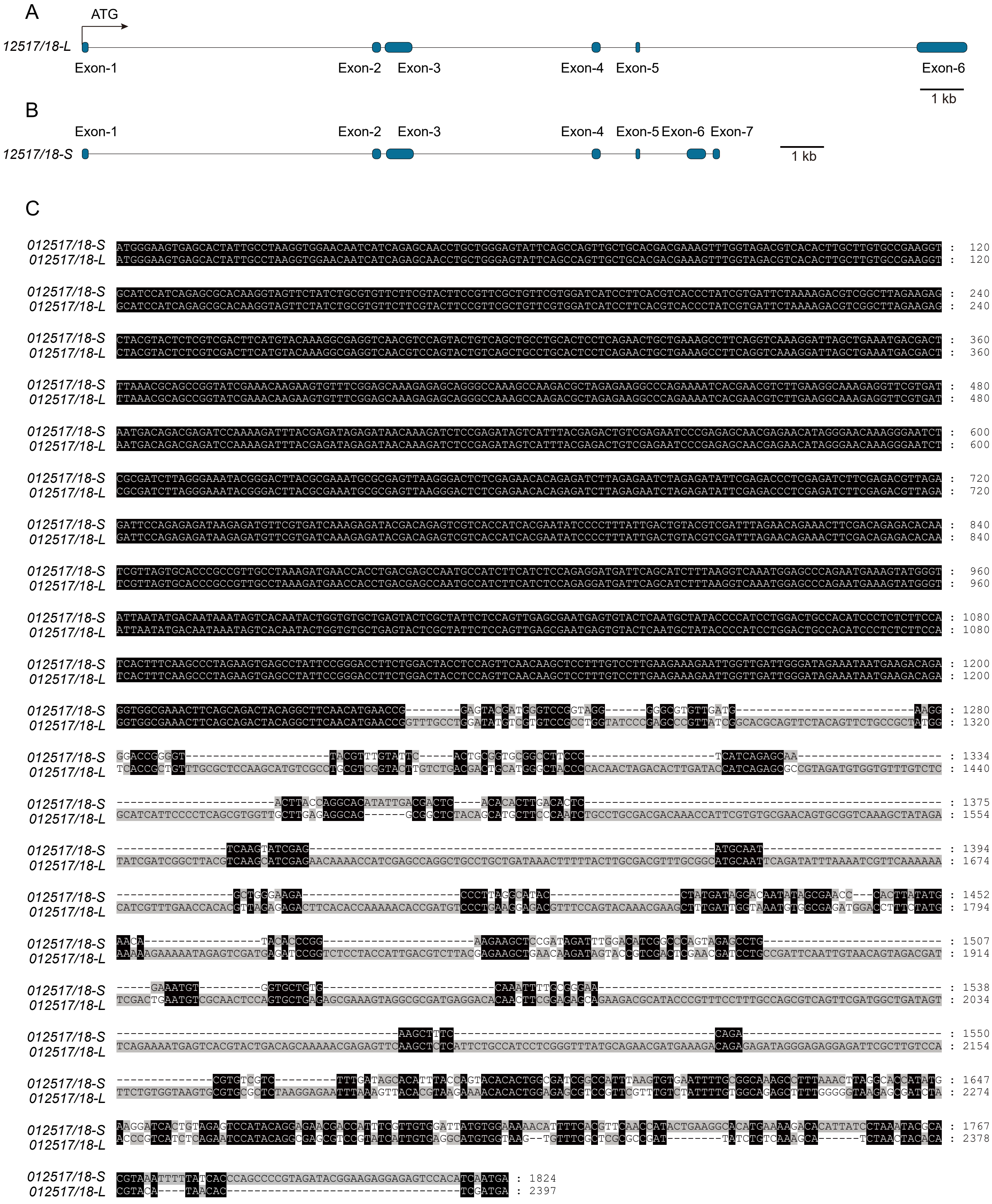


Fig.S2. Sequence analysis of the *12517/18-L* and *12517/18-S* transcripts.
(A)The gene structure of *12517/18-L*; (B) the gene structure of *12517/18-S*; (C) the sequence alignment of *12517/18-L*and *12517/18-S*.

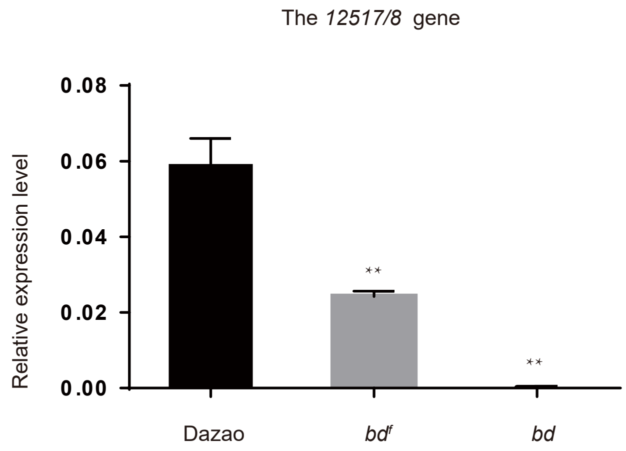


Fig.S3.The expression level of *12517/18* in wildtype Dazao strain and *bd* allele mutants.

```
Dazao: TGTCAGTGGAAAGTAATCGTAGCAAAACACTTCCAGATAATACAACATGAATGCCGCCATCCACCTTGAGACATCCGATC 7485394
   bd: TGTCAGTTGAAAGTAATCGTAGCAAAACACTTCCAGATAATACAACATGAATGCCGCCATCCACCTTGAGACATCCGATC

Dazao: ATAAGTACATACTAGTTCTATTACATAATGGATTTTTCACCCTTCAAAACAAAATACTTTACTGTTGCGCAGCAGAAATA 7485474
   bd: ATAAGTACATACTAGTTCTATTACATAATGGATTTTTCACCCTTCAAAACAAAATACTTTACTGTTGCGCAGCAGAAATA

Dazao: GATAGCAT---------------------AATGGGACCTATCAATGTAGGCTTAGAATACGTCACCCTAGCACGAACCAA 7485533
   bd: GATAGCATCCGGTCGCTCAACAGACGAGAGGTGGGACCTATCAATGTGGGCTTAGAATACGTCACCCTAGCACGAACCAA

Dazao: CTTCGTAAATTTTGTGGGCTTAGTTATTAGTACCTGATATTATAACTTCATCGTGAAGTAGTTTTGTAGTCACGTGTGAA 7485613
   bd: CTTCGTAACTTTTGTGGGCTTAGTTATTAGTACCTGATATTATAACTTCATCGTGAAGTAGTTTTGTAGTCACGTGTGAA

Dazao: GTACGTAT 7485621
   bd: GTACGTAT

Dazao: --------------------------------------------------------------------------------
   bd: TTCCTACCTGCTGTTGAAAAAATTTGTCATGGTACGTTTACTTAAAATATGGTATACTGAAAATAATGCAATTTACAAAA 80
Dazao: --------------------------------------------------------------------------------
   bd: AAAAAAAAAAAAAACATCGGTCGGGTTATGCAGGAGCTCTGTGTAAATGGCGTGAAATACTTTCTAGTTAGCTGGCTGAC 160
Dazao: --------------------------------------------------------------------------------
   bd: TTGAATTTGGAATGATTGATTACTATTCGAAAAAGCTCTATCGTATTATTTTTAACCAAGAAATAAAATAAAAAAGAGAA 240
Dazao: --------------------------------------------------------------------------------
   bd: ACTGAATTCCTTGAGTTCTTCAATTTAAATTTTCTGGATAGGTTCGTCAATCCAAATAATTTAGCATATTATATTAGATA 320
Dazao: --------------------------------------------------------------------------------
   bd: AAATAAAACTGTTAAAAGTCAATTTAGTATAGAACGGACGCCAACTTATACTTAATATTTATTTCAACAAATTTCCATAT 400
Dazao: --------------------------------------------------------------------------------
   bd: GCATCGCAAAGATTTAACTATTGTTAATCAACCAGGAGATATCTTTGTCACATTCGAGAAAACTATTATATTTTGTCGAT 480
Dazao: --------------------------------------------------------------------------------
   bd: TGATTCTTTACTTTCTATGCTAATATTGTTCACTGAGTATAACCTGTCCGTTTTGGAATCCTCATTTCCTGGCACAATAT 560
Dazao: --------------------------------------------------------------------------------
   bd: TATTGGCCCAGAATTATTGTATCCGGAATTAAGTATTGTTGGTGATCCAAAAATAATTCATGGTGGTGGTTGGTTGGCTG 640
Dazao: --------------------------------------------------------------------------------
   bd: GTTTTACGCGTGGAGGACGAAATTTCGAATTCTATACTTTTACACATTCGTTGTTATTACGCATGAAAATACTGTTCGTA 720
Dazao: --------------------------------------------------------------------------------
   bd: AATTTATTACTACTTATATAAACCCTATGAGATACCTTCGTTAAATTGTAATATTTGGTTTCAGCGCTTAACATGACAAA 800
Dazao: --------------------------------------------------------------------------------
   bd: CGATGTGGCTGGAGCGACTTTCATGGCCGCAGCTACTTCTGCACCAGAATTATTCGTGAACGTCATAGGTACCTTCATTA 880
Dazao: --------------------------------------------------------------------------------
   bd: CGGAGGGTGATATCGGAGTGGGGACCATAGTGGGTTCGGCTGTCTTCAATATATTAGCTGTGGCGGCTTGTTGTGGTATT 960
Dazao: --------------------------------------------------------------------------------
   bd: GGTGCGGGAATGGTTTGTATTTCTGATGTCCTTCGAAGATCAATTTTATTTTCAGGTTTTTAATAAAAGTCATTAGTATT 1040
Dazao: --------------------------------------------------------------------------------
   bd: AGTAGCTATACTTGAGACCTTAGAACTTATATCTCAAGGTGGGTGGCGCATTTACGTTGTAGATGGCTATGGGCTCCAGT 1120
Dazao: --------------------------------------------------------------------------------
   bd: AACCACTTAACACCAGGTGGGCTGTGAGCTGGTCCAACCATATAAGCAATAAACAAATAAGAATAATAATAAATATGATC 1200
Dazao: --------------------------------------------------------------------------------
   bd: ATTAGTAAAACAGTTACATATATGTTAAAAATCATCACGAGGCCTTAACTCTTGCCTACAATACGCATACATGCCGCATT 1280
Dazao: --------------------------------------------------------------------------------
   bd: ACAAGGATTAATAGTAATGTCTCATTAAGATACTCGCGGAATTAACTCAATACGCACGAATACCAGGATGTAAAGTGTTC 1360
Dazao: --------------------------------------------------------------------------------
   bd: GTTCGGGTGTATTCAGTATTAGCCACGAATACTGGTCCGTTAAGTGCACGTTGTCCGCTTTTTGGCACGATTCAAAAGGC 1440
Dazao: --------------------------------------------------------------------------------
   bd: GTTTGTGCGCGGTTTTTACGTATTCGTCATTGTTTAATTGACTTGTGAACTTTGTCGCAAGCGGTTTATTATCTACGATG 1520
Dazao: --------------------------------------------------------------------------------
   bd: GATTGGACGAACGATTACTTTAAAGTGTTTCACAAAATTTTAAAGGGTGTAAAGTCGTATTGTTTTTTATTTTATAATAT 1600
Dazao: --------------------------------------------------------------------------------
   bd: TTTATGATACCCATTAAAATATATGCATGAAGTCATAGCAACGATTCTAATGTCATCCATTTTGAAATTTAAGTTCGCTG 1680
Dazao: --------------------------------------------------------------------------------
   bd: AACATAGAAAAATTTATAATTGATTCATGATGTATTCTTGTACAAGATATGAGGCAAATTGTTCGGCGATACTTTGGATG 1760
Dazao: --------------------------------------------------------------------------------
   bd: AGGCAAAACAATGGTTTTAATTTATTAAAATAGTACCTAGCTAAAATATGATTGGTAGTTGAGAATTTAAAGTTTTCTAA 1840
```

```
Dazao: --------------------------------------------------------------------------------
   bd: TTTTTTAACTTTTCAGGTTGTTCCTTGGACTGGTGGCCTCTTACTAGAGATTGTCTGGCTTATGGGATTACAGTTTCAAT 1920

Dazao: --------------------------------------------------------------------------------
   bd: ACTTATTTGTATTATACACGATGAGTATGTTCAGTGGTATGAAGCATTTCTTCTAGTTTCATTGTACGGGGTTTACATTT 2000

Dazao: --------------------------------------------------------------------------------
   bd: GCATAATGTATTATGACAAGCCTATACAGAATTTTGCGAAAGGTGAGTTTCGGAAATAAAGGTTATTTATTAATTGACGT 2080

Dazao: --------------------------------------------------------------------------------
   bd: ATAGATGAATGTTTTAAATTTCACAGGGAGCTGGAAATGGGTATCAAACATGTTCAAAGATAATGAAGAGAAACAAGATG 2160

Dazao: --------------------------------------------------------------------------------
   bd: CAGAAAATGGTAAATATACGTTAATTACTTATTTTTATGCCACATTAGAATAGATAAATTTAGTTTGGATGTTTCGATAT 2240

Dazao: --------------------------------------------------------------------------------
   bd: TTCACACGATTCTTTTAAACCTCACATAGCTTTATTTTATTATCTCATAAGTTAGATAAAACATGAGAAGTATAGTGAAA 2320

Dazao: --------------------------------------------------------------------------------
   bd: TAATCATTACACACTTATAAAAACGATATTAATCCTATGAATTTTCAGGTAAAGATTTAAAGTTATCGACCCATATACTA 2400

Dazao: --------------------------------------------------------------------------------
   bd: AAAATAGATGTACCTATAGAACAACGCCATCAACATAACGGTAATATTAAAACATCACCCCTTCGTCAAGATGCGAAAGA 2480

Dazao: --------------------------------------------------------------------------------
   bd: AAAGATTTTAGAAAGTGTCGATGTTTCGGATAAAGTTTCCGCCAGTGAAGAAGGAAAAGATAAAAATGTAAATATTGACA 2560

Dazao: --------------------------------------------------------------------------------
   bd: TGGATTCTAAGCCTAATGAAATGGAAATCGCTAGCCCAGAGGAAAACTTATATCAGCCAACTGCAGAAGATAATAAAGAT 2640

Dazao: --------------------------------------------------------------------------------
   bd: GTTGAAGATAGATCCGAGAACTCACTGTGGAAATGGCCTAGCAACAAGAGTTGCCTAGCTAAGGTATTTTAATTTTGATT 2720

Dazao: --------------------------------------------------------------------------------
   bd: TCTATATTTCACAAATAATATTTTATTCCAATTTTTATAAAGCTTAATTATTTTCCTTTCAGATAACCAAAATTGTAACA 2800

Dazao: --------------------------------------------------------------------------------
   bd: TGGCCCATTCACTTGGTGTTCCTGTTTACAATTCCAGATTGTGAGAAACCACGATTTAAAAACTGGTTTCCGCTTACTTT 2880

Dazao: --------------------------------------------------------------------------------
   bd: CATTATGTGTATTGTTTGGATCGGTTCACTGTCTTATATAGTTGCATGGATGATAACAATAATAGGTAAGAAAAAAATAAG 2960

Dazao: --------------------------------------------------------------------------------
   bd: CCATTTAAGTTAAAAATTTACATATATTGAATGAATATTTTAAACATGTATATTTTTAGGTGATACGCTGATGATCCCAG 3040

Dazao: --------------------------------------------------------------------------------
   bd: ATTCAGTTATGGGTATTACCTTTCTTGCTGCTGGCACTTCAGTTCCCGAAGCTGTTTCCAGCGTTATAGTAGTTAAGCAA 3120

Dazao: --------------------------------------------------------------------------------
   bd: GGTAGTATTTACAAGAAACCAAGCTACAACTAAGGGTTTTAGAAAAGATAACTAAAATTGATTGTTTTTTGACAGGGCAT 3200

Dazao: --------------------------------------------------------------------------------
   bd: GGTTCGATGGGAATCAGTAACTCAATAGGATCGAATACATTTGATATTTTACTTTGCCTTGGCTTACCGTGACTTATAAA 3380

Dazao: --------------------------------------------------------------------------------
   bd: AGCATCGCTAACTCCAGCCGAGCCAGGACATTATTGGGTCAGATTTTGTTTTATAACTTATTTTTCTGTATGCGAATTCC 3460

Dazao: --------------------------------------------------------------------------------
   bd: TTGTGTACTTTTCCATACCTTGCGAATGTATGACTTTGCGATTCTTTGTTTGCCTTTAGACTTGCTCTGCGAAATTTAAT 3520

Dazao: ----------------------------------------
   bd: AAGTTAATTTGAAACGAATTACAATTATTGTGTACCACTA 3560

Dazao: AGGTGTGTTGCTGTTGCTGAGTTGCTATACGTTTTCTTATAATTAAAGATTATCAGTAAATCGTTAATTCCCATAAATGA 7653983
   bd: AGGTGTGTTGCTGTTGCTGAGTTGCTATACGTTTT-TTATAATTAAAGATTATCAGTAAATCGTTAATTCCCATAAATGA

Dazao: GATAAAGCCTGTATGTAACCTAAATTGATGTGGCAAGATGAACAACAGTTTCTTTATATTATATTTATCGGCGCAACGTC 7654063
   bd: GATAAAGCCTGTATGTAACCTAAATTGATGTGGCAAGATGAACAACAGTTTCTTTATATTATATTTATCGGCGCAACGTC

Dazao: ACTATTTAGTTAACTATAAATAATTAAAGTTAACTAAGACAGTATTCAAGAGAATTCATGACTAAGCTGCCCTCACTTTC 7654143
   bd: ACTATTTAGTTAACTATAAATAATTAAAGTTAACTAAGACAGTATTCAAGAGAATTCATGACTAAGCTGCCCTCACTTTC

Dazao: TACTATAAATACCTGTGATTATACGAGTGACTATTGTCGCAAAAACATTCCGGCACGAGCGCTCAAACGGAGCGGAGCTC 7654223
   bd: TACTATAAATACCTGTGATTATACGAGTGACTATTGTCGCAAAAACATTCCGGCACGAGCGCTCAAACGGAGCGGAGCTC

Dazao: TCTGTTGTTATGGTCGTACATTTAAAATGTAATAAATGGAATTCATTGGAAAAGTGAGTCTTAATTACTGTGACTATGGA 7654303
   bd: TCTGTTGTTATGGTCGTACATTTAAAATGTAATAAATGGAATTCATTGGAAAAGTGAGTCTTAATTACTGTGACTATGGA

Dazao: AGGCAATAATAATAAAGGAGAATGGCGTCAGCAACGCTGGTGTCATCATATTAAAAACACCATGGAATCCGCCTCTCCGC 7654383
   bd: AGGCAATAATAATAAAGGAGAATGGCGTCAGCAACGCTGGTGTCATCATATTAAAAACACCATGGAATCCGCCTCTCCGC

Dazao: CATACTATATTTTTAGAAAACT 7654405
   bd: CATACTATATTTTTAGAAAACT
```

Fig.S4. The responsible sequence of *bd* mutant.
In the *bd* mutant, an over 168kb genome sequence was absented and a 3560bp sequence was inserted.

A

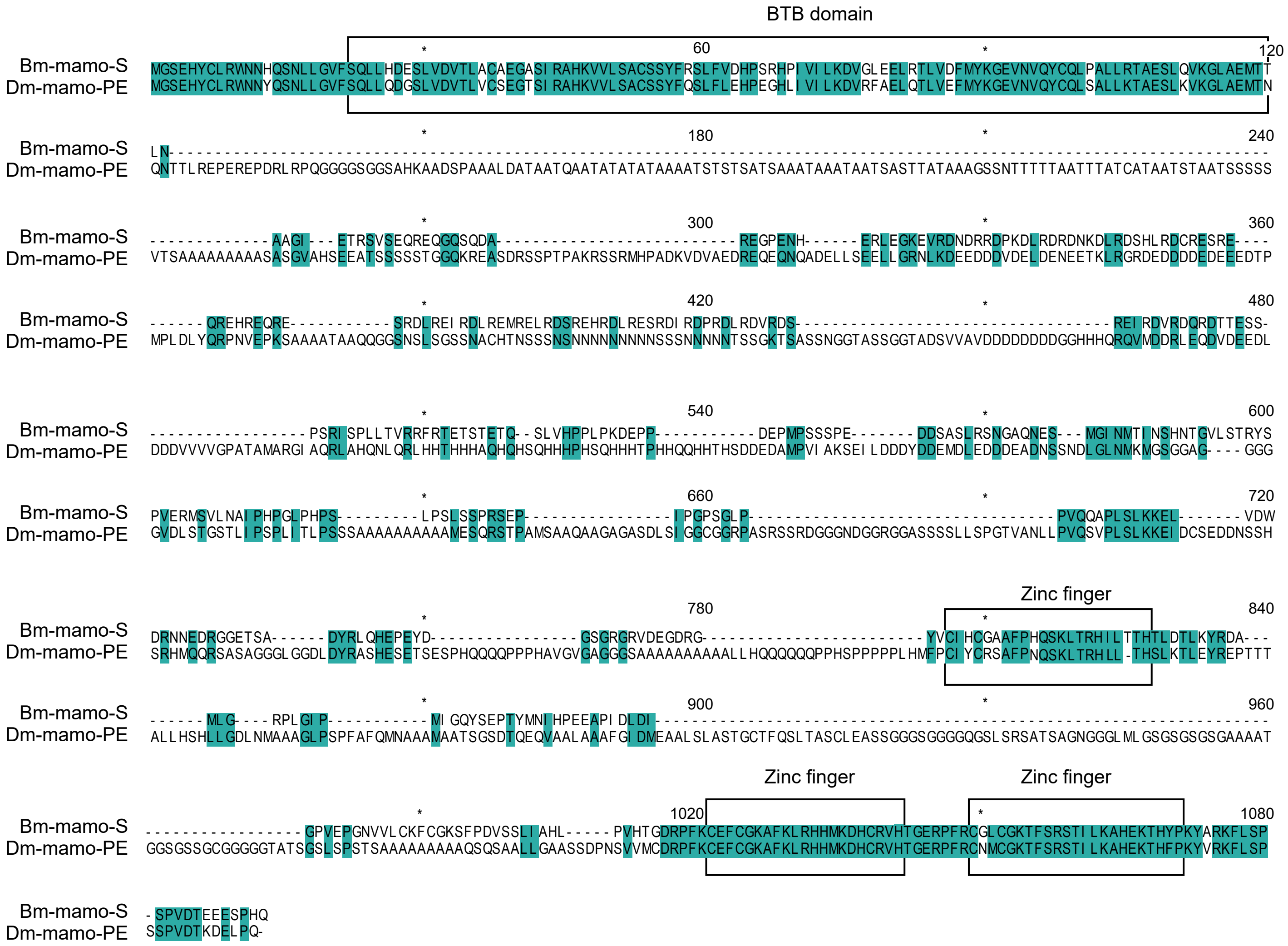
BTB domain
Zinc finger
Zinc finger
Zinc finger
Bm-mamo-S
Dm-mamo-PE

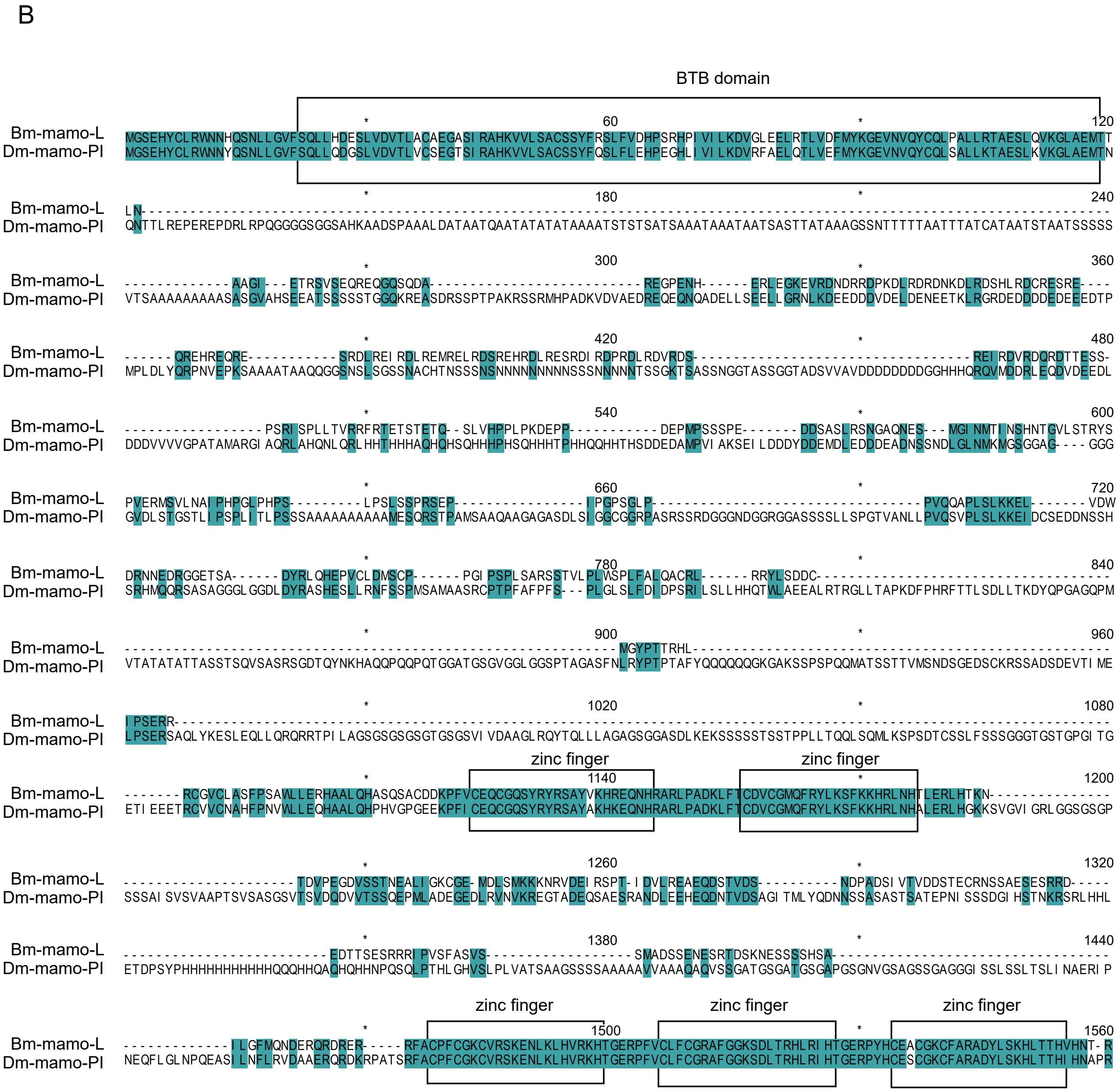


Fig.S5. The sequence alignment between the mamo protein of *Drosophila melanogaster* and the amino acid sequence encoded by the *012517/18* gene.

(A) The sequence alignment of Bm-mamo-S and Dm-mamo-PE; (B) the sequence alignment of Bm-mamo-L and Dm-mamo-PI.

A

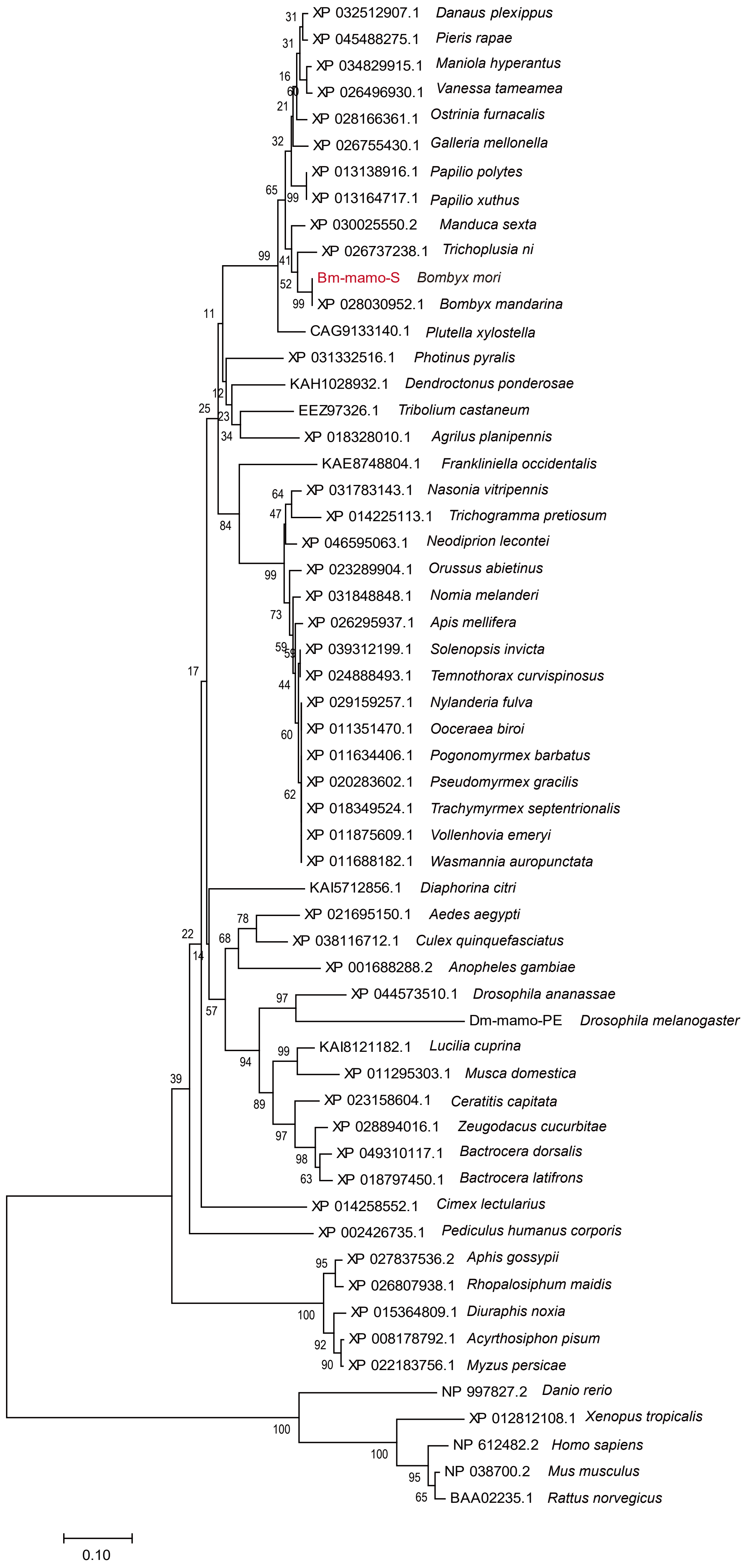

XP 032512907.1 Danaus plexippus
XP 045488275.1 Pieris rapae
XP 034829915.1 Maniola hyperantus
XP 026496930.1 Vanessa tameamea
XP 028166361.1 Ostrinia furnacalis
XP 026755430.1 Galleria mellonella
XP 013138916.1 Papilio polytes
XP 013164717.1 Papilio xuthus
XP 030025550.2 Manduca sexta
XP 026737238.1 Trichoplusia ni
Bm-mamo-S Bombyx mori
XP 028030952.1 Bombyx mandarina
CAG9133140.1 Plutella xylostella
XP 031332516.1 Photinus pyralis
KAH1028932.1 Dendroctonus ponderosae
EEZ97326.1 Tribolium castaneum
XP 018328010.1 Agrilus planipennis
KAE8748804.1 Frankliniella occidentalis
XP 031783143.1 Nasonia vitripennis
XP 014225113.1 Trichogramma pretiosum
XP 046595063.1 Neodiprion lecontei
XP 023289904.1 Orussus abietinus
XP 031848848.1 Nomia melanderi
XP 026295937.1 Apis mellifera
XP 039312199.1 Solenopsis invicta
XP 024888493.1 Temnothorax curvispinosus
XP 029159257.1 Nylanderia fulva
XP 011351470.1 Ooceraea biroi
XP 011634406.1 Pogonomyrmex barbatus
XP 020283602.1 Pseudomyrmex gracilis
XP 018349524.1 Trachymyrmex septentrionalis
XP 011875609.1 Vollenhovia emeryi
XP 011688182.1 Wasmannia auropunctata
KAI5712856.1 Diaphorina citri
XP 021695150.1 Aedes aegypti
XP 038116712.1 Culex quinquefasciatus
XP 001688288.2 Anopheles gambiae
XP 044573510.1 Drosophila ananassae
Dm-mamo-PE Drosophila melanogaster
KAI8121182.1 Lucilia cuprina
XP 011295303.1 Musca domestica
XP 023158604.1 Ceratitis capitata
XP 028894016.1 Zeugodacus cucurbitae
XP 049310117.1 Bactrocera dorsalis
XP 018797450.1 Bactrocera latifrons
XP 014258552.1 Cimex lectularius
XP 002426735.1 Pediculus humanus corporis
XP 027837536.2 Aphis gossypii
XP 026807938.1 Rhopalosiphum maidis
XP 015364809.1 Diuraphis noxia
XP 008178792.1 Acyrthosiphon pisum
XP 022183756.1 Myzus persicae
NP 997827.2 Danio rerio
XP 012812108.1 Xenopus tropicalis
NP 612482.2 Homo sapiens
NP 038700.2 Mus musculus
BAA02235.1 Rattus norvegicus
0.10

B

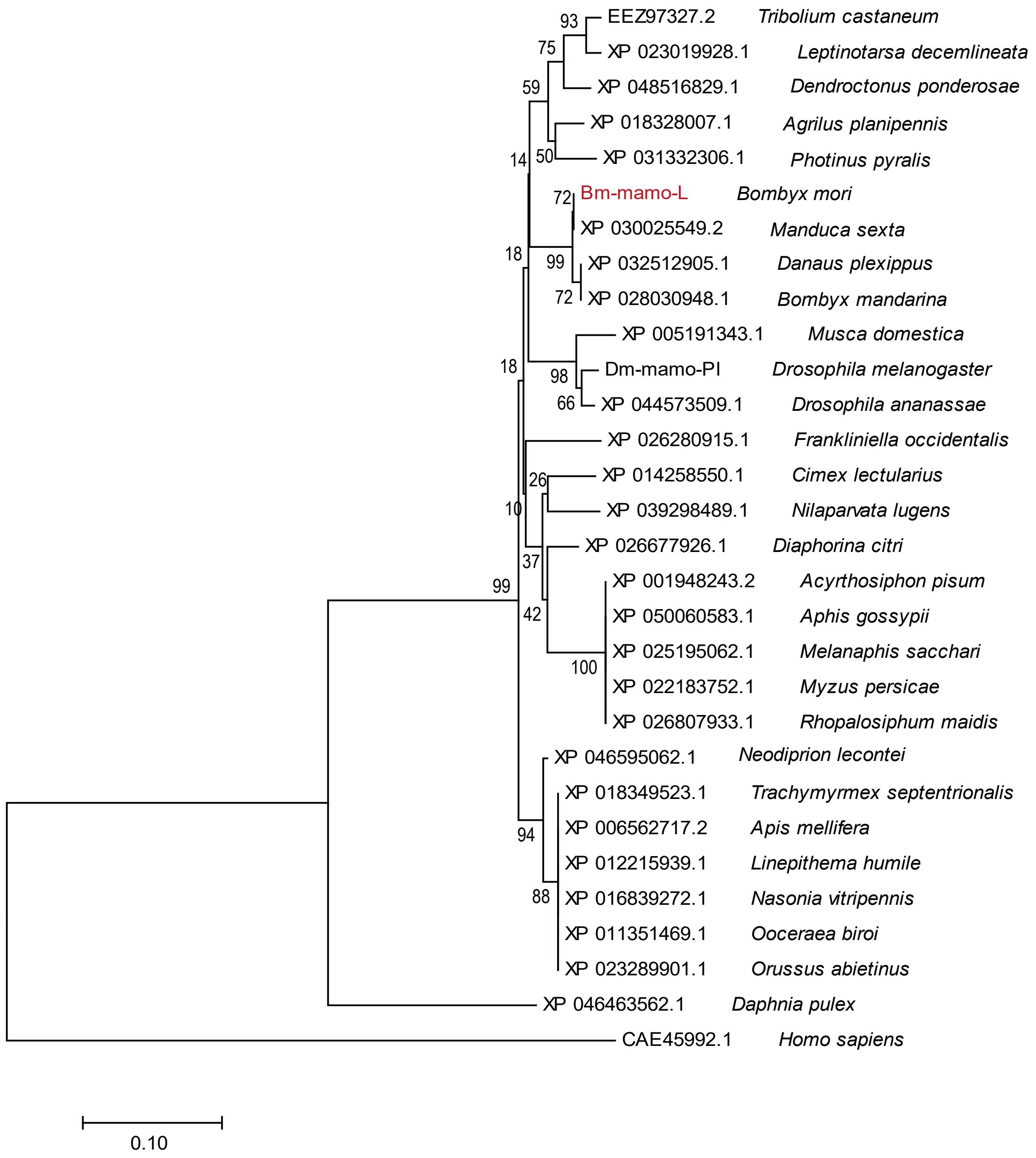


Fig.S6. The phylogenetic analysis.
(A)The phylogenetic analysis of orthologs of Bm-mamo-S; (B) the phylogenetic analysis of orthologs of Bm-mamo-L.

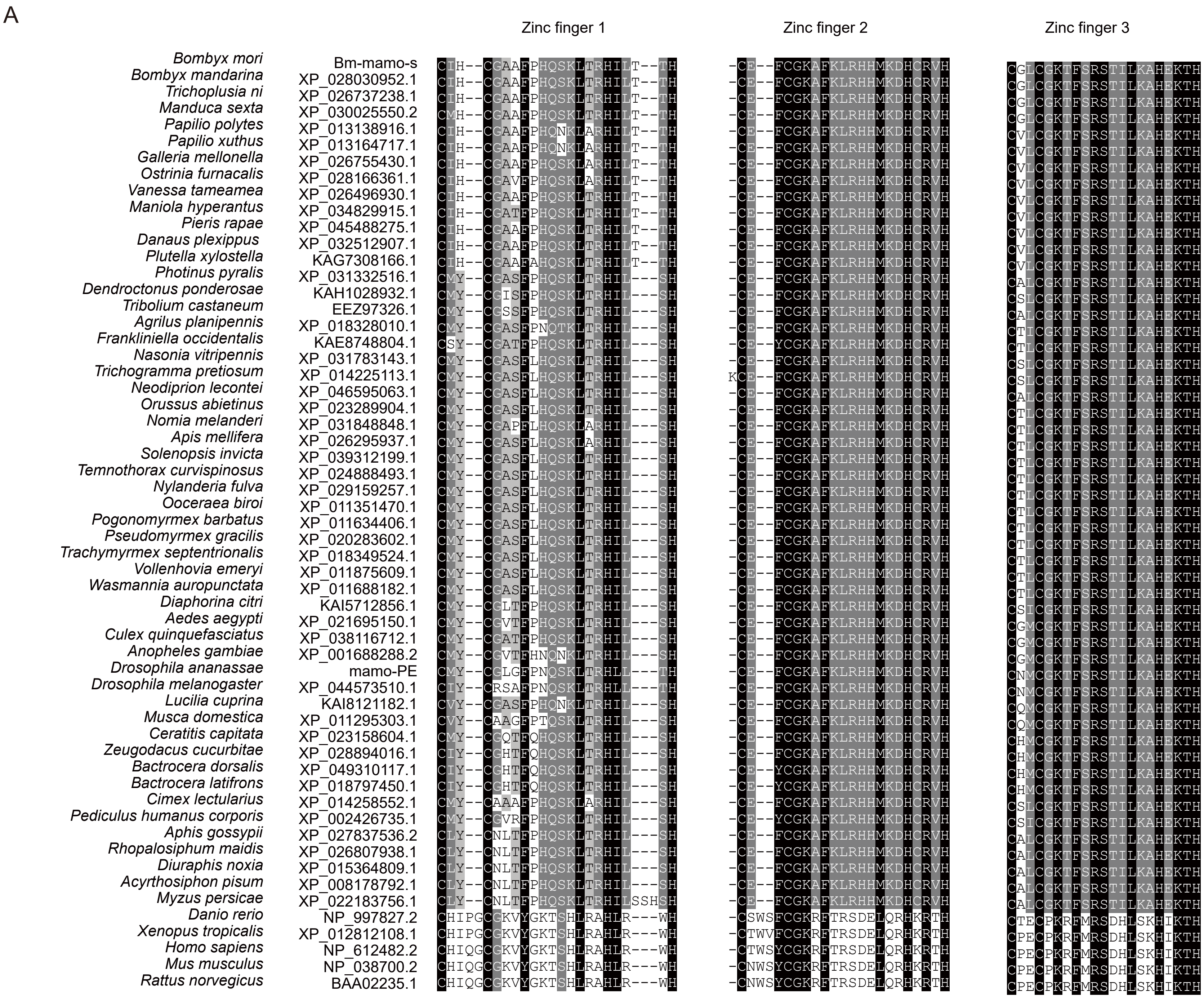
A
Zinc finger 1
Zinc finger 2
Zinc finger 3
Bombyx mori Bm-mamo-s CIH--CGAAFPHQSKLTRHILT--TH -CE--FCGKAFKLRHHMKDHCRVH CGLCGKTFSRSTILKAHEKTH
Bombyx mandarina XP_028030952.1 CIH--CGAAFPHQSKLTRHILT--TH -CE--FCGKAFKLRHHMKDHCRVH CGLCGKTFSRSTILKAHEKTH
Trichoplusia ni XP_026737238.1 CIH--CGAAFPHQSKLTRHILT--TH -CE--FCGKAFKLRHHMKDHCRVH CGLCGKTFSRSTILKAHEKTH
Manduca sexta XP_030025550.2 CMH--CGAAFPHQSKLTRHILT--TH -CE--FCGKAFKLRHHMKDHCRVH CGLCGKTFSRSTILKAHEKTH
Papilio polytes XP_013138916.1 CIH--CGAAFPHQNKLARHILT--TH -CE--FCGKAFKLRHHMKDHCRVH CVLCGKTFSRSTILKAHEKTH
Papilio xuthus XP_013164717.1 CIH--CGAAFPHQNKLARHILT--TH -CE--FCGKAFKLRHHMKDHCRVH CVLCGKTFSRSTILKAHEKTH
Galleria mellonella XP_026755430.1 CIH--CGAAFPHQSKLARHILT--TH -CE--FCGKAFKLRHHMKDHCRVH CVLCGKTFSRSTILKAHEKTH
Ostrinia furnacalis XP_028166361.1 CIH--CGAVFPHQSKLARHILT--TH -CE--FCGKAFKLRHHMKDHCRVH CVLCGKTFSRSTILKAHEKTH
Vanessa tameamea XP_026496930.1 CIH--CGAAFPHQSKLTRHILT--TH -CE--FCGKAFKLRHHMKDHCRVH CVLCGKTFSRSTILKAHEKTH
Maniola hyperantus XP_034829915.1 CIH--CGATFPHQSKLTRHILT--TH -CE--FCGKAFKLRHHMKDHCRVH CVLCGKTFSRSTILKAHEKTH
Pieris rapae XP_045488275.1 CIH--CGAAFPHQSKLTRHILT--TH -CE--FCGKAFKLRHHMKDHCRVH CVLCGKTFSRSTILKAHEKTH
Danaus plexippus XP_032512907.1 CIH--CGAAFPHQSKLTRHILT--TH -CE--FCGKAFKLRHHMKDHCRVH CVLCGKTFSRSTILKAHEKTH
Plutella xylostella KAG7308166.1 CIH--CGAAFAHQSKLTRHILT--TH -CE--FCGKAFKLRHHMKDHCRVH CVLCGKTFSRSTILKAHEKTH
Photinus pyralis XP_031332516.1 CMY--CGASFPHQSKLTRHIL---SH -CE--FCGKAFKLRHHMKDHCRVH CALCGKTFSRSTILKAHEKTH
Dendroctonus ponderosae KAH1028932.1 CMY--CGISFPHQSKLTRHIL---SH -CE--FCGKAFKLRHHMKDHCRVH CSLCGKTFSRSTILKAHEKTH
Tribolium castaneum EEZ97326.1 CMY--CGSSFPHQSKLTRHIL---SH -CE--FCGKAFKLRHHMKDHCRVH CALCGKTFSRSTILKAHEKTH
Agrilus planipennis XP_018328010.1 CMY--CGASFPNQTKLTRHIL---SH -CE--FCGKAFKLRHHMKDHCRVH CTICGKTFSRSTILKAHEKTH
Frankliniella occidentalis KAE8748804.1 CSY--CGATFPHQSKLTRHIL---SH -CE--YCGKAFKLRHHMKDHCRVH CTLCGKTFSRSTILKAHEKTH
Nasonia vitripennis XP_031783143.1 CMY--CGASFLHQSKLTRHIL---SH -CE--FCGKAFKLRHHMKDHCRVH CSLCGKTFSRSTILKAHEKTH
Trichogramma pretiosum XP_014225113.1 CMY--CGASFLHQSKLTRHIL---SH KCE--FCGKAFKLRHHMKDHCRVH CSLCGKTFSRSTILKAHEKTH
Neodiprion lecontei XP_046595063.1 CMY--CGASFLHQSKLTRHIL---SH -CE--FCGKAFKLRHHMKDHCRVH CALCGKTFSRSTILKAHEKTH
Orussus abietinus XP_023289904.1 CMY--CGASFLHQSKLTRHIL---SH -CE--FCGKAFKLRHHMKDHCRVH CTLCGKTFSRSTILKAHEKTH
Nomia melanderi XP_031848848.1 CMY--CGAPFLHQSKLARHIL---SH -CE--FCGKAFKLRHHMKDHCRVH CTLCGKTFSRSTILKAHEKTH
Apis mellifera XP_026295937.1 CMY--CGASFLHQSKLARHIL---SH -CE--FCGKAFKLRHHMKDHCRVH CTLCGKTFSRSTILKAHEKTH
Solenopsis invicta XP_039312199.1 CMY--CGASFLHQSKLTRHIL---SH -CE--FCGKAFKLRHHMKDHCRVH CTLCGKTFSRSTILKAHEKTH
Temnothorax curvispinosus XP_024888493.1 CMY--CGASFLHQSKLTRHIL---SH -CE--FCGKAFKLRHHMKDHCRVH CTLCGKTFSRSTILKAHEKTH
Nylanderia fulva XP_029159257.1 CMY--CGASFLHQSKLTRHIL---SH -CE--FCGKAFKLRHHMKDHCRVH CTLCGKTFSRSTILKAHEKTH
Ooceraea biroi XP_011351470.1 CMY--CGASFLHQSKLTRHIL---SH -CE--FCGKAFKLRHHMKDHCRVH CTLCGKTFSRSTILKAHEKTH
Pogonomyrmex barbatus XP_011634406.1 CMY--CGASFLHQSKLTRHIL---SH -CE--FCGKAFKLRHHMKDHCRVH CTLCGKTFSRSTILKAHEKTH
Pseudomyrmex gracilis XP_020283602.1 CMY--CGASFLHQSKLTRHIL---SH -CE--FCGKAFKLRHHMKDHCRVH CTLCGKTFSRSTILKAHEKTH
Trachymyrmex septentrionalis XP_018349524.1 CMY--CGASFLHQSKLTRHIL---SH -CE--FCGKAFKLRHHMKDHCRVH CTLCGKTFSRSTILKAHEKTH
Vollenhovia emeryi XP_011875609.1 CMY--CGASFLHQSKLTRHIL---SH -CE--FCGKAFKLRHHMKDHCRVH CTLCGKTFSRSTILKAHEKTH
Wasmannia auropunctata XP_011688182.1 CMY--CGASFLHQSKLTRHIL---SH -CE--FCGKAFKLRHHMKDHCRVH CTLCGKTFSRSTILKAHEKTH
Diaphorina citri KAI5712856.1 CMY--CGLTFPHQSKLTRHIL---SH -CE--FCGKAFKLRHHMKDHCRVH CSICGKTFSRSTILKAHEKTH
Aedes aegypti XP_021695150.1 CMY--CGVTFPHQSKLTRHIL---SH -CE--FCGKAFKLRHHMKDHCRVH CGMCGKTFSRSTILKAHEKTH
Culex quinquefasciatus XP_038116712.1 CMY--CGATFPHQSKLTRHIL---SH -CE--FCGKAFKLRHHMKDHCRVH CGMCGKTFSRSTILKAHEKTH
Anopheles gambiae XP_001688288.2 CMY--CGVTFHNQNKLTRHIL---SH -CE--FCGKAFKLRHHMKDHCRVH CGMCGKTFSRSTILKAHEKTH
Drosophila ananassae mamo-PE CMY--CGLGFPNQSKLTRHLL---TH -CE--FCGKAFKLRHHMKDHCRVH CNMCGKTFSRSTILKAHEKTH
Drosophila melanogaster XP_044573510.1 CIY--CRSAFPNQSKLTRHLL---TH -CE--FCGKAFKLRHHMKDHCRVH CNMCGKTFSRSTILKAHEKTH
Lucilia cuprina KAI8121182.1 CVY--CGASFPHQNKLTRHIL---SH -CE--FCGKAFKLRHHMKDHCRVH CQMCGKTFSRSTILKAHEKTH
Musca domestica XP_011295303.1 CVY--CAAGFPTQSKLTRHIL---SH -CE--FCGKAFKLRHHMKDHCRVH CQMCGKTFSRSTILKAHEKTH
Ceratitis capitata XP_023158604.1 CMY--CGQTFQHQSKLTRHIL---SH -CE--FCGKAFKLRHHMKDHCRVH CHMCGKTFSRSTILKAHEKTH
Zeugodacus cucurbitae XP_028894016.1 CIY--CGHTFQHQSKLTRHIL---SH -CE--FCGKAFKLRHHMKDHCRVH CHMCGKTFSRSTILKAHEKTH
Bactrocera dorsalis XP_049310117.1 CIY--CGHTFQHQSKLTRHIL---SH -CE--YCGKAFKLRHHMKDHCRVH CHMCGKTFSRSTILKAHEKTH
Bactrocera latifrons XP_018797450.1 CIY--CGHTFQHQSKLTRHIL---SH -CE--YCGKAFKLRHHMKDHCRVH CHMCGKTFSRSTILKAHEKTH
Cimex lectularius XP_014258552.1 CMY--CAAAFPHQSKLARHIL---SH -CE--FCGKAFKLRHHMKDHCRVH CSLCGKTFSRSTILKAHEKTH
Pediculus humanus corporis XP_002426735.1 CMY--CGVRFPHQSKLTRHIL---SH -CE--YCGKAFKLRHHMKDHCRVH CSICGKTFSRSTILKAHEKTH
Aphis gossypii XP_027837536.2 CLY--CNLTFPHQSKLTRHIL---SH -CE--FCGKAFKLRHHMKDHCRVH CALCGKTFSRSTILKAHEKTH
Rhopalosiphum maidis XP_026807938.1 CLY--CNLTFPHQSKLTRHIL---SH -CE--FCGKAFKLRHHMKDHCRVH CALCGKTFSRSTILKAHEKTH
Diuraphis noxia XP_015364809.1 CLY--CNLTFPHQSKLTRHIL---SH -CE--FCGKAFKLRHHMKDHCRVH CALCGKTFSRSTILKAHEKTH
Acyrthosiphon pisum XP_008178792.1 CLY--CNLTFPHQSKLTRHIL---SH -CE--FCGKAFKLRHHMKDHCRVH CALCGKTFSRSTILKAHEKTH
Myzus persicae XP_022183756.1 CLY--CNLTFPHQSKLTRHILSSHSH -CE--FCGKAFKLRHHMKDHCRVH CALCGKTFSRSTILKAHEKTH
Danio rerio NP_997827.2 CHIPGCGKVYGKTSHLRAHLR---WH -CSWSFCGKRFTRSDELQRHKRTH CTECPKRFMRSDHLSKHIKTH
Xenopus tropicalis XP_012812108.1 CHIPGCGKVYGKTSHLRAHLR---WH -CTWVFCGKRFTRSDELQRHKRTH CPECPKRFMRSDHLSKHIKTH
Homo sapiens NP_612482.2 CHIQGCGKVYGKTSHLRAHLR---WH -CTWSYCGKRFTRSDELQRHKRTH CPECPKRFMRSDHLSKHIKTH
Mus musculus NP_038700.2 CHIQGCGKVYGKTSHLRAHLR---WH -CNWSYCGKRFTRSDELQRHKRTH CPECPKRFMRSDHLSKHIKTH
Rattus norvegicus BAA02235.1 CHIQGCGKVYGKTSHLRAHLR---WH -CNWSYCGKRFTRSDELQRHKRTH CPECPKRFMRSDHLSKHIKTH

B

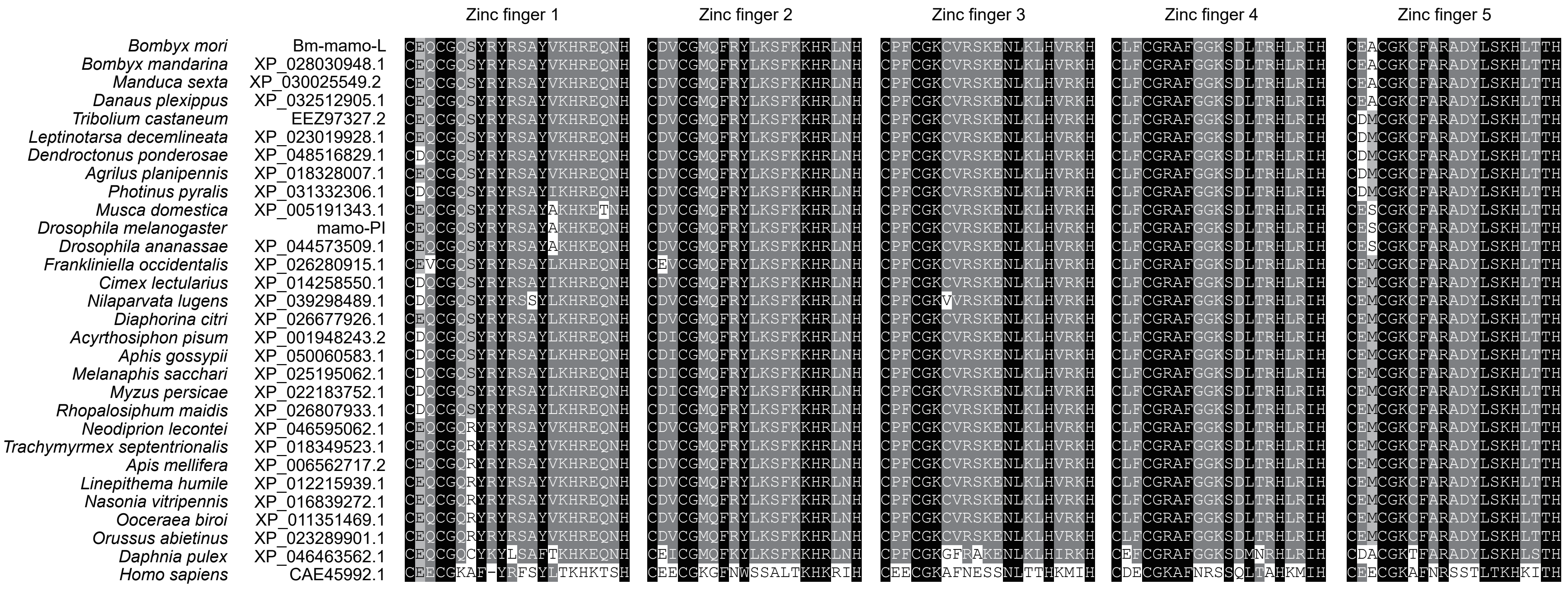


Fig.S7. The sequence alignment of zinc finger motif of orthologs of mamo in multiple species.

(A) The sequence alignment of zinc finger of orthologs of Bm-mamo-S; (B) the sequence alignment of zinc finger of orthologs of Bm-mamo-L.

A

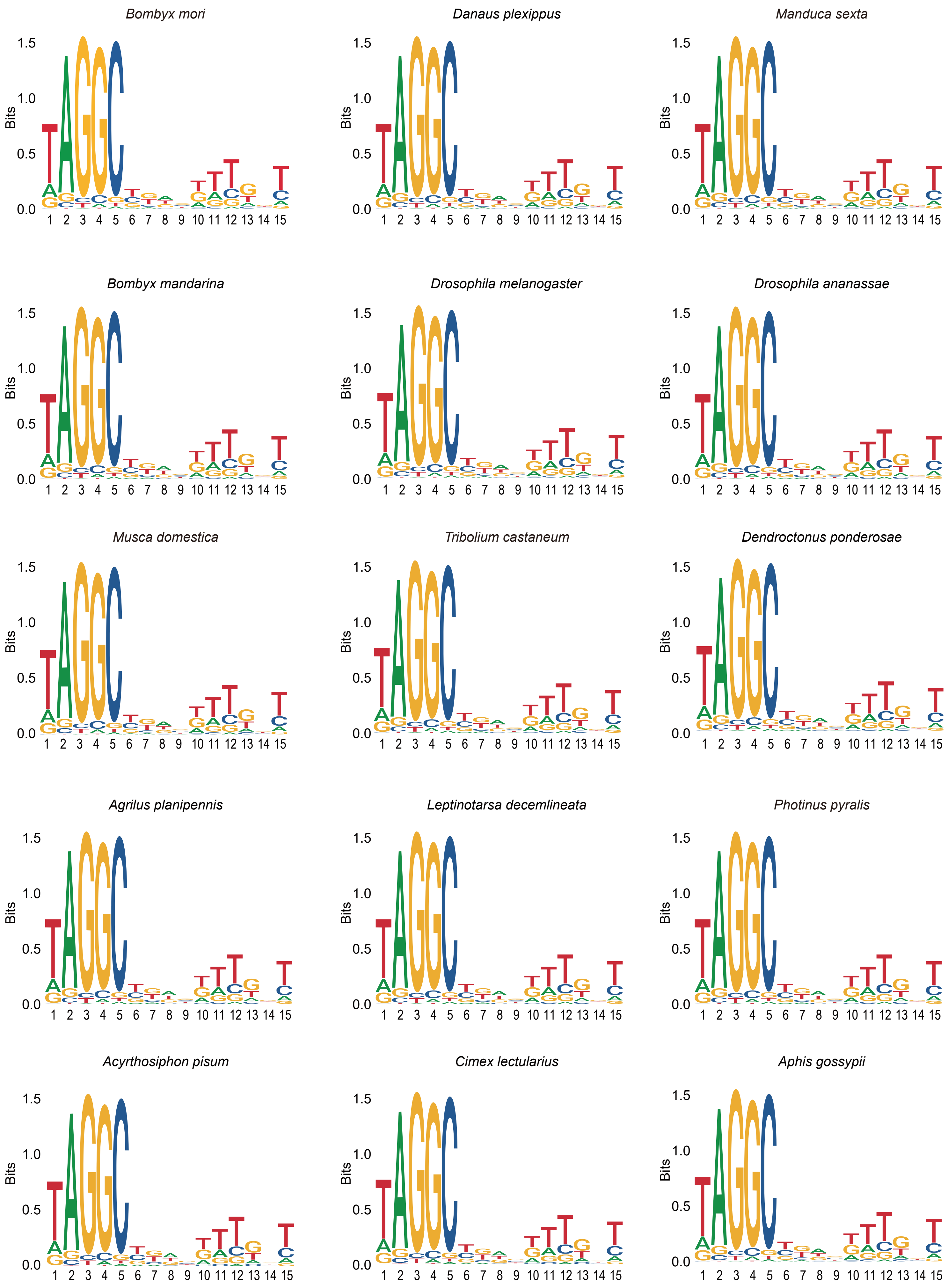
Bombyx mori
Danaus plexippus
Manduca sexta
Bombyx mandarina
Drosophila melanogaster
Drosophila ananassae
Musca domestica
Tribolium castaneum
Dendroctonus ponderosae
Agrilus planipennis
Leptinotarsa decemlineata
Photinus pyralis
Acyrthosiphon pisum
Cimex lectularius
Aphis gossypii
Bits
0.0
0.5
1.0
1.5
1 2 3 4 5 6 7 8 9 10 11 12 13 14 15

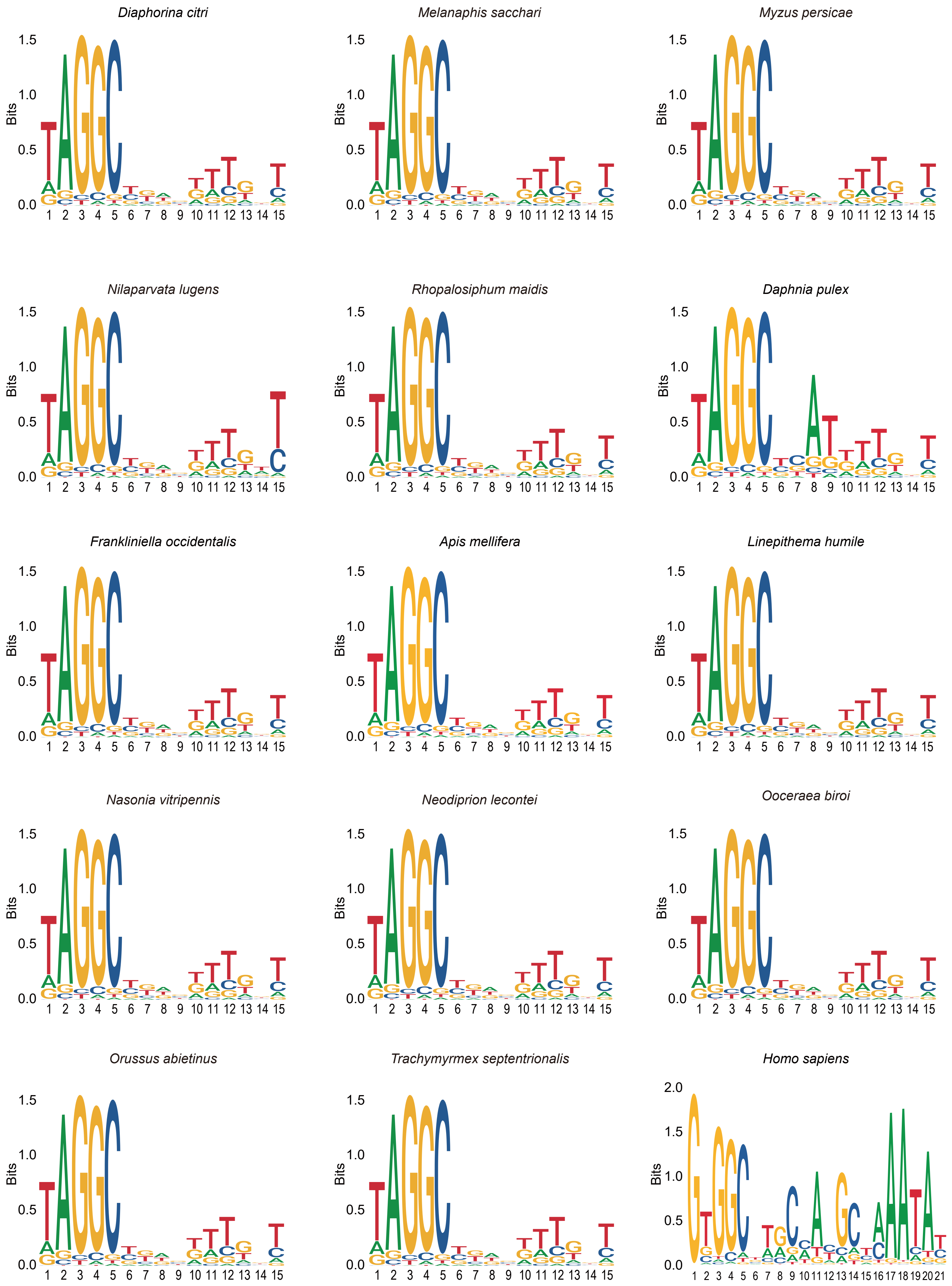
Diaphorina citri
Melanaphis sacchari
Myzus persicae
Nilaparvata lugens
Rhopalosiphum maidis
Daphnia pulex
Frankliniella occidentalis
Apis mellifera
Linepithema humile
Nasonia vitripennis
Neodiprion lecontei
Ooceraea biroi
Orussus abietinus
Trachymyrmex septentrionalis
Homo sapiens
Bits

B

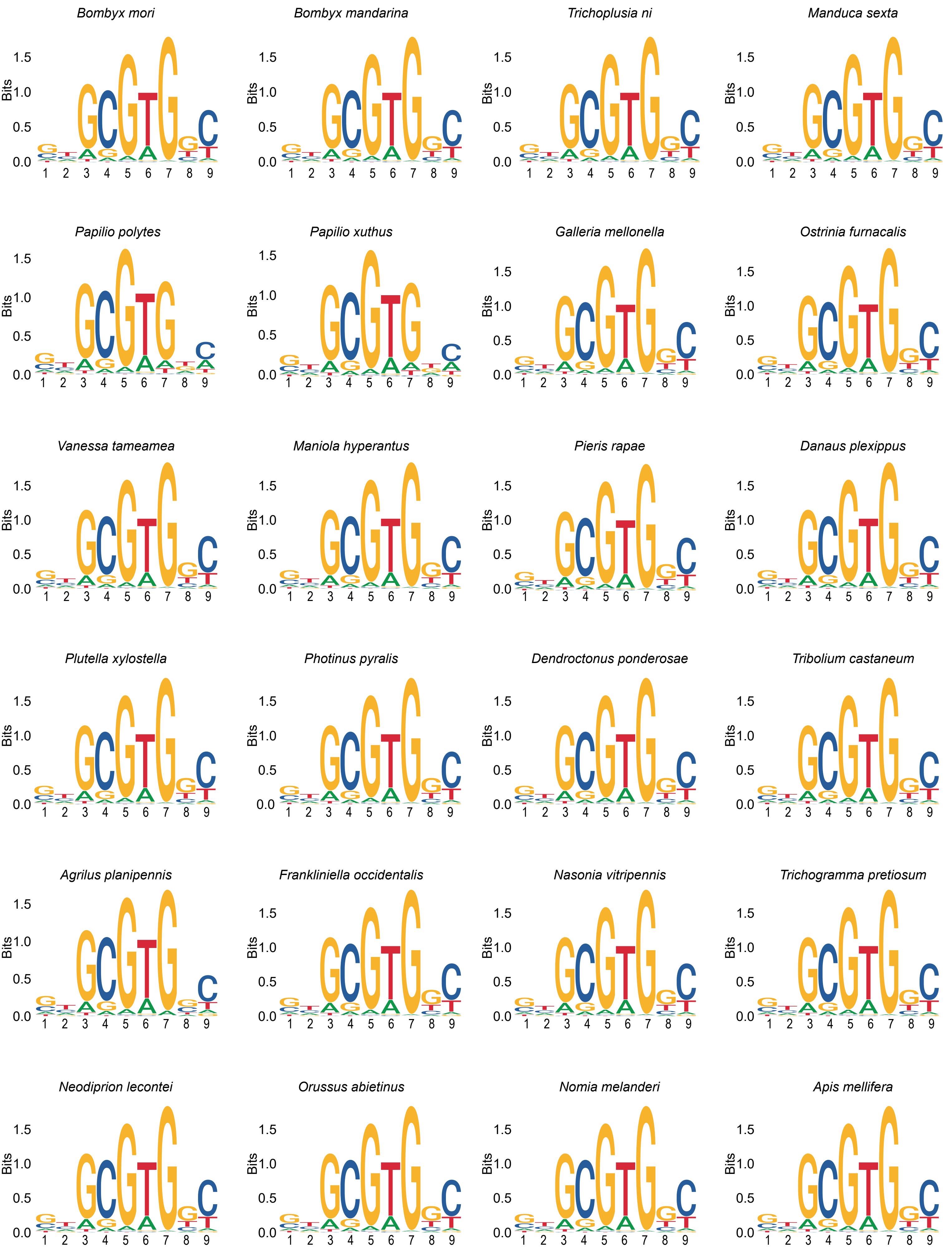

Bombyx mori
Bombyx mandarina
Trichoplusia ni
Manduca sexta
Papilio polytes
Papilio xuthus
Galleria mellonella
Ostrinia furnacalis
Vanessa tameamea
Maniola hyperantus
Pieris rapae
Danaus plexippus
Plutella xylostella
Photinus pyralis
Dendroctonus ponderosae
Tribolium castaneum
Agrilus planipennis
Frankliniella occidentalis
Nasonia vitripennis
Trichogramma pretiosum
Neodiprion lecontei
Orussus abietinus
Nomia melanderi
Apis mellifera
Bits
0.0
0.5
1.0
1.5
1 2 3 4 5 6 7 8 9

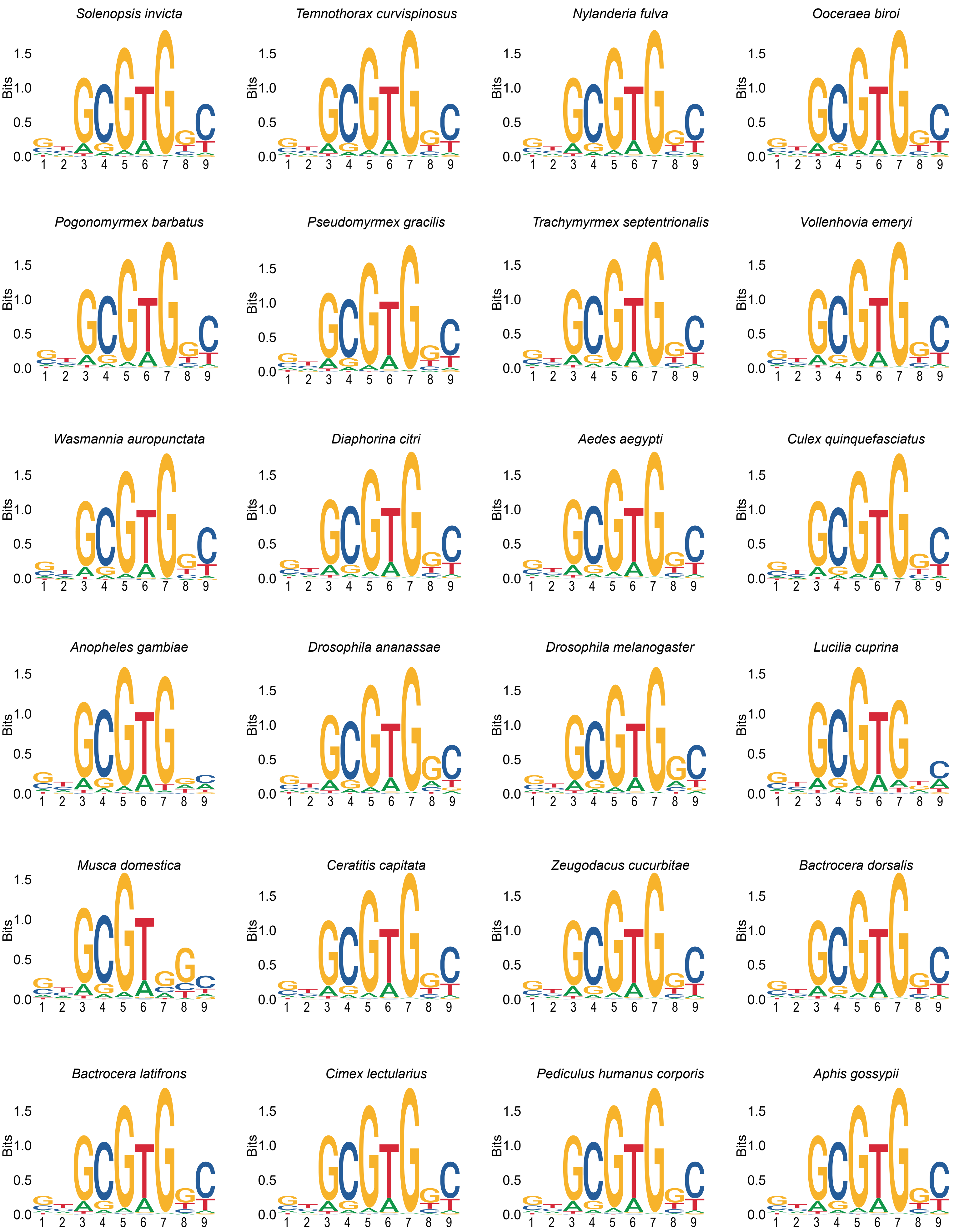
Solenopsis invicta
Temnothorax curvispinosus
Nylanderia fulva
Ooceraea biroi
Pogonomyrmex barbatus
Pseudomyrmex gracilis
Trachymyrmex septentrionalis
Vollenhovia emeryi
Wasmannia auropunctata
Diaphorina citri
Aedes aegypti
Culex quinquefasciatus
Anopheles gambiae
Drosophila ananassae
Drosophila melanogaster
Lucilia cuprina
Musca domestica
Ceratitis capitata
Zeugodacus cucurbitae
Bactrocera dorsalis
Bactrocera latifrons
Cimex lectularius
Pediculus humanus corporis
Aphis gossypii
Bits

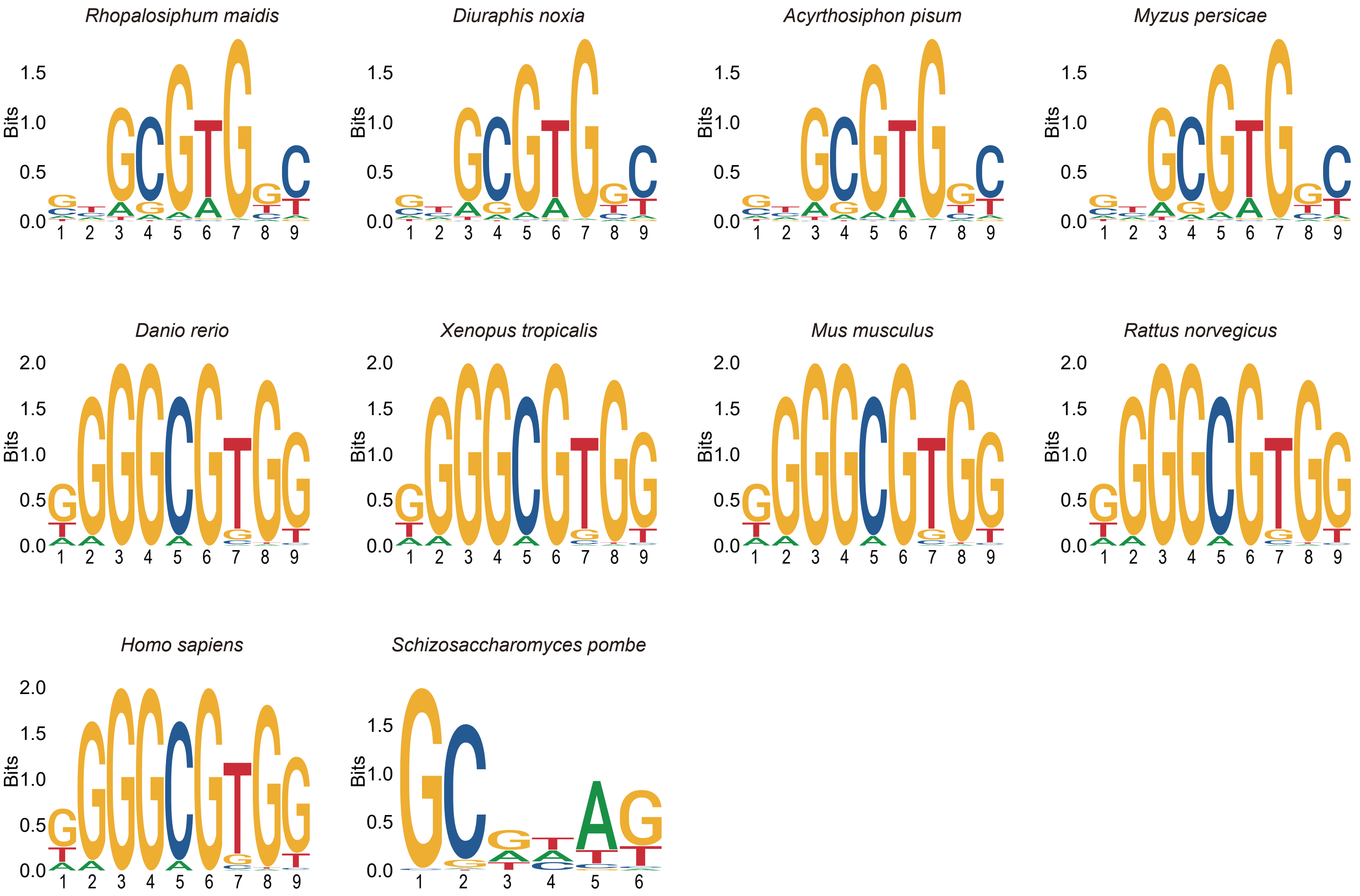


Fig.S8. The sequence logo of DNA binding site of orthologs of mamo in multiple species.
(A) The sequence logo of DNA binding site of orthologs of Bm-mamo-L; (B) the sequence logo of DNA binding site of orthologs of Bm-mamo-S.

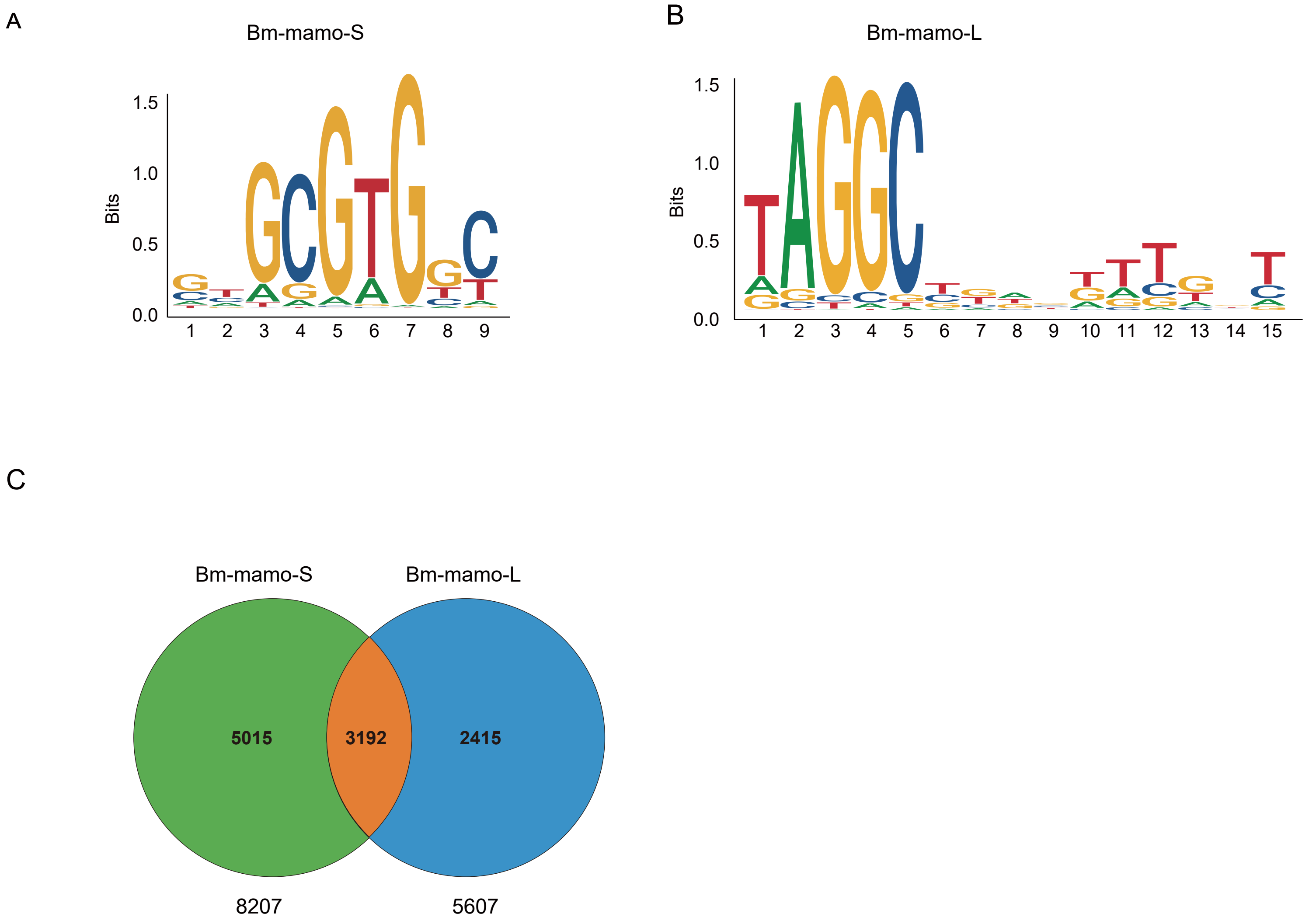


Fig.S9. The DNA-binding site sequence logo of Bm-mamo protein and the analysis of downstream target genes.
(A) The DNA-binding site sequence logo of Bm-mamo-s protein.
(B) The DNA-binding site sequence logo of Bm-mamo-L protein.
(C) According to MEME's FIMO program, potential downstream target genes of the Bm-mamo protein in the silkworm genome were identified.

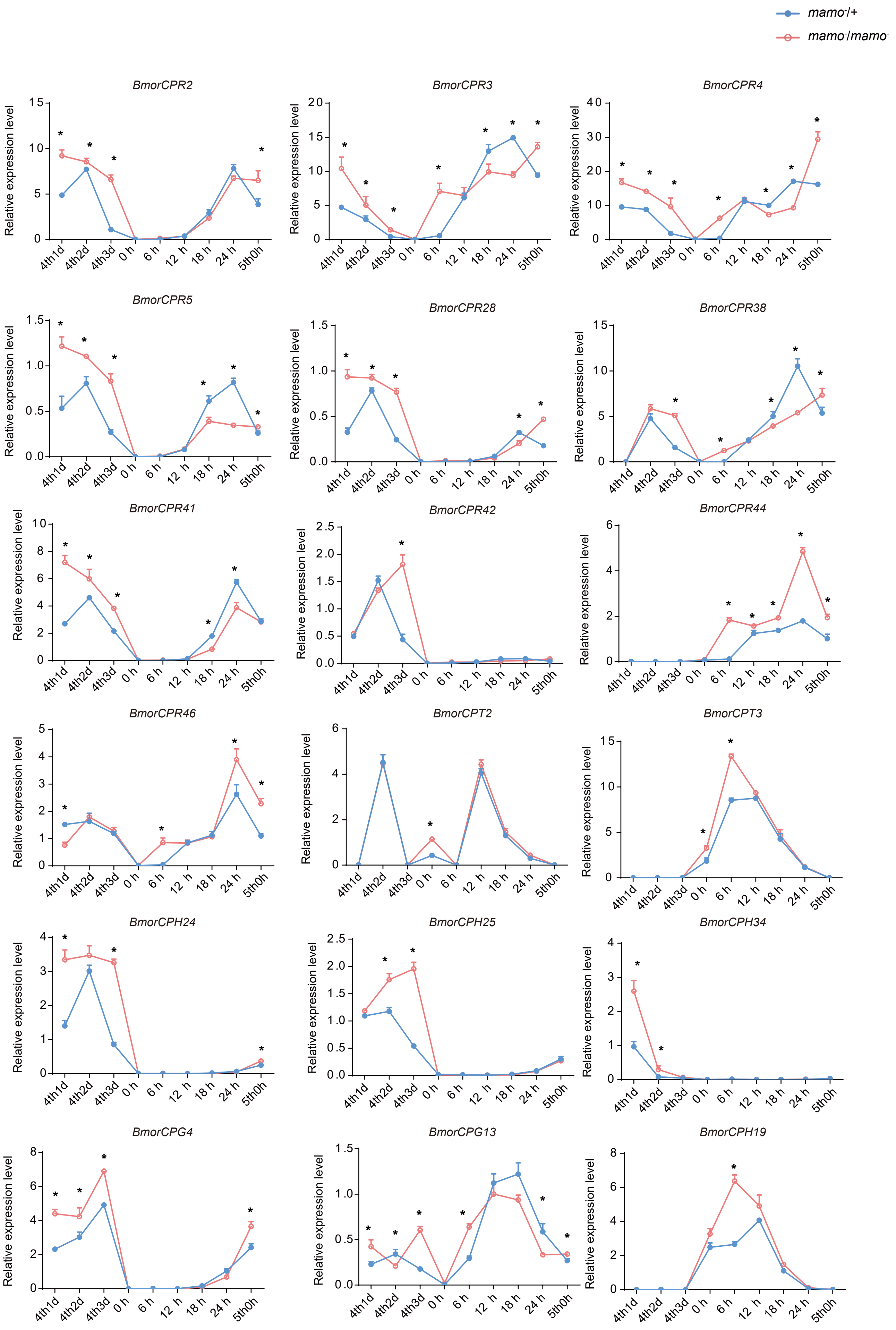


Fig.S10. The expression levels of cuticular protein genes were measured by qRT–PCR. The red line indicates the homozygote of *Bm-mamo* knockout individuals, and the blue line indicates the heterozygote of *Bm-mamo* knockout individuals. The means ± s.d.s. * P<0.05, paired Student's t test.

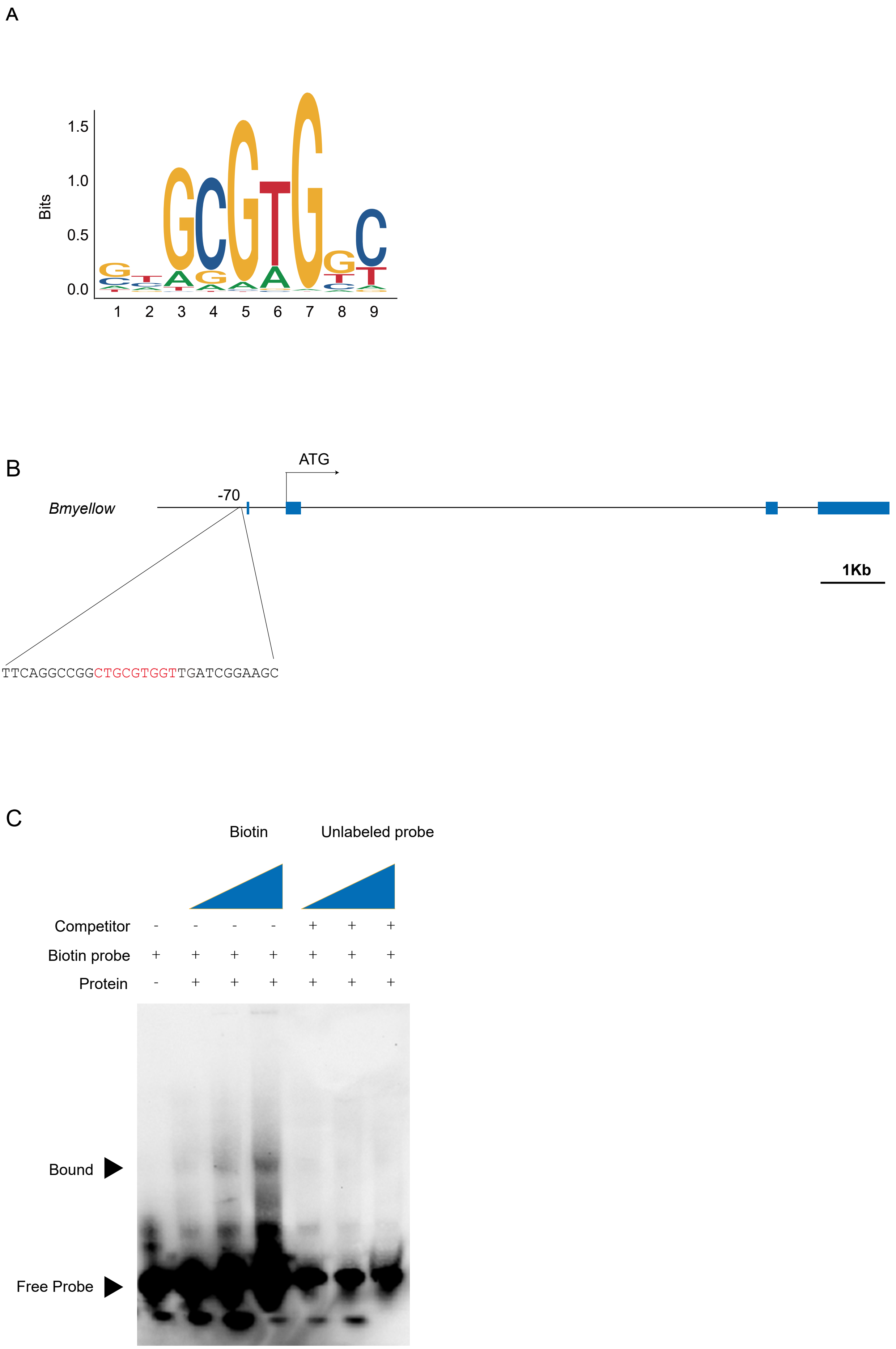


Fig.S11. Bm-mamo directly binds *yellow*.(A)The sequence logo of Bm-mamo-S, (B) the gene structureof *Bm-yellow*, which contains a Bm-mamo-S binding consensus sequence (red), (C) the EMSA analyses of Bm-mamo-S.

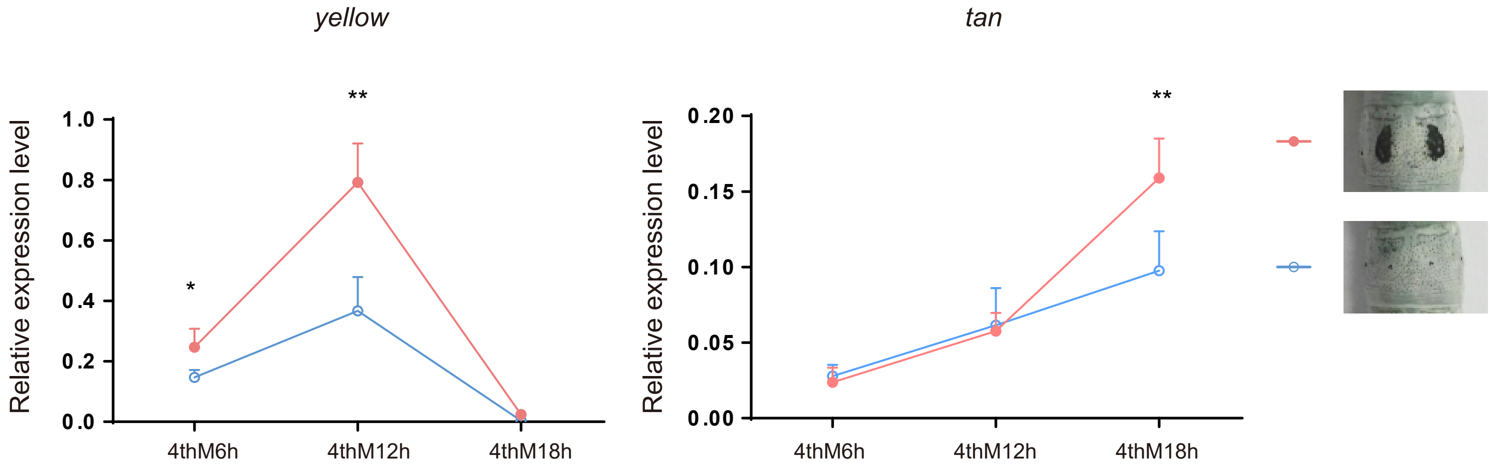


Fig.S12. The expression level of *yellow* and *tan* in the epidermis of pigmentation and non-pigmentation areas in Dazao strain.

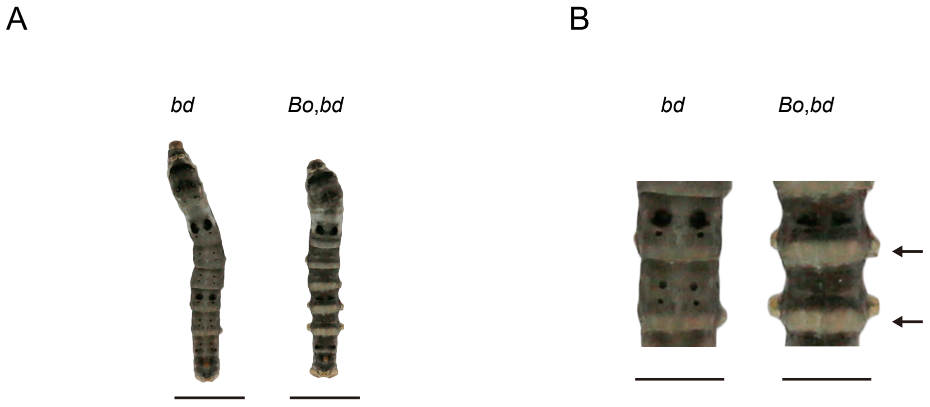


Fig.S13. Phenotypes of offspring from hybridization between *Bo* and *bd.*
(A) The phenotype of 5th instar and  day 3. the bars indicate 1 centimeter.
(B) The magnification of body segments. The bars indicate 0.5 cm. The indicated by the arrow is the area where the pigment is significantly reduced.

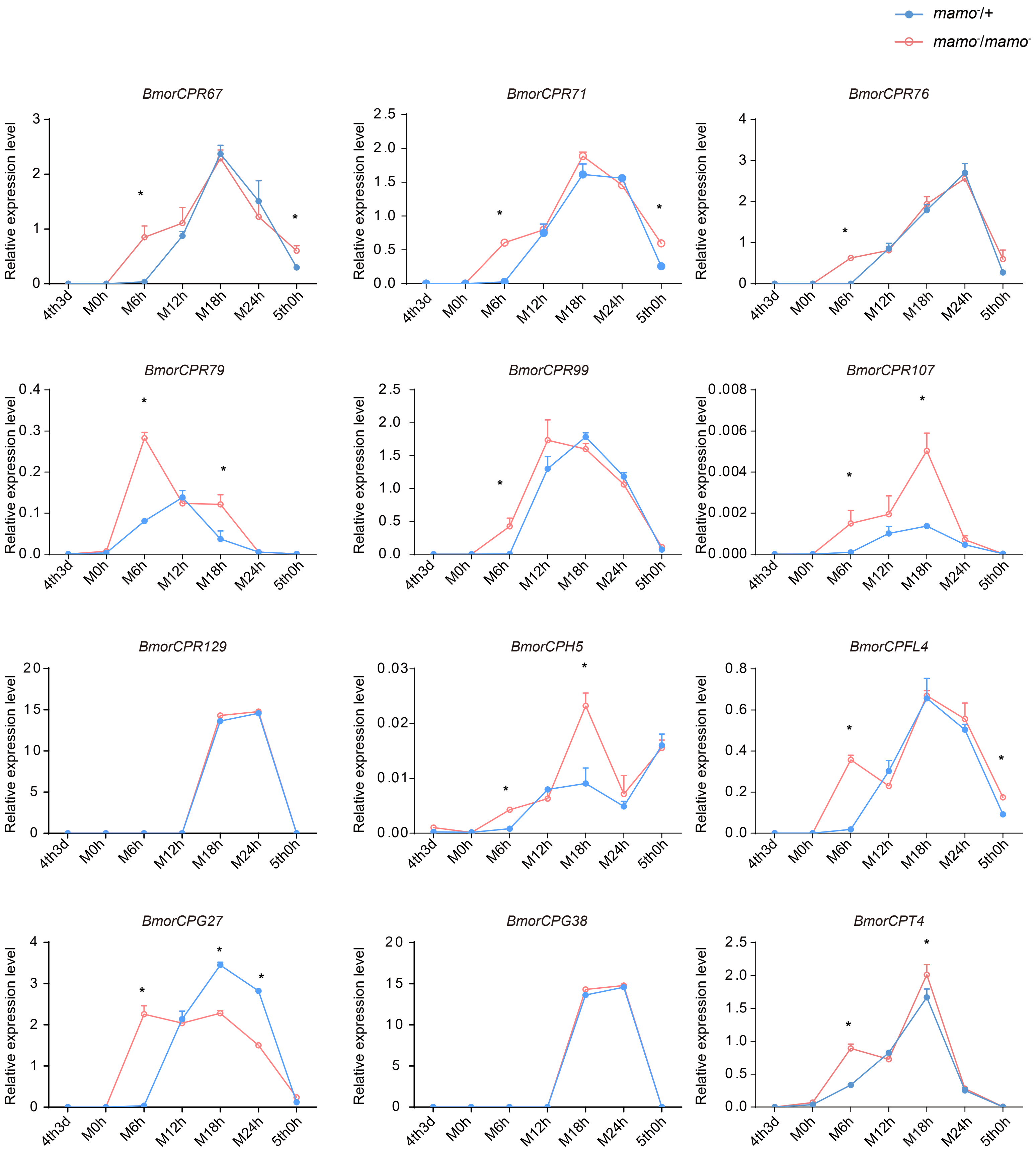


Fig.S14. Detection of the expression level of the orthologous genes of the cuticular protein genes highly expressed in the black marking of the *Papilio xuthus* larvae in the silkworm larvae.

The blue indicates heterozygous *Bm-mamo* gene knockout individuals, and the red indicates homozygous individulas.

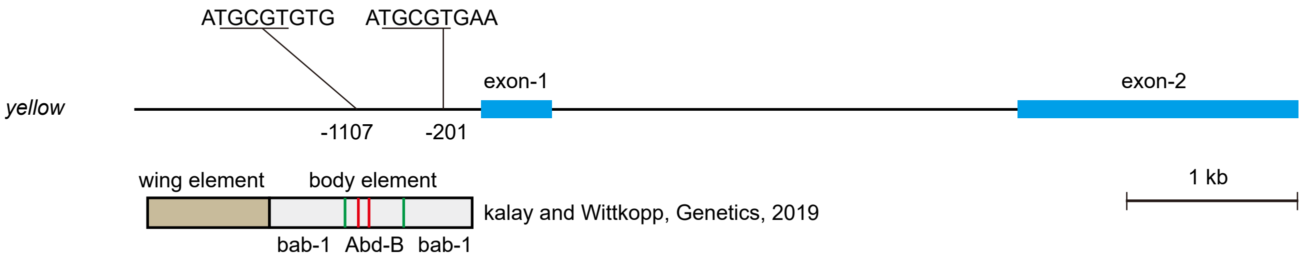


Fig.S15. The *Cis*-regulatory element model of *yellow* in *D.melanogaster*.

Blue blocks indicate exons, black lines indicate introns, brown block indicates wings element, gray block indicates body element, green lines indicate the binding site of bab-1, and the red lines indicate the binding site of Abd-B.

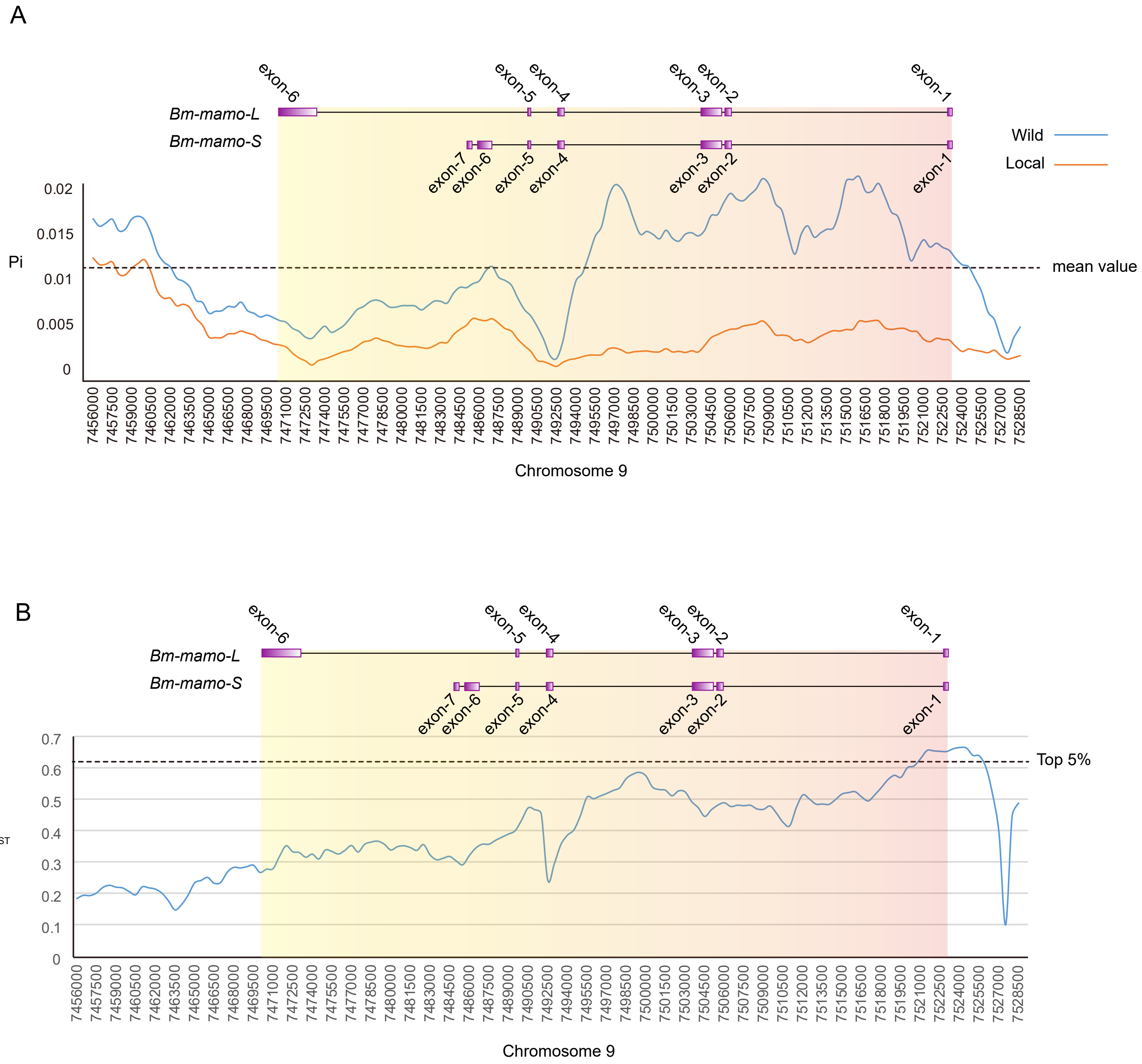


Fig. S16. Investigation of nucleotide diversity and fixation index in the genomic region of the *Bm-mamo* using 51 wild and 171 domestic silkworm strains.

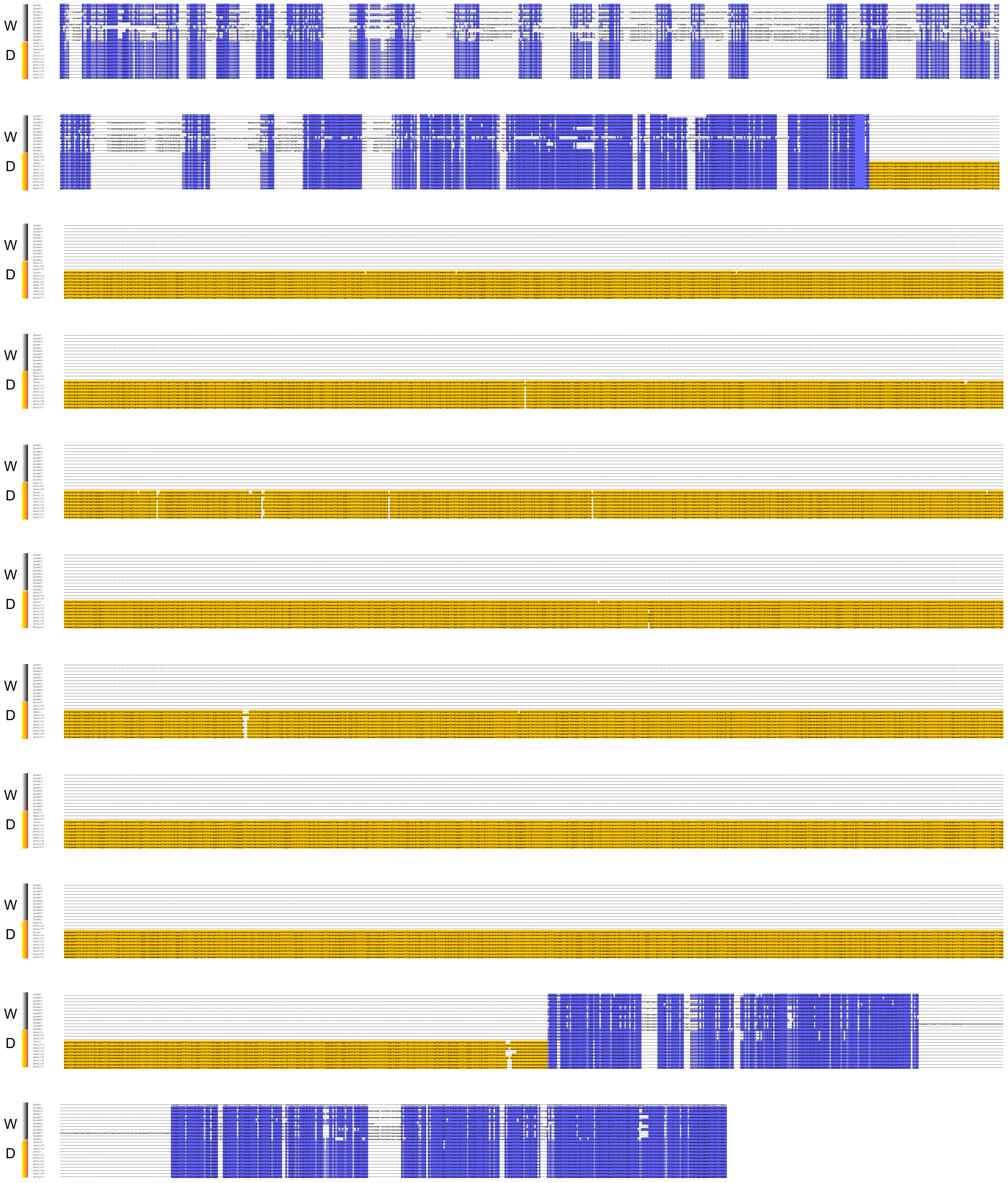


Fig. S17. The multiple sequence alignment of upstream of *Bm-mamo* among 12 wild silkworms and 12 domesticated silkworms. W indicates wild strains. D indicates domesticated strains. The yellow indicats long interspersed nuclear element (LINE).
