## Supplemental Table for "The BTB-ZF gene *Bm-mamo* regulates pigmentation in silkworm caterpillars"

Supplementary Table S1. *bd*$^f$ linkage analysis of BC1 population.

| Phenotype | BC1 number | DNA marker a | b | c | d | e | f | k | g | h | l | m | i | j |
|---|---|---|---|---|---|---|---|---|---|---|---|---|---|---|
| *bd*$^f$ | 71045 | H | H | H | A | A | A | A | A | A | A | A | A | A |
| *bd*$^f$ | 71115 | H | H | A | A | A | A | A | A | A | A | A | A | A |
| *bd*$^f$ | 91051 | H | A | A | A | A | A | A | A | A | A | A | A | A |
| *bd*$^f$ | 51068 | A | A | A | A | A | A | A | A | A | H | H | H | H |
| *bd*$^f$ | 61052 | A | A | A | A | A | A | A | A | A | A | A | A | H |
| *bd*$^f$ | 81077 | A | A | A | A | A | A | A | A | A | A | A | H | H |
| *bd*$^f$ | 91032 | A | A | A | A | A | A | A | A | A | A | A | H | H |
| *bd*$^f$ | 91074 | A | A | A | A | A | A | A | A | A | A | A | H | H |
| WT | 72017 | A | A | A | A | H | H | H | H | H | H | H | H | H |
| WT | 82013 | A | A | A | A | A | A | H | H | H | H | H | H | H |

The "A" indicates homozygous, and the "H" indicates heterozygous type.

Supplementary Table S2. The indel and snp in responsible region of *bd*$^{f}$

| number | Position | Reference sequence | ALT sequence |
|---|---|---|---|
| 1 | 7316807 | A | G |
| 2 | 7320156 | CAATAATT | C |
| 3 | 7320199 | T | C |
| 4 | 7320204 | TTCTTAATTGAGGGCAAC | T |
| 5 | 7320230 | T | A |
| 6 | 7324797 | G | A |
| 7 | 7324801 | T | C |
| 8 | 7324808 | G | A |
| 9 | 7324814 | A | C |
| 10 | 7324817 | C | T |
| 11 | 7324819 | C | A |
| 12 | 7325061 | T | C |
| 13 | 7332513 | GGATAGTTTTTTTT | G |
| 14 | 7332530 | T | TC |
| 15 | 7332555 | G | GT |
| 16 | 7332795 | G | T |
| 17 | 7332808 | A | G |
| 18 | 7332810 | C | G |
| 19 | 7332831 | G | A |
| 20 | 7333293 | C | T |
| 21 | 7333297 | A | G |
| 22 | 7333303 | C | A |
| 23 | 7333500 | C | CG |
| 24 | 7333639 | A | G |
| 25 | 7333677 | G | A |
| 26 | 7333822 | G | A |
| 27 | 7333831 | C | T |
| 28 | 7333851 | A | AAGG |
| 29 | 7334520 | T | A |
| 30 | 7334544 | C | T |
| 31 | 7334546 | G | T |
| 32 | 7334552 | A | C |
| 33 | 7334555 | A | T |
| 34 | 7334569 | G | C |
| 35 | 7334570 | A | C |
| 36 | 7334576 | A | G |
| 37 | 7334597 | G | C |
| 38 | 7334756 | A | G |
| 39 | 7335114 | AG | A |
| 40 | 7336847 | C | G |
| 41 | 7336848 | G | A |
| 42 | 7337265 | T | A |
| 43 | 7353264 | C | A |
| 44 | 7353464 | G | A |
| 45 | 7356217 | G | C |
| 46 | 7361617 | C | T |
| 47 | 7361980 | C | A |
| 48 | 7362171 | C | T |
| 49 | 7370432 | T | A |
| 50 | 7382697 | G | GT |
| 51 | 7396071 | A | C |
| 52 | 7415771 | A | G |
| 53 | 7417110 | AAT | A |
| 54 | 7439577 | A | T |
| 55 | 7439974 | T | A |

| | | | |
|---|---|---|---|
| 56 | 7440103 | A | T |
| 57 | 7440348 | AT | A |
| 58 | 7440485 | C | T |
| 59 | 7441732 | A | T |
| 60 | 7441917 | T | TA |
| 61 | 7441926 | T | G |
| 62 | 7442075 | A | G |
| 63 | 7446204 | C | CTTAGAGCGATCCCTATGGA |
| 64 | 7446212 | T | TCTCCGTAGACCC |
| 65 | 7446367 | G | A |
| 66 | 7446425 | G | C |
| 67 | 7446464 | A | T |
| 68 | 7446537 | T | TTTTTTTTTTTTTTTTTTTTTATTG CTTTGATGGA |
| 69 | 7446596 | G | C |
| 70 | 7446950 | GAAAAAAAA | G |
| 71 | 7446959 | A | ATTTTTTTTTT |
| 72 | 7446962 | C | T |
| 73 | 7446991 | A | T |
| 74 | 7447827 | G | GAATATATACACCTACCAAGCTT CATCTCTGCACTCCCCTAAGGTA GACTGTTAGAGAATGCCTATGGC ATTAAGTCCGCCTTTATACTTTAT AGTATAAGAAGTTATAAATAAAT AAATAAAAAATAAA |
| 75 | 7447880 | T | G |
| 76 | 7448447 | G | T |
| 77 | 7448454 | T | A |
| 78 | 7456427 | T | C |
| 79 | 7458064 | A | ATG |
| 80 | 7458724 | C | T |
| 81 | 7458737 | C | G |
| 82 | 7458763 | C | A |
| 83 | 7458829 | A | T |
| 84 | 7458839 | T | A |
| 85 | 7458850 | C | G |
| 86 | 7460891 | A | G |
| 87 | 7461384 | G | GTTTAA |
| 88 | 7461965 | T | TAA |
| 89 | 7462106 | T | TGAAGTGAGAAGAGTGGATATTG CACCACTTTCCCGA |
| 90 | 7462111 | T | TAGTAATGCG |
| 91 | 7462115 | T | C |
| 92 | 7462761 | G | GA |
| 93 | 7464027 | TAA | T |
| 94 | 7464129 | T | C |
| 95 | 7464549 | A | T |
| 96 | 7464694 | A | G |
| 97 | 7464873 | G | A |
| 98 | 7465153 | T | C |
| 99 | 7465197 | T | G |
| 100 | 7465649 | ATTACGATT | A |
| 101 | 7466488 | T | G |
| 102 | 7469552 | A | G |
| 103 | 7471051 | G | A |
| 104 | 7473092 | T | A |
| 105 | 7474382 | T | TA |
| 106 | 7474509 | A | AAAAAG |
| 107 | 7475952 | T | A |

| | | | |
|---|---|---|---|
| 108 | 7479011 | T | A |
| 109 | 7479141 | TCCATC | T |
| 110 | 7481220 | T | C |
| 111 | 7481380 | T | A |
| 112 | 7481382 | TATA | T |
| 113 | 7481715 | T | A |
| 114 | 7483572 | G | T |
| 115 | 7484658 | T | C |
| 116 | 7486058 | A | C |
| 117 | 7486171 | T | G |
| 118 | 7486292 | C | A |
| 119 | 7486372 | G | T |
| 120 | 7486383 | T | C |
| 121 | 7486407 | A | T |
| 122 | 7486408 | A | T |
| 123 | 7486518 | A | C |
| 124 | 7486540 | G | C |
| 125 | 7487789 | G | A |
| 126 | 7487794 | A | T |
| 127 | 7487825 | C | G |
| 128 | 7487880 | C | CT |
| 129 | 7487891 | T | C |
| 130 | 7487893 | TAA | T |
| 131 | 7487904 | T | C |
| 132 | 7487909 | T | A |
| 133 | 7487911 | G | A |
| 134 | 7487913 | T | C |
| 135 | 7487968 | A | G |
| 136 | 7487974 | T | C |
| 137 | 7487995 | A | G |
| 138 | 7487996 | G | A |
| 139 | 7487999 | T | C |
| 140 | 7488015 | A | T |
| 141 | 7488025 | C | T |
| 142 | 7488035 | G | A |
| 143 | 7488058 | G | A |
| 144 | 7488077 | TAA | T |
| 145 | 7488082 | A | ACC |
| 146 | 7488088 | A | T |
| 147 | 7488478 | T | TC |
| 148 | 7489305 | G | C |
| 149 | 7490124 | A | AC |
| 150 | 7491068 | A | C |
| 151 | 7491681 | T | TAA |
| 152 | 7491684 | ACT | A |
| 153 | 7498645 | T | TA |
| 154 | 7498775 | T | TA |
| 155 | 7508717 | T | C |
| 156 | 7508719 | C | G |
| 157 | 7508735 | A | G |
| 158 | 7508871 | T | TCGTC |
| 159 | 7509233 | G | A |
| 160 | 7509234 | A | AC |
| 161 | 7509361 | G | A |
| 162 | 7509385 | T | C |
| 163 | 7509386 | T | C |
| 164 | 7509399 | G | A |
| 165 | 7509408 | A | G |

| | | | |
|---|---|---|---|
| 166 | 7509418 | A | C |
| 167 | 7509448 | TGA | T |
| 168 | 7509461 | TGCACGG | T |
| 169 | 7509470 | A | T |
| 170 | 7509482 | G | C |
| 171 | 7509503 | T | C |
| 172 | 7509512 | A | T |
| 173 | 7509584 | A | G |
| 174 | 7509711 | A | AT |
| 175 | 7509743 | T | A |
| 176 | 7509767 | G | C |
| 177 | 7511669 | GTT | G |
| 178 | 7512904 | G | C |
| 179 | 7513238 | T | G |
| 180 | 7513253 | A | G |
| 181 | 7513277 | T | C |
| 182 | 7513312 | G | T |
| 183 | 7513327 | C | G |
| 184 | 7514492 | A | C |
| 185 | 7514783 | C | A |
| 186 | 7517289 | G | T |
| 187 | 7517316 | G | *,A |
| 188 | 7517318 | G | *,A |
| 189 | 7517320 | G | A |
| 190 | 7517374 | T | A |
| 191 | 7518059 | G | C |
| 192 | 7518388 | T | TCG |
| 193 | 7518970 | G | C |
| 194 | 7519653 | AT | A |
| 195 | 7521134 | T | G |
| 196 | 7523470 | G | GA |
| 197 | 7525558 | T | C |
| 198 | 7526835 | C | T |
| 199 | 7527254 | C | A |
| 200 | 7527362 | C | A |
| 201 | 7527386 | A | T |
| 202 | 7527429 | A | T |
| 203 | 7527489 | GA | G |
| 204 | 7527619 | T | C |
| 205 | 7527625 | GC | G |
| 206 | 7527627 | T | A |
| 207 | 7527635 | A | T |
| 208 | 7527649 | A | T |
| 209 | 7527675 | G | C |
| 210 | 7527682 | C | T |
| 211 | 7527697 | C | G |
| 212 | 7527709 | T | A |
| 213 | 7527713 | G | A |
| 214 | 7527720 | G | C |
| 215 | 7527737 | T | C |
| 216 | 7527769 | C | T |
| 217 | 7527794 | C | T |
| 218 | 7527806 | A | G |
| 219 | 7527828 | A | T |
| 220 | 7527872 | C | T |
| 221 | 7527879 | T | G |
| 222 | 7527925 | T | C |
| 223 | 7527976 | A | C |

| | | | |
|---|---|---|---|
| 224 | 7527982 | T | A |
| 225 | 7527989 | C | T |
| 226 | 7528045 | T | C |
| 227 | 7528054 | CAG | C |
| 228 | 7528089 | T | C |
| 229 | 7528137 | G | A |
| 230 | 7528351 | A | G |
| 231 | 7528354 | CG | C |
| 232 | 7528356 | T | C |
| 233 | 7528361 | G | A |
| 234 | 7528379 | G | A |
| 235 | 7528386 | T | A |
| 236 | 7528391 | T | A |
| 237 | 7528393 | A | T |
| 238 | 7528395 | A | C |
| 239 | 7528837 | A | G |
| 240 | 7529117 | C | T |
| 241 | 7529238 | C | A |
| 242 | 7529301 | C | T |
| 243 | 7529303 | C | T |
| 244 | 7529305 | C | T |
| 245 | 7529329 | C | T |
| 246 | 7529362 | G | C |
| 247 | 7529426 | C | A |
| 248 | 7529524 | GGCCCCCTTTGCCGGTTCCCA A | G |
| 249 | 7529601 | CG | C |
| 250 | 7529609 | TG | T |
| 251 | 7529700 | T | C |
| 252 | 7529714 | T | C |
| 253 | 7529715 | A | G |
| 254 | 7529732 | G | C |
| 255 | 7529831 | C | T |
| 256 | 7529901 | G | A |
| 257 | 7529938 | G | A |
| 258 | 7530529 | C | T |
| 259 | 7530679 | C | CTGTGTG |
| 260 | 7530906 | A | AT |
| 261 | 7531022 | C | T |
| 262 | 7531195 | G | C |
| 263 | 7531469 | C | T |
| 264 | 7531515 | A | G |
| 265 | 7531540 | A | G |
| 266 | 7532630 | AT | A |
| 267 | 7533035 | G | A |
| 268 | 7533101 | T | A |
| 269 | 7533249 | A | AT |
| 270 | 7535118 | G | GA |
| 271 | 7535150 | G | T |
| 272 | 7535674 | T | A |
| 273 | 7536199 | A | C |
| 274 | 7536497 | G | A |
| 275 | 7536518 | C | A |
| 276 | 7536558 | G | T |
| 277 | 7536993 | T | A |
| 278 | 7537411 | C | CG |
| 279 | 7537677 | T | C |
| 280 | 7538389 | A | T |

| | | | |
|---|---|---|---|
| 281 | 7540238 | T | C |
| 282 | 7540739 | ATTTTT | A |
| 283 | 7542588 | TACATGA | T |
| 284 | 7545321 | T | TTC |
| 285 | 7546576 | A | ATGT |
| 286 | 7547703 | T | A |
| 287 | 7547933 | A | G |
| 288 | 7547936 | GA | G |
| 289 | 7548447 | T | G |
| 290 | 7548448 | G | A |
| 291 | 7548461 | C | G |
| 292 | 7548464 | C | T |
| 293 | 7548465 | C | T |
| 294 | 7548491 | A | G |
| 295 | 7548498 | G | A |
| 296 | 7548519 | C | T |
| 297 | 7548524 | A | G |
| 298 | 7548724 | A | G |
| 299 | 7548748 | A | C |
| 300 | 7548800 | G | A |
| 301 | 7548838 | G | T |
| 302 | 7548843 | G | A |
| 303 | 7548850 | C | T |
| 304 | 7548912 | A | G |
| 305 | 7549670 | T | C |
| 306 | 7549826 | C | CATACATCGTATGCCATATTCGTTACTCAAGACGGTTA |
| 307 | 7550504 | A | ATT |
| 308 | 7550950 | A | AT |
| 309 | 7551972 | A | AT |
| 310 | 7553642 | T | A |
| 311 | 7554703 | CT | C |
| 312 | 7555964 | C | A |
| 313 | 7555999 | A | ATTTT |
| 314 | 7557362 | A | G |
| 315 | 7557393 | G | A |
| 316 | 7557406 | T | TGAG |
| 317 | 7557435 | A | G |
| 318 | 7557480 | C | T |
| 319 | 7557486 | A | T |
| 320 | 7564403 | AGACCAT | A |
| 321 | 7564413 | CAGATTTTGCGACCTGATCA | C |
| 322 | 7564483 | G | A |
| 323 | 7564505 | G | A |
| 324 | 7564513 | C | G |
| 325 | 7564519 | CACGA | C |
| 326 | 7564524 | T | TCCC |
| 327 | 7564526 | C | T |
| 328 | 7564588 | G | A |
| 329 | 7564625 | T | TG |
| 330 | 7564628 | CCG | C |
| 331 | 7564633 | C | G |
| 332 | 7564634 | ACG | A |
| 333 | 7564674 | G | A |
| 334 | 7569062 | T | C |
| 335 | 7569185 | G | A |
| 336 | 7569233 | G | A |
| 337 | 7569252 | T | G |

| | | | |
|---|---|---|---|
| 338 | 7569813 | G | T |
| 339 | 7569869 | T | C |
| 340 | 7569963 | T | TTGCAAGTATCATCATCATCATCATCATCATCAACAGCCCACAGTCGTCCACTGCTGGACATAGGCCTCTCCCAATCCACACCACAATAAGTGTAATATAATATAA |
| 341 | 7570882 | GATATGCT | G |
| 342 | 7570997 | A | ACTATTC |
| 343 | 7571002 | A | T |
| 344 | 7571007 | A | ATT |
| 345 | 7571192 | A | AT |
| 346 | 7571212 | T | TCA |
| 347 | 7571424 | A | G |
| 348 | 7571519 | A | T |
| 349 | 7571521 | C | T |
| 350 | 7571524 | T | TTTTTTTTAAGATTGAAGGATTACTGGTGG |
| 351 | 7571566 | G | A |
| 352 | 7571593 | C | T |
| 353 | 7571646 | C | A |
| 354 | 7571656 | A | T |
| 355 | 7571679 | G | A |
| 356 | 7571786 | T | C |
| 357 | 7574636 | T | C |
| 358 | 7574772 | C | T |
| 359 | 7575846 | A | AC |
| 360 | 7575848 | A | C |
| 361 | 7579171 | AAATAATTAGTTTAATTAAAAATTTTAGATCGCAGAATACTAAACAGTTATGT | A |
| 362 | 7585542 | A | T |
| 363 | 7586186 | G | A |
| 364 | 7588585 | T | TC |
| 365 | 7588586 | A | C |
| 366 | 7588589 | T | A |
| 367 | 7588590 | A | ACTCTGAACC |
| 368 | 7588593 | A | G |
| 369 | 7588597 | A | T |
| 370 | 7589395 | T | A |
| 371 | 7589396 | T | A |
| 372 | 7590492 | G | A |
| 373 | 7590507 | G | C |
| 374 | 7590529 | G | C |
| 375 | 7590531 | C | T |
| 376 | 7590535 | C | A |
| 377 | 7590702 | A | G |
| 378 | 7590713 | G | GATTTTA |
| 379 | 7590718 | G | C |
| 380 | 7590725 | A | C |
| 381 | 7590759 | A | G |
| 382 | 7591612 | G | T |
| 383 | 7591661 | T | C |
| 384 | 7591731 | A | T |
| 385 | 7592491 | A | G |
| 386 | 7592567 | A | T |
| 387 | 7630004 | A | AG |
| 388 | 7643232 | A | T |
| 389 | 7643238 | T | TTTTA |

| | | | |
|---|---|---|---|
| 390 | 7643289 | T | G |
| 391 | 7644620 | A | AG |
| 392 | 7645583 | G | A |
| 393 | 7646672 | G | T |
| 394 | 7646843 | C | CATAAA |
| 395 | 7646863 | G | GT |
| 396 | 7647457 | T | G |
| 397 | 7647973 | C | T |
| 398 | 7648842 | G | A |
| 399 | 7648851 | G | C |
| 400 | 7648854 | A | C |
| 401 | 7648866 | A | C |
| 402 | 7648918 | C | T |
| 403 | 7648951 | C | T |
| 404 | 7648974 | G | A |
| 405 | 7648977 | A | T |
| 406 | 7648992 | TC | T |
| 407 | 7649163 | A | G |
| 408 | 7649304 | A | G |
| 409 | 7649305 | G | A |
| 410 | 7649310 | T | A |
| 411 | 7649329 | C | T |
| 412 | 7649330 | A | G |
| 413 | 7649332 | T | C |
| 414 | 7649337 | G | A |
| 415 | 7649338 | C | A |
| 416 | 7649357 | G | A |
| 417 | 7649397 | T | A |
| 418 | 7649441 | A | T |
| 419 | 7649442 | C | A |
| 420 | 7650312 | A | AT |
| 421 | 7650330 | C | T |
| 422 | 7651156 | AT | A |
| 423 | 7651581 | T | A |
| 424 | 7652532 | C | T |
| 425 | 7652973 | GT | G |
| 426 | 7655115 | TA | T |
| 427 | 7656544 | G | A |
| 428 | 7656656 | T | C |
| 429 | 7656696 | C | T |
| 430 | 7656699 | G | A |
| 431 | 7656701 | G | A |
| 432 | 7656705 | C | T |
| 433 | 7656831 | C | T |
| 434 | 7657063 | A | T |
| 435 | 7657162 | A | C |
| 436 | 7657332 | C | T |
| 437 | 7657387 | C | A |
| 438 | 7657413 | A | T |
| 439 | 7665074 | A | G |
| 440 | 7665086 | T | G |
| 441 | 7665097 | C | G |
| 442 | 7665109 | G | T |
| 443 | 7665165 | G | A |
| 444 | 7665172 | G | C |
| 445 | 7665188 | G | T |
| 446 | 7672196 | A | T |
| 447 | 7673986 | G | A |

| | | | |
|---|---|---|---|
| 448 | 7674015 | T | G |
| 449 | 7674053 | T | G |
| 450 | 7675625 | C | T |
| 451 | 7675680 | T | C |
| 452 | 7676125 | A | C |
| 453 | 7676821 | G | A |
| 454 | 7676973 | T | TA |
| 455 | 7677441 | T | A |
| 456 | 7679225 | TA | T |
| 457 | 7679784 | T | C |
| 458 | 7681014 | A | AACGT |
| 459 | 7681058 | G | A |
| 460 | 7681606 | T | C |
| 461 | 7681610 | G | A |
| 462 | 7681612 | G | A |
| 463 | 7681624 | C | T |
| 464 | 7681625 | T | G |
| 465 | 7681629 | A | G |
| 466 | 7681642 | G | A |
| 467 | 7681658 | C | T |
| 468 | 7681665 | T | C |
| 469 | 7681778 | G | A |
| 470 | 7681780 | A | T |
| 471 | 7681781 | T | C |
| 472 | 7681785 | C | CGT |
| 473 | 7681788 | T | C |
| 474 | 7681789 | T | TAACTCCGCAC |
| 475 | 7681792 | G | A |
| 476 | 7681794 | T | C |
| 477 | 7681797 | T | C |
| 478 | 7681804 | G | C |
| 479 | 7681811 | T | TCG |
| 480 | 7681817 | C | G |
| 481 | 7681854 | A | G |
| 482 | 7682567 | G | A |
| 483 | 7683565 | G | T |
| 484 | 7684449 | A | ACT |
| 485 | 7684935 | T | A |
| 486 | 7684936 | G | A |
| 487 | 7684944 | T | C |
| 488 | 7684945 | G | A |
| 489 | 7684968 | A | T |
| 490 | 7686030 | C | CA |
| 491 | 7689020 | A | G |
| 492 | 7690131 | A | G |
| 493 | 7690724 | T | TGAAC |
| 494 | 7691590 | T | TA |

**Supplementary Table S3 The binding site analysis of Bm-mamo-L**

Using the position weight matrix of mamo-L protein to search the 2000 base pair regions upstream and downstream of the silkworm prediction gene.

| Number | motif_id | gene ID | up/down stream | start | stop | strand | score | p-value | q-value | matched_sequence |
|---|---|---|---|---|---|---|---|---|---|---|
| 1 | mamo-L | *BGIBMGA000004* | down | 131 | 145 | - | 10.8226 | 7.58E-05 | 1 | AAGGCGGGCGATTTT |
| 2 | mamo-L | *BGIBMGA000005* | up | 1470 | 1484 | - | 10.5726 | 9.33E-05 | 1 | TAGGCTAATATTTAT |
| 3 | mamo-L | *BGIBMGA000006* | up | 1289 | 1303 | + | 11.4435 | 4.31E-05 | 1 | GAGGCCCGGTTTTTT |
| 4 | mamo-L | *BGIBMGA000011* | down | 1676 | 1690 | - | 10.8629 | 7.33E-05 | 1 | TAGGCTCTGATTAGC |
| 5 | mamo-L | *BGIBMGA000012* | down | 515 | 529 | - | 12.5323 | 1.39E-05 | 1 | TAGGCCTTTTATGAT |
| 6 | mamo-L | *BGIBMGA000013* | up | 950 | 964 | - | 11.2177 | 5.34E-05 | 1 | TAGGCTCAGTCTTTT |
| 7 | mamo-L | *BGIBMGA000014* | up | 625 | 639 | + | 13.2258 | 6.08E-06 | 1 | TAGGCTTACAATGCT |
| 8 | mamo-L | *BGIBMGA000014* | up | 733 | 747 | - | 11.5323 | 3.96E-05 | 1 | TAGGCGAACATTTGT |
| 9 | mamo-L | *BGIBMGA000014* | up | 1311 | 1325 | - | 10.5726 | 9.33E-05 | 1 | TAGGCTAATATTTAT |
| 10 | mamo-L | *BGIBMGA000015* | up | 1355 | 1369 | + | 10.5726 | 9.33E-05 | 1 | TAGGCTAATATTTAT |
| 11 | mamo-L | *BGIBMGA000016* | up | 1911 | 1925 | - | 13.1532 | 6.65E-06 | 1 | AAGGCTGTTGTTGCT |
| 12 | mamo-L | *BGIBMGA000017* | up | 611 | 625 | - | 11.4194 | 4.42E-05 | 1 | TAGGCTGAATTCATT |
| 13 | mamo-L | *BGIBMGA000021* | down | 938 | 952 | + | 12.1935 | 2.05E-05 | 1 | TAGGCGGGCTTTGAA |
| 14 | mamo-L | *BGIBMGA000022* | up | 209 | 223 | + | 12.1935 | 2.05E-05 | 1 | TAGGCGGGCTTTGAA |
| 15 | mamo-L | *BGIBMGA000026* | down | 1663 | 1677 | - | 10.6613 | 8.67E-05 | 1 | TAGGCTTAGAATAGC |
| 16 | mamo-L | *BGIBMGA000027* | down | 63 | 77 | + | 13.8871 | 2.34E-06 | 1 | TAGGCCGTTTTTTTT |
| 17 | mamo-L | *BGIBMGA000030* | down | 1167 | 1181 | + | 11.0081 | 6.45E-05 | 1 | TAGGCTCTCAATTCT |
| 18 | mamo-L | *BGIBMGA000031* | down | 1825 | 1839 | - | 11.0081 | 6.45E-05 | 1 | TAGGCTCTCAATTCT |
| 19 | mamo-L | *BGIBMGA000033* | down | 1329 | 1343 | - | 12.7097 | 1.13E-05 | 1 | TAGGCTGCTGATGTT |
| 20 | mamo-L | *BGIBMGA000035* | up | 412 | 426 | + | 11.9274 | 2.69E-05 | 1 | TAGGCTAGGTTTCTT |
| 21 | mamo-L | *BGIBMGA000036* | down | 1030 | 1044 | - | 11.75 | 3.21E-05 | 1 | TAGGCTCGAATTGGC |
| 22 | mamo-L | *BGIBMGA000038* | down | 1280 | 1294 | - | 10.75 | 8.06E-05 | 1 | GAGGCCGATTGTGCC |
| 23 | mamo-L | *BGIBMGA000042* | down | 1316 | 1330 | - | 11.7742 | 3.14E-05 | 1 | TAGGCTGATTTGTTC |
| 24 | mamo-L | *BGIBMGA000046* | down | 178 | 192 | - | 11.4113 | 4.45E-05 | 1 | TAGGCTTTTTTCCCT |
| 25 | mamo-L | *BGIBMGA000047* | up | 814 | 828 | - | 10.9839 | 6.59E-05 | 1 | TAGGCTTTTTTTAGA |
| 26 | mamo-L | *BGIBMGA000047* | up | 1530 | 1544 | - | 10.6452 | 8.79E-05 | 1 | TAGCCTCTGGTTGTT |
| 27 | mamo-L | *BGIBMGA000048* | up | 129 | 143 | - | 10.7419 | 8.11E-05 | 1 | TAGGGCGTTGTTGGT |
| 28 | mamo-L | *BGIBMGA000049* | up | 1518 | 1532 | + | 12.25 | 1.92E-05 | 1 | TAGGCTCAGGTTAAT |
| 29 | mamo-L | *BGIBMGA000050* | up | 697 | 711 | - | 11.0806 | 6.05E-05 | 1 | TAGGCTTATGTGGTG |
| 30 | mamo-L | *BGIBMGA000051* | up | 1744 | 1758 | - | 11.4516 | 4.28E-05 | 1 | AAGGCTCACTATGTT |
| 31 | mamo-L | *BGIBMGA000052* | up | 1147 | 1161 | + | 10.5806 | 9.27E-05 | 1 | TAGGGCGACGTTGGC |
| 32 | mamo-L | *BGIBMGA000054* | up | 869 | 883 | + | 10.8065 | 7.69E-05 | 1 | TAGGCCACCGTGGTT |
| 33 | mamo-L | *BGIBMGA000055* | down | 1800 | 1814 | - | 11.4435 | 4.31E-05 | 1 | GAGGCCCGGTTTTTT |
| 34 | mamo-L | *BGIBMGA000055* | down | 671 | 685 | - | 11.4032 | 4.49E-05 | 1 | GAGGCTGAAGTCGTT |
| 35 | mamo-L | *BGIBMGA000060* | up | 911 | 925 | - | 11.0161 | 6.40E-05 | 1 | TCGGCGGGGTTTGTT |
| 36 | mamo-L | *BGIBMGA000062* | down | 988 | 1002 | - | 11.9274 | 2.69E-05 | 1 | TAGGCTCCGTGTGCT |
| 37 | mamo-L | *BGIBMGA000063* | up | 655 | 669 | - | 13.3468 | 5.19E-06 | 1 | TAGGCCCGTTTTTGT |
| 38 | mamo-L | *BGIBMGA000065* | up | 1299 | 1313 | + | 10.9194 | 6.98E-05 | 1 | TAGGCTACTTTTTTT |
| 39 | mamo-L | *BGIBMGA000065* | up | 9 | 23 | + | 10.5726 | 9.33E-05 | 1 | TAGGCTAATATTTAT |
| 40 | mamo-L | *BGIBMGA000066* | up | 308 | 322 | + | 14.5161 | 8.88E-07 | 1 | TAGGCCGATATTGGT |
| 41 | mamo-L | *BGIBMGA000067* | up | 1602 | 1616 | + | 10.5726 | 9.33E-05 | 1 | TAGGCTAATATTTAT |
| 42 | mamo-L | *BGIBMGA000070* | up | 190 | 204 | + | 11.8871 | 2.81E-05 | 1 | TAGGCCGGTATTGCA |
| 43 | mamo-L | *BGIBMGA000073* | down | 1869 | 1883 | + | 12.0645 | 2.34E-05 | 1 | TAGGCGGGCTTCGGA |
| 44 | mamo-L | *BGIBMGA000074* | down | 1458 | 1472 | - | 12.4677 | 1.49E-05 | 1 | TAGGCTTCTATTGCT |
| 45 | mamo-L | *BGIBMGA000076* | up | 845 | 859 | - | 10.5726 | 9.33E-05 | 1 | TAGGCTAATATTTAT |
| 46 | mamo-L | *BGIBMGA000077* | down | 1615 | 1629 | + | 10.9194 | 6.98E-05 | 1 | TAGGCTACTTTTTTT |
| 47 | mamo-L | *BGIBMGA000077* | up | 693 | 707 | - | 10.5726 | 9.33E-05 | 1 | TAGGCTAATATTTAT |
| 48 | mamo-L | *BGIBMGA000079* | down | 1214 | 1228 | - | 11.1855 | 5.50E-05 | 1 | TAGGCTGGGTGTTGG |
| 49 | mamo-L | *BGIBMGA000080* | down | 946 | 960 | - | 11.1129 | 5.88E-05 | 1 | TAGGCTGCGGGTTTC |
| 50 | mamo-L | *BGIBMGA000080* | up | 640 | 654 | - | 11 | 6.50E-05 | 1 | TAGGCATGCGTGTGT |
| 51 | mamo-L | *BGIBMGA000083* | up | 1110 | 1124 | - | 10.9435 | 6.83E-05 | 1 | TAGGCTTAGTGGTGC |
| 52 | mamo-L | *BGIBMGA000086* | up | 1558 | 1572 | + | 11.621 | 3.63E-05 | 1 | TAGGCAGGCGTTTCC |
| 53 | mamo-L | *BGIBMGA000086* | up | 1283 | 1297 | + | 10.879 | 7.22E-05 | 1 | TAGGCCAATAGTGCT |
| 54 | mamo-L | *BGIBMGA000088* | down | 921 | 935 | + | 12.8306 | 9.76E-06 | 1 | AAGGCTGTGTTTTGT |
| 55 | mamo-L | *BGIBMGA000090* | down | 1635 | 1649 | - | 11.7903 | 3.08E-05 | 1 | TAGGCCTGGTTTTTG |
| 56 | mamo-L | *BGIBMGA000091* | down | 456 | 470 | - | 12.4113 | 1.60E-05 | 1 | AAGGCCGATTTTTGT |
| 57 | mamo-L | *BGIBMGA000091* | down | 392 | 406 | - | 10.621 | 8.97E-05 | 1 | TAGGCCTCATTTATT |
| 58 | mamo-L | *BGIBMGA000093* | up | 1465 | 1479 | + | 11.2016 | 5.41E-05 | 1 | TAGGCTGACTTTCCG |
| 59 | mamo-L | *BGIBMGA000093* | down | 820 | 834 | - | 10.5726 | 9.33E-05 | 1 | TAGGCTAATATTTAT |
| 60 | mamo-L | *BGIBMGA000096* | up | 900 | 914 | + | 11.7581 | 3.18E-05 | 1 | TAGGCCGTAAATGTT |
| 61 | mamo-L | *BGIBMGA000097* | up | 1643 | 1657 | + | 12.4919 | 1.46E-05 | 1 | TAGGCCGTCTTTTGA |
| 62 | mamo-L | *BGIBMGA000100* | up | 1486 | 1500 | - | 11.7984 | 3.06E-05 | 1 | TAGGCTTTGCTTTTT |
| 63 | mamo-L | *BGIBMGA000103* | up | 373 | 387 | - | 14.2016 | 1.47E-06 | 1 | TAGGCCGTTTTTTGT |
| 64 | mamo-L | *BGIBMGA000105* | up | 1862 | 1876 | - | 10.8468 | 7.42E-05 | 1 | TAGGCCAGCGTCCGT |
| 65 | mamo-L | *BGIBMGA000105* | up | 744 | 758 | - | 10.5726 | 9.33E-05 | 1 | TAGGCTAATATTTAT |
| 66 | mamo-L | *BGIBMGA000108* | down | 1639 | 1653 | + | 11.6048 | 3.70E-05 | 1 | TAGGCCTGCAGCGCT |
| 67 | mamo-L | *BGIBMGA000108* | down | 803 | 817 | - | 11.4597 | 4.25E-05 | 1 | TAGGCGGGGTGGTGT |
| 68 | mamo-L | *BGIBMGA000112* | up | 384 | 398 | - | 13.1129 | 6.98E-06 | 1 | TAGGCTCTTGGTGGT |
| 69 | mamo-L | *BGIBMGA000115* | up | 349 | 363 | + | 12.0645 | 2.34E-05 | 1 | TAGGCGGGCTTCGGA |
| 70 | mamo-L | *BGIBMGA000121* | up | 534 | 548 | + | 10.9194 | 6.98E-05 | 1 | TAGGCTACTTTTTTT |
| 71 | mamo-L | *BGIBMGA000122* | down | 600 | 614 | + | 10.8468 | 7.42E-05 | 1 | TAGGCCAGCGTCCGT |
| 72 | mamo-L | *BGIBMGA000122* | down | 1718 | 1732 | + | 10.5726 | 9.33E-05 | 1 | TAGGCTAATATTTAT |
| 73 | mamo-L | *BGIBMGA000124* | up | 1935 | 1949 | + | 14.2016 | 1.47E-06 | 1 | TAGGCCGTTTTTTGT |
| 74 | mamo-L | *BGIBMGA000125* | up | 862 | 876 | + | 10.8306 | 7.52E-05 | 1 | GAGGCTCTTTTTTTT |
| 75 | mamo-L | *BGIBMGA000126* | down | 1545 | 1559 | - | 11.371 | 4.62E-05 | 1 | TAGGCTGTATAGGCT |
| 76 | mamo-L | *BGIBMGA000131* | up | 1674 | 1688 | + | 12.2661 | 1.89E-05 | 1 | TAGGCTGTGTGCTGT |
| 77 | mamo-L | *BGIBMGA000131* | up | 1506 | 1520 | + | 11.4435 | 4.31E-05 | 1 | TAGGCTTGAGAGGCT |
| 78 | mamo-L | *BGIBMGA000136* | down | 641 | 655 | - | 12.2742 | 1.87E-05 | 1 | TAGGCTTTTTTTGTG |
| 79 | mamo-L | *BGIBMGA000136* | up | 1874 | 1888 | - | 10.5645 | 9.40E-05 | 1 | TAGGCTTACTAGCTT |
| 80 | mamo-L | *BGIBMGA000138* | up | 624 | 638 | + | 10.5726 | 9.33E-05 | 1 | TAGGCTAATATTTAT |
| 81 | mamo-L | *BGIBMGA000139* | down | 146 | 160 | + | 13.1935 | 6.33E-06 | 1 | TAGGCCGGTTTTACT |
| 82 | mamo-L | *BGIBMGA000139* | down | 1489 | 1503 | + | 12.5242 | 1.40E-05 | 1 | TAGGCAGATTTTGGC |
| 83 | mamo-L | *BGIBMGA000142* | up | 327 | 341 | - | 12.7984 | 1.02E-05 | 1 | TAGGCCTTTTTTGGA |
| 84 | mamo-L | *BGIBMGA000143* | up | 1665 | 1679 | + | 10.5806 | 9.27E-05 | 1 | GAGGCGCGCTGTGTT |
| 85 | mamo-L | *BGIBMGA000153* | up | 1679 | 1693 | + | 13.0403 | 7.65E-06 | 1 | TAGGCCTTGATTTTT |

| | | | | | | | | | | |
|---|---|---|---|---|---|---|---|---|---|---|
| 86 | mamo-L | *BGIBMGA000153* | up | 1043 | 1057 | + | 10.5887 | 9.21E-05 | 1 | AAGGCCGGTGTGGCC |
| 87 | mamo-L | *BGIBMGA000153* | down | 1516 | 1530 | - | 10.5726 | 9.33E-05 | 1 | TAGGCTAATATTTAT |
| 88 | mamo-L | *BGIBMGA000156* | up | 365 | 379 | - | 10.4919 | 9.96E-05 | 1 | TAGGCTGTGTCTGTG |
| 89 | mamo-L | *BGIBMGA000161* | down | 928 | 942 | - | 13.2903 | 5.60E-06 | 1 | TAGGCTCTCGGTGTT |
| 90 | mamo-L | *BGIBMGA000161* | down | 1262 | 1276 | - | 11.3952 | 4.52E-05 | 1 | TAGGCTCATTAGGCT |
| 91 | mamo-L | *BGIBMGA000161* | down | 1253 | 1267 | - | 11.2339 | 5.26E-05 | 1 | TAGGCTCATTTTCCC |
| 92 | mamo-L | *BGIBMGA000161* | up | 1146 | 1160 | - | 10.9194 | 6.98E-05 | 1 | TAGGCTACTTTTTTT |
| 93 | mamo-L | *BGIBMGA000162* | up | 89 | 103 | + | 10.9597 | 6.73E-05 | 1 | TAGGCTGATAACGAT |
| 94 | mamo-L | *BGIBMGA000164* | up | 1438 | 1452 | - | 10.5 | 9.90E-05 | 1 | AAGGCTGGGTATACT |
| 95 | mamo-L | *BGIBMGA000166* | up | 1813 | 1827 | + | 12.6855 | 1.16E-05 | 1 | TAGGCCTTTTTTATT |
| 96 | mamo-L | *BGIBMGA000166* | down | 1847 | 1861 | - | 10.9194 | 6.98E-05 | 1 | TAGGCTACTTTTTTT |
| 97 | mamo-L | *BGIBMGA000167* | down | 550 | 564 | - | 11.2984 | 4.95E-05 | 1 | TAGGCTTACTTTCCA |
| 98 | mamo-L | *BGIBMGA000168* | up | 492 | 506 | + | 11.0887 | 6.01E-05 | 1 | TAGCCTGAGGTTGTC |
| 99 | mamo-L | *BGIBMGA000174* | down | 200 | 214 | - | 11.9677 | 2.59E-05 | 1 | TAGGCTTTATATTGT |
| 100 | mamo-L | *BGIBMGA000174* | down | 1643 | 1657 | - | 10.5726 | 9.33E-05 | 1 | TAGGCTAATATTTAT |
| 101 | mamo-L | *BGIBMGA000176* | down | 1874 | 1888 | + | 11.871 | 2.85E-05 | 1 | TAGGCTTTATTTGTA |
| 102 | mamo-L | *BGIBMGA000177* | up | 222 | 236 | + | 10.7984 | 7.74E-05 | 1 | TAGGCTCATGTTTCG |
| 103 | mamo-L | *BGIBMGA000177* | up | 908 | 922 | + | 10.5403 | 9.58E-05 | 1 | TAGGCTGTATGGTGT |
| 104 | mamo-L | *BGIBMGA000178* | down | 817 | 831 | - | 11.1048 | 5.92E-05 | 1 | TAGGCCAGTTTTGCA |
| 105 | mamo-L | *BGIBMGA000178* | up | 857 | 871 | - | 10.5968 | 9.15E-05 | 1 | GAGGCTATGTTTTTT |
| 106 | mamo-L | *BGIBMGA000182* | down | 862 | 876 | + | 14.6532 | 6.86E-07 | 1 | TAGGCTTAGTTTTTT |
| 107 | mamo-L | *BGIBMGA000188* | up | 1186 | 1200 | + | 10.5887 | 9.21E-05 | 1 | TAGGCGGAGTGTTGG |
| 108 | mamo-L | *BGIBMGA000190* | down | 1836 | 1850 | + | 12.1129 | 2.22E-05 | 1 | AAGGCCTATTTTTGT |
| 109 | mamo-L | *BGIBMGA000191* | up | 528 | 542 | + | 12.1129 | 2.22E-05 | 1 | AAGGCCTATTTTTGT |
| 110 | mamo-L | *BGIBMGA000192* | up | 981 | 995 | - | 10.9113 | 7.02E-05 | 1 | TAGGCTCGTTTGTGC |
| 111 | mamo-L | *BGIBMGA000196* | down | 1092 | 1106 | + | 10.9355 | 6.88E-05 | 1 | TAGGCTTCCGGTGGA |
| 112 | mamo-L | *BGIBMGA000197* | up | 1927 | 1941 | - | 12.1129 | 2.22E-05 | 1 | AAGGCCTATTTTTGT |
| 113 | mamo-L | *BGIBMGA000198* | up | 1962 | 1976 | - | 11.2581 | 5.14E-05 | 1 | TAGGCTATTGTTCTT |
| 114 | mamo-L | *BGIBMGA000198* | down | 1123 | 1137 | - | 11.0806 | 6.05E-05 | 1 | AAGGCTGTGCTTGTT |
| 115 | mamo-L | *BGIBMGA000201* | down | 743 | 757 | + | 14.9677 | 3.56E-07 | 1 | TAGGCTTAGTTTTGT |
| 116 | mamo-L | *BGIBMGA000201* | up | 560 | 574 | - | 10.5887 | 9.21E-05 | 1 | TAGGCGGAGTGTTGG |
| 117 | mamo-L | *BGIBMGA000206* | down | 109 | 123 | + | 10.5726 | 9.33E-05 | 1 | TAGGCTAATATTTAT |
| 118 | mamo-L | *BGIBMGA000207* | down | 1936 | 1950 | + | 12.3145 | 1.79E-05 | 1 | TAGGCGGGTTATTTT |
| 119 | mamo-L | *BGIBMGA000207* | up | 224 | 238 | + | 10.5726 | 9.33E-05 | 1 | TAGGCTAATATTTAT |
| 120 | mamo-L | *BGIBMGA000209* | up | 1052 | 1066 | + | 12.3145 | 1.79E-05 | 1 | TAGGCGGGTTATTTT |
| 121 | mamo-L | *BGIBMGA000210* | down | 1117 | 1131 | - | 12.8548 | 9.51E-06 | 1 | TAGGCGGGGATGGGT |
| 122 | mamo-L | *BGIBMGA000211* | up | 963 | 977 | + | 14.7258 | 5.83E-07 | 1 | TAGGCTTGCGTTGCC |
| 123 | mamo-L | *BGIBMGA000216* | up | 1266 | 1280 | + | 10.5161 | 9.77E-05 | 1 | TAGCCTGTGTTTTTT |
| 124 | mamo-L | *BGIBMGA000219* | up | 878 | 892 | - | 11.5323 | 3.96E-05 | 1 | TAGGCGTGAGGTGAT |
| 125 | mamo-L | *BGIBMGA000219* | down | 1190 | 1204 | + | 11.0645 | 6.14E-05 | 1 | GAGGCTGGTGATGCC |
| 126 | mamo-L | *BGIBMGA000219* | up | 179 | 193 | - | 11.0403 | 6.28E-05 | 1 | TAGGCTGTATGTTAT |
| 127 | mamo-L | *BGIBMGA000222* | down | 511 | 525 | + | 10.5726 | 9.33E-05 | 1 | TAGGCTAATATTTAT |
| 128 | mamo-L | *BGIBMGA000223* | up | 1579 | 1593 | - | 11.2823 | 5.03E-05 | 1 | AAGGCGGTTTTTTTT |
| 129 | mamo-L | *BGIBMGA000225* | down | 1209 | 1223 | - | 10.6129 | 9.03E-05 | 1 | TAGGCCTGAAGTGCC |
| 130 | mamo-L | *BGIBMGA000228* | down | 1292 | 1306 | - | 11.129 | 5.79E-05 | 1 | GAGGCTGTGTATTCT |
| 131 | mamo-L | *BGIBMGA000229* | down | 88 | 102 | + | 10.4919 | 9.96E-05 | 1 | TAGGCCTGTAGTTAT |
| 132 | mamo-L | *BGIBMGA000238* | up | 1170 | 1184 | + | 10.5645 | 9.40E-05 | 1 | TAGGCTTATTACACT |
| 133 | mamo-L | *BGIBMGA000239* | up | 791 | 805 | + | 11.8306 | 2.97E-05 | 1 | TAGGCTGTTGCTGAT |
| 134 | mamo-L | *BGIBMGA000239* | up | 785 | 799 | + | 10.9113 | 7.02E-05 | 1 | GAGGCCTAGGCTGTT |
| 135 | mamo-L | *BGIBMGA000240* | down | 1572 | 1586 | + | 11.2661 | 5.10E-05 | 1 | GAGGCTTGCAGTGGT |
| 136 | mamo-L | *BGIBMGA000240* | down | 1601 | 1615 | + | 11.1774 | 5.54E-05 | 1 | TAGGCTTAGGACTCC |
| 137 | mamo-L | *BGIBMGA000242* | up | 62 | 76 | - | 10.9919 | 6.55E-05 | 1 | TAGGCTTAAAATGTC |
| 138 | mamo-L | *BGIBMGA000244* | down | 642 | 656 | + | 10.6935 | 8.45E-05 | 1 | AAGGCCTTCTTCTTT |
| 139 | mamo-L | *BGIBMGA000247* | up | 1473 | 1487 | + | 11.2097 | 5.38E-05 | 1 | TAGGCCGTTTGCTTT |
| 140 | mamo-L | *BGIBMGA000250* | up | 955 | 969 | + | 13.1048 | 7.05E-06 | 1 | TAGGCCTTCGTTGTA |
| 141 | mamo-L | *BGIBMGA000250* | down | 1487 | 1501 | + | 11.5242 | 3.99E-05 | 1 | TAGGCAGACGTTGGG |
| 142 | mamo-L | *BGIBMGA000251* | up | 1125 | 1139 | + | 11.5726 | 3.81E-05 | 1 | TAGGCTGTTGTTTCG |
| 143 | mamo-L | *BGIBMGA000251* | up | 135 | 149 | + | 11.5242 | 3.99E-05 | 1 | TAGGCAGACGTTGGG |
| 144 | mamo-L | *BGIBMGA000253* | down | 136 | 150 | + | 10.5726 | 9.33E-05 | 1 | AAGGCTGACTTTTTA |
| 145 | mamo-L | *BGIBMGA000257* | up | 1058 | 1072 | + | 11.7903 | 3.08E-05 | 1 | TAGGCCTCATTTGAT |
| 146 | mamo-L | *BGIBMGA000260* | up | 493 | 507 | - | 10.5726 | 9.33E-05 | 1 | TAGGCTAATATTTAT |
| 147 | mamo-L | *BGIBMGA000261* | down | 1668 | 1682 | - | 11.5 | 4.08E-05 | 1 | TAGGCAGATTTTAGT |
| 148 | mamo-L | *BGIBMGA000262* | up | 1457 | 1471 | - | 11.5 | 4.08E-05 | 1 | TAGGCAGATTTTAGT |
| 149 | mamo-L | *BGIBMGA000274* | down | 1698 | 1712 | - | 10.5 | 9.90E-05 | 1 | AAGGCTTTGTTTCGC |
| 150 | mamo-L | *BGIBMGA000275* | up | 1578 | 1592 | + | 11.5484 | 3.90E-05 | 1 | TAGGCTGTGGTCGAA |
| 151 | mamo-L | *BGIBMGA000280* | down | 412 | 426 | + | 11.371 | 4.62E-05 | 1 | GAGGCTTAGATTTCT |
| 152 | mamo-L | *BGIBMGA000284* | up | 1963 | 1977 | - | 12.6129 | 1.26E-05 | 1 | TAGGCTGATGATAGT |
| 153 | mamo-L | *BGIBMGA000285* | down | 12 | 26 | - | 11.1129 | 5.88E-05 | 1 | AAGGCCTGCAGTGGT |
| 154 | mamo-L | *BGIBMGA000287* | down | 1565 | 1579 | + | 11.5323 | 3.96E-05 | 1 | TAGGCGTGAGGTGAT |
| 155 | mamo-L | *BGIBMGA000287* | down | 175 | 189 | - | 10.9435 | 6.83E-05 | 1 | TAGGCCGGTTACAGT |
| 156 | mamo-L | *BGIBMGA000288* | up | 1229 | 1243 | + | 11.5323 | 3.96E-05 | 1 | TAGGCGTGAGGTGAT |
| 157 | mamo-L | *BGIBMGA000290* | down | 621 | 635 | + | 11.4597 | 4.25E-05 | 1 | TAGGCGGGGTGGTGT |
| 158 | mamo-L | *BGIBMGA000291* | down | 125 | 139 | - | 13.7823 | 2.76E-06 | 1 | TAGGCTTTTTGTGGT |
| 159 | mamo-L | *BGIBMGA000293* | down | 1921 | 1935 | - | 10.5726 | 9.33E-05 | 1 | TAGGCTAATATTTAT |
| 160 | mamo-L | *BGIBMGA000294* | down | 1905 | 1919 | - | 10.7419 | 8.11E-05 | 1 | TCGGCGGTGTTTGCT |
| 161 | mamo-L | *BGIBMGA000294* | up | 514 | 528 | + | 10.5887 | 9.21E-05 | 1 | TAGGCTCTGTTTCGA |
| 162 | mamo-L | *BGIBMGA000297* | up | 600 | 614 | + | 13.2177 | 6.14E-06 | 1 | TAGGCCTACTACGGT |
| 163 | mamo-L | *BGIBMGA000301* | down | 1066 | 1080 | + | 12.1613 | 2.11E-05 | 1 | TAGGCCCTCCTTGGT |
| 164 | mamo-L | *BGIBMGA000301* | down | 900 | 914 | + | 10.6371 | 8.85E-05 | 1 | TAGGCAGGCGGTAGT |
| 165 | mamo-L | *BGIBMGA000304* | down | 1276 | 1290 | - | 12.0403 | 2.40E-05 | 1 | TAGGCGTAGATTTAT |
| 166 | mamo-L | *BGIBMGA000305* | down | 1554 | 1568 | - | 10.629 | 8.91E-05 | 1 | TAGGCGGTCTACTGC |
| 167 | mamo-L | *BGIBMGA000311* | down | 1724 | 1738 | + | 10.7661 | 7.95E-05 | 1 | TAGGCCCTAGATTCT |
| 168 | mamo-L | *BGIBMGA000312* | down | 1951 | 1965 | - | 10.8065 | 7.69E-05 | 1 | AAGGCCTTAGTTTTT |
| 169 | mamo-L | *BGIBMGA000324* | up | 1155 | 1169 | + | 11.1935 | 5.45E-05 | 1 | TAGGCCACGGATGTT |
| 170 | mamo-L | *BGIBMGA000326* | down | 1463 | 1477 | + | 12.4032 | 1.61E-05 | 1 | TAGGCATGGGGTGGT |
| 171 | mamo-L | *BGIBMGA000327* | up | 336 | 350 | - | 11.1613 | 5.63E-05 | 1 | TAGGCTCTCTGTATT |
| 172 | mamo-L | *BGIBMGA000329* | up | 134 | 148 | + | 11.7581 | 3.18E-05 | 1 | TAGGCCGTATTGTGT |
| 173 | mamo-L | *BGIBMGA000329* | down | 1093 | 1107 | - | 10.8629 | 7.33E-05 | 1 | TAGGCCCCCTTCGTC |
| 174 | mamo-L | *BGIBMGA000330* | down | 1103 | 1117 | - | 13.1129 | 6.98E-06 | 1 | TAGGCGTATTGTGTT |
| 175 | mamo-L | *BGIBMGA000330* | down | 1407 | 1421 | + | 12.4355 | 1.55E-05 | 1 | GAGGCTTACATTGTT |
| 176 | mamo-L | *BGIBMGA000332* | up | 1327 | 1341 | - | 11.9839 | 2.54E-05 | 1 | GAGGCCTGATTTGTT |
| 177 | mamo-L | *BGIBMGA000332* | up | 699 | 713 | - | 10.6371 | 8.85E-05 | 1 | TAGGCATTTGTCTGT |

| | | | | | | | | | | |
|---|---|---|---|---|---|---|---|---|---|---|
| 178 | mamo-L | *BGIBMGA000334* | up | 1925 | 1939 | - | 10.7097 | 8.33E-05 | 1 | TAGGCGCCGGATTGT |
| 179 | mamo-L | *BGIBMGA000335* | down | 884 | 898 | - | 12.8065 | 1.01E-05 | 1 | TAGGCTGCTTTTTTT |
| 180 | mamo-L | *BGIBMGA000335* | up | 1621 | 1635 | + | 10.5161 | 9.77E-05 | 1 | TAGGCCAACGATGTA |
| 181 | mamo-L | *BGIBMGA000337* | down | 1645 | 1659 | + | 12.3226 | 1.77E-05 | 1 | TAGGCCTCTTTTTTT |
| 182 | mamo-L | *BGIBMGA000337* | down | 1181 | 1195 | - | 11.1613 | 5.63E-05 | 1 | TAGGCTCCGATTTTT |
| 183 | mamo-L | *BGIBMGA000340* | up | 309 | 323 | - | 11.5645 | 3.84E-05 | 1 | TAGGCTGGGGGTCTC |
| 184 | mamo-L | *BGIBMGA000341* | down | 427 | 441 | - | 10.6048 | 9.09E-05 | 1 | TAGGCCGTCACTTCT |
| 185 | mamo-L | *BGIBMGA000345* | down | 1685 | 1699 | - | 12.3629 | 1.69E-05 | 1 | TAGGCGAGATTTGGT |
| 186 | mamo-L | *BGIBMGA000345* | down | 320 | 334 | - | 10.5726 | 9.33E-05 | 1 | TAGGCTAATATTTAT |
| 187 | mamo-L | *BGIBMGA000346* | up | 355 | 369 | + | 14.3226 | 1.22E-06 | 1 | TAGGCTTTGATTGTT |
| 188 | mamo-L | *BGIBMGA000348* | up | 173 | 187 | + | 11.6048 | 3.70E-05 | 1 | TAGGCTCATATTAGT |
| 189 | mamo-L | *BGIBMGA000354* | down | 1914 | 1928 | - | 11.1935 | 5.45E-05 | 1 | TAGGCTTAAGTGAGT |
| 190 | mamo-L | *BGIBMGA000355* | up | 889 | 903 | - | 11.1935 | 5.45E-05 | 1 | TAGGCTTAAGTGAGT |
| 191 | mamo-L | *BGIBMGA000355* | down | 1492 | 1506 | - | 10.5726 | 9.33E-05 | 1 | TAGGCTTCTATTTAT |
| 192 | mamo-L | *BGIBMGA000356* | up | 653 | 667 | - | 11.4516 | 4.28E-05 | 1 | TAGGCGTTGGTTGAG |
| 193 | mamo-L | *BGIBMGA000358* | up | 548 | 562 | + | 12.8145 | 9.96E-06 | 1 | GAGGCTGATGTTGGC |
| 194 | mamo-L | *BGIBMGA000364* | up | 1774 | 1788 | - | 12.3065 | 1.80E-05 | 1 | TAGGCGTTTATTTGT |
| 195 | mamo-L | *BGIBMGA000364* | down | 404 | 418 | + | 10.5726 | 9.33E-05 | 1 | TAGGCTTCTATTTAT |
| 196 | mamo-L | *BGIBMGA000366* | up | 400 | 414 | - | 13.4677 | 4.39E-06 | 1 | TAGGCTGATGATGAT |
| 197 | mamo-L | *BGIBMGA000369* | down | 1180 | 1194 | + | 12.3629 | 1.69E-05 | 1 | TAGGCGAGATTTGGT |
| 198 | mamo-L | *BGIBMGA000370* | down | 679 | 693 | - | 10.9194 | 6.98E-05 | 1 | TAGGCTACTTTTTTT |
| 199 | mamo-L | *BGIBMGA000370* | up | 1079 | 1093 | - | 10.75 | 8.06E-05 | 1 | TAGGCTTATGTCTTA |
| 200 | mamo-L | *BGIBMGA000370* | up | 1667 | 1681 | - | 10.5726 | 9.33E-05 | 1 | TAGGCTAATATTTAT |
| 201 | mamo-L | *BGIBMGA000371* | up | 100 | 114 | - | 10.5161 | 9.77E-05 | 1 | TAGGCCAACGATGTA |
| 202 | mamo-L | *BGIBMGA000378* | down | 1099 | 1113 | + | 13.3387 | 5.24E-06 | 1 | TAGGCGGGCTTTGGA |
| 203 | mamo-L | *BGIBMGA000378* | up | 610 | 624 | + | 10.9194 | 6.98E-05 | 1 | AAGGCTTGTATTGTC |
| 204 | mamo-L | *BGIBMGA000378* | down | 946 | 960 | - | 10.8306 | 7.52E-05 | 1 | TAGGCCCCCTTCGCC |
| 205 | mamo-L | *BGIBMGA000383* | up | 889 | 903 | + | 10.9113 | 7.02E-05 | 1 | TAGGCTGATGGCACT |
| 206 | mamo-L | *BGIBMGA000383* | down | 98 | 112 | - | 10.5726 | 9.33E-05 | 1 | TAGGCTAATATTTAT |
| 207 | mamo-L | *BGIBMGA000385* | down | 1150 | 1164 | + | 11.3871 | 4.55E-05 | 1 | GAGGCTCGCGGTGTT |
| 208 | mamo-L | *BGIBMGA000390* | down | 812 | 826 | - | 10.7661 | 7.95E-05 | 1 | TAGGCCCTAGATTCT |
| 209 | mamo-L | *BGIBMGA000394* | up | 430 | 444 | - | 11.9113 | 2.74E-05 | 1 | TAGGCGTAGTTCGGA |
| 210 | mamo-L | *BGIBMGA000394* | down | 541 | 555 | - | 11.2581 | 5.14E-05 | 1 | TAGGCCGTTTACTCT |
| 211 | mamo-L | *BGIBMGA000394* | up | 272 | 286 | + | 10.5403 | 9.58E-05 | 1 | TAGGCTGTATGGTGT |
| 212 | mamo-L | *BGIBMGA000398* | up | 1245 | 1259 | + | 12.5081 | 1.43E-05 | 1 | TAGGCTAATTTTTTT |
| 213 | mamo-L | *BGIBMGA000400* | up | 1370 | 1384 | + | 11.1694 | 5.59E-05 | 1 | TAGGCCATATTTTCT |
| 214 | mamo-L | *BGIBMGA000401* | down | 1175 | 1189 | - | 11.3468 | 4.73E-05 | 1 | TAGGCCTTTATTGCA |
| 215 | mamo-L | *BGIBMGA000406* | down | 1411 | 1425 | - | 12.0161 | 2.46E-05 | 1 | TAGGCCGTGAATTTT |
| 216 | mamo-L | *BGIBMGA000408* | up | 1958 | 1972 | + | 10.9839 | 6.59E-05 | 1 | TAGGCCCTTGTTGAA |
| 217 | mamo-L | *BGIBMGA000416* | up | 771 | 785 | - | 10.6129 | 9.03E-05 | 1 | TAGGCCCGCATTTAC |
| 218 | mamo-L | *BGIBMGA000426* | up | 1259 | 1273 | + | 11.379 | 4.58E-05 | 1 | TAGGCAGACGTGTGT |
| 219 | mamo-L | *BGIBMGA000431* | up | 1518 | 1532 | + | 10.5726 | 9.33E-05 | 1 | TAGGCTAATATTTAT |
| 220 | mamo-L | *BGIBMGA000434* | up | 838 | 852 | + | 12.1452 | 2.15E-05 | 1 | TAGGCGGACTTCGGA |
| 221 | mamo-L | *BGIBMGA000435* | down | 275 | 289 | - | 10.8468 | 7.42E-05 | 1 | AAGGCTTATTGTGAT |
| 222 | mamo-L | *BGIBMGA000435* | down | 1221 | 1235 | - | 10.8306 | 7.52E-05 | 1 | TAGGCCCCCTTCGCC |
| 223 | mamo-L | *BGIBMGA000437* | down | 1783 | 1797 | + | 11.5887 | 3.75E-05 | 1 | AAGGCCCACGTTTCT |
| 224 | mamo-L | *BGIBMGA000438* | up | 902 | 916 | - | 13.4677 | 4.39E-06 | 1 | TAGGCTGATGATGAT |
| 225 | mamo-L | *BGIBMGA000438* | up | 1786 | 1800 | + | 11.2097 | 5.38E-05 | 1 | TAGGCTGGTTTTATA |
| 226 | mamo-L | *BGIBMGA000439* | up | 1838 | 1852 | + | 10.5726 | 9.33E-05 | 1 | TAGGCTAATATTTAT |
| 227 | mamo-L | *BGIBMGA000440* | up | 1777 | 1791 | + | 10.5403 | 9.58E-05 | 1 | TAGGCTGTATGGTGT |
| 228 | mamo-L | *BGIBMGA000444* | down | 1072 | 1086 | - | 11.4919 | 4.11E-05 | 1 | TAGGCGGTAATTTTT |
| 229 | mamo-L | *BGIBMGA000446* | up | 1147 | 1161 | + | 12.0645 | 2.34E-05 | 1 | TAGGCGGGCTTCGGA |
| 230 | mamo-L | *BGIBMGA000446* | up | 994 | 1008 | - | 10.8306 | 7.52E-05 | 1 | TAGGCCCCCTTCGCC |
| 231 | mamo-L | *BGIBMGA000449* | up | 1196 | 1210 | - | 10.9677 | 6.68E-05 | 1 | TAGGCTTAGGGCGCA |
| 232 | mamo-L | *BGIBMGA000451* | down | 1567 | 1581 | - | 10.9919 | 6.55E-05 | 1 | AAGGCTGTTTGTGGC |
| 233 | mamo-L | *BGIBMGA000451* | up | 1141 | 1155 | + | 10.5726 | 9.33E-05 | 1 | TAGGCTAATATTTAT |
| 234 | mamo-L | *BGIBMGA000456* | down | 1512 | 1526 | + | 12.5968 | 1.28E-05 | 1 | TAGGCTCTGTTTACT |
| 235 | mamo-L | *BGIBMGA000459* | down | 423 | 437 | + | 12.1613 | 2.11E-05 | 1 | TAGGCCTACATTGCA |
| 236 | mamo-L | *BGIBMGA000464* | up | 1414 | 1428 | - | 11.4435 | 4.31E-05 | 1 | GAGGCCTTTTTTTTT |
| 237 | mamo-L | *BGIBMGA000464* | up | 1627 | 1641 | + | 10.5887 | 9.21E-05 | 1 | TAGGCTTTGACTTTT |
| 238 | mamo-L | *BGIBMGA000467* | down | 812 | 826 | + | 10.5403 | 9.58E-05 | 1 | TAGGCCGATCTTGCA |
| 239 | mamo-L | *BGIBMGA000470* | down | 1479 | 1493 | - | 10.621 | 8.97E-05 | 1 | TAGGCTTGTGATTTA |
| 240 | mamo-L | *BGIBMGA000473* | up | 617 | 631 | + | 11.0403 | 6.28E-05 | 1 | TAGGCTGTATGTTAT |
| 241 | mamo-L | *BGIBMGA000473* | down | 1912 | 1926 | - | 11.0081 | 6.45E-05 | 1 | TAGGCCAACTTTTAC |
| 242 | mamo-L | *BGIBMGA000476* | up | 99 | 113 | + | 12.0645 | 2.34E-05 | 1 | TAGGCGGGCTTCGGA |
| 243 | mamo-L | *BGIBMGA000479* | up | 1681 | 1695 | - | 10.7984 | 7.74E-05 | 1 | TAGGCCTACGAAGCT |
| 244 | mamo-L | *BGIBMGA000482* | down | 1693 | 1707 | + | 11.3306 | 4.80E-05 | 1 | GAGGCTTTCTTTGGA |
| 245 | mamo-L | *BGIBMGA000484* | down | 1239 | 1253 | - | 10.5645 | 9.40E-05 | 1 | TAGGCTTTTTATCAT |
| 246 | mamo-L | *BGIBMGA000490* | down | 517 | 531 | - | 11.9839 | 2.54E-05 | 1 | TAGGCCTACAGTGCC |
| 247 | mamo-L | *BGIBMGA000494* | down | 1205 | 1219 | - | 13.2419 | 5.96E-06 | 1 | TAGGCCCATGTTTTT |
| 248 | mamo-L | *BGIBMGA000495* | down | 1718 | 1732 | + | 13.2016 | 6.26E-06 | 1 | TAGGCGGACGGTGAT |
| 249 | mamo-L | *BGIBMGA000496* | up | 133 | 147 | + | 10.5726 | 9.33E-05 | 1 | TAGCCTGTATTTGGT |
| 250 | mamo-L | *BGIBMGA000499* | down | 208 | 222 | - | 11.3548 | 4.69E-05 | 1 | TAGGCCACCGTTTTT |
| 251 | mamo-L | *BGIBMGA000499* | up | 1365 | 1379 | - | 10.5323 | 9.63E-05 | 1 | AAGGCGTTGTTCGAT |
| 252 | mamo-L | *BGIBMGA000501* | down | 924 | 938 | - | 10.9194 | 6.98E-05 | 1 | TAGGCTACTTTTTTT |
| 253 | mamo-L | *BGIBMGA000502* | down | 1509 | 1523 | - | 10.5645 | 9.40E-05 | 1 | TAGGCGGAGTGGTGC |
| 254 | mamo-L | *BGIBMGA000503* | up | 1937 | 1951 | + | 11.9677 | 2.59E-05 | 1 | TAGGCTTTGTGCTGT |
| 255 | mamo-L | *BGIBMGA000504* | up | 1698 | 1712 | - | 10.6129 | 9.03E-05 | 1 | TAGGCATGTGCTGGT |
| 256 | mamo-L | *BGIBMGA000510* | down | 336 | 350 | + | 10.9194 | 6.98E-05 | 1 | TAGGCTACTTTTTTT |
| 257 | mamo-L | *BGIBMGA000513* | up | 1493 | 1507 | + | 11.7661 | 3.16E-05 | 1 | TAGGCGTTGTTTTGA |
| 258 | mamo-L | *BGIBMGA000515* | up | 1000 | 1014 | + | 13.8629 | 2.44E-06 | 1 | TAGGCTCGGTATGGT |
| 259 | mamo-L | *BGIBMGA000515* | up | 981 | 995 | + | 11.5161 | 4.02E-05 | 1 | TAGGCTCTAGTTGGA |
| 260 | mamo-L | *BGIBMGA000516* | down | 612 | 626 | + | 11.8871 | 2.81E-05 | 1 | TAGGCTCGTAATGTT |
| 261 | mamo-L | *BGIBMGA000517* | up | 1418 | 1432 | - | 11.4032 | 4.49E-05 | 1 | GAGGCCGGGATTTCT |
| 262 | mamo-L | *BGIBMGA000519* | down | 872 | 886 | + | 11.3548 | 4.69E-05 | 1 | AAGGCTGATGATGAT |
| 263 | mamo-L | *BGIBMGA000519* | down | 1936 | 1950 | - | 10.6855 | 8.51E-05 | 1 | TAGGCCCTGATGGCC |
| 264 | mamo-L | *BGIBMGA000525* | up | 1313 | 1327 | - | 13.7016 | 3.11E-06 | 1 | TAGGCTTCTGTTGCT |
| 265 | mamo-L | *BGIBMGA000527* | up | 781 | 795 | + | 10.5806 | 9.27E-05 | 1 | TAGGCGTTCGTTACA |
| 266 | mamo-L | *BGIBMGA000529* | down | 1955 | 1969 | + | 11.8065 | 3.03E-05 | 1 | TAGGCTGACGATCAT |
| 267 | mamo-L | *BGIBMGA000536* | up | 163 | 177 | + | 11.5726 | 3.81E-05 | 1 | TAGGCAGACGGTGAT |
| 268 | mamo-L | *BGIBMGA000537* | down | 418 | 432 | + | 11.2823 | 5.03E-05 | 1 | TAGGCCTGGTTTATA |
| 269 | mamo-L | *BGIBMGA000538* | up | 882 | 896 | - | 10.5403 | 9.58E-05 | 1 | AAGGCTCTGGATTGT |

| | | | | | | | | | | |
|---|---|---|---|---|---|---|---|---|---|---|
| 270 | mamo-L | *BGIBMGA000541* | up | 1861 | 1875 | + | 11.2581 | 5.14E-05 | 1 | TAGGCTGAGGCTGGG |
| 271 | mamo-L | *BGIBMGA000542* | up | 1793 | 1807 | + | 10.5403 | 9.58E-05 | 1 | TAGGCTGTATGGTGT |
| 272 | mamo-L | *BGIBMGA000546* | up | 954 | 968 | + | 11.7984 | 3.06E-05 | 1 | TAGGCACGCGTTGAT |
| 273 | mamo-L | *BGIBMGA000551* | up | 119 | 133 | - | 12.1694 | 2.09E-05 | 1 | TAGGCCGACTAGGGC |
| 274 | mamo-L | *BGIBMGA000551* | down | 718 | 732 | - | 12.129 | 2.19E-05 | 1 | TAGGCTTTTTTGTTT |
| 275 | mamo-L | *BGIBMGA000551* | up | 1938 | 1952 | + | 10.6855 | 8.51E-05 | 1 | TAGGCTTGCGATTCG |
| 276 | mamo-L | *BGIBMGA000552* | up | 1844 | 1858 | + | 10.7823 | 7.84E-05 | 1 | AAGGCCGACTTTTAC |
| 277 | mamo-L | *BGIBMGA000553* | up | 1645 | 1659 | + | 10.5726 | 9.33E-05 | 1 | TAGGCTAATATTTAT |
| 278 | mamo-L | *BGIBMGA000553* | down | 506 | 520 | + | 10.5081 | 9.83E-05 | 1 | TAGGCCTATAATTTC |
| 279 | mamo-L | *BGIBMGA000555* | down | 1203 | 1217 | + | 10.5726 | 9.33E-05 | 1 | TAGGCTAATATTTAT |
| 280 | mamo-L | *BGIBMGA000560* | down | 793 | 807 | + | 11.75 | 3.21E-05 | 1 | TAGGCACACTTTGAT |
| 281 | mamo-L | *BGIBMGA000561* | down | 1860 | 1874 | + | 10.6855 | 8.51E-05 | 1 | TAGGCCGCAATTTCT |
| 282 | mamo-L | *BGIBMGA000562* | up | 1856 | 1870 | + | 10.7581 | 8.00E-05 | 1 | TAGGCCTTATTTCTC |
| 283 | mamo-L | *BGIBMGA000564* | up | 539 | 553 | - | 10.7177 | 8.28E-05 | 1 | AAGGCCGAGGATCGT |
| 284 | mamo-L | *BGIBMGA000570* | down | 796 | 810 | + | 13.5968 | 3.65E-06 | 1 | TAGGCCGTGGTTTTC |
| 285 | mamo-L | *BGIBMGA000573* | up | 1684 | 1698 | - | 11.0242 | 6.36E-05 | 1 | TAGGCTGATGACATT |
| 286 | mamo-L | *BGIBMGA000574* | down | 1336 | 1350 | + | 11.8065 | 3.03E-05 | 1 | TAGGCTAATGTTTAT |
| 287 | mamo-L | *BGIBMGA000575* | up | 1369 | 1383 | - | 12.4919 | 1.46E-05 | 1 | TAGGCTGAAATTTTT |
| 288 | mamo-L | *BGIBMGA000577* | up | 213 | 227 | - | 10.7581 | 8.00E-05 | 1 | TACGCTTGGTTTGTT |
| 289 | mamo-L | *BGIBMGA000582* | up | 117 | 131 | + | 11.9919 | 2.52E-05 | 1 | TAGGCGTGATTGGTT |
| 290 | mamo-L | *BGIBMGA000582* | up | 161 | 175 | - | 10.5806 | 9.27E-05 | 1 | TAGGCTGGGACGGTT |
| 291 | mamo-L | *BGIBMGA000583* | up | 1211 | 1225 | - | 12.0726 | 2.32E-05 | 1 | TAGGCAGTTATTGGT |
| 292 | mamo-L | *BGIBMGA000585* | down | 489 | 503 | - | 11.7016 | 3.37E-05 | 1 | TAGGCCCGGTTTCAT |
| 293 | mamo-L | *BGIBMGA000585* | up | 489 | 503 | + | 11.3548 | 4.69E-05 | 1 | AAGGCTGATGATGAT |
| 294 | mamo-L | *BGIBMGA000590* | down | 950 | 964 | - | 10.5081 | 9.83E-05 | 1 | TAGGCGGGGTGTTGG |
| 295 | mamo-L | *BGIBMGA000594* | up | 1455 | 1469 | - | 11.7339 | 3.26E-05 | 1 | TAGGCTATGTTTTCC |
| 296 | mamo-L | *BGIBMGA000595* | down | 293 | 307 | + | 14.4597 | 9.75E-07 | 1 | TAGGCTTGTGTTTGT |
| 297 | mamo-L | *BGIBMGA000598* | down | 258 | 272 | - | 10.5484 | 9.52E-05 | 1 | TAGGCGCCATTTTGT |
| 298 | mamo-L | *BGIBMGA000603* | down | 659 | 673 | + | 10.5484 | 9.52E-05 | 1 | TAGGCATCGTTCGCT |
| 299 | mamo-L | *BGIBMGA000604* | down | 835 | 849 | + | 12.5726 | 1.33E-05 | 1 | AAGGCTTACGTTTCT |
| 300 | mamo-L | *BGIBMGA000605* | up | 1971 | 1985 | + | 12.5726 | 1.33E-05 | 1 | AAGGCTTACGTTTCT |
| 301 | mamo-L | *BGIBMGA000605* | up | 819 | 833 | + | 11.5968 | 3.73E-05 | 1 | AAGGCTTACGTTTCC |
| 302 | mamo-L | *BGIBMGA000612* | up | 88 | 102 | - | 11.3548 | 4.69E-05 | 1 | AAGGCTGATGATGAT |
| 303 | mamo-L | *BGIBMGA000613* | down | 1872 | 1886 | + | 11.3629 | 4.66E-05 | 1 | TAGGCTAATGTCTTT |
| 304 | mamo-L | *BGIBMGA000616* | up | 1927 | 1941 | + | 11.5242 | 3.99E-05 | 1 | TAGGCGTGGTCTGAT |
| 305 | mamo-L | *BGIBMGA000616* | down | 1786 | 1800 | - | 11.2661 | 5.10E-05 | 1 | TAGGCTGAAGATGCA |
| 306 | mamo-L | *BGIBMGA000629* | down | 499 | 513 | - | 11.1935 | 5.45E-05 | 1 | TAGGCAGGGGTTGTG |
| 307 | mamo-L | *BGIBMGA000632* | up | 1653 | 1667 | - | 11.1048 | 5.92E-05 | 1 | TAGGCTAGTTATTTT |
| 308 | mamo-L | *BGIBMGA000637* | up | 378 | 392 | + | 10.7742 | 7.89E-05 | 1 | TAGGCCAAGGTTTCA |
| 309 | mamo-L | *BGIBMGA000637* | up | 1462 | 1476 | - | 10.5726 | 9.33E-05 | 1 | TAGGCTAATATTTAT |
| 310 | mamo-L | *BGIBMGA000637* | down | 1109 | 1123 | + | 10.5403 | 9.58E-05 | 1 | AAGGCTTTTCTTGGT |
| 311 | mamo-L | *BGIBMGA000638* | down | 1333 | 1347 | - | 12.7984 | 1.02E-05 | 1 | TAGGCTTGGGAGGCT |
| 312 | mamo-L | *BGIBMGA000640* | down | 524 | 538 | - | 10.879 | 7.22E-05 | 1 | TAGGCTAAATTTTAT |
| 313 | mamo-L | *BGIBMGA000644* | up | 1862 | 1876 | + | 11.0968 | 5.97E-05 | 1 | TAGGCGTATTCTTGT |
| 314 | mamo-L | *BGIBMGA000644* | up | 747 | 761 | + | 10.5726 | 9.33E-05 | 1 | TAGGCTAATATTTAT |
| 315 | mamo-L | *BGIBMGA000645* | up | 1028 | 1042 | - | 11.2581 | 5.14E-05 | 1 | AAGGCCGAAATTGCT |
| 316 | mamo-L | *BGIBMGA000647* | up | 18 | 32 | + | 10.7339 | 8.17E-05 | 1 | TAGGCCACTTTTTTT |
| 317 | mamo-L | *BGIBMGA000649* | down | 977 | 991 | - | 14.3306 | 1.21E-06 | 1 | TAGGCCGTCGTCGTT |
| 318 | mamo-L | *BGIBMGA000651* | down | 1533 | 1547 | - | 10.7581 | 8.00E-05 | 1 | TAGGCCGTGCTCGTC |
| 319 | mamo-L | *BGIBMGA000652* | up | 1480 | 1494 | + | 10.8548 | 7.38E-05 | 1 | TAGGCATACGTGGCC |
| 320 | mamo-L | *BGIBMGA000657* | down | 1389 | 1403 | - | 14.8548 | 4.50E-07 | 1 | TAGGCTTACTTTGAT |
| 321 | mamo-L | *BGIBMGA000657* | down | 1741 | 1755 | + | 11.1774 | 5.54E-05 | 1 | TAGGCTAAGTTTTGA |
| 322 | mamo-L | *BGIBMGA000657* | up | 23 | 37 | + | 11.1048 | 5.92E-05 | 1 | TAGGCGTTTAATGGC |
| 323 | mamo-L | *BGIBMGA000662* | up | 588 | 602 | + | 10.629 | 8.91E-05 | 1 | TAGGCTTTGTACGTA |
| 324 | mamo-L | *BGIBMGA000664* | down | 1067 | 1081 | + | 10.7823 | 7.84E-05 | 1 | TAGGCTATTTGTTTT |
| 325 | mamo-L | *BGIBMGA000665* | up | 201 | 215 | - | 13.4435 | 4.51E-06 | 1 | TAGGCTTAGTTGGGC |
| 326 | mamo-L | *BGIBMGA000665* | down | 1682 | 1696 | + | 11.4435 | 4.31E-05 | 1 | TAGGCTTAGTTGAGC |
| 327 | mamo-L | *BGIBMGA000667* | up | 1370 | 1384 | - | 10.7258 | 8.23E-05 | 1 | TAGGCTTAGGAGCCT |
| 328 | mamo-L | *BGIBMGA000668* | down | 1035 | 1049 | - | 10.5081 | 9.83E-05 | 1 | TAGGCCTATAATTTC |
| 329 | mamo-L | *BGIBMGA000669* | up | 1078 | 1092 | + | 12.2097 | 2.01E-05 | 1 | TAGGCTTGAGGTGAT |
| 330 | mamo-L | *BGIBMGA000674* | up | 426 | 440 | - | 11.3548 | 4.69E-05 | 1 | AAGGCTGATGATGAT |
| 331 | mamo-L | *BGIBMGA000677* | up | 260 | 274 | - | 10.6129 | 9.03E-05 | 1 | TAGGCACTCTTGGTT |
| 332 | mamo-L | *BGIBMGA000678* | down | 899 | 913 | - | 10.8871 | 7.17E-05 | 1 | TAGGCTGTGTCTGTA |
| 333 | mamo-L | *BGIBMGA000680* | up | 1152 | 1166 | + | 11.9677 | 2.59E-05 | 1 | AAGGCTTTTATTGGT |
| 334 | mamo-L | *BGIBMGA000683* | up | 263 | 277 | - | 11.2581 | 5.14E-05 | 1 | TAGGCTGAGGCTGGG |
| 335 | mamo-L | *BGIBMGA000690* | down | 1484 | 1498 | + | 12.5887 | 1.30E-05 | 1 | TAGGCAGACTGTGGT |
| 336 | mamo-L | *BGIBMGA000690* | down | 246 | 260 | - | 10.7016 | 8.39E-05 | 1 | TAGGCCTATTTGAGC |
| 337 | mamo-L | *BGIBMGA000691* | up | 176 | 190 | + | 12.5887 | 1.30E-05 | 1 | TAGGCAGACTGTGGT |
| 338 | mamo-L | *BGIBMGA000696* | down | 1158 | 1172 | + | 12.25 | 1.92E-05 | 1 | TAGGCGATAGTTGGT |
| 339 | mamo-L | *BGIBMGA000697* | down | 1715 | 1729 | + | 11.8952 | 2.79E-05 | 1 | TAGGCCATGTTTTGC |
| 340 | mamo-L | *BGIBMGA000698* | down | 306 | 320 | - | 10.5565 | 9.46E-05 | 1 | TAGGCAGACGATTAT |
| 341 | mamo-L | *BGIBMGA000701* | up | 904 | 918 | + | 12.0161 | 2.46E-05 | 1 | AAGGCTTACGTTAGT |
| 342 | mamo-L | *BGIBMGA000703* | up | 883 | 897 | - | 11.5887 | 3.75E-05 | 1 | GAGGCCCACGTTTTT |
| 343 | mamo-L | *BGIBMGA000704* | down | 987 | 1001 | - | 10.9194 | 6.98E-05 | 1 | AAGGCCTATGTTTCC |
| 344 | mamo-L | *BGIBMGA000706* | down | 1801 | 1815 | + | 12.1855 | 2.06E-05 | 1 | GAGGCCGTCGTCGTT |
| 345 | mamo-L | *BGIBMGA000708* | up | 587 | 601 | + | 10.6694 | 8.62E-05 | 1 | TAGGCGACAGTTGTT |
| 346 | mamo-L | *BGIBMGA000711* | up | 1388 | 1402 | + | 11.25 | 5.18E-05 | 1 | TAGGCGCTCTCTGTT |
| 347 | mamo-L | *BGIBMGA000713* | down | 17 | 31 | - | 14.5 | 9.07E-07 | 1 | TAGGCTGTAGTTGTT |
| 348 | mamo-L | *BGIBMGA000715* | up | 364 | 378 | + | 10.7984 | 7.74E-05 | 1 | TAGGCGTGGGCTAGT |
| 349 | mamo-L | *BGIBMGA000717* | down | 1425 | 1439 | - | 10.6371 | 8.85E-05 | 1 | TAGGCTTTTTTTACA |
| 350 | mamo-L | *BGIBMGA000719* | down | 922 | 936 | - | 11.1935 | 5.45E-05 | 1 | TAGGCGCATGAGGGT |
| 351 | mamo-L | *BGIBMGA000720* | up | 590 | 604 | - | 11.1935 | 5.45E-05 | 1 | TAGGCGCATGAGGGT |
| 352 | mamo-L | *BGIBMGA000725* | up | 300 | 314 | + | 11.621 | 3.63E-05 | 1 | AAGGCTTTTATTGCT |
| 353 | mamo-L | *BGIBMGA000726* | down | 945 | 959 | - | 10.9194 | 6.98E-05 | 1 | TAGGCTACTTTTTTT |
| 354 | mamo-L | *BGIBMGA000729* | up | 1663 | 1677 | - | 11.9919 | 2.52E-05 | 1 | AAGGCCGTTGATGGT |
| 355 | mamo-L | *BGIBMGA000730* | down | 1752 | 1766 | + | 13.2419 | 5.96E-06 | 1 | TAGGCCCATGTTTTT |
| 356 | mamo-L | *BGIBMGA000731* | up | 1704 | 1718 | + | 11.8387 | 2.94E-05 | 1 | TAGGCATGTGTTTTT |
| 357 | mamo-L | *BGIBMGA000732* | down | 927 | 941 | - | 10.9194 | 6.98E-05 | 1 | TAGGCTACTTTTTTT |
| 358 | mamo-L | *BGIBMGA000732* | down | 1946 | 1960 | + | 10.5 | 9.90E-05 | 1 | GAGGCCCAGGTCGTC |
| 359 | mamo-L | *BGIBMGA000734* | up | 355 | 369 | - | 12.2016 | 2.03E-05 | 1 | TAGGCTGTAGATGTC |
| 360 | mamo-L | *BGIBMGA000734* | down | 645 | 659 | + | 10.9194 | 6.98E-05 | 1 | TAGGCTACTTTTTTT |
| 361 | mamo-L | *BGIBMGA000735* | down | 1983 | 1997 | + | 10.5484 | 9.52E-05 | 1 | TAGGCGGGCTTGGAA |

| | | | | | | | | | | |
|---|---|---|---|---|---|---|---|---|---|---|
| 362 | mamo-L | *BGIBMGA000737* | down | 1594 | 1608 | - | 10.5 | 9.90E-05 | 1 | GAGGCCCAGGTCGTC |
| 363 | mamo-L | *BGIBMGA000741* | up | 1518 | 1532 | + | 11.0887 | 6.01E-05 | 1 | TAGGCGAGTGTTGGA |
| 364 | mamo-L | *BGIBMGA000742* | down | 1904 | 1918 | + | 11.8548 | 2.89E-05 | 1 | AAGGCTGTTGTGGGT |
| 365 | mamo-L | *BGIBMGA000742* | up | 1632 | 1646 | - | 11.0968 | 5.97E-05 | 1 | TAGGCCGAGCTTGCA |
| 366 | mamo-L | *BGIBMGA000745* | down | 1461 | 1475 | + | 12.1935 | 2.05E-05 | 1 | TAGGCTTAAATTTTT |
| 367 | mamo-L | *BGIBMGA000749* | up | 1393 | 1407 | - | 11.9677 | 2.59E-05 | 1 | TAGGCCGTCTGCGAT |
| 368 | mamo-L | *BGIBMGA000752* | up | 1095 | 1109 | + | 11.6129 | 3.66E-05 | 1 | TAGGCTTTTAATGAT |
| 369 | mamo-L | *BGIBMGA000752* | up | 755 | 769 | - | 10.621 | 8.97E-05 | 1 | TAGGCTTTTGGTATC |
| 370 | mamo-L | *BGIBMGA000753* | up | 435 | 449 | - | 10.8952 | 7.12E-05 | 1 | TAGGCTGTATGTTTC |
| 371 | mamo-L | *BGIBMGA000753* | down | 368 | 382 | - | 10.5968 | 9.15E-05 | 1 | GAGGCTAGTTTTTGT |
| 372 | mamo-L | *BGIBMGA000758* | down | 1314 | 1328 | + | 12.7742 | 1.04E-05 | 1 | TAGGCCTATTGTGAT |
| 373 | mamo-L | *BGIBMGA000759* | down | 961 | 975 | + | 11.879 | 2.83E-05 | 1 | TAGGCCCGTGGTGTC |
| 374 | mamo-L | *BGIBMGA000760* | down | 149 | 163 | - | 10.5726 | 9.33E-05 | 1 | TAGGCTAATATTTAT |
| 375 | mamo-L | *BGIBMGA000761* | up | 205 | 219 | - | 11.5484 | 3.90E-05 | 1 | AAGGCGGAGGTTCGT |
| 376 | mamo-L | *BGIBMGA000763* | down | 525 | 539 | - | 12.5968 | 1.28E-05 | 1 | AAGGCTTAATTTGGT |
| 377 | mamo-L | *BGIBMGA000764* | up | 1822 | 1836 | + | 12.0806 | 2.31E-05 | 1 | TGGGCTTGGTTTGTT |
| 378 | mamo-L | *BGIBMGA000767* | up | 1809 | 1823 | - | 11.9274 | 2.69E-05 | 1 | GAGGCTTTGTATGCT |
| 379 | mamo-L | *BGIBMGA000767* | up | 492 | 506 | - | 11.6774 | 3.44E-05 | 1 | TCGGCTGACTTTGCT |
| 380 | mamo-L | *BGIBMGA000769* | up | 316 | 330 | + | 11.0484 | 6.23E-05 | 1 | TAGGCGGTCATTAAT |
| 381 | mamo-L | *BGIBMGA000770* | up | 15 | 29 | + | 11.5323 | 3.96E-05 | 1 | TAGGCGTGAGGTGAT |
| 382 | mamo-L | *BGIBMGA000770* | down | 1445 | 1459 | - | 11.0887 | 6.01E-05 | 1 | TAGGCCGCTTTTGCG |
| 383 | mamo-L | *BGIBMGA000773* | up | 964 | 978 | + | 12.25 | 1.92E-05 | 1 | TAGGCCCACTATTCT |
| 384 | mamo-L | *BGIBMGA000775* | down | 689 | 703 | - | 12.6855 | 1.16E-05 | 1 | TAGGCGTTCTTTATT |
| 385 | mamo-L | *BGIBMGA000775* | up | 282 | 296 | - | 10.5484 | 9.52E-05 | 1 | GAGGCGTTAGTTGAT |
| 386 | mamo-L | *BGIBMGA000776* | down | 1375 | 1389 | - | 11.3306 | 4.80E-05 | 1 | TAGGCTTACTATGAA |
| 387 | mamo-L | *BGIBMGA000776* | down | 1772 | 1786 | - | 10.9194 | 6.98E-05 | 1 | TAGGCTACTTTTTTT |
| 388 | mamo-L | *BGIBMGA000777* | up | 89 | 103 | - | 11.3306 | 4.80E-05 | 1 | TAGGCTTACTATGAA |
| 389 | mamo-L | *BGIBMGA000777* | up | 486 | 500 | - | 10.9194 | 6.98E-05 | 1 | TAGGCTACTTTTTTT |
| 390 | mamo-L | *BGIBMGA000778* | up | 1495 | 1509 | + | 10.9194 | 6.98E-05 | 1 | TAGGCTACTTTTTTT |
| 391 | mamo-L | *BGIBMGA000780* | up | 91 | 105 | + | 10.9355 | 6.88E-05 | 1 | TAGGCCGCCTTCATT |
| 392 | mamo-L | *BGIBMGA000781* | up | 1547 | 1561 | - | 11.3065 | 4.91E-05 | 1 | GAGGCTGTGGTCTCT |
| 393 | mamo-L | *BGIBMGA000785* | down | 741 | 755 | + | 10.6694 | 8.62E-05 | 1 | TAGGCCCTGTGCTTT |
| 394 | mamo-L | *BGIBMGA000788* | up | 1624 | 1638 | + | 11.4194 | 4.42E-05 | 1 | AAGGCTTGCTTTTTC |
| 395 | mamo-L | *BGIBMGA000790* | down | 1723 | 1737 | + | 11.6855 | 3.42E-05 | 1 | TAGGCGCTCTTTGTA |
| 396 | mamo-L | *BGIBMGA000791* | up | 41 | 55 | + | 11.2016 | 5.41E-05 | 1 | TAGGCAAAGGTTTGT |
| 397 | mamo-L | *BGIBMGA000792* | down | 597 | 611 | - | 12.6371 | 1.23E-05 | 1 | TAGGCTCGTTTGGCT |
| 398 | mamo-L | *BGIBMGA000796* | up | 1358 | 1372 | + | 10.7177 | 8.28E-05 | 1 | TAGGCTTAGTTTAAA |
| 399 | mamo-L | *BGIBMGA000798* | down | 643 | 657 | - | 10.8387 | 7.47E-05 | 1 | TAGGCTTGAGACGCC |
| 400 | mamo-L | *BGIBMGA000799* | up | 71 | 85 | - | 11.5323 | 3.96E-05 | 1 | TAGGCGTGAGGTGAT |
| 401 | mamo-L | *BGIBMGA000801* | down | 189 | 203 | + | 11.3871 | 4.55E-05 | 1 | GAGGCTCGTTTTTGT |
| 402 | mamo-L | *BGIBMGA000809* | up | 617 | 631 | - | 11.0242 | 6.36E-05 | 1 | TAGGCTTGAATTCCT |
| 403 | mamo-L | *BGIBMGA000810* | down | 1246 | 1260 | + | 10.5726 | 9.33E-05 | 1 | TAGGCTAATATTTAT |
| 404 | mamo-L | *BGIBMGA000813* | down | 46 | 60 | + | 11.0806 | 6.05E-05 | 1 | AAGGCTTTTGTTGAC |
| 405 | mamo-L | *BGIBMGA000821* | down | 1370 | 1384 | + | 10.5968 | 9.15E-05 | 1 | GAGGCTAGTTTTTGT |
| 406 | mamo-L | *BGIBMGA000822* | up | 1735 | 1749 | + | 10.8952 | 7.12E-05 | 1 | TAGGCTGTATGTTTC |
| 407 | mamo-L | *BGIBMGA000825* | down | 359 | 373 | - | 11.8145 | 3.01E-05 | 1 | TAGGCGGAAGATGCC |
| 408 | mamo-L | *BGIBMGA000826* | up | 1130 | 1144 | - | 12.1048 | 2.24E-05 | 1 | TAGGCATATTTTTGT |
| 409 | mamo-L | *BGIBMGA000826* | up | 743 | 757 | + | 11.6532 | 3.53E-05 | 1 | AAGGCGTGGGATGCT |
| 410 | mamo-L | *BGIBMGA000830* | down | 76 | 90 | + | 11.6935 | 3.39E-05 | 1 | GAGGCTATGTTTGTT |
| 411 | mamo-L | *BGIBMGA000830* | down | 1402 | 1416 | - | 11.0887 | 6.01E-05 | 1 | AAGGCCGCCTATGGT |
| 412 | mamo-L | *BGIBMGA000831* | up | 922 | 936 | + | 11.9435 | 2.65E-05 | 1 | GAGGCTTTTTTTTGT |
| 413 | mamo-L | *BGIBMGA000836* | down | 618 | 632 | - | 11.4758 | 4.18E-05 | 1 | TAGGCCTTATGTGCC |
| 414 | mamo-L | *BGIBMGA000836* | up | 67 | 81 | - | 11.2016 | 5.41E-05 | 1 | AAGGCTTAGATGGGT |
| 415 | mamo-L | *BGIBMGA000838* | down | 896 | 910 | + | 10.6452 | 8.79E-05 | 1 | TAGGCTGATGTGTCA |
| 416 | mamo-L | *BGIBMGA000840* | up | 1902 | 1916 | + | 12.5161 | 1.42E-05 | 1 | TAGGCGCATGTGGGT |
| 417 | mamo-L | *BGIBMGA000842* | up | 982 | 996 | + | 11.75 | 3.21E-05 | 1 | TAGGCTTACATTAAT |
| 418 | mamo-L | *BGIBMGA000842* | down | 740 | 754 | - | 10.75 | 8.06E-05 | 1 | GAGGCCGATTGTGCC |
| 419 | mamo-L | *BGIBMGA000843* | down | 194 | 208 | + | 12.629 | 1.24E-05 | 1 | TAGGCTAATTTTGTC |
| 420 | mamo-L | *BGIBMGA000843* | down | 1500 | 1514 | - | 11.5968 | 3.73E-05 | 1 | GAGGCCGTGGTTTAT |
| 421 | mamo-L | *BGIBMGA000846* | down | 1790 | 1804 | - | 12.6371 | 1.23E-05 | 1 | TAGGCGGAGTGGGGT |
| 422 | mamo-L | *BGIBMGA000851* | up | 138 | 152 | + | 11.5323 | 3.96E-05 | 1 | TAGGCGTGAGGTGAT |
| 423 | mamo-L | *BGIBMGA000852* | up | 1497 | 1511 | + | 12.621 | 1.25E-05 | 1 | TAGGCCGACATTTTC |
| 424 | mamo-L | *BGIBMGA000853* | up | 1180 | 1194 | + | 11.7581 | 3.18E-05 | 1 | TAGGCGGGGTTGGGA |
| 425 | mamo-L | *BGIBMGA000859* | down | 829 | 843 | - | 11.3145 | 4.87E-05 | 1 | TAGGCAGGCTTTGGG |
| 426 | mamo-L | *BGIBMGA000860* | down | 617 | 631 | + | 10.75 | 8.06E-05 | 1 | GAGGCCGATTGTGCC |
| 427 | mamo-L | *BGIBMGA000863* | down | 118 | 132 | - | 11.6855 | 3.42E-05 | 1 | TAGGCTGATTGCTCT |
| 428 | mamo-L | *BGIBMGA000863* | up | 1523 | 1537 | - | 10.7177 | 8.28E-05 | 1 | TAGGCTTATTTGAAT |
| 429 | mamo-L | *BGIBMGA000864* | up | 997 | 1011 | - | 11.6855 | 3.42E-05 | 1 | TAGGCTGATTGCTCT |
| 430 | mamo-L | *BGIBMGA000866* | up | 364 | 378 | + | 10.9194 | 6.98E-05 | 1 | TAGGCTACTTTTTTT |
| 431 | mamo-L | *BGIBMGA000866* | down | 1447 | 1461 | + | 10.5 | 9.90E-05 | 1 | TAGCCTGATGTTGCC |
| 432 | mamo-L | *BGIBMGA000867* | up | 300 | 314 | + | 10.5 | 9.90E-05 | 1 | TAGCCTGATGTTGCC |
| 433 | mamo-L | *BGIBMGA000870* | down | 1003 | 1017 | - | 10.5403 | 9.58E-05 | 1 | TAGGCTTTAATGTGT |
| 434 | mamo-L | *BGIBMGA000871* | up | 1566 | 1580 | - | 12.0161 | 2.46E-05 | 1 | GAGGCGGATGTTTGT |
| 435 | mamo-L | *BGIBMGA000871* | up | 1427 | 1441 | - | 10.9839 | 6.59E-05 | 1 | TAGGCCGACGTGTTA |
| 436 | mamo-L | *BGIBMGA000872* | down | 1320 | 1334 | - | 13.8548 | 2.48E-06 | 1 | TAGGCGGATGGTGGT |
| 437 | mamo-L | *BGIBMGA000874* | up | 1883 | 1897 | + | 10.5323 | 9.63E-05 | 1 | TAGGCTCTGTAGTCT |
| 438 | mamo-L | *BGIBMGA000879* | down | 1453 | 1467 | + | 13.5242 | 4.05E-06 | 1 | TAGGCTGTTTTGGTT |
| 439 | mamo-L | *BGIBMGA000882* | down | 1155 | 1169 | + | 10.75 | 8.06E-05 | 1 | GAGGCCGATTGTGCC |
| 440 | mamo-L | *BGIBMGA000885* | up | 1819 | 1833 | - | 10.8871 | 7.17E-05 | 1 | TAGGCCTACATAGCT |
| 441 | mamo-L | *BGIBMGA000885* | up | 1826 | 1840 | + | 10.5806 | 9.27E-05 | 1 | TAGGCCTACGATTCG |
| 442 | mamo-L | *BGIBMGA000886* | up | 828 | 842 | - | 11.9355 | 2.67E-05 | 1 | TAGGCTCACGGGGCT |
| 443 | mamo-L | *BGIBMGA000886* | up | 1821 | 1835 | - | 10.8871 | 7.17E-05 | 1 | TAGGCCTACATAGCT |
| 444 | mamo-L | *BGIBMGA000886* | up | 1828 | 1842 | + | 10.5806 | 9.27E-05 | 1 | TAGGCCTACGATTCG |
| 445 | mamo-L | *BGIBMGA000887* | up | 485 | 499 | + | 11.4597 | 4.25E-05 | 1 | TAGGCGCTTATTGAT |
| 446 | mamo-L | *BGIBMGA000889* | down | 1277 | 1291 | - | 11.4274 | 4.38E-05 | 1 | TAGGCTTGCGTAGAT |
| 447 | mamo-L | *BGIBMGA000892* | up | 1089 | 1103 | + | 11.6452 | 3.56E-05 | 1 | TAGGCCTTGGTTTCG |
| 448 | mamo-L | *BGIBMGA000892* | up | 966 | 980 | - | 10.6855 | 8.51E-05 | 1 | TAGGCCGGTAGTACT |
| 449 | mamo-L | *BGIBMGA000893* | down | 919 | 933 | - | 11.4435 | 4.31E-05 | 1 | TAGGCGCGTAGTGGT |
| 450 | mamo-L | *BGIBMGA000895* | down | 431 | 445 | - | 12.0645 | 2.34E-05 | 1 | TAGGCGGGCTTCGGA |
| 451 | mamo-L | *BGIBMGA000895* | up | 644 | 658 | - | 11.7903 | 3.08E-05 | 1 | TAGGCTTCGTTCTTT |
| 452 | mamo-L | *BGIBMGA000899* | down | 1594 | 1608 | - | 11.1694 | 5.59E-05 | 1 | AAGGCTTTAATTGGT |
| 453 | mamo-L | *BGIBMGA000904* | down | 361 | 375 | + | 10.9194 | 6.98E-05 | 1 | TAGGCTACTTTTTTT |

| | | | | | | | | | | |
|---|---|---|---|---|---|---|---|---|---|---|
| 454 | mamo-L | *BGIBMGA000909* | up | 1194 | 1208 | - | 11.621 | 3.63E-05 | 1 | TAGGCCTTCTTCGGG |
| 455 | mamo-L | *BGIBMGA000913* | down | 994 | 1008 | + | 10.5726 | 9.33E-05 | 1 | TAGGCTAATATTTAT |
| 456 | mamo-L | *BGIBMGA000914* | up | 996 | 1010 | + | 10.5726 | 9.33E-05 | 1 | TAGGCTAATATTTAT |
| 457 | mamo-L | *BGIBMGA000919* | up | 125 | 139 | + | 12.5081 | 1.43E-05 | 1 | TAGGCTTCTTTTTTT |
| 458 | mamo-L | *BGIBMGA000919* | down | 1037 | 1051 | - | 12.2419 | 1.94E-05 | 1 | TAGGCTCATTATGAT |
| 459 | mamo-L | *BGIBMGA000919* | down | 1916 | 1930 | + | 12.1371 | 2.17E-05 | 1 | GAGGCTTGATTTGCT |
| 460 | mamo-L | *BGIBMGA000924* | down | 734 | 748 | - | 11.2742 | 5.06E-05 | 1 | TAGGCATGCGTTCTT |
| 461 | mamo-L | *BGIBMGA000924* | down | 908 | 922 | + | 10.7097 | 8.33E-05 | 1 | TAGGCCGAATTGCTT |
| 462 | mamo-L | *BGIBMGA000926* | down | 1339 | 1353 | - | 11.4113 | 4.45E-05 | 1 | TAGGCTGAGCTTTCC |
| 463 | mamo-L | *BGIBMGA000927* | up | 504 | 518 | - | 11.4113 | 4.45E-05 | 1 | TAGGCTGAGCTTTCC |
| 464 | mamo-L | *BGIBMGA000928* | down | 536 | 550 | - | 13.1452 | 6.69E-06 | 1 | TAGGCTGTTGATGAT |
| 465 | mamo-L | *BGIBMGA000928* | up | 1063 | 1077 | + | 10.5081 | 9.83E-05 | 1 | GAGGCCCGGTTTACT |
| 466 | mamo-L | *BGIBMGA000936* | down | 1370 | 1384 | + | 13.9113 | 2.26E-06 | 1 | TAGGCCTATTTTTTT |
| 467 | mamo-L | *BGIBMGA000942* | down | 1561 | 1575 | - | 10.5806 | 9.27E-05 | 1 | TAGGCCTCCTTTTCA |
| 468 | mamo-L | *BGIBMGA000943* | up | 1172 | 1186 | - | 11.0806 | 6.05E-05 | 1 | GAGGCTTTGATTTTT |
| 469 | mamo-L | *BGIBMGA000947* | up | 412 | 426 | - | 10.9194 | 6.98E-05 | 1 | AAGGCGCGCTTTTTT |
| 470 | mamo-L | *BGIBMGA000958* | down | 1853 | 1867 | - | 13.0887 | 7.18E-06 | 1 | TAGGCTTGAGATGCT |
| 471 | mamo-L | *BGIBMGA000959* | up | 1460 | 1474 | - | 14 | 1.98E-06 | 1 | TAGGCTGCTGTTGCT |
| 472 | mamo-L | *BGIBMGA000967* | down | 299 | 313 | + | 11.7177 | 3.32E-05 | 1 | TAGGCTTGTTATTTC |
| 473 | mamo-L | *BGIBMGA000968* | up | 1192 | 1206 | + | 11.7177 | 3.32E-05 | 1 | TAGGCTTGTTATTTC |
| 474 | mamo-L | *BGIBMGA000969* | down | 1009 | 1023 | - | 11.7339 | 3.26E-05 | 1 | TAGGCTGAAGGTAGT |
| 475 | mamo-L | *BGIBMGA000969* | down | 65 | 79 | - | 10.7581 | 8.00E-05 | 1 | TAGGCTTAGATTAGA |
| 476 | mamo-L | *BGIBMGA000978* | down | 478 | 492 | - | 11.0726 | 6.09E-05 | 1 | TAGGCTGCTTTTAAT |
| 477 | mamo-L | *BGIBMGA000983* | down | 566 | 580 | - | 11.1935 | 5.45E-05 | 1 | TAGGCGCATGAGGGT |
| 478 | mamo-L | *BGIBMGA000984* | down | 932 | 946 | - | 10.5161 | 9.77E-05 | 1 | TAGGCCGCCTTTTTG |
| 479 | mamo-L | *BGIBMGA000985* | down | 1510 | 1524 | + | 10.879 | 7.22E-05 | 1 | TAGGCCGCCGGCTGT |
| 480 | mamo-L | *BGIBMGA000985* | up | 1039 | 1053 | - | 10.5161 | 9.77E-05 | 1 | TAGGCCGCCTTTTTG |
| 481 | mamo-L | *BGIBMGA000986* | up | 1590 | 1604 | - | 10.9113 | 7.02E-05 | 1 | AAGGCTCAGTTTTAT |
| 482 | mamo-L | *BGIBMGA000988* | up | 1825 | 1839 | - | 12.9597 | 8.38E-06 | 1 | TAGGCTGATCTTGTT |
| 483 | mamo-L | *BGIBMGA000989* | up | 554 | 568 | - | 10.5726 | 9.33E-05 | 1 | TAGGCTAATATTTAT |
| 484 | mamo-L | *BGIBMGA000991* | up | 256 | 270 | - | 13.6371 | 3.42E-06 | 1 | AAGGCTTAGTTTGTT |
| 485 | mamo-L | *BGIBMGA000991* | down | 1613 | 1627 | + | 13.0565 | 7.51E-06 | 1 | TAGGCGTGGGGTGGC |
| 486 | mamo-L | *BGIBMGA000992* | up | 125 | 139 | + | 13.0565 | 7.51E-06 | 1 | TAGGCGTGGGGTGGC |
| 487 | mamo-L | *BGIBMGA000994* | up | 1027 | 1041 | - | 12.0645 | 2.34E-05 | 1 | TAGGCGGGCTTCGGA |
| 488 | mamo-L | *BGIBMGA000995* | up | 932 | 946 | - | 14.3952 | 1.09E-06 | 1 | TAGGCCTTCTTTTGT |
| 489 | mamo-L | *BGIBMGA000997* | down | 1436 | 1450 | - | 11.1452 | 5.70E-05 | 1 | TAGGCTTCTAATGCT |
| 490 | mamo-L | *BGIBMGA000998* | up | 1432 | 1446 | + | 10.5403 | 9.58E-05 | 1 | TAGGCTGTATGGTGT |
| 491 | mamo-L | *BGIBMGA000999* | up | 12 | 26 | + | 11.7097 | 3.34E-05 | 1 | TAGGCAGTCTTGGTT |
| 492 | mamo-L | *BGIBMGA001001* | down | 893 | 907 | - | 11.2097 | 5.38E-05 | 1 | TAGGCCCAAATTTTT |
| 493 | mamo-L | *BGIBMGA001004* | up | 1958 | 1972 | + | 11.1129 | 5.88E-05 | 1 | AAGGCTTTGTTTGTA |
| 494 | mamo-L | *BGIBMGA001004* | up | 1127 | 1141 | - | 10.5 | 9.90E-05 | 1 | GAGGCCCAGGTCGTC |
| 495 | mamo-L | *BGIBMGA001006* | up | 542 | 556 | + | 11.5484 | 3.90E-05 | 1 | GAGGCTTGGGTCGAT |
| 496 | mamo-L | *BGIBMGA001008* | down | 1222 | 1236 | + | 11.5887 | 3.75E-05 | 1 | TAGGCGTTGTTCGGA |
| 497 | mamo-L | *BGIBMGA001013* | down | 1575 | 1589 | - | 11.6935 | 3.39E-05 | 1 | TAGGCGTGTTTGTTT |
| 498 | mamo-L | *BGIBMGA001016* | up | 421 | 435 | - | 10.5403 | 9.58E-05 | 1 | TAGGCTGTATGGTGT |
| 499 | mamo-L | *BGIBMGA001018* | up | 137 | 151 | - | 10.9194 | 6.98E-05 | 1 | TAGGCTACTTTTTTT |
| 500 | mamo-L | *BGIBMGA001018* | up | 1265 | 1279 | - | 10.5565 | 9.46E-05 | 1 | AAGGCTTTTATTTTT |
| 501 | mamo-L | *BGIBMGA001026* | up | 1913 | 1927 | + | 10.5726 | 9.33E-05 | 1 | TAGGCTAATATTTAT |
| 502 | mamo-L | *BGIBMGA001029* | up | 1029 | 1043 | - | 11.371 | 4.62E-05 | 1 | TAGGCTGTATAGGCT |
| 503 | mamo-L | *BGIBMGA001033* | down | 1714 | 1728 | + | 10.6371 | 8.85E-05 | 1 | TAGGCATATTTCGTC |
| 504 | mamo-L | *BGIBMGA001034* | down | 470 | 484 | - | 11.2177 | 5.34E-05 | 1 | AAGGCCGATGTCGAT |
| 505 | mamo-L | *BGIBMGA001036* | down | 1747 | 1761 | - | 11.8548 | 2.89E-05 | 1 | TAGGCTGGGGTGGTG |
| 506 | mamo-L | *BGIBMGA001042* | up | 1269 | 1283 | - | 10.5323 | 9.63E-05 | 1 | TCGGCTGGCTTTTTT |
| 507 | mamo-L | *BGIBMGA001043* | down | 1850 | 1864 | - | 10.9839 | 6.59E-05 | 1 | TAGGCATTATTTTGT |
| 508 | mamo-L | *BGIBMGA001044* | down | 1195 | 1209 | + | 11.371 | 4.62E-05 | 1 | TAGGCTTAGATCATT |
| 509 | mamo-L | *BGIBMGA001044* | up | 803 | 817 | + | 11.0323 | 6.32E-05 | 1 | TAGGCTGGGAGGTGT |
| 510 | mamo-L | *BGIBMGA001046* | up | 415 | 429 | - | 12.4113 | 1.60E-05 | 1 | TAGGCTCGAATTGTT |
| 511 | mamo-L | *BGIBMGA001047* | down | 1117 | 1131 | - | 10.7581 | 8.00E-05 | 1 | TAGGCTAACAGTGAT |
| 512 | mamo-L | *BGIBMGA001049* | down | 423 | 437 | + | 12.0968 | 2.26E-05 | 1 | TAGGCTTAAGTGTGT |
| 513 | mamo-L | *BGIBMGA001050* | down | 1755 | 1769 | + | 10.6532 | 8.73E-05 | 1 | GAGGCTTTTTTTTTC |
| 514 | mamo-L | *BGIBMGA001052* | up | 1790 | 1804 | - | 12.8952 | 9.05E-06 | 1 | TAGGCTTGTCTTGGT |
| 515 | mamo-L | *BGIBMGA001053* | down | 1717 | 1731 | - | 13.4677 | 4.39E-06 | 1 | TAGGCTGATGATGAT |
| 516 | mamo-L | *BGIBMGA001056* | up | 1346 | 1360 | - | 11.4435 | 4.31E-05 | 1 | TAGGCTTGAGAGGCT |
| 517 | mamo-L | *BGIBMGA001059* | up | 1284 | 1298 | + | 13.0968 | 7.11E-06 | 1 | TAGGCTGTTTTTTTC |
| 518 | mamo-L | *BGIBMGA001062* | down | 255 | 269 | + | 12.7177 | 1.12E-05 | 1 | AAGGCTGATTTTGGC |
| 519 | mamo-L | *BGIBMGA001064* | up | 1224 | 1238 | - | 10.9435 | 6.83E-05 | 1 | TAGGCGTATTTGTAT |
| 520 | mamo-L | *BGIBMGA001065* | down | 915 | 929 | - | 12.1613 | 2.11E-05 | 1 | TAGGCTGAATGTTCT |
| 521 | mamo-L | *BGIBMGA001067* | up | 1405 | 1419 | + | 11.2258 | 5.30E-05 | 1 | TAGGCCTTTTTCCCT |
| 522 | mamo-L | *BGIBMGA001067* | up | 205 | 219 | - | 10.9194 | 6.98E-05 | 1 | TAGGCTACTTTTTTT |
| 523 | mamo-L | *BGIBMGA001074* | up | 1493 | 1507 | - | 10.5726 | 9.33E-05 | 1 | TAGGCTAATATTTAT |
| 524 | mamo-L | *BGIBMGA001076* | down | 1523 | 1537 | - | 11.9274 | 2.69E-05 | 1 | GAGGCCTTCATTGTT |
| 525 | mamo-L | *BGIBMGA001076* | up | 1785 | 1799 | - | 10.7661 | 7.95E-05 | 1 | AAGGCTGGTGTTAGC |
| 526 | mamo-L | *BGIBMGA001086* | down | 1619 | 1633 | + | 12.5806 | 1.31E-05 | 1 | AAGGCGGGCGTTGGC |
| 527 | mamo-L | *BGIBMGA001088* | up | 530 | 544 | + | 12.3468 | 1.72E-05 | 1 | TAGGCTGAGAACGTT |
| 528 | mamo-L | *BGIBMGA001088* | up | 511 | 525 | + | 10.5726 | 9.33E-05 | 1 | TAGGCTGACAATTGA |
| 529 | mamo-L | *BGIBMGA001089* | up | 487 | 501 | - | 11.7177 | 3.32E-05 | 1 | TCGGCCGAGGTTGTT |
| 530 | mamo-L | *BGIBMGA001091* | down | 420 | 434 | - | 13.3226 | 5.34E-06 | 1 | TAGGCTCTGGGTGCT |
| 531 | mamo-L | *BGIBMGA001091* | up | 1649 | 1663 | + | 10.9355 | 6.88E-05 | 1 | TAGGCGGTGGTGGTG |
| 532 | mamo-L | *BGIBMGA001096* | up | 1334 | 1348 | - | 12.1613 | 2.11E-05 | 1 | TAGGCCTTTGTGGCC |
| 533 | mamo-L | *BGIBMGA001097* | up | 259 | 273 | - | 11.5887 | 3.75E-05 | 1 | TAGGCATGCGTTCGT |
| 534 | mamo-L | *BGIBMGA001103* | up | 78 | 92 | - | 12.3548 | 1.70E-05 | 1 | TAGGCTTTCTTTACC |
| 535 | mamo-L | *BGIBMGA001105* | up | 151 | 165 | + | 11.4435 | 4.31E-05 | 1 | TAGGCTTGAGAGGCT |
| 536 | mamo-L | *BGIBMGA001106* | up | 240 | 254 | + | 11.1694 | 5.59E-05 | 1 | TAGGCTGAAAATTTT |
| 537 | mamo-L | *BGIBMGA001110* | down | 700 | 714 | + | 13.9758 | 2.07E-06 | 1 | TAGGCGTAGTTTTTT |
| 538 | mamo-L | *BGIBMGA001110* | up | 1043 | 1057 | + | 12.5726 | 1.33E-05 | 1 | TAGGCCCGGATTGCC |
| 539 | mamo-L | *BGIBMGA001110* | down | 517 | 531 | + | 11.0887 | 6.01E-05 | 1 | TAGGCTTATTTCAAT |
| 540 | mamo-L | *BGIBMGA001112* | up | 852 | 866 | - | 10.9194 | 6.98E-05 | 1 | TAGGCGGGCTTCGAA |
| 541 | mamo-L | *BGIBMGA001113* | down | 13 | 27 | - | 10.7581 | 8.00E-05 | 1 | AAGGCTTTTTTTATT |
| 542 | mamo-L | *BGIBMGA001116* | up | 618 | 632 | - | 11.0323 | 6.32E-05 | 1 | TAGGCTGGGAGGTGT |
| 543 | mamo-L | *BGIBMGA001116* | down | 433 | 447 | + | 10.9839 | 6.59E-05 | 1 | TAGGCATTATTTTGT |
| 544 | mamo-L | *BGIBMGA001116* | down | 25 | 39 | - | 10.9032 | 7.07E-05 | 1 | TAGGCCTTATTTCAT |
| 545 | mamo-L | *BGIBMGA001117* | down | 280 | 294 | + | 12.0565 | 2.36E-05 | 1 | TAGGCTGTCTATGGG |

| | | | | | | | | | | |
|---|---|---|---|---|---|---|---|---|---|---|
| 546 | mamo-L | *BGIBMGA001117* | down | 27 | 41 | + | 11.3468 | 4.73E-05 | 1 | TAGGCGAAATTTTGT |
| 547 | mamo-L | *BGIBMGA001118* | down | 640 | 654 | - | 10.5081 | 9.83E-05 | 1 | TAGGCCAGTTTTAAT |
| 548 | mamo-L | *BGIBMGA001124* | down | 609 | 623 | - | 12.0161 | 2.46E-05 | 1 | TAGGCTGGTTATTTC |
| 549 | mamo-L | *BGIBMGA001124* | down | 1268 | 1282 | + | 11.8548 | 2.89E-05 | 1 | TAGGCTGGGGTGGTG |
| 550 | mamo-L | *BGIBMGA001133* | down | 1519 | 1533 | - | 11.3306 | 4.80E-05 | 1 | AAGGCCGTCTTTACT |
| 551 | mamo-L | *BGIBMGA001133* | up | 151 | 165 | + | 10.5403 | 9.58E-05 | 1 | TAGGCTTTATTGGGA |
| 552 | mamo-L | *BGIBMGA001134* | down | 340 | 354 | - | 11.3306 | 4.80E-05 | 1 | AAGGCCGTCTTTACT |
| 553 | mamo-L | *BGIBMGA001137* | up | 1537 | 1551 | + | 10.5403 | 9.58E-05 | 1 | TAGGCTGTATGGTGT |
| 554 | mamo-L | *BGIBMGA001138* | up | 1606 | 1620 | + | 11.3548 | 4.69E-05 | 1 | AAGGCTGATGATGAT |
| 555 | mamo-L | *BGIBMGA001141* | down | 1633 | 1647 | + | 11.6935 | 3.39E-05 | 1 | TAGGCGTGTTTGTTT |
| 556 | mamo-L | *BGIBMGA001142* | up | 1512 | 1526 | + | 10.9032 | 7.07E-05 | 1 | TAGGCTCTATTCTTT |
| 557 | mamo-L | *BGIBMGA001145* | up | 519 | 533 | - | 11.0323 | 6.32E-05 | 1 | TAGGCTTACGACCCT |
| 558 | mamo-L | *BGIBMGA001145* | up | 950 | 964 | - | 10.9194 | 6.98E-05 | 1 | TAGGCTACTTTTTTT |
| 559 | mamo-L | *BGIBMGA001145* | down | 1338 | 1352 | - | 10.5726 | 9.33E-05 | 1 | TAGGCTAATATTTAT |
| 560 | mamo-L | *BGIBMGA001147* | down | 1804 | 1818 | + | 11.1371 | 5.75E-05 | 1 | AAGGCTGCTGTTTGT |
| 561 | mamo-L | *BGIBMGA001149* | up | 1395 | 1409 | + | 13.8387 | 2.54E-06 | 1 | TAGGCTAGTTTTGGT |
| 562 | mamo-L | *BGIBMGA001150* | up | 1287 | 1301 | + | 13.2742 | 5.70E-06 | 1 | TAGGCGTAGGTTTAT |
| 563 | mamo-L | *BGIBMGA001150* | up | 781 | 795 | - | 10.75 | 8.06E-05 | 1 | GAGGCCGATTGTGCC |
| 564 | mamo-L | *BGIBMGA001151* | up | 711 | 725 | + | 13.0565 | 7.51E-06 | 1 | TAGGCTTACTACGCT |
| 565 | mamo-L | *BGIBMGA001152* | down | 546 | 560 | - | 10.8871 | 7.17E-05 | 1 | TAGGCATTTTTGGCT |
| 566 | mamo-L | *BGIBMGA001154* | down | 353 | 367 | - | 12.9919 | 8.10E-06 | 1 | TAGGCCGGTTTTGCA |
| 567 | mamo-L | *BGIBMGA001154* | up | 1329 | 1343 | + | 10.9839 | 6.59E-05 | 1 | AAGGCCATTTTTGTT |
| 568 | mamo-L | *BGIBMGA001155* | up | 1203 | 1217 | + | 11.5323 | 3.96E-05 | 1 | TAGGCGTGAGGTGAT |
| 569 | mamo-L | *BGIBMGA001156* | up | 1177 | 1191 | + | 12.0645 | 2.34E-05 | 1 | TAGGCGGGCTTCGGA |
| 570 | mamo-L | *BGIBMGA001160* | down | 1136 | 1150 | - | 10.5726 | 9.33E-05 | 1 | TAGGCTAATATTTAT |
| 571 | mamo-L | *BGIBMGA001162* | up | 621 | 635 | + | 10.9194 | 6.98E-05 | 1 | TAGGCTACTTTTTTT |
| 572 | mamo-L | *BGIBMGA001164* | up | 1210 | 1224 | + | 11.1935 | 5.45E-05 | 1 | AAGGCCGTGATTTCT |
| 573 | mamo-L | *BGIBMGA001168* | down | 465 | 479 | - | 11 | 6.50E-05 | 1 | TAGGCATATTTTGGA |
| 574 | mamo-L | *BGIBMGA001177* | down | 1181 | 1195 | - | 11.121 | 5.84E-05 | 1 | TAGGCTTAGTTGTGA |
| 575 | mamo-L | *BGIBMGA001179* | up | 907 | 921 | - | 10.9194 | 6.98E-05 | 1 | TAGGCTACTTTTTTT |
| 576 | mamo-L | *BGIBMGA001181* | down | 925 | 939 | - | 10.8226 | 7.58E-05 | 1 | AAGGCTGGTTTGGAT |
| 577 | mamo-L | *BGIBMGA001185* | up | 1539 | 1553 | - | 10.9194 | 6.98E-05 | 1 | TAGGCTACTTTTTTT |
| 578 | mamo-L | *BGIBMGA001188* | up | 1415 | 1429 | + | 14 | 1.98E-06 | 1 | TAGGCTGCTGTTGCT |
| 579 | mamo-L | *BGIBMGA001188* | down | 1054 | 1068 | + | 13.0887 | 7.18E-06 | 1 | TAGGCTTGAGATGCT |
| 580 | mamo-L | *BGIBMGA001191* | down | 1231 | 1245 | + | 11.6532 | 3.53E-05 | 1 | AAGGCTTTTATTGTT |
| 581 | mamo-L | *BGIBMGA001201* | up | 38 | 52 | + | 11.3065 | 4.91E-05 | 1 | TAGGCTGACGAGCGT |
| 582 | mamo-L | *BGIBMGA001201* | up | 1168 | 1182 | + | 11.1774 | 5.54E-05 | 1 | TGGGCCGAGTTTTTT |
| 583 | mamo-L | *BGIBMGA001202* | down | 1034 | 1048 | + | 10.9194 | 6.98E-05 | 1 | AAGGCGCGCTTTTTT |
| 584 | mamo-L | *BGIBMGA001204* | down | 460 | 474 | - | 12.7742 | 1.04E-05 | 1 | TAGGCCTCGGTGGGT |
| 585 | mamo-L | *BGIBMGA001204* | up | 180 | 194 | - | 11.5323 | 3.96E-05 | 1 | TAGGCTAATGTTGTA |
| 586 | mamo-L | *BGIBMGA001204* | up | 1524 | 1538 | - | 10.7742 | 7.89E-05 | 1 | TAGGCTCGATTGTTT |
| 587 | mamo-L | *BGIBMGA001205* | down | 1768 | 1782 | - | 10.6048 | 9.09E-05 | 1 | TAGGCGCACAGTTTT |
| 588 | mamo-L | *BGIBMGA001206* | up | 1766 | 1780 | + | 11.0806 | 6.05E-05 | 1 | GAGGCTTTGATTTTT |
| 589 | mamo-L | *BGIBMGA001209* | up | 1290 | 1304 | + | 12.3952 | 1.63E-05 | 1 | AAGGCCGAATTTGTT |
| 590 | mamo-L | *BGIBMGA001209* | up | 1271 | 1285 | + | 11.1774 | 5.54E-05 | 1 | AAGGCCGGAATTGCT |
| 591 | mamo-L | *BGIBMGA001211* | down | 883 | 897 | + | 11 | 6.50E-05 | 1 | AAGGCTAGGGTTTTT |
| 592 | mamo-L | *BGIBMGA001212* | down | 275 | 289 | - | 13.9113 | 2.26E-06 | 1 | TAGGCCTATTTTTTT |
| 593 | mamo-L | *BGIBMGA001212* | up | 651 | 665 | + | 11 | 6.50E-05 | 1 | AAGGCTAGGGTTTTT |
| 594 | mamo-L | *BGIBMGA001215* | down | 1168 | 1182 | - | 10.9839 | 6.59E-05 | 1 | TAGGCGTGGATTTAC |
| 595 | mamo-L | *BGIBMGA001217* | up | 1636 | 1650 | + | 11.8952 | 2.79E-05 | 1 | TAGGCTGTGTGGTGT |
| 596 | mamo-L | *BGIBMGA001218* | down | 32 | 46 | - | 10.75 | 8.06E-05 | 1 | AAGGCGGGTGTTATT |
| 597 | mamo-L | *BGIBMGA001219* | up | 1724 | 1738 | + | 12.4355 | 1.55E-05 | 1 | TAGGCTCACTATTCT |
| 598 | mamo-L | *BGIBMGA001224* | down | 1474 | 1488 | + | 12.8226 | 9.86E-06 | 1 | TAGGCTTCCTTCGTT |
| 599 | mamo-L | *BGIBMGA001226* | down | 114 | 128 | - | 12.121 | 2.20E-05 | 1 | TAGGCAGACGTTAGT |
| 600 | mamo-L | *BGIBMGA001231* | down | 1367 | 1381 | - | 10.5968 | 9.15E-05 | 1 | TAGGCTTAATTGCTT |
| 601 | mamo-L | *BGIBMGA001232* | down | 608 | 622 | + | 12.2419 | 1.94E-05 | 1 | TAGGCTCATTATGAT |
| 602 | mamo-L | *BGIBMGA001233* | up | 289 | 303 | - | 12.5081 | 1.43E-05 | 1 | TAGGCTTCTTTTTTT |
| 603 | mamo-L | *BGIBMGA001234* | down | 807 | 821 | + | 11.7823 | 3.11E-05 | 1 | TAGGCTGTGTAGGTC |
| 604 | mamo-L | *BGIBMGA001235* | down | 890 | 904 | + | 13.0081 | 7.94E-06 | 1 | TAGGCCTGCTGTGCC |
| 605 | mamo-L | *BGIBMGA001236* | down | 340 | 354 | - | 11.3952 | 4.52E-05 | 1 | TAGGCTTTATGCGTT |
| 606 | mamo-L | *BGIBMGA001237* | down | 30 | 44 | - | 12.5081 | 1.43E-05 | 1 | TAGGCCGGCGGCGGC |
| 607 | mamo-L | *BGIBMGA001237* | up | 1021 | 1035 | - | 11.3952 | 4.52E-05 | 1 | TAGGCTTTATGCGTT |
| 608 | mamo-L | *BGIBMGA001240* | down | 973 | 987 | - | 12.3065 | 1.80E-05 | 1 | TAGGCAGGATTTGTT |
| 609 | mamo-L | *BGIBMGA001240* | up | 370 | 384 | + | 10.629 | 8.91E-05 | 1 | TAGGCCGTGAGCTCT |
| 610 | mamo-L | *BGIBMGA001242* | down | 1505 | 1519 | + | 11.1694 | 5.59E-05 | 1 | AAGGCTTTAATTGGT |
| 611 | mamo-L | *BGIBMGA001244* | down | 1005 | 1019 | - | 10.5887 | 9.21E-05 | 1 | TAGGCGCATGACGGC |
| 612 | mamo-L | *BGIBMGA001245* | up | 562 | 576 | - | 10.9435 | 6.83E-05 | 1 | TAGGCTGTGATGACT |
| 613 | mamo-L | *BGIBMGA001247* | down | 1862 | 1876 | + | 12.0645 | 2.34E-05 | 1 | TAGGCGGGCTTCGGA |
| 614 | mamo-L | *BGIBMGA001247* | up | 1055 | 1069 | + | 11.2016 | 5.41E-05 | 1 | TAGGCTTAAGATATT |
| 615 | mamo-L | *BGIBMGA001248* | up | 1273 | 1287 | + | 11.7903 | 3.08E-05 | 1 | TAGGCTTCGTTCTTT |
| 616 | mamo-L | *BGIBMGA001249* | down | 1418 | 1432 | - | 13.5887 | 3.68E-06 | 1 | TAGGCCTTTTTTTTT |
| 617 | mamo-L | *BGIBMGA001253* | up | 46 | 60 | - | 11.6452 | 3.56E-05 | 1 | TAGGCCTTGGTTTCG |
| 618 | mamo-L | *BGIBMGA001253* | up | 169 | 183 | + | 10.6855 | 8.51E-05 | 1 | TAGGCCGGTAGTACT |
| 619 | mamo-L | *BGIBMGA001256* | down | 1695 | 1709 | - | 12.9919 | 8.10E-06 | 1 | TAGGCCGGTTTTGCA |
| 620 | mamo-L | *BGIBMGA001256* | down | 427 | 441 | + | 10.9032 | 7.07E-05 | 1 | AAGGCGTAGTTCTGT |
| 621 | mamo-L | *BGIBMGA001256* | down | 1847 | 1861 | + | 10.8629 | 7.33E-05 | 1 | TAGGCCATTTTTGCA |
| 622 | mamo-L | *BGIBMGA001257* | down | 1339 | 1353 | - | 11.4597 | 4.25E-05 | 1 | TAGGCGGGGTGGTGT |
| 623 | mamo-L | *BGIBMGA001257* | down | 1718 | 1732 | + | 10.5 | 9.90E-05 | 1 | GAGGCCCAGGTCGTC |
| 624 | mamo-L | *BGIBMGA001258* | up | 103 | 117 | - | 11.4597 | 4.25E-05 | 1 | TAGGCGGGGTGGTGT |
| 625 | mamo-L | *BGIBMGA001258* | up | 482 | 496 | + | 10.5 | 9.90E-05 | 1 | GAGGCCCAGGTCGTC |
| 626 | mamo-L | *BGIBMGA001262* | down | 829 | 843 | - | 12.9435 | 8.54E-06 | 1 | TAGGCTTTTTTTTAT |
| 627 | mamo-L | *BGIBMGA001262* | up | 1506 | 1520 | + | 11.7581 | 3.18E-05 | 1 | TAGGCTAAGTTCTCT |
| 628 | mamo-L | *BGIBMGA001266* | up | 294 | 308 | - | 14.7177 | 5.94E-07 | 1 | TAGGCTTACGTTTTT |
| 629 | mamo-L | *BGIBMGA001267* | down | 801 | 815 | - | 10.5726 | 9.33E-05 | 1 | TAGGCTAATATTTAT |
| 630 | mamo-L | *BGIBMGA001274* | down | 681 | 695 | + | 11.8065 | 3.03E-05 | 1 | TAGGCTGACGATCAT |
| 631 | mamo-L | *BGIBMGA001277* | down | 480 | 494 | + | 10.75 | 8.06E-05 | 1 | GAGGCCGATTGTGCC |
| 632 | mamo-L | *BGIBMGA001283* | down | 285 | 299 | - | 11.4274 | 4.38E-05 | 1 | TAGGCATTTATTGCT |
| 633 | mamo-L | *BGIBMGA001283* | down | 1835 | 1849 | - | 11.2742 | 5.06E-05 | 1 | AAGGCTTTCGTCGAT |
| 634 | mamo-L | *BGIBMGA001284* | up | 1496 | 1510 | + | 11.4516 | 4.28E-05 | 1 | TAGGCTTCTTTTCTT |
| 635 | mamo-L | *BGIBMGA001284* | down | 1386 | 1400 | - | 10.75 | 8.06E-05 | 1 | GAGGCCGATTGTGCC |
| 636 | mamo-L | *BGIBMGA001284* | up | 1798 | 1812 | + | 10.5726 | 9.33E-05 | 1 | TAGGCTTCATGTGAT |
| 637 | mamo-L | *BGIBMGA001284* | down | 273 | 287 | - | 10.5565 | 9.46E-05 | 1 | TAGGCTTATCATTGT |

| | | | | | | | | | | |
|---|---|---|---|---|---|---|---|---|---|---|
| 638 | mamo-L | *BGIBMGA001286* | up | 1523 | 1537 | + | 14.2339 | 1.38E-06 | 1 | TAGGCTTTCTTTTCT |
| 639 | mamo-L | *BGIBMGA001290* | up | 1462 | 1476 | - | 13.129 | 6.83E-06 | 1 | TAGGCTTTCATTTCT |
| 640 | mamo-L | *BGIBMGA001290* | down | 800 | 814 | - | 12.2984 | 1.81E-05 | 1 | TAGGCCTGTTTTGCG |
| 641 | mamo-L | *BGIBMGA001293* | down | 102 | 116 | - | 10.5968 | 9.15E-05 | 1 | GAGGCTAGTTTTTGT |
| 642 | mamo-L | *BGIBMGA001294* | up | 586 | 600 | - | 10.5968 | 9.15E-05 | 1 | GAGGCTAGTTTTTGT |
| 643 | mamo-L | *BGIBMGA001295* | down | 1235 | 1249 | - | 11.0887 | 6.01E-05 | 1 | AAGGCTTAGGGCGTT |
| 644 | mamo-L | *BGIBMGA001297* | down | 1762 | 1776 | + | 13.1048 | 7.05E-06 | 1 | TAGGCTCTTGTTTTT |
| 645 | mamo-L | *BGIBMGA001298* | up | 855 | 869 | + | 13.2823 | 5.66E-06 | 1 | TAGGCTGTTTTTGGA |
| 646 | mamo-L | *BGIBMGA001298* | down | 1328 | 1342 | + | 10.75 | 8.06E-05 | 1 | GAGGCCGATTGTGCC |
| 647 | mamo-L | *BGIBMGA001302* | down | 1526 | 1540 | - | 11.4516 | 4.28E-05 | 1 | TAGGCTTGCGTCACC |
| 648 | mamo-L | *BGIBMGA001303* | up | 880 | 894 | - | 12.7258 | 1.11E-05 | 1 | TAGGCGTGCTTTGTA |
| 649 | mamo-L | *BGIBMGA001304* | up | 1979 | 1993 | - | 11.3548 | 4.69E-05 | 1 | TCGGCTGTCTTTGCT |
| 650 | mamo-L | *BGIBMGA001305* | up | 1752 | 1766 | - | 11.3629 | 4.66E-05 | 1 | TAGGCTTAATTTGAA |
| 651 | mamo-L | *BGIBMGA001306* | down | 656 | 670 | - | 13.1048 | 7.05E-06 | 1 | TAGGCTGACGATAGT |
| 652 | mamo-L | *BGIBMGA001306* | up | 1929 | 1943 | - | 11.0726 | 6.09E-05 | 1 | TAGGCGGCCGTGTCT |
| 653 | mamo-L | *BGIBMGA001310* | up | 1489 | 1503 | - | 10.9919 | 6.55E-05 | 1 | AAGGCTAGGTTTGTC |
| 654 | mamo-L | *BGIBMGA001310* | up | 1633 | 1647 | - | 10.7984 | 7.74E-05 | 1 | AAGGCTTGTATTTTT |
| 655 | mamo-L | *BGIBMGA001318* | up | 1193 | 1207 | + | 11.0403 | 6.28E-05 | 1 | TAGGCCTACAATACT |
| 656 | mamo-L | *BGIBMGA001320* | up | 1006 | 1020 | + | 12.121 | 2.20E-05 | 1 | TAGGCTTTGATTGTA |
| 657 | mamo-L | *BGIBMGA001321* | up | 834 | 848 | + | 10.9919 | 6.55E-05 | 1 | AAGGCTAGGTTTGTC |
| 658 | mamo-L | *BGIBMGA001321* | up | 690 | 704 | + | 10.7984 | 7.74E-05 | 1 | AAGGCTTGTATTTTT |
| 659 | mamo-L | *BGIBMGA001325* | up | 941 | 955 | - | 12.7258 | 1.11E-05 | 1 | AAGGCTTGGTTTGAT |
| 660 | mamo-L | *BGIBMGA001325* | up | 1221 | 1235 | + | 10.8145 | 7.63E-05 | 1 | TAGGCTTGCAGCGCC |
| 661 | mamo-L | *BGIBMGA001335* | down | 406 | 420 | + | 11.8065 | 3.03E-05 | 1 | TAGGCTGACGTCGAA |
| 662 | mamo-L | *BGIBMGA001335* | down | 1079 | 1093 | - | 11.3387 | 4.77E-05 | 1 | TAGGCTGCTCTTGCT |
| 663 | mamo-L | *BGIBMGA001336* | up | 1395 | 1409 | - | 10.6371 | 8.85E-05 | 1 | TAGGCACTTTTTTCT |
| 664 | mamo-L | *BGIBMGA001337* | up | 1008 | 1022 | - | 10.5968 | 9.15E-05 | 1 | GAGGCTAGTTTTTGT |
| 665 | mamo-L | *BGIBMGA001338* | down | 1590 | 1604 | - | 10.4919 | 9.96E-05 | 1 | TAGGCTGTGTCTGTG |
| 666 | mamo-L | *BGIBMGA001340* | up | 734 | 748 | + | 12.2016 | 2.03E-05 | 1 | TAGGCTCCCTTTTTT |
| 667 | mamo-L | *BGIBMGA001345* | down | 1063 | 1077 | + | 11.2742 | 5.06E-05 | 1 | TAGGCGGCGGCTGTT |
| 668 | mamo-L | *BGIBMGA001345* | up | 461 | 475 | - | 10.9194 | 6.98E-05 | 1 | TAGGCTACTTTTTTT |
| 669 | mamo-L | *BGIBMGA001347* | down | 817 | 831 | - | 10.9194 | 6.98E-05 | 1 | TAGGCTACTTTTTTT |
| 670 | mamo-L | *BGIBMGA001347* | up | 1145 | 1159 | + | 10.6694 | 8.62E-05 | 1 | TAGGCGTATTTGGTA |
| 671 | mamo-L | *BGIBMGA001353* | up | 1416 | 1430 | + | 10.9839 | 6.59E-05 | 1 | GAGGCTGTCGTTAGC |
| 672 | mamo-L | *BGIBMGA001355* | down | 51 | 65 | + | 11.9113 | 2.74E-05 | 1 | AAGGCTTTGTGTGTT |
| 673 | mamo-L | *BGIBMGA001356* | up | 309 | 323 | + | 11.2984 | 4.95E-05 | 1 | AAGGCGTATATTGTT |
| 674 | mamo-L | *BGIBMGA001357* | up | 1248 | 1262 | - | 11.4677 | 4.22E-05 | 1 | TAGGCTGATGATAAT |
| 675 | mamo-L | *BGIBMGA001361* | down | 1086 | 1100 | + | 11.3548 | 4.69E-05 | 1 | AAGGCTGATGATGAT |
| 676 | mamo-L | *BGIBMGA001361* | down | 1804 | 1818 | - | 10.5081 | 9.83E-05 | 1 | TAGGCGGGGTGTTGG |
| 677 | mamo-L | *BGIBMGA001365* | down | 782 | 796 | - | 11.1935 | 5.45E-05 | 1 | TAGGCGCATGAGGGT |
| 678 | mamo-L | *BGIBMGA001369* | up | 64 | 78 | + | 13.1935 | 6.33E-06 | 1 | TAGGCTTTAGATGGT |
| 679 | mamo-L | *BGIBMGA001370* | up | 945 | 959 | - | 10.621 | 8.97E-05 | 1 | AAGGCGGGTTTTATT |
| 680 | mamo-L | *BGIBMGA001372* | up | 46 | 60 | + | 10.5968 | 9.15E-05 | 1 | AAGGCTTGGGGTAGT |
| 681 | mamo-L | *BGIBMGA001376* | up | 1759 | 1773 | + | 10.8226 | 7.58E-05 | 1 | AAGGCTCTTATTGCT |
| 682 | mamo-L | *BGIBMGA001377* | down | 1495 | 1509 | + | 11.2903 | 4.99E-05 | 1 | TAGGCGCGGTTTTAC |
| 683 | mamo-L | *BGIBMGA001379* | down | 1918 | 1932 | - | 11.1935 | 5.45E-05 | 1 | AAGGCTGAGTTGTTT |
| 684 | mamo-L | *BGIBMGA001381* | down | 911 | 925 | - | 11.2742 | 5.06E-05 | 1 | TAGGCGGCGGCTGTT |
| 685 | mamo-L | *BGIBMGA001386* | down | 115 | 129 | + | 13.3871 | 4.90E-06 | 1 | TAGGCTTAATATGGT |
| 686 | mamo-L | *BGIBMGA001392* | down | 1343 | 1357 | - | 11.1774 | 5.54E-05 | 1 | AAGGCTGATTTTGTA |
| 687 | mamo-L | *BGIBMGA001392* | up | 1425 | 1439 | - | 10.7419 | 8.11E-05 | 1 | TCGGCTGACGTTTTT |
| 688 | mamo-L | *BGIBMGA001394* | down | 387 | 401 | - | 11.5726 | 3.81E-05 | 1 | TAGGCGTGGTCTTGT |
| 689 | mamo-L | *BGIBMGA001398* | down | 1200 | 1214 | - | 12.7258 | 1.11E-05 | 1 | TAGGCTTTGGTTAAT |
| 690 | mamo-L | *BGIBMGA001399* | down | 1151 | 1165 | + | 11.7419 | 3.23E-05 | 1 | TAGGCCTAAGACGTT |
| 691 | mamo-L | *BGIBMGA001401* | up | 581 | 595 | + | 11.5323 | 3.96E-05 | 1 | TAGGCGTGAGGTGAT |
| 692 | mamo-L | *BGIBMGA001401* | down | 1644 | 1658 | - | 10.9194 | 6.98E-05 | 1 | TAGGCTTAGCATGTC |
| 693 | mamo-L | *BGIBMGA001406* | down | 1542 | 1556 | - | 12.5 | 1.44E-05 | 1 | TAGGCTAATATTGTT |
| 694 | mamo-L | *BGIBMGA001407* | up | 34 | 48 | + | 13.5565 | 3.86E-06 | 1 | TAGGCTTGTGTTAGT |
| 695 | mamo-L | *BGIBMGA001407* | down | 619 | 633 | + | 10.8145 | 7.63E-05 | 1 | GAGGCTTCGGTGGGT |
| 696 | mamo-L | *BGIBMGA001414* | down | 1747 | 1761 | + | 11.4597 | 4.25E-05 | 1 | TAGGCGGCTATTGGC |
| 697 | mamo-L | *BGIBMGA001415* | up | 42 | 56 | + | 12.3306 | 1.75E-05 | 1 | TAGGCTTACTATACT |
| 698 | mamo-L | *BGIBMGA001415* | down | 731 | 745 | + | 11.1935 | 5.45E-05 | 1 | TAGGCCGTGGCTCGT |
| 699 | mamo-L | *BGIBMGA001420* | down | 132 | 146 | - | 11.6129 | 3.66E-05 | 1 | TAGGCTTTTAATGAT |
| 700 | mamo-L | *BGIBMGA001423* | down | 141 | 155 | + | 13.6613 | 3.32E-06 | 1 | TAGGCTCGTGTTTGT |
| 701 | mamo-L | *BGIBMGA001428* | down | 395 | 409 | + | 11.121 | 5.84E-05 | 1 | TAGGCTGAATTGTAT |
| 702 | mamo-L | *BGIBMGA001429* | down | 249 | 263 | + | 11.5323 | 3.96E-05 | 1 | TAGGCGGGATTCTCT |
| 703 | mamo-L | *BGIBMGA001433* | down | 853 | 867 | - | 10.5726 | 9.33E-05 | 1 | TAGGCTAATATTTAT |
| 704 | mamo-L | *BGIBMGA001435* | up | 597 | 611 | - | 10.8629 | 7.33E-05 | 1 | TAGGCTGTCTAAGTT |
| 705 | mamo-L | *BGIBMGA001437* | up | 1802 | 1816 | - | 11.9194 | 2.72E-05 | 1 | TAGGCTTCTTTTAGT |
| 706 | mamo-L | *BGIBMGA001438* | down | 108 | 122 | + | 13.2823 | 5.66E-06 | 1 | TAGGCTGTCATCGTT |
| 707 | mamo-L | *BGIBMGA001439* | up | 1094 | 1108 | + | 12.4677 | 1.49E-05 | 1 | TAGGCTTACGTCTTC |
| 708 | mamo-L | *BGIBMGA001439* | up | 448 | 462 | - | 11.2177 | 5.34E-05 | 1 | TAGGCCGATTTCCGC |
| 709 | mamo-L | *BGIBMGA001440* | up | 81 | 95 | + | 11.7661 | 3.16E-05 | 1 | TAGGCAGTTTTTTTT |
| 710 | mamo-L | *BGIBMGA001440* | down | 743 | 757 | + | 10.8871 | 7.17E-05 | 1 | AAGGCTGTTGATGTC |
| 711 | mamo-L | *BGIBMGA001440* | up | 1949 | 1963 | + | 10.5726 | 9.33E-05 | 1 | AAGGCCTACTTTCGC |
| 712 | mamo-L | *BGIBMGA001441* | up | 517 | 531 | - | 10.6371 | 8.85E-05 | 1 | TAGGCTTTTTTTACA |
| 713 | mamo-L | *BGIBMGA001444* | up | 1033 | 1047 | - | 12.2258 | 1.97E-05 | 1 | AAGGCTGTTTTTGAT |
| 714 | mamo-L | *BGIBMGA001446* | down | 1290 | 1304 | + | 12.7581 | 1.07E-05 | 1 | TAGGCTGTCTTTTAC |
| 715 | mamo-L | *BGIBMGA001446* | down | 153 | 167 | - | 11.6129 | 3.66E-05 | 1 | TAGGCTTTTAATGAT |
| 716 | mamo-L | *BGIBMGA001451* | down | 36 | 50 | + | 13.3306 | 5.30E-06 | 1 | TAGGCTTAGTATTTT |
| 717 | mamo-L | *BGIBMGA001451* | down | 389 | 403 | + | 10.7258 | 8.23E-05 | 1 | TAGGCTTTGTGTTTA |
| 718 | mamo-L | *BGIBMGA001453* | up | 649 | 663 | + | 10.8387 | 7.47E-05 | 1 | TAGGCTTATTATGAA |
| 719 | mamo-L | *BGIBMGA001463* | up | 1684 | 1698 | + | 10.7984 | 7.74E-05 | 1 | TAGGCTGTATTGTAT |
| 720 | mamo-L | *BGIBMGA001465* | up | 886 | 900 | - | 10.9194 | 6.98E-05 | 1 | TAGGCTACTTTTTTT |
| 721 | mamo-L | *BGIBMGA001471* | up | 651 | 665 | + | 10.7823 | 7.84E-05 | 1 | TAGGCGGTGAGTTGC |
| 722 | mamo-L | *BGIBMGA001474* | up | 780 | 794 | - | 13.3145 | 5.41E-06 | 1 | TAGGCGTTGTGTGCT |
| 723 | mamo-L | *BGIBMGA001478* | down | 1433 | 1447 | - | 12.6129 | 1.26E-05 | 1 | TAGGCCTTTTTTTTC |
| 724 | mamo-L | *BGIBMGA001480* | up | 889 | 903 | - | 13.4677 | 4.39E-06 | 1 | TAGGCTGATGATGAT |
| 725 | mamo-L | *BGIBMGA001481* | up | 1823 | 1837 | + | 11.1774 | 5.54E-05 | 1 | TAGGCTTAGTTCTTA |
| 726 | mamo-L | *BGIBMGA001482* | down | 11 | 25 | - | 11.621 | 3.63E-05 | 1 | TAGGCCTTGGAGTGT |
| 727 | mamo-L | *BGIBMGA001482* | down | 18 | 32 | + | 10.6452 | 8.79E-05 | 1 | AAGGCCTACTTGTTT |
| 728 | mamo-L | *BGIBMGA001484* | down | 444 | 458 | - | 10.4919 | 9.96E-05 | 1 | TAGGCTTAGAATAAT |
| 729 | mamo-L | *BGIBMGA001488* | down | 862 | 876 | + | 11.8387 | 2.94E-05 | 1 | TAGGCTATGTTTATT |

| | | | | | | | | | | |
|---|---|---|---|---|---|---|---|---|---|---|
| 730 | mamo-L | *BGIBMGA001490* | up | 625 | 639 | - | 12.6935 | 1.15E-05 | 1 | TAGGCTGTATTGGCT |
| 731 | mamo-L | *BGIBMGA001490* | down | 1203 | 1217 | + | 11.621 | 3.63E-05 | 1 | TAGGCCTTGGAGTGT |
| 732 | mamo-L | *BGIBMGA001490* | down | 1196 | 1210 | - | 10.6452 | 8.79E-05 | 1 | AAGGCCTACTTGTTT |
| 733 | mamo-L | *BGIBMGA001491* | up | 1242 | 1256 | + | 12.879 | 9.26E-06 | 1 | TAGGCCTCGTTTTTT |
| 734 | mamo-L | *BGIBMGA001493* | up | 1282 | 1296 | + | 13.4677 | 4.39E-06 | 1 | TAGGCTGATGATGAT |
| 735 | mamo-L | *BGIBMGA001495* | down | 1010 | 1024 | - | 11.629 | 3.61E-05 | 1 | TAGGCCTATGGCGAT |
| 736 | mamo-L | *BGIBMGA001497* | up | 1196 | 1210 | + | 13.3065 | 5.48E-06 | 1 | GAGGCGTACGTTGGT |
| 737 | mamo-L | *BGIBMGA001499* | down | 1876 | 1890 | - | 10.5968 | 9.15E-05 | 1 | GAGGCTAGTTTTTGT |
| 738 | mamo-L | *BGIBMGA001500* | down | 89 | 103 | - | 10.629 | 8.91E-05 | 1 | TAGGCCTGCAGCGCC |
| 739 | mamo-L | *BGIBMGA001501* | down | 1129 | 1143 | - | 11.3145 | 4.87E-05 | 1 | TAGGCTTAGGCTTAT |
| 740 | mamo-L | *BGIBMGA001503* | up | 1103 | 1117 | - | 11.4274 | 4.38E-05 | 1 | AAGGCCGAGGATTCT |
| 741 | mamo-L | *BGIBMGA001504* | down | 1353 | 1367 | + | 11.2984 | 4.95E-05 | 1 | TAGGCTACGGGTGTT |
| 742 | mamo-L | *BGIBMGA001509* | up | 1333 | 1347 | - | 12.7258 | 1.11E-05 | 1 | TAGGCCTAGGAGGTT |
| 743 | mamo-L | *BGIBMGA001509* | up | 931 | 945 | - | 12.0887 | 2.29E-05 | 1 | GAGGCTTATGGTGGT |
| 744 | mamo-L | *BGIBMGA001509* | up | 1340 | 1354 | + | 11.8387 | 2.94E-05 | 1 | TAGGCCTATGCTGGC |
| 745 | mamo-L | *BGIBMGA001511* | up | 339 | 353 | - | 11.7984 | 3.06E-05 | 1 | GAGGCTGAGGTTGCA |
| 746 | mamo-L | *BGIBMGA001513* | up | 872 | 886 | + | 11.0323 | 6.32E-05 | 1 | TAGGCAGACGTGTCT |
| 747 | mamo-L | *BGIBMGA001513* | up | 1665 | 1679 | - | 10.5726 | 9.33E-05 | 1 | TAGGCTAATATTTAT |
| 748 | mamo-L | *BGIBMGA001514* | down | 1789 | 1803 | + | 10.5161 | 9.77E-05 | 1 | TAGGCTTTTTTTCTA |
| 749 | mamo-L | *BGIBMGA001521* | down | 646 | 660 | + | 11.1048 | 5.92E-05 | 1 | TAGGCAGGTTTTATT |
| 750 | mamo-L | *BGIBMGA001522* | down | 1864 | 1878 | + | 12.1774 | 2.08E-05 | 1 | TAGGCTTGCACTGGT |
| 751 | mamo-L | *BGIBMGA001525* | down | 1813 | 1827 | - | 12.7581 | 1.07E-05 | 1 | TAGGCTGTCTTTTAC |
| 752 | mamo-L | *BGIBMGA001532* | up | 879 | 893 | - | 11.25 | 5.18E-05 | 1 | TAGGCCCATTTCAGT |
| 753 | mamo-L | *BGIBMGA001535* | down | 198 | 212 | + | 11.2177 | 5.34E-05 | 1 | TAGGCCGATTTCCGC |
| 754 | mamo-L | *BGIBMGA001535* | down | 708 | 722 | + | 10.8145 | 7.63E-05 | 1 | TGGGCGCAGGTTGTT |
| 755 | mamo-L | *BGIBMGA001536* | down | 1673 | 1687 | + | 11.9194 | 2.72E-05 | 1 | TAGGCTTCTTTTAGT |
| 756 | mamo-L | *BGIBMGA001537* | up | 1074 | 1088 | + | 11.9194 | 2.72E-05 | 1 | TAGGCTTCTTTTAGT |
| 757 | mamo-L | *BGIBMGA001538* | up | 1186 | 1200 | + | 12.9355 | 8.65E-06 | 1 | TAGGCTGTATGTGCT |
| 758 | mamo-L | *BGIBMGA001541* | down | 519 | 533 | - | 12.5242 | 1.40E-05 | 1 | TAGGCCCATTTTAGT |
| 759 | mamo-L | *BGIBMGA001547* | down | 951 | 965 | - | 11.121 | 5.84E-05 | 1 | TAGGCTGAATTGTAT |
| 760 | mamo-L | *BGIBMGA001548* | down | 1457 | 1471 | - | 11.7016 | 3.37E-05 | 1 | TAGGCAGACGATTGT |
| 761 | mamo-L | *BGIBMGA001548* | down | 59 | 73 | + | 11.4758 | 4.18E-05 | 1 | TAGGCTTAGTTGGCG |
| 762 | mamo-L | *BGIBMGA001551* | down | 1082 | 1096 | + | 11.6774 | 3.44E-05 | 1 | GAGGCTTAGTTTTAT |
| 763 | mamo-L | *BGIBMGA001555* | down | 278 | 292 | + | 11.0968 | 5.97E-05 | 1 | GAGGCCGATTGTGGC |
| 764 | mamo-L | *BGIBMGA001560* | down | 1413 | 1427 | + | 11.0726 | 6.09E-05 | 1 | TAGGCTTGTTTTCGA |
| 765 | mamo-L | *BGIBMGA001561* | up | 1341 | 1355 | + | 11.5323 | 3.96E-05 | 1 | TAGGCGTGAGGTGAT |
| 766 | mamo-L | *BGIBMGA001562* | down | 570 | 584 | + | 12.4516 | 1.53E-05 | 1 | TGGGCTGTGTTTGGT |
| 767 | mamo-L | *BGIBMGA001563* | up | 886 | 900 | + | 11.5726 | 3.81E-05 | 1 | TAGGCGGGTTTCGGA |
| 768 | mamo-L | *BGIBMGA001565* | down | 219 | 233 | - | 11.4597 | 4.25E-05 | 1 | TAGGCGGCTATTGGC |
| 769 | mamo-L | *BGIBMGA001568* | up | 1070 | 1084 | + | 10.5 | 9.90E-05 | 1 | GAGGCCCAGGTCGTC |
| 770 | mamo-L | *BGIBMGA001570* | down | 1882 | 1896 | + | 11.5968 | 3.73E-05 | 1 | GAGGCTTTTTTTTCT |
| 771 | mamo-L | *BGIBMGA001571* | down | 1958 | 1972 | + | 10.5726 | 9.33E-05 | 1 | TAGGCTAATATTTAT |
| 772 | mamo-L | *BGIBMGA001573* | up | 567 | 581 | - | 11.2742 | 5.06E-05 | 1 | AAGGCCGGTTTTCGT |
| 773 | mamo-L | *BGIBMGA001574* | down | 1761 | 1775 | - | 10.9839 | 6.59E-05 | 1 | TAGGCAGTGATGGGT |
| 774 | mamo-L | *BGIBMGA001578* | up | 497 | 511 | + | 10.7823 | 7.84E-05 | 1 | TAGGCTGGATGGTGT |
| 775 | mamo-L | *BGIBMGA001580* | up | 1805 | 1819 | + | 12.3952 | 1.63E-05 | 1 | AAGGCGGGGTTTTGT |
| 776 | mamo-L | *BGIBMGA001581* | down | 965 | 979 | + | 11.9274 | 2.69E-05 | 1 | TAGGCCACTGTTGCT |
| 777 | mamo-L | *BGIBMGA001583* | up | 1144 | 1158 | - | 10.5726 | 9.33E-05 | 1 | TAGGCTAATATTTAT |
| 778 | mamo-L | *BGIBMGA001586* | up | 41 | 55 | + | 12.0645 | 2.34E-05 | 1 | TAGGCGGGCTTCGGA |
| 779 | mamo-L | *BGIBMGA001588* | up | 1961 | 1975 | - | 10.5726 | 9.33E-05 | 1 | TAGGCTAATATTTAT |
| 780 | mamo-L | *BGIBMGA001596* | down | 1458 | 1472 | - | 10.7258 | 8.23E-05 | 1 | TAGGCTGGGATAGGC |
| 781 | mamo-L | *BGIBMGA001598* | down | 353 | 367 | + | 12.8387 | 9.67E-06 | 1 | TAGGCTCGGTTTACT |
| 782 | mamo-L | *BGIBMGA001607* | up | 755 | 769 | + | 11.0323 | 6.32E-05 | 1 | TAGGCGGTATTTAGC |
| 783 | mamo-L | *BGIBMGA001607* | up | 1767 | 1781 | + | 10.9194 | 6.98E-05 | 1 | TAGGCTACTTTTTTT |
| 784 | mamo-L | *BGIBMGA001608* | up | 606 | 620 | + | 10.7258 | 8.23E-05 | 1 | TAGGCTTCTGTTACC |
| 785 | mamo-L | *BGIBMGA001612* | down | 1319 | 1333 | - | 10.8871 | 7.17E-05 | 1 | TAGGCTAGGCTTGGC |
| 786 | mamo-L | *BGIBMGA001613* | down | 645 | 659 | + | 12.0565 | 2.36E-05 | 1 | AAGGCTCTTGTTGCT |
| 787 | mamo-L | *BGIBMGA001614* | up | 1759 | 1773 | + | 12.0565 | 2.36E-05 | 1 | AAGGCTCTTGTTGCT |
| 788 | mamo-L | *BGIBMGA001614* | up | 394 | 408 | + | 10.8306 | 7.52E-05 | 1 | AAGGCCTCCGTTTTT |
| 789 | mamo-L | *BGIBMGA001616* | down | 220 | 234 | - | 11.6855 | 3.42E-05 | 1 | TAGGCGGAGTGTGGG |
| 790 | mamo-L | *BGIBMGA001616* | down | 1611 | 1625 | - | 10.8306 | 7.52E-05 | 1 | AAGGCTTTATTTTCT |
| 791 | mamo-L | *BGIBMGA001618* | up | 418 | 432 | - | 12.0887 | 2.29E-05 | 1 | TAGGCAGATTTTTTT |
| 792 | mamo-L | *BGIBMGA001619* | up | 428 | 442 | - | 11 | 6.50E-05 | 1 | TAGGCATGCGTGTGT |
| 793 | mamo-L | *BGIBMGA001620* | up | 750 | 764 | - | 10.75 | 8.06E-05 | 1 | GAGGCCGATTGTGCC |
| 794 | mamo-L | *BGIBMGA001621* | up | 1437 | 1451 | + | 11.2339 | 5.26E-05 | 1 | TAGGCAGGGTTGTGT |
| 795 | mamo-L | *BGIBMGA001625* | down | 699 | 713 | + | 10.8387 | 7.47E-05 | 1 | TAGGCGGACCTCGGC |
| 796 | mamo-L | *BGIBMGA001628* | down | 636 | 650 | - | 12.5323 | 1.39E-05 | 1 | TAGGCCTAGTTCTGC |
| 797 | mamo-L | *BGIBMGA001630* | down | 1958 | 1972 | + | 10.7984 | 7.74E-05 | 1 | AAGGCGGTATTTTGT |
| 798 | mamo-L | *BGIBMGA001634* | down | 1170 | 1184 | + | 10.7258 | 8.23E-05 | 1 | TAGCCTTTTTTTGCT |
| 799 | mamo-L | *BGIBMGA001634* | up | 1576 | 1590 | + | 10.5161 | 9.77E-05 | 1 | TAGGCCTATGATTTA |
| 800 | mamo-L | *BGIBMGA001635* | down | 1674 | 1688 | - | 12.1371 | 2.17E-05 | 1 | GAGGCCGTTGTTGAT |
| 801 | mamo-L | *BGIBMGA001636* | up | 1764 | 1778 | - | 12.1371 | 2.17E-05 | 1 | GAGGCCGTTGTTGAT |
| 802 | mamo-L | *BGIBMGA001637* | down | 1013 | 1027 | + | 11.1129 | 5.88E-05 | 1 | AAGGCTTTGTTTGTA |
| 803 | mamo-L | *BGIBMGA001639* | up | 348 | 362 | - | 11.3629 | 4.66E-05 | 1 | TAGGCAGTGGGTTGT |
| 804 | mamo-L | *BGIBMGA001641* | up | 516 | 530 | - | 11.4194 | 4.42E-05 | 1 | TAGGCAGTCATTGAT |
| 805 | mamo-L | *BGIBMGA001641* | down | 793 | 807 | - | 10.9274 | 6.92E-05 | 1 | TAGGCCGGCAGCGCC |
| 806 | mamo-L | *BGIBMGA001642* | down | 1076 | 1090 | + | 10.75 | 8.06E-05 | 1 | TAGGCGTATATTAGC |
| 807 | mamo-L | *BGIBMGA001654* | up | 58 | 72 | + | 12.7581 | 1.07E-05 | 1 | TAGGCGGGGGTTAGC |
| 808 | mamo-L | *BGIBMGA001655* | down | 264 | 278 | + | 10.9758 | 6.64E-05 | 1 | AAGGCCGTATTTTTT |
| 809 | mamo-L | *BGIBMGA001655* | up | 1256 | 1270 | + | 10.5726 | 9.33E-05 | 1 | TAGGCTAATATTTAT |
| 810 | mamo-L | *BGIBMGA001660* | up | 1858 | 1872 | - | 10.7984 | 7.74E-05 | 1 | TAGGCACTTGTTTTT |
| 811 | mamo-L | *BGIBMGA001661* | down | 493 | 507 | + | 13.1048 | 7.05E-06 | 1 | TAGGCCGATATTTTT |
| 812 | mamo-L | *BGIBMGA001662* | up | 1073 | 1087 | + | 13.1048 | 7.05E-06 | 1 | TAGGCCGATATTTTT |
| 813 | mamo-L | *BGIBMGA001662* | down | 564 | 578 | + | 12.2903 | 1.83E-05 | 1 | TAGGCGTTGAATGCT |
| 814 | mamo-L | *BGIBMGA001664* | down | 1230 | 1244 | - | 12.4597 | 1.51E-05 | 1 | AAGGCTTGGTTTTTT |
| 815 | mamo-L | *BGIBMGA001664* | down | 855 | 869 | + | 10.5726 | 9.33E-05 | 1 | TAGGCTAATATTTAT |
| 816 | mamo-L | *BGIBMGA001665* | down | 1697 | 1711 | - | 11.1048 | 5.92E-05 | 1 | GAGGCCTGTTTGGCT |
| 817 | mamo-L | *BGIBMGA001666* | down | 1335 | 1349 | + | 10.5565 | 9.46E-05 | 1 | TAGGCGGGGTGGAGT |
| 818 | mamo-L | *BGIBMGA001668* | down | 545 | 559 | - | 10.9677 | 6.68E-05 | 1 | TAGGCAGTATTTTTT |
| 819 | mamo-L | *BGIBMGA001674* | down | 581 | 595 | - | 11.0887 | 6.01E-05 | 1 | AAGGCCTATGTCGGC |
| 820 | mamo-L | *BGIBMGA001674* | down | 726 | 740 | + | 10.9758 | 6.64E-05 | 1 | TAGGCTGTATATTTC |
| 821 | mamo-L | *BGIBMGA001675* | up | 1339 | 1353 | - | 11.0887 | 6.01E-05 | 1 | AAGGCCTATGTCGGC |

| | | | | | | | | | | |
|---|---|---|---|---|---|---|---|---|---|---|
| 822 | mamo-L | *BGIBMGA001675* | up | 1484 | 1498 | + | 10.9758 | 6.64E-05 | 1 | TAGGCTGTATATTTC |
| 823 | mamo-L | *BGIBMGA001678* | up | 1683 | 1697 | + | 12.3226 | 1.77E-05 | 1 | TAGGCCAATTTTTTT |
| 824 | mamo-L | *BGIBMGA001679* | down | 953 | 967 | + | 12.4032 | 1.61E-05 | 1 | AAGGCGTATTTTGTT |
| 825 | mamo-L | *BGIBMGA001688* | down | 229 | 243 | + | 12.6855 | 1.16E-05 | 1 | TAGGCTTTTTGTTGT |
| 826 | mamo-L | *BGIBMGA001688* | up | 760 | 774 | - | 11.2661 | 5.10E-05 | 1 | TAGGCCAGTTTTTTC |
| 827 | mamo-L | *BGIBMGA001688* | up | 925 | 939 | + | 10.5081 | 9.83E-05 | 1 | AAGGCTCGCTTTCCT |
| 828 | mamo-L | *BGIBMGA001693* | up | 1572 | 1586 | - | 12.7177 | 1.12E-05 | 1 | TAGGCTTTTTATGAT |
| 829 | mamo-L | *BGIBMGA001693* | down | 965 | 979 | + | 11.2258 | 5.30E-05 | 1 | TAGGCCTTTTTCCCT |
| 830 | mamo-L | *BGIBMGA001693* | up | 1587 | 1601 | + | 11.2258 | 5.30E-05 | 1 | TAGGCTTTCATTGAA |
| 831 | mamo-L | *BGIBMGA001695* | up | 1733 | 1747 | + | 11.3629 | 4.66E-05 | 1 | TAGGCTTTTTTAGCT |
| 832 | mamo-L | *BGIBMGA001696* | up | 787 | 801 | + | 10.5726 | 9.33E-05 | 1 | TAGGCTAATATTTAT |
| 833 | mamo-L | *BGIBMGA001698* | down | 1716 | 1730 | - | 11.4919 | 4.11E-05 | 1 | TAGGCATACATTTGT |
| 834 | mamo-L | *BGIBMGA001702* | up | 1698 | 1712 | + | 13.3548 | 5.14E-06 | 1 | TAGGCAGTCTTTGTT |
| 835 | mamo-L | *BGIBMGA001702* | down | 29 | 43 | + | 11.7823 | 3.11E-05 | 1 | TAGGCTTGGCATGCT |
| 836 | mamo-L | *BGIBMGA001704* | up | 1724 | 1738 | - | 10.4919 | 9.96E-05 | 1 | TAGGCTGTGGTAGGA |
| 837 | mamo-L | *BGIBMGA001706* | up | 775 | 789 | - | 11.6613 | 3.50E-05 | 1 | AAGGCTTTTTTTTTT |
| 838 | mamo-L | *BGIBMGA001709* | up | 1119 | 1133 | + | 12.5726 | 1.33E-05 | 1 | TAGGCAGGGTGTGGT |
| 839 | mamo-L | *BGIBMGA001713* | up | 499 | 513 | - | 11.4355 | 4.34E-05 | 1 | AAGGCTTATTTGGTT |
| 840 | mamo-L | *BGIBMGA001717* | down | 1948 | 1962 | - | 10.7581 | 8.00E-05 | 1 | AAGGCGTTTTATGTT |
| 841 | mamo-L | *BGIBMGA001718* | up | 1931 | 1945 | - | 10.7581 | 8.00E-05 | 1 | AAGGCGTTTTATGTT |
| 842 | mamo-L | *BGIBMGA001720* | down | 895 | 909 | + | 13.5403 | 3.97E-06 | 1 | TAGGCTGTATTTGAT |
| 843 | mamo-L | *BGIBMGA001720* | down | 987 | 1001 | + | 11.1129 | 5.88E-05 | 1 | TAGGCCTTGTTTTAA |
| 844 | mamo-L | *BGIBMGA001722* | down | 665 | 679 | - | 11.4113 | 4.45E-05 | 1 | TAGGCTACCTTTTTT |
| 845 | mamo-L | *BGIBMGA001724* | up | 1177 | 1191 | - | 13.3548 | 5.14E-06 | 1 | TAGGCAGTCTTTGTT |
| 846 | mamo-L | *BGIBMGA001724* | up | 492 | 506 | - | 11.7823 | 3.11E-05 | 1 | TAGGCTTGGCATGCT |
| 847 | mamo-L | *BGIBMGA001727* | down | 1529 | 1543 | - | 11.129 | 5.79E-05 | 1 | TAGGCTGGTGGTACC |
| 848 | mamo-L | *BGIBMGA001728* | up | 1459 | 1473 | - | 10.5726 | 9.33E-05 | 1 | TAGGCTAATATTTAT |
| 849 | mamo-L | *BGIBMGA001729* | up | 293 | 307 | - | 10.5726 | 9.33E-05 | 1 | TAGGCTAATATTTAT |
| 850 | mamo-L | *BGIBMGA001731* | down | 1228 | 1242 | - | 11.2258 | 5.30E-05 | 1 | TAGGCCTTTTTCCCT |
| 851 | mamo-L | *BGIBMGA001733* | up | 155 | 169 | + | 11.5242 | 3.99E-05 | 1 | TAGGCTATTTTTTGC |
| 852 | mamo-L | *BGIBMGA001733* | up | 372 | 386 | + | 11.2258 | 5.30E-05 | 1 | TAGGCCTTTTTCCCT |
| 853 | mamo-L | *BGIBMGA001734* | down | 512 | 526 | + | 10.8306 | 7.52E-05 | 1 | AAGGCTTTCTATTTT |
| 854 | mamo-L | *BGIBMGA001736* | down | 1623 | 1637 | + | 10.9597 | 6.73E-05 | 1 | TAGGCGCTTATGGGT |
| 855 | mamo-L | *BGIBMGA001738* | up | 920 | 934 | + | 11.2661 | 5.10E-05 | 1 | TAGGCCAGTTTTTTC |
| 856 | mamo-L | *BGIBMGA001738* | down | 1061 | 1075 | + | 10.621 | 8.97E-05 | 1 | TAGGCAGAAATTGGC |
| 857 | mamo-L | *BGIBMGA001738* | up | 755 | 769 | - | 10.5081 | 9.83E-05 | 1 | AAGGCTCGCTTTCCT |
| 858 | mamo-L | *BGIBMGA001746* | down | 1021 | 1035 | + | 10.8952 | 7.12E-05 | 1 | TAGGCTCTTGTCACT |
| 859 | mamo-L | *BGIBMGA001746* | down | 1751 | 1765 | + | 10.629 | 8.91E-05 | 1 | AAGGCCTTAGTCGTT |
| 860 | mamo-L | *BGIBMGA001748* | down | 50 | 64 | + | 10.7984 | 7.74E-05 | 1 | AAGGCGGAGAATGCT |
| 861 | mamo-L | *BGIBMGA001761* | down | 937 | 951 | + | 11.5484 | 3.90E-05 | 1 | TAGGCAGAGAGTGGT |
| 862 | mamo-L | *BGIBMGA001763* | down | 1660 | 1674 | - | 11.3548 | 4.69E-05 | 1 | TAGGCGGGGCGTGTT |
| 863 | mamo-L | *BGIBMGA001770* | down | 1069 | 1083 | - | 12.2581 | 1.90E-05 | 1 | TAGGCTTGTGTTTGA |
| 864 | mamo-L | *BGIBMGA001771* | up | 1399 | 1413 | - | 12.2581 | 1.90E-05 | 1 | TAGGCTTGTGTTTGA |
| 865 | mamo-L | *BGIBMGA001771* | down | 1359 | 1373 | - | 10.9194 | 6.98E-05 | 1 | TAGGCTACTTTTTTT |
| 866 | mamo-L | *BGIBMGA001772* | down | 575 | 589 | - | 10.5565 | 9.46E-05 | 1 | TAGGCGGGGTGGAGT |
| 867 | mamo-L | *BGIBMGA001774* | up | 1559 | 1573 | + | 10.5968 | 9.15E-05 | 1 | GAGGCTAGTTTTTGT |
| 868 | mamo-L | *BGIBMGA001775* | down | 1815 | 1829 | - | 10.5726 | 9.33E-05 | 1 | TAGGCTAATATTTAT |
| 869 | mamo-L | *BGIBMGA001777* | up | 1362 | 1376 | - | 10.5806 | 9.27E-05 | 1 | AAGGCTGAATTTATT |
| 870 | mamo-L | *BGIBMGA001779* | up | 1564 | 1578 | - | 11.9597 | 2.61E-05 | 1 | TAGGCGCATTTTTGC |
| 871 | mamo-L | *BGIBMGA001783* | up | 1334 | 1348 | + | 10.8629 | 7.33E-05 | 1 | GAGGCGTTCTTGGCT |
| 872 | mamo-L | *BGIBMGA001784* | up | 1764 | 1778 | - | 11.5403 | 3.93E-05 | 1 | TAGGCTTACTGGTTT |
| 873 | mamo-L | *BGIBMGA001784* | down | 1575 | 1589 | + | 11.1613 | 5.63E-05 | 1 | TAGGCGGACGTGGCG |
| 874 | mamo-L | *BGIBMGA001785* | down | 1573 | 1587 | - | 12.7419 | 1.09E-05 | 1 | TAGGCGGAGTTTGCG |
| 875 | mamo-L | *BGIBMGA001785* | up | 336 | 350 | + | 11.1613 | 5.63E-05 | 1 | TAGGCGGACGTGGCG |
| 876 | mamo-L | *BGIBMGA001788* | up | 18 | 32 | + | 12.3629 | 1.69E-05 | 1 | TAGGCTAAGGTTTAT |
| 877 | mamo-L | *BGIBMGA001790* | down | 1708 | 1722 | + | 11.1129 | 5.88E-05 | 1 | TAGGCGTTTTTCGAC |
| 878 | mamo-L | *BGIBMGA001793* | up | 1381 | 1395 | - | 14.5806 | 7.86E-07 | 1 | TAGGCTGTGGTCGTT |
| 879 | mamo-L | *BGIBMGA001793* | down | 474 | 488 | + | 10.5806 | 9.27E-05 | 1 | TAGGCTACCTTTTAT |
| 880 | mamo-L | *BGIBMGA001794* | up | 77 | 91 | + | 10.5726 | 9.33E-05 | 1 | TAGGCTAATATTTAT |
| 881 | mamo-L | *BGIBMGA001797* | down | 586 | 600 | + | 12.5968 | 1.28E-05 | 1 | AAGGCTTACTTTGTC |
| 882 | mamo-L | *BGIBMGA001798* | down | 410 | 424 | - | 11.0403 | 6.28E-05 | 1 | TAGGCTTAAATCGTC |
| 883 | mamo-L | *BGIBMGA001806* | down | 174 | 188 | + | 12.0887 | 2.29E-05 | 1 | TAGGCTTTCTTCATT |
| 884 | mamo-L | *BGIBMGA001807* | down | 1454 | 1468 | + | 11.3145 | 4.87E-05 | 1 | AAGGCTGTCTGTGAT |
| 885 | mamo-L | *BGIBMGA001812* | down | 898 | 912 | + | 11.4194 | 4.42E-05 | 1 | TAGGCCTTTGATTTC |
| 886 | mamo-L | *BGIBMGA001814* | up | 1710 | 1724 | + | 12.2984 | 1.81E-05 | 1 | TAGGCGCTTTTTTTT |
| 887 | mamo-L | *BGIBMGA001814* | down | 1953 | 1967 | - | 10.7581 | 8.00E-05 | 1 | TAGGCGGGAATTTTC |
| 888 | mamo-L | *BGIBMGA001815* | down | 39 | 53 | + | 11.3952 | 4.52E-05 | 1 | TAGGCGTTCATTCCT |
| 889 | mamo-L | *BGIBMGA001816* | down | 1843 | 1857 | + | 11.5403 | 3.93E-05 | 1 | TAGGCGGAGTGGTGT |
| 890 | mamo-L | *BGIBMGA001818* | up | 1396 | 1410 | + | 12.871 | 9.35E-06 | 1 | TAGGCTTTTTTTATT |
| 891 | mamo-L | *BGIBMGA001818* | down | 1221 | 1235 | + | 11.3952 | 4.52E-05 | 1 | GAGGCGTTTGTTTGT |
| 892 | mamo-L | *BGIBMGA001819* | up | 524 | 538 | + | 13.6452 | 3.39E-06 | 1 | TAGGCCGACGACGGT |
| 893 | mamo-L | *BGIBMGA001822* | down | 1320 | 1334 | + | 11.2823 | 5.03E-05 | 1 | AAGGCGGTTTTTTTT |
| 894 | mamo-L | *BGIBMGA001824* | up | 585 | 599 | - | 11.1048 | 5.92E-05 | 1 | TAGGCCGTGATTTCA |
| 895 | mamo-L | *BGIBMGA001830* | down | 409 | 423 | + | 12.6371 | 1.23E-05 | 1 | TAGGCGGAGTGGGGT |
| 896 | mamo-L | *BGIBMGA001833* | up | 410 | 424 | - | 11.5887 | 3.75E-05 | 1 | TAGGCTGACGACTAT |
| 897 | mamo-L | *BGIBMGA001840* | down | 906 | 920 | - | 10.6371 | 8.85E-05 | 1 | TAGGCATTTTTTTAT |
| 898 | mamo-L | *BGIBMGA001840* | up | 1089 | 1103 | + | 10.5726 | 9.33E-05 | 1 | TAGGCTAATATTTAT |
| 899 | mamo-L | *BGIBMGA001843* | down | 1090 | 1104 | - | 10.5081 | 9.83E-05 | 1 | TGGGCCTGTTTTGAT |
| 900 | mamo-L | *BGIBMGA001845* | up | 1742 | 1756 | + | 11.1855 | 5.50E-05 | 1 | TAGGCTGGGTGTTGG |
| 901 | mamo-L | *BGIBMGA001845* | down | 229 | 243 | - | 10.7742 | 7.89E-05 | 1 | AAGGCTCTCGTTGAC |
| 902 | mamo-L | *BGIBMGA001850* | down | 1145 | 1159 | + | 12.7177 | 1.12E-05 | 1 | TAGGCGTTGTTTACT |
| 903 | mamo-L | *BGIBMGA001853* | down | 910 | 924 | + | 11.7339 | 3.26E-05 | 1 | TAGGCTAATGTTATT |
| 904 | mamo-L | *BGIBMGA001861* | up | 1407 | 1421 | - | 10.9274 | 6.92E-05 | 1 | GAGGCTTTAGTTTCT |
| 905 | mamo-L | *BGIBMGA001864* | down | 1978 | 1992 | - | 10.7661 | 7.95E-05 | 1 | GAGGCGGTCGTTGTA |
| 906 | mamo-L | *BGIBMGA001865* | down | 330 | 344 | - | 11.2177 | 5.34E-05 | 1 | TAGGCGTCTGTTCGT |
| 907 | mamo-L | *BGIBMGA001872* | up | 722 | 736 | + | 11.2903 | 4.99E-05 | 1 | TAGGCTGTATGGGCT |
| 908 | mamo-L | *BGIBMGA001874* | down | 1326 | 1340 | - | 11.3145 | 4.87E-05 | 1 | TAGGCGGTGTCTTTT |
| 909 | mamo-L | *BGIBMGA001875* | up | 978 | 992 | - | 12.9839 | 8.17E-06 | 1 | TAGGCTTTTTTTGGA |
| 910 | mamo-L | *BGIBMGA001876* | up | 158 | 172 | + | 11.3145 | 4.87E-05 | 1 | TAGGCGGTCGAGGAT |
| 911 | mamo-L | *BGIBMGA001877* | up | 1687 | 1701 | - | 11 | 6.50E-05 | 1 | AAGGCCTAATTTTTT |
| 912 | mamo-L | *BGIBMGA001878* | up | 1585 | 1599 | - | 10.7016 | 8.39E-05 | 1 | TAGGCGCACTGTTCC |
| 913 | mamo-L | *BGIBMGA001881* | up | 639 | 653 | - | 11.2258 | 5.30E-05 | 1 | TAGGCCTTTTTCCCT |

| | | | | | | | | | | |
|---|---|---|---|---|---|---|---|---|---|---|
| 914 | mamo-L | *BGIBMGA001882* | down | 1684 | 1698 | - | 13.3629 | 5.09E-06 | 1 | TAGGCTGCGTTTTTT |
| 915 | mamo-L | *BGIBMGA001883* | up | 477 | 491 | + | 13.8548 | 2.48E-06 | 1 | TAGGCCTACTTGGTT |
| 916 | mamo-L | *BGIBMGA001887* | down | 702 | 716 | + | 10.5726 | 9.33E-05 | 1 | TAGGCTAATATTTAT |
| 917 | mamo-L | *BGIBMGA001891* | up | 282 | 296 | - | 10.8145 | 7.63E-05 | 1 | TAGGCTGTTAATTAT |
| 918 | mamo-L | *BGIBMGA001892* | down | 1545 | 1559 | + | 11.1129 | 5.88E-05 | 1 | TAGGCTTTTAAGGGT |
| 919 | mamo-L | *BGIBMGA001893* | up | 1625 | 1639 | - | 11.9597 | 2.61E-05 | 1 | GAGGCCGTTGTTGCC |
| 920 | mamo-L | *BGIBMGA001894* | up | 232 | 246 | + | 14.6855 | 6.44E-07 | 1 | TAGGCGTTCTTTGTT |
| 921 | mamo-L | *BGIBMGA001894* | up | 1678 | 1692 | + | 10.5403 | 9.58E-05 | 1 | TAGGCTGTATGGTGT |
| 922 | mamo-L | *BGIBMGA001896* | up | 1407 | 1421 | - | 10.9194 | 6.98E-05 | 1 | TAGGCTACTTTTTTT |
| 923 | mamo-L | *BGIBMGA001897* | down | 185 | 199 | - | 11.8065 | 3.03E-05 | 1 | TAGGCTAATGTTTAT |
| 924 | mamo-L | *BGIBMGA001898* | down | 1812 | 1826 | + | 10.9274 | 6.92E-05 | 1 | TAGGCTCCCGGTTTT |
| 925 | mamo-L | *BGIBMGA001900* | down | 286 | 300 | + | 13.0484 | 7.57E-06 | 1 | TAGGCTGTATATGTT |
| 926 | mamo-L | *BGIBMGA001901* | down | 690 | 704 | - | 12.9355 | 8.65E-06 | 1 | GAGGCTGAGGTTTTT |
| 927 | mamo-L | *BGIBMGA001901* | up | 587 | 601 | - | 10.8468 | 7.42E-05 | 1 | TAGGCTATGATTGGA |
| 928 | mamo-L | *BGIBMGA001902* | up | 341 | 355 | - | 11.0645 | 6.14E-05 | 1 | TAGGCCCTAGATGAT |
| 929 | mamo-L | *BGIBMGA001903* | up | 1921 | 1935 | - | 11.75 | 3.21E-05 | 1 | TAGGCTGACATTCTC |
| 930 | mamo-L | *BGIBMGA001907* | up | 1800 | 1814 | - | 11.25 | 5.18E-05 | 1 | AAGGCTTACGATTCT |
| 931 | mamo-L | *BGIBMGA001907* | down | 415 | 429 | + | 10.9194 | 6.98E-05 | 1 | TAGGCTACTTTTTTT |
| 932 | mamo-L | *BGIBMGA001907* | up | 182 | 196 | - | 10.5 | 9.90E-05 | 1 | GAGGCCCAGGTCGTC |
| 933 | mamo-L | *BGIBMGA001910* | up | 95 | 109 | - | 11.0242 | 6.36E-05 | 1 | AAGGCCTTGTTTTCC |
| 934 | mamo-L | *BGIBMGA001910* | down | 1714 | 1728 | + | 10.5968 | 9.15E-05 | 1 | GAGGCTAGTTTTTGT |
| 935 | mamo-L | *BGIBMGA001910* | down | 1940 | 1954 | - | 10.5806 | 9.27E-05 | 1 | GAGGCTGAGGTGGAC |
| 936 | mamo-L | *BGIBMGA001911* | down | 1232 | 1246 | - | 10.5726 | 9.33E-05 | 1 | TAGGCGATTTTTACT |
| 937 | mamo-L | *BGIBMGA001912* | down | 382 | 396 | - | 13.4113 | 4.75E-06 | 1 | TAGGCCGAATTTTTT |
| 938 | mamo-L | *BGIBMGA001912* | down | 688 | 702 | - | 10.9919 | 6.55E-05 | 1 | TAGGCGTATTATGTA |
| 939 | mamo-L | *BGIBMGA001917* | up | 351 | 365 | + | 10.5645 | 9.40E-05 | 1 | AAGGCTTACTATAGT |
| 940 | mamo-L | *BGIBMGA001918* | down | 1144 | 1158 | - | 12.5645 | 1.34E-05 | 1 | TAGGCGGTTGTCTGT |
| 941 | mamo-L | *BGIBMGA001918* | up | 27 | 41 | + | 11.75 | 3.21E-05 | 1 | TAGGCTGACATTCTC |
| 942 | mamo-L | *BGIBMGA001919* | up | 1749 | 1763 | - | 12.7258 | 1.11E-05 | 1 | TAGGCGTGCTTTGTA |
| 943 | mamo-L | *BGIBMGA001920* | down | 493 | 507 | - | 11.0403 | 6.28E-05 | 1 | TAGGCCAGCATTAGT |
| 944 | mamo-L | *BGIBMGA001923* | down | 1512 | 1526 | + | 10.7258 | 8.23E-05 | 1 | GAGGCTTTTTTTATT |
| 945 | mamo-L | *BGIBMGA001929* | up | 1068 | 1082 | - | 10.5403 | 9.58E-05 | 1 | TAGGCTGTATGGTGT |
| 946 | mamo-L | *BGIBMGA001930* | up | 1857 | 1871 | + | 10.5726 | 9.33E-05 | 1 | TAGGCTAATATTTAT |
| 947 | mamo-L | *BGIBMGA001931* | down | 773 | 787 | + | 11.6613 | 3.50E-05 | 1 | GAGGCCTAGGTTGGA |
| 948 | mamo-L | *BGIBMGA001933* | up | 1302 | 1316 | + | 12.7581 | 1.07E-05 | 1 | TAGGCGTTTTGTGCT |
| 949 | mamo-L | *BGIBMGA001933* | up | 83 | 97 | + | 11.1694 | 5.59E-05 | 1 | TAGGCCGTCGACCGT |
| 950 | mamo-L | *BGIBMGA001934* | up | 1166 | 1180 | + | 10.6694 | 8.62E-05 | 1 | AAGGCGGGGAGTGTT |
| 951 | mamo-L | *BGIBMGA001936* | down | 538 | 552 | - | 13.0565 | 7.51E-06 | 1 | TAGGCGGGGGGCGGT |
| 952 | mamo-L | *BGIBMGA001937* | up | 1511 | 1525 | + | 10.7097 | 8.33E-05 | 1 | TAGGCCTGGTAGGAC |
| 953 | mamo-L | *BGIBMGA001941* | down | 178 | 192 | - | 12.2581 | 1.90E-05 | 1 | TAGGCTAGCTTTTGC |
| 954 | mamo-L | *BGIBMGA001941* | down | 319 | 333 | - | 11.0242 | 6.36E-05 | 1 | AAGGCTTATTTCTGT |
| 955 | mamo-L | *BGIBMGA001949* | up | 859 | 873 | - | 11.2903 | 4.99E-05 | 1 | TAGGCTGTATGGGCT |
| 956 | mamo-L | *BGIBMGA001953* | up | 791 | 805 | + | 11.2419 | 5.21E-05 | 1 | TAGGCATAGATTTTT |
| 957 | mamo-L | *BGIBMGA001954* | up | 1085 | 1099 | + | 10.5726 | 9.33E-05 | 1 | TAGGCTAATATTTAT |
| 958 | mamo-L | *BGIBMGA001955* | down | 1217 | 1231 | + | 11.2177 | 5.34E-05 | 1 | TAGGCGTCTGTTCGT |
| 959 | mamo-L | *BGIBMGA001957* | up | 1893 | 1907 | - | 10.9355 | 6.88E-05 | 1 | TAGGCCTCATTTAGT |
| 960 | mamo-L | *BGIBMGA001958* | down | 442 | 456 | - | 10.9194 | 6.98E-05 | 1 | TAGGCTACTTTTTTT |
| 961 | mamo-L | *BGIBMGA001959* | up | 20 | 34 | - | 10.9194 | 6.98E-05 | 1 | TAGGCTACTTTTTTT |
| 962 | mamo-L | *BGIBMGA001963* | up | 615 | 629 | - | 10.5726 | 9.33E-05 | 1 | TAGGCTAATATTTAT |
| 963 | mamo-L | *BGIBMGA001964* | down | 1401 | 1415 | + | 10.9194 | 6.98E-05 | 1 | TAGGCTACTTTTTTT |
| 964 | mamo-L | *BGIBMGA001964* | down | 1617 | 1631 | + | 10.9194 | 6.98E-05 | 1 | TAGGCTACTTTTTTT |
| 965 | mamo-L | *BGIBMGA001964* | down | 1829 | 1843 | + | 10.9194 | 6.98E-05 | 1 | TAGGCTACTTTTTTT |
| 966 | mamo-L | *BGIBMGA001971* | up | 1191 | 1205 | + | 11.7661 | 3.16E-05 | 1 | TAGGCAGTTTTTTTT |
| 967 | mamo-L | *BGIBMGA001974* | down | 1642 | 1656 | - | 11.5726 | 3.81E-05 | 1 | TAGGCTTAGGTAGAT |
| 968 | mamo-L | *BGIBMGA001974* | down | 1556 | 1570 | - | 10.7258 | 8.23E-05 | 1 | TAGGCAGACGATTGC |
| 969 | mamo-L | *BGIBMGA001976* | down | 1605 | 1619 | - | 11.7823 | 3.11E-05 | 1 | GAGGCTCACTTCGGT |
| 970 | mamo-L | *BGIBMGA001977* | down | 1018 | 1032 | - | 11.3548 | 4.69E-05 | 1 | AAGGCTGATGATGAT |
| 971 | mamo-L | *BGIBMGA001977* | down | 1671 | 1685 | - | 10.4919 | 9.96E-05 | 1 | TAGGCGTTTATTGTG |
| 972 | mamo-L | *BGIBMGA001983* | up | 891 | 905 | - | 10.9274 | 6.92E-05 | 1 | TAGGCTTAATCTGAT |
| 973 | mamo-L | *BGIBMGA001983* | up | 406 | 420 | + | 10.5726 | 9.33E-05 | 1 | TAGGCTAATATTTAT |
| 974 | mamo-L | *BGIBMGA001988* | up | 1153 | 1167 | - | 11.6613 | 3.50E-05 | 1 | AAGGCTCGGTTTTTT |
| 975 | mamo-L | *BGIBMGA001988* | down | 1901 | 1915 | + | 11.4677 | 4.22E-05 | 1 | TAGGCCCTGTTTATC |
| 976 | mamo-L | *BGIBMGA001993* | down | 743 | 757 | - | 10.9194 | 6.98E-05 | 1 | TAGGCTACTTTTTTT |
| 977 | mamo-L | *BGIBMGA001993* | down | 456 | 470 | + | 10.5726 | 9.33E-05 | 1 | TAGGCTAATATTTAT |
| 978 | mamo-L | *BGIBMGA001998* | down | 672 | 686 | + | 12.0484 | 2.38E-05 | 1 | TAGGCGTATGACGTT |
| 979 | mamo-L | *BGIBMGA002000* | up | 819 | 833 | - | 12.9919 | 8.10E-06 | 1 | TAGGCCGGTTTTGCA |
| 980 | mamo-L | *BGIBMGA002001* | down | 115 | 129 | + | 10.9435 | 6.83E-05 | 1 | AAGGCTGCGTGTGTT |
| 981 | mamo-L | *BGIBMGA002003* | up | 797 | 811 | + | 12.1129 | 2.22E-05 | 1 | TAGGCTGGTTCTGGC |
| 982 | mamo-L | *BGIBMGA002004* | down | 585 | 599 | + | 10.6452 | 8.79E-05 | 1 | TAGGCTGTTACTTGT |
| 983 | mamo-L | *BGIBMGA002005* | up | 1085 | 1099 | + | 12.0242 | 2.44E-05 | 1 | TAGGCCTCCTTTGGA |
| 984 | mamo-L | *BGIBMGA002010* | down | 1819 | 1833 | - | 11.2016 | 5.41E-05 | 1 | TAGGCTGGCTGTTTA |
| 985 | mamo-L | *BGIBMGA002010* | up | 1810 | 1824 | - | 10.7823 | 7.84E-05 | 1 | TAGGCATATGGTGAT |
| 986 | mamo-L | *BGIBMGA002012* | up | 1487 | 1501 | + | 10.5726 | 9.33E-05 | 1 | TAGGCTAATATTTAT |
| 987 | mamo-L | *BGIBMGA002014* | up | 209 | 223 | + | 12.3629 | 1.69E-05 | 1 | AAGGCTAAGTTTGGT |
| 988 | mamo-L | *BGIBMGA002015* | up | 883 | 897 | + | 11.2823 | 5.03E-05 | 1 | AAGGCGGTTTTTTTT |
| 989 | mamo-L | *BGIBMGA002015* | down | 792 | 806 | + | 10.9677 | 6.68E-05 | 1 | TAGGCCAGTTTCTTT |
| 990 | mamo-L | *BGIBMGA002017* | up | 220 | 234 | - | 10.5726 | 9.33E-05 | 1 | TAGGCTAATATTTAT |
| 991 | mamo-L | *BGIBMGA002019* | down | 1794 | 1808 | + | 11.2177 | 5.34E-05 | 1 | TAGGCGAGATTTGAT |
| 992 | mamo-L | *BGIBMGA002023* | down | 656 | 670 | - | 10.6371 | 8.85E-05 | 1 | TAGGCCGTGCTCTTT |
| 993 | mamo-L | *BGIBMGA002026* | up | 1696 | 1710 | + | 10.5726 | 9.33E-05 | 1 | TAGGCTAATATTTAT |
| 994 | mamo-L | *BGIBMGA002030* | down | 197 | 211 | - | 13.0887 | 7.18E-06 | 1 | TAGGCCTTGTTTCTT |
| 995 | mamo-L | *BGIBMGA002031* | up | 1724 | 1738 | + | 10.5726 | 9.33E-05 | 1 | TAGGCTAATATTTAT |
| 996 | mamo-L | *BGIBMGA002032* | up | 1268 | 1282 | - | 11.1452 | 5.70E-05 | 1 | TAGGCCAAGTTTAAT |
| 997 | mamo-L | *BGIBMGA002033* | up | 1544 | 1558 | - | 10.8145 | 7.63E-05 | 1 | AAGGCTTTGTGTTTT |
| 998 | mamo-L | *BGIBMGA002037* | down | 504 | 518 | + | 10.7823 | 7.84E-05 | 1 | TAGGCCCGTGATCTT |
| 999 | mamo-L | *BGIBMGA002040* | up | 471 | 485 | + | 11.8548 | 2.89E-05 | 1 | AAGGCTGTGTTTTGC |
| 1000 | mamo-L | *BGIBMGA002042* | up | 904 | 918 | - | 10.629 | 8.91E-05 | 1 | TAGGCCTGCAGCGCC |
| 1001 | mamo-L | *BGIBMGA002049* | up | 1143 | 1157 | - | 10.4919 | 9.96E-05 | 1 | TAGGCTTCCTGTGTA |
| 1002 | mamo-L | *BGIBMGA002050* | down | 1165 | 1179 | - | 12.3468 | 1.72E-05 | 1 | TAGGCTGTGGGCGAT |
| 1003 | mamo-L | *BGIBMGA002050* | down | 1799 | 1813 | - | 11.2984 | 4.95E-05 | 1 | TAGGCCAATGTCGTC |
| 1004 | mamo-L | *BGIBMGA002050* | down | 441 | 455 | + | 10.5081 | 9.83E-05 | 1 | TAGGCTCGTCTCGTT |
| 1005 | mamo-L | *BGIBMGA002051* | up | 1638 | 1652 | - | 11.8065 | 3.03E-05 | 1 | TAGGCTAATGTTTAT |

| | | | | | | | | | | |
|---|---|---|---|---|---|---|---|---|---|---|
| 1006 | mamo-L | *BGIBMGA002053* | down | 205 | 219 | - | 11.4758 | 4.18E-05 | 1 | TAGGCACTCTTTTGT |
| 1007 | mamo-L | *BGIBMGA002054* | down | 1908 | 1922 | + | 11.9516 | 2.63E-05 | 1 | TAGGCTTATTGTCGT |
| 1008 | mamo-L | *BGIBMGA002060* | down | 231 | 245 | + | 11.0484 | 6.23E-05 | 1 | TAGGCGCTCGAGGTT |
| 1009 | mamo-L | *BGIBMGA002061* | up | 1735 | 1749 | + | 11.0484 | 6.23E-05 | 1 | TAGGCGCTCGAGGTT |
| 1010 | mamo-L | *BGIBMGA002061* | up | 1408 | 1422 | - | 10.6452 | 8.79E-05 | 1 | GAGGCTGAAGTTACT |
| 1011 | mamo-L | *BGIBMGA002064* | up | 1370 | 1384 | - | 12.7419 | 1.09E-05 | 1 | GAGGCTCACTTTGTT |
| 1012 | mamo-L | *BGIBMGA002064* | down | 39 | 53 | - | 10.871 | 7.28E-05 | 1 | TAGGCCCGGAGCGCT |
| 1013 | mamo-L | *BGIBMGA002065* | up | 671 | 685 | - | 10.871 | 7.28E-05 | 1 | TAGGCCCGGAGCGCT |
| 1014 | mamo-L | *BGIBMGA002069* | up | 165 | 179 | + | 12.8952 | 9.05E-06 | 1 | TAGGCTGTGTTTAAT |
| 1015 | mamo-L | *BGIBMGA002070* | down | 285 | 299 | + | 11.1129 | 5.88E-05 | 1 | AAGGCGGCTTTTGTT |
| 1016 | mamo-L | *BGIBMGA002073* | down | 1332 | 1346 | - | 11.9113 | 2.74E-05 | 1 | AAGGCTTGGTTGGTT |
| 1017 | mamo-L | *BGIBMGA002073* | down | 113 | 127 | + | 10.629 | 8.91E-05 | 1 | AAGGCGGATGTTGTA |
| 1018 | mamo-L | *BGIBMGA002074* | up | 1637 | 1651 | - | 11.9113 | 2.74E-05 | 1 | AAGGCTTGGTTGGTT |
| 1019 | mamo-L | *BGIBMGA002074* | down | 647 | 661 | + | 11.4113 | 4.45E-05 | 1 | TAGGCCTAGTATAAT |
| 1020 | mamo-L | *BGIBMGA002074* | up | 418 | 432 | + | 10.629 | 8.91E-05 | 1 | AAGGCGGATGTTGTA |
| 1021 | mamo-L | *BGIBMGA002076* | up | 1588 | 1602 | - | 12.121 | 2.20E-05 | 1 | TAGGCGTTTGTTGTA |
| 1022 | mamo-L | *BGIBMGA002078* | down | 1486 | 1500 | + | 12.629 | 1.24E-05 | 1 | TAGGCCGATTGCGTT |
| 1023 | mamo-L | *BGIBMGA002080* | down | 147 | 161 | + | 10.5726 | 9.33E-05 | 1 | TAGGCTAATATTTAT |
| 1024 | mamo-L | *BGIBMGA002085* | down | 893 | 907 | - | 14.7258 | 5.83E-07 | 1 | TAGGCTTGCGTTGCC |
| 1025 | mamo-L | *BGIBMGA002090* | up | 1580 | 1594 | + | 11.2097 | 5.38E-05 | 1 | TAGGCAGGATTTTTT |
| 1026 | mamo-L | *BGIBMGA002091* | down | 1001 | 1015 | - | 10.5726 | 9.33E-05 | 1 | TAGGCTAATATTTAT |
| 1027 | mamo-L | *BGIBMGA002094* | up | 1426 | 1440 | + | 14.3306 | 1.21E-06 | 1 | TAGGCGGGGGGTGGT |
| 1028 | mamo-L | *BGIBMGA002097* | down | 783 | 797 | + | 11.129 | 5.79E-05 | 1 | TAGGCCTAAGATGGA |
| 1029 | mamo-L | *BGIBMGA002097* | down | 113 | 127 | - | 10.75 | 8.06E-05 | 1 | GAGGCCGATTGTGCC |
| 1030 | mamo-L | *BGIBMGA002100* | up | 964 | 978 | - | 11.25 | 5.18E-05 | 1 | TAGGCTTAGGTGTGA |
| 1031 | mamo-L | *BGIBMGA002102* | down | 256 | 270 | + | 11.0645 | 6.14E-05 | 1 | TAGGCGCTTTATGCC |
| 1032 | mamo-L | *BGIBMGA002108* | up | 922 | 936 | - | 13.7742 | 2.79E-06 | 1 | TAGGCGGGTGGTGGT |
| 1033 | mamo-L | *BGIBMGA002108* | down | 841 | 855 | - | 10.5726 | 9.33E-05 | 1 | TAGGCTAATATTTAT |
| 1034 | mamo-L | *BGIBMGA002110* | up | 786 | 800 | + | 11.371 | 4.62E-05 | 1 | TAGGCTGTATAGGCT |
| 1035 | mamo-L | *BGIBMGA002112* | up | 1424 | 1438 | + | 13.1935 | 6.33E-06 | 1 | TAGGCTTTAGTTGCC |
| 1036 | mamo-L | *BGIBMGA002113* | up | 1977 | 1991 | - | 10.621 | 8.97E-05 | 1 | AAGGCTGGTTGCGTT |
| 1037 | mamo-L | *BGIBMGA002114* | up | 485 | 499 | + | 11.4516 | 4.28E-05 | 1 | AAGGCTTAGTGTTGT |
| 1038 | mamo-L | *BGIBMGA002116* | up | 1380 | 1394 | - | 11.3548 | 4.69E-05 | 1 | AAGGCTGATGATGAT |
| 1039 | mamo-L | *BGIBMGA002126* | up | 146 | 160 | - | 10.7661 | 7.95E-05 | 1 | GAGGCCGTATTCGTT |
| 1040 | mamo-L | *BGIBMGA002127* | down | 43 | 57 | - | 12.3548 | 1.70E-05 | 1 | TAGGCTGTCATTGTA |
| 1041 | mamo-L | *BGIBMGA002128* | down | 1226 | 1240 | - | 11.3629 | 4.66E-05 | 1 | TAGGCTCATATTTAT |
| 1042 | mamo-L | *BGIBMGA002129* | up | 1905 | 1919 | - | 10.9677 | 6.68E-05 | 1 | TAGGCTGAATTTTCG |
| 1043 | mamo-L | *BGIBMGA002130* | down | 1277 | 1291 | + | 12.0323 | 2.42E-05 | 1 | AAGGCCTTGTTTTTT |
| 1044 | mamo-L | *BGIBMGA002130* | down | 216 | 230 | + | 10.629 | 8.91E-05 | 1 | TAGGCCTGCAGCGCC |
| 1045 | mamo-L | *BGIBMGA002130* | up | 1932 | 1946 | + | 10.5726 | 9.33E-05 | 1 | TAGGCTAATATTTAT |
| 1046 | mamo-L | *BGIBMGA002135* | up | 685 | 699 | - | 10.6048 | 9.09E-05 | 1 | TAGGCTTCAATTTTT |
| 1047 | mamo-L | *BGIBMGA002135* | up | 841 | 855 | - | 10.5081 | 9.83E-05 | 1 | TAGGCTGCATTTGTG |
| 1048 | mamo-L | *BGIBMGA002138* | down | 1213 | 1227 | - | 10.5726 | 9.33E-05 | 1 | TAGGCTAATATTTAT |
| 1049 | mamo-L | *BGIBMGA002153* | down | 757 | 771 | - | 11.7419 | 3.23E-05 | 1 | TAGGCGGGAAGTGGT |
| 1050 | mamo-L | *BGIBMGA002155* | up | 1195 | 1209 | - | 12.3548 | 1.70E-05 | 1 | TAGGCTGCGGTTGCA |
| 1051 | mamo-L | *BGIBMGA002156* | down | 1311 | 1325 | + | 12.25 | 1.92E-05 | 1 | TAGGCTGTGGACGCC |
| 1052 | mamo-L | *BGIBMGA002156* | up | 340 | 354 | + | 11.9758 | 2.57E-05 | 1 | TAGGCTGTCTGTGGG |
| 1053 | mamo-L | *BGIBMGA002158* | up | 1312 | 1326 | - | 10.75 | 8.06E-05 | 1 | AAGGCCTGTTGTGGC |
| 1054 | mamo-L | *BGIBMGA002159* | up | 1647 | 1661 | - | 12.4194 | 1.58E-05 | 1 | TAGGCGTTATTTGTC |
| 1055 | mamo-L | *BGIBMGA002169* | down | 292 | 306 | - | 12.2258 | 1.97E-05 | 1 | TAGGCTTTGGGTTAT |
| 1056 | mamo-L | *BGIBMGA002171* | down | 1442 | 1456 | + | 12.75 | 1.08E-05 | 1 | TAGGCCAAAGTTGTT |
| 1057 | mamo-L | *BGIBMGA002171* | up | 1906 | 1920 | + | 12.4919 | 1.46E-05 | 1 | TAGGCTATTATTGGT |
| 1058 | mamo-L | *BGIBMGA002174* | up | 281 | 295 | + | 11.121 | 5.84E-05 | 1 | TAGGCGAAATTTGCC |
| 1059 | mamo-L | *BGIBMGA002175* | up | 820 | 834 | + | 12.4113 | 1.60E-05 | 1 | TAGGCTTGTGTTAAT |
| 1060 | mamo-L | *BGIBMGA002177* | down | 191 | 205 | + | 10.629 | 8.91E-05 | 1 | TAGGCAGGGTTTGAA |
| 1061 | mamo-L | *BGIBMGA002179* | up | 1041 | 1055 | + | 12.5565 | 1.35E-05 | 1 | TAGGCCAGTTTTTGT |
| 1062 | mamo-L | *BGIBMGA002182* | down | 1345 | 1359 | - | 11.7823 | 3.11E-05 | 1 | TAGGCGTGGGGCGGC |
| 1063 | mamo-L | *BGIBMGA002182* | up | 1582 | 1596 | - | 10.75 | 8.06E-05 | 1 | GAGGCCGATTGTGCC |
| 1064 | mamo-L | *BGIBMGA002183* | up | 1913 | 1927 | + | 10.5726 | 9.33E-05 | 1 | TAGGCTAATATTTAT |
| 1065 | mamo-L | *BGIBMGA002193* | down | 277 | 291 | - | 12.25 | 1.92E-05 | 1 | TAGGCTGTGGACGCC |
| 1066 | mamo-L | *BGIBMGA002193* | up | 1253 | 1267 | + | 11.5806 | 3.78E-05 | 1 | TAGGCTGACATTTTA |
| 1067 | mamo-L | *BGIBMGA002194* | down | 113 | 127 | + | 11.3952 | 4.52E-05 | 1 | GAGGCTGTGTTTCCT |
| 1068 | mamo-L | *BGIBMGA002195* | down | 929 | 943 | - | 11.6129 | 3.66E-05 | 1 | TAGGCTTTTAATGAT |
| 1069 | mamo-L | *BGIBMGA002196* | down | 128 | 142 | - | 11.6129 | 3.66E-05 | 1 | TAGGCTTTTAATGAT |
| 1070 | mamo-L | *BGIBMGA002198* | down | 642 | 656 | - | 10.5726 | 9.33E-05 | 1 | TAGGCTAATATTTAT |
| 1071 | mamo-L | *BGIBMGA002199* | up | 183 | 197 | - | 10.4919 | 9.96E-05 | 1 | AAGGCCGATGTTAAT |
| 1072 | mamo-L | *BGIBMGA002200* | down | 819 | 833 | + | 10.8145 | 7.63E-05 | 1 | TAGGCTTCTATTAGT |
| 1073 | mamo-L | *BGIBMGA002200* | up | 743 | 757 | - | 10.5403 | 9.58E-05 | 1 | TAGGCCAATGATAGT |
| 1074 | mamo-L | *BGIBMGA002204* | up | 283 | 297 | - | 11.1935 | 5.45E-05 | 1 | TAGGCTTTGTTTACA |
| 1075 | mamo-L | *BGIBMGA002205* | up | 1864 | 1878 | + | 10.5726 | 9.33E-05 | 1 | TAGGCTAATATTTAT |
| 1076 | mamo-L | *BGIBMGA002209* | up | 801 | 815 | + | 11.1613 | 5.63E-05 | 1 | TAGGCATGGATTTTT |
| 1077 | mamo-L | *BGIBMGA002213* | up | 1709 | 1723 | + | 12.1855 | 2.06E-05 | 1 | AAGGCTTTGTTTTCT |
| 1078 | mamo-L | *BGIBMGA002213* | up | 340 | 354 | - | 10.7742 | 7.89E-05 | 1 | TAGGCATATAATGGT |
| 1079 | mamo-L | *BGIBMGA002214* | down | 515 | 529 | + | 10.629 | 8.91E-05 | 1 | TAGGCCTGCAGCGCC |
| 1080 | mamo-L | *BGIBMGA002215* | down | 1131 | 1145 | + | 12.1532 | 2.13E-05 | 1 | TAGGCTGATGCTGAT |
| 1081 | mamo-L | *BGIBMGA002217* | down | 1533 | 1547 | + | 12.9919 | 8.10E-06 | 1 | TAGGCCGGTTTTGCA |
| 1082 | mamo-L | *BGIBMGA002217* | up | 1934 | 1948 | + | 10.5726 | 9.33E-05 | 1 | TAGGCTAATATTTAT |
| 1083 | mamo-L | *BGIBMGA002218* | down | 1153 | 1167 | - | 12.1532 | 2.13E-05 | 1 | TAGGCTGATGCTGAT |
| 1084 | mamo-L | *BGIBMGA002223* | up | 639 | 653 | + | 12.0323 | 2.42E-05 | 1 | GAGGCGGAGGATGTT |
| 1085 | mamo-L | *BGIBMGA002224* | up | 1068 | 1082 | + | 11.7984 | 3.06E-05 | 1 | AAGGCCTATTTTTTT |
| 1086 | mamo-L | *BGIBMGA002224* | up | 1493 | 1507 | - | 10.9032 | 7.07E-05 | 1 | TAGGCCTTGCTTGAC |
| 1087 | mamo-L | *BGIBMGA002226* | down | 875 | 889 | - | 10.9113 | 7.02E-05 | 1 | GAGGCGCATTTTGGC |
| 1088 | mamo-L | *BGIBMGA002227* | up | 433 | 447 | + | 12.0645 | 2.34E-05 | 1 | TAGGCGGGCTTCGGA |
| 1089 | mamo-L | *BGIBMGA002227* | up | 280 | 294 | - | 10.8306 | 7.52E-05 | 1 | TAGGCCCCCTTCGCC |
| 1090 | mamo-L | *BGIBMGA002230* | down | 239 | 253 | - | 13.3065 | 5.48E-06 | 1 | TAGGCTTATATTTGT |
| 1091 | mamo-L | *BGIBMGA002232* | down | 150 | 164 | - | 12.3226 | 1.77E-05 | 1 | TAGGCCAATTTTTTT |
| 1092 | mamo-L | *BGIBMGA002232* | up | 1344 | 1358 | - | 10.5726 | 9.33E-05 | 1 | TAGGCTAATATTTAT |
| 1093 | mamo-L | *BGIBMGA002234* | down | 647 | 661 | - | 11.5726 | 3.81E-05 | 1 | TAGGCTTTTTTTTTA |
| 1094 | mamo-L | *BGIBMGA002239* | down | 719 | 733 | + | 10.8871 | 7.17E-05 | 1 | TAGGCAAAGGTTTTT |
| 1095 | mamo-L | *BGIBMGA002240* | up | 401 | 415 | + | 12.2581 | 1.90E-05 | 1 | AAGGCTGAGGTTGAC |
| 1096 | mamo-L | *BGIBMGA002244* | up | 286 | 300 | - | 11.4919 | 4.11E-05 | 1 | TAGGCTTGGTGGTCT |
| 1097 | mamo-L | *BGIBMGA002245* | down | 1408 | 1422 | - | 12.7984 | 1.02E-05 | 1 | TAGGCTTGAGTGGTT |

| | | | | | | | | | | |
|---|---|---|---|---|---|---|---|---|---|---|
| 1098 | mamo-L | *BGIBMGA002248* | up | 680 | 694 | + | 10.9194 | 6.98E-05 | 1 | TAGGCTACTTTTTTT |
| 1099 | mamo-L | *BGIBMGA002248* | down | 1496 | 1510 | + | 10.5726 | 9.33E-05 | 1 | TAGGCTAATATTTAT |
| 1100 | mamo-L | *BGIBMGA002252* | up | 1079 | 1093 | + | 12.3306 | 1.75E-05 | 1 | TAGGCTTACTATACT |
| 1101 | mamo-L | *BGIBMGA002269* | up | 1254 | 1268 | - | 11.5968 | 3.73E-05 | 1 | TAGGCGGCGTGTTGT |
| 1102 | mamo-L | *BGIBMGA002273* | up | 549 | 563 | - | 10.5484 | 9.52E-05 | 1 | TAGGCCTTCTTGTGA |
| 1103 | mamo-L | *BGIBMGA002274* | down | 1180 | 1194 | - | 11.0968 | 5.97E-05 | 1 | TAGGCTGAGATGGTA |
| 1104 | mamo-L | *BGIBMGA002277* | up | 603 | 617 | - | 11.629 | 3.61E-05 | 1 | GAGGCTTTTTTTTTT |
| 1105 | mamo-L | *BGIBMGA002280* | up | 1846 | 1860 | + | 11.2097 | 5.38E-05 | 1 | TAGGCTGCGGTTTGG |
| 1106 | mamo-L | *BGIBMGA002282* | down | 1983 | 1997 | - | 13.6452 | 3.39E-06 | 1 | TAGGCTTAGTATTGT |
| 1107 | mamo-L | *BGIBMGA002284* | down | 1229 | 1243 | - | 11.3548 | 4.69E-05 | 1 | AAGGCTGATGATGAT |
| 1108 | mamo-L | *BGIBMGA002285* | down | 287 | 301 | + | 11.4758 | 4.18E-05 | 1 | TAGGCTTATTTGTTC |
| 1109 | mamo-L | *BGIBMGA002286* | up | 1455 | 1469 | - | 11.9355 | 2.67E-05 | 1 | TAGGCTCACGGGGCT |
| 1110 | mamo-L | *BGIBMGA002288* | down | 794 | 808 | - | 11.7903 | 3.08E-05 | 1 | TAGGCTGAGGGTCAT |
| 1111 | mamo-L | *BGIBMGA002289* | down | 406 | 420 | + | 10.5726 | 9.33E-05 | 1 | TAGGCTAATATTTAT |
| 1112 | mamo-L | *BGIBMGA002291* | down | 1294 | 1308 | + | 10.7581 | 8.00E-05 | 1 | TAGGCGGGGAACGAT |
| 1113 | mamo-L | *BGIBMGA002301* | up | 978 | 992 | - | 10.5726 | 9.33E-05 | 1 | TAGGCTAATATTTAT |
| 1114 | mamo-L | *BGIBMGA002302* | up | 1704 | 1718 | + | 11.5323 | 3.96E-05 | 1 | TAGGCAGTGATTTGT |
| 1115 | mamo-L | *BGIBMGA002302* | up | 511 | 525 | - | 10.9516 | 6.78E-05 | 1 | TAGGCTTTGGTATCT |
| 1116 | mamo-L | *BGIBMGA002302* | up | 1580 | 1594 | + | 10.5806 | 9.27E-05 | 1 | TAGGCTTTTCTTTGC |
| 1117 | mamo-L | *BGIBMGA002305* | up | 1616 | 1630 | + | 11.5968 | 3.73E-05 | 1 | TAGGCGGCGTGTTGT |
| 1118 | mamo-L | *BGIBMGA002305* | down | 1923 | 1937 | + | 10.879 | 7.22E-05 | 1 | TAGGCAGTGGCTGCT |
| 1119 | mamo-L | *BGIBMGA002306* | up | 709 | 723 | - | 12.5161 | 1.42E-05 | 1 | AAGGCTGTGTTTTTT |
| 1120 | mamo-L | *BGIBMGA002306* | down | 320 | 334 | - | 10.7903 | 7.79E-05 | 1 | AAGGCTTATGATTTT |
| 1121 | mamo-L | *BGIBMGA002306* | down | 190 | 204 | + | 10.5968 | 9.15E-05 | 1 | GAGGCTAGTTTTTGT |
| 1122 | mamo-L | *BGIBMGA002308* | down | 598 | 612 | - | 11.7903 | 3.08E-05 | 1 | TAGGCTCAGATCTGT |
| 1123 | mamo-L | *BGIBMGA002314* | down | 1137 | 1151 | + | 13.3145 | 5.41E-06 | 1 | TAGGCTTAGATGGGT |
| 1124 | mamo-L | *BGIBMGA002314* | down | 555 | 569 | + | 12.1855 | 2.06E-05 | 1 | TAGGCTATTTTTTTT |
| 1125 | mamo-L | *BGIBMGA002315* | up | 1131 | 1145 | - | 10.9919 | 6.55E-05 | 1 | TAGGCTTTAGATGGA |
| 1126 | mamo-L | *BGIBMGA002315* | down | 1888 | 1902 | - | 10.9194 | 6.98E-05 | 1 | TAGGCTACTTTTTTT |
| 1127 | mamo-L | *BGIBMGA002316* | down | 754 | 768 | + | 11.629 | 3.61E-05 | 1 | GAGGCTTTTTTTTTT |
| 1128 | mamo-L | *BGIBMGA002316* | up | 1096 | 1110 | + | 10.4919 | 9.96E-05 | 1 | TAGGCTGTGTCTGTG |
| 1129 | mamo-L | *BGIBMGA002318* | up | 539 | 553 | - | 12.871 | 9.35E-06 | 1 | TAGGCTTTTTTTATT |
| 1130 | mamo-L | *BGIBMGA002322* | up | 1769 | 1783 | - | 10.4919 | 9.96E-05 | 1 | TAGGCTAGTATTTAT |
| 1131 | mamo-L | *BGIBMGA002330* | up | 510 | 524 | - | 12.6935 | 1.15E-05 | 1 | TAGGCTGAGGTGCGT |
| 1132 | mamo-L | *BGIBMGA002331* | down | 727 | 741 | + | 13.2984 | 5.54E-06 | 1 | TAGGCGTACGTTCGT |
| 1133 | mamo-L | *BGIBMGA002331* | down | 934 | 948 | - | 12.1129 | 2.22E-05 | 1 | TAGGCCGATGATATT |
| 1134 | mamo-L | *BGIBMGA002331* | down | 1444 | 1458 | + | 11.4677 | 4.22E-05 | 1 | AAGGCTTACTATTGT |
| 1135 | mamo-L | *BGIBMGA002331* | down | 548 | 562 | - | 10.629 | 8.91E-05 | 1 | GAGGCTAATTTTGAT |
| 1136 | mamo-L | *BGIBMGA002332* | up | 793 | 807 | - | 10.7581 | 8.00E-05 | 1 | TAGGCCGTGTTTCCG |
| 1137 | mamo-L | *BGIBMGA002334* | down | 1161 | 1175 | + | 11.3065 | 4.91E-05 | 1 | AAGGCGTATTTTTTT |
| 1138 | mamo-L | *BGIBMGA002338* | up | 436 | 450 | - | 13.3226 | 5.34E-06 | 1 | TAGGCCGTCGACGGT |
| 1139 | mamo-L | *BGIBMGA002339* | down | 1470 | 1484 | - | 10.7097 | 8.33E-05 | 1 | TAGGCTTGGCTGTGT |
| 1140 | mamo-L | *BGIBMGA002342* | down | 187 | 201 | - | 14.4032 | 1.06E-06 | 1 | TAGGCTGATTGTGGT |
| 1141 | mamo-L | *BGIBMGA002344* | up | 9 | 23 | + | 12.3468 | 1.72E-05 | 1 | AAGGCTTTGGTTTTT |
| 1142 | mamo-L | *BGIBMGA002355* | down | 604 | 618 | + | 11.4758 | 4.18E-05 | 1 | TAGGCTGTTTTTTTG |
| 1143 | mamo-L | *BGIBMGA002358* | down | 642 | 656 | - | 11.2823 | 5.03E-05 | 1 | AAGGCTGGCAGTGTT |
| 1144 | mamo-L | *BGIBMGA002362* | down | 1889 | 1903 | + | 12.25 | 1.92E-05 | 1 | TAGGCCCACTATTCT |
| 1145 | mamo-L | *BGIBMGA002375* | down | 1783 | 1797 | - | 10.6129 | 9.03E-05 | 1 | TAGGCTTTCATGATT |
| 1146 | mamo-L | *BGIBMGA002377* | down | 856 | 870 | + | 10.629 | 8.91E-05 | 1 | TAGGCCTGCAGCGCC |
| 1147 | mamo-L | *BGIBMGA002380* | up | 1727 | 1741 | - | 12.1774 | 2.08E-05 | 1 | TAGGCTTACGGTGCA |
| 1148 | mamo-L | *BGIBMGA002381* | up | 65 | 79 | - | 13.3548 | 5.14E-06 | 1 | TAGGCTTATTTTCGT |
| 1149 | mamo-L | *BGIBMGA002384* | up | 1865 | 1879 | - | 11.9113 | 2.74E-05 | 1 | TAGGCCGTCGTTTTG |
| 1150 | mamo-L | *BGIBMGA002388* | down | 1711 | 1725 | - | 11.371 | 4.62E-05 | 1 | TAGGCTGTATAGGCT |
| 1151 | mamo-L | *BGIBMGA002391* | up | 1573 | 1587 | - | 10.9839 | 6.59E-05 | 1 | GAGGCGATGTTTGCT |
| 1152 | mamo-L | *BGIBMGA002392* | up | 1134 | 1148 | + | 10.5726 | 9.33E-05 | 1 | TAGGCTAATATTTAT |
| 1153 | mamo-L | *BGIBMGA002394* | up | 1833 | 1847 | + | 12.7742 | 1.04E-05 | 1 | TAGGCTTATTATTTT |
| 1154 | mamo-L | *BGIBMGA002396* | up | 1648 | 1662 | + | 11.5242 | 3.99E-05 | 1 | TAGGCTGTGTTTATA |
| 1155 | mamo-L | *BGIBMGA002399* | down | 1441 | 1455 | + | 12.0645 | 2.34E-05 | 1 | TAGGCGGGCTTCGGA |
| 1156 | mamo-L | *BGIBMGA002399* | up | 667 | 681 | + | 10.9919 | 6.55E-05 | 1 | TAGGCTTTAGATGGA |
| 1157 | mamo-L | *BGIBMGA002401* | down | 1950 | 1964 | - | 11.1935 | 5.45E-05 | 1 | TAGGCGCATGAGGGT |
| 1158 | mamo-L | *BGIBMGA002403* | up | 259 | 273 | + | 11.0323 | 6.32E-05 | 1 | TAGGCTGTCTTGCAT |
| 1159 | mamo-L | *BGIBMGA002404* | down | 1534 | 1548 | + | 11.0484 | 6.23E-05 | 1 | TAGGCCTTTAGTTCT |
| 1160 | mamo-L | *BGIBMGA002405* | down | 1267 | 1281 | - | 11.6855 | 3.42E-05 | 1 | TAGGCTGCTCTTGGT |
| 1161 | mamo-L | *BGIBMGA002407* | up | 1640 | 1654 | - | 13.4194 | 4.70E-06 | 1 | TAGGCCAATTTTGTT |
| 1162 | mamo-L | *BGIBMGA002411* | up | 1869 | 1883 | - | 10.8468 | 7.42E-05 | 1 | GAGGCTTATTTTGTA |
| 1163 | mamo-L | *BGIBMGA002420* | down | 584 | 598 | - | 13.9597 | 2.11E-06 | 1 | TAGGCCCGCTTTGGC |
| 1164 | mamo-L | *BGIBMGA002421* | up | 1685 | 1699 | - | 13.9597 | 2.11E-06 | 1 | TAGGCCCGCTTTGGC |
| 1165 | mamo-L | *BGIBMGA002425* | up | 938 | 952 | + | 11.7823 | 3.11E-05 | 1 | TAGGCGTGGGGCGGC |
| 1166 | mamo-L | *BGIBMGA002428* | down | 1879 | 1893 | - | 10.6613 | 8.67E-05 | 1 | GAGGCTAAATTTGTT |
| 1167 | mamo-L | *BGIBMGA002429* | down | 692 | 706 | + | 10.9194 | 6.98E-05 | 1 | TAGGCTACTTTTTTT |
| 1168 | mamo-L | *BGIBMGA002430* | up | 1563 | 1577 | - | 11.0968 | 5.97E-05 | 1 | TAGGCAGTAGTTTTT |
| 1169 | mamo-L | *BGIBMGA002432* | down | 1120 | 1134 | - | 12.9919 | 8.10E-06 | 1 | TAGGCCGGTTTTGCA |
| 1170 | mamo-L | *BGIBMGA002433* | down | 1666 | 1680 | + | 12.2177 | 1.99E-05 | 1 | AAGGCTTTGTTTTTT |
| 1171 | mamo-L | *BGIBMGA002434* | up | 1872 | 1886 | + | 11.4516 | 4.28E-05 | 1 | TAGGCACAGGATGTT |
| 1172 | mamo-L | *BGIBMGA002434* | up | 1147 | 1161 | + | 11.4113 | 4.45E-05 | 1 | TAGGCGTGGGGGGGC |
| 1173 | mamo-L | *BGIBMGA002435* | up | 605 | 619 | + | 10.5968 | 9.15E-05 | 1 | TAGGCTATAATTTGT |
| 1174 | mamo-L | *BGIBMGA002436* | up | 1117 | 1131 | - | 12.0645 | 2.34E-05 | 1 | TAGGCGGGCTTCGGA |
| 1175 | mamo-L | *BGIBMGA002440* | up | 1166 | 1180 | + | 10.5968 | 9.15E-05 | 1 | TAGGCTTTAATCTTT |
| 1176 | mamo-L | *BGIBMGA002441* | up | 513 | 527 | + | 11.7339 | 3.26E-05 | 1 | TAGGCTTCGAATGTT |
| 1177 | mamo-L | *BGIBMGA002442* | down | 1115 | 1129 | + | 12.3065 | 1.80E-05 | 1 | TAGGCCCTCGTTGTA |
| 1178 | mamo-L | *BGIBMGA002442* | up | 783 | 797 | - | 10.879 | 7.22E-05 | 1 | AAGGCTGGTTATTTT |
| 1179 | mamo-L | *BGIBMGA002448* | down | 1753 | 1767 | + | 10.5081 | 9.83E-05 | 1 | TAGGCGGGGTGTTGG |
| 1180 | mamo-L | *BGIBMGA002449* | down | 1653 | 1667 | - | 11.7661 | 3.16E-05 | 1 | TAGGCAGTTTTTTTT |
| 1181 | mamo-L | *BGIBMGA002449* | up | 997 | 1011 | + | 10.5645 | 9.40E-05 | 1 | AAGGCTTACTATAGT |
| 1182 | mamo-L | *BGIBMGA002449* | up | 1551 | 1565 | + | 10.5645 | 9.40E-05 | 1 | AAGGCTTACTATAGT |
| 1183 | mamo-L | *BGIBMGA002450* | down | 1053 | 1067 | + | 12.0161 | 2.46E-05 | 1 | TAGGCGTGTTATTTT |
| 1184 | mamo-L | *BGIBMGA002452* | up | 756 | 770 | - | 12.0887 | 2.29E-05 | 1 | TAGGCTGTTGGTGAC |
| 1185 | mamo-L | *BGIBMGA002452* | down | 577 | 591 | + | 10.879 | 7.22E-05 | 1 | TAGGCCTCCTTCTGC |
| 1186 | mamo-L | *BGIBMGA002453* | up | 303 | 317 | + | 10.75 | 8.06E-05 | 1 | TAGGCTATGTGTAGT |
| 1187 | mamo-L | *BGIBMGA002454* | down | 137 | 151 | + | 11.0242 | 6.36E-05 | 1 | TAGGCTGTTTGGTTT |
| 1188 | mamo-L | *BGIBMGA002455* | down | 271 | 285 | + | 12.0081 | 2.48E-05 | 1 | TAGGCTATTGGTGTT |
| 1189 | mamo-L | *BGIBMGA002455* | down | 930 | 944 | + | 11.0081 | 6.45E-05 | 1 | TAGGCAGGGTAGGGT |

| | | | | | | | | | | |
|---|---|---|---|---|---|---|---|---|---|---|
| 1190 | mamo-L | *BGIBMGA002456* | down | 123 | 137 | + | 12.871 | 9.35E-06 | 1 | TAGGCGTAGATTTTT |
| 1191 | mamo-L | *BGIBMGA002460* | down | 1517 | 1531 | - | 13.3387 | 5.24E-06 | 1 | TAGGCGGGCTTTGGA |
| 1192 | mamo-L | *BGIBMGA002460* | down | 557 | 571 | - | 12.7661 | 1.05E-05 | 1 | TAGGCCGACGATCGT |
| 1193 | mamo-L | *BGIBMGA002460* | down | 1670 | 1684 | + | 12.0968 | 2.26E-05 | 1 | TAGGCCCTCTTCGCC |
| 1194 | mamo-L | *BGIBMGA002462* | down | 75 | 89 | - | 13.2258 | 6.08E-06 | 1 | TAGGCGTTTGTTTTT |
| 1195 | mamo-L | *BGIBMGA002466* | down | 766 | 780 | - | 12.2177 | 1.99E-05 | 1 | AAGGCTTTGTTTTTT |
| 1196 | mamo-L | *BGIBMGA002466* | up | 667 | 681 | - | 11.4516 | 4.28E-05 | 1 | TAGGCACAGGATGTT |
| 1197 | mamo-L | *BGIBMGA002466* | up | 1392 | 1406 | - | 11.4113 | 4.45E-05 | 1 | TAGGCGTGGGGGGGC |
| 1198 | mamo-L | *BGIBMGA002469* | down | 802 | 816 | + | 12.9919 | 8.10E-06 | 1 | TAGGCCGGTTTTGCA |
| 1199 | mamo-L | *BGIBMGA002470* | up | 301 | 315 | + | 12.9194 | 8.81E-06 | 1 | TAGGCCTTAGTTTTT |
| 1200 | mamo-L | *BGIBMGA002472* | down | 1707 | 1721 | + | 10.7419 | 8.11E-05 | 1 | AAGGCTCACTTTACT |
| 1201 | mamo-L | *BGIBMGA002473* | up | 566 | 580 | + | 12.9919 | 8.10E-06 | 1 | TAGGCCGGTTTTGCA |
| 1202 | mamo-L | *BGIBMGA002475* | up | 291 | 305 | + | 11.9758 | 2.57E-05 | 1 | AAGGCTTTTTTTTGT |
| 1203 | mamo-L | *BGIBMGA002475* | up | 2 | 16 | - | 10.9839 | 6.59E-05 | 1 | AAGGCGTTTTTTTTT |
| 1204 | mamo-L | *BGIBMGA002477* | up | 1440 | 1454 | + | 10.7823 | 7.84E-05 | 1 | TAGGCCCGCGCGGGT |
| 1205 | mamo-L | *BGIBMGA002489* | up | 1014 | 1028 | + | 11.2823 | 5.03E-05 | 1 | AAGGCTTTGTTTACT |
| 1206 | mamo-L | *BGIBMGA002490* | down | 810 | 824 | + | 11.5726 | 3.81E-05 | 1 | TAGCCTTACTTTGTT |
| 1207 | mamo-L | *BGIBMGA002490* | up | 134 | 148 | - | 11.5323 | 3.96E-05 | 1 | TAGGCGTGAGGTGAT |
| 1208 | mamo-L | *BGIBMGA002490* | down | 1440 | 1454 | - | 11.3387 | 4.77E-05 | 1 | TAGGCTGGTGTTATA |
| 1209 | mamo-L | *BGIBMGA002491* | up | 804 | 818 | - | 12.5242 | 1.40E-05 | 1 | TGGGCCGACTTTGGT |
| 1210 | mamo-L | *BGIBMGA002493* | up | 984 | 998 | - | 13.4194 | 4.70E-06 | 1 | TAGGCGTATTTTTTT |
| 1211 | mamo-L | *BGIBMGA002493* | up | 1074 | 1088 | + | 11.5323 | 3.96E-05 | 1 | TAGGCGTGAGGTGAT |
| 1212 | mamo-L | *BGIBMGA002500* | up | 313 | 327 | + | 12.4355 | 1.55E-05 | 1 | TAGGCTAGTTGTGGT |
| 1213 | mamo-L | *BGIBMGA002505* | down | 1394 | 1408 | - | 10.5081 | 9.83E-05 | 1 | TGGGCCTGTTTTGAT |
| 1214 | mamo-L | *BGIBMGA002512* | down | 947 | 961 | + | 10.7177 | 8.28E-05 | 1 | TAGGCGGGCAGTATT |
| 1215 | mamo-L | *BGIBMGA002515* | down | 554 | 568 | - | 10.5 | 9.90E-05 | 1 | AAGGCGGGCGTGTTT |
| 1216 | mamo-L | *BGIBMGA002516* | up | 351 | 365 | - | 10.5 | 9.90E-05 | 1 | AAGGCGGGCGTGTTT |
| 1217 | mamo-L | *BGIBMGA002517* | up | 402 | 416 | - | 12.1048 | 2.24E-05 | 1 | AAGGCTTATTTTGTC |
| 1218 | mamo-L | *BGIBMGA002517* | up | 89 | 103 | + | 10.6048 | 9.09E-05 | 1 | TAGGCCCTCTGCTTT |
| 1219 | mamo-L | *BGIBMGA002518* | down | 436 | 450 | - | 12.4839 | 1.47E-05 | 1 | TAGGCATGGGTTGCC |
| 1220 | mamo-L | *BGIBMGA002518* | down | 335 | 349 | + | 11.4032 | 4.49E-05 | 1 | TAGGCTTCTTTTGTA |
| 1221 | mamo-L | *BGIBMGA002521* | up | 1727 | 1741 | - | 12.8065 | 1.01E-05 | 1 | TAGGCTGCTTTTTTT |
| 1222 | mamo-L | *BGIBMGA002521* | up | 1062 | 1076 | + | 11.621 | 3.63E-05 | 1 | TAGGCTTTATATTCT |
| 1223 | mamo-L | *BGIBMGA002521* | down | 434 | 448 | - | 10.9194 | 6.98E-05 | 1 | TAGGCTACTTTTTTT |
| 1224 | mamo-L | *BGIBMGA002522* | down | 1467 | 1481 | + | 10.9516 | 6.78E-05 | 1 | AAGGCTGGTGTGGAT |
| 1225 | mamo-L | *BGIBMGA002525* | up | 1637 | 1651 | - | 11.2177 | 5.34E-05 | 1 | TAGGCTAGGATTTGC |
| 1226 | mamo-L | *BGIBMGA002525* | up | 677 | 691 | - | 10.7742 | 7.89E-05 | 1 | TAGGCTCTAGACGCT |
| 1227 | mamo-L | *BGIBMGA002529* | up | 1850 | 1864 | + | 10.6694 | 8.62E-05 | 1 | TAGGCCGGTATGGAC |
| 1228 | mamo-L | *BGIBMGA002531* | up | 324 | 338 | - | 10.871 | 7.28E-05 | 1 | AAGGCTGTTTGTTGT |
| 1229 | mamo-L | *BGIBMGA002535* | up | 369 | 383 | - | 10.7016 | 8.39E-05 | 1 | TAGGCCTAATATCCT |
| 1230 | mamo-L | *BGIBMGA002544* | down | 472 | 486 | - | 11.0484 | 6.23E-05 | 1 | TAGGCCTGGTGTGAA |
| 1231 | mamo-L | *BGIBMGA002546* | down | 43 | 57 | + | 10.6129 | 9.03E-05 | 1 | TAGGCTTTCATGATT |
| 1232 | mamo-L | *BGIBMGA002555* | down | 791 | 805 | + | 10.5 | 9.90E-05 | 1 | GAGGCCCAGGTCGTC |
| 1233 | mamo-L | *BGIBMGA002558* | down | 160 | 174 | + | 11.2097 | 5.38E-05 | 1 | TAGGCGGACGATTAC |
| 1234 | mamo-L | *BGIBMGA002561* | down | 1783 | 1797 | - | 11.121 | 5.84E-05 | 1 | TAGGCTTTGTGTTAC |
| 1235 | mamo-L | *BGIBMGA002562* | up | 168 | 182 | - | 11.121 | 5.84E-05 | 1 | TAGGCTTTGTGTTAC |
| 1236 | mamo-L | *BGIBMGA002565* | up | 383 | 397 | - | 10.5726 | 9.33E-05 | 1 | TAGGCTAATATTTAT |
| 1237 | mamo-L | *BGIBMGA002570* | down | 1173 | 1187 | - | 11.0565 | 6.18E-05 | 1 | TAGGGTTTGTTTGGT |
| 1238 | mamo-L | *BGIBMGA002573* | down | 673 | 687 | + | 10.9194 | 6.98E-05 | 1 | TAGGCTACTTTTTTT |
| 1239 | mamo-L | *BGIBMGA002574* | down | 337 | 351 | - | 10.5726 | 9.33E-05 | 1 | TAGGCTAATATTTAT |
| 1240 | mamo-L | *BGIBMGA002582* | up | 1441 | 1455 | - | 10.7823 | 7.84E-05 | 1 | TAGGCCCGCAGGGGT |
| 1241 | mamo-L | *BGIBMGA002583* | up | 1565 | 1579 | - | 13.7419 | 2.91E-06 | 1 | TAGGCTGCCGTTTGT |
| 1242 | mamo-L | *BGIBMGA002584* | up | 1233 | 1247 | + | 11.6855 | 3.42E-05 | 1 | TAGGCGTATTTTAAT |
| 1243 | mamo-L | *BGIBMGA002587* | up | 1522 | 1536 | - | 12.2097 | 2.01E-05 | 1 | TAGGCCTAATTTATT |
| 1244 | mamo-L | *BGIBMGA002588* | up | 1458 | 1472 | + | 10.9194 | 6.98E-05 | 1 | TAGGCTACTTTTTTT |
| 1245 | mamo-L | *BGIBMGA002588* | down | 1097 | 1111 | + | 10.7661 | 7.95E-05 | 1 | AAGGCGGGTGATGGC |
| 1246 | mamo-L | *BGIBMGA002592* | up | 615 | 629 | + | 10.9194 | 6.98E-05 | 1 | TAGGCTACTTTTTTT |
| 1247 | mamo-L | *BGIBMGA002599* | up | 1513 | 1527 | - | 10.75 | 8.06E-05 | 1 | GAGGCCGATTGTGCC |
| 1248 | mamo-L | *BGIBMGA002603* | up | 1557 | 1571 | + | 11.2903 | 4.99E-05 | 1 | TAGGCCACGTTTTTT |
| 1249 | mamo-L | *BGIBMGA002603* | down | 890 | 904 | + | 11.0726 | 6.09E-05 | 1 | AAGGCTTGATTTTCT |
| 1250 | mamo-L | *BGIBMGA002604* | down | 1792 | 1806 | - | 10.7581 | 8.00E-05 | 1 | AAGGCTTTTTTTATT |
| 1251 | mamo-L | *BGIBMGA002605* | down | 1854 | 1868 | + | 13.3548 | 5.14E-06 | 1 | TAGGCTGTGGGTTTT |
| 1252 | mamo-L | *BGIBMGA002605* | up | 62 | 76 | - | 10.7581 | 8.00E-05 | 1 | AAGGCTTTTTTTATT |
| 1253 | mamo-L | *BGIBMGA002607* | up | 215 | 229 | + | 10.5565 | 9.46E-05 | 1 | TAGGCTGAAATCTGC |
| 1254 | mamo-L | *BGIBMGA002610* | down | 1113 | 1127 | + | 10.7258 | 8.23E-05 | 1 | TAGGCGGGGTGCGGA |
| 1255 | mamo-L | *BGIBMGA002611* | up | 1236 | 1250 | + | 10.5645 | 9.40E-05 | 1 | TAGGCTTCGGTTTCG |
| 1256 | mamo-L | *BGIBMGA002616* | down | 1976 | 1990 | - | 12.379 | 1.66E-05 | 1 | AAGGCGGTTTTTGTT |
| 1257 | mamo-L | *BGIBMGA002616* | up | 1073 | 1087 | - | 12.2419 | 1.94E-05 | 1 | TAGGCTTAGTGCGAT |
| 1258 | mamo-L | *BGIBMGA002617* | up | 1236 | 1250 | - | 12.379 | 1.66E-05 | 1 | AAGGCGGTTTTTGTT |
| 1259 | mamo-L | *BGIBMGA002621* | down | 644 | 658 | - | 13.6774 | 3.24E-06 | 1 | TAGGCAGACTTTGTT |
| 1260 | mamo-L | *BGIBMGA002622* | up | 171 | 185 | - | 13.6774 | 3.24E-06 | 1 | TAGGCAGACTTTGTT |
| 1261 | mamo-L | *BGIBMGA002624* | up | 1850 | 1864 | - | 10.7581 | 8.00E-05 | 1 | AAGGCCTGTTTCTGT |
| 1262 | mamo-L | *BGIBMGA002625* | down | 197 | 211 | + | 12.4919 | 1.46E-05 | 1 | TAGGCTGAGGGCGCC |
| 1263 | mamo-L | *BGIBMGA002627* | down | 632 | 646 | + | 12.7177 | 1.12E-05 | 1 | TAGGCTTAGTTCTGC |
| 1264 | mamo-L | *BGIBMGA002628* | down | 1419 | 1433 | - | 11.3387 | 4.77E-05 | 1 | TAGGCCTTTGTAGTT |
| 1265 | mamo-L | *BGIBMGA002635* | down | 1176 | 1190 | + | 11.2016 | 5.41E-05 | 1 | TAGGCCAGCTTTGCG |
| 1266 | mamo-L | *BGIBMGA002644* | up | 751 | 765 | + | 11.7097 | 3.34E-05 | 1 | TAGGCATGTTTTTTT |
| 1267 | mamo-L | *BGIBMGA002644* | up | 813 | 827 | - | 11.6048 | 3.70E-05 | 1 | TAGGCTCAGGTAGTT |
| 1268 | mamo-L | *BGIBMGA002646* | down | 86 | 100 | + | 12.2177 | 1.99E-05 | 1 | TAGGCTTAGATGTGT |
| 1269 | mamo-L | *BGIBMGA002646* | up | 1786 | 1800 | + | 10.5726 | 9.33E-05 | 1 | TAGGCTAATATTTAT |
| 1270 | mamo-L | *BGIBMGA002647* | up | 1607 | 1621 | - | 11.1129 | 5.88E-05 | 1 | TAGGCCCGCTGTTCC |
| 1271 | mamo-L | *BGIBMGA002647* | up | 1835 | 1849 | - | 10.5968 | 9.15E-05 | 1 | TAGGCTCTTATCTTT |
| 1272 | mamo-L | *BGIBMGA002650* | up | 231 | 245 | + | 10.5484 | 9.52E-05 | 1 | GAGGCTGGCTATGGA |
| 1273 | mamo-L | *BGIBMGA002651* | down | 1861 | 1875 | - | 10.5645 | 9.40E-05 | 1 | AAGGCTGGGTGTGGA |
| 1274 | mamo-L | *BGIBMGA002656* | up | 1265 | 1279 | - | 12.9032 | 8.96E-06 | 1 | TAGGCCGGGGATAGT |
| 1275 | mamo-L | *BGIBMGA002660* | up | 99 | 113 | + | 11.5806 | 3.78E-05 | 1 | TAGGCTGTAATTAGT |
| 1276 | mamo-L | *BGIBMGA002664* | down | 1081 | 1095 | + | 10.9597 | 6.73E-05 | 1 | AAGGCTAATGTTGGC |
| 1277 | mamo-L | *BGIBMGA002666* | up | 1312 | 1326 | + | 10.6452 | 8.79E-05 | 1 | TAGGCTTCTTTCAGT |
| 1278 | mamo-L | *BGIBMGA002667* | down | 985 | 999 | - | 12.4919 | 1.46E-05 | 1 | TAGGCTGAGGGCGCC |
| 1279 | mamo-L | *BGIBMGA002669* | up | 1480 | 1494 | + | 11.8871 | 2.81E-05 | 1 | AAGGCTTATGATGTT |
| 1280 | mamo-L | *BGIBMGA002672* | up | 93 | 107 | - | 10.7823 | 7.84E-05 | 1 | AAGGCCGATTGTGCC |
| 1281 | mamo-L | *BGIBMGA002673* | up | 1512 | 1526 | - | 12.4113 | 1.60E-05 | 1 | TAGGCATACGTTTTT |

| | | | | | | | | | | |
|---|---|---|---|---|---|---|---|---|---|---|
| 1282 | mamo-L | *BGIBMGA002676* | up | 859 | 873 | + | 10.5081 | 9.83E-05 | 1 | TAGGCATGTGTGTGT |
| 1283 | mamo-L | *BGIBMGA002677* | down | 520 | 534 | + | 13.1048 | 7.05E-06 | 1 | TAGGCCTAGAATGCT |
| 1284 | mamo-L | *BGIBMGA002677* | down | 1958 | 1972 | + | 11.871 | 2.85E-05 | 1 | TAGGCCGTCAGTTTT |
| 1285 | mamo-L | *BGIBMGA002679* | up | 1248 | 1262 | - | 10.5887 | 9.21E-05 | 1 | TAGGCTTGTTTTTAG |
| 1286 | mamo-L | *BGIBMGA002680* | down | 1830 | 1844 | - | 11.3387 | 4.77E-05 | 1 | GAGGCTTACATTTTT |
| 1287 | mamo-L | *BGIBMGA002682* | up | 114 | 128 | - | 11.0403 | 6.28E-05 | 1 | GAGGCCGGGTTTGTG |
| 1288 | mamo-L | *BGIBMGA002682* | up | 10 | 24 | + | 10.9194 | 6.98E-05 | 1 | TAGGCTACTTTTTTT |
| 1289 | mamo-L | *BGIBMGA002687* | down | 261 | 275 | - | 11.0726 | 6.09E-05 | 1 | AAGGCTTGATTTTCT |
| 1290 | mamo-L | *BGIBMGA002687* | up | 1224 | 1238 | + | 10.9758 | 6.64E-05 | 1 | AAGGCCCTGTATGCT |
| 1291 | mamo-L | *BGIBMGA002689* | down | 1827 | 1841 | + | 11.8548 | 2.89E-05 | 1 | TAGGCCAACGTTCCT |
| 1292 | mamo-L | *BGIBMGA002690* | down | 1548 | 1562 | + | 12.5645 | 1.34E-05 | 1 | TAGGCTGTATTTGAC |
| 1293 | mamo-L | *BGIBMGA002706* | up | 182 | 196 | - | 10.5726 | 9.33E-05 | 1 | TAGGCTAATATTTAT |
| 1294 | mamo-L | *BGIBMGA002708* | up | 1364 | 1378 | + | 10.5403 | 9.58E-05 | 1 | TAGGCTGTATGGTGT |
| 1295 | mamo-L | *BGIBMGA002709* | down | 1024 | 1038 | + | 12.1129 | 2.22E-05 | 1 | TAGGCTTTGCTTTGT |
| 1296 | mamo-L | *BGIBMGA002709* | up | 1329 | 1343 | - | 11.0726 | 6.09E-05 | 1 | TAGGCCAAGTTTTAC |
| 1297 | mamo-L | *BGIBMGA002710* | up | 138 | 152 | + | 12.1129 | 2.22E-05 | 1 | TAGGCTTTGCTTTGT |
| 1298 | mamo-L | *BGIBMGA002712* | up | 1646 | 1660 | - | 11.4113 | 4.45E-05 | 1 | AAGGCTGTGATTTTT |
| 1299 | mamo-L | *BGIBMGA002713* | down | 1102 | 1116 | + | 10.5565 | 9.46E-05 | 1 | TAGGCGCGGAGTTCT |
| 1300 | mamo-L | *BGIBMGA002718* | down | 1655 | 1669 | - | 11.371 | 4.62E-05 | 1 | GAGGCTGATGATTGT |
| 1301 | mamo-L | *BGIBMGA002719* | down | 1153 | 1167 | + | 11.0565 | 6.18E-05 | 1 | TAGGGTTTGTTTGGT |
| 1302 | mamo-L | *BGIBMGA002719* | up | 924 | 938 | - | 10.6129 | 9.03E-05 | 1 | AAGGCCAGCATTGTT |
| 1303 | mamo-L | *BGIBMGA002721* | up | 864 | 878 | - | 10.629 | 8.91E-05 | 1 | TAGGCCTGCAGCGCC |
| 1304 | mamo-L | *BGIBMGA002724* | up | 1898 | 1912 | - | 10.7661 | 7.95E-05 | 1 | TAGGCTGGAATGTTT |
| 1305 | mamo-L | *BGIBMGA002725* | down | 762 | 776 | - | 11.4194 | 4.42E-05 | 1 | GAGGCCTACGGTGGC |
| 1306 | mamo-L | *BGIBMGA002727* | up | 1433 | 1447 | - | 11.25 | 5.18E-05 | 1 | TAGGCTATTTTTACT |
| 1307 | mamo-L | *BGIBMGA002727* | down | 1594 | 1608 | - | 11.0403 | 6.28E-05 | 1 | TAGGCTTGCTGCGGA |
| 1308 | mamo-L | *BGIBMGA002728* | up | 1538 | 1552 | - | 11.0403 | 6.28E-05 | 1 | TAGGCTTGCTGCGGA |
| 1309 | mamo-L | *BGIBMGA002731* | up | 1637 | 1651 | + | 12.2823 | 1.85E-05 | 1 | TAGGCGTTGGTTGTG |
| 1310 | mamo-L | *BGIBMGA002731* | down | 1618 | 1632 | - | 10.9194 | 6.98E-05 | 1 | TAGGCTACTTTTTTT |
| 1311 | mamo-L | *BGIBMGA002731* | down | 595 | 609 | - | 10.879 | 7.22E-05 | 1 | TAGGCTCACATTTAC |
| 1312 | mamo-L | *BGIBMGA002732* | up | 298 | 312 | + | 11.1774 | 5.54E-05 | 1 | GAGGCTTAGAATGTT |
| 1313 | mamo-L | *BGIBMGA002732* | up | 824 | 838 | + | 11.0645 | 6.14E-05 | 1 | AAGGCTGGTGGCGGT |
| 1314 | mamo-L | *BGIBMGA002733* | up | 1110 | 1124 | - | 11.3468 | 4.73E-05 | 1 | AAGGCGGGTGGTGTT |
| 1315 | mamo-L | *BGIBMGA002735* | up | 1658 | 1672 | + | 12.5726 | 1.33E-05 | 1 | TAGGCAGGGTGTGGT |
| 1316 | mamo-L | *BGIBMGA002735* | up | 1459 | 1473 | + | 12.3145 | 1.79E-05 | 1 | TAGGCTATTGTTTTT |
| 1317 | mamo-L | *BGIBMGA002737* | down | 102 | 116 | - | 11.0242 | 6.36E-05 | 1 | AAGGCCTTGTTTTCC |
| 1318 | mamo-L | *BGIBMGA002738* | down | 354 | 368 | - | 11.6694 | 3.47E-05 | 1 | TAGGCTGATGATGCG |
| 1319 | mamo-L | *BGIBMGA002738* | down | 1427 | 1441 | - | 11.4032 | 4.49E-05 | 1 | TAGGCTTTTAGCGGT |
| 1320 | mamo-L | *BGIBMGA002738* | up | 1552 | 1566 | - | 10.9516 | 6.78E-05 | 1 | TGGGCTTATGTTTGT |
| 1321 | mamo-L | *BGIBMGA002746* | down | 358 | 372 | - | 11.5403 | 3.93E-05 | 1 | TAGGCTGAGAGTATT |
| 1322 | mamo-L | *BGIBMGA002746* | down | 1970 | 1984 | - | 10.8952 | 7.12E-05 | 1 | TAGGCGCTTTGTTTT |
| 1323 | mamo-L | *BGIBMGA002748* | up | 1607 | 1621 | + | 12.3065 | 1.80E-05 | 1 | TAGGCTTGCGTTGAG |
| 1324 | mamo-L | *BGIBMGA002748* | up | 1771 | 1785 | + | 11.9435 | 2.65E-05 | 1 | TAGGCGGAGACTGGT |
| 1325 | mamo-L | *BGIBMGA002750* | down | 1750 | 1764 | - | 12.7581 | 1.07E-05 | 1 | TAGGCTGTGTAGGTT |
| 1326 | mamo-L | *BGIBMGA002752* | up | 571 | 585 | - | 10.8548 | 7.38E-05 | 1 | GAGGCCTGTTTTTAT |
| 1327 | mamo-L | *BGIBMGA002754* | up | 627 | 641 | + | 12.7258 | 1.11E-05 | 1 | TAGGCGTGCTTTGTA |
| 1328 | mamo-L | *BGIBMGA002758* | up | 1751 | 1765 | + | 12.4597 | 1.51E-05 | 1 | TAGGCTGATATTTAT |
| 1329 | mamo-L | *BGIBMGA002759* | down | 1537 | 1551 | - | 11.371 | 4.62E-05 | 1 | TAGGCATAGTTTTTC |
| 1330 | mamo-L | *BGIBMGA002759* | up | 124 | 138 | - | 11.0968 | 5.97E-05 | 1 | TAGGCTGAGATGGTA |
| 1331 | mamo-L | *BGIBMGA002766* | up | 1735 | 1749 | - | 12.7258 | 1.11E-05 | 1 | TAGGCGTGCTTTGTA |
| 1332 | mamo-L | *BGIBMGA002773* | up | 1200 | 1214 | + | 10.8952 | 7.12E-05 | 1 | TAGGCTTACGCTGGG |
| 1333 | mamo-L | *BGIBMGA002774* | down | 1844 | 1858 | - | 10.4919 | 9.96E-05 | 1 | TAGGCTGTGTCTGTG |
| 1334 | mamo-L | *BGIBMGA002775* | down | 538 | 552 | - | 10.8468 | 7.42E-05 | 1 | TAGGCTGATGAGGAC |
| 1335 | mamo-L | *BGIBMGA002782* | up | 189 | 203 | - | 10.5403 | 9.58E-05 | 1 | TAGGCGGGATATTTC |
| 1336 | mamo-L | *BGIBMGA002785* | up | 39 | 53 | + | 11.25 | 5.18E-05 | 1 | TAGGCCCATTTCAGT |
| 1337 | mamo-L | *BGIBMGA002786* | up | 358 | 372 | - | 12.9677 | 8.30E-06 | 1 | TAGGCGTTTGATGCT |
| 1338 | mamo-L | *BGIBMGA002787* | up | 974 | 988 | + | 12.0645 | 2.34E-05 | 1 | TAGGCGGGCTTCGGA |
| 1339 | mamo-L | *BGIBMGA002788* | up | 887 | 901 | - | 10.8306 | 7.52E-05 | 1 | TGGGCTGTGTTCGCT |
| 1340 | mamo-L | *BGIBMGA002789* | down | 1432 | 1446 | - | 11.3548 | 4.69E-05 | 1 | AAGGCTGATGATGAT |
| 1341 | mamo-L | *BGIBMGA002790* | up | 121 | 135 | + | 12.6855 | 1.16E-05 | 1 | TAGGCTTTTTTTCCT |
| 1342 | mamo-L | *BGIBMGA002791* | down | 1850 | 1864 | + | 11.0242 | 6.36E-05 | 1 | TAGGCTGGATGTCCT |
| 1343 | mamo-L | *BGIBMGA002794* | down | 1441 | 1455 | - | 12.5887 | 1.30E-05 | 1 | TAGGCCAATTTTGAT |
| 1344 | mamo-L | *BGIBMGA002795* | down | 689 | 703 | - | 12.5887 | 1.30E-05 | 1 | TAGGCCAATTTTGAT |
| 1345 | mamo-L | *BGIBMGA002798* | up | 1632 | 1646 | - | 13.0242 | 7.77E-06 | 1 | TAGGCAGACGTTTGT |
| 1346 | mamo-L | *BGIBMGA002800* | up | 908 | 922 | + | 10.629 | 8.91E-05 | 1 | GAGGCTTCCTATGTT |
| 1347 | mamo-L | *BGIBMGA002800* | up | 1217 | 1231 | + | 10.629 | 8.91E-05 | 1 | GAGGCTTCCTATGTT |
| 1348 | mamo-L | *BGIBMGA002800* | down | 1204 | 1218 | - | 10.5726 | 9.33E-05 | 1 | TAGGCTAATATTTAT |
| 1349 | mamo-L | *BGIBMGA002804* | down | 1546 | 1560 | - | 11.1613 | 5.63E-05 | 1 | TAGGCGTTTATTTAT |
| 1350 | mamo-L | *BGIBMGA002805* | down | 188 | 202 | - | 10.9758 | 6.64E-05 | 1 | TAGGCCTGCAGCGGC |
| 1351 | mamo-L | *BGIBMGA002807* | up | 468 | 482 | - | 11.4435 | 4.31E-05 | 1 | AAGGCGTGTGATGGT |
| 1352 | mamo-L | *BGIBMGA002810* | up | 1863 | 1877 | + | 10.8226 | 7.58E-05 | 1 | AAGGCGTAATTTTGT |
| 1353 | mamo-L | *BGIBMGA002813* | down | 729 | 743 | + | 10.9274 | 6.92E-05 | 1 | TAGGCTGTGATTTTG |
| 1354 | mamo-L | *BGIBMGA002815* | up | 1249 | 1263 | + | 10.5726 | 9.33E-05 | 1 | TAGGCTAATATTTAT |
| 1355 | mamo-L | *BGIBMGA002820* | up | 815 | 829 | - | 10.8387 | 7.47E-05 | 1 | TAGGCTGAGATCAAT |
| 1356 | mamo-L | *BGIBMGA002820* | down | 379 | 393 | + | 10.8226 | 7.58E-05 | 1 | AAGGCTGGTTTGGAT |
| 1357 | mamo-L | *BGIBMGA002820* | down | 571 | 585 | - | 10.75 | 8.06E-05 | 1 | GAGGCCGATTGTGCC |
| 1358 | mamo-L | *BGIBMGA002822* | up | 198 | 212 | - | 12.1452 | 2.15E-05 | 1 | TAGGCAGGTGGTGGT |
| 1359 | mamo-L | *BGIBMGA002822* | down | 1855 | 1869 | - | 12.0323 | 2.42E-05 | 1 | TAGGCTTTGTATTTC |
| 1360 | mamo-L | *BGIBMGA002824* | down | 1826 | 1840 | + | 11.1371 | 5.75E-05 | 1 | AAGGCTATTTTTGCT |
| 1361 | mamo-L | *BGIBMGA002824* | down | 66 | 80 | - | 10.7097 | 8.33E-05 | 1 | TAGGCAGAGTTCTGC |
| 1362 | mamo-L | *BGIBMGA002824* | down | 787 | 801 | - | 10.5565 | 9.46E-05 | 1 | AAGGCTGTTTGTTTT |
| 1363 | mamo-L | *BGIBMGA002828* | up | 1605 | 1619 | + | 11.9435 | 2.65E-05 | 1 | GAGGCTTATATTGTT |
| 1364 | mamo-L | *BGIBMGA002833* | down | 1768 | 1782 | - | 10.5726 | 9.33E-05 | 1 | TAGGCTAATATTTAT |
| 1365 | mamo-L | *BGIBMGA002834* | up | 1848 | 1862 | - | 10.5726 | 9.33E-05 | 1 | TAGGCTAATATTTAT |
| 1366 | mamo-L | *BGIBMGA002837* | up | 209 | 223 | - | 12.0645 | 2.34E-05 | 1 | TAGGCGGGCTTCGGA |
| 1367 | mamo-L | *BGIBMGA002841* | up | 1867 | 1881 | - | 12.1935 | 2.05E-05 | 1 | TAGGCTTAAATTTTT |
| 1368 | mamo-L | *BGIBMGA002842* | down | 1945 | 1959 | + | 11.4758 | 4.18E-05 | 1 | GAGGCTAGTGTTGCT |
| 1369 | mamo-L | *BGIBMGA002843* | up | 102 | 116 | + | 12.6935 | 1.15E-05 | 1 | TAGGCCGACATTATT |
| 1370 | mamo-L | *BGIBMGA002844* | down | 1104 | 1118 | - | 10.6129 | 9.03E-05 | 1 | TAGGCTTAAAGCGTT |
| 1371 | mamo-L | *BGIBMGA002845* | down | 1016 | 1030 | - | 12.7258 | 1.11E-05 | 1 | TAGGCCCACGATTGT |
| 1372 | mamo-L | *BGIBMGA002845* | up | 606 | 620 | - | 11.7016 | 3.37E-05 | 1 | GAGGCGCATGTTGTT |
| 1373 | mamo-L | *BGIBMGA002845* | up | 1814 | 1828 | - | 10.6129 | 9.03E-05 | 1 | TAGGCTTAAAGCGTT |

| | | | | | | | | | | |
|---|---|---|---|---|---|---|---|---|---|---|
| 1374 | mamo-L | *BGIBMGA002846* | up | 1930 | 1944 | + | 11.7016 | 3.37E-05 | 1 | TAGGCAGGTTGTGTT |
| 1375 | mamo-L | *BGIBMGA002847* | down | 1684 | 1698 | - | 11.75 | 3.21E-05 | 1 | TAGGCATATATTGCT |
| 1376 | mamo-L | *BGIBMGA002848* | down | 848 | 862 | - | 10.8306 | 7.52E-05 | 1 | AAGGCTTTCTATTTT |
| 1377 | mamo-L | *BGIBMGA002849* | up | 1364 | 1378 | + | 11.5645 | 3.84E-05 | 1 | GAGGCTTGTTGTGTT |
| 1378 | mamo-L | *BGIBMGA002850* | up | 909 | 923 | - | 13.2984 | 5.54E-06 | 1 | TAGGCGCGCGTCGGT |
| 1379 | mamo-L | *BGIBMGA002851* | up | 406 | 420 | + | 10.7823 | 7.84E-05 | 1 | TAGGCCGTTTTTATA |
| 1380 | mamo-L | *BGIBMGA002851* | up | 1621 | 1635 | - | 10.5403 | 9.58E-05 | 1 | GAGGCTTGTTTGTGT |
| 1381 | mamo-L | *BGIBMGA002857* | down | 1868 | 1882 | - | 11.371 | 4.62E-05 | 1 | TAGGCGTGCGGTGGG |
| 1382 | mamo-L | *BGIBMGA002864* | down | 1860 | 1874 | + | 12.5968 | 1.28E-05 | 1 | AAGGCTTTCGTTTGT |
| 1383 | mamo-L | *BGIBMGA002864* | up | 608 | 622 | - | 10.8548 | 7.38E-05 | 1 | GAGGCCTGTTTTTAT |
| 1384 | mamo-L | *BGIBMGA002865* | down | 1182 | 1196 | + | 12.0726 | 2.32E-05 | 1 | TAGGCTTATGATTAT |
| 1385 | mamo-L | *BGIBMGA002867* | down | 470 | 484 | + | 10.6855 | 8.51E-05 | 1 | AAGGCGGGTGGTGGC |
| 1386 | mamo-L | *BGIBMGA002871* | up | 539 | 553 | + | 10.6935 | 8.45E-05 | 1 | TAGGCAAACTTTTTT |
| 1387 | mamo-L | *BGIBMGA002872* | up | 308 | 322 | + | 11.1048 | 5.92E-05 | 1 | TAGGCTTTTATTAGC |
| 1388 | mamo-L | *BGIBMGA002873* | up | 1237 | 1251 | + | 12.379 | 1.66E-05 | 1 | TAGGCATACGTTTCT |
| 1389 | mamo-L | *BGIBMGA002876* | up | 1283 | 1297 | - | 14.7258 | 5.83E-07 | 1 | TAGGCTTGCGTTGCC |
| 1390 | mamo-L | *BGIBMGA002877* | up | 1652 | 1666 | - | 11.9274 | 2.69E-05 | 1 | TAGGCTGTGGTGGGG |
| 1391 | mamo-L | *BGIBMGA002877* | up | 1509 | 1523 | - | 11.7258 | 3.29E-05 | 1 | AAGGCCGGGGTTTTC |
| 1392 | mamo-L | *BGIBMGA002881* | down | 907 | 921 | + | 12.2177 | 1.99E-05 | 1 | TAGGCTGTGTGTTCC |
| 1393 | mamo-L | *BGIBMGA002884* | up | 899 | 913 | + | 10.8387 | 7.47E-05 | 1 | TAGGCTGAGATCAAT |
| 1394 | mamo-L | *BGIBMGA002884* | down | 709 | 723 | + | 10.4919 | 9.96E-05 | 1 | TAGGCACGTGTGGTT |
| 1395 | mamo-L | *BGIBMGA002885* | down | 50 | 64 | - | 10.8548 | 7.38E-05 | 1 | AAGGCGGGTATTGGC |
| 1396 | mamo-L | *BGIBMGA002887* | up | 769 | 783 | - | 13.7742 | 2.79E-06 | 1 | TAGGCTTTTTTTTTT |
| 1397 | mamo-L | *BGIBMGA002887* | up | 317 | 331 | + | 11.7823 | 3.11E-05 | 1 | GAGGCTCACTTCGGT |
| 1398 | mamo-L | *BGIBMGA002889* | down | 1744 | 1758 | + | 11.9435 | 2.65E-05 | 1 | AAGGCTGGTTATGCT |
| 1399 | mamo-L | *BGIBMGA002897* | down | 900 | 914 | - | 10.5484 | 9.52E-05 | 1 | GAGGCTTATTGTTTT |
| 1400 | mamo-L | *BGIBMGA002898* | down | 409 | 423 | + | 11.9274 | 2.69E-05 | 1 | TAGGCATTCTTTTCT |
| 1401 | mamo-L | *BGIBMGA002898* | down | 685 | 699 | - | 11.6774 | 3.44E-05 | 1 | AAGGCTCACTTTTTT |
| 1402 | mamo-L | *BGIBMGA002901* | up | 1226 | 1240 | + | 10.629 | 8.91E-05 | 1 | TAGGCCTGCAGCGCC |
| 1403 | mamo-L | *BGIBMGA002902* | up | 631 | 645 | - | 12.0806 | 2.31E-05 | 1 | TAGGCTTGTGTAGGT |
| 1404 | mamo-L | *BGIBMGA002904* | down | 280 | 294 | + | 11.7419 | 3.23E-05 | 1 | AAGGCGGATGGTGGT |
| 1405 | mamo-L | *BGIBMGA002907* | down | 1942 | 1956 | - | 10.5968 | 9.15E-05 | 1 | TAGGCTAATTTTACC |
| 1406 | mamo-L | *BGIBMGA002908* | down | 1761 | 1775 | + | 11.1452 | 5.70E-05 | 1 | TAGGCCGGCTTATTT |
| 1407 | mamo-L | *BGIBMGA002908* | up | 17 | 31 | - | 10.5968 | 9.15E-05 | 1 | TAGGCTAATTTTACC |
| 1408 | mamo-L | *BGIBMGA002911* | up | 1147 | 1161 | - | 12.9919 | 8.10E-06 | 1 | TAGGCCGGTTTTGCA |
| 1409 | mamo-L | *BGIBMGA002911* | up | 1280 | 1294 | - | 12.6774 | 1.17E-05 | 1 | TAGGCTTACTATAGT |
| 1410 | mamo-L | *BGIBMGA002914* | up | 409 | 423 | + | 10.8065 | 7.69E-05 | 1 | TAGGCCTGCAGCGAT |
| 1411 | mamo-L | *BGIBMGA002915* | up | 1285 | 1299 | - | 12.6855 | 1.16E-05 | 1 | TAGGCTTTTTTTCCT |
| 1412 | mamo-L | *BGIBMGA002918* | up | 1978 | 1992 | + | 12.9677 | 8.30E-06 | 1 | TAGGCGTTTGATGCT |
| 1413 | mamo-L | *BGIBMGA002923* | up | 309 | 323 | + | 12.0645 | 2.34E-05 | 1 | TAGGCTGTTATTATT |
| 1414 | mamo-L | *BGIBMGA002924* | up | 1580 | 1594 | - | 11.4032 | 4.49E-05 | 1 | TAGGCTAATATTTTT |
| 1415 | mamo-L | *BGIBMGA002924* | up | 1409 | 1423 | + | 11.1613 | 5.63E-05 | 1 | GAGGCTGTGTATTTT |
| 1416 | mamo-L | *BGIBMGA002924* | up | 1936 | 1950 | + | 10.7903 | 7.79E-05 | 1 | TAGGCCTCTTTTGCG |
| 1417 | mamo-L | *BGIBMGA002927* | down | 1695 | 1709 | + | 10.9516 | 6.78E-05 | 1 | TAGGCTTTGTTGAAT |
| 1418 | mamo-L | *BGIBMGA002931* | up | 712 | 726 | - | 10.9194 | 6.98E-05 | 1 | TAGGCTACTTTTTTT |
| 1419 | mamo-L | *BGIBMGA002933* | up | 223 | 237 | + | 13.0887 | 7.18E-06 | 1 | TAGGCTTAGTTTAGC |
| 1420 | mamo-L | *BGIBMGA002933* | down | 1749 | 1763 | + | 10.8145 | 7.63E-05 | 1 | TAGGCTGTTAATTAT |
| 1421 | mamo-L | *BGIBMGA002934* | up | 1223 | 1237 | + | 11.2581 | 5.14E-05 | 1 | TAGGCTCCGGTTGCA |
| 1422 | mamo-L | *BGIBMGA002938* | down | 1254 | 1268 | + | 10.6048 | 9.09E-05 | 1 | TAGGCCTGAAATTTT |
| 1423 | mamo-L | *BGIBMGA002943* | up | 1192 | 1206 | - | 11.1935 | 5.45E-05 | 1 | TAGGCTTTGTTTACA |
| 1424 | mamo-L | *BGIBMGA002945* | down | 1456 | 1470 | - | 13.9677 | 2.09E-06 | 1 | TAGGCTATGGTTGTT |
| 1425 | mamo-L | *BGIBMGA002954* | down | 1947 | 1961 | - | 13.4274 | 4.64E-06 | 1 | TAGGCTCTGGTGGGT |
| 1426 | mamo-L | *BGIBMGA002956* | down | 608 | 622 | - | 11.8065 | 3.03E-05 | 1 | TAGGCGCTAATTGGT |
| 1427 | mamo-L | *BGIBMGA002958* | down | 311 | 325 | - | 12.4032 | 1.61E-05 | 1 | TAGGCTTTTGTTGTG |
| 1428 | mamo-L | *BGIBMGA002961* | down | 1092 | 1106 | - | 10.7177 | 8.28E-05 | 1 | TCGGCTGGGTTTGTC |
| 1429 | mamo-L | *BGIBMGA002962* | up | 1058 | 1072 | + | 10.7823 | 7.84E-05 | 1 | TAGGCTAGGGACGAT |
| 1430 | mamo-L | *BGIBMGA002965* | up | 222 | 236 | - | 11.0161 | 6.40E-05 | 1 | TAGGCCGCAGTTACT |
| 1431 | mamo-L | *BGIBMGA002967* | up | 917 | 931 | - | 12.0484 | 2.38E-05 | 1 | TAGGCCGATAATGAT |
| 1432 | mamo-L | *BGIBMGA002970* | down | 1142 | 1156 | - | 10.5645 | 9.40E-05 | 1 | GAGGCCTTTGTCGAT |
| 1433 | mamo-L | *BGIBMGA002972* | down | 238 | 252 | - | 10.9194 | 6.98E-05 | 1 | TAGGCTACTTTTTTT |
| 1434 | mamo-L | *BGIBMGA002977* | up | 570 | 584 | - | 11.1048 | 5.92E-05 | 1 | AAGGCCCAGGTGGCT |
| 1435 | mamo-L | *BGIBMGA002978* | up | 599 | 613 | - | 10.5403 | 9.58E-05 | 1 | TAGGCTGTATGGTGT |
| 1436 | mamo-L | *BGIBMGA002979* | up | 256 | 270 | - | 10.5726 | 9.33E-05 | 1 | TAGGCTAATATTTAT |
| 1437 | mamo-L | *BGIBMGA002980* | up | 1222 | 1236 | - | 12.1452 | 2.15E-05 | 1 | TAGGCCGGGGCTTCT |
| 1438 | mamo-L | *BGIBMGA002980* | up | 1621 | 1635 | + | 11.3468 | 4.73E-05 | 1 | GAGGCGGTGGTTAGT |
| 1439 | mamo-L | *BGIBMGA002980* | down | 1546 | 1560 | + | 10.8548 | 7.38E-05 | 1 | TAGGCATTCATTTTT |
| 1440 | mamo-L | *BGIBMGA002981* | down | 1585 | 1599 | + | 14.2258 | 1.41E-06 | 1 | TAGGCTTTGGTGGGT |
| 1441 | mamo-L | *BGIBMGA002984* | down | 232 | 246 | + | 11.7661 | 3.16E-05 | 1 | TAGGCCCTCGTGTTT |
| 1442 | mamo-L | *BGIBMGA002986* | down | 1738 | 1752 | - | 11.6935 | 3.39E-05 | 1 | TAGGCTGTTTGTTTC |
| 1443 | mamo-L | *BGIBMGA002988* | up | 1219 | 1233 | - | 13.4274 | 4.64E-06 | 1 | TAGGCTTGTTTTAGT |
| 1444 | mamo-L | *BGIBMGA002992* | down | 1486 | 1500 | + | 13.2258 | 6.08E-06 | 1 | TAGGCCATTGTTGTT |
| 1445 | mamo-L | *BGIBMGA002992* | down | 1473 | 1487 | - | 11.2097 | 5.38E-05 | 1 | TAGGCGTGTGATACT |
| 1446 | mamo-L | *BGIBMGA002996* | up | 182 | 196 | - | 12.1855 | 2.06E-05 | 1 | TAGGCTTTAATTTGT |
| 1447 | mamo-L | *BGIBMGA002996* | down | 1650 | 1664 | + | 12.1048 | 2.24E-05 | 1 | TAGGCATGCGATGTT |
| 1448 | mamo-L | *BGIBMGA002997* | down | 791 | 805 | + | 10.9355 | 6.88E-05 | 1 | TCGGCTTGTGTTGCT |
| 1449 | mamo-L | *BGIBMGA002997* | up | 46 | 60 | - | 10.7742 | 7.89E-05 | 1 | TAGGCTTGGTAGTAT |
| 1450 | mamo-L | *BGIBMGA002999* | up | 1710 | 1724 | + | 10.6532 | 8.73E-05 | 1 | TAGGCTTTCAGTGTA |
| 1451 | mamo-L | *BGIBMGA002999* | up | 1048 | 1062 | - | 10.5323 | 9.63E-05 | 1 | TAGGCGGCGGTTTGG |
| 1452 | mamo-L | *BGIBMGA003000* | down | 1074 | 1088 | + | 10.9194 | 6.98E-05 | 1 | TAGGCTACTTTTTTT |
| 1453 | mamo-L | *BGIBMGA003002* | down | 1738 | 1752 | - | 12.2742 | 1.87E-05 | 1 | TAGGCTCAATATGTT |
| 1454 | mamo-L | *BGIBMGA003003* | down | 751 | 765 | + | 11.3871 | 4.55E-05 | 1 | TAGGCCGATTATGTG |
| 1455 | mamo-L | *BGIBMGA003005* | up | 573 | 587 | - | 10.8306 | 7.52E-05 | 1 | TAGGCCCCCTTCGCC |
| 1456 | mamo-L | *BGIBMGA003007* | up | 434 | 448 | + | 12.7258 | 1.11E-05 | 1 | TAGGCGTGCTTTGTA |
| 1457 | mamo-L | *BGIBMGA003007* | up | 882 | 896 | + | 12.7258 | 1.11E-05 | 1 | TAGGCGTGCTTTGTA |
| 1458 | mamo-L | *BGIBMGA003007* | down | 670 | 684 | - | 10.5 | 9.90E-05 | 1 | GAGGCCCAGGTCGTC |
| 1459 | mamo-L | *BGIBMGA003015* | up | 247 | 261 | - | 11.4355 | 4.34E-05 | 1 | GAGGCTTGCGTTCTT |
| 1460 | mamo-L | *BGIBMGA003024* | up | 1247 | 1261 | + | 12.2258 | 1.97E-05 | 1 | TAGGCTGCGTTTGCA |
| 1461 | mamo-L | *BGIBMGA003029* | up | 1647 | 1661 | + | 12.1371 | 2.17E-05 | 1 | TAGGCTGTATGTGAT |
| 1462 | mamo-L | *BGIBMGA003031* | up | 1492 | 1506 | + | 12.129 | 2.19E-05 | 1 | TAGGCTGTGGACTCT |
| 1463 | mamo-L | *BGIBMGA003033* | down | 498 | 512 | + | 10.7581 | 8.00E-05 | 1 | TAGGCGGTTTCTTTT |
| 1464 | mamo-L | *BGIBMGA003034* | up | 1006 | 1020 | - | 11.0081 | 6.45E-05 | 1 | GAGGCGTGGTTTCGT |
| 1465 | mamo-L | *BGIBMGA003035* | up | 1666 | 1680 | - | 11.3548 | 4.69E-05 | 1 | TAGGCGGAATTTAGC |

| | | | | | | | | | | |
|---|---|---|---|---|---|---|---|---|---|---|
| 1466 | mamo-L | *BGIBMGA003037* | up | 895 | 909 | - | 10.8145 | 7.63E-05 | 1 | AAGGCTTGGTTGTTT |
| 1467 | mamo-L | *BGIBMGA003040* | up | 6 | 20 | - | 11.1855 | 5.50E-05 | 1 | TAGGCGGAGGCGGCT |
| 1468 | mamo-L | *BGIBMGA003041* | up | 1979 | 1993 | - | 10.5403 | 9.58E-05 | 1 | TAGGCTTCCAATTCT |
| 1469 | mamo-L | *BGIBMGA003043* | up | 1556 | 1570 | + | 13.7742 | 2.79E-06 | 1 | TAGGCGGGTGGTGGT |
| 1470 | mamo-L | *BGIBMGA003044* | up | 762 | 776 | + | 10.5726 | 9.33E-05 | 1 | TAGGCTAATATTTAT |
| 1471 | mamo-L | *BGIBMGA003045* | up | 1071 | 1085 | - | 13.3871 | 4.90E-06 | 1 | TAGGCTGATGGTGAT |
| 1472 | mamo-L | *BGIBMGA003045* | up | 1746 | 1760 | + | 12.75 | 1.08E-05 | 1 | TAGGCGGTCGGCGGT |
| 1473 | mamo-L | *BGIBMGA003046* | up | 74 | 88 | - | 11.8468 | 2.92E-05 | 1 | TAGGCCGGGATTAAT |
| 1474 | mamo-L | *BGIBMGA003046* | down | 1704 | 1718 | - | 10.9355 | 6.88E-05 | 1 | GAGGCTTGTTTTACT |
| 1475 | mamo-L | *BGIBMGA003047* | up | 1589 | 1603 | - | 12.6371 | 1.23E-05 | 1 | TAGGCTGTTCTTGTT |
| 1476 | mamo-L | *BGIBMGA003054* | up | 1666 | 1680 | + | 11.0081 | 6.45E-05 | 1 | AAGGCCGGTTTCGAT |
| 1477 | mamo-L | *BGIBMGA003055* | up | 378 | 392 | - | 11.2984 | 4.95E-05 | 1 | TAGGCGTAATATTTT |
| 1478 | mamo-L | *BGIBMGA003060* | up | 807 | 821 | + | 10.6452 | 8.79E-05 | 1 | TAGGCTCACTTGGTG |
| 1479 | mamo-L | *BGIBMGA003061* | up | 366 | 380 | - | 12.7339 | 1.10E-05 | 1 | TAGGCCTTGGCTGTT |
| 1480 | mamo-L | *BGIBMGA003061* | down | 671 | 685 | + | 11.4435 | 4.31E-05 | 1 | TAGGCTCATGGGGCT |
| 1481 | mamo-L | *BGIBMGA003066* | down | 464 | 478 | - | 11.3145 | 4.87E-05 | 1 | AAGGCCTAATTTTGT |
| 1482 | mamo-L | *BGIBMGA003073* | down | 988 | 1002 | + | 10.5968 | 9.15E-05 | 1 | TGGGCCGTGTATGCT |
| 1483 | mamo-L | *BGIBMGA003074* | up | 943 | 957 | - | 10.6532 | 8.73E-05 | 1 | GAGGCGGGTGGTGGC |
| 1484 | mamo-L | *BGIBMGA003076* | up | 1160 | 1174 | - | 11.1935 | 5.45E-05 | 1 | GAGGCTGATGTCGCC |
| 1485 | mamo-L | *BGIBMGA003080* | up | 1909 | 1923 | + | 12.5242 | 1.40E-05 | 1 | GAGGCTTGGGTTTCT |
| 1486 | mamo-L | *BGIBMGA003080* | down | 193 | 207 | - | 10.8871 | 7.17E-05 | 1 | TAGGGTTGCTTTGCT |
| 1487 | mamo-L | *BGIBMGA003081* | up | 229 | 243 | + | 11.6774 | 3.44E-05 | 1 | TAGGCGGCGTTTTCC |
| 1488 | mamo-L | *BGIBMGA003081* | down | 1808 | 1822 | - | 11.379 | 4.58E-05 | 1 | TAGGCGTAGTTTTTG |
| 1489 | mamo-L | *BGIBMGA003083* | up | 1113 | 1127 | + | 10.8871 | 7.17E-05 | 1 | TAGGCCTGCTGTCTC |
| 1490 | mamo-L | *BGIBMGA003084* | up | 13 | 27 | + | 11.9839 | 2.54E-05 | 1 | AAGGCTTATTTTTTT |
| 1491 | mamo-L | *BGIBMGA003085* | down | 639 | 653 | + | 11.8306 | 2.97E-05 | 1 | TAGGCTTGGAGGGGT |
| 1492 | mamo-L | *BGIBMGA003089* | down | 1276 | 1290 | - | 12.7339 | 1.10E-05 | 1 | TAGGCAGAGTATGGT |
| 1493 | mamo-L | *BGIBMGA003089* | down | 1806 | 1820 | + | 12.129 | 2.19E-05 | 1 | TAGGCCGACGAGGAT |
| 1494 | mamo-L | *BGIBMGA003089* | down | 1153 | 1167 | + | 10.6532 | 8.73E-05 | 1 | TAGGCTGTGACGGGT |
| 1495 | mamo-L | *BGIBMGA003090* | up | 154 | 168 | + | 12.129 | 2.19E-05 | 1 | TAGGCCGACGAGGAT |
| 1496 | mamo-L | *BGIBMGA003090* | down | 578 | 592 | - | 11.379 | 4.58E-05 | 1 | TAGGCTGTAATTGGA |
| 1497 | mamo-L | *BGIBMGA003095* | up | 679 | 693 | + | 10.8306 | 7.52E-05 | 1 | AAGGCGGACTGCGGT |
| 1498 | mamo-L | *BGIBMGA003097* | up | 118 | 132 | + | 10.5161 | 9.77E-05 | 1 | TAGGCTATCAATGAT |
| 1499 | mamo-L | *BGIBMGA003098* | up | 968 | 982 | + | 12.9919 | 8.10E-06 | 1 | TAGGCCGGTTTTGCA |
| 1500 | mamo-L | *BGIBMGA003098* | down | 280 | 294 | - | 11.3145 | 4.87E-05 | 1 | TAGGCTTGGGAAGGT |
| 1501 | mamo-L | *BGIBMGA003099* | down | 1491 | 1505 | + | 11.3145 | 4.87E-05 | 1 | GAGGCCTAGTTTTCC |
| 1502 | mamo-L | *BGIBMGA003099* | up | 882 | 896 | + | 10.6855 | 8.51E-05 | 1 | TAGGCGTACTCTAGT |
| 1503 | mamo-L | *BGIBMGA003100* | down | 419 | 433 | + | 11.6613 | 3.50E-05 | 1 | TAGGCTGGTTAGTGT |
| 1504 | mamo-L | *BGIBMGA003101* | down | 1518 | 1532 | + | 11.9355 | 2.67E-05 | 1 | GAGGCGTGAGTTGGT |
| 1505 | mamo-L | *BGIBMGA003101* | up | 1248 | 1262 | + | 11.6613 | 3.50E-05 | 1 | TAGGCTGGTTAGTGT |
| 1506 | mamo-L | *BGIBMGA003102* | up | 1274 | 1288 | - | 11.6048 | 3.70E-05 | 1 | TAGGCACGGGGTGGT |
| 1507 | mamo-L | *BGIBMGA003103* | up | 77 | 91 | + | 12.871 | 9.35E-06 | 1 | AAGGCTTACGTTGAT |
| 1508 | mamo-L | *BGIBMGA003103* | down | 731 | 745 | + | 10.8306 | 7.52E-05 | 1 | TAGGCCCCCTTCGCC |
| 1509 | mamo-L | *BGIBMGA003108* | down | 968 | 982 | - | 12.0323 | 2.42E-05 | 1 | GAGGCCTAATTTGCT |
| 1510 | mamo-L | *BGIBMGA003108* | down | 975 | 989 | + | 11.6129 | 3.66E-05 | 1 | TAGGCCTCGTGCGGT |
| 1511 | mamo-L | *BGIBMGA003108* | up | 522 | 536 | - | 10.5968 | 9.15E-05 | 1 | GAGGCTAGTTTTTGT |
| 1512 | mamo-L | *BGIBMGA003112* | up | 714 | 728 | + | 12.0806 | 2.31E-05 | 1 | TAGGCCTTGTTCGGA |
| 1513 | mamo-L | *BGIBMGA003113* | up | 1971 | 1985 | + | 13.3871 | 4.90E-06 | 1 | TAGGCTGATGGTGAT |
| 1514 | mamo-L | *BGIBMGA003113* | up | 1296 | 1310 | - | 12.75 | 1.08E-05 | 1 | TAGGCGGTCGGCGGT |
| 1515 | mamo-L | *BGIBMGA003115* | up | 376 | 390 | + | 11.5806 | 3.78E-05 | 1 | GAGGCTGTGTTTATT |
| 1516 | mamo-L | *BGIBMGA003116* | down | 1672 | 1686 | - | 10.629 | 8.91E-05 | 1 | TAGGCCTGCAGCGCC |
| 1517 | mamo-L | *BGIBMGA003121* | down | 1195 | 1209 | + | 10.5726 | 9.33E-05 | 1 | TAGGCTAATATTTAT |
| 1518 | mamo-L | *BGIBMGA003123* | up | 929 | 943 | + | 11.0081 | 6.45E-05 | 1 | GAGGCGTGGTTTCGT |
| 1519 | mamo-L | *BGIBMGA003124* | up | 1276 | 1290 | + | 11.5323 | 3.96E-05 | 1 | TAGGCGTGAGGTGAT |
| 1520 | mamo-L | *BGIBMGA003126* | down | 1063 | 1077 | - | 10.9597 | 6.73E-05 | 1 | TAGGCTAAGGTTTCA |
| 1521 | mamo-L | *BGIBMGA003127* | up | 1819 | 1833 | - | 10.9597 | 6.73E-05 | 1 | TAGGCTAAGGTTTCA |
| 1522 | mamo-L | *BGIBMGA003129* | up | 1097 | 1111 | + | 12.8306 | 9.76E-06 | 1 | TAGGCAGTTTTTGCT |
| 1523 | mamo-L | *BGIBMGA003134* | down | 1469 | 1483 | - | 11.3548 | 4.69E-05 | 1 | GAGGCTTGCGTTGCA |
| 1524 | mamo-L | *BGIBMGA003135* | up | 1880 | 1894 | - | 11.8065 | 3.03E-05 | 1 | TAGGCGGTGTGTCGT |
| 1525 | mamo-L | *BGIBMGA003137* | down | 1885 | 1899 | + | 11.4032 | 4.49E-05 | 1 | AAGGCGGTTTTTGTC |
| 1526 | mamo-L | *BGIBMGA003137* | down | 1864 | 1878 | - | 10.5323 | 9.63E-05 | 1 | AAGGCTTGCGGTAGT |
| 1527 | mamo-L | *BGIBMGA003138* | up | 794 | 808 | - | 11.4032 | 4.49E-05 | 1 | TAGGCGGTTTGTAGT |
| 1528 | mamo-L | *BGIBMGA003141* | down | 983 | 997 | + | 14.75 | 5.63E-07 | 1 | TAGGCGTTGTTTGTT |
| 1529 | mamo-L | *BGIBMGA003143* | up | 456 | 470 | + | 12.2258 | 1.97E-05 | 1 | TAGGCTGCGTGTGAT |
| 1530 | mamo-L | *BGIBMGA003149* | down | 1855 | 1869 | - | 10.8952 | 7.12E-05 | 1 | AAGGCCGTTGATTGT |
| 1531 | mamo-L | *BGIBMGA003150* | down | 1187 | 1201 | - | 13.3871 | 4.90E-06 | 1 | TAGGCCGTGTATGAT |
| 1532 | mamo-L | *BGIBMGA003151* | down | 1872 | 1886 | - | 12.0242 | 2.44E-05 | 1 | TAGGCTTATGTTTTA |
| 1533 | mamo-L | *BGIBMGA003155* | up | 1261 | 1275 | - | 12.7258 | 1.11E-05 | 1 | TAGGCGTGCTTTGTA |
| 1534 | mamo-L | *BGIBMGA003155* | up | 1709 | 1723 | - | 12.7258 | 1.11E-05 | 1 | TAGGCGTGCTTTGTA |
| 1535 | mamo-L | *BGIBMGA003156* | up | 293 | 307 | + | 11.8548 | 2.89E-05 | 1 | TAGGCCCAGATTGAC |
| 1536 | mamo-L | *BGIBMGA003159* | up | 158 | 172 | + | 10.8145 | 7.63E-05 | 1 | TAGGCCGAATACTTT |
| 1537 | mamo-L | *BGIBMGA003161* | down | 400 | 414 | + | 12.2742 | 1.87E-05 | 1 | TAGGCTCAATATGTT |
| 1538 | mamo-L | *BGIBMGA003161* | up | 1524 | 1538 | + | 12.2661 | 1.89E-05 | 1 | TAGGCTGATGATACT |
| 1539 | mamo-L | *BGIBMGA003165* | down | 760 | 774 | - | 10.9194 | 6.98E-05 | 1 | TAGGCTACTTTTTTT |
| 1540 | mamo-L | *BGIBMGA003166* | up | 1035 | 1049 | + | 12.4677 | 1.49E-05 | 1 | TAGGCTTCTATTGCT |
| 1541 | mamo-L | *BGIBMGA003168* | up | 969 | 983 | - | 10.5726 | 9.33E-05 | 1 | TAGGCTAATATTTAT |
| 1542 | mamo-L | *BGIBMGA003169* | up | 1551 | 1565 | + | 10.7742 | 7.89E-05 | 1 | TAGGCTTGGTAGTAT |
| 1543 | mamo-L | *BGIBMGA003176* | up | 773 | 787 | + | 12.4919 | 1.46E-05 | 1 | TAGGCTGGGTATCTT |
| 1544 | mamo-L | *BGIBMGA003176* | down | 868 | 882 | + | 10.621 | 8.97E-05 | 1 | TAGGCCTGATTCGCA |
| 1545 | mamo-L | *BGIBMGA003181* | down | 359 | 373 | - | 11.8065 | 3.03E-05 | 1 | AAGGCGCTGTTTGCT |
| 1546 | mamo-L | *BGIBMGA003184* | down | 1321 | 1335 | + | 11.5242 | 3.99E-05 | 1 | TAGGCAGACGACGGT |
| 1547 | mamo-L | *BGIBMGA003185* | up | 1062 | 1076 | + | 12.7258 | 1.11E-05 | 1 | TAGGCTTGCGTTACC |
| 1548 | mamo-L | *BGIBMGA003191* | up | 1620 | 1634 | + | 10.5403 | 9.58E-05 | 1 | TAGGCTGTATGGTGT |
| 1549 | mamo-L | *BGIBMGA003192* | up | 142 | 156 | + | 11.1048 | 5.92E-05 | 1 | AAGGCCCAGGTGGCT |
| 1550 | mamo-L | *BGIBMGA003198* | down | 1811 | 1825 | + | 11.2258 | 5.30E-05 | 1 | TAGGCTATTTTCTGT |
| 1551 | mamo-L | *BGIBMGA003199* | up | 1854 | 1868 | + | 11.9435 | 2.65E-05 | 1 | TAGGCCTTGTTTTTA |
| 1552 | mamo-L | *BGIBMGA003199* | up | 1798 | 1812 | + | 10.7339 | 8.17E-05 | 1 | TAGGCTTATAATACT |
| 1553 | mamo-L | *BGIBMGA003199* | up | 1210 | 1224 | + | 10.6613 | 8.67E-05 | 1 | AAGGCTGGCGACGGC |
| 1554 | mamo-L | *BGIBMGA003206* | down | 1516 | 1530 | - | 10.8387 | 7.47E-05 | 1 | TAGGCCTTTTTGGTA |
| 1555 | mamo-L | *BGIBMGA003208* | up | 991 | 1005 | + | 11.371 | 4.62E-05 | 1 | TAGGCTGTATAGGCT |
| 1556 | mamo-L | *BGIBMGA003214* | down | 349 | 363 | + | 11.2419 | 5.21E-05 | 1 | AAGGCGGCTGTTGTT |
| 1557 | mamo-L | *BGIBMGA003216* | down | 200 | 214 | - | 11.1935 | 5.45E-05 | 1 | TAGGCTAACTTTTAC |

| | | | | | | | | | | |
|---|---|---|---|---|---|---|---|---|---|---|
| 1558 | mamo-L | *BGIBMGA003217* | down | 1520 | 1534 | - | 11.0484 | 6.23E-05 | 1 | TAGGCACGTGGTGGT |
| 1559 | mamo-L | *BGIBMGA003219* | up | 890 | 904 | - | 10.5565 | 9.46E-05 | 1 | TAGGCTATAGTCTGT |
| 1560 | mamo-L | *BGIBMGA003222* | down | 324 | 338 | + | 11.3952 | 4.52E-05 | 1 | TAGGCCTTTAGTTGT |
| 1561 | mamo-L | *BGIBMGA003224* | up | 266 | 280 | + | 12.2419 | 1.94E-05 | 1 | TAGGCTTAATTTCTT |
| 1562 | mamo-L | *BGIBMGA003226* | down | 1311 | 1325 | - | 10.8387 | 7.47E-05 | 1 | TAGGCCCTCCTTGCC |
| 1563 | mamo-L | *BGIBMGA003227* | up | 461 | 475 | - | 10.8387 | 7.47E-05 | 1 | TAGGCCCTCCTTGCC |
| 1564 | mamo-L | *BGIBMGA003230* | down | 235 | 249 | - | 12.0887 | 2.29E-05 | 1 | TAGGCCGGCTAGGGC |
| 1565 | mamo-L | *BGIBMGA003231* | down | 1689 | 1703 | + | 10.75 | 8.06E-05 | 1 | GAGGCCGATTGTGCC |
| 1566 | mamo-L | *BGIBMGA003231* | down | 34 | 48 | - | 10.6371 | 8.85E-05 | 1 | TAGGCTTTAAATGCC |
| 1567 | mamo-L | *BGIBMGA003233* | up | 682 | 696 | + | 10.7581 | 8.00E-05 | 1 | TCGGCTTACGTTTGT |
| 1568 | mamo-L | *BGIBMGA003233* | up | 1538 | 1552 | + | 10.6774 | 8.56E-05 | 1 | GAGGCTGGCTTCGAC |
| 1569 | mamo-L | *BGIBMGA003235* | down | 228 | 242 | - | 10.5726 | 9.33E-05 | 1 | TAGGCTAATATTTAT |
| 1570 | mamo-L | *BGIBMGA003243* | up | 716 | 730 | - | 11.129 | 5.79E-05 | 1 | GAGGCTTACTGTGCC |
| 1571 | mamo-L | *BGIBMGA003244* | up | 1234 | 1248 | - | 10.9194 | 6.98E-05 | 1 | TAGGCTACTTTTTTT |
| 1572 | mamo-L | *BGIBMGA003245* | up | 429 | 443 | + | 11.371 | 4.62E-05 | 1 | TAGGCTGTATAGGCT |
| 1573 | mamo-L | *BGIBMGA003245* | up | 1876 | 1890 | + | 11.371 | 4.62E-05 | 1 | TAGGCTGTATAGGCT |
| 1574 | mamo-L | *BGIBMGA003252* | up | 1182 | 1196 | - | 11.2823 | 5.03E-05 | 1 | AAGGCGGTTTTTTTT |
| 1575 | mamo-L | *BGIBMGA003253* | down | 1138 | 1152 | - | 13.4113 | 4.75E-06 | 1 | GAGGCTTGTGTTGGT |
| 1576 | mamo-L | *BGIBMGA003253* | down | 515 | 529 | + | 11.3145 | 4.87E-05 | 1 | AAGGCTGGGGCTGCT |
| 1577 | mamo-L | *BGIBMGA003254* | down | 1343 | 1357 | - | 11.2258 | 5.30E-05 | 1 | TAGGCCTTTTTCCCT |
| 1578 | mamo-L | *BGIBMGA003255* | up | 1519 | 1533 | - | 10.8306 | 7.52E-05 | 1 | TAGGCCCCCTTCGCC |
| 1579 | mamo-L | *BGIBMGA003256* | down | 71 | 85 | + | 12.2742 | 1.87E-05 | 1 | TAGGCGAATGTTTGT |
| 1580 | mamo-L | *BGIBMGA003257* | down | 1479 | 1493 | + | 11.1371 | 5.75E-05 | 1 | AAGGCTTCAGTTGGT |
| 1581 | mamo-L | *BGIBMGA003258* | down | 6 | 20 | + | 11.2177 | 5.34E-05 | 1 | TAGGCTGGTAGTAGT |
| 1582 | mamo-L | *BGIBMGA003258* | down | 1438 | 1452 | - | 10.5403 | 9.58E-05 | 1 | TAGGCTGTATGGTGT |
| 1583 | mamo-L | *BGIBMGA003259* | down | 1189 | 1203 | - | 13.1774 | 6.47E-06 | 1 | TAGGCTTTGGGTGTC |
| 1584 | mamo-L | *BGIBMGA003262* | down | 365 | 379 | - | 11.371 | 4.62E-05 | 1 | TAGGCTAATATTTCT |
| 1585 | mamo-L | *BGIBMGA003265* | down | 416 | 430 | + | 10.9597 | 6.73E-05 | 1 | AAGGCTTTTGTTTAT |
| 1586 | mamo-L | *BGIBMGA003266* | up | 1009 | 1023 | + | 10.5726 | 9.33E-05 | 1 | TAGGCTAATATTTAT |
| 1587 | mamo-L | *BGIBMGA003268* | down | 1230 | 1244 | + | 10.9194 | 6.98E-05 | 1 | TAGGCTACTTTTTTT |
| 1588 | mamo-L | *BGIBMGA003268* | down | 711 | 725 | + | 10.6048 | 9.09E-05 | 1 | GAGGCTGAGTCTGGC |
| 1589 | mamo-L | *BGIBMGA003272* | down | 624 | 638 | - | 11.1935 | 5.45E-05 | 1 | TAGGCGCATGAGGGT |
| 1590 | mamo-L | *BGIBMGA003272* | up | 1595 | 1609 | - | 10.8468 | 7.42E-05 | 1 | AAGGCTTGTTTTCTT |
| 1591 | mamo-L | *BGIBMGA003274* | up | 1886 | 1900 | + | 11 | 6.50E-05 | 1 | TAGGCAGGTGGTGAT |
| 1592 | mamo-L | *BGIBMGA003274* | up | 564 | 578 | - | 10.9032 | 7.07E-05 | 1 | TAGGCGTTGATGTTT |
| 1593 | mamo-L | *BGIBMGA003276* | down | 778 | 792 | - | 13.5887 | 3.68E-06 | 1 | TAGGCCTTTTTTTTT |
| 1594 | mamo-L | *BGIBMGA003276* | down | 59 | 73 | + | 12.871 | 9.35E-06 | 1 | TAGGCTTTTTTTATT |
| 1595 | mamo-L | *BGIBMGA003276* | up | 1140 | 1154 | + | 12.0645 | 2.34E-05 | 1 | TAGGCGGGCTTCGGA |
| 1596 | mamo-L | *BGIBMGA003281* | up | 965 | 979 | + | 13.7742 | 2.79E-06 | 1 | TAGGCGGGTGGTGGT |
| 1597 | mamo-L | *BGIBMGA003282* | down | 593 | 607 | + | 10.5726 | 9.33E-05 | 1 | TAGGCCTAAAGTTCT |
| 1598 | mamo-L | *BGIBMGA003284* | down | 1349 | 1363 | + | 12.2016 | 2.03E-05 | 1 | TAGGCGCTCTTTAGT |
| 1599 | mamo-L | *BGIBMGA003284* | down | 853 | 867 | + | 11.371 | 4.62E-05 | 1 | TAGGCTAGTTTTCTT |
| 1600 | mamo-L | *BGIBMGA003287* | down | 996 | 1010 | + | 10.9194 | 6.98E-05 | 1 | TAGGCTACTTTTTTT |
| 1601 | mamo-L | *BGIBMGA003287* | up | 1807 | 1821 | - | 10.7419 | 8.11E-05 | 1 | AAGGCGCTGTTTTTT |
| 1602 | mamo-L | *BGIBMGA003289* | up | 1608 | 1622 | + | 13.3145 | 5.41E-06 | 1 | TAGGCTTAGATGGGT |
| 1603 | mamo-L | *BGIBMGA003291* | down | 1028 | 1042 | - | 10.5726 | 9.33E-05 | 1 | TAGGCTGATGGGATT |
| 1604 | mamo-L | *BGIBMGA003293* | down | 185 | 199 | + | 11.3548 | 4.69E-05 | 1 | AAGGCTGATGATGAT |
| 1605 | mamo-L | *BGIBMGA003293* | up | 1515 | 1529 | + | 10.9194 | 6.98E-05 | 1 | TAGGCTACTTTTTTT |
| 1606 | mamo-L | *BGIBMGA003295* | up | 648 | 662 | + | 13.4677 | 4.39E-06 | 1 | TAGGCTGATGATGAT |
| 1607 | mamo-L | *BGIBMGA003296* | down | 139 | 153 | + | 11.5726 | 3.81E-05 | 1 | TAGGCTGAGGTCTCA |
| 1608 | mamo-L | *BGIBMGA003297* | down | 1851 | 1865 | - | 10.9516 | 6.78E-05 | 1 | AAGGCTGAGTTTCAT |
| 1609 | mamo-L | *BGIBMGA003302* | up | 6 | 20 | + | 12.3306 | 1.75E-05 | 1 | TAGGCCGTGATTTCC |
| 1610 | mamo-L | *BGIBMGA003302* | up | 1373 | 1387 | + | 11.9032 | 2.76E-05 | 1 | TAGGCCGATTGTATT |
| 1611 | mamo-L | *BGIBMGA003302* | up | 1576 | 1590 | + | 11.9032 | 2.76E-05 | 1 | TAGGCCGATTGTATT |
| 1612 | mamo-L | *BGIBMGA003303* | up | 214 | 228 | - | 11.7661 | 3.16E-05 | 1 | TAGGCCGATCTTGCC |
| 1613 | mamo-L | *BGIBMGA003303* | up | 1939 | 1953 | + | 10.9919 | 6.55E-05 | 1 | TAGGCCGGTTTTACA |
| 1614 | mamo-L | *BGIBMGA003304* | up | 1657 | 1671 | - | 11.8065 | 3.03E-05 | 1 | TAGGCTAATGTTTAT |
| 1615 | mamo-L | *BGIBMGA003305* | down | 1470 | 1484 | - | 11.4355 | 4.34E-05 | 1 | AAGGCTTATTTGGTT |
| 1616 | mamo-L | *BGIBMGA003307* | down | 144 | 158 | - | 13.0887 | 7.18E-06 | 1 | TAGGCTGTGTCTGTT |
| 1617 | mamo-L | *BGIBMGA003308* | down | 1658 | 1672 | + | 12.0323 | 2.42E-05 | 1 | TAGGCACGGGTTGGC |
| 1618 | mamo-L | *BGIBMGA003309* | up | 1210 | 1224 | - | 11.6371 | 3.58E-05 | 1 | TAGGCGCTCGGTGTC |
| 1619 | mamo-L | *BGIBMGA003309* | down | 574 | 588 | + | 11.4355 | 4.34E-05 | 1 | AAGGCTTGCGGTTGT |
| 1620 | mamo-L | *BGIBMGA003309* | up | 1560 | 1574 | - | 10.9435 | 6.83E-05 | 1 | TAGGCCGGTTACAGT |
| 1621 | mamo-L | *BGIBMGA003314* | up | 1635 | 1649 | + | 10.5968 | 9.15E-05 | 1 | GAGGCTAGTTTTTGT |
| 1622 | mamo-L | *BGIBMGA003315* | up | 1322 | 1336 | - | 10.9274 | 6.92E-05 | 1 | TAGGCTGTGAACTTT |
| 1623 | mamo-L | *BGIBMGA003319* | down | 1504 | 1518 | - | 12.6613 | 1.19E-05 | 1 | TAGGCATAGTTTTGT |
| 1624 | mamo-L | *BGIBMGA003320* | down | 1121 | 1135 | + | 13.0806 | 7.24E-06 | 1 | TAGGCTTTGGTGGAT |
| 1625 | mamo-L | *BGIBMGA003330* | down | 476 | 490 | - | 13.0968 | 7.11E-06 | 1 | TAGGCTGAGGTCGAC |
| 1626 | mamo-L | *BGIBMGA003330* | up | 735 | 749 | - | 11.1935 | 5.45E-05 | 1 | TAGGCCGACGGTTCA |
| 1627 | mamo-L | *BGIBMGA003332* | up | 166 | 180 | - | 11.1371 | 5.75E-05 | 1 | AAGGCCTTCTTTTAT |
| 1628 | mamo-L | *BGIBMGA003332* | up | 1884 | 1898 | + | 11 | 6.50E-05 | 1 | TAGGCCGCGTTCATT |
| 1629 | mamo-L | *BGIBMGA003334* | up | 483 | 497 | - | 11.5806 | 3.78E-05 | 1 | TAGGCCCACGATTAT |
| 1630 | mamo-L | *BGIBMGA003334* | up | 1211 | 1225 | + | 11.5161 | 4.02E-05 | 1 | TAGCCCTACGTTGTT |
| 1631 | mamo-L | *BGIBMGA003335* | up | 1860 | 1874 | - | 11.9516 | 2.63E-05 | 1 | AAGGCTTAGTTTAGT |
| 1632 | mamo-L | *BGIBMGA003336* | down | 1932 | 1946 | - | 10.9919 | 6.55E-05 | 1 | GAGGCGGAGGGCGGT |
| 1633 | mamo-L | *BGIBMGA003348* | down | 1004 | 1018 | + | 11 | 6.50E-05 | 1 | TAGGCATGCGTGTGT |
| 1634 | mamo-L | *BGIBMGA003349* | up | 1617 | 1631 | - | 10.5726 | 9.33E-05 | 1 | TAGGCTAATTTCTGC |
| 1635 | mamo-L | *BGIBMGA003351* | down | 1537 | 1551 | + | 10.75 | 8.06E-05 | 1 | GAGGCCGATTGTGCC |
| 1636 | mamo-L | *BGIBMGA003355* | up | 1801 | 1815 | - | 10.8226 | 7.58E-05 | 1 | GAGGCGTAGTTTTCC |
| 1637 | mamo-L | *BGIBMGA003356* | down | 1352 | 1366 | - | 11.7258 | 3.29E-05 | 1 | TAGGCTTATTCTGAT |
| 1638 | mamo-L | *BGIBMGA003356* | up | 24 | 38 | - | 10.8226 | 7.58E-05 | 1 | AAGGCTGGTTTGGAT |
| 1639 | mamo-L | *BGIBMGA003359* | up | 378 | 392 | - | 11.371 | 4.62E-05 | 1 | TAGGCGGACTTCAGC |
| 1640 | mamo-L | *BGIBMGA003360* | up | 202 | 216 | + | 10.75 | 8.06E-05 | 1 | GAGGCTGAAGTTTAT |
| 1641 | mamo-L | *BGIBMGA003362* | up | 1490 | 1504 | - | 11.9032 | 2.76E-05 | 1 | AAGGCTTGTTTTTTT |
| 1642 | mamo-L | *BGIBMGA003363* | down | 324 | 338 | - | 10.9113 | 7.02E-05 | 1 | TAGGCTGTTGGTTGA |
| 1643 | mamo-L | *BGIBMGA003364* | down | 1263 | 1277 | + | 12.7903 | 1.02E-05 | 1 | TAGGCTTGTTCTGGT |
| 1644 | mamo-L | *BGIBMGA003364* | up | 590 | 604 | + | 10.9194 | 6.98E-05 | 1 | TAGGCTACTTTTTTT |
| 1645 | mamo-L | *BGIBMGA003365* | down | 911 | 925 | + | 10.629 | 8.91E-05 | 1 | TAGGCCTGCAGCGCC |
| 1646 | mamo-L | *BGIBMGA003366* | down | 834 | 848 | + | 11.9032 | 2.76E-05 | 1 | TAGGCGGTGGTCATT |
| 1647 | mamo-L | *BGIBMGA003366* | up | 1653 | 1667 | + | 10.629 | 8.91E-05 | 1 | TAGGCCTGCAGCGCC |
| 1648 | mamo-L | *BGIBMGA003366* | down | 1870 | 1884 | + | 10.5726 | 9.33E-05 | 1 | TCGGCGTTGGTTGCT |
| 1649 | mamo-L | *BGIBMGA003368* | up | 1080 | 1094 | + | 11.7097 | 3.34E-05 | 1 | TAGGCTAAATTTTTT |

| | | | | | | | | | | |
|---|---|---|---|---|---|---|---|---|---|---|
| 1650 | mamo-L | *BGIBMGA003372* | down | 605 | 619 | + | 10.5726 | 9.33E-05 | 1 | TAGGCTAATATTTAT |
| 1651 | mamo-L | *BGIBMGA003374* | up | 1729 | 1743 | + | 12.2742 | 1.87E-05 | 1 | TAGGCATTCTTTTGT |
| 1652 | mamo-L | *BGIBMGA003376* | down | 590 | 604 | - | 12.7823 | 1.03E-05 | 1 | TAGGCCTTTGTTACT |
| 1653 | mamo-L | *BGIBMGA003379* | up | 1360 | 1374 | + | 11.1855 | 5.50E-05 | 1 | TAGGCTTATATTTAC |
| 1654 | mamo-L | *BGIBMGA003381* | down | 1663 | 1677 | + | 11.2339 | 5.26E-05 | 1 | TAGGCGGATATGGAT |
| 1655 | mamo-L | *BGIBMGA003382* | up | 1478 | 1492 | + | 11.2339 | 5.26E-05 | 1 | TAGGCGGATATGGAT |
| 1656 | mamo-L | *BGIBMGA003382* | down | 535 | 549 | - | 10.621 | 8.97E-05 | 1 | TCGGCGTGCTTTGCT |
| 1657 | mamo-L | *BGIBMGA003383* | up | 339 | 353 | + | 11.2339 | 5.26E-05 | 1 | TAGGCGGATATGGAT |
| 1658 | mamo-L | *BGIBMGA003384* | up | 1909 | 1923 | - | 11.9597 | 2.61E-05 | 1 | TAGGCTTCGATTTTT |
| 1659 | mamo-L | *BGIBMGA003384* | down | 1442 | 1456 | - | 10.9435 | 6.83E-05 | 1 | TAGGCGTTTTATTAT |
| 1660 | mamo-L | *BGIBMGA003388* | up | 1159 | 1173 | - | 11.7258 | 3.29E-05 | 1 | AAGGCTATGTTTGTT |
| 1661 | mamo-L | *BGIBMGA003388* | up | 1742 | 1756 | - | 11.6532 | 3.53E-05 | 1 | TAGGCAGACGATGAT |
| 1662 | mamo-L | *BGIBMGA003388* | up | 69 | 83 | + | 11.25 | 5.18E-05 | 1 | TAGGCCCATTTCAGT |
| 1663 | mamo-L | *BGIBMGA003388* | up | 1167 | 1181 | + | 10.6129 | 9.03E-05 | 1 | TAGCCTTGCGTTGCC |
| 1664 | mamo-L | *BGIBMGA003389* | up | 1608 | 1622 | - | 10.8468 | 7.42E-05 | 1 | TAGGCTTGTATTCCC |
| 1665 | mamo-L | *BGIBMGA003392* | up | 282 | 296 | + | 11.0968 | 5.97E-05 | 1 | TAGGCCATTTTTATT |
| 1666 | mamo-L | *BGIBMGA003394* | down | 245 | 259 | + | 11.7258 | 3.29E-05 | 1 | TAGGCTTATTCTGAT |
| 1667 | mamo-L | *BGIBMGA003400* | down | 122 | 136 | + | 13.1048 | 7.05E-06 | 1 | TAGGCTTAGGTGTCT |
| 1668 | mamo-L | *BGIBMGA003402* | down | 676 | 690 | + | 12.6855 | 1.16E-05 | 1 | TAGGCTTTTTTTCCT |
| 1669 | mamo-L | *BGIBMGA003403* | down | 1975 | 1989 | + | 10.8306 | 7.52E-05 | 1 | TAGGCCCCCTTCGCC |
| 1670 | mamo-L | *BGIBMGA003407* | up | 1844 | 1858 | + | 11.6371 | 3.58E-05 | 1 | TAGGCTTATTGTCTT |
| 1671 | mamo-L | *BGIBMGA003407* | down | 1114 | 1128 | + | 11.2258 | 5.30E-05 | 1 | TAGGCTTACAATACT |
| 1672 | mamo-L | *BGIBMGA003407* | down | 190 | 204 | + | 10.9194 | 6.98E-05 | 1 | TAGGCTACTTTTTTT |
| 1673 | mamo-L | *BGIBMGA003409* | down | 381 | 395 | + | 11.7177 | 3.32E-05 | 1 | AAGGCGTGCTTTTTT |
| 1674 | mamo-L | *BGIBMGA003410* | down | 198 | 212 | + | 10.9194 | 6.98E-05 | 1 | TAGGCTACTTTTTTT |
| 1675 | mamo-L | *BGIBMGA003411* | down | 1800 | 1814 | - | 11.3226 | 4.83E-05 | 1 | AAGGCCGAGAATGTT |
| 1676 | mamo-L | *BGIBMGA003412* | up | 323 | 337 | - | 10.5 | 9.90E-05 | 1 | GAGGCCCAGGTCGTC |
| 1677 | mamo-L | *BGIBMGA003414* | down | 205 | 219 | - | 12.5081 | 1.43E-05 | 1 | TAGGCTAATTTTTTT |
| 1678 | mamo-L | *BGIBMGA003415* | up | 1079 | 1093 | - | 12.5081 | 1.43E-05 | 1 | TAGGCTAATTTTTTT |
| 1679 | mamo-L | *BGIBMGA003416* | up | 831 | 845 | - | 11.4597 | 4.25E-05 | 1 | TAGGCTTACTATTAC |
| 1680 | mamo-L | *BGIBMGA003419* | up | 413 | 427 | - | 11.371 | 4.62E-05 | 1 | TAGGCGCATGTGGAT |
| 1681 | mamo-L | *BGIBMGA003421* | down | 49 | 63 | - | 12.2097 | 2.01E-05 | 1 | TAGGCTTGTTTTTAC |
| 1682 | mamo-L | *BGIBMGA003423* | down | 1003 | 1017 | - | 12.5806 | 1.31E-05 | 1 | TAGGCTTACATTATT |
| 1683 | mamo-L | *BGIBMGA003425* | up | 27 | 41 | + | 11.5484 | 3.90E-05 | 1 | GAGGCTTAAGTTGAT |
| 1684 | mamo-L | *BGIBMGA003426* | up | 937 | 951 | - | 11.2258 | 5.30E-05 | 1 | TAGGCCTTTTTCCCT |
| 1685 | mamo-L | *BGIBMGA003426* | up | 1944 | 1958 | + | 10.5726 | 9.33E-05 | 1 | TAGGCTAATATTTAT |
| 1686 | mamo-L | *BGIBMGA003427* | up | 1605 | 1619 | - | 11.4516 | 4.28E-05 | 1 | AAGGCCGGGCTTGGT |
| 1687 | mamo-L | *BGIBMGA003429* | down | 1828 | 1842 | + | 11.121 | 5.84E-05 | 1 | GAGGCTAACTTTGAT |
| 1688 | mamo-L | *BGIBMGA003431* | down | 857 | 871 | + | 10.5968 | 9.15E-05 | 1 | GAGGCTAGTTTTTGT |
| 1689 | mamo-L | *BGIBMGA003432* | up | 1743 | 1757 | + | 10.5968 | 9.15E-05 | 1 | GAGGCTAGTTTTTGT |
| 1690 | mamo-L | *BGIBMGA003435* | down | 489 | 503 | - | 12.7016 | 1.14E-05 | 1 | AAGGCGGATTTTGTT |
| 1691 | mamo-L | *BGIBMGA003435* | up | 425 | 439 | + | 11.6371 | 3.58E-05 | 1 | TAGGCGCTCGGTGTC |
| 1692 | mamo-L | *BGIBMGA003435* | up | 75 | 89 | + | 10.9435 | 6.83E-05 | 1 | TAGGCCGGTTACAGT |
| 1693 | mamo-L | *BGIBMGA003436* | up | 1833 | 1847 | - | 12.7016 | 1.14E-05 | 1 | AAGGCGGATTTTGTT |
| 1694 | mamo-L | *BGIBMGA003436* | down | 1878 | 1892 | - | 11.3548 | 4.69E-05 | 1 | AAGGCTGATGATGAT |
| 1695 | mamo-L | *BGIBMGA003437* | down | 1031 | 1045 | + | 11.4355 | 4.34E-05 | 1 | AAGGCTTATTTGGTT |
| 1696 | mamo-L | *BGIBMGA003439* | up | 775 | 789 | + | 12.7258 | 1.11E-05 | 1 | TAGGCGTGCTTTGTA |
| 1697 | mamo-L | *BGIBMGA003439* | down | 1828 | 1842 | - | 10.621 | 8.97E-05 | 1 | GAGGCTCAATTTGAT |
| 1698 | mamo-L | *BGIBMGA003442* | down | 749 | 763 | - | 11.5726 | 3.81E-05 | 1 | TAGGCTGAGGTCTCA |
| 1699 | mamo-L | *BGIBMGA003442* | down | 1867 | 1881 | - | 11.2097 | 5.38E-05 | 1 | AAGGCTGCGATTGCT |
| 1700 | mamo-L | *BGIBMGA003444* | down | 1575 | 1589 | + | 10.9194 | 6.98E-05 | 1 | TAGGCTACTTTTTTT |
| 1701 | mamo-L | *BGIBMGA003452* | down | 681 | 695 | + | 10.5968 | 9.15E-05 | 1 | TAGCCTGATTTTTGT |
| 1702 | mamo-L | *BGIBMGA003453* | up | 646 | 660 | + | 10.5968 | 9.15E-05 | 1 | TAGCCTGATTTTTGT |
| 1703 | mamo-L | *BGIBMGA003456* | up | 890 | 904 | + | 12.3306 | 1.75E-05 | 1 | TAGGCTTACTATACT |
| 1704 | mamo-L | *BGIBMGA003457* | down | 48 | 62 | + | 12.2339 | 1.95E-05 | 1 | GAGGCGGGAGTTGGT |
| 1705 | mamo-L | *BGIBMGA003457* | up | 463 | 477 | + | 12.2097 | 2.01E-05 | 1 | TAGGCCGTGTTGAGT |
| 1706 | mamo-L | *BGIBMGA003462* | down | 1525 | 1539 | + | 10.6452 | 8.79E-05 | 1 | GAGGCTGAGAATGAT |
| 1707 | mamo-L | *BGIBMGA003463* | up | 531 | 545 | - | 10.6371 | 8.85E-05 | 1 | TAGGCTAATTAGGTT |
| 1708 | mamo-L | *BGIBMGA003465* | down | 1560 | 1574 | - | 11.4274 | 4.38E-05 | 1 | TAGGCGCAGGTTATC |
| 1709 | mamo-L | *BGIBMGA003466* | up | 590 | 604 | + | 12.7258 | 1.11E-05 | 1 | TAGGCGTGCTTTGTA |
| 1710 | mamo-L | *BGIBMGA003466* | down | 836 | 850 | + | 11.0968 | 5.97E-05 | 1 | TAGGCTGAGATGGTA |
| 1711 | mamo-L | *BGIBMGA003473* | down | 366 | 380 | + | 12.75 | 1.08E-05 | 1 | TAGGCCGAGGGTCGT |
| 1712 | mamo-L | *BGIBMGA003473* | down | 320 | 334 | - | 10.871 | 7.28E-05 | 1 | TAGGGCTTGTTTGGT |
| 1713 | mamo-L | *BGIBMGA003475* | up | 1 | 15 | - | 12.0484 | 2.38E-05 | 1 | TAGGCTGACGGTAGC |
| 1714 | mamo-L | *BGIBMGA003475* | up | 272 | 286 | + | 10.6371 | 8.85E-05 | 1 | TAGGCGTTTGGTGGG |
| 1715 | mamo-L | *BGIBMGA003477* | down | 282 | 296 | - | 11.25 | 5.18E-05 | 1 | TAGGCTCTGGTAGCT |
| 1716 | mamo-L | *BGIBMGA003478* | up | 241 | 255 | - | 11.25 | 5.18E-05 | 1 | TAGGCTCTGGTAGCT |
| 1717 | mamo-L | *BGIBMGA003480* | up | 545 | 559 | - | 14.621 | 7.29E-07 | 1 | TAGGCTTAGTTTTCT |
| 1718 | mamo-L | *BGIBMGA003480* | down | 1934 | 1948 | - | 11.2016 | 5.41E-05 | 1 | GAGGCTGACTCTGTT |
| 1719 | mamo-L | *BGIBMGA003488* | down | 1214 | 1228 | + | 10.6694 | 8.62E-05 | 1 | TAGGCTGTTAATTTC |
| 1720 | mamo-L | *BGIBMGA003507* | up | 1391 | 1405 | + | 11.3145 | 4.87E-05 | 1 | TAGGCGTACTTTTTG |
| 1721 | mamo-L | *BGIBMGA003516* | down | 351 | 365 | + | 14.2177 | 1.43E-06 | 1 | TAGGCTGATTTCGTT |
| 1722 | mamo-L | *BGIBMGA003517* | up | 1077 | 1091 | + | 10.5645 | 9.40E-05 | 1 | TCGGCTTGGTTTGAT |
| 1723 | mamo-L | *BGIBMGA003521* | up | 1599 | 1613 | + | 12.2903 | 1.83E-05 | 1 | TAGGCTCCCTTTGCC |
| 1724 | mamo-L | *BGIBMGA003522* | up | 582 | 596 | - | 10.5 | 9.90E-05 | 1 | GAGGCCCAGGTCGTC |
| 1725 | mamo-L | *BGIBMGA003527* | down | 1163 | 1177 | - | 11.7984 | 3.06E-05 | 1 | TAGGCTTTGCTTTTT |
| 1726 | mamo-L | *BGIBMGA003530* | down | 623 | 637 | + | 14.7177 | 5.94E-07 | 1 | TAGGCGCAGGTTGGT |
| 1727 | mamo-L | *BGIBMGA003531* | down | 1667 | 1681 | + | 12.5968 | 1.28E-05 | 1 | AAGGCTTTCGTTTGT |
| 1728 | mamo-L | *BGIBMGA003531* | down | 1779 | 1793 | - | 10.9597 | 6.73E-05 | 1 | TAGGCGTGGTGTGCG |
| 1729 | mamo-L | *BGIBMGA003535* | up | 128 | 142 | + | 13.4677 | 4.39E-06 | 1 | TAGGCTTGGATTTTT |
| 1730 | mamo-L | *BGIBMGA003536* | up | 657 | 671 | + | 10.8548 | 7.38E-05 | 1 | GAGGCCTGTTTTTAT |
| 1731 | mamo-L | *BGIBMGA003538* | down | 819 | 833 | - | 11.7984 | 3.06E-05 | 1 | TAGGCCACTTTTGCT |
| 1732 | mamo-L | *BGIBMGA003538* | down | 1044 | 1058 | + | 11.0887 | 6.01E-05 | 1 | TAGGCCGCTTTTGCG |
| 1733 | mamo-L | *BGIBMGA003539* | down | 561 | 575 | - | 10.4919 | 9.96E-05 | 1 | TAGGCCGACATTATA |
| 1734 | mamo-L | *BGIBMGA003541* | down | 792 | 806 | + | 10.5645 | 9.40E-05 | 1 | TAGGCCTACTGGGGA |
| 1735 | mamo-L | *BGIBMGA003543* | down | 1312 | 1326 | + | 10.7097 | 8.33E-05 | 1 | GAGGCATATTTTGCT |
| 1736 | mamo-L | *BGIBMGA003543* | up | 435 | 449 | - | 10.5645 | 9.40E-05 | 1 | TAGGCGGGGTAGTGC |
| 1737 | mamo-L | *BGIBMGA003550* | down | 1752 | 1766 | + | 12.2984 | 1.81E-05 | 1 | TAGGCTGTGACTGGT |
| 1738 | mamo-L | *BGIBMGA003550* | down | 819 | 833 | + | 11.5484 | 3.90E-05 | 1 | TAGGCTTTTTATATT |
| 1739 | mamo-L | *BGIBMGA003550* | up | 856 | 870 | - | 10.871 | 7.28E-05 | 1 | AAGGCTGTTTGTTGT |
| 1740 | mamo-L | *BGIBMGA003552* | up | 1747 | 1761 | - | 11.3952 | 4.52E-05 | 1 | TAGGCTCATGATGAC |
| 1741 | mamo-L | *BGIBMGA003552* | up | 1073 | 1087 | + | 10.8065 | 7.69E-05 | 1 | TAGGCCGACATTAGA |

| | | | | | | | | | | |
|---|---|---|---|---|---|---|---|---|---|---|
| 1742 | mamo-L | *BGIBMGA003559* | up | 532 | 546 | - | 11.9919 | 2.52E-05 | 1 | AAGGCTTAGTTGGTT |
| 1743 | mamo-L | *BGIBMGA003562* | down | 1715 | 1729 | - | 11.5887 | 3.75E-05 | 1 | GAGGCTGGTGTTGAC |
| 1744 | mamo-L | *BGIBMGA003563* | up | 1786 | 1800 | - | 11.5887 | 3.75E-05 | 1 | GAGGCTGGTGTTGAC |
| 1745 | mamo-L | *BGIBMGA003567* | down | 915 | 929 | + | 12.3952 | 1.63E-05 | 1 | TAGGCCGACTGTATT |
| 1746 | mamo-L | *BGIBMGA003567* | up | 1706 | 1720 | - | 11.9355 | 2.67E-05 | 1 | TAGGCGATAGTTGTT |
| 1747 | mamo-L | *BGIBMGA003567* | up | 1564 | 1578 | - | 11.6532 | 3.53E-05 | 1 | AAGGCTTTTATTGTT |
| 1748 | mamo-L | *BGIBMGA003567* | down | 1801 | 1815 | - | 10.9194 | 6.98E-05 | 1 | TAGGCTACTTTTTTT |
| 1749 | mamo-L | *BGIBMGA003570* | down | 1095 | 1109 | - | 12.8065 | 1.01E-05 | 1 | TAGGCTTACGTTACC |
| 1750 | mamo-L | *BGIBMGA003576* | up | 1980 | 1994 | - | 10.5726 | 9.33E-05 | 1 | TAGGCTAATATTTAT |
| 1751 | mamo-L | *BGIBMGA003579* | up | 1225 | 1239 | + | 12.2419 | 1.94E-05 | 1 | AAGGCCTGGTTTTCT |
| 1752 | mamo-L | *BGIBMGA003580* | down | 1836 | 1850 | + | 10.621 | 8.97E-05 | 1 | GAGGCGAACGTTTGT |
| 1753 | mamo-L | *BGIBMGA003581* | down | 741 | 755 | - | 11.4597 | 4.25E-05 | 1 | TAGGCGGGGTGGTGT |
| 1754 | mamo-L | *BGIBMGA003582* | up | 1455 | 1469 | + | 10.7581 | 8.00E-05 | 1 | TAGGCTGAGTGTAGA |
| 1755 | mamo-L | *BGIBMGA003582* | down | 915 | 929 | + | 10.7419 | 8.11E-05 | 1 | TAGGCTTTTTTTTAA |
| 1756 | mamo-L | *BGIBMGA003591* | down | 270 | 284 | - | 14.2016 | 1.47E-06 | 1 | TAGGCGGGGTGTGGT |
| 1757 | mamo-L | *BGIBMGA003592* | up | 993 | 1007 | - | 14.2016 | 1.47E-06 | 1 | TAGGCGGGGTGTGGT |
| 1758 | mamo-L | *BGIBMGA003598* | up | 791 | 805 | - | 12.9919 | 8.10E-06 | 1 | TAGGCCGGTTTTGCA |
| 1759 | mamo-L | *BGIBMGA003598* | down | 1448 | 1462 | + | 11.4758 | 4.18E-05 | 1 | AAGGCGGATGATGCT |
| 1760 | mamo-L | *BGIBMGA003603* | down | 586 | 600 | - | 11.75 | 3.21E-05 | 1 | AAGGCTCTCGTTGAT |
| 1761 | mamo-L | *BGIBMGA003609* | down | 824 | 838 | - | 12.7419 | 1.09E-05 | 1 | TAGGCGTGCTGTTGT |
| 1762 | mamo-L | *BGIBMGA003609* | up | 1425 | 1439 | + | 12.4516 | 1.53E-05 | 1 | TAGGCTGAAGTCTTT |
| 1763 | mamo-L | *BGIBMGA003609* | down | 1093 | 1107 | - | 10.9194 | 6.98E-05 | 1 | TAGGCTACTTTTTTT |
| 1764 | mamo-L | *BGIBMGA003610* | up | 1563 | 1577 | + | 11.8306 | 2.97E-05 | 1 | TAGGCTGTCTGGTGT |
| 1765 | mamo-L | *BGIBMGA003612* | up | 1529 | 1543 | + | 11.5242 | 3.99E-05 | 1 | TAGGCTGGCATCATT |
| 1766 | mamo-L | *BGIBMGA003615* | up | 798 | 812 | - | 10.5968 | 9.15E-05 | 1 | GAGGCTAGTTTTTGT |
| 1767 | mamo-L | *BGIBMGA003617* | down | 404 | 418 | + | 13.3387 | 5.24E-06 | 1 | TAGGCGGGCTTTGGA |
| 1768 | mamo-L | *BGIBMGA003622* | up | 385 | 399 | + | 11.9032 | 2.76E-05 | 1 | TAGGCCTGCAGTGCC |
| 1769 | mamo-L | *BGIBMGA003623* | up | 1046 | 1060 | + | 11.8387 | 2.94E-05 | 1 | AAGGCCGATTATGCT |
| 1770 | mamo-L | *BGIBMGA003624* | up | 1969 | 1983 | + | 11.621 | 3.63E-05 | 1 | AAGGCTTTCGTTTGC |
| 1771 | mamo-L | *BGIBMGA003626* | down | 327 | 341 | + | 12.4274 | 1.57E-05 | 1 | TAGGCTAGTTTTTTT |
| 1772 | mamo-L | *BGIBMGA003636* | up | 254 | 268 | - | 10.9839 | 6.59E-05 | 1 | TCGGCGGGGTTTGCT |
| 1773 | mamo-L | *BGIBMGA003638* | up | 1472 | 1486 | - | 12.7177 | 1.12E-05 | 1 | TAGGCTTAGTTCTGC |
| 1774 | mamo-L | *BGIBMGA003641* | up | 426 | 440 | + | 11.3548 | 4.69E-05 | 1 | AAGGCTGATGATGAT |
| 1775 | mamo-L | *BGIBMGA003643* | up | 1014 | 1028 | + | 12.4113 | 1.60E-05 | 1 | TAGGCCTATTACGTT |
| 1776 | mamo-L | *BGIBMGA003643* | up | 1599 | 1613 | - | 12.0645 | 2.34E-05 | 1 | TAGGCGGGCTTCGGA |
| 1777 | mamo-L | *BGIBMGA003643* | up | 1749 | 1763 | - | 10.5645 | 9.40E-05 | 1 | TAGGCTTATTACACT |
| 1778 | mamo-L | *BGIBMGA003647* | down | 293 | 307 | - | 10.7097 | 8.33E-05 | 1 | GAGGCATATTTTGCT |
| 1779 | mamo-L | *BGIBMGA003649* | down | 658 | 672 | - | 12.2984 | 1.81E-05 | 1 | TAGGCGTTGTATTCT |
| 1780 | mamo-L | *BGIBMGA003651* | down | 704 | 718 | + | 10.4919 | 9.96E-05 | 1 | TAGGCCGACATTATA |
| 1781 | mamo-L | *BGIBMGA003661* | up | 1600 | 1614 | - | 11.4919 | 4.11E-05 | 1 | TAGGCCTCTTTTTAT |
| 1782 | mamo-L | *BGIBMGA003661* | up | 1606 | 1620 | - | 10.8387 | 7.47E-05 | 1 | TAGGCTTAGGCCTCT |
| 1783 | mamo-L | *BGIBMGA003678* | up | 1606 | 1620 | - | 10.5645 | 9.40E-05 | 1 | TCGGCTTGGTTTGAT |
| 1784 | mamo-L | *BGIBMGA003679* | down | 1609 | 1623 | - | 14.2177 | 1.43E-06 | 1 | TAGGCTGATTTCGTT |
| 1785 | mamo-L | *BGIBMGA003681* | up | 1066 | 1080 | + | 10.7016 | 8.39E-05 | 1 | TAGGCCTAATTAGCT |
| 1786 | mamo-L | *BGIBMGA003686* | up | 83 | 97 | - | 11.4758 | 4.18E-05 | 1 | GAGGCCCAGGTCGTT |
| 1787 | mamo-L | *BGIBMGA003688* | up | 545 | 559 | - | 11.0726 | 6.09E-05 | 1 | TAGGCGTATGACGTC |
| 1788 | mamo-L | *BGIBMGA003689* | up | 1420 | 1434 | - | 11.3145 | 4.87E-05 | 1 | TAGGCGTACTTTTTG |
| 1789 | mamo-L | *BGIBMGA003690* | up | 1961 | 1975 | - | 12.1371 | 2.17E-05 | 1 | TAGGCTTTAATTGAT |
| 1790 | mamo-L | *BGIBMGA003692* | up | 884 | 898 | + | 10.8226 | 7.58E-05 | 1 | TAGGCAGACGTTCAT |
| 1791 | mamo-L | *BGIBMGA003693* | down | 1736 | 1750 | + | 11.2097 | 5.38E-05 | 1 | TAGGCTCGGCTTTCT |
| 1792 | mamo-L | *BGIBMGA003696* | up | 1357 | 1371 | + | 12.1935 | 2.05E-05 | 1 | TAGGCTTAGTGTCTT |
| 1793 | mamo-L | *BGIBMGA003699* | up | 806 | 820 | + | 12.2661 | 1.89E-05 | 1 | TAGGCCTAGGTCCTT |
| 1794 | mamo-L | *BGIBMGA003699* | down | 462 | 476 | - | 10.7823 | 7.84E-05 | 1 | GAGGCTTTTGTTTTC |
| 1795 | mamo-L | *BGIBMGA003700* | down | 439 | 453 | + | 12.3145 | 1.79E-05 | 1 | TAGGCTATTGTTTTT |
| 1796 | mamo-L | *BGIBMGA003700* | up | 1785 | 1799 | + | 10.5403 | 9.58E-05 | 1 | TAGGCTGTATGGTGT |
| 1797 | mamo-L | *BGIBMGA003703* | down | 162 | 176 | + | 11.5323 | 3.96E-05 | 1 | TAGGCTTAAATTTGC |
| 1798 | mamo-L | *BGIBMGA003703* | down | 1174 | 1188 | + | 10.5 | 9.90E-05 | 1 | TAGGCAGGGTGTTGC |
| 1799 | mamo-L | *BGIBMGA003706* | down | 1357 | 1371 | + | 10.9677 | 6.68E-05 | 1 | TAGGCTGGTTCTGAC |
| 1800 | mamo-L | *BGIBMGA003708* | up | 1621 | 1635 | - | 10.8468 | 7.42E-05 | 1 | TAGGCATCGTTTGCC |
| 1801 | mamo-L | *BGIBMGA003708* | up | 1020 | 1034 | + | 10.629 | 8.91E-05 | 1 | TAGGCCGTGAGCTCT |
| 1802 | mamo-L | *BGIBMGA003714* | down | 1750 | 1764 | + | 10.5726 | 9.33E-05 | 1 | TAGGCTAATATTTAT |
| 1803 | mamo-L | *BGIBMGA003715* | up | 1905 | 1919 | + | 12.3145 | 1.79E-05 | 1 | GAGGCGGGGGTCGGT |
| 1804 | mamo-L | *BGIBMGA003715* | down | 101 | 115 | - | 11.4677 | 4.22E-05 | 1 | GAGGCCGGCGGTGAT |
| 1805 | mamo-L | *BGIBMGA003715* | up | 1029 | 1043 | + | 10.9194 | 6.98E-05 | 1 | GAGGCTGTAGGTGCT |
| 1806 | mamo-L | *BGIBMGA003716* | up | 68 | 82 | - | 11.4032 | 4.49E-05 | 1 | TAGGCCGAGCTTTAT |
| 1807 | mamo-L | *BGIBMGA003720* | up | 1601 | 1615 | + | 13.2177 | 6.14E-06 | 1 | TAGGCTTCCGTTGCC |
| 1808 | mamo-L | *BGIBMGA003720* | up | 1158 | 1172 | + | 11.5242 | 3.99E-05 | 1 | TAGGCTTTTGTAGTT |
| 1809 | mamo-L | *BGIBMGA003721* | up | 1860 | 1874 | + | 12.2661 | 1.89E-05 | 1 | AAGGCTGTTATTGGT |
| 1810 | mamo-L | *BGIBMGA003722* | up | 1744 | 1758 | - | 14.6371 | 7.08E-07 | 1 | TAGGCGGGGGTTTGT |
| 1811 | mamo-L | *BGIBMGA003723* | up | 186 | 200 | - | 14.6371 | 7.08E-07 | 1 | TAGGCGGGGGTTTGT |
| 1812 | mamo-L | *BGIBMGA003729* | up | 1637 | 1651 | - | 12.7258 | 1.11E-05 | 1 | TAGGCGTGCTTTGTA |
| 1813 | mamo-L | *BGIBMGA003730* | down | 1215 | 1229 | + | 12.1452 | 2.15E-05 | 1 | TAGGCAGGTGGTGGT |
| 1814 | mamo-L | *BGIBMGA003735* | down | 884 | 898 | - | 11.4032 | 4.49E-05 | 1 | TAGGCGGTTTGTAGT |
| 1815 | mamo-L | *BGIBMGA003736* | up | 1244 | 1258 | + | 12.0565 | 2.36E-05 | 1 | TAGGCAGATTTTTCT |
| 1816 | mamo-L | *BGIBMGA003736* | down | 1387 | 1401 | + | 11.6855 | 3.42E-05 | 1 | TAGGCTTGTTTCCTT |
| 1817 | mamo-L | *BGIBMGA003737* | up | 459 | 473 | + | 11.6855 | 3.42E-05 | 1 | TAGGCTTGTTTCCTT |
| 1818 | mamo-L | *BGIBMGA003739* | down | 1360 | 1374 | - | 11.2661 | 5.10E-05 | 1 | TAGGCAGGTTTTCGT |
| 1819 | mamo-L | *BGIBMGA003739* | down | 1705 | 1719 | + | 10.9194 | 6.98E-05 | 1 | TAGGCTACTTTTTTT |
| 1820 | mamo-L | *BGIBMGA003743* | down | 1275 | 1289 | + | 10.7742 | 7.89E-05 | 1 | TAGGCCCTCGCTTTT |
| 1821 | mamo-L | *BGIBMGA003743* | up | 665 | 679 | + | 10.5726 | 9.33E-05 | 1 | TAGGCTAATATTTAT |
| 1822 | mamo-L | *BGIBMGA003747* | down | 1966 | 1980 | + | 10.5 | 9.90E-05 | 1 | GAGGCCCAGGTCGTC |
| 1823 | mamo-L | *BGIBMGA003749* | down | 231 | 245 | + | 10.9516 | 6.78E-05 | 1 | TAGGCATTCTATTGT |
| 1824 | mamo-L | *BGIBMGA003754* | down | 1285 | 1299 | + | 10.5 | 9.90E-05 | 1 | GAGGCCCAGGTCGTC |
| 1825 | mamo-L | *BGIBMGA003755* | up | 149 | 163 | - | 14.2661 | 1.33E-06 | 1 | TAGGCTTGTGTTGTC |
| 1826 | mamo-L | *BGIBMGA003755* | down | 122 | 136 | + | 11.129 | 5.79E-05 | 1 | TAGGCTGCAGATTGT |
| 1827 | mamo-L | *BGIBMGA003756* | down | 585 | 599 | + | 10.5484 | 9.52E-05 | 1 | TGGGCGGGCTGTGGT |
| 1828 | mamo-L | *BGIBMGA003757* | down | 602 | 616 | + | 11.4839 | 4.15E-05 | 1 | GAGGCTTACGGTTGT |
| 1829 | mamo-L | *BGIBMGA003757* | down | 1675 | 1689 | + | 10.5968 | 9.15E-05 | 1 | AAGGCTGTTAATGCT |
| 1830 | mamo-L | *BGIBMGA003767* | up | 254 | 268 | - | 12.7258 | 1.11E-05 | 1 | TAGGCGTGCTTTGTA |
| 1831 | mamo-L | *BGIBMGA003768* | up | 1358 | 1372 | - | 11.4516 | 4.28E-05 | 1 | TAGGCTAATGTTGGG |
| 1832 | mamo-L | *BGIBMGA003775* | up | 648 | 662 | + | 10.6129 | 9.03E-05 | 1 | TAGGCGTTCGTTATA |
| 1833 | mamo-L | *BGIBMGA003777* | down | 208 | 222 | + | 12.7016 | 1.14E-05 | 1 | TAGGCTTGCTTTTAC |

| | | | | | | | | | | |
|---|---|---|---|---|---|---|---|---|---|---|
| 1834 | mamo-L | *BGIBMGA003779* | up | 916 | 930 | - | 11.7258 | 3.29E-05 | 1 | AAGGCTATGTTTGTT |
| 1835 | mamo-L | *BGIBMGA003784* | up | 1098 | 1112 | - | 11.1371 | 5.75E-05 | 1 | AAGGCCGGTGTTTCC |
| 1836 | mamo-L | *BGIBMGA003786* | up | 1920 | 1934 | - | 11.7661 | 3.16E-05 | 1 | TAGGCGTATATGGTT |
| 1837 | mamo-L | *BGIBMGA003786* | down | 1307 | 1321 | + | 11.7097 | 3.34E-05 | 1 | TAGGCGGTGATTGCA |
| 1838 | mamo-L | *BGIBMGA003791* | up | 49 | 63 | + | 12.8065 | 1.01E-05 | 1 | TAGGCGGGTTTTTAT |
| 1839 | mamo-L | *BGIBMGA003793* | down | 1502 | 1516 | - | 10.5726 | 9.33E-05 | 1 | TAGGCTAATATTTAT |
| 1840 | mamo-L | *BGIBMGA003795* | up | 1584 | 1598 | + | 11.2097 | 5.38E-05 | 1 | TAGGCCTGTTTGTTC |
| 1841 | mamo-L | *BGIBMGA003798* | up | 354 | 368 | - | 10.7661 | 7.95E-05 | 1 | AAGGCGGAAATTGCT |
| 1842 | mamo-L | *BGIBMGA003802* | up | 460 | 474 | + | 11.2177 | 5.34E-05 | 1 | GAGGCGCTTTTTGCT |
| 1843 | mamo-L | *BGIBMGA003804* | up | 681 | 695 | - | 11.1935 | 5.45E-05 | 1 | GAGGCCGAGGTGGCC |
| 1844 | mamo-L | *BGIBMGA003807* | down | 996 | 1010 | - | 10.5726 | 9.33E-05 | 1 | TAGGCTAATATTTAT |
| 1845 | mamo-L | *BGIBMGA003811* | up | 494 | 508 | + | 11.3548 | 4.69E-05 | 1 | AAGGCTGATGATGAT |
| 1846 | mamo-L | *BGIBMGA003813* | down | 623 | 637 | + | 11.5645 | 3.84E-05 | 1 | TAGGCGCATGACGGT |
| 1847 | mamo-L | *BGIBMGA003813* | up | 1945 | 1959 | - | 10.9274 | 6.92E-05 | 1 | TAGGCTAGTTACGTT |
| 1848 | mamo-L | *BGIBMGA003815* | down | 677 | 691 | + | 12.9839 | 8.17E-06 | 1 | TAGGCTTTTTTTGGA |
| 1849 | mamo-L | *BGIBMGA003818* | up | 706 | 720 | - | 13.3226 | 5.34E-06 | 1 | TAGGCTTAGAATGTT |
| 1850 | mamo-L | *BGIBMGA003818* | down | 1322 | 1336 | + | 10.9194 | 6.98E-05 | 1 | TAGGCTACTTTTTTT |
| 1851 | mamo-L | *BGIBMGA003818* | up | 634 | 648 | - | 10.8065 | 7.69E-05 | 1 | TAGGCCGTATTTTGG |
| 1852 | mamo-L | *BGIBMGA003821* | down | 1051 | 1065 | - | 11 | 6.50E-05 | 1 | TAGGCATGCGTGTGT |
| 1853 | mamo-L | *BGIBMGA003822* | up | 1085 | 1099 | - | 10.9516 | 6.78E-05 | 1 | AAGGCTTCGTTTTTT |
| 1854 | mamo-L | *BGIBMGA003823* | up | 188 | 202 | - | 10.5968 | 9.15E-05 | 1 | GAGGCTAGTTTTTGT |
| 1855 | mamo-L | *BGIBMGA003824* | down | 1349 | 1363 | + | 10.629 | 8.91E-05 | 1 | TAGGCCTGCAGCGCC |
| 1856 | mamo-L | *BGIBMGA003828* | down | 1387 | 1401 | + | 10.9194 | 6.98E-05 | 1 | TAGGCTACTTTTTTT |
| 1857 | mamo-L | *BGIBMGA003834* | down | 1445 | 1459 | + | 11.7016 | 3.37E-05 | 1 | TAGGCGGGTTTTGAA |
| 1858 | mamo-L | *BGIBMGA003839* | down | 773 | 787 | - | 10.5645 | 9.40E-05 | 1 | TAGGCTGTATGCTCT |
| 1859 | mamo-L | *BGIBMGA003840* | down | 634 | 648 | + | 10.5726 | 9.33E-05 | 1 | TAGGCTAATATTTAT |
| 1860 | mamo-L | *BGIBMGA003841* | down | 1113 | 1127 | + | 11 | 6.50E-05 | 1 | AAGGCTTGTTTTATT |
| 1861 | mamo-L | *BGIBMGA003842* | up | 220 | 234 | - | 12.0161 | 2.46E-05 | 1 | TAGGCTTTCTTCTTC |
| 1862 | mamo-L | *BGIBMGA003843* | up | 68 | 82 | - | 11.4113 | 4.45E-05 | 1 | TAGGCGGGGTTGGCA |
| 1863 | mamo-L | *BGIBMGA003843* | up | 1815 | 1829 | - | 10.5726 | 9.33E-05 | 1 | TAGGCTAATATTTAT |
| 1864 | mamo-L | *BGIBMGA003845* | down | 1473 | 1487 | + | 13.0726 | 7.34E-06 | 1 | TAGGCTGGCTTTAAT |
| 1865 | mamo-L | *BGIBMGA003850* | down | 1079 | 1093 | + | 10.6532 | 8.73E-05 | 1 | TAGGCGAAGTTTAAT |
| 1866 | mamo-L | *BGIBMGA003855* | down | 847 | 861 | + | 13.3952 | 4.84E-06 | 1 | TAGGCGTAGGTTGAC |
| 1867 | mamo-L | *BGIBMGA003861* | up | 129 | 143 | + | 10.9839 | 6.59E-05 | 1 | AAGGCGTTTTTTTTT |
| 1868 | mamo-L | *BGIBMGA003864* | down | 270 | 284 | + | 12.9919 | 8.10E-06 | 1 | TAGGCCGGTTTTGCA |
| 1869 | mamo-L | *BGIBMGA003864* | down | 353 | 367 | - | 10.8065 | 7.69E-05 | 1 | GAGGCCGTCGTGGCC |
| 1870 | mamo-L | *BGIBMGA003867* | up | 1124 | 1138 | - | 11.9597 | 2.61E-05 | 1 | AAGGCTGTTTTTTTT |
| 1871 | mamo-L | *BGIBMGA003869* | down | 129 | 143 | - | 11.621 | 3.63E-05 | 1 | TAGGCTGTGAAGGCT |
| 1872 | mamo-L | *BGIBMGA003874* | down | 1156 | 1170 | + | 10.8871 | 7.17E-05 | 1 | TAGGCTACTTTTTCT |
| 1873 | mamo-L | *BGIBMGA003877* | down | 1402 | 1416 | + | 10.5968 | 9.15E-05 | 1 | TAGGCGAGTGGTGTC |
| 1874 | mamo-L | *BGIBMGA003877* | down | 1523 | 1537 | - | 10.5 | 9.90E-05 | 1 | GAGGCCCAGGTCGTC |
| 1875 | mamo-L | *BGIBMGA003878* | up | 645 | 659 | + | 11.0323 | 6.32E-05 | 1 | TAGGCGTGTAATGTC |
| 1876 | mamo-L | *BGIBMGA003878* | down | 1129 | 1143 | - | 10.75 | 8.06E-05 | 1 | GAGGCCGATTGTGCC |
| 1877 | mamo-L | *BGIBMGA003878* | down | 1693 | 1707 | + | 10.6532 | 8.73E-05 | 1 | AAGGCTTTTTTCGAT |
| 1878 | mamo-L | *BGIBMGA003881* | up | 290 | 304 | + | 11.5645 | 3.84E-05 | 1 | TAGGCCCAGGGTTAT |
| 1879 | mamo-L | *BGIBMGA003883* | down | 170 | 184 | + | 11.371 | 4.62E-05 | 1 | TAGGCTGTATAGGCT |
| 1880 | mamo-L | *BGIBMGA003883* | down | 179 | 193 | + | 11.0242 | 6.36E-05 | 1 | TAGGCTTACGCTCTT |
| 1881 | mamo-L | *BGIBMGA003884* | down | 1554 | 1568 | - | 10.4919 | 9.96E-05 | 1 | TAGGCTGTGTCTGTG |
| 1882 | mamo-L | *BGIBMGA003885* | up | 493 | 507 | + | 10.8548 | 7.38E-05 | 1 | GAGGCCTGTTTTTAT |
| 1883 | mamo-L | *BGIBMGA003886* | down | 846 | 860 | + | 10.9113 | 7.02E-05 | 1 | TAGGCTTTGTGCCGT |
| 1884 | mamo-L | *BGIBMGA003886* | down | 1582 | 1596 | + | 10.7581 | 8.00E-05 | 1 | TAGGCAGATAATGTT |
| 1885 | mamo-L | *BGIBMGA003886* | up | 1674 | 1688 | - | 10.5242 | 9.70E-05 | 1 | AAGGCCTGTGATTTT |
| 1886 | mamo-L | *BGIBMGA003888* | up | 814 | 828 | + | 10.7258 | 8.23E-05 | 1 | TAGGCCGTTTTATGT |
| 1887 | mamo-L | *BGIBMGA003904* | down | 182 | 196 | + | 11.8871 | 2.81E-05 | 1 | TAGGCCTTATTTATT |
| 1888 | mamo-L | *BGIBMGA003904* | down | 540 | 554 | - | 10.9839 | 6.59E-05 | 1 | TAGGCGAAGTGTTTT |
| 1889 | mamo-L | *BGIBMGA003905* | up | 602 | 616 | - | 11.379 | 4.58E-05 | 1 | TAGGCGGATTGTACT |
| 1890 | mamo-L | *BGIBMGA003906* | up | 1434 | 1448 | - | 10.5645 | 9.40E-05 | 1 | TGGGCCTTGTGTGGT |
| 1891 | mamo-L | *BGIBMGA003908* | up | 1870 | 1884 | - | 11.3065 | 4.91E-05 | 1 | TAGGCTGACGAGCGT |
| 1892 | mamo-L | *BGIBMGA003913* | down | 761 | 775 | - | 11.1452 | 5.70E-05 | 1 | TAGGCTGTGATAGTT |
| 1893 | mamo-L | *BGIBMGA003913* | down | 718 | 732 | - | 10.8468 | 7.42E-05 | 1 | TAGGCTGGTAGCTGT |
| 1894 | mamo-L | *BGIBMGA003914* | up | 657 | 671 | + | 11.25 | 5.18E-05 | 1 | GAGGCGGTTTTTTTT |
| 1895 | mamo-L | *BGIBMGA003914* | down | 1165 | 1179 | + | 11.1613 | 5.63E-05 | 1 | TAGGCTAAGTAGGCT |
| 1896 | mamo-L | *BGIBMGA003916* | down | 1838 | 1852 | - | 11.25 | 5.18E-05 | 1 | TAGGCCCATTTCAGT |
| 1897 | mamo-L | *BGIBMGA003917* | up | 1981 | 1995 | + | 12.5323 | 1.39E-05 | 1 | AAGGCTGATGTTGTC |
| 1898 | mamo-L | *BGIBMGA003919* | down | 1607 | 1621 | + | 11.2258 | 5.30E-05 | 1 | TAGGCCTTTTTCCCT |
| 1899 | mamo-L | *BGIBMGA003920* | up | 1680 | 1694 | - | 10.5726 | 9.33E-05 | 1 | TAGGCTAATATTTAT |
| 1900 | mamo-L | *BGIBMGA003922* | down | 1401 | 1415 | + | 11.25 | 5.18E-05 | 1 | TAGGCCCATTTCAGT |
| 1901 | mamo-L | *BGIBMGA003923* | down | 1358 | 1372 | + | 11.1452 | 5.70E-05 | 1 | TAGGCTGTGATAGTT |
| 1902 | mamo-L | *BGIBMGA003923* | down | 1401 | 1415 | + | 10.8468 | 7.42E-05 | 1 | TAGGCTGGTAGCTGT |
| 1903 | mamo-L | *BGIBMGA003924* | up | 962 | 976 | - | 10.5081 | 9.83E-05 | 1 | AAGGCTCTCGTTTTC |
| 1904 | mamo-L | *BGIBMGA003926* | down | 1431 | 1445 | - | 11.8952 | 2.79E-05 | 1 | AAGGCCGTTTTTGTC |
| 1905 | mamo-L | *BGIBMGA003927* | up | 105 | 119 | + | 10.5726 | 9.33E-05 | 1 | TAGGCTAATATTTAT |
| 1906 | mamo-L | *BGIBMGA003929* | up | 1823 | 1837 | + | 12.0484 | 2.38E-05 | 1 | AAGGCTAAGTTTGTT |
| 1907 | mamo-L | *BGIBMGA003930* | down | 1729 | 1743 | + | 12.0484 | 2.38E-05 | 1 | AAGGCTAAGTTTGTT |
| 1908 | mamo-L | *BGIBMGA003932* | down | 1048 | 1062 | - | 12.7258 | 1.11E-05 | 1 | AAGGCTGATGTTTGT |
| 1909 | mamo-L | *BGIBMGA003932* | down | 1482 | 1496 | - | 10.7903 | 7.79E-05 | 1 | TAGGCGATGTTCTTT |
| 1910 | mamo-L | *BGIBMGA003937* | up | 1553 | 1567 | - | 11.9355 | 2.67E-05 | 1 | TAGGCGCGCAGTGGT |
| 1911 | mamo-L | *BGIBMGA003938* | up | 1934 | 1948 | + | 10.879 | 7.22E-05 | 1 | TAGGCGCGCGACTGT |
| 1912 | mamo-L | *BGIBMGA003940* | up | 623 | 637 | + | 12.9839 | 8.17E-06 | 1 | TAGGCTAGGTTTTTT |
| 1913 | mamo-L | *BGIBMGA003941* | down | 1608 | 1622 | - | 11.129 | 5.79E-05 | 1 | TAGGCTGGTGGTACC |
| 1914 | mamo-L | *BGIBMGA003947* | down | 740 | 754 | - | 11.371 | 4.62E-05 | 1 | TAGGCTGTATAGGCT |
| 1915 | mamo-L | *BGIBMGA003951* | down | 1933 | 1947 | + | 11.121 | 5.84E-05 | 1 | TAGGCTTAATTCATT |
| 1916 | mamo-L | *BGIBMGA003952* | down | 1484 | 1498 | - | 12.1855 | 2.06E-05 | 1 | TAGGCTATTTTTTTT |
| 1917 | mamo-L | *BGIBMGA003954* | down | 1307 | 1321 | + | 11.371 | 4.62E-05 | 1 | TAGGCTGTATAGGCT |
| 1918 | mamo-L | *BGIBMGA003956* | up | 819 | 833 | + | 11.5726 | 3.81E-05 | 1 | TAGGCTTTTTTTTTA |
| 1919 | mamo-L | *BGIBMGA003956* | up | 1659 | 1673 | + | 10.9194 | 6.98E-05 | 1 | TAGGCTACTTTTTTT |
| 1920 | mamo-L | *BGIBMGA003956* | up | 1388 | 1402 | + | 10.6935 | 8.45E-05 | 1 | AAGGCTGCTTTTTTT |
| 1921 | mamo-L | *BGIBMGA003957* | down | 124 | 138 | - | 12.0968 | 2.26E-05 | 1 | TAGGCTTGGTTGTAT |
| 1922 | mamo-L | *BGIBMGA003959* | down | 1850 | 1864 | + | 10.5968 | 9.15E-05 | 1 | GAGGCTAGTTTTTGT |
| 1923 | mamo-L | *BGIBMGA003960* | down | 1935 | 1949 | - | 11.25 | 5.18E-05 | 1 | AAGGCTGCGTTTTTT |
| 1924 | mamo-L | *BGIBMGA003960* | up | 1949 | 1963 | + | 10.9032 | 7.07E-05 | 1 | AAGGCTGTTTTTCTT |
| 1925 | mamo-L | *BGIBMGA003961* | down | 1171 | 1185 | - | 12.5 | 1.44E-05 | 1 | TAGGCTTAGGACTGT |

| | | | | | | | | | | |
|---|---|---|---|---|---|---|---|---|---|---|
| 1926 | mamo-L | *BGIBMGA003962* | down | 1861 | 1875 | - | 12.7258 | 1.11E-05 | 1 | TAGGCTGTGTAGGCT |
| 1927 | mamo-L | *BGIBMGA003962* | down | 1236 | 1250 | - | 11.2177 | 5.34E-05 | 1 | TAGGCCTCATGTGTT |
| 1928 | mamo-L | *BGIBMGA003964* | up | 1333 | 1347 | - | 13.4677 | 4.39E-06 | 1 | TAGGCTGATGATGAT |
| 1929 | mamo-L | *BGIBMGA003965* | down | 854 | 868 | - | 11.2903 | 4.99E-05 | 1 | TAGGCATAGGTTGGG |
| 1930 | mamo-L | *BGIBMGA003965* | up | 707 | 721 | - | 10.5323 | 9.63E-05 | 1 | AAGGCCCAGGTCGTC |
| 1931 | mamo-L | *BGIBMGA003966* | up | 1101 | 1115 | + | 11.6774 | 3.44E-05 | 1 | TAGGCATGTTTTTCT |
| 1932 | mamo-L | *BGIBMGA003969* | down | 1479 | 1493 | + | 13.7742 | 2.79E-06 | 1 | TAGGCGGGTGGTGGT |
| 1933 | mamo-L | *BGIBMGA003970* | down | 571 | 585 | - | 10.9194 | 6.98E-05 | 1 | TAGGCTACTTTTTTT |
| 1934 | mamo-L | *BGIBMGA003971* | down | 78 | 92 | - | 13.129 | 6.83E-06 | 1 | TAGGCTGTCCTTGTT |
| 1935 | mamo-L | *BGIBMGA003972* | down | 1762 | 1776 | - | 10.9597 | 6.73E-05 | 1 | AAGGCTTTCGATTTT |
| 1936 | mamo-L | *BGIBMGA003972* | down | 866 | 880 | + | 10.8548 | 7.38E-05 | 1 | TAGGCTTACTACGCA |
| 1937 | mamo-L | *BGIBMGA003974* | up | 424 | 438 | - | 10.6048 | 9.09E-05 | 1 | AAGGCTGAAGTTTCC |
| 1938 | mamo-L | *BGIBMGA003975* | up | 86 | 100 | + | 11 | 6.50E-05 | 1 | TAGGCTTATAGTAGT |
| 1939 | mamo-L | *BGIBMGA003976* | up | 244 | 258 | + | 10.7823 | 7.84E-05 | 1 | AAGGCTGACAATGGC |
| 1940 | mamo-L | *BGIBMGA003980* | up | 217 | 231 | - | 10.8226 | 7.58E-05 | 1 | TAGGCAGTCTGTTCT |
| 1941 | mamo-L | *BGIBMGA003980* | down | 62 | 76 | + | 10.5726 | 9.33E-05 | 1 | TAGGCTAATATTTAT |
| 1942 | mamo-L | *BGIBMGA003981* | up | 1875 | 1889 | + | 12.7581 | 1.07E-05 | 1 | AAGGCTTTTTTTGTT |
| 1943 | mamo-L | *BGIBMGA003981* | down | 160 | 174 | + | 10.5242 | 9.70E-05 | 1 | TAGGCGTTGTATTAC |
| 1944 | mamo-L | *BGIBMGA003982* | down | 1824 | 1838 | + | 11.1855 | 5.50E-05 | 1 | TGGGCTTAGTTTGTC |
| 1945 | mamo-L | *BGIBMGA003983* | up | 696 | 710 | + | 10.5968 | 9.15E-05 | 1 | GAGGCCGTTGTCTTT |
| 1946 | mamo-L | *BGIBMGA003990* | down | 1896 | 1910 | + | 11.5726 | 3.81E-05 | 1 | TAGGCTTTAGTTGCG |
| 1947 | mamo-L | *BGIBMGA003996* | up | 824 | 838 | + | 11.1613 | 5.63E-05 | 1 | TAGGCGTTCAATTTT |
| 1948 | mamo-L | *BGIBMGA004001* | down | 1588 | 1602 | + | 12.1371 | 2.17E-05 | 1 | TAGGCTGTTGTAGGT |
| 1949 | mamo-L | *BGIBMGA004001* | down | 27 | 41 | + | 11.1371 | 5.75E-05 | 1 | TAGGCTGTCTTTTAG |
| 1950 | mamo-L | *BGIBMGA004003* | up | 1059 | 1073 | + | 10.7823 | 7.84E-05 | 1 | TAGGCCCGTGATCTT |
| 1951 | mamo-L | *BGIBMGA004004* | up | 1603 | 1617 | + | 13.7742 | 2.79E-06 | 1 | TAGGCTTTTTTTTTT |
| 1952 | mamo-L | *BGIBMGA004005* | down | 1649 | 1663 | - | 10.5726 | 9.33E-05 | 1 | TAGGCTAATATTTAT |
| 1953 | mamo-L | *BGIBMGA004007* | up | 828 | 842 | - | 10.9194 | 6.98E-05 | 1 | AAGGCTTGTATTGTC |
| 1954 | mamo-L | *BGIBMGA004012* | up | 825 | 839 | + | 10.7177 | 8.28E-05 | 1 | TAGGCTTAATTGACT |
| 1955 | mamo-L | *BGIBMGA004013* | up | 95 | 109 | - | 10.9194 | 6.98E-05 | 1 | TAGGCTACTTTTTTT |
| 1956 | mamo-L | *BGIBMGA004016* | down | 1372 | 1386 | - | 12.1452 | 2.15E-05 | 1 | TAGGCAGGTGGTGGT |
| 1957 | mamo-L | *BGIBMGA004020* | up | 963 | 977 | + | 11.371 | 4.62E-05 | 1 | TAGGCCTATAGTTCT |
| 1958 | mamo-L | *BGIBMGA004021* | down | 1797 | 1811 | - | 10.6371 | 8.85E-05 | 1 | AAGGCGTAAGTTTTT |
| 1959 | mamo-L | *BGIBMGA004022* | down | 877 | 891 | - | 10.5726 | 9.33E-05 | 1 | GAGGCCGACGGTGAC |
| 1960 | mamo-L | *BGIBMGA004024* | up | 1727 | 1741 | - | 11.7339 | 3.26E-05 | 1 | TAGGCCGTCGTCGTG |
| 1961 | mamo-L | *BGIBMGA004028* | up | 383 | 397 | - | 11.0403 | 6.28E-05 | 1 | AAGGCCCGTTTTGTC |
| 1962 | mamo-L | *BGIBMGA004029* | up | 708 | 722 | + | 11.5403 | 3.93E-05 | 1 | GAGGCGGATTTTTCT |
| 1963 | mamo-L | *BGIBMGA004029* | down | 1143 | 1157 | - | 10.5 | 9.90E-05 | 1 | AAGGCCATATTTGGT |
| 1964 | mamo-L | *BGIBMGA004030* | down | 1819 | 1833 | - | 12.4677 | 1.49E-05 | 1 | TAGGCTTCTATTGCT |
| 1965 | mamo-L | *BGIBMGA004030* | down | 1552 | 1566 | - | 12.0645 | 2.34E-05 | 1 | TAGGCGGGCTTCGGA |
| 1966 | mamo-L | *BGIBMGA004032* | up | 1432 | 1446 | - | 11.3548 | 4.69E-05 | 1 | AAGGCTGATGATGAT |
| 1967 | mamo-L | *BGIBMGA004035* | down | 1073 | 1087 | + | 11.7903 | 3.08E-05 | 1 | TAGGCTGAATTTTAC |
| 1968 | mamo-L | *BGIBMGA004038* | up | 1032 | 1046 | + | 11.5323 | 3.96E-05 | 1 | TAGGCGTGAGGTGAT |
| 1969 | mamo-L | *BGIBMGA004039* | up | 862 | 876 | + | 11.0484 | 6.23E-05 | 1 | AAGGCGGGCTGTGGC |
| 1970 | mamo-L | *BGIBMGA004040* | down | 1043 | 1057 | - | 12.1935 | 2.05E-05 | 1 | GAGGCTTTGTGTGGT |
| 1971 | mamo-L | *BGIBMGA004044* | down | 738 | 752 | - | 10.7097 | 8.33E-05 | 1 | TAGGCAAAGGTCGTT |
| 1972 | mamo-L | *BGIBMGA004046* | up | 1533 | 1547 | + | 10.5242 | 9.70E-05 | 1 | TAGGCTGCTTACTGT |
| 1973 | mamo-L | *BGIBMGA004048* | up | 1272 | 1286 | + | 11.3065 | 4.91E-05 | 1 | AAGGCGCACATTGGT |
| 1974 | mamo-L | *BGIBMGA004048* | down | 883 | 897 | - | 10.8548 | 7.38E-05 | 1 | GAGGCCTGTTTTTAT |
| 1975 | mamo-L | *BGIBMGA004050* | up | 1524 | 1538 | + | 11.2984 | 4.95E-05 | 1 | GAGGCCTAGGTCGTC |
| 1976 | mamo-L | *BGIBMGA004051* | down | 1550 | 1564 | - | 10.8226 | 7.58E-05 | 1 | TAGGCGTGATTTCGC |
| 1977 | mamo-L | *BGIBMGA004065* | down | 140 | 154 | + | 11.8065 | 3.03E-05 | 1 | AAGGCTTGTGATGTT |
| 1978 | mamo-L | *BGIBMGA004070* | up | 910 | 924 | - | 11.2823 | 5.03E-05 | 1 | AAGGCTTATGTTTAT |
| 1979 | mamo-L | *BGIBMGA004072* | down | 1265 | 1279 | - | 10.6613 | 8.67E-05 | 1 | AAGGCCGGTTATTCT |
| 1980 | mamo-L | *BGIBMGA004073* | up | 1166 | 1180 | - | 10.6613 | 8.67E-05 | 1 | AAGGCCGGTTATTCT |
| 1981 | mamo-L | *BGIBMGA004076* | down | 1803 | 1817 | - | 10.8952 | 7.12E-05 | 1 | TAGGCTGTAATCTTT |
| 1982 | mamo-L | *BGIBMGA004087* | up | 1399 | 1413 | + | 12.6532 | 1.20E-05 | 1 | TAGGCAGGGTATGGT |
| 1983 | mamo-L | *BGIBMGA004087* | down | 1535 | 1549 | - | 12.5726 | 1.33E-05 | 1 | TAGGCCCGGATTGCC |
| 1984 | mamo-L | *BGIBMGA004087* | down | 195 | 209 | - | 12.4597 | 1.51E-05 | 1 | TAGGCTGTCGGTTAT |
| 1985 | mamo-L | *BGIBMGA004088* | up | 1395 | 1409 | - | 11.5484 | 3.90E-05 | 1 | TAGGCCAATGTTATT |
| 1986 | mamo-L | *BGIBMGA004090* | down | 1191 | 1205 | + | 11.3306 | 4.80E-05 | 1 | TAGGCCTAGTTTACA |
| 1987 | mamo-L | *BGIBMGA004093* | up | 1692 | 1706 | - | 12.2742 | 1.87E-05 | 1 | TAGGCGAATGTTTGT |
| 1988 | mamo-L | *BGIBMGA004096* | up | 394 | 408 | + | 11.0887 | 6.01E-05 | 1 | TAGGCTTTATTTCAT |
| 1989 | mamo-L | *BGIBMGA004098* | up | 268 | 282 | - | 11.371 | 4.62E-05 | 1 | TAGGCCTATAGTTCT |
| 1990 | mamo-L | *BGIBMGA004100* | down | 1264 | 1278 | + | 12.0645 | 2.34E-05 | 1 | TAGGCGGGCTTCGGA |
| 1991 | mamo-L | *BGIBMGA004104* | down | 1386 | 1400 | + | 11.0968 | 5.97E-05 | 1 | TAGGCTGTAGGTATT |
| 1992 | mamo-L | *BGIBMGA004108* | up | 1515 | 1529 | + | 11.6532 | 3.53E-05 | 1 | TAGGCCTGCATTAGC |
| 1993 | mamo-L | *BGIBMGA004112* | up | 1482 | 1496 | - | 12.0081 | 2.48E-05 | 1 | TAGGCAGGTTTTTTT |
| 1994 | mamo-L | *BGIBMGA004122* | up | 1149 | 1163 | - | 11 | 6.50E-05 | 1 | TAGGCATGCGTGTGT |
| 1995 | mamo-L | *BGIBMGA004127* | up | 453 | 467 | - | 11.1613 | 5.63E-05 | 1 | TAGGCGTTCAATTTT |
| 1996 | mamo-L | *BGIBMGA004128* | down | 832 | 846 | - | 10.5081 | 9.83E-05 | 1 | TAGGCACAATTTTGT |
| 1997 | mamo-L | *BGIBMGA004129* | up | 851 | 865 | - | 10.6855 | 8.51E-05 | 1 | TCGGCGTGGTTTGCT |
| 1998 | mamo-L | *BGIBMGA004132* | up | 331 | 345 | + | 10.5726 | 9.33E-05 | 1 | TAGGCTAATATTTAT |
| 1999 | mamo-L | *BGIBMGA004134* | down | 1826 | 1840 | - | 10.5 | 9.90E-05 | 1 | GAGGCCCAGGTCGTC |
| 2000 | mamo-L | *BGIBMGA004138* | up | 1460 | 1474 | + | 10.6048 | 9.09E-05 | 1 | AAGGCTGAAGTTTCC |
| 2001 | mamo-L | *BGIBMGA004142* | up | 373 | 387 | - | 11.8387 | 2.94E-05 | 1 | GAGGCTGACTTTATT |
| 2002 | mamo-L | *BGIBMGA004144* | down | 1887 | 1901 | - | 10.9194 | 6.98E-05 | 1 | TAGGCTACTTTTTTT |
| 2003 | mamo-L | *BGIBMGA004145* | up | 1510 | 1524 | - | 10.629 | 8.91E-05 | 1 | TAGGCCTGCAGCGCC |
| 2004 | mamo-L | *BGIBMGA004149* | up | 492 | 506 | + | 11.1774 | 5.54E-05 | 1 | TAGGCACTGGTCGTT |
| 2005 | mamo-L | *BGIBMGA004151* | down | 1801 | 1815 | + | 10.8226 | 7.58E-05 | 1 | TAGGCTAGGATTCTT |
| 2006 | mamo-L | *BGIBMGA004152* | down | 1604 | 1618 | + | 11.3548 | 4.69E-05 | 1 | AAGGCTGATGATGAT |
| 2007 | mamo-L | *BGIBMGA004152* | up | 76 | 90 | + | 10.9194 | 6.98E-05 | 1 | TAGGCTACTTTTTTT |
| 2008 | mamo-L | *BGIBMGA004152* | down | 1833 | 1847 | + | 10.9194 | 6.98E-05 | 1 | TAGGCTACTTTTTTT |
| 2009 | mamo-L | *BGIBMGA004152* | down | 1260 | 1274 | - | 10.7258 | 8.23E-05 | 1 | TAGGCCCTTTTTCCC |
| 2010 | mamo-L | *BGIBMGA004157* | up | 484 | 498 | + | 12.8871 | 9.15E-06 | 1 | TAGGCATATTTTGTT |
| 2011 | mamo-L | *BGIBMGA004161* | down | 873 | 887 | + | 13.7258 | 3.01E-06 | 1 | TAGGCTGGTTTTAGT |
| 2012 | mamo-L | *BGIBMGA004161* | down | 1564 | 1578 | - | 11.6129 | 3.66E-05 | 1 | AAGGCTGGTTTTAGT |
| 2013 | mamo-L | *BGIBMGA004165* | down | 1886 | 1900 | - | 11.0323 | 6.32E-05 | 1 | AAGGCGTAGTTTTAT |
| 2014 | mamo-L | *BGIBMGA004170* | up | 978 | 992 | - | 15.0726 | 2.89E-07 | 1 | TAGGCTGGCTTTGAT |
| 2015 | mamo-L | *BGIBMGA004172* | up | 1349 | 1363 | - | 11.7339 | 3.26E-05 | 1 | TAGGCTATGTTTTCC |
| 2016 | mamo-L | *BGIBMGA004173* | down | 640 | 654 | - | 11.2177 | 5.34E-05 | 1 | TAGGCCGGAAATTGT |
| 2017 | mamo-L | *BGIBMGA004174* | up | 1602 | 1616 | + | 12.4758 | 1.48E-05 | 1 | TAGGCTTGTTCTGTT |

| | | | | | | | | | | |
|---|---|---|---|---|---|---|---|---|---|---|
| 2018 | mamo-L | *BGIBMGA004179* | up | 175 | 189 | - | 11.371 | 4.62E-05 | 1 | TAGGCTGTATAGGCT |
| 2019 | mamo-L | *BGIBMGA004181* | up | 644 | 658 | - | 10.5323 | 9.63E-05 | 1 | TAGGCATGGTTTAAT |
| 2020 | mamo-L | *BGIBMGA004182* | up | 1878 | 1892 | - | 13.0726 | 7.34E-06 | 1 | TAGGCCGATTTTGCA |
| 2021 | mamo-L | *BGIBMGA004182* | down | 191 | 205 | + | 11.2742 | 5.06E-05 | 1 | TAGGCTTTGTATAAT |
| 2022 | mamo-L | *BGIBMGA004183* | up | 1925 | 1939 | - | 11.0323 | 6.32E-05 | 1 | TAGGCAGGTGTTGTA |
| 2023 | mamo-L | *BGIBMGA004189* | up | 271 | 285 | - | 10.6774 | 8.56E-05 | 1 | TAGGCCTTTATTTAC |
| 2024 | mamo-L | *BGIBMGA004189* | up | 433 | 447 | - | 10.5726 | 9.33E-05 | 1 | TAGGCTAATATTTAT |
| 2025 | mamo-L | *BGIBMGA004191* | up | 1086 | 1100 | - | 10.5645 | 9.40E-05 | 1 | AAGGCCTTGCTTGCT |
| 2026 | mamo-L | *BGIBMGA004192* | down | 809 | 823 | - | 10.9758 | 6.64E-05 | 1 | GAGGCTTCATTTGGT |
| 2027 | mamo-L | *BGIBMGA004192* | up | 589 | 603 | - | 10.8871 | 7.17E-05 | 1 | AAGGCTCTGGTTTGC |
| 2028 | mamo-L | *BGIBMGA004193* | up | 1707 | 1721 | + | 11.7581 | 3.18E-05 | 1 | GAGGCTGTCTTTTGC |
| 2029 | mamo-L | *BGIBMGA004194* | down | 149 | 163 | + | 10.8387 | 7.47E-05 | 1 | TAGGCTTATGAAGGT |
| 2030 | mamo-L | *BGIBMGA004199* | down | 1746 | 1760 | - | 12.4919 | 1.46E-05 | 1 | TAGGCCGTCTTTTGA |
| 2031 | mamo-L | *BGIBMGA004206* | down | 310 | 324 | - | 11.129 | 5.79E-05 | 1 | AAGGCTCACTTGGTT |
| 2032 | mamo-L | *BGIBMGA004207* | down | 352 | 366 | + | 10.7984 | 7.74E-05 | 1 | TAGGCCATGGTTTGA |
| 2033 | mamo-L | *BGIBMGA004207* | down | 297 | 311 | - | 10.7097 | 8.33E-05 | 1 | TAGGCTGTTCTTTAT |
| 2034 | mamo-L | *BGIBMGA004208* | up | 494 | 508 | + | 13.129 | 6.83E-06 | 1 | TAGGCTTTCTTTGCA |
| 2035 | mamo-L | *BGIBMGA004214* | up | 879 | 893 | + | 10.6935 | 8.45E-05 | 1 | TGGGCGTAGATTGGT |
| 2036 | mamo-L | *BGIBMGA004217* | down | 1731 | 1745 | - | 10.8629 | 7.33E-05 | 1 | AAGGCCTATTTTACT |
| 2037 | mamo-L | *BGIBMGA004220* | up | 37 | 51 | + | 10.7097 | 8.33E-05 | 1 | TAGCCTTGCTTTTGT |
| 2038 | mamo-L | *BGIBMGA004220* | up | 819 | 833 | - | 10.5968 | 9.15E-05 | 1 | GAGGCTAGTTTTTGT |
| 2039 | mamo-L | *BGIBMGA004223* | up | 1846 | 1860 | - | 10.7258 | 8.23E-05 | 1 | AAGGCGTGGTTTCTT |
| 2040 | mamo-L | *BGIBMGA004224* | down | 302 | 316 | + | 13.2419 | 5.96E-06 | 1 | TAGGCTGTTTTTTAT |
| 2041 | mamo-L | *BGIBMGA004224* | down | 149 | 163 | + | 12.1452 | 2.15E-05 | 1 | TAGGCTCTTTTTTAT |
| 2042 | mamo-L | *BGIBMGA004224* | down | 1084 | 1098 | - | 10.7339 | 8.17E-05 | 1 | TAGGCTTTTATTGAA |
| 2043 | mamo-L | *BGIBMGA004225* | down | 1555 | 1569 | - | 10.5 | 9.90E-05 | 1 | GAGGCCCAGGTCGTC |
| 2044 | mamo-L | *BGIBMGA004226* | down | 530 | 544 | - | 10.5726 | 9.33E-05 | 1 | TAGGCTAATATTTAT |
| 2045 | mamo-L | *BGIBMGA004232* | down | 1003 | 1017 | + | 12.1452 | 2.15E-05 | 1 | TAGGCCTAAATCGGT |
| 2046 | mamo-L | *BGIBMGA004233* | up | 1032 | 1046 | + | 12.1452 | 2.15E-05 | 1 | TAGGCCTAAATCGGT |
| 2047 | mamo-L | *BGIBMGA004237* | down | 1184 | 1198 | - | 11.2177 | 5.34E-05 | 1 | TAGGCGTCCATTTTT |
| 2048 | mamo-L | *BGIBMGA004239* | up | 1479 | 1493 | + | 13.3629 | 5.09E-06 | 1 | TAGGCTTTCTTTATT |
| 2049 | mamo-L | *BGIBMGA004244* | down | 250 | 264 | - | 10.5726 | 9.33E-05 | 1 | TAGGCTAATATTTAT |
| 2050 | mamo-L | *BGIBMGA004246* | up | 1015 | 1029 | + | 13.3387 | 5.24E-06 | 1 | TAGGCTGATTATGAT |
| 2051 | mamo-L | *BGIBMGA004252* | down | 1166 | 1180 | + | 11.1613 | 5.63E-05 | 1 | TAGGCCATTTTCGGC |
| 2052 | mamo-L | *BGIBMGA004254* | down | 1915 | 1929 | + | 11.3548 | 4.69E-05 | 1 | AAGGCTGATGATGAT |
| 2053 | mamo-L | *BGIBMGA004255* | down | 678 | 692 | - | 12.0645 | 2.34E-05 | 1 | TAGGCGGGCTTCGGA |
| 2054 | mamo-L | *BGIBMGA004255* | up | 1364 | 1378 | + | 11.3548 | 4.69E-05 | 1 | AAGGCTGATGATGAT |
| 2055 | mamo-L | *BGIBMGA004255* | down | 831 | 845 | + | 10.8306 | 7.52E-05 | 1 | TAGGCCCCCTTCGCC |
| 2056 | mamo-L | *BGIBMGA004256* | up | 860 | 874 | - | 12.0645 | 2.34E-05 | 1 | TAGGCGGGCTTCGGA |
| 2057 | mamo-L | *BGIBMGA004256* | up | 1013 | 1027 | + | 10.8306 | 7.52E-05 | 1 | TAGGCCCCCTTCGCC |
| 2058 | mamo-L | *BGIBMGA004258* | down | 284 | 298 | - | 12.0645 | 2.34E-05 | 1 | TAGGCGGGCTTCGGA |
| 2059 | mamo-L | *BGIBMGA004258* | up | 1010 | 1024 | + | 10.7903 | 7.79E-05 | 1 | TAGGCTATCTTTCAT |
| 2060 | mamo-L | *BGIBMGA004258* | up | 1607 | 1621 | - | 10.6935 | 8.45E-05 | 1 | TAGGCTAAGTCTGAT |
| 2061 | mamo-L | *BGIBMGA004259* | up | 1782 | 1796 | + | 12.1532 | 2.13E-05 | 1 | AAGGCTTGTGTTGTC |
| 2062 | mamo-L | *BGIBMGA004260* | down | 817 | 831 | + | 12.4032 | 1.61E-05 | 1 | TAGGCTTGCGTGAGT |
| 2063 | mamo-L | *BGIBMGA004263* | up | 691 | 705 | + | 11.3871 | 4.55E-05 | 1 | AAGGCTTGCTTCGAT |
| 2064 | mamo-L | *BGIBMGA004264* | up | 1969 | 1983 | + | 10.9919 | 6.55E-05 | 1 | TAGGCTGTGAAGGGC |
| 2065 | mamo-L | *BGIBMGA004265* | down | 773 | 787 | - | 11.0968 | 5.97E-05 | 1 | AAGGCGCACGTTTCT |
| 2066 | mamo-L | *BGIBMGA004265* | up | 36 | 50 | - | 10.5726 | 9.33E-05 | 1 | TAGGCTAATATTTAT |
| 2067 | mamo-L | *BGIBMGA004269* | down | 803 | 817 | + | 11.6774 | 3.44E-05 | 1 | AAGGCTTATTGTGTT |
| 2068 | mamo-L | *BGIBMGA004269* | up | 1295 | 1309 | - | 10.7823 | 7.84E-05 | 1 | GAGGCGAATTTTGTT |
| 2069 | mamo-L | *BGIBMGA004281* | down | 193 | 207 | - | 11.7984 | 3.06E-05 | 1 | TAGGCACACTTTTGT |
| 2070 | mamo-L | *BGIBMGA004284* | down | 272 | 286 | + | 13.7742 | 2.79E-06 | 1 | TAGGCTTTTTTTTTT |
| 2071 | mamo-L | *BGIBMGA004285* | down | 1015 | 1029 | + | 11.3548 | 4.69E-05 | 1 | AAGGCTGATGATGAT |
| 2072 | mamo-L | *BGIBMGA004286* | down | 1633 | 1647 | + | 11.5726 | 3.81E-05 | 1 | TAGGCTTTTTTTTTA |
| 2073 | mamo-L | *BGIBMGA004286* | up | 1425 | 1439 | - | 10.5968 | 9.15E-05 | 1 | GAGGCTAGTTTTTGT |
| 2074 | mamo-L | *BGIBMGA004296* | down | 150 | 164 | + | 10.5726 | 9.33E-05 | 1 | TAGGCTAATATTTAT |
| 2075 | mamo-L | *BGIBMGA004297* | down | 188 | 202 | + | 14.1855 | 1.50E-06 | 1 | TAGGCTGACGTTTAT |
| 2076 | mamo-L | *BGIBMGA004304* | up | 180 | 194 | + | 10.8145 | 7.63E-05 | 1 | AAGGCTTTGTGTTTT |
| 2077 | mamo-L | *BGIBMGA004304* | up | 766 | 780 | - | 10.7177 | 8.28E-05 | 1 | GAGGCTCGCGTTTTC |
| 2078 | mamo-L | *BGIBMGA004305* | up | 353 | 367 | + | 10.8548 | 7.38E-05 | 1 | GAGGCCTGTTTTTAT |
| 2079 | mamo-L | *BGIBMGA004310* | down | 355 | 369 | - | 11.4597 | 4.25E-05 | 1 | TAGGCGGGGTGGTGT |
| 2080 | mamo-L | *BGIBMGA004310* | down | 451 | 465 | + | 10.6855 | 8.51E-05 | 1 | AAGGCGGGTGGTGGC |
| 2081 | mamo-L | *BGIBMGA004312* | down | 693 | 707 | - | 12.2823 | 1.85E-05 | 1 | TAGGCCTTTGGTTCT |
| 2082 | mamo-L | *BGIBMGA004312* | down | 1346 | 1360 | + | 11.6694 | 3.47E-05 | 1 | AAGGCGTGTGTTTGT |
| 2083 | mamo-L | *BGIBMGA004314* | down | 871 | 885 | + | 11.4597 | 4.25E-05 | 1 | TAGGCGGGGTGGTGT |
| 2084 | mamo-L | *BGIBMGA004314* | down | 775 | 789 | - | 10.6855 | 8.51E-05 | 1 | AAGGCGGGTGGTGGC |
| 2085 | mamo-L | *BGIBMGA004315* | up | 153 | 167 | - | 11.4032 | 4.49E-05 | 1 | GAGGCCTTTATTGCT |
| 2086 | mamo-L | *BGIBMGA004315* | down | 539 | 553 | - | 10.9194 | 6.98E-05 | 1 | TAGGCTACTTTTTTT |
| 2087 | mamo-L | *BGIBMGA004317* | down | 207 | 221 | + | 12.7419 | 1.09E-05 | 1 | TAGGCGGATTTTTTC |
| 2088 | mamo-L | *BGIBMGA004317* | up | 320 | 334 | - | 10.5 | 9.90E-05 | 1 | GAGGCCCAGGTCGTC |
| 2089 | mamo-L | *BGIBMGA004319* | down | 171 | 185 | - | 10.9194 | 6.98E-05 | 1 | TAGGCTACTTTTTTT |
| 2090 | mamo-L | *BGIBMGA004322* | up | 1794 | 1808 | + | 11.4919 | 4.11E-05 | 1 | AAGGCTTCTTTTGTT |
| 2091 | mamo-L | *BGIBMGA004322* | up | 1672 | 1686 | - | 11.2177 | 5.34E-05 | 1 | TAGGCCGGCGACACT |
| 2092 | mamo-L | *BGIBMGA004323* | down | 475 | 489 | - | 11.7661 | 3.16E-05 | 1 | TAGGCTTATTTCCTT |
| 2093 | mamo-L | *BGIBMGA004324* | down | 1353 | 1367 | + | 10.5726 | 9.33E-05 | 1 | TAGGCTAATATTTAT |
| 2094 | mamo-L | *BGIBMGA004325* | up | 1638 | 1652 | + | 11.4919 | 4.11E-05 | 1 | AAGGCTTCTTTTGTT |
| 2095 | mamo-L | *BGIBMGA004325* | up | 1525 | 1539 | - | 11.1694 | 5.59E-05 | 1 | TAGGCCGGCGGCATT |
| 2096 | mamo-L | *BGIBMGA004325* | down | 1964 | 1978 | + | 10.5484 | 9.52E-05 | 1 | TAGGCGGGCGAGCGT |
| 2097 | mamo-L | *BGIBMGA004328* | down | 617 | 631 | - | 14.9677 | 3.56E-07 | 1 | TAGGCTCGGGTTGCT |
| 2098 | mamo-L | *BGIBMGA004329* | down | 122 | 136 | - | 10.629 | 8.91E-05 | 1 | TAGGCCTGCAGCGCC |
| 2099 | mamo-L | *BGIBMGA004330* | up | 979 | 993 | + | 10.5403 | 9.58E-05 | 1 | GAGGCTTACGATGAC |
| 2100 | mamo-L | *BGIBMGA004333* | down | 1644 | 1658 | + | 11.7984 | 3.06E-05 | 1 | TAGGCACACTTTTGT |
| 2101 | mamo-L | *BGIBMGA004335* | down | 80 | 94 | + | 11.9839 | 2.54E-05 | 1 | TAGGCCGGTTGTCGT |
| 2102 | mamo-L | *BGIBMGA004341* | up | 835 | 849 | - | 10.6532 | 8.73E-05 | 1 | TAGGCATCCATTGCT |
| 2103 | mamo-L | *BGIBMGA004341* | up | 1217 | 1231 | - | 10.5726 | 9.33E-05 | 1 | TAGGCTAATATTTAT |
| 2104 | mamo-L | *BGIBMGA004344* | up | 1843 | 1857 | + | 13.1452 | 6.69E-06 | 1 | TAGGCAGACGTTGGC |
| 2105 | mamo-L | *BGIBMGA004344* | up | 1846 | 1860 | - | 11.0081 | 6.45E-05 | 1 | TAGGCCAACGTCTGC |
| 2106 | mamo-L | *BGIBMGA004347* | down | 98 | 112 | - | 11.871 | 2.85E-05 | 1 | TAGGCTTTAATTTTT |
| 2107 | mamo-L | *BGIBMGA004350* | down | 916 | 930 | + | 11.1613 | 5.63E-05 | 1 | AAGGCCCTCATTGTT |
| 2108 | mamo-L | *BGIBMGA004353* | up | 39 | 53 | - | 10.5726 | 9.33E-05 | 1 | TAGGCTAATATTTAT |
| 2109 | mamo-L | *BGIBMGA004354* | up | 683 | 697 | - | 12.8145 | 9.96E-06 | 1 | TAGGCCTTTGTTATT |

| | | | | | | | | | | |
|---|---|---|---|---|---|---|---|---|---|---|
| 2110 | mamo-L | *BGIBMGA004354* | down | 1412 | 1426 | + | 11.5726 | 3.81E-05 | 1 | TAGGCTTAGTTCTAC |
| 2111 | mamo-L | *BGIBMGA004357* | up | 256 | 270 | - | 10.5968 | 9.15E-05 | 1 | GAGGCTAGTTTTTGT |
| 2112 | mamo-L | *BGIBMGA004363* | up | 759 | 773 | - | 10.5968 | 9.15E-05 | 1 | TAGGCTTATTTTATG |
| 2113 | mamo-L | *BGIBMGA004366* | down | 10 | 24 | + | 10.9435 | 6.83E-05 | 1 | TAGGCTGCTTTCAGT |
| 2114 | mamo-L | *BGIBMGA004368* | down | 1467 | 1481 | - | 12.9919 | 8.10E-06 | 1 | TAGGCCGGTTTTGCA |
| 2115 | mamo-L | *BGIBMGA004368* | down | 692 | 706 | + | 12.7097 | 1.13E-05 | 1 | TAGGCTTAATTTAGT |
| 2116 | mamo-L | *BGIBMGA004369* | down | 1073 | 1087 | - | 12.0968 | 2.26E-05 | 1 | TAGGCAGTGTATGTT |
| 2117 | mamo-L | *BGIBMGA004369* | down | 1052 | 1066 | + | 11.0484 | 6.23E-05 | 1 | TAGGCCGCCTTAGGT |
| 2118 | mamo-L | *BGIBMGA004374* | down | 370 | 384 | + | 11.7661 | 3.16E-05 | 1 | TAGGCTTTTATTATT |
| 2119 | mamo-L | *BGIBMGA004375* | up | 329 | 343 | + | 10.8306 | 7.52E-05 | 1 | TAGGCTTTCATTGAG |
| 2120 | mamo-L | *BGIBMGA004377* | up | 1394 | 1408 | + | 11.5323 | 3.96E-05 | 1 | TAGGCGTGAGGTGAT |
| 2121 | mamo-L | *BGIBMGA004378* | up | 1424 | 1438 | + | 13.621 | 3.50E-06 | 1 | TAGGCGTTGTTTTCT |
| 2122 | mamo-L | *BGIBMGA004379* | down | 548 | 562 | - | 10.6613 | 8.67E-05 | 1 | TAGGCGTGTGTCGCG |
| 2123 | mamo-L | *BGIBMGA004382* | down | 1279 | 1293 | - | 11.3871 | 4.55E-05 | 1 | TAGGCTATATTTTTT |
| 2124 | mamo-L | *BGIBMGA004386* | up | 89 | 103 | - | 11.4194 | 4.42E-05 | 1 | TAGGCTGTTTTGTCC |
| 2125 | mamo-L | *BGIBMGA004386* | down | 1373 | 1387 | + | 10.8145 | 7.63E-05 | 1 | TAGGCCCTGCTTTTT |
| 2126 | mamo-L | *BGIBMGA004388* | down | 755 | 769 | - | 10.6855 | 8.51E-05 | 1 | AAGGCGGGTGGTGGC |
| 2127 | mamo-L | *BGIBMGA004389* | up | 199 | 213 | - | 12.2984 | 1.81E-05 | 1 | TAGGCTGAGTAGTGT |
| 2128 | mamo-L | *BGIBMGA004389* | down | 1369 | 1383 | - | 10.9194 | 6.98E-05 | 1 | TAGGCTACTTTTTTT |
| 2129 | mamo-L | *BGIBMGA004393* | down | 234 | 248 | + | 11.1855 | 5.50E-05 | 1 | TAGGCGTATTGTTAT |
| 2130 | mamo-L | *BGIBMGA004395* | up | 490 | 504 | + | 12.0645 | 2.34E-05 | 1 | TAGGCGGGCTTCGGA |
| 2131 | mamo-L | *BGIBMGA004395* | up | 1689 | 1703 | + | 10.8548 | 7.38E-05 | 1 | TAGGCTAGCTTAGGT |
| 2132 | mamo-L | *BGIBMGA004395* | down | 588 | 602 | + | 10.5968 | 9.15E-05 | 1 | GAGGCTAGTTTTTGT |
| 2133 | mamo-L | *BGIBMGA004396* | down | 842 | 856 | + | 10.6048 | 9.09E-05 | 1 | TAGGCTAAATTTGTA |
| 2134 | mamo-L | *BGIBMGA004398* | down | 1330 | 1344 | - | 10.5 | 9.90E-05 | 1 | GAGGCCCAGGTCGTC |
| 2135 | mamo-L | *BGIBMGA004404* | down | 655 | 669 | + | 13.129 | 6.83E-06 | 1 | TAGGCTGTGTTTGTG |
| 2136 | mamo-L | *BGIBMGA004408* | down | 756 | 770 | - | 10.5726 | 9.33E-05 | 1 | TAGGCTAATATTTAT |
| 2137 | mamo-L | *BGIBMGA004410* | down | 1851 | 1865 | + | 10.6774 | 8.56E-05 | 1 | TAGCCCGACTATGGT |
| 2138 | mamo-L | *BGIBMGA004412* | down | 1901 | 1915 | - | 11.5565 | 3.87E-05 | 1 | TAGGCGGTCGGTCTT |
| 2139 | mamo-L | *BGIBMGA004414* | down | 1890 | 1904 | + | 10.7661 | 7.95E-05 | 1 | TAGGCTTCCTTTTCA |
| 2140 | mamo-L | *BGIBMGA004415* | up | 115 | 129 | - | 11.9597 | 2.61E-05 | 1 | TAGGCTAAGATTTTT |
| 2141 | mamo-L | *BGIBMGA004420* | down | 693 | 707 | - | 10.5403 | 9.58E-05 | 1 | TAGGCGGGAGATGTA |
| 2142 | mamo-L | *BGIBMGA004420* | up | 549 | 563 | - | 10.4919 | 9.96E-05 | 1 | TAGGCTGTGTCTGTG |
| 2143 | mamo-L | *BGIBMGA004422* | up | 1425 | 1439 | + | 11.2177 | 5.34E-05 | 1 | TAGGCCACCATTGTT |
| 2144 | mamo-L | *BGIBMGA004427* | up | 1667 | 1681 | - | 10.5726 | 9.33E-05 | 1 | TAGGCTAATATTTAT |
| 2145 | mamo-L | *BGIBMGA004428* | up | 97 | 111 | + | 11.3548 | 4.69E-05 | 1 | AAGGCTGATGATGAT |
| 2146 | mamo-L | *BGIBMGA004429* | up | 933 | 947 | - | 10.9194 | 6.98E-05 | 1 | TAGGCTACTTTTTTT |
| 2147 | mamo-L | *BGIBMGA004432* | down | 543 | 557 | - | 12.5242 | 1.40E-05 | 1 | TAGGCTGGCGATCCT |
| 2148 | mamo-L | *BGIBMGA004434* | up | 321 | 335 | - | 11.3548 | 4.69E-05 | 1 | AAGGCTGATGATGAT |
| 2149 | mamo-L | *BGIBMGA004437* | down | 1657 | 1671 | + | 12.1452 | 2.15E-05 | 1 | TAGGCTCTTTTTTAT |
| 2150 | mamo-L | *BGIBMGA004440* | down | 1085 | 1099 | - | 10.9435 | 6.83E-05 | 1 | TAGGCCTGGAGTACT |
| 2151 | mamo-L | *BGIBMGA004441* | up | 580 | 594 | + | 10.9597 | 6.73E-05 | 1 | TAGGCTGAGAATAGC |
| 2152 | mamo-L | *BGIBMGA004443* | up | 104 | 118 | + | 10.629 | 8.91E-05 | 1 | TAGGCCTGCAGCGCC |
| 2153 | mamo-L | *BGIBMGA004447* | up | 1612 | 1626 | + | 10.7258 | 8.23E-05 | 1 | AAGGCCCAGTTTTAT |
| 2154 | mamo-L | *BGIBMGA004448* | up | 1074 | 1088 | + | 10.9194 | 6.98E-05 | 1 | TAGGCTACTTTTTTT |
| 2155 | mamo-L | *BGIBMGA004449* | down | 1866 | 1880 | + | 10.6532 | 8.73E-05 | 1 | GAGGCTTTTTTTTTC |
| 2156 | mamo-L | *BGIBMGA004450* | down | 1284 | 1298 | + | 14.121 | 1.65E-06 | 1 | TAGGCTTAGGTTTGC |
| 2157 | mamo-L | *BGIBMGA004451* | up | 1139 | 1153 | + | 11.3065 | 4.91E-05 | 1 | AAGGCGTATTTTTTT |
| 2158 | mamo-L | *BGIBMGA004457* | up | 207 | 221 | + | 11.3387 | 4.77E-05 | 1 | TAGGCTCTAGGTGGC |
| 2159 | mamo-L | *BGIBMGA004458* | down | 870 | 884 | + | 10.5726 | 9.33E-05 | 1 | TAGGCTAATATTTAT |
| 2160 | mamo-L | *BGIBMGA004466* | up | 582 | 596 | + | 10.9194 | 6.98E-05 | 1 | TAGGCTACTTTTTTT |
| 2161 | mamo-L | *BGIBMGA004466* | down | 226 | 240 | - | 10.629 | 8.91E-05 | 1 | TAGGCCTGCAGCGCC |
| 2162 | mamo-L | *BGIBMGA004470* | down | 671 | 685 | - | 11.2581 | 5.14E-05 | 1 | TAGGCATAATTTGAT |
| 2163 | mamo-L | *BGIBMGA004471* | up | 59 | 73 | - | 10.5968 | 9.15E-05 | 1 | AAGGCTGTTAATGCT |
| 2164 | mamo-L | *BGIBMGA004473* | down | 1752 | 1766 | - | 10.629 | 8.91E-05 | 1 | TAGGCCTGCAGCGCC |
| 2165 | mamo-L | *BGIBMGA004474* | down | 505 | 519 | + | 12.0161 | 2.46E-05 | 1 | TAGGCAGGTTGTGGT |
| 2166 | mamo-L | *BGIBMGA004475* | up | 959 | 973 | + | 11.1048 | 5.92E-05 | 1 | TAGGCCGTGTTCCCC |
| 2167 | mamo-L | *BGIBMGA004477* | down | 79 | 93 | + | 10.5645 | 9.40E-05 | 1 | TAGGCTGCTTGCGGC |
| 2168 | mamo-L | *BGIBMGA004480* | down | 126 | 140 | + | 12.9919 | 8.10E-06 | 1 | TAGGCCGGTTTTGCA |
| 2169 | mamo-L | *BGIBMGA004483* | down | 588 | 602 | - | 10.7419 | 8.11E-05 | 1 | AAGGCGCTGTTTTTT |
| 2170 | mamo-L | *BGIBMGA004484* | up | 802 | 816 | - | 10.7419 | 8.11E-05 | 1 | AAGGCGCTGTTTTTT |
| 2171 | mamo-L | *BGIBMGA004487* | down | 28 | 42 | + | 10.7258 | 8.23E-05 | 1 | TAGGCCCTTTATCGT |
| 2172 | mamo-L | *BGIBMGA004489* | down | 829 | 843 | - | 11.0242 | 6.36E-05 | 1 | TAGGCTGCTGTTACC |
| 2173 | mamo-L | *BGIBMGA004491* | down | 804 | 818 | - | 13.4274 | 4.64E-06 | 1 | TAGGCCGGTGTTTAT |
| 2174 | mamo-L | *BGIBMGA004491* | down | 1131 | 1145 | + | 11.7661 | 3.16E-05 | 1 | TAGGCGTTAGTTTGC |
| 2175 | mamo-L | *BGIBMGA004492* | down | 130 | 144 | - | 11.7661 | 3.16E-05 | 1 | TAGGCTTTTATTATT |
| 2176 | mamo-L | *BGIBMGA004497* | down | 1339 | 1353 | + | 11.4919 | 4.11E-05 | 1 | TAGGCCGCGATGGCT |
| 2177 | mamo-L | *BGIBMGA004498* | up | 638 | 652 | + | 10.5645 | 9.40E-05 | 1 | TAGGCTTATTACACT |
| 2178 | mamo-L | *BGIBMGA004499* | down | 669 | 683 | + | 13.1048 | 7.05E-06 | 1 | TAGGCCTTATTTTGT |
| 2179 | mamo-L | *BGIBMGA004499* | up | 345 | 359 | - | 12.2984 | 1.81E-05 | 1 | TAGGCGCTTTTTTTT |
| 2180 | mamo-L | *BGIBMGA004500* | up | 69 | 83 | + | 11.7984 | 3.06E-05 | 1 | TAGGCTTTGCTTTTT |
| 2181 | mamo-L | *BGIBMGA004502* | up | 687 | 701 | + | 13.1774 | 6.47E-06 | 1 | TAGGCCGAATTGGGT |
| 2182 | mamo-L | *BGIBMGA004502* | up | 657 | 671 | - | 10.7661 | 7.95E-05 | 1 | TAGGCTTTTGTTACA |
| 2183 | mamo-L | *BGIBMGA004503* | up | 349 | 363 | + | 10.9839 | 6.59E-05 | 1 | AAGGCGTTTTTTTTT |
| 2184 | mamo-L | *BGIBMGA004516* | up | 860 | 874 | - | 11.0645 | 6.14E-05 | 1 | TAGGCCGCGAATTGT |
| 2185 | mamo-L | *BGIBMGA004521* | up | 1578 | 1592 | - | 12.0161 | 2.46E-05 | 1 | TAGGCCGCCGCTGGT |
| 2186 | mamo-L | *BGIBMGA004522* | up | 1398 | 1412 | + | 11.3548 | 4.69E-05 | 1 | TAGGCCTAAGTTTGA |
| 2187 | mamo-L | *BGIBMGA004522* | up | 1616 | 1630 | + | 11.3548 | 4.69E-05 | 1 | TAGGCCTAAGTTTGA |
| 2188 | mamo-L | *BGIBMGA004522* | up | 1582 | 1596 | + | 10.7258 | 8.23E-05 | 1 | TAGGCTTGGTTGTTA |
| 2189 | mamo-L | *BGIBMGA004522* | down | 1771 | 1785 | + | 10.5806 | 9.27E-05 | 1 | TAGGCGTACAATATT |
| 2190 | mamo-L | *BGIBMGA004523* | up | 772 | 786 | + | 10.7258 | 8.23E-05 | 1 | GAGGCTTTTTTTATT |
| 2191 | mamo-L | *BGIBMGA004524* | down | 158 | 172 | - | 12.2258 | 1.97E-05 | 1 | TAGGCTGGCTTGACT |
| 2192 | mamo-L | *BGIBMGA004525* | down | 1014 | 1028 | + | 12.4435 | 1.54E-05 | 1 | TAGGCTTTTAATGTT |
| 2193 | mamo-L | *BGIBMGA004526* | up | 1887 | 1901 | + | 10.9194 | 6.98E-05 | 1 | TAGGCTACTTTTTTT |
| 2194 | mamo-L | *BGIBMGA004527* | up | 78 | 92 | - | 10.6452 | 8.79E-05 | 1 | TAGGCGGTGTTCCTC |
| 2195 | mamo-L | *BGIBMGA004529* | down | 1097 | 1111 | - | 14.7258 | 5.83E-07 | 1 | TAGGCTTGCGTTGCC |
| 2196 | mamo-L | *BGIBMGA004529* | down | 1883 | 1897 | + | 11.6855 | 3.42E-05 | 1 | TAGGCGGAGTGTGGG |
| 2197 | mamo-L | *BGIBMGA004529* | down | 1352 | 1366 | + | 10.4919 | 9.96E-05 | 1 | TAGGCTAGTATTTAT |
| 2198 | mamo-L | *BGIBMGA004532* | up | 127 | 141 | + | 12.5323 | 1.39E-05 | 1 | TAGGCAGGGTTTTCT |
| 2199 | mamo-L | *BGIBMGA004533* | up | 1025 | 1039 | + | 11.3548 | 4.69E-05 | 1 | AAGGCTGATGATGAT |
| 2200 | mamo-L | *BGIBMGA004535* | down | 1778 | 1792 | - | 12.6532 | 1.20E-05 | 1 | TAGGCAGAGTGTGGT |
| 2201 | mamo-L | *BGIBMGA004538* | up | 1016 | 1030 | + | 10.5726 | 9.33E-05 | 1 | TAGGCTAATATTTAT |

| | | | | | | | | | | |
|---|---|---|---|---|---|---|---|---|---|---|
| 2202 | mamo-L | *BGIBMGA004539* | up | 174 | 188 | - | 12.1774 | 2.08E-05 | 1 | TAGGCTTTGTATTAT |
| 2203 | mamo-L | *BGIBMGA004542* | up | 835 | 849 | + | 11.8548 | 2.89E-05 | 1 | GAGGCCTTCGTCGCT |
| 2204 | mamo-L | *BGIBMGA004545* | up | 475 | 489 | + | 13.1613 | 6.59E-06 | 1 | TAGGCTTATTTTACT |
| 2205 | mamo-L | *BGIBMGA004546* | down | 1115 | 1129 | - | 11.4355 | 4.34E-05 | 1 | TAGGCCTTTTATTAT |
| 2206 | mamo-L | *BGIBMGA004547* | up | 851 | 865 | - | 11.3468 | 4.73E-05 | 1 | TAGGCCGAGCATGGC |
| 2207 | mamo-L | *BGIBMGA004550* | down | 101 | 115 | - | 10.5726 | 9.33E-05 | 1 | TAGGCTAATATTTAT |
| 2208 | mamo-L | *BGIBMGA004551* | down | 394 | 408 | - | 11.4032 | 4.49E-05 | 1 | TAGGCTTAGGTGAAT |
| 2209 | mamo-L | *BGIBMGA004554* | down | 848 | 862 | + | 11.2823 | 5.03E-05 | 1 | TCGGCTTGTGTTGGT |
| 2210 | mamo-L | *BGIBMGA004555* | up | 1153 | 1167 | + | 11.2823 | 5.03E-05 | 1 | TCGGCTTGTGTTGGT |
| 2211 | mamo-L | *BGIBMGA004557* | down | 1806 | 1820 | + | 12.3952 | 1.63E-05 | 1 | TAGGCGTACGTGTTT |
| 2212 | mamo-L | *BGIBMGA004562* | down | 1696 | 1710 | + | 12.6774 | 1.17E-05 | 1 | TAGGCTATCTTTTTT |
| 2213 | mamo-L | *BGIBMGA004565* | up | 1202 | 1216 | + | 11.0565 | 6.18E-05 | 1 | TAGGCTTAAAGTGAT |
| 2214 | mamo-L | *BGIBMGA004566* | up | 519 | 533 | - | 14.0887 | 1.74E-06 | 1 | TAGGCTGATTGTGTT |
| 2215 | mamo-L | *BGIBMGA004566* | up | 690 | 704 | + | 13.3387 | 5.24E-06 | 1 | TAGGCTGATTTTCTT |
| 2216 | mamo-L | *BGIBMGA004568* | up | 97 | 111 | - | 13.3548 | 5.14E-06 | 1 | TAGGCTTGTTTTTGC |
| 2217 | mamo-L | *BGIBMGA004571* | down | 647 | 661 | - | 14.1371 | 1.62E-06 | 1 | TAGGCTTTCGATGCT |
| 2218 | mamo-L | *BGIBMGA004573* | down | 1770 | 1784 | + | 12.7097 | 1.13E-05 | 1 | TAGGCCTTCGTTGTG |
| 2219 | mamo-L | *BGIBMGA004573* | down | 1763 | 1777 | - | 11.5242 | 3.99E-05 | 1 | AAGGCCTAGTTTTAT |
| 2220 | mamo-L | *BGIBMGA004573* | down | 313 | 327 | - | 10.9194 | 6.98E-05 | 1 | TAGGCTACTTTTTTT |
| 2221 | mamo-L | *BGIBMGA004575* | up | 813 | 827 | - | 11.1129 | 5.88E-05 | 1 | AAGGCGGCTTTTGTT |
| 2222 | mamo-L | *BGIBMGA004576* | up | 1363 | 1377 | - | 12.5968 | 1.28E-05 | 1 | TAGGCCCACTTCGAT |
| 2223 | mamo-L | *BGIBMGA004577* | down | 362 | 376 | - | 11.7661 | 3.16E-05 | 1 | TAGGCAGTTTTTTTT |
| 2224 | mamo-L | *BGIBMGA004578* | up | 428 | 442 | + | 10.9194 | 6.98E-05 | 1 | TAGGCTACTTTTTTT |
| 2225 | mamo-L | *BGIBMGA004579* | up | 1488 | 1502 | - | 10.6371 | 8.85E-05 | 1 | AAGGCCGTATGTGCT |
| 2226 | mamo-L | *BGIBMGA004580* | up | 318 | 332 | + | 14.2177 | 1.43E-06 | 1 | TAGGCCGGTTTTGCC |
| 2227 | mamo-L | *BGIBMGA004580* | down | 1152 | 1166 | + | 10.5806 | 9.27E-05 | 1 | TAGGCGTCTGTGGAT |
| 2228 | mamo-L | *BGIBMGA004587* | up | 34 | 48 | - | 10.5726 | 9.33E-05 | 1 | TAGGCTAATATTTAT |
| 2229 | mamo-L | *BGIBMGA004588* | up | 1425 | 1439 | + | 11.3065 | 4.91E-05 | 1 | AAGGCGTATTTTTTT |
| 2230 | mamo-L | *BGIBMGA004589* | up | 1808 | 1822 | - | 10.5726 | 9.33E-05 | 1 | TAGGCTAATATTTAT |
| 2231 | mamo-L | *BGIBMGA004590* | down | 1544 | 1558 | + | 10.9194 | 6.98E-05 | 1 | TAGGCTACTTTTTTT |
| 2232 | mamo-L | *BGIBMGA004592* | down | 857 | 871 | + | 11.5968 | 3.73E-05 | 1 | TAGGCTGTGGTATGT |
| 2233 | mamo-L | *BGIBMGA004594* | down | 1664 | 1678 | - | 10.5726 | 9.33E-05 | 1 | TAGGCTAATATTTAT |
| 2234 | mamo-L | *BGIBMGA004595* | up | 553 | 567 | + | 12.0323 | 2.42E-05 | 1 | AAGGCTTGTGTTTTT |
| 2235 | mamo-L | *BGIBMGA004596* | up | 1868 | 1882 | - | 13.6774 | 3.24E-06 | 1 | AAGGCTGTCGTTGTT |
| 2236 | mamo-L | *BGIBMGA004598* | up | 1282 | 1296 | + | 10.5403 | 9.58E-05 | 1 | TAGGCTGTATGGTGT |
| 2237 | mamo-L | *BGIBMGA004599* | down | 91 | 105 | - | 10.8145 | 7.63E-05 | 1 | TAGGCATCCTTTGTC |
| 2238 | mamo-L | *BGIBMGA004603* | up | 887 | 901 | - | 11.8548 | 2.89E-05 | 1 | GAGGCCTTCGTCGCT |
| 2239 | mamo-L | *BGIBMGA004603* | down | 826 | 840 | - | 10.5726 | 9.33E-05 | 1 | TAGGCTAATATTTAT |
| 2240 | mamo-L | *BGIBMGA004606* | up | 516 | 530 | - | 10.9194 | 6.98E-05 | 1 | TAGGCTACTTTTTTT |
| 2241 | mamo-L | *BGIBMGA004610* | down | 1890 | 1904 | + | 11.7016 | 3.37E-05 | 1 | TAGGCTCAGTATTAT |
| 2242 | mamo-L | *BGIBMGA004612* | down | 1417 | 1431 | + | 10.9758 | 6.64E-05 | 1 | TGGGCTGTCTTTTTT |
| 2243 | mamo-L | *BGIBMGA004617* | up | 750 | 764 | + | 12.25 | 1.92E-05 | 1 | TAGGCCCACTATTCT |
| 2244 | mamo-L | *BGIBMGA004617* | up | 1851 | 1865 | - | 10.9839 | 6.59E-05 | 1 | GAGGCGTGTGGTGCT |
| 2245 | mamo-L | *BGIBMGA004618* | down | 643 | 657 | - | 10.7419 | 8.11E-05 | 1 | TAGGCTCTTTTGAGT |
| 2246 | mamo-L | *BGIBMGA004620* | up | 1154 | 1168 | - | 11.4758 | 4.18E-05 | 1 | TAGGCCAATGTTTTC |
| 2247 | mamo-L | *BGIBMGA004621* | down | 1442 | 1456 | - | 11.3548 | 4.69E-05 | 1 | AAGGCTGATGATGAT |
| 2248 | mamo-L | *BGIBMGA004623* | up | 389 | 403 | - | 13.6774 | 3.24E-06 | 1 | TAGGCCTTTTATGGT |
| 2249 | mamo-L | *BGIBMGA004626* | down | 765 | 779 | - | 11.2581 | 5.14E-05 | 1 | GAGGCTGGCGACGCT |
| 2250 | mamo-L | *BGIBMGA004628* | down | 159 | 173 | - | 14.3468 | 1.17E-06 | 1 | TAGGCGGACGGTGGT |
| 2251 | mamo-L | *BGIBMGA004636* | up | 42 | 56 | - | 12.6452 | 1.21E-05 | 1 | TAGGCTTATGGCGTT |
| 2252 | mamo-L | *BGIBMGA004637* | up | 1482 | 1496 | + | 12.6452 | 1.21E-05 | 1 | TAGGCTTATGGCGTT |
| 2253 | mamo-L | *BGIBMGA004638* | down | 1600 | 1614 | - | 12.3306 | 1.75E-05 | 1 | TAGGCTTACTATACT |
| 2254 | mamo-L | *BGIBMGA004642* | down | 1056 | 1070 | - | 11.879 | 2.83E-05 | 1 | TAGGCTTTAAGTGGT |
| 2255 | mamo-L | *BGIBMGA004644* | up | 987 | 1001 | + | 11.9435 | 2.65E-05 | 1 | TAGGCCTGTTTTCAT |
| 2256 | mamo-L | *BGIBMGA004645* | up | 465 | 479 | + | 13.2016 | 6.26E-06 | 1 | TAGGCCGATGGTGAT |
| 2257 | mamo-L | *BGIBMGA004645* | down | 674 | 688 | + | 12.9435 | 8.54E-06 | 1 | TAGGCTTTTTTTTAT |
| 2258 | mamo-L | *BGIBMGA004651* | down | 1829 | 1843 | + | 10.9919 | 6.55E-05 | 1 | TAGGCAGTTGTTATT |
| 2259 | mamo-L | *BGIBMGA004654* | down | 1614 | 1628 | + | 11.5403 | 3.93E-05 | 1 | TAGGCGGGGGGGGAT |
| 2260 | mamo-L | *BGIBMGA004655* | up | 1428 | 1442 | + | 10.9194 | 6.98E-05 | 1 | TAGGCTACTTTTTTT |
| 2261 | mamo-L | *BGIBMGA004659* | down | 69 | 83 | - | 13.4677 | 4.39E-06 | 1 | TAGGCTGATGATGAT |
| 2262 | mamo-L | *BGIBMGA004664* | up | 1124 | 1138 | + | 11.1774 | 5.54E-05 | 1 | TAGGCGGGGGTGGTG |
| 2263 | mamo-L | *BGIBMGA004669* | down | 1936 | 1950 | + | 11.1935 | 5.45E-05 | 1 | TAGGCGCATGTGGCC |
| 2264 | mamo-L | *BGIBMGA004669* | up | 1087 | 1101 | - | 10.6855 | 8.51E-05 | 1 | AAGGCTTTTTTTTTC |
| 2265 | mamo-L | *BGIBMGA004671* | down | 1881 | 1895 | - | 10.7258 | 8.23E-05 | 1 | TAGGCGTGCTTTATA |
| 2266 | mamo-L | *BGIBMGA004671* | down | 1707 | 1721 | + | 10.6613 | 8.67E-05 | 1 | TAGGCGCTGAGTTGT |
| 2267 | mamo-L | *BGIBMGA004674* | up | 1435 | 1449 | - | 12.1855 | 2.06E-05 | 1 | TAGCCTGACTTTGGT |
| 2268 | mamo-L | *BGIBMGA004676* | down | 1549 | 1563 | + | 10.9194 | 6.98E-05 | 1 | TAGGCTACTTTTTTT |
| 2269 | mamo-L | *BGIBMGA004677* | up | 241 | 255 | + | 11.7097 | 3.34E-05 | 1 | GAGGCCGTGTTTAGT |
| 2270 | mamo-L | *BGIBMGA004678* | down | 1755 | 1769 | - | 11.6371 | 3.58E-05 | 1 | TAGGCGCTCGGTGTC |
| 2271 | mamo-L | *BGIBMGA004681* | down | 1342 | 1356 | + | 11.4194 | 4.42E-05 | 1 | TAGGCCATGATTTCT |
| 2272 | mamo-L | *BGIBMGA004688* | down | 1102 | 1116 | + | 11.5565 | 3.87E-05 | 1 | TAGGCTTTTCTTTGT |
| 2273 | mamo-L | *BGIBMGA004693* | up | 319 | 333 | - | 10.8145 | 7.63E-05 | 1 | TAGGCGGACGTGATC |
| 2274 | mamo-L | *BGIBMGA004697* | down | 657 | 671 | - | 11.1694 | 5.59E-05 | 1 | TAGGCATTCTTTGGA |
| 2275 | mamo-L | *BGIBMGA004698* | down | 1466 | 1480 | - | 11.0403 | 6.28E-05 | 1 | TAGGCCGTGGACACT |
| 2276 | mamo-L | *BGIBMGA004699* | down | 1522 | 1536 | + | 11.0403 | 6.28E-05 | 1 | TAGGCCGTGGACACT |
| 2277 | mamo-L | *BGIBMGA004700* | down | 1886 | 1900 | - | 11.4516 | 4.28E-05 | 1 | TAGGCTTCTTTTCTT |
| 2278 | mamo-L | *BGIBMGA004704* | down | 842 | 856 | - | 11.0968 | 5.97E-05 | 1 | TAGGCTCAATGTTTT |
| 2279 | mamo-L | *BGIBMGA004705* | down | 640 | 654 | - | 10.8306 | 7.52E-05 | 1 | TAGGCCTAAGTAGCT |
| 2280 | mamo-L | *BGIBMGA004706* | up | 1378 | 1392 | - | 10.8306 | 7.52E-05 | 1 | TAGGCCTAAGTAGCT |
| 2281 | mamo-L | *BGIBMGA004707* | down | 466 | 480 | + | 10.8306 | 7.52E-05 | 1 | TAGGCCCCCTTCGCC |
| 2282 | mamo-L | *BGIBMGA004709* | down | 415 | 429 | + | 11.4274 | 4.38E-05 | 1 | TAGGCCGAGAGTTAT |
| 2283 | mamo-L | *BGIBMGA004710* | up | 1584 | 1598 | + | 11.4274 | 4.38E-05 | 1 | TAGGCCGAGAGTTAT |
| 2284 | mamo-L | *BGIBMGA004712* | up | 1211 | 1225 | + | 11.7177 | 3.32E-05 | 1 | TAGGCTGTATTGGCC |
| 2285 | mamo-L | *BGIBMGA004712* | down | 928 | 942 | - | 11.5565 | 3.87E-05 | 1 | TAGGCTTTTCTTTGT |
| 2286 | mamo-L | *BGIBMGA004721* | up | 1849 | 1863 | + | 10.5 | 9.90E-05 | 1 | AAGGCCGCTATTGTT |
| 2287 | mamo-L | *BGIBMGA004729* | down | 655 | 669 | - | 10.8226 | 7.58E-05 | 1 | AAGGCTGTTTTTGCA |
| 2288 | mamo-L | *BGIBMGA004736* | down | 1217 | 1231 | + | 10.621 | 8.97E-05 | 1 | AAGGCTTACGATTGC |
| 2289 | mamo-L | *BGIBMGA004737* | down | 1131 | 1145 | + | 12.4274 | 1.57E-05 | 1 | TAGGCTTTGGTTCTC |
| 2290 | mamo-L | *BGIBMGA004737* | down | 865 | 879 | - | 11.3548 | 4.69E-05 | 1 | AAGGCTGATGATGAT |
| 2291 | mamo-L | *BGIBMGA004738* | up | 1802 | 1816 | - | 10.5565 | 9.46E-05 | 1 | TAGGCGTCTGGTTTT |
| 2292 | mamo-L | *BGIBMGA004740* | down | 1141 | 1155 | + | 12.25 | 1.92E-05 | 1 | TAGGCCCACTATTCT |
| 2293 | mamo-L | *BGIBMGA004740* | down | 1840 | 1854 | - | 11.6048 | 3.70E-05 | 1 | TAGGCTGCGGTTTGA |

| | | | | | | | | | | |
|---|---|---|---|---|---|---|---|---|---|---|
| 2294 | mamo-L | *BGIBMGA004741* | down | 422 | 436 | + | 10.6532 | 8.73E-05 | 1 | TAGGCGCGTTTTCAT |
| 2295 | mamo-L | *BGIBMGA004742* | down | 1120 | 1134 | - | 11.1935 | 5.45E-05 | 1 | TAGGCTTATTGTGTG |
| 2296 | mamo-L | *BGIBMGA004742* | up | 1260 | 1274 | + | 10.6532 | 8.73E-05 | 1 | TAGGCGCGTTTTCAT |
| 2297 | mamo-L | *BGIBMGA004747* | down | 1300 | 1314 | - | 10.5887 | 9.21E-05 | 1 | TAGGCCTTGTATTCA |
| 2298 | mamo-L | *BGIBMGA004751* | down | 995 | 1009 | - | 11.5323 | 3.96E-05 | 1 | TAGGCCGATTGCTTT |
| 2299 | mamo-L | *BGIBMGA004751* | down | 1819 | 1833 | - | 10.75 | 8.06E-05 | 1 | GAGGCCGATTGTGCC |
| 2300 | mamo-L | *BGIBMGA004752* | down | 1311 | 1325 | - | 11.6613 | 3.50E-05 | 1 | TAGGCAGTGGTTGGA |
| 2301 | mamo-L | *BGIBMGA004752* | up | 1127 | 1141 | + | 10.9194 | 6.98E-05 | 1 | TAGGCTACTTTTTTT |
| 2302 | mamo-L | *BGIBMGA004753* | up | 785 | 799 | + | 11.4274 | 4.38E-05 | 1 | AAGGCCTAGGTTCTT |
| 2303 | mamo-L | *BGIBMGA004759* | down | 1906 | 1920 | + | 13.5645 | 3.81E-06 | 1 | TAGGCCTAGTTTATT |
| 2304 | mamo-L | *BGIBMGA004759* | up | 1228 | 1242 | - | 12.5726 | 1.33E-05 | 1 | GAGGCCTAGGTTGTC |
| 2305 | mamo-L | *BGIBMGA004760* | up | 1655 | 1669 | - | 13.5645 | 3.81E-06 | 1 | TAGGCCTAGTTTATT |
| 2306 | mamo-L | *BGIBMGA004761* | up | 219 | 233 | + | 10.7258 | 8.23E-05 | 1 | TAGGCTGGTCTTCTT |
| 2307 | mamo-L | *BGIBMGA004764* | up | 1112 | 1126 | - | 10.7823 | 7.84E-05 | 1 | GAGGCCTGAGTGGGT |
| 2308 | mamo-L | *BGIBMGA004764* | up | 1084 | 1098 | - | 10.6048 | 9.09E-05 | 1 | TAGGCTACCGTTACT |
| 2309 | mamo-L | *BGIBMGA004765* | down | 505 | 519 | - | 10.5726 | 9.33E-05 | 1 | TAGGCTAATATTTAT |
| 2310 | mamo-L | *BGIBMGA004769* | up | 619 | 633 | + | 11.9194 | 2.72E-05 | 1 | AAGGCGGACGGTGTT |
| 2311 | mamo-L | *BGIBMGA004775* | down | 951 | 965 | + | 13.5081 | 4.15E-06 | 1 | TAGGCTGATTATGGC |
| 2312 | mamo-L | *BGIBMGA004775* | down | 546 | 560 | + | 11.4355 | 4.34E-05 | 1 | AAGGCCGAAGGTGGT |
| 2313 | mamo-L | *BGIBMGA004779* | down | 822 | 836 | + | 11.4919 | 4.11E-05 | 1 | AAGGCCTGTTATGTT |
| 2314 | mamo-L | *BGIBMGA004780* | down | 1594 | 1608 | + | 10.5726 | 9.33E-05 | 1 | TAGGCTAATATTTAT |
| 2315 | mamo-L | *BGIBMGA004781* | down | 1478 | 1492 | - | 11.1048 | 5.92E-05 | 1 | TAGGCCTGGTGTATC |
| 2316 | mamo-L | *BGIBMGA004781* | down | 472 | 486 | - | 10.7581 | 8.00E-05 | 1 | TAGGCAAAGTTTTTT |
| 2317 | mamo-L | *BGIBMGA004783* | down | 440 | 454 | - | 11.6129 | 3.66E-05 | 1 | TAGGCGTATTTTTAC |
| 2318 | mamo-L | *BGIBMGA004784* | down | 217 | 231 | - | 12.6613 | 1.19E-05 | 1 | AAGGCCGGGTTTGCC |
| 2319 | mamo-L | *BGIBMGA004784* | down | 116 | 130 | - | 10.5968 | 9.15E-05 | 1 | GAGGCTAGTTTTTGT |
| 2320 | mamo-L | *BGIBMGA004785* | down | 1690 | 1704 | - | 11.1613 | 5.63E-05 | 1 | TAGGCGGCGTTGGTC |
| 2321 | mamo-L | *BGIBMGA004788* | down | 230 | 244 | + | 10.9194 | 6.98E-05 | 1 | TAGGCTACTTTTTTT |
| 2322 | mamo-L | *BGIBMGA004790* | down | 676 | 690 | - | 13.6452 | 3.39E-06 | 1 | TAGGCTGAGTTCTCT |
| 2323 | mamo-L | *BGIBMGA004790* | down | 1929 | 1943 | + | 11.1855 | 5.50E-05 | 1 | TAGGCGCAGTGTAGT |
| 2324 | mamo-L | *BGIBMGA004793* | down | 416 | 430 | + | 10.6694 | 8.62E-05 | 1 | TAGGCTGGCTTCCAC |
| 2325 | mamo-L | *BGIBMGA004794* | down | 1634 | 1648 | + | 10.8952 | 7.12E-05 | 1 | TAGGCCATTTTTGTA |
| 2326 | mamo-L | *BGIBMGA004797* | down | 1931 | 1945 | + | 10.5161 | 9.77E-05 | 1 | GAGGCGGATGTCGCC |
| 2327 | mamo-L | *BGIBMGA004800* | up | 518 | 532 | + | 10.5726 | 9.33E-05 | 1 | TAGGCTAATATTTAT |
| 2328 | mamo-L | *BGIBMGA004800* | down | 1676 | 1690 | - | 10.5403 | 9.58E-05 | 1 | TAGGCTCTCGATAAT |
| 2329 | mamo-L | *BGIBMGA004801* | up | 987 | 1001 | - | 11.5081 | 4.05E-05 | 1 | AAGGCTGAGTTGTGT |
| 2330 | mamo-L | *BGIBMGA004803* | down | 363 | 377 | + | 10.5806 | 9.27E-05 | 1 | TAGGCTTAGAAGTTT |
| 2331 | mamo-L | *BGIBMGA004805* | up | 212 | 226 | - | 11.3548 | 4.69E-05 | 1 | AAGGCTGATGATGAT |
| 2332 | mamo-L | *BGIBMGA004805* | down | 792 | 806 | + | 10.5726 | 9.33E-05 | 1 | TAGGCTAATATTTAT |
| 2333 | mamo-L | *BGIBMGA004806* | down | 1355 | 1369 | + | 10.75 | 8.06E-05 | 1 | TAGGCGAAGTGGGGT |
| 2334 | mamo-L | *BGIBMGA004807* | up | 1240 | 1254 | + | 10.75 | 8.06E-05 | 1 | TAGGCGAAGTGGGGT |
| 2335 | mamo-L | *BGIBMGA004812* | down | 866 | 880 | - | 11.4516 | 4.28E-05 | 1 | TAGGCTGGATTTCCC |
| 2336 | mamo-L | *BGIBMGA004813* | up | 369 | 383 | + | 11.6129 | 3.66E-05 | 1 | TAGGCTTTTAATGAT |
| 2337 | mamo-L | *BGIBMGA004814* | down | 43 | 57 | - | 12.5726 | 1.33E-05 | 1 | TAGGCTTAGGTCACT |
| 2338 | mamo-L | *BGIBMGA004814* | down | 194 | 208 | + | 12.1935 | 2.05E-05 | 1 | TAGGCCGGATTTGCA |
| 2339 | mamo-L | *BGIBMGA004814* | up | 192 | 206 | + | 12.0806 | 2.31E-05 | 1 | TAGGCTTGTATTTAT |
| 2340 | mamo-L | *BGIBMGA004815* | down | 315 | 329 | - | 14.0887 | 1.74E-06 | 1 | TAGGCTTTTTTTTGT |
| 2341 | mamo-L | *BGIBMGA004815* | down | 385 | 399 | - | 10.7097 | 8.33E-05 | 1 | TAGGCCGGTTGCCGT |
| 2342 | mamo-L | *BGIBMGA004826* | down | 598 | 612 | + | 13.871 | 2.40E-06 | 1 | AAGGCTGACTTTGTT |
| 2343 | mamo-L | *BGIBMGA004834* | down | 1423 | 1437 | + | 12.7258 | 1.11E-05 | 1 | TAGGCGTGCTTTGTA |
| 2344 | mamo-L | *BGIBMGA004835* | down | 1319 | 1333 | - | 12.7258 | 1.11E-05 | 1 | TAGGCGTGCTTTGTA |
| 2345 | mamo-L | *BGIBMGA004837* | down | 1340 | 1354 | + | 11.9435 | 2.65E-05 | 1 | TAGGCTGCCTATTCT |
| 2346 | mamo-L | *BGIBMGA004839* | up | 589 | 603 | - | 11.8226 | 2.99E-05 | 1 | TAGGCGTAGGTTTGG |
| 2347 | mamo-L | *BGIBMGA004840* | up | 686 | 700 | + | 12 | 2.50E-05 | 1 | TAGGCCTATTATAGT |
| 2348 | mamo-L | *BGIBMGA004840* | down | 1077 | 1091 | + | 11.5968 | 3.73E-05 | 1 | AAGGCCGAGGACGGT |
| 2349 | mamo-L | *BGIBMGA004841* | up | 888 | 902 | + | 12.5645 | 1.34E-05 | 1 | TAGGCAGGGTTTTTT |
| 2350 | mamo-L | *BGIBMGA004842* | up | 794 | 808 | + | 10.8226 | 7.58E-05 | 1 | GAGGCTGAGTATTGC |
| 2351 | mamo-L | *BGIBMGA004844* | down | 10 | 24 | + | 11.9194 | 2.72E-05 | 1 | GAGGCTTAGTTTAGT |
| 2352 | mamo-L | *BGIBMGA004848* | down | 648 | 662 | - | 11.4677 | 4.22E-05 | 1 | TAGGCTTAGGGTACC |
| 2353 | mamo-L | *BGIBMGA004849* | up | 1270 | 1284 | - | 11.4677 | 4.22E-05 | 1 | TAGGCTTAGGGTACC |
| 2354 | mamo-L | *BGIBMGA004852* | down | 1300 | 1314 | - | 11 | 6.50E-05 | 1 | TAGGCGTACATTTAC |
| 2355 | mamo-L | *BGIBMGA004859* | down | 1828 | 1842 | - | 12.5081 | 1.43E-05 | 1 | TAGGCTACCTTTGTT |
| 2356 | mamo-L | *BGIBMGA004861* | down | 869 | 883 | + | 16.1694 | 1.16E-08 | 1 | TAGGCTGTGGTTGGT |
| 2357 | mamo-L | *BGIBMGA004861* | up | 976 | 990 | - | 10.5726 | 9.33E-05 | 1 | TAGGCTAATATTTAT |
| 2358 | mamo-L | *BGIBMGA004863* | up | 650 | 664 | - | 10.5 | 9.90E-05 | 1 | GAGGCCCAGGTCGTC |
| 2359 | mamo-L | *BGIBMGA004865* | down | 106 | 120 | - | 12.6935 | 1.15E-05 | 1 | TAGGCTGGCTATGGA |
| 2360 | mamo-L | *BGIBMGA004866* | up | 1537 | 1551 | + | 11.3548 | 4.69E-05 | 1 | AAGGCTGATGATGAT |
| 2361 | mamo-L | *BGIBMGA004867* | up | 1771 | 1785 | + | 11.0242 | 6.36E-05 | 1 | AAGGCTTGGGTTAGC |
| 2362 | mamo-L | *BGIBMGA004870* | down | 1727 | 1741 | - | 12.9758 | 8.23E-06 | 1 | TAGGCCGGCTTGTTT |
| 2363 | mamo-L | *BGIBMGA004870* | down | 1305 | 1319 | - | 10.5726 | 9.33E-05 | 1 | TAGGCTAATATTTAT |
| 2364 | mamo-L | *BGIBMGA004871* | down | 215 | 229 | + | 10.5726 | 9.33E-05 | 1 | TAGGCTAATATTTAT |
| 2365 | mamo-L | *BGIBMGA004873* | down | 483 | 497 | - | 10.9194 | 6.98E-05 | 1 | TAGGCTCTGTGGGGC |
| 2366 | mamo-L | *BGIBMGA004882* | up | 486 | 500 | + | 12.4435 | 1.54E-05 | 1 | TAGGCTTTGTTTCAT |
| 2367 | mamo-L | *BGIBMGA004883* | up | 666 | 680 | - | 10.9194 | 6.98E-05 | 1 | TAGGCTACTTTTTTT |
| 2368 | mamo-L | *BGIBMGA004884* | down | 879 | 893 | + | 10.7581 | 8.00E-05 | 1 | TAGGCAAAGTTTTTT |
| 2369 | mamo-L | *BGIBMGA004887* | up | 1436 | 1450 | + | 10.5726 | 9.33E-05 | 1 | TAGGCTAATATTTAT |
| 2370 | mamo-L | *BGIBMGA004892* | down | 1562 | 1576 | - | 10.6532 | 8.73E-05 | 1 | TAGGCCCGCTCGGGT |
| 2371 | mamo-L | *BGIBMGA004893* | up | 1356 | 1370 | - | 10.6532 | 8.73E-05 | 1 | TAGGCCCGCTCGGGT |
| 2372 | mamo-L | *BGIBMGA004895* | up | 1515 | 1529 | + | 13.0242 | 7.77E-06 | 1 | GAGGCTGTTTTTGTT |
| 2373 | mamo-L | *BGIBMGA004899* | down | 873 | 887 | - | 10.9919 | 6.55E-05 | 1 | TAGGCATAATTTTTT |
| 2374 | mamo-L | *BGIBMGA004899* | up | 879 | 893 | - | 10.5323 | 9.63E-05 | 1 | TAGGCGGCGGTTTGG |
| 2375 | mamo-L | *BGIBMGA004900* | down | 1047 | 1061 | - | 11.6694 | 3.47E-05 | 1 | TAGGCCTTCCTTGTC |
| 2376 | mamo-L | *BGIBMGA004902* | up | 291 | 305 | - | 11.3065 | 4.91E-05 | 1 | AAGGCGGACATTTGT |
| 2377 | mamo-L | *BGIBMGA004909* | up | 1879 | 1893 | - | 11.5968 | 3.73E-05 | 1 | TAGGCGTAGTATCTT |
| 2378 | mamo-L | *BGIBMGA004910* | down | 1103 | 1117 | + | 12.379 | 1.66E-05 | 1 | TAGGCTTTGTGGGTT |
| 2379 | mamo-L | *BGIBMGA004910* | up | 805 | 819 | + | 10.75 | 8.06E-05 | 1 | TAGGCGTTGGTTTAA |
| 2380 | mamo-L | *BGIBMGA004910* | up | 661 | 675 | + | 10.6774 | 8.56E-05 | 1 | TAGGCGTAGGTTTAG |
| 2381 | mamo-L | *BGIBMGA004918* | down | 275 | 289 | - | 10.7581 | 8.00E-05 | 1 | AAGGCCAGCTTCGGT |
| 2382 | mamo-L | *BGIBMGA004919* | up | 1602 | 1616 | - | 13.9758 | 2.07E-06 | 1 | TAGGCTGGTTGTGCT |
| 2383 | mamo-L | *BGIBMGA004923* | up | 254 | 268 | + | 11.2339 | 5.26E-05 | 1 | TAGGCAGTCGTCGTC |
| 2384 | mamo-L | *BGIBMGA004928* | up | 1792 | 1806 | + | 10.5403 | 9.58E-05 | 1 | TAGGCTGTATGGTGT |
| 2385 | mamo-L | *BGIBMGA004931* | down | 1552 | 1566 | - | 11.4435 | 4.31E-05 | 1 | TAGGCTTGAGAGGCT |

| | | | | | | | | | | |
|---|---|---|---|---|---|---|---|---|---|---|
| 2386 | mamo-L | *BGIBMGA004931* | up | 425 | 439 | - | 10.9194 | 6.98E-05 | 1 | TAGGCTACTTTTTTT |
| 2387 | mamo-L | *BGIBMGA004932* | up | 1366 | 1380 | + | 13.5726 | 3.75E-06 | 1 | TAGGCTTACATTGCC |
| 2388 | mamo-L | *BGIBMGA004932* | up | 593 | 607 | + | 11.8387 | 2.94E-05 | 1 | TAGGCCTATGTTTTA |
| 2389 | mamo-L | *BGIBMGA004932* | down | 380 | 394 | - | 10.5726 | 9.33E-05 | 1 | TAGGCTAATATTTAT |
| 2390 | mamo-L | *BGIBMGA004934* | down | 1876 | 1890 | - | 11.0081 | 6.45E-05 | 1 | AAGGCTTACCTTGCT |
| 2391 | mamo-L | *BGIBMGA004938* | down | 517 | 531 | + | 10.9919 | 6.55E-05 | 1 | TAGGCAGATAGTGGT |
| 2392 | mamo-L | *BGIBMGA004941* | down | 1824 | 1838 | - | 10.8387 | 7.47E-05 | 1 | GAGGCTTACGTTAAT |
| 2393 | mamo-L | *BGIBMGA004951* | up | 456 | 470 | + | 12.4677 | 1.49E-05 | 1 | TAGGCTTCTATTGCT |
| 2394 | mamo-L | *BGIBMGA004953* | down | 1555 | 1569 | - | 12.871 | 9.35E-06 | 1 | TAGGCTTTTTTTATT |
| 2395 | mamo-L | *BGIBMGA004953* | up | 1397 | 1411 | - | 12.1855 | 2.06E-05 | 1 | TAGGCTGTATGTTGT |
| 2396 | mamo-L | *BGIBMGA004953* | up | 716 | 730 | + | 11.7097 | 3.34E-05 | 1 | TAGGCTGTGGGGTTT |
| 2397 | mamo-L | *BGIBMGA004953* | up | 121 | 135 | - | 11.0645 | 6.14E-05 | 1 | GAGGCTGACTTGTCT |
| 2398 | mamo-L | *BGIBMGA004959* | down | 1006 | 1020 | - | 10.629 | 8.91E-05 | 1 | TAGGCTCAATAGGTT |
| 2399 | mamo-L | *BGIBMGA004959* | down | 553 | 567 | - | 10.621 | 8.97E-05 | 1 | TAGGCGGGTGTGGTG |
| 2400 | mamo-L | *BGIBMGA004960* | up | 1685 | 1699 | - | 10.621 | 8.97E-05 | 1 | TAGGCGGGTGTGGTG |
| 2401 | mamo-L | *BGIBMGA004962* | up | 1330 | 1344 | + | 12.1855 | 2.06E-05 | 1 | TAGGCTGTATGTTGT |
| 2402 | mamo-L | *BGIBMGA004965* | up | 53 | 67 | - | 11.9839 | 2.54E-05 | 1 | AAGGCGGCGTTTGGT |
| 2403 | mamo-L | *BGIBMGA004967* | down | 707 | 721 | + | 12.5242 | 1.40E-05 | 1 | AAGGCTTGCGTTTTT |
| 2404 | mamo-L | *BGIBMGA004969* | up | 1829 | 1843 | + | 10.5726 | 9.33E-05 | 1 | TAGGCTAATATTTAT |
| 2405 | mamo-L | *BGIBMGA004973* | up | 1652 | 1666 | - | 10.5726 | 9.33E-05 | 1 | GAGGCCGACTATTGC |
| 2406 | mamo-L | *BGIBMGA004977* | down | 307 | 321 | - | 10.9919 | 6.55E-05 | 1 | TAGGCAGATAGTGGT |
| 2407 | mamo-L | *BGIBMGA004979* | up | 619 | 633 | + | 11.3387 | 4.77E-05 | 1 | TAGGCGTATATTTTC |
| 2408 | mamo-L | *BGIBMGA004980* | down | 867 | 881 | - | 11.3306 | 4.80E-05 | 1 | AAGGCCCGCGTCGCT |
| 2409 | mamo-L | *BGIBMGA004981* | up | 908 | 922 | - | 11.3306 | 4.80E-05 | 1 | AAGGCCCGCGTCGCT |
| 2410 | mamo-L | *BGIBMGA004982* | up | 1230 | 1244 | + | 11.1774 | 5.54E-05 | 1 | TAGGCTTAGTTCTTA |
| 2411 | mamo-L | *BGIBMGA004988* | down | 453 | 467 | - | 11.2903 | 4.99E-05 | 1 | TAGGCATTTTTCGTT |
| 2412 | mamo-L | *BGIBMGA004997* | up | 1119 | 1133 | - | 11.4758 | 4.18E-05 | 1 | AAGGCTGTCGTTGTA |
| 2413 | mamo-L | *BGIBMGA004998* | up | 438 | 452 | + | 10.5 | 9.90E-05 | 1 | GAGGCCCAGGTCGTC |
| 2414 | mamo-L | *BGIBMGA004999* | up | 213 | 227 | + | 10.879 | 7.22E-05 | 1 | AAGGCTCATTGTGTT |
| 2415 | mamo-L | *BGIBMGA005000* | down | 122 | 136 | - | 11.6129 | 3.66E-05 | 1 | TAGGCTTTTAATGAT |
| 2416 | mamo-L | *BGIBMGA005008* | down | 302 | 316 | + | 10.7177 | 8.28E-05 | 1 | TAGGCTTAGTTTAAA |
| 2417 | mamo-L | *BGIBMGA005008* | up | 1877 | 1891 | - | 10.6855 | 8.51E-05 | 1 | AAGGCTCTTTTCGTT |
| 2418 | mamo-L | *BGIBMGA005010* | up | 1412 | 1426 | - | 12.1694 | 2.09E-05 | 1 | TAGGCGTTTGTTCTT |
| 2419 | mamo-L | *BGIBMGA005010* | down | 175 | 189 | + | 11.7016 | 3.37E-05 | 1 | AAGGCTTAGATTGAT |
| 2420 | mamo-L | *BGIBMGA005011* | down | 1732 | 1746 | + | 12.3871 | 1.65E-05 | 1 | TAGGCGGAAGTTTGC |
| 2421 | mamo-L | *BGIBMGA005012* | up | 1139 | 1153 | + | 12.3871 | 1.65E-05 | 1 | TAGGCGGAAGTTTGC |
| 2422 | mamo-L | *BGIBMGA005013* | down | 591 | 605 | - | 11.0484 | 6.23E-05 | 1 | TAGGCTTGTTAGTTT |
| 2423 | mamo-L | *BGIBMGA005017* | up | 1817 | 1831 | - | 10.8629 | 7.33E-05 | 1 | AAGGCTCTTTTTTTT |
| 2424 | mamo-L | *BGIBMGA005019* | up | 1184 | 1198 | - | 13.0806 | 7.24E-06 | 1 | TAGGCTGTGGTGTCT |
| 2425 | mamo-L | *BGIBMGA005019* | down | 1016 | 1030 | + | 10.5645 | 9.40E-05 | 1 | TAGGCTCTTTGCGAT |
| 2426 | mamo-L | *BGIBMGA005023* | down | 685 | 699 | - | 10.5968 | 9.15E-05 | 1 | GAGGCTAGTTTTTGT |
| 2427 | mamo-L | *BGIBMGA005025* | up | 9 | 23 | + | 11.6935 | 3.39E-05 | 1 | AAGGCCGAAGTTGAT |
| 2428 | mamo-L | *BGIBMGA005026* | down | 940 | 954 | + | 10.7016 | 8.39E-05 | 1 | TGGGCGTTTGTTGCT |
| 2429 | mamo-L | *BGIBMGA005028* | down | 442 | 456 | + | 10.75 | 8.06E-05 | 1 | AAGGCAGAGGATGGT |
| 2430 | mamo-L | *BGIBMGA005029* | up | 1659 | 1673 | - | 10.9194 | 6.98E-05 | 1 | TAGGCTACTTTTTTT |
| 2431 | mamo-L | *BGIBMGA005032* | down | 537 | 551 | - | 12.0645 | 2.34E-05 | 1 | TAGGCGGGCTTCGGA |
| 2432 | mamo-L | *BGIBMGA005036* | down | 1861 | 1875 | - | 13.2339 | 6.02E-06 | 1 | TAGGCTGGTGTGGGC |
| 2433 | mamo-L | *BGIBMGA005038* | up | 1070 | 1084 | + | 11.5323 | 3.96E-05 | 1 | TAGGCGTGAGGTGAT |
| 2434 | mamo-L | *BGIBMGA005039* | down | 835 | 849 | + | 11.9839 | 2.54E-05 | 1 | TAGGCTTAGTCTTCT |
| 2435 | mamo-L | *BGIBMGA005039* | down | 1281 | 1295 | - | 11.1935 | 5.45E-05 | 1 | AAGGCTCTGTATGTT |
| 2436 | mamo-L | *BGIBMGA005040* | up | 1451 | 1465 | - | 12.4194 | 1.58E-05 | 1 | TAGGCCGACGGTTCC |
| 2437 | mamo-L | *BGIBMGA005040* | up | 129 | 143 | + | 11.9839 | 2.54E-05 | 1 | TAGGCTTAGTCTTCT |
| 2438 | mamo-L | *BGIBMGA005040* | up | 575 | 589 | - | 11.1935 | 5.45E-05 | 1 | AAGGCTCTGTATGTT |
| 2439 | mamo-L | *BGIBMGA005043* | up | 604 | 618 | - | 12.4677 | 1.49E-05 | 1 | TAGGCTTCTATTGCT |
| 2440 | mamo-L | *BGIBMGA005048* | down | 1352 | 1366 | + | 14.3145 | 1.24E-06 | 1 | TAGGCTTGATTTGTT |
| 2441 | mamo-L | *BGIBMGA005048* | up | 324 | 338 | - | 12.3629 | 1.69E-05 | 1 | TAGGCTTAATTTACT |
| 2442 | mamo-L | *BGIBMGA005048* | up | 1579 | 1593 | + | 11.8226 | 2.99E-05 | 1 | TAGGCCTGTATTATT |
| 2443 | mamo-L | *BGIBMGA005049* | down | 804 | 818 | + | 11.3387 | 4.77E-05 | 1 | AAGGCTTTGGATTGT |
| 2444 | mamo-L | *BGIBMGA005050* | up | 686 | 700 | + | 11.3387 | 4.77E-05 | 1 | AAGGCTTTGGATTGT |
| 2445 | mamo-L | *BGIBMGA005051* | down | 376 | 390 | + | 10.5 | 9.90E-05 | 1 | GAGGCCCAGGTCGTC |
| 2446 | mamo-L | *BGIBMGA005054* | up | 713 | 727 | + | 10.9839 | 6.59E-05 | 1 | AAGGCCTAGGACGTT |
| 2447 | mamo-L | *BGIBMGA005060* | up | 446 | 460 | - | 12.2339 | 1.95E-05 | 1 | AAGGCTTATGTTGTC |
| 2448 | mamo-L | *BGIBMGA005060* | up | 60 | 74 | + | 12.1452 | 2.15E-05 | 1 | TAGGCCTCCATTGGC |
| 2449 | mamo-L | *BGIBMGA005061* | down | 1284 | 1298 | + | 12.0403 | 2.40E-05 | 1 | TAGGCTTTTGTCAGT |
| 2450 | mamo-L | *BGIBMGA005063* | down | 950 | 964 | - | 11.4435 | 4.31E-05 | 1 | TAGGCTACGTTTTCT |
| 2451 | mamo-L | *BGIBMGA005063* | down | 1982 | 1996 | - | 11.0887 | 6.01E-05 | 1 | TAGGCTCTCTGTTTC |
| 2452 | mamo-L | *BGIBMGA005064* | down | 124 | 138 | + | 13.0887 | 7.18E-06 | 1 | TAGGCTTAGTTTAGC |
| 2453 | mamo-L | *BGIBMGA005065* | up | 26 | 40 | + | 11.5968 | 3.73E-05 | 1 | TAGGCATTTGTTTTT |
| 2454 | mamo-L | *BGIBMGA005067* | up | 1881 | 1895 | - | 11.1774 | 5.54E-05 | 1 | TAGGCCTCTGTCTTT |
| 2455 | mamo-L | *BGIBMGA005067* | down | 1577 | 1591 | - | 10.629 | 8.91E-05 | 1 | TAGGCCTGCAGCGCC |
| 2456 | mamo-L | *BGIBMGA005068* | down | 1487 | 1501 | + | 12.5081 | 1.43E-05 | 1 | TAGGCTTCTTTTTTT |
| 2457 | mamo-L | *BGIBMGA005068* | up | 502 | 516 | - | 10.629 | 8.91E-05 | 1 | TAGGCCTGCAGCGCC |
| 2458 | mamo-L | *BGIBMGA005069* | up | 373 | 387 | - | 12.7903 | 1.02E-05 | 1 | TAGGCCTTATTTTTT |
| 2459 | mamo-L | *BGIBMGA005069* | down | 1962 | 1976 | + | 10.7742 | 7.89E-05 | 1 | TAGGCCGAAAACGCT |
| 2460 | mamo-L | *BGIBMGA005071* | down | 1070 | 1084 | + | 13.7984 | 2.69E-06 | 1 | TAGGCTGTGTTTTAT |
| 2461 | mamo-L | *BGIBMGA005071* | up | 1103 | 1117 | - | 11.4194 | 4.42E-05 | 1 | TAGGCGTGCTGCGAT |
| 2462 | mamo-L | *BGIBMGA005072* | down | 1519 | 1533 | + | 10.621 | 8.97E-05 | 1 | TAGGCAAGCTGTGGT |
| 2463 | mamo-L | *BGIBMGA005073* | up | 1494 | 1508 | - | 11.1371 | 5.75E-05 | 1 | TAGGCCGCGATTACT |
| 2464 | mamo-L | *BGIBMGA005074* | down | 1056 | 1070 | + | 13.3871 | 4.90E-06 | 1 | TAGGCCAATTTTGCT |
| 2465 | mamo-L | *BGIBMGA005074* | up | 825 | 839 | - | 11.2903 | 4.99E-05 | 1 | TAGGCTGTATGGGCT |
| 2466 | mamo-L | *BGIBMGA005075* | up | 346 | 360 | + | 13.3871 | 4.90E-06 | 1 | TAGGCCAATTTTGCT |
| 2467 | mamo-L | *BGIBMGA005075* | up | 1057 | 1071 | - | 12.4355 | 1.55E-05 | 1 | TAGGCGTGTTTTATT |
| 2468 | mamo-L | *BGIBMGA005079* | down | 656 | 670 | + | 10.7097 | 8.33E-05 | 1 | TAGGCAGATTTGGAT |
| 2469 | mamo-L | *BGIBMGA005082* | up | 1567 | 1581 | + | 11.9516 | 2.63E-05 | 1 | TAGGCGCTGTTTATT |
| 2470 | mamo-L | *BGIBMGA005082* | up | 1704 | 1718 | - | 11.4032 | 4.49E-05 | 1 | TAGGCTAATATTTTT |
| 2471 | mamo-L | *BGIBMGA005084* | down | 117 | 131 | + | 10.5726 | 9.33E-05 | 1 | TAGGCTAATATTTAT |
| 2472 | mamo-L | *BGIBMGA005089* | up | 449 | 463 | + | 11.5323 | 3.96E-05 | 1 | TAGGCGTGAGGTGAT |
| 2473 | mamo-L | *BGIBMGA005091* | down | 259 | 273 | + | 12.7661 | 1.05E-05 | 1 | TAGGCTTTTTTTTCC |
| 2474 | mamo-L | *BGIBMGA005092* | up | 1550 | 1564 | - | 10.5726 | 9.33E-05 | 1 | TAGGCTAATATTTAT |
| 2475 | mamo-L | *BGIBMGA005093* | up | 1842 | 1856 | + | 13.0887 | 7.18E-06 | 1 | TAGGCTTCCTTTGCC |
| 2476 | mamo-L | *BGIBMGA005093* | up | 184 | 198 | - | 10.9194 | 6.98E-05 | 1 | AAGGCGCGCTTTTTT |
| 2477 | mamo-L | *BGIBMGA005094* | up | 978 | 992 | + | 13.9194 | 2.23E-06 | 1 | TAGGCTAATTTTGGT |

| | | | | | | | | | | |
|---|---|---|---|---|---|---|---|---|---|---|
| 2478 | mamo-L | *BGIBMGA005095* | down | 4 | 18 | + | 14.5242 | 8.74E-07 | 1 | TAGGCTTAAGTTGTT |
| 2479 | mamo-L | *BGIBMGA005095* | up | 715 | 729 | + | 10.5726 | 9.33E-05 | 1 | TAGGCTAATATTTAT |
| 2480 | mamo-L | *BGIBMGA005104* | down | 263 | 277 | - | 10.9194 | 6.98E-05 | 1 | TAGGCTACTTTTTTT |
| 2481 | mamo-L | *BGIBMGA005109* | up | 1207 | 1221 | - | 10.9032 | 7.07E-05 | 1 | TAGGCTTTAGTTTTA |
| 2482 | mamo-L | *BGIBMGA005110* | up | 287 | 301 | - | 11.3548 | 4.69E-05 | 1 | AAGGCTGATGATGAT |
| 2483 | mamo-L | *BGIBMGA005111* | down | 1838 | 1852 | + | 14.7258 | 5.83E-07 | 1 | TAGGCTTGCGTTGCC |
| 2484 | mamo-L | *BGIBMGA005116* | up | 1343 | 1357 | + | 11.121 | 5.84E-05 | 1 | AAGGCTGCAGTTGTT |
| 2485 | mamo-L | *BGIBMGA005117* | down | 958 | 972 | - | 10.7097 | 8.33E-05 | 1 | TAGGCAGATTTGGAT |
| 2486 | mamo-L | *BGIBMGA005117* | up | 33 | 47 | + | 10.6855 | 8.51E-05 | 1 | TAGGCGCCGTTTATT |
| 2487 | mamo-L | *BGIBMGA005119* | up | 1386 | 1400 | - | 10.5242 | 9.70E-05 | 1 | TAGGCGTGAGGTCGT |
| 2488 | mamo-L | *BGIBMGA005120* | up | 1432 | 1446 | - | 12.1452 | 2.15E-05 | 1 | TAGGCGTATGGTTTT |
| 2489 | mamo-L | *BGIBMGA005121* | down | 370 | 384 | + | 11.4435 | 4.31E-05 | 1 | TAGGCTTGAGAGGCT |
| 2490 | mamo-L | *BGIBMGA005122* | up | 1023 | 1037 | + | 11.4435 | 4.31E-05 | 1 | TAGGCTTGAGAGGCT |
| 2491 | mamo-L | *BGIBMGA005123* | up | 1648 | 1662 | + | 11.379 | 4.58E-05 | 1 | TAGGCCTCGTTTGTG |
| 2492 | mamo-L | *BGIBMGA005124* | up | 1119 | 1133 | + | 10.5242 | 9.70E-05 | 1 | AAGGCCTTCAGTGCT |
| 2493 | mamo-L | *BGIBMGA005126* | down | 1245 | 1259 | - | 10.8387 | 7.47E-05 | 1 | GAGGCTGTTTGTTGT |
| 2494 | mamo-L | *BGIBMGA005126* | down | 1692 | 1706 | - | 10.5726 | 9.33E-05 | 1 | TAGGCTCGAATTCGT |
| 2495 | mamo-L | *BGIBMGA005127* | up | 1801 | 1815 | - | 10.8387 | 7.47E-05 | 1 | GAGGCTGTTTGTTGT |
| 2496 | mamo-L | *BGIBMGA005128* | up | 1902 | 1916 | - | 11.5081 | 4.05E-05 | 1 | TAGGCGATTTTTTTT |
| 2497 | mamo-L | *BGIBMGA005129* | down | 258 | 272 | + | 12.4355 | 1.55E-05 | 1 | TAGGCTATTGTTGTC |
| 2498 | mamo-L | *BGIBMGA005129* | up | 1787 | 1801 | + | 11.0403 | 6.28E-05 | 1 | TAGGCCCGCGGGGGC |
| 2499 | mamo-L | *BGIBMGA005130* | down | 1813 | 1827 | - | 13.7984 | 2.69E-06 | 1 | TAGGCTGTGTTTTAT |
| 2500 | mamo-L | *BGIBMGA005130* | up | 1583 | 1597 | + | 12.4355 | 1.55E-05 | 1 | TAGGCTATTGTTGTC |
| 2501 | mamo-L | *BGIBMGA005132* | down | 1832 | 1846 | + | 12.1048 | 2.24E-05 | 1 | TAGGCCGAGATGGCC |
| 2502 | mamo-L | *BGIBMGA005132* | up | 1650 | 1664 | + | 10.5806 | 9.27E-05 | 1 | TAGGCTCACGACCGT |
| 2503 | mamo-L | *BGIBMGA005133* | down | 492 | 506 | - | 11.0323 | 6.32E-05 | 1 | TAGGCTTGCGATTGG |
| 2504 | mamo-L | *BGIBMGA005134* | down | 1748 | 1762 | + | 10.8952 | 7.12E-05 | 1 | TAGGCTATCGTTACC |
| 2505 | mamo-L | *BGIBMGA005134* | down | 1146 | 1160 | - | 10.75 | 8.06E-05 | 1 | TAGGCCTTTATTAAT |
| 2506 | mamo-L | *BGIBMGA005135* | down | 1230 | 1244 | - | 12.5081 | 1.43E-05 | 1 | TAGGCTTCTTTTTTT |
| 2507 | mamo-L | *BGIBMGA005140* | down | 252 | 266 | + | 11.1935 | 5.45E-05 | 1 | GAGGCTGGATTCGTT |
| 2508 | mamo-L | *BGIBMGA005142* | down | 632 | 646 | + | 10.9194 | 6.98E-05 | 1 | TAGGCTACTTTTTTT |
| 2509 | mamo-L | *BGIBMGA005142* | down | 773 | 787 | - | 10.8145 | 7.63E-05 | 1 | AAGGCCTGTTTTATT |
| 2510 | mamo-L | *BGIBMGA005148* | up | 1778 | 1792 | + | 11.7742 | 3.14E-05 | 1 | TAGGCCTTCAATGTC |
| 2511 | mamo-L | *BGIBMGA005151* | up | 362 | 376 | + | 12.3629 | 1.69E-05 | 1 | TAGGCTTAATTTACT |
| 2512 | mamo-L | *BGIBMGA005153* | up | 282 | 296 | + | 10.6613 | 8.67E-05 | 1 | TAGGCTTAGAATAGC |
| 2513 | mamo-L | *BGIBMGA005158* | up | 21 | 35 | + | 12.4677 | 1.49E-05 | 1 | TAGGCTTCTATTGCT |
| 2514 | mamo-L | *BGIBMGA005163* | down | 791 | 805 | - | 11.3548 | 4.69E-05 | 1 | AAGGCTGATGATGAT |
| 2515 | mamo-L | *BGIBMGA005164* | down | 661 | 675 | - | 11.1694 | 5.59E-05 | 1 | GAGGCTTTGATTGCC |
| 2516 | mamo-L | *BGIBMGA005170* | up | 34 | 48 | + | 12.1694 | 2.09E-05 | 1 | TAGGCGTTTGTTCTT |
| 2517 | mamo-L | *BGIBMGA005171* | down | 454 | 468 | - | 10.7177 | 8.28E-05 | 1 | TAGGCTTAGTTTAAA |
| 2518 | mamo-L | *BGIBMGA005172* | down | 896 | 910 | + | 11.4032 | 4.49E-05 | 1 | TAGGCTTTCTTTCAC |
| 2519 | mamo-L | *BGIBMGA005174* | up | 441 | 455 | - | 14.7661 | 5.44E-07 | 1 | TAGGCCGAGTTTTTT |
| 2520 | mamo-L | *BGIBMGA005176* | up | 152 | 166 | + | 10.629 | 8.91E-05 | 1 | GAGGCCGGCATTCGT |
| 2521 | mamo-L | *BGIBMGA005177* | down | 245 | 259 | - | 11.1613 | 5.63E-05 | 1 | TAGGCATGGTTTGTA |
| 2522 | mamo-L | *BGIBMGA005179* | down | 483 | 497 | + | 11.3065 | 4.91E-05 | 1 | AAGGCCAATTTTGTT |
| 2523 | mamo-L | *BGIBMGA005179* | up | 1861 | 1875 | + | 10.8145 | 7.63E-05 | 1 | TAGGCTGAGGATCGA |
| 2524 | mamo-L | *BGIBMGA005181* | down | 771 | 785 | - | 11.371 | 4.62E-05 | 1 | TAGGCTGTATAGGCT |
| 2525 | mamo-L | *BGIBMGA005182* | down | 1881 | 1895 | + | 12.1048 | 2.24E-05 | 1 | TAGGCCCATTATTGT |
| 2526 | mamo-L | *BGIBMGA005182* | up | 580 | 594 | + | 11.2661 | 5.10E-05 | 1 | TAGGCAAATTTTGCT |
| 2527 | mamo-L | *BGIBMGA005182* | up | 1474 | 1488 | - | 10.9274 | 6.92E-05 | 1 | AAGGCCTTGATTTTT |
| 2528 | mamo-L | *BGIBMGA005183* | up | 923 | 937 | + | 10.9194 | 6.98E-05 | 1 | TAGGCTACTTTTTTT |
| 2529 | mamo-L | *BGIBMGA005183* | up | 1119 | 1133 | - | 10.9194 | 6.98E-05 | 1 | TAGGCTACTTTTTTT |
| 2530 | mamo-L | *BGIBMGA005185* | down | 1577 | 1591 | - | 10.7903 | 7.79E-05 | 1 | AAGGCCTTCGTCTCT |
| 2531 | mamo-L | *BGIBMGA005186* | up | 323 | 337 | - | 11.7984 | 3.06E-05 | 1 | TAGGCTTTGCTTTTT |
| 2532 | mamo-L | *BGIBMGA005188* | down | 1380 | 1394 | + | 10.6694 | 8.62E-05 | 1 | TAGGCTTATTTTTAG |
| 2533 | mamo-L | *BGIBMGA005190* | up | 1564 | 1578 | + | 10.9194 | 6.98E-05 | 1 | TAGGCTACTTTTTTT |
| 2534 | mamo-L | *BGIBMGA005194* | down | 984 | 998 | + | 11.1694 | 5.59E-05 | 1 | TAGGCTTTTAACGTT |
| 2535 | mamo-L | *BGIBMGA005195* | up | 16 | 30 | - | 11.0403 | 6.28E-05 | 1 | TAGGCGTATATCTTT |
| 2536 | mamo-L | *BGIBMGA005195* | down | 1483 | 1497 | - | 10.6935 | 8.45E-05 | 1 | AAGGCTTTTATCGGT |
| 2537 | mamo-L | *BGIBMGA005200* | up | 1977 | 1991 | + | 10.9435 | 6.83E-05 | 1 | TAGGCTAGGATTACT |
| 2538 | mamo-L | *BGIBMGA005202* | up | 1159 | 1173 | + | 11.7661 | 3.16E-05 | 1 | GAGGCCTATTTTTTT |
| 2539 | mamo-L | *BGIBMGA005203* | down | 966 | 980 | + | 11.3548 | 4.69E-05 | 1 | AAGGCTGATGATGAT |
| 2540 | mamo-L | *BGIBMGA005204* | up | 1222 | 1236 | - | 11.5081 | 4.05E-05 | 1 | GAGGCGTTGTTTTTT |
| 2541 | mamo-L | *BGIBMGA005204* | up | 1007 | 1021 | - | 11.1774 | 5.54E-05 | 1 | TAGGCGGGGGTGGTG |
| 2542 | mamo-L | *BGIBMGA005209* | up | 684 | 698 | - | 10.7419 | 8.11E-05 | 1 | TAGGCAGTATATGTT |
| 2543 | mamo-L | *BGIBMGA005209* | down | 1788 | 1802 | - | 10.6371 | 8.85E-05 | 1 | TGGGCTTCCGTTGTT |
| 2544 | mamo-L | *BGIBMGA005210* | down | 1514 | 1528 | - | 10.6694 | 8.62E-05 | 1 | TAGGCTTATTTTTAG |
| 2545 | mamo-L | *BGIBMGA005215* | down | 475 | 489 | - | 12.6774 | 1.17E-05 | 1 | TAGGCTGGGGCTTGT |
| 2546 | mamo-L | *BGIBMGA005215* | down | 307 | 321 | + | 10.621 | 8.97E-05 | 1 | TAGGCTTTCTCTTCC |
| 2547 | mamo-L | *BGIBMGA005216* | down | 1849 | 1863 | - | 12.1048 | 2.24E-05 | 1 | TAGGCCCATTATTGT |
| 2548 | mamo-L | *BGIBMGA005219* | up | 1939 | 1953 | - | 11.379 | 4.58E-05 | 1 | AAGGCCTTCTTTAGT |
| 2549 | mamo-L | *BGIBMGA005221* | down | 743 | 757 | - | 10.7258 | 8.23E-05 | 1 | TAGGCAAAGTTTTCT |
| 2550 | mamo-L | *BGIBMGA005222* | down | 637 | 651 | + | 11.5968 | 3.73E-05 | 1 | TAGGCTTAGTATAAT |
| 2551 | mamo-L | *BGIBMGA005222* | up | 1709 | 1723 | + | 11.1048 | 5.92E-05 | 1 | TAGGCTTAAAGTTGT |
| 2552 | mamo-L | *BGIBMGA005223* | down | 563 | 577 | - | 12.0161 | 2.46E-05 | 1 | TAGGCCGTCGGTCCT |
| 2553 | mamo-L | *BGIBMGA005223* | up | 1474 | 1488 | - | 11.5887 | 3.75E-05 | 1 | GAGGCGTATTTTTGT |
| 2554 | mamo-L | *BGIBMGA005224* | up | 845 | 859 | - | 10.5403 | 9.58E-05 | 1 | TAGGCCGTTGTATTT |
| 2555 | mamo-L | *BGIBMGA005225* | up | 61 | 75 | + | 10.8629 | 7.33E-05 | 1 | TAGGCCTTCAGTGAC |
| 2556 | mamo-L | *BGIBMGA005226* | up | 1969 | 1983 | - | 12.3065 | 1.80E-05 | 1 | TAGGCCGGAGTCGTC |
| 2557 | mamo-L | *BGIBMGA005226* | up | 1558 | 1572 | - | 10.7097 | 8.33E-05 | 1 | TAGGCCTACAGCGCC |
| 2558 | mamo-L | *BGIBMGA005228* | up | 1625 | 1639 | - | 12.0645 | 2.34E-05 | 1 | TAGGCGGGCTTCGGA |
| 2559 | mamo-L | *BGIBMGA005228* | up | 505 | 519 | + | 10.9516 | 6.78E-05 | 1 | TAGGCAGATATTTCT |
| 2560 | mamo-L | *BGIBMGA005228* | up | 312 | 326 | + | 10.9194 | 6.98E-05 | 1 | TAGGCTACTTTTTTT |
| 2561 | mamo-L | *BGIBMGA005229* | up | 682 | 696 | + | 11.6452 | 3.56E-05 | 1 | GAGGCTCACTTTTTT |
| 2562 | mamo-L | *BGIBMGA005230* | down | 1507 | 1521 | - | 11.5968 | 3.73E-05 | 1 | TAGGCTTAGTATAAT |
| 2563 | mamo-L | *BGIBMGA005231* | down | 162 | 176 | - | 10.7339 | 8.17E-05 | 1 | GAGGCTGGTCTTGTT |
| 2564 | mamo-L | *BGIBMGA005232* | down | 1795 | 1809 | - | 11.3548 | 4.69E-05 | 1 | AAGGCTGATGATGAT |
| 2565 | mamo-L | *BGIBMGA005232* | down | 1629 | 1643 | - | 10.9194 | 6.98E-05 | 1 | TAGGCTACTTTTTTT |
| 2566 | mamo-L | *BGIBMGA005233* | up | 144 | 158 | + | 12.1613 | 2.11E-05 | 1 | TAGGCTTGTGTCGAC |
| 2567 | mamo-L | *BGIBMGA005233* | up | 927 | 941 | - | 11.6694 | 3.47E-05 | 1 | TAGGCCGGTGGCGCC |
| 2568 | mamo-L | *BGIBMGA005234* | down | 952 | 966 | + | 11.0806 | 6.05E-05 | 1 | AAGGCTTATTTTATT |
| 2569 | mamo-L | *BGIBMGA005234* | down | 1330 | 1344 | - | 10.5403 | 9.58E-05 | 1 | TAGGCTCTGATTTGA |

| | | | | | | | | | | |
|---|---|---|---|---|---|---|---|---|---|---|
| 2570 | mamo-L | *BGIBMGA005235* | up | 1054 | 1068 | - | 10.6048 | 9.09E-05 | 1 | AAGGCTTATTTGGAT |
| 2571 | mamo-L | *BGIBMGA005242* | up | 852 | 866 | + | 10.7258 | 8.23E-05 | 1 | TAGGCCAAGGACTGT |
| 2572 | mamo-L | *BGIBMGA005245* | down | 1769 | 1783 | + | 10.8871 | 7.17E-05 | 1 | AAGGCCTTGGTCTTT |
| 2573 | mamo-L | *BGIBMGA005245* | down | 1836 | 1850 | + | 10.7581 | 8.00E-05 | 1 | AAGGCTTTTTTTATT |
| 2574 | mamo-L | *BGIBMGA005247* | down | 389 | 403 | - | 10.5161 | 9.77E-05 | 1 | AAGGCTGGGGCTGAT |
| 2575 | mamo-L | *BGIBMGA005253* | up | 314 | 328 | - | 11.0242 | 6.36E-05 | 1 | AAGGCCTTGTTTTCC |
| 2576 | mamo-L | *BGIBMGA005257* | up | 360 | 374 | - | 10.6371 | 8.85E-05 | 1 | TAGGCTTTTAATGAC |
| 2577 | mamo-L | *BGIBMGA005258* | up | 1922 | 1936 | - | 11.3065 | 4.91E-05 | 1 | TAGGCTTTTGTTTTG |
| 2578 | mamo-L | *BGIBMGA005259* | up | 217 | 231 | - | 10.5726 | 9.33E-05 | 1 | TAGGCTAATATTTAT |
| 2579 | mamo-L | *BGIBMGA005260* | up | 488 | 502 | - | 10.9435 | 6.83E-05 | 1 | AAGGCGGATTTTTGC |
| 2580 | mamo-L | *BGIBMGA005261* | down | 1306 | 1320 | - | 11 | 6.50E-05 | 1 | TAGGCTGGCCTCTTT |
| 2581 | mamo-L | *BGIBMGA005262* | down | 1362 | 1376 | - | 11.8065 | 3.03E-05 | 1 | TAGGCCTATGTTTCA |
| 2582 | mamo-L | *BGIBMGA005262* | up | 660 | 674 | + | 11.0484 | 6.23E-05 | 1 | TAGGCCAATGGTTTT |
| 2583 | mamo-L | *BGIBMGA005263* | up | 147 | 161 | + | 11.3871 | 4.55E-05 | 1 | TAGGCTATATTTTTT |
| 2584 | mamo-L | *BGIBMGA005267* | down | 1370 | 1384 | - | 10.6129 | 9.03E-05 | 1 | TAGGCGTTCAAGGTT |
| 2585 | mamo-L | *BGIBMGA005268* | down | 1092 | 1106 | + | 11.6452 | 3.56E-05 | 1 | TAGGCTGTTAATTTT |
| 2586 | mamo-L | *BGIBMGA005269* | up | 197 | 211 | - | 10.5726 | 9.33E-05 | 1 | TAGGCTAATATTTAT |
| 2587 | mamo-L | *BGIBMGA005277* | up | 666 | 680 | + | 12.4194 | 1.58E-05 | 1 | AAGGCTTGGATTGCT |
| 2588 | mamo-L | *BGIBMGA005277* | up | 1863 | 1877 | - | 11.7581 | 3.18E-05 | 1 | TAGGCCTGATTCTTT |
| 2589 | mamo-L | *BGIBMGA005278* | up | 1900 | 1914 | + | 10.5726 | 9.33E-05 | 1 | TAGGCTAATATTTAT |
| 2590 | mamo-L | *BGIBMGA005280* | up | 160 | 174 | + | 11.6452 | 3.56E-05 | 1 | AAGGCTTACTTTTAT |
| 2591 | mamo-L | *BGIBMGA005286* | down | 293 | 307 | + | 10.9194 | 6.98E-05 | 1 | TAGGCCTTATAGGTT |
| 2592 | mamo-L | *BGIBMGA005289* | up | 1678 | 1692 | - | 11.4919 | 4.11E-05 | 1 | TAGGCTTGGGTCGAA |
| 2593 | mamo-L | *BGIBMGA005292* | down | 1806 | 1820 | - | 11.2742 | 5.06E-05 | 1 | TAGGCCCCGGTTACT |
| 2594 | mamo-L | *BGIBMGA005296* | up | 567 | 581 | - | 13.1129 | 6.98E-06 | 1 | TAGGCTTGTTTTATT |
| 2595 | mamo-L | *BGIBMGA005296* | up | 1338 | 1352 | - | 11.629 | 3.61E-05 | 1 | TAGGCGAGCGTTCGT |
| 2596 | mamo-L | *BGIBMGA005297* | down | 278 | 292 | + | 11.6129 | 3.66E-05 | 1 | TAGGCTTAGATTGAA |
| 2597 | mamo-L | *BGIBMGA005298* | up | 739 | 753 | - | 12.3226 | 1.77E-05 | 1 | TAGGCAGGTTTTTGT |
| 2598 | mamo-L | *BGIBMGA005299* | down | 133 | 147 | + | 12.4355 | 1.55E-05 | 1 | TAGGCGTTTTTTTGC |
| 2599 | mamo-L | *BGIBMGA005307* | down | 767 | 781 | + | 10.8952 | 7.12E-05 | 1 | TAGGCTGTGTAAGCT |
| 2600 | mamo-L | *BGIBMGA005308* | up | 1687 | 1701 | + | 10.8952 | 7.12E-05 | 1 | TAGGCTGTGTAAGCT |
| 2601 | mamo-L | *BGIBMGA005308* | down | 974 | 988 | - | 10.5645 | 9.40E-05 | 1 | TAGGCTGAAGTCCAT |
| 2602 | mamo-L | *BGIBMGA005308* | down | 1260 | 1274 | - | 10.5645 | 9.40E-05 | 1 | TAGGCTGAAGTCCAT |
| 2603 | mamo-L | *BGIBMGA005310* | up | 297 | 311 | + | 12.9516 | 8.46E-06 | 1 | TAGGCTTCTGTTTGT |
| 2604 | mamo-L | *BGIBMGA005311* | down | 1246 | 1260 | - | 10.8387 | 7.47E-05 | 1 | TAGGCGTAGTTTACA |
| 2605 | mamo-L | *BGIBMGA005312* | up | 464 | 478 | + | 10.6129 | 9.03E-05 | 1 | TAGGCATATGTCTCT |
| 2606 | mamo-L | *BGIBMGA005318* | down | 1689 | 1703 | - | 13.3145 | 5.41E-06 | 1 | AAGGCGGTCGTTGGT |
| 2607 | mamo-L | *BGIBMGA005321* | down | 468 | 482 | + | 15.129 | 2.58E-07 | 1 | TAGGCGGATTTTGGT |
| 2608 | mamo-L | *BGIBMGA005326* | down | 1787 | 1801 | - | 11.4032 | 4.49E-05 | 1 | TAGGCTAATATTTTT |
| 2609 | mamo-L | *BGIBMGA005330* | up | 1947 | 1961 | + | 11.4758 | 4.18E-05 | 1 | TAGGCCTTTAATTGT |
| 2610 | mamo-L | *BGIBMGA005330* | up | 1157 | 1171 | - | 10.7177 | 8.28E-05 | 1 | TGGGCTGTCTATGCT |
| 2611 | mamo-L | *BGIBMGA005331* | down | 1129 | 1143 | + | 11.4194 | 4.42E-05 | 1 | TAGGCGATCGTTGAC |
| 2612 | mamo-L | *BGIBMGA005331* | up | 1662 | 1676 | + | 10.9032 | 7.07E-05 | 1 | TAGGCGAGGTGTTTT |
| 2613 | mamo-L | *BGIBMGA005331* | down | 1552 | 1566 | - | 10.5726 | 9.33E-05 | 1 | TAGGCTAATATTTAT |
| 2614 | mamo-L | *BGIBMGA005337* | up | 150 | 164 | - | 11.3306 | 4.80E-05 | 1 | TAGGCCGTCGTGGCG |
| 2615 | mamo-L | *BGIBMGA005342* | down | 1928 | 1942 | - | 15.8065 | 6.11E-08 | 1 | TAGGCTGATTTTGGT |
| 2616 | mamo-L | *BGIBMGA005343* | down | 431 | 445 | - | 12.2419 | 1.94E-05 | 1 | TAGGCTTGATTTTTC |
| 2617 | mamo-L | *BGIBMGA005347* | down | 627 | 641 | + | 10.629 | 8.91E-05 | 1 | TAGGCTGTTAGTACT |
| 2618 | mamo-L | *BGIBMGA005348* | down | 521 | 535 | + | 11.9597 | 2.61E-05 | 1 | TAGGCGCATTTTTGC |
| 2619 | mamo-L | *BGIBMGA005350* | up | 610 | 624 | + | 11.2339 | 5.26E-05 | 1 | TAGGCTACTTTTTGT |
| 2620 | mamo-L | *BGIBMGA005352* | up | 692 | 706 | - | 10.75 | 8.06E-05 | 1 | AAGGCTTGCTTGTTT |
| 2621 | mamo-L | *BGIBMGA005353* | up | 229 | 243 | - | 12.9919 | 8.10E-06 | 1 | TAGGCCGGTTTTGCA |
| 2622 | mamo-L | *BGIBMGA005353* | up | 376 | 390 | + | 10.8226 | 7.58E-05 | 1 | TAGGCCACTTTTGCC |
| 2623 | mamo-L | *BGIBMGA005356* | down | 1198 | 1212 | - | 11.6935 | 3.39E-05 | 1 | GAGGCTATGTTTGTT |
| 2624 | mamo-L | *BGIBMGA005357* | up | 1345 | 1359 | - | 10.8548 | 7.38E-05 | 1 | AAGGCTGTTATTTTT |
| 2625 | mamo-L | *BGIBMGA005357* | down | 1658 | 1672 | + | 10.7097 | 8.33E-05 | 1 | TAGGCGCAGGTTTTG |
| 2626 | mamo-L | *BGIBMGA005362* | down | 181 | 195 | + | 12.9919 | 8.10E-06 | 1 | TAGGCCGGTTTTGCA |
| 2627 | mamo-L | *BGIBMGA005362* | down | 494 | 508 | + | 11.1855 | 5.50E-05 | 1 | TAGGCCTCTTTTGCA |
| 2628 | mamo-L | *BGIBMGA005363* | up | 1232 | 1246 | + | 11.4194 | 4.42E-05 | 1 | TAGGCTGAGTTTACG |
| 2629 | mamo-L | *BGIBMGA005364* | up | 1962 | 1976 | - | 11.0403 | 6.28E-05 | 1 | AAGGCTAAGTTTGCC |
| 2630 | mamo-L | *BGIBMGA005367* | up | 452 | 466 | - | 12.4677 | 1.49E-05 | 1 | TAGGCTTCTATTGCT |
| 2631 | mamo-L | *BGIBMGA005370* | down | 1841 | 1855 | + | 14.7258 | 5.83E-07 | 1 | TAGGCTTGCGTTGCC |
| 2632 | mamo-L | *BGIBMGA005370* | up | 163 | 177 | - | 10.8387 | 7.47E-05 | 1 | TAGGGTTTGGTTGCT |
| 2633 | mamo-L | *BGIBMGA005371* | down | 1530 | 1544 | - | 11.3065 | 4.91E-05 | 1 | TAGGCTTTTGTTTTG |
| 2634 | mamo-L | *BGIBMGA005371* | up | 1832 | 1846 | + | 10.879 | 7.22E-05 | 1 | TAGGCTTAGGTCTCG |
| 2635 | mamo-L | *BGIBMGA005373* | up | 488 | 502 | + | 12.6532 | 1.20E-05 | 1 | AAGGCTTTGTTTGGC |
| 2636 | mamo-L | *BGIBMGA005373* | up | 859 | 873 | - | 10.5403 | 9.58E-05 | 1 | GAGGCTTTTTTTCCT |
| 2637 | mamo-L | *BGIBMGA005375* | down | 1678 | 1692 | - | 10.6371 | 8.85E-05 | 1 | GAGGCTTTGGTGTCT |
| 2638 | mamo-L | *BGIBMGA005378* | down | 1205 | 1219 | + | 14.0403 | 1.87E-06 | 1 | TAGGCCGACTTTTGC |
| 2639 | mamo-L | *BGIBMGA005378* | down | 1087 | 1101 | + | 11.1855 | 5.50E-05 | 1 | AAGGCTGTCTGCGGT |
| 2640 | mamo-L | *BGIBMGA005379* | down | 1375 | 1389 | - | 13.4597 | 4.43E-06 | 1 | TAGGCCGTGATTGTC |
| 2641 | mamo-L | *BGIBMGA005381* | down | 1202 | 1216 | + | 13.1371 | 6.76E-06 | 1 | TAGGCCTAGAATGTT |
| 2642 | mamo-L | *BGIBMGA005386* | up | 1476 | 1490 | + | 10.9194 | 6.98E-05 | 1 | TAGGCTACTTTTTTT |
| 2643 | mamo-L | *BGIBMGA005389* | up | 1945 | 1959 | + | 10.9355 | 6.88E-05 | 1 | GAGGCTGAGTAGGTT |
| 2644 | mamo-L | *BGIBMGA005391* | down | 416 | 430 | - | 10.4919 | 9.96E-05 | 1 | TAGGCATTTTTTTTC |
| 2645 | mamo-L | *BGIBMGA005399* | up | 1704 | 1718 | + | 13.7742 | 2.79E-06 | 1 | TAGGCGGGTGGTGGT |
| 2646 | mamo-L | *BGIBMGA005401* | up | 1315 | 1329 | + | 11.121 | 5.84E-05 | 1 | TAGGCGGCTTATTGT |
| 2647 | mamo-L | *BGIBMGA005404* | down | 1056 | 1070 | + | 11.9435 | 2.65E-05 | 1 | TAGGCTGGTTCTGAT |
| 2648 | mamo-L | *BGIBMGA005408* | down | 83 | 97 | + | 10.4919 | 9.96E-05 | 1 | TAGGCATTTTTTTTC |
| 2649 | mamo-L | *BGIBMGA005411* | up | 180 | 194 | - | 10.9194 | 6.98E-05 | 1 | TAGGCTACTTTTTTT |
| 2650 | mamo-L | *BGIBMGA005417* | down | 169 | 183 | - | 12.1855 | 2.06E-05 | 1 | TAGGCTATTTTTTTT |
| 2651 | mamo-L | *BGIBMGA005423* | up | 894 | 908 | + | 13.9113 | 2.26E-06 | 1 | TAGGCGTACTTTTTT |
| 2652 | mamo-L | *BGIBMGA005427* | down | 1986 | 2000 | + | 10.5161 | 9.77E-05 | 1 | AAGGCGTGGTGCGGT |
| 2653 | mamo-L | *BGIBMGA005428* | up | 1968 | 1982 | + | 10.5161 | 9.77E-05 | 1 | AAGGCGTGGTGCGGT |
| 2654 | mamo-L | *BGIBMGA005429* | up | 671 | 685 | + | 10.6371 | 8.85E-05 | 1 | GAGGCTTTGGTGTCT |
| 2655 | mamo-L | *BGIBMGA005432* | up | 416 | 430 | - | 10.7742 | 7.89E-05 | 1 | TGGGCTGATTTTTCT |
| 2656 | mamo-L | *BGIBMGA005433* | down | 1143 | 1157 | + | 10.7984 | 7.74E-05 | 1 | TAGGCCGTAATTGGG |
| 2657 | mamo-L | *BGIBMGA005434* | up | 1384 | 1398 | + | 12.7258 | 1.11E-05 | 1 | TAGGCGTGCTTTGTA |
| 2658 | mamo-L | *BGIBMGA005434* | up | 115 | 129 | - | 12.6532 | 1.20E-05 | 1 | AAGGCTTTGTTTGGC |
| 2659 | mamo-L | *BGIBMGA005436* | down | 1633 | 1647 | + | 11.3065 | 4.91E-05 | 1 | TAGGCTTTTGTTTTG |
| 2660 | mamo-L | *BGIBMGA005439* | up | 1598 | 1612 | + | 12.4677 | 1.49E-05 | 1 | TAGGCTTCTATTGCT |
| 2661 | mamo-L | *BGIBMGA005441* | down | 1259 | 1273 | - | 13.7258 | 3.01E-06 | 1 | TAGGCCTAGTTTCGT |

| | | | | | | | | | | |
|---|---|---|---|---|---|---|---|---|---|---|
| 2662 | mamo-L | *BGIBMGA005442* | down | 813 | 827 | + | 13.4597 | 4.43E-06 | 1 | TAGGCTGACGAGGGT |
| 2663 | mamo-L | *BGIBMGA005442* | down | 156 | 170 | + | 10.9194 | 6.98E-05 | 1 | TAGGCAGTGTGTTTT |
| 2664 | mamo-L | *BGIBMGA005444* | up | 902 | 916 | - | 10.871 | 7.28E-05 | 1 | TAGGCGATGGATTTT |
| 2665 | mamo-L | *BGIBMGA005444* | down | 1661 | 1675 | - | 10.7823 | 7.84E-05 | 1 | TAGGCCCGTGATCTT |
| 2666 | mamo-L | *BGIBMGA005445* | up | 1472 | 1486 | + | 10.9194 | 6.98E-05 | 1 | AAGGCTTTTTTTCGT |
| 2667 | mamo-L | *BGIBMGA005447* | up | 832 | 846 | - | 10.5726 | 9.33E-05 | 1 | TAGGCTAATATTTAT |
| 2668 | mamo-L | *BGIBMGA005448* | down | 787 | 801 | + | 12.7339 | 1.10E-05 | 1 | TAGGCTGTTATGGGT |
| 2669 | mamo-L | *BGIBMGA005448* | down | 586 | 600 | - | 11.8548 | 2.89E-05 | 1 | TAGGCTGGGGTGGTG |
| 2670 | mamo-L | *BGIBMGA005450* | down | 668 | 682 | + | 11.6694 | 3.47E-05 | 1 | AAGGCGTTGGTTTTT |
| 2671 | mamo-L | *BGIBMGA005450* | down | 1562 | 1576 | - | 11.5403 | 3.93E-05 | 1 | TAGGCTGCCGTTTGA |
| 2672 | mamo-L | *BGIBMGA005451* | down | 465 | 479 | - | 10.9597 | 6.73E-05 | 1 | TAGGCGATGATTTTT |
| 2673 | mamo-L | *BGIBMGA005455* | up | 452 | 466 | + | 10.75 | 8.06E-05 | 1 | AAGGCTTGCTTGTTT |
| 2674 | mamo-L | *BGIBMGA005457* | up | 1930 | 1944 | - | 13.9758 | 2.07E-06 | 1 | TAGGCCTCGTTTGTT |
| 2675 | mamo-L | *BGIBMGA005457* | down | 337 | 351 | + | 11.5726 | 3.81E-05 | 1 | TAGGCTGTGAGGGTT |
| 2676 | mamo-L | *BGIBMGA005457* | up | 801 | 815 | + | 11.1855 | 5.50E-05 | 1 | TAGGCTTACGTTACG |
| 2677 | mamo-L | *BGIBMGA005457* | up | 93 | 107 | - | 10.8226 | 7.58E-05 | 1 | TAGGCTTTATATTAT |
| 2678 | mamo-L | *BGIBMGA005463* | up | 258 | 272 | - | 10.7661 | 7.95E-05 | 1 | AAGGCGGGTGATGGC |
| 2679 | mamo-L | *BGIBMGA005464* | down | 325 | 339 | + | 11.1694 | 5.59E-05 | 1 | TAGGCTGATGATCTC |
| 2680 | mamo-L | *BGIBMGA005469* | down | 941 | 955 | + | 11.2742 | 5.06E-05 | 1 | TAGGCATGCGTTCTT |
| 2681 | mamo-L | *BGIBMGA005473* | down | 1303 | 1317 | - | 11.4194 | 4.42E-05 | 1 | TAGGCGATCGTTGAC |
| 2682 | mamo-L | *BGIBMGA005473* | down | 880 | 894 | + | 10.5726 | 9.33E-05 | 1 | TAGGCTAATATTTAT |
| 2683 | mamo-L | *BGIBMGA005474* | down | 238 | 252 | + | 10.5726 | 9.33E-05 | 1 | TAGGCTAATATTTAT |
| 2684 | mamo-L | *BGIBMGA005475* | up | 1889 | 1903 | + | 10.9355 | 6.88E-05 | 1 | GAGGCTTGTTTTACT |
| 2685 | mamo-L | *BGIBMGA005479* | down | 1736 | 1750 | - | 11.8306 | 2.97E-05 | 1 | TAGGCCTTATTCTGT |
| 2686 | mamo-L | *BGIBMGA005480* | up | 209 | 223 | - | 11.4758 | 4.18E-05 | 1 | TAGGCCTTTAATTGT |
| 2687 | mamo-L | *BGIBMGA005480* | up | 146 | 160 | + | 10.9435 | 6.83E-05 | 1 | AAGGCGGGGATTTCT |
| 2688 | mamo-L | *BGIBMGA005480* | up | 999 | 1013 | + | 10.7177 | 8.28E-05 | 1 | TGGGCTGTCTATGCT |
| 2689 | mamo-L | *BGIBMGA005487* | up | 619 | 633 | + | 11.2742 | 5.06E-05 | 1 | TAGGCATGCGTTCTT |
| 2690 | mamo-L | *BGIBMGA005490* | up | 557 | 571 | - | 12.2016 | 2.03E-05 | 1 | AAGGCCGTAGTTGTT |
| 2691 | mamo-L | *BGIBMGA005493* | up | 1310 | 1324 | + | 11.9516 | 2.63E-05 | 1 | TAGGCTGTGTGCTTT |
| 2692 | mamo-L | *BGIBMGA005496* | up | 1064 | 1078 | + | 11.1613 | 5.63E-05 | 1 | TAGGCCGGTATCAGT |
| 2693 | mamo-L | *BGIBMGA005496* | up | 1920 | 1934 | - | 11.0403 | 6.28E-05 | 1 | TAGGCGTATTTAGTT |
| 2694 | mamo-L | *BGIBMGA005498* | up | 1726 | 1740 | - | 11.2419 | 5.21E-05 | 1 | TAGGCTTTTCTTTTT |
| 2695 | mamo-L | *BGIBMGA005498* | up | 1310 | 1324 | + | 10.9194 | 6.98E-05 | 1 | TAGGCCAACATTGGA |
| 2696 | mamo-L | *BGIBMGA005501* | down | 1372 | 1386 | + | 10.5645 | 9.40E-05 | 1 | TAGGCTGAAGTCCAT |
| 2697 | mamo-L | *BGIBMGA005501* | down | 1658 | 1672 | + | 10.5645 | 9.40E-05 | 1 | TAGGCTGAAGTCCAT |
| 2698 | mamo-L | *BGIBMGA005503* | down | 1173 | 1187 | - | 11.2581 | 5.14E-05 | 1 | TAGGCATTCGTTTAT |
| 2699 | mamo-L | *BGIBMGA005506* | down | 1446 | 1460 | + | 12.5968 | 1.28E-05 | 1 | TAGGCATACTTTTGT |
| 2700 | mamo-L | *BGIBMGA005510* | up | 385 | 399 | + | 13.3952 | 4.84E-06 | 1 | TAGGCTTTGTTTACT |
| 2701 | mamo-L | *BGIBMGA005517* | down | 849 | 863 | - | 13.9435 | 2.15E-06 | 1 | TAGGCCAAGTTTGCT |
| 2702 | mamo-L | *BGIBMGA005518* | down | 678 | 692 | + | 12.7984 | 1.02E-05 | 1 | TAGGCTTGAGTGGTT |
| 2703 | mamo-L | *BGIBMGA005521* | up | 412 | 426 | - | 10.9194 | 6.98E-05 | 1 | TAGGCTACTTTTTTT |
| 2704 | mamo-L | *BGIBMGA005525* | up | 1146 | 1160 | - | 10.5726 | 9.33E-05 | 1 | TAGGCTAATATTTAT |
| 2705 | mamo-L | *BGIBMGA005527* | down | 312 | 326 | - | 10.75 | 8.06E-05 | 1 | GAGGCCGATTGTGCC |
| 2706 | mamo-L | *BGIBMGA005529* | up | 1796 | 1810 | + | 12.6532 | 1.20E-05 | 1 | TAGGCAGAGTGTGGT |
| 2707 | mamo-L | *BGIBMGA005531* | down | 827 | 841 | + | 10.8629 | 7.33E-05 | 1 | AAGGCTTAGGTTTAC |
| 2708 | mamo-L | *BGIBMGA005536* | down | 90 | 104 | + | 12.4516 | 1.53E-05 | 1 | TAGGCTTAGGTCCTT |
| 2709 | mamo-L | *BGIBMGA005537* | up | 754 | 768 | - | 12.5565 | 1.35E-05 | 1 | TAGGCTGAGCTCGGT |
| 2710 | mamo-L | *BGIBMGA005541* | down | 655 | 669 | + | 11.0968 | 5.97E-05 | 1 | GAGGCCGATTGTGGC |
| 2711 | mamo-L | *BGIBMGA005543* | up | 1799 | 1813 | - | 11.3548 | 4.69E-05 | 1 | AAGGCTGATGATGAT |
| 2712 | mamo-L | *BGIBMGA005543* | down | 206 | 220 | - | 10.6694 | 8.62E-05 | 1 | TAGGCTGACTTACGT |
| 2713 | mamo-L | *BGIBMGA005548* | up | 732 | 746 | - | 10.621 | 8.97E-05 | 1 | TAGGCTACCATTTGT |
| 2714 | mamo-L | *BGIBMGA005553* | down | 355 | 369 | - | 10.7339 | 8.17E-05 | 1 | TAGGCTGATGGGCGT |
| 2715 | mamo-L | *BGIBMGA005554* | up | 84 | 98 | - | 10.7339 | 8.17E-05 | 1 | TAGGCTGATGGGCGT |
| 2716 | mamo-L | *BGIBMGA005557* | down | 1797 | 1811 | + | 10.6855 | 8.51E-05 | 1 | TAGGCTGAGTTAGGA |
| 2717 | mamo-L | *BGIBMGA005560* | down | 274 | 288 | - | 12.9274 | 8.74E-06 | 1 | GAGGCTGGCTTTGAT |
| 2718 | mamo-L | *BGIBMGA005563* | up | 634 | 648 | - | 11.75 | 3.21E-05 | 1 | GAGGCCAGGTTTGTT |
| 2719 | mamo-L | *BGIBMGA005564* | down | 473 | 487 | + | 11.9194 | 2.72E-05 | 1 | TAGGCCGTCTGTCTT |
| 2720 | mamo-L | *BGIBMGA005564* | down | 101 | 115 | + | 11.4355 | 4.34E-05 | 1 | GAGGCGTTCATTGTT |
| 2721 | mamo-L | *BGIBMGA005566* | down | 359 | 373 | + | 10.5403 | 9.58E-05 | 1 | TAGGCCATATTTTGC |
| 2722 | mamo-L | *BGIBMGA005567* | up | 578 | 592 | + | 13.0081 | 7.94E-06 | 1 | TAGGCTTGCTACGTT |
| 2723 | mamo-L | *BGIBMGA005567* | down | 241 | 255 | + | 10.5726 | 9.33E-05 | 1 | TAGGCTAATATTTAT |
| 2724 | mamo-L | *BGIBMGA005569* | up | 807 | 821 | + | 10.8629 | 7.33E-05 | 1 | GAGGCTAGCTTTGCC |
| 2725 | mamo-L | *BGIBMGA005572* | up | 790 | 804 | + | 10.5 | 9.90E-05 | 1 | GAGGCCCAGGTCGTC |
| 2726 | mamo-L | *BGIBMGA005574* | down | 650 | 664 | + | 12.1855 | 2.06E-05 | 1 | TAGGCTATTTTTTTT |
| 2727 | mamo-L | *BGIBMGA005574* | down | 1126 | 1140 | + | 11.5968 | 3.73E-05 | 1 | TAGGCTAGTTTTTAT |
| 2728 | mamo-L | *BGIBMGA005575* | up | 613 | 627 | + | 12.1855 | 2.06E-05 | 1 | TAGGCTATTTTTTTT |
| 2729 | mamo-L | *BGIBMGA005575* | up | 1089 | 1103 | + | 11.5968 | 3.73E-05 | 1 | TAGGCTAGTTTTTAT |
| 2730 | mamo-L | *BGIBMGA005576* | up | 388 | 402 | - | 10.621 | 8.97E-05 | 1 | AAGGCGATTGTTGTT |
| 2731 | mamo-L | *BGIBMGA005578* | down | 818 | 832 | + | 12.6129 | 1.26E-05 | 1 | TAGGCTTTCATGGTT |
| 2732 | mamo-L | *BGIBMGA005579* | up | 1392 | 1406 | + | 12.6129 | 1.26E-05 | 1 | TAGGCTTTCATGGTT |
| 2733 | mamo-L | *BGIBMGA005579* | down | 1750 | 1764 | + | 11.4032 | 4.49E-05 | 1 | GAGGCAGTGGTTGTT |
| 2734 | mamo-L | *BGIBMGA005580* | up | 1791 | 1805 | + | 11.4032 | 4.49E-05 | 1 | GAGGCAGTGGTTGTT |
| 2735 | mamo-L | *BGIBMGA005581* | up | 976 | 990 | + | 13.3145 | 5.41E-06 | 1 | AAGGCTTTGTTTGTT |
| 2736 | mamo-L | *BGIBMGA005584* | up | 296 | 310 | - | 10.5968 | 9.15E-05 | 1 | GAGGCTAGTTTTTGT |
| 2737 | mamo-L | *BGIBMGA005585* | down | 1855 | 1869 | + | 12.8306 | 9.76E-06 | 1 | TAGGCGGTCTTTCTT |
| 2738 | mamo-L | *BGIBMGA005587* | up | 635 | 649 | + | 10.7823 | 7.84E-05 | 1 | TAGGCCCCCTATGCC |
| 2739 | mamo-L | *BGIBMGA005587* | up | 315 | 329 | + | 10.7581 | 8.00E-05 | 1 | TAGGCGACGATTGCT |
| 2740 | mamo-L | *BGIBMGA005587* | up | 298 | 312 | + | 10.5161 | 9.77E-05 | 1 | TAGCCTGGCTTCGTT |
| 2741 | mamo-L | *BGIBMGA005588* | down | 1290 | 1304 | + | 11.5806 | 3.78E-05 | 1 | AAGGCTTTTGTCGCT |
| 2742 | mamo-L | *BGIBMGA005589* | down | 896 | 910 | + | 10.5645 | 9.40E-05 | 1 | TAGGCTCAATTTGAA |
| 2743 | mamo-L | *BGIBMGA005591* | down | 908 | 922 | + | 11.5968 | 3.73E-05 | 1 | TAGGCGGAGTGCTTT |
| 2744 | mamo-L | *BGIBMGA005600* | up | 1791 | 1805 | - | 11.0968 | 5.97E-05 | 1 | TGGGCGTTCTTTGTT |
| 2745 | mamo-L | *BGIBMGA005600* | down | 519 | 533 | + | 10.6371 | 8.85E-05 | 1 | TAGGCCTTATATTAT |
| 2746 | mamo-L | *BGIBMGA005600* | up | 1613 | 1627 | - | 10.5645 | 9.40E-05 | 1 | TAGGCTTATTACACT |
| 2747 | mamo-L | *BGIBMGA005601* | down | 1428 | 1442 | + | 10.629 | 8.91E-05 | 1 | TAGGCCTGCAGCGCC |
| 2748 | mamo-L | *BGIBMGA005602* | up | 1163 | 1177 | + | 10.629 | 8.91E-05 | 1 | TAGGCCTGCAGCGCC |
| 2749 | mamo-L | *BGIBMGA005604* | down | 250 | 264 | + | 12.0887 | 2.29E-05 | 1 | TAGGCTCAGATTTGC |
| 2750 | mamo-L | *BGIBMGA005605* | down | 1956 | 1970 | + | 11.0887 | 6.01E-05 | 1 | AAGGCTTAAGATGTT |
| 2751 | mamo-L | *BGIBMGA005610* | up | 137 | 151 | + | 12.2016 | 2.03E-05 | 1 | TAGGCTGTAGATGTC |
| 2752 | mamo-L | *BGIBMGA005611* | down | 1911 | 1925 | + | 11.0565 | 6.18E-05 | 1 | TAGGCTTTGCTTCGT |
| 2753 | mamo-L | *BGIBMGA005612* | up | 1270 | 1284 | - | 11.7097 | 3.34E-05 | 1 | TAGGCTTTGTTGTTC |

| | | | | | | | | | | |
|---|---|---|---|---|---|---|---|---|---|---|
| 2754 | mamo-L | *BGIBMGA005612* | up | 1213 | 1227 | + | 10.5565 | 9.46E-05 | 1 | GAGGCAGAATTTGGT |
| 2755 | mamo-L | *BGIBMGA005612* | up | 55 | 69 | - | 10.5 | 9.90E-05 | 1 | AAGGCGTAAATTGTT |
| 2756 | mamo-L | *BGIBMGA005614* | up | 807 | 821 | - | 10.7823 | 7.84E-05 | 1 | TAGGCCCCCTATGCC |
| 2757 | mamo-L | *BGIBMGA005614* | up | 1127 | 1141 | - | 10.7581 | 8.00E-05 | 1 | TAGGCGACGATTGCT |
| 2758 | mamo-L | *BGIBMGA005614* | up | 1144 | 1158 | - | 10.5161 | 9.77E-05 | 1 | TAGCCTGGCTTCGTT |
| 2759 | mamo-L | *BGIBMGA005617* | down | 985 | 999 | - | 11.0645 | 6.14E-05 | 1 | AAGGCTTTGTTCGTC |
| 2760 | mamo-L | *BGIBMGA005618* | down | 364 | 378 | - | 11.0645 | 6.14E-05 | 1 | AAGGCTTTGTTCGTC |
| 2761 | mamo-L | *BGIBMGA005619* | down | 1790 | 1804 | + | 12.0968 | 2.26E-05 | 1 | TAGGCGGCTTTTTCT |
| 2762 | mamo-L | *BGIBMGA005623* | down | 140 | 154 | + | 10.8145 | 7.63E-05 | 1 | AAGGCTTGGTTGTTT |
| 2763 | mamo-L | *BGIBMGA005625* | up | 1294 | 1308 | - | 13.7177 | 3.05E-06 | 1 | TAGGCGGATTTTTTT |
| 2764 | mamo-L | *BGIBMGA005627* | up | 1174 | 1188 | + | 11.0645 | 6.14E-05 | 1 | TAGGCTAACATTTAT |
| 2765 | mamo-L | *BGIBMGA005629* | up | 725 | 739 | - | 11.3871 | 4.55E-05 | 1 | TAGGCGCGTTTCGTC |
| 2766 | mamo-L | *BGIBMGA005630* | down | 1481 | 1495 | + | 11.8952 | 2.79E-05 | 1 | TAGGCCCGTTTTGCA |
| 2767 | mamo-L | *BGIBMGA005630* | down | 1270 | 1284 | + | 10.7419 | 8.11E-05 | 1 | TAGGCGGTGTTCGAA |
| 2768 | mamo-L | *BGIBMGA005633* | down | 1028 | 1042 | + | 10.5726 | 9.33E-05 | 1 | TAGGCTAATATTTAT |
| 2769 | mamo-L | *BGIBMGA005641* | down | 1972 | 1986 | + | 10.9677 | 6.68E-05 | 1 | TAGGCTGGTTCTGAC |
| 2770 | mamo-L | *BGIBMGA005642* | down | 595 | 609 | - | 11.3871 | 4.55E-05 | 1 | TAGGCACATGTTGAT |
| 2771 | mamo-L | *BGIBMGA005642* | up | 1563 | 1577 | + | 10.6129 | 9.03E-05 | 1 | TAGGCGTAAATTATT |
| 2772 | mamo-L | *BGIBMGA005644* | down | 188 | 202 | - | 12.9919 | 8.10E-06 | 1 | TAGGCCGGTTTTGCA |
| 2773 | mamo-L | *BGIBMGA005647* | down | 87 | 101 | - | 10.6371 | 8.85E-05 | 1 | TAGGCCAAATATGGC |
| 2774 | mamo-L | *BGIBMGA005650* | down | 834 | 848 | - | 10.9113 | 7.02E-05 | 1 | TAGGCTATTGGTTTT |
| 2775 | mamo-L | *BGIBMGA005652* | up | 1560 | 1574 | - | 10.9032 | 7.07E-05 | 1 | TAGGCTTTGGTGACC |
| 2776 | mamo-L | *BGIBMGA005655* | down | 1574 | 1588 | - | 10.6855 | 8.51E-05 | 1 | TAGGCTGAGTTAGGA |
| 2777 | mamo-L | *BGIBMGA005659* | up | 818 | 832 | - | 10.9435 | 6.83E-05 | 1 | AAGGCCTAGAGTGTT |
| 2778 | mamo-L | *BGIBMGA005661* | up | 1115 | 1129 | + | 11.0968 | 5.97E-05 | 1 | TAGGCTGAGATGGTA |
| 2779 | mamo-L | *BGIBMGA005663* | up | 1470 | 1484 | + | 11.1532 | 5.67E-05 | 1 | TAGGCTAGCATTTGC |
| 2780 | mamo-L | *BGIBMGA005663* | up | 575 | 589 | + | 10.621 | 8.97E-05 | 1 | TAGGCTACCATTTGT |
| 2781 | mamo-L | *BGIBMGA005669* | up | 86 | 100 | - | 10.9194 | 6.98E-05 | 1 | TAGGCTTAAATCTTT |
| 2782 | mamo-L | *BGIBMGA005676* | down | 989 | 1003 | + | 10.75 | 8.06E-05 | 1 | GAGGCCGATTGTGCC |
| 2783 | mamo-L | *BGIBMGA005677* | down | 501 | 515 | + | 13.7177 | 3.05E-06 | 1 | TAGGCTTTCTTGGTT |
| 2784 | mamo-L | *BGIBMGA005678* | down | 1532 | 1546 | + | 10.9194 | 6.98E-05 | 1 | TAGGCTACTTTTTTT |
| 2785 | mamo-L | *BGIBMGA005679* | down | 1048 | 1062 | - | 11.5645 | 3.84E-05 | 1 | AAGGCCTAAGTTGGC |
| 2786 | mamo-L | *BGIBMGA005680* | down | 159 | 173 | - | 11.4113 | 4.45E-05 | 1 | TAGGCCTGCATTTAC |
| 2787 | mamo-L | *BGIBMGA005682* | down | 846 | 860 | + | 10.8548 | 7.38E-05 | 1 | GAGGCCTGTTTTTAT |
| 2788 | mamo-L | *BGIBMGA005685* | up | 758 | 772 | - | 12.371 | 1.68E-05 | 1 | TAGGCTGATGATTAT |
| 2789 | mamo-L | *BGIBMGA005687* | up | 117 | 131 | - | 12.6532 | 1.20E-05 | 1 | TAGGCAGAGTGTGGT |
| 2790 | mamo-L | *BGIBMGA005689* | up | 1703 | 1717 | + | 11.6613 | 3.50E-05 | 1 | TAGGCTATCATTGCC |
| 2791 | mamo-L | *BGIBMGA005690* | up | 1723 | 1737 | - | 11.121 | 5.84E-05 | 1 | AAGGCTGTAATTGCT |
| 2792 | mamo-L | *BGIBMGA005690* | up | 511 | 525 | + | 10.5726 | 9.33E-05 | 1 | TAGGCTAATATTTAT |
| 2793 | mamo-L | *BGIBMGA005692* | up | 624 | 638 | + | 11.1452 | 5.70E-05 | 1 | TAGGCTCAGTTTAAC |
| 2794 | mamo-L | *BGIBMGA005693* | up | 885 | 899 | + | 11.4435 | 4.31E-05 | 1 | TAGGCTTGAGAGGCT |
| 2795 | mamo-L | *BGIBMGA005693* | down | 1764 | 1778 | - | 10.5726 | 9.33E-05 | 1 | TAGGCTAATATTTAT |
| 2796 | mamo-L | *BGIBMGA005694* | down | 265 | 279 | + | 10.7984 | 7.74E-05 | 1 | TAGGCGGGTCGTGTT |
| 2797 | mamo-L | *BGIBMGA005696* | down | 802 | 816 | - | 10.6371 | 8.85E-05 | 1 | AAGGCTGGTTTTAGC |
| 2798 | mamo-L | *BGIBMGA005696* | down | 117 | 131 | + | 10.5081 | 9.83E-05 | 1 | TAGGCTGCATACGTT |
| 2799 | mamo-L | *BGIBMGA005696* | up | 1166 | 1180 | - | 10.5 | 9.90E-05 | 1 | GAGGCCCAGGTCGTC |
| 2800 | mamo-L | *BGIBMGA005704* | up | 341 | 355 | - | 11.4677 | 4.22E-05 | 1 | GAGGCCCTGTTTGAT |
| 2801 | mamo-L | *BGIBMGA005706* | up | 1888 | 1902 | + | 12.6613 | 1.19E-05 | 1 | TAGGCCGTTGATTCT |
| 2802 | mamo-L | *BGIBMGA005709* | up | 340 | 354 | + | 12.2339 | 1.95E-05 | 1 | GAGGCCCTCTTTGTT |
| 2803 | mamo-L | *BGIBMGA005717* | down | 35 | 49 | + | 11.8548 | 2.89E-05 | 1 | TAGGCCGAATAGGGT |
| 2804 | mamo-L | *BGIBMGA005718* | up | 629 | 643 | + | 11.8548 | 2.89E-05 | 1 | TAGGCCGAATAGGGT |
| 2805 | mamo-L | *BGIBMGA005719* | up | 1905 | 1919 | + | 12.4113 | 1.60E-05 | 1 | TAGGCTTACTTCATT |
| 2806 | mamo-L | *BGIBMGA005719* | up | 66 | 80 | + | 11.6613 | 3.50E-05 | 1 | AAGGCTTTTTTTTTT |
| 2807 | mamo-L | *BGIBMGA005719* | up | 452 | 466 | - | 11.629 | 3.61E-05 | 1 | GAGGCTTTTTTTTTT |
| 2808 | mamo-L | *BGIBMGA005720* | up | 299 | 313 | + | 10.7016 | 8.39E-05 | 1 | AAGGCCGGTTGTGCC |
| 2809 | mamo-L | *BGIBMGA005721* | down | 253 | 267 | + | 12.9919 | 8.10E-06 | 1 | TAGGCCGGTTTTGCA |
| 2810 | mamo-L | *BGIBMGA005721* | down | 1087 | 1101 | - | 11.8952 | 2.79E-05 | 1 | TAGGCCGGTTGTTAT |
| 2811 | mamo-L | *BGIBMGA005721* | down | 110 | 124 | - | 10.7097 | 8.33E-05 | 1 | TAGGCCAGTTTTGCG |
| 2812 | mamo-L | *BGIBMGA005726* | up | 1794 | 1808 | - | 10.7661 | 7.95E-05 | 1 | TAGGCTCAATTTAAT |
| 2813 | mamo-L | *BGIBMGA005727* | up | 1467 | 1481 | + | 12.4919 | 1.46E-05 | 1 | TAGGCTGATGATGAC |
| 2814 | mamo-L | *BGIBMGA005728* | down | 758 | 772 | + | 10.5726 | 9.33E-05 | 1 | TAGGCTAATATTTAT |
| 2815 | mamo-L | *BGIBMGA005729* | up | 434 | 448 | + | 10.5726 | 9.33E-05 | 1 | TAGGCTAATATTTAT |
| 2816 | mamo-L | *BGIBMGA005732* | down | 1660 | 1674 | + | 10.6774 | 8.56E-05 | 1 | AAGGCCCTTTTTTTT |
| 2817 | mamo-L | *BGIBMGA005732* | down | 1008 | 1022 | - | 10.5887 | 9.21E-05 | 1 | TAGGCGTTTGCTTTT |
| 2818 | mamo-L | *BGIBMGA005734* | up | 755 | 769 | - | 10.5726 | 9.33E-05 | 1 | TAGGCTAATATTTAT |
| 2819 | mamo-L | *BGIBMGA005735* | down | 1372 | 1386 | + | 11.9839 | 2.54E-05 | 1 | TAGGCCGAAGAGGGT |
| 2820 | mamo-L | *BGIBMGA005735* | down | 692 | 706 | + | 10.5 | 9.90E-05 | 1 | GAGGCCCAGGTCGTC |
| 2821 | mamo-L | *BGIBMGA005736* | up | 620 | 634 | + | 11.0242 | 6.36E-05 | 1 | TAGGCCGCGGTTTGG |
| 2822 | mamo-L | *BGIBMGA005738* | up | 1670 | 1684 | + | 10.5726 | 9.33E-05 | 1 | TAGGCTAATATTTAT |
| 2823 | mamo-L | *BGIBMGA005739* | up | 1019 | 1033 | - | 11.3952 | 4.52E-05 | 1 | TAGGCTGTCAGTTGC |
| 2824 | mamo-L | *BGIBMGA005740* | up | 1727 | 1741 | - | 11.8468 | 2.92E-05 | 1 | TAGGCGTAAGTTATT |
| 2825 | mamo-L | *BGIBMGA005744* | up | 294 | 308 | - | 10.8065 | 7.69E-05 | 1 | TGGGCTTTCGTTTTT |
| 2826 | mamo-L | *BGIBMGA005744* | down | 443 | 457 | + | 10.5726 | 9.33E-05 | 1 | TAGGCTAATATTTAT |
| 2827 | mamo-L | *BGIBMGA005745* | up | 338 | 352 | - | 10.5 | 9.90E-05 | 1 | GAGGCCCAGGTCGTC |
| 2828 | mamo-L | *BGIBMGA005747* | up | 1270 | 1284 | + | 13.0081 | 7.94E-06 | 1 | TAGGCTTATTGTTGT |
| 2829 | mamo-L | *BGIBMGA005747* | up | 688 | 702 | - | 10.5323 | 9.63E-05 | 1 | AAGGCCGAGGTCTTC |
| 2830 | mamo-L | *BGIBMGA005751* | down | 307 | 321 | - | 12.7258 | 1.11E-05 | 1 | TAGGCGTGCTTTGTA |
| 2831 | mamo-L | *BGIBMGA005754* | up | 1362 | 1376 | - | 13.8548 | 2.48E-06 | 1 | TAGGCCTTTTTTGAT |
| 2832 | mamo-L | *BGIBMGA005754* | up | 1369 | 1383 | + | 11.3629 | 4.66E-05 | 1 | AAGGCCTACGTTCTT |
| 2833 | mamo-L | *BGIBMGA005755* | up | 687 | 701 | + | 10.8306 | 7.52E-05 | 1 | TAGGCCCCCTTTGGG |
| 2834 | mamo-L | *BGIBMGA005756* | up | 12 | 26 | + | 11.3548 | 4.69E-05 | 1 | TAGGCTGATTATCGC |
| 2835 | mamo-L | *BGIBMGA005759* | up | 1788 | 1802 | + | 11.7984 | 3.06E-05 | 1 | TAGGCTTTGCTTTTT |
| 2836 | mamo-L | *BGIBMGA005760* | down | 1789 | 1803 | - | 10.5887 | 9.21E-05 | 1 | TAGGCCAGGTTCACT |
| 2837 | mamo-L | *BGIBMGA005763* | down | 1766 | 1780 | - | 10.8548 | 7.38E-05 | 1 | AAGGCTGTTATTTTT |
| 2838 | mamo-L | *BGIBMGA005764* | up | 1174 | 1188 | - | 11.2419 | 5.21E-05 | 1 | GAGGCCGACTGTGCC |
| 2839 | mamo-L | *BGIBMGA005764* | down | 323 | 337 | + | 10.5726 | 9.33E-05 | 1 | TAGGCTAATATTTAT |
| 2840 | mamo-L | *BGIBMGA005765* | up | 1867 | 1881 | + | 10.5726 | 9.33E-05 | 1 | TAGGCTAATATTTAT |
| 2841 | mamo-L | *BGIBMGA005766* | up | 1512 | 1526 | + | 13.2419 | 5.96E-06 | 1 | TAGGCTTGGTTGTGT |
| 2842 | mamo-L | *BGIBMGA005766* | down | 369 | 383 | + | 10.9194 | 6.98E-05 | 1 | TAGGCTACTTTTTTT |
| 2843 | mamo-L | *BGIBMGA005767* | down | 414 | 428 | - | 12.2742 | 1.87E-05 | 1 | TAGGCGTACTGGGGT |
| 2844 | mamo-L | *BGIBMGA005769* | up | 255 | 269 | - | 11.4274 | 4.38E-05 | 1 | GAGGCTGTGTATGAT |
| 2845 | mamo-L | *BGIBMGA005776* | up | 463 | 477 | - | 13.2016 | 6.26E-06 | 1 | TAGGCTGATGATTTT |

| | | | | | | | | | | |
|---|---|---|---|---|---|---|---|---|---|---|
| 2846 | mamo-L | *BGIBMGA005778* | down | 882 | 896 | - | 12.2419 | 1.94E-05 | 1 | TAGGCCTTATTGGTT |
| 2847 | mamo-L | *BGIBMGA005782* | down | 1122 | 1136 | - | 11.129 | 5.79E-05 | 1 | AAGGCCTTCATTGAT |
| 2848 | mamo-L | *BGIBMGA005784* | up | 494 | 508 | + | 11.7258 | 3.29E-05 | 1 | TAGGCTCACTTTCCC |
| 2849 | mamo-L | *BGIBMGA005787* | down | 776 | 790 | + | 11.0968 | 5.97E-05 | 1 | TAGGCTGAGATGGTA |
| 2850 | mamo-L | *BGIBMGA005793* | up | 1802 | 1816 | - | 14.1935 | 1.49E-06 | 1 | TAGGCGTTTTTTGTT |
| 2851 | mamo-L | *BGIBMGA005795* | up | 631 | 645 | - | 13.5403 | 3.97E-06 | 1 | TAGGCTGTGTTTCCT |
| 2852 | mamo-L | *BGIBMGA005796* | up | 1896 | 1910 | - | 10.7661 | 7.95E-05 | 1 | TAGGCTCGAATGGTT |
| 2853 | mamo-L | *BGIBMGA005805* | up | 1908 | 1922 | - | 12.6452 | 1.21E-05 | 1 | TAGGCGTTGTTCGAT |
| 2854 | mamo-L | *BGIBMGA005805* | down | 1411 | 1425 | - | 11.371 | 4.62E-05 | 1 | TAGGCTGTATAGGCT |
| 2855 | mamo-L | *BGIBMGA005806* | up | 1612 | 1626 | + | 11.2419 | 5.21E-05 | 1 | GAGGCCGACTGTGCC |
| 2856 | mamo-L | *BGIBMGA005811* | up | 913 | 927 | + | 12.9677 | 8.30E-06 | 1 | TAGGCCGACTTTAAT |
| 2857 | mamo-L | *BGIBMGA005812* | up | 1714 | 1728 | + | 10.9355 | 6.88E-05 | 1 | TAGGCCGTGGGTTCA |
| 2858 | mamo-L | *BGIBMGA005812* | up | 832 | 846 | + | 10.8226 | 7.58E-05 | 1 | AAGGCTGGTTTGGAT |
| 2859 | mamo-L | *BGIBMGA005813* | down | 1892 | 1906 | - | 10.6532 | 8.73E-05 | 1 | AAGGCTTTTTATTGT |
| 2860 | mamo-L | *BGIBMGA005818* | up | 1261 | 1275 | - | 10.9758 | 6.64E-05 | 1 | TAGGCTTATCTTAGT |
| 2861 | mamo-L | *BGIBMGA005818* | down | 1049 | 1063 | + | 10.7016 | 8.39E-05 | 1 | TAGGCGACGGATGTT |
| 2862 | mamo-L | *BGIBMGA005822* | down | 189 | 203 | - | 11.2984 | 4.95E-05 | 1 | TAGGCTGCGTGTTGC |
| 2863 | mamo-L | *BGIBMGA005822* | up | 1510 | 1524 | + | 11.1452 | 5.70E-05 | 1 | TAGGCCCGTTTTCAT |
| 2864 | mamo-L | *BGIBMGA005822* | down | 1475 | 1489 | + | 11.0242 | 6.36E-05 | 1 | TAGGCTGCAATTGTC |
| 2865 | mamo-L | *BGIBMGA005822* | down | 1712 | 1726 | - | 10.8065 | 7.69E-05 | 1 | AAGGCCTTAGTTTTT |
| 2866 | mamo-L | *BGIBMGA005832* | down | 309 | 323 | - | 10.5726 | 9.33E-05 | 1 | TAGGCTAATATTTAT |
| 2867 | mamo-L | *BGIBMGA005838* | up | 920 | 934 | - | 12.3387 | 1.73E-05 | 1 | TAGGCCGACGGTCCT |
| 2868 | mamo-L | *BGIBMGA005842* | down | 1722 | 1736 | + | 13.0726 | 7.34E-06 | 1 | TAGGCCGATTTTGCA |
| 2869 | mamo-L | *BGIBMGA005844* | down | 471 | 485 | - | 10.9194 | 6.98E-05 | 1 | TAGGCTACTTTTTTT |
| 2870 | mamo-L | *BGIBMGA005850* | up | 1215 | 1229 | - | 11.3548 | 4.69E-05 | 1 | AAGGCTGATGATGAT |
| 2871 | mamo-L | *BGIBMGA005851* | up | 959 | 973 | + | 10.8548 | 7.38E-05 | 1 | AAGGCTCGCTTTCGT |
| 2872 | mamo-L | *BGIBMGA005852* | down | 1814 | 1828 | - | 14.621 | 7.29E-07 | 1 | TAGGCTTAGTTTTCT |
| 2873 | mamo-L | *BGIBMGA005853* | up | 1535 | 1549 | - | 11.8871 | 2.81E-05 | 1 | TAGGCTTTTTTTTGA |
| 2874 | mamo-L | *BGIBMGA005853* | down | 320 | 334 | - | 10.7984 | 7.74E-05 | 1 | AAGGCTTATTTTGGG |
| 2875 | mamo-L | *BGIBMGA005854* | up | 969 | 983 | - | 10.7984 | 7.74E-05 | 1 | AAGGCTTATTTTGGG |
| 2876 | mamo-L | *BGIBMGA005855* | up | 1834 | 1848 | + | 10.5403 | 9.58E-05 | 1 | TAGGCTGTATGGTGT |
| 2877 | mamo-L | *BGIBMGA005856* | up | 245 | 259 | + | 10.5403 | 9.58E-05 | 1 | TAGGCTGTATGGTGT |
| 2878 | mamo-L | *BGIBMGA005860* | down | 1407 | 1421 | + | 11.1855 | 5.50E-05 | 1 | TAGGCTGTCTCTCGT |
| 2879 | mamo-L | *BGIBMGA005861* | up | 1820 | 1834 | - | 11.4113 | 4.45E-05 | 1 | TAGGCAGTAGTTTGT |
| 2880 | mamo-L | *BGIBMGA005863* | up | 1093 | 1107 | - | 11.7661 | 3.16E-05 | 1 | TAGGCCCGTTGCGGT |
| 2881 | mamo-L | *BGIBMGA005864* | up | 1628 | 1642 | - | 12.5081 | 1.43E-05 | 1 | TAGGCGTTTTTTAGT |
| 2882 | mamo-L | *BGIBMGA005874* | up | 1424 | 1438 | - | 10.8306 | 7.52E-05 | 1 | GAGGCCGATTATGCC |
| 2883 | mamo-L | *BGIBMGA005875* | up | 91 | 105 | + | 10.5565 | 9.46E-05 | 1 | TAGGCGGTTGAGTTT |
| 2884 | mamo-L | *BGIBMGA005880* | up | 1248 | 1262 | + | 12.1774 | 2.08E-05 | 1 | TAGGCCCTCATTTTT |
| 2885 | mamo-L | *BGIBMGA005881* | down | 1548 | 1562 | + | 10.629 | 8.91E-05 | 1 | TAGGCCTGCAGCGCC |
| 2886 | mamo-L | *BGIBMGA005883* | up | 30 | 44 | - | 11.7984 | 3.06E-05 | 1 | TAGGCTTTGCTTTTT |
| 2887 | mamo-L | *BGIBMGA005885* | up | 1120 | 1134 | + | 10.8548 | 7.38E-05 | 1 | GAGGCCTGTTTTTAT |
| 2888 | mamo-L | *BGIBMGA005888* | down | 1181 | 1195 | + | 12.5323 | 1.39E-05 | 1 | TAGGCTTATTTTAGC |
| 2889 | mamo-L | *BGIBMGA005890* | down | 1874 | 1888 | + | 11.3548 | 4.69E-05 | 1 | AAGGCTGATGATGAT |
| 2890 | mamo-L | *BGIBMGA005893* | up | 505 | 519 | - | 12.25 | 1.92E-05 | 1 | TAGGCCCACTATTCT |
| 2891 | mamo-L | *BGIBMGA005894* | up | 997 | 1011 | - | 10.8548 | 7.38E-05 | 1 | GAGGCCTGTTTTTAT |
| 2892 | mamo-L | *BGIBMGA005895* | up | 1448 | 1462 | + | 11.7984 | 3.06E-05 | 1 | TAGGCTTTGCTTTTT |
| 2893 | mamo-L | *BGIBMGA005896* | up | 1168 | 1182 | - | 11.9839 | 2.54E-05 | 1 | TAGGCCGAAGAGGGT |
| 2894 | mamo-L | *BGIBMGA005898* | down | 482 | 496 | + | 10.879 | 7.22E-05 | 1 | TAGGCTTTCATTTGG |
| 2895 | mamo-L | *BGIBMGA005902* | down | 103 | 117 | + | 10.5726 | 9.33E-05 | 1 | TAGGCTAATATTTAT |
| 2896 | mamo-L | *BGIBMGA005903* | up | 1012 | 1026 | - | 10.9194 | 6.98E-05 | 1 | TAGGCTACTTTTTTT |
| 2897 | mamo-L | *BGIBMGA005906* | down | 412 | 426 | + | 12.2903 | 1.83E-05 | 1 | TAGGCTGGTGATTAT |
| 2898 | mamo-L | *BGIBMGA005908* | up | 1116 | 1130 | - | 10.8629 | 7.33E-05 | 1 | TAGGCTAAGTTTTTA |
| 2899 | mamo-L | *BGIBMGA005909* | down | 1393 | 1407 | + | 11.4435 | 4.31E-05 | 1 | TAGGCTTGAGAGGCT |
| 2900 | mamo-L | *BGIBMGA005909* | down | 1464 | 1478 | - | 11.0323 | 6.32E-05 | 1 | TAGGCTTGTATTATC |
| 2901 | mamo-L | *BGIBMGA005914* | up | 1115 | 1129 | + | 12.2016 | 2.03E-05 | 1 | TAGGCTGTAGATGTC |
| 2902 | mamo-L | *BGIBMGA005916* | down | 779 | 793 | + | 11.0242 | 6.36E-05 | 1 | TAGGCTTACAATGCA |
| 2903 | mamo-L | *BGIBMGA005919* | down | 1964 | 1978 | + | 10.8306 | 7.52E-05 | 1 | AAGGCTTTTTTTTAT |
| 2904 | mamo-L | *BGIBMGA005920* | down | 1776 | 1790 | - | 13.5968 | 3.65E-06 | 1 | TAGGCTTGTGTGGTT |
| 2905 | mamo-L | *BGIBMGA005923* | up | 446 | 460 | - | 11.2097 | 5.38E-05 | 1 | AAGGCTTGCTTCGCC |
| 2906 | mamo-L | *BGIBMGA005926* | down | 1448 | 1462 | - | 12.879 | 9.26E-06 | 1 | AAGGCGTGGTTTGTT |
| 2907 | mamo-L | *BGIBMGA005926* | down | 918 | 932 | + | 12.5081 | 1.43E-05 | 1 | TAGGCTAATTTTTTT |
| 2908 | mamo-L | *BGIBMGA005930* | up | 1934 | 1948 | + | 11.3145 | 4.87E-05 | 1 | GAGGCTGTCTTTGTA |
| 2909 | mamo-L | *BGIBMGA005932* | up | 1167 | 1181 | + | 10.6613 | 8.67E-05 | 1 | TAGGCGGACGAGGAC |
| 2910 | mamo-L | *BGIBMGA005933* | up | 1935 | 1949 | - | 12.5242 | 1.40E-05 | 1 | TAGGCTGTGGTTTCA |
| 2911 | mamo-L | *BGIBMGA005935* | down | 867 | 881 | + | 12.5484 | 1.36E-05 | 1 | AAGGCTGATTTTGAT |
| 2912 | mamo-L | *BGIBMGA005935* | up | 66 | 80 | + | 11.6129 | 3.66E-05 | 1 | TAGGCTTTTAATGAT |
| 2913 | mamo-L | *BGIBMGA005944* | down | 1053 | 1067 | - | 10.4919 | 9.96E-05 | 1 | TAGGCCATGTTCGGA |
| 2914 | mamo-L | *BGIBMGA005949* | up | 1576 | 1590 | + | 12.4677 | 1.49E-05 | 1 | TAGGCTTCTATTGCT |
| 2915 | mamo-L | *BGIBMGA005953* | down | 1119 | 1133 | + | 10.75 | 8.06E-05 | 1 | AAGGCGGTGTTTCCT |
| 2916 | mamo-L | *BGIBMGA005956* | down | 1102 | 1116 | + | 11.9919 | 2.52E-05 | 1 | TAGGCGTTTATTTTT |
| 2917 | mamo-L | *BGIBMGA005957* | down | 1911 | 1925 | + | 12.2661 | 1.89E-05 | 1 | TAGGCTCGCAGTGCT |
| 2918 | mamo-L | *BGIBMGA005959* | down | 1662 | 1676 | + | 13.2016 | 6.26E-06 | 1 | TAGGCTTGGGTTGTG |
| 2919 | mamo-L | *BGIBMGA005960* | up | 209 | 223 | + | 13.9194 | 2.23E-06 | 1 | TAGGCCTAGTTGGTT |
| 2920 | mamo-L | *BGIBMGA005960* | down | 671 | 685 | - | 11.7339 | 3.26E-05 | 1 | TAGGCTAGCTTCGCC |
| 2921 | mamo-L | *BGIBMGA005960* | down | 685 | 699 | - | 10.9194 | 6.98E-05 | 1 | TAGGCTACTTTTTTT |
| 2922 | mamo-L | *BGIBMGA005967* | down | 422 | 436 | - | 10.5968 | 9.15E-05 | 1 | GAGGCGGATTTTTTC |
| 2923 | mamo-L | *BGIBMGA005973* | up | 980 | 994 | - | 11.9194 | 2.72E-05 | 1 | TAGGCTTTTGTCGAC |
| 2924 | mamo-L | *BGIBMGA005974* | up | 1944 | 1958 | + | 10.5726 | 9.33E-05 | 1 | TAGGCTAATATTTAT |
| 2925 | mamo-L | *BGIBMGA005978* | up | 1655 | 1669 | + | 10.7016 | 8.39E-05 | 1 | TAGACTGTCGTTGGT |
| 2926 | mamo-L | *BGIBMGA005980* | up | 1793 | 1807 | - | 10.7661 | 7.95E-05 | 1 | TAGGCGCGGGTCGGG |
| 2927 | mamo-L | *BGIBMGA005980* | up | 839 | 853 | + | 10.5726 | 9.33E-05 | 1 | TAGGCGTGCAATTAT |
| 2928 | mamo-L | *BGIBMGA005981* | up | 173 | 187 | + | 10.629 | 8.91E-05 | 1 | TAGGCTATCGTCATT |
| 2929 | mamo-L | *BGIBMGA005988* | down | 1401 | 1415 | - | 10.8226 | 7.58E-05 | 1 | TAGGCGGGCATTTTA |
| 2930 | mamo-L | *BGIBMGA005989* | down | 932 | 946 | + | 12.129 | 2.19E-05 | 1 | TAGGCTTTTTTGTTT |
| 2931 | mamo-L | *BGIBMGA005989* | up | 36 | 50 | - | 10.5726 | 9.33E-05 | 1 | AAGGCGGGCGCTGCT |
| 2932 | mamo-L | *BGIBMGA005990* | up | 1983 | 1997 | - | 12.0081 | 2.48E-05 | 1 | TAGGCCTGGTTAGTT |
| 2933 | mamo-L | *BGIBMGA005991* | up | 480 | 494 | - | 11.1694 | 5.59E-05 | 1 | GAGGCTGATGTGGGC |
| 2934 | mamo-L | *BGIBMGA005992* | up | 576 | 590 | - | 11.9919 | 2.52E-05 | 1 | TAGGCACAGGTTTGT |
| 2935 | mamo-L | *BGIBMGA005994* | up | 438 | 452 | + | 13.2903 | 5.60E-06 | 1 | TAGGCTGATATTTTT |
| 2936 | mamo-L | *BGIBMGA005994* | up | 667 | 681 | + | 12.4677 | 1.49E-05 | 1 | TAGGCTTCTATTGCT |
| 2937 | mamo-L | *BGIBMGA006001* | up | 849 | 863 | - | 11.2258 | 5.30E-05 | 1 | TAGGCGAACAGTGGT |

| | | | | | | | | | | |
|---|---|---|---|---|---|---|---|---|---|---|
| 2938 | mamo-L | *BGIBMGA006002* | down | 373 | 387 | + | 11.2016 | 5.41E-05 | 1 | TAGGCCGGGGGCACT |
| 2939 | mamo-L | *BGIBMGA006004* | down | 258 | 272 | + | 14.5242 | 8.74E-07 | 1 | TAGGCTTAAGTTGTT |
| 2940 | mamo-L | *BGIBMGA006005* | up | 31 | 45 | + | 10.9194 | 6.98E-05 | 1 | TAGGCTACTTTTTTT |
| 2941 | mamo-L | *BGIBMGA006015* | down | 800 | 814 | + | 11.3952 | 4.52E-05 | 1 | AAGGCGTTCGTCGCT |
| 2942 | mamo-L | *BGIBMGA006018* | down | 1719 | 1733 | - | 10.6048 | 9.09E-05 | 1 | TAGGCCCGTAATTTT |
| 2943 | mamo-L | *BGIBMGA006021* | down | 1974 | 1988 | + | 11.1935 | 5.45E-05 | 1 | TAGGCTAACTTTTAC |
| 2944 | mamo-L | *BGIBMGA006025* | down | 1359 | 1373 | - | 10.5242 | 9.70E-05 | 1 | TAGGCGACCGGTGCT |
| 2945 | mamo-L | *BGIBMGA006027* | up | 1858 | 1872 | + | 10.5 | 9.90E-05 | 1 | TAGGCTCGCGTTCGG |
| 2946 | mamo-L | *BGIBMGA006031* | down | 684 | 698 | - | 11.6855 | 3.42E-05 | 1 | TAGGCTTATTGTTCC |
| 2947 | mamo-L | *BGIBMGA006032* | up | 1212 | 1226 | - | 10.5968 | 9.15E-05 | 1 | TAGGCCGTATGTCCT |
| 2948 | mamo-L | *BGIBMGA006034* | down | 98 | 112 | + | 12.4435 | 1.54E-05 | 1 | TAGGCCTAAATTGGC |
| 2949 | mamo-L | *BGIBMGA006034* | down | 58 | 72 | - | 10.6452 | 8.79E-05 | 1 | TAGGCTGAGTTATAT |
| 2950 | mamo-L | *BGIBMGA006037* | up | 1887 | 1901 | - | 10.6129 | 9.03E-05 | 1 | GAGGCTGATTGGGGT |
| 2951 | mamo-L | *BGIBMGA006037* | down | 1308 | 1322 | - | 10.5484 | 9.52E-05 | 1 | TAGGCTGTTTATTTA |
| 2952 | mamo-L | *BGIBMGA006038* | up | 662 | 676 | + | 11.4194 | 4.42E-05 | 1 | TAGGCTTGGTTCATC |
| 2953 | mamo-L | *BGIBMGA006039* | down | 501 | 515 | - | 12.2984 | 1.81E-05 | 1 | TAGGCGTTATTTTTT |
| 2954 | mamo-L | *BGIBMGA006040* | up | 654 | 668 | - | 12.2984 | 1.81E-05 | 1 | TAGGCGTTATTTTTT |
| 2955 | mamo-L | *BGIBMGA006041* | down | 132 | 146 | + | 11.2984 | 4.95E-05 | 1 | AAGGCCCAATTTGTT |
| 2956 | mamo-L | *BGIBMGA006042* | up | 1781 | 1795 | + | 13.0403 | 7.65E-06 | 1 | TAGGCTTATTTTCTT |
| 2957 | mamo-L | *BGIBMGA006043* | up | 1484 | 1498 | - | 11.6452 | 3.56E-05 | 1 | TAGGCCGTCTTGCCT |
| 2958 | mamo-L | *BGIBMGA006045* | down | 643 | 657 | - | 12.8871 | 9.15E-06 | 1 | TAGGCTGAGTATCGT |
| 2959 | mamo-L | *BGIBMGA006046* | up | 1877 | 1891 | - | 11.3871 | 4.55E-05 | 1 | TAGGCGCGTTTCGTC |
| 2960 | mamo-L | *BGIBMGA006047* | down | 1833 | 1847 | + | 11.1532 | 5.67E-05 | 1 | AAGGCTTATTTTTAT |
| 2961 | mamo-L | *BGIBMGA006051* | down | 203 | 217 | + | 10.5726 | 9.33E-05 | 1 | TAGGCTAATATTTAT |
| 2962 | mamo-L | *BGIBMGA006054* | up | 141 | 155 | + | 13.5887 | 3.68E-06 | 1 | TAGGCTGAAATTGTT |
| 2963 | mamo-L | *BGIBMGA006061* | up | 1394 | 1408 | + | 11.1371 | 5.75E-05 | 1 | TAGGCCGTGATTTTA |
| 2964 | mamo-L | *BGIBMGA006063* | down | 663 | 677 | - | 12.0565 | 2.36E-05 | 1 | GAGGCTGTTGTTTTT |
| 2965 | mamo-L | *BGIBMGA006065* | down | 1487 | 1501 | + | 11.1129 | 5.88E-05 | 1 | TAGGCGCATGGGGGT |
| 2966 | mamo-L | *BGIBMGA006066* | up | 1805 | 1819 | - | 10.7581 | 8.00E-05 | 1 | TAGGCGCTATTTCGT |
| 2967 | mamo-L | *BGIBMGA006071* | up | 296 | 310 | + | 12.0645 | 2.34E-05 | 1 | TAGGCGGGCTTCGGA |
| 2968 | mamo-L | *BGIBMGA006072* | down | 79 | 93 | + | 11.371 | 4.62E-05 | 1 | TAGGCTGTATAGGCT |
| 2969 | mamo-L | *BGIBMGA006073* | up | 1176 | 1190 | + | 11.371 | 4.62E-05 | 1 | TAGGCTGTATAGGCT |
| 2970 | mamo-L | *BGIBMGA006073* | down | 1536 | 1550 | + | 11.0323 | 6.32E-05 | 1 | TAGGCCGCGTGTCGT |
| 2971 | mamo-L | *BGIBMGA006074* | up | 132 | 146 | - | 14.7258 | 5.83E-07 | 1 | TAGGCTTGCGTTGCC |
| 2972 | mamo-L | *BGIBMGA006079* | down | 313 | 327 | - | 10.8306 | 7.52E-05 | 1 | AAGGCTTTTTTTTAT |
| 2973 | mamo-L | *BGIBMGA006080* | up | 28 | 42 | - | 10.8306 | 7.52E-05 | 1 | AAGGCTTTTTTTTAT |
| 2974 | mamo-L | *BGIBMGA006082* | down | 339 | 353 | + | 11.0323 | 6.32E-05 | 1 | TAGGCCGAAGGTGTA |
| 2975 | mamo-L | *BGIBMGA006083* | down | 1800 | 1814 | - | 11.5806 | 3.78E-05 | 1 | TAGGCGTTGGCTGGC |
| 2976 | mamo-L | *BGIBMGA006084* | up | 669 | 683 | - | 10.621 | 8.97E-05 | 1 | AAGGCTATGATTGTT |
| 2977 | mamo-L | *BGIBMGA006084* | up | 1043 | 1057 | - | 10.5484 | 9.52E-05 | 1 | TAGGCGCGTAATGGC |
| 2978 | mamo-L | *BGIBMGA006086* | down | 1763 | 1777 | + | 10.5161 | 9.77E-05 | 1 | AAGGCCCGGTTCTGT |
| 2979 | mamo-L | *BGIBMGA006089* | up | 1821 | 1835 | - | 11.4516 | 4.28E-05 | 1 | TAGGCGAGCATTTGT |
| 2980 | mamo-L | *BGIBMGA006091* | down | 120 | 134 | + | 10.5645 | 9.40E-05 | 1 | TAGGCCTACTGGGGA |
| 2981 | mamo-L | *BGIBMGA006095* | down | 890 | 904 | - | 12.4435 | 1.54E-05 | 1 | TAGGCTTTGTTTTGA |
| 2982 | mamo-L | *BGIBMGA006095* | down | 1682 | 1696 | - | 11.6935 | 3.39E-05 | 1 | TAGGCGGTGATCGTC |
| 2983 | mamo-L | *BGIBMGA006101* | down | 1503 | 1517 | - | 10.7742 | 7.89E-05 | 1 | AAGGCAGACATTGGT |
| 2984 | mamo-L | *BGIBMGA006102* | down | 622 | 636 | + | 10.9194 | 6.98E-05 | 1 | TAGGCTACTTTTTTT |
| 2985 | mamo-L | *BGIBMGA006103* | down | 546 | 560 | - | 10.7258 | 8.23E-05 | 1 | TAGGCTCATGAGGAT |
| 2986 | mamo-L | *BGIBMGA006104* | up | 1405 | 1419 | + | 11.7742 | 3.14E-05 | 1 | GAGGCCCAGGTTGTC |
| 2987 | mamo-L | *BGIBMGA006109* | down | 1706 | 1720 | - | 10.7742 | 7.89E-05 | 1 | AAGGCTGAGTATCGT |
| 2988 | mamo-L | *BGIBMGA006111* | up | 978 | 992 | + | 13.5403 | 3.97E-06 | 1 | TAGGCTCTTGTTGGC |
| 2989 | mamo-L | *BGIBMGA006113* | down | 994 | 1008 | + | 10.9677 | 6.68E-05 | 1 | TAGGCTTAATTTGAG |
| 2990 | mamo-L | *BGIBMGA006115* | down | 1894 | 1908 | + | 14.6452 | 6.98E-07 | 1 | TAGGCTGATGTTGTC |
| 2991 | mamo-L | *BGIBMGA006119* | down | 479 | 493 | + | 12.6694 | 1.18E-05 | 1 | TAGGCCGAATTTCGT |
| 2992 | mamo-L | *BGIBMGA006122* | up | 1782 | 1796 | + | 11.1048 | 5.92E-05 | 1 | TAGGCTTGAATCGAT |
| 2993 | mamo-L | *BGIBMGA006122* | down | 184 | 198 | - | 10.8387 | 7.47E-05 | 1 | TAGGCCGGTTTTCCA |
| 2994 | mamo-L | *BGIBMGA006123* | up | 403 | 417 | + | 11.2903 | 4.99E-05 | 1 | GAGGCTCATTTCGGT |
| 2995 | mamo-L | *BGIBMGA006123* | down | 508 | 522 | - | 11.0403 | 6.28E-05 | 1 | TAGGCCGTCTACCGT |
| 2996 | mamo-L | *BGIBMGA006127* | up | 321 | 335 | - | 11.4274 | 4.38E-05 | 1 | TAGGCAGGAGATGGT |
| 2997 | mamo-L | *BGIBMGA006129* | down | 575 | 589 | - | 11.1935 | 5.45E-05 | 1 | TAGGCTAACTTTTAC |
| 2998 | mamo-L | *BGIBMGA006135* | up | 3 | 17 | - | 12.0645 | 2.34E-05 | 1 | TAGGCTTTCGACGAT |
| 2999 | mamo-L | *BGIBMGA006145* | down | 1383 | 1397 | - | 11.4758 | 4.18E-05 | 1 | TAGGCTTGAGAGGTT |
| 3000 | mamo-L | *BGIBMGA006145* | up | 1546 | 1560 | - | 10.9597 | 6.73E-05 | 1 | TAGGCTGACGAGCCT |
| 3001 | mamo-L | *BGIBMGA006147* | up | 139 | 153 | - | 11.379 | 4.58E-05 | 1 | TAGGCCTCCCTTGTT |
| 3002 | mamo-L | *BGIBMGA006147* | down | 117 | 131 | - | 10.7903 | 7.79E-05 | 1 | TAGGCCAACGATTAT |
| 3003 | mamo-L | *BGIBMGA006149* | down | 239 | 253 | + | 10.9194 | 6.98E-05 | 1 | TAGGCTACTTTTTTT |
| 3004 | mamo-L | *BGIBMGA006151* | up | 1073 | 1087 | + | 11.3629 | 4.66E-05 | 1 | TAGGCTGATTTTTAA |
| 3005 | mamo-L | *BGIBMGA006152* | up | 518 | 532 | - | 13.0242 | 7.77E-06 | 1 | TAGGCGGGCTTTGTA |
| 3006 | mamo-L | *BGIBMGA006152* | up | 1453 | 1467 | - | 10.8871 | 7.17E-05 | 1 | TAGGCCATCGTTAAT |
| 3007 | mamo-L | *BGIBMGA006161* | up | 81 | 95 | + | 11.9919 | 2.52E-05 | 1 | TAGGCACAGGTTTGT |
| 3008 | mamo-L | *BGIBMGA006162* | down | 1523 | 1537 | + | 12.0081 | 2.48E-05 | 1 | TAGGCCTGGTTAGTT |
| 3009 | mamo-L | *BGIBMGA006162* | up | 1722 | 1736 | + | 11.1694 | 5.59E-05 | 1 | GAGGCTGATGTGGGC |
| 3010 | mamo-L | *BGIBMGA006163* | down | 1070 | 1084 | - | 13.9597 | 2.11E-06 | 1 | TAGGCGGTGTGTGGT |
| 3011 | mamo-L | *BGIBMGA006163* | up | 43 | 57 | - | 11.2581 | 5.14E-05 | 1 | AAGGCGGGCGATGGC |
| 3012 | mamo-L | *BGIBMGA006166* | up | 300 | 314 | + | 11.7339 | 3.26E-05 | 1 | TAGGCCTCTTTTAGT |
| 3013 | mamo-L | *BGIBMGA006168* | down | 1193 | 1207 | - | 12.25 | 1.92E-05 | 1 | TAGGCCCACTATTCT |
| 3014 | mamo-L | *BGIBMGA006168* | up | 1068 | 1082 | - | 11.3065 | 4.91E-05 | 1 | GAGGCGGGGTTTCGT |
| 3015 | mamo-L | *BGIBMGA006172* | up | 513 | 527 | + | 11.871 | 2.85E-05 | 1 | TAGGCAAGGGTTGCT |
| 3016 | mamo-L | *BGIBMGA006176* | up | 1603 | 1617 | - | 10.5726 | 9.33E-05 | 1 | TAGGCTAATATTTAT |
| 3017 | mamo-L | *BGIBMGA006177* | up | 1154 | 1168 | - | 10.6371 | 8.85E-05 | 1 | TAGGCAAGGATTGCT |
| 3018 | mamo-L | *BGIBMGA006177* | up | 1168 | 1182 | - | 10.6048 | 9.09E-05 | 1 | AAGGCTTAGGTAGGT |
| 3019 | mamo-L | *BGIBMGA006178* | down | 514 | 528 | + | 10.9194 | 6.98E-05 | 1 | TAGGCTACTTTTTTT |
| 3020 | mamo-L | *BGIBMGA006179* | down | 1803 | 1817 | + | 10.9194 | 6.98E-05 | 1 | TAGGCTACTTTTTTT |
| 3021 | mamo-L | *BGIBMGA006181* | down | 312 | 326 | - | 11.5403 | 3.93E-05 | 1 | TAGGCGAGTGGTGCT |
| 3022 | mamo-L | *BGIBMGA006183* | up | 270 | 284 | - | 11.1129 | 5.88E-05 | 1 | TAGGCGTTTGATGGA |
| 3023 | mamo-L | *BGIBMGA006185* | down | 1078 | 1092 | - | 11.7419 | 3.23E-05 | 1 | AAGGCTGGTGTTAGT |
| 3024 | mamo-L | *BGIBMGA006189* | up | 1246 | 1260 | - | 11.5806 | 3.78E-05 | 1 | AAGGCGGTGTATGCT |
| 3025 | mamo-L | *BGIBMGA006190* | down | 904 | 918 | + | 10.9194 | 6.98E-05 | 1 | TAGGCTTAAATCTTT |
| 3026 | mamo-L | *BGIBMGA006192* | up | 164 | 178 | - | 12.4677 | 1.49E-05 | 1 | TAGGCTTCTATTGCT |
| 3027 | mamo-L | *BGIBMGA006192* | down | 1595 | 1609 | - | 11.1774 | 5.54E-05 | 1 | AAGGCTGAATGTGTT |
| 3028 | mamo-L | *BGIBMGA006193* | down | 136 | 150 | - | 13.4677 | 4.39E-06 | 1 | TAGGCTGATGATGAT |
| 3029 | mamo-L | *BGIBMGA006196* | down | 781 | 795 | - | 11.2016 | 5.41E-05 | 1 | TAGGCAGACTTGGAT |

| | | | | | | | | | | |
|---|---|---|---|---|---|---|---|---|---|---|
| 3030 | mamo-L | *BGIBMGA006198* | up | 495 | 509 | + | 10.8306 | 7.52E-05 | 1 | AAGGCTTTTTTTTAT |
| 3031 | mamo-L | *BGIBMGA006200* | up | 1694 | 1708 | - | 12.7258 | 1.11E-05 | 1 | TAGGCGTGCTTTGTA |
| 3032 | mamo-L | *BGIBMGA006201* | down | 284 | 298 | - | 10.5968 | 9.15E-05 | 1 | GAGGCTAGTTTTTGT |
| 3033 | mamo-L | *BGIBMGA006203* | down | 338 | 352 | + | 12.6532 | 1.20E-05 | 1 | TAGGCCGATTTGGCC |
| 3034 | mamo-L | *BGIBMGA006205* | down | 1455 | 1469 | - | 12.9516 | 8.46E-06 | 1 | TAGGCCGTCGAGGGT |
| 3035 | mamo-L | *BGIBMGA006205* | down | 1313 | 1327 | - | 11.7742 | 3.14E-05 | 1 | GAGGCCCAGGTTGTC |
| 3036 | mamo-L | *BGIBMGA006205* | up | 1331 | 1345 | + | 10.5726 | 9.33E-05 | 1 | TAGGCTAATATTTAT |
| 3037 | mamo-L | *BGIBMGA006206* | down | 779 | 793 | + | 12.0645 | 2.34E-05 | 1 | TAGGCGGGCTTCGGA |
| 3038 | mamo-L | *BGIBMGA006206* | up | 195 | 209 | + | 10.5968 | 9.15E-05 | 1 | TAGGCATTGGAGGGT |
| 3039 | mamo-L | *BGIBMGA006208* | down | 1715 | 1729 | + | 11.6855 | 3.42E-05 | 1 | TAGGCCTTGAATTCT |
| 3040 | mamo-L | *BGIBMGA006211* | up | 668 | 682 | - | 11.4435 | 4.31E-05 | 1 | TAGGCTTGAGAGGCT |
| 3041 | mamo-L | *BGIBMGA006211* | down | 94 | 108 | + | 10.5645 | 9.40E-05 | 1 | TAGGCTGCTGCTGAT |
| 3042 | mamo-L | *BGIBMGA006215* | up | 1162 | 1176 | + | 12.0645 | 2.34E-05 | 1 | TAGGCGGGCTTCGGA |
| 3043 | mamo-L | *BGIBMGA006217* | down | 458 | 472 | + | 12.9355 | 8.65E-06 | 1 | TAGGCGTGAGTTGAT |
| 3044 | mamo-L | *BGIBMGA006218* | up | 1825 | 1839 | - | 10.5726 | 9.33E-05 | 1 | TAGGCTAATATTTAT |
| 3045 | mamo-L | *BGIBMGA006223* | down | 882 | 896 | + | 11.6129 | 3.66E-05 | 1 | TAGGCATATTTCGTT |
| 3046 | mamo-L | *BGIBMGA006223* | down | 541 | 555 | + | 10.7339 | 8.17E-05 | 1 | TAGGCTAAGTTCCTT |
| 3047 | mamo-L | *BGIBMGA006224* | down | 324 | 338 | + | 10.5726 | 9.33E-05 | 1 | GAGGCCTATGATTTT |
| 3048 | mamo-L | *BGIBMGA006231* | down | 1026 | 1040 | + | 11.129 | 5.79E-05 | 1 | GAGGCTCAGTTGGCT |
| 3049 | mamo-L | *BGIBMGA006232* | up | 1800 | 1814 | - | 13.1935 | 6.33E-06 | 1 | TAGGCTTTGTTTGCA |
| 3050 | mamo-L | *BGIBMGA006235* | down | 1489 | 1503 | + | 10.879 | 7.22E-05 | 1 | TGGGCTGTTGTTGAT |
| 3051 | mamo-L | *BGIBMGA006238* | up | 664 | 678 | - | 11.0806 | 6.05E-05 | 1 | AAGGCTTATTTTATT |
| 3052 | mamo-L | *BGIBMGA006239* | up | 1428 | 1442 | + | 10.9194 | 6.98E-05 | 1 | TAGGCTACTTTTTTT |
| 3053 | mamo-L | *BGIBMGA006242* | down | 1261 | 1275 | + | 13.4677 | 4.39E-06 | 1 | TAGGCTGATGATGAT |
| 3054 | mamo-L | *BGIBMGA006243* | down | 985 | 999 | - | 10.9516 | 6.78E-05 | 1 | TAGGCACACGTTTGC |
| 3055 | mamo-L | *BGIBMGA006246* | up | 1743 | 1757 | - | 10.879 | 7.22E-05 | 1 | AAGGCCGGTTTTGCA |
| 3056 | mamo-L | *BGIBMGA006247* | up | 449 | 463 | - | 12.1048 | 2.24E-05 | 1 | TAGGCCGACTTTTTG |
| 3057 | mamo-L | *BGIBMGA006248* | up | 1115 | 1129 | + | 14.4677 | 9.58E-07 | 1 | TAGGCCTAGTTTTTT |
| 3058 | mamo-L | *BGIBMGA006252* | down | 1577 | 1591 | + | 12.9919 | 8.10E-06 | 1 | TAGGCTTTCTTCTTT |
| 3059 | mamo-L | *BGIBMGA006255* | up | 425 | 439 | + | 11.1532 | 5.67E-05 | 1 | TAGGCTCTGTGTTTC |
| 3060 | mamo-L | *BGIBMGA006256* | down | 768 | 782 | - | 10.75 | 8.06E-05 | 1 | GAGGCCGATTGTGCC |
| 3061 | mamo-L | *BGIBMGA006258* | up | 330 | 344 | + | 10.9194 | 6.98E-05 | 1 | TAGGCTACTTTTTTT |
| 3062 | mamo-L | *BGIBMGA006268* | down | 1451 | 1465 | + | 10.5726 | 9.33E-05 | 1 | TAGGCTAATATTTAT |
| 3063 | mamo-L | *BGIBMGA006270* | up | 1409 | 1423 | - | 11.129 | 5.79E-05 | 1 | AAGGCTTTATTTGAT |
| 3064 | mamo-L | *BGIBMGA006272* | down | 231 | 245 | + | 11.5403 | 3.93E-05 | 1 | TAGGCGTATTTTATC |
| 3065 | mamo-L | *BGIBMGA006275* | down | 193 | 207 | + | 10.6774 | 8.56E-05 | 1 | TAGGCGATATTTTCT |
| 3066 | mamo-L | *BGIBMGA006277* | down | 284 | 298 | - | 11.3548 | 4.69E-05 | 1 | TAGGCTCGGGTGATT |
| 3067 | mamo-L | *BGIBMGA006278* | up | 1645 | 1659 | + | 12.4516 | 1.53E-05 | 1 | TAGGCCTGGGTAGGT |
| 3068 | mamo-L | *BGIBMGA006278* | up | 113 | 127 | + | 11.6532 | 3.53E-05 | 1 | TAGGCCGTTTTTTCA |
| 3069 | mamo-L | *BGIBMGA006280* | down | 956 | 970 | + | 10.5565 | 9.46E-05 | 1 | TAGGCCGGCTTCAGA |
| 3070 | mamo-L | *BGIBMGA006281* | down | 1315 | 1329 | + | 11.2258 | 5.30E-05 | 1 | TAGGCCTTTTTCCCT |
| 3071 | mamo-L | *BGIBMGA006287* | down | 1309 | 1323 | - | 11.7984 | 3.06E-05 | 1 | TAGGCTTTGCTTTTT |
| 3072 | mamo-L | *BGIBMGA006288* | down | 1183 | 1197 | + | 11.3548 | 4.69E-05 | 1 | AAGGCTGATGATGAT |
| 3073 | mamo-L | *BGIBMGA006289* | up | 1153 | 1167 | - | 13.7258 | 3.01E-06 | 1 | TAGGCTGTGTTTATT |
| 3074 | mamo-L | *BGIBMGA006289* | down | 1332 | 1346 | + | 11.4839 | 4.15E-05 | 1 | TAGGCCGCTTTTGCA |
| 3075 | mamo-L | *BGIBMGA006290* | up | 759 | 773 | + | 14.2661 | 1.33E-06 | 1 | TAGGCGGGCGGTGGT |
| 3076 | mamo-L | *BGIBMGA006291* | down | 1434 | 1448 | - | 11.4839 | 4.15E-05 | 1 | TAGGCCGCTTTTGCA |
| 3077 | mamo-L | *BGIBMGA006291* | down | 384 | 398 | + | 10.5403 | 9.58E-05 | 1 | TAGGCGTAAGTCACT |
| 3078 | mamo-L | *BGIBMGA006292* | down | 1607 | 1621 | - | 11.7823 | 3.11E-05 | 1 | TAGGCATTTTTTTGT |
| 3079 | mamo-L | *BGIBMGA006293* | down | 951 | 965 | - | 12.129 | 2.19E-05 | 1 | TAGGCTGTGGTTTCG |
| 3080 | mamo-L | *BGIBMGA006294* | up | 680 | 694 | + | 11.9839 | 2.54E-05 | 1 | TAGGCAGAGTTTTGC |
| 3081 | mamo-L | *BGIBMGA006299* | up | 1712 | 1726 | + | 10.5726 | 9.33E-05 | 1 | TAGGCTAATATTTAT |
| 3082 | mamo-L | *BGIBMGA006303* | down | 362 | 376 | - | 10.75 | 8.06E-05 | 1 | GAGGCCGATTGTGCC |
| 3083 | mamo-L | *BGIBMGA006304* | down | 108 | 122 | + | 13.0242 | 7.77E-06 | 1 | TAGGCCGGTTTTGTA |
| 3084 | mamo-L | *BGIBMGA006305* | up | 1900 | 1914 | + | 10.5726 | 9.33E-05 | 1 | TAGGCTAATATTTAT |
| 3085 | mamo-L | *BGIBMGA006306* | down | 898 | 912 | + | 13.4919 | 4.25E-06 | 1 | TAGGCTGTGTGTGAT |
| 3086 | mamo-L | *BGIBMGA006306* | up | 421 | 435 | - | 11.1613 | 5.63E-05 | 1 | TAGGCGTTTTTTGAA |
| 3087 | mamo-L | *BGIBMGA006307* | down | 148 | 162 | + | 11.2903 | 4.99E-05 | 1 | TAGGCTTAAATTATT |
| 3088 | mamo-L | *BGIBMGA006309* | down | 1164 | 1178 | + | 13.0645 | 7.42E-06 | 1 | TAGGCTAAGTTTTTT |
| 3089 | mamo-L | *BGIBMGA006315* | up | 1550 | 1564 | + | 11.2097 | 5.38E-05 | 1 | TAGGCTCTCATCGTC |
| 3090 | mamo-L | *BGIBMGA006323* | up | 1621 | 1635 | + | 11.5968 | 3.73E-05 | 1 | TAGGCTTGTGTGATT |
| 3091 | mamo-L | *BGIBMGA006324* | down | 1651 | 1665 | - | 12.5323 | 1.39E-05 | 1 | TAGGCCCAAGTTGCC |
| 3092 | mamo-L | *BGIBMGA006324* | down | 1658 | 1672 | + | 11.879 | 2.83E-05 | 1 | TGGGCCTACTTTGCT |
| 3093 | mamo-L | *BGIBMGA006325* | up | 1712 | 1726 | - | 12.5323 | 1.39E-05 | 1 | TAGGCCCAAGTTGCC |
| 3094 | mamo-L | *BGIBMGA006325* | up | 1719 | 1733 | + | 11.879 | 2.83E-05 | 1 | TGGGCCTACTTTGCT |
| 3095 | mamo-L | *BGIBMGA006326* | up | 369 | 383 | - | 12.9194 | 8.81E-06 | 1 | TAGGCGGAATTTTTT |
| 3096 | mamo-L | *BGIBMGA006329* | down | 1352 | 1366 | + | 10.9113 | 7.02E-05 | 1 | TAGGCCAGGTTTTGA |
| 3097 | mamo-L | *BGIBMGA006331* | down | 1576 | 1590 | - | 10.9194 | 6.98E-05 | 1 | TAGGCTACTTTTTTT |
| 3098 | mamo-L | *BGIBMGA006333* | up | 1268 | 1282 | + | 11.0806 | 6.05E-05 | 1 | GAGGCGGTCTTTTGC |
| 3099 | mamo-L | *BGIBMGA006333* | down | 1939 | 1953 | + | 10.8065 | 7.69E-05 | 1 | TAGGCCCATTGTATT |
| 3100 | mamo-L | *BGIBMGA006335* | down | 1391 | 1405 | - | 10.9194 | 6.98E-05 | 1 | TAGGCTACTTTTTTT |
| 3101 | mamo-L | *BGIBMGA006340* | up | 1717 | 1731 | + | 12.0161 | 2.46E-05 | 1 | GAGGCGTTTTTTGCT |
| 3102 | mamo-L | *BGIBMGA006343* | down | 534 | 548 | + | 11.0887 | 6.01E-05 | 1 | TAGGCATATGTTTAT |
| 3103 | mamo-L | *BGIBMGA006343* | up | 1109 | 1123 | + | 10.8306 | 7.52E-05 | 1 | AAGGCCTCCGTTTTT |
| 3104 | mamo-L | *BGIBMGA006343* | up | 82 | 96 | + | 10.6048 | 9.09E-05 | 1 | TCGGCCGTGTTTGGC |
| 3105 | mamo-L | *BGIBMGA006344* | down | 310 | 324 | + | 11.1613 | 5.63E-05 | 1 | TAGGCCTTTAATTTT |
| 3106 | mamo-L | *BGIBMGA006348* | up | 75 | 89 | - | 10.5242 | 9.70E-05 | 1 | GAGGCTTGGGATGAC |
| 3107 | mamo-L | *BGIBMGA006349* | up | 1932 | 1946 | + | 11.7177 | 3.32E-05 | 1 | TAGGCTAATATTTGT |
| 3108 | mamo-L | *BGIBMGA006353* | up | 1652 | 1666 | - | 10.7661 | 7.95E-05 | 1 | TAGGCGTGGGATGAA |
| 3109 | mamo-L | *BGIBMGA006357* | down | 1320 | 1334 | + | 10.5565 | 9.46E-05 | 1 | AAGGCTTTTGATGCC |
| 3110 | mamo-L | *BGIBMGA006361* | down | 1578 | 1592 | - | 11.379 | 4.58E-05 | 1 | TAGGCCCTCTAGGCT |
| 3111 | mamo-L | *BGIBMGA006367* | up | 1622 | 1636 | - | 12.0968 | 2.26E-05 | 1 | AAGGCTGGGGTTGGA |
| 3112 | mamo-L | *BGIBMGA006367* | down | 1926 | 1940 | + | 10.5726 | 9.33E-05 | 1 | TAGGCTAATATTTAT |
| 3113 | mamo-L | *BGIBMGA006368* | up | 466 | 480 | + | 12.4435 | 1.54E-05 | 1 | GAGGCTTACTTTTTT |
| 3114 | mamo-L | *BGIBMGA006368* | up | 87 | 101 | + | 12.129 | 2.19E-05 | 1 | TAGGCTTTGTTTTTA |
| 3115 | mamo-L | *BGIBMGA006376* | up | 1277 | 1291 | + | 10.6694 | 8.62E-05 | 1 | TAGGCTATTGTGTTT |
| 3116 | mamo-L | *BGIBMGA006378* | up | 51 | 65 | - | 13.621 | 3.50E-06 | 1 | TAGGCTAGTGTTGCT |
| 3117 | mamo-L | *BGIBMGA006381* | down | 1570 | 1584 | + | 13.621 | 3.50E-06 | 1 | TAGGCTGATGTTATT |
| 3118 | mamo-L | *BGIBMGA006383* | down | 1593 | 1607 | - | 12.5403 | 1.38E-05 | 1 | TAGGCTTCCGTTAGT |
| 3119 | mamo-L | *BGIBMGA006383* | down | 1485 | 1499 | - | 12.4919 | 1.46E-05 | 1 | AAGGCCGGGGTCGCT |
| 3120 | mamo-L | *BGIBMGA006384* | down | 1598 | 1612 | + | 11.4597 | 4.25E-05 | 1 | TAGGCTTTTATGGGC |
| 3121 | mamo-L | *BGIBMGA006387* | up | 1920 | 1934 | - | 10.7258 | 8.23E-05 | 1 | TAGGCCGTTTTATGT |

| | | | | | | | | | | |
|---|---|---|---|---|---|---|---|---|---|---|
| 3122 | mamo-L | *BGIBMGA006391* | down | 58 | 72 | + | 12.2581 | 1.90E-05 | 1 | TAGGCTCTGTTCTTT |
| 3123 | mamo-L | *BGIBMGA006391* | down | 1359 | 1373 | - | 11.4516 | 4.28E-05 | 1 | TAGGCTTACATTCTC |
| 3124 | mamo-L | *BGIBMGA006392* | down | 157 | 171 | + | 10.4919 | 9.96E-05 | 1 | AAGGCCGAAGTTACT |
| 3125 | mamo-L | *BGIBMGA006393* | up | 645 | 659 | + | 12.5806 | 1.31E-05 | 1 | AAGGCTGGGGGTGTT |
| 3126 | mamo-L | *BGIBMGA006396* | down | 1966 | 1980 | + | 11.6613 | 3.50E-05 | 1 | AAGGCGGGTGGTGGT |
| 3127 | mamo-L | *BGIBMGA006401* | up | 955 | 969 | - | 12.3871 | 1.65E-05 | 1 | TAGGCAGAATTTGTT |
| 3128 | mamo-L | *BGIBMGA006401* | down | 1380 | 1394 | - | 12.2581 | 1.90E-05 | 1 | TAGGCTTGCGTAGTT |
| 3129 | mamo-L | *BGIBMGA006407* | down | 794 | 808 | + | 11.3629 | 4.66E-05 | 1 | TAGGCTGAGCATGAT |
| 3130 | mamo-L | *BGIBMGA006410* | down | 9 | 23 | + | 11.1935 | 5.45E-05 | 1 | TAGGCGCATGAGGGT |
| 3131 | mamo-L | *BGIBMGA006411* | down | 989 | 1003 | - | 13.5081 | 4.15E-06 | 1 | TAGGCTGTCGTCGCC |
| 3132 | mamo-L | *BGIBMGA006414* | down | 1056 | 1070 | + | 11.4274 | 4.38E-05 | 1 | TAGGCGTCGTTCTGT |
| 3133 | mamo-L | *BGIBMGA006417* | up | 1428 | 1442 | - | 10.8065 | 7.69E-05 | 1 | TAGGCCGGCTGAGCT |
| 3134 | mamo-L | *BGIBMGA006421* | up | 1962 | 1976 | + | 10.6048 | 9.09E-05 | 1 | TAGGCATGTATTTTT |
| 3135 | mamo-L | *BGIBMGA006422* | up | 1365 | 1379 | + | 11.2823 | 5.03E-05 | 1 | TAGGCAGTCGTTGTA |
| 3136 | mamo-L | *BGIBMGA006422* | down | 234 | 248 | + | 10.5484 | 9.52E-05 | 1 | TAGGCTTAGAAGTCT |
| 3137 | mamo-L | *BGIBMGA006426* | down | 774 | 788 | + | 12.121 | 2.20E-05 | 1 | TAGGCGTTTGTTGTA |
| 3138 | mamo-L | *BGIBMGA006427* | up | 1050 | 1064 | - | 12.379 | 1.66E-05 | 1 | TAGGCGATGTTTTGT |
| 3139 | mamo-L | *BGIBMGA006433* | down | 990 | 1004 | + | 11.9758 | 2.57E-05 | 1 | TAGGCCTATTTCTGC |
| 3140 | mamo-L | *BGIBMGA006435* | up | 727 | 741 | + | 10.8871 | 7.17E-05 | 1 | TAGGCTTCCTATGGA |
| 3141 | mamo-L | *BGIBMGA006435* | down | 530 | 544 | - | 10.7339 | 8.17E-05 | 1 | TAGGCTTGAATGGAT |
| 3142 | mamo-L | *BGIBMGA006436* | down | 666 | 680 | + | 12.4597 | 1.51E-05 | 1 | TAGGCTGATATTTAT |
| 3143 | mamo-L | *BGIBMGA006436* | up | 862 | 876 | - | 11.9597 | 2.61E-05 | 1 | AAGGCTGTTTTTTTT |
| 3144 | mamo-L | *BGIBMGA006436* | up | 1366 | 1380 | - | 10.7097 | 8.33E-05 | 1 | TAGGCGATATTTTTT |
| 3145 | mamo-L | *BGIBMGA006437* | up | 579 | 593 | - | 13.9516 | 2.13E-06 | 1 | TAGGCGGTGTTTTTT |
| 3146 | mamo-L | *BGIBMGA006439* | down | 1094 | 1108 | - | 11 | 6.50E-05 | 1 | AAGGCTGGGGTTTGA |
| 3147 | mamo-L | *BGIBMGA006443* | up | 255 | 269 | + | 12.25 | 1.92E-05 | 1 | TAGGCCCACTATTCT |
| 3148 | mamo-L | *BGIBMGA006446* | up | 1730 | 1744 | - | 11.3629 | 4.66E-05 | 1 | TAGGCTCATATTTAT |
| 3149 | mamo-L | *BGIBMGA006448* | up | 628 | 642 | + | 12.0484 | 2.38E-05 | 1 | TAGGCATTCTTTGCC |
| 3150 | mamo-L | *BGIBMGA006449* | down | 1612 | 1626 | + | 10.5 | 9.90E-05 | 1 | GAGGCCCAGGTCGTC |
| 3151 | mamo-L | *BGIBMGA006450* | up | 275 | 289 | + | 11.7984 | 3.06E-05 | 1 | TAGGCCACTTTTGCT |
| 3152 | mamo-L | *BGIBMGA006450* | up | 1463 | 1477 | + | 11.7016 | 3.37E-05 | 1 | AAGGCTGGGGTCGCC |
| 3153 | mamo-L | *BGIBMGA006453* | down | 54 | 68 | - | 11.7742 | 3.14E-05 | 1 | AAGGCCGTTTTTTTT |
| 3154 | mamo-L | *BGIBMGA006459* | down | 1366 | 1380 | + | 13.0726 | 7.34E-06 | 1 | TAGGCGTACATTGAT |
| 3155 | mamo-L | *BGIBMGA006460* | up | 1212 | 1226 | + | 13.0726 | 7.34E-06 | 1 | TAGGCGTACATTGAT |
| 3156 | mamo-L | *BGIBMGA006463* | up | 1981 | 1995 | - | 12.0968 | 2.26E-05 | 1 | AAGGCTGGGGTTGGA |
| 3157 | mamo-L | *BGIBMGA006464* | down | 244 | 258 | - | 11.1452 | 5.70E-05 | 1 | TAGGCGGGGTGGTTT |
| 3158 | mamo-L | *BGIBMGA006465* | up | 1859 | 1873 | + | 11 | 6.50E-05 | 1 | AAGGCTGTGGTGTTT |
| 3159 | mamo-L | *BGIBMGA006468* | down | 1404 | 1418 | - | 10.5726 | 9.33E-05 | 1 | TAGGCTAATATTTAT |
| 3160 | mamo-L | *BGIBMGA006470* | up | 523 | 537 | - | 12.0645 | 2.34E-05 | 1 | TAGGCGGGCTTCGGA |
| 3161 | mamo-L | *BGIBMGA006472* | down | 914 | 928 | - | 11.7419 | 3.23E-05 | 1 | TAGGCGTGGGTTTGG |
| 3162 | mamo-L | *BGIBMGA006472* | down | 122 | 136 | - | 11.6694 | 3.47E-05 | 1 | GAGGCTTAGGGTGGC |
| 3163 | mamo-L | *BGIBMGA006473* | up | 584 | 598 | - | 11.7419 | 3.23E-05 | 1 | TAGGCGTGGGTTTGG |
| 3164 | mamo-L | *BGIBMGA006477* | up | 1789 | 1803 | + | 12.3871 | 1.65E-05 | 1 | TAGGCAGAATTTGTT |
| 3165 | mamo-L | *BGIBMGA006483* | up | 554 | 568 | + | 11.5806 | 3.78E-05 | 1 | TCGGCCGTGTTTGGT |
| 3166 | mamo-L | *BGIBMGA006484* | down | 568 | 582 | - | 11.6613 | 3.50E-05 | 1 | AAGGCGGGTGGTGGT |
| 3167 | mamo-L | *BGIBMGA006485* | down | 1064 | 1078 | + | 13.7823 | 2.76E-06 | 1 | TAGGCTCGTGTTGGC |
| 3168 | mamo-L | *BGIBMGA006485* | up | 1772 | 1786 | + | 11.8065 | 3.03E-05 | 1 | AAGGCTTATTTCGTT |
| 3169 | mamo-L | *BGIBMGA006489* | up | 706 | 720 | - | 11.4435 | 4.31E-05 | 1 | TAGGCTGGCAATACT |
| 3170 | mamo-L | *BGIBMGA006491* | down | 1815 | 1829 | + | 12.5403 | 1.38E-05 | 1 | TAGGCTTCCGTTAGT |
| 3171 | mamo-L | *BGIBMGA006491* | down | 1923 | 1937 | + | 12.4919 | 1.46E-05 | 1 | AAGGCCGGGGTCGCT |
| 3172 | mamo-L | *BGIBMGA006495* | up | 1874 | 1888 | + | 11.371 | 4.62E-05 | 1 | TAGGCTGTATAGGCT |
| 3173 | mamo-L | *BGIBMGA006497* | down | 151 | 165 | - | 10.75 | 8.06E-05 | 1 | TAGGCCCGCTTCGTG |
| 3174 | mamo-L | *BGIBMGA006498* | up | 1278 | 1292 | + | 13.621 | 3.50E-06 | 1 | TAGGCTAGTGTTGCT |
| 3175 | mamo-L | *BGIBMGA006499* | down | 1135 | 1149 | + | 10.8871 | 7.17E-05 | 1 | TAGGCCCTTGTTCTC |
| 3176 | mamo-L | *BGIBMGA006500* | down | 423 | 437 | + | 10.9113 | 7.02E-05 | 1 | TAGGCTTTCTTAGTC |
| 3177 | mamo-L | *BGIBMGA006502* | down | 1102 | 1116 | - | 11.2903 | 4.99E-05 | 1 | TAGGCGTTATTTTCC |
| 3178 | mamo-L | *BGIBMGA006503* | down | 973 | 987 | - | 10.75 | 8.06E-05 | 1 | GAGGCCGATTGTGCC |
| 3179 | mamo-L | *BGIBMGA006504* | down | 548 | 562 | + | 11.5323 | 3.96E-05 | 1 | TAGGCTAAATTCGTT |
| 3180 | mamo-L | *BGIBMGA006510* | up | 315 | 329 | + | 13.0565 | 7.51E-06 | 1 | AAGGCGTAGGTTGCT |
| 3181 | mamo-L | *BGIBMGA006513* | down | 729 | 743 | - | 13.4677 | 4.39E-06 | 1 | TAGGCTGATGATGAT |
| 3182 | mamo-L | *BGIBMGA006513* | down | 1175 | 1189 | + | 10.5161 | 9.77E-05 | 1 | AAGGCTTTTGTCTTT |
| 3183 | mamo-L | *BGIBMGA006514* | down | 978 | 992 | - | 10.9194 | 6.98E-05 | 1 | TAGGCTGACGACGTG |
| 3184 | mamo-L | *BGIBMGA006515* | down | 274 | 288 | - | 11.0484 | 6.23E-05 | 1 | TAGGCGGTAGTGTCT |
| 3185 | mamo-L | *BGIBMGA006517* | down | 1938 | 1952 | + | 13.7742 | 2.79E-06 | 1 | TAGGCTTTTTTTTTT |
| 3186 | mamo-L | *BGIBMGA006521* | up | 1644 | 1658 | + | 12.4516 | 1.53E-05 | 1 | TAGGCTTATTTGTTT |
| 3187 | mamo-L | *BGIBMGA006524* | up | 1512 | 1526 | + | 13.1129 | 6.98E-06 | 1 | TAGGCCGCCTTTTTT |
| 3188 | mamo-L | *BGIBMGA006525* | down | 1821 | 1835 | - | 10.5565 | 9.46E-05 | 1 | AAGGCTTTTGATGCC |
| 3189 | mamo-L | *BGIBMGA006529* | down | 1728 | 1742 | - | 10.8145 | 7.63E-05 | 1 | TAGGCCGCCTATTTC |
| 3190 | mamo-L | *BGIBMGA006530* | down | 1070 | 1084 | + | 12.5081 | 1.43E-05 | 1 | TAGGCTGAATTTCCT |
| 3191 | mamo-L | *BGIBMGA006531* | up | 1099 | 1113 | + | 13.4597 | 4.43E-06 | 1 | TAGGCTCTCATTGTT |
| 3192 | mamo-L | *BGIBMGA006531* | up | 1375 | 1389 | - | 11.7016 | 3.37E-05 | 1 | TAGGCGGTTATTAGT |
| 3193 | mamo-L | *BGIBMGA006531* | up | 496 | 510 | + | 10.9194 | 6.98E-05 | 1 | TAGGCTACTTTTTTT |
| 3194 | mamo-L | *BGIBMGA006534* | up | 278 | 292 | - | 11.0726 | 6.09E-05 | 1 | TAGGCTCTTTAGGCT |
| 3195 | mamo-L | *BGIBMGA006535* | up | 557 | 571 | - | 10.7177 | 8.28E-05 | 1 | TAGGCATATGATGTC |
| 3196 | mamo-L | *BGIBMGA006538* | down | 392 | 406 | + | 11.4435 | 4.31E-05 | 1 | TAGGCTTGAGAGGCT |
| 3197 | mamo-L | *BGIBMGA006540* | down | 709 | 723 | - | 11.5968 | 3.73E-05 | 1 | TAGGCTGTATTGTCT |
| 3198 | mamo-L | *BGIBMGA006541* | down | 1136 | 1150 | - | 12.1452 | 2.15E-05 | 1 | TAGGCAGGTGGTGGT |
| 3199 | mamo-L | *BGIBMGA006543* | up | 954 | 968 | + | 12.5161 | 1.42E-05 | 1 | TAGGCGGACATTAGT |
| 3200 | mamo-L | *BGIBMGA006550* | down | 1805 | 1819 | + | 10.6371 | 8.85E-05 | 1 | AAGGCGCCCGTTGTT |
| 3201 | mamo-L | *BGIBMGA006550* | down | 1577 | 1591 | - | 10.5726 | 9.33E-05 | 1 | TAGGCTAATATTTAT |
| 3202 | mamo-L | *BGIBMGA006554* | up | 1981 | 1995 | + | 12.2984 | 1.81E-05 | 1 | GAGGCTCGTGTTGTT |
| 3203 | mamo-L | *BGIBMGA006557* | down | 1472 | 1486 | + | 11.371 | 4.62E-05 | 1 | TAGGCTGTATAGGCT |
| 3204 | mamo-L | *BGIBMGA006558* | up | 1015 | 1029 | + | 11.371 | 4.62E-05 | 1 | TAGGCTGTATAGGCT |
| 3205 | mamo-L | *BGIBMGA006559* | up | 1394 | 1408 | - | 14.3145 | 1.24E-06 | 1 | TAGGCTCGTTTTGTT |
| 3206 | mamo-L | *BGIBMGA006560* | up | 1114 | 1128 | + | 11.1048 | 5.92E-05 | 1 | AAGGCTGAGGTCAGT |
| 3207 | mamo-L | *BGIBMGA006561* | down | 472 | 486 | - | 11.371 | 4.62E-05 | 1 | TAGGCTGTGGATCCC |
| 3208 | mamo-L | *BGIBMGA006561* | up | 62 | 76 | - | 10.9194 | 6.98E-05 | 1 | TAGGCTACTTTTTTT |
| 3209 | mamo-L | *BGIBMGA006565* | down | 745 | 759 | - | 11.8065 | 3.03E-05 | 1 | TAGGCTAATGTTTAT |
| 3210 | mamo-L | *BGIBMGA006565* | down | 1416 | 1430 | + | 11.7177 | 3.32E-05 | 1 | TAGGCGGGCTGCGAT |
| 3211 | mamo-L | *BGIBMGA006566* | down | 1337 | 1351 | + | 11.2823 | 5.03E-05 | 1 | TAGGCTGGATTTTCA |
| 3212 | mamo-L | *BGIBMGA006566* | down | 563 | 577 | - | 11.0161 | 6.40E-05 | 1 | TAGGCTTTGATCACT |
| 3213 | mamo-L | *BGIBMGA006566* | down | 1087 | 1101 | - | 10.7742 | 7.89E-05 | 1 | TGGGCTTAATTTGCT |

| | | | | | | | | | | |
|---|---|---|---|---|---|---|---|---|---|---|
| 3214 | mamo-L | *BGIBMGA006568* | down | 1872 | 1886 | - | 10.9597 | 6.73E-05 | 1 | AAGGCTTGCTGTTCT |
| 3215 | mamo-L | *BGIBMGA006568* | up | 121 | 135 | + | 10.5403 | 9.58E-05 | 1 | TAGGCTGTATGGTGT |
| 3216 | mamo-L | *BGIBMGA006569* | up | 1795 | 1809 | - | 11.2258 | 5.30E-05 | 1 | TAGGCTGGTAGGGCT |
| 3217 | mamo-L | *BGIBMGA006572* | down | 588 | 602 | + | 12.2984 | 1.81E-05 | 1 | AAGGCTTGAGTTGCT |
| 3218 | mamo-L | *BGIBMGA006580* | up | 565 | 579 | + | 11.2661 | 5.10E-05 | 1 | TAGGCTTTTTATGGG |
| 3219 | mamo-L | *BGIBMGA006581* | down | 1412 | 1426 | - | 11.371 | 4.62E-05 | 1 | TAGGCTGTATAGGCT |
| 3220 | mamo-L | *BGIBMGA006581* | up | 1102 | 1116 | - | 11.0887 | 6.01E-05 | 1 | TAGGCTGGAAATTTT |
| 3221 | mamo-L | *BGIBMGA006581* | up | 724 | 738 | + | 11.0484 | 6.23E-05 | 1 | TAGGCTATTGGCGGT |
| 3222 | mamo-L | *BGIBMGA006588* | up | 122 | 136 | + | 10.5645 | 9.40E-05 | 1 | GAGGCTGGTGTTAAT |
| 3223 | mamo-L | *BGIBMGA006592* | up | 1497 | 1511 | - | 12.9355 | 8.65E-06 | 1 | TAGGCCAAATTTGGT |
| 3224 | mamo-L | *BGIBMGA006596* | down | 605 | 619 | + | 11.3468 | 4.73E-05 | 1 | TAGGCGAAATTTTGT |
| 3225 | mamo-L | *BGIBMGA006599* | up | 1610 | 1624 | + | 10.7742 | 7.89E-05 | 1 | AAGGCGTAGTGTTGT |
| 3226 | mamo-L | *BGIBMGA006605* | down | 781 | 795 | - | 10.5 | 9.90E-05 | 1 | GAGGCCCAGGTCGTC |
| 3227 | mamo-L | *BGIBMGA006607* | up | 1144 | 1158 | + | 10.7016 | 8.39E-05 | 1 | AAGGCCAACTTTTTT |
| 3228 | mamo-L | *BGIBMGA006611* | down | 1091 | 1105 | + | 11.0565 | 6.18E-05 | 1 | AAGGCTTATGTTCTT |
| 3229 | mamo-L | *BGIBMGA006611* | up | 1410 | 1424 | + | 10.5323 | 9.63E-05 | 1 | TAGGCAGCCGTTAGT |
| 3230 | mamo-L | *BGIBMGA006615* | down | 521 | 535 | - | 10.5726 | 9.33E-05 | 1 | TAGGCTAATATTTAT |
| 3231 | mamo-L | *BGIBMGA006618* | down | 1205 | 1219 | - | 13.7661 | 2.82E-06 | 1 | TAGGCTTTTATTGTT |
| 3232 | mamo-L | *BGIBMGA006618* | down | 738 | 752 | - | 10.8387 | 7.47E-05 | 1 | AAGGCGTACGGTTGT |
| 3233 | mamo-L | *BGIBMGA006620* | up | 542 | 556 | + | 11.2823 | 5.03E-05 | 1 | AAGGCGGTTTTTTTT |
| 3234 | mamo-L | *BGIBMGA006620* | up | 949 | 963 | + | 10.8306 | 7.52E-05 | 1 | TAGGCCCCCTTCGCC |
| 3235 | mamo-L | *BGIBMGA006623* | down | 923 | 937 | + | 12.1371 | 2.17E-05 | 1 | TAGGCCGATGTTTTA |
| 3236 | mamo-L | *BGIBMGA006624* | up | 1492 | 1506 | + | 13.0726 | 7.34E-06 | 1 | TAGGCTCTGGTTAGT |
| 3237 | mamo-L | *BGIBMGA006628* | up | 1726 | 1740 | + | 11.6532 | 3.53E-05 | 1 | TAGGCTATGTTTCCT |
| 3238 | mamo-L | *BGIBMGA006633* | down | 198 | 212 | - | 10.5726 | 9.33E-05 | 1 | TAGGCTAATATTTAT |
| 3239 | mamo-L | *BGIBMGA006634* | up | 1909 | 1923 | - | 10.9597 | 6.73E-05 | 1 | TAGGCTTATTGTAAT |
| 3240 | mamo-L | *BGIBMGA006644* | up | 1584 | 1598 | - | 11.9032 | 2.76E-05 | 1 | TAGGCTCGCTTTTAC |
| 3241 | mamo-L | *BGIBMGA006646* | up | 1162 | 1176 | + | 10.5968 | 9.15E-05 | 1 | TACGCTTATTTTGGT |
| 3242 | mamo-L | *BGIBMGA006650* | up | 660 | 674 | - | 10.7177 | 8.28E-05 | 1 | AAGGCGGGGAATGCT |
| 3243 | mamo-L | *BGIBMGA006651* | down | 115 | 129 | - | 12.629 | 1.24E-05 | 1 | GAGGCTGAGTTCGTT |
| 3244 | mamo-L | *BGIBMGA006652* | down | 793 | 807 | - | 11.5242 | 3.99E-05 | 1 | TCGGCTTGGGTTGTT |
| 3245 | mamo-L | *BGIBMGA006653* | up | 1594 | 1608 | - | 11.5242 | 3.99E-05 | 1 | TCGGCTTGGGTTGTT |
| 3246 | mamo-L | *BGIBMGA006659* | down | 1174 | 1188 | + | 10.8548 | 7.38E-05 | 1 | TACGCTTGGGTTGCT |
| 3247 | mamo-L | *BGIBMGA006662* | up | 1483 | 1497 | - | 11.9355 | 2.67E-05 | 1 | TAGGCCTATGTCTAT |
| 3248 | mamo-L | *BGIBMGA006668* | up | 1229 | 1243 | + | 14.2016 | 1.47E-06 | 1 | TAGGCGGGGTGTGGT |
| 3249 | mamo-L | *BGIBMGA006670* | up | 1603 | 1617 | + | 10.7177 | 8.28E-05 | 1 | GAGGCCTACTCTGTT |
| 3250 | mamo-L | *BGIBMGA006674* | up | 259 | 273 | - | 10.5161 | 9.77E-05 | 1 | TAGGCGTATTGTGTG |
| 3251 | mamo-L | *BGIBMGA006679* | up | 1465 | 1479 | - | 11.7097 | 3.34E-05 | 1 | TAGGCTGTATTTAGC |
| 3252 | mamo-L | *BGIBMGA006679* | down | 1191 | 1205 | + | 10.9194 | 6.98E-05 | 1 | TAGGCTACTTTTTTT |
| 3253 | mamo-L | *BGIBMGA006682* | up | 1422 | 1436 | - | 11.7823 | 3.11E-05 | 1 | TAGGCGACTGTTGGT |
| 3254 | mamo-L | *BGIBMGA006686* | down | 1791 | 1805 | + | 11.4435 | 4.31E-05 | 1 | AAGGCCTTTTTTTCT |
| 3255 | mamo-L | *BGIBMGA006690* | up | 864 | 878 | - | 10.5968 | 9.15E-05 | 1 | TACGCTTATTTTGGT |
| 3256 | mamo-L | *BGIBMGA006702* | up | 1566 | 1580 | - | 11.9839 | 2.54E-05 | 1 | TAGGCTGGTACTGGT |
| 3257 | mamo-L | *BGIBMGA006703* | down | 556 | 570 | - | 10.7339 | 8.17E-05 | 1 | TAGGCGCGGGGTCCT |
| 3258 | mamo-L | *BGIBMGA006706* | down | 1452 | 1466 | + | 10.5726 | 9.33E-05 | 1 | TAGGCTAATATTTAT |
| 3259 | mamo-L | *BGIBMGA006718* | down | 873 | 887 | - | 12.1371 | 2.17E-05 | 1 | TAGGCCGATGTTTTA |
| 3260 | mamo-L | *BGIBMGA006720* | down | 288 | 302 | + | 13.7661 | 2.82E-06 | 1 | TAGGCTTTTATTGTT |
| 3261 | mamo-L | *BGIBMGA006720* | down | 755 | 769 | + | 10.8387 | 7.47E-05 | 1 | AAGGCGTACGGTTGT |
| 3262 | mamo-L | *BGIBMGA006724* | down | 967 | 981 | - | 11.0806 | 6.05E-05 | 1 | TAGGCTATTATTTTT |
| 3263 | mamo-L | *BGIBMGA006724* | up | 1107 | 1121 | - | 11.0323 | 6.32E-05 | 1 | TAGGCCGACGAGTAT |
| 3264 | mamo-L | *BGIBMGA006725* | down | 801 | 815 | - | 11.5323 | 3.96E-05 | 1 | TAGGCTCTCGATCGT |
| 3265 | mamo-L | *BGIBMGA006726* | down | 389 | 403 | - | 12.0645 | 2.34E-05 | 1 | TAGGCGGGCTTCGGA |
| 3266 | mamo-L | *BGIBMGA006729* | up | 755 | 769 | - | 10.5806 | 9.27E-05 | 1 | TAGGCTCACTATCAT |
| 3267 | mamo-L | *BGIBMGA006730* | down | 477 | 491 | - | 11.3145 | 4.87E-05 | 1 | GAGGCGTGTTTTGTC |
| 3268 | mamo-L | *BGIBMGA006735* | down | 65 | 79 | + | 11.7097 | 3.34E-05 | 1 | TAGGCTGAATTTCAT |
| 3269 | mamo-L | *BGIBMGA006736* | down | 1391 | 1405 | - | 10.5645 | 9.40E-05 | 1 | TAGGCCAAATTCTGT |
| 3270 | mamo-L | *BGIBMGA006739* | down | 1823 | 1837 | + | 13.7984 | 2.69E-06 | 1 | TAGGCGAAGTTTGGT |
| 3271 | mamo-L | *BGIBMGA006739* | down | 1560 | 1574 | - | 10.871 | 7.28E-05 | 1 | TAGGCTTGGTAAGTT |
| 3272 | mamo-L | *BGIBMGA006741* | down | 409 | 423 | + | 10.7339 | 8.17E-05 | 1 | AAGGCTTAGTTTTAC |
| 3273 | mamo-L | *BGIBMGA006743* | up | 643 | 657 | - | 10.8306 | 7.52E-05 | 1 | TAGGCCCCCTTCGCC |
| 3274 | mamo-L | *BGIBMGA006746* | up | 101 | 115 | - | 10.5081 | 9.83E-05 | 1 | AAGGCGGTGAATGTT |
| 3275 | mamo-L | *BGIBMGA006747* | down | 1906 | 1920 | + | 10.9032 | 7.07E-05 | 1 | TAGGCGGGGTGTTGA |
| 3276 | mamo-L | *BGIBMGA006748* | down | 762 | 776 | - | 10.8629 | 7.33E-05 | 1 | AAGGCTTTATTTTTT |
| 3277 | mamo-L | *BGIBMGA006749* | up | 1531 | 1545 | - | 10.8629 | 7.33E-05 | 1 | AAGGCTTTATTTTTT |
| 3278 | mamo-L | *BGIBMGA006753* | down | 667 | 681 | + | 11.1129 | 5.88E-05 | 1 | TAGGCGTTTTATTGC |
| 3279 | mamo-L | *BGIBMGA006753* | up | 762 | 776 | + | 10.5726 | 9.33E-05 | 1 | TAGGCTAATATTTAT |
| 3280 | mamo-L | *BGIBMGA006760* | up | 1952 | 1966 | - | 11.2661 | 5.10E-05 | 1 | TAGGCTTTTTATGGG |
| 3281 | mamo-L | *BGIBMGA006763* | down | 448 | 462 | - | 10.629 | 8.91E-05 | 1 | TAGGCCTGCAGCGCC |
| 3282 | mamo-L | *BGIBMGA006764* | down | 835 | 849 | + | 12.0968 | 2.26E-05 | 1 | TGGGCGGAGTTTGGT |
| 3283 | mamo-L | *BGIBMGA006766* | up | 922 | 936 | + | 13.5968 | 3.65E-06 | 1 | TAGGCTATTTTTGGT |
| 3284 | mamo-L | *BGIBMGA006767* | down | 250 | 264 | - | 11.6935 | 3.39E-05 | 1 | TAGGCTTCCGGTTCT |
| 3285 | mamo-L | *BGIBMGA006770* | up | 114 | 128 | - | 10.9194 | 6.98E-05 | 1 | TAGGCTACTTTTTTT |
| 3286 | mamo-L | *BGIBMGA006770* | up | 808 | 822 | + | 10.879 | 7.22E-05 | 1 | TAGGCTAAATTTTAT |
| 3287 | mamo-L | *BGIBMGA006771* | up | 1028 | 1042 | + | 10.9597 | 6.73E-05 | 1 | AAGGCTTGCTGTTCT |
| 3288 | mamo-L | *BGIBMGA006780* | down | 100 | 114 | + | 14.2258 | 1.41E-06 | 1 | TAGGCTTATGTTTTT |
| 3289 | mamo-L | *BGIBMGA006780* | up | 941 | 955 | - | 11.1048 | 5.92E-05 | 1 | AAGGCTGAGGTCAGT |
| 3290 | mamo-L | *BGIBMGA006783* | up | 1648 | 1662 | + | 10.5726 | 9.33E-05 | 1 | TAGGCTAATATTTAT |
| 3291 | mamo-L | *BGIBMGA006786* | up | 924 | 938 | + | 10.5726 | 9.33E-05 | 1 | TAGGCTAATATTTAT |
| 3292 | mamo-L | *BGIBMGA006787* | up | 183 | 197 | + | 10.9194 | 6.98E-05 | 1 | TAGGCTACTTTTTTT |
| 3293 | mamo-L | *BGIBMGA006788* | down | 400 | 414 | + | 11.2903 | 4.99E-05 | 1 | TAGGCTGAATGTATT |
| 3294 | mamo-L | *BGIBMGA006789* | down | 1119 | 1133 | - | 12.0645 | 2.34E-05 | 1 | TAGGCGGGCTTCGGA |
| 3295 | mamo-L | *BGIBMGA006789* | down | 1889 | 1903 | - | 10.5645 | 9.40E-05 | 1 | GAGGCGCTCGTCGCT |
| 3296 | mamo-L | *BGIBMGA006790* | up | 1417 | 1431 | + | 10.5726 | 9.33E-05 | 1 | TAGGCTAATATTTAT |
| 3297 | mamo-L | *BGIBMGA006798* | down | 1582 | 1596 | + | 12.0645 | 2.34E-05 | 1 | TAGGCGGGCTTCGGA |
| 3298 | mamo-L | *BGIBMGA006800* | down | 1129 | 1143 | - | 11.9032 | 2.76E-05 | 1 | TAGGCTTTCTTCCCT |
| 3299 | mamo-L | *BGIBMGA006801* | down | 898 | 912 | + | 11.8306 | 2.97E-05 | 1 | TAGGCTTGGAGGGGT |
| 3300 | mamo-L | *BGIBMGA006801* | up | 1449 | 1463 | - | 11.621 | 3.63E-05 | 1 | TAGGCTTAGGTCTGA |
| 3301 | mamo-L | *BGIBMGA006802* | down | 1342 | 1356 | + | 10.7903 | 7.79E-05 | 1 | TAGGCTCCCAGTGTT |
| 3302 | mamo-L | *BGIBMGA006803* | up | 883 | 897 | + | 10.7661 | 7.95E-05 | 1 | TAGGCGGAATATTAT |
| 3303 | mamo-L | *BGIBMGA006806* | down | 1172 | 1186 | + | 12.2742 | 1.87E-05 | 1 | TAGGCTTTAGTTTAT |
| 3304 | mamo-L | *BGIBMGA006809* | down | 165 | 179 | + | 10.9194 | 6.98E-05 | 1 | TAGGCTTAAATCTTT |
| 3305 | mamo-L | *BGIBMGA006810* | down | 610 | 624 | - | 10.8226 | 7.58E-05 | 1 | AAGGCTTTTTTCGGC |

| | | | | | | | | | | |
|---|---|---|---|---|---|---|---|---|---|---|
| 3306 | mamo-L | *BGIBMGA006811* | up | 837 | 851 | + | 13.4194 | 4.70E-06 | 1 | TAGGCCGGTTTTGAC |
| 3307 | mamo-L | *BGIBMGA006812* | up | 559 | 573 | + | 12.2823 | 1.85E-05 | 1 | TAGGCCCACTATTTT |
| 3308 | mamo-L | *BGIBMGA006813* | up | 80 | 94 | + | 11.5726 | 3.81E-05 | 1 | GAGGCTGAATTTGTC |
| 3309 | mamo-L | *BGIBMGA006813* | down | 1180 | 1194 | + | 10.5726 | 9.33E-05 | 1 | TAGGCTAATATTTAT |
| 3310 | mamo-L | *BGIBMGA006814* | up | 1251 | 1265 | + | 10.5726 | 9.33E-05 | 1 | TAGGCTAATATTTAT |
| 3311 | mamo-L | *BGIBMGA006816* | down | 971 | 985 | + | 11.2097 | 5.38E-05 | 1 | TAGGCTTATTATAGC |
| 3312 | mamo-L | *BGIBMGA006820* | down | 584 | 598 | - | 13.1371 | 6.76E-06 | 1 | TAGGCTAGCGTTGCC |
| 3313 | mamo-L | *BGIBMGA006824* | up | 1295 | 1309 | + | 11.6371 | 3.58E-05 | 1 | TAGGCTTGCCGTGCT |
| 3314 | mamo-L | *BGIBMGA006825* | down | 1057 | 1071 | - | 12.879 | 9.26E-06 | 1 | TAGGCACGGTTTGGT |
| 3315 | mamo-L | *BGIBMGA006829* | down | 678 | 692 | + | 10.8306 | 7.52E-05 | 1 | TAGGCTGTGTAGTAT |
| 3316 | mamo-L | *BGIBMGA006835* | down | 1683 | 1697 | - | 11.3548 | 4.69E-05 | 1 | AAGGCTGATGATGAT |
| 3317 | mamo-L | *BGIBMGA006836* | up | 4 | 18 | + | 11.4194 | 4.42E-05 | 1 | TAGGCGTTGTTGAGT |
| 3318 | mamo-L | *BGIBMGA006837* | up | 1887 | 1901 | + | 10.9839 | 6.59E-05 | 1 | TAGGCCTATGATAAT |
| 3319 | mamo-L | *BGIBMGA006837* | down | 1343 | 1357 | + | 10.9355 | 6.88E-05 | 1 | TAGGCGCAGGCTGAT |
| 3320 | mamo-L | *BGIBMGA006837* | up | 1880 | 1894 | - | 10.7419 | 8.11E-05 | 1 | TAGGCCTATATAGGT |
| 3321 | mamo-L | *BGIBMGA006837* | down | 1643 | 1657 | + | 10.5726 | 9.33E-05 | 1 | TAGGCAGGGAATGGC |
| 3322 | mamo-L | *BGIBMGA006838* | up | 112 | 126 | + | 12.7823 | 1.03E-05 | 1 | TAGGCAGGGGATGGT |
| 3323 | mamo-L | *BGIBMGA006840* | up | 1636 | 1650 | - | 12.5403 | 1.38E-05 | 1 | AAGGCTTAGTTTTTT |
| 3324 | mamo-L | *BGIBMGA006842* | down | 976 | 990 | - | 12.0484 | 2.38E-05 | 1 | TAGGCTAGGTTTACT |
| 3325 | mamo-L | *BGIBMGA006844* | up | 1305 | 1319 | + | 11.5806 | 3.78E-05 | 1 | TAGGCCATTGTGGTT |
| 3326 | mamo-L | *BGIBMGA006852* | down | 1804 | 1818 | - | 11.5081 | 4.05E-05 | 1 | TAGGCTTGTAGTTTT |
| 3327 | mamo-L | *BGIBMGA006855* | up | 1777 | 1791 | + | 10.5403 | 9.58E-05 | 1 | TAGGCTGTATGGTGT |
| 3328 | mamo-L | *BGIBMGA006860* | down | 1695 | 1709 | + | 12.879 | 9.26E-06 | 1 | TAGGCACGGTTTGGT |
| 3329 | mamo-L | *BGIBMGA006861* | down | 1923 | 1937 | - | 10.9919 | 6.55E-05 | 1 | TAGGCATAATTTTTT |
| 3330 | mamo-L | *BGIBMGA006862* | down | 124 | 138 | - | 10.9919 | 6.55E-05 | 1 | TAGGCATAATTTTTT |
| 3331 | mamo-L | *BGIBMGA006863* | down | 158 | 172 | - | 12.7097 | 1.13E-05 | 1 | TAGGCGTAATATGGT |
| 3332 | mamo-L | *BGIBMGA006864* | down | 1925 | 1939 | + | 13.1371 | 6.76E-06 | 1 | TAGGCTAGCGTTGCC |
| 3333 | mamo-L | *BGIBMGA006864* | up | 1025 | 1039 | + | 11.7823 | 3.11E-05 | 1 | TAGGCCGATTATGTA |
| 3334 | mamo-L | *BGIBMGA006864* | up | 1780 | 1794 | + | 10.9194 | 6.98E-05 | 1 | TAGGCTACTTTTTTT |
| 3335 | mamo-L | *BGIBMGA006871* | up | 938 | 952 | + | 11.7903 | 3.08E-05 | 1 | TAGGCTCTTTTCGCC |
| 3336 | mamo-L | *BGIBMGA006871* | down | 1439 | 1453 | - | 10.5968 | 9.15E-05 | 1 | GAGGCTAGTTTTTGT |
| 3337 | mamo-L | *BGIBMGA006872* | up | 861 | 875 | - | 10.5968 | 9.15E-05 | 1 | GAGGCTAGTTTTTGT |
| 3338 | mamo-L | *BGIBMGA006874* | down | 1136 | 1150 | - | 11.1532 | 5.67E-05 | 1 | AAGGCTGTAATTGTT |
| 3339 | mamo-L | *BGIBMGA006876* | up | 810 | 824 | - | 13.4194 | 4.70E-06 | 1 | TAGGCCGGTTTTGAC |
| 3340 | mamo-L | *BGIBMGA006878* | up | 1816 | 1830 | - | 10.9516 | 6.78E-05 | 1 | GAGGCGTTTTTTTTT |
| 3341 | mamo-L | *BGIBMGA006880* | down | 525 | 539 | + | 12.9919 | 8.10E-06 | 1 | TAGGCCGGTTTTGCA |
| 3342 | mamo-L | *BGIBMGA006880* | up | 1105 | 1119 | - | 11.4516 | 4.28E-05 | 1 | GAGGCTGAATTTTTT |
| 3343 | mamo-L | *BGIBMGA006891* | up | 414 | 428 | - | 12.4758 | 1.48E-05 | 1 | AAGGCCGAGTTCGTT |
| 3344 | mamo-L | *BGIBMGA006899* | down | 1465 | 1479 | + | 12.879 | 9.26E-06 | 1 | TAGGCCAAGTTTTTT |
| 3345 | mamo-L | *BGIBMGA006901* | down | 1779 | 1793 | - | 12.4032 | 1.61E-05 | 1 | TAGGCTTAGGTAGTT |
| 3346 | mamo-L | *BGIBMGA006903* | down | 671 | 685 | + | 10.5645 | 9.40E-05 | 1 | TCGGCTTGGTTTGAT |
| 3347 | mamo-L | *BGIBMGA006911* | up | 318 | 332 | + | 12.5161 | 1.42E-05 | 1 | TAGGCGCATGTGGGT |
| 3348 | mamo-L | *BGIBMGA006917* | down | 544 | 558 | + | 10.9194 | 6.98E-05 | 1 | TAGGCTACTTTTTTT |
| 3349 | mamo-L | *BGIBMGA006918* | up | 1386 | 1400 | - | 15.6613 | 9.32E-08 | 1 | TAGGCTGTCTTTGTT |
| 3350 | mamo-L | *BGIBMGA006920* | down | 939 | 953 | + | 10.5645 | 9.40E-05 | 1 | TAGGCTTATTACACT |
| 3351 | mamo-L | *BGIBMGA006921* | up | 227 | 241 | + | 11.379 | 4.58E-05 | 1 | TAGGCGAGGGTTCTT |
| 3352 | mamo-L | *BGIBMGA006923* | down | 1102 | 1116 | - | 10.7016 | 8.39E-05 | 1 | AAGGCGGATTTTATT |
| 3353 | mamo-L | *BGIBMGA006924* | down | 1063 | 1077 | - | 11 | 6.50E-05 | 1 | TAGGCAATGGTTGGC |
| 3354 | mamo-L | *BGIBMGA006926* | up | 769 | 783 | - | 10.6613 | 8.67E-05 | 1 | TAGGCAGAGGGTGAC |
| 3355 | mamo-L | *BGIBMGA006932* | down | 137 | 151 | - | 11.4758 | 4.18E-05 | 1 | TAGGCAGGGTGTTGT |
| 3356 | mamo-L | *BGIBMGA006934* | up | 438 | 452 | - | 11.1129 | 5.88E-05 | 1 | TAGGCGCATGGGGGT |
| 3357 | mamo-L | *BGIBMGA006938* | down | 350 | 364 | + | 10.9355 | 6.88E-05 | 1 | TAGGCCGAATTGTAT |
| 3358 | mamo-L | *BGIBMGA006943* | up | 1557 | 1571 | + | 11.129 | 5.79E-05 | 1 | AAGGCGTATTTCGTT |
| 3359 | mamo-L | *BGIBMGA006947* | down | 233 | 247 | + | 13.4113 | 4.75E-06 | 1 | TAGGCCTTTTTCGTT |
| 3360 | mamo-L | *BGIBMGA006949* | up | 166 | 180 | - | 12.4194 | 1.58E-05 | 1 | TAGGCTGAGCTTTTT |
| 3361 | mamo-L | *BGIBMGA006950* | up | 502 | 516 | + | 11.9435 | 2.65E-05 | 1 | TAGGCCAAGTTTACT |
| 3362 | mamo-L | *BGIBMGA006952* | up | 777 | 791 | + | 13.5403 | 3.97E-06 | 1 | TAGGCTGTATTTGAT |
| 3363 | mamo-L | *BGIBMGA006956* | up | 983 | 997 | + | 10.629 | 8.91E-05 | 1 | TAGGCAAAAGTTGTT |
| 3364 | mamo-L | *BGIBMGA006958* | down | 628 | 642 | + | 13.1855 | 6.42E-06 | 1 | TAGGCTAAGTTTGTC |
| 3365 | mamo-L | *BGIBMGA006958* | down | 1945 | 1959 | - | 11.0484 | 6.23E-05 | 1 | AAGGCTTTCATTTTT |
| 3366 | mamo-L | *BGIBMGA006965* | up | 861 | 875 | - | 11.4194 | 4.42E-05 | 1 | TAGGCCCTTGTTGTG |
| 3367 | mamo-L | *BGIBMGA006965* | up | 104 | 118 | - | 10.7097 | 8.33E-05 | 1 | TAGCCTGAAGTTGTT |
| 3368 | mamo-L | *BGIBMGA006966* | down | 1097 | 1111 | + | 10.5403 | 9.58E-05 | 1 | TAGGCTTAGGTGTTG |
| 3369 | mamo-L | *BGIBMGA006967* | down | 1694 | 1708 | + | 11.2823 | 5.03E-05 | 1 | GAGGCCCGCGATGTT |
| 3370 | mamo-L | *BGIBMGA006974* | up | 723 | 737 | - | 10.5 | 9.90E-05 | 1 | GAGGCTGTGTATTGC |
| 3371 | mamo-L | *BGIBMGA006978* | up | 292 | 306 | - | 13.121 | 6.92E-06 | 1 | TAGGCTAAATTTGGT |
| 3372 | mamo-L | *BGIBMGA006984* | down | 1026 | 1040 | + | 11.2097 | 5.38E-05 | 1 | TAGGCAGATGATTGT |
| 3373 | mamo-L | *BGIBMGA006989* | up | 1758 | 1772 | + | 11.0968 | 5.97E-05 | 1 | AAGGCTTTCTTTCTT |
| 3374 | mamo-L | *BGIBMGA006990* | up | 1609 | 1623 | + | 12.0726 | 2.32E-05 | 1 | TAGGCTCTTTTTATT |
| 3375 | mamo-L | *BGIBMGA006996* | up | 418 | 432 | - | 11.379 | 4.58E-05 | 1 | TAGGCTTTTCTCGGT |
| 3376 | mamo-L | *BGIBMGA006999* | up | 35 | 49 | - | 11.0645 | 6.14E-05 | 1 | TAGGCGGATGTGGCA |
| 3377 | mamo-L | *BGIBMGA007001* | up | 1605 | 1619 | - | 10.5726 | 9.33E-05 | 1 | AAGGCCTTTTTTATT |
| 3378 | mamo-L | *BGIBMGA007002* | up | 1003 | 1017 | - | 11.4194 | 4.42E-05 | 1 | TAGGCTATGTATTTT |
| 3379 | mamo-L | *BGIBMGA007003* | up | 1064 | 1078 | - | 11.371 | 4.62E-05 | 1 | AAGGCTGGTTTTTAT |
| 3380 | mamo-L | *BGIBMGA007003* | up | 919 | 933 | + | 10.8065 | 7.69E-05 | 1 | TAGGCTTTTTAGTTT |
| 3381 | mamo-L | *BGIBMGA007003* | down | 1714 | 1728 | + | 10.7742 | 7.89E-05 | 1 | TGGGCTGATTTTTCT |
| 3382 | mamo-L | *BGIBMGA007004* | up | 873 | 887 | - | 10.5726 | 9.33E-05 | 1 | TAGGCTAATATTTAT |
| 3383 | mamo-L | *BGIBMGA007005* | down | 1743 | 1757 | + | 11.129 | 5.79E-05 | 1 | TAGGCTGGTGGTACC |
| 3384 | mamo-L | *BGIBMGA007008* | up | 1215 | 1229 | + | 11.9839 | 2.54E-05 | 1 | TAGGCTTATAATTGT |
| 3385 | mamo-L | *BGIBMGA007009* | up | 775 | 789 | - | 13.4274 | 4.64E-06 | 1 | TAGGCTTAAGTTTTT |
| 3386 | mamo-L | *BGIBMGA007013* | down | 1824 | 1838 | + | 13.2177 | 6.14E-06 | 1 | TAGGCTCGTTTTTTT |
| 3387 | mamo-L | *BGIBMGA007013* | down | 1890 | 1904 | + | 11.9355 | 2.67E-05 | 1 | TAGGCTCACGGGGCT |
| 3388 | mamo-L | *BGIBMGA007013* | down | 765 | 779 | + | 10.75 | 8.06E-05 | 1 | GAGGCCGATTGTGCC |
| 3389 | mamo-L | *BGIBMGA007013* | down | 1357 | 1371 | + | 10.5323 | 9.63E-05 | 1 | TAGGCTAAGCTTTTT |
| 3390 | mamo-L | *BGIBMGA007014* | down | 858 | 872 | - | 12.4839 | 1.47E-05 | 1 | GAGGCTGGTTTTTGT |
| 3391 | mamo-L | *BGIBMGA007017* | down | 1908 | 1922 | - | 12.2823 | 1.85E-05 | 1 | TAGGCCTTTTTCTCT |
| 3392 | mamo-L | *BGIBMGA007018* | up | 1315 | 1329 | - | 12.2823 | 1.85E-05 | 1 | TAGGCCTTTTTCTCT |
| 3393 | mamo-L | *BGIBMGA007018* | up | 1431 | 1445 | + | 10.871 | 7.28E-05 | 1 | AAGGCTTTCATCGTT |
| 3394 | mamo-L | *BGIBMGA007019* | down | 1578 | 1592 | + | 11.4274 | 4.38E-05 | 1 | AAGGCCTAGGTTCTT |
| 3395 | mamo-L | *BGIBMGA007020* | down | 1913 | 1927 | + | 10.5806 | 9.27E-05 | 1 | TAGGCTACCTTTTAT |
| 3396 | mamo-L | *BGIBMGA007021* | up | 1977 | 1991 | + | 11.4677 | 4.22E-05 | 1 | TAGGCGAGCGTTATT |
| 3397 | mamo-L | *BGIBMGA007023* | down | 683 | 697 | - | 10.5726 | 9.33E-05 | 1 | TAGGCTAATATTTAT |

| | | | | | | | | | | |
|---|---|---|---|---|---|---|---|---|---|---|
| 3398 | mamo-L | *BGIBMGA007024* | down | 303 | 317 | + | 12.5 | 1.44E-05 | 1 | TAGGCGTACAGTGTT |
| 3399 | mamo-L | *BGIBMGA007024* | down | 1878 | 1892 | + | 12.3629 | 1.69E-05 | 1 | TAGGCGTATTTTCTT |
| 3400 | mamo-L | *BGIBMGA007025* | down | 1213 | 1227 | + | 12.7984 | 1.02E-05 | 1 | TAGGCGGTCTTTCCT |
| 3401 | mamo-L | *BGIBMGA007025* | up | 1464 | 1478 | - | 12.0645 | 2.34E-05 | 1 | TAGGCGGGCTTCGGA |
| 3402 | mamo-L | *BGIBMGA007025* | up | 1687 | 1701 | - | 10.5161 | 9.77E-05 | 1 | GAGGCGGATGTCGCC |
| 3403 | mamo-L | *BGIBMGA007027* | down | 516 | 530 | - | 10.5968 | 9.15E-05 | 1 | TAGGCGTTCATTCAT |
| 3404 | mamo-L | *BGIBMGA007028* | down | 1962 | 1976 | - | 13.6048 | 3.60E-06 | 1 | TAGGCGTGTTTTGAT |
| 3405 | mamo-L | *BGIBMGA007028* | up | 1289 | 1303 | - | 10.5968 | 9.15E-05 | 1 | TAGGCGTTCATTCAT |
| 3406 | mamo-L | *BGIBMGA007029* | down | 1694 | 1708 | + | 11.1129 | 5.88E-05 | 1 | TAGGCCGAGGACGGG |
| 3407 | mamo-L | *BGIBMGA007033* | down | 862 | 876 | - | 12.2903 | 1.83E-05 | 1 | TAGGCCGGCGTCGGG |
| 3408 | mamo-L | *BGIBMGA007037* | up | 1827 | 1841 | + | 10.9194 | 6.98E-05 | 1 | TAGGCTACTTTTTTT |
| 3409 | mamo-L | *BGIBMGA007039* | up | 1383 | 1397 | - | 11.0806 | 6.05E-05 | 1 | GAGGCTTTTTTGGTT |
| 3410 | mamo-L | *BGIBMGA007040* | down | 1726 | 1740 | + | 11.7823 | 3.11E-05 | 1 | TAGGCGTGGGGCGGC |
| 3411 | mamo-L | *BGIBMGA007044* | up | 878 | 892 | - | 10.6129 | 9.03E-05 | 1 | TAGGCTTTTGGTCAT |
| 3412 | mamo-L | *BGIBMGA007044* | down | 1052 | 1066 | + | 10.5484 | 9.52E-05 | 1 | GAGGCTGTGATTTAT |
| 3413 | mamo-L | *BGIBMGA007046* | up | 880 | 894 | - | 11.7823 | 3.11E-05 | 1 | TAGGCCCAGTTTTGA |
| 3414 | mamo-L | *BGIBMGA007047* | down | 812 | 826 | - | 11.8387 | 2.94E-05 | 1 | TAGGCTGAAGTTTGA |
| 3415 | mamo-L | *BGIBMGA007048* | down | 1688 | 1702 | - | 11.3468 | 4.73E-05 | 1 | TAGGCGGGTAACGGT |
| 3416 | mamo-L | *BGIBMGA007049* | up | 313 | 327 | + | 12.0645 | 2.34E-05 | 1 | TAGGCGGGCTTCGGA |
| 3417 | mamo-L | *BGIBMGA007049* | up | 160 | 174 | - | 10.8306 | 7.52E-05 | 1 | TAGGCCCCCTTCGCC |
| 3418 | mamo-L | *BGIBMGA007049* | up | 1359 | 1373 | + | 10.5726 | 9.33E-05 | 1 | TAGGCTAATATTTAT |
| 3419 | mamo-L | *BGIBMGA007051* | up | 1553 | 1567 | + | 10.8871 | 7.17E-05 | 1 | TGGGCCTTCGTTGAT |
| 3420 | mamo-L | *BGIBMGA007052* | up | 644 | 658 | - | 11.25 | 5.18E-05 | 1 | TAGGCTTTGCTGGTT |
| 3421 | mamo-L | *BGIBMGA007056* | down | 121 | 135 | - | 11.0887 | 6.01E-05 | 1 | TAGGCTCTCATCTTT |
| 3422 | mamo-L | *BGIBMGA007059* | down | 1687 | 1701 | - | 11.0242 | 6.36E-05 | 1 | AAGGCCTTGTTTTCC |
| 3423 | mamo-L | *BGIBMGA007059* | up | 785 | 799 | - | 10.5 | 9.90E-05 | 1 | GAGGCCCAGGTCGTC |
| 3424 | mamo-L | *BGIBMGA007066* | up | 1919 | 1933 | + | 11.0968 | 5.97E-05 | 1 | TAGGCTTCCGATGAC |
| 3425 | mamo-L | *BGIBMGA007069* | down | 736 | 750 | + | 10.9435 | 6.83E-05 | 1 | AAGGCTGCTGTTGTC |
| 3426 | mamo-L | *BGIBMGA007076* | down | 1533 | 1547 | + | 11.5484 | 3.90E-05 | 1 | TAGGCCATGTTTTCC |
| 3427 | mamo-L | *BGIBMGA007076* | down | 67 | 81 | - | 11.0161 | 6.40E-05 | 1 | TAGGCCACCTTCGCT |
| 3428 | mamo-L | *BGIBMGA007081* | up | 66 | 80 | - | 11.7258 | 3.29E-05 | 1 | AAGGCTTATTATGCT |
| 3429 | mamo-L | *BGIBMGA007081* | up | 382 | 396 | - | 11.4677 | 4.22E-05 | 1 | TAGGCGAGCGTTATT |
| 3430 | mamo-L | *BGIBMGA007083* | up | 1859 | 1873 | + | 12.2823 | 1.85E-05 | 1 | TAGGCCTTTTTCTCT |
| 3431 | mamo-L | *BGIBMGA007083* | up | 1743 | 1757 | - | 10.871 | 7.28E-05 | 1 | AAGGCTTTCATCGTT |
| 3432 | mamo-L | *BGIBMGA007086* | up | 1944 | 1958 | + | 10.5726 | 9.33E-05 | 1 | TAGGCTAATATTTAT |
| 3433 | mamo-L | *BGIBMGA007089* | down | 658 | 672 | + | 12.4597 | 1.51E-05 | 1 | TAGGCTGATTTTGAA |
| 3434 | mamo-L | *BGIBMGA007090* | up | 1060 | 1074 | + | 11.7419 | 3.23E-05 | 1 | TCGGCTGAGTTTGCT |
| 3435 | mamo-L | *BGIBMGA007097* | down | 1686 | 1700 | - | 11.4758 | 4.18E-05 | 1 | TAGGCGATGTTTAGT |
| 3436 | mamo-L | *BGIBMGA007099* | up | 1800 | 1814 | + | 13.7742 | 2.79E-06 | 1 | TAGGCGGGTGGTGGT |
| 3437 | mamo-L | *BGIBMGA007100* | down | 1667 | 1681 | + | 10.5726 | 9.33E-05 | 1 | AAGGCCTTTTTTATT |
| 3438 | mamo-L | *BGIBMGA007101* | up | 43 | 57 | + | 10.9274 | 6.92E-05 | 1 | TAGGCTAGTTACGTT |
| 3439 | mamo-L | *BGIBMGA007105* | up | 131 | 145 | + | 11.3548 | 4.69E-05 | 1 | AAGGCTGATGATGAT |
| 3440 | mamo-L | *BGIBMGA007107* | down | 1597 | 1611 | - | 13.8629 | 2.44E-06 | 1 | TAGGCGTATGTTTGT |
| 3441 | mamo-L | *BGIBMGA007108* | up | 1289 | 1303 | - | 10.9194 | 6.98E-05 | 1 | TAGGCTACTTTTTTT |
| 3442 | mamo-L | *BGIBMGA007111* | down | 175 | 189 | + | 11.5161 | 4.02E-05 | 1 | TAGGCTTAGGGCAGT |
| 3443 | mamo-L | *BGIBMGA007112* | down | 90 | 104 | + | 11.9597 | 2.61E-05 | 1 | TAGGCCTTATTTTAT |
| 3444 | mamo-L | *BGIBMGA007112* | down | 83 | 97 | - | 10.7097 | 8.33E-05 | 1 | AAGGCCTATTTTCCT |
| 3445 | mamo-L | *BGIBMGA007113* | down | 669 | 683 | - | 12.0645 | 2.34E-05 | 1 | TAGGCGGGCTTCGGA |
| 3446 | mamo-L | *BGIBMGA007124* | up | 1839 | 1853 | + | 11.3226 | 4.83E-05 | 1 | TAGGCGTTGTTTGAG |
| 3447 | mamo-L | *BGIBMGA007125* | down | 1461 | 1475 | - | 11.2097 | 5.38E-05 | 1 | TAGGCAGATGATTGT |
| 3448 | mamo-L | *BGIBMGA007127* | up | 1702 | 1716 | - | 13.1935 | 6.33E-06 | 1 | TAGGCTTATTTTATT |
| 3449 | mamo-L | *BGIBMGA007129* | down | 724 | 738 | + | 11.8226 | 2.99E-05 | 1 | TAGGCGTTTTTCTTT |
| 3450 | mamo-L | *BGIBMGA007129* | down | 368 | 382 | + | 11.0565 | 6.18E-05 | 1 | TAGGCCGTATATGAC |
| 3451 | mamo-L | *BGIBMGA007130* | up | 1980 | 1994 | + | 11.0565 | 6.18E-05 | 1 | TAGGCCGTATATGAC |
| 3452 | mamo-L | *BGIBMGA007134* | up | 635 | 649 | + | 13.9194 | 2.23E-06 | 1 | TAGGCGGGCGGTGCT |
| 3453 | mamo-L | *BGIBMGA007134* | down | 463 | 477 | + | 11.7339 | 3.26E-05 | 1 | TAGGCTTTGTACTTT |
| 3454 | mamo-L | *BGIBMGA007135* | up | 1231 | 1245 | + | 11.7339 | 3.26E-05 | 1 | TAGGCTTTGTACTTT |
| 3455 | mamo-L | *BGIBMGA007140* | up | 794 | 808 | + | 10.9194 | 6.98E-05 | 1 | TAGGCCGGTTATCAT |
| 3456 | mamo-L | *BGIBMGA007142* | down | 1299 | 1313 | - | 10.5726 | 9.33E-05 | 1 | GAGGCTGTGTATAGT |
| 3457 | mamo-L | *BGIBMGA007143* | up | 1139 | 1153 | - | 11.7581 | 3.18E-05 | 1 | TAGGCCCGTGGTTTT |
| 3458 | mamo-L | *BGIBMGA007144* | up | 433 | 447 | - | 11.0726 | 6.09E-05 | 1 | TAGGCGTAGGTTTAA |
| 3459 | mamo-L | *BGIBMGA007144* | up | 286 | 300 | + | 10.6935 | 8.45E-05 | 1 | TAGGCGAATATTTCT |
| 3460 | mamo-L | *BGIBMGA007150* | up | 204 | 218 | + | 11.371 | 4.62E-05 | 1 | TAGGCTGTGTTTCTA |
| 3461 | mamo-L | *BGIBMGA007150* | down | 1917 | 1931 | - | 10.5 | 9.90E-05 | 1 | GAGGCCCAGGTCGTC |
| 3462 | mamo-L | *BGIBMGA007153* | up | 1566 | 1580 | + | 10.7258 | 8.23E-05 | 1 | GAGGCTTTTTTTATT |
| 3463 | mamo-L | *BGIBMGA007154* | down | 192 | 206 | + | 11.6855 | 3.42E-05 | 1 | TAGGCCGACAATAGT |
| 3464 | mamo-L | *BGIBMGA007156* | up | 1364 | 1378 | + | 11 | 6.50E-05 | 1 | AAGGCGTGTTATGTT |
| 3465 | mamo-L | *BGIBMGA007160* | up | 1804 | 1818 | - | 13.4677 | 4.39E-06 | 1 | TAGGCTGATGATGAT |
| 3466 | mamo-L | *BGIBMGA007160* | up | 1964 | 1978 | + | 11.1129 | 5.88E-05 | 1 | AAGGCTGTGTGTTTT |
| 3467 | mamo-L | *BGIBMGA007163* | up | 303 | 317 | - | 10.7097 | 8.33E-05 | 1 | TAGGCTGTGTATTTG |
| 3468 | mamo-L | *BGIBMGA007164* | down | 1339 | 1353 | + | 11.9597 | 2.61E-05 | 1 | TAGGCCAGGGGTGGC |
| 3469 | mamo-L | *BGIBMGA007166* | down | 190 | 204 | - | 11.2016 | 5.41E-05 | 1 | TAGGCGGACAAGGCT |
| 3470 | mamo-L | *BGIBMGA007172* | down | 1490 | 1504 | - | 11.6855 | 3.42E-05 | 1 | TAGGCCTTGTATGCA |
| 3471 | mamo-L | *BGIBMGA007178* | up | 815 | 829 | - | 11.9758 | 2.57E-05 | 1 | AAGGCTTTTTTTTGT |
| 3472 | mamo-L | *BGIBMGA007178* | up | 497 | 511 | - | 11.6452 | 3.56E-05 | 1 | TAGGCTGGGGTAGTC |
| 3473 | mamo-L | *BGIBMGA007178* | down | 261 | 275 | + | 10.9194 | 6.98E-05 | 1 | TAGGCTACTTTTTTT |
| 3474 | mamo-L | *BGIBMGA007180* | down | 1935 | 1949 | + | 11.2742 | 5.06E-05 | 1 | AAGGCTGATTTCGAT |
| 3475 | mamo-L | *BGIBMGA007180* | down | 1232 | 1246 | + | 11.0968 | 5.97E-05 | 1 | TAGGCTGAGATGGTA |
| 3476 | mamo-L | *BGIBMGA007181* | up | 1126 | 1140 | + | 11.2742 | 5.06E-05 | 1 | AAGGCTGATTTCGAT |
| 3477 | mamo-L | *BGIBMGA007181* | up | 423 | 437 | + | 11.0968 | 5.97E-05 | 1 | TAGGCTGAGATGGTA |
| 3478 | mamo-L | *BGIBMGA007182* | up | 90 | 104 | + | 11.2742 | 5.06E-05 | 1 | AAGGCTGATTTCGAT |
| 3479 | mamo-L | *BGIBMGA007183* | down | 921 | 935 | + | 11.3548 | 4.69E-05 | 1 | AAGGCTGATGATGAT |
| 3480 | mamo-L | *BGIBMGA007184* | up | 830 | 844 | - | 12.4194 | 1.58E-05 | 1 | TAGGCGGTTTTTTTC |
| 3481 | mamo-L | *BGIBMGA007187* | up | 1028 | 1042 | + | 11.3548 | 4.69E-05 | 1 | AAGGCTGATGATGAT |
| 3482 | mamo-L | *BGIBMGA007189* | down | 854 | 868 | - | 11.4758 | 4.18E-05 | 1 | GAGGCCCAGGTCGTT |
| 3483 | mamo-L | *BGIBMGA007193* | down | 1304 | 1318 | + | 11.6855 | 3.42E-05 | 1 | TAGGCCTTGTATGCA |
| 3484 | mamo-L | *BGIBMGA007196* | down | 1676 | 1690 | - | 11.9597 | 2.61E-05 | 1 | TAGGCCAGGGGTGGC |
| 3485 | mamo-L | *BGIBMGA007204* | down | 5 | 19 | - | 11.6048 | 3.70E-05 | 1 | TAGGCTCGGGATTTC |
| 3486 | mamo-L | *BGIBMGA007204* | down | 474 | 488 | - | 10.9194 | 6.98E-05 | 1 | TAGGCTACTTTTTTT |
| 3487 | mamo-L | *BGIBMGA007207* | down | 1500 | 1514 | + | 10.9597 | 6.73E-05 | 1 | GAGGCTTGCGTTGCG |
| 3488 | mamo-L | *BGIBMGA007210* | up | 777 | 791 | + | 10.4919 | 9.96E-05 | 1 | TAGGCTAGTATTTAT |
| 3489 | mamo-L | *BGIBMGA007211* | up | 975 | 989 | - | 10.5323 | 9.63E-05 | 1 | AAGGCCCAGGTCGTC |

| | | | | | | | | | | |
|---|---|---|---|---|---|---|---|---|---|---|
| 3490 | mamo-L | *BGIBMGA007213* | down | 883 | 897 | - | 12.25 | 1.92E-05 | 1 | TAGGCCTACTATTAT |
| 3491 | mamo-L | *BGIBMGA007218* | down | 557 | 571 | - | 11.6613 | 3.50E-05 | 1 | GAGGCTATGTTTGCT |
| 3492 | mamo-L | *BGIBMGA007222* | down | 675 | 689 | - | 10.7016 | 8.39E-05 | 1 | TAGGCCCAGTAGTTT |
| 3493 | mamo-L | *BGIBMGA007224* | up | 35 | 49 | - | 11.4677 | 4.22E-05 | 1 | TAGGCCCTGTTTATC |
| 3494 | mamo-L | *BGIBMGA007227* | down | 967 | 981 | + | 10.9194 | 6.98E-05 | 1 | TAGGCTACTTTTTTT |
| 3495 | mamo-L | *BGIBMGA007228* | up | 1125 | 1139 | + | 10.9355 | 6.88E-05 | 1 | GAGGCCTATGTCTGT |
| 3496 | mamo-L | *BGIBMGA007230* | down | 1469 | 1483 | - | 10.6532 | 8.73E-05 | 1 | TAGCCCCAGTTTGTT |
| 3497 | mamo-L | *BGIBMGA007233* | down | 376 | 390 | + | 10.6694 | 8.62E-05 | 1 | TATGCTGGTGTTGGT |
| 3498 | mamo-L | *BGIBMGA007237* | up | 1527 | 1541 | + | 11.25 | 5.18E-05 | 1 | TAGGCGAAGGGCGGT |
| 3499 | mamo-L | *BGIBMGA007237* | down | 104 | 118 | - | 11.0968 | 5.97E-05 | 1 | TGGGCCCGGTTTGTT |
| 3500 | mamo-L | *BGIBMGA007242* | down | 392 | 406 | - | 10.8145 | 7.63E-05 | 1 | AAGGCGAATTTTGTT |
| 3501 | mamo-L | *BGIBMGA007243* | down | 1453 | 1467 | + | 11.0887 | 6.01E-05 | 1 | AAGGCAGGTGTTGCT |
| 3502 | mamo-L | *BGIBMGA007243* | up | 515 | 529 | - | 10.6694 | 8.62E-05 | 1 | AAGGCGAAGGTTGAT |
| 3503 | mamo-L | *BGIBMGA007250* | down | 974 | 988 | + | 11.4597 | 4.25E-05 | 1 | TAGGCTTACGATGAA |
| 3504 | mamo-L | *BGIBMGA007253* | up | 787 | 801 | + | 11.8065 | 3.03E-05 | 1 | TAGGCTTTCAATTCT |
| 3505 | mamo-L | *BGIBMGA007253* | down | 1874 | 1888 | + | 11.3065 | 4.91E-05 | 1 | TAGGCTATGTGTTCT |
| 3506 | mamo-L | *BGIBMGA007254* | down | 1263 | 1277 | - | 11.7903 | 3.08E-05 | 1 | TAGGCGTAATTTTAT |
| 3507 | mamo-L | *BGIBMGA007257* | up | 1483 | 1497 | + | 13.0484 | 7.57E-06 | 1 | TAGGCTCACTTTCGT |
| 3508 | mamo-L | *BGIBMGA007258* | up | 635 | 649 | - | 12.6371 | 1.23E-05 | 1 | TAGGCTTAATTTTGC |
| 3509 | mamo-L | *BGIBMGA007258* | down | 1314 | 1328 | + | 10.5 | 9.90E-05 | 1 | GAGGCCCAGGTCGTC |
| 3510 | mamo-L | *BGIBMGA007259* | up | 985 | 999 | + | 10.5 | 9.90E-05 | 1 | GAGGCCCAGGTCGTC |
| 3511 | mamo-L | *BGIBMGA007260* | down | 1458 | 1472 | - | 11.3548 | 4.69E-05 | 1 | AAGGCTGATGATGAT |
| 3512 | mamo-L | *BGIBMGA007261* | up | 1080 | 1094 | + | 11.7258 | 3.29E-05 | 1 | TAGGCCATGTTTTAT |
| 3513 | mamo-L | *BGIBMGA007261* | down | 995 | 1009 | + | 10.5968 | 9.15E-05 | 1 | TAGGCTGTCCGTTCT |
| 3514 | mamo-L | *BGIBMGA007262* | down | 1459 | 1473 | - | 11.3387 | 4.77E-05 | 1 | TAGGCGGATTTAGTT |
| 3515 | mamo-L | *BGIBMGA007266* | up | 937 | 951 | + | 11.5323 | 3.96E-05 | 1 | TAGGCGTGAGGTGAT |
| 3516 | mamo-L | *BGIBMGA007269* | up | 405 | 419 | + | 12.2339 | 1.95E-05 | 1 | TAGGCTGATAATGAT |
| 3517 | mamo-L | *BGIBMGA007270* | down | 28 | 42 | + | 13.1532 | 6.65E-06 | 1 | GAGGCTGTTGTTGTT |
| 3518 | mamo-L | *BGIBMGA007273* | down | 1262 | 1276 | + | 11.1855 | 5.50E-05 | 1 | TAGGCCAGTTTTCTT |
| 3519 | mamo-L | *BGIBMGA007274* | up | 962 | 976 | + | 11.1855 | 5.50E-05 | 1 | TAGGCCAGTTTTCTT |
| 3520 | mamo-L | *BGIBMGA007276* | down | 1373 | 1387 | - | 10.5968 | 9.15E-05 | 1 | TAGGCTTATTTTATG |
| 3521 | mamo-L | *BGIBMGA007280* | down | 1659 | 1673 | + | 10.5726 | 9.33E-05 | 1 | TAGGCTAATATTTAT |
| 3522 | mamo-L | *BGIBMGA007281* | up | 1051 | 1065 | - | 12.8306 | 9.76E-06 | 1 | TAGGCCCAGATTGAT |
| 3523 | mamo-L | *BGIBMGA007282* | up | 1848 | 1862 | - | 11.0323 | 6.32E-05 | 1 | TAGGCTGACATGGTA |
| 3524 | mamo-L | *BGIBMGA007282* | up | 189 | 203 | + | 10.5968 | 9.15E-05 | 1 | TAGGCTTTAATCTTT |
| 3525 | mamo-L | *BGIBMGA007282* | up | 435 | 449 | - | 10.5806 | 9.27E-05 | 1 | AAGGCTTGCATTGAC |
| 3526 | mamo-L | *BGIBMGA007285* | up | 1857 | 1871 | + | 13 | 8.01E-06 | 1 | TAGGCTAACTTTTTT |
| 3527 | mamo-L | *BGIBMGA007286* | up | 1855 | 1869 | + | 13 | 8.01E-06 | 1 | TAGGCTAACTTTTTT |
| 3528 | mamo-L | *BGIBMGA007290* | down | 550 | 564 | + | 10.9919 | 6.55E-05 | 1 | TAGGCCTTTTACTTT |
| 3529 | mamo-L | *BGIBMGA007296* | down | 1731 | 1745 | - | 10.9194 | 6.98E-05 | 1 | TAGGCTACTTTTTTT |
| 3530 | mamo-L | *BGIBMGA007296* | up | 895 | 909 | + | 10.8226 | 7.58E-05 | 1 | TAGGCAGTCGTTAGC |
| 3531 | mamo-L | *BGIBMGA007298* | up | 220 | 234 | - | 11.5323 | 3.96E-05 | 1 | TAGGCGTGAGGTGAT |
| 3532 | mamo-L | *BGIBMGA007299* | down | 1959 | 1973 | - | 12.9274 | 8.74E-06 | 1 | TAGGCCGGCATTAGT |
| 3533 | mamo-L | *BGIBMGA007300* | down | 458 | 472 | - | 10.5968 | 9.15E-05 | 1 | TAGGCTGTCCGTTCT |
| 3534 | mamo-L | *BGIBMGA007301* | down | 1926 | 1940 | + | 13.0565 | 7.51E-06 | 1 | TAGGCGGGGATTTCT |
| 3535 | mamo-L | *BGIBMGA007302* | down | 141 | 155 | - | 10.5403 | 9.58E-05 | 1 | TAGGCCTGTTTTCCA |
| 3536 | mamo-L | *BGIBMGA007307* | up | 1402 | 1416 | + | 11.7903 | 3.08E-05 | 1 | TAGGCGTAATTTTAT |
| 3537 | mamo-L | *BGIBMGA007308* | down | 1791 | 1805 | + | 14.6371 | 7.08E-07 | 1 | TAGGCCTGCTTTTGT |
| 3538 | mamo-L | *BGIBMGA007308* | up | 635 | 649 | - | 11.4355 | 4.34E-05 | 1 | TAGGCGGATTACTGT |
| 3539 | mamo-L | *BGIBMGA007309* | down | 406 | 420 | + | 14.6371 | 7.08E-07 | 1 | TAGGCCTGCTTTTGT |
| 3540 | mamo-L | *BGIBMGA007311* | up | 805 | 819 | - | 14.3952 | 1.09E-06 | 1 | TAGGCGGGCTTTGAT |
| 3541 | mamo-L | *BGIBMGA007313* | down | 1829 | 1843 | - | 13.4677 | 4.39E-06 | 1 | TAGGCTGATGATGAT |
| 3542 | mamo-L | *BGIBMGA007314* | down | 370 | 384 | - | 10.5726 | 9.33E-05 | 1 | TAGGCTAATATTTAT |
| 3543 | mamo-L | *BGIBMGA007316* | down | 854 | 868 | - | 11.6452 | 3.56E-05 | 1 | TAGGCCGTCTGGTGT |
| 3544 | mamo-L | *BGIBMGA007316* | up | 373 | 387 | + | 10.8145 | 7.63E-05 | 1 | TAGGCTCTTAATGAT |
| 3545 | mamo-L | *BGIBMGA007317* | up | 284 | 298 | - | 13.7661 | 2.82E-06 | 1 | TAGGCTGTTTGTGTT |
| 3546 | mamo-L | *BGIBMGA007320* | down | 1386 | 1400 | - | 10.6452 | 8.79E-05 | 1 | AAGGCTTGGATTGAC |
| 3547 | mamo-L | *BGIBMGA007327* | down | 645 | 659 | + | 10.6452 | 8.79E-05 | 1 | AAGGCTTGGATTGAC |
| 3548 | mamo-L | *BGIBMGA007329* | up | 831 | 845 | - | 11.3145 | 4.87E-05 | 1 | TAGGCTGACGAAGTT |
| 3549 | mamo-L | *BGIBMGA007331* | up | 1958 | 1972 | - | 11.1774 | 5.54E-05 | 1 | TAGGCTAAGTTTCAT |
| 3550 | mamo-L | *BGIBMGA007331* | up | 725 | 739 | + | 10.9839 | 6.59E-05 | 1 | TAGGCTTTTTTTAGA |
| 3551 | mamo-L | *BGIBMGA007332* | down | 1466 | 1480 | - | 10.5726 | 9.33E-05 | 1 | TAGGCTAATATTTAT |
| 3552 | mamo-L | *BGIBMGA007334* | up | 1512 | 1526 | + | 11.2258 | 5.30E-05 | 1 | TAGGCGCATGTGGTC |
| 3553 | mamo-L | *BGIBMGA007339* | up | 1252 | 1266 | + | 12.4032 | 1.61E-05 | 1 | TAGGCTGATTGTAGT |
| 3554 | mamo-L | *BGIBMGA007339* | up | 506 | 520 | - | 10.7984 | 7.74E-05 | 1 | GAGGCCTATTGTGGC |
| 3555 | mamo-L | *BGIBMGA007341* | up | 254 | 268 | - | 11.7258 | 3.29E-05 | 1 | AAGGCCTGGTTGGTT |
| 3556 | mamo-L | *BGIBMGA007341* | down | 233 | 247 | - | 11.3387 | 4.77E-05 | 1 | TAGGCTGACGTGCTC |
| 3557 | mamo-L | *BGIBMGA007345* | down | 106 | 120 | - | 10.5726 | 9.33E-05 | 1 | TAGGCTAATATTTAT |
| 3558 | mamo-L | *BGIBMGA007348* | down | 1718 | 1732 | + | 11.4435 | 4.31E-05 | 1 | TAGGCTGGCAATACT |
| 3559 | mamo-L | *BGIBMGA007350* | up | 91 | 105 | - | 10.5726 | 9.33E-05 | 1 | TAGGCTAATATTTAT |
| 3560 | mamo-L | *BGIBMGA007352* | down | 258 | 272 | + | 13.0242 | 7.77E-06 | 1 | TAGGCCGGTTTTGTA |
| 3561 | mamo-L | *BGIBMGA007357* | up | 1201 | 1215 | - | 10.629 | 8.91E-05 | 1 | TAGGCCAAGGATCTT |
| 3562 | mamo-L | *BGIBMGA007364* | up | 1493 | 1507 | + | 10.8387 | 7.47E-05 | 1 | TGGGCTTTGGTTTCT |
| 3563 | mamo-L | *BGIBMGA007365* | up | 49 | 63 | + | 10.5726 | 9.33E-05 | 1 | TAGGCTAATATTTAT |
| 3564 | mamo-L | *BGIBMGA007366* | up | 488 | 502 | - | 11.5323 | 3.96E-05 | 1 | TAGGCGTGAGGTGAT |
| 3565 | mamo-L | *BGIBMGA007370* | up | 1488 | 1502 | + | 11.9032 | 2.76E-05 | 1 | TAGGCCGATTGTATT |
| 3566 | mamo-L | *BGIBMGA007370* | up | 1691 | 1705 | + | 10.7984 | 7.74E-05 | 1 | TAGGCCGATAGTATT |
| 3567 | mamo-L | *BGIBMGA007374* | down | 1966 | 1980 | + | 10.9194 | 6.98E-05 | 1 | TAGGCTACTTTTTTT |
| 3568 | mamo-L | *BGIBMGA007375* | up | 672 | 686 | - | 10.621 | 8.97E-05 | 1 | TAGGCTGTATAGTGT |
| 3569 | mamo-L | *BGIBMGA007377* | down | 428 | 442 | - | 10.5 | 9.90E-05 | 1 | GAGGCCCAGGTCGTC |
| 3570 | mamo-L | *BGIBMGA007380* | down | 202 | 216 | + | 13.6048 | 3.60E-06 | 1 | TAGGCTGGTTTTGAC |
| 3571 | mamo-L | *BGIBMGA007382* | down | 70 | 84 | + | 10.4919 | 9.96E-05 | 1 | TAGGCTGGCGGAGGC |
| 3572 | mamo-L | *BGIBMGA007384* | down | 1621 | 1635 | - | 13.7097 | 3.08E-06 | 1 | TAGGCTGACGTCTCT |
| 3573 | mamo-L | *BGIBMGA007385* | down | 1552 | 1566 | - | 11.9032 | 2.76E-05 | 1 | TAGGCCGATTGTATT |
| 3574 | mamo-L | *BGIBMGA007385* | down | 1755 | 1769 | - | 11.9032 | 2.76E-05 | 1 | TAGGCCGATTGTATT |
| 3575 | mamo-L | *BGIBMGA007387* | down | 460 | 474 | + | 13.6129 | 3.55E-06 | 1 | TAGGCCGACTGTTGT |
| 3576 | mamo-L | *BGIBMGA007387* | up | 1857 | 1871 | - | 11.6694 | 3.47E-05 | 1 | TAGGCTGACAGTGAC |
| 3577 | mamo-L | *BGIBMGA007388* | up | 868 | 882 | + | 11.0968 | 5.97E-05 | 1 | TGGGCCCGGTTTGTT |
| 3578 | mamo-L | *BGIBMGA007389* | down | 1009 | 1023 | + | 10.5 | 9.90E-05 | 1 | GAGGCCCAGGTCGTC |
| 3579 | mamo-L | *BGIBMGA007392* | down | 60 | 74 | - | 11.0242 | 6.36E-05 | 1 | GAGGCGCTGTTTTGT |
| 3580 | mamo-L | *BGIBMGA007393* | down | 328 | 342 | - | 11.5403 | 3.93E-05 | 1 | TAGGCTCGGTTTTCA |
| 3581 | mamo-L | *BGIBMGA007393* | down | 1755 | 1769 | + | 10.7984 | 7.74E-05 | 1 | TAGGCCCTAGATTTT |

| | | | | | | | | | | |
|---|---|---|---|---|---|---|---|---|---|---|
| 3582 | mamo-L | *BGIBMGA007394* | up | 1396 | 1410 | - | 11.5403 | 3.93E-05 | 1 | TAGGCTCGGTTTTCA |
| 3583 | mamo-L | *BGIBMGA007395* | down | 270 | 284 | - | 10.9274 | 6.92E-05 | 1 | TAGGCAGGGTGGGGT |
| 3584 | mamo-L | *BGIBMGA007400* | up | 1228 | 1242 | - | 14.121 | 1.65E-06 | 1 | TAGGCTGAGGTCTGT |
| 3585 | mamo-L | *BGIBMGA007405* | down | 1119 | 1133 | - | 12.1048 | 2.24E-05 | 1 | TAGGCTGGTGGTACT |
| 3586 | mamo-L | *BGIBMGA007405* | down | 1185 | 1199 | - | 11.4839 | 4.15E-05 | 1 | TAGGCCTCAATTGCT |
| 3587 | mamo-L | *BGIBMGA007405* | up | 122 | 136 | - | 10.8387 | 7.47E-05 | 1 | TGGGCTTTGGTTTCT |
| 3588 | mamo-L | *BGIBMGA007406* | up | 650 | 664 | + | 13.2581 | 5.81E-06 | 1 | TAGGCTGATATTTCT |
| 3589 | mamo-L | *BGIBMGA007406* | up | 390 | 404 | + | 10.5161 | 9.77E-05 | 1 | AAGGCTTTTGTCTTT |
| 3590 | mamo-L | *BGIBMGA007409* | up | 1502 | 1516 | - | 12.8468 | 9.58E-06 | 1 | TAGGCCGACGTCGAC |
| 3591 | mamo-L | *BGIBMGA007413* | down | 1388 | 1402 | - | 13.0242 | 7.77E-06 | 1 | TAGGCCGGTTTTGTA |
| 3592 | mamo-L | *BGIBMGA007417* | down | 626 | 640 | - | 11.1855 | 5.50E-05 | 1 | TAGGCCAGATATGCT |
| 3593 | mamo-L | *BGIBMGA007417* | up | 807 | 821 | + | 10.5726 | 9.33E-05 | 1 | TAGGCTAATATTTAT |
| 3594 | mamo-L | *BGIBMGA007420* | down | 568 | 582 | - | 11.4597 | 4.25E-05 | 1 | TAGGCTGACGAGAGT |
| 3595 | mamo-L | *BGIBMGA007420* | down | 1291 | 1305 | - | 11.4435 | 4.31E-05 | 1 | TAGGCTGGCAATACT |
| 3596 | mamo-L | *BGIBMGA007423* | down | 802 | 816 | - | 12.379 | 1.66E-05 | 1 | TAGGCTTACGGTACT |
| 3597 | mamo-L | *BGIBMGA007424* | up | 1282 | 1296 | + | 11.7097 | 3.34E-05 | 1 | TAGGCTTTTTTTCCC |
| 3598 | mamo-L | *BGIBMGA007427* | down | 1770 | 1784 | + | 10.9194 | 6.98E-05 | 1 | TAGGCTACTTTTTTT |
| 3599 | mamo-L | *BGIBMGA007428* | down | 1699 | 1713 | - | 12.621 | 1.25E-05 | 1 | TAGGCCAAATTTGTT |
| 3600 | mamo-L | *BGIBMGA007429* | down | 1116 | 1130 | - | 12.9919 | 8.10E-06 | 1 | TAGGCCGGTTTTGCA |
| 3601 | mamo-L | *BGIBMGA007431* | down | 1475 | 1489 | - | 10.8387 | 7.47E-05 | 1 | AAGGCTAATGTTTGT |
| 3602 | mamo-L | *BGIBMGA007434* | up | 447 | 461 | - | 12.1048 | 2.24E-05 | 1 | TAGGCCTAGTGCTGT |
| 3603 | mamo-L | *BGIBMGA007436* | up | 1709 | 1723 | - | 10.5726 | 9.33E-05 | 1 | TAGGCTAATATTTAT |
| 3604 | mamo-L | *BGIBMGA007437* | up | 716 | 730 | - | 10.9839 | 6.59E-05 | 1 | TAGGCTTTTTTTAGA |
| 3605 | mamo-L | *BGIBMGA007438* | down | 1872 | 1886 | + | 10.9194 | 6.98E-05 | 1 | TAGGCTACTTTTTTT |
| 3606 | mamo-L | *BGIBMGA007441* | up | 542 | 556 | + | 12.4677 | 1.49E-05 | 1 | TAGGCTTCTATTGCT |
| 3607 | mamo-L | *BGIBMGA007442* | down | 1551 | 1565 | + | 11.4758 | 4.18E-05 | 1 | TAGGCCCACGATACT |
| 3608 | mamo-L | *BGIBMGA007443* | up | 1119 | 1133 | - | 12.5645 | 1.34E-05 | 1 | TAGGCTAGTTTCGGT |
| 3609 | mamo-L | *BGIBMGA007443* | down | 1863 | 1877 | + | 12.1694 | 2.09E-05 | 1 | GAGGCTTGATTTGTT |
| 3610 | mamo-L | *BGIBMGA007444* | up | 368 | 382 | - | 10.5645 | 9.40E-05 | 1 | TAGGCTTAGTTTCAA |
| 3611 | mamo-L | *BGIBMGA007446* | up | 1282 | 1296 | + | 11.1694 | 5.59E-05 | 1 | GAGGCTTGTGTTTAT |
| 3612 | mamo-L | *BGIBMGA007447* | up | 466 | 480 | + | 12.0968 | 2.26E-05 | 1 | TAGGCTGTCGTTTTG |
| 3613 | mamo-L | *BGIBMGA007448* | up | 958 | 972 | + | 11.7661 | 3.16E-05 | 1 | TAGGCCATGATTTGT |
| 3614 | mamo-L | *BGIBMGA007449* | up | 80 | 94 | - | 10.9194 | 6.98E-05 | 1 | TAGGCTACTTTTTTT |
| 3615 | mamo-L | *BGIBMGA007451* | down | 1943 | 1957 | - | 11.8145 | 3.01E-05 | 1 | AAGGCTTGCTTGGCT |
| 3616 | mamo-L | *BGIBMGA007452* | up | 199 | 213 | - | 10.5403 | 9.58E-05 | 1 | TAGGCTGTATGGTGT |
| 3617 | mamo-L | *BGIBMGA007455* | down | 126 | 140 | - | 10.5726 | 9.33E-05 | 1 | TAGGCTAATATTTAT |
| 3618 | mamo-L | *BGIBMGA007456* | up | 1055 | 1069 | - | 10.5726 | 9.33E-05 | 1 | TAGGCTAATATTTAT |
| 3619 | mamo-L | *BGIBMGA007457* | down | 935 | 949 | + | 12.9597 | 8.38E-06 | 1 | TAGGCTTATATTTCT |
| 3620 | mamo-L | *BGIBMGA007458* | up | 480 | 494 | - | 10.7903 | 7.79E-05 | 1 | TAGGCTATCTTTCAT |
| 3621 | mamo-L | *BGIBMGA007458* | up | 1332 | 1346 | + | 10.5403 | 9.58E-05 | 1 | GAGGCGTACGATTCT |
| 3622 | mamo-L | *BGIBMGA007459* | down | 1813 | 1827 | + | 10.7258 | 8.23E-05 | 1 | TAGGCTCATGAGGAT |
| 3623 | mamo-L | *BGIBMGA007460* | up | 1278 | 1292 | + | 11.1129 | 5.88E-05 | 1 | TAGGCAGAGTACGCT |
| 3624 | mamo-L | *BGIBMGA007462* | up | 577 | 591 | - | 11.9032 | 2.76E-05 | 1 | TAGGCCGATTGTATT |
| 3625 | mamo-L | *BGIBMGA007463* | down | 1504 | 1518 | - | 11.5806 | 3.78E-05 | 1 | TAGGCATACGTTTAT |
| 3626 | mamo-L | *BGIBMGA007463* | up | 1822 | 1836 | - | 11.371 | 4.62E-05 | 1 | AAGGCTTACATTTTT |
| 3627 | mamo-L | *BGIBMGA007463* | down | 1317 | 1331 | + | 10.5968 | 9.15E-05 | 1 | AAGGCTGAGGCTGAT |
| 3628 | mamo-L | *BGIBMGA007464* | up | 1707 | 1721 | + | 10.5161 | 9.77E-05 | 1 | AAGGCTGGCTTCATT |
| 3629 | mamo-L | *BGIBMGA007466* | down | 1912 | 1926 | - | 11.621 | 3.63E-05 | 1 | AAGGCGTATTTTTGT |
| 3630 | mamo-L | *BGIBMGA007468* | up | 1424 | 1438 | - | 13.5565 | 3.86E-06 | 1 | TAGGCGTATGGTGGT |
| 3631 | mamo-L | *BGIBMGA007470* | down | 1359 | 1373 | - | 10.5726 | 9.33E-05 | 1 | TAGGCTCCTATTTCT |
| 3632 | mamo-L | *BGIBMGA007471* | up | 1536 | 1550 | + | 13.7177 | 3.05E-06 | 1 | TAGGCTGTAGTTTGT |
| 3633 | mamo-L | *BGIBMGA007471* | down | 1903 | 1917 | - | 11.7823 | 3.11E-05 | 1 | GAGGCTTACTTTTGC |
| 3634 | mamo-L | *BGIBMGA007472* | up | 156 | 170 | + | 11.0887 | 6.01E-05 | 1 | GAGGCTTGCGGTTTT |
| 3635 | mamo-L | *BGIBMGA007474* | up | 1834 | 1848 | + | 13.4032 | 4.80E-06 | 1 | TAGGCGGTTTGTGGT |
| 3636 | mamo-L | *BGIBMGA007477* | up | 894 | 908 | - | 11.1129 | 5.88E-05 | 1 | TAGGCAGAGTACGCT |
| 3637 | mamo-L | *BGIBMGA007481* | up | 1813 | 1827 | + | 11.0806 | 6.05E-05 | 1 | TAGGCCTGTGAGGCC |
| 3638 | mamo-L | *BGIBMGA007481* | down | 1603 | 1617 | - | 10.9194 | 6.98E-05 | 1 | TAGGCTACTTTTTTT |
| 3639 | mamo-L | *BGIBMGA007482* | down | 1566 | 1580 | - | 12.9597 | 8.38E-06 | 1 | TAGGCTTATATTTCT |
| 3640 | mamo-L | *BGIBMGA007484* | down | 1473 | 1487 | + | 10.5726 | 9.33E-05 | 1 | TAGGCTAATATTTAT |
| 3641 | mamo-L | *BGIBMGA007487* | down | 369 | 383 | - | 12.3629 | 1.69E-05 | 1 | TAGGCTGTCGACGAT |
| 3642 | mamo-L | *BGIBMGA007488* | up | 1803 | 1817 | + | 10.5403 | 9.58E-05 | 1 | TAGGCTGTATGGTGT |
| 3643 | mamo-L | *BGIBMGA007489* | up | 203 | 217 | - | 10.5161 | 9.77E-05 | 1 | TAGGCTTATTTGGAA |
| 3644 | mamo-L | *BGIBMGA007498* | up | 1779 | 1793 | - | 15.129 | 2.58E-07 | 1 | TAGGCCGGGGTTTGT |
| 3645 | mamo-L | *BGIBMGA007499* | down | 605 | 619 | + | 11.1452 | 5.70E-05 | 1 | TAGGCTGTTGATAAT |
| 3646 | mamo-L | *BGIBMGA007502* | up | 288 | 302 | + | 10.5081 | 9.83E-05 | 1 | TAGGCGGATGGTGAA |
| 3647 | mamo-L | *BGIBMGA007503* | down | 863 | 877 | - | 11.2742 | 5.06E-05 | 1 | TAGGCATGCGTTCTT |
| 3648 | mamo-L | *BGIBMGA007505* | down | 342 | 356 | - | 12.1532 | 2.13E-05 | 1 | TAGGCTCATGTCTTT |
| 3649 | mamo-L | *BGIBMGA007505* | up | 1715 | 1729 | - | 10.5484 | 9.52E-05 | 1 | TAGGCGCGTTGTAGT |
| 3650 | mamo-L | *BGIBMGA007506* | down | 510 | 524 | + | 12.7419 | 1.09E-05 | 1 | TAGGCTATGTTTTTT |
| 3651 | mamo-L | *BGIBMGA007510* | up | 1177 | 1191 | + | 11.0242 | 6.36E-05 | 1 | AAGGCCTTGTTTTCC |
| 3652 | mamo-L | *BGIBMGA007512* | down | 320 | 334 | + | 10.9194 | 6.98E-05 | 1 | TAGGCTACTTTTTTT |
| 3653 | mamo-L | *BGIBMGA007517* | up | 1911 | 1925 | + | 10.75 | 8.06E-05 | 1 | TAGGCCTTCAATATT |
| 3654 | mamo-L | *BGIBMGA007527* | down | 1393 | 1407 | + | 11.5968 | 3.73E-05 | 1 | TAGGCTTTTTTCATT |
| 3655 | mamo-L | *BGIBMGA007527* | up | 1065 | 1079 | + | 10.5726 | 9.33E-05 | 1 | TAGGCTAATATTTAT |
| 3656 | mamo-L | *BGIBMGA007528* | down | 836 | 850 | + | 10.9758 | 6.64E-05 | 1 | TAGGCCTAGCTCTGT |
| 3657 | mamo-L | *BGIBMGA007529* | up | 552 | 566 | - | 11.4435 | 4.31E-05 | 1 | TAGGCTTGAGAGGCT |
| 3658 | mamo-L | *BGIBMGA007531* | up | 1841 | 1855 | - | 10.8065 | 7.69E-05 | 1 | GAGGCCTGAGTCGCT |
| 3659 | mamo-L | *BGIBMGA007532* | down | 1019 | 1033 | + | 11.5887 | 3.75E-05 | 1 | TAGGCTTTCGTCGCG |
| 3660 | mamo-L | *BGIBMGA007534* | down | 1375 | 1389 | - | 12.4355 | 1.55E-05 | 1 | GAGGCTTACATTGTT |
| 3661 | mamo-L | *BGIBMGA007534* | up | 925 | 939 | - | 11.3952 | 4.52E-05 | 1 | TAGGCCTGATGGGGT |
| 3662 | mamo-L | *BGIBMGA007536* | up | 1564 | 1578 | - | 11.3548 | 4.69E-05 | 1 | AAGGCTTTTTGTGTT |
| 3663 | mamo-L | *BGIBMGA007536* | down | 36 | 50 | + | 10.5161 | 9.77E-05 | 1 | GAGGCCTAAGTGGCT |
| 3664 | mamo-L | *BGIBMGA007540* | down | 1634 | 1648 | - | 13.4839 | 4.31E-06 | 1 | TAGGCTTACTTTGTA |
| 3665 | mamo-L | *BGIBMGA007541* | down | 1884 | 1898 | + | 13.7984 | 2.69E-06 | 1 | TAGGCTTACTTTGGA |
| 3666 | mamo-L | *BGIBMGA007546* | up | 656 | 670 | - | 10.9194 | 6.98E-05 | 1 | TAGGCTTAATGTTTC |
| 3667 | mamo-L | *BGIBMGA007546* | up | 7 | 21 | + | 10.8306 | 7.52E-05 | 1 | TAGGCCCCCTTCGCC |
| 3668 | mamo-L | *BGIBMGA007548* | down | 215 | 229 | - | 13.8871 | 2.34E-06 | 1 | TAGGCCTTATTTGTT |
| 3669 | mamo-L | *BGIBMGA007548* | down | 580 | 594 | - | 13.379 | 4.96E-06 | 1 | TAGGCCTAAGTCGGT |
| 3670 | mamo-L | *BGIBMGA007548* | down | 465 | 479 | - | 12.6694 | 1.18E-05 | 1 | TAGGCTTATGAGGGT |
| 3671 | mamo-L | *BGIBMGA007548* | down | 1868 | 1882 | + | 11.2984 | 4.95E-05 | 1 | TAGGCGAAGTGTTGT |
| 3672 | mamo-L | *BGIBMGA007548* | up | 1947 | 1961 | + | 11.2984 | 4.95E-05 | 1 | TAGGCGAAGTGTTGT |
| 3673 | mamo-L | *BGIBMGA007549* | up | 60 | 74 | - | 13.379 | 4.96E-06 | 1 | TAGGCTGATAATGGT |

| | | | | | | | | | | |
|---|---|---|---|---|---|---|---|---|---|---|
| 3674 | mamo-L | *BGIBMGA007551* | up | 771 | 785 | + | 13.379 | 4.96E-06 | 1 | TAGGCTGATAATGGT |
| 3675 | mamo-L | *BGIBMGA007552* | down | 1478 | 1492 | - | 11.2984 | 4.95E-05 | 1 | TAGGCGAAGTGTTGT |
| 3676 | mamo-L | *BGIBMGA007552* | up | 916 | 930 | - | 10.6371 | 8.85E-05 | 1 | TAGGCATTTTTTTAT |
| 3677 | mamo-L | *BGIBMGA007553* | up | 734 | 748 | + | 13.3145 | 5.41E-06 | 1 | AAGGCTTTGTTTGTT |
| 3678 | mamo-L | *BGIBMGA007556* | down | 158 | 172 | - | 13.7984 | 2.69E-06 | 1 | TAGGCTTACTTTGGA |
| 3679 | mamo-L | *BGIBMGA007557* | up | 599 | 613 | + | 11.5323 | 3.96E-05 | 1 | TAGGCGTGAGGTGAT |
| 3680 | mamo-L | *BGIBMGA007558* | down | 1309 | 1323 | + | 10.5323 | 9.63E-05 | 1 | TAGGCTGACGACCAT |
| 3681 | mamo-L | *BGIBMGA007560* | up | 1597 | 1611 | - | 10.9677 | 6.68E-05 | 1 | AAGGCCGCCTTTTCT |
| 3682 | mamo-L | *BGIBMGA007561* | down | 613 | 627 | - | 12.7016 | 1.14E-05 | 1 | TAGGCTGTTGTTGTG |
| 3683 | mamo-L | *BGIBMGA007562* | up | 749 | 763 | - | 12.7016 | 1.14E-05 | 1 | TAGGCTGTTGTTGTG |
| 3684 | mamo-L | *BGIBMGA007566* | down | 594 | 608 | + | 12.4355 | 1.55E-05 | 1 | GAGGCTTACATTGTT |
| 3685 | mamo-L | *BGIBMGA007567* | up | 488 | 502 | + | 11.0806 | 6.05E-05 | 1 | TAGGCGCTGGAGGCT |
| 3686 | mamo-L | *BGIBMGA007569* | down | 1531 | 1545 | - | 11.8952 | 2.79E-05 | 1 | TAGGCTGTTGGTATT |
| 3687 | mamo-L | *BGIBMGA007569* | up | 764 | 778 | - | 11.4758 | 4.18E-05 | 1 | TAGGCTGCGTTTCAT |
| 3688 | mamo-L | *BGIBMGA007570* | up | 399 | 413 | - | 12.2581 | 1.90E-05 | 1 | AAGGCTGTATTTGTT |
| 3689 | mamo-L | *BGIBMGA007572* | down | 1153 | 1167 | - | 11.5968 | 3.73E-05 | 1 | TAGGCTTTTTTCATT |
| 3690 | mamo-L | *BGIBMGA007574* | up | 440 | 454 | + | 11.3548 | 4.69E-05 | 1 | TAGGCTGACAATGTA |
| 3691 | mamo-L | *BGIBMGA007578* | down | 363 | 377 | + | 10.7742 | 7.89E-05 | 1 | TAGGCTTATCTTGGA |
| 3692 | mamo-L | *BGIBMGA007583* | down | 1658 | 1672 | + | 13.379 | 4.96E-06 | 1 | AAGGCTGATTTTGTT |
| 3693 | mamo-L | *BGIBMGA007583* | down | 619 | 633 | - | 12.7419 | 1.09E-05 | 1 | TAGGCTATGTTTTTT |
| 3694 | mamo-L | *BGIBMGA007585* | down | 1180 | 1194 | + | 11.2742 | 5.06E-05 | 1 | TAGGCATGCGTTCTT |
| 3695 | mamo-L | *BGIBMGA007592* | down | 1090 | 1104 | + | 11.0242 | 6.36E-05 | 1 | GAGGCCGGCGGCGTT |
| 3696 | mamo-L | *BGIBMGA007593* | down | 789 | 803 | - | 12.0645 | 2.34E-05 | 1 | TAGGCGGGCTTCGGA |
| 3697 | mamo-L | *BGIBMGA007595* | up | 711 | 725 | + | 10.9839 | 6.59E-05 | 1 | TAGGCTCAAGTGTTT |
| 3698 | mamo-L | *BGIBMGA007596* | up | 466 | 480 | - | 10.5968 | 9.15E-05 | 1 | AAGGCTGTTAATGCT |
| 3699 | mamo-L | *BGIBMGA007599* | up | 1222 | 1236 | - | 10.5645 | 9.40E-05 | 1 | TAGGCTTTCATTTTG |
| 3700 | mamo-L | *BGIBMGA007601* | up | 146 | 160 | - | 10.9194 | 6.98E-05 | 1 | TAGGCTTAAATCTTT |
| 3701 | mamo-L | *BGIBMGA007603* | up | 1341 | 1355 | - | 12.7258 | 1.11E-05 | 1 | TAGGCGTGCTTTGTA |
| 3702 | mamo-L | *BGIBMGA007607* | down | 1533 | 1547 | - | 11.371 | 4.62E-05 | 1 | TAGGCTGTATAGGCT |
| 3703 | mamo-L | *BGIBMGA007609* | up | 889 | 903 | + | 12.8145 | 9.96E-06 | 1 | TAGGCCCACATTTGT |
| 3704 | mamo-L | *BGIBMGA007610* | up | 1039 | 1053 | - | 11.0565 | 6.18E-05 | 1 | TAGGCTTGAAATGAT |
| 3705 | mamo-L | *BGIBMGA007611* | down | 887 | 901 | + | 10.7097 | 8.33E-05 | 1 | AAGGCGTTGGTCTGT |
| 3706 | mamo-L | *BGIBMGA007614* | down | 682 | 696 | + | 10.8629 | 7.33E-05 | 1 | TAGGCAGTTTTTATT |
| 3707 | mamo-L | *BGIBMGA007615* | down | 1537 | 1551 | + | 10.7016 | 8.39E-05 | 1 | TAGGCCCGTGATGCA |
| 3708 | mamo-L | *BGIBMGA007617* | up | 673 | 687 | - | 10.4919 | 9.96E-05 | 1 | TAGGCTGTGTCTGTG |
| 3709 | mamo-L | *BGIBMGA007624* | down | 449 | 463 | + | 11.0968 | 5.97E-05 | 1 | TAGGCAGGCTGTTTT |
| 3710 | mamo-L | *BGIBMGA007628* | down | 633 | 647 | - | 11.2742 | 5.06E-05 | 1 | TAGGCATACTATTGT |
| 3711 | mamo-L | *BGIBMGA007628* | up | 878 | 892 | - | 10.7097 | 8.33E-05 | 1 | TGGGCTGATGATGTT |
| 3712 | mamo-L | *BGIBMGA007631* | down | 794 | 808 | - | 10.5403 | 9.58E-05 | 1 | TAGGCTGTATGGTGT |
| 3713 | mamo-L | *BGIBMGA007632* | up | 1184 | 1198 | + | 10.6129 | 9.03E-05 | 1 | TAGGCTTATAGTTTC |
| 3714 | mamo-L | *BGIBMGA007636* | up | 469 | 483 | + | 10.9516 | 6.78E-05 | 1 | GAGGCGCGGTTTTTT |
| 3715 | mamo-L | *BGIBMGA007638* | down | 320 | 334 | + | 13.6855 | 3.18E-06 | 1 | TAGGCTTACTTTATT |
| 3716 | mamo-L | *BGIBMGA007642* | down | 954 | 968 | + | 12.1935 | 2.05E-05 | 1 | TAGGCTCATATTTTT |
| 3717 | mamo-L | *BGIBMGA007646* | down | 870 | 884 | + | 10.5726 | 9.33E-05 | 1 | TAGGCTAATATTTAT |
| 3718 | mamo-L | *BGIBMGA007648* | down | 1379 | 1393 | + | 11.3871 | 4.55E-05 | 1 | TAGGCTGCGGTCTAT |
| 3719 | mamo-L | *BGIBMGA007648* | down | 455 | 469 | + | 11.25 | 5.18E-05 | 1 | TAGGCTTGCAATTAT |
| 3720 | mamo-L | *BGIBMGA007648* | up | 1236 | 1250 | + | 10.4919 | 9.96E-05 | 1 | TAGGCTTATTGTTTA |
| 3721 | mamo-L | *BGIBMGA007650* | up | 1356 | 1370 | + | 10.8387 | 7.47E-05 | 1 | GAGGCTGTTTTTCCT |
| 3722 | mamo-L | *BGIBMGA007658* | down | 1677 | 1691 | + | 11.3548 | 4.69E-05 | 1 | AAGGCTGATGATGAT |
| 3723 | mamo-L | *BGIBMGA007659* | down | 452 | 466 | + | 11.0403 | 6.28E-05 | 1 | TAGGCCGTCGCTTAT |
| 3724 | mamo-L | *BGIBMGA007661* | up | 1320 | 1334 | - | 10.7661 | 7.95E-05 | 1 | AAGGCTGGTGTTAGC |
| 3725 | mamo-L | *BGIBMGA007663* | up | 1824 | 1838 | - | 12.8387 | 9.67E-06 | 1 | TAGGCTCGGTTTACT |
| 3726 | mamo-L | *BGIBMGA007666* | up | 538 | 552 | + | 11 | 6.50E-05 | 1 | AAGGCTGGTGTGTGT |
| 3727 | mamo-L | *BGIBMGA007668* | up | 1363 | 1377 | - | 10.9194 | 6.98E-05 | 1 | TAGGCTACTTTTTTT |
| 3728 | mamo-L | *BGIBMGA007669* | down | 816 | 830 | - | 10.5726 | 9.33E-05 | 1 | TAGGCTAATATTTAT |
| 3729 | mamo-L | *BGIBMGA007670* | up | 719 | 733 | - | 11.3548 | 4.69E-05 | 1 | AAGGCTGATGATGAT |
| 3730 | mamo-L | *BGIBMGA007670* | up | 11 | 25 | - | 11.3145 | 4.87E-05 | 1 | AAGGCCCATTTTTGT |
| 3731 | mamo-L | *BGIBMGA007672* | up | 518 | 532 | + | 10.9194 | 6.98E-05 | 1 | TAGGCTACTTTTTTT |
| 3732 | mamo-L | *BGIBMGA007675* | up | 1302 | 1316 | + | 10.9839 | 6.59E-05 | 1 | AAGGCGTTTTTTTTT |
| 3733 | mamo-L | *BGIBMGA007679* | down | 645 | 659 | - | 15.4516 | 1.42E-07 | 1 | TAGGCCTGGTTTGCT |
| 3734 | mamo-L | *BGIBMGA007680* | up | 1078 | 1092 | - | 10.9194 | 6.98E-05 | 1 | TAGGCTACTTTTTTT |
| 3735 | mamo-L | *BGIBMGA007681* | down | 498 | 512 | - | 10.6694 | 8.62E-05 | 1 | TAGGCACTTTTTTTT |
| 3736 | mamo-L | *BGIBMGA007690* | down | 634 | 648 | + | 10.7984 | 7.74E-05 | 1 | GAGGCTTTATTTTCT |
| 3737 | mamo-L | *BGIBMGA007693* | up | 568 | 582 | - | 10.5806 | 9.27E-05 | 1 | TAGGCAGTCTTGTCT |
| 3738 | mamo-L | *BGIBMGA007697* | down | 1286 | 1300 | + | 10.621 | 8.97E-05 | 1 | TAGGCTTGATACTTT |
| 3739 | mamo-L | *BGIBMGA007698* | up | 1817 | 1831 | - | 12.7258 | 1.11E-05 | 1 | TAGGCGTGCTTTGTA |
| 3740 | mamo-L | *BGIBMGA007699* | down | 1115 | 1129 | + | 11.3871 | 4.55E-05 | 1 | TAGGCTATATTTTTT |
| 3741 | mamo-L | *BGIBMGA007701* | down | 788 | 802 | + | 10.5726 | 9.33E-05 | 1 | TAGGCTAATATTTAT |
| 3742 | mamo-L | *BGIBMGA007701* | down | 1026 | 1040 | + | 10.5565 | 9.46E-05 | 1 | TAGGCGACTTTTTGT |
| 3743 | mamo-L | *BGIBMGA007705* | down | 120 | 134 | + | 10.5726 | 9.33E-05 | 1 | TAGGCTAATATTTAT |
| 3744 | mamo-L | *BGIBMGA007706* | down | 378 | 392 | + | 11.7823 | 3.11E-05 | 1 | TAGGCTTTGTTGATT |
| 3745 | mamo-L | *BGIBMGA007706* | down | 1416 | 1430 | - | 11.4355 | 4.34E-05 | 1 | AAGGCTTTTTATGTT |
| 3746 | mamo-L | *BGIBMGA007709* | down | 632 | 646 | - | 10.9355 | 6.88E-05 | 1 | TAGGCCCAGTGGTGT |
| 3747 | mamo-L | *BGIBMGA007710* | up | 1477 | 1491 | - | 13.4677 | 4.39E-06 | 1 | TAGGCTGATGATGAT |
| 3748 | mamo-L | *BGIBMGA007713* | down | 69 | 83 | + | 10.7823 | 7.84E-05 | 1 | TAGGCCTTAATTATT |
| 3749 | mamo-L | *BGIBMGA007715* | down | 15 | 29 | + | 10.7016 | 8.39E-05 | 1 | TAGGCATTGTATTTT |
| 3750 | mamo-L | *BGIBMGA007717* | down | 1103 | 1117 | + | 11.6371 | 3.58E-05 | 1 | TAGGCTTGTATCTTT |
| 3751 | mamo-L | *BGIBMGA007720* | down | 1193 | 1207 | - | 11.3871 | 4.55E-05 | 1 | TAGGCTTGCCTTAGT |
| 3752 | mamo-L | *BGIBMGA007721* | up | 1472 | 1486 | - | 12.0403 | 2.40E-05 | 1 | TAGGCTTTCTATATT |
| 3753 | mamo-L | *BGIBMGA007722* | up | 1017 | 1031 | + | 10.629 | 8.91E-05 | 1 | TAGGCAGCTGTTTTT |
| 3754 | mamo-L | *BGIBMGA007725* | up | 14 | 28 | - | 10.5484 | 9.52E-05 | 1 | GAGGCGGCTGTTGGC |
| 3755 | mamo-L | *BGIBMGA007726* | down | 63 | 77 | + | 10.871 | 7.28E-05 | 1 | AAGGCTCTTTGTGGT |
| 3756 | mamo-L | *BGIBMGA007731* | down | 50 | 64 | + | 11.4919 | 4.11E-05 | 1 | TAGGCTGAATTCTAT |
| 3757 | mamo-L | *BGIBMGA007731* | up | 1326 | 1340 | - | 11.2419 | 5.21E-05 | 1 | TAGGCCAGGTTGGCC |
| 3758 | mamo-L | *BGIBMGA007732* | up | 673 | 687 | + | 11.2823 | 5.03E-05 | 1 | TAGGCTAATTCTGGT |
| 3759 | mamo-L | *BGIBMGA007734* | up | 503 | 517 | + | 12.7339 | 1.10E-05 | 1 | TAGGCTTATAATGCT |
| 3760 | mamo-L | *BGIBMGA007734* | up | 1763 | 1777 | + | 12.6613 | 1.19E-05 | 1 | TAGGCTGTCTAGGCT |
| 3761 | mamo-L | *BGIBMGA007734* | up | 1772 | 1786 | + | 10.871 | 7.28E-05 | 1 | TAGGCTATGTAGGTT |
| 3762 | mamo-L | *BGIBMGA007736* | up | 1490 | 1504 | + | 11.3871 | 4.55E-05 | 1 | TAGGCCTTTTTTTTA |
| 3763 | mamo-L | *BGIBMGA007736* | up | 1559 | 1573 | - | 10.7258 | 8.23E-05 | 1 | TAGGCCTATTTTAGG |
| 3764 | mamo-L | *BGIBMGA007744* | down | 372 | 386 | + | 10.9194 | 6.98E-05 | 1 | TAGGCTAGGTTAGGT |
| 3765 | mamo-L | *BGIBMGA007747* | up | 1433 | 1447 | + | 12.1855 | 2.06E-05 | 1 | TAGGCTATTTTTTTT |

| | | | | | | | | | | |
|---|---|---|---|---|---|---|---|---|---|---|
| 3766 | mamo-L | *BGIBMGA007752* | up | 1699 | 1713 | - | 10.8952 | 7.12E-05 | 1 | TAGGCAGCTGTTGAT |
| 3767 | mamo-L | *BGIBMGA007753* | up | 1466 | 1480 | - | 11.1774 | 5.54E-05 | 1 | TAGGCTGCATTTTAT |
| 3768 | mamo-L | *BGIBMGA007754* | up | 657 | 671 | + | 13.2903 | 5.60E-06 | 1 | TAGGCTTTATTTTGT |
| 3769 | mamo-L | *BGIBMGA007757* | down | 1894 | 1908 | - | 11.4919 | 4.11E-05 | 1 | GAGGCTGAGATCGCT |
| 3770 | mamo-L | *BGIBMGA007760* | up | 464 | 478 | - | 10.8306 | 7.52E-05 | 1 | TAGGCCCCCTTCGCC |
| 3771 | mamo-L | *BGIBMGA007762* | down | 91 | 105 | - | 10.5726 | 9.33E-05 | 1 | TAGGCTAATATTTAT |
| 3772 | mamo-L | *BGIBMGA007762* | up | 1974 | 1988 | - | 10.5726 | 9.33E-05 | 1 | TAGGCTAATATTTAT |
| 3773 | mamo-L | *BGIBMGA007763* | down | 750 | 764 | - | 10.5968 | 9.15E-05 | 1 | GAGGCTAGTTTTTGT |
| 3774 | mamo-L | *BGIBMGA007768* | up | 1331 | 1345 | + | 14.0403 | 1.87E-06 | 1 | TAGGCCGGCTTGGCT |
| 3775 | mamo-L | *BGIBMGA007773* | up | 1601 | 1615 | - | 10.7097 | 8.33E-05 | 1 | TAGGCCGAATTGCTT |
| 3776 | mamo-L | *BGIBMGA007774* | up | 1440 | 1454 | + | 11.1774 | 5.54E-05 | 1 | TAGGCCTAAGTAGGT |
| 3777 | mamo-L | *BGIBMGA007777* | up | 1329 | 1343 | + | 11.1935 | 5.45E-05 | 1 | AAGGCTGTGTATTTT |
| 3778 | mamo-L | *BGIBMGA007779* | up | 839 | 853 | + | 11.3629 | 4.66E-05 | 1 | TAGGCTTTCGTTTAA |
| 3779 | mamo-L | *BGIBMGA007779* | up | 1113 | 1127 | + | 11.2581 | 5.14E-05 | 1 | TAGGCTAAAGTGGCT |
| 3780 | mamo-L | *BGIBMGA007781* | up | 1264 | 1278 | + | 10.5 | 9.90E-05 | 1 | GAGGCCCAGGTCGTC |
| 3781 | mamo-L | *BGIBMGA007785* | down | 1013 | 1027 | - | 10.9194 | 6.98E-05 | 1 | TAGGCTAGGTTAGGT |
| 3782 | mamo-L | *BGIBMGA007786* | up | 545 | 559 | + | 11.5403 | 3.93E-05 | 1 | TAGGCTTTTGGCTGT |
| 3783 | mamo-L | *BGIBMGA007786* | up | 1737 | 1751 | - | 10.5242 | 9.70E-05 | 1 | GAGGCCGTGGAGGCT |
| 3784 | mamo-L | *BGIBMGA007787* | up | 1568 | 1582 | - | 10.5645 | 9.40E-05 | 1 | AAGGCGGTGTTCTTT |
| 3785 | mamo-L | *BGIBMGA007789* | up | 1799 | 1813 | + | 11.4113 | 4.45E-05 | 1 | TAGGCTACCTTTTTT |
| 3786 | mamo-L | *BGIBMGA007794* | up | 1539 | 1553 | - | 10.8629 | 7.33E-05 | 1 | GAGGCCGAGGTTCAT |
| 3787 | mamo-L | *BGIBMGA007795* | up | 852 | 866 | - | 11.2823 | 5.03E-05 | 1 | TAGGCTAATTCTGGT |
| 3788 | mamo-L | *BGIBMGA007795* | down | 1852 | 1866 | + | 11.1694 | 5.59E-05 | 1 | TAGGCGAGGTGTGAT |
| 3789 | mamo-L | *BGIBMGA007798* | down | 1508 | 1522 | + | 13 | 8.01E-06 | 1 | TAGGCTTTGTTGTGT |
| 3790 | mamo-L | *BGIBMGA007798* | up | 724 | 738 | + | 11.2419 | 5.21E-05 | 1 | TAGGCCCACGGTCCT |
| 3791 | mamo-L | *BGIBMGA007804* | down | 959 | 973 | - | 11.4032 | 4.49E-05 | 1 | TAGGCCCTCTTTATC |
| 3792 | mamo-L | *BGIBMGA007805* | down | 794 | 808 | - | 12.0403 | 2.40E-05 | 1 | TAGGCGTTTTATGAT |
| 3793 | mamo-L | *BGIBMGA007809* | down | 443 | 457 | - | 10.7581 | 8.00E-05 | 1 | TAGGCTAACATTGCA |
| 3794 | mamo-L | *BGIBMGA007816* | down | 1714 | 1728 | - | 11.6371 | 3.58E-05 | 1 | TAGGCTTGTATCTTT |
| 3795 | mamo-L | *BGIBMGA007817* | down | 252 | 266 | - | 13.6694 | 3.27E-06 | 1 | TAGGCTTTGTTTTGC |
| 3796 | mamo-L | *BGIBMGA007819* | up | 727 | 741 | - | 11.0806 | 6.05E-05 | 1 | TCGGCCGGTGTTGTT |
| 3797 | mamo-L | *BGIBMGA007820* | down | 526 | 540 | - | 14.0726 | 1.78E-06 | 1 | TAGGCTCTTTTTGTT |
| 3798 | mamo-L | *BGIBMGA007822* | down | 244 | 258 | + | 10.5726 | 9.33E-05 | 1 | TAGGCTAATATTTAT |
| 3799 | mamo-L | *BGIBMGA007826* | up | 264 | 278 | + | 13.3226 | 5.34E-06 | 1 | AAGGCCGATGTTGTT |
| 3800 | mamo-L | *BGIBMGA007829* | down | 159 | 173 | - | 11.5806 | 3.78E-05 | 1 | TAGGCTAGGGACGCT |
| 3801 | mamo-L | *BGIBMGA007829* | down | 1608 | 1622 | - | 10.5726 | 9.33E-05 | 1 | TAGGCTAATATTTAT |
| 3802 | mamo-L | *BGIBMGA007831* | up | 443 | 457 | - | 14.2177 | 1.43E-06 | 1 | TAGGCTGACATTGGC |
| 3803 | mamo-L | *BGIBMGA007831* | down | 1727 | 1741 | - | 13.8952 | 2.32E-06 | 1 | TAGGCGTTCATTGGT |
| 3804 | mamo-L | *BGIBMGA007834* | down | 165 | 179 | - | 10.5726 | 9.33E-05 | 1 | TAGGCTAATATTTAT |
| 3805 | mamo-L | *BGIBMGA007835* | down | 1200 | 1214 | - | 11.0484 | 6.23E-05 | 1 | TAGGCACTGTTCGTT |
| 3806 | mamo-L | *BGIBMGA007837* | up | 1752 | 1766 | - | 10.5726 | 9.33E-05 | 1 | TAGGCTAATATTTAT |
| 3807 | mamo-L | *BGIBMGA007843* | down | 165 | 179 | + | 12.5161 | 1.42E-05 | 1 | TAGGCTTTAGTTAGT |
| 3808 | mamo-L | *BGIBMGA007844* | down | 384 | 398 | + | 10.8306 | 7.52E-05 | 1 | GAGGCTCTTTTTTTT |
| 3809 | mamo-L | *BGIBMGA007847* | up | 1102 | 1116 | + | 10.7661 | 7.95E-05 | 1 | TAGGCTTACGATTCG |
| 3810 | mamo-L | *BGIBMGA007851* | down | 302 | 316 | - | 12.1935 | 2.05E-05 | 1 | TAGGCCGATTTCCGT |
| 3811 | mamo-L | *BGIBMGA007851* | up | 1908 | 1922 | - | 11.0161 | 6.40E-05 | 1 | TAGGCCTACATTTGG |
| 3812 | mamo-L | *BGIBMGA007853* | up | 109 | 123 | - | 13.9435 | 2.15E-06 | 1 | TAGGCTGTTGATGCT |
| 3813 | mamo-L | *BGIBMGA007861* | up | 1420 | 1434 | - | 13.5726 | 3.75E-06 | 1 | TAGGCTTAGGTGGGC |
| 3814 | mamo-L | *BGIBMGA007864* | down | 1486 | 1500 | - | 12.0242 | 2.44E-05 | 1 | AAGGCTTTCGATGCT |
| 3815 | mamo-L | *BGIBMGA007870* | up | 1656 | 1670 | - | 12.25 | 1.92E-05 | 1 | GAGGCTGATTTTTTT |
| 3816 | mamo-L | *BGIBMGA007871* | down | 840 | 854 | - | 10.9758 | 6.64E-05 | 1 | TAGGCATCGGTTGCC |
| 3817 | mamo-L | *BGIBMGA007873* | up | 1651 | 1665 | + | 11.8306 | 2.97E-05 | 1 | TAGGCTTATCTTGAT |
| 3818 | mamo-L | *BGIBMGA007875* | down | 1518 | 1532 | + | 13.5403 | 3.97E-06 | 1 | TAGGCTTTGATTTGT |
| 3819 | mamo-L | *BGIBMGA007876* | down | 1014 | 1028 | - | 12.2903 | 1.83E-05 | 1 | TAGGCCTCGTTTAGT |
| 3820 | mamo-L | *BGIBMGA007877* | up | 962 | 976 | + | 12.7258 | 1.11E-05 | 1 | TAGGCGTGCTTTGTA |
| 3821 | mamo-L | *BGIBMGA007880* | up | 1369 | 1383 | - | 12.4677 | 1.49E-05 | 1 | TAGGCTTCTATTGCT |
| 3822 | mamo-L | *BGIBMGA007880* | up | 371 | 385 | + | 10.7581 | 8.00E-05 | 1 | TAGGCTATGTTCGAC |
| 3823 | mamo-L | *BGIBMGA007883* | down | 1282 | 1296 | + | 11.5806 | 3.78E-05 | 1 | GAGGCTGTGTTTATT |
| 3824 | mamo-L | *BGIBMGA007884* | up | 1466 | 1480 | + | 10.6774 | 8.56E-05 | 1 | TAGGCGGGCGGCATT |
| 3825 | mamo-L | *BGIBMGA007888* | up | 663 | 677 | - | 10.5645 | 9.40E-05 | 1 | GAGGCCTTCGACGTT |
| 3826 | mamo-L | *BGIBMGA007889* | down | 37 | 51 | - | 10.7984 | 7.74E-05 | 1 | GAGGCTTTATTTTCT |
| 3827 | mamo-L | *BGIBMGA007889* | up | 722 | 736 | - | 10.7097 | 8.33E-05 | 1 | TAGGCATTTGATGGC |
| 3828 | mamo-L | *BGIBMGA007891* | down | 753 | 767 | - | 11.129 | 5.79E-05 | 1 | GAGGCTGAGTTGTCT |
| 3829 | mamo-L | *BGIBMGA007891* | down | 437 | 451 | + | 10.7258 | 8.23E-05 | 1 | TAGGCGGGGTGCGGA |
| 3830 | mamo-L | *BGIBMGA007894* | down | 758 | 772 | - | 10.9516 | 6.78E-05 | 1 | TAGGCGGTTCATGGT |
| 3831 | mamo-L | *BGIBMGA007898* | down | 928 | 942 | + | 10.7823 | 7.84E-05 | 1 | AAGGCTGCTTTTGCC |
| 3832 | mamo-L | *BGIBMGA007899* | up | 1449 | 1463 | - | 12.4435 | 1.54E-05 | 1 | TAGGCTTTGTTTCAT |
| 3833 | mamo-L | *BGIBMGA007900* | up | 1007 | 1021 | - | 13.25 | 5.89E-06 | 1 | TAGGCGTACGTTGGA |
| 3834 | mamo-L | *BGIBMGA007903* | up | 278 | 292 | + | 12.3306 | 1.75E-05 | 1 | TAGGCTTACTATACT |
| 3835 | mamo-L | *BGIBMGA007906* | up | 1838 | 1852 | + | 12.6371 | 1.23E-05 | 1 | TAGGCAGGCGGTGGT |
| 3836 | mamo-L | *BGIBMGA007906* | up | 1608 | 1622 | - | 11.7419 | 3.23E-05 | 1 | TAGGCCAACGATGTC |
| 3837 | mamo-L | *BGIBMGA007909* | down | 627 | 641 | + | 11.5081 | 4.05E-05 | 1 | TAGCCTGATGTTGTT |
| 3838 | mamo-L | *BGIBMGA007910* | up | 1371 | 1385 | + | 11.379 | 4.58E-05 | 1 | TAGGCGTTCATTGTA |
| 3839 | mamo-L | *BGIBMGA007910* | down | 1523 | 1537 | + | 10.5726 | 9.33E-05 | 1 | TAGGCTTCCAATTTT |
| 3840 | mamo-L | *BGIBMGA007915* | down | 992 | 1006 | - | 10.5726 | 9.33E-05 | 1 | TAGGCTAATATTTAT |
| 3841 | mamo-L | *BGIBMGA007917* | down | 314 | 328 | + | 12.1532 | 2.13E-05 | 1 | TAGGCTAGGTTTTAT |
| 3842 | mamo-L | *BGIBMGA007918* | down | 589 | 603 | + | 10.5565 | 9.46E-05 | 1 | TAGGCATTCTGTTTT |
| 3843 | mamo-L | *BGIBMGA007919* | down | 1460 | 1474 | + | 10.5726 | 9.33E-05 | 1 | TAGGCTAATATTTAT |
| 3844 | mamo-L | *BGIBMGA007921* | up | 637 | 651 | - | 12.4597 | 1.51E-05 | 1 | AAGGCTTGGTTTTTT |
| 3845 | mamo-L | *BGIBMGA007923* | down | 1578 | 1592 | - | 11.4032 | 4.49E-05 | 1 | TAGGCTAATATTTTT |
| 3846 | mamo-L | *BGIBMGA007931* | down | 511 | 525 | + | 14.3629 | 1.14E-06 | 1 | TAGGCTTTCGTTTCT |
| 3847 | mamo-L | *BGIBMGA007931* | up | 186 | 200 | - | 11.3468 | 4.73E-05 | 1 | TAGGCGTAATTCTTT |
| 3848 | mamo-L | *BGIBMGA007933* | down | 475 | 489 | - | 12.8871 | 9.15E-06 | 1 | TAGGCTGTCTTGTCT |
| 3849 | mamo-L | *BGIBMGA007935* | up | 1912 | 1926 | + | 11.129 | 5.79E-05 | 1 | AAGGCCCTCATTGCT |
| 3850 | mamo-L | *BGIBMGA007937* | down | 114 | 128 | + | 11.7339 | 3.26E-05 | 1 | TAGGCTTCTGTTATT |
| 3851 | mamo-L | *BGIBMGA007938* | up | 1194 | 1208 | + | 11.7339 | 3.26E-05 | 1 | TAGGCTTCTGTTATT |
| 3852 | mamo-L | *BGIBMGA007942* | down | 1363 | 1377 | - | 12.1935 | 2.05E-05 | 1 | TAGGCTCATATTTTT |
| 3853 | mamo-L | *BGIBMGA007942* | up | 790 | 804 | + | 10.4919 | 9.96E-05 | 1 | TAGGCTGTGTCTGTG |
| 3854 | mamo-L | *BGIBMGA007951* | down | 1433 | 1447 | - | 12.0484 | 2.38E-05 | 1 | AAGGCGTTTTTTGCT |
| 3855 | mamo-L | *BGIBMGA007955* | down | 1539 | 1553 | - | 10.5726 | 9.33E-05 | 1 | TAGGCTAATATTTAT |
| 3856 | mamo-L | *BGIBMGA007957* | down | 1110 | 1124 | - | 11.1532 | 5.67E-05 | 1 | TAGGCAGCGGTTTCT |
| 3857 | mamo-L | *BGIBMGA007969* | up | 431 | 445 | + | 10.9194 | 6.98E-05 | 1 | TAGGCTACTTTTTTT |

| | | | | | | | | | | |
|---|---|---|---|---|---|---|---|---|---|---|
| 3858 | mamo-L | *BGIBMGA007969* | down | 171 | 185 | - | 10.8629 | 7.33E-05 | 1 | TAGGCAGTTTTTATT |
| 3859 | mamo-L | *BGIBMGA007971* | down | 449 | 463 | - | 11.5726 | 3.81E-05 | 1 | TAGGCGGTGGACGCC |
| 3860 | mamo-L | *BGIBMGA007975* | up | 1566 | 1580 | - | 10.5726 | 9.33E-05 | 1 | TAGGCTAATATTTAT |
| 3861 | mamo-L | *BGIBMGA007978* | up | 542 | 556 | + | 11.3548 | 4.69E-05 | 1 | AAGGCTGATGATGAT |
| 3862 | mamo-L | *BGIBMGA007979* | up | 699 | 713 | + | 10.6452 | 8.79E-05 | 1 | AAGGCCTTTTTTTAT |
| 3863 | mamo-L | *BGIBMGA007980* | up | 1956 | 1970 | + | 13.2823 | 5.66E-06 | 1 | TAGGCGGACGATGAT |
| 3864 | mamo-L | *BGIBMGA007982* | up | 1625 | 1639 | - | 10.9194 | 6.98E-05 | 1 | TAGGCTACTTTTTTT |
| 3865 | mamo-L | *BGIBMGA007984* | up | 810 | 824 | + | 10.621 | 8.97E-05 | 1 | TAGGCTGAAAAGGTT |
| 3866 | mamo-L | *BGIBMGA007985* | down | 116 | 130 | - | 10.9194 | 6.98E-05 | 1 | TAGGCTACTTTTTTT |
| 3867 | mamo-L | *BGIBMGA007988* | down | 1248 | 1262 | + | 13.9194 | 2.23E-06 | 1 | TAGGCTAATTTTGGT |
| 3868 | mamo-L | *BGIBMGA007993* | down | 1306 | 1320 | + | 11.9274 | 2.69E-05 | 1 | GAGGCCGGGGTTCGT |
| 3869 | mamo-L | *BGIBMGA007994* | up | 1917 | 1931 | + | 10.8387 | 7.47E-05 | 1 | TAGGCTCGTTTAGTT |
| 3870 | mamo-L | *BGIBMGA007994* | down | 1341 | 1355 | - | 10.4919 | 9.96E-05 | 1 | TAGGCTGTGTCTGTG |
| 3871 | mamo-L | *BGIBMGA007996* | down | 774 | 788 | + | 11.2903 | 4.99E-05 | 1 | TAGGCAATTTTTGGT |
| 3872 | mamo-L | *BGIBMGA007997* | up | 307 | 321 | - | 10.9194 | 6.98E-05 | 1 | TAGGCTACTTTTTTT |
| 3873 | mamo-L | *BGIBMGA007998* | up | 950 | 964 | + | 11.7661 | 3.16E-05 | 1 | TAGGCAGTTTTTTTT |
| 3874 | mamo-L | *BGIBMGA007998* | up | 557 | 571 | + | 11.1694 | 5.59E-05 | 1 | TAGGCTTATGATAAT |
| 3875 | mamo-L | *BGIBMGA007999* | up | 618 | 632 | + | 12.1532 | 2.13E-05 | 1 | GAGGCTGATGATGTT |
| 3876 | mamo-L | *BGIBMGA007999* | down | 1719 | 1733 | + | 11.7258 | 3.29E-05 | 1 | AAGGCTATGTTTGTT |
| 3877 | mamo-L | *BGIBMGA008000* | up | 527 | 541 | - | 13.0242 | 7.77E-06 | 1 | TAGGCAGACGTTTGT |
| 3878 | mamo-L | *BGIBMGA008000* | up | 1005 | 1019 | + | 11.8065 | 3.03E-05 | 1 | TAGGCTTTCTTCGGG |
| 3879 | mamo-L | *BGIBMGA008001* | up | 589 | 603 | - | 11.4758 | 4.18E-05 | 1 | TAGGCTGTTTTTTTG |
| 3880 | mamo-L | *BGIBMGA008004* | down | 727 | 741 | - | 11.2419 | 5.21E-05 | 1 | TAGGCTTTATTTAAT |
| 3881 | mamo-L | *BGIBMGA008004* | down | 1927 | 1941 | - | 10.621 | 8.97E-05 | 1 | TAGGCCTCATTTATT |
| 3882 | mamo-L | *BGIBMGA008005* | up | 1092 | 1106 | + | 10.9194 | 6.98E-05 | 1 | TAGGCTACTTTTTTT |
| 3883 | mamo-L | *BGIBMGA008007* | down | 667 | 681 | + | 10.75 | 8.06E-05 | 1 | GAGGCCGATTGTGCC |
| 3884 | mamo-L | *BGIBMGA008008* | down | 1191 | 1205 | - | 11.1935 | 5.45E-05 | 1 | TAGGCGTATTATATT |
| 3885 | mamo-L | *BGIBMGA008008* | down | 1263 | 1277 | + | 10.5081 | 9.83E-05 | 1 | AAGGCGTGGTTCTTT |
| 3886 | mamo-L | *BGIBMGA008012* | up | 1338 | 1352 | + | 13.5242 | 4.05E-06 | 1 | TAGGCCTACGTTTCC |
| 3887 | mamo-L | *BGIBMGA008012* | down | 923 | 937 | + | 11.8226 | 2.99E-05 | 1 | TAGGCGTAGGTTTGG |
| 3888 | mamo-L | *BGIBMGA008012* | down | 1143 | 1157 | + | 11.6048 | 3.70E-05 | 1 | AAGGCTGGCTGTTGT |
| 3889 | mamo-L | *BGIBMGA008013* | up | 805 | 819 | + | 11.8226 | 2.99E-05 | 1 | TAGGCGTAGGTTTGG |
| 3890 | mamo-L | *BGIBMGA008013* | up | 1025 | 1039 | + | 11.6048 | 3.70E-05 | 1 | AAGGCTGGCTGTTGT |
| 3891 | mamo-L | *BGIBMGA008013* | down | 229 | 243 | + | 10.9194 | 6.98E-05 | 1 | TAGGCTACTTTTTTT |
| 3892 | mamo-L | *BGIBMGA008015* | up | 1761 | 1775 | - | 10.5726 | 9.33E-05 | 1 | TAGGCTAATATTTAT |
| 3893 | mamo-L | *BGIBMGA008016* | down | 316 | 330 | + | 11.5806 | 3.78E-05 | 1 | AAGGCTCAAGTTGCT |
| 3894 | mamo-L | *BGIBMGA008017* | up | 1349 | 1363 | + | 11.9516 | 2.63E-05 | 1 | TAGGCTGTATATTTT |
| 3895 | mamo-L | *BGIBMGA008023* | down | 1601 | 1615 | + | 11.379 | 4.58E-05 | 1 | GAGGCCTGTTGTGTT |
| 3896 | mamo-L | *BGIBMGA008024* | down | 35 | 49 | + | 11.379 | 4.58E-05 | 1 | GAGGCCTGTTGTGTT |
| 3897 | mamo-L | *BGIBMGA008025* | down | 1236 | 1250 | + | 11.4597 | 4.25E-05 | 1 | TAGGCGGGGTGGTGT |
| 3898 | mamo-L | *BGIBMGA008026* | down | 1956 | 1970 | - | 10.5 | 9.90E-05 | 1 | GAGGCCCAGGTCGTC |
| 3899 | mamo-L | *BGIBMGA008027* | up | 117 | 131 | - | 11.3065 | 4.91E-05 | 1 | TAGGCCCTGATTACT |
| 3900 | mamo-L | *BGIBMGA008028* | down | 702 | 716 | + | 10.5968 | 9.15E-05 | 1 | GAGGCTAGTTTTTGT |
| 3901 | mamo-L | *BGIBMGA008029* | down | 900 | 914 | - | 11.8145 | 3.01E-05 | 1 | TAGGCTTGATGTTTT |
| 3902 | mamo-L | *BGIBMGA008030* | up | 1409 | 1423 | + | 11 | 6.50E-05 | 1 | TAGGCAATGGTTGGC |
| 3903 | mamo-L | *BGIBMGA008030* | down | 682 | 696 | - | 10.9194 | 6.98E-05 | 1 | TAGGCTACTTTTTTT |
| 3904 | mamo-L | *BGIBMGA008035* | down | 376 | 390 | - | 12.0403 | 2.40E-05 | 1 | TAGGCCTAGTATGTA |
| 3905 | mamo-L | *BGIBMGA008035* | down | 78 | 92 | - | 11.6694 | 3.47E-05 | 1 | TAGGCTAATATTGAT |
| 3906 | mamo-L | *BGIBMGA008036* | down | 968 | 982 | - | 12.9919 | 8.10E-06 | 1 | TAGGCAGTTGTTGTT |
| 3907 | mamo-L | *BGIBMGA008038* | down | 1133 | 1147 | + | 11.5726 | 3.81E-05 | 1 | AAGGCGTGGGGTGCT |
| 3908 | mamo-L | *BGIBMGA008041* | down | 375 | 389 | - | 11.8145 | 3.01E-05 | 1 | AAGGCTGCGGTTGGC |
| 3909 | mamo-L | *BGIBMGA008044* | up | 365 | 379 | - | 11.4758 | 4.18E-05 | 1 | TAGGCATTTTGTGGT |
| 3910 | mamo-L | *BGIBMGA008046* | down | 146 | 160 | - | 12.4919 | 1.46E-05 | 1 | TAGGCTGATGATGAC |
| 3911 | mamo-L | *BGIBMGA008048* | down | 429 | 443 | + | 11.9597 | 2.61E-05 | 1 | AAGGCTGTTTTTTTT |
| 3912 | mamo-L | *BGIBMGA008053* | up | 1199 | 1213 | + | 11.7258 | 3.29E-05 | 1 | TAGGCGAAGGTTGGA |
| 3913 | mamo-L | *BGIBMGA008053* | down | 458 | 472 | - | 10.9113 | 7.02E-05 | 1 | AAGGCTCAGTTTTAT |
| 3914 | mamo-L | *BGIBMGA008056* | down | 1699 | 1713 | - | 10.5726 | 9.33E-05 | 1 | TAGGCTAATATTTAT |
| 3915 | mamo-L | *BGIBMGA008057* | up | 811 | 825 | + | 11.0565 | 6.18E-05 | 1 | TAGGCGGAATTCAGT |
| 3916 | mamo-L | *BGIBMGA008057* | down | 262 | 276 | + | 10.621 | 8.97E-05 | 1 | TAGGCGTGCTATCAT |
| 3917 | mamo-L | *BGIBMGA008058* | up | 964 | 978 | + | 10.621 | 8.97E-05 | 1 | TAGGCGTGCTATCAT |
| 3918 | mamo-L | *BGIBMGA008060* | down | 1666 | 1680 | + | 10.7661 | 7.95E-05 | 1 | TAGGCGCAGTGTTCC |
| 3919 | mamo-L | *BGIBMGA008061* | up | 379 | 393 | + | 10.7661 | 7.95E-05 | 1 | TAGGCGCAGTGTTCC |
| 3920 | mamo-L | *BGIBMGA008066* | down | 500 | 514 | - | 12.6935 | 1.15E-05 | 1 | AAGGCGGTTTTTGGT |
| 3921 | mamo-L | *BGIBMGA008067* | up | 1878 | 1892 | - | 11.3871 | 4.55E-05 | 1 | TAGGCGCGTTTCGTC |
| 3922 | mamo-L | *BGIBMGA008068* | down | 1441 | 1455 | - | 10.9677 | 6.68E-05 | 1 | AAGGCCAACTTTGAT |
| 3923 | mamo-L | *BGIBMGA008069* | down | 1279 | 1293 | + | 12.3306 | 1.75E-05 | 1 | TAGGCAGTTGTTGGC |
| 3924 | mamo-L | *BGIBMGA008071* | down | 1590 | 1604 | - | 10.9194 | 6.98E-05 | 1 | TAGGCTACTTTTTTT |
| 3925 | mamo-L | *BGIBMGA008076* | down | 911 | 925 | + | 10.621 | 8.97E-05 | 1 | TAGGCCTCATTTATT |
| 3926 | mamo-L | *BGIBMGA008079* | up | 1850 | 1864 | - | 11.7258 | 3.29E-05 | 1 | AAGGCTATGTTTGTT |
| 3927 | mamo-L | *BGIBMGA008081* | down | 1632 | 1646 | + | 10.5645 | 9.40E-05 | 1 | AAGGCCGATTACGCT |
| 3928 | mamo-L | *BGIBMGA008085* | up | 1333 | 1347 | - | 10.871 | 7.28E-05 | 1 | TAGGCTAGAGATGGC |
| 3929 | mamo-L | *BGIBMGA008091* | up | 652 | 666 | + | 10.5726 | 9.33E-05 | 1 | TAGGCTAATATTTAT |
| 3930 | mamo-L | *BGIBMGA008097* | up | 406 | 420 | + | 11.3871 | 4.55E-05 | 1 | AAGGCGGACTTTGAC |
| 3931 | mamo-L | *BGIBMGA008097* | down | 1351 | 1365 | + | 10.5887 | 9.21E-05 | 1 | TAGGCGTGCCTTGAC |
| 3932 | mamo-L | *BGIBMGA008102* | up | 1115 | 1129 | - | 11.6694 | 3.47E-05 | 1 | TAGGCGGTGTTTTGG |
| 3933 | mamo-L | *BGIBMGA008104* | down | 230 | 244 | + | 10.9194 | 6.98E-05 | 1 | TAGGCTACTTTTTTT |
| 3934 | mamo-L | *BGIBMGA008105* | down | 1252 | 1266 | + | 14.1935 | 1.49E-06 | 1 | TAGGCTTAGGTTAGT |
| 3935 | mamo-L | *BGIBMGA008111* | down | 1532 | 1546 | - | 12.4274 | 1.57E-05 | 1 | TAGGCTAGTTTTTTT |
| 3936 | mamo-L | *BGIBMGA008111* | down | 1162 | 1176 | - | 12.0645 | 2.34E-05 | 1 | TAGGCGGGCTTCGGA |
| 3937 | mamo-L | *BGIBMGA008112* | up | 1083 | 1097 | + | 11.4435 | 4.31E-05 | 1 | TAGGCTTGAGAGGCT |
| 3938 | mamo-L | *BGIBMGA008112* | down | 1591 | 1605 | - | 10.7661 | 7.95E-05 | 1 | TAGGCACTTGTTTCT |
| 3939 | mamo-L | *BGIBMGA008112* | up | 1849 | 1863 | - | 10.629 | 8.91E-05 | 1 | TAGGCCTGCAGCGCC |
| 3940 | mamo-L | *BGIBMGA008114* | down | 742 | 756 | - | 10.5 | 9.90E-05 | 1 | GAGGCCCAGGTCGTC |
| 3941 | mamo-L | *BGIBMGA008123* | down | 1603 | 1617 | - | 13.1774 | 6.47E-06 | 1 | TAGGCTGGTTTTGCA |
| 3942 | mamo-L | *BGIBMGA008124* | down | 636 | 650 | - | 10.6855 | 8.51E-05 | 1 | TAGGCCTGCTGGAGT |
| 3943 | mamo-L | *BGIBMGA008127* | down | 260 | 274 | - | 12.9919 | 8.10E-06 | 1 | TAGGCCGGTTTTGCA |
| 3944 | mamo-L | *BGIBMGA008130* | down | 320 | 334 | - | 10.75 | 8.06E-05 | 1 | GAGGCCGATTGTGCC |
| 3945 | mamo-L | *BGIBMGA008132* | down | 1687 | 1701 | - | 10.5726 | 9.33E-05 | 1 | TAGGCTAATATTTAT |
| 3946 | mamo-L | *BGIBMGA008133* | down | 7 | 21 | - | 11.5726 | 3.81E-05 | 1 | TAGGCTAGGGCTGTT |
| 3947 | mamo-L | *BGIBMGA008134* | down | 60 | 74 | - | 13.0726 | 7.34E-06 | 1 | TAGGCCGATTTTGCA |
| 3948 | mamo-L | *BGIBMGA008135* | down | 1491 | 1505 | - | 12.3306 | 1.75E-05 | 1 | TAGGCTTACTATACT |
| 3949 | mamo-L | *BGIBMGA008138* | up | 1721 | 1735 | - | 15.6613 | 9.32E-08 | 1 | TAGGCTGTCTTTGTT |

| | | | | | | | | | | |
|---|---|---|---|---|---|---|---|---|---|---|
| 3950 | mamo-L | *BGIBMGA008143* | down | 1257 | 1271 | + | 12.0645 | 2.34E-05 | 1 | TAGGCGGGCTTCGGA |
| 3951 | mamo-L | *BGIBMGA008143* | up | 1723 | 1737 | + | 10.9194 | 6.98E-05 | 1 | TAGGCTACTTTTTTT |
| 3952 | mamo-L | *BGIBMGA008143* | down | 1104 | 1118 | - | 10.8306 | 7.52E-05 | 1 | TAGGCCCCCTTCGCC |
| 3953 | mamo-L | *BGIBMGA008143* | down | 1502 | 1516 | + | 10.8065 | 7.69E-05 | 1 | TAGGCGGAGTGAGGT |
| 3954 | mamo-L | *BGIBMGA008148* | up | 697 | 711 | + | 10.7177 | 8.28E-05 | 1 | AAGGCCTGGTTGGCC |
| 3955 | mamo-L | *BGIBMGA008148* | down | 1050 | 1064 | - | 10.5726 | 9.33E-05 | 1 | TAGGCTAATATTTAT |
| 3956 | mamo-L | *BGIBMGA008149* | down | 1299 | 1313 | - | 10.9194 | 6.98E-05 | 1 | TAGGCTACTTTTTTT |
| 3957 | mamo-L | *BGIBMGA008152* | down | 1032 | 1046 | + | 13.0726 | 7.34E-06 | 1 | TAGGCCGATTTTGCA |
| 3958 | mamo-L | *BGIBMGA008157* | down | 132 | 146 | - | 11.1048 | 5.92E-05 | 1 | TAGGCCAGTTTTGCA |
| 3959 | mamo-L | *BGIBMGA008158* | down | 538 | 552 | - | 10.9919 | 6.55E-05 | 1 | GAGGCGGGGGTTGGG |
| 3960 | mamo-L | *BGIBMGA008161* | up | 1721 | 1735 | - | 11.1855 | 5.50E-05 | 1 | TAGGCGCAGTGTAGT |
| 3961 | mamo-L | *BGIBMGA008164* | up | 1589 | 1603 | + | 12.3226 | 1.77E-05 | 1 | TAGGCCGATTTTTGA |
| 3962 | mamo-L | *BGIBMGA008167* | down | 1838 | 1852 | - | 12.0645 | 2.34E-05 | 1 | TAGGCGGGCTTCGGA |
| 3963 | mamo-L | *BGIBMGA008168* | up | 1321 | 1335 | + | 11.5323 | 3.96E-05 | 1 | TAGGCGTGAGGTGAT |
| 3964 | mamo-L | *BGIBMGA008168* | down | 1202 | 1216 | + | 11.0887 | 6.01E-05 | 1 | TAGGCCTGGTCTTGC |
| 3965 | mamo-L | *BGIBMGA008169* | up | 652 | 666 | + | 11.0887 | 6.01E-05 | 1 | TAGGCCTGGTCTTGC |
| 3966 | mamo-L | *BGIBMGA008171* | down | 623 | 637 | + | 11.2742 | 5.06E-05 | 1 | TAGGCATGCGTTCTT |
| 3967 | mamo-L | *BGIBMGA008173* | down | 660 | 674 | + | 10.5726 | 9.33E-05 | 1 | TAGGCTAATATTTAT |
| 3968 | mamo-L | *BGIBMGA008176* | down | 1632 | 1646 | - | 13.6048 | 3.60E-06 | 1 | TAGGCGTGTTTTGAT |
| 3969 | mamo-L | *BGIBMGA008177* | up | 1416 | 1430 | - | 13.6048 | 3.60E-06 | 1 | TAGGCGTGTTTTGAT |
| 3970 | mamo-L | *BGIBMGA008178* | up | 535 | 549 | + | 11.4355 | 4.34E-05 | 1 | AAGGCTTATTTGGTT |
| 3971 | mamo-L | *BGIBMGA008183* | up | 1944 | 1958 | + | 10.5726 | 9.33E-05 | 1 | TAGGCTAATATTTAT |
| 3972 | mamo-L | *BGIBMGA008184* | up | 1258 | 1272 | - | 11.4194 | 4.42E-05 | 1 | TAGGCCCTTGACGTT |
| 3973 | mamo-L | *BGIBMGA008188* | down | 494 | 508 | + | 13.3871 | 4.90E-06 | 1 | TAGGCTGGTGATGAT |
| 3974 | mamo-L | *BGIBMGA008189* | down | 1049 | 1063 | + | 10.9194 | 6.98E-05 | 1 | TAGGCTACTTTTTTT |
| 3975 | mamo-L | *BGIBMGA008191* | up | 1560 | 1574 | + | 11.8387 | 2.94E-05 | 1 | TAGGCTATGTTTATT |
| 3976 | mamo-L | *BGIBMGA008191* | down | 502 | 516 | + | 11.0242 | 6.36E-05 | 1 | GAGGCCCGTTTCGGT |
| 3977 | mamo-L | *BGIBMGA008194* | up | 1484 | 1498 | - | 10.75 | 8.06E-05 | 1 | GAGGCCGATTGTGCC |
| 3978 | mamo-L | *BGIBMGA008197* | up | 1468 | 1482 | + | 10.9194 | 6.98E-05 | 1 | TAGGCTACTTTTTTT |
| 3979 | mamo-L | *BGIBMGA008197* | down | 1067 | 1081 | + | 10.6371 | 8.85E-05 | 1 | TAGGCCTCATATGGC |
| 3980 | mamo-L | *BGIBMGA008198* | down | 1812 | 1826 | + | 11.2016 | 5.41E-05 | 1 | GAGGCTTTGATTGTC |
| 3981 | mamo-L | *BGIBMGA008201* | down | 1136 | 1150 | + | 11.8145 | 3.01E-05 | 1 | TAGGCCTACTGCGCC |
| 3982 | mamo-L | *BGIBMGA008204* | up | 408 | 422 | - | 10.75 | 8.06E-05 | 1 | TAGGCCTATTTCTGA |
| 3983 | mamo-L | *BGIBMGA008204* | up | 415 | 429 | + | 10.7258 | 8.23E-05 | 1 | TAGGCCTATGGTACC |
| 3984 | mamo-L | *BGIBMGA008211* | down | 1385 | 1399 | + | 11.7097 | 3.34E-05 | 1 | TAGGCTCCTTTTTTT |
| 3985 | mamo-L | *BGIBMGA008215* | down | 1148 | 1162 | - | 10.5726 | 9.33E-05 | 1 | TAGGCTAATATTTAT |
| 3986 | mamo-L | *BGIBMGA008216* | up | 1430 | 1444 | - | 10.5726 | 9.33E-05 | 1 | TAGGCTAATATTTAT |
| 3987 | mamo-L | *BGIBMGA008217* | down | 130 | 144 | + | 10.5968 | 9.15E-05 | 1 | GAGGCTAGTTTTTGT |
| 3988 | mamo-L | *BGIBMGA008218* | down | 771 | 785 | - | 12.7581 | 1.07E-05 | 1 | AAGGCTTTTTTTGTT |
| 3989 | mamo-L | *BGIBMGA008219* | up | 240 | 254 | + | 11.6694 | 3.47E-05 | 1 | TAGGCCGGCGTCTAC |
| 3990 | mamo-L | *BGIBMGA008221* | up | 1720 | 1734 | - | 11.0323 | 6.32E-05 | 1 | TAGGCCCAAATCGTT |
| 3991 | mamo-L | *BGIBMGA008223* | down | 559 | 573 | + | 12.8952 | 9.05E-06 | 1 | AAGGCCTATTTTGTT |
| 3992 | mamo-L | *BGIBMGA008223* | down | 1754 | 1768 | + | 10.5726 | 9.33E-05 | 1 | TAGGCTAATATTTAT |
| 3993 | mamo-L | *BGIBMGA008224* | down | 1771 | 1785 | - | 13.9919 | 2.01E-06 | 1 | TAGGCTGAAGTTGAT |
| 3994 | mamo-L | *BGIBMGA008225* | up | 58 | 72 | + | 13.3226 | 5.34E-06 | 1 | TAGGCTGATGATGTC |
| 3995 | mamo-L | *BGIBMGA008225* | down | 57 | 71 | + | 11.0323 | 6.32E-05 | 1 | TAGGCCGCGTGTCGT |
| 3996 | mamo-L | *BGIBMGA008229* | down | 835 | 849 | + | 12.6129 | 1.26E-05 | 1 | GAGGCTGGTGTTTGT |
| 3997 | mamo-L | *BGIBMGA008229* | down | 1762 | 1776 | - | 10.9194 | 6.98E-05 | 1 | TAGGCTACTTTTTTT |
| 3998 | mamo-L | *BGIBMGA008231* | down | 361 | 375 | - | 12.629 | 1.24E-05 | 1 | TAGGCGGGTGGTGAT |
| 3999 | mamo-L | *BGIBMGA008236* | down | 804 | 818 | + | 10.9194 | 6.98E-05 | 1 | TAGGCTACTTTTTTT |
| 4000 | mamo-L | *BGIBMGA008238* | up | 1333 | 1347 | + | 11.4113 | 4.45E-05 | 1 | GAGGCTTTCGTCGGC |
| 4001 | mamo-L | *BGIBMGA008239* | down | 870 | 884 | + | 12.0161 | 2.46E-05 | 1 | TAGGCTTAGAACGCT |
| 4002 | mamo-L | *BGIBMGA008239* | up | 469 | 483 | + | 11.4839 | 4.15E-05 | 1 | AAGGCTTGTGTGGTT |
| 4003 | mamo-L | *BGIBMGA008240* | down | 1008 | 1022 | + | 11.4435 | 4.31E-05 | 1 | TAGGCTTAGTTAGAT |
| 4004 | mamo-L | *BGIBMGA008240* | down | 781 | 795 | + | 10.6129 | 9.03E-05 | 1 | AAGGCGGTAGTTTTT |
| 4005 | mamo-L | *BGIBMGA008240* | up | 611 | 625 | - | 10.5645 | 9.40E-05 | 1 | TAGGCTGTATGCTCT |
| 4006 | mamo-L | *BGIBMGA008244* | down | 200 | 214 | + | 10.9194 | 6.98E-05 | 1 | TAGGCTACTTTTTTT |
| 4007 | mamo-L | *BGIBMGA008245* | up | 1659 | 1673 | - | 12.5887 | 1.30E-05 | 1 | TAGGCTGAATATTGT |
| 4008 | mamo-L | *BGIBMGA008245* | down | 448 | 462 | + | 11.3226 | 4.83E-05 | 1 | TAGGCTTTGGGTAAT |
| 4009 | mamo-L | *BGIBMGA008246* | up | 229 | 243 | - | 10.6613 | 8.67E-05 | 1 | TAGGCCTGTAGTTGC |
| 4010 | mamo-L | *BGIBMGA008247* | up | 795 | 809 | - | 14.7742 | 5.37E-07 | 1 | TAGGCTTTGGTTTGT |
| 4011 | mamo-L | *BGIBMGA008248* | down | 1319 | 1333 | - | 11.2097 | 5.38E-05 | 1 | TAGGCTCGTATTATT |
| 4012 | mamo-L | *BGIBMGA008250* | up | 58 | 72 | + | 11.6935 | 3.39E-05 | 1 | TAGGCTCAGAATGAT |
| 4013 | mamo-L | *BGIBMGA008253* | down | 1789 | 1803 | - | 11.2581 | 5.14E-05 | 1 | GAGGCCGGGTGTGTC |
| 4014 | mamo-L | *BGIBMGA008260* | up | 433 | 447 | - | 10.9919 | 6.55E-05 | 1 | TCGGCTGAGTTTTGT |
| 4015 | mamo-L | *BGIBMGA008262* | down | 16 | 30 | - | 11.9758 | 2.57E-05 | 1 | TAGGCTGGTTGTACT |
| 4016 | mamo-L | *BGIBMGA008266* | up | 1485 | 1499 | - | 12.4032 | 1.61E-05 | 1 | TAGGCTGAAGATTTT |
| 4017 | mamo-L | *BGIBMGA008268* | up | 68 | 82 | + | 11.8387 | 2.94E-05 | 1 | TAGGCTATGTTTATT |
| 4018 | mamo-L | *BGIBMGA008271* | up | 509 | 523 | - | 11.25 | 5.18E-05 | 1 | AAGGCGTTTTTTGAT |
| 4019 | mamo-L | *BGIBMGA008274* | up | 1895 | 1909 | + | 10.7258 | 8.23E-05 | 1 | TAGGCTGGTCTTCTT |
| 4020 | mamo-L | *BGIBMGA008278* | down | 622 | 636 | - | 13.7742 | 2.79E-06 | 1 | TAGGCGGGTGGTGGT |
| 4021 | mamo-L | *BGIBMGA008278* | down | 1169 | 1183 | + | 12.4516 | 1.53E-05 | 1 | TAGGCTTTTTATTTT |
| 4022 | mamo-L | *BGIBMGA008279* | up | 97 | 111 | + | 12.4516 | 1.53E-05 | 1 | TAGGCTTTTTATTTT |
| 4023 | mamo-L | *BGIBMGA008280* | down | 1405 | 1419 | - | 12.6371 | 1.23E-05 | 1 | TAGGCGGAGTGGGGT |
| 4024 | mamo-L | *BGIBMGA008287* | up | 542 | 556 | + | 10.9194 | 6.98E-05 | 1 | TAGGCTACTTTTTTT |
| 4025 | mamo-L | *BGIBMGA008293* | up | 1400 | 1414 | + | 10.9758 | 6.64E-05 | 1 | TGGGCTGTCTTTTTT |
| 4026 | mamo-L | *BGIBMGA008294* | up | 1062 | 1076 | - | 11.2016 | 5.41E-05 | 1 | TAGGCGGTGATGTTT |
| 4027 | mamo-L | *BGIBMGA008297* | up | 1760 | 1774 | + | 10.5726 | 9.33E-05 | 1 | TAGGCTAATATTTAT |
| 4028 | mamo-L | *BGIBMGA008301* | down | 324 | 338 | + | 10.9194 | 6.98E-05 | 1 | TAGGCTACTTTTTTT |
| 4029 | mamo-L | *BGIBMGA008302* | up | 1773 | 1787 | + | 11.8387 | 2.94E-05 | 1 | TAGGCTTTTGGTTGC |
| 4030 | mamo-L | *BGIBMGA008303* | down | 127 | 141 | - | 11.6129 | 3.66E-05 | 1 | TAGGCTTTTAATGAT |
| 4031 | mamo-L | *BGIBMGA008304* | up | 949 | 963 | - | 11.6129 | 3.66E-05 | 1 | TAGGCTTTTAATGAT |
| 4032 | mamo-L | *BGIBMGA008305* | down | 305 | 319 | + | 11.6694 | 3.47E-05 | 1 | TAGGCAGTCTTTAGT |
| 4033 | mamo-L | *BGIBMGA008305* | down | 1619 | 1633 | + | 11.2419 | 5.21E-05 | 1 | AAGGCTGGTGGTTGT |
| 4034 | mamo-L | *BGIBMGA008309* | up | 1592 | 1606 | + | 10.4919 | 9.96E-05 | 1 | TAGGCGGTCGTGATC |
| 4035 | mamo-L | *BGIBMGA008311* | down | 1655 | 1669 | - | 11.0887 | 6.01E-05 | 1 | AAGGCCTACGTGTGT |
| 4036 | mamo-L | *BGIBMGA008315* | down | 578 | 592 | - | 10.6855 | 8.51E-05 | 1 | TAGGCCATTTGTGCC |
| 4037 | mamo-L | *BGIBMGA008316* | down | 1287 | 1301 | + | 11.379 | 4.58E-05 | 1 | GAGGCTGTGATTTTT |
| 4038 | mamo-L | *BGIBMGA008318* | down | 462 | 476 | - | 12.5161 | 1.42E-05 | 1 | TAGGCCGAGGATCTT |
| 4039 | mamo-L | *BGIBMGA008319* | down | 845 | 859 | - | 12.7661 | 1.05E-05 | 1 | TAGGCTTTAGGTGCT |
| 4040 | mamo-L | *BGIBMGA008321* | up | 1982 | 1996 | + | 10.7823 | 7.84E-05 | 1 | GAGGCTATAGTTGGT |
| 4041 | mamo-L | *BGIBMGA008322* | up | 1940 | 1954 | - | 11.4677 | 4.22E-05 | 1 | GAGGCTGGCGATTTT |

| | | | | | | | | | | |
|---|---|---|---|---|---|---|---|---|---|---|
| 4042 | mamo-L | *BGIBMGA008322* | up | 1682 | 1696 | - | 11.3468 | 4.73E-05 | 1 | AAGGCTGTCGATGCC |
| 4043 | mamo-L | *BGIBMGA008322* | up | 1063 | 1077 | - | 11.0645 | 6.14E-05 | 1 | TAGGCTAACATTTAT |
| 4044 | mamo-L | *BGIBMGA008325* | down | 1865 | 1879 | + | 11.3629 | 4.66E-05 | 1 | TAGGCTTTCGTTTAA |
| 4045 | mamo-L | *BGIBMGA008327* | up | 1752 | 1766 | - | 12.3145 | 1.79E-05 | 1 | TAGGCCATTTTTTGT |
| 4046 | mamo-L | *BGIBMGA008331* | up | 321 | 335 | + | 10.4919 | 9.96E-05 | 1 | TAGGCTGTGTCTGTG |
| 4047 | mamo-L | *BGIBMGA008334* | down | 1421 | 1435 | - | 11.3468 | 4.73E-05 | 1 | TAGGCAGGAGGTGGT |
| 4048 | mamo-L | *BGIBMGA008334* | down | 1693 | 1707 | + | 10.5726 | 9.33E-05 | 1 | TAGGCTAATATTTAT |
| 4049 | mamo-L | *BGIBMGA008336* | up | 38 | 52 | + | 11.9032 | 2.76E-05 | 1 | TAGGCTATCGTTATT |
| 4050 | mamo-L | *BGIBMGA008337* | down | 562 | 576 | + | 12.0968 | 2.26E-05 | 1 | GAGGCCTGCGTCGCT |
| 4051 | mamo-L | *BGIBMGA008343* | down | 1932 | 1946 | - | 10.8629 | 7.33E-05 | 1 | TAGGCTTTAAATTGT |
| 4052 | mamo-L | *BGIBMGA008344* | down | 1608 | 1622 | - | 13.5323 | 4.01E-06 | 1 | TAGGCTGTGGCTGGT |
| 4053 | mamo-L | *BGIBMGA008344* | down | 1009 | 1023 | - | 10.9435 | 6.83E-05 | 1 | GAGGCCGGCTTTGCG |
| 4054 | mamo-L | *BGIBMGA008346* | down | 134 | 148 | - | 14.1613 | 1.57E-06 | 1 | TAGGCGTAAGTTGGT |
| 4055 | mamo-L | *BGIBMGA008350* | down | 343 | 357 | + | 12.0081 | 2.48E-05 | 1 | TAGGCGTATGCTGTT |
| 4056 | mamo-L | *BGIBMGA008351* | up | 141 | 155 | + | 11.1048 | 5.92E-05 | 1 | TAGGCTTGATGTGAC |
| 4057 | mamo-L | *BGIBMGA008352* | down | 610 | 624 | + | 10.5 | 9.90E-05 | 1 | GAGGCCCAGGTCGTC |
| 4058 | mamo-L | *BGIBMGA008353* | up | 1216 | 1230 | + | 11.8871 | 2.81E-05 | 1 | TAGGCTTTCTATCTT |
| 4059 | mamo-L | *BGIBMGA008356* | up | 1953 | 1967 | - | 11.8145 | 3.01E-05 | 1 | TAGGCCTTGACTGGT |
| 4060 | mamo-L | *BGIBMGA008361* | up | 1332 | 1346 | - | 14.0403 | 1.87E-06 | 1 | TAGGCAGGCGTTGGT |
| 4061 | mamo-L | *BGIBMGA008362* | up | 1907 | 1921 | + | 11.2258 | 5.30E-05 | 1 | TAGGCCTTTTTCCCT |
| 4062 | mamo-L | *BGIBMGA008363* | up | 648 | 662 | - | 12.4677 | 1.49E-05 | 1 | TAGGCTTCTATTGCT |
| 4063 | mamo-L | *BGIBMGA008364* | up | 1497 | 1511 | + | 11.7984 | 3.06E-05 | 1 | TAGGCTTTGCTTTTT |
| 4064 | mamo-L | *BGIBMGA008369* | down | 1417 | 1431 | + | 10.5968 | 9.15E-05 | 1 | GAGGCGCTTGTTTGT |
| 4065 | mamo-L | *BGIBMGA008371* | up | 1829 | 1843 | + | 12.1855 | 2.06E-05 | 1 | TAGGCTGTATGTTGT |
| 4066 | mamo-L | *BGIBMGA008372* | down | 1022 | 1036 | + | 11.7339 | 3.26E-05 | 1 | TAGGCCTAGTTGCCT |
| 4067 | mamo-L | *BGIBMGA008373* | down | 1819 | 1833 | - | 11.2339 | 5.26E-05 | 1 | TAGGCAGGGTTGTGT |
| 4068 | mamo-L | *BGIBMGA008375* | up | 1262 | 1276 | + | 10.9677 | 6.68E-05 | 1 | TAGGCCTTTTTGTTC |
| 4069 | mamo-L | *BGIBMGA008377* | up | 435 | 449 | + | 11.1532 | 5.67E-05 | 1 | TAGGCCAGTTGTTGT |
| 4070 | mamo-L | *BGIBMGA008383* | down | 176 | 190 | - | 12.0484 | 2.38E-05 | 1 | AAGGCTTCGTTTGTT |
| 4071 | mamo-L | *BGIBMGA008385* | up | 1199 | 1213 | - | 11.371 | 4.62E-05 | 1 | TAGGCTGTATAGGCT |
| 4072 | mamo-L | *BGIBMGA008385* | up | 1513 | 1527 | - | 10.9435 | 6.83E-05 | 1 | GAGGCTTGGGGCGCT |
| 4073 | mamo-L | *BGIBMGA008391* | up | 693 | 707 | - | 11.0161 | 6.40E-05 | 1 | AAGGCTTATTGTGGC |
| 4074 | mamo-L | *BGIBMGA008392* | up | 1391 | 1405 | + | 13.4274 | 4.64E-06 | 1 | TAGGCTCATGTTTTT |
| 4075 | mamo-L | *BGIBMGA008392* | down | 1876 | 1890 | + | 10.5726 | 9.33E-05 | 1 | TAGGCTAATATTTAT |
| 4076 | mamo-L | *BGIBMGA008396* | up | 1370 | 1384 | - | 14.2097 | 1.45E-06 | 1 | TAGGCTGTGGTGGTT |
| 4077 | mamo-L | *BGIBMGA008397* | up | 1177 | 1191 | - | 10.8065 | 7.69E-05 | 1 | TAGGCCGACGAGCTT |
| 4078 | mamo-L | *BGIBMGA008404* | up | 1000 | 1014 | + | 11.5645 | 3.84E-05 | 1 | TAGGCTTAATTTAAT |
| 4079 | mamo-L | *BGIBMGA008405* | down | 1685 | 1699 | + | 10.9355 | 6.88E-05 | 1 | TAGGCGGTTTGTCTT |
| 4080 | mamo-L | *BGIBMGA008405* | up | 1509 | 1523 | + | 10.9194 | 6.98E-05 | 1 | TAGGCTACTTTTTTT |
| 4081 | mamo-L | *BGIBMGA008406* | up | 1130 | 1144 | - | 14.0968 | 1.72E-06 | 1 | TAGGCTTATTTTTTT |
| 4082 | mamo-L | *BGIBMGA008406* | up | 278 | 292 | - | 10.5726 | 9.33E-05 | 1 | TAGGCTAATATTTAT |
| 4083 | mamo-L | *BGIBMGA008407* | down | 847 | 861 | - | 12.7742 | 1.04E-05 | 1 | TAGGCTTTGTTGGCC |
| 4084 | mamo-L | *BGIBMGA008409* | up | 1218 | 1232 | + | 13.5 | 4.19E-06 | 1 | TAGGCCGATTTTGAC |
| 4085 | mamo-L | *BGIBMGA008409* | up | 1626 | 1640 | - | 10.8065 | 7.69E-05 | 1 | TGGGCGGAGTTTGTC |
| 4086 | mamo-L | *BGIBMGA008412* | up | 254 | 268 | - | 13.1855 | 6.42E-06 | 1 | TAGGCGGAGTGTTGT |
| 4087 | mamo-L | *BGIBMGA008413* | up | 569 | 583 | - | 12.9758 | 8.23E-06 | 1 | TAGGCGGGCTTCGTC |
| 4088 | mamo-L | *BGIBMGA008416* | down | 1922 | 1936 | - | 12.9919 | 8.10E-06 | 1 | TAGGCTTATATTTTT |
| 4089 | mamo-L | *BGIBMGA008416* | down | 1562 | 1576 | - | 10.5806 | 9.27E-05 | 1 | TAGGCTACCTTTTAT |
| 4090 | mamo-L | *BGIBMGA008418* | down | 322 | 336 | - | 13.5968 | 3.65E-06 | 1 | TAGGCCGGAGTCGGT |
| 4091 | mamo-L | *BGIBMGA008419* | up | 1193 | 1207 | + | 13.621 | 3.50E-06 | 1 | TAGGCTTACATCGGT |
| 4092 | mamo-L | *BGIBMGA008422* | up | 120 | 134 | + | 11.1129 | 5.88E-05 | 1 | TAGGCGCATGGGGGT |
| 4093 | mamo-L | *BGIBMGA008422* | down | 1866 | 1880 | + | 10.9597 | 6.73E-05 | 1 | TAGGCTTTCACTGGC |
| 4094 | mamo-L | *BGIBMGA008422* | up | 688 | 702 | - | 10.879 | 7.22E-05 | 1 | TAGGCCAACGTTCCC |
| 4095 | mamo-L | *BGIBMGA008423* | up | 818 | 832 | + | 10.5726 | 9.33E-05 | 1 | TAGGCTAATATTTAT |
| 4096 | mamo-L | *BGIBMGA008425* | up | 958 | 972 | - | 13.4274 | 4.64E-06 | 1 | TAGGCTCATGTTTTT |
| 4097 | mamo-L | *BGIBMGA008430* | down | 813 | 827 | - | 11.3548 | 4.69E-05 | 1 | AAGGCTGATGATGAT |
| 4098 | mamo-L | *BGIBMGA008430* | up | 228 | 242 | - | 10.6855 | 8.51E-05 | 1 | AAGGCGGGTGGTGGC |
| 4099 | mamo-L | *BGIBMGA008432* | down | 1541 | 1555 | + | 10.9919 | 6.55E-05 | 1 | AAGGCCTATGTTACT |
| 4100 | mamo-L | *BGIBMGA008435* | up | 1239 | 1253 | - | 11.7097 | 3.34E-05 | 1 | TAGGCGGTGATTGCA |
| 4101 | mamo-L | *BGIBMGA008440* | up | 296 | 310 | + | 10.5806 | 9.27E-05 | 1 | AAGGCTTGTTATTTT |
| 4102 | mamo-L | *BGIBMGA008441* | up | 1175 | 1189 | + | 12.0645 | 2.34E-05 | 1 | TAGGCGGGCTTCGGA |
| 4103 | mamo-L | *BGIBMGA008441* | up | 1022 | 1036 | - | 10.8306 | 7.52E-05 | 1 | TAGGCCCCCTTCGCC |
| 4104 | mamo-L | *BGIBMGA008445* | up | 1923 | 1937 | - | 14.0161 | 1.95E-06 | 1 | TAGGCTTGTTTTTTT |
| 4105 | mamo-L | *BGIBMGA008448* | down | 1940 | 1954 | + | 11.9274 | 2.69E-05 | 1 | TAGGCCGGTTTTTTA |
| 4106 | mamo-L | *BGIBMGA008448* | down | 1202 | 1216 | - | 11.629 | 3.61E-05 | 1 | AAGGCTTGGTTTTAT |
| 4107 | mamo-L | *BGIBMGA008448* | down | 1818 | 1832 | + | 10.629 | 8.91E-05 | 1 | TAGGCCTGCAGCGCC |
| 4108 | mamo-L | *BGIBMGA008453* | down | 1788 | 1802 | + | 13 | 8.01E-06 | 1 | TAGGCCGTGGCTGCT |
| 4109 | mamo-L | *BGIBMGA008456* | down | 1620 | 1634 | + | 11 | 6.50E-05 | 1 | TAGGCGAATTTTTAT |
| 4110 | mamo-L | *BGIBMGA008456* | up | 1221 | 1235 | - | 10.5726 | 9.33E-05 | 1 | TAGGCTAATATTTAT |
| 4111 | mamo-L | *BGIBMGA008461* | down | 878 | 892 | - | 10.9194 | 6.98E-05 | 1 | TAGGCTACTTTTTTT |
| 4112 | mamo-L | *BGIBMGA008462* | down | 1121 | 1135 | + | 10.5645 | 9.40E-05 | 1 | TAGGCTCAAATTTAT |
| 4113 | mamo-L | *BGIBMGA008468* | down | 729 | 743 | + | 11.6935 | 3.39E-05 | 1 | TGGGCTGACGTTGAT |
| 4114 | mamo-L | *BGIBMGA008468* | up | 16 | 30 | - | 10.9194 | 6.98E-05 | 1 | TAGGCTACTTTTTTT |
| 4115 | mamo-L | *BGIBMGA008470* | down | 1897 | 1911 | + | 12.0645 | 2.34E-05 | 1 | TAGGCGGGCTTCGGA |
| 4116 | mamo-L | *BGIBMGA008470* | down | 1744 | 1758 | - | 10.8306 | 7.52E-05 | 1 | TAGGCCCCCTTCGCC |
| 4117 | mamo-L | *BGIBMGA008471* | down | 1020 | 1034 | + | 11.6048 | 3.70E-05 | 1 | TAGGCCTGGATGTCT |
| 4118 | mamo-L | *BGIBMGA008472* | up | 1208 | 1222 | + | 10.7016 | 8.39E-05 | 1 | TAGGCTGGTAGTGTA |
| 4119 | mamo-L | *BGIBMGA008473* | up | 1934 | 1948 | - | 12.4597 | 1.51E-05 | 1 | TAGGCTGATATTTAT |
| 4120 | mamo-L | *BGIBMGA008474* | up | 1190 | 1204 | + | 10.8226 | 7.58E-05 | 1 | TAGGCATTGGTGTGT |
| 4121 | mamo-L | *BGIBMGA008475* | up | 822 | 836 | - | 11.2742 | 5.06E-05 | 1 | AAGGCCGTGGTTGGG |
| 4122 | mamo-L | *BGIBMGA008475* | up | 181 | 195 | + | 10.4919 | 9.96E-05 | 1 | TAGGCCTGTGCTTAT |
| 4123 | mamo-L | *BGIBMGA008478* | up | 1360 | 1374 | + | 11.7984 | 3.06E-05 | 1 | TAGGCTTTGCTTTTT |
| 4124 | mamo-L | *BGIBMGA008479* | down | 1333 | 1347 | + | 14.1613 | 1.57E-06 | 1 | TAGGCGTAAGTTGGT |
| 4125 | mamo-L | *BGIBMGA008480* | up | 1617 | 1631 | - | 13.8952 | 2.32E-06 | 1 | TAGGCTGTCATTGGC |
| 4126 | mamo-L | *BGIBMGA008480* | up | 1236 | 1250 | - | 13.5323 | 4.01E-06 | 1 | TAGGCTGTGGCTGGT |
| 4127 | mamo-L | *BGIBMGA008481* | up | 1214 | 1228 | - | 11 | 6.50E-05 | 1 | TAGGCCGATTTAGAT |
| 4128 | mamo-L | *BGIBMGA008482* | down | 1439 | 1453 | + | 13.5323 | 4.01E-06 | 1 | TAGGCTGTGGCTGGT |
| 4129 | mamo-L | *BGIBMGA008483* | up | 1356 | 1370 | - | 10.5565 | 9.46E-05 | 1 | TAGGGTGGCTTTGGC |
| 4130 | mamo-L | *BGIBMGA008484* | down | 463 | 477 | + | 11.7097 | 3.34E-05 | 1 | TAGGCCAACGATGCC |
| 4131 | mamo-L | *BGIBMGA008484* | down | 408 | 422 | - | 11.4032 | 4.49E-05 | 1 | TAGGCTATCTTCTTT |
| 4132 | mamo-L | *BGIBMGA008486* | down | 1428 | 1442 | + | 11.1048 | 5.92E-05 | 1 | AAGGCTGGGATGGTT |
| 4133 | mamo-L | *BGIBMGA008490* | down | 80 | 94 | - | 10.5161 | 9.77E-05 | 1 | TAGGCTACAGTTGAT |

| | | | | | | | | | | |
|---|---|---|---|---|---|---|---|---|---|---|
| 4134 | mamo-L | *BGIBMGA008494* | down | 1629 | 1643 | + | 11.1129 | 5.88E-05 | 1 | TAGGCGCAGATCTGT |
| 4135 | mamo-L | *BGIBMGA008495* | up | 78 | 92 | + | 11.1129 | 5.88E-05 | 1 | TAGGCGCAGATCTGT |
| 4136 | mamo-L | *BGIBMGA008496* | up | 1433 | 1447 | - | 11.25 | 5.18E-05 | 1 | TAGGCGTCTGTTGAC |
| 4137 | mamo-L | *BGIBMGA008497* | up | 1732 | 1746 | + | 11.3952 | 4.52E-05 | 1 | TAGGCCGGTTGGTGT |
| 4138 | mamo-L | *BGIBMGA008501* | up | 1873 | 1887 | - | 10.7581 | 8.00E-05 | 1 | TAGCCTTTTTTTGTT |
| 4139 | mamo-L | *BGIBMGA008505* | down | 1528 | 1542 | + | 11.0565 | 6.18E-05 | 1 | TAGGCATGCGTCTTT |
| 4140 | mamo-L | *BGIBMGA008507* | down | 1641 | 1655 | + | 10.8226 | 7.58E-05 | 1 | AAGGCTGGTTTGGAT |
| 4141 | mamo-L | *BGIBMGA008508* | up | 1648 | 1662 | + | 10.6935 | 8.45E-05 | 1 | TAGGCTCGTGTGTCC |
| 4142 | mamo-L | *BGIBMGA008512* | down | 428 | 442 | + | 10.5161 | 9.77E-05 | 1 | TACGCTTTGTTTGTT |
| 4143 | mamo-L | *BGIBMGA008513* | down | 630 | 644 | - | 10.9677 | 6.68E-05 | 1 | TAGGCTCATTATTCC |
| 4144 | mamo-L | *BGIBMGA008516* | up | 1981 | 1995 | - | 12.5645 | 1.34E-05 | 1 | TAGGCTATGTTCGTT |
| 4145 | mamo-L | *BGIBMGA008517* | down | 1229 | 1243 | + | 10.9032 | 7.07E-05 | 1 | AAGGCCTGCTTCTCT |
| 4146 | mamo-L | *BGIBMGA008522* | up | 1332 | 1346 | - | 11.3145 | 4.87E-05 | 1 | AAGGCTGTGGTGTGT |
| 4147 | mamo-L | *BGIBMGA008527* | up | 733 | 747 | + | 10.9194 | 6.98E-05 | 1 | TAGGCTACTTTTTTT |
| 4148 | mamo-L | *BGIBMGA008531* | down | 1967 | 1981 | + | 10.4919 | 9.96E-05 | 1 | TAGGCTTTTTGTATC |
| 4149 | mamo-L | *BGIBMGA008535* | down | 605 | 619 | + | 12.7258 | 1.11E-05 | 1 | TAGGCGTGCTTTGTA |
| 4150 | mamo-L | *BGIBMGA008537* | down | 1640 | 1654 | - | 10.4919 | 9.96E-05 | 1 | TAGGCTGTGTCTGTG |
| 4151 | mamo-L | *BGIBMGA008539* | down | 1018 | 1032 | + | 13.3871 | 4.90E-06 | 1 | TAGGCTGGTGATGAT |
| 4152 | mamo-L | *BGIBMGA008543* | up | 1151 | 1165 | + | 12.3226 | 1.77E-05 | 1 | GAGGCTGTTGTTGAT |
| 4153 | mamo-L | *BGIBMGA008550* | down | 416 | 430 | + | 10.9919 | 6.55E-05 | 1 | TAGGCCTTTTACTTT |
| 4154 | mamo-L | *BGIBMGA008551* | up | 1505 | 1519 | + | 10.9919 | 6.55E-05 | 1 | TAGGCCTTTTACTTT |
| 4155 | mamo-L | *BGIBMGA008552* | down | 1410 | 1424 | - | 11.9839 | 2.54E-05 | 1 | TAGGCATTGATTGCT |
| 4156 | mamo-L | *BGIBMGA008553* | up | 1629 | 1643 | - | 10.75 | 8.06E-05 | 1 | TAGGCGTATATTAGC |
| 4157 | mamo-L | *BGIBMGA008554* | up | 1336 | 1350 | - | 11.2984 | 4.95E-05 | 1 | AAGGCCTAGTTTCTT |
| 4158 | mamo-L | *BGIBMGA008558* | up | 161 | 175 | - | 12.0645 | 2.34E-05 | 1 | TAGGCGGGCTTCGGA |
| 4159 | mamo-L | *BGIBMGA008560* | down | 95 | 109 | - | 11.6532 | 3.53E-05 | 1 | TAGGCTGCTGGTGTC |
| 4160 | mamo-L | *BGIBMGA008560* | down | 1601 | 1615 | + | 11.0806 | 6.05E-05 | 1 | GAGGCTTTGTTTGTA |
| 4161 | mamo-L | *BGIBMGA008563* | up | 1686 | 1700 | - | 12.4113 | 1.60E-05 | 1 | TAGGCATACGTTTTT |
| 4162 | mamo-L | *BGIBMGA008563* | down | 1558 | 1572 | + | 10.5161 | 9.77E-05 | 1 | TAGGGTGGCGTTGAT |
| 4163 | mamo-L | *BGIBMGA008564* | up | 1531 | 1545 | + | 11.3952 | 4.52E-05 | 1 | AAGGCTTATTTTAGT |
| 4164 | mamo-L | *BGIBMGA008564* | up | 1442 | 1456 | + | 11.0968 | 5.97E-05 | 1 | TAGGCTAATGCTGTT |
| 4165 | mamo-L | *BGIBMGA008567* | down | 1528 | 1542 | + | 12.8871 | 9.15E-06 | 1 | AAGGCTTTTGTTGTT |
| 4166 | mamo-L | *BGIBMGA008579* | down | 903 | 917 | + | 11.3306 | 4.80E-05 | 1 | TAGGCTGCGTATGAC |
| 4167 | mamo-L | *BGIBMGA008582* | up | 1862 | 1876 | + | 10.5726 | 9.33E-05 | 1 | TAGGCTAATATTTAT |
| 4168 | mamo-L | *BGIBMGA008583* | down | 816 | 830 | + | 10.9194 | 6.98E-05 | 1 | TAGGCTACTTTTTTT |
| 4169 | mamo-L | *BGIBMGA008584* | up | 491 | 505 | + | 10.8065 | 7.69E-05 | 1 | TAGGCCCTTTATTGC |
| 4170 | mamo-L | *BGIBMGA008589* | down | 884 | 898 | + | 11.6613 | 3.50E-05 | 1 | AAGGCGGGTGGTGGT |
| 4171 | mamo-L | *BGIBMGA008592* | down | 1817 | 1831 | + | 11.3629 | 4.66E-05 | 1 | GAGGCCGATGTTTAT |
| 4172 | mamo-L | *BGIBMGA008596* | down | 1693 | 1707 | + | 10.5726 | 9.33E-05 | 1 | TAGGCCGGGCGCGTT |
| 4173 | mamo-L | *BGIBMGA008600* | up | 1253 | 1267 | + | 10.8548 | 7.38E-05 | 1 | GAGGCCGTGAGTGCT |
| 4174 | mamo-L | *BGIBMGA008601* | down | 857 | 871 | + | 10.7742 | 7.89E-05 | 1 | TAGGCGGACATTGAG |
| 4175 | mamo-L | *BGIBMGA008602* | down | 297 | 311 | + | 11.1694 | 5.59E-05 | 1 | AAGGCTATTTTTGTT |
| 4176 | mamo-L | *BGIBMGA008602* | down | 1344 | 1358 | + | 10.5968 | 9.15E-05 | 1 | GAGGCTAGTTTTTGT |
| 4177 | mamo-L | *BGIBMGA008603* | down | 1456 | 1470 | + | 11.6048 | 3.70E-05 | 1 | TAGGCCTTGAGTTCT |
| 4178 | mamo-L | *BGIBMGA008604* | up | 1174 | 1188 | + | 11.6048 | 3.70E-05 | 1 | TAGGCCTTGAGTTCT |
| 4179 | mamo-L | *BGIBMGA008605* | down | 1215 | 1229 | - | 10.9194 | 6.98E-05 | 1 | TAGGCTACTTTTTTT |
| 4180 | mamo-L | *BGIBMGA008616* | up | 964 | 978 | + | 10.9677 | 6.68E-05 | 1 | TAGGCATGTTTTCGT |
| 4181 | mamo-L | *BGIBMGA008617* | down | 429 | 443 | - | 11.3629 | 4.66E-05 | 1 | AAGGCCGTCTTTATT |
| 4182 | mamo-L | *BGIBMGA008617* | up | 622 | 636 | - | 10.8065 | 7.69E-05 | 1 | TAGGCGTTCTGGGAT |
| 4183 | mamo-L | *BGIBMGA008622* | down | 1318 | 1332 | - | 11.6532 | 3.53E-05 | 1 | TAGGCTTAATTGTTT |
| 4184 | mamo-L | *BGIBMGA008626* | down | 1477 | 1491 | - | 10.7823 | 7.84E-05 | 1 | TGGGCTGTGTATGCT |
| 4185 | mamo-L | *BGIBMGA008630* | up | 1846 | 1860 | - | 10.5726 | 9.33E-05 | 1 | TAGGCTAATATTTAT |
| 4186 | mamo-L | *BGIBMGA008632* | down | 445 | 459 | + | 11.6371 | 3.58E-05 | 1 | TAGGCTTTCTTTTCG |
| 4187 | mamo-L | *BGIBMGA008638* | down | 1214 | 1228 | - | 10.9516 | 6.78E-05 | 1 | AAGGCTAAGTTTTTT |
| 4188 | mamo-L | *BGIBMGA008642* | up | 955 | 969 | + | 13.371 | 5.03E-06 | 1 | GAGGCTGAGGTTGGC |
| 4189 | mamo-L | *BGIBMGA008643* | up | 1291 | 1305 | + | 10.5968 | 9.15E-05 | 1 | TAGGCTTATCGTGGC |
| 4190 | mamo-L | *BGIBMGA008647* | down | 1644 | 1658 | + | 11.3226 | 4.83E-05 | 1 | AAGGCTGGGCTTGTT |
| 4191 | mamo-L | *BGIBMGA008648* | down | 1654 | 1668 | - | 11.1855 | 5.50E-05 | 1 | TAGGCATTGATTGAT |
| 4192 | mamo-L | *BGIBMGA008651* | down | 1750 | 1764 | + | 12.1935 | 2.05E-05 | 1 | TAGGCTGATTTTTTA |
| 4193 | mamo-L | *BGIBMGA008652* | up | 1393 | 1407 | + | 12.1935 | 2.05E-05 | 1 | TAGGCTGATTTTTTA |
| 4194 | mamo-L | *BGIBMGA008658* | down | 140 | 154 | + | 10.5806 | 9.27E-05 | 1 | AAGGCGGCGTGTGGT |
| 4195 | mamo-L | *BGIBMGA008665* | up | 989 | 1003 | + | 10.9435 | 6.83E-05 | 1 | TAGGCATCAGTTGGT |
| 4196 | mamo-L | *BGIBMGA008666* | up | 1747 | 1761 | - | 10.5726 | 9.33E-05 | 1 | TAGGCTAATATTTAT |
| 4197 | mamo-L | *BGIBMGA008668* | down | 1615 | 1629 | + | 11.3548 | 4.69E-05 | 1 | AAGGCTGATGATGAT |
| 4198 | mamo-L | *BGIBMGA008669* | up | 72 | 86 | + | 11.3548 | 4.69E-05 | 1 | AAGGCTGATGATGAT |
| 4199 | mamo-L | *BGIBMGA008671* | down | 1112 | 1126 | + | 10.8306 | 7.52E-05 | 1 | TGGGCTGTGTTCGCT |
| 4200 | mamo-L | *BGIBMGA008672* | down | 1937 | 1951 | - | 11.2258 | 5.30E-05 | 1 | AAGGCGTGTTTTTTT |
| 4201 | mamo-L | *BGIBMGA008686* | up | 1531 | 1545 | - | 11.7742 | 3.14E-05 | 1 | TAGGCGGCAGTTTGT |
| 4202 | mamo-L | *BGIBMGA008688* | up | 570 | 584 | + | 12.5081 | 1.43E-05 | 1 | TAGGCTTTCGTTTGA |
| 4203 | mamo-L | *BGIBMGA008691* | up | 1309 | 1323 | + | 11.4839 | 4.15E-05 | 1 | TAGGCTGGGATTTGG |
| 4204 | mamo-L | *BGIBMGA008691* | up | 154 | 168 | + | 10.879 | 7.22E-05 | 1 | TAGGCCTATGTCCAT |
| 4205 | mamo-L | *BGIBMGA008692* | down | 1712 | 1726 | - | 10.9032 | 7.07E-05 | 1 | TGGGCGGTTTTTGTT |
| 4206 | mamo-L | *BGIBMGA008693* | down | 1377 | 1391 | + | 10.9839 | 6.59E-05 | 1 | TAGGCGAAGTGTTTT |
| 4207 | mamo-L | *BGIBMGA008693* | up | 445 | 459 | - | 10.7177 | 8.28E-05 | 1 | AAGGCTGGGATTACT |
| 4208 | mamo-L | *BGIBMGA008695* | down | 171 | 185 | - | 10.6048 | 9.09E-05 | 1 | AAGGCCAACGATGTT |
| 4209 | mamo-L | *BGIBMGA008698* | up | 209 | 223 | - | 10.8306 | 7.52E-05 | 1 | TAGGCCCCCTTCGCC |
| 4210 | mamo-L | *BGIBMGA008699* | down | 502 | 516 | - | 12.8145 | 9.96E-06 | 1 | TAGGCTGGCAATGGC |
| 4211 | mamo-L | *BGIBMGA008700* | up | 767 | 781 | + | 12.7661 | 1.05E-05 | 1 | TAGGCTGGTGTGTCT |
| 4212 | mamo-L | *BGIBMGA008700* | up | 885 | 899 | + | 12.6371 | 1.23E-05 | 1 | TAGGCGGCGGTCGTT |
| 4213 | mamo-L | *BGIBMGA008702* | up | 1111 | 1125 | + | 11.0565 | 6.18E-05 | 1 | GAGGCTGATGATTTT |
| 4214 | mamo-L | *BGIBMGA008703* | up | 430 | 444 | + | 10.5968 | 9.15E-05 | 1 | AAGGCGTATTTTGAC |
| 4215 | mamo-L | *BGIBMGA008705* | down | 894 | 908 | + | 11.0403 | 6.28E-05 | 1 | TGGGCTCTGTTTGTT |
| 4216 | mamo-L | *BGIBMGA008706* | up | 756 | 770 | - | 12.5081 | 1.43E-05 | 1 | TAGGCTTTCGTTTGA |
| 4217 | mamo-L | *BGIBMGA008706* | up | 1634 | 1648 | - | 10.6129 | 9.03E-05 | 1 | TAGGCGGTATGGGCT |
| 4218 | mamo-L | *BGIBMGA008707* | down | 277 | 291 | + | 10.9194 | 6.98E-05 | 1 | TAGGCTACTTTTTTT |
| 4219 | mamo-L | *BGIBMGA008709* | down | 232 | 246 | - | 13.4677 | 4.39E-06 | 1 | TAGGCTGATGATGAT |
| 4220 | mamo-L | *BGIBMGA008715* | down | 283 | 297 | + | 10.9194 | 6.98E-05 | 1 | TAGGCTACTTTTTTT |
| 4221 | mamo-L | *BGIBMGA008721* | down | 1247 | 1261 | + | 11.4435 | 4.31E-05 | 1 | TAGGCTTAGTTAGAT |
| 4222 | mamo-L | *BGIBMGA008722* | down | 1300 | 1314 | - | 10.8306 | 7.52E-05 | 1 | TGGGCTGTGTTCGCT |
| 4223 | mamo-L | *BGIBMGA008727* | up | 1288 | 1302 | - | 11.7177 | 3.32E-05 | 1 | TAGGCTAATATTTGT |
| 4224 | mamo-L | *BGIBMGA008727* | up | 1654 | 1668 | + | 11.0565 | 6.18E-05 | 1 | TAGGCTAGTGACGTT |
| 4225 | mamo-L | *BGIBMGA008729* | down | 760 | 774 | - | 10.7903 | 7.79E-05 | 1 | GAGGCCTATTTTTTC |

| | | | | | | | | | | |
|---|---|---|---|---|---|---|---|---|---|---|
| 4226 | mamo-L | *BGIBMGA008729* | down | 1138 | 1152 | - | 10.7823 | 7.84E-05 | 1 | TAGGCCTAGATGACT |
| 4227 | mamo-L | *BGIBMGA008730* | up | 352 | 366 | - | 11.0242 | 6.36E-05 | 1 | TAGGCTGCGTGTACT |
| 4228 | mamo-L | *BGIBMGA008730* | up | 1308 | 1322 | - | 10.5726 | 9.33E-05 | 1 | TAGGCTAATATTTAT |
| 4229 | mamo-L | *BGIBMGA008734* | down | 1354 | 1368 | - | 10.9194 | 6.98E-05 | 1 | TAGGCTACTTTTTTT |
| 4230 | mamo-L | *BGIBMGA008734* | up | 904 | 918 | - | 10.4919 | 9.96E-05 | 1 | TAGGCTCTGTTAGGC |
| 4231 | mamo-L | *BGIBMGA008736* | up | 158 | 172 | - | 10.5806 | 9.27E-05 | 1 | TAGGCTACCTTTTAT |
| 4232 | mamo-L | *BGIBMGA008738* | down | 1916 | 1930 | - | 10.8226 | 7.58E-05 | 1 | TAGGCTCTTTATTAT |
| 4233 | mamo-L | *BGIBMGA008739* | down | 1223 | 1237 | - | 10.8226 | 7.58E-05 | 1 | TAGGCTCTTTATTAT |
| 4234 | mamo-L | *BGIBMGA008740* | up | 1367 | 1381 | - | 10.8226 | 7.58E-05 | 1 | TAGGCTCTTTATTAT |
| 4235 | mamo-L | *BGIBMGA008744* | down | 1548 | 1562 | + | 11.6048 | 3.70E-05 | 1 | TAGGCGATTGTTTCT |
| 4236 | mamo-L | *BGIBMGA008745* | up | 782 | 796 | + | 11.6048 | 3.70E-05 | 1 | TAGGCGATTGTTTCT |
| 4237 | mamo-L | *BGIBMGA008748* | down | 1683 | 1697 | + | 13.2177 | 6.14E-06 | 1 | TAGGCGGAGTTTCTT |
| 4238 | mamo-L | *BGIBMGA008751* | up | 1277 | 1291 | - | 12.9435 | 8.54E-06 | 1 | TAGGCTTTTTTTTAT |
| 4239 | mamo-L | *BGIBMGA008752* | up | 564 | 578 | - | 10.75 | 8.06E-05 | 1 | GAGGCCGATTGTGCC |
| 4240 | mamo-L | *BGIBMGA008752* | up | 104 | 118 | - | 10.5726 | 9.33E-05 | 1 | TAGGCTAATATTTAT |
| 4241 | mamo-L | *BGIBMGA008753* | down | 159 | 173 | - | 12.6613 | 1.19E-05 | 1 | AAGGCCGATGTTGGC |
| 4242 | mamo-L | *BGIBMGA008754* | up | 1745 | 1759 | - | 10.9194 | 6.98E-05 | 1 | TAGGCTACTTTTTTT |
| 4243 | mamo-L | *BGIBMGA008755* | down | 852 | 866 | + | 10.7984 | 7.74E-05 | 1 | TAGGCGTGCTTTTAA |
| 4244 | mamo-L | *BGIBMGA008763* | up | 1540 | 1554 | + | 10.9355 | 6.88E-05 | 1 | TAGGCCTTAGATTGC |
| 4245 | mamo-L | *BGIBMGA008763* | up | 1798 | 1812 | + | 10.5726 | 9.33E-05 | 1 | TAGGCTAATATTTAT |
| 4246 | mamo-L | *BGIBMGA008766* | up | 1607 | 1621 | + | 10.7339 | 8.17E-05 | 1 | AAGGCGTGAATTGGT |
| 4247 | mamo-L | *BGIBMGA008772* | down | 1392 | 1406 | + | 13.2097 | 6.20E-06 | 1 | TAGGCCTATGTTTAT |
| 4248 | mamo-L | *BGIBMGA008774* | down | 800 | 814 | - | 12.7823 | 1.03E-05 | 1 | TAGGCCTTAATTGTT |
| 4249 | mamo-L | *BGIBMGA008777* | up | 199 | 213 | - | 12.1532 | 2.13E-05 | 1 | TAGGCCAACTTTTGC |
| 4250 | mamo-L | *BGIBMGA008778* | up | 117 | 131 | + | 10.5323 | 9.63E-05 | 1 | AAGGCCGCGGTTTGC |
| 4251 | mamo-L | *BGIBMGA008779* | up | 352 | 366 | + | 13.9435 | 2.15E-06 | 1 | TAGGCCAAGTTTGCT |
| 4252 | mamo-L | *BGIBMGA008781* | down | 249 | 263 | - | 10.6935 | 8.45E-05 | 1 | TAGGCTGCAATCGCT |
| 4253 | mamo-L | *BGIBMGA008782* | down | 705 | 719 | + | 11.629 | 3.61E-05 | 1 | GAGGCTTTTTTTTTT |
| 4254 | mamo-L | *BGIBMGA008783* | up | 929 | 943 | + | 10.9194 | 6.98E-05 | 1 | TAGGCTACTTTTTTT |
| 4255 | mamo-L | *BGIBMGA008787* | up | 932 | 946 | + | 10.7581 | 8.00E-05 | 1 | AAGGCGTATTTGGTT |
| 4256 | mamo-L | *BGIBMGA008788* | up | 265 | 279 | - | 13.2177 | 6.14E-06 | 1 | TAGGCGGAGTTTCTT |
| 4257 | mamo-L | *BGIBMGA008789* | up | 1256 | 1270 | + | 12.25 | 1.92E-05 | 1 | AAGGCTTATTTTGAT |
| 4258 | mamo-L | *BGIBMGA008791* | up | 818 | 832 | + | 13.1855 | 6.42E-06 | 1 | TAGGCGGGGGGTGAT |
| 4259 | mamo-L | *BGIBMGA008791* | down | 61 | 75 | + | 11.4113 | 4.45E-05 | 1 | TAGGCTTGAGTTGAA |
| 4260 | mamo-L | *BGIBMGA008791* | down | 289 | 303 | + | 11.4113 | 4.45E-05 | 1 | AAGGCTGTTTTGGTT |
| 4261 | mamo-L | *BGIBMGA008791* | down | 1630 | 1644 | + | 11.121 | 5.84E-05 | 1 | TAGGCTCATTTCATT |
| 4262 | mamo-L | *BGIBMGA008791* | down | 1124 | 1138 | + | 10.6371 | 8.85E-05 | 1 | TAGGCTGAGGCTTAC |
| 4263 | mamo-L | *BGIBMGA008792* | down | 1851 | 1865 | - | 10.9435 | 6.83E-05 | 1 | AAGGCGAATGTTGTT |
| 4264 | mamo-L | *BGIBMGA008793* | down | 1763 | 1777 | - | 14.5726 | 7.98E-07 | 1 | TAGGCCTGCGTTGTC |
| 4265 | mamo-L | *BGIBMGA008794* | down | 1430 | 1444 | + | 12.0806 | 2.31E-05 | 1 | TAGGCCTTGTTCGGA |
| 4266 | mamo-L | *BGIBMGA008803* | up | 867 | 881 | - | 11.9516 | 2.63E-05 | 1 | TAGGCCTGCTCTGAT |
| 4267 | mamo-L | *BGIBMGA008807* | down | 40 | 54 | - | 10.629 | 8.91E-05 | 1 | TAGGCCTGCAGCGCC |
| 4268 | mamo-L | *BGIBMGA008810* | up | 1284 | 1298 | + | 13.5887 | 3.68E-06 | 1 | TAGGCCTTTTTTTTT |
| 4269 | mamo-L | *BGIBMGA008810* | up | 1560 | 1574 | + | 12.0645 | 2.34E-05 | 1 | TAGGCGGGCTTCGGA |
| 4270 | mamo-L | *BGIBMGA008810* | up | 1554 | 1568 | + | 10.75 | 8.06E-05 | 1 | TAGGCCTAGGCGGGC |
| 4271 | mamo-L | *BGIBMGA008815* | down | 174 | 188 | + | 10.9919 | 6.55E-05 | 1 | TAGGCTCTTGATGGA |
| 4272 | mamo-L | *BGIBMGA008816* | up | 1923 | 1937 | - | 12.1371 | 2.17E-05 | 1 | TAGGCTGTTATTTAT |
| 4273 | mamo-L | *BGIBMGA008817* | down | 1812 | 1826 | + | 11.5806 | 3.78E-05 | 1 | TAGGCTCGGATTCCT |
| 4274 | mamo-L | *BGIBMGA008819* | down | 165 | 179 | + | 12.7581 | 1.07E-05 | 1 | TAGGCCTTTTTTTAT |
| 4275 | mamo-L | *BGIBMGA008819* | down | 524 | 538 | + | 10.8145 | 7.63E-05 | 1 | AAGGCGCATATTGGT |
| 4276 | mamo-L | *BGIBMGA008820* | up | 1780 | 1794 | - | 11.4194 | 4.42E-05 | 1 | TAGGCCTGGTGTAGC |
| 4277 | mamo-L | *BGIBMGA008824* | down | 1374 | 1388 | - | 12.4194 | 1.58E-05 | 1 | TAGGCTTAAGATTGT |
| 4278 | mamo-L | *BGIBMGA008824* | up | 1600 | 1614 | + | 11.5323 | 3.96E-05 | 1 | GAGGCCTTTTTTGCC |
| 4279 | mamo-L | *BGIBMGA008826* | down | 1228 | 1242 | + | 10.9839 | 6.59E-05 | 1 | TAGGCTTTTTTTAGA |
| 4280 | mamo-L | *BGIBMGA008832* | up | 714 | 728 | + | 10.5968 | 9.15E-05 | 1 | GAGGCTAGTTTTTGT |
| 4281 | mamo-L | *BGIBMGA008833* | down | 874 | 888 | - | 12.6855 | 1.16E-05 | 1 | TAGGCGAGCGTTTGT |
| 4282 | mamo-L | *BGIBMGA008833* | down | 90 | 104 | + | 10.6371 | 8.85E-05 | 1 | TAGGCATTATTTTCT |
| 4283 | mamo-L | *BGIBMGA008836* | down | 544 | 558 | + | 12.6694 | 1.18E-05 | 1 | GAGGCCTTTGTTGTT |
| 4284 | mamo-L | *BGIBMGA008836* | up | 1686 | 1700 | - | 11.7339 | 3.26E-05 | 1 | TAGGCTAGCTTCGCC |
| 4285 | mamo-L | *BGIBMGA008846* | up | 474 | 488 | + | 10.8468 | 7.42E-05 | 1 | AAGGCTTATATTTCT |
| 4286 | mamo-L | *BGIBMGA008848* | up | 472 | 486 | + | 10.9194 | 6.98E-05 | 1 | TAGGCTACTTTTTTT |
| 4287 | mamo-L | *BGIBMGA008849* | down | 274 | 288 | - | 11 | 6.50E-05 | 1 | TAGGCTCGTCTTTGT |
| 4288 | mamo-L | *BGIBMGA008850* | down | 1964 | 1978 | + | 10.9677 | 6.68E-05 | 1 | TAGGCTGAAATGGTC |
| 4289 | mamo-L | *BGIBMGA008851* | down | 593 | 607 | - | 12.0806 | 2.31E-05 | 1 | TAGGCCTTGTTCGGA |
| 4290 | mamo-L | *BGIBMGA008851* | down | 1400 | 1414 | - | 10.5484 | 9.52E-05 | 1 | GAGGCTGTGGTTCCC |
| 4291 | mamo-L | *BGIBMGA008853* | up | 140 | 154 | + | 10.5726 | 9.33E-05 | 1 | AAGGCTTTTTTTCCT |
| 4292 | mamo-L | *BGIBMGA008855* | down | 691 | 705 | - | 12.6371 | 1.23E-05 | 1 | TAGGCCGACTTTCCC |
| 4293 | mamo-L | *BGIBMGA008856* | up | 69 | 83 | - | 11.3548 | 4.69E-05 | 1 | AAGGCTGATGATGAT |
| 4294 | mamo-L | *BGIBMGA008860* | down | 750 | 764 | + | 11.6129 | 3.66E-05 | 1 | TAGGCTTTAAATGCT |
| 4295 | mamo-L | *BGIBMGA008860* | down | 1717 | 1731 | - | 10.5 | 9.90E-05 | 1 | GAGGCCCAGGTCGTC |
| 4296 | mamo-L | *BGIBMGA008861* | down | 1836 | 1850 | - | 10.7581 | 8.00E-05 | 1 | TAGGCCGGGTTGTGG |
| 4297 | mamo-L | *BGIBMGA008862* | down | 256 | 270 | + | 11.1613 | 5.63E-05 | 1 | TAGGCTGGGTAGCGT |
| 4298 | mamo-L | *BGIBMGA008862* | up | 1512 | 1526 | - | 10.7581 | 8.00E-05 | 1 | TAGGCCGGGTTGTGG |
| 4299 | mamo-L | *BGIBMGA008863* | down | 683 | 697 | - | 14.7258 | 5.83E-07 | 1 | TAGGCTTGCGTTGCC |
| 4300 | mamo-L | *BGIBMGA008864* | up | 1830 | 1844 | + | 10.5403 | 9.58E-05 | 1 | TAGGCTGTATGGTGT |
| 4301 | mamo-L | *BGIBMGA008865* | down | 1250 | 1264 | + | 12.0645 | 2.34E-05 | 1 | TAGGCGGGCTTCGGA |
| 4302 | mamo-L | *BGIBMGA008865* | down | 1104 | 1118 | - | 10.8306 | 7.52E-05 | 1 | TAGGCCCCCTTCGCC |
| 4303 | mamo-L | *BGIBMGA008866* | up | 1288 | 1302 | - | 10.8226 | 7.58E-05 | 1 | AAGGCTGGTTTGGAT |
| 4304 | mamo-L | *BGIBMGA008867* | up | 1923 | 1937 | - | 11.7823 | 3.11E-05 | 1 | GAGGCTGCGGTTGGC |
| 4305 | mamo-L | *BGIBMGA008867* | down | 377 | 391 | - | 11.0968 | 5.97E-05 | 1 | TAGGCTTTCGCTTGC |
| 4306 | mamo-L | *BGIBMGA008870* | up | 937 | 951 | - | 11.0887 | 6.01E-05 | 1 | TAGGCTGATATTTTA |
| 4307 | mamo-L | *BGIBMGA008871* | up | 823 | 837 | + | 10.9194 | 6.98E-05 | 1 | TAGGCTACTTTTTTT |
| 4308 | mamo-L | *BGIBMGA008871* | down | 1826 | 1840 | - | 10.8468 | 7.42E-05 | 1 | TAGGCCGTCTTGTGA |
| 4309 | mamo-L | *BGIBMGA008872* | down | 1448 | 1462 | - | 10.9597 | 6.73E-05 | 1 | TAGGCATATTTTTAT |
| 4310 | mamo-L | *BGIBMGA008872* | down | 500 | 514 | - | 10.5726 | 9.33E-05 | 1 | TAGGCTAATATTTAT |
| 4311 | mamo-L | *BGIBMGA008875* | up | 537 | 551 | + | 10.7258 | 8.23E-05 | 1 | TAGGCTGACAATAAT |
| 4312 | mamo-L | *BGIBMGA008876* | up | 972 | 986 | - | 10.6694 | 8.62E-05 | 1 | TAGGCCTGCTGCTTC |
| 4313 | mamo-L | *BGIBMGA008877* | up | 888 | 902 | + | 10.5726 | 9.33E-05 | 1 | TAGGCTAATATTTAT |
| 4314 | mamo-L | *BGIBMGA008880* | up | 1559 | 1573 | + | 11.7984 | 3.06E-05 | 1 | TAGGCTTTGCTTTTT |
| 4315 | mamo-L | *BGIBMGA008886* | up | 1627 | 1641 | - | 10.5645 | 9.40E-05 | 1 | TAGGCTCGAGGGGCT |
| 4316 | mamo-L | *BGIBMGA008887* | up | 1310 | 1324 | + | 11.0161 | 6.40E-05 | 1 | TAGGCTGAGCGTTTT |
| 4317 | mamo-L | *BGIBMGA008888* | up | 1835 | 1849 | + | 11.5726 | 3.81E-05 | 1 | TAGGCTTTTTTTTTA |

| | | | | | | | | | | |
|---|---|---|---|---|---|---|---|---|---|---|
| 4318 | mamo-L | *BGIBMGA008889* | down | 623 | 637 | - | 13.3145 | 5.41E-06 | 1 | AAGGCTTTGTTTGTT |
| 4319 | mamo-L | *BGIBMGA008889* | down | 285 | 299 | + | 10.75 | 8.06E-05 | 1 | GAGGCCGATTGTGCC |
| 4320 | mamo-L | *BGIBMGA008891* | up | 274 | 288 | - | 12.1371 | 2.17E-05 | 1 | GAGGCTCGTTTTGCT |
| 4321 | mamo-L | *BGIBMGA008891* | down | 767 | 781 | + | 10.6048 | 9.09E-05 | 1 | TAGGCTTGTATGTGC |
| 4322 | mamo-L | *BGIBMGA008897* | up | 695 | 709 | + | 12.2661 | 1.89E-05 | 1 | TAGGCCTAGGTCCTT |
| 4323 | mamo-L | *BGIBMGA008897* | up | 911 | 925 | - | 10.4919 | 9.96E-05 | 1 | GAGGCTGGAGTCGAT |
| 4324 | mamo-L | *BGIBMGA008900* | down | 1526 | 1540 | + | 13.5161 | 4.10E-06 | 1 | TAGGCGTAGGTTAGT |
| 4325 | mamo-L | *BGIBMGA008901* | down | 1938 | 1952 | - | 10.9677 | 6.68E-05 | 1 | TAGGCGGGCTTCTGA |
| 4326 | mamo-L | *BGIBMGA008902* | up | 642 | 656 | + | 11.4032 | 4.49E-05 | 1 | TAGGCTAATATTTTT |
| 4327 | mamo-L | *BGIBMGA008902* | down | 220 | 234 | - | 11.3468 | 4.73E-05 | 1 | AAGGCTAAGGTTGAT |
| 4328 | mamo-L | *BGIBMGA008905* | down | 836 | 850 | - | 10.5726 | 9.33E-05 | 1 | TAGGCTGTTTTTATG |
| 4329 | mamo-L | *BGIBMGA008907* | up | 917 | 931 | - | 10.8468 | 7.42E-05 | 1 | TAGGCCGAGTATTTG |
| 4330 | mamo-L | *BGIBMGA008907* | up | 1055 | 1069 | + | 10.5806 | 9.27E-05 | 1 | TAGGCTGAAAATAGT |
| 4331 | mamo-L | *BGIBMGA008909* | down | 633 | 647 | - | 11.3629 | 4.66E-05 | 1 | TAGGCTTGGTGTTAC |
| 4332 | mamo-L | *BGIBMGA008910* | down | 1333 | 1347 | + | 10.7823 | 7.84E-05 | 1 | TAGGCTTGTTATACC |
| 4333 | mamo-L | *BGIBMGA008911* | up | 1667 | 1681 | + | 10.7823 | 7.84E-05 | 1 | TAGGCTTGTTATACC |
| 4334 | mamo-L | *BGIBMGA008913* | down | 1273 | 1287 | + | 12.1774 | 2.08E-05 | 1 | TAGGCCAAGGTTTAT |
| 4335 | mamo-L | *BGIBMGA008915* | down | 600 | 614 | + | 11.5403 | 3.93E-05 | 1 | TAGGCGGATTTCATT |
| 4336 | mamo-L | *BGIBMGA008916* | up | 360 | 374 | - | 10.5968 | 9.15E-05 | 1 | TAGGCCCGTATGTGT |
| 4337 | mamo-L | *BGIBMGA008919* | down | 1081 | 1095 | - | 11.3871 | 4.55E-05 | 1 | TAGGCTGAAATTGTA |
| 4338 | mamo-L | *BGIBMGA008920* | up | 440 | 454 | - | 10.75 | 8.06E-05 | 1 | GAGGCCGATTGTGCC |
| 4339 | mamo-L | *BGIBMGA008926* | down | 666 | 680 | + | 12.0161 | 2.46E-05 | 1 | TAGGCCGAGATGTTT |
| 4340 | mamo-L | *BGIBMGA008927* | up | 1446 | 1460 | + | 11.7419 | 3.23E-05 | 1 | TAGGCGTAGTTTTCA |
| 4341 | mamo-L | *BGIBMGA008928* | up | 1238 | 1252 | + | 11.2742 | 5.06E-05 | 1 | AAGGCTGATTTTTCC |
| 4342 | mamo-L | *BGIBMGA008928* | down | 1873 | 1887 | - | 10.9032 | 7.07E-05 | 1 | TAGGCGGAGCTCGGC |
| 4343 | mamo-L | *BGIBMGA008929* | down | 1359 | 1373 | - | 10.5968 | 9.15E-05 | 1 | GAGGCTAGTTTTTGT |
| 4344 | mamo-L | *BGIBMGA008932* | up | 1902 | 1916 | + | 10.7258 | 8.23E-05 | 1 | TAGGCCGGGGTTCAA |
| 4345 | mamo-L | *BGIBMGA008933* | up | 377 | 391 | + | 12.2419 | 1.94E-05 | 1 | TAGGCCGACGGCGCC |
| 4346 | mamo-L | *BGIBMGA008934* | up | 1895 | 1909 | + | 11.5565 | 3.87E-05 | 1 | TAGGCCGTTACTGGT |
| 4347 | mamo-L | *BGIBMGA008934* | down | 1170 | 1184 | + | 10.8145 | 7.63E-05 | 1 | AAGGCTATAGTTGGT |
| 4348 | mamo-L | *BGIBMGA008938* | down | 1525 | 1539 | + | 12.5403 | 1.38E-05 | 1 | AAGGCCTTTTTTGCT |
| 4349 | mamo-L | *BGIBMGA008938* | down | 214 | 228 | - | 10.8145 | 7.63E-05 | 1 | AAGGCTATAGTTGGT |
| 4350 | mamo-L | *BGIBMGA008939* | up | 483 | 497 | - | 13.7742 | 2.79E-06 | 1 | TAGGCGGGTGGTGGT |
| 4351 | mamo-L | *BGIBMGA008941* | up | 1947 | 1961 | - | 11.9435 | 2.65E-05 | 1 | TAGGCGTGTGTGGTC |
| 4352 | mamo-L | *BGIBMGA008942* | up | 1280 | 1294 | + | 11.6855 | 3.42E-05 | 1 | TAGGCGTATTTTAAT |
| 4353 | mamo-L | *BGIBMGA008942* | up | 1679 | 1693 | + | 10.75 | 8.06E-05 | 1 | AAGGCTCGCGTTTTC |
| 4354 | mamo-L | *BGIBMGA008943* | down | 1963 | 1977 | - | 12.25 | 1.92E-05 | 1 | TAGGCCCACTATTCT |
| 4355 | mamo-L | *BGIBMGA008945* | down | 1669 | 1683 | - | 12.871 | 9.35E-06 | 1 | TAGGCGTAGATTTTT |
| 4356 | mamo-L | *BGIBMGA008949* | up | 137 | 151 | - | 10.9919 | 6.55E-05 | 1 | TAGGCCGGTTGTAAT |
| 4357 | mamo-L | *BGIBMGA008950* | up | 655 | 669 | - | 11.7984 | 3.06E-05 | 1 | TAGGCGGTGGTTTGG |
| 4358 | mamo-L | *BGIBMGA008950* | up | 73 | 87 | - | 10.5726 | 9.33E-05 | 1 | TAGGCTAATATTTAT |
| 4359 | mamo-L | *BGIBMGA008951* | down | 1901 | 1915 | - | 12.0161 | 2.46E-05 | 1 | TAGGCCGAGATGTTT |
| 4360 | mamo-L | *BGIBMGA008951* | up | 891 | 905 | - | 11.7419 | 3.23E-05 | 1 | TAGGCGTAGTTTTCA |
| 4361 | mamo-L | *BGIBMGA008962* | down | 1875 | 1889 | + | 10.5726 | 9.33E-05 | 1 | TAGGCTAATATTTAT |
| 4362 | mamo-L | *BGIBMGA008963* | down | 1118 | 1132 | - | 13.0887 | 7.18E-06 | 1 | TAGGCCGTATTTTTT |
| 4363 | mamo-L | *BGIBMGA008963* | down | 1497 | 1511 | + | 11.2339 | 5.26E-05 | 1 | TAGGCCGATAAGGTT |
| 4364 | mamo-L | *BGIBMGA008966* | down | 650 | 664 | - | 12.1774 | 2.08E-05 | 1 | TAGGCCAAGGTTTAT |
| 4365 | mamo-L | *BGIBMGA008973* | up | 1100 | 1114 | - | 10.75 | 8.06E-05 | 1 | GAGGCCGATTGTGCC |
| 4366 | mamo-L | *BGIBMGA008988* | up | 1501 | 1515 | - | 11.0806 | 6.05E-05 | 1 | TAGGCGCACTTTCTC |
| 4367 | mamo-L | *BGIBMGA008991* | down | 1606 | 1620 | - | 10.9194 | 6.98E-05 | 1 | TAGGCTACTTTTTTT |
| 4368 | mamo-L | *BGIBMGA008992* | up | 660 | 674 | - | 12.2661 | 1.89E-05 | 1 | TAGGCCTAGGTCCTT |
| 4369 | mamo-L | *BGIBMGA008992* | up | 444 | 458 | + | 10.4919 | 9.96E-05 | 1 | GAGGCTGGAGTCGAT |
| 4370 | mamo-L | *BGIBMGA008993* | up | 429 | 443 | + | 11.7661 | 3.16E-05 | 1 | TAGGCTTTTATTATT |
| 4371 | mamo-L | *BGIBMGA008994* | down | 1922 | 1936 | - | 11.6613 | 3.50E-05 | 1 | AAGGCTTTTTTTTTT |
| 4372 | mamo-L | *BGIBMGA008994* | down | 1414 | 1428 | - | 11.3145 | 4.87E-05 | 1 | TAGGCAGGCGGTGCC |
| 4373 | mamo-L | *BGIBMGA008998* | down | 1869 | 1883 | + | 11.2016 | 5.41E-05 | 1 | GAGGCCTGATTTTGT |
| 4374 | mamo-L | *BGIBMGA009005* | up | 1448 | 1462 | + | 10.6129 | 9.03E-05 | 1 | TAGGCGTGGTGTATC |
| 4375 | mamo-L | *BGIBMGA009006* | up | 1927 | 1941 | + | 10.9597 | 6.73E-05 | 1 | AAGGCCGATATTTCT |
| 4376 | mamo-L | *BGIBMGA009007* | down | 1920 | 1934 | - | 11.4435 | 4.31E-05 | 1 | AAGGCTTGTGTTAGT |
| 4377 | mamo-L | *BGIBMGA009008* | down | 448 | 462 | + | 12.7984 | 1.02E-05 | 1 | TAGGCTTTTTTTTTC |
| 4378 | mamo-L | *BGIBMGA009008* | up | 905 | 919 | - | 11.5242 | 3.99E-05 | 1 | TAGGCTAGTTTTATT |
| 4379 | mamo-L | *BGIBMGA009008* | up | 961 | 975 | - | 11.5242 | 3.99E-05 | 1 | TAGGCTAGTTTTATT |
| 4380 | mamo-L | *BGIBMGA009008* | up | 1017 | 1031 | - | 11.5242 | 3.99E-05 | 1 | TAGGCTAGTTTTATT |
| 4381 | mamo-L | *BGIBMGA009008* | up | 1073 | 1087 | - | 11.5242 | 3.99E-05 | 1 | TAGGCTAGTTTTATT |
| 4382 | mamo-L | *BGIBMGA009008* | up | 1129 | 1143 | - | 11.5242 | 3.99E-05 | 1 | TAGGCTAGTTTTATT |
| 4383 | mamo-L | *BGIBMGA009008* | up | 1185 | 1199 | - | 11.5242 | 3.99E-05 | 1 | TAGGCTAGTTTTATT |
| 4384 | mamo-L | *BGIBMGA009008* | up | 1241 | 1255 | - | 11.5242 | 3.99E-05 | 1 | TAGGCTAGTTTTATT |
| 4385 | mamo-L | *BGIBMGA009008* | up | 1297 | 1311 | - | 11.5242 | 3.99E-05 | 1 | TAGGCTAGTTTTATT |
| 4386 | mamo-L | *BGIBMGA009008* | up | 1353 | 1367 | - | 11.5242 | 3.99E-05 | 1 | TAGGCTAGTTTTATT |
| 4387 | mamo-L | *BGIBMGA009008* | up | 1409 | 1423 | - | 11.5242 | 3.99E-05 | 1 | TAGGCTAGTTTTATT |
| 4388 | mamo-L | *BGIBMGA009008* | up | 1465 | 1479 | - | 11.5242 | 3.99E-05 | 1 | TAGGCTAGTTTTATT |
| 4389 | mamo-L | *BGIBMGA009008* | up | 1521 | 1535 | - | 11.5242 | 3.99E-05 | 1 | TAGGCTAGTTTTATT |
| 4390 | mamo-L | *BGIBMGA009009* | down | 1481 | 1495 | - | 11.2742 | 5.06E-05 | 1 | TAGGCCTACGTTCTA |
| 4391 | mamo-L | *BGIBMGA009009* | down | 1488 | 1502 | + | 10.6855 | 8.51E-05 | 1 | TAGGCCTAAAATTTT |
| 4392 | mamo-L | *BGIBMGA009010* | up | 1200 | 1214 | + | 12.0645 | 2.34E-05 | 1 | TAGGCGGGCTTCGGA |
| 4393 | mamo-L | *BGIBMGA009017* | down | 826 | 840 | + | 10.9597 | 6.73E-05 | 1 | TAGGCATATTTTTAT |
| 4394 | mamo-L | *BGIBMGA009017* | down | 1774 | 1788 | + | 10.5726 | 9.33E-05 | 1 | TAGGCTAATATTTAT |
| 4395 | mamo-L | *BGIBMGA009025* | up | 1174 | 1188 | - | 11.3629 | 4.66E-05 | 1 | TAGGCTCACAATTTT |
| 4396 | mamo-L | *BGIBMGA009026* | up | 91 | 105 | + | 11.8548 | 2.89E-05 | 1 | TAGGCTTTGTTGTAT |
| 4397 | mamo-L | *BGIBMGA009026* | down | 384 | 398 | - | 11.2823 | 5.03E-05 | 1 | TAGGCTTTAATTAGT |
| 4398 | mamo-L | *BGIBMGA009028* | down | 1558 | 1572 | - | 11.1613 | 5.63E-05 | 1 | TAGGCTGGGTAGCGT |
| 4399 | mamo-L | *BGIBMGA009030* | up | 520 | 534 | + | 11.7661 | 3.16E-05 | 1 | AAGGCTTAGGTTATT |
| 4400 | mamo-L | *BGIBMGA009031* | down | 174 | 188 | - | 11.6129 | 3.66E-05 | 1 | TAGGCTTTTAATGAT |
| 4401 | mamo-L | *BGIBMGA009037* | up | 1479 | 1493 | - | 10.8468 | 7.42E-05 | 1 | AAGGCTTATATTTCT |
| 4402 | mamo-L | *BGIBMGA009040* | up | 427 | 441 | - | 12.2339 | 1.95E-05 | 1 | AAGGCGGGCGTTGCC |
| 4403 | mamo-L | *BGIBMGA009040* | up | 632 | 646 | + | 10.5726 | 9.33E-05 | 1 | TAGGCTAATATTTAT |
| 4404 | mamo-L | *BGIBMGA009041* | up | 1808 | 1822 | - | 12.5484 | 1.36E-05 | 1 | TAGGCCCACTTTCTT |
| 4405 | mamo-L | *BGIBMGA009041* | up | 941 | 955 | - | 11.3548 | 4.69E-05 | 1 | AAGGCTGATGATGAT |
| 4406 | mamo-L | *BGIBMGA009041* | down | 1585 | 1599 | - | 10.9194 | 6.98E-05 | 1 | GAGGCGTTTTTTTCT |
| 4407 | mamo-L | *BGIBMGA009046* | down | 1767 | 1781 | - | 12.3306 | 1.75E-05 | 1 | TAGGCTTACTATACT |
| 4408 | mamo-L | *BGIBMGA009048* | down | 1213 | 1227 | + | 11.1129 | 5.88E-05 | 1 | TAGGCTTCCTTTTGA |
| 4409 | mamo-L | *BGIBMGA009048* | up | 1432 | 1446 | + | 10.75 | 8.06E-05 | 1 | GAGGCCGATTGTGCC |

| | | | | | | | | | | |
|---|---|---|---|---|---|---|---|---|---|---|
| 4410 | mamo-L | *BGIBMGA009049* | down | 1010 | 1024 | + | 12.2823 | 1.85E-05 | 1 | AAGGCTCATTTTGTT |
| 4411 | mamo-L | *BGIBMGA009050* | down | 347 | 361 | - | 12.6048 | 1.27E-05 | 1 | TAGGCTCACGGTGCC |
| 4412 | mamo-L | *BGIBMGA009054* | down | 1646 | 1660 | + | 10.5161 | 9.77E-05 | 1 | TAGGCCTGCGTTAAA |
| 4413 | mamo-L | *BGIBMGA009055* | down | 1871 | 1885 | + | 11.2016 | 5.41E-05 | 1 | TAGGCGTATCTTTGT |
| 4414 | mamo-L | *BGIBMGA009059* | down | 1788 | 1802 | + | 12.0565 | 2.36E-05 | 1 | TAGGCAGATTTTTCT |
| 4415 | mamo-L | *BGIBMGA009060* | up | 1284 | 1298 | + | 10.8306 | 7.52E-05 | 1 | AAGGCTTTCTATTTT |
| 4416 | mamo-L | *BGIBMGA009061* | up | 1254 | 1268 | - | 11.8306 | 2.97E-05 | 1 | AAGGCGTAGTTTTCT |
| 4417 | mamo-L | *BGIBMGA009062* | up | 1318 | 1332 | - | 10.6371 | 8.85E-05 | 1 | TAGGCAGGCGGTAGT |
| 4418 | mamo-L | *BGIBMGA009063* | down | 699 | 713 | + | 12.6129 | 1.26E-05 | 1 | TAGGCCCGGTTTTTC |
| 4419 | mamo-L | *BGIBMGA009063* | up | 1364 | 1378 | + | 12.129 | 2.19E-05 | 1 | TAGGCGCCTTTTGTT |
| 4420 | mamo-L | *BGIBMGA009063* | down | 778 | 792 | + | 10.7097 | 8.33E-05 | 1 | TAGGCCAGTTTTGCG |
| 4421 | mamo-L | *BGIBMGA009064* | up | 1156 | 1170 | + | 12.6129 | 1.26E-05 | 1 | TAGGCCCGGTTTTTC |
| 4422 | mamo-L | *BGIBMGA009064* | up | 1235 | 1249 | + | 10.7097 | 8.33E-05 | 1 | TAGGCCAGTTTTGCG |
| 4423 | mamo-L | *BGIBMGA009066* | down | 141 | 155 | + | 12.5645 | 1.34E-05 | 1 | GAGGCTGATTTTTGT |
| 4424 | mamo-L | *BGIBMGA009068* | down | 114 | 128 | + | 12.5242 | 1.40E-05 | 1 | TAGGCTTGCTATTGC |
| 4425 | mamo-L | *BGIBMGA009075* | up | 495 | 509 | - | 11.4355 | 4.34E-05 | 1 | TAGGCTACGATTGCT |
| 4426 | mamo-L | *BGIBMGA009076* | down | 209 | 223 | - | 10.6694 | 8.62E-05 | 1 | GAGGCCGGTTGTGCC |
| 4427 | mamo-L | *BGIBMGA009077* | up | 757 | 771 | + | 10.5323 | 9.63E-05 | 1 | AAGGCAGAGTTTTTT |
| 4428 | mamo-L | *BGIBMGA009081* | down | 323 | 337 | - | 10.7339 | 8.17E-05 | 1 | TAGGCATATTATGAT |
| 4429 | mamo-L | *BGIBMGA009084* | down | 699 | 713 | - | 10.7984 | 7.74E-05 | 1 | TAGGCTTTAGGTATT |
| 4430 | mamo-L | *BGIBMGA009085* | up | 1113 | 1127 | - | 10.7984 | 7.74E-05 | 1 | TAGGCTTTAGGTATT |
| 4431 | mamo-L | *BGIBMGA009088* | up | 593 | 607 | - | 10.5242 | 9.70E-05 | 1 | TAGGCGTTGTATTAC |
| 4432 | mamo-L | *BGIBMGA009095* | up | 1385 | 1399 | - | 10.5081 | 9.83E-05 | 1 | TAGGCGGGGTGTTGG |
| 4433 | mamo-L | *BGIBMGA009099* | down | 1067 | 1081 | + | 10.5726 | 9.33E-05 | 1 | TAGGCTAATATTTAT |
| 4434 | mamo-L | *BGIBMGA009100* | down | 1974 | 1988 | - | 10.9194 | 6.98E-05 | 1 | TAGGCTACTTTTTTT |
| 4435 | mamo-L | *BGIBMGA009102* | down | 960 | 974 | - | 12.7419 | 1.09E-05 | 1 | TAGGCCGATCTTGCT |
| 4436 | mamo-L | *BGIBMGA009103* | up | 1154 | 1168 | + | 11.8306 | 2.97E-05 | 1 | AAGGCGTAGTTTTCT |
| 4437 | mamo-L | *BGIBMGA009107* | down | 764 | 778 | - | 12.4677 | 1.49E-05 | 1 | TAGGCTTCTATTGCT |
| 4438 | mamo-L | *BGIBMGA009107* | down | 1372 | 1386 | - | 11.2016 | 5.41E-05 | 1 | TAGGCGTATCTTTGT |
| 4439 | mamo-L | *BGIBMGA009107* | up | 1805 | 1819 | + | 10.5726 | 9.33E-05 | 1 | TAGGCTAATATTTAT |
| 4440 | mamo-L | *BGIBMGA009108* | up | 839 | 853 | + | 12.1048 | 2.24E-05 | 1 | TAGGCCCCCTTTGCC |
| 4441 | mamo-L | *BGIBMGA009108* | up | 909 | 923 | - | 10.879 | 7.22E-05 | 1 | GAGGCGGACTTCGCC |
| 4442 | mamo-L | *BGIBMGA009109* | down | 1683 | 1697 | - | 11.3065 | 4.91E-05 | 1 | TAGGCGTATAATTGT |
| 4443 | mamo-L | *BGIBMGA009110* | down | 1140 | 1154 | - | 11.0403 | 6.28E-05 | 1 | TAGGCTTTTTGGTGT |
| 4444 | mamo-L | *BGIBMGA009110* | down | 959 | 973 | - | 10.5726 | 9.33E-05 | 1 | TAGGCTAATATTTAT |
| 4445 | mamo-L | *BGIBMGA009111* | down | 257 | 271 | - | 11.1129 | 5.88E-05 | 1 | TAGGCTTCCTTTTGA |
| 4446 | mamo-L | *BGIBMGA009112* | up | 119 | 133 | - | 10.75 | 8.06E-05 | 1 | GAGGCCGATTGTGCC |
| 4447 | mamo-L | *BGIBMGA009113* | up | 562 | 576 | + | 10.5645 | 9.40E-05 | 1 | TAGGCTGCTGCTGAT |
| 4448 | mamo-L | *BGIBMGA009114* | up | 283 | 297 | + | 10.7258 | 8.23E-05 | 1 | TAGGCGTCTATTTTT |
| 4449 | mamo-L | *BGIBMGA009116* | up | 1544 | 1558 | + | 11.0403 | 6.28E-05 | 1 | TAGGCGGGCAGGGCT |
| 4450 | mamo-L | *BGIBMGA009118* | down | 551 | 565 | - | 10.5726 | 9.33E-05 | 1 | GAGGCGTACGATTTT |
| 4451 | mamo-L | *BGIBMGA009120* | down | 112 | 126 | + | 10.5726 | 9.33E-05 | 1 | TAGGCTAATATTTAT |
| 4452 | mamo-L | *BGIBMGA009121* | up | 287 | 301 | + | 10.9194 | 6.98E-05 | 1 | TAGGCTACTTTTTTT |
| 4453 | mamo-L | *BGIBMGA009125* | up | 1473 | 1487 | + | 10.9194 | 6.98E-05 | 1 | TAGGCTACTTTTTTT |
| 4454 | mamo-L | *BGIBMGA009125* | down | 1693 | 1707 | - | 10.9113 | 7.02E-05 | 1 | TAGGCCGTATTCATT |
| 4455 | mamo-L | *BGIBMGA009129* | down | 499 | 513 | + | 10.8306 | 7.52E-05 | 1 | TAGGCGTACGATCAT |
| 4456 | mamo-L | *BGIBMGA009130* | down | 570 | 584 | - | 13.3226 | 5.34E-06 | 1 | TAGGCCGTCGACGGT |
| 4457 | mamo-L | *BGIBMGA009130* | up | 1008 | 1022 | + | 10.8306 | 7.52E-05 | 1 | TAGGCGTACGATCAT |
| 4458 | mamo-L | *BGIBMGA009133* | down | 1143 | 1157 | + | 11.1935 | 5.45E-05 | 1 | TAGGCCTGCGACTGC |
| 4459 | mamo-L | *BGIBMGA009138* | down | 1097 | 1111 | + | 10.6452 | 8.79E-05 | 1 | TCGGCTGAGTTTTCT |
| 4460 | mamo-L | *BGIBMGA009140* | down | 280 | 294 | + | 14.7258 | 5.83E-07 | 1 | TAGGCTTGCGTTGCC |
| 4461 | mamo-L | *BGIBMGA009140* | down | 1218 | 1232 | - | 13.3871 | 4.90E-06 | 1 | TAGGCTTAATATGGT |
| 4462 | mamo-L | *BGIBMGA009140* | down | 5 | 19 | + | 11.0565 | 6.18E-05 | 1 | AAGGCGGGCGGTTGT |
| 4463 | mamo-L | *BGIBMGA009141* | up | 1410 | 1424 | - | 10.7097 | 8.33E-05 | 1 | TAGGCCTACAGCGCC |
| 4464 | mamo-L | *BGIBMGA009148* | down | 166 | 180 | - | 10.9194 | 6.98E-05 | 1 | TAGGCTACTTTTTTT |
| 4465 | mamo-L | *BGIBMGA009153* | up | 1454 | 1468 | - | 12.129 | 2.19E-05 | 1 | TAGGCCGTCGGTTTC |
| 4466 | mamo-L | *BGIBMGA009154* | up | 131 | 145 | + | 13.0726 | 7.34E-06 | 1 | TAGGCCGATTTTGCA |
| 4467 | mamo-L | *BGIBMGA009155* | up | 1293 | 1307 | + | 13.2177 | 6.14E-06 | 1 | TAGGCTGAGTTTAAT |
| 4468 | mamo-L | *BGIBMGA009157* | down | 122 | 136 | + | 10.9113 | 7.02E-05 | 1 | TAGGCCGTATTCATT |
| 4469 | mamo-L | *BGIBMGA009163* | down | 167 | 181 | - | 10.5726 | 9.33E-05 | 1 | TAGGCTAATATTTAT |
| 4470 | mamo-L | *BGIBMGA009163* | up | 1932 | 1946 | + | 10.5242 | 9.70E-05 | 1 | TAGGCAGCGGTTTGC |
| 4471 | mamo-L | *BGIBMGA009165* | up | 573 | 587 | + | 12.879 | 9.26E-06 | 1 | TAGGCGGAAATTGCT |
| 4472 | mamo-L | *BGIBMGA009166* | down | 1542 | 1556 | + | 13.3145 | 5.41E-06 | 1 | TAGGCTTAGATGGGT |
| 4473 | mamo-L | *BGIBMGA009172* | up | 935 | 949 | - | 12.9516 | 8.46E-06 | 1 | TAGGCTGACGATCGT |
| 4474 | mamo-L | *BGIBMGA009175* | up | 1232 | 1246 | + | 10.5726 | 9.33E-05 | 1 | TAGGCTAATATTTAT |
| 4475 | mamo-L | *BGIBMGA009176* | down | 1096 | 1110 | - | 10.6855 | 8.51E-05 | 1 | TAGGCGACCGTCGTT |
| 4476 | mamo-L | *BGIBMGA009178* | up | 617 | 631 | + | 11.7903 | 3.08E-05 | 1 | TAGGCCATCGTTTAT |
| 4477 | mamo-L | *BGIBMGA009180* | down | 353 | 367 | + | 11.7823 | 3.11E-05 | 1 | TAGGCGTGGATCGAT |
| 4478 | mamo-L | *BGIBMGA009185* | up | 1770 | 1784 | + | 12.2823 | 1.85E-05 | 1 | GAGGCTGCGTTTGCT |
| 4479 | mamo-L | *BGIBMGA009185* | down | 42 | 56 | + | 11.2661 | 5.10E-05 | 1 | GAGGCTGGCGTGGTC |
| 4480 | mamo-L | *BGIBMGA009192* | down | 1329 | 1343 | - | 10.5 | 9.90E-05 | 1 | GAGGCCCAGGTCGTC |
| 4481 | mamo-L | *BGIBMGA009194* | down | 1571 | 1585 | + | 11.2823 | 5.03E-05 | 1 | TAGGCTCGCGGCGTC |
| 4482 | mamo-L | *BGIBMGA009197* | down | 648 | 662 | - | 11.5403 | 3.93E-05 | 1 | TAGGCGGAGTTGCCT |
| 4483 | mamo-L | *BGIBMGA009198* | up | 361 | 375 | - | 11.1694 | 5.59E-05 | 1 | TAGGCCACTTTTGGC |
| 4484 | mamo-L | *BGIBMGA009199* | up | 1538 | 1552 | + | 12.5726 | 1.33E-05 | 1 | TAGGCCCGGATTGCC |
| 4485 | mamo-L | *BGIBMGA009201* | up | 878 | 892 | + | 12.2984 | 1.81E-05 | 1 | AAGGCTTGCGATGTT |
| 4486 | mamo-L | *BGIBMGA009202* | up | 1143 | 1157 | + | 11.8145 | 3.01E-05 | 1 | TAGGCTTGTTCTGGC |
| 4487 | mamo-L | *BGIBMGA009202* | down | 1645 | 1659 | + | 11.5403 | 3.93E-05 | 1 | TAGGCGGAGTTGCCT |
| 4488 | mamo-L | *BGIBMGA009203* | down | 835 | 849 | - | 11.6613 | 3.50E-05 | 1 | AAGGCTTTTTTTTTT |
| 4489 | mamo-L | *BGIBMGA009205* | up | 241 | 255 | + | 11.379 | 4.58E-05 | 1 | TAGGCGAAGTTCGAT |
| 4490 | mamo-L | *BGIBMGA009205* | up | 1611 | 1625 | + | 10.6371 | 8.85E-05 | 1 | GAGGCTGGCGATTAT |
| 4491 | mamo-L | *BGIBMGA009206* | down | 790 | 804 | + | 11.1774 | 5.54E-05 | 1 | TAGGCTTTTTTTTTG |
| 4492 | mamo-L | *BGIBMGA009207* | down | 172 | 186 | + | 10.5726 | 9.33E-05 | 1 | TAGGCTAATATTTAT |
| 4493 | mamo-L | *BGIBMGA009212* | down | 458 | 472 | + | 10.8629 | 7.33E-05 | 1 | TAGGCAGTTTTTATT |
| 4494 | mamo-L | *BGIBMGA009216* | up | 1263 | 1277 | - | 10.5323 | 9.63E-05 | 1 | AAGGCCTTCTGTTCT |
| 4495 | mamo-L | *BGIBMGA009221* | down | 1761 | 1775 | + | 11.5645 | 3.84E-05 | 1 | AAGGCCTAGTTTGGA |
| 4496 | mamo-L | *BGIBMGA009223* | down | 1890 | 1904 | + | 12.5887 | 1.30E-05 | 1 | TAGGCCAAATTTGCT |
| 4497 | mamo-L | *BGIBMGA009227* | down | 1724 | 1738 | + | 12.9435 | 8.54E-06 | 1 | TAGGCTTTTTTTTAT |
| 4498 | mamo-L | *BGIBMGA009228* | down | 1807 | 1821 | - | 11.371 | 4.62E-05 | 1 | AAGGCTTGCGTCGTC |
| 4499 | mamo-L | *BGIBMGA009228* | down | 1750 | 1764 | - | 11.0323 | 6.32E-05 | 1 | TAGGCTGTGGCTCCT |
| 4500 | mamo-L | *BGIBMGA009230* | down | 902 | 916 | + | 12.2984 | 1.81E-05 | 1 | AAGGCTTATTTTTGT |
| 4501 | mamo-L | *BGIBMGA009231* | up | 307 | 321 | + | 12.2984 | 1.81E-05 | 1 | AAGGCTTATTTTTGT |

| | | | | | | | | | | |
|---|---|---|---|---|---|---|---|---|---|---|
| 4502 | mamo-L | *BGIBMGA009236* | up | 48 | 62 | - | 11.3548 | 4.69E-05 | 1 | TAGGCTGTCTATTGA |
| 4503 | mamo-L | *BGIBMGA009241* | up | 293 | 307 | - | 11.9677 | 2.59E-05 | 1 | TAGGCTAAGAGTGGT |
| 4504 | mamo-L | *BGIBMGA009245* | up | 772 | 786 | - | 13.1935 | 6.33E-06 | 1 | TAGGCTTTGATTTCT |
| 4505 | mamo-L | *BGIBMGA009245* | up | 1201 | 1215 | + | 12.0242 | 2.44E-05 | 1 | TAGGCTAAATTTTGT |
| 4506 | mamo-L | *BGIBMGA009246* | down | 300 | 314 | - | 11.121 | 5.84E-05 | 1 | TAGGCCGACGAGCGT |
| 4507 | mamo-L | *BGIBMGA009256* | down | 568 | 582 | - | 10.6532 | 8.73E-05 | 1 | TAGGCCGGTTTATTT |
| 4508 | mamo-L | *BGIBMGA009259* | up | 1385 | 1399 | - | 11.6452 | 3.56E-05 | 1 | TAGGCGTAGTATTCC |
| 4509 | mamo-L | *BGIBMGA009261* | up | 1568 | 1582 | - | 12.9919 | 8.10E-06 | 1 | TAGGCCGGTTTTGCA |
| 4510 | mamo-L | *BGIBMGA009262* | down | 679 | 693 | - | 11.4758 | 4.18E-05 | 1 | TAGGCAGACTTTGTA |
| 4511 | mamo-L | *BGIBMGA009264* | up | 1689 | 1703 | - | 10.5323 | 9.63E-05 | 1 | TAGGCCAACGTCGCA |
| 4512 | mamo-L | *BGIBMGA009267* | down | 18 | 32 | - | 11.371 | 4.62E-05 | 1 | TGGGCTGAGTGTGGT |
| 4513 | mamo-L | *BGIBMGA009270* | down | 377 | 391 | - | 11 | 6.50E-05 | 1 | TAGGCCTAGCTTACT |
| 4514 | mamo-L | *BGIBMGA009272* | up | 286 | 300 | + | 11.25 | 5.18E-05 | 1 | TAGGCTTTGGATTGA |
| 4515 | mamo-L | *BGIBMGA009277* | down | 1087 | 1101 | + | 11.371 | 4.62E-05 | 1 | AAGGCTTGCGTCGTC |
| 4516 | mamo-L | *BGIBMGA009277* | down | 1144 | 1158 | + | 11.0323 | 6.32E-05 | 1 | TAGGCTGTGGCTCCT |
| 4517 | mamo-L | *BGIBMGA009278* | down | 1354 | 1368 | - | 11.4758 | 4.18E-05 | 1 | AAGGCCATCTTTGTT |
| 4518 | mamo-L | *BGIBMGA009278* | up | 1276 | 1290 | - | 10.9194 | 6.98E-05 | 1 | TAGGCATTGATTTTT |
| 4519 | mamo-L | *BGIBMGA009279* | down | 875 | 889 | - | 14.2258 | 1.41E-06 | 1 | TAGGCTTATGTTTTT |
| 4520 | mamo-L | *BGIBMGA009279* | down | 318 | 332 | + | 10.5726 | 9.33E-05 | 1 | TAGGCTAATATTTAT |
| 4521 | mamo-L | *BGIBMGA009285* | up | 313 | 327 | - | 11.5323 | 3.96E-05 | 1 | TAGGCGTGAGGTGAT |
| 4522 | mamo-L | *BGIBMGA009287* | up | 156 | 170 | - | 13.4919 | 4.25E-06 | 1 | TAGGCCTAGGTTGTA |
| 4523 | mamo-L | *BGIBMGA009288* | down | 435 | 449 | - | 10.9516 | 6.78E-05 | 1 | AAGGCTTTTTTTGAC |
| 4524 | mamo-L | *BGIBMGA009288* | up | 214 | 228 | + | 10.5645 | 9.40E-05 | 1 | TGGGCCGTCTATGTT |
| 4525 | mamo-L | *BGIBMGA009292* | down | 863 | 877 | - | 10.6129 | 9.03E-05 | 1 | GAGGCCTTTTTTTAT |
| 4526 | mamo-L | *BGIBMGA009295* | down | 157 | 171 | + | 12.2016 | 2.03E-05 | 1 | TAGGCATGCTTTTTT |
| 4527 | mamo-L | *BGIBMGA009299* | down | 1848 | 1862 | - | 13.129 | 6.83E-06 | 1 | TAGGCCTACTTCTTT |
| 4528 | mamo-L | *BGIBMGA009306* | up | 1901 | 1915 | + | 10.5726 | 9.33E-05 | 1 | TAGGCTAATATTTAT |
| 4529 | mamo-L | *BGIBMGA009312* | up | 1471 | 1485 | + | 11 | 6.50E-05 | 1 | TAGGCTTGCGGTTCA |
| 4530 | mamo-L | *BGIBMGA009314* | up | 718 | 732 | - | 11.2177 | 5.34E-05 | 1 | TAGGCCGGTATTTAC |
| 4531 | mamo-L | *BGIBMGA009316* | down | 48 | 62 | + | 10.5726 | 9.33E-05 | 1 | TAGGCTAATATTTAT |
| 4532 | mamo-L | *BGIBMGA009319* | down | 656 | 670 | - | 11.3548 | 4.69E-05 | 1 | AAGGCTGATGATGAT |
| 4533 | mamo-L | *BGIBMGA009319* | down | 173 | 187 | + | 10.5726 | 9.33E-05 | 1 | TAGGCTAATATTTAT |
| 4534 | mamo-L | *BGIBMGA009320* | up | 674 | 688 | - | 11.7339 | 3.26E-05 | 1 | TAGGCTAGCTTCGCC |
| 4535 | mamo-L | *BGIBMGA009322* | down | 1518 | 1532 | + | 12.3468 | 1.72E-05 | 1 | TAGGCGGCGTGTGCT |
| 4536 | mamo-L | *BGIBMGA009325* | down | 1109 | 1123 | - | 10.9194 | 6.98E-05 | 1 | TAGGCTACTTTTTTT |
| 4537 | mamo-L | *BGIBMGA009326* | down | 1771 | 1785 | + | 12.0645 | 2.34E-05 | 1 | TAGGCGGGCTTCGGA |
| 4538 | mamo-L | *BGIBMGA009326* | up | 1317 | 1331 | - | 10.9677 | 6.68E-05 | 1 | TAGGCCGTGTTATTT |
| 4539 | mamo-L | *BGIBMGA009326* | down | 1618 | 1632 | - | 10.8629 | 7.33E-05 | 1 | TAGGCCCCCTTCGTC |
| 4540 | mamo-L | *BGIBMGA009327* | up | 1149 | 1163 | + | 12.1452 | 2.15E-05 | 1 | TAGGCAGGTGGTGGT |
| 4541 | mamo-L | *BGIBMGA009327* | down | 376 | 390 | + | 11.3387 | 4.77E-05 | 1 | GAGGCTTACATTTTT |
| 4542 | mamo-L | *BGIBMGA009327* | down | 1212 | 1226 | - | 10.4919 | 9.96E-05 | 1 | TAGGCTGTGTCTGTG |
| 4543 | mamo-L | *BGIBMGA009328* | down | 1176 | 1190 | - | 10.75 | 8.06E-05 | 1 | TGGGCTGACTTGGTT |
| 4544 | mamo-L | *BGIBMGA009328* | up | 423 | 437 | - | 10.4919 | 9.96E-05 | 1 | TAGGCTGTGTCTGTG |
| 4545 | mamo-L | *BGIBMGA009333* | up | 920 | 934 | - | 12.8065 | 1.01E-05 | 1 | TAGGCGTGGTGTTGT |
| 4546 | mamo-L | *BGIBMGA009335* | down | 644 | 658 | - | 11.6855 | 3.42E-05 | 1 | TAGGCTTACGTTTAA |
| 4547 | mamo-L | *BGIBMGA009337* | down | 1203 | 1217 | + | 11.9597 | 2.61E-05 | 1 | AAGGCTGTTTTTTTT |
| 4548 | mamo-L | *BGIBMGA009337* | up | 959 | 973 | + | 11.8871 | 2.81E-05 | 1 | TAGGCGAGTGGTGGT |
| 4549 | mamo-L | *BGIBMGA009337* | up | 1643 | 1657 | + | 11.129 | 5.79E-05 | 1 | AAGGCGAATTTTGGT |
| 4550 | mamo-L | *BGIBMGA009340* | down | 1240 | 1254 | - | 11.4758 | 4.18E-05 | 1 | AAGGCTGAAATTGTT |
| 4551 | mamo-L | *BGIBMGA009341* | down | 4 | 18 | + | 11.0081 | 6.45E-05 | 1 | GAGGCTGCGGGTGCT |
| 4552 | mamo-L | *BGIBMGA009352* | down | 694 | 708 | + | 10.5887 | 9.21E-05 | 1 | TAGGCCTACTGAGCT |
| 4553 | mamo-L | *BGIBMGA009353* | down | 1916 | 1930 | + | 13.7742 | 2.79E-06 | 1 | TAGGCGGGTGGTGGT |
| 4554 | mamo-L | *BGIBMGA009354* | up | 1922 | 1936 | + | 13.7742 | 2.79E-06 | 1 | TAGGCGGGTGGTGGT |
| 4555 | mamo-L | *BGIBMGA009359* | up | 1263 | 1277 | - | 12.6532 | 1.20E-05 | 1 | TAGGCTGGCGTCTCC |
| 4556 | mamo-L | *BGIBMGA009360* | down | 1483 | 1497 | - | 10.9597 | 6.73E-05 | 1 | AAGGCTGTGTTGGCC |
| 4557 | mamo-L | *BGIBMGA009362* | down | 315 | 329 | - | 10.8145 | 7.63E-05 | 1 | TAGGCCTCATTTGAC |
| 4558 | mamo-L | *BGIBMGA009363* | down | 1794 | 1808 | - | 11.3548 | 4.69E-05 | 1 | AAGGCTGATGATGAT |
| 4559 | mamo-L | *BGIBMGA009364* | down | 1390 | 1404 | + | 11.4113 | 4.45E-05 | 1 | TAGGCTACCTTTTTT |
| 4560 | mamo-L | *BGIBMGA009364* | down | 1727 | 1741 | + | 10.7984 | 7.74E-05 | 1 | TAGGCTTAGGTCAAC |
| 4561 | mamo-L | *BGIBMGA009365* | down | 1111 | 1125 | - | 11.6048 | 3.70E-05 | 1 | TAGGCGGGGTGTGGG |
| 4562 | mamo-L | *BGIBMGA009368* | up | 1938 | 1952 | + | 11.1452 | 5.70E-05 | 1 | AAGGCGATGGTTGCT |
| 4563 | mamo-L | *BGIBMGA009368* | down | 1132 | 1146 | - | 10.7823 | 7.84E-05 | 1 | TAGGCCTGAGGTTTC |
| 4564 | mamo-L | *BGIBMGA009369* | down | 258 | 272 | + | 10.75 | 8.06E-05 | 1 | TGGGCTGACTTGGTT |
| 4565 | mamo-L | *BGIBMGA009370* | down | 1955 | 1969 | - | 11.75 | 3.21E-05 | 1 | TAGGCCGGTTGTTTC |
| 4566 | mamo-L | *BGIBMGA009371* | down | 14 | 28 | - | 12.3468 | 1.72E-05 | 1 | TAGGCGGCGTGTGCT |
| 4567 | mamo-L | *BGIBMGA009379* | down | 1871 | 1885 | + | 12.5726 | 1.33E-05 | 1 | AAGGCCTTTTTTGTT |
| 4568 | mamo-L | *BGIBMGA009382* | down | 1702 | 1716 | + | 11.1129 | 5.88E-05 | 1 | TAGGCTGTGATAGCT |
| 4569 | mamo-L | *BGIBMGA009385* | down | 1851 | 1865 | + | 11.0242 | 6.36E-05 | 1 | TAGGCCTTGCTTAGT |
| 4570 | mamo-L | *BGIBMGA009388* | down | 669 | 683 | - | 11.1613 | 5.63E-05 | 1 | TAGGCAGATGATGAT |
| 4571 | mamo-L | *BGIBMGA009391* | down | 1487 | 1501 | - | 12.3226 | 1.77E-05 | 1 | TAGGCTGTCAGTGAT |
| 4572 | mamo-L | *BGIBMGA009391* | down | 1797 | 1811 | + | 12.0081 | 2.48E-05 | 1 | TAGGCCGATTCTGGC |
| 4573 | mamo-L | *BGIBMGA009396* | down | 146 | 160 | + | 10.7016 | 8.39E-05 | 1 | TAGGCGGTGGTGAAT |
| 4574 | mamo-L | *BGIBMGA009397* | up | 567 | 581 | - | 13.4677 | 4.39E-06 | 1 | TAGGCTGATGATGAT |
| 4575 | mamo-L | *BGIBMGA009399* | down | 1473 | 1487 | - | 12.4194 | 1.58E-05 | 1 | TAGGCCGTAATTGGC |
| 4576 | mamo-L | *BGIBMGA009399* | up | 732 | 746 | + | 12.1048 | 2.24E-05 | 1 | TAGGCTTTGTATATT |
| 4577 | mamo-L | *BGIBMGA009400* | down | 178 | 192 | + | 11.0484 | 6.23E-05 | 1 | AAGGCTTATTTTACT |
| 4578 | mamo-L | *BGIBMGA009401* | up | 1705 | 1719 | + | 11.7984 | 3.06E-05 | 1 | TAGGCTTCTGTCGGC |
| 4579 | mamo-L | *BGIBMGA009401* | up | 1387 | 1401 | - | 10.6371 | 8.85E-05 | 1 | TAGGCTGTTCTTATT |
| 4580 | mamo-L | *BGIBMGA009402* | down | 482 | 496 | + | 10.7581 | 8.00E-05 | 1 | TAGGCCATCTTTAAT |
| 4581 | mamo-L | *BGIBMGA009405* | up | 1295 | 1309 | - | 10.5726 | 9.33E-05 | 1 | TAGGCTAATATTTAT |
| 4582 | mamo-L | *BGIBMGA009406* | down | 960 | 974 | - | 10.8306 | 7.52E-05 | 1 | AAGGCACGCGTTGGT |
| 4583 | mamo-L | *BGIBMGA009406* | up | 1425 | 1439 | - | 10.4919 | 9.96E-05 | 1 | TAGGCTGTGTCTGTG |
| 4584 | mamo-L | *BGIBMGA009407* | up | 21 | 35 | + | 11.6613 | 3.50E-05 | 1 | TAGGCTAGGTATTTT |
| 4585 | mamo-L | *BGIBMGA009411* | down | 253 | 267 | - | 12.0242 | 2.44E-05 | 1 | TAGGCCTCCTTTGGA |
| 4586 | mamo-L | *BGIBMGA009411* | down | 825 | 839 | + | 11.7177 | 3.32E-05 | 1 | TAGGCCTTGAATTTT |
| 4587 | mamo-L | *BGIBMGA009413* | down | 124 | 138 | + | 12.1613 | 2.11E-05 | 1 | TAGGCTAACATTGAT |
| 4588 | mamo-L | *BGIBMGA009414* | up | 404 | 418 | + | 12.7258 | 1.11E-05 | 1 | TAGGCCGGTGACGCT |
| 4589 | mamo-L | *BGIBMGA009419* | down | 1141 | 1155 | - | 11.9032 | 2.76E-05 | 1 | TAGGCTTTAGATGTC |
| 4590 | mamo-L | *BGIBMGA009420* | up | 1134 | 1148 | - | 11.9032 | 2.76E-05 | 1 | TAGGCTTTAGATGTC |
| 4591 | mamo-L | *BGIBMGA009422* | down | 354 | 368 | - | 10.5726 | 9.33E-05 | 1 | TAGGCTAATATTTAT |
| 4592 | mamo-L | *BGIBMGA009424* | down | 426 | 440 | - | 10.621 | 8.97E-05 | 1 | TGGGCTTTTTTTGGC |
| 4593 | mamo-L | *BGIBMGA009430* | down | 632 | 646 | - | 11.1855 | 5.50E-05 | 1 | TAGGCCTTGTGGGCC |

| | | | | | | | | | | |
|---|---|---|---|---|---|---|---|---|---|---|
| 4594 | mamo-L | *BGIBMGA009431* | up | 130 | 144 | + | 13.4677 | 4.39E-06 | 1 | TAGGCTGATGATGAT |
| 4595 | mamo-L | *BGIBMGA009437* | up | 1720 | 1734 | + | 12.4919 | 1.46E-05 | 1 | TAGGCGGGTGTCTTT |
| 4596 | mamo-L | *BGIBMGA009441* | up | 607 | 621 | - | 12.3548 | 1.70E-05 | 1 | TAGGCCGTTTTTGCG |
| 4597 | mamo-L | *BGIBMGA009441* | down | 1448 | 1462 | + | 11.1935 | 5.45E-05 | 1 | TAGGCGCATGAGGGT |
| 4598 | mamo-L | *BGIBMGA009443* | up | 508 | 522 | - | 12.2581 | 1.90E-05 | 1 | TAGGCGCTTATTGCT |
| 4599 | mamo-L | *BGIBMGA009445* | up | 943 | 957 | + | 12.0323 | 2.42E-05 | 1 | TAGGCCGATTTCATT |
| 4600 | mamo-L | *BGIBMGA009447* | up | 1085 | 1099 | - | 13.0806 | 7.24E-06 | 1 | TAGGCCTATTTTTAT |
| 4601 | mamo-L | *BGIBMGA009448* | up | 1578 | 1592 | - | 11.0403 | 6.28E-05 | 1 | TAGGCTTTTTGGTGT |
| 4602 | mamo-L | *BGIBMGA009448* | down | 1858 | 1872 | - | 10.9435 | 6.83E-05 | 1 | TAGGCCTTAATTCGT |
| 4603 | mamo-L | *BGIBMGA009453* | up | 910 | 924 | - | 11.7984 | 3.06E-05 | 1 | TAGGCTGATTACTTT |
| 4604 | mamo-L | *BGIBMGA009456* | down | 457 | 471 | + | 12.5645 | 1.34E-05 | 1 | GAGGCTTTCGTTTGT |
| 4605 | mamo-L | *BGIBMGA009456* | up | 294 | 308 | + | 12.0081 | 2.48E-05 | 1 | TAGGCCTAATTTGTA |
| 4606 | mamo-L | *BGIBMGA009462* | down | 712 | 726 | + | 11.0806 | 6.05E-05 | 1 | TAGGCCGGCCATTGT |
| 4607 | mamo-L | *BGIBMGA009466* | up | 764 | 778 | - | 11.3548 | 4.69E-05 | 1 | AAGGCTGATGATGAT |
| 4608 | mamo-L | *BGIBMGA009467* | up | 1201 | 1215 | + | 11.4758 | 4.18E-05 | 1 | TAGGCTACGTTTTTT |
| 4609 | mamo-L | *BGIBMGA009469* | down | 1783 | 1797 | + | 10.6774 | 8.56E-05 | 1 | TAGGCGGTCGGTGAA |
| 4610 | mamo-L | *BGIBMGA009469* | up | 492 | 506 | + | 10.5403 | 9.58E-05 | 1 | GAGGCTGACGACGAT |
| 4611 | mamo-L | *BGIBMGA009470* | up | 1931 | 1945 | + | 10.6774 | 8.56E-05 | 1 | TAGGCGGTCGGTGAA |
| 4612 | mamo-L | *BGIBMGA009476* | up | 83 | 97 | - | 10.8548 | 7.38E-05 | 1 | TAGGCCTGCAGCTGT |
| 4613 | mamo-L | *BGIBMGA009484* | up | 1008 | 1022 | - | 13.6855 | 3.18E-06 | 1 | TAGGCCATCGTTGCT |
| 4614 | mamo-L | *BGIBMGA009484* | up | 1592 | 1606 | - | 12.0565 | 2.36E-05 | 1 | TAGGCTGTGGTGCTT |
| 4615 | mamo-L | *BGIBMGA009485* | up | 523 | 537 | - | 13.6855 | 3.18E-06 | 1 | TAGGCCATCGTTGCT |
| 4616 | mamo-L | *BGIBMGA009485* | up | 1107 | 1121 | - | 12.0565 | 2.36E-05 | 1 | TAGGCTGTGGTGCTT |
| 4617 | mamo-L | *BGIBMGA009488* | up | 1184 | 1198 | - | 11.6129 | 3.66E-05 | 1 | TAGGCTCTCGGGGCT |
| 4618 | mamo-L | *BGIBMGA009490* | down | 559 | 573 | - | 13.129 | 6.83E-06 | 1 | AAGGCCTGTTTTGGT |
| 4619 | mamo-L | *BGIBMGA009491* | up | 423 | 437 | - | 13.129 | 6.83E-06 | 1 | AAGGCCTGTTTTGGT |
| 4620 | mamo-L | *BGIBMGA009492* | down | 1081 | 1095 | + | 12.0161 | 2.46E-05 | 1 | TAGGCCCCCTTTTTT |
| 4621 | mamo-L | *BGIBMGA009496* | up | 1566 | 1580 | + | 11.3548 | 4.69E-05 | 1 | AAGGCTGATGATGAT |
| 4622 | mamo-L | *BGIBMGA009497* | down | 1503 | 1517 | - | 12.7339 | 1.10E-05 | 1 | GAGGCTGGTGTTGGC |
| 4623 | mamo-L | *BGIBMGA009497* | down | 741 | 755 | - | 10.5726 | 9.33E-05 | 1 | TAGGCTAATATTTAT |
| 4624 | mamo-L | *BGIBMGA009497* | down | 984 | 998 | + | 10.5726 | 9.33E-05 | 1 | TAGGCTAATATTTAT |
| 4625 | mamo-L | *BGIBMGA009505* | up | 1283 | 1297 | - | 11.2339 | 5.26E-05 | 1 | TAGGCTGATTTATGT |
| 4626 | mamo-L | *BGIBMGA009506* | down | 1446 | 1460 | - | 12.1129 | 2.22E-05 | 1 | TAGGCTGAGCGTGTT |
| 4627 | mamo-L | *BGIBMGA009506* | up | 717 | 731 | + | 10.7984 | 7.74E-05 | 1 | TAGGCGGCTTTGTGT |
| 4628 | mamo-L | *BGIBMGA009506* | up | 840 | 854 | - | 10.5403 | 9.58E-05 | 1 | TAGGCTATGTTTTTA |
| 4629 | mamo-L | *BGIBMGA009508* | up | 1283 | 1297 | - | 12.0081 | 2.48E-05 | 1 | TAGGCCTAATTTGTA |
| 4630 | mamo-L | *BGIBMGA009513* | up | 229 | 243 | + | 11.7984 | 3.06E-05 | 1 | TAGGCCTGGTGCGCC |
| 4631 | mamo-L | *BGIBMGA009518* | down | 1459 | 1473 | - | 10.5 | 9.90E-05 | 1 | GAGGCCCAGGTCGTC |
| 4632 | mamo-L | *BGIBMGA009521* | up | 1739 | 1753 | - | 11.4839 | 4.15E-05 | 1 | TAGGCTTTTATTGGG |
| 4633 | mamo-L | *BGIBMGA009525* | down | 1401 | 1415 | - | 10.5806 | 9.27E-05 | 1 | TAGGCGGGTGACAGT |
| 4634 | mamo-L | *BGIBMGA009525* | up | 501 | 515 | + | 10.5484 | 9.52E-05 | 1 | TAGGCGTACAATACT |
| 4635 | mamo-L | *BGIBMGA009529* | down | 741 | 755 | - | 11.0565 | 6.18E-05 | 1 | TAGGCTTGAAATGAT |
| 4636 | mamo-L | *BGIBMGA009530* | down | 1208 | 1222 | - | 13.4677 | 4.39E-06 | 1 | TAGGCTGATGATGAT |
| 4637 | mamo-L | *BGIBMGA009532* | up | 1444 | 1458 | + | 11.2581 | 5.14E-05 | 1 | TAGCCCTTGGTTGTT |
| 4638 | mamo-L | *BGIBMGA009532* | down | 445 | 459 | + | 11.0968 | 5.97E-05 | 1 | TAGGCTCTTTTTATC |
| 4639 | mamo-L | *BGIBMGA009532* | up | 1134 | 1148 | - | 10.5726 | 9.33E-05 | 1 | TAGGCTAATATTTAT |
| 4640 | mamo-L | *BGIBMGA009536* | up | 1851 | 1865 | + | 13.2823 | 5.66E-06 | 1 | TAGGCCGATGATGAT |
| 4641 | mamo-L | *BGIBMGA009537* | up | 1281 | 1295 | + | 11.4032 | 4.49E-05 | 1 | TAGGCTTGAATTGAC |
| 4642 | mamo-L | *BGIBMGA009538* | down | 1436 | 1450 | + | 13.0565 | 7.51E-06 | 1 | TAGGCCTGGAATGTT |
| 4643 | mamo-L | *BGIBMGA009539* | down | 1083 | 1097 | + | 12.9919 | 8.10E-06 | 1 | TAGGCCGGTTTTGCA |
| 4644 | mamo-L | *BGIBMGA009543* | down | 1440 | 1454 | + | 13.4839 | 4.31E-06 | 1 | TAGGCTCGTTTTGAT |
| 4645 | mamo-L | *BGIBMGA009546* | up | 1457 | 1471 | + | 11.0242 | 6.36E-05 | 1 | TAGGCTGTAGTCGTA |
| 4646 | mamo-L | *BGIBMGA009550* | down | 505 | 519 | - | 11.2742 | 5.06E-05 | 1 | AAGGCTTTCGTCGAT |
| 4647 | mamo-L | *BGIBMGA009551* | up | 1576 | 1590 | + | 10.75 | 8.06E-05 | 1 | GAGGCCGATTGTGCC |
| 4648 | mamo-L | *BGIBMGA009552* | down | 1574 | 1588 | - | 10.8387 | 7.47E-05 | 1 | TAGGCCGGTTTTCCA |
| 4649 | mamo-L | *BGIBMGA009554* | down | 449 | 463 | - | 11.0403 | 6.28E-05 | 1 | TAGGCTTTTTGGTGT |
| 4650 | mamo-L | *BGIBMGA009554* | down | 1844 | 1858 | - | 10.5726 | 9.33E-05 | 1 | TAGGCTAATATTTAT |
| 4651 | mamo-L | *BGIBMGA009555* | down | 1306 | 1320 | - | 12.5242 | 1.40E-05 | 1 | TAGGCTTACTTCGGA |
| 4652 | mamo-L | *BGIBMGA009558* | up | 1552 | 1566 | - | 11.2016 | 5.41E-05 | 1 | AAGGCTAAGGTTGTC |
| 4653 | mamo-L | *BGIBMGA009559* | up | 1981 | 1995 | + | 11.3871 | 4.55E-05 | 1 | TAGGCCTAGCTGGTT |
| 4654 | mamo-L | *BGIBMGA009560* | down | 81 | 95 | + | 10.9597 | 6.73E-05 | 1 | GAGGCCGATATTTTT |
| 4655 | mamo-L | *BGIBMGA009560* | down | 264 | 278 | - | 10.5968 | 9.15E-05 | 1 | GAGGCTAGTTTTTGT |
| 4656 | mamo-L | *BGIBMGA009561* | up | 1192 | 1206 | + | 11.371 | 4.62E-05 | 1 | AAGGCGTTCGTGGGT |
| 4657 | mamo-L | *BGIBMGA009562* | up | 911 | 925 | + | 13.6613 | 3.32E-06 | 1 | TAGGCTTACGATGAT |
| 4658 | mamo-L | *BGIBMGA009569* | down | 138 | 152 | + | 10.9194 | 6.98E-05 | 1 | TAGGCTACTTTTTTT |
| 4659 | mamo-L | *BGIBMGA009571* | down | 257 | 271 | + | 10.5081 | 9.83E-05 | 1 | TGGGCTGACATTTGT |
| 4660 | mamo-L | *BGIBMGA009572* | down | 42 | 56 | - | 12.8226 | 9.86E-06 | 1 | AAGGCGGTTGTTGGT |
| 4661 | mamo-L | *BGIBMGA009572* | up | 218 | 232 | + | 10.5081 | 9.83E-05 | 1 | TGGGCTGACATTTGT |
| 4662 | mamo-L | *BGIBMGA009573* | down | 1463 | 1477 | + | 12.0645 | 2.34E-05 | 1 | TAGGCGGGCTTCGGA |
| 4663 | mamo-L | *BGIBMGA009574* | up | 122 | 136 | + | 12.0645 | 2.34E-05 | 1 | TAGGCGGGCTTCGGA |
| 4664 | mamo-L | *BGIBMGA009575* | down | 1833 | 1847 | + | 10.4919 | 9.96E-05 | 1 | TAGGCTTAGAATAAT |
| 4665 | mamo-L | *BGIBMGA009576* | up | 1703 | 1717 | - | 10.5323 | 9.63E-05 | 1 | TAGGCTGACGGCTAC |
| 4666 | mamo-L | *BGIBMGA009577* | up | 398 | 412 | - | 12.7581 | 1.07E-05 | 1 | TAGGCTTTTATTGCC |
| 4667 | mamo-L | *BGIBMGA009580* | down | 1225 | 1239 | + | 12.0645 | 2.34E-05 | 1 | TAGGCGGGCTTCGGA |
| 4668 | mamo-L | *BGIBMGA009580* | up | 983 | 997 | - | 11.3145 | 4.87E-05 | 1 | AAGGCTGTCATTTCT |
| 4669 | mamo-L | *BGIBMGA009580* | up | 1468 | 1482 | - | 10.5081 | 9.83E-05 | 1 | TAGGCGGGGTGTTGG |
| 4670 | mamo-L | *BGIBMGA009583* | up | 187 | 201 | + | 10.5726 | 9.33E-05 | 1 | TAGGCTAATATTTAT |
| 4671 | mamo-L | *BGIBMGA009585* | up | 737 | 751 | - | 10.5726 | 9.33E-05 | 1 | TAGGCTAATATTTAT |
| 4672 | mamo-L | *BGIBMGA009591* | down | 1149 | 1163 | + | 10.9194 | 6.98E-05 | 1 | TAGGCTGCATGTTGT |
| 4673 | mamo-L | *BGIBMGA009601* | up | 1386 | 1400 | + | 10.9194 | 6.98E-05 | 1 | TAGGCTACTTTTTTT |
| 4674 | mamo-L | *BGIBMGA009603* | up | 449 | 463 | + | 11.4032 | 4.49E-05 | 1 | TAGGCTGTATAGGTT |
| 4675 | mamo-L | *BGIBMGA009605* | down | 1387 | 1401 | - | 11.8145 | 3.01E-05 | 1 | GAGGCGGTGTGTGGT |
| 4676 | mamo-L | *BGIBMGA009609* | down | 1943 | 1957 | + | 12.1935 | 2.05E-05 | 1 | TAGGCCGGTTTTGAA |
| 4677 | mamo-L | *BGIBMGA009609* | down | 206 | 220 | + | 10.5968 | 9.15E-05 | 1 | TCGGCTTTTTTTGTT |
| 4678 | mamo-L | *BGIBMGA009610* | up | 308 | 322 | + | 13.1452 | 6.69E-06 | 1 | TAGGCCGTTTTTCGT |
| 4679 | mamo-L | *BGIBMGA009610* | up | 368 | 382 | + | 12.8145 | 9.96E-06 | 1 | TAGGCCGTCTTTAGC |
| 4680 | mamo-L | *BGIBMGA009611* | up | 1305 | 1319 | - | 13.6774 | 3.24E-06 | 1 | TAGGCCTTTTATGGT |
| 4681 | mamo-L | *BGIBMGA009611* | down | 727 | 741 | - | 11.7742 | 3.14E-05 | 1 | TAGGCCTTCTGTATT |
| 4682 | mamo-L | *BGIBMGA009611* | down | 1252 | 1266 | - | 10.8226 | 7.58E-05 | 1 | TGGGCGGAGTTCGGT |
| 4683 | mamo-L | *BGIBMGA009611* | down | 1473 | 1487 | - | 10.5242 | 9.70E-05 | 1 | TAGGCGTTGTATTAC |
| 4684 | mamo-L | *BGIBMGA009623* | up | 639 | 653 | + | 10.9194 | 6.98E-05 | 1 | TAGGCTTCAATTTGT |
| 4685 | mamo-L | *BGIBMGA009628* | down | 1891 | 1905 | + | 10.5726 | 9.33E-05 | 1 | TAGGCTAATATTTAT |

| | | | | | | | | | | |
|---|---|---|---|---|---|---|---|---|---|---|
| 4686 | mamo-L | *BGIBMGA009633* | up | 1472 | 1486 | - | 10.8226 | 7.58E-05 | 1 | AAGGCTGGTTTGGAT |
| 4687 | mamo-L | *BGIBMGA009634* | up | 200 | 214 | - | 11.0403 | 6.28E-05 | 1 | TAGGCTTTTTGGTGT |
| 4688 | mamo-L | *BGIBMGA009634* | up | 1826 | 1840 | + | 10.9194 | 6.98E-05 | 1 | TAGGCTACTTTTTTT |
| 4689 | mamo-L | *BGIBMGA009639* | up | 1723 | 1737 | + | 13.1048 | 7.05E-06 | 1 | TAGGCTTTAGTTTTT |
| 4690 | mamo-L | *BGIBMGA009640* | up | 336 | 350 | - | 11.25 | 5.18E-05 | 1 | TAGGCCCATTTCAGT |
| 4691 | mamo-L | *BGIBMGA009640* | down | 1004 | 1018 | - | 10.5726 | 9.33E-05 | 1 | TAGGCTAATATTTAT |
| 4692 | mamo-L | *BGIBMGA009640* | down | 1154 | 1168 | + | 10.5726 | 9.33E-05 | 1 | AAGGCTTTGTTGTTT |
| 4693 | mamo-L | *BGIBMGA009642* | up | 971 | 985 | + | 10.5565 | 9.46E-05 | 1 | AAGGCTTTTTTTGTA |
| 4694 | mamo-L | *BGIBMGA009644* | down | 899 | 913 | - | 11.1129 | 5.88E-05 | 1 | TAGGCGCATGGGGGT |
| 4695 | mamo-L | *BGIBMGA009648* | down | 1542 | 1556 | - | 11.8387 | 2.94E-05 | 1 | TAGGCGTGCAATGGC |
| 4696 | mamo-L | *BGIBMGA009649* | up | 1894 | 1908 | - | 11.8387 | 2.94E-05 | 1 | TAGGCGTGCAATGGC |
| 4697 | mamo-L | *BGIBMGA009650* | up | 1304 | 1318 | - | 11.5081 | 4.05E-05 | 1 | TAGGCGGGATTGTGT |
| 4698 | mamo-L | *BGIBMGA009652* | up | 257 | 271 | + | 11.0403 | 6.28E-05 | 1 | TAGGCCAATTGTGTC |
| 4699 | mamo-L | *BGIBMGA009652* | down | 209 | 223 | - | 10.8226 | 7.58E-05 | 1 | AAGGCTTTTATTGAT |
| 4700 | mamo-L | *BGIBMGA009653* | down | 402 | 416 | + | 12.1613 | 2.11E-05 | 1 | TAGGCTTATATTTAT |
| 4701 | mamo-L | *BGIBMGA009656* | down | 165 | 179 | - | 11.2419 | 5.21E-05 | 1 | TAGGCTCTTTTTAAT |
| 4702 | mamo-L | *BGIBMGA009657* | up | 1920 | 1934 | + | 11.3548 | 4.69E-05 | 1 | AAGGCTGATGATGAT |
| 4703 | mamo-L | *BGIBMGA009661* | up | 983 | 997 | + | 10.9194 | 6.98E-05 | 1 | TAGGCTACTTTTTTT |
| 4704 | mamo-L | *BGIBMGA009670* | down | 1895 | 1909 | - | 13.2661 | 5.76E-06 | 1 | TAGGCTGTAATTGTT |
| 4705 | mamo-L | *BGIBMGA009670* | down | 385 | 399 | - | 11.0968 | 5.97E-05 | 1 | TAGGCTCGATATTTT |
| 4706 | mamo-L | *BGIBMGA009676* | down | 133 | 147 | - | 11.129 | 5.79E-05 | 1 | TAGGCTGTGTTTATG |
| 4707 | mamo-L | *BGIBMGA009677* | up | 616 | 630 | - | 10.9194 | 6.98E-05 | 1 | TAGGCTACTTTTTTT |
| 4708 | mamo-L | *BGIBMGA009677* | down | 123 | 137 | + | 10.5726 | 9.33E-05 | 1 | TAGGCTAATATTTAT |
| 4709 | mamo-L | *BGIBMGA009678* | up | 1565 | 1579 | - | 10.7581 | 8.00E-05 | 1 | TAGGCATTCGTGTGT |
| 4710 | mamo-L | *BGIBMGA009679* | down | 120 | 134 | + | 11.4355 | 4.34E-05 | 1 | AAGGCTTTTTATGTT |
| 4711 | mamo-L | *BGIBMGA009681* | down | 504 | 518 | - | 12.3871 | 1.65E-05 | 1 | TAGGCGTCGTTTTTT |
| 4712 | mamo-L | *BGIBMGA009689* | down | 19 | 33 | + | 12.0887 | 2.29E-05 | 1 | TAGGCGTATTTGTGT |
| 4713 | mamo-L | *BGIBMGA009689* | up | 1574 | 1588 | - | 10.9194 | 6.98E-05 | 1 | TAGGCTACTTTTTTT |
| 4714 | mamo-L | *BGIBMGA009694* | down | 1647 | 1661 | - | 11.6532 | 3.53E-05 | 1 | TAGGCTGTCATTTAC |
| 4715 | mamo-L | *BGIBMGA009696* | up | 747 | 761 | - | 11.0887 | 6.01E-05 | 1 | TAGGCTGTGTCTATT |
| 4716 | mamo-L | *BGIBMGA009698* | down | 135 | 149 | + | 10.5323 | 9.63E-05 | 1 | TAGGCTAACGTTTTG |
| 4717 | mamo-L | *BGIBMGA009699* | up | 352 | 366 | + | 10.5323 | 9.63E-05 | 1 | TAGGCTAACGTTTTG |
| 4718 | mamo-L | *BGIBMGA009700* | down | 650 | 664 | - | 11.1129 | 5.88E-05 | 1 | AAGGCTCTGTGTGTT |
| 4719 | mamo-L | *BGIBMGA009700* | up | 432 | 446 | - | 10.9194 | 6.98E-05 | 1 | TAGGCTACTTTTTTT |
| 4720 | mamo-L | *BGIBMGA009701* | up | 1306 | 1320 | - | 11.1129 | 5.88E-05 | 1 | AAGGCTCTGTGTGTT |
| 4721 | mamo-L | *BGIBMGA009702* | up | 682 | 696 | + | 13.121 | 6.92E-06 | 1 | TAGGCTAAATTTGGT |
| 4722 | mamo-L | *BGIBMGA009702* | up | 588 | 602 | + | 11.629 | 3.61E-05 | 1 | AAGGCTTAAGTTTGT |
| 4723 | mamo-L | *BGIBMGA009703* | up | 575 | 589 | - | 12.1452 | 2.15E-05 | 1 | TAGGCGGACTTCGGA |
| 4724 | mamo-L | *BGIBMGA009703* | up | 724 | 738 | + | 10.8306 | 7.52E-05 | 1 | TAGGCCCCCTTCGCC |
| 4725 | mamo-L | *BGIBMGA009705* | up | 1556 | 1570 | - | 10.5081 | 9.83E-05 | 1 | TAGGCGGGGTGTTGG |
| 4726 | mamo-L | *BGIBMGA009709* | up | 109 | 123 | - | 11.8065 | 3.03E-05 | 1 | TAGGCTAATGTTTAT |
| 4727 | mamo-L | *BGIBMGA009709* | down | 759 | 773 | + | 10.629 | 8.91E-05 | 1 | TAGGCCTGCAGCGCC |
| 4728 | mamo-L | *BGIBMGA009711* | down | 571 | 585 | + | 12.3468 | 1.72E-05 | 1 | TAGGCTGTCCTTTGT |
| 4729 | mamo-L | *BGIBMGA009712* | down | 302 | 316 | - | 10.9839 | 6.59E-05 | 1 | AAGGCTGTTTTTTTC |
| 4730 | mamo-L | *BGIBMGA009715* | up | 623 | 637 | - | 10.5968 | 9.15E-05 | 1 | AAGGCTGTTAATGCT |
| 4731 | mamo-L | *BGIBMGA009715* | down | 1212 | 1226 | - | 10.5726 | 9.33E-05 | 1 | TAGGCTAATATTTAT |
| 4732 | mamo-L | *BGIBMGA009718* | down | 1372 | 1386 | - | 10.5726 | 9.33E-05 | 1 | TAGGCTAATATTTAT |
| 4733 | mamo-L | *BGIBMGA009719* | down | 1603 | 1617 | + | 12.25 | 1.92E-05 | 1 | TAGGCTTCTTATGCT |
| 4734 | mamo-L | *BGIBMGA009719* | down | 614 | 628 | - | 10.6855 | 8.51E-05 | 1 | AAGGCTTTTTTTTTC |
| 4735 | mamo-L | *BGIBMGA009720* | down | 348 | 362 | - | 10.9919 | 6.55E-05 | 1 | TAGGCTAGTGTTAGC |
| 4736 | mamo-L | *BGIBMGA009721* | up | 30 | 44 | - | 10.75 | 8.06E-05 | 1 | GAGGCCGATTGTGCC |
| 4737 | mamo-L | *BGIBMGA009722* | down | 748 | 762 | + | 14.2661 | 1.33E-06 | 1 | TAGGCGGGCGGTGGT |
| 4738 | mamo-L | *BGIBMGA009724* | up | 80 | 94 | + | 12.7903 | 1.02E-05 | 1 | TAGGCCCTTTTTTTT |
| 4739 | mamo-L | *BGIBMGA009725* | up | 1455 | 1469 | - | 11.3548 | 4.69E-05 | 1 | AAGGCTGATGATGAT |
| 4740 | mamo-L | *BGIBMGA009727* | up | 1847 | 1861 | + | 11.8226 | 2.99E-05 | 1 | TAGGCTTTGTATGGG |
| 4741 | mamo-L | *BGIBMGA009729* | up | 129 | 143 | - | 11.5323 | 3.96E-05 | 1 | TAGGCTGAGTGCCGT |
| 4742 | mamo-L | *BGIBMGA009729* | down | 197 | 211 | - | 11.3548 | 4.69E-05 | 1 | AAGGCTGATGATGAT |
| 4743 | mamo-L | *BGIBMGA009730* | down | 1426 | 1440 | + | 10.9194 | 6.98E-05 | 1 | TAGGCTACTTTTTTT |
| 4744 | mamo-L | *BGIBMGA009732* | down | 755 | 769 | - | 13.7742 | 2.79E-06 | 1 | TAGGCGGGTGGTGGT |
| 4745 | mamo-L | *BGIBMGA009733* | up | 1357 | 1371 | - | 13.7742 | 2.79E-06 | 1 | TAGGCGGGTGGTGGT |
| 4746 | mamo-L | *BGIBMGA009734* | down | 1551 | 1565 | - | 12.2742 | 1.87E-05 | 1 | AAGGCTTTATTTGGT |
| 4747 | mamo-L | *BGIBMGA009738* | up | 1650 | 1664 | + | 12.2823 | 1.85E-05 | 1 | AAGGCGGAGTTTGTC |
| 4748 | mamo-L | *BGIBMGA009739* | up | 1227 | 1241 | - | 11.3871 | 4.55E-05 | 1 | TAGGCGCGTTTCGTC |
| 4749 | mamo-L | *BGIBMGA009741* | up | 308 | 322 | - | 10.4919 | 9.96E-05 | 1 | TAGGCTCACAGTGGA |
| 4750 | mamo-L | *BGIBMGA009746* | up | 512 | 526 | - | 11.5968 | 3.73E-05 | 1 | TAGGCTAGTTTTTAT |
| 4751 | mamo-L | *BGIBMGA009751* | down | 578 | 592 | + | 11.0887 | 6.01E-05 | 1 | AAGGCCTAATATGGT |
| 4752 | mamo-L | *BGIBMGA009752* | up | 823 | 837 | + | 12.3871 | 1.65E-05 | 1 | TAGGCGTTTTTTGAC |
| 4753 | mamo-L | *BGIBMGA009752* | up | 547 | 561 | + | 11.5242 | 3.99E-05 | 1 | TAGGCTTTTTTCTTC |
| 4754 | mamo-L | *BGIBMGA009754* | down | 1864 | 1878 | - | 10.9677 | 6.68E-05 | 1 | AAGGCCCATTTTTCT |
| 4755 | mamo-L | *BGIBMGA009755* | up | 393 | 407 | + | 10.5726 | 9.33E-05 | 1 | TAGGCTAATATTTAT |
| 4756 | mamo-L | *BGIBMGA009756* | down | 256 | 270 | - | 11.1935 | 5.45E-05 | 1 | TAGGCTGTGTGTCTC |
| 4757 | mamo-L | *BGIBMGA009756* | down | 359 | 373 | - | 10.621 | 8.97E-05 | 1 | TAGGCTGTATAGTGT |
| 4758 | mamo-L | *BGIBMGA009760* | down | 938 | 952 | + | 11.4516 | 4.28E-05 | 1 | TAGGCTTTTTCTTGT |
| 4759 | mamo-L | *BGIBMGA009761* | down | 1829 | 1843 | + | 11.75 | 3.21E-05 | 1 | TAGGCTGCTGACGGT |
| 4760 | mamo-L | *BGIBMGA009761* | up | 435 | 449 | + | 10.629 | 8.91E-05 | 1 | TAGGCTTCTTTTATC |
| 4761 | mamo-L | *BGIBMGA009762* | up | 1856 | 1870 | + | 11.3871 | 4.55E-05 | 1 | TAGGCCTCTTTTACT |
| 4762 | mamo-L | *BGIBMGA009764* | up | 1981 | 1995 | - | 10.5323 | 9.63E-05 | 1 | TAGGCCTAATTGACT |
| 4763 | mamo-L | *BGIBMGA009773* | up | 414 | 428 | + | 10.9113 | 7.02E-05 | 1 | TAGGCTGTCGGAGTT |
| 4764 | mamo-L | *BGIBMGA009774* | up | 1973 | 1987 | + | 11.2097 | 5.38E-05 | 1 | TAGGCTTGAATTATT |
| 4765 | mamo-L | *BGIBMGA009776* | down | 1013 | 1027 | - | 11.8065 | 3.03E-05 | 1 | TAGGCTAATGTTTAT |
| 4766 | mamo-L | *BGIBMGA009778* | down | 1573 | 1587 | + | 10.9032 | 7.07E-05 | 1 | TAGGCTCTTAGTGGC |
| 4767 | mamo-L | *BGIBMGA009778* | down | 109 | 123 | - | 10.5726 | 9.33E-05 | 1 | AAGGCCGGCATTTAT |
| 4768 | mamo-L | *BGIBMGA009780* | down | 1265 | 1279 | + | 11.0403 | 6.28E-05 | 1 | TAGGCTTTTTGGTGT |
| 4769 | mamo-L | *BGIBMGA009782* | down | 1282 | 1296 | + | 10.7177 | 8.28E-05 | 1 | TAGGCTTAGTTTAAA |
| 4770 | mamo-L | *BGIBMGA009782* | down | 290 | 304 | - | 10.5726 | 9.33E-05 | 1 | TAGGCTAATATTTAT |
| 4771 | mamo-L | *BGIBMGA009783* | up | 359 | 373 | + | 11.5081 | 4.05E-05 | 1 | TAGGCGGGATTGTGT |
| 4772 | mamo-L | *BGIBMGA009791* | down | 1378 | 1392 | - | 11.6774 | 3.44E-05 | 1 | TAGGCCTCGTATGTC |
| 4773 | mamo-L | *BGIBMGA009792* | up | 477 | 491 | + | 11.4274 | 4.38E-05 | 1 | TAGGCTGTCAATGAC |
| 4774 | mamo-L | *BGIBMGA009793* | down | 617 | 631 | - | 10.9194 | 6.98E-05 | 1 | TAGGCTCTCGGCTTT |
| 4775 | mamo-L | *BGIBMGA009796* | down | 652 | 666 | - | 12.4597 | 1.51E-05 | 1 | TAGGCTGATATTTAT |
| 4776 | mamo-L | *BGIBMGA009803* | down | 235 | 249 | + | 11.4274 | 4.38E-05 | 1 | AAGGCTGGGAATGTT |
| 4777 | mamo-L | *BGIBMGA009803* | down | 1872 | 1886 | - | 11.0403 | 6.28E-05 | 1 | TAGGCTTTTTGGTGT |

| | | | | | | | | | | |
|---|---|---|---|---|---|---|---|---|---|---|
| 4778 | mamo-L | *BGIBMGA009811* | down | 434 | 448 | - | 10.6452 | 8.79E-05 | 1 | TAGGCGGAGGGGGAC |
| 4779 | mamo-L | *BGIBMGA009815* | up | 711 | 725 | - | 12.4839 | 1.47E-05 | 1 | GAGGCTGGTTTTTGT |
| 4780 | mamo-L | *BGIBMGA009819* | up | 1859 | 1873 | - | 11.871 | 2.85E-05 | 1 | AAGGCTGACTTTATT |
| 4781 | mamo-L | *BGIBMGA009819* | down | 1024 | 1038 | + | 10.4919 | 9.96E-05 | 1 | TAGGCACAGTATGAT |
| 4782 | mamo-L | *BGIBMGA009821* | down | 170 | 184 | - | 10.7016 | 8.39E-05 | 1 | TAGGCGGGTTGTGCG |
| 4783 | mamo-L | *BGIBMGA009825* | down | 636 | 650 | + | 10.4919 | 9.96E-05 | 1 | TAGGCATTTTTTTTC |
| 4784 | mamo-L | *BGIBMGA009826* | up | 648 | 662 | + | 10.9597 | 6.73E-05 | 1 | TAGGCCAACGGGGCT |
| 4785 | mamo-L | *BGIBMGA009833* | up | 1968 | 1982 | - | 10.6855 | 8.51E-05 | 1 | AAGGCTGGTGTGTTT |
| 4786 | mamo-L | *BGIBMGA009833* | down | 68 | 82 | + | 10.6048 | 9.09E-05 | 1 | AAGGCCTTCAATGCT |
| 4787 | mamo-L | *BGIBMGA009836* | up | 825 | 839 | + | 12.9839 | 8.17E-06 | 1 | TAGGCCTGTGTTTTC |
| 4788 | mamo-L | *BGIBMGA009839* | down | 160 | 174 | + | 15.7419 | 7.43E-08 | 1 | TAGGCTTTGTTTGGT |
| 4789 | mamo-L | *BGIBMGA009839* | down | 96 | 110 | - | 10.9839 | 6.59E-05 | 1 | TAGGCTTCGATTTTC |
| 4790 | mamo-L | *BGIBMGA009842* | down | 1963 | 1977 | + | 11.7903 | 3.08E-05 | 1 | AAGGCCCGGTTTTGT |
| 4791 | mamo-L | *BGIBMGA009850* | up | 1777 | 1791 | - | 11.1129 | 5.88E-05 | 1 | AAGGCTGTGTGTTTT |
| 4792 | mamo-L | *BGIBMGA009852* | down | 241 | 255 | + | 11.129 | 5.79E-05 | 1 | TAGGCTTAGGTTCTG |
| 4793 | mamo-L | *BGIBMGA009853* | down | 1846 | 1860 | - | 13.1774 | 6.47E-06 | 1 | TAGGCTGGTTTTGCA |
| 4794 | mamo-L | *BGIBMGA009857* | up | 699 | 713 | + | 10.5726 | 9.33E-05 | 1 | TAGGCTAATATTTAT |
| 4795 | mamo-L | *BGIBMGA009858* | up | 1160 | 1174 | - | 11.4597 | 4.25E-05 | 1 | TAGGCAGACCTTGGT |
| 4796 | mamo-L | *BGIBMGA009860* | up | 716 | 730 | + | 10.9113 | 7.02E-05 | 1 | TAGGCTGATGGCACT |
| 4797 | mamo-L | *BGIBMGA009861* | up | 1461 | 1475 | + | 11.4677 | 4.22E-05 | 1 | TAGGCCCTGTTTATC |
| 4798 | mamo-L | *BGIBMGA009867* | up | 327 | 341 | + | 11.9758 | 2.57E-05 | 1 | AAGGCTTTTTTTTGT |
| 4799 | mamo-L | *BGIBMGA009872* | down | 667 | 681 | + | 11.0806 | 6.05E-05 | 1 | TAGGCCGGCCATTGT |
| 4800 | mamo-L | *BGIBMGA009877* | up | 1346 | 1360 | + | 11.3387 | 4.77E-05 | 1 | AAGGCTTACTGTGAT |
| 4801 | mamo-L | *BGIBMGA009878* | down | 927 | 941 | - | 10.6048 | 9.09E-05 | 1 | TAGGCTGGGCTTAAT |
| 4802 | mamo-L | *BGIBMGA009878* | down | 1624 | 1638 | - | 10.5968 | 9.15E-05 | 1 | AAGGCGTTCGATTGT |
| 4803 | mamo-L | *BGIBMGA009879* | up | 920 | 934 | - | 12.0645 | 2.34E-05 | 1 | TAGGCGGGCTTCGGA |
| 4804 | mamo-L | *BGIBMGA009880* | up | 645 | 659 | - | 10.9113 | 7.02E-05 | 1 | TAGGCTGATGGCACT |
| 4805 | mamo-L | *BGIBMGA009884* | down | 1016 | 1030 | + | 12.9919 | 8.10E-06 | 1 | TAGGCCGGTTTTGCA |
| 4806 | mamo-L | *BGIBMGA009886* | down | 1325 | 1339 | + | 13.1774 | 6.47E-06 | 1 | TAGGCTGGTTTTGCA |
| 4807 | mamo-L | *BGIBMGA009887* | down | 1772 | 1786 | - | 11.129 | 5.79E-05 | 1 | TAGGCTTAGGTTCTG |
| 4808 | mamo-L | *BGIBMGA009889* | down | 1909 | 1923 | - | 12.1371 | 2.17E-05 | 1 | TAGGCGTGTGTGTGT |
| 4809 | mamo-L | *BGIBMGA009889* | down | 1543 | 1557 | - | 10.75 | 8.06E-05 | 1 | TAGGCTTAATTGATT |
| 4810 | mamo-L | *BGIBMGA009890* | down | 1075 | 1089 | + | 11.3548 | 4.69E-05 | 1 | AAGGCTGATGATGAT |
| 4811 | mamo-L | *BGIBMGA009897* | up | 723 | 737 | - | 10.5 | 9.90E-05 | 1 | TAGGCTTAAGGGGTC |
| 4812 | mamo-L | *BGIBMGA009899* | down | 338 | 352 | - | 10.8145 | 7.63E-05 | 1 | TAGACTGGCTTTGGT |
| 4813 | mamo-L | *BGIBMGA009902* | up | 295 | 309 | + | 12.0887 | 2.29E-05 | 1 | TAGGCTATTGATGTT |
| 4814 | mamo-L | *BGIBMGA009906* | up | 650 | 664 | + | 11.2016 | 5.41E-05 | 1 | TAGGCGTTAAGTGGT |
| 4815 | mamo-L | *BGIBMGA009907* | up | 521 | 535 | - | 13.7742 | 2.79E-06 | 1 | TAGGCTTTTTTTTTT |
| 4816 | mamo-L | *BGIBMGA009908* | down | 405 | 419 | + | 11.3629 | 4.66E-05 | 1 | TAGGCTTAAATTTAT |
| 4817 | mamo-L | *BGIBMGA009908* | down | 1628 | 1642 | + | 10.6371 | 8.85E-05 | 1 | TAGGCTAATTAGGTT |
| 4818 | mamo-L | *BGIBMGA009909* | up | 339 | 353 | + | 10.9919 | 6.55E-05 | 1 | TAGGCGTATTATGTA |
| 4819 | mamo-L | *BGIBMGA009911* | down | 774 | 788 | + | 12.2258 | 1.97E-05 | 1 | GAGGCTGTATTTGTT |
| 4820 | mamo-L | *BGIBMGA009917* | down | 1880 | 1894 | - | 10.7903 | 7.79E-05 | 1 | TAGGCTTTCTTATTT |
| 4821 | mamo-L | *BGIBMGA009918* | down | 66 | 80 | - | 11.6613 | 3.50E-05 | 1 | GAGGCCGAAGTTGAT |
| 4822 | mamo-L | *BGIBMGA009918* | up | 505 | 519 | + | 11.0806 | 6.05E-05 | 1 | TAGGCTGTGGTGCTC |
| 4823 | mamo-L | *BGIBMGA009921* | up | 578 | 592 | - | 11.7823 | 3.11E-05 | 1 | TAGGCAATCTTTGGT |
| 4824 | mamo-L | *BGIBMGA009922* | up | 587 | 601 | - | 10.9597 | 6.73E-05 | 1 | TAGGCCAACGGGGCT |
| 4825 | mamo-L | *BGIBMGA009923* | up | 605 | 619 | - | 12.9113 | 8.87E-06 | 1 | TAGGCTTACTTGTCT |
| 4826 | mamo-L | *BGIBMGA009924* | down | 1272 | 1286 | - | 12.0645 | 2.34E-05 | 1 | TAGGCGGGCTTCGGA |
| 4827 | mamo-L | *BGIBMGA009925* | up | 1169 | 1183 | - | 11.5081 | 4.05E-05 | 1 | AAGGCTGGCGGTGCC |
| 4828 | mamo-L | *BGIBMGA009926* | down | 791 | 805 | + | 11.4274 | 4.38E-05 | 1 | TAGGCTCAGTATGTA |
| 4829 | mamo-L | *BGIBMGA009927* | down | 490 | 504 | + | 10.7016 | 8.39E-05 | 1 | TAGGCGGGTTGTGCG |
| 4830 | mamo-L | *BGIBMGA009928* | up | 744 | 758 | + | 12.0645 | 2.34E-05 | 1 | TAGGCGGGCTTCGGA |
| 4831 | mamo-L | *BGIBMGA009928* | up | 591 | 605 | - | 10.8306 | 7.52E-05 | 1 | TAGGCCCCCTTCGCC |
| 4832 | mamo-L | *BGIBMGA009928* | up | 1495 | 1509 | - | 10.629 | 8.91E-05 | 1 | TAGGCCTGCAGCGCC |
| 4833 | mamo-L | *BGIBMGA009930* | down | 1171 | 1185 | + | 11.0484 | 6.23E-05 | 1 | TAGGCTCATGGTTTC |
| 4834 | mamo-L | *BGIBMGA009931* | up | 1234 | 1248 | + | 11.0484 | 6.23E-05 | 1 | TAGGCTCATGGTTTC |
| 4835 | mamo-L | *BGIBMGA009933* | down | 1563 | 1577 | - | 12.6371 | 1.23E-05 | 1 | TAGGCAGGCGGTGGT |
| 4836 | mamo-L | *BGIBMGA009934* | up | 225 | 239 | - | 12.6371 | 1.23E-05 | 1 | TAGGCAGGCGGTGGT |
| 4837 | mamo-L | *BGIBMGA009934* | down | 703 | 717 | - | 11.3548 | 4.69E-05 | 1 | AAGGCTGATGATGAT |
| 4838 | mamo-L | *BGIBMGA009936* | down | 1885 | 1899 | + | 10.9194 | 6.98E-05 | 1 | TAGGCTACTTTTTTT |
| 4839 | mamo-L | *BGIBMGA009936* | down | 1254 | 1268 | + | 10.8145 | 7.63E-05 | 1 | GAGGCTTTGGTTGTG |
| 4840 | mamo-L | *BGIBMGA009941* | down | 820 | 834 | + | 11.3548 | 4.69E-05 | 1 | AAGGCTGATGATGAT |
| 4841 | mamo-L | *BGIBMGA009945* | up | 1098 | 1112 | - | 12.1774 | 2.08E-05 | 1 | TAGGCCTTTAGTGTT |
| 4842 | mamo-L | *BGIBMGA009948* | down | 786 | 800 | - | 10.8629 | 7.33E-05 | 1 | TAGGCCACTGTTTTT |
| 4843 | mamo-L | *BGIBMGA009957* | up | 392 | 406 | - | 12.0645 | 2.34E-05 | 1 | TAGGCGGGCTTCGGA |
| 4844 | mamo-L | *BGIBMGA009959* | up | 541 | 555 | + | 10.5726 | 9.33E-05 | 1 | TAGGCTAATATTTAT |
| 4845 | mamo-L | *BGIBMGA009962* | down | 1582 | 1596 | + | 12.7016 | 1.14E-05 | 1 | TAGGCCGGGTATTGC |
| 4846 | mamo-L | *BGIBMGA009962* | up | 81 | 95 | - | 11.1774 | 5.54E-05 | 1 | TAGGCTTTTTTTTTG |
| 4847 | mamo-L | *BGIBMGA009964* | down | 1807 | 1821 | - | 11.9274 | 2.69E-05 | 1 | AAGGCTGTTTTTTCT |
| 4848 | mamo-L | *BGIBMGA009964* | up | 1014 | 1028 | - | 11.2258 | 5.30E-05 | 1 | TAGGCCTTTTTCCCT |
| 4849 | mamo-L | *BGIBMGA009966* | up | 1116 | 1130 | + | 14.5 | 9.07E-07 | 1 | TAGGCCTACGTTTCT |
| 4850 | mamo-L | *BGIBMGA009968* | down | 1641 | 1655 | + | 12.3226 | 1.77E-05 | 1 | TAGGCTGTGTGTATT |
| 4851 | mamo-L | *BGIBMGA009973* | up | 1913 | 1927 | - | 14.3952 | 1.09E-06 | 1 | TAGGCTTTCGTTTTT |
| 4852 | mamo-L | *BGIBMGA009975* | up | 385 | 399 | + | 14.3952 | 1.09E-06 | 1 | TAGGCTTTCGTTTTT |
| 4853 | mamo-L | *BGIBMGA009976* | down | 477 | 491 | - | 12.5968 | 1.28E-05 | 1 | TAGGCCGGTTACGCT |
| 4854 | mamo-L | *BGIBMGA009976* | down | 1006 | 1020 | - | 10.8468 | 7.42E-05 | 1 | TAGGCCGATAATACT |
| 4855 | mamo-L | *BGIBMGA009976* | down | 144 | 158 | + | 10.5726 | 9.33E-05 | 1 | TAGGCTAATATTTAT |
| 4856 | mamo-L | *BGIBMGA009980* | up | 611 | 625 | + | 11.2419 | 5.21E-05 | 1 | AAGGCTGCGATTGTT |
| 4857 | mamo-L | *BGIBMGA009981* | up | 556 | 570 | + | 11.7984 | 3.06E-05 | 1 | TAGGCTTTGCTTTTT |
| 4858 | mamo-L | *BGIBMGA009982* | down | 1206 | 1220 | - | 11.0081 | 6.45E-05 | 1 | AAGGCCGGTTATTGT |
| 4859 | mamo-L | *BGIBMGA009983* | up | 1501 | 1515 | - | 11.0081 | 6.45E-05 | 1 | AAGGCCGGTTATTGT |
| 4860 | mamo-L | *BGIBMGA009985* | down | 1897 | 1911 | - | 12.0323 | 2.42E-05 | 1 | AAGGCTGGCGTTGGA |
| 4861 | mamo-L | *BGIBMGA009986* | up | 1244 | 1258 | + | 11.2258 | 5.30E-05 | 1 | TAGGCCTTTTTCCCT |
| 4862 | mamo-L | *BGIBMGA009987* | down | 1418 | 1432 | - | 11 | 6.50E-05 | 1 | AAGGCCCATTTTTTT |
| 4863 | mamo-L | *BGIBMGA009989* | down | 796 | 810 | - | 10.4919 | 9.96E-05 | 1 | TAGGCTATTATTAGT |
| 4864 | mamo-L | *BGIBMGA009990* | up | 855 | 869 | - | 13.4435 | 4.51E-06 | 1 | TAGGCTTCCGTTTGT |
| 4865 | mamo-L | *BGIBMGA009992* | up | 190 | 204 | - | 11.2984 | 4.95E-05 | 1 | TGGGCCGTGGTTTGT |
| 4866 | mamo-L | *BGIBMGA009993* | down | 1442 | 1456 | + | 10.879 | 7.22E-05 | 1 | TAGGCGGACAATATT |
| 4867 | mamo-L | *BGIBMGA009994* | up | 1258 | 1272 | + | 10.879 | 7.22E-05 | 1 | TAGGCGGACAATATT |
| 4868 | mamo-L | *BGIBMGA009994* | down | 739 | 753 | - | 10.5242 | 9.70E-05 | 1 | GAGGCTGTTTGTTTT |
| 4869 | mamo-L | *BGIBMGA009999* | down | 1333 | 1347 | + | 11.5161 | 4.02E-05 | 1 | AAGGCTTTGGTTTAT |

| | | | | | | | | | | |
|---|---|---|---|---|---|---|---|---|---|---|
| 4870 | mamo-L | *BGIBMGA010000* | up | 1596 | 1610 | + | 11.5161 | 4.02E-05 | 1 | AAGGCTTTGGTTTAT |
| 4871 | mamo-L | *BGIBMGA010000* | down | 581 | 595 | - | 10.9919 | 6.55E-05 | 1 | TAGGCTTTGATTTCA |
| 4872 | mamo-L | *BGIBMGA010001* | up | 515 | 529 | - | 10.9839 | 6.59E-05 | 1 | AAGGCGTTTTTTTTT |
| 4873 | mamo-L | *BGIBMGA010008* | up | 796 | 810 | + | 11.2742 | 5.06E-05 | 1 | GAGGCGTATTTTTTT |
| 4874 | mamo-L | *BGIBMGA010008* | up | 1692 | 1706 | - | 11.2419 | 5.21E-05 | 1 | TAGGCTTTGAATTGC |
| 4875 | mamo-L | *BGIBMGA010013* | down | 771 | 785 | + | 11.6613 | 3.50E-05 | 1 | AAGGCTATCTTTGTT |
| 4876 | mamo-L | *BGIBMGA010014* | up | 1368 | 1382 | - | 11.2016 | 5.41E-05 | 1 | AAGGCGGTTGTCGCT |
| 4877 | mamo-L | *BGIBMGA010017* | up | 815 | 829 | - | 11.9758 | 2.57E-05 | 1 | TAGGCCTATTTTGAA |
| 4878 | mamo-L | *BGIBMGA010021* | up | 1576 | 1590 | + | 13.0806 | 7.24E-06 | 1 | TAGGCTTTCTTTGGG |
| 4879 | mamo-L | *BGIBMGA010023* | up | 1661 | 1675 | - | 10.9113 | 7.02E-05 | 1 | TAGGCCAACGTTCTC |
| 4880 | mamo-L | *BGIBMGA010024* | up | 1836 | 1850 | - | 11.621 | 3.63E-05 | 1 | TAGGCGGTAGTTGTA |
| 4881 | mamo-L | *BGIBMGA010032* | up | 1440 | 1454 | + | 10.6613 | 8.67E-05 | 1 | TAGGCCGAGCCTGCT |
| 4882 | mamo-L | *BGIBMGA010032* | up | 518 | 532 | - | 10.629 | 8.91E-05 | 1 | TAGGCCTGCAGCGCC |
| 4883 | mamo-L | *BGIBMGA010034* | up | 498 | 512 | + | 12.9758 | 8.23E-06 | 1 | TAGGCTTTATTTTTT |
| 4884 | mamo-L | *BGIBMGA010034* | up | 512 | 526 | + | 11.5484 | 3.90E-05 | 1 | TAGGCTTTTTATATT |
| 4885 | mamo-L | *BGIBMGA010044* | up | 260 | 274 | + | 12.9435 | 8.54E-06 | 1 | TAGGCTTTTTTTTAT |
| 4886 | mamo-L | *BGIBMGA010044* | up | 777 | 791 | + | 10.5 | 9.90E-05 | 1 | GAGGCCCAGGTCGTC |
| 4887 | mamo-L | *BGIBMGA010045* | down | 462 | 476 | - | 12.0645 | 2.34E-05 | 1 | TAGGCGGGCTTCGGA |
| 4888 | mamo-L | *BGIBMGA010046* | down | 86 | 100 | - | 11.6129 | 3.66E-05 | 1 | TAGGCTTTTAATGAT |
| 4889 | mamo-L | *BGIBMGA010050* | up | 267 | 281 | + | 10.6452 | 8.79E-05 | 1 | TAGGCAGACGATCGT |
| 4890 | mamo-L | *BGIBMGA010055* | down | 756 | 770 | + | 12.2097 | 2.01E-05 | 1 | TAGGCTTGTGTTGAA |
| 4891 | mamo-L | *BGIBMGA010060* | up | 99 | 113 | + | 11.6855 | 3.42E-05 | 1 | TAGGCAGAGTTCTGT |
| 4892 | mamo-L | *BGIBMGA010060* | down | 553 | 567 | - | 10.9194 | 6.98E-05 | 1 | TAGGCTACTTTTTTT |
| 4893 | mamo-L | *BGIBMGA010061* | up | 1365 | 1379 | - | 12.5081 | 1.43E-05 | 1 | TAGGCTTCTTTTTTT |
| 4894 | mamo-L | *BGIBMGA010061* | up | 1576 | 1590 | - | 10.9194 | 6.98E-05 | 1 | TAGGCTACTTTTTTT |
| 4895 | mamo-L | *BGIBMGA010062* | up | 1836 | 1850 | - | 11.621 | 3.63E-05 | 1 | TAGGCGGTAGTTGTA |
| 4896 | mamo-L | *BGIBMGA010064* | up | 964 | 978 | + | 13.3952 | 4.84E-06 | 1 | TAGGCGGTTTTTTTT |
| 4897 | mamo-L | *BGIBMGA010065* | up | 1492 | 1506 | + | 11.1774 | 5.54E-05 | 1 | TAGGCCGGATATTAT |
| 4898 | mamo-L | *BGIBMGA010066* | down | 336 | 350 | + | 10.5726 | 9.33E-05 | 1 | TAGGCTAATATTTAT |
| 4899 | mamo-L | *BGIBMGA010067* | down | 252 | 266 | - | 10.5726 | 9.33E-05 | 1 | TAGGCTAATATTTAT |
| 4900 | mamo-L | *BGIBMGA010067* | up | 1860 | 1874 | + | 10.5726 | 9.33E-05 | 1 | TAGGCTAATATTTAT |
| 4901 | mamo-L | *BGIBMGA010068* | up | 1052 | 1066 | - | 11.0242 | 6.36E-05 | 1 | TAGGCTGTAGTAGTT |
| 4902 | mamo-L | *BGIBMGA010073* | up | 746 | 760 | - | 11.0887 | 6.01E-05 | 1 | TAGGCTGAATATGGG |
| 4903 | mamo-L | *BGIBMGA010076* | down | 996 | 1010 | + | 10.5 | 9.90E-05 | 1 | GAGGCCCAGGTCGTC |
| 4904 | mamo-L | *BGIBMGA010079* | down | 1501 | 1515 | + | 11.1935 | 5.45E-05 | 1 | TAGGCGCATGAGGGT |
| 4905 | mamo-L | *BGIBMGA010080* | down | 1655 | 1669 | + | 11.9032 | 2.76E-05 | 1 | TAGGCGGGTTTTAAT |
| 4906 | mamo-L | *BGIBMGA010080* | down | 37 | 51 | + | 11.1935 | 5.45E-05 | 1 | TAGGCGCATGAGGGT |
| 4907 | mamo-L | *BGIBMGA010081* | down | 1449 | 1463 | - | 10.9516 | 6.78E-05 | 1 | GAGGCGTTTTTTTTT |
| 4908 | mamo-L | *BGIBMGA010085* | down | 46 | 60 | - | 13.1855 | 6.42E-06 | 1 | TAGGCTTGCTATTTT |
| 4909 | mamo-L | *BGIBMGA010085* | up | 662 | 676 | + | 12.5323 | 1.39E-05 | 1 | TAGGCCTTTGACGGT |
| 4910 | mamo-L | *BGIBMGA010091* | down | 451 | 465 | - | 11 | 6.50E-05 | 1 | TAGGCGGTGGTCTAC |
| 4911 | mamo-L | *BGIBMGA010093* | up | 69 | 83 | - | 13.2339 | 6.02E-06 | 1 | TAGGCCGATTTTTTC |
| 4912 | mamo-L | *BGIBMGA010096* | down | 793 | 807 | + | 11.8145 | 3.01E-05 | 1 | TAGGCTTGATGTTTT |
| 4913 | mamo-L | *BGIBMGA010098* | up | 292 | 306 | - | 13.2339 | 6.02E-06 | 1 | TAGGCCTGCTTTCCT |
| 4914 | mamo-L | *BGIBMGA010102* | up | 1676 | 1690 | - | 11.4758 | 4.18E-05 | 1 | AAGGCTTTGTTTCGT |
| 4915 | mamo-L | *BGIBMGA010103* | down | 207 | 221 | - | 11.9516 | 2.63E-05 | 1 | TAGGCTGTATATTTT |
| 4916 | mamo-L | *BGIBMGA010103* | down | 1133 | 1147 | + | 11.8226 | 2.99E-05 | 1 | TAGGCTGAAGAGGCT |
| 4917 | mamo-L | *BGIBMGA010103* | down | 696 | 710 | + | 11.1855 | 5.50E-05 | 1 | TAGGCAGTAGATGGT |
| 4918 | mamo-L | *BGIBMGA010106* | down | 1088 | 1102 | + | 10.5968 | 9.15E-05 | 1 | GAGGCTAGTTTTTGT |
| 4919 | mamo-L | *BGIBMGA010111* | down | 1513 | 1527 | + | 10.5726 | 9.33E-05 | 1 | TAGGCCAAGTGTATT |
| 4920 | mamo-L | *BGIBMGA010112* | down | 516 | 530 | + | 10.7419 | 8.11E-05 | 1 | GAGGCGCAATTTGCT |
| 4921 | mamo-L | *BGIBMGA010118* | up | 564 | 578 | + | 10.7661 | 7.95E-05 | 1 | AAGGCTGGTGTTAGC |
| 4922 | mamo-L | *BGIBMGA010119* | up | 1570 | 1584 | - | 13.5484 | 3.92E-06 | 1 | TAGGCTGAGTGTTTT |
| 4923 | mamo-L | *BGIBMGA010122* | up | 212 | 226 | + | 10.621 | 8.97E-05 | 1 | TAGGCTGTGGCTGTG |
| 4924 | mamo-L | *BGIBMGA010123* | up | 1036 | 1050 | + | 12.0726 | 2.32E-05 | 1 | TAGGCTGATTCTTGT |
| 4925 | mamo-L | *BGIBMGA010125* | up | 1909 | 1923 | - | 10.6855 | 8.51E-05 | 1 | TAGGCTATTTTTGTG |
| 4926 | mamo-L | *BGIBMGA010126* | down | 710 | 724 | - | 13.0484 | 7.57E-06 | 1 | TAGGCTTACTCTGTT |
| 4927 | mamo-L | *BGIBMGA010128* | up | 690 | 704 | + | 10.6371 | 8.85E-05 | 1 | TAGGCCTGTCTTTGC |
| 4928 | mamo-L | *BGIBMGA010132* | down | 1837 | 1851 | - | 11.2581 | 5.14E-05 | 1 | TAGGCACATTTTGAT |
| 4929 | mamo-L | *BGIBMGA010133* | down | 222 | 236 | - | 11.3387 | 4.77E-05 | 1 | GAGGCTTACATTTTT |
| 4930 | mamo-L | *BGIBMGA010136* | up | 113 | 127 | + | 11.9839 | 2.54E-05 | 1 | AAGGCTTATTTTTTT |
| 4931 | mamo-L | *BGIBMGA010137* | up | 745 | 759 | + | 12.2016 | 2.03E-05 | 1 | TAGGCGCTCTTTAGT |
| 4932 | mamo-L | *BGIBMGA010137* | down | 1207 | 1221 | + | 10.8871 | 7.17E-05 | 1 | TAGGCACTGTGTGCT |
| 4933 | mamo-L | *BGIBMGA010137* | down | 853 | 867 | - | 10.871 | 7.28E-05 | 1 | GAGGCGCACGATGTT |
| 4934 | mamo-L | *BGIBMGA010141* | up | 745 | 759 | - | 11.5081 | 4.05E-05 | 1 | TAGGCGGTCTATCTT |
| 4935 | mamo-L | *BGIBMGA010141* | up | 362 | 376 | + | 10.5242 | 9.70E-05 | 1 | AAGGCGTAGATGGGT |
| 4936 | mamo-L | *BGIBMGA010142* | down | 1388 | 1402 | + | 10.9194 | 6.98E-05 | 1 | TAGGCTACTTTTTTT |
| 4937 | mamo-L | *BGIBMGA010143* | up | 383 | 397 | + | 10.9194 | 6.98E-05 | 1 | TAGGCTACTTTTTTT |
| 4938 | mamo-L | *BGIBMGA010143* | down | 746 | 760 | + | 10.9194 | 6.98E-05 | 1 | TAGGCTACTTTTTTT |
| 4939 | mamo-L | *BGIBMGA010144* | down | 473 | 487 | + | 11.4677 | 4.22E-05 | 1 | TAGGCATTTTTTTTT |
| 4940 | mamo-L | *BGIBMGA010144* | up | 1361 | 1375 | + | 10.9194 | 6.98E-05 | 1 | TAGGCTACTTTTTTT |
| 4941 | mamo-L | *BGIBMGA010145* | up | 490 | 504 | - | 11.3548 | 4.69E-05 | 1 | AAGGCTGATGATGAT |
| 4942 | mamo-L | *BGIBMGA010149* | up | 422 | 436 | + | 11.129 | 5.79E-05 | 1 | TAGGCGAAAGTTTCT |
| 4943 | mamo-L | *BGIBMGA010150* | down | 636 | 650 | + | 11.4758 | 4.18E-05 | 1 | TAGGCACACATTGTT |
| 4944 | mamo-L | *BGIBMGA010151* | down | 688 | 702 | - | 11.621 | 3.63E-05 | 1 | AAGGCTTGCGTTATT |
| 4945 | mamo-L | *BGIBMGA010153* | down | 551 | 565 | - | 10.8468 | 7.42E-05 | 1 | AAGGCTTATATTTCT |
| 4946 | mamo-L | *BGIBMGA010155* | up | 1047 | 1061 | - | 12.5 | 1.44E-05 | 1 | TAGGCTTTTTTCTTT |
| 4947 | mamo-L | *BGIBMGA010157* | down | 1583 | 1597 | + | 12.5645 | 1.34E-05 | 1 | TAGGCGTGTGTTATT |
| 4948 | mamo-L | *BGIBMGA010158* | up | 926 | 940 | + | 11.5403 | 3.93E-05 | 1 | AAGGCTGGATTCGGT |
| 4949 | mamo-L | *BGIBMGA010159* | up | 1382 | 1396 | + | 14.371 | 1.12E-06 | 1 | TAGGCGTAGGTTGAT |
| 4950 | mamo-L | *BGIBMGA010159* | up | 1915 | 1929 | + | 12.0645 | 2.34E-05 | 1 | GAGGCTGAGTTTCGT |
| 4951 | mamo-L | *BGIBMGA010159* | up | 22 | 36 | + | 10.6694 | 8.62E-05 | 1 | TAGGCCTACAACTCT |
| 4952 | mamo-L | *BGIBMGA010163* | down | 1012 | 1026 | + | 13.8548 | 2.48E-06 | 1 | TAGGCGGATGGTGGT |
| 4953 | mamo-L | *BGIBMGA010164* | up | 340 | 354 | + | 10.5726 | 9.33E-05 | 1 | TAGGCTAATATTTAT |
| 4954 | mamo-L | *BGIBMGA010165* | up | 790 | 804 | - | 11.5323 | 3.96E-05 | 1 | TAGGCGTGAGGTGAT |
| 4955 | mamo-L | *BGIBMGA010166* | down | 1736 | 1750 | - | 11.6129 | 3.66E-05 | 1 | TAGGCTGTTAATTCT |
| 4956 | mamo-L | *BGIBMGA010169* | down | 941 | 955 | + | 10.5968 | 9.15E-05 | 1 | TAGGCTTAATTGCTT |
| 4957 | mamo-L | *BGIBMGA010171* | up | 1327 | 1341 | + | 12.2097 | 2.01E-05 | 1 | TAGGCTTTTTTTAGC |
| 4958 | mamo-L | *BGIBMGA010176* | up | 135 | 149 | + | 11.5242 | 3.99E-05 | 1 | AAGGCTTATGTTAGT |
| 4959 | mamo-L | *BGIBMGA010179* | down | 1818 | 1832 | - | 11.7177 | 3.32E-05 | 1 | TAGGCTAATATTTGT |
| 4960 | mamo-L | *BGIBMGA010179* | down | 972 | 986 | + | 10.9194 | 6.98E-05 | 1 | TAGGCTACTTTTTTT |
| 4961 | mamo-L | *BGIBMGA010179* | up | 866 | 880 | + | 10.6532 | 8.73E-05 | 1 | GAGGCTGCTATTGTT |

| | | | | | | | | | | |
|---|---|---|---|---|---|---|---|---|---|---|
| 4962 | mamo-L | *BGIBMGA010179* | down | 1456 | 1470 | + | 10.5726 | 9.33E-05 | 1 | TAGGCTAATATTTAT |
| 4963 | mamo-L | *BGIBMGA010182* | up | 1438 | 1452 | + | 10.5726 | 9.33E-05 | 1 | TAGGCTAATATTTAT |
| 4964 | mamo-L | *BGIBMGA010185* | up | 1173 | 1187 | - | 10.5726 | 9.33E-05 | 1 | TAGGCTAATATTTAT |
| 4965 | mamo-L | *BGIBMGA010189* | up | 1281 | 1295 | - | 10.6371 | 8.85E-05 | 1 | TAGGCCCAGTTTTAA |
| 4966 | mamo-L | *BGIBMGA010190* | up | 336 | 350 | + | 10.5 | 9.90E-05 | 1 | GAGGCCCAGGTCGTC |
| 4967 | mamo-L | *BGIBMGA010194* | up | 1528 | 1542 | - | 10.5645 | 9.40E-05 | 1 | TAGGCTTATTACACT |
| 4968 | mamo-L | *BGIBMGA010196* | down | 1438 | 1452 | + | 12.0726 | 2.32E-05 | 1 | GAGGCTGATTTCGTT |
| 4969 | mamo-L | *BGIBMGA010197* | up | 1582 | 1596 | + | 11.0806 | 6.05E-05 | 1 | AAGGCTTATTTTATT |
| 4970 | mamo-L | *BGIBMGA010197* | up | 1463 | 1477 | - | 11.0565 | 6.18E-05 | 1 | GAGGCCTGCCTTGGT |
| 4971 | mamo-L | *BGIBMGA010201* | up | 130 | 144 | + | 11.8226 | 2.99E-05 | 1 | TAGGCTCGTGTGGTC |
| 4972 | mamo-L | *BGIBMGA010202* | down | 128 | 142 | - | 12.9355 | 8.65E-06 | 1 | TAGGCTTTCAATGTT |
| 4973 | mamo-L | *BGIBMGA010202* | down | 1606 | 1620 | - | 10.5968 | 9.15E-05 | 1 | TAGGCTTAATTGCTT |
| 4974 | mamo-L | *BGIBMGA010203* | down | 525 | 539 | + | 12.7823 | 1.03E-05 | 1 | TAGGCGGTCTTTGTA |
| 4975 | mamo-L | *BGIBMGA010205* | down | 1507 | 1521 | + | 11.8065 | 3.03E-05 | 1 | TAGGCTGACTTCTAC |
| 4976 | mamo-L | *BGIBMGA010205* | down | 1620 | 1634 | - | 11.3306 | 4.80E-05 | 1 | TAGGCGGTGGTGGTA |
| 4977 | mamo-L | *BGIBMGA010206* | down | 1118 | 1132 | + | 11.6129 | 3.66E-05 | 1 | TAGGCTGTTAATTCT |
| 4978 | mamo-L | *BGIBMGA010206* | down | 805 | 819 | - | 10.5645 | 9.40E-05 | 1 | TAGGCAATGGTTTTT |
| 4979 | mamo-L | *BGIBMGA010210* | down | 405 | 419 | + | 12.1371 | 2.17E-05 | 1 | TGGGCTGTGTTTGTT |
| 4980 | mamo-L | *BGIBMGA010213* | down | 1508 | 1522 | + | 10.8306 | 7.52E-05 | 1 | TAGGCCACTGTTTCT |
| 4981 | mamo-L | *BGIBMGA010214* | down | 178 | 192 | - | 11.25 | 5.18E-05 | 1 | AAGGCGTTTTTTGAT |
| 4982 | mamo-L | *BGIBMGA010215* | up | 115 | 129 | - | 12.6694 | 1.18E-05 | 1 | AAGGCTTAGGTTTTT |
| 4983 | mamo-L | *BGIBMGA010215* | up | 318 | 332 | - | 11.25 | 5.18E-05 | 1 | AAGGCGTTTTTTGAT |
| 4984 | mamo-L | *BGIBMGA010217* | down | 296 | 310 | + | 14.8065 | 5.03E-07 | 1 | TAGGCTTACGTTGCC |
| 4985 | mamo-L | *BGIBMGA010217* | up | 284 | 298 | - | 10.7742 | 7.89E-05 | 1 | TAGGCTGGCCACGTT |
| 4986 | mamo-L | *BGIBMGA010218* | down | 902 | 916 | - | 11.2258 | 5.30E-05 | 1 | TAGGCCTTTTTCCCT |
| 4987 | mamo-L | *BGIBMGA010219* | up | 1192 | 1206 | - | 11.3548 | 4.69E-05 | 1 | TAGGCTGTCGACCGT |
| 4988 | mamo-L | *BGIBMGA010220* | down | 1365 | 1379 | - | 12.5645 | 1.34E-05 | 1 | TAGGCGTGTGTTATT |
| 4989 | mamo-L | *BGIBMGA010223* | up | 370 | 384 | - | 11.5323 | 3.96E-05 | 1 | TAGGCGTGAGGTGAT |
| 4990 | mamo-L | *BGIBMGA010223* | up | 55 | 69 | - | 10.9194 | 6.98E-05 | 1 | TAGGCTACTTTTTTT |
| 4991 | mamo-L | *BGIBMGA010225* | down | 1744 | 1758 | + | 12.0645 | 2.34E-05 | 1 | TAGGCGGGCTTCGGA |
| 4992 | mamo-L | *BGIBMGA010229* | up | 480 | 494 | - | 11.129 | 5.79E-05 | 1 | TAGGCGAAAGTTTCT |
| 4993 | mamo-L | *BGIBMGA010231* | up | 1779 | 1793 | + | 12.5726 | 1.33E-05 | 1 | TAGGCCAGGTATGTT |
| 4994 | mamo-L | *BGIBMGA010235* | up | 816 | 830 | - | 10.9919 | 6.55E-05 | 1 | TAGGCTTTAGATGGA |
| 4995 | mamo-L | *BGIBMGA010237* | down | 674 | 688 | + | 10.5726 | 9.33E-05 | 1 | TAGGCTAATATTTAT |
| 4996 | mamo-L | *BGIBMGA010238* | up | 690 | 704 | + | 10.5726 | 9.33E-05 | 1 | TAGGCTAATATTTAT |
| 4997 | mamo-L | *BGIBMGA010239* | up | 520 | 534 | - | 12.0645 | 2.34E-05 | 1 | TAGGCGGGCTTCGGA |
| 4998 | mamo-L | *BGIBMGA010240* | up | 187 | 201 | - | 11.9113 | 2.74E-05 | 1 | TAGGCATTTGTTTGT |
| 4999 | mamo-L | *BGIBMGA010240* | up | 365 | 379 | - | 11.2016 | 5.41E-05 | 1 | AAGGCTTTGATTGCC |
| 5000 | mamo-L | *BGIBMGA010242* | down | 720 | 734 | - | 10.629 | 8.91E-05 | 1 | TAGGCAGTTGGCGGT |
| 5001 | mamo-L | *BGIBMGA010242* | down | 1206 | 1220 | - | 10.5565 | 9.46E-05 | 1 | TAGGCGGGATACTGT |
| 5002 | mamo-L | *BGIBMGA010247* | down | 629 | 643 | + | 12.3145 | 1.79E-05 | 1 | TAGGCAGTGATTGTT |
| 5003 | mamo-L | *BGIBMGA010249* | up | 1233 | 1247 | - | 10.8226 | 7.58E-05 | 1 | TAGGCTATTAATGCT |
| 5004 | mamo-L | *BGIBMGA010252* | down | 1130 | 1144 | - | 11.5161 | 4.02E-05 | 1 | AAGGCGGTCTATGCT |
| 5005 | mamo-L | *BGIBMGA010253* | down | 1403 | 1417 | - | 11.6048 | 3.70E-05 | 1 | TAGGCAGGGGTTCCT |
| 5006 | mamo-L | *BGIBMGA010253* | up | 39 | 53 | + | 10.6935 | 8.45E-05 | 1 | TAGGCTGGGGTCTAA |
| 5007 | mamo-L | *BGIBMGA010255* | down | 307 | 321 | + | 10.8145 | 7.63E-05 | 1 | TAGGCTGTGAGCTCT |
| 5008 | mamo-L | *BGIBMGA010261* | down | 1006 | 1020 | - | 13.2823 | 5.66E-06 | 1 | GAGGCTTGTTTTGGT |
| 5009 | mamo-L | *BGIBMGA010262* | up | 225 | 239 | + | 13.5484 | 3.92E-06 | 1 | TAGGCTGAGTGTTTT |
| 5010 | mamo-L | *BGIBMGA010268* | down | 1594 | 1608 | + | 11.5887 | 3.75E-05 | 1 | TAGGCTTATAGTTTT |
| 5011 | mamo-L | *BGIBMGA010270* | up | 33 | 47 | + | 10.7903 | 7.79E-05 | 1 | TAGGCCAATTTTGCG |
| 5012 | mamo-L | *BGIBMGA010273* | down | 1813 | 1827 | + | 12.5323 | 1.39E-05 | 1 | AAGGCTTTGTTTTGT |
| 5013 | mamo-L | *BGIBMGA010273* | down | 1957 | 1971 | - | 12.4758 | 1.48E-05 | 1 | TAGGCTTGGCTTGGC |
| 5014 | mamo-L | *BGIBMGA010278* | down | 1493 | 1507 | + | 11.1613 | 5.63E-05 | 1 | TAGGCGTGGTGGTGT |
| 5015 | mamo-L | *BGIBMGA010281* | down | 165 | 179 | + | 11.9516 | 2.63E-05 | 1 | TAGGCTGTATATTTT |
| 5016 | mamo-L | *BGIBMGA010283* | up | 446 | 460 | + | 11.8952 | 2.79E-05 | 1 | TAGGCCAGCATTGAT |
| 5017 | mamo-L | *BGIBMGA010284* | up | 1453 | 1467 | + | 11.4758 | 4.18E-05 | 1 | AAGGCTTTGTTTCGT |
| 5018 | mamo-L | *BGIBMGA010285* | down | 1806 | 1820 | - | 11.0968 | 5.97E-05 | 1 | TAGGCATGTGTTCGT |
| 5019 | mamo-L | *BGIBMGA010287* | down | 476 | 490 | - | 11.3548 | 4.69E-05 | 1 | AAGGCTGATGATGAT |
| 5020 | mamo-L | *BGIBMGA010287* | down | 984 | 998 | - | 11.2742 | 5.06E-05 | 1 | TAGGCATGCGTTCTT |
| 5021 | mamo-L | *BGIBMGA010290* | up | 587 | 601 | - | 11.8306 | 2.97E-05 | 1 | TAGGCTGTTGCTGAT |
| 5022 | mamo-L | *BGIBMGA010290* | up | 593 | 607 | - | 10.9113 | 7.02E-05 | 1 | GAGGCCTAGGCTGTT |
| 5023 | mamo-L | *BGIBMGA010293* | up | 871 | 885 | - | 11.4435 | 4.31E-05 | 1 | TAGGCTTGAGAGGCT |
| 5024 | mamo-L | *BGIBMGA010293* | down | 1909 | 1923 | + | 11.1774 | 5.54E-05 | 1 | TAGGCTGGTTGTAAT |
| 5025 | mamo-L | *BGIBMGA010296* | up | 1162 | 1176 | + | 10.9194 | 6.98E-05 | 1 | TAGGCTACTTTTTTT |
| 5026 | mamo-L | *BGIBMGA010298* | up | 871 | 885 | - | 11.1129 | 5.88E-05 | 1 | TCGGCTGAGTTTGGC |
| 5027 | mamo-L | *BGIBMGA010301* | down | 1935 | 1949 | - | 10.9194 | 6.98E-05 | 1 | TAGGCTACTTTTTTT |
| 5028 | mamo-L | *BGIBMGA010302* | up | 1240 | 1254 | + | 10.9194 | 6.98E-05 | 1 | TAGGCTACTTTTTTT |
| 5029 | mamo-L | *BGIBMGA010311* | down | 1465 | 1479 | - | 12.3306 | 1.75E-05 | 1 | GAGGCTTGCTTTTCT |
| 5030 | mamo-L | *BGIBMGA010314* | down | 525 | 539 | - | 10.8306 | 7.52E-05 | 1 | AAGGCTTTTTTTTAT |
| 5031 | mamo-L | *BGIBMGA010316* | up | 468 | 482 | - | 10.8065 | 7.69E-05 | 1 | TGGGCGGTGGTTTGT |
| 5032 | mamo-L | *BGIBMGA010317* | up | 1649 | 1663 | + | 11.0726 | 6.09E-05 | 1 | TAGGCTAGTTATTCT |
| 5033 | mamo-L | *BGIBMGA010318* | down | 1865 | 1879 | + | 10.8065 | 7.69E-05 | 1 | TAGGCAGTTGGTTGT |
| 5034 | mamo-L | *BGIBMGA010322* | up | 143 | 157 | - | 10.6613 | 8.67E-05 | 1 | TAGGCCTGTAGTTGC |
| 5035 | mamo-L | *BGIBMGA010326* | up | 420 | 434 | - | 12.2742 | 1.87E-05 | 1 | TAGGCTGAATATTTT |
| 5036 | mamo-L | *BGIBMGA010327* | up | 1441 | 1455 | - | 10.6613 | 8.67E-05 | 1 | TAGGCACGGGTTACT |
| 5037 | mamo-L | *BGIBMGA010329* | up | 1014 | 1028 | - | 11.3226 | 4.83E-05 | 1 | AAGGCTCAGGTGGTT |
| 5038 | mamo-L | *BGIBMGA010330* | down | 1869 | 1883 | - | 10.6129 | 9.03E-05 | 1 | TAGGCTCGTATTGTG |
| 5039 | mamo-L | *BGIBMGA010332* | down | 402 | 416 | + | 12.7258 | 1.11E-05 | 1 | TAGGCGTGCTTTGTA |
| 5040 | mamo-L | *BGIBMGA010332* | up | 1110 | 1124 | + | 11.1774 | 5.54E-05 | 1 | TAGGCAGGCTATTTT |
| 5041 | mamo-L | *BGIBMGA010334* | down | 791 | 805 | - | 11.0645 | 6.14E-05 | 1 | AAGGCCGAGGTTCGC |
| 5042 | mamo-L | *BGIBMGA010336* | up | 1152 | 1166 | - | 11.7742 | 3.14E-05 | 1 | TAGGCCGTGATTAGC |
| 5043 | mamo-L | *BGIBMGA010338* | up | 608 | 622 | - | 10.5806 | 9.27E-05 | 1 | AAGGCTAGTTTTGAT |
| 5044 | mamo-L | *BGIBMGA010340* | down | 1612 | 1626 | + | 13.5403 | 3.97E-06 | 1 | TAGGCGTATTTTGTC |
| 5045 | mamo-L | *BGIBMGA010342* | up | 1805 | 1819 | + | 10.6774 | 8.56E-05 | 1 | GAGGCCTGTGTTGCA |
| 5046 | mamo-L | *BGIBMGA010345* | up | 1803 | 1817 | - | 11.4113 | 4.45E-05 | 1 | TAGGCAGTAGTTTGT |
| 5047 | mamo-L | *BGIBMGA010345* | up | 1677 | 1691 | + | 10.7258 | 8.23E-05 | 1 | AAGGCTCAGAATGGT |
| 5048 | mamo-L | *BGIBMGA010352* | down | 1247 | 1261 | + | 11.3629 | 4.66E-05 | 1 | TAGGCCGATTAGGTC |
| 5049 | mamo-L | *BGIBMGA010352* | down | 811 | 825 | - | 10.8306 | 7.52E-05 | 1 | AAGGCTTTTTTTTAT |
| 5050 | mamo-L | *BGIBMGA010356* | down | 1883 | 1897 | + | 12.6694 | 1.18E-05 | 1 | AAGGCTTAGGTTTTT |
| 5051 | mamo-L | *BGIBMGA010356* | up | 220 | 234 | + | 11.4677 | 4.22E-05 | 1 | TAGGCCTTGACTGCT |
| 5052 | mamo-L | *BGIBMGA010358* | down | 553 | 567 | + | 10.5726 | 9.33E-05 | 1 | TAGGCTAATATTTAT |
| 5053 | mamo-L | *BGIBMGA010359* | up | 735 | 749 | - | 11.9597 | 2.61E-05 | 1 | AAGGCTGTTTTTTTT |

| | | | | | | | | | | |
|---|---|---|---|---|---|---|---|---|---|---|
| 5054 | mamo-L | *BGIBMGA010359* | up | 757 | 771 | - | 10.75 | 8.06E-05 | 1 | TAGGCCGGCGGTTTG |
| 5055 | mamo-L | *BGIBMGA010359* | up | 1744 | 1758 | + | 10.5726 | 9.33E-05 | 1 | TAGGCTAATATTTAT |
| 5056 | mamo-L | *BGIBMGA010365* | down | 1589 | 1603 | - | 11.4516 | 4.28E-05 | 1 | TAGGCGAAGTTTACT |
| 5057 | mamo-L | *BGIBMGA010367* | down | 1135 | 1149 | + | 12.8952 | 9.05E-06 | 1 | TAGGCTTTTGATTGT |
| 5058 | mamo-L | *BGIBMGA010369* | down | 730 | 744 | - | 10.9194 | 6.98E-05 | 1 | TAGGCTACTTTTTTT |
| 5059 | mamo-L | *BGIBMGA010374* | up | 1714 | 1728 | - | 10.5968 | 9.15E-05 | 1 | GAGGCTAGTTTTTGT |
| 5060 | mamo-L | *BGIBMGA010375* | up | 1245 | 1259 | - | 10.5403 | 9.58E-05 | 1 | TATGCTGTGTTTGTT |
| 5061 | mamo-L | *BGIBMGA010379* | up | 1448 | 1462 | - | 10.8306 | 7.52E-05 | 1 | TAGGCAGAGATTGAC |
| 5062 | mamo-L | *BGIBMGA010382* | down | 1538 | 1552 | + | 11.0161 | 6.40E-05 | 1 | TAGGCGCTTTGTGTC |
| 5063 | mamo-L | *BGIBMGA010382* | up | 1513 | 1527 | + | 10.7984 | 7.74E-05 | 1 | GAGGCTATCTTTGAT |
| 5064 | mamo-L | *BGIBMGA010385* | up | 1857 | 1871 | - | 10.9597 | 6.73E-05 | 1 | AAGGCTGATTATTTT |
| 5065 | mamo-L | *BGIBMGA010386* | down | 1161 | 1175 | - | 13.5565 | 3.86E-06 | 1 | TAGGCCTGGTTTTAT |
| 5066 | mamo-L | *BGIBMGA010389* | down | 567 | 581 | + | 12.629 | 1.24E-05 | 1 | TAGGCCTATATCGTT |
| 5067 | mamo-L | *BGIBMGA010391* | down | 676 | 690 | - | 12.621 | 1.25E-05 | 1 | TAGGCTTTCTTGTTT |
| 5068 | mamo-L | *BGIBMGA010397* | up | 819 | 833 | + | 10.9516 | 6.78E-05 | 1 | AAGGCTTCGTTTTTT |
| 5069 | mamo-L | *BGIBMGA010398* | down | 1382 | 1396 | + | 10.7097 | 8.33E-05 | 1 | TAGGCCGTCGAGTAT |
| 5070 | mamo-L | *BGIBMGA010406* | up | 1865 | 1879 | + | 12.3306 | 1.75E-05 | 1 | TAGGCTCTCATTTCT |
| 5071 | mamo-L | *BGIBMGA010409* | down | 1901 | 1915 | + | 10.7581 | 8.00E-05 | 1 | TAGGCCGTGTTTCCG |
| 5072 | mamo-L | *BGIBMGA010409* | down | 386 | 400 | - | 10.5726 | 9.33E-05 | 1 | TAGGCTAATATTTAT |
| 5073 | mamo-L | *BGIBMGA010415* | up | 1628 | 1642 | - | 10.6855 | 8.51E-05 | 1 | TAGGCCTTTTGTGTG |
| 5074 | mamo-L | *BGIBMGA010422* | up | 1296 | 1310 | - | 12.8952 | 9.05E-06 | 1 | TAGGCCCAGGTTATT |
| 5075 | mamo-L | *BGIBMGA010422* | up | 1331 | 1345 | + | 11.75 | 3.21E-05 | 1 | GAGGCTGAGTTTCTT |
| 5076 | mamo-L | *BGIBMGA010424* | up | 75 | 89 | + | 11.8145 | 3.01E-05 | 1 | GAGGCTTTCTGTGTT |
| 5077 | mamo-L | *BGIBMGA010429* | down | 1299 | 1313 | - | 11.2097 | 5.38E-05 | 1 | TAGGCAGAAGTCGCT |
| 5078 | mamo-L | *BGIBMGA010430* | down | 715 | 729 | - | 12.0242 | 2.44E-05 | 1 | TAGGCTTATGTTTTA |
| 5079 | mamo-L | *BGIBMGA010432* | down | 1276 | 1290 | + | 11.379 | 4.58E-05 | 1 | TAGGCTTGGCTTTGC |
| 5080 | mamo-L | *BGIBMGA010433* | up | 1742 | 1756 | - | 10.8629 | 7.33E-05 | 1 | TAGGCTATTTATTTT |
| 5081 | mamo-L | *BGIBMGA010436* | up | 652 | 666 | + | 12.9597 | 8.38E-06 | 1 | AAGGCTGGCTTTGAT |
| 5082 | mamo-L | *BGIBMGA010436* | down | 1027 | 1041 | - | 11.2903 | 4.99E-05 | 1 | TAGGCCTGTAGTTCT |
| 5083 | mamo-L | *BGIBMGA010437* | down | 1476 | 1490 | + | 10.75 | 8.06E-05 | 1 | GAGGCCGATTGTGCC |
| 5084 | mamo-L | *BGIBMGA010440* | down | 307 | 321 | + | 11.0645 | 6.14E-05 | 1 | TAGGCTAACATTTAT |
| 5085 | mamo-L | *BGIBMGA010442* | down | 682 | 696 | + | 10.75 | 8.06E-05 | 1 | AAGGCTGGGATTATT |
| 5086 | mamo-L | *BGIBMGA010444* | up | 1682 | 1696 | - | 12 | 2.50E-05 | 1 | TAGGCTTACTGTGGG |
| 5087 | mamo-L | *BGIBMGA010448* | down | 761 | 775 | - | 11.4758 | 4.18E-05 | 1 | AAGGCCTTTTTTTTT |
| 5088 | mamo-L | *BGIBMGA010449* | up | 1099 | 1113 | - | 12.7258 | 1.11E-05 | 1 | TAGGCGTGCTTTGTA |
| 5089 | mamo-L | *BGIBMGA010450* | down | 1375 | 1389 | + | 11.2742 | 5.06E-05 | 1 | TAGGCCTTAGTGTTT |
| 5090 | mamo-L | *BGIBMGA010453* | up | 1837 | 1851 | + | 11.0565 | 6.18E-05 | 1 | AAGGCGGGCTTCTGT |
| 5091 | mamo-L | *BGIBMGA010454* | up | 951 | 965 | + | 10.7581 | 8.00E-05 | 1 | TAGGCTTGATATGCA |
| 5092 | mamo-L | *BGIBMGA010458* | up | 465 | 479 | + | 13.0565 | 7.51E-06 | 1 | TAGGCGTGGGGTGGC |
| 5093 | mamo-L | *BGIBMGA010459* | up | 1569 | 1583 | + | 11.5806 | 3.78E-05 | 1 | TAGGCCTTTATTATT |
| 5094 | mamo-L | *BGIBMGA010465* | down | 491 | 505 | + | 12.5887 | 1.30E-05 | 1 | TAGGCCAATTTTGAT |
| 5095 | mamo-L | *BGIBMGA010471* | down | 1516 | 1530 | - | 12.5 | 1.44E-05 | 1 | TAGGCGGGTTTTGCA |
| 5096 | mamo-L | *BGIBMGA010471* | up | 1095 | 1109 | - | 10.5806 | 9.27E-05 | 1 | TAGGCTGATGAGTTC |
| 5097 | mamo-L | *BGIBMGA010477* | up | 923 | 937 | + | 13.3548 | 5.14E-06 | 1 | TAGGCTTGTGTTGGA |
| 5098 | mamo-L | *BGIBMGA010477* | up | 1312 | 1326 | - | 11.0242 | 6.36E-05 | 1 | AAGGCCTTGTTTTCC |
| 5099 | mamo-L | *BGIBMGA010478* | down | 278 | 292 | + | 10.75 | 8.06E-05 | 1 | TGGGCTGAGGTTCGT |
| 5100 | mamo-L | *BGIBMGA010480* | down | 1822 | 1836 | + | 11.1774 | 5.54E-05 | 1 | TAGGCTTTTTTTTTG |
| 5101 | mamo-L | *BGIBMGA010480* | down | 1338 | 1352 | - | 11.0645 | 6.14E-05 | 1 | TAGGCTGTTAAGGCT |
| 5102 | mamo-L | *BGIBMGA010482* | up | 111 | 125 | + | 10.75 | 8.06E-05 | 1 | GAGGCCGATTGTGCC |
| 5103 | mamo-L | *BGIBMGA010483* | up | 866 | 880 | + | 11.5242 | 3.99E-05 | 1 | TAGGCCGAATTTCAT |
| 5104 | mamo-L | *BGIBMGA010483* | down | 1798 | 1812 | + | 11.2339 | 5.26E-05 | 1 | TAGGCTACTTTTTGT |
| 5105 | mamo-L | *BGIBMGA010484* | up | 1829 | 1843 | - | 10.8226 | 7.58E-05 | 1 | AAGGCTGTTTTTGCA |
| 5106 | mamo-L | *BGIBMGA010485* | down | 1868 | 1882 | - | 12.1129 | 2.22E-05 | 1 | TAGGCTGGTTCTGGC |
| 5107 | mamo-L | *BGIBMGA010486* | up | 1147 | 1161 | - | 10.6371 | 8.85E-05 | 1 | TAGGCTTTTATTCTC |
| 5108 | mamo-L | *BGIBMGA010488* | down | 1552 | 1566 | + | 12.9274 | 8.74E-06 | 1 | TAGGCCTTTGTTGGA |
| 5109 | mamo-L | *BGIBMGA010489* | up | 968 | 982 | + | 12.9274 | 8.74E-06 | 1 | TAGGCCTTTGTTGGA |
| 5110 | mamo-L | *BGIBMGA010496* | up | 660 | 674 | + | 10.5726 | 9.33E-05 | 1 | TAGGCTAATATTTAT |
| 5111 | mamo-L | *BGIBMGA010498* | down | 611 | 625 | + | 12.1129 | 2.22E-05 | 1 | TAGGCTGGTTCTGGC |
| 5112 | mamo-L | *BGIBMGA010501* | up | 982 | 996 | - | 11.0968 | 5.97E-05 | 1 | TAGGCTGAGATGGTA |
| 5113 | mamo-L | *BGIBMGA010503* | up | 1399 | 1413 | + | 10.9113 | 7.02E-05 | 1 | TAGGCTATTTTCTTT |
| 5114 | mamo-L | *BGIBMGA010515* | down | 950 | 964 | - | 11.2823 | 5.03E-05 | 1 | TAGGCTGTGTCTGAC |
| 5115 | mamo-L | *BGIBMGA010517* | down | 1343 | 1357 | - | 11.0726 | 6.09E-05 | 1 | TAGGCAAAGTTTTGT |
| 5116 | mamo-L | *BGIBMGA010521* | up | 603 | 617 | + | 11.371 | 4.62E-05 | 1 | TAGGCTGTATAGGCT |
| 5117 | mamo-L | *BGIBMGA010523* | down | 165 | 179 | - | 12.1935 | 2.05E-05 | 1 | TAGGCCGGATTTGCA |
| 5118 | mamo-L | *BGIBMGA010523* | down | 313 | 327 | + | 12.0887 | 2.29E-05 | 1 | TAGGCCATTTTTGCC |
| 5119 | mamo-L | *BGIBMGA010528* | down | 817 | 831 | + | 10.7903 | 7.79E-05 | 1 | TAGGCGGATGATAAT |
| 5120 | mamo-L | *BGIBMGA010530* | down | 440 | 454 | - | 11.9032 | 2.76E-05 | 1 | TAGGCAGGGGTTGGA |
| 5121 | mamo-L | *BGIBMGA010531* | up | 770 | 784 | + | 12.0645 | 2.34E-05 | 1 | TAGGCGGGCTTCGGA |
| 5122 | mamo-L | *BGIBMGA010533* | down | 541 | 555 | + | 10.871 | 7.28E-05 | 1 | TAGGCTCAGATTATC |
| 5123 | mamo-L | *BGIBMGA010534* | down | 1968 | 1982 | - | 10.5484 | 9.52E-05 | 1 | AAGGCCTAGGTTGAA |
| 5124 | mamo-L | *BGIBMGA010535* | up | 315 | 329 | - | 12.7823 | 1.03E-05 | 1 | TAGGCAGGGGATGGT |
| 5125 | mamo-L | *BGIBMGA010539* | up | 298 | 312 | + | 11.3629 | 4.66E-05 | 1 | TAGGCTTTTTTAGCT |
| 5126 | mamo-L | *BGIBMGA010540* | down | 903 | 917 | - | 12.9919 | 8.10E-06 | 1 | TAGGCCGGTTTTGCA |
| 5127 | mamo-L | *BGIBMGA010542* | up | 995 | 1009 | + | 11.7823 | 3.11E-05 | 1 | TAGGCGTGGGGCGGC |
| 5128 | mamo-L | *BGIBMGA010545* | up | 1152 | 1166 | - | 12.3387 | 1.73E-05 | 1 | TAGGCCGTGGTTTCA |
| 5129 | mamo-L | *BGIBMGA010545* | up | 1446 | 1460 | - | 11.0242 | 6.36E-05 | 1 | TGGGCTGATGATGGT |
| 5130 | mamo-L | *BGIBMGA010546* | down | 1017 | 1031 | + | 10.879 | 7.22E-05 | 1 | TAGGCCAGTAATGCT |
| 5131 | mamo-L | *BGIBMGA010547* | down | 709 | 723 | + | 10.6452 | 8.79E-05 | 1 | AAGGCCGAGATTATT |
| 5132 | mamo-L | *BGIBMGA010548* | down | 21 | 35 | + | 11.0645 | 6.14E-05 | 1 | AAGGCAGTTTTTGGT |
| 5133 | mamo-L | *BGIBMGA010549* | down | 117 | 131 | + | 10.75 | 8.06E-05 | 1 | AAGGCAGTTTTTGTT |
| 5134 | mamo-L | *BGIBMGA010549* | up | 982 | 996 | + | 10.5887 | 9.21E-05 | 1 | TAGGCATATGTGTGT |
| 5135 | mamo-L | *BGIBMGA010556* | up | 586 | 600 | + | 11.871 | 2.85E-05 | 1 | TAGGCTGTGGATAGC |
| 5136 | mamo-L | *BGIBMGA010557* | up | 769 | 783 | + | 11.8306 | 2.97E-05 | 1 | TAGGCTGTTGCTGAT |
| 5137 | mamo-L | *BGIBMGA010557* | up | 763 | 777 | + | 10.9113 | 7.02E-05 | 1 | GAGGCCTAGGCTGTT |
| 5138 | mamo-L | *BGIBMGA010557* | up | 1129 | 1143 | - | 10.7823 | 7.84E-05 | 1 | AAGGCTTTTGATTGT |
| 5139 | mamo-L | *BGIBMGA010558* | down | 1259 | 1273 | - | 10.7097 | 8.33E-05 | 1 | TAGGCCGTCGAGTAT |
| 5140 | mamo-L | *BGIBMGA010559* | up | 1453 | 1467 | - | 10.9194 | 6.98E-05 | 1 | TAGGCTACTTTTTTT |
| 5141 | mamo-L | *BGIBMGA010561* | up | 156 | 170 | - | 10.9516 | 6.78E-05 | 1 | AAGGCTTCGTTTTTT |
| 5142 | mamo-L | *BGIBMGA010563* | up | 1324 | 1338 | - | 11.25 | 5.18E-05 | 1 | TAGGCCCATTTCAGT |
| 5143 | mamo-L | *BGIBMGA010564* | down | 1642 | 1656 | - | 11.6774 | 3.44E-05 | 1 | TAGGCTAATTTTTAT |
| 5144 | mamo-L | *BGIBMGA010567* | up | 108 | 122 | - | 11.0484 | 6.23E-05 | 1 | AAGGCTTTCTTTGTA |
| 5145 | mamo-L | *BGIBMGA010568* | down | 648 | 662 | + | 13.5565 | 3.86E-06 | 1 | TAGGCCTGGTTTTAT |

| | | | | | | | | | | |
|---|---|---|---|---|---|---|---|---|---|---|
| 5146 | mamo-L | *BGIBMGA010569* | down | 1504 | 1518 | + | 12.7742 | 1.04E-05 | 1 | TAGGCTGACGGCGGC |
| 5147 | mamo-L | *BGIBMGA010569* | down | 142 | 156 | + | 10.7016 | 8.39E-05 | 1 | TAGGCCTCCAATTGT |
| 5148 | mamo-L | *BGIBMGA010571* | down | 653 | 667 | - | 10.7419 | 8.11E-05 | 1 | TAGGCGTTGATTTAC |
| 5149 | mamo-L | *BGIBMGA010572* | down | 972 | 986 | - | 10.5081 | 9.83E-05 | 1 | TAGGCTAAATATGTC |
| 5150 | mamo-L | *BGIBMGA010573* | up | 1552 | 1566 | - | 11.5968 | 3.73E-05 | 1 | TAGGCACGGGTTTTT |
| 5151 | mamo-L | *BGIBMGA010577* | down | 1919 | 1933 | + | 11.9839 | 2.54E-05 | 1 | TAGGCTCACGTCTGC |
| 5152 | mamo-L | *BGIBMGA010578* | up | 200 | 214 | + | 12.4677 | 1.49E-05 | 1 | TAGGCTTCTATTGCT |
| 5153 | mamo-L | *BGIBMGA010578* | up | 1362 | 1376 | - | 11.5968 | 3.73E-05 | 1 | TAGGCCATGGTTCCT |
| 5154 | mamo-L | *BGIBMGA010578* | up | 1414 | 1428 | - | 10.9758 | 6.64E-05 | 1 | TAGGCTGGTGGCGGA |
| 5155 | mamo-L | *BGIBMGA010579* | down | 357 | 371 | + | 10.8871 | 7.17E-05 | 1 | TAGGCGTAATTTAAT |
| 5156 | mamo-L | *BGIBMGA010580* | down | 466 | 480 | - | 12.0645 | 2.34E-05 | 1 | TAGGCGGGCTTCGGA |
| 5157 | mamo-L | *BGIBMGA010587* | down | 1584 | 1598 | + | 12.8468 | 9.58E-06 | 1 | AAGGCTTAGATTGGT |
| 5158 | mamo-L | *BGIBMGA010587* | up | 945 | 959 | - | 10.9194 | 6.98E-05 | 1 | TAGGCTACTTTTTTT |
| 5159 | mamo-L | *BGIBMGA010588* | up | 1941 | 1955 | + | 10.9839 | 6.59E-05 | 1 | TAGGCATAAATTGTT |
| 5160 | mamo-L | *BGIBMGA010589* | up | 1621 | 1635 | + | 11.0161 | 6.40E-05 | 1 | TAGGCTGGGATGGTA |
| 5161 | mamo-L | *BGIBMGA010601* | up | 1808 | 1822 | - | 11 | 6.50E-05 | 1 | AAGGCTGTTTTCTGT |
| 5162 | mamo-L | *BGIBMGA010602* | up | 959 | 973 | - | 11.8952 | 2.79E-05 | 1 | TAGGCTTCCATTTTT |
| 5163 | mamo-L | *BGIBMGA010604* | down | 1041 | 1055 | - | 13.3306 | 5.30E-06 | 1 | TAGGCTTAGTATTTT |
| 5164 | mamo-L | *BGIBMGA010604* | down | 402 | 416 | - | 11.4113 | 4.45E-05 | 1 | TAGGCTTAATTTTGA |
| 5165 | mamo-L | *BGIBMGA010607* | down | 1604 | 1618 | - | 11.8629 | 2.87E-05 | 1 | TAGGCTTCCGGCGGT |
| 5166 | mamo-L | *BGIBMGA010607* | up | 1393 | 1407 | + | 11.3952 | 4.52E-05 | 1 | TAGGCCGAAGGTCGT |
| 5167 | mamo-L | *BGIBMGA010609* | down | 1491 | 1505 | + | 10.75 | 8.06E-05 | 1 | GAGGCCGATTGTGCC |
| 5168 | mamo-L | *BGIBMGA010611* | up | 591 | 605 | - | 10.7419 | 8.11E-05 | 1 | TACGCTTACTTTGCT |
| 5169 | mamo-L | *BGIBMGA010616* | up | 866 | 880 | - | 13.5 | 4.19E-06 | 1 | TAGGCTTTGTTTTAT |
| 5170 | mamo-L | *BGIBMGA010619* | up | 1666 | 1680 | + | 10.629 | 8.91E-05 | 1 | TAGGCTGAATAGTTT |
| 5171 | mamo-L | *BGIBMGA010620* | down | 183 | 197 | - | 11.2823 | 5.03E-05 | 1 | TAGGCTCTTATTAGT |
| 5172 | mamo-L | *BGIBMGA010626* | up | 267 | 281 | - | 13.121 | 6.92E-06 | 1 | TAGGCCGGTTTTTCC |
| 5173 | mamo-L | *BGIBMGA010627* | down | 1869 | 1883 | + | 10.871 | 7.28E-05 | 1 | TGGGCCTTGTTTTGT |
| 5174 | mamo-L | *BGIBMGA010629* | down | 700 | 714 | + | 12.5161 | 1.42E-05 | 1 | GAGGCTGATTTTGAT |
| 5175 | mamo-L | *BGIBMGA010630* | up | 37 | 51 | + | 12.5161 | 1.42E-05 | 1 | GAGGCTGATTTTGAT |
| 5176 | mamo-L | *BGIBMGA010631* | down | 1789 | 1803 | + | 12.2258 | 1.97E-05 | 1 | TAGGCAGGTGATGGT |
| 5177 | mamo-L | *BGIBMGA010639* | down | 590 | 604 | - | 10.9194 | 6.98E-05 | 1 | TAGGCTACTTTTTTT |
| 5178 | mamo-L | *BGIBMGA010641* | down | 884 | 898 | + | 12.0403 | 2.40E-05 | 1 | TAGGCCTTTGTGTCT |
| 5179 | mamo-L | *BGIBMGA010641* | up | 963 | 977 | - | 10.5 | 9.90E-05 | 1 | GAGGCCCAGGTCGTC |
| 5180 | mamo-L | *BGIBMGA010642* | down | 621 | 635 | - | 12.4355 | 1.55E-05 | 1 | TAGGCCTTCTTGTTT |
| 5181 | mamo-L | *BGIBMGA010642* | up | 329 | 343 | + | 10.9194 | 6.98E-05 | 1 | TAGGCTACTTTTTTT |
| 5182 | mamo-L | *BGIBMGA010644* | up | 498 | 512 | + | 10.7661 | 7.95E-05 | 1 | TAGGCCGTCTGCACT |
| 5183 | mamo-L | *BGIBMGA010645* | up | 1019 | 1033 | + | 12.0081 | 2.48E-05 | 1 | TAGGCAGGTTTTTTT |
| 5184 | mamo-L | *BGIBMGA010645* | down | 530 | 544 | + | 10.879 | 7.22E-05 | 1 | AAGGCTTGCGTGTTT |
| 5185 | mamo-L | *BGIBMGA010646* | up | 935 | 949 | + | 10.879 | 7.22E-05 | 1 | AAGGCTTGCGTGTTT |
| 5186 | mamo-L | *BGIBMGA010650* | up | 1523 | 1537 | + | 12.6774 | 1.17E-05 | 1 | TAGGCGTGAGGTGGT |
| 5187 | mamo-L | *BGIBMGA010655* | up | 929 | 943 | - | 11.5323 | 3.96E-05 | 1 | TAGGCGTGAGGTGAT |
| 5188 | mamo-L | *BGIBMGA010666* | up | 1410 | 1424 | - | 11.2742 | 5.06E-05 | 1 | GAGGCGTATTTTTTT |
| 5189 | mamo-L | *BGIBMGA010668* | down | 912 | 926 | + | 11.2823 | 5.03E-05 | 1 | TAGGCTCTTATTAGT |
| 5190 | mamo-L | *BGIBMGA010674* | up | 777 | 791 | + | 10.5 | 9.90E-05 | 1 | GAGGCCCAGGTCGTC |
| 5191 | mamo-L | *BGIBMGA010676* | down | 1113 | 1127 | + | 14.2016 | 1.47E-06 | 1 | TAGGCTCTTGTTGTT |
| 5192 | mamo-L | *BGIBMGA010676* | up | 1762 | 1776 | + | 12.75 | 1.08E-05 | 1 | TAGGCCTCGGATGCT |
| 5193 | mamo-L | *BGIBMGA010678* | down | 1883 | 1897 | + | 10.871 | 7.28E-05 | 1 | TAGGCCAAGATTATT |
| 5194 | mamo-L | *BGIBMGA010680* | down | 1326 | 1340 | + | 11.5081 | 4.05E-05 | 1 | AAGGCTGGCGGTGCC |
| 5195 | mamo-L | *BGIBMGA010683* | down | 1137 | 1151 | + | 13.3306 | 5.30E-06 | 1 | TAGGCTTAGTATTTT |
| 5196 | mamo-L | *BGIBMGA010683* | down | 1776 | 1790 | + | 11.4113 | 4.45E-05 | 1 | TAGGCTTAATTTTGA |
| 5197 | mamo-L | *BGIBMGA010683* | up | 1986 | 2000 | - | 11.3306 | 4.80E-05 | 1 | TGGGCTCAGTTTGCT |
| 5198 | mamo-L | *BGIBMGA010685* | down | 1369 | 1383 | - | 11.5565 | 3.87E-05 | 1 | AAGGCGTAGTGTGTT |
| 5199 | mamo-L | *BGIBMGA010686* | down | 258 | 272 | - | 13.7742 | 2.79E-06 | 1 | TAGGCGGGTGGTGGT |
| 5200 | mamo-L | *BGIBMGA010688* | down | 62 | 76 | + | 10.5726 | 9.33E-05 | 1 | TAGGCTAATATTTAT |
| 5201 | mamo-L | *BGIBMGA010693* | up | 1235 | 1249 | + | 13.2903 | 5.60E-06 | 1 | TAGGCTTAAATTGTT |
| 5202 | mamo-L | *BGIBMGA010693* | up | 990 | 1004 | + | 10.5726 | 9.33E-05 | 1 | AAGGCCAGCGTCGTT |
| 5203 | mamo-L | *BGIBMGA010694* | down | 391 | 405 | - | 10.879 | 7.22E-05 | 1 | AAGGCTTATATTTTT |
| 5204 | mamo-L | *BGIBMGA010696* | up | 1009 | 1023 | - | 11.1774 | 5.54E-05 | 1 | TAGGCGTACTGGTGT |
| 5205 | mamo-L | *BGIBMGA010697* | up | 531 | 545 | - | 10.9435 | 6.83E-05 | 1 | TAGGCCGGTGGTACC |
| 5206 | mamo-L | *BGIBMGA010703* | up | 467 | 481 | - | 13.1774 | 6.47E-06 | 1 | TAGGCTGTAGATGTT |
| 5207 | mamo-L | *BGIBMGA010704* | down | 1294 | 1308 | + | 11.2742 | 5.06E-05 | 1 | AAGGCTTTCGTCGAT |
| 5208 | mamo-L | *BGIBMGA010704* | down | 783 | 797 | + | 10.871 | 7.28E-05 | 1 | TAGGCTGCGTGTCCT |
| 5209 | mamo-L | *BGIBMGA010707* | down | 1163 | 1177 | + | 11.629 | 3.61E-05 | 1 | TAGGCTTTTTGTCGT |
| 5210 | mamo-L | *BGIBMGA010708* | down | 423 | 437 | - | 12.5484 | 1.36E-05 | 1 | TAGGCCTACGATTGC |
| 5211 | mamo-L | *BGIBMGA010713* | down | 1856 | 1870 | + | 10.5726 | 9.33E-05 | 1 | TAGGCTAATATTTAT |
| 5212 | mamo-L | *BGIBMGA010715* | down | 729 | 743 | - | 10.4919 | 9.96E-05 | 1 | AAGGCGGTTTTTGGA |
| 5213 | mamo-L | *BGIBMGA010716* | up | 1120 | 1134 | - | 11.5968 | 3.73E-05 | 1 | TAGGCCTTTTGTAGT |
| 5214 | mamo-L | *BGIBMGA010720* | up | 589 | 603 | - | 12.629 | 1.24E-05 | 1 | TAGGCGGGTGGTGAT |
| 5215 | mamo-L | *BGIBMGA010720* | up | 949 | 963 | - | 11.4274 | 4.38E-05 | 1 | TAGGCAGTCGTCTGT |
| 5216 | mamo-L | *BGIBMGA010721* | down | 373 | 387 | - | 10.5726 | 9.33E-05 | 1 | TAGGCTAATATTTAT |
| 5217 | mamo-L | *BGIBMGA010727* | up | 1135 | 1149 | + | 12.0484 | 2.38E-05 | 1 | TAGGCCGATAATGAT |
| 5218 | mamo-L | *BGIBMGA010727* | down | 378 | 392 | + | 10.5726 | 9.33E-05 | 1 | TAGGCTAATATTTAT |
| 5219 | mamo-L | *BGIBMGA010730* | down | 1512 | 1526 | + | 10.871 | 7.28E-05 | 1 | TAGGCTCTTGTGAGT |
| 5220 | mamo-L | *BGIBMGA010731* | up | 195 | 209 | + | 12.1855 | 2.06E-05 | 1 | TAGGCTGTATTTCCT |
| 5221 | mamo-L | *BGIBMGA010731* | up | 1824 | 1838 | + | 10.5161 | 9.77E-05 | 1 | AAGGCTGTCTTTGAA |
| 5222 | mamo-L | *BGIBMGA010733* | up | 1200 | 1214 | + | 11.379 | 4.58E-05 | 1 | TAGGCTATAATTGTT |
| 5223 | mamo-L | *BGIBMGA010734* | up | 1386 | 1400 | - | 10.5 | 9.90E-05 | 1 | AAGGCATAGTTTGAT |
| 5224 | mamo-L | *BGIBMGA010737* | down | 301 | 315 | + | 11.2016 | 5.41E-05 | 1 | TAGGCAGTCTTTCTT |
| 5225 | mamo-L | *BGIBMGA010738* | up | 423 | 437 | + | 11.0484 | 6.23E-05 | 1 | TAGGCTGTATATATT |
| 5226 | mamo-L | *BGIBMGA010738* | down | 751 | 765 | + | 10.6613 | 8.67E-05 | 1 | AAGGCAGAGGTTTTT |
| 5227 | mamo-L | *BGIBMGA010740* | up | 250 | 264 | + | 10.5968 | 9.15E-05 | 1 | GAGGCTAGTTTTTGT |
| 5228 | mamo-L | *BGIBMGA010742* | down | 843 | 857 | - | 10.9194 | 6.98E-05 | 1 | TAGGCTACTTTTTTT |
| 5229 | mamo-L | *BGIBMGA010743* | up | 894 | 908 | + | 10.9194 | 6.98E-05 | 1 | TAGGCTACTTTTTTT |
| 5230 | mamo-L | *BGIBMGA010746* | up | 286 | 300 | + | 12.4032 | 1.61E-05 | 1 | TAGGCTCAAGATGTT |
| 5231 | mamo-L | *BGIBMGA010753* | up | 131 | 145 | + | 11.6129 | 3.66E-05 | 1 | TAGGCTCACGGTATT |
| 5232 | mamo-L | *BGIBMGA010755* | up | 1226 | 1240 | + | 10.5 | 9.90E-05 | 1 | GAGGCCCAGGTCGTC |
| 5233 | mamo-L | *BGIBMGA010760* | down | 441 | 455 | + | 10.7339 | 8.17E-05 | 1 | AAGGCGGTGTTTGTA |
| 5234 | mamo-L | *BGIBMGA010761* | down | 152 | 166 | + | 10.9194 | 6.98E-05 | 1 | GAGGCCGCTGTTTGT |
| 5235 | mamo-L | *BGIBMGA010765* | up | 8 | 22 | + | 12.0161 | 2.46E-05 | 1 | TAGGCGGGCTGTGAC |
| 5236 | mamo-L | *BGIBMGA010768* | down | 1168 | 1182 | + | 11.121 | 5.84E-05 | 1 | TAGGCGAGGTTCGCC |
| 5237 | mamo-L | *BGIBMGA010773* | up | 535 | 549 | - | 10.5403 | 9.58E-05 | 1 | GAGGCGGGCGCTGCT |

| | | | | | | | | | | |
|---|---|---|---|---|---|---|---|---|---|---|
| 5238 | mamo-L | *BGIBMGA010775* | down | 1380 | 1394 | - | 11.379 | 4.58E-05 | 1 | TAGGCCAACGTTAGC |
| 5239 | mamo-L | *BGIBMGA010776* | up | 1772 | 1786 | + | 11.379 | 4.58E-05 | 1 | TAGGCCAACGTTAGC |
| 5240 | mamo-L | *BGIBMGA010778* | up | 1322 | 1336 | - | 11.2581 | 5.14E-05 | 1 | TAGGCCTGAGAGGCT |
| 5241 | mamo-L | *BGIBMGA010779* | up | 1515 | 1529 | + | 10.5645 | 9.40E-05 | 1 | TAGGCTTCGTCCGGT |
| 5242 | mamo-L | *BGIBMGA010781* | down | 1205 | 1219 | - | 12.0645 | 2.34E-05 | 1 | TAGGCTCTCGACGCT |
| 5243 | mamo-L | *BGIBMGA010781* | down | 647 | 661 | + | 11.0403 | 6.28E-05 | 1 | TAGGCCTGAAATGGC |
| 5244 | mamo-L | *BGIBMGA010781* | up | 468 | 482 | + | 10.5726 | 9.33E-05 | 1 | TAGGCTAATATTTAT |
| 5245 | mamo-L | *BGIBMGA010782* | up | 934 | 948 | - | 11.6048 | 3.70E-05 | 1 | TAGGCTTGTTGCGAT |
| 5246 | mamo-L | *BGIBMGA010782* | down | 1237 | 1251 | - | 10.5726 | 9.33E-05 | 1 | TAGGCTAATATTTAT |
| 5247 | mamo-L | *BGIBMGA010783* | down | 890 | 904 | - | 14.5887 | 7.72E-07 | 1 | TAGGCTTACTTTTTT |
| 5248 | mamo-L | *BGIBMGA010783* | down | 1274 | 1288 | - | 11.0242 | 6.36E-05 | 1 | GAGGCTGTTTTTATT |
| 5249 | mamo-L | *BGIBMGA010783* | up | 1856 | 1870 | - | 10.5726 | 9.33E-05 | 1 | TAGGCTAATATTTAT |
| 5250 | mamo-L | *BGIBMGA010784* | down | 1707 | 1721 | - | 11.4677 | 4.22E-05 | 1 | TAGGCCGCTGGTGTC |
| 5251 | mamo-L | *BGIBMGA010788* | up | 1360 | 1374 | + | 12.5726 | 1.33E-05 | 1 | TAGGCAGGGTGTGGT |
| 5252 | mamo-L | *BGIBMGA010795* | down | 1310 | 1324 | - | 10.9194 | 6.98E-05 | 1 | TAGGCTACTTTTTTT |
| 5253 | mamo-L | *BGIBMGA010795* | down | 1844 | 1858 | + | 10.5726 | 9.33E-05 | 1 | TAGGCTAATATTTAT |
| 5254 | mamo-L | *BGIBMGA010795* | down | 1543 | 1557 | + | 10.4919 | 9.96E-05 | 1 | TAGCCTAAGGTTGGT |
| 5255 | mamo-L | *BGIBMGA010796* | down | 1785 | 1799 | + | 11.0403 | 6.28E-05 | 1 | TAGGCTTTTTGGTGT |
| 5256 | mamo-L | *BGIBMGA010797* | down | 1976 | 1990 | - | 12.7258 | 1.11E-05 | 1 | TAGGCGTGCTTTGTA |
| 5257 | mamo-L | *BGIBMGA010797* | down | 485 | 499 | + | 10.9758 | 6.64E-05 | 1 | GAGGCTTTCGTCTTT |
| 5258 | mamo-L | *BGIBMGA010797* | up | 865 | 879 | + | 10.7419 | 8.11E-05 | 1 | TAGGCTTATTATCTC |
| 5259 | mamo-L | *BGIBMGA010797* | down | 1351 | 1365 | - | 10.5726 | 9.33E-05 | 1 | TAGGCTAATATTTAT |
| 5260 | mamo-L | *BGIBMGA010798* | up | 474 | 488 | - | 10.9677 | 6.68E-05 | 1 | TAGGCAGTATTTTTT |
| 5261 | mamo-L | *BGIBMGA010801* | up | 1392 | 1406 | - | 10.5726 | 9.33E-05 | 1 | TAGGCTAATATTTAT |
| 5262 | mamo-L | *BGIBMGA010802* | down | 1264 | 1278 | + | 10.9194 | 6.98E-05 | 1 | TAGGCTACTTTTTTT |
| 5263 | mamo-L | *BGIBMGA010804* | down | 152 | 166 | + | 11.0968 | 5.97E-05 | 1 | TAGGCGGGCGTATGT |
| 5264 | mamo-L | *BGIBMGA010805* | up | 1426 | 1440 | + | 11.0968 | 5.97E-05 | 1 | TAGGCGGGCGTATGT |
| 5265 | mamo-L | *BGIBMGA010806* | down | 1597 | 1611 | - | 10.6613 | 8.67E-05 | 1 | AAGGCAGAGGTTTTT |
| 5266 | mamo-L | *BGIBMGA010806* | down | 192 | 206 | - | 10.5726 | 9.33E-05 | 1 | TAGGCTAATATTTAT |
| 5267 | mamo-L | *BGIBMGA010808* | down | 1008 | 1022 | - | 12.7823 | 1.03E-05 | 1 | TAGGCCGTTATTTTT |
| 5268 | mamo-L | *BGIBMGA010809* | down | 1852 | 1866 | + | 11.5968 | 3.73E-05 | 1 | TAGGCGCAATATGTT |
| 5269 | mamo-L | *BGIBMGA010812* | up | 1403 | 1417 | + | 10.5565 | 9.46E-05 | 1 | TAGGCGTGGTTCCCC |
| 5270 | mamo-L | *BGIBMGA010813* | down | 932 | 946 | - | 11.0484 | 6.23E-05 | 1 | TAGGCCTAATATCGT |
| 5271 | mamo-L | *BGIBMGA010815* | down | 1730 | 1744 | + | 11.2177 | 5.34E-05 | 1 | TAGGCGTATTTTTTA |
| 5272 | mamo-L | *BGIBMGA010816* | down | 1574 | 1588 | + | 10.7097 | 8.33E-05 | 1 | GAGGCGTACTTTCTT |
| 5273 | mamo-L | *BGIBMGA010817* | down | 505 | 519 | + | 10.5726 | 9.33E-05 | 1 | TAGGCTAATATTTAT |
| 5274 | mamo-L | *BGIBMGA010818* | up | 1647 | 1661 | - | 12.1855 | 2.06E-05 | 1 | TAGGCTGTATTTCCT |
| 5275 | mamo-L | *BGIBMGA010818* | up | 18 | 32 | - | 10.5161 | 9.77E-05 | 1 | AAGGCTGTCTTTGAA |
| 5276 | mamo-L | *BGIBMGA010822* | down | 149 | 163 | - | 11.5484 | 3.90E-05 | 1 | TAGGCCTGTTTTTGG |
| 5277 | mamo-L | *BGIBMGA010823* | up | 771 | 785 | + | 10.9194 | 6.98E-05 | 1 | TAGGCTACTTTTTTT |
| 5278 | mamo-L | *BGIBMGA010824* | down | 1832 | 1846 | - | 10.5968 | 9.15E-05 | 1 | GAGGCTAGTTTTTGT |
| 5279 | mamo-L | *BGIBMGA010825* | up | 1225 | 1239 | - | 10.9194 | 6.98E-05 | 1 | TAGGCTACTTTTTTT |
| 5280 | mamo-L | *BGIBMGA010826* | up | 1466 | 1480 | - | 12.1855 | 2.06E-05 | 1 | TAGGCCTAGGTCGCA |
| 5281 | mamo-L | *BGIBMGA010828* | up | 1152 | 1166 | + | 12.3629 | 1.69E-05 | 1 | TAGGCCTACATTACT |
| 5282 | mamo-L | *BGIBMGA010834* | down | 573 | 587 | + | 10.7016 | 8.39E-05 | 1 | TAGGCTGAGGTAATT |
| 5283 | mamo-L | *BGIBMGA010836* | down | 1042 | 1056 | - | 11.1774 | 5.54E-05 | 1 | AAGGCTGATTTTGTA |
| 5284 | mamo-L | *BGIBMGA010841* | down | 1891 | 1905 | + | 10.7016 | 8.39E-05 | 1 | AAGGCTTTTTTCTGT |
| 5285 | mamo-L | *BGIBMGA010845* | down | 1307 | 1321 | + | 13.5887 | 3.68E-06 | 1 | TAGGCCTTTTTTTTT |
| 5286 | mamo-L | *BGIBMGA010845* | up | 1384 | 1398 | - | 11.4194 | 4.42E-05 | 1 | TAGGCGTTGTTTTCA |
| 5287 | mamo-L | *BGIBMGA010846* | down | 795 | 809 | - | 12.1694 | 2.09E-05 | 1 | TAGGCTAACTTTTAT |
| 5288 | mamo-L | *BGIBMGA010847* | up | 780 | 794 | - | 12.1694 | 2.09E-05 | 1 | TAGGCTAACTTTTAT |
| 5289 | mamo-L | *BGIBMGA010853* | down | 439 | 453 | + | 12.2177 | 1.99E-05 | 1 | TAGGCGATCGTTGCC |
| 5290 | mamo-L | *BGIBMGA010854* | up | 313 | 327 | - | 11.9194 | 2.72E-05 | 1 | TAGGCCGATATTGGG |
| 5291 | mamo-L | *BGIBMGA010856* | up | 200 | 214 | - | 11.2258 | 5.30E-05 | 1 | TAGGCCGCCGTCCGT |
| 5292 | mamo-L | *BGIBMGA010857* | down | 858 | 872 | + | 13.7742 | 2.79E-06 | 1 | TAGGCTTTTTTTTTT |
| 5293 | mamo-L | *BGIBMGA010857* | down | 199 | 213 | + | 11.3629 | 4.66E-05 | 1 | TAGGCCGACTGTGAA |
| 5294 | mamo-L | *BGIBMGA010857* | down | 77 | 91 | + | 10.9919 | 6.55E-05 | 1 | TAGGCATAATTTTTT |
| 5295 | mamo-L | *BGIBMGA010859* | up | 924 | 938 | + | 10.5726 | 9.33E-05 | 1 | AAGGCTCACGATGCC |
| 5296 | mamo-L | *BGIBMGA010861* | down | 223 | 237 | - | 10.6532 | 8.73E-05 | 1 | TAGGCTAGGTTTGAG |
| 5297 | mamo-L | *BGIBMGA010862* | down | 355 | 369 | - | 12.5081 | 1.43E-05 | 1 | GAGGCTGACTTGGGT |
| 5298 | mamo-L | *BGIBMGA010863* | up | 1172 | 1186 | - | 12.7258 | 1.11E-05 | 1 | TAGGCGTGCTTTGTA |
| 5299 | mamo-L | *BGIBMGA010866* | up | 1686 | 1700 | + | 11.3548 | 4.69E-05 | 1 | AAGGCTGATGATGAT |
| 5300 | mamo-L | *BGIBMGA010869* | up | 1798 | 1812 | + | 11.1855 | 5.50E-05 | 1 | TAGGCTGAAATCTCT |
| 5301 | mamo-L | *BGIBMGA010871* | up | 134 | 148 | + | 11.1129 | 5.88E-05 | 1 | TGGGCCCACTTTGTT |
| 5302 | mamo-L | *BGIBMGA010871* | down | 658 | 672 | + | 11.0645 | 6.14E-05 | 1 | TAGGCTGTAGGTACT |
| 5303 | mamo-L | *BGIBMGA010875* | down | 850 | 864 | - | 10.9194 | 6.98E-05 | 1 | TAGGCTACTTTTTTT |
| 5304 | mamo-L | *BGIBMGA010876* | down | 1888 | 1902 | + | 12.5 | 1.44E-05 | 1 | TAGGCTTAGGTTTGG |
| 5305 | mamo-L | *BGIBMGA010880* | up | 1716 | 1730 | + | 13.5323 | 4.01E-06 | 1 | TAGGCTTGATTTTGT |
| 5306 | mamo-L | *BGIBMGA010880* | down | 422 | 436 | - | 11.0645 | 6.14E-05 | 1 | TAGGCTGTAGGTACT |
| 5307 | mamo-L | *BGIBMGA010881* | up | 208 | 222 | - | 11.1129 | 5.88E-05 | 1 | TGGGCCCACTTTGTT |
| 5308 | mamo-L | *BGIBMGA010887* | up | 704 | 718 | + | 12.4677 | 1.49E-05 | 1 | TAGGCTTCTATTGCT |
| 5309 | mamo-L | *BGIBMGA010896* | down | 873 | 887 | + | 13.0726 | 7.34E-06 | 1 | TAGGCCGATTTTGCA |
| 5310 | mamo-L | *BGIBMGA010897* | down | 1458 | 1472 | + | 10.5726 | 9.33E-05 | 1 | TAGGCTAATATTTAT |
| 5311 | mamo-L | *BGIBMGA010898* | down | 1985 | 1999 | + | 10.6532 | 8.73E-05 | 1 | TAGGCCAGCATTTTC |
| 5312 | mamo-L | *BGIBMGA010900* | up | 241 | 255 | + | 10.9113 | 7.02E-05 | 1 | GAGGCTTGCGGCGTT |
| 5313 | mamo-L | *BGIBMGA010902* | down | 311 | 325 | + | 12.8629 | 9.42E-06 | 1 | TAGGCGGGGTTGTGT |
| 5314 | mamo-L | *BGIBMGA010904* | up | 703 | 717 | - | 11.6371 | 3.58E-05 | 1 | GAGGCTCACATTGTT |
| 5315 | mamo-L | *BGIBMGA010909* | up | 1314 | 1328 | - | 12.8387 | 9.67E-06 | 1 | AAGGCTGAGTTTTTT |
| 5316 | mamo-L | *BGIBMGA010911* | up | 276 | 290 | - | 11.8065 | 3.03E-05 | 1 | TAGGCGTACGGTTAT |
| 5317 | mamo-L | *BGIBMGA010913* | up | 104 | 118 | - | 10.9355 | 6.88E-05 | 1 | GAGGCTGATTGTGCC |
| 5318 | mamo-L | *BGIBMGA010916* | down | 1119 | 1133 | + | 10.5161 | 9.77E-05 | 1 | TAGGCACGCTATGGC |
| 5319 | mamo-L | *BGIBMGA010927* | down | 426 | 440 | - | 11.1774 | 5.54E-05 | 1 | TAGGCTTTTTTTTTG |
| 5320 | mamo-L | *BGIBMGA010928* | up | 1980 | 1994 | - | 11.5161 | 4.02E-05 | 1 | GAGGCCGAAGTTGTC |
| 5321 | mamo-L | *BGIBMGA010930* | down | 1496 | 1510 | + | 12.3226 | 1.77E-05 | 1 | TAGGCGTGGTCTGCT |
| 5322 | mamo-L | *BGIBMGA010930* | down | 337 | 351 | + | 10.5726 | 9.33E-05 | 1 | TAGGCTAATATTTAT |
| 5323 | mamo-L | *BGIBMGA010931* | up | 607 | 621 | + | 12.5 | 1.44E-05 | 1 | TAGGCTCAGTATTCT |
| 5324 | mamo-L | *BGIBMGA010931* | down | 1257 | 1271 | + | 12.4032 | 1.61E-05 | 1 | AAGGCGTATTTTGTT |
| 5325 | mamo-L | *BGIBMGA010935* | down | 1300 | 1314 | - | 12.25 | 1.92E-05 | 1 | TAGGCCCACTATTCT |
| 5326 | mamo-L | *BGIBMGA010941* | down | 1038 | 1052 | - | 10.5726 | 9.33E-05 | 1 | TAGGCTAATATTTAT |
| 5327 | mamo-L | *BGIBMGA010943* | up | 1406 | 1420 | + | 10.7903 | 7.79E-05 | 1 | TAGGCGCTTGGGGGT |
| 5328 | mamo-L | *BGIBMGA010944* | down | 1653 | 1667 | + | 10.9194 | 6.98E-05 | 1 | TAGGCTACTTTTTTT |
| 5329 | mamo-L | *BGIBMGA010952* | up | 1819 | 1833 | - | 10.5403 | 9.58E-05 | 1 | GAGGCTTGTTTGTGT |

| | | | | | | | | | | |
|---|---|---|---|---|---|---|---|---|---|---|
| 5330 | mamo-L | *BGIBMGA010954* | up | 1039 | 1053 | + | 10.8306 | 7.52E-05 | 1 | TAGGCCCCCTTCGCC |
| 5331 | mamo-L | *BGIBMGA010959* | up | 785 | 799 | - | 12.9597 | 8.38E-06 | 1 | TAGGCTTAGATTAGT |
| 5332 | mamo-L | *BGIBMGA010962* | down | 1485 | 1499 | + | 11.0081 | 6.45E-05 | 1 | TAGGCTGTGCGTTGT |
| 5333 | mamo-L | *BGIBMGA010966* | down | 368 | 382 | - | 10.5565 | 9.46E-05 | 1 | AAGGCTGTTTGTTTT |
| 5334 | mamo-L | *BGIBMGA010967* | up | 1471 | 1485 | + | 11.7419 | 3.23E-05 | 1 | TAGGCTAAGTATTTT |
| 5335 | mamo-L | *BGIBMGA010968* | down | 350 | 364 | - | 13.1855 | 6.42E-06 | 1 | AAGGCTGTTGTTGTT |
| 5336 | mamo-L | *BGIBMGA010971* | down | 133 | 147 | - | 11.3952 | 4.52E-05 | 1 | TAGGCTTAGAATTAT |
| 5337 | mamo-L | *BGIBMGA010972* | down | 1277 | 1291 | + | 11.7984 | 3.06E-05 | 1 | TAGGCCACTTTTGCT |
| 5338 | mamo-L | *BGIBMGA010975* | up | 1420 | 1434 | + | 11.5323 | 3.96E-05 | 1 | TAGGCGTGAGGTGAT |
| 5339 | mamo-L | *BGIBMGA010975* | down | 37 | 51 | - | 10.6613 | 8.67E-05 | 1 | GAGGCTGCTTTTTTT |
| 5340 | mamo-L | *BGIBMGA010977* | up | 1075 | 1089 | - | 10.5323 | 9.63E-05 | 1 | TGGGCTTTGTTCGCT |
| 5341 | mamo-L | *BGIBMGA010982* | up | 143 | 157 | - | 10.9435 | 6.83E-05 | 1 | AAGGCTTTGTTCTTT |
| 5342 | mamo-L | *BGIBMGA010987* | down | 2 | 16 | - | 11.1774 | 5.54E-05 | 1 | TAGGCCCAAATTTCT |
| 5343 | mamo-L | *BGIBMGA010990* | down | 1265 | 1279 | + | 11.8387 | 2.94E-05 | 1 | TAGGCTGTTTGTTAT |
| 5344 | mamo-L | *BGIBMGA010991* | down | 103 | 117 | + | 10.5726 | 9.33E-05 | 1 | TAGGCTAATATTTAT |
| 5345 | mamo-L | *BGIBMGA010992* | up | 1063 | 1077 | + | 10.5726 | 9.33E-05 | 1 | TAGGCTAATATTTAT |
| 5346 | mamo-L | *BGIBMGA010994* | down | 40 | 54 | + | 10.5726 | 9.33E-05 | 1 | TAGGCTAATATTTAT |
| 5347 | mamo-L | *BGIBMGA010997* | down | 739 | 753 | + | 12.0565 | 2.36E-05 | 1 | TAGGCCTGAGTTGTA |
| 5348 | mamo-L | *BGIBMGA010997* | down | 197 | 211 | - | 11.0887 | 6.01E-05 | 1 | TAGGCCGCTTTTGCG |
| 5349 | mamo-L | *BGIBMGA011001* | up | 1333 | 1347 | - | 11 | 6.50E-05 | 1 | AAGGCTGTTGTTCCT |
| 5350 | mamo-L | *BGIBMGA011005* | up | 532 | 546 | - | 12.4597 | 1.51E-05 | 1 | TAGGCTTGCGTCATT |
| 5351 | mamo-L | *BGIBMGA011007* | down | 871 | 885 | - | 11.371 | 4.62E-05 | 1 | TAGGCTGTATAGGCT |
| 5352 | mamo-L | *BGIBMGA011016* | down | 370 | 384 | + | 12.3629 | 1.69E-05 | 1 | TAGGCTGTGCTTGAT |
| 5353 | mamo-L | *BGIBMGA011016* | down | 22 | 36 | - | 10.6532 | 8.73E-05 | 1 | TAGCCTTTGTTTGGC |
| 5354 | mamo-L | *BGIBMGA011019* | up | 759 | 773 | + | 11.5323 | 3.96E-05 | 1 | TAGGCGTGAGGTGAT |
| 5355 | mamo-L | *BGIBMGA011023* | down | 1000 | 1014 | + | 10.629 | 8.91E-05 | 1 | TAGGCCTGCAGCGCC |
| 5356 | mamo-L | *BGIBMGA011024* | up | 338 | 352 | - | 12.3548 | 1.70E-05 | 1 | TAGGCTGAGTTTTTG |
| 5357 | mamo-L | *BGIBMGA011024* | up | 1318 | 1332 | + | 11.3629 | 4.66E-05 | 1 | GAGGCGGATGGTGCT |
| 5358 | mamo-L | *BGIBMGA011025* | up | 1182 | 1196 | - | 11.7742 | 3.14E-05 | 1 | TAGGCCTGCTTGATT |
| 5359 | mamo-L | *BGIBMGA011029* | up | 548 | 562 | + | 10.7177 | 8.28E-05 | 1 | TAGCCTGATTTTGGC |
| 5360 | mamo-L | *BGIBMGA011030* | up | 1776 | 1790 | + | 11.0887 | 6.01E-05 | 1 | TAGGCTCTATTTCCT |
| 5361 | mamo-L | *BGIBMGA011031* | up | 776 | 790 | + | 11.3548 | 4.69E-05 | 1 | GAGGCCTACTTTATT |
| 5362 | mamo-L | *BGIBMGA011031* | up | 1062 | 1076 | + | 10.621 | 8.97E-05 | 1 | GAGGCTTTTTATTGT |
| 5363 | mamo-L | *BGIBMGA011036* | down | 1276 | 1290 | - | 14.5806 | 7.86E-07 | 1 | TAGGCTTGTGTTGGC |
| 5364 | mamo-L | *BGIBMGA011037* | up | 1100 | 1114 | - | 11.3548 | 4.69E-05 | 1 | AAGGCGTGTGTTTTT |
| 5365 | mamo-L | *BGIBMGA011044* | down | 1504 | 1518 | + | 13 | 8.01E-06 | 1 | TAGGCGTTCTTTAGT |
| 5366 | mamo-L | *BGIBMGA011045* | down | 1794 | 1808 | - | 13.4839 | 4.31E-06 | 1 | TAGGCCGGCTTTGCA |
| 5367 | mamo-L | *BGIBMGA011046* | up | 1115 | 1129 | + | 12.5968 | 1.28E-05 | 1 | TAGGCCGGTTTTGCG |
| 5368 | mamo-L | *BGIBMGA011048* | up | 1951 | 1965 | - | 12.7016 | 1.14E-05 | 1 | AAGGCGGATTTTGTT |
| 5369 | mamo-L | *BGIBMGA011049* | down | 1160 | 1174 | - | 10.5161 | 9.77E-05 | 1 | AAGGCCTTTGGTTGT |
| 5370 | mamo-L | *BGIBMGA011053* | down | 1391 | 1405 | - | 10.8226 | 7.58E-05 | 1 | TAGGCCGATATCTCC |
| 5371 | mamo-L | *BGIBMGA011054* | up | 589 | 603 | - | 13.7742 | 2.79E-06 | 1 | TAGGCGGGTGGTGGT |
| 5372 | mamo-L | *BGIBMGA011057* | up | 449 | 463 | + | 11.8871 | 2.81E-05 | 1 | TAGGCGGTGTTCGGA |
| 5373 | mamo-L | *BGIBMGA011064* | up | 663 | 677 | + | 11.9677 | 2.59E-05 | 1 | TAGGCTGTTTTCTAT |
| 5374 | mamo-L | *BGIBMGA011070* | down | 552 | 566 | + | 10.5726 | 9.33E-05 | 1 | TAGGCTTTGCCTGGT |
| 5375 | mamo-L | *BGIBMGA011074* | up | 641 | 655 | + | 11.4113 | 4.45E-05 | 1 | TAGGCTCGTTTTTAC |
| 5376 | mamo-L | *BGIBMGA011076* | up | 1444 | 1458 | + | 10.9113 | 7.02E-05 | 1 | TAGGCTGTATGCTGT |
| 5377 | mamo-L | *BGIBMGA011077* | up | 1050 | 1064 | + | 10.5968 | 9.15E-05 | 1 | AAGGCTGTTAATGCT |
| 5378 | mamo-L | *BGIBMGA011077* | up | 1781 | 1795 | - | 10.5726 | 9.33E-05 | 1 | TAGGCTAATATTTAT |
| 5379 | mamo-L | *BGIBMGA011079* | down | 596 | 610 | + | 10.9194 | 6.98E-05 | 1 | TAGGCTACTTTTTTT |
| 5380 | mamo-L | *BGIBMGA011079* | down | 1836 | 1850 | + | 10.5 | 9.90E-05 | 1 | GAGGCCCAGGTCGTC |
| 5381 | mamo-L | *BGIBMGA011084* | up | 1941 | 1955 | - | 10.5645 | 9.40E-05 | 1 | TAGGCTCAGTGTCAT |
| 5382 | mamo-L | *BGIBMGA011085* | up | 418 | 432 | + | 13.0726 | 7.34E-06 | 1 | TAGGCCGATTTTGCA |
| 5383 | mamo-L | *BGIBMGA011089* | down | 1556 | 1570 | + | 12.0484 | 2.38E-05 | 1 | TAGGCTGGACTTGCT |
| 5384 | mamo-L | *BGIBMGA011089* | up | 1350 | 1364 | - | 11.5323 | 3.96E-05 | 1 | TAGGCGTGAGGTGAT |
| 5385 | mamo-L | *BGIBMGA011092* | down | 1235 | 1249 | - | 12.1452 | 2.15E-05 | 1 | TAGGCGTATTTCTTT |
| 5386 | mamo-L | *BGIBMGA011098* | down | 1652 | 1666 | - | 11.4677 | 4.22E-05 | 1 | TAGGCTGATGATAAT |
| 5387 | mamo-L | *BGIBMGA011100* | down | 26 | 40 | - | 10.5726 | 9.33E-05 | 1 | TAGGCTTCAGGCGGT |
| 5388 | mamo-L | *BGIBMGA011101* | up | 426 | 440 | - | 12.4032 | 1.61E-05 | 1 | AAGGCGTATTTTGTT |
| 5389 | mamo-L | *BGIBMGA011103* | up | 1896 | 1910 | + | 11.8952 | 2.79E-05 | 1 | TAGGCTGTGTGGTGT |
| 5390 | mamo-L | *BGIBMGA011108* | down | 1849 | 1863 | - | 11.1935 | 5.45E-05 | 1 | TAGGCGCATGAGGGT |
| 5391 | mamo-L | *BGIBMGA011112* | down | 1642 | 1656 | + | 11.1452 | 5.70E-05 | 1 | AAGGCTTAAATTGCT |
| 5392 | mamo-L | *BGIBMGA011114* | up | 926 | 940 | - | 11.25 | 5.18E-05 | 1 | TAGGCAGTCTTTTCC |
| 5393 | mamo-L | *BGIBMGA011117* | down | 1771 | 1785 | - | 12.871 | 9.35E-06 | 1 | TAGGCTTTTTTTATT |
| 5394 | mamo-L | *BGIBMGA011118* | down | 1796 | 1810 | + | 11.25 | 5.18E-05 | 1 | TAGGCCTCTGATGTC |
| 5395 | mamo-L | *BGIBMGA011119* | down | 1286 | 1300 | + | 12.1532 | 2.13E-05 | 1 | AAGGCTTTCTTTTTT |
| 5396 | mamo-L | *BGIBMGA011120* | up | 893 | 907 | + | 11.0161 | 6.40E-05 | 1 | AAGGCTCTCTGTGCT |
| 5397 | mamo-L | *BGIBMGA011124* | up | 1206 | 1220 | + | 13.5403 | 3.97E-06 | 1 | TAGGCTGTTGTTTGC |
| 5398 | mamo-L | *BGIBMGA011125* | down | 273 | 287 | - | 10.8871 | 7.17E-05 | 1 | TAGGCATATTTTATT |
| 5399 | mamo-L | *BGIBMGA011129* | up | 1420 | 1434 | + | 12.1129 | 2.22E-05 | 1 | TAGGCTTAATTCGCC |
| 5400 | mamo-L | *BGIBMGA011136* | up | 1322 | 1336 | + | 10.8065 | 7.69E-05 | 1 | TAGGCGGAAAATTGT |
| 5401 | mamo-L | *BGIBMGA011137* | down | 255 | 269 | - | 11.0242 | 6.36E-05 | 1 | AAGGCTTCCTTCGGT |
| 5402 | mamo-L | *BGIBMGA011137* | up | 711 | 725 | + | 10.7419 | 8.11E-05 | 1 | AAGGCTTCGTTCGCT |
| 5403 | mamo-L | *BGIBMGA011144* | down | 664 | 678 | - | 10.6694 | 8.62E-05 | 1 | TAGGCTAGTGTTCAT |
| 5404 | mamo-L | *BGIBMGA011145* | down | 310 | 324 | + | 10.5565 | 9.46E-05 | 1 | AAGGCTGTTTGTTTT |
| 5405 | mamo-L | *BGIBMGA011147* | down | 952 | 966 | + | 11.4839 | 4.15E-05 | 1 | TAGGCAGACAGTGGT |
| 5406 | mamo-L | *BGIBMGA011153* | down | 959 | 973 | + | 11.9355 | 2.67E-05 | 1 | GAGGCCTTCTTTTTT |
| 5407 | mamo-L | *BGIBMGA011154* | up | 842 | 856 | + | 11.2984 | 4.95E-05 | 1 | TAGGCTGAATATTTC |
| 5408 | mamo-L | *BGIBMGA011156* | down | 1248 | 1262 | + | 10.8548 | 7.38E-05 | 1 | GAGGCCTGTTTTTAT |
| 5409 | mamo-L | *BGIBMGA011157* | up | 87 | 101 | + | 10.8548 | 7.38E-05 | 1 | GAGGCCTGTTTTTAT |
| 5410 | mamo-L | *BGIBMGA011163* | up | 780 | 794 | - | 10.5323 | 9.63E-05 | 1 | TAGGCTGACGACCAT |
| 5411 | mamo-L | *BGIBMGA011164* | down | 1354 | 1368 | - | 11.1935 | 5.45E-05 | 1 | TAGGCGCATGAGGGT |
| 5412 | mamo-L | *BGIBMGA011166* | up | 1267 | 1281 | - | 10.5565 | 9.46E-05 | 1 | TAGGCCGGCTTCAGA |
| 5413 | mamo-L | *BGIBMGA011171* | up | 1776 | 1790 | - | 10.7097 | 8.33E-05 | 1 | TAGGCTTGAAGTTTT |
| 5414 | mamo-L | *BGIBMGA011172* | up | 113 | 127 | + | 11.0161 | 6.40E-05 | 1 | TAGGCCAGTGATTCT |
| 5415 | mamo-L | *BGIBMGA011176* | down | 1532 | 1546 | + | 10.5403 | 9.58E-05 | 1 | TAGGCTGTATGGTGT |
| 5416 | mamo-L | *BGIBMGA011189* | down | 1046 | 1060 | + | 13.4113 | 4.75E-06 | 1 | TAGGCGCGGTTTTGT |
| 5417 | mamo-L | *BGIBMGA011191* | down | 706 | 720 | - | 11.9516 | 2.63E-05 | 1 | GAGGCTTATTTTTTT |
| 5418 | mamo-L | *BGIBMGA011192* | up | 1695 | 1709 | - | 10.8952 | 7.12E-05 | 1 | TAGGCTGGGACGGGT |
| 5419 | mamo-L | *BGIBMGA011193* | up | 1295 | 1309 | - | 12.8145 | 9.96E-06 | 1 | TAGGCCTGGTCTGCT |
| 5420 | mamo-L | *BGIBMGA011195* | down | 381 | 395 | - | 11.7339 | 3.26E-05 | 1 | TAGGCTAGCTTCGCC |
| 5421 | mamo-L | *BGIBMGA011195* | up | 1667 | 1681 | - | 11.5242 | 3.99E-05 | 1 | AAGGCTTATGTTAGT |

| | | | | | | | | | | |
|---|---|---|---|---|---|---|---|---|---|---|
| 5422 | mamo-L | *BGIBMGA011199* | up | 1794 | 1808 | + | 11.121 | 5.84E-05 | 1 | TAGGCATAAGTTTTT |
| 5423 | mamo-L | *BGIBMGA011205* | down | 773 | 787 | - | 11.5726 | 3.81E-05 | 1 | AAGGCGTGGGGTGCT |
| 5424 | mamo-L | *BGIBMGA011210* | down | 732 | 746 | + | 12.2016 | 2.03E-05 | 1 | TAGGCTGTAGATGTC |
| 5425 | mamo-L | *BGIBMGA011211* | up | 1271 | 1285 | + | 11.1694 | 5.59E-05 | 1 | TAGGCGGAAGTTATC |
| 5426 | mamo-L | *BGIBMGA011212* | down | 1154 | 1168 | - | 10.5726 | 9.33E-05 | 1 | TAGGCTAATATTTAT |
| 5427 | mamo-L | *BGIBMGA011213* | down | 305 | 319 | - | 10.9194 | 6.98E-05 | 1 | TAGGCTACTTTTTTT |
| 5428 | mamo-L | *BGIBMGA011215* | down | 449 | 463 | + | 13.0565 | 7.51E-06 | 1 | AAGGCTGTTTTTGTT |
| 5429 | mamo-L | *BGIBMGA011220* | down | 1194 | 1208 | - | 10.5726 | 9.33E-05 | 1 | TAGGCTAATATTTAT |
| 5430 | mamo-L | *BGIBMGA011222* | up | 1275 | 1289 | + | 10.4919 | 9.96E-05 | 1 | GAGGCTTGGTTCTGC |
| 5431 | mamo-L | *BGIBMGA011223* | up | 226 | 240 | + | 14.879 | 4.31E-07 | 1 | TAGGCTGGGTGTGGT |
| 5432 | mamo-L | *BGIBMGA011223* | up | 1803 | 1817 | - | 11.0484 | 6.23E-05 | 1 | TAGGCTGTCAGCGAT |
| 5433 | mamo-L | *BGIBMGA011224* | up | 1592 | 1606 | + | 10.5726 | 9.33E-05 | 1 | TAGGCTAATATTTAT |
| 5434 | mamo-L | *BGIBMGA011226* | down | 1676 | 1690 | + | 12.9919 | 8.10E-06 | 1 | TAGGCCGGTTTTGCA |
| 5435 | mamo-L | *BGIBMGA011226* | down | 786 | 800 | + | 12.3226 | 1.77E-05 | 1 | AAGGCTGGTTTTGTC |
| 5436 | mamo-L | *BGIBMGA011226* | down | 1539 | 1553 | - | 10.9758 | 6.64E-05 | 1 | TAGGCTTCTTTTGCG |
| 5437 | mamo-L | *BGIBMGA011229* | up | 356 | 370 | + | 12.8145 | 9.96E-06 | 1 | TAGGCGGCGGTTTTT |
| 5438 | mamo-L | *BGIBMGA011231* | down | 401 | 415 | + | 12.9919 | 8.10E-06 | 1 | TAGGCCGGTTTTGCA |
| 5439 | mamo-L | *BGIBMGA011231* | up | 148 | 162 | - | 10.7581 | 8.00E-05 | 1 | TAGGCAGTTGGTGAT |
| 5440 | mamo-L | *BGIBMGA011235* | down | 1295 | 1309 | - | 13.1935 | 6.33E-06 | 1 | TAGGCTTGCATCGCT |
| 5441 | mamo-L | *BGIBMGA011235* | down | 1955 | 1969 | - | 11.4355 | 4.34E-05 | 1 | AAGGCCGCCTTTGGC |
| 5442 | mamo-L | *BGIBMGA011235* | up | 417 | 431 | - | 10.8226 | 7.58E-05 | 1 | TAGGCCGTCGACCCT |
| 5443 | mamo-L | *BGIBMGA011235* | up | 894 | 908 | + | 10.7742 | 7.89E-05 | 1 | AAGGCTGTTTACGGT |
| 5444 | mamo-L | *BGIBMGA011236* | down | 1366 | 1380 | - | 12.9919 | 8.10E-06 | 1 | TAGGCCGGTTTTGCA |
| 5445 | mamo-L | *BGIBMGA011238* | down | 1255 | 1269 | - | 11.8629 | 2.87E-05 | 1 | AAGGCCAAGTTTGTT |
| 5446 | mamo-L | *BGIBMGA011238* | down | 1052 | 1066 | - | 11.4194 | 4.42E-05 | 1 | GAGGCTCATTTTGAT |
| 5447 | mamo-L | *BGIBMGA011238* | up | 766 | 780 | - | 11.0081 | 6.45E-05 | 1 | TAGGCAGGGTAGGGT |
| 5448 | mamo-L | *BGIBMGA011241* | up | 95 | 109 | - | 10.5726 | 9.33E-05 | 1 | TAGGCTAATATTTAT |
| 5449 | mamo-L | *BGIBMGA011246* | down | 1765 | 1779 | + | 11.2097 | 5.38E-05 | 1 | TAGGCAGGATTTTTT |
| 5450 | mamo-L | *BGIBMGA011248* | up | 671 | 685 | - | 11.8629 | 2.87E-05 | 1 | TAGGCCTTAGATGAT |
| 5451 | mamo-L | *BGIBMGA011248* | up | 1931 | 1945 | + | 10.7661 | 7.95E-05 | 1 | TAGGCTTGTAGTCGT |
| 5452 | mamo-L | *BGIBMGA011249* | up | 1437 | 1451 | + | 12.6694 | 1.18E-05 | 1 | TAGGCCGACGACGGC |
| 5453 | mamo-L | *BGIBMGA011250* | up | 929 | 943 | + | 11.7016 | 3.37E-05 | 1 | TAGGCTTTAGGTTTT |
| 5454 | mamo-L | *BGIBMGA011255* | down | 917 | 931 | + | 10.629 | 8.91E-05 | 1 | TAGGCCTGCAGCGCC |
| 5455 | mamo-L | *BGIBMGA011262* | up | 1776 | 1790 | + | 13.629 | 3.46E-06 | 1 | AAGGCTTTGTTTGGT |
| 5456 | mamo-L | *BGIBMGA011262* | up | 1851 | 1865 | + | 11.2742 | 5.06E-05 | 1 | TAGGCCCACGATGCA |
| 5457 | mamo-L | *BGIBMGA011266* | up | 1115 | 1129 | - | 11.6935 | 3.39E-05 | 1 | GAGGCGTGCTGTGGT |
| 5458 | mamo-L | *BGIBMGA011267* | down | 952 | 966 | - | 10.9194 | 6.98E-05 | 1 | TAGGCTACTTTTTTT |
| 5459 | mamo-L | *BGIBMGA011267* | up | 415 | 429 | - | 10.7823 | 7.84E-05 | 1 | AAGGCCGATTGTGCC |
| 5460 | mamo-L | *BGIBMGA011272* | up | 1704 | 1718 | - | 10.7339 | 8.17E-05 | 1 | TAGGCCTAAATCTTT |
| 5461 | mamo-L | *BGIBMGA011273* | down | 404 | 418 | + | 10.621 | 8.97E-05 | 1 | GAGGCTGAATTTTAT |
| 5462 | mamo-L | *BGIBMGA011274* | up | 1046 | 1060 | - | 10.5887 | 9.21E-05 | 1 | TAGGCGGATGATGAA |
| 5463 | mamo-L | *BGIBMGA011275* | down | 531 | 545 | + | 11.9274 | 2.69E-05 | 1 | TAGGCCTAGTGTGCA |
| 5464 | mamo-L | *BGIBMGA011276* | up | 813 | 827 | + | 11.9274 | 2.69E-05 | 1 | TAGGCCTAGTGTGCA |
| 5465 | mamo-L | *BGIBMGA011282* | up | 283 | 297 | - | 11.9435 | 2.65E-05 | 1 | GAGGCTGGTTATGTT |
| 5466 | mamo-L | *BGIBMGA011282* | down | 1104 | 1118 | + | 11.0645 | 6.14E-05 | 1 | TAGGCTGTTGGTAAT |
| 5467 | mamo-L | *BGIBMGA011285* | down | 1154 | 1168 | + | 10.9919 | 6.55E-05 | 1 | TAGGCCAGGTTTTAC |
| 5468 | mamo-L | *BGIBMGA011286* | up | 801 | 815 | + | 11.5565 | 3.87E-05 | 1 | TAGGCTGTCGGTAAT |
| 5469 | mamo-L | *BGIBMGA011286* | up | 1826 | 1840 | + | 11.2742 | 5.06E-05 | 1 | TAGGCAGACGGTTCT |
| 5470 | mamo-L | *BGIBMGA011288* | up | 1952 | 1966 | - | 11.4677 | 4.22E-05 | 1 | TAGGCCCTGTTTATC |
| 5471 | mamo-L | *BGIBMGA011288* | up | 993 | 1007 | - | 11.1613 | 5.63E-05 | 1 | AAGGCTCTGTATGCT |
| 5472 | mamo-L | *BGIBMGA011288* | down | 813 | 827 | - | 10.9194 | 6.98E-05 | 1 | TAGGCGACGTTTGTC |
| 5473 | mamo-L | *BGIBMGA011292* | up | 948 | 962 | + | 13.6774 | 3.24E-06 | 1 | TAGGCCTAGTTTGGA |
| 5474 | mamo-L | *BGIBMGA011292* | up | 941 | 955 | - | 10.6371 | 8.85E-05 | 1 | TAGGCCTAATTGTAT |
| 5475 | mamo-L | *BGIBMGA011294* | down | 62 | 76 | + | 10.5726 | 9.33E-05 | 1 | TAGGCTAATATTTAT |
| 5476 | mamo-L | *BGIBMGA011295* | up | 1197 | 1211 | + | 10.5726 | 9.33E-05 | 1 | TAGGCTAATATTTAT |
| 5477 | mamo-L | *BGIBMGA011301* | up | 1883 | 1897 | - | 11.3629 | 4.66E-05 | 1 | AAGGCTGACAGTGTT |
| 5478 | mamo-L | *BGIBMGA011301* | down | 992 | 1006 | - | 10.7258 | 8.23E-05 | 1 | TAGGCCTGTTTTATA |
| 5479 | mamo-L | *BGIBMGA011303* | up | 1973 | 1987 | + | 11.129 | 5.79E-05 | 1 | AAGGCGAATTTTGGT |
| 5480 | mamo-L | *BGIBMGA011303* | down | 246 | 260 | - | 10.5806 | 9.27E-05 | 1 | TAGGCCTCCTTTTCA |
| 5481 | mamo-L | *BGIBMGA011305* | down | 1109 | 1123 | - | 11.75 | 3.21E-05 | 1 | TAGGCTAGATTTGTC |
| 5482 | mamo-L | *BGIBMGA011306* | up | 716 | 730 | - | 10.5726 | 9.33E-05 | 1 | TAGGCTAATATTTAT |
| 5483 | mamo-L | *BGIBMGA011312* | up | 1233 | 1247 | - | 11.5645 | 3.84E-05 | 1 | TAGGCCGCTTTTCTT |
| 5484 | mamo-L | *BGIBMGA011316* | up | 610 | 624 | + | 10.8629 | 7.33E-05 | 1 | GAGGCGTTCTTGGCT |
| 5485 | mamo-L | *BGIBMGA011320* | down | 493 | 507 | + | 13.0968 | 7.11E-06 | 1 | TAGGCTGTTTTTTTC |
| 5486 | mamo-L | *BGIBMGA011322* | down | 1775 | 1789 | - | 11.4597 | 4.25E-05 | 1 | TAGGCGGGGTGGTGT |
| 5487 | mamo-L | *BGIBMGA011322* | down | 1496 | 1510 | + | 11.1129 | 5.88E-05 | 1 | TAGGCTTAGCATTGT |
| 5488 | mamo-L | *BGIBMGA011323* | down | 337 | 351 | + | 11.9435 | 2.65E-05 | 1 | GAGGCTGGTTATGTT |
| 5489 | mamo-L | *BGIBMGA011324* | up | 1346 | 1360 | - | 11.4677 | 4.22E-05 | 1 | TAGGCCCTGTTTATC |
| 5490 | mamo-L | *BGIBMGA011327* | down | 765 | 779 | + | 10.6371 | 8.85E-05 | 1 | TAGGCTTTGTCTCTT |
| 5491 | mamo-L | *BGIBMGA011332* | up | 1715 | 1729 | - | 10.9194 | 6.98E-05 | 1 | TAGGCTACTTTTTTT |
| 5492 | mamo-L | *BGIBMGA011339* | down | 1000 | 1014 | - | 12.0645 | 2.34E-05 | 1 | TAGGCGGGCTTCGGA |
| 5493 | mamo-L | *BGIBMGA011343* | up | 1139 | 1153 | + | 11.5806 | 3.78E-05 | 1 | TAGGCTGGGTGTTGA |
| 5494 | mamo-L | *BGIBMGA011344* | down | 1868 | 1882 | + | 12.1371 | 2.17E-05 | 1 | TAGGCATGGGATGCT |
| 5495 | mamo-L | *BGIBMGA011344* | down | 1702 | 1716 | + | 11.3306 | 4.80E-05 | 1 | TAGGCGGCATTTTTT |
| 5496 | mamo-L | *BGIBMGA011347* | up | 1314 | 1328 | + | 10.4919 | 9.96E-05 | 1 | TAGGCCGACATTATA |
| 5497 | mamo-L | *BGIBMGA011352* | up | 200 | 214 | + | 10.5887 | 9.21E-05 | 1 | TAGGCTGGGGTCACA |
| 5498 | mamo-L | *BGIBMGA011353* | down | 1825 | 1839 | + | 11.9274 | 2.69E-05 | 1 | GAGGCTTTATTTGTT |
| 5499 | mamo-L | *BGIBMGA011355* | down | 592 | 606 | + | 13.5161 | 4.10E-06 | 1 | TAGGCTGGATTTTTT |
| 5500 | mamo-L | *BGIBMGA011355* | down | 1264 | 1278 | - | 10.5726 | 9.33E-05 | 1 | TAGGCTAATATTTAT |
| 5501 | mamo-L | *BGIBMGA011355* | up | 1932 | 1946 | + | 10.5726 | 9.33E-05 | 1 | TAGGCTAATATTTAT |
| 5502 | mamo-L | *BGIBMGA011357* | up | 1424 | 1438 | + | 11.8629 | 2.87E-05 | 1 | TAGGCCTTAGATGAT |
| 5503 | mamo-L | *BGIBMGA011357* | up | 164 | 178 | - | 10.7661 | 7.95E-05 | 1 | TAGGCTTGTAGTCGT |
| 5504 | mamo-L | *BGIBMGA011359* | up | 441 | 455 | + | 10.5161 | 9.77E-05 | 1 | TAGGCCGGTTCTTTC |
| 5505 | mamo-L | *BGIBMGA011363* | up | 1475 | 1489 | - | 10.5726 | 9.33E-05 | 1 | TAGGCTAATATTTAT |
| 5506 | mamo-L | *BGIBMGA011368* | up | 1461 | 1475 | - | 12.2258 | 1.97E-05 | 1 | TAGGCCGGTGATGAC |
| 5507 | mamo-L | *BGIBMGA011368* | up | 635 | 649 | - | 11.121 | 5.84E-05 | 1 | TAGGCTCAGTGGTGT |
| 5508 | mamo-L | *BGIBMGA011369* | down | 1973 | 1987 | - | 10.5726 | 9.33E-05 | 1 | TAGGCTAATATTTAT |
| 5509 | mamo-L | *BGIBMGA011370* | down | 1438 | 1452 | - | 11.8226 | 2.99E-05 | 1 | TAGGCTGAGGACTGC |
| 5510 | mamo-L | *BGIBMGA011374* | down | 112 | 126 | - | 13.0726 | 7.34E-06 | 1 | TAGGCCGATTTTGCA |
| 5511 | mamo-L | *BGIBMGA011374* | down | 37 | 51 | - | 11.1855 | 5.50E-05 | 1 | TAGGCCAATTTTGCA |
| 5512 | mamo-L | *BGIBMGA011374* | down | 1635 | 1649 | - | 11.0726 | 6.09E-05 | 1 | TAGGCCGATTTTACA |
| 5513 | mamo-L | *BGIBMGA011374* | up | 665 | 679 | + | 10.7581 | 8.00E-05 | 1 | TAGGCAGTTGGTGAT |

| | | | | | | | | | | |
|---|---|---|---|---|---|---|---|---|---|---|
| 5514 | mamo-L | *BGIBMGA011374* | down | 659 | 673 | + | 10.5726 | 9.33E-05 | 1 | TAGGCTAATATTTAT |
| 5515 | mamo-L | *BGIBMGA011376* | down | 1840 | 1854 | + | 10.5726 | 9.33E-05 | 1 | TAGGCTAATATTTAT |
| 5516 | mamo-L | *BGIBMGA011379* | up | 1610 | 1624 | - | 14.879 | 4.31E-07 | 1 | TAGGCTGGGTGTGGT |
| 5517 | mamo-L | *BGIBMGA011379* | up | 33 | 47 | + | 11.0484 | 6.23E-05 | 1 | TAGGCTGTCAGCGAT |
| 5518 | mamo-L | *BGIBMGA011381* | up | 54 | 68 | + | 10.6694 | 8.62E-05 | 1 | AAGGCTCACTATTGT |
| 5519 | mamo-L | *BGIBMGA011390* | down | 533 | 547 | + | 11.3871 | 4.55E-05 | 1 | AAGGCTTTGTTTTAT |
| 5520 | mamo-L | *BGIBMGA011399* | down | 1141 | 1155 | - | 13.2177 | 6.14E-06 | 1 | TAGGCCTGTGTTCGT |
| 5521 | mamo-L | *BGIBMGA011399* | up | 237 | 251 | + | 10.9194 | 6.98E-05 | 1 | TAGGCTACTTTTTTT |
| 5522 | mamo-L | *BGIBMGA011401* | down | 1230 | 1244 | - | 11.2823 | 5.03E-05 | 1 | TAGGCGCATATGGGT |
| 5523 | mamo-L | *BGIBMGA011406* | down | 482 | 496 | - | 10.5161 | 9.77E-05 | 1 | TAGGCCCGCGTTACA |
| 5524 | mamo-L | *BGIBMGA011410* | up | 464 | 478 | - | 13.0887 | 7.18E-06 | 1 | TAGGCTGGTTCTGGT |
| 5525 | mamo-L | *BGIBMGA011412* | down | 1775 | 1789 | + | 10.5565 | 9.46E-05 | 1 | GAGGCGTACTTGGGC |
| 5526 | mamo-L | *BGIBMGA011413* | up | 571 | 585 | + | 11.1129 | 5.88E-05 | 1 | AAGGCGTTTGTTTTT |
| 5527 | mamo-L | *BGIBMGA011415* | down | 1921 | 1935 | + | 11.6613 | 3.50E-05 | 1 | AAGGCTTTTTTTTTT |
| 5528 | mamo-L | *BGIBMGA011418* | down | 1775 | 1789 | - | 12.6371 | 1.23E-05 | 1 | TAGGCAGGCGGTGGT |
| 5529 | mamo-L | *BGIBMGA011418* | up | 626 | 640 | + | 11.2097 | 5.38E-05 | 1 | TAGGCCTTTATCTTT |
| 5530 | mamo-L | *BGIBMGA011425* | up | 792 | 806 | + | 10.6855 | 8.51E-05 | 1 | TAGGCCGTGTAGGGA |
| 5531 | mamo-L | *BGIBMGA011431* | up | 1862 | 1876 | - | 13.1452 | 6.69E-06 | 1 | TAGGCCTAGTATTTT |
| 5532 | mamo-L | *BGIBMGA011432* | down | 1332 | 1346 | + | 13.7419 | 2.91E-06 | 1 | TAGGCTTTTTTTTCT |
| 5533 | mamo-L | *BGIBMGA011432* | up | 1335 | 1349 | + | 11.5726 | 3.81E-05 | 1 | TAGGCTGGGGTTTAG |
| 5534 | mamo-L | *BGIBMGA011434* | down | 1591 | 1605 | - | 11.8952 | 2.79E-05 | 1 | TAGGCTTTTTTTATC |
| 5535 | mamo-L | *BGIBMGA011439* | down | 1157 | 1171 | + | 10.9677 | 6.68E-05 | 1 | TAGGCAGTATTTTTT |
| 5536 | mamo-L | *BGIBMGA011445* | down | 1772 | 1786 | + | 11.2581 | 5.14E-05 | 1 | TAGGCCTAGTATTGA |
| 5537 | mamo-L | *BGIBMGA011448* | down | 724 | 738 | - | 13.5645 | 3.81E-06 | 1 | TAGGCCTACATTGAT |
| 5538 | mamo-L | *BGIBMGA011449* | up | 579 | 593 | - | 13.5645 | 3.81E-06 | 1 | TAGGCCTACATTGAT |
| 5539 | mamo-L | *BGIBMGA011449* | down | 649 | 663 | + | 10.8871 | 7.17E-05 | 1 | AAGGCTTTGAATGTT |
| 5540 | mamo-L | *BGIBMGA011451* | up | 1657 | 1671 | - | 11.621 | 3.63E-05 | 1 | AAGGCTAATGTTGTT |
| 5541 | mamo-L | *BGIBMGA011451* | down | 576 | 590 | + | 11.4919 | 4.11E-05 | 1 | TAGGCGTTCGATATT |
| 5542 | mamo-L | *BGIBMGA011452* | up | 1838 | 1852 | - | 12.0645 | 2.34E-05 | 1 | TAGGCAGTATTTGTT |
| 5543 | mamo-L | *BGIBMGA011455* | up | 342 | 356 | + | 11.0081 | 6.45E-05 | 1 | TAGGCTATGTTTAAT |
| 5544 | mamo-L | *BGIBMGA011458* | down | 1443 | 1457 | - | 12.0887 | 2.29E-05 | 1 | AAGGCTTTAGTTGTT |
| 5545 | mamo-L | *BGIBMGA011459* | down | 1479 | 1493 | + | 10.6613 | 8.67E-05 | 1 | GAGGCGTACATTTTT |
| 5546 | mamo-L | *BGIBMGA011460* | down | 1496 | 1510 | - | 10.5645 | 9.40E-05 | 1 | TAGGCTTATTACACT |
| 5547 | mamo-L | *BGIBMGA011461* | up | 72 | 86 | - | 11.3629 | 4.66E-05 | 1 | TAGGCTGGTGTCTAC |
| 5548 | mamo-L | *BGIBMGA011461* | down | 1740 | 1754 | + | 10.5 | 9.90E-05 | 1 | GAGGCCCAGGTCGTC |
| 5549 | mamo-L | *BGIBMGA011462* | up | 1811 | 1825 | + | 11.5645 | 3.84E-05 | 1 | TAGGCCATCGATGAT |
| 5550 | mamo-L | *BGIBMGA011463* | down | 1432 | 1446 | + | 10.5 | 9.90E-05 | 1 | GAGGCCCAGGTCGTC |
| 5551 | mamo-L | *BGIBMGA011465* | down | 1044 | 1058 | + | 10.9919 | 6.55E-05 | 1 | TAGGCTGCTATTGAC |
| 5552 | mamo-L | *BGIBMGA011466* | up | 326 | 340 | + | 11.6694 | 3.47E-05 | 1 | AAGGCTGGATTTGAT |
| 5553 | mamo-L | *BGIBMGA011468* | down | 888 | 902 | - | 10.8065 | 7.69E-05 | 1 | TAGGCCTAATGTATT |
| 5554 | mamo-L | *BGIBMGA011469* | up | 550 | 564 | + | 12.5 | 1.44E-05 | 1 | TAGGCTATTTTTTGT |
| 5555 | mamo-L | *BGIBMGA011471* | down | 1647 | 1661 | + | 10.6129 | 9.03E-05 | 1 | TAGGCCGTTCTTCGT |
| 5556 | mamo-L | *BGIBMGA011474* | up | 796 | 810 | + | 10.5726 | 9.33E-05 | 1 | TAGGCTAATATTTAT |
| 5557 | mamo-L | *BGIBMGA011477* | down | 123 | 137 | + | 10.8065 | 7.69E-05 | 1 | TAGGCCAACATTATT |
| 5558 | mamo-L | *BGIBMGA011478* | up | 1778 | 1792 | - | 10.5726 | 9.33E-05 | 1 | TAGGCTAATATTTAT |
| 5559 | mamo-L | *BGIBMGA011481* | down | 177 | 191 | + | 12.2984 | 1.81E-05 | 1 | TAGGCTGTAGTTGTA |
| 5560 | mamo-L | *BGIBMGA011483* | down | 789 | 803 | + | 10.5968 | 9.15E-05 | 1 | AAGGCGTGGTTTGGG |
| 5561 | mamo-L | *BGIBMGA011484* | down | 848 | 862 | + | 10.6048 | 9.09E-05 | 1 | TAGGCAGTCATGGTT |
| 5562 | mamo-L | *BGIBMGA011486* | up | 1268 | 1282 | - | 10.5 | 9.90E-05 | 1 | GAGGCCCAGGTCGTC |
| 5563 | mamo-L | *BGIBMGA011490* | down | 329 | 343 | + | 12.2097 | 2.01E-05 | 1 | TAGGCCGGCTATCCT |
| 5564 | mamo-L | *BGIBMGA011492* | up | 1539 | 1553 | - | 11.2581 | 5.14E-05 | 1 | AAGGCGAATGTTGGT |
| 5565 | mamo-L | *BGIBMGA011496* | up | 1597 | 1611 | + | 11.7742 | 3.14E-05 | 1 | TAGGCAAGGTTTGTT |
| 5566 | mamo-L | *BGIBMGA011499* | down | 638 | 652 | + | 11.0161 | 6.40E-05 | 1 | TAGGCTGTGAGTGTA |
| 5567 | mamo-L | *BGIBMGA011500* | up | 306 | 320 | - | 11.2177 | 5.34E-05 | 1 | GAGGCCTAGATTTTT |
| 5568 | mamo-L | *BGIBMGA011500* | up | 1951 | 1965 | + | 11.0161 | 6.40E-05 | 1 | TAGGCTGTGAGTGTA |
| 5569 | mamo-L | *BGIBMGA011503* | down | 1890 | 1904 | - | 12.1774 | 2.08E-05 | 1 | TAGGCTTTGTATTAT |
| 5570 | mamo-L | *BGIBMGA011505* | down | 787 | 801 | + | 12.3952 | 1.63E-05 | 1 | TAGGCTGTAGATTGT |
| 5571 | mamo-L | *BGIBMGA011508* | down | 1836 | 1850 | - | 12.3952 | 1.63E-05 | 1 | TAGGCTGTAGATTGT |
| 5572 | mamo-L | *BGIBMGA011509* | down | 477 | 491 | + | 11.629 | 3.61E-05 | 1 | TAGGCCTATTTTTGG |
| 5573 | mamo-L | *BGIBMGA011511* | down | 1870 | 1884 | - | 12.5726 | 1.33E-05 | 1 | TAGGCCCGGATTGCC |
| 5574 | mamo-L | *BGIBMGA011511* | down | 890 | 904 | - | 11.5242 | 3.99E-05 | 1 | TAGGCGGGTGGCGGC |
| 5575 | mamo-L | *BGIBMGA011514* | up | 354 | 368 | - | 10.9194 | 6.98E-05 | 1 | TAGGCTACTTTTTTT |
| 5576 | mamo-L | *BGIBMGA011514* | down | 1087 | 1101 | + | 10.5 | 9.90E-05 | 1 | GAGGCCCAGGTCGTC |
| 5577 | mamo-L | *BGIBMGA011517* | down | 1688 | 1702 | - | 11.0968 | 5.97E-05 | 1 | GAGGCTTTAGTCGGT |
| 5578 | mamo-L | *BGIBMGA011529* | up | 376 | 390 | - | 11.1371 | 5.75E-05 | 1 | AAGGCCGGTGTTTCC |
| 5579 | mamo-L | *BGIBMGA011535* | up | 1296 | 1310 | + | 12.6371 | 1.23E-05 | 1 | TAGGCAGGCGGTGGT |
| 5580 | mamo-L | *BGIBMGA011539* | up | 1024 | 1038 | - | 11.4113 | 4.45E-05 | 1 | TAGGCTACCTTTTTT |
| 5581 | mamo-L | *BGIBMGA011542* | up | 199 | 213 | - | 12.5 | 1.44E-05 | 1 | TAGGCTATTTTTTGT |
| 5582 | mamo-L | *BGIBMGA011543* | up | 1333 | 1347 | + | 12.0484 | 2.38E-05 | 1 | GAGGCGGGGTTTTTT |
| 5583 | mamo-L | *BGIBMGA011544* | down | 315 | 329 | - | 11.4516 | 4.28E-05 | 1 | TAGGCTTGTCTTTCT |
| 5584 | mamo-L | *BGIBMGA011545* | down | 1448 | 1462 | + | 10.6935 | 8.45E-05 | 1 | AAGGCGTGTGTTTGC |
| 5585 | mamo-L | *BGIBMGA011546* | up | 908 | 922 | + | 10.6935 | 8.45E-05 | 1 | AAGGCGTGTGTTTGC |
| 5586 | mamo-L | *BGIBMGA011549* | up | 109 | 123 | + | 10.5403 | 9.58E-05 | 1 | TAGGCTGTATGGTGT |
| 5587 | mamo-L | *BGIBMGA011550* | down | 461 | 475 | - | 10.5 | 9.90E-05 | 1 | GAGGCCCAGGTCGTC |
| 5588 | mamo-L | *BGIBMGA011555* | down | 1675 | 1689 | - | 11.2903 | 4.99E-05 | 1 | TAGGCCTGTAGTTCT |
| 5589 | mamo-L | *BGIBMGA011556* | up | 1261 | 1275 | + | 13 | 8.01E-06 | 1 | TAGGCTGTTGATGTC |
| 5590 | mamo-L | *BGIBMGA011558* | up | 543 | 557 | - | 10.7339 | 8.17E-05 | 1 | TAGGCCGCTTTTTGA |
| 5591 | mamo-L | *BGIBMGA011559* | down | 1684 | 1698 | - | 11.3468 | 4.73E-05 | 1 | GAGGCGGTTGTTTCT |
| 5592 | mamo-L | *BGIBMGA011560* | up | 236 | 250 | - | 11.6129 | 3.66E-05 | 1 | TAGGCTGAAGATGGA |
| 5593 | mamo-L | *BGIBMGA011561* | up | 1214 | 1228 | + | 10.9516 | 6.78E-05 | 1 | TAGGCTCTGGTTAAC |
| 5594 | mamo-L | *BGIBMGA011567* | down | 82 | 96 | - | 12.1935 | 2.05E-05 | 1 | TAGGCGGATTTGGTC |
| 5595 | mamo-L | *BGIBMGA011568* | down | 1101 | 1115 | - | 10.8871 | 7.17E-05 | 1 | AAGGCTTTGAATGTT |
| 5596 | mamo-L | *BGIBMGA011572* | up | 1341 | 1355 | + | 12 | 2.50E-05 | 1 | TAGGCCAAGGTCGAT |
| 5597 | mamo-L | *BGIBMGA011575* | down | 721 | 735 | + | 11.2419 | 5.21E-05 | 1 | AAGGCTTTGTTTTTC |
| 5598 | mamo-L | *BGIBMGA011578* | down | 25 | 39 | - | 11.2581 | 5.14E-05 | 1 | TAGGCCTAGTATTGA |
| 5599 | mamo-L | *BGIBMGA011579* | down | 1967 | 1981 | + | 12.2742 | 1.87E-05 | 1 | TAGGCTTTTTTTGTG |
| 5600 | mamo-L | *BGIBMGA011580* | down | 1467 | 1481 | - | 11.8468 | 2.92E-05 | 1 | TAGGCTTATGTCGTA |
| 5601 | mamo-L | *BGIBMGA011581* | down | 311 | 325 | - | 11.8468 | 2.92E-05 | 1 | TAGGCTAGGTTTGCA |
| 5602 | mamo-L | *BGIBMGA011581* | down | 1570 | 1584 | - | 10.879 | 7.22E-05 | 1 | TAGGCTGACAGTGTG |
| 5603 | mamo-L | *BGIBMGA011582* | up | 930 | 944 | - | 10.879 | 7.22E-05 | 1 | TAGGCTGACAGTGTG |
| 5604 | mamo-L | *BGIBMGA011584* | down | 185 | 199 | + | 12.1613 | 2.11E-05 | 1 | TAGGCTTATTTTGAA |
| 5605 | mamo-L | *BGIBMGA011584* | up | 1232 | 1246 | - | 11.0484 | 6.23E-05 | 1 | GAGGCTTATTTTATT |

| | | | | | | | | | | |
|---|---|---|---|---|---|---|---|---|---|---|
| 5606 | mamo-L | *BGIBMGA011587* | up | 753 | 767 | - | 12.7258 | 1.11E-05 | 1 | GAGGCTGGGTTTTTT |
| 5607 | mamo-L | *BGIBMGA011587* | up | 167 | 181 | - | 12.0242 | 2.44E-05 | 1 | AAGGCGGTCGTTGTC |
| 5608 | mamo-L | *BGIBMGA011591* | up | 1088 | 1102 | - | 13.4355 | 4.58E-06 | 1 | TAGGCTTATTTTTGC |
| 5609 | mamo-L | *BGIBMGA011591* | up | 768 | 782 | + | 11.3145 | 4.87E-05 | 1 | TAGGCTTAGATGAGT |
| 5610 | mamo-L | *BGIBMGA011592* | down | 143 | 157 | + | 12.7581 | 1.07E-05 | 1 | AAGGCTTTTTTTGTT |
| 5611 | mamo-L | *BGIBMGA011596* | up | 1690 | 1704 | - | 11.3468 | 4.73E-05 | 1 | TAGGCTATCAATGTT |
| 5612 | mamo-L | *BGIBMGA011597* | up | 1563 | 1577 | - | 11.5806 | 3.78E-05 | 1 | TAGGCATACGTTTAT |
| 5613 | mamo-L | *BGIBMGA011599* | down | 1446 | 1460 | + | 10.5 | 9.90E-05 | 1 | GAGGCCCAGGTCGTC |
| 5614 | mamo-L | *BGIBMGA011601* | up | 1748 | 1762 | - | 11.7903 | 3.08E-05 | 1 | TAGGCTAAGTTCTTT |
| 5615 | mamo-L | *BGIBMGA011602* | up | 247 | 261 | + | 10.7258 | 8.23E-05 | 1 | AAGGCTCATGTTAGT |
| 5616 | mamo-L | *BGIBMGA011603* | up | 505 | 519 | + | 11 | 6.50E-05 | 1 | TAGGCCACTTTTGAT |
| 5617 | mamo-L | *BGIBMGA011606* | up | 637 | 651 | - | 11.2097 | 5.38E-05 | 1 | TAGGCTATTTTTTTC |
| 5618 | mamo-L | *BGIBMGA011606* | down | 1455 | 1469 | + | 11.1935 | 5.45E-05 | 1 | TAGGCGCATGAGGGT |
| 5619 | mamo-L | *BGIBMGA011611* | up | 1532 | 1546 | - | 11.7823 | 3.11E-05 | 1 | TAGGCGGGGCTTGTC |
| 5620 | mamo-L | *BGIBMGA011613* | up | 1545 | 1559 | - | 11.0806 | 6.05E-05 | 1 | AAGGCTTATTTTATT |
| 5621 | mamo-L | *BGIBMGA011614* | up | 82 | 96 | + | 11.5081 | 4.05E-05 | 1 | TAGGCGAAGGTCGAT |
| 5622 | mamo-L | *BGIBMGA011618* | up | 371 | 385 | + | 12.4194 | 1.58E-05 | 1 | TAGGCTTAGTGTTAT |
| 5623 | mamo-L | *BGIBMGA011623* | up | 1248 | 1262 | - | 10.5726 | 9.33E-05 | 1 | TAGGCCGAAATTAAT |
| 5624 | mamo-L | *BGIBMGA011624* | up | 1869 | 1883 | - | 11.9516 | 2.63E-05 | 1 | GAGGCCGGTTTTTCT |
| 5625 | mamo-L | *BGIBMGA011626* | down | 443 | 457 | - | 12.5887 | 1.30E-05 | 1 | TAGGCTTAGTTAGGT |
| 5626 | mamo-L | *BGIBMGA011628* | up | 1 | 15 | - | 10.9194 | 6.98E-05 | 1 | TAGGCTACTTTTTTT |
| 5627 | mamo-L | *BGIBMGA011630* | up | 7 | 21 | - | 11.5161 | 4.02E-05 | 1 | AAGGCTTACTTCTGT |
| 5628 | mamo-L | *BGIBMGA011635* | down | 270 | 284 | + | 11.2258 | 5.30E-05 | 1 | AAGGCTGGTTTTTTC |
| 5629 | mamo-L | *BGIBMGA011635* | down | 704 | 718 | - | 10.75 | 8.06E-05 | 1 | TAGGCCGTAAATGCC |
| 5630 | mamo-L | *BGIBMGA011636* | down | 953 | 967 | - | 13.0242 | 7.77E-06 | 1 | TAGGCTAAGATTGCT |
| 5631 | mamo-L | *BGIBMGA011637* | up | 1688 | 1702 | + | 11.6532 | 3.53E-05 | 1 | TAGGCTAGGTTGTGT |
| 5632 | mamo-L | *BGIBMGA011638* | up | 1636 | 1650 | - | 11.3629 | 4.66E-05 | 1 | TAGGCTCATATTTAT |
| 5633 | mamo-L | *BGIBMGA011639* | down | 639 | 653 | - | 11.2661 | 5.10E-05 | 1 | TAGGCGAGATTTTGT |
| 5634 | mamo-L | *BGIBMGA011640* | up | 722 | 736 | - | 10.9194 | 6.98E-05 | 1 | TAGGCTACTTTTTTT |
| 5635 | mamo-L | *BGIBMGA011642* | down | 1821 | 1835 | + | 13.379 | 4.96E-06 | 1 | TAGGCTTCGTTTTGT |
| 5636 | mamo-L | *BGIBMGA011644* | down | 1001 | 1015 | + | 10.5726 | 9.33E-05 | 1 | TAGGCTAATATTTAT |
| 5637 | mamo-L | *BGIBMGA011646* | up | 955 | 969 | - | 11.6371 | 3.58E-05 | 1 | GAGGCTGACATTTTT |
| 5638 | mamo-L | *BGIBMGA011647* | down | 1438 | 1452 | + | 10.6774 | 8.56E-05 | 1 | TGGGCTGATGATGCT |
| 5639 | mamo-L | *BGIBMGA011648* | up | 1314 | 1328 | - | 13.4677 | 4.39E-06 | 1 | TAGGCTGATGATGAT |
| 5640 | mamo-L | *BGIBMGA011651* | up | 991 | 1005 | - | 10.6532 | 8.73E-05 | 1 | GAGGCTTTCCTTGCT |
| 5641 | mamo-L | *BGIBMGA011652* | down | 1446 | 1460 | - | 11.371 | 4.62E-05 | 1 | TAGGCTGAGAACGTC |
| 5642 | mamo-L | *BGIBMGA011653* | up | 1593 | 1607 | + | 10.7016 | 8.39E-05 | 1 | TAGGCCTACATTTTG |
| 5643 | mamo-L | *BGIBMGA011661* | up | 1326 | 1340 | - | 11.2097 | 5.38E-05 | 1 | TAGGCGCTTTGTTGT |
| 5644 | mamo-L | *BGIBMGA011662* | up | 1887 | 1901 | + | 11.4032 | 4.49E-05 | 1 | AAGGCTTAAGATGGT |
| 5645 | mamo-L | *BGIBMGA011670* | down | 292 | 306 | - | 14.2661 | 1.33E-06 | 1 | TAGGCTTTCTTTTTT |
| 5646 | mamo-L | *BGIBMGA011675* | down | 1963 | 1977 | + | 12.629 | 1.24E-05 | 1 | TAGGCGTATTTTGGA |
| 5647 | mamo-L | *BGIBMGA011675* | down | 362 | 376 | - | 10.9274 | 6.92E-05 | 1 | TAGGCCAACTTTCAT |
| 5648 | mamo-L | *BGIBMGA011676* | up | 312 | 326 | + | 11.5323 | 3.96E-05 | 1 | TAGGCGTGAGGTGAT |
| 5649 | mamo-L | *BGIBMGA011678* | down | 1822 | 1836 | + | 14.6855 | 6.44E-07 | 1 | TAGGCTGTATTTGGT |
| 5650 | mamo-L | *BGIBMGA011678* | up | 395 | 409 | + | 11.5968 | 3.73E-05 | 1 | TAGGCTTTATTGGAT |
| 5651 | mamo-L | *BGIBMGA011679* | up | 1816 | 1830 | - | 11.629 | 3.61E-05 | 1 | TAGGCTTCGCTTGTT |
| 5652 | mamo-L | *BGIBMGA011680* | down | 172 | 186 | + | 10.5726 | 9.33E-05 | 1 | TAGGCTAATATTTAT |
| 5653 | mamo-L | *BGIBMGA011683* | down | 1473 | 1487 | - | 10.9194 | 6.98E-05 | 1 | TAGGCTACTTTTTTT |
| 5654 | mamo-L | *BGIBMGA011684* | down | 566 | 580 | - | 11.7661 | 3.16E-05 | 1 | TAGGCTGACCTTAGT |
| 5655 | mamo-L | *BGIBMGA011686* | up | 1516 | 1530 | + | 11.0242 | 6.36E-05 | 1 | AAGGCCTTGTTTTCC |
| 5656 | mamo-L | *BGIBMGA011687* | down | 37 | 51 | + | 11.5081 | 4.05E-05 | 1 | TAGGCTTGTTGTGTA |
| 5657 | mamo-L | *BGIBMGA011691* | up | 956 | 970 | + | 11.2177 | 5.34E-05 | 1 | GAGGCTTTCTTTATT |
| 5658 | mamo-L | *BGIBMGA011692* | down | 793 | 807 | + | 10.5968 | 9.15E-05 | 1 | AAGGCTGTTAATGCT |
| 5659 | mamo-L | *BGIBMGA011695* | down | 139 | 153 | - | 11.129 | 5.79E-05 | 1 | TAGGCGGGCGATTAC |
| 5660 | mamo-L | *BGIBMGA011697* | up | 1910 | 1924 | - | 11.4194 | 4.42E-05 | 1 | GAGGCTTAATTTGAT |
| 5661 | mamo-L | *BGIBMGA011697* | down | 1401 | 1415 | + | 10.5 | 9.90E-05 | 1 | GAGGCCCAGGTCGTC |
| 5662 | mamo-L | *BGIBMGA011701* | up | 274 | 288 | - | 10.7339 | 8.17E-05 | 1 | TAGGCTAAGTTCCTT |
| 5663 | mamo-L | *BGIBMGA011703* | down | 456 | 470 | - | 10.9194 | 6.98E-05 | 1 | TAGGCTACTTTTTTT |
| 5664 | mamo-L | *BGIBMGA011704* | down | 1085 | 1099 | + | 11.4194 | 4.42E-05 | 1 | GAGGCGGAGGTCGGC |
| 5665 | mamo-L | *BGIBMGA011707* | up | 1117 | 1131 | + | 11.2177 | 5.34E-05 | 1 | TAGGCTTAGATCCTT |
| 5666 | mamo-L | *BGIBMGA011710* | down | 1144 | 1158 | - | 10.9435 | 6.83E-05 | 1 | TGGGCTGGTGATGGT |
| 5667 | mamo-L | *BGIBMGA011711* | up | 732 | 746 | - | 10.5403 | 9.58E-05 | 1 | TAGGCTGTATGGTGT |
| 5668 | mamo-L | *BGIBMGA011716* | down | 1792 | 1806 | - | 10.7097 | 8.33E-05 | 1 | TAGGCAGTTGTTGGG |
| 5669 | mamo-L | *BGIBMGA011717* | up | 206 | 220 | + | 10.9194 | 6.98E-05 | 1 | TAGGCTACTTTTTTT |
| 5670 | mamo-L | *BGIBMGA011718* | down | 1362 | 1376 | - | 12.7823 | 1.03E-05 | 1 | TAGGCTCACTATTGT |
| 5671 | mamo-L | *BGIBMGA011720* | up | 104 | 118 | + | 12.0242 | 2.44E-05 | 1 | TAGGCCTCCTTTGGA |
| 5672 | mamo-L | *BGIBMGA011720* | down | 1358 | 1372 | - | 10.9194 | 6.98E-05 | 1 | TAGGCTACTTTTTTT |
| 5673 | mamo-L | *BGIBMGA011721* | up | 1585 | 1599 | - | 10.9194 | 6.98E-05 | 1 | TAGGCTACTTTTTTT |
| 5674 | mamo-L | *BGIBMGA011722* | down | 1366 | 1380 | + | 10.5968 | 9.15E-05 | 1 | AAGGCCTTGTGTTCT |
| 5675 | mamo-L | *BGIBMGA011722* | down | 1663 | 1677 | + | 10.5726 | 9.33E-05 | 1 | AAGGCCGGTGCTGCT |
| 5676 | mamo-L | *BGIBMGA011723* | up | 508 | 522 | + | 10.5968 | 9.15E-05 | 1 | AAGGCCTTGTGTTCT |
| 5677 | mamo-L | *BGIBMGA011723* | up | 805 | 819 | + | 10.5726 | 9.33E-05 | 1 | AAGGCCGGTGCTGCT |
| 5678 | mamo-L | *BGIBMGA011724* | up | 1862 | 1876 | + | 11.6613 | 3.50E-05 | 1 | TAGGCTTGCTAGGTC |
| 5679 | mamo-L | *BGIBMGA011725* | up | 505 | 519 | - | 10.6855 | 8.51E-05 | 1 | TAGGCAAGGTGTGGT |
| 5680 | mamo-L | *BGIBMGA011727* | down | 748 | 762 | + | 10.8145 | 7.63E-05 | 1 | TAGGCTCTTATTCTT |
| 5681 | mamo-L | *BGIBMGA011731* | down | 1493 | 1507 | + | 11.0565 | 6.18E-05 | 1 | TAGGCTTATTTAGGC |
| 5682 | mamo-L | *BGIBMGA011731* | down | 1503 | 1517 | + | 11 | 6.50E-05 | 1 | TAGGCCGACAACTTT |
| 5683 | mamo-L | *BGIBMGA011738* | up | 1854 | 1868 | - | 12.2258 | 1.97E-05 | 1 | TAGGCCCACTTGGAT |
| 5684 | mamo-L | *BGIBMGA011739* | down | 999 | 1013 | + | 11.3387 | 4.77E-05 | 1 | TAGGCCGTAGTTATC |
| 5685 | mamo-L | *BGIBMGA011740* | down | 43 | 57 | - | 14.2742 | 1.32E-06 | 1 | TAGGCCGTCGTGGGT |
| 5686 | mamo-L | *BGIBMGA011740* | down | 1868 | 1882 | - | 12.0161 | 2.46E-05 | 1 | TAGGCCGGTTGTGAC |
| 5687 | mamo-L | *BGIBMGA011740* | down | 136 | 150 | - | 11.3548 | 4.69E-05 | 1 | AAGGCTGATGATGAT |
| 5688 | mamo-L | *BGIBMGA011740* | up | 1394 | 1408 | + | 11.3387 | 4.77E-05 | 1 | TAGGCCGTAGTTATC |
| 5689 | mamo-L | *BGIBMGA011741* | down | 1285 | 1299 | - | 11.3548 | 4.69E-05 | 1 | AAGGCTGATGATGAT |
| 5690 | mamo-L | *BGIBMGA011742* | up | 669 | 683 | - | 10.5968 | 9.15E-05 | 1 | TAGGCTTATTTTATG |
| 5691 | mamo-L | *BGIBMGA011743* | up | 568 | 582 | - | 10.5565 | 9.46E-05 | 1 | AAGGCTTTTATTTTT |
| 5692 | mamo-L | *BGIBMGA011751* | down | 1666 | 1680 | + | 11.3871 | 4.55E-05 | 1 | TAGGCGTAGTGTGGG |
| 5693 | mamo-L | *BGIBMGA011751* | up | 539 | 553 | + | 10.9194 | 6.98E-05 | 1 | TAGGCTACTTTTTTT |
| 5694 | mamo-L | *BGIBMGA011754* | down | 1090 | 1104 | + | 10.6452 | 8.79E-05 | 1 | TAGGCTGTTGAGAGT |
| 5695 | mamo-L | *BGIBMGA011758* | up | 1466 | 1480 | + | 10.5726 | 9.33E-05 | 1 | TAGGCTAATATTTAT |
| 5696 | mamo-L | *BGIBMGA011761* | down | 9 | 23 | - | 10.5968 | 9.15E-05 | 1 | AAGGCCTTGTGTTCT |
| 5697 | mamo-L | *BGIBMGA011762* | down | 352 | 366 | - | 10.6855 | 8.51E-05 | 1 | AAGGCTTTATTCGTT |

| | | | | | | | | | | |
|---|---|---|---|---|---|---|---|---|---|---|
| 5698 | mamo-L | *BGIBMGA011764* | up | 668 | 682 | + | 12.621 | 1.25E-05 | 1 | TAGGCTTGCTTTTGA |
| 5699 | mamo-L | *BGIBMGA011764* | up | 717 | 731 | - | 10.5806 | 9.27E-05 | 1 | AAGGCTAGTTTTGAT |
| 5700 | mamo-L | *BGIBMGA011767* | up | 1053 | 1067 | + | 10.9194 | 6.98E-05 | 1 | TAGGCATTTTTGGTT |
| 5701 | mamo-L | *BGIBMGA011768* | up | 385 | 399 | - | 11.5161 | 4.02E-05 | 1 | TAGGCCTCAATTGTT |
| 5702 | mamo-L | *BGIBMGA011770* | up | 1973 | 1987 | - | 11.8548 | 2.89E-05 | 1 | TAGGCCTTTTTTAAT |
| 5703 | mamo-L | *BGIBMGA011771* | up | 1706 | 1720 | + | 10.5403 | 9.58E-05 | 1 | TAGGCTGTATGGTGT |
| 5704 | mamo-L | *BGIBMGA011773* | up | 1201 | 1215 | - | 12.5323 | 1.39E-05 | 1 | AAGGCTTTGTTTTGT |
| 5705 | mamo-L | *BGIBMGA011773* | down | 35 | 49 | - | 10.5161 | 9.77E-05 | 1 | TAGGCGGGCGCTGTA |
| 5706 | mamo-L | *BGIBMGA011775* | down | 1561 | 1575 | + | 14.7258 | 5.83E-07 | 1 | TAGGCTTGCGTTGCC |
| 5707 | mamo-L | *BGIBMGA011777* | down | 382 | 396 | + | 11.3145 | 4.87E-05 | 1 | TAGGCTTGTTTGCTT |
| 5708 | mamo-L | *BGIBMGA011778* | up | 1506 | 1520 | - | 14.9435 | 3.78E-07 | 1 | TAGGCTGTGTTTTGT |
| 5709 | mamo-L | *BGIBMGA011779* | up | 1870 | 1884 | + | 10.7339 | 8.17E-05 | 1 | TAGGCTAAGTTCCTT |
| 5710 | mamo-L | *BGIBMGA011788* | down | 1819 | 1833 | - | 11.8548 | 2.89E-05 | 1 | TAGGCTGGGGTGGTG |
| 5711 | mamo-L | *BGIBMGA011788* | down | 850 | 864 | - | 11.1774 | 5.54E-05 | 1 | TAGGCGTGGGAGGTC |
| 5712 | mamo-L | *BGIBMGA011791* | up | 1548 | 1562 | - | 10.8306 | 7.52E-05 | 1 | TAGGCCCCCTTCGCC |
| 5713 | mamo-L | *BGIBMGA011795* | down | 1944 | 1958 | - | 10.9194 | 6.98E-05 | 1 | TAGGCTACTTTTTTT |
| 5714 | mamo-L | *BGIBMGA011797* | down | 1757 | 1771 | - | 11.5968 | 3.73E-05 | 1 | TAGGCGTAGTTAGTT |
| 5715 | mamo-L | *BGIBMGA011798* | up | 1198 | 1212 | + | 10.8548 | 7.38E-05 | 1 | GAGGCCTGTTTTTAT |
| 5716 | mamo-L | *BGIBMGA011799* | up | 1952 | 1966 | - | 14.1694 | 1.54E-06 | 1 | TAGGCTGATTATGTT |
| 5717 | mamo-L | *BGIBMGA011800* | down | 511 | 525 | - | 12.629 | 1.24E-05 | 1 | TAGGCGTATTTTGGA |
| 5718 | mamo-L | *BGIBMGA011800* | down | 282 | 296 | + | 11.5726 | 3.81E-05 | 1 | TAGGCTTTGGATAGC |
| 5719 | mamo-L | *BGIBMGA011807* | down | 1652 | 1666 | - | 11.5565 | 3.87E-05 | 1 | AAGGCTGGCATTTCT |
| 5720 | mamo-L | *BGIBMGA011809* | up | 33 | 47 | + | 12.9758 | 8.23E-06 | 1 | TAGGCTCTTTTTTTT |
| 5721 | mamo-L | *BGIBMGA011810* | up | 19 | 33 | - | 13.0081 | 7.94E-06 | 1 | TAGGCTAGCTATGGT |
| 5722 | mamo-L | *BGIBMGA011811* | up | 1881 | 1895 | + | 11.4435 | 4.31E-05 | 1 | AAGGCTCTCTTTGCC |
| 5723 | mamo-L | *BGIBMGA011815* | up | 1014 | 1028 | + | 10.5403 | 9.58E-05 | 1 | TAGGCTGTATGGTGT |
| 5724 | mamo-L | *BGIBMGA011822* | down | 1349 | 1363 | + | 10.7419 | 8.11E-05 | 1 | AAGGCCCGTTTCGTT |
| 5725 | mamo-L | *BGIBMGA011825* | down | 1081 | 1095 | - | 10.7419 | 8.11E-05 | 1 | TAGGCTTTTTTTTAA |
| 5726 | mamo-L | *BGIBMGA011830* | up | 1371 | 1385 | + | 10.9194 | 6.98E-05 | 1 | TAGGCTACTTTTTTT |
| 5727 | mamo-L | *BGIBMGA011831* | down | 446 | 460 | + | 13.1694 | 6.54E-06 | 1 | TAGGCTGTTTTTATT |
| 5728 | mamo-L | *BGIBMGA011831* | down | 1114 | 1128 | + | 11.5645 | 3.84E-05 | 1 | TAGGCTGTTAGTTTT |
| 5729 | mamo-L | *BGIBMGA011832* | up | 1311 | 1325 | + | 13.1694 | 6.54E-06 | 1 | TAGGCTGTTTTTATT |
| 5730 | mamo-L | *BGIBMGA011832* | up | 1979 | 1993 | + | 11.5645 | 3.84E-05 | 1 | TAGGCTGTTAGTTTT |
| 5731 | mamo-L | *BGIBMGA011835* | down | 541 | 555 | - | 10.6774 | 8.56E-05 | 1 | TGGGCTGATGATGCT |
| 5732 | mamo-L | *BGIBMGA011839* | up | 1003 | 1017 | - | 11.2016 | 5.41E-05 | 1 | TAGGCCATATTTTTT |
| 5733 | mamo-L | *BGIBMGA011840* | up | 1916 | 1930 | - | 11.4839 | 4.15E-05 | 1 | AAGGCTATTTTTGGT |
| 5734 | mamo-L | *BGIBMGA011840* | down | 1119 | 1133 | - | 10.5726 | 9.33E-05 | 1 | TAGGCTAATATTTAT |
| 5735 | mamo-L | *BGIBMGA011841* | down | 1769 | 1783 | - | 12.5968 | 1.28E-05 | 1 | AAGGCTGATTTTTGT |
| 5736 | mamo-L | *BGIBMGA011841* | down | 1643 | 1657 | + | 11.2581 | 5.14E-05 | 1 | TAGGCACAATTTGCT |
| 5737 | mamo-L | *BGIBMGA011845* | down | 1479 | 1493 | + | 11.2661 | 5.10E-05 | 1 | TAGGCGAGATTTTGT |
| 5738 | mamo-L | *BGIBMGA011848* | up | 165 | 179 | + | 10.75 | 8.06E-05 | 1 | GAGGCCGATTGTGCC |
| 5739 | mamo-L | *BGIBMGA011849* | up | 1721 | 1735 | + | 11.2097 | 5.38E-05 | 1 | TAGGCTTTTCTTTCT |
| 5740 | mamo-L | *BGIBMGA011850* | down | 465 | 479 | + | 11.8871 | 2.81E-05 | 1 | TAGGCCGGTATTGCA |
| 5741 | mamo-L | *BGIBMGA011853* | up | 1469 | 1483 | + | 10.4919 | 9.96E-05 | 1 | TAGGCTGTGTCTGTG |
| 5742 | mamo-L | *BGIBMGA011855* | down | 500 | 514 | - | 10.9194 | 6.98E-05 | 1 | TAGGCTACTTTTTTT |
| 5743 | mamo-L | *BGIBMGA011856* | up | 1549 | 1563 | + | 10.5726 | 9.33E-05 | 1 | TAGGCCGAAATTAAT |
| 5744 | mamo-L | *BGIBMGA011860* | up | 1971 | 1985 | + | 11.629 | 3.61E-05 | 1 | TAGGCGCGCTGTTTT |
| 5745 | mamo-L | *BGIBMGA011860* | up | 1780 | 1794 | - | 10.8387 | 7.47E-05 | 1 | TCGGCTTGTTTTGTT |
| 5746 | mamo-L | *BGIBMGA011861* | up | 255 | 269 | + | 13.3065 | 5.48E-06 | 1 | TAGGCTTATTTTGGA |
| 5747 | mamo-L | *BGIBMGA011864* | up | 1691 | 1705 | - | 10.5968 | 9.15E-05 | 1 | TAGGCTTAATTGCTT |
| 5748 | mamo-L | *BGIBMGA011867* | down | 871 | 885 | - | 13.0161 | 7.84E-06 | 1 | TAGGCGTAAGTTGAT |
| 5749 | mamo-L | *BGIBMGA011869* | up | 556 | 570 | - | 10.5323 | 9.63E-05 | 1 | AAGGCCGATTTTAGC |
| 5750 | mamo-L | *BGIBMGA011870* | up | 668 | 682 | + | 11.6129 | 3.66E-05 | 1 | TAGGCTTTTAATGAT |
| 5751 | mamo-L | *BGIBMGA011873* | down | 1126 | 1140 | - | 13.7742 | 2.79E-06 | 1 | TAGGCGGGTGGTGGT |
| 5752 | mamo-L | *BGIBMGA011875* | up | 1025 | 1039 | - | 10.879 | 7.22E-05 | 1 | AAGGCTAGTGTTGGC |
| 5753 | mamo-L | *BGIBMGA011880* | down | 132 | 146 | - | 12.4516 | 1.53E-05 | 1 | TAGGCTTTTTATTTT |
| 5754 | mamo-L | *BGIBMGA011881* | up | 1467 | 1481 | - | 11.3952 | 4.52E-05 | 1 | TAGGCAGAGGTGGAT |
| 5755 | mamo-L | *BGIBMGA011882* | up | 450 | 464 | + | 13.5645 | 3.81E-06 | 1 | TAGGCTGAATTTTCT |
| 5756 | mamo-L | *BGIBMGA011887* | up | 299 | 313 | - | 10.9032 | 7.07E-05 | 1 | TAGGCTTGCTGTTTA |
| 5757 | mamo-L | *BGIBMGA011888* | up | 390 | 404 | - | 11.1935 | 5.45E-05 | 1 | AAGGCCGATTTTATT |
| 5758 | mamo-L | *BGIBMGA011888* | up | 1533 | 1547 | - | 10.75 | 8.06E-05 | 1 | GAGGCCGATTGTGCC |
| 5759 | mamo-L | *BGIBMGA011892* | down | 793 | 807 | + | 10.5565 | 9.46E-05 | 1 | TAGGCCGGCTTCAGA |
| 5760 | mamo-L | *BGIBMGA011894* | up | 234 | 248 | - | 12.5968 | 1.28E-05 | 1 | TAGGCTTATTTTGTG |
| 5761 | mamo-L | *BGIBMGA011894* | down | 167 | 181 | - | 11.6613 | 3.50E-05 | 1 | TAGGCCGTGATGTCT |
| 5762 | mamo-L | *BGIBMGA011894* | down | 385 | 399 | + | 10.7984 | 7.74E-05 | 1 | GAGGCTTTTTTTTAT |
| 5763 | mamo-L | *BGIBMGA011895* | up | 570 | 584 | - | 11.6613 | 3.50E-05 | 1 | TAGGCCGTGATGTCT |
| 5764 | mamo-L | *BGIBMGA011895* | up | 788 | 802 | + | 10.7984 | 7.74E-05 | 1 | GAGGCTTTTTTTTAT |
| 5765 | mamo-L | *BGIBMGA011896* | up | 1886 | 1900 | - | 11.0887 | 6.01E-05 | 1 | TAGGCGATGTTTTTC |
| 5766 | mamo-L | *BGIBMGA011897* | up | 350 | 364 | + | 11.2581 | 5.14E-05 | 1 | TAGGCGTGTATTTTC |
| 5767 | mamo-L | *BGIBMGA011900* | down | 450 | 464 | + | 13.5645 | 3.81E-06 | 1 | TAGGCTGATTTTTAT |
| 5768 | mamo-L | *BGIBMGA011901* | up | 287 | 301 | + | 13.5645 | 3.81E-06 | 1 | TAGGCTGATTTTTAT |
| 5769 | mamo-L | *BGIBMGA011901* | down | 1607 | 1621 | - | 10.9194 | 6.98E-05 | 1 | TAGGCTACTTTTTTT |
| 5770 | mamo-L | *BGIBMGA011909* | down | 936 | 950 | + | 10.9194 | 6.98E-05 | 1 | TAGGCTACTTTTTTT |
| 5771 | mamo-L | *BGIBMGA011910* | down | 1106 | 1120 | + | 12.6371 | 1.23E-05 | 1 | TAGGCGGAGTGGGGT |
| 5772 | mamo-L | *BGIBMGA011915* | up | 690 | 704 | + | 12.4194 | 1.58E-05 | 1 | TAGGCTTAGTGTTAT |
| 5773 | mamo-L | *BGIBMGA011916* | up | 1554 | 1568 | - | 11.7419 | 3.23E-05 | 1 | AAGGCCTTTGTCGGT |
| 5774 | mamo-L | *BGIBMGA011918* | down | 1824 | 1838 | + | 12.7258 | 1.11E-05 | 1 | TAGGCGTGTGTTCGT |
| 5775 | mamo-L | *BGIBMGA011920* | down | 1823 | 1837 | + | 10.5726 | 9.33E-05 | 1 | TAGGCTAATATTTAT |
| 5776 | mamo-L | *BGIBMGA011920* | up | 1812 | 1826 | - | 10.5242 | 9.70E-05 | 1 | TAGGCTTATGAAGTT |
| 5777 | mamo-L | *BGIBMGA011923* | up | 399 | 413 | - | 14.7661 | 5.44E-07 | 1 | TAGGCTTTGTTTGGC |
| 5778 | mamo-L | *BGIBMGA011923* | up | 71 | 85 | - | 10.4919 | 9.96E-05 | 1 | GAGGCTTGGTTCTGC |
| 5779 | mamo-L | *BGIBMGA011925* | down | 443 | 457 | - | 10.9194 | 6.98E-05 | 1 | TAGGCTACTTTTTTT |
| 5780 | mamo-L | *BGIBMGA011926* | up | 104 | 118 | + | 14.4597 | 9.75E-07 | 1 | TAGGCTTTGGTTTTT |
| 5781 | mamo-L | *BGIBMGA011931* | up | 1267 | 1281 | - | 11.7016 | 3.37E-05 | 1 | TAGGCCTTTTTTCAT |
| 5782 | mamo-L | *BGIBMGA011934* | up | 1361 | 1375 | + | 11.6774 | 3.44E-05 | 1 | TAGGCATGTTTTTCT |
| 5783 | mamo-L | *BGIBMGA011934* | up | 762 | 776 | - | 10.5 | 9.90E-05 | 1 | TAGGCGATTTTCGAT |
| 5784 | mamo-L | *BGIBMGA011935* | up | 1815 | 1829 | - | 10.5806 | 9.27E-05 | 1 | TAGGCCTTTGGTAAT |
| 5785 | mamo-L | *BGIBMGA011937* | up | 1766 | 1780 | + | 10.7177 | 8.28E-05 | 1 | TAGGCTTAAGTTAAC |
| 5786 | mamo-L | *BGIBMGA011938* | down | 1201 | 1215 | - | 12.9194 | 8.81E-06 | 1 | TAGGCGGAATTTTTT |
| 5787 | mamo-L | *BGIBMGA011938* | up | 816 | 830 | + | 10.8871 | 7.17E-05 | 1 | TGGGCTTCGTTTGGT |
| 5788 | mamo-L | *BGIBMGA011939* | up | 748 | 762 | - | 12.9194 | 8.81E-06 | 1 | TAGGCGGAATTTTTT |
| 5789 | mamo-L | *BGIBMGA011940* | down | 1854 | 1868 | - | 14.2823 | 1.31E-06 | 1 | TAGGCGTAGATTGGT |

| | | | | | | | | | | |
|---|---|---|---|---|---|---|---|---|---|---|
| 5790 | mamo-L | *BGIBMGA011941* | up | 1370 | 1384 | + | 12.7258 | 1.11E-05 | 1 | TAGGCGTGCTTTGTA |
| 5791 | mamo-L | *BGIBMGA011943* | up | 1648 | 1662 | + | 10.8065 | 7.69E-05 | 1 | TAGGCCGTTTTCTAC |
| 5792 | mamo-L | *BGIBMGA011946* | down | 1810 | 1824 | + | 10.5726 | 9.33E-05 | 1 | TAGGCTAATATTTAT |
| 5793 | mamo-L | *BGIBMGA011947* | down | 1327 | 1341 | + | 11.7016 | 3.37E-05 | 1 | TAGGCTGCTATTTTT |
| 5794 | mamo-L | *BGIBMGA011958* | up | 373 | 387 | + | 11.9597 | 2.61E-05 | 1 | TAGGCATTCTTTTTT |
| 5795 | mamo-L | *BGIBMGA011963* | up | 1886 | 1900 | - | 10.8548 | 7.38E-05 | 1 | TAGGCTCTATATTTT |
| 5796 | mamo-L | *BGIBMGA011965* | up | 1480 | 1494 | - | 10.7177 | 8.28E-05 | 1 | TAGGCTTAAGTTAAC |
| 5797 | mamo-L | *BGIBMGA011966* | down | 497 | 511 | + | 11.75 | 3.21E-05 | 1 | TAGGCTCTGGTTACC |
| 5798 | mamo-L | *BGIBMGA011967* | up | 1721 | 1735 | + | 11.75 | 3.21E-05 | 1 | TAGGCTCTGGTTACC |
| 5799 | mamo-L | *BGIBMGA011970* | down | 1929 | 1943 | - | 11.6774 | 3.44E-05 | 1 | TAGGCTAATTTTTAT |
| 5800 | mamo-L | *BGIBMGA011975* | down | 1771 | 1785 | + | 12.2823 | 1.85E-05 | 1 | AAGGCTCATTTTGTT |
| 5801 | mamo-L | *BGIBMGA011976* | up | 381 | 395 | + | 12.2823 | 1.85E-05 | 1 | AAGGCTCATTTTGTT |
| 5802 | mamo-L | *BGIBMGA011977* | up | 1380 | 1394 | + | 10.5242 | 9.70E-05 | 1 | TAGGCTTATGAAGTT |
| 5803 | mamo-L | *BGIBMGA011977* | up | 537 | 551 | + | 10.5081 | 9.83E-05 | 1 | TAGGCTATGTTTTCA |
| 5804 | mamo-L | *BGIBMGA011978* | down | 653 | 667 | - | 10.621 | 8.97E-05 | 1 | TAGGCTAAGCATGGT |
| 5805 | mamo-L | *BGIBMGA011980* | up | 156 | 170 | + | 11.7419 | 3.23E-05 | 1 | AAGGCCTTTGTCGGT |
| 5806 | mamo-L | *BGIBMGA011982* | up | 840 | 854 | + | 14.3306 | 1.21E-06 | 1 | TAGGCGGGGGGTGGT |
| 5807 | mamo-L | *BGIBMGA011983* | up | 1914 | 1928 | + | 10.5726 | 9.33E-05 | 1 | TAGGCTAATATTTAT |
| 5808 | mamo-L | *BGIBMGA011984* | up | 212 | 226 | + | 10.5726 | 9.33E-05 | 1 | TAGGCTAATATTTAT |
| 5809 | mamo-L | *BGIBMGA011986* | down | 524 | 538 | + | 12.0242 | 2.44E-05 | 1 | TAGGCCTAATTTCCT |
| 5810 | mamo-L | *BGIBMGA011990* | down | 752 | 766 | + | 12.5081 | 1.43E-05 | 1 | TAGGCTAATTTTTTT |
| 5811 | mamo-L | *BGIBMGA011990* | down | 163 | 177 | + | 10.9113 | 7.02E-05 | 1 | GAGGCTTTCTTGGGC |
| 5812 | mamo-L | *BGIBMGA011991* | up | 1512 | 1526 | + | 12.5081 | 1.43E-05 | 1 | TAGGCTAATTTTTTT |
| 5813 | mamo-L | *BGIBMGA011991* | up | 923 | 937 | + | 10.9113 | 7.02E-05 | 1 | GAGGCTTTCTTGGGC |
| 5814 | mamo-L | *BGIBMGA011996* | up | 461 | 475 | + | 12.5968 | 1.28E-05 | 1 | TAGGCTTATTTTGTG |
| 5815 | mamo-L | *BGIBMGA011997* | up | 474 | 488 | - | 10.9597 | 6.73E-05 | 1 | TAGGCTTATTGTAAT |
| 5816 | mamo-L | *BGIBMGA011998* | up | 1283 | 1297 | - | 12.7581 | 1.07E-05 | 1 | TAGGCGTTCTTTTAT |
| 5817 | mamo-L | *BGIBMGA011999* | up | 387 | 401 | + | 11.1935 | 5.45E-05 | 1 | AAGGCCGATTTTATT |
| 5818 | mamo-L | *BGIBMGA011999* | down | 1552 | 1566 | + | 11.0081 | 6.45E-05 | 1 | TAGGCTTACATGTAT |
| 5819 | mamo-L | *BGIBMGA012003* | down | 159 | 173 | - | 11.0403 | 6.28E-05 | 1 | TAGGCTTTTTGGTGT |
| 5820 | mamo-L | *BGIBMGA012005* | up | 991 | 1005 | - | 10.5806 | 9.27E-05 | 1 | TAGGCTATCGATATT |
| 5821 | mamo-L | *BGIBMGA012010* | up | 812 | 826 | - | 13.5806 | 3.72E-06 | 1 | TAGGCGTTCATTGTT |
| 5822 | mamo-L | *BGIBMGA012010* | down | 448 | 462 | + | 10.7097 | 8.33E-05 | 1 | AAGGCTGGCTGGGCT |
| 5823 | mamo-L | *BGIBMGA012013* | up | 360 | 374 | - | 12.9274 | 8.74E-06 | 1 | TAGGCCGTTTTCTGT |
| 5824 | mamo-L | *BGIBMGA012027* | down | 1482 | 1496 | - | 11.2742 | 5.06E-05 | 1 | TAGGCATGCGTTCTT |
| 5825 | mamo-L | *BGIBMGA012027* | down | 1321 | 1335 | - | 10.7419 | 8.11E-05 | 1 | TAGGCTAAATGTGGC |
| 5826 | mamo-L | *BGIBMGA012031* | up | 607 | 621 | - | 10.6694 | 8.62E-05 | 1 | TAGGCATTATTTTTT |
| 5827 | mamo-L | *BGIBMGA012032* | down | 1013 | 1027 | + | 11.3065 | 4.91E-05 | 1 | TAGGCTCAAGATTTT |
| 5828 | mamo-L | *BGIBMGA012032* | up | 1075 | 1089 | - | 10.8548 | 7.38E-05 | 1 | GAGGCCTGTTTTTAT |
| 5829 | mamo-L | *BGIBMGA012033* | up | 18 | 32 | - | 11.9758 | 2.57E-05 | 1 | TAGGCTCATTATTTT |
| 5830 | mamo-L | *BGIBMGA012034* | up | 347 | 361 | + | 10.7661 | 7.95E-05 | 1 | AAGGCTTGTGGCGGT |
| 5831 | mamo-L | *BGIBMGA012035* | up | 807 | 821 | - | 11.5645 | 3.84E-05 | 1 | TAGGCTAAGTTTGTG |
| 5832 | mamo-L | *BGIBMGA012035* | up | 186 | 200 | - | 11.121 | 5.84E-05 | 1 | TAGGCTTGAGATATT |
| 5833 | mamo-L | *BGIBMGA012036* | up | 1105 | 1119 | + | 11.2903 | 4.99E-05 | 1 | TAGGCTGTATGGGCT |
| 5834 | mamo-L | *BGIBMGA012042* | down | 1028 | 1042 | + | 12.9919 | 8.10E-06 | 1 | TAGGCCGGTTTTGCA |
| 5835 | mamo-L | *BGIBMGA012044* | up | 1222 | 1236 | - | 11.4435 | 4.31E-05 | 1 | TAGGCTTGAGAGGCT |
| 5836 | mamo-L | *BGIBMGA012045* | down | 1701 | 1715 | - | 11.121 | 5.84E-05 | 1 | TAGGCGGATTTTTTG |
| 5837 | mamo-L | *BGIBMGA012048* | up | 731 | 745 | - | 11.4677 | 4.22E-05 | 1 | TAGGCCCTGTTTATC |
| 5838 | mamo-L | *BGIBMGA012053* | up | 1522 | 1536 | - | 12.2581 | 1.90E-05 | 1 | GAGGCTGTGTATGTT |
| 5839 | mamo-L | *BGIBMGA012053* | up | 1549 | 1563 | + | 11.0323 | 6.32E-05 | 1 | TAGGCAGATGTTGGG |
| 5840 | mamo-L | *BGIBMGA012053* | down | 885 | 899 | + | 10.8871 | 7.17E-05 | 1 | TAGGCTGGTTTTTAG |
| 5841 | mamo-L | *BGIBMGA012054* | down | 1193 | 1207 | - | 11.129 | 5.79E-05 | 1 | TAGGCCGGAGACGAT |
| 5842 | mamo-L | *BGIBMGA012054* | down | 796 | 810 | + | 10.8145 | 7.63E-05 | 1 | AAGGCGTTCGTTGGA |
| 5843 | mamo-L | *BGIBMGA012057* | down | 1582 | 1596 | + | 10.6855 | 8.51E-05 | 1 | AAGGCGGGTGGTGGC |
| 5844 | mamo-L | *BGIBMGA012059* | down | 1666 | 1680 | - | 11.1694 | 5.59E-05 | 1 | AAGGCTTCGGATGGT |
| 5845 | mamo-L | *BGIBMGA012060* | up | 69 | 83 | - | 11.6129 | 3.66E-05 | 1 | AAGGCTGTGTTTATT |
| 5846 | mamo-L | *BGIBMGA012062* | down | 1982 | 1996 | + | 11.7177 | 3.32E-05 | 1 | TAGGCCCACTTTTGA |
| 5847 | mamo-L | *BGIBMGA012062* | down | 576 | 590 | + | 11.629 | 3.61E-05 | 1 | GAGGCTTTTTTTTTT |
| 5848 | mamo-L | *BGIBMGA012064* | up | 589 | 603 | - | 11.6613 | 3.50E-05 | 1 | GAGGCCGGGGATTGT |
| 5849 | mamo-L | *BGIBMGA012065* | up | 687 | 701 | - | 11.9758 | 2.57E-05 | 1 | TAGGCTCATTATTTT |
| 5850 | mamo-L | *BGIBMGA012066* | down | 251 | 265 | - | 11.7661 | 3.16E-05 | 1 | TAGGCAGTTTTTTTT |
| 5851 | mamo-L | *BGIBMGA012067* | down | 1 | 15 | - | 11.1371 | 5.75E-05 | 1 | AAGGCTTCAGTTGGT |
| 5852 | mamo-L | *BGIBMGA012068* | up | 1159 | 1173 | + | 11.9274 | 2.69E-05 | 1 | TAGGCTTCTTTGGCT |
| 5853 | mamo-L | *BGIBMGA012072* | down | 260 | 274 | + | 12.2581 | 1.90E-05 | 1 | TAGGCAGTCTTTTTT |
| 5854 | mamo-L | *BGIBMGA012085* | down | 184 | 198 | + | 10.8548 | 7.38E-05 | 1 | AAGGCTTAGATGGCT |
| 5855 | mamo-L | *BGIBMGA012088* | up | 1496 | 1510 | - | 13.1774 | 6.47E-06 | 1 | TAGGCTTACAGTGTT |
| 5856 | mamo-L | *BGIBMGA012092* | up | 1541 | 1555 | - | 10.5968 | 9.15E-05 | 1 | GAGGCTAGTTTTTGT |
| 5857 | mamo-L | *BGIBMGA012093* | up | 1350 | 1364 | + | 11.9274 | 2.69E-05 | 1 | TAGGCTGTCTCTTTT |
| 5858 | mamo-L | *BGIBMGA012095* | up | 173 | 187 | + | 11.6935 | 3.39E-05 | 1 | AAGGCTGATTTTAGT |
| 5859 | mamo-L | *BGIBMGA012096* | down | 992 | 1006 | + | 12.7258 | 1.11E-05 | 1 | TAGGCGTGCTTTGTA |
| 5860 | mamo-L | *BGIBMGA012096* | up | 1671 | 1685 | - | 10.9194 | 6.98E-05 | 1 | TAGGCTACTTTTTTT |
| 5861 | mamo-L | *BGIBMGA012097* | up | 1018 | 1032 | - | 10.5726 | 9.33E-05 | 1 | TAGGCTAATATTTAT |
| 5862 | mamo-L | *BGIBMGA012101* | up | 1299 | 1313 | - | 10.9194 | 6.98E-05 | 1 | TAGGCTACTTTTTTT |
| 5863 | mamo-L | *BGIBMGA012104* | down | 159 | 173 | + | 10.5726 | 9.33E-05 | 1 | TAGGCTAATATTTAT |
| 5864 | mamo-L | *BGIBMGA012106* | up | 418 | 432 | - | 11.5242 | 3.99E-05 | 1 | GAGGCTTTGGTTGGA |
| 5865 | mamo-L | *BGIBMGA012107* | down | 631 | 645 | + | 10.8468 | 7.42E-05 | 1 | TAGGCCTGCACTGAT |
| 5866 | mamo-L | *BGIBMGA012108* | down | 788 | 802 | + | 11.121 | 5.84E-05 | 1 | TAGGCGGATTTTTTG |
| 5867 | mamo-L | *BGIBMGA012110* | down | 13 | 27 | - | 10.6371 | 8.85E-05 | 1 | TAGGCGGGGGGTTGG |
| 5868 | mamo-L | *BGIBMGA012111* | down | 124 | 138 | - | 11.4516 | 4.28E-05 | 1 | GAGGCTGAATTTTTT |
| 5869 | mamo-L | *BGIBMGA012112* | up | 903 | 917 | - | 11.6532 | 3.53E-05 | 1 | TAGGCAGACGATGAT |
| 5870 | mamo-L | *BGIBMGA012115* | down | 1295 | 1309 | - | 11.0161 | 6.40E-05 | 1 | GAGGCTTTCATTTTT |
| 5871 | mamo-L | *BGIBMGA012117* | up | 1738 | 1752 | - | 12.7258 | 1.11E-05 | 1 | TAGGCGTGCTTTGTA |
| 5872 | mamo-L | *BGIBMGA012118* | down | 1767 | 1781 | - | 12.7258 | 1.11E-05 | 1 | TAGGCGTGCTTTGTA |
| 5873 | mamo-L | *BGIBMGA012118* | up | 489 | 503 | - | 11.5887 | 3.75E-05 | 1 | TAGGCAGTTTTCGTT |
| 5874 | mamo-L | *BGIBMGA012120* | up | 1705 | 1719 | + | 10.5726 | 9.33E-05 | 1 | TAGGCTAATATTTAT |
| 5875 | mamo-L | *BGIBMGA012121* | down | 1961 | 1975 | + | 11.8871 | 2.81E-05 | 1 | AAGGCTTACTTTAGT |
| 5876 | mamo-L | *BGIBMGA012121* | down | 868 | 882 | - | 11.3065 | 4.91E-05 | 1 | TAGGCTCAAGATTTT |
| 5877 | mamo-L | *BGIBMGA012121* | up | 441 | 455 | - | 10.5 | 9.90E-05 | 1 | GAGGCCCAGGTCGTC |
| 5878 | mamo-L | *BGIBMGA012123* | down | 613 | 627 | + | 13.621 | 3.50E-06 | 1 | TAGGCTGACTGCGGT |
| 5879 | mamo-L | *BGIBMGA012135* | up | 1125 | 1139 | + | 11.6452 | 3.56E-05 | 1 | TAGGCTCTTAATGTT |
| 5880 | mamo-L | *BGIBMGA012138* | down | 1657 | 1671 | + | 10.879 | 7.22E-05 | 1 | TAGGCTTGTATTCTC |
| 5881 | mamo-L | *BGIBMGA012142* | up | 487 | 501 | - | 11.1935 | 5.45E-05 | 1 | TAGGCGCATGAGGGT |

| | | | | | | | | | | |
|---|---|---|---|---|---|---|---|---|---|---|
| 5882 | mamo-L | *BGIBMGA012143* | down | 1804 | 1818 | + | 11.1532 | 5.67E-05 | 1 | AAGGCGGATGTGGCT |
| 5883 | mamo-L | *BGIBMGA012146* | down | 199 | 213 | - | 10.9194 | 6.98E-05 | 1 | TAGGCTACTTTTTTT |
| 5884 | mamo-L | *BGIBMGA012147* | up | 577 | 591 | - | 11.3548 | 4.69E-05 | 1 | AAGGCTGATGATGAT |
| 5885 | mamo-L | *BGIBMGA012147* | down | 780 | 794 | + | 10.5242 | 9.70E-05 | 1 | GAGGCTGTTTGTTTT |
| 5886 | mamo-L | *BGIBMGA012157* | up | 318 | 332 | + | 11.2177 | 5.34E-05 | 1 | AAGGCCTAGTTTGCA |
| 5887 | mamo-L | *BGIBMGA012160* | down | 215 | 229 | - | 12.1129 | 2.22E-05 | 1 | TAGGCTTAGGCTTCT |
| 5888 | mamo-L | *BGIBMGA012164* | up | 162 | 176 | + | 11.5403 | 3.93E-05 | 1 | TAGGCCATGATTGCC |
| 5889 | mamo-L | *BGIBMGA012166* | down | 197 | 211 | + | 10.879 | 7.22E-05 | 1 | AAGGCTGATTGTTTT |
| 5890 | mamo-L | *BGIBMGA012170* | down | 1879 | 1893 | + | 10.8387 | 7.47E-05 | 1 | TAGGCCGGTTTTCCA |
| 5891 | mamo-L | *BGIBMGA012171* | up | 449 | 463 | - | 10.9194 | 6.98E-05 | 1 | TAGGCTACTTTTTTT |
| 5892 | mamo-L | *BGIBMGA012173* | up | 255 | 269 | + | 11.5484 | 3.90E-05 | 1 | TAGGCTGGTCTGGGT |
| 5893 | mamo-L | *BGIBMGA012173* | up | 1599 | 1613 | + | 10.8145 | 7.63E-05 | 1 | TAGGCATATTTTTTC |
| 5894 | mamo-L | *BGIBMGA012173* | down | 881 | 895 | - | 10.629 | 8.91E-05 | 1 | TAGGCCTGCAGCGCC |
| 5895 | mamo-L | *BGIBMGA012174* | up | 1442 | 1456 | - | 10.9194 | 6.98E-05 | 1 | TAGGCTACTTTTTTT |
| 5896 | mamo-L | *BGIBMGA012174* | down | 788 | 802 | - | 10.5726 | 9.33E-05 | 1 | TAGGCTAATATTTAT |
| 5897 | mamo-L | *BGIBMGA012175* | up | 1064 | 1078 | - | 10.5726 | 9.33E-05 | 1 | TAGGCTAATATTTAT |
| 5898 | mamo-L | *BGIBMGA012177* | down | 425 | 439 | + | 10.9435 | 6.83E-05 | 1 | AAGGCCGGTGATGTC |
| 5899 | mamo-L | *BGIBMGA012177* | down | 563 | 577 | - | 10.5 | 9.90E-05 | 1 | GAGGCCCAGGTCGTC |
| 5900 | mamo-L | *BGIBMGA012178* | up | 1504 | 1518 | - | 10.8145 | 7.63E-05 | 1 | TAGGCATATTTTTTC |
| 5901 | mamo-L | *BGIBMGA012179* | up | 1970 | 1984 | + | 10.7984 | 7.74E-05 | 1 | TAGGCGCAGTTCGTA |
| 5902 | mamo-L | *BGIBMGA012188* | down | 756 | 770 | + | 10.9194 | 6.98E-05 | 1 | TAGGCTACTTTTTTT |
| 5903 | mamo-L | *BGIBMGA012189* | down | 406 | 420 | + | 10.5726 | 9.33E-05 | 1 | TAGGCTAATATTTAT |
| 5904 | mamo-L | *BGIBMGA012190* | up | 1079 | 1093 | + | 10.5726 | 9.33E-05 | 1 | TAGGCTAATATTTAT |
| 5905 | mamo-L | *BGIBMGA012191* | down | 1021 | 1035 | - | 10.5726 | 9.33E-05 | 1 | TAGGCTAATATTTAT |
| 5906 | mamo-L | *BGIBMGA012192* | up | 817 | 831 | - | 10.6694 | 8.62E-05 | 1 | TGGGCCTGGGATGCT |
| 5907 | mamo-L | *BGIBMGA012193* | down | 1525 | 1539 | - | 11.1935 | 5.45E-05 | 1 | TAGGCGCATGAGGGT |
| 5908 | mamo-L | *BGIBMGA012193* | up | 1334 | 1348 | + | 10.5161 | 9.77E-05 | 1 | TAGGCGTTTGTTAAC |
| 5909 | mamo-L | *BGIBMGA012194* | up | 577 | 591 | - | 11.8065 | 3.03E-05 | 1 | AAGGCGGGCTTCGCT |
| 5910 | mamo-L | *BGIBMGA012195* | down | 1502 | 1516 | + | 11.8952 | 2.79E-05 | 1 | TAGGCTTATTTTTTA |
| 5911 | mamo-L | *BGIBMGA012198* | up | 1317 | 1331 | + | 11.4758 | 4.18E-05 | 1 | GAGGCTGTGTTTTCC |
| 5912 | mamo-L | *BGIBMGA012201* | up | 1760 | 1774 | + | 10.5081 | 9.83E-05 | 1 | TAGGCGGGGTGTTGG |
| 5913 | mamo-L | *BGIBMGA012202* | up | 1411 | 1425 | - | 11.4516 | 4.28E-05 | 1 | TCGGCTGTGTTTGTT |
| 5914 | mamo-L | *BGIBMGA012206* | up | 70 | 84 | + | 10.7177 | 8.28E-05 | 1 | TAGGCTTAGTTTAAA |
| 5915 | mamo-L | *BGIBMGA012211* | up | 94 | 108 | + | 12.8145 | 9.96E-06 | 1 | TAGGCCTCCTTTTTT |
| 5916 | mamo-L | *BGIBMGA012211* | up | 916 | 930 | + | 11.1532 | 5.67E-05 | 1 | TAGGCGAAATTTGTC |
| 5917 | mamo-L | *BGIBMGA012214* | up | 303 | 317 | + | 12.3629 | 1.69E-05 | 1 | TAGGCTGATTATGAC |
| 5918 | mamo-L | *BGIBMGA012215* | up | 1879 | 1893 | - | 11.4113 | 4.45E-05 | 1 | TAGGCCTGTAGTGCC |
| 5919 | mamo-L | *BGIBMGA012215* | down | 1371 | 1385 | - | 11.0161 | 6.40E-05 | 1 | TAGGCTGTTTATAAT |
| 5920 | mamo-L | *BGIBMGA012216* | up | 396 | 410 | - | 11.0161 | 6.40E-05 | 1 | TAGGCTGTTTATAAT |
| 5921 | mamo-L | *BGIBMGA012219* | down | 1866 | 1880 | - | 12.5081 | 1.43E-05 | 1 | TAGGCTAATTTTTTT |
| 5922 | mamo-L | *BGIBMGA012221* | up | 1842 | 1856 | - | 10.5726 | 9.33E-05 | 1 | TAGGCTAATATTTAT |
| 5923 | mamo-L | *BGIBMGA012224* | up | 1820 | 1834 | + | 10.5726 | 9.33E-05 | 1 | TAGGCTAATATTTAT |
| 5924 | mamo-L | *BGIBMGA012225* | up | 153 | 167 | + | 10.8306 | 7.52E-05 | 1 | TAGCCTGAGATTGTT |
| 5925 | mamo-L | *BGIBMGA012227* | down | 1695 | 1709 | - | 10.7016 | 8.39E-05 | 1 | GAGGCTTTTGTTCTT |
| 5926 | mamo-L | *BGIBMGA012228* | up | 1512 | 1526 | - | 13.379 | 4.96E-06 | 1 | AAGGCTGATTTTGTT |
| 5927 | mamo-L | *BGIBMGA012228* | down | 804 | 818 | - | 12.4839 | 1.47E-05 | 1 | TAGGCCTTTGATGCC |
| 5928 | mamo-L | *BGIBMGA012231* | down | 1346 | 1360 | + | 11.0161 | 6.40E-05 | 1 | TAGGCTCATTACTGT |
| 5929 | mamo-L | *BGIBMGA012231* | up | 1725 | 1739 | + | 11.0081 | 6.45E-05 | 1 | AAGGCCTTGGTCGTC |
| 5930 | mamo-L | *BGIBMGA012237* | down | 1699 | 1713 | + | 14.7258 | 5.83E-07 | 1 | TAGGCTTGCGTTGCC |
| 5931 | mamo-L | *BGIBMGA012238* | down | 1942 | 1956 | + | 12.0645 | 2.34E-05 | 1 | TAGGCCAGTTTCGTT |
| 5932 | mamo-L | *BGIBMGA012242* | up | 444 | 458 | + | 11.0081 | 6.45E-05 | 1 | GAGGCGTGGTTTCGT |
| 5933 | mamo-L | *BGIBMGA012242* | down | 1960 | 1974 | + | 10.9194 | 6.98E-05 | 1 | TAGGCTTAGTGTGAG |
| 5934 | mamo-L | *BGIBMGA012243* | down | 184 | 198 | - | 11.1694 | 5.59E-05 | 1 | TAGGCTTGCAGTTAT |
| 5935 | mamo-L | *BGIBMGA012250* | down | 1603 | 1617 | + | 10.9516 | 6.78E-05 | 1 | TAGGCTTAAGTGTAT |
| 5936 | mamo-L | *BGIBMGA012251* | down | 1336 | 1350 | + | 10.8387 | 7.47E-05 | 1 | TAGGCTCATAATTCT |
| 5937 | mamo-L | *BGIBMGA012251* | down | 945 | 959 | + | 10.5726 | 9.33E-05 | 1 | TAGGCTAATATTTAT |
| 5938 | mamo-L | *BGIBMGA012256* | down | 1479 | 1493 | - | 12.2742 | 1.87E-05 | 1 | AAGGCTGTCTTCGTT |
| 5939 | mamo-L | *BGIBMGA012256* | down | 657 | 671 | + | 10.5403 | 9.58E-05 | 1 | TAGGCTAGCTTAGTT |
| 5940 | mamo-L | *BGIBMGA012257* | down | 1578 | 1592 | - | 12.7258 | 1.11E-05 | 1 | TAGGCGTGCTTTGTA |
| 5941 | mamo-L | *BGIBMGA012257* | up | 1126 | 1140 | - | 10.6855 | 8.51E-05 | 1 | AAGGCTATATTTGGT |
| 5942 | mamo-L | *BGIBMGA012259* | down | 186 | 200 | - | 10.5726 | 9.33E-05 | 1 | TAGGCTAATATTTAT |
| 5943 | mamo-L | *BGIBMGA012260* | down | 1973 | 1987 | - | 10.7016 | 8.39E-05 | 1 | GAGGCTCGGGATGAT |
| 5944 | mamo-L | *BGIBMGA012264* | down | 1461 | 1475 | + | 13.371 | 5.03E-06 | 1 | TAGGCTGTTGTTTAT |
| 5945 | mamo-L | *BGIBMGA012264* | up | 24 | 38 | + | 12.0645 | 2.34E-05 | 1 | TAGGCGGGCTTCGGA |
| 5946 | mamo-L | *BGIBMGA012265* | up | 377 | 391 | - | 11.8065 | 3.03E-05 | 1 | TAGGCTATGTTTACT |
| 5947 | mamo-L | *BGIBMGA012265* | up | 75 | 89 | - | 11.0161 | 6.40E-05 | 1 | TAGGCGTTTATTTTC |
| 5948 | mamo-L | *BGIBMGA012271* | up | 1985 | 1999 | - | 14.9597 | 3.63E-07 | 1 | TAGGCTGTCGTTGAT |
| 5949 | mamo-L | *BGIBMGA012273* | down | 900 | 914 | + | 12.1452 | 2.15E-05 | 1 | TAGGCTTATTGGGTT |
| 5950 | mamo-L | *BGIBMGA012276* | up | 128 | 142 | + | 10.7661 | 7.95E-05 | 1 | AAGGCTGTTGATTTT |
| 5951 | mamo-L | *BGIBMGA012276* | down | 675 | 689 | + | 10.5726 | 9.33E-05 | 1 | TAGGCTAATATTTAT |
| 5952 | mamo-L | *BGIBMGA012277* | up | 1461 | 1475 | - | 10.5403 | 9.58E-05 | 1 | TAGGCTGTATGGTGT |
| 5953 | mamo-L | *BGIBMGA012278* | up | 1435 | 1449 | - | 11.9274 | 2.69E-05 | 1 | TAGGCCAGCGTTACT |
| 5954 | mamo-L | *BGIBMGA012278* | down | 1877 | 1891 | - | 11.0645 | 6.14E-05 | 1 | TAGGCTGGCACTTTT |
| 5955 | mamo-L | *BGIBMGA012280* | up | 51 | 65 | + | 13.4677 | 4.39E-06 | 1 | TAGGCTGATGATGAT |
| 5956 | mamo-L | *BGIBMGA012281* | up | 1819 | 1833 | + | 10.6532 | 8.73E-05 | 1 | AAGGCTTAGTTTTGA |
| 5957 | mamo-L | *BGIBMGA012283* | down | 1194 | 1208 | + | 10.9274 | 6.92E-05 | 1 | TAGGCTAGTTACGTT |
| 5958 | mamo-L | *BGIBMGA012285* | up | 1801 | 1815 | + | 11.7661 | 3.16E-05 | 1 | GAGGCTGAATTTTGT |
| 5959 | mamo-L | *BGIBMGA012286* | up | 1016 | 1030 | + | 13.4274 | 4.64E-06 | 1 | TAGGCTTGTTTTAGT |
| 5960 | mamo-L | *BGIBMGA012287* | up | 1684 | 1698 | + | 11.8065 | 3.03E-05 | 1 | TAGGCCCCCTTCGCT |
| 5961 | mamo-L | *BGIBMGA012289* | up | 593 | 607 | + | 11.9194 | 2.72E-05 | 1 | TAGGCATATGTTTTT |
| 5962 | mamo-L | *BGIBMGA012294* | up | 1278 | 1292 | - | 10.9194 | 6.98E-05 | 1 | TAGGCTACTTTTTTT |
| 5963 | mamo-L | *BGIBMGA012299* | up | 1426 | 1440 | + | 10.5403 | 9.58E-05 | 1 | TAGGCTATTTTGTTT |
| 5964 | mamo-L | *BGIBMGA012300* | down | 1448 | 1462 | + | 11.7339 | 3.26E-05 | 1 | TAGGCTTCTGTTATT |
| 5965 | mamo-L | *BGIBMGA012301* | down | 1793 | 1807 | - | 13.0726 | 7.34E-06 | 1 | TAGGCCGATTTTGCA |
| 5966 | mamo-L | *BGIBMGA012302* | down | 1885 | 1899 | + | 10.5484 | 9.52E-05 | 1 | GAGGCGCACGTGGTT |
| 5967 | mamo-L | *BGIBMGA012303* | up | 1198 | 1212 | + | 10.5484 | 9.52E-05 | 1 | GAGGCGCACGTGGTT |
| 5968 | mamo-L | *BGIBMGA012304* | down | 1263 | 1277 | + | 10.6613 | 8.67E-05 | 1 | TAGGCTTAGAATAGC |
| 5969 | mamo-L | *BGIBMGA012307* | down | 725 | 739 | + | 12.5323 | 1.39E-05 | 1 | TAGGCCCGGTTTCTT |
| 5970 | mamo-L | *BGIBMGA012308* | up | 1572 | 1586 | - | 11.129 | 5.79E-05 | 1 | TAGGCCCGAATTTTT |
| 5971 | mamo-L | *BGIBMGA012309* | up | 90 | 104 | - | 12.0403 | 2.40E-05 | 1 | TAGGCGGGGTATTAT |
| 5972 | mamo-L | *BGIBMGA012311* | down | 1684 | 1698 | + | 12.7258 | 1.11E-05 | 1 | TAGGCGTGCTTTGTA |
| 5973 | mamo-L | *BGIBMGA012314* | up | 1215 | 1229 | - | 11.2258 | 5.30E-05 | 1 | TAGGCGTTCTTCCCT |

| | | | | | | | | | | |
|---|---|---|---|---|---|---|---|---|---|---|
| 5974 | mamo-L | *BGIBMGA012314* | up | 794 | 808 | + | 10.5161 | 9.77E-05 | 1 | TGGGCCGAGGTTGGA |
| 5975 | mamo-L | *BGIBMGA012315* | down | 494 | 508 | + | 13.0323 | 7.70E-06 | 1 | TAGGCGGGCAGTGGT |
| 5976 | mamo-L | *BGIBMGA012317* | up | 476 | 490 | + | 12.6613 | 1.19E-05 | 1 | TAGGCTTATTGTTCT |
| 5977 | mamo-L | *BGIBMGA012317* | down | 1330 | 1344 | + | 12.25 | 1.92E-05 | 1 | TAGGCTTGCTATACT |
| 5978 | mamo-L | *BGIBMGA012317* | up | 1544 | 1558 | + | 10.871 | 7.28E-05 | 1 | TAGGCTAAAATTGAT |
| 5979 | mamo-L | *BGIBMGA012320* | up | 1620 | 1634 | - | 11 | 6.50E-05 | 1 | AAGGCTTTTTTTTGC |
| 5980 | mamo-L | *BGIBMGA012321* | down | 1412 | 1426 | - | 10.5 | 9.90E-05 | 1 | TAGGCTGGCTTATAT |
| 5981 | mamo-L | *BGIBMGA012322* | down | 51 | 65 | + | 10.7419 | 8.11E-05 | 1 | GAGGCGTTCGTTTAT |
| 5982 | mamo-L | *BGIBMGA012326* | up | 1651 | 1665 | - | 11.2258 | 5.30E-05 | 1 | TAGGCTAGTGTGTGT |
| 5983 | mamo-L | *BGIBMGA012327* | down | 1484 | 1498 | - | 11.0645 | 6.14E-05 | 1 | TAGGCATGGGTGTGT |
| 5984 | mamo-L | *BGIBMGA012330* | up | 543 | 557 | + | 11.371 | 4.62E-05 | 1 | TAGGCTGTATAGGCT |
| 5985 | mamo-L | *BGIBMGA012332* | down | 1316 | 1330 | + | 12.0484 | 2.38E-05 | 1 | TAGGCCGATAATGAT |
| 5986 | mamo-L | *BGIBMGA012332* | up | 85 | 99 | + | 10.5726 | 9.33E-05 | 1 | TAGGCTAATATTTAT |
| 5987 | mamo-L | *BGIBMGA012334* | down | 365 | 379 | + | 10.7258 | 8.23E-05 | 1 | AAGGCTTTTTTTACT |
| 5988 | mamo-L | *BGIBMGA012334* | up | 715 | 729 | + | 10.7177 | 8.28E-05 | 1 | GAGGCTTTCTGTTTT |
| 5989 | mamo-L | *BGIBMGA012335* | down | 1540 | 1554 | + | 10.7823 | 7.84E-05 | 1 | TATGCTGGGTTTGTT |
| 5990 | mamo-L | *BGIBMGA012337* | up | 1726 | 1740 | - | 11.2339 | 5.26E-05 | 1 | TAGGCTACTTTTTGT |
| 5991 | mamo-L | *BGIBMGA012337* | down | 1751 | 1765 | - | 10.75 | 8.06E-05 | 1 | TAGGCTTACGCGTGT |
| 5992 | mamo-L | *BGIBMGA012340* | down | 194 | 208 | - | 10.5726 | 9.33E-05 | 1 | TAGGCTAATATTTAT |
| 5993 | mamo-L | *BGIBMGA012345* | down | 172 | 186 | + | 11.3306 | 4.80E-05 | 1 | TAGGCTTTATTGTTT |
| 5994 | mamo-L | *BGIBMGA012346* | up | 1241 | 1255 | + | 10.5645 | 9.40E-05 | 1 | TAGGCTTTTTTAGAT |
| 5995 | mamo-L | *BGIBMGA012349* | down | 583 | 597 | + | 10.6613 | 8.67E-05 | 1 | TACGCTTGCTTTGCT |
| 5996 | mamo-L | *BGIBMGA012353* | down | 463 | 477 | + | 12.5565 | 1.35E-05 | 1 | TAGGCTTATTCTGTT |
| 5997 | mamo-L | *BGIBMGA012361* | up | 199 | 213 | + | 11.1048 | 5.92E-05 | 1 | TAGGCCAGTTTTGCA |
| 5998 | mamo-L | *BGIBMGA012364* | up | 1600 | 1614 | + | 10.5242 | 9.70E-05 | 1 | AAGGCTTTTTTTGCA |
| 5999 | mamo-L | *BGIBMGA012367* | down | 662 | 676 | + | 12.25 | 1.92E-05 | 1 | TAGGCCCACTATTCT |
| 6000 | mamo-L | *BGIBMGA012368* | up | 946 | 960 | + | 12.25 | 1.92E-05 | 1 | TAGGCCCACTATTCT |
| 6001 | mamo-L | *BGIBMGA012368* | down | 1135 | 1149 | - | 10.6532 | 8.73E-05 | 1 | TAGGCGTACAATTAT |
| 6002 | mamo-L | *BGIBMGA012370* | up | 740 | 754 | - | 11.1452 | 5.70E-05 | 1 | TAGGCTGGCGGGCGT |
| 6003 | mamo-L | *BGIBMGA012378* | up | 778 | 792 | + | 10.5968 | 9.15E-05 | 1 | GAGGCTAGTTTTTGT |
| 6004 | mamo-L | *BGIBMGA012379* | down | 704 | 718 | + | 10.629 | 8.91E-05 | 1 | TAGGCCTGCAGCGCC |
| 6005 | mamo-L | *BGIBMGA012380* | up | 803 | 817 | + | 10.629 | 8.91E-05 | 1 | TAGGCCTGCAGCGCC |
| 6006 | mamo-L | *BGIBMGA012381* | up | 1184 | 1198 | + | 12.1935 | 2.05E-05 | 1 | TAGGCCTATTTTCGC |
| 6007 | mamo-L | *BGIBMGA012384* | up | 101 | 115 | + | 10.5484 | 9.52E-05 | 1 | TAGCCCGTGGATGGT |
| 6008 | mamo-L | *BGIBMGA012387* | down | 1398 | 1412 | + | 12.0645 | 2.34E-05 | 1 | TAGGCGGGCTTCGGA |
| 6009 | mamo-L | *BGIBMGA012387* | up | 210 | 224 | - | 11.0968 | 5.97E-05 | 1 | TAGGCTATGTTGTTT |
| 6010 | mamo-L | *BGIBMGA012389* | down | 145 | 159 | + | 11.0161 | 6.40E-05 | 1 | GAGGCTTTCATTTTT |
| 6011 | mamo-L | *BGIBMGA012390* | up | 1764 | 1778 | + | 11.5484 | 3.90E-05 | 1 | TAGGCCGGCGATATC |
| 6012 | mamo-L | *BGIBMGA012392* | down | 150 | 164 | + | 10.7177 | 8.28E-05 | 1 | TAGGCTTAGTTTAAA |
| 6013 | mamo-L | *BGIBMGA012397* | up | 329 | 343 | - | 10.8145 | 7.63E-05 | 1 | TAGGCCGCCTATTTC |
| 6014 | mamo-L | *BGIBMGA012400* | up | 824 | 838 | - | 12.2742 | 1.87E-05 | 1 | TAGGCTTAGTGTTTC |
| 6015 | mamo-L | *BGIBMGA012401* | up | 1744 | 1758 | - | 10.5726 | 9.33E-05 | 1 | TAGGCTAATATTTAT |
| 6016 | mamo-L | *BGIBMGA012402* | up | 14 | 28 | + | 12.9435 | 8.54E-06 | 1 | TAGGCTTTTTTTTAT |
| 6017 | mamo-L | *BGIBMGA012403* | down | 1225 | 1239 | - | 10.5323 | 9.63E-05 | 1 | GAGGCGTTGGTTGTA |
| 6018 | mamo-L | *BGIBMGA012406* | up | 1249 | 1263 | + | 11.3952 | 4.52E-05 | 1 | TAGGCGTTATTTATT |
| 6019 | mamo-L | *BGIBMGA012408* | up | 26 | 40 | - | 11.8306 | 2.97E-05 | 1 | TAGGCTGTTGCTGAT |
| 6020 | mamo-L | *BGIBMGA012408* | up | 32 | 46 | - | 10.9113 | 7.02E-05 | 1 | GAGGCCTAGGCTGTT |
| 6021 | mamo-L | *BGIBMGA012409* | up | 900 | 914 | + | 10.5726 | 9.33E-05 | 1 | TAGGCTAATATTTAT |
| 6022 | mamo-L | *BGIBMGA012413* | down | 1116 | 1130 | - | 11.1855 | 5.50E-05 | 1 | GAGGCTCCGTTTGCT |
| 6023 | mamo-L | *BGIBMGA012413* | up | 623 | 637 | + | 10.7016 | 8.39E-05 | 1 | TAGGCATATTGTTGT |
| 6024 | mamo-L | *BGIBMGA012415* | up | 498 | 512 | + | 12.2661 | 1.89E-05 | 1 | TAGGCGGCTGGTGGT |
| 6025 | mamo-L | *BGIBMGA012415* | down | 839 | 853 | - | 10.9758 | 6.64E-05 | 1 | TAGGCAGTTTTTGGA |
| 6026 | mamo-L | *BGIBMGA012423* | down | 859 | 873 | - | 11.0323 | 6.32E-05 | 1 | TAGGCGCCTTTTTTT |
| 6027 | mamo-L | *BGIBMGA012424* | up | 1535 | 1549 | + | 12.1129 | 2.22E-05 | 1 | TAGGCTTAGTGTGCA |
| 6028 | mamo-L | *BGIBMGA012425* | up | 246 | 260 | + | 10.629 | 8.91E-05 | 1 | AAGGCTAGTTTTTGT |
| 6029 | mamo-L | *BGIBMGA012425* | up | 1601 | 1615 | + | 10.6048 | 9.09E-05 | 1 | TAGGCCTAATGTGTA |
| 6030 | mamo-L | *BGIBMGA012426* | down | 259 | 273 | + | 13 | 8.01E-06 | 1 | AAGGCTTGTTTTGTT |
| 6031 | mamo-L | *BGIBMGA012435* | up | 1726 | 1740 | + | 10.5806 | 9.27E-05 | 1 | TACGCTTTCGTTGTT |
| 6032 | mamo-L | *BGIBMGA012437* | up | 241 | 255 | - | 13.0242 | 7.77E-06 | 1 | TAGGCTTGCGGCGCT |
| 6033 | mamo-L | *BGIBMGA012437* | down | 1014 | 1028 | + | 11.6694 | 3.47E-05 | 1 | TAGGCTTATAATTTT |
| 6034 | mamo-L | *BGIBMGA012438* | down | 1357 | 1371 | + | 11.7903 | 3.08E-05 | 1 | TAGGCTTGAGAGGGT |
| 6035 | mamo-L | *BGIBMGA012442* | up | 760 | 774 | - | 11.4274 | 4.38E-05 | 1 | AAGGCGTTTGTTTGT |
| 6036 | mamo-L | *BGIBMGA012442* | down | 81 | 95 | - | 11.4032 | 4.49E-05 | 1 | TAGGCTAATATTTTT |
| 6037 | mamo-L | *BGIBMGA012443* | down | 1330 | 1344 | - | 11.9274 | 2.69E-05 | 1 | TAGGCTGTGTAGGAT |
| 6038 | mamo-L | *BGIBMGA012444* | down | 1013 | 1027 | + | 11.0565 | 6.18E-05 | 1 | TAGCCGTAGGTTGCT |
| 6039 | mamo-L | *BGIBMGA012445* | up | 968 | 982 | - | 11.7419 | 3.23E-05 | 1 | AAGGCCGGGGTCTGT |
| 6040 | mamo-L | *BGIBMGA012445* | up | 1515 | 1529 | - | 10.5726 | 9.33E-05 | 1 | TAGGCTAATATTTAT |
| 6041 | mamo-L | *BGIBMGA012451* | up | 1743 | 1757 | + | 10.9194 | 6.98E-05 | 1 | TAGGCTACTTTTTTT |
| 6042 | mamo-L | *BGIBMGA012452* | up | 785 | 799 | + | 13 | 8.01E-06 | 1 | TAGGCGGAGTTCTTT |
| 6043 | mamo-L | *BGIBMGA012452* | down | 665 | 679 | + | 11.129 | 5.79E-05 | 1 | TAGGCTGTGTTTATG |
| 6044 | mamo-L | *BGIBMGA012452* | down | 195 | 209 | + | 10.7419 | 8.11E-05 | 1 | GAGGCGTTCGTTTAT |
| 6045 | mamo-L | *BGIBMGA012456* | up | 1102 | 1116 | - | 10.5968 | 9.15E-05 | 1 | GAGGCTAGTTTTTGT |
| 6046 | mamo-L | *BGIBMGA012465* | down | 1402 | 1416 | - | 10.5968 | 9.15E-05 | 1 | AAGGCTCATTTTAGT |
| 6047 | mamo-L | *BGIBMGA012466* | down | 206 | 220 | - | 11.6129 | 3.66E-05 | 1 | AAGGCTATTGTTGGT |
| 6048 | mamo-L | *BGIBMGA012467* | up | 1470 | 1484 | - | 11.6129 | 3.66E-05 | 1 | AAGGCTATTGTTGGT |
| 6049 | mamo-L | *BGIBMGA012469* | up | 1357 | 1371 | + | 11.3548 | 4.69E-05 | 1 | TAGGCTTTCATTTAC |
| 6050 | mamo-L | *BGIBMGA012470* | down | 1046 | 1060 | - | 11.621 | 3.63E-05 | 1 | TAGGCTTTTTATTAT |
| 6051 | mamo-L | *BGIBMGA012471* | down | 549 | 563 | + | 11.4597 | 4.25E-05 | 1 | TAGGCTGGCAGCGGC |
| 6052 | mamo-L | *BGIBMGA012476* | up | 1704 | 1718 | + | 12.0403 | 2.40E-05 | 1 | TAGGCGTTTTATGAT |
| 6053 | mamo-L | *BGIBMGA012477* | down | 1943 | 1957 | - | 11.4113 | 4.45E-05 | 1 | TAGGCGGGTGTGGAC |
| 6054 | mamo-L | *BGIBMGA012478* | up | 672 | 686 | - | 10.5726 | 9.33E-05 | 1 | TACGCTGTTTTTGGT |
| 6055 | mamo-L | *BGIBMGA012481* | up | 800 | 814 | + | 10.5726 | 9.33E-05 | 1 | TAGGCTAATATTTAT |
| 6056 | mamo-L | *BGIBMGA012485* | down | 1793 | 1807 | + | 11.4435 | 4.31E-05 | 1 | TAGGCTGAGTTATCT |
| 6057 | mamo-L | *BGIBMGA012485* | down | 569 | 583 | + | 10.5726 | 9.33E-05 | 1 | TAGGCTAATATTTAT |
| 6058 | mamo-L | *BGIBMGA012486* | down | 405 | 419 | - | 13.2903 | 5.60E-06 | 1 | TAGGCTGATATTTTT |
| 6059 | mamo-L | *BGIBMGA012487* | up | 1344 | 1358 | - | 10.5726 | 9.33E-05 | 1 | TAGGCTAATATTTAT |
| 6060 | mamo-L | *BGIBMGA012490* | up | 1053 | 1067 | + | 13.2742 | 5.70E-06 | 1 | TAGGCTGTATTTTTT |
| 6061 | mamo-L | *BGIBMGA012492* | up | 1353 | 1367 | + | 11.0887 | 6.01E-05 | 1 | GAGGCGGACGTTGTA |
| 6062 | mamo-L | *BGIBMGA012495* | up | 1155 | 1169 | - | 11.9839 | 2.54E-05 | 1 | TAGGCGTAGAATGGC |
| 6063 | mamo-L | *BGIBMGA012498* | up | 1027 | 1041 | + | 12.5726 | 1.33E-05 | 1 | TAGGCTGTTTACGTT |
| 6064 | mamo-L | *BGIBMGA012499* | down | 685 | 699 | - | 11.25 | 5.18E-05 | 1 | TAGGCGGTCATTGCG |
| 6065 | mamo-L | *BGIBMGA012500* | down | 811 | 825 | + | 11.621 | 3.63E-05 | 1 | TAGGCTTTTTATTAT |

| | | | | | | | | | | |
|---|---|---|---|---|---|---|---|---|---|---|
| 6066 | mamo-L | *BGIBMGA012502* | down | 191 | 205 | - | 12.6371 | 1.23E-05 | 1 | TAGGCGGAGTGGGGT |
| 6067 | mamo-L | *BGIBMGA012503* | down | 1062 | 1076 | + | 10.9194 | 6.98E-05 | 1 | TAGGCTACTTTTTTT |
| 6068 | mamo-L | *BGIBMGA012507* | down | 1186 | 1200 | - | 13.4919 | 4.25E-06 | 1 | TAGGCCGTCTATGGC |
| 6069 | mamo-L | *BGIBMGA012507* | up | 1604 | 1618 | - | 11.4839 | 4.15E-05 | 1 | TAGGCTTACTTTATA |
| 6070 | mamo-L | *BGIBMGA012515* | down | 1960 | 1974 | - | 11.129 | 5.79E-05 | 1 | TAGGCTGTGTTTATG |
| 6071 | mamo-L | *BGIBMGA012518* | down | 209 | 223 | - | 11.5081 | 4.05E-05 | 1 | TAGGCTTTGTATGTG |
| 6072 | mamo-L | *BGIBMGA012521* | down | 1370 | 1384 | + | 10.7661 | 7.95E-05 | 1 | GAGGCGGTCGTTGTA |
| 6073 | mamo-L | *BGIBMGA012522* | up | 1863 | 1877 | + | 10.5726 | 9.33E-05 | 1 | TAGGCTAATATTTAT |
| 6074 | mamo-L | *BGIBMGA012523* | up | 1571 | 1585 | + | 10.6129 | 9.03E-05 | 1 | TAGGCTGATTGCGTA |
| 6075 | mamo-L | *BGIBMGA012524* | down | 218 | 232 | + | 10.4919 | 9.96E-05 | 1 | TAGGCTTTAATGGAT |
| 6076 | mamo-L | *BGIBMGA012525* | down | 1412 | 1426 | - | 10.4919 | 9.96E-05 | 1 | GAGGCGTACGGTTTT |
| 6077 | mamo-L | *BGIBMGA012526* | up | 1272 | 1286 | + | 12.1694 | 2.09E-05 | 1 | TAGGCTGTGTGTCTT |
| 6078 | mamo-L | *BGIBMGA012528* | up | 1894 | 1908 | + | 13.4677 | 4.39E-06 | 1 | TAGGCTGATGATGAT |
| 6079 | mamo-L | *BGIBMGA012528* | up | 1551 | 1565 | - | 10.7097 | 8.33E-05 | 1 | TAGGCTCGTTGTGTA |
| 6080 | mamo-L | *BGIBMGA012529* | up | 1328 | 1342 | + | 12.3468 | 1.72E-05 | 1 | TAGGCTGACAGTTCT |
| 6081 | mamo-L | *BGIBMGA012530* | down | 1146 | 1160 | + | 12.1048 | 2.24E-05 | 1 | TAGGCTTATTGTAGT |
| 6082 | mamo-L | *BGIBMGA012531* | up | 1642 | 1656 | + | 12.4435 | 1.54E-05 | 1 | GAGGCTTACTTTTTT |
| 6083 | mamo-L | *BGIBMGA012532* | down | 1810 | 1824 | + | 11.2258 | 5.30E-05 | 1 | TAGGCTTGTTTTAGA |
| 6084 | mamo-L | *BGIBMGA012534* | up | 1204 | 1218 | - | 11.1935 | 5.45E-05 | 1 | TAGGCTTGTTATGTG |
| 6085 | mamo-L | *BGIBMGA012534* | up | 1486 | 1500 | - | 10.9194 | 6.98E-05 | 1 | TAGGCTACTTTTTTT |
| 6086 | mamo-L | *BGIBMGA012543* | up | 392 | 406 | - | 11.1694 | 5.59E-05 | 1 | GAGGCGCAGGGTGGT |
| 6087 | mamo-L | *BGIBMGA012546* | up | 649 | 663 | - | 10.9194 | 6.98E-05 | 1 | TAGGCTACTTTTTTT |
| 6088 | mamo-L | *BGIBMGA012547* | down | 841 | 855 | - | 10.9274 | 6.92E-05 | 1 | TAGGCAGCAGTTGTT |
| 6089 | mamo-L | *BGIBMGA012547* | down | 890 | 904 | + | 10.75 | 8.06E-05 | 1 | TAGGCAGCGTGTGTT |
| 6090 | mamo-L | *BGIBMGA012548* | down | 1465 | 1479 | - | 15.2016 | 2.22E-07 | 1 | TAGGCGTAGGTTGTT |
| 6091 | mamo-L | *BGIBMGA012549* | down | 461 | 475 | + | 11.3145 | 4.87E-05 | 1 | TAGGCTTGATAGGCT |
| 6092 | mamo-L | *BGIBMGA012549* | up | 662 | 676 | - | 11.2016 | 5.41E-05 | 1 | TAGGCGGTGATGTTT |
| 6093 | mamo-L | *BGIBMGA012551* | down | 311 | 325 | - | 10.5323 | 9.63E-05 | 1 | TGGGCTGAGTTTTAT |
| 6094 | mamo-L | *BGIBMGA012562* | up | 1197 | 1211 | - | 13.5887 | 3.68E-06 | 1 | TAGGCCTTTTTTTTT |
| 6095 | mamo-L | *BGIBMGA012566* | up | 445 | 459 | + | 12.6694 | 1.18E-05 | 1 | TAGGCTTTTATTTTT |
| 6096 | mamo-L | *BGIBMGA012569* | up | 925 | 939 | - | 12.8548 | 9.51E-06 | 1 | TAGGCCGGTTTCTTT |
| 6097 | mamo-L | *BGIBMGA012571* | down | 1039 | 1053 | + | 12.621 | 1.25E-05 | 1 | TAGGCTTGCTTTTGA |
| 6098 | mamo-L | *BGIBMGA012575* | up | 1110 | 1124 | + | 10.871 | 7.28E-05 | 1 | TAGGCTTAAAATTTT |
| 6099 | mamo-L | *BGIBMGA012576* | down | 1217 | 1231 | + | 10.6613 | 8.67E-05 | 1 | GAGGCTTCATTTGTT |
| 6100 | mamo-L | *BGIBMGA012576* | down | 1843 | 1857 | - | 10.5887 | 9.21E-05 | 1 | AAGGCGGATAATGGT |
| 6101 | mamo-L | *BGIBMGA012576* | up | 796 | 810 | + | 10.5403 | 9.58E-05 | 1 | TAGGCTGTATGGTGT |
| 6102 | mamo-L | *BGIBMGA012577* | down | 660 | 674 | - | 10.9355 | 6.88E-05 | 1 | GAGGCTTATAATGGT |
| 6103 | mamo-L | *BGIBMGA012579* | up | 714 | 728 | + | 10.5 | 9.90E-05 | 1 | GAGGCCCAGGTCGTC |
| 6104 | mamo-L | *BGIBMGA012581* | down | 874 | 888 | - | 11.5323 | 3.96E-05 | 1 | TAGGCGTGAGGTGAT |
| 6105 | mamo-L | *BGIBMGA012582* | down | 282 | 296 | - | 10.6774 | 8.56E-05 | 1 | GAGGCCATGTTTGAT |
| 6106 | mamo-L | *BGIBMGA012583* | up | 1305 | 1319 | - | 11.1129 | 5.88E-05 | 1 | GAGGCGTGTGTCGCT |
| 6107 | mamo-L | *BGIBMGA012584* | down | 801 | 815 | + | 10.7097 | 8.33E-05 | 1 | AAGGCTTCTTTTTGT |
| 6108 | mamo-L | *BGIBMGA012585* | up | 1964 | 1978 | - | 11.3145 | 4.87E-05 | 1 | TGGGCTTACTTTTGT |
| 6109 | mamo-L | *BGIBMGA012593* | down | 635 | 649 | - | 11.8548 | 2.89E-05 | 1 | TAGGCTGTTCTTTGT |
| 6110 | mamo-L | *BGIBMGA012594* | up | 1949 | 1963 | + | 12.8629 | 9.42E-06 | 1 | GAGGCTTGGTTTGGC |
| 6111 | mamo-L | *BGIBMGA012594* | down | 886 | 900 | - | 10.5726 | 9.33E-05 | 1 | TAGGCTAATATTTAT |
| 6112 | mamo-L | *BGIBMGA012595* | up | 1292 | 1306 | + | 12.0645 | 2.34E-05 | 1 | TAGGCGGGCTTCGGA |
| 6113 | mamo-L | *BGIBMGA012603* | down | 907 | 921 | + | 14.7258 | 5.83E-07 | 1 | TAGGCTTGCGTTGCC |
| 6114 | mamo-L | *BGIBMGA012607* | up | 422 | 436 | + | 10.9194 | 6.98E-05 | 1 | TAGGCTACTTTTTTT |
| 6115 | mamo-L | *BGIBMGA012612* | up | 1353 | 1367 | + | 10.6694 | 8.62E-05 | 1 | TAGGCGCTAGTCTGT |
| 6116 | mamo-L | *BGIBMGA012615* | up | 684 | 698 | - | 10.9194 | 6.98E-05 | 1 | TAGGCTACTTTTTTT |
| 6117 | mamo-L | *BGIBMGA012616* | down | 380 | 394 | - | 10.621 | 8.97E-05 | 1 | TAGGCTTCTTTTCAT |
| 6118 | mamo-L | *BGIBMGA012619* | down | 597 | 611 | + | 12.3226 | 1.77E-05 | 1 | TAGGCGGTTGATGTC |
| 6119 | mamo-L | *BGIBMGA012624* | up | 1040 | 1054 | + | 11.1129 | 5.88E-05 | 1 | GAGGCGTGTGTCGCT |
| 6120 | mamo-L | *BGIBMGA012626* | up | 1337 | 1351 | - | 10.9194 | 6.98E-05 | 1 | TAGGCTACTTTTTTT |
| 6121 | mamo-L | *BGIBMGA012628* | down | 332 | 346 | - | 11.371 | 4.62E-05 | 1 | TAGGCTGTATAGGCT |
| 6122 | mamo-L | *BGIBMGA012628* | down | 1034 | 1048 | + | 11.0484 | 6.23E-05 | 1 | GAGGCGGAGGTGGGC |
| 6123 | mamo-L | *BGIBMGA012629* | up | 1368 | 1382 | - | 11.371 | 4.62E-05 | 1 | TAGGCTGTATAGGCT |
| 6124 | mamo-L | *BGIBMGA012631* | down | 1915 | 1929 | - | 11.0645 | 6.14E-05 | 1 | TAGGCTTCCGTAGGT |
| 6125 | mamo-L | *BGIBMGA012636* | down | 1961 | 1975 | - | 11.0806 | 6.05E-05 | 1 | TAGGCTGTAAGTTGT |
| 6126 | mamo-L | *BGIBMGA012637* | up | 1271 | 1285 | + | 10.7742 | 7.89E-05 | 1 | TAGGCTATTAGTGTT |
| 6127 | mamo-L | *BGIBMGA012639* | down | 1723 | 1737 | - | 13.3952 | 4.84E-06 | 1 | AAGGCTGGTGTTGCT |
| 6128 | mamo-L | *BGIBMGA012639* | down | 1433 | 1447 | + | 11.2339 | 5.26E-05 | 1 | TAGGCTACTTTTTGT |
| 6129 | mamo-L | *BGIBMGA012639* | down | 1156 | 1170 | + | 10.9194 | 6.98E-05 | 1 | TAGGCTACTTTTTTT |
| 6130 | mamo-L | *BGIBMGA012648* | down | 130 | 144 | - | 11.1935 | 5.45E-05 | 1 | TAGGCTAACTTTTAC |
| 6131 | mamo-L | *BGIBMGA012654* | down | 1889 | 1903 | - | 13.1935 | 6.33E-06 | 1 | TAGGCTCTTGATGGT |
| 6132 | mamo-L | *BGIBMGA012655* | down | 547 | 561 | - | 12.8145 | 9.96E-06 | 1 | GAGGCTGATGTTGGC |
| 6133 | mamo-L | *BGIBMGA012659* | up | 1095 | 1109 | - | 11.9194 | 2.72E-05 | 1 | TAGGCATATGTTTTT |
| 6134 | mamo-L | *BGIBMGA012660* | up | 1570 | 1584 | - | 11.6129 | 3.66E-05 | 1 | TAGGCTTTTAATGAT |
| 6135 | mamo-L | *BGIBMGA012664* | up | 95 | 109 | - | 10.5968 | 9.15E-05 | 1 | AAGGCTGTTAATGCT |
| 6136 | mamo-L | *BGIBMGA012667* | up | 1456 | 1470 | - | 10.9274 | 6.92E-05 | 1 | AAGGCTGGTTTCTTT |
| 6137 | mamo-L | *BGIBMGA012668* | down | 1290 | 1304 | - | 11.3548 | 4.69E-05 | 1 | TAGGCTATATTTTCT |
| 6138 | mamo-L | *BGIBMGA012670* | down | 915 | 929 | + | 13.9839 | 2.04E-06 | 1 | TAGGCGGATGTCGGT |
| 6139 | mamo-L | *BGIBMGA012670* | up | 205 | 219 | + | 11.7177 | 3.32E-05 | 1 | TAGGCTTGTTATTTC |
| 6140 | mamo-L | *BGIBMGA012671* | up | 1210 | 1224 | - | 12.25 | 1.92E-05 | 1 | AAGGCTTTCGTTTCT |
| 6141 | mamo-L | *BGIBMGA012671* | down | 1266 | 1280 | + | 10.9839 | 6.59E-05 | 1 | TAGGCGAGGTATTTT |
| 6142 | mamo-L | *BGIBMGA012672* | up | 477 | 491 | - | 11.2016 | 5.41E-05 | 1 | GAGGCTCGTGTTTTT |
| 6143 | mamo-L | *BGIBMGA012674* | up | 1170 | 1184 | + | 11.0161 | 6.40E-05 | 1 | AAGGCCTGTGTTTAT |
| 6144 | mamo-L | *BGIBMGA012676* | up | 851 | 865 | + | 10.8306 | 7.52E-05 | 1 | AAGGCTTTTTTTTAT |
| 6145 | mamo-L | *BGIBMGA012676* | up | 1580 | 1594 | + | 10.5726 | 9.33E-05 | 1 | TAGGCTAATATTTAT |
| 6146 | mamo-L | *BGIBMGA012678* | up | 1938 | 1952 | - | 12.7419 | 1.09E-05 | 1 | TAGGCTGTTAATGTT |
| 6147 | mamo-L | *BGIBMGA012679* | up | 1783 | 1797 | - | 12.7581 | 1.07E-05 | 1 | AAGGCTGAGGTCGCT |
| 6148 | mamo-L | *BGIBMGA012680* | up | 1608 | 1622 | + | 12.2823 | 1.85E-05 | 1 | TAGGCTCGTTTTACT |
| 6149 | mamo-L | *BGIBMGA012682* | up | 1559 | 1573 | + | 11.3548 | 4.69E-05 | 1 | TAGGCTTAAATCGGC |
| 6150 | mamo-L | *BGIBMGA012686* | up | 1041 | 1055 | + | 11.0565 | 6.18E-05 | 1 | TAGGCGGGGTGTAAT |
| 6151 | mamo-L | *BGIBMGA012695* | up | 1966 | 1980 | + | 11.3629 | 4.66E-05 | 1 | TAGGCTAATGTCTTT |
| 6152 | mamo-L | *BGIBMGA012697* | down | 1310 | 1324 | + | 10.6371 | 8.85E-05 | 1 | TAGGCTTCGAATTTT |
| 6153 | mamo-L | *BGIBMGA012697* | up | 524 | 538 | + | 10.5726 | 9.33E-05 | 1 | TAGGCAGATGTGTTT |
| 6154 | mamo-L | *BGIBMGA012700* | down | 915 | 929 | + | 12.5806 | 1.31E-05 | 1 | TAGGCCTATTTGTGT |
| 6155 | mamo-L | *BGIBMGA012704* | down | 1905 | 1919 | - | 10.6855 | 8.51E-05 | 1 | AAGGCTATATTTGGT |
| 6156 | mamo-L | *BGIBMGA012707* | up | 1149 | 1163 | - | 11.0161 | 6.40E-05 | 1 | AAGGCCTGTGTTTAT |
| 6157 | mamo-L | *BGIBMGA012707* | down | 644 | 658 | - | 10.9839 | 6.59E-05 | 1 | TAGGCTTTTTTTAGA |

| | | | | | | | | | | |
|---|---|---|---|---|---|---|---|---|---|---|
| 6158 | mamo-L | *BGIBMGA012708* | down | 1103 | 1117 | - | 12.9435 | 8.54E-06 | 1 | TAGGCTTTTTTTTAT |
| 6159 | mamo-L | *BGIBMGA012708* | up | 1134 | 1148 | + | 10.5726 | 9.33E-05 | 1 | TAGGCTAATATTTAT |
| 6160 | mamo-L | *BGIBMGA012711* | up | 526 | 540 | + | 11.2016 | 5.41E-05 | 1 | GAGGCTCGTGTTTTT |
| 6161 | mamo-L | *BGIBMGA012711* | down | 1067 | 1081 | - | 10.9839 | 6.59E-05 | 1 | TAGGCGAGGTATTTT |
| 6162 | mamo-L | *BGIBMGA012714* | up | 894 | 908 | - | 12.6452 | 1.21E-05 | 1 | TAGGCCCAGGTCGTC |
| 6163 | mamo-L | *BGIBMGA012717* | up | 1117 | 1131 | - | 11.1855 | 5.50E-05 | 1 | TAGGCCCTAGTTACT |
| 6164 | mamo-L | *BGIBMGA012725* | down | 199 | 213 | - | 11 | 6.50E-05 | 1 | TAGGCATATATTTGT |
| 6165 | mamo-L | *BGIBMGA012726* | up | 1335 | 1349 | - | 11 | 6.50E-05 | 1 | TAGGCATATATTTGT |
| 6166 | mamo-L | *BGIBMGA012732* | down | 1848 | 1862 | - | 10.5968 | 9.15E-05 | 1 | GAGGCTAGTTTTTGT |
| 6167 | mamo-L | *BGIBMGA012734* | down | 1557 | 1571 | + | 13.4194 | 4.70E-06 | 1 | TAGGCCGTGGTCGTC |
| 6168 | mamo-L | *BGIBMGA012734* | down | 353 | 367 | + | 12.3065 | 1.80E-05 | 1 | TAGGCCATTATTGGT |
| 6169 | mamo-L | *BGIBMGA012737* | down | 512 | 526 | - | 11.4919 | 4.11E-05 | 1 | GAGGCTTATGTTAGT |
| 6170 | mamo-L | *BGIBMGA012741* | up | 1198 | 1212 | + | 11.4032 | 4.49E-05 | 1 | TAGGCCGGCAATGGA |
| 6171 | mamo-L | *BGIBMGA012742* | up | 432 | 446 | - | 11.9435 | 2.65E-05 | 1 | GAGGCTTTTTTTTGT |
| 6172 | mamo-L | *BGIBMGA012743* | up | 1222 | 1236 | + | 11.7339 | 3.26E-05 | 1 | TAGGCTAGCTTCGCC |
| 6173 | mamo-L | *BGIBMGA012751* | down | 1500 | 1514 | + | 10.5726 | 9.33E-05 | 1 | TAGGCTAATATTTAT |
| 6174 | mamo-L | *BGIBMGA012756* | down | 289 | 303 | - | 11.3871 | 4.55E-05 | 1 | TAGGCAGAAGTTTCT |
| 6175 | mamo-L | *BGIBMGA012757* | down | 278 | 292 | - | 12.8226 | 9.86E-06 | 1 | TAGGCCATGTTTGAT |
| 6176 | mamo-L | *BGIBMGA012757* | down | 1937 | 1951 | + | 11.121 | 5.84E-05 | 1 | TAGGCTTACTGCGGA |
| 6177 | mamo-L | *BGIBMGA012758* | down | 511 | 525 | + | 10.9113 | 7.02E-05 | 1 | TAGGCTGTATGCTGT |
| 6178 | mamo-L | *BGIBMGA012761* | up | 329 | 343 | - | 11.379 | 4.58E-05 | 1 | AAGGCTTACAGTGGT |
| 6179 | mamo-L | *BGIBMGA012761* | down | 485 | 499 | + | 10.9355 | 6.88E-05 | 1 | TAGGCCTACGCCTGT |
| 6180 | mamo-L | *BGIBMGA012766* | up | 1828 | 1842 | - | 11.0806 | 6.05E-05 | 1 | GAGGCTCGTTGTGGT |
| 6181 | mamo-L | *BGIBMGA012769* | down | 1167 | 1181 | - | 10.5242 | 9.70E-05 | 1 | TAGGCAGCGGTTTGC |
| 6182 | mamo-L | *BGIBMGA012772* | down | 1206 | 1220 | - | 11.1774 | 5.54E-05 | 1 | TAGGCTGTGGTGGAA |
| 6183 | mamo-L | *BGIBMGA012774* | up | 159 | 173 | + | 11.4677 | 4.22E-05 | 1 | TAGGCCCTGTTTATC |
| 6184 | mamo-L | *BGIBMGA012775* | down | 81 | 95 | + | 10.629 | 8.91E-05 | 1 | GAGGCGTTGGTTTCC |
| 6185 | mamo-L | *BGIBMGA012776* | down | 31 | 45 | + | 11.7097 | 3.34E-05 | 1 | AAGGCTTAGTTTTAT |
| 6186 | mamo-L | *BGIBMGA012781* | up | 1289 | 1303 | - | 11.9677 | 2.59E-05 | 1 | AAGGCCTTCTTTTTT |
| 6187 | mamo-L | *BGIBMGA012781* | up | 1330 | 1344 | - | 11.2823 | 5.03E-05 | 1 | AAGGCGGTTTTTTTT |
| 6188 | mamo-L | *BGIBMGA012782* | up | 1691 | 1705 | - | 10.5645 | 9.40E-05 | 1 | TAGGCTTATTACACT |
| 6189 | mamo-L | *BGIBMGA012783* | up | 70 | 84 | + | 10.5726 | 9.33E-05 | 1 | TAGGCTAATATTTAT |
| 6190 | mamo-L | *BGIBMGA012784* | up | 686 | 700 | + | 12.8306 | 9.76E-06 | 1 | TAGGCAGGGGTCGGT |
| 6191 | mamo-L | *BGIBMGA012787* | up | 1738 | 1752 | + | 11.5403 | 3.93E-05 | 1 | AAGGCGTTGTTTTTT |
| 6192 | mamo-L | *BGIBMGA012789* | down | 1289 | 1303 | - | 10.5081 | 9.83E-05 | 1 | TAGGCTTTCAAGTGT |
| 6193 | mamo-L | *BGIBMGA012791* | down | 1429 | 1443 | + | 10.9194 | 6.98E-05 | 1 | TAGGCTACTTTTTTT |
| 6194 | mamo-L | *BGIBMGA012803* | up | 1268 | 1282 | - | 12.0645 | 2.34E-05 | 1 | TAGGCAGTATTTGTT |
| 6195 | mamo-L | *BGIBMGA012803* | up | 1315 | 1329 | - | 11.7016 | 3.37E-05 | 1 | TAGGCTTTGTTTTCG |
| 6196 | mamo-L | *BGIBMGA012803* | up | 1065 | 1079 | - | 11.4597 | 4.25E-05 | 1 | TAGGCGGGGTGGTGT |
| 6197 | mamo-L | *BGIBMGA012805* | up | 781 | 795 | - | 11.7661 | 3.16E-05 | 1 | TAGGCACTTTTTGTT |
| 6198 | mamo-L | *BGIBMGA012806* | down | 18 | 32 | + | 12.3387 | 1.73E-05 | 1 | AAGGCTGGTGGTGGT |
| 6199 | mamo-L | *BGIBMGA012806* | down | 1185 | 1199 | - | 10.5565 | 9.46E-05 | 1 | AAGGCTGTTTGTTTT |
| 6200 | mamo-L | *BGIBMGA012807* | up | 1518 | 1532 | + | 12.3387 | 1.73E-05 | 1 | AAGGCTGGTGGTGGT |
| 6201 | mamo-L | *BGIBMGA012808* | down | 1620 | 1634 | + | 13.3145 | 5.41E-06 | 1 | TAGGCAGATGTTGTT |
| 6202 | mamo-L | *BGIBMGA012810* | up | 1268 | 1282 | + | 10.6532 | 8.73E-05 | 1 | TAGGCTTTCTCTTTC |
| 6203 | mamo-L | *BGIBMGA012810* | down | 1755 | 1769 | - | 10.5403 | 9.58E-05 | 1 | TAGGCTGTATGGTGT |
| 6204 | mamo-L | *BGIBMGA012813* | down | 1757 | 1771 | - | 12.0161 | 2.46E-05 | 1 | TAGGCAGTGTGTGTT |
| 6205 | mamo-L | *BGIBMGA012815* | down | 1299 | 1313 | - | 13.7339 | 2.97E-06 | 1 | TAGGCTTTTATTGCT |
| 6206 | mamo-L | *BGIBMGA012815* | down | 1119 | 1133 | + | 11.7742 | 3.14E-05 | 1 | TAGGCGGTCAATTGT |
| 6207 | mamo-L | *BGIBMGA012816* | up | 1602 | 1616 | - | 11.5968 | 3.73E-05 | 1 | AAGGCTCGCTTTTTT |
| 6208 | mamo-L | *BGIBMGA012817* | down | 1982 | 1996 | - | 14.121 | 1.65E-06 | 1 | TAGGCTGAGTTTTAT |
| 6209 | mamo-L | *BGIBMGA012817* | down | 1558 | 1572 | + | 11.75 | 3.21E-05 | 1 | TAGGCTCTGGTTACC |
| 6210 | mamo-L | *BGIBMGA012817* | down | 1657 | 1671 | - | 10.7903 | 7.79E-05 | 1 | TAGGCCTCTTTTGCG |
| 6211 | mamo-L | *BGIBMGA012818* | up | 383 | 397 | - | 12.5081 | 1.43E-05 | 1 | TAGGCTGGCTATTTC |
| 6212 | mamo-L | *BGIBMGA012820* | up | 1663 | 1677 | + | 10.5726 | 9.33E-05 | 1 | TAGGCTAATATTTAT |
| 6213 | mamo-L | *BGIBMGA012824* | down | 933 | 947 | - | 14.2742 | 1.32E-06 | 1 | TAGGCGGAGTTTTTT |
| 6214 | mamo-L | *BGIBMGA012831* | up | 803 | 817 | + | 13.7742 | 2.79E-06 | 1 | TAGGCGGGTGGTGGT |
| 6215 | mamo-L | *BGIBMGA012838* | down | 857 | 871 | - | 11.9194 | 2.72E-05 | 1 | TAGGCTTATTTCATT |
| 6216 | mamo-L | *BGIBMGA012841* | up | 1490 | 1504 | - | 12.9919 | 8.10E-06 | 1 | TAGGCTTATATTTTT |
| 6217 | mamo-L | *BGIBMGA012846* | down | 1492 | 1506 | + | 11.2177 | 5.34E-05 | 1 | TAGCCTGACGTTTGT |
| 6218 | mamo-L | *BGIBMGA012847* | down | 1082 | 1096 | + | 12.0968 | 2.26E-05 | 1 | TAGGCTCTGTGTTCT |
| 6219 | mamo-L | *BGIBMGA012847* | up | 1733 | 1747 | + | 11.2177 | 5.34E-05 | 1 | TAGCCTGACGTTTGT |
| 6220 | mamo-L | *BGIBMGA012847* | down | 237 | 251 | + | 10.5726 | 9.33E-05 | 1 | TAGGCTAATATTTAT |
| 6221 | mamo-L | *BGIBMGA012848* | down | 1763 | 1777 | - | 12.3306 | 1.75E-05 | 1 | TAGGCTTACTATACT |
| 6222 | mamo-L | *BGIBMGA012850* | up | 1448 | 1462 | + | 11.5323 | 3.96E-05 | 1 | TAGGCGTGAGGTGAT |
| 6223 | mamo-L | *BGIBMGA012859* | down | 1360 | 1374 | + | 10.5161 | 9.77E-05 | 1 | TAGGCGTATGTTTAA |
| 6224 | mamo-L | *BGIBMGA012862* | down | 171 | 185 | - | 11.371 | 4.62E-05 | 1 | TAGGCTGTATAGGCT |
| 6225 | mamo-L | *BGIBMGA012864* | down | 44 | 58 | + | 11.2984 | 4.95E-05 | 1 | AAGGCTATTGTTGTT |
| 6226 | mamo-L | *BGIBMGA012865* | down | 1000 | 1014 | - | 11.8306 | 2.97E-05 | 1 | TAGGCTTGGAGGGGT |
| 6227 | mamo-L | *BGIBMGA012865* | down | 35 | 49 | + | 11.2984 | 4.95E-05 | 1 | AAGGCTATTGTTGTT |
| 6228 | mamo-L | *BGIBMGA012867* | down | 1913 | 1927 | - | 11.4919 | 4.11E-05 | 1 | TAGGCTATCTTCGCC |
| 6229 | mamo-L | *BGIBMGA012868* | up | 1351 | 1365 | - | 11.4758 | 4.18E-05 | 1 | TAGGCCGTGATCAGT |
| 6230 | mamo-L | *BGIBMGA012871* | down | 1633 | 1647 | - | 10.6855 | 8.51E-05 | 1 | AAGGCCTTTATTTGT |
| 6231 | mamo-L | *BGIBMGA012880* | up | 368 | 382 | - | 12.2097 | 2.01E-05 | 1 | TAGGCCTTGTTTGAA |
| 6232 | mamo-L | *BGIBMGA012884* | down | 142 | 156 | + | 11.4758 | 4.18E-05 | 1 | AAGGCCTTTTTTTTT |
| 6233 | mamo-L | *BGIBMGA012888* | up | 1144 | 1158 | + | 10.5968 | 9.15E-05 | 1 | GAGGCTAGTTTTTGT |
| 6234 | mamo-L | *BGIBMGA012889* | down | 1257 | 1271 | - | 13.0887 | 7.18E-06 | 1 | TAGGCTTGTGATGAT |
| 6235 | mamo-L | *BGIBMGA012889* | up | 220 | 234 | - | 10.8226 | 7.58E-05 | 1 | TAGGCCGTCGACCCT |
| 6236 | mamo-L | *BGIBMGA012895* | down | 1462 | 1476 | + | 12.4274 | 1.57E-05 | 1 | TAGGCTGGCTTAGTT |
| 6237 | mamo-L | *BGIBMGA012895* | down | 739 | 753 | + | 11.3548 | 4.69E-05 | 1 | AAGGCTGATGATGAT |
| 6238 | mamo-L | *BGIBMGA012895* | up | 1461 | 1475 | + | 10.9194 | 6.98E-05 | 1 | TAGGCTACTTTTTTT |
| 6239 | mamo-L | *BGIBMGA012896* | down | 506 | 520 | + | 11.6452 | 3.56E-05 | 1 | TAGGCGCATTTTTTC |
| 6240 | mamo-L | *BGIBMGA012896* | down | 329 | 343 | + | 11.1855 | 5.50E-05 | 1 | TAGGCTTAAAATTGT |
| 6241 | mamo-L | *BGIBMGA012898* | down | 1242 | 1256 | - | 10.8871 | 7.17E-05 | 1 | TAGGCTTATTATTGA |
| 6242 | mamo-L | *BGIBMGA012899* | down | 78 | 92 | + | 11.0161 | 6.40E-05 | 1 | TAGGCATTGGTTGCA |
| 6243 | mamo-L | *BGIBMGA012902* | down | 1595 | 1609 | - | 10.5726 | 9.33E-05 | 1 | GAGGCTGATGTTTAC |
| 6244 | mamo-L | *BGIBMGA012907* | up | 1020 | 1034 | - | 11.2823 | 5.03E-05 | 1 | TGGGCTTTTTTTTGTT |
| 6245 | mamo-L | *BGIBMGA012908* | down | 825 | 839 | + | 12.3306 | 1.75E-05 | 1 | AAGGCTGGTGTTTTT |
| 6246 | mamo-L | *BGIBMGA012908* | down | 216 | 230 | + | 11.7339 | 3.26E-05 | 1 | TAGGCTAGCTTCGCC |
| 6247 | mamo-L | *BGIBMGA012908* | down | 206 | 220 | + | 10.7339 | 8.17E-05 | 1 | TAGGCTTTTTTAGGC |
| 6248 | mamo-L | *BGIBMGA012909* | up | 1227 | 1241 | + | 12.3306 | 1.75E-05 | 1 | AAGGCTGGTGTTTTT |
| 6249 | mamo-L | *BGIBMGA012909* | up | 618 | 632 | + | 11.7339 | 3.26E-05 | 1 | TAGGCTAGCTTCGCC |

| | | | | | | | | | | |
|---|---|---|---|---|---|---|---|---|---|---|
| 6250 | mamo-L | *BGIBMGA012909* | up | 608 | 622 | + | 10.7339 | 8.17E-05 | 1 | TAGGCTTTTTTAGGC |
| 6251 | mamo-L | *BGIBMGA012910* | down | 1936 | 1950 | + | 11.1774 | 5.54E-05 | 1 | GAGGCCGTGCTTGGT |
| 6252 | mamo-L | *BGIBMGA012912* | up | 226 | 240 | + | 11.1774 | 5.54E-05 | 1 | GAGGCCGTGCTTGGT |
| 6253 | mamo-L | *BGIBMGA012913* | down | 343 | 357 | + | 10.5726 | 9.33E-05 | 1 | TAGGCTAATATTTAT |
| 6254 | mamo-L | *BGIBMGA012918* | down | 1387 | 1401 | + | 11.2177 | 5.34E-05 | 1 | TAGGCGTATGACGAT |
| 6255 | mamo-L | *BGIBMGA012921* | up | 461 | 475 | - | 11.5242 | 3.99E-05 | 1 | TAGGCTTGGTGGTTT |
| 6256 | mamo-L | *BGIBMGA012921* | up | 1095 | 1109 | + | 10.5242 | 9.70E-05 | 1 | TAGGCGTTTGAGGAT |
| 6257 | mamo-L | *BGIBMGA012924* | up | 1152 | 1166 | + | 12.0645 | 2.34E-05 | 1 | TAGGCGGGCTTCGGA |
| 6258 | mamo-L | *BGIBMGA012927* | up | 80 | 94 | - | 11.1774 | 5.54E-05 | 1 | GAGGCCGTGCTTGGT |
| 6259 | mamo-L | *BGIBMGA012928* | down | 711 | 725 | + | 10.9194 | 6.98E-05 | 1 | TAGGCTACTTTTTTT |
| 6260 | mamo-L | *BGIBMGA012928* | up | 641 | 655 | - | 10.879 | 7.22E-05 | 1 | AAGGCTGTCTCTGCT |
| 6261 | mamo-L | *BGIBMGA012928* | down | 564 | 578 | - | 10.7984 | 7.74E-05 | 1 | AAGGCTTATTTCGCC |
| 6262 | mamo-L | *BGIBMGA012932* | down | 839 | 853 | - | 11.7903 | 3.08E-05 | 1 | TAGGCGTAATTTTAT |
| 6263 | mamo-L | *BGIBMGA012934* | up | 692 | 706 | - | 11.7984 | 3.06E-05 | 1 | TAGGCTGAGGGTATC |
| 6264 | mamo-L | *BGIBMGA012935* | up | 1197 | 1211 | + | 12.0645 | 2.34E-05 | 1 | TAGGCGGGCTTCGGA |
| 6265 | mamo-L | *BGIBMGA012935* | up | 1044 | 1058 | - | 10.8306 | 7.52E-05 | 1 | TAGGCCCCCTTCGCC |
| 6266 | mamo-L | *BGIBMGA012936* | down | 1186 | 1200 | + | 10.5726 | 9.33E-05 | 1 | GAGGCTGATGTTTAC |
| 6267 | mamo-L | *BGIBMGA012938* | up | 1349 | 1363 | + | 10.6855 | 8.51E-05 | 1 | TAGGCGTTTATCTCT |
| 6268 | mamo-L | *BGIBMGA012940* | down | 448 | 462 | + | 10.9597 | 6.73E-05 | 1 | AAGGCCTACAATGTT |
| 6269 | mamo-L | *BGIBMGA012941* | up | 358 | 372 | - | 12.0645 | 2.34E-05 | 1 | TAGGCGGGCTTCGGA |
| 6270 | mamo-L | *BGIBMGA012941* | up | 511 | 525 | + | 11.1774 | 5.54E-05 | 1 | TAGGCCCCCTTCGGC |
| 6271 | mamo-L | *BGIBMGA012947* | down | 680 | 694 | - | 10.8871 | 7.17E-05 | 1 | TAGGCCAATGTTAGC |
| 6272 | mamo-L | *BGIBMGA012951* | up | 1448 | 1462 | + | 11.9194 | 2.72E-05 | 1 | AAGGCTGGCGTTATT |
| 6273 | mamo-L | *BGIBMGA012951* | up | 1094 | 1108 | - | 10.7258 | 8.23E-05 | 1 | TAGGCTGCAATCGTT |
| 6274 | mamo-L | *BGIBMGA012953* | down | 694 | 708 | - | 10.5726 | 9.33E-05 | 1 | TAGGCTAATATTTAT |
| 6275 | mamo-L | *BGIBMGA012955* | down | 1390 | 1404 | + | 12.8871 | 9.15E-06 | 1 | TAGGCGTAATTTGAT |
| 6276 | mamo-L | *BGIBMGA012955* | down | 1572 | 1586 | - | 11.8387 | 2.94E-05 | 1 | GAGGCTTGTTTTTCT |
| 6277 | mamo-L | *BGIBMGA012956* | up | 1373 | 1387 | - | 11.0323 | 6.32E-05 | 1 | AAGGCTGTTGATGAT |
| 6278 | mamo-L | *BGIBMGA012957* | up | 1234 | 1248 | + | 10.5726 | 9.33E-05 | 1 | TAGGCTAATATTTAT |
| 6279 | mamo-L | *BGIBMGA012959* | down | 460 | 474 | - | 13.4677 | 4.39E-06 | 1 | TAGGCTGATGATGAT |
| 6280 | mamo-L | *BGIBMGA012960* | up | 1747 | 1761 | - | 14.2661 | 1.33E-06 | 1 | TAGGCGGGCGGTGGT |
| 6281 | mamo-L | *BGIBMGA012960* | down | 722 | 736 | - | 10.5 | 9.90E-05 | 1 | GAGGCCCAGGTCGTC |
| 6282 | mamo-L | *BGIBMGA012964* | up | 217 | 231 | + | 11.5403 | 3.93E-05 | 1 | TAGGCTTTCGATGCG |
| 6283 | mamo-L | *BGIBMGA012965* | down | 221 | 235 | + | 10.6855 | 8.51E-05 | 1 | AAGGCCTTTATTTGT |
| 6284 | mamo-L | *BGIBMGA012967* | up | 834 | 848 | + | 12.4194 | 1.58E-05 | 1 | TAGGCCCGCATTTTT |
| 6285 | mamo-L | *BGIBMGA012970* | down | 920 | 934 | - | 12.0403 | 2.40E-05 | 1 | TAGGCCTTGTTCTAT |
| 6286 | mamo-L | *BGIBMGA012973* | up | 283 | 297 | - | 11.9597 | 2.61E-05 | 1 | TAGGCGACCGTTGTT |
| 6287 | mamo-L | *BGIBMGA012974* | up | 1181 | 1195 | - | 11.0806 | 6.05E-05 | 1 | TAGGCTCTGGTGACT |
| 6288 | mamo-L | *BGIBMGA012975* | up | 1328 | 1342 | - | 11.379 | 4.58E-05 | 1 | TAGGCGCTCAGTGTT |
| 6289 | mamo-L | *BGIBMGA012975* | up | 971 | 985 | - | 10.8306 | 7.52E-05 | 1 | AAGGCTTTTTTTTAT |
| 6290 | mamo-L | *BGIBMGA012979* | up | 777 | 791 | + | 12.0645 | 2.34E-05 | 1 | TAGGCGGGCTTCGGA |
| 6291 | mamo-L | *BGIBMGA012984* | down | 1242 | 1256 | - | 11.6774 | 3.44E-05 | 1 | TAGGCCTAATGTTCT |
| 6292 | mamo-L | *BGIBMGA012984* | down | 1037 | 1051 | + | 11.0242 | 6.36E-05 | 1 | TAGGCCGTTTTTCAC |
| 6293 | mamo-L | *BGIBMGA012984* | down | 1249 | 1263 | + | 10.6855 | 8.51E-05 | 1 | TAGGCCTATGTAGTC |
| 6294 | mamo-L | *BGIBMGA012987* | down | 151 | 165 | - | 11.5645 | 3.84E-05 | 1 | TAGGCCTACCGTGTT |
| 6295 | mamo-L | *BGIBMGA012991* | down | 193 | 207 | + | 12.9919 | 8.10E-06 | 1 | TAGGCCGGTTTTGCA |
| 6296 | mamo-L | *BGIBMGA012991* | down | 1088 | 1102 | - | 11.7097 | 3.34E-05 | 1 | TAGGCATGTTTTTTT |
| 6297 | mamo-L | *BGIBMGA012991* | up | 1647 | 1661 | + | 10.5081 | 9.83E-05 | 1 | TAGGCGGGGTGTTGG |
| 6298 | mamo-L | *BGIBMGA012992* | down | 1516 | 1530 | + | 11.8629 | 2.87E-05 | 1 | TAGGCCTGCGGCGCC |
| 6299 | mamo-L | *BGIBMGA012993* | up | 838 | 852 | + | 11.2903 | 4.99E-05 | 1 | TAGGCCTGTAGTTCT |
| 6300 | mamo-L | *BGIBMGA012994* | down | 46 | 60 | - | 11.7823 | 3.11E-05 | 1 | TAGGCTTGTTTTTCA |
| 6301 | mamo-L | *BGIBMGA012994* | down | 359 | 373 | - | 10.8145 | 7.63E-05 | 1 | TAGGCGCAATTTTCC |
| 6302 | mamo-L | *BGIBMGA012996* | down | 688 | 702 | - | 11.7823 | 3.11E-05 | 1 | TAGGCGTGGGGCGGC |
| 6303 | mamo-L | *BGIBMGA012998* | up | 1001 | 1015 | - | 10.5968 | 9.15E-05 | 1 | GAGGCTAGTTTTTGT |
| 6304 | mamo-L | *BGIBMGA013001* | down | 1282 | 1296 | - | 11.8468 | 2.92E-05 | 1 | TAGGCTCCTGGTGGT |
| 6305 | mamo-L | *BGIBMGA013002* | down | 1346 | 1360 | - | 10.9194 | 6.98E-05 | 1 | TAGGCTACTTTTTTT |
| 6306 | mamo-L | *BGIBMGA013003* | up | 1428 | 1442 | - | 10.9194 | 6.98E-05 | 1 | TAGGCTACTTTTTTT |
| 6307 | mamo-L | *BGIBMGA013004* | up | 436 | 450 | + | 12.0645 | 2.34E-05 | 1 | TAGGCGGGCTTCGGA |
| 6308 | mamo-L | *BGIBMGA013004* | up | 1606 | 1620 | - | 11.871 | 2.85E-05 | 1 | AAGGCGGATGTCGGT |
| 6309 | mamo-L | *BGIBMGA013004* | down | 1057 | 1071 | + | 11.4032 | 4.49E-05 | 1 | TAGGCGTACGCTTTT |
| 6310 | mamo-L | *BGIBMGA013005* | down | 85 | 99 | + | 11.3145 | 4.87E-05 | 1 | TAGGCCTATTTTTTG |
| 6311 | mamo-L | *BGIBMGA013009* | up | 1235 | 1249 | + | 12.7097 | 1.13E-05 | 1 | TAGGCTGCTGATGTT |
| 6312 | mamo-L | *BGIBMGA013010* | down | 1438 | 1452 | + | 13.2903 | 5.60E-06 | 1 | TAGGCCCGCTTGGGT |
| 6313 | mamo-L | *BGIBMGA013010* | down | 1299 | 1313 | + | 11.7661 | 3.16E-05 | 1 | TAGGCAGTTTTTTTT |
| 6314 | mamo-L | *BGIBMGA013014* | down | 1634 | 1648 | - | 10.9194 | 6.98E-05 | 1 | TAGGCTACTTTTTTT |
| 6315 | mamo-L | *BGIBMGA013014* | up | 61 | 75 | - | 10.621 | 8.97E-05 | 1 | AAGGCTGGTTGCGTT |
| 6316 | mamo-L | *BGIBMGA013021* | down | 1190 | 1204 | + | 12.2016 | 2.03E-05 | 1 | TAGGCTGAAGTGGTC |
| 6317 | mamo-L | *BGIBMGA013021* | down | 1418 | 1432 | + | 10.5484 | 9.52E-05 | 1 | TAGGCGCCATTTTGT |
| 6318 | mamo-L | *BGIBMGA013024* | down | 588 | 602 | + | 11.8468 | 2.92E-05 | 1 | AAGGCTGACGATGAT |
| 6319 | mamo-L | *BGIBMGA013024* | up | 240 | 254 | - | 11.3548 | 4.69E-05 | 1 | AAGGCTGATGATGAT |
| 6320 | mamo-L | *BGIBMGA013026* | down | 697 | 711 | + | 14.121 | 1.65E-06 | 1 | TAGGCTGAGTTTTAT |
| 6321 | mamo-L | *BGIBMGA013026* | down | 1121 | 1135 | - | 11.75 | 3.21E-05 | 1 | TAGGCTCTGGTTACC |
| 6322 | mamo-L | *BGIBMGA013026* | down | 1022 | 1036 | + | 10.7903 | 7.79E-05 | 1 | TAGGCCTCTTTTGCG |
| 6323 | mamo-L | *BGIBMGA013031* | up | 1635 | 1649 | + | 10.5726 | 9.33E-05 | 1 | TAGGCTAATATTTAT |
| 6324 | mamo-L | *BGIBMGA013035* | down | 1261 | 1275 | + | 10.621 | 8.97E-05 | 1 | TAGGCGTTTTTGTAT |
| 6325 | mamo-L | *BGIBMGA013036* | up | 222 | 236 | + | 10.7581 | 8.00E-05 | 1 | AAGGCGGAAGTCGTT |
| 6326 | mamo-L | *BGIBMGA013036* | up | 1166 | 1180 | - | 10.4919 | 9.96E-05 | 1 | TGGGCGGGCGTCGTT |
| 6327 | mamo-L | *BGIBMGA013038* | up | 1379 | 1393 | - | 13.0968 | 7.11E-06 | 1 | TAGGCGTTTTTTTTT |
| 6328 | mamo-L | *BGIBMGA013047* | down | 1214 | 1228 | - | 11.6694 | 3.47E-05 | 1 | TAGGCCGATTGTGCA |
| 6329 | mamo-L | *BGIBMGA013047* | up | 1304 | 1318 | + | 11.0081 | 6.45E-05 | 1 | TAGGCATACGGTTTT |
| 6330 | mamo-L | *BGIBMGA013047* | down | 903 | 917 | - | 10.9677 | 6.68E-05 | 1 | GAGGCTCGTGTGGGT |
| 6331 | mamo-L | *BGIBMGA013048* | down | 214 | 228 | - | 10.5726 | 9.33E-05 | 1 | TAGGCTAATATTTAT |
| 6332 | mamo-L | *BGIBMGA013052* | down | 1872 | 1886 | - | 11.5403 | 3.93E-05 | 1 | TAGGCTATCTGTGAT |
| 6333 | mamo-L | *BGIBMGA013054* | down | 1578 | 1592 | - | 10.6532 | 8.73E-05 | 1 | TAGGCTTTCTCTTTC |
| 6334 | mamo-L | *BGIBMGA013054* | up | 1764 | 1778 | + | 10.5403 | 9.58E-05 | 1 | TAGGCTGTATGGTGT |
| 6335 | mamo-L | *BGIBMGA013056* | down | 698 | 712 | - | 11.1613 | 5.63E-05 | 1 | TAGGCGGTTTGTTAT |
| 6336 | mamo-L | *BGIBMGA013056* | down | 764 | 778 | + | 10.629 | 8.91E-05 | 1 | TAGGCCTGCAGCGCC |
| 6337 | mamo-L | *BGIBMGA013057* | down | 1004 | 1018 | - | 12.5081 | 1.43E-05 | 1 | TAGGCTGATTTTTGA |
| 6338 | mamo-L | *BGIBMGA013061* | up | 1555 | 1569 | + | 12.5081 | 1.43E-05 | 1 | TAGGCTTCTTTTTTT |
| 6339 | mamo-L | *BGIBMGA013061* | down | 1485 | 1499 | - | 10.5726 | 9.33E-05 | 1 | TAGGCTAATATTTAT |
| 6340 | mamo-L | *BGIBMGA013063* | down | 262 | 276 | - | 10.9194 | 6.98E-05 | 1 | TAGGCTACTTTTTTT |
| 6341 | mamo-L | *BGIBMGA013064* | down | 1689 | 1703 | - | 14.6371 | 7.08E-07 | 1 | TAGGCTTGCGTTTTT |

| | | | | | | | | | | |
|---|---|---|---|---|---|---|---|---|---|---|
| 6342 | mamo-L | *BGIBMGA013065* | up | 1457 | 1471 | - | 12.7661 | 1.05E-05 | 1 | TAGGCTTTTTTTTCC |
| 6343 | mamo-L | *BGIBMGA013069* | up | 691 | 705 | - | 10.6694 | 8.62E-05 | 1 | TAGGCGGATTGGTTT |
| 6344 | mamo-L | *BGIBMGA013071* | down | 1250 | 1264 | - | 10.5565 | 9.46E-05 | 1 | TAGGCAGACGATTAT |
| 6345 | mamo-L | *BGIBMGA013079* | down | 131 | 145 | + | 12.3871 | 1.65E-05 | 1 | TAGGCTCCGTTTGTC |
| 6346 | mamo-L | *BGIBMGA013083* | up | 1133 | 1147 | - | 11.4677 | 4.22E-05 | 1 | TAGGCCCTGTTTATC |
| 6347 | mamo-L | *BGIBMGA013084* | up | 445 | 459 | - | 11.0081 | 6.45E-05 | 1 | AAGGCTGATTTCTTT |
| 6348 | mamo-L | *BGIBMGA013085* | down | 379 | 393 | + | 11.1774 | 5.54E-05 | 1 | TAGGCTGTGGTGGAA |
| 6349 | mamo-L | *BGIBMGA013088* | up | 951 | 965 | - | 11.2339 | 5.26E-05 | 1 | AAGGCTTAGTTCTCT |
| 6350 | mamo-L | *BGIBMGA013089* | up | 1639 | 1653 | + | 10.6694 | 8.62E-05 | 1 | AAGGCCGTTATTTTT |
| 6351 | mamo-L | *BGIBMGA013090* | up | 386 | 400 | - | 10.5726 | 9.33E-05 | 1 | TAGGCTAATATTTAT |
| 6352 | mamo-L | *BGIBMGA013091* | down | 17 | 31 | - | 10.5726 | 9.33E-05 | 1 | TAGGCTAATATTTAT |
| 6353 | mamo-L | *BGIBMGA013092* | down | 1211 | 1225 | + | 13.1048 | 7.05E-06 | 1 | TAGGCCGATTTTGTA |
| 6354 | mamo-L | *BGIBMGA013092* | up | 367 | 381 | - | 11.2419 | 5.21E-05 | 1 | TAGGCGTATTTCATT |
| 6355 | mamo-L | *BGIBMGA013092* | down | 1343 | 1357 | - | 10.5081 | 9.83E-05 | 1 | TGGGCTTATTTTTTT |
| 6356 | mamo-L | *BGIBMGA013097* | up | 1444 | 1458 | + | 11.379 | 4.58E-05 | 1 | AAGGCTTACAGTGGT |
| 6357 | mamo-L | *BGIBMGA013099* | down | 270 | 284 | + | 12.8226 | 9.86E-06 | 1 | TAGGCCATGTTTGAT |
| 6358 | mamo-L | *BGIBMGA013099* | up | 234 | 248 | - | 12.0645 | 2.34E-05 | 1 | TAGGCGGGCTTCGGA |
| 6359 | mamo-L | *BGIBMGA013101* | up | 753 | 767 | - | 11.9677 | 2.59E-05 | 1 | AAGGCGGTGGTTTTT |
| 6360 | mamo-L | *BGIBMGA013108* | down | 1368 | 1382 | - | 10.5968 | 9.15E-05 | 1 | AAGGCTGTTAATGCT |
| 6361 | mamo-L | *BGIBMGA013111* | up | 380 | 394 | + | 10.9194 | 6.98E-05 | 1 | TAGGCTACTTTTTTT |
| 6362 | mamo-L | *BGIBMGA013113* | up | 959 | 973 | - | 12.5484 | 1.36E-05 | 1 | TAGGCATACTTTGAT |
| 6363 | mamo-L | *BGIBMGA013115* | down | 383 | 397 | - | 10.8226 | 7.58E-05 | 1 | TAGGCCTGATTCACT |
| 6364 | mamo-L | *BGIBMGA013119* | down | 1897 | 1911 | + | 11.2097 | 5.38E-05 | 1 | TAGGCTTTTCTTTCT |
| 6365 | mamo-L | *BGIBMGA013119* | up | 166 | 180 | + | 10.7419 | 8.11E-05 | 1 | GAGGCTTGGATGGCT |
| 6366 | mamo-L | *BGIBMGA013119* | down | 1315 | 1329 | - | 10.7097 | 8.33E-05 | 1 | TAGGCGGATGTTACA |
| 6367 | mamo-L | *BGIBMGA013123* | down | 1642 | 1656 | + | 11.4274 | 4.38E-05 | 1 | AAGGCCTAGGTTCTT |
| 6368 | mamo-L | *BGIBMGA013124* | up | 957 | 971 | + | 10.9194 | 6.98E-05 | 1 | TAGGCTACTTTTTTT |
| 6369 | mamo-L | *BGIBMGA013124* | down | 391 | 405 | + | 10.879 | 7.22E-05 | 1 | TAGGCTTGTTTTACA |
| 6370 | mamo-L | *BGIBMGA013125* | down | 1372 | 1386 | + | 12.129 | 2.19E-05 | 1 | TAGGCCCGTTTTATT |
| 6371 | mamo-L | *BGIBMGA013125* | down | 615 | 629 | + | 11.9597 | 2.61E-05 | 1 | TAGGCCTTCGTCTTC |
| 6372 | mamo-L | *BGIBMGA013126* | up | 1147 | 1161 | + | 12.129 | 2.19E-05 | 1 | TAGGCCCGTTTTATT |
| 6373 | mamo-L | *BGIBMGA013126* | up | 390 | 404 | + | 11.9597 | 2.61E-05 | 1 | TAGGCCTTCGTCTTC |
| 6374 | mamo-L | *BGIBMGA013127* | up | 888 | 902 | + | 13.5484 | 3.92E-06 | 1 | TAGGCTTAGATTTTT |
| 6375 | mamo-L | *BGIBMGA013127* | down | 597 | 611 | - | 11.2339 | 5.26E-05 | 1 | TAGGCTGGCTCTACT |
| 6376 | mamo-L | *BGIBMGA013131* | up | 907 | 921 | - | 10.5 | 9.90E-05 | 1 | TAGGCGTTTTTTTTG |
| 6377 | mamo-L | *BGIBMGA013134* | up | 1193 | 1207 | - | 13.5484 | 3.92E-06 | 1 | TAGGCTTAGATTTTT |
| 6378 | mamo-L | *BGIBMGA013137* | up | 124 | 138 | - | 11.2742 | 5.06E-05 | 1 | TAGGCATGCGTTCTT |
| 6379 | mamo-L | *BGIBMGA013137* | down | 312 | 326 | - | 10.9194 | 6.98E-05 | 1 | TAGGCTACTTTTTTT |
| 6380 | mamo-L | *BGIBMGA013138* | down | 1481 | 1495 | - | 11.4274 | 4.38E-05 | 1 | AAGGCCTAGGTTCTT |
| 6381 | mamo-L | *BGIBMGA013138* | up | 957 | 971 | - | 10.9194 | 6.98E-05 | 1 | TAGGCTACTTTTTTT |
| 6382 | mamo-L | *BGIBMGA013141* | down | 703 | 717 | - | 11.2097 | 5.38E-05 | 1 | TAGGCTTTTCTTTCT |
| 6383 | mamo-L | *BGIBMGA013141* | down | 1285 | 1299 | + | 10.7097 | 8.33E-05 | 1 | TAGGCGGATGTTACA |
| 6384 | mamo-L | *BGIBMGA013142* | up | 1528 | 1542 | + | 11.6774 | 3.44E-05 | 1 | TAGGCCTAACTTGTT |
| 6385 | mamo-L | *BGIBMGA013142* | up | 1727 | 1741 | + | 11.0242 | 6.36E-05 | 1 | AAGGCCTGGGTGGAT |
| 6386 | mamo-L | *BGIBMGA013146* | down | 543 | 557 | - | 11.3629 | 4.66E-05 | 1 | TAGGCTCGTGGGGCT |
| 6387 | mamo-L | *BGIBMGA013147* | up | 1092 | 1106 | - | 11.3629 | 4.66E-05 | 1 | TAGGCTCGTGGGGCT |
| 6388 | mamo-L | *BGIBMGA013148* | down | 1148 | 1162 | - | 11.0726 | 6.09E-05 | 1 | GAGGCGTTTTTTGTC |
| 6389 | mamo-L | *BGIBMGA013154* | up | 1769 | 1783 | - | 11.5645 | 3.84E-05 | 1 | TAGGCATTGGTTAGT |
| 6390 | mamo-L | *BGIBMGA013154* | up | 1188 | 1202 | - | 10.5887 | 9.21E-05 | 1 | TAGGCATTGGTTAGC |
| 6391 | mamo-L | *BGIBMGA013157* | up | 510 | 524 | - | 10.6613 | 8.67E-05 | 1 | TAGGCCTTTGATAAT |
| 6392 | mamo-L | *BGIBMGA013158* | down | 333 | 347 | + | 10.9194 | 6.98E-05 | 1 | TAGGCTACTTTTTTT |
| 6393 | mamo-L | *BGIBMGA013160* | up | 231 | 245 | - | 11.1048 | 5.92E-05 | 1 | TAGGCACTTTTTGGC |
| 6394 | mamo-L | *BGIBMGA013160* | up | 1933 | 1947 | + | 11.0323 | 6.32E-05 | 1 | TAGGCACGTTTTGTC |
| 6395 | mamo-L | *BGIBMGA013162* | up | 1240 | 1254 | - | 10.7419 | 8.11E-05 | 1 | TAGGCTAGCGGTATT |
| 6396 | mamo-L | *BGIBMGA013166* | up | 1534 | 1548 | - | 10.9919 | 6.55E-05 | 1 | GAGGCCTATGTTATT |
| 6397 | mamo-L | *BGIBMGA013167* | up | 1114 | 1128 | - | 11.5806 | 3.78E-05 | 1 | AAGGCATACTTTGGT |
| 6398 | mamo-L | *BGIBMGA013170* | up | 355 | 369 | - | 12.0645 | 2.34E-05 | 1 | TAGGCGGGCTTCGGA |
| 6399 | mamo-L | *BGIBMGA013172* | down | 17 | 31 | - | 11.7419 | 3.23E-05 | 1 | AAGGCGGTCTTTTCT |
| 6400 | mamo-L | *BGIBMGA013174* | down | 455 | 469 | + | 11.4194 | 4.42E-05 | 1 | GAGGCTTAAGTCGGT |
| 6401 | mamo-L | *BGIBMGA013177* | down | 1907 | 1921 | + | 10.5726 | 9.33E-05 | 1 | TAGGCTAATATTTAT |
| 6402 | mamo-L | *BGIBMGA013178* | up | 1530 | 1544 | + | 10.5726 | 9.33E-05 | 1 | TAGGCTAATATTTAT |
| 6403 | mamo-L | *BGIBMGA013181* | up | 354 | 368 | - | 13.0726 | 7.34E-06 | 1 | TAGGCCGATTTTGCA |
| 6404 | mamo-L | *BGIBMGA013183* | up | 1848 | 1862 | - | 10.5968 | 9.15E-05 | 1 | GAGGCTGTGTTTCAT |
| 6405 | mamo-L | *BGIBMGA013185* | down | 1196 | 1210 | + | 12.7903 | 1.02E-05 | 1 | TAGGCCCTTTTTTTT |
| 6406 | mamo-L | *BGIBMGA013188* | up | 891 | 905 | + | 10.4919 | 9.96E-05 | 1 | TAGGCTAGTATTTAT |
| 6407 | mamo-L | *BGIBMGA013190* | up | 1377 | 1391 | + | 11.3548 | 4.69E-05 | 1 | TAGGCTATATTTTCT |
| 6408 | mamo-L | *BGIBMGA013191* | up | 1586 | 1600 | + | 12 | 2.50E-05 | 1 | TAGGCTTAGTAGTGT |
| 6409 | mamo-L | *BGIBMGA013192* | up | 1577 | 1591 | + | 13.7742 | 2.79E-06 | 1 | TAGGCCGTGATTGGC |
| 6410 | mamo-L | *BGIBMGA013192* | up | 313 | 327 | - | 10.5 | 9.90E-05 | 1 | GAGGCCCAGGTCGTC |
| 6411 | mamo-L | *BGIBMGA013193* | down | 1690 | 1704 | - | 11.7177 | 3.32E-05 | 1 | TAGGCGTTAGTTGAC |
| 6412 | mamo-L | *BGIBMGA013194* | up | 935 | 949 | - | 10.6371 | 8.85E-05 | 1 | TAGGCTCAGAATGGA |
| 6413 | mamo-L | *BGIBMGA013196* | up | 391 | 405 | + | 11.6935 | 3.39E-05 | 1 | TAGGCTGTATGCGTT |
| 6414 | mamo-L | *BGIBMGA013197* | down | 446 | 460 | + | 11.75 | 3.21E-05 | 1 | TAGGCGCTGTTTGTA |
| 6415 | mamo-L | *BGIBMGA013197* | up | 1195 | 1209 | + | 11.1855 | 5.50E-05 | 1 | AAGGCTGGTAATGGT |
| 6416 | mamo-L | *BGIBMGA013200* | down | 1607 | 1621 | - | 13.2903 | 5.60E-06 | 1 | TAGGCTGATATTTTT |
| 6417 | mamo-L | *BGIBMGA013200* | up | 505 | 519 | + | 11.9919 | 2.52E-05 | 1 | TAGGCACAGGTTTGT |
| 6418 | mamo-L | *BGIBMGA013200* | down | 741 | 755 | + | 10.5726 | 9.33E-05 | 1 | TAGGCTAATATTTAT |
| 6419 | mamo-L | *BGIBMGA013205* | up | 369 | 383 | + | 12.2823 | 1.85E-05 | 1 | TAGGCCTTTTTCTCT |
| 6420 | mamo-L | *BGIBMGA013207* | down | 576 | 590 | - | 10.5161 | 9.77E-05 | 1 | TGGGCTTTGTATGTT |
| 6421 | mamo-L | *BGIBMGA013212* | down | 200 | 214 | + | 11.4919 | 4.11E-05 | 1 | TAGGCTGTTTTGACT |
| 6422 | mamo-L | *BGIBMGA013213* | up | 1524 | 1538 | + | 11.2258 | 5.30E-05 | 1 | TAGGCTTGGGTATTT |
| 6423 | mamo-L | *BGIBMGA013214* | up | 1656 | 1670 | + | 12.1532 | 2.13E-05 | 1 | TAGGCTAGGTTTTAT |
| 6424 | mamo-L | *BGIBMGA013214* | down | 1790 | 1804 | + | 11.4919 | 4.11E-05 | 1 | AAGGCGTACTGTGTT |
| 6425 | mamo-L | *BGIBMGA013218* | up | 1365 | 1379 | + | 11.3387 | 4.77E-05 | 1 | GAGGCTCGTTTTGAT |
| 6426 | mamo-L | *BGIBMGA013219* | up | 1435 | 1449 | - | 11.6935 | 3.39E-05 | 1 | TAGGCTGTATGCGTT |
| 6427 | mamo-L | *BGIBMGA013219* | down | 517 | 531 | - | 11.1129 | 5.88E-05 | 1 | TAGGCGTATGTGGGA |
| 6428 | mamo-L | *BGIBMGA013221* | down | 829 | 843 | + | 11.7177 | 3.32E-05 | 1 | TAGGCGTTAGTTGAC |
| 6429 | mamo-L | *BGIBMGA013226* | up | 207 | 221 | - | 11.3548 | 4.69E-05 | 1 | TAGGCTATATTTTCT |
| 6430 | mamo-L | *BGIBMGA013227* | up | 1451 | 1465 | - | 11.4597 | 4.25E-05 | 1 | TAGGCGGGGTGGTGT |
| 6431 | mamo-L | *BGIBMGA013231* | up | 821 | 835 | - | 11 | 6.50E-05 | 1 | TAGGCGTCTTTTTAT |
| 6432 | mamo-L | *BGIBMGA013236* | up | 547 | 561 | + | 13.7177 | 3.05E-06 | 1 | TAGGCGGATTTTTTT |
| 6433 | mamo-L | *BGIBMGA013239* | down | 906 | 920 | + | 11.5565 | 3.87E-05 | 1 | TAGGCGTGGTGTACT |

| | | | | | | | | | | |
|---|---|---|---|---|---|---|---|---|---|---|
| 6434 | mamo-L | *BGIBMGA013241* | down | 1183 | 1197 | + | 14.0645 | 1.80E-06 | 1 | TAGGCTTAGTTTAGT |
| 6435 | mamo-L | *BGIBMGA013243* | down | 1343 | 1357 | - | 11.7581 | 3.18E-05 | 1 | TAGGCGCGGATGGGT |
| 6436 | mamo-L | *BGIBMGA013243* | down | 540 | 554 | - | 11.1129 | 5.88E-05 | 1 | TAGGCGTATTGTATT |
| 6437 | mamo-L | *BGIBMGA013245* | down | 1843 | 1857 | + | 11.5484 | 3.90E-05 | 1 | TAGGCCGGTTTGACT |
| 6438 | mamo-L | *BGIBMGA013247* | down | 1785 | 1799 | - | 11.4194 | 4.42E-05 | 1 | GAGGCTTAAGTCGGT |
| 6439 | mamo-L | *BGIBMGA013248* | down | 884 | 898 | - | 10.4919 | 9.96E-05 | 1 | TAGGCGACTGTTGTC |
| 6440 | mamo-L | *BGIBMGA013249* | up | 423 | 437 | + | 11.5484 | 3.90E-05 | 1 | GAGGCTGTGTTTACT |
| 6441 | mamo-L | *BGIBMGA013249* | down | 1662 | 1676 | - | 11.1774 | 5.54E-05 | 1 | GAGGCCGTCTTTCTT |
| 6442 | mamo-L | *BGIBMGA013250* | down | 803 | 817 | - | 12.6452 | 1.21E-05 | 1 | GAGGCTGGGGTCGCT |
| 6443 | mamo-L | *BGIBMGA013253* | down | 334 | 348 | + | 10.8065 | 7.69E-05 | 1 | TAGGCTATGATTTAT |
| 6444 | mamo-L | *BGIBMGA013253* | down | 630 | 644 | + | 10.6129 | 9.03E-05 | 1 | AAGGCTAGATTTGTT |
| 6445 | mamo-L | *BGIBMGA013253* | down | 814 | 828 | - | 10.5726 | 9.33E-05 | 1 | TAGGCTAATATTTAT |
| 6446 | mamo-L | *BGIBMGA013254* | up | 286 | 300 | - | 10.5887 | 9.21E-05 | 1 | TAGGCTTAGTTAAGT |
| 6447 | mamo-L | *BGIBMGA013256* | up | 955 | 969 | - | 10.7903 | 7.79E-05 | 1 | GAGGCCGCCTTCGTT |
| 6448 | mamo-L | *BGIBMGA013261* | down | 930 | 944 | - | 10.6935 | 8.45E-05 | 1 | AAGGCTTCGTATGCT |
| 6449 | mamo-L | *BGIBMGA013262* | up | 134 | 148 | - | 10.6935 | 8.45E-05 | 1 | AAGGCTTCGTATGCT |
| 6450 | mamo-L | *BGIBMGA013268* | down | 504 | 518 | - | 11.3548 | 4.69E-05 | 1 | AAGGCTGATGATGAT |
| 6451 | mamo-L | *BGIBMGA013270* | down | 227 | 241 | - | 11.1935 | 5.45E-05 | 1 | TAGGCCACGGATGTT |
| 6452 | mamo-L | *BGIBMGA013270* | up | 526 | 540 | - | 10.7419 | 8.11E-05 | 1 | TAGGCTGGCCTTGCG |
| 6453 | mamo-L | *BGIBMGA013272* | down | 1541 | 1555 | + | 10.7742 | 7.89E-05 | 1 | TAGGCTCTTGTTGAG |
| 6454 | mamo-L | *BGIBMGA013273* | up | 110 | 124 | + | 10.7742 | 7.89E-05 | 1 | TAGGCGGGATTCGGA |
| 6455 | mamo-L | *BGIBMGA013276* | down | 1362 | 1376 | - | 10.8145 | 7.63E-05 | 1 | TACGCTGTGTTTGTT |
| 6456 | mamo-L | *BGIBMGA013277* | down | 1264 | 1278 | + | 10.7661 | 7.95E-05 | 1 | TAGGCCTCGAATTGT |
| 6457 | mamo-L | *BGIBMGA013280* | up | 15 | 29 | + | 12.0081 | 2.48E-05 | 1 | TAGGCCGAAGACGCT |
| 6458 | mamo-L | *BGIBMGA013282* | up | 1477 | 1491 | + | 11.0565 | 6.18E-05 | 1 | TAGGCGTGTTACTGT |
| 6459 | mamo-L | *BGIBMGA013292* | down | 775 | 789 | + | 10.9194 | 6.98E-05 | 1 | TAGGCTACTTTTTTT |
| 6460 | mamo-L | *BGIBMGA013293* | down | 619 | 633 | - | 10.5726 | 9.33E-05 | 1 | TAGGCTAATATTTAT |
| 6461 | mamo-L | *BGIBMGA013294* | up | 1152 | 1166 | + | 13.121 | 6.92E-06 | 1 | GAGGCTGTTGTTGCT |
| 6462 | mamo-L | *BGIBMGA013299* | up | 583 | 597 | - | 10.5887 | 9.21E-05 | 1 | TAGGCCTCATTTACT |
| 6463 | mamo-L | *BGIBMGA013302* | up | 1387 | 1401 | - | 10.7339 | 8.17E-05 | 1 | GAGGCTGAGGTTTTA |
| 6464 | mamo-L | *BGIBMGA013304* | up | 1532 | 1546 | + | 10.5726 | 9.33E-05 | 1 | TAGGCTCCGATTAGT |
| 6465 | mamo-L | *BGIBMGA013307* | down | 1869 | 1883 | + | 10.5726 | 9.33E-05 | 1 | TAGGCTAATATTTAT |
| 6466 | mamo-L | *BGIBMGA013308* | down | 652 | 666 | + | 11.2258 | 5.30E-05 | 1 | TAGGCCTTTTTCCCT |
| 6467 | mamo-L | *BGIBMGA013309* | up | 1721 | 1735 | + | 11.7903 | 3.08E-05 | 1 | AAGGCCTATATTGTT |
| 6468 | mamo-L | *BGIBMGA013309* | down | 782 | 796 | - | 10.5565 | 9.46E-05 | 1 | TAGGCGAATTTCTTT |
| 6469 | mamo-L | *BGIBMGA013314* | down | 378 | 392 | + | 10.75 | 8.06E-05 | 1 | GAGGCCGATTGTGCC |
| 6470 | mamo-L | *BGIBMGA013319* | up | 851 | 865 | + | 12.5242 | 1.40E-05 | 1 | TAGGCCGTCGATTGC |
| 6471 | mamo-L | *BGIBMGA013322* | down | 214 | 228 | + | 12.0081 | 2.48E-05 | 1 | TAGGCCCATATTTTT |
| 6472 | mamo-L | *BGIBMGA013326* | down | 13 | 27 | + | 11.0726 | 6.09E-05 | 1 | GAGGCCGGGCTTGCT |
| 6473 | mamo-L | *BGIBMGA013329* | up | 1396 | 1410 | - | 10.5565 | 9.46E-05 | 1 | TAGGTTGGCTTTGGT |
| 6474 | mamo-L | *BGIBMGA013333* | up | 363 | 377 | - | 13.4274 | 4.64E-06 | 1 | TAGGCATGCGTTGTT |
| 6475 | mamo-L | *BGIBMGA013333* | down | 88 | 102 | - | 11.1613 | 5.63E-05 | 1 | TAGGCAGATGATGAT |
| 6476 | mamo-L | *BGIBMGA013336* | down | 416 | 430 | + | 12.9919 | 8.10E-06 | 1 | TAGGCCGGTTTTGCA |
| 6477 | mamo-L | *BGIBMGA013340* | up | 539 | 553 | - | 11.9758 | 2.57E-05 | 1 | AAGGCTTTTTTTTGT |
| 6478 | mamo-L | *BGIBMGA013340* | down | 1769 | 1783 | - | 11.0323 | 6.32E-05 | 1 | TAGGCAGGTTTTTTC |
| 6479 | mamo-L | *BGIBMGA013348* | up | 1812 | 1826 | + | 10.8226 | 7.58E-05 | 1 | TAGGCTTTATATTAT |
| 6480 | mamo-L | *BGIBMGA013349* | down | 834 | 848 | + | 10.5887 | 9.21E-05 | 1 | TAGGCTTGTACTTGT |
| 6481 | mamo-L | *BGIBMGA013353* | down | 143 | 157 | + | 10.6452 | 8.79E-05 | 1 | TAGGCTGTTGATTCA |
| 6482 | mamo-L | *BGIBMGA013356* | down | 330 | 344 | + | 11.4032 | 4.49E-05 | 1 | GAGGCTGAAGTCGTT |
| 6483 | mamo-L | *BGIBMGA013359* | down | 1429 | 1443 | + | 10.5968 | 9.15E-05 | 1 | AAGGCTGTTAATGCT |
| 6484 | mamo-L | *BGIBMGA013362* | up | 1759 | 1773 | + | 12.4032 | 1.61E-05 | 1 | TAGGCTGAGGGCTTT |
| 6485 | mamo-L | *BGIBMGA013362* | down | 441 | 455 | - | 10.5726 | 9.33E-05 | 1 | TAGGCTAATATTTAT |
| 6486 | mamo-L | *BGIBMGA013364* | down | 1204 | 1218 | - | 11.5403 | 3.93E-05 | 1 | TAGGCAGGCGTTCCT |
| 6487 | mamo-L | *BGIBMGA013365* | down | 996 | 1010 | - | 10.5565 | 9.46E-05 | 1 | AAGGCCGTAGTGGTT |
| 6488 | mamo-L | *BGIBMGA013366* | up | 1368 | 1382 | - | 10.8226 | 7.58E-05 | 1 | TAGGCTTTATATTAT |
| 6489 | mamo-L | *BGIBMGA013368* | down | 258 | 272 | - | 12.8145 | 9.96E-06 | 1 | TAGGCTTGCATTAGT |
| 6490 | mamo-L | *BGIBMGA013369* | up | 1869 | 1883 | - | 10.5403 | 9.58E-05 | 1 | TAGCCTTAGTTTTTT |
| 6491 | mamo-L | *BGIBMGA013371* | down | 1144 | 1158 | + | 12.4194 | 1.58E-05 | 1 | TAGGCCCGCATTTTT |
| 6492 | mamo-L | *BGIBMGA013371* | down | 178 | 192 | + | 11.4677 | 4.22E-05 | 1 | TAGGCCCTGTTTATC |
| 6493 | mamo-L | *BGIBMGA013372* | up | 771 | 785 | + | 12.4194 | 1.58E-05 | 1 | TAGGCCCGCATTTTT |
| 6494 | mamo-L | *BGIBMGA013372* | down | 1368 | 1382 | + | 10.5 | 9.90E-05 | 1 | GAGGCCCAGGTCGTC |
| 6495 | mamo-L | *BGIBMGA013378* | up | 1354 | 1368 | + | 11.6774 | 3.44E-05 | 1 | TAGGCGGTCTTCACT |
| 6496 | mamo-L | *BGIBMGA013382* | up | 1610 | 1624 | - | 13 | 8.01E-06 | 1 | TAGGCTTTGAATGTT |
| 6497 | mamo-L | *BGIBMGA013383* | up | 834 | 848 | + | 10.9194 | 6.98E-05 | 1 | TAGGCTACTTTTTTT |
| 6498 | mamo-L | *BGIBMGA013392* | down | 1554 | 1568 | + | 10.4919 | 9.96E-05 | 1 | GAGGCAGAGATTGTT |
| 6499 | mamo-L | *BGIBMGA013393* | up | 1516 | 1530 | + | 11.4435 | 4.31E-05 | 1 | TAGGCTTGAGAGGCT |
| 6500 | mamo-L | *BGIBMGA013395* | up | 1086 | 1100 | + | 12.3468 | 1.72E-05 | 1 | TAGGCCATGTTCGCT |
| 6501 | mamo-L | *BGIBMGA013396* | down | 827 | 841 | - | 10.6371 | 8.85E-05 | 1 | TAGGCTTGGTTTAAA |
| 6502 | mamo-L | *BGIBMGA013397* | down | 1652 | 1666 | + | 12.1855 | 2.06E-05 | 1 | TAGGCTATTTTTTTT |
| 6503 | mamo-L | *BGIBMGA013399* | down | 845 | 859 | + | 11.8468 | 2.92E-05 | 1 | TAGGCTATCTTTTAT |
| 6504 | mamo-L | *BGIBMGA013399* | down | 1939 | 1953 | - | 10.9194 | 6.98E-05 | 1 | TAGGCTACTTTTTTT |
| 6505 | mamo-L | *BGIBMGA013404* | up | 910 | 924 | - | 11.0806 | 6.05E-05 | 1 | TAGGCGTCGTTCTCT |
| 6506 | mamo-L | *BGIBMGA013406* | up | 729 | 743 | - | 10.9355 | 6.88E-05 | 1 | TAGGCAGGCTTTAGC |
| 6507 | mamo-L | *BGIBMGA013407* | down | 579 | 593 | + | 12.7258 | 1.11E-05 | 1 | TAGGCGTGCTTTGTA |
| 6508 | mamo-L | *BGIBMGA013410* | up | 1977 | 1991 | + | 10.7984 | 7.74E-05 | 1 | TAGGCCGATGCTATT |
| 6509 | mamo-L | *BGIBMGA013411* | up | 312 | 326 | + | 10.7581 | 8.00E-05 | 1 | TAGGCTTTCTGCTAT |
| 6510 | mamo-L | *BGIBMGA013414* | up | 45 | 59 | - | 10.9355 | 6.88E-05 | 1 | TAGGCCTAAGTGTGC |
| 6511 | mamo-L | *BGIBMGA013414* | down | 288 | 302 | + | 10.5968 | 9.15E-05 | 1 | GAGGCTAGTTTTTGT |
| 6512 | mamo-L | *BGIBMGA013416* | down | 248 | 262 | + | 10.5726 | 9.33E-05 | 1 | TAGGCTAATATTTAT |
| 6513 | mamo-L | *BGIBMGA013416* | down | 1432 | 1446 | + | 10.5726 | 9.33E-05 | 1 | TAGGCTAATATTTAT |
| 6514 | mamo-L | *BGIBMGA013417* | up | 1900 | 1914 | + | 10.5726 | 9.33E-05 | 1 | TAGGCTAATATTTAT |
| 6515 | mamo-L | *BGIBMGA013419* | up | 1490 | 1504 | - | 12.2258 | 1.97E-05 | 1 | TAGGCAGGGTGTGCT |
| 6516 | mamo-L | *BGIBMGA013419* | up | 943 | 957 | + | 12.0645 | 2.34E-05 | 1 | TAGGCGGGCTTCGGA |
| 6517 | mamo-L | *BGIBMGA013420* | up | 168 | 182 | + | 12.4597 | 1.51E-05 | 1 | TAGGCCGATGTTATC |
| 6518 | mamo-L | *BGIBMGA013421* | down | 1737 | 1751 | - | 10.8871 | 7.17E-05 | 1 | GAGGCCTAGTGTTCT |
| 6519 | mamo-L | *BGIBMGA013423* | up | 867 | 881 | - | 10.8065 | 7.69E-05 | 1 | AAGGCCTTAGTTTTT |
| 6520 | mamo-L | *BGIBMGA013425* | down | 988 | 1002 | + | 10.5726 | 9.33E-05 | 1 | TAGGCTAATATTTAT |
| 6521 | mamo-L | *BGIBMGA013429* | down | 796 | 810 | - | 11.2016 | 5.41E-05 | 1 | AAGGCGGAGGGTTGT |
| 6522 | mamo-L | *BGIBMGA013437* | down | 1679 | 1693 | + | 12.0645 | 2.34E-05 | 1 | TAGGCGGGCTTCGGA |
| 6523 | mamo-L | *BGIBMGA013437* | down | 1537 | 1551 | - | 10.8306 | 7.52E-05 | 1 | TAGGCCCCCTTCGCC |
| 6524 | mamo-L | *BGIBMGA013443* | down | 1405 | 1419 | + | 11.621 | 3.63E-05 | 1 | TAGGCTGTGAAGGCT |
| 6525 | mamo-L | *BGIBMGA013444* | down | 139 | 153 | - | 11.6129 | 3.66E-05 | 1 | TAGGCTTTTAATGAT |

| | | | | | | | | | | |
|---|---|---|---|---|---|---|---|---|---|---|
| 6526 | mamo-L | *BGIBMGA013444* | down | 36 | 50 | - | 10.8226 | 7.58E-05 | 1 | TAGGCATAGTTGGTC |
| 6527 | mamo-L | *BGIBMGA013447* | up | 1465 | 1479 | - | 11.0726 | 6.09E-05 | 1 | AAGGCTTGTTTTTAT |
| 6528 | mamo-L | *BGIBMGA013448* | down | 1287 | 1301 | + | 10.5968 | 9.15E-05 | 1 | AAGGCTGTTAATGCT |
| 6529 | mamo-L | *BGIBMGA013449* | up | 507 | 521 | + | 10.5968 | 9.15E-05 | 1 | AAGGCTGTTAATGCT |
| 6530 | mamo-L | *BGIBMGA013452* | down | 964 | 978 | + | 11.6129 | 3.66E-05 | 1 | TAGGCTTTTAATGAT |
| 6531 | mamo-L | *BGIBMGA013454* | up | 1677 | 1691 | - | 10.7823 | 7.84E-05 | 1 | TAGGCCTTAATTATT |
| 6532 | mamo-L | *BGIBMGA013456* | down | 250 | 264 | - | 11.3629 | 4.66E-05 | 1 | TAGGCCAGGTGTTCT |
| 6533 | mamo-L | *BGIBMGA013458* | up | 1443 | 1457 | - | 10.7258 | 8.23E-05 | 1 | TAGGCTGCAATCGTT |
| 6534 | mamo-L | *BGIBMGA013460* | down | 655 | 669 | + | 10.5968 | 9.15E-05 | 1 | GAGGCTAGTTTTTGT |
| 6535 | mamo-L | *BGIBMGA013461* | down | 128 | 142 | - | 11.6129 | 3.66E-05 | 1 | TAGGCTTTTAATGAT |
| 6536 | mamo-L | *BGIBMGA013463* | down | 17 | 31 | - | 14.379 | 1.11E-06 | 1 | TAGGCCGGGTGTGTT |
| 6537 | mamo-L | *BGIBMGA013464* | down | 420 | 434 | - | 10.7661 | 7.95E-05 | 1 | TAGGCTGAGTGGGCA |
| 6538 | mamo-L | *BGIBMGA013465* | up | 1681 | 1695 | - | 11.2661 | 5.10E-05 | 1 | TAGGCTTATGGGGCC |
| 6539 | mamo-L | *BGIBMGA013466* | down | 1059 | 1073 | - | 11.3226 | 4.83E-05 | 1 | AAGGCCTTGTTTGAC |
| 6540 | mamo-L | *BGIBMGA013466* | up | 597 | 611 | + | 10.5806 | 9.27E-05 | 1 | TAGGCGCAGTACTTT |
| 6541 | mamo-L | *BGIBMGA013467* | up | 710 | 724 | - | 10.9194 | 6.98E-05 | 1 | TAGGCTACTTTTTTT |
| 6542 | mamo-L | *BGIBMGA013468* | up | 729 | 743 | - | 12.0242 | 2.44E-05 | 1 | TAGGCTAACTTTTTC |
| 6543 | mamo-L | *BGIBMGA013472* | up | 25 | 39 | - | 11.3387 | 4.77E-05 | 1 | TAGGCTTTGATTTGA |
| 6544 | mamo-L | *BGIBMGA013473* | down | 818 | 832 | + | 11.1935 | 5.45E-05 | 1 | AAGGCTTTCTTCTGT |
| 6545 | mamo-L | *BGIBMGA013478* | down | 1537 | 1551 | + | 10.6129 | 9.03E-05 | 1 | GAGGCGTGCGATGTC |
| 6546 | mamo-L | *BGIBMGA013481* | down | 399 | 413 | - | 10.621 | 8.97E-05 | 1 | GAGGCTGACGTCTTC |
| 6547 | mamo-L | *BGIBMGA013482* | up | 1549 | 1563 | + | 12.0323 | 2.42E-05 | 1 | TAGGCCTCGGTTTTC |
| 6548 | mamo-L | *BGIBMGA013485* | up | 735 | 749 | + | 10.5726 | 9.33E-05 | 1 | TAGGCTAATATTTAT |
| 6549 | mamo-L | *BGIBMGA013488* | down | 255 | 269 | - | 10.4919 | 9.96E-05 | 1 | TAGGCTAGTATTTAT |
| 6550 | mamo-L | *BGIBMGA013489* | up | 582 | 596 | + | 11.4435 | 4.31E-05 | 1 | TAGGCTTGAGAGGCT |
| 6551 | mamo-L | *BGIBMGA013492* | down | 1636 | 1650 | + | 11.75 | 3.21E-05 | 1 | TAGGCTGTTTCTTGT |
| 6552 | mamo-L | *BGIBMGA013496* | up | 1207 | 1221 | + | 12.4677 | 1.49E-05 | 1 | TAGGCTTCTATTGCT |
| 6553 | mamo-L | *BGIBMGA013499* | down | 1730 | 1744 | - | 10.8629 | 7.33E-05 | 1 | AAGGCTCAGGATTGT |
| 6554 | mamo-L | *BGIBMGA013503* | down | 100 | 114 | - | 10.8065 | 7.69E-05 | 1 | TAGGCTGCATATGTC |
| 6555 | mamo-L | *BGIBMGA013505* | down | 459 | 473 | + | 10.8065 | 7.69E-05 | 1 | TAGGCTGCATATGTC |
| 6556 | mamo-L | *BGIBMGA013505* | up | 394 | 408 | - | 10.5 | 9.90E-05 | 1 | GAGGCCCAGGTCGTC |
| 6557 | mamo-L | *BGIBMGA013506* | down | 1459 | 1473 | + | 12.7984 | 1.02E-05 | 1 | GAGGCTTGCTTTGGC |
| 6558 | mamo-L | *BGIBMGA013509* | down | 1129 | 1143 | - | 11.2258 | 5.30E-05 | 1 | AAGGCGTGTTTTTTT |
| 6559 | mamo-L | *BGIBMGA013512* | up | 1628 | 1642 | - | 12.4677 | 1.49E-05 | 1 | TAGGCTTCTATTGCT |
| 6560 | mamo-L | *BGIBMGA013514* | down | 1442 | 1456 | - | 10.9839 | 6.59E-05 | 1 | TAGGCTCTCGGGGGC |
| 6561 | mamo-L | *BGIBMGA013516* | down | 994 | 1008 | - | 11.75 | 3.21E-05 | 1 | TAGGCTGTTTCTTGT |
| 6562 | mamo-L | *BGIBMGA013519* | down | 726 | 740 | + | 11.5161 | 4.02E-05 | 1 | TAGGCCATATTTTGT |
| 6563 | mamo-L | *BGIBMGA013520* | up | 373 | 387 | + | 12.5968 | 1.28E-05 | 1 | TAGGCAGCGGTTGGT |
| 6564 | mamo-L | *BGIBMGA013520* | up | 736 | 750 | + | 10.5806 | 9.27E-05 | 1 | AAGGCTGTGGTAGGT |
| 6565 | mamo-L | *BGIBMGA013530* | down | 927 | 941 | + | 11.3952 | 4.52E-05 | 1 | GAGGCCGTGTTTATT |
| 6566 | mamo-L | *BGIBMGA013530* | up | 43 | 57 | + | 11.25 | 5.18E-05 | 1 | GAGGCGGTTTTTTTT |
| 6567 | mamo-L | *BGIBMGA013532* | down | 73 | 87 | - | 12.2903 | 1.83E-05 | 1 | GAGGCTGACGTGGCT |
| 6568 | mamo-L | *BGIBMGA013539* | down | 546 | 560 | + | 11.1935 | 5.45E-05 | 1 | TAGGCGCATGAGGGT |
| 6569 | mamo-L | *BGIBMGA013541* | up | 1965 | 1979 | + | 12.2177 | 1.99E-05 | 1 | TAGGCCTGCGTGAGT |
| 6570 | mamo-L | *BGIBMGA013542* | up | 1142 | 1156 | + | 12.0645 | 2.34E-05 | 1 | TAGGCGGGCTTCGGA |
| 6571 | mamo-L | *BGIBMGA013542* | up | 989 | 1003 | - | 10.8306 | 7.52E-05 | 1 | TAGGCCCCCTTCGCC |
| 6572 | mamo-L | *BGIBMGA013546* | down | 1942 | 1956 | - | 14.7258 | 5.83E-07 | 1 | TAGGCTTGCGTTGCC |
| 6573 | mamo-L | *BGIBMGA013547* | down | 816 | 830 | - | 12.2177 | 1.99E-05 | 1 | TAGGCCTGCGTGAGT |
| 6574 | mamo-L | *BGIBMGA013547* | up | 505 | 519 | + | 11.0403 | 6.28E-05 | 1 | TAGGCTTTTTGGTGT |
| 6575 | mamo-L | *BGIBMGA013549* | down | 768 | 782 | + | 11.6129 | 3.66E-05 | 1 | TAGGCCCATTTTGTG |
| 6576 | mamo-L | *BGIBMGA013556* | up | 714 | 728 | + | 10.9839 | 6.59E-05 | 1 | AAGGCGTTTTTTTTT |
| 6577 | mamo-L | *BGIBMGA013557* | up | 598 | 612 | - | 10.9194 | 6.98E-05 | 1 | TAGGCTACTTTTTTT |
| 6578 | mamo-L | *BGIBMGA013560* | up | 346 | 360 | + | 10.6129 | 9.03E-05 | 1 | TAGGCTTTTTTCCAT |
| 6579 | mamo-L | *BGIBMGA013563* | up | 368 | 382 | + | 11.4839 | 4.15E-05 | 1 | TAGGCCGCTTTTGCA |
| 6580 | mamo-L | *BGIBMGA013565* | down | 98 | 112 | - | 11.7016 | 3.37E-05 | 1 | AAGGCTTACGTTATT |
| 6581 | mamo-L | *BGIBMGA013570* | up | 850 | 864 | + | 10.9194 | 6.98E-05 | 1 | TAGGCTACTTTTTTT |
| 6582 | mamo-L | *BGIBMGA013573* | down | 906 | 920 | - | 12.0323 | 2.42E-05 | 1 | AAGGCCTTGTTTTTT |
| 6583 | mamo-L | *BGIBMGA013574* | down | 1196 | 1210 | - | 11.8145 | 3.01E-05 | 1 | TAGGCTAGCTTTGTA |
| 6584 | mamo-L | *BGIBMGA013574* | down | 1419 | 1433 | - | 10.8952 | 7.12E-05 | 1 | TAGGCTGTTTTGGCG |
| 6585 | mamo-L | *BGIBMGA013575* | up | 821 | 835 | - | 12.0161 | 2.46E-05 | 1 | TAGGCTGCTTTTGGA |
| 6586 | mamo-L | *BGIBMGA013578* | up | 1259 | 1273 | - | 13.4677 | 4.39E-06 | 1 | TAGGCTGATGATGAT |
| 6587 | mamo-L | *BGIBMGA013578* | up | 1220 | 1234 | + | 10.7742 | 7.89E-05 | 1 | TAGGCTTACATTAAC |
| 6588 | mamo-L | *BGIBMGA013579* | down | 1218 | 1232 | - | 13.8952 | 2.32E-06 | 1 | TAGGCGTGGTTTTTT |
| 6589 | mamo-L | *BGIBMGA013582* | down | 76 | 90 | - | 11.0484 | 6.23E-05 | 1 | TAGGCCTATTTTCAC |
| 6590 | mamo-L | *BGIBMGA013586* | down | 1235 | 1249 | - | 10.9355 | 6.88E-05 | 1 | TAGGCTTCCTTCGGA |
| 6591 | mamo-L | *BGIBMGA013587* | down | 1261 | 1275 | + | 12.9919 | 8.10E-06 | 1 | TAGGCCGGTTTTGCA |
| 6592 | mamo-L | *BGIBMGA013587* | up | 1008 | 1022 | + | 12.7903 | 1.02E-05 | 1 | TAGGCTTTTATTGTC |
| 6593 | mamo-L | *BGIBMGA013591* | down | 225 | 239 | + | 10.9032 | 7.07E-05 | 1 | TAGGCGAGTGTTTTC |
| 6594 | mamo-L | *BGIBMGA013594* | down | 1798 | 1812 | - | 12.0484 | 2.38E-05 | 1 | AAGGCCGATGTCGTT |
| 6595 | mamo-L | *BGIBMGA013602* | down | 1495 | 1509 | - | 11.371 | 4.62E-05 | 1 | TAGGCTGTATAGGCT |
| 6596 | mamo-L | *BGIBMGA013603* | up | 1252 | 1266 | - | 11.371 | 4.62E-05 | 1 | TAGGCTGTATAGGCT |
| 6597 | mamo-L | *BGIBMGA013604* | down | 1544 | 1558 | + | 11.6694 | 3.47E-05 | 1 | TAGGCGGGCTGTCTT |
| 6598 | mamo-L | *BGIBMGA013605* | up | 1500 | 1514 | - | 11.7419 | 3.23E-05 | 1 | TAGGCCATTTATGCT |
| 6599 | mamo-L | *BGIBMGA013605* | up | 1681 | 1695 | - | 11.6694 | 3.47E-05 | 1 | AAGGCCGTGTTTTGC |
| 6600 | mamo-L | *BGIBMGA013606* | up | 249 | 263 | - | 11.6129 | 3.66E-05 | 1 | TAGGCTTTTAATGAT |
| 6601 | mamo-L | *BGIBMGA013609* | up | 1613 | 1627 | + | 11.7419 | 3.23E-05 | 1 | TAGGCCATTTATGCT |
| 6602 | mamo-L | *BGIBMGA013609* | up | 1432 | 1446 | + | 11.6694 | 3.47E-05 | 1 | AAGGCCGTGTTTTGC |
| 6603 | mamo-L | *BGIBMGA013610* | down | 1687 | 1701 | + | 11.3548 | 4.69E-05 | 1 | AAGGCTGATGATGAT |
| 6604 | mamo-L | *BGIBMGA013610* | down | 471 | 485 | - | 10.9435 | 6.83E-05 | 1 | AAGGCTTTGTTCTTT |
| 6605 | mamo-L | *BGIBMGA013611* | up | 295 | 309 | + | 11.3548 | 4.69E-05 | 1 | AAGGCTGATGATGAT |
| 6606 | mamo-L | *BGIBMGA013613* | up | 275 | 289 | + | 13.7339 | 2.97E-06 | 1 | TAGGCTTTTATTGCT |
| 6607 | mamo-L | *BGIBMGA013619* | down | 1159 | 1173 | - | 12.0161 | 2.46E-05 | 1 | TAGGCCGTAGATGTC |
| 6608 | mamo-L | *BGIBMGA013619* | down | 1114 | 1128 | - | 10.8145 | 7.63E-05 | 1 | TAGGCTGTGAGCTCT |
| 6609 | mamo-L | *BGIBMGA013624* | up | 527 | 541 | + | 10.8145 | 7.63E-05 | 1 | TAGGCGCAGTGCTGT |
| 6610 | mamo-L | *BGIBMGA013624* | up | 1781 | 1795 | + | 10.5403 | 9.58E-05 | 1 | TAGGCTGTATGGTGT |
| 6611 | mamo-L | *BGIBMGA013625* | down | 1384 | 1398 | + | 11.7742 | 3.14E-05 | 1 | TCGGCTGAGTTTGTT |
| 6612 | mamo-L | *BGIBMGA013634* | up | 1510 | 1524 | - | 12.8145 | 9.96E-06 | 1 | TAGGCTCACGTTGTA |
| 6613 | mamo-L | *BGIBMGA013635* | down | 131 | 145 | - | 12.121 | 2.20E-05 | 1 | TAGGCCGACGTTAAC |
| 6614 | mamo-L | *BGIBMGA013637* | up | 419 | 433 | + | 12.2984 | 1.81E-05 | 1 | TAGGCCTCGTTGGCT |
| 6615 | mamo-L | *BGIBMGA013637* | down | 321 | 335 | - | 11.6694 | 3.47E-05 | 1 | AAGGCTGACTTTGTA |
| 6616 | mamo-L | *BGIBMGA013641* | down | 1368 | 1382 | - | 12.3952 | 1.63E-05 | 1 | TAGGCATGGGTTTTT |
| 6617 | mamo-L | *BGIBMGA013643* | up | 1536 | 1550 | - | 10.8387 | 7.47E-05 | 1 | AAGGCCTTGGATTTT |

| | | | | | | | | | | |
|---|---|---|---|---|---|---|---|---|---|---|
| 6618 | mamo-L | *BGIBMGA013644* | down | 416 | 430 | - | 10.8226 | 7.58E-05 | 1 | TGGGCTAACTTTGGT |
| 6619 | mamo-L | *BGIBMGA013645* | up | 886 | 900 | + | 11.4839 | 4.15E-05 | 1 | GAGGCTTGCGTCGAT |
| 6620 | mamo-L | *BGIBMGA013652* | up | 1057 | 1071 | + | 14.2258 | 1.41E-06 | 1 | TAGGCTTATGTTTTT |
| 6621 | mamo-L | *BGIBMGA013660* | down | 83 | 97 | + | 11.9677 | 2.59E-05 | 1 | TAGGCTTTATATTGT |
| 6622 | mamo-L | *BGIBMGA013661* | up | 1374 | 1388 | + | 11.7903 | 3.08E-05 | 1 | AAGGCGGGCTATGTT |
| 6623 | mamo-L | *BGIBMGA013661* | down | 1858 | 1872 | + | 10.9274 | 6.92E-05 | 1 | TAGGCAGGTGTGGGC |
| 6624 | mamo-L | *BGIBMGA013664* | up | 1404 | 1418 | + | 14.7903 | 5.14E-07 | 1 | TAGGCTGATGTTGAT |
| 6625 | mamo-L | *BGIBMGA013665* | up | 915 | 929 | - | 13.6855 | 3.18E-06 | 1 | TAGGCTGTCGATTGT |
| 6626 | mamo-L | *BGIBMGA013665* | down | 833 | 847 | + | 10.7581 | 8.00E-05 | 1 | TAGGCGGAAAATGAT |
| 6627 | mamo-L | *BGIBMGA013668* | down | 1178 | 1192 | - | 10.9516 | 6.78E-05 | 1 | TAGGCAGGTTTTCTT |
| 6628 | mamo-L | *BGIBMGA013671* | up | 96 | 110 | - | 12.621 | 1.25E-05 | 1 | TAGGCCTTTTGTGGC |
| 6629 | mamo-L | *BGIBMGA013674* | down | 288 | 302 | - | 11.2177 | 5.34E-05 | 1 | AAGGCCGGATTTTTT |
| 6630 | mamo-L | *BGIBMGA013674* | down | 1340 | 1354 | - | 11.1452 | 5.70E-05 | 1 | AAGGCTTAAATTGCT |
| 6631 | mamo-L | *BGIBMGA013678* | down | 111 | 125 | + | 10.9516 | 6.78E-05 | 1 | TAGGCAGGTTTTCTT |
| 6632 | mamo-L | *BGIBMGA013679* | up | 1462 | 1476 | + | 13.6855 | 3.18E-06 | 1 | TAGGCTGTCGATTGT |
| 6633 | mamo-L | *BGIBMGA013679* | down | 246 | 260 | - | 11.2016 | 5.41E-05 | 1 | TAGGCTGACCTCGTC |
| 6634 | mamo-L | *BGIBMGA013681* | down | 508 | 522 | - | 10.7742 | 7.89E-05 | 1 | TGGGCTTATTTTGAT |
| 6635 | mamo-L | *BGIBMGA013682* | down | 321 | 335 | + | 12.8387 | 9.67E-06 | 1 | TAGGCTCGGTTTACT |
| 6636 | mamo-L | *BGIBMGA013682* | up | 877 | 891 | + | 10.5 | 9.90E-05 | 1 | GAGGCCCAGGTCGTC |
| 6637 | mamo-L | *BGIBMGA013683* | up | 1029 | 1043 | + | 14.25 | 1.36E-06 | 1 | TAGGCCGTCGATGCT |
| 6638 | mamo-L | *BGIBMGA013683* | up | 1475 | 1489 | + | 11.1532 | 5.67E-05 | 1 | TAGGCTTAGTTTATG |
| 6639 | mamo-L | *BGIBMGA013685* | up | 609 | 623 | + | 11.0806 | 6.05E-05 | 1 | TAGGCTTGGCTCTGT |
| 6640 | mamo-L | *BGIBMGA013689* | up | 1042 | 1056 | + | 11.9839 | 2.54E-05 | 1 | TAGGCCTTTGTTAAT |
| 6641 | mamo-L | *BGIBMGA013691* | up | 468 | 482 | + | 10.5645 | 9.40E-05 | 1 | TAGGCTTGTTTGTAC |
| 6642 | mamo-L | *BGIBMGA013693* | down | 950 | 964 | + | 11.3548 | 4.69E-05 | 1 | AAGGCTGATGATGAT |
| 6643 | mamo-L | *BGIBMGA013696* | up | 1095 | 1109 | + | 11.2016 | 5.41E-05 | 1 | AAGGCTGTGATCGCT |
| 6644 | mamo-L | *BGIBMGA013696* | down | 413 | 427 | - | 11 | 6.50E-05 | 1 | AAGGCTTGTTTTATT |
| 6645 | mamo-L | *BGIBMGA013700* | up | 727 | 741 | + | 12.1613 | 2.11E-05 | 1 | TAGGCTAAGTTTATT |
| 6646 | mamo-L | *BGIBMGA013701* | down | 1328 | 1342 | - | 11.1532 | 5.67E-05 | 1 | AAGGCTTATTTTTAT |
| 6647 | mamo-L | *BGIBMGA013702* | up | 1024 | 1038 | - | 10.5968 | 9.15E-05 | 1 | GAGGCTAGTTTTTGT |
| 6648 | mamo-L | *BGIBMGA013703* | up | 241 | 255 | - | 11.7984 | 3.06E-05 | 1 | TAGGCTTTGCTTTTT |
| 6649 | mamo-L | *BGIBMGA013703* | down | 1320 | 1334 | - | 10.8871 | 7.17E-05 | 1 | GAGGCTGGCGAGGCT |
| 6650 | mamo-L | *BGIBMGA013704* | down | 1207 | 1221 | - | 10.5726 | 9.33E-05 | 1 | TAGGCTAATATTTAT |
| 6651 | mamo-L | *BGIBMGA013704* | down | 564 | 578 | + | 10.5161 | 9.77E-05 | 1 | TAGGCGAACTTTTAC |
| 6652 | mamo-L | *BGIBMGA013705* | up | 564 | 578 | + | 11.0242 | 6.36E-05 | 1 | TGGGCTGATGATGGT |
| 6653 | mamo-L | *BGIBMGA013706* | up | 1690 | 1704 | + | 11.4435 | 4.31E-05 | 1 | TAGGCTTAGGACCGT |
| 6654 | mamo-L | *BGIBMGA013708* | up | 1343 | 1357 | - | 11.3952 | 4.52E-05 | 1 | TAGGCGAACGATGAT |
| 6655 | mamo-L | *BGIBMGA013708* | up | 1136 | 1150 | + | 11.1855 | 5.50E-05 | 1 | TAGGCTTCTTATTTT |
| 6656 | mamo-L | *BGIBMGA013711* | down | 684 | 698 | - | 12.3952 | 1.63E-05 | 1 | TAGGCCTTTGATTTT |
| 6657 | mamo-L | *BGIBMGA013711* | down | 1000 | 1014 | - | 11.1532 | 5.67E-05 | 1 | TAGGCCAGTTGTTGT |
| 6658 | mamo-L | *BGIBMGA013712* | down | 1102 | 1116 | + | 11.4194 | 4.42E-05 | 1 | GAGGCCGGCGTCGCC |
| 6659 | mamo-L | *BGIBMGA013714* | down | 1705 | 1719 | - | 11.8952 | 2.79E-05 | 1 | TAGGCCAAGATTGTC |
| 6660 | mamo-L | *BGIBMGA013714* | down | 45 | 59 | + | 10.8871 | 7.17E-05 | 1 | GAGGCTGGCGAGGCT |
| 6661 | mamo-L | *BGIBMGA013715* | down | 45 | 59 | + | 11.1532 | 5.67E-05 | 1 | AAGGCTTATTTTTAT |
| 6662 | mamo-L | *BGIBMGA013716* | up | 1602 | 1616 | + | 11.1532 | 5.67E-05 | 1 | AAGGCTTATTTTTAT |
| 6663 | mamo-L | *BGIBMGA013720* | up | 1221 | 1235 | - | 13.5081 | 4.15E-06 | 1 | TAGGCTGTCTATGAT |
| 6664 | mamo-L | *BGIBMGA013721* | down | 475 | 489 | + | 10.8145 | 7.63E-05 | 1 | TAGGCCTTAGTCTAT |
| 6665 | mamo-L | *BGIBMGA013726* | down | 1956 | 1970 | - | 12.4919 | 1.46E-05 | 1 | TAGGCGGAGGATAGT |
| 6666 | mamo-L | *BGIBMGA013726* | up | 1347 | 1361 | - | 11.2016 | 5.41E-05 | 1 | AAGGCTGTGATCGCT |
| 6667 | mamo-L | *BGIBMGA013729* | up | 1853 | 1867 | - | 13.8065 | 2.66E-06 | 1 | TAGGCTATGTTTGCT |
| 6668 | mamo-L | *BGIBMGA013730* | up | 33 | 47 | + | 10.9355 | 6.88E-05 | 1 | AAGGCCGCGGATGCT |
| 6669 | mamo-L | *BGIBMGA013731* | up | 357 | 371 | + | 13.7823 | 2.76E-06 | 1 | TAGGCTTGTTTGGGT |
| 6670 | mamo-L | *BGIBMGA013731* | up | 1556 | 1570 | + | 12.1935 | 2.05E-05 | 1 | TAGGCGGGCGTCGGA |
| 6671 | mamo-L | *BGIBMGA013732* | up | 632 | 646 | + | 10.9435 | 6.83E-05 | 1 | TAGGCCGACAAGTGT |
| 6672 | mamo-L | *BGIBMGA013735* | down | 679 | 693 | - | 10.6129 | 9.03E-05 | 1 | TAGGCTTCGGTGACT |
| 6673 | mamo-L | *BGIBMGA013739* | down | 1974 | 1988 | + | 11.3065 | 4.91E-05 | 1 | TAGGCTGACGAGCGT |
| 6674 | mamo-L | *BGIBMGA013739* | down | 1257 | 1271 | + | 11.1935 | 5.45E-05 | 1 | TAGGCGCATGAGGGT |
| 6675 | mamo-L | *BGIBMGA013740* | down | 720 | 734 | + | 13.2581 | 5.81E-06 | 1 | TAGGCTGGTTTTCTT |
| 6676 | mamo-L | *BGIBMGA013741* | up | 1286 | 1300 | + | 10.5726 | 9.33E-05 | 1 | TAGGCTAATATTTAT |
| 6677 | mamo-L | *BGIBMGA013742* | down | 644 | 658 | + | 11.2258 | 5.30E-05 | 1 | TAGGCTATTGTTCCT |
| 6678 | mamo-L | *BGIBMGA013746* | up | 839 | 853 | + | 11.621 | 3.63E-05 | 1 | AAGGCGTATTTTTGT |
| 6679 | mamo-L | *BGIBMGA013747* | up | 613 | 627 | + | 12.0484 | 2.38E-05 | 1 | AAGGCTAAGTTTGTT |
| 6680 | mamo-L | *BGIBMGA013747* | up | 1401 | 1415 | + | 10.7742 | 7.89E-05 | 1 | AAGGCTAAGTTCGTT |
| 6681 | mamo-L | *BGIBMGA013748* | down | 1415 | 1429 | - | 11.4194 | 4.42E-05 | 1 | GAGGCCGACATTTCT |
| 6682 | mamo-L | *BGIBMGA013748* | down | 287 | 301 | - | 10.75 | 8.06E-05 | 1 | GAGGCCGATTGTGCC |
| 6683 | mamo-L | *BGIBMGA013748* | down | 1575 | 1589 | + | 10.5726 | 9.33E-05 | 1 | TAGGCTAATATTTAT |
| 6684 | mamo-L | *BGIBMGA013749* | up | 1230 | 1244 | - | 11.5645 | 3.84E-05 | 1 | AAGGCTTGTTGTGCT |
| 6685 | mamo-L | *BGIBMGA013749* | down | 513 | 527 | - | 11.4194 | 4.42E-05 | 1 | GAGGCCGACATTTCT |
| 6686 | mamo-L | *BGIBMGA013749* | up | 1556 | 1570 | - | 10.75 | 8.06E-05 | 1 | GAGGCCGATTGTGCC |
| 6687 | mamo-L | *BGIBMGA013749* | down | 673 | 687 | + | 10.5726 | 9.33E-05 | 1 | TAGGCTAATATTTAT |
| 6688 | mamo-L | *BGIBMGA013749* | down | 1880 | 1894 | + | 10.5645 | 9.40E-05 | 1 | GAGGCTGTCGATTGC |
| 6689 | mamo-L | *BGIBMGA013750* | down | 462 | 476 | + | 10.9194 | 6.98E-05 | 1 | TAGGCTACTTTTTTT |
| 6690 | mamo-L | *BGIBMGA013752* | down | 1985 | 1999 | - | 10.7742 | 7.89E-05 | 1 | TAGGCGCACAATGCC |
| 6691 | mamo-L | *BGIBMGA013752* | up | 1757 | 1771 | + | 10.5403 | 9.58E-05 | 1 | TAGGCTGTATGGTGT |
| 6692 | mamo-L | *BGIBMGA013755* | up | 601 | 615 | + | 11.4597 | 4.25E-05 | 1 | TAGGCGCTGGTGGTC |
| 6693 | mamo-L | *BGIBMGA013757* | down | 246 | 260 | + | 11.6371 | 3.58E-05 | 1 | TAGGCTTGTGGTGTA |
| 6694 | mamo-L | *BGIBMGA013759* | up | 416 | 430 | - | 11.0403 | 6.28E-05 | 1 | TAGGCATTGATTGTC |
| 6695 | mamo-L | *BGIBMGA013762* | up | 472 | 486 | + | 10.9194 | 6.98E-05 | 1 | TAGGCTACTTTTTTT |
| 6696 | mamo-L | *BGIBMGA013763* | up | 1305 | 1319 | + | 11.379 | 4.58E-05 | 1 | TAGGCCAGCTATTCT |
| 6697 | mamo-L | *BGIBMGA013764* | up | 1189 | 1203 | - | 11.4355 | 4.34E-05 | 1 | TAGGCGGGGCATGTT |
| 6698 | mamo-L | *BGIBMGA013764* | down | 302 | 316 | - | 10.9516 | 6.78E-05 | 1 | TAGGCTGAAGTCGTG |
| 6699 | mamo-L | *BGIBMGA013766* | up | 905 | 919 | - | 10.5726 | 9.33E-05 | 1 | TAGGCTAATATTTAT |
| 6700 | mamo-L | *BGIBMGA013767* | down | 925 | 939 | + | 11.4355 | 4.34E-05 | 1 | AAGGCTTAGATTTTT |
| 6701 | mamo-L | *BGIBMGA013769* | up | 438 | 452 | - | 10.5323 | 9.63E-05 | 1 | AAGGCGTTGTTCGAT |
| 6702 | mamo-L | *BGIBMGA013772* | down | 135 | 149 | - | 11.129 | 5.79E-05 | 1 | TAGGCTAAGATTTAT |
| 6703 | mamo-L | *BGIBMGA013775* | down | 1454 | 1468 | + | 11.9677 | 2.59E-05 | 1 | TAGGCCGTGTACGTC |
| 6704 | mamo-L | *BGIBMGA013778* | up | 1757 | 1771 | + | 14.3306 | 1.21E-06 | 1 | TAGGCGGGGGGTGGT |
| 6705 | mamo-L | *BGIBMGA013781* | up | 1772 | 1786 | - | 10.9677 | 6.68E-05 | 1 | TAGGCAGTATTTTTT |
| 6706 | mamo-L | *BGIBMGA013782* | down | 588 | 602 | - | 14 | 1.98E-06 | 1 | TAGGCCGGTGATGCT |
| 6707 | mamo-L | *BGIBMGA013785* | down | 696 | 710 | + | 11.3387 | 4.77E-05 | 1 | AAGGCCGACGTGGAT |
| 6708 | mamo-L | *BGIBMGA013792* | down | 1544 | 1558 | + | 12.0645 | 2.34E-05 | 1 | TAGGCGGGCTTCGGA |
| 6709 | mamo-L | *BGIBMGA013792* | up | 198 | 212 | + | 11.371 | 4.62E-05 | 1 | TAGGCTGTATAGGCT |

| | | | | | | | | | | |
|---|---|---|---|---|---|---|---|---|---|---|
| 6710 | mamo-L | *BGIBMGA013792* | down | 1167 | 1181 | + | 11.3468 | 4.73E-05 | 1 | TAGGCTTGCTTATGT |
| 6711 | mamo-L | *BGIBMGA013792* | down | 243 | 257 | - | 10.75 | 8.06E-05 | 1 | TAGGCCTCCTTCGGA |
| 6712 | mamo-L | *BGIBMGA013792* | up | 1957 | 1971 | + | 10.6129 | 9.03E-05 | 1 | TAGGCCTTCATCCCT |
| 6713 | mamo-L | *BGIBMGA013794* | up | 1741 | 1755 | + | 10.9194 | 6.98E-05 | 1 | TAGGCTACTTTTTTT |
| 6714 | mamo-L | *BGIBMGA013795* | down | 1400 | 1414 | + | 10.9677 | 6.68E-05 | 1 | TAGGCCACCTATGCT |
| 6715 | mamo-L | *BGIBMGA013796* | down | 677 | 691 | + | 11.4194 | 4.42E-05 | 1 | GAGGCCGACATTTCT |
| 6716 | mamo-L | *BGIBMGA013796* | down | 1805 | 1819 | + | 10.75 | 8.06E-05 | 1 | GAGGCCGATTGTGCC |
| 6717 | mamo-L | *BGIBMGA013796* | down | 517 | 531 | - | 10.5726 | 9.33E-05 | 1 | TAGGCTAATATTTAT |
| 6718 | mamo-L | *BGIBMGA013808* | down | 978 | 992 | - | 10.9274 | 6.92E-05 | 1 | AAGGCCCTTGTTGTC |
| 6719 | mamo-L | *BGIBMGA013808* | down | 1662 | 1676 | + | 10.8468 | 7.42E-05 | 1 | TAGGCGGGGTCTGAC |
| 6720 | mamo-L | *BGIBMGA013809* | up | 306 | 320 | + | 11.6613 | 3.50E-05 | 1 | TAGGCCGTTGTGGAC |
| 6721 | mamo-L | *BGIBMGA013811* | up | 673 | 687 | - | 11.4677 | 4.22E-05 | 1 | TAGGCGTAGAGTTTT |
| 6722 | mamo-L | *BGIBMGA013813* | down | 255 | 269 | - | 12.9194 | 8.81E-06 | 1 | TAGGCGGAATTTTTT |
| 6723 | mamo-L | *BGIBMGA013815* | up | 227 | 241 | + | 10.9194 | 6.98E-05 | 1 | TAGGCTACTTTTTTT |
| 6724 | mamo-L | *BGIBMGA013815* | down | 1125 | 1139 | + | 10.5726 | 9.33E-05 | 1 | TAGGCTAATATTTAT |
| 6725 | mamo-L | *BGIBMGA013818* | down | 1547 | 1561 | + | 12.9919 | 8.10E-06 | 1 | TAGGCGGTGGTTCCT |
| 6726 | mamo-L | *BGIBMGA013822* | down | 1571 | 1585 | + | 11.5 | 4.08E-05 | 1 | TAGGCTAATTATTGT |
| 6727 | mamo-L | *BGIBMGA013823* | down | 1043 | 1057 | + | 11.9919 | 2.52E-05 | 1 | TAGGCTTTAATTGTC |
| 6728 | mamo-L | *BGIBMGA013823* | up | 674 | 688 | + | 10.5726 | 9.33E-05 | 1 | TAGGCTAATATTTAT |
| 6729 | mamo-L | *BGIBMGA013828* | up | 560 | 574 | - | 11.0242 | 6.36E-05 | 1 | AAGGCCTTGTTTTCC |
| 6730 | mamo-L | *BGIBMGA013829* | up | 1956 | 1970 | + | 11.5323 | 3.96E-05 | 1 | TAGGCTGTGGCTAGT |
| 6731 | mamo-L | *BGIBMGA013832* | down | 321 | 335 | - | 10.7984 | 7.74E-05 | 1 | TAGGCCGATAGTATT |
| 6732 | mamo-L | *BGIBMGA013838* | down | 927 | 941 | + | 12.7903 | 1.02E-05 | 1 | TAGGCTGTTTGTGTC |
| 6733 | mamo-L | *BGIBMGA013839* | down | 491 | 505 | + | 11.8952 | 2.79E-05 | 1 | GAGGCCTATGTTTTT |
| 6734 | mamo-L | *BGIBMGA013840* | up | 1033 | 1047 | - | 12.2661 | 1.89E-05 | 1 | TAGGCCTTTTATTTT |
| 6735 | mamo-L | *BGIBMGA013840* | up | 1040 | 1054 | + | 11.7984 | 3.06E-05 | 1 | AAGGCCTATTTTTTT |
| 6736 | mamo-L | *BGIBMGA013843* | up | 51 | 65 | + | 11.6613 | 3.50E-05 | 1 | TAGGCACTGTTTGGC |
| 6737 | mamo-L | *BGIBMGA013843* | up | 74 | 88 | + | 10.8952 | 7.12E-05 | 1 | TAGGCTGTAATCTTT |
| 6738 | mamo-L | *BGIBMGA013845* | down | 143 | 157 | - | 12.1129 | 2.22E-05 | 1 | TAGGCTTAATTTGGG |
| 6739 | mamo-L | *BGIBMGA013846* | down | 498 | 512 | - | 10.5726 | 9.33E-05 | 1 | TAGGCTAATATTTAT |
| 6740 | mamo-L | *BGIBMGA013850* | down | 552 | 566 | - | 10.9194 | 6.98E-05 | 1 | TAGGCTACTTTTTTT |
| 6741 | mamo-L | *BGIBMGA013852* | down | 1165 | 1179 | + | 11.4516 | 4.28E-05 | 1 | TAGGCTGAGTACATT |
| 6742 | mamo-L | *BGIBMGA013855* | down | 767 | 781 | - | 12.3468 | 1.72E-05 | 1 | TAGGCCGTATTTCGT |
| 6743 | mamo-L | *BGIBMGA013856* | up | 949 | 963 | - | 12.0887 | 2.29E-05 | 1 | TAGGCTCGGATGGCT |
| 6744 | mamo-L | *BGIBMGA013856* | up | 1172 | 1186 | - | 12.0806 | 2.31E-05 | 1 | TAGGCGAGGTATGTT |
| 6745 | mamo-L | *BGIBMGA013856* | down | 1919 | 1933 | + | 11.0806 | 6.05E-05 | 1 | TAGGCTTGCTTTTAG |
| 6746 | mamo-L | *BGIBMGA013858* | up | 1085 | 1099 | + | 10.8629 | 7.33E-05 | 1 | TGGGCGGTCGTTGGC |
| 6747 | mamo-L | *BGIBMGA013860* | down | 316 | 330 | + | 10.5968 | 9.15E-05 | 1 | GAGGCTAGTTTTTGT |
| 6748 | mamo-L | *BGIBMGA013861* | down | 1116 | 1130 | + | 10.5968 | 9.15E-05 | 1 | AAGGCGGAGTTTAGC |
| 6749 | mamo-L | *BGIBMGA013861* | down | 1500 | 1514 | + | 10.5968 | 9.15E-05 | 1 | AAGGCGGAGTTTAGC |
| 6750 | mamo-L | *BGIBMGA013862* | down | 39 | 53 | - | 11.1129 | 5.88E-05 | 1 | TAGGCTATTGATGTC |
| 6751 | mamo-L | *BGIBMGA013863* | up | 1639 | 1653 | + | 11.871 | 2.85E-05 | 1 | TAGGCTTAGTGGGAT |
| 6752 | mamo-L | *BGIBMGA013863* | down | 1267 | 1281 | - | 11.2742 | 5.06E-05 | 1 | GAGGCGTATTTTTTT |
| 6753 | mamo-L | *BGIBMGA013863* | down | 339 | 353 | - | 10.6129 | 9.03E-05 | 1 | TAGGCTTACGAGGCA |
| 6754 | mamo-L | *BGIBMGA013864* | up | 1804 | 1818 | + | 10.75 | 8.06E-05 | 1 | GAGGCCGATTGTGCC |
| 6755 | mamo-L | *BGIBMGA013869* | down | 336 | 350 | - | 11.4516 | 4.28E-05 | 1 | TAGGCCTGCTCGGGT |
| 6756 | mamo-L | *BGIBMGA013871* | down | 1539 | 1553 | - | 12.25 | 1.92E-05 | 1 | TAGGCCCACTATTCT |
| 6757 | mamo-L | *BGIBMGA013872* | up | 1182 | 1196 | + | 10.9516 | 6.78E-05 | 1 | AAGGCTTAATTGGGT |
| 6758 | mamo-L | *BGIBMGA013873* | down | 1087 | 1101 | + | 12.0887 | 2.29E-05 | 1 | TAGGCCGGCTAGGGC |
| 6759 | mamo-L | *BGIBMGA013873* | up | 359 | 373 | - | 11.25 | 5.18E-05 | 1 | AAGGCTGCGTTTTTT |
| 6760 | mamo-L | *BGIBMGA013873* | up | 860 | 874 | - | 11.1613 | 5.63E-05 | 1 | TAGGCAGGGTGTTTT |
| 6761 | mamo-L | *BGIBMGA013874* | up | 1359 | 1373 | + | 12.0887 | 2.29E-05 | 1 | TAGGCCGGCTAGGGC |
| 6762 | mamo-L | *BGIBMGA013874* | down | 123 | 137 | + | 11.621 | 3.63E-05 | 1 | AAGGCGGGTGTTTCT |
| 6763 | mamo-L | *BGIBMGA013876* | up | 1239 | 1253 | + | 12.9839 | 8.17E-06 | 1 | TAGGCCGTTTTTATT |
| 6764 | mamo-L | *BGIBMGA013877* | down | 50 | 64 | + | 11.0806 | 6.05E-05 | 1 | TAGCCTTATTTTGTT |
| 6765 | mamo-L | *BGIBMGA013881* | down | 334 | 348 | - | 12.3226 | 1.77E-05 | 1 | TAGGCTGTTGTTATC |
| 6766 | mamo-L | *BGIBMGA013884* | up | 1249 | 1263 | + | 13.0242 | 7.77E-06 | 1 | TAGGCTGATGTTGTG |
| 6767 | mamo-L | *BGIBMGA013884* | up | 850 | 864 | - | 11.6613 | 3.50E-05 | 1 | TAGGCCGTTGTGGAC |
| 6768 | mamo-L | *BGIBMGA013886* | up | 491 | 505 | - | 11.1935 | 5.45E-05 | 1 | TAGGCGCATGAGGGT |
| 6769 | mamo-L | *BGIBMGA013888* | up | 1375 | 1389 | + | 11.4435 | 4.31E-05 | 1 | TAGGCTTTTACTGGT |
| 6770 | mamo-L | *BGIBMGA013889* | down | 284 | 298 | + | 10.7258 | 8.23E-05 | 1 | AAGGCTCATGTTAGT |
| 6771 | mamo-L | *BGIBMGA013891* | up | 1897 | 1911 | + | 10.5565 | 9.46E-05 | 1 | TAGGCCGTACTTTTT |
| 6772 | mamo-L | *BGIBMGA013893* | down | 1685 | 1699 | - | 10.8145 | 7.63E-05 | 1 | TAGGCATATGTTGTA |
| 6773 | mamo-L | *BGIBMGA013897* | down | 348 | 362 | - | 11.9435 | 2.65E-05 | 1 | TAGGCTGACTTTCGA |
| 6774 | mamo-L | *BGIBMGA013898* | up | 1493 | 1507 | + | 11.7661 | 3.16E-05 | 1 | TAGGCGTAGGGTACT |
| 6775 | mamo-L | *BGIBMGA013901* | up | 239 | 253 | - | 11.7903 | 3.08E-05 | 1 | TAGGCTGTTTTTTGG |
| 6776 | mamo-L | *BGIBMGA013902* | up | 1553 | 1567 | - | 11.0242 | 6.36E-05 | 1 | AAGGCTGTTTTTACT |
| 6777 | mamo-L | *BGIBMGA013903* | down | 953 | 967 | + | 10.9919 | 6.55E-05 | 1 | TAGGCCAGGTTTTAC |
| 6778 | mamo-L | *BGIBMGA013904* | up | 758 | 772 | + | 10.9919 | 6.55E-05 | 1 | TAGGCCAGGTTTTAC |
| 6779 | mamo-L | *BGIBMGA013906* | up | 1297 | 1311 | - | 10.8629 | 7.33E-05 | 1 | TAGGCTAATTTGTTT |
| 6780 | mamo-L | *BGIBMGA013909* | down | 217 | 231 | - | 10.5968 | 9.15E-05 | 1 | GAGGCTAGTTTTTGT |
| 6781 | mamo-L | *BGIBMGA013911* | down | 1801 | 1815 | - | 12.9919 | 8.10E-06 | 1 | TAGGCTATCTTTTGT |
| 6782 | mamo-L | *BGIBMGA013911* | up | 1732 | 1746 | - | 10.6774 | 8.56E-05 | 1 | AAGGCGTTTTGTGTT |
| 6783 | mamo-L | *BGIBMGA013912* | up | 1669 | 1683 | - | 12.2177 | 1.99E-05 | 1 | TAGGCCATTGTTGCC |
| 6784 | mamo-L | *BGIBMGA013914* | down | 1815 | 1829 | - | 12.4032 | 1.61E-05 | 1 | AAGGCGTATTTTGTT |
| 6785 | mamo-L | *BGIBMGA013914* | down | 1091 | 1105 | + | 11.2097 | 5.38E-05 | 1 | TAGGCCTTTTGTTTC |
| 6786 | mamo-L | *BGIBMGA013915* | down | 124 | 138 | - | 10.5968 | 9.15E-05 | 1 | GAGGCTAGTTTTTGT |
| 6787 | mamo-L | *BGIBMGA013916* | down | 979 | 993 | + | 13.3952 | 4.84E-06 | 1 | TAGGCTTTGTTTACT |
| 6788 | mamo-L | *BGIBMGA013917* | down | 158 | 172 | - | 11.2339 | 5.26E-05 | 1 | GAGGCGTTCTTCGCT |
| 6789 | mamo-L | *BGIBMGA013918* | up | 1084 | 1098 | + | 12.5081 | 1.43E-05 | 1 | TAGGCTAATTTTTTT |
| 6790 | mamo-L | *BGIBMGA013922* | down | 156 | 170 | - | 15.5323 | 1.20E-07 | 1 | TAGGCCTAGTTTGCT |
| 6791 | mamo-L | *BGIBMGA013922* | down | 163 | 177 | + | 10.8065 | 7.69E-05 | 1 | TAGGCCTAGGGGCGT |
| 6792 | mamo-L | *BGIBMGA013924* | down | 254 | 268 | + | 11.7581 | 3.18E-05 | 1 | AAGGCTTTTGTTTCT |
| 6793 | mamo-L | *BGIBMGA013929* | down | 1694 | 1708 | - | 10.75 | 8.06E-05 | 1 | GAGGCCGATTGTGCC |
| 6794 | mamo-L | *BGIBMGA013929* | up | 1215 | 1229 | - | 10.6774 | 8.56E-05 | 1 | AAGGCTTTTATTGTC |
| 6795 | mamo-L | *BGIBMGA013930* | down | 244 | 258 | - | 10.5968 | 9.15E-05 | 1 | GAGGCTAGTTTTTGT |
| 6796 | mamo-L | *BGIBMGA013931* | up | 1382 | 1396 | + | 12.1774 | 2.08E-05 | 1 | TAGGCTTAGTTGTAT |
| 6797 | mamo-L | *BGIBMGA013935* | down | 1585 | 1599 | - | 11.5 | 4.08E-05 | 1 | TAGGCAGCGGTTTGT |
| 6798 | mamo-L | *BGIBMGA013936* | up | 583 | 597 | - | 11.5 | 4.08E-05 | 1 | TAGGCAGCGGTTTGT |
| 6799 | mamo-L | *BGIBMGA013939* | up | 1014 | 1028 | - | 10.6855 | 8.51E-05 | 1 | TAGGCTTTGTCCGAT |
| 6800 | mamo-L | *BGIBMGA013941* | up | 1235 | 1249 | + | 10.6613 | 8.67E-05 | 1 | TAGGCCTGCAGCGTC |
| 6801 | mamo-L | *BGIBMGA013942* | up | 883 | 897 | + | 14.2339 | 1.38E-06 | 1 | TAGGCCTAGTTGGGT |

| | | | | | | | | | | |
|---|---|---|---|---|---|---|---|---|---|---|
| 6802 | mamo-L | *BGIBMGA013942* | down | 1157 | 1171 | + | 10.5887 | 9.21E-05 | 1 | AAGGCTGAGGTAGTT |
| 6803 | mamo-L | *BGIBMGA013944* | down | 1267 | 1281 | - | 10.8306 | 7.52E-05 | 1 | AAGGCCGGTTTCGCC |
| 6804 | mamo-L | *BGIBMGA013947* | up | 1003 | 1017 | - | 10.9032 | 7.07E-05 | 1 | TAGGCGTTTAGTTGT |
| 6805 | mamo-L | *BGIBMGA013947* | down | 1022 | 1036 | + | 10.5726 | 9.33E-05 | 1 | TAGGCTAATATTTAT |
| 6806 | mamo-L | *BGIBMGA013949* | down | 175 | 189 | + | 11.1935 | 5.45E-05 | 1 | TAGGCGCATGAGGGT |
| 6807 | mamo-L | *BGIBMGA013952* | up | 492 | 506 | + | 10.5806 | 9.27E-05 | 1 | TAGGCAAAGGGTGTT |
| 6808 | mamo-L | *BGIBMGA013953* | down | 1497 | 1511 | - | 10.9113 | 7.02E-05 | 1 | TAGGCTGTATGCTGT |
| 6809 | mamo-L | *BGIBMGA013954* | up | 637 | 651 | - | 12.8387 | 9.67E-06 | 1 | TAGGCTCGGTTTACT |
| 6810 | mamo-L | *BGIBMGA013956* | down | 181 | 195 | - | 10.9194 | 6.98E-05 | 1 | TAGGCTACTTTTTTT |
| 6811 | mamo-L | *BGIBMGA013958* | up | 1861 | 1875 | - | 10.5 | 9.90E-05 | 1 | TAGGCAGCTTTTTTT |
| 6812 | mamo-L | *BGIBMGA013961* | down | 334 | 348 | - | 10.9032 | 7.07E-05 | 1 | AAGGCTGACTAGGTT |
| 6813 | mamo-L | *BGIBMGA013963* | up | 1871 | 1885 | - | 11.1694 | 5.59E-05 | 1 | AAGGCTTTCATTGTC |
| 6814 | mamo-L | *BGIBMGA013964* | down | 1984 | 1998 | - | 11.4435 | 4.31E-05 | 1 | AAGGCCTTTTTTTCT |
| 6815 | mamo-L | *BGIBMGA013967* | down | 859 | 873 | + | 10.879 | 7.22E-05 | 1 | TGGGCTGTTGTTGAT |
| 6816 | mamo-L | *BGIBMGA013971* | down | 1765 | 1779 | + | 12.0968 | 2.26E-05 | 1 | TAGGCCGTCGTCGCA |
| 6817 | mamo-L | *BGIBMGA013971* | down | 29 | 43 | + | 11.5484 | 3.90E-05 | 1 | TAGGCCGGTGGCTCT |
| 6818 | mamo-L | *BGIBMGA013976* | up | 95 | 109 | - | 10.9194 | 6.98E-05 | 1 | TAGGCTACTTTTTTT |
| 6819 | mamo-L | *BGIBMGA013976* | up | 373 | 387 | - | 10.5081 | 9.83E-05 | 1 | AAGGCGTGGGGTTTT |
| 6820 | mamo-L | *BGIBMGA013978* | down | 1044 | 1058 | + | 10.6694 | 8.62E-05 | 1 | TAGGCATTGTATTCT |
| 6821 | mamo-L | *BGIBMGA013978* | up | 1529 | 1543 | - | 10.6532 | 8.73E-05 | 1 | TAGGCCGCGCGTGGT |
| 6822 | mamo-L | *BGIBMGA013980* | up | 1708 | 1722 | + | 12.7177 | 1.12E-05 | 1 | TAGGCTTAGTTCTGC |
| 6823 | mamo-L | *BGIBMGA013981* | up | 1658 | 1672 | + | 10.5726 | 9.33E-05 | 1 | TAGGCTAATATTTAT |
| 6824 | mamo-L | *BGIBMGA013982* | up | 111 | 125 | - | 12.25 | 1.92E-05 | 1 | AAGGCTTATTTTGAT |
| 6825 | mamo-L | *BGIBMGA013982* | up | 439 | 453 | + | 10.5323 | 9.63E-05 | 1 | TAGGCATTTTTTACT |
| 6826 | mamo-L | *BGIBMGA013990* | up | 1661 | 1675 | + | 10.5726 | 9.33E-05 | 1 | TAGGCTAATATTTAT |
| 6827 | mamo-L | *BGIBMGA013999* | down | 729 | 743 | - | 11.3387 | 4.77E-05 | 1 | TAGGCAGGAGTTTTT |
| 6828 | mamo-L | *BGIBMGA014000* | up | 749 | 763 | + | 12.6371 | 1.23E-05 | 1 | TAGGCAGGCGGTGGT |
| 6829 | mamo-L | *BGIBMGA014002* | up | 585 | 599 | - | 11.2661 | 5.10E-05 | 1 | AAGGCTGATATTGCC |
| 6830 | mamo-L | *BGIBMGA014004* | down | 988 | 1002 | + | 11.1935 | 5.45E-05 | 1 | TAGGCGCATGAGGGT |
| 6831 | mamo-L | *BGIBMGA014008* | up | 705 | 719 | + | 11.5323 | 3.96E-05 | 1 | TAGGCGTGAGGTGAT |
| 6832 | mamo-L | *BGIBMGA014009* | up | 1626 | 1640 | + | 10.8306 | 7.52E-05 | 1 | TAGGCCTGATTTTTA |
| 6833 | mamo-L | *BGIBMGA014010* | down | 1217 | 1231 | - | 10.5726 | 9.33E-05 | 1 | TAGGCTAATATTTAT |
| 6834 | mamo-L | *BGIBMGA014011* | down | 35 | 49 | + | 12.9758 | 8.23E-06 | 1 | AAGGCTTAGTTTGGC |
| 6835 | mamo-L | *BGIBMGA014011* | up | 1705 | 1719 | - | 10.5726 | 9.33E-05 | 1 | TAGGCTAATATTTAT |
| 6836 | mamo-L | *BGIBMGA014014* | down | 1876 | 1890 | + | 10.9194 | 6.98E-05 | 1 | TAGGCTACTTTTTTT |
| 6837 | mamo-L | *BGIBMGA014014* | down | 281 | 295 | - | 10.75 | 8.06E-05 | 1 | GAGGCCGATTGTGCC |
| 6838 | mamo-L | *BGIBMGA014015* | down | 888 | 902 | - | 10.9194 | 6.98E-05 | 1 | TAGGCTACTTTTTTT |
| 6839 | mamo-L | *BGIBMGA014015* | down | 1100 | 1114 | - | 10.9194 | 6.98E-05 | 1 | TAGGCTACTTTTTTT |
| 6840 | mamo-L | *BGIBMGA014018* | up | 835 | 849 | + | 12.0645 | 2.34E-05 | 1 | TAGGCGGGCTTCGGA |
| 6841 | mamo-L | *BGIBMGA014020* | up | 207 | 221 | + | 13.0565 | 7.51E-06 | 1 | TAGGCGGGGGGCGGT |
| 6842 | mamo-L | *BGIBMGA014020* | down | 739 | 753 | - | 12.5403 | 1.38E-05 | 1 | TAGGCGCGTTTTTTT |
| 6843 | mamo-L | *BGIBMGA014020* | down | 714 | 728 | + | 10.5645 | 9.40E-05 | 1 | TAGGCCTACTGGGGA |
| 6844 | mamo-L | *BGIBMGA014027* | down | 1448 | 1462 | - | 11.621 | 3.63E-05 | 1 | TAGGCAGACGGTTGT |
| 6845 | mamo-L | *BGIBMGA014032* | down | 1640 | 1654 | - | 11.121 | 5.84E-05 | 1 | TAGGCCTATTTTAGA |
| 6846 | mamo-L | *BGIBMGA014033* | up | 459 | 473 | + | 11.4435 | 4.31E-05 | 1 | TAGGCTTGAGAGGCT |
| 6847 | mamo-L | *BGIBMGA014033* | up | 489 | 503 | + | 10.7661 | 7.95E-05 | 1 | TAGGCCCACGAGTTT |
| 6848 | mamo-L | *BGIBMGA014035* | down | 810 | 824 | - | 11.2823 | 5.03E-05 | 1 | TAGGCGGGGATTTAC |
| 6849 | mamo-L | *BGIBMGA014036* | down | 1318 | 1332 | - | 11.4113 | 4.45E-05 | 1 | TAGGCTTCTAGTGGT |
| 6850 | mamo-L | *BGIBMGA014036* | up | 882 | 896 | - | 11.1613 | 5.63E-05 | 1 | GAGGCTGGCTACGTT |
| 6851 | mamo-L | *BGIBMGA014038* | down | 1413 | 1427 | + | 10.7581 | 8.00E-05 | 1 | AAGGCTTTTTTTATT |
| 6852 | mamo-L | *BGIBMGA014039* | down | 207 | 221 | + | 12.8387 | 9.67E-06 | 1 | TAGGCTTAAGTTAGT |
| 6853 | mamo-L | *BGIBMGA014045* | up | 999 | 1013 | + | 10.5323 | 9.63E-05 | 1 | AAGGCCCAGGTCGTC |
| 6854 | mamo-L | *BGIBMGA014046* | down | 850 | 864 | + | 12.5726 | 1.33E-05 | 1 | TAGGCTCTGTTCTGT |
| 6855 | mamo-L | *BGIBMGA014046* | down | 1641 | 1655 | + | 10.5726 | 9.33E-05 | 1 | TAGGCTAATATTTAT |
| 6856 | mamo-L | *BGIBMGA014047* | up | 1013 | 1027 | - | 11.2984 | 4.95E-05 | 1 | TAGGCTGATAGTAGT |
| 6857 | mamo-L | *BGIBMGA014047* | down | 961 | 975 | + | 10.5645 | 9.40E-05 | 1 | TGGGCTTGTTTCGGT |
| 6858 | mamo-L | *BGIBMGA014048* | down | 1446 | 1460 | + | 13.4194 | 4.70E-06 | 1 | TAGGCCAATTTTGTT |
| 6859 | mamo-L | *BGIBMGA014049* | down | 1757 | 1771 | - | 13.4194 | 4.70E-06 | 1 | TAGGCCAATTTTGTT |
| 6860 | mamo-L | *BGIBMGA014053* | up | 1341 | 1355 | + | 10.6371 | 8.85E-05 | 1 | TCGGCCTAGGTTTGT |
| 6861 | mamo-L | *BGIBMGA014054* | down | 522 | 536 | - | 11.3548 | 4.69E-05 | 1 | AAGGCTGATGATGAT |
| 6862 | mamo-L | *BGIBMGA014055* | down | 769 | 783 | + | 10.621 | 8.97E-05 | 1 | TGGGCTTTTTTTGGC |
| 6863 | mamo-L | *BGIBMGA014056* | up | 553 | 567 | - | 10.75 | 8.06E-05 | 1 | GAGGCCGATTGTGCC |
| 6864 | mamo-L | *BGIBMGA014059* | up | 640 | 654 | + | 11.129 | 5.79E-05 | 1 | TAGGCTTCTTTGGAT |
| 6865 | mamo-L | *BGIBMGA014063* | down | 149 | 163 | - | 10.5726 | 9.33E-05 | 1 | TAGGCTAATATTTAT |
| 6866 | mamo-L | *BGIBMGA014065* | down | 991 | 1005 | + | 10.9677 | 6.68E-05 | 1 | TAGGCCCTTGGGGTT |
| 6867 | mamo-L | *BGIBMGA014065* | down | 17 | 31 | - | 10.5242 | 9.70E-05 | 1 | TAGGCTAATGATGGA |
| 6868 | mamo-L | *BGIBMGA014067* | up | 99 | 113 | + | 12.5242 | 1.40E-05 | 1 | TAGGCCCATTTTAGT |
| 6869 | mamo-L | *BGIBMGA014068* | down | 14 | 28 | - | 14.5887 | 7.72E-07 | 1 | TAGGCGGAGTTTTGT |
| 6870 | mamo-L | *BGIBMGA014070* | up | 1601 | 1615 | - | 11.4435 | 4.31E-05 | 1 | AAGGCCTTTTTTTCT |
| 6871 | mamo-L | *BGIBMGA014071* | down | 1129 | 1143 | + | 11.2742 | 5.06E-05 | 1 | AAGGCTTTCGTCGAT |
| 6872 | mamo-L | *BGIBMGA014071* | down | 1953 | 1967 | - | 10.9194 | 6.98E-05 | 1 | TAGGCTACTTTTTTT |
| 6873 | mamo-L | *BGIBMGA014074* | up | 992 | 1006 | + | 11.25 | 5.18E-05 | 1 | TAGGCAGATGGTGGC |
| 6874 | mamo-L | *BGIBMGA014074* | up | 1011 | 1025 | - | 11.0161 | 6.40E-05 | 1 | TAGGCCTGCACTGGC |
| 6875 | mamo-L | *BGIBMGA014078* | down | 1267 | 1281 | + | 11.2016 | 5.41E-05 | 1 | TAGGCATATTTTAGT |
| 6876 | mamo-L | *BGIBMGA014090* | down | 276 | 290 | + | 10.629 | 8.91E-05 | 1 | TAGGCCTGCAGCGCC |
| 6877 | mamo-L | *BGIBMGA014093* | up | 167 | 181 | + | 11.3548 | 4.69E-05 | 1 | AAGGCTGATGATGAT |
| 6878 | mamo-L | *BGIBMGA014093* | down | 614 | 628 | + | 10.5242 | 9.70E-05 | 1 | TAGGCTTTTGTGAAT |
| 6879 | mamo-L | *BGIBMGA014094* | down | 951 | 965 | + | 12.8468 | 9.58E-06 | 1 | AAGGCTGATGTTGGC |
| 6880 | mamo-L | *BGIBMGA014096* | up | 672 | 686 | + | 10.5726 | 9.33E-05 | 1 | TAGGCTAATATTTAT |
| 6881 | mamo-L | *BGIBMGA014097* | down | 1522 | 1536 | - | 11.0081 | 6.45E-05 | 1 | TAGGCCGCCGTTTCA |
| 6882 | mamo-L | *BGIBMGA014107* | up | 1185 | 1199 | - | 11.0242 | 6.36E-05 | 1 | AAGGCCTTGTTTTCC |
| 6883 | mamo-L | *BGIBMGA014110* | up | 1333 | 1347 | + | 13.1452 | 6.69E-06 | 1 | TAGGCTGTTGATGAT |
| 6884 | mamo-L | *BGIBMGA014113* | up | 546 | 560 | - | 11.9113 | 2.74E-05 | 1 | GAGGCTGACTTTTAT |
| 6885 | mamo-L | *BGIBMGA014114* | up | 196 | 210 | + | 10.5887 | 9.21E-05 | 1 | TAGGCTTTGTCTGTA |
| 6886 | mamo-L | *BGIBMGA014114* | up | 961 | 975 | - | 10.5726 | 9.33E-05 | 1 | TAGGCTAATATTTAT |
| 6887 | mamo-L | *BGIBMGA014116* | up | 1075 | 1089 | + | 12.0645 | 2.34E-05 | 1 | TAGGCGGGCTTCGGA |
| 6888 | mamo-L | *BGIBMGA014116* | down | 1783 | 1797 | + | 11.9758 | 2.57E-05 | 1 | AAGGCTTTTTTTTGT |
| 6889 | mamo-L | *BGIBMGA014123* | down | 1431 | 1445 | - | 12.0887 | 2.29E-05 | 1 | TAGGCCGGCTAGGGC |
| 6890 | mamo-L | *BGIBMGA014123* | up | 1138 | 1152 | - | 11.129 | 5.79E-05 | 1 | TAGGCTTTTACTGTT |
| 6891 | mamo-L | *BGIBMGA014123* | up | 1013 | 1027 | - | 10.629 | 8.91E-05 | 1 | TAGGCTGCTGTGTGC |
| 6892 | mamo-L | *BGIBMGA014124* | up | 1966 | 1980 | + | 12.6613 | 1.19E-05 | 1 | TAGGCGTTCGATGAT |
| 6893 | mamo-L | *BGIBMGA014125* | up | 1168 | 1182 | + | 10.5887 | 9.21E-05 | 1 | AAGGCACTCGTTGGT |

| | | | | | | | | | | |
|---|---|---|---|---|---|---|---|---|---|---|
| 6894 | mamo-L | *BGIBMGA014126* | down | 649 | 663 | + | 10.871 | 7.28E-05 | 1 | TAGGCTCTTTTCTAT |
| 6895 | mamo-L | *BGIBMGA014128* | up | 1805 | 1819 | - | 10.9194 | 6.98E-05 | 1 | TAGGCTACTTTTTTT |
| 6896 | mamo-L | *BGIBMGA014130* | down | 1656 | 1670 | + | 11.4919 | 4.11E-05 | 1 | TAGGCGGTAATTTTT |
| 6897 | mamo-L | *BGIBMGA014132* | down | 519 | 533 | + | 10.9032 | 7.07E-05 | 1 | TAGGCTTTGTACTAT |
| 6898 | mamo-L | *BGIBMGA014133* | down | 1835 | 1849 | + | 10.5726 | 9.33E-05 | 1 | TAGGCTAATATTTAT |
| 6899 | mamo-L | *BGIBMGA014135* | up | 1760 | 1774 | + | 10.8387 | 7.47E-05 | 1 | TAGGCTCCCAATGCT |
| 6900 | mamo-L | *BGIBMGA014138* | up | 271 | 285 | - | 11.6371 | 3.58E-05 | 1 | TAGGCCTGCAGCGTT |
| 6901 | mamo-L | *BGIBMGA014139* | up | 582 | 596 | + | 10.75 | 8.06E-05 | 1 | GAGGCCGATTGTGCC |
| 6902 | mamo-L | *BGIBMGA014148* | up | 1374 | 1388 | + | 12.1613 | 2.11E-05 | 1 | TAGGCTTATTTCTGC |
| 6903 | mamo-L | *BGIBMGA014150* | up | 1056 | 1070 | - | 11.1935 | 5.45E-05 | 1 | TAGGCGCATGAGGGT |
| 6904 | mamo-L | *BGIBMGA014150* | down | 1457 | 1471 | + | 10.5887 | 9.21E-05 | 1 | TAGGCTTGTTTTTAG |
| 6905 | mamo-L | *BGIBMGA014150* | up | 1728 | 1742 | + | 10.5726 | 9.33E-05 | 1 | TAGGCTAATATTTAT |
| 6906 | mamo-L | *BGIBMGA014151* | down | 1340 | 1354 | + | 11.3871 | 4.55E-05 | 1 | TAGGCCGTGGTTAGG |
| 6907 | mamo-L | *BGIBMGA014152* | up | 1731 | 1745 | - | 10.5726 | 9.33E-05 | 1 | TAGGCTAATATTTAT |
| 6908 | mamo-L | *BGIBMGA014153* | down | 413 | 427 | + | 10.9194 | 6.98E-05 | 1 | TAGGCTACTTTTTTT |
| 6909 | mamo-L | *BGIBMGA014156* | up | 1759 | 1773 | + | 11.2661 | 5.10E-05 | 1 | AAGGCTTGGGATTTT |
| 6910 | mamo-L | *BGIBMGA014156* | down | 1648 | 1662 | - | 10.5726 | 9.33E-05 | 1 | TAGGCTAATATTTAT |
| 6911 | mamo-L | *BGIBMGA014158* | down | 725 | 739 | - | 11.3226 | 4.83E-05 | 1 | AAGGCTGGTGTTTCC |
| 6912 | mamo-L | *BGIBMGA014158* | up | 1118 | 1132 | - | 10.6452 | 8.79E-05 | 1 | TAGGCCAAGTGTTAT |
| 6913 | mamo-L | *BGIBMGA014160* | down | 1878 | 1892 | - | 10.5726 | 9.33E-05 | 1 | TAGGCTAATATTTAT |
| 6914 | mamo-L | *BGIBMGA014161* | up | 951 | 965 | - | 10.5726 | 9.33E-05 | 1 | TAGGCTAATATTTAT |
| 6915 | mamo-L | *BGIBMGA014168* | down | 1824 | 1838 | - | 12.2016 | 2.03E-05 | 1 | TAGGCTGTAGATGTC |
| 6916 | mamo-L | *BGIBMGA014169* | up | 836 | 850 | - | 12.2016 | 2.03E-05 | 1 | TAGGCTGTAGATGTC |
| 6917 | mamo-L | *BGIBMGA014170* | up | 999 | 1013 | - | 13.7742 | 2.79E-06 | 1 | TAGGCTTAGGTTTCC |
| 6918 | mamo-L | *BGIBMGA014170* | up | 356 | 370 | + | 12.6774 | 1.17E-05 | 1 | TAGGCTGGGGCTTGT |
| 6919 | mamo-L | *BGIBMGA014170* | up | 424 | 438 | - | 10.8387 | 7.47E-05 | 1 | TAGGCAGACGAGGTT |
| 6920 | mamo-L | *BGIBMGA014172* | down | 300 | 314 | - | 10.75 | 8.06E-05 | 1 | GAGGCCGATTGTGCC |
| 6921 | mamo-L | *BGIBMGA014175* | up | 522 | 536 | - | 12.4355 | 1.55E-05 | 1 | TAGGCTGTGTGTGGA |
| 6922 | mamo-L | *BGIBMGA014179* | up | 565 | 579 | - | 11.3871 | 4.55E-05 | 1 | GAGGCCGAATTTGTC |
| 6923 | mamo-L | *BGIBMGA014180* | down | 313 | 327 | + | 10.5726 | 9.33E-05 | 1 | TAGGCTAATATTTAT |
| 6924 | mamo-L | *BGIBMGA014181* | up | 83 | 97 | + | 11.5726 | 3.81E-05 | 1 | TAGGCTTTTTTTTTA |
| 6925 | mamo-L | *BGIBMGA014184* | up | 449 | 463 | - | 10.5968 | 9.15E-05 | 1 | AAGGCTTAATTTAGT |
| 6926 | mamo-L | *BGIBMGA014185* | up | 1495 | 1509 | + | 12.3306 | 1.75E-05 | 1 | TAGGCTTACTATACT |
| 6927 | mamo-L | *BGIBMGA014186* | up | 247 | 261 | + | 10.629 | 8.91E-05 | 1 | AAGGCTAGTTTTTGT |
| 6928 | mamo-L | *BGIBMGA014186* | up | 56 | 70 | - | 10.6129 | 9.03E-05 | 1 | TAGGCTAATATTGGA |
| 6929 | mamo-L | *BGIBMGA014188* | down | 1198 | 1212 | + | 10.5726 | 9.33E-05 | 1 | TAGGCTAATATTTAT |
| 6930 | mamo-L | *BGIBMGA014198* | down | 1489 | 1503 | + | 12.1935 | 2.05E-05 | 1 | TAGGCCGGATTTGCA |
| 6931 | mamo-L | *BGIBMGA014199* | up | 1368 | 1382 | + | 10.7661 | 7.95E-05 | 1 | GAGGCTCACGATTGT |
| 6932 | mamo-L | *BGIBMGA014201* | up | 1357 | 1371 | + | 11.5242 | 3.99E-05 | 1 | AAGGCGTATGATGGT |
| 6933 | mamo-L | *BGIBMGA014201* | up | 800 | 814 | + | 10.5726 | 9.33E-05 | 1 | TAGGCTAATATTTAT |
| 6934 | mamo-L | *BGIBMGA014205* | down | 432 | 446 | + | 11.3468 | 4.73E-05 | 1 | TAGGCTTTTAATTTT |
| 6935 | mamo-L | *BGIBMGA014205* | up | 653 | 667 | + | 10.9194 | 6.98E-05 | 1 | TAGGCTACTTTTTTT |
| 6936 | mamo-L | *BGIBMGA014205* | down | 1050 | 1064 | - | 10.4919 | 9.96E-05 | 1 | TAGGCTTACATCCGC |
| 6937 | mamo-L | *BGIBMGA014205* | up | 1206 | 1220 | - | 10.4919 | 9.96E-05 | 1 | TAGGCTTACATCCGC |
| 6938 | mamo-L | *BGIBMGA014207* | up | 134 | 148 | + | 11.5968 | 3.73E-05 | 1 | AAGGCGGTTTTTTGT |
| 6939 | mamo-L | *BGIBMGA014209* | up | 469 | 483 | - | 11.75 | 3.21E-05 | 1 | TAGGCTCTCTTTCGC |
| 6940 | mamo-L | *BGIBMGA014210* | down | 774 | 788 | - | 12.5726 | 1.33E-05 | 1 | TAGGCAGGGTGTGGT |
| 6941 | mamo-L | *BGIBMGA014210* | down | 1513 | 1527 | + | 10.9194 | 6.98E-05 | 1 | TAGGCTACTTTTTTT |
| 6942 | mamo-L | *BGIBMGA014221* | down | 1014 | 1028 | - | 12.7742 | 1.04E-05 | 1 | TAGGCTGAGTTCATT |
| 6943 | mamo-L | *BGIBMGA014222* | up | 1478 | 1492 | + | 10.9194 | 6.98E-05 | 1 | TAGGCTACTTTTTTT |
| 6944 | mamo-L | *BGIBMGA014224* | up | 1573 | 1587 | + | 10.5968 | 9.15E-05 | 1 | GAGGCTAGTTTTTGT |
| 6945 | mamo-L | *BGIBMGA014226* | down | 942 | 956 | - | 11.5081 | 4.05E-05 | 1 | GAGGCGGTCTTGGGT |
| 6946 | mamo-L | *BGIBMGA014228* | up | 1011 | 1025 | - | 12.9113 | 8.87E-06 | 1 | TAGGCTTATTTCGCC |
| 6947 | mamo-L | *BGIBMGA014231* | up | 1489 | 1503 | - | 11.6694 | 3.47E-05 | 1 | GAGGCTTTCGTGGCT |
| 6948 | mamo-L | *BGIBMGA014231* | down | 980 | 994 | - | 10.5968 | 9.15E-05 | 1 | GAGGCTAGTTTTTGT |
| 6949 | mamo-L | *BGIBMGA014235* | up | 1167 | 1181 | - | 11.1452 | 5.70E-05 | 1 | TAGGCTCATTATTAT |
| 6950 | mamo-L | *BGIBMGA014237* | down | 613 | 627 | + | 11.9032 | 2.76E-05 | 1 | GAGGCCTGGTGTGCT |
| 6951 | mamo-L | *BGIBMGA014238* | up | 384 | 398 | - | 10.5726 | 9.33E-05 | 1 | TAGGCTAATATTTAT |
| 6952 | mamo-L | *BGIBMGA014243* | down | 11 | 25 | - | 10.8065 | 7.69E-05 | 1 | AAGGCGTTTTTCGTT |
| 6953 | mamo-L | *BGIBMGA014243* | down | 226 | 240 | - | 10.5726 | 9.33E-05 | 1 | TAGGCTAATATTTAT |
| 6954 | mamo-L | *BGIBMGA014244* | down | 280 | 294 | - | 10.75 | 8.06E-05 | 1 | GAGGCCGATTGTGCC |
| 6955 | mamo-L | *BGIBMGA014246* | down | 539 | 553 | + | 11.7823 | 3.11E-05 | 1 | TAGGCGTGGGGCGGC |
| 6956 | mamo-L | *BGIBMGA014247* | down | 1234 | 1248 | - | 12.2097 | 2.01E-05 | 1 | TAGGCCTAGGTGCGT |
| 6957 | mamo-L | *BGIBMGA014247* | up | 204 | 218 | + | 11.8871 | 2.81E-05 | 1 | TAGGCTTTCTATCTT |
| 6958 | mamo-L | *BGIBMGA014248* | up | 1924 | 1938 | - | 12.2097 | 2.01E-05 | 1 | TAGGCCTAGGTGCGT |
| 6959 | mamo-L | *BGIBMGA014250* | up | 1430 | 1444 | - | 11.4435 | 4.31E-05 | 1 | TAGGCTTTTTTCCTT |
| 6960 | mamo-L | *BGIBMGA014253* | down | 1030 | 1044 | + | 10.621 | 8.97E-05 | 1 | TAGGCTAGATATTGT |
| 6961 | mamo-L | *BGIBMGA014258* | down | 1951 | 1965 | - | 10.6371 | 8.85E-05 | 1 | AAGGCCTTTATTGAT |
| 6962 | mamo-L | *BGIBMGA014260* | down | 487 | 501 | + | 12.6935 | 1.15E-05 | 1 | AAGGCTGGCTTTTTT |
| 6963 | mamo-L | *BGIBMGA014263* | up | 55 | 69 | - | 12.879 | 9.26E-06 | 1 | TAGGCTCAGTTTTTC |
| 6964 | mamo-L | *BGIBMGA014263* | down | 1867 | 1881 | + | 11.2339 | 5.26E-05 | 1 | TAGGCTACTTTTTGT |
| 6965 | mamo-L | *BGIBMGA014264* | up | 329 | 343 | + | 11.2339 | 5.26E-05 | 1 | TAGGCTACTTTTTGT |
| 6966 | mamo-L | *BGIBMGA014265* | down | 1091 | 1105 | + | 10.7419 | 8.11E-05 | 1 | TAGGCAGTCGTGTTT |
| 6967 | mamo-L | *BGIBMGA014266* | down | 729 | 743 | + | 11.3387 | 4.77E-05 | 1 | TAGGCGTAATGTGTC |
| 6968 | mamo-L | *BGIBMGA014270* | up | 1547 | 1561 | + | 11.3629 | 4.66E-05 | 1 | TAGGCTTTTTGTTCC |
| 6969 | mamo-L | *BGIBMGA014270* | up | 425 | 439 | - | 11.2177 | 5.34E-05 | 1 | TAGGCAGTGTTTGTA |
| 6970 | mamo-L | *BGIBMGA014272* | down | 1090 | 1104 | + | 10.5968 | 9.15E-05 | 1 | GAGGCTAGTTTTTGT |
| 6971 | mamo-L | *BGIBMGA014275* | up | 60 | 74 | + | 10.629 | 8.91E-05 | 1 | TAGGCCTGCAGCGCC |
| 6972 | mamo-L | *BGIBMGA014278* | down | 59 | 73 | - | 13.621 | 3.50E-06 | 1 | TAGGCTGTGTATTGT |
| 6973 | mamo-L | *BGIBMGA014290* | down | 908 | 922 | - | 10.8387 | 7.47E-05 | 1 | TAGGCTTTCACTTGT |
| 6974 | mamo-L | *BGIBMGA014292* | down | 1839 | 1853 | + | 11.6613 | 3.50E-05 | 1 | TAGGCTTAGATTCAT |
| 6975 | mamo-L | *BGIBMGA014293* | up | 799 | 813 | - | 13.8387 | 2.54E-06 | 1 | TAGGCGTTAGTTGGT |
| 6976 | mamo-L | *BGIBMGA014293* | up | 1016 | 1030 | + | 10.9194 | 6.98E-05 | 1 | TAGGCATTTTTGGTT |
| 6977 | mamo-L | *BGIBMGA014296* | down | 1944 | 1958 | + | 14.2823 | 1.31E-06 | 1 | TAGGCTTACTGTGTT |
| 6978 | mamo-L | *BGIBMGA014296* | down | 815 | 829 | + | 12.5726 | 1.33E-05 | 1 | TAGGCCCGGATTGCC |
| 6979 | mamo-L | *BGIBMGA014296* | up | 544 | 558 | + | 11.371 | 4.62E-05 | 1 | TAGGCTGTATAGGCT |
| 6980 | mamo-L | *BGIBMGA014296* | down | 746 | 760 | + | 10.9032 | 7.07E-05 | 1 | TAGGCGGGATTTGAA |
| 6981 | mamo-L | *BGIBMGA014297* | up | 1806 | 1820 | - | 10.5726 | 9.33E-05 | 1 | TAGGCTAATATTTAT |
| 6982 | mamo-L | *BGIBMGA014299* | up | 1211 | 1225 | + | 10.75 | 8.06E-05 | 1 | GAGGCAGACTTTTGT |
| 6983 | mamo-L | *BGIBMGA014300* | up | 899 | 913 | + | 14.7258 | 5.83E-07 | 1 | TAGGCTTGCGTTGCC |
| 6984 | mamo-L | *BGIBMGA014305* | up | 1662 | 1676 | + | 10.5726 | 9.33E-05 | 1 | TAGGCTAATATTTAT |
| 6985 | mamo-L | *BGIBMGA014306* | down | 565 | 579 | - | 12.5081 | 1.43E-05 | 1 | TAGGCCCACAGTGGT |

| | | | | | | | | | | |
|---|---|---|---|---|---|---|---|---|---|---|
| 6986 | mamo-L | *BGIBMGA014307* | up | 14 | 28 | + | 12.5081 | 1.43E-05 | 1 | TAGGCCCACAGTGGT |
| 6987 | mamo-L | *BGIBMGA014307* | down | 358 | 372 | - | 10.9194 | 6.98E-05 | 1 | TAGGCTACTTTTTTT |
| 6988 | mamo-L | *BGIBMGA014308* | down | 183 | 197 | - | 11.9032 | 2.76E-05 | 1 | TAGGCCTGCAGTGCC |
| 6989 | mamo-L | *BGIBMGA014312* | down | 633 | 647 | - | 10.5726 | 9.33E-05 | 1 | TAGGCTAATATTTAT |
| 6990 | mamo-L | *BGIBMGA014314* | down | 1643 | 1657 | - | 12.9758 | 8.23E-06 | 1 | TAGGCTTTATTTTTT |
| 6991 | mamo-L | *BGIBMGA014316* | down | 1674 | 1688 | - | 12.6371 | 1.23E-05 | 1 | TAGGCAGGCGGTGGT |
| 6992 | mamo-L | *BGIBMGA014317* | down | 828 | 842 | + | 10.8306 | 7.52E-05 | 1 | TAGGCCCCCTTCGCC |
| 6993 | mamo-L | *BGIBMGA014324* | up | 1272 | 1286 | - | 13.0081 | 7.94E-06 | 1 | TAGGCCTATTTTATT |
| 6994 | mamo-L | *BGIBMGA014325* | up | 929 | 943 | + | 10.5726 | 9.33E-05 | 1 | TAGGCTAATATTTAT |
| 6995 | mamo-L | *BGIBMGA014327* | up | 422 | 436 | + | 10.5726 | 9.33E-05 | 1 | TAGGCTAATATTTAT |
| 6996 | mamo-L | *BGIBMGA014336* | down | 48 | 62 | + | 10.8306 | 7.52E-05 | 1 | TAGGCGGGCAGTGGA |
| 6997 | mamo-L | *BGIBMGA014338* | down | 540 | 554 | + | 11.5968 | 3.73E-05 | 1 | TAGGCTTAGGTTAGG |
| 6998 | mamo-L | *BGIBMGA014338* | down | 551 | 565 | + | 11.5968 | 3.73E-05 | 1 | TAGGCTTAGGTTAGG |
| 6999 | mamo-L | *BGIBMGA014338* | down | 562 | 576 | + | 11.5968 | 3.73E-05 | 1 | TAGGCTTAGGTTAGG |
| 7000 | mamo-L | *BGIBMGA014338* | down | 595 | 609 | + | 11.5968 | 3.73E-05 | 1 | TAGGCTTAGGTTAGG |
| 7001 | mamo-L | *BGIBMGA014338* | down | 661 | 675 | + | 11.5968 | 3.73E-05 | 1 | TAGGCTTAGGTTAGG |
| 7002 | mamo-L | *BGIBMGA014338* | down | 672 | 686 | + | 11.5968 | 3.73E-05 | 1 | TAGGCTTAGGTTAGG |
| 7003 | mamo-L | *BGIBMGA014338* | down | 705 | 719 | + | 11.5968 | 3.73E-05 | 1 | TAGGCTTAGGTTAGG |
| 7004 | mamo-L | *BGIBMGA014338* | down | 716 | 730 | + | 11.5968 | 3.73E-05 | 1 | TAGGCTTAGGTTAGG |
| 7005 | mamo-L | *BGIBMGA014338* | down | 738 | 752 | + | 11.5968 | 3.73E-05 | 1 | TAGGCTTAGGTTAGG |
| 7006 | mamo-L | *BGIBMGA014358* | down | 527 | 541 | - | 10.7661 | 7.95E-05 | 1 | GAGGCTTGTATTTTT |
| 7007 | mamo-L | *BGIBMGA014358* | up | 223 | 237 | - | 10.5 | 9.90E-05 | 1 | TAGGCGTTTTACTTT |
| 7008 | mamo-L | *BGIBMGA014363* | down | 1360 | 1374 | - | 12.8145 | 9.96E-06 | 1 | TAGGCGGCGGTTTTT |
| 7009 | mamo-L | *BGIBMGA014363* | up | 1810 | 1824 | - | 10.5726 | 9.33E-05 | 1 | TAGGCTAATATTTAT |
| 7010 | mamo-L | *BGIBMGA014372* | up | 611 | 625 | + | 12.5565 | 1.35E-05 | 1 | TAGGCCGAGATTGTA |
| 7011 | mamo-L | *BGIBMGA014373* | down | 1598 | 1612 | + | 11.4032 | 4.49E-05 | 1 | TAGGCTGTATAGGTT |
| 7012 | mamo-L | *BGIBMGA014373* | down | 1122 | 1136 | - | 10.9194 | 6.98E-05 | 1 | TAGGCTACTTTTTTT |
| 7013 | mamo-L | *BGIBMGA014375* | up | 432 | 446 | + | 10.5887 | 9.21E-05 | 1 | GAGGCCGGCGTCCGT |
| 7014 | mamo-L | *BGIBMGA014377* | up | 304 | 318 | - | 10.5726 | 9.33E-05 | 1 | TAGGCTAATATTTAT |
| 7015 | mamo-L | *BGIBMGA014378* | up | 960 | 974 | - | 11.25 | 5.18E-05 | 1 | TAGGCTCTGGGTTCC |
| 7016 | mamo-L | *BGIBMGA014381* | down | 747 | 761 | - | 11.371 | 4.62E-05 | 1 | TAGGCTGTATAGGCT |
| 7017 | mamo-L | *BGIBMGA014382* | down | 1390 | 1404 | + | 12.5 | 1.44E-05 | 1 | TAGGCTATTTTTTGT |
| 7018 | mamo-L | *BGIBMGA014382* | down | 137 | 151 | - | 11.8468 | 2.92E-05 | 1 | TAGGCGGTGTTCTAT |
| 7019 | mamo-L | *BGIBMGA014385* | up | 1558 | 1572 | - | 10.7823 | 7.84E-05 | 1 | TAGGCCCGTGATCTT |
| 7020 | mamo-L | *BGIBMGA014392* | down | 1607 | 1621 | + | 12.2097 | 2.01E-05 | 1 | TAGGCCGGCTATCCT |
| 7021 | mamo-L | *BGIBMGA014392* | down | 1691 | 1705 | + | 12.0645 | 2.34E-05 | 1 | TAGGCGGGCTTCGGA |
| 7022 | mamo-L | *BGIBMGA014396* | down | 544 | 558 | + | 13.3952 | 4.84E-06 | 1 | TAGGCCTGCGTTCTT |
| 7023 | mamo-L | *BGIBMGA014396* | up | 74 | 88 | + | 11.7984 | 3.06E-05 | 1 | TAGGCTTTGCTTTTT |
| 7024 | mamo-L | *BGIBMGA014398* | down | 1387 | 1401 | + | 10.9919 | 6.55E-05 | 1 | TAGGCGTATAATTTT |
| 7025 | mamo-L | *BGIBMGA014401* | up | 1411 | 1425 | - | 10.9194 | 6.98E-05 | 1 | TAGGCTACTTTTTTT |
| 7026 | mamo-L | *BGIBMGA014402* | up | 1777 | 1791 | - | 14.3952 | 1.09E-06 | 1 | TAGGCTGATTTTTTT |
| 7027 | mamo-L | *BGIBMGA014402* | down | 1889 | 1903 | + | 12.25 | 1.92E-05 | 1 | TAGGCCCACTATTCT |
| 7028 | mamo-L | *BGIBMGA014405* | down | 575 | 589 | + | 12.25 | 1.92E-05 | 1 | TAGGCCCACTATTCT |
| 7029 | mamo-L | *BGIBMGA014405* | up | 1583 | 1597 | - | 10.8065 | 7.69E-05 | 1 | TAGGGTGATTTTGTT |
| 7030 | mamo-L | *BGIBMGA014405* | up | 1722 | 1736 | - | 10.5726 | 9.33E-05 | 1 | TAGGCTAATATTTAT |
| 7031 | mamo-L | *BGIBMGA014407* | up | 1656 | 1670 | + | 10.9194 | 6.98E-05 | 1 | TAGGCTACTTTTTTT |
| 7032 | mamo-L | *BGIBMGA014412* | up | 1156 | 1170 | - | 10.8548 | 7.38E-05 | 1 | GAGGCCTGTTTTTAT |
| 7033 | mamo-L | *BGIBMGA014420* | down | 230 | 244 | + | 11.1855 | 5.50E-05 | 1 | TAGGCTGTGGACTTC |
| 7034 | mamo-L | *BGIBMGA014420* | up | 969 | 983 | + | 10.6855 | 8.51E-05 | 1 | TAGGCTTACTTCTCG |
| 7035 | mamo-L | *BGIBMGA014422* | down | 1094 | 1108 | + | 12.9516 | 8.46E-06 | 1 | GAGGCCGACGTTGAT |
| 7036 | mamo-L | *BGIBMGA014422* | down | 631 | 645 | + | 11.7177 | 3.32E-05 | 1 | GAGGCTGACGTCGTC |
| 7037 | mamo-L | *BGIBMGA014423* | down | 1 | 15 | - | 12.0323 | 2.42E-05 | 1 | TAGGCGGACGTGTGC |
| 7038 | mamo-L | *BGIBMGA014426* | up | 1748 | 1762 | - | 12.9516 | 8.46E-06 | 1 | GAGGCCGACGTTGAT |
| 7039 | mamo-L | *BGIBMGA014426* | up | 473 | 487 | + | 11.3548 | 4.69E-05 | 1 | AAGGCTGATGATGAT |
| 7040 | mamo-L | *BGIBMGA014427* | down | 1963 | 1977 | + | 10.7823 | 7.84E-05 | 1 | AAGGCTTTTGATTGT |
| 7041 | mamo-L | *BGIBMGA014427* | down | 336 | 350 | - | 10.75 | 8.06E-05 | 1 | GAGGCCGATTGTGCC |
| 7042 | mamo-L | *BGIBMGA014429* | up | 1500 | 1514 | + | 11.8871 | 2.81E-05 | 1 | TAGGCGTGTGGCGTT |
| 7043 | mamo-L | *BGIBMGA014438* | up | 794 | 808 | + | 11.2097 | 5.38E-05 | 1 | TAGGCGCGCGGGGTT |
| 7044 | mamo-L | *BGIBMGA014441* | up | 534 | 548 | + | 10.9919 | 6.55E-05 | 1 | TAGGCTGCAAATGGT |
| 7045 | mamo-L | *BGIBMGA014444* | up | 1961 | 1975 | - | 10.5726 | 9.33E-05 | 1 | TAGGCTAATATTTAT |
| 7046 | mamo-L | *BGIBMGA014448* | down | 341 | 355 | + | 14.3226 | 1.22E-06 | 1 | TAGGCTGTGTGTGTT |
| 7047 | mamo-L | *BGIBMGA014450* | up | 134 | 148 | + | 11.1855 | 5.50E-05 | 1 | TAGGCTAATTATTTT |
| 7048 | mamo-L | *BGIBMGA014453* | down | 710 | 724 | - | 11.2339 | 5.26E-05 | 1 | TAGGCCAGTTTTTCC |
| 7049 | mamo-L | *BGIBMGA014453* | down | 688 | 702 | + | 10.629 | 8.91E-05 | 1 | TAGGCCTGCAGCGCC |
| 7050 | mamo-L | *BGIBMGA014454* | up | 311 | 325 | - | 11.2339 | 5.26E-05 | 1 | TAGGCCAGTTTTTCC |
| 7051 | mamo-L | *BGIBMGA014454* | up | 289 | 303 | + | 10.629 | 8.91E-05 | 1 | TAGGCCTGCAGCGCC |
| 7052 | mamo-L | *BGIBMGA014462* | up | 1851 | 1865 | - | 15.5081 | 1.26E-07 | 1 | TAGGCTGGTGTTGCT |
| 7053 | mamo-L | *BGIBMGA014470* | down | 1804 | 1818 | - | 10.9839 | 6.59E-05 | 1 | TAGGCTTTTTTTAGA |
| 7054 | mamo-L | *BGIBMGA014470* | down | 1032 | 1046 | - | 10.6048 | 9.09E-05 | 1 | TAGGCGGACGGTGAG |
| 7055 | mamo-L | *BGIBMGA014471* | up | 577 | 591 | - | 10.9839 | 6.59E-05 | 1 | TAGGCTTTTTTTAGA |
| 7056 | mamo-L | *BGIBMGA014473* | down | 1625 | 1639 | + | 11.2742 | 5.06E-05 | 1 | TAGGCTCTGGTGCGT |
| 7057 | mamo-L | *BGIBMGA014474* | up | 1969 | 1983 | + | 10.5242 | 9.70E-05 | 1 | TAGGCCGTTAATACT |
| 7058 | mamo-L | *BGIBMGA014475* | up | 1667 | 1681 | + | 11.1694 | 5.59E-05 | 1 | TAGGCTGAAAATTTT |
| 7059 | mamo-L | *BGIBMGA014477* | up | 1071 | 1085 | - | 10.9032 | 7.07E-05 | 1 | TAGGCTTTTCGTGCT |
| 7060 | mamo-L | *BGIBMGA014477* | up | 1084 | 1098 | + | 10.7823 | 7.84E-05 | 1 | TAGGCTAAGTTTTGG |
| 7061 | mamo-L | *BGIBMGA014477* | up | 1975 | 1989 | - | 10.7016 | 8.39E-05 | 1 | TAGGCTGCGATGGCC |
| 7062 | mamo-L | *BGIBMGA014478* | up | 35 | 49 | - | 10.9194 | 6.98E-05 | 1 | TAGGCTACTTTTTTT |
| 7063 | mamo-L | *BGIBMGA014481* | down | 395 | 409 | + | 13.4677 | 4.39E-06 | 1 | TAGGCTGATGATGAT |
| 7064 | mamo-L | *BGIBMGA014483* | down | 927 | 941 | - | 10.6774 | 8.56E-05 | 1 | TAGGCCGTAATCTCT |
| 7065 | mamo-L | *BGIBMGA014485* | up | 717 | 731 | - | 10.75 | 8.06E-05 | 1 | GAGGCCGATTGTGCC |
| 7066 | mamo-L | *BGIBMGA014489* | down | 1322 | 1336 | - | 10.5726 | 9.33E-05 | 1 | TAGGCTAATATTTAT |
| 7067 | mamo-L | *BGIBMGA014494* | down | 665 | 679 | + | 11.3468 | 4.73E-05 | 1 | TAGGCTTTTAATTTT |
| 7068 | mamo-L | *BGIBMGA014494* | down | 705 | 719 | - | 10.7823 | 7.84E-05 | 1 | TAGGCTGTTGGCCGT |
| 7069 | mamo-L | *BGIBMGA014495* | down | 1573 | 1587 | - | 11.5242 | 3.99E-05 | 1 | TAGGCTGTTGGCTTT |
| 7070 | mamo-L | *BGIBMGA014495* | down | 897 | 911 | - | 10.7823 | 7.84E-05 | 1 | TAGGCTGTTGGCCGT |
| 7071 | mamo-L | *BGIBMGA014497* | down | 1839 | 1853 | + | 11.5242 | 3.99E-05 | 1 | TAGGCTGTTGGCTTT |
| 7072 | mamo-L | *BGIBMGA014500* | up | 202 | 216 | + | 12.4355 | 1.55E-05 | 1 | TAGGCTGGTTTTATC |
| 7073 | mamo-L | *BGIBMGA014500* | up | 697 | 711 | + | 11.0726 | 6.09E-05 | 1 | TAGGCTGGCTAAGCT |
| 7074 | mamo-L | *BGIBMGA014500* | up | 1840 | 1854 | - | 10.5726 | 9.33E-05 | 1 | TAGGCTAATATTTAT |
| 7075 | mamo-L | *BGIBMGA014505* | up | 1957 | 1971 | + | 11.5726 | 3.81E-05 | 1 | GAGGCGTTCGTTTTT |
| 7076 | mamo-L | *BGIBMGA014508* | down | 1620 | 1634 | + | 10.9194 | 6.98E-05 | 1 | TAGGCTACTTTTTTT |
| 7077 | mamo-L | *BGIBMGA014513* | up | 1319 | 1333 | - | 11.1613 | 5.63E-05 | 1 | TAGGCCCAGTATGGG |

| | | | | | | | | | | |
|---|---|---|---|---|---|---|---|---|---|---|
| 7078 | mamo-L | *BGIBMGA014517* | up | 587 | 601 | + | 11.9516 | 2.63E-05 | 1 | TAGGCCGACATGTTT |
| 7079 | mamo-L | *BGIBMGA014517* | up | 279 | 293 | + | 10.5645 | 9.40E-05 | 1 | TAGGCTTATTACACT |
| 7080 | mamo-L | *BGIBMGA014518* | up | 474 | 488 | - | 10.7581 | 8.00E-05 | 1 | TAGGCATCGTTTTTT |
| 7081 | mamo-L | *BGIBMGA014520* | up | 655 | 669 | + | 11.5403 | 3.93E-05 | 1 | TAGGCTTTCGATGCG |
| 7082 | mamo-L | *BGIBMGA014520* | down | 1335 | 1349 | - | 10.9194 | 6.98E-05 | 1 | TAGGCTACTTTTTTT |
| 7083 | mamo-L | *BGIBMGA014522* | up | 236 | 250 | + | 10.9194 | 6.98E-05 | 1 | AAGGCCTGATTTTTT |
| 7084 | mamo-L | *BGIBMGA014529* | down | 1297 | 1311 | - | 12.0645 | 2.34E-05 | 1 | TAGGCGGTTTTGTGT |
| 7085 | mamo-L | *BGIBMGA014529* | down | 1168 | 1182 | + | 10.5403 | 9.58E-05 | 1 | TAGGCTATGTTTTTA |
| 7086 | mamo-L | *BGIBMGA014531* | down | 1006 | 1020 | - | 13.2419 | 5.96E-06 | 1 | TAGGCCTTCGGTGGC |
| 7087 | mamo-L | *BGIBMGA014531* | down | 1041 | 1055 | + | 11.3548 | 4.69E-05 | 1 | AAGGCTGATGATGAT |
| 7088 | mamo-L | *BGIBMGA014532* | up | 1116 | 1130 | - | 13.2419 | 5.96E-06 | 1 | TAGGCCTTCGGTGGC |
| 7089 | mamo-L | *BGIBMGA014532* | up | 1151 | 1165 | + | 11.3548 | 4.69E-05 | 1 | AAGGCTGATGATGAT |
| 7090 | mamo-L | *BGIBMGA014534* | down | 272 | 286 | - | 11.0645 | 6.14E-05 | 1 | TAGGCTTCCGTCGGA |
| 7091 | mamo-L | *BGIBMGA014536* | up | 666 | 680 | + | 11.5403 | 3.93E-05 | 1 | AAGGCTTACTTTACT |
| 7092 | mamo-L | *BGIBMGA014536* | up | 1317 | 1331 | - | 10.8629 | 7.33E-05 | 1 | TAGGCTATTTATTTT |
| 7093 | mamo-L | *BGIBMGA014539* | up | 1634 | 1648 | + | 10.7177 | 8.28E-05 | 1 | TAGGCTTAGTTTAAA |
| 7094 | mamo-L | *BGIBMGA014544* | up | 1755 | 1769 | - | 11.629 | 3.61E-05 | 1 | TAGGCTGACGACGGA |
| 7095 | mamo-L | *BGIBMGA014545* | up | 188 | 202 | + | 10.8629 | 7.33E-05 | 1 | AAGGCTTGCTTTGCG |
| 7096 | mamo-L | *BGIBMGA014547* | up | 527 | 541 | + | 11 | 6.50E-05 | 1 | TAGGCATGCGTGTGT |
| 7097 | mamo-L | *BGIBMGA014548* | down | 661 | 675 | - | 10.5726 | 9.33E-05 | 1 | TAGGCTAATATTTAT |
| 7098 | mamo-L | *BGIBMGA014552* | down | 1183 | 1197 | - | 11.9274 | 2.69E-05 | 1 | TAGGCTCTGGTTAAT |
| 7099 | mamo-L | *BGIBMGA014553* | down | 1198 | 1212 | + | 11.5484 | 3.90E-05 | 1 | TAGGCTCTGGATGGA |
| 7100 | mamo-L | *BGIBMGA014557* | down | 344 | 358 | - | 11.3871 | 4.55E-05 | 1 | TAGGCGTAGTGTGGG |
| 7101 | mamo-L | *BGIBMGA014559* | down | 85 | 99 | + | 10.5 | 9.90E-05 | 1 | TAGGCTTATTCCTGT |
| 7102 | mamo-L | *BGIBMGA014565* | down | 204 | 218 | + | 10.871 | 7.28E-05 | 1 | TAGGCTGCTATTTAT |
| 7103 | mamo-L | *BGIBMGA014566* | up | 1641 | 1655 | + | 10.871 | 7.28E-05 | 1 | TAGGCTGCTATTTAT |
| 7104 | mamo-L | *BGIBMGA014571* | up | 1192 | 1206 | + | 13.0726 | 7.34E-06 | 1 | TAGGCCGGCGTGTCT |
| 7105 | mamo-L | *BGIBMGA014571* | down | 515 | 529 | + | 10.5 | 9.90E-05 | 1 | GAGGCCCAGGTCGTC |
| 7106 | mamo-L | *BGIBMGA014574* | up | 934 | 948 | - | 12.7258 | 1.11E-05 | 1 | TAGGCGTGCTTTGTA |
| 7107 | mamo-L | *BGIBMGA014574* | up | 1567 | 1581 | - | 10.7177 | 8.28E-05 | 1 | TAGGCTATTCTTGCT |
| 7108 | mamo-L | *BGIBMGA014576* | down | 1588 | 1602 | - | 11.9113 | 2.74E-05 | 1 | TAGGCCTCGATCGGT |
| 7109 | mamo-L | *BGIBMGA014594* | down | 854 | 868 | + | 11.3306 | 4.80E-05 | 1 | TAGGCCCGGTATACT |
| 7110 | mamo-L | *BGIBMGA014594* | up | 1270 | 1284 | + | 10.4919 | 9.96E-05 | 1 | TAGGCTAGTATTTAT |
| 7111 | mamo-L | *BGIBMGA014599* | up | 1540 | 1554 | + | 10.9758 | 6.64E-05 | 1 | AAGGCTGGTTCTGGT |
| 7112 | mamo-L | *BGIBMGA014600* | down | 550 | 564 | + | 12.0484 | 2.38E-05 | 1 | AAGGCTAAGTTTGTT |
| 7113 | mamo-L | *BGIBMGA014606* | down | 1409 | 1423 | - | 12.4032 | 1.61E-05 | 1 | AAGGCGTATTTTGTT |
| 7114 | mamo-L | *BGIBMGA014607* | up | 16 | 30 | - | 12.4032 | 1.61E-05 | 1 | AAGGCGTATTTTGTT |
| 7115 | mamo-L | *BGIBMGA014610* | down | 608 | 622 | - | 12.6129 | 1.26E-05 | 1 | TAGGCCATCTTTGTC |
| 7116 | mamo-L | *BGIBMGA014611* | down | 1620 | 1634 | + | 11.0726 | 6.09E-05 | 1 | TAGGCGGGGCTTAGT |
| 7117 | mamo-L | *BGIBMGA014612* | up | 950 | 964 | + | 11.0726 | 6.09E-05 | 1 | TAGGCGGGGCTTAGT |
| 7118 | mamo-L | *BGIBMGA014612* | down | 131 | 145 | - | 10.6774 | 8.56E-05 | 1 | TAGGCCAGTTTTAGC |
| 7119 | mamo-L | *BGIBMGA014613* | up | 220 | 234 | - | 10.6774 | 8.56E-05 | 1 | TAGGCCAGTTTTAGC |
| 7120 | mamo-L | *BGIBMGA014616* | down | 46 | 60 | - | 10.8226 | 7.58E-05 | 1 | TAGGCATAGTTGGTC |
| 7121 | mamo-L | *BGIBMGA014617* | down | 1472 | 1486 | + | 10.9194 | 6.98E-05 | 1 | TAGGCTACTTTTTTT |
| 7122 | mamo-L | *BGIBMGA014617* | up | 935 | 949 | - | 10.8629 | 7.33E-05 | 1 | TAGGCAGAGGATAGT |
| 7123 | mamo-L | *BGIBMGA014618* | down | 1175 | 1189 | + | 10.75 | 8.06E-05 | 1 | TAGGCGTATATTAGC |
| 7124 | mamo-L | *BGIBMGA014619* | up | 1210 | 1224 | + | 10.5 | 9.90E-05 | 1 | GAGGCCCAGGTCGTC |
| 7125 | mamo-L | *BGIBMGA014621* | down | 59 | 73 | - | 11.4113 | 4.45E-05 | 1 | TAGGCTATTGTTATT |

**Supplementary Table S4 The binding site analysis of Bm-mamo-s**

Using the position weight matrix of mamo-s protein to search the 2000 base pair regions upstream and downstream of the silkworm prediction gene.

| Number | motif_id | gene ID | up/down stream | start | stop | strand | score | p-value | q-value | matched_sequence |
|---|---|---|---|---|---|---|---|---|---|---|
| 1 | mamo-s | *BGIBMGA000001* | down | 684 | 692 | - | 11.3303 | 4.96E-05 | 0.615 | GTGGGTGGC |
| 2 | mamo-s | *BGIBMGA000004* | down | 235 | 243 | + | 11.3303 | 4.96E-05 | 0.615 | GTGGGTGGC |
| 3 | mamo-s | *BGIBMGA000006* | up | 300 | 308 | + | 10.9358 | 7.30E-05 | 0.728 | GCGGGTGGC |
| 4 | mamo-s | *BGIBMGA000007* | down | 1840 | 1848 | + | 10.9358 | 7.30E-05 | 0.728 | GCGGGTGGC |
| 5 | mamo-s | *BGIBMGA000009* | down | 1398 | 1406 | - | 11.3303 | 4.96E-05 | 0.615 | GTGGGTGGC |
| 6 | mamo-s | *BGIBMGA000012* | up | 1435 | 1443 | + | 11.3303 | 4.96E-05 | 0.615 | GTGGGTGGC |
| 7 | mamo-s | *BGIBMGA000013* | up | 484 | 492 | + | 11.3303 | 4.96E-05 | 0.615 | GTGGGTGGC |
| 8 | mamo-s | *BGIBMGA000013* | down | 829 | 837 | + | 11.3303 | 4.96E-05 | 0.615 | GTGGGTGGC |
| 9 | mamo-s | *BGIBMGA000013* | down | 1549 | 1557 | + | 11.3303 | 4.96E-05 | 0.615 | GTGGGTGGC |
| 10 | mamo-s | *BGIBMGA000014* | down | 1388 | 1396 | + | 11.3303 | 4.96E-05 | 0.615 | GTGGGTGGC |
| 11 | mamo-s | *BGIBMGA000014* | up | 1872 | 1880 | + | 10.5413 | 9.42E-05 | 0.857 | CAGCGTGTC |
| 12 | mamo-s | *BGIBMGA000016* | down | 511 | 519 | + | 11.2844 | 5.18E-05 | 0.63 | CTGCGTGCC |
| 13 | mamo-s | *BGIBMGA000018* | down | 1291 | 1299 | + | 11.3303 | 4.96E-05 | 0.615 | GTGGGTGGC |
| 14 | mamo-s | *BGIBMGA000019* | up | 1023 | 1031 | - | 11.3303 | 4.96E-05 | 0.615 | GTGGGTGGC |
| 15 | mamo-s | *BGIBMGA000020* | up | 368 | 376 | + | 14.055 | 2.23E-06 | 0.615 | GTGCGTGGC |
| 16 | mamo-s | *BGIBMGA000020* | down | 251 | 259 | + | 11.3303 | 4.96E-05 | 0.615 | GTGGGTGGC |
| 17 | mamo-s | *BGIBMGA000021* | up | 1499 | 1507 | + | 11.3303 | 4.96E-05 | 0.615 | GTGGGTGGC |
| 18 | mamo-s | *BGIBMGA000024* | up | 1873 | 1881 | - | 11.1376 | 6.35E-05 | 0.709 | TCGCGTGGC |
| 19 | mamo-s | *BGIBMGA000025* | up | 1450 | 1458 | + | 14.055 | 2.23E-06 | 0.615 | GTGCGTGGC |
| 20 | mamo-s | *BGIBMGA000025* | up | 513 | 521 | - | 12.1284 | 2.16E-05 | 0.615 | CAGCGTGGC |
| 21 | mamo-s | *BGIBMGA000025* | down | 148 | 156 | + | 11.3303 | 4.96E-05 | 0.615 | GTGGGTGGC |
| 22 | mamo-s | *BGIBMGA000026* | down | 936 | 944 | - | 11.3303 | 4.96E-05 | 0.615 | GTGGGTGGC |
| 23 | mamo-s | *BGIBMGA000026* | up | 1571 | 1579 | - | 11.0826 | 6.84E-05 | 0.709 | GTACGTGGC |
| 24 | mamo-s | *BGIBMGA000027* | down | 1480 | 1488 | - | 10.5688 | 8.88E-05 | 0.824 | GTGCGTGAC |
| 25 | mamo-s | *BGIBMGA000028* | down | 312 | 320 | - | 11.3303 | 4.96E-05 | 0.615 | GTGGGTGGC |
| 26 | mamo-s | *BGIBMGA000028* | up | 591 | 599 | - | 11.3303 | 4.96E-05 | 0.615 | GTGGGTGGC |
| 27 | mamo-s | *BGIBMGA000030* | up | 487 | 495 | - | 11.3303 | 4.96E-05 | 0.615 | GTGGGTGGC |
| 28 | mamo-s | *BGIBMGA000033* | down | 391 | 399 | - | 12.0734 | 2.66E-05 | 0.615 | GCGCGTGTC |
| 29 | mamo-s | *BGIBMGA000036* | up | 1509 | 1517 | + | 11.3303 | 4.96E-05 | 0.615 | GTGGGTGGC |
| 30 | mamo-s | *BGIBMGA000036* | up | 1873 | 1881 | - | 11.1376 | 6.35E-05 | 0.709 | TCGCGTGGC |
| 31 | mamo-s | *BGIBMGA000038* | up | 1503 | 1511 | + | 11.3303 | 4.96E-05 | 0.615 | GTGGGTGGC |
| 32 | mamo-s | *BGIBMGA000039* | up | 285 | 293 | + | 11.3303 | 4.96E-05 | 0.615 | GTGGGTGGC |
| 33 | mamo-s | *BGIBMGA000041* | down | 1974 | 1982 | - | 11.7064 | 3.52E-05 | 0.615 | GCGCGTGGT |
| 34 | mamo-s | *BGIBMGA000041* | up | 42 | 50 | - | 11.3303 | 4.96E-05 | 0.615 | GTGGGTGGC |
| 35 | mamo-s | *BGIBMGA000044* | up | 1337 | 1345 | - | 10.5413 | 9.42E-05 | 0.857 | CAGCGTGTC |
| 36 | mamo-s | *BGIBMGA000045* | up | 1927 | 1935 | - | 11.1376 | 6.35E-05 | 0.709 | TCGCGTGGC |
| 37 | mamo-s | *BGIBMGA000048* | down | 1646 | 1654 | + | 12.1009 | 2.43E-05 | 0.615 | GTGCGTGGT |
| 38 | mamo-s | *BGIBMGA000048* | up | 1729 | 1737 | + | 11.7064 | 3.52E-05 | 0.615 | GCGCGTGGT |
| 39 | mamo-s | *BGIBMGA000048* | up | 994 | 1002 | + | 11.3303 | 4.96E-05 | 0.615 | GTGGGTGGC |
| 40 | mamo-s | *BGIBMGA000049* | up | 762 | 770 | + | 12.1009 | 2.43E-05 | 0.615 | GTGCGTGGT |
| 41 | mamo-s | *BGIBMGA000049* | down | 198 | 206 | - | 10.9358 | 7.30E-05 | 0.728 | GCGGGTGGC |
| 42 | mamo-s | *BGIBMGA000050* | up | 410 | 418 | + | 11.3303 | 4.96E-05 | 0.615 | GTGGGTGGC |
| 43 | mamo-s | *BGIBMGA000050* | up | 595 | 603 | - | 11.156 | 6.13E-05 | 0.705 | GAGCGTGCC |
| 44 | mamo-s | *BGIBMGA000051* | up | 811 | 819 | + | 11.3303 | 4.96E-05 | 0.615 | GTGGGTGGC |
| 45 | mamo-s | *BGIBMGA000051* | up | 1175 | 1183 | + | 11.3303 | 4.96E-05 | 0.615 | GTGGGTGGC |
| 46 | mamo-s | *BGIBMGA000054* | down | 1865 | 1873 | - | 10.7523 | 8.38E-05 | 0.797 | GGGCGTGTC |
| 47 | mamo-s | *BGIBMGA000059* | down | 1486 | 1494 | + | 11.3303 | 4.96E-05 | 0.615 | GTGGGTGGC |
| 48 | mamo-s | *BGIBMGA000062* | down | 667 | 675 | + | 11.3303 | 4.96E-05 | 0.615 | GTGGGTGGC |
| 49 | mamo-s | *BGIBMGA000062* | down | 250 | 258 | - | 10.5413 | 9.42E-05 | 0.857 | GTGCGAGTC |
| 50 | mamo-s | *BGIBMGA000064* | down | 1792 | 1800 | + | 13.6606 | 4.06E-06 | 0.615 | GCGCGTGGC |
| 51 | mamo-s | *BGIBMGA000064* | down | 1383 | 1391 | - | 11.3303 | 4.96E-05 | 0.615 | GTGGGTGGC |
| 52 | mamo-s | *BGIBMGA000065* | up | 827 | 835 | + | 11.0826 | 6.84E-05 | 0.709 | GTACGTGGC |
| 53 | mamo-s | *BGIBMGA000066* | up | 1508 | 1516 | + | 10.8073 | 7.89E-05 | 0.766 | CCGCGTGGT |
| 54 | mamo-s | *BGIBMGA000068* | up | 218 | 226 | - | 11.3303 | 4.96E-05 | 0.615 | GTGGGTGGC |
| 55 | mamo-s | *BGIBMGA000068* | up | 1204 | 1212 | + | 11.1743 | 5.90E-05 | 0.684 | CCGCGTGTC |
| 56 | mamo-s | *BGIBMGA000071* | down | 1845 | 1853 | + | 11.7064 | 3.52E-05 | 0.615 | GCGCGTGGT |
| 57 | mamo-s | *BGIBMGA000071* | up | 614 | 622 | - | 10.5413 | 9.42E-05 | 0.857 | CAGCGTGTC |
| 58 | mamo-s | *BGIBMGA000072* | down | 884 | 892 | - | 12.1835 | 1.71E-05 | 0.615 | GTGCGTGCC |
| 59 | mamo-s | *BGIBMGA000073* | down | 217 | 225 | + | 11.3303 | 4.96E-05 | 0.615 | GTGGGTGGC |
| 60 | mamo-s | *BGIBMGA000073* | up | 1812 | 1820 | - | 11.3303 | 4.96E-05 | 0.615 | GTGGGTGGC |
| 61 | mamo-s | *BGIBMGA000073* | up | 408 | 416 | - | 11.1743 | 5.90E-05 | 0.684 | CCGCGTGTC |
| 62 | mamo-s | *BGIBMGA000078* | up | 1694 | 1702 | - | 11.3303 | 4.96E-05 | 0.615 | GTGGGTGGC |
| 63 | mamo-s | *BGIBMGA000079* | down | 777 | 785 | - | 11.3303 | 4.96E-05 | 0.615 | GTGGGTGGC |
| 64 | mamo-s | *BGIBMGA000079* | down | 1317 | 1325 | + | 11.3303 | 4.96E-05 | 0.615 | GTGGGTGGC |
| 65 | mamo-s | *BGIBMGA000082* | down | 1971 | 1979 | - | 11.3303 | 4.96E-05 | 0.615 | GTGGGTGGC |
| 66 | mamo-s | *BGIBMGA000083* | up | 1214 | 1222 | + | 14.055 | 2.23E-06 | 0.615 | GTGCGTGGC |
| 67 | mamo-s | *BGIBMGA000083* | down | 147 | 155 | - | 11.0826 | 6.84E-05 | 0.709 | GTACGTGGC |
| 68 | mamo-s | *BGIBMGA000085* | up | 1828 | 1836 | - | 10.8073 | 7.89E-05 | 0.766 | CCGCGTGGT |
| 69 | mamo-s | *BGIBMGA000089* | up | 282 | 290 | - | 11.3303 | 4.96E-05 | 0.615 | GTGGGTGGC |
| 70 | mamo-s | *BGIBMGA000089* | down | 647 | 655 | + | 10.9358 | 7.30E-05 | 0.728 | GCGGGTGGC |
| 71 | mamo-s | *BGIBMGA000090* | up | 1530 | 1538 | + | 11.3303 | 4.96E-05 | 0.615 | GTGGGTGGC |
| 72 | mamo-s | *BGIBMGA000091* | up | 497 | 505 | + | 11.0826 | 6.84E-05 | 0.709 | GTACGTGGC |
| 73 | mamo-s | *BGIBMGA000092* | down | 1928 | 1936 | - | 10.8073 | 7.89E-05 | 0.766 | CCGCGTGGT |
| 74 | mamo-s | *BGIBMGA000093* | up | 1340 | 1348 | + | 11.3303 | 4.96E-05 | 0.615 | GTGGGTGGC |
| 75 | mamo-s | *BGIBMGA000094* | down | 1139 | 1147 | - | 11.3303 | 4.96E-05 | 0.615 | GTGGGTGGC |
| 76 | mamo-s | *BGIBMGA000094* | up | 1389 | 1397 | + | 11.3303 | 4.96E-05 | 0.615 | GTGGGTGGC |
| 77 | mamo-s | *BGIBMGA000095* | up | 220 | 228 | - | 12.3394 | 1.49E-05 | 0.615 | GGGCGTGGC |
| 78 | mamo-s | *BGIBMGA000095* | down | 875 | 883 | - | 10.9358 | 7.30E-05 | 0.728 | GCGGGTGGC |
| 79 | mamo-s | *BGIBMGA000097* | down | 386 | 394 | - | 13.0275 | 8.53E-06 | 0.615 | GAGCGTGGC |
| 80 | mamo-s | *BGIBMGA000098* | down | 1849 | 1857 | + | 12.1009 | 2.43E-05 | 0.615 | GTGCGTGGT |
| 81 | mamo-s | *BGIBMGA000104* | down | 1180 | 1188 | + | 11.3303 | 4.96E-05 | 0.615 | GTGGGTGGC |

| | | | | | | | | | | |
|---|---|---|---|---|---|---|---|---|---|---|
| 82 | mamo-s | *BGIBMGA000106* | up | 1814 | 1822 | + | 11.3303 | 4.96E-05 | 0.615 | GTGGGTGGC |
| 83 | mamo-s | *BGIBMGA000108* | down | 900 | 908 | + | 10.9358 | 7.30E-05 | 0.728 | GCGGGTGGC |
| 84 | mamo-s | *BGIBMGA000109* | down | 1887 | 1895 | - | 11.3303 | 4.96E-05 | 0.615 | GTGGGTGGC |
| 85 | mamo-s | *BGIBMGA000110* | up | 1514 | 1522 | + | 11.3303 | 4.96E-05 | 0.615 | GTGGGTGGC |
| 86 | mamo-s | *BGIBMGA000113* | down | 294 | 302 | - | 11.3303 | 4.96E-05 | 0.615 | GTGGGTGGC |
| 87 | mamo-s | *BGIBMGA000114* | down | 1349 | 1357 | - | 11.7798 | 3.11E-05 | 0.615 | ATGCGTGGC |
| 88 | mamo-s | *BGIBMGA000115* | down | 167 | 175 | - | 11.3303 | 4.96E-05 | 0.615 | GTGGGTGGC |
| 89 | mamo-s | *BGIBMGA000115* | down | 1185 | 1193 | - | 10.7523 | 8.38E-05 | 0.797 | AAGCGTGGC |
| 90 | mamo-s | *BGIBMGA000116* | down | 191 | 199 | + | 11.3303 | 4.96E-05 | 0.615 | GTGGGTGGC |
| 91 | mamo-s | *BGIBMGA000116* | down | 484 | 492 | - | 10.5138 | 9.75E-05 | 0.865 | GTGCGTGTT |
| 92 | mamo-s | *BGIBMGA000117* | up | 1667 | 1675 | + | 12.3394 | 1.49E-05 | 0.615 | GGGCGTGGC |
| 93 | mamo-s | *BGIBMGA000117* | up | 1719 | 1727 | + | 11.3303 | 4.96E-05 | 0.615 | GTGGGTGGC |
| 94 | mamo-s | *BGIBMGA000119* | down | 542 | 550 | - | 11.3303 | 4.96E-05 | 0.615 | GTGGGTGGC |
| 95 | mamo-s | *BGIBMGA000119* | up | 379 | 387 | + | 11.0826 | 6.84E-05 | 0.709 | GTACGTGGC |
| 96 | mamo-s | *BGIBMGA000121* | up | 1198 | 1206 | - | 11.3303 | 4.96E-05 | 0.615 | GTGGGTGGC |
| 97 | mamo-s | *BGIBMGA000121* | up | 560 | 568 | - | 10.8349 | 7.66E-05 | 0.751 | CCGCGAGGC |
| 98 | mamo-s | *BGIBMGA000123* | up | 1786 | 1794 | - | 11.3303 | 4.96E-05 | 0.615 | GTGGGTGGC |
| 99 | mamo-s | *BGIBMGA000125* | up | 1718 | 1726 | + | 11.3303 | 4.96E-05 | 0.615 | GTGGGTGGC |
| 100 | mamo-s | *BGIBMGA000126* | up | 244 | 252 | - | 11.3303 | 4.96E-05 | 0.615 | GTGGGTGGC |
| 101 | mamo-s | *BGIBMGA000129* | up | 101 | 109 | + | 11.3303 | 4.96E-05 | 0.615 | GTGGGTGGC |
| 102 | mamo-s | *BGIBMGA000129* | up | 931 | 939 | - | 11.3303 | 4.96E-05 | 0.615 | GTGGGTGGC |
| 103 | mamo-s | *BGIBMGA000130* | down | 242 | 250 | + | 10.9358 | 7.30E-05 | 0.728 | GCGGGTGGC |
| 104 | mamo-s | *BGIBMGA000136* | down | 1327 | 1335 | + | 10.5688 | 8.88E-05 | 0.824 | GTGCGTGAC |
| 105 | mamo-s | *BGIBMGA000137* | up | 453 | 461 | - | 11.3303 | 4.96E-05 | 0.615 | GTGGGTGGC |
| 106 | mamo-s | *BGIBMGA000138* | down | 156 | 164 | - | 11.3303 | 4.96E-05 | 0.615 | GTGGGTGGC |
| 107 | mamo-s | *BGIBMGA000140* | down | 1577 | 1585 | + | 11.3303 | 4.96E-05 | 0.615 | GTGGGTGGC |
| 108 | mamo-s | *BGIBMGA000142* | down | 872 | 880 | + | 11.3303 | 4.96E-05 | 0.615 | GTGGGTGGC |
| 109 | mamo-s | *BGIBMGA000142* | up | 1004 | 1012 | - | 11.3303 | 4.96E-05 | 0.615 | GTGGGTGGC |
| 110 | mamo-s | *BGIBMGA000143* | up | 1902 | 1910 | + | 11.3303 | 4.96E-05 | 0.615 | GTGGGTGGC |
| 111 | mamo-s | *BGIBMGA000144* | down | 1674 | 1682 | + | 12.3394 | 1.49E-05 | 0.615 | GGGCGTGGC |
| 112 | mamo-s | *BGIBMGA000147* | down | 84 | 92 | - | 11.1009 | 6.57E-05 | 0.709 | GAGCGAGGC |
| 113 | mamo-s | *BGIBMGA000148* | up | 103 | 111 | - | 11.1009 | 6.57E-05 | 0.709 | GAGCGAGGC |
| 114 | mamo-s | *BGIBMGA000149* | up | 1176 | 1184 | - | 11.4404 | 4.51E-05 | 0.615 | CGGCGTGGC |
| 115 | mamo-s | *BGIBMGA000149* | down | 1286 | 1294 | + | 11.3303 | 4.96E-05 | 0.615 | GTGGGTGGC |
| 116 | mamo-s | *BGIBMGA000150* | up | 14 | 22 | + | 11.3303 | 4.96E-05 | 0.615 | GTGGGTGGC |
| 117 | mamo-s | *BGIBMGA000150* | up | 1294 | 1302 | - | 11.3303 | 4.96E-05 | 0.615 | GTGGGTGGC |
| 118 | mamo-s | *BGIBMGA000152* | up | 751 | 759 | + | 11.2844 | 5.18E-05 | 0.63 | CTGCGTGCC |
| 119 | mamo-s | *BGIBMGA000153* | down | 1184 | 1192 | + | 11.3303 | 4.96E-05 | 0.615 | GTGGGTGGC |
| 120 | mamo-s | *BGIBMGA000154* | down | 904 | 912 | + | 11.3303 | 4.96E-05 | 0.615 | GTGGGTGGC |
| 121 | mamo-s | *BGIBMGA000155* | down | 1733 | 1741 | - | 11.3303 | 4.96E-05 | 0.615 | GTGGGTGGC |
| 122 | mamo-s | *BGIBMGA000157* | down | 1258 | 1266 | + | 11.3303 | 4.96E-05 | 0.615 | GTGGGTGGC |
| 123 | mamo-s | *BGIBMGA000161* | up | 685 | 693 | + | 11.3303 | 4.96E-05 | 0.615 | GTGGGTGGC |
| 124 | mamo-s | *BGIBMGA000161* | up | 1074 | 1082 | - | 10.9358 | 7.30E-05 | 0.728 | GCGGGTGGC |
| 125 | mamo-s | *BGIBMGA000163* | down | 1258 | 1266 | - | 11.3303 | 4.96E-05 | 0.615 | GTGGGTGGC |
| 126 | mamo-s | *BGIBMGA000164* | down | 845 | 853 | - | 11.7798 | 3.11E-05 | 0.615 | ATGCGTGGC |
| 127 | mamo-s | *BGIBMGA000164* | up | 1257 | 1265 | - | 11.3303 | 4.96E-05 | 0.615 | GTGGGTGGC |
| 128 | mamo-s | *BGIBMGA000164* | up | 775 | 783 | + | 11.0734 | 7.12E-05 | 0.726 | GAGCGTGGT |
| 129 | mamo-s | *BGIBMGA000165* | up | 1585 | 1593 | - | 11.7798 | 3.11E-05 | 0.615 | ATGCGTGGC |
| 130 | mamo-s | *BGIBMGA000166* | up | 231 | 239 | + | 11.3303 | 4.96E-05 | 0.615 | GTGGGTGGC |
| 131 | mamo-s | *BGIBMGA000166* | down | 1349 | 1357 | + | 11.3303 | 4.96E-05 | 0.615 | GTGGGTGGC |
| 132 | mamo-s | *BGIBMGA000166* | down | 648 | 656 | - | 10.5688 | 8.88E-05 | 0.824 | GTGCGTGAC |
| 133 | mamo-s | *BGIBMGA000168* | up | 957 | 965 | - | 13.6606 | 4.06E-06 | 0.615 | GCGCGTGGC |
| 134 | mamo-s | *BGIBMGA000169* | up | 1432 | 1440 | + | 12.1284 | 2.16E-05 | 0.615 | GTGCGAGGC |
| 135 | mamo-s | *BGIBMGA000169* | down | 1971 | 1979 | - | 11.3303 | 4.96E-05 | 0.615 | GTGGGTGGC |
| 136 | mamo-s | *BGIBMGA000169* | down | 777 | 785 | - | 10.5688 | 8.88E-05 | 0.824 | GTGCGTGAC |
| 137 | mamo-s | *BGIBMGA000169* | up | 1237 | 1245 | + | 10.5138 | 9.75E-05 | 0.865 | GTGCGTGTT |
| 138 | mamo-s | *BGIBMGA000170* | down | 1647 | 1655 | + | 11.3303 | 4.96E-05 | 0.615 | GTGGGTGGC |
| 139 | mamo-s | *BGIBMGA000171* | up | 1408 | 1416 | - | 11.3303 | 4.96E-05 | 0.615 | GTGGGTGGC |
| 140 | mamo-s | *BGIBMGA000172* | down | 690 | 698 | + | 11.3303 | 4.96E-05 | 0.615 | GTGGGTGGC |
| 141 | mamo-s | *BGIBMGA000174* | down | 958 | 966 | + | 11.3303 | 4.96E-05 | 0.615 | GTGGGTGGC |
| 142 | mamo-s | *BGIBMGA000175* | up | 153 | 161 | + | 11.4404 | 4.51E-05 | 0.615 | GAGCGTGTC |
| 143 | mamo-s | *BGIBMGA000179* | down | 1816 | 1824 | + | 11.3303 | 4.96E-05 | 0.615 | GTGGGTGGC |
| 144 | mamo-s | *BGIBMGA000179* | up | 1988 | 1996 | + | 10.5413 | 9.42E-05 | 0.857 | CAGCGTGTC |
| 145 | mamo-s | *BGIBMGA000181* | up | 574 | 582 | + | 11.3303 | 4.96E-05 | 0.615 | GTGGGTGGC |
| 146 | mamo-s | *BGIBMGA000181* | down | 1597 | 1605 | - | 11.3303 | 4.96E-05 | 0.615 | GTGGGTGGC |
| 147 | mamo-s | *BGIBMGA000182* | down | 508 | 516 | - | 11.1376 | 6.35E-05 | 0.709 | TCGCGTGGC |
| 148 | mamo-s | *BGIBMGA000183* | up | 251 | 259 | + | 11.3303 | 4.96E-05 | 0.615 | GTGGGTGGC |
| 149 | mamo-s | *BGIBMGA000183* | up | 1236 | 1244 | - | 11.3303 | 4.96E-05 | 0.615 | GTGGGTGGC |
| 150 | mamo-s | *BGIBMGA000183* | down | 1374 | 1382 | - | 11.3303 | 4.96E-05 | 0.615 | GTGGGTGGC |
| 151 | mamo-s | *BGIBMGA000184* | up | 704 | 712 | + | 11.3303 | 4.96E-05 | 0.615 | GTGGGTGGC |
| 152 | mamo-s | *BGIBMGA000184* | down | 1106 | 1114 | - | 11.3303 | 4.96E-05 | 0.615 | GTGGGTGGC |
| 153 | mamo-s | *BGIBMGA000188* | up | 1089 | 1097 | - | 11.3303 | 4.96E-05 | 0.615 | GTGGGTGGC |
| 154 | mamo-s | *BGIBMGA000188* | down | 1769 | 1777 | + | 11.2018 | 5.68E-05 | 0.674 | CTGCGTGGT |
| 155 | mamo-s | *BGIBMGA000190* | up | 420 | 428 | + | 11.3303 | 4.96E-05 | 0.615 | GTGGGTGGC |
| 156 | mamo-s | *BGIBMGA000191* | down | 349 | 357 | + | 11.3303 | 4.96E-05 | 0.615 | GTGGGTGGC |
| 157 | mamo-s | *BGIBMGA000191* | down | 1959 | 1967 | - | 11.2844 | 5.18E-05 | 0.63 | CTGCGTGCC |
| 158 | mamo-s | *BGIBMGA000193* | up | 202 | 210 | - | 11.3303 | 4.96E-05 | 0.615 | GTGGGTGGC |
| 159 | mamo-s | *BGIBMGA000193* | down | 1257 | 1265 | + | 11.3303 | 4.96E-05 | 0.615 | GTGGGTGGC |
| 160 | mamo-s | *BGIBMGA000193* | up | 1611 | 1619 | + | 11.2844 | 5.18E-05 | 0.63 | CTGCGTGCC |
| 161 | mamo-s | *BGIBMGA000194* | up | 241 | 249 | - | 13.0275 | 8.53E-06 | 0.615 | GAGCGTGGC |
| 162 | mamo-s | *BGIBMGA000196* | down | 878 | 886 | - | 11.2018 | 5.68E-05 | 0.674 | CTGCGTGGT |
| 163 | mamo-s | *BGIBMGA000197* | down | 1387 | 1395 | + | 11.3303 | 4.96E-05 | 0.615 | GTGGGTGGC |
| 164 | mamo-s | *BGIBMGA000199* | up | 19 | 27 | + | 11.3303 | 4.96E-05 | 0.615 | GTGGGTGGC |
| 165 | mamo-s | *BGIBMGA000200* | up | 780 | 788 | + | 11.3853 | 4.74E-05 | 0.615 | ACGCGTGGC |
| 166 | mamo-s | *BGIBMGA000201* | up | 663 | 671 | + | 11.3303 | 4.96E-05 | 0.615 | GTGGGTGGC |
| 167 | mamo-s | *BGIBMGA000202* | up | 141 | 149 | + | 11.3303 | 4.96E-05 | 0.615 | GTGGGTGGC |
| 168 | mamo-s | *BGIBMGA000203* | up | 206 | 214 | + | 11.1009 | 6.57E-05 | 0.709 | GAGCGAGGC |
| 169 | mamo-s | *BGIBMGA000204* | up | 1035 | 1043 | - | 12.3394 | 1.49E-05 | 0.615 | GGGCGTGGC |

| | | | | | | | | | | |
|---|---|---|---|---|---|---|---|---|---|---|
| 170 | mamo-s | *BGIBMGA000204* | up | 494 | 502 | + | 11.3303 | 4.96E-05 | 0.615 | GTGGGTGGC |
| 171 | mamo-s | *BGIBMGA000205* | up | 604 | 612 | + | 11.3303 | 4.96E-05 | 0.615 | GTGGGTGGC |
| 172 | mamo-s | *BGIBMGA000206* | down | 340 | 348 | - | 11.3303 | 4.96E-05 | 0.615 | GTGGGTGGC |
| 173 | mamo-s | *BGIBMGA000206* | up | 341 | 349 | - | 11.3303 | 4.96E-05 | 0.615 | GTGGGTGGC |
| 174 | mamo-s | *BGIBMGA000206* | down | 1814 | 1822 | + | 10.5138 | 9.75E-05 | 0.865 | GTGCGTGTT |
| 175 | mamo-s | *BGIBMGA000207* | up | 455 | 463 | - | 11.3303 | 4.96E-05 | 0.615 | GTGGGTGGC |
| 176 | mamo-s | *BGIBMGA000207* | up | 1929 | 1937 | + | 10.5138 | 9.75E-05 | 0.865 | GTGCGTGTT |
| 177 | mamo-s | *BGIBMGA000208* | down | 371 | 379 | + | 12.1284 | 2.16E-05 | 0.615 | GTGCGAGGC |
| 178 | mamo-s | *BGIBMGA000208* | up | 733 | 741 | - | 10.9358 | 7.30E-05 | 0.728 | GCGGGTGGC |
| 179 | mamo-s | *BGIBMGA000208* | down | 522 | 530 | - | 10.7523 | 8.38E-05 | 0.797 | GGGCGTGTC |
| 180 | mamo-s | *BGIBMGA000208* | up | 687 | 695 | - | 10.6881 | 8.61E-05 | 0.81 | GCACGTGGC |
| 181 | mamo-s | *BGIBMGA000210* | up | 21 | 29 | - | 11.3303 | 4.96E-05 | 0.615 | GTGGGTGGC |
| 182 | mamo-s | *BGIBMGA000212* | down | 831 | 839 | + | 10.5138 | 9.75E-05 | 0.865 | GTGCGTGTT |
| 183 | mamo-s | *BGIBMGA000215* | up | 193 | 201 | - | 11.3303 | 4.96E-05 | 0.615 | GTGGGTGGC |
| 184 | mamo-s | *BGIBMGA000219* | up | 1205 | 1213 | - | 10.9358 | 7.30E-05 | 0.728 | GCGGGTGGC |
| 185 | mamo-s | *BGIBMGA000222* | down | 868 | 876 | + | 11.7798 | 3.11E-05 | 0.615 | ATGCGTGGC |
| 186 | mamo-s | *BGIBMGA000223* | up | 1015 | 1023 | + | 11.3303 | 4.96E-05 | 0.615 | GTGGGTGGC |
| 187 | mamo-s | *BGIBMGA000223* | up | 1121 | 1129 | - | 10.5688 | 8.88E-05 | 0.824 | GTGCGTGAC |
| 188 | mamo-s | *BGIBMGA000224* | up | 1394 | 1402 | - | 11.1376 | 6.35E-05 | 0.709 | TCGCGTGGC |
| 189 | mamo-s | *BGIBMGA000225* | up | 69 | 77 | + | 11.3303 | 4.96E-05 | 0.615 | GTGGGTGGC |
| 190 | mamo-s | *BGIBMGA000225* | down | 922 | 930 | + | 11.3303 | 4.96E-05 | 0.615 | GTGGGTGGC |
| 191 | mamo-s | *BGIBMGA000228* | up | 1425 | 1433 | - | 11.3303 | 4.96E-05 | 0.615 | GTGGGTGGC |
| 192 | mamo-s | *BGIBMGA000232* | down | 658 | 666 | - | 13.6606 | 4.06E-06 | 0.615 | GCGCGTGGC |
| 193 | mamo-s | *BGIBMGA000232* | up | 1984 | 1992 | + | 11.5321 | 4.06E-05 | 0.615 | TTGCGTGGC |
| 194 | mamo-s | *BGIBMGA000232* | down | 1083 | 1091 | + | 11.3303 | 4.96E-05 | 0.615 | GTGGGTGGC |
| 195 | mamo-s | *BGIBMGA000234* | down | 911 | 919 | + | 11.3303 | 4.96E-05 | 0.615 | GTGGGTGGC |
| 196 | mamo-s | *BGIBMGA000234* | up | 1705 | 1713 | - | 11.2018 | 5.68E-05 | 0.674 | CTGCGTGGT |
| 197 | mamo-s | *BGIBMGA000235* | up | 920 | 928 | + | 11.3303 | 4.96E-05 | 0.615 | GTGGGTGGC |
| 198 | mamo-s | *BGIBMGA000235* | down | 1717 | 1725 | + | 11.3303 | 4.96E-05 | 0.615 | GTGGGTGGC |
| 199 | mamo-s | *BGIBMGA000237* | up | 355 | 363 | - | 11.2018 | 5.68E-05 | 0.674 | CTGCGTGGT |
| 200 | mamo-s | *BGIBMGA000241* | down | 1599 | 1607 | + | 11.3303 | 4.96E-05 | 0.615 | GTGGGTGGC |
| 201 | mamo-s | *BGIBMGA000241* | down | 1968 | 1976 | + | 11.3303 | 4.96E-05 | 0.615 | GTGGGTGGC |
| 202 | mamo-s | *BGIBMGA000242* | up | 729 | 737 | - | 14.055 | 2.23E-06 | 0.615 | GTGCGTGGC |
| 203 | mamo-s | *BGIBMGA000242* | up | 656 | 664 | + | 10.7523 | 8.38E-05 | 0.797 | GGGCGTGTC |
| 204 | mamo-s | *BGIBMGA000244* | down | 1248 | 1256 | + | 11.4404 | 4.51E-05 | 0.615 | GAGCGTGTC |
| 205 | mamo-s | *BGIBMGA000245* | down | 434 | 442 | + | 11.3303 | 4.96E-05 | 0.615 | GTGGGTGGC |
| 206 | mamo-s | *BGIBMGA000247* | up | 1096 | 1104 | + | 11.3303 | 4.96E-05 | 0.615 | GTGGGTGGC |
| 207 | mamo-s | *BGIBMGA000249* | up | 1335 | 1343 | - | 11.3303 | 4.96E-05 | 0.615 | GTGGGTGGC |
| 208 | mamo-s | *BGIBMGA000249* | up | 1854 | 1862 | + | 11.3303 | 4.96E-05 | 0.615 | GTGGGTGGC |
| 209 | mamo-s | *BGIBMGA000251* | down | 852 | 860 | + | 11.0826 | 6.84E-05 | 0.709 | GTACGTGGC |
| 210 | mamo-s | *BGIBMGA000252* | down | 708 | 716 | - | 11.3303 | 4.96E-05 | 0.615 | GTGGGTGGC |
| 211 | mamo-s | *BGIBMGA000254* | down | 401 | 409 | - | 11.3303 | 4.96E-05 | 0.615 | GTGGGTGGC |
| 212 | mamo-s | *BGIBMGA000255* | up | 1121 | 1129 | - | 11.3303 | 4.96E-05 | 0.615 | GTGGGTGGC |
| 213 | mamo-s | *BGIBMGA000257* | down | 748 | 756 | + | 11.3303 | 4.96E-05 | 0.615 | GTGGGTGGC |
| 214 | mamo-s | *BGIBMGA000259* | down | 576 | 584 | + | 11.3303 | 4.96E-05 | 0.615 | GTGGGTGGC |
| 215 | mamo-s | *BGIBMGA000260* | down | 215 | 223 | - | 11.3303 | 4.96E-05 | 0.615 | GTGGGTGGC |
| 216 | mamo-s | *BGIBMGA000260* | down | 1814 | 1822 | - | 10.5413 | 9.42E-05 | 0.857 | CAGCGTGTC |
| 217 | mamo-s | *BGIBMGA000266* | down | 780 | 788 | + | 11.3303 | 4.96E-05 | 0.615 | GTGGGTGGC |
| 218 | mamo-s | *BGIBMGA000268* | down | 391 | 399 | - | 11.3303 | 4.96E-05 | 0.615 | GTGGGTGGC |
| 219 | mamo-s | *BGIBMGA000268* | down | 1584 | 1592 | + | 11.3303 | 4.96E-05 | 0.615 | GTGGGTGGC |
| 220 | mamo-s | *BGIBMGA000271* | up | 877 | 885 | - | 11.7064 | 3.52E-05 | 0.615 | GCGCGTGGT |
| 221 | mamo-s | *BGIBMGA000273* | up | 715 | 723 | + | 11.3303 | 4.96E-05 | 0.615 | GTGGGTGGC |
| 222 | mamo-s | *BGIBMGA000275* | up | 1199 | 1207 | + | 11.1743 | 5.90E-05 | 0.684 | CCGCGTGTC |
| 223 | mamo-s | *BGIBMGA000278* | up | 138 | 146 | + | 11.3303 | 4.96E-05 | 0.615 | GTGGGTGGC |
| 224 | mamo-s | *BGIBMGA000279* | down | 1850 | 1858 | - | 11.3303 | 4.96E-05 | 0.615 | GTGGGTGGC |
| 225 | mamo-s | *BGIBMGA000280* | down | 1283 | 1291 | - | 12.3394 | 1.49E-05 | 0.615 | GGGCGTGGC |
| 226 | mamo-s | *BGIBMGA000280* | down | 1220 | 1228 | + | 11.3303 | 4.96E-05 | 0.615 | GTGGGTGGC |
| 227 | mamo-s | *BGIBMGA000280* | up | 1261 | 1269 | - | 11.3303 | 4.96E-05 | 0.615 | GTGGGTGGC |
| 228 | mamo-s | *BGIBMGA000280* | down | 1546 | 1554 | + | 11.0826 | 6.84E-05 | 0.709 | GTACGTGGC |
| 229 | mamo-s | *BGIBMGA000281* | up | 11 | 19 | - | 11.1743 | 5.90E-05 | 0.684 | CCGCGTGTC |
| 230 | mamo-s | *BGIBMGA000283* | up | 1189 | 1197 | + | 11.3303 | 4.96E-05 | 0.615 | GTGGGTGGC |
| 231 | mamo-s | *BGIBMGA000286* | up | 375 | 383 | - | 11.1743 | 5.90E-05 | 0.684 | CCGCGTGTC |
| 232 | mamo-s | *BGIBMGA000290* | down | 528 | 536 | - | 11.3303 | 4.96E-05 | 0.615 | GTGGGTGGC |
| 233 | mamo-s | *BGIBMGA000292* | down | 111 | 119 | - | 11.0826 | 6.84E-05 | 0.709 | GTACGTGGC |
| 234 | mamo-s | *BGIBMGA000294* | down | 647 | 655 | - | 11.3303 | 4.96E-05 | 0.615 | GTGGGTGGC |
| 235 | mamo-s | *BGIBMGA000296* | up | 1111 | 1119 | - | 13.156 | 6.30E-06 | 0.615 | CTGCGTGGC |
| 236 | mamo-s | *BGIBMGA000296* | down | 1752 | 1760 | - | 12.1284 | 2.16E-05 | 0.615 | GTGCGAGGC |
| 237 | mamo-s | *BGIBMGA000297* | up | 220 | 228 | + | 11.3303 | 4.96E-05 | 0.615 | GTGGGTGGC |
| 238 | mamo-s | *BGIBMGA000298* | down | 1344 | 1352 | - | 10.6881 | 8.61E-05 | 0.81 | GCACGTGGC |
| 239 | mamo-s | *BGIBMGA000301* | up | 732 | 740 | - | 10.7523 | 8.38E-05 | 0.797 | AAGCGTGGC |
| 240 | mamo-s | *BGIBMGA000302* | up | 391 | 399 | - | 12.1284 | 2.16E-05 | 0.615 | CAGCGTGGC |
| 241 | mamo-s | *BGIBMGA000305* | down | 1270 | 1278 | - | 11.3303 | 4.96E-05 | 0.615 | GTGGGTGGC |
| 242 | mamo-s | *BGIBMGA000306* | down | 1645 | 1653 | - | 11.3303 | 4.96E-05 | 0.615 | GTGGGTGGC |
| 243 | mamo-s | *BGIBMGA000308* | up | 459 | 467 | + | 11.3303 | 4.96E-05 | 0.615 | GTGGGTGGC |
| 244 | mamo-s | *BGIBMGA000313* | down | 773 | 781 | - | 11.3303 | 4.96E-05 | 0.615 | GTGGGTGGC |
| 245 | mamo-s | *BGIBMGA000315* | down | 861 | 869 | + | 11.3303 | 4.96E-05 | 0.615 | GTGGGTGGC |
| 246 | mamo-s | *BGIBMGA000320* | down | 1491 | 1499 | - | 11.7339 | 3.29E-05 | 0.615 | GCGCGAGGC |
| 247 | mamo-s | *BGIBMGA000320* | up | 1836 | 1844 | + | 10.5138 | 9.75E-05 | 0.865 | GTGCGTGTT |
| 248 | mamo-s | *BGIBMGA000321* | up | 472 | 480 | - | 12.1835 | 1.71E-05 | 0.615 | GTGCGTGCC |
| 249 | mamo-s | *BGIBMGA000321* | up | 1145 | 1153 | - | 11.0826 | 6.84E-05 | 0.709 | GTACGTGGC |
| 250 | mamo-s | *BGIBMGA000322* | up | 1328 | 1336 | + | 11.3303 | 4.96E-05 | 0.615 | GTGGGTGGC |
| 251 | mamo-s | *BGIBMGA000323* | down | 1778 | 1786 | - | 11.3303 | 4.96E-05 | 0.615 | GTGGGTGGC |
| 252 | mamo-s | *BGIBMGA000328* | down | 1606 | 1614 | - | 11.3303 | 4.96E-05 | 0.615 | GTGGGTGGC |
| 253 | mamo-s | *BGIBMGA000330* | down | 594 | 602 | - | 10.8899 | 7.48E-05 | 0.739 | CCGCGTGCC |
| 254 | mamo-s | *BGIBMGA000333* | up | 1740 | 1748 | + | 11.0826 | 6.84E-05 | 0.709 | GTACGTGGC |
| 255 | mamo-s | *BGIBMGA000336* | down | 1509 | 1517 | + | 11.3303 | 4.96E-05 | 0.615 | GTGGGTGGC |
| 256 | mamo-s | *BGIBMGA000336* | up | 53 | 61 | + | 10.5413 | 9.42E-05 | 0.857 | CAGCGTGTC |
| 257 | mamo-s | *BGIBMGA000339* | up | 483 | 491 | + | 11.3303 | 4.96E-05 | 0.615 | GTGGGTGGC |

| | | | | | | | | | | |
|---|---|---|---|---|---|---|---|---|---|---|
| 258 | mamo-s | *BGIBMGA000341* | down | 779 | 787 | - | 12.1835 | 1.71E-05 | 0.615 | GTGCGTGCC |
| 259 | mamo-s | *BGIBMGA000348* | up | 1591 | 1599 | + | 11.3303 | 4.96E-05 | 0.615 | GTGGGTGGC |
| 260 | mamo-s | *BGIBMGA000353* | up | 1753 | 1761 | - | 11.0826 | 6.84E-05 | 0.709 | GTACGTGGC |
| 261 | mamo-s | *BGIBMGA000355* | down | 804 | 812 | + | 11.3303 | 4.96E-05 | 0.615 | GTGGGTGGC |
| 262 | mamo-s | *BGIBMGA000358* | up | 695 | 703 | + | 10.8899 | 7.48E-05 | 0.739 | CCGCGTGCC |
| 263 | mamo-s | *BGIBMGA000361* | down | 663 | 671 | - | 11.3303 | 4.96E-05 | 0.615 | GTGGGTGGC |
| 264 | mamo-s | *BGIBMGA000364* | down | 1098 | 1106 | - | 11.3303 | 4.96E-05 | 0.615 | GTGGGTGGC |
| 265 | mamo-s | *BGIBMGA000371* | down | 594 | 602 | + | 14.055 | 2.23E-06 | 0.615 | GTGCGTGGC |
| 266 | mamo-s | *BGIBMGA000375* | up | 234 | 242 | - | 10.5138 | 9.75E-05 | 0.865 | GTGCGTGTT |
| 267 | mamo-s | *BGIBMGA000377* | up | 723 | 731 | - | 11.3303 | 4.96E-05 | 0.615 | GTGGGTGGC |
| 268 | mamo-s | *BGIBMGA000377* | down | 1834 | 1842 | - | 10.8349 | 7.66E-05 | 0.751 | CCGCGAGGC |
| 269 | mamo-s | *BGIBMGA000384* | up | 1339 | 1347 | + | 11.3303 | 4.96E-05 | 0.615 | GTGGGTGGC |
| 270 | mamo-s | *BGIBMGA000384* | down | 1577 | 1585 | + | 11.1743 | 5.90E-05 | 0.684 | CCGCGTGTC |
| 271 | mamo-s | *BGIBMGA000384* | down | 1987 | 1995 | - | 11.0826 | 6.84E-05 | 0.709 | GTACGTGGC |
| 272 | mamo-s | *BGIBMGA000384* | down | 1747 | 1755 | + | 10.8899 | 7.48E-05 | 0.739 | CCGCGTGCC |
| 273 | mamo-s | *BGIBMGA000386* | up | 1035 | 1043 | + | 11.4404 | 4.51E-05 | 0.615 | CGGCGTGGC |
| 274 | mamo-s | *BGIBMGA000388* | up | 1725 | 1733 | + | 10.5688 | 8.88E-05 | 0.824 | GTGCGTGAC |
| 275 | mamo-s | *BGIBMGA000398* | up | 315 | 323 | - | 12.3394 | 1.49E-05 | 0.615 | GGGCGTGGC |
| 276 | mamo-s | *BGIBMGA000400* | up | 990 | 998 | + | 11.3303 | 4.96E-05 | 0.615 | GTGGGTGGC |
| 277 | mamo-s | *BGIBMGA000403* | down | 1547 | 1555 | - | 11.3303 | 4.96E-05 | 0.615 | GTGGGTGGC |
| 278 | mamo-s | *BGIBMGA000405* | down | 1060 | 1068 | + | 11.156 | 6.13E-05 | 0.705 | GAGCGTGCC |
| 279 | mamo-s | *BGIBMGA000409* | up | 739 | 747 | + | 11.3303 | 4.96E-05 | 0.615 | GTGGGTGGC |
| 280 | mamo-s | *BGIBMGA000411* | up | 294 | 302 | - | 10.7523 | 8.38E-05 | 0.797 | AAGCGTGGC |
| 281 | mamo-s | *BGIBMGA000411* | down | 179 | 187 | - | 10.5138 | 9.75E-05 | 0.865 | GTGCGTGTT |
| 282 | mamo-s | *BGIBMGA000412* | up | 38 | 46 | - | 10.5138 | 9.75E-05 | 0.865 | GTGCGTGTT |
| 283 | mamo-s | *BGIBMGA000413* | down | 286 | 294 | + | 11.5688 | 3.79E-05 | 0.615 | CTGCGTGTC |
| 284 | mamo-s | *BGIBMGA000417* | down | 726 | 734 | - | 11.3303 | 4.96E-05 | 0.615 | GTGGGTGGC |
| 285 | mamo-s | *BGIBMGA000419* | up | 1596 | 1604 | + | 11.3303 | 4.96E-05 | 0.615 | GTGGGTGGC |
| 286 | mamo-s | *BGIBMGA000419* | down | 1361 | 1369 | - | 10.9358 | 7.30E-05 | 0.728 | GCGGGTGGC |
| 287 | mamo-s | *BGIBMGA000420* | down | 1661 | 1669 | + | 11.3303 | 4.96E-05 | 0.615 | GTGGGTGGC |
| 288 | mamo-s | *BGIBMGA000420* | up | 634 | 642 | - | 10.9358 | 7.30E-05 | 0.728 | GCGGGTGGC |
| 289 | mamo-s | *BGIBMGA000421* | up | 99 | 107 | - | 11.0826 | 6.84E-05 | 0.709 | GTACGTGGC |
| 290 | mamo-s | *BGIBMGA000423* | up | 592 | 600 | + | 11.1376 | 6.35E-05 | 0.709 | TCGCGTGGC |
| 291 | mamo-s | *BGIBMGA000425* | down | 1207 | 1215 | - | 12.4679 | 1.31E-05 | 0.615 | GTGCGTGTC |
| 292 | mamo-s | *BGIBMGA000428* | down | 1611 | 1619 | - | 10.7523 | 8.38E-05 | 0.797 | GGGCGTGTC |
| 293 | mamo-s | *BGIBMGA000429* | down | 389 | 397 | - | 11.3303 | 4.96E-05 | 0.615 | GTGGGTGGC |
| 294 | mamo-s | *BGIBMGA000429* | down | 1235 | 1243 | - | 11.3303 | 4.96E-05 | 0.615 | GTGGGTGGC |
| 295 | mamo-s | *BGIBMGA000429* | up | 571 | 579 | - | 10.7523 | 8.38E-05 | 0.797 | GGGCGTGTC |
| 296 | mamo-s | *BGIBMGA000433* | down | 1592 | 1600 | - | 11.7798 | 3.11E-05 | 0.615 | ATGCGTGGC |
| 297 | mamo-s | *BGIBMGA000433* | down | 547 | 555 | + | 11.0734 | 7.12E-05 | 0.726 | GAGCGTGGT |
| 298 | mamo-s | *BGIBMGA000436* | up | 1316 | 1324 | + | 11.3303 | 4.96E-05 | 0.615 | GTGGGTGGC |
| 299 | mamo-s | *BGIBMGA000438* | up | 247 | 255 | - | 11.2844 | 5.18E-05 | 0.63 | CTGCGTGCC |
| 300 | mamo-s | *BGIBMGA000439* | down | 451 | 459 | - | 11.3303 | 4.96E-05 | 0.615 | GTGGGTGGC |
| 301 | mamo-s | *BGIBMGA000440* | down | 1886 | 1894 | - | 12.1284 | 2.16E-05 | 0.615 | CAGCGTGGC |
| 302 | mamo-s | *BGIBMGA000441* | up | 505 | 513 | - | 12.1284 | 2.16E-05 | 0.615 | CAGCGTGGC |
| 303 | mamo-s | *BGIBMGA000442* | up | 1087 | 1095 | + | 11.3303 | 4.96E-05 | 0.615 | GTGGGTGGC |
| 304 | mamo-s | *BGIBMGA000446* | up | 272 | 280 | + | 12.3394 | 1.49E-05 | 0.615 | GGGCGTGGC |
| 305 | mamo-s | *BGIBMGA000448* | up | 760 | 768 | + | 11.3303 | 4.96E-05 | 0.615 | GTGGGTGGC |
| 306 | mamo-s | *BGIBMGA000449* | down | 1092 | 1100 | - | 11.3303 | 4.96E-05 | 0.615 | GTGGGTGGC |
| 307 | mamo-s | *BGIBMGA000450* | up | 632 | 640 | + | 12.0734 | 2.66E-05 | 0.615 | GCGCGTGTC |
| 308 | mamo-s | *BGIBMGA000453* | up | 1322 | 1330 | + | 10.9358 | 7.30E-05 | 0.728 | GCGGGTGGC |
| 309 | mamo-s | *BGIBMGA000453* | up | 1038 | 1046 | + | 10.5413 | 9.42E-05 | 0.857 | GTGCGAGTC |
| 310 | mamo-s | *BGIBMGA000455* | down | 1647 | 1655 | + | 11.3303 | 4.96E-05 | 0.615 | GTGGGTGGC |
| 311 | mamo-s | *BGIBMGA000456* | down | 1885 | 1893 | - | 11.3303 | 4.96E-05 | 0.615 | GTGGGTGGC |
| 312 | mamo-s | *BGIBMGA000458* | down | 407 | 415 | + | 11.2018 | 5.68E-05 | 0.674 | CTGCGTGGT |
| 313 | mamo-s | *BGIBMGA000459* | up | 1179 | 1187 | + | 11.2018 | 5.68E-05 | 0.674 | CTGCGTGGT |
| 314 | mamo-s | *BGIBMGA000460* | up | 1243 | 1251 | + | 11.2018 | 5.68E-05 | 0.674 | CTGCGTGGT |
| 315 | mamo-s | *BGIBMGA000462* | up | 1957 | 1965 | + | 12.4679 | 1.31E-05 | 0.615 | GTGCGTGTC |
| 316 | mamo-s | *BGIBMGA000464* | up | 1390 | 1398 | + | 14.055 | 2.23E-06 | 0.615 | GTGCGTGGC |
| 317 | mamo-s | *BGIBMGA000465* | down | 1835 | 1843 | + | 11.3303 | 4.96E-05 | 0.615 | GTGGGTGGC |
| 318 | mamo-s | *BGIBMGA000466* | down | 1740 | 1748 | + | 14.055 | 2.23E-06 | 0.615 | GTGCGTGGC |
| 319 | mamo-s | *BGIBMGA000466* | up | 725 | 733 | - | 11.3303 | 4.96E-05 | 0.615 | GTGGGTGGC |
| 320 | mamo-s | *BGIBMGA000471* | down | 1570 | 1578 | - | 11.0826 | 6.84E-05 | 0.709 | GTACGTGGC |
| 321 | mamo-s | *BGIBMGA000472* | up | 1240 | 1248 | - | 11.3303 | 4.96E-05 | 0.615 | GTGGGTGGC |
| 322 | mamo-s | *BGIBMGA000473* | down | 518 | 526 | - | 11.3303 | 4.96E-05 | 0.615 | GTGGGTGGC |
| 323 | mamo-s | *BGIBMGA000475* | up | 647 | 655 | + | 11.0734 | 7.12E-05 | 0.726 | GAGCGTGGT |
| 324 | mamo-s | *BGIBMGA000476* | up | 824 | 832 | + | 11.3303 | 4.96E-05 | 0.615 | GTGGGTGGC |
| 325 | mamo-s | *BGIBMGA000478* | down | 187 | 195 | + | 11.2018 | 5.68E-05 | 0.674 | CTGCGTGGT |
| 326 | mamo-s | *BGIBMGA000484* | up | 1848 | 1856 | - | 11.2018 | 5.68E-05 | 0.674 | CTGCGTGGT |
| 327 | mamo-s | *BGIBMGA000487* | up | 742 | 750 | + | 13.6606 | 4.06E-06 | 0.615 | GCGCGTGGC |
| 328 | mamo-s | *BGIBMGA000487* | up | 1716 | 1724 | + | 11.7798 | 3.11E-05 | 0.615 | ATGCGTGGC |
| 329 | mamo-s | *BGIBMGA000488* | down | 585 | 593 | + | 12.4679 | 1.31E-05 | 0.615 | GTGCGTGTC |
| 330 | mamo-s | *BGIBMGA000489* | down | 8 | 16 | + | 11.789 | 2.84E-05 | 0.615 | GCGCGTGCC |
| 331 | mamo-s | *BGIBMGA000489* | down | 919 | 927 | - | 11.3303 | 4.96E-05 | 0.615 | GTGGGTGGC |
| 332 | mamo-s | *BGIBMGA000490* | up | 1543 | 1551 | - | 11.3303 | 4.96E-05 | 0.615 | GTGGGTGGC |
| 333 | mamo-s | *BGIBMGA000491* | down | 1763 | 1771 | + | 11.3303 | 4.96E-05 | 0.615 | GTGGGTGGC |
| 334 | mamo-s | *BGIBMGA000492* | down | 224 | 232 | - | 11.789 | 2.84E-05 | 0.615 | GCGCGTGCC |
| 335 | mamo-s | *BGIBMGA000495* | down | 1081 | 1089 | - | 11.3303 | 4.96E-05 | 0.615 | GTGGGTGGC |
| 336 | mamo-s | *BGIBMGA000495* | down | 1398 | 1406 | + | 11.3303 | 4.96E-05 | 0.615 | GTGGGTGGC |
| 337 | mamo-s | *BGIBMGA000496* | down | 1122 | 1130 | - | 13.156 | 6.30E-06 | 0.615 | CTGCGTGGC |
| 338 | mamo-s | *BGIBMGA000496* | up | 1500 | 1508 | - | 11.2294 | 5.41E-05 | 0.65 | CTGCGAGGC |
| 339 | mamo-s | *BGIBMGA000497* | up | 1786 | 1794 | - | 12.4679 | 1.31E-05 | 0.615 | GTGCGTGTC |
| 340 | mamo-s | *BGIBMGA000497* | down | 196 | 204 | + | 12.1835 | 1.71E-05 | 0.615 | GTGCGTGCC |
| 341 | mamo-s | *BGIBMGA000499* | down | 1120 | 1128 | - | 11.3303 | 4.96E-05 | 0.615 | GTGGGTGGC |
| 342 | mamo-s | *BGIBMGA000499* | up | 635 | 643 | + | 10.5138 | 9.75E-05 | 0.865 | GTGCGTGTT |
| 343 | mamo-s | *BGIBMGA000502* | up | 1232 | 1240 | - | 11.3303 | 4.96E-05 | 0.615 | GTGGGTGGC |
| 344 | mamo-s | *BGIBMGA000505* | down | 1370 | 1378 | - | 11.0734 | 7.12E-05 | 0.726 | GAGCGTGGT |
| 345 | mamo-s | *BGIBMGA000506* | down | 1132 | 1140 | - | 11.3853 | 4.74E-05 | 0.615 | ACGCGTGGC |

| | | | | | | | | | | |
|---|---|---|---|---|---|---|---|---|---|---|
| 346 | mamo-s | *BGIBMGA000508* | down | 1909 | 1917 | + | 11.0826 | 6.84E-05 | 0.709 | GTACGTGGC |
| 347 | mamo-s | *BGIBMGA000510* | up | 386 | 394 | + | 11.3303 | 4.96E-05 | 0.615 | GTGGGTGGC |
| 348 | mamo-s | *BGIBMGA000513* | down | 473 | 481 | - | 11.3303 | 4.96E-05 | 0.615 | GTGGGTGGC |
| 349 | mamo-s | *BGIBMGA000513* | down | 793 | 801 | - | 11.3303 | 4.96E-05 | 0.615 | GTGGGTGGC |
| 350 | mamo-s | *BGIBMGA000517* | down | 995 | 1003 | + | 10.5138 | 9.75E-05 | 0.865 | GTGCGTGTT |
| 351 | mamo-s | *BGIBMGA000520* | down | 615 | 623 | - | 12.1284 | 2.16E-05 | 0.615 | CAGCGTGGC |
| 352 | mamo-s | *BGIBMGA000520* | down | 580 | 588 | - | 10.5688 | 8.88E-05 | 0.824 | GTGCGTGAC |
| 353 | mamo-s | *BGIBMGA000522* | down | 1388 | 1396 | + | 11.0826 | 6.84E-05 | 0.709 | GTACGTGGC |
| 354 | mamo-s | *BGIBMGA000526* | down | 1935 | 1943 | + | 11.7064 | 3.52E-05 | 0.615 | GCGCGTGGT |
| 355 | mamo-s | *BGIBMGA000527* | down | 1806 | 1814 | - | 12.1284 | 2.16E-05 | 0.615 | CAGCGTGGC |
| 356 | mamo-s | *BGIBMGA000528* | up | 285 | 293 | + | 11.3303 | 4.96E-05 | 0.615 | GTGGGTGGC |
| 357 | mamo-s | *BGIBMGA000535* | down | 1121 | 1129 | - | 11.3303 | 4.96E-05 | 0.615 | GTGGGTGGC |
| 358 | mamo-s | *BGIBMGA000536* | down | 1742 | 1750 | - | 10.6881 | 8.61E-05 | 0.81 | GCACGTGGC |
| 359 | mamo-s | *BGIBMGA000542* | up | 741 | 749 | + | 11.1743 | 5.90E-05 | 0.684 | CCGCGTGTC |
| 360 | mamo-s | *BGIBMGA000551* | up | 1826 | 1834 | + | 13.6606 | 4.06E-06 | 0.615 | GCGCGTGGC |
| 361 | mamo-s | *BGIBMGA000553* | down | 777 | 785 | - | 11.0826 | 6.84E-05 | 0.709 | GTACGTGGC |
| 362 | mamo-s | *BGIBMGA000554* | down | 179 | 187 | - | 11.1376 | 6.35E-05 | 0.709 | TCGCGTGGC |
| 363 | mamo-s | *BGIBMGA000556* | down | 910 | 918 | + | 12.1835 | 1.71E-05 | 0.615 | GTGCGTGCC |
| 364 | mamo-s | *BGIBMGA000556* | down | 1858 | 1866 | + | 11.3303 | 4.96E-05 | 0.615 | GTGGGTGGC |
| 365 | mamo-s | *BGIBMGA000559* | up | 411 | 419 | - | 10.5413 | 9.42E-05 | 0.857 | GTGCGAGTC |
| 366 | mamo-s | *BGIBMGA000561* | down | 899 | 907 | + | 11.1376 | 6.35E-05 | 0.709 | TCGCGTGGC |
| 367 | mamo-s | *BGIBMGA000562* | down | 103 | 111 | + | 11.7798 | 3.11E-05 | 0.615 | ATGCGTGGC |
| 368 | mamo-s | *BGIBMGA000566* | down | 72 | 80 | + | 11.3853 | 4.74E-05 | 0.615 | ACGCGTGGC |
| 369 | mamo-s | *BGIBMGA000571* | up | 1874 | 1882 | - | 11.789 | 2.84E-05 | 0.615 | GCGCGTGCC |
| 370 | mamo-s | *BGIBMGA000571* | down | 1071 | 1079 | + | 11.3303 | 4.96E-05 | 0.615 | GTGGGTGGC |
| 371 | mamo-s | *BGIBMGA000572* | up | 768 | 776 | - | 12.1835 | 1.71E-05 | 0.615 | GTGCGTGCC |
| 372 | mamo-s | *BGIBMGA000575* | up | 1727 | 1735 | - | 11.3303 | 4.96E-05 | 0.615 | GTGGGTGGC |
| 373 | mamo-s | *BGIBMGA000576* | down | 923 | 931 | + | 10.5138 | 9.75E-05 | 0.865 | GTGCGTGTT |
| 374 | mamo-s | *BGIBMGA000579* | up | 1559 | 1567 | + | 11.3303 | 4.96E-05 | 0.615 | GTGGGTGGC |
| 375 | mamo-s | *BGIBMGA000580* | up | 1246 | 1254 | + | 11.3303 | 4.96E-05 | 0.615 | GTGGGTGGC |
| 376 | mamo-s | *BGIBMGA000582* | up | 391 | 399 | + | 11.3303 | 4.96E-05 | 0.615 | GTGGGTGGC |
| 377 | mamo-s | *BGIBMGA000584* | down | 1892 | 1900 | - | 11.3303 | 4.96E-05 | 0.615 | GTGGGTGGC |
| 378 | mamo-s | *BGIBMGA000585* | up | 1081 | 1089 | + | 11.1376 | 6.35E-05 | 0.709 | TCGCGTGGC |
| 379 | mamo-s | *BGIBMGA000587* | up | 1840 | 1848 | + | 11.3303 | 4.96E-05 | 0.615 | GTGGGTGGC |
| 380 | mamo-s | *BGIBMGA000591* | up | 553 | 561 | - | 11.3303 | 4.96E-05 | 0.615 | GTGGGTGGC |
| 381 | mamo-s | *BGIBMGA000592* | up | 334 | 342 | - | 11.1743 | 5.90E-05 | 0.684 | CCGCGTGTC |
| 382 | mamo-s | *BGIBMGA000593* | up | 1524 | 1532 | - | 11.3303 | 4.96E-05 | 0.615 | GTGGGTGGC |
| 383 | mamo-s | *BGIBMGA000594* | up | 104 | 112 | - | 11.3303 | 4.96E-05 | 0.615 | GTGGGTGGC |
| 384 | mamo-s | *BGIBMGA000594* | down | 488 | 496 | + | 11.3303 | 4.96E-05 | 0.615 | GTGGGTGGC |
| 385 | mamo-s | *BGIBMGA000595* | up | 190 | 198 | - | 10.9358 | 7.30E-05 | 0.728 | GCGGGTGGC |
| 386 | mamo-s | *BGIBMGA000597* | down | 1648 | 1656 | + | 11.4404 | 4.51E-05 | 0.615 | GAGCGTGTC |
| 387 | mamo-s | *BGIBMGA000597* | up | 686 | 694 | + | 11.3303 | 4.96E-05 | 0.615 | GTGGGTGGC |
| 388 | mamo-s | *BGIBMGA000598* | down | 581 | 589 | + | 12.1284 | 2.16E-05 | 0.615 | CAGCGTGGC |
| 389 | mamo-s | *BGIBMGA000600* | up | 676 | 684 | + | 12.4679 | 1.31E-05 | 0.615 | GTGCGTGTC |
| 390 | mamo-s | *BGIBMGA000601* | up | 506 | 514 | + | 11.3303 | 4.96E-05 | 0.615 | GTGGGTGGC |
| 391 | mamo-s | *BGIBMGA000604* | up | 411 | 419 | + | 11.3303 | 4.96E-05 | 0.615 | GTGGGTGGC |
| 392 | mamo-s | *BGIBMGA000604* | down | 937 | 945 | + | 11.2294 | 5.41E-05 | 0.65 | CTGCGAGGC |
| 393 | mamo-s | *BGIBMGA000604* | down | 944 | 952 | - | 10.6881 | 8.61E-05 | 0.81 | GCACGTGGC |
| 394 | mamo-s | *BGIBMGA000605* | up | 921 | 929 | + | 11.2294 | 5.41E-05 | 0.65 | CTGCGAGGC |
| 395 | mamo-s | *BGIBMGA000605* | up | 928 | 936 | - | 10.6881 | 8.61E-05 | 0.81 | GCACGTGGC |
| 396 | mamo-s | *BGIBMGA000608* | down | 1663 | 1671 | - | 10.5138 | 9.75E-05 | 0.865 | GTGCGTGTT |
| 397 | mamo-s | *BGIBMGA000610* | up | 1209 | 1217 | + | 11.4404 | 4.51E-05 | 0.615 | GAGCGTGTC |
| 398 | mamo-s | *BGIBMGA000610* | down | 915 | 923 | + | 10.5688 | 8.88E-05 | 0.824 | GTGCGTGAC |
| 399 | mamo-s | *BGIBMGA000611* | down | 1663 | 1671 | + | 11.3303 | 4.96E-05 | 0.615 | GTGGGTGGC |
| 400 | mamo-s | *BGIBMGA000611* | down | 685 | 693 | + | 11.1009 | 6.57E-05 | 0.709 | GAGCGAGGC |
| 401 | mamo-s | *BGIBMGA000612* | up | 1803 | 1811 | + | 10.7523 | 8.38E-05 | 0.797 | GGGCGTGTC |
| 402 | mamo-s | *BGIBMGA000613* | down | 1717 | 1725 | - | 12.7615 | 1.04E-05 | 0.615 | CCGCGTGGC |
| 403 | mamo-s | *BGIBMGA000615* | up | 1778 | 1786 | - | 11.3303 | 4.96E-05 | 0.615 | GTGGGTGGC |
| 404 | mamo-s | *BGIBMGA000618* | up | 1737 | 1745 | + | 11.1376 | 6.35E-05 | 0.709 | TCGCGTGGC |
| 405 | mamo-s | *BGIBMGA000619* | up | 1580 | 1588 | - | 11.3303 | 4.96E-05 | 0.615 | GTGGGTGGC |
| 406 | mamo-s | *BGIBMGA000620* | up | 45 | 53 | - | 12.1009 | 2.43E-05 | 0.615 | GTGCGTGGT |
| 407 | mamo-s | *BGIBMGA000622* | up | 1664 | 1672 | - | 10.8899 | 7.48E-05 | 0.739 | CCGCGTGCC |
| 408 | mamo-s | *BGIBMGA000628* | up | 583 | 591 | - | 10.8073 | 7.89E-05 | 0.766 | CCGCGTGGT |
| 409 | mamo-s | *BGIBMGA000630* | up | 465 | 473 | - | 10.5688 | 8.88E-05 | 0.824 | GTGCGTGAC |
| 410 | mamo-s | *BGIBMGA000631* | up | 108 | 116 | + | 13.0275 | 8.53E-06 | 0.615 | GAGCGTGGC |
| 411 | mamo-s | *BGIBMGA000631* | down | 508 | 516 | - | 11.3303 | 4.96E-05 | 0.615 | GTGGGTGGC |
| 412 | mamo-s | *BGIBMGA000631* | down | 1519 | 1527 | - | 11.3303 | 4.96E-05 | 0.615 | GTGGGTGGC |
| 413 | mamo-s | *BGIBMGA000632* | up | 1259 | 1267 | - | 11.3303 | 4.96E-05 | 0.615 | GTGGGTGGC |
| 414 | mamo-s | *BGIBMGA000634* | up | 682 | 690 | + | 14.055 | 2.23E-06 | 0.615 | GTGCGTGGC |
| 415 | mamo-s | *BGIBMGA000635* | down | 1491 | 1499 | + | 11.3303 | 4.96E-05 | 0.615 | GTGGGTGGC |
| 416 | mamo-s | *BGIBMGA000636* | up | 267 | 275 | - | 11.3303 | 4.96E-05 | 0.615 | GTGGGTGGC |
| 417 | mamo-s | *BGIBMGA000637* | down | 1791 | 1799 | - | 11.4404 | 4.51E-05 | 0.615 | GAGCGTGTC |
| 418 | mamo-s | *BGIBMGA000640* | down | 1386 | 1394 | - | 11.4404 | 4.51E-05 | 0.615 | GAGCGTGTC |
| 419 | mamo-s | *BGIBMGA000640* | up | 538 | 546 | - | 11.3303 | 4.96E-05 | 0.615 | GTGGGTGGC |
| 420 | mamo-s | *BGIBMGA000641* | down | 1852 | 1860 | + | 12.0734 | 2.66E-05 | 0.615 | GCGCGTGTC |
| 421 | mamo-s | *BGIBMGA000644* | up | 1947 | 1955 | + | 10.6881 | 8.61E-05 | 0.81 | GCACGTGGC |
| 422 | mamo-s | *BGIBMGA000648* | up | 1686 | 1694 | - | 11.3303 | 4.96E-05 | 0.615 | GTGGGTGGC |
| 423 | mamo-s | *BGIBMGA000648* | down | 1757 | 1765 | + | 11.0826 | 6.84E-05 | 0.709 | GTACGTGGC |
| 424 | mamo-s | *BGIBMGA000649* | down | 1040 | 1048 | - | 11.3303 | 4.96E-05 | 0.615 | GTGGGTGGC |
| 425 | mamo-s | *BGIBMGA000649* | down | 564 | 572 | + | 11.0826 | 6.84E-05 | 0.709 | GTACGTGGC |
| 426 | mamo-s | *BGIBMGA000650* | up | 296 | 304 | + | 13.6606 | 4.06E-06 | 0.615 | GCGCGTGGC |
| 427 | mamo-s | *BGIBMGA000650* | down | 1657 | 1665 | + | 10.7523 | 8.38E-05 | 0.797 | AAGCGTGGC |
| 428 | mamo-s | *BGIBMGA000653* | down | 1122 | 1130 | + | 11.3303 | 4.96E-05 | 0.615 | GTGGGTGGC |
| 429 | mamo-s | *BGIBMGA000653* | down | 1748 | 1756 | - | 11.3303 | 4.96E-05 | 0.615 | GTGGGTGGC |
| 430 | mamo-s | *BGIBMGA000655* | down | 1945 | 1953 | + | 11.2018 | 5.68E-05 | 0.674 | CTGCGTGGT |
| 431 | mamo-s | *BGIBMGA000656* | down | 1212 | 1220 | + | 10.8899 | 7.48E-05 | 0.739 | CCGCGTGCC |
| 432 | mamo-s | *BGIBMGA000658* | down | 1847 | 1855 | - | 11.3303 | 4.96E-05 | 0.615 | GTGGGTGGC |
| 433 | mamo-s | *BGIBMGA000659* | up | 236 | 244 | - | 11.0826 | 6.84E-05 | 0.709 | GTACGTGGC |

| | | | | | | | | | | |
|---|---|---|---|---|---|---|---|---|---|---|
| 434 | mamo-s | *BGIBMGA000660* | down | 1646 | 1654 | + | 11.0734 | 7.12E-05 | 0.726 | GAGCGTGGT |
| 435 | mamo-s | *BGIBMGA000660* | up | 1625 | 1633 | + | 10.5413 | 9.42E-05 | 0.857 | GTGCGAGTC |
| 436 | mamo-s | *BGIBMGA000662* | down | 1779 | 1787 | + | 11.0826 | 6.84E-05 | 0.709 | GTACGTGGC |
| 437 | mamo-s | *BGIBMGA000666* | up | 578 | 586 | - | 11.3303 | 4.96E-05 | 0.615 | GTGGGTGGC |
| 438 | mamo-s | *BGIBMGA000667* | down | 493 | 501 | + | 11.1376 | 6.35E-05 | 0.709 | TCGCGTGGC |
| 439 | mamo-s | *BGIBMGA000668* | down | 770 | 778 | + | 11.0826 | 6.84E-05 | 0.709 | GTACGTGGC |
| 440 | mamo-s | *BGIBMGA000674* | up | 689 | 697 | + | 14.055 | 2.23E-06 | 0.615 | GTGCGTGGC |
| 441 | mamo-s | *BGIBMGA000675* | up | 821 | 829 | - | 10.7523 | 8.38E-05 | 0.797 | AAGCGTGGC |
| 442 | mamo-s | *BGIBMGA000676* | down | 1404 | 1412 | + | 13.0275 | 8.53E-06 | 0.615 | GAGCGTGGC |
| 443 | mamo-s | *BGIBMGA000676* | up | 614 | 622 | + | 12.1835 | 1.71E-05 | 0.615 | GTGCGTGCC |
| 444 | mamo-s | *BGIBMGA000686* | up | 587 | 595 | - | 10.5138 | 9.75E-05 | 0.865 | GTGCGTGTT |
| 445 | mamo-s | *BGIBMGA000687* | down | 1360 | 1368 | - | 11.3303 | 4.96E-05 | 0.615 | GTGGGTGGC |
| 446 | mamo-s | *BGIBMGA000688* | up | 202 | 210 | - | 11.3303 | 4.96E-05 | 0.615 | GTGGGTGGC |
| 447 | mamo-s | *BGIBMGA000688* | down | 1377 | 1385 | - | 11.3303 | 4.96E-05 | 0.615 | GTGGGTGGC |
| 448 | mamo-s | *BGIBMGA000689* | down | 1147 | 1155 | + | 10.6881 | 8.61E-05 | 0.81 | GCACGTGGC |
| 449 | mamo-s | *BGIBMGA000691* | up | 1054 | 1062 | - | 10.5138 | 9.75E-05 | 0.865 | GTGCGTGTT |
| 450 | mamo-s | *BGIBMGA000692* | down | 1269 | 1277 | + | 12.1835 | 1.71E-05 | 0.615 | GTGCGTGCC |
| 451 | mamo-s | *BGIBMGA000692* | down | 696 | 704 | + | 11.3303 | 4.96E-05 | 0.615 | GTGGGTGGC |
| 452 | mamo-s | *BGIBMGA000693* | up | 1183 | 1191 | + | 12.1835 | 1.71E-05 | 0.615 | GTGCGTGCC |
| 453 | mamo-s | *BGIBMGA000693* | up | 610 | 618 | + | 11.3303 | 4.96E-05 | 0.615 | GTGGGTGGC |
| 454 | mamo-s | *BGIBMGA000695* | down | 1317 | 1325 | - | 11.3303 | 4.96E-05 | 0.615 | GTGGGTGGC |
| 455 | mamo-s | *BGIBMGA000696* | down | 1097 | 1105 | + | 10.5413 | 9.42E-05 | 0.857 | CAGCGTGTC |
| 456 | mamo-s | *BGIBMGA000697* | up | 1513 | 1521 | - | 11.0826 | 6.84E-05 | 0.709 | GTACGTGGC |
| 457 | mamo-s | *BGIBMGA000698* | up | 74 | 82 | + | 11.5321 | 4.06E-05 | 0.615 | TTGCGTGGC |
| 458 | mamo-s | *BGIBMGA000698* | up | 194 | 202 | - | 11.3303 | 4.96E-05 | 0.615 | GTGGGTGGC |
| 459 | mamo-s | *BGIBMGA000698* | down | 1429 | 1437 | + | 11.3303 | 4.96E-05 | 0.615 | GTGGGTGGC |
| 460 | mamo-s | *BGIBMGA000699* | up | 440 | 448 | + | 12.1009 | 2.43E-05 | 0.615 | GTGCGTGGT |
| 461 | mamo-s | *BGIBMGA000699* | up | 349 | 357 | + | 11.5321 | 4.06E-05 | 0.615 | TTGCGTGGC |
| 462 | mamo-s | *BGIBMGA000701* | up | 384 | 392 | - | 11.2018 | 5.68E-05 | 0.674 | CTGCGTGGT |
| 463 | mamo-s | *BGIBMGA000702* | down | 426 | 434 | - | 11.3303 | 4.96E-05 | 0.615 | GTGGGTGGC |
| 464 | mamo-s | *BGIBMGA000703* | up | 1124 | 1132 | - | 11.3303 | 4.96E-05 | 0.615 | GTGGGTGGC |
| 465 | mamo-s | *BGIBMGA000704* | down | 1467 | 1475 | - | 14.055 | 2.23E-06 | 0.615 | GTGCGTGGC |
| 466 | mamo-s | *BGIBMGA000705* | down | 1318 | 1326 | - | 12.3394 | 1.49E-05 | 0.615 | GGGCGTGGC |
| 467 | mamo-s | *BGIBMGA000705* | up | 1372 | 1380 | - | 11.2018 | 5.68E-05 | 0.674 | CTGCGTGGT |
| 468 | mamo-s | *BGIBMGA000708* | up | 391 | 399 | - | 11.3303 | 4.96E-05 | 0.615 | GTGGGTGGC |
| 469 | mamo-s | *BGIBMGA000714* | down | 61 | 69 | + | 10.7523 | 8.38E-05 | 0.797 | AAGCGTGGC |
| 470 | mamo-s | *BGIBMGA000715* | down | 1216 | 1224 | + | 11.3303 | 4.96E-05 | 0.615 | GTGGGTGGC |
| 471 | mamo-s | *BGIBMGA000716* | up | 1623 | 1631 | - | 11.789 | 2.84E-05 | 0.615 | GCGCGTGCC |
| 472 | mamo-s | *BGIBMGA000718* | down | 1700 | 1708 | - | 11.3303 | 4.96E-05 | 0.615 | GTGGGTGGC |
| 473 | mamo-s | *BGIBMGA000718* | up | 1316 | 1324 | + | 10.5413 | 9.42E-05 | 0.857 | GTGCGAGTC |
| 474 | mamo-s | *BGIBMGA000720* | down | 1857 | 1865 | - | 10.5688 | 8.88E-05 | 0.824 | GTGCGTGAC |
| 475 | mamo-s | *BGIBMGA000721* | up | 810 | 818 | + | 11.0826 | 6.84E-05 | 0.709 | GTACGTGGC |
| 476 | mamo-s | *BGIBMGA000721* | down | 564 | 572 | + | 11.0734 | 7.12E-05 | 0.726 | GAGCGTGGT |
| 477 | mamo-s | *BGIBMGA000722* | down | 288 | 296 | - | 11.3303 | 4.96E-05 | 0.615 | GTGGGTGGC |
| 478 | mamo-s | *BGIBMGA000725* | up | 1983 | 1991 | - | 14.055 | 2.23E-06 | 0.615 | GTGCGTGGC |
| 479 | mamo-s | *BGIBMGA000726* | up | 1972 | 1980 | + | 11.7064 | 3.52E-05 | 0.615 | GCGCGTGGT |
| 480 | mamo-s | *BGIBMGA000726* | up | 1208 | 1216 | + | 11.3303 | 4.96E-05 | 0.615 | GTGGGTGGC |
| 481 | mamo-s | *BGIBMGA000726* | down | 1398 | 1406 | - | 11.3303 | 4.96E-05 | 0.615 | GTGGGTGGC |
| 482 | mamo-s | *BGIBMGA000728* | down | 1107 | 1115 | + | 11.3303 | 4.96E-05 | 0.615 | GTGGGTGGC |
| 483 | mamo-s | *BGIBMGA000729* | up | 1052 | 1060 | + | 11.789 | 2.84E-05 | 0.615 | GCGCGTGCC |
| 484 | mamo-s | *BGIBMGA000729* | down | 290 | 298 | - | 11.0826 | 6.84E-05 | 0.709 | GTACGTGGC |
| 485 | mamo-s | *BGIBMGA000734* | up | 375 | 383 | - | 11.3303 | 4.96E-05 | 0.615 | GTGGGTGGC |
| 486 | mamo-s | *BGIBMGA000734* | down | 381 | 389 | - | 11.3303 | 4.96E-05 | 0.615 | GTGGGTGGC |
| 487 | mamo-s | *BGIBMGA000734* | down | 1157 | 1165 | + | 11.3303 | 4.96E-05 | 0.615 | GTGGGTGGC |
| 488 | mamo-s | *BGIBMGA000734* | up | 1429 | 1437 | - | 11.2018 | 5.68E-05 | 0.674 | CTGCGTGGT |
| 489 | mamo-s | *BGIBMGA000737* | up | 916 | 924 | + | 13.156 | 6.30E-06 | 0.615 | CTGCGTGGC |
| 490 | mamo-s | *BGIBMGA000737* | down | 401 | 409 | - | 11.3303 | 4.96E-05 | 0.615 | GTGGGTGGC |
| 491 | mamo-s | *BGIBMGA000737* | up | 815 | 823 | - | 11.3303 | 4.96E-05 | 0.615 | GTGGGTGGC |
| 492 | mamo-s | *BGIBMGA000739* | down | 1762 | 1770 | + | 11.3303 | 4.96E-05 | 0.615 | GTGGGTGGC |
| 493 | mamo-s | *BGIBMGA000740* | up | 1680 | 1688 | - | 11.0826 | 6.84E-05 | 0.709 | GTACGTGGC |
| 494 | mamo-s | *BGIBMGA000741* | up | 213 | 221 | - | 11.3303 | 4.96E-05 | 0.615 | GTGGGTGGC |
| 495 | mamo-s | *BGIBMGA000741* | down | 1720 | 1728 | - | 11.3303 | 4.96E-05 | 0.615 | GTGGGTGGC |
| 496 | mamo-s | *BGIBMGA000741* | up | 823 | 831 | - | 11.0734 | 7.12E-05 | 0.726 | GAGCGTGGT |
| 497 | mamo-s | *BGIBMGA000741* | down | 1544 | 1552 | - | 10.7523 | 8.38E-05 | 0.797 | AAGCGTGGC |
| 498 | mamo-s | *BGIBMGA000743* | down | 1173 | 1181 | + | 11.3303 | 4.96E-05 | 0.615 | GTGGGTGGC |
| 499 | mamo-s | *BGIBMGA000745* | down | 254 | 262 | + | 11.0826 | 6.84E-05 | 0.709 | GTACGTGGC |
| 500 | mamo-s | *BGIBMGA000746* | up | 1550 | 1558 | + | 11.3303 | 4.96E-05 | 0.615 | GTGGGTGGC |
| 501 | mamo-s | *BGIBMGA000748* | down | 1029 | 1037 | + | 10.8073 | 7.89E-05 | 0.766 | CCGCGTGGT |
| 502 | mamo-s | *BGIBMGA000749* | up | 1728 | 1736 | + | 11.3303 | 4.96E-05 | 0.615 | GTGGGTGGC |
| 503 | mamo-s | *BGIBMGA000749* | down | 1696 | 1704 | + | 11.2294 | 5.41E-05 | 0.65 | CTGCGAGGC |
| 504 | mamo-s | *BGIBMGA000750* | down | 65 | 73 | + | 12.1835 | 1.71E-05 | 0.615 | GTGCGTGCC |
| 505 | mamo-s | *BGIBMGA000750* | down | 888 | 896 | - | 11.3303 | 4.96E-05 | 0.615 | GTGGGTGGC |
| 506 | mamo-s | *BGIBMGA000750* | down | 1603 | 1611 | - | 11.3303 | 4.96E-05 | 0.615 | GTGGGTGGC |
| 507 | mamo-s | *BGIBMGA000750* | down | 557 | 565 | - | 10.9358 | 7.30E-05 | 0.728 | GCGGGTGGC |
| 508 | mamo-s | *BGIBMGA000751* | up | 1935 | 1943 | + | 12.4679 | 1.31E-05 | 0.615 | GTGCGTGTC |
| 509 | mamo-s | *BGIBMGA000751* | down | 297 | 305 | + | 11.3303 | 4.96E-05 | 0.615 | GTGGGTGGC |
| 510 | mamo-s | *BGIBMGA000751* | up | 595 | 603 | + | 11.3303 | 4.96E-05 | 0.615 | GTGGGTGGC |
| 511 | mamo-s | *BGIBMGA000754* | up | 627 | 635 | - | 11.0826 | 6.84E-05 | 0.709 | GTACGTGGC |
| 512 | mamo-s | *BGIBMGA000756* | down | 1530 | 1538 | + | 10.5413 | 9.42E-05 | 0.857 | GTGCGAGTC |
| 513 | mamo-s | *BGIBMGA000758* | down | 1956 | 1964 | + | 11.3853 | 4.74E-05 | 0.615 | ACGCGTGGC |
| 514 | mamo-s | *BGIBMGA000760* | down | 948 | 956 | + | 11.3303 | 4.96E-05 | 0.615 | GTGGGTGGC |
| 515 | mamo-s | *BGIBMGA000762* | up | 652 | 660 | + | 11.3303 | 4.96E-05 | 0.615 | GTGGGTGGC |
| 516 | mamo-s | *BGIBMGA000762* | up | 1819 | 1827 | + | 11.3303 | 4.96E-05 | 0.615 | GTGGGTGGC |
| 517 | mamo-s | *BGIBMGA000763* | up | 1877 | 1885 | + | 11.3303 | 4.96E-05 | 0.615 | GTGGGTGGC |
| 518 | mamo-s | *BGIBMGA000763* | up | 1506 | 1514 | + | 10.8073 | 7.89E-05 | 0.766 | CCGCGTGGT |
| 519 | mamo-s | *BGIBMGA000764* | down | 1377 | 1385 | - | 12.7615 | 1.04E-05 | 0.615 | CCGCGTGGC |
| 520 | mamo-s | *BGIBMGA000764* | down | 801 | 809 | - | 11.3303 | 4.96E-05 | 0.615 | GTGGGTGGC |
| 521 | mamo-s | *BGIBMGA000764* | up | 1340 | 1348 | + | 11.3303 | 4.96E-05 | 0.615 | GTGGGTGGC |

| | | | | | | | | | | |
|---|---|---|---|---|---|---|---|---|---|---|
| 522 | mamo-s | *BGIBMGA000764* | up | 1552 | 1560 | - | 11.3303 | 4.96E-05 | 0.615 | GTGGGTGGC |
| 523 | mamo-s | *BGIBMGA000764* | down | 1556 | 1564 | + | 10.5413 | 9.42E-05 | 0.857 | GTGCGAGTC |
| 524 | mamo-s | *BGIBMGA000765* | down | 1595 | 1603 | - | 10.5138 | 9.75E-05 | 0.865 | GTGCGTGTT |
| 525 | mamo-s | *BGIBMGA000767* | down | 1926 | 1934 | + | 14.055 | 2.23E-06 | 0.615 | GTGCGTGGC |
| 526 | mamo-s | *BGIBMGA000767* | down | 1694 | 1702 | - | 11.1009 | 6.57E-05 | 0.709 | GAGCGAGGC |
| 527 | mamo-s | *BGIBMGA000767* | up | 142 | 150 | - | 10.7523 | 8.38E-05 | 0.797 | AAGCGTGGC |
| 528 | mamo-s | *BGIBMGA000769* | up | 1619 | 1627 | - | 11.3303 | 4.96E-05 | 0.615 | GTGGGTGGC |
| 529 | mamo-s | *BGIBMGA000771* | down | 1039 | 1047 | - | 11.2294 | 5.41E-05 | 0.65 | CTGCGAGGC |
| 530 | mamo-s | *BGIBMGA000775* | up | 552 | 560 | + | 11.3303 | 4.96E-05 | 0.615 | GTGGGTGGC |
| 531 | mamo-s | *BGIBMGA000776* | down | 362 | 370 | - | 11.3303 | 4.96E-05 | 0.615 | GTGGGTGGC |
| 532 | mamo-s | *BGIBMGA000777* | up | 938 | 946 | + | 11.3303 | 4.96E-05 | 0.615 | GTGGGTGGC |
| 533 | mamo-s | *BGIBMGA000778* | down | 654 | 662 | + | 11.3303 | 4.96E-05 | 0.615 | GTGGGTGGC |
| 534 | mamo-s | *BGIBMGA000779* | down | 58 | 66 | - | 13.6606 | 4.06E-06 | 0.615 | GCGCGTGGC |
| 535 | mamo-s | *BGIBMGA000779* | down | 321 | 329 | - | 11.7339 | 3.29E-05 | 0.615 | GCGCGAGGC |
| 536 | mamo-s | *BGIBMGA000781* | down | 990 | 998 | - | 10.9358 | 7.30E-05 | 0.728 | GCGGGTGGC |
| 537 | mamo-s | *BGIBMGA000782* | down | 1858 | 1866 | + | 11.3303 | 4.96E-05 | 0.615 | GTGGGTGGC |
| 538 | mamo-s | *BGIBMGA000783* | down | 719 | 727 | - | 11.1009 | 6.57E-05 | 0.709 | GAGCGAGGC |
| 539 | mamo-s | *BGIBMGA000783* | up | 1674 | 1682 | + | 10.5138 | 9.75E-05 | 0.865 | GTGCGTGTT |
| 540 | mamo-s | *BGIBMGA000784* | down | 541 | 549 | + | 11.789 | 2.84E-05 | 0.615 | GCGCGTGCC |
| 541 | mamo-s | *BGIBMGA000784* | down | 952 | 960 | + | 11.5321 | 4.06E-05 | 0.615 | TTGCGTGGC |
| 542 | mamo-s | *BGIBMGA000784* | up | 241 | 249 | - | 10.5138 | 9.75E-05 | 0.865 | GTGCGTGTT |
| 543 | mamo-s | *BGIBMGA000786* | down | 1526 | 1534 | - | 11.3303 | 4.96E-05 | 0.615 | GTGGGTGGC |
| 544 | mamo-s | *BGIBMGA000787* | up | 231 | 239 | + | 12.4679 | 1.31E-05 | 0.615 | GTGCGTGTC |
| 545 | mamo-s | *BGIBMGA000787* | down | 768 | 776 | - | 11.3303 | 4.96E-05 | 0.615 | GTGGGTGGC |
| 546 | mamo-s | *BGIBMGA000787* | up | 1740 | 1748 | - | 11.3303 | 4.96E-05 | 0.615 | GTGGGTGGC |
| 547 | mamo-s | *BGIBMGA000787* | down | 338 | 346 | + | 11.2844 | 5.18E-05 | 0.63 | CTGCGTGCC |
| 548 | mamo-s | *BGIBMGA000788* | up | 632 | 640 | + | 11.3303 | 4.96E-05 | 0.615 | GTGGGTGGC |
| 549 | mamo-s | *BGIBMGA000790* | up | 1752 | 1760 | + | 11.0826 | 6.84E-05 | 0.709 | GTACGTGGC |
| 550 | mamo-s | *BGIBMGA000791* | down | 197 | 205 | + | 12.1284 | 2.16E-05 | 0.615 | CAGCGTGGC |
| 551 | mamo-s | *BGIBMGA000791* | down | 1639 | 1647 | - | 11.156 | 6.13E-05 | 0.705 | GAGCGTGCC |
| 552 | mamo-s | *BGIBMGA000797* | up | 1361 | 1369 | + | 11.3853 | 4.74E-05 | 0.615 | ACGCGTGGC |
| 553 | mamo-s | *BGIBMGA000798* | down | 34 | 42 | - | 13.6606 | 4.06E-06 | 0.615 | GCGCGTGGC |
| 554 | mamo-s | *BGIBMGA000799* | up | 1306 | 1314 | + | 11.3303 | 4.96E-05 | 0.615 | GTGGGTGGC |
| 555 | mamo-s | *BGIBMGA000800* | down | 1389 | 1397 | + | 11.7339 | 3.29E-05 | 0.615 | GCGCGAGGC |
| 556 | mamo-s | *BGIBMGA000800* | up | 959 | 967 | + | 11.3303 | 4.96E-05 | 0.615 | GTGGGTGGC |
| 557 | mamo-s | *BGIBMGA000800* | up | 143 | 151 | - | 11.0826 | 6.84E-05 | 0.709 | GTACGTGGC |
| 558 | mamo-s | *BGIBMGA000801* | down | 501 | 509 | + | 11.0826 | 6.84E-05 | 0.709 | GTACGTGGC |
| 559 | mamo-s | *BGIBMGA000802* | down | 129 | 137 | + | 11.3303 | 4.96E-05 | 0.615 | GTGGGTGGC |
| 560 | mamo-s | *BGIBMGA000802* | down | 1543 | 1551 | + | 11.3303 | 4.96E-05 | 0.615 | GTGGGTGGC |
| 561 | mamo-s | *BGIBMGA000806* | up | 1844 | 1852 | + | 11.3303 | 4.96E-05 | 0.615 | GTGGGTGGC |
| 562 | mamo-s | *BGIBMGA000807* | down | 1254 | 1262 | + | 11.0734 | 7.12E-05 | 0.726 | GAGCGTGGT |
| 563 | mamo-s | *BGIBMGA000809* | down | 354 | 362 | - | 11.3303 | 4.96E-05 | 0.615 | GTGGGTGGC |
| 564 | mamo-s | *BGIBMGA000810* | up | 1359 | 1367 | + | 11.789 | 2.84E-05 | 0.615 | GCGCGTGCC |
| 565 | mamo-s | *BGIBMGA000811* | up | 1767 | 1775 | + | 12.0734 | 2.66E-05 | 0.615 | GCGCGTGTC |
| 566 | mamo-s | *BGIBMGA000811* | up | 141 | 149 | - | 11.3303 | 4.96E-05 | 0.615 | GTGGGTGGC |
| 567 | mamo-s | *BGIBMGA000811* | up | 1584 | 1592 | + | 11.3303 | 4.96E-05 | 0.615 | GTGGGTGGC |
| 568 | mamo-s | *BGIBMGA000813* | down | 1847 | 1855 | + | 13.156 | 6.30E-06 | 0.615 | CTGCGTGGC |
| 569 | mamo-s | *BGIBMGA000813* | down | 648 | 656 | - | 11.3303 | 4.96E-05 | 0.615 | GTGGGTGGC |
| 570 | mamo-s | *BGIBMGA000813* | down | 1941 | 1949 | - | 10.7523 | 8.38E-05 | 0.797 | GGGCGTGTC |
| 571 | mamo-s | *BGIBMGA000814* | down | 1254 | 1262 | - | 11.3303 | 4.96E-05 | 0.615 | GTGGGTGGC |
| 572 | mamo-s | *BGIBMGA000815* | up | 800 | 808 | + | 11.3303 | 4.96E-05 | 0.615 | GTGGGTGGC |
| 573 | mamo-s | *BGIBMGA000815* | down | 1065 | 1073 | - | 10.8899 | 7.48E-05 | 0.739 | CCGCGTGCC |
| 574 | mamo-s | *BGIBMGA000816* | down | 1348 | 1356 | + | 11.3303 | 4.96E-05 | 0.615 | GTGGGTGGC |
| 575 | mamo-s | *BGIBMGA000820* | up | 1849 | 1857 | - | 11.0734 | 7.12E-05 | 0.726 | GAGCGTGGT |
| 576 | mamo-s | *BGIBMGA000821* | up | 1742 | 1750 | + | 11.3303 | 4.96E-05 | 0.615 | GTGGGTGGC |
| 577 | mamo-s | *BGIBMGA000821* | up | 1162 | 1170 | + | 10.5688 | 8.88E-05 | 0.824 | GTGCGTGAC |
| 578 | mamo-s | *BGIBMGA000823* | down | 358 | 366 | - | 14.055 | 2.23E-06 | 0.615 | GTGCGTGGC |
| 579 | mamo-s | *BGIBMGA000824* | down | 1042 | 1050 | + | 11.3303 | 4.96E-05 | 0.615 | GTGGGTGGC |
| 580 | mamo-s | *BGIBMGA000824* | down | 1757 | 1765 | + | 11.3303 | 4.96E-05 | 0.615 | GTGGGTGGC |
| 581 | mamo-s | *BGIBMGA000824* | down | 648 | 656 | + | 11.2018 | 5.68E-05 | 0.674 | CTGCGTGGT |
| 582 | mamo-s | *BGIBMGA000825* | down | 1888 | 1896 | + | 12.1009 | 2.43E-05 | 0.615 | GTGCGTGGT |
| 583 | mamo-s | *BGIBMGA000827* | down | 1079 | 1087 | - | 11.3303 | 4.96E-05 | 0.615 | GTGGGTGGC |
| 584 | mamo-s | *BGIBMGA000828* | down | 1562 | 1570 | - | 11.5321 | 4.06E-05 | 0.615 | TTGCGTGGC |
| 585 | mamo-s | *BGIBMGA000828* | down | 1063 | 1071 | - | 11.4404 | 4.51E-05 | 0.615 | GAGCGTGTC |
| 586 | mamo-s | *BGIBMGA000828* | down | 612 | 620 | + | 11.3303 | 4.96E-05 | 0.615 | GTGGGTGGC |
| 587 | mamo-s | *BGIBMGA000828* | down | 1421 | 1429 | + | 11.3303 | 4.96E-05 | 0.615 | GTGGGTGGC |
| 588 | mamo-s | *BGIBMGA000830* | down | 1969 | 1977 | + | 11.7339 | 3.29E-05 | 0.615 | GCGCGAGGC |
| 589 | mamo-s | *BGIBMGA000830* | down | 1992 | 2000 | - | 11.2018 | 5.68E-05 | 0.674 | CTGCGTGGT |
| 590 | mamo-s | *BGIBMGA000831* | up | 152 | 160 | + | 14.055 | 2.23E-06 | 0.615 | GTGCGTGGC |
| 591 | mamo-s | *BGIBMGA000831* | down | 125 | 133 | + | 11.1009 | 6.57E-05 | 0.709 | GAGCGAGGC |
| 592 | mamo-s | *BGIBMGA000833* | down | 1399 | 1407 | - | 10.5138 | 9.75E-05 | 0.865 | GTGCGTGTT |
| 593 | mamo-s | *BGIBMGA000835* | up | 1254 | 1262 | + | 11.3303 | 4.96E-05 | 0.615 | GTGGGTGGC |
| 594 | mamo-s | *BGIBMGA000835* | up | 445 | 453 | - | 11.0826 | 6.84E-05 | 0.709 | GTACGTGGC |
| 595 | mamo-s | *BGIBMGA000835* | up | 1729 | 1737 | - | 11.0826 | 6.84E-05 | 0.709 | GTACGTGGC |
| 596 | mamo-s | *BGIBMGA000836* | up | 1017 | 1025 | - | 11.3303 | 4.96E-05 | 0.615 | GTGGGTGGC |
| 597 | mamo-s | *BGIBMGA000836* | up | 542 | 550 | + | 11.0826 | 6.84E-05 | 0.709 | GTACGTGGC |
| 598 | mamo-s | *BGIBMGA000836* | up | 1826 | 1834 | + | 11.0826 | 6.84E-05 | 0.709 | GTACGTGGC |
| 599 | mamo-s | *BGIBMGA000837* | up | 471 | 479 | - | 13.156 | 6.30E-06 | 0.615 | CTGCGTGGC |
| 600 | mamo-s | *BGIBMGA000837* | up | 1114 | 1122 | + | 11.3303 | 4.96E-05 | 0.615 | GTGGGTGGC |
| 601 | mamo-s | *BGIBMGA000838* | down | 1971 | 1979 | - | 11.7339 | 3.29E-05 | 0.615 | GCGCGAGGC |
| 602 | mamo-s | *BGIBMGA000838* | down | 1948 | 1956 | + | 11.5321 | 4.06E-05 | 0.615 | TTGCGTGGC |
| 603 | mamo-s | *BGIBMGA000838* | up | 10 | 18 | + | 11.3303 | 4.96E-05 | 0.615 | GTGGGTGGC |
| 604 | mamo-s | *BGIBMGA000838* | up | 1918 | 1926 | - | 11.156 | 6.13E-05 | 0.705 | GAGCGTGCC |
| 605 | mamo-s | *BGIBMGA000839* | down | 498 | 506 | + | 11.3303 | 4.96E-05 | 0.615 | GTGGGTGGC |
| 606 | mamo-s | *BGIBMGA000840* | down | 1628 | 1636 | + | 11.3303 | 4.96E-05 | 0.615 | GTGGGTGGC |
| 607 | mamo-s | *BGIBMGA000841* | up | 454 | 462 | + | 11.3303 | 4.96E-05 | 0.615 | GTGGGTGGC |
| 608 | mamo-s | *BGIBMGA000841* | up | 1661 | 1669 | - | 11.3303 | 4.96E-05 | 0.615 | GTGGGTGGC |
| 609 | mamo-s | *BGIBMGA000841* | up | 210 | 218 | + | 10.9358 | 7.30E-05 | 0.728 | GCGGGTGGC |

| | | | | | | | | | | |
|---|---|---|---|---|---|---|---|---|---|---|
| 610 | mamo-s | *BGIBMGA000842* | down | 458 | 466 | + | 11.0826 | 6.84E-05 | 0.709 | GTACGTGGC |
| 611 | mamo-s | *BGIBMGA000844* | up | 387 | 395 | + | 11.3303 | 4.96E-05 | 0.615 | GTGGGTGGC |
| 612 | mamo-s | *BGIBMGA000849* | down | 893 | 901 | - | 11.3303 | 4.96E-05 | 0.615 | GTGGGTGGC |
| 613 | mamo-s | *BGIBMGA000849* | up | 1266 | 1274 | + | 11.3303 | 4.96E-05 | 0.615 | GTGGGTGGC |
| 614 | mamo-s | *BGIBMGA000850* | up | 1852 | 1860 | - | 11.3303 | 4.96E-05 | 0.615 | GTGGGTGGC |
| 615 | mamo-s | *BGIBMGA000852* | down | 405 | 413 | + | 11.4404 | 4.51E-05 | 0.615 | CGGCGTGGC |
| 616 | mamo-s | *BGIBMGA000852* | down | 421 | 429 | + | 11.3303 | 4.96E-05 | 0.615 | GTGGGTGGC |
| 617 | mamo-s | *BGIBMGA000853* | up | 1958 | 1966 | + | 11.4404 | 4.51E-05 | 0.615 | CGGCGTGGC |
| 618 | mamo-s | *BGIBMGA000853* | up | 1974 | 1982 | + | 11.3303 | 4.96E-05 | 0.615 | GTGGGTGGC |
| 619 | mamo-s | *BGIBMGA000854* | up | 631 | 639 | + | 11.3303 | 4.96E-05 | 0.615 | GTGGGTGGC |
| 620 | mamo-s | *BGIBMGA000859* | up | 1646 | 1654 | - | 12.4679 | 1.31E-05 | 0.615 | GTGCGTGTC |
| 621 | mamo-s | *BGIBMGA000860* | up | 1707 | 1715 | + | 11.0826 | 6.84E-05 | 0.709 | GTACGTGGC |
| 622 | mamo-s | *BGIBMGA000863* | up | 1376 | 1384 | + | 11.3303 | 4.96E-05 | 0.615 | GTGGGTGGC |
| 623 | mamo-s | *BGIBMGA000865* | up | 260 | 268 | + | 14.055 | 2.23E-06 | 0.615 | GTGCGTGGC |
| 624 | mamo-s | *BGIBMGA000865* | down | 486 | 494 | + | 11.3303 | 4.96E-05 | 0.615 | GTGGGTGGC |
| 625 | mamo-s | *BGIBMGA000866* | down | 820 | 828 | - | 11.3303 | 4.96E-05 | 0.615 | GTGGGTGGC |
| 626 | mamo-s | *BGIBMGA000867* | down | 381 | 389 | - | 11.3303 | 4.96E-05 | 0.615 | GTGGGTGGC |
| 627 | mamo-s | *BGIBMGA000868* | up | 1776 | 1784 | - | 11.3303 | 4.96E-05 | 0.615 | GTGGGTGGC |
| 628 | mamo-s | *BGIBMGA000869* | down | 1050 | 1058 | + | 11.3303 | 4.96E-05 | 0.615 | GTGGGTGGC |
| 629 | mamo-s | *BGIBMGA000870* | down | 1229 | 1237 | - | 11.3303 | 4.96E-05 | 0.615 | GTGGGTGGC |
| 630 | mamo-s | *BGIBMGA000871* | up | 927 | 935 | - | 11.1009 | 6.57E-05 | 0.709 | GAGCGAGGC |
| 631 | mamo-s | *BGIBMGA000871* | down | 661 | 669 | + | 10.5688 | 8.88E-05 | 0.824 | GTGCGTGAC |
| 632 | mamo-s | *BGIBMGA000872* | down | 1711 | 1719 | - | 11.3303 | 4.96E-05 | 0.615 | GTGGGTGGC |
| 633 | mamo-s | *BGIBMGA000873* | down | 363 | 371 | - | 11.3303 | 4.96E-05 | 0.615 | GTGGGTGGC |
| 634 | mamo-s | *BGIBMGA000875* | up | 1229 | 1237 | + | 11.3303 | 4.96E-05 | 0.615 | GTGGGTGGC |
| 635 | mamo-s | *BGIBMGA000876* | down | 1646 | 1654 | - | 11.3303 | 4.96E-05 | 0.615 | GTGGGTGGC |
| 636 | mamo-s | *BGIBMGA000877* | up | 1893 | 1901 | + | 11.7339 | 3.29E-05 | 0.615 | GCGCGAGGC |
| 637 | mamo-s | *BGIBMGA000877* | down | 818 | 826 | - | 11.3303 | 4.96E-05 | 0.615 | GTGGGTGGC |
| 638 | mamo-s | *BGIBMGA000878* | down | 1902 | 1910 | - | 11.789 | 2.84E-05 | 0.615 | GCGCGTGCC |
| 639 | mamo-s | *BGIBMGA000881* | up | 1216 | 1224 | - | 11.3303 | 4.96E-05 | 0.615 | GTGGGTGGC |
| 640 | mamo-s | *BGIBMGA000883* | up | 1348 | 1356 | - | 11.0826 | 6.84E-05 | 0.709 | GTACGTGGC |
| 641 | mamo-s | *BGIBMGA000885* | down | 1236 | 1244 | + | 11.3303 | 4.96E-05 | 0.615 | GTGGGTGGC |
| 642 | mamo-s | *BGIBMGA000886* | down | 1050 | 1058 | + | 11.3303 | 4.96E-05 | 0.615 | GTGGGTGGC |
| 643 | mamo-s | *BGIBMGA000889* | down | 1151 | 1159 | + | 11.3303 | 4.96E-05 | 0.615 | GTGGGTGGC |
| 644 | mamo-s | *BGIBMGA000894* | down | 964 | 972 | + | 11.3303 | 4.96E-05 | 0.615 | GTGGGTGGC |
| 645 | mamo-s | *BGIBMGA000894* | down | 604 | 612 | + | 10.7523 | 8.38E-05 | 0.797 | GGGCGTGTC |
| 646 | mamo-s | *BGIBMGA000895* | down | 570 | 578 | + | 11.7339 | 3.29E-05 | 0.615 | GCGCGAGGC |
| 647 | mamo-s | *BGIBMGA000898* | down | 1702 | 1710 | - | 13.156 | 6.30E-06 | 0.615 | CTGCGTGGC |
| 648 | mamo-s | *BGIBMGA000898* | down | 1850 | 1858 | + | 13.156 | 6.30E-06 | 0.615 | CTGCGTGGC |
| 649 | mamo-s | *BGIBMGA000899* | up | 1509 | 1517 | - | 13.156 | 6.30E-06 | 0.615 | CTGCGTGGC |
| 650 | mamo-s | *BGIBMGA000899* | up | 1657 | 1665 | + | 13.156 | 6.30E-06 | 0.615 | CTGCGTGGC |
| 651 | mamo-s | *BGIBMGA000899* | down | 1343 | 1351 | + | 11.3303 | 4.96E-05 | 0.615 | GTGGGTGGC |
| 652 | mamo-s | *BGIBMGA000901* | up | 566 | 574 | + | 12.1009 | 2.43E-05 | 0.615 | GTGCGTGGT |
| 653 | mamo-s | *BGIBMGA000902* | up | 121 | 129 | + | 11.3303 | 4.96E-05 | 0.615 | GTGGGTGGC |
| 654 | mamo-s | *BGIBMGA000902* | down | 8 | 16 | + | 10.8899 | 7.48E-05 | 0.739 | CCGCGTGCC |
| 655 | mamo-s | *BGIBMGA000904* | up | 948 | 956 | + | 11.1743 | 5.90E-05 | 0.684 | CCGCGTGTC |
| 656 | mamo-s | *BGIBMGA000904* | down | 1299 | 1307 | - | 11.0826 | 6.84E-05 | 0.709 | GTACGTGGC |
| 657 | mamo-s | *BGIBMGA000906* | up | 663 | 671 | - | 11.3303 | 4.96E-05 | 0.615 | GTGGGTGGC |
| 658 | mamo-s | *BGIBMGA000906* | down | 33 | 41 | + | 10.5413 | 9.42E-05 | 0.857 | GTGCGAGTC |
| 659 | mamo-s | *BGIBMGA000908* | down | 531 | 539 | - | 10.5138 | 9.75E-05 | 0.865 | GTGCGTGTT |
| 660 | mamo-s | *BGIBMGA000912* | up | 833 | 841 | + | 11.3303 | 4.96E-05 | 0.615 | GTGGGTGGC |
| 661 | mamo-s | *BGIBMGA000912* | down | 1757 | 1765 | + | 11.1009 | 6.57E-05 | 0.709 | GAGCGAGGC |
| 662 | mamo-s | *BGIBMGA000913* | up | 900 | 908 | + | 10.7523 | 8.38E-05 | 0.797 | GGGCGTGTC |
| 663 | mamo-s | *BGIBMGA000917* | down | 1639 | 1647 | - | 11.3303 | 4.96E-05 | 0.615 | GTGGGTGGC |
| 664 | mamo-s | *BGIBMGA000918* | down | 1872 | 1880 | + | 10.5138 | 9.75E-05 | 0.865 | GTGCGTGTT |
| 665 | mamo-s | *BGIBMGA000920* | down | 776 | 784 | - | 11.3303 | 4.96E-05 | 0.615 | GTGGGTGGC |
| 666 | mamo-s | *BGIBMGA000924* | up | 729 | 737 | - | 11.0826 | 6.84E-05 | 0.709 | GTACGTGGC |
| 667 | mamo-s | *BGIBMGA000924* | up | 45 | 53 | + | 11.0734 | 7.12E-05 | 0.726 | GAGCGTGGT |
| 668 | mamo-s | *BGIBMGA000926* | up | 1291 | 1299 | - | 11.3853 | 4.74E-05 | 0.615 | ACGCGTGGC |
| 669 | mamo-s | *BGIBMGA000929* | up | 365 | 373 | + | 11.3303 | 4.96E-05 | 0.615 | GTGGGTGGC |
| 670 | mamo-s | *BGIBMGA000931* | down | 1806 | 1814 | - | 11.1009 | 6.57E-05 | 0.709 | GAGCGAGGC |
| 671 | mamo-s | *BGIBMGA000934* | down | 1376 | 1384 | + | 11.3303 | 4.96E-05 | 0.615 | GTGGGTGGC |
| 672 | mamo-s | *BGIBMGA000935* | down | 1529 | 1537 | + | 11.0826 | 6.84E-05 | 0.709 | GTACGTGGC |
| 673 | mamo-s | *BGIBMGA000937* | up | 744 | 752 | - | 11.3303 | 4.96E-05 | 0.615 | GTGGGTGGC |
| 674 | mamo-s | *BGIBMGA000948* | up | 687 | 695 | - | 11.3303 | 4.96E-05 | 0.615 | GTGGGTGGC |
| 675 | mamo-s | *BGIBMGA000949* | down | 1073 | 1081 | - | 12.1284 | 2.16E-05 | 0.615 | GTGCGAGGC |
| 676 | mamo-s | *BGIBMGA000950* | up | 371 | 379 | - | 11.3303 | 4.96E-05 | 0.615 | GTGGGTGGC |
| 677 | mamo-s | *BGIBMGA000951* | up | 196 | 204 | + | 10.8349 | 7.66E-05 | 0.751 | CCGCGAGGC |
| 678 | mamo-s | *BGIBMGA000952* | down | 1683 | 1691 | + | 12.1009 | 2.43E-05 | 0.615 | GTGCGTGGT |
| 679 | mamo-s | *BGIBMGA000955* | up | 391 | 399 | - | 14.055 | 2.23E-06 | 0.615 | GTGCGTGGC |
| 680 | mamo-s | *BGIBMGA000960* | down | 761 | 769 | - | 11.3303 | 4.96E-05 | 0.615 | GTGGGTGGC |
| 681 | mamo-s | *BGIBMGA000963* | up | 1220 | 1228 | + | 11.2844 | 5.18E-05 | 0.63 | CTGCGTGCC |
| 682 | mamo-s | *BGIBMGA000966* | down | 580 | 588 | + | 11.2294 | 5.41E-05 | 0.65 | CTGCGAGGC |
| 683 | mamo-s | *BGIBMGA000967* | up | 845 | 853 | - | 11.5321 | 4.06E-05 | 0.615 | TTGCGTGGC |
| 684 | mamo-s | *BGIBMGA000969* | down | 1 | 9 | - | 11.7798 | 3.11E-05 | 0.615 | ATGCGTGGC |
| 685 | mamo-s | *BGIBMGA000969* | down | 1427 | 1435 | + | 10.7523 | 8.38E-05 | 0.797 | AAGCGTGGC |
| 686 | mamo-s | *BGIBMGA000969* | up | 180 | 188 | + | 10.5138 | 9.75E-05 | 0.865 | GTGCGTGTT |
| 687 | mamo-s | *BGIBMGA000971* | down | 1362 | 1370 | + | 11.3303 | 4.96E-05 | 0.615 | GTGGGTGGC |
| 688 | mamo-s | *BGIBMGA000971* | down | 342 | 350 | - | 11.2018 | 5.68E-05 | 0.674 | CTGCGTGGT |
| 689 | mamo-s | *BGIBMGA000972* | up | 1668 | 1676 | - | 11.3303 | 4.96E-05 | 0.615 | GTGGGTGGC |
| 690 | mamo-s | *BGIBMGA000984* | up | 1710 | 1718 | - | 11.3303 | 4.96E-05 | 0.615 | GTGGGTGGC |
| 691 | mamo-s | *BGIBMGA000986* | down | 334 | 342 | - | 11.3303 | 4.96E-05 | 0.615 | GTGGGTGGC |
| 692 | mamo-s | *BGIBMGA000987* | up | 1405 | 1413 | + | 11.3303 | 4.96E-05 | 0.615 | GTGGGTGGC |
| 693 | mamo-s | *BGIBMGA000988* | up | 140 | 148 | + | 12.1835 | 1.71E-05 | 0.615 | GTGCGTGCC |
| 694 | mamo-s | *BGIBMGA000988* | down | 1744 | 1752 | - | 10.5413 | 9.42E-05 | 0.857 | GTGCGAGTC |
| 695 | mamo-s | *BGIBMGA000989* | up | 96 | 104 | - | 11.3303 | 4.96E-05 | 0.615 | GTGGGTGGC |
| 696 | mamo-s | *BGIBMGA000990* | down | 1179 | 1187 | - | 12.1284 | 2.16E-05 | 0.615 | CAGCGTGGC |
| 697 | mamo-s | *BGIBMGA000993* | down | 1087 | 1095 | - | 11.3303 | 4.96E-05 | 0.615 | GTGGGTGGC |

| | | | | | | | | | | |
|---|---|---|---|---|---|---|---|---|---|---|
| 698 | mamo-s | *BGIBMGA000994* | down | 1492 | 1500 | - | 11.3303 | 4.96E-05 | 0.615 | GTGGGTGGC |
| 699 | mamo-s | *BGIBMGA000995* | down | 1980 | 1988 | + | 11.0826 | 6.84E-05 | 0.709 | GTACGTGGC |
| 700 | mamo-s | *BGIBMGA000995* | down | 1416 | 1424 | - | 10.7523 | 8.38E-05 | 0.797 | GGGCGTGTC |
| 701 | mamo-s | *BGIBMGA000999* | up | 1558 | 1566 | - | 11.3303 | 4.96E-05 | 0.615 | GTGGGTGGC |
| 702 | mamo-s | *BGIBMGA001000* | up | 1704 | 1712 | - | 11.789 | 2.84E-05 | 0.615 | GCGCGTGCC |
| 703 | mamo-s | *BGIBMGA001001* | up | 1862 | 1870 | - | 11.789 | 2.84E-05 | 0.615 | GCGCGTGCC |
| 704 | mamo-s | *BGIBMGA001002* | up | 1835 | 1843 | - | 11.789 | 2.84E-05 | 0.615 | GCGCGTGCC |
| 705 | mamo-s | *BGIBMGA001003* | up | 1518 | 1526 | - | 11.5321 | 4.06E-05 | 0.615 | TTGCGTGGC |
| 706 | mamo-s | *BGIBMGA001003* | down | 745 | 753 | - | 11.3303 | 4.96E-05 | 0.615 | GTGGGTGGC |
| 707 | mamo-s | *BGIBMGA001006* | up | 1377 | 1385 | + | 11.3303 | 4.96E-05 | 0.615 | GTGGGTGGC |
| 708 | mamo-s | *BGIBMGA001006* | up | 2 | 10 | - | 11.2018 | 5.68E-05 | 0.674 | CTGCGTGGT |
| 709 | mamo-s | *BGIBMGA001006* | up | 670 | 678 | - | 10.8899 | 7.48E-05 | 0.739 | CCGCGTGCC |
| 710 | mamo-s | *BGIBMGA001007* | up | 446 | 454 | - | 11.789 | 2.84E-05 | 0.615 | GCGCGTGCC |
| 711 | mamo-s | *BGIBMGA001007* | down | 1441 | 1449 | + | 10.5688 | 8.88E-05 | 0.824 | GTGCGTGAC |
| 712 | mamo-s | *BGIBMGA001008* | up | 139 | 147 | + | 10.5688 | 8.88E-05 | 0.824 | GTGCGTGAC |
| 713 | mamo-s | *BGIBMGA001009* | up | 1578 | 1586 | + | 11.3303 | 4.96E-05 | 0.615 | GTGGGTGGC |
| 714 | mamo-s | *BGIBMGA001009* | down | 964 | 972 | + | 10.9358 | 7.30E-05 | 0.728 | GCGGGTGGC |
| 715 | mamo-s | *BGIBMGA001011* | down | 1768 | 1776 | + | 12.4679 | 1.31E-05 | 0.615 | GTGCGTGTC |
| 716 | mamo-s | *BGIBMGA001011* | down | 1624 | 1632 | - | 11.1376 | 6.35E-05 | 0.709 | TCGCGTGGC |
| 717 | mamo-s | *BGIBMGA001011* | up | 443 | 451 | - | 11.0734 | 7.12E-05 | 0.726 | GAGCGTGGT |
| 718 | mamo-s | *BGIBMGA001015* | up | 398 | 406 | + | 11.1009 | 6.57E-05 | 0.709 | GAGCGAGGC |
| 719 | mamo-s | *BGIBMGA001016* | down | 1675 | 1683 | + | 11.0734 | 7.12E-05 | 0.726 | GAGCGTGGT |
| 720 | mamo-s | *BGIBMGA001017* | up | 783 | 791 | + | 11.3303 | 4.96E-05 | 0.615 | GTGGGTGGC |
| 721 | mamo-s | *BGIBMGA001017* | down | 1432 | 1440 | - | 10.7523 | 8.38E-05 | 0.797 | GGGCGTGTC |
| 722 | mamo-s | *BGIBMGA001018* | up | 1435 | 1443 | - | 11.3303 | 4.96E-05 | 0.615 | GTGGGTGGC |
| 723 | mamo-s | *BGIBMGA001029* | down | 1818 | 1826 | + | 12.0734 | 2.66E-05 | 0.615 | GCGCGTGTC |
| 724 | mamo-s | *BGIBMGA001032* | down | 1958 | 1966 | - | 11.3303 | 4.96E-05 | 0.615 | GTGGGTGGC |
| 725 | mamo-s | *BGIBMGA001034* | down | 1394 | 1402 | + | 11.0826 | 6.84E-05 | 0.709 | GTACGTGGC |
| 726 | mamo-s | *BGIBMGA001034* | down | 779 | 787 | - | 10.6881 | 8.61E-05 | 0.81 | GCACGTGGC |
| 727 | mamo-s | *BGIBMGA001036* | down | 1027 | 1035 | + | 11.3303 | 4.96E-05 | 0.615 | GTGGGTGGC |
| 728 | mamo-s | *BGIBMGA001036* | down | 1850 | 1858 | + | 11.3303 | 4.96E-05 | 0.615 | GTGGGTGGC |
| 729 | mamo-s | *BGIBMGA001042* | up | 1282 | 1290 | - | 11.5688 | 3.79E-05 | 0.615 | CTGCGTGTC |
| 730 | mamo-s | *BGIBMGA001043* | down | 217 | 225 | - | 11.3303 | 4.96E-05 | 0.615 | GTGGGTGGC |
| 731 | mamo-s | *BGIBMGA001043* | up | 953 | 961 | + | 11.3303 | 4.96E-05 | 0.615 | GTGGGTGGC |
| 732 | mamo-s | *BGIBMGA001044* | up | 1028 | 1036 | - | 11.3303 | 4.96E-05 | 0.615 | GTGGGTGGC |
| 733 | mamo-s | *BGIBMGA001046* | up | 70 | 78 | - | 11.7339 | 3.29E-05 | 0.615 | GCGCGAGGC |
| 734 | mamo-s | *BGIBMGA001046* | up | 1112 | 1120 | - | 11.3303 | 4.96E-05 | 0.615 | GTGGGTGGC |
| 735 | mamo-s | *BGIBMGA001046* | down | 1298 | 1306 | - | 11.3303 | 4.96E-05 | 0.615 | GTGGGTGGC |
| 736 | mamo-s | *BGIBMGA001046* | up | 47 | 55 | + | 11.2018 | 5.68E-05 | 0.674 | CTGCGTGGT |
| 737 | mamo-s | *BGIBMGA001047* | up | 658 | 666 | + | 11.3303 | 4.96E-05 | 0.615 | GTGGGTGGC |
| 738 | mamo-s | *BGIBMGA001048* | down | 395 | 403 | - | 11.7064 | 3.52E-05 | 0.615 | GCGCGTGGT |
| 739 | mamo-s | *BGIBMGA001051* | down | 413 | 421 | - | 11.3303 | 4.96E-05 | 0.615 | GTGGGTGGC |
| 740 | mamo-s | *BGIBMGA001051* | down | 798 | 806 | - | 11.1743 | 5.90E-05 | 0.684 | CCGCGTGTC |
| 741 | mamo-s | *BGIBMGA001051* | up | 743 | 751 | + | 11.0826 | 6.84E-05 | 0.709 | GTACGTGGC |
| 742 | mamo-s | *BGIBMGA001052* | down | 573 | 581 | + | 12.0734 | 2.66E-05 | 0.615 | GCGCGTGTC |
| 743 | mamo-s | *BGIBMGA001052* | down | 1678 | 1686 | + | 11.1743 | 5.90E-05 | 0.684 | CCGCGTGTC |
| 744 | mamo-s | *BGIBMGA001053* | down | 1526 | 1534 | + | 12.1009 | 2.43E-05 | 0.615 | GTGCGTGGT |
| 745 | mamo-s | *BGIBMGA001054* | down | 961 | 969 | - | 11.0826 | 6.84E-05 | 0.709 | GTACGTGGC |
| 746 | mamo-s | *BGIBMGA001056* | down | 119 | 127 | - | 11.3303 | 4.96E-05 | 0.615 | GTGGGTGGC |
| 747 | mamo-s | *BGIBMGA001056* | up | 1331 | 1339 | + | 10.8349 | 7.66E-05 | 0.751 | CCGCGAGGC |
| 748 | mamo-s | *BGIBMGA001057* | up | 1780 | 1788 | - | 11.3303 | 4.96E-05 | 0.615 | GTGGGTGGC |
| 749 | mamo-s | *BGIBMGA001058* | up | 1581 | 1589 | - | 13.156 | 6.30E-06 | 0.615 | CTGCGTGGC |
| 750 | mamo-s | *BGIBMGA001058* | up | 138 | 146 | + | 11.3303 | 4.96E-05 | 0.615 | GTGGGTGGC |
| 751 | mamo-s | *BGIBMGA001058* | up | 1699 | 1707 | + | 10.9358 | 7.30E-05 | 0.728 | GCGGGTGGC |
| 752 | mamo-s | *BGIBMGA001058* | down | 1211 | 1219 | - | 10.7523 | 8.38E-05 | 0.797 | AAGCGTGGC |
| 753 | mamo-s | *BGIBMGA001059* | down | 756 | 764 | + | 14.055 | 2.23E-06 | 0.615 | GTGCGTGGC |
| 754 | mamo-s | *BGIBMGA001059* | down | 1832 | 1840 | + | 11.1009 | 6.57E-05 | 0.709 | GAGCGAGGC |
| 755 | mamo-s | *BGIBMGA001062* | down | 1067 | 1075 | + | 11.3303 | 4.96E-05 | 0.615 | GTGGGTGGC |
| 756 | mamo-s | *BGIBMGA001062* | down | 1396 | 1404 | - | 11.2844 | 5.18E-05 | 0.63 | CTGCGTGCC |
| 757 | mamo-s | *BGIBMGA001062* | down | 1796 | 1804 | - | 10.9358 | 7.30E-05 | 0.728 | GCGGGTGGC |
| 758 | mamo-s | *BGIBMGA001066* | down | 929 | 937 | + | 11.3303 | 4.96E-05 | 0.615 | GTGGGTGGC |
| 759 | mamo-s | *BGIBMGA001066* | up | 1107 | 1115 | - | 11.3303 | 4.96E-05 | 0.615 | GTGGGTGGC |
| 760 | mamo-s | *BGIBMGA001066* | up | 1565 | 1573 | - | 11.3303 | 4.96E-05 | 0.615 | GTGGGTGGC |
| 761 | mamo-s | *BGIBMGA001067* | down | 1012 | 1020 | + | 12.3394 | 1.49E-05 | 0.615 | GGGCGTGGC |
| 762 | mamo-s | *BGIBMGA001067* | up | 692 | 700 | + | 11.3853 | 4.74E-05 | 0.615 | ACGCGTGGC |
| 763 | mamo-s | *BGIBMGA001068* | down | 407 | 415 | + | 12.4679 | 1.31E-05 | 0.615 | GTGCGTGTC |
| 764 | mamo-s | *BGIBMGA001068* | up | 53 | 61 | + | 12.3394 | 1.49E-05 | 0.615 | GGGCGTGGC |
| 765 | mamo-s | *BGIBMGA001068* | down | 1276 | 1284 | + | 10.5688 | 8.88E-05 | 0.824 | GTGCGTGAC |
| 766 | mamo-s | *BGIBMGA001072* | down | 137 | 145 | + | 10.8349 | 7.66E-05 | 0.751 | CCGCGAGGC |
| 767 | mamo-s | *BGIBMGA001073* | up | 126 | 134 | + | 10.8349 | 7.66E-05 | 0.751 | CCGCGAGGC |
| 768 | mamo-s | *BGIBMGA001074* | down | 1553 | 1561 | - | 10.6881 | 8.61E-05 | 0.81 | GCACGTGGC |
| 769 | mamo-s | *BGIBMGA001075* | up | 1341 | 1349 | - | 13.6606 | 4.06E-06 | 0.615 | GCGCGTGGC |
| 770 | mamo-s | *BGIBMGA001075* | down | 592 | 600 | + | 11.2294 | 5.41E-05 | 0.65 | CTGCGAGGC |
| 771 | mamo-s | *BGIBMGA001079* | down | 67 | 75 | + | 11.7339 | 3.29E-05 | 0.615 | GCGCGAGGC |
| 772 | mamo-s | *BGIBMGA001079* | down | 1377 | 1385 | - | 10.9358 | 7.30E-05 | 0.728 | GCGGGTGGC |
| 773 | mamo-s | *BGIBMGA001079* | up | 1761 | 1769 | + | 10.8073 | 7.89E-05 | 0.766 | CCGCGTGGT |
| 774 | mamo-s | *BGIBMGA001083* | up | 1721 | 1729 | + | 11.5321 | 4.06E-05 | 0.615 | TTGCGTGGC |
| 775 | mamo-s | *BGIBMGA001085* | down | 181 | 189 | + | 10.7523 | 8.38E-05 | 0.797 | AAGCGTGGC |
| 776 | mamo-s | *BGIBMGA001086* | up | 960 | 968 | - | 11.3303 | 4.96E-05 | 0.615 | GTGGGTGGC |
| 777 | mamo-s | *BGIBMGA001087* | up | 39 | 47 | + | 13.0275 | 8.53E-06 | 0.615 | GAGCGTGGC |
| 778 | mamo-s | *BGIBMGA001087* | down | 773 | 781 | - | 11.2294 | 5.41E-05 | 0.65 | CTGCGAGGC |
| 779 | mamo-s | *BGIBMGA001088* | up | 1440 | 1448 | - | 11.0826 | 6.84E-05 | 0.709 | GTACGTGGC |
| 780 | mamo-s | *BGIBMGA001090* | down | 1498 | 1506 | - | 13.0275 | 8.53E-06 | 0.615 | GAGCGTGGC |
| 781 | mamo-s | *BGIBMGA001090* | down | 1560 | 1568 | - | 12.1284 | 2.16E-05 | 0.615 | GTGCGAGGC |
| 782 | mamo-s | *BGIBMGA001092* | down | 721 | 729 | - | 11.3303 | 4.96E-05 | 0.615 | GTGGGTGGC |
| 783 | mamo-s | *BGIBMGA001093* | down | 1449 | 1457 | - | 10.5688 | 8.88E-05 | 0.824 | GTGCGTGAC |
| 784 | mamo-s | *BGIBMGA001094* | down | 290 | 298 | + | 11.3303 | 4.96E-05 | 0.615 | GTGGGTGGC |
| 785 | mamo-s | *BGIBMGA001094* | down | 1718 | 1726 | + | 11.3303 | 4.96E-05 | 0.615 | GTGGGTGGC |

| | | | | | | | | | | |
|---|---|---|---|---|---|---|---|---|---|---|
| 786 | mamo-s | *BGIBMGA001095* | up | 1061 | 1069 | + | 11.4404 | 4.51E-05 | 0.615 | GAGCGTGTC |
| 787 | mamo-s | *BGIBMGA001100* | down | 1314 | 1322 | + | 11.0826 | 6.84E-05 | 0.709 | GTACGTGGC |
| 788 | mamo-s | *BGIBMGA001103* | down | 724 | 732 | - | 11.3303 | 4.96E-05 | 0.615 | GTGGGTGGC |
| 789 | mamo-s | *BGIBMGA001103* | down | 1070 | 1078 | - | 11.1009 | 6.57E-05 | 0.709 | GAGCGAGGC |
| 790 | mamo-s | *BGIBMGA001105* | up | 172 | 180 | - | 10.8349 | 7.66E-05 | 0.751 | CCGCGAGGC |
| 791 | mamo-s | *BGIBMGA001106* | down | 149 | 157 | + | 11.0826 | 6.84E-05 | 0.709 | GTACGTGGC |
| 792 | mamo-s | *BGIBMGA001107* | down | 50 | 58 | - | 10.5138 | 9.75E-05 | 0.865 | GTGCGTGTT |
| 793 | mamo-s | *BGIBMGA001108* | down | 794 | 802 | + | 11.3303 | 4.96E-05 | 0.615 | GTGGGTGGC |
| 794 | mamo-s | *BGIBMGA001108* | down | 1779 | 1787 | + | 11.3303 | 4.96E-05 | 0.615 | GTGGGTGGC |
| 795 | mamo-s | *BGIBMGA001108* | up | 601 | 609 | + | 10.8073 | 7.89E-05 | 0.766 | CCGCGTGGT |
| 796 | mamo-s | *BGIBMGA001111* | down | 1420 | 1428 | + | 11.7064 | 3.52E-05 | 0.615 | GCGCGTGGT |
| 797 | mamo-s | *BGIBMGA001112* | down | 404 | 412 | + | 14.055 | 2.23E-06 | 0.615 | GTGCGTGGC |
| 798 | mamo-s | *BGIBMGA001112* | down | 707 | 715 | + | 12.7615 | 1.04E-05 | 0.615 | CCGCGTGGC |
| 799 | mamo-s | *BGIBMGA001112* | up | 1349 | 1357 | + | 11.4404 | 4.51E-05 | 0.615 | CGGCGTGGC |
| 800 | mamo-s | *BGIBMGA001112* | up | 1889 | 1897 | + | 11.3303 | 4.96E-05 | 0.615 | GTGGGTGGC |
| 801 | mamo-s | *BGIBMGA001115* | up | 880 | 888 | - | 11.4404 | 4.51E-05 | 0.615 | GAGCGTGTC |
| 802 | mamo-s | *BGIBMGA001116* | up | 399 | 407 | + | 11.3303 | 4.96E-05 | 0.615 | GTGGGTGGC |
| 803 | mamo-s | *BGIBMGA001117* | up | 1347 | 1355 | - | 10.6881 | 8.61E-05 | 0.81 | GCACGTGGC |
| 804 | mamo-s | *BGIBMGA001120* | up | 1045 | 1053 | + | 11.3303 | 4.96E-05 | 0.615 | GTGGGTGGC |
| 805 | mamo-s | *BGIBMGA001121* | up | 1479 | 1487 | - | 10.8073 | 7.89E-05 | 0.766 | CCGCGTGGT |
| 806 | mamo-s | *BGIBMGA001122* | down | 1471 | 1479 | + | 11.5688 | 3.79E-05 | 0.615 | CTGCGTGTC |
| 807 | mamo-s | *BGIBMGA001122* | up | 1213 | 1221 | + | 11.3303 | 4.96E-05 | 0.615 | GTGGGTGGC |
| 808 | mamo-s | *BGIBMGA001123* | down | 800 | 808 | + | 11.4404 | 4.51E-05 | 0.615 | CGGCGTGGC |
| 809 | mamo-s | *BGIBMGA001124* | down | 1171 | 1179 | - | 11.3303 | 4.96E-05 | 0.615 | GTGGGTGGC |
| 810 | mamo-s | *BGIBMGA001125* | down | 1207 | 1215 | - | 11.4404 | 4.51E-05 | 0.615 | CGGCGTGGC |
| 811 | mamo-s | *BGIBMGA001129* | up | 112 | 120 | - | 11.3303 | 4.96E-05 | 0.615 | GTGGGTGGC |
| 812 | mamo-s | *BGIBMGA001133* | down | 971 | 979 | - | 12.1835 | 1.71E-05 | 0.615 | GTGCGTGCC |
| 813 | mamo-s | *BGIBMGA001134* | down | 1490 | 1498 | + | 11.3303 | 4.96E-05 | 0.615 | GTGGGTGGC |
| 814 | mamo-s | *BGIBMGA001136* | down | 1322 | 1330 | + | 11.3303 | 4.96E-05 | 0.615 | GTGGGTGGC |
| 815 | mamo-s | *BGIBMGA001141* | up | 707 | 715 | + | 11.3303 | 4.96E-05 | 0.615 | GTGGGTGGC |
| 816 | mamo-s | *BGIBMGA001142* | up | 1252 | 1260 | - | 11.0826 | 6.84E-05 | 0.709 | GTACGTGGC |
| 817 | mamo-s | *BGIBMGA001142* | up | 585 | 593 | + | 11.0734 | 7.12E-05 | 0.726 | GAGCGTGGT |
| 818 | mamo-s | *BGIBMGA001145* | up | 397 | 405 | + | 10.7523 | 8.38E-05 | 0.797 | GGGCGTGTC |
| 819 | mamo-s | *BGIBMGA001146* | down | 1844 | 1852 | - | 11.3303 | 4.96E-05 | 0.615 | GTGGGTGGC |
| 820 | mamo-s | *BGIBMGA001149* | up | 1288 | 1296 | + | 11.2018 | 5.68E-05 | 0.674 | CTGCGTGGT |
| 821 | mamo-s | *BGIBMGA001150* | up | 1494 | 1502 | - | 11.3303 | 4.96E-05 | 0.615 | GTGGGTGGC |
| 822 | mamo-s | *BGIBMGA001150* | up | 1691 | 1699 | - | 11.3303 | 4.96E-05 | 0.615 | GTGGGTGGC |
| 823 | mamo-s | *BGIBMGA001151* | down | 576 | 584 | + | 11.3303 | 4.96E-05 | 0.615 | GTGGGTGGC |
| 824 | mamo-s | *BGIBMGA001152* | up | 1106 | 1114 | + | 11.3303 | 4.96E-05 | 0.615 | GTGGGTGGC |
| 825 | mamo-s | *BGIBMGA001153* | up | 834 | 842 | + | 12.1835 | 1.71E-05 | 0.615 | GTGCGTGCC |
| 826 | mamo-s | *BGIBMGA001154* | down | 1061 | 1069 | - | 11.0826 | 6.84E-05 | 0.709 | GTACGTGGC |
| 827 | mamo-s | *BGIBMGA001154* | down | 1625 | 1633 | + | 10.7523 | 8.38E-05 | 0.797 | GGGCGTGTC |
| 828 | mamo-s | *BGIBMGA001157* | up | 778 | 786 | - | 12.1284 | 2.16E-05 | 0.615 | CAGCGTGGC |
| 829 | mamo-s | *BGIBMGA001159* | down | 1390 | 1398 | + | 12.1284 | 2.16E-05 | 0.615 | CAGCGTGGC |
| 830 | mamo-s | *BGIBMGA001160* | up | 419 | 427 | + | 11.3303 | 4.96E-05 | 0.615 | GTGGGTGGC |
| 831 | mamo-s | *BGIBMGA001160* | down | 1652 | 1660 | + | 11.3303 | 4.96E-05 | 0.615 | GTGGGTGGC |
| 832 | mamo-s | *BGIBMGA001162* | down | 1445 | 1453 | + | 11.3303 | 4.96E-05 | 0.615 | GTGGGTGGC |
| 833 | mamo-s | *BGIBMGA001163* | down | 751 | 759 | - | 11.3303 | 4.96E-05 | 0.615 | GTGGGTGGC |
| 834 | mamo-s | *BGIBMGA001167* | down | 1301 | 1309 | - | 11.3303 | 4.96E-05 | 0.615 | GTGGGTGGC |
| 835 | mamo-s | *BGIBMGA001167* | up | 194 | 202 | + | 11.2844 | 5.18E-05 | 0.63 | CTGCGTGCC |
| 836 | mamo-s | *BGIBMGA001168* | down | 1257 | 1265 | - | 11.2294 | 5.41E-05 | 0.65 | CTGCGAGGC |
| 837 | mamo-s | *BGIBMGA001170* | up | 400 | 408 | - | 11.3303 | 4.96E-05 | 0.615 | GTGGGTGGC |
| 838 | mamo-s | *BGIBMGA001171* | down | 1964 | 1972 | + | 11.3303 | 4.96E-05 | 0.615 | GTGGGTGGC |
| 839 | mamo-s | *BGIBMGA001176* | up | 149 | 157 | + | 12.1284 | 2.16E-05 | 0.615 | GTGCGAGGC |
| 840 | mamo-s | *BGIBMGA001176* | down | 397 | 405 | + | 11.2018 | 5.68E-05 | 0.674 | CTGCGTGGT |
| 841 | mamo-s | *BGIBMGA001176* | down | 1382 | 1390 | + | 11.1009 | 6.57E-05 | 0.709 | GAGCGAGGC |
| 842 | mamo-s | *BGIBMGA001178* | down | 93 | 101 | - | 11.3303 | 4.96E-05 | 0.615 | GTGGGTGGC |
| 843 | mamo-s | *BGIBMGA001179* | down | 945 | 953 | - | 11.3303 | 4.96E-05 | 0.615 | GTGGGTGGC |
| 844 | mamo-s | *BGIBMGA001180* | up | 1786 | 1794 | - | 13.6606 | 4.06E-06 | 0.615 | GCGCGTGGC |
| 845 | mamo-s | *BGIBMGA001180* | up | 679 | 687 | - | 11.3303 | 4.96E-05 | 0.615 | GTGGGTGGC |
| 846 | mamo-s | *BGIBMGA001181* | down | 1691 | 1699 | + | 11.4404 | 4.51E-05 | 0.615 | CGGCGTGGC |
| 847 | mamo-s | *BGIBMGA001182* | up | 1427 | 1435 | - | 11.0734 | 7.12E-05 | 0.726 | GAGCGTGGT |
| 848 | mamo-s | *BGIBMGA001182* | up | 258 | 266 | - | 10.5138 | 9.75E-05 | 0.865 | GTGCGTGTT |
| 849 | mamo-s | *BGIBMGA001184* | up | 1553 | 1561 | - | 11.3303 | 4.96E-05 | 0.615 | GTGGGTGGC |
| 850 | mamo-s | *BGIBMGA001186* | down | 998 | 1006 | - | 11.4404 | 4.51E-05 | 0.615 | CGGCGTGGC |
| 851 | mamo-s | *BGIBMGA001186* | down | 1695 | 1703 | + | 11.0734 | 7.12E-05 | 0.726 | GAGCGTGGT |
| 852 | mamo-s | *BGIBMGA001186* | down | 182 | 190 | - | 10.6881 | 8.61E-05 | 0.81 | GCACGTGGC |
| 853 | mamo-s | *BGIBMGA001192* | up | 1345 | 1353 | - | 14.055 | 2.23E-06 | 0.615 | GTGCGTGGC |
| 854 | mamo-s | *BGIBMGA001193* | down | 1459 | 1467 | + | 11.1009 | 6.57E-05 | 0.709 | GAGCGAGGC |
| 855 | mamo-s | *BGIBMGA001195* | down | 1781 | 1789 | + | 12.1284 | 2.16E-05 | 0.615 | GTGCGAGGC |
| 856 | mamo-s | *BGIBMGA001195* | up | 1648 | 1656 | + | 11.3303 | 4.96E-05 | 0.615 | GTGGGTGGC |
| 857 | mamo-s | *BGIBMGA001196* | down | 884 | 892 | + | 12.1284 | 2.16E-05 | 0.615 | GTGCGAGGC |
| 858 | mamo-s | *BGIBMGA001196* | up | 1015 | 1023 | + | 11.3303 | 4.96E-05 | 0.615 | GTGGGTGGC |
| 859 | mamo-s | *BGIBMGA001198* | down | 1504 | 1512 | - | 11.0734 | 7.12E-05 | 0.726 | GAGCGTGGT |
| 860 | mamo-s | *BGIBMGA001200* | up | 1160 | 1168 | + | 11.3303 | 4.96E-05 | 0.615 | GTGGGTGGC |
| 861 | mamo-s | *BGIBMGA001203* | down | 1877 | 1885 | + | 11.0826 | 6.84E-05 | 0.709 | GTACGTGGC |
| 862 | mamo-s | *BGIBMGA001204* | down | 232 | 240 | + | 11.3303 | 4.96E-05 | 0.615 | GTGGGTGGC |
| 863 | mamo-s | *BGIBMGA001206* | down | 980 | 988 | + | 11.1009 | 6.57E-05 | 0.709 | GAGCGAGGC |
| 864 | mamo-s | *BGIBMGA001213* | down | 1779 | 1787 | - | 11.3303 | 4.96E-05 | 0.615 | GTGGGTGGC |
| 865 | mamo-s | *BGIBMGA001217* | up | 121 | 129 | + | 11.3303 | 4.96E-05 | 0.615 | GTGGGTGGC |
| 866 | mamo-s | *BGIBMGA001219* | down | 867 | 875 | - | 11.3303 | 4.96E-05 | 0.615 | GTGGGTGGC |
| 867 | mamo-s | *BGIBMGA001219* | down | 116 | 124 | + | 11.1009 | 6.57E-05 | 0.709 | GAGCGAGGC |
| 868 | mamo-s | *BGIBMGA001221* | up | 1691 | 1699 | - | 11.3303 | 4.96E-05 | 0.615 | GTGGGTGGC |
| 869 | mamo-s | *BGIBMGA001221* | down | 1303 | 1311 | + | 10.5138 | 9.75E-05 | 0.865 | GTGCGTGTT |
| 870 | mamo-s | *BGIBMGA001222* | down | 1091 | 1099 | - | 13.0275 | 8.53E-06 | 0.615 | GAGCGTGGC |
| 871 | mamo-s | *BGIBMGA001222* | down | 763 | 771 | - | 10.8349 | 7.66E-05 | 0.751 | CCGCGAGGC |
| 872 | mamo-s | *BGIBMGA001224* | down | 1342 | 1350 | - | 14.055 | 2.23E-06 | 0.615 | GTGCGTGGC |
| 873 | mamo-s | *BGIBMGA001224* | up | 492 | 500 | - | 11.3303 | 4.96E-05 | 0.615 | GTGGGTGGC |

| | | | | | | | | | | |
|---|---|---|---|---|---|---|---|---|---|---|
| 874 | mamo-s | *BGIBMGA001224* | up | 1262 | 1270 | - | 11.3303 | 4.96E-05 | 0.615 | GTGGGTGGC |
| 875 | mamo-s | *BGIBMGA001225* | down | 393 | 401 | - | 11.3303 | 4.96E-05 | 0.615 | GTGGGTGGC |
| 876 | mamo-s | *BGIBMGA001226* | up | 558 | 566 | + | 11.3303 | 4.96E-05 | 0.615 | GTGGGTGGC |
| 877 | mamo-s | *BGIBMGA001226* | up | 1301 | 1309 | - | 11.3303 | 4.96E-05 | 0.615 | GTGGGTGGC |
| 878 | mamo-s | *BGIBMGA001226* | down | 44 | 52 | - | 11.2294 | 5.41E-05 | 0.65 | CTGCGAGGC |
| 879 | mamo-s | *BGIBMGA001227* | down | 424 | 432 | + | 11.3303 | 4.96E-05 | 0.615 | GTGGGTGGC |
| 880 | mamo-s | *BGIBMGA001228* | down | 493 | 501 | - | 11.3303 | 4.96E-05 | 0.615 | GTGGGTGGC |
| 881 | mamo-s | *BGIBMGA001229* | down | 669 | 677 | - | 11.3303 | 4.96E-05 | 0.615 | GTGGGTGGC |
| 882 | mamo-s | *BGIBMGA001230* | up | 1958 | 1966 | + | 12.1284 | 2.16E-05 | 0.615 | GTGCGAGGC |
| 883 | mamo-s | *BGIBMGA001231* | down | 1803 | 1811 | + | 12.1835 | 1.71E-05 | 0.615 | GTGCGTGCC |
| 884 | mamo-s | *BGIBMGA001231* | down | 1243 | 1251 | + | 11.3303 | 4.96E-05 | 0.615 | GTGGGTGGC |
| 885 | mamo-s | *BGIBMGA001232* | up | 269 | 277 | + | 12.1835 | 1.71E-05 | 0.615 | GTGCGTGCC |
| 886 | mamo-s | *BGIBMGA001234* | up | 1265 | 1273 | - | 11.5688 | 3.79E-05 | 0.615 | CTGCGTGTC |
| 887 | mamo-s | *BGIBMGA001234* | up | 1045 | 1053 | - | 11.3303 | 4.96E-05 | 0.615 | GTGGGTGGC |
| 888 | mamo-s | *BGIBMGA001235* | up | 1348 | 1356 | - | 11.3303 | 4.96E-05 | 0.615 | GTGGGTGGC |
| 889 | mamo-s | *BGIBMGA001238* | down | 1269 | 1277 | - | 11.3303 | 4.96E-05 | 0.615 | GTGGGTGGC |
| 890 | mamo-s | *BGIBMGA001239* | down | 635 | 643 | + | 11.1743 | 5.90E-05 | 0.684 | CCGCGTGTC |
| 891 | mamo-s | *BGIBMGA001240* | up | 1195 | 1203 | + | 11.1743 | 5.90E-05 | 0.684 | CCGCGTGTC |
| 892 | mamo-s | *BGIBMGA001242* | down | 1762 | 1770 | - | 11.3303 | 4.96E-05 | 0.615 | GTGGGTGGC |
| 893 | mamo-s | *BGIBMGA001244* | down | 160 | 168 | + | 14.055 | 2.23E-06 | 0.615 | GTGCGTGGC |
| 894 | mamo-s | *BGIBMGA001244* | down | 937 | 945 | + | 11.3303 | 4.96E-05 | 0.615 | GTGGGTGGC |
| 895 | mamo-s | *BGIBMGA001245* | down | 362 | 370 | + | 11.3303 | 4.96E-05 | 0.615 | GTGGGTGGC |
| 896 | mamo-s | *BGIBMGA001247* | down | 1729 | 1737 | - | 11.7339 | 3.29E-05 | 0.615 | GCGCGAGGC |
| 897 | mamo-s | *BGIBMGA001248* | down | 530 | 538 | - | 11.5321 | 4.06E-05 | 0.615 | TTGCGTGGC |
| 898 | mamo-s | *BGIBMGA001249* | up | 117 | 125 | - | 11.5321 | 4.06E-05 | 0.615 | TTGCGTGGC |
| 899 | mamo-s | *BGIBMGA001250* | up | 788 | 796 | + | 10.5413 | 9.42E-05 | 0.857 | CAGCGTGTC |
| 900 | mamo-s | *BGIBMGA001251* | up | 1958 | 1966 | + | 11.4404 | 4.51E-05 | 0.615 | CGGCGTGGC |
| 901 | mamo-s | *BGIBMGA001251* | up | 1968 | 1976 | - | 11.1376 | 6.35E-05 | 0.709 | TCGCGTGGC |
| 902 | mamo-s | *BGIBMGA001254* | up | 930 | 938 | + | 10.5138 | 9.75E-05 | 0.865 | GTGCGTGTT |
| 903 | mamo-s | *BGIBMGA001257* | down | 1441 | 1449 | + | 11.3303 | 4.96E-05 | 0.615 | GTGGGTGGC |
| 904 | mamo-s | *BGIBMGA001258* | down | 1341 | 1349 | + | 12.4679 | 1.31E-05 | 0.615 | GTGCGTGTC |
| 905 | mamo-s | *BGIBMGA001258* | up | 205 | 213 | + | 11.3303 | 4.96E-05 | 0.615 | GTGGGTGGC |
| 906 | mamo-s | *BGIBMGA001260* | up | 831 | 839 | + | 11.3853 | 4.74E-05 | 0.615 | ACGCGTGGC |
| 907 | mamo-s | *BGIBMGA001260* | down | 1213 | 1221 | - | 11.3303 | 4.96E-05 | 0.615 | GTGGGTGGC |
| 908 | mamo-s | *BGIBMGA001261* | up | 1372 | 1380 | + | 10.8899 | 7.48E-05 | 0.739 | CCGCGTGCC |
| 909 | mamo-s | *BGIBMGA001264* | down | 1096 | 1104 | + | 11.3303 | 4.96E-05 | 0.615 | GTGGGTGGC |
| 910 | mamo-s | *BGIBMGA001265* | down | 795 | 803 | + | 11.0826 | 6.84E-05 | 0.709 | GTACGTGGC |
| 911 | mamo-s | *BGIBMGA001266* | down | 1921 | 1929 | - | 11.7339 | 3.29E-05 | 0.615 | GCGCGAGGC |
| 912 | mamo-s | *BGIBMGA001266* | down | 1727 | 1735 | - | 11.5321 | 4.06E-05 | 0.615 | TTGCGTGGC |
| 913 | mamo-s | *BGIBMGA001266* | up | 486 | 494 | + | 11.3303 | 4.96E-05 | 0.615 | GTGGGTGGC |
| 914 | mamo-s | *BGIBMGA001268* | down | 1454 | 1462 | + | 11.3303 | 4.96E-05 | 0.615 | GTGGGTGGC |
| 915 | mamo-s | *BGIBMGA001269* | up | 832 | 840 | - | 11.7339 | 3.29E-05 | 0.615 | GCGCGAGGC |
| 916 | mamo-s | *BGIBMGA001269* | up | 475 | 483 | + | 11.3303 | 4.96E-05 | 0.615 | GTGGGTGGC |
| 917 | mamo-s | *BGIBMGA001269* | down | 1451 | 1459 | + | 11.3303 | 4.96E-05 | 0.615 | GTGGGTGGC |
| 918 | mamo-s | *BGIBMGA001270* | up | 574 | 582 | - | 11.3303 | 4.96E-05 | 0.615 | GTGGGTGGC |
| 919 | mamo-s | *BGIBMGA001271* | down | 1735 | 1743 | - | 10.5688 | 8.88E-05 | 0.824 | GTGCGTGAC |
| 920 | mamo-s | *BGIBMGA001272* | down | 570 | 578 | + | 11.3303 | 4.96E-05 | 0.615 | GTGGGTGGC |
| 921 | mamo-s | *BGIBMGA001272* | up | 1083 | 1091 | - | 11.3303 | 4.96E-05 | 0.615 | GTGGGTGGC |
| 922 | mamo-s | *BGIBMGA001273* | down | 641 | 649 | - | 11.3303 | 4.96E-05 | 0.615 | GTGGGTGGC |
| 923 | mamo-s | *BGIBMGA001273* | up | 1770 | 1778 | + | 11.3303 | 4.96E-05 | 0.615 | GTGGGTGGC |
| 924 | mamo-s | *BGIBMGA001276* | down | 1926 | 1934 | - | 11.3303 | 4.96E-05 | 0.615 | GTGGGTGGC |
| 925 | mamo-s | *BGIBMGA001277* | down | 1457 | 1465 | + | 11.4404 | 4.51E-05 | 0.615 | GAGCGTGTC |
| 926 | mamo-s | *BGIBMGA001277* | up | 369 | 377 | - | 11.3303 | 4.96E-05 | 0.615 | GTGGGTGGC |
| 927 | mamo-s | *BGIBMGA001278* | down | 1171 | 1179 | - | 12.1835 | 1.71E-05 | 0.615 | GTGCGTGCC |
| 928 | mamo-s | *BGIBMGA001278* | down | 1536 | 1544 | - | 11.3303 | 4.96E-05 | 0.615 | GTGGGTGGC |
| 929 | mamo-s | *BGIBMGA001278* | down | 1040 | 1048 | - | 11.1009 | 6.57E-05 | 0.709 | GAGCGAGGC |
| 930 | mamo-s | *BGIBMGA001278* | down | 449 | 457 | + | 11.0826 | 6.84E-05 | 0.709 | GTACGTGGC |
| 931 | mamo-s | *BGIBMGA001279* | down | 693 | 701 | - | 11.3303 | 4.96E-05 | 0.615 | GTGGGTGGC |
| 932 | mamo-s | *BGIBMGA001279* | down | 1833 | 1841 | + | 11.1743 | 5.90E-05 | 0.684 | CCGCGTGTC |
| 933 | mamo-s | *BGIBMGA001280* | down | 1784 | 1792 | - | 11.3303 | 4.96E-05 | 0.615 | GTGGGTGGC |
| 934 | mamo-s | *BGIBMGA001280* | up | 1117 | 1125 | + | 11.1743 | 5.90E-05 | 0.684 | CCGCGTGTC |
| 935 | mamo-s | *BGIBMGA001281* | up | 632 | 640 | - | 14.055 | 2.23E-06 | 0.615 | GTGCGTGGC |
| 936 | mamo-s | *BGIBMGA001282* | up | 692 | 700 | + | 11.3303 | 4.96E-05 | 0.615 | GTGGGTGGC |
| 937 | mamo-s | *BGIBMGA001283* | down | 1241 | 1249 | + | 13.6606 | 4.06E-06 | 0.615 | GCGCGTGGC |
| 938 | mamo-s | *BGIBMGA001283* | down | 1488 | 1496 | + | 13.6606 | 4.06E-06 | 0.615 | GCGCGTGGC |
| 939 | mamo-s | *BGIBMGA001283* | down | 797 | 805 | - | 11.3303 | 4.96E-05 | 0.615 | GTGGGTGGC |
| 940 | mamo-s | *BGIBMGA001289* | down | 1403 | 1411 | - | 11.3303 | 4.96E-05 | 0.615 | GTGGGTGGC |
| 941 | mamo-s | *BGIBMGA001292* | up | 1073 | 1081 | + | 11.3303 | 4.96E-05 | 0.615 | GTGGGTGGC |
| 942 | mamo-s | *BGIBMGA001293* | down | 1327 | 1335 | + | 11.3303 | 4.96E-05 | 0.615 | GTGGGTGGC |
| 943 | mamo-s | *BGIBMGA001293* | up | 1790 | 1798 | + | 11.3303 | 4.96E-05 | 0.615 | GTGGGTGGC |
| 944 | mamo-s | *BGIBMGA001294* | up | 1811 | 1819 | + | 11.3303 | 4.96E-05 | 0.615 | GTGGGTGGC |
| 945 | mamo-s | *BGIBMGA001295* | down | 1422 | 1430 | - | 10.5688 | 8.88E-05 | 0.824 | GTGCGTGAC |
| 946 | mamo-s | *BGIBMGA001296* | down | 1628 | 1636 | - | 11.3303 | 4.96E-05 | 0.615 | GTGGGTGGC |
| 947 | mamo-s | *BGIBMGA001298* | up | 732 | 740 | + | 11.3303 | 4.96E-05 | 0.615 | GTGGGTGGC |
| 948 | mamo-s | *BGIBMGA001298* | down | 1616 | 1624 | - | 11.0826 | 6.84E-05 | 0.709 | GTACGTGGC |
| 949 | mamo-s | *BGIBMGA001299* | up | 1803 | 1811 | + | 11.0826 | 6.84E-05 | 0.709 | GTACGTGGC |
| 950 | mamo-s | *BGIBMGA001300* | up | 1585 | 1593 | - | 11.3303 | 4.96E-05 | 0.615 | GTGGGTGGC |
| 951 | mamo-s | *BGIBMGA001301* | up | 438 | 446 | + | 13.6606 | 4.06E-06 | 0.615 | GCGCGTGGC |
| 952 | mamo-s | *BGIBMGA001301* | up | 394 | 402 | + | 11.3853 | 4.74E-05 | 0.615 | ACGCGTGGC |
| 953 | mamo-s | *BGIBMGA001302* | up | 333 | 341 | + | 11.0826 | 6.84E-05 | 0.709 | GTACGTGGC |
| 954 | mamo-s | *BGIBMGA001304* | down | 153 | 161 | + | 13.0275 | 8.53E-06 | 0.615 | GAGCGTGGC |
| 955 | mamo-s | *BGIBMGA001304* | up | 669 | 677 | + | 11.3303 | 4.96E-05 | 0.615 | GTGGGTGGC |
| 956 | mamo-s | *BGIBMGA001305* | down | 230 | 238 | + | 11.3303 | 4.96E-05 | 0.615 | GTGGGTGGC |
| 957 | mamo-s | *BGIBMGA001305* | down | 1116 | 1124 | + | 11.3303 | 4.96E-05 | 0.615 | GTGGGTGGC |
| 958 | mamo-s | *BGIBMGA001305* | up | 1221 | 1229 | + | 11.3303 | 4.96E-05 | 0.615 | GTGGGTGGC |
| 959 | mamo-s | *BGIBMGA001307* | up | 1511 | 1519 | + | 11.3303 | 4.96E-05 | 0.615 | GTGGGTGGC |
| 960 | mamo-s | *BGIBMGA001308* | up | 681 | 689 | + | 10.9358 | 7.30E-05 | 0.728 | GCGGGTGGC |
| 961 | mamo-s | *BGIBMGA001312* | up | 873 | 881 | + | 11.3303 | 4.96E-05 | 0.615 | GTGGGTGGC |

| | | | | | | | | | | |
|---|---|---|---|---|---|---|---|---|---|---|
| 962 | mamo-s | *BGIBMGA001313* | down | 852 | 860 | + | 10.5413 | 9.42E-05 | 0.857 | CAGCGTGTC |
| 963 | mamo-s | *BGIBMGA001314* | up | 200 | 208 | + | 10.5413 | 9.42E-05 | 0.857 | CAGCGTGTC |
| 964 | mamo-s | *BGIBMGA001315* | down | 741 | 749 | - | 11.3303 | 4.96E-05 | 0.615 | GTGGGTGGC |
| 965 | mamo-s | *BGIBMGA001317* | up | 358 | 366 | - | 11.3303 | 4.96E-05 | 0.615 | GTGGGTGGC |
| 966 | mamo-s | *BGIBMGA001317* | down | 1268 | 1276 | - | 11.3303 | 4.96E-05 | 0.615 | GTGGGTGGC |
| 967 | mamo-s | *BGIBMGA001318* | up | 715 | 723 | - | 11.3303 | 4.96E-05 | 0.615 | GTGGGTGGC |
| 968 | mamo-s | *BGIBMGA001320* | up | 1254 | 1262 | - | 11.3303 | 4.96E-05 | 0.615 | GTGGGTGGC |
| 969 | mamo-s | *BGIBMGA001322* | down | 1805 | 1813 | + | 11.3303 | 4.96E-05 | 0.615 | GTGGGTGGC |
| 970 | mamo-s | *BGIBMGA001323* | up | 1307 | 1315 | - | 11.156 | 6.13E-05 | 0.705 | GAGCGTGCC |
| 971 | mamo-s | *BGIBMGA001327* | up | 201 | 209 | - | 11.3303 | 4.96E-05 | 0.615 | GTGGGTGGC |
| 972 | mamo-s | *BGIBMGA001328* | up | 887 | 895 | + | 11.3303 | 4.96E-05 | 0.615 | GTGGGTGGC |
| 973 | mamo-s | *BGIBMGA001333* | up | 486 | 494 | - | 11.3303 | 4.96E-05 | 0.615 | GTGGGTGGC |
| 974 | mamo-s | *BGIBMGA001334* | up | 1878 | 1886 | - | 11.7798 | 3.11E-05 | 0.615 | ATGCGTGGC |
| 975 | mamo-s | *BGIBMGA001335* | down | 1253 | 1261 | - | 11.3303 | 4.96E-05 | 0.615 | GTGGGTGGC |
| 976 | mamo-s | *BGIBMGA001338* | up | 1837 | 1845 | + | 11.3303 | 4.96E-05 | 0.615 | GTGGGTGGC |
| 977 | mamo-s | *BGIBMGA001338* | up | 1653 | 1661 | - | 11.0734 | 7.12E-05 | 0.726 | GAGCGTGGT |
| 978 | mamo-s | *BGIBMGA001338* | down | 1894 | 1902 | + | 10.8349 | 7.66E-05 | 0.751 | CCGCGAGGC |
| 979 | mamo-s | *BGIBMGA001339* | down | 1100 | 1108 | - | 11.0826 | 6.84E-05 | 0.709 | GTACGTGGC |
| 980 | mamo-s | *BGIBMGA001342* | up | 1933 | 1941 | - | 11.789 | 2.84E-05 | 0.615 | GCGCGTGCC |
| 981 | mamo-s | *BGIBMGA001342* | up | 362 | 370 | - | 11.3303 | 4.96E-05 | 0.615 | GTGGGTGGC |
| 982 | mamo-s | *BGIBMGA001344* | down | 97 | 105 | - | 11.3303 | 4.96E-05 | 0.615 | GTGGGTGGC |
| 983 | mamo-s | *BGIBMGA001345* | up | 153 | 161 | + | 11.789 | 2.84E-05 | 0.615 | GCGCGTGCC |
| 984 | mamo-s | *BGIBMGA001345* | down | 1746 | 1754 | + | 10.7523 | 8.38E-05 | 0.797 | GGGCGTGTC |
| 985 | mamo-s | *BGIBMGA001346* | up | 1873 | 1881 | + | 11.7798 | 3.11E-05 | 0.615 | ATGCGTGGC |
| 986 | mamo-s | *BGIBMGA001347* | down | 796 | 804 | + | 10.8349 | 7.66E-05 | 0.751 | CCGCGAGGC |
| 987 | mamo-s | *BGIBMGA001348* | down | 933 | 941 | - | 13.6606 | 4.06E-06 | 0.615 | GCGCGTGGC |
| 988 | mamo-s | *BGIBMGA001348* | down | 1763 | 1771 | - | 13.0275 | 8.53E-06 | 0.615 | GAGCGTGGC |
| 989 | mamo-s | *BGIBMGA001348* | down | 137 | 145 | - | 11.5688 | 3.79E-05 | 0.615 | CTGCGTGTC |
| 990 | mamo-s | *BGIBMGA001348* | down | 1173 | 1181 | - | 11.0826 | 6.84E-05 | 0.709 | GTACGTGGC |
| 991 | mamo-s | *BGIBMGA001349* | up | 1345 | 1353 | - | 13.6606 | 4.06E-06 | 0.615 | GCGCGTGGC |
| 992 | mamo-s | *BGIBMGA001349* | up | 549 | 557 | - | 11.5688 | 3.79E-05 | 0.615 | CTGCGTGTC |
| 993 | mamo-s | *BGIBMGA001349* | up | 1585 | 1593 | - | 11.0826 | 6.84E-05 | 0.709 | GTACGTGGC |
| 994 | mamo-s | *BGIBMGA001350* | down | 332 | 340 | + | 10.5413 | 9.42E-05 | 0.857 | CAGCGTGTC |
| 995 | mamo-s | *BGIBMGA001354* | down | 1712 | 1720 | - | 12.0734 | 2.66E-05 | 0.615 | GCGCGTGTC |
| 996 | mamo-s | *BGIBMGA001356* | down | 767 | 775 | + | 11.3303 | 4.96E-05 | 0.615 | GTGGGTGGC |
| 997 | mamo-s | *BGIBMGA001356* | down | 1762 | 1770 | + | 11.3303 | 4.96E-05 | 0.615 | GTGGGTGGC |
| 998 | mamo-s | *BGIBMGA001356* | up | 1761 | 1769 | - | 11.3303 | 4.96E-05 | 0.615 | GTGGGTGGC |
| 999 | mamo-s | *BGIBMGA001357* | down | 1832 | 1840 | + | 11.3303 | 4.96E-05 | 0.615 | GTGGGTGGC |
| 1000 | mamo-s | *BGIBMGA001357* | up | 1879 | 1887 | + | 11.3303 | 4.96E-05 | 0.615 | GTGGGTGGC |
| 1001 | mamo-s | *BGIBMGA001357* | down | 1286 | 1294 | + | 11.2844 | 5.18E-05 | 0.63 | CTGCGTGCC |
| 1002 | mamo-s | *BGIBMGA001361* | up | 886 | 894 | - | 11.1743 | 5.90E-05 | 0.684 | CCGCGTGTC |
| 1003 | mamo-s | *BGIBMGA001364* | down | 1145 | 1153 | + | 11.3303 | 4.96E-05 | 0.615 | GTGGGTGGC |
| 1004 | mamo-s | *BGIBMGA001365* | up | 60 | 68 | - | 12.4679 | 1.31E-05 | 0.615 | GTGCGTGTC |
| 1005 | mamo-s | *BGIBMGA001365* | down | 1712 | 1720 | + | 12.0734 | 2.66E-05 | 0.615 | GCGCGTGTC |
| 1006 | mamo-s | *BGIBMGA001365* | up | 1522 | 1530 | + | 11.3303 | 4.96E-05 | 0.615 | GTGGGTGGC |
| 1007 | mamo-s | *BGIBMGA001367* | up | 623 | 631 | - | 11.3303 | 4.96E-05 | 0.615 | GTGGGTGGC |
| 1008 | mamo-s | *BGIBMGA001368* | down | 1713 | 1721 | - | 11.3303 | 4.96E-05 | 0.615 | GTGGGTGGC |
| 1009 | mamo-s | *BGIBMGA001368* | up | 759 | 767 | - | 11.2018 | 5.68E-05 | 0.674 | CTGCGTGGT |
| 1010 | mamo-s | *BGIBMGA001368* | down | 295 | 303 | + | 11.0826 | 6.84E-05 | 0.709 | GTACGTGGC |
| 1011 | mamo-s | *BGIBMGA001370* | up | 1698 | 1706 | - | 11.3853 | 4.74E-05 | 0.615 | ACGCGTGGC |
| 1012 | mamo-s | *BGIBMGA001370* | down | 990 | 998 | - | 11.1376 | 6.35E-05 | 0.709 | TCGCGTGGC |
| 1013 | mamo-s | *BGIBMGA001371* | down | 1123 | 1131 | + | 12.1284 | 2.16E-05 | 0.615 | CAGCGTGGC |
| 1014 | mamo-s | *BGIBMGA001372* | down | 1680 | 1688 | - | 11.5321 | 4.06E-05 | 0.615 | TTGCGTGGC |
| 1015 | mamo-s | *BGIBMGA001373* | up | 336 | 344 | + | 11.3303 | 4.96E-05 | 0.615 | GTGGGTGGC |
| 1016 | mamo-s | *BGIBMGA001373* | up | 481 | 489 | - | 10.8899 | 7.48E-05 | 0.739 | CCGCGTGCC |
| 1017 | mamo-s | *BGIBMGA001375* | down | 1550 | 1558 | + | 11.3303 | 4.96E-05 | 0.615 | GTGGGTGGC |
| 1018 | mamo-s | *BGIBMGA001376* | up | 167 | 175 | - | 11.3303 | 4.96E-05 | 0.615 | GTGGGTGGC |
| 1019 | mamo-s | *BGIBMGA001377* | up | 24 | 32 | + | 11.3303 | 4.96E-05 | 0.615 | GTGGGTGGC |
| 1020 | mamo-s | *BGIBMGA001378* | down | 334 | 342 | - | 11.3303 | 4.96E-05 | 0.615 | GTGGGTGGC |
| 1021 | mamo-s | *BGIBMGA001379* | up | 374 | 382 | - | 11.3303 | 4.96E-05 | 0.615 | GTGGGTGGC |
| 1022 | mamo-s | *BGIBMGA001379* | down | 283 | 291 | - | 10.9358 | 7.30E-05 | 0.728 | GCGGGTGGC |
| 1023 | mamo-s | *BGIBMGA001380* | down | 519 | 527 | - | 11.4404 | 4.51E-05 | 0.615 | GAGCGTGTC |
| 1024 | mamo-s | *BGIBMGA001380* | up | 1933 | 1941 | + | 11.4404 | 4.51E-05 | 0.615 | GAGCGTGTC |
| 1025 | mamo-s | *BGIBMGA001381* | up | 1820 | 1828 | - | 12.0734 | 2.66E-05 | 0.615 | GCGCGTGTC |
| 1026 | mamo-s | *BGIBMGA001381* | down | 234 | 242 | - | 10.7523 | 8.38E-05 | 0.797 | GGGCGTGTC |
| 1027 | mamo-s | *BGIBMGA001383* | up | 556 | 564 | - | 11.1376 | 6.35E-05 | 0.709 | TCGCGTGGC |
| 1028 | mamo-s | *BGIBMGA001384* | down | 276 | 284 | - | 12.4679 | 1.31E-05 | 0.615 | GTGCGTGTC |
| 1029 | mamo-s | *BGIBMGA001388* | up | 790 | 798 | + | 14.055 | 2.23E-06 | 0.615 | GTGCGTGGC |
| 1030 | mamo-s | *BGIBMGA001389* | down | 163 | 171 | - | 11.3303 | 4.96E-05 | 0.615 | GTGGGTGGC |
| 1031 | mamo-s | *BGIBMGA001389* | down | 1286 | 1294 | + | 11.3303 | 4.96E-05 | 0.615 | GTGGGTGGC |
| 1032 | mamo-s | *BGIBMGA001391* | up | 285 | 293 | + | 11.3853 | 4.74E-05 | 0.615 | ACGCGTGGC |
| 1033 | mamo-s | *BGIBMGA001391* | up | 1012 | 1020 | - | 11.3303 | 4.96E-05 | 0.615 | GTGGGTGGC |
| 1034 | mamo-s | *BGIBMGA001392* | up | 360 | 368 | + | 11.3303 | 4.96E-05 | 0.615 | GTGGGTGGC |
| 1035 | mamo-s | *BGIBMGA001392* | up | 1698 | 1706 | - | 11.3303 | 4.96E-05 | 0.615 | GTGGGTGGC |
| 1036 | mamo-s | *BGIBMGA001392* | up | 242 | 250 | + | 10.8073 | 7.89E-05 | 0.766 | CCGCGTGGT |
| 1037 | mamo-s | *BGIBMGA001393* | down | 307 | 315 | + | 11.3303 | 4.96E-05 | 0.615 | GTGGGTGGC |
| 1038 | mamo-s | *BGIBMGA001394* | up | 52 | 60 | + | 11.3303 | 4.96E-05 | 0.615 | GTGGGTGGC |
| 1039 | mamo-s | *BGIBMGA001394* | down | 985 | 993 | + | 11.3303 | 4.96E-05 | 0.615 | GTGGGTGGC |
| 1040 | mamo-s | *BGIBMGA001395* | up | 759 | 767 | - | 13.0275 | 8.53E-06 | 0.615 | GAGCGTGGC |
| 1041 | mamo-s | *BGIBMGA001395* | down | 219 | 227 | - | 11.3303 | 4.96E-05 | 0.615 | GTGGGTGGC |
| 1042 | mamo-s | *BGIBMGA001396* | up | 499 | 507 | + | 11.5321 | 4.06E-05 | 0.615 | TTGCGTGGC |
| 1043 | mamo-s | *BGIBMGA001396* | down | 1063 | 1071 | + | 11.3303 | 4.96E-05 | 0.615 | GTGGGTGGC |
| 1044 | mamo-s | *BGIBMGA001402* | down | 1348 | 1356 | + | 11.5688 | 3.79E-05 | 0.615 | CTGCGTGTC |
| 1045 | mamo-s | *BGIBMGA001402* | down | 1538 | 1546 | + | 11.5688 | 3.79E-05 | 0.615 | CTGCGTGTC |
| 1046 | mamo-s | *BGIBMGA001402* | down | 1918 | 1926 | + | 11.5688 | 3.79E-05 | 0.615 | CTGCGTGTC |
| 1047 | mamo-s | *BGIBMGA001403* | down | 484 | 492 | + | 11.3303 | 4.96E-05 | 0.615 | GTGGGTGGC |
| 1048 | mamo-s | *BGIBMGA001404* | up | 1109 | 1117 | - | 11.5321 | 4.06E-05 | 0.615 | TTGCGTGGC |
| 1049 | mamo-s | *BGIBMGA001404* | down | 1419 | 1427 | - | 11.3303 | 4.96E-05 | 0.615 | GTGGGTGGC |

| | | | | | | | | | | |
|---|---|---|---|---|---|---|---|---|---|---|
| 1050 | mamo-s | *BGIBMGA001405* | down | 985 | 993 | - | 10.8899 | 7.48E-05 | 0.739 | CCGCGTGCC |
| 1051 | mamo-s | *BGIBMGA001407* | up | 1844 | 1852 | + | 11.3303 | 4.96E-05 | 0.615 | GTGGGTGGC |
| 1052 | mamo-s | *BGIBMGA001408* | up | 1141 | 1149 | + | 10.8899 | 7.48E-05 | 0.739 | CCGCGTGCC |
| 1053 | mamo-s | *BGIBMGA001409* | down | 1805 | 1813 | - | 12.4679 | 1.31E-05 | 0.615 | GTGCGTGTC |
| 1054 | mamo-s | *BGIBMGA001409* | up | 887 | 895 | + | 11.0826 | 6.84E-05 | 0.709 | GTACGTGGC |
| 1055 | mamo-s | *BGIBMGA001410* | up | 1636 | 1644 | - | 11.3303 | 4.96E-05 | 0.615 | GTGGGTGGC |
| 1056 | mamo-s | *BGIBMGA001410* | down | 654 | 662 | - | 11.0826 | 6.84E-05 | 0.709 | GTACGTGGC |
| 1057 | mamo-s | *BGIBMGA001412* | down | 497 | 505 | + | 11.3303 | 4.96E-05 | 0.615 | GTGGGTGGC |
| 1058 | mamo-s | *BGIBMGA001412* | down | 1266 | 1274 | + | 11.3303 | 4.96E-05 | 0.615 | GTGGGTGGC |
| 1059 | mamo-s | *BGIBMGA001413* | down | 1263 | 1271 | - | 11.3303 | 4.96E-05 | 0.615 | GTGGGTGGC |
| 1060 | mamo-s | *BGIBMGA001415* | down | 1500 | 1508 | - | 11.3303 | 4.96E-05 | 0.615 | GTGGGTGGC |
| 1061 | mamo-s | *BGIBMGA001415* | down | 1839 | 1847 | + | 10.5138 | 9.75E-05 | 0.865 | GTGCGTGTT |
| 1062 | mamo-s | *BGIBMGA001416* | up | 930 | 938 | + | 11.3303 | 4.96E-05 | 0.615 | GTGGGTGGC |
| 1063 | mamo-s | *BGIBMGA001417* | up | 873 | 881 | + | 11.3303 | 4.96E-05 | 0.615 | GTGGGTGGC |
| 1064 | mamo-s | *BGIBMGA001419* | up | 244 | 252 | - | 11.3303 | 4.96E-05 | 0.615 | GTGGGTGGC |
| 1065 | mamo-s | *BGIBMGA001419* | down | 1362 | 1370 | - | 11.3303 | 4.96E-05 | 0.615 | GTGGGTGGC |
| 1066 | mamo-s | *BGIBMGA001419* | down | 1469 | 1477 | - | 10.9358 | 7.30E-05 | 0.728 | GCGGGTGGC |
| 1067 | mamo-s | *BGIBMGA001420* | down | 1832 | 1840 | - | 11.3303 | 4.96E-05 | 0.615 | GTGGGTGGC |
| 1068 | mamo-s | *BGIBMGA001421* | down | 1434 | 1442 | - | 11.4404 | 4.51E-05 | 0.615 | CGGCGTGGC |
| 1069 | mamo-s | *BGIBMGA001424* | down | 1780 | 1788 | - | 11.3303 | 4.96E-05 | 0.615 | GTGGGTGGC |
| 1070 | mamo-s | *BGIBMGA001432* | up | 236 | 244 | + | 10.7523 | 8.38E-05 | 0.797 | AAGCGTGGC |
| 1071 | mamo-s | *BGIBMGA001438* | down | 878 | 886 | - | 11.3303 | 4.96E-05 | 0.615 | GTGGGTGGC |
| 1072 | mamo-s | *BGIBMGA001438* | up | 1107 | 1115 | + | 11.3303 | 4.96E-05 | 0.615 | GTGGGTGGC |
| 1073 | mamo-s | *BGIBMGA001438* | down | 1897 | 1905 | + | 10.7523 | 8.38E-05 | 0.797 | GGGCGTGTC |
| 1074 | mamo-s | *BGIBMGA001439* | down | 1408 | 1416 | + | 11.3303 | 4.96E-05 | 0.615 | GTGGGTGGC |
| 1075 | mamo-s | *BGIBMGA001440* | down | 201 | 209 | - | 11.7339 | 3.29E-05 | 0.615 | GCGCGAGGC |
| 1076 | mamo-s | *BGIBMGA001440* | down | 279 | 287 | - | 11.7339 | 3.29E-05 | 0.615 | GCGCGAGGC |
| 1077 | mamo-s | *BGIBMGA001440* | up | 1932 | 1940 | + | 11.5688 | 3.79E-05 | 0.615 | CTGCGTGTC |
| 1078 | mamo-s | *BGIBMGA001440* | up | 1215 | 1223 | - | 11.3303 | 4.96E-05 | 0.615 | GTGGGTGGC |
| 1079 | mamo-s | *BGIBMGA001440* | down | 1886 | 1894 | + | 11.3303 | 4.96E-05 | 0.615 | GTGGGTGGC |
| 1080 | mamo-s | *BGIBMGA001443* | up | 36 | 44 | + | 11.3303 | 4.96E-05 | 0.615 | GTGGGTGGC |
| 1081 | mamo-s | *BGIBMGA001444* | up | 850 | 858 | - | 11.3303 | 4.96E-05 | 0.615 | GTGGGTGGC |
| 1082 | mamo-s | *BGIBMGA001444* | up | 1331 | 1339 | + | 11.3303 | 4.96E-05 | 0.615 | GTGGGTGGC |
| 1083 | mamo-s | *BGIBMGA001445* | down | 1546 | 1554 | - | 11.3303 | 4.96E-05 | 0.615 | GTGGGTGGC |
| 1084 | mamo-s | *BGIBMGA001445* | down | 1290 | 1298 | + | 10.5138 | 9.75E-05 | 0.865 | GTGCGTGTT |
| 1085 | mamo-s | *BGIBMGA001446* | up | 90 | 98 | - | 11.3303 | 4.96E-05 | 0.615 | GTGGGTGGC |
| 1086 | mamo-s | *BGIBMGA001446* | down | 1656 | 1664 | + | 11.3303 | 4.96E-05 | 0.615 | GTGGGTGGC |
| 1087 | mamo-s | *BGIBMGA001447* | down | 1146 | 1154 | - | 11.1743 | 5.90E-05 | 0.684 | CCGCGTGTC |
| 1088 | mamo-s | *BGIBMGA001447* | up | 1745 | 1753 | - | 10.8899 | 7.48E-05 | 0.739 | CCGCGTGCC |
| 1089 | mamo-s | *BGIBMGA001448* | up | 300 | 308 | + | 11.3303 | 4.96E-05 | 0.615 | GTGGGTGGC |
| 1090 | mamo-s | *BGIBMGA001448* | down | 1691 | 1699 | - | 11.3303 | 4.96E-05 | 0.615 | GTGGGTGGC |
| 1091 | mamo-s | *BGIBMGA001449* | up | 483 | 491 | - | 11.3303 | 4.96E-05 | 0.615 | GTGGGTGGC |
| 1092 | mamo-s | *BGIBMGA001449* | up | 719 | 727 | + | 11.3303 | 4.96E-05 | 0.615 | GTGGGTGGC |
| 1093 | mamo-s | *BGIBMGA001451* | down | 216 | 224 | - | 11.3303 | 4.96E-05 | 0.615 | GTGGGTGGC |
| 1094 | mamo-s | *BGIBMGA001451* | up | 268 | 276 | - | 10.6881 | 8.61E-05 | 0.81 | GCACGTGGC |
| 1095 | mamo-s | *BGIBMGA001451* | down | 592 | 600 | - | 10.5138 | 9.75E-05 | 0.865 | GTGCGTGTT |
| 1096 | mamo-s | *BGIBMGA001453* | down | 669 | 677 | + | 11.3303 | 4.96E-05 | 0.615 | GTGGGTGGC |
| 1097 | mamo-s | *BGIBMGA001453* | up | 1470 | 1478 | + | 11.3303 | 4.96E-05 | 0.615 | GTGGGTGGC |
| 1098 | mamo-s | *BGIBMGA001455* | up | 1146 | 1154 | - | 11.0826 | 6.84E-05 | 0.709 | GTACGTGGC |
| 1099 | mamo-s | *BGIBMGA001456* | up | 715 | 723 | - | 12.4679 | 1.31E-05 | 0.615 | GTGCGTGTC |
| 1100 | mamo-s | *BGIBMGA001456* | up | 834 | 842 | - | 12.4679 | 1.31E-05 | 0.615 | GTGCGTGTC |
| 1101 | mamo-s | *BGIBMGA001456* | up | 680 | 688 | + | 11.4404 | 4.51E-05 | 0.615 | CGGCGTGGC |
| 1102 | mamo-s | *BGIBMGA001456* | down | 1459 | 1467 | - | 11.3303 | 4.96E-05 | 0.615 | GTGGGTGGC |
| 1103 | mamo-s | *BGIBMGA001456* | down | 1597 | 1605 | - | 11.1009 | 6.57E-05 | 0.709 | GAGCGAGGC |
| 1104 | mamo-s | *BGIBMGA001456* | down | 1730 | 1738 | - | 11.1009 | 6.57E-05 | 0.709 | GAGCGAGGC |
| 1105 | mamo-s | *BGIBMGA001456* | down | 1834 | 1842 | + | 10.5138 | 9.75E-05 | 0.865 | GTGCGTGTT |
| 1106 | mamo-s | *BGIBMGA001457* | up | 1232 | 1240 | + | 11.3303 | 4.96E-05 | 0.615 | GTGGGTGGC |
| 1107 | mamo-s | *BGIBMGA001461* | down | 1774 | 1782 | - | 11.3303 | 4.96E-05 | 0.615 | GTGGGTGGC |
| 1108 | mamo-s | *BGIBMGA001462* | down | 920 | 928 | - | 11.3303 | 4.96E-05 | 0.615 | GTGGGTGGC |
| 1109 | mamo-s | *BGIBMGA001464* | down | 1203 | 1211 | - | 11.0826 | 6.84E-05 | 0.709 | GTACGTGGC |
| 1110 | mamo-s | *BGIBMGA001466* | up | 1412 | 1420 | + | 11.3303 | 4.96E-05 | 0.615 | GTGGGTGGC |
| 1111 | mamo-s | *BGIBMGA001469* | up | 860 | 868 | - | 11.3303 | 4.96E-05 | 0.615 | GTGGGTGGC |
| 1112 | mamo-s | *BGIBMGA001470* | up | 555 | 563 | - | 11.3303 | 4.96E-05 | 0.615 | GTGGGTGGC |
| 1113 | mamo-s | *BGIBMGA001472* | up | 1081 | 1089 | + | 11.1743 | 5.90E-05 | 0.684 | CCGCGTGTC |
| 1114 | mamo-s | *BGIBMGA001473* | up | 285 | 293 | - | 11.3303 | 4.96E-05 | 0.615 | GTGGGTGGC |
| 1115 | mamo-s | *BGIBMGA001474* | down | 951 | 959 | + | 11.0826 | 6.84E-05 | 0.709 | GTACGTGGC |
| 1116 | mamo-s | *BGIBMGA001477* | down | 625 | 633 | + | 10.5688 | 8.88E-05 | 0.824 | GTGCGTGAC |
| 1117 | mamo-s | *BGIBMGA001479* | down | 1819 | 1827 | + | 11.3303 | 4.96E-05 | 0.615 | GTGGGTGGC |
| 1118 | mamo-s | *BGIBMGA001479* | down | 1167 | 1175 | - | 11.2844 | 5.18E-05 | 0.63 | CTGCGTGCC |
| 1119 | mamo-s | *BGIBMGA001479* | down | 1382 | 1390 | - | 10.7523 | 8.38E-05 | 0.797 | AAGCGTGGC |
| 1120 | mamo-s | *BGIBMGA001480* | down | 389 | 397 | + | 13.0275 | 8.53E-06 | 0.615 | GAGCGTGGC |
| 1121 | mamo-s | *BGIBMGA001480* | down | 951 | 959 | + | 11.3303 | 4.96E-05 | 0.615 | GTGGGTGGC |
| 1122 | mamo-s | *BGIBMGA001480* | up | 1249 | 1257 | - | 11.3303 | 4.96E-05 | 0.615 | GTGGGTGGC |
| 1123 | mamo-s | *BGIBMGA001482* | down | 1691 | 1699 | - | 11.3303 | 4.96E-05 | 0.615 | GTGGGTGGC |
| 1124 | mamo-s | *BGIBMGA001484* | down | 1414 | 1422 | + | 10.5138 | 9.75E-05 | 0.865 | GTGCGTGTT |
| 1125 | mamo-s | *BGIBMGA001485* | down | 170 | 178 | + | 12.1009 | 2.43E-05 | 0.615 | GTGCGTGGT |
| 1126 | mamo-s | *BGIBMGA001485* | up | 1787 | 1795 | - | 12.1009 | 2.43E-05 | 0.615 | GTGCGTGGT |
| 1127 | mamo-s | *BGIBMGA001485* | down | 1798 | 1806 | - | 11.5321 | 4.06E-05 | 0.615 | TTGCGTGGC |
| 1128 | mamo-s | *BGIBMGA001485* | up | 1172 | 1180 | + | 11.3303 | 4.96E-05 | 0.615 | GTGGGTGGC |
| 1129 | mamo-s | *BGIBMGA001486* | down | 1858 | 1866 | - | 14.055 | 2.23E-06 | 0.615 | GTGCGTGGC |
| 1130 | mamo-s | *BGIBMGA001486* | up | 675 | 683 | + | 11.3303 | 4.96E-05 | 0.615 | GTGGGTGGC |
| 1131 | mamo-s | *BGIBMGA001486* | up | 1279 | 1287 | + | 11.3303 | 4.96E-05 | 0.615 | GTGGGTGGC |
| 1132 | mamo-s | *BGIBMGA001487* | up | 406 | 414 | + | 11.3303 | 4.96E-05 | 0.615 | GTGGGTGGC |
| 1133 | mamo-s | *BGIBMGA001493* | up | 928 | 936 | + | 11.3303 | 4.96E-05 | 0.615 | GTGGGTGGC |
| 1134 | mamo-s | *BGIBMGA001495* | down | 497 | 505 | - | 11.0734 | 7.12E-05 | 0.726 | GAGCGTGGT |
| 1135 | mamo-s | *BGIBMGA001496* | up | 1242 | 1250 | - | 10.8073 | 7.89E-05 | 0.766 | CCGCGTGGT |
| 1136 | mamo-s | *BGIBMGA001498* | down | 1751 | 1759 | - | 10.8349 | 7.66E-05 | 0.751 | CCGCGAGGC |
| 1137 | mamo-s | *BGIBMGA001499* | up | 853 | 861 | - | 11.3303 | 4.96E-05 | 0.615 | GTGGGTGGC |

| | | | | | | | | | | |
|---|---|---|---|---|---|---|---|---|---|---|
| 1138 | mamo-s | *BGIBMGA001502* | up | 340 | 348 | - | 11.3303 | 4.96E-05 | 0.615 | GTGGGTGGC |
| 1139 | mamo-s | *BGIBMGA001502* | down | 383 | 391 | + | 11.3303 | 4.96E-05 | 0.615 | GTGGGTGGC |
| 1140 | mamo-s | *BGIBMGA001502* | up | 1024 | 1032 | - | 11.3303 | 4.96E-05 | 0.615 | GTGGGTGGC |
| 1141 | mamo-s | *BGIBMGA001504* | down | 435 | 443 | - | 11.7064 | 3.52E-05 | 0.615 | GCGCGTGGT |
| 1142 | mamo-s | *BGIBMGA001504* | down | 1755 | 1763 | - | 10.5413 | 9.42E-05 | 0.857 | CAGCGTGTC |
| 1143 | mamo-s | *BGIBMGA001506* | up | 1624 | 1632 | + | 11.0826 | 6.84E-05 | 0.709 | GTACGTGGC |
| 1144 | mamo-s | *BGIBMGA001508* | down | 299 | 307 | + | 12.3394 | 1.49E-05 | 0.615 | GGGCGTGGC |
| 1145 | mamo-s | *BGIBMGA001510* | up | 1095 | 1103 | + | 13.156 | 6.30E-06 | 0.615 | CTGCGTGGC |
| 1146 | mamo-s | *BGIBMGA001510* | up | 699 | 707 | - | 11.3303 | 4.96E-05 | 0.615 | GTGGGTGGC |
| 1147 | mamo-s | *BGIBMGA001510* | up | 1237 | 1245 | + | 11.3303 | 4.96E-05 | 0.615 | GTGGGTGGC |
| 1148 | mamo-s | *BGIBMGA001511* | up | 976 | 984 | + | 11.3303 | 4.96E-05 | 0.615 | GTGGGTGGC |
| 1149 | mamo-s | *BGIBMGA001512* | up | 1740 | 1748 | - | 11.3303 | 4.96E-05 | 0.615 | GTGGGTGGC |
| 1150 | mamo-s | *BGIBMGA001513* | down | 1123 | 1131 | - | 11.3303 | 4.96E-05 | 0.615 | GTGGGTGGC |
| 1151 | mamo-s | *BGIBMGA001515* | up | 1360 | 1368 | - | 11.3303 | 4.96E-05 | 0.615 | GTGGGTGGC |
| 1152 | mamo-s | *BGIBMGA001516* | up | 621 | 629 | + | 11.3303 | 4.96E-05 | 0.615 | GTGGGTGGC |
| 1153 | mamo-s | *BGIBMGA001518* | up | 384 | 392 | + | 11.3303 | 4.96E-05 | 0.615 | GTGGGTGGC |
| 1154 | mamo-s | *BGIBMGA001519* | down | 1553 | 1561 | - | 11.4404 | 4.51E-05 | 0.615 | CGGCGTGGC |
| 1155 | mamo-s | *BGIBMGA001519* | down | 746 | 754 | + | 11.3303 | 4.96E-05 | 0.615 | GTGGGTGGC |
| 1156 | mamo-s | *BGIBMGA001519* | down | 1143 | 1151 | + | 11.2294 | 5.41E-05 | 0.65 | CTGCGAGGC |
| 1157 | mamo-s | *BGIBMGA001522* | down | 1055 | 1063 | + | 11.3303 | 4.96E-05 | 0.615 | GTGGGTGGC |
| 1158 | mamo-s | *BGIBMGA001522* | up | 411 | 419 | - | 11.0734 | 7.12E-05 | 0.726 | GAGCGTGGT |
| 1159 | mamo-s | *BGIBMGA001524* | up | 1055 | 1063 | - | 11.3303 | 4.96E-05 | 0.615 | GTGGGTGGC |
| 1160 | mamo-s | *BGIBMGA001524* | up | 217 | 225 | - | 10.9358 | 7.30E-05 | 0.728 | GCGGGTGGC |
| 1161 | mamo-s | *BGIBMGA001525* | down | 1453 | 1461 | - | 11.3303 | 4.96E-05 | 0.615 | GTGGGTGGC |
| 1162 | mamo-s | *BGIBMGA001526* | down | 74 | 82 | + | 11.0734 | 7.12E-05 | 0.726 | GAGCGTGGT |
| 1163 | mamo-s | *BGIBMGA001527* | up | 1686 | 1694 | + | 11.2018 | 5.68E-05 | 0.674 | CTGCGTGGT |
| 1164 | mamo-s | *BGIBMGA001529* | up | 1542 | 1550 | - | 11.7064 | 3.52E-05 | 0.615 | GCGCGTGGT |
| 1165 | mamo-s | *BGIBMGA001530* | up | 1714 | 1722 | - | 11.3303 | 4.96E-05 | 0.615 | GTGGGTGGC |
| 1166 | mamo-s | *BGIBMGA001531* | up | 1615 | 1623 | + | 11.3303 | 4.96E-05 | 0.615 | GTGGGTGGC |
| 1167 | mamo-s | *BGIBMGA001531* | up | 276 | 284 | - | 11.0826 | 6.84E-05 | 0.709 | GTACGTGGC |
| 1168 | mamo-s | *BGIBMGA001534* | up | 206 | 214 | + | 11.3303 | 4.96E-05 | 0.615 | GTGGGTGGC |
| 1169 | mamo-s | *BGIBMGA001534* | up | 1395 | 1403 | - | 11.3303 | 4.96E-05 | 0.615 | GTGGGTGGC |
| 1170 | mamo-s | *BGIBMGA001541* | down | 1019 | 1027 | - | 11.3303 | 4.96E-05 | 0.615 | GTGGGTGGC |
| 1171 | mamo-s | *BGIBMGA001544* | up | 1505 | 1513 | - | 12.1284 | 2.16E-05 | 0.615 | GTGCGAGGC |
| 1172 | mamo-s | *BGIBMGA001545* | up | 1742 | 1750 | + | 11.3303 | 4.96E-05 | 0.615 | GTGGGTGGC |
| 1173 | mamo-s | *BGIBMGA001552* | up | 716 | 724 | + | 11.3303 | 4.96E-05 | 0.615 | GTGGGTGGC |
| 1174 | mamo-s | *BGIBMGA001552* | up | 1610 | 1618 | + | 11.3303 | 4.96E-05 | 0.615 | GTGGGTGGC |
| 1175 | mamo-s | *BGIBMGA001554* | down | 852 | 860 | + | 11.2844 | 5.18E-05 | 0.63 | CTGCGTGCC |
| 1176 | mamo-s | *BGIBMGA001555* | down | 341 | 349 | + | 11.0734 | 7.12E-05 | 0.726 | GAGCGTGGT |
| 1177 | mamo-s | *BGIBMGA001556* | down | 1854 | 1862 | - | 11.3303 | 4.96E-05 | 0.615 | GTGGGTGGC |
| 1178 | mamo-s | *BGIBMGA001557* | down | 280 | 288 | + | 11.1376 | 6.35E-05 | 0.709 | TCGCGTGGC |
| 1179 | mamo-s | *BGIBMGA001559* | up | 330 | 338 | - | 12.7615 | 1.04E-05 | 0.615 | CCGCGTGGC |
| 1180 | mamo-s | *BGIBMGA001559* | up | 757 | 765 | + | 10.5138 | 9.75E-05 | 0.865 | GTGCGTGTT |
| 1181 | mamo-s | *BGIBMGA001561* | down | 1524 | 1532 | + | 11.3303 | 4.96E-05 | 0.615 | GTGGGTGGC |
| 1182 | mamo-s | *BGIBMGA001561* | down | 731 | 739 | + | 11.0826 | 6.84E-05 | 0.709 | GTACGTGGC |
| 1183 | mamo-s | *BGIBMGA001562* | down | 348 | 356 | - | 11.7798 | 3.11E-05 | 0.615 | ATGCGTGGC |
| 1184 | mamo-s | *BGIBMGA001562* | up | 569 | 577 | + | 11.3303 | 4.96E-05 | 0.615 | GTGGGTGGC |
| 1185 | mamo-s | *BGIBMGA001562* | up | 904 | 912 | - | 11.3303 | 4.96E-05 | 0.615 | GTGGGTGGC |
| 1186 | mamo-s | *BGIBMGA001562* | up | 1090 | 1098 | - | 11.3303 | 4.96E-05 | 0.615 | GTGGGTGGC |
| 1187 | mamo-s | *BGIBMGA001562* | down | 1918 | 1926 | + | 11.3303 | 4.96E-05 | 0.615 | GTGGGTGGC |
| 1188 | mamo-s | *BGIBMGA001564* | down | 1600 | 1608 | - | 11.0826 | 6.84E-05 | 0.709 | GTACGTGGC |
| 1189 | mamo-s | *BGIBMGA001565* | up | 1857 | 1865 | + | 11.7339 | 3.29E-05 | 0.615 | GCGCGAGGC |
| 1190 | mamo-s | *BGIBMGA001565* | up | 257 | 265 | + | 11.3303 | 4.96E-05 | 0.615 | GTGGGTGGC |
| 1191 | mamo-s | *BGIBMGA001566* | down | 1664 | 1672 | - | 12.0734 | 2.66E-05 | 0.615 | GCGCGTGTC |
| 1192 | mamo-s | *BGIBMGA001566* | down | 264 | 272 | - | 11.3303 | 4.96E-05 | 0.615 | GTGGGTGGC |
| 1193 | mamo-s | *BGIBMGA001568* | up | 78 | 86 | + | 11.3303 | 4.96E-05 | 0.615 | GTGGGTGGC |
| 1194 | mamo-s | *BGIBMGA001573* | down | 1106 | 1114 | + | 11.3303 | 4.96E-05 | 0.615 | GTGGGTGGC |
| 1195 | mamo-s | *BGIBMGA001574* | down | 820 | 828 | - | 11.3303 | 4.96E-05 | 0.615 | GTGGGTGGC |
| 1196 | mamo-s | *BGIBMGA001577* | down | 646 | 654 | - | 11.5688 | 3.79E-05 | 0.615 | CTGCGTGTC |
| 1197 | mamo-s | *BGIBMGA001577* | down | 1216 | 1224 | - | 11.5688 | 3.79E-05 | 0.615 | CTGCGTGTC |
| 1198 | mamo-s | *BGIBMGA001578* | down | 1572 | 1580 | - | 11.3303 | 4.96E-05 | 0.615 | GTGGGTGGC |
| 1199 | mamo-s | *BGIBMGA001579* | down | 1448 | 1456 | + | 10.5138 | 9.75E-05 | 0.865 | GTGCGTGTT |
| 1200 | mamo-s | *BGIBMGA001580* | up | 1014 | 1022 | - | 11.3303 | 4.96E-05 | 0.615 | GTGGGTGGC |
| 1201 | mamo-s | *BGIBMGA001580* | down | 1066 | 1074 | - | 11.3303 | 4.96E-05 | 0.615 | GTGGGTGGC |
| 1202 | mamo-s | *BGIBMGA001581* | down | 609 | 617 | - | 11.3303 | 4.96E-05 | 0.615 | GTGGGTGGC |
| 1203 | mamo-s | *BGIBMGA001582* | up | 1820 | 1828 | + | 11.3303 | 4.96E-05 | 0.615 | GTGGGTGGC |
| 1204 | mamo-s | *BGIBMGA001583* | down | 1621 | 1629 | + | 14.055 | 2.23E-06 | 0.615 | GTGCGTGGC |
| 1205 | mamo-s | *BGIBMGA001583* | down | 1174 | 1182 | + | 11.3303 | 4.96E-05 | 0.615 | GTGGGTGGC |
| 1206 | mamo-s | *BGIBMGA001584* | up | 1905 | 1913 | + | 11.1009 | 6.57E-05 | 0.709 | GAGCGAGGC |
| 1207 | mamo-s | *BGIBMGA001584* | up | 1061 | 1069 | + | 10.8073 | 7.89E-05 | 0.766 | CCGCGTGGT |
| 1208 | mamo-s | *BGIBMGA001584* | down | 1551 | 1559 | + | 10.5413 | 9.42E-05 | 0.857 | GTGCGAGTC |
| 1209 | mamo-s | *BGIBMGA001585* | down | 1221 | 1229 | - | 11.3303 | 4.96E-05 | 0.615 | GTGGGTGGC |
| 1210 | mamo-s | *BGIBMGA001585* | down | 1440 | 1448 | + | 11.3303 | 4.96E-05 | 0.615 | GTGGGTGGC |
| 1211 | mamo-s | *BGIBMGA001586* | down | 1705 | 1713 | - | 11.3303 | 4.96E-05 | 0.615 | GTGGGTGGC |
| 1212 | mamo-s | *BGIBMGA001587* | down | 1080 | 1088 | - | 11.7339 | 3.29E-05 | 0.615 | GCGCGAGGC |
| 1213 | mamo-s | *BGIBMGA001587* | down | 430 | 438 | - | 11.3303 | 4.96E-05 | 0.615 | GTGGGTGGC |
| 1214 | mamo-s | *BGIBMGA001589* | down | 925 | 933 | + | 11.3303 | 4.96E-05 | 0.615 | GTGGGTGGC |
| 1215 | mamo-s | *BGIBMGA001592* | up | 872 | 880 | - | 10.5688 | 8.88E-05 | 0.824 | GTGCGTGAC |
| 1216 | mamo-s | *BGIBMGA001594* | up | 1504 | 1512 | - | 11.3303 | 4.96E-05 | 0.615 | GTGGGTGGC |
| 1217 | mamo-s | *BGIBMGA001595* | up | 54 | 62 | + | 13.0275 | 8.53E-06 | 0.615 | GAGCGTGGC |
| 1218 | mamo-s | *BGIBMGA001595* | down | 1934 | 1942 | - | 11.7339 | 3.29E-05 | 0.615 | GCGCGAGGC |
| 1219 | mamo-s | *BGIBMGA001595* | down | 1911 | 1919 | + | 11.5321 | 4.06E-05 | 0.615 | TTGCGTGGC |
| 1220 | mamo-s | *BGIBMGA001595* | down | 873 | 881 | - | 11.3303 | 4.96E-05 | 0.615 | GTGGGTGGC |
| 1221 | mamo-s | *BGIBMGA001596* | down | 978 | 986 | - | 11.3303 | 4.96E-05 | 0.615 | GTGGGTGGC |
| 1222 | mamo-s | *BGIBMGA001597* | down | 735 | 743 | + | 11.3303 | 4.96E-05 | 0.615 | GTGGGTGGC |
| 1223 | mamo-s | *BGIBMGA001598* | up | 302 | 310 | + | 11.5321 | 4.06E-05 | 0.615 | TTGCGTGGC |
| 1224 | mamo-s | *BGIBMGA001598* | up | 1552 | 1560 | + | 11.0826 | 6.84E-05 | 0.709 | GTACGTGGC |
| 1225 | mamo-s | *BGIBMGA001599* | up | 1533 | 1541 | + | 11.3303 | 4.96E-05 | 0.615 | GTGGGTGGC |

| | | | | | | | | | | |
|---|---|---|---|---|---|---|---|---|---|---|
| 1226 | mamo-s | *BGIBMGA001600* | up | 198 | 206 | + | 11.3303 | 4.96E-05 | 0.615 | GTGGGTGGC |
| 1227 | mamo-s | *BGIBMGA001600* | down | 510 | 518 | + | 11.3303 | 4.96E-05 | 0.615 | GTGGGTGGC |
| 1228 | mamo-s | *BGIBMGA001600* | down | 1260 | 1268 | + | 11.3303 | 4.96E-05 | 0.615 | GTGGGTGGC |
| 1229 | mamo-s | *BGIBMGA001602* | up | 413 | 421 | - | 14.055 | 2.23E-06 | 0.615 | GTGCGTGGC |
| 1230 | mamo-s | *BGIBMGA001606* | down | 636 | 644 | - | 11.3303 | 4.96E-05 | 0.615 | GTGGGTGGC |
| 1231 | mamo-s | *BGIBMGA001606* | down | 1835 | 1843 | + | 11.3303 | 4.96E-05 | 0.615 | GTGGGTGGC |
| 1232 | mamo-s | *BGIBMGA001607* | up | 137 | 145 | + | 11.3303 | 4.96E-05 | 0.615 | GTGGGTGGC |
| 1233 | mamo-s | *BGIBMGA001610* | up | 783 | 791 | + | 13.0275 | 8.53E-06 | 0.615 | GAGCGTGGC |
| 1234 | mamo-s | *BGIBMGA001613* | up | 155 | 163 | + | 11.3303 | 4.96E-05 | 0.615 | GTGGGTGGC |
| 1235 | mamo-s | *BGIBMGA001613* | up | 663 | 671 | - | 11.3303 | 4.96E-05 | 0.615 | GTGGGTGGC |
| 1236 | mamo-s | *BGIBMGA001614* | down | 1370 | 1378 | - | 11.3303 | 4.96E-05 | 0.615 | GTGGGTGGC |
| 1237 | mamo-s | *BGIBMGA001615* | up | 1984 | 1992 | - | 10.5138 | 9.75E-05 | 0.865 | GTGCGTGTT |
| 1238 | mamo-s | *BGIBMGA001617* | down | 820 | 828 | - | 11.3303 | 4.96E-05 | 0.615 | GTGGGTGGC |
| 1239 | mamo-s | *BGIBMGA001618* | up | 1059 | 1067 | + | 11.3303 | 4.96E-05 | 0.615 | GTGGGTGGC |
| 1240 | mamo-s | *BGIBMGA001618* | down | 1405 | 1413 | + | 10.5138 | 9.75E-05 | 0.865 | GTGCGTGTT |
| 1241 | mamo-s | *BGIBMGA001619* | up | 670 | 678 | + | 11.789 | 2.84E-05 | 0.615 | GCGCGTGCC |
| 1242 | mamo-s | *BGIBMGA001624* | up | 1107 | 1115 | - | 11.7064 | 3.52E-05 | 0.615 | GCGCGTGGT |
| 1243 | mamo-s | *BGIBMGA001627* | up | 1959 | 1967 | + | 13.6606 | 4.06E-06 | 0.615 | GCGCGTGGC |
| 1244 | mamo-s | *BGIBMGA001631* | down | 1096 | 1104 | + | 11.3303 | 4.96E-05 | 0.615 | GTGGGTGGC |
| 1245 | mamo-s | *BGIBMGA001631* | up | 1800 | 1808 | - | 11.3303 | 4.96E-05 | 0.615 | GTGGGTGGC |
| 1246 | mamo-s | *BGIBMGA001631* | down | 848 | 856 | - | 11.1009 | 6.57E-05 | 0.709 | GAGCGAGGC |
| 1247 | mamo-s | *BGIBMGA001634* | up | 212 | 220 | + | 11.3303 | 4.96E-05 | 0.615 | GTGGGTGGC |
| 1248 | mamo-s | *BGIBMGA001635* | down | 987 | 995 | - | 11.3303 | 4.96E-05 | 0.615 | GTGGGTGGC |
| 1249 | mamo-s | *BGIBMGA001636* | up | 1077 | 1085 | - | 11.3303 | 4.96E-05 | 0.615 | GTGGGTGGC |
| 1250 | mamo-s | *BGIBMGA001637* | up | 961 | 969 | - | 11.3303 | 4.96E-05 | 0.615 | GTGGGTGGC |
| 1251 | mamo-s | *BGIBMGA001640* | up | 938 | 946 | - | 11.3303 | 4.96E-05 | 0.615 | GTGGGTGGC |
| 1252 | mamo-s | *BGIBMGA001640* | down | 1178 | 1186 | + | 11.3303 | 4.96E-05 | 0.615 | GTGGGTGGC |
| 1253 | mamo-s | *BGIBMGA001641* | down | 465 | 473 | + | 11.3303 | 4.96E-05 | 0.615 | GTGGGTGGC |
| 1254 | mamo-s | *BGIBMGA001641* | up | 1261 | 1269 | - | 11.2844 | 5.18E-05 | 0.63 | CTGCGTGCC |
| 1255 | mamo-s | *BGIBMGA001641* | up | 918 | 926 | + | 11.0826 | 6.84E-05 | 0.709 | GTACGTGGC |
| 1256 | mamo-s | *BGIBMGA001642* | up | 1888 | 1896 | + | 11.3303 | 4.96E-05 | 0.615 | GTGGGTGGC |
| 1257 | mamo-s | *BGIBMGA001644* | up | 1704 | 1712 | + | 13.156 | 6.30E-06 | 0.615 | CTGCGTGGC |
| 1258 | mamo-s | *BGIBMGA001645* | down | 1905 | 1913 | - | 11.3303 | 4.96E-05 | 0.615 | GTGGGTGGC |
| 1259 | mamo-s | *BGIBMGA001646* | up | 596 | 604 | - | 11.3303 | 4.96E-05 | 0.615 | GTGGGTGGC |
| 1260 | mamo-s | *BGIBMGA001647* | up | 804 | 812 | + | 11.3303 | 4.96E-05 | 0.615 | GTGGGTGGC |
| 1261 | mamo-s | *BGIBMGA001648* | up | 1226 | 1234 | - | 11.3303 | 4.96E-05 | 0.615 | GTGGGTGGC |
| 1262 | mamo-s | *BGIBMGA001649* | down | 1250 | 1258 | - | 11.789 | 2.84E-05 | 0.615 | GCGCGTGCC |
| 1263 | mamo-s | *BGIBMGA001649* | down | 1965 | 1973 | - | 11.3853 | 4.74E-05 | 0.615 | ACGCGTGGC |
| 1264 | mamo-s | *BGIBMGA001650* | up | 1489 | 1497 | + | 11.3303 | 4.96E-05 | 0.615 | GTGGGTGGC |
| 1265 | mamo-s | *BGIBMGA001651* | up | 564 | 572 | - | 11.3303 | 4.96E-05 | 0.615 | GTGGGTGGC |
| 1266 | mamo-s | *BGIBMGA001651* | up | 1530 | 1538 | + | 11.3303 | 4.96E-05 | 0.615 | GTGGGTGGC |
| 1267 | mamo-s | *BGIBMGA001651* | up | 199 | 207 | + | 10.7523 | 8.38E-05 | 0.797 | AAGCGTGGC |
| 1268 | mamo-s | *BGIBMGA001652* | up | 1225 | 1233 | + | 11.3303 | 4.96E-05 | 0.615 | GTGGGTGGC |
| 1269 | mamo-s | *BGIBMGA001652* | up | 684 | 692 | + | 11.0826 | 6.84E-05 | 0.709 | GTACGTGGC |
| 1270 | mamo-s | *BGIBMGA001655* | up | 784 | 792 | - | 11.3303 | 4.96E-05 | 0.615 | GTGGGTGGC |
| 1271 | mamo-s | *BGIBMGA001656* | up | 1989 | 1997 | - | 10.5413 | 9.42E-05 | 0.857 | GTGCGAGTC |
| 1272 | mamo-s | *BGIBMGA001657* | down | 1419 | 1427 | + | 11.3303 | 4.96E-05 | 0.615 | GTGGGTGGC |
| 1273 | mamo-s | *BGIBMGA001658* | down | 688 | 696 | - | 11.3303 | 4.96E-05 | 0.615 | GTGGGTGGC |
| 1274 | mamo-s | *BGIBMGA001659* | down | 1219 | 1227 | + | 14.055 | 2.23E-06 | 0.615 | GTGCGTGGC |
| 1275 | mamo-s | *BGIBMGA001659* | down | 479 | 487 | + | 11.3303 | 4.96E-05 | 0.615 | GTGGGTGGC |
| 1276 | mamo-s | *BGIBMGA001659* | down | 1563 | 1571 | + | 10.5138 | 9.75E-05 | 0.865 | GTGCGTGTT |
| 1277 | mamo-s | *BGIBMGA001660* | down | 586 | 594 | - | 11.0826 | 6.84E-05 | 0.709 | GTACGTGGC |
| 1278 | mamo-s | *BGIBMGA001661* | up | 1013 | 1021 | - | 11.3303 | 4.96E-05 | 0.615 | GTGGGTGGC |
| 1279 | mamo-s | *BGIBMGA001662* | up | 502 | 510 | + | 12.1284 | 2.16E-05 | 0.615 | GTGCGAGGC |
| 1280 | mamo-s | *BGIBMGA001663* | down | 798 | 806 | - | 11.7798 | 3.11E-05 | 0.615 | ATGCGTGGC |
| 1281 | mamo-s | *BGIBMGA001664* | down | 1835 | 1843 | + | 11.0826 | 6.84E-05 | 0.709 | GTACGTGGC |
| 1282 | mamo-s | *BGIBMGA001665* | down | 445 | 453 | - | 11.3303 | 4.96E-05 | 0.615 | GTGGGTGGC |
| 1283 | mamo-s | *BGIBMGA001665* | down | 913 | 921 | + | 11.3303 | 4.96E-05 | 0.615 | GTGGGTGGC |
| 1284 | mamo-s | *BGIBMGA001666* | up | 897 | 905 | - | 11.3303 | 4.96E-05 | 0.615 | GTGGGTGGC |
| 1285 | mamo-s | *BGIBMGA001666* | down | 1246 | 1254 | - | 11.3303 | 4.96E-05 | 0.615 | GTGGGTGGC |
| 1286 | mamo-s | *BGIBMGA001666* | up | 1365 | 1373 | + | 11.3303 | 4.96E-05 | 0.615 | GTGGGTGGC |
| 1287 | mamo-s | *BGIBMGA001668* | up | 1429 | 1437 | + | 11.156 | 6.13E-05 | 0.705 | GAGCGTGCC |
| 1288 | mamo-s | *BGIBMGA001670* | down | 389 | 397 | - | 11.3303 | 4.96E-05 | 0.615 | GTGGGTGGC |
| 1289 | mamo-s | *BGIBMGA001670* | up | 929 | 937 | + | 10.8073 | 7.89E-05 | 0.766 | CCGCGTGGT |
| 1290 | mamo-s | *BGIBMGA001670* | up | 206 | 214 | + | 10.5138 | 9.75E-05 | 0.865 | GTGCGTGTT |
| 1291 | mamo-s | *BGIBMGA001671* | down | 289 | 297 | - | 10.9358 | 7.30E-05 | 0.728 | GCGGGTGGC |
| 1292 | mamo-s | *BGIBMGA001674* | up | 1023 | 1031 | - | 11.7798 | 3.11E-05 | 0.615 | ATGCGTGGC |
| 1293 | mamo-s | *BGIBMGA001675* | down | 1332 | 1340 | - | 12.1284 | 2.16E-05 | 0.615 | CAGCGTGGC |
| 1294 | mamo-s | *BGIBMGA001676* | up | 940 | 948 | - | 11.3303 | 4.96E-05 | 0.615 | GTGGGTGGC |
| 1295 | mamo-s | *BGIBMGA001678* | down | 1337 | 1345 | - | 14.055 | 2.23E-06 | 0.615 | GTGCGTGGC |
| 1296 | mamo-s | *BGIBMGA001679* | down | 1551 | 1559 | + | 11.3303 | 4.96E-05 | 0.615 | GTGGGTGGC |
| 1297 | mamo-s | *BGIBMGA001680* | down | 1688 | 1696 | - | 11.3303 | 4.96E-05 | 0.615 | GTGGGTGGC |
| 1298 | mamo-s | *BGIBMGA001683* | down | 1342 | 1350 | - | 11.3303 | 4.96E-05 | 0.615 | GTGGGTGGC |
| 1299 | mamo-s | *BGIBMGA001683* | up | 1734 | 1742 | + | 11.3303 | 4.96E-05 | 0.615 | GTGGGTGGC |
| 1300 | mamo-s | *BGIBMGA001683* | down | 1544 | 1552 | + | 10.8899 | 7.48E-05 | 0.739 | CCGCGTGCC |
| 1301 | mamo-s | *BGIBMGA001685* | down | 1632 | 1640 | + | 11.3303 | 4.96E-05 | 0.615 | GTGGGTGGC |
| 1302 | mamo-s | *BGIBMGA001685* | down | 456 | 464 | + | 11.0826 | 6.84E-05 | 0.709 | GTACGTGGC |
| 1303 | mamo-s | *BGIBMGA001686* | down | 1644 | 1652 | + | 11.0826 | 6.84E-05 | 0.709 | GTACGTGGC |
| 1304 | mamo-s | *BGIBMGA001686* | down | 791 | 799 | + | 11.0734 | 7.12E-05 | 0.726 | GAGCGTGGT |
| 1305 | mamo-s | *BGIBMGA001687* | down | 464 | 472 | + | 11.4404 | 4.51E-05 | 0.615 | CGGCGTGGC |
| 1306 | mamo-s | *BGIBMGA001687* | up | 1685 | 1693 | + | 11.3303 | 4.96E-05 | 0.615 | GTGGGTGGC |
| 1307 | mamo-s | *BGIBMGA001687* | down | 453 | 461 | - | 11.2018 | 5.68E-05 | 0.674 | CTGCGTGGT |
| 1308 | mamo-s | *BGIBMGA001688* | down | 1764 | 1772 | + | 11.3303 | 4.96E-05 | 0.615 | GTGGGTGGC |
| 1309 | mamo-s | *BGIBMGA001689* | up | 1062 | 1070 | + | 11.3303 | 4.96E-05 | 0.615 | GTGGGTGGC |
| 1310 | mamo-s | *BGIBMGA001691* | down | 1182 | 1190 | - | 11.3303 | 4.96E-05 | 0.615 | GTGGGTGGC |
| 1311 | mamo-s | *BGIBMGA001691* | up | 1263 | 1271 | - | 11.3303 | 4.96E-05 | 0.615 | GTGGGTGGC |
| 1312 | mamo-s | *BGIBMGA001693* | down | 590 | 598 | - | 11.789 | 2.84E-05 | 0.615 | GCGCGTGCC |
| 1313 | mamo-s | *BGIBMGA001693* | down | 1686 | 1694 | - | 11.3303 | 4.96E-05 | 0.615 | GTGGGTGGC |

| | | | | | | | | | | |
|---|---|---|---|---|---|---|---|---|---|---|
| 1314 | mamo-s | *BGIBMGA001695* | up | 538 | 546 | - | 11.3303 | 4.96E-05 | 0.615 | GTGGGTGGC |
| 1315 | mamo-s | *BGIBMGA001697* | down | 1814 | 1822 | - | 11.0826 | 6.84E-05 | 0.709 | GTACGTGGC |
| 1316 | mamo-s | *BGIBMGA001700* | up | 64 | 72 | + | 12.0734 | 2.66E-05 | 0.615 | GCGCGTGTC |
| 1317 | mamo-s | *BGIBMGA001700* | down | 1123 | 1131 | + | 11.3303 | 4.96E-05 | 0.615 | GTGGGTGGC |
| 1318 | mamo-s | *BGIBMGA001701* | up | 52 | 60 | + | 11.3303 | 4.96E-05 | 0.615 | GTGGGTGGC |
| 1319 | mamo-s | *BGIBMGA001703* | down | 1276 | 1284 | - | 11.5688 | 3.79E-05 | 0.615 | CTGCGTGTC |
| 1320 | mamo-s | *BGIBMGA001703* | up | 564 | 572 | - | 10.8899 | 7.48E-05 | 0.739 | CCGCGTGCC |
| 1321 | mamo-s | *BGIBMGA001705* | down | 480 | 488 | - | 11.3303 | 4.96E-05 | 0.615 | GTGGGTGGC |
| 1322 | mamo-s | *BGIBMGA001705* | down | 141 | 149 | - | 11.0826 | 6.84E-05 | 0.709 | GTACGTGGC |
| 1323 | mamo-s | *BGIBMGA001707* | down | 1428 | 1436 | + | 11.4404 | 4.51E-05 | 0.615 | CGGCGTGGC |
| 1324 | mamo-s | *BGIBMGA001707* | up | 120 | 128 | + | 11.3303 | 4.96E-05 | 0.615 | GTGGGTGGC |
| 1325 | mamo-s | *BGIBMGA001708* | up | 881 | 889 | + | 13.6606 | 4.06E-06 | 0.615 | GCGCGTGGC |
| 1326 | mamo-s | *BGIBMGA001710* | up | 1545 | 1553 | + | 11.1743 | 5.90E-05 | 0.684 | CCGCGTGTC |
| 1327 | mamo-s | *BGIBMGA001712* | up | 1263 | 1271 | + | 11.3303 | 4.96E-05 | 0.615 | GTGGGTGGC |
| 1328 | mamo-s | *BGIBMGA001712* | down | 1931 | 1939 | + | 11.3303 | 4.96E-05 | 0.615 | GTGGGTGGC |
| 1329 | mamo-s | *BGIBMGA001712* | down | 1417 | 1425 | - | 11.0826 | 6.84E-05 | 0.709 | GTACGTGGC |
| 1330 | mamo-s | *BGIBMGA001713* | down | 661 | 669 | - | 11.4404 | 4.51E-05 | 0.615 | GAGCGTGTC |
| 1331 | mamo-s | *BGIBMGA001713* | up | 1226 | 1234 | + | 11.4404 | 4.51E-05 | 0.615 | GAGCGTGTC |
| 1332 | mamo-s | *BGIBMGA001714* | down | 41 | 49 | - | 10.7523 | 8.38E-05 | 0.797 | GGGCGTGTC |
| 1333 | mamo-s | *BGIBMGA001720* | up | 166 | 174 | - | 11.3303 | 4.96E-05 | 0.615 | GTGGGTGGC |
| 1334 | mamo-s | *BGIBMGA001721* | up | 1488 | 1496 | + | 13.0275 | 8.53E-06 | 0.615 | GAGCGTGGC |
| 1335 | mamo-s | *BGIBMGA001721* | up | 158 | 166 | + | 11.3303 | 4.96E-05 | 0.615 | GTGGGTGGC |
| 1336 | mamo-s | *BGIBMGA001722* | down | 1986 | 1994 | - | 11.3303 | 4.96E-05 | 0.615 | GTGGGTGGC |
| 1337 | mamo-s | *BGIBMGA001723* | down | 1844 | 1852 | - | 11.3303 | 4.96E-05 | 0.615 | GTGGGTGGC |
| 1338 | mamo-s | *BGIBMGA001727* | down | 358 | 366 | + | 11.3303 | 4.96E-05 | 0.615 | GTGGGTGGC |
| 1339 | mamo-s | *BGIBMGA001727* | down | 691 | 699 | - | 11.0826 | 6.84E-05 | 0.709 | GTACGTGGC |
| 1340 | mamo-s | *BGIBMGA001727* | up | 1576 | 1584 | + | 11.0826 | 6.84E-05 | 0.709 | GTACGTGGC |
| 1341 | mamo-s | *BGIBMGA001727* | up | 1104 | 1112 | + | 10.5138 | 9.75E-05 | 0.865 | GTGCGTGTT |
| 1342 | mamo-s | *BGIBMGA001728* | down | 1821 | 1829 | - | 11.3303 | 4.96E-05 | 0.615 | GTGGGTGGC |
| 1343 | mamo-s | *BGIBMGA001729* | down | 1696 | 1704 | + | 12.1009 | 2.43E-05 | 0.615 | GTGCGTGGT |
| 1344 | mamo-s | *BGIBMGA001729* | down | 306 | 314 | + | 11.3303 | 4.96E-05 | 0.615 | GTGGGTGGC |
| 1345 | mamo-s | *BGIBMGA001730* | up | 1322 | 1330 | - | 11.2844 | 5.18E-05 | 0.63 | CTGCGTGCC |
| 1346 | mamo-s | *BGIBMGA001731* | down | 1609 | 1617 | + | 11.789 | 2.84E-05 | 0.615 | GCGCGTGCC |
| 1347 | mamo-s | *BGIBMGA001731* | down | 62 | 70 | + | 11.3303 | 4.96E-05 | 0.615 | GTGGGTGGC |
| 1348 | mamo-s | *BGIBMGA001731* | down | 513 | 521 | + | 11.3303 | 4.96E-05 | 0.615 | GTGGGTGGC |
| 1349 | mamo-s | *BGIBMGA001732* | down | 478 | 486 | - | 11.3303 | 4.96E-05 | 0.615 | GTGGGTGGC |
| 1350 | mamo-s | *BGIBMGA001733* | down | 1524 | 1532 | - | 12.0734 | 2.66E-05 | 0.615 | GCGCGTGTC |
| 1351 | mamo-s | *BGIBMGA001736* | up | 287 | 295 | - | 11.3303 | 4.96E-05 | 0.615 | GTGGGTGGC |
| 1352 | mamo-s | *BGIBMGA001736* | down | 1166 | 1174 | + | 11.3303 | 4.96E-05 | 0.615 | GTGGGTGGC |
| 1353 | mamo-s | *BGIBMGA001737* | down | 636 | 644 | - | 11.3303 | 4.96E-05 | 0.615 | GTGGGTGGC |
| 1354 | mamo-s | *BGIBMGA001737* | down | 965 | 973 | - | 11.3303 | 4.96E-05 | 0.615 | GTGGGTGGC |
| 1355 | mamo-s | *BGIBMGA001740* | up | 1280 | 1288 | + | 11.3303 | 4.96E-05 | 0.615 | GTGGGTGGC |
| 1356 | mamo-s | *BGIBMGA001741* | down | 956 | 964 | - | 11.3303 | 4.96E-05 | 0.615 | GTGGGTGGC |
| 1357 | mamo-s | *BGIBMGA001742* | up | 879 | 887 | + | 10.5138 | 9.75E-05 | 0.865 | GTGCGTGTT |
| 1358 | mamo-s | *BGIBMGA001743* | up | 1778 | 1786 | + | 11.2844 | 5.18E-05 | 0.63 | CTGCGTGCC |
| 1359 | mamo-s | *BGIBMGA001743* | down | 1826 | 1834 | + | 10.5138 | 9.75E-05 | 0.865 | GTGCGTGTT |
| 1360 | mamo-s | *BGIBMGA001744* | down | 387 | 395 | + | 11.3303 | 4.96E-05 | 0.615 | GTGGGTGGC |
| 1361 | mamo-s | *BGIBMGA001744* | up | 1185 | 1193 | + | 11.0826 | 6.84E-05 | 0.709 | GTACGTGGC |
| 1362 | mamo-s | *BGIBMGA001744* | down | 185 | 193 | - | 10.8899 | 7.48E-05 | 0.739 | CCGCGTGCC |
| 1363 | mamo-s | *BGIBMGA001744* | up | 896 | 904 | + | 10.5413 | 9.42E-05 | 0.857 | CAGCGTGTC |
| 1364 | mamo-s | *BGIBMGA001745* | down | 307 | 315 | + | 11.3303 | 4.96E-05 | 0.615 | GTGGGTGGC |
| 1365 | mamo-s | *BGIBMGA001745* | up | 1807 | 1815 | - | 11.3303 | 4.96E-05 | 0.615 | GTGGGTGGC |
| 1366 | mamo-s | *BGIBMGA001745* | up | 884 | 892 | + | 11.0734 | 7.12E-05 | 0.726 | GAGCGTGGT |
| 1367 | mamo-s | *BGIBMGA001748* | down | 757 | 765 | + | 11.3303 | 4.96E-05 | 0.615 | GTGGGTGGC |
| 1368 | mamo-s | *BGIBMGA001748* | down | 1730 | 1738 | + | 11.3303 | 4.96E-05 | 0.615 | GTGGGTGGC |
| 1369 | mamo-s | *BGIBMGA001750* | down | 933 | 941 | + | 11.2844 | 5.18E-05 | 0.63 | CTGCGTGCC |
| 1370 | mamo-s | *BGIBMGA001752* | down | 1595 | 1603 | + | 14.055 | 2.23E-06 | 0.615 | GTGCGTGGC |
| 1371 | mamo-s | *BGIBMGA001754* | up | 483 | 491 | - | 11.5688 | 3.79E-05 | 0.615 | CTGCGTGTC |
| 1372 | mamo-s | *BGIBMGA001755* | down | 1175 | 1183 | + | 11.3303 | 4.96E-05 | 0.615 | GTGGGTGGC |
| 1373 | mamo-s | *BGIBMGA001756* | down | 971 | 979 | - | 13.6606 | 4.06E-06 | 0.615 | GCGCGTGGC |
| 1374 | mamo-s | *BGIBMGA001757* | up | 1601 | 1609 | + | 11.3303 | 4.96E-05 | 0.615 | GTGGGTGGC |
| 1375 | mamo-s | *BGIBMGA001758* | up | 1525 | 1533 | - | 11.3303 | 4.96E-05 | 0.615 | GTGGGTGGC |
| 1376 | mamo-s | *BGIBMGA001760* | down | 446 | 454 | - | 11.0826 | 6.84E-05 | 0.709 | GTACGTGGC |
| 1377 | mamo-s | *BGIBMGA001761* | down | 138 | 146 | - | 11.3303 | 4.96E-05 | 0.615 | GTGGGTGGC |
| 1378 | mamo-s | *BGIBMGA001761* | up | 1123 | 1131 | + | 11.3303 | 4.96E-05 | 0.615 | GTGGGTGGC |
| 1379 | mamo-s | *BGIBMGA001761* | down | 1226 | 1234 | - | 11.3303 | 4.96E-05 | 0.615 | GTGGGTGGC |
| 1380 | mamo-s | *BGIBMGA001761* | up | 1710 | 1718 | + | 11.0826 | 6.84E-05 | 0.709 | GTACGTGGC |
| 1381 | mamo-s | *BGIBMGA001762* | down | 1129 | 1137 | - | 11.3303 | 4.96E-05 | 0.615 | GTGGGTGGC |
| 1382 | mamo-s | *BGIBMGA001763* | up | 1471 | 1479 | + | 12.0734 | 2.66E-05 | 0.615 | GCGCGTGTC |
| 1383 | mamo-s | *BGIBMGA001764* | down | 1719 | 1727 | + | 11.3303 | 4.96E-05 | 0.615 | GTGGGTGGC |
| 1384 | mamo-s | *BGIBMGA001765* | up | 1780 | 1788 | + | 11.7064 | 3.52E-05 | 0.615 | GCGCGTGGT |
| 1385 | mamo-s | *BGIBMGA001766* | down | 524 | 532 | - | 11.3303 | 4.96E-05 | 0.615 | GTGGGTGGC |
| 1386 | mamo-s | *BGIBMGA001766* | up | 957 | 965 | - | 10.8073 | 7.89E-05 | 0.766 | CCGCGTGGT |
| 1387 | mamo-s | *BGIBMGA001766* | up | 1680 | 1688 | - | 10.5138 | 9.75E-05 | 0.865 | GTGCGTGTT |
| 1388 | mamo-s | *BGIBMGA001767* | up | 1228 | 1236 | - | 11.4404 | 4.51E-05 | 0.615 | CGGCGTGGC |
| 1389 | mamo-s | *BGIBMGA001768* | up | 191 | 199 | - | 11.156 | 6.13E-05 | 0.705 | GAGCGTGCC |
| 1390 | mamo-s | *BGIBMGA001769* | down | 1471 | 1479 | - | 12.4679 | 1.31E-05 | 0.615 | GTGCGTGTC |
| 1391 | mamo-s | *BGIBMGA001769* | up | 362 | 370 | - | 11.3303 | 4.96E-05 | 0.615 | GTGGGTGGC |
| 1392 | mamo-s | *BGIBMGA001770* | down | 1733 | 1741 | + | 10.5413 | 9.42E-05 | 0.857 | CAGCGTGTC |
| 1393 | mamo-s | *BGIBMGA001771* | down | 350 | 358 | - | 11.5688 | 3.79E-05 | 0.615 | CTGCGTGTC |
| 1394 | mamo-s | *BGIBMGA001771* | down | 1654 | 1662 | - | 11.3303 | 4.96E-05 | 0.615 | GTGGGTGGC |
| 1395 | mamo-s | *BGIBMGA001772* | up | 454 | 462 | - | 11.3303 | 4.96E-05 | 0.615 | GTGGGTGGC |
| 1396 | mamo-s | *BGIBMGA001772* | down | 670 | 678 | + | 11.3303 | 4.96E-05 | 0.615 | GTGGGTGGC |
| 1397 | mamo-s | *BGIBMGA001773* | down | 1801 | 1809 | + | 11.3303 | 4.96E-05 | 0.615 | GTGGGTGGC |
| 1398 | mamo-s | *BGIBMGA001777* | up | 195 | 203 | + | 12.3394 | 1.49E-05 | 0.615 | GGGCGTGGC |
| 1399 | mamo-s | *BGIBMGA001777* | up | 864 | 872 | - | 11.3303 | 4.96E-05 | 0.615 | GTGGGTGGC |
| 1400 | mamo-s | *BGIBMGA001778* | down | 167 | 175 | - | 11.2294 | 5.41E-05 | 0.65 | CTGCGAGGC |
| 1401 | mamo-s | *BGIBMGA001779* | down | 1067 | 1075 | - | 11.3303 | 4.96E-05 | 0.615 | GTGGGTGGC |

| | | | | | | | | | | |
|---|---|---|---|---|---|---|---|---|---|---|
| 1402 | mamo-s | *BGIBMGA001781* | up | 154 | 162 | + | 11.3303 | 4.96E-05 | 0.615 | GTGGGTGGC |
| 1403 | mamo-s | *BGIBMGA001781* | down | 1971 | 1979 | + | 11.3303 | 4.96E-05 | 0.615 | GTGGGTGGC |
| 1404 | mamo-s | *BGIBMGA001782* | down | 876 | 884 | + | 11.3303 | 4.96E-05 | 0.615 | GTGGGTGGC |
| 1405 | mamo-s | *BGIBMGA001782* | down | 498 | 506 | - | 11.1009 | 6.57E-05 | 0.709 | GAGCGAGGC |
| 1406 | mamo-s | *BGIBMGA001784* | up | 1725 | 1733 | + | 10.8349 | 7.66E-05 | 0.751 | CCGCGAGGC |
| 1407 | mamo-s | *BGIBMGA001785* | up | 974 | 982 | - | 12.4679 | 1.31E-05 | 0.615 | GTGCGTGTC |
| 1408 | mamo-s | *BGIBMGA001785* | down | 1020 | 1028 | + | 11.3303 | 4.96E-05 | 0.615 | GTGGGTGGC |
| 1409 | mamo-s | *BGIBMGA001789* | down | 514 | 522 | + | 12.7615 | 1.04E-05 | 0.615 | CCGCGTGGC |
| 1410 | mamo-s | *BGIBMGA001790* | up | 1599 | 1607 | - | 11.1009 | 6.57E-05 | 0.709 | GAGCGAGGC |
| 1411 | mamo-s | *BGIBMGA001791* | up | 1259 | 1267 | + | 11.3303 | 4.96E-05 | 0.615 | GTGGGTGGC |
| 1412 | mamo-s | *BGIBMGA001792* | up | 38 | 46 | - | 11.7798 | 3.11E-05 | 0.615 | ATGCGTGGC |
| 1413 | mamo-s | *BGIBMGA001793* | up | 194 | 202 | + | 10.5413 | 9.42E-05 | 0.857 | CAGCGTGTC |
| 1414 | mamo-s | *BGIBMGA001795* | down | 1317 | 1325 | - | 11.3303 | 4.96E-05 | 0.615 | GTGGGTGGC |
| 1415 | mamo-s | *BGIBMGA001797* | down | 1387 | 1395 | - | 11.3303 | 4.96E-05 | 0.615 | GTGGGTGGC |
| 1416 | mamo-s | *BGIBMGA001799* | down | 526 | 534 | + | 11.3303 | 4.96E-05 | 0.615 | GTGGGTGGC |
| 1417 | mamo-s | *BGIBMGA001803* | up | 1318 | 1326 | - | 11.3303 | 4.96E-05 | 0.615 | GTGGGTGGC |
| 1418 | mamo-s | *BGIBMGA001805* | down | 524 | 532 | + | 11.3303 | 4.96E-05 | 0.615 | GTGGGTGGC |
| 1419 | mamo-s | *BGIBMGA001808* | down | 1798 | 1806 | - | 11.3303 | 4.96E-05 | 0.615 | GTGGGTGGC |
| 1420 | mamo-s | *BGIBMGA001808* | down | 1712 | 1720 | + | 10.5413 | 9.42E-05 | 0.857 | CAGCGTGTC |
| 1421 | mamo-s | *BGIBMGA001809* | up | 1650 | 1658 | - | 11.3303 | 4.96E-05 | 0.615 | GTGGGTGGC |
| 1422 | mamo-s | *BGIBMGA001809* | up | 1564 | 1572 | + | 10.5413 | 9.42E-05 | 0.857 | CAGCGTGTC |
| 1423 | mamo-s | *BGIBMGA001810* | down | 1350 | 1358 | + | 11.7064 | 3.52E-05 | 0.615 | GCGCGTGGT |
| 1424 | mamo-s | *BGIBMGA001810* | up | 406 | 414 | - | 11.3303 | 4.96E-05 | 0.615 | GTGGGTGGC |
| 1425 | mamo-s | *BGIBMGA001813* | up | 55 | 63 | + | 11.3303 | 4.96E-05 | 0.615 | GTGGGTGGC |
| 1426 | mamo-s | *BGIBMGA001814* | up | 920 | 928 | + | 11.3303 | 4.96E-05 | 0.615 | GTGGGTGGC |
| 1427 | mamo-s | *BGIBMGA001814* | down | 378 | 386 | + | 11.0826 | 6.84E-05 | 0.709 | GTACGTGGC |
| 1428 | mamo-s | *BGIBMGA001815* | down | 1238 | 1246 | + | 11.3303 | 4.96E-05 | 0.615 | GTGGGTGGC |
| 1429 | mamo-s | *BGIBMGA001816* | down | 933 | 941 | + | 11.0734 | 7.12E-05 | 0.726 | GAGCGTGGT |
| 1430 | mamo-s | *BGIBMGA001816* | up | 1214 | 1222 | + | 11.0734 | 7.12E-05 | 0.726 | GAGCGTGGT |
| 1431 | mamo-s | *BGIBMGA001817* | up | 1130 | 1138 | - | 11.0734 | 7.12E-05 | 0.726 | GAGCGTGGT |
| 1432 | mamo-s | *BGIBMGA001818* | up | 1495 | 1503 | - | 11.3303 | 4.96E-05 | 0.615 | GTGGGTGGC |
| 1433 | mamo-s | *BGIBMGA001819* | down | 630 | 638 | + | 11.3303 | 4.96E-05 | 0.615 | GTGGGTGGC |
| 1434 | mamo-s | *BGIBMGA001819* | down | 843 | 851 | - | 11.3303 | 4.96E-05 | 0.615 | GTGGGTGGC |
| 1435 | mamo-s | *BGIBMGA001819* | up | 241 | 249 | - | 11.0826 | 6.84E-05 | 0.709 | GTACGTGGC |
| 1436 | mamo-s | *BGIBMGA001820* | down | 1522 | 1530 | - | 11.3303 | 4.96E-05 | 0.615 | GTGGGTGGC |
| 1437 | mamo-s | *BGIBMGA001824* | up | 750 | 758 | + | 11.3303 | 4.96E-05 | 0.615 | GTGGGTGGC |
| 1438 | mamo-s | *BGIBMGA001826* | up | 930 | 938 | - | 11.3303 | 4.96E-05 | 0.615 | GTGGGTGGC |
| 1439 | mamo-s | *BGIBMGA001827* | up | 142 | 150 | - | 11.3303 | 4.96E-05 | 0.615 | GTGGGTGGC |
| 1440 | mamo-s | *BGIBMGA001829* | down | 213 | 221 | - | 11.3303 | 4.96E-05 | 0.615 | GTGGGTGGC |
| 1441 | mamo-s | *BGIBMGA001836* | up | 493 | 501 | - | 11.3303 | 4.96E-05 | 0.615 | GTGGGTGGC |
| 1442 | mamo-s | *BGIBMGA001836* | down | 746 | 754 | - | 11.3303 | 4.96E-05 | 0.615 | GTGGGTGGC |
| 1443 | mamo-s | *BGIBMGA001837* | up | 878 | 886 | - | 11.3303 | 4.96E-05 | 0.615 | GTGGGTGGC |
| 1444 | mamo-s | *BGIBMGA001839* | up | 1049 | 1057 | - | 12.1009 | 2.43E-05 | 0.615 | GTGCGTGGT |
| 1445 | mamo-s | *BGIBMGA001841* | up | 372 | 380 | + | 13.6606 | 4.06E-06 | 0.615 | GCGCGTGGC |
| 1446 | mamo-s | *BGIBMGA001841* | down | 1729 | 1737 | - | 11.3303 | 4.96E-05 | 0.615 | GTGGGTGGC |
| 1447 | mamo-s | *BGIBMGA001842* | down | 1827 | 1835 | - | 11.3303 | 4.96E-05 | 0.615 | GTGGGTGGC |
| 1448 | mamo-s | *BGIBMGA001843* | up | 1756 | 1764 | + | 11.0826 | 6.84E-05 | 0.709 | GTACGTGGC |
| 1449 | mamo-s | *BGIBMGA001845* | up | 1651 | 1659 | - | 11.3303 | 4.96E-05 | 0.615 | GTGGGTGGC |
| 1450 | mamo-s | *BGIBMGA001846* | up | 1090 | 1098 | + | 11.3303 | 4.96E-05 | 0.615 | GTGGGTGGC |
| 1451 | mamo-s | *BGIBMGA001849* | down | 300 | 308 | - | 11.7064 | 3.52E-05 | 0.615 | GCGCGTGGT |
| 1452 | mamo-s | *BGIBMGA001849* | up | 468 | 476 | + | 11.2018 | 5.68E-05 | 0.674 | CTGCGTGGT |
| 1453 | mamo-s | *BGIBMGA001850* | up | 1610 | 1618 | + | 11.3303 | 4.96E-05 | 0.615 | GTGGGTGGC |
| 1454 | mamo-s | *BGIBMGA001851* | up | 698 | 706 | - | 11.3303 | 4.96E-05 | 0.615 | GTGGGTGGC |
| 1455 | mamo-s | *BGIBMGA001851* | down | 1518 | 1526 | - | 11.3303 | 4.96E-05 | 0.615 | GTGGGTGGC |
| 1456 | mamo-s | *BGIBMGA001858* | up | 1081 | 1089 | + | 12.7615 | 1.04E-05 | 0.615 | CCGCGTGGC |
| 1457 | mamo-s | *BGIBMGA001859* | down | 848 | 856 | + | 11.3303 | 4.96E-05 | 0.615 | GTGGGTGGC |
| 1458 | mamo-s | *BGIBMGA001863* | up | 1126 | 1134 | - | 11.3303 | 4.96E-05 | 0.615 | GTGGGTGGC |
| 1459 | mamo-s | *BGIBMGA001863* | down | 1708 | 1716 | + | 11.3303 | 4.96E-05 | 0.615 | GTGGGTGGC |
| 1460 | mamo-s | *BGIBMGA001864* | down | 1936 | 1944 | - | 11.3303 | 4.96E-05 | 0.615 | GTGGGTGGC |
| 1461 | mamo-s | *BGIBMGA001866* | up | 609 | 617 | + | 12.1009 | 2.43E-05 | 0.615 | GTGCGTGGT |
| 1462 | mamo-s | *BGIBMGA001870* | down | 878 | 886 | + | 11.3303 | 4.96E-05 | 0.615 | GTGGGTGGC |
| 1463 | mamo-s | *BGIBMGA001873* | up | 302 | 310 | + | 11.3303 | 4.96E-05 | 0.615 | GTGGGTGGC |
| 1464 | mamo-s | *BGIBMGA001874* | up | 1682 | 1690 | - | 11.3303 | 4.96E-05 | 0.615 | GTGGGTGGC |
| 1465 | mamo-s | *BGIBMGA001875* | down | 291 | 299 | - | 11.3303 | 4.96E-05 | 0.615 | GTGGGTGGC |
| 1466 | mamo-s | *BGIBMGA001876* | down | 334 | 342 | - | 11.7798 | 3.11E-05 | 0.615 | ATGCGTGGC |
| 1467 | mamo-s | *BGIBMGA001876* | down | 1588 | 1596 | + | 11.3303 | 4.96E-05 | 0.615 | GTGGGTGGC |
| 1468 | mamo-s | *BGIBMGA001878* | up | 971 | 979 | - | 11.0826 | 6.84E-05 | 0.709 | GTACGTGGC |
| 1469 | mamo-s | *BGIBMGA001880* | up | 796 | 804 | - | 11.3303 | 4.96E-05 | 0.615 | GTGGGTGGC |
| 1470 | mamo-s | *BGIBMGA001882* | up | 1590 | 1598 | - | 11.3303 | 4.96E-05 | 0.615 | GTGGGTGGC |
| 1471 | mamo-s | *BGIBMGA001883* | down | 260 | 268 | + | 13.0275 | 8.53E-06 | 0.615 | GAGCGTGGC |
| 1472 | mamo-s | *BGIBMGA001884* | up | 766 | 774 | + | 13.0275 | 8.53E-06 | 0.615 | GAGCGTGGC |
| 1473 | mamo-s | *BGIBMGA001886* | down | 1618 | 1626 | + | 11.2294 | 5.41E-05 | 0.65 | CTGCGAGGC |
| 1474 | mamo-s | *BGIBMGA001888* | up | 765 | 773 | + | 11.7798 | 3.11E-05 | 0.615 | ATGCGTGGC |
| 1475 | mamo-s | *BGIBMGA001888* | up | 461 | 469 | + | 11.3303 | 4.96E-05 | 0.615 | GTGGGTGGC |
| 1476 | mamo-s | *BGIBMGA001890* | up | 1572 | 1580 | - | 10.5138 | 9.75E-05 | 0.865 | GTGCGTGTT |
| 1477 | mamo-s | *BGIBMGA001891* | down | 940 | 948 | - | 11.3303 | 4.96E-05 | 0.615 | GTGGGTGGC |
| 1478 | mamo-s | *BGIBMGA001893* | up | 124 | 132 | + | 11.3303 | 4.96E-05 | 0.615 | GTGGGTGGC |
| 1479 | mamo-s | *BGIBMGA001896* | up | 1670 | 1678 | - | 11.3303 | 4.96E-05 | 0.615 | GTGGGTGGC |
| 1480 | mamo-s | *BGIBMGA001899* | up | 802 | 810 | + | 14.055 | 2.23E-06 | 0.615 | GTGCGTGGC |
| 1481 | mamo-s | *BGIBMGA001901* | up | 1496 | 1504 | + | 11.3303 | 4.96E-05 | 0.615 | GTGGGTGGC |
| 1482 | mamo-s | *BGIBMGA001901* | up | 863 | 871 | + | 10.9358 | 7.30E-05 | 0.728 | GCGGGTGGC |
| 1483 | mamo-s | *BGIBMGA001902* | down | 1140 | 1148 | + | 11.3303 | 4.96E-05 | 0.615 | GTGGGTGGC |
| 1484 | mamo-s | *BGIBMGA001905* | down | 1621 | 1629 | - | 11.3303 | 4.96E-05 | 0.615 | GTGGGTGGC |
| 1485 | mamo-s | *BGIBMGA001908* | up | 144 | 152 | - | 13.6606 | 4.06E-06 | 0.615 | GCGCGTGGC |
| 1486 | mamo-s | *BGIBMGA001911* | up | 817 | 825 | - | 10.5413 | 9.42E-05 | 0.857 | GTGCGAGTC |
| 1487 | mamo-s | *BGIBMGA001912* | down | 1692 | 1700 | - | 11.4404 | 4.51E-05 | 0.615 | CGGCGTGGC |
| 1488 | mamo-s | *BGIBMGA001912* | down | 1801 | 1809 | + | 11.0826 | 6.84E-05 | 0.709 | GTACGTGGC |
| 1489 | mamo-s | *BGIBMGA001913* | up | 743 | 751 | + | 11.3303 | 4.96E-05 | 0.615 | GTGGGTGGC |

| | | | | | | | | | | |
|---|---|---|---|---|---|---|---|---|---|---|
| 1490 | mamo-s | *BGIBMGA001913* | up | 1008 | 1016 | + | 11.3303 | 4.96E-05 | 0.615 | GTGGGTGGC |
| 1491 | mamo-s | *BGIBMGA001915* | up | 353 | 361 | - | 11.4404 | 4.51E-05 | 0.615 | CGGCGTGGC |
| 1492 | mamo-s | *BGIBMGA001915* | up | 521 | 529 | - | 11.4404 | 4.51E-05 | 0.615 | CGGCGTGGC |
| 1493 | mamo-s | *BGIBMGA001916* | down | 1088 | 1096 | + | 11.3303 | 4.96E-05 | 0.615 | GTGGGTGGC |
| 1494 | mamo-s | *BGIBMGA001917* | down | 1737 | 1745 | + | 11.3303 | 4.96E-05 | 0.615 | GTGGGTGGC |
| 1495 | mamo-s | *BGIBMGA001923* | up | 1825 | 1833 | + | 11.3303 | 4.96E-05 | 0.615 | GTGGGTGGC |
| 1496 | mamo-s | *BGIBMGA001926* | up | 1312 | 1320 | + | 12.7615 | 1.04E-05 | 0.615 | CCGCGTGGC |
| 1497 | mamo-s | *BGIBMGA001930* | down | 274 | 282 | - | 11.4404 | 4.51E-05 | 0.615 | CGGCGTGGC |
| 1498 | mamo-s | *BGIBMGA001930* | down | 795 | 803 | + | 11.3303 | 4.96E-05 | 0.615 | GTGGGTGGC |
| 1499 | mamo-s | *BGIBMGA001930* | down | 1146 | 1154 | - | 10.6881 | 8.61E-05 | 0.81 | GCACGTGGC |
| 1500 | mamo-s | *BGIBMGA001933* | up | 1129 | 1137 | - | 10.5138 | 9.75E-05 | 0.865 | GTGCGTGTT |
| 1501 | mamo-s | *BGIBMGA001934* | up | 1219 | 1227 | - | 13.6606 | 4.06E-06 | 0.615 | GCGCGTGGC |
| 1502 | mamo-s | *BGIBMGA001935* | up | 1793 | 1801 | - | 11.7798 | 3.11E-05 | 0.615 | ATGCGTGGC |
| 1503 | mamo-s | *BGIBMGA001936* | down | 1874 | 1882 | + | 11.1743 | 5.90E-05 | 0.684 | CCGCGTGTC |
| 1504 | mamo-s | *BGIBMGA001938* | down | 1991 | 1999 | - | 11.2294 | 5.41E-05 | 0.65 | CTGCGAGGC |
| 1505 | mamo-s | *BGIBMGA001938* | down | 424 | 432 | - | 10.5138 | 9.75E-05 | 0.865 | GTGCGTGTT |
| 1506 | mamo-s | *BGIBMGA001941* | down | 1936 | 1944 | + | 12.0734 | 2.66E-05 | 0.615 | GCGCGTGTC |
| 1507 | mamo-s | *BGIBMGA001941* | down | 849 | 857 | - | 11.3303 | 4.96E-05 | 0.615 | GTGGGTGGC |
| 1508 | mamo-s | *BGIBMGA001942* | up | 1701 | 1709 | - | 11.1009 | 6.57E-05 | 0.709 | GAGCGAGGC |
| 1509 | mamo-s | *BGIBMGA001943* | down | 1608 | 1616 | + | 12.1284 | 2.16E-05 | 0.615 | CAGCGTGGC |
| 1510 | mamo-s | *BGIBMGA001943* | up | 1565 | 1573 | + | 11.3303 | 4.96E-05 | 0.615 | GTGGGTGGC |
| 1511 | mamo-s | *BGIBMGA001944* | down | 537 | 545 | + | 11.3303 | 4.96E-05 | 0.615 | GTGGGTGGC |
| 1512 | mamo-s | *BGIBMGA001944* | down | 698 | 706 | + | 11.3303 | 4.96E-05 | 0.615 | GTGGGTGGC |
| 1513 | mamo-s | *BGIBMGA001944* | down | 1054 | 1062 | - | 11.0826 | 6.84E-05 | 0.709 | GTACGTGGC |
| 1514 | mamo-s | *BGIBMGA001945* | up | 263 | 271 | - | 11.3303 | 4.96E-05 | 0.615 | GTGGGTGGC |
| 1515 | mamo-s | *BGIBMGA001946* | up | 1300 | 1308 | + | 11.3853 | 4.74E-05 | 0.615 | ACGCGTGGC |
| 1516 | mamo-s | *BGIBMGA001948* | down | 1972 | 1980 | - | 11.156 | 6.13E-05 | 0.705 | GAGCGTGCC |
| 1517 | mamo-s | *BGIBMGA001950* | down | 803 | 811 | + | 12.7615 | 1.04E-05 | 0.615 | CCGCGTGGC |
| 1518 | mamo-s | *BGIBMGA001952* | down | 243 | 251 | + | 11.3303 | 4.96E-05 | 0.615 | GTGGGTGGC |
| 1519 | mamo-s | *BGIBMGA001953* | down | 1454 | 1462 | + | 11.7339 | 3.29E-05 | 0.615 | GCGCGAGGC |
| 1520 | mamo-s | *BGIBMGA001953* | up | 56 | 64 | + | 11.3303 | 4.96E-05 | 0.615 | GTGGGTGGC |
| 1521 | mamo-s | *BGIBMGA001953* | down | 326 | 334 | + | 11.156 | 6.13E-05 | 0.705 | GAGCGTGCC |
| 1522 | mamo-s | *BGIBMGA001957* | down | 53 | 61 | + | 11.3303 | 4.96E-05 | 0.615 | GTGGGTGGC |
| 1523 | mamo-s | *BGIBMGA001958* | down | 1323 | 1331 | + | 11.3303 | 4.96E-05 | 0.615 | GTGGGTGGC |
| 1524 | mamo-s | *BGIBMGA001959* | up | 901 | 909 | + | 11.3303 | 4.96E-05 | 0.615 | GTGGGTGGC |
| 1525 | mamo-s | *BGIBMGA001960* | down | 473 | 481 | - | 11.3303 | 4.96E-05 | 0.615 | GTGGGTGGC |
| 1526 | mamo-s | *BGIBMGA001960* | down | 1576 | 1584 | - | 11.3303 | 4.96E-05 | 0.615 | GTGGGTGGC |
| 1527 | mamo-s | *BGIBMGA001960* | down | 1950 | 1958 | + | 11.3303 | 4.96E-05 | 0.615 | GTGGGTGGC |
| 1528 | mamo-s | *BGIBMGA001961* | up | 1939 | 1947 | + | 12.4679 | 1.31E-05 | 0.615 | GTGCGTGTC |
| 1529 | mamo-s | *BGIBMGA001961* | down | 274 | 282 | + | 11.3303 | 4.96E-05 | 0.615 | GTGGGTGGC |
| 1530 | mamo-s | *BGIBMGA001961* | down | 1709 | 1717 | + | 11.3303 | 4.96E-05 | 0.615 | GTGGGTGGC |
| 1531 | mamo-s | *BGIBMGA001961* | up | 283 | 291 | + | 11.0826 | 6.84E-05 | 0.709 | GTACGTGGC |
| 1532 | mamo-s | *BGIBMGA001964* | up | 212 | 220 | + | 10.5138 | 9.75E-05 | 0.865 | GTGCGTGTT |
| 1533 | mamo-s | *BGIBMGA001968* | up | 1297 | 1305 | + | 11.3303 | 4.96E-05 | 0.615 | GTGGGTGGC |
| 1534 | mamo-s | *BGIBMGA001970* | down | 698 | 706 | - | 13.6606 | 4.06E-06 | 0.615 | GCGCGTGGC |
| 1535 | mamo-s | *BGIBMGA001970* | up | 304 | 312 | - | 11.3303 | 4.96E-05 | 0.615 | GTGGGTGGC |
| 1536 | mamo-s | *BGIBMGA001972* | up | 101 | 109 | + | 11.3303 | 4.96E-05 | 0.615 | GTGGGTGGC |
| 1537 | mamo-s | *BGIBMGA001972* | down | 1788 | 1796 | - | 10.8073 | 7.89E-05 | 0.766 | CCGCGTGGT |
| 1538 | mamo-s | *BGIBMGA001973* | up | 419 | 427 | + | 11.3303 | 4.96E-05 | 0.615 | GTGGGTGGC |
| 1539 | mamo-s | *BGIBMGA001974* | down | 1297 | 1305 | - | 14.055 | 2.23E-06 | 0.615 | GTGCGTGGC |
| 1540 | mamo-s | *BGIBMGA001974* | up | 572 | 580 | + | 10.9358 | 7.30E-05 | 0.728 | GCGGGTGGC |
| 1541 | mamo-s | *BGIBMGA001975* | up | 1505 | 1513 | - | 11.2018 | 5.68E-05 | 0.674 | CTGCGTGGT |
| 1542 | mamo-s | *BGIBMGA001977* | down | 1545 | 1553 | + | 11.3303 | 4.96E-05 | 0.615 | GTGGGTGGC |
| 1543 | mamo-s | *BGIBMGA001978* | up | 1498 | 1506 | - | 11.3303 | 4.96E-05 | 0.615 | GTGGGTGGC |
| 1544 | mamo-s | *BGIBMGA001980* | up | 1139 | 1147 | + | 11.2018 | 5.68E-05 | 0.674 | CTGCGTGGT |
| 1545 | mamo-s | *BGIBMGA001982* | down | 1131 | 1139 | - | 11.7798 | 3.11E-05 | 0.615 | ATGCGTGGC |
| 1546 | mamo-s | *BGIBMGA001983* | up | 998 | 1006 | + | 10.5138 | 9.75E-05 | 0.865 | GTGCGTGTT |
| 1547 | mamo-s | *BGIBMGA001984* | down | 1455 | 1463 | - | 12.1284 | 2.16E-05 | 0.615 | GTGCGAGGC |
| 1548 | mamo-s | *BGIBMGA001984* | down | 739 | 747 | - | 11.3303 | 4.96E-05 | 0.615 | GTGGGTGGC |
| 1549 | mamo-s | *BGIBMGA001985* | down | 198 | 206 | + | 12.4679 | 1.31E-05 | 0.615 | GTGCGTGTC |
| 1550 | mamo-s | *BGIBMGA001986* | up | 660 | 668 | + | 11.3303 | 4.96E-05 | 0.615 | GTGGGTGGC |
| 1551 | mamo-s | *BGIBMGA001987* | up | 1327 | 1335 | + | 12.0734 | 2.66E-05 | 0.615 | GCGCGTGTC |
| 1552 | mamo-s | *BGIBMGA001988* | down | 207 | 215 | + | 11.3303 | 4.96E-05 | 0.615 | GTGGGTGGC |
| 1553 | mamo-s | *BGIBMGA001988* | up | 805 | 813 | + | 11.3303 | 4.96E-05 | 0.615 | GTGGGTGGC |
| 1554 | mamo-s | *BGIBMGA001988* | up | 1828 | 1836 | + | 11.2844 | 5.18E-05 | 0.63 | CTGCGTGCC |
| 1555 | mamo-s | *BGIBMGA001988* | up | 359 | 367 | - | 11.0826 | 6.84E-05 | 0.709 | GTACGTGGC |
| 1556 | mamo-s | *BGIBMGA001990* | up | 435 | 443 | - | 11.3303 | 4.96E-05 | 0.615 | GTGGGTGGC |
| 1557 | mamo-s | *BGIBMGA001990* | down | 841 | 849 | - | 11.3303 | 4.96E-05 | 0.615 | GTGGGTGGC |
| 1558 | mamo-s | *BGIBMGA001990* | up | 954 | 962 | - | 10.8073 | 7.89E-05 | 0.766 | CCGCGTGGT |
| 1559 | mamo-s | *BGIBMGA001991* | up | 492 | 500 | + | 11.3303 | 4.96E-05 | 0.615 | GTGGGTGGC |
| 1560 | mamo-s | *BGIBMGA001992* | up | 1397 | 1405 | - | 11.3303 | 4.96E-05 | 0.615 | GTGGGTGGC |
| 1561 | mamo-s | *BGIBMGA001994* | down | 452 | 460 | + | 12.1009 | 2.43E-05 | 0.615 | GTGCGTGGT |
| 1562 | mamo-s | *BGIBMGA001994* | down | 445 | 453 | + | 11.2018 | 5.68E-05 | 0.674 | CTGCGTGGT |
| 1563 | mamo-s | *BGIBMGA001994* | up | 481 | 489 | + | 10.5138 | 9.75E-05 | 0.865 | GTGCGTGTT |
| 1564 | mamo-s | *BGIBMGA001996* | up | 230 | 238 | - | 11.3303 | 4.96E-05 | 0.615 | GTGGGTGGC |
| 1565 | mamo-s | *BGIBMGA001996* | up | 1802 | 1810 | + | 11.3303 | 4.96E-05 | 0.615 | GTGGGTGGC |
| 1566 | mamo-s | *BGIBMGA001996* | down | 1128 | 1136 | + | 10.9358 | 7.30E-05 | 0.728 | GCGGGTGGC |
| 1567 | mamo-s | *BGIBMGA002002* | down | 41 | 49 | - | 11.1743 | 5.90E-05 | 0.684 | CCGCGTGTC |
| 1568 | mamo-s | *BGIBMGA002003* | up | 985 | 993 | + | 11.1743 | 5.90E-05 | 0.684 | CCGCGTGTC |
| 1569 | mamo-s | *BGIBMGA002004* | up | 681 | 689 | - | 12.4679 | 1.31E-05 | 0.615 | GTGCGTGTC |
| 1570 | mamo-s | *BGIBMGA002004* | up | 646 | 654 | + | 11.4404 | 4.51E-05 | 0.615 | CGGCGTGGC |
| 1571 | mamo-s | *BGIBMGA002004* | down | 1952 | 1960 | - | 10.7523 | 8.38E-05 | 0.797 | GGGCGTGTC |
| 1572 | mamo-s | *BGIBMGA002005* | up | 535 | 543 | - | 10.7523 | 8.38E-05 | 0.797 | GGGCGTGTC |
| 1573 | mamo-s | *BGIBMGA002007* | up | 738 | 746 | + | 11.3303 | 4.96E-05 | 0.615 | GTGGGTGGC |
| 1574 | mamo-s | *BGIBMGA002009* | up | 1760 | 1768 | - | 13.156 | 6.30E-06 | 0.615 | CTGCGTGGC |
| 1575 | mamo-s | *BGIBMGA002011* | up | 570 | 578 | - | 11.0826 | 6.84E-05 | 0.709 | GTACGTGGC |
| 1576 | mamo-s | *BGIBMGA002012* | down | 1503 | 1511 | - | 11.1376 | 6.35E-05 | 0.709 | TCGCGTGGC |
| 1577 | mamo-s | *BGIBMGA002016* | up | 1195 | 1203 | + | 12.1835 | 1.71E-05 | 0.615 | GTGCGTGCC |

| | | | | | | | | | | |
|---|---|---|---|---|---|---|---|---|---|---|
| 1578 | mamo-s | *BGIBMGA002016* | up | 812 | 820 | + | 11.3303 | 4.96E-05 | 0.615 | GTGGGTGGC |
| 1579 | mamo-s | *BGIBMGA002017* | down | 706 | 714 | - | 13.0275 | 8.53E-06 | 0.615 | GAGCGTGGC |
| 1580 | mamo-s | *BGIBMGA002017* | down | 673 | 681 | + | 12.1835 | 1.71E-05 | 0.615 | GTGCGTGCC |
| 1581 | mamo-s | *BGIBMGA002022* | up | 1837 | 1845 | - | 13.156 | 6.30E-06 | 0.615 | CTGCGTGGC |
| 1582 | mamo-s | *BGIBMGA002022* | down | 13 | 21 | - | 10.7523 | 8.38E-05 | 0.797 | AAGCGTGGC |
| 1583 | mamo-s | *BGIBMGA002022* | up | 728 | 736 | - | 10.7523 | 8.38E-05 | 0.797 | AAGCGTGGC |
| 1584 | mamo-s | *BGIBMGA002025* | up | 1723 | 1731 | + | 11.3303 | 4.96E-05 | 0.615 | GTGGGTGGC |
| 1585 | mamo-s | *BGIBMGA002026* | up | 281 | 289 | + | 11.5321 | 4.06E-05 | 0.615 | TTGCGTGGC |
| 1586 | mamo-s | *BGIBMGA002029* | up | 719 | 727 | - | 11.3303 | 4.96E-05 | 0.615 | GTGGGTGGC |
| 1587 | mamo-s | *BGIBMGA002030* | up | 316 | 324 | + | 11.7798 | 3.11E-05 | 0.615 | ATGCGTGGC |
| 1588 | mamo-s | *BGIBMGA002030* | down | 803 | 811 | - | 11.3303 | 4.96E-05 | 0.615 | GTGGGTGGC |
| 1589 | mamo-s | *BGIBMGA002030* | down | 1605 | 1613 | - | 11.3303 | 4.96E-05 | 0.615 | GTGGGTGGC |
| 1590 | mamo-s | *BGIBMGA002031* | up | 1063 | 1071 | + | 14.055 | 2.23E-06 | 0.615 | GTGCGTGGC |
| 1591 | mamo-s | *BGIBMGA002031* | down | 624 | 632 | + | 10.8073 | 7.89E-05 | 0.766 | CCGCGTGGT |
| 1592 | mamo-s | *BGIBMGA002034* | up | 512 | 520 | - | 12.1009 | 2.43E-05 | 0.615 | GTGCGTGGT |
| 1593 | mamo-s | *BGIBMGA002037* | up | 1629 | 1637 | - | 11.3303 | 4.96E-05 | 0.615 | GTGGGTGGC |
| 1594 | mamo-s | *BGIBMGA002038* | down | 509 | 517 | - | 12.1835 | 1.71E-05 | 0.615 | GTGCGTGCC |
| 1595 | mamo-s | *BGIBMGA002038* | up | 114 | 122 | + | 11.3303 | 4.96E-05 | 0.615 | GTGGGTGGC |
| 1596 | mamo-s | *BGIBMGA002038* | down | 1312 | 1320 | + | 11.3303 | 4.96E-05 | 0.615 | GTGGGTGGC |
| 1597 | mamo-s | *BGIBMGA002038* | down | 1633 | 1641 | + | 11.3303 | 4.96E-05 | 0.615 | GTGGGTGGC |
| 1598 | mamo-s | *BGIBMGA002039* | up | 477 | 485 | + | 11.7064 | 3.52E-05 | 0.615 | GCGCGTGGT |
| 1599 | mamo-s | *BGIBMGA002039* | down | 1368 | 1376 | - | 10.5138 | 9.75E-05 | 0.865 | GTGCGTGTT |
| 1600 | mamo-s | *BGIBMGA002044* | down | 1803 | 1811 | + | 11.2294 | 5.41E-05 | 0.65 | CTGCGAGGC |
| 1601 | mamo-s | *BGIBMGA002044* | up | 106 | 114 | - | 11.0826 | 6.84E-05 | 0.709 | GTACGTGGC |
| 1602 | mamo-s | *BGIBMGA002047* | down | 8 | 16 | + | 11.7339 | 3.29E-05 | 0.615 | GCGCGAGGC |
| 1603 | mamo-s | *BGIBMGA002047* | up | 378 | 386 | + | 11.1743 | 5.90E-05 | 0.684 | CCGCGTGTC |
| 1604 | mamo-s | *BGIBMGA002049* | down | 1277 | 1285 | + | 11.3303 | 4.96E-05 | 0.615 | GTGGGTGGC |
| 1605 | mamo-s | *BGIBMGA002049* | down | 1340 | 1348 | - | 10.9358 | 7.30E-05 | 0.728 | GCGGGTGGC |
| 1606 | mamo-s | *BGIBMGA002050* | up | 879 | 887 | - | 10.7523 | 8.38E-05 | 0.797 | AAGCGTGGC |
| 1607 | mamo-s | *BGIBMGA002051* | down | 185 | 193 | + | 12.1284 | 2.16E-05 | 0.615 | GTGCGAGGC |
| 1608 | mamo-s | *BGIBMGA002053* | up | 158 | 166 | - | 10.9358 | 7.30E-05 | 0.728 | GCGGGTGGC |
| 1609 | mamo-s | *BGIBMGA002055* | down | 1441 | 1449 | - | 11.3303 | 4.96E-05 | 0.615 | GTGGGTGGC |
| 1610 | mamo-s | *BGIBMGA002056* | down | 1069 | 1077 | + | 10.8349 | 7.66E-05 | 0.751 | CCGCGAGGC |
| 1611 | mamo-s | *BGIBMGA002056* | up | 1067 | 1075 | - | 10.6881 | 8.61E-05 | 0.81 | GCACGTGGC |
| 1612 | mamo-s | *BGIBMGA002057* | down | 1369 | 1377 | - | 13.0275 | 8.53E-06 | 0.615 | GAGCGTGGC |
| 1613 | mamo-s | *BGIBMGA002060* | down | 205 | 213 | + | 12.1284 | 2.16E-05 | 0.615 | CAGCGTGGC |
| 1614 | mamo-s | *BGIBMGA002061* | up | 1709 | 1717 | + | 12.1284 | 2.16E-05 | 0.615 | CAGCGTGGC |
| 1615 | mamo-s | *BGIBMGA002063* | up | 1015 | 1023 | + | 12.3394 | 1.49E-05 | 0.615 | GGGCGTGGC |
| 1616 | mamo-s | *BGIBMGA002064* | up | 1853 | 1861 | - | 11.4404 | 4.51E-05 | 0.615 | GAGCGTGTC |
| 1617 | mamo-s | *BGIBMGA002065* | up | 4 | 12 | + | 10.5138 | 9.75E-05 | 0.865 | GTGCGTGTT |
| 1618 | mamo-s | *BGIBMGA002068* | up | 822 | 830 | - | 11.3303 | 4.96E-05 | 0.615 | GTGGGTGGC |
| 1619 | mamo-s | *BGIBMGA002069* | down | 542 | 550 | - | 12.1835 | 1.71E-05 | 0.615 | GTGCGTGCC |
| 1620 | mamo-s | *BGIBMGA002070* | up | 486 | 494 | - | 12.1835 | 1.71E-05 | 0.615 | GTGCGTGCC |
| 1621 | mamo-s | *BGIBMGA002071* | down | 1038 | 1046 | + | 11.0826 | 6.84E-05 | 0.709 | GTACGTGGC |
| 1622 | mamo-s | *BGIBMGA002071* | down | 1802 | 1810 | - | 10.5138 | 9.75E-05 | 0.865 | GTGCGTGTT |
| 1623 | mamo-s | *BGIBMGA002072* | up | 1787 | 1795 | - | 11.3303 | 4.96E-05 | 0.615 | GTGGGTGGC |
| 1624 | mamo-s | *BGIBMGA002075* | down | 776 | 784 | - | 12.1835 | 1.71E-05 | 0.615 | GTGCGTGCC |
| 1625 | mamo-s | *BGIBMGA002075* | up | 930 | 938 | + | 11.7064 | 3.52E-05 | 0.615 | GCGCGTGGT |
| 1626 | mamo-s | *BGIBMGA002075* | down | 475 | 483 | + | 11.0826 | 6.84E-05 | 0.709 | GTACGTGGC |
| 1627 | mamo-s | *BGIBMGA002075* | up | 452 | 460 | + | 10.5688 | 8.88E-05 | 0.824 | GTGCGTGAC |
| 1628 | mamo-s | *BGIBMGA002077* | down | 1580 | 1588 | - | 12.7615 | 1.04E-05 | 0.615 | CCGCGTGGC |
| 1629 | mamo-s | *BGIBMGA002077* | down | 225 | 233 | + | 11.3303 | 4.96E-05 | 0.615 | GTGGGTGGC |
| 1630 | mamo-s | *BGIBMGA002077* | up | 1479 | 1487 | + | 11.3303 | 4.96E-05 | 0.615 | GTGGGTGGC |
| 1631 | mamo-s | *BGIBMGA002079* | up | 979 | 987 | + | 13.6606 | 4.06E-06 | 0.615 | GCGCGTGGC |
| 1632 | mamo-s | *BGIBMGA002080* | down | 1354 | 1362 | + | 11.3303 | 4.96E-05 | 0.615 | GTGGGTGGC |
| 1633 | mamo-s | *BGIBMGA002082* | down | 1531 | 1539 | + | 12.0734 | 2.66E-05 | 0.615 | GCGCGTGTC |
| 1634 | mamo-s | *BGIBMGA002082* | down | 840 | 848 | - | 11.3303 | 4.96E-05 | 0.615 | GTGGGTGGC |
| 1635 | mamo-s | *BGIBMGA002083* | up | 148 | 156 | + | 11.3303 | 4.96E-05 | 0.615 | GTGGGTGGC |
| 1636 | mamo-s | *BGIBMGA002083* | up | 1644 | 1652 | - | 11.3303 | 4.96E-05 | 0.615 | GTGGGTGGC |
| 1637 | mamo-s | *BGIBMGA002084* | up | 622 | 630 | + | 12.1284 | 2.16E-05 | 0.615 | GTGCGAGGC |
| 1638 | mamo-s | *BGIBMGA002085* | down | 1441 | 1449 | - | 11.7064 | 3.52E-05 | 0.615 | GCGCGTGGT |
| 1639 | mamo-s | *BGIBMGA002085* | down | 216 | 224 | - | 11.3303 | 4.96E-05 | 0.615 | GTGGGTGGC |
| 1640 | mamo-s | *BGIBMGA002085* | up | 642 | 650 | + | 11.3303 | 4.96E-05 | 0.615 | GTGGGTGGC |
| 1641 | mamo-s | *BGIBMGA002085* | up | 1038 | 1046 | + | 11.3303 | 4.96E-05 | 0.615 | GTGGGTGGC |
| 1642 | mamo-s | *BGIBMGA002086* | down | 430 | 438 | - | 11.3303 | 4.96E-05 | 0.615 | GTGGGTGGC |
| 1643 | mamo-s | *BGIBMGA002086* | up | 663 | 671 | - | 10.8073 | 7.89E-05 | 0.766 | CCGCGTGGT |
| 1644 | mamo-s | *BGIBMGA002090* | up | 1925 | 1933 | - | 12.1284 | 2.16E-05 | 0.615 | CAGCGTGGC |
| 1645 | mamo-s | *BGIBMGA002090* | up | 1787 | 1795 | + | 11.3303 | 4.96E-05 | 0.615 | GTGGGTGGC |
| 1646 | mamo-s | *BGIBMGA002092* | down | 1992 | 2000 | - | 11.0734 | 7.12E-05 | 0.726 | GAGCGTGGT |
| 1647 | mamo-s | *BGIBMGA002092* | up | 304 | 312 | - | 10.8899 | 7.48E-05 | 0.739 | CCGCGTGCC |
| 1648 | mamo-s | *BGIBMGA002093* | up | 212 | 220 | - | 10.5138 | 9.75E-05 | 0.865 | GTGCGTGTT |
| 1649 | mamo-s | *BGIBMGA002096* | up | 767 | 775 | - | 11.3303 | 4.96E-05 | 0.615 | GTGGGTGGC |
| 1650 | mamo-s | *BGIBMGA002097* | down | 1603 | 1611 | - | 11.3303 | 4.96E-05 | 0.615 | GTGGGTGGC |
| 1651 | mamo-s | *BGIBMGA002100* | up | 1914 | 1922 | + | 11.4404 | 4.51E-05 | 0.615 | CGGCGTGGC |
| 1652 | mamo-s | *BGIBMGA002101* | up | 462 | 470 | - | 11.3303 | 4.96E-05 | 0.615 | GTGGGTGGC |
| 1653 | mamo-s | *BGIBMGA002101* | down | 475 | 483 | + | 11.3303 | 4.96E-05 | 0.615 | GTGGGTGGC |
| 1654 | mamo-s | *BGIBMGA002102* | down | 196 | 204 | - | 11.3303 | 4.96E-05 | 0.615 | GTGGGTGGC |
| 1655 | mamo-s | *BGIBMGA002102* | up | 783 | 791 | + | 11.3303 | 4.96E-05 | 0.615 | GTGGGTGGC |
| 1656 | mamo-s | *BGIBMGA002103* | up | 127 | 135 | + | 11.3303 | 4.96E-05 | 0.615 | GTGGGTGGC |
| 1657 | mamo-s | *BGIBMGA002106* | up | 1117 | 1125 | - | 11.3303 | 4.96E-05 | 0.615 | GTGGGTGGC |
| 1658 | mamo-s | *BGIBMGA002110* | down | 295 | 303 | - | 11.3303 | 4.96E-05 | 0.615 | GTGGGTGGC |
| 1659 | mamo-s | *BGIBMGA002110* | up | 1856 | 1864 | - | 11.3303 | 4.96E-05 | 0.615 | GTGGGTGGC |
| 1660 | mamo-s | *BGIBMGA002111* | down | 1697 | 1705 | - | 11.3303 | 4.96E-05 | 0.615 | GTGGGTGGC |
| 1661 | mamo-s | *BGIBMGA002111* | down | 1966 | 1974 | - | 11.3303 | 4.96E-05 | 0.615 | GTGGGTGGC |
| 1662 | mamo-s | *BGIBMGA002115* | down | 774 | 782 | - | 10.5138 | 9.75E-05 | 0.865 | GTGCGTGTT |
| 1663 | mamo-s | *BGIBMGA002119* | up | 1897 | 1905 | - | 11.3303 | 4.96E-05 | 0.615 | GTGGGTGGC |
| 1664 | mamo-s | *BGIBMGA002119* | down | 346 | 354 | + | 10.5413 | 9.42E-05 | 0.857 | CAGCGTGTC |
| 1665 | mamo-s | *BGIBMGA002124* | down | 353 | 361 | + | 11.3303 | 4.96E-05 | 0.615 | GTGGGTGGC |

| | | | | | | | | | | |
|---|---|---|---|---|---|---|---|---|---|---|
| 1666 | mamo-s | *BGIBMGA002124* | down | 1214 | 1222 | + | 11.3303 | 4.96E-05 | 0.615 | GTGGGTGGC |
| 1667 | mamo-s | *BGIBMGA002124* | up | 1231 | 1239 | + | 11.3303 | 4.96E-05 | 0.615 | GTGGGTGGC |
| 1668 | mamo-s | *BGIBMGA002126* | up | 1004 | 1012 | - | 11.3303 | 4.96E-05 | 0.615 | GTGGGTGGC |
| 1669 | mamo-s | *BGIBMGA002126* | down | 1898 | 1906 | - | 11.3303 | 4.96E-05 | 0.615 | GTGGGTGGC |
| 1670 | mamo-s | *BGIBMGA002127* | up | 966 | 974 | + | 11.0826 | 6.84E-05 | 0.709 | GTACGTGGC |
| 1671 | mamo-s | *BGIBMGA002132* | up | 1370 | 1378 | + | 11.3303 | 4.96E-05 | 0.615 | GTGGGTGGC |
| 1672 | mamo-s | *BGIBMGA002132* | down | 1875 | 1883 | + | 10.7523 | 8.38E-05 | 0.797 | GGGCGTGTC |
| 1673 | mamo-s | *BGIBMGA002133* | down | 298 | 306 | + | 11.3303 | 4.96E-05 | 0.615 | GTGGGTGGC |
| 1674 | mamo-s | *BGIBMGA002137* | up | 47 | 55 | + | 11.3303 | 4.96E-05 | 0.615 | GTGGGTGGC |
| 1675 | mamo-s | *BGIBMGA002138* | up | 93 | 101 | - | 11.3303 | 4.96E-05 | 0.615 | GTGGGTGGC |
| 1676 | mamo-s | *BGIBMGA002139* | down | 1015 | 1023 | + | 11.3303 | 4.96E-05 | 0.615 | GTGGGTGGC |
| 1677 | mamo-s | *BGIBMGA002139* | down | 832 | 840 | + | 10.7523 | 8.38E-05 | 0.797 | GGGCGTGTC |
| 1678 | mamo-s | *BGIBMGA002141* | up | 25 | 33 | + | 12.1009 | 2.43E-05 | 0.615 | GTGCGTGGT |
| 1679 | mamo-s | *BGIBMGA002141* | up | 299 | 307 | - | 11.3303 | 4.96E-05 | 0.615 | GTGGGTGGC |
| 1680 | mamo-s | *BGIBMGA002142* | down | 1489 | 1497 | - | 10.5688 | 8.88E-05 | 0.824 | GTGCGTGAC |
| 1681 | mamo-s | *BGIBMGA002144* | down | 1155 | 1163 | - | 13.0275 | 8.53E-06 | 0.615 | GAGCGTGGC |
| 1682 | mamo-s | *BGIBMGA002147* | up | 1001 | 1009 | + | 11.3303 | 4.96E-05 | 0.615 | GTGGGTGGC |
| 1683 | mamo-s | *BGIBMGA002149* | down | 380 | 388 | + | 11.3303 | 4.96E-05 | 0.615 | GTGGGTGGC |
| 1684 | mamo-s | *BGIBMGA002150* | up | 746 | 754 | + | 11.4404 | 4.51E-05 | 0.615 | GAGCGTGTC |
| 1685 | mamo-s | *BGIBMGA002150* | down | 555 | 563 | + | 11.3303 | 4.96E-05 | 0.615 | GTGGGTGGC |
| 1686 | mamo-s | *BGIBMGA002151* | down | 1144 | 1152 | + | 11.0826 | 6.84E-05 | 0.709 | GTACGTGGC |
| 1687 | mamo-s | *BGIBMGA002153* | down | 239 | 247 | - | 11.3303 | 4.96E-05 | 0.615 | GTGGGTGGC |
| 1688 | mamo-s | *BGIBMGA002153* | down | 860 | 868 | + | 11.3303 | 4.96E-05 | 0.615 | GTGGGTGGC |
| 1689 | mamo-s | *BGIBMGA002154* | up | 1614 | 1622 | + | 11.4404 | 4.51E-05 | 0.615 | CGGCGTGGC |
| 1690 | mamo-s | *BGIBMGA002156* | up | 1493 | 1501 | - | 11.3303 | 4.96E-05 | 0.615 | GTGGGTGGC |
| 1691 | mamo-s | *BGIBMGA002156* | down | 725 | 733 | - | 11.0734 | 7.12E-05 | 0.726 | GAGCGTGGT |
| 1692 | mamo-s | *BGIBMGA002157* | up | 1761 | 1769 | - | 11.789 | 2.84E-05 | 0.615 | GCGCGTGCC |
| 1693 | mamo-s | *BGIBMGA002157* | down | 456 | 464 | + | 11.4404 | 4.51E-05 | 0.615 | CGGCGTGGC |
| 1694 | mamo-s | *BGIBMGA002158* | up | 392 | 400 | + | 11.4404 | 4.51E-05 | 0.615 | CGGCGTGGC |
| 1695 | mamo-s | *BGIBMGA002158* | up | 1464 | 1472 | - | 11.0734 | 7.12E-05 | 0.726 | GAGCGTGGT |
| 1696 | mamo-s | *BGIBMGA002161* | up | 596 | 604 | + | 11.1743 | 5.90E-05 | 0.684 | CCGCGTGTC |
| 1697 | mamo-s | *BGIBMGA002163* | down | 1457 | 1465 | + | 12.1009 | 2.43E-05 | 0.615 | GTGCGTGGT |
| 1698 | mamo-s | *BGIBMGA002163* | down | 625 | 633 | + | 11.4404 | 4.51E-05 | 0.615 | CGGCGTGGC |
| 1699 | mamo-s | *BGIBMGA002163* | up | 355 | 363 | - | 11.3303 | 4.96E-05 | 0.615 | GTGGGTGGC |
| 1700 | mamo-s | *BGIBMGA002163* | up | 1016 | 1024 | + | 11.3303 | 4.96E-05 | 0.615 | GTGGGTGGC |
| 1701 | mamo-s | *BGIBMGA002165* | up | 1 | 9 | - | 11.3303 | 4.96E-05 | 0.615 | GTGGGTGGC |
| 1702 | mamo-s | *BGIBMGA002166* | up | 1145 | 1153 | + | 11.3303 | 4.96E-05 | 0.615 | GTGGGTGGC |
| 1703 | mamo-s | *BGIBMGA002167* | up | 592 | 600 | + | 11.3303 | 4.96E-05 | 0.615 | GTGGGTGGC |
| 1704 | mamo-s | *BGIBMGA002168* | down | 42 | 50 | + | 12.1835 | 1.71E-05 | 0.615 | GTGCGTGCC |
| 1705 | mamo-s | *BGIBMGA002168* | up | 1234 | 1242 | + | 11.789 | 2.84E-05 | 0.615 | GCGCGTGCC |
| 1706 | mamo-s | *BGIBMGA002169* | down | 686 | 694 | + | 11.3303 | 4.96E-05 | 0.615 | GTGGGTGGC |
| 1707 | mamo-s | *BGIBMGA002169* | down | 1421 | 1429 | + | 11.3303 | 4.96E-05 | 0.615 | GTGGGTGGC |
| 1708 | mamo-s | *BGIBMGA002170* | up | 291 | 299 | - | 10.7523 | 8.38E-05 | 0.797 | GGGCGTGTC |
| 1709 | mamo-s | *BGIBMGA002171* | down | 877 | 885 | - | 11.3303 | 4.96E-05 | 0.615 | GTGGGTGGC |
| 1710 | mamo-s | *BGIBMGA002173* | up | 1528 | 1536 | + | 10.9358 | 7.30E-05 | 0.728 | GCGGGTGGC |
| 1711 | mamo-s | *BGIBMGA002174* | up | 71 | 79 | + | 11.3303 | 4.96E-05 | 0.615 | GTGGGTGGC |
| 1712 | mamo-s | *BGIBMGA002174* | down | 1368 | 1376 | - | 11.3303 | 4.96E-05 | 0.615 | GTGGGTGGC |
| 1713 | mamo-s | *BGIBMGA002175* | up | 1793 | 1801 | - | 11.4404 | 4.51E-05 | 0.615 | CGGCGTGGC |
| 1714 | mamo-s | *BGIBMGA002175* | up | 1468 | 1476 | - | 11.3303 | 4.96E-05 | 0.615 | GTGGGTGGC |
| 1715 | mamo-s | *BGIBMGA002177* | down | 1714 | 1722 | + | 11.3303 | 4.96E-05 | 0.615 | GTGGGTGGC |
| 1716 | mamo-s | *BGIBMGA002179* | down | 5 | 13 | + | 13.0275 | 8.53E-06 | 0.615 | GAGCGTGGC |
| 1717 | mamo-s | *BGIBMGA002180* | down | 1311 | 1319 | + | 11.4404 | 4.51E-05 | 0.615 | GAGCGTGTC |
| 1718 | mamo-s | *BGIBMGA002180* | down | 348 | 356 | + | 11.3303 | 4.96E-05 | 0.615 | GTGGGTGGC |
| 1719 | mamo-s | *BGIBMGA002181* | down | 380 | 388 | - | 11.3303 | 4.96E-05 | 0.615 | GTGGGTGGC |
| 1720 | mamo-s | *BGIBMGA002181* | up | 1414 | 1422 | - | 11.3303 | 4.96E-05 | 0.615 | GTGGGTGGC |
| 1721 | mamo-s | *BGIBMGA002182* | down | 382 | 390 | - | 11.3303 | 4.96E-05 | 0.615 | GTGGGTGGC |
| 1722 | mamo-s | *BGIBMGA002182* | up | 297 | 305 | + | 11.0826 | 6.84E-05 | 0.709 | GTACGTGGC |
| 1723 | mamo-s | *BGIBMGA002183* | up | 1416 | 1424 | - | 12.1009 | 2.43E-05 | 0.615 | GTGCGTGGT |
| 1724 | mamo-s | *BGIBMGA002183* | down | 397 | 405 | - | 10.5138 | 9.75E-05 | 0.865 | GTGCGTGTT |
| 1725 | mamo-s | *BGIBMGA002184* | up | 962 | 970 | + | 11.3303 | 4.96E-05 | 0.615 | GTGGGTGGC |
| 1726 | mamo-s | *BGIBMGA002184* | down | 1433 | 1441 | + | 11.3303 | 4.96E-05 | 0.615 | GTGGGTGGC |
| 1727 | mamo-s | *BGIBMGA002185* | down | 1897 | 1905 | + | 12.7615 | 1.04E-05 | 0.615 | CCGCGTGGC |
| 1728 | mamo-s | *BGIBMGA002185* | up | 1088 | 1096 | + | 11.3303 | 4.96E-05 | 0.615 | GTGGGTGGC |
| 1729 | mamo-s | *BGIBMGA002186* | down | 379 | 387 | - | 11.2018 | 5.68E-05 | 0.674 | CTGCGTGGT |
| 1730 | mamo-s | *BGIBMGA002187* | up | 108 | 116 | - | 11.0826 | 6.84E-05 | 0.709 | GTACGTGGC |
| 1731 | mamo-s | *BGIBMGA002188* | down | 44 | 52 | - | 12.1009 | 2.43E-05 | 0.615 | GTGCGTGGT |
| 1732 | mamo-s | *BGIBMGA002188* | down | 876 | 884 | - | 11.4404 | 4.51E-05 | 0.615 | CGGCGTGGC |
| 1733 | mamo-s | *BGIBMGA002189* | up | 599 | 607 | - | 11.3303 | 4.96E-05 | 0.615 | GTGGGTGGC |
| 1734 | mamo-s | *BGIBMGA002191* | down | 1388 | 1396 | - | 13.6606 | 4.06E-06 | 0.615 | GCGCGTGGC |
| 1735 | mamo-s | *BGIBMGA002193* | down | 1976 | 1984 | + | 13.0275 | 8.53E-06 | 0.615 | GAGCGTGGC |
| 1736 | mamo-s | *BGIBMGA002193* | up | 1497 | 1505 | + | 11.0826 | 6.84E-05 | 0.709 | GTACGTGGC |
| 1737 | mamo-s | *BGIBMGA002193* | down | 869 | 877 | + | 11.0734 | 7.12E-05 | 0.726 | GAGCGTGGT |
| 1738 | mamo-s | *BGIBMGA002194* | down | 867 | 875 | - | 11.1009 | 6.57E-05 | 0.709 | GAGCGAGGC |
| 1739 | mamo-s | *BGIBMGA002196* | up | 1172 | 1180 | + | 11.4404 | 4.51E-05 | 0.615 | CGGCGTGGC |
| 1740 | mamo-s | *BGIBMGA002197* | down | 1475 | 1483 | - | 11.789 | 2.84E-05 | 0.615 | GCGCGTGCC |
| 1741 | mamo-s | *BGIBMGA002197* | up | 1158 | 1166 | - | 11.3303 | 4.96E-05 | 0.615 | GTGGGTGGC |
| 1742 | mamo-s | *BGIBMGA002197* | down | 781 | 789 | + | 11.2844 | 5.18E-05 | 0.63 | CTGCGTGCC |
| 1743 | mamo-s | *BGIBMGA002197* | up | 1966 | 1974 | + | 11.156 | 6.13E-05 | 0.705 | GAGCGTGCC |
| 1744 | mamo-s | *BGIBMGA002198* | down | 1760 | 1768 | - | 11.3303 | 4.96E-05 | 0.615 | GTGGGTGGC |
| 1745 | mamo-s | *BGIBMGA002199* | down | 190 | 198 | + | 11.3303 | 4.96E-05 | 0.615 | GTGGGTGGC |
| 1746 | mamo-s | *BGIBMGA002199* | up | 313 | 321 | - | 11.3303 | 4.96E-05 | 0.615 | GTGGGTGGC |
| 1747 | mamo-s | *BGIBMGA002201* | down | 513 | 521 | - | 13.0275 | 8.53E-06 | 0.615 | GAGCGTGGC |
| 1748 | mamo-s | *BGIBMGA002204* | up | 1213 | 1221 | - | 12.0734 | 2.66E-05 | 0.615 | GCGCGTGTC |
| 1749 | mamo-s | *BGIBMGA002205* | up | 1142 | 1150 | + | 11.3303 | 4.96E-05 | 0.615 | GTGGGTGGC |
| 1750 | mamo-s | *BGIBMGA002205* | down | 1127 | 1135 | - | 11.0826 | 6.84E-05 | 0.709 | GTACGTGGC |
| 1751 | mamo-s | *BGIBMGA002206* | down | 1829 | 1837 | - | 11.0826 | 6.84E-05 | 0.709 | GTACGTGGC |
| 1752 | mamo-s | *BGIBMGA002209* | down | 479 | 487 | - | 11.3303 | 4.96E-05 | 0.615 | GTGGGTGGC |
| 1753 | mamo-s | *BGIBMGA002210* | down | 1600 | 1608 | - | 11.3303 | 4.96E-05 | 0.615 | GTGGGTGGC |

| | | | | | | | | | | |
|---|---|---|---|---|---|---|---|---|---|---|
| 1754 | mamo-s | *BGIBMGA002211* | down | 1613 | 1621 | - | 11.3303 | 4.96E-05 | 0.615 | GTGGGTGGC |
| 1755 | mamo-s | *BGIBMGA002212* | down | 1822 | 1830 | + | 11.3303 | 4.96E-05 | 0.615 | GTGGGTGGC |
| 1756 | mamo-s | *BGIBMGA002213* | down | 1930 | 1938 | - | 12.4679 | 1.31E-05 | 0.615 | GTGCGTGTC |
| 1757 | mamo-s | *BGIBMGA002213* | down | 798 | 806 | + | 12.3394 | 1.49E-05 | 0.615 | GGGCGTGGC |
| 1758 | mamo-s | *BGIBMGA002213* | up | 108 | 116 | + | 11.3303 | 4.96E-05 | 0.615 | GTGGGTGGC |
| 1759 | mamo-s | *BGIBMGA002213* | down | 23 | 31 | - | 10.6881 | 8.61E-05 | 0.81 | GCACGTGGC |
| 1760 | mamo-s | *BGIBMGA002214* | up | 1498 | 1506 | + | 11.3303 | 4.96E-05 | 0.615 | GTGGGTGGC |
| 1761 | mamo-s | *BGIBMGA002216* | up | 1286 | 1294 | + | 11.789 | 2.84E-05 | 0.615 | GCGCGTGCC |
| 1762 | mamo-s | *BGIBMGA002220* | down | 1211 | 1219 | - | 11.7339 | 3.29E-05 | 0.615 | GCGCGAGGC |
| 1763 | mamo-s | *BGIBMGA002220* | down | 835 | 843 | + | 11.0734 | 7.12E-05 | 0.726 | GAGCGTGGT |
| 1764 | mamo-s | *BGIBMGA002221* | up | 724 | 732 | - | 11.789 | 2.84E-05 | 0.615 | GCGCGTGCC |
| 1765 | mamo-s | *BGIBMGA002221* | up | 1647 | 1655 | + | 11.0734 | 7.12E-05 | 0.726 | GAGCGTGGT |
| 1766 | mamo-s | *BGIBMGA002222* | up | 1784 | 1792 | - | 11.3303 | 4.96E-05 | 0.615 | GTGGGTGGC |
| 1767 | mamo-s | *BGIBMGA002222* | down | 1871 | 1879 | + | 11.0826 | 6.84E-05 | 0.709 | GTACGTGGC |
| 1768 | mamo-s | *BGIBMGA002226* | down | 907 | 915 | + | 13.0275 | 8.53E-06 | 0.615 | GAGCGTGGC |
| 1769 | mamo-s | *BGIBMGA002228* | down | 1801 | 1809 | - | 11.3303 | 4.96E-05 | 0.615 | GTGGGTGGC |
| 1770 | mamo-s | *BGIBMGA002229* | down | 219 | 227 | + | 11.0826 | 6.84E-05 | 0.709 | GTACGTGGC |
| 1771 | mamo-s | *BGIBMGA002231* | up | 1664 | 1672 | - | 10.5413 | 9.42E-05 | 0.857 | CAGCGTGTC |
| 1772 | mamo-s | *BGIBMGA002232* | down | 1239 | 1247 | - | 11.3303 | 4.96E-05 | 0.615 | GTGGGTGGC |
| 1773 | mamo-s | *BGIBMGA002232* | down | 130 | 138 | + | 10.8349 | 7.66E-05 | 0.751 | CCGCGAGGC |
| 1774 | mamo-s | *BGIBMGA002233* | down | 1855 | 1863 | + | 12.1009 | 2.43E-05 | 0.615 | GTGCGTGGT |
| 1775 | mamo-s | *BGIBMGA002233* | down | 752 | 760 | - | 11.3303 | 4.96E-05 | 0.615 | GTGGGTGGC |
| 1776 | mamo-s | *BGIBMGA002234* | up | 197 | 205 | + | 11.3303 | 4.96E-05 | 0.615 | GTGGGTGGC |
| 1777 | mamo-s | *BGIBMGA002238* | up | 512 | 520 | - | 10.8899 | 7.48E-05 | 0.739 | CCGCGTGCC |
| 1778 | mamo-s | *BGIBMGA002239* | up | 1769 | 1777 | - | 11.3303 | 4.96E-05 | 0.615 | GTGGGTGGC |
| 1779 | mamo-s | *BGIBMGA002240* | down | 230 | 238 | + | 10.7523 | 8.38E-05 | 0.797 | AAGCGTGGC |
| 1780 | mamo-s | *BGIBMGA002240* | up | 1719 | 1727 | + | 10.7523 | 8.38E-05 | 0.797 | AAGCGTGGC |
| 1781 | mamo-s | *BGIBMGA002244* | up | 621 | 629 | - | 11.2844 | 5.18E-05 | 0.63 | CTGCGTGCC |
| 1782 | mamo-s | *BGIBMGA002248* | down | 334 | 342 | + | 11.3303 | 4.96E-05 | 0.615 | GTGGGTGGC |
| 1783 | mamo-s | *BGIBMGA002249* | down | 256 | 264 | - | 11.3303 | 4.96E-05 | 0.615 | GTGGGTGGC |
| 1784 | mamo-s | *BGIBMGA002251* | down | 242 | 250 | + | 11.0826 | 6.84E-05 | 0.709 | GTACGTGGC |
| 1785 | mamo-s | *BGIBMGA002254* | up | 1474 | 1482 | - | 11.3303 | 4.96E-05 | 0.615 | GTGGGTGGC |
| 1786 | mamo-s | *BGIBMGA002254* | up | 471 | 479 | - | 11.2844 | 5.18E-05 | 0.63 | CTGCGTGCC |
| 1787 | mamo-s | *BGIBMGA002254* | down | 627 | 635 | - | 11.1743 | 5.90E-05 | 0.684 | CCGCGTGTC |
| 1788 | mamo-s | *BGIBMGA002254* | up | 52 | 60 | - | 11.0826 | 6.84E-05 | 0.709 | GTACGTGGC |
| 1789 | mamo-s | *BGIBMGA002255* | down | 1358 | 1366 | + | 11.3303 | 4.96E-05 | 0.615 | GTGGGTGGC |
| 1790 | mamo-s | *BGIBMGA002255* | up | 1424 | 1432 | - | 11.3303 | 4.96E-05 | 0.615 | GTGGGTGGC |
| 1791 | mamo-s | *BGIBMGA002255* | down | 1901 | 1909 | - | 11.3303 | 4.96E-05 | 0.615 | GTGGGTGGC |
| 1792 | mamo-s | *BGIBMGA002255* | down | 1010 | 1018 | + | 11.0826 | 6.84E-05 | 0.709 | GTACGTGGC |
| 1793 | mamo-s | *BGIBMGA002255* | up | 1664 | 1672 | - | 10.5413 | 9.42E-05 | 0.857 | CAGCGTGTC |
| 1794 | mamo-s | *BGIBMGA002255* | up | 1034 | 1042 | + | 10.5138 | 9.75E-05 | 0.865 | GTGCGTGTT |
| 1795 | mamo-s | *BGIBMGA002256* | down | 320 | 328 | + | 11.3303 | 4.96E-05 | 0.615 | GTGGGTGGC |
| 1796 | mamo-s | *BGIBMGA002256* | down | 1509 | 1517 | + | 11.3303 | 4.96E-05 | 0.615 | GTGGGTGGC |
| 1797 | mamo-s | *BGIBMGA002257* | up | 1599 | 1607 | + | 11.3303 | 4.96E-05 | 0.615 | GTGGGTGGC |
| 1798 | mamo-s | *BGIBMGA002263* | up | 1599 | 1607 | + | 13.0275 | 8.53E-06 | 0.615 | GAGCGTGGC |
| 1799 | mamo-s | *BGIBMGA002264* | up | 563 | 571 | + | 10.8349 | 7.66E-05 | 0.751 | CCGCGAGGC |
| 1800 | mamo-s | *BGIBMGA002265* | down | 1294 | 1302 | + | 12.1835 | 1.71E-05 | 0.615 | GTGCGTGCC |
| 1801 | mamo-s | *BGIBMGA002266* | up | 96 | 104 | - | 13.6606 | 4.06E-06 | 0.615 | GCGCGTGGC |
| 1802 | mamo-s | *BGIBMGA002266* | up | 437 | 445 | - | 11.3303 | 4.96E-05 | 0.615 | GTGGGTGGC |
| 1803 | mamo-s | *BGIBMGA002266* | up | 1637 | 1645 | - | 11.0826 | 6.84E-05 | 0.709 | GTACGTGGC |
| 1804 | mamo-s | *BGIBMGA002267* | down | 1351 | 1359 | - | 11.1009 | 6.57E-05 | 0.709 | GAGCGAGGC |
| 1805 | mamo-s | *BGIBMGA002269* | up | 87 | 95 | + | 11.2844 | 5.18E-05 | 0.63 | CTGCGTGCC |
| 1806 | mamo-s | *BGIBMGA002269* | up | 1172 | 1180 | - | 10.5688 | 8.88E-05 | 0.824 | GTGCGTGAC |
| 1807 | mamo-s | *BGIBMGA002270* | down | 1913 | 1921 | - | 11.3303 | 4.96E-05 | 0.615 | GTGGGTGGC |
| 1808 | mamo-s | *BGIBMGA002271* | down | 1663 | 1671 | + | 11.3303 | 4.96E-05 | 0.615 | GTGGGTGGC |
| 1809 | mamo-s | *BGIBMGA002271* | up | 1754 | 1762 | + | 11.3303 | 4.96E-05 | 0.615 | GTGGGTGGC |
| 1810 | mamo-s | *BGIBMGA002272* | down | 896 | 904 | + | 11.4404 | 4.51E-05 | 0.615 | CGGCGTGGC |
| 1811 | mamo-s | *BGIBMGA002274* | up | 1632 | 1640 | + | 11.3303 | 4.96E-05 | 0.615 | GTGGGTGGC |
| 1812 | mamo-s | *BGIBMGA002275* | up | 1520 | 1528 | + | 14.055 | 2.23E-06 | 0.615 | GTGCGTGGC |
| 1813 | mamo-s | *BGIBMGA002276* | down | 1408 | 1416 | + | 11.3303 | 4.96E-05 | 0.615 | GTGGGTGGC |
| 1814 | mamo-s | *BGIBMGA002276* | down | 955 | 963 | - | 10.5138 | 9.75E-05 | 0.865 | GTGCGTGTT |
| 1815 | mamo-s | *BGIBMGA002277* | up | 509 | 517 | + | 11.3303 | 4.96E-05 | 0.615 | GTGGGTGGC |
| 1816 | mamo-s | *BGIBMGA002278* | up | 820 | 828 | - | 11.3303 | 4.96E-05 | 0.615 | GTGGGTGGC |
| 1817 | mamo-s | *BGIBMGA002278* | up | 664 | 672 | + | 10.5413 | 9.42E-05 | 0.857 | CAGCGTGTC |
| 1818 | mamo-s | *BGIBMGA002279* | up | 342 | 350 | - | 11.0826 | 6.84E-05 | 0.709 | GTACGTGGC |
| 1819 | mamo-s | *BGIBMGA002280* | down | 1364 | 1372 | + | 11.3303 | 4.96E-05 | 0.615 | GTGGGTGGC |
| 1820 | mamo-s | *BGIBMGA002281* | down | 1184 | 1192 | - | 11.3303 | 4.96E-05 | 0.615 | GTGGGTGGC |
| 1821 | mamo-s | *BGIBMGA002281* | up | 1750 | 1758 | - | 11.3303 | 4.96E-05 | 0.615 | GTGGGTGGC |
| 1822 | mamo-s | *BGIBMGA002282* | down | 1510 | 1518 | - | 10.9358 | 7.30E-05 | 0.728 | GCGGGTGGC |
| 1823 | mamo-s | *BGIBMGA002283* | down | 1044 | 1052 | + | 11.3303 | 4.96E-05 | 0.615 | GTGGGTGGC |
| 1824 | mamo-s | *BGIBMGA002284* | up | 1598 | 1606 | - | 11.7798 | 3.11E-05 | 0.615 | ATGCGTGGC |
| 1825 | mamo-s | *BGIBMGA002284* | up | 248 | 256 | + | 11.3303 | 4.96E-05 | 0.615 | GTGGGTGGC |
| 1826 | mamo-s | *BGIBMGA002285* | down | 1454 | 1462 | - | 11.3303 | 4.96E-05 | 0.615 | GTGGGTGGC |
| 1827 | mamo-s | *BGIBMGA002287* | up | 341 | 349 | + | 11.3303 | 4.96E-05 | 0.615 | GTGGGTGGC |
| 1828 | mamo-s | *BGIBMGA002287* | down | 1385 | 1393 | - | 10.7523 | 8.38E-05 | 0.797 | AAGCGTGGC |
| 1829 | mamo-s | *BGIBMGA002289* | up | 1280 | 1288 | + | 11.2844 | 5.18E-05 | 0.63 | CTGCGTGCC |
| 1830 | mamo-s | *BGIBMGA002290* | up | 404 | 412 | + | 11.3303 | 4.96E-05 | 0.615 | GTGGGTGGC |
| 1831 | mamo-s | *BGIBMGA002290* | down | 926 | 934 | + | 11.3303 | 4.96E-05 | 0.615 | GTGGGTGGC |
| 1832 | mamo-s | *BGIBMGA002290* | up | 1678 | 1686 | - | 11.3303 | 4.96E-05 | 0.615 | GTGGGTGGC |
| 1833 | mamo-s | *BGIBMGA002290* | down | 1897 | 1905 | + | 11.3303 | 4.96E-05 | 0.615 | GTGGGTGGC |
| 1834 | mamo-s | *BGIBMGA002291* | down | 1144 | 1152 | - | 11.3303 | 4.96E-05 | 0.615 | GTGGGTGGC |
| 1835 | mamo-s | *BGIBMGA002292* | up | 1898 | 1906 | + | 11.4404 | 4.51E-05 | 0.615 | CGGCGTGGC |
| 1836 | mamo-s | *BGIBMGA002298* | up | 1722 | 1730 | + | 11.3303 | 4.96E-05 | 0.615 | GTGGGTGGC |
| 1837 | mamo-s | *BGIBMGA002301* | up | 1846 | 1854 | - | 11.7064 | 3.52E-05 | 0.615 | GCGCGTGGT |
| 1838 | mamo-s | *BGIBMGA002301* | up | 159 | 167 | + | 11.5688 | 3.79E-05 | 0.615 | CTGCGTGTC |
| 1839 | mamo-s | *BGIBMGA002301* | down | 149 | 157 | + | 11.3303 | 4.96E-05 | 0.615 | GTGGGTGGC |
| 1840 | mamo-s | *BGIBMGA002301* | up | 436 | 444 | - | 11.3303 | 4.96E-05 | 0.615 | GTGGGTGGC |
| 1841 | mamo-s | *BGIBMGA002303* | down | 1842 | 1850 | + | 12.7615 | 1.04E-05 | 0.615 | CCGCGTGGC |

| | | | | | | | | | | |
|---|---|---|---|---|---|---|---|---|---|---|
| 1842 | mamo-s | *BGIBMGA002304* | down | 1570 | 1578 | - | 11.4404 | 4.51E-05 | 0.615 | CGGCGTGGC |
| 1843 | mamo-s | *BGIBMGA002305* | down | 1474 | 1482 | - | 11.3303 | 4.96E-05 | 0.615 | GTGGGTGGC |
| 1844 | mamo-s | *BGIBMGA002305* | down | 243 | 251 | - | 11.2844 | 5.18E-05 | 0.63 | CTGCGTGCC |
| 1845 | mamo-s | *BGIBMGA002305* | up | 1704 | 1712 | + | 10.5688 | 8.88E-05 | 0.824 | GTGCGTGAC |
| 1846 | mamo-s | *BGIBMGA002306* | up | 467 | 475 | + | 11.3303 | 4.96E-05 | 0.615 | GTGGGTGGC |
| 1847 | mamo-s | *BGIBMGA002306* | down | 1080 | 1088 | + | 11.3303 | 4.96E-05 | 0.615 | GTGGGTGGC |
| 1848 | mamo-s | *BGIBMGA002306* | down | 624 | 632 | - | 11.1743 | 5.90E-05 | 0.684 | CCGCGTGTC |
| 1849 | mamo-s | *BGIBMGA002307* | up | 1043 | 1051 | + | 11.2294 | 5.41E-05 | 0.65 | CTGCGAGGC |
| 1850 | mamo-s | *BGIBMGA002307* | down | 1797 | 1805 | - | 11.0826 | 6.84E-05 | 0.709 | GTACGTGGC |
| 1851 | mamo-s | *BGIBMGA002308* | up | 743 | 751 | + | 11.3303 | 4.96E-05 | 0.615 | GTGGGTGGC |
| 1852 | mamo-s | *BGIBMGA002308* | down | 1932 | 1940 | + | 11.3303 | 4.96E-05 | 0.615 | GTGGGTGGC |
| 1853 | mamo-s | *BGIBMGA002309* | up | 743 | 751 | + | 11.0826 | 6.84E-05 | 0.709 | GTACGTGGC |
| 1854 | mamo-s | *BGIBMGA002311* | up | 337 | 345 | + | 11.3303 | 4.96E-05 | 0.615 | GTGGGTGGC |
| 1855 | mamo-s | *BGIBMGA002311* | down | 1425 | 1433 | - | 11.3303 | 4.96E-05 | 0.615 | GTGGGTGGC |
| 1856 | mamo-s | *BGIBMGA002312* | up | 988 | 996 | - | 11.3303 | 4.96E-05 | 0.615 | GTGGGTGGC |
| 1857 | mamo-s | *BGIBMGA002313* | up | 1575 | 1583 | + | 14.055 | 2.23E-06 | 0.615 | GTGCGTGGC |
| 1858 | mamo-s | *BGIBMGA002313* | up | 684 | 692 | + | 11.3303 | 4.96E-05 | 0.615 | GTGGGTGGC |
| 1859 | mamo-s | *BGIBMGA002313* | up | 1076 | 1084 | - | 11.3303 | 4.96E-05 | 0.615 | GTGGGTGGC |
| 1860 | mamo-s | *BGIBMGA002314* | down | 1220 | 1228 | - | 11.3303 | 4.96E-05 | 0.615 | GTGGGTGGC |
| 1861 | mamo-s | *BGIBMGA002316* | up | 1828 | 1836 | - | 11.3303 | 4.96E-05 | 0.615 | GTGGGTGGC |
| 1862 | mamo-s | *BGIBMGA002317* | up | 351 | 359 | + | 11.3303 | 4.96E-05 | 0.615 | GTGGGTGGC |
| 1863 | mamo-s | *BGIBMGA002318* | down | 1299 | 1307 | - | 11.3303 | 4.96E-05 | 0.615 | GTGGGTGGC |
| 1864 | mamo-s | *BGIBMGA002319* | down | 578 | 586 | - | 11.0826 | 6.84E-05 | 0.709 | GTACGTGGC |
| 1865 | mamo-s | *BGIBMGA002320* | up | 654 | 662 | - | 11.3303 | 4.96E-05 | 0.615 | GTGGGTGGC |
| 1866 | mamo-s | *BGIBMGA002320* | up | 1475 | 1483 | - | 11.3303 | 4.96E-05 | 0.615 | GTGGGTGGC |
| 1867 | mamo-s | *BGIBMGA002321* | up | 372 | 380 | + | 11.3303 | 4.96E-05 | 0.615 | GTGGGTGGC |
| 1868 | mamo-s | *BGIBMGA002321* | up | 1406 | 1414 | - | 10.5688 | 8.88E-05 | 0.824 | GTGCGTGAC |
| 1869 | mamo-s | *BGIBMGA002323* | up | 1708 | 1716 | - | 11.0826 | 6.84E-05 | 0.709 | GTACGTGGC |
| 1870 | mamo-s | *BGIBMGA002324* | up | 684 | 692 | + | 11.3303 | 4.96E-05 | 0.615 | GTGGGTGGC |
| 1871 | mamo-s | *BGIBMGA002325* | up | 672 | 680 | + | 12.7615 | 1.04E-05 | 0.615 | CCGCGTGGC |
| 1872 | mamo-s | *BGIBMGA002325* | up | 435 | 443 | - | 11.3853 | 4.74E-05 | 0.615 | ACGCGTGGC |
| 1873 | mamo-s | *BGIBMGA002326* | up | 1590 | 1598 | - | 11.3303 | 4.96E-05 | 0.615 | GTGGGTGGC |
| 1874 | mamo-s | *BGIBMGA002328* | down | 712 | 720 | + | 11.3303 | 4.96E-05 | 0.615 | GTGGGTGGC |
| 1875 | mamo-s | *BGIBMGA002329* | down | 1036 | 1044 | - | 11.7064 | 3.52E-05 | 0.615 | GCGCGTGGT |
| 1876 | mamo-s | *BGIBMGA002329* | down | 626 | 634 | - | 11.3303 | 4.96E-05 | 0.615 | GTGGGTGGC |
| 1877 | mamo-s | *BGIBMGA002331* | down | 1737 | 1745 | - | 11.4404 | 4.51E-05 | 0.615 | GAGCGTGTC |
| 1878 | mamo-s | *BGIBMGA002331* | up | 635 | 643 | - | 11.0826 | 6.84E-05 | 0.709 | GTACGTGGC |
| 1879 | mamo-s | *BGIBMGA002332* | down | 1575 | 1583 | - | 12.1009 | 2.43E-05 | 0.615 | GTGCGTGGT |
| 1880 | mamo-s | *BGIBMGA002332* | down | 512 | 520 | + | 11.3303 | 4.96E-05 | 0.615 | GTGGGTGGC |
| 1881 | mamo-s | *BGIBMGA002333* | down | 191 | 199 | - | 11.3303 | 4.96E-05 | 0.615 | GTGGGTGGC |
| 1882 | mamo-s | *BGIBMGA002333* | up | 973 | 981 | - | 11.3303 | 4.96E-05 | 0.615 | GTGGGTGGC |
| 1883 | mamo-s | *BGIBMGA002333* | up | 1458 | 1466 | + | 11.3303 | 4.96E-05 | 0.615 | GTGGGTGGC |
| 1884 | mamo-s | *BGIBMGA002334* | down | 152 | 160 | - | 11.3303 | 4.96E-05 | 0.615 | GTGGGTGGC |
| 1885 | mamo-s | *BGIBMGA002334* | up | 1013 | 1021 | - | 11.3303 | 4.96E-05 | 0.615 | GTGGGTGGC |
| 1886 | mamo-s | *BGIBMGA002334* | up | 1718 | 1726 | + | 11.3303 | 4.96E-05 | 0.615 | GTGGGTGGC |
| 1887 | mamo-s | *BGIBMGA002335* | up | 1341 | 1349 | - | 11.0826 | 6.84E-05 | 0.709 | GTACGTGGC |
| 1888 | mamo-s | *BGIBMGA002336* | down | 571 | 579 | + | 11.3303 | 4.96E-05 | 0.615 | GTGGGTGGC |
| 1889 | mamo-s | *BGIBMGA002337* | down | 1598 | 1606 | + | 11.3303 | 4.96E-05 | 0.615 | GTGGGTGGC |
| 1890 | mamo-s | *BGIBMGA002338* | down | 181 | 189 | + | 11.3303 | 4.96E-05 | 0.615 | GTGGGTGGC |
| 1891 | mamo-s | *BGIBMGA002338* | down | 1748 | 1756 | + | 10.7523 | 8.38E-05 | 0.797 | AAGCGTGGC |
| 1892 | mamo-s | *BGIBMGA002339* | down | 680 | 688 | - | 14.055 | 2.23E-06 | 0.615 | GTGCGTGGC |
| 1893 | mamo-s | *BGIBMGA002339* | up | 1899 | 1907 | + | 10.8899 | 7.48E-05 | 0.739 | CCGCGTGCC |
| 1894 | mamo-s | *BGIBMGA002343* | up | 598 | 606 | + | 11.3303 | 4.96E-05 | 0.615 | GTGGGTGGC |
| 1895 | mamo-s | *BGIBMGA002344* | up | 126 | 134 | + | 11.3303 | 4.96E-05 | 0.615 | GTGGGTGGC |
| 1896 | mamo-s | *BGIBMGA002344* | down | 1260 | 1268 | + | 11.3303 | 4.96E-05 | 0.615 | GTGGGTGGC |
| 1897 | mamo-s | *BGIBMGA002345* | up | 1651 | 1659 | - | 12.1009 | 2.43E-05 | 0.615 | GTGCGTGGT |
| 1898 | mamo-s | *BGIBMGA002348* | down | 416 | 424 | - | 11.2844 | 5.18E-05 | 0.63 | CTGCGTGCC |
| 1899 | mamo-s | *BGIBMGA002350* | down | 1063 | 1071 | + | 10.7523 | 8.38E-05 | 0.797 | GGGCGTGTC |
| 1900 | mamo-s | *BGIBMGA002351* | down | 986 | 994 | + | 11.3303 | 4.96E-05 | 0.615 | GTGGGTGGC |
| 1901 | mamo-s | *BGIBMGA002354* | up | 100 | 108 | - | 10.7523 | 8.38E-05 | 0.797 | AAGCGTGGC |
| 1902 | mamo-s | *BGIBMGA002355* | down | 1521 | 1529 | - | 11.3303 | 4.96E-05 | 0.615 | GTGGGTGGC |
| 1903 | mamo-s | *BGIBMGA002356* | down | 292 | 300 | - | 12.3394 | 1.49E-05 | 0.615 | GGGCGTGGC |
| 1904 | mamo-s | *BGIBMGA002356* | up | 263 | 271 | - | 10.5413 | 9.42E-05 | 0.857 | GTGCGAGTC |
| 1905 | mamo-s | *BGIBMGA002357* | down | 111 | 119 | - | 11.3303 | 4.96E-05 | 0.615 | GTGGGTGGC |
| 1906 | mamo-s | *BGIBMGA002358* | up | 577 | 585 | - | 11.1743 | 5.90E-05 | 0.684 | CCGCGTGTC |
| 1907 | mamo-s | *BGIBMGA002358* | up | 1622 | 1630 | + | 11.1376 | 6.35E-05 | 0.709 | TCGCGTGGC |
| 1908 | mamo-s | *BGIBMGA002359* | down | 1661 | 1669 | - | 11.3303 | 4.96E-05 | 0.615 | GTGGGTGGC |
| 1909 | mamo-s | *BGIBMGA002360* | up | 592 | 600 | + | 11.3303 | 4.96E-05 | 0.615 | GTGGGTGGC |
| 1910 | mamo-s | *BGIBMGA002362* | down | 662 | 670 | + | 13.156 | 6.30E-06 | 0.615 | CTGCGTGGC |
| 1911 | mamo-s | *BGIBMGA002363* | up | 896 | 904 | - | 11.3303 | 4.96E-05 | 0.615 | GTGGGTGGC |
| 1912 | mamo-s | *BGIBMGA002363* | down | 1792 | 1800 | + | 11.3303 | 4.96E-05 | 0.615 | GTGGGTGGC |
| 1913 | mamo-s | *BGIBMGA002364* | up | 1979 | 1987 | + | 10.5688 | 8.88E-05 | 0.824 | GTGCGTGAC |
| 1914 | mamo-s | *BGIBMGA002365* | down | 1488 | 1496 | + | 11.3303 | 4.96E-05 | 0.615 | GTGGGTGGC |
| 1915 | mamo-s | *BGIBMGA002366* | up | 810 | 818 | + | 11.3303 | 4.96E-05 | 0.615 | GTGGGTGGC |
| 1916 | mamo-s | *BGIBMGA002367* | down | 279 | 287 | + | 11.2294 | 5.41E-05 | 0.65 | CTGCGAGGC |
| 1917 | mamo-s | *BGIBMGA002368* | up | 131 | 139 | + | 11.1743 | 5.90E-05 | 0.684 | CCGCGTGTC |
| 1918 | mamo-s | *BGIBMGA002368* | down | 1104 | 1112 | + | 11.0826 | 6.84E-05 | 0.709 | GTACGTGGC |
| 1919 | mamo-s | *BGIBMGA002369* | up | 346 | 354 | - | 11.3303 | 4.96E-05 | 0.615 | GTGGGTGGC |
| 1920 | mamo-s | *BGIBMGA002369* | down | 378 | 386 | - | 11.0826 | 6.84E-05 | 0.709 | GTACGTGGC |
| 1921 | mamo-s | *BGIBMGA002369* | down | 1228 | 1236 | + | 10.5413 | 9.42E-05 | 0.857 | CAGCGTGTC |
| 1922 | mamo-s | *BGIBMGA002371* | down | 1728 | 1736 | - | 13.6606 | 4.06E-06 | 0.615 | GCGCGTGGC |
| 1923 | mamo-s | *BGIBMGA002375* | up | 1435 | 1443 | + | 11.3303 | 4.96E-05 | 0.615 | GTGGGTGGC |
| 1924 | mamo-s | *BGIBMGA002376* | up | 1191 | 1199 | - | 11.3303 | 4.96E-05 | 0.615 | GTGGGTGGC |
| 1925 | mamo-s | *BGIBMGA002377* | down | 174 | 182 | + | 11.5688 | 3.79E-05 | 0.615 | CTGCGTGTC |
| 1926 | mamo-s | *BGIBMGA002380* | down | 1207 | 1215 | - | 11.3303 | 4.96E-05 | 0.615 | GTGGGTGGC |
| 1927 | mamo-s | *BGIBMGA002382* | up | 1133 | 1141 | + | 11.0734 | 7.12E-05 | 0.726 | GAGCGTGGT |
| 1928 | mamo-s | *BGIBMGA002385* | up | 396 | 404 | - | 11.3303 | 4.96E-05 | 0.615 | GTGGGTGGC |
| 1929 | mamo-s | *BGIBMGA002385* | up | 1099 | 1107 | - | 11.3303 | 4.96E-05 | 0.615 | GTGGGTGGC |

| | | | | | | | | | | |
|---|---|---|---|---|---|---|---|---|---|---|
| 1930 | mamo-s | *BGIBMGA002387* | up | 512 | 520 | - | 13.6606 | 4.06E-06 | 0.615 | GCGCGTGGC |
| 1931 | mamo-s | *BGIBMGA002388* | down | 875 | 883 | + | 11.3303 | 4.96E-05 | 0.615 | GTGGGTGGC |
| 1932 | mamo-s | *BGIBMGA002389* | down | 1105 | 1113 | - | 11.0826 | 6.84E-05 | 0.709 | GTACGTGGC |
| 1933 | mamo-s | *BGIBMGA002390* | up | 692 | 700 | - | 11.3303 | 4.96E-05 | 0.615 | GTGGGTGGC |
| 1934 | mamo-s | *BGIBMGA002390* | down | 1075 | 1083 | + | 11.3303 | 4.96E-05 | 0.615 | GTGGGTGGC |
| 1935 | mamo-s | *BGIBMGA002391* | up | 1018 | 1026 | - | 11.3303 | 4.96E-05 | 0.615 | GTGGGTGGC |
| 1936 | mamo-s | *BGIBMGA002391* | up | 1765 | 1773 | + | 11.3303 | 4.96E-05 | 0.615 | GTGGGTGGC |
| 1937 | mamo-s | *BGIBMGA002394* | up | 411 | 419 | + | 12.1009 | 2.43E-05 | 0.615 | GTGCGTGGT |
| 1938 | mamo-s | *BGIBMGA002394* | down | 1206 | 1214 | + | 11.789 | 2.84E-05 | 0.615 | GCGCGTGCC |
| 1939 | mamo-s | *BGIBMGA002395* | up | 1340 | 1348 | - | 11.3303 | 4.96E-05 | 0.615 | GTGGGTGGC |
| 1940 | mamo-s | *BGIBMGA002396* | down | 1205 | 1213 | + | 11.3303 | 4.96E-05 | 0.615 | GTGGGTGGC |
| 1941 | mamo-s | *BGIBMGA002397* | up | 1782 | 1790 | + | 11.3303 | 4.96E-05 | 0.615 | GTGGGTGGC |
| 1942 | mamo-s | *BGIBMGA002399* | up | 1983 | 1991 | - | 11.7339 | 3.29E-05 | 0.615 | GCGCGAGGC |
| 1943 | mamo-s | *BGIBMGA002400* | up | 211 | 219 | + | 11.0734 | 7.12E-05 | 0.726 | GAGCGTGGT |
| 1944 | mamo-s | *BGIBMGA002401* | up | 98 | 106 | - | 11.3303 | 4.96E-05 | 0.615 | GTGGGTGGC |
| 1945 | mamo-s | *BGIBMGA002401* | down | 306 | 314 | + | 11.3303 | 4.96E-05 | 0.615 | GTGGGTGGC |
| 1946 | mamo-s | *BGIBMGA002401* | up | 460 | 468 | + | 11.3303 | 4.96E-05 | 0.615 | GTGGGTGGC |
| 1947 | mamo-s | *BGIBMGA002403* | down | 1668 | 1676 | - | 12.0734 | 2.66E-05 | 0.615 | GCGCGTGTC |
| 1948 | mamo-s | *BGIBMGA002404* | up | 145 | 153 | - | 11.4404 | 4.51E-05 | 0.615 | CGGCGTGGC |
| 1949 | mamo-s | *BGIBMGA002407* | up | 256 | 264 | + | 11.3303 | 4.96E-05 | 0.615 | GTGGGTGGC |
| 1950 | mamo-s | *BGIBMGA002411* | up | 1135 | 1143 | - | 11.3303 | 4.96E-05 | 0.615 | GTGGGTGGC |
| 1951 | mamo-s | *BGIBMGA002415* | down | 1155 | 1163 | - | 11.3303 | 4.96E-05 | 0.615 | GTGGGTGGC |
| 1952 | mamo-s | *BGIBMGA002418* | up | 1354 | 1362 | - | 11.3303 | 4.96E-05 | 0.615 | GTGGGTGGC |
| 1953 | mamo-s | *BGIBMGA002418* | down | 1667 | 1675 | - | 11.3303 | 4.96E-05 | 0.615 | GTGGGTGGC |
| 1954 | mamo-s | *BGIBMGA002419* | up | 464 | 472 | + | 11.3303 | 4.96E-05 | 0.615 | GTGGGTGGC |
| 1955 | mamo-s | *BGIBMGA002420* | up | 1576 | 1584 | - | 11.3303 | 4.96E-05 | 0.615 | GTGGGTGGC |
| 1956 | mamo-s | *BGIBMGA002421* | up | 434 | 442 | - | 11.3303 | 4.96E-05 | 0.615 | GTGGGTGGC |
| 1957 | mamo-s | *BGIBMGA002422* | up | 1615 | 1623 | - | 11.3303 | 4.96E-05 | 0.615 | GTGGGTGGC |
| 1958 | mamo-s | *BGIBMGA002422* | down | 1546 | 1554 | - | 11.1376 | 6.35E-05 | 0.709 | TCGCGTGGC |
| 1959 | mamo-s | *BGIBMGA002425* | down | 486 | 494 | - | 11.4404 | 4.51E-05 | 0.615 | CGGCGTGGC |
| 1960 | mamo-s | *BGIBMGA002426* | down | 1826 | 1834 | - | 11.5321 | 4.06E-05 | 0.615 | TTGCGTGGC |
| 1961 | mamo-s | *BGIBMGA002426* | down | 1267 | 1275 | - | 10.5138 | 9.75E-05 | 0.865 | GTGCGTGTT |
| 1962 | mamo-s | *BGIBMGA002427* | down | 825 | 833 | - | 10.5138 | 9.75E-05 | 0.865 | GTGCGTGTT |
| 1963 | mamo-s | *BGIBMGA002430* | up | 848 | 856 | - | 11.5321 | 4.06E-05 | 0.615 | TTGCGTGGC |
| 1964 | mamo-s | *BGIBMGA002430* | down | 1567 | 1575 | + | 11.3303 | 4.96E-05 | 0.615 | GTGGGTGGC |
| 1965 | mamo-s | *BGIBMGA002432* | up | 867 | 875 | - | 11.3303 | 4.96E-05 | 0.615 | GTGGGTGGC |
| 1966 | mamo-s | *BGIBMGA002432* | down | 1660 | 1668 | + | 11.3303 | 4.96E-05 | 0.615 | GTGGGTGGC |
| 1967 | mamo-s | *BGIBMGA002432* | up | 1868 | 1876 | + | 11.156 | 6.13E-05 | 0.705 | GAGCGTGCC |
| 1968 | mamo-s | *BGIBMGA002433* | up | 994 | 1002 | + | 11.2294 | 5.41E-05 | 0.65 | CTGCGAGGC |
| 1969 | mamo-s | *BGIBMGA002433* | down | 519 | 527 | - | 11.1009 | 6.57E-05 | 0.709 | GAGCGAGGC |
| 1970 | mamo-s | *BGIBMGA002435* | up | 1267 | 1275 | - | 12.0734 | 2.66E-05 | 0.615 | GCGCGTGTC |
| 1971 | mamo-s | *BGIBMGA002435* | down | 1518 | 1526 | - | 11.3303 | 4.96E-05 | 0.615 | GTGGGTGGC |
| 1972 | mamo-s | *BGIBMGA002436* | down | 1505 | 1513 | + | 11.3853 | 4.74E-05 | 0.615 | ACGCGTGGC |
| 1973 | mamo-s | *BGIBMGA002437* | up | 1213 | 1221 | + | 11.3853 | 4.74E-05 | 0.615 | ACGCGTGGC |
| 1974 | mamo-s | *BGIBMGA002438* | up | 966 | 974 | + | 11.3303 | 4.96E-05 | 0.615 | GTGGGTGGC |
| 1975 | mamo-s | *BGIBMGA002441* | down | 1501 | 1509 | - | 11.7798 | 3.11E-05 | 0.615 | ATGCGTGGC |
| 1976 | mamo-s | *BGIBMGA002441* | up | 1708 | 1716 | - | 11.3303 | 4.96E-05 | 0.615 | GTGGGTGGC |
| 1977 | mamo-s | *BGIBMGA002442* | up | 261 | 269 | + | 11.3303 | 4.96E-05 | 0.615 | GTGGGTGGC |
| 1978 | mamo-s | *BGIBMGA002445* | up | 1470 | 1478 | - | 12.1835 | 1.71E-05 | 0.615 | GTGCGTGCC |
| 1979 | mamo-s | *BGIBMGA002446* | up | 1401 | 1409 | + | 11.3303 | 4.96E-05 | 0.615 | GTGGGTGGC |
| 1980 | mamo-s | *BGIBMGA002447* | up | 625 | 633 | - | 12.1284 | 2.16E-05 | 0.615 | CAGCGTGGC |
| 1981 | mamo-s | *BGIBMGA002448* | up | 963 | 971 | + | 11.3303 | 4.96E-05 | 0.615 | GTGGGTGGC |
| 1982 | mamo-s | *BGIBMGA002448* | down | 1656 | 1664 | - | 11.3303 | 4.96E-05 | 0.615 | GTGGGTGGC |
| 1983 | mamo-s | *BGIBMGA002450* | up | 883 | 891 | - | 10.5413 | 9.42E-05 | 0.857 | CAGCGTGTC |
| 1984 | mamo-s | *BGIBMGA002452* | up | 398 | 406 | + | 11.5688 | 3.79E-05 | 0.615 | CTGCGTGTC |
| 1985 | mamo-s | *BGIBMGA002455* | up | 1482 | 1490 | + | 10.9358 | 7.30E-05 | 0.728 | GCGGGTGGC |
| 1986 | mamo-s | *BGIBMGA002456* | down | 1550 | 1558 | - | 10.5413 | 9.42E-05 | 0.857 | GTGCGAGTC |
| 1987 | mamo-s | *BGIBMGA002458* | up | 433 | 441 | + | 11.1743 | 5.90E-05 | 0.684 | CCGCGTGTC |
| 1988 | mamo-s | *BGIBMGA002461* | up | 743 | 751 | - | 11.1743 | 5.90E-05 | 0.684 | CCGCGTGTC |
| 1989 | mamo-s | *BGIBMGA002462* | up | 1607 | 1615 | - | 11.3303 | 4.96E-05 | 0.615 | GTGGGTGGC |
| 1990 | mamo-s | *BGIBMGA002463* | down | 129 | 137 | - | 11.3303 | 4.96E-05 | 0.615 | GTGGGTGGC |
| 1991 | mamo-s | *BGIBMGA002463* | down | 1246 | 1254 | - | 11.0826 | 6.84E-05 | 0.709 | GTACGTGGC |
| 1992 | mamo-s | *BGIBMGA002464* | down | 606 | 614 | - | 11.3303 | 4.96E-05 | 0.615 | GTGGGTGGC |
| 1993 | mamo-s | *BGIBMGA002465* | up | 1136 | 1144 | - | 11.0826 | 6.84E-05 | 0.709 | GTACGTGGC |
| 1994 | mamo-s | *BGIBMGA002466* | down | 1919 | 1927 | + | 11.1009 | 6.57E-05 | 0.709 | GAGCGAGGC |
| 1995 | mamo-s | *BGIBMGA002467* | down | 477 | 485 | - | 11.0826 | 6.84E-05 | 0.709 | GTACGTGGC |
| 1996 | mamo-s | *BGIBMGA002468* | up | 447 | 455 | + | 12.1284 | 2.16E-05 | 0.615 | GTGCGAGGC |
| 1997 | mamo-s | *BGIBMGA002468* | up | 849 | 857 | + | 11.1376 | 6.35E-05 | 0.709 | TCGCGTGGC |
| 1998 | mamo-s | *BGIBMGA002469* | down | 268 | 276 | - | 11.3303 | 4.96E-05 | 0.615 | GTGGGTGGC |
| 1999 | mamo-s | *BGIBMGA002469* | up | 44 | 52 | + | 11.1376 | 6.35E-05 | 0.709 | TCGCGTGGC |
| 2000 | mamo-s | *BGIBMGA002470* | down | 1755 | 1763 | - | 11.3303 | 4.96E-05 | 0.615 | GTGGGTGGC |
| 2001 | mamo-s | *BGIBMGA002472* | up | 332 | 340 | - | 12.3394 | 1.49E-05 | 0.615 | GGGCGTGGC |
| 2002 | mamo-s | *BGIBMGA002472* | down | 435 | 443 | + | 11.3303 | 4.96E-05 | 0.615 | GTGGGTGGC |
| 2003 | mamo-s | *BGIBMGA002472* | up | 1144 | 1152 | - | 11.3303 | 4.96E-05 | 0.615 | GTGGGTGGC |
| 2004 | mamo-s | *BGIBMGA002473* | down | 1983 | 1991 | - | 12.4679 | 1.31E-05 | 0.615 | GTGCGTGTC |
| 2005 | mamo-s | *BGIBMGA002474* | up | 95 | 103 | - | 11.3303 | 4.96E-05 | 0.615 | GTGGGTGGC |
| 2006 | mamo-s | *BGIBMGA002474* | up | 708 | 716 | + | 11.3303 | 4.96E-05 | 0.615 | GTGGGTGGC |
| 2007 | mamo-s | *BGIBMGA002475* | down | 255 | 263 | + | 11.0826 | 6.84E-05 | 0.709 | GTACGTGGC |
| 2008 | mamo-s | *BGIBMGA002478* | down | 516 | 524 | + | 11.3303 | 4.96E-05 | 0.615 | GTGGGTGGC |
| 2009 | mamo-s | *BGIBMGA002478* | down | 905 | 913 | - | 11.1743 | 5.90E-05 | 0.684 | CCGCGTGTC |
| 2010 | mamo-s | *BGIBMGA002480* | down | 319 | 327 | + | 11.3303 | 4.96E-05 | 0.615 | GTGGGTGGC |
| 2011 | mamo-s | *BGIBMGA002480* | up | 369 | 377 | - | 11.3303 | 4.96E-05 | 0.615 | GTGGGTGGC |
| 2012 | mamo-s | *BGIBMGA002481* | up | 1030 | 1038 | - | 11.3303 | 4.96E-05 | 0.615 | GTGGGTGGC |
| 2013 | mamo-s | *BGIBMGA002481* | down | 361 | 369 | + | 10.5138 | 9.75E-05 | 0.865 | GTGCGTGTT |
| 2014 | mamo-s | *BGIBMGA002487* | down | 1468 | 1476 | + | 12.3394 | 1.49E-05 | 0.615 | GGGCGTGGC |
| 2015 | mamo-s | *BGIBMGA002489* | down | 1014 | 1022 | + | 11.3303 | 4.96E-05 | 0.615 | GTGGGTGGC |
| 2016 | mamo-s | *BGIBMGA002490* | down | 1140 | 1148 | - | 11.0826 | 6.84E-05 | 0.709 | GTACGTGGC |
| 2017 | mamo-s | *BGIBMGA002491* | down | 46 | 54 | - | 13.0275 | 8.53E-06 | 0.615 | GAGCGTGGC |

| | | | | | | | | | | |
|---|---|---|---|---|---|---|---|---|---|---|
| 2018 | mamo-s | *BGIBMGA002491* | up | 1206 | 1214 | - | 11.3303 | 4.96E-05 | 0.615 | GTGGGTGGC |
| 2019 | mamo-s | *BGIBMGA002491* | down | 810 | 818 | + | 11.1376 | 6.35E-05 | 0.709 | TCGCGTGGC |
| 2020 | mamo-s | *BGIBMGA002495* | up | 909 | 917 | - | 11.3303 | 4.96E-05 | 0.615 | GTGGGTGGC |
| 2021 | mamo-s | *BGIBMGA002498* | up | 959 | 967 | - | 13.0275 | 8.53E-06 | 0.615 | GAGCGTGGC |
| 2022 | mamo-s | *BGIBMGA002498* | up | 474 | 482 | + | 11.3303 | 4.96E-05 | 0.615 | GTGGGTGGC |
| 2023 | mamo-s | *BGIBMGA002499* | up | 129 | 137 | + | 11.2294 | 5.41E-05 | 0.65 | CTGCGAGGC |
| 2024 | mamo-s | *BGIBMGA002500* | up | 36 | 44 | - | 11.3303 | 4.96E-05 | 0.615 | GTGGGTGGC |
| 2025 | mamo-s | *BGIBMGA002500* | up | 1579 | 1587 | - | 11.3303 | 4.96E-05 | 0.615 | GTGGGTGGC |
| 2026 | mamo-s | *BGIBMGA002500* | down | 1851 | 1859 | - | 11.3303 | 4.96E-05 | 0.615 | GTGGGTGGC |
| 2027 | mamo-s | *BGIBMGA002505* | up | 1035 | 1043 | + | 13.6606 | 4.06E-06 | 0.615 | GCGCGTGGC |
| 2028 | mamo-s | *BGIBMGA002506* | up | 649 | 657 | - | 11.3303 | 4.96E-05 | 0.615 | GTGGGTGGC |
| 2029 | mamo-s | *BGIBMGA002506* | up | 1411 | 1419 | + | 11.3303 | 4.96E-05 | 0.615 | GTGGGTGGC |
| 2030 | mamo-s | *BGIBMGA002509* | up | 1849 | 1857 | - | 11.3303 | 4.96E-05 | 0.615 | GTGGGTGGC |
| 2031 | mamo-s | *BGIBMGA002510* | down | 1589 | 1597 | + | 12.0734 | 2.66E-05 | 0.615 | GCGCGTGTC |
| 2032 | mamo-s | *BGIBMGA002510* | up | 32 | 40 | + | 11.3303 | 4.96E-05 | 0.615 | GTGGGTGGC |
| 2033 | mamo-s | *BGIBMGA002513* | up | 1985 | 1993 | + | 14.055 | 2.23E-06 | 0.615 | GTGCGTGGC |
| 2034 | mamo-s | *BGIBMGA002513* | up | 1220 | 1228 | + | 11.3303 | 4.96E-05 | 0.615 | GTGGGTGGC |
| 2035 | mamo-s | *BGIBMGA002513* | up | 187 | 195 | - | 11.1743 | 5.90E-05 | 0.684 | CCGCGTGTC |
| 2036 | mamo-s | *BGIBMGA002516* | down | 719 | 727 | - | 11.3303 | 4.96E-05 | 0.615 | GTGGGTGGC |
| 2037 | mamo-s | *BGIBMGA002517* | down | 1257 | 1265 | - | 11.789 | 2.84E-05 | 0.615 | GCGCGTGCC |
| 2038 | mamo-s | *BGIBMGA002518* | up | 424 | 432 | - | 11.3303 | 4.96E-05 | 0.615 | GTGGGTGGC |
| 2039 | mamo-s | *BGIBMGA002519* | down | 363 | 371 | + | 11.3303 | 4.96E-05 | 0.615 | GTGGGTGGC |
| 2040 | mamo-s | *BGIBMGA002520* | up | 1002 | 1010 | + | 11.3303 | 4.96E-05 | 0.615 | GTGGGTGGC |
| 2041 | mamo-s | *BGIBMGA002522* | down | 1710 | 1718 | + | 11.2844 | 5.18E-05 | 0.63 | CTGCGTGCC |
| 2042 | mamo-s | *BGIBMGA002523* | down | 937 | 945 | - | 13.6606 | 4.06E-06 | 0.615 | GCGCGTGGC |
| 2043 | mamo-s | *BGIBMGA002523* | down | 599 | 607 | + | 11.7798 | 3.11E-05 | 0.615 | ATGCGTGGC |
| 2044 | mamo-s | *BGIBMGA002525* | up | 393 | 401 | - | 11.3303 | 4.96E-05 | 0.615 | GTGGGTGGC |
| 2045 | mamo-s | *BGIBMGA002526* | up | 720 | 728 | - | 11.0826 | 6.84E-05 | 0.709 | GTACGTGGC |
| 2046 | mamo-s | *BGIBMGA002527* | up | 1667 | 1675 | - | 11.3303 | 4.96E-05 | 0.615 | GTGGGTGGC |
| 2047 | mamo-s | *BGIBMGA002527* | down | 666 | 674 | - | 11.0826 | 6.84E-05 | 0.709 | GTACGTGGC |
| 2048 | mamo-s | *BGIBMGA002528* | down | 1478 | 1486 | - | 14.055 | 2.23E-06 | 0.615 | GTGCGTGGC |
| 2049 | mamo-s | *BGIBMGA002529* | down | 1330 | 1338 | + | 11.3303 | 4.96E-05 | 0.615 | GTGGGTGGC |
| 2050 | mamo-s | *BGIBMGA002529* | down | 1804 | 1812 | - | 11.3303 | 4.96E-05 | 0.615 | GTGGGTGGC |
| 2051 | mamo-s | *BGIBMGA002532* | up | 582 | 590 | - | 11.0826 | 6.84E-05 | 0.709 | GTACGTGGC |
| 2052 | mamo-s | *BGIBMGA002533* | down | 658 | 666 | + | 11.3303 | 4.96E-05 | 0.615 | GTGGGTGGC |
| 2053 | mamo-s | *BGIBMGA002533* | down | 1919 | 1927 | + | 11.1009 | 6.57E-05 | 0.709 | GAGCGAGGC |
| 2054 | mamo-s | *BGIBMGA002534* | up | 523 | 531 | - | 11.3303 | 4.96E-05 | 0.615 | GTGGGTGGC |
| 2055 | mamo-s | *BGIBMGA002534* | up | 1775 | 1783 | - | 11.3303 | 4.96E-05 | 0.615 | GTGGGTGGC |
| 2056 | mamo-s | *BGIBMGA002535* | up | 1168 | 1176 | + | 11.3303 | 4.96E-05 | 0.615 | GTGGGTGGC |
| 2057 | mamo-s | *BGIBMGA002536* | up | 679 | 687 | + | 11.3303 | 4.96E-05 | 0.615 | GTGGGTGGC |
| 2058 | mamo-s | *BGIBMGA002536* | down | 860 | 868 | + | 11.3303 | 4.96E-05 | 0.615 | GTGGGTGGC |
| 2059 | mamo-s | *BGIBMGA002536* | up | 1739 | 1747 | - | 11.3303 | 4.96E-05 | 0.615 | GTGGGTGGC |
| 2060 | mamo-s | *BGIBMGA002536* | down | 1821 | 1829 | - | 11.3303 | 4.96E-05 | 0.615 | GTGGGTGGC |
| 2061 | mamo-s | *BGIBMGA002537* | up | 117 | 125 | + | 11.3303 | 4.96E-05 | 0.615 | GTGGGTGGC |
| 2062 | mamo-s | *BGIBMGA002538* | down | 1208 | 1216 | + | 11.3303 | 4.96E-05 | 0.615 | GTGGGTGGC |
| 2063 | mamo-s | *BGIBMGA002539* | up | 56 | 64 | - | 12.4679 | 1.31E-05 | 0.615 | GTGCGTGTC |
| 2064 | mamo-s | *BGIBMGA002539* | down | 1389 | 1397 | - | 11.3303 | 4.96E-05 | 0.615 | GTGGGTGGC |
| 2065 | mamo-s | *BGIBMGA002541* | up | 780 | 788 | + | 11.3303 | 4.96E-05 | 0.615 | GTGGGTGGC |
| 2066 | mamo-s | *BGIBMGA002542* | up | 1167 | 1175 | - | 11.3303 | 4.96E-05 | 0.615 | GTGGGTGGC |
| 2067 | mamo-s | *BGIBMGA002542* | down | 924 | 932 | - | 11.1009 | 6.57E-05 | 0.709 | GAGCGAGGC |
| 2068 | mamo-s | *BGIBMGA002543* | down | 755 | 763 | + | 11.2844 | 5.18E-05 | 0.63 | CTGCGTGCC |
| 2069 | mamo-s | *BGIBMGA002544* | up | 694 | 702 | - | 11.3303 | 4.96E-05 | 0.615 | GTGGGTGGC |
| 2070 | mamo-s | *BGIBMGA002544* | down | 1527 | 1535 | + | 11.0826 | 6.84E-05 | 0.709 | GTACGTGGC |
| 2071 | mamo-s | *BGIBMGA002545* | down | 1838 | 1846 | - | 11.3303 | 4.96E-05 | 0.615 | GTGGGTGGC |
| 2072 | mamo-s | *BGIBMGA002548* | down | 570 | 578 | + | 11.3303 | 4.96E-05 | 0.615 | GTGGGTGGC |
| 2073 | mamo-s | *BGIBMGA002548* | down | 1286 | 1294 | - | 11.3303 | 4.96E-05 | 0.615 | GTGGGTGGC |
| 2074 | mamo-s | *BGIBMGA002549* | down | 795 | 803 | + | 11.3303 | 4.96E-05 | 0.615 | GTGGGTGGC |
| 2075 | mamo-s | *BGIBMGA002549* | up | 122 | 130 | + | 11.1009 | 6.57E-05 | 0.709 | GAGCGAGGC |
| 2076 | mamo-s | *BGIBMGA002550* | up | 812 | 820 | - | 11.3303 | 4.96E-05 | 0.615 | GTGGGTGGC |
| 2077 | mamo-s | *BGIBMGA002552* | up | 198 | 206 | + | 11.5321 | 4.06E-05 | 0.615 | TTGCGTGGC |
| 2078 | mamo-s | *BGIBMGA002554* | down | 1150 | 1158 | - | 11.3303 | 4.96E-05 | 0.615 | GTGGGTGGC |
| 2079 | mamo-s | *BGIBMGA002557* | up | 243 | 251 | + | 14.055 | 2.23E-06 | 0.615 | GTGCGTGGC |
| 2080 | mamo-s | *BGIBMGA002562* | down | 1083 | 1091 | + | 11.3303 | 4.96E-05 | 0.615 | GTGGGTGGC |
| 2081 | mamo-s | *BGIBMGA002562* | down | 1579 | 1587 | - | 11.3303 | 4.96E-05 | 0.615 | GTGGGTGGC |
| 2082 | mamo-s | *BGIBMGA002563* | down | 467 | 475 | + | 11.3303 | 4.96E-05 | 0.615 | GTGGGTGGC |
| 2083 | mamo-s | *BGIBMGA002563* | up | 593 | 601 | - | 11.3303 | 4.96E-05 | 0.615 | GTGGGTGGC |
| 2084 | mamo-s | *BGIBMGA002563* | down | 698 | 706 | + | 11.3303 | 4.96E-05 | 0.615 | GTGGGTGGC |
| 2085 | mamo-s | *BGIBMGA002563* | down | 931 | 939 | + | 11.3303 | 4.96E-05 | 0.615 | GTGGGTGGC |
| 2086 | mamo-s | *BGIBMGA002564* | up | 118 | 126 | + | 11.3303 | 4.96E-05 | 0.615 | GTGGGTGGC |
| 2087 | mamo-s | *BGIBMGA002565* | down | 1942 | 1950 | - | 11.3303 | 4.96E-05 | 0.615 | GTGGGTGGC |
| 2088 | mamo-s | *BGIBMGA002566* | down | 670 | 678 | - | 11.3303 | 4.96E-05 | 0.615 | GTGGGTGGC |
| 2089 | mamo-s | *BGIBMGA002567* | down | 1459 | 1467 | + | 11.3303 | 4.96E-05 | 0.615 | GTGGGTGGC |
| 2090 | mamo-s | *BGIBMGA002568* | up | 309 | 317 | + | 11.3303 | 4.96E-05 | 0.615 | GTGGGTGGC |
| 2091 | mamo-s | *BGIBMGA002569* | up | 684 | 692 | + | 11.3303 | 4.96E-05 | 0.615 | GTGGGTGGC |
| 2092 | mamo-s | *BGIBMGA002571* | down | 1472 | 1480 | - | 10.6881 | 8.61E-05 | 0.81 | GCACGTGGC |
| 2093 | mamo-s | *BGIBMGA002574* | down | 1106 | 1114 | - | 11.3303 | 4.96E-05 | 0.615 | GTGGGTGGC |
| 2094 | mamo-s | *BGIBMGA002576* | down | 1257 | 1265 | + | 11.3303 | 4.96E-05 | 0.615 | GTGGGTGGC |
| 2095 | mamo-s | *BGIBMGA002576* | down | 1911 | 1919 | - | 11.3303 | 4.96E-05 | 0.615 | GTGGGTGGC |
| 2096 | mamo-s | *BGIBMGA002578* | up | 165 | 173 | + | 11.3303 | 4.96E-05 | 0.615 | GTGGGTGGC |
| 2097 | mamo-s | *BGIBMGA002580* | down | 1510 | 1518 | - | 12.3394 | 1.49E-05 | 0.615 | GGGCGTGGC |
| 2098 | mamo-s | *BGIBMGA002581* | up | 1833 | 1841 | + | 11.3303 | 4.96E-05 | 0.615 | GTGGGTGGC |
| 2099 | mamo-s | *BGIBMGA002582* | down | 523 | 531 | - | 11.3303 | 4.96E-05 | 0.615 | GTGGGTGGC |
| 2100 | mamo-s | *BGIBMGA002583* | up | 1961 | 1969 | + | 10.5138 | 9.75E-05 | 0.865 | GTGCGTGTT |
| 2101 | mamo-s | *BGIBMGA002585* | down | 1418 | 1426 | - | 10.5138 | 9.75E-05 | 0.865 | GTGCGTGTT |
| 2102 | mamo-s | *BGIBMGA002588* | up | 504 | 512 | + | 11.3303 | 4.96E-05 | 0.615 | GTGGGTGGC |
| 2103 | mamo-s | *BGIBMGA002588* | up | 1799 | 1807 | - | 11.3303 | 4.96E-05 | 0.615 | GTGGGTGGC |
| 2104 | mamo-s | *BGIBMGA002590* | down | 1575 | 1583 | + | 11.3303 | 4.96E-05 | 0.615 | GTGGGTGGC |
| 2105 | mamo-s | *BGIBMGA002593* | up | 265 | 273 | + | 11.4404 | 4.51E-05 | 0.615 | GAGCGTGTC |

| | | | | | | | | | | |
|---|---|---|---|---|---|---|---|---|---|---|
| 2106 | mamo-s | *BGIBMGA002593* | down | 1223 | 1231 | - | 10.5413 | 9.42E-05 | 0.857 | GTGCGAGTC |
| 2107 | mamo-s | *BGIBMGA002595* | down | 1481 | 1489 | + | 10.5138 | 9.75E-05 | 0.865 | GTGCGTGTT |
| 2108 | mamo-s | *BGIBMGA002597* | down | 863 | 871 | - | 11.3303 | 4.96E-05 | 0.615 | GTGGGTGGC |
| 2109 | mamo-s | *BGIBMGA002598* | up | 1193 | 1201 | + | 10.9358 | 7.30E-05 | 0.728 | GCGGGTGGC |
| 2110 | mamo-s | *BGIBMGA002599* | down | 978 | 986 | + | 11.3303 | 4.96E-05 | 0.615 | GTGGGTGGC |
| 2111 | mamo-s | *BGIBMGA002599* | up | 1252 | 1260 | - | 10.5413 | 9.42E-05 | 0.857 | CAGCGTGTC |
| 2112 | mamo-s | *BGIBMGA002600* | up | 1907 | 1915 | + | 11.789 | 2.84E-05 | 0.615 | GCGCGTGCC |
| 2113 | mamo-s | *BGIBMGA002604* | up | 406 | 414 | + | 10.5138 | 9.75E-05 | 0.865 | GTGCGTGTT |
| 2114 | mamo-s | *BGIBMGA002606* | up | 1709 | 1717 | - | 11.3303 | 4.96E-05 | 0.615 | GTGGGTGGC |
| 2115 | mamo-s | *BGIBMGA002606* | up | 781 | 789 | - | 10.8349 | 7.66E-05 | 0.751 | CCGCGAGGC |
| 2116 | mamo-s | *BGIBMGA002608* | down | 1326 | 1334 | + | 11.3303 | 4.96E-05 | 0.615 | GTGGGTGGC |
| 2117 | mamo-s | *BGIBMGA002612* | up | 747 | 755 | - | 11.3303 | 4.96E-05 | 0.615 | GTGGGTGGC |
| 2118 | mamo-s | *BGIBMGA002614* | up | 824 | 832 | - | 11.3303 | 4.96E-05 | 0.615 | GTGGGTGGC |
| 2119 | mamo-s | *BGIBMGA002615* | up | 1734 | 1742 | - | 13.0275 | 8.53E-06 | 0.615 | GAGCGTGGC |
| 2120 | mamo-s | *BGIBMGA002617* | down | 1697 | 1705 | + | 11.3303 | 4.96E-05 | 0.615 | GTGGGTGGC |
| 2121 | mamo-s | *BGIBMGA002618* | up | 1161 | 1169 | + | 11.3303 | 4.96E-05 | 0.615 | GTGGGTGGC |
| 2122 | mamo-s | *BGIBMGA002618* | down | 1807 | 1815 | - | 11.2018 | 5.68E-05 | 0.674 | CTGCGTGGT |
| 2123 | mamo-s | *BGIBMGA002620* | up | 895 | 903 | + | 11.3303 | 4.96E-05 | 0.615 | GTGGGTGGC |
| 2124 | mamo-s | *BGIBMGA002620* | down | 1502 | 1510 | - | 11.3303 | 4.96E-05 | 0.615 | GTGGGTGGC |
| 2125 | mamo-s | *BGIBMGA002621* | down | 1637 | 1645 | - | 13.6606 | 4.06E-06 | 0.615 | GCGCGTGGC |
| 2126 | mamo-s | *BGIBMGA002621* | up | 829 | 837 | + | 11.3303 | 4.96E-05 | 0.615 | GTGGGTGGC |
| 2127 | mamo-s | *BGIBMGA002622* | up | 1164 | 1172 | - | 13.6606 | 4.06E-06 | 0.615 | GCGCGTGGC |
| 2128 | mamo-s | *BGIBMGA002627* | down | 422 | 430 | + | 11.7064 | 3.52E-05 | 0.615 | GCGCGTGGT |
| 2129 | mamo-s | *BGIBMGA002627* | down | 1881 | 1889 | + | 11.3303 | 4.96E-05 | 0.615 | GTGGGTGGC |
| 2130 | mamo-s | *BGIBMGA002628* | down | 1209 | 1217 | + | 11.2018 | 5.68E-05 | 0.674 | CTGCGTGGT |
| 2131 | mamo-s | *BGIBMGA002628* | down | 1965 | 1973 | + | 11.1743 | 5.90E-05 | 0.684 | CCGCGTGTC |
| 2132 | mamo-s | *BGIBMGA002629* | up | 235 | 243 | + | 11.3303 | 4.96E-05 | 0.615 | GTGGGTGGC |
| 2133 | mamo-s | *BGIBMGA002629* | down | 1689 | 1697 | + | 11.3303 | 4.96E-05 | 0.615 | GTGGGTGGC |
| 2134 | mamo-s | *BGIBMGA002630* | up | 1364 | 1372 | + | 11.2844 | 5.18E-05 | 0.63 | CTGCGTGCC |
| 2135 | mamo-s | *BGIBMGA002630* | up | 1022 | 1030 | + | 11.1009 | 6.57E-05 | 0.709 | GAGCGAGGC |
| 2136 | mamo-s | *BGIBMGA002631* | up | 1406 | 1414 | - | 10.9358 | 7.30E-05 | 0.728 | GCGGGTGGC |
| 2137 | mamo-s | *BGIBMGA002633* | down | 767 | 775 | - | 11.3303 | 4.96E-05 | 0.615 | GTGGGTGGC |
| 2138 | mamo-s | *BGIBMGA002634* | up | 1170 | 1178 | + | 11.5321 | 4.06E-05 | 0.615 | TTGCGTGGC |
| 2139 | mamo-s | *BGIBMGA002635* | up | 779 | 787 | - | 11.3303 | 4.96E-05 | 0.615 | GTGGGTGGC |
| 2140 | mamo-s | *BGIBMGA002635* | down | 1619 | 1627 | + | 11.0826 | 6.84E-05 | 0.709 | GTACGTGGC |
| 2141 | mamo-s | *BGIBMGA002639* | down | 1568 | 1576 | + | 11.4404 | 4.51E-05 | 0.615 | CGGCGTGGC |
| 2142 | mamo-s | *BGIBMGA002640* | down | 1037 | 1045 | + | 11.3303 | 4.96E-05 | 0.615 | GTGGGTGGC |
| 2143 | mamo-s | *BGIBMGA002640* | up | 1162 | 1170 | + | 11.0826 | 6.84E-05 | 0.709 | GTACGTGGC |
| 2144 | mamo-s | *BGIBMGA002640* | down | 1646 | 1654 | - | 10.5413 | 9.42E-05 | 0.857 | CAGCGTGTC |
| 2145 | mamo-s | *BGIBMGA002641* | down | 1223 | 1231 | - | 11.3853 | 4.74E-05 | 0.615 | ACGCGTGGC |
| 2146 | mamo-s | *BGIBMGA002641* | down | 493 | 501 | - | 11.1743 | 5.90E-05 | 0.684 | CCGCGTGTC |
| 2147 | mamo-s | *BGIBMGA002644* | up | 1665 | 1673 | + | 11.789 | 2.84E-05 | 0.615 | GCGCGTGCC |
| 2148 | mamo-s | *BGIBMGA002645* | down | 1148 | 1156 | + | 12.4679 | 1.31E-05 | 0.615 | GTGCGTGTC |
| 2149 | mamo-s | *BGIBMGA002645* | down | 1379 | 1387 | - | 11.3303 | 4.96E-05 | 0.615 | GTGGGTGGC |
| 2150 | mamo-s | *BGIBMGA002647* | up | 1890 | 1898 | - | 12.1284 | 2.16E-05 | 0.615 | CAGCGTGGC |
| 2151 | mamo-s | *BGIBMGA002650* | up | 314 | 322 | + | 11.1376 | 6.35E-05 | 0.709 | TCGCGTGGC |
| 2152 | mamo-s | *BGIBMGA002651* | down | 1187 | 1195 | + | 12.1835 | 1.71E-05 | 0.615 | GTGCGTGCC |
| 2153 | mamo-s | *BGIBMGA002651* | down | 1079 | 1087 | + | 11.7064 | 3.52E-05 | 0.615 | GCGCGTGGT |
| 2154 | mamo-s | *BGIBMGA002651* | up | 287 | 295 | + | 11.3303 | 4.96E-05 | 0.615 | GTGGGTGGC |
| 2155 | mamo-s | *BGIBMGA002651* | down | 774 | 782 | + | 11.3303 | 4.96E-05 | 0.615 | GTGGGTGGC |
| 2156 | mamo-s | *BGIBMGA002652* | up | 1477 | 1485 | + | 12.1835 | 1.71E-05 | 0.615 | GTGCGTGCC |
| 2157 | mamo-s | *BGIBMGA002652* | up | 1369 | 1377 | + | 11.7064 | 3.52E-05 | 0.615 | GCGCGTGGT |
| 2158 | mamo-s | *BGIBMGA002652* | up | 1064 | 1072 | + | 11.3303 | 4.96E-05 | 0.615 | GTGGGTGGC |
| 2159 | mamo-s | *BGIBMGA002656* | down | 1347 | 1355 | - | 11.2844 | 5.18E-05 | 0.63 | CTGCGTGCC |
| 2160 | mamo-s | *BGIBMGA002658* | up | 421 | 429 | - | 11.0826 | 6.84E-05 | 0.709 | GTACGTGGC |
| 2161 | mamo-s | *BGIBMGA002660* | down | 582 | 590 | + | 11.3303 | 4.96E-05 | 0.615 | GTGGGTGGC |
| 2162 | mamo-s | *BGIBMGA002661* | up | 1176 | 1184 | + | 11.3853 | 4.74E-05 | 0.615 | ACGCGTGGC |
| 2163 | mamo-s | *BGIBMGA002661* | down | 843 | 851 | - | 11.3303 | 4.96E-05 | 0.615 | GTGGGTGGC |
| 2164 | mamo-s | *BGIBMGA002662* | up | 816 | 824 | - | 11.4404 | 4.51E-05 | 0.615 | CGGCGTGGC |
| 2165 | mamo-s | *BGIBMGA002663* | down | 1249 | 1257 | - | 11.3303 | 4.96E-05 | 0.615 | GTGGGTGGC |
| 2166 | mamo-s | *BGIBMGA002664* | down | 1980 | 1988 | + | 12.1009 | 2.43E-05 | 0.615 | GTGCGTGGT |
| 2167 | mamo-s | *BGIBMGA002664* | up | 1589 | 1597 | + | 11.3303 | 4.96E-05 | 0.615 | GTGGGTGGC |
| 2168 | mamo-s | *BGIBMGA002664* | up | 1803 | 1811 | + | 10.7523 | 8.38E-05 | 0.797 | AAGCGTGGC |
| 2169 | mamo-s | *BGIBMGA002665* | up | 119 | 127 | + | 11.3303 | 4.96E-05 | 0.615 | GTGGGTGGC |
| 2170 | mamo-s | *BGIBMGA002665* | up | 626 | 634 | - | 11.3303 | 4.96E-05 | 0.615 | GTGGGTGGC |
| 2171 | mamo-s | *BGIBMGA002668* | down | 742 | 750 | - | 10.8073 | 7.89E-05 | 0.766 | CCGCGTGGT |
| 2172 | mamo-s | *BGIBMGA002669* | down | 1847 | 1855 | - | 11.3303 | 4.96E-05 | 0.615 | GTGGGTGGC |
| 2173 | mamo-s | *BGIBMGA002670* | down | 1668 | 1676 | + | 11.0826 | 6.84E-05 | 0.709 | GTACGTGGC |
| 2174 | mamo-s | *BGIBMGA002671* | up | 1065 | 1073 | - | 11.3303 | 4.96E-05 | 0.615 | GTGGGTGGC |
| 2175 | mamo-s | *BGIBMGA002671* | down | 1099 | 1107 | + | 11.3303 | 4.96E-05 | 0.615 | GTGGGTGGC |
| 2176 | mamo-s | *BGIBMGA002672* | up | 1702 | 1710 | + | 11.3303 | 4.96E-05 | 0.615 | GTGGGTGGC |
| 2177 | mamo-s | *BGIBMGA002673* | up | 1027 | 1035 | - | 10.5413 | 9.42E-05 | 0.857 | CAGCGTGTC |
| 2178 | mamo-s | *BGIBMGA002674* | down | 673 | 681 | + | 11.2018 | 5.68E-05 | 0.674 | CTGCGTGGT |
| 2179 | mamo-s | *BGIBMGA002678* | up | 734 | 742 | + | 11.3303 | 4.96E-05 | 0.615 | GTGGGTGGC |
| 2180 | mamo-s | *BGIBMGA002679* | up | 1777 | 1785 | + | 10.5413 | 9.42E-05 | 0.857 | CAGCGTGTC |
| 2181 | mamo-s | *BGIBMGA002682* | down | 247 | 255 | + | 11.0826 | 6.84E-05 | 0.709 | GTACGTGGC |
| 2182 | mamo-s | *BGIBMGA002685* | down | 974 | 982 | - | 10.7523 | 8.38E-05 | 0.797 | AAGCGTGGC |
| 2183 | mamo-s | *BGIBMGA002685* | up | 1792 | 1800 | - | 10.6881 | 8.61E-05 | 0.81 | GCACGTGGC |
| 2184 | mamo-s | *BGIBMGA002687* | down | 1324 | 1332 | + | 10.7523 | 8.38E-05 | 0.797 | GGGCGTGTC |
| 2185 | mamo-s | *BGIBMGA002688* | down | 499 | 507 | + | 11.3303 | 4.96E-05 | 0.615 | GTGGGTGGC |
| 2186 | mamo-s | *BGIBMGA002692* | down | 454 | 462 | - | 11.3303 | 4.96E-05 | 0.615 | GTGGGTGGC |
| 2187 | mamo-s | *BGIBMGA002692* | down | 617 | 625 | - | 11.3303 | 4.96E-05 | 0.615 | GTGGGTGGC |
| 2188 | mamo-s | *BGIBMGA002695* | down | 805 | 813 | + | 10.5413 | 9.42E-05 | 0.857 | GTGCGAGTC |
| 2189 | mamo-s | *BGIBMGA002696* | up | 978 | 986 | + | 11.3303 | 4.96E-05 | 0.615 | GTGGGTGGC |
| 2190 | mamo-s | *BGIBMGA002697* | down | 425 | 433 | + | 11.3303 | 4.96E-05 | 0.615 | GTGGGTGGC |
| 2191 | mamo-s | *BGIBMGA002699* | down | 140 | 148 | + | 11.0826 | 6.84E-05 | 0.709 | GTACGTGGC |
| 2192 | mamo-s | *BGIBMGA002700* | down | 799 | 807 | - | 11.3303 | 4.96E-05 | 0.615 | GTGGGTGGC |
| 2193 | mamo-s | *BGIBMGA002701* | up | 1035 | 1043 | + | 12.1009 | 2.43E-05 | 0.615 | GTGCGTGGT |

| | | | | | | | | | | |
|---|---|---|---|---|---|---|---|---|---|---|
| 2194 | mamo-s | *BGIBMGA002701* | down | 1084 | 1092 | + | 11.3303 | 4.96E-05 | 0.615 | GTGGGTGGC |
| 2195 | mamo-s | *BGIBMGA002702* | up | 1850 | 1858 | + | 12.1835 | 1.71E-05 | 0.615 | GTGCGTGCC |
| 2196 | mamo-s | *BGIBMGA002702* | down | 1881 | 1889 | + | 11.4404 | 4.51E-05 | 0.615 | GAGCGTGTC |
| 2197 | mamo-s | *BGIBMGA002702* | up | 1083 | 1091 | - | 11.3303 | 4.96E-05 | 0.615 | GTGGGTGGC |
| 2198 | mamo-s | *BGIBMGA002703* | down | 1032 | 1040 | - | 13.0275 | 8.53E-06 | 0.615 | GAGCGTGGC |
| 2199 | mamo-s | *BGIBMGA002703* | up | 1533 | 1541 | + | 11.3303 | 4.96E-05 | 0.615 | GTGGGTGGC |
| 2200 | mamo-s | *BGIBMGA002703* | up | 594 | 602 | + | 11.0826 | 6.84E-05 | 0.709 | GTACGTGGC |
| 2201 | mamo-s | *BGIBMGA002704* | up | 863 | 871 | + | 11.3303 | 4.96E-05 | 0.615 | GTGGGTGGC |
| 2202 | mamo-s | *BGIBMGA002705* | down | 1509 | 1517 | + | 11.3303 | 4.96E-05 | 0.615 | GTGGGTGGC |
| 2203 | mamo-s | *BGIBMGA002707* | up | 1787 | 1795 | + | 11.3853 | 4.74E-05 | 0.615 | ACGCGTGGC |
| 2204 | mamo-s | *BGIBMGA002708* | down | 1176 | 1184 | + | 11.3303 | 4.96E-05 | 0.615 | GTGGGTGGC |
| 2205 | mamo-s | *BGIBMGA002709* | down | 1605 | 1613 | + | 13.156 | 6.30E-06 | 0.615 | CTGCGTGGC |
| 2206 | mamo-s | *BGIBMGA002710* | up | 719 | 727 | + | 13.156 | 6.30E-06 | 0.615 | CTGCGTGGC |
| 2207 | mamo-s | *BGIBMGA002710* | down | 1393 | 1401 | + | 11.3303 | 4.96E-05 | 0.615 | GTGGGTGGC |
| 2208 | mamo-s | *BGIBMGA002711* | up | 1854 | 1862 | + | 11.3303 | 4.96E-05 | 0.615 | GTGGGTGGC |
| 2209 | mamo-s | *BGIBMGA002711* | down | 1381 | 1389 | - | 11.0826 | 6.84E-05 | 0.709 | GTACGTGGC |
| 2210 | mamo-s | *BGIBMGA002713* | up | 726 | 734 | + | 11.3303 | 4.96E-05 | 0.615 | GTGGGTGGC |
| 2211 | mamo-s | *BGIBMGA002714* | up | 1727 | 1735 | - | 11.3303 | 4.96E-05 | 0.615 | GTGGGTGGC |
| 2212 | mamo-s | *BGIBMGA002714* | up | 85 | 93 | - | 11.1009 | 6.57E-05 | 0.709 | GAGCGAGGC |
| 2213 | mamo-s | *BGIBMGA002714* | down | 21 | 29 | - | 10.6881 | 8.61E-05 | 0.81 | GCACGTGGC |
| 2214 | mamo-s | *BGIBMGA002715* | up | 1313 | 1321 | + | 11.3303 | 4.96E-05 | 0.615 | GTGGGTGGC |
| 2215 | mamo-s | *BGIBMGA002716* | down | 271 | 279 | + | 11.3303 | 4.96E-05 | 0.615 | GTGGGTGGC |
| 2216 | mamo-s | *BGIBMGA002717* | down | 1532 | 1540 | - | 11.3303 | 4.96E-05 | 0.615 | GTGGGTGGC |
| 2217 | mamo-s | *BGIBMGA002721* | up | 157 | 165 | - | 11.3303 | 4.96E-05 | 0.615 | GTGGGTGGC |
| 2218 | mamo-s | *BGIBMGA002721* | down | 1309 | 1317 | - | 11.3303 | 4.96E-05 | 0.615 | GTGGGTGGC |
| 2219 | mamo-s | *BGIBMGA002722* | up | 1981 | 1989 | + | 11.5688 | 3.79E-05 | 0.615 | CTGCGTGTC |
| 2220 | mamo-s | *BGIBMGA002722* | up | 455 | 463 | + | 11.3303 | 4.96E-05 | 0.615 | GTGGGTGGC |
| 2221 | mamo-s | *BGIBMGA002722* | up | 844 | 852 | + | 11.3303 | 4.96E-05 | 0.615 | GTGGGTGGC |
| 2222 | mamo-s | *BGIBMGA002723* | up | 1058 | 1066 | + | 13.156 | 6.30E-06 | 0.615 | CTGCGTGGC |
| 2223 | mamo-s | *BGIBMGA002723* | up | 1372 | 1380 | - | 12.1284 | 2.16E-05 | 0.615 | CAGCGTGGC |
| 2224 | mamo-s | *BGIBMGA002723* | down | 1808 | 1816 | + | 11.3303 | 4.96E-05 | 0.615 | GTGGGTGGC |
| 2225 | mamo-s | *BGIBMGA002724* | up | 1988 | 1996 | + | 12.4679 | 1.31E-05 | 0.615 | GTGCGTGTC |
| 2226 | mamo-s | *BGIBMGA002724* | down | 572 | 580 | + | 11.3303 | 4.96E-05 | 0.615 | GTGGGTGGC |
| 2227 | mamo-s | *BGIBMGA002724* | up | 1416 | 1424 | - | 11.3303 | 4.96E-05 | 0.615 | GTGGGTGGC |
| 2228 | mamo-s | *BGIBMGA002724* | down | 1440 | 1448 | - | 11.3303 | 4.96E-05 | 0.615 | GTGGGTGGC |
| 2229 | mamo-s | *BGIBMGA002728* | down | 1004 | 1012 | - | 11.3303 | 4.96E-05 | 0.615 | GTGGGTGGC |
| 2230 | mamo-s | *BGIBMGA002729* | up | 401 | 409 | - | 11.3303 | 4.96E-05 | 0.615 | GTGGGTGGC |
| 2231 | mamo-s | *BGIBMGA002730* | up | 68 | 76 | - | 11.3303 | 4.96E-05 | 0.615 | GTGGGTGGC |
| 2232 | mamo-s | *BGIBMGA002732* | down | 1680 | 1688 | - | 11.3303 | 4.96E-05 | 0.615 | GTGGGTGGC |
| 2233 | mamo-s | *BGIBMGA002733* | up | 242 | 250 | + | 12.0734 | 2.66E-05 | 0.615 | GCGCGTGTC |
| 2234 | mamo-s | *BGIBMGA002733* | up | 403 | 411 | + | 11.3303 | 4.96E-05 | 0.615 | GTGGGTGGC |
| 2235 | mamo-s | *BGIBMGA002733* | down | 1973 | 1981 | + | 11.3303 | 4.96E-05 | 0.615 | GTGGGTGGC |
| 2236 | mamo-s | *BGIBMGA002735* | up | 352 | 360 | - | 11.3303 | 4.96E-05 | 0.615 | GTGGGTGGC |
| 2237 | mamo-s | *BGIBMGA002735* | down | 1499 | 1507 | - | 11.3303 | 4.96E-05 | 0.615 | GTGGGTGGC |
| 2238 | mamo-s | *BGIBMGA002738* | up | 1135 | 1143 | + | 11.3303 | 4.96E-05 | 0.615 | GTGGGTGGC |
| 2239 | mamo-s | *BGIBMGA002743* | up | 681 | 689 | - | 11.4404 | 4.51E-05 | 0.615 | CGGCGTGGC |
| 2240 | mamo-s | *BGIBMGA002743* | down | 807 | 815 | - | 11.3303 | 4.96E-05 | 0.615 | GTGGGTGGC |
| 2241 | mamo-s | *BGIBMGA002743* | up | 1907 | 1915 | + | 11.1009 | 6.57E-05 | 0.709 | GAGCGAGGC |
| 2242 | mamo-s | *BGIBMGA002744* | down | 702 | 710 | - | 11.3303 | 4.96E-05 | 0.615 | GTGGGTGGC |
| 2243 | mamo-s | *BGIBMGA002744* | up | 1218 | 1226 | + | 11.3303 | 4.96E-05 | 0.615 | GTGGGTGGC |
| 2244 | mamo-s | *BGIBMGA002745* | up | 1526 | 1534 | - | 12.0734 | 2.66E-05 | 0.615 | GCGCGTGTC |
| 2245 | mamo-s | *BGIBMGA002747* | up | 1833 | 1841 | + | 11.789 | 2.84E-05 | 0.615 | GCGCGTGCC |
| 2246 | mamo-s | *BGIBMGA002747* | up | 1741 | 1749 | + | 11.7064 | 3.52E-05 | 0.615 | GCGCGTGGT |
| 2247 | mamo-s | *BGIBMGA002747* | up | 1650 | 1658 | + | 11.3303 | 4.96E-05 | 0.615 | GTGGGTGGC |
| 2248 | mamo-s | *BGIBMGA002748* | down | 528 | 536 | - | 11.5688 | 3.79E-05 | 0.615 | CTGCGTGTC |
| 2249 | mamo-s | *BGIBMGA002748* | up | 192 | 200 | + | 11.3303 | 4.96E-05 | 0.615 | GTGGGTGGC |
| 2250 | mamo-s | *BGIBMGA002749* | up | 1494 | 1502 | + | 11.3303 | 4.96E-05 | 0.615 | GTGGGTGGC |
| 2251 | mamo-s | *BGIBMGA002750* | up | 1152 | 1160 | - | 12.1284 | 2.16E-05 | 0.615 | CAGCGTGGC |
| 2252 | mamo-s | *BGIBMGA002750* | up | 1195 | 1203 | - | 11.3853 | 4.74E-05 | 0.615 | ACGCGTGGC |
| 2253 | mamo-s | *BGIBMGA002752* | down | 1173 | 1181 | + | 11.1743 | 5.90E-05 | 0.684 | CCGCGTGTC |
| 2254 | mamo-s | *BGIBMGA002753* | up | 415 | 423 | + | 11.3303 | 4.96E-05 | 0.615 | GTGGGTGGC |
| 2255 | mamo-s | *BGIBMGA002753* | up | 1405 | 1413 | - | 11.3303 | 4.96E-05 | 0.615 | GTGGGTGGC |
| 2256 | mamo-s | *BGIBMGA002755* | down | 1099 | 1107 | - | 11.3303 | 4.96E-05 | 0.615 | GTGGGTGGC |
| 2257 | mamo-s | *BGIBMGA002755* | up | 926 | 934 | + | 11.0826 | 6.84E-05 | 0.709 | GTACGTGGC |
| 2258 | mamo-s | *BGIBMGA002756* | down | 1835 | 1843 | - | 11.3303 | 4.96E-05 | 0.615 | GTGGGTGGC |
| 2259 | mamo-s | *BGIBMGA002758* | down | 817 | 825 | + | 12.1835 | 1.71E-05 | 0.615 | GTGCGTGCC |
| 2260 | mamo-s | *BGIBMGA002758* | down | 1239 | 1247 | - | 11.0826 | 6.84E-05 | 0.709 | GTACGTGGC |
| 2261 | mamo-s | *BGIBMGA002759* | up | 407 | 415 | - | 12.1284 | 2.16E-05 | 0.615 | CAGCGTGGC |
| 2262 | mamo-s | *BGIBMGA002759* | down | 701 | 709 | + | 11.3303 | 4.96E-05 | 0.615 | GTGGGTGGC |
| 2263 | mamo-s | *BGIBMGA002759* | down | 1051 | 1059 | - | 11.3303 | 4.96E-05 | 0.615 | GTGGGTGGC |
| 2264 | mamo-s | *BGIBMGA002759* | up | 1289 | 1297 | - | 11.3303 | 4.96E-05 | 0.615 | GTGGGTGGC |
| 2265 | mamo-s | *BGIBMGA002761* | up | 1877 | 1885 | + | 11.2844 | 5.18E-05 | 0.63 | CTGCGTGCC |
| 2266 | mamo-s | *BGIBMGA002764* | down | 370 | 378 | + | 11.3303 | 4.96E-05 | 0.615 | GTGGGTGGC |
| 2267 | mamo-s | *BGIBMGA002764* | up | 1828 | 1836 | + | 11.3303 | 4.96E-05 | 0.615 | GTGGGTGGC |
| 2268 | mamo-s | *BGIBMGA002767* | up | 986 | 994 | - | 11.3303 | 4.96E-05 | 0.615 | GTGGGTGGC |
| 2269 | mamo-s | *BGIBMGA002768* | down | 639 | 647 | - | 11.3303 | 4.96E-05 | 0.615 | GTGGGTGGC |
| 2270 | mamo-s | *BGIBMGA002769* | down | 1231 | 1239 | + | 11.3303 | 4.96E-05 | 0.615 | GTGGGTGGC |
| 2271 | mamo-s | *BGIBMGA002769* | down | 1714 | 1722 | - | 10.5138 | 9.75E-05 | 0.865 | GTGCGTGTT |
| 2272 | mamo-s | *BGIBMGA002770* | down | 983 | 991 | + | 11.3303 | 4.96E-05 | 0.615 | GTGGGTGGC |
| 2273 | mamo-s | *BGIBMGA002771* | down | 750 | 758 | + | 11.4404 | 4.51E-05 | 0.615 | CGGCGTGGC |
| 2274 | mamo-s | *BGIBMGA002772* | down | 997 | 1005 | + | 10.5413 | 9.42E-05 | 0.857 | GTGCGAGTC |
| 2275 | mamo-s | *BGIBMGA002773* | down | 534 | 542 | - | 13.6606 | 4.06E-06 | 0.615 | GCGCGTGGC |
| 2276 | mamo-s | *BGIBMGA002774* | up | 50 | 58 | - | 11.3303 | 4.96E-05 | 0.615 | GTGGGTGGC |
| 2277 | mamo-s | *BGIBMGA002775* | up | 213 | 221 | + | 11.3303 | 4.96E-05 | 0.615 | GTGGGTGGC |
| 2278 | mamo-s | *BGIBMGA002775* | down | 676 | 684 | - | 11.3303 | 4.96E-05 | 0.615 | GTGGGTGGC |
| 2279 | mamo-s | *BGIBMGA002775* | down | 296 | 304 | - | 11.0826 | 6.84E-05 | 0.709 | GTACGTGGC |
| 2280 | mamo-s | *BGIBMGA002775* | up | 1412 | 1420 | + | 11.0826 | 6.84E-05 | 0.709 | GTACGTGGC |
| 2281 | mamo-s | *BGIBMGA002776* | up | 897 | 905 | + | 11.3303 | 4.96E-05 | 0.615 | GTGGGTGGC |

| | | | | | | | | | | |
|---|---|---|---|---|---|---|---|---|---|---|
| 2282 | mamo-s | *BGIBMGA002777* | down | 240 | 248 | - | 11.3303 | 4.96E-05 | 0.615 | GTGGGTGGC |
| 2283 | mamo-s | *BGIBMGA002777* | down | 1378 | 1386 | - | 11.1743 | 5.90E-05 | 0.684 | CCGCGTGTC |
| 2284 | mamo-s | *BGIBMGA002779* | up | 933 | 941 | + | 14.055 | 2.23E-06 | 0.615 | GTGCGTGGC |
| 2285 | mamo-s | *BGIBMGA002782* | down | 831 | 839 | - | 11.3303 | 4.96E-05 | 0.615 | GTGGGTGGC |
| 2286 | mamo-s | *BGIBMGA002784* | up | 1900 | 1908 | + | 10.6881 | 8.61E-05 | 0.81 | GCACGTGGC |
| 2287 | mamo-s | *BGIBMGA002785* | up | 1763 | 1771 | - | 11.3303 | 4.96E-05 | 0.615 | GTGGGTGGC |
| 2288 | mamo-s | *BGIBMGA002787* | down | 547 | 555 | + | 11.3303 | 4.96E-05 | 0.615 | GTGGGTGGC |
| 2289 | mamo-s | *BGIBMGA002787* | up | 1642 | 1650 | + | 11.3303 | 4.96E-05 | 0.615 | GTGGGTGGC |
| 2290 | mamo-s | *BGIBMGA002797* | down | 1962 | 1970 | - | 11.3303 | 4.96E-05 | 0.615 | GTGGGTGGC |
| 2291 | mamo-s | *BGIBMGA002798* | down | 1033 | 1041 | - | 11.4404 | 4.51E-05 | 0.615 | GAGCGTGTC |
| 2292 | mamo-s | *BGIBMGA002798* | down | 1653 | 1661 | + | 11.3303 | 4.96E-05 | 0.615 | GTGGGTGGC |
| 2293 | mamo-s | *BGIBMGA002802* | down | 1912 | 1920 | - | 11.1009 | 6.57E-05 | 0.709 | GAGCGAGGC |
| 2294 | mamo-s | *BGIBMGA002803* | down | 779 | 787 | + | 11.3303 | 4.96E-05 | 0.615 | GTGGGTGGC |
| 2295 | mamo-s | *BGIBMGA002804* | up | 1160 | 1168 | - | 11.3303 | 4.96E-05 | 0.615 | GTGGGTGGC |
| 2296 | mamo-s | *BGIBMGA002807* | up | 1087 | 1095 | + | 11.3303 | 4.96E-05 | 0.615 | GTGGGTGGC |
| 2297 | mamo-s | *BGIBMGA002808* | down | 736 | 744 | - | 13.6606 | 4.06E-06 | 0.615 | GCGCGTGGC |
| 2298 | mamo-s | *BGIBMGA002809* | down | 1838 | 1846 | + | 11.2018 | 5.68E-05 | 0.674 | CTGCGTGGT |
| 2299 | mamo-s | *BGIBMGA002812* | down | 1339 | 1347 | + | 10.5688 | 8.88E-05 | 0.824 | GTGCGTGAC |
| 2300 | mamo-s | *BGIBMGA002813* | up | 1158 | 1166 | - | 12.3394 | 1.49E-05 | 0.615 | GGGCGTGGC |
| 2301 | mamo-s | *BGIBMGA002813* | up | 431 | 439 | - | 11.3303 | 4.96E-05 | 0.615 | GTGGGTGGC |
| 2302 | mamo-s | *BGIBMGA002814* | down | 1286 | 1294 | + | 11.1743 | 5.90E-05 | 0.684 | CCGCGTGTC |
| 2303 | mamo-s | *BGIBMGA002814* | down | 1920 | 1928 | + | 11.1009 | 6.57E-05 | 0.709 | GAGCGAGGC |
| 2304 | mamo-s | *BGIBMGA002817* | up | 244 | 252 | - | 10.5138 | 9.75E-05 | 0.865 | GTGCGTGTT |
| 2305 | mamo-s | *BGIBMGA002818* | down | 458 | 466 | + | 11.3303 | 4.96E-05 | 0.615 | GTGGGTGGC |
| 2306 | mamo-s | *BGIBMGA002818* | up | 1509 | 1517 | - | 11.3303 | 4.96E-05 | 0.615 | GTGGGTGGC |
| 2307 | mamo-s | *BGIBMGA002819* | up | 456 | 464 | + | 13.6606 | 4.06E-06 | 0.615 | GCGCGTGGC |
| 2308 | mamo-s | *BGIBMGA002820* | down | 66 | 74 | + | 11.3303 | 4.96E-05 | 0.615 | GTGGGTGGC |
| 2309 | mamo-s | *BGIBMGA002821* | down | 1634 | 1642 | - | 13.6606 | 4.06E-06 | 0.615 | GCGCGTGGC |
| 2310 | mamo-s | *BGIBMGA002821* | up | 1023 | 1031 | - | 11.3303 | 4.96E-05 | 0.615 | GTGGGTGGC |
| 2311 | mamo-s | *BGIBMGA002823* | down | 824 | 832 | + | 10.5138 | 9.75E-05 | 0.865 | GTGCGTGTT |
| 2312 | mamo-s | *BGIBMGA002825* | down | 1725 | 1733 | + | 11.3303 | 4.96E-05 | 0.615 | GTGGGTGGC |
| 2313 | mamo-s | *BGIBMGA002826* | down | 1897 | 1905 | + | 11.5321 | 4.06E-05 | 0.615 | TTGCGTGGC |
| 2314 | mamo-s | *BGIBMGA002827* | down | 616 | 624 | - | 11.3303 | 4.96E-05 | 0.615 | GTGGGTGGC |
| 2315 | mamo-s | *BGIBMGA002827* | down | 1848 | 1856 | + | 11.3303 | 4.96E-05 | 0.615 | GTGGGTGGC |
| 2316 | mamo-s | *BGIBMGA002831* | up | 1769 | 1777 | + | 11.0826 | 6.84E-05 | 0.709 | GTACGTGGC |
| 2317 | mamo-s | *BGIBMGA002832* | down | 142 | 150 | - | 11.7339 | 3.29E-05 | 0.615 | GCGCGAGGC |
| 2318 | mamo-s | *BGIBMGA002832* | up | 458 | 466 | - | 11.3303 | 4.96E-05 | 0.615 | GTGGGTGGC |
| 2319 | mamo-s | *BGIBMGA002832* | down | 729 | 737 | + | 11.1743 | 5.90E-05 | 0.684 | CCGCGTGTC |
| 2320 | mamo-s | *BGIBMGA002833* | up | 127 | 135 | + | 11.3303 | 4.96E-05 | 0.615 | GTGGGTGGC |
| 2321 | mamo-s | *BGIBMGA002834* | down | 492 | 500 | + | 12.4679 | 1.31E-05 | 0.615 | GTGCGTGTC |
| 2322 | mamo-s | *BGIBMGA002834* | down | 1896 | 1904 | - | 11.3303 | 4.96E-05 | 0.615 | GTGGGTGGC |
| 2323 | mamo-s | *BGIBMGA002836* | down | 1920 | 1928 | - | 12.3394 | 1.49E-05 | 0.615 | GGGCGTGGC |
| 2324 | mamo-s | *BGIBMGA002837* | up | 949 | 957 | - | 12.3394 | 1.49E-05 | 0.615 | GGGCGTGGC |
| 2325 | mamo-s | *BGIBMGA002838* | down | 1778 | 1786 | - | 11.3303 | 4.96E-05 | 0.615 | GTGGGTGGC |
| 2326 | mamo-s | *BGIBMGA002845* | up | 76 | 84 | - | 12.0734 | 2.66E-05 | 0.615 | GCGCGTGTC |
| 2327 | mamo-s | *BGIBMGA002845* | down | 1793 | 1801 | + | 11.7798 | 3.11E-05 | 0.615 | ATGCGTGGC |
| 2328 | mamo-s | *BGIBMGA002846* | up | 7 | 15 | + | 11.7798 | 3.11E-05 | 0.615 | ATGCGTGGC |
| 2329 | mamo-s | *BGIBMGA002846* | down | 288 | 296 | + | 11.3303 | 4.96E-05 | 0.615 | GTGGGTGGC |
| 2330 | mamo-s | *BGIBMGA002847* | down | 1899 | 1907 | - | 11.3303 | 4.96E-05 | 0.615 | GTGGGTGGC |
| 2331 | mamo-s | *BGIBMGA002850* | down | 1571 | 1579 | + | 11.7064 | 3.52E-05 | 0.615 | GCGCGTGGT |
| 2332 | mamo-s | *BGIBMGA002850* | down | 1893 | 1901 | - | 11.5321 | 4.06E-05 | 0.615 | TTGCGTGGC |
| 2333 | mamo-s | *BGIBMGA002850* | down | 1138 | 1146 | + | 11.2018 | 5.68E-05 | 0.674 | CTGCGTGGT |
| 2334 | mamo-s | *BGIBMGA002851* | up | 1778 | 1786 | - | 10.6881 | 8.61E-05 | 0.81 | GCACGTGGC |
| 2335 | mamo-s | *BGIBMGA002851* | up | 1783 | 1791 | + | 10.5688 | 8.88E-05 | 0.824 | GTGCGTGAC |
| 2336 | mamo-s | *BGIBMGA002852* | down | 822 | 830 | - | 11.0826 | 6.84E-05 | 0.709 | GTACGTGGC |
| 2337 | mamo-s | *BGIBMGA002853* | up | 1957 | 1965 | - | 12.4679 | 1.31E-05 | 0.615 | GTGCGTGTC |
| 2338 | mamo-s | *BGIBMGA002854* | down | 551 | 559 | - | 12.1009 | 2.43E-05 | 0.615 | GTGCGTGGT |
| 2339 | mamo-s | *BGIBMGA002856* | down | 1728 | 1736 | - | 11.3303 | 4.96E-05 | 0.615 | GTGGGTGGC |
| 2340 | mamo-s | *BGIBMGA002860* | up | 594 | 602 | + | 13.6606 | 4.06E-06 | 0.615 | GCGCGTGGC |
| 2341 | mamo-s | *BGIBMGA002860* | up | 1019 | 1027 | + | 10.9358 | 7.30E-05 | 0.728 | GCGGGTGGC |
| 2342 | mamo-s | *BGIBMGA002864* | down | 818 | 826 | - | 12.4679 | 1.31E-05 | 0.615 | GTGCGTGTC |
| 2343 | mamo-s | *BGIBMGA002865* | down | 478 | 486 | - | 11.3303 | 4.96E-05 | 0.615 | GTGGGTGGC |
| 2344 | mamo-s | *BGIBMGA002866* | down | 498 | 506 | + | 11.7339 | 3.29E-05 | 0.615 | GCGCGAGGC |
| 2345 | mamo-s | *BGIBMGA002867* | up | 319 | 327 | - | 11.3303 | 4.96E-05 | 0.615 | GTGGGTGGC |
| 2346 | mamo-s | *BGIBMGA002869* | down | 1128 | 1136 | + | 11.3303 | 4.96E-05 | 0.615 | GTGGGTGGC |
| 2347 | mamo-s | *BGIBMGA002870* | up | 860 | 868 | + | 11.3303 | 4.96E-05 | 0.615 | GTGGGTGGC |
| 2348 | mamo-s | *BGIBMGA002872* | down | 1328 | 1336 | + | 11.3853 | 4.74E-05 | 0.615 | ACGCGTGGC |
| 2349 | mamo-s | *BGIBMGA002872* | down | 432 | 440 | + | 11.3303 | 4.96E-05 | 0.615 | GTGGGTGGC |
| 2350 | mamo-s | *BGIBMGA002873* | down | 1978 | 1986 | + | 11.7798 | 3.11E-05 | 0.615 | ATGCGTGGC |
| 2351 | mamo-s | *BGIBMGA002873* | up | 515 | 523 | + | 11.3853 | 4.74E-05 | 0.615 | ACGCGTGGC |
| 2352 | mamo-s | *BGIBMGA002873* | down | 1273 | 1281 | - | 11.3303 | 4.96E-05 | 0.615 | GTGGGTGGC |
| 2353 | mamo-s | *BGIBMGA002874* | down | 1239 | 1247 | + | 11.3303 | 4.96E-05 | 0.615 | GTGGGTGGC |
| 2354 | mamo-s | *BGIBMGA002876* | down | 149 | 157 | + | 10.9358 | 7.30E-05 | 0.728 | GCGGGTGGC |
| 2355 | mamo-s | *BGIBMGA002877* | up | 1397 | 1405 | + | 11.3303 | 4.96E-05 | 0.615 | GTGGGTGGC |
| 2356 | mamo-s | *BGIBMGA002878* | up | 1926 | 1934 | + | 10.5138 | 9.75E-05 | 0.865 | GTGCGTGTT |
| 2357 | mamo-s | *BGIBMGA002879* | up | 1292 | 1300 | + | 12.4679 | 1.31E-05 | 0.615 | GTGCGTGTC |
| 2358 | mamo-s | *BGIBMGA002879* | down | 849 | 857 | + | 11.3303 | 4.96E-05 | 0.615 | GTGGGTGGC |
| 2359 | mamo-s | *BGIBMGA002879* | up | 1640 | 1648 | + | 11.3303 | 4.96E-05 | 0.615 | GTGGGTGGC |
| 2360 | mamo-s | *BGIBMGA002880* | down | 448 | 456 | - | 11.3303 | 4.96E-05 | 0.615 | GTGGGTGGC |
| 2361 | mamo-s | *BGIBMGA002881* | up | 232 | 240 | + | 11.3303 | 4.96E-05 | 0.615 | GTGGGTGGC |
| 2362 | mamo-s | *BGIBMGA002882* | down | 1807 | 1815 | - | 11.3303 | 4.96E-05 | 0.615 | GTGGGTGGC |
| 2363 | mamo-s | *BGIBMGA002883* | up | 1182 | 1190 | - | 11.3303 | 4.96E-05 | 0.615 | GTGGGTGGC |
| 2364 | mamo-s | *BGIBMGA002883* | up | 1757 | 1765 | - | 11.0826 | 6.84E-05 | 0.709 | GTACGTGGC |
| 2365 | mamo-s | *BGIBMGA002885* | up | 1652 | 1660 | - | 13.6606 | 4.06E-06 | 0.615 | GCGCGTGGC |
| 2366 | mamo-s | *BGIBMGA002886* | up | 122 | 130 | + | 11.3303 | 4.96E-05 | 0.615 | GTGGGTGGC |
| 2367 | mamo-s | *BGIBMGA002889* | up | 931 | 939 | - | 11.4404 | 4.51E-05 | 0.615 | CGGCGTGGC |
| 2368 | mamo-s | *BGIBMGA002889* | up | 1147 | 1155 | + | 11.4404 | 4.51E-05 | 0.615 | GAGCGTGTC |
| 2369 | mamo-s | *BGIBMGA002889* | down | 1963 | 1971 | - | 11.1009 | 6.57E-05 | 0.709 | GAGCGAGGC |

| | | | | | | | | | | |
|---|---|---|---|---|---|---|---|---|---|---|
| 2370 | mamo-s | *BGIBMGA002892* | up | 506 | 514 | + | 11.2294 | 5.41E-05 | 0.65 | CTGCGAGGC |
| 2371 | mamo-s | *BGIBMGA002896* | down | 878 | 886 | + | 13.6606 | 4.06E-06 | 0.615 | GCGCGTGGC |
| 2372 | mamo-s | *BGIBMGA002898* | down | 1784 | 1792 | + | 11.3303 | 4.96E-05 | 0.615 | GTGGGTGGC |
| 2373 | mamo-s | *BGIBMGA002899* | up | 1874 | 1882 | + | 10.5688 | 8.88E-05 | 0.824 | GTGCGTGAC |
| 2374 | mamo-s | *BGIBMGA002902* | down | 1204 | 1212 | - | 11.3303 | 4.96E-05 | 0.615 | GTGGGTGGC |
| 2375 | mamo-s | *BGIBMGA002902* | up | 1689 | 1697 | + | 11.3303 | 4.96E-05 | 0.615 | GTGGGTGGC |
| 2376 | mamo-s | *BGIBMGA002903* | down | 161 | 169 | + | 11.3303 | 4.96E-05 | 0.615 | GTGGGTGGC |
| 2377 | mamo-s | *BGIBMGA002906* | down | 1298 | 1306 | + | 11.4404 | 4.51E-05 | 0.615 | GAGCGTGTC |
| 2378 | mamo-s | *BGIBMGA002906* | down | 678 | 686 | - | 11.3303 | 4.96E-05 | 0.615 | GTGGGTGGC |
| 2379 | mamo-s | *BGIBMGA002907* | down | 483 | 491 | - | 11.3303 | 4.96E-05 | 0.615 | GTGGGTGGC |
| 2380 | mamo-s | *BGIBMGA002908* | down | 244 | 252 | - | 12.1009 | 2.43E-05 | 0.615 | GTGCGTGGT |
| 2381 | mamo-s | *BGIBMGA002908* | down | 618 | 626 | + | 11.3303 | 4.96E-05 | 0.615 | GTGGGTGGC |
| 2382 | mamo-s | *BGIBMGA002909* | up | 1700 | 1708 | - | 11.3303 | 4.96E-05 | 0.615 | GTGGGTGGC |
| 2383 | mamo-s | *BGIBMGA002909* | up | 1216 | 1224 | + | 11.0826 | 6.84E-05 | 0.709 | GTACGTGGC |
| 2384 | mamo-s | *BGIBMGA002911* | down | 1394 | 1402 | + | 11.3303 | 4.96E-05 | 0.615 | GTGGGTGGC |
| 2385 | mamo-s | *BGIBMGA002911* | up | 1704 | 1712 | - | 11.3303 | 4.96E-05 | 0.615 | GTGGGTGGC |
| 2386 | mamo-s | *BGIBMGA002913* | down | 797 | 805 | + | 10.5413 | 9.42E-05 | 0.857 | GTGCGAGTC |
| 2387 | mamo-s | *BGIBMGA002914* | down | 7 | 15 | - | 12.3394 | 1.49E-05 | 0.615 | GGGCGTGGC |
| 2388 | mamo-s | *BGIBMGA002916* | up | 1269 | 1277 | + | 11.1376 | 6.35E-05 | 0.709 | TCGCGTGGC |
| 2389 | mamo-s | *BGIBMGA002916* | up | 977 | 985 | - | 10.5413 | 9.42E-05 | 0.857 | CAGCGTGTC |
| 2390 | mamo-s | *BGIBMGA002918* | down | 1362 | 1370 | - | 11.7064 | 3.52E-05 | 0.615 | GCGCGTGGT |
| 2391 | mamo-s | *BGIBMGA002919* | down | 355 | 363 | + | 11.3303 | 4.96E-05 | 0.615 | GTGGGTGGC |
| 2392 | mamo-s | *BGIBMGA002919* | up | 1037 | 1045 | - | 11.3303 | 4.96E-05 | 0.615 | GTGGGTGGC |
| 2393 | mamo-s | *BGIBMGA002920* | down | 1973 | 1981 | - | 10.9358 | 7.30E-05 | 0.728 | GCGGGTGGC |
| 2394 | mamo-s | *BGIBMGA002920* | down | 81 | 89 | + | 10.8899 | 7.48E-05 | 0.739 | CCGCGTGCC |
| 2395 | mamo-s | *BGIBMGA002922* | up | 1827 | 1835 | - | 11.3303 | 4.96E-05 | 0.615 | GTGGGTGGC |
| 2396 | mamo-s | *BGIBMGA002922* | up | 1449 | 1457 | - | 11.0734 | 7.12E-05 | 0.726 | GAGCGTGGT |
| 2397 | mamo-s | *BGIBMGA002923* | down | 1372 | 1380 | + | 11.4404 | 4.51E-05 | 0.615 | GAGCGTGTC |
| 2398 | mamo-s | *BGIBMGA002923* | up | 214 | 222 | - | 11.3303 | 4.96E-05 | 0.615 | GTGGGTGGC |
| 2399 | mamo-s | *BGIBMGA002924* | down | 1147 | 1155 | - | 14.055 | 2.23E-06 | 0.615 | GTGCGTGGC |
| 2400 | mamo-s | *BGIBMGA002924* | down | 475 | 483 | + | 11.0734 | 7.12E-05 | 0.726 | GAGCGTGGT |
| 2401 | mamo-s | *BGIBMGA002925* | down | 166 | 174 | - | 11.3303 | 4.96E-05 | 0.615 | GTGGGTGGC |
| 2402 | mamo-s | *BGIBMGA002925* | down | 789 | 797 | + | 11.3303 | 4.96E-05 | 0.615 | GTGGGTGGC |
| 2403 | mamo-s | *BGIBMGA002925* | up | 796 | 804 | + | 11.3303 | 4.96E-05 | 0.615 | GTGGGTGGC |
| 2404 | mamo-s | *BGIBMGA002930* | up | 406 | 414 | - | 12.4679 | 1.31E-05 | 0.615 | GTGCGTGTC |
| 2405 | mamo-s | *BGIBMGA002938* | down | 1702 | 1710 | - | 11.0826 | 6.84E-05 | 0.709 | GTACGTGGC |
| 2406 | mamo-s | *BGIBMGA002939* | up | 415 | 423 | + | 11.3303 | 4.96E-05 | 0.615 | GTGGGTGGC |
| 2407 | mamo-s | *BGIBMGA002942* | up | 214 | 222 | - | 10.5413 | 9.42E-05 | 0.857 | GTGCGAGTC |
| 2408 | mamo-s | *BGIBMGA002943* | down | 435 | 443 | - | 11.3303 | 4.96E-05 | 0.615 | GTGGGTGGC |
| 2409 | mamo-s | *BGIBMGA002943* | down | 1806 | 1814 | - | 11.3303 | 4.96E-05 | 0.615 | GTGGGTGGC |
| 2410 | mamo-s | *BGIBMGA002943* | up | 981 | 989 | - | 11.0734 | 7.12E-05 | 0.726 | GAGCGTGGT |
| 2411 | mamo-s | *BGIBMGA002944* | up | 20 | 28 | - | 11.3303 | 4.96E-05 | 0.615 | GTGGGTGGC |
| 2412 | mamo-s | *BGIBMGA002944* | up | 1391 | 1399 | - | 11.3303 | 4.96E-05 | 0.615 | GTGGGTGGC |
| 2413 | mamo-s | *BGIBMGA002945* | up | 1855 | 1863 | + | 10.7523 | 8.38E-05 | 0.797 | GGGCGTGTC |
| 2414 | mamo-s | *BGIBMGA002946* | up | 194 | 202 | + | 11.3303 | 4.96E-05 | 0.615 | GTGGGTGGC |
| 2415 | mamo-s | *BGIBMGA002948* | down | 1175 | 1183 | - | 11.3303 | 4.96E-05 | 0.615 | GTGGGTGGC |
| 2416 | mamo-s | *BGIBMGA002949* | up | 935 | 943 | - | 13.0275 | 8.53E-06 | 0.615 | GAGCGTGGC |
| 2417 | mamo-s | *BGIBMGA002950* | down | 1116 | 1124 | - | 11.0826 | 6.84E-05 | 0.709 | GTACGTGGC |
| 2418 | mamo-s | *BGIBMGA002951* | up | 1078 | 1086 | - | 12.1284 | 2.16E-05 | 0.615 | GTGCGAGGC |
| 2419 | mamo-s | *BGIBMGA002951* | down | 1444 | 1452 | + | 12.1009 | 2.43E-05 | 0.615 | GTGCGTGGT |
| 2420 | mamo-s | *BGIBMGA002951* | up | 1516 | 1524 | + | 11.3303 | 4.96E-05 | 0.615 | GTGGGTGGC |
| 2421 | mamo-s | *BGIBMGA002953* | down | 329 | 337 | + | 11.3303 | 4.96E-05 | 0.615 | GTGGGTGGC |
| 2422 | mamo-s | *BGIBMGA002954* | up | 218 | 226 | + | 11.3303 | 4.96E-05 | 0.615 | GTGGGTGGC |
| 2423 | mamo-s | *BGIBMGA002956* | down | 791 | 799 | - | 12.7615 | 1.04E-05 | 0.615 | CCGCGTGGC |
| 2424 | mamo-s | *BGIBMGA002957* | up | 894 | 902 | - | 11.789 | 2.84E-05 | 0.615 | GCGCGTGCC |
| 2425 | mamo-s | *BGIBMGA002957* | up | 1175 | 1183 | - | 11.3303 | 4.96E-05 | 0.615 | GTGGGTGGC |
| 2426 | mamo-s | *BGIBMGA002959* | down | 1502 | 1510 | + | 11.3303 | 4.96E-05 | 0.615 | GTGGGTGGC |
| 2427 | mamo-s | *BGIBMGA002961* | down | 161 | 169 | + | 11.3303 | 4.96E-05 | 0.615 | GTGGGTGGC |
| 2428 | mamo-s | *BGIBMGA002962* | up | 1604 | 1612 | + | 11.3303 | 4.96E-05 | 0.615 | GTGGGTGGC |
| 2429 | mamo-s | *BGIBMGA002963* | up | 1805 | 1813 | + | 11.3303 | 4.96E-05 | 0.615 | GTGGGTGGC |
| 2430 | mamo-s | *BGIBMGA002964* | up | 1537 | 1545 | + | 12.3394 | 1.49E-05 | 0.615 | GGGCGTGGC |
| 2431 | mamo-s | *BGIBMGA002967* | up | 1861 | 1869 | - | 11.3303 | 4.96E-05 | 0.615 | GTGGGTGGC |
| 2432 | mamo-s | *BGIBMGA002968* | down | 1732 | 1740 | - | 11.3303 | 4.96E-05 | 0.615 | GTGGGTGGC |
| 2433 | mamo-s | *BGIBMGA002968* | up | 1950 | 1958 | + | 11.0826 | 6.84E-05 | 0.709 | GTACGTGGC |
| 2434 | mamo-s | *BGIBMGA002971* | down | 1367 | 1375 | - | 12.1009 | 2.43E-05 | 0.615 | GTGCGTGGT |
| 2435 | mamo-s | *BGIBMGA002972* | down | 218 | 226 | + | 10.8349 | 7.66E-05 | 0.751 | CCGCGAGGC |
| 2436 | mamo-s | *BGIBMGA002973* | down | 977 | 985 | - | 10.9358 | 7.30E-05 | 0.728 | GCGGGTGGC |
| 2437 | mamo-s | *BGIBMGA002974* | down | 718 | 726 | + | 11.3303 | 4.96E-05 | 0.615 | GTGGGTGGC |
| 2438 | mamo-s | *BGIBMGA002974* | up | 31 | 39 | + | 11.0826 | 6.84E-05 | 0.709 | GTACGTGGC |
| 2439 | mamo-s | *BGIBMGA002975* | up | 825 | 833 | + | 11.3303 | 4.96E-05 | 0.615 | GTGGGTGGC |
| 2440 | mamo-s | *BGIBMGA002976* | down | 913 | 921 | - | 11.3303 | 4.96E-05 | 0.615 | GTGGGTGGC |
| 2441 | mamo-s | *BGIBMGA002977* | up | 211 | 219 | - | 11.0734 | 7.12E-05 | 0.726 | GAGCGTGGT |
| 2442 | mamo-s | *BGIBMGA002983* | up | 1628 | 1636 | - | 11.2844 | 5.18E-05 | 0.63 | CTGCGTGCC |
| 2443 | mamo-s | *BGIBMGA002984* | down | 711 | 719 | - | 10.7523 | 8.38E-05 | 0.797 | AAGCGTGGC |
| 2444 | mamo-s | *BGIBMGA002985* | up | 294 | 302 | + | 11.3303 | 4.96E-05 | 0.615 | GTGGGTGGC |
| 2445 | mamo-s | *BGIBMGA002986* | down | 1929 | 1937 | - | 11.3303 | 4.96E-05 | 0.615 | GTGGGTGGC |
| 2446 | mamo-s | *BGIBMGA002988* | down | 138 | 146 | + | 11.3303 | 4.96E-05 | 0.615 | GTGGGTGGC |
| 2447 | mamo-s | *BGIBMGA002989* | up | 398 | 406 | - | 11.0826 | 6.84E-05 | 0.709 | GTACGTGGC |
| 2448 | mamo-s | *BGIBMGA002990* | down | 1312 | 1320 | + | 11.3303 | 4.96E-05 | 0.615 | GTGGGTGGC |
| 2449 | mamo-s | *BGIBMGA002992* | down | 252 | 260 | + | 11.3303 | 4.96E-05 | 0.615 | GTGGGTGGC |
| 2450 | mamo-s | *BGIBMGA002993* | up | 1525 | 1533 | + | 12.1009 | 2.43E-05 | 0.615 | GTGCGTGGT |
| 2451 | mamo-s | *BGIBMGA002993* | down | 1374 | 1382 | + | 11.7339 | 3.29E-05 | 0.615 | GCGCGAGGC |
| 2452 | mamo-s | *BGIBMGA002994* | down | 552 | 560 | - | 13.6606 | 4.06E-06 | 0.615 | GCGCGTGGC |
| 2453 | mamo-s | *BGIBMGA002995* | up | 1599 | 1607 | + | 10.9358 | 7.30E-05 | 0.728 | GCGGGTGGC |
| 2454 | mamo-s | *BGIBMGA002996* | up | 1149 | 1157 | - | 12.1284 | 2.16E-05 | 0.615 | GTGCGAGGC |
| 2455 | mamo-s | *BGIBMGA002996* | down | 158 | 166 | + | 11.3303 | 4.96E-05 | 0.615 | GTGGGTGGC |
| 2456 | mamo-s | *BGIBMGA002996* | up | 1733 | 1741 | - | 11.1743 | 5.90E-05 | 0.684 | CCGCGTGTC |
| 2457 | mamo-s | *BGIBMGA002999* | down | 1803 | 1811 | + | 11.3303 | 4.96E-05 | 0.615 | GTGGGTGGC |

| | | | | | | | | | | |
|---|---|---|---|---|---|---|---|---|---|---|
| 2458 | mamo-s | *BGIBMGA002999* | up | 1156 | 1164 | + | 11.0826 | 6.84E-05 | 0.709 | GTACGTGGC |
| 2459 | mamo-s | *BGIBMGA003001* | down | 483 | 491 | - | 12.4679 | 1.31E-05 | 0.615 | GTGCGTGTC |
| 2460 | mamo-s | *BGIBMGA003001* | down | 1443 | 1451 | - | 11.3303 | 4.96E-05 | 0.615 | GTGGGTGGC |
| 2461 | mamo-s | *BGIBMGA003001* | down | 334 | 342 | - | 11.1743 | 5.90E-05 | 0.684 | CCGCGTGTC |
| 2462 | mamo-s | *BGIBMGA003006* | down | 1098 | 1106 | + | 11.3303 | 4.96E-05 | 0.615 | GTGGGTGGC |
| 2463 | mamo-s | *BGIBMGA003006* | up | 165 | 173 | - | 10.8349 | 7.66E-05 | 0.751 | CCGCGAGGC |
| 2464 | mamo-s | *BGIBMGA003007* | down | 367 | 375 | - | 14.055 | 2.23E-06 | 0.615 | GTGCGTGGC |
| 2465 | mamo-s | *BGIBMGA003007* | up | 670 | 678 | - | 11.3303 | 4.96E-05 | 0.615 | GTGGGTGGC |
| 2466 | mamo-s | *BGIBMGA003008* | down | 679 | 687 | - | 11.3303 | 4.96E-05 | 0.615 | GTGGGTGGC |
| 2467 | mamo-s | *BGIBMGA003011* | up | 1801 | 1809 | - | 10.5688 | 8.88E-05 | 0.824 | GTGCGTGAC |
| 2468 | mamo-s | *BGIBMGA003013* | up | 1662 | 1670 | + | 10.7523 | 8.38E-05 | 0.797 | GGGCGTGTC |
| 2469 | mamo-s | *BGIBMGA003014* | up | 896 | 904 | + | 11.789 | 2.84E-05 | 0.615 | GCGCGTGCC |
| 2470 | mamo-s | *BGIBMGA003015* | down | 105 | 113 | - | 12.1284 | 2.16E-05 | 0.615 | GTGCGAGGC |
| 2471 | mamo-s | *BGIBMGA003015* | down | 1963 | 1971 | + | 11.3303 | 4.96E-05 | 0.615 | GTGGGTGGC |
| 2472 | mamo-s | *BGIBMGA003016* | down | 1377 | 1385 | + | 11.3303 | 4.96E-05 | 0.615 | GTGGGTGGC |
| 2473 | mamo-s | *BGIBMGA003017* | down | 1782 | 1790 | - | 12.4679 | 1.31E-05 | 0.615 | GTGCGTGTC |
| 2474 | mamo-s | *BGIBMGA003017* | up | 822 | 830 | + | 11.3303 | 4.96E-05 | 0.615 | GTGGGTGGC |
| 2475 | mamo-s | *BGIBMGA003025* | up | 1250 | 1258 | - | 12.1284 | 2.16E-05 | 0.615 | GTGCGAGGC |
| 2476 | mamo-s | *BGIBMGA003026* | down | 741 | 749 | - | 11.3303 | 4.96E-05 | 0.615 | GTGGGTGGC |
| 2477 | mamo-s | *BGIBMGA003028* | up | 670 | 678 | - | 11.3303 | 4.96E-05 | 0.615 | GTGGGTGGC |
| 2478 | mamo-s | *BGIBMGA003029* | down | 1321 | 1329 | + | 11.3303 | 4.96E-05 | 0.615 | GTGGGTGGC |
| 2479 | mamo-s | *BGIBMGA003030* | down | 1786 | 1794 | - | 11.3303 | 4.96E-05 | 0.615 | GTGGGTGGC |
| 2480 | mamo-s | *BGIBMGA003031* | up | 245 | 253 | - | 11.3303 | 4.96E-05 | 0.615 | GTGGGTGGC |
| 2481 | mamo-s | *BGIBMGA003031* | up | 1218 | 1226 | - | 11.0826 | 6.84E-05 | 0.709 | GTACGTGGC |
| 2482 | mamo-s | *BGIBMGA003033* | up | 1225 | 1233 | + | 13.6606 | 4.06E-06 | 0.615 | GCGCGTGGC |
| 2483 | mamo-s | *BGIBMGA003034* | down | 399 | 407 | + | 11.3303 | 4.96E-05 | 0.615 | GTGGGTGGC |
| 2484 | mamo-s | *BGIBMGA003035* | down | 278 | 286 | + | 11.3303 | 4.96E-05 | 0.615 | GTGGGTGGC |
| 2485 | mamo-s | *BGIBMGA003035* | down | 902 | 910 | - | 10.8899 | 7.48E-05 | 0.739 | CCGCGTGCC |
| 2486 | mamo-s | *BGIBMGA003036* | down | 167 | 175 | - | 11.3303 | 4.96E-05 | 0.615 | GTGGGTGGC |
| 2487 | mamo-s | *BGIBMGA003036* | up | 1274 | 1282 | - | 11.3303 | 4.96E-05 | 0.615 | GTGGGTGGC |
| 2488 | mamo-s | *BGIBMGA003036* | down | 1977 | 1985 | - | 11.3303 | 4.96E-05 | 0.615 | GTGGGTGGC |
| 2489 | mamo-s | *BGIBMGA003036* | up | 590 | 598 | + | 11.0826 | 6.84E-05 | 0.709 | GTACGTGGC |
| 2490 | mamo-s | *BGIBMGA003038* | up | 350 | 358 | + | 12.1009 | 2.43E-05 | 0.615 | GTGCGTGGT |
| 2491 | mamo-s | *BGIBMGA003038* | down | 1199 | 1207 | - | 11.3303 | 4.96E-05 | 0.615 | GTGGGTGGC |
| 2492 | mamo-s | *BGIBMGA003038* | down | 1884 | 1892 | + | 11.3303 | 4.96E-05 | 0.615 | GTGGGTGGC |
| 2493 | mamo-s | *BGIBMGA003042* | down | 226 | 234 | + | 12.7615 | 1.04E-05 | 0.615 | CCGCGTGGC |
| 2494 | mamo-s | *BGIBMGA003042* | down | 1626 | 1634 | - | 11.3303 | 4.96E-05 | 0.615 | GTGGGTGGC |
| 2495 | mamo-s | *BGIBMGA003049* | down | 812 | 820 | + | 11.3303 | 4.96E-05 | 0.615 | GTGGGTGGC |
| 2496 | mamo-s | *BGIBMGA003049* | down | 265 | 273 | + | 11.2844 | 5.18E-05 | 0.63 | CTGCGTGCC |
| 2497 | mamo-s | *BGIBMGA003054* | up | 342 | 350 | - | 11.3303 | 4.96E-05 | 0.615 | GTGGGTGGC |
| 2498 | mamo-s | *BGIBMGA003054* | down | 1404 | 1412 | + | 11.0826 | 6.84E-05 | 0.709 | GTACGTGGC |
| 2499 | mamo-s | *BGIBMGA003057* | down | 1423 | 1431 | - | 11.3303 | 4.96E-05 | 0.615 | GTGGGTGGC |
| 2500 | mamo-s | *BGIBMGA003057* | up | 1425 | 1433 | - | 10.5138 | 9.75E-05 | 0.865 | GTGCGTGTT |
| 2501 | mamo-s | *BGIBMGA003059* | up | 1922 | 1930 | - | 11.7064 | 3.52E-05 | 0.615 | GCGCGTGGT |
| 2502 | mamo-s | *BGIBMGA003059* | up | 84 | 92 | + | 11.3303 | 4.96E-05 | 0.615 | GTGGGTGGC |
| 2503 | mamo-s | *BGIBMGA003059* | up | 825 | 833 | - | 11.0826 | 6.84E-05 | 0.709 | GTACGTGGC |
| 2504 | mamo-s | *BGIBMGA003060* | down | 122 | 130 | - | 14.055 | 2.23E-06 | 0.615 | GTGCGTGGC |
| 2505 | mamo-s | *BGIBMGA003061* | up | 664 | 672 | - | 14.055 | 2.23E-06 | 0.615 | GTGCGTGGC |
| 2506 | mamo-s | *BGIBMGA003062* | down | 1916 | 1924 | + | 12.1284 | 2.16E-05 | 0.615 | CAGCGTGGC |
| 2507 | mamo-s | *BGIBMGA003062* | up | 1284 | 1292 | + | 11.3303 | 4.96E-05 | 0.615 | GTGGGTGGC |
| 2508 | mamo-s | *BGIBMGA003062* | down | 867 | 875 | + | 10.5413 | 9.42E-05 | 0.857 | CAGCGTGTC |
| 2509 | mamo-s | *BGIBMGA003064* | down | 1720 | 1728 | + | 11.3303 | 4.96E-05 | 0.615 | GTGGGTGGC |
| 2510 | mamo-s | *BGIBMGA003065* | up | 1913 | 1921 | - | 12.3394 | 1.49E-05 | 0.615 | GGGCGTGGC |
| 2511 | mamo-s | *BGIBMGA003065* | up | 760 | 768 | + | 11.5688 | 3.79E-05 | 0.615 | CTGCGTGTC |
| 2512 | mamo-s | *BGIBMGA003068* | up | 767 | 775 | + | 11.0826 | 6.84E-05 | 0.709 | GTACGTGGC |
| 2513 | mamo-s | *BGIBMGA003071* | up | 1347 | 1355 | + | 12.0734 | 2.66E-05 | 0.615 | GCGCGTGTC |
| 2514 | mamo-s | *BGIBMGA003071* | down | 75 | 83 | + | 11.3303 | 4.96E-05 | 0.615 | GTGGGTGGC |
| 2515 | mamo-s | *BGIBMGA003071* | up | 600 | 608 | + | 10.8349 | 7.66E-05 | 0.751 | CCGCGAGGC |
| 2516 | mamo-s | *BGIBMGA003074* | down | 828 | 836 | + | 11.3303 | 4.96E-05 | 0.615 | GTGGGTGGC |
| 2517 | mamo-s | *BGIBMGA003074* | up | 1653 | 1661 | - | 11.0826 | 6.84E-05 | 0.709 | GTACGTGGC |
| 2518 | mamo-s | *BGIBMGA003075* | down | 1062 | 1070 | + | 11.3303 | 4.96E-05 | 0.615 | GTGGGTGGC |
| 2519 | mamo-s | *BGIBMGA003076* | up | 867 | 875 | + | 11.3303 | 4.96E-05 | 0.615 | GTGGGTGGC |
| 2520 | mamo-s | *BGIBMGA003078* | up | 1546 | 1554 | - | 12.1284 | 2.16E-05 | 0.615 | GTGCGAGGC |
| 2521 | mamo-s | *BGIBMGA003078* | up | 237 | 245 | - | 11.3303 | 4.96E-05 | 0.615 | GTGGGTGGC |
| 2522 | mamo-s | *BGIBMGA003079* | up | 32 | 40 | - | 11.3303 | 4.96E-05 | 0.615 | GTGGGTGGC |
| 2523 | mamo-s | *BGIBMGA003082* | down | 1131 | 1139 | + | 11.3303 | 4.96E-05 | 0.615 | GTGGGTGGC |
| 2524 | mamo-s | *BGIBMGA003083* | down | 1700 | 1708 | + | 11.3303 | 4.96E-05 | 0.615 | GTGGGTGGC |
| 2525 | mamo-s | *BGIBMGA003083* | up | 1355 | 1363 | - | 10.8073 | 7.89E-05 | 0.766 | CCGCGTGGT |
| 2526 | mamo-s | *BGIBMGA003084* | down | 1118 | 1126 | - | 11.2844 | 5.18E-05 | 0.63 | CTGCGTGCC |
| 2527 | mamo-s | *BGIBMGA003084* | up | 1481 | 1489 | - | 11.2844 | 5.18E-05 | 0.63 | CTGCGTGCC |
| 2528 | mamo-s | *BGIBMGA003085* | down | 445 | 453 | - | 11.7339 | 3.29E-05 | 0.615 | GCGCGAGGC |
| 2529 | mamo-s | *BGIBMGA003086* | up | 1836 | 1844 | - | 11.3303 | 4.96E-05 | 0.615 | GTGGGTGGC |
| 2530 | mamo-s | *BGIBMGA003091* | up | 1028 | 1036 | + | 11.3303 | 4.96E-05 | 0.615 | GTGGGTGGC |
| 2531 | mamo-s | *BGIBMGA003091* | up | 828 | 836 | - | 11.0826 | 6.84E-05 | 0.709 | GTACGTGGC |
| 2532 | mamo-s | *BGIBMGA003094* | down | 411 | 419 | + | 11.3303 | 4.96E-05 | 0.615 | GTGGGTGGC |
| 2533 | mamo-s | *BGIBMGA003095* | up | 191 | 199 | - | 11.5688 | 3.79E-05 | 0.615 | CTGCGTGTC |
| 2534 | mamo-s | *BGIBMGA003095* | up | 1524 | 1532 | + | 11.3303 | 4.96E-05 | 0.615 | GTGGGTGGC |
| 2535 | mamo-s | *BGIBMGA003096* | down | 487 | 495 | - | 10.5413 | 9.42E-05 | 0.857 | CAGCGTGTC |
| 2536 | mamo-s | *BGIBMGA003098* | up | 1416 | 1424 | + | 11.3303 | 4.96E-05 | 0.615 | GTGGGTGGC |
| 2537 | mamo-s | *BGIBMGA003102* | up | 1378 | 1386 | + | 11.3303 | 4.96E-05 | 0.615 | GTGGGTGGC |
| 2538 | mamo-s | *BGIBMGA003102* | down | 1457 | 1465 | + | 11.3303 | 4.96E-05 | 0.615 | GTGGGTGGC |
| 2539 | mamo-s | *BGIBMGA003104* | down | 702 | 710 | - | 11.3303 | 4.96E-05 | 0.615 | GTGGGTGGC |
| 2540 | mamo-s | *BGIBMGA003104* | down | 277 | 285 | - | 10.9358 | 7.30E-05 | 0.728 | GCGGGTGGC |
| 2541 | mamo-s | *BGIBMGA003105* | up | 984 | 992 | - | 11.3303 | 4.96E-05 | 0.615 | GTGGGTGGC |
| 2542 | mamo-s | *BGIBMGA003105* | down | 1018 | 1026 | - | 11.3303 | 4.96E-05 | 0.615 | GTGGGTGGC |
| 2543 | mamo-s | *BGIBMGA003105* | up | 559 | 567 | - | 10.9358 | 7.30E-05 | 0.728 | GCGGGTGGC |
| 2544 | mamo-s | *BGIBMGA003106* | up | 635 | 643 | - | 12.1009 | 2.43E-05 | 0.615 | GTGCGTGGT |
| 2545 | mamo-s | *BGIBMGA003106* | up | 70 | 78 | + | 11.3303 | 4.96E-05 | 0.615 | GTGGGTGGC |

| | | | | | | | | | | |
|---|---|---|---|---|---|---|---|---|---|---|
| 2546 | mamo-s | *BGIBMGA003106* | down | 718 | 726 | - | 11.3303 | 4.96E-05 | 0.615 | GTGGGTGGC |
| 2547 | mamo-s | *BGIBMGA003107* | up | 1159 | 1167 | - | 11.3303 | 4.96E-05 | 0.615 | GTGGGTGGC |
| 2548 | mamo-s | *BGIBMGA003107* | up | 330 | 338 | + | 11.2844 | 5.18E-05 | 0.63 | CTGCGTGCC |
| 2549 | mamo-s | *BGIBMGA003108* | down | 1325 | 1333 | - | 11.3303 | 4.96E-05 | 0.615 | GTGGGTGGC |
| 2550 | mamo-s | *BGIBMGA003110* | up | 1332 | 1340 | - | 11.3303 | 4.96E-05 | 0.615 | GTGGGTGGC |
| 2551 | mamo-s | *BGIBMGA003114* | down | 1245 | 1253 | - | 11.3303 | 4.96E-05 | 0.615 | GTGGGTGGC |
| 2552 | mamo-s | *BGIBMGA003117* | up | 613 | 621 | - | 11.3303 | 4.96E-05 | 0.615 | GTGGGTGGC |
| 2553 | mamo-s | *BGIBMGA003117* | down | 679 | 687 | + | 11.3303 | 4.96E-05 | 0.615 | GTGGGTGGC |
| 2554 | mamo-s | *BGIBMGA003117* | down | 415 | 423 | - | 11.0826 | 6.84E-05 | 0.709 | GTACGTGGC |
| 2555 | mamo-s | *BGIBMGA003119* | down | 820 | 828 | - | 11.3303 | 4.96E-05 | 0.615 | GTGGGTGGC |
| 2556 | mamo-s | *BGIBMGA003120* | up | 1846 | 1854 | - | 11.3303 | 4.96E-05 | 0.615 | GTGGGTGGC |
| 2557 | mamo-s | *BGIBMGA003121* | up | 160 | 168 | - | 12.1009 | 2.43E-05 | 0.615 | GTGCGTGGT |
| 2558 | mamo-s | *BGIBMGA003122* | up | 842 | 850 | + | 11.5321 | 4.06E-05 | 0.615 | TTGCGTGGC |
| 2559 | mamo-s | *BGIBMGA003122* | down | 1794 | 1802 | - | 11.3303 | 4.96E-05 | 0.615 | GTGGGTGGC |
| 2560 | mamo-s | *BGIBMGA003124* | down | 1397 | 1405 | - | 11.3303 | 4.96E-05 | 0.615 | GTGGGTGGC |
| 2561 | mamo-s | *BGIBMGA003127* | up | 178 | 186 | + | 13.6606 | 4.06E-06 | 0.615 | GCGCGTGGC |
| 2562 | mamo-s | *BGIBMGA003127* | down | 1890 | 1898 | + | 11.5688 | 3.79E-05 | 0.615 | CTGCGTGTC |
| 2563 | mamo-s | *BGIBMGA003128* | up | 1528 | 1536 | + | 11.1376 | 6.35E-05 | 0.709 | TCGCGTGGC |
| 2564 | mamo-s | *BGIBMGA003130* | down | 1353 | 1361 | + | 11.7339 | 3.29E-05 | 0.615 | GCGCGAGGC |
| 2565 | mamo-s | *BGIBMGA003130* | up | 1179 | 1187 | - | 11.3303 | 4.96E-05 | 0.615 | GTGGGTGGC |
| 2566 | mamo-s | *BGIBMGA003130* | down | 1376 | 1384 | - | 11.2018 | 5.68E-05 | 0.674 | CTGCGTGGT |
| 2567 | mamo-s | *BGIBMGA003134* | up | 1859 | 1867 | + | 13.156 | 6.30E-06 | 0.615 | CTGCGTGGC |
| 2568 | mamo-s | *BGIBMGA003135* | up | 1598 | 1606 | + | 11.5321 | 4.06E-05 | 0.615 | TTGCGTGGC |
| 2569 | mamo-s | *BGIBMGA003136* | up | 1536 | 1544 | + | 11.3303 | 4.96E-05 | 0.615 | GTGGGTGGC |
| 2570 | mamo-s | *BGIBMGA003137* | up | 515 | 523 | - | 11.3303 | 4.96E-05 | 0.615 | GTGGGTGGC |
| 2571 | mamo-s | *BGIBMGA003138* | down | 1427 | 1435 | - | 11.3303 | 4.96E-05 | 0.615 | GTGGGTGGC |
| 2572 | mamo-s | *BGIBMGA003140* | down | 1282 | 1290 | + | 11.3303 | 4.96E-05 | 0.615 | GTGGGTGGC |
| 2573 | mamo-s | *BGIBMGA003147* | down | 484 | 492 | + | 11.3303 | 4.96E-05 | 0.615 | GTGGGTGGC |
| 2574 | mamo-s | *BGIBMGA003148* | up | 73 | 81 | + | 11.3303 | 4.96E-05 | 0.615 | GTGGGTGGC |
| 2575 | mamo-s | *BGIBMGA003149* | up | 329 | 337 | + | 10.5138 | 9.75E-05 | 0.865 | GTGCGTGTT |
| 2576 | mamo-s | *BGIBMGA003153* | down | 1860 | 1868 | + | 11.3303 | 4.96E-05 | 0.615 | GTGGGTGGC |
| 2577 | mamo-s | *BGIBMGA003155* | up | 1479 | 1487 | + | 11.3303 | 4.96E-05 | 0.615 | GTGGGTGGC |
| 2578 | mamo-s | *BGIBMGA003158* | down | 342 | 350 | - | 11.7339 | 3.29E-05 | 0.615 | GCGCGAGGC |
| 2579 | mamo-s | *BGIBMGA003159* | up | 392 | 400 | + | 12.0734 | 2.66E-05 | 0.615 | GCGCGTGTC |
| 2580 | mamo-s | *BGIBMGA003159* | up | 818 | 826 | - | 11.7339 | 3.29E-05 | 0.615 | GCGCGAGGC |
| 2581 | mamo-s | *BGIBMGA003159* | up | 347 | 355 | + | 10.8899 | 7.48E-05 | 0.739 | CCGCGTGCC |
| 2582 | mamo-s | *BGIBMGA003160* | up | 1823 | 1831 | - | 13.6606 | 4.06E-06 | 0.615 | GCGCGTGGC |
| 2583 | mamo-s | *BGIBMGA003161* | up | 1188 | 1196 | - | 11.3303 | 4.96E-05 | 0.615 | GTGGGTGGC |
| 2584 | mamo-s | *BGIBMGA003163* | down | 1676 | 1684 | - | 12.7615 | 1.04E-05 | 0.615 | CCGCGTGGC |
| 2585 | mamo-s | *BGIBMGA003163* | up | 41 | 49 | + | 11.3303 | 4.96E-05 | 0.615 | GTGGGTGGC |
| 2586 | mamo-s | *BGIBMGA003163* | up | 282 | 290 | + | 11.3303 | 4.96E-05 | 0.615 | GTGGGTGGC |
| 2587 | mamo-s | *BGIBMGA003164* | up | 673 | 681 | + | 10.9358 | 7.30E-05 | 0.728 | GCGGGTGGC |
| 2588 | mamo-s | *BGIBMGA003168* | up | 1654 | 1662 | - | 12.1835 | 1.71E-05 | 0.615 | GTGCGTGCC |
| 2589 | mamo-s | *BGIBMGA003171* | down | 403 | 411 | - | 11.3303 | 4.96E-05 | 0.615 | GTGGGTGGC |
| 2590 | mamo-s | *BGIBMGA003173* | down | 1703 | 1711 | + | 11.3303 | 4.96E-05 | 0.615 | GTGGGTGGC |
| 2591 | mamo-s | *BGIBMGA003174* | up | 1800 | 1808 | + | 11.0826 | 6.84E-05 | 0.709 | GTACGTGGC |
| 2592 | mamo-s | *BGIBMGA003177* | down | 1375 | 1383 | - | 12.4679 | 1.31E-05 | 0.615 | GTGCGTGTC |
| 2593 | mamo-s | *BGIBMGA003183* | up | 530 | 538 | + | 11.1376 | 6.35E-05 | 0.709 | TCGCGTGGC |
| 2594 | mamo-s | *BGIBMGA003183* | down | 1806 | 1814 | + | 10.7523 | 8.38E-05 | 0.797 | AAGCGTGGC |
| 2595 | mamo-s | *BGIBMGA003185* | down | 959 | 967 | - | 11.3303 | 4.96E-05 | 0.615 | GTGGGTGGC |
| 2596 | mamo-s | *BGIBMGA003185* | down | 1888 | 1896 | - | 11.3303 | 4.96E-05 | 0.615 | GTGGGTGGC |
| 2597 | mamo-s | *BGIBMGA003189* | down | 1904 | 1912 | - | 14.055 | 2.23E-06 | 0.615 | GTGCGTGGC |
| 2598 | mamo-s | *BGIBMGA003189* | up | 153 | 161 | - | 12.7615 | 1.04E-05 | 0.615 | CCGCGTGGC |
| 2599 | mamo-s | *BGIBMGA003190* | down | 1190 | 1198 | - | 11.3303 | 4.96E-05 | 0.615 | GTGGGTGGC |
| 2600 | mamo-s | *BGIBMGA003192* | up | 1161 | 1169 | - | 11.0826 | 6.84E-05 | 0.709 | GTACGTGGC |
| 2601 | mamo-s | *BGIBMGA003192* | up | 507 | 515 | + | 11.0734 | 7.12E-05 | 0.726 | GAGCGTGGT |
| 2602 | mamo-s | *BGIBMGA003193* | down | 387 | 395 | - | 11.3303 | 4.96E-05 | 0.615 | GTGGGTGGC |
| 2603 | mamo-s | *BGIBMGA003193* | up | 934 | 942 | - | 11.1743 | 5.90E-05 | 0.684 | CCGCGTGTC |
| 2604 | mamo-s | *BGIBMGA003196* | up | 427 | 435 | - | 11.0826 | 6.84E-05 | 0.709 | GTACGTGGC |
| 2605 | mamo-s | *BGIBMGA003202* | up | 10 | 18 | + | 11.3303 | 4.96E-05 | 0.615 | GTGGGTGGC |
| 2606 | mamo-s | *BGIBMGA003202* | down | 1894 | 1902 | + | 11.3303 | 4.96E-05 | 0.615 | GTGGGTGGC |
| 2607 | mamo-s | *BGIBMGA003203* | down | 673 | 681 | + | 12.1009 | 2.43E-05 | 0.615 | GTGCGTGGT |
| 2608 | mamo-s | *BGIBMGA003210* | up | 1764 | 1772 | + | 11.0734 | 7.12E-05 | 0.726 | GAGCGTGGT |
| 2609 | mamo-s | *BGIBMGA003212* | down | 426 | 434 | + | 10.9358 | 7.30E-05 | 0.728 | GCGGGTGGC |
| 2610 | mamo-s | *BGIBMGA003213* | up | 1564 | 1572 | - | 12.0734 | 2.66E-05 | 0.615 | GCGCGTGTC |
| 2611 | mamo-s | *BGIBMGA003213* | up | 1584 | 1592 | - | 11.5688 | 3.79E-05 | 0.615 | CTGCGTGTC |
| 2612 | mamo-s | *BGIBMGA003215* | down | 1635 | 1643 | + | 14.055 | 2.23E-06 | 0.615 | GTGCGTGGC |
| 2613 | mamo-s | *BGIBMGA003215* | down | 1971 | 1979 | - | 12.0734 | 2.66E-05 | 0.615 | GCGCGTGTC |
| 2614 | mamo-s | *BGIBMGA003215* | down | 572 | 580 | - | 11.4404 | 4.51E-05 | 0.615 | CGGCGTGGC |
| 2615 | mamo-s | *BGIBMGA003216* | up | 214 | 222 | + | 12.4679 | 1.31E-05 | 0.615 | GTGCGTGTC |
| 2616 | mamo-s | *BGIBMGA003216* | down | 1162 | 1170 | + | 11.3303 | 4.96E-05 | 0.615 | GTGGGTGGC |
| 2617 | mamo-s | *BGIBMGA003218* | up | 1281 | 1289 | + | 11.3303 | 4.96E-05 | 0.615 | GTGGGTGGC |
| 2618 | mamo-s | *BGIBMGA003219* | up | 1489 | 1497 | + | 11.3303 | 4.96E-05 | 0.615 | GTGGGTGGC |
| 2619 | mamo-s | *BGIBMGA003220* | up | 1813 | 1821 | - | 11.1376 | 6.35E-05 | 0.709 | TCGCGTGGC |
| 2620 | mamo-s | *BGIBMGA003221* | up | 1465 | 1473 | + | 12.7615 | 1.04E-05 | 0.615 | CCGCGTGGC |
| 2621 | mamo-s | *BGIBMGA003221* | down | 1825 | 1833 | - | 11.0826 | 6.84E-05 | 0.709 | GTACGTGGC |
| 2622 | mamo-s | *BGIBMGA003223* | down | 1246 | 1254 | + | 11.3303 | 4.96E-05 | 0.615 | GTGGGTGGC |
| 2623 | mamo-s | *BGIBMGA003223* | up | 190 | 198 | + | 10.9358 | 7.30E-05 | 0.728 | GCGGGTGGC |
| 2624 | mamo-s | *BGIBMGA003224* | up | 874 | 882 | + | 11.3303 | 4.96E-05 | 0.615 | GTGGGTGGC |
| 2625 | mamo-s | *BGIBMGA003225* | up | 1147 | 1155 | + | 11.3303 | 4.96E-05 | 0.615 | GTGGGTGGC |
| 2626 | mamo-s | *BGIBMGA003227* | down | 883 | 891 | - | 11.7798 | 3.11E-05 | 0.615 | ATGCGTGGC |
| 2627 | mamo-s | *BGIBMGA003228* | down | 447 | 455 | - | 11.3303 | 4.96E-05 | 0.615 | GTGGGTGGC |
| 2628 | mamo-s | *BGIBMGA003229* | up | 1326 | 1334 | - | 11.3303 | 4.96E-05 | 0.615 | GTGGGTGGC |
| 2629 | mamo-s | *BGIBMGA003229* | up | 1731 | 1739 | + | 11.0826 | 6.84E-05 | 0.709 | GTACGTGGC |
| 2630 | mamo-s | *BGIBMGA003230* | up | 622 | 630 | + | 11.3303 | 4.96E-05 | 0.615 | GTGGGTGGC |
| 2631 | mamo-s | *BGIBMGA003231* | up | 1246 | 1254 | + | 14.055 | 2.23E-06 | 0.615 | GTGCGTGGC |
| 2632 | mamo-s | *BGIBMGA003232* | down | 1461 | 1469 | + | 11.3303 | 4.96E-05 | 0.615 | GTGGGTGGC |
| 2633 | mamo-s | *BGIBMGA003232* | up | 1473 | 1481 | + | 11.3303 | 4.96E-05 | 0.615 | GTGGGTGGC |

| | | | | | | | | | | |
|---|---|---|---|---|---|---|---|---|---|---|
| 2634 | mamo-s | *BGIBMGA003233* | up | 1742 | 1750 | + | 12.3394 | 1.49E-05 | 0.615 | GGGCGTGGC |
| 2635 | mamo-s | *BGIBMGA003234* | up | 368 | 376 | - | 11.3303 | 4.96E-05 | 0.615 | GTGGGTGGC |
| 2636 | mamo-s | *BGIBMGA003234* | down | 464 | 472 | + | 11.3303 | 4.96E-05 | 0.615 | GTGGGTGGC |
| 2637 | mamo-s | *BGIBMGA003234* | down | 1399 | 1407 | - | 11.3303 | 4.96E-05 | 0.615 | GTGGGTGGC |
| 2638 | mamo-s | *BGIBMGA003235* | down | 1891 | 1899 | - | 11.3303 | 4.96E-05 | 0.615 | GTGGGTGGC |
| 2639 | mamo-s | *BGIBMGA003235* | up | 828 | 836 | - | 10.5138 | 9.75E-05 | 0.865 | GTGCGTGTT |
| 2640 | mamo-s | *BGIBMGA003239* | down | 1807 | 1815 | + | 11.0826 | 6.84E-05 | 0.709 | GTACGTGGC |
| 2641 | mamo-s | *BGIBMGA003241* | down | 1960 | 1968 | + | 11.3303 | 4.96E-05 | 0.615 | GTGGGTGGC |
| 2642 | mamo-s | *BGIBMGA003244* | down | 1199 | 1207 | + | 11.3303 | 4.96E-05 | 0.615 | GTGGGTGGC |
| 2643 | mamo-s | *BGIBMGA003245* | down | 697 | 705 | + | 11.1743 | 5.90E-05 | 0.684 | CCGCGTGTC |
| 2644 | mamo-s | *BGIBMGA003246* | down | 1921 | 1929 | + | 11.3303 | 4.96E-05 | 0.615 | GTGGGTGGC |
| 2645 | mamo-s | *BGIBMGA003248* | up | 269 | 277 | + | 12.7615 | 1.04E-05 | 0.615 | CCGCGTGGC |
| 2646 | mamo-s | *BGIBMGA003248* | up | 1895 | 1903 | - | 11.5688 | 3.79E-05 | 0.615 | CTGCGTGTC |
| 2647 | mamo-s | *BGIBMGA003252* | down | 1484 | 1492 | - | 11.3303 | 4.96E-05 | 0.615 | GTGGGTGGC |
| 2648 | mamo-s | *BGIBMGA003252* | up | 1507 | 1515 | + | 11.3303 | 4.96E-05 | 0.615 | GTGGGTGGC |
| 2649 | mamo-s | *BGIBMGA003252* | down | 440 | 448 | + | 10.8899 | 7.48E-05 | 0.739 | CCGCGTGCC |
| 2650 | mamo-s | *BGIBMGA003253* | down | 377 | 385 | + | 11.3303 | 4.96E-05 | 0.615 | GTGGGTGGC |
| 2651 | mamo-s | *BGIBMGA003253* | down | 1684 | 1692 | - | 11.3303 | 4.96E-05 | 0.615 | GTGGGTGGC |
| 2652 | mamo-s | *BGIBMGA003253* | up | 1860 | 1868 | + | 11.3303 | 4.96E-05 | 0.615 | GTGGGTGGC |
| 2653 | mamo-s | *BGIBMGA003253* | up | 794 | 802 | + | 10.7523 | 8.38E-05 | 0.797 | GGGCGTGTC |
| 2654 | mamo-s | *BGIBMGA003254* | up | 1828 | 1836 | + | 14.055 | 2.23E-06 | 0.615 | GTGCGTGGC |
| 2655 | mamo-s | *BGIBMGA003254* | up | 113 | 121 | - | 11.3303 | 4.96E-05 | 0.615 | GTGGGTGGC |
| 2656 | mamo-s | *BGIBMGA003255* | down | 1078 | 1086 | - | 11.3303 | 4.96E-05 | 0.615 | GTGGGTGGC |
| 2657 | mamo-s | *BGIBMGA003255* | up | 1409 | 1417 | + | 11.3303 | 4.96E-05 | 0.615 | GTGGGTGGC |
| 2658 | mamo-s | *BGIBMGA003255* | up | 216 | 224 | - | 11.0826 | 6.84E-05 | 0.709 | GTACGTGGC |
| 2659 | mamo-s | *BGIBMGA003256* | down | 333 | 341 | + | 11.3303 | 4.96E-05 | 0.615 | GTGGGTGGC |
| 2660 | mamo-s | *BGIBMGA003256* | down | 722 | 730 | - | 11.0826 | 6.84E-05 | 0.709 | GTACGTGGC |
| 2661 | mamo-s | *BGIBMGA003257* | down | 770 | 778 | + | 11.3303 | 4.96E-05 | 0.615 | GTGGGTGGC |
| 2662 | mamo-s | *BGIBMGA003257* | up | 837 | 845 | - | 11.3303 | 4.96E-05 | 0.615 | GTGGGTGGC |
| 2663 | mamo-s | *BGIBMGA003258* | up | 843 | 851 | - | 11.3303 | 4.96E-05 | 0.615 | GTGGGTGGC |
| 2664 | mamo-s | *BGIBMGA003259* | up | 921 | 929 | - | 11.3303 | 4.96E-05 | 0.615 | GTGGGTGGC |
| 2665 | mamo-s | *BGIBMGA003260* | up | 794 | 802 | + | 14.055 | 2.23E-06 | 0.615 | GTGCGTGGC |
| 2666 | mamo-s | *BGIBMGA003260* | up | 157 | 165 | - | 11.3303 | 4.96E-05 | 0.615 | GTGGGTGGC |
| 2667 | mamo-s | *BGIBMGA003261* | up | 175 | 183 | + | 11.3303 | 4.96E-05 | 0.615 | GTGGGTGGC |
| 2668 | mamo-s | *BGIBMGA003262* | up | 1254 | 1262 | - | 14.055 | 2.23E-06 | 0.615 | GTGCGTGGC |
| 2669 | mamo-s | *BGIBMGA003263* | up | 1772 | 1780 | - | 11.3303 | 4.96E-05 | 0.615 | GTGGGTGGC |
| 2670 | mamo-s | *BGIBMGA003264* | up | 475 | 483 | - | 11.3303 | 4.96E-05 | 0.615 | GTGGGTGGC |
| 2671 | mamo-s | *BGIBMGA003264* | up | 1642 | 1650 | + | 11.3303 | 4.96E-05 | 0.615 | GTGGGTGGC |
| 2672 | mamo-s | *BGIBMGA003265* | down | 724 | 732 | + | 11.3303 | 4.96E-05 | 0.615 | GTGGGTGGC |
| 2673 | mamo-s | *BGIBMGA003265* | up | 962 | 970 | - | 11.3303 | 4.96E-05 | 0.615 | GTGGGTGGC |
| 2674 | mamo-s | *BGIBMGA003265* | up | 1268 | 1276 | - | 10.5413 | 9.42E-05 | 0.857 | CAGCGTGTC |
| 2675 | mamo-s | *BGIBMGA003266* | up | 1303 | 1311 | - | 11.3303 | 4.96E-05 | 0.615 | GTGGGTGGC |
| 2676 | mamo-s | *BGIBMGA003267* | down | 1162 | 1170 | - | 11.0826 | 6.84E-05 | 0.709 | GTACGTGGC |
| 2677 | mamo-s | *BGIBMGA003268* | down | 673 | 681 | - | 10.5688 | 8.88E-05 | 0.824 | GTGCGTGAC |
| 2678 | mamo-s | *BGIBMGA003269* | up | 547 | 555 | - | 10.5413 | 9.42E-05 | 0.857 | GTGCGAGTC |
| 2679 | mamo-s | *BGIBMGA003270* | up | 1884 | 1892 | + | 11.3303 | 4.96E-05 | 0.615 | GTGGGTGGC |
| 2680 | mamo-s | *BGIBMGA003271* | down | 1298 | 1306 | - | 11.3303 | 4.96E-05 | 0.615 | GTGGGTGGC |
| 2681 | mamo-s | *BGIBMGA003271* | down | 744 | 752 | + | 11.0826 | 6.84E-05 | 0.709 | GTACGTGGC |
| 2682 | mamo-s | *BGIBMGA003272* | up | 226 | 234 | + | 11.3303 | 4.96E-05 | 0.615 | GTGGGTGGC |
| 2683 | mamo-s | *BGIBMGA003273* | up | 1474 | 1482 | + | 11.4404 | 4.51E-05 | 0.615 | CGGCGTGGC |
| 2684 | mamo-s | *BGIBMGA003273* | up | 392 | 400 | + | 11.3303 | 4.96E-05 | 0.615 | GTGGGTGGC |
| 2685 | mamo-s | *BGIBMGA003273* | up | 915 | 923 | - | 11.3303 | 4.96E-05 | 0.615 | GTGGGTGGC |
| 2686 | mamo-s | *BGIBMGA003274* | down | 1039 | 1047 | - | 11.3303 | 4.96E-05 | 0.615 | GTGGGTGGC |
| 2687 | mamo-s | *BGIBMGA003274* | up | 1111 | 1119 | - | 11.3303 | 4.96E-05 | 0.615 | GTGGGTGGC |
| 2688 | mamo-s | *BGIBMGA003275* | down | 656 | 664 | - | 11.3303 | 4.96E-05 | 0.615 | GTGGGTGGC |
| 2689 | mamo-s | *BGIBMGA003276* | down | 1305 | 1313 | + | 11.3303 | 4.96E-05 | 0.615 | GTGGGTGGC |
| 2690 | mamo-s | *BGIBMGA003277* | up | 497 | 505 | + | 11.3303 | 4.96E-05 | 0.615 | GTGGGTGGC |
| 2691 | mamo-s | *BGIBMGA003277* | down | 398 | 406 | - | 11.0826 | 6.84E-05 | 0.709 | GTACGTGGC |
| 2692 | mamo-s | *BGIBMGA003278* | down | 1690 | 1698 | + | 11.0826 | 6.84E-05 | 0.709 | GTACGTGGC |
| 2693 | mamo-s | *BGIBMGA003279* | up | 1251 | 1259 | - | 12.7615 | 1.04E-05 | 0.615 | CCGCGTGGC |
| 2694 | mamo-s | *BGIBMGA003279* | down | 764 | 772 | + | 11.3303 | 4.96E-05 | 0.615 | GTGGGTGGC |
| 2695 | mamo-s | *BGIBMGA003280* | up | 1559 | 1567 | + | 10.5413 | 9.42E-05 | 0.857 | GTGCGAGTC |
| 2696 | mamo-s | *BGIBMGA003281* | up | 1171 | 1179 | - | 13.6606 | 4.06E-06 | 0.615 | GCGCGTGGC |
| 2697 | mamo-s | *BGIBMGA003282* | up | 270 | 278 | - | 11.7339 | 3.29E-05 | 0.615 | GCGCGAGGC |
| 2698 | mamo-s | *BGIBMGA003283* | down | 1559 | 1567 | - | 11.3303 | 4.96E-05 | 0.615 | GTGGGTGGC |
| 2699 | mamo-s | *BGIBMGA003284* | up | 1225 | 1233 | - | 12.1009 | 2.43E-05 | 0.615 | GTGCGTGGT |
| 2700 | mamo-s | *BGIBMGA003284* | down | 1395 | 1403 | + | 10.5138 | 9.75E-05 | 0.865 | GTGCGTGTT |
| 2701 | mamo-s | *BGIBMGA003285* | down | 1229 | 1237 | + | 11.789 | 2.84E-05 | 0.615 | GCGCGTGCC |
| 2702 | mamo-s | *BGIBMGA003286* | up | 1 | 9 | + | 13.6606 | 4.06E-06 | 0.615 | GCGCGTGGC |
| 2703 | mamo-s | *BGIBMGA003286* | down | 634 | 642 | + | 12.7615 | 1.04E-05 | 0.615 | CCGCGTGGC |
| 2704 | mamo-s | *BGIBMGA003286* | up | 392 | 400 | + | 11.5688 | 3.79E-05 | 0.615 | CTGCGTGTC |
| 2705 | mamo-s | *BGIBMGA003286* | up | 490 | 498 | - | 11.1009 | 6.57E-05 | 0.709 | GAGCGAGGC |
| 2706 | mamo-s | *BGIBMGA003286* | down | 1055 | 1063 | + | 10.8073 | 7.89E-05 | 0.766 | CCGCGTGGT |
| 2707 | mamo-s | *BGIBMGA003287* | up | 46 | 54 | + | 12.7615 | 1.04E-05 | 0.615 | CCGCGTGGC |
| 2708 | mamo-s | *BGIBMGA003287* | down | 225 | 233 | + | 12.0734 | 2.66E-05 | 0.615 | GCGCGTGTC |
| 2709 | mamo-s | *BGIBMGA003287* | up | 467 | 475 | + | 10.8073 | 7.89E-05 | 0.766 | CCGCGTGGT |
| 2710 | mamo-s | *BGIBMGA003289* | down | 1859 | 1867 | - | 10.8349 | 7.66E-05 | 0.751 | CCGCGAGGC |
| 2711 | mamo-s | *BGIBMGA003290* | down | 24 | 32 | + | 12.0734 | 2.66E-05 | 0.615 | GCGCGTGTC |
| 2712 | mamo-s | *BGIBMGA003290* | up | 1274 | 1282 | - | 10.8349 | 7.66E-05 | 0.751 | CCGCGAGGC |
| 2713 | mamo-s | *BGIBMGA003291* | up | 1477 | 1485 | - | 11.3303 | 4.96E-05 | 0.615 | GTGGGTGGC |
| 2714 | mamo-s | *BGIBMGA003293* | down | 1228 | 1236 | + | 12.7615 | 1.04E-05 | 0.615 | CCGCGTGGC |
| 2715 | mamo-s | *BGIBMGA003293* | up | 828 | 836 | - | 11.3303 | 4.96E-05 | 0.615 | GTGGGTGGC |
| 2716 | mamo-s | *BGIBMGA003294* | up | 753 | 761 | - | 11.3303 | 4.96E-05 | 0.615 | GTGGGTGGC |
| 2717 | mamo-s | *BGIBMGA003294* | down | 1117 | 1125 | - | 11.3303 | 4.96E-05 | 0.615 | GTGGGTGGC |
| 2718 | mamo-s | *BGIBMGA003297* | down | 578 | 586 | + | 11.7339 | 3.29E-05 | 0.615 | GCGCGAGGC |
| 2719 | mamo-s | *BGIBMGA003298* | down | 1895 | 1903 | - | 11.789 | 2.84E-05 | 0.615 | GCGCGTGCC |
| 2720 | mamo-s | *BGIBMGA003298* | down | 806 | 814 | + | 11.3303 | 4.96E-05 | 0.615 | GTGGGTGGC |
| 2721 | mamo-s | *BGIBMGA003301* | down | 1861 | 1869 | + | 11.3303 | 4.96E-05 | 0.615 | GTGGGTGGC |

| | | | | | | | | | | |
|---|---|---|---|---|---|---|---|---|---|---|
| 2722 | mamo-s | *BGIBMGA003303* | down | 238 | 246 | + | 11.3303 | 4.96E-05 | 0.615 | GTGGGTGGC |
| 2723 | mamo-s | *BGIBMGA003303* | up | 1357 | 1365 | + | 11.3303 | 4.96E-05 | 0.615 | GTGGGTGGC |
| 2724 | mamo-s | *BGIBMGA003304* | up | 1064 | 1072 | + | 11.3303 | 4.96E-05 | 0.615 | GTGGGTGGC |
| 2725 | mamo-s | *BGIBMGA003308* | up | 379 | 387 | - | 12.7615 | 1.04E-05 | 0.615 | CCGCGTGGC |
| 2726 | mamo-s | *BGIBMGA003309* | up | 619 | 627 | - | 11.2294 | 5.41E-05 | 0.65 | CTGCGAGGC |
| 2727 | mamo-s | *BGIBMGA003310* | up | 396 | 404 | - | 11.3303 | 4.96E-05 | 0.615 | GTGGGTGGC |
| 2728 | mamo-s | *BGIBMGA003310* | up | 971 | 979 | + | 11.0826 | 6.84E-05 | 0.709 | GTACGTGGC |
| 2729 | mamo-s | *BGIBMGA003311* | up | 722 | 730 | - | 11.1009 | 6.57E-05 | 0.709 | GAGCGAGGC |
| 2730 | mamo-s | *BGIBMGA003312* | down | 1683 | 1691 | - | 11.3303 | 4.96E-05 | 0.615 | GTGGGTGGC |
| 2731 | mamo-s | *BGIBMGA003312* | down | 209 | 217 | + | 11.0826 | 6.84E-05 | 0.709 | GTACGTGGC |
| 2732 | mamo-s | *BGIBMGA003314* | down | 2 | 10 | - | 12.1284 | 2.16E-05 | 0.615 | CAGCGTGGC |
| 2733 | mamo-s | *BGIBMGA003314* | up | 392 | 400 | - | 11.3303 | 4.96E-05 | 0.615 | GTGGGTGGC |
| 2734 | mamo-s | *BGIBMGA003315* | down | 680 | 688 | - | 11.7798 | 3.11E-05 | 0.615 | ATGCGTGGC |
| 2735 | mamo-s | *BGIBMGA003315* | down | 1145 | 1153 | - | 10.5138 | 9.75E-05 | 0.865 | GTGCGTGTT |
| 2736 | mamo-s | *BGIBMGA003315* | up | 1654 | 1662 | - | 10.5138 | 9.75E-05 | 0.865 | GTGCGTGTT |
| 2737 | mamo-s | *BGIBMGA003316* | down | 796 | 804 | + | 12.1835 | 1.71E-05 | 0.615 | GTGCGTGCC |
| 2738 | mamo-s | *BGIBMGA003317* | down | 1183 | 1191 | + | 11.3303 | 4.96E-05 | 0.615 | GTGGGTGGC |
| 2739 | mamo-s | *BGIBMGA003319* | up | 1175 | 1183 | - | 14.055 | 2.23E-06 | 0.615 | GTGCGTGGC |
| 2740 | mamo-s | *BGIBMGA003323* | up | 566 | 574 | + | 11.3303 | 4.96E-05 | 0.615 | GTGGGTGGC |
| 2741 | mamo-s | *BGIBMGA003326* | up | 1400 | 1408 | - | 11.1376 | 6.35E-05 | 0.709 | TCGCGTGGC |
| 2742 | mamo-s | *BGIBMGA003327* | up | 82 | 90 | + | 11.3303 | 4.96E-05 | 0.615 | GTGGGTGGC |
| 2743 | mamo-s | *BGIBMGA003329* | down | 1424 | 1432 | - | 11.1743 | 5.90E-05 | 0.684 | CCGCGTGTC |
| 2744 | mamo-s | *BGIBMGA003330* | up | 601 | 609 | - | 11.3303 | 4.96E-05 | 0.615 | GTGGGTGGC |
| 2745 | mamo-s | *BGIBMGA003330* | down | 1525 | 1533 | - | 11.3303 | 4.96E-05 | 0.615 | GTGGGTGGC |
| 2746 | mamo-s | *BGIBMGA003343* | up | 1275 | 1283 | - | 11.3303 | 4.96E-05 | 0.615 | GTGGGTGGC |
| 2747 | mamo-s | *BGIBMGA003344* | up | 367 | 375 | - | 11.3303 | 4.96E-05 | 0.615 | GTGGGTGGC |
| 2748 | mamo-s | *BGIBMGA003347* | down | 225 | 233 | - | 14.055 | 2.23E-06 | 0.615 | GTGCGTGGC |
| 2749 | mamo-s | *BGIBMGA003348* | up | 166 | 174 | - | 14.055 | 2.23E-06 | 0.615 | GTGCGTGGC |
| 2750 | mamo-s | *BGIBMGA003350* | down | 1321 | 1329 | + | 12.1009 | 2.43E-05 | 0.615 | GTGCGTGGT |
| 2751 | mamo-s | *BGIBMGA003350* | down | 1763 | 1771 | + | 11.5321 | 4.06E-05 | 0.615 | TTGCGTGGC |
| 2752 | mamo-s | *BGIBMGA003352* | down | 1740 | 1748 | - | 11.3303 | 4.96E-05 | 0.615 | GTGGGTGGC |
| 2753 | mamo-s | *BGIBMGA003353* | up | 280 | 288 | + | 10.7523 | 8.38E-05 | 0.797 | AAGCGTGGC |
| 2754 | mamo-s | *BGIBMGA003354* | down | 1637 | 1645 | - | 11.3303 | 4.96E-05 | 0.615 | GTGGGTGGC |
| 2755 | mamo-s | *BGIBMGA003357* | down | 660 | 668 | + | 10.5138 | 9.75E-05 | 0.865 | GTGCGTGTT |
| 2756 | mamo-s | *BGIBMGA003358* | up | 608 | 616 | + | 10.5138 | 9.75E-05 | 0.865 | GTGCGTGTT |
| 2757 | mamo-s | *BGIBMGA003359* | up | 1194 | 1202 | - | 11.7798 | 3.11E-05 | 0.615 | ATGCGTGGC |
| 2758 | mamo-s | *BGIBMGA003360* | up | 324 | 332 | - | 11.3303 | 4.96E-05 | 0.615 | GTGGGTGGC |
| 2759 | mamo-s | *BGIBMGA003362* | up | 1764 | 1772 | - | 11.3303 | 4.96E-05 | 0.615 | GTGGGTGGC |
| 2760 | mamo-s | *BGIBMGA003362* | up | 1972 | 1980 | - | 11.1743 | 5.90E-05 | 0.684 | CCGCGTGTC |
| 2761 | mamo-s | *BGIBMGA003363* | up | 881 | 889 | + | 12.4679 | 1.31E-05 | 0.615 | GTGCGTGTC |
| 2762 | mamo-s | *BGIBMGA003363* | up | 941 | 949 | + | 12.4679 | 1.31E-05 | 0.615 | GTGCGTGTC |
| 2763 | mamo-s | *BGIBMGA003363* | up | 1001 | 1009 | + | 12.4679 | 1.31E-05 | 0.615 | GTGCGTGTC |
| 2764 | mamo-s | *BGIBMGA003363* | up | 1035 | 1043 | - | 11.4404 | 4.51E-05 | 0.615 | CGGCGTGGC |
| 2765 | mamo-s | *BGIBMGA003363* | down | 1302 | 1310 | + | 11.3853 | 4.74E-05 | 0.615 | ACGCGTGGC |
| 2766 | mamo-s | *BGIBMGA003365* | up | 1220 | 1228 | + | 11.1743 | 5.90E-05 | 0.684 | CCGCGTGTC |
| 2767 | mamo-s | *BGIBMGA003366* | up | 415 | 423 | - | 11.3303 | 4.96E-05 | 0.615 | GTGGGTGGC |
| 2768 | mamo-s | *BGIBMGA003368* | down | 942 | 950 | + | 11.3303 | 4.96E-05 | 0.615 | GTGGGTGGC |
| 2769 | mamo-s | *BGIBMGA003370* | up | 1478 | 1486 | + | 12.1835 | 1.71E-05 | 0.615 | GTGCGTGCC |
| 2770 | mamo-s | *BGIBMGA003373* | up | 414 | 422 | + | 11.5321 | 4.06E-05 | 0.615 | TTGCGTGGC |
| 2771 | mamo-s | *BGIBMGA003373* | down | 1504 | 1512 | + | 11.3303 | 4.96E-05 | 0.615 | GTGGGTGGC |
| 2772 | mamo-s | *BGIBMGA003378* | up | 781 | 789 | + | 11.3853 | 4.74E-05 | 0.615 | ACGCGTGGC |
| 2773 | mamo-s | *BGIBMGA003378* | up | 479 | 487 | - | 11.2844 | 5.18E-05 | 0.63 | CTGCGTGCC |
| 2774 | mamo-s | *BGIBMGA003379* | down | 1377 | 1385 | - | 10.8073 | 7.89E-05 | 0.766 | CCGCGTGGT |
| 2775 | mamo-s | *BGIBMGA003380* | down | 538 | 546 | - | 10.5688 | 8.88E-05 | 0.824 | GTGCGTGAC |
| 2776 | mamo-s | *BGIBMGA003381* | up | 911 | 919 | - | 10.5688 | 8.88E-05 | 0.824 | GTGCGTGAC |
| 2777 | mamo-s | *BGIBMGA003382* | down | 1891 | 1899 | + | 10.5413 | 9.42E-05 | 0.857 | CAGCGTGTC |
| 2778 | mamo-s | *BGIBMGA003387* | down | 1142 | 1150 | + | 11.3303 | 4.96E-05 | 0.615 | GTGGGTGGC |
| 2779 | mamo-s | *BGIBMGA003387* | down | 1003 | 1011 | + | 11.2018 | 5.68E-05 | 0.674 | CTGCGTGGT |
| 2780 | mamo-s | *BGIBMGA003390* | up | 16 | 24 | - | 12.4679 | 1.31E-05 | 0.615 | GTGCGTGTC |
| 2781 | mamo-s | *BGIBMGA003390* | up | 76 | 84 | - | 12.4679 | 1.31E-05 | 0.615 | GTGCGTGTC |
| 2782 | mamo-s | *BGIBMGA003391* | down | 546 | 554 | + | 11.2294 | 5.41E-05 | 0.65 | CTGCGAGGC |
| 2783 | mamo-s | *BGIBMGA003393* | down | 1431 | 1439 | + | 11.1009 | 6.57E-05 | 0.709 | GAGCGAGGC |
| 2784 | mamo-s | *BGIBMGA003393* | down | 1620 | 1628 | - | 11.1009 | 6.57E-05 | 0.709 | GAGCGAGGC |
| 2785 | mamo-s | *BGIBMGA003397* | up | 1116 | 1124 | + | 11.0826 | 6.84E-05 | 0.709 | GTACGTGGC |
| 2786 | mamo-s | *BGIBMGA003401* | down | 1229 | 1237 | + | 13.6606 | 4.06E-06 | 0.615 | GCGCGTGGC |
| 2787 | mamo-s | *BGIBMGA003401* | down | 1839 | 1847 | - | 11.3303 | 4.96E-05 | 0.615 | GTGGGTGGC |
| 2788 | mamo-s | *BGIBMGA003402* | down | 329 | 337 | + | 11.3303 | 4.96E-05 | 0.615 | GTGGGTGGC |
| 2789 | mamo-s | *BGIBMGA003402* | down | 630 | 638 | + | 11.3303 | 4.96E-05 | 0.615 | GTGGGTGGC |
| 2790 | mamo-s | *BGIBMGA003403* | down | 1574 | 1582 | + | 13.6606 | 4.06E-06 | 0.615 | GCGCGTGGC |
| 2791 | mamo-s | *BGIBMGA003405* | down | 412 | 420 | - | 13.6606 | 4.06E-06 | 0.615 | GCGCGTGGC |
| 2792 | mamo-s | *BGIBMGA003405* | up | 382 | 390 | + | 11.3303 | 4.96E-05 | 0.615 | GTGGGTGGC |
| 2793 | mamo-s | *BGIBMGA003407* | down | 486 | 494 | + | 12.7615 | 1.04E-05 | 0.615 | CCGCGTGGC |
| 2794 | mamo-s | *BGIBMGA003407* | up | 505 | 513 | - | 11.3303 | 4.96E-05 | 0.615 | GTGGGTGGC |
| 2795 | mamo-s | *BGIBMGA003407* | down | 1837 | 1845 | + | 11.2294 | 5.41E-05 | 0.65 | CTGCGAGGC |
| 2796 | mamo-s | *BGIBMGA003409* | down | 1928 | 1936 | + | 11.3303 | 4.96E-05 | 0.615 | GTGGGTGGC |
| 2797 | mamo-s | *BGIBMGA003409* | down | 352 | 360 | - | 11.1743 | 5.90E-05 | 0.684 | CCGCGTGTC |
| 2798 | mamo-s | *BGIBMGA003411* | up | 944 | 952 | - | 10.8073 | 7.89E-05 | 0.766 | CCGCGTGGT |
| 2799 | mamo-s | *BGIBMGA003416* | up | 440 | 448 | + | 14.055 | 2.23E-06 | 0.615 | GTGCGTGGC |
| 2800 | mamo-s | *BGIBMGA003419* | up | 1509 | 1517 | - | 11.4404 | 4.51E-05 | 0.615 | CGGCGTGGC |
| 2801 | mamo-s | *BGIBMGA003419* | up | 137 | 145 | + | 11.3303 | 4.96E-05 | 0.615 | GTGGGTGGC |
| 2802 | mamo-s | *BGIBMGA003419* | down | 1565 | 1573 | - | 11.0826 | 6.84E-05 | 0.709 | GTACGTGGC |
| 2803 | mamo-s | *BGIBMGA003420* | down | 1958 | 1966 | + | 11.5321 | 4.06E-05 | 0.615 | TTGCGTGGC |
| 2804 | mamo-s | *BGIBMGA003421* | down | 320 | 328 | - | 11.3303 | 4.96E-05 | 0.615 | GTGGGTGGC |
| 2805 | mamo-s | *BGIBMGA003425* | up | 863 | 871 | + | 12.4679 | 1.31E-05 | 0.615 | GTGCGTGTC |
| 2806 | mamo-s | *BGIBMGA003426* | down | 1842 | 1850 | + | 11.3303 | 4.96E-05 | 0.615 | GTGGGTGGC |
| 2807 | mamo-s | *BGIBMGA003426* | up | 904 | 912 | - | 11.2844 | 5.18E-05 | 0.63 | CTGCGTGCC |
| 2808 | mamo-s | *BGIBMGA003427* | up | 1353 | 1361 | - | 11.3303 | 4.96E-05 | 0.615 | GTGGGTGGC |
| 2809 | mamo-s | *BGIBMGA003428* | up | 1857 | 1865 | - | 12.1284 | 2.16E-05 | 0.615 | CAGCGTGGC |

| | | | | | | | | | | |
|---|---|---|---|---|---|---|---|---|---|---|
| 2810 | mamo-s | *BGIBMGA003430* | down | 408 | 416 | - | 11.3303 | 4.96E-05 | 0.615 | GTGGGTGGC |
| 2811 | mamo-s | *BGIBMGA003431* | down | 1357 | 1365 | - | 11.3303 | 4.96E-05 | 0.615 | GTGGGTGGC |
| 2812 | mamo-s | *BGIBMGA003431* | down | 1969 | 1977 | - | 11.3303 | 4.96E-05 | 0.615 | GTGGGTGGC |
| 2813 | mamo-s | *BGIBMGA003432* | up | 841 | 849 | - | 11.0826 | 6.84E-05 | 0.709 | GTACGTGGC |
| 2814 | mamo-s | *BGIBMGA003432* | down | 1036 | 1044 | + | 10.5413 | 9.42E-05 | 0.857 | GTGCGAGTC |
| 2815 | mamo-s | *BGIBMGA003435* | up | 1022 | 1030 | + | 11.2294 | 5.41E-05 | 0.65 | CTGCGAGGC |
| 2816 | mamo-s | *BGIBMGA003435* | down | 1951 | 1959 | + | 11.0826 | 6.84E-05 | 0.709 | GTACGTGGC |
| 2817 | mamo-s | *BGIBMGA003436* | up | 1171 | 1179 | - | 11.1743 | 5.90E-05 | 0.684 | CCGCGTGTC |
| 2818 | mamo-s | *BGIBMGA003441* | down | 1480 | 1488 | - | 13.6606 | 4.06E-06 | 0.615 | GCGCGTGGC |
| 2819 | mamo-s | *BGIBMGA003441* | up | 966 | 974 | + | 11.3303 | 4.96E-05 | 0.615 | GTGGGTGGC |
| 2820 | mamo-s | *BGIBMGA003443* | up | 906 | 914 | + | 11.3303 | 4.96E-05 | 0.615 | GTGGGTGGC |
| 2821 | mamo-s | *BGIBMGA003444* | up | 326 | 334 | + | 11.3303 | 4.96E-05 | 0.615 | GTGGGTGGC |
| 2822 | mamo-s | *BGIBMGA003444* | down | 686 | 694 | - | 11.3303 | 4.96E-05 | 0.615 | GTGGGTGGC |
| 2823 | mamo-s | *BGIBMGA003444* | down | 1903 | 1911 | + | 11.3303 | 4.96E-05 | 0.615 | GTGGGTGGC |
| 2824 | mamo-s | *BGIBMGA003445* | down | 1196 | 1204 | - | 11.7339 | 3.29E-05 | 0.615 | GCGCGAGGC |
| 2825 | mamo-s | *BGIBMGA003445* | up | 610 | 618 | - | 11.3303 | 4.96E-05 | 0.615 | GTGGGTGGC |
| 2826 | mamo-s | *BGIBMGA003445* | down | 1173 | 1181 | + | 11.2018 | 5.68E-05 | 0.674 | CTGCGTGGT |
| 2827 | mamo-s | *BGIBMGA003447* | down | 1136 | 1144 | + | 11.3303 | 4.96E-05 | 0.615 | GTGGGTGGC |
| 2828 | mamo-s | *BGIBMGA003448* | down | 1093 | 1101 | - | 11.3303 | 4.96E-05 | 0.615 | GTGGGTGGC |
| 2829 | mamo-s | *BGIBMGA003448* | up | 815 | 823 | + | 10.5688 | 8.88E-05 | 0.824 | GTGCGTGAC |
| 2830 | mamo-s | *BGIBMGA003452* | down | 502 | 510 | - | 10.5688 | 8.88E-05 | 0.824 | GTGCGTGAC |
| 2831 | mamo-s | *BGIBMGA003453* | up | 467 | 475 | - | 10.5688 | 8.88E-05 | 0.824 | GTGCGTGAC |
| 2832 | mamo-s | *BGIBMGA003455* | up | 1764 | 1772 | - | 11.7798 | 3.11E-05 | 0.615 | ATGCGTGGC |
| 2833 | mamo-s | *BGIBMGA003456* | down | 487 | 495 | - | 11.3303 | 4.96E-05 | 0.615 | GTGGGTGGC |
| 2834 | mamo-s | *BGIBMGA003457* | down | 44 | 52 | + | 11.1009 | 6.57E-05 | 0.709 | GAGCGAGGC |
| 2835 | mamo-s | *BGIBMGA003458* | up | 1800 | 1808 | - | 11.3303 | 4.96E-05 | 0.615 | GTGGGTGGC |
| 2836 | mamo-s | *BGIBMGA003458* | down | 219 | 227 | + | 10.7523 | 8.38E-05 | 0.797 | AAGCGTGGC |
| 2837 | mamo-s | *BGIBMGA003459* | up | 482 | 490 | + | 10.7523 | 8.38E-05 | 0.797 | AAGCGTGGC |
| 2838 | mamo-s | *BGIBMGA003460* | down | 1424 | 1432 | + | 11.3303 | 4.96E-05 | 0.615 | GTGGGTGGC |
| 2839 | mamo-s | *BGIBMGA003467* | down | 1911 | 1919 | - | 11.3303 | 4.96E-05 | 0.615 | GTGGGTGGC |
| 2840 | mamo-s | *BGIBMGA003467* | up | 556 | 564 | - | 11.1743 | 5.90E-05 | 0.684 | CCGCGTGTC |
| 2841 | mamo-s | *BGIBMGA003468* | down | 1140 | 1148 | - | 11.5688 | 3.79E-05 | 0.615 | CTGCGTGTC |
| 2842 | mamo-s | *BGIBMGA003469* | down | 1123 | 1131 | + | 11.3303 | 4.96E-05 | 0.615 | GTGGGTGGC |
| 2843 | mamo-s | *BGIBMGA003471* | down | 1722 | 1730 | + | 11.0826 | 6.84E-05 | 0.709 | GTACGTGGC |
| 2844 | mamo-s | *BGIBMGA003473* | up | 491 | 499 | + | 11.0826 | 6.84E-05 | 0.709 | GTACGTGGC |
| 2845 | mamo-s | *BGIBMGA003475* | up | 282 | 290 | + | 11.3303 | 4.96E-05 | 0.615 | GTGGGTGGC |
| 2846 | mamo-s | *BGIBMGA003476* | up | 1922 | 1930 | - | 12.1835 | 1.71E-05 | 0.615 | GTGCGTGCC |
| 2847 | mamo-s | *BGIBMGA003477* | down | 999 | 1007 | + | 12.7615 | 1.04E-05 | 0.615 | CCGCGTGGC |
| 2848 | mamo-s | *BGIBMGA003477* | up | 1765 | 1773 | + | 11.4404 | 4.51E-05 | 0.615 | GAGCGTGTC |
| 2849 | mamo-s | *BGIBMGA003478* | up | 958 | 966 | + | 12.7615 | 1.04E-05 | 0.615 | CCGCGTGGC |
| 2850 | mamo-s | *BGIBMGA003480* | down | 224 | 232 | + | 11.3303 | 4.96E-05 | 0.615 | GTGGGTGGC |
| 2851 | mamo-s | *BGIBMGA003480* | up | 886 | 894 | - | 11.3303 | 4.96E-05 | 0.615 | GTGGGTGGC |
| 2852 | mamo-s | *BGIBMGA003480* | up | 1869 | 1877 | + | 10.5138 | 9.75E-05 | 0.865 | GTGCGTGTT |
| 2853 | mamo-s | *BGIBMGA003481* | down | 669 | 677 | - | 14.055 | 2.23E-06 | 0.615 | GTGCGTGGC |
| 2854 | mamo-s | *BGIBMGA003481* | up | 1225 | 1233 | + | 11.3303 | 4.96E-05 | 0.615 | GTGGGTGGC |
| 2855 | mamo-s | *BGIBMGA003485* | down | 167 | 175 | - | 11.3303 | 4.96E-05 | 0.615 | GTGGGTGGC |
| 2856 | mamo-s | *BGIBMGA003485* | down | 1653 | 1661 | + | 11.3303 | 4.96E-05 | 0.615 | GTGGGTGGC |
| 2857 | mamo-s | *BGIBMGA003486* | down | 1244 | 1252 | - | 11.3303 | 4.96E-05 | 0.615 | GTGGGTGGC |
| 2858 | mamo-s | *BGIBMGA003489* | down | 527 | 535 | + | 14.055 | 2.23E-06 | 0.615 | GTGCGTGGC |
| 2859 | mamo-s | *BGIBMGA003489* | up | 576 | 584 | + | 11.3303 | 4.96E-05 | 0.615 | GTGGGTGGC |
| 2860 | mamo-s | *BGIBMGA003497* | down | 1906 | 1914 | + | 11.3303 | 4.96E-05 | 0.615 | GTGGGTGGC |
| 2861 | mamo-s | *BGIBMGA003498* | up | 1328 | 1336 | - | 11.7339 | 3.29E-05 | 0.615 | GCGCGAGGC |
| 2862 | mamo-s | *BGIBMGA003498* | down | 207 | 215 | + | 11.7064 | 3.52E-05 | 0.615 | GCGCGTGGT |
| 2863 | mamo-s | *BGIBMGA003499* | down | 86 | 94 | + | 12.0734 | 2.66E-05 | 0.615 | GCGCGTGTC |
| 2864 | mamo-s | *BGIBMGA003499* | down | 699 | 707 | + | 11.3303 | 4.96E-05 | 0.615 | GTGGGTGGC |
| 2865 | mamo-s | *BGIBMGA003500* | down | 1489 | 1497 | - | 11.3303 | 4.96E-05 | 0.615 | GTGGGTGGC |
| 2866 | mamo-s | *BGIBMGA003501* | down | 1437 | 1445 | + | 10.9358 | 7.30E-05 | 0.728 | GCGGGTGGC |
| 2867 | mamo-s | *BGIBMGA003503* | down | 543 | 551 | - | 11.3303 | 4.96E-05 | 0.615 | GTGGGTGGC |
| 2868 | mamo-s | *BGIBMGA003503* | down | 1801 | 1809 | + | 11.3303 | 4.96E-05 | 0.615 | GTGGGTGGC |
| 2869 | mamo-s | *BGIBMGA003503* | up | 1002 | 1010 | - | 11.1009 | 6.57E-05 | 0.709 | GAGCGAGGC |
| 2870 | mamo-s | *BGIBMGA003506* | up | 49 | 57 | + | 13.156 | 6.30E-06 | 0.615 | CTGCGTGGC |
| 2871 | mamo-s | *BGIBMGA003508* | down | 569 | 577 | - | 14.055 | 2.23E-06 | 0.615 | GTGCGTGGC |
| 2872 | mamo-s | *BGIBMGA003508* | down | 943 | 951 | + | 11.0734 | 7.12E-05 | 0.726 | GAGCGTGGT |
| 2873 | mamo-s | *BGIBMGA003508* | up | 1558 | 1566 | - | 10.9358 | 7.30E-05 | 0.728 | GCGGGTGGC |
| 2874 | mamo-s | *BGIBMGA003509* | down | 1265 | 1273 | - | 12.1009 | 2.43E-05 | 0.615 | GTGCGTGGT |
| 2875 | mamo-s | *BGIBMGA003514* | up | 674 | 682 | - | 11.3303 | 4.96E-05 | 0.615 | GTGGGTGGC |
| 2876 | mamo-s | *BGIBMGA003516* | down | 508 | 516 | - | 11.4404 | 4.51E-05 | 0.615 | CGGCGTGGC |
| 2877 | mamo-s | *BGIBMGA003516* | up | 596 | 604 | - | 10.5688 | 8.88E-05 | 0.824 | GTGCGTGAC |
| 2878 | mamo-s | *BGIBMGA003517* | up | 1592 | 1600 | + | 11.3303 | 4.96E-05 | 0.615 | GTGGGTGGC |
| 2879 | mamo-s | *BGIBMGA003518* | up | 1164 | 1172 | + | 13.0275 | 8.53E-06 | 0.615 | GAGCGTGGC |
| 2880 | mamo-s | *BGIBMGA003519* | down | 74 | 82 | + | 10.8349 | 7.66E-05 | 0.751 | CCGCGAGGC |
| 2881 | mamo-s | *BGIBMGA003520* | up | 1238 | 1246 | + | 11.4404 | 4.51E-05 | 0.615 | CGGCGTGGC |
| 2882 | mamo-s | *BGIBMGA003520* | down | 1239 | 1247 | + | 11.0826 | 6.84E-05 | 0.709 | GTACGTGGC |
| 2883 | mamo-s | *BGIBMGA003522* | up | 1297 | 1305 | - | 11.3303 | 4.96E-05 | 0.615 | GTGGGTGGC |
| 2884 | mamo-s | *BGIBMGA003525* | up | 1950 | 1958 | + | 11.4404 | 4.51E-05 | 0.615 | CGGCGTGGC |
| 2885 | mamo-s | *BGIBMGA003526* | up | 677 | 685 | - | 13.0275 | 8.53E-06 | 0.615 | GAGCGTGGC |
| 2886 | mamo-s | *BGIBMGA003528* | up | 1229 | 1237 | + | 10.8899 | 7.48E-05 | 0.739 | CCGCGTGCC |
| 2887 | mamo-s | *BGIBMGA003529* | down | 1099 | 1107 | - | 11.3303 | 4.96E-05 | 0.615 | GTGGGTGGC |
| 2888 | mamo-s | *BGIBMGA003530* | up | 1760 | 1768 | - | 10.8899 | 7.48E-05 | 0.739 | CCGCGTGCC |
| 2889 | mamo-s | *BGIBMGA003531* | down | 1450 | 1458 | - | 11.3303 | 4.96E-05 | 0.615 | GTGGGTGGC |
| 2890 | mamo-s | *BGIBMGA003531* | up | 1735 | 1743 | - | 11.3303 | 4.96E-05 | 0.615 | GTGGGTGGC |
| 2891 | mamo-s | *BGIBMGA003532* | up | 702 | 710 | - | 11.3303 | 4.96E-05 | 0.615 | GTGGGTGGC |
| 2892 | mamo-s | *BGIBMGA003533* | up | 493 | 501 | - | 11.3303 | 4.96E-05 | 0.615 | GTGGGTGGC |
| 2893 | mamo-s | *BGIBMGA003535* | down | 655 | 663 | + | 11.4404 | 4.51E-05 | 0.615 | GAGCGTGTC |
| 2894 | mamo-s | *BGIBMGA003535* | up | 1448 | 1456 | - | 10.9358 | 7.30E-05 | 0.728 | GCGGGTGGC |
| 2895 | mamo-s | *BGIBMGA003538* | down | 418 | 426 | + | 12.1284 | 2.16E-05 | 0.615 | GTGCGAGGC |
| 2896 | mamo-s | *BGIBMGA003538* | down | 1139 | 1147 | + | 11.2294 | 5.41E-05 | 0.65 | CTGCGAGGC |
| 2897 | mamo-s | *BGIBMGA003548* | down | 1156 | 1164 | - | 11.3303 | 4.96E-05 | 0.615 | GTGGGTGGC |

| | | | | | | | | | | |
|---|---|---|---|---|---|---|---|---|---|---|
| 2898 | mamo-s | *BGIBMGA003548* | up | 1466 | 1474 | - | 11.3303 | 4.96E-05 | 0.615 | GTGGGTGGC |
| 2899 | mamo-s | *BGIBMGA003550* | up | 1136 | 1144 | - | 11.3303 | 4.96E-05 | 0.615 | GTGGGTGGC |
| 2900 | mamo-s | *BGIBMGA003552* | up | 1405 | 1413 | - | 10.5138 | 9.75E-05 | 0.865 | GTGCGTGTT |
| 2901 | mamo-s | *BGIBMGA003554* | down | 1565 | 1573 | - | 11.3303 | 4.96E-05 | 0.615 | GTGGGTGGC |
| 2902 | mamo-s | *BGIBMGA003555* | down | 1112 | 1120 | - | 13.0275 | 8.53E-06 | 0.615 | GAGCGTGGC |
| 2903 | mamo-s | *BGIBMGA003555* | down | 1320 | 1328 | + | 12.1284 | 2.16E-05 | 0.615 | GTGCGAGGC |
| 2904 | mamo-s | *BGIBMGA003555* | up | 1441 | 1449 | + | 11.3303 | 4.96E-05 | 0.615 | GTGGGTGGC |
| 2905 | mamo-s | *BGIBMGA003555* | down | 1134 | 1142 | - | 11.2844 | 5.18E-05 | 0.63 | CTGCGTGCC |
| 2906 | mamo-s | *BGIBMGA003555* | down | 1016 | 1024 | + | 10.8899 | 7.48E-05 | 0.739 | CCGCGTGCC |
| 2907 | mamo-s | *BGIBMGA003558* | down | 757 | 765 | + | 11.0734 | 7.12E-05 | 0.726 | GAGCGTGGT |
| 2908 | mamo-s | *BGIBMGA003560* | up | 1397 | 1405 | - | 11.3303 | 4.96E-05 | 0.615 | GTGGGTGGC |
| 2909 | mamo-s | *BGIBMGA003561* | down | 1313 | 1321 | + | 11.3303 | 4.96E-05 | 0.615 | GTGGGTGGC |
| 2910 | mamo-s | *BGIBMGA003568* | down | 860 | 868 | + | 11.3303 | 4.96E-05 | 0.615 | GTGGGTGGC |
| 2911 | mamo-s | *BGIBMGA003569* | down | 514 | 522 | + | 11.7798 | 3.11E-05 | 0.615 | ATGCGTGGC |
| 2912 | mamo-s | *BGIBMGA003569* | up | 545 | 553 | + | 11.3303 | 4.96E-05 | 0.615 | GTGGGTGGC |
| 2913 | mamo-s | *BGIBMGA003579* | up | 548 | 556 | - | 11.3303 | 4.96E-05 | 0.615 | GTGGGTGGC |
| 2914 | mamo-s | *BGIBMGA003579* | up | 766 | 774 | + | 11.3303 | 4.96E-05 | 0.615 | GTGGGTGGC |
| 2915 | mamo-s | *BGIBMGA003580* | up | 102 | 110 | - | 11.0826 | 6.84E-05 | 0.709 | GTACGTGGC |
| 2916 | mamo-s | *BGIBMGA003581* | down | 1657 | 1665 | - | 11.4404 | 4.51E-05 | 0.615 | GAGCGTGTC |
| 2917 | mamo-s | *BGIBMGA003581* | down | 253 | 261 | + | 11.3303 | 4.96E-05 | 0.615 | GTGGGTGGC |
| 2918 | mamo-s | *BGIBMGA003582* | down | 264 | 272 | + | 11.5688 | 3.79E-05 | 0.615 | CTGCGTGTC |
| 2919 | mamo-s | *BGIBMGA003582* | up | 1585 | 1593 | - | 11.4404 | 4.51E-05 | 0.615 | CGGCGTGGC |
| 2920 | mamo-s | *BGIBMGA003584* | up | 1970 | 1978 | + | 13.156 | 6.30E-06 | 0.615 | CTGCGTGGC |
| 2921 | mamo-s | *BGIBMGA003586* | down | 1926 | 1934 | - | 10.5413 | 9.42E-05 | 0.857 | GTGCGAGTC |
| 2922 | mamo-s | *BGIBMGA003590* | down | 1386 | 1394 | - | 11.4404 | 4.51E-05 | 0.615 | CGGCGTGGC |
| 2923 | mamo-s | *BGIBMGA003591* | up | 329 | 337 | - | 11.3303 | 4.96E-05 | 0.615 | GTGGGTGGC |
| 2924 | mamo-s | *BGIBMGA003591* | up | 489 | 497 | - | 11.3303 | 4.96E-05 | 0.615 | GTGGGTGGC |
| 2925 | mamo-s | *BGIBMGA003591* | down | 1638 | 1646 | + | 11.3303 | 4.96E-05 | 0.615 | GTGGGTGGC |
| 2926 | mamo-s | *BGIBMGA003592* | down | 11 | 19 | - | 12.1009 | 2.43E-05 | 0.615 | GTGCGTGGT |
| 2927 | mamo-s | *BGIBMGA003592* | up | 648 | 656 | - | 11.789 | 2.84E-05 | 0.615 | GCGCGTGCC |
| 2928 | mamo-s | *BGIBMGA003594* | down | 531 | 539 | + | 14.055 | 2.23E-06 | 0.615 | GTGCGTGGC |
| 2929 | mamo-s | *BGIBMGA003595* | down | 450 | 458 | + | 11.3303 | 4.96E-05 | 0.615 | GTGGGTGGC |
| 2930 | mamo-s | *BGIBMGA003598* | down | 1682 | 1690 | + | 11.0826 | 6.84E-05 | 0.709 | GTACGTGGC |
| 2931 | mamo-s | *BGIBMGA003598* | up | 1117 | 1125 | + | 10.7523 | 8.38E-05 | 0.797 | AAGCGTGGC |
| 2932 | mamo-s | *BGIBMGA003599* | down | 1074 | 1082 | + | 13.156 | 6.30E-06 | 0.615 | CTGCGTGGC |
| 2933 | mamo-s | *BGIBMGA003599* | up | 683 | 691 | - | 11.3303 | 4.96E-05 | 0.615 | GTGGGTGGC |
| 2934 | mamo-s | *BGIBMGA003600* | down | 75 | 83 | - | 13.156 | 6.30E-06 | 0.615 | CTGCGTGGC |
| 2935 | mamo-s | *BGIBMGA003601* | up | 307 | 315 | + | 11.3303 | 4.96E-05 | 0.615 | GTGGGTGGC |
| 2936 | mamo-s | *BGIBMGA003604* | down | 205 | 213 | + | 11.3303 | 4.96E-05 | 0.615 | GTGGGTGGC |
| 2937 | mamo-s | *BGIBMGA003605* | up | 256 | 264 | - | 12.3394 | 1.49E-05 | 0.615 | GGGCGTGGC |
| 2938 | mamo-s | *BGIBMGA003605* | down | 1790 | 1798 | - | 11.2844 | 5.18E-05 | 0.63 | CTGCGTGCC |
| 2939 | mamo-s | *BGIBMGA003607* | up | 1211 | 1219 | - | 11.3303 | 4.96E-05 | 0.615 | GTGGGTGGC |
| 2940 | mamo-s | *BGIBMGA003609* | up | 97 | 105 | - | 13.6606 | 4.06E-06 | 0.615 | GCGCGTGGC |
| 2941 | mamo-s | *BGIBMGA003609* | up | 586 | 594 | + | 10.8899 | 7.48E-05 | 0.739 | CCGCGTGCC |
| 2942 | mamo-s | *BGIBMGA003610* | up | 1542 | 1550 | + | 11.3303 | 4.96E-05 | 0.615 | GTGGGTGGC |
| 2943 | mamo-s | *BGIBMGA003610* | down | 1981 | 1989 | + | 10.5413 | 9.42E-05 | 0.857 | GTGCGAGTC |
| 2944 | mamo-s | *BGIBMGA003612* | up | 1486 | 1494 | + | 14.055 | 2.23E-06 | 0.615 | GTGCGTGGC |
| 2945 | mamo-s | *BGIBMGA003612* | up | 97 | 105 | - | 11.3303 | 4.96E-05 | 0.615 | GTGGGTGGC |
| 2946 | mamo-s | *BGIBMGA003613* | up | 1747 | 1755 | - | 11.0826 | 6.84E-05 | 0.709 | GTACGTGGC |
| 2947 | mamo-s | *BGIBMGA003613* | up | 735 | 743 | - | 10.5688 | 8.88E-05 | 0.824 | GTGCGTGAC |
| 2948 | mamo-s | *BGIBMGA003614* | up | 1521 | 1529 | + | 11.3303 | 4.96E-05 | 0.615 | GTGGGTGGC |
| 2949 | mamo-s | *BGIBMGA003614* | up | 788 | 796 | + | 11.0826 | 6.84E-05 | 0.709 | GTACGTGGC |
| 2950 | mamo-s | *BGIBMGA003615* | up | 844 | 852 | + | 10.7523 | 8.38E-05 | 0.797 | GGGCGTGTC |
| 2951 | mamo-s | *BGIBMGA003617* | down | 1421 | 1429 | + | 11.3303 | 4.96E-05 | 0.615 | GTGGGTGGC |
| 2952 | mamo-s | *BGIBMGA003619* | down | 1567 | 1575 | - | 11.0826 | 6.84E-05 | 0.709 | GTACGTGGC |
| 2953 | mamo-s | *BGIBMGA003621* | up | 1418 | 1426 | + | 11.7064 | 3.52E-05 | 0.615 | GCGCGTGGT |
| 2954 | mamo-s | *BGIBMGA003621* | down | 1609 | 1617 | + | 11.3303 | 4.96E-05 | 0.615 | GTGGGTGGC |
| 2955 | mamo-s | *BGIBMGA003622* | down | 718 | 726 | - | 11.3303 | 4.96E-05 | 0.615 | GTGGGTGGC |
| 2956 | mamo-s | *BGIBMGA003623* | down | 688 | 696 | + | 10.6881 | 8.61E-05 | 0.81 | GCACGTGGC |
| 2957 | mamo-s | *BGIBMGA003624* | down | 967 | 975 | - | 12.4679 | 1.31E-05 | 0.615 | GTGCGTGTC |
| 2958 | mamo-s | *BGIBMGA003624* | up | 115 | 123 | - | 11.0826 | 6.84E-05 | 0.709 | GTACGTGGC |
| 2959 | mamo-s | *BGIBMGA003624* | down | 1191 | 1199 | + | 10.6881 | 8.61E-05 | 0.81 | GCACGTGGC |
| 2960 | mamo-s | *BGIBMGA003626* | up | 1623 | 1631 | - | 14.055 | 2.23E-06 | 0.615 | GTGCGTGGC |
| 2961 | mamo-s | *BGIBMGA003627* | up | 1560 | 1568 | - | 11.3303 | 4.96E-05 | 0.615 | GTGGGTGGC |
| 2962 | mamo-s | *BGIBMGA003628* | up | 612 | 620 | - | 14.055 | 2.23E-06 | 0.615 | GTGCGTGGC |
| 2963 | mamo-s | *BGIBMGA003628* | down | 314 | 322 | - | 13.6606 | 4.06E-06 | 0.615 | GCGCGTGGC |
| 2964 | mamo-s | *BGIBMGA003629* | down | 870 | 878 | - | 11.1743 | 5.90E-05 | 0.684 | CCGCGTGTC |
| 2965 | mamo-s | *BGIBMGA003630* | up | 1571 | 1579 | - | 11.1743 | 5.90E-05 | 0.684 | CCGCGTGTC |
| 2966 | mamo-s | *BGIBMGA003635* | up | 1157 | 1165 | + | 11.3303 | 4.96E-05 | 0.615 | GTGGGTGGC |
| 2967 | mamo-s | *BGIBMGA003638* | down | 1044 | 1052 | + | 14.055 | 2.23E-06 | 0.615 | GTGCGTGGC |
| 2968 | mamo-s | *BGIBMGA003638* | up | 1688 | 1696 | - | 11.7064 | 3.52E-05 | 0.615 | GCGCGTGGT |
| 2969 | mamo-s | *BGIBMGA003639* | up | 67 | 75 | + | 11.3303 | 4.96E-05 | 0.615 | GTGGGTGGC |
| 2970 | mamo-s | *BGIBMGA003639* | down | 290 | 298 | + | 11.3303 | 4.96E-05 | 0.615 | GTGGGTGGC |
| 2971 | mamo-s | *BGIBMGA003639* | down | 1403 | 1411 | + | 11.3303 | 4.96E-05 | 0.615 | GTGGGTGGC |
| 2972 | mamo-s | *BGIBMGA003640* | down | 84 | 92 | + | 11.4404 | 4.51E-05 | 0.615 | CGGCGTGGC |
| 2973 | mamo-s | *BGIBMGA003640* | up | 1488 | 1496 | - | 11.3303 | 4.96E-05 | 0.615 | GTGGGTGGC |
| 2974 | mamo-s | *BGIBMGA003641* | up | 1377 | 1385 | - | 11.3303 | 4.96E-05 | 0.615 | GTGGGTGGC |
| 2975 | mamo-s | *BGIBMGA003642* | down | 1020 | 1028 | - | 11.3303 | 4.96E-05 | 0.615 | GTGGGTGGC |
| 2976 | mamo-s | *BGIBMGA003646* | down | 124 | 132 | - | 10.5138 | 9.75E-05 | 0.865 | GTGCGTGTT |
| 2977 | mamo-s | *BGIBMGA003647* | down | 1690 | 1698 | + | 10.5413 | 9.42E-05 | 0.857 | CAGCGTGTC |
| 2978 | mamo-s | *BGIBMGA003648* | down | 767 | 775 | + | 11.1743 | 5.90E-05 | 0.684 | CCGCGTGTC |
| 2979 | mamo-s | *BGIBMGA003648* | down | 1190 | 1198 | - | 10.5413 | 9.42E-05 | 0.857 | CAGCGTGTC |
| 2980 | mamo-s | *BGIBMGA003649* | up | 1625 | 1633 | - | 11.2294 | 5.41E-05 | 0.65 | CTGCGAGGC |
| 2981 | mamo-s | *BGIBMGA003649* | up | 219 | 227 | + | 11.0826 | 6.84E-05 | 0.709 | GTACGTGGC |
| 2982 | mamo-s | *BGIBMGA003649* | up | 944 | 952 | - | 11.0734 | 7.12E-05 | 0.726 | GAGCGTGGT |
| 2983 | mamo-s | *BGIBMGA003650* | down | 111 | 119 | - | 11.3303 | 4.96E-05 | 0.615 | GTGGGTGGC |
| 2984 | mamo-s | *BGIBMGA003653* | down | 578 | 586 | - | 11.3303 | 4.96E-05 | 0.615 | GTGGGTGGC |
| 2985 | mamo-s | *BGIBMGA003654* | down | 598 | 606 | + | 11.156 | 6.13E-05 | 0.705 | GAGCGTGCC |

| | | | | | | | | | | |
|---|---|---|---|---|---|---|---|---|---|---|
| 2986 | mamo-s | *BGIBMGA003655* | up | 883 | 891 | + | 10.9358 | 7.30E-05 | 0.728 | GCGGGTGGC |
| 2987 | mamo-s | *BGIBMGA003656* | down | 75 | 83 | - | 11.7798 | 3.11E-05 | 0.615 | ATGCGTGGC |
| 2988 | mamo-s | *BGIBMGA003665* | down | 21 | 29 | + | 11.5321 | 4.06E-05 | 0.615 | TTGCGTGGC |
| 2989 | mamo-s | *BGIBMGA003670* | up | 270 | 278 | - | 11.0826 | 6.84E-05 | 0.709 | GTACGTGGC |
| 2990 | mamo-s | *BGIBMGA003671* | down | 1466 | 1474 | - | 11.1743 | 5.90E-05 | 0.684 | CCGCGTGTC |
| 2991 | mamo-s | *BGIBMGA003671* | down | 1883 | 1891 | - | 10.5138 | 9.75E-05 | 0.865 | GTGCGTGTT |
| 2992 | mamo-s | *BGIBMGA003672* | down | 489 | 497 | + | 11.3303 | 4.96E-05 | 0.615 | GTGGGTGGC |
| 2993 | mamo-s | *BGIBMGA003673* | up | 1626 | 1634 | + | 11.3303 | 4.96E-05 | 0.615 | GTGGGTGGC |
| 2994 | mamo-s | *BGIBMGA003673* | up | 287 | 295 | - | 10.5413 | 9.42E-05 | 0.857 | CAGCGTGTC |
| 2995 | mamo-s | *BGIBMGA003678* | down | 1833 | 1841 | + | 12.1009 | 2.43E-05 | 0.615 | GTGCGTGGT |
| 2996 | mamo-s | *BGIBMGA003678* | up | 1097 | 1105 | - | 11.3303 | 4.96E-05 | 0.615 | GTGGGTGGC |
| 2997 | mamo-s | *BGIBMGA003679* | down | 1458 | 1466 | + | 11.4404 | 4.51E-05 | 0.615 | CGGCGTGGC |
| 2998 | mamo-s | *BGIBMGA003679* | up | 1890 | 1898 | + | 11.1743 | 5.90E-05 | 0.684 | CCGCGTGTC |
| 2999 | mamo-s | *BGIBMGA003680* | up | 1027 | 1035 | + | 11.3303 | 4.96E-05 | 0.615 | GTGGGTGGC |
| 3000 | mamo-s | *BGIBMGA003681* | down | 1670 | 1678 | + | 10.5138 | 9.75E-05 | 0.865 | GTGCGTGTT |
| 3001 | mamo-s | *BGIBMGA003683* | up | 90 | 98 | - | 11.3303 | 4.96E-05 | 0.615 | GTGGGTGGC |
| 3002 | mamo-s | *BGIBMGA003683* | up | 744 | 752 | + | 11.3303 | 4.96E-05 | 0.615 | GTGGGTGGC |
| 3003 | mamo-s | *BGIBMGA003688* | up | 429 | 437 | - | 11.1009 | 6.57E-05 | 0.709 | GAGCGAGGC |
| 3004 | mamo-s | *BGIBMGA003688* | up | 1297 | 1305 | + | 10.9358 | 7.30E-05 | 0.728 | GCGGGTGGC |
| 3005 | mamo-s | *BGIBMGA003689* | down | 1700 | 1708 | - | 10.5138 | 9.75E-05 | 0.865 | GTGCGTGTT |
| 3006 | mamo-s | *BGIBMGA003691* | down | 1211 | 1219 | - | 11.3303 | 4.96E-05 | 0.615 | GTGGGTGGC |
| 3007 | mamo-s | *BGIBMGA003692* | down | 476 | 484 | - | 11.3303 | 4.96E-05 | 0.615 | GTGGGTGGC |
| 3008 | mamo-s | *BGIBMGA003692* | down | 1416 | 1424 | - | 11.3303 | 4.96E-05 | 0.615 | GTGGGTGGC |
| 3009 | mamo-s | *BGIBMGA003695* | up | 1035 | 1043 | + | 11.3303 | 4.96E-05 | 0.615 | GTGGGTGGC |
| 3010 | mamo-s | *BGIBMGA003695* | down | 1697 | 1705 | - | 11.3303 | 4.96E-05 | 0.615 | GTGGGTGGC |
| 3011 | mamo-s | *BGIBMGA003697* | down | 1502 | 1510 | - | 11.3303 | 4.96E-05 | 0.615 | GTGGGTGGC |
| 3012 | mamo-s | *BGIBMGA003698* | up | 421 | 429 | + | 11.5688 | 3.79E-05 | 0.615 | CTGCGTGTC |
| 3013 | mamo-s | *BGIBMGA003701* | down | 1343 | 1351 | + | 11.2018 | 5.68E-05 | 0.674 | CTGCGTGGT |
| 3014 | mamo-s | *BGIBMGA003709* | up | 323 | 331 | - | 11.3303 | 4.96E-05 | 0.615 | GTGGGTGGC |
| 3015 | mamo-s | *BGIBMGA003709* | down | 796 | 804 | - | 11.3303 | 4.96E-05 | 0.615 | GTGGGTGGC |
| 3016 | mamo-s | *BGIBMGA003709* | up | 813 | 821 | + | 11.1743 | 5.90E-05 | 0.684 | CCGCGTGTC |
| 3017 | mamo-s | *BGIBMGA003711* | down | 721 | 729 | - | 11.5688 | 3.79E-05 | 0.615 | CTGCGTGTC |
| 3018 | mamo-s | *BGIBMGA003711* | up | 1370 | 1378 | - | 11.0826 | 6.84E-05 | 0.709 | GTACGTGGC |
| 3019 | mamo-s | *BGIBMGA003712* | down | 392 | 400 | + | 11.3303 | 4.96E-05 | 0.615 | GTGGGTGGC |
| 3020 | mamo-s | *BGIBMGA003712* | up | 1596 | 1604 | + | 11.3303 | 4.96E-05 | 0.615 | GTGGGTGGC |
| 3021 | mamo-s | *BGIBMGA003714* | down | 167 | 175 | - | 11.3303 | 4.96E-05 | 0.615 | GTGGGTGGC |
| 3022 | mamo-s | *BGIBMGA003714* | down | 690 | 698 | + | 11.3303 | 4.96E-05 | 0.615 | GTGGGTGGC |
| 3023 | mamo-s | *BGIBMGA003716* | down | 1275 | 1283 | + | 12.1009 | 2.43E-05 | 0.615 | GTGCGTGGT |
| 3024 | mamo-s | *BGIBMGA003722* | up | 464 | 472 | - | 10.7523 | 8.38E-05 | 0.797 | GGGCGTGTC |
| 3025 | mamo-s | *BGIBMGA003727* | down | 1510 | 1518 | - | 11.3303 | 4.96E-05 | 0.615 | GTGGGTGGC |
| 3026 | mamo-s | *BGIBMGA003728* | down | 948 | 956 | + | 11.3303 | 4.96E-05 | 0.615 | GTGGGTGGC |
| 3027 | mamo-s | *BGIBMGA003729* | down | 1545 | 1553 | + | 11.3303 | 4.96E-05 | 0.615 | GTGGGTGGC |
| 3028 | mamo-s | *BGIBMGA003730* | down | 1989 | 1997 | + | 12.4679 | 1.31E-05 | 0.615 | GTGCGTGTC |
| 3029 | mamo-s | *BGIBMGA003733* | up | 296 | 304 | - | 11.1009 | 6.57E-05 | 0.709 | GAGCGAGGC |
| 3030 | mamo-s | *BGIBMGA003739* | down | 1190 | 1198 | - | 14.055 | 2.23E-06 | 0.615 | GTGCGTGGC |
| 3031 | mamo-s | *BGIBMGA003742* | down | 919 | 927 | - | 11.0826 | 6.84E-05 | 0.709 | GTACGTGGC |
| 3032 | mamo-s | *BGIBMGA003744* | down | 536 | 544 | - | 12.1284 | 2.16E-05 | 0.615 | GTGCGAGGC |
| 3033 | mamo-s | *BGIBMGA003745* | down | 1454 | 1462 | + | 11.7798 | 3.11E-05 | 0.615 | ATGCGTGGC |
| 3034 | mamo-s | *BGIBMGA003746* | down | 1324 | 1332 | - | 12.3394 | 1.49E-05 | 0.615 | GGGCGTGGC |
| 3035 | mamo-s | *BGIBMGA003746* | down | 1486 | 1494 | + | 11.3853 | 4.74E-05 | 0.615 | ACGCGTGGC |
| 3036 | mamo-s | *BGIBMGA003749* | up | 1248 | 1256 | - | 11.3303 | 4.96E-05 | 0.615 | GTGGGTGGC |
| 3037 | mamo-s | *BGIBMGA003750* | up | 1486 | 1494 | - | 11.3303 | 4.96E-05 | 0.615 | GTGGGTGGC |
| 3038 | mamo-s | *BGIBMGA003751* | down | 1750 | 1758 | + | 11.7339 | 3.29E-05 | 0.615 | GCGCGAGGC |
| 3039 | mamo-s | *BGIBMGA003751* | down | 17 | 25 | + | 11.2294 | 5.41E-05 | 0.65 | CTGCGAGGC |
| 3040 | mamo-s | *BGIBMGA003751* | down | 1773 | 1781 | - | 11.2018 | 5.68E-05 | 0.674 | CTGCGTGGT |
| 3041 | mamo-s | *BGIBMGA003754* | down | 1884 | 1892 | - | 11.7064 | 3.52E-05 | 0.615 | GCGCGTGGT |
| 3042 | mamo-s | *BGIBMGA003755* | up | 1819 | 1827 | - | 14.055 | 2.23E-06 | 0.615 | GTGCGTGGC |
| 3043 | mamo-s | *BGIBMGA003759* | down | 978 | 986 | - | 11.3303 | 4.96E-05 | 0.615 | GTGGGTGGC |
| 3044 | mamo-s | *BGIBMGA003762* | down | 1282 | 1290 | + | 11.3303 | 4.96E-05 | 0.615 | GTGGGTGGC |
| 3045 | mamo-s | *BGIBMGA003764* | down | 97 | 105 | - | 11.3303 | 4.96E-05 | 0.615 | GTGGGTGGC |
| 3046 | mamo-s | *BGIBMGA003764* | down | 1039 | 1047 | + | 11.3303 | 4.96E-05 | 0.615 | GTGGGTGGC |
| 3047 | mamo-s | *BGIBMGA003766* | up | 1785 | 1793 | + | 11.3303 | 4.96E-05 | 0.615 | GTGGGTGGC |
| 3048 | mamo-s | *BGIBMGA003767* | down | 1501 | 1509 | + | 13.156 | 6.30E-06 | 0.615 | CTGCGTGGC |
| 3049 | mamo-s | *BGIBMGA003768* | down | 827 | 835 | + | 11.7064 | 3.52E-05 | 0.615 | GCGCGTGGT |
| 3050 | mamo-s | *BGIBMGA003768* | down | 218 | 226 | + | 11.3303 | 4.96E-05 | 0.615 | GTGGGTGGC |
| 3051 | mamo-s | *BGIBMGA003768* | down | 1067 | 1075 | - | 11.3303 | 4.96E-05 | 0.615 | GTGGGTGGC |
| 3052 | mamo-s | *BGIBMGA003768* | down | 772 | 780 | - | 11.2294 | 5.41E-05 | 0.65 | CTGCGAGGC |
| 3053 | mamo-s | *BGIBMGA003768* | down | 862 | 870 | + | 10.5413 | 9.42E-05 | 0.857 | GTGCGAGTC |
| 3054 | mamo-s | *BGIBMGA003769* | up | 1124 | 1132 | + | 11.7064 | 3.52E-05 | 0.615 | GCGCGTGGT |
| 3055 | mamo-s | *BGIBMGA003769* | up | 515 | 523 | + | 11.3303 | 4.96E-05 | 0.615 | GTGGGTGGC |
| 3056 | mamo-s | *BGIBMGA003769* | up | 1364 | 1372 | - | 11.3303 | 4.96E-05 | 0.615 | GTGGGTGGC |
| 3057 | mamo-s | *BGIBMGA003769* | up | 1069 | 1077 | - | 11.2294 | 5.41E-05 | 0.65 | CTGCGAGGC |
| 3058 | mamo-s | *BGIBMGA003769* | up | 1159 | 1167 | + | 10.5413 | 9.42E-05 | 0.857 | GTGCGAGTC |
| 3059 | mamo-s | *BGIBMGA003772* | down | 1217 | 1225 | + | 10.5688 | 8.88E-05 | 0.824 | GTGCGTGAC |
| 3060 | mamo-s | *BGIBMGA003773* | up | 226 | 234 | - | 11.0826 | 6.84E-05 | 0.709 | GTACGTGGC |
| 3061 | mamo-s | *BGIBMGA003774* | up | 552 | 560 | + | 11.0826 | 6.84E-05 | 0.709 | GTACGTGGC |
| 3062 | mamo-s | *BGIBMGA003775* | down | 1447 | 1455 | - | 11.0826 | 6.84E-05 | 0.709 | GTACGTGGC |
| 3063 | mamo-s | *BGIBMGA003778* | up | 301 | 309 | - | 11.3303 | 4.96E-05 | 0.615 | GTGGGTGGC |
| 3064 | mamo-s | *BGIBMGA003780* | down | 1118 | 1126 | + | 11.0826 | 6.84E-05 | 0.709 | GTACGTGGC |
| 3065 | mamo-s | *BGIBMGA003781* | up | 714 | 722 | + | 11.0826 | 6.84E-05 | 0.709 | GTACGTGGC |
| 3066 | mamo-s | *BGIBMGA003782* | up | 958 | 966 | - | 11.3303 | 4.96E-05 | 0.615 | GTGGGTGGC |
| 3067 | mamo-s | *BGIBMGA003783* | down | 735 | 743 | - | 11.5688 | 3.79E-05 | 0.615 | CTGCGTGTC |
| 3068 | mamo-s | *BGIBMGA003784* | down | 1094 | 1102 | + | 14.055 | 2.23E-06 | 0.615 | GTGCGTGGC |
| 3069 | mamo-s | *BGIBMGA003784* | up | 583 | 591 | + | 12.3394 | 1.49E-05 | 0.615 | GGGCGTGGC |
| 3070 | mamo-s | *BGIBMGA003784* | up | 57 | 65 | - | 11.3303 | 4.96E-05 | 0.615 | GTGGGTGGC |
| 3071 | mamo-s | *BGIBMGA003788* | up | 786 | 794 | - | 11.3303 | 4.96E-05 | 0.615 | GTGGGTGGC |
| 3072 | mamo-s | *BGIBMGA003788* | up | 367 | 375 | - | 11.1009 | 6.57E-05 | 0.709 | GAGCGAGGC |
| 3073 | mamo-s | *BGIBMGA003788* | down | 160 | 168 | + | 10.8349 | 7.66E-05 | 0.751 | CCGCGAGGC |

| | | | | | | | | | | |
|---|---|---|---|---|---|---|---|---|---|---|
| 3074 | mamo-s | *BGIBMGA003788* | down | 1352 | 1360 | + | 10.8349 | 7.66E-05 | 0.751 | CCGCGAGGC |
| 3075 | mamo-s | *BGIBMGA003791* | down | 1443 | 1451 | + | 11.3303 | 4.96E-05 | 0.615 | GTGGGTGGC |
| 3076 | mamo-s | *BGIBMGA003792* | up | 509 | 517 | + | 11.156 | 6.13E-05 | 0.705 | GAGCGTGCC |
| 3077 | mamo-s | *BGIBMGA003794* | up | 45 | 53 | + | 12.7615 | 1.04E-05 | 0.615 | CCGCGTGGC |
| 3078 | mamo-s | *BGIBMGA003794* | up | 521 | 529 | - | 11.3303 | 4.96E-05 | 0.615 | GTGGGTGGC |
| 3079 | mamo-s | *BGIBMGA003794* | down | 835 | 843 | - | 11.3303 | 4.96E-05 | 0.615 | GTGGGTGGC |
| 3080 | mamo-s | *BGIBMGA003794* | up | 1520 | 1528 | - | 11.0826 | 6.84E-05 | 0.709 | GTACGTGGC |
| 3081 | mamo-s | *BGIBMGA003796* | down | 1080 | 1088 | - | 11.1009 | 6.57E-05 | 0.709 | GAGCGAGGC |
| 3082 | mamo-s | *BGIBMGA003798* | up | 117 | 125 | + | 13.156 | 6.30E-06 | 0.615 | CTGCGTGGC |
| 3083 | mamo-s | *BGIBMGA003800* | down | 1002 | 1010 | - | 11.5688 | 3.79E-05 | 0.615 | CTGCGTGTC |
| 3084 | mamo-s | *BGIBMGA003800* | down | 1365 | 1373 | - | 11.5321 | 4.06E-05 | 0.615 | TTGCGTGGC |
| 3085 | mamo-s | *BGIBMGA003801* | up | 416 | 424 | - | 11.5688 | 3.79E-05 | 0.615 | CTGCGTGTC |
| 3086 | mamo-s | *BGIBMGA003801* | up | 779 | 787 | - | 11.5321 | 4.06E-05 | 0.615 | TTGCGTGGC |
| 3087 | mamo-s | *BGIBMGA003801* | down | 1844 | 1852 | + | 11.1743 | 5.90E-05 | 0.684 | CCGCGTGTC |
| 3088 | mamo-s | *BGIBMGA003802* | down | 1503 | 1511 | - | 11.3303 | 4.96E-05 | 0.615 | GTGGGTGGC |
| 3089 | mamo-s | *BGIBMGA003803* | up | 411 | 419 | - | 11.3303 | 4.96E-05 | 0.615 | GTGGGTGGC |
| 3090 | mamo-s | *BGIBMGA003807* | up | 477 | 485 | + | 11.3303 | 4.96E-05 | 0.615 | GTGGGTGGC |
| 3091 | mamo-s | *BGIBMGA003807* | up | 1274 | 1282 | - | 11.3303 | 4.96E-05 | 0.615 | GTGGGTGGC |
| 3092 | mamo-s | *BGIBMGA003808* | up | 1679 | 1687 | - | 11.3303 | 4.96E-05 | 0.615 | GTGGGTGGC |
| 3093 | mamo-s | *BGIBMGA003808* | up | 357 | 365 | + | 10.6881 | 8.61E-05 | 0.81 | GCACGTGGC |
| 3094 | mamo-s | *BGIBMGA003809* | up | 1272 | 1280 | - | 11.3303 | 4.96E-05 | 0.615 | GTGGGTGGC |
| 3095 | mamo-s | *BGIBMGA003810* | up | 1666 | 1674 | - | 10.7523 | 8.38E-05 | 0.797 | GGGCGTGTC |
| 3096 | mamo-s | *BGIBMGA003811* | up | 302 | 310 | + | 10.9358 | 7.30E-05 | 0.728 | GCGGGTGGC |
| 3097 | mamo-s | *BGIBMGA003813* | down | 219 | 227 | + | 11.3303 | 4.96E-05 | 0.615 | GTGGGTGGC |
| 3098 | mamo-s | *BGIBMGA003815* | down | 570 | 578 | - | 11.0826 | 6.84E-05 | 0.709 | GTACGTGGC |
| 3099 | mamo-s | *BGIBMGA003816* | up | 1573 | 1581 | - | 11.3303 | 4.96E-05 | 0.615 | GTGGGTGGC |
| 3100 | mamo-s | *BGIBMGA003816* | up | 1341 | 1349 | + | 10.7523 | 8.38E-05 | 0.797 | GGGCGTGTC |
| 3101 | mamo-s | *BGIBMGA003817* | down | 1716 | 1724 | - | 10.8349 | 7.66E-05 | 0.751 | CCGCGAGGC |
| 3102 | mamo-s | *BGIBMGA003818* | up | 261 | 269 | - | 11.3303 | 4.96E-05 | 0.615 | GTGGGTGGC |
| 3103 | mamo-s | *BGIBMGA003818* | down | 1683 | 1691 | - | 11.3303 | 4.96E-05 | 0.615 | GTGGGTGGC |
| 3104 | mamo-s | *BGIBMGA003820* | up | 483 | 491 | + | 11.156 | 6.13E-05 | 0.705 | GAGCGTGCC |
| 3105 | mamo-s | *BGIBMGA003821* | up | 1618 | 1626 | - | 13.6606 | 4.06E-06 | 0.615 | GCGCGTGGC |
| 3106 | mamo-s | *BGIBMGA003821* | down | 72 | 80 | + | 11.3303 | 4.96E-05 | 0.615 | GTGGGTGGC |
| 3107 | mamo-s | *BGIBMGA003821* | up | 218 | 226 | + | 11.3303 | 4.96E-05 | 0.615 | GTGGGTGGC |
| 3108 | mamo-s | *BGIBMGA003821* | up | 780 | 788 | - | 11.3303 | 4.96E-05 | 0.615 | GTGGGTGGC |
| 3109 | mamo-s | *BGIBMGA003822* | down | 1373 | 1381 | + | 11.3303 | 4.96E-05 | 0.615 | GTGGGTGGC |
| 3110 | mamo-s | *BGIBMGA003823* | down | 354 | 362 | - | 11.3303 | 4.96E-05 | 0.615 | GTGGGTGGC |
| 3111 | mamo-s | *BGIBMGA003824* | up | 800 | 808 | - | 11.3303 | 4.96E-05 | 0.615 | GTGGGTGGC |
| 3112 | mamo-s | *BGIBMGA003824* | down | 1884 | 1892 | - | 11.0826 | 6.84E-05 | 0.709 | GTACGTGGC |
| 3113 | mamo-s | *BGIBMGA003825* | down | 103 | 111 | - | 11.3303 | 4.96E-05 | 0.615 | GTGGGTGGC |
| 3114 | mamo-s | *BGIBMGA003825* | down | 1344 | 1352 | - | 11.3303 | 4.96E-05 | 0.615 | GTGGGTGGC |
| 3115 | mamo-s | *BGIBMGA003827* | down | 479 | 487 | + | 12.1284 | 2.16E-05 | 0.615 | CAGCGTGGC |
| 3116 | mamo-s | *BGIBMGA003827* | down | 1147 | 1155 | - | 11.3303 | 4.96E-05 | 0.615 | GTGGGTGGC |
| 3117 | mamo-s | *BGIBMGA003827* | up | 1343 | 1351 | - | 11.3303 | 4.96E-05 | 0.615 | GTGGGTGGC |
| 3118 | mamo-s | *BGIBMGA003828* | down | 247 | 255 | - | 11.3303 | 4.96E-05 | 0.615 | GTGGGTGGC |
| 3119 | mamo-s | *BGIBMGA003828* | down | 1723 | 1731 | + | 11.3303 | 4.96E-05 | 0.615 | GTGGGTGGC |
| 3120 | mamo-s | *BGIBMGA003831* | up | 296 | 304 | - | 11.3303 | 4.96E-05 | 0.615 | GTGGGTGGC |
| 3121 | mamo-s | *BGIBMGA003831* | down | 94 | 102 | - | 10.9358 | 7.30E-05 | 0.728 | GCGGGTGGC |
| 3122 | mamo-s | *BGIBMGA003832* | up | 372 | 380 | - | 11.3303 | 4.96E-05 | 0.615 | GTGGGTGGC |
| 3123 | mamo-s | *BGIBMGA003832* | down | 446 | 454 | - | 11.3303 | 4.96E-05 | 0.615 | GTGGGTGGC |
| 3124 | mamo-s | *BGIBMGA003832* | down | 1007 | 1015 | + | 11.3303 | 4.96E-05 | 0.615 | GTGGGTGGC |
| 3125 | mamo-s | *BGIBMGA003832* | up | 1190 | 1198 | + | 10.5688 | 8.88E-05 | 0.824 | GTGCGTGAC |
| 3126 | mamo-s | *BGIBMGA003836* | up | 633 | 641 | - | 11.3303 | 4.96E-05 | 0.615 | GTGGGTGGC |
| 3127 | mamo-s | *BGIBMGA003837* | up | 230 | 238 | + | 11.3303 | 4.96E-05 | 0.615 | GTGGGTGGC |
| 3128 | mamo-s | *BGIBMGA003839* | down | 1627 | 1635 | + | 11.3303 | 4.96E-05 | 0.615 | GTGGGTGGC |
| 3129 | mamo-s | *BGIBMGA003839* | up | 649 | 657 | - | 10.6881 | 8.61E-05 | 0.81 | GCACGTGGC |
| 3130 | mamo-s | *BGIBMGA003840* | down | 1963 | 1971 | - | 11.3303 | 4.96E-05 | 0.615 | GTGGGTGGC |
| 3131 | mamo-s | *BGIBMGA003841* | down | 887 | 895 | - | 11.3303 | 4.96E-05 | 0.615 | GTGGGTGGC |
| 3132 | mamo-s | *BGIBMGA003843* | up | 459 | 467 | - | 11.3303 | 4.96E-05 | 0.615 | GTGGGTGGC |
| 3133 | mamo-s | *BGIBMGA003843* | down | 1946 | 1954 | + | 11.3303 | 4.96E-05 | 0.615 | GTGGGTGGC |
| 3134 | mamo-s | *BGIBMGA003846* | down | 164 | 172 | + | 11.3303 | 4.96E-05 | 0.615 | GTGGGTGGC |
| 3135 | mamo-s | *BGIBMGA003847* | up | 1828 | 1836 | + | 11.7798 | 3.11E-05 | 0.615 | ATGCGTGGC |
| 3136 | mamo-s | *BGIBMGA003847* | down | 1919 | 1927 | + | 11.7798 | 3.11E-05 | 0.615 | ATGCGTGGC |
| 3137 | mamo-s | *BGIBMGA003848* | up | 1828 | 1836 | + | 11.7798 | 3.11E-05 | 0.615 | ATGCGTGGC |
| 3138 | mamo-s | *BGIBMGA003848* | down | 703 | 711 | - | 10.8073 | 7.89E-05 | 0.766 | CCGCGTGGT |
| 3139 | mamo-s | *BGIBMGA003849* | up | 1931 | 1939 | + | 11.5321 | 4.06E-05 | 0.615 | TTGCGTGGC |
| 3140 | mamo-s | *BGIBMGA003849* | up | 25 | 33 | + | 11.3303 | 4.96E-05 | 0.615 | GTGGGTGGC |
| 3141 | mamo-s | *BGIBMGA003849* | down | 754 | 762 | - | 10.8349 | 7.66E-05 | 0.751 | CCGCGAGGC |
| 3142 | mamo-s | *BGIBMGA003851* | up | 345 | 353 | + | 11.3303 | 4.96E-05 | 0.615 | GTGGGTGGC |
| 3143 | mamo-s | *BGIBMGA003854* | down | 571 | 579 | + | 11.1009 | 6.57E-05 | 0.709 | GAGCGAGGC |
| 3144 | mamo-s | *BGIBMGA003863* | down | 559 | 567 | - | 12.4679 | 1.31E-05 | 0.615 | GTGCGTGTC |
| 3145 | mamo-s | *BGIBMGA003865* | up | 615 | 623 | + | 11.5321 | 4.06E-05 | 0.615 | TTGCGTGGC |
| 3146 | mamo-s | *BGIBMGA003865* | up | 1806 | 1814 | - | 11.3303 | 4.96E-05 | 0.615 | GTGGGTGGC |
| 3147 | mamo-s | *BGIBMGA003869* | up | 1932 | 1940 | - | 11.7339 | 3.29E-05 | 0.615 | GCGCGAGGC |
| 3148 | mamo-s | *BGIBMGA003871* | up | 1278 | 1286 | - | 12.3394 | 1.49E-05 | 0.615 | GGGCGTGGC |
| 3149 | mamo-s | *BGIBMGA003872* | down | 417 | 425 | + | 11.789 | 2.84E-05 | 0.615 | GCGCGTGCC |
| 3150 | mamo-s | *BGIBMGA003873* | up | 1068 | 1076 | - | 13.0275 | 8.53E-06 | 0.615 | GAGCGTGGC |
| 3151 | mamo-s | *BGIBMGA003873* | up | 1839 | 1847 | + | 12.1009 | 2.43E-05 | 0.615 | GTGCGTGGT |
| 3152 | mamo-s | *BGIBMGA003873* | up | 513 | 521 | + | 11.789 | 2.84E-05 | 0.615 | GCGCGTGCC |
| 3153 | mamo-s | *BGIBMGA003873* | down | 53 | 61 | - | 11.5688 | 3.79E-05 | 0.615 | CTGCGTGTC |
| 3154 | mamo-s | *BGIBMGA003874* | up | 613 | 621 | + | 12.1284 | 2.16E-05 | 0.615 | CAGCGTGGC |
| 3155 | mamo-s | *BGIBMGA003874* | down | 32 | 40 | - | 11.1376 | 6.35E-05 | 0.709 | TCGCGTGGC |
| 3156 | mamo-s | *BGIBMGA003875* | down | 285 | 293 | + | 12.7615 | 1.04E-05 | 0.615 | CCGCGTGGC |
| 3157 | mamo-s | *BGIBMGA003875* | down | 441 | 449 | + | 11.4404 | 4.51E-05 | 0.615 | CGGCGTGGC |
| 3158 | mamo-s | *BGIBMGA003875* | down | 1817 | 1825 | + | 11.1743 | 5.90E-05 | 0.684 | CCGCGTGTC |
| 3159 | mamo-s | *BGIBMGA003875* | down | 1918 | 1926 | + | 11.1009 | 6.57E-05 | 0.709 | GAGCGAGGC |
| 3160 | mamo-s | *BGIBMGA003876* | down | 7 | 15 | + | 10.5413 | 9.42E-05 | 0.857 | CAGCGTGTC |
| 3161 | mamo-s | *BGIBMGA003879* | down | 701 | 709 | + | 10.5688 | 8.88E-05 | 0.824 | GTGCGTGAC |

| | | | | | | | | | | |
|---|---|---|---|---|---|---|---|---|---|---|
| 3162 | mamo-s | *BGIBMGA003880* | up | 1462 | 1470 | + | 11.3303 | 4.96E-05 | 0.615 | GTGGGTGGC |
| 3163 | mamo-s | *BGIBMGA003881* | down | 1648 | 1656 | + | 11.3303 | 4.96E-05 | 0.615 | GTGGGTGGC |
| 3164 | mamo-s | *BGIBMGA003882* | down | 1579 | 1587 | - | 11.3303 | 4.96E-05 | 0.615 | GTGGGTGGC |
| 3165 | mamo-s | *BGIBMGA003883* | up | 164 | 172 | + | 13.6606 | 4.06E-06 | 0.615 | GCGCGTGGC |
| 3166 | mamo-s | *BGIBMGA003883* | up | 721 | 729 | + | 12.4679 | 1.31E-05 | 0.615 | GTGCGTGTC |
| 3167 | mamo-s | *BGIBMGA003887* | up | 324 | 332 | + | 13.156 | 6.30E-06 | 0.615 | CTGCGTGGC |
| 3168 | mamo-s | *BGIBMGA003887* | up | 824 | 832 | - | 11.7064 | 3.52E-05 | 0.615 | GCGCGTGGT |
| 3169 | mamo-s | *BGIBMGA003893* | down | 243 | 251 | - | 12.4679 | 1.31E-05 | 0.615 | GTGCGTGTC |
| 3170 | mamo-s | *BGIBMGA003894* | down | 890 | 898 | + | 11.1009 | 6.57E-05 | 0.709 | GAGCGAGGC |
| 3171 | mamo-s | *BGIBMGA003897* | up | 807 | 815 | - | 11.4404 | 4.51E-05 | 0.615 | CGGCGTGGC |
| 3172 | mamo-s | *BGIBMGA003897* | up | 116 | 124 | - | 11.3303 | 4.96E-05 | 0.615 | GTGGGTGGC |
| 3173 | mamo-s | *BGIBMGA003898* | down | 1892 | 1900 | - | 10.8899 | 7.48E-05 | 0.739 | CCGCGTGCC |
| 3174 | mamo-s | *BGIBMGA003900* | up | 1816 | 1824 | + | 11.3303 | 4.96E-05 | 0.615 | GTGGGTGGC |
| 3175 | mamo-s | *BGIBMGA003901* | down | 1113 | 1121 | - | 10.8899 | 7.48E-05 | 0.739 | CCGCGTGCC |
| 3176 | mamo-s | *BGIBMGA003902* | up | 346 | 354 | + | 11.7064 | 3.52E-05 | 0.615 | GCGCGTGGT |
| 3177 | mamo-s | *BGIBMGA003902* | up | 1005 | 1013 | - | 11.0826 | 6.84E-05 | 0.709 | GTACGTGGC |
| 3178 | mamo-s | *BGIBMGA003903* | down | 847 | 855 | + | 11.3303 | 4.96E-05 | 0.615 | GTGGGTGGC |
| 3179 | mamo-s | *BGIBMGA003904* | down | 1600 | 1608 | - | 11.156 | 6.13E-05 | 0.705 | GAGCGTGCC |
| 3180 | mamo-s | *BGIBMGA003905* | up | 1597 | 1605 | - | 10.5138 | 9.75E-05 | 0.865 | GTGCGTGTT |
| 3181 | mamo-s | *BGIBMGA003906* | up | 147 | 155 | + | 11.3303 | 4.96E-05 | 0.615 | GTGGGTGGC |
| 3182 | mamo-s | *BGIBMGA003906* | down | 349 | 357 | - | 11.3303 | 4.96E-05 | 0.615 | GTGGGTGGC |
| 3183 | mamo-s | *BGIBMGA003906* | up | 369 | 377 | - | 10.7523 | 8.38E-05 | 0.797 | AAGCGTGGC |
| 3184 | mamo-s | *BGIBMGA003908* | down | 602 | 610 | + | 12.1835 | 1.71E-05 | 0.615 | GTGCGTGCC |
| 3185 | mamo-s | *BGIBMGA003908* | down | 175 | 183 | - | 11.3303 | 4.96E-05 | 0.615 | GTGGGTGGC |
| 3186 | mamo-s | *BGIBMGA003908* | down | 1090 | 1098 | + | 11.3303 | 4.96E-05 | 0.615 | GTGGGTGGC |
| 3187 | mamo-s | *BGIBMGA003908* | down | 760 | 768 | - | 11.0826 | 6.84E-05 | 0.709 | GTACGTGGC |
| 3188 | mamo-s | *BGIBMGA003910* | up | 1991 | 1999 | - | 13.156 | 6.30E-06 | 0.615 | CTGCGTGGC |
| 3189 | mamo-s | *BGIBMGA003910* | down | 1425 | 1433 | + | 11.3303 | 4.96E-05 | 0.615 | GTGGGTGGC |
| 3190 | mamo-s | *BGIBMGA003911* | up | 417 | 425 | - | 11.3303 | 4.96E-05 | 0.615 | GTGGGTGGC |
| 3191 | mamo-s | *BGIBMGA003912* | down | 1680 | 1688 | - | 11.0826 | 6.84E-05 | 0.709 | GTACGTGGC |
| 3192 | mamo-s | *BGIBMGA003915* | down | 153 | 161 | - | 11.3303 | 4.96E-05 | 0.615 | GTGGGTGGC |
| 3193 | mamo-s | *BGIBMGA003916* | down | 206 | 214 | + | 11.3303 | 4.96E-05 | 0.615 | GTGGGTGGC |
| 3194 | mamo-s | *BGIBMGA003917* | down | 1581 | 1589 | - | 11.3303 | 4.96E-05 | 0.615 | GTGGGTGGC |
| 3195 | mamo-s | *BGIBMGA003917* | up | 1697 | 1705 | - | 11.3303 | 4.96E-05 | 0.615 | GTGGGTGGC |
| 3196 | mamo-s | *BGIBMGA003919* | up | 523 | 531 | + | 11.3303 | 4.96E-05 | 0.615 | GTGGGTGGC |
| 3197 | mamo-s | *BGIBMGA003920* | down | 1136 | 1144 | + | 11.3303 | 4.96E-05 | 0.615 | GTGGGTGGC |
| 3198 | mamo-s | *BGIBMGA003921* | down | 1186 | 1194 | + | 10.5138 | 9.75E-05 | 0.865 | GTGCGTGTT |
| 3199 | mamo-s | *BGIBMGA003922* | up | 846 | 854 | + | 10.5138 | 9.75E-05 | 0.865 | GTGCGTGTT |
| 3200 | mamo-s | *BGIBMGA003925* | up | 595 | 603 | - | 11.3303 | 4.96E-05 | 0.615 | GTGGGTGGC |
| 3201 | mamo-s | *BGIBMGA003925* | down | 1424 | 1432 | - | 11.0734 | 7.12E-05 | 0.726 | GAGCGTGGT |
| 3202 | mamo-s | *BGIBMGA003925* | up | 1468 | 1476 | - | 11.0734 | 7.12E-05 | 0.726 | GAGCGTGGT |
| 3203 | mamo-s | *BGIBMGA003926* | down | 1911 | 1919 | - | 11.3303 | 4.96E-05 | 0.615 | GTGGGTGGC |
| 3204 | mamo-s | *BGIBMGA003929* | down | 929 | 937 | + | 11.1743 | 5.90E-05 | 0.684 | CCGCGTGTC |
| 3205 | mamo-s | *BGIBMGA003931* | down | 629 | 637 | + | 10.6881 | 8.61E-05 | 0.81 | GCACGTGGC |
| 3206 | mamo-s | *BGIBMGA003932* | up | 1232 | 1240 | - | 11.789 | 2.84E-05 | 0.615 | GCGCGTGCC |
| 3207 | mamo-s | *BGIBMGA003932* | down | 178 | 186 | + | 11.0734 | 7.12E-05 | 0.726 | GAGCGTGGT |
| 3208 | mamo-s | *BGIBMGA003933* | down | 613 | 621 | + | 11.7064 | 3.52E-05 | 0.615 | GCGCGTGGT |
| 3209 | mamo-s | *BGIBMGA003933* | up | 474 | 482 | - | 11.3303 | 4.96E-05 | 0.615 | GTGGGTGGC |
| 3210 | mamo-s | *BGIBMGA003934* | up | 1014 | 1022 | + | 11.7064 | 3.52E-05 | 0.615 | GCGCGTGGT |
| 3211 | mamo-s | *BGIBMGA003935* | up | 827 | 835 | - | 12.3394 | 1.49E-05 | 0.615 | GGGCGTGGC |
| 3212 | mamo-s | *BGIBMGA003935* | up | 170 | 178 | + | 11.3303 | 4.96E-05 | 0.615 | GTGGGTGGC |
| 3213 | mamo-s | *BGIBMGA003936* | down | 209 | 217 | + | 11.3303 | 4.96E-05 | 0.615 | GTGGGTGGC |
| 3214 | mamo-s | *BGIBMGA003939* | down | 1800 | 1808 | + | 10.5138 | 9.75E-05 | 0.865 | GTGCGTGTT |
| 3215 | mamo-s | *BGIBMGA003941* | up | 1681 | 1689 | - | 11.3303 | 4.96E-05 | 0.615 | GTGGGTGGC |
| 3216 | mamo-s | *BGIBMGA003941* | down | 1708 | 1716 | + | 11.3303 | 4.96E-05 | 0.615 | GTGGGTGGC |
| 3217 | mamo-s | *BGIBMGA003943* | up | 590 | 598 | - | 11.3303 | 4.96E-05 | 0.615 | GTGGGTGGC |
| 3218 | mamo-s | *BGIBMGA003944* | up | 1817 | 1825 | - | 11.3303 | 4.96E-05 | 0.615 | GTGGGTGGC |
| 3219 | mamo-s | *BGIBMGA003945* | up | 954 | 962 | + | 11.3303 | 4.96E-05 | 0.615 | GTGGGTGGC |
| 3220 | mamo-s | *BGIBMGA003949* | down | 173 | 181 | + | 11.3303 | 4.96E-05 | 0.615 | GTGGGTGGC |
| 3221 | mamo-s | *BGIBMGA003954* | up | 20 | 28 | + | 11.1743 | 5.90E-05 | 0.684 | CCGCGTGTC |
| 3222 | mamo-s | *BGIBMGA003955* | up | 1335 | 1343 | + | 11.3303 | 4.96E-05 | 0.615 | GTGGGTGGC |
| 3223 | mamo-s | *BGIBMGA003957* | up | 1081 | 1089 | + | 11.3303 | 4.96E-05 | 0.615 | GTGGGTGGC |
| 3224 | mamo-s | *BGIBMGA003960* | up | 526 | 534 | + | 11.3303 | 4.96E-05 | 0.615 | GTGGGTGGC |
| 3225 | mamo-s | *BGIBMGA003960* | up | 907 | 915 | + | 11.3303 | 4.96E-05 | 0.615 | GTGGGTGGC |
| 3226 | mamo-s | *BGIBMGA003961* | down | 787 | 795 | - | 12.3394 | 1.49E-05 | 0.615 | GGGCGTGGC |
| 3227 | mamo-s | *BGIBMGA003961* | up | 1048 | 1056 | + | 11.3303 | 4.96E-05 | 0.615 | GTGGGTGGC |
| 3228 | mamo-s | *BGIBMGA003961* | down | 851 | 859 | + | 10.8349 | 7.66E-05 | 0.751 | CCGCGAGGC |
| 3229 | mamo-s | *BGIBMGA003963* | down | 1842 | 1850 | - | 11.3303 | 4.96E-05 | 0.615 | GTGGGTGGC |
| 3230 | mamo-s | *BGIBMGA003963* | down | 305 | 313 | - | 11.2294 | 5.41E-05 | 0.65 | CTGCGAGGC |
| 3231 | mamo-s | *BGIBMGA003964* | up | 809 | 817 | + | 10.5413 | 9.42E-05 | 0.857 | GTGCGAGTC |
| 3232 | mamo-s | *BGIBMGA003965* | up | 1628 | 1636 | - | 11.2844 | 5.18E-05 | 0.63 | CTGCGTGCC |
| 3233 | mamo-s | *BGIBMGA003966* | down | 165 | 173 | - | 11.3303 | 4.96E-05 | 0.615 | GTGGGTGGC |
| 3234 | mamo-s | *BGIBMGA003966* | up | 1831 | 1839 | - | 11.3303 | 4.96E-05 | 0.615 | GTGGGTGGC |
| 3235 | mamo-s | *BGIBMGA003967* | up | 210 | 218 | + | 11.3303 | 4.96E-05 | 0.615 | GTGGGTGGC |
| 3236 | mamo-s | *BGIBMGA003968* | up | 682 | 690 | + | 11.3303 | 4.96E-05 | 0.615 | GTGGGTGGC |
| 3237 | mamo-s | *BGIBMGA003968* | up | 1665 | 1673 | - | 11.3303 | 4.96E-05 | 0.615 | GTGGGTGGC |
| 3238 | mamo-s | *BGIBMGA003969* | down | 1028 | 1036 | + | 11.3303 | 4.96E-05 | 0.615 | GTGGGTGGC |
| 3239 | mamo-s | *BGIBMGA003970* | up | 64 | 72 | - | 11.3303 | 4.96E-05 | 0.615 | GTGGGTGGC |
| 3240 | mamo-s | *BGIBMGA003970* | up | 861 | 869 | + | 11.3303 | 4.96E-05 | 0.615 | GTGGGTGGC |
| 3241 | mamo-s | *BGIBMGA003970* | down | 1579 | 1587 | + | 10.7523 | 8.38E-05 | 0.797 | GGGCGTGTC |
| 3242 | mamo-s | *BGIBMGA003971* | up | 974 | 982 | + | 11.1376 | 6.35E-05 | 0.709 | TCGCGTGGC |
| 3243 | mamo-s | *BGIBMGA003973* | down | 1689 | 1697 | + | 11.3303 | 4.96E-05 | 0.615 | GTGGGTGGC |
| 3244 | mamo-s | *BGIBMGA003977* | down | 1053 | 1061 | - | 11.3303 | 4.96E-05 | 0.615 | GTGGGTGGC |
| 3245 | mamo-s | *BGIBMGA003980* | down | 337 | 345 | - | 11.0826 | 6.84E-05 | 0.709 | GTACGTGGC |
| 3246 | mamo-s | *BGIBMGA003982* | up | 1142 | 1150 | - | 10.5138 | 9.75E-05 | 0.865 | GTGCGTGTT |
| 3247 | mamo-s | *BGIBMGA003982* | down | 1208 | 1216 | - | 10.5138 | 9.75E-05 | 0.865 | GTGCGTGTT |
| 3248 | mamo-s | *BGIBMGA003985* | down | 75 | 83 | + | 11.5688 | 3.79E-05 | 0.615 | CTGCGTGTC |
| 3249 | mamo-s | *BGIBMGA003985* | down | 1332 | 1340 | - | 11.3303 | 4.96E-05 | 0.615 | GTGGGTGGC |

| | | | | | | | | | | |
|---|---|---|---|---|---|---|---|---|---|---|
| 3250 | mamo-s | *BGIBMGA003986* | down | 1142 | 1150 | + | 11.3303 | 4.96E-05 | 0.615 | GTGGGTGGC |
| 3251 | mamo-s | *BGIBMGA003987* | down | 742 | 750 | - | 11.7339 | 3.29E-05 | 0.615 | GCGCGAGGC |
| 3252 | mamo-s | *BGIBMGA003991* | down | 1067 | 1075 | - | 11.1743 | 5.90E-05 | 0.684 | CCGCGTGTC |
| 3253 | mamo-s | *BGIBMGA003993* | up | 408 | 416 | - | 11.3303 | 4.96E-05 | 0.615 | GTGGGTGGC |
| 3254 | mamo-s | *BGIBMGA003995* | down | 583 | 591 | + | 12.4679 | 1.31E-05 | 0.615 | GTGCGTGTC |
| 3255 | mamo-s | *BGIBMGA003995* | up | 1963 | 1971 | + | 10.9358 | 7.30E-05 | 0.728 | GCGGGTGGC |
| 3256 | mamo-s | *BGIBMGA003997* | up | 641 | 649 | - | 12.1835 | 1.71E-05 | 0.615 | GTGCGTGCC |
| 3257 | mamo-s | *BGIBMGA003999* | up | 1716 | 1724 | - | 11.3303 | 4.96E-05 | 0.615 | GTGGGTGGC |
| 3258 | mamo-s | *BGIBMGA004000* | up | 1519 | 1527 | + | 11.3303 | 4.96E-05 | 0.615 | GTGGGTGGC |
| 3259 | mamo-s | *BGIBMGA004001* | up | 1836 | 1844 | + | 12.4679 | 1.31E-05 | 0.615 | GTGCGTGTC |
| 3260 | mamo-s | *BGIBMGA004002* | up | 909 | 917 | + | 11.3303 | 4.96E-05 | 0.615 | GTGGGTGGC |
| 3261 | mamo-s | *BGIBMGA004002* | down | 1582 | 1590 | - | 11.0826 | 6.84E-05 | 0.709 | GTACGTGGC |
| 3262 | mamo-s | *BGIBMGA004004* | up | 585 | 593 | - | 11.3303 | 4.96E-05 | 0.615 | GTGGGTGGC |
| 3263 | mamo-s | *BGIBMGA004005* | down | 1345 | 1353 | - | 11.0826 | 6.84E-05 | 0.709 | GTACGTGGC |
| 3264 | mamo-s | *BGIBMGA004005* | down | 681 | 689 | + | 11.0734 | 7.12E-05 | 0.726 | GAGCGTGGT |
| 3265 | mamo-s | *BGIBMGA004008* | down | 1637 | 1645 | + | 11.3303 | 4.96E-05 | 0.615 | GTGGGTGGC |
| 3266 | mamo-s | *BGIBMGA004008* | up | 1685 | 1693 | + | 11.3303 | 4.96E-05 | 0.615 | GTGGGTGGC |
| 3267 | mamo-s | *BGIBMGA004009* | down | 944 | 952 | - | 11.3303 | 4.96E-05 | 0.615 | GTGGGTGGC |
| 3268 | mamo-s | *BGIBMGA004009* | down | 1192 | 1200 | - | 11.3303 | 4.96E-05 | 0.615 | GTGGGTGGC |
| 3269 | mamo-s | *BGIBMGA004010* | down | 1083 | 1091 | + | 11.3853 | 4.74E-05 | 0.615 | ACGCGTGGC |
| 3270 | mamo-s | *BGIBMGA004013* | up | 470 | 478 | - | 11.0826 | 6.84E-05 | 0.709 | GTACGTGGC |
| 3271 | mamo-s | *BGIBMGA004014* | down | 1137 | 1145 | - | 11.3303 | 4.96E-05 | 0.615 | GTGGGTGGC |
| 3272 | mamo-s | *BGIBMGA004015* | up | 493 | 501 | - | 11.3303 | 4.96E-05 | 0.615 | GTGGGTGGC |
| 3273 | mamo-s | *BGIBMGA004015* | down | 534 | 542 | + | 11.3303 | 4.96E-05 | 0.615 | GTGGGTGGC |
| 3274 | mamo-s | *BGIBMGA004016* | down | 1476 | 1484 | + | 11.3303 | 4.96E-05 | 0.615 | GTGGGTGGC |
| 3275 | mamo-s | *BGIBMGA004017* | down | 801 | 809 | + | 11.3303 | 4.96E-05 | 0.615 | GTGGGTGGC |
| 3276 | mamo-s | *BGIBMGA004022* | down | 3 | 11 | - | 12.1284 | 2.16E-05 | 0.615 | GTGCGAGGC |
| 3277 | mamo-s | *BGIBMGA004022* | down | 887 | 895 | - | 11.1009 | 6.57E-05 | 0.709 | GAGCGAGGC |
| 3278 | mamo-s | *BGIBMGA004023* | up | 747 | 755 | - | 11.7339 | 3.29E-05 | 0.615 | GCGCGAGGC |
| 3279 | mamo-s | *BGIBMGA004023* | down | 1660 | 1668 | + | 11.3303 | 4.96E-05 | 0.615 | GTGGGTGGC |
| 3280 | mamo-s | *BGIBMGA004030* | up | 928 | 936 | - | 11.4404 | 4.51E-05 | 0.615 | CGGCGTGGC |
| 3281 | mamo-s | *BGIBMGA004030* | down | 1895 | 1903 | - | 11.0826 | 6.84E-05 | 0.709 | GTACGTGGC |
| 3282 | mamo-s | *BGIBMGA004031* | up | 1044 | 1052 | + | 11.4404 | 4.51E-05 | 0.615 | CGGCGTGGC |
| 3283 | mamo-s | *BGIBMGA004032* | down | 1845 | 1853 | + | 12.3394 | 1.49E-05 | 0.615 | GGGCGTGGC |
| 3284 | mamo-s | *BGIBMGA004032* | down | 1897 | 1905 | + | 11.3303 | 4.96E-05 | 0.615 | GTGGGTGGC |
| 3285 | mamo-s | *BGIBMGA004036* | up | 1033 | 1041 | - | 12.4679 | 1.31E-05 | 0.615 | GTGCGTGTC |
| 3286 | mamo-s | *BGIBMGA004040* | down | 1259 | 1267 | + | 11.3303 | 4.96E-05 | 0.615 | GTGGGTGGC |
| 3287 | mamo-s | *BGIBMGA004041* | up | 1535 | 1543 | - | 11.3303 | 4.96E-05 | 0.615 | GTGGGTGGC |
| 3288 | mamo-s | *BGIBMGA004043* | up | 518 | 526 | + | 11.3303 | 4.96E-05 | 0.615 | GTGGGTGGC |
| 3289 | mamo-s | *BGIBMGA004043* | up | 1783 | 1791 | - | 11.3303 | 4.96E-05 | 0.615 | GTGGGTGGC |
| 3290 | mamo-s | *BGIBMGA004044* | up | 1283 | 1291 | + | 11.1009 | 6.57E-05 | 0.709 | GAGCGAGGC |
| 3291 | mamo-s | *BGIBMGA004047* | down | 367 | 375 | + | 11.1743 | 5.90E-05 | 0.684 | CCGCGTGTC |
| 3292 | mamo-s | *BGIBMGA004049* | up | 563 | 571 | + | 10.8899 | 7.48E-05 | 0.739 | CCGCGTGCC |
| 3293 | mamo-s | *BGIBMGA004050* | down | 1406 | 1414 | + | 11.3303 | 4.96E-05 | 0.615 | GTGGGTGGC |
| 3294 | mamo-s | *BGIBMGA004051* | up | 1051 | 1059 | - | 11.3303 | 4.96E-05 | 0.615 | GTGGGTGGC |
| 3295 | mamo-s | *BGIBMGA004052* | down | 1405 | 1413 | - | 11.3303 | 4.96E-05 | 0.615 | GTGGGTGGC |
| 3296 | mamo-s | *BGIBMGA004053* | up | 1330 | 1338 | + | 11.0734 | 7.12E-05 | 0.726 | GAGCGTGGT |
| 3297 | mamo-s | *BGIBMGA004054* | up | 635 | 643 | - | 11.3303 | 4.96E-05 | 0.615 | GTGGGTGGC |
| 3298 | mamo-s | *BGIBMGA004054* | down | 1479 | 1487 | - | 11.3303 | 4.96E-05 | 0.615 | GTGGGTGGC |
| 3299 | mamo-s | *BGIBMGA004060* | up | 436 | 444 | + | 11.3303 | 4.96E-05 | 0.615 | GTGGGTGGC |
| 3300 | mamo-s | *BGIBMGA004061* | up | 1952 | 1960 | + | 11.0734 | 7.12E-05 | 0.726 | GAGCGTGGT |
| 3301 | mamo-s | *BGIBMGA004063* | up | 88 | 96 | - | 10.8899 | 7.48E-05 | 0.739 | CCGCGTGCC |
| 3302 | mamo-s | *BGIBMGA004064* | down | 1585 | 1593 | + | 11.3303 | 4.96E-05 | 0.615 | GTGGGTGGC |
| 3303 | mamo-s | *BGIBMGA004069* | up | 21 | 29 | - | 11.1009 | 6.57E-05 | 0.709 | GAGCGAGGC |
| 3304 | mamo-s | *BGIBMGA004070* | down | 606 | 614 | + | 11.3303 | 4.96E-05 | 0.615 | GTGGGTGGC |
| 3305 | mamo-s | *BGIBMGA004070* | up | 1863 | 1871 | + | 11.3303 | 4.96E-05 | 0.615 | GTGGGTGGC |
| 3306 | mamo-s | *BGIBMGA004071* | down | 825 | 833 | - | 10.9358 | 7.30E-05 | 0.728 | GCGGGTGGC |
| 3307 | mamo-s | *BGIBMGA004072* | down | 1981 | 1989 | + | 11.3303 | 4.96E-05 | 0.615 | GTGGGTGGC |
| 3308 | mamo-s | *BGIBMGA004073* | up | 1882 | 1890 | + | 11.3303 | 4.96E-05 | 0.615 | GTGGGTGGC |
| 3309 | mamo-s | *BGIBMGA004074* | down | 1018 | 1026 | - | 11.3303 | 4.96E-05 | 0.615 | GTGGGTGGC |
| 3310 | mamo-s | *BGIBMGA004074* | down | 616 | 624 | - | 11.1009 | 6.57E-05 | 0.709 | GAGCGAGGC |
| 3311 | mamo-s | *BGIBMGA004075* | down | 867 | 875 | - | 10.5413 | 9.42E-05 | 0.857 | GTGCGAGTC |
| 3312 | mamo-s | *BGIBMGA004079* | up | 224 | 232 | + | 14.055 | 2.23E-06 | 0.615 | GTGCGTGGC |
| 3313 | mamo-s | *BGIBMGA004080* | down | 492 | 500 | - | 11.3303 | 4.96E-05 | 0.615 | GTGGGTGGC |
| 3314 | mamo-s | *BGIBMGA004080* | up | 845 | 853 | - | 11.3303 | 4.96E-05 | 0.615 | GTGGGTGGC |
| 3315 | mamo-s | *BGIBMGA004080* | up | 1810 | 1818 | + | 11.2294 | 5.41E-05 | 0.65 | CTGCGAGGC |
| 3316 | mamo-s | *BGIBMGA004080* | down | 1636 | 1644 | - | 10.5413 | 9.42E-05 | 0.857 | CAGCGTGTC |
| 3317 | mamo-s | *BGIBMGA004081* | up | 597 | 605 | + | 11.3303 | 4.96E-05 | 0.615 | GTGGGTGGC |
| 3318 | mamo-s | *BGIBMGA004081* | up | 1666 | 1674 | + | 11.3303 | 4.96E-05 | 0.615 | GTGGGTGGC |
| 3319 | mamo-s | *BGIBMGA004081* | up | 800 | 808 | - | 11.0826 | 6.84E-05 | 0.709 | GTACGTGGC |
| 3320 | mamo-s | *BGIBMGA004082* | up | 507 | 515 | + | 12.4679 | 1.31E-05 | 0.615 | GTGCGTGTC |
| 3321 | mamo-s | *BGIBMGA004082* | down | 1255 | 1263 | - | 12.1284 | 2.16E-05 | 0.615 | GTGCGAGGC |
| 3322 | mamo-s | *BGIBMGA004084* | up | 44 | 52 | - | 11.3303 | 4.96E-05 | 0.615 | GTGGGTGGC |
| 3323 | mamo-s | *BGIBMGA004084* | down | 165 | 173 | + | 11.3303 | 4.96E-05 | 0.615 | GTGGGTGGC |
| 3324 | mamo-s | *BGIBMGA004085* | up | 1721 | 1729 | - | 11.3303 | 4.96E-05 | 0.615 | GTGGGTGGC |
| 3325 | mamo-s | *BGIBMGA004087* | up | 1299 | 1307 | - | 11.3303 | 4.96E-05 | 0.615 | GTGGGTGGC |
| 3326 | mamo-s | *BGIBMGA004093* | down | 409 | 417 | - | 10.9358 | 7.30E-05 | 0.728 | GCGGGTGGC |
| 3327 | mamo-s | *BGIBMGA004094* | up | 1310 | 1318 | + | 11.3303 | 4.96E-05 | 0.615 | GTGGGTGGC |
| 3328 | mamo-s | *BGIBMGA004097* | up | 1148 | 1156 | + | 12.0734 | 2.66E-05 | 0.615 | GCGCGTGTC |
| 3329 | mamo-s | *BGIBMGA004097* | up | 944 | 952 | - | 11.3303 | 4.96E-05 | 0.615 | GTGGGTGGC |
| 3330 | mamo-s | *BGIBMGA004107* | up | 791 | 799 | - | 11.3303 | 4.96E-05 | 0.615 | GTGGGTGGC |
| 3331 | mamo-s | *BGIBMGA004107* | up | 1869 | 1877 | + | 11.3303 | 4.96E-05 | 0.615 | GTGGGTGGC |
| 3332 | mamo-s | *BGIBMGA004107* | down | 1507 | 1515 | + | 10.5138 | 9.75E-05 | 0.865 | GTGCGTGTT |
| 3333 | mamo-s | *BGIBMGA004109* | up | 1978 | 1986 | - | 11.5688 | 3.79E-05 | 0.615 | CTGCGTGTC |
| 3334 | mamo-s | *BGIBMGA004110* | down | 264 | 272 | - | 11.0826 | 6.84E-05 | 0.709 | GTACGTGGC |
| 3335 | mamo-s | *BGIBMGA004111* | up | 709 | 717 | + | 11.3303 | 4.96E-05 | 0.615 | GTGGGTGGC |
| 3336 | mamo-s | *BGIBMGA004111* | down | 1784 | 1792 | + | 11.3303 | 4.96E-05 | 0.615 | GTGGGTGGC |
| 3337 | mamo-s | *BGIBMGA004111* | down | 148 | 156 | - | 10.9358 | 7.30E-05 | 0.728 | GCGGGTGGC |

| | | | | | | | | | | |
|---|---|---|---|---|---|---|---|---|---|---|
| 3338 | mamo-s | *BGIBMGA004112* | up | 1586 | 1594 | + | 11.3303 | 4.96E-05 | 0.615 | GTGGGTGGC |
| 3339 | mamo-s | *BGIBMGA004115* | up | 1429 | 1437 | + | 11.3303 | 4.96E-05 | 0.615 | GTGGGTGGC |
| 3340 | mamo-s | *BGIBMGA004116* | down | 1002 | 1010 | - | 11.3303 | 4.96E-05 | 0.615 | GTGGGTGGC |
| 3341 | mamo-s | *BGIBMGA004116* | down | 656 | 664 | - | 10.5138 | 9.75E-05 | 0.865 | GTGCGTGTT |
| 3342 | mamo-s | *BGIBMGA004118* | up | 726 | 734 | - | 11.3303 | 4.96E-05 | 0.615 | GTGGGTGGC |
| 3343 | mamo-s | *BGIBMGA004120* | up | 429 | 437 | - | 11.3303 | 4.96E-05 | 0.615 | GTGGGTGGC |
| 3344 | mamo-s | *BGIBMGA004122* | up | 469 | 477 | - | 13.6606 | 4.06E-06 | 0.615 | GCGCGTGGC |
| 3345 | mamo-s | *BGIBMGA004122* | up | 458 | 466 | - | 10.8073 | 7.89E-05 | 0.766 | CCGCGTGGT |
| 3346 | mamo-s | *BGIBMGA004122* | down | 120 | 128 | + | 10.5138 | 9.75E-05 | 0.865 | GTGCGTGTT |
| 3347 | mamo-s | *BGIBMGA004123* | down | 1440 | 1448 | + | 11.3303 | 4.96E-05 | 0.615 | GTGGGTGGC |
| 3348 | mamo-s | *BGIBMGA004123* | down | 1895 | 1903 | + | 11.1009 | 6.57E-05 | 0.709 | GAGCGAGGC |
| 3349 | mamo-s | *BGIBMGA004124* | down | 1349 | 1357 | - | 11.0826 | 6.84E-05 | 0.709 | GTACGTGGC |
| 3350 | mamo-s | *BGIBMGA004125* | down | 607 | 615 | + | 11.3303 | 4.96E-05 | 0.615 | GTGGGTGGC |
| 3351 | mamo-s | *BGIBMGA004126* | up | 1755 | 1763 | - | 11.3303 | 4.96E-05 | 0.615 | GTGGGTGGC |
| 3352 | mamo-s | *BGIBMGA004128* | down | 1534 | 1542 | + | 11.7064 | 3.52E-05 | 0.615 | GCGCGTGGT |
| 3353 | mamo-s | *BGIBMGA004129* | down | 1115 | 1123 | + | 11.3303 | 4.96E-05 | 0.615 | GTGGGTGGC |
| 3354 | mamo-s | *BGIBMGA004129* | down | 493 | 501 | + | 11.0826 | 6.84E-05 | 0.709 | GTACGTGGC |
| 3355 | mamo-s | *BGIBMGA004129* | up | 516 | 524 | - | 11.0826 | 6.84E-05 | 0.709 | GTACGTGGC |
| 3356 | mamo-s | *BGIBMGA004130* | down | 1854 | 1862 | + | 11.3303 | 4.96E-05 | 0.615 | GTGGGTGGC |
| 3357 | mamo-s | *BGIBMGA004130* | up | 134 | 142 | - | 11.1743 | 5.90E-05 | 0.684 | CCGCGTGTC |
| 3358 | mamo-s | *BGIBMGA004130* | up | 279 | 287 | + | 10.7523 | 8.38E-05 | 0.797 | AAGCGTGGC |
| 3359 | mamo-s | *BGIBMGA004132* | up | 562 | 570 | - | 11.7339 | 3.29E-05 | 0.615 | GCGCGAGGC |
| 3360 | mamo-s | *BGIBMGA004133* | down | 1424 | 1432 | + | 11.3303 | 4.96E-05 | 0.615 | GTGGGTGGC |
| 3361 | mamo-s | *BGIBMGA004136* | down | 1233 | 1241 | + | 11.3303 | 4.96E-05 | 0.615 | GTGGGTGGC |
| 3362 | mamo-s | *BGIBMGA004141* | down | 15 | 23 | + | 12.4679 | 1.31E-05 | 0.615 | GTGCGTGTC |
| 3363 | mamo-s | *BGIBMGA004141* | up | 1102 | 1110 | - | 12.3394 | 1.49E-05 | 0.615 | GGGCGTGGC |
| 3364 | mamo-s | *BGIBMGA004141* | up | 1960 | 1968 | - | 11.1376 | 6.35E-05 | 0.709 | TCGCGTGGC |
| 3365 | mamo-s | *BGIBMGA004144* | down | 243 | 251 | + | 12.7615 | 1.04E-05 | 0.615 | CCGCGTGGC |
| 3366 | mamo-s | *BGIBMGA004144* | down | 253 | 261 | + | 12.3394 | 1.49E-05 | 0.615 | GGGCGTGGC |
| 3367 | mamo-s | *BGIBMGA004144* | down | 268 | 276 | + | 12.3394 | 1.49E-05 | 0.615 | GGGCGTGGC |
| 3368 | mamo-s | *BGIBMGA004144* | up | 1032 | 1040 | - | 11.3303 | 4.96E-05 | 0.615 | GTGGGTGGC |
| 3369 | mamo-s | *BGIBMGA004144* | down | 1868 | 1876 | + | 10.8349 | 7.66E-05 | 0.751 | CCGCGAGGC |
| 3370 | mamo-s | *BGIBMGA004144* | up | 731 | 739 | + | 10.7523 | 8.38E-05 | 0.797 | AAGCGTGGC |
| 3371 | mamo-s | *BGIBMGA004146* | down | 519 | 527 | + | 12.4679 | 1.31E-05 | 0.615 | GTGCGTGTC |
| 3372 | mamo-s | *BGIBMGA004146* | down | 564 | 572 | + | 11.1376 | 6.35E-05 | 0.709 | TCGCGTGGC |
| 3373 | mamo-s | *BGIBMGA004147* | down | 520 | 528 | + | 11.3303 | 4.96E-05 | 0.615 | GTGGGTGGC |
| 3374 | mamo-s | *BGIBMGA004148* | up | 1535 | 1543 | - | 11.3303 | 4.96E-05 | 0.615 | GTGGGTGGC |
| 3375 | mamo-s | *BGIBMGA004149* | up | 1048 | 1056 | + | 11.1376 | 6.35E-05 | 0.709 | TCGCGTGGC |
| 3376 | mamo-s | *BGIBMGA004150* | up | 555 | 563 | + | 11.1376 | 6.35E-05 | 0.709 | TCGCGTGGC |
| 3377 | mamo-s | *BGIBMGA004151* | up | 1339 | 1347 | + | 11.3303 | 4.96E-05 | 0.615 | GTGGGTGGC |
| 3378 | mamo-s | *BGIBMGA004151* | up | 285 | 293 | + | 10.7523 | 8.38E-05 | 0.797 | AAGCGTGGC |
| 3379 | mamo-s | *BGIBMGA004152* | up | 627 | 635 | - | 13.6606 | 4.06E-06 | 0.615 | GCGCGTGGC |
| 3380 | mamo-s | *BGIBMGA004152* | up | 1367 | 1375 | - | 11.3303 | 4.96E-05 | 0.615 | GTGGGTGGC |
| 3381 | mamo-s | *BGIBMGA004154* | up | 983 | 991 | + | 12.4679 | 1.31E-05 | 0.615 | GTGCGTGTC |
| 3382 | mamo-s | *BGIBMGA004154* | down | 305 | 313 | + | 11.3303 | 4.96E-05 | 0.615 | GTGGGTGGC |
| 3383 | mamo-s | *BGIBMGA004154* | down | 1465 | 1473 | - | 11.3303 | 4.96E-05 | 0.615 | GTGGGTGGC |
| 3384 | mamo-s | *BGIBMGA004154* | up | 1761 | 1769 | - | 11.0826 | 6.84E-05 | 0.709 | GTACGTGGC |
| 3385 | mamo-s | *BGIBMGA004155* | down | 688 | 696 | + | 12.1009 | 2.43E-05 | 0.615 | GTGCGTGGT |
| 3386 | mamo-s | *BGIBMGA004155* | up | 1039 | 1047 | - | 11.3303 | 4.96E-05 | 0.615 | GTGGGTGGC |
| 3387 | mamo-s | *BGIBMGA004155* | down | 954 | 962 | + | 10.9358 | 7.30E-05 | 0.728 | GCGGGTGGC |
| 3388 | mamo-s | *BGIBMGA004156* | up | 509 | 517 | + | 12.1009 | 2.43E-05 | 0.615 | GTGCGTGGT |
| 3389 | mamo-s | *BGIBMGA004156* | down | 673 | 681 | + | 11.3303 | 4.96E-05 | 0.615 | GTGGGTGGC |
| 3390 | mamo-s | *BGIBMGA004156* | up | 775 | 783 | + | 10.9358 | 7.30E-05 | 0.728 | GCGGGTGGC |
| 3391 | mamo-s | *BGIBMGA004158* | down | 926 | 934 | + | 12.3394 | 1.49E-05 | 0.615 | GGGCGTGGC |
| 3392 | mamo-s | *BGIBMGA004158* | down | 1329 | 1337 | + | 11.3303 | 4.96E-05 | 0.615 | GTGGGTGGC |
| 3393 | mamo-s | *BGIBMGA004159* | down | 1657 | 1665 | + | 11.7339 | 3.29E-05 | 0.615 | GCGCGAGGC |
| 3394 | mamo-s | *BGIBMGA004161* | up | 302 | 310 | - | 10.5138 | 9.75E-05 | 0.865 | GTGCGTGTT |
| 3395 | mamo-s | *BGIBMGA004164* | up | 1847 | 1855 | - | 12.1284 | 2.16E-05 | 0.615 | GTGCGAGGC |
| 3396 | mamo-s | *BGIBMGA004166* | up | 1559 | 1567 | - | 10.5138 | 9.75E-05 | 0.865 | GTGCGTGTT |
| 3397 | mamo-s | *BGIBMGA004167* | down | 684 | 692 | - | 11.3303 | 4.96E-05 | 0.615 | GTGGGTGGC |
| 3398 | mamo-s | *BGIBMGA004167* | down | 1580 | 1588 | - | 10.8349 | 7.66E-05 | 0.751 | CCGCGAGGC |
| 3399 | mamo-s | *BGIBMGA004168* | up | 648 | 656 | - | 11.3303 | 4.96E-05 | 0.615 | GTGGGTGGC |
| 3400 | mamo-s | *BGIBMGA004169* | up | 1382 | 1390 | + | 11.3303 | 4.96E-05 | 0.615 | GTGGGTGGC |
| 3401 | mamo-s | *BGIBMGA004170* | up | 22 | 30 | - | 11.3303 | 4.96E-05 | 0.615 | GTGGGTGGC |
| 3402 | mamo-s | *BGIBMGA004170* | down | 1218 | 1226 | - | 10.6881 | 8.61E-05 | 0.81 | GCACGTGGC |
| 3403 | mamo-s | *BGIBMGA004171* | down | 299 | 307 | + | 11.3303 | 4.96E-05 | 0.615 | GTGGGTGGC |
| 3404 | mamo-s | *BGIBMGA004172* | down | 858 | 866 | + | 11.3303 | 4.96E-05 | 0.615 | GTGGGTGGC |
| 3405 | mamo-s | *BGIBMGA004173* | down | 1115 | 1123 | - | 11.3303 | 4.96E-05 | 0.615 | GTGGGTGGC |
| 3406 | mamo-s | *BGIBMGA004173* | down | 1160 | 1168 | - | 11.1009 | 6.57E-05 | 0.709 | GAGCGAGGC |
| 3407 | mamo-s | *BGIBMGA004175* | down | 1672 | 1680 | - | 11.3303 | 4.96E-05 | 0.615 | GTGGGTGGC |
| 3408 | mamo-s | *BGIBMGA004176* | down | 588 | 596 | - | 11.5321 | 4.06E-05 | 0.615 | TTGCGTGGC |
| 3409 | mamo-s | *BGIBMGA004177* | down | 1445 | 1453 | - | 11.3303 | 4.96E-05 | 0.615 | GTGGGTGGC |
| 3410 | mamo-s | *BGIBMGA004178* | up | 110 | 118 | - | 12.1284 | 2.16E-05 | 0.615 | GTGCGAGGC |
| 3411 | mamo-s | *BGIBMGA004178* | down | 1395 | 1403 | + | 11.0826 | 6.84E-05 | 0.709 | GTACGTGGC |
| 3412 | mamo-s | *BGIBMGA004179* | up | 1051 | 1059 | - | 11.3303 | 4.96E-05 | 0.615 | GTGGGTGGC |
| 3413 | mamo-s | *BGIBMGA004180* | down | 1958 | 1966 | + | 10.7523 | 8.38E-05 | 0.797 | AAGCGTGGC |
| 3414 | mamo-s | *BGIBMGA004181* | down | 752 | 760 | + | 11.3303 | 4.96E-05 | 0.615 | GTGGGTGGC |
| 3415 | mamo-s | *BGIBMGA004182* | up | 1342 | 1350 | + | 11.3303 | 4.96E-05 | 0.615 | GTGGGTGGC |
| 3416 | mamo-s | *BGIBMGA004182* | up | 1511 | 1519 | + | 11.3303 | 4.96E-05 | 0.615 | GTGGGTGGC |
| 3417 | mamo-s | *BGIBMGA004183* | up | 495 | 503 | + | 11.3303 | 4.96E-05 | 0.615 | GTGGGTGGC |
| 3418 | mamo-s | *BGIBMGA004183* | down | 1433 | 1441 | - | 11.3303 | 4.96E-05 | 0.615 | GTGGGTGGC |
| 3419 | mamo-s | *BGIBMGA004183* | up | 1757 | 1765 | + | 11.3303 | 4.96E-05 | 0.615 | GTGGGTGGC |
| 3420 | mamo-s | *BGIBMGA004184* | down | 600 | 608 | - | 11.3303 | 4.96E-05 | 0.615 | GTGGGTGGC |
| 3421 | mamo-s | *BGIBMGA004185* | up | 1022 | 1030 | - | 11.3303 | 4.96E-05 | 0.615 | GTGGGTGGC |
| 3422 | mamo-s | *BGIBMGA004189* | up | 1660 | 1668 | + | 11.3303 | 4.96E-05 | 0.615 | GTGGGTGGC |
| 3423 | mamo-s | *BGIBMGA004190* | up | 226 | 234 | + | 11.3303 | 4.96E-05 | 0.615 | GTGGGTGGC |
| 3424 | mamo-s | *BGIBMGA004191* | down | 47 | 55 | - | 14.055 | 2.23E-06 | 0.615 | GTGCGTGGC |
| 3425 | mamo-s | *BGIBMGA004191* | down | 74 | 82 | - | 13.6606 | 4.06E-06 | 0.615 | GCGCGTGGC |

| | | | | | | | | | | |
|---|---|---|---|---|---|---|---|---|---|---|
| 3426 | mamo-s | *BGIBMGA004191* | down | 1946 | 1954 | - | 11.7339 | 3.29E-05 | 0.615 | GCGCGAGGC |
| 3427 | mamo-s | *BGIBMGA004194* | down | 1211 | 1219 | - | 10.5138 | 9.75E-05 | 0.865 | GTGCGTGTT |
| 3428 | mamo-s | *BGIBMGA004195* | down | 1130 | 1138 | + | 12.0734 | 2.66E-05 | 0.615 | GCGCGTGTC |
| 3429 | mamo-s | *BGIBMGA004195* | down | 1329 | 1337 | + | 12.0734 | 2.66E-05 | 0.615 | GCGCGTGTC |
| 3430 | mamo-s | *BGIBMGA004195* | down | 832 | 840 | + | 11.3303 | 4.96E-05 | 0.615 | GTGGGTGGC |
| 3431 | mamo-s | *BGIBMGA004195* | up | 152 | 160 | - | 10.5138 | 9.75E-05 | 0.865 | GTGCGTGTT |
| 3432 | mamo-s | *BGIBMGA004196* | down | 1860 | 1868 | - | 11.7339 | 3.29E-05 | 0.615 | GCGCGAGGC |
| 3433 | mamo-s | *BGIBMGA004197* | up | 1058 | 1066 | + | 11.3303 | 4.96E-05 | 0.615 | GTGGGTGGC |
| 3434 | mamo-s | *BGIBMGA004197* | down | 713 | 721 | - | 11.0826 | 6.84E-05 | 0.709 | GTACGTGGC |
| 3435 | mamo-s | *BGIBMGA004198* | down | 1048 | 1056 | - | 11.3303 | 4.96E-05 | 0.615 | GTGGGTGGC |
| 3436 | mamo-s | *BGIBMGA004200* | down | 1383 | 1391 | + | 13.6606 | 4.06E-06 | 0.615 | GCGCGTGGC |
| 3437 | mamo-s | *BGIBMGA004202* | down | 477 | 485 | + | 11.2294 | 5.41E-05 | 0.65 | CTGCGAGGC |
| 3438 | mamo-s | *BGIBMGA004203* | down | 1597 | 1605 | + | 11.7339 | 3.29E-05 | 0.615 | GCGCGAGGC |
| 3439 | mamo-s | *BGIBMGA004203* | up | 1308 | 1316 | - | 11.3303 | 4.96E-05 | 0.615 | GTGGGTGGC |
| 3440 | mamo-s | *BGIBMGA004203* | down | 1620 | 1628 | - | 11.2018 | 5.68E-05 | 0.674 | CTGCGTGGT |
| 3441 | mamo-s | *BGIBMGA004204* | up | 638 | 646 | + | 11.3303 | 4.96E-05 | 0.615 | GTGGGTGGC |
| 3442 | mamo-s | *BGIBMGA004205* | up | 1917 | 1925 | - | 11.5321 | 4.06E-05 | 0.615 | TTGCGTGGC |
| 3443 | mamo-s | *BGIBMGA004206* | down | 602 | 610 | - | 11.3303 | 4.96E-05 | 0.615 | GTGGGTGGC |
| 3444 | mamo-s | *BGIBMGA004210* | down | 199 | 207 | - | 12.7615 | 1.04E-05 | 0.615 | CCGCGTGGC |
| 3445 | mamo-s | *BGIBMGA004210* | down | 174 | 182 | - | 12.3394 | 1.49E-05 | 0.615 | GGGCGTGGC |
| 3446 | mamo-s | *BGIBMGA004210* | down | 189 | 197 | - | 12.3394 | 1.49E-05 | 0.615 | GGGCGTGGC |
| 3447 | mamo-s | *BGIBMGA004210* | down | 1607 | 1615 | + | 11.3303 | 4.96E-05 | 0.615 | GTGGGTGGC |
| 3448 | mamo-s | *BGIBMGA004213* | down | 751 | 759 | + | 11.1743 | 5.90E-05 | 0.684 | CCGCGTGTC |
| 3449 | mamo-s | *BGIBMGA004214* | up | 1592 | 1600 | - | 11.3303 | 4.96E-05 | 0.615 | GTGGGTGGC |
| 3450 | mamo-s | *BGIBMGA004215* | up | 346 | 354 | + | 11.7798 | 3.11E-05 | 0.615 | ATGCGTGGC |
| 3451 | mamo-s | *BGIBMGA004215* | up | 456 | 464 | - | 11.3303 | 4.96E-05 | 0.615 | GTGGGTGGC |
| 3452 | mamo-s | *BGIBMGA004217* | up | 1798 | 1806 | + | 11.3303 | 4.96E-05 | 0.615 | GTGGGTGGC |
| 3453 | mamo-s | *BGIBMGA004218* | down | 545 | 553 | - | 11.3303 | 4.96E-05 | 0.615 | GTGGGTGGC |
| 3454 | mamo-s | *BGIBMGA004218* | down | 1991 | 1999 | + | 11.3303 | 4.96E-05 | 0.615 | GTGGGTGGC |
| 3455 | mamo-s | *BGIBMGA004219* | up | 1188 | 1196 | - | 11.3303 | 4.96E-05 | 0.615 | GTGGGTGGC |
| 3456 | mamo-s | *BGIBMGA004219* | up | 507 | 515 | + | 11.1009 | 6.57E-05 | 0.709 | GAGCGAGGC |
| 3457 | mamo-s | *BGIBMGA004220* | up | 1755 | 1763 | - | 11.3303 | 4.96E-05 | 0.615 | GTGGGTGGC |
| 3458 | mamo-s | *BGIBMGA004222* | up | 93 | 101 | - | 11.3303 | 4.96E-05 | 0.615 | GTGGGTGGC |
| 3459 | mamo-s | *BGIBMGA004222* | up | 1176 | 1184 | + | 11.3303 | 4.96E-05 | 0.615 | GTGGGTGGC |
| 3460 | mamo-s | *BGIBMGA004223* | up | 534 | 542 | - | 12.7615 | 1.04E-05 | 0.615 | CCGCGTGGC |
| 3461 | mamo-s | *BGIBMGA004223* | up | 265 | 273 | - | 11.3303 | 4.96E-05 | 0.615 | GTGGGTGGC |
| 3462 | mamo-s | *BGIBMGA004224* | up | 677 | 685 | - | 11.3303 | 4.96E-05 | 0.615 | GTGGGTGGC |
| 3463 | mamo-s | *BGIBMGA004224* | up | 1216 | 1224 | + | 11.3303 | 4.96E-05 | 0.615 | GTGGGTGGC |
| 3464 | mamo-s | *BGIBMGA004225* | down | 174 | 182 | - | 11.3303 | 4.96E-05 | 0.615 | GTGGGTGGC |
| 3465 | mamo-s | *BGIBMGA004225* | up | 888 | 896 | - | 11.3303 | 4.96E-05 | 0.615 | GTGGGTGGC |
| 3466 | mamo-s | *BGIBMGA004226* | up | 182 | 190 | - | 11.3303 | 4.96E-05 | 0.615 | GTGGGTGGC |
| 3467 | mamo-s | *BGIBMGA004228* | down | 1955 | 1963 | + | 11.3303 | 4.96E-05 | 0.615 | GTGGGTGGC |
| 3468 | mamo-s | *BGIBMGA004229* | up | 742 | 750 | - | 11.3303 | 4.96E-05 | 0.615 | GTGGGTGGC |
| 3469 | mamo-s | *BGIBMGA004230* | up | 736 | 744 | - | 14.055 | 2.23E-06 | 0.615 | GTGCGTGGC |
| 3470 | mamo-s | *BGIBMGA004232* | up | 2 | 10 | - | 11.2294 | 5.41E-05 | 0.65 | CTGCGAGGC |
| 3471 | mamo-s | *BGIBMGA004234* | up | 1992 | 2000 | + | 11.789 | 2.84E-05 | 0.615 | GCGCGTGCC |
| 3472 | mamo-s | *BGIBMGA004235* | up | 393 | 401 | + | 11.0734 | 7.12E-05 | 0.726 | GAGCGTGGT |
| 3473 | mamo-s | *BGIBMGA004236* | down | 1201 | 1209 | - | 11.3303 | 4.96E-05 | 0.615 | GTGGGTGGC |
| 3474 | mamo-s | *BGIBMGA004236* | up | 1180 | 1188 | + | 10.7523 | 8.38E-05 | 0.797 | AAGCGTGGC |
| 3475 | mamo-s | *BGIBMGA004237* | down | 810 | 818 | + | 13.6606 | 4.06E-06 | 0.615 | GCGCGTGGC |
| 3476 | mamo-s | *BGIBMGA004237* | up | 238 | 246 | - | 11.3303 | 4.96E-05 | 0.615 | GTGGGTGGC |
| 3477 | mamo-s | *BGIBMGA004238* | up | 1565 | 1573 | - | 11.0734 | 7.12E-05 | 0.726 | GAGCGTGGT |
| 3478 | mamo-s | *BGIBMGA004238* | up | 1738 | 1746 | + | 10.6881 | 8.61E-05 | 0.81 | GCACGTGGC |
| 3479 | mamo-s | *BGIBMGA004239* | down | 207 | 215 | + | 11.2294 | 5.41E-05 | 0.65 | CTGCGAGGC |
| 3480 | mamo-s | *BGIBMGA004241* | down | 223 | 231 | + | 11.3303 | 4.96E-05 | 0.615 | GTGGGTGGC |
| 3481 | mamo-s | *BGIBMGA004242* | up | 1669 | 1677 | - | 11.3303 | 4.96E-05 | 0.615 | GTGGGTGGC |
| 3482 | mamo-s | *BGIBMGA004242* | down | 31 | 39 | - | 11.0826 | 6.84E-05 | 0.709 | GTACGTGGC |
| 3483 | mamo-s | *BGIBMGA004243* | down | 1083 | 1091 | - | 11.3303 | 4.96E-05 | 0.615 | GTGGGTGGC |
| 3484 | mamo-s | *BGIBMGA004243* | down | 1435 | 1443 | + | 11.3303 | 4.96E-05 | 0.615 | GTGGGTGGC |
| 3485 | mamo-s | *BGIBMGA004244* | down | 1501 | 1509 | + | 13.156 | 6.30E-06 | 0.615 | CTGCGTGGC |
| 3486 | mamo-s | *BGIBMGA004245* | up | 703 | 711 | + | 13.156 | 6.30E-06 | 0.615 | CTGCGTGGC |
| 3487 | mamo-s | *BGIBMGA004245* | down | 247 | 255 | - | 11.3303 | 4.96E-05 | 0.615 | GTGGGTGGC |
| 3488 | mamo-s | *BGIBMGA004248* | down | 937 | 945 | + | 11.3303 | 4.96E-05 | 0.615 | GTGGGTGGC |
| 3489 | mamo-s | *BGIBMGA004248* | up | 1763 | 1771 | - | 11.3303 | 4.96E-05 | 0.615 | GTGGGTGGC |
| 3490 | mamo-s | *BGIBMGA004248* | down | 1359 | 1367 | - | 10.5138 | 9.75E-05 | 0.865 | GTGCGTGTT |
| 3491 | mamo-s | *BGIBMGA004249* | up | 961 | 969 | + | 11.3303 | 4.96E-05 | 0.615 | GTGGGTGGC |
| 3492 | mamo-s | *BGIBMGA004249* | up | 1383 | 1391 | - | 10.5138 | 9.75E-05 | 0.865 | GTGCGTGTT |
| 3493 | mamo-s | *BGIBMGA004250* | up | 451 | 459 | + | 11.3303 | 4.96E-05 | 0.615 | GTGGGTGGC |
| 3494 | mamo-s | *BGIBMGA004251* | down | 6 | 14 | + | 11.789 | 2.84E-05 | 0.615 | GCGCGTGCC |
| 3495 | mamo-s | *BGIBMGA004252* | up | 1900 | 1908 | - | 11.0734 | 7.12E-05 | 0.726 | GAGCGTGGT |
| 3496 | mamo-s | *BGIBMGA004255* | down | 1927 | 1935 | + | 10.5688 | 8.88E-05 | 0.824 | GTGCGTGAC |
| 3497 | mamo-s | *BGIBMGA004257* | up | 1240 | 1248 | - | 11.5321 | 4.06E-05 | 0.615 | TTGCGTGGC |
| 3498 | mamo-s | *BGIBMGA004259* | down | 1318 | 1326 | - | 12.3394 | 1.49E-05 | 0.615 | GGGCGTGGC |
| 3499 | mamo-s | *BGIBMGA004259* | up | 222 | 230 | + | 12.1284 | 2.16E-05 | 0.615 | CAGCGTGGC |
| 3500 | mamo-s | *BGIBMGA004259* | down | 1249 | 1257 | - | 12.0734 | 2.66E-05 | 0.615 | GCGCGTGTC |
| 3501 | mamo-s | *BGIBMGA004260* | up | 99 | 107 | - | 11.3303 | 4.96E-05 | 0.615 | GTGGGTGGC |
| 3502 | mamo-s | *BGIBMGA004262* | down | 645 | 653 | - | 12.1284 | 2.16E-05 | 0.615 | GTGCGAGGC |
| 3503 | mamo-s | *BGIBMGA004263* | down | 1032 | 1040 | + | 11.1743 | 5.90E-05 | 0.684 | CCGCGTGTC |
| 3504 | mamo-s | *BGIBMGA004263* | up | 1170 | 1178 | - | 10.5413 | 9.42E-05 | 0.857 | CAGCGTGTC |
| 3505 | mamo-s | *BGIBMGA004264* | down | 1907 | 1915 | - | 11.156 | 6.13E-05 | 0.705 | GAGCGTGCC |
| 3506 | mamo-s | *BGIBMGA004266* | down | 897 | 905 | - | 11.3303 | 4.96E-05 | 0.615 | GTGGGTGGC |
| 3507 | mamo-s | *BGIBMGA004267* | down | 926 | 934 | + | 11.0826 | 6.84E-05 | 0.709 | GTACGTGGC |
| 3508 | mamo-s | *BGIBMGA004268* | up | 1861 | 1869 | - | 10.5688 | 8.88E-05 | 0.824 | GTGCGTGAC |
| 3509 | mamo-s | *BGIBMGA004269* | up | 81 | 89 | - | 11.3303 | 4.96E-05 | 0.615 | GTGGGTGGC |
| 3510 | mamo-s | *BGIBMGA004270* | down | 1873 | 1881 | - | 11.3303 | 4.96E-05 | 0.615 | GTGGGTGGC |
| 3511 | mamo-s | *BGIBMGA004271* | up | 1364 | 1372 | - | 11.3303 | 4.96E-05 | 0.615 | GTGGGTGGC |
| 3512 | mamo-s | *BGIBMGA004271* | down | 1643 | 1651 | + | 11.3303 | 4.96E-05 | 0.615 | GTGGGTGGC |
| 3513 | mamo-s | *BGIBMGA004271* | down | 207 | 215 | - | 11.0826 | 6.84E-05 | 0.709 | GTACGTGGC |

| | | | | | | | | | | |
|---|---|---|---|---|---|---|---|---|---|---|
| 3514 | mamo-s | *BGIBMGA004272* | up | 662 | 670 | - | 14.055 | 2.23E-06 | 0.615 | GTGCGTGGC |
| 3515 | mamo-s | *BGIBMGA004272* | up | 1177 | 1185 | - | 11.0826 | 6.84E-05 | 0.709 | GTACGTGGC |
| 3516 | mamo-s | *BGIBMGA004272* | down | 360 | 368 | + | 10.8073 | 7.89E-05 | 0.766 | CCGCGTGGT |
| 3517 | mamo-s | *BGIBMGA004274* | up | 1059 | 1067 | + | 11.3303 | 4.96E-05 | 0.615 | GTGGGTGGC |
| 3518 | mamo-s | *BGIBMGA004274* | down | 1460 | 1468 | - | 11.3303 | 4.96E-05 | 0.615 | GTGGGTGGC |
| 3519 | mamo-s | *BGIBMGA004275* | up | 1699 | 1707 | - | 11.3303 | 4.96E-05 | 0.615 | GTGGGTGGC |
| 3520 | mamo-s | *BGIBMGA004276* | up | 1718 | 1726 | + | 11.5321 | 4.06E-05 | 0.615 | TTGCGTGGC |
| 3521 | mamo-s | *BGIBMGA004276* | up | 160 | 168 | - | 11.3303 | 4.96E-05 | 0.615 | GTGGGTGGC |
| 3522 | mamo-s | *BGIBMGA004277* | up | 798 | 806 | + | 11.3303 | 4.96E-05 | 0.615 | GTGGGTGGC |
| 3523 | mamo-s | *BGIBMGA004277* | down | 1803 | 1811 | + | 11.3303 | 4.96E-05 | 0.615 | GTGGGTGGC |
| 3524 | mamo-s | *BGIBMGA004278* | up | 1102 | 1110 | - | 11.2294 | 5.41E-05 | 0.65 | CTGCGAGGC |
| 3525 | mamo-s | *BGIBMGA004280* | up | 831 | 839 | - | 11.3303 | 4.96E-05 | 0.615 | GTGGGTGGC |
| 3526 | mamo-s | *BGIBMGA004280* | up | 1008 | 1016 | + | 11.3303 | 4.96E-05 | 0.615 | GTGGGTGGC |
| 3527 | mamo-s | *BGIBMGA004281* | down | 1410 | 1418 | + | 11.3303 | 4.96E-05 | 0.615 | GTGGGTGGC |
| 3528 | mamo-s | *BGIBMGA004283* | down | 1502 | 1510 | + | 11.3303 | 4.96E-05 | 0.615 | GTGGGTGGC |
| 3529 | mamo-s | *BGIBMGA004285* | up | 1904 | 1912 | + | 11.3303 | 4.96E-05 | 0.615 | GTGGGTGGC |
| 3530 | mamo-s | *BGIBMGA004286* | up | 221 | 229 | + | 13.6606 | 4.06E-06 | 0.615 | GCGCGTGGC |
| 3531 | mamo-s | *BGIBMGA004286* | down | 1712 | 1720 | - | 11.2844 | 5.18E-05 | 0.63 | CTGCGTGCC |
| 3532 | mamo-s | *BGIBMGA004287* | up | 507 | 515 | + | 11.3303 | 4.96E-05 | 0.615 | GTGGGTGGC |
| 3533 | mamo-s | *BGIBMGA004287* | down | 879 | 887 | + | 11.3303 | 4.96E-05 | 0.615 | GTGGGTGGC |
| 3534 | mamo-s | *BGIBMGA004291* | down | 724 | 732 | - | 12.0734 | 2.66E-05 | 0.615 | GCGCGTGTC |
| 3535 | mamo-s | *BGIBMGA004293* | down | 1453 | 1461 | - | 11.3303 | 4.96E-05 | 0.615 | GTGGGTGGC |
| 3536 | mamo-s | *BGIBMGA004294* | down | 1823 | 1831 | + | 11.3303 | 4.96E-05 | 0.615 | GTGGGTGGC |
| 3537 | mamo-s | *BGIBMGA004295* | up | 1195 | 1203 | - | 14.055 | 2.23E-06 | 0.615 | GTGCGTGGC |
| 3538 | mamo-s | *BGIBMGA004295* | up | 169 | 177 | + | 11.3303 | 4.96E-05 | 0.615 | GTGGGTGGC |
| 3539 | mamo-s | *BGIBMGA004297* | down | 331 | 339 | + | 11.7339 | 3.29E-05 | 0.615 | GCGCGAGGC |
| 3540 | mamo-s | *BGIBMGA004298* | down | 1285 | 1293 | - | 11.3303 | 4.96E-05 | 0.615 | GTGGGTGGC |
| 3541 | mamo-s | *BGIBMGA004299* | down | 1767 | 1775 | + | 11.3303 | 4.96E-05 | 0.615 | GTGGGTGGC |
| 3542 | mamo-s | *BGIBMGA004300* | down | 1173 | 1181 | - | 12.1009 | 2.43E-05 | 0.615 | GTGCGTGGT |
| 3543 | mamo-s | *BGIBMGA004300* | up | 1425 | 1433 | + | 11.789 | 2.84E-05 | 0.615 | GCGCGTGCC |
| 3544 | mamo-s | *BGIBMGA004300* | down | 115 | 123 | - | 11.3303 | 4.96E-05 | 0.615 | GTGGGTGGC |
| 3545 | mamo-s | *BGIBMGA004302* | up | 446 | 454 | - | 11.3303 | 4.96E-05 | 0.615 | GTGGGTGGC |
| 3546 | mamo-s | *BGIBMGA004303* | down | 424 | 432 | + | 11.3303 | 4.96E-05 | 0.615 | GTGGGTGGC |
| 3547 | mamo-s | *BGIBMGA004303* | down | 1156 | 1164 | + | 10.9358 | 7.30E-05 | 0.728 | GCGGGTGGC |
| 3548 | mamo-s | *BGIBMGA004304* | up | 1318 | 1326 | - | 11.2018 | 5.68E-05 | 0.674 | CTGCGTGGT |
| 3549 | mamo-s | *BGIBMGA004304* | up | 727 | 735 | + | 11.1009 | 6.57E-05 | 0.709 | GAGCGAGGC |
| 3550 | mamo-s | *BGIBMGA004305* | down | 1361 | 1369 | + | 11.7064 | 3.52E-05 | 0.615 | GCGCGTGGT |
| 3551 | mamo-s | *BGIBMGA004306* | down | 1352 | 1360 | - | 13.156 | 6.30E-06 | 0.615 | CTGCGTGGC |
| 3552 | mamo-s | *BGIBMGA004306* | up | 921 | 929 | + | 11.7064 | 3.52E-05 | 0.615 | GCGCGTGGT |
| 3553 | mamo-s | *BGIBMGA004310* | up | 266 | 274 | - | 11.0734 | 7.12E-05 | 0.726 | GAGCGTGGT |
| 3554 | mamo-s | *BGIBMGA004312* | down | 942 | 950 | + | 11.3303 | 4.96E-05 | 0.615 | GTGGGTGGC |
| 3555 | mamo-s | *BGIBMGA004312* | down | 1680 | 1688 | - | 11.3303 | 4.96E-05 | 0.615 | GTGGGTGGC |
| 3556 | mamo-s | *BGIBMGA004313* | up | 208 | 216 | - | 10.5138 | 9.75E-05 | 0.865 | GTGCGTGTT |
| 3557 | mamo-s | *BGIBMGA004318* | down | 1562 | 1570 | + | 12.3394 | 1.49E-05 | 0.615 | GGGCGTGGC |
| 3558 | mamo-s | *BGIBMGA004319* | down | 1097 | 1105 | + | 11.3303 | 4.96E-05 | 0.615 | GTGGGTGGC |
| 3559 | mamo-s | *BGIBMGA004320* | up | 1225 | 1233 | + | 11.3303 | 4.96E-05 | 0.615 | GTGGGTGGC |
| 3560 | mamo-s | *BGIBMGA004320* | down | 1703 | 1711 | - | 11.3303 | 4.96E-05 | 0.615 | GTGGGTGGC |
| 3561 | mamo-s | *BGIBMGA004320* | up | 235 | 243 | - | 11.1743 | 5.90E-05 | 0.684 | CCGCGTGTC |
| 3562 | mamo-s | *BGIBMGA004320* | up | 702 | 710 | + | 11.0826 | 6.84E-05 | 0.709 | GTACGTGGC |
| 3563 | mamo-s | *BGIBMGA004321* | down | 1395 | 1403 | + | 11.0826 | 6.84E-05 | 0.709 | GTACGTGGC |
| 3564 | mamo-s | *BGIBMGA004325* | down | 385 | 393 | - | 13.0275 | 8.53E-06 | 0.615 | GAGCGTGGC |
| 3565 | mamo-s | *BGIBMGA004325* | down | 496 | 504 | + | 10.9358 | 7.30E-05 | 0.728 | GCGGGTGGC |
| 3566 | mamo-s | *BGIBMGA004327* | up | 1783 | 1791 | + | 11.5321 | 4.06E-05 | 0.615 | TTGCGTGGC |
| 3567 | mamo-s | *BGIBMGA004329* | up | 1030 | 1038 | + | 11.3303 | 4.96E-05 | 0.615 | GTGGGTGGC |
| 3568 | mamo-s | *BGIBMGA004330* | down | 627 | 635 | - | 11.3303 | 4.96E-05 | 0.615 | GTGGGTGGC |
| 3569 | mamo-s | *BGIBMGA004331* | up | 1623 | 1631 | + | 11.3303 | 4.96E-05 | 0.615 | GTGGGTGGC |
| 3570 | mamo-s | *BGIBMGA004331* | down | 1931 | 1939 | + | 11.3303 | 4.96E-05 | 0.615 | GTGGGTGGC |
| 3571 | mamo-s | *BGIBMGA004333* | down | 433 | 441 | - | 11.3303 | 4.96E-05 | 0.615 | GTGGGTGGC |
| 3572 | mamo-s | *BGIBMGA004334* | up | 1597 | 1605 | + | 11.2844 | 5.18E-05 | 0.63 | CTGCGTGCC |
| 3573 | mamo-s | *BGIBMGA004335* | up | 379 | 387 | - | 11.3303 | 4.96E-05 | 0.615 | GTGGGTGGC |
| 3574 | mamo-s | *BGIBMGA004339* | down | 235 | 243 | - | 11.3303 | 4.96E-05 | 0.615 | GTGGGTGGC |
| 3575 | mamo-s | *BGIBMGA004339* | up | 1361 | 1369 | + | 11.2294 | 5.41E-05 | 0.65 | CTGCGAGGC |
| 3576 | mamo-s | *BGIBMGA004341* | up | 652 | 660 | + | 11.3303 | 4.96E-05 | 0.615 | GTGGGTGGC |
| 3577 | mamo-s | *BGIBMGA004341* | down | 1322 | 1330 | + | 11.3303 | 4.96E-05 | 0.615 | GTGGGTGGC |
| 3578 | mamo-s | *BGIBMGA004341* | up | 1675 | 1683 | - | 11.3303 | 4.96E-05 | 0.615 | GTGGGTGGC |
| 3579 | mamo-s | *BGIBMGA004342* | up | 936 | 944 | - | 11.3303 | 4.96E-05 | 0.615 | GTGGGTGGC |
| 3580 | mamo-s | *BGIBMGA004342* | down | 1657 | 1665 | + | 11.0826 | 6.84E-05 | 0.709 | GTACGTGGC |
| 3581 | mamo-s | *BGIBMGA004343* | up | 469 | 477 | + | 11.0826 | 6.84E-05 | 0.709 | GTACGTGGC |
| 3582 | mamo-s | *BGIBMGA004344* | down | 1194 | 1202 | - | 11.3303 | 4.96E-05 | 0.615 | GTGGGTGGC |
| 3583 | mamo-s | *BGIBMGA004345* | up | 1272 | 1280 | - | 11.1009 | 6.57E-05 | 0.709 | GAGCGAGGC |
| 3584 | mamo-s | *BGIBMGA004348* | up | 209 | 217 | + | 11.3303 | 4.96E-05 | 0.615 | GTGGGTGGC |
| 3585 | mamo-s | *BGIBMGA004353* | up | 1385 | 1393 | - | 11.3303 | 4.96E-05 | 0.615 | GTGGGTGGC |
| 3586 | mamo-s | *BGIBMGA004354* | down | 992 | 1000 | - | 11.3303 | 4.96E-05 | 0.615 | GTGGGTGGC |
| 3587 | mamo-s | *BGIBMGA004354* | up | 1301 | 1309 | + | 11.3303 | 4.96E-05 | 0.615 | GTGGGTGGC |
| 3588 | mamo-s | *BGIBMGA004354* | down | 1982 | 1990 | - | 11.3303 | 4.96E-05 | 0.615 | GTGGGTGGC |
| 3589 | mamo-s | *BGIBMGA004355* | up | 951 | 959 | - | 11.3303 | 4.96E-05 | 0.615 | GTGGGTGGC |
| 3590 | mamo-s | *BGIBMGA004355* | up | 1142 | 1150 | + | 10.9358 | 7.30E-05 | 0.728 | GCGGGTGGC |
| 3591 | mamo-s | *BGIBMGA004358* | down | 1545 | 1553 | + | 11.3853 | 4.74E-05 | 0.615 | ACGCGTGGC |
| 3592 | mamo-s | *BGIBMGA004358* | down | 1124 | 1132 | + | 11.0826 | 6.84E-05 | 0.709 | GTACGTGGC |
| 3593 | mamo-s | *BGIBMGA004359* | up | 1666 | 1674 | + | 11.0826 | 6.84E-05 | 0.709 | GTACGTGGC |
| 3594 | mamo-s | *BGIBMGA004361* | up | 291 | 299 | + | 11.0734 | 7.12E-05 | 0.726 | GAGCGTGGT |
| 3595 | mamo-s | *BGIBMGA004362* | down | 1182 | 1190 | + | 11.3303 | 4.96E-05 | 0.615 | GTGGGTGGC |
| 3596 | mamo-s | *BGIBMGA004364* | down | 1687 | 1695 | + | 11.3303 | 4.96E-05 | 0.615 | GTGGGTGGC |
| 3597 | mamo-s | *BGIBMGA004365* | up | 596 | 604 | + | 11.3303 | 4.96E-05 | 0.615 | GTGGGTGGC |
| 3598 | mamo-s | *BGIBMGA004366* | down | 464 | 472 | + | 11.3303 | 4.96E-05 | 0.615 | GTGGGTGGC |
| 3599 | mamo-s | *BGIBMGA004367* | up | 1271 | 1279 | - | 11.3303 | 4.96E-05 | 0.615 | GTGGGTGGC |
| 3600 | mamo-s | *BGIBMGA004367* | up | 712 | 720 | + | 11.1743 | 5.90E-05 | 0.684 | CCGCGTGTC |
| 3601 | mamo-s | *BGIBMGA004369* | down | 323 | 331 | + | 11.5688 | 3.79E-05 | 0.615 | CTGCGTGTC |

| | | | | | | | | | | |
|---|---|---|---|---|---|---|---|---|---|---|
| 3602 | mamo-s | *BGIBMGA004370* | up | 1506 | 1514 | - | 14.055 | 2.23E-06 | 0.615 | GTGCGTGGC |
| 3603 | mamo-s | *BGIBMGA004370* | up | 265 | 273 | - | 11.3303 | 4.96E-05 | 0.615 | GTGGGTGGC |
| 3604 | mamo-s | *BGIBMGA004371* | up | 831 | 839 | - | 11.3303 | 4.96E-05 | 0.615 | GTGGGTGGC |
| 3605 | mamo-s | *BGIBMGA004372* | up | 1641 | 1649 | - | 10.7523 | 8.38E-05 | 0.797 | AAGCGTGGC |
| 3606 | mamo-s | *BGIBMGA004375* | down | 239 | 247 | - | 11.3303 | 4.96E-05 | 0.615 | GTGGGTGGC |
| 3607 | mamo-s | *BGIBMGA004375* | up | 1332 | 1340 | - | 11.2294 | 5.41E-05 | 0.65 | CTGCGAGGC |
| 3608 | mamo-s | *BGIBMGA004375* | down | 689 | 697 | + | 11.156 | 6.13E-05 | 0.705 | GAGCGTGCC |
| 3609 | mamo-s | *BGIBMGA004375* | up | 1489 | 1497 | - | 11.1009 | 6.57E-05 | 0.709 | GAGCGAGGC |
| 3610 | mamo-s | *BGIBMGA004378* | up | 54 | 62 | + | 12.1009 | 2.43E-05 | 0.615 | GTGCGTGGT |
| 3611 | mamo-s | *BGIBMGA004379* | up | 224 | 232 | - | 11.3303 | 4.96E-05 | 0.615 | GTGGGTGGC |
| 3612 | mamo-s | *BGIBMGA004381* | up | 1228 | 1236 | - | 10.5138 | 9.75E-05 | 0.865 | GTGCGTGTT |
| 3613 | mamo-s | *BGIBMGA004383* | down | 1885 | 1893 | - | 12.1835 | 1.71E-05 | 0.615 | GTGCGTGCC |
| 3614 | mamo-s | *BGIBMGA004384* | down | 1281 | 1289 | - | 12.1835 | 1.71E-05 | 0.615 | GTGCGTGCC |
| 3615 | mamo-s | *BGIBMGA004386* | down | 1662 | 1670 | - | 12.3394 | 1.49E-05 | 0.615 | GGGCGTGGC |
| 3616 | mamo-s | *BGIBMGA004386* | down | 167 | 175 | - | 11.3303 | 4.96E-05 | 0.615 | GTGGGTGGC |
| 3617 | mamo-s | *BGIBMGA004391* | up | 1082 | 1090 | + | 11.3303 | 4.96E-05 | 0.615 | GTGGGTGGC |
| 3618 | mamo-s | *BGIBMGA004392* | down | 1080 | 1088 | + | 11.3303 | 4.96E-05 | 0.615 | GTGGGTGGC |
| 3619 | mamo-s | *BGIBMGA004393* | up | 1379 | 1387 | + | 11.3303 | 4.96E-05 | 0.615 | GTGGGTGGC |
| 3620 | mamo-s | *BGIBMGA004393* | up | 332 | 340 | - | 11.0826 | 6.84E-05 | 0.709 | GTACGTGGC |
| 3621 | mamo-s | *BGIBMGA004396* | up | 434 | 442 | - | 11.3303 | 4.96E-05 | 0.615 | GTGGGTGGC |
| 3622 | mamo-s | *BGIBMGA004397* | down | 1304 | 1312 | + | 11.3303 | 4.96E-05 | 0.615 | GTGGGTGGC |
| 3623 | mamo-s | *BGIBMGA004397* | down | 1582 | 1590 | - | 11.3303 | 4.96E-05 | 0.615 | GTGGGTGGC |
| 3624 | mamo-s | *BGIBMGA004399* | down | 253 | 261 | - | 11.3853 | 4.74E-05 | 0.615 | ACGCGTGGC |
| 3625 | mamo-s | *BGIBMGA004399* | down | 1731 | 1739 | + | 11.3303 | 4.96E-05 | 0.615 | GTGGGTGGC |
| 3626 | mamo-s | *BGIBMGA004401* | up | 272 | 280 | - | 11.3303 | 4.96E-05 | 0.615 | GTGGGTGGC |
| 3627 | mamo-s | *BGIBMGA004402* | up | 768 | 776 | - | 11.1743 | 5.90E-05 | 0.684 | CCGCGTGTC |
| 3628 | mamo-s | *BGIBMGA004405* | up | 1116 | 1124 | + | 12.1009 | 2.43E-05 | 0.615 | GTGCGTGGT |
| 3629 | mamo-s | *BGIBMGA004405* | down | 1302 | 1310 | + | 11.3303 | 4.96E-05 | 0.615 | GTGGGTGGC |
| 3630 | mamo-s | *BGIBMGA004407* | down | 1311 | 1319 | + | 11.3303 | 4.96E-05 | 0.615 | GTGGGTGGC |
| 3631 | mamo-s | *BGIBMGA004408* | up | 1094 | 1102 | - | 11.3303 | 4.96E-05 | 0.615 | GTGGGTGGC |
| 3632 | mamo-s | *BGIBMGA004408* | down | 1301 | 1309 | - | 11.3303 | 4.96E-05 | 0.615 | GTGGGTGGC |
| 3633 | mamo-s | *BGIBMGA004408* | down | 307 | 315 | - | 11.0826 | 6.84E-05 | 0.709 | GTACGTGGC |
| 3634 | mamo-s | *BGIBMGA004410* | up | 322 | 330 | - | 11.3303 | 4.96E-05 | 0.615 | GTGGGTGGC |
| 3635 | mamo-s | *BGIBMGA004418* | down | 1851 | 1859 | - | 12.3394 | 1.49E-05 | 0.615 | GGGCGTGGC |
| 3636 | mamo-s | *BGIBMGA004418* | up | 457 | 465 | + | 12.1284 | 2.16E-05 | 0.615 | CAGCGTGGC |
| 3637 | mamo-s | *BGIBMGA004419* | up | 417 | 425 | + | 11.3303 | 4.96E-05 | 0.615 | GTGGGTGGC |
| 3638 | mamo-s | *BGIBMGA004420* | down | 1137 | 1145 | + | 12.4679 | 1.31E-05 | 0.615 | GTGCGTGTC |
| 3639 | mamo-s | *BGIBMGA004421* | down | 654 | 662 | - | 11.3303 | 4.96E-05 | 0.615 | GTGGGTGGC |
| 3640 | mamo-s | *BGIBMGA004425* | down | 341 | 349 | - | 11.3303 | 4.96E-05 | 0.615 | GTGGGTGGC |
| 3641 | mamo-s | *BGIBMGA004425* | up | 1459 | 1467 | - | 11.3303 | 4.96E-05 | 0.615 | GTGGGTGGC |
| 3642 | mamo-s | *BGIBMGA004425* | down | 1796 | 1804 | + | 11.2294 | 5.41E-05 | 0.65 | CTGCGAGGC |
| 3643 | mamo-s | *BGIBMGA004427* | up | 615 | 623 | + | 11.0826 | 6.84E-05 | 0.709 | GTACGTGGC |
| 3644 | mamo-s | *BGIBMGA004428* | down | 796 | 804 | + | 11.3303 | 4.96E-05 | 0.615 | GTGGGTGGC |
| 3645 | mamo-s | *BGIBMGA004431* | up | 988 | 996 | - | 13.6606 | 4.06E-06 | 0.615 | GCGCGTGGC |
| 3646 | mamo-s | *BGIBMGA004431* | up | 127 | 135 | - | 11.3303 | 4.96E-05 | 0.615 | GTGGGTGGC |
| 3647 | mamo-s | *BGIBMGA004431* | down | 135 | 143 | - | 11.1743 | 5.90E-05 | 0.684 | CCGCGTGTC |
| 3648 | mamo-s | *BGIBMGA004433* | down | 1256 | 1264 | - | 11.3303 | 4.96E-05 | 0.615 | GTGGGTGGC |
| 3649 | mamo-s | *BGIBMGA004434* | up | 1918 | 1926 | + | 11.7064 | 3.52E-05 | 0.615 | GCGCGTGGT |
| 3650 | mamo-s | *BGIBMGA004434* | down | 923 | 931 | - | 11.3303 | 4.96E-05 | 0.615 | GTGGGTGGC |
| 3651 | mamo-s | *BGIBMGA004435* | up | 1792 | 1800 | - | 13.6606 | 4.06E-06 | 0.615 | GCGCGTGGC |
| 3652 | mamo-s | *BGIBMGA004435* | down | 347 | 355 | - | 11.3303 | 4.96E-05 | 0.615 | GTGGGTGGC |
| 3653 | mamo-s | *BGIBMGA004437* | down | 188 | 196 | - | 11.3303 | 4.96E-05 | 0.615 | GTGGGTGGC |
| 3654 | mamo-s | *BGIBMGA004439* | down | 387 | 395 | - | 11.3303 | 4.96E-05 | 0.615 | GTGGGTGGC |
| 3655 | mamo-s | *BGIBMGA004442* | up | 80 | 88 | + | 11.1009 | 6.57E-05 | 0.709 | GAGCGAGGC |
| 3656 | mamo-s | *BGIBMGA004443* | down | 872 | 880 | + | 11.5688 | 3.79E-05 | 0.615 | CTGCGTGTC |
| 3657 | mamo-s | *BGIBMGA004444* | down | 511 | 519 | - | 11.789 | 2.84E-05 | 0.615 | GCGCGTGCC |
| 3658 | mamo-s | *BGIBMGA004444* | up | 302 | 310 | + | 11.3303 | 4.96E-05 | 0.615 | GTGGGTGGC |
| 3659 | mamo-s | *BGIBMGA004445* | up | 1065 | 1073 | - | 11.3303 | 4.96E-05 | 0.615 | GTGGGTGGC |
| 3660 | mamo-s | *BGIBMGA004446* | down | 911 | 919 | + | 11.3303 | 4.96E-05 | 0.615 | GTGGGTGGC |
| 3661 | mamo-s | *BGIBMGA004446* | down | 1366 | 1374 | - | 11.2294 | 5.41E-05 | 0.65 | CTGCGAGGC |
| 3662 | mamo-s | *BGIBMGA004446* | up | 1512 | 1520 | + | 11.1743 | 5.90E-05 | 0.684 | CCGCGTGTC |
| 3663 | mamo-s | *BGIBMGA004446* | up | 1981 | 1989 | - | 10.8899 | 7.48E-05 | 0.739 | CCGCGTGCC |
| 3664 | mamo-s | *BGIBMGA004446* | down | 424 | 432 | + | 10.7523 | 8.38E-05 | 0.797 | AAGCGTGGC |
| 3665 | mamo-s | *BGIBMGA004447* | down | 980 | 988 | - | 11.3303 | 4.96E-05 | 0.615 | GTGGGTGGC |
| 3666 | mamo-s | *BGIBMGA004449* | up | 1209 | 1217 | + | 11.3303 | 4.96E-05 | 0.615 | GTGGGTGGC |
| 3667 | mamo-s | *BGIBMGA004449* | down | 279 | 287 | + | 11.0826 | 6.84E-05 | 0.709 | GTACGTGGC |
| 3668 | mamo-s | *BGIBMGA004451* | down | 365 | 373 | - | 11.7798 | 3.11E-05 | 0.615 | ATGCGTGGC |
| 3669 | mamo-s | *BGIBMGA004451* | down | 311 | 319 | - | 10.6881 | 8.61E-05 | 0.81 | GCACGTGGC |
| 3670 | mamo-s | *BGIBMGA004452* | up | 229 | 237 | + | 11.7339 | 3.29E-05 | 0.615 | GCGCGAGGC |
| 3671 | mamo-s | *BGIBMGA004452* | down | 1087 | 1095 | - | 11.3303 | 4.96E-05 | 0.615 | GTGGGTGGC |
| 3672 | mamo-s | *BGIBMGA004452* | up | 252 | 260 | - | 11.2018 | 5.68E-05 | 0.674 | CTGCGTGGT |
| 3673 | mamo-s | *BGIBMGA004454* | down | 1173 | 1181 | - | 11.156 | 6.13E-05 | 0.705 | GAGCGTGCC |
| 3674 | mamo-s | *BGIBMGA004454* | up | 1005 | 1013 | + | 11.1376 | 6.35E-05 | 0.709 | TCGCGTGGC |
| 3675 | mamo-s | *BGIBMGA004456* | down | 1434 | 1442 | - | 10.8073 | 7.89E-05 | 0.766 | CCGCGTGGT |
| 3676 | mamo-s | *BGIBMGA004457* | up | 1803 | 1811 | + | 11.3303 | 4.96E-05 | 0.615 | GTGGGTGGC |
| 3677 | mamo-s | *BGIBMGA004458* | down | 1627 | 1635 | - | 13.6606 | 4.06E-06 | 0.615 | GCGCGTGGC |
| 3678 | mamo-s | *BGIBMGA004460* | up | 366 | 374 | + | 11.3303 | 4.96E-05 | 0.615 | GTGGGTGGC |
| 3679 | mamo-s | *BGIBMGA004461* | up | 267 | 275 | + | 11.3303 | 4.96E-05 | 0.615 | GTGGGTGGC |
| 3680 | mamo-s | *BGIBMGA004461* | down | 1743 | 1751 | + | 10.9358 | 7.30E-05 | 0.728 | GCGGGTGGC |
| 3681 | mamo-s | *BGIBMGA004463* | down | 950 | 958 | + | 11.3303 | 4.96E-05 | 0.615 | GTGGGTGGC |
| 3682 | mamo-s | *BGIBMGA004464* | up | 1670 | 1678 | - | 11.3303 | 4.96E-05 | 0.615 | GTGGGTGGC |
| 3683 | mamo-s | *BGIBMGA004465* | down | 831 | 839 | + | 11.2844 | 5.18E-05 | 0.63 | CTGCGTGCC |
| 3684 | mamo-s | *BGIBMGA004467* | up | 1336 | 1344 | + | 11.3303 | 4.96E-05 | 0.615 | GTGGGTGGC |
| 3685 | mamo-s | *BGIBMGA004473* | down | 539 | 547 | + | 11.3303 | 4.96E-05 | 0.615 | GTGGGTGGC |
| 3686 | mamo-s | *BGIBMGA004474* | down | 408 | 416 | - | 11.3303 | 4.96E-05 | 0.615 | GTGGGTGGC |
| 3687 | mamo-s | *BGIBMGA004474* | down | 824 | 832 | - | 11.3303 | 4.96E-05 | 0.615 | GTGGGTGGC |
| 3688 | mamo-s | *BGIBMGA004475* | down | 577 | 585 | + | 11.3303 | 4.96E-05 | 0.615 | GTGGGTGGC |
| 3689 | mamo-s | *BGIBMGA004479* | up | 586 | 594 | - | 11.3303 | 4.96E-05 | 0.615 | GTGGGTGGC |

| | | | | | | | | | | |
|---|---|---|---|---|---|---|---|---|---|---|
| 3690 | mamo-s | *BGIBMGA004479* | down | 1799 | 1807 | - | 11.3303 | 4.96E-05 | 0.615 | GTGGGTGGC |
| 3691 | mamo-s | *BGIBMGA004481* | up | 328 | 336 | + | 11.3853 | 4.74E-05 | 0.615 | ACGCGTGGC |
| 3692 | mamo-s | *BGIBMGA004481* | up | 1269 | 1277 | + | 11.3303 | 4.96E-05 | 0.615 | GTGGGTGGC |
| 3693 | mamo-s | *BGIBMGA004482* | down | 941 | 949 | + | 11.3303 | 4.96E-05 | 0.615 | GTGGGTGGC |
| 3694 | mamo-s | *BGIBMGA004482* | up | 1107 | 1115 | + | 11.3303 | 4.96E-05 | 0.615 | GTGGGTGGC |
| 3695 | mamo-s | *BGIBMGA004483* | up | 398 | 406 | + | 11.3303 | 4.96E-05 | 0.615 | GTGGGTGGC |
| 3696 | mamo-s | *BGIBMGA004485* | up | 1084 | 1092 | - | 11.3303 | 4.96E-05 | 0.615 | GTGGGTGGC |
| 3697 | mamo-s | *BGIBMGA004485* | up | 640 | 648 | + | 11.0826 | 6.84E-05 | 0.709 | GTACGTGGC |
| 3698 | mamo-s | *BGIBMGA004486* | up | 1840 | 1848 | + | 11.3303 | 4.96E-05 | 0.615 | GTGGGTGGC |
| 3699 | mamo-s | *BGIBMGA004488* | down | 1626 | 1634 | - | 11.3303 | 4.96E-05 | 0.615 | GTGGGTGGC |
| 3700 | mamo-s | *BGIBMGA004488* | up | 1536 | 1544 | + | 10.9358 | 7.30E-05 | 0.728 | GCGGGTGGC |
| 3701 | mamo-s | *BGIBMGA004490* | up | 1125 | 1133 | + | 11.2294 | 5.41E-05 | 0.65 | CTGCGAGGC |
| 3702 | mamo-s | *BGIBMGA004490* | up | 459 | 467 | - | 10.5688 | 8.88E-05 | 0.824 | GTGCGTGAC |
| 3703 | mamo-s | *BGIBMGA004492* | down | 1015 | 1023 | + | 12.1284 | 2.16E-05 | 0.615 | GTGCGAGGC |
| 3704 | mamo-s | *BGIBMGA004492* | up | 1107 | 1115 | - | 11.7798 | 3.11E-05 | 0.615 | ATGCGTGGC |
| 3705 | mamo-s | *BGIBMGA004497* | up | 100 | 108 | + | 11.3303 | 4.96E-05 | 0.615 | GTGGGTGGC |
| 3706 | mamo-s | *BGIBMGA004497* | up | 659 | 667 | - | 11.1743 | 5.90E-05 | 0.684 | CCGCGTGTC |
| 3707 | mamo-s | *BGIBMGA004498* | down | 1047 | 1055 | + | 11.3303 | 4.96E-05 | 0.615 | GTGGGTGGC |
| 3708 | mamo-s | *BGIBMGA004498* | down | 830 | 838 | - | 11.0826 | 6.84E-05 | 0.709 | GTACGTGGC |
| 3709 | mamo-s | *BGIBMGA004502* | up | 856 | 864 | - | 11.3303 | 4.96E-05 | 0.615 | GTGGGTGGC |
| 3710 | mamo-s | *BGIBMGA004502* | down | 590 | 598 | + | 11.0734 | 7.12E-05 | 0.726 | GAGCGTGGT |
| 3711 | mamo-s | *BGIBMGA004503* | down | 288 | 296 | - | 11.3303 | 4.96E-05 | 0.615 | GTGGGTGGC |
| 3712 | mamo-s | *BGIBMGA004504* | down | 1357 | 1365 | + | 11.4404 | 4.51E-05 | 0.615 | CGGCGTGGC |
| 3713 | mamo-s | *BGIBMGA004504* | up | 561 | 569 | - | 11.3303 | 4.96E-05 | 0.615 | GTGGGTGGC |
| 3714 | mamo-s | *BGIBMGA004505* | up | 57 | 65 | + | 11.3303 | 4.96E-05 | 0.615 | GTGGGTGGC |
| 3715 | mamo-s | *BGIBMGA004505* | up | 1088 | 1096 | - | 11.3303 | 4.96E-05 | 0.615 | GTGGGTGGC |
| 3716 | mamo-s | *BGIBMGA004506* | up | 932 | 940 | - | 14.055 | 2.23E-06 | 0.615 | GTGCGTGGC |
| 3717 | mamo-s | *BGIBMGA004506* | down | 956 | 964 | - | 13.0275 | 8.53E-06 | 0.615 | GAGCGTGGC |
| 3718 | mamo-s | *BGIBMGA004506* | up | 1849 | 1857 | + | 11.3303 | 4.96E-05 | 0.615 | GTGGGTGGC |
| 3719 | mamo-s | *BGIBMGA004507* | up | 1415 | 1423 | - | 10.5138 | 9.75E-05 | 0.865 | GTGCGTGTT |
| 3720 | mamo-s | *BGIBMGA004508* | up | 452 | 460 | - | 11.3303 | 4.96E-05 | 0.615 | GTGGGTGGC |
| 3721 | mamo-s | *BGIBMGA004508* | up | 1902 | 1910 | - | 11.1009 | 6.57E-05 | 0.709 | GAGCGAGGC |
| 3722 | mamo-s | *BGIBMGA004511* | down | 527 | 535 | + | 11.3303 | 4.96E-05 | 0.615 | GTGGGTGGC |
| 3723 | mamo-s | *BGIBMGA004511* | up | 1466 | 1474 | + | 11.3303 | 4.96E-05 | 0.615 | GTGGGTGGC |
| 3724 | mamo-s | *BGIBMGA004511* | down | 157 | 165 | - | 11.1743 | 5.90E-05 | 0.684 | CCGCGTGTC |
| 3725 | mamo-s | *BGIBMGA004511* | down | 893 | 901 | + | 11.0826 | 6.84E-05 | 0.709 | GTACGTGGC |
| 3726 | mamo-s | *BGIBMGA004512* | up | 1063 | 1071 | - | 11.1743 | 5.90E-05 | 0.684 | CCGCGTGTC |
| 3727 | mamo-s | *BGIBMGA004512* | up | 1900 | 1908 | + | 11.0826 | 6.84E-05 | 0.709 | GTACGTGGC |
| 3728 | mamo-s | *BGIBMGA004514* | up | 1831 | 1839 | - | 11.7339 | 3.29E-05 | 0.615 | GCGCGAGGC |
| 3729 | mamo-s | *BGIBMGA004515* | down | 699 | 707 | - | 11.3303 | 4.96E-05 | 0.615 | GTGGGTGGC |
| 3730 | mamo-s | *BGIBMGA004518* | down | 256 | 264 | - | 11.3303 | 4.96E-05 | 0.615 | GTGGGTGGC |
| 3731 | mamo-s | *BGIBMGA004518* | up | 700 | 708 | - | 11.3303 | 4.96E-05 | 0.615 | GTGGGTGGC |
| 3732 | mamo-s | *BGIBMGA004518* | down | 1721 | 1729 | - | 11.3303 | 4.96E-05 | 0.615 | GTGGGTGGC |
| 3733 | mamo-s | *BGIBMGA004519* | down | 1503 | 1511 | - | 11.3303 | 4.96E-05 | 0.615 | GTGGGTGGC |
| 3734 | mamo-s | *BGIBMGA004520* | up | 606 | 614 | - | 11.3303 | 4.96E-05 | 0.615 | GTGGGTGGC |
| 3735 | mamo-s | *BGIBMGA004520* | down | 1833 | 1841 | - | 11.3303 | 4.96E-05 | 0.615 | GTGGGTGGC |
| 3736 | mamo-s | *BGIBMGA004520* | down | 390 | 398 | - | 11.2844 | 5.18E-05 | 0.63 | CTGCGTGCC |
| 3737 | mamo-s | *BGIBMGA004521* | up | 1233 | 1241 | - | 10.9358 | 7.30E-05 | 0.728 | GCGGGTGGC |
| 3738 | mamo-s | *BGIBMGA004524* | down | 707 | 715 | + | 10.8073 | 7.89E-05 | 0.766 | CCGCGTGGT |
| 3739 | mamo-s | *BGIBMGA004526* | up | 1136 | 1144 | + | 11.3303 | 4.96E-05 | 0.615 | GTGGGTGGC |
| 3740 | mamo-s | *BGIBMGA004527* | down | 456 | 464 | + | 11.3303 | 4.96E-05 | 0.615 | GTGGGTGGC |
| 3741 | mamo-s | *BGIBMGA004527* | down | 1190 | 1198 | - | 11.3303 | 4.96E-05 | 0.615 | GTGGGTGGC |
| 3742 | mamo-s | *BGIBMGA004527* | down | 900 | 908 | - | 11.2844 | 5.18E-05 | 0.63 | CTGCGTGCC |
| 3743 | mamo-s | *BGIBMGA004529* | down | 1809 | 1817 | - | 14.055 | 2.23E-06 | 0.615 | GTGCGTGGC |
| 3744 | mamo-s | *BGIBMGA004529* | up | 1944 | 1952 | + | 11.4404 | 4.51E-05 | 0.615 | GAGCGTGTC |
| 3745 | mamo-s | *BGIBMGA004530* | up | 1734 | 1742 | - | 11.3303 | 4.96E-05 | 0.615 | GTGGGTGGC |
| 3746 | mamo-s | *BGIBMGA004531* | down | 1557 | 1565 | + | 11.3303 | 4.96E-05 | 0.615 | GTGGGTGGC |
| 3747 | mamo-s | *BGIBMGA004532* | up | 32 | 40 | - | 11.3303 | 4.96E-05 | 0.615 | GTGGGTGGC |
| 3748 | mamo-s | *BGIBMGA004533* | down | 1938 | 1946 | - | 11.7064 | 3.52E-05 | 0.615 | GCGCGTGGT |
| 3749 | mamo-s | *BGIBMGA004533* | down | 1959 | 1967 | - | 11.7064 | 3.52E-05 | 0.615 | GCGCGTGGT |
| 3750 | mamo-s | *BGIBMGA004535* | down | 1884 | 1892 | + | 11.3303 | 4.96E-05 | 0.615 | GTGGGTGGC |
| 3751 | mamo-s | *BGIBMGA004535* | up | 571 | 579 | - | 10.6881 | 8.61E-05 | 0.81 | GCACGTGGC |
| 3752 | mamo-s | *BGIBMGA004536* | down | 704 | 712 | - | 11.3303 | 4.96E-05 | 0.615 | GTGGGTGGC |
| 3753 | mamo-s | *BGIBMGA004537* | up | 1171 | 1179 | - | 11.3303 | 4.96E-05 | 0.615 | GTGGGTGGC |
| 3754 | mamo-s | *BGIBMGA004537* | down | 1745 | 1753 | - | 11.0826 | 6.84E-05 | 0.709 | GTACGTGGC |
| 3755 | mamo-s | *BGIBMGA004538* | down | 253 | 261 | + | 12.1835 | 1.71E-05 | 0.615 | GTGCGTGCC |
| 3756 | mamo-s | *BGIBMGA004538* | down | 964 | 972 | - | 11.3303 | 4.96E-05 | 0.615 | GTGGGTGGC |
| 3757 | mamo-s | *BGIBMGA004540* | down | 1140 | 1148 | + | 11.3303 | 4.96E-05 | 0.615 | GTGGGTGGC |
| 3758 | mamo-s | *BGIBMGA004541* | up | 749 | 757 | - | 12.1009 | 2.43E-05 | 0.615 | GTGCGTGGT |
| 3759 | mamo-s | *BGIBMGA004541* | down | 1069 | 1077 | + | 10.9358 | 7.30E-05 | 0.728 | GCGGGTGGC |
| 3760 | mamo-s | *BGIBMGA004542* | down | 787 | 795 | - | 11.0826 | 6.84E-05 | 0.709 | GTACGTGGC |
| 3761 | mamo-s | *BGIBMGA004544* | down | 319 | 327 | - | 11.3303 | 4.96E-05 | 0.615 | GTGGGTGGC |
| 3762 | mamo-s | *BGIBMGA004545* | up | 1409 | 1417 | - | 11.3303 | 4.96E-05 | 0.615 | GTGGGTGGC |
| 3763 | mamo-s | *BGIBMGA004545* | up | 56 | 64 | - | 10.8899 | 7.48E-05 | 0.739 | CCGCGTGCC |
| 3764 | mamo-s | *BGIBMGA004547* | down | 1266 | 1274 | - | 11.3303 | 4.96E-05 | 0.615 | GTGGGTGGC |
| 3765 | mamo-s | *BGIBMGA004547* | down | 324 | 332 | - | 11.0826 | 6.84E-05 | 0.709 | GTACGTGGC |
| 3766 | mamo-s | *BGIBMGA004548* | down | 1203 | 1211 | + | 11.3303 | 4.96E-05 | 0.615 | GTGGGTGGC |
| 3767 | mamo-s | *BGIBMGA004548* | down | 1625 | 1633 | + | 11.0826 | 6.84E-05 | 0.709 | GTACGTGGC |
| 3768 | mamo-s | *BGIBMGA004550* | up | 1658 | 1666 | + | 11.3303 | 4.96E-05 | 0.615 | GTGGGTGGC |
| 3769 | mamo-s | *BGIBMGA004550* | down | 265 | 273 | + | 10.8073 | 7.89E-05 | 0.766 | CCGCGTGGT |
| 3770 | mamo-s | *BGIBMGA004552* | down | 90 | 98 | - | 12.4679 | 1.31E-05 | 0.615 | GTGCGTGTC |
| 3771 | mamo-s | *BGIBMGA004552* | down | 1843 | 1851 | - | 11.3303 | 4.96E-05 | 0.615 | GTGGGTGGC |
| 3772 | mamo-s | *BGIBMGA004553* | down | 412 | 420 | - | 12.4679 | 1.31E-05 | 0.615 | GTGCGTGTC |
| 3773 | mamo-s | *BGIBMGA004553* | up | 765 | 773 | + | 11.1743 | 5.90E-05 | 0.684 | CCGCGTGTC |
| 3774 | mamo-s | *BGIBMGA004554* | up | 693 | 701 | + | 11.3303 | 4.96E-05 | 0.615 | GTGGGTGGC |
| 3775 | mamo-s | *BGIBMGA004556* | down | 1940 | 1948 | + | 10.8899 | 7.48E-05 | 0.739 | CCGCGTGCC |
| 3776 | mamo-s | *BGIBMGA004557* | up | 1554 | 1562 | - | 11.3303 | 4.96E-05 | 0.615 | GTGGGTGGC |
| 3777 | mamo-s | *BGIBMGA004561* | down | 957 | 965 | + | 11.3303 | 4.96E-05 | 0.615 | GTGGGTGGC |

| | | | | | | | | | | |
|---|---|---|---|---|---|---|---|---|---|---|
| 3778 | mamo-s | *BGIBMGA004561* | up | 1690 | 1698 | - | 11.0734 | 7.12E-05 | 0.726 | GAGCGTGGT |
| 3779 | mamo-s | *BGIBMGA004563* | down | 283 | 291 | - | 11.3303 | 4.96E-05 | 0.615 | GTGGGTGGC |
| 3780 | mamo-s | *BGIBMGA004563* | down | 1551 | 1559 | + | 11.3303 | 4.96E-05 | 0.615 | GTGGGTGGC |
| 3781 | mamo-s | *BGIBMGA004563* | up | 1844 | 1852 | + | 11.3303 | 4.96E-05 | 0.615 | GTGGGTGGC |
| 3782 | mamo-s | *BGIBMGA004564* | up | 642 | 650 | + | 11.3303 | 4.96E-05 | 0.615 | GTGGGTGGC |
| 3783 | mamo-s | *BGIBMGA004564* | down | 1963 | 1971 | - | 11.3303 | 4.96E-05 | 0.615 | GTGGGTGGC |
| 3784 | mamo-s | *BGIBMGA004570* | up | 98 | 106 | + | 10.8899 | 7.48E-05 | 0.739 | CCGCGTGCC |
| 3785 | mamo-s | *BGIBMGA004571* | up | 396 | 404 | + | 12.1835 | 1.71E-05 | 0.615 | GTGCGTGCC |
| 3786 | mamo-s | *BGIBMGA004571* | down | 555 | 563 | + | 10.7523 | 8.38E-05 | 0.797 | AAGCGTGGC |
| 3787 | mamo-s | *BGIBMGA004572* | up | 995 | 1003 | + | 11.3303 | 4.96E-05 | 0.615 | GTGGGTGGC |
| 3788 | mamo-s | *BGIBMGA004574* | down | 112 | 120 | - | 14.055 | 2.23E-06 | 0.615 | GTGCGTGGC |
| 3789 | mamo-s | *BGIBMGA004574* | down | 900 | 908 | - | 11.3853 | 4.74E-05 | 0.615 | ACGCGTGGC |
| 3790 | mamo-s | *BGIBMGA004574* | up | 443 | 451 | + | 11.3303 | 4.96E-05 | 0.615 | GTGGGTGGC |
| 3791 | mamo-s | *BGIBMGA004574* | down | 791 | 799 | + | 11.3303 | 4.96E-05 | 0.615 | GTGGGTGGC |
| 3792 | mamo-s | *BGIBMGA004574* | up | 351 | 359 | - | 10.9358 | 7.30E-05 | 0.728 | GCGGGTGGC |
| 3793 | mamo-s | *BGIBMGA004575* | down | 1455 | 1463 | - | 11.3303 | 4.96E-05 | 0.615 | GTGGGTGGC |
| 3794 | mamo-s | *BGIBMGA004576* | up | 401 | 409 | - | 11.7064 | 3.52E-05 | 0.615 | GCGCGTGGT |
| 3795 | mamo-s | *BGIBMGA004576* | up | 585 | 593 | - | 10.6881 | 8.61E-05 | 0.81 | GCACGTGGC |
| 3796 | mamo-s | *BGIBMGA004577* | down | 240 | 248 | + | 11.3303 | 4.96E-05 | 0.615 | GTGGGTGGC |
| 3797 | mamo-s | *BGIBMGA004577* | up | 1541 | 1549 | - | 11.3303 | 4.96E-05 | 0.615 | GTGGGTGGC |
| 3798 | mamo-s | *BGIBMGA004578* | up | 1335 | 1343 | + | 11.3303 | 4.96E-05 | 0.615 | GTGGGTGGC |
| 3799 | mamo-s | *BGIBMGA004579* | up | 36 | 44 | - | 11.3303 | 4.96E-05 | 0.615 | GTGGGTGGC |
| 3800 | mamo-s | *BGIBMGA004580* | up | 171 | 179 | - | 11.3303 | 4.96E-05 | 0.615 | GTGGGTGGC |
| 3801 | mamo-s | *BGIBMGA004580* | down | 1283 | 1291 | - | 11.3303 | 4.96E-05 | 0.615 | GTGGGTGGC |
| 3802 | mamo-s | *BGIBMGA004582* | up | 9 | 17 | + | 11.3303 | 4.96E-05 | 0.615 | GTGGGTGGC |
| 3803 | mamo-s | *BGIBMGA004582* | up | 1833 | 1841 | + | 10.6881 | 8.61E-05 | 0.81 | GCACGTGGC |
| 3804 | mamo-s | *BGIBMGA004584* | up | 1853 | 1861 | - | 11.3853 | 4.74E-05 | 0.615 | ACGCGTGGC |
| 3805 | mamo-s | *BGIBMGA004584* | up | 882 | 890 | + | 11.3303 | 4.96E-05 | 0.615 | GTGGGTGGC |
| 3806 | mamo-s | *BGIBMGA004584* | down | 1200 | 1208 | - | 11.3303 | 4.96E-05 | 0.615 | GTGGGTGGC |
| 3807 | mamo-s | *BGIBMGA004584* | up | 1214 | 1222 | + | 10.5688 | 8.88E-05 | 0.824 | GTGCGTGAC |
| 3808 | mamo-s | *BGIBMGA004585* | down | 868 | 876 | - | 11.3303 | 4.96E-05 | 0.615 | GTGGGTGGC |
| 3809 | mamo-s | *BGIBMGA004586* | up | 665 | 673 | - | 10.5413 | 9.42E-05 | 0.857 | GTGCGAGTC |
| 3810 | mamo-s | *BGIBMGA004587* | down | 408 | 416 | - | 11.3303 | 4.96E-05 | 0.615 | GTGGGTGGC |
| 3811 | mamo-s | *BGIBMGA004589* | down | 372 | 380 | - | 11.3303 | 4.96E-05 | 0.615 | GTGGGTGGC |
| 3812 | mamo-s | *BGIBMGA004589* | up | 1475 | 1483 | + | 10.9358 | 7.30E-05 | 0.728 | GCGGGTGGC |
| 3813 | mamo-s | *BGIBMGA004590* | down | 1963 | 1971 | - | 11.3303 | 4.96E-05 | 0.615 | GTGGGTGGC |
| 3814 | mamo-s | *BGIBMGA004591* | up | 1152 | 1160 | - | 11.3303 | 4.96E-05 | 0.615 | GTGGGTGGC |
| 3815 | mamo-s | *BGIBMGA004594* | down | 614 | 622 | - | 11.3303 | 4.96E-05 | 0.615 | GTGGGTGGC |
| 3816 | mamo-s | *BGIBMGA004599* | up | 404 | 412 | - | 11.0826 | 6.84E-05 | 0.709 | GTACGTGGC |
| 3817 | mamo-s | *BGIBMGA004602* | up | 1589 | 1597 | + | 11.3303 | 4.96E-05 | 0.615 | GTGGGTGGC |
| 3818 | mamo-s | *BGIBMGA004603* | down | 1338 | 1346 | + | 11.3303 | 4.96E-05 | 0.615 | GTGGGTGGC |
| 3819 | mamo-s | *BGIBMGA004604* | up | 352 | 360 | + | 11.3303 | 4.96E-05 | 0.615 | GTGGGTGGC |
| 3820 | mamo-s | *BGIBMGA004605* | up | 1788 | 1796 | + | 12.1835 | 1.71E-05 | 0.615 | GTGCGTGCC |
| 3821 | mamo-s | *BGIBMGA004606* | down | 530 | 538 | - | 11.3303 | 4.96E-05 | 0.615 | GTGGGTGGC |
| 3822 | mamo-s | *BGIBMGA004606* | down | 1888 | 1896 | - | 11.3303 | 4.96E-05 | 0.615 | GTGGGTGGC |
| 3823 | mamo-s | *BGIBMGA004607* | up | 660 | 668 | - | 11.3303 | 4.96E-05 | 0.615 | GTGGGTGGC |
| 3824 | mamo-s | *BGIBMGA004609* | down | 1676 | 1684 | + | 11.0734 | 7.12E-05 | 0.726 | GAGCGTGGT |
| 3825 | mamo-s | *BGIBMGA004610* | down | 513 | 521 | + | 11.0734 | 7.12E-05 | 0.726 | GAGCGTGGT |
| 3826 | mamo-s | *BGIBMGA004611* | down | 1015 | 1023 | - | 11.3303 | 4.96E-05 | 0.615 | GTGGGTGGC |
| 3827 | mamo-s | *BGIBMGA004613* | up | 971 | 979 | + | 11.3303 | 4.96E-05 | 0.615 | GTGGGTGGC |
| 3828 | mamo-s | *BGIBMGA004613* | down | 1966 | 1974 | + | 11.3303 | 4.96E-05 | 0.615 | GTGGGTGGC |
| 3829 | mamo-s | *BGIBMGA004615* | down | 561 | 569 | + | 11.0826 | 6.84E-05 | 0.709 | GTACGTGGC |
| 3830 | mamo-s | *BGIBMGA004616* | up | 470 | 478 | - | 10.5138 | 9.75E-05 | 0.865 | GTGCGTGTT |
| 3831 | mamo-s | *BGIBMGA004617* | up | 1787 | 1795 | - | 12.1284 | 2.16E-05 | 0.615 | CAGCGTGGC |
| 3832 | mamo-s | *BGIBMGA004618* | up | 999 | 1007 | + | 11.3303 | 4.96E-05 | 0.615 | GTGGGTGGC |
| 3833 | mamo-s | *BGIBMGA004618* | down | 1828 | 1836 | - | 11.0734 | 7.12E-05 | 0.726 | GAGCGTGGT |
| 3834 | mamo-s | *BGIBMGA004619* | down | 1948 | 1956 | - | 11.3303 | 4.96E-05 | 0.615 | GTGGGTGGC |
| 3835 | mamo-s | *BGIBMGA004620* | up | 27 | 35 | - | 11.3303 | 4.96E-05 | 0.615 | GTGGGTGGC |
| 3836 | mamo-s | *BGIBMGA004620* | down | 1898 | 1906 | + | 11.3303 | 4.96E-05 | 0.615 | GTGGGTGGC |
| 3837 | mamo-s | *BGIBMGA004620* | down | 485 | 493 | + | 10.5138 | 9.75E-05 | 0.865 | GTGCGTGTT |
| 3838 | mamo-s | *BGIBMGA004621* | down | 1358 | 1366 | + | 11.3303 | 4.96E-05 | 0.615 | GTGGGTGGC |
| 3839 | mamo-s | *BGIBMGA004622* | down | 1525 | 1533 | - | 13.6606 | 4.06E-06 | 0.615 | GCGCGTGGC |
| 3840 | mamo-s | *BGIBMGA004622* | up | 1404 | 1412 | - | 11.3303 | 4.96E-05 | 0.615 | GTGGGTGGC |
| 3841 | mamo-s | *BGIBMGA004623* | up | 54 | 62 | - | 11.3303 | 4.96E-05 | 0.615 | GTGGGTGGC |
| 3842 | mamo-s | *BGIBMGA004623* | down | 1049 | 1057 | + | 11.3303 | 4.96E-05 | 0.615 | GTGGGTGGC |
| 3843 | mamo-s | *BGIBMGA004624* | down | 1946 | 1954 | - | 11.0826 | 6.84E-05 | 0.709 | GTACGTGGC |
| 3844 | mamo-s | *BGIBMGA004627* | up | 847 | 855 | - | 12.1009 | 2.43E-05 | 0.615 | GTGCGTGGT |
| 3845 | mamo-s | *BGIBMGA004627* | up | 1911 | 1919 | - | 10.5138 | 9.75E-05 | 0.865 | GTGCGTGTT |
| 3846 | mamo-s | *BGIBMGA004628* | down | 435 | 443 | + | 11.3303 | 4.96E-05 | 0.615 | GTGGGTGGC |
| 3847 | mamo-s | *BGIBMGA004628* | up | 221 | 229 | + | 11.2844 | 5.18E-05 | 0.63 | CTGCGTGCC |
| 3848 | mamo-s | *BGIBMGA004629* | down | 47 | 55 | - | 11.0826 | 6.84E-05 | 0.709 | GTACGTGGC |
| 3849 | mamo-s | *BGIBMGA004629* | down | 971 | 979 | - | 10.7523 | 8.38E-05 | 0.797 | GGGCGTGTC |
| 3850 | mamo-s | *BGIBMGA004631* | down | 263 | 271 | + | 11.3303 | 4.96E-05 | 0.615 | GTGGGTGGC |
| 3851 | mamo-s | *BGIBMGA004631* | up | 1928 | 1936 | - | 11.2844 | 5.18E-05 | 0.63 | CTGCGTGCC |
| 3852 | mamo-s | *BGIBMGA004631* | down | 1455 | 1463 | - | 11.0826 | 6.84E-05 | 0.709 | GTACGTGGC |
| 3853 | mamo-s | *BGIBMGA004633* | down | 1947 | 1955 | - | 11.5321 | 4.06E-05 | 0.615 | TTGCGTGGC |
| 3854 | mamo-s | *BGIBMGA004634* | down | 422 | 430 | - | 11.7339 | 3.29E-05 | 0.615 | GCGCGAGGC |
| 3855 | mamo-s | *BGIBMGA004636* | up | 620 | 628 | + | 12.1284 | 2.16E-05 | 0.615 | GTGCGAGGC |
| 3856 | mamo-s | *BGIBMGA004636* | down | 497 | 505 | - | 10.5138 | 9.75E-05 | 0.865 | GTGCGTGTT |
| 3857 | mamo-s | *BGIBMGA004637* | up | 910 | 918 | - | 12.1284 | 2.16E-05 | 0.615 | GTGCGAGGC |
| 3858 | mamo-s | *BGIBMGA004639* | down | 992 | 1000 | - | 10.9358 | 7.30E-05 | 0.728 | GCGGGTGGC |
| 3859 | mamo-s | *BGIBMGA004641* | up | 728 | 736 | - | 11.3303 | 4.96E-05 | 0.615 | GTGGGTGGC |
| 3860 | mamo-s | *BGIBMGA004641* | down | 202 | 210 | - | 11.2844 | 5.18E-05 | 0.63 | CTGCGTGCC |
| 3861 | mamo-s | *BGIBMGA004641* | up | 238 | 246 | - | 11.1009 | 6.57E-05 | 0.709 | GAGCGAGGC |
| 3862 | mamo-s | *BGIBMGA004641* | down | 673 | 681 | - | 11.1009 | 6.57E-05 | 0.709 | GAGCGAGGC |
| 3863 | mamo-s | *BGIBMGA004642* | down | 1821 | 1829 | - | 11.0826 | 6.84E-05 | 0.709 | GTACGTGGC |
| 3864 | mamo-s | *BGIBMGA004644* | down | 691 | 699 | + | 11.3303 | 4.96E-05 | 0.615 | GTGGGTGGC |
| 3865 | mamo-s | *BGIBMGA004645* | down | 1939 | 1947 | - | 11.2294 | 5.41E-05 | 0.65 | CTGCGAGGC |

| | | | | | | | | | | |
|---|---|---|---|---|---|---|---|---|---|---|
| 3866 | mamo-s | *BGIBMGA004648* | down | 776 | 784 | - | 11.3303 | 4.96E-05 | 0.615 | GTGGGTGGC |
| 3867 | mamo-s | *BGIBMGA004650* | up | 850 | 858 | - | 13.0275 | 8.53E-06 | 0.615 | GAGCGTGGC |
| 3868 | mamo-s | *BGIBMGA004651* | up | 1022 | 1030 | - | 12.3394 | 1.49E-05 | 0.615 | GGGCGTGGC |
| 3869 | mamo-s | *BGIBMGA004651* | up | 892 | 900 | + | 11.5688 | 3.79E-05 | 0.615 | CTGCGTGTC |
| 3870 | mamo-s | *BGIBMGA004652* | up | 1417 | 1425 | - | 13.6606 | 4.06E-06 | 0.615 | GCGCGTGGC |
| 3871 | mamo-s | *BGIBMGA004652* | down | 1518 | 1526 | - | 10.5138 | 9.75E-05 | 0.865 | GTGCGTGTT |
| 3872 | mamo-s | *BGIBMGA004653* | up | 705 | 713 | - | 11.7798 | 3.11E-05 | 0.615 | ATGCGTGGC |
| 3873 | mamo-s | *BGIBMGA004656* | up | 463 | 471 | - | 11.3303 | 4.96E-05 | 0.615 | GTGGGTGGC |
| 3874 | mamo-s | *BGIBMGA004656* | down | 1746 | 1754 | - | 10.9358 | 7.30E-05 | 0.728 | GCGGGTGGC |
| 3875 | mamo-s | *BGIBMGA004657* | down | 559 | 567 | - | 13.0275 | 8.53E-06 | 0.615 | GAGCGTGGC |
| 3876 | mamo-s | *BGIBMGA004657* | down | 465 | 473 | + | 12.1835 | 1.71E-05 | 0.615 | GTGCGTGCC |
| 3877 | mamo-s | *BGIBMGA004657* | up | 293 | 301 | + | 11.3303 | 4.96E-05 | 0.615 | GTGGGTGGC |
| 3878 | mamo-s | *BGIBMGA004660* | down | 798 | 806 | - | 12.1284 | 2.16E-05 | 0.615 | CAGCGTGGC |
| 3879 | mamo-s | *BGIBMGA004660* | up | 353 | 361 | - | 11.3303 | 4.96E-05 | 0.615 | GTGGGTGGC |
| 3880 | mamo-s | *BGIBMGA004661* | up | 1652 | 1660 | - | 12.1284 | 2.16E-05 | 0.615 | CAGCGTGGC |
| 3881 | mamo-s | *BGIBMGA004662* | up | 900 | 908 | - | 12.1284 | 2.16E-05 | 0.615 | CAGCGTGGC |
| 3882 | mamo-s | *BGIBMGA004664* | down | 921 | 929 | + | 11.3303 | 4.96E-05 | 0.615 | GTGGGTGGC |
| 3883 | mamo-s | *BGIBMGA004664* | up | 1026 | 1034 | - | 11.3303 | 4.96E-05 | 0.615 | GTGGGTGGC |
| 3884 | mamo-s | *BGIBMGA004667* | up | 571 | 579 | + | 11.0826 | 6.84E-05 | 0.709 | GTACGTGGC |
| 3885 | mamo-s | *BGIBMGA004671* | up | 1402 | 1410 | - | 12.1009 | 2.43E-05 | 0.615 | GTGCGTGGT |
| 3886 | mamo-s | *BGIBMGA004672* | down | 302 | 310 | - | 12.7615 | 1.04E-05 | 0.615 | CCGCGTGGC |
| 3887 | mamo-s | *BGIBMGA004672* | down | 722 | 730 | + | 12.0734 | 2.66E-05 | 0.615 | GCGCGTGTC |
| 3888 | mamo-s | *BGIBMGA004672* | up | 1401 | 1409 | - | 11.3303 | 4.96E-05 | 0.615 | GTGGGTGGC |
| 3889 | mamo-s | *BGIBMGA004672* | down | 1639 | 1647 | - | 11.3303 | 4.96E-05 | 0.615 | GTGGGTGGC |
| 3890 | mamo-s | *BGIBMGA004674* | down | 682 | 690 | - | 12.1009 | 2.43E-05 | 0.615 | GTGCGTGGT |
| 3891 | mamo-s | *BGIBMGA004675* | down | 1508 | 1516 | - | 11.3303 | 4.96E-05 | 0.615 | GTGGGTGGC |
| 3892 | mamo-s | *BGIBMGA004677* | down | 1050 | 1058 | + | 11.3303 | 4.96E-05 | 0.615 | GTGGGTGGC |
| 3893 | mamo-s | *BGIBMGA004677* | down | 1767 | 1775 | + | 11.3303 | 4.96E-05 | 0.615 | GTGGGTGGC |
| 3894 | mamo-s | *BGIBMGA004677* | down | 1967 | 1975 | - | 11.3303 | 4.96E-05 | 0.615 | GTGGGTGGC |
| 3895 | mamo-s | *BGIBMGA004677* | up | 1568 | 1576 | - | 11.0826 | 6.84E-05 | 0.709 | GTACGTGGC |
| 3896 | mamo-s | *BGIBMGA004679* | down | 808 | 816 | - | 11.3303 | 4.96E-05 | 0.615 | GTGGGTGGC |
| 3897 | mamo-s | *BGIBMGA004680* | up | 784 | 792 | - | 12.1009 | 2.43E-05 | 0.615 | GTGCGTGGT |
| 3898 | mamo-s | *BGIBMGA004680* | up | 682 | 690 | - | 10.7523 | 8.38E-05 | 0.797 | AAGCGTGGC |
| 3899 | mamo-s | *BGIBMGA004681* | down | 1630 | 1638 | - | 11.3303 | 4.96E-05 | 0.615 | GTGGGTGGC |
| 3900 | mamo-s | *BGIBMGA004685* | down | 135 | 143 | - | 11.3303 | 4.96E-05 | 0.615 | GTGGGTGGC |
| 3901 | mamo-s | *BGIBMGA004688* | up | 32 | 40 | - | 11.3303 | 4.96E-05 | 0.615 | GTGGGTGGC |
| 3902 | mamo-s | *BGIBMGA004689* | up | 1260 | 1268 | + | 11.3303 | 4.96E-05 | 0.615 | GTGGGTGGC |
| 3903 | mamo-s | *BGIBMGA004689* | down | 1417 | 1425 | + | 11.3303 | 4.96E-05 | 0.615 | GTGGGTGGC |
| 3904 | mamo-s | *BGIBMGA004689* | up | 969 | 977 | + | 11.0826 | 6.84E-05 | 0.709 | GTACGTGGC |
| 3905 | mamo-s | *BGIBMGA004692* | up | 397 | 405 | + | 12.3394 | 1.49E-05 | 0.615 | GGGCGTGGC |
| 3906 | mamo-s | *BGIBMGA004692* | down | 1034 | 1042 | - | 11.3303 | 4.96E-05 | 0.615 | GTGGGTGGC |
| 3907 | mamo-s | *BGIBMGA004693* | up | 1799 | 1807 | - | 13.6606 | 4.06E-06 | 0.615 | GCGCGTGGC |
| 3908 | mamo-s | *BGIBMGA004693* | down | 1983 | 1991 | - | 11.7064 | 3.52E-05 | 0.615 | GCGCGTGGT |
| 3909 | mamo-s | *BGIBMGA004693* | up | 1417 | 1425 | + | 10.7523 | 8.38E-05 | 0.797 | GGGCGTGTC |
| 3910 | mamo-s | *BGIBMGA004694* | down | 1733 | 1741 | + | 11.3303 | 4.96E-05 | 0.615 | GTGGGTGGC |
| 3911 | mamo-s | *BGIBMGA004696* | up | 682 | 690 | - | 11.3303 | 4.96E-05 | 0.615 | GTGGGTGGC |
| 3912 | mamo-s | *BGIBMGA004696* | up | 1343 | 1351 | - | 11.3303 | 4.96E-05 | 0.615 | GTGGGTGGC |
| 3913 | mamo-s | *BGIBMGA004697* | up | 1044 | 1052 | - | 11.7339 | 3.29E-05 | 0.615 | GCGCGAGGC |
| 3914 | mamo-s | *BGIBMGA004701* | up | 1481 | 1489 | - | 11.3303 | 4.96E-05 | 0.615 | GTGGGTGGC |
| 3915 | mamo-s | *BGIBMGA004703* | down | 400 | 408 | + | 11.1376 | 6.35E-05 | 0.709 | TCGCGTGGC |
| 3916 | mamo-s | *BGIBMGA004706* | down | 958 | 966 | + | 11.3303 | 4.96E-05 | 0.615 | GTGGGTGGC |
| 3917 | mamo-s | *BGIBMGA004707* | up | 492 | 500 | - | 12.3394 | 1.49E-05 | 0.615 | GGGCGTGGC |
| 3918 | mamo-s | *BGIBMGA004708* | up | 892 | 900 | - | 11.3303 | 4.96E-05 | 0.615 | GTGGGTGGC |
| 3919 | mamo-s | *BGIBMGA004710* | up | 127 | 135 | + | 11.5688 | 3.79E-05 | 0.615 | CTGCGTGTC |
| 3920 | mamo-s | *BGIBMGA004711* | up | 142 | 150 | - | 11.7798 | 3.11E-05 | 0.615 | ATGCGTGGC |
| 3921 | mamo-s | *BGIBMGA004713* | up | 1258 | 1266 | + | 11.3303 | 4.96E-05 | 0.615 | GTGGGTGGC |
| 3922 | mamo-s | *BGIBMGA004714* | down | 941 | 949 | + | 11.3303 | 4.96E-05 | 0.615 | GTGGGTGGC |
| 3923 | mamo-s | *BGIBMGA004714* | down | 838 | 846 | + | 10.5413 | 9.42E-05 | 0.857 | GTGCGAGTC |
| 3924 | mamo-s | *BGIBMGA004715* | down | 1024 | 1032 | - | 11.3303 | 4.96E-05 | 0.615 | GTGGGTGGC |
| 3925 | mamo-s | *BGIBMGA004715* | up | 1189 | 1197 | + | 11.3303 | 4.96E-05 | 0.615 | GTGGGTGGC |
| 3926 | mamo-s | *BGIBMGA004715* | up | 1086 | 1094 | + | 10.5413 | 9.42E-05 | 0.857 | GTGCGAGTC |
| 3927 | mamo-s | *BGIBMGA004718* | up | 36 | 44 | + | 11.3303 | 4.96E-05 | 0.615 | GTGGGTGGC |
| 3928 | mamo-s | *BGIBMGA004720* | down | 851 | 859 | - | 11.1743 | 5.90E-05 | 0.684 | CCGCGTGTC |
| 3929 | mamo-s | *BGIBMGA004721* | up | 1245 | 1253 | - | 12.1835 | 1.71E-05 | 0.615 | GTGCGTGCC |
| 3930 | mamo-s | *BGIBMGA004722* | up | 1191 | 1199 | - | 11.0826 | 6.84E-05 | 0.709 | GTACGTGGC |
| 3931 | mamo-s | *BGIBMGA004724* | down | 304 | 312 | - | 12.3394 | 1.49E-05 | 0.615 | GGGCGTGGC |
| 3932 | mamo-s | *BGIBMGA004725* | up | 1509 | 1517 | - | 11.3303 | 4.96E-05 | 0.615 | GTGGGTGGC |
| 3933 | mamo-s | *BGIBMGA004725* | up | 905 | 913 | + | 11.0826 | 6.84E-05 | 0.709 | GTACGTGGC |
| 3934 | mamo-s | *BGIBMGA004726* | up | 988 | 996 | - | 11.3303 | 4.96E-05 | 0.615 | GTGGGTGGC |
| 3935 | mamo-s | *BGIBMGA004730* | up | 1240 | 1248 | + | 11.3303 | 4.96E-05 | 0.615 | GTGGGTGGC |
| 3936 | mamo-s | *BGIBMGA004732* | down | 157 | 165 | + | 12.4679 | 1.31E-05 | 0.615 | GTGCGTGTC |
| 3937 | mamo-s | *BGIBMGA004732* | up | 1751 | 1759 | + | 11.3303 | 4.96E-05 | 0.615 | GTGGGTGGC |
| 3938 | mamo-s | *BGIBMGA004733* | down | 633 | 641 | - | 10.7523 | 8.38E-05 | 0.797 | AAGCGTGGC |
| 3939 | mamo-s | *BGIBMGA004733* | down | 1146 | 1154 | + | 10.7523 | 8.38E-05 | 0.797 | AAGCGTGGC |
| 3940 | mamo-s | *BGIBMGA004734* | up | 1006 | 1014 | - | 14.055 | 2.23E-06 | 0.615 | GTGCGTGGC |
| 3941 | mamo-s | *BGIBMGA004735* | down | 1283 | 1291 | + | 12.1009 | 2.43E-05 | 0.615 | GTGCGTGGT |
| 3942 | mamo-s | *BGIBMGA004735* | down | 230 | 238 | - | 11.3303 | 4.96E-05 | 0.615 | GTGGGTGGC |
| 3943 | mamo-s | *BGIBMGA004735* | up | 1495 | 1503 | + | 11.3303 | 4.96E-05 | 0.615 | GTGGGTGGC |
| 3944 | mamo-s | *BGIBMGA004737* | down | 1420 | 1428 | - | 11.3303 | 4.96E-05 | 0.615 | GTGGGTGGC |
| 3945 | mamo-s | *BGIBMGA004737* | up | 821 | 829 | + | 10.9358 | 7.30E-05 | 0.728 | GCGGGTGGC |
| 3946 | mamo-s | *BGIBMGA004739* | down | 1707 | 1715 | - | 11.3303 | 4.96E-05 | 0.615 | GTGGGTGGC |
| 3947 | mamo-s | *BGIBMGA004740* | up | 1987 | 1995 | + | 11.7064 | 3.52E-05 | 0.615 | GCGCGTGGT |
| 3948 | mamo-s | *BGIBMGA004740* | up | 336 | 344 | - | 11.3303 | 4.96E-05 | 0.615 | GTGGGTGGC |
| 3949 | mamo-s | *BGIBMGA004741* | down | 1309 | 1317 | + | 11.1743 | 5.90E-05 | 0.684 | CCGCGTGTC |
| 3950 | mamo-s | *BGIBMGA004741* | up | 1269 | 1277 | - | 10.5688 | 8.88E-05 | 0.824 | GTGCGTGAC |
| 3951 | mamo-s | *BGIBMGA004744* | up | 1297 | 1305 | - | 11.3303 | 4.96E-05 | 0.615 | GTGGGTGGC |
| 3952 | mamo-s | *BGIBMGA004747* | down | 1650 | 1658 | + | 11.3303 | 4.96E-05 | 0.615 | GTGGGTGGC |
| 3953 | mamo-s | *BGIBMGA004747* | up | 1542 | 1550 | - | 11.0826 | 6.84E-05 | 0.709 | GTACGTGGC |

| | | | | | | | | | | |
|---|---|---|---|---|---|---|---|---|---|---|
| 3954 | mamo-s | *BGIBMGA004748* | down | 947 | 955 | - | 12.1284 | 2.16E-05 | 0.615 | CAGCGTGGC |
| 3955 | mamo-s | *BGIBMGA004748* | up | 1514 | 1522 | + | 11.3303 | 4.96E-05 | 0.615 | GTGGGTGGC |
| 3956 | mamo-s | *BGIBMGA004749* | up | 162 | 170 | - | 11.3303 | 4.96E-05 | 0.615 | GTGGGTGGC |
| 3957 | mamo-s | *BGIBMGA004749* | down | 1251 | 1259 | + | 10.8899 | 7.48E-05 | 0.739 | CCGCGTGCC |
| 3958 | mamo-s | *BGIBMGA004752* | down | 1502 | 1510 | + | 11.3303 | 4.96E-05 | 0.615 | GTGGGTGGC |
| 3959 | mamo-s | *BGIBMGA004755* | up | 1505 | 1513 | - | 11.0826 | 6.84E-05 | 0.709 | GTACGTGGC |
| 3960 | mamo-s | *BGIBMGA004755* | down | 21 | 29 | - | 10.9358 | 7.30E-05 | 0.728 | GCGGGTGGC |
| 3961 | mamo-s | *BGIBMGA004758* | down | 561 | 569 | + | 11.0826 | 6.84E-05 | 0.709 | GTACGTGGC |
| 3962 | mamo-s | *BGIBMGA004759* | down | 1263 | 1271 | - | 10.5413 | 9.42E-05 | 0.857 | CAGCGTGTC |
| 3963 | mamo-s | *BGIBMGA004760* | down | 1508 | 1516 | - | 10.7523 | 8.38E-05 | 0.797 | GGGCGTGTC |
| 3964 | mamo-s | *BGIBMGA004762* | down | 727 | 735 | + | 11.3303 | 4.96E-05 | 0.615 | GTGGGTGGC |
| 3965 | mamo-s | *BGIBMGA004763* | down | 572 | 580 | + | 11.3303 | 4.96E-05 | 0.615 | GTGGGTGGC |
| 3966 | mamo-s | *BGIBMGA004763* | down | 253 | 261 | - | 10.8073 | 7.89E-05 | 0.766 | CCGCGTGGT |
| 3967 | mamo-s | *BGIBMGA004765* | down | 757 | 765 | - | 11.1743 | 5.90E-05 | 0.684 | CCGCGTGTC |
| 3968 | mamo-s | *BGIBMGA004765* | up | 379 | 387 | + | 11.0734 | 7.12E-05 | 0.726 | GAGCGTGGT |
| 3969 | mamo-s | *BGIBMGA004766* | up | 338 | 346 | - | 11.3303 | 4.96E-05 | 0.615 | GTGGGTGGC |
| 3970 | mamo-s | *BGIBMGA004766* | down | 810 | 818 | + | 11.3303 | 4.96E-05 | 0.615 | GTGGGTGGC |
| 3971 | mamo-s | *BGIBMGA004768* | up | 762 | 770 | - | 11.3303 | 4.96E-05 | 0.615 | GTGGGTGGC |
| 3972 | mamo-s | *BGIBMGA004768* | up | 1778 | 1786 | + | 11.0826 | 6.84E-05 | 0.709 | GTACGTGGC |
| 3973 | mamo-s | *BGIBMGA004768* | down | 1931 | 1939 | - | 11.0826 | 6.84E-05 | 0.709 | GTACGTGGC |
| 3974 | mamo-s | *BGIBMGA004771* | up | 1450 | 1458 | - | 11.3303 | 4.96E-05 | 0.615 | GTGGGTGGC |
| 3975 | mamo-s | *BGIBMGA004772* | up | 97 | 105 | - | 11.3303 | 4.96E-05 | 0.615 | GTGGGTGGC |
| 3976 | mamo-s | *BGIBMGA004773* | down | 1522 | 1530 | - | 11.3303 | 4.96E-05 | 0.615 | GTGGGTGGC |
| 3977 | mamo-s | *BGIBMGA004774* | down | 1717 | 1725 | - | 11.3303 | 4.96E-05 | 0.615 | GTGGGTGGC |
| 3978 | mamo-s | *BGIBMGA004776* | down | 1109 | 1117 | + | 11.2844 | 5.18E-05 | 0.63 | CTGCGTGCC |
| 3979 | mamo-s | *BGIBMGA004777* | up | 690 | 698 | + | 11.5688 | 3.79E-05 | 0.615 | CTGCGTGTC |
| 3980 | mamo-s | *BGIBMGA004779* | down | 693 | 701 | - | 11.3303 | 4.96E-05 | 0.615 | GTGGGTGGC |
| 3981 | mamo-s | *BGIBMGA004780* | up | 373 | 381 | - | 10.7523 | 8.38E-05 | 0.797 | AAGCGTGGC |
| 3982 | mamo-s | *BGIBMGA004781* | up | 1944 | 1952 | + | 10.5688 | 8.88E-05 | 0.824 | GTGCGTGAC |
| 3983 | mamo-s | *BGIBMGA004783* | up | 905 | 913 | + | 11.3303 | 4.96E-05 | 0.615 | GTGGGTGGC |
| 3984 | mamo-s | *BGIBMGA004784* | down | 1231 | 1239 | + | 13.0275 | 8.53E-06 | 0.615 | GAGCGTGGC |
| 3985 | mamo-s | *BGIBMGA004785* | down | 1611 | 1619 | + | 13.6606 | 4.06E-06 | 0.615 | GCGCGTGGC |
| 3986 | mamo-s | *BGIBMGA004786* | up | 472 | 480 | + | 12.7615 | 1.04E-05 | 0.615 | CCGCGTGGC |
| 3987 | mamo-s | *BGIBMGA004786* | up | 103 | 111 | - | 11.3303 | 4.96E-05 | 0.615 | GTGGGTGGC |
| 3988 | mamo-s | *BGIBMGA004786* | up | 369 | 377 | - | 10.6881 | 8.61E-05 | 0.81 | GCACGTGGC |
| 3989 | mamo-s | *BGIBMGA004786* | up | 278 | 286 | + | 10.5688 | 8.88E-05 | 0.824 | GTGCGTGAC |
| 3990 | mamo-s | *BGIBMGA004788* | down | 460 | 468 | - | 11.3303 | 4.96E-05 | 0.615 | GTGGGTGGC |
| 3991 | mamo-s | *BGIBMGA004792* | down | 1359 | 1367 | - | 11.3303 | 4.96E-05 | 0.615 | GTGGGTGGC |
| 3992 | mamo-s | *BGIBMGA004793* | up | 360 | 368 | - | 12.3394 | 1.49E-05 | 0.615 | GGGCGTGGC |
| 3993 | mamo-s | *BGIBMGA004793* | up | 1421 | 1429 | - | 10.7523 | 8.38E-05 | 0.797 | AAGCGTGGC |
| 3994 | mamo-s | *BGIBMGA004794* | down | 919 | 927 | + | 11.5688 | 3.79E-05 | 0.615 | CTGCGTGTC |
| 3995 | mamo-s | *BGIBMGA004794* | up | 748 | 756 | - | 11.3303 | 4.96E-05 | 0.615 | GTGGGTGGC |
| 3996 | mamo-s | *BGIBMGA004794* | down | 1758 | 1766 | + | 11.0826 | 6.84E-05 | 0.709 | GTACGTGGC |
| 3997 | mamo-s | *BGIBMGA004795* | down | 1351 | 1359 | + | 11.3303 | 4.96E-05 | 0.615 | GTGGGTGGC |
| 3998 | mamo-s | *BGIBMGA004795* | down | 1880 | 1888 | - | 11.0826 | 6.84E-05 | 0.709 | GTACGTGGC |
| 3999 | mamo-s | *BGIBMGA004796* | up | 550 | 558 | + | 11.3303 | 4.96E-05 | 0.615 | GTGGGTGGC |
| 4000 | mamo-s | *BGIBMGA004796* | down | 871 | 879 | - | 11.3303 | 4.96E-05 | 0.615 | GTGGGTGGC |
| 4001 | mamo-s | *BGIBMGA004796* | up | 1271 | 1279 | - | 11.3303 | 4.96E-05 | 0.615 | GTGGGTGGC |
| 4002 | mamo-s | *BGIBMGA004797* | up | 894 | 902 | + | 12.4679 | 1.31E-05 | 0.615 | GTGCGTGTC |
| 4003 | mamo-s | *BGIBMGA004798* | down | 1398 | 1406 | + | 10.8899 | 7.48E-05 | 0.739 | CCGCGTGCC |
| 4004 | mamo-s | *BGIBMGA004799* | up | 350 | 358 | - | 11.3303 | 4.96E-05 | 0.615 | GTGGGTGGC |
| 4005 | mamo-s | *BGIBMGA004799* | down | 765 | 773 | - | 11.3303 | 4.96E-05 | 0.615 | GTGGGTGGC |
| 4006 | mamo-s | *BGIBMGA004799* | down | 1863 | 1871 | + | 11.2844 | 5.18E-05 | 0.63 | CTGCGTGCC |
| 4007 | mamo-s | *BGIBMGA004802* | up | 203 | 211 | + | 11.3303 | 4.96E-05 | 0.615 | GTGGGTGGC |
| 4008 | mamo-s | *BGIBMGA004802* | up | 960 | 968 | - | 11.0826 | 6.84E-05 | 0.709 | GTACGTGGC |
| 4009 | mamo-s | *BGIBMGA004803* | down | 1830 | 1838 | - | 11.3303 | 4.96E-05 | 0.615 | GTGGGTGGC |
| 4010 | mamo-s | *BGIBMGA004804* | up | 387 | 395 | - | 12.1009 | 2.43E-05 | 0.615 | GTGCGTGGT |
| 4011 | mamo-s | *BGIBMGA004804* | up | 712 | 720 | + | 11.3303 | 4.96E-05 | 0.615 | GTGGGTGGC |
| 4012 | mamo-s | *BGIBMGA004806* | down | 1288 | 1296 | - | 11.7339 | 3.29E-05 | 0.615 | GCGCGAGGC |
| 4013 | mamo-s | *BGIBMGA004807* | up | 1173 | 1181 | - | 11.7339 | 3.29E-05 | 0.615 | GCGCGAGGC |
| 4014 | mamo-s | *BGIBMGA004809* | up | 226 | 234 | + | 11.3303 | 4.96E-05 | 0.615 | GTGGGTGGC |
| 4015 | mamo-s | *BGIBMGA004809* | down | 687 | 695 | + | 11.3303 | 4.96E-05 | 0.615 | GTGGGTGGC |
| 4016 | mamo-s | *BGIBMGA004809* | down | 1982 | 1990 | + | 11.0734 | 7.12E-05 | 0.726 | GAGCGTGGT |
| 4017 | mamo-s | *BGIBMGA004810* | down | 1544 | 1552 | + | 11.3303 | 4.96E-05 | 0.615 | GTGGGTGGC |
| 4018 | mamo-s | *BGIBMGA004810* | down | 273 | 281 | + | 11.0826 | 6.84E-05 | 0.709 | GTACGTGGC |
| 4019 | mamo-s | *BGIBMGA004810* | up | 638 | 646 | + | 11.0734 | 7.12E-05 | 0.726 | GAGCGTGGT |
| 4020 | mamo-s | *BGIBMGA004813* | down | 988 | 996 | - | 11.1743 | 5.90E-05 | 0.684 | CCGCGTGTC |
| 4021 | mamo-s | *BGIBMGA004814* | up | 1070 | 1078 | + | 11.0826 | 6.84E-05 | 0.709 | GTACGTGGC |
| 4022 | mamo-s | *BGIBMGA004816* | up | 348 | 356 | + | 11.3303 | 4.96E-05 | 0.615 | GTGGGTGGC |
| 4023 | mamo-s | *BGIBMGA004816* | up | 758 | 766 | - | 11.3303 | 4.96E-05 | 0.615 | GTGGGTGGC |
| 4024 | mamo-s | *BGIBMGA004817* | up | 841 | 849 | + | 14.055 | 2.23E-06 | 0.615 | GTGCGTGGC |
| 4025 | mamo-s | *BGIBMGA004817* | down | 1591 | 1599 | - | 11.7064 | 3.52E-05 | 0.615 | GCGCGTGGT |
| 4026 | mamo-s | *BGIBMGA004820* | up | 1193 | 1201 | - | 11.3303 | 4.96E-05 | 0.615 | GTGGGTGGC |
| 4027 | mamo-s | *BGIBMGA004821* | down | 504 | 512 | - | 11.3303 | 4.96E-05 | 0.615 | GTGGGTGGC |
| 4028 | mamo-s | *BGIBMGA004821* | up | 1876 | 1884 | + | 11.3303 | 4.96E-05 | 0.615 | GTGGGTGGC |
| 4029 | mamo-s | *BGIBMGA004827* | down | 1524 | 1532 | + | 13.0275 | 8.53E-06 | 0.615 | GAGCGTGGC |
| 4030 | mamo-s | *BGIBMGA004829* | up | 994 | 1002 | - | 11.3303 | 4.96E-05 | 0.615 | GTGGGTGGC |
| 4031 | mamo-s | *BGIBMGA004831* | down | 1170 | 1178 | + | 11.3303 | 4.96E-05 | 0.615 | GTGGGTGGC |
| 4032 | mamo-s | *BGIBMGA004832* | up | 1363 | 1371 | + | 11.3303 | 4.96E-05 | 0.615 | GTGGGTGGC |
| 4033 | mamo-s | *BGIBMGA004832* | down | 420 | 428 | - | 10.5138 | 9.75E-05 | 0.865 | GTGCGTGTT |
| 4034 | mamo-s | *BGIBMGA004834* | down | 450 | 458 | - | 11.7339 | 3.29E-05 | 0.615 | GCGCGAGGC |
| 4035 | mamo-s | *BGIBMGA004834* | up | 841 | 849 | - | 11.3303 | 4.96E-05 | 0.615 | GTGGGTGGC |
| 4036 | mamo-s | *BGIBMGA004836* | up | 1307 | 1315 | + | 11.3303 | 4.96E-05 | 0.615 | GTGGGTGGC |
| 4037 | mamo-s | *BGIBMGA004838* | down | 451 | 459 | - | 10.6881 | 8.61E-05 | 0.81 | GCACGTGGC |
| 4038 | mamo-s | *BGIBMGA004839* | down | 459 | 467 | + | 11.3303 | 4.96E-05 | 0.615 | GTGGGTGGC |
| 4039 | mamo-s | *BGIBMGA004841* | up | 791 | 799 | - | 11.3303 | 4.96E-05 | 0.615 | GTGGGTGGC |
| 4040 | mamo-s | *BGIBMGA004841* | up | 992 | 1000 | - | 11.3303 | 4.96E-05 | 0.615 | GTGGGTGGC |
| 4041 | mamo-s | *BGIBMGA004842* | down | 1250 | 1258 | + | 14.055 | 2.23E-06 | 0.615 | GTGCGTGGC |

| | | | | | | | | | | |
|---|---|---|---|---|---|---|---|---|---|---|
| 4042 | mamo-s | *BGIBMGA004842* | down | 1036 | 1044 | + | 11.0826 | 6.84E-05 | 0.709 | GTACGTGGC |
| 4043 | mamo-s | *BGIBMGA004843* | down | 1950 | 1958 | - | 11.3303 | 4.96E-05 | 0.615 | GTGGGTGGC |
| 4044 | mamo-s | *BGIBMGA004844* | down | 962 | 970 | + | 10.5138 | 9.75E-05 | 0.865 | GTGCGTGTT |
| 4045 | mamo-s | *BGIBMGA004846* | down | 1485 | 1493 | - | 11.3303 | 4.96E-05 | 0.615 | GTGGGTGGC |
| 4046 | mamo-s | *BGIBMGA004851* | down | 248 | 256 | + | 12.0734 | 2.66E-05 | 0.615 | GCGCGTGTC |
| 4047 | mamo-s | *BGIBMGA004851* | down | 1172 | 1180 | - | 11.7339 | 3.29E-05 | 0.615 | GCGCGAGGC |
| 4048 | mamo-s | *BGIBMGA004854* | down | 1707 | 1715 | + | 11.3303 | 4.96E-05 | 0.615 | GTGGGTGGC |
| 4049 | mamo-s | *BGIBMGA004855* | down | 1034 | 1042 | + | 11.3303 | 4.96E-05 | 0.615 | GTGGGTGGC |
| 4050 | mamo-s | *BGIBMGA004856* | down | 890 | 898 | + | 11.7798 | 3.11E-05 | 0.615 | ATGCGTGGC |
| 4051 | mamo-s | *BGIBMGA004856* | up | 872 | 880 | + | 11.3303 | 4.96E-05 | 0.615 | GTGGGTGGC |
| 4052 | mamo-s | *BGIBMGA004856* | up | 1831 | 1839 | - | 11.3303 | 4.96E-05 | 0.615 | GTGGGTGGC |
| 4053 | mamo-s | *BGIBMGA004858* | up | 223 | 231 | + | 11.3303 | 4.96E-05 | 0.615 | GTGGGTGGC |
| 4054 | mamo-s | *BGIBMGA004858* | down | 1102 | 1110 | - | 11.3303 | 4.96E-05 | 0.615 | GTGGGTGGC |
| 4055 | mamo-s | *BGIBMGA004859* | down | 884 | 892 | - | 11.0826 | 6.84E-05 | 0.709 | GTACGTGGC |
| 4056 | mamo-s | *BGIBMGA004860* | up | 240 | 248 | - | 11.3303 | 4.96E-05 | 0.615 | GTGGGTGGC |
| 4057 | mamo-s | *BGIBMGA004861* | down | 1065 | 1073 | + | 11.3303 | 4.96E-05 | 0.615 | GTGGGTGGC |
| 4058 | mamo-s | *BGIBMGA004861* | down | 1597 | 1605 | + | 11.3303 | 4.96E-05 | 0.615 | GTGGGTGGC |
| 4059 | mamo-s | *BGIBMGA004863* | up | 1583 | 1591 | + | 11.3303 | 4.96E-05 | 0.615 | GTGGGTGGC |
| 4060 | mamo-s | *BGIBMGA004864* | down | 410 | 418 | + | 10.5413 | 9.42E-05 | 0.857 | GTGCGAGTC |
| 4061 | mamo-s | *BGIBMGA004868* | down | 509 | 517 | + | 11.3303 | 4.96E-05 | 0.615 | GTGGGTGGC |
| 4062 | mamo-s | *BGIBMGA004869* | down | 292 | 300 | - | 11.7064 | 3.52E-05 | 0.615 | GCGCGTGGT |
| 4063 | mamo-s | *BGIBMGA004869* | down | 913 | 921 | + | 11.3303 | 4.96E-05 | 0.615 | GTGGGTGGC |
| 4064 | mamo-s | *BGIBMGA004871* | down | 1943 | 1951 | + | 11.3303 | 4.96E-05 | 0.615 | GTGGGTGGC |
| 4065 | mamo-s | *BGIBMGA004871* | up | 1987 | 1995 | - | 11.2018 | 5.68E-05 | 0.674 | CTGCGTGGT |
| 4066 | mamo-s | *BGIBMGA004872* | down | 393 | 401 | + | 11.7798 | 3.11E-05 | 0.615 | ATGCGTGGC |
| 4067 | mamo-s | *BGIBMGA004872* | up | 175 | 183 | + | 11.3303 | 4.96E-05 | 0.615 | GTGGGTGGC |
| 4068 | mamo-s | *BGIBMGA004872* | down | 1903 | 1911 | + | 11.0826 | 6.84E-05 | 0.709 | GTACGTGGC |
| 4069 | mamo-s | *BGIBMGA004874* | down | 129 | 137 | - | 11.3303 | 4.96E-05 | 0.615 | GTGGGTGGC |
| 4070 | mamo-s | *BGIBMGA004874* | down | 1963 | 1971 | - | 11.3303 | 4.96E-05 | 0.615 | GTGGGTGGC |
| 4071 | mamo-s | *BGIBMGA004874* | down | 497 | 505 | - | 11.0826 | 6.84E-05 | 0.709 | GTACGTGGC |
| 4072 | mamo-s | *BGIBMGA004875* | down | 1716 | 1724 | + | 11.3303 | 4.96E-05 | 0.615 | GTGGGTGGC |
| 4073 | mamo-s | *BGIBMGA004876* | up | 778 | 786 | + | 11.3303 | 4.96E-05 | 0.615 | GTGGGTGGC |
| 4074 | mamo-s | *BGIBMGA004877* | down | 1353 | 1361 | + | 11.5688 | 3.79E-05 | 0.615 | CTGCGTGTC |
| 4075 | mamo-s | *BGIBMGA004877* | down | 216 | 224 | + | 11.3303 | 4.96E-05 | 0.615 | GTGGGTGGC |
| 4076 | mamo-s | *BGIBMGA004878* | up | 202 | 210 | + | 11.3303 | 4.96E-05 | 0.615 | GTGGGTGGC |
| 4077 | mamo-s | *BGIBMGA004878* | up | 983 | 991 | + | 11.3303 | 4.96E-05 | 0.615 | GTGGGTGGC |
| 4078 | mamo-s | *BGIBMGA004878* | down | 1536 | 1544 | + | 11.1743 | 5.90E-05 | 0.684 | CCGCGTGTC |
| 4079 | mamo-s | *BGIBMGA004879* | up | 1530 | 1538 | + | 12.1835 | 1.71E-05 | 0.615 | GTGCGTGCC |
| 4080 | mamo-s | *BGIBMGA004880* | down | 828 | 836 | + | 11.3303 | 4.96E-05 | 0.615 | GTGGGTGGC |
| 4081 | mamo-s | *BGIBMGA004880* | up | 1853 | 1861 | + | 11.3303 | 4.96E-05 | 0.615 | GTGGGTGGC |
| 4082 | mamo-s | *BGIBMGA004883* | down | 31 | 39 | - | 12.1835 | 1.71E-05 | 0.615 | GTGCGTGCC |
| 4083 | mamo-s | *BGIBMGA004884* | up | 1243 | 1251 | - | 12.1835 | 1.71E-05 | 0.615 | GTGCGTGCC |
| 4084 | mamo-s | *BGIBMGA004887* | up | 1676 | 1684 | - | 11.3303 | 4.96E-05 | 0.615 | GTGGGTGGC |
| 4085 | mamo-s | *BGIBMGA004888* | up | 19 | 27 | + | 11.3303 | 4.96E-05 | 0.615 | GTGGGTGGC |
| 4086 | mamo-s | *BGIBMGA004895* | down | 1260 | 1268 | + | 11.3303 | 4.96E-05 | 0.615 | GTGGGTGGC |
| 4087 | mamo-s | *BGIBMGA004899* | down | 159 | 167 | - | 11.3303 | 4.96E-05 | 0.615 | GTGGGTGGC |
| 4088 | mamo-s | *BGIBMGA004899* | down | 1762 | 1770 | + | 11.3303 | 4.96E-05 | 0.615 | GTGGGTGGC |
| 4089 | mamo-s | *BGIBMGA004900* | down | 733 | 741 | + | 12.1284 | 2.16E-05 | 0.615 | CAGCGTGGC |
| 4090 | mamo-s | *BGIBMGA004901* | up | 639 | 647 | - | 11.0826 | 6.84E-05 | 0.709 | GTACGTGGC |
| 4091 | mamo-s | *BGIBMGA004903* | up | 465 | 473 | - | 11.3303 | 4.96E-05 | 0.615 | GTGGGTGGC |
| 4092 | mamo-s | *BGIBMGA004905* | up | 551 | 559 | - | 11.3303 | 4.96E-05 | 0.615 | GTGGGTGGC |
| 4093 | mamo-s | *BGIBMGA004907* | down | 691 | 699 | - | 11.3303 | 4.96E-05 | 0.615 | GTGGGTGGC |
| 4094 | mamo-s | *BGIBMGA004908* | down | 225 | 233 | + | 13.0275 | 8.53E-06 | 0.615 | GAGCGTGGC |
| 4095 | mamo-s | *BGIBMGA004909* | down | 706 | 714 | - | 10.7523 | 8.38E-05 | 0.797 | AAGCGTGGC |
| 4096 | mamo-s | *BGIBMGA004912* | down | 962 | 970 | - | 10.8349 | 7.66E-05 | 0.751 | CCGCGAGGC |
| 4097 | mamo-s | *BGIBMGA004913* | down | 661 | 669 | + | 11.3303 | 4.96E-05 | 0.615 | GTGGGTGGC |
| 4098 | mamo-s | *BGIBMGA004914* | down | 78 | 86 | + | 12.0734 | 2.66E-05 | 0.615 | GCGCGTGTC |
| 4099 | mamo-s | *BGIBMGA004914* | down | 537 | 545 | + | 12.0734 | 2.66E-05 | 0.615 | GCGCGTGTC |
| 4100 | mamo-s | *BGIBMGA004914* | down | 219 | 227 | + | 11.3303 | 4.96E-05 | 0.615 | GTGGGTGGC |
| 4101 | mamo-s | *BGIBMGA004915* | down | 78 | 86 | + | 12.0734 | 2.66E-05 | 0.615 | GCGCGTGTC |
| 4102 | mamo-s | *BGIBMGA004919* | down | 1621 | 1629 | + | 11.3303 | 4.96E-05 | 0.615 | GTGGGTGGC |
| 4103 | mamo-s | *BGIBMGA004920* | down | 1520 | 1528 | - | 11.3303 | 4.96E-05 | 0.615 | GTGGGTGGC |
| 4104 | mamo-s | *BGIBMGA004921* | down | 683 | 691 | + | 11.3303 | 4.96E-05 | 0.615 | GTGGGTGGC |
| 4105 | mamo-s | *BGIBMGA004922* | up | 507 | 515 | + | 10.8073 | 7.89E-05 | 0.766 | CCGCGTGGT |
| 4106 | mamo-s | *BGIBMGA004923* | down | 1412 | 1420 | - | 11.3303 | 4.96E-05 | 0.615 | GTGGGTGGC |
| 4107 | mamo-s | *BGIBMGA004925* | up | 1378 | 1386 | - | 13.6606 | 4.06E-06 | 0.615 | GCGCGTGGC |
| 4108 | mamo-s | *BGIBMGA004925* | up | 1148 | 1156 | + | 11.7798 | 3.11E-05 | 0.615 | ATGCGTGGC |
| 4109 | mamo-s | *BGIBMGA004926* | down | 877 | 885 | - | 11.3303 | 4.96E-05 | 0.615 | GTGGGTGGC |
| 4110 | mamo-s | *BGIBMGA004927* | down | 1774 | 1782 | + | 11.3303 | 4.96E-05 | 0.615 | GTGGGTGGC |
| 4111 | mamo-s | *BGIBMGA004928* | up | 880 | 888 | + | 11.3303 | 4.96E-05 | 0.615 | GTGGGTGGC |
| 4112 | mamo-s | *BGIBMGA004928* | down | 1790 | 1798 | - | 10.9358 | 7.30E-05 | 0.728 | GCGGGTGGC |
| 4113 | mamo-s | *BGIBMGA004929* | up | 1049 | 1057 | - | 11.7339 | 3.29E-05 | 0.615 | GCGCGAGGC |
| 4114 | mamo-s | *BGIBMGA004929* | down | 895 | 903 | + | 11.3303 | 4.96E-05 | 0.615 | GTGGGTGGC |
| 4115 | mamo-s | *BGIBMGA004929* | up | 319 | 327 | - | 10.9358 | 7.30E-05 | 0.728 | GCGGGTGGC |
| 4116 | mamo-s | *BGIBMGA004935* | down | 147 | 155 | - | 11.3303 | 4.96E-05 | 0.615 | GTGGGTGGC |
| 4117 | mamo-s | *BGIBMGA004940* | up | 1346 | 1354 | - | 11.3303 | 4.96E-05 | 0.615 | GTGGGTGGC |
| 4118 | mamo-s | *BGIBMGA004942* | up | 1289 | 1297 | - | 11.3303 | 4.96E-05 | 0.615 | GTGGGTGGC |
| 4119 | mamo-s | *BGIBMGA004943* | up | 1245 | 1253 | - | 10.5688 | 8.88E-05 | 0.824 | GTGCGTGAC |
| 4120 | mamo-s | *BGIBMGA004944* | down | 1131 | 1139 | + | 11.2018 | 5.68E-05 | 0.674 | CTGCGTGGT |
| 4121 | mamo-s | *BGIBMGA004945* | down | 1718 | 1726 | + | 14.055 | 2.23E-06 | 0.615 | GTGCGTGGC |
| 4122 | mamo-s | *BGIBMGA004946* | up | 947 | 955 | - | 11.2844 | 5.18E-05 | 0.63 | CTGCGTGCC |
| 4123 | mamo-s | *BGIBMGA004950* | down | 321 | 329 | + | 10.5138 | 9.75E-05 | 0.865 | GTGCGTGTT |
| 4124 | mamo-s | *BGIBMGA004953* | up | 1682 | 1690 | - | 11.3303 | 4.96E-05 | 0.615 | GTGGGTGGC |
| 4125 | mamo-s | *BGIBMGA004954* | up | 307 | 315 | + | 12.7615 | 1.04E-05 | 0.615 | CCGCGTGGC |
| 4126 | mamo-s | *BGIBMGA004954* | down | 938 | 946 | - | 11.3303 | 4.96E-05 | 0.615 | GTGGGTGGC |
| 4127 | mamo-s | *BGIBMGA004955* | down | 1472 | 1480 | + | 11.3303 | 4.96E-05 | 0.615 | GTGGGTGGC |
| 4128 | mamo-s | *BGIBMGA004956* | down | 426 | 434 | + | 12.4679 | 1.31E-05 | 0.615 | GTGCGTGTC |
| 4129 | mamo-s | *BGIBMGA004956* | down | 1135 | 1143 | + | 12.1835 | 1.71E-05 | 0.615 | GTGCGTGCC |

| | | | | | | | | | | |
|---|---|---|---|---|---|---|---|---|---|---|
| 4130 | mamo-s | *BGIBMGA004956* | down | 1623 | 1631 | - | 11.5321 | 4.06E-05 | 0.615 | TTGCGTGGC |
| 4131 | mamo-s | *BGIBMGA004957* | down | 417 | 425 | - | 12.4679 | 1.31E-05 | 0.615 | GTGCGTGTC |
| 4132 | mamo-s | *BGIBMGA004959* | up | 1577 | 1585 | + | 11.7339 | 3.29E-05 | 0.615 | GCGCGAGGC |
| 4133 | mamo-s | *BGIBMGA004961* | up | 537 | 545 | + | 12.1835 | 1.71E-05 | 0.615 | GTGCGTGCC |
| 4134 | mamo-s | *BGIBMGA004961* | down | 1356 | 1364 | - | 10.8073 | 7.89E-05 | 0.766 | CCGCGTGGT |
| 4135 | mamo-s | *BGIBMGA004962* | up | 1051 | 1059 | + | 11.3303 | 4.96E-05 | 0.615 | GTGGGTGGC |
| 4136 | mamo-s | *BGIBMGA004963* | down | 1376 | 1384 | + | 11.3303 | 4.96E-05 | 0.615 | GTGGGTGGC |
| 4137 | mamo-s | *BGIBMGA004965* | up | 1912 | 1920 | + | 11.2844 | 5.18E-05 | 0.63 | CTGCGTGCC |
| 4138 | mamo-s | *BGIBMGA004965* | down | 1239 | 1247 | + | 10.7523 | 8.38E-05 | 0.797 | GGGCGTGTC |
| 4139 | mamo-s | *BGIBMGA004967* | up | 1759 | 1767 | + | 11.3303 | 4.96E-05 | 0.615 | GTGGGTGGC |
| 4140 | mamo-s | *BGIBMGA004967* | up | 52 | 60 | + | 11.1743 | 5.90E-05 | 0.684 | CCGCGTGTC |
| 4141 | mamo-s | *BGIBMGA004968* | up | 1805 | 1813 | - | 11.3303 | 4.96E-05 | 0.615 | GTGGGTGGC |
| 4142 | mamo-s | *BGIBMGA004969* | up | 782 | 790 | - | 11.3303 | 4.96E-05 | 0.615 | GTGGGTGGC |
| 4143 | mamo-s | *BGIBMGA004970* | up | 646 | 654 | + | 11.3303 | 4.96E-05 | 0.615 | GTGGGTGGC |
| 4144 | mamo-s | *BGIBMGA004970* | down | 651 | 659 | - | 11.3303 | 4.96E-05 | 0.615 | GTGGGTGGC |
| 4145 | mamo-s | *BGIBMGA004972* | up | 1812 | 1820 | + | 12.7615 | 1.04E-05 | 0.615 | CCGCGTGGC |
| 4146 | mamo-s | *BGIBMGA004976* | up | 1311 | 1319 | + | 11.3303 | 4.96E-05 | 0.615 | GTGGGTGGC |
| 4147 | mamo-s | *BGIBMGA004979* | up | 634 | 642 | + | 11.4404 | 4.51E-05 | 0.615 | CGGCGTGGC |
| 4148 | mamo-s | *BGIBMGA004980* | down | 718 | 726 | - | 10.5688 | 8.88E-05 | 0.824 | GTGCGTGAC |
| 4149 | mamo-s | *BGIBMGA004981* | down | 1875 | 1883 | - | 11.7798 | 3.11E-05 | 0.615 | ATGCGTGGC |
| 4150 | mamo-s | *BGIBMGA004981* | down | 1759 | 1767 | + | 11.3303 | 4.96E-05 | 0.615 | GTGGGTGGC |
| 4151 | mamo-s | *BGIBMGA004981* | up | 759 | 767 | - | 10.5688 | 8.88E-05 | 0.824 | GTGCGTGAC |
| 4152 | mamo-s | *BGIBMGA004982* | up | 559 | 567 | + | 11.3303 | 4.96E-05 | 0.615 | GTGGGTGGC |
| 4153 | mamo-s | *BGIBMGA004982* | down | 1661 | 1669 | + | 11.3303 | 4.96E-05 | 0.615 | GTGGGTGGC |
| 4154 | mamo-s | *BGIBMGA004984* | down | 1292 | 1300 | - | 11.3303 | 4.96E-05 | 0.615 | GTGGGTGGC |
| 4155 | mamo-s | *BGIBMGA004984* | down | 1474 | 1482 | - | 11.3303 | 4.96E-05 | 0.615 | GTGGGTGGC |
| 4156 | mamo-s | *BGIBMGA004986* | down | 287 | 295 | + | 11.3303 | 4.96E-05 | 0.615 | GTGGGTGGC |
| 4157 | mamo-s | *BGIBMGA004987* | down | 618 | 626 | + | 11.3303 | 4.96E-05 | 0.615 | GTGGGTGGC |
| 4158 | mamo-s | *BGIBMGA004989* | up | 67 | 75 | + | 10.8899 | 7.48E-05 | 0.739 | CCGCGTGCC |
| 4159 | mamo-s | *BGIBMGA004992* | down | 1364 | 1372 | + | 10.7523 | 8.38E-05 | 0.797 | AAGCGTGGC |
| 4160 | mamo-s | *BGIBMGA004994* | up | 36 | 44 | - | 12.1009 | 2.43E-05 | 0.615 | GTGCGTGGT |
| 4161 | mamo-s | *BGIBMGA004996* | up | 704 | 712 | + | 11.3853 | 4.74E-05 | 0.615 | ACGCGTGGC |
| 4162 | mamo-s | *BGIBMGA004996* | down | 164 | 172 | - | 11.3303 | 4.96E-05 | 0.615 | GTGGGTGGC |
| 4163 | mamo-s | *BGIBMGA004999* | up | 993 | 1001 | - | 11.0826 | 6.84E-05 | 0.709 | GTACGTGGC |
| 4164 | mamo-s | *BGIBMGA005000* | up | 214 | 222 | - | 11.3303 | 4.96E-05 | 0.615 | GTGGGTGGC |
| 4165 | mamo-s | *BGIBMGA005000* | down | 1715 | 1723 | - | 11.0826 | 6.84E-05 | 0.709 | GTACGTGGC |
| 4166 | mamo-s | *BGIBMGA005005* | down | 1230 | 1238 | + | 11.3303 | 4.96E-05 | 0.615 | GTGGGTGGC |
| 4167 | mamo-s | *BGIBMGA005007* | up | 1422 | 1430 | + | 11.4404 | 4.51E-05 | 0.615 | CGGCGTGGC |
| 4168 | mamo-s | *BGIBMGA005007* | up | 960 | 968 | - | 11.3303 | 4.96E-05 | 0.615 | GTGGGTGGC |
| 4169 | mamo-s | *BGIBMGA005009* | down | 1168 | 1176 | + | 11.0826 | 6.84E-05 | 0.709 | GTACGTGGC |
| 4170 | mamo-s | *BGIBMGA005010* | up | 1802 | 1810 | - | 11.7064 | 3.52E-05 | 0.615 | GCGCGTGGT |
| 4171 | mamo-s | *BGIBMGA005014* | up | 1497 | 1505 | + | 11.0734 | 7.12E-05 | 0.726 | GAGCGTGGT |
| 4172 | mamo-s | *BGIBMGA005017* | up | 845 | 853 | - | 11.3303 | 4.96E-05 | 0.615 | GTGGGTGGC |
| 4173 | mamo-s | *BGIBMGA005018* | up | 1793 | 1801 | + | 11.3303 | 4.96E-05 | 0.615 | GTGGGTGGC |
| 4174 | mamo-s | *BGIBMGA005019* | down | 1545 | 1553 | - | 11.7339 | 3.29E-05 | 0.615 | GCGCGAGGC |
| 4175 | mamo-s | *BGIBMGA005019* | up | 1452 | 1460 | - | 11.3303 | 4.96E-05 | 0.615 | GTGGGTGGC |
| 4176 | mamo-s | *BGIBMGA005019* | down | 1522 | 1530 | + | 11.2018 | 5.68E-05 | 0.674 | CTGCGTGGT |
| 4177 | mamo-s | *BGIBMGA005020* | up | 1608 | 1616 | + | 11.789 | 2.84E-05 | 0.615 | GCGCGTGCC |
| 4178 | mamo-s | *BGIBMGA005020* | up | 1120 | 1128 | + | 10.7523 | 8.38E-05 | 0.797 | AAGCGTGGC |
| 4179 | mamo-s | *BGIBMGA005021* | up | 860 | 868 | - | 11.3303 | 4.96E-05 | 0.615 | GTGGGTGGC |
| 4180 | mamo-s | *BGIBMGA005021* | down | 674 | 682 | + | 11.0734 | 7.12E-05 | 0.726 | GAGCGTGGT |
| 4181 | mamo-s | *BGIBMGA005022* | up | 918 | 926 | + | 10.8073 | 7.89E-05 | 0.766 | CCGCGTGGT |
| 4182 | mamo-s | *BGIBMGA005024* | down | 139 | 147 | - | 11.789 | 2.84E-05 | 0.615 | GCGCGTGCC |
| 4183 | mamo-s | *BGIBMGA005025* | up | 234 | 242 | - | 11.789 | 2.84E-05 | 0.615 | GCGCGTGCC |
| 4184 | mamo-s | *BGIBMGA005030* | down | 1756 | 1764 | + | 14.055 | 2.23E-06 | 0.615 | GTGCGTGGC |
| 4185 | mamo-s | *BGIBMGA005030* | down | 142 | 150 | - | 11.3303 | 4.96E-05 | 0.615 | GTGGGTGGC |
| 4186 | mamo-s | *BGIBMGA005031* | up | 512 | 520 | - | 11.3303 | 4.96E-05 | 0.615 | GTGGGTGGC |
| 4187 | mamo-s | *BGIBMGA005032* | up | 700 | 708 | + | 11.3303 | 4.96E-05 | 0.615 | GTGGGTGGC |
| 4188 | mamo-s | *BGIBMGA005033* | up | 1443 | 1451 | - | 10.5138 | 9.75E-05 | 0.865 | GTGCGTGTT |
| 4189 | mamo-s | *BGIBMGA005034* | up | 41 | 49 | + | 11.3303 | 4.96E-05 | 0.615 | GTGGGTGGC |
| 4190 | mamo-s | *BGIBMGA005034* | down | 1424 | 1432 | - | 11.3303 | 4.96E-05 | 0.615 | GTGGGTGGC |
| 4191 | mamo-s | *BGIBMGA005035* | up | 1978 | 1986 | + | 11.4404 | 4.51E-05 | 0.615 | GAGCGTGTC |
| 4192 | mamo-s | *BGIBMGA005038* | down | 895 | 903 | + | 11.4404 | 4.51E-05 | 0.615 | GAGCGTGTC |
| 4193 | mamo-s | *BGIBMGA005039* | up | 881 | 889 | + | 11.4404 | 4.51E-05 | 0.615 | GAGCGTGTC |
| 4194 | mamo-s | *BGIBMGA005042* | up | 1670 | 1678 | + | 11.1376 | 6.35E-05 | 0.709 | TCGCGTGGC |
| 4195 | mamo-s | *BGIBMGA005044* | down | 666 | 674 | - | 11.3303 | 4.96E-05 | 0.615 | GTGGGTGGC |
| 4196 | mamo-s | *BGIBMGA005045* | down | 445 | 453 | - | 11.3303 | 4.96E-05 | 0.615 | GTGGGTGGC |
| 4197 | mamo-s | *BGIBMGA005047* | up | 1156 | 1164 | - | 11.3303 | 4.96E-05 | 0.615 | GTGGGTGGC |
| 4198 | mamo-s | *BGIBMGA005054* | up | 693 | 701 | - | 11.7798 | 3.11E-05 | 0.615 | ATGCGTGGC |
| 4199 | mamo-s | *BGIBMGA005055* | up | 1151 | 1159 | + | 11.0826 | 6.84E-05 | 0.709 | GTACGTGGC |
| 4200 | mamo-s | *BGIBMGA005056* | up | 813 | 821 | + | 11.3303 | 4.96E-05 | 0.615 | GTGGGTGGC |
| 4201 | mamo-s | *BGIBMGA005057* | down | 195 | 203 | + | 11.3303 | 4.96E-05 | 0.615 | GTGGGTGGC |
| 4202 | mamo-s | *BGIBMGA005058* | up | 842 | 850 | - | 13.156 | 6.30E-06 | 0.615 | CTGCGTGGC |
| 4203 | mamo-s | *BGIBMGA005058* | down | 1892 | 1900 | + | 11.3303 | 4.96E-05 | 0.615 | GTGGGTGGC |
| 4204 | mamo-s | *BGIBMGA005058* | down | 974 | 982 | - | 11.2844 | 5.18E-05 | 0.63 | CTGCGTGCC |
| 4205 | mamo-s | *BGIBMGA005060* | down | 36 | 44 | - | 10.5138 | 9.75E-05 | 0.865 | GTGCGTGTT |
| 4206 | mamo-s | *BGIBMGA005062* | up | 1067 | 1075 | + | 11.3303 | 4.96E-05 | 0.615 | GTGGGTGGC |
| 4207 | mamo-s | *BGIBMGA005062* | down | 1256 | 1264 | + | 11.3303 | 4.96E-05 | 0.615 | GTGGGTGGC |
| 4208 | mamo-s | *BGIBMGA005063* | up | 715 | 723 | + | 11.3303 | 4.96E-05 | 0.615 | GTGGGTGGC |
| 4209 | mamo-s | *BGIBMGA005063* | up | 147 | 155 | - | 11.0826 | 6.84E-05 | 0.709 | GTACGTGGC |
| 4210 | mamo-s | *BGIBMGA005064* | down | 1038 | 1046 | + | 10.9358 | 7.30E-05 | 0.728 | GCGGGTGGC |
| 4211 | mamo-s | *BGIBMGA005065* | up | 1230 | 1238 | - | 11.3303 | 4.96E-05 | 0.615 | GTGGGTGGC |
| 4212 | mamo-s | *BGIBMGA005070* | up | 1793 | 1801 | - | 11.3303 | 4.96E-05 | 0.615 | GTGGGTGGC |
| 4213 | mamo-s | *BGIBMGA005070* | down | 762 | 770 | - | 11.0826 | 6.84E-05 | 0.709 | GTACGTGGC |
| 4214 | mamo-s | *BGIBMGA005070* | up | 1404 | 1412 | + | 11.0826 | 6.84E-05 | 0.709 | GTACGTGGC |
| 4215 | mamo-s | *BGIBMGA005071* | up | 1010 | 1018 | + | 11.3303 | 4.96E-05 | 0.615 | GTGGGTGGC |
| 4216 | mamo-s | *BGIBMGA005072* | up | 1709 | 1717 | - | 12.4679 | 1.31E-05 | 0.615 | GTGCGTGTC |
| 4217 | mamo-s | *BGIBMGA005072* | up | 1828 | 1836 | - | 12.4679 | 1.31E-05 | 0.615 | GTGCGTGTC |

| | | | | | | | | | | |
|---|---|---|---|---|---|---|---|---|---|---|
| 4218 | mamo-s | *BGIBMGA005072* | up | 1674 | 1682 | + | 11.4404 | 4.51E-05 | 0.615 | CGGCGTGGC |
| 4219 | mamo-s | *BGIBMGA005073* | up | 474 | 482 | - | 11.3303 | 4.96E-05 | 0.615 | GTGGGTGGC |
| 4220 | mamo-s | *BGIBMGA005075* | down | 1030 | 1038 | - | 10.8073 | 7.89E-05 | 0.766 | CCGCGTGGT |
| 4221 | mamo-s | *BGIBMGA005076* | down | 463 | 471 | - | 11.3303 | 4.96E-05 | 0.615 | GTGGGTGGC |
| 4222 | mamo-s | *BGIBMGA005077* | down | 1159 | 1167 | + | 11.3303 | 4.96E-05 | 0.615 | GTGGGTGGC |
| 4223 | mamo-s | *BGIBMGA005078* | up | 1815 | 1823 | - | 11.3303 | 4.96E-05 | 0.615 | GTGGGTGGC |
| 4224 | mamo-s | *BGIBMGA005082* | down | 1953 | 1961 | - | 13.0275 | 8.53E-06 | 0.615 | GAGCGTGGC |
| 4225 | mamo-s | *BGIBMGA005082* | up | 1188 | 1196 | + | 11.3303 | 4.96E-05 | 0.615 | GTGGGTGGC |
| 4226 | mamo-s | *BGIBMGA005082* | down | 1709 | 1717 | - | 11.0826 | 6.84E-05 | 0.709 | GTACGTGGC |
| 4227 | mamo-s | *BGIBMGA005083* | up | 1369 | 1377 | + | 11.3303 | 4.96E-05 | 0.615 | GTGGGTGGC |
| 4228 | mamo-s | *BGIBMGA005085* | down | 1680 | 1688 | - | 10.8073 | 7.89E-05 | 0.766 | CCGCGTGGT |
| 4229 | mamo-s | *BGIBMGA005090* | up | 1108 | 1116 | + | 11.3303 | 4.96E-05 | 0.615 | GTGGGTGGC |
| 4230 | mamo-s | *BGIBMGA005090* | down | 1522 | 1530 | - | 11.1009 | 6.57E-05 | 0.709 | GAGCGAGGC |
| 4231 | mamo-s | *BGIBMGA005091* | up | 1598 | 1606 | - | 11.3303 | 4.96E-05 | 0.615 | GTGGGTGGC |
| 4232 | mamo-s | *BGIBMGA005091* | down | 492 | 500 | + | 11.0826 | 6.84E-05 | 0.709 | GTACGTGGC |
| 4233 | mamo-s | *BGIBMGA005095* | up | 1828 | 1836 | + | 11.3303 | 4.96E-05 | 0.615 | GTGGGTGGC |
| 4234 | mamo-s | *BGIBMGA005100* | up | 1463 | 1471 | - | 11.3303 | 4.96E-05 | 0.615 | GTGGGTGGC |
| 4235 | mamo-s | *BGIBMGA005100* | up | 599 | 607 | - | 11.0826 | 6.84E-05 | 0.709 | GTACGTGGC |
| 4236 | mamo-s | *BGIBMGA005101* | down | 42 | 50 | + | 12.1835 | 1.71E-05 | 0.615 | GTGCGTGCC |
| 4237 | mamo-s | *BGIBMGA005101* | up | 1207 | 1215 | - | 11.3303 | 4.96E-05 | 0.615 | GTGGGTGGC |
| 4238 | mamo-s | *BGIBMGA005102* | down | 1234 | 1242 | - | 13.156 | 6.30E-06 | 0.615 | CTGCGTGGC |
| 4239 | mamo-s | *BGIBMGA005102* | down | 1306 | 1314 | - | 11.4404 | 4.51E-05 | 0.615 | GAGCGTGTC |
| 4240 | mamo-s | *BGIBMGA005103* | up | 418 | 426 | - | 10.9358 | 7.30E-05 | 0.728 | GCGGGTGGC |
| 4241 | mamo-s | *BGIBMGA005104* | up | 1793 | 1801 | - | 11.3303 | 4.96E-05 | 0.615 | GTGGGTGGC |
| 4242 | mamo-s | *BGIBMGA005105* | down | 771 | 779 | + | 13.0275 | 8.53E-06 | 0.615 | GAGCGTGGC |
| 4243 | mamo-s | *BGIBMGA005105* | down | 613 | 621 | + | 11.3303 | 4.96E-05 | 0.615 | GTGGGTGGC |
| 4244 | mamo-s | *BGIBMGA005105* | up | 555 | 563 | - | 11.2294 | 5.41E-05 | 0.65 | CTGCGAGGC |
| 4245 | mamo-s | *BGIBMGA005106* | up | 668 | 676 | + | 11.3303 | 4.96E-05 | 0.615 | GTGGGTGGC |
| 4246 | mamo-s | *BGIBMGA005108* | down | 1674 | 1682 | - | 11.3303 | 4.96E-05 | 0.615 | GTGGGTGGC |
| 4247 | mamo-s | *BGIBMGA005110* | down | 1438 | 1446 | - | 11.3303 | 4.96E-05 | 0.615 | GTGGGTGGC |
| 4248 | mamo-s | *BGIBMGA005111* | down | 1054 | 1062 | + | 11.3303 | 4.96E-05 | 0.615 | GTGGGTGGC |
| 4249 | mamo-s | *BGIBMGA005113* | up | 844 | 852 | + | 11.3303 | 4.96E-05 | 0.615 | GTGGGTGGC |
| 4250 | mamo-s | *BGIBMGA005113* | up | 1746 | 1754 | - | 11.3303 | 4.96E-05 | 0.615 | GTGGGTGGC |
| 4251 | mamo-s | *BGIBMGA005114* | up | 1394 | 1402 | + | 11.7339 | 3.29E-05 | 0.615 | GCGCGAGGC |
| 4252 | mamo-s | *BGIBMGA005114* | down | 1339 | 1347 | - | 11.3303 | 4.96E-05 | 0.615 | GTGGGTGGC |
| 4253 | mamo-s | *BGIBMGA005114* | down | 317 | 325 | - | 10.6881 | 8.61E-05 | 0.81 | GCACGTGGC |
| 4254 | mamo-s | *BGIBMGA005115* | down | 573 | 581 | + | 11.3303 | 4.96E-05 | 0.615 | GTGGGTGGC |
| 4255 | mamo-s | *BGIBMGA005115* | down | 795 | 803 | + | 11.3303 | 4.96E-05 | 0.615 | GTGGGTGGC |
| 4256 | mamo-s | *BGIBMGA005115* | up | 1116 | 1124 | - | 11.3303 | 4.96E-05 | 0.615 | GTGGGTGGC |
| 4257 | mamo-s | *BGIBMGA005115* | up | 94 | 102 | - | 10.6881 | 8.61E-05 | 0.81 | GCACGTGGC |
| 4258 | mamo-s | *BGIBMGA005115* | down | 958 | 966 | + | 10.5688 | 8.88E-05 | 0.824 | GTGCGTGAC |
| 4259 | mamo-s | *BGIBMGA005117* | up | 1613 | 1621 | + | 11.3303 | 4.96E-05 | 0.615 | GTGGGTGGC |
| 4260 | mamo-s | *BGIBMGA005118* | down | 1270 | 1278 | + | 10.7523 | 8.38E-05 | 0.797 | AAGCGTGGC |
| 4261 | mamo-s | *BGIBMGA005120* | down | 755 | 763 | - | 11.3303 | 4.96E-05 | 0.615 | GTGGGTGGC |
| 4262 | mamo-s | *BGIBMGA005121* | down | 690 | 698 | + | 11.3303 | 4.96E-05 | 0.615 | GTGGGTGGC |
| 4263 | mamo-s | *BGIBMGA005121* | up | 1063 | 1071 | - | 11.3303 | 4.96E-05 | 0.615 | GTGGGTGGC |
| 4264 | mamo-s | *BGIBMGA005122* | down | 574 | 582 | - | 11.3303 | 4.96E-05 | 0.615 | GTGGGTGGC |
| 4265 | mamo-s | *BGIBMGA005122* | up | 1343 | 1351 | + | 11.3303 | 4.96E-05 | 0.615 | GTGGGTGGC |
| 4266 | mamo-s | *BGIBMGA005122* | down | 1275 | 1283 | + | 11.156 | 6.13E-05 | 0.705 | GAGCGTGCC |
| 4267 | mamo-s | *BGIBMGA005123* | up | 932 | 940 | - | 11.3303 | 4.96E-05 | 0.615 | GTGGGTGGC |
| 4268 | mamo-s | *BGIBMGA005123* | up | 1318 | 1326 | + | 11.2018 | 5.68E-05 | 0.674 | CTGCGTGGT |
| 4269 | mamo-s | *BGIBMGA005123* | up | 655 | 663 | - | 11.0826 | 6.84E-05 | 0.709 | GTACGTGGC |
| 4270 | mamo-s | *BGIBMGA005125* | down | 462 | 470 | + | 11.3303 | 4.96E-05 | 0.615 | GTGGGTGGC |
| 4271 | mamo-s | *BGIBMGA005126* | down | 281 | 289 | - | 11.3303 | 4.96E-05 | 0.615 | GTGGGTGGC |
| 4272 | mamo-s | *BGIBMGA005127* | up | 837 | 845 | - | 11.3303 | 4.96E-05 | 0.615 | GTGGGTGGC |
| 4273 | mamo-s | *BGIBMGA005127* | down | 996 | 1004 | - | 11.3303 | 4.96E-05 | 0.615 | GTGGGTGGC |
| 4274 | mamo-s | *BGIBMGA005128* | down | 232 | 240 | + | 11.3303 | 4.96E-05 | 0.615 | GTGGGTGGC |
| 4275 | mamo-s | *BGIBMGA005128* | down | 840 | 848 | + | 11.3303 | 4.96E-05 | 0.615 | GTGGGTGGC |
| 4276 | mamo-s | *BGIBMGA005128* | up | 1151 | 1159 | + | 11.3303 | 4.96E-05 | 0.615 | GTGGGTGGC |
| 4277 | mamo-s | *BGIBMGA005128* | down | 1490 | 1498 | - | 11.3303 | 4.96E-05 | 0.615 | GTGGGTGGC |
| 4278 | mamo-s | *BGIBMGA005129* | up | 1419 | 1427 | + | 11.3303 | 4.96E-05 | 0.615 | GTGGGTGGC |
| 4279 | mamo-s | *BGIBMGA005129* | down | 572 | 580 | + | 11.2844 | 5.18E-05 | 0.63 | CTGCGTGCC |
| 4280 | mamo-s | *BGIBMGA005130* | up | 759 | 767 | - | 11.3303 | 4.96E-05 | 0.615 | GTGGGTGGC |
| 4281 | mamo-s | *BGIBMGA005130* | up | 1897 | 1905 | + | 11.2844 | 5.18E-05 | 0.63 | CTGCGTGCC |
| 4282 | mamo-s | *BGIBMGA005131* | up | 869 | 877 | - | 11.1743 | 5.90E-05 | 0.684 | CCGCGTGTC |
| 4283 | mamo-s | *BGIBMGA005135* | up | 408 | 416 | + | 11.789 | 2.84E-05 | 0.615 | GCGCGTGCC |
| 4284 | mamo-s | *BGIBMGA005135* | up | 1098 | 1106 | - | 11.3303 | 4.96E-05 | 0.615 | GTGGGTGGC |
| 4285 | mamo-s | *BGIBMGA005135* | up | 1675 | 1683 | + | 11.3303 | 4.96E-05 | 0.615 | GTGGGTGGC |
| 4286 | mamo-s | *BGIBMGA005135* | up | 1298 | 1306 | + | 11.1376 | 6.35E-05 | 0.709 | TCGCGTGGC |
| 4287 | mamo-s | *BGIBMGA005138* | down | 651 | 659 | + | 11.3303 | 4.96E-05 | 0.615 | GTGGGTGGC |
| 4288 | mamo-s | *BGIBMGA005139* | down | 1944 | 1952 | - | 10.6881 | 8.61E-05 | 0.81 | GCACGTGGC |
| 4289 | mamo-s | *BGIBMGA005140* | down | 621 | 629 | - | 11.1009 | 6.57E-05 | 0.709 | GAGCGAGGC |
| 4290 | mamo-s | *BGIBMGA005141* | up | 282 | 290 | + | 13.156 | 6.30E-06 | 0.615 | CTGCGTGGC |
| 4291 | mamo-s | *BGIBMGA005143* | up | 1685 | 1693 | + | 11.7064 | 3.52E-05 | 0.615 | GCGCGTGGT |
| 4292 | mamo-s | *BGIBMGA005143* | down | 1171 | 1179 | + | 11.3303 | 4.96E-05 | 0.615 | GTGGGTGGC |
| 4293 | mamo-s | *BGIBMGA005148* | up | 986 | 994 | - | 12.0734 | 2.66E-05 | 0.615 | GCGCGTGTC |
| 4294 | mamo-s | *BGIBMGA005148* | down | 1053 | 1061 | - | 11.1743 | 5.90E-05 | 0.684 | CCGCGTGTC |
| 4295 | mamo-s | *BGIBMGA005148* | up | 420 | 428 | + | 11.1009 | 6.57E-05 | 0.709 | GAGCGAGGC |
| 4296 | mamo-s | *BGIBMGA005148* | up | 83 | 91 | + | 10.9358 | 7.30E-05 | 0.728 | GCGGGTGGC |
| 4297 | mamo-s | *BGIBMGA005150* | up | 1825 | 1833 | + | 11.5321 | 4.06E-05 | 0.615 | TTGCGTGGC |
| 4298 | mamo-s | *BGIBMGA005150* | up | 823 | 831 | + | 11.3303 | 4.96E-05 | 0.615 | GTGGGTGGC |
| 4299 | mamo-s | *BGIBMGA005150* | down | 1113 | 1121 | + | 11.3303 | 4.96E-05 | 0.615 | GTGGGTGGC |
| 4300 | mamo-s | *BGIBMGA005152* | down | 713 | 721 | - | 11.3303 | 4.96E-05 | 0.615 | GTGGGTGGC |
| 4301 | mamo-s | *BGIBMGA005153* | up | 1645 | 1653 | - | 10.9358 | 7.30E-05 | 0.728 | GCGGGTGGC |
| 4302 | mamo-s | *BGIBMGA005156* | down | 913 | 921 | + | 11.3303 | 4.96E-05 | 0.615 | GTGGGTGGC |
| 4303 | mamo-s | *BGIBMGA005160* | up | 1008 | 1016 | - | 11.3303 | 4.96E-05 | 0.615 | GTGGGTGGC |
| 4304 | mamo-s | *BGIBMGA005164* | down | 365 | 373 | + | 11.3303 | 4.96E-05 | 0.615 | GTGGGTGGC |
| 4305 | mamo-s | *BGIBMGA005166* | up | 687 | 695 | + | 10.5138 | 9.75E-05 | 0.865 | GTGCGTGTT |

| | | | | | | | | | | |
|---|---|---|---|---|---|---|---|---|---|---|
| 4306 | mamo-s | *BGIBMGA005169* | up | 459 | 467 | - | 11.0734 | 7.12E-05 | 0.726 | GAGCGTGGT |
| 4307 | mamo-s | *BGIBMGA005171* | down | 1837 | 1845 | - | 11.3303 | 4.96E-05 | 0.615 | GTGGGTGGC |
| 4308 | mamo-s | *BGIBMGA005172* | up | 473 | 481 | - | 11.4404 | 4.51E-05 | 0.615 | CGGCGTGGC |
| 4309 | mamo-s | *BGIBMGA005172* | up | 935 | 943 | + | 11.3303 | 4.96E-05 | 0.615 | GTGGGTGGC |
| 4310 | mamo-s | *BGIBMGA005174* | up | 1025 | 1033 | + | 11.0826 | 6.84E-05 | 0.709 | GTACGTGGC |
| 4311 | mamo-s | *BGIBMGA005175* | up | 24 | 32 | + | 11.3303 | 4.96E-05 | 0.615 | GTGGGTGGC |
| 4312 | mamo-s | *BGIBMGA005176* | up | 909 | 917 | - | 12.3394 | 1.49E-05 | 0.615 | GGGCGTGGC |
| 4313 | mamo-s | *BGIBMGA005176* | up | 973 | 981 | + | 12.1835 | 1.71E-05 | 0.615 | GTGCGTGCC |
| 4314 | mamo-s | *BGIBMGA005176* | down | 1925 | 1933 | + | 11.3303 | 4.96E-05 | 0.615 | GTGGGTGGC |
| 4315 | mamo-s | *BGIBMGA005176* | up | 1808 | 1816 | + | 10.5138 | 9.75E-05 | 0.865 | GTGCGTGTT |
| 4316 | mamo-s | *BGIBMGA005177* | down | 1726 | 1734 | - | 11.3303 | 4.96E-05 | 0.615 | GTGGGTGGC |
| 4317 | mamo-s | *BGIBMGA005180* | up | 1755 | 1763 | - | 13.6606 | 4.06E-06 | 0.615 | GCGCGTGGC |
| 4318 | mamo-s | *BGIBMGA005181* | up | 924 | 932 | - | 11.3303 | 4.96E-05 | 0.615 | GTGGGTGGC |
| 4319 | mamo-s | *BGIBMGA005183* | down | 565 | 573 | - | 11.0826 | 6.84E-05 | 0.709 | GTACGTGGC |
| 4320 | mamo-s | *BGIBMGA005184* | up | 1473 | 1481 | + | 11.3303 | 4.96E-05 | 0.615 | GTGGGTGGC |
| 4321 | mamo-s | *BGIBMGA005185* | up | 1085 | 1093 | + | 14.055 | 2.23E-06 | 0.615 | GTGCGTGGC |
| 4322 | mamo-s | *BGIBMGA005185* | up | 199 | 207 | + | 11.3303 | 4.96E-05 | 0.615 | GTGGGTGGC |
| 4323 | mamo-s | *BGIBMGA005185* | up | 1336 | 1344 | + | 11.1009 | 6.57E-05 | 0.709 | GAGCGAGGC |
| 4324 | mamo-s | *BGIBMGA005185* | down | 1739 | 1747 | - | 10.5138 | 9.75E-05 | 0.865 | GTGCGTGTT |
| 4325 | mamo-s | *BGIBMGA005186* | down | 1475 | 1483 | + | 11.3303 | 4.96E-05 | 0.615 | GTGGGTGGC |
| 4326 | mamo-s | *BGIBMGA005187* | down | 313 | 321 | - | 11.3303 | 4.96E-05 | 0.615 | GTGGGTGGC |
| 4327 | mamo-s | *BGIBMGA005190* | up | 1805 | 1813 | - | 11.3303 | 4.96E-05 | 0.615 | GTGGGTGGC |
| 4328 | mamo-s | *BGIBMGA005190* | up | 542 | 550 | + | 11.2018 | 5.68E-05 | 0.674 | CTGCGTGGT |
| 4329 | mamo-s | *BGIBMGA005193* | up | 534 | 542 | - | 11.3303 | 4.96E-05 | 0.615 | GTGGGTGGC |
| 4330 | mamo-s | *BGIBMGA005194* | up | 42 | 50 | + | 13.6606 | 4.06E-06 | 0.615 | GCGCGTGGC |
| 4331 | mamo-s | *BGIBMGA005194* | up | 1883 | 1891 | - | 13.156 | 6.30E-06 | 0.615 | CTGCGTGGC |
| 4332 | mamo-s | *BGIBMGA005194* | down | 204 | 212 | + | 11.3303 | 4.96E-05 | 0.615 | GTGGGTGGC |
| 4333 | mamo-s | *BGIBMGA005195* | up | 204 | 212 | + | 11.3303 | 4.96E-05 | 0.615 | GTGGGTGGC |
| 4334 | mamo-s | *BGIBMGA005195* | up | 1284 | 1292 | - | 11.3303 | 4.96E-05 | 0.615 | GTGGGTGGC |
| 4335 | mamo-s | *BGIBMGA005195* | down | 1306 | 1314 | + | 10.5138 | 9.75E-05 | 0.865 | GTGCGTGTT |
| 4336 | mamo-s | *BGIBMGA005199* | down | 454 | 462 | + | 11.3303 | 4.96E-05 | 0.615 | GTGGGTGGC |
| 4337 | mamo-s | *BGIBMGA005201* | down | 42 | 50 | - | 11.2844 | 5.18E-05 | 0.63 | CTGCGTGCC |
| 4338 | mamo-s | *BGIBMGA005204* | up | 1110 | 1118 | + | 11.3303 | 4.96E-05 | 0.615 | GTGGGTGGC |
| 4339 | mamo-s | *BGIBMGA005210* | down | 363 | 371 | + | 11.7798 | 3.11E-05 | 0.615 | ATGCGTGGC |
| 4340 | mamo-s | *BGIBMGA005211* | up | 1907 | 1915 | + | 11.4404 | 4.51E-05 | 0.615 | CGGCGTGGC |
| 4341 | mamo-s | *BGIBMGA005213* | down | 1462 | 1470 | + | 11.3303 | 4.96E-05 | 0.615 | GTGGGTGGC |
| 4342 | mamo-s | *BGIBMGA005214* | up | 1625 | 1633 | - | 12.0734 | 2.66E-05 | 0.615 | GCGCGTGTC |
| 4343 | mamo-s | *BGIBMGA005214* | up | 1535 | 1543 | + | 10.5688 | 8.88E-05 | 0.824 | GTGCGTGAC |
| 4344 | mamo-s | *BGIBMGA005215* | down | 1024 | 1032 | + | 11.3303 | 4.96E-05 | 0.615 | GTGGGTGGC |
| 4345 | mamo-s | *BGIBMGA005215* | down | 1671 | 1679 | - | 11.3303 | 4.96E-05 | 0.615 | GTGGGTGGC |
| 4346 | mamo-s | *BGIBMGA005218* | down | 1553 | 1561 | + | 12.3394 | 1.49E-05 | 0.615 | GGGCGTGGC |
| 4347 | mamo-s | *BGIBMGA005218* | up | 1571 | 1579 | - | 11.3303 | 4.96E-05 | 0.615 | GTGGGTGGC |
| 4348 | mamo-s | *BGIBMGA005219* | down | 1432 | 1440 | + | 10.7523 | 8.38E-05 | 0.797 | GGGCGTGTC |
| 4349 | mamo-s | *BGIBMGA005220* | up | 1785 | 1793 | + | 11.3303 | 4.96E-05 | 0.615 | GTGGGTGGC |
| 4350 | mamo-s | *BGIBMGA005221* | up | 402 | 410 | + | 11.2844 | 5.18E-05 | 0.63 | CTGCGTGCC |
| 4351 | mamo-s | *BGIBMGA005224* | up | 1737 | 1745 | - | 11.3303 | 4.96E-05 | 0.615 | GTGGGTGGC |
| 4352 | mamo-s | *BGIBMGA005225* | down | 1568 | 1576 | + | 11.1376 | 6.35E-05 | 0.709 | TCGCGTGGC |
| 4353 | mamo-s | *BGIBMGA005225* | up | 795 | 803 | - | 11.0826 | 6.84E-05 | 0.709 | GTACGTGGC |
| 4354 | mamo-s | *BGIBMGA005226* | up | 848 | 856 | - | 11.3303 | 4.96E-05 | 0.615 | GTGGGTGGC |
| 4355 | mamo-s | *BGIBMGA005227* | up | 1197 | 1205 | - | 11.0826 | 6.84E-05 | 0.709 | GTACGTGGC |
| 4356 | mamo-s | *BGIBMGA005232* | down | 1374 | 1382 | - | 11.3303 | 4.96E-05 | 0.615 | GTGGGTGGC |
| 4357 | mamo-s | *BGIBMGA005234* | up | 239 | 247 | - | 11.3303 | 4.96E-05 | 0.615 | GTGGGTGGC |
| 4358 | mamo-s | *BGIBMGA005236* | down | 1754 | 1762 | - | 11.3303 | 4.96E-05 | 0.615 | GTGGGTGGC |
| 4359 | mamo-s | *BGIBMGA005236* | up | 265 | 273 | - | 11.0826 | 6.84E-05 | 0.709 | GTACGTGGC |
| 4360 | mamo-s | *BGIBMGA005237* | up | 731 | 739 | - | 11.3303 | 4.96E-05 | 0.615 | GTGGGTGGC |
| 4361 | mamo-s | *BGIBMGA005237* | up | 1862 | 1870 | + | 11.3303 | 4.96E-05 | 0.615 | GTGGGTGGC |
| 4362 | mamo-s | *BGIBMGA005238* | down | 973 | 981 | - | 11.3303 | 4.96E-05 | 0.615 | GTGGGTGGC |
| 4363 | mamo-s | *BGIBMGA005239* | down | 97 | 105 | + | 12.1835 | 1.71E-05 | 0.615 | GTGCGTGCC |
| 4364 | mamo-s | *BGIBMGA005239* | up | 321 | 329 | + | 11.3303 | 4.96E-05 | 0.615 | GTGGGTGGC |
| 4365 | mamo-s | *BGIBMGA005239* | down | 390 | 398 | - | 11.3303 | 4.96E-05 | 0.615 | GTGGGTGGC |
| 4366 | mamo-s | *BGIBMGA005240* | down | 810 | 818 | + | 11.3853 | 4.74E-05 | 0.615 | ACGCGTGGC |
| 4367 | mamo-s | *BGIBMGA005240* | down | 1872 | 1880 | + | 11.1743 | 5.90E-05 | 0.684 | CCGCGTGTC |
| 4368 | mamo-s | *BGIBMGA005241* | down | 302 | 310 | + | 11.4404 | 4.51E-05 | 0.615 | GAGCGTGTC |
| 4369 | mamo-s | *BGIBMGA005243* | down | 706 | 714 | - | 14.055 | 2.23E-06 | 0.615 | GTGCGTGGC |
| 4370 | mamo-s | *BGIBMGA005243* | down | 1095 | 1103 | - | 11.3303 | 4.96E-05 | 0.615 | GTGGGTGGC |
| 4371 | mamo-s | *BGIBMGA005244* | up | 617 | 625 | + | 11.156 | 6.13E-05 | 0.705 | GAGCGTGCC |
| 4372 | mamo-s | *BGIBMGA005245* | down | 623 | 631 | + | 11.3303 | 4.96E-05 | 0.615 | GTGGGTGGC |
| 4373 | mamo-s | *BGIBMGA005246* | down | 1226 | 1234 | + | 13.6606 | 4.06E-06 | 0.615 | GCGCGTGGC |
| 4374 | mamo-s | *BGIBMGA005251* | up | 520 | 528 | - | 11.0826 | 6.84E-05 | 0.709 | GTACGTGGC |
| 4375 | mamo-s | *BGIBMGA005252* | down | 182 | 190 | + | 11.5321 | 4.06E-05 | 0.615 | TTGCGTGGC |
| 4376 | mamo-s | *BGIBMGA005259* | down | 1474 | 1482 | - | 11.3303 | 4.96E-05 | 0.615 | GTGGGTGGC |
| 4377 | mamo-s | *BGIBMGA005260* | down | 216 | 224 | + | 11.3303 | 4.96E-05 | 0.615 | GTGGGTGGC |
| 4378 | mamo-s | *BGIBMGA005262* | up | 237 | 245 | - | 11.3303 | 4.96E-05 | 0.615 | GTGGGTGGC |
| 4379 | mamo-s | *BGIBMGA005262* | up | 1962 | 1970 | - | 10.5413 | 9.42E-05 | 0.857 | GTGCGAGTC |
| 4380 | mamo-s | *BGIBMGA005263* | up | 1704 | 1712 | + | 11.3303 | 4.96E-05 | 0.615 | GTGGGTGGC |
| 4381 | mamo-s | *BGIBMGA005265* | down | 1466 | 1474 | + | 11.3303 | 4.96E-05 | 0.615 | GTGGGTGGC |
| 4382 | mamo-s | *BGIBMGA005265* | down | 675 | 683 | + | 11.0734 | 7.12E-05 | 0.726 | GAGCGTGGT |
| 4383 | mamo-s | *BGIBMGA005265* | up | 1721 | 1729 | + | 11.0734 | 7.12E-05 | 0.726 | GAGCGTGGT |
| 4384 | mamo-s | *BGIBMGA005267* | down | 1069 | 1077 | - | 14.055 | 2.23E-06 | 0.615 | GTGCGTGGC |
| 4385 | mamo-s | *BGIBMGA005267* | up | 1018 | 1026 | - | 12.4679 | 1.31E-05 | 0.615 | GTGCGTGTC |
| 4386 | mamo-s | *BGIBMGA005268* | up | 1224 | 1232 | + | 12.4679 | 1.31E-05 | 0.615 | GTGCGTGTC |
| 4387 | mamo-s | *BGIBMGA005268* | down | 1801 | 1809 | - | 11.3303 | 4.96E-05 | 0.615 | GTGGGTGGC |
| 4388 | mamo-s | *BGIBMGA005269* | up | 745 | 753 | + | 11.3303 | 4.96E-05 | 0.615 | GTGGGTGGC |
| 4389 | mamo-s | *BGIBMGA005269* | up | 1229 | 1237 | - | 11.3303 | 4.96E-05 | 0.615 | GTGGGTGGC |
| 4390 | mamo-s | *BGIBMGA005269* | up | 810 | 818 | + | 10.5413 | 9.42E-05 | 0.857 | GTGCGAGTC |
| 4391 | mamo-s | *BGIBMGA005271* | up | 521 | 529 | + | 10.5138 | 9.75E-05 | 0.865 | GTGCGTGTT |
| 4392 | mamo-s | *BGIBMGA005272* | up | 905 | 913 | + | 11.3303 | 4.96E-05 | 0.615 | GTGGGTGGC |
| 4393 | mamo-s | *BGIBMGA005272* | down | 1764 | 1772 | + | 11.3303 | 4.96E-05 | 0.615 | GTGGGTGGC |

| | | | | | | | | | | |
|---|---|---|---|---|---|---|---|---|---|---|
| 4394 | mamo-s | *BGIBMGA005276* | up | 530 | 538 | - | 11.7064 | 3.52E-05 | 0.615 | GCGCGTGGT |
| 4395 | mamo-s | *BGIBMGA005278* | down | 406 | 414 | + | 11.3303 | 4.96E-05 | 0.615 | GTGGGTGGC |
| 4396 | mamo-s | *BGIBMGA005278* | up | 1355 | 1363 | + | 11.3303 | 4.96E-05 | 0.615 | GTGGGTGGC |
| 4397 | mamo-s | *BGIBMGA005283* | down | 855 | 863 | + | 11.3303 | 4.96E-05 | 0.615 | GTGGGTGGC |
| 4398 | mamo-s | *BGIBMGA005283* | up | 1089 | 1097 | + | 10.7523 | 8.38E-05 | 0.797 | GGGCGTGTC |
| 4399 | mamo-s | *BGIBMGA005286* | down | 1103 | 1111 | - | 12.1835 | 1.71E-05 | 0.615 | GTGCGTGCC |
| 4400 | mamo-s | *BGIBMGA005286* | up | 325 | 333 | - | 11.1743 | 5.90E-05 | 0.684 | CCGCGTGTC |
| 4401 | mamo-s | *BGIBMGA005287* | up | 349 | 357 | - | 12.1835 | 1.71E-05 | 0.615 | GTGCGTGCC |
| 4402 | mamo-s | *BGIBMGA005289* | up | 68 | 76 | - | 12.1009 | 2.43E-05 | 0.615 | GTGCGTGGT |
| 4403 | mamo-s | *BGIBMGA005289* | down | 1598 | 1606 | - | 10.5413 | 9.42E-05 | 0.857 | GTGCGAGTC |
| 4404 | mamo-s | *BGIBMGA005292* | down | 166 | 174 | - | 11.1009 | 6.57E-05 | 0.709 | GAGCGAGGC |
| 4405 | mamo-s | *BGIBMGA005292* | up | 12 | 20 | + | 10.9358 | 7.30E-05 | 0.728 | GCGGGTGGC |
| 4406 | mamo-s | *BGIBMGA005293* | down | 1586 | 1594 | - | 11.3303 | 4.96E-05 | 0.615 | GTGGGTGGC |
| 4407 | mamo-s | *BGIBMGA005293* | down | 1907 | 1915 | + | 11.3303 | 4.96E-05 | 0.615 | GTGGGTGGC |
| 4408 | mamo-s | *BGIBMGA005293* | up | 728 | 736 | - | 11.1009 | 6.57E-05 | 0.709 | GAGCGAGGC |
| 4409 | mamo-s | *BGIBMGA005293* | down | 1454 | 1462 | + | 10.5138 | 9.75E-05 | 0.865 | GTGCGTGTT |
| 4410 | mamo-s | *BGIBMGA005295* | up | 1617 | 1625 | + | 12.1835 | 1.71E-05 | 0.615 | GTGCGTGCC |
| 4411 | mamo-s | *BGIBMGA005296* | up | 1765 | 1773 | + | 13.0275 | 8.53E-06 | 0.615 | GAGCGTGGC |
| 4412 | mamo-s | *BGIBMGA005296* | down | 1457 | 1465 | - | 11.3303 | 4.96E-05 | 0.615 | GTGGGTGGC |
| 4413 | mamo-s | *BGIBMGA005296* | up | 1189 | 1197 | - | 11.1743 | 5.90E-05 | 0.684 | CCGCGTGTC |
| 4414 | mamo-s | *BGIBMGA005297* | down | 30 | 38 | + | 10.5138 | 9.75E-05 | 0.865 | GTGCGTGTT |
| 4415 | mamo-s | *BGIBMGA005301* | down | 1953 | 1961 | + | 11.3303 | 4.96E-05 | 0.615 | GTGGGTGGC |
| 4416 | mamo-s | *BGIBMGA005305* | up | 735 | 743 | - | 11.3303 | 4.96E-05 | 0.615 | GTGGGTGGC |
| 4417 | mamo-s | *BGIBMGA005310* | up | 1872 | 1880 | - | 11.3303 | 4.96E-05 | 0.615 | GTGGGTGGC |
| 4418 | mamo-s | *BGIBMGA005311* | down | 1967 | 1975 | + | 12.4679 | 1.31E-05 | 0.615 | GTGCGTGTC |
| 4419 | mamo-s | *BGIBMGA005314* | down | 208 | 216 | + | 11.3303 | 4.96E-05 | 0.615 | GTGGGTGGC |
| 4420 | mamo-s | *BGIBMGA005315* | up | 73 | 81 | + | 11.3303 | 4.96E-05 | 0.615 | GTGGGTGGC |
| 4421 | mamo-s | *BGIBMGA005318* | down | 1061 | 1069 | - | 11.3303 | 4.96E-05 | 0.615 | GTGGGTGGC |
| 4422 | mamo-s | *BGIBMGA005319* | down | 1333 | 1341 | - | 11.3303 | 4.96E-05 | 0.615 | GTGGGTGGC |
| 4423 | mamo-s | *BGIBMGA005320* | up | 763 | 771 | - | 11.0826 | 6.84E-05 | 0.709 | GTACGTGGC |
| 4424 | mamo-s | *BGIBMGA005325* | down | 1386 | 1394 | - | 11.2294 | 5.41E-05 | 0.65 | CTGCGAGGC |
| 4425 | mamo-s | *BGIBMGA005326* | down | 901 | 909 | + | 11.3303 | 4.96E-05 | 0.615 | GTGGGTGGC |
| 4426 | mamo-s | *BGIBMGA005327* | up | 1341 | 1349 | + | 12.0734 | 2.66E-05 | 0.615 | GCGCGTGTC |
| 4427 | mamo-s | *BGIBMGA005327* | up | 75 | 83 | + | 11.4404 | 4.51E-05 | 0.615 | CGGCGTGGC |
| 4428 | mamo-s | *BGIBMGA005329* | down | 721 | 729 | + | 11.3303 | 4.96E-05 | 0.615 | GTGGGTGGC |
| 4429 | mamo-s | *BGIBMGA005331* | up | 702 | 710 | + | 11.3303 | 4.96E-05 | 0.615 | GTGGGTGGC |
| 4430 | mamo-s | *BGIBMGA005333* | down | 258 | 266 | + | 11.0826 | 6.84E-05 | 0.709 | GTACGTGGC |
| 4431 | mamo-s | *BGIBMGA005340* | up | 1453 | 1461 | - | 10.5138 | 9.75E-05 | 0.865 | GTGCGTGTT |
| 4432 | mamo-s | *BGIBMGA005340* | down | 1711 | 1719 | + | 10.5138 | 9.75E-05 | 0.865 | GTGCGTGTT |
| 4433 | mamo-s | *BGIBMGA005341* | down | 1631 | 1639 | + | 12.1835 | 1.71E-05 | 0.615 | GTGCGTGCC |
| 4434 | mamo-s | *BGIBMGA005342* | down | 1270 | 1278 | + | 10.6881 | 8.61E-05 | 0.81 | GCACGTGGC |
| 4435 | mamo-s | *BGIBMGA005345* | up | 591 | 599 | + | 11.3303 | 4.96E-05 | 0.615 | GTGGGTGGC |
| 4436 | mamo-s | *BGIBMGA005345* | up | 1418 | 1426 | + | 11.0826 | 6.84E-05 | 0.709 | GTACGTGGC |
| 4437 | mamo-s | *BGIBMGA005346* | up | 62 | 70 | - | 12.1284 | 2.16E-05 | 0.615 | CAGCGTGGC |
| 4438 | mamo-s | *BGIBMGA005348* | up | 99 | 107 | + | 11.156 | 6.13E-05 | 0.705 | GAGCGTGCC |
| 4439 | mamo-s | *BGIBMGA005348* | down | 698 | 706 | + | 10.5413 | 9.42E-05 | 0.857 | GTGCGAGTC |
| 4440 | mamo-s | *BGIBMGA005349* | up | 1084 | 1092 | - | 11.0826 | 6.84E-05 | 0.709 | GTACGTGGC |
| 4441 | mamo-s | *BGIBMGA005349* | down | 1714 | 1722 | + | 11.0826 | 6.84E-05 | 0.709 | GTACGTGGC |
| 4442 | mamo-s | *BGIBMGA005350* | up | 1892 | 1900 | - | 10.9358 | 7.30E-05 | 0.728 | GCGGGTGGC |
| 4443 | mamo-s | *BGIBMGA005351* | up | 878 | 886 | + | 14.055 | 2.23E-06 | 0.615 | GTGCGTGGC |
| 4444 | mamo-s | *BGIBMGA005351* | up | 1028 | 1036 | + | 10.5688 | 8.88E-05 | 0.824 | GTGCGTGAC |
| 4445 | mamo-s | *BGIBMGA005351* | up | 1190 | 1198 | + | 10.5688 | 8.88E-05 | 0.824 | GTGCGTGAC |
| 4446 | mamo-s | *BGIBMGA005353* | up | 39 | 47 | + | 11.4404 | 4.51E-05 | 0.615 | CGGCGTGGC |
| 4447 | mamo-s | *BGIBMGA005354* | down | 349 | 357 | - | 11.3303 | 4.96E-05 | 0.615 | GTGGGTGGC |
| 4448 | mamo-s | *BGIBMGA005355* | up | 131 | 139 | + | 11.3303 | 4.96E-05 | 0.615 | GTGGGTGGC |
| 4449 | mamo-s | *BGIBMGA005357* | down | 950 | 958 | - | 11.7798 | 3.11E-05 | 0.615 | ATGCGTGGC |
| 4450 | mamo-s | *BGIBMGA005357* | up | 1861 | 1869 | + | 10.9358 | 7.30E-05 | 0.728 | GCGGGTGGC |
| 4451 | mamo-s | *BGIBMGA005358* | down | 199 | 207 | + | 12.4679 | 1.31E-05 | 0.615 | GTGCGTGTC |
| 4452 | mamo-s | *BGIBMGA005359* | up | 1793 | 1801 | + | 12.4679 | 1.31E-05 | 0.615 | GTGCGTGTC |
| 4453 | mamo-s | *BGIBMGA005362* | down | 1478 | 1486 | - | 11.4404 | 4.51E-05 | 0.615 | CGGCGTGGC |
| 4454 | mamo-s | *BGIBMGA005362* | up | 550 | 558 | - | 11.3303 | 4.96E-05 | 0.615 | GTGGGTGGC |
| 4455 | mamo-s | *BGIBMGA005363* | up | 707 | 715 | - | 11.4404 | 4.51E-05 | 0.615 | CGGCGTGGC |
| 4456 | mamo-s | *BGIBMGA005364* | down | 127 | 135 | - | 11.3303 | 4.96E-05 | 0.615 | GTGGGTGGC |
| 4457 | mamo-s | *BGIBMGA005365* | down | 1965 | 1973 | - | 11.3303 | 4.96E-05 | 0.615 | GTGGGTGGC |
| 4458 | mamo-s | *BGIBMGA005366* | up | 185 | 193 | - | 11.3303 | 4.96E-05 | 0.615 | GTGGGTGGC |
| 4459 | mamo-s | *BGIBMGA005369* | up | 27 | 35 | - | 12.7615 | 1.04E-05 | 0.615 | CCGCGTGGC |
| 4460 | mamo-s | *BGIBMGA005369* | up | 1086 | 1094 | + | 11.0826 | 6.84E-05 | 0.709 | GTACGTGGC |
| 4461 | mamo-s | *BGIBMGA005370* | up | 483 | 491 | + | 10.5138 | 9.75E-05 | 0.865 | GTGCGTGTT |
| 4462 | mamo-s | *BGIBMGA005371* | up | 1705 | 1713 | + | 11.2844 | 5.18E-05 | 0.63 | CTGCGTGCC |
| 4463 | mamo-s | *BGIBMGA005372* | up | 390 | 398 | - | 12.1835 | 1.71E-05 | 0.615 | GTGCGTGCC |
| 4464 | mamo-s | *BGIBMGA005372* | up | 162 | 170 | - | 11.0734 | 7.12E-05 | 0.726 | GAGCGTGGT |
| 4465 | mamo-s | *BGIBMGA005373* | down | 1641 | 1649 | + | 10.5413 | 9.42E-05 | 0.857 | GTGCGAGTC |
| 4466 | mamo-s | *BGIBMGA005374* | up | 1596 | 1604 | + | 11.3303 | 4.96E-05 | 0.615 | GTGGGTGGC |
| 4467 | mamo-s | *BGIBMGA005377* | down | 1921 | 1929 | + | 14.055 | 2.23E-06 | 0.615 | GTGCGTGGC |
| 4468 | mamo-s | *BGIBMGA005377* | down | 939 | 947 | - | 11.3303 | 4.96E-05 | 0.615 | GTGGGTGGC |
| 4469 | mamo-s | *BGIBMGA005378* | up | 303 | 311 | + | 12.7615 | 1.04E-05 | 0.615 | CCGCGTGGC |
| 4470 | mamo-s | *BGIBMGA005378* | down | 577 | 585 | - | 11.2294 | 5.41E-05 | 0.65 | CTGCGAGGC |
| 4471 | mamo-s | *BGIBMGA005386* | down | 227 | 235 | - | 11.3303 | 4.96E-05 | 0.615 | GTGGGTGGC |
| 4472 | mamo-s | *BGIBMGA005386* | up | 116 | 124 | - | 11.0734 | 7.12E-05 | 0.726 | GAGCGTGGT |
| 4473 | mamo-s | *BGIBMGA005387* | up | 1584 | 1592 | - | 11.3303 | 4.96E-05 | 0.615 | GTGGGTGGC |
| 4474 | mamo-s | *BGIBMGA005389* | up | 1562 | 1570 | - | 12.1835 | 1.71E-05 | 0.615 | GTGCGTGCC |
| 4475 | mamo-s | *BGIBMGA005390* | up | 819 | 827 | - | 11.4404 | 4.51E-05 | 0.615 | GAGCGTGTC |
| 4476 | mamo-s | *BGIBMGA005394* | up | 1209 | 1217 | - | 10.9358 | 7.30E-05 | 0.728 | GCGGGTGGC |
| 4477 | mamo-s | *BGIBMGA005395* | up | 434 | 442 | + | 11.3303 | 4.96E-05 | 0.615 | GTGGGTGGC |
| 4478 | mamo-s | *BGIBMGA005396* | up | 216 | 224 | - | 11.7798 | 3.11E-05 | 0.615 | ATGCGTGGC |
| 4479 | mamo-s | *BGIBMGA005396* | up | 95 | 103 | + | 10.8073 | 7.89E-05 | 0.766 | CCGCGTGGT |
| 4480 | mamo-s | *BGIBMGA005397* | up | 809 | 817 | + | 12.7615 | 1.04E-05 | 0.615 | CCGCGTGGC |
| 4481 | mamo-s | *BGIBMGA005397* | up | 367 | 375 | + | 11.2294 | 5.41E-05 | 0.65 | CTGCGAGGC |

| | | | | | | | | | | |
|---|---|---|---|---|---|---|---|---|---|---|
| 4482 | mamo-s | *BGIBMGA005399* | up | 915 | 923 | - | 11.3303 | 4.96E-05 | 0.615 | GTGGGTGGC |
| 4483 | mamo-s | *BGIBMGA005400* | up | 1813 | 1821 | - | 11.0826 | 6.84E-05 | 0.709 | GTACGTGGC |
| 4484 | mamo-s | *BGIBMGA005403* | up | 621 | 629 | - | 11.3303 | 4.96E-05 | 0.615 | GTGGGTGGC |
| 4485 | mamo-s | *BGIBMGA005403* | up | 1731 | 1739 | - | 11.3303 | 4.96E-05 | 0.615 | GTGGGTGGC |
| 4486 | mamo-s | *BGIBMGA005404* | down | 1227 | 1235 | - | 11.7064 | 3.52E-05 | 0.615 | GCGCGTGGT |
| 4487 | mamo-s | *BGIBMGA005404* | down | 341 | 349 | - | 11.3303 | 4.96E-05 | 0.615 | GTGGGTGGC |
| 4488 | mamo-s | *BGIBMGA005405* | down | 729 | 737 | + | 11.3303 | 4.96E-05 | 0.615 | GTGGGTGGC |
| 4489 | mamo-s | *BGIBMGA005406* | down | 1241 | 1249 | - | 10.6881 | 8.61E-05 | 0.81 | GCACGTGGC |
| 4490 | mamo-s | *BGIBMGA005407* | up | 1810 | 1818 | - | 10.6881 | 8.61E-05 | 0.81 | GCACGTGGC |
| 4491 | mamo-s | *BGIBMGA005409* | up | 882 | 890 | - | 12.7615 | 1.04E-05 | 0.615 | CCGCGTGGC |
| 4492 | mamo-s | *BGIBMGA005409* | down | 1507 | 1515 | + | 11.3303 | 4.96E-05 | 0.615 | GTGGGTGGC |
| 4493 | mamo-s | *BGIBMGA005410* | down | 175 | 183 | + | 11.3303 | 4.96E-05 | 0.615 | GTGGGTGGC |
| 4494 | mamo-s | *BGIBMGA005411* | up | 1546 | 1554 | + | 11.0734 | 7.12E-05 | 0.726 | GAGCGTGGT |
| 4495 | mamo-s | *BGIBMGA005417* | up | 1233 | 1241 | + | 11.3303 | 4.96E-05 | 0.615 | GTGGGTGGC |
| 4496 | mamo-s | *BGIBMGA005417* | up | 1659 | 1667 | - | 11.3303 | 4.96E-05 | 0.615 | GTGGGTGGC |
| 4497 | mamo-s | *BGIBMGA005417* | up | 600 | 608 | + | 10.5413 | 9.42E-05 | 0.857 | GTGCGAGTC |
| 4498 | mamo-s | *BGIBMGA005419* | down | 1863 | 1871 | + | 11.3303 | 4.96E-05 | 0.615 | GTGGGTGGC |
| 4499 | mamo-s | *BGIBMGA005420* | up | 182 | 190 | + | 11.3303 | 4.96E-05 | 0.615 | GTGGGTGGC |
| 4500 | mamo-s | *BGIBMGA005423* | down | 773 | 781 | + | 11.2844 | 5.18E-05 | 0.63 | CTGCGTGCC |
| 4501 | mamo-s | *BGIBMGA005423* | down | 70 | 78 | + | 11.2294 | 5.41E-05 | 0.65 | CTGCGAGGC |
| 4502 | mamo-s | *BGIBMGA005424* | down | 1586 | 1594 | - | 14.055 | 2.23E-06 | 0.615 | GTGCGTGGC |
| 4503 | mamo-s | *BGIBMGA005424* | up | 1877 | 1885 | + | 11.3303 | 4.96E-05 | 0.615 | GTGGGTGGC |
| 4504 | mamo-s | *BGIBMGA005426* | up | 1872 | 1880 | + | 10.7523 | 8.38E-05 | 0.797 | AAGCGTGGC |
| 4505 | mamo-s | *BGIBMGA005430* | up | 1764 | 1772 | + | 11.3303 | 4.96E-05 | 0.615 | GTGGGTGGC |
| 4506 | mamo-s | *BGIBMGA005433* | up | 1335 | 1343 | - | 11.3303 | 4.96E-05 | 0.615 | GTGGGTGGC |
| 4507 | mamo-s | *BGIBMGA005433* | down | 1243 | 1251 | + | 10.5413 | 9.42E-05 | 0.857 | GTGCGAGTC |
| 4508 | mamo-s | *BGIBMGA005436* | up | 1007 | 1015 | + | 13.6606 | 4.06E-06 | 0.615 | GCGCGTGGC |
| 4509 | mamo-s | *BGIBMGA005436* | down | 555 | 563 | - | 11.3303 | 4.96E-05 | 0.615 | GTGGGTGGC |
| 4510 | mamo-s | *BGIBMGA005437* | up | 1238 | 1246 | - | 11.3303 | 4.96E-05 | 0.615 | GTGGGTGGC |
| 4511 | mamo-s | *BGIBMGA005437* | down | 1834 | 1842 | + | 11.3303 | 4.96E-05 | 0.615 | GTGGGTGGC |
| 4512 | mamo-s | *BGIBMGA005439* | down | 1241 | 1249 | + | 11.3303 | 4.96E-05 | 0.615 | GTGGGTGGC |
| 4513 | mamo-s | *BGIBMGA005439* | down | 1359 | 1367 | - | 11.3303 | 4.96E-05 | 0.615 | GTGGGTGGC |
| 4514 | mamo-s | *BGIBMGA005439* | down | 1719 | 1727 | - | 11.3303 | 4.96E-05 | 0.615 | GTGGGTGGC |
| 4515 | mamo-s | *BGIBMGA005441* | up | 1331 | 1339 | - | 11.3303 | 4.96E-05 | 0.615 | GTGGGTGGC |
| 4516 | mamo-s | *BGIBMGA005442* | down | 1911 | 1919 | + | 11.3303 | 4.96E-05 | 0.615 | GTGGGTGGC |
| 4517 | mamo-s | *BGIBMGA005442* | up | 1130 | 1138 | - | 11.0826 | 6.84E-05 | 0.709 | GTACGTGGC |
| 4518 | mamo-s | *BGIBMGA005444* | down | 934 | 942 | + | 11.5688 | 3.79E-05 | 0.615 | CTGCGTGTC |
| 4519 | mamo-s | *BGIBMGA005444* | up | 165 | 173 | + | 11.3303 | 4.96E-05 | 0.615 | GTGGGTGGC |
| 4520 | mamo-s | *BGIBMGA005446* | up | 529 | 537 | + | 11.3303 | 4.96E-05 | 0.615 | GTGGGTGGC |
| 4521 | mamo-s | *BGIBMGA005446* | down | 1219 | 1227 | + | 11.3303 | 4.96E-05 | 0.615 | GTGGGTGGC |
| 4522 | mamo-s | *BGIBMGA005450* | down | 785 | 793 | + | 10.5138 | 9.75E-05 | 0.865 | GTGCGTGTT |
| 4523 | mamo-s | *BGIBMGA005459* | down | 1612 | 1620 | + | 11.3303 | 4.96E-05 | 0.615 | GTGGGTGGC |
| 4524 | mamo-s | *BGIBMGA005460* | up | 1233 | 1241 | - | 11.3303 | 4.96E-05 | 0.615 | GTGGGTGGC |
| 4525 | mamo-s | *BGIBMGA005461* | down | 596 | 604 | + | 11.3303 | 4.96E-05 | 0.615 | GTGGGTGGC |
| 4526 | mamo-s | *BGIBMGA005461* | down | 654 | 662 | + | 11.3303 | 4.96E-05 | 0.615 | GTGGGTGGC |
| 4527 | mamo-s | *BGIBMGA005463* | up | 648 | 656 | + | 11.3303 | 4.96E-05 | 0.615 | GTGGGTGGC |
| 4528 | mamo-s | *BGIBMGA005466* | up | 890 | 898 | + | 11.3303 | 4.96E-05 | 0.615 | GTGGGTGGC |
| 4529 | mamo-s | *BGIBMGA005467* | up | 1248 | 1256 | - | 11.3303 | 4.96E-05 | 0.615 | GTGGGTGGC |
| 4530 | mamo-s | *BGIBMGA005468* | down | 250 | 258 | - | 10.6881 | 8.61E-05 | 0.81 | GCACGTGGC |
| 4531 | mamo-s | *BGIBMGA005470* | up | 282 | 290 | - | 11.3303 | 4.96E-05 | 0.615 | GTGGGTGGC |
| 4532 | mamo-s | *BGIBMGA005474* | down | 506 | 514 | + | 11.3303 | 4.96E-05 | 0.615 | GTGGGTGGC |
| 4533 | mamo-s | *BGIBMGA005474* | down | 1422 | 1430 | + | 11.3303 | 4.96E-05 | 0.615 | GTGGGTGGC |
| 4534 | mamo-s | *BGIBMGA005475* | down | 1861 | 1869 | + | 11.3303 | 4.96E-05 | 0.615 | GTGGGTGGC |
| 4535 | mamo-s | *BGIBMGA005476* | up | 477 | 485 | + | 11.3303 | 4.96E-05 | 0.615 | GTGGGTGGC |
| 4536 | mamo-s | *BGIBMGA005476* | down | 1558 | 1566 | + | 11.3303 | 4.96E-05 | 0.615 | GTGGGTGGC |
| 4537 | mamo-s | *BGIBMGA005477* | up | 870 | 878 | - | 11.3303 | 4.96E-05 | 0.615 | GTGGGTGGC |
| 4538 | mamo-s | *BGIBMGA005478* | down | 981 | 989 | - | 13.6606 | 4.06E-06 | 0.615 | GCGCGTGGC |
| 4539 | mamo-s | *BGIBMGA005478* | down | 1460 | 1468 | - | 11.3303 | 4.96E-05 | 0.615 | GTGGGTGGC |
| 4540 | mamo-s | *BGIBMGA005481* | up | 394 | 402 | + | 11.3303 | 4.96E-05 | 0.615 | GTGGGTGGC |
| 4541 | mamo-s | *BGIBMGA005482* | up | 1135 | 1143 | + | 11.3303 | 4.96E-05 | 0.615 | GTGGGTGGC |
| 4542 | mamo-s | *BGIBMGA005483* | down | 1458 | 1466 | - | 11.3303 | 4.96E-05 | 0.615 | GTGGGTGGC |
| 4543 | mamo-s | *BGIBMGA005484* | up | 1315 | 1323 | + | 10.5413 | 9.42E-05 | 0.857 | CAGCGTGTC |
| 4544 | mamo-s | *BGIBMGA005489* | down | 1751 | 1759 | - | 11.3303 | 4.96E-05 | 0.615 | GTGGGTGGC |
| 4545 | mamo-s | *BGIBMGA005493* | down | 139 | 147 | + | 10.5138 | 9.75E-05 | 0.865 | GTGCGTGTT |
| 4546 | mamo-s | *BGIBMGA005493* | down | 818 | 826 | - | 10.5138 | 9.75E-05 | 0.865 | GTGCGTGTT |
| 4547 | mamo-s | *BGIBMGA005494* | up | 191 | 199 | + | 10.5138 | 9.75E-05 | 0.865 | GTGCGTGTT |
| 4548 | mamo-s | *BGIBMGA005494* | up | 870 | 878 | - | 10.5138 | 9.75E-05 | 0.865 | GTGCGTGTT |
| 4549 | mamo-s | *BGIBMGA005502* | down | 367 | 375 | - | 10.8899 | 7.48E-05 | 0.739 | CCGCGTGCC |
| 4550 | mamo-s | *BGIBMGA005502* | down | 262 | 270 | - | 10.8073 | 7.89E-05 | 0.766 | CCGCGTGGT |
| 4551 | mamo-s | *BGIBMGA005503* | up | 1319 | 1327 | + | 11.3303 | 4.96E-05 | 0.615 | GTGGGTGGC |
| 4552 | mamo-s | *BGIBMGA005504* | up | 691 | 699 | + | 11.3303 | 4.96E-05 | 0.615 | GTGGGTGGC |
| 4553 | mamo-s | *BGIBMGA005504* | down | 712 | 720 | + | 11.3303 | 4.96E-05 | 0.615 | GTGGGTGGC |
| 4554 | mamo-s | *BGIBMGA005505* | up | 1551 | 1559 | + | 11.3303 | 4.96E-05 | 0.615 | GTGGGTGGC |
| 4555 | mamo-s | *BGIBMGA005507* | up | 1132 | 1140 | + | 11.3303 | 4.96E-05 | 0.615 | GTGGGTGGC |
| 4556 | mamo-s | *BGIBMGA005507* | down | 1255 | 1263 | + | 11.3303 | 4.96E-05 | 0.615 | GTGGGTGGC |
| 4557 | mamo-s | *BGIBMGA005510* | up | 488 | 496 | - | 11.3303 | 4.96E-05 | 0.615 | GTGGGTGGC |
| 4558 | mamo-s | *BGIBMGA005510* | down | 413 | 421 | + | 11.1743 | 5.90E-05 | 0.684 | CCGCGTGTC |
| 4559 | mamo-s | *BGIBMGA005512* | down | 515 | 523 | - | 11.3303 | 4.96E-05 | 0.615 | GTGGGTGGC |
| 4560 | mamo-s | *BGIBMGA005512* | down | 1238 | 1246 | + | 11.0826 | 6.84E-05 | 0.709 | GTACGTGGC |
| 4561 | mamo-s | *BGIBMGA005513* | down | 1650 | 1658 | - | 10.7523 | 8.38E-05 | 0.797 | AAGCGTGGC |
| 4562 | mamo-s | *BGIBMGA005514* | up | 1472 | 1480 | + | 11.3303 | 4.96E-05 | 0.615 | GTGGGTGGC |
| 4563 | mamo-s | *BGIBMGA005515* | down | 1353 | 1361 | - | 10.8899 | 7.48E-05 | 0.739 | CCGCGTGCC |
| 4564 | mamo-s | *BGIBMGA005516* | down | 462 | 470 | + | 11.3303 | 4.96E-05 | 0.615 | GTGGGTGGC |
| 4565 | mamo-s | *BGIBMGA005520* | down | 1514 | 1522 | - | 13.0275 | 8.53E-06 | 0.615 | GAGCGTGGC |
| 4566 | mamo-s | *BGIBMGA005520* | down | 1694 | 1702 | + | 11.5688 | 3.79E-05 | 0.615 | CTGCGTGTC |
| 4567 | mamo-s | *BGIBMGA005522* | down | 1071 | 1079 | + | 11.3303 | 4.96E-05 | 0.615 | GTGGGTGGC |
| 4568 | mamo-s | *BGIBMGA005523* | up | 1869 | 1877 | + | 11.3303 | 4.96E-05 | 0.615 | GTGGGTGGC |
| 4569 | mamo-s | *BGIBMGA005524* | down | 233 | 241 | - | 11.3303 | 4.96E-05 | 0.615 | GTGGGTGGC |

| | | | | | | | | | | |
|---|---|---|---|---|---|---|---|---|---|---|
| 4570 | mamo-s | *BGIBMGA005524* | up | 442 | 450 | - | 11.3303 | 4.96E-05 | 0.615 | GTGGGTGGC |
| 4571 | mamo-s | *BGIBMGA005524* | up | 1652 | 1660 | - | 11.0826 | 6.84E-05 | 0.709 | GTACGTGGC |
| 4572 | mamo-s | *BGIBMGA005525* | down | 1450 | 1458 | + | 11.3303 | 4.96E-05 | 0.615 | GTGGGTGGC |
| 4573 | mamo-s | *BGIBMGA005526* | down | 586 | 594 | - | 11.2294 | 5.41E-05 | 0.65 | CTGCGAGGC |
| 4574 | mamo-s | *BGIBMGA005527* | up | 1001 | 1009 | - | 11.3303 | 4.96E-05 | 0.615 | GTGGGTGGC |
| 4575 | mamo-s | *BGIBMGA005529* | down | 1411 | 1419 | + | 10.5138 | 9.75E-05 | 0.865 | GTGCGTGTT |
| 4576 | mamo-s | *BGIBMGA005530* | up | 1391 | 1399 | + | 10.5138 | 9.75E-05 | 0.865 | GTGCGTGTT |
| 4577 | mamo-s | *BGIBMGA005536* | up | 1522 | 1530 | - | 11.3303 | 4.96E-05 | 0.615 | GTGGGTGGC |
| 4578 | mamo-s | *BGIBMGA005536* | up | 443 | 451 | + | 10.8349 | 7.66E-05 | 0.751 | CCGCGAGGC |
| 4579 | mamo-s | *BGIBMGA005537* | up | 690 | 698 | + | 11.1743 | 5.90E-05 | 0.684 | CCGCGTGTC |
| 4580 | mamo-s | *BGIBMGA005537* | down | 830 | 838 | + | 11.0826 | 6.84E-05 | 0.709 | GTACGTGGC |
| 4581 | mamo-s | *BGIBMGA005538* | down | 1837 | 1845 | + | 11.3303 | 4.96E-05 | 0.615 | GTGGGTGGC |
| 4582 | mamo-s | *BGIBMGA005539* | down | 620 | 628 | - | 11.3303 | 4.96E-05 | 0.615 | GTGGGTGGC |
| 4583 | mamo-s | *BGIBMGA005541* | down | 1255 | 1263 | - | 11.3303 | 4.96E-05 | 0.615 | GTGGGTGGC |
| 4584 | mamo-s | *BGIBMGA005542* | up | 499 | 507 | - | 11.3303 | 4.96E-05 | 0.615 | GTGGGTGGC |
| 4585 | mamo-s | *BGIBMGA005542* | down | 685 | 693 | - | 11.3303 | 4.96E-05 | 0.615 | GTGGGTGGC |
| 4586 | mamo-s | *BGIBMGA005544* | up | 933 | 941 | + | 11.3303 | 4.96E-05 | 0.615 | GTGGGTGGC |
| 4587 | mamo-s | *BGIBMGA005544* | up | 1037 | 1045 | + | 11.3303 | 4.96E-05 | 0.615 | GTGGGTGGC |
| 4588 | mamo-s | *BGIBMGA005548* | up | 432 | 440 | + | 12.4679 | 1.31E-05 | 0.615 | GTGCGTGTC |
| 4589 | mamo-s | *BGIBMGA005549* | up | 785 | 793 | - | 14.055 | 2.23E-06 | 0.615 | GTGCGTGGC |
| 4590 | mamo-s | *BGIBMGA005550* | up | 863 | 871 | - | 11.3303 | 4.96E-05 | 0.615 | GTGGGTGGC |
| 4591 | mamo-s | *BGIBMGA005555* | down | 689 | 697 | + | 11.3303 | 4.96E-05 | 0.615 | GTGGGTGGC |
| 4592 | mamo-s | *BGIBMGA005556* | down | 1908 | 1916 | + | 13.156 | 6.30E-06 | 0.615 | CTGCGTGGC |
| 4593 | mamo-s | *BGIBMGA005556* | down | 1325 | 1333 | - | 11.5321 | 4.06E-05 | 0.615 | TTGCGTGGC |
| 4594 | mamo-s | *BGIBMGA005556* | down | 1852 | 1860 | - | 11.3303 | 4.96E-05 | 0.615 | GTGGGTGGC |
| 4595 | mamo-s | *BGIBMGA005557* | down | 1565 | 1573 | + | 11.3303 | 4.96E-05 | 0.615 | GTGGGTGGC |
| 4596 | mamo-s | *BGIBMGA005559* | down | 1275 | 1283 | + | 11.3303 | 4.96E-05 | 0.615 | GTGGGTGGC |
| 4597 | mamo-s | *BGIBMGA005560* | down | 1809 | 1817 | - | 11.3303 | 4.96E-05 | 0.615 | GTGGGTGGC |
| 4598 | mamo-s | *BGIBMGA005561* | down | 1279 | 1287 | - | 12.4679 | 1.31E-05 | 0.615 | GTGCGTGTC |
| 4599 | mamo-s | *BGIBMGA005566* | down | 738 | 746 | + | 11.3303 | 4.96E-05 | 0.615 | GTGGGTGGC |
| 4600 | mamo-s | *BGIBMGA005567* | up | 1289 | 1297 | + | 11.3303 | 4.96E-05 | 0.615 | GTGGGTGGC |
| 4601 | mamo-s | *BGIBMGA005569* | up | 1063 | 1071 | - | 10.5138 | 9.75E-05 | 0.865 | GTGCGTGTT |
| 4602 | mamo-s | *BGIBMGA005570* | up | 1516 | 1524 | - | 11.3303 | 4.96E-05 | 0.615 | GTGGGTGGC |
| 4603 | mamo-s | *BGIBMGA005571* | up | 1011 | 1019 | - | 10.8073 | 7.89E-05 | 0.766 | CCGCGTGGT |
| 4604 | mamo-s | *BGIBMGA005572* | down | 796 | 804 | + | 14.055 | 2.23E-06 | 0.615 | GTGCGTGGC |
| 4605 | mamo-s | *BGIBMGA005573* | up | 936 | 944 | + | 11.3303 | 4.96E-05 | 0.615 | GTGGGTGGC |
| 4606 | mamo-s | *BGIBMGA005574* | down | 993 | 1001 | - | 11.4404 | 4.51E-05 | 0.615 | GAGCGTGTC |
| 4607 | mamo-s | *BGIBMGA005574* | down | 1867 | 1875 | + | 11.3303 | 4.96E-05 | 0.615 | GTGGGTGGC |
| 4608 | mamo-s | *BGIBMGA005575* | up | 956 | 964 | - | 11.4404 | 4.51E-05 | 0.615 | GAGCGTGTC |
| 4609 | mamo-s | *BGIBMGA005575* | down | 825 | 833 | + | 11.3303 | 4.96E-05 | 0.615 | GTGGGTGGC |
| 4610 | mamo-s | *BGIBMGA005575* | up | 1830 | 1838 | + | 11.3303 | 4.96E-05 | 0.615 | GTGGGTGGC |
| 4611 | mamo-s | *BGIBMGA005579* | down | 746 | 754 | - | 14.055 | 2.23E-06 | 0.615 | GTGCGTGGC |
| 4612 | mamo-s | *BGIBMGA005580* | up | 787 | 795 | - | 14.055 | 2.23E-06 | 0.615 | GTGCGTGGC |
| 4613 | mamo-s | *BGIBMGA005582* | up | 1665 | 1673 | + | 10.5688 | 8.88E-05 | 0.824 | GTGCGTGAC |
| 4614 | mamo-s | *BGIBMGA005585* | down | 1098 | 1106 | - | 11.3303 | 4.96E-05 | 0.615 | GTGGGTGGC |
| 4615 | mamo-s | *BGIBMGA005586* | up | 1332 | 1340 | + | 11.3303 | 4.96E-05 | 0.615 | GTGGGTGGC |
| 4616 | mamo-s | *BGIBMGA005588* | down | 201 | 209 | + | 11.5321 | 4.06E-05 | 0.615 | TTGCGTGGC |
| 4617 | mamo-s | *BGIBMGA005589* | down | 1525 | 1533 | - | 11.789 | 2.84E-05 | 0.615 | GCGCGTGCC |
| 4618 | mamo-s | *BGIBMGA005589* | down | 1876 | 1884 | + | 10.7523 | 8.38E-05 | 0.797 | GGGCGTGTC |
| 4619 | mamo-s | *BGIBMGA005591* | up | 350 | 358 | - | 11.3303 | 4.96E-05 | 0.615 | GTGGGTGGC |
| 4620 | mamo-s | *BGIBMGA005592* | down | 804 | 812 | + | 11.3303 | 4.96E-05 | 0.615 | GTGGGTGGC |
| 4621 | mamo-s | *BGIBMGA005593* | up | 1514 | 1522 | + | 11.3303 | 4.96E-05 | 0.615 | GTGGGTGGC |
| 4622 | mamo-s | *BGIBMGA005593* | down | 1731 | 1739 | + | 10.9358 | 7.30E-05 | 0.728 | GCGGGTGGC |
| 4623 | mamo-s | *BGIBMGA005595* | up | 933 | 941 | - | 11.3303 | 4.96E-05 | 0.615 | GTGGGTGGC |
| 4624 | mamo-s | *BGIBMGA005596* | up | 1213 | 1221 | - | 11.3303 | 4.96E-05 | 0.615 | GTGGGTGGC |
| 4625 | mamo-s | *BGIBMGA005598* | down | 125 | 133 | - | 11.0826 | 6.84E-05 | 0.709 | GTACGTGGC |
| 4626 | mamo-s | *BGIBMGA005599* | up | 1514 | 1522 | + | 11.3303 | 4.96E-05 | 0.615 | GTGGGTGGC |
| 4627 | mamo-s | *BGIBMGA005600* | down | 1402 | 1410 | - | 11.3303 | 4.96E-05 | 0.615 | GTGGGTGGC |
| 4628 | mamo-s | *BGIBMGA005602* | down | 1736 | 1744 | + | 11.3303 | 4.96E-05 | 0.615 | GTGGGTGGC |
| 4629 | mamo-s | *BGIBMGA005603* | down | 1849 | 1857 | - | 11.3303 | 4.96E-05 | 0.615 | GTGGGTGGC |
| 4630 | mamo-s | *BGIBMGA005604* | down | 1142 | 1150 | - | 11.3303 | 4.96E-05 | 0.615 | GTGGGTGGC |
| 4631 | mamo-s | *BGIBMGA005604* | up | 1242 | 1250 | - | 11.3303 | 4.96E-05 | 0.615 | GTGGGTGGC |
| 4632 | mamo-s | *BGIBMGA005604* | down | 1284 | 1292 | + | 10.5688 | 8.88E-05 | 0.824 | GTGCGTGAC |
| 4633 | mamo-s | *BGIBMGA005605* | up | 1799 | 1807 | - | 11.3303 | 4.96E-05 | 0.615 | GTGGGTGGC |
| 4634 | mamo-s | *BGIBMGA005607* | down | 1618 | 1626 | + | 11.3303 | 4.96E-05 | 0.615 | GTGGGTGGC |
| 4635 | mamo-s | *BGIBMGA005609* | down | 121 | 129 | + | 11.3303 | 4.96E-05 | 0.615 | GTGGGTGGC |
| 4636 | mamo-s | *BGIBMGA005609* | up | 1290 | 1298 | - | 11.3303 | 4.96E-05 | 0.615 | GTGGGTGGC |
| 4637 | mamo-s | *BGIBMGA005610* | up | 123 | 131 | + | 11.3303 | 4.96E-05 | 0.615 | GTGGGTGGC |
| 4638 | mamo-s | *BGIBMGA005611* | down | 1386 | 1394 | + | 11.3303 | 4.96E-05 | 0.615 | GTGGGTGGC |
| 4639 | mamo-s | *BGIBMGA005611* | up | 1765 | 1773 | + | 11.3303 | 4.96E-05 | 0.615 | GTGGGTGGC |
| 4640 | mamo-s | *BGIBMGA005613* | up | 549 | 557 | + | 11.3303 | 4.96E-05 | 0.615 | GTGGGTGGC |
| 4641 | mamo-s | *BGIBMGA005616* | down | 447 | 455 | + | 11.3303 | 4.96E-05 | 0.615 | GTGGGTGGC |
| 4642 | mamo-s | *BGIBMGA005616* | up | 1375 | 1383 | + | 10.5138 | 9.75E-05 | 0.865 | GTGCGTGTT |
| 4643 | mamo-s | *BGIBMGA005617* | down | 333 | 341 | + | 11.156 | 6.13E-05 | 0.705 | GAGCGTGCC |
| 4644 | mamo-s | *BGIBMGA005621* | down | 600 | 608 | + | 14.055 | 2.23E-06 | 0.615 | GTGCGTGGC |
| 4645 | mamo-s | *BGIBMGA005622* | up | 345 | 353 | + | 10.5688 | 8.88E-05 | 0.824 | GTGCGTGAC |
| 4646 | mamo-s | *BGIBMGA005622* | up | 1596 | 1604 | - | 10.5138 | 9.75E-05 | 0.865 | GTGCGTGTT |
| 4647 | mamo-s | *BGIBMGA005623* | up | 872 | 880 | + | 11.3303 | 4.96E-05 | 0.615 | GTGGGTGGC |
| 4648 | mamo-s | *BGIBMGA005624* | up | 706 | 714 | - | 11.0826 | 6.84E-05 | 0.709 | GTACGTGGC |
| 4649 | mamo-s | *BGIBMGA005626* | up | 355 | 363 | + | 11.0826 | 6.84E-05 | 0.709 | GTACGTGGC |
| 4650 | mamo-s | *BGIBMGA005628* | up | 1376 | 1384 | + | 11.3303 | 4.96E-05 | 0.615 | GTGGGTGGC |
| 4651 | mamo-s | *BGIBMGA005629* | down | 162 | 170 | + | 11.3303 | 4.96E-05 | 0.615 | GTGGGTGGC |
| 4652 | mamo-s | *BGIBMGA005630* | down | 814 | 822 | + | 11.3303 | 4.96E-05 | 0.615 | GTGGGTGGC |
| 4653 | mamo-s | *BGIBMGA005633* | up | 65 | 73 | - | 11.3303 | 4.96E-05 | 0.615 | GTGGGTGGC |
| 4654 | mamo-s | *BGIBMGA005633* | up | 711 | 719 | - | 11.1376 | 6.35E-05 | 0.709 | TCGCGTGGC |
| 4655 | mamo-s | *BGIBMGA005634* | down | 351 | 359 | - | 12.7615 | 1.04E-05 | 0.615 | CCGCGTGGC |
| 4656 | mamo-s | *BGIBMGA005635* | up | 924 | 932 | + | 11.3303 | 4.96E-05 | 0.615 | GTGGGTGGC |
| 4657 | mamo-s | *BGIBMGA005639* | down | 396 | 404 | - | 11.1743 | 5.90E-05 | 0.684 | CCGCGTGTC |

| | | | | | | | | | | |
|---|---|---|---|---|---|---|---|---|---|---|
| 4658 | mamo-s | *BGIBMGA005643* | down | 1759 | 1767 | + | 11.3303 | 4.96E-05 | 0.615 | GTGGGTGGC |
| 4659 | mamo-s | *BGIBMGA005644* | up | 1484 | 1492 | + | 11.1009 | 6.57E-05 | 0.709 | GAGCGAGGC |
| 4660 | mamo-s | *BGIBMGA005644* | up | 1230 | 1238 | + | 10.7523 | 8.38E-05 | 0.797 | AAGCGTGGC |
| 4661 | mamo-s | *BGIBMGA005645* | up | 729 | 737 | - | 12.1835 | 1.71E-05 | 0.615 | GTGCGTGCC |
| 4662 | mamo-s | *BGIBMGA005645* | up | 814 | 822 | + | 11.4404 | 4.51E-05 | 0.615 | CGGCGTGGC |
| 4663 | mamo-s | *BGIBMGA005646* | down | 744 | 752 | + | 11.2844 | 5.18E-05 | 0.63 | CTGCGTGCC |
| 4664 | mamo-s | *BGIBMGA005647* | up | 1097 | 1105 | + | 11.3303 | 4.96E-05 | 0.615 | GTGGGTGGC |
| 4665 | mamo-s | *BGIBMGA005652* | down | 1440 | 1448 | + | 11.3303 | 4.96E-05 | 0.615 | GTGGGTGGC |
| 4666 | mamo-s | *BGIBMGA005652* | up | 412 | 420 | + | 11.2844 | 5.18E-05 | 0.63 | CTGCGTGCC |
| 4667 | mamo-s | *BGIBMGA005653* | up | 1841 | 1849 | + | 11.0826 | 6.84E-05 | 0.709 | GTACGTGGC |
| 4668 | mamo-s | *BGIBMGA005654* | down | 296 | 304 | - | 11.3303 | 4.96E-05 | 0.615 | GTGGGTGGC |
| 4669 | mamo-s | *BGIBMGA005654* | down | 630 | 638 | - | 11.0826 | 6.84E-05 | 0.709 | GTACGTGGC |
| 4670 | mamo-s | *BGIBMGA005655* | down | 516 | 524 | + | 11.3303 | 4.96E-05 | 0.615 | GTGGGTGGC |
| 4671 | mamo-s | *BGIBMGA005655* | down | 1812 | 1820 | - | 11.3303 | 4.96E-05 | 0.615 | GTGGGTGGC |
| 4672 | mamo-s | *BGIBMGA005658* | up | 1711 | 1719 | - | 11.3303 | 4.96E-05 | 0.615 | GTGGGTGGC |
| 4673 | mamo-s | *BGIBMGA005660* | up | 1063 | 1071 | + | 11.3303 | 4.96E-05 | 0.615 | GTGGGTGGC |
| 4674 | mamo-s | *BGIBMGA005661* | down | 239 | 247 | - | 11.789 | 2.84E-05 | 0.615 | GCGCGTGCC |
| 4675 | mamo-s | *BGIBMGA005662* | up | 1396 | 1404 | + | 11.1376 | 6.35E-05 | 0.709 | TCGCGTGGC |
| 4676 | mamo-s | *BGIBMGA005663* | up | 881 | 889 | - | 12.4679 | 1.31E-05 | 0.615 | GTGCGTGTC |
| 4677 | mamo-s | *BGIBMGA005663* | up | 1824 | 1832 | - | 10.5688 | 8.88E-05 | 0.824 | GTGCGTGAC |
| 4678 | mamo-s | *BGIBMGA005665* | down | 1135 | 1143 | - | 11.7798 | 3.11E-05 | 0.615 | ATGCGTGGC |
| 4679 | mamo-s | *BGIBMGA005665* | up | 1017 | 1025 | + | 11.3303 | 4.96E-05 | 0.615 | GTGGGTGGC |
| 4680 | mamo-s | *BGIBMGA005665* | down | 1847 | 1855 | - | 11.3303 | 4.96E-05 | 0.615 | GTGGGTGGC |
| 4681 | mamo-s | *BGIBMGA005666* | up | 487 | 495 | - | 11.7798 | 3.11E-05 | 0.615 | ATGCGTGGC |
| 4682 | mamo-s | *BGIBMGA005666* | up | 1199 | 1207 | - | 11.3303 | 4.96E-05 | 0.615 | GTGGGTGGC |
| 4683 | mamo-s | *BGIBMGA005667* | down | 778 | 786 | - | 12.1835 | 1.71E-05 | 0.615 | GTGCGTGCC |
| 4684 | mamo-s | *BGIBMGA005668* | up | 1833 | 1841 | - | 12.1835 | 1.71E-05 | 0.615 | GTGCGTGCC |
| 4685 | mamo-s | *BGIBMGA005672* | down | 225 | 233 | + | 11.3303 | 4.96E-05 | 0.615 | GTGGGTGGC |
| 4686 | mamo-s | *BGIBMGA005673* | down | 1943 | 1951 | + | 11.1743 | 5.90E-05 | 0.684 | CCGCGTGTC |
| 4687 | mamo-s | *BGIBMGA005674* | down | 933 | 941 | - | 11.3303 | 4.96E-05 | 0.615 | GTGGGTGGC |
| 4688 | mamo-s | *BGIBMGA005675* | down | 1745 | 1753 | - | 11.3303 | 4.96E-05 | 0.615 | GTGGGTGGC |
| 4689 | mamo-s | *BGIBMGA005676* | down | 519 | 527 | - | 11.5321 | 4.06E-05 | 0.615 | TTGCGTGGC |
| 4690 | mamo-s | *BGIBMGA005676* | down | 1594 | 1602 | - | 11.3303 | 4.96E-05 | 0.615 | GTGGGTGGC |
| 4691 | mamo-s | *BGIBMGA005677* | up | 977 | 985 | - | 11.3303 | 4.96E-05 | 0.615 | GTGGGTGGC |
| 4692 | mamo-s | *BGIBMGA005677* | down | 1600 | 1608 | - | 11.3303 | 4.96E-05 | 0.615 | GTGGGTGGC |
| 4693 | mamo-s | *BGIBMGA005677* | down | 193 | 201 | - | 11.0826 | 6.84E-05 | 0.709 | GTACGTGGC |
| 4694 | mamo-s | *BGIBMGA005678* | down | 972 | 980 | + | 11.3303 | 4.96E-05 | 0.615 | GTGGGTGGC |
| 4695 | mamo-s | *BGIBMGA005678* | up | 1023 | 1031 | - | 11.3303 | 4.96E-05 | 0.615 | GTGGGTGGC |
| 4696 | mamo-s | *BGIBMGA005678* | up | 1843 | 1851 | + | 11.3303 | 4.96E-05 | 0.615 | GTGGGTGGC |
| 4697 | mamo-s | *BGIBMGA005678* | down | 1558 | 1566 | - | 10.8349 | 7.66E-05 | 0.751 | CCGCGAGGC |
| 4698 | mamo-s | *BGIBMGA005679* | up | 615 | 623 | + | 11.3303 | 4.96E-05 | 0.615 | GTGGGTGGC |
| 4699 | mamo-s | *BGIBMGA005680* | up | 1480 | 1488 | + | 11.3303 | 4.96E-05 | 0.615 | GTGGGTGGC |
| 4700 | mamo-s | *BGIBMGA005686* | down | 169 | 177 | + | 11.3303 | 4.96E-05 | 0.615 | GTGGGTGGC |
| 4701 | mamo-s | *BGIBMGA005691* | down | 1150 | 1158 | - | 13.6606 | 4.06E-06 | 0.615 | GCGCGTGGC |
| 4702 | mamo-s | *BGIBMGA005691* | down | 1383 | 1391 | + | 12.1009 | 2.43E-05 | 0.615 | GTGCGTGGT |
| 4703 | mamo-s | *BGIBMGA005691* | down | 617 | 625 | + | 11.3853 | 4.74E-05 | 0.615 | ACGCGTGGC |
| 4704 | mamo-s | *BGIBMGA005692* | up | 906 | 914 | + | 11.0826 | 6.84E-05 | 0.709 | GTACGTGGC |
| 4705 | mamo-s | *BGIBMGA005693* | up | 1657 | 1665 | - | 11.3303 | 4.96E-05 | 0.615 | GTGGGTGGC |
| 4706 | mamo-s | *BGIBMGA005694* | down | 1171 | 1179 | - | 13.0275 | 8.53E-06 | 0.615 | GAGCGTGGC |
| 4707 | mamo-s | *BGIBMGA005694* | down | 1379 | 1387 | - | 11.3303 | 4.96E-05 | 0.615 | GTGGGTGGC |
| 4708 | mamo-s | *BGIBMGA005694* | up | 1469 | 1477 | - | 11.3303 | 4.96E-05 | 0.615 | GTGGGTGGC |
| 4709 | mamo-s | *BGIBMGA005695* | down | 811 | 819 | - | 12.1284 | 2.16E-05 | 0.615 | GTGCGAGGC |
| 4710 | mamo-s | *BGIBMGA005696* | up | 1873 | 1881 | - | 10.5688 | 8.88E-05 | 0.824 | GTGCGTGAC |
| 4711 | mamo-s | *BGIBMGA005697* | down | 1839 | 1847 | - | 11.3853 | 4.74E-05 | 0.615 | ACGCGTGGC |
| 4712 | mamo-s | *BGIBMGA005697* | down | 836 | 844 | - | 11.1743 | 5.90E-05 | 0.684 | CCGCGTGTC |
| 4713 | mamo-s | *BGIBMGA005698* | up | 607 | 615 | + | 11.3303 | 4.96E-05 | 0.615 | GTGGGTGGC |
| 4714 | mamo-s | *BGIBMGA005699* | down | 951 | 959 | - | 11.3303 | 4.96E-05 | 0.615 | GTGGGTGGC |
| 4715 | mamo-s | *BGIBMGA005700* | down | 1729 | 1737 | + | 12.0734 | 2.66E-05 | 0.615 | GCGCGTGTC |
| 4716 | mamo-s | *BGIBMGA005701* | down | 846 | 854 | - | 12.0734 | 2.66E-05 | 0.615 | GCGCGTGTC |
| 4717 | mamo-s | *BGIBMGA005701* | up | 1873 | 1881 | + | 11.3303 | 4.96E-05 | 0.615 | GTGGGTGGC |
| 4718 | mamo-s | *BGIBMGA005702* | down | 1325 | 1333 | + | 11.3303 | 4.96E-05 | 0.615 | GTGGGTGGC |
| 4719 | mamo-s | *BGIBMGA005703* | up | 666 | 674 | + | 11.0826 | 6.84E-05 | 0.709 | GTACGTGGC |
| 4720 | mamo-s | *BGIBMGA005704* | down | 658 | 666 | - | 10.5413 | 9.42E-05 | 0.857 | CAGCGTGTC |
| 4721 | mamo-s | *BGIBMGA005705* | down | 78 | 86 | - | 12.4679 | 1.31E-05 | 0.615 | GTGCGTGTC |
| 4722 | mamo-s | *BGIBMGA005705* | up | 1052 | 1060 | - | 11.3303 | 4.96E-05 | 0.615 | GTGGGTGGC |
| 4723 | mamo-s | *BGIBMGA005706* | up | 50 | 58 | - | 11.1009 | 6.57E-05 | 0.709 | GAGCGAGGC |
| 4724 | mamo-s | *BGIBMGA005707* | down | 1317 | 1325 | + | 10.5688 | 8.88E-05 | 0.824 | GTGCGTGAC |
| 4725 | mamo-s | *BGIBMGA005708* | up | 1189 | 1197 | + | 12.1835 | 1.71E-05 | 0.615 | GTGCGTGCC |
| 4726 | mamo-s | *BGIBMGA005708* | up | 41 | 49 | - | 11.3303 | 4.96E-05 | 0.615 | GTGGGTGGC |
| 4727 | mamo-s | *BGIBMGA005709* | up | 1340 | 1348 | + | 11.3303 | 4.96E-05 | 0.615 | GTGGGTGGC |
| 4728 | mamo-s | *BGIBMGA005709* | up | 570 | 578 | + | 11.2018 | 5.68E-05 | 0.674 | CTGCGTGGT |
| 4729 | mamo-s | *BGIBMGA005709* | up | 702 | 710 | - | 11.2018 | 5.68E-05 | 0.674 | CTGCGTGGT |
| 4730 | mamo-s | *BGIBMGA005710* | up | 904 | 912 | - | 12.3394 | 1.49E-05 | 0.615 | GGGCGTGGC |
| 4731 | mamo-s | *BGIBMGA005710* | down | 492 | 500 | + | 11.3303 | 4.96E-05 | 0.615 | GTGGGTGGC |
| 4732 | mamo-s | *BGIBMGA005710* | up | 1524 | 1532 | - | 10.9358 | 7.30E-05 | 0.728 | GCGGGTGGC |
| 4733 | mamo-s | *BGIBMGA005711* | up | 622 | 630 | - | 10.7523 | 8.38E-05 | 0.797 | AAGCGTGGC |
| 4734 | mamo-s | *BGIBMGA005712* | up | 628 | 636 | + | 10.5138 | 9.75E-05 | 0.865 | GTGCGTGTT |
| 4735 | mamo-s | *BGIBMGA005715* | up | 1956 | 1964 | - | 10.8073 | 7.89E-05 | 0.766 | CCGCGTGGT |
| 4736 | mamo-s | *BGIBMGA005715* | down | 435 | 443 | + | 10.5413 | 9.42E-05 | 0.857 | CAGCGTGTC |
| 4737 | mamo-s | *BGIBMGA005717* | up | 18 | 26 | + | 11.3303 | 4.96E-05 | 0.615 | GTGGGTGGC |
| 4738 | mamo-s | *BGIBMGA005717* | down | 151 | 159 | - | 11.3303 | 4.96E-05 | 0.615 | GTGGGTGGC |
| 4739 | mamo-s | *BGIBMGA005717* | up | 1490 | 1498 | + | 11.3303 | 4.96E-05 | 0.615 | GTGGGTGGC |
| 4740 | mamo-s | *BGIBMGA005717* | down | 1202 | 1210 | + | 11.2844 | 5.18E-05 | 0.63 | CTGCGTGCC |
| 4741 | mamo-s | *BGIBMGA005718* | up | 474 | 482 | + | 13.6606 | 4.06E-06 | 0.615 | GCGCGTGGC |
| 4742 | mamo-s | *BGIBMGA005718* | up | 745 | 753 | - | 11.3303 | 4.96E-05 | 0.615 | GTGGGTGGC |
| 4743 | mamo-s | *BGIBMGA005718* | up | 1796 | 1804 | + | 11.2844 | 5.18E-05 | 0.63 | CTGCGTGCC |
| 4744 | mamo-s | *BGIBMGA005719* | down | 1304 | 1312 | - | 10.5688 | 8.88E-05 | 0.824 | GTGCGTGAC |
| 4745 | mamo-s | *BGIBMGA005720* | up | 1017 | 1025 | + | 11.3303 | 4.96E-05 | 0.615 | GTGGGTGGC |

| | | | | | | | | | | |
|---|---|---|---|---|---|---|---|---|---|---|
| 4746 | mamo-s | *BGIBMGA005721* | up | 1742 | 1750 | - | 12.1009 | 2.43E-05 | 0.615 | GTGCGTGGT |
| 4747 | mamo-s | *BGIBMGA005721* | up | 1845 | 1853 | + | 12.1009 | 2.43E-05 | 0.615 | GTGCGTGGT |
| 4748 | mamo-s | *BGIBMGA005725* | down | 420 | 428 | + | 11.789 | 2.84E-05 | 0.615 | GCGCGTGCC |
| 4749 | mamo-s | *BGIBMGA005725* | up | 896 | 904 | - | 11.3303 | 4.96E-05 | 0.615 | GTGGGTGGC |
| 4750 | mamo-s | *BGIBMGA005725* | up | 1799 | 1807 | + | 11.3303 | 4.96E-05 | 0.615 | GTGGGTGGC |
| 4751 | mamo-s | *BGIBMGA005725* | down | 137 | 145 | + | 10.7523 | 8.38E-05 | 0.797 | AAGCGTGGC |
| 4752 | mamo-s | *BGIBMGA005728* | down | 356 | 364 | + | 11.3303 | 4.96E-05 | 0.615 | GTGGGTGGC |
| 4753 | mamo-s | *BGIBMGA005728* | up | 868 | 876 | + | 11.3303 | 4.96E-05 | 0.615 | GTGGGTGGC |
| 4754 | mamo-s | *BGIBMGA005729* | up | 32 | 40 | + | 11.3303 | 4.96E-05 | 0.615 | GTGGGTGGC |
| 4755 | mamo-s | *BGIBMGA005730* | down | 1276 | 1284 | + | 11.1743 | 5.90E-05 | 0.684 | CCGCGTGTC |
| 4756 | mamo-s | *BGIBMGA005730* | up | 419 | 427 | - | 11.0734 | 7.12E-05 | 0.726 | GAGCGTGGT |
| 4757 | mamo-s | *BGIBMGA005731* | down | 226 | 234 | + | 11.3303 | 4.96E-05 | 0.615 | GTGGGTGGC |
| 4758 | mamo-s | *BGIBMGA005731* | down | 1410 | 1418 | - | 11.0826 | 6.84E-05 | 0.709 | GTACGTGGC |
| 4759 | mamo-s | *BGIBMGA005732* | up | 1530 | 1538 | + | 12.1009 | 2.43E-05 | 0.615 | GTGCGTGGT |
| 4760 | mamo-s | *BGIBMGA005733* | down | 1719 | 1727 | - | 11.3303 | 4.96E-05 | 0.615 | GTGGGTGGC |
| 4761 | mamo-s | *BGIBMGA005734* | down | 704 | 712 | - | 11.1743 | 5.90E-05 | 0.684 | CCGCGTGTC |
| 4762 | mamo-s | *BGIBMGA005736* | up | 518 | 526 | - | 11.0826 | 6.84E-05 | 0.709 | GTACGTGGC |
| 4763 | mamo-s | *BGIBMGA005740* | up | 143 | 151 | - | 11.3303 | 4.96E-05 | 0.615 | GTGGGTGGC |
| 4764 | mamo-s | *BGIBMGA005746* | down | 150 | 158 | - | 11.3303 | 4.96E-05 | 0.615 | GTGGGTGGC |
| 4765 | mamo-s | *BGIBMGA005746* | up | 959 | 967 | + | 11.3303 | 4.96E-05 | 0.615 | GTGGGTGGC |
| 4766 | mamo-s | *BGIBMGA005748* | down | 1378 | 1386 | + | 11.0826 | 6.84E-05 | 0.709 | GTACGTGGC |
| 4767 | mamo-s | *BGIBMGA005750* | down | 244 | 252 | - | 11.3303 | 4.96E-05 | 0.615 | GTGGGTGGC |
| 4768 | mamo-s | *BGIBMGA005750* | up | 1305 | 1313 | - | 11.0826 | 6.84E-05 | 0.709 | GTACGTGGC |
| 4769 | mamo-s | *BGIBMGA005751* | up | 1514 | 1522 | + | 12.1009 | 2.43E-05 | 0.615 | GTGCGTGGT |
| 4770 | mamo-s | *BGIBMGA005751* | down | 1446 | 1454 | - | 11.3303 | 4.96E-05 | 0.615 | GTGGGTGGC |
| 4771 | mamo-s | *BGIBMGA005753* | down | 1252 | 1260 | - | 11.3303 | 4.96E-05 | 0.615 | GTGGGTGGC |
| 4772 | mamo-s | *BGIBMGA005754* | up | 1245 | 1253 | + | 11.3303 | 4.96E-05 | 0.615 | GTGGGTGGC |
| 4773 | mamo-s | *BGIBMGA005755* | down | 125 | 133 | + | 13.6606 | 4.06E-06 | 0.615 | GCGCGTGGC |
| 4774 | mamo-s | *BGIBMGA005756* | down | 926 | 934 | - | 11.3303 | 4.96E-05 | 0.615 | GTGGGTGGC |
| 4775 | mamo-s | *BGIBMGA005758* | up | 1120 | 1128 | - | 11.3303 | 4.96E-05 | 0.615 | GTGGGTGGC |
| 4776 | mamo-s | *BGIBMGA005758* | down | 94 | 102 | - | 10.9358 | 7.30E-05 | 0.728 | GCGGGTGGC |
| 4777 | mamo-s | *BGIBMGA005758* | up | 921 | 929 | - | 10.9358 | 7.30E-05 | 0.728 | GCGGGTGGC |
| 4778 | mamo-s | *BGIBMGA005761* | up | 240 | 248 | + | 11.3303 | 4.96E-05 | 0.615 | GTGGGTGGC |
| 4779 | mamo-s | *BGIBMGA005762* | down | 494 | 502 | - | 11.3303 | 4.96E-05 | 0.615 | GTGGGTGGC |
| 4780 | mamo-s | *BGIBMGA005762* | up | 766 | 774 | + | 11.3303 | 4.96E-05 | 0.615 | GTGGGTGGC |
| 4781 | mamo-s | *BGIBMGA005764* | up | 1558 | 1566 | - | 11.0734 | 7.12E-05 | 0.726 | GAGCGTGGT |
| 4782 | mamo-s | *BGIBMGA005764* | down | 829 | 837 | - | 10.7523 | 8.38E-05 | 0.797 | GGGCGTGTC |
| 4783 | mamo-s | *BGIBMGA005765* | up | 368 | 376 | + | 11.3303 | 4.96E-05 | 0.615 | GTGGGTGGC |
| 4784 | mamo-s | *BGIBMGA005765* | up | 732 | 740 | + | 11.3303 | 4.96E-05 | 0.615 | GTGGGTGGC |
| 4785 | mamo-s | *BGIBMGA005766* | up | 1040 | 1048 | - | 11.7339 | 3.29E-05 | 0.615 | GCGCGAGGC |
| 4786 | mamo-s | *BGIBMGA005766* | down | 720 | 728 | - | 11.3303 | 4.96E-05 | 0.615 | GTGGGTGGC |
| 4787 | mamo-s | *BGIBMGA005767* | down | 523 | 531 | - | 11.789 | 2.84E-05 | 0.615 | GCGCGTGCC |
| 4788 | mamo-s | *BGIBMGA005767* | up | 1588 | 1596 | + | 11.3303 | 4.96E-05 | 0.615 | GTGGGTGGC |
| 4789 | mamo-s | *BGIBMGA005767* | down | 642 | 650 | + | 11.0826 | 6.84E-05 | 0.709 | GTACGTGGC |
| 4790 | mamo-s | *BGIBMGA005768* | up | 291 | 299 | + | 11.3303 | 4.96E-05 | 0.615 | GTGGGTGGC |
| 4791 | mamo-s | *BGIBMGA005769* | down | 1644 | 1652 | + | 12.3394 | 1.49E-05 | 0.615 | GGGCGTGGC |
| 4792 | mamo-s | *BGIBMGA005769* | up | 1640 | 1648 | - | 11.0826 | 6.84E-05 | 0.709 | GTACGTGGC |
| 4793 | mamo-s | *BGIBMGA005771* | down | 1327 | 1335 | - | 11.3303 | 4.96E-05 | 0.615 | GTGGGTGGC |
| 4794 | mamo-s | *BGIBMGA005772* | down | 1060 | 1068 | + | 10.8349 | 7.66E-05 | 0.751 | CCGCGAGGC |
| 4795 | mamo-s | *BGIBMGA005773* | down | 869 | 877 | - | 11.3303 | 4.96E-05 | 0.615 | GTGGGTGGC |
| 4796 | mamo-s | *BGIBMGA005773* | up | 1650 | 1658 | + | 10.6881 | 8.61E-05 | 0.81 | GCACGTGGC |
| 4797 | mamo-s | *BGIBMGA005776* | down | 741 | 749 | + | 12.1009 | 2.43E-05 | 0.615 | GTGCGTGGT |
| 4798 | mamo-s | *BGIBMGA005776* | up | 71 | 79 | + | 11.3303 | 4.96E-05 | 0.615 | GTGGGTGGC |
| 4799 | mamo-s | *BGIBMGA005776* | down | 982 | 990 | - | 11.156 | 6.13E-05 | 0.705 | GAGCGTGCC |
| 4800 | mamo-s | *BGIBMGA005776* | down | 535 | 543 | + | 10.5413 | 9.42E-05 | 0.857 | GTGCGAGTC |
| 4801 | mamo-s | *BGIBMGA005779* | down | 580 | 588 | - | 11.4404 | 4.51E-05 | 0.615 | GAGCGTGTC |
| 4802 | mamo-s | *BGIBMGA005779* | down | 617 | 625 | + | 11.1743 | 5.90E-05 | 0.684 | CCGCGTGTC |
| 4803 | mamo-s | *BGIBMGA005779* | down | 1260 | 1268 | - | 11.1009 | 6.57E-05 | 0.709 | GAGCGAGGC |
| 4804 | mamo-s | *BGIBMGA005779* | down | 1424 | 1432 | - | 10.5413 | 9.42E-05 | 0.857 | GTGCGAGTC |
| 4805 | mamo-s | *BGIBMGA005781* | down | 1054 | 1062 | - | 11.3303 | 4.96E-05 | 0.615 | GTGGGTGGC |
| 4806 | mamo-s | *BGIBMGA005783* | up | 1456 | 1464 | - | 12.1835 | 1.71E-05 | 0.615 | GTGCGTGCC |
| 4807 | mamo-s | *BGIBMGA005784* | up | 1910 | 1918 | + | 11.156 | 6.13E-05 | 0.705 | GAGCGTGCC |
| 4808 | mamo-s | *BGIBMGA005785* | down | 387 | 395 | + | 11.4404 | 4.51E-05 | 0.615 | GAGCGTGTC |
| 4809 | mamo-s | *BGIBMGA005785* | down | 350 | 358 | - | 11.1743 | 5.90E-05 | 0.684 | CCGCGTGTC |
| 4810 | mamo-s | *BGIBMGA005786* | down | 1010 | 1018 | - | 11.3303 | 4.96E-05 | 0.615 | GTGGGTGGC |
| 4811 | mamo-s | *BGIBMGA005787* | down | 500 | 508 | + | 12.1284 | 2.16E-05 | 0.615 | CAGCGTGGC |
| 4812 | mamo-s | *BGIBMGA005787* | down | 1764 | 1772 | - | 10.8349 | 7.66E-05 | 0.751 | CCGCGAGGC |
| 4813 | mamo-s | *BGIBMGA005788* | down | 1583 | 1591 | - | 11.3303 | 4.96E-05 | 0.615 | GTGGGTGGC |
| 4814 | mamo-s | *BGIBMGA005793* | up | 1710 | 1718 | + | 11.3303 | 4.96E-05 | 0.615 | GTGGGTGGC |
| 4815 | mamo-s | *BGIBMGA005794* | up | 1109 | 1117 | + | 13.0275 | 8.53E-06 | 0.615 | GAGCGTGGC |
| 4816 | mamo-s | *BGIBMGA005794* | down | 961 | 969 | - | 11.3853 | 4.74E-05 | 0.615 | ACGCGTGGC |
| 4817 | mamo-s | *BGIBMGA005794* | down | 1908 | 1916 | - | 10.5413 | 9.42E-05 | 0.857 | GTGCGAGTC |
| 4818 | mamo-s | *BGIBMGA005795* | down | 856 | 864 | + | 13.6606 | 4.06E-06 | 0.615 | GCGCGTGGC |
| 4819 | mamo-s | *BGIBMGA005798* | down | 1649 | 1657 | + | 11.3303 | 4.96E-05 | 0.615 | GTGGGTGGC |
| 4820 | mamo-s | *BGIBMGA005798* | up | 1519 | 1527 | + | 11.0826 | 6.84E-05 | 0.709 | GTACGTGGC |
| 4821 | mamo-s | *BGIBMGA005803* | up | 726 | 734 | - | 11.0826 | 6.84E-05 | 0.709 | GTACGTGGC |
| 4822 | mamo-s | *BGIBMGA005804* | up | 444 | 452 | + | 11.3303 | 4.96E-05 | 0.615 | GTGGGTGGC |
| 4823 | mamo-s | *BGIBMGA005804* | down | 1114 | 1122 | - | 11.3303 | 4.96E-05 | 0.615 | GTGGGTGGC |
| 4824 | mamo-s | *BGIBMGA005806* | up | 1234 | 1242 | + | 11.0734 | 7.12E-05 | 0.726 | GAGCGTGGT |
| 4825 | mamo-s | *BGIBMGA005806* | down | 418 | 426 | - | 10.5413 | 9.42E-05 | 0.857 | GTGCGAGTC |
| 4826 | mamo-s | *BGIBMGA005807* | down | 1553 | 1561 | + | 10.5413 | 9.42E-05 | 0.857 | GTGCGAGTC |
| 4827 | mamo-s | *BGIBMGA005808* | down | 1046 | 1054 | + | 11.3303 | 4.96E-05 | 0.615 | GTGGGTGGC |
| 4828 | mamo-s | *BGIBMGA005810* | down | 1477 | 1485 | - | 11.3303 | 4.96E-05 | 0.615 | GTGGGTGGC |
| 4829 | mamo-s | *BGIBMGA005812* | up | 393 | 401 | - | 11.3303 | 4.96E-05 | 0.615 | GTGGGTGGC |
| 4830 | mamo-s | *BGIBMGA005814* | up | 1610 | 1618 | - | 11.3303 | 4.96E-05 | 0.615 | GTGGGTGGC |
| 4831 | mamo-s | *BGIBMGA005815* | down | 215 | 223 | - | 12.3394 | 1.49E-05 | 0.615 | GGGCGTGGC |
| 4832 | mamo-s | *BGIBMGA005815* | down | 163 | 171 | - | 11.3303 | 4.96E-05 | 0.615 | GTGGGTGGC |
| 4833 | mamo-s | *BGIBMGA005816* | down | 1718 | 1726 | + | 10.9358 | 7.30E-05 | 0.728 | GCGGGTGGC |

| | | | | | | | | | | |
|---|---|---|---|---|---|---|---|---|---|---|
| 4834 | mamo-s | *BGIBMGA005818* | up | 1017 | 1025 | - | 11.3303 | 4.96E-05 | 0.615 | GTGGGTGGC |
| 4835 | mamo-s | *BGIBMGA005819* | down | 1559 | 1567 | + | 11.3303 | 4.96E-05 | 0.615 | GTGGGTGGC |
| 4836 | mamo-s | *BGIBMGA005820* | up | 353 | 361 | + | 11.3303 | 4.96E-05 | 0.615 | GTGGGTGGC |
| 4837 | mamo-s | *BGIBMGA005820* | down | 1685 | 1693 | - | 11.1743 | 5.90E-05 | 0.684 | CCGCGTGTC |
| 4838 | mamo-s | *BGIBMGA005822* | down | 1744 | 1752 | + | 11.3303 | 4.96E-05 | 0.615 | GTGGGTGGC |
| 4839 | mamo-s | *BGIBMGA005824* | down | 922 | 930 | + | 10.6881 | 8.61E-05 | 0.81 | GCACGTGGC |
| 4840 | mamo-s | *BGIBMGA005828* | down | 1180 | 1188 | + | 11.1743 | 5.90E-05 | 0.684 | CCGCGTGTC |
| 4841 | mamo-s | *BGIBMGA005830* | down | 684 | 692 | + | 11.0826 | 6.84E-05 | 0.709 | GTACGTGGC |
| 4842 | mamo-s | *BGIBMGA005830* | down | 1801 | 1809 | - | 11.0734 | 7.12E-05 | 0.726 | GAGCGTGGT |
| 4843 | mamo-s | *BGIBMGA005831* | down | 1631 | 1639 | + | 12.1284 | 2.16E-05 | 0.615 | GTGCGAGGC |
| 4844 | mamo-s | *BGIBMGA005831* | down | 476 | 484 | - | 10.9358 | 7.30E-05 | 0.728 | GCGGGTGGC |
| 4845 | mamo-s | *BGIBMGA005832* | up | 1448 | 1456 | + | 12.1284 | 2.16E-05 | 0.615 | GTGCGAGGC |
| 4846 | mamo-s | *BGIBMGA005832* | down | 1594 | 1602 | - | 11.3303 | 4.96E-05 | 0.615 | GTGGGTGGC |
| 4847 | mamo-s | *BGIBMGA005832* | up | 293 | 301 | - | 10.9358 | 7.30E-05 | 0.728 | GCGGGTGGC |
| 4848 | mamo-s | *BGIBMGA005834* | up | 1293 | 1301 | - | 11.0826 | 6.84E-05 | 0.709 | GTACGTGGC |
| 4849 | mamo-s | *BGIBMGA005835* | down | 672 | 680 | + | 11.3303 | 4.96E-05 | 0.615 | GTGGGTGGC |
| 4850 | mamo-s | *BGIBMGA005837* | up | 135 | 143 | - | 12.1009 | 2.43E-05 | 0.615 | GTGCGTGGT |
| 4851 | mamo-s | *BGIBMGA005837* | up | 861 | 869 | - | 11.7798 | 3.11E-05 | 0.615 | ATGCGTGGC |
| 4852 | mamo-s | *BGIBMGA005837* | up | 1028 | 1036 | - | 11.3303 | 4.96E-05 | 0.615 | GTGGGTGGC |
| 4853 | mamo-s | *BGIBMGA005837* | down | 1761 | 1769 | - | 10.5688 | 8.88E-05 | 0.824 | GTGCGTGAC |
| 4854 | mamo-s | *BGIBMGA005839* | down | 649 | 657 | + | 12.1284 | 2.16E-05 | 0.615 | CAGCGTGGC |
| 4855 | mamo-s | *BGIBMGA005841* | up | 299 | 307 | - | 10.6881 | 8.61E-05 | 0.81 | GCACGTGGC |
| 4856 | mamo-s | *BGIBMGA005842* | down | 718 | 726 | - | 11.1743 | 5.90E-05 | 0.684 | CCGCGTGTC |
| 4857 | mamo-s | *BGIBMGA005843* | up | 1213 | 1221 | + | 13.6606 | 4.06E-06 | 0.615 | GCGCGTGGC |
| 4858 | mamo-s | *BGIBMGA005843* | up | 1544 | 1552 | - | 11.3303 | 4.96E-05 | 0.615 | GTGGGTGGC |
| 4859 | mamo-s | *BGIBMGA005845* | down | 695 | 703 | - | 11.4404 | 4.51E-05 | 0.615 | CGGCGTGGC |
| 4860 | mamo-s | *BGIBMGA005846* | up | 1046 | 1054 | - | 11.4404 | 4.51E-05 | 0.615 | CGGCGTGGC |
| 4861 | mamo-s | *BGIBMGA005847* | up | 1050 | 1058 | + | 11.3853 | 4.74E-05 | 0.615 | ACGCGTGGC |
| 4862 | mamo-s | *BGIBMGA005847* | up | 1724 | 1732 | + | 10.8349 | 7.66E-05 | 0.751 | CCGCGAGGC |
| 4863 | mamo-s | *BGIBMGA005848* | down | 1866 | 1874 | - | 10.8073 | 7.89E-05 | 0.766 | CCGCGTGGT |
| 4864 | mamo-s | *BGIBMGA005848* | down | 1442 | 1450 | - | 10.5413 | 9.42E-05 | 0.857 | GTGCGAGTC |
| 4865 | mamo-s | *BGIBMGA005849* | down | 305 | 313 | - | 10.8073 | 7.89E-05 | 0.766 | CCGCGTGGT |
| 4866 | mamo-s | *BGIBMGA005850* | down | 110 | 118 | + | 11.0734 | 7.12E-05 | 0.726 | GAGCGTGGT |
| 4867 | mamo-s | *BGIBMGA005851* | down | 770 | 778 | - | 11.4404 | 4.51E-05 | 0.615 | CGGCGTGGC |
| 4868 | mamo-s | *BGIBMGA005856* | down | 905 | 913 | - | 13.0275 | 8.53E-06 | 0.615 | GAGCGTGGC |
| 4869 | mamo-s | *BGIBMGA005856* | down | 504 | 512 | - | 11.3303 | 4.96E-05 | 0.615 | GTGGGTGGC |
| 4870 | mamo-s | *BGIBMGA005857* | down | 1060 | 1068 | + | 11.3303 | 4.96E-05 | 0.615 | GTGGGTGGC |
| 4871 | mamo-s | *BGIBMGA005858* | up | 223 | 231 | + | 12.1835 | 1.71E-05 | 0.615 | GTGCGTGCC |
| 4872 | mamo-s | *BGIBMGA005858* | down | 713 | 721 | + | 11.4404 | 4.51E-05 | 0.615 | CGGCGTGGC |
| 4873 | mamo-s | *BGIBMGA005859* | up | 33 | 41 | - | 11.156 | 6.13E-05 | 0.705 | GAGCGTGCC |
| 4874 | mamo-s | *BGIBMGA005860* | down | 1985 | 1993 | + | 11.3303 | 4.96E-05 | 0.615 | GTGGGTGGC |
| 4875 | mamo-s | *BGIBMGA005861* | up | 1514 | 1522 | + | 11.5321 | 4.06E-05 | 0.615 | TTGCGTGGC |
| 4876 | mamo-s | *BGIBMGA005861* | up | 1565 | 1573 | - | 11.4404 | 4.51E-05 | 0.615 | CGGCGTGGC |
| 4877 | mamo-s | *BGIBMGA005861* | down | 1947 | 1955 | + | 11.4404 | 4.51E-05 | 0.615 | GAGCGTGTC |
| 4878 | mamo-s | *BGIBMGA005861* | up | 505 | 513 | - | 11.156 | 6.13E-05 | 0.705 | GAGCGTGCC |
| 4879 | mamo-s | *BGIBMGA005861* | up | 1599 | 1607 | - | 11.156 | 6.13E-05 | 0.705 | GAGCGTGCC |
| 4880 | mamo-s | *BGIBMGA005861* | up | 14 | 22 | + | 10.5138 | 9.75E-05 | 0.865 | GTGCGTGTT |
| 4881 | mamo-s | *BGIBMGA005862* | down | 363 | 371 | + | 11.3303 | 4.96E-05 | 0.615 | GTGGGTGGC |
| 4882 | mamo-s | *BGIBMGA005863* | down | 1071 | 1079 | - | 11.7064 | 3.52E-05 | 0.615 | GCGCGTGGT |
| 4883 | mamo-s | *BGIBMGA005863* | down | 1292 | 1300 | - | 11.2294 | 5.41E-05 | 0.65 | CTGCGAGGC |
| 4884 | mamo-s | *BGIBMGA005867* | down | 1407 | 1415 | - | 12.1284 | 2.16E-05 | 0.615 | CAGCGTGGC |
| 4885 | mamo-s | *BGIBMGA005870* | down | 1585 | 1593 | - | 12.4679 | 1.31E-05 | 0.615 | GTGCGTGTC |
| 4886 | mamo-s | *BGIBMGA005870* | down | 1645 | 1653 | - | 12.4679 | 1.31E-05 | 0.615 | GTGCGTGTC |
| 4887 | mamo-s | *BGIBMGA005870* | down | 1551 | 1559 | + | 11.4404 | 4.51E-05 | 0.615 | CGGCGTGGC |
| 4888 | mamo-s | *BGIBMGA005870* | up | 1980 | 1988 | + | 11.4404 | 4.51E-05 | 0.615 | CGGCGTGGC |
| 4889 | mamo-s | *BGIBMGA005871* | up | 818 | 826 | - | 12.4679 | 1.31E-05 | 0.615 | GTGCGTGTC |
| 4890 | mamo-s | *BGIBMGA005871* | up | 878 | 886 | - | 12.4679 | 1.31E-05 | 0.615 | GTGCGTGTC |
| 4891 | mamo-s | *BGIBMGA005871* | up | 784 | 792 | + | 11.4404 | 4.51E-05 | 0.615 | CGGCGTGGC |
| 4892 | mamo-s | *BGIBMGA005872* | up | 700 | 708 | - | 11.3303 | 4.96E-05 | 0.615 | GTGGGTGGC |
| 4893 | mamo-s | *BGIBMGA005873* | up | 1742 | 1750 | - | 10.9358 | 7.30E-05 | 0.728 | GCGGGTGGC |
| 4894 | mamo-s | *BGIBMGA005874* | up | 465 | 473 | + | 11.5688 | 3.79E-05 | 0.615 | CTGCGTGTC |
| 4895 | mamo-s | *BGIBMGA005875* | down | 1935 | 1943 | - | 11.3303 | 4.96E-05 | 0.615 | GTGGGTGGC |
| 4896 | mamo-s | *BGIBMGA005879* | up | 1804 | 1812 | + | 11.3303 | 4.96E-05 | 0.615 | GTGGGTGGC |
| 4897 | mamo-s | *BGIBMGA005880* | down | 1748 | 1756 | - | 13.6606 | 4.06E-06 | 0.615 | GCGCGTGGC |
| 4898 | mamo-s | *BGIBMGA005880* | down | 41 | 49 | + | 13.0275 | 8.53E-06 | 0.615 | GAGCGTGGC |
| 4899 | mamo-s | *BGIBMGA005881* | up | 1539 | 1547 | + | 13.0275 | 8.53E-06 | 0.615 | GAGCGTGGC |
| 4900 | mamo-s | *BGIBMGA005882* | up | 1363 | 1371 | + | 11.3303 | 4.96E-05 | 0.615 | GTGGGTGGC |
| 4901 | mamo-s | *BGIBMGA005883* | down | 1258 | 1266 | - | 11.3303 | 4.96E-05 | 0.615 | GTGGGTGGC |
| 4902 | mamo-s | *BGIBMGA005885* | down | 851 | 859 | + | 11.1009 | 6.57E-05 | 0.709 | GAGCGAGGC |
| 4903 | mamo-s | *BGIBMGA005886* | down | 1296 | 1304 | - | 11.7798 | 3.11E-05 | 0.615 | ATGCGTGGC |
| 4904 | mamo-s | *BGIBMGA005886* | down | 965 | 973 | + | 11.3303 | 4.96E-05 | 0.615 | GTGGGTGGC |
| 4905 | mamo-s | *BGIBMGA005895* | down | 1673 | 1681 | - | 11.0826 | 6.84E-05 | 0.709 | GTACGTGGC |
| 4906 | mamo-s | *BGIBMGA005896* | down | 1128 | 1136 | + | 14.055 | 2.23E-06 | 0.615 | GTGCGTGGC |
| 4907 | mamo-s | *BGIBMGA005896* | down | 1897 | 1905 | + | 11.3303 | 4.96E-05 | 0.615 | GTGGGTGGC |
| 4908 | mamo-s | *BGIBMGA005898* | down | 1989 | 1997 | + | 12.1284 | 2.16E-05 | 0.615 | CAGCGTGGC |
| 4909 | mamo-s | *BGIBMGA005907* | up | 1126 | 1134 | - | 11.3303 | 4.96E-05 | 0.615 | GTGGGTGGC |
| 4910 | mamo-s | *BGIBMGA005909* | up | 12 | 20 | - | 12.0734 | 2.66E-05 | 0.615 | GCGCGTGTC |
| 4911 | mamo-s | *BGIBMGA005909* | up | 221 | 229 | + | 11.0734 | 7.12E-05 | 0.726 | GAGCGTGGT |
| 4912 | mamo-s | *BGIBMGA005914* | down | 395 | 403 | + | 11.2844 | 5.18E-05 | 0.63 | CTGCGTGCC |
| 4913 | mamo-s | *BGIBMGA005914* | down | 189 | 197 | + | 11.2294 | 5.41E-05 | 0.65 | CTGCGAGGC |
| 4914 | mamo-s | *BGIBMGA005915* | up | 1372 | 1380 | + | 11.2844 | 5.18E-05 | 0.63 | CTGCGTGCC |
| 4915 | mamo-s | *BGIBMGA005915* | up | 1166 | 1174 | + | 11.2294 | 5.41E-05 | 0.65 | CTGCGAGGC |
| 4916 | mamo-s | *BGIBMGA005917* | down | 408 | 416 | - | 11.7339 | 3.29E-05 | 0.615 | GCGCGAGGC |
| 4917 | mamo-s | *BGIBMGA005919* | down | 1878 | 1886 | + | 10.5138 | 9.75E-05 | 0.865 | GTGCGTGTT |
| 4918 | mamo-s | *BGIBMGA005920* | down | 1805 | 1813 | + | 11.5688 | 3.79E-05 | 0.615 | CTGCGTGTC |
| 4919 | mamo-s | *BGIBMGA005920* | up | 1547 | 1555 | - | 11.3303 | 4.96E-05 | 0.615 | GTGGGTGGC |
| 4920 | mamo-s | *BGIBMGA005921* | down | 1353 | 1361 | + | 11.3303 | 4.96E-05 | 0.615 | GTGGGTGGC |
| 4921 | mamo-s | *BGIBMGA005922* | down | 1469 | 1477 | + | 11.3303 | 4.96E-05 | 0.615 | GTGGGTGGC |

| | | | | | | | | | | |
|---|---|---|---|---|---|---|---|---|---|---|
| 4922 | mamo-s | *BGIBMGA005923* | up | 575 | 583 | - | 11.3303 | 4.96E-05 | 0.615 | GTGGGTGGC |
| 4923 | mamo-s | *BGIBMGA005927* | up | 550 | 558 | + | 11.3853 | 4.74E-05 | 0.615 | ACGCGTGGC |
| 4924 | mamo-s | *BGIBMGA005928* | down | 1444 | 1452 | + | 11.1743 | 5.90E-05 | 0.684 | CCGCGTGTC |
| 4925 | mamo-s | *BGIBMGA005930* | up | 1812 | 1820 | + | 12.4679 | 1.31E-05 | 0.615 | GTGCGTGTC |
| 4926 | mamo-s | *BGIBMGA005931* | up | 411 | 419 | - | 12.3394 | 1.49E-05 | 0.615 | GGGCGTGGC |
| 4927 | mamo-s | *BGIBMGA005931* | up | 1481 | 1489 | + | 10.8899 | 7.48E-05 | 0.739 | CCGCGTGCC |
| 4928 | mamo-s | *BGIBMGA005932* | up | 103 | 111 | + | 11.3303 | 4.96E-05 | 0.615 | GTGGGTGGC |
| 4929 | mamo-s | *BGIBMGA005932* | down | 905 | 913 | + | 11.2844 | 5.18E-05 | 0.63 | CTGCGTGCC |
| 4930 | mamo-s | *BGIBMGA005932* | down | 153 | 161 | + | 11.0734 | 7.12E-05 | 0.726 | GAGCGTGGT |
| 4931 | mamo-s | *BGIBMGA005933* | up | 996 | 1004 | + | 10.5688 | 8.88E-05 | 0.824 | GTGCGTGAC |
| 4932 | mamo-s | *BGIBMGA005934* | down | 227 | 235 | - | 12.1009 | 2.43E-05 | 0.615 | GTGCGTGGT |
| 4933 | mamo-s | *BGIBMGA005934* | up | 269 | 277 | + | 11.2018 | 5.68E-05 | 0.674 | CTGCGTGGT |
| 4934 | mamo-s | *BGIBMGA005935* | down | 1183 | 1191 | - | 11.3303 | 4.96E-05 | 0.615 | GTGGGTGGC |
| 4935 | mamo-s | *BGIBMGA005936* | down | 966 | 974 | - | 11.3303 | 4.96E-05 | 0.615 | GTGGGTGGC |
| 4936 | mamo-s | *BGIBMGA005939* | down | 817 | 825 | + | 10.5138 | 9.75E-05 | 0.865 | GTGCGTGTT |
| 4937 | mamo-s | *BGIBMGA005940* | up | 925 | 933 | + | 11.3303 | 4.96E-05 | 0.615 | GTGGGTGGC |
| 4938 | mamo-s | *BGIBMGA005940* | down | 1034 | 1042 | - | 11.1009 | 6.57E-05 | 0.709 | GAGCGAGGC |
| 4939 | mamo-s | *BGIBMGA005940* | down | 1331 | 1339 | - | 11.1009 | 6.57E-05 | 0.709 | GAGCGAGGC |
| 4940 | mamo-s | *BGIBMGA005941* | down | 939 | 947 | - | 11.3303 | 4.96E-05 | 0.615 | GTGGGTGGC |
| 4941 | mamo-s | *BGIBMGA005942* | up | 303 | 311 | + | 11.3303 | 4.96E-05 | 0.615 | GTGGGTGGC |
| 4942 | mamo-s | *BGIBMGA005942* | up | 902 | 910 | + | 11.3303 | 4.96E-05 | 0.615 | GTGGGTGGC |
| 4943 | mamo-s | *BGIBMGA005942* | up | 1425 | 1433 | - | 11.3303 | 4.96E-05 | 0.615 | GTGGGTGGC |
| 4944 | mamo-s | *BGIBMGA005943* | up | 879 | 887 | - | 12.1284 | 2.16E-05 | 0.615 | GTGCGAGGC |
| 4945 | mamo-s | *BGIBMGA005944* | up | 909 | 917 | - | 13.156 | 6.30E-06 | 0.615 | CTGCGTGGC |
| 4946 | mamo-s | *BGIBMGA005944* | up | 592 | 600 | - | 11.3303 | 4.96E-05 | 0.615 | GTGGGTGGC |
| 4947 | mamo-s | *BGIBMGA005946* | up | 729 | 737 | + | 11.3303 | 4.96E-05 | 0.615 | GTGGGTGGC |
| 4948 | mamo-s | *BGIBMGA005946* | up | 47 | 55 | - | 10.8073 | 7.89E-05 | 0.766 | CCGCGTGGT |
| 4949 | mamo-s | *BGIBMGA005948* | down | 1203 | 1211 | + | 11.1376 | 6.35E-05 | 0.709 | TCGCGTGGC |
| 4950 | mamo-s | *BGIBMGA005949* | down | 1713 | 1721 | - | 11.3303 | 4.96E-05 | 0.615 | GTGGGTGGC |
| 4951 | mamo-s | *BGIBMGA005950* | down | 1802 | 1810 | - | 11.3303 | 4.96E-05 | 0.615 | GTGGGTGGC |
| 4952 | mamo-s | *BGIBMGA005951* | up | 1428 | 1436 | - | 11.3303 | 4.96E-05 | 0.615 | GTGGGTGGC |
| 4953 | mamo-s | *BGIBMGA005955* | down | 860 | 868 | + | 12.4679 | 1.31E-05 | 0.615 | GTGCGTGTC |
| 4954 | mamo-s | *BGIBMGA005955* | up | 656 | 664 | - | 10.9358 | 7.30E-05 | 0.728 | GCGGGTGGC |
| 4955 | mamo-s | *BGIBMGA005957* | down | 906 | 914 | + | 12.1009 | 2.43E-05 | 0.615 | GTGCGTGGT |
| 4956 | mamo-s | *BGIBMGA005958* | down | 439 | 447 | + | 12.1009 | 2.43E-05 | 0.615 | GTGCGTGGT |
| 4957 | mamo-s | *BGIBMGA005958* | up | 1531 | 1539 | + | 10.5138 | 9.75E-05 | 0.865 | GTGCGTGTT |
| 4958 | mamo-s | *BGIBMGA005961* | down | 1720 | 1728 | + | 11.3303 | 4.96E-05 | 0.615 | GTGGGTGGC |
| 4959 | mamo-s | *BGIBMGA005962* | down | 825 | 833 | + | 11.1743 | 5.90E-05 | 0.684 | CCGCGTGTC |
| 4960 | mamo-s | *BGIBMGA005968* | up | 1209 | 1217 | - | 10.5138 | 9.75E-05 | 0.865 | GTGCGTGTT |
| 4961 | mamo-s | *BGIBMGA005973* | down | 1209 | 1217 | + | 12.4679 | 1.31E-05 | 0.615 | GTGCGTGTC |
| 4962 | mamo-s | *BGIBMGA005974* | up | 1386 | 1394 | + | 11.7798 | 3.11E-05 | 0.615 | ATGCGTGGC |
| 4963 | mamo-s | *BGIBMGA005977* | down | 1632 | 1640 | - | 12.1835 | 1.71E-05 | 0.615 | GTGCGTGCC |
| 4964 | mamo-s | *BGIBMGA005977* | up | 1732 | 1740 | + | 11.3303 | 4.96E-05 | 0.615 | GTGGGTGGC |
| 4965 | mamo-s | *BGIBMGA005978* | down | 1403 | 1411 | + | 11.3303 | 4.96E-05 | 0.615 | GTGGGTGGC |
| 4966 | mamo-s | *BGIBMGA005980* | up | 1440 | 1448 | - | 10.7523 | 8.38E-05 | 0.797 | AAGCGTGGC |
| 4967 | mamo-s | *BGIBMGA005981* | up | 323 | 331 | + | 11.2018 | 5.68E-05 | 0.674 | CTGCGTGGT |
| 4968 | mamo-s | *BGIBMGA005982* | up | 148 | 156 | + | 12.1835 | 1.71E-05 | 0.615 | GTGCGTGCC |
| 4969 | mamo-s | *BGIBMGA005982* | up | 544 | 552 | + | 11.3303 | 4.96E-05 | 0.615 | GTGGGTGGC |
| 4970 | mamo-s | *BGIBMGA005982* | down | 926 | 934 | - | 11.0826 | 6.84E-05 | 0.709 | GTACGTGGC |
| 4971 | mamo-s | *BGIBMGA005983* | down | 987 | 995 | - | 11.7339 | 3.29E-05 | 0.615 | GCGCGAGGC |
| 4972 | mamo-s | *BGIBMGA005983* | down | 1065 | 1073 | - | 11.7339 | 3.29E-05 | 0.615 | GCGCGAGGC |
| 4973 | mamo-s | *BGIBMGA005983* | up | 824 | 832 | - | 11.0826 | 6.84E-05 | 0.709 | GTACGTGGC |
| 4974 | mamo-s | *BGIBMGA005984* | up | 7 | 15 | - | 12.1284 | 2.16E-05 | 0.615 | GTGCGAGGC |
| 4975 | mamo-s | *BGIBMGA005986* | down | 1832 | 1840 | + | 11.3303 | 4.96E-05 | 0.615 | GTGGGTGGC |
| 4976 | mamo-s | *BGIBMGA005986* | down | 1941 | 1949 | - | 11.3303 | 4.96E-05 | 0.615 | GTGGGTGGC |
| 4977 | mamo-s | *BGIBMGA005990* | up | 1906 | 1914 | + | 10.7523 | 8.38E-05 | 0.797 | AAGCGTGGC |
| 4978 | mamo-s | *BGIBMGA005993* | up | 1588 | 1596 | + | 11.2018 | 5.68E-05 | 0.674 | CTGCGTGGT |
| 4979 | mamo-s | *BGIBMGA005994* | down | 1397 | 1405 | + | 12.4679 | 1.31E-05 | 0.615 | GTGCGTGTC |
| 4980 | mamo-s | *BGIBMGA005994* | down | 942 | 950 | + | 11.3303 | 4.96E-05 | 0.615 | GTGGGTGGC |
| 4981 | mamo-s | *BGIBMGA005995* | up | 1612 | 1620 | + | 12.4679 | 1.31E-05 | 0.615 | GTGCGTGTC |
| 4982 | mamo-s | *BGIBMGA005995* | up | 1157 | 1165 | + | 11.3303 | 4.96E-05 | 0.615 | GTGGGTGGC |
| 4983 | mamo-s | *BGIBMGA005996* | down | 625 | 633 | - | 11.5321 | 4.06E-05 | 0.615 | TTGCGTGGC |
| 4984 | mamo-s | *BGIBMGA005997* | down | 1662 | 1670 | + | 12.4679 | 1.31E-05 | 0.615 | GTGCGTGTC |
| 4985 | mamo-s | *BGIBMGA005998* | up | 1287 | 1295 | + | 11.3303 | 4.96E-05 | 0.615 | GTGGGTGGC |
| 4986 | mamo-s | *BGIBMGA005998* | down | 614 | 622 | + | 11.1009 | 6.57E-05 | 0.709 | GAGCGAGGC |
| 4987 | mamo-s | *BGIBMGA006000* | up | 523 | 531 | - | 11.3303 | 4.96E-05 | 0.615 | GTGGGTGGC |
| 4988 | mamo-s | *BGIBMGA006000* | up | 1087 | 1095 | - | 11.3303 | 4.96E-05 | 0.615 | GTGGGTGGC |
| 4989 | mamo-s | *BGIBMGA006002* | up | 1523 | 1531 | - | 11.3303 | 4.96E-05 | 0.615 | GTGGGTGGC |
| 4990 | mamo-s | *BGIBMGA006004* | up | 644 | 652 | + | 11.4404 | 4.51E-05 | 0.615 | GAGCGTGTC |
| 4991 | mamo-s | *BGIBMGA006007* | down | 1557 | 1565 | - | 11.3303 | 4.96E-05 | 0.615 | GTGGGTGGC |
| 4992 | mamo-s | *BGIBMGA006009* | up | 1806 | 1814 | - | 12.4679 | 1.31E-05 | 0.615 | GTGCGTGTC |
| 4993 | mamo-s | *BGIBMGA006009* | up | 513 | 521 | - | 11.0826 | 6.84E-05 | 0.709 | GTACGTGGC |
| 4994 | mamo-s | *BGIBMGA006010* | down | 661 | 669 | + | 11.5321 | 4.06E-05 | 0.615 | TTGCGTGGC |
| 4995 | mamo-s | *BGIBMGA006010* | down | 1147 | 1155 | - | 11.3303 | 4.96E-05 | 0.615 | GTGGGTGGC |
| 4996 | mamo-s | *BGIBMGA006011* | down | 1831 | 1839 | + | 12.7615 | 1.04E-05 | 0.615 | CCGCGTGGC |
| 4997 | mamo-s | *BGIBMGA006011* | up | 109 | 117 | + | 11.5321 | 4.06E-05 | 0.615 | TTGCGTGGC |
| 4998 | mamo-s | *BGIBMGA006011* | up | 595 | 603 | - | 11.3303 | 4.96E-05 | 0.615 | GTGGGTGGC |
| 4999 | mamo-s | *BGIBMGA006013* | down | 1772 | 1780 | - | 11.3303 | 4.96E-05 | 0.615 | GTGGGTGGC |
| 5000 | mamo-s | *BGIBMGA006015* | down | 1954 | 1962 | + | 11.5688 | 3.79E-05 | 0.615 | CTGCGTGTC |
| 5001 | mamo-s | *BGIBMGA006015* | down | 1636 | 1644 | + | 11.3303 | 4.96E-05 | 0.615 | GTGGGTGGC |
| 5002 | mamo-s | *BGIBMGA006019* | down | 1372 | 1380 | + | 11.3303 | 4.96E-05 | 0.615 | GTGGGTGGC |
| 5003 | mamo-s | *BGIBMGA006029* | down | 573 | 581 | - | 12.1009 | 2.43E-05 | 0.615 | GTGCGTGGT |
| 5004 | mamo-s | *BGIBMGA006031* | up | 497 | 505 | - | 11.3303 | 4.96E-05 | 0.615 | GTGGGTGGC |
| 5005 | mamo-s | *BGIBMGA006032* | up | 956 | 964 | + | 11.4404 | 4.51E-05 | 0.615 | GAGCGTGTC |
| 5006 | mamo-s | *BGIBMGA006033* | up | 1147 | 1155 | + | 11.3303 | 4.96E-05 | 0.615 | GTGGGTGGC |
| 5007 | mamo-s | *BGIBMGA006035* | up | 896 | 904 | - | 11.3303 | 4.96E-05 | 0.615 | GTGGGTGGC |
| 5008 | mamo-s | *BGIBMGA006035* | down | 1400 | 1408 | + | 10.7523 | 8.38E-05 | 0.797 | GGGCGTGTC |
| 5009 | mamo-s | *BGIBMGA006041* | up | 1172 | 1180 | - | 14.055 | 2.23E-06 | 0.615 | GTGCGTGGC |

| | | | | | | | | | | |
|---|---|---|---|---|---|---|---|---|---|---|
| 5010 | mamo-s | *BGIBMGA006042* | up | 1319 | 1327 | + | 12.1284 | 2.16E-05 | 0.615 | CAGCGTGGC |
| 5011 | mamo-s | *BGIBMGA006042* | up | 1349 | 1357 | + | 10.9358 | 7.30E-05 | 0.728 | GCGGGTGGC |
| 5012 | mamo-s | *BGIBMGA006044* | down | 1093 | 1101 | + | 11.3303 | 4.96E-05 | 0.615 | GTGGGTGGC |
| 5013 | mamo-s | *BGIBMGA006045* | down | 400 | 408 | - | 12.4679 | 1.31E-05 | 0.615 | GTGCGTGTC |
| 5014 | mamo-s | *BGIBMGA006045* | down | 369 | 377 | - | 10.5138 | 9.75E-05 | 0.865 | GTGCGTGTT |
| 5015 | mamo-s | *BGIBMGA006050* | up | 478 | 486 | + | 12.1835 | 1.71E-05 | 0.615 | GTGCGTGCC |
| 5016 | mamo-s | *BGIBMGA006050* | down | 684 | 692 | + | 11.3303 | 4.96E-05 | 0.615 | GTGGGTGGC |
| 5017 | mamo-s | *BGIBMGA006051* | down | 1018 | 1026 | + | 13.6606 | 4.06E-06 | 0.615 | GCGCGTGGC |
| 5018 | mamo-s | *BGIBMGA006052* | up | 1763 | 1771 | - | 11.3303 | 4.96E-05 | 0.615 | GTGGGTGGC |
| 5019 | mamo-s | *BGIBMGA006052* | down | 1578 | 1586 | + | 11.0826 | 6.84E-05 | 0.709 | GTACGTGGC |
| 5020 | mamo-s | *BGIBMGA006053* | down | 691 | 699 | - | 11.3303 | 4.96E-05 | 0.615 | GTGGGTGGC |
| 5021 | mamo-s | *BGIBMGA006054* | up | 1051 | 1059 | - | 11.3303 | 4.96E-05 | 0.615 | GTGGGTGGC |
| 5022 | mamo-s | *BGIBMGA006058* | down | 1934 | 1942 | + | 11.3303 | 4.96E-05 | 0.615 | GTGGGTGGC |
| 5023 | mamo-s | *BGIBMGA006058* | down | 1183 | 1191 | + | 11.2844 | 5.18E-05 | 0.63 | CTGCGTGCC |
| 5024 | mamo-s | *BGIBMGA006059* | up | 771 | 779 | + | 11.3303 | 4.96E-05 | 0.615 | GTGGGTGGC |
| 5025 | mamo-s | *BGIBMGA006059* | down | 1393 | 1401 | - | 11.3303 | 4.96E-05 | 0.615 | GTGGGTGGC |
| 5026 | mamo-s | *BGIBMGA006059* | down | 1784 | 1792 | - | 11.3303 | 4.96E-05 | 0.615 | GTGGGTGGC |
| 5027 | mamo-s | *BGIBMGA006059* | up | 20 | 28 | + | 11.2844 | 5.18E-05 | 0.63 | CTGCGTGCC |
| 5028 | mamo-s | *BGIBMGA006060* | up | 380 | 388 | + | 11.3303 | 4.96E-05 | 0.615 | GTGGGTGGC |
| 5029 | mamo-s | *BGIBMGA006061* | down | 1283 | 1291 | + | 11.3303 | 4.96E-05 | 0.615 | GTGGGTGGC |
| 5030 | mamo-s | *BGIBMGA006061* | up | 819 | 827 | + | 10.5413 | 9.42E-05 | 0.857 | CAGCGTGTC |
| 5031 | mamo-s | *BGIBMGA006062* | down | 1001 | 1009 | - | 12.1009 | 2.43E-05 | 0.615 | GTGCGTGGT |
| 5032 | mamo-s | *BGIBMGA006062* | up | 268 | 276 | + | 11.3303 | 4.96E-05 | 0.615 | GTGGGTGGC |
| 5033 | mamo-s | *BGIBMGA006062* | down | 1290 | 1298 | - | 11.3303 | 4.96E-05 | 0.615 | GTGGGTGGC |
| 5034 | mamo-s | *BGIBMGA006063* | up | 1083 | 1091 | - | 11.3303 | 4.96E-05 | 0.615 | GTGGGTGGC |
| 5035 | mamo-s | *BGIBMGA006064* | down | 1202 | 1210 | - | 11.3303 | 4.96E-05 | 0.615 | GTGGGTGGC |
| 5036 | mamo-s | *BGIBMGA006065* | up | 1760 | 1768 | - | 11.3303 | 4.96E-05 | 0.615 | GTGGGTGGC |
| 5037 | mamo-s | *BGIBMGA006067* | down | 697 | 705 | - | 11.3303 | 4.96E-05 | 0.615 | GTGGGTGGC |
| 5038 | mamo-s | *BGIBMGA006068* | down | 1236 | 1244 | + | 11.0826 | 6.84E-05 | 0.709 | GTACGTGGC |
| 5039 | mamo-s | *BGIBMGA006069* | up | 89 | 97 | - | 11.3303 | 4.96E-05 | 0.615 | GTGGGTGGC |
| 5040 | mamo-s | *BGIBMGA006069* | up | 1214 | 1222 | + | 11.3303 | 4.96E-05 | 0.615 | GTGGGTGGC |
| 5041 | mamo-s | *BGIBMGA006069* | down | 1321 | 1329 | - | 10.5413 | 9.42E-05 | 0.857 | CAGCGTGTC |
| 5042 | mamo-s | *BGIBMGA006070* | down | 458 | 466 | + | 13.0275 | 8.53E-06 | 0.615 | GAGCGTGGC |
| 5043 | mamo-s | *BGIBMGA006070* | up | 816 | 824 | - | 11.3303 | 4.96E-05 | 0.615 | GTGGGTGGC |
| 5044 | mamo-s | *BGIBMGA006071* | down | 1646 | 1654 | + | 12.4679 | 1.31E-05 | 0.615 | GTGCGTGTC |
| 5045 | mamo-s | *BGIBMGA006071* | up | 1238 | 1246 | + | 11.3303 | 4.96E-05 | 0.615 | GTGGGTGGC |
| 5046 | mamo-s | *BGIBMGA006072* | down | 1141 | 1149 | - | 12.7615 | 1.04E-05 | 0.615 | CCGCGTGGC |
| 5047 | mamo-s | *BGIBMGA006073* | down | 1540 | 1548 | + | 11.1743 | 5.90E-05 | 0.684 | CCGCGTGTC |
| 5048 | mamo-s | *BGIBMGA006075* | down | 327 | 335 | + | 12.3394 | 1.49E-05 | 0.615 | GGGCGTGGC |
| 5049 | mamo-s | *BGIBMGA006076* | up | 704 | 712 | + | 12.3394 | 1.49E-05 | 0.615 | GGGCGTGGC |
| 5050 | mamo-s | *BGIBMGA006078* | down | 405 | 413 | - | 11.3303 | 4.96E-05 | 0.615 | GTGGGTGGC |
| 5051 | mamo-s | *BGIBMGA006078* | down | 1302 | 1310 | - | 11.0826 | 6.84E-05 | 0.709 | GTACGTGGC |
| 5052 | mamo-s | *BGIBMGA006079* | up | 861 | 869 | - | 11.0826 | 6.84E-05 | 0.709 | GTACGTGGC |
| 5053 | mamo-s | *BGIBMGA006082* | down | 1627 | 1635 | + | 11.3303 | 4.96E-05 | 0.615 | GTGGGTGGC |
| 5054 | mamo-s | *BGIBMGA006082* | up | 1705 | 1713 | - | 11.3303 | 4.96E-05 | 0.615 | GTGGGTGGC |
| 5055 | mamo-s | *BGIBMGA006083* | up | 273 | 281 | + | 11.0826 | 6.84E-05 | 0.709 | GTACGTGGC |
| 5056 | mamo-s | *BGIBMGA006084* | up | 993 | 1001 | + | 11.7339 | 3.29E-05 | 0.615 | GCGCGAGGC |
| 5057 | mamo-s | *BGIBMGA006084* | up | 148 | 156 | + | 11.3303 | 4.96E-05 | 0.615 | GTGGGTGGC |
| 5058 | mamo-s | *BGIBMGA006084* | up | 1686 | 1694 | - | 11.1376 | 6.35E-05 | 0.709 | TCGCGTGGC |
| 5059 | mamo-s | *BGIBMGA006085* | up | 323 | 331 | - | 11.3303 | 4.96E-05 | 0.615 | GTGGGTGGC |
| 5060 | mamo-s | *BGIBMGA006086* | up | 253 | 261 | - | 12.4679 | 1.31E-05 | 0.615 | GTGCGTGTC |
| 5061 | mamo-s | *BGIBMGA006087* | down | 838 | 846 | - | 11.3303 | 4.96E-05 | 0.615 | GTGGGTGGC |
| 5062 | mamo-s | *BGIBMGA006087* | down | 864 | 872 | - | 11.3303 | 4.96E-05 | 0.615 | GTGGGTGGC |
| 5063 | mamo-s | *BGIBMGA006087* | up | 213 | 221 | - | 11.1009 | 6.57E-05 | 0.709 | GAGCGAGGC |
| 5064 | mamo-s | *BGIBMGA006088* | down | 775 | 783 | - | 11.3303 | 4.96E-05 | 0.615 | GTGGGTGGC |
| 5065 | mamo-s | *BGIBMGA006088* | up | 1479 | 1487 | - | 11.3303 | 4.96E-05 | 0.615 | GTGGGTGGC |
| 5066 | mamo-s | *BGIBMGA006088* | up | 1505 | 1513 | - | 11.3303 | 4.96E-05 | 0.615 | GTGGGTGGC |
| 5067 | mamo-s | *BGIBMGA006088* | down | 426 | 434 | - | 11.0826 | 6.84E-05 | 0.709 | GTACGTGGC |
| 5068 | mamo-s | *BGIBMGA006090* | down | 531 | 539 | - | 13.0275 | 8.53E-06 | 0.615 | GAGCGTGGC |
| 5069 | mamo-s | *BGIBMGA006091* | down | 1208 | 1216 | + | 11.7798 | 3.11E-05 | 0.615 | ATGCGTGGC |
| 5070 | mamo-s | *BGIBMGA006092* | up | 1639 | 1647 | - | 11.2844 | 5.18E-05 | 0.63 | CTGCGTGCC |
| 5071 | mamo-s | *BGIBMGA006092* | up | 161 | 169 | - | 10.7523 | 8.38E-05 | 0.797 | AAGCGTGGC |
| 5072 | mamo-s | *BGIBMGA006094* | down | 353 | 361 | - | 11.3303 | 4.96E-05 | 0.615 | GTGGGTGGC |
| 5073 | mamo-s | *BGIBMGA006094* | up | 710 | 718 | - | 11.3303 | 4.96E-05 | 0.615 | GTGGGTGGC |
| 5074 | mamo-s | *BGIBMGA006095* | down | 1750 | 1758 | + | 10.9358 | 7.30E-05 | 0.728 | GCGGGTGGC |
| 5075 | mamo-s | *BGIBMGA006097* | down | 441 | 449 | - | 11.3303 | 4.96E-05 | 0.615 | GTGGGTGGC |
| 5076 | mamo-s | *BGIBMGA006098* | up | 133 | 141 | - | 11.3303 | 4.96E-05 | 0.615 | GTGGGTGGC |
| 5077 | mamo-s | *BGIBMGA006098* | up | 1094 | 1102 | - | 11.3303 | 4.96E-05 | 0.615 | GTGGGTGGC |
| 5078 | mamo-s | *BGIBMGA006098* | up | 1395 | 1403 | - | 11.0734 | 7.12E-05 | 0.726 | GAGCGTGGT |
| 5079 | mamo-s | *BGIBMGA006099* | down | 16 | 24 | - | 11.5688 | 3.79E-05 | 0.615 | CTGCGTGTC |
| 5080 | mamo-s | *BGIBMGA006099* | down | 247 | 255 | + | 11.3303 | 4.96E-05 | 0.615 | GTGGGTGGC |
| 5081 | mamo-s | *BGIBMGA006099* | down | 286 | 294 | - | 11.0826 | 6.84E-05 | 0.709 | GTACGTGGC |
| 5082 | mamo-s | *BGIBMGA006100* | up | 138 | 146 | - | 12.3394 | 1.49E-05 | 0.615 | GGGCGTGGC |
| 5083 | mamo-s | *BGIBMGA006100* | up | 910 | 918 | - | 11.3303 | 4.96E-05 | 0.615 | GTGGGTGGC |
| 5084 | mamo-s | *BGIBMGA006104* | up | 1087 | 1095 | - | 10.9358 | 7.30E-05 | 0.728 | GCGGGTGGC |
| 5085 | mamo-s | *BGIBMGA006108* | down | 878 | 886 | - | 11.7339 | 3.29E-05 | 0.615 | GCGCGAGGC |
| 5086 | mamo-s | *BGIBMGA006108* | up | 1201 | 1209 | - | 11.3303 | 4.96E-05 | 0.615 | GTGGGTGGC |
| 5087 | mamo-s | *BGIBMGA006108* | down | 1056 | 1064 | - | 10.8899 | 7.48E-05 | 0.739 | CCGCGTGCC |
| 5088 | mamo-s | *BGIBMGA006108* | down | 1050 | 1058 | + | 10.6881 | 8.61E-05 | 0.81 | GCACGTGGC |
| 5089 | mamo-s | *BGIBMGA006109* | down | 1631 | 1639 | - | 14.055 | 2.23E-06 | 0.615 | GTGCGTGGC |
| 5090 | mamo-s | *BGIBMGA006110* | up | 1330 | 1338 | + | 11.3303 | 4.96E-05 | 0.615 | GTGGGTGGC |
| 5091 | mamo-s | *BGIBMGA006112* | down | 991 | 999 | + | 11.3303 | 4.96E-05 | 0.615 | GTGGGTGGC |
| 5092 | mamo-s | *BGIBMGA006112* | down | 1464 | 1472 | + | 11.0826 | 6.84E-05 | 0.709 | GTACGTGGC |
| 5093 | mamo-s | *BGIBMGA006113* | down | 652 | 660 | + | 10.5413 | 9.42E-05 | 0.857 | GTGCGAGTC |
| 5094 | mamo-s | *BGIBMGA006114* | down | 1164 | 1172 | - | 11.0826 | 6.84E-05 | 0.709 | GTACGTGGC |
| 5095 | mamo-s | *BGIBMGA006115* | up | 869 | 877 | + | 11.3303 | 4.96E-05 | 0.615 | GTGGGTGGC |
| 5096 | mamo-s | *BGIBMGA006116* | up | 7 | 15 | - | 11.3303 | 4.96E-05 | 0.615 | GTGGGTGGC |
| 5097 | mamo-s | *BGIBMGA006117* | down | 954 | 962 | - | 11.3303 | 4.96E-05 | 0.615 | GTGGGTGGC |

| | | | | | | | | | | |
|---|---|---|---|---|---|---|---|---|---|---|
| 5098 | mamo-s | *BGIBMGA006117* | down | 1935 | 1943 | - | 11.3303 | 4.96E-05 | 0.615 | GTGGGTGGC |
| 5099 | mamo-s | *BGIBMGA006119* | up | 1091 | 1099 | - | 13.0275 | 8.53E-06 | 0.615 | GAGCGTGGC |
| 5100 | mamo-s | *BGIBMGA006119* | down | 326 | 334 | - | 11.3303 | 4.96E-05 | 0.615 | GTGGGTGGC |
| 5101 | mamo-s | *BGIBMGA006119* | down | 1676 | 1684 | + | 11.3303 | 4.96E-05 | 0.615 | GTGGGTGGC |
| 5102 | mamo-s | *BGIBMGA006120* | up | 525 | 533 | - | 11.3303 | 4.96E-05 | 0.615 | GTGGGTGGC |
| 5103 | mamo-s | *BGIBMGA006122* | up | 491 | 499 | - | 11.3303 | 4.96E-05 | 0.615 | GTGGGTGGC |
| 5104 | mamo-s | *BGIBMGA006122* | down | 1892 | 1900 | + | 11.3303 | 4.96E-05 | 0.615 | GTGGGTGGC |
| 5105 | mamo-s | *BGIBMGA006125* | up | 690 | 698 | + | 11.3303 | 4.96E-05 | 0.615 | GTGGGTGGC |
| 5106 | mamo-s | *BGIBMGA006125* | down | 1452 | 1460 | + | 11.3303 | 4.96E-05 | 0.615 | GTGGGTGGC |
| 5107 | mamo-s | *BGIBMGA006126* | up | 1829 | 1837 | + | 11.789 | 2.84E-05 | 0.615 | GCGCGTGCC |
| 5108 | mamo-s | *BGIBMGA006127* | down | 1529 | 1537 | - | 11.3303 | 4.96E-05 | 0.615 | GTGGGTGGC |
| 5109 | mamo-s | *BGIBMGA006127* | up | 1843 | 1851 | + | 11.3303 | 4.96E-05 | 0.615 | GTGGGTGGC |
| 5110 | mamo-s | *BGIBMGA006130* | down | 565 | 573 | + | 10.5138 | 9.75E-05 | 0.865 | GTGCGTGTT |
| 5111 | mamo-s | *BGIBMGA006132* | up | 866 | 874 | - | 11.3303 | 4.96E-05 | 0.615 | GTGGGTGGC |
| 5112 | mamo-s | *BGIBMGA006132* | down | 1696 | 1704 | - | 11.156 | 6.13E-05 | 0.705 | GAGCGTGCC |
| 5113 | mamo-s | *BGIBMGA006135* | up | 761 | 769 | - | 11.3303 | 4.96E-05 | 0.615 | GTGGGTGGC |
| 5114 | mamo-s | *BGIBMGA006136* | up | 1525 | 1533 | + | 11.7798 | 3.11E-05 | 0.615 | ATGCGTGGC |
| 5115 | mamo-s | *BGIBMGA006137* | down | 859 | 867 | - | 11.4404 | 4.51E-05 | 0.615 | GAGCGTGTC |
| 5116 | mamo-s | *BGIBMGA006138* | down | 1520 | 1528 | + | 11.3303 | 4.96E-05 | 0.615 | GTGGGTGGC |
| 5117 | mamo-s | *BGIBMGA006140* | up | 1409 | 1417 | - | 11.1376 | 6.35E-05 | 0.709 | TCGCGTGGC |
| 5118 | mamo-s | *BGIBMGA006141* | down | 315 | 323 | - | 11.3303 | 4.96E-05 | 0.615 | GTGGGTGGC |
| 5119 | mamo-s | *BGIBMGA006141* | up | 840 | 848 | + | 11.0826 | 6.84E-05 | 0.709 | GTACGTGGC |
| 5120 | mamo-s | *BGIBMGA006141* | down | 674 | 682 | - | 10.6881 | 8.61E-05 | 0.81 | GCACGTGGC |
| 5121 | mamo-s | *BGIBMGA006142* | up | 1104 | 1112 | - | 10.7523 | 8.38E-05 | 0.797 | GGGCGTGTC |
| 5122 | mamo-s | *BGIBMGA006145* | down | 180 | 188 | - | 12.1284 | 2.16E-05 | 0.615 | CAGCGTGGC |
| 5123 | mamo-s | *BGIBMGA006146* | up | 395 | 403 | + | 10.8349 | 7.66E-05 | 0.751 | CCGCGAGGC |
| 5124 | mamo-s | *BGIBMGA006147* | down | 971 | 979 | + | 11.3303 | 4.96E-05 | 0.615 | GTGGGTGGC |
| 5125 | mamo-s | *BGIBMGA006147* | up | 1229 | 1237 | + | 11.3303 | 4.96E-05 | 0.615 | GTGGGTGGC |
| 5126 | mamo-s | *BGIBMGA006149* | up | 1622 | 1630 | + | 11.3303 | 4.96E-05 | 0.615 | GTGGGTGGC |
| 5127 | mamo-s | *BGIBMGA006155* | down | 272 | 280 | + | 11.3303 | 4.96E-05 | 0.615 | GTGGGTGGC |
| 5128 | mamo-s | *BGIBMGA006156* | up | 1692 | 1700 | - | 11.1376 | 6.35E-05 | 0.709 | TCGCGTGGC |
| 5129 | mamo-s | *BGIBMGA006157* | up | 48 | 56 | + | 11.3303 | 4.96E-05 | 0.615 | GTGGGTGGC |
| 5130 | mamo-s | *BGIBMGA006157* | down | 374 | 382 | - | 11.3303 | 4.96E-05 | 0.615 | GTGGGTGGC |
| 5131 | mamo-s | *BGIBMGA006157* | up | 1872 | 1880 | + | 11.3303 | 4.96E-05 | 0.615 | GTGGGTGGC |
| 5132 | mamo-s | *BGIBMGA006158* | down | 1111 | 1119 | + | 11.5321 | 4.06E-05 | 0.615 | TTGCGTGGC |
| 5133 | mamo-s | *BGIBMGA006162* | down | 575 | 583 | + | 11.5688 | 3.79E-05 | 0.615 | CTGCGTGTC |
| 5134 | mamo-s | *BGIBMGA006162* | down | 1606 | 1614 | - | 10.7523 | 8.38E-05 | 0.797 | AAGCGTGGC |
| 5135 | mamo-s | *BGIBMGA006164* | down | 1909 | 1917 | + | 10.8073 | 7.89E-05 | 0.766 | CCGCGTGGT |
| 5136 | mamo-s | *BGIBMGA006169* | down | 955 | 963 | - | 11.3303 | 4.96E-05 | 0.615 | GTGGGTGGC |
| 5137 | mamo-s | *BGIBMGA006169* | down | 1492 | 1500 | - | 11.2844 | 5.18E-05 | 0.63 | CTGCGTGCC |
| 5138 | mamo-s | *BGIBMGA006170* | up | 1929 | 1937 | + | 12.1009 | 2.43E-05 | 0.615 | GTGCGTGGT |
| 5139 | mamo-s | *BGIBMGA006171* | down | 1525 | 1533 | + | 10.5138 | 9.75E-05 | 0.865 | GTGCGTGTT |
| 5140 | mamo-s | *BGIBMGA006176* | up | 613 | 621 | - | 12.1835 | 1.71E-05 | 0.615 | GTGCGTGCC |
| 5141 | mamo-s | *BGIBMGA006176* | up | 36 | 44 | - | 11.3303 | 4.96E-05 | 0.615 | GTGGGTGGC |
| 5142 | mamo-s | *BGIBMGA006180* | down | 580 | 588 | - | 11.3303 | 4.96E-05 | 0.615 | GTGGGTGGC |
| 5143 | mamo-s | *BGIBMGA006182* | down | 394 | 402 | - | 12.4679 | 1.31E-05 | 0.615 | GTGCGTGTC |
| 5144 | mamo-s | *BGIBMGA006186* | down | 1280 | 1288 | - | 11.3303 | 4.96E-05 | 0.615 | GTGGGTGGC |
| 5145 | mamo-s | *BGIBMGA006187* | down | 485 | 493 | - | 13.156 | 6.30E-06 | 0.615 | CTGCGTGGC |
| 5146 | mamo-s | *BGIBMGA006188* | down | 1079 | 1087 | + | 11.3303 | 4.96E-05 | 0.615 | GTGGGTGGC |
| 5147 | mamo-s | *BGIBMGA006188* | up | 1918 | 1926 | + | 10.6881 | 8.61E-05 | 0.81 | GCACGTGGC |
| 5148 | mamo-s | *BGIBMGA006189* | up | 1497 | 1505 | + | 11.0826 | 6.84E-05 | 0.709 | GTACGTGGC |
| 5149 | mamo-s | *BGIBMGA006193* | down | 399 | 407 | - | 13.156 | 6.30E-06 | 0.615 | CTGCGTGGC |
| 5150 | mamo-s | *BGIBMGA006195* | down | 1009 | 1017 | - | 11.1376 | 6.35E-05 | 0.709 | TCGCGTGGC |
| 5151 | mamo-s | *BGIBMGA006197* | down | 159 | 167 | - | 11.3303 | 4.96E-05 | 0.615 | GTGGGTGGC |
| 5152 | mamo-s | *BGIBMGA006199* | down | 411 | 419 | - | 11.3303 | 4.96E-05 | 0.615 | GTGGGTGGC |
| 5153 | mamo-s | *BGIBMGA006202* | up | 1964 | 1972 | + | 12.0734 | 2.66E-05 | 0.615 | GCGCGTGTC |
| 5154 | mamo-s | *BGIBMGA006203* | down | 408 | 416 | + | 11.1743 | 5.90E-05 | 0.684 | CCGCGTGTC |
| 5155 | mamo-s | *BGIBMGA006203* | down | 978 | 986 | + | 10.5138 | 9.75E-05 | 0.865 | GTGCGTGTT |
| 5156 | mamo-s | *BGIBMGA006204* | down | 230 | 238 | - | 11.3303 | 4.96E-05 | 0.615 | GTGGGTGGC |
| 5157 | mamo-s | *BGIBMGA006204* | up | 1108 | 1116 | + | 11.3303 | 4.96E-05 | 0.615 | GTGGGTGGC |
| 5158 | mamo-s | *BGIBMGA006204* | down | 1124 | 1132 | + | 11.3303 | 4.96E-05 | 0.615 | GTGGGTGGC |
| 5159 | mamo-s | *BGIBMGA006204* | up | 1914 | 1922 | + | 10.8899 | 7.48E-05 | 0.739 | CCGCGTGCC |
| 5160 | mamo-s | *BGIBMGA006205* | down | 534 | 542 | - | 12.4679 | 1.31E-05 | 0.615 | GTGCGTGTC |
| 5161 | mamo-s | *BGIBMGA006205* | down | 1992 | 2000 | + | 11.3303 | 4.96E-05 | 0.615 | GTGGGTGGC |
| 5162 | mamo-s | *BGIBMGA006207* | up | 666 | 674 | - | 12.1284 | 2.16E-05 | 0.615 | GTGCGAGGC |
| 5163 | mamo-s | *BGIBMGA006207* | up | 1511 | 1519 | + | 11.3303 | 4.96E-05 | 0.615 | GTGGGTGGC |
| 5164 | mamo-s | *BGIBMGA006207* | up | 1261 | 1269 | + | 10.8073 | 7.89E-05 | 0.766 | CCGCGTGGT |
| 5165 | mamo-s | *BGIBMGA006209* | down | 1251 | 1259 | - | 11.0826 | 6.84E-05 | 0.709 | GTACGTGGC |
| 5166 | mamo-s | *BGIBMGA006210* | up | 471 | 479 | - | 12.1835 | 1.71E-05 | 0.615 | GTGCGTGCC |
| 5167 | mamo-s | *BGIBMGA006210* | down | 710 | 718 | + | 12.1009 | 2.43E-05 | 0.615 | GTGCGTGGT |
| 5168 | mamo-s | *BGIBMGA006211* | up | 725 | 733 | + | 11.3303 | 4.96E-05 | 0.615 | GTGGGTGGC |
| 5169 | mamo-s | *BGIBMGA006211* | down | 852 | 860 | + | 11.3303 | 4.96E-05 | 0.615 | GTGGGTGGC |
| 5170 | mamo-s | *BGIBMGA006211* | up | 1453 | 1461 | - | 11.1009 | 6.57E-05 | 0.709 | GAGCGAGGC |
| 5171 | mamo-s | *BGIBMGA006212* | down | 1889 | 1897 | - | 10.5413 | 9.42E-05 | 0.857 | GTGCGAGTC |
| 5172 | mamo-s | *BGIBMGA006213* | up | 626 | 634 | + | 11.3303 | 4.96E-05 | 0.615 | GTGGGTGGC |
| 5173 | mamo-s | *BGIBMGA006213* | down | 806 | 814 | + | 11.3303 | 4.96E-05 | 0.615 | GTGGGTGGC |
| 5174 | mamo-s | *BGIBMGA006217* | up | 1570 | 1578 | - | 11.3303 | 4.96E-05 | 0.615 | GTGGGTGGC |
| 5175 | mamo-s | *BGIBMGA006218* | down | 1374 | 1382 | - | 11.2844 | 5.18E-05 | 0.63 | CTGCGTGCC |
| 5176 | mamo-s | *BGIBMGA006219* | down | 1308 | 1316 | + | 11.3303 | 4.96E-05 | 0.615 | GTGGGTGGC |
| 5177 | mamo-s | *BGIBMGA006222* | up | 445 | 453 | + | 11.2018 | 5.68E-05 | 0.674 | CTGCGTGGT |
| 5178 | mamo-s | *BGIBMGA006223* | up | 132 | 140 | + | 11.3303 | 4.96E-05 | 0.615 | GTGGGTGGC |
| 5179 | mamo-s | *BGIBMGA006223* | up | 567 | 575 | + | 11.3303 | 4.96E-05 | 0.615 | GTGGGTGGC |
| 5180 | mamo-s | *BGIBMGA006223* | up | 1544 | 1552 | - | 11.3303 | 4.96E-05 | 0.615 | GTGGGTGGC |
| 5181 | mamo-s | *BGIBMGA006224* | up | 838 | 846 | - | 11.3303 | 4.96E-05 | 0.615 | GTGGGTGGC |
| 5182 | mamo-s | *BGIBMGA006224* | down | 1224 | 1232 | + | 11.3303 | 4.96E-05 | 0.615 | GTGGGTGGC |
| 5183 | mamo-s | *BGIBMGA006226* | up | 1576 | 1584 | - | 11.2294 | 5.41E-05 | 0.65 | CTGCGAGGC |
| 5184 | mamo-s | *BGIBMGA006226* | up | 769 | 777 | + | 10.9358 | 7.30E-05 | 0.728 | GCGGGTGGC |
| 5185 | mamo-s | *BGIBMGA006227* | up | 1811 | 1819 | + | 11.3303 | 4.96E-05 | 0.615 | GTGGGTGGC |

| | | | | | | | | | | |
|---|---|---|---|---|---|---|---|---|---|---|
| 5186 | mamo-s | *BGIBMGA006230* | up | 1725 | 1733 | + | 12.1284 | 2.16E-05 | 0.615 | GTGCGAGGC |
| 5187 | mamo-s | *BGIBMGA006230* | up | 470 | 478 | + | 11.3303 | 4.96E-05 | 0.615 | GTGGGTGGC |
| 5188 | mamo-s | *BGIBMGA006235* | up | 816 | 824 | - | 11.3303 | 4.96E-05 | 0.615 | GTGGGTGGC |
| 5189 | mamo-s | *BGIBMGA006236* | down | 1587 | 1595 | - | 11.0826 | 6.84E-05 | 0.709 | GTACGTGGC |
| 5190 | mamo-s | *BGIBMGA006237* | up | 1625 | 1633 | + | 11.7798 | 3.11E-05 | 0.615 | ATGCGTGGC |
| 5191 | mamo-s | *BGIBMGA006237* | down | 1723 | 1731 | + | 11.3303 | 4.96E-05 | 0.615 | GTGGGTGGC |
| 5192 | mamo-s | *BGIBMGA006237* | up | 658 | 666 | + | 11.0826 | 6.84E-05 | 0.709 | GTACGTGGC |
| 5193 | mamo-s | *BGIBMGA006239* | up | 121 | 129 | - | 11.3303 | 4.96E-05 | 0.615 | GTGGGTGGC |
| 5194 | mamo-s | *BGIBMGA006240* | down | 1342 | 1350 | + | 10.7523 | 8.38E-05 | 0.797 | AAGCGTGGC |
| 5195 | mamo-s | *BGIBMGA006244* | down | 855 | 863 | + | 11.3303 | 4.96E-05 | 0.615 | GTGGGTGGC |
| 5196 | mamo-s | *BGIBMGA006246* | up | 909 | 917 | + | 11.3303 | 4.96E-05 | 0.615 | GTGGGTGGC |
| 5197 | mamo-s | *BGIBMGA006246* | up | 1161 | 1169 | + | 11.3303 | 4.96E-05 | 0.615 | GTGGGTGGC |
| 5198 | mamo-s | *BGIBMGA006248* | down | 1522 | 1530 | + | 11.3303 | 4.96E-05 | 0.615 | GTGGGTGGC |
| 5199 | mamo-s | *BGIBMGA006248* | up | 1808 | 1816 | - | 11.3303 | 4.96E-05 | 0.615 | GTGGGTGGC |
| 5200 | mamo-s | *BGIBMGA006249* | up | 270 | 278 | - | 11.7798 | 3.11E-05 | 0.615 | ATGCGTGGC |
| 5201 | mamo-s | *BGIBMGA006249* | up | 1237 | 1245 | - | 11.0826 | 6.84E-05 | 0.709 | GTACGTGGC |
| 5202 | mamo-s | *BGIBMGA006251* | up | 1787 | 1795 | + | 12.1284 | 2.16E-05 | 0.615 | CAGCGTGGC |
| 5203 | mamo-s | *BGIBMGA006251* | up | 650 | 658 | - | 11.0826 | 6.84E-05 | 0.709 | GTACGTGGC |
| 5204 | mamo-s | *BGIBMGA006253* | up | 795 | 803 | - | 11.3303 | 4.96E-05 | 0.615 | GTGGGTGGC |
| 5205 | mamo-s | *BGIBMGA006254* | up | 1740 | 1748 | + | 10.9358 | 7.30E-05 | 0.728 | GCGGGTGGC |
| 5206 | mamo-s | *BGIBMGA006255* | up | 1624 | 1632 | - | 11.3303 | 4.96E-05 | 0.615 | GTGGGTGGC |
| 5207 | mamo-s | *BGIBMGA006256* | up | 578 | 586 | - | 11.3303 | 4.96E-05 | 0.615 | GTGGGTGGC |
| 5208 | mamo-s | *BGIBMGA006256* | down | 1128 | 1136 | - | 11.3303 | 4.96E-05 | 0.615 | GTGGGTGGC |
| 5209 | mamo-s | *BGIBMGA006257* | up | 681 | 689 | - | 11.0826 | 6.84E-05 | 0.709 | GTACGTGGC |
| 5210 | mamo-s | *BGIBMGA006258* | down | 529 | 537 | - | 11.3303 | 4.96E-05 | 0.615 | GTGGGTGGC |
| 5211 | mamo-s | *BGIBMGA006259* | up | 1756 | 1764 | + | 12.0734 | 2.66E-05 | 0.615 | GCGCGTGTC |
| 5212 | mamo-s | *BGIBMGA006259* | down | 1176 | 1184 | - | 11.0826 | 6.84E-05 | 0.709 | GTACGTGGC |
| 5213 | mamo-s | *BGIBMGA006260* | up | 844 | 852 | - | 10.5413 | 9.42E-05 | 0.857 | GTGCGAGTC |
| 5214 | mamo-s | *BGIBMGA006264* | down | 128 | 136 | - | 11.3303 | 4.96E-05 | 0.615 | GTGGGTGGC |
| 5215 | mamo-s | *BGIBMGA006266* | down | 1001 | 1009 | - | 12.4679 | 1.31E-05 | 0.615 | GTGCGTGTC |
| 5216 | mamo-s | *BGIBMGA006267* | up | 1709 | 1717 | - | 12.4679 | 1.31E-05 | 0.615 | GTGCGTGTC |
| 5217 | mamo-s | *BGIBMGA006268* | down | 1884 | 1892 | - | 11.3303 | 4.96E-05 | 0.615 | GTGGGTGGC |
| 5218 | mamo-s | *BGIBMGA006269* | up | 954 | 962 | - | 11.3303 | 4.96E-05 | 0.615 | GTGGGTGGC |
| 5219 | mamo-s | *BGIBMGA006269* | up | 1145 | 1153 | - | 11.3303 | 4.96E-05 | 0.615 | GTGGGTGGC |
| 5220 | mamo-s | *BGIBMGA006270* | up | 626 | 634 | - | 10.5138 | 9.75E-05 | 0.865 | GTGCGTGTT |
| 5221 | mamo-s | *BGIBMGA006271* | down | 740 | 748 | - | 11.7798 | 3.11E-05 | 0.615 | ATGCGTGGC |
| 5222 | mamo-s | *BGIBMGA006271* | up | 439 | 447 | - | 11.3303 | 4.96E-05 | 0.615 | GTGGGTGGC |
| 5223 | mamo-s | *BGIBMGA006273* | up | 538 | 546 | + | 11.3303 | 4.96E-05 | 0.615 | GTGGGTGGC |
| 5224 | mamo-s | *BGIBMGA006274* | down | 1509 | 1517 | - | 11.4404 | 4.51E-05 | 0.615 | GAGCGTGTC |
| 5225 | mamo-s | *BGIBMGA006274* | up | 1059 | 1067 | - | 11.3303 | 4.96E-05 | 0.615 | GTGGGTGGC |
| 5226 | mamo-s | *BGIBMGA006274* | down | 1577 | 1585 | - | 11.3303 | 4.96E-05 | 0.615 | GTGGGTGGC |
| 5227 | mamo-s | *BGIBMGA006276* | up | 1372 | 1380 | - | 11.0826 | 6.84E-05 | 0.709 | GTACGTGGC |
| 5228 | mamo-s | *BGIBMGA006278* | down | 1657 | 1665 | - | 10.8073 | 7.89E-05 | 0.766 | CCGCGTGGT |
| 5229 | mamo-s | *BGIBMGA006279* | up | 1169 | 1177 | + | 11.3303 | 4.96E-05 | 0.615 | GTGGGTGGC |
| 5230 | mamo-s | *BGIBMGA006279* | down | 967 | 975 | - | 10.9358 | 7.30E-05 | 0.728 | GCGGGTGGC |
| 5231 | mamo-s | *BGIBMGA006280* | down | 1075 | 1083 | - | 12.1009 | 2.43E-05 | 0.615 | GTGCGTGGT |
| 5232 | mamo-s | *BGIBMGA006280* | up | 1353 | 1361 | - | 11.3303 | 4.96E-05 | 0.615 | GTGGGTGGC |
| 5233 | mamo-s | *BGIBMGA006281* | up | 428 | 436 | - | 10.8899 | 7.48E-05 | 0.739 | CCGCGTGCC |
| 5234 | mamo-s | *BGIBMGA006282* | down | 1413 | 1421 | + | 11.3303 | 4.96E-05 | 0.615 | GTGGGTGGC |
| 5235 | mamo-s | *BGIBMGA006282* | up | 1656 | 1664 | - | 11.3303 | 4.96E-05 | 0.615 | GTGGGTGGC |
| 5236 | mamo-s | *BGIBMGA006283* | down | 1656 | 1664 | - | 11.2844 | 5.18E-05 | 0.63 | CTGCGTGCC |
| 5237 | mamo-s | *BGIBMGA006288* | up | 603 | 611 | - | 10.8899 | 7.48E-05 | 0.739 | CCGCGTGCC |
| 5238 | mamo-s | *BGIBMGA006289* | down | 368 | 376 | - | 12.1009 | 2.43E-05 | 0.615 | GTGCGTGGT |
| 5239 | mamo-s | *BGIBMGA006293* | down | 1624 | 1632 | + | 11.3303 | 4.96E-05 | 0.615 | GTGGGTGGC |
| 5240 | mamo-s | *BGIBMGA006295* | up | 1901 | 1909 | - | 12.1284 | 2.16E-05 | 0.615 | GTGCGAGGC |
| 5241 | mamo-s | *BGIBMGA006295* | up | 17 | 25 | + | 11.3303 | 4.96E-05 | 0.615 | GTGGGTGGC |
| 5242 | mamo-s | *BGIBMGA006295* | down | 441 | 449 | - | 11.3303 | 4.96E-05 | 0.615 | GTGGGTGGC |
| 5243 | mamo-s | *BGIBMGA006296* | down | 630 | 638 | + | 11.3303 | 4.96E-05 | 0.615 | GTGGGTGGC |
| 5244 | mamo-s | *BGIBMGA006299* | up | 1215 | 1223 | - | 11.3303 | 4.96E-05 | 0.615 | GTGGGTGGC |
| 5245 | mamo-s | *BGIBMGA006302* | down | 172 | 180 | - | 11.3303 | 4.96E-05 | 0.615 | GTGGGTGGC |
| 5246 | mamo-s | *BGIBMGA006302* | up | 1644 | 1652 | - | 11.3303 | 4.96E-05 | 0.615 | GTGGGTGGC |
| 5247 | mamo-s | *BGIBMGA006303* | up | 698 | 706 | + | 11.3303 | 4.96E-05 | 0.615 | GTGGGTGGC |
| 5248 | mamo-s | *BGIBMGA006303* | up | 1805 | 1813 | - | 11.3303 | 4.96E-05 | 0.615 | GTGGGTGGC |
| 5249 | mamo-s | *BGIBMGA006304* | up | 1902 | 1910 | + | 11.3303 | 4.96E-05 | 0.615 | GTGGGTGGC |
| 5250 | mamo-s | *BGIBMGA006306* | up | 1898 | 1906 | + | 11.3303 | 4.96E-05 | 0.615 | GTGGGTGGC |
| 5251 | mamo-s | *BGIBMGA006307* | down | 1072 | 1080 | - | 11.3303 | 4.96E-05 | 0.615 | GTGGGTGGC |
| 5252 | mamo-s | *BGIBMGA006308* | up | 950 | 958 | - | 11.3303 | 4.96E-05 | 0.615 | GTGGGTGGC |
| 5253 | mamo-s | *BGIBMGA006308* | up | 1660 | 1668 | + | 11.3303 | 4.96E-05 | 0.615 | GTGGGTGGC |
| 5254 | mamo-s | *BGIBMGA006311* | up | 572 | 580 | - | 11.3303 | 4.96E-05 | 0.615 | GTGGGTGGC |
| 5255 | mamo-s | *BGIBMGA006313* | down | 1479 | 1487 | + | 13.0275 | 8.53E-06 | 0.615 | GAGCGTGGC |
| 5256 | mamo-s | *BGIBMGA006314* | up | 609 | 617 | - | 11.3303 | 4.96E-05 | 0.615 | GTGGGTGGC |
| 5257 | mamo-s | *BGIBMGA006315* | up | 1777 | 1785 | + | 13.6606 | 4.06E-06 | 0.615 | GCGCGTGGC |
| 5258 | mamo-s | *BGIBMGA006317* | down | 1066 | 1074 | + | 11.2844 | 5.18E-05 | 0.63 | CTGCGTGCC |
| 5259 | mamo-s | *BGIBMGA006317* | up | 1148 | 1156 | - | 11.1376 | 6.35E-05 | 0.709 | TCGCGTGGC |
| 5260 | mamo-s | *BGIBMGA006319* | up | 1894 | 1902 | - | 12.4679 | 1.31E-05 | 0.615 | GTGCGTGTC |
| 5261 | mamo-s | *BGIBMGA006320* | up | 423 | 431 | - | 11.3303 | 4.96E-05 | 0.615 | GTGGGTGGC |
| 5262 | mamo-s | *BGIBMGA006322* | down | 1418 | 1426 | - | 11.3303 | 4.96E-05 | 0.615 | GTGGGTGGC |
| 5263 | mamo-s | *BGIBMGA006322* | down | 1992 | 2000 | - | 11.3303 | 4.96E-05 | 0.615 | GTGGGTGGC |
| 5264 | mamo-s | *BGIBMGA006322* | down | 248 | 256 | + | 11.2844 | 5.18E-05 | 0.63 | CTGCGTGCC |
| 5265 | mamo-s | *BGIBMGA006324* | up | 1214 | 1222 | - | 11.3303 | 4.96E-05 | 0.615 | GTGGGTGGC |
| 5266 | mamo-s | *BGIBMGA006325* | down | 1317 | 1325 | + | 11.0826 | 6.84E-05 | 0.709 | GTACGTGGC |
| 5267 | mamo-s | *BGIBMGA006327* | up | 1884 | 1892 | + | 11.3303 | 4.96E-05 | 0.615 | GTGGGTGGC |
| 5268 | mamo-s | *BGIBMGA006330* | down | 1647 | 1655 | + | 11.3303 | 4.96E-05 | 0.615 | GTGGGTGGC |
| 5269 | mamo-s | *BGIBMGA006331* | down | 585 | 593 | + | 10.7523 | 8.38E-05 | 0.797 | AAGCGTGGC |
| 5270 | mamo-s | *BGIBMGA006332* | down | 3 | 11 | - | 12.7615 | 1.04E-05 | 0.615 | CCGCGTGGC |
| 5271 | mamo-s | *BGIBMGA006334* | up | 1539 | 1547 | + | 11.3303 | 4.96E-05 | 0.615 | GTGGGTGGC |
| 5272 | mamo-s | *BGIBMGA006335* | down | 80 | 88 | + | 11.789 | 2.84E-05 | 0.615 | GCGCGTGCC |
| 5273 | mamo-s | *BGIBMGA006335* | down | 1641 | 1649 | + | 10.5413 | 9.42E-05 | 0.857 | CAGCGTGTC |

| | | | | | | | | | | |
|---|---|---|---|---|---|---|---|---|---|---|
| 5274 | mamo-s | *BGIBMGA006337* | down | 1122 | 1130 | - | 12.1284 | 2.16E-05 | 0.615 | GTGCGAGGC |
| 5275 | mamo-s | *BGIBMGA006337* | down | 1856 | 1864 | + | 11.0826 | 6.84E-05 | 0.709 | GTACGTGGC |
| 5276 | mamo-s | *BGIBMGA006338* | up | 696 | 704 | + | 11.3303 | 4.96E-05 | 0.615 | GTGGGTGGC |
| 5277 | mamo-s | *BGIBMGA006339* | up | 1930 | 1938 | - | 10.5413 | 9.42E-05 | 0.857 | GTGCGAGTC |
| 5278 | mamo-s | *BGIBMGA006340* | down | 327 | 335 | + | 11.3303 | 4.96E-05 | 0.615 | GTGGGTGGC |
| 5279 | mamo-s | *BGIBMGA006341* | up | 1813 | 1821 | - | 11.3303 | 4.96E-05 | 0.615 | GTGGGTGGC |
| 5280 | mamo-s | *BGIBMGA006342* | up | 1859 | 1867 | + | 11.3303 | 4.96E-05 | 0.615 | GTGGGTGGC |
| 5281 | mamo-s | *BGIBMGA006343* | up | 384 | 392 | - | 11.3303 | 4.96E-05 | 0.615 | GTGGGTGGC |
| 5282 | mamo-s | *BGIBMGA006343* | down | 1506 | 1514 | + | 11.3303 | 4.96E-05 | 0.615 | GTGGGTGGC |
| 5283 | mamo-s | *BGIBMGA006343* | up | 1084 | 1092 | - | 10.9358 | 7.30E-05 | 0.728 | GCGGGTGGC |
| 5284 | mamo-s | *BGIBMGA006349* | down | 1553 | 1561 | - | 11.3303 | 4.96E-05 | 0.615 | GTGGGTGGC |
| 5285 | mamo-s | *BGIBMGA006350* | down | 1216 | 1224 | - | 10.9358 | 7.30E-05 | 0.728 | GCGGGTGGC |
| 5286 | mamo-s | *BGIBMGA006351* | up | 176 | 184 | + | 12.4679 | 1.31E-05 | 0.615 | GTGCGTGTC |
| 5287 | mamo-s | *BGIBMGA006352* | up | 1547 | 1555 | + | 11.2018 | 5.68E-05 | 0.674 | CTGCGTGGT |
| 5288 | mamo-s | *BGIBMGA006354* | up | 1613 | 1621 | + | 11.1743 | 5.90E-05 | 0.684 | CCGCGTGTC |
| 5289 | mamo-s | *BGIBMGA006359* | down | 1145 | 1153 | + | 11.1743 | 5.90E-05 | 0.684 | CCGCGTGTC |
| 5290 | mamo-s | *BGIBMGA006360* | up | 965 | 973 | + | 11.3303 | 4.96E-05 | 0.615 | GTGGGTGGC |
| 5291 | mamo-s | *BGIBMGA006361* | down | 1624 | 1632 | - | 12.1009 | 2.43E-05 | 0.615 | GTGCGTGGT |
| 5292 | mamo-s | *BGIBMGA006362* | down | 762 | 770 | + | 11.3303 | 4.96E-05 | 0.615 | GTGGGTGGC |
| 5293 | mamo-s | *BGIBMGA006362* | down | 1576 | 1584 | - | 10.9358 | 7.30E-05 | 0.728 | GCGGGTGGC |
| 5294 | mamo-s | *BGIBMGA006363* | up | 838 | 846 | + | 11.3303 | 4.96E-05 | 0.615 | GTGGGTGGC |
| 5295 | mamo-s | *BGIBMGA006363* | up | 1652 | 1660 | - | 10.9358 | 7.30E-05 | 0.728 | GCGGGTGGC |
| 5296 | mamo-s | *BGIBMGA006365* | up | 1859 | 1867 | + | 11.4404 | 4.51E-05 | 0.615 | CGGCGTGGC |
| 5297 | mamo-s | *BGIBMGA006365* | up | 1681 | 1689 | + | 11.3303 | 4.96E-05 | 0.615 | GTGGGTGGC |
| 5298 | mamo-s | *BGIBMGA006366* | up | 558 | 566 | + | 11.3303 | 4.96E-05 | 0.615 | GTGGGTGGC |
| 5299 | mamo-s | *BGIBMGA006367* | down | 490 | 498 | + | 11.3303 | 4.96E-05 | 0.615 | GTGGGTGGC |
| 5300 | mamo-s | *BGIBMGA006369* | down | 767 | 775 | - | 11.3303 | 4.96E-05 | 0.615 | GTGGGTGGC |
| 5301 | mamo-s | *BGIBMGA006372* | down | 492 | 500 | - | 10.6881 | 8.61E-05 | 0.81 | GCACGTGGC |
| 5302 | mamo-s | *BGIBMGA006375* | up | 1054 | 1062 | - | 11.0734 | 7.12E-05 | 0.726 | GAGCGTGGT |
| 5303 | mamo-s | *BGIBMGA006377* | up | 860 | 868 | + | 11.3303 | 4.96E-05 | 0.615 | GTGGGTGGC |
| 5304 | mamo-s | *BGIBMGA006379* | up | 249 | 257 | + | 11.3303 | 4.96E-05 | 0.615 | GTGGGTGGC |
| 5305 | mamo-s | *BGIBMGA006382* | up | 1073 | 1081 | - | 11.3303 | 4.96E-05 | 0.615 | GTGGGTGGC |
| 5306 | mamo-s | *BGIBMGA006383* | down | 1937 | 1945 | + | 11.156 | 6.13E-05 | 0.705 | GAGCGTGCC |
| 5307 | mamo-s | *BGIBMGA006384* | down | 1924 | 1932 | + | 11.3303 | 4.96E-05 | 0.615 | GTGGGTGGC |
| 5308 | mamo-s | *BGIBMGA006385* | up | 874 | 882 | - | 12.1835 | 1.71E-05 | 0.615 | GTGCGTGCC |
| 5309 | mamo-s | *BGIBMGA006386* | up | 1715 | 1723 | - | 13.156 | 6.30E-06 | 0.615 | CTGCGTGGC |
| 5310 | mamo-s | *BGIBMGA006386* | up | 1962 | 1970 | + | 10.8349 | 7.66E-05 | 0.751 | CCGCGAGGC |
| 5311 | mamo-s | *BGIBMGA006387* | up | 1851 | 1859 | - | 10.5413 | 9.42E-05 | 0.857 | CAGCGTGTC |
| 5312 | mamo-s | *BGIBMGA006389* | up | 1525 | 1533 | + | 12.0734 | 2.66E-05 | 0.615 | GCGCGTGTC |
| 5313 | mamo-s | *BGIBMGA006389* | up | 1762 | 1770 | - | 11.789 | 2.84E-05 | 0.615 | GCGCGTGCC |
| 5314 | mamo-s | *BGIBMGA006389* | up | 1395 | 1403 | - | 10.6881 | 8.61E-05 | 0.81 | GCACGTGGC |
| 5315 | mamo-s | *BGIBMGA006391* | down | 651 | 659 | - | 10.5688 | 8.88E-05 | 0.824 | GTGCGTGAC |
| 5316 | mamo-s | *BGIBMGA006393* | up | 766 | 774 | + | 11.3303 | 4.96E-05 | 0.615 | GTGGGTGGC |
| 5317 | mamo-s | *BGIBMGA006395* | down | 1076 | 1084 | + | 11.0734 | 7.12E-05 | 0.726 | GAGCGTGGT |
| 5318 | mamo-s | *BGIBMGA006396* | up | 817 | 825 | - | 11.3303 | 4.96E-05 | 0.615 | GTGGGTGGC |
| 5319 | mamo-s | *BGIBMGA006396* | down | 857 | 865 | - | 10.7523 | 8.38E-05 | 0.797 | AAGCGTGGC |
| 5320 | mamo-s | *BGIBMGA006397* | up | 119 | 127 | + | 11.3303 | 4.96E-05 | 0.615 | GTGGGTGGC |
| 5321 | mamo-s | *BGIBMGA006400* | up | 508 | 516 | - | 11.3303 | 4.96E-05 | 0.615 | GTGGGTGGC |
| 5322 | mamo-s | *BGIBMGA006401* | up | 91 | 99 | + | 11.3303 | 4.96E-05 | 0.615 | GTGGGTGGC |
| 5323 | mamo-s | *BGIBMGA006404* | down | 1184 | 1192 | + | 11.3303 | 4.96E-05 | 0.615 | GTGGGTGGC |
| 5324 | mamo-s | *BGIBMGA006406* | down | 804 | 812 | + | 11.3303 | 4.96E-05 | 0.615 | GTGGGTGGC |
| 5325 | mamo-s | *BGIBMGA006407* | up | 1565 | 1573 | + | 11.3303 | 4.96E-05 | 0.615 | GTGGGTGGC |
| 5326 | mamo-s | *BGIBMGA006408* | down | 68 | 76 | - | 12.4679 | 1.31E-05 | 0.615 | GTGCGTGTC |
| 5327 | mamo-s | *BGIBMGA006408* | down | 1517 | 1525 | - | 10.5138 | 9.75E-05 | 0.865 | GTGCGTGTT |
| 5328 | mamo-s | *BGIBMGA006409* | down | 1436 | 1444 | + | 10.7523 | 8.38E-05 | 0.797 | AAGCGTGGC |
| 5329 | mamo-s | *BGIBMGA006410* | up | 599 | 607 | - | 11.3303 | 4.96E-05 | 0.615 | GTGGGTGGC |
| 5330 | mamo-s | *BGIBMGA006411* | down | 129 | 137 | + | 11.2018 | 5.68E-05 | 0.674 | CTGCGTGGT |
| 5331 | mamo-s | *BGIBMGA006413* | up | 936 | 944 | + | 12.4679 | 1.31E-05 | 0.615 | GTGCGTGTC |
| 5332 | mamo-s | *BGIBMGA006415* | up | 1556 | 1564 | + | 11.7798 | 3.11E-05 | 0.615 | ATGCGTGGC |
| 5333 | mamo-s | *BGIBMGA006417* | up | 1581 | 1589 | - | 11.3303 | 4.96E-05 | 0.615 | GTGGGTGGC |
| 5334 | mamo-s | *BGIBMGA006419* | down | 64 | 72 | + | 10.5138 | 9.75E-05 | 0.865 | GTGCGTGTT |
| 5335 | mamo-s | *BGIBMGA006420* | up | 1662 | 1670 | + | 12.1284 | 2.16E-05 | 0.615 | GTGCGAGGC |
| 5336 | mamo-s | *BGIBMGA006420* | up | 1729 | 1737 | - | 11.7064 | 3.52E-05 | 0.615 | GCGCGTGGT |
| 5337 | mamo-s | *BGIBMGA006420* | up | 469 | 477 | + | 11.3303 | 4.96E-05 | 0.615 | GTGGGTGGC |
| 5338 | mamo-s | *BGIBMGA006432* | up | 316 | 324 | + | 13.0275 | 8.53E-06 | 0.615 | GAGCGTGGC |
| 5339 | mamo-s | *BGIBMGA006432* | down | 345 | 353 | + | 11.3303 | 4.96E-05 | 0.615 | GTGGGTGGC |
| 5340 | mamo-s | *BGIBMGA006433* | down | 1079 | 1087 | + | 11.3303 | 4.96E-05 | 0.615 | GTGGGTGGC |
| 5341 | mamo-s | *BGIBMGA006434* | down | 981 | 989 | - | 13.0275 | 8.53E-06 | 0.615 | GAGCGTGGC |
| 5342 | mamo-s | *BGIBMGA006434* | down | 542 | 550 | + | 11.0826 | 6.84E-05 | 0.709 | GTACGTGGC |
| 5343 | mamo-s | *BGIBMGA006436* | up | 47 | 55 | - | 11.3303 | 4.96E-05 | 0.615 | GTGGGTGGC |
| 5344 | mamo-s | *BGIBMGA006437* | down | 663 | 671 | + | 11.0826 | 6.84E-05 | 0.709 | GTACGTGGC |
| 5345 | mamo-s | *BGIBMGA006440* | down | 1066 | 1074 | + | 11.3303 | 4.96E-05 | 0.615 | GTGGGTGGC |
| 5346 | mamo-s | *BGIBMGA006441* | up | 601 | 609 | - | 10.8073 | 7.89E-05 | 0.766 | CCGCGTGGT |
| 5347 | mamo-s | *BGIBMGA006441* | down | 1114 | 1122 | + | 10.5688 | 8.88E-05 | 0.824 | GTGCGTGAC |
| 5348 | mamo-s | *BGIBMGA006442* | up | 125 | 133 | + | 11.3303 | 4.96E-05 | 0.615 | GTGGGTGGC |
| 5349 | mamo-s | *BGIBMGA006443* | down | 696 | 704 | - | 11.5688 | 3.79E-05 | 0.615 | CTGCGTGTC |
| 5350 | mamo-s | *BGIBMGA006443* | up | 1318 | 1326 | + | 11.3303 | 4.96E-05 | 0.615 | GTGGGTGGC |
| 5351 | mamo-s | *BGIBMGA006445* | down | 1150 | 1158 | - | 12.7615 | 1.04E-05 | 0.615 | CCGCGTGGC |
| 5352 | mamo-s | *BGIBMGA006446* | down | 362 | 370 | + | 11.3303 | 4.96E-05 | 0.615 | GTGGGTGGC |
| 5353 | mamo-s | *BGIBMGA006452* | up | 576 | 584 | + | 11.3303 | 4.96E-05 | 0.615 | GTGGGTGGC |
| 5354 | mamo-s | *BGIBMGA006452* | up | 1270 | 1278 | - | 11.0826 | 6.84E-05 | 0.709 | GTACGTGGC |
| 5355 | mamo-s | *BGIBMGA006452* | down | 260 | 268 | + | 10.7523 | 8.38E-05 | 0.797 | AAGCGTGGC |
| 5356 | mamo-s | *BGIBMGA006453* | up | 1539 | 1547 | - | 11.3303 | 4.96E-05 | 0.615 | GTGGGTGGC |
| 5357 | mamo-s | *BGIBMGA006454* | up | 1550 | 1558 | - | 11.3303 | 4.96E-05 | 0.615 | GTGGGTGGC |
| 5358 | mamo-s | *BGIBMGA006454* | down | 1769 | 1777 | + | 11.3303 | 4.96E-05 | 0.615 | GTGGGTGGC |
| 5359 | mamo-s | *BGIBMGA006458* | down | 107 | 115 | + | 11.0734 | 7.12E-05 | 0.726 | GAGCGTGGT |
| 5360 | mamo-s | *BGIBMGA006461* | down | 171 | 179 | + | 11.3303 | 4.96E-05 | 0.615 | GTGGGTGGC |
| 5361 | mamo-s | *BGIBMGA006461* | down | 56 | 64 | + | 10.9358 | 7.30E-05 | 0.728 | GCGGGTGGC |

| | | | | | | | | | | |
|---|---|---|---|---|---|---|---|---|---|---|
| 5362 | mamo-s | *BGIBMGA006462* | down | 859 | 867 | + | 11.3303 | 4.96E-05 | 0.615 | GTGGGTGGC |
| 5363 | mamo-s | *BGIBMGA006463* | down | 96 | 104 | + | 11.0826 | 6.84E-05 | 0.709 | GTACGTGGC |
| 5364 | mamo-s | *BGIBMGA006464* | up | 193 | 201 | + | 11.5321 | 4.06E-05 | 0.615 | TTGCGTGGC |
| 5365 | mamo-s | *BGIBMGA006465* | down | 1774 | 1782 | + | 11.3303 | 4.96E-05 | 0.615 | GTGGGTGGC |
| 5366 | mamo-s | *BGIBMGA006467* | down | 1395 | 1403 | - | 11.3303 | 4.96E-05 | 0.615 | GTGGGTGGC |
| 5367 | mamo-s | *BGIBMGA006469* | up | 1829 | 1837 | - | 11.3303 | 4.96E-05 | 0.615 | GTGGGTGGC |
| 5368 | mamo-s | *BGIBMGA006469* | up | 698 | 706 | - | 10.8899 | 7.48E-05 | 0.739 | CCGCGTGCC |
| 5369 | mamo-s | *BGIBMGA006471* | down | 1257 | 1265 | - | 11.3303 | 4.96E-05 | 0.615 | GTGGGTGGC |
| 5370 | mamo-s | *BGIBMGA006474* | down | 1975 | 1983 | - | 11.3303 | 4.96E-05 | 0.615 | GTGGGTGGC |
| 5371 | mamo-s | *BGIBMGA006475* | up | 1533 | 1541 | - | 11.0826 | 6.84E-05 | 0.709 | GTACGTGGC |
| 5372 | mamo-s | *BGIBMGA006482* | up | 1602 | 1610 | - | 11.0826 | 6.84E-05 | 0.709 | GTACGTGGC |
| 5373 | mamo-s | *BGIBMGA006483* | up | 378 | 386 | + | 11.3303 | 4.96E-05 | 0.615 | GTGGGTGGC |
| 5374 | mamo-s | *BGIBMGA006483* | down | 1672 | 1680 | + | 11.3303 | 4.96E-05 | 0.615 | GTGGGTGGC |
| 5375 | mamo-s | *BGIBMGA006483* | down | 538 | 546 | - | 10.5688 | 8.88E-05 | 0.824 | GTGCGTGAC |
| 5376 | mamo-s | *BGIBMGA006484* | down | 1683 | 1691 | + | 10.7523 | 8.38E-05 | 0.797 | AAGCGTGGC |
| 5377 | mamo-s | *BGIBMGA006485* | down | 1258 | 1266 | + | 11.3303 | 4.96E-05 | 0.615 | GTGGGTGGC |
| 5378 | mamo-s | *BGIBMGA006486* | up | 1275 | 1283 | + | 12.1009 | 2.43E-05 | 0.615 | GTGCGTGGT |
| 5379 | mamo-s | *BGIBMGA006487* | up | 201 | 209 | + | 11.3303 | 4.96E-05 | 0.615 | GTGGGTGGC |
| 5380 | mamo-s | *BGIBMGA006488* | down | 1140 | 1148 | - | 11.0734 | 7.12E-05 | 0.726 | GAGCGTGGT |
| 5381 | mamo-s | *BGIBMGA006489* | down | 903 | 911 | + | 11.3303 | 4.96E-05 | 0.615 | GTGGGTGGC |
| 5382 | mamo-s | *BGIBMGA006490* | down | 784 | 792 | - | 10.5413 | 9.42E-05 | 0.857 | GTGCGAGTC |
| 5383 | mamo-s | *BGIBMGA006491* | down | 1477 | 1485 | - | 11.156 | 6.13E-05 | 0.705 | GAGCGTGCC |
| 5384 | mamo-s | *BGIBMGA006492* | down | 1968 | 1976 | - | 12.0734 | 2.66E-05 | 0.615 | GCGCGTGTC |
| 5385 | mamo-s | *BGIBMGA006492* | down | 1902 | 1910 | - | 11.0826 | 6.84E-05 | 0.709 | GTACGTGGC |
| 5386 | mamo-s | *BGIBMGA006492* | up | 1354 | 1362 | - | 10.8899 | 7.48E-05 | 0.739 | CCGCGTGCC |
| 5387 | mamo-s | *BGIBMGA006493* | down | 1590 | 1598 | + | 11.4404 | 4.51E-05 | 0.615 | GAGCGTGTC |
| 5388 | mamo-s | *BGIBMGA006494* | up | 148 | 156 | + | 11.4404 | 4.51E-05 | 0.615 | GAGCGTGTC |
| 5389 | mamo-s | *BGIBMGA006497* | up | 1240 | 1248 | - | 11.3303 | 4.96E-05 | 0.615 | GTGGGTGGC |
| 5390 | mamo-s | *BGIBMGA006498* | down | 1614 | 1622 | - | 11.4404 | 4.51E-05 | 0.615 | CGGCGTGGC |
| 5391 | mamo-s | *BGIBMGA006499* | down | 1451 | 1459 | + | 11.789 | 2.84E-05 | 0.615 | GCGCGTGCC |
| 5392 | mamo-s | *BGIBMGA006499* | down | 1378 | 1386 | + | 11.7064 | 3.52E-05 | 0.615 | GCGCGTGGT |
| 5393 | mamo-s | *BGIBMGA006500* | down | 970 | 978 | + | 11.3303 | 4.96E-05 | 0.615 | GTGGGTGGC |
| 5394 | mamo-s | *BGIBMGA006500* | up | 1183 | 1191 | + | 11.3303 | 4.96E-05 | 0.615 | GTGGGTGGC |
| 5395 | mamo-s | *BGIBMGA006502* | up | 284 | 292 | + | 11.3303 | 4.96E-05 | 0.615 | GTGGGTGGC |
| 5396 | mamo-s | *BGIBMGA006504* | up | 1663 | 1671 | - | 10.6881 | 8.61E-05 | 0.81 | GCACGTGGC |
| 5397 | mamo-s | *BGIBMGA006505* | down | 1737 | 1745 | - | 11.3303 | 4.96E-05 | 0.615 | GTGGGTGGC |
| 5398 | mamo-s | *BGIBMGA006510* | down | 184 | 192 | + | 10.5138 | 9.75E-05 | 0.865 | GTGCGTGTT |
| 5399 | mamo-s | *BGIBMGA006513* | up | 689 | 697 | + | 11.3303 | 4.96E-05 | 0.615 | GTGGGTGGC |
| 5400 | mamo-s | *BGIBMGA006516* | up | 385 | 393 | - | 12.3394 | 1.49E-05 | 0.615 | GGGCGTGGC |
| 5401 | mamo-s | *BGIBMGA006518* | down | 188 | 196 | + | 11.4404 | 4.51E-05 | 0.615 | GAGCGTGTC |
| 5402 | mamo-s | *BGIBMGA006518* | up | 1681 | 1689 | + | 10.7523 | 8.38E-05 | 0.797 | GGGCGTGTC |
| 5403 | mamo-s | *BGIBMGA006519* | up | 607 | 615 | + | 11.3303 | 4.96E-05 | 0.615 | GTGGGTGGC |
| 5404 | mamo-s | *BGIBMGA006521* | down | 1376 | 1384 | - | 13.6606 | 4.06E-06 | 0.615 | GCGCGTGGC |
| 5405 | mamo-s | *BGIBMGA006521* | up | 1403 | 1411 | - | 11.3303 | 4.96E-05 | 0.615 | GTGGGTGGC |
| 5406 | mamo-s | *BGIBMGA006521* | down | 1923 | 1931 | + | 11.3303 | 4.96E-05 | 0.615 | GTGGGTGGC |
| 5407 | mamo-s | *BGIBMGA006521* | up | 734 | 742 | - | 10.7523 | 8.38E-05 | 0.797 | AAGCGTGGC |
| 5408 | mamo-s | *BGIBMGA006522* | up | 1837 | 1845 | + | 11.3303 | 4.96E-05 | 0.615 | GTGGGTGGC |
| 5409 | mamo-s | *BGIBMGA006523* | up | 383 | 391 | + | 11.3303 | 4.96E-05 | 0.615 | GTGGGTGGC |
| 5410 | mamo-s | *BGIBMGA006523* | down | 1542 | 1550 | + | 10.8899 | 7.48E-05 | 0.739 | CCGCGTGCC |
| 5411 | mamo-s | *BGIBMGA006524* | up | 1606 | 1614 | - | 11.3303 | 4.96E-05 | 0.615 | GTGGGTGGC |
| 5412 | mamo-s | *BGIBMGA006524* | up | 905 | 913 | + | 10.8899 | 7.48E-05 | 0.739 | CCGCGTGCC |
| 5413 | mamo-s | *BGIBMGA006525* | up | 248 | 256 | + | 11.3303 | 4.96E-05 | 0.615 | GTGGGTGGC |
| 5414 | mamo-s | *BGIBMGA006525* | up | 436 | 444 | + | 11.3303 | 4.96E-05 | 0.615 | GTGGGTGGC |
| 5415 | mamo-s | *BGIBMGA006527* | down | 804 | 812 | + | 14.055 | 2.23E-06 | 0.615 | GTGCGTGGC |
| 5416 | mamo-s | *BGIBMGA006527* | down | 988 | 996 | + | 11.3303 | 4.96E-05 | 0.615 | GTGGGTGGC |
| 5417 | mamo-s | *BGIBMGA006527* | up | 1163 | 1171 | - | 11.3303 | 4.96E-05 | 0.615 | GTGGGTGGC |
| 5418 | mamo-s | *BGIBMGA006528* | up | 613 | 621 | - | 11.3303 | 4.96E-05 | 0.615 | GTGGGTGGC |
| 5419 | mamo-s | *BGIBMGA006528* | up | 1111 | 1119 | + | 11.3303 | 4.96E-05 | 0.615 | GTGGGTGGC |
| 5420 | mamo-s | *BGIBMGA006529* | up | 1016 | 1024 | + | 11.5321 | 4.06E-05 | 0.615 | TTGCGTGGC |
| 5421 | mamo-s | *BGIBMGA006530* | down | 1303 | 1311 | - | 11.0826 | 6.84E-05 | 0.709 | GTACGTGGC |
| 5422 | mamo-s | *BGIBMGA006532* | down | 1165 | 1173 | - | 11.3303 | 4.96E-05 | 0.615 | GTGGGTGGC |
| 5423 | mamo-s | *BGIBMGA006533* | up | 299 | 307 | - | 11.0826 | 6.84E-05 | 0.709 | GTACGTGGC |
| 5424 | mamo-s | *BGIBMGA006535* | up | 989 | 997 | - | 11.3303 | 4.96E-05 | 0.615 | GTGGGTGGC |
| 5425 | mamo-s | *BGIBMGA006536* | up | 414 | 422 | + | 11.3303 | 4.96E-05 | 0.615 | GTGGGTGGC |
| 5426 | mamo-s | *BGIBMGA006537* | up | 851 | 859 | - | 11.3303 | 4.96E-05 | 0.615 | GTGGGTGGC |
| 5427 | mamo-s | *BGIBMGA006539* | up | 1371 | 1379 | + | 12.1009 | 2.43E-05 | 0.615 | GTGCGTGGT |
| 5428 | mamo-s | *BGIBMGA006539* | up | 1555 | 1563 | - | 11.3303 | 4.96E-05 | 0.615 | GTGGGTGGC |
| 5429 | mamo-s | *BGIBMGA006541* | down | 1239 | 1247 | + | 11.3303 | 4.96E-05 | 0.615 | GTGGGTGGC |
| 5430 | mamo-s | *BGIBMGA006542* | up | 441 | 449 | - | 11.3303 | 4.96E-05 | 0.615 | GTGGGTGGC |
| 5431 | mamo-s | *BGIBMGA006544* | up | 1094 | 1102 | + | 14.055 | 2.23E-06 | 0.615 | GTGCGTGGC |
| 5432 | mamo-s | *BGIBMGA006544* | down | 1611 | 1619 | - | 11.4404 | 4.51E-05 | 0.615 | GAGCGTGTC |
| 5433 | mamo-s | *BGIBMGA006545* | up | 1200 | 1208 | + | 10.5413 | 9.42E-05 | 0.857 | CAGCGTGTC |
| 5434 | mamo-s | *BGIBMGA006547* | up | 1013 | 1021 | + | 11.3303 | 4.96E-05 | 0.615 | GTGGGTGGC |
| 5435 | mamo-s | *BGIBMGA006550* | up | 900 | 908 | + | 11.3303 | 4.96E-05 | 0.615 | GTGGGTGGC |
| 5436 | mamo-s | *BGIBMGA006552* | up | 990 | 998 | - | 11.3303 | 4.96E-05 | 0.615 | GTGGGTGGC |
| 5437 | mamo-s | *BGIBMGA006552* | down | 1719 | 1727 | - | 10.8349 | 7.66E-05 | 0.751 | CCGCGAGGC |
| 5438 | mamo-s | *BGIBMGA006553* | up | 799 | 807 | + | 11.3303 | 4.96E-05 | 0.615 | GTGGGTGGC |
| 5439 | mamo-s | *BGIBMGA006553* | up | 1709 | 1717 | + | 11.3303 | 4.96E-05 | 0.615 | GTGGGTGGC |
| 5440 | mamo-s | *BGIBMGA006553* | down | 1071 | 1079 | + | 11.0826 | 6.84E-05 | 0.709 | GTACGTGGC |
| 5441 | mamo-s | *BGIBMGA006554* | down | 462 | 470 | + | 11.3303 | 4.96E-05 | 0.615 | GTGGGTGGC |
| 5442 | mamo-s | *BGIBMGA006554* | down | 1092 | 1100 | - | 11.3303 | 4.96E-05 | 0.615 | GTGGGTGGC |
| 5443 | mamo-s | *BGIBMGA006554* | up | 1934 | 1942 | - | 11.0826 | 6.84E-05 | 0.709 | GTACGTGGC |
| 5444 | mamo-s | *BGIBMGA006555* | up | 1702 | 1710 | - | 11.3303 | 4.96E-05 | 0.615 | GTGGGTGGC |
| 5445 | mamo-s | *BGIBMGA006557* | up | 428 | 436 | - | 12.3394 | 1.49E-05 | 0.615 | GGGCGTGGC |
| 5446 | mamo-s | *BGIBMGA006557* | down | 88 | 96 | + | 11.7798 | 3.11E-05 | 0.615 | ATGCGTGGC |
| 5447 | mamo-s | *BGIBMGA006558* | down | 762 | 770 | + | 11.7339 | 3.29E-05 | 0.615 | GCGCGAGGC |
| 5448 | mamo-s | *BGIBMGA006558* | down | 1006 | 1014 | + | 11.3303 | 4.96E-05 | 0.615 | GTGGGTGGC |
| 5449 | mamo-s | *BGIBMGA006562* | up | 701 | 709 | + | 14.055 | 2.23E-06 | 0.615 | GTGCGTGGC |

| | | | | | | | | | | |
|---|---|---|---|---|---|---|---|---|---|---|
| 5450 | mamo-s | *BGIBMGA006566* | up | 370 | 378 | + | 11.3303 | 4.96E-05 | 0.615 | GTGGGTGGC |
| 5451 | mamo-s | *BGIBMGA006569* | down | 136 | 144 | + | 11.3303 | 4.96E-05 | 0.615 | GTGGGTGGC |
| 5452 | mamo-s | *BGIBMGA006570* | up | 1204 | 1212 | - | 14.055 | 2.23E-06 | 0.615 | GTGCGTGGC |
| 5453 | mamo-s | *BGIBMGA006570* | down | 185 | 193 | - | 11.5321 | 4.06E-05 | 0.615 | TTGCGTGGC |
| 5454 | mamo-s | *BGIBMGA006571* | down | 1332 | 1340 | - | 13.0275 | 8.53E-06 | 0.615 | GAGCGTGGC |
| 5455 | mamo-s | *BGIBMGA006571* | down | 318 | 326 | - | 11.4404 | 4.51E-05 | 0.615 | CGGCGTGGC |
| 5456 | mamo-s | *BGIBMGA006573* | up | 64 | 72 | + | 11.3303 | 4.96E-05 | 0.615 | GTGGGTGGC |
| 5457 | mamo-s | *BGIBMGA006574* | down | 1953 | 1961 | - | 10.5688 | 8.88E-05 | 0.824 | GTGCGTGAC |
| 5458 | mamo-s | *BGIBMGA006575* | up | 982 | 990 | - | 10.5688 | 8.88E-05 | 0.824 | GTGCGTGAC |
| 5459 | mamo-s | *BGIBMGA006577* | up | 173 | 181 | - | 11.1743 | 5.90E-05 | 0.684 | CCGCGTGTC |
| 5460 | mamo-s | *BGIBMGA006585* | down | 831 | 839 | - | 11.3303 | 4.96E-05 | 0.615 | GTGGGTGGC |
| 5461 | mamo-s | *BGIBMGA006585* | down | 1409 | 1417 | - | 11.3303 | 4.96E-05 | 0.615 | GTGGGTGGC |
| 5462 | mamo-s | *BGIBMGA006587* | up | 1232 | 1240 | + | 11.3303 | 4.96E-05 | 0.615 | GTGGGTGGC |
| 5463 | mamo-s | *BGIBMGA006589* | up | 650 | 658 | + | 11.3303 | 4.96E-05 | 0.615 | GTGGGTGGC |
| 5464 | mamo-s | *BGIBMGA006589* | down | 1201 | 1209 | + | 11.3303 | 4.96E-05 | 0.615 | GTGGGTGGC |
| 5465 | mamo-s | *BGIBMGA006590* | down | 1083 | 1091 | + | 11.3303 | 4.96E-05 | 0.615 | GTGGGTGGC |
| 5466 | mamo-s | *BGIBMGA006591* | down | 903 | 911 | - | 11.2844 | 5.18E-05 | 0.63 | CTGCGTGCC |
| 5467 | mamo-s | *BGIBMGA006591* | down | 1206 | 1214 | - | 10.5138 | 9.75E-05 | 0.865 | GTGCGTGTT |
| 5468 | mamo-s | *BGIBMGA006593* | down | 453 | 461 | - | 11.0826 | 6.84E-05 | 0.709 | GTACGTGGC |
| 5469 | mamo-s | *BGIBMGA006595* | up | 981 | 989 | - | 12.3394 | 1.49E-05 | 0.615 | GGGCGTGGC |
| 5470 | mamo-s | *BGIBMGA006595* | down | 1843 | 1851 | - | 11.2844 | 5.18E-05 | 0.63 | CTGCGTGCC |
| 5471 | mamo-s | *BGIBMGA006597* | down | 1590 | 1598 | - | 11.3303 | 4.96E-05 | 0.615 | GTGGGTGGC |
| 5472 | mamo-s | *BGIBMGA006598* | up | 217 | 225 | + | 11.0826 | 6.84E-05 | 0.709 | GTACGTGGC |
| 5473 | mamo-s | *BGIBMGA006599* | down | 145 | 153 | + | 11.3303 | 4.96E-05 | 0.615 | GTGGGTGGC |
| 5474 | mamo-s | *BGIBMGA006600* | down | 1262 | 1270 | - | 11.3303 | 4.96E-05 | 0.615 | GTGGGTGGC |
| 5475 | mamo-s | *BGIBMGA006600* | up | 1637 | 1645 | + | 11.3303 | 4.96E-05 | 0.615 | GTGGGTGGC |
| 5476 | mamo-s | *BGIBMGA006600* | up | 660 | 668 | - | 11.0826 | 6.84E-05 | 0.709 | GTACGTGGC |
| 5477 | mamo-s | *BGIBMGA006603* | up | 1327 | 1335 | + | 11.3303 | 4.96E-05 | 0.615 | GTGGGTGGC |
| 5478 | mamo-s | *BGIBMGA006603* | down | 11 | 19 | + | 11.1743 | 5.90E-05 | 0.684 | CCGCGTGTC |
| 5479 | mamo-s | *BGIBMGA006603* | down | 151 | 159 | + | 10.9358 | 7.30E-05 | 0.728 | GCGGGTGGC |
| 5480 | mamo-s | *BGIBMGA006603* | down | 662 | 670 | - | 10.5413 | 9.42E-05 | 0.857 | GTGCGAGTC |
| 5481 | mamo-s | *BGIBMGA006603* | up | 931 | 939 | - | 10.5138 | 9.75E-05 | 0.865 | GTGCGTGTT |
| 5482 | mamo-s | *BGIBMGA006605* | down | 1384 | 1392 | + | 11.3303 | 4.96E-05 | 0.615 | GTGGGTGGC |
| 5483 | mamo-s | *BGIBMGA006606* | up | 356 | 364 | + | 11.3303 | 4.96E-05 | 0.615 | GTGGGTGGC |
| 5484 | mamo-s | *BGIBMGA006606* | up | 1304 | 1312 | - | 11.0826 | 6.84E-05 | 0.709 | GTACGTGGC |
| 5485 | mamo-s | *BGIBMGA006610* | up | 580 | 588 | + | 11.3303 | 4.96E-05 | 0.615 | GTGGGTGGC |
| 5486 | mamo-s | *BGIBMGA006611* | up | 1327 | 1335 | + | 12.1284 | 2.16E-05 | 0.615 | GTGCGAGGC |
| 5487 | mamo-s | *BGIBMGA006613* | down | 830 | 838 | - | 10.6881 | 8.61E-05 | 0.81 | GCACGTGGC |
| 5488 | mamo-s | *BGIBMGA006614* | up | 280 | 288 | - | 11.1009 | 6.57E-05 | 0.709 | GAGCGAGGC |
| 5489 | mamo-s | *BGIBMGA006614* | down | 1343 | 1351 | - | 10.5413 | 9.42E-05 | 0.857 | GTGCGAGTC |
| 5490 | mamo-s | *BGIBMGA006621* | down | 1045 | 1053 | - | 11.3303 | 4.96E-05 | 0.615 | GTGGGTGGC |
| 5491 | mamo-s | *BGIBMGA006621* | down | 33 | 41 | - | 10.9358 | 7.30E-05 | 0.728 | GCGGGTGGC |
| 5492 | mamo-s | *BGIBMGA006622* | up | 1925 | 1933 | + | 10.9358 | 7.30E-05 | 0.728 | GCGGGTGGC |
| 5493 | mamo-s | *BGIBMGA006623* | down | 1859 | 1867 | + | 11.7339 | 3.29E-05 | 0.615 | GCGCGAGGC |
| 5494 | mamo-s | *BGIBMGA006624* | up | 237 | 245 | + | 11.3303 | 4.96E-05 | 0.615 | GTGGGTGGC |
| 5495 | mamo-s | *BGIBMGA006624* | up | 360 | 368 | + | 11.0826 | 6.84E-05 | 0.709 | GTACGTGGC |
| 5496 | mamo-s | *BGIBMGA006625* | down | 365 | 373 | + | 11.3303 | 4.96E-05 | 0.615 | GTGGGTGGC |
| 5497 | mamo-s | *BGIBMGA006626* | down | 213 | 221 | - | 11.3303 | 4.96E-05 | 0.615 | GTGGGTGGC |
| 5498 | mamo-s | *BGIBMGA006628* | down | 1049 | 1057 | - | 11.3303 | 4.96E-05 | 0.615 | GTGGGTGGC |
| 5499 | mamo-s | *BGIBMGA006629* | up | 182 | 190 | - | 11.3303 | 4.96E-05 | 0.615 | GTGGGTGGC |
| 5500 | mamo-s | *BGIBMGA006630* | down | 1656 | 1664 | + | 11.3303 | 4.96E-05 | 0.615 | GTGGGTGGC |
| 5501 | mamo-s | *BGIBMGA006632* | down | 1640 | 1648 | - | 11.789 | 2.84E-05 | 0.615 | GCGCGTGCC |
| 5502 | mamo-s | *BGIBMGA006632* | down | 633 | 641 | - | 11.3303 | 4.96E-05 | 0.615 | GTGGGTGGC |
| 5503 | mamo-s | *BGIBMGA006634* | down | 1562 | 1570 | + | 11.3303 | 4.96E-05 | 0.615 | GTGGGTGGC |
| 5504 | mamo-s | *BGIBMGA006635* | down | 1760 | 1768 | - | 11.3303 | 4.96E-05 | 0.615 | GTGGGTGGC |
| 5505 | mamo-s | *BGIBMGA006636* | up | 974 | 982 | + | 11.789 | 2.84E-05 | 0.615 | GCGCGTGCC |
| 5506 | mamo-s | *BGIBMGA006637* | up | 984 | 992 | + | 11.3303 | 4.96E-05 | 0.615 | GTGGGTGGC |
| 5507 | mamo-s | *BGIBMGA006638* | down | 1390 | 1398 | + | 11.3303 | 4.96E-05 | 0.615 | GTGGGTGGC |
| 5508 | mamo-s | *BGIBMGA006639* | up | 1819 | 1827 | + | 11.0826 | 6.84E-05 | 0.709 | GTACGTGGC |
| 5509 | mamo-s | *BGIBMGA006639* | up | 433 | 441 | - | 10.6881 | 8.61E-05 | 0.81 | GCACGTGGC |
| 5510 | mamo-s | *BGIBMGA006640* | down | 1130 | 1138 | + | 12.1284 | 2.16E-05 | 0.615 | GTGCGAGGC |
| 5511 | mamo-s | *BGIBMGA006641* | down | 1677 | 1685 | - | 10.5138 | 9.75E-05 | 0.865 | GTGCGTGTT |
| 5512 | mamo-s | *BGIBMGA006644* | down | 704 | 712 | + | 11.3303 | 4.96E-05 | 0.615 | GTGGGTGGC |
| 5513 | mamo-s | *BGIBMGA006644* | up | 1433 | 1441 | - | 11.3303 | 4.96E-05 | 0.615 | GTGGGTGGC |
| 5514 | mamo-s | *BGIBMGA006644* | up | 116 | 124 | - | 10.9358 | 7.30E-05 | 0.728 | GCGGGTGGC |
| 5515 | mamo-s | *BGIBMGA006644* | up | 1868 | 1876 | + | 10.5688 | 8.88E-05 | 0.824 | GTGCGTGAC |
| 5516 | mamo-s | *BGIBMGA006645* | down | 1019 | 1027 | + | 11.4404 | 4.51E-05 | 0.615 | CGGCGTGGC |
| 5517 | mamo-s | *BGIBMGA006646* | down | 1361 | 1369 | + | 10.5688 | 8.88E-05 | 0.824 | GTGCGTGAC |
| 5518 | mamo-s | *BGIBMGA006646* | up | 1880 | 1888 | + | 10.5688 | 8.88E-05 | 0.824 | GTGCGTGAC |
| 5519 | mamo-s | *BGIBMGA006647* | up | 849 | 857 | - | 11.3303 | 4.96E-05 | 0.615 | GTGGGTGGC |
| 5520 | mamo-s | *BGIBMGA006649* | up | 1877 | 1885 | - | 10.5413 | 9.42E-05 | 0.857 | CAGCGTGTC |
| 5521 | mamo-s | *BGIBMGA006650* | down | 1098 | 1106 | - | 11.7064 | 3.52E-05 | 0.615 | GCGCGTGGT |
| 5522 | mamo-s | *BGIBMGA006650* | up | 1637 | 1645 | - | 11.3303 | 4.96E-05 | 0.615 | GTGGGTGGC |
| 5523 | mamo-s | *BGIBMGA006651* | down | 279 | 287 | - | 11.0734 | 7.12E-05 | 0.726 | GAGCGTGGT |
| 5524 | mamo-s | *BGIBMGA006655* | down | 1826 | 1834 | - | 10.7523 | 8.38E-05 | 0.797 | GGGCGTGTC |
| 5525 | mamo-s | *BGIBMGA006656* | down | 1477 | 1485 | - | 10.5413 | 9.42E-05 | 0.857 | CAGCGTGTC |
| 5526 | mamo-s | *BGIBMGA006657* | down | 1563 | 1571 | - | 11.3303 | 4.96E-05 | 0.615 | GTGGGTGGC |
| 5527 | mamo-s | *BGIBMGA006658* | up | 83 | 91 | - | 11.3303 | 4.96E-05 | 0.615 | GTGGGTGGC |
| 5528 | mamo-s | *BGIBMGA006659* | down | 1884 | 1892 | - | 12.4679 | 1.31E-05 | 0.615 | GTGCGTGTC |
| 5529 | mamo-s | *BGIBMGA006659* | up | 142 | 150 | + | 11.3303 | 4.96E-05 | 0.615 | GTGGGTGGC |
| 5530 | mamo-s | *BGIBMGA006662* | up | 206 | 214 | - | 11.3303 | 4.96E-05 | 0.615 | GTGGGTGGC |
| 5531 | mamo-s | *BGIBMGA006663* | down | 407 | 415 | - | 12.1284 | 2.16E-05 | 0.615 | CAGCGTGGC |
| 5532 | mamo-s | *BGIBMGA006667* | down | 1001 | 1009 | + | 11.3303 | 4.96E-05 | 0.615 | GTGGGTGGC |
| 5533 | mamo-s | *BGIBMGA006667* | up | 922 | 930 | + | 11.0826 | 6.84E-05 | 0.709 | GTACGTGGC |
| 5534 | mamo-s | *BGIBMGA006668* | down | 1122 | 1130 | - | 11.3303 | 4.96E-05 | 0.615 | GTGGGTGGC |
| 5535 | mamo-s | *BGIBMGA006668* | up | 1132 | 1140 | - | 11.3303 | 4.96E-05 | 0.615 | GTGGGTGGC |
| 5536 | mamo-s | *BGIBMGA006668* | up | 1806 | 1814 | + | 11.3303 | 4.96E-05 | 0.615 | GTGGGTGGC |
| 5537 | mamo-s | *BGIBMGA006671* | up | 259 | 267 | - | 12.4679 | 1.31E-05 | 0.615 | GTGCGTGTC |

| | | | | | | | | | | |
|---|---|---|---|---|---|---|---|---|---|---|
| 5538 | mamo-s | *BGIBMGA006671* | up | 1198 | 1206 | + | 10.7523 | 8.38E-05 | 0.797 | GGGCGTGTC |
| 5539 | mamo-s | *BGIBMGA006671* | up | 141 | 149 | + | 10.6881 | 8.61E-05 | 0.81 | GCACGTGGC |
| 5540 | mamo-s | *BGIBMGA006673* | down | 1216 | 1224 | - | 11.3303 | 4.96E-05 | 0.615 | GTGGGTGGC |
| 5541 | mamo-s | *BGIBMGA006674* | down | 561 | 569 | - | 11.4404 | 4.51E-05 | 0.615 | CGGCGTGGC |
| 5542 | mamo-s | *BGIBMGA006674* | up | 1821 | 1829 | - | 10.7523 | 8.38E-05 | 0.797 | AAGCGTGGC |
| 5543 | mamo-s | *BGIBMGA006675* | up | 438 | 446 | - | 11.0826 | 6.84E-05 | 0.709 | GTACGTGGC |
| 5544 | mamo-s | *BGIBMGA006676* | down | 34 | 42 | - | 11.7339 | 3.29E-05 | 0.615 | GCGCGAGGC |
| 5545 | mamo-s | *BGIBMGA006676* | down | 807 | 815 | + | 11.7339 | 3.29E-05 | 0.615 | GCGCGAGGC |
| 5546 | mamo-s | *BGIBMGA006679* | down | 814 | 822 | + | 11.3303 | 4.96E-05 | 0.615 | GTGGGTGGC |
| 5547 | mamo-s | *BGIBMGA006679* | up | 1042 | 1050 | - | 11.3303 | 4.96E-05 | 0.615 | GTGGGTGGC |
| 5548 | mamo-s | *BGIBMGA006681* | up | 1648 | 1656 | + | 11.3303 | 4.96E-05 | 0.615 | GTGGGTGGC |
| 5549 | mamo-s | *BGIBMGA006683* | up | 330 | 338 | - | 11.3303 | 4.96E-05 | 0.615 | GTGGGTGGC |
| 5550 | mamo-s | *BGIBMGA006684* | down | 1758 | 1766 | - | 11.3303 | 4.96E-05 | 0.615 | GTGGGTGGC |
| 5551 | mamo-s | *BGIBMGA006685* | down | 245 | 253 | + | 11.3853 | 4.74E-05 | 0.615 | ACGCGTGGC |
| 5552 | mamo-s | *BGIBMGA006685* | up | 1679 | 1687 | - | 10.9358 | 7.30E-05 | 0.728 | GCGGGTGGC |
| 5553 | mamo-s | *BGIBMGA006686* | up | 1565 | 1573 | - | 13.6606 | 4.06E-06 | 0.615 | GCGCGTGGC |
| 5554 | mamo-s | *BGIBMGA006687* | up | 1141 | 1149 | - | 12.1284 | 2.16E-05 | 0.615 | GTGCGAGGC |
| 5555 | mamo-s | *BGIBMGA006687* | up | 1007 | 1015 | + | 12.0734 | 2.66E-05 | 0.615 | GCGCGTGTC |
| 5556 | mamo-s | *BGIBMGA006688* | down | 1759 | 1767 | + | 12.1835 | 1.71E-05 | 0.615 | GTGCGTGCC |
| 5557 | mamo-s | *BGIBMGA006688* | up | 1800 | 1808 | + | 12.1835 | 1.71E-05 | 0.615 | GTGCGTGCC |
| 5558 | mamo-s | *BGIBMGA006690* | up | 152 | 160 | - | 10.5688 | 8.88E-05 | 0.824 | GTGCGTGAC |
| 5559 | mamo-s | *BGIBMGA006692* | down | 165 | 173 | + | 11.3303 | 4.96E-05 | 0.615 | GTGGGTGGC |
| 5560 | mamo-s | *BGIBMGA006693* | down | 1529 | 1537 | + | 11.2844 | 5.18E-05 | 0.63 | CTGCGTGCC |
| 5561 | mamo-s | *BGIBMGA006694* | up | 1971 | 1979 | + | 12.0734 | 2.66E-05 | 0.615 | GCGCGTGTC |
| 5562 | mamo-s | *BGIBMGA006695* | up | 1640 | 1648 | + | 11.3303 | 4.96E-05 | 0.615 | GTGGGTGGC |
| 5563 | mamo-s | *BGIBMGA006695* | up | 57 | 65 | + | 10.7523 | 8.38E-05 | 0.797 | GGGCGTGTC |
| 5564 | mamo-s | *BGIBMGA006696* | down | 550 | 558 | + | 10.5413 | 9.42E-05 | 0.857 | CAGCGTGTC |
| 5565 | mamo-s | *BGIBMGA006697* | up | 567 | 575 | + | 12.1284 | 2.16E-05 | 0.615 | GTGCGAGGC |
| 5566 | mamo-s | *BGIBMGA006697* | up | 1295 | 1303 | + | 10.5413 | 9.42E-05 | 0.857 | CAGCGTGTC |
| 5567 | mamo-s | *BGIBMGA006700* | down | 1966 | 1974 | - | 11.2018 | 5.68E-05 | 0.674 | CTGCGTGGT |
| 5568 | mamo-s | *BGIBMGA006706* | up | 71 | 79 | - | 11.4404 | 4.51E-05 | 0.615 | GAGCGTGTC |
| 5569 | mamo-s | *BGIBMGA006707* | down | 909 | 917 | + | 11.3303 | 4.96E-05 | 0.615 | GTGGGTGGC |
| 5570 | mamo-s | *BGIBMGA006707* | down | 1822 | 1830 | - | 11.3303 | 4.96E-05 | 0.615 | GTGGGTGGC |
| 5571 | mamo-s | *BGIBMGA006711* | up | 1429 | 1437 | - | 11.3303 | 4.96E-05 | 0.615 | GTGGGTGGC |
| 5572 | mamo-s | *BGIBMGA006713* | down | 399 | 407 | + | 11.0826 | 6.84E-05 | 0.709 | GTACGTGGC |
| 5573 | mamo-s | *BGIBMGA006714* | up | 1847 | 1855 | - | 11.1009 | 6.57E-05 | 0.709 | GAGCGAGGC |
| 5574 | mamo-s | *BGIBMGA006715* | down | 1547 | 1555 | + | 11.3303 | 4.96E-05 | 0.615 | GTGGGTGGC |
| 5575 | mamo-s | *BGIBMGA006716* | down | 1413 | 1421 | - | 11.3303 | 4.96E-05 | 0.615 | GTGGGTGGC |
| 5576 | mamo-s | *BGIBMGA006717* | up | 1767 | 1775 | - | 11.3303 | 4.96E-05 | 0.615 | GTGGGTGGC |
| 5577 | mamo-s | *BGIBMGA006718* | down | 1891 | 1899 | + | 11.156 | 6.13E-05 | 0.705 | GAGCGTGCC |
| 5578 | mamo-s | *BGIBMGA006721* | up | 761 | 769 | + | 11.3303 | 4.96E-05 | 0.615 | GTGGGTGGC |
| 5579 | mamo-s | *BGIBMGA006721* | up | 1539 | 1547 | + | 11.0826 | 6.84E-05 | 0.709 | GTACGTGGC |
| 5580 | mamo-s | *BGIBMGA006725* | up | 1137 | 1145 | - | 10.9358 | 7.30E-05 | 0.728 | GCGGGTGGC |
| 5581 | mamo-s | *BGIBMGA006726* | down | 1176 | 1184 | - | 10.9358 | 7.30E-05 | 0.728 | GCGGGTGGC |
| 5582 | mamo-s | *BGIBMGA006728* | down | 1975 | 1983 | + | 11.4404 | 4.51E-05 | 0.615 | GAGCGTGTC |
| 5583 | mamo-s | *BGIBMGA006729* | down | 1496 | 1504 | + | 10.5138 | 9.75E-05 | 0.865 | GTGCGTGTT |
| 5584 | mamo-s | *BGIBMGA006730* | up | 263 | 271 | - | 11.3303 | 4.96E-05 | 0.615 | GTGGGTGGC |
| 5585 | mamo-s | *BGIBMGA006732* | down | 1480 | 1488 | + | 10.6881 | 8.61E-05 | 0.81 | GCACGTGGC |
| 5586 | mamo-s | *BGIBMGA006736* | down | 169 | 177 | + | 11.1376 | 6.35E-05 | 0.709 | TCGCGTGGC |
| 5587 | mamo-s | *BGIBMGA006736* | up | 1448 | 1456 | + | 10.7523 | 8.38E-05 | 0.797 | AAGCGTGGC |
| 5588 | mamo-s | *BGIBMGA006738* | up | 165 | 173 | + | 12.1284 | 2.16E-05 | 0.615 | CAGCGTGGC |
| 5589 | mamo-s | *BGIBMGA006739* | down | 1459 | 1467 | + | 11.3303 | 4.96E-05 | 0.615 | GTGGGTGGC |
| 5590 | mamo-s | *BGIBMGA006740* | up | 360 | 368 | + | 11.3303 | 4.96E-05 | 0.615 | GTGGGTGGC |
| 5591 | mamo-s | *BGIBMGA006740* | up | 310 | 318 | - | 11.2844 | 5.18E-05 | 0.63 | CTGCGTGCC |
| 5592 | mamo-s | *BGIBMGA006741* | up | 1033 | 1041 | + | 11.3303 | 4.96E-05 | 0.615 | GTGGGTGGC |
| 5593 | mamo-s | *BGIBMGA006742* | up | 1656 | 1664 | - | 11.3303 | 4.96E-05 | 0.615 | GTGGGTGGC |
| 5594 | mamo-s | *BGIBMGA006742* | up | 142 | 150 | - | 11.0826 | 6.84E-05 | 0.709 | GTACGTGGC |
| 5595 | mamo-s | *BGIBMGA006746* | up | 481 | 489 | - | 11.2844 | 5.18E-05 | 0.63 | CTGCGTGCC |
| 5596 | mamo-s | *BGIBMGA006747* | down | 1042 | 1050 | - | 11.3303 | 4.96E-05 | 0.615 | GTGGGTGGC |
| 5597 | mamo-s | *BGIBMGA006747* | down | 1813 | 1821 | - | 11.3303 | 4.96E-05 | 0.615 | GTGGGTGGC |
| 5598 | mamo-s | *BGIBMGA006747* | down | 1705 | 1713 | + | 10.5413 | 9.42E-05 | 0.857 | CAGCGTGTC |
| 5599 | mamo-s | *BGIBMGA006750* | down | 1930 | 1938 | - | 11.1743 | 5.90E-05 | 0.684 | CCGCGTGTC |
| 5600 | mamo-s | *BGIBMGA006751* | down | 1156 | 1164 | + | 11.5321 | 4.06E-05 | 0.615 | TTGCGTGGC |
| 5601 | mamo-s | *BGIBMGA006752* | up | 202 | 210 | + | 11.5321 | 4.06E-05 | 0.615 | TTGCGTGGC |
| 5602 | mamo-s | *BGIBMGA006755* | up | 582 | 590 | + | 11.3303 | 4.96E-05 | 0.615 | GTGGGTGGC |
| 5603 | mamo-s | *BGIBMGA006755* | down | 650 | 658 | + | 11.3303 | 4.96E-05 | 0.615 | GTGGGTGGC |
| 5604 | mamo-s | *BGIBMGA006756* | up | 834 | 842 | + | 11.3303 | 4.96E-05 | 0.615 | GTGGGTGGC |
| 5605 | mamo-s | *BGIBMGA006759* | up | 1444 | 1452 | + | 11.3303 | 4.96E-05 | 0.615 | GTGGGTGGC |
| 5606 | mamo-s | *BGIBMGA006763* | up | 789 | 797 | - | 13.6606 | 4.06E-06 | 0.615 | GCGCGTGGC |
| 5607 | mamo-s | *BGIBMGA006763* | up | 650 | 658 | + | 13.6606 | 4.06E-06 | 0.615 | GCGCGTGGC |
| 5608 | mamo-s | *BGIBMGA006763* | up | 70 | 78 | - | 11.3303 | 4.96E-05 | 0.615 | GTGGGTGGC |
| 5609 | mamo-s | *BGIBMGA006764* | down | 42 | 50 | + | 11.7339 | 3.29E-05 | 0.615 | GCGCGAGGC |
| 5610 | mamo-s | *BGIBMGA006764* | down | 1580 | 1588 | + | 11.3303 | 4.96E-05 | 0.615 | GTGGGTGGC |
| 5611 | mamo-s | *BGIBMGA006765* | up | 258 | 266 | - | 11.3303 | 4.96E-05 | 0.615 | GTGGGTGGC |
| 5612 | mamo-s | *BGIBMGA006765* | up | 1047 | 1055 | - | 11.3303 | 4.96E-05 | 0.615 | GTGGGTGGC |
| 5613 | mamo-s | *BGIBMGA006765* | down | 1659 | 1667 | + | 11.2294 | 5.41E-05 | 0.65 | CTGCGAGGC |
| 5614 | mamo-s | *BGIBMGA006765* | up | 48 | 56 | - | 10.5413 | 9.42E-05 | 0.857 | GTGCGAGTC |
| 5615 | mamo-s | *BGIBMGA006766* | up | 516 | 524 | + | 11.3303 | 4.96E-05 | 0.615 | GTGGGTGGC |
| 5616 | mamo-s | *BGIBMGA006767* | up | 1157 | 1165 | + | 10.5138 | 9.75E-05 | 0.865 | GTGCGTGTT |
| 5617 | mamo-s | *BGIBMGA006768* | up | 1850 | 1858 | - | 13.0275 | 8.53E-06 | 0.615 | GAGCGTGGC |
| 5618 | mamo-s | *BGIBMGA006768* | down | 1132 | 1140 | - | 11.3303 | 4.96E-05 | 0.615 | GTGGGTGGC |
| 5619 | mamo-s | *BGIBMGA006768* | up | 749 | 757 | - | 10.5413 | 9.42E-05 | 0.857 | GTGCGAGTC |
| 5620 | mamo-s | *BGIBMGA006770* | down | 1110 | 1118 | - | 11.5321 | 4.06E-05 | 0.615 | TTGCGTGGC |
| 5621 | mamo-s | *BGIBMGA006770* | up | 1006 | 1014 | + | 11.1743 | 5.90E-05 | 0.684 | CCGCGTGTC |
| 5622 | mamo-s | *BGIBMGA006770* | up | 1770 | 1778 | - | 10.5688 | 8.88E-05 | 0.824 | GTGCGTGAC |
| 5623 | mamo-s | *BGIBMGA006772* | down | 1762 | 1770 | - | 11.3303 | 4.96E-05 | 0.615 | GTGGGTGGC |
| 5624 | mamo-s | *BGIBMGA006773* | up | 1846 | 1854 | + | 12.1835 | 1.71E-05 | 0.615 | GTGCGTGCC |
| 5625 | mamo-s | *BGIBMGA006774* | down | 1695 | 1703 | + | 12.1009 | 2.43E-05 | 0.615 | GTGCGTGGT |

| | | | | | | | | | | |
|---|---|---|---|---|---|---|---|---|---|---|
| 5626 | mamo-s | *BGIBMGA006774* | up | 1823 | 1831 | - | 11.3303 | 4.96E-05 | 0.615 | GTGGGTGGC |
| 5627 | mamo-s | *BGIBMGA006774* | up | 1180 | 1188 | - | 10.7523 | 8.38E-05 | 0.797 | GGGCGTGTC |
| 5628 | mamo-s | *BGIBMGA006777* | down | 1091 | 1099 | + | 12.1284 | 2.16E-05 | 0.615 | GTGCGAGGC |
| 5629 | mamo-s | *BGIBMGA006780* | down | 1695 | 1703 | - | 11.3303 | 4.96E-05 | 0.615 | GTGGGTGGC |
| 5630 | mamo-s | *BGIBMGA006780* | down | 1929 | 1937 | - | 11.1009 | 6.57E-05 | 0.709 | GAGCGAGGC |
| 5631 | mamo-s | *BGIBMGA006781* | down | 1590 | 1598 | + | 11.3303 | 4.96E-05 | 0.615 | GTGGGTGGC |
| 5632 | mamo-s | *BGIBMGA006784* | down | 169 | 177 | + | 10.5138 | 9.75E-05 | 0.865 | GTGCGTGTT |
| 5633 | mamo-s | *BGIBMGA006785* | down | 912 | 920 | + | 11.0734 | 7.12E-05 | 0.726 | GAGCGTGGT |
| 5634 | mamo-s | *BGIBMGA006786* | up | 1551 | 1559 | - | 11.3303 | 4.96E-05 | 0.615 | GTGGGTGGC |
| 5635 | mamo-s | *BGIBMGA006790* | up | 574 | 582 | + | 11.3303 | 4.96E-05 | 0.615 | GTGGGTGGC |
| 5636 | mamo-s | *BGIBMGA006791* | up | 375 | 383 | + | 11.3303 | 4.96E-05 | 0.615 | GTGGGTGGC |
| 5637 | mamo-s | *BGIBMGA006793* | up | 1874 | 1882 | + | 11.3303 | 4.96E-05 | 0.615 | GTGGGTGGC |
| 5638 | mamo-s | *BGIBMGA006796* | up | 1745 | 1753 | + | 14.055 | 2.23E-06 | 0.615 | GTGCGTGGC |
| 5639 | mamo-s | *BGIBMGA006796* | up | 1447 | 1455 | - | 11.3303 | 4.96E-05 | 0.615 | GTGGGTGGC |
| 5640 | mamo-s | *BGIBMGA006801* | down | 705 | 713 | - | 11.7339 | 3.29E-05 | 0.615 | GCGCGAGGC |
| 5641 | mamo-s | *BGIBMGA006803* | down | 1291 | 1299 | - | 11.1009 | 6.57E-05 | 0.709 | GAGCGAGGC |
| 5642 | mamo-s | *BGIBMGA006804* | up | 330 | 338 | + | 11.7798 | 3.11E-05 | 0.615 | ATGCGTGGC |
| 5643 | mamo-s | *BGIBMGA006804* | up | 591 | 599 | + | 11.3853 | 4.74E-05 | 0.615 | ACGCGTGGC |
| 5644 | mamo-s | *BGIBMGA006804* | down | 1253 | 1261 | + | 11.3303 | 4.96E-05 | 0.615 | GTGGGTGGC |
| 5645 | mamo-s | *BGIBMGA006805* | down | 1309 | 1317 | + | 10.7523 | 8.38E-05 | 0.797 | AAGCGTGGC |
| 5646 | mamo-s | *BGIBMGA006808* | up | 494 | 502 | - | 11.3303 | 4.96E-05 | 0.615 | GTGGGTGGC |
| 5647 | mamo-s | *BGIBMGA006808* | up | 1390 | 1398 | - | 11.0734 | 7.12E-05 | 0.726 | GAGCGTGGT |
| 5648 | mamo-s | *BGIBMGA006809* | up | 1389 | 1397 | + | 10.9358 | 7.30E-05 | 0.728 | GCGGGTGGC |
| 5649 | mamo-s | *BGIBMGA006811* | up | 283 | 291 | + | 12.4679 | 1.31E-05 | 0.615 | GTGCGTGTC |
| 5650 | mamo-s | *BGIBMGA006819* | up | 864 | 872 | + | 11.3303 | 4.96E-05 | 0.615 | GTGGGTGGC |
| 5651 | mamo-s | *BGIBMGA006822* | up | 1257 | 1265 | - | 11.3303 | 4.96E-05 | 0.615 | GTGGGTGGC |
| 5652 | mamo-s | *BGIBMGA006823* | up | 669 | 677 | + | 11.0826 | 6.84E-05 | 0.709 | GTACGTGGC |
| 5653 | mamo-s | *BGIBMGA006827* | up | 1912 | 1920 | + | 13.6606 | 4.06E-06 | 0.615 | GCGCGTGGC |
| 5654 | mamo-s | *BGIBMGA006827* | up | 1827 | 1835 | - | 11.789 | 2.84E-05 | 0.615 | GCGCGTGCC |
| 5655 | mamo-s | *BGIBMGA006827* | up | 1378 | 1386 | - | 11.3303 | 4.96E-05 | 0.615 | GTGGGTGGC |
| 5656 | mamo-s | *BGIBMGA006828* | down | 337 | 345 | + | 11.3303 | 4.96E-05 | 0.615 | GTGGGTGGC |
| 5657 | mamo-s | *BGIBMGA006830* | down | 1034 | 1042 | + | 11.3303 | 4.96E-05 | 0.615 | GTGGGTGGC |
| 5658 | mamo-s | *BGIBMGA006833* | down | 1578 | 1586 | - | 11.5688 | 3.79E-05 | 0.615 | CTGCGTGTC |
| 5659 | mamo-s | *BGIBMGA006834* | down | 1290 | 1298 | - | 11.3303 | 4.96E-05 | 0.615 | GTGGGTGGC |
| 5660 | mamo-s | *BGIBMGA006835* | down | 954 | 962 | - | 11.3303 | 4.96E-05 | 0.615 | GTGGGTGGC |
| 5661 | mamo-s | *BGIBMGA006835* | up | 844 | 852 | + | 11.1376 | 6.35E-05 | 0.709 | TCGCGTGGC |
| 5662 | mamo-s | *BGIBMGA006835* | up | 863 | 871 | + | 10.5413 | 9.42E-05 | 0.857 | CAGCGTGTC |
| 5663 | mamo-s | *BGIBMGA006836* | up | 834 | 842 | + | 11.3303 | 4.96E-05 | 0.615 | GTGGGTGGC |
| 5664 | mamo-s | *BGIBMGA006839* | down | 132 | 140 | + | 14.055 | 2.23E-06 | 0.615 | GTGCGTGGC |
| 5665 | mamo-s | *BGIBMGA006842* | up | 1070 | 1078 | + | 11.3303 | 4.96E-05 | 0.615 | GTGGGTGGC |
| 5666 | mamo-s | *BGIBMGA006843* | down | 883 | 891 | + | 11.3303 | 4.96E-05 | 0.615 | GTGGGTGGC |
| 5667 | mamo-s | *BGIBMGA006844* | down | 702 | 710 | - | 10.6881 | 8.61E-05 | 0.81 | GCACGTGGC |
| 5668 | mamo-s | *BGIBMGA006845* | up | 525 | 533 | + | 13.6606 | 4.06E-06 | 0.615 | GCGCGTGGC |
| 5669 | mamo-s | *BGIBMGA006845* | up | 1600 | 1608 | + | 10.5413 | 9.42E-05 | 0.857 | CAGCGTGTC |
| 5670 | mamo-s | *BGIBMGA006847* | up | 1317 | 1325 | - | 10.5413 | 9.42E-05 | 0.857 | CAGCGTGTC |
| 5671 | mamo-s | *BGIBMGA006850* | up | 982 | 990 | - | 12.7615 | 1.04E-05 | 0.615 | CCGCGTGGC |
| 5672 | mamo-s | *BGIBMGA006850* | down | 586 | 594 | + | 10.7523 | 8.38E-05 | 0.797 | GGGCGTGTC |
| 5673 | mamo-s | *BGIBMGA006852* | up | 1484 | 1492 | + | 11.3303 | 4.96E-05 | 0.615 | GTGGGTGGC |
| 5674 | mamo-s | *BGIBMGA006853* | down | 61 | 69 | + | 11.4404 | 4.51E-05 | 0.615 | GAGCGTGTC |
| 5675 | mamo-s | *BGIBMGA006856* | down | 6 | 14 | - | 14.055 | 2.23E-06 | 0.615 | GTGCGTGGC |
| 5676 | mamo-s | *BGIBMGA006856* | down | 1933 | 1941 | - | 13.156 | 6.30E-06 | 0.615 | CTGCGTGGC |
| 5677 | mamo-s | *BGIBMGA006856* | down | 813 | 821 | + | 11.3303 | 4.96E-05 | 0.615 | GTGGGTGGC |
| 5678 | mamo-s | *BGIBMGA006857* | up | 1200 | 1208 | - | 13.156 | 6.30E-06 | 0.615 | CTGCGTGGC |
| 5679 | mamo-s | *BGIBMGA006857* | up | 80 | 88 | + | 11.3303 | 4.96E-05 | 0.615 | GTGGGTGGC |
| 5680 | mamo-s | *BGIBMGA006858* | down | 1477 | 1485 | - | 11.3303 | 4.96E-05 | 0.615 | GTGGGTGGC |
| 5681 | mamo-s | *BGIBMGA006861* | down | 358 | 366 | + | 11.3303 | 4.96E-05 | 0.615 | GTGGGTGGC |
| 5682 | mamo-s | *BGIBMGA006862* | up | 715 | 723 | + | 11.3303 | 4.96E-05 | 0.615 | GTGGGTGGC |
| 5683 | mamo-s | *BGIBMGA006863* | up | 1149 | 1157 | + | 11.3303 | 4.96E-05 | 0.615 | GTGGGTGGC |
| 5684 | mamo-s | *BGIBMGA006864* | down | 55 | 63 | + | 12.0734 | 2.66E-05 | 0.615 | GCGCGTGTC |
| 5685 | mamo-s | *BGIBMGA006864* | up | 674 | 682 | + | 11.3303 | 4.96E-05 | 0.615 | GTGGGTGGC |
| 5686 | mamo-s | *BGIBMGA006866* | down | 436 | 444 | + | 11.3303 | 4.96E-05 | 0.615 | GTGGGTGGC |
| 5687 | mamo-s | *BGIBMGA006870* | up | 1887 | 1895 | + | 10.5688 | 8.88E-05 | 0.824 | GTGCGTGAC |
| 5688 | mamo-s | *BGIBMGA006871* | up | 671 | 679 | + | 11.3303 | 4.96E-05 | 0.615 | GTGGGTGGC |
| 5689 | mamo-s | *BGIBMGA006872* | down | 913 | 921 | + | 11.3303 | 4.96E-05 | 0.615 | GTGGGTGGC |
| 5690 | mamo-s | *BGIBMGA006873* | up | 1321 | 1329 | - | 11.3303 | 4.96E-05 | 0.615 | GTGGGTGGC |
| 5691 | mamo-s | *BGIBMGA006874* | down | 1390 | 1398 | + | 11.4404 | 4.51E-05 | 0.615 | CGGCGTGGC |
| 5692 | mamo-s | *BGIBMGA006875* | up | 697 | 705 | - | 11.2018 | 5.68E-05 | 0.674 | CTGCGTGGT |
| 5693 | mamo-s | *BGIBMGA006876* | up | 1370 | 1378 | - | 12.4679 | 1.31E-05 | 0.615 | GTGCGTGTC |
| 5694 | mamo-s | *BGIBMGA006882* | up | 934 | 942 | - | 12.1009 | 2.43E-05 | 0.615 | GTGCGTGGT |
| 5695 | mamo-s | *BGIBMGA006883* | up | 83 | 91 | + | 11.3303 | 4.96E-05 | 0.615 | GTGGGTGGC |
| 5696 | mamo-s | *BGIBMGA006885* | up | 1669 | 1677 | - | 11.3303 | 4.96E-05 | 0.615 | GTGGGTGGC |
| 5697 | mamo-s | *BGIBMGA006885* | up | 1846 | 1854 | - | 10.5138 | 9.75E-05 | 0.865 | GTGCGTGTT |
| 5698 | mamo-s | *BGIBMGA006886* | up | 1103 | 1111 | - | 11.4404 | 4.51E-05 | 0.615 | GAGCGTGTC |
| 5699 | mamo-s | *BGIBMGA006886* | down | 922 | 930 | + | 11.3303 | 4.96E-05 | 0.615 | GTGGGTGGC |
| 5700 | mamo-s | *BGIBMGA006887* | down | 819 | 827 | - | 11.3303 | 4.96E-05 | 0.615 | GTGGGTGGC |
| 5701 | mamo-s | *BGIBMGA006888* | up | 704 | 712 | + | 11.3303 | 4.96E-05 | 0.615 | GTGGGTGGC |
| 5702 | mamo-s | *BGIBMGA006889* | down | 1056 | 1064 | + | 11.3303 | 4.96E-05 | 0.615 | GTGGGTGGC |
| 5703 | mamo-s | *BGIBMGA006892* | up | 892 | 900 | + | 11.3303 | 4.96E-05 | 0.615 | GTGGGTGGC |
| 5704 | mamo-s | *BGIBMGA006894* | down | 576 | 584 | + | 11.3303 | 4.96E-05 | 0.615 | GTGGGTGGC |
| 5705 | mamo-s | *BGIBMGA006897* | down | 1103 | 1111 | - | 10.8073 | 7.89E-05 | 0.766 | CCGCGTGGT |
| 5706 | mamo-s | *BGIBMGA006898* | up | 407 | 415 | - | 11.3303 | 4.96E-05 | 0.615 | GTGGGTGGC |
| 5707 | mamo-s | *BGIBMGA006898* | up | 1095 | 1103 | + | 11.3303 | 4.96E-05 | 0.615 | GTGGGTGGC |
| 5708 | mamo-s | *BGIBMGA006899* | up | 1480 | 1488 | + | 13.6606 | 4.06E-06 | 0.615 | GCGCGTGGC |
| 5709 | mamo-s | *BGIBMGA006900* | up | 800 | 808 | - | 11.2844 | 5.18E-05 | 0.63 | CTGCGTGCC |
| 5710 | mamo-s | *BGIBMGA006902* | down | 1055 | 1063 | - | 11.3303 | 4.96E-05 | 0.615 | GTGGGTGGC |
| 5711 | mamo-s | *BGIBMGA006903* | up | 100 | 108 | + | 11.3303 | 4.96E-05 | 0.615 | GTGGGTGGC |
| 5712 | mamo-s | *BGIBMGA006904* | up | 230 | 238 | + | 14.055 | 2.23E-06 | 0.615 | GTGCGTGGC |
| 5713 | mamo-s | *BGIBMGA006907* | up | 950 | 958 | - | 11.3303 | 4.96E-05 | 0.615 | GTGGGTGGC |

| | | | | | | | | | | |
|---|---|---|---|---|---|---|---|---|---|---|
| 5714 | mamo-s | *BGIBMGA006907* | up | 1375 | 1383 | + | 11.3303 | 4.96E-05 | 0.615 | GTGGGTGGC |
| 5715 | mamo-s | *BGIBMGA006907* | down | 1796 | 1804 | - | 11.3303 | 4.96E-05 | 0.615 | GTGGGTGGC |
| 5716 | mamo-s | *BGIBMGA006908* | up | 369 | 377 | + | 11.3303 | 4.96E-05 | 0.615 | GTGGGTGGC |
| 5717 | mamo-s | *BGIBMGA006908* | down | 1460 | 1468 | - | 11.3303 | 4.96E-05 | 0.615 | GTGGGTGGC |
| 5718 | mamo-s | *BGIBMGA006909* | down | 518 | 526 | - | 14.055 | 2.23E-06 | 0.615 | GTGCGTGGC |
| 5719 | mamo-s | *BGIBMGA006909* | down | 149 | 157 | + | 11.3303 | 4.96E-05 | 0.615 | GTGGGTGGC |
| 5720 | mamo-s | *BGIBMGA006912* | down | 243 | 251 | + | 10.5138 | 9.75E-05 | 0.865 | GTGCGTGTT |
| 5721 | mamo-s | *BGIBMGA006913* | up | 1833 | 1841 | + | 11.3303 | 4.96E-05 | 0.615 | GTGGGTGGC |
| 5722 | mamo-s | *BGIBMGA006914* | down | 1385 | 1393 | - | 11.3303 | 4.96E-05 | 0.615 | GTGGGTGGC |
| 5723 | mamo-s | *BGIBMGA006915* | down | 579 | 587 | - | 11.0826 | 6.84E-05 | 0.709 | GTACGTGGC |
| 5724 | mamo-s | *BGIBMGA006921* | down | 1658 | 1666 | + | 11.3303 | 4.96E-05 | 0.615 | GTGGGTGGC |
| 5725 | mamo-s | *BGIBMGA006927* | down | 756 | 764 | + | 11.3303 | 4.96E-05 | 0.615 | GTGGGTGGC |
| 5726 | mamo-s | *BGIBMGA006928* | down | 594 | 602 | + | 11.0826 | 6.84E-05 | 0.709 | GTACGTGGC |
| 5727 | mamo-s | *BGIBMGA006929* | up | 169 | 177 | + | 10.8349 | 7.66E-05 | 0.751 | CCGCGAGGC |
| 5728 | mamo-s | *BGIBMGA006930* | up | 1218 | 1226 | + | 11.3303 | 4.96E-05 | 0.615 | GTGGGTGGC |
| 5729 | mamo-s | *BGIBMGA006931* | up | 96 | 104 | - | 11.0826 | 6.84E-05 | 0.709 | GTACGTGGC |
| 5730 | mamo-s | *BGIBMGA006932* | up | 416 | 424 | - | 11.7064 | 3.52E-05 | 0.615 | GCGCGTGGT |
| 5731 | mamo-s | *BGIBMGA006932* | down | 244 | 252 | + | 11.3303 | 4.96E-05 | 0.615 | GTGGGTGGC |
| 5732 | mamo-s | *BGIBMGA006932* | down | 1541 | 1549 | + | 11.3303 | 4.96E-05 | 0.615 | GTGGGTGGC |
| 5733 | mamo-s | *BGIBMGA006933* | down | 1272 | 1280 | - | 12.7615 | 1.04E-05 | 0.615 | CCGCGTGGC |
| 5734 | mamo-s | *BGIBMGA006936* | up | 316 | 324 | - | 11.0826 | 6.84E-05 | 0.709 | GTACGTGGC |
| 5735 | mamo-s | *BGIBMGA006937* | up | 754 | 762 | - | 10.5413 | 9.42E-05 | 0.857 | CAGCGTGTC |
| 5736 | mamo-s | *BGIBMGA006937* | down | 1750 | 1758 | - | 10.5413 | 9.42E-05 | 0.857 | GTGCGAGTC |
| 5737 | mamo-s | *BGIBMGA006938* | up | 1231 | 1239 | + | 11.3303 | 4.96E-05 | 0.615 | GTGGGTGGC |
| 5738 | mamo-s | *BGIBMGA006940* | down | 1043 | 1051 | - | 10.5138 | 9.75E-05 | 0.865 | GTGCGTGTT |
| 5739 | mamo-s | *BGIBMGA006944* | up | 900 | 908 | + | 11.5321 | 4.06E-05 | 0.615 | TTGCGTGGC |
| 5740 | mamo-s | *BGIBMGA006949* | down | 1372 | 1380 | + | 11.1743 | 5.90E-05 | 0.684 | CCGCGTGTC |
| 5741 | mamo-s | *BGIBMGA006949* | down | 1018 | 1026 | - | 10.8899 | 7.48E-05 | 0.739 | CCGCGTGCC |
| 5742 | mamo-s | *BGIBMGA006949* | up | 1038 | 1046 | - | 10.8349 | 7.66E-05 | 0.751 | CCGCGAGGC |
| 5743 | mamo-s | *BGIBMGA006950* | up | 669 | 677 | + | 11.3303 | 4.96E-05 | 0.615 | GTGGGTGGC |
| 5744 | mamo-s | *BGIBMGA006950* | up | 1870 | 1878 | + | 11.3303 | 4.96E-05 | 0.615 | GTGGGTGGC |
| 5745 | mamo-s | *BGIBMGA006950* | up | 152 | 160 | + | 10.8899 | 7.48E-05 | 0.739 | CCGCGTGCC |
| 5746 | mamo-s | *BGIBMGA006951* | down | 1541 | 1549 | - | 11.156 | 6.13E-05 | 0.705 | GAGCGTGCC |
| 5747 | mamo-s | *BGIBMGA006952* | down | 1041 | 1049 | + | 11.7798 | 3.11E-05 | 0.615 | ATGCGTGGC |
| 5748 | mamo-s | *BGIBMGA006953* | up | 1316 | 1324 | + | 11.7798 | 3.11E-05 | 0.615 | ATGCGTGGC |
| 5749 | mamo-s | *BGIBMGA006955* | up | 347 | 355 | - | 11.3303 | 4.96E-05 | 0.615 | GTGGGTGGC |
| 5750 | mamo-s | *BGIBMGA006956* | down | 1652 | 1660 | - | 11.3303 | 4.96E-05 | 0.615 | GTGGGTGGC |
| 5751 | mamo-s | *BGIBMGA006958* | up | 736 | 744 | + | 12.0734 | 2.66E-05 | 0.615 | GCGCGTGTC |
| 5752 | mamo-s | *BGIBMGA006959* | up | 656 | 664 | + | 11.2844 | 5.18E-05 | 0.63 | CTGCGTGCC |
| 5753 | mamo-s | *BGIBMGA006961* | up | 1374 | 1382 | - | 10.9358 | 7.30E-05 | 0.728 | GCGGGTGGC |
| 5754 | mamo-s | *BGIBMGA006964* | down | 839 | 847 | + | 11.2294 | 5.41E-05 | 0.65 | CTGCGAGGC |
| 5755 | mamo-s | *BGIBMGA006965* | up | 730 | 738 | - | 11.3303 | 4.96E-05 | 0.615 | GTGGGTGGC |
| 5756 | mamo-s | *BGIBMGA006968* | down | 1157 | 1165 | + | 11.3303 | 4.96E-05 | 0.615 | GTGGGTGGC |
| 5757 | mamo-s | *BGIBMGA006968* | down | 448 | 456 | + | 11.0826 | 6.84E-05 | 0.709 | GTACGTGGC |
| 5758 | mamo-s | *BGIBMGA006969* | down | 1157 | 1165 | - | 11.4404 | 4.51E-05 | 0.615 | CGGCGTGGC |
| 5759 | mamo-s | *BGIBMGA006969* | down | 329 | 337 | + | 11.3303 | 4.96E-05 | 0.615 | GTGGGTGGC |
| 5760 | mamo-s | *BGIBMGA006969* | up | 782 | 790 | - | 11.3303 | 4.96E-05 | 0.615 | GTGGGTGGC |
| 5761 | mamo-s | *BGIBMGA006969* | up | 1008 | 1016 | + | 11.3303 | 4.96E-05 | 0.615 | GTGGGTGGC |
| 5762 | mamo-s | *BGIBMGA006971* | down | 1438 | 1446 | + | 11.3303 | 4.96E-05 | 0.615 | GTGGGTGGC |
| 5763 | mamo-s | *BGIBMGA006971* | down | 1917 | 1925 | + | 11.0826 | 6.84E-05 | 0.709 | GTACGTGGC |
| 5764 | mamo-s | *BGIBMGA006972* | down | 1342 | 1350 | + | 11.3303 | 4.96E-05 | 0.615 | GTGGGTGGC |
| 5765 | mamo-s | *BGIBMGA006973* | down | 1707 | 1715 | - | 11.3303 | 4.96E-05 | 0.615 | GTGGGTGGC |
| 5766 | mamo-s | *BGIBMGA006976* | down | 979 | 987 | - | 11.3303 | 4.96E-05 | 0.615 | GTGGGTGGC |
| 5767 | mamo-s | *BGIBMGA006976* | down | 1596 | 1604 | + | 11.1743 | 5.90E-05 | 0.684 | CCGCGTGTC |
| 5768 | mamo-s | *BGIBMGA006978* | down | 491 | 499 | - | 11.0734 | 7.12E-05 | 0.726 | GAGCGTGGT |
| 5769 | mamo-s | *BGIBMGA006981* | down | 280 | 288 | + | 11.3303 | 4.96E-05 | 0.615 | GTGGGTGGC |
| 5770 | mamo-s | *BGIBMGA006984* | up | 511 | 519 | + | 11.3303 | 4.96E-05 | 0.615 | GTGGGTGGC |
| 5771 | mamo-s | *BGIBMGA006984* | down | 1407 | 1415 | + | 11.3303 | 4.96E-05 | 0.615 | GTGGGTGGC |
| 5772 | mamo-s | *BGIBMGA006986* | down | 1567 | 1575 | - | 11.3303 | 4.96E-05 | 0.615 | GTGGGTGGC |
| 5773 | mamo-s | *BGIBMGA006986* | down | 1881 | 1889 | + | 11.1009 | 6.57E-05 | 0.709 | GAGCGAGGC |
| 5774 | mamo-s | *BGIBMGA006987* | down | 93 | 101 | + | 12.1835 | 1.71E-05 | 0.615 | GTGCGTGCC |
| 5775 | mamo-s | *BGIBMGA006987* | down | 787 | 795 | - | 10.7523 | 8.38E-05 | 0.797 | GGGCGTGTC |
| 5776 | mamo-s | *BGIBMGA006988* | up | 737 | 745 | + | 11.3303 | 4.96E-05 | 0.615 | GTGGGTGGC |
| 5777 | mamo-s | *BGIBMGA006988* | up | 1801 | 1809 | + | 11.156 | 6.13E-05 | 0.705 | GAGCGTGCC |
| 5778 | mamo-s | *BGIBMGA006989* | down | 1807 | 1815 | - | 11.3303 | 4.96E-05 | 0.615 | GTGGGTGGC |
| 5779 | mamo-s | *BGIBMGA006992* | down | 1684 | 1692 | + | 11.3303 | 4.96E-05 | 0.615 | GTGGGTGGC |
| 5780 | mamo-s | *BGIBMGA006992* | down | 1404 | 1412 | - | 11.0734 | 7.12E-05 | 0.726 | GAGCGTGGT |
| 5781 | mamo-s | *BGIBMGA006995* | up | 1438 | 1446 | + | 11.3303 | 4.96E-05 | 0.615 | GTGGGTGGC |
| 5782 | mamo-s | *BGIBMGA006996* | up | 1338 | 1346 | - | 11.2844 | 5.18E-05 | 0.63 | CTGCGTGCC |
| 5783 | mamo-s | *BGIBMGA007002* | up | 1791 | 1799 | + | 11.2844 | 5.18E-05 | 0.63 | CTGCGTGCC |
| 5784 | mamo-s | *BGIBMGA007003* | down | 137 | 145 | - | 10.9358 | 7.30E-05 | 0.728 | GCGGGTGGC |
| 5785 | mamo-s | *BGIBMGA007004* | up | 682 | 690 | + | 11.3303 | 4.96E-05 | 0.615 | GTGGGTGGC |
| 5786 | mamo-s | *BGIBMGA007005* | down | 1645 | 1653 | - | 11.3303 | 4.96E-05 | 0.615 | GTGGGTGGC |
| 5787 | mamo-s | *BGIBMGA007007* | up | 861 | 869 | + | 12.1284 | 2.16E-05 | 0.615 | CAGCGTGGC |
| 5788 | mamo-s | *BGIBMGA007008* | down | 1151 | 1159 | + | 10.7523 | 8.38E-05 | 0.797 | AAGCGTGGC |
| 5789 | mamo-s | *BGIBMGA007010* | down | 1178 | 1186 | + | 11.0826 | 6.84E-05 | 0.709 | GTACGTGGC |
| 5790 | mamo-s | *BGIBMGA007011* | up | 1868 | 1876 | - | 11.789 | 2.84E-05 | 0.615 | GCGCGTGCC |
| 5791 | mamo-s | *BGIBMGA007011* | up | 1612 | 1620 | - | 11.3303 | 4.96E-05 | 0.615 | GTGGGTGGC |
| 5792 | mamo-s | *BGIBMGA007011* | up | 1145 | 1153 | + | 11.1376 | 6.35E-05 | 0.709 | TCGCGTGGC |
| 5793 | mamo-s | *BGIBMGA007012* | up | 713 | 721 | - | 11.3303 | 4.96E-05 | 0.615 | GTGGGTGGC |
| 5794 | mamo-s | *BGIBMGA007015* | down | 1134 | 1142 | + | 11.3303 | 4.96E-05 | 0.615 | GTGGGTGGC |
| 5795 | mamo-s | *BGIBMGA007015* | up | 828 | 836 | - | 11.1009 | 6.57E-05 | 0.709 | GAGCGAGGC |
| 5796 | mamo-s | *BGIBMGA007015* | up | 1638 | 1646 | - | 10.7523 | 8.38E-05 | 0.797 | GGGCGTGTC |
| 5797 | mamo-s | *BGIBMGA007018* | down | 558 | 566 | + | 11.3303 | 4.96E-05 | 0.615 | GTGGGTGGC |
| 5798 | mamo-s | *BGIBMGA007021* | down | 1009 | 1017 | - | 11.3303 | 4.96E-05 | 0.615 | GTGGGTGGC |
| 5799 | mamo-s | *BGIBMGA007022* | up | 1183 | 1191 | + | 11.3303 | 4.96E-05 | 0.615 | GTGGGTGGC |
| 5800 | mamo-s | *BGIBMGA007023* | down | 976 | 984 | - | 11.0826 | 6.84E-05 | 0.709 | GTACGTGGC |
| 5801 | mamo-s | *BGIBMGA007025* | up | 1532 | 1540 | + | 12.1284 | 2.16E-05 | 0.615 | CAGCGTGGC |

| | | | | | | | | | | |
|---|---|---|---|---|---|---|---|---|---|---|
| 5802 | mamo-s | *BGIBMGA007027* | down | 1350 | 1358 | + | 14.055 | 2.23E-06 | 0.615 | GTGCGTGGC |
| 5803 | mamo-s | *BGIBMGA007027* | down | 1649 | 1657 | + | 11.3303 | 4.96E-05 | 0.615 | GTGGGTGGC |
| 5804 | mamo-s | *BGIBMGA007027* | down | 1369 | 1377 | - | 10.7523 | 8.38E-05 | 0.797 | GGGCGTGTC |
| 5805 | mamo-s | *BGIBMGA007030* | down | 1435 | 1443 | + | 14.055 | 2.23E-06 | 0.615 | GTGCGTGGC |
| 5806 | mamo-s | *BGIBMGA007030* | up | 1279 | 1287 | - | 11.3303 | 4.96E-05 | 0.615 | GTGGGTGGC |
| 5807 | mamo-s | *BGIBMGA007030* | down | 1561 | 1569 | + | 11.3303 | 4.96E-05 | 0.615 | GTGGGTGGC |
| 5808 | mamo-s | *BGIBMGA007030* | down | 1454 | 1462 | - | 10.7523 | 8.38E-05 | 0.797 | GGGCGTGTC |
| 5809 | mamo-s | *BGIBMGA007033* | up | 1235 | 1243 | - | 11.3303 | 4.96E-05 | 0.615 | GTGGGTGGC |
| 5810 | mamo-s | *BGIBMGA007036* | up | 1053 | 1061 | + | 11.3303 | 4.96E-05 | 0.615 | GTGGGTGGC |
| 5811 | mamo-s | *BGIBMGA007037* | down | 1470 | 1478 | + | 11.3303 | 4.96E-05 | 0.615 | GTGGGTGGC |
| 5812 | mamo-s | *BGIBMGA007039* | down | 1872 | 1880 | + | 11.3303 | 4.96E-05 | 0.615 | GTGGGTGGC |
| 5813 | mamo-s | *BGIBMGA007041* | up | 303 | 311 | - | 11.3303 | 4.96E-05 | 0.615 | GTGGGTGGC |
| 5814 | mamo-s | *BGIBMGA007044* | down | 1628 | 1636 | + | 11.3303 | 4.96E-05 | 0.615 | GTGGGTGGC |
| 5815 | mamo-s | *BGIBMGA007046* | up | 1470 | 1478 | - | 11.7798 | 3.11E-05 | 0.615 | ATGCGTGGC |
| 5816 | mamo-s | *BGIBMGA007047* | up | 1369 | 1377 | - | 11.3303 | 4.96E-05 | 0.615 | GTGGGTGGC |
| 5817 | mamo-s | *BGIBMGA007049* | down | 208 | 216 | + | 11.3303 | 4.96E-05 | 0.615 | GTGGGTGGC |
| 5818 | mamo-s | *BGIBMGA007049* | up | 696 | 704 | + | 10.9358 | 7.30E-05 | 0.728 | GCGGGTGGC |
| 5819 | mamo-s | *BGIBMGA007051* | down | 1883 | 1891 | + | 11.3303 | 4.96E-05 | 0.615 | GTGGGTGGC |
| 5820 | mamo-s | *BGIBMGA007053* | down | 1473 | 1481 | + | 14.055 | 2.23E-06 | 0.615 | GTGCGTGGC |
| 5821 | mamo-s | *BGIBMGA007054* | up | 838 | 846 | + | 11.3303 | 4.96E-05 | 0.615 | GTGGGTGGC |
| 5822 | mamo-s | *BGIBMGA007055* | down | 1597 | 1605 | + | 11.3303 | 4.96E-05 | 0.615 | GTGGGTGGC |
| 5823 | mamo-s | *BGIBMGA007055* | down | 1736 | 1744 | - | 11.3303 | 4.96E-05 | 0.615 | GTGGGTGGC |
| 5824 | mamo-s | *BGIBMGA007059* | down | 1485 | 1493 | + | 11.3303 | 4.96E-05 | 0.615 | GTGGGTGGC |
| 5825 | mamo-s | *BGIBMGA007061* | down | 518 | 526 | + | 11.3303 | 4.96E-05 | 0.615 | GTGGGTGGC |
| 5826 | mamo-s | *BGIBMGA007064* | down | 366 | 374 | + | 11.3303 | 4.96E-05 | 0.615 | GTGGGTGGC |
| 5827 | mamo-s | *BGIBMGA007067* | down | 380 | 388 | + | 11.3303 | 4.96E-05 | 0.615 | GTGGGTGGC |
| 5828 | mamo-s | *BGIBMGA007067* | down | 1007 | 1015 | + | 11.3303 | 4.96E-05 | 0.615 | GTGGGTGGC |
| 5829 | mamo-s | *BGIBMGA007067* | up | 1721 | 1729 | + | 11.3303 | 4.96E-05 | 0.615 | GTGGGTGGC |
| 5830 | mamo-s | *BGIBMGA007070* | up | 219 | 227 | + | 11.3303 | 4.96E-05 | 0.615 | GTGGGTGGC |
| 5831 | mamo-s | *BGIBMGA007070* | down | 334 | 342 | - | 10.5413 | 9.42E-05 | 0.857 | GTGCGAGTC |
| 5832 | mamo-s | *BGIBMGA007071* | up | 1723 | 1731 | + | 11.3303 | 4.96E-05 | 0.615 | GTGGGTGGC |
| 5833 | mamo-s | *BGIBMGA007072* | down | 1028 | 1036 | + | 11.0826 | 6.84E-05 | 0.709 | GTACGTGGC |
| 5834 | mamo-s | *BGIBMGA007075* | up | 207 | 215 | + | 11.3303 | 4.96E-05 | 0.615 | GTGGGTGGC |
| 5835 | mamo-s | *BGIBMGA007076* | up | 649 | 657 | + | 11.0826 | 6.84E-05 | 0.709 | GTACGTGGC |
| 5836 | mamo-s | *BGIBMGA007079* | down | 218 | 226 | + | 11.5688 | 3.79E-05 | 0.615 | CTGCGTGTC |
| 5837 | mamo-s | *BGIBMGA007079* | up | 727 | 735 | + | 11.3303 | 4.96E-05 | 0.615 | GTGGGTGGC |
| 5838 | mamo-s | *BGIBMGA007080* | down | 66 | 74 | + | 10.8073 | 7.89E-05 | 0.766 | CCGCGTGGT |
| 5839 | mamo-s | *BGIBMGA007081* | up | 278 | 286 | + | 11.1009 | 6.57E-05 | 0.709 | GAGCGAGGC |
| 5840 | mamo-s | *BGIBMGA007086* | up | 538 | 546 | - | 11.3303 | 4.96E-05 | 0.615 | GTGGGTGGC |
| 5841 | mamo-s | *BGIBMGA007087* | up | 722 | 730 | + | 14.055 | 2.23E-06 | 0.615 | GTGCGTGGC |
| 5842 | mamo-s | *BGIBMGA007087* | up | 1794 | 1802 | - | 11.3303 | 4.96E-05 | 0.615 | GTGGGTGGC |
| 5843 | mamo-s | *BGIBMGA007090* | down | 459 | 467 | + | 11.1009 | 6.57E-05 | 0.709 | GAGCGAGGC |
| 5844 | mamo-s | *BGIBMGA007090* | down | 755 | 763 | + | 10.5688 | 8.88E-05 | 0.824 | GTGCGTGAC |
| 5845 | mamo-s | *BGIBMGA007090* | down | 1571 | 1579 | + | 10.5413 | 9.42E-05 | 0.857 | CAGCGTGTC |
| 5846 | mamo-s | *BGIBMGA007091* | up | 1625 | 1633 | + | 11.3303 | 4.96E-05 | 0.615 | GTGGGTGGC |
| 5847 | mamo-s | *BGIBMGA007092* | up | 444 | 452 | - | 11.3303 | 4.96E-05 | 0.615 | GTGGGTGGC |
| 5848 | mamo-s | *BGIBMGA007092* | up | 1330 | 1338 | - | 11.3303 | 4.96E-05 | 0.615 | GTGGGTGGC |
| 5849 | mamo-s | *BGIBMGA007095* | up | 1272 | 1280 | - | 13.156 | 6.30E-06 | 0.615 | CTGCGTGGC |
| 5850 | mamo-s | *BGIBMGA007095* | down | 1545 | 1553 | - | 11.7798 | 3.11E-05 | 0.615 | ATGCGTGGC |
| 5851 | mamo-s | *BGIBMGA007096* | up | 1584 | 1592 | + | 12.4679 | 1.31E-05 | 0.615 | GTGCGTGTC |
| 5852 | mamo-s | *BGIBMGA007096* | up | 676 | 684 | + | 11.3303 | 4.96E-05 | 0.615 | GTGGGTGGC |
| 5853 | mamo-s | *BGIBMGA007097* | down | 183 | 191 | - | 11.3303 | 4.96E-05 | 0.615 | GTGGGTGGC |
| 5854 | mamo-s | *BGIBMGA007097* | down | 522 | 530 | - | 11.3303 | 4.96E-05 | 0.615 | GTGGGTGGC |
| 5855 | mamo-s | *BGIBMGA007097* | up | 734 | 742 | + | 11.3303 | 4.96E-05 | 0.615 | GTGGGTGGC |
| 5856 | mamo-s | *BGIBMGA007098* | up | 1517 | 1525 | - | 11.3303 | 4.96E-05 | 0.615 | GTGGGTGGC |
| 5857 | mamo-s | *BGIBMGA007099* | down | 177 | 185 | - | 10.5413 | 9.42E-05 | 0.857 | GTGCGAGTC |
| 5858 | mamo-s | *BGIBMGA007101* | down | 990 | 998 | + | 11.3303 | 4.96E-05 | 0.615 | GTGGGTGGC |
| 5859 | mamo-s | *BGIBMGA007102* | up | 184 | 192 | - | 11.3303 | 4.96E-05 | 0.615 | GTGGGTGGC |
| 5860 | mamo-s | *BGIBMGA007102* | up | 1053 | 1061 | - | 11.3303 | 4.96E-05 | 0.615 | GTGGGTGGC |
| 5861 | mamo-s | *BGIBMGA007105* | up | 1687 | 1695 | - | 11.789 | 2.84E-05 | 0.615 | GCGCGTGCC |
| 5862 | mamo-s | *BGIBMGA007106* | down | 339 | 347 | - | 12.1284 | 2.16E-05 | 0.615 | CAGCGTGGC |
| 5863 | mamo-s | *BGIBMGA007106* | up | 1069 | 1077 | + | 10.5688 | 8.88E-05 | 0.824 | GTGCGTGAC |
| 5864 | mamo-s | *BGIBMGA007108* | up | 510 | 518 | + | 11.3303 | 4.96E-05 | 0.615 | GTGGGTGGC |
| 5865 | mamo-s | *BGIBMGA007110* | down | 186 | 194 | + | 11.3303 | 4.96E-05 | 0.615 | GTGGGTGGC |
| 5866 | mamo-s | *BGIBMGA007110* | up | 283 | 291 | + | 11.3303 | 4.96E-05 | 0.615 | GTGGGTGGC |
| 5867 | mamo-s | *BGIBMGA007113* | down | 82 | 90 | + | 11.3303 | 4.96E-05 | 0.615 | GTGGGTGGC |
| 5868 | mamo-s | *BGIBMGA007116* | down | 680 | 688 | - | 11.3303 | 4.96E-05 | 0.615 | GTGGGTGGC |
| 5869 | mamo-s | *BGIBMGA007116* | down | 792 | 800 | - | 11.3303 | 4.96E-05 | 0.615 | GTGGGTGGC |
| 5870 | mamo-s | *BGIBMGA007116* | up | 1350 | 1358 | + | 11.0826 | 6.84E-05 | 0.709 | GTACGTGGC |
| 5871 | mamo-s | *BGIBMGA007117* | up | 894 | 902 | - | 11.3303 | 4.96E-05 | 0.615 | GTGGGTGGC |
| 5872 | mamo-s | *BGIBMGA007119* | up | 1781 | 1789 | + | 11.3303 | 4.96E-05 | 0.615 | GTGGGTGGC |
| 5873 | mamo-s | *BGIBMGA007120* | down | 1025 | 1033 | - | 11.3303 | 4.96E-05 | 0.615 | GTGGGTGGC |
| 5874 | mamo-s | *BGIBMGA007121* | up | 328 | 336 | - | 11.3303 | 4.96E-05 | 0.615 | GTGGGTGGC |
| 5875 | mamo-s | *BGIBMGA007121* | up | 1754 | 1762 | - | 10.5413 | 9.42E-05 | 0.857 | CAGCGTGTC |
| 5876 | mamo-s | *BGIBMGA007124* | down | 911 | 919 | + | 10.5138 | 9.75E-05 | 0.865 | GTGCGTGTT |
| 5877 | mamo-s | *BGIBMGA007125* | down | 1086 | 1094 | - | 11.3303 | 4.96E-05 | 0.615 | GTGGGTGGC |
| 5878 | mamo-s | *BGIBMGA007125* | up | 1242 | 1250 | + | 11.0826 | 6.84E-05 | 0.709 | GTACGTGGC |
| 5879 | mamo-s | *BGIBMGA007127* | down | 860 | 868 | + | 11.3303 | 4.96E-05 | 0.615 | GTGGGTGGC |
| 5880 | mamo-s | *BGIBMGA007128* | down | 1901 | 1909 | + | 11.3303 | 4.96E-05 | 0.615 | GTGGGTGGC |
| 5881 | mamo-s | *BGIBMGA007131* | up | 497 | 505 | - | 11.3303 | 4.96E-05 | 0.615 | GTGGGTGGC |
| 5882 | mamo-s | *BGIBMGA007131* | up | 1963 | 1971 | + | 11.2018 | 5.68E-05 | 0.674 | CTGCGTGGT |
| 5883 | mamo-s | *BGIBMGA007132* | down | 729 | 737 | - | 11.3303 | 4.96E-05 | 0.615 | GTGGGTGGC |
| 5884 | mamo-s | *BGIBMGA007134* | down | 1747 | 1755 | - | 13.156 | 6.30E-06 | 0.615 | CTGCGTGGC |
| 5885 | mamo-s | *BGIBMGA007134* | up | 537 | 545 | - | 11.3303 | 4.96E-05 | 0.615 | GTGGGTGGC |
| 5886 | mamo-s | *BGIBMGA007134* | down | 1785 | 1793 | - | 10.9358 | 7.30E-05 | 0.728 | GCGGGTGGC |
| 5887 | mamo-s | *BGIBMGA007136* | down | 1861 | 1869 | + | 11.0734 | 7.12E-05 | 0.726 | GAGCGTGGT |
| 5888 | mamo-s | *BGIBMGA007137* | down | 173 | 181 | + | 11.3303 | 4.96E-05 | 0.615 | GTGGGTGGC |
| 5889 | mamo-s | *BGIBMGA007137* | up | 777 | 785 | + | 11.0826 | 6.84E-05 | 0.709 | GTACGTGGC |

| | | | | | | | | | | |
|---|---|---|---|---|---|---|---|---|---|---|
| 5890 | mamo-s | *BGIBMGA007140* | down | 237 | 245 | - | 12.7615 | 1.04E-05 | 0.615 | CCGCGTGGC |
| 5891 | mamo-s | *BGIBMGA007141* | up | 1028 | 1036 | + | 11.0826 | 6.84E-05 | 0.709 | GTACGTGGC |
| 5892 | mamo-s | *BGIBMGA007142* | up | 814 | 822 | - | 13.6606 | 4.06E-06 | 0.615 | GCGCGTGGC |
| 5893 | mamo-s | *BGIBMGA007142* | up | 907 | 915 | - | 10.5413 | 9.42E-05 | 0.857 | GTGCGAGTC |
| 5894 | mamo-s | *BGIBMGA007143* | up | 1648 | 1656 | - | 11.3303 | 4.96E-05 | 0.615 | GTGGGTGGC |
| 5895 | mamo-s | *BGIBMGA007145* | down | 694 | 702 | - | 12.7615 | 1.04E-05 | 0.615 | CCGCGTGGC |
| 5896 | mamo-s | *BGIBMGA007145* | down | 730 | 738 | + | 12.0734 | 2.66E-05 | 0.615 | GCGCGTGTC |
| 5897 | mamo-s | *BGIBMGA007145* | down | 1328 | 1336 | - | 10.5688 | 8.88E-05 | 0.824 | GTGCGTGAC |
| 5898 | mamo-s | *BGIBMGA007145* | down | 1444 | 1452 | + | 10.5688 | 8.88E-05 | 0.824 | GTGCGTGAC |
| 5899 | mamo-s | *BGIBMGA007145* | down | 1467 | 1475 | - | 10.5688 | 8.88E-05 | 0.824 | GTGCGTGAC |
| 5900 | mamo-s | *BGIBMGA007146* | down | 1624 | 1632 | - | 13.6606 | 4.06E-06 | 0.615 | GCGCGTGGC |
| 5901 | mamo-s | *BGIBMGA007146* | down | 1831 | 1839 | - | 11.789 | 2.84E-05 | 0.615 | GCGCGTGCC |
| 5902 | mamo-s | *BGIBMGA007146* | up | 156 | 164 | + | 11.3303 | 4.96E-05 | 0.615 | GTGGGTGGC |
| 5903 | mamo-s | *BGIBMGA007146* | up | 1034 | 1042 | - | 11.3303 | 4.96E-05 | 0.615 | GTGGGTGGC |
| 5904 | mamo-s | *BGIBMGA007147* | up | 1226 | 1234 | + | 10.6881 | 8.61E-05 | 0.81 | GCACGTGGC |
| 5905 | mamo-s | *BGIBMGA007148* | up | 519 | 527 | - | 11.0826 | 6.84E-05 | 0.709 | GTACGTGGC |
| 5906 | mamo-s | *BGIBMGA007150* | up | 1383 | 1391 | + | 11.0826 | 6.84E-05 | 0.709 | GTACGTGGC |
| 5907 | mamo-s | *BGIBMGA007151* | up | 1347 | 1355 | + | 11.3303 | 4.96E-05 | 0.615 | GTGGGTGGC |
| 5908 | mamo-s | *BGIBMGA007151* | down | 1717 | 1725 | + | 11.3303 | 4.96E-05 | 0.615 | GTGGGTGGC |
| 5909 | mamo-s | *BGIBMGA007152* | up | 1218 | 1226 | + | 12.1835 | 1.71E-05 | 0.615 | GTGCGTGCC |
| 5910 | mamo-s | *BGIBMGA007153* | down | 1940 | 1948 | - | 11.3303 | 4.96E-05 | 0.615 | GTGGGTGGC |
| 5911 | mamo-s | *BGIBMGA007156* | up | 1656 | 1664 | - | 11.3303 | 4.96E-05 | 0.615 | GTGGGTGGC |
| 5912 | mamo-s | *BGIBMGA007156* | up | 1432 | 1440 | - | 10.5413 | 9.42E-05 | 0.857 | CAGCGTGTC |
| 5913 | mamo-s | *BGIBMGA007157* | down | 920 | 928 | - | 11.3303 | 4.96E-05 | 0.615 | GTGGGTGGC |
| 5914 | mamo-s | *BGIBMGA007158* | down | 1172 | 1180 | - | 12.1009 | 2.43E-05 | 0.615 | GTGCGTGGT |
| 5915 | mamo-s | *BGIBMGA007158* | up | 1445 | 1453 | + | 11.3303 | 4.96E-05 | 0.615 | GTGGGTGGC |
| 5916 | mamo-s | *BGIBMGA007162* | down | 245 | 253 | + | 14.055 | 2.23E-06 | 0.615 | GTGCGTGGC |
| 5917 | mamo-s | *BGIBMGA007162* | down | 1947 | 1955 | - | 11.3303 | 4.96E-05 | 0.615 | GTGGGTGGC |
| 5918 | mamo-s | *BGIBMGA007163* | up | 591 | 599 | + | 12.4679 | 1.31E-05 | 0.615 | GTGCGTGTC |
| 5919 | mamo-s | *BGIBMGA007163* | down | 1346 | 1354 | + | 11.3303 | 4.96E-05 | 0.615 | GTGGGTGGC |
| 5920 | mamo-s | *BGIBMGA007165* | down | 762 | 770 | + | 14.055 | 2.23E-06 | 0.615 | GTGCGTGGC |
| 5921 | mamo-s | *BGIBMGA007167* | down | 856 | 864 | + | 11.3303 | 4.96E-05 | 0.615 | GTGGGTGGC |
| 5922 | mamo-s | *BGIBMGA007167* | up | 1031 | 1039 | + | 11.2844 | 5.18E-05 | 0.63 | CTGCGTGCC |
| 5923 | mamo-s | *BGIBMGA007168* | up | 310 | 318 | + | 11.3303 | 4.96E-05 | 0.615 | GTGGGTGGC |
| 5924 | mamo-s | *BGIBMGA007169* | up | 1232 | 1240 | + | 11.3303 | 4.96E-05 | 0.615 | GTGGGTGGC |
| 5925 | mamo-s | *BGIBMGA007171* | down | 1553 | 1561 | - | 11.3303 | 4.96E-05 | 0.615 | GTGGGTGGC |
| 5926 | mamo-s | *BGIBMGA007171* | down | 1293 | 1301 | + | 11.0826 | 6.84E-05 | 0.709 | GTACGTGGC |
| 5927 | mamo-s | *BGIBMGA007172* | up | 997 | 1005 | + | 11.789 | 2.84E-05 | 0.615 | GCGCGTGCC |
| 5928 | mamo-s | *BGIBMGA007172* | down | 1120 | 1128 | + | 11.4404 | 4.51E-05 | 0.615 | CGGCGTGGC |
| 5929 | mamo-s | *BGIBMGA007172* | up | 193 | 201 | - | 11.0826 | 6.84E-05 | 0.709 | GTACGTGGC |
| 5930 | mamo-s | *BGIBMGA007173* | down | 528 | 536 | - | 12.1835 | 1.71E-05 | 0.615 | GTGCGTGCC |
| 5931 | mamo-s | *BGIBMGA007173* | up | 1055 | 1063 | - | 11.0734 | 7.12E-05 | 0.726 | GAGCGTGGT |
| 5932 | mamo-s | *BGIBMGA007174* | down | 265 | 273 | - | 10.7523 | 8.38E-05 | 0.797 | AAGCGTGGC |
| 5933 | mamo-s | *BGIBMGA007175* | up | 799 | 807 | + | 11.0826 | 6.84E-05 | 0.709 | GTACGTGGC |
| 5934 | mamo-s | *BGIBMGA007176* | up | 705 | 713 | - | 10.9358 | 7.30E-05 | 0.728 | GCGGGTGGC |
| 5935 | mamo-s | *BGIBMGA007177* | up | 111 | 119 | - | 13.0275 | 8.53E-06 | 0.615 | GAGCGTGGC |
| 5936 | mamo-s | *BGIBMGA007177* | down | 214 | 222 | + | 11.3303 | 4.96E-05 | 0.615 | GTGGGTGGC |
| 5937 | mamo-s | *BGIBMGA007179* | down | 239 | 247 | + | 11.3303 | 4.96E-05 | 0.615 | GTGGGTGGC |
| 5938 | mamo-s | *BGIBMGA007183* | up | 1144 | 1152 | - | 11.0826 | 6.84E-05 | 0.709 | GTACGTGGC |
| 5939 | mamo-s | *BGIBMGA007183* | down | 1253 | 1261 | + | 11.0826 | 6.84E-05 | 0.709 | GTACGTGGC |
| 5940 | mamo-s | *BGIBMGA007184* | up | 903 | 911 | + | 13.6606 | 4.06E-06 | 0.615 | GCGCGTGGC |
| 5941 | mamo-s | *BGIBMGA007185* | up | 1846 | 1854 | + | 11.7339 | 3.29E-05 | 0.615 | GCGCGAGGC |
| 5942 | mamo-s | *BGIBMGA007185* | up | 1869 | 1877 | - | 11.5321 | 4.06E-05 | 0.615 | TTGCGTGGC |
| 5943 | mamo-s | *BGIBMGA007187* | down | 307 | 315 | + | 11.3303 | 4.96E-05 | 0.615 | GTGGGTGGC |
| 5944 | mamo-s | *BGIBMGA007188* | down | 1566 | 1574 | + | 11.3303 | 4.96E-05 | 0.615 | GTGGGTGGC |
| 5945 | mamo-s | *BGIBMGA007188* | down | 1717 | 1725 | - | 11.0826 | 6.84E-05 | 0.709 | GTACGTGGC |
| 5946 | mamo-s | *BGIBMGA007189* | down | 222 | 230 | + | 11.3303 | 4.96E-05 | 0.615 | GTGGGTGGC |
| 5947 | mamo-s | *BGIBMGA007189* | up | 1839 | 1847 | + | 11.3303 | 4.96E-05 | 0.615 | GTGGGTGGC |
| 5948 | mamo-s | *BGIBMGA007189* | up | 1990 | 1998 | - | 11.0826 | 6.84E-05 | 0.709 | GTACGTGGC |
| 5949 | mamo-s | *BGIBMGA007189* | down | 454 | 462 | - | 10.9358 | 7.30E-05 | 0.728 | GCGGGTGGC |
| 5950 | mamo-s | *BGIBMGA007190* | down | 808 | 816 | + | 11.3303 | 4.96E-05 | 0.615 | GTGGGTGGC |
| 5951 | mamo-s | *BGIBMGA007190* | down | 1661 | 1669 | + | 11.3303 | 4.96E-05 | 0.615 | GTGGGTGGC |
| 5952 | mamo-s | *BGIBMGA007190* | down | 1406 | 1414 | - | 10.5138 | 9.75E-05 | 0.865 | GTGCGTGTT |
| 5953 | mamo-s | *BGIBMGA007191* | down | 1408 | 1416 | - | 11.3303 | 4.96E-05 | 0.615 | GTGGGTGGC |
| 5954 | mamo-s | *BGIBMGA007191* | up | 1499 | 1507 | + | 10.9358 | 7.30E-05 | 0.728 | GCGGGTGGC |
| 5955 | mamo-s | *BGIBMGA007193* | down | 1680 | 1688 | - | 11.4404 | 4.51E-05 | 0.615 | CGGCGTGGC |
| 5956 | mamo-s | *BGIBMGA007193* | up | 1029 | 1037 | + | 11.0734 | 7.12E-05 | 0.726 | GAGCGTGGT |
| 5957 | mamo-s | *BGIBMGA007195* | down | 407 | 415 | + | 11.3303 | 4.96E-05 | 0.615 | GTGGGTGGC |
| 5958 | mamo-s | *BGIBMGA007198* | up | 642 | 650 | + | 11.3303 | 4.96E-05 | 0.615 | GTGGGTGGC |
| 5959 | mamo-s | *BGIBMGA007198* | up | 1168 | 1176 | + | 11.3303 | 4.96E-05 | 0.615 | GTGGGTGGC |
| 5960 | mamo-s | *BGIBMGA007199* | down | 1225 | 1233 | - | 14.055 | 2.23E-06 | 0.615 | GTGCGTGGC |
| 5961 | mamo-s | *BGIBMGA007199* | down | 1843 | 1851 | - | 11.5321 | 4.06E-05 | 0.615 | TTGCGTGGC |
| 5962 | mamo-s | *BGIBMGA007199* | up | 1243 | 1251 | + | 11.2844 | 5.18E-05 | 0.63 | CTGCGTGCC |
| 5963 | mamo-s | *BGIBMGA007200* | down | 1453 | 1461 | + | 12.1009 | 2.43E-05 | 0.615 | GTGCGTGGT |
| 5964 | mamo-s | *BGIBMGA007203* | down | 13 | 21 | - | 11.1009 | 6.57E-05 | 0.709 | GAGCGAGGC |
| 5965 | mamo-s | *BGIBMGA007204* | down | 1444 | 1452 | + | 11.3303 | 4.96E-05 | 0.615 | GTGGGTGGC |
| 5966 | mamo-s | *BGIBMGA007206* | down | 1898 | 1906 | + | 12.3394 | 1.49E-05 | 0.615 | GGGCGTGGC |
| 5967 | mamo-s | *BGIBMGA007207* | up | 1974 | 1982 | + | 11.5688 | 3.79E-05 | 0.615 | CTGCGTGTC |
| 5968 | mamo-s | *BGIBMGA007207* | down | 517 | 525 | - | 11.4404 | 4.51E-05 | 0.615 | GAGCGTGTC |
| 5969 | mamo-s | *BGIBMGA007210* | down | 138 | 146 | + | 13.6606 | 4.06E-06 | 0.615 | GCGCGTGGC |
| 5970 | mamo-s | *BGIBMGA007211* | down | 1552 | 1560 | + | 11.3303 | 4.96E-05 | 0.615 | GTGGGTGGC |
| 5971 | mamo-s | *BGIBMGA007212* | down | 1608 | 1616 | - | 12.1835 | 1.71E-05 | 0.615 | GTGCGTGCC |
| 5972 | mamo-s | *BGIBMGA007213* | up | 142 | 150 | - | 11.3303 | 4.96E-05 | 0.615 | GTGGGTGGC |
| 5973 | mamo-s | *BGIBMGA007213* | down | 1377 | 1385 | - | 11.3303 | 4.96E-05 | 0.615 | GTGGGTGGC |
| 5974 | mamo-s | *BGIBMGA007215* | up | 1795 | 1803 | - | 11.3853 | 4.74E-05 | 0.615 | ACGCGTGGC |
| 5975 | mamo-s | *BGIBMGA007215* | down | 1066 | 1074 | + | 11.3303 | 4.96E-05 | 0.615 | GTGGGTGGC |
| 5976 | mamo-s | *BGIBMGA007215* | up | 1026 | 1034 | + | 11.1009 | 6.57E-05 | 0.709 | GAGCGAGGC |
| 5977 | mamo-s | *BGIBMGA007215* | up | 587 | 595 | - | 10.5413 | 9.42E-05 | 0.857 | CAGCGTGTC |

| | | | | | | | | | | |
|---|---|---|---|---|---|---|---|---|---|---|
| 5978 | mamo-s | *BGIBMGA007216* | down | 1008 | 1016 | + | 12.7615 | 1.04E-05 | 0.615 | CCGCGTGGC |
| 5979 | mamo-s | *BGIBMGA007216* | down | 972 | 980 | - | 12.0734 | 2.66E-05 | 0.615 | GCGCGTGTC |
| 5980 | mamo-s | *BGIBMGA007216* | up | 23 | 31 | - | 10.8349 | 7.66E-05 | 0.751 | CCGCGAGGC |
| 5981 | mamo-s | *BGIBMGA007216* | down | 235 | 243 | + | 10.5688 | 8.88E-05 | 0.824 | GTGCGTGAC |
| 5982 | mamo-s | *BGIBMGA007216* | down | 258 | 266 | - | 10.5688 | 8.88E-05 | 0.824 | GTGCGTGAC |
| 5983 | mamo-s | *BGIBMGA007216* | down | 374 | 382 | + | 10.5688 | 8.88E-05 | 0.824 | GTGCGTGAC |
| 5984 | mamo-s | *BGIBMGA007219* | down | 1749 | 1757 | - | 10.5688 | 8.88E-05 | 0.824 | GTGCGTGAC |
| 5985 | mamo-s | *BGIBMGA007222* | down | 1213 | 1221 | - | 11.3303 | 4.96E-05 | 0.615 | GTGGGTGGC |
| 5986 | mamo-s | *BGIBMGA007222* | up | 1865 | 1873 | + | 10.5413 | 9.42E-05 | 0.857 | GTGCGAGTC |
| 5987 | mamo-s | *BGIBMGA007224* | down | 201 | 209 | + | 11.2844 | 5.18E-05 | 0.63 | CTGCGTGCC |
| 5988 | mamo-s | *BGIBMGA007225* | down | 1409 | 1417 | - | 11.3303 | 4.96E-05 | 0.615 | GTGGGTGGC |
| 5989 | mamo-s | *BGIBMGA007226* | down | 595 | 603 | - | 11.0734 | 7.12E-05 | 0.726 | GAGCGTGGT |
| 5990 | mamo-s | *BGIBMGA007229* | down | 298 | 306 | + | 11.5688 | 3.79E-05 | 0.615 | CTGCGTGTC |
| 5991 | mamo-s | *BGIBMGA007229* | up | 1657 | 1665 | - | 11.3303 | 4.96E-05 | 0.615 | GTGGGTGGC |
| 5992 | mamo-s | *BGIBMGA007230* | up | 1292 | 1300 | + | 11.3303 | 4.96E-05 | 0.615 | GTGGGTGGC |
| 5993 | mamo-s | *BGIBMGA007232* | up | 1208 | 1216 | - | 11.3303 | 4.96E-05 | 0.615 | GTGGGTGGC |
| 5994 | mamo-s | *BGIBMGA007232* | up | 76 | 84 | + | 10.7523 | 8.38E-05 | 0.797 | AAGCGTGGC |
| 5995 | mamo-s | *BGIBMGA007238* | down | 27 | 35 | + | 10.5688 | 8.88E-05 | 0.824 | GTGCGTGAC |
| 5996 | mamo-s | *BGIBMGA007244* | up | 251 | 259 | - | 11.3303 | 4.96E-05 | 0.615 | GTGGGTGGC |
| 5997 | mamo-s | *BGIBMGA007245* | down | 602 | 610 | - | 13.6606 | 4.06E-06 | 0.615 | GCGCGTGGC |
| 5998 | mamo-s | *BGIBMGA007246* | down | 957 | 965 | - | 11.3303 | 4.96E-05 | 0.615 | GTGGGTGGC |
| 5999 | mamo-s | *BGIBMGA007249* | down | 1601 | 1609 | - | 11.3303 | 4.96E-05 | 0.615 | GTGGGTGGC |
| 6000 | mamo-s | *BGIBMGA007249* | up | 1109 | 1117 | - | 11.0826 | 6.84E-05 | 0.709 | GTACGTGGC |
| 6001 | mamo-s | *BGIBMGA007252* | down | 141 | 149 | - | 11.3303 | 4.96E-05 | 0.615 | GTGGGTGGC |
| 6002 | mamo-s | *BGIBMGA007252* | up | 894 | 902 | - | 11.3303 | 4.96E-05 | 0.615 | GTGGGTGGC |
| 6003 | mamo-s | *BGIBMGA007253* | up | 364 | 372 | - | 10.8349 | 7.66E-05 | 0.751 | CCGCGAGGC |
| 6004 | mamo-s | *BGIBMGA007255* | up | 1075 | 1083 | + | 11.3303 | 4.96E-05 | 0.615 | GTGGGTGGC |
| 6005 | mamo-s | *BGIBMGA007255* | up | 1596 | 1604 | - | 11.3303 | 4.96E-05 | 0.615 | GTGGGTGGC |
| 6006 | mamo-s | *BGIBMGA007255* | down | 1619 | 1627 | + | 11.3303 | 4.96E-05 | 0.615 | GTGGGTGGC |
| 6007 | mamo-s | *BGIBMGA007256* | up | 1566 | 1574 | - | 10.9358 | 7.30E-05 | 0.728 | GCGGGTGGC |
| 6008 | mamo-s | *BGIBMGA007261* | up | 1287 | 1295 | - | 11.3303 | 4.96E-05 | 0.615 | GTGGGTGGC |
| 6009 | mamo-s | *BGIBMGA007261* | up | 18 | 26 | + | 11.2294 | 5.41E-05 | 0.65 | CTGCGAGGC |
| 6010 | mamo-s | *BGIBMGA007262* | up | 664 | 672 | - | 11.3303 | 4.96E-05 | 0.615 | GTGGGTGGC |
| 6011 | mamo-s | *BGIBMGA007264* | up | 227 | 235 | - | 11.4404 | 4.51E-05 | 0.615 | CGGCGTGGC |
| 6012 | mamo-s | *BGIBMGA007264* | down | 241 | 249 | + | 11.3303 | 4.96E-05 | 0.615 | GTGGGTGGC |
| 6013 | mamo-s | *BGIBMGA007266* | up | 1680 | 1688 | - | 11.0826 | 6.84E-05 | 0.709 | GTACGTGGC |
| 6014 | mamo-s | *BGIBMGA007267* | down | 730 | 738 | + | 11.1743 | 5.90E-05 | 0.684 | CCGCGTGTC |
| 6015 | mamo-s | *BGIBMGA007270* | up | 872 | 880 | - | 11.3303 | 4.96E-05 | 0.615 | GTGGGTGGC |
| 6016 | mamo-s | *BGIBMGA007273* | up | 814 | 822 | + | 13.0275 | 8.53E-06 | 0.615 | GAGCGTGGC |
| 6017 | mamo-s | *BGIBMGA007274* | down | 1112 | 1120 | + | 11.3853 | 4.74E-05 | 0.615 | ACGCGTGGC |
| 6018 | mamo-s | *BGIBMGA007277* | down | 87 | 95 | - | 11.7339 | 3.29E-05 | 0.615 | GCGCGAGGC |
| 6019 | mamo-s | *BGIBMGA007277* | down | 549 | 557 | + | 10.5138 | 9.75E-05 | 0.865 | GTGCGTGTT |
| 6020 | mamo-s | *BGIBMGA007278* | up | 46 | 54 | + | 12.0734 | 2.66E-05 | 0.615 | GCGCGTGTC |
| 6021 | mamo-s | *BGIBMGA007278* | down | 1710 | 1718 | + | 11.789 | 2.84E-05 | 0.615 | GCGCGTGCC |
| 6022 | mamo-s | *BGIBMGA007278* | down | 1462 | 1470 | + | 11.3303 | 4.96E-05 | 0.615 | GTGGGTGGC |
| 6023 | mamo-s | *BGIBMGA007280* | up | 777 | 785 | - | 13.156 | 6.30E-06 | 0.615 | CTGCGTGGC |
| 6024 | mamo-s | *BGIBMGA007282* | down | 1033 | 1041 | - | 14.055 | 2.23E-06 | 0.615 | GTGCGTGGC |
| 6025 | mamo-s | *BGIBMGA007283* | up | 338 | 346 | + | 14.055 | 2.23E-06 | 0.615 | GTGCGTGGC |
| 6026 | mamo-s | *BGIBMGA007283* | up | 1742 | 1750 | + | 11.0826 | 6.84E-05 | 0.709 | GTACGTGGC |
| 6027 | mamo-s | *BGIBMGA007285* | down | 378 | 386 | + | 12.7615 | 1.04E-05 | 0.615 | CCGCGTGGC |
| 6028 | mamo-s | *BGIBMGA007285* | up | 703 | 711 | - | 10.5413 | 9.42E-05 | 0.857 | CAGCGTGTC |
| 6029 | mamo-s | *BGIBMGA007286* | down | 522 | 530 | + | 11.0826 | 6.84E-05 | 0.709 | GTACGTGGC |
| 6030 | mamo-s | *BGIBMGA007286* | up | 1881 | 1889 | - | 10.8349 | 7.66E-05 | 0.751 | CCGCGAGGC |
| 6031 | mamo-s | *BGIBMGA007288* | down | 1735 | 1743 | + | 13.6606 | 4.06E-06 | 0.615 | GCGCGTGGC |
| 6032 | mamo-s | *BGIBMGA007288* | down | 193 | 201 | - | 11.3303 | 4.96E-05 | 0.615 | GTGGGTGGC |
| 6033 | mamo-s | *BGIBMGA007299* | down | 732 | 740 | - | 11.7339 | 3.29E-05 | 0.615 | GCGCGAGGC |
| 6034 | mamo-s | *BGIBMGA007299* | down | 994 | 1002 | + | 11.3303 | 4.96E-05 | 0.615 | GTGGGTGGC |
| 6035 | mamo-s | *BGIBMGA007300* | up | 844 | 852 | + | 10.7523 | 8.38E-05 | 0.797 | AAGCGTGGC |
| 6036 | mamo-s | *BGIBMGA007301* | down | 1002 | 1010 | - | 11.3303 | 4.96E-05 | 0.615 | GTGGGTGGC |
| 6037 | mamo-s | *BGIBMGA007303* | down | 1815 | 1823 | + | 11.3303 | 4.96E-05 | 0.615 | GTGGGTGGC |
| 6038 | mamo-s | *BGIBMGA007308* | down | 843 | 851 | + | 12.1284 | 2.16E-05 | 0.615 | GTGCGAGGC |
| 6039 | mamo-s | *BGIBMGA007308* | up | 97 | 105 | + | 11.2844 | 5.18E-05 | 0.63 | CTGCGTGCC |
| 6040 | mamo-s | *BGIBMGA007309* | down | 669 | 677 | + | 11.3303 | 4.96E-05 | 0.615 | GTGGGTGGC |
| 6041 | mamo-s | *BGIBMGA007311* | down | 1927 | 1935 | + | 12.4679 | 1.31E-05 | 0.615 | GTGCGTGTC |
| 6042 | mamo-s | *BGIBMGA007312* | down | 648 | 656 | - | 11.3303 | 4.96E-05 | 0.615 | GTGGGTGGC |
| 6043 | mamo-s | *BGIBMGA007314* | up | 845 | 853 | - | 11.3303 | 4.96E-05 | 0.615 | GTGGGTGGC |
| 6044 | mamo-s | *BGIBMGA007315* | up | 548 | 556 | + | 11.3303 | 4.96E-05 | 0.615 | GTGGGTGGC |
| 6045 | mamo-s | *BGIBMGA007315* | up | 144 | 152 | - | 10.7523 | 8.38E-05 | 0.797 | AAGCGTGGC |
| 6046 | mamo-s | *BGIBMGA007316* | down | 1633 | 1641 | - | 11.5688 | 3.79E-05 | 0.615 | CTGCGTGTC |
| 6047 | mamo-s | *BGIBMGA007316* | down | 1437 | 1445 | - | 11.3303 | 4.96E-05 | 0.615 | GTGGGTGGC |
| 6048 | mamo-s | *BGIBMGA007318* | up | 204 | 212 | + | 11.3303 | 4.96E-05 | 0.615 | GTGGGTGGC |
| 6049 | mamo-s | *BGIBMGA007318* | up | 1667 | 1675 | - | 11.1376 | 6.35E-05 | 0.709 | TCGCGTGGC |
| 6050 | mamo-s | *BGIBMGA007320* | down | 1133 | 1141 | - | 10.9358 | 7.30E-05 | 0.728 | GCGGGTGGC |
| 6051 | mamo-s | *BGIBMGA007321* | up | 1828 | 1836 | + | 12.1835 | 1.71E-05 | 0.615 | GTGCGTGCC |
| 6052 | mamo-s | *BGIBMGA007321* | down | 154 | 162 | + | 12.1009 | 2.43E-05 | 0.615 | GTGCGTGGT |
| 6053 | mamo-s | *BGIBMGA007321* | up | 441 | 449 | + | 11.3303 | 4.96E-05 | 0.615 | GTGGGTGGC |
| 6054 | mamo-s | *BGIBMGA007323* | up | 582 | 590 | + | 11.1743 | 5.90E-05 | 0.684 | CCGCGTGTC |
| 6055 | mamo-s | *BGIBMGA007327* | down | 904 | 912 | + | 10.9358 | 7.30E-05 | 0.728 | GCGGGTGGC |
| 6056 | mamo-s | *BGIBMGA007328* | up | 1340 | 1348 | - | 11.3303 | 4.96E-05 | 0.615 | GTGGGTGGC |
| 6057 | mamo-s | *BGIBMGA007329* | down | 1489 | 1497 | + | 11.7798 | 3.11E-05 | 0.615 | ATGCGTGGC |
| 6058 | mamo-s | *BGIBMGA007329* | down | 1774 | 1782 | + | 11.3303 | 4.96E-05 | 0.615 | GTGGGTGGC |
| 6059 | mamo-s | *BGIBMGA007332* | up | 875 | 883 | - | 11.1009 | 6.57E-05 | 0.709 | GAGCGAGGC |
| 6060 | mamo-s | *BGIBMGA007334* | up | 6 | 14 | - | 11.789 | 2.84E-05 | 0.615 | GCGCGTGCC |
| 6061 | mamo-s | *BGIBMGA007334* | up | 402 | 410 | + | 11.3303 | 4.96E-05 | 0.615 | GTGGGTGGC |
| 6062 | mamo-s | *BGIBMGA007335* | up | 1720 | 1728 | - | 11.3303 | 4.96E-05 | 0.615 | GTGGGTGGC |
| 6063 | mamo-s | *BGIBMGA007338* | down | 930 | 938 | + | 11.3303 | 4.96E-05 | 0.615 | GTGGGTGGC |
| 6064 | mamo-s | *BGIBMGA007338* | down | 1849 | 1857 | - | 11.3303 | 4.96E-05 | 0.615 | GTGGGTGGC |
| 6065 | mamo-s | *BGIBMGA007340* | down | 1010 | 1018 | - | 12.7615 | 1.04E-05 | 0.615 | CCGCGTGGC |

| | | | | | | | | | | |
|---|---|---|---|---|---|---|---|---|---|---|
| 6066 | mamo-s | *BGIBMGA007342* | up | 238 | 246 | + | 11.3303 | 4.96E-05 | 0.615 | GTGGGTGGC |
| 6067 | mamo-s | *BGIBMGA007344* | down | 1384 | 1392 | + | 12.1284 | 2.16E-05 | 0.615 | GTGCGAGGC |
| 6068 | mamo-s | *BGIBMGA007344* | up | 941 | 949 | + | 11.1009 | 6.57E-05 | 0.709 | GAGCGAGGC |
| 6069 | mamo-s | *BGIBMGA007346* | up | 1392 | 1400 | + | 11.3303 | 4.96E-05 | 0.615 | GTGGGTGGC |
| 6070 | mamo-s | *BGIBMGA007348* | up | 91 | 99 | - | 11.3303 | 4.96E-05 | 0.615 | GTGGGTGGC |
| 6071 | mamo-s | *BGIBMGA007348* | down | 167 | 175 | + | 11.3303 | 4.96E-05 | 0.615 | GTGGGTGGC |
| 6072 | mamo-s | *BGIBMGA007348* | down | 892 | 900 | + | 10.5138 | 9.75E-05 | 0.865 | GTGCGTGTT |
| 6073 | mamo-s | *BGIBMGA007351* | down | 1568 | 1576 | + | 13.6606 | 4.06E-06 | 0.615 | GCGCGTGGC |
| 6074 | mamo-s | *BGIBMGA007351* | down | 981 | 989 | - | 11.3303 | 4.96E-05 | 0.615 | GTGGGTGGC |
| 6075 | mamo-s | *BGIBMGA007352* | up | 1912 | 1920 | + | 13.6606 | 4.06E-06 | 0.615 | GCGCGTGGC |
| 6076 | mamo-s | *BGIBMGA007352* | down | 1110 | 1118 | - | 11.3303 | 4.96E-05 | 0.615 | GTGGGTGGC |
| 6077 | mamo-s | *BGIBMGA007352* | up | 1325 | 1333 | - | 11.3303 | 4.96E-05 | 0.615 | GTGGGTGGC |
| 6078 | mamo-s | *BGIBMGA007353* | up | 1770 | 1778 | + | 11.3853 | 4.74E-05 | 0.615 | ACGCGTGGC |
| 6079 | mamo-s | *BGIBMGA007353* | down | 261 | 269 | - | 10.5138 | 9.75E-05 | 0.865 | GTGCGTGTT |
| 6080 | mamo-s | *BGIBMGA007354* | down | 261 | 269 | - | 10.5138 | 9.75E-05 | 0.865 | GTGCGTGTT |
| 6081 | mamo-s | *BGIBMGA007357* | down | 1298 | 1306 | - | 11.7064 | 3.52E-05 | 0.615 | GCGCGTGGT |
| 6082 | mamo-s | *BGIBMGA007358* | up | 843 | 851 | - | 11.7064 | 3.52E-05 | 0.615 | GCGCGTGGT |
| 6083 | mamo-s | *BGIBMGA007367* | up | 1315 | 1323 | - | 11.3303 | 4.96E-05 | 0.615 | GTGGGTGGC |
| 6084 | mamo-s | *BGIBMGA007367* | up | 426 | 434 | - | 11.1743 | 5.90E-05 | 0.684 | CCGCGTGTC |
| 6085 | mamo-s | *BGIBMGA007368* | down | 1228 | 1236 | + | 11.1009 | 6.57E-05 | 0.709 | GAGCGAGGC |
| 6086 | mamo-s | *BGIBMGA007369* | up | 799 | 807 | + | 11.3303 | 4.96E-05 | 0.615 | GTGGGTGGC |
| 6087 | mamo-s | *BGIBMGA007369* | down | 1558 | 1566 | + | 10.8899 | 7.48E-05 | 0.739 | CCGCGTGCC |
| 6088 | mamo-s | *BGIBMGA007370* | down | 489 | 497 | + | 11.2844 | 5.18E-05 | 0.63 | CTGCGTGCC |
| 6089 | mamo-s | *BGIBMGA007371* | up | 1364 | 1372 | - | 12.1284 | 2.16E-05 | 0.615 | CAGCGTGGC |
| 6090 | mamo-s | *BGIBMGA007372* | down | 1764 | 1772 | + | 12.1009 | 2.43E-05 | 0.615 | GTGCGTGGT |
| 6091 | mamo-s | *BGIBMGA007372* | down | 641 | 649 | - | 11.4404 | 4.51E-05 | 0.615 | GAGCGTGTC |
| 6092 | mamo-s | *BGIBMGA007372* | down | 136 | 144 | + | 10.5138 | 9.75E-05 | 0.865 | GTGCGTGTT |
| 6093 | mamo-s | *BGIBMGA007373* | up | 308 | 316 | + | 11.4404 | 4.51E-05 | 0.615 | CGGCGTGGC |
| 6094 | mamo-s | *BGIBMGA007373* | up | 979 | 987 | - | 11.4404 | 4.51E-05 | 0.615 | GAGCGTGTC |
| 6095 | mamo-s | *BGIBMGA007373* | up | 474 | 482 | + | 10.5138 | 9.75E-05 | 0.865 | GTGCGTGTT |
| 6096 | mamo-s | *BGIBMGA007374* | down | 1515 | 1523 | + | 11.5321 | 4.06E-05 | 0.615 | TTGCGTGGC |
| 6097 | mamo-s | *BGIBMGA007374* | down | 483 | 491 | - | 11.3303 | 4.96E-05 | 0.615 | GTGGGTGGC |
| 6098 | mamo-s | *BGIBMGA007374* | up | 1169 | 1177 | + | 11.3303 | 4.96E-05 | 0.615 | GTGGGTGGC |
| 6099 | mamo-s | *BGIBMGA007374* | up | 902 | 910 | - | 10.8073 | 7.89E-05 | 0.766 | CCGCGTGGT |
| 6100 | mamo-s | *BGIBMGA007375* | down | 1721 | 1729 | + | 12.1835 | 1.71E-05 | 0.615 | GTGCGTGCC |
| 6101 | mamo-s | *BGIBMGA007375* | down | 934 | 942 | - | 11.3303 | 4.96E-05 | 0.615 | GTGGGTGGC |
| 6102 | mamo-s | *BGIBMGA007376* | up | 1760 | 1768 | - | 11.1009 | 6.57E-05 | 0.709 | GAGCGAGGC |
| 6103 | mamo-s | *BGIBMGA007377* | up | 218 | 226 | - | 11.7339 | 3.29E-05 | 0.615 | GCGCGAGGC |
| 6104 | mamo-s | *BGIBMGA007377* | up | 1297 | 1305 | + | 11.3303 | 4.96E-05 | 0.615 | GTGGGTGGC |
| 6105 | mamo-s | *BGIBMGA007377* | up | 1062 | 1070 | + | 10.7523 | 8.38E-05 | 0.797 | AAGCGTGGC |
| 6106 | mamo-s | *BGIBMGA007378* | up | 1526 | 1534 | + | 11.3303 | 4.96E-05 | 0.615 | GTGGGTGGC |
| 6107 | mamo-s | *BGIBMGA007378* | down | 1954 | 1962 | - | 11.1743 | 5.90E-05 | 0.684 | CCGCGTGTC |
| 6108 | mamo-s | *BGIBMGA007379* | up | 16 | 24 | - | 11.0734 | 7.12E-05 | 0.726 | GAGCGTGGT |
| 6109 | mamo-s | *BGIBMGA007381* | up | 1730 | 1738 | - | 11.1743 | 5.90E-05 | 0.684 | CCGCGTGTC |
| 6110 | mamo-s | *BGIBMGA007383* | up | 207 | 215 | + | 12.0734 | 2.66E-05 | 0.615 | GCGCGTGTC |
| 6111 | mamo-s | *BGIBMGA007383* | down | 1207 | 1215 | - | 11.5321 | 4.06E-05 | 0.615 | TTGCGTGGC |
| 6112 | mamo-s | *BGIBMGA007383* | down | 1353 | 1361 | + | 10.9358 | 7.30E-05 | 0.728 | GCGGGTGGC |
| 6113 | mamo-s | *BGIBMGA007383* | up | 1769 | 1777 | - | 10.5413 | 9.42E-05 | 0.857 | GTGCGAGTC |
| 6114 | mamo-s | *BGIBMGA007384* | down | 423 | 431 | - | 11.7339 | 3.29E-05 | 0.615 | GCGCGAGGC |
| 6115 | mamo-s | *BGIBMGA007384* | down | 1184 | 1192 | - | 11.3303 | 4.96E-05 | 0.615 | GTGGGTGGC |
| 6116 | mamo-s | *BGIBMGA007385* | down | 39 | 47 | + | 11.7798 | 3.11E-05 | 0.615 | ATGCGTGGC |
| 6117 | mamo-s | *BGIBMGA007385* | up | 1487 | 1495 | - | 11.2844 | 5.18E-05 | 0.63 | CTGCGTGCC |
| 6118 | mamo-s | *BGIBMGA007385* | down | 762 | 770 | + | 10.5413 | 9.42E-05 | 0.857 | GTGCGAGTC |
| 6119 | mamo-s | *BGIBMGA007386* | up | 872 | 880 | + | 12.3394 | 1.49E-05 | 0.615 | GGGCGTGGC |
| 6120 | mamo-s | *BGIBMGA007386* | up | 941 | 949 | + | 12.0734 | 2.66E-05 | 0.615 | GCGCGTGTC |
| 6121 | mamo-s | *BGIBMGA007386* | up | 1147 | 1155 | - | 11.2844 | 5.18E-05 | 0.63 | CTGCGTGCC |
| 6122 | mamo-s | *BGIBMGA007388* | down | 228 | 236 | + | 12.0734 | 2.66E-05 | 0.615 | GCGCGTGTC |
| 6123 | mamo-s | *BGIBMGA007388* | down | 291 | 299 | - | 11.2018 | 5.68E-05 | 0.674 | CTGCGTGGT |
| 6124 | mamo-s | *BGIBMGA007388* | up | 1535 | 1543 | + | 11.1376 | 6.35E-05 | 0.709 | TCGCGTGGC |
| 6125 | mamo-s | *BGIBMGA007389* | up | 937 | 945 | - | 13.156 | 6.30E-06 | 0.615 | CTGCGTGGC |
| 6126 | mamo-s | *BGIBMGA007389* | up | 1248 | 1256 | - | 11.3303 | 4.96E-05 | 0.615 | GTGGGTGGC |
| 6127 | mamo-s | *BGIBMGA007389* | up | 1693 | 1701 | - | 11.0826 | 6.84E-05 | 0.709 | GTACGTGGC |
| 6128 | mamo-s | *BGIBMGA007391* | up | 790 | 798 | - | 11.0826 | 6.84E-05 | 0.709 | GTACGTGGC |
| 6129 | mamo-s | *BGIBMGA007392* | up | 788 | 796 | - | 11.3303 | 4.96E-05 | 0.615 | GTGGGTGGC |
| 6130 | mamo-s | *BGIBMGA007393* | down | 853 | 861 | + | 12.0734 | 2.66E-05 | 0.615 | GCGCGTGTC |
| 6131 | mamo-s | *BGIBMGA007393* | down | 661 | 669 | + | 11.7339 | 3.29E-05 | 0.615 | GCGCGAGGC |
| 6132 | mamo-s | *BGIBMGA007394* | up | 1921 | 1929 | + | 12.0734 | 2.66E-05 | 0.615 | GCGCGTGTC |
| 6133 | mamo-s | *BGIBMGA007394* | up | 1729 | 1737 | + | 11.7339 | 3.29E-05 | 0.615 | GCGCGAGGC |
| 6134 | mamo-s | *BGIBMGA007395* | up | 623 | 631 | - | 10.5413 | 9.42E-05 | 0.857 | CAGCGTGTC |
| 6135 | mamo-s | *BGIBMGA007396* | down | 51 | 59 | + | 13.6606 | 4.06E-06 | 0.615 | GCGCGTGGC |
| 6136 | mamo-s | *BGIBMGA007396* | down | 1196 | 1204 | + | 11.3303 | 4.96E-05 | 0.615 | GTGGGTGGC |
| 6137 | mamo-s | *BGIBMGA007396* | up | 989 | 997 | + | 10.9358 | 7.30E-05 | 0.728 | GCGGGTGGC |
| 6138 | mamo-s | *BGIBMGA007397* | down | 623 | 631 | + | 10.5413 | 9.42E-05 | 0.857 | GTGCGAGTC |
| 6139 | mamo-s | *BGIBMGA007399* | up | 534 | 542 | + | 11.3303 | 4.96E-05 | 0.615 | GTGGGTGGC |
| 6140 | mamo-s | *BGIBMGA007399* | up | 1423 | 1431 | + | 11.1743 | 5.90E-05 | 0.684 | CCGCGTGTC |
| 6141 | mamo-s | *BGIBMGA007400* | down | 1342 | 1350 | - | 11.3303 | 4.96E-05 | 0.615 | GTGGGTGGC |
| 6142 | mamo-s | *BGIBMGA007400* | down | 22 | 30 | + | 10.8899 | 7.48E-05 | 0.739 | CCGCGTGCC |
| 6143 | mamo-s | *BGIBMGA007401* | down | 251 | 259 | - | 11.3303 | 4.96E-05 | 0.615 | GTGGGTGGC |
| 6144 | mamo-s | *BGIBMGA007401* | down | 922 | 930 | - | 11.3303 | 4.96E-05 | 0.615 | GTGGGTGGC |
| 6145 | mamo-s | *BGIBMGA007402* | up | 878 | 886 | - | 10.7523 | 8.38E-05 | 0.797 | AAGCGTGGC |
| 6146 | mamo-s | *BGIBMGA007404* | up | 694 | 702 | + | 11.3303 | 4.96E-05 | 0.615 | GTGGGTGGC |
| 6147 | mamo-s | *BGIBMGA007405* | up | 1676 | 1684 | + | 10.5688 | 8.88E-05 | 0.824 | GTGCGTGAC |
| 6148 | mamo-s | *BGIBMGA007406* | up | 931 | 939 | - | 12.1835 | 1.71E-05 | 0.615 | GTGCGTGCC |
| 6149 | mamo-s | *BGIBMGA007406* | up | 98 | 106 | - | 11.5688 | 3.79E-05 | 0.615 | CTGCGTGTC |
| 6150 | mamo-s | *BGIBMGA007407* | down | 464 | 472 | + | 10.6881 | 8.61E-05 | 0.81 | GCACGTGGC |
| 6151 | mamo-s | *BGIBMGA007413* | down | 542 | 550 | + | 11.3303 | 4.96E-05 | 0.615 | GTGGGTGGC |
| 6152 | mamo-s | *BGIBMGA007413* | up | 1390 | 1398 | + | 10.5138 | 9.75E-05 | 0.865 | GTGCGTGTT |
| 6153 | mamo-s | *BGIBMGA007414* | down | 1410 | 1418 | + | 11.4404 | 4.51E-05 | 0.615 | GAGCGTGTC |

| | | | | | | | | | | |
|---|---|---|---|---|---|---|---|---|---|---|
| 6154 | mamo-s | *BGIBMGA007415* | up | 852 | 860 | + | 13.6606 | 4.06E-06 | 0.615 | GCGCGTGGC |
| 6155 | mamo-s | *BGIBMGA007417* | down | 1500 | 1508 | + | 14.055 | 2.23E-06 | 0.615 | GTGCGTGGC |
| 6156 | mamo-s | *BGIBMGA007419* | down | 1340 | 1348 | - | 14.055 | 2.23E-06 | 0.615 | GTGCGTGGC |
| 6157 | mamo-s | *BGIBMGA007419* | down | 1303 | 1311 | - | 10.9358 | 7.30E-05 | 0.728 | GCGGGTGGC |
| 6158 | mamo-s | *BGIBMGA007420* | up | 1986 | 1994 | - | 14.055 | 2.23E-06 | 0.615 | GTGCGTGGC |
| 6159 | mamo-s | *BGIBMGA007420* | up | 198 | 206 | - | 11.7064 | 3.52E-05 | 0.615 | GCGCGTGGT |
| 6160 | mamo-s | *BGIBMGA007420* | down | 263 | 271 | + | 11.3303 | 4.96E-05 | 0.615 | GTGGGTGGC |
| 6161 | mamo-s | *BGIBMGA007420* | up | 1949 | 1957 | - | 10.9358 | 7.30E-05 | 0.728 | GCGGGTGGC |
| 6162 | mamo-s | *BGIBMGA007423* | up | 1293 | 1301 | + | 14.055 | 2.23E-06 | 0.615 | GTGCGTGGC |
| 6163 | mamo-s | *BGIBMGA007423* | up | 1154 | 1162 | + | 10.8073 | 7.89E-05 | 0.766 | CCGCGTGGT |
| 6164 | mamo-s | *BGIBMGA007424* | down | 18 | 26 | - | 11.7798 | 3.11E-05 | 0.615 | ATGCGTGGC |
| 6165 | mamo-s | *BGIBMGA007424* | down | 1863 | 1871 | - | 10.5413 | 9.42E-05 | 0.857 | GTGCGAGTC |
| 6166 | mamo-s | *BGIBMGA007426* | down | 114 | 122 | + | 11.1743 | 5.90E-05 | 0.684 | CCGCGTGTC |
| 6167 | mamo-s | *BGIBMGA007428* | up | 646 | 654 | + | 10.9358 | 7.30E-05 | 0.728 | GCGGGTGGC |
| 6168 | mamo-s | *BGIBMGA007428* | down | 416 | 424 | - | 10.5688 | 8.88E-05 | 0.824 | GTGCGTGAC |
| 6169 | mamo-s | *BGIBMGA007429* | down | 1341 | 1349 | - | 11.3303 | 4.96E-05 | 0.615 | GTGGGTGGC |
| 6170 | mamo-s | *BGIBMGA007430* | up | 102 | 110 | - | 12.1009 | 2.43E-05 | 0.615 | GTGCGTGGT |
| 6171 | mamo-s | *BGIBMGA007430* | up | 1014 | 1022 | - | 11.0826 | 6.84E-05 | 0.709 | GTACGTGGC |
| 6172 | mamo-s | *BGIBMGA007431* | up | 1230 | 1238 | + | 11.3303 | 4.96E-05 | 0.615 | GTGGGTGGC |
| 6173 | mamo-s | *BGIBMGA007431* | down | 1260 | 1268 | + | 10.5413 | 9.42E-05 | 0.857 | GTGCGAGTC |
| 6174 | mamo-s | *BGIBMGA007432* | down | 1333 | 1341 | + | 12.4679 | 1.31E-05 | 0.615 | GTGCGTGTC |
| 6175 | mamo-s | *BGIBMGA007432* | down | 1992 | 2000 | + | 11.3303 | 4.96E-05 | 0.615 | GTGGGTGGC |
| 6176 | mamo-s | *BGIBMGA007433* | up | 147 | 155 | + | 11.3303 | 4.96E-05 | 0.615 | GTGGGTGGC |
| 6177 | mamo-s | *BGIBMGA007433* | up | 549 | 557 | - | 11.3303 | 4.96E-05 | 0.615 | GTGGGTGGC |
| 6178 | mamo-s | *BGIBMGA007434* | down | 972 | 980 | - | 11.3303 | 4.96E-05 | 0.615 | GTGGGTGGC |
| 6179 | mamo-s | *BGIBMGA007437* | down | 1294 | 1302 | + | 11.3303 | 4.96E-05 | 0.615 | GTGGGTGGC |
| 6180 | mamo-s | *BGIBMGA007438* | up | 623 | 631 | - | 11.3303 | 4.96E-05 | 0.615 | GTGGGTGGC |
| 6181 | mamo-s | *BGIBMGA007440* | up | 225 | 233 | - | 12.1835 | 1.71E-05 | 0.615 | GTGCGTGCC |
| 6182 | mamo-s | *BGIBMGA007440* | up | 396 | 404 | - | 12.1835 | 1.71E-05 | 0.615 | GTGCGTGCC |
| 6183 | mamo-s | *BGIBMGA007440* | up | 149 | 157 | + | 10.5138 | 9.75E-05 | 0.865 | GTGCGTGTT |
| 6184 | mamo-s | *BGIBMGA007440* | up | 320 | 328 | + | 10.5138 | 9.75E-05 | 0.865 | GTGCGTGTT |
| 6185 | mamo-s | *BGIBMGA007440* | up | 491 | 499 | + | 10.5138 | 9.75E-05 | 0.865 | GTGCGTGTT |
| 6186 | mamo-s | *BGIBMGA007442* | up | 1499 | 1507 | + | 11.3303 | 4.96E-05 | 0.615 | GTGGGTGGC |
| 6187 | mamo-s | *BGIBMGA007443* | down | 1292 | 1300 | + | 11.7798 | 3.11E-05 | 0.615 | ATGCGTGGC |
| 6188 | mamo-s | *BGIBMGA007444* | up | 566 | 574 | - | 11.3303 | 4.96E-05 | 0.615 | GTGGGTGGC |
| 6189 | mamo-s | *BGIBMGA007445* | down | 1827 | 1835 | - | 11.3303 | 4.96E-05 | 0.615 | GTGGGTGGC |
| 6190 | mamo-s | *BGIBMGA007446* | up | 194 | 202 | - | 13.0275 | 8.53E-06 | 0.615 | GAGCGTGGC |
| 6191 | mamo-s | *BGIBMGA007448* | down | 1411 | 1419 | + | 11.3303 | 4.96E-05 | 0.615 | GTGGGTGGC |
| 6192 | mamo-s | *BGIBMGA007448* | up | 1559 | 1567 | - | 11.3303 | 4.96E-05 | 0.615 | GTGGGTGGC |
| 6193 | mamo-s | *BGIBMGA007449* | up | 885 | 893 | + | 11.3303 | 4.96E-05 | 0.615 | GTGGGTGGC |
| 6194 | mamo-s | *BGIBMGA007452* | down | 1783 | 1791 | + | 11.1743 | 5.90E-05 | 0.684 | CCGCGTGTC |
| 6195 | mamo-s | *BGIBMGA007453* | down | 1417 | 1425 | - | 11.3303 | 4.96E-05 | 0.615 | GTGGGTGGC |
| 6196 | mamo-s | *BGIBMGA007453* | down | 1825 | 1833 | - | 11.3303 | 4.96E-05 | 0.615 | GTGGGTGGC |
| 6197 | mamo-s | *BGIBMGA007454* | up | 1616 | 1624 | - | 11.0826 | 6.84E-05 | 0.709 | GTACGTGGC |
| 6198 | mamo-s | *BGIBMGA007455* | up | 490 | 498 | - | 14.055 | 2.23E-06 | 0.615 | GTGCGTGGC |
| 6199 | mamo-s | *BGIBMGA007455* | down | 872 | 880 | - | 11.3303 | 4.96E-05 | 0.615 | GTGGGTGGC |
| 6200 | mamo-s | *BGIBMGA007456* | up | 1801 | 1809 | - | 11.3303 | 4.96E-05 | 0.615 | GTGGGTGGC |
| 6201 | mamo-s | *BGIBMGA007457* | down | 1807 | 1815 | - | 11.0826 | 6.84E-05 | 0.709 | GTACGTGGC |
| 6202 | mamo-s | *BGIBMGA007460* | down | 789 | 797 | + | 11.4404 | 4.51E-05 | 0.615 | GAGCGTGTC |
| 6203 | mamo-s | *BGIBMGA007461* | down | 145 | 153 | - | 11.3303 | 4.96E-05 | 0.615 | GTGGGTGGC |
| 6204 | mamo-s | *BGIBMGA007462* | up | 67 | 75 | + | 11.7798 | 3.11E-05 | 0.615 | ATGCGTGGC |
| 6205 | mamo-s | *BGIBMGA007465* | up | 1421 | 1429 | - | 11.3303 | 4.96E-05 | 0.615 | GTGGGTGGC |
| 6206 | mamo-s | *BGIBMGA007465* | up | 688 | 696 | + | 11.2294 | 5.41E-05 | 0.65 | CTGCGAGGC |
| 6207 | mamo-s | *BGIBMGA007467* | up | 192 | 200 | - | 10.9358 | 7.30E-05 | 0.728 | GCGGGTGGC |
| 6208 | mamo-s | *BGIBMGA007467* | up | 33 | 41 | - | 10.5688 | 8.88E-05 | 0.824 | GTGCGTGAC |
| 6209 | mamo-s | *BGIBMGA007469* | down | 642 | 650 | + | 11.1009 | 6.57E-05 | 0.709 | GAGCGAGGC |
| 6210 | mamo-s | *BGIBMGA007471* | down | 1665 | 1673 | + | 11.3303 | 4.96E-05 | 0.615 | GTGGGTGGC |
| 6211 | mamo-s | *BGIBMGA007472* | up | 1157 | 1165 | + | 11.3303 | 4.96E-05 | 0.615 | GTGGGTGGC |
| 6212 | mamo-s | *BGIBMGA007474* | up | 1236 | 1244 | - | 11.3303 | 4.96E-05 | 0.615 | GTGGGTGGC |
| 6213 | mamo-s | *BGIBMGA007478* | down | 1489 | 1497 | - | 10.7523 | 8.38E-05 | 0.797 | GGGCGTGTC |
| 6214 | mamo-s | *BGIBMGA007479* | up | 953 | 961 | - | 11.3303 | 4.96E-05 | 0.615 | GTGGGTGGC |
| 6215 | mamo-s | *BGIBMGA007479* | up | 1583 | 1591 | + | 10.9358 | 7.30E-05 | 0.728 | GCGGGTGGC |
| 6216 | mamo-s | *BGIBMGA007481* | up | 611 | 619 | - | 14.055 | 2.23E-06 | 0.615 | GTGCGTGGC |
| 6217 | mamo-s | *BGIBMGA007482* | down | 700 | 708 | + | 11.0826 | 6.84E-05 | 0.709 | GTACGTGGC |
| 6218 | mamo-s | *BGIBMGA007484* | up | 936 | 944 | + | 11.2844 | 5.18E-05 | 0.63 | CTGCGTGCC |
| 6219 | mamo-s | *BGIBMGA007485* | down | 226 | 234 | + | 11.3303 | 4.96E-05 | 0.615 | GTGGGTGGC |
| 6220 | mamo-s | *BGIBMGA007486* | up | 1850 | 1858 | + | 10.5138 | 9.75E-05 | 0.865 | GTGCGTGTT |
| 6221 | mamo-s | *BGIBMGA007487* | down | 1783 | 1791 | - | 11.1743 | 5.90E-05 | 0.684 | CCGCGTGTC |
| 6222 | mamo-s | *BGIBMGA007487* | up | 111 | 119 | + | 10.5413 | 9.42E-05 | 0.857 | GTGCGAGTC |
| 6223 | mamo-s | *BGIBMGA007491* | down | 837 | 845 | - | 11.4404 | 4.51E-05 | 0.615 | CGGCGTGGC |
| 6224 | mamo-s | *BGIBMGA007493* | down | 1109 | 1117 | - | 12.3394 | 1.49E-05 | 0.615 | GGGCGTGGC |
| 6225 | mamo-s | *BGIBMGA007493* | down | 927 | 935 | - | 11.3303 | 4.96E-05 | 0.615 | GTGGGTGGC |
| 6226 | mamo-s | *BGIBMGA007495* | up | 523 | 531 | - | 10.9358 | 7.30E-05 | 0.728 | GCGGGTGGC |
| 6227 | mamo-s | *BGIBMGA007498* | up | 1213 | 1221 | + | 11.5321 | 4.06E-05 | 0.615 | TTGCGTGGC |
| 6228 | mamo-s | *BGIBMGA007498* | up | 141 | 149 | + | 10.8899 | 7.48E-05 | 0.739 | CCGCGTGCC |
| 6229 | mamo-s | *BGIBMGA007499* | down | 73 | 81 | - | 11.1009 | 6.57E-05 | 0.709 | GAGCGAGGC |
| 6230 | mamo-s | *BGIBMGA007500* | down | 1369 | 1377 | + | 11.0826 | 6.84E-05 | 0.709 | GTACGTGGC |
| 6231 | mamo-s | *BGIBMGA007502* | up | 1423 | 1431 | + | 12.7615 | 1.04E-05 | 0.615 | CCGCGTGGC |
| 6232 | mamo-s | *BGIBMGA007503* | up | 652 | 660 | - | 11.0734 | 7.12E-05 | 0.726 | GAGCGTGGT |
| 6233 | mamo-s | *BGIBMGA007503* | up | 700 | 708 | + | 10.5413 | 9.42E-05 | 0.857 | GTGCGAGTC |
| 6234 | mamo-s | *BGIBMGA007505* | up | 1991 | 1999 | - | 12.1835 | 1.71E-05 | 0.615 | GTGCGTGCC |
| 6235 | mamo-s | *BGIBMGA007505* | down | 1733 | 1741 | - | 11.3303 | 4.96E-05 | 0.615 | GTGGGTGGC |
| 6236 | mamo-s | *BGIBMGA007510* | down | 1237 | 1245 | + | 11.3303 | 4.96E-05 | 0.615 | GTGGGTGGC |
| 6237 | mamo-s | *BGIBMGA007511* | down | 183 | 191 | + | 12.0734 | 2.66E-05 | 0.615 | GCGCGTGTC |
| 6238 | mamo-s | *BGIBMGA007511* | up | 555 | 563 | - | 11.4404 | 4.51E-05 | 0.615 | CGGCGTGGC |
| 6239 | mamo-s | *BGIBMGA007511* | up | 215 | 223 | - | 11.3303 | 4.96E-05 | 0.615 | GTGGGTGGC |
| 6240 | mamo-s | *BGIBMGA007511* | up | 1878 | 1886 | + | 11.3303 | 4.96E-05 | 0.615 | GTGGGTGGC |
| 6241 | mamo-s | *BGIBMGA007512* | up | 297 | 305 | + | 10.5688 | 8.88E-05 | 0.824 | GTGCGTGAC |

| | | | | | | | | | | |
|---|---|---|---|---|---|---|---|---|---|---|
| 6242 | mamo-s | *BGIBMGA007515* | down | 1413 | 1421 | - | 11.3303 | 4.96E-05 | 0.615 | GTGGGTGGC |
| 6243 | mamo-s | *BGIBMGA007516* | down | 298 | 306 | - | 11.4404 | 4.51E-05 | 0.615 | GAGCGTGTC |
| 6244 | mamo-s | *BGIBMGA007519* | up | 82 | 90 | - | 12.1835 | 1.71E-05 | 0.615 | GTGCGTGCC |
| 6245 | mamo-s | *BGIBMGA007520* | down | 1472 | 1480 | + | 11.3303 | 4.96E-05 | 0.615 | GTGGGTGGC |
| 6246 | mamo-s | *BGIBMGA007520* | down | 1812 | 1820 | - | 11.3303 | 4.96E-05 | 0.615 | GTGGGTGGC |
| 6247 | mamo-s | *BGIBMGA007524* | up | 1835 | 1843 | + | 11.0734 | 7.12E-05 | 0.726 | GAGCGTGGT |
| 6248 | mamo-s | *BGIBMGA007526* | down | 1593 | 1601 | - | 11.3303 | 4.96E-05 | 0.615 | GTGGGTGGC |
| 6249 | mamo-s | *BGIBMGA007527* | up | 180 | 188 | - | 11.5321 | 4.06E-05 | 0.615 | TTGCGTGGC |
| 6250 | mamo-s | *BGIBMGA007527* | up | 346 | 354 | - | 11.3303 | 4.96E-05 | 0.615 | GTGGGTGGC |
| 6251 | mamo-s | *BGIBMGA007528* | down | 1169 | 1177 | + | 11.2844 | 5.18E-05 | 0.63 | CTGCGTGCC |
| 6252 | mamo-s | *BGIBMGA007529* | up | 177 | 185 | - | 11.3303 | 4.96E-05 | 0.615 | GTGGGTGGC |
| 6253 | mamo-s | *BGIBMGA007531* | up | 1752 | 1760 | + | 12.3394 | 1.49E-05 | 0.615 | GGGCGTGGC |
| 6254 | mamo-s | *BGIBMGA007531* | down | 1410 | 1418 | - | 12.1284 | 2.16E-05 | 0.615 | CAGCGTGGC |
| 6255 | mamo-s | *BGIBMGA007533* | up | 1040 | 1048 | + | 11.3303 | 4.96E-05 | 0.615 | GTGGGTGGC |
| 6256 | mamo-s | *BGIBMGA007534* | down | 1825 | 1833 | + | 14.055 | 2.23E-06 | 0.615 | GTGCGTGGC |
| 6257 | mamo-s | *BGIBMGA007534* | up | 1770 | 1778 | - | 11.7339 | 3.29E-05 | 0.615 | GCGCGAGGC |
| 6258 | mamo-s | *BGIBMGA007537* | up | 1360 | 1368 | + | 11.3303 | 4.96E-05 | 0.615 | GTGGGTGGC |
| 6259 | mamo-s | *BGIBMGA007541* | up | 1043 | 1051 | + | 11.3303 | 4.96E-05 | 0.615 | GTGGGTGGC |
| 6260 | mamo-s | *BGIBMGA007541* | down | 501 | 509 | - | 10.6881 | 8.61E-05 | 0.81 | GCACGTGGC |
| 6261 | mamo-s | *BGIBMGA007542* | up | 1422 | 1430 | - | 11.3303 | 4.96E-05 | 0.615 | GTGGGTGGC |
| 6262 | mamo-s | *BGIBMGA007545* | up | 106 | 114 | + | 11.3303 | 4.96E-05 | 0.615 | GTGGGTGGC |
| 6263 | mamo-s | *BGIBMGA007546* | up | 544 | 552 | - | 12.3394 | 1.49E-05 | 0.615 | GGGCGTGGC |
| 6264 | mamo-s | *BGIBMGA007549* | down | 1235 | 1243 | - | 11.3303 | 4.96E-05 | 0.615 | GTGGGTGGC |
| 6265 | mamo-s | *BGIBMGA007552* | up | 534 | 542 | - | 11.3303 | 4.96E-05 | 0.615 | GTGGGTGGC |
| 6266 | mamo-s | *BGIBMGA007552* | up | 1849 | 1857 | + | 11.3303 | 4.96E-05 | 0.615 | GTGGGTGGC |
| 6267 | mamo-s | *BGIBMGA007553* | up | 1940 | 1948 | - | 14.055 | 2.23E-06 | 0.615 | GTGCGTGGC |
| 6268 | mamo-s | *BGIBMGA007556* | up | 78 | 86 | + | 11.3303 | 4.96E-05 | 0.615 | GTGGGTGGC |
| 6269 | mamo-s | *BGIBMGA007556* | down | 1547 | 1555 | + | 10.6881 | 8.61E-05 | 0.81 | GCACGTGGC |
| 6270 | mamo-s | *BGIBMGA007557* | down | 1612 | 1620 | + | 11.3303 | 4.96E-05 | 0.615 | GTGGGTGGC |
| 6271 | mamo-s | *BGIBMGA007558* | down | 1136 | 1144 | + | 13.156 | 6.30E-06 | 0.615 | CTGCGTGGC |
| 6272 | mamo-s | *BGIBMGA007558* | down | 1471 | 1479 | + | 11.0826 | 6.84E-05 | 0.709 | GTACGTGGC |
| 6273 | mamo-s | *BGIBMGA007560* | up | 783 | 791 | + | 11.3303 | 4.96E-05 | 0.615 | GTGGGTGGC |
| 6274 | mamo-s | *BGIBMGA007560* | down | 1132 | 1140 | - | 11.3303 | 4.96E-05 | 0.615 | GTGGGTGGC |
| 6275 | mamo-s | *BGIBMGA007560* | down | 429 | 437 | - | 10.5138 | 9.75E-05 | 0.865 | GTGCGTGTT |
| 6276 | mamo-s | *BGIBMGA007561* | down | 1089 | 1097 | + | 11.789 | 2.84E-05 | 0.615 | GCGCGTGCC |
| 6277 | mamo-s | *BGIBMGA007561* | up | 1177 | 1185 | - | 11.3303 | 4.96E-05 | 0.615 | GTGGGTGGC |
| 6278 | mamo-s | *BGIBMGA007561* | up | 474 | 482 | - | 10.5138 | 9.75E-05 | 0.865 | GTGCGTGTT |
| 6279 | mamo-s | *BGIBMGA007562* | up | 1225 | 1233 | + | 11.789 | 2.84E-05 | 0.615 | GCGCGTGCC |
| 6280 | mamo-s | *BGIBMGA007566* | down | 150 | 158 | - | 14.055 | 2.23E-06 | 0.615 | GTGCGTGGC |
| 6281 | mamo-s | *BGIBMGA007567* | up | 1251 | 1259 | + | 11.7339 | 3.29E-05 | 0.615 | GCGCGAGGC |
| 6282 | mamo-s | *BGIBMGA007568* | up | 1277 | 1285 | + | 11.3303 | 4.96E-05 | 0.615 | GTGGGTGGC |
| 6283 | mamo-s | *BGIBMGA007570* | up | 930 | 938 | - | 10.5138 | 9.75E-05 | 0.865 | GTGCGTGTT |
| 6284 | mamo-s | *BGIBMGA007571* | up | 1532 | 1540 | - | 11.3303 | 4.96E-05 | 0.615 | GTGGGTGGC |
| 6285 | mamo-s | *BGIBMGA007572* | down | 215 | 223 | - | 11.3303 | 4.96E-05 | 0.615 | GTGGGTGGC |
| 6286 | mamo-s | *BGIBMGA007574* | down | 1016 | 1024 | - | 11.3303 | 4.96E-05 | 0.615 | GTGGGTGGC |
| 6287 | mamo-s | *BGIBMGA007575* | up | 1164 | 1172 | - | 12.1284 | 2.16E-05 | 0.615 | CAGCGTGGC |
| 6288 | mamo-s | *BGIBMGA007580* | down | 1659 | 1667 | + | 12.3394 | 1.49E-05 | 0.615 | GGGCGTGGC |
| 6289 | mamo-s | *BGIBMGA007580* | up | 1316 | 1324 | + | 12.1284 | 2.16E-05 | 0.615 | GTGCGAGGC |
| 6290 | mamo-s | *BGIBMGA007581* | down | 1113 | 1121 | - | 11.3303 | 4.96E-05 | 0.615 | GTGGGTGGC |
| 6291 | mamo-s | *BGIBMGA007581* | up | 1000 | 1008 | - | 10.5688 | 8.88E-05 | 0.824 | GTGCGTGAC |
| 6292 | mamo-s | *BGIBMGA007584* | down | 1856 | 1864 | - | 11.3303 | 4.96E-05 | 0.615 | GTGGGTGGC |
| 6293 | mamo-s | *BGIBMGA007587* | down | 1516 | 1524 | - | 11.3303 | 4.96E-05 | 0.615 | GTGGGTGGC |
| 6294 | mamo-s | *BGIBMGA007588* | up | 560 | 568 | + | 11.3303 | 4.96E-05 | 0.615 | GTGGGTGGC |
| 6295 | mamo-s | *BGIBMGA007588* | up | 1773 | 1781 | - | 11.3303 | 4.96E-05 | 0.615 | GTGGGTGGC |
| 6296 | mamo-s | *BGIBMGA007590* | up | 371 | 379 | + | 11.3303 | 4.96E-05 | 0.615 | GTGGGTGGC |
| 6297 | mamo-s | *BGIBMGA007593* | up | 336 | 344 | + | 12.7615 | 1.04E-05 | 0.615 | CCGCGTGGC |
| 6298 | mamo-s | *BGIBMGA007593* | down | 1525 | 1533 | - | 12.3394 | 1.49E-05 | 0.615 | GGGCGTGGC |
| 6299 | mamo-s | *BGIBMGA007595* | up | 622 | 630 | - | 11.4404 | 4.51E-05 | 0.615 | CGGCGTGGC |
| 6300 | mamo-s | *BGIBMGA007595* | up | 637 | 645 | - | 11.2294 | 5.41E-05 | 0.65 | CTGCGAGGC |
| 6301 | mamo-s | *BGIBMGA007595* | down | 1957 | 1965 | + | 11.0826 | 6.84E-05 | 0.709 | GTACGTGGC |
| 6302 | mamo-s | *BGIBMGA007596* | down | 1910 | 1918 | + | 14.055 | 2.23E-06 | 0.615 | GTGCGTGGC |
| 6303 | mamo-s | *BGIBMGA007596* | down | 1012 | 1020 | - | 11.7798 | 3.11E-05 | 0.615 | ATGCGTGGC |
| 6304 | mamo-s | *BGIBMGA007596* | down | 1478 | 1486 | + | 11.4404 | 4.51E-05 | 0.615 | CGGCGTGGC |
| 6305 | mamo-s | *BGIBMGA007597* | down | 1270 | 1278 | - | 11.3303 | 4.96E-05 | 0.615 | GTGGGTGGC |
| 6306 | mamo-s | *BGIBMGA007598* | down | 75 | 83 | - | 11.3303 | 4.96E-05 | 0.615 | GTGGGTGGC |
| 6307 | mamo-s | *BGIBMGA007600* | down | 1239 | 1247 | - | 11.3303 | 4.96E-05 | 0.615 | GTGGGTGGC |
| 6308 | mamo-s | *BGIBMGA007601* | down | 1821 | 1829 | + | 11.5321 | 4.06E-05 | 0.615 | TTGCGTGGC |
| 6309 | mamo-s | *BGIBMGA007601* | up | 265 | 273 | - | 11.3303 | 4.96E-05 | 0.615 | GTGGGTGGC |
| 6310 | mamo-s | *BGIBMGA007601* | down | 1101 | 1109 | - | 11.3303 | 4.96E-05 | 0.615 | GTGGGTGGC |
| 6311 | mamo-s | *BGIBMGA007602* | up | 921 | 929 | + | 11.0826 | 6.84E-05 | 0.709 | GTACGTGGC |
| 6312 | mamo-s | *BGIBMGA007602* | up | 1601 | 1609 | - | 11.0734 | 7.12E-05 | 0.726 | GAGCGTGGT |
| 6313 | mamo-s | *BGIBMGA007605* | up | 84 | 92 | - | 12.0734 | 2.66E-05 | 0.615 | GCGCGTGTC |
| 6314 | mamo-s | *BGIBMGA007606* | up | 1474 | 1482 | - | 12.4679 | 1.31E-05 | 0.615 | GTGCGTGTC |
| 6315 | mamo-s | *BGIBMGA007606* | up | 1918 | 1926 | + | 12.1835 | 1.71E-05 | 0.615 | GTGCGTGCC |
| 6316 | mamo-s | *BGIBMGA007609* | down | 1948 | 1956 | - | 12.1835 | 1.71E-05 | 0.615 | GTGCGTGCC |
| 6317 | mamo-s | *BGIBMGA007610* | down | 468 | 476 | - | 11.0826 | 6.84E-05 | 0.709 | GTACGTGGC |
| 6318 | mamo-s | *BGIBMGA007610* | down | 1371 | 1379 | + | 11.0826 | 6.84E-05 | 0.709 | GTACGTGGC |
| 6319 | mamo-s | *BGIBMGA007611* | up | 290 | 298 | - | 11.3303 | 4.96E-05 | 0.615 | GTGGGTGGC |
| 6320 | mamo-s | *BGIBMGA007611* | up | 1692 | 1700 | - | 11.0826 | 6.84E-05 | 0.709 | GTACGTGGC |
| 6321 | mamo-s | *BGIBMGA007613* | up | 862 | 870 | - | 11.3303 | 4.96E-05 | 0.615 | GTGGGTGGC |
| 6322 | mamo-s | *BGIBMGA007614* | up | 748 | 756 | - | 11.3303 | 4.96E-05 | 0.615 | GTGGGTGGC |
| 6323 | mamo-s | *BGIBMGA007616* | down | 854 | 862 | + | 10.5688 | 8.88E-05 | 0.824 | GTGCGTGAC |
| 6324 | mamo-s | *BGIBMGA007621* | down | 1076 | 1084 | + | 11.3303 | 4.96E-05 | 0.615 | GTGGGTGGC |
| 6325 | mamo-s | *BGIBMGA007621* | up | 296 | 304 | + | 11.0826 | 6.84E-05 | 0.709 | GTACGTGGC |
| 6326 | mamo-s | *BGIBMGA007621* | down | 950 | 958 | - | 10.9358 | 7.30E-05 | 0.728 | GCGGGTGGC |
| 6327 | mamo-s | *BGIBMGA007622* | up | 822 | 830 | + | 11.3303 | 4.96E-05 | 0.615 | GTGGGTGGC |
| 6328 | mamo-s | *BGIBMGA007622* | down | 1393 | 1401 | - | 11.0826 | 6.84E-05 | 0.709 | GTACGTGGC |
| 6329 | mamo-s | *BGIBMGA007622* | up | 696 | 704 | - | 10.9358 | 7.30E-05 | 0.728 | GCGGGTGGC |

| | | | | | | | | | | |
|---|---|---|---|---|---|---|---|---|---|---|
| 6330 | mamo-s | *BGIBMGA007623* | down | 465 | 473 | + | 11.3303 | 4.96E-05 | 0.615 | GTGGGTGGC |
| 6331 | mamo-s | *BGIBMGA007624* | down | 204 | 212 | + | 11.3303 | 4.96E-05 | 0.615 | GTGGGTGGC |
| 6332 | mamo-s | *BGIBMGA007625* | up | 1023 | 1031 | + | 11.3303 | 4.96E-05 | 0.615 | GTGGGTGGC |
| 6333 | mamo-s | *BGIBMGA007626* | down | 1054 | 1062 | - | 11.3303 | 4.96E-05 | 0.615 | GTGGGTGGC |
| 6334 | mamo-s | *BGIBMGA007627* | down | 1719 | 1727 | - | 12.3394 | 1.49E-05 | 0.615 | GGGCGTGGC |
| 6335 | mamo-s | *BGIBMGA007627* | up | 718 | 726 | - | 10.8899 | 7.48E-05 | 0.739 | CCGCGTGCC |
| 6336 | mamo-s | *BGIBMGA007629* | up | 662 | 670 | - | 11.3303 | 4.96E-05 | 0.615 | GTGGGTGGC |
| 6337 | mamo-s | *BGIBMGA007633* | up | 1461 | 1469 | + | 11.3303 | 4.96E-05 | 0.615 | GTGGGTGGC |
| 6338 | mamo-s | *BGIBMGA007634* | down | 1839 | 1847 | - | 12.0734 | 2.66E-05 | 0.615 | GCGCGTGTC |
| 6339 | mamo-s | *BGIBMGA007634* | up | 1868 | 1876 | - | 11.5688 | 3.79E-05 | 0.615 | CTGCGTGTC |
| 6340 | mamo-s | *BGIBMGA007634* | down | 1338 | 1346 | + | 11.3303 | 4.96E-05 | 0.615 | GTGGGTGGC |
| 6341 | mamo-s | *BGIBMGA007641* | up | 1682 | 1690 | - | 12.4679 | 1.31E-05 | 0.615 | GTGCGTGTC |
| 6342 | mamo-s | *BGIBMGA007644* | down | 1522 | 1530 | + | 13.6606 | 4.06E-06 | 0.615 | GCGCGTGGC |
| 6343 | mamo-s | *BGIBMGA007644* | up | 1687 | 1695 | + | 11.3303 | 4.96E-05 | 0.615 | GTGGGTGGC |
| 6344 | mamo-s | *BGIBMGA007647* | up | 108 | 116 | - | 11.3303 | 4.96E-05 | 0.615 | GTGGGTGGC |
| 6345 | mamo-s | *BGIBMGA007647* | up | 1487 | 1495 | - | 11.3303 | 4.96E-05 | 0.615 | GTGGGTGGC |
| 6346 | mamo-s | *BGIBMGA007650* | up | 622 | 630 | + | 11.0826 | 6.84E-05 | 0.709 | GTACGTGGC |
| 6347 | mamo-s | *BGIBMGA007651* | up | 1371 | 1379 | - | 11.3303 | 4.96E-05 | 0.615 | GTGGGTGGC |
| 6348 | mamo-s | *BGIBMGA007651* | up | 28 | 36 | - | 11.0826 | 6.84E-05 | 0.709 | GTACGTGGC |
| 6349 | mamo-s | *BGIBMGA007651* | down | 1760 | 1768 | - | 10.8073 | 7.89E-05 | 0.766 | CCGCGTGGT |
| 6350 | mamo-s | *BGIBMGA007653* | up | 1586 | 1594 | + | 11.2844 | 5.18E-05 | 0.63 | CTGCGTGCC |
| 6351 | mamo-s | *BGIBMGA007654* | down | 1595 | 1603 | - | 11.3303 | 4.96E-05 | 0.615 | GTGGGTGGC |
| 6352 | mamo-s | *BGIBMGA007654* | up | 1715 | 1723 | + | 11.3303 | 4.96E-05 | 0.615 | GTGGGTGGC |
| 6353 | mamo-s | *BGIBMGA007655* | up | 1472 | 1480 | + | 11.3303 | 4.96E-05 | 0.615 | GTGGGTGGC |
| 6354 | mamo-s | *BGIBMGA007659* | up | 1771 | 1779 | + | 11.5321 | 4.06E-05 | 0.615 | TTGCGTGGC |
| 6355 | mamo-s | *BGIBMGA007660* | up | 1808 | 1816 | + | 11.5688 | 3.79E-05 | 0.615 | CTGCGTGTC |
| 6356 | mamo-s | *BGIBMGA007660* | up | 1551 | 1559 | - | 11.3303 | 4.96E-05 | 0.615 | GTGGGTGGC |
| 6357 | mamo-s | *BGIBMGA007661* | down | 252 | 260 | - | 11.0826 | 6.84E-05 | 0.709 | GTACGTGGC |
| 6358 | mamo-s | *BGIBMGA007664* | up | 1422 | 1430 | - | 11.2844 | 5.18E-05 | 0.63 | CTGCGTGCC |
| 6359 | mamo-s | *BGIBMGA007664* | up | 1358 | 1366 | - | 10.5138 | 9.75E-05 | 0.865 | GTGCGTGTT |
| 6360 | mamo-s | *BGIBMGA007670* | up | 1942 | 1950 | - | 11.0826 | 6.84E-05 | 0.709 | GTACGTGGC |
| 6361 | mamo-s | *BGIBMGA007672* | up | 203 | 211 | + | 11.3303 | 4.96E-05 | 0.615 | GTGGGTGGC |
| 6362 | mamo-s | *BGIBMGA007675* | up | 1808 | 1816 | + | 11.3303 | 4.96E-05 | 0.615 | GTGGGTGGC |
| 6363 | mamo-s | *BGIBMGA007676* | down | 1988 | 1996 | - | 11.3303 | 4.96E-05 | 0.615 | GTGGGTGGC |
| 6364 | mamo-s | *BGIBMGA007677* | up | 311 | 319 | + | 11.3303 | 4.96E-05 | 0.615 | GTGGGTGGC |
| 6365 | mamo-s | *BGIBMGA007677* | down | 1211 | 1219 | + | 10.5138 | 9.75E-05 | 0.865 | GTGCGTGTT |
| 6366 | mamo-s | *BGIBMGA007678* | down | 637 | 645 | + | 11.3303 | 4.96E-05 | 0.615 | GTGGGTGGC |
| 6367 | mamo-s | *BGIBMGA007679* | down | 530 | 538 | + | 11.0734 | 7.12E-05 | 0.726 | GAGCGTGGT |
| 6368 | mamo-s | *BGIBMGA007681* | down | 405 | 413 | + | 11.3303 | 4.96E-05 | 0.615 | GTGGGTGGC |
| 6369 | mamo-s | *BGIBMGA007681* | up | 455 | 463 | - | 11.3303 | 4.96E-05 | 0.615 | GTGGGTGGC |
| 6370 | mamo-s | *BGIBMGA007682* | down | 1282 | 1290 | - | 11.3303 | 4.96E-05 | 0.615 | GTGGGTGGC |
| 6371 | mamo-s | *BGIBMGA007683* | up | 314 | 322 | - | 11.0826 | 6.84E-05 | 0.709 | GTACGTGGC |
| 6372 | mamo-s | *BGIBMGA007683* | up | 233 | 241 | + | 10.8899 | 7.48E-05 | 0.739 | CCGCGTGCC |
| 6373 | mamo-s | *BGIBMGA007685* | down | 735 | 743 | - | 10.8349 | 7.66E-05 | 0.751 | CCGCGAGGC |
| 6374 | mamo-s | *BGIBMGA007686* | up | 765 | 773 | - | 10.8349 | 7.66E-05 | 0.751 | CCGCGAGGC |
| 6375 | mamo-s | *BGIBMGA007687* | up | 799 | 807 | - | 11.1743 | 5.90E-05 | 0.684 | CCGCGTGTC |
| 6376 | mamo-s | *BGIBMGA007687* | up | 1886 | 1894 | + | 11.156 | 6.13E-05 | 0.705 | GAGCGTGCC |
| 6377 | mamo-s | *BGIBMGA007688* | down | 528 | 536 | - | 12.1284 | 2.16E-05 | 0.615 | GTGCGAGGC |
| 6378 | mamo-s | *BGIBMGA007688* | up | 203 | 211 | + | 11.3303 | 4.96E-05 | 0.615 | GTGGGTGGC |
| 6379 | mamo-s | *BGIBMGA007691* | up | 393 | 401 | + | 11.3303 | 4.96E-05 | 0.615 | GTGGGTGGC |
| 6380 | mamo-s | *BGIBMGA007691* | up | 538 | 546 | - | 11.3303 | 4.96E-05 | 0.615 | GTGGGTGGC |
| 6381 | mamo-s | *BGIBMGA007694* | up | 1453 | 1461 | + | 11.3303 | 4.96E-05 | 0.615 | GTGGGTGGC |
| 6382 | mamo-s | *BGIBMGA007700* | up | 1337 | 1345 | + | 12.1835 | 1.71E-05 | 0.615 | GTGCGTGCC |
| 6383 | mamo-s | *BGIBMGA007700* | down | 1948 | 1956 | - | 11.0826 | 6.84E-05 | 0.709 | GTACGTGGC |
| 6384 | mamo-s | *BGIBMGA007703* | down | 151 | 159 | + | 11.3303 | 4.96E-05 | 0.615 | GTGGGTGGC |
| 6385 | mamo-s | *BGIBMGA007703* | up | 1244 | 1252 | + | 11.3303 | 4.96E-05 | 0.615 | GTGGGTGGC |
| 6386 | mamo-s | *BGIBMGA007705* | up | 1240 | 1248 | - | 11.3303 | 4.96E-05 | 0.615 | GTGGGTGGC |
| 6387 | mamo-s | *BGIBMGA007706* | up | 1121 | 1129 | - | 11.3853 | 4.74E-05 | 0.615 | ACGCGTGGC |
| 6388 | mamo-s | *BGIBMGA007706* | up | 1215 | 1223 | + | 11.3303 | 4.96E-05 | 0.615 | GTGGGTGGC |
| 6389 | mamo-s | *BGIBMGA007708* | down | 399 | 407 | - | 11.3303 | 4.96E-05 | 0.615 | GTGGGTGGC |
| 6390 | mamo-s | *BGIBMGA007708* | down | 1147 | 1155 | - | 11.0826 | 6.84E-05 | 0.709 | GTACGTGGC |
| 6391 | mamo-s | *BGIBMGA007710* | down | 1707 | 1715 | + | 11.3303 | 4.96E-05 | 0.615 | GTGGGTGGC |
| 6392 | mamo-s | *BGIBMGA007711* | up | 1456 | 1464 | + | 14.055 | 2.23E-06 | 0.615 | GTGCGTGGC |
| 6393 | mamo-s | *BGIBMGA007713* | up | 482 | 490 | + | 11.3303 | 4.96E-05 | 0.615 | GTGGGTGGC |
| 6394 | mamo-s | *BGIBMGA007714* | up | 669 | 677 | + | 11.3303 | 4.96E-05 | 0.615 | GTGGGTGGC |
| 6395 | mamo-s | *BGIBMGA007719* | down | 304 | 312 | - | 11.0826 | 6.84E-05 | 0.709 | GTACGTGGC |
| 6396 | mamo-s | *BGIBMGA007722* | down | 957 | 965 | - | 11.5321 | 4.06E-05 | 0.615 | TTGCGTGGC |
| 6397 | mamo-s | *BGIBMGA007722* | down | 1289 | 1297 | - | 11.0734 | 7.12E-05 | 0.726 | GAGCGTGGT |
| 6398 | mamo-s | *BGIBMGA007723* | up | 1655 | 1663 | - | 12.1009 | 2.43E-05 | 0.615 | GTGCGTGGT |
| 6399 | mamo-s | *BGIBMGA007723* | up | 56 | 64 | + | 11.1743 | 5.90E-05 | 0.684 | CCGCGTGTC |
| 6400 | mamo-s | *BGIBMGA007724* | up | 696 | 704 | + | 11.156 | 6.13E-05 | 0.705 | GAGCGTGCC |
| 6401 | mamo-s | *BGIBMGA007726* | down | 1220 | 1228 | - | 14.055 | 2.23E-06 | 0.615 | GTGCGTGGC |
| 6402 | mamo-s | *BGIBMGA007728* | down | 1790 | 1798 | - | 12.3394 | 1.49E-05 | 0.615 | GGGCGTGGC |
| 6403 | mamo-s | *BGIBMGA007728* | down | 1854 | 1862 | - | 11.5688 | 3.79E-05 | 0.615 | CTGCGTGTC |
| 6404 | mamo-s | *BGIBMGA007729* | up | 1814 | 1822 | + | 11.3303 | 4.96E-05 | 0.615 | GTGGGTGGC |
| 6405 | mamo-s | *BGIBMGA007730* | down | 1615 | 1623 | - | 11.3303 | 4.96E-05 | 0.615 | GTGGGTGGC |
| 6406 | mamo-s | *BGIBMGA007733* | up | 1723 | 1731 | - | 11.3303 | 4.96E-05 | 0.615 | GTGGGTGGC |
| 6407 | mamo-s | *BGIBMGA007735* | down | 1047 | 1055 | - | 11.3303 | 4.96E-05 | 0.615 | GTGGGTGGC |
| 6408 | mamo-s | *BGIBMGA007737* | up | 1765 | 1773 | - | 13.156 | 6.30E-06 | 0.615 | CTGCGTGGC |
| 6409 | mamo-s | *BGIBMGA007738* | up | 1300 | 1308 | - | 10.7523 | 8.38E-05 | 0.797 | GGGCGTGTC |
| 6410 | mamo-s | *BGIBMGA007741* | up | 1007 | 1015 | + | 12.0734 | 2.66E-05 | 0.615 | GCGCGTGTC |
| 6411 | mamo-s | *BGIBMGA007741* | up | 901 | 909 | - | 10.6881 | 8.61E-05 | 0.81 | GCACGTGGC |
| 6412 | mamo-s | *BGIBMGA007742* | up | 1734 | 1742 | - | 11.4404 | 4.51E-05 | 0.615 | GAGCGTGTC |
| 6413 | mamo-s | *BGIBMGA007742* | up | 581 | 589 | + | 11.3303 | 4.96E-05 | 0.615 | GTGGGTGGC |
| 6414 | mamo-s | *BGIBMGA007742* | up | 1713 | 1721 | + | 11.3303 | 4.96E-05 | 0.615 | GTGGGTGGC |
| 6415 | mamo-s | *BGIBMGA007745* | down | 1048 | 1056 | + | 10.9358 | 7.30E-05 | 0.728 | GCGGGTGGC |
| 6416 | mamo-s | *BGIBMGA007746* | up | 872 | 880 | + | 11.3303 | 4.96E-05 | 0.615 | GTGGGTGGC |
| 6417 | mamo-s | *BGIBMGA007746* | down | 1752 | 1760 | + | 11.1743 | 5.90E-05 | 0.684 | CCGCGTGTC |

| | | | | | | | | | | |
|---|---|---|---|---|---|---|---|---|---|---|
| 6418 | mamo-s | *BGIBMGA007747* | up | 744 | 752 | + | 11.1743 | 5.90E-05 | 0.684 | CCGCGTGTC |
| 6419 | mamo-s | *BGIBMGA007749* | down | 285 | 293 | + | 11.3303 | 4.96E-05 | 0.615 | GTGGGTGGC |
| 6420 | mamo-s | *BGIBMGA007749* | up | 1239 | 1247 | - | 11.3303 | 4.96E-05 | 0.615 | GTGGGTGGC |
| 6421 | mamo-s | *BGIBMGA007751* | up | 1948 | 1956 | + | 13.6606 | 4.06E-06 | 0.615 | GCGCGTGGC |
| 6422 | mamo-s | *BGIBMGA007751* | down | 99 | 107 | + | 11.7798 | 3.11E-05 | 0.615 | ATGCGTGGC |
| 6423 | mamo-s | *BGIBMGA007752* | up | 627 | 635 | - | 12.4679 | 1.31E-05 | 0.615 | GTGCGTGTC |
| 6424 | mamo-s | *BGIBMGA007753* | down | 1986 | 1994 | + | 11.3303 | 4.96E-05 | 0.615 | GTGGGTGGC |
| 6425 | mamo-s | *BGIBMGA007754* | down | 674 | 682 | - | 11.3303 | 4.96E-05 | 0.615 | GTGGGTGGC |
| 6426 | mamo-s | *BGIBMGA007758* | down | 205 | 213 | - | 11.3303 | 4.96E-05 | 0.615 | GTGGGTGGC |
| 6427 | mamo-s | *BGIBMGA007759* | up | 113 | 121 | - | 11.3303 | 4.96E-05 | 0.615 | GTGGGTGGC |
| 6428 | mamo-s | *BGIBMGA007764* | up | 1958 | 1966 | + | 11.5321 | 4.06E-05 | 0.615 | TTGCGTGGC |
| 6429 | mamo-s | *BGIBMGA007766* | down | 360 | 368 | + | 10.8073 | 7.89E-05 | 0.766 | CCGCGTGGT |
| 6430 | mamo-s | *BGIBMGA007767* | down | 1487 | 1495 | - | 11.0734 | 7.12E-05 | 0.726 | GAGCGTGGT |
| 6431 | mamo-s | *BGIBMGA007768* | down | 796 | 804 | + | 11.0826 | 6.84E-05 | 0.709 | GTACGTGGC |
| 6432 | mamo-s | *BGIBMGA007769* | up | 1966 | 1974 | - | 11.7798 | 3.11E-05 | 0.615 | ATGCGTGGC |
| 6433 | mamo-s | *BGIBMGA007770* | up | 1831 | 1839 | + | 11.3303 | 4.96E-05 | 0.615 | GTGGGTGGC |
| 6434 | mamo-s | *BGIBMGA007770* | down | 473 | 481 | + | 10.9358 | 7.30E-05 | 0.728 | GCGGGTGGC |
| 6435 | mamo-s | *BGIBMGA007774* | down | 332 | 340 | - | 12.1835 | 1.71E-05 | 0.615 | GTGCGTGCC |
| 6436 | mamo-s | *BGIBMGA007775* | up | 1359 | 1367 | + | 11.0826 | 6.84E-05 | 0.709 | GTACGTGGC |
| 6437 | mamo-s | *BGIBMGA007778* | down | 1125 | 1133 | + | 11.3303 | 4.96E-05 | 0.615 | GTGGGTGGC |
| 6438 | mamo-s | *BGIBMGA007779* | down | 938 | 946 | + | 11.3303 | 4.96E-05 | 0.615 | GTGGGTGGC |
| 6439 | mamo-s | *BGIBMGA007780* | up | 1040 | 1048 | + | 12.4679 | 1.31E-05 | 0.615 | GTGCGTGTC |
| 6440 | mamo-s | *BGIBMGA007781* | down | 1858 | 1866 | + | 11.3303 | 4.96E-05 | 0.615 | GTGGGTGGC |
| 6441 | mamo-s | *BGIBMGA007783* | down | 1290 | 1298 | - | 11.0826 | 6.84E-05 | 0.709 | GTACGTGGC |
| 6442 | mamo-s | *BGIBMGA007784* | up | 120 | 128 | - | 12.1284 | 2.16E-05 | 0.615 | GTGCGAGGC |
| 6443 | mamo-s | *BGIBMGA007786* | up | 630 | 638 | - | 11.4404 | 4.51E-05 | 0.615 | GAGCGTGTC |
| 6444 | mamo-s | *BGIBMGA007786* | up | 933 | 941 | - | 10.5413 | 9.42E-05 | 0.857 | CAGCGTGTC |
| 6445 | mamo-s | *BGIBMGA007787* | up | 1788 | 1796 | - | 14.055 | 2.23E-06 | 0.615 | GTGCGTGGC |
| 6446 | mamo-s | *BGIBMGA007789* | up | 908 | 916 | - | 11.3303 | 4.96E-05 | 0.615 | GTGGGTGGC |
| 6447 | mamo-s | *BGIBMGA007789* | down | 1761 | 1769 | - | 11.1009 | 6.57E-05 | 0.709 | GAGCGAGGC |
| 6448 | mamo-s | *BGIBMGA007790* | down | 1978 | 1986 | + | 11.3303 | 4.96E-05 | 0.615 | GTGGGTGGC |
| 6449 | mamo-s | *BGIBMGA007791* | down | 1019 | 1027 | - | 12.1284 | 2.16E-05 | 0.615 | CAGCGTGGC |
| 6450 | mamo-s | *BGIBMGA007792* | up | 316 | 324 | - | 12.1284 | 2.16E-05 | 0.615 | CAGCGTGGC |
| 6451 | mamo-s | *BGIBMGA007792* | up | 1426 | 1434 | - | 11.789 | 2.84E-05 | 0.615 | GCGCGTGCC |
| 6452 | mamo-s | *BGIBMGA007792* | down | 824 | 832 | - | 11.3303 | 4.96E-05 | 0.615 | GTGGGTGGC |
| 6453 | mamo-s | *BGIBMGA007801* | down | 1307 | 1315 | - | 11.3303 | 4.96E-05 | 0.615 | GTGGGTGGC |
| 6454 | mamo-s | *BGIBMGA007804* | up | 1822 | 1830 | - | 11.7798 | 3.11E-05 | 0.615 | ATGCGTGGC |
| 6455 | mamo-s | *BGIBMGA007805* | down | 1970 | 1978 | - | 10.5413 | 9.42E-05 | 0.857 | CAGCGTGTC |
| 6456 | mamo-s | *BGIBMGA007806* | up | 886 | 894 | + | 11.7798 | 3.11E-05 | 0.615 | ATGCGTGGC |
| 6457 | mamo-s | *BGIBMGA007810* | up | 952 | 960 | + | 11.7339 | 3.29E-05 | 0.615 | GCGCGAGGC |
| 6458 | mamo-s | *BGIBMGA007812* | down | 806 | 814 | - | 11.3303 | 4.96E-05 | 0.615 | GTGGGTGGC |
| 6459 | mamo-s | *BGIBMGA007815* | up | 1134 | 1142 | + | 10.7523 | 8.38E-05 | 0.797 | AAGCGTGGC |
| 6460 | mamo-s | *BGIBMGA007816* | up | 397 | 405 | - | 11.3303 | 4.96E-05 | 0.615 | GTGGGTGGC |
| 6461 | mamo-s | *BGIBMGA007818* | down | 1767 | 1775 | + | 11.0826 | 6.84E-05 | 0.709 | GTACGTGGC |
| 6462 | mamo-s | *BGIBMGA007821* | up | 530 | 538 | - | 11.3303 | 4.96E-05 | 0.615 | GTGGGTGGC |
| 6463 | mamo-s | *BGIBMGA007824* | down | 1715 | 1723 | + | 11.0734 | 7.12E-05 | 0.726 | GAGCGTGGT |
| 6464 | mamo-s | *BGIBMGA007824* | down | 1441 | 1449 | + | 10.7523 | 8.38E-05 | 0.797 | AAGCGTGGC |
| 6465 | mamo-s | *BGIBMGA007825* | down | 477 | 485 | - | 11.4404 | 4.51E-05 | 0.615 | CGGCGTGGC |
| 6466 | mamo-s | *BGIBMGA007826* | down | 705 | 713 | + | 11.3303 | 4.96E-05 | 0.615 | GTGGGTGGC |
| 6467 | mamo-s | *BGIBMGA007827* | down | 324 | 332 | + | 10.5413 | 9.42E-05 | 0.857 | CAGCGTGTC |
| 6468 | mamo-s | *BGIBMGA007829* | down | 263 | 271 | + | 11.3303 | 4.96E-05 | 0.615 | GTGGGTGGC |
| 6469 | mamo-s | *BGIBMGA007830* | up | 185 | 193 | - | 11.3303 | 4.96E-05 | 0.615 | GTGGGTGGC |
| 6470 | mamo-s | *BGIBMGA007831* | up | 1037 | 1045 | + | 12.4679 | 1.31E-05 | 0.615 | GTGCGTGTC |
| 6471 | mamo-s | *BGIBMGA007835* | down | 1646 | 1654 | + | 12.1835 | 1.71E-05 | 0.615 | GTGCGTGCC |
| 6472 | mamo-s | *BGIBMGA007839* | up | 754 | 762 | - | 11.3303 | 4.96E-05 | 0.615 | GTGGGTGGC |
| 6473 | mamo-s | *BGIBMGA007840* | down | 1747 | 1755 | - | 11.5688 | 3.79E-05 | 0.615 | CTGCGTGTC |
| 6474 | mamo-s | *BGIBMGA007846* | down | 779 | 787 | + | 13.6606 | 4.06E-06 | 0.615 | GCGCGTGGC |
| 6475 | mamo-s | *BGIBMGA007846* | down | 1955 | 1963 | + | 11.0826 | 6.84E-05 | 0.709 | GTACGTGGC |
| 6476 | mamo-s | *BGIBMGA007849* | up | 510 | 518 | - | 10.5138 | 9.75E-05 | 0.865 | GTGCGTGTT |
| 6477 | mamo-s | *BGIBMGA007851* | down | 276 | 284 | + | 10.7523 | 8.38E-05 | 0.797 | AAGCGTGGC |
| 6478 | mamo-s | *BGIBMGA007852* | up | 1026 | 1034 | + | 11.3303 | 4.96E-05 | 0.615 | GTGGGTGGC |
| 6479 | mamo-s | *BGIBMGA007853* | down | 500 | 508 | + | 11.3303 | 4.96E-05 | 0.615 | GTGGGTGGC |
| 6480 | mamo-s | *BGIBMGA007854* | up | 1009 | 1017 | + | 11.5321 | 4.06E-05 | 0.615 | TTGCGTGGC |
| 6481 | mamo-s | *BGIBMGA007856* | up | 709 | 717 | - | 11.3303 | 4.96E-05 | 0.615 | GTGGGTGGC |
| 6482 | mamo-s | *BGIBMGA007856* | down | 538 | 546 | + | 11.0826 | 6.84E-05 | 0.709 | GTACGTGGC |
| 6483 | mamo-s | *BGIBMGA007857* | up | 699 | 707 | + | 11.2018 | 5.68E-05 | 0.674 | CTGCGTGGT |
| 6484 | mamo-s | *BGIBMGA007857* | up | 528 | 536 | + | 11.0826 | 6.84E-05 | 0.709 | GTACGTGGC |
| 6485 | mamo-s | *BGIBMGA007859* | down | 116 | 124 | + | 10.7523 | 8.38E-05 | 0.797 | AAGCGTGGC |
| 6486 | mamo-s | *BGIBMGA007860* | down | 1541 | 1549 | + | 11.3303 | 4.96E-05 | 0.615 | GTGGGTGGC |
| 6487 | mamo-s | *BGIBMGA007861* | up | 495 | 503 | + | 11.3303 | 4.96E-05 | 0.615 | GTGGGTGGC |
| 6488 | mamo-s | *BGIBMGA007862* | up | 186 | 194 | + | 11.3303 | 4.96E-05 | 0.615 | GTGGGTGGC |
| 6489 | mamo-s | *BGIBMGA007864* | up | 734 | 742 | + | 12.4679 | 1.31E-05 | 0.615 | GTGCGTGTC |
| 6490 | mamo-s | *BGIBMGA007864* | down | 1002 | 1010 | + | 11.3303 | 4.96E-05 | 0.615 | GTGGGTGGC |
| 6491 | mamo-s | *BGIBMGA007867* | down | 1015 | 1023 | - | 11.3303 | 4.96E-05 | 0.615 | GTGGGTGGC |
| 6492 | mamo-s | *BGIBMGA007868* | down | 608 | 616 | + | 11.3303 | 4.96E-05 | 0.615 | GTGGGTGGC |
| 6493 | mamo-s | *BGIBMGA007868* | down | 1320 | 1328 | - | 10.8073 | 7.89E-05 | 0.766 | CCGCGTGGT |
| 6494 | mamo-s | *BGIBMGA007869* | down | 173 | 181 | + | 12.4679 | 1.31E-05 | 0.615 | GTGCGTGTC |
| 6495 | mamo-s | *BGIBMGA007870* | down | 143 | 151 | - | 11.3303 | 4.96E-05 | 0.615 | GTGGGTGGC |
| 6496 | mamo-s | *BGIBMGA007870* | up | 340 | 348 | - | 11.156 | 6.13E-05 | 0.705 | GAGCGTGCC |
| 6497 | mamo-s | *BGIBMGA007873* | down | 510 | 518 | - | 13.0275 | 8.53E-06 | 0.615 | GAGCGTGGC |
| 6498 | mamo-s | *BGIBMGA007873* | up | 308 | 316 | - | 11.3303 | 4.96E-05 | 0.615 | GTGGGTGGC |
| 6499 | mamo-s | *BGIBMGA007875* | up | 1026 | 1034 | - | 11.3303 | 4.96E-05 | 0.615 | GTGGGTGGC |
| 6500 | mamo-s | *BGIBMGA007875* | down | 1824 | 1832 | - | 10.5138 | 9.75E-05 | 0.865 | GTGCGTGTT |
| 6501 | mamo-s | *BGIBMGA007879* | down | 650 | 658 | + | 11.3303 | 4.96E-05 | 0.615 | GTGGGTGGC |
| 6502 | mamo-s | *BGIBMGA007881* | down | 1804 | 1812 | + | 11.0734 | 7.12E-05 | 0.726 | GAGCGTGGT |
| 6503 | mamo-s | *BGIBMGA007882* | down | 764 | 772 | + | 11.0826 | 6.84E-05 | 0.709 | GTACGTGGC |
| 6504 | mamo-s | *BGIBMGA007882* | up | 1266 | 1274 | - | 11.0826 | 6.84E-05 | 0.709 | GTACGTGGC |
| 6505 | mamo-s | *BGIBMGA007882* | up | 960 | 968 | + | 11.0734 | 7.12E-05 | 0.726 | GAGCGTGGT |

| | | | | | | | | | | |
|---|---|---|---|---|---|---|---|---|---|---|
| 6506 | mamo-s | *BGIBMGA007883* | up | 199 | 207 | - | 11.156 | 6.13E-05 | 0.705 | GAGCGTGCC |
| 6507 | mamo-s | *BGIBMGA007884* | up | 154 | 162 | - | 11.3303 | 4.96E-05 | 0.615 | GTGGGTGGC |
| 6508 | mamo-s | *BGIBMGA007884* | up | 1142 | 1150 | - | 11.3303 | 4.96E-05 | 0.615 | GTGGGTGGC |
| 6509 | mamo-s | *BGIBMGA007884* | down | 1690 | 1698 | + | 11.3303 | 4.96E-05 | 0.615 | GTGGGTGGC |
| 6510 | mamo-s | *BGIBMGA007885* | up | 820 | 828 | + | 11.3303 | 4.96E-05 | 0.615 | GTGGGTGGC |
| 6511 | mamo-s | *BGIBMGA007886* | up | 857 | 865 | - | 13.0275 | 8.53E-06 | 0.615 | GAGCGTGGC |
| 6512 | mamo-s | *BGIBMGA007890* | up | 1093 | 1101 | - | 11.4404 | 4.51E-05 | 0.615 | CGGCGTGGC |
| 6513 | mamo-s | *BGIBMGA007890* | down | 1795 | 1803 | + | 11.3303 | 4.96E-05 | 0.615 | GTGGGTGGC |
| 6514 | mamo-s | *BGIBMGA007891* | up | 1536 | 1544 | + | 11.3303 | 4.96E-05 | 0.615 | GTGGGTGGC |
| 6515 | mamo-s | *BGIBMGA007891* | up | 699 | 707 | + | 11.1009 | 6.57E-05 | 0.709 | GAGCGAGGC |
| 6516 | mamo-s | *BGIBMGA007892* | up | 931 | 939 | + | 11.1743 | 5.90E-05 | 0.684 | CCGCGTGTC |
| 6517 | mamo-s | *BGIBMGA007893* | down | 648 | 656 | + | 10.6881 | 8.61E-05 | 0.81 | GCACGTGGC |
| 6518 | mamo-s | *BGIBMGA007896* | up | 201 | 209 | + | 11.3303 | 4.96E-05 | 0.615 | GTGGGTGGC |
| 6519 | mamo-s | *BGIBMGA007896* | down | 1332 | 1340 | - | 11.3303 | 4.96E-05 | 0.615 | GTGGGTGGC |
| 6520 | mamo-s | *BGIBMGA007896* | down | 1070 | 1078 | - | 10.8073 | 7.89E-05 | 0.766 | CCGCGTGGT |
| 6521 | mamo-s | *BGIBMGA007897* | up | 925 | 933 | - | 10.7523 | 8.38E-05 | 0.797 | AAGCGTGGC |
| 6522 | mamo-s | *BGIBMGA007898* | up | 916 | 924 | + | 11.7064 | 3.52E-05 | 0.615 | GCGCGTGGT |
| 6523 | mamo-s | *BGIBMGA007899* | up | 1054 | 1062 | - | 11.3303 | 4.96E-05 | 0.615 | GTGGGTGGC |
| 6524 | mamo-s | *BGIBMGA007899* | down | 1464 | 1472 | + | 11.3303 | 4.96E-05 | 0.615 | GTGGGTGGC |
| 6525 | mamo-s | *BGIBMGA007901* | down | 1099 | 1107 | - | 11.3303 | 4.96E-05 | 0.615 | GTGGGTGGC |
| 6526 | mamo-s | *BGIBMGA007902* | down | 505 | 513 | - | 11.3303 | 4.96E-05 | 0.615 | GTGGGTGGC |
| 6527 | mamo-s | *BGIBMGA007902* | down | 1873 | 1881 | - | 10.5138 | 9.75E-05 | 0.865 | GTGCGTGTT |
| 6528 | mamo-s | *BGIBMGA007903* | up | 870 | 878 | + | 11.0826 | 6.84E-05 | 0.709 | GTACGTGGC |
| 6529 | mamo-s | *BGIBMGA007906* | up | 200 | 208 | + | 11.3303 | 4.96E-05 | 0.615 | GTGGGTGGC |
| 6530 | mamo-s | *BGIBMGA007911* | up | 1081 | 1089 | - | 12.0734 | 2.66E-05 | 0.615 | GCGCGTGTC |
| 6531 | mamo-s | *BGIBMGA007912* | up | 581 | 589 | - | 12.1009 | 2.43E-05 | 0.615 | GTGCGTGGT |
| 6532 | mamo-s | *BGIBMGA007912* | up | 748 | 756 | - | 11.3303 | 4.96E-05 | 0.615 | GTGGGTGGC |
| 6533 | mamo-s | *BGIBMGA007912* | up | 401 | 409 | + | 11.1743 | 5.90E-05 | 0.684 | CCGCGTGTC |
| 6534 | mamo-s | *BGIBMGA007913* | up | 1057 | 1065 | - | 11.3853 | 4.74E-05 | 0.615 | ACGCGTGGC |
| 6535 | mamo-s | *BGIBMGA007913* | down | 464 | 472 | + | 11.3303 | 4.96E-05 | 0.615 | GTGGGTGGC |
| 6536 | mamo-s | *BGIBMGA007913* | up | 491 | 499 | - | 11.3303 | 4.96E-05 | 0.615 | GTGGGTGGC |
| 6537 | mamo-s | *BGIBMGA007913* | down | 1485 | 1493 | + | 11.3303 | 4.96E-05 | 0.615 | GTGGGTGGC |
| 6538 | mamo-s | *BGIBMGA007913* | down | 1864 | 1872 | - | 11.3303 | 4.96E-05 | 0.615 | GTGGGTGGC |
| 6539 | mamo-s | *BGIBMGA007913* | down | 1276 | 1284 | + | 10.5138 | 9.75E-05 | 0.865 | GTGCGTGTT |
| 6540 | mamo-s | *BGIBMGA007914* | up | 96 | 104 | - | 11.3303 | 4.96E-05 | 0.615 | GTGGGTGGC |
| 6541 | mamo-s | *BGIBMGA007915* | down | 323 | 331 | - | 11.3303 | 4.96E-05 | 0.615 | GTGGGTGGC |
| 6542 | mamo-s | *BGIBMGA007915* | up | 1792 | 1800 | + | 10.8349 | 7.66E-05 | 0.751 | CCGCGAGGC |
| 6543 | mamo-s | *BGIBMGA007920* | up | 597 | 605 | + | 11.3303 | 4.96E-05 | 0.615 | GTGGGTGGC |
| 6544 | mamo-s | *BGIBMGA007920* | up | 1580 | 1588 | + | 10.5413 | 9.42E-05 | 0.857 | CAGCGTGTC |
| 6545 | mamo-s | *BGIBMGA007922* | up | 767 | 775 | - | 11.3303 | 4.96E-05 | 0.615 | GTGGGTGGC |
| 6546 | mamo-s | *BGIBMGA007923* | down | 552 | 560 | + | 11.3853 | 4.74E-05 | 0.615 | ACGCGTGGC |
| 6547 | mamo-s | *BGIBMGA007925* | up | 1700 | 1708 | + | 11.3303 | 4.96E-05 | 0.615 | GTGGGTGGC |
| 6548 | mamo-s | *BGIBMGA007928* | up | 650 | 658 | - | 11.3303 | 4.96E-05 | 0.615 | GTGGGTGGC |
| 6549 | mamo-s | *BGIBMGA007930* | down | 1758 | 1766 | - | 11.3303 | 4.96E-05 | 0.615 | GTGGGTGGC |
| 6550 | mamo-s | *BGIBMGA007931* | down | 1332 | 1340 | - | 10.5138 | 9.75E-05 | 0.865 | GTGCGTGTT |
| 6551 | mamo-s | *BGIBMGA007932* | up | 847 | 855 | - | 11.3303 | 4.96E-05 | 0.615 | GTGGGTGGC |
| 6552 | mamo-s | *BGIBMGA007933* | down | 764 | 772 | + | 11.5688 | 3.79E-05 | 0.615 | CTGCGTGTC |
| 6553 | mamo-s | *BGIBMGA007933* | down | 831 | 839 | + | 11.1376 | 6.35E-05 | 0.709 | TCGCGTGGC |
| 6554 | mamo-s | *BGIBMGA007934* | down | 152 | 160 | + | 11.3303 | 4.96E-05 | 0.615 | GTGGGTGGC |
| 6555 | mamo-s | *BGIBMGA007934* | down | 570 | 578 | + | 10.7523 | 8.38E-05 | 0.797 | AAGCGTGGC |
| 6556 | mamo-s | *BGIBMGA007936* | down | 490 | 498 | - | 11.3303 | 4.96E-05 | 0.615 | GTGGGTGGC |
| 6557 | mamo-s | *BGIBMGA007937* | down | 859 | 867 | + | 12.4679 | 1.31E-05 | 0.615 | GTGCGTGTC |
| 6558 | mamo-s | *BGIBMGA007937* | down | 1027 | 1035 | - | 11.7064 | 3.52E-05 | 0.615 | GCGCGTGGT |
| 6559 | mamo-s | *BGIBMGA007937* | down | 749 | 757 | + | 11.3303 | 4.96E-05 | 0.615 | GTGGGTGGC |
| 6560 | mamo-s | *BGIBMGA007938* | up | 1939 | 1947 | + | 12.4679 | 1.31E-05 | 0.615 | GTGCGTGTC |
| 6561 | mamo-s | *BGIBMGA007938* | down | 699 | 707 | + | 11.3303 | 4.96E-05 | 0.615 | GTGGGTGGC |
| 6562 | mamo-s | *BGIBMGA007938* | up | 1829 | 1837 | + | 11.3303 | 4.96E-05 | 0.615 | GTGGGTGGC |
| 6563 | mamo-s | *BGIBMGA007938* | up | 847 | 855 | + | 10.8899 | 7.48E-05 | 0.739 | CCGCGTGCC |
| 6564 | mamo-s | *BGIBMGA007939* | up | 326 | 334 | + | 11.3303 | 4.96E-05 | 0.615 | GTGGGTGGC |
| 6565 | mamo-s | *BGIBMGA007941* | up | 1216 | 1224 | - | 12.1009 | 2.43E-05 | 0.615 | GTGCGTGGT |
| 6566 | mamo-s | *BGIBMGA007942* | up | 356 | 364 | + | 11.3303 | 4.96E-05 | 0.615 | GTGGGTGGC |
| 6567 | mamo-s | *BGIBMGA007942* | up | 1720 | 1728 | + | 10.7523 | 8.38E-05 | 0.797 | AAGCGTGGC |
| 6568 | mamo-s | *BGIBMGA007943* | down | 31 | 39 | + | 14.055 | 2.23E-06 | 0.615 | GTGCGTGGC |
| 6569 | mamo-s | *BGIBMGA007947* | down | 652 | 660 | + | 10.5688 | 8.88E-05 | 0.824 | GTGCGTGAC |
| 6570 | mamo-s | *BGIBMGA007948* | down | 277 | 285 | + | 12.0734 | 2.66E-05 | 0.615 | GCGCGTGTC |
| 6571 | mamo-s | *BGIBMGA007948* | down | 778 | 786 | - | 11.3303 | 4.96E-05 | 0.615 | GTGGGTGGC |
| 6572 | mamo-s | *BGIBMGA007949* | up | 111 | 119 | + | 11.5688 | 3.79E-05 | 0.615 | CTGCGTGTC |
| 6573 | mamo-s | *BGIBMGA007949* | down | 1411 | 1419 | + | 10.5413 | 9.42E-05 | 0.857 | CAGCGTGTC |
| 6574 | mamo-s | *BGIBMGA007950* | up | 1304 | 1312 | + | 10.5413 | 9.42E-05 | 0.857 | CAGCGTGTC |
| 6575 | mamo-s | *BGIBMGA007953* | up | 1260 | 1268 | + | 11.2294 | 5.41E-05 | 0.65 | CTGCGAGGC |
| 6576 | mamo-s | *BGIBMGA007956* | up | 101 | 109 | - | 11.3303 | 4.96E-05 | 0.615 | GTGGGTGGC |
| 6577 | mamo-s | *BGIBMGA007956* | down | 416 | 424 | + | 11.3303 | 4.96E-05 | 0.615 | GTGGGTGGC |
| 6578 | mamo-s | *BGIBMGA007956* | up | 1666 | 1674 | + | 11.3303 | 4.96E-05 | 0.615 | GTGGGTGGC |
| 6579 | mamo-s | *BGIBMGA007957* | down | 1700 | 1708 | - | 12.1009 | 2.43E-05 | 0.615 | GTGCGTGGT |
| 6580 | mamo-s | *BGIBMGA007957* | up | 1452 | 1460 | - | 11.3303 | 4.96E-05 | 0.615 | GTGGGTGGC |
| 6581 | mamo-s | *BGIBMGA007957* | up | 1165 | 1173 | + | 10.5688 | 8.88E-05 | 0.824 | GTGCGTGAC |
| 6582 | mamo-s | *BGIBMGA007959* | up | 1204 | 1212 | + | 11.3303 | 4.96E-05 | 0.615 | GTGGGTGGC |
| 6583 | mamo-s | *BGIBMGA007960* | down | 1921 | 1929 | - | 11.3303 | 4.96E-05 | 0.615 | GTGGGTGGC |
| 6584 | mamo-s | *BGIBMGA007961* | up | 1448 | 1456 | - | 11.0826 | 6.84E-05 | 0.709 | GTACGTGGC |
| 6585 | mamo-s | *BGIBMGA007964* | up | 440 | 448 | + | 11.3303 | 4.96E-05 | 0.615 | GTGGGTGGC |
| 6586 | mamo-s | *BGIBMGA007964* | up | 1734 | 1742 | + | 10.5138 | 9.75E-05 | 0.865 | GTGCGTGTT |
| 6587 | mamo-s | *BGIBMGA007965* | up | 1915 | 1923 | + | 11.3303 | 4.96E-05 | 0.615 | GTGGGTGGC |
| 6588 | mamo-s | *BGIBMGA007967* | down | 409 | 417 | - | 11.3303 | 4.96E-05 | 0.615 | GTGGGTGGC |
| 6589 | mamo-s | *BGIBMGA007967* | down | 1795 | 1803 | - | 10.5688 | 8.88E-05 | 0.824 | GTGCGTGAC |
| 6590 | mamo-s | *BGIBMGA007973* | up | 1407 | 1415 | + | 11.3303 | 4.96E-05 | 0.615 | GTGGGTGGC |
| 6591 | mamo-s | *BGIBMGA007975* | down | 985 | 993 | + | 11.789 | 2.84E-05 | 0.615 | GCGCGTGCC |
| 6592 | mamo-s | *BGIBMGA007975* | down | 121 | 129 | - | 10.7523 | 8.38E-05 | 0.797 | GGGCGTGTC |
| 6593 | mamo-s | *BGIBMGA007976* | up | 636 | 644 | - | 12.1284 | 2.16E-05 | 0.615 | GTGCGAGGC |

| | | | | | | | | | | |
|---|---|---|---|---|---|---|---|---|---|---|
| 6594 | mamo-s | *BGIBMGA007976* | up | 594 | 602 | - | 11.4404 | 4.51E-05 | 0.615 | CGGCGTGGC |
| 6595 | mamo-s | *BGIBMGA007977* | up | 653 | 661 | + | 11.3303 | 4.96E-05 | 0.615 | GTGGGTGGC |
| 6596 | mamo-s | *BGIBMGA007977* | down | 1148 | 1156 | - | 11.3303 | 4.96E-05 | 0.615 | GTGGGTGGC |
| 6597 | mamo-s | *BGIBMGA007977* | down | 1619 | 1627 | + | 11.0826 | 6.84E-05 | 0.709 | GTACGTGGC |
| 6598 | mamo-s | *BGIBMGA007980* | up | 1722 | 1730 | + | 11.3303 | 4.96E-05 | 0.615 | GTGGGTGGC |
| 6599 | mamo-s | *BGIBMGA007980* | up | 1490 | 1498 | - | 10.8349 | 7.66E-05 | 0.751 | CCGCGAGGC |
| 6600 | mamo-s | *BGIBMGA007981* | up | 1040 | 1048 | - | 11.3303 | 4.96E-05 | 0.615 | GTGGGTGGC |
| 6601 | mamo-s | *BGIBMGA007981* | down | 1866 | 1874 | - | 11.1743 | 5.90E-05 | 0.684 | CCGCGTGTC |
| 6602 | mamo-s | *BGIBMGA007983* | down | 780 | 788 | - | 11.3303 | 4.96E-05 | 0.615 | GTGGGTGGC |
| 6603 | mamo-s | *BGIBMGA007983* | up | 1312 | 1320 | - | 11.3303 | 4.96E-05 | 0.615 | GTGGGTGGC |
| 6604 | mamo-s | *BGIBMGA007986* | up | 1228 | 1236 | + | 11.3303 | 4.96E-05 | 0.615 | GTGGGTGGC |
| 6605 | mamo-s | *BGIBMGA007986* | down | 406 | 414 | - | 11.2294 | 5.41E-05 | 0.65 | CTGCGAGGC |
| 6606 | mamo-s | *BGIBMGA007986* | up | 1739 | 1747 | + | 11.0826 | 6.84E-05 | 0.709 | GTACGTGGC |
| 6607 | mamo-s | *BGIBMGA007986* | down | 6 | 14 | - | 10.9358 | 7.30E-05 | 0.728 | GCGGGTGGC |
| 6608 | mamo-s | *BGIBMGA007987* | up | 417 | 425 | - | 11.3303 | 4.96E-05 | 0.615 | GTGGGTGGC |
| 6609 | mamo-s | *BGIBMGA007988* | up | 992 | 1000 | + | 11.3303 | 4.96E-05 | 0.615 | GTGGGTGGC |
| 6610 | mamo-s | *BGIBMGA007991* | up | 981 | 989 | - | 11.3303 | 4.96E-05 | 0.615 | GTGGGTGGC |
| 6611 | mamo-s | *BGIBMGA007992* | up | 797 | 805 | - | 11.2018 | 5.68E-05 | 0.674 | CTGCGTGGT |
| 6612 | mamo-s | *BGIBMGA007993* | up | 1423 | 1431 | + | 11.2844 | 5.18E-05 | 0.63 | CTGCGTGCC |
| 6613 | mamo-s | *BGIBMGA007993* | up | 1602 | 1610 | - | 11.0826 | 6.84E-05 | 0.709 | GTACGTGGC |
| 6614 | mamo-s | *BGIBMGA007996* | down | 376 | 384 | - | 11.0826 | 6.84E-05 | 0.709 | GTACGTGGC |
| 6615 | mamo-s | *BGIBMGA007997* | up | 617 | 625 | - | 11.3303 | 4.96E-05 | 0.615 | GTGGGTGGC |
| 6616 | mamo-s | *BGIBMGA007997* | up | 46 | 54 | + | 10.5413 | 9.42E-05 | 0.857 | CAGCGTGTC |
| 6617 | mamo-s | *BGIBMGA007998* | up | 1667 | 1675 | + | 11.4404 | 4.51E-05 | 0.615 | CGGCGTGGC |
| 6618 | mamo-s | *BGIBMGA007998* | down | 798 | 806 | - | 11.3303 | 4.96E-05 | 0.615 | GTGGGTGGC |
| 6619 | mamo-s | *BGIBMGA007998* | up | 1048 | 1056 | - | 11.3303 | 4.96E-05 | 0.615 | GTGGGTGGC |
| 6620 | mamo-s | *BGIBMGA007999* | up | 1806 | 1814 | + | 11.3303 | 4.96E-05 | 0.615 | GTGGGTGGC |
| 6621 | mamo-s | *BGIBMGA008000* | up | 333 | 341 | - | 11.3303 | 4.96E-05 | 0.615 | GTGGGTGGC |
| 6622 | mamo-s | *BGIBMGA008001* | up | 843 | 851 | + | 11.3303 | 4.96E-05 | 0.615 | GTGGGTGGC |
| 6623 | mamo-s | *BGIBMGA008002* | down | 939 | 947 | - | 13.6606 | 4.06E-06 | 0.615 | GCGCGTGGC |
| 6624 | mamo-s | *BGIBMGA008004* | up | 732 | 740 | + | 11.3303 | 4.96E-05 | 0.615 | GTGGGTGGC |
| 6625 | mamo-s | *BGIBMGA008005* | up | 462 | 470 | + | 10.8073 | 7.89E-05 | 0.766 | CCGCGTGGT |
| 6626 | mamo-s | *BGIBMGA008006* | down | 323 | 331 | + | 11.3303 | 4.96E-05 | 0.615 | GTGGGTGGC |
| 6627 | mamo-s | *BGIBMGA008007* | down | 1544 | 1552 | - | 11.3303 | 4.96E-05 | 0.615 | GTGGGTGGC |
| 6628 | mamo-s | *BGIBMGA008008* | down | 1479 | 1487 | - | 11.3303 | 4.96E-05 | 0.615 | GTGGGTGGC |
| 6629 | mamo-s | *BGIBMGA008008* | up | 1386 | 1394 | - | 11.0826 | 6.84E-05 | 0.709 | GTACGTGGC |
| 6630 | mamo-s | *BGIBMGA008008* | down | 828 | 836 | - | 10.8349 | 7.66E-05 | 0.751 | CCGCGAGGC |
| 6631 | mamo-s | *BGIBMGA008010* | down | 544 | 552 | - | 11.3303 | 4.96E-05 | 0.615 | GTGGGTGGC |
| 6632 | mamo-s | *BGIBMGA008015* | down | 1584 | 1592 | - | 12.1835 | 1.71E-05 | 0.615 | GTGCGTGCC |
| 6633 | mamo-s | *BGIBMGA008016* | up | 658 | 666 | - | 12.1835 | 1.71E-05 | 0.615 | GTGCGTGCC |
| 6634 | mamo-s | *BGIBMGA008016* | down | 441 | 449 | - | 11.3303 | 4.96E-05 | 0.615 | GTGGGTGGC |
| 6635 | mamo-s | *BGIBMGA008016* | up | 1378 | 1386 | - | 10.9358 | 7.30E-05 | 0.728 | GCGGGTGGC |
| 6636 | mamo-s | *BGIBMGA008020* | down | 1800 | 1808 | + | 11.2844 | 5.18E-05 | 0.63 | CTGCGTGCC |
| 6637 | mamo-s | *BGIBMGA008022* | up | 1074 | 1082 | - | 11.0826 | 6.84E-05 | 0.709 | GTACGTGGC |
| 6638 | mamo-s | *BGIBMGA008024* | down | 1197 | 1205 | + | 11.5688 | 3.79E-05 | 0.615 | CTGCGTGTC |
| 6639 | mamo-s | *BGIBMGA008026* | up | 1620 | 1628 | - | 11.3303 | 4.96E-05 | 0.615 | GTGGGTGGC |
| 6640 | mamo-s | *BGIBMGA008027* | up | 1830 | 1838 | + | 11.3303 | 4.96E-05 | 0.615 | GTGGGTGGC |
| 6641 | mamo-s | *BGIBMGA008028* | down | 662 | 670 | - | 10.7523 | 8.38E-05 | 0.797 | GGGCGTGTC |
| 6642 | mamo-s | *BGIBMGA008031* | down | 106 | 114 | + | 11.3303 | 4.96E-05 | 0.615 | GTGGGTGGC |
| 6643 | mamo-s | *BGIBMGA008033* | up | 1334 | 1342 | + | 12.1284 | 2.16E-05 | 0.615 | CAGCGTGGC |
| 6644 | mamo-s | *BGIBMGA008034* | down | 672 | 680 | + | 11.3303 | 4.96E-05 | 0.615 | GTGGGTGGC |
| 6645 | mamo-s | *BGIBMGA008035* | up | 692 | 700 | + | 11.3303 | 4.96E-05 | 0.615 | GTGGGTGGC |
| 6646 | mamo-s | *BGIBMGA008036* | up | 623 | 631 | - | 11.0826 | 6.84E-05 | 0.709 | GTACGTGGC |
| 6647 | mamo-s | *BGIBMGA008037* | down | 1478 | 1486 | - | 11.0734 | 7.12E-05 | 0.726 | GAGCGTGGT |
| 6648 | mamo-s | *BGIBMGA008038* | down | 1980 | 1988 | + | 11.0734 | 7.12E-05 | 0.726 | GAGCGTGGT |
| 6649 | mamo-s | *BGIBMGA008039* | down | 1826 | 1834 | + | 11.1376 | 6.35E-05 | 0.709 | TCGCGTGGC |
| 6650 | mamo-s | *BGIBMGA008040* | down | 136 | 144 | + | 11.5321 | 4.06E-05 | 0.615 | TTGCGTGGC |
| 6651 | mamo-s | *BGIBMGA008041* | up | 305 | 313 | + | 11.3303 | 4.96E-05 | 0.615 | GTGGGTGGC |
| 6652 | mamo-s | *BGIBMGA008041* | up | 789 | 797 | - | 11.3303 | 4.96E-05 | 0.615 | GTGGGTGGC |
| 6653 | mamo-s | *BGIBMGA008043* | up | 1092 | 1100 | - | 12.1284 | 2.16E-05 | 0.615 | CAGCGTGGC |
| 6654 | mamo-s | *BGIBMGA008045* | down | 1359 | 1367 | - | 11.3303 | 4.96E-05 | 0.615 | GTGGGTGGC |
| 6655 | mamo-s | *BGIBMGA008046* | up | 955 | 963 | - | 14.055 | 2.23E-06 | 0.615 | GTGCGTGGC |
| 6656 | mamo-s | *BGIBMGA008048* | down | 798 | 806 | + | 12.0734 | 2.66E-05 | 0.615 | GCGCGTGTC |
| 6657 | mamo-s | *BGIBMGA008049* | up | 1821 | 1829 | - | 11.3303 | 4.96E-05 | 0.615 | GTGGGTGGC |
| 6658 | mamo-s | *BGIBMGA008051* | up | 623 | 631 | + | 11.3303 | 4.96E-05 | 0.615 | GTGGGTGGC |
| 6659 | mamo-s | *BGIBMGA008052* | up | 757 | 765 | + | 13.6606 | 4.06E-06 | 0.615 | GCGCGTGGC |
| 6660 | mamo-s | *BGIBMGA008052* | down | 60 | 68 | - | 11.2844 | 5.18E-05 | 0.63 | CTGCGTGCC |
| 6661 | mamo-s | *BGIBMGA008053* | down | 641 | 649 | + | 11.4404 | 4.51E-05 | 0.615 | CGGCGTGGC |
| 6662 | mamo-s | *BGIBMGA008055* | down | 774 | 782 | + | 11.3303 | 4.96E-05 | 0.615 | GTGGGTGGC |
| 6663 | mamo-s | *BGIBMGA008058* | down | 1160 | 1168 | + | 11.0826 | 6.84E-05 | 0.709 | GTACGTGGC |
| 6664 | mamo-s | *BGIBMGA008059* | up | 464 | 472 | - | 14.055 | 2.23E-06 | 0.615 | GTGCGTGGC |
| 6665 | mamo-s | *BGIBMGA008060* | down | 1005 | 1013 | + | 11.3303 | 4.96E-05 | 0.615 | GTGGGTGGC |
| 6666 | mamo-s | *BGIBMGA008060* | up | 986 | 994 | + | 11.1743 | 5.90E-05 | 0.684 | CCGCGTGTC |
| 6667 | mamo-s | *BGIBMGA008062* | down | 795 | 803 | + | 11.3303 | 4.96E-05 | 0.615 | GTGGGTGGC |
| 6668 | mamo-s | *BGIBMGA008062* | down | 1199 | 1207 | - | 11.3303 | 4.96E-05 | 0.615 | GTGGGTGGC |
| 6669 | mamo-s | *BGIBMGA008063* | up | 492 | 500 | - | 11.3303 | 4.96E-05 | 0.615 | GTGGGTGGC |
| 6670 | mamo-s | *BGIBMGA008063* | down | 539 | 547 | + | 11.3303 | 4.96E-05 | 0.615 | GTGGGTGGC |
| 6671 | mamo-s | *BGIBMGA008064* | up | 765 | 773 | + | 11.3303 | 4.96E-05 | 0.615 | GTGGGTGGC |
| 6672 | mamo-s | *BGIBMGA008064* | up | 1357 | 1365 | - | 11.3303 | 4.96E-05 | 0.615 | GTGGGTGGC |
| 6673 | mamo-s | *BGIBMGA008068* | down | 1017 | 1025 | + | 11.7798 | 3.11E-05 | 0.615 | ATGCGTGGC |
| 6674 | mamo-s | *BGIBMGA008069* | down | 1188 | 1196 | - | 14.055 | 2.23E-06 | 0.615 | GTGCGTGGC |
| 6675 | mamo-s | *BGIBMGA008069* | down | 953 | 961 | - | 11.3303 | 4.96E-05 | 0.615 | GTGGGTGGC |
| 6676 | mamo-s | *BGIBMGA008070* | up | 1847 | 1855 | - | 11.3303 | 4.96E-05 | 0.615 | GTGGGTGGC |
| 6677 | mamo-s | *BGIBMGA008076* | down | 132 | 140 | - | 10.5413 | 9.42E-05 | 0.857 | GTGCGAGTC |
| 6678 | mamo-s | *BGIBMGA008079* | down | 1898 | 1906 | - | 11.3303 | 4.96E-05 | 0.615 | GTGGGTGGC |
| 6679 | mamo-s | *BGIBMGA008080* | up | 297 | 305 | - | 11.3303 | 4.96E-05 | 0.615 | GTGGGTGGC |
| 6680 | mamo-s | *BGIBMGA008081* | down | 134 | 142 | - | 11.3303 | 4.96E-05 | 0.615 | GTGGGTGGC |
| 6681 | mamo-s | *BGIBMGA008083* | up | 744 | 752 | + | 11.3303 | 4.96E-05 | 0.615 | GTGGGTGGC |

| | | | | | | | | | | |
|---|---|---|---|---|---|---|---|---|---|---|
| 6682 | mamo-s | *BGIBMGA008087* | up | 310 | 318 | + | 11.3303 | 4.96E-05 | 0.615 | GTGGGTGGC |
| 6683 | mamo-s | *BGIBMGA008090* | up | 1073 | 1081 | - | 11.0826 | 6.84E-05 | 0.709 | GTACGTGGC |
| 6684 | mamo-s | *BGIBMGA008092* | up | 1930 | 1938 | - | 11.789 | 2.84E-05 | 0.615 | GCGCGTGCC |
| 6685 | mamo-s | *BGIBMGA008093* | up | 1031 | 1039 | - | 11.3303 | 4.96E-05 | 0.615 | GTGGGTGGC |
| 6686 | mamo-s | *BGIBMGA008093* | up | 1507 | 1515 | + | 11.3303 | 4.96E-05 | 0.615 | GTGGGTGGC |
| 6687 | mamo-s | *BGIBMGA008094* | up | 1017 | 1025 | + | 11.0826 | 6.84E-05 | 0.709 | GTACGTGGC |
| 6688 | mamo-s | *BGIBMGA008097* | down | 1702 | 1710 | + | 11.3303 | 4.96E-05 | 0.615 | GTGGGTGGC |
| 6689 | mamo-s | *BGIBMGA008098* | up | 401 | 409 | + | 11.3303 | 4.96E-05 | 0.615 | GTGGGTGGC |
| 6690 | mamo-s | *BGIBMGA008099* | up | 1852 | 1860 | + | 11.7339 | 3.29E-05 | 0.615 | GCGCGAGGC |
| 6691 | mamo-s | *BGIBMGA008099* | up | 1839 | 1847 | - | 11.3303 | 4.96E-05 | 0.615 | GTGGGTGGC |
| 6692 | mamo-s | *BGIBMGA008100* | up | 519 | 527 | + | 12.4679 | 1.31E-05 | 0.615 | GTGCGTGTC |
| 6693 | mamo-s | *BGIBMGA008104* | down | 852 | 860 | - | 11.3303 | 4.96E-05 | 0.615 | GTGGGTGGC |
| 6694 | mamo-s | *BGIBMGA008105* | up | 64 | 72 | + | 11.3303 | 4.96E-05 | 0.615 | GTGGGTGGC |
| 6695 | mamo-s | *BGIBMGA008105* | down | 288 | 296 | + | 11.3303 | 4.96E-05 | 0.615 | GTGGGTGGC |
| 6696 | mamo-s | *BGIBMGA008106* | down | 1131 | 1139 | + | 14.055 | 2.23E-06 | 0.615 | GTGCGTGGC |
| 6697 | mamo-s | *BGIBMGA008106* | down | 243 | 251 | + | 11.3303 | 4.96E-05 | 0.615 | GTGGGTGGC |
| 6698 | mamo-s | *BGIBMGA008106* | up | 1065 | 1073 | - | 11.3303 | 4.96E-05 | 0.615 | GTGGGTGGC |
| 6699 | mamo-s | *BGIBMGA008106* | down | 558 | 566 | - | 11.2844 | 5.18E-05 | 0.63 | CTGCGTGCC |
| 6700 | mamo-s | *BGIBMGA008108* | down | 1009 | 1017 | - | 11.3303 | 4.96E-05 | 0.615 | GTGGGTGGC |
| 6701 | mamo-s | *BGIBMGA008108* | up | 1585 | 1593 | - | 11.1743 | 5.90E-05 | 0.684 | CCGCGTGTC |
| 6702 | mamo-s | *BGIBMGA008108* | up | 1116 | 1124 | + | 10.8899 | 7.48E-05 | 0.739 | CCGCGTGCC |
| 6703 | mamo-s | *BGIBMGA008109* | down | 1438 | 1446 | - | 11.3303 | 4.96E-05 | 0.615 | GTGGGTGGC |
| 6704 | mamo-s | *BGIBMGA008112* | up | 1158 | 1166 | - | 10.5138 | 9.75E-05 | 0.865 | GTGCGTGTT |
| 6705 | mamo-s | *BGIBMGA008113* | up | 1711 | 1719 | + | 10.5138 | 9.75E-05 | 0.865 | GTGCGTGTT |
| 6706 | mamo-s | *BGIBMGA008114* | down | 1431 | 1439 | + | 11.3303 | 4.96E-05 | 0.615 | GTGGGTGGC |
| 6707 | mamo-s | *BGIBMGA008115* | down | 247 | 255 | - | 11.0734 | 7.12E-05 | 0.726 | GAGCGTGGT |
| 6708 | mamo-s | *BGIBMGA008116* | up | 1239 | 1247 | - | 11.0734 | 7.12E-05 | 0.726 | GAGCGTGGT |
| 6709 | mamo-s | *BGIBMGA008117* | up | 17 | 25 | + | 11.3303 | 4.96E-05 | 0.615 | GTGGGTGGC |
| 6710 | mamo-s | *BGIBMGA008117* | up | 835 | 843 | - | 11.3303 | 4.96E-05 | 0.615 | GTGGGTGGC |
| 6711 | mamo-s | *BGIBMGA008120* | down | 496 | 504 | + | 11.3303 | 4.96E-05 | 0.615 | GTGGGTGGC |
| 6712 | mamo-s | *BGIBMGA008120* | down | 1876 | 1884 | + | 11.3303 | 4.96E-05 | 0.615 | GTGGGTGGC |
| 6713 | mamo-s | *BGIBMGA008120* | up | 1200 | 1208 | + | 11.1743 | 5.90E-05 | 0.684 | CCGCGTGTC |
| 6714 | mamo-s | *BGIBMGA008121* | down | 733 | 741 | - | 12.3394 | 1.49E-05 | 0.615 | GGGCGTGGC |
| 6715 | mamo-s | *BGIBMGA008121* | down | 229 | 237 | + | 11.3303 | 4.96E-05 | 0.615 | GTGGGTGGC |
| 6716 | mamo-s | *BGIBMGA008122* | down | 235 | 243 | - | 12.1009 | 2.43E-05 | 0.615 | GTGCGTGGT |
| 6717 | mamo-s | *BGIBMGA008123* | down | 822 | 830 | + | 12.3394 | 1.49E-05 | 0.615 | GGGCGTGGC |
| 6718 | mamo-s | *BGIBMGA008123* | down | 1785 | 1793 | + | 11.3303 | 4.96E-05 | 0.615 | GTGGGTGGC |
| 6719 | mamo-s | *BGIBMGA008124* | down | 1452 | 1460 | - | 11.3303 | 4.96E-05 | 0.615 | GTGGGTGGC |
| 6720 | mamo-s | *BGIBMGA008125* | up | 449 | 457 | - | 11.3303 | 4.96E-05 | 0.615 | GTGGGTGGC |
| 6721 | mamo-s | *BGIBMGA008125* | up | 1280 | 1288 | + | 10.6881 | 8.61E-05 | 0.81 | GCACGTGGC |
| 6722 | mamo-s | *BGIBMGA008126* | up | 1085 | 1093 | + | 11.156 | 6.13E-05 | 0.705 | GAGCGTGCC |
| 6723 | mamo-s | *BGIBMGA008129* | up | 95 | 103 | + | 11.3303 | 4.96E-05 | 0.615 | GTGGGTGGC |
| 6724 | mamo-s | *BGIBMGA008129* | down | 1642 | 1650 | - | 11.3303 | 4.96E-05 | 0.615 | GTGGGTGGC |
| 6725 | mamo-s | *BGIBMGA008130* | up | 715 | 723 | + | 11.3303 | 4.96E-05 | 0.615 | GTGGGTGGC |
| 6726 | mamo-s | *BGIBMGA008131* | up | 485 | 493 | - | 11.3303 | 4.96E-05 | 0.615 | GTGGGTGGC |
| 6727 | mamo-s | *BGIBMGA008131* | up | 1350 | 1358 | - | 11.3303 | 4.96E-05 | 0.615 | GTGGGTGGC |
| 6728 | mamo-s | *BGIBMGA008132* | down | 725 | 733 | + | 11.3303 | 4.96E-05 | 0.615 | GTGGGTGGC |
| 6729 | mamo-s | *BGIBMGA008133* | up | 1723 | 1731 | + | 11.3303 | 4.96E-05 | 0.615 | GTGGGTGGC |
| 6730 | mamo-s | *BGIBMGA008133* | down | 1105 | 1113 | + | 10.5413 | 9.42E-05 | 0.857 | CAGCGTGTC |
| 6731 | mamo-s | *BGIBMGA008134* | up | 1095 | 1103 | + | 11.3303 | 4.96E-05 | 0.615 | GTGGGTGGC |
| 6732 | mamo-s | *BGIBMGA008135* | up | 1199 | 1207 | + | 11.3303 | 4.96E-05 | 0.615 | GTGGGTGGC |
| 6733 | mamo-s | *BGIBMGA008136* | up | 845 | 853 | - | 11.0826 | 6.84E-05 | 0.709 | GTACGTGGC |
| 6734 | mamo-s | *BGIBMGA008141* | down | 1274 | 1282 | + | 11.3303 | 4.96E-05 | 0.615 | GTGGGTGGC |
| 6735 | mamo-s | *BGIBMGA008142* | down | 599 | 607 | - | 11.3303 | 4.96E-05 | 0.615 | GTGGGTGGC |
| 6736 | mamo-s | *BGIBMGA008143* | down | 215 | 223 | + | 12.3394 | 1.49E-05 | 0.615 | GGGCGTGGC |
| 6737 | mamo-s | *BGIBMGA008143* | down | 524 | 532 | + | 12.3394 | 1.49E-05 | 0.615 | GGGCGTGGC |
| 6738 | mamo-s | *BGIBMGA008143* | down | 988 | 996 | + | 11.7064 | 3.52E-05 | 0.615 | GCGCGTGGT |
| 6739 | mamo-s | *BGIBMGA008144* | up | 482 | 490 | + | 11.3303 | 4.96E-05 | 0.615 | GTGGGTGGC |
| 6740 | mamo-s | *BGIBMGA008144* | down | 861 | 869 | + | 11.3303 | 4.96E-05 | 0.615 | GTGGGTGGC |
| 6741 | mamo-s | *BGIBMGA008144* | up | 1122 | 1130 | - | 11.0826 | 6.84E-05 | 0.709 | GTACGTGGC |
| 6742 | mamo-s | *BGIBMGA008145* | down | 159 | 167 | - | 14.055 | 2.23E-06 | 0.615 | GTGCGTGGC |
| 6743 | mamo-s | *BGIBMGA008145* | up | 718 | 726 | + | 11.3303 | 4.96E-05 | 0.615 | GTGGGTGGC |
| 6744 | mamo-s | *BGIBMGA008147* | up | 384 | 392 | + | 11.3303 | 4.96E-05 | 0.615 | GTGGGTGGC |
| 6745 | mamo-s | *BGIBMGA008150* | up | 410 | 418 | - | 11.7064 | 3.52E-05 | 0.615 | GCGCGTGGT |
| 6746 | mamo-s | *BGIBMGA008151* | up | 210 | 218 | - | 11.3303 | 4.96E-05 | 0.615 | GTGGGTGGC |
| 6747 | mamo-s | *BGIBMGA008152* | up | 387 | 395 | - | 11.7798 | 3.11E-05 | 0.615 | ATGCGTGGC |
| 6748 | mamo-s | *BGIBMGA008152* | up | 168 | 176 | - | 11.3303 | 4.96E-05 | 0.615 | GTGGGTGGC |
| 6749 | mamo-s | *BGIBMGA008153* | up | 378 | 386 | - | 11.3303 | 4.96E-05 | 0.615 | GTGGGTGGC |
| 6750 | mamo-s | *BGIBMGA008153* | up | 1562 | 1570 | - | 11.3303 | 4.96E-05 | 0.615 | GTGGGTGGC |
| 6751 | mamo-s | *BGIBMGA008153* | down | 1886 | 1894 | - | 11.0826 | 6.84E-05 | 0.709 | GTACGTGGC |
| 6752 | mamo-s | *BGIBMGA008154* | down | 403 | 411 | - | 12.0734 | 2.66E-05 | 0.615 | GCGCGTGTC |
| 6753 | mamo-s | *BGIBMGA008154* | down | 431 | 439 | - | 12.0734 | 2.66E-05 | 0.615 | GCGCGTGTC |
| 6754 | mamo-s | *BGIBMGA008154* | down | 463 | 471 | - | 12.0734 | 2.66E-05 | 0.615 | GCGCGTGTC |
| 6755 | mamo-s | *BGIBMGA008154* | down | 483 | 491 | - | 12.0734 | 2.66E-05 | 0.615 | GCGCGTGTC |
| 6756 | mamo-s | *BGIBMGA008154* | down | 515 | 523 | - | 12.0734 | 2.66E-05 | 0.615 | GCGCGTGTC |
| 6757 | mamo-s | *BGIBMGA008154* | down | 575 | 583 | - | 12.0734 | 2.66E-05 | 0.615 | GCGCGTGTC |
| 6758 | mamo-s | *BGIBMGA008154* | down | 595 | 603 | - | 12.0734 | 2.66E-05 | 0.615 | GCGCGTGTC |
| 6759 | mamo-s | *BGIBMGA008154* | down | 615 | 623 | - | 12.0734 | 2.66E-05 | 0.615 | GCGCGTGTC |
| 6760 | mamo-s | *BGIBMGA008154* | down | 647 | 655 | - | 12.0734 | 2.66E-05 | 0.615 | GCGCGTGTC |
| 6761 | mamo-s | *BGIBMGA008154* | down | 687 | 695 | - | 12.0734 | 2.66E-05 | 0.615 | GCGCGTGTC |
| 6762 | mamo-s | *BGIBMGA008154* | down | 719 | 727 | - | 12.0734 | 2.66E-05 | 0.615 | GCGCGTGTC |
| 6763 | mamo-s | *BGIBMGA008154* | down | 835 | 843 | - | 12.0734 | 2.66E-05 | 0.615 | GCGCGTGTC |
| 6764 | mamo-s | *BGIBMGA008154* | down | 886 | 894 | - | 12.0734 | 2.66E-05 | 0.615 | GCGCGTGTC |
| 6765 | mamo-s | *BGIBMGA008154* | down | 906 | 914 | - | 12.0734 | 2.66E-05 | 0.615 | GCGCGTGTC |
| 6766 | mamo-s | *BGIBMGA008154* | down | 926 | 934 | - | 12.0734 | 2.66E-05 | 0.615 | GCGCGTGTC |
| 6767 | mamo-s | *BGIBMGA008154* | down | 958 | 966 | - | 12.0734 | 2.66E-05 | 0.615 | GCGCGTGTC |
| 6768 | mamo-s | *BGIBMGA008154* | down | 970 | 978 | - | 12.0734 | 2.66E-05 | 0.615 | GCGCGTGTC |
| 6769 | mamo-s | *BGIBMGA008154* | down | 990 | 998 | - | 12.0734 | 2.66E-05 | 0.615 | GCGCGTGTC |

| | | | | | | | | | | |
|---|---|---|---|---|---|---|---|---|---|---|
| 6770 | mamo-s | *BGIBMGA008154* | down | 1042 | 1050 | - | 12.0734 | 2.66E-05 | 0.615 | GCGCGTGTC |
| 6771 | mamo-s | *BGIBMGA008154* | down | 1062 | 1070 | - | 12.0734 | 2.66E-05 | 0.615 | GCGCGTGTC |
| 6772 | mamo-s | *BGIBMGA008154* | down | 1114 | 1122 | - | 12.0734 | 2.66E-05 | 0.615 | GCGCGTGTC |
| 6773 | mamo-s | *BGIBMGA008154* | down | 1134 | 1142 | - | 12.0734 | 2.66E-05 | 0.615 | GCGCGTGTC |
| 6774 | mamo-s | *BGIBMGA008154* | down | 1186 | 1194 | - | 12.0734 | 2.66E-05 | 0.615 | GCGCGTGTC |
| 6775 | mamo-s | *BGIBMGA008154* | down | 1198 | 1206 | - | 12.0734 | 2.66E-05 | 0.615 | GCGCGTGTC |
| 6776 | mamo-s | *BGIBMGA008154* | down | 1218 | 1226 | - | 12.0734 | 2.66E-05 | 0.615 | GCGCGTGTC |
| 6777 | mamo-s | *BGIBMGA008154* | down | 1250 | 1258 | - | 12.0734 | 2.66E-05 | 0.615 | GCGCGTGTC |
| 6778 | mamo-s | *BGIBMGA008154* | down | 395 | 403 | - | 11.5688 | 3.79E-05 | 0.615 | CTGCGTGTC |
| 6779 | mamo-s | *BGIBMGA008154* | down | 415 | 423 | - | 11.5688 | 3.79E-05 | 0.615 | CTGCGTGTC |
| 6780 | mamo-s | *BGIBMGA008154* | down | 423 | 431 | - | 11.5688 | 3.79E-05 | 0.615 | CTGCGTGTC |
| 6781 | mamo-s | *BGIBMGA008154* | down | 475 | 483 | - | 11.5688 | 3.79E-05 | 0.615 | CTGCGTGTC |
| 6782 | mamo-s | *BGIBMGA008154* | down | 507 | 515 | - | 11.5688 | 3.79E-05 | 0.615 | CTGCGTGTC |
| 6783 | mamo-s | *BGIBMGA008154* | down | 535 | 543 | - | 11.5688 | 3.79E-05 | 0.615 | CTGCGTGTC |
| 6784 | mamo-s | *BGIBMGA008154* | down | 567 | 575 | - | 11.5688 | 3.79E-05 | 0.615 | CTGCGTGTC |
| 6785 | mamo-s | *BGIBMGA008154* | down | 607 | 615 | - | 11.5688 | 3.79E-05 | 0.615 | CTGCGTGTC |
| 6786 | mamo-s | *BGIBMGA008154* | down | 639 | 647 | - | 11.5688 | 3.79E-05 | 0.615 | CTGCGTGTC |
| 6787 | mamo-s | *BGIBMGA008154* | down | 679 | 687 | - | 11.5688 | 3.79E-05 | 0.615 | CTGCGTGTC |
| 6788 | mamo-s | *BGIBMGA008154* | down | 763 | 771 | - | 11.5688 | 3.79E-05 | 0.615 | CTGCGTGTC |
| 6789 | mamo-s | *BGIBMGA008154* | down | 847 | 855 | - | 11.5688 | 3.79E-05 | 0.615 | CTGCGTGTC |
| 6790 | mamo-s | *BGIBMGA008154* | down | 878 | 886 | - | 11.5688 | 3.79E-05 | 0.615 | CTGCGTGTC |
| 6791 | mamo-s | *BGIBMGA008154* | down | 918 | 926 | - | 11.5688 | 3.79E-05 | 0.615 | CTGCGTGTC |
| 6792 | mamo-s | *BGIBMGA008154* | down | 1002 | 1010 | - | 11.5688 | 3.79E-05 | 0.615 | CTGCGTGTC |
| 6793 | mamo-s | *BGIBMGA008154* | down | 1034 | 1042 | - | 11.5688 | 3.79E-05 | 0.615 | CTGCGTGTC |
| 6794 | mamo-s | *BGIBMGA008154* | down | 1054 | 1062 | - | 11.5688 | 3.79E-05 | 0.615 | CTGCGTGTC |
| 6795 | mamo-s | *BGIBMGA008154* | down | 1074 | 1082 | - | 11.5688 | 3.79E-05 | 0.615 | CTGCGTGTC |
| 6796 | mamo-s | *BGIBMGA008154* | down | 1106 | 1114 | - | 11.5688 | 3.79E-05 | 0.615 | CTGCGTGTC |
| 6797 | mamo-s | *BGIBMGA008154* | down | 1126 | 1134 | - | 11.5688 | 3.79E-05 | 0.615 | CTGCGTGTC |
| 6798 | mamo-s | *BGIBMGA008154* | down | 1146 | 1154 | - | 11.5688 | 3.79E-05 | 0.615 | CTGCGTGTC |
| 6799 | mamo-s | *BGIBMGA008154* | down | 1178 | 1186 | - | 11.5688 | 3.79E-05 | 0.615 | CTGCGTGTC |
| 6800 | mamo-s | *BGIBMGA008154* | down | 1242 | 1250 | - | 11.5688 | 3.79E-05 | 0.615 | CTGCGTGTC |
| 6801 | mamo-s | *BGIBMGA008156* | down | 954 | 962 | + | 14.055 | 2.23E-06 | 0.615 | GTGCGTGGC |
| 6802 | mamo-s | *BGIBMGA008156* | up | 728 | 736 | + | 11.3303 | 4.96E-05 | 0.615 | GTGGGTGGC |
| 6803 | mamo-s | *BGIBMGA008156* | down | 1336 | 1344 | - | 11.3303 | 4.96E-05 | 0.615 | GTGGGTGGC |
| 6804 | mamo-s | *BGIBMGA008156* | up | 1444 | 1452 | + | 10.6881 | 8.61E-05 | 0.81 | GCACGTGGC |
| 6805 | mamo-s | *BGIBMGA008157* | down | 1351 | 1359 | + | 11.3303 | 4.96E-05 | 0.615 | GTGGGTGGC |
| 6806 | mamo-s | *BGIBMGA008157* | up | 1913 | 1921 | + | 11.1009 | 6.57E-05 | 0.709 | GAGCGAGGC |
| 6807 | mamo-s | *BGIBMGA008158* | up | 1030 | 1038 | + | 11.3303 | 4.96E-05 | 0.615 | GTGGGTGGC |
| 6808 | mamo-s | *BGIBMGA008158* | up | 982 | 990 | - | 10.8073 | 7.89E-05 | 0.766 | CCGCGTGGT |
| 6809 | mamo-s | *BGIBMGA008158* | down | 1441 | 1449 | + | 10.5413 | 9.42E-05 | 0.857 | CAGCGTGTC |
| 6810 | mamo-s | *BGIBMGA008159* | up | 647 | 655 | - | 11.7339 | 3.29E-05 | 0.615 | GCGCGAGGC |
| 6811 | mamo-s | *BGIBMGA008159* | up | 347 | 355 | - | 11.3303 | 4.96E-05 | 0.615 | GTGGGTGGC |
| 6812 | mamo-s | *BGIBMGA008159* | down | 1032 | 1040 | + | 11.3303 | 4.96E-05 | 0.615 | GTGGGTGGC |
| 6813 | mamo-s | *BGIBMGA008159* | up | 1164 | 1172 | + | 10.8899 | 7.48E-05 | 0.739 | CCGCGTGCC |
| 6814 | mamo-s | *BGIBMGA008160* | up | 27 | 35 | - | 11.3303 | 4.96E-05 | 0.615 | GTGGGTGGC |
| 6815 | mamo-s | *BGIBMGA008160* | down | 1939 | 1947 | - | 11.3303 | 4.96E-05 | 0.615 | GTGGGTGGC |
| 6816 | mamo-s | *BGIBMGA008161* | up | 1904 | 1912 | - | 11.3853 | 4.74E-05 | 0.615 | ACGCGTGGC |
| 6817 | mamo-s | *BGIBMGA008162* | up | 243 | 251 | - | 11.3303 | 4.96E-05 | 0.615 | GTGGGTGGC |
| 6818 | mamo-s | *BGIBMGA008164* | up | 1220 | 1228 | + | 10.5688 | 8.88E-05 | 0.824 | GTGCGTGAC |
| 6819 | mamo-s | *BGIBMGA008165* | down | 438 | 446 | - | 12.1284 | 2.16E-05 | 0.615 | CAGCGTGGC |
| 6820 | mamo-s | *BGIBMGA008165* | up | 75 | 83 | - | 10.5688 | 8.88E-05 | 0.824 | GTGCGTGAC |
| 6821 | mamo-s | *BGIBMGA008166* | up | 1563 | 1571 | - | 11.7064 | 3.52E-05 | 0.615 | GCGCGTGGT |
| 6822 | mamo-s | *BGIBMGA008166* | down | 1612 | 1620 | + | 11.3303 | 4.96E-05 | 0.615 | GTGGGTGGC |
| 6823 | mamo-s | *BGIBMGA008169* | up | 1495 | 1503 | + | 12.0734 | 2.66E-05 | 0.615 | GCGCGTGTC |
| 6824 | mamo-s | *BGIBMGA008169* | up | 1527 | 1535 | + | 12.0734 | 2.66E-05 | 0.615 | GCGCGTGTC |
| 6825 | mamo-s | *BGIBMGA008169* | up | 1547 | 1555 | + | 12.0734 | 2.66E-05 | 0.615 | GCGCGTGTC |
| 6826 | mamo-s | *BGIBMGA008169* | up | 1559 | 1567 | + | 12.0734 | 2.66E-05 | 0.615 | GCGCGTGTC |
| 6827 | mamo-s | *BGIBMGA008169* | up | 1611 | 1619 | + | 12.0734 | 2.66E-05 | 0.615 | GCGCGTGTC |
| 6828 | mamo-s | *BGIBMGA008169* | up | 1631 | 1639 | + | 12.0734 | 2.66E-05 | 0.615 | GCGCGTGTC |
| 6829 | mamo-s | *BGIBMGA008169* | up | 1683 | 1691 | + | 12.0734 | 2.66E-05 | 0.615 | GCGCGTGTC |
| 6830 | mamo-s | *BGIBMGA008169* | up | 1703 | 1711 | + | 12.0734 | 2.66E-05 | 0.615 | GCGCGTGTC |
| 6831 | mamo-s | *BGIBMGA008169* | up | 1755 | 1763 | + | 12.0734 | 2.66E-05 | 0.615 | GCGCGTGTC |
| 6832 | mamo-s | *BGIBMGA008169* | up | 1775 | 1783 | + | 12.0734 | 2.66E-05 | 0.615 | GCGCGTGTC |
| 6833 | mamo-s | *BGIBMGA008169* | up | 1787 | 1795 | + | 12.0734 | 2.66E-05 | 0.615 | GCGCGTGTC |
| 6834 | mamo-s | *BGIBMGA008169* | up | 1819 | 1827 | + | 12.0734 | 2.66E-05 | 0.615 | GCGCGTGTC |
| 6835 | mamo-s | *BGIBMGA008169* | up | 1839 | 1847 | + | 12.0734 | 2.66E-05 | 0.615 | GCGCGTGTC |
| 6836 | mamo-s | *BGIBMGA008169* | up | 1859 | 1867 | + | 12.0734 | 2.66E-05 | 0.615 | GCGCGTGTC |
| 6837 | mamo-s | *BGIBMGA008169* | up | 1910 | 1918 | + | 12.0734 | 2.66E-05 | 0.615 | GCGCGTGTC |
| 6838 | mamo-s | *BGIBMGA008169* | up | 1503 | 1511 | + | 11.5688 | 3.79E-05 | 0.615 | CTGCGTGTC |
| 6839 | mamo-s | *BGIBMGA008169* | up | 1567 | 1575 | + | 11.5688 | 3.79E-05 | 0.615 | CTGCGTGTC |
| 6840 | mamo-s | *BGIBMGA008169* | up | 1599 | 1607 | + | 11.5688 | 3.79E-05 | 0.615 | CTGCGTGTC |
| 6841 | mamo-s | *BGIBMGA008169* | up | 1619 | 1627 | + | 11.5688 | 3.79E-05 | 0.615 | CTGCGTGTC |
| 6842 | mamo-s | *BGIBMGA008169* | up | 1639 | 1647 | + | 11.5688 | 3.79E-05 | 0.615 | CTGCGTGTC |
| 6843 | mamo-s | *BGIBMGA008169* | up | 1671 | 1679 | + | 11.5688 | 3.79E-05 | 0.615 | CTGCGTGTC |
| 6844 | mamo-s | *BGIBMGA008169* | up | 1691 | 1699 | + | 11.5688 | 3.79E-05 | 0.615 | CTGCGTGTC |
| 6845 | mamo-s | *BGIBMGA008169* | up | 1711 | 1719 | + | 11.5688 | 3.79E-05 | 0.615 | CTGCGTGTC |
| 6846 | mamo-s | *BGIBMGA008169* | up | 1743 | 1751 | + | 11.5688 | 3.79E-05 | 0.615 | CTGCGTGTC |
| 6847 | mamo-s | *BGIBMGA008169* | up | 1827 | 1835 | + | 11.5688 | 3.79E-05 | 0.615 | CTGCGTGTC |
| 6848 | mamo-s | *BGIBMGA008169* | up | 1867 | 1875 | + | 11.5688 | 3.79E-05 | 0.615 | CTGCGTGTC |
| 6849 | mamo-s | *BGIBMGA008169* | up | 1898 | 1906 | + | 11.5688 | 3.79E-05 | 0.615 | CTGCGTGTC |
| 6850 | mamo-s | *BGIBMGA008169* | up | 1982 | 1990 | + | 11.5688 | 3.79E-05 | 0.615 | CTGCGTGTC |
| 6851 | mamo-s | *BGIBMGA008169* | down | 1066 | 1074 | - | 11.3303 | 4.96E-05 | 0.615 | GTGGGTGGC |
| 6852 | mamo-s | *BGIBMGA008170* | down | 798 | 806 | + | 13.6606 | 4.06E-06 | 0.615 | GCGCGTGGC |
| 6853 | mamo-s | *BGIBMGA008172* | down | 58 | 66 | - | 12.1835 | 1.71E-05 | 0.615 | GTGCGTGCC |
| 6854 | mamo-s | *BGIBMGA008173* | up | 1390 | 1398 | - | 11.0734 | 7.12E-05 | 0.726 | GAGCGTGGT |
| 6855 | mamo-s | *BGIBMGA008174* | up | 1159 | 1167 | + | 11.3303 | 4.96E-05 | 0.615 | GTGGGTGGC |
| 6856 | mamo-s | *BGIBMGA008174* | up | 1698 | 1706 | + | 11.3303 | 4.96E-05 | 0.615 | GTGGGTGGC |
| 6857 | mamo-s | *BGIBMGA008175* | down | 544 | 552 | + | 14.055 | 2.23E-06 | 0.615 | GTGCGTGGC |

| | | | | | | | | | | |
|---|---|---|---|---|---|---|---|---|---|---|
| 6858 | mamo-s | *BGIBMGA008175* | down | 885 | 893 | + | 11.1009 | 6.57E-05 | 0.709 | GAGCGAGGC |
| 6859 | mamo-s | *BGIBMGA008176* | down | 1783 | 1791 | + | 11.3303 | 4.96E-05 | 0.615 | GTGGGTGGC |
| 6860 | mamo-s | *BGIBMGA008176* | down | 1957 | 1965 | - | 11.3303 | 4.96E-05 | 0.615 | GTGGGTGGC |
| 6861 | mamo-s | *BGIBMGA008177* | up | 1567 | 1575 | + | 11.3303 | 4.96E-05 | 0.615 | GTGGGTGGC |
| 6862 | mamo-s | *BGIBMGA008177* | up | 1741 | 1749 | - | 11.3303 | 4.96E-05 | 0.615 | GTGGGTGGC |
| 6863 | mamo-s | *BGIBMGA008177* | down | 383 | 391 | + | 10.6881 | 8.61E-05 | 0.81 | GCACGTGGC |
| 6864 | mamo-s | *BGIBMGA008178* | down | 1582 | 1590 | + | 13.0275 | 8.53E-06 | 0.615 | GAGCGTGGC |
| 6865 | mamo-s | *BGIBMGA008178* | up | 1547 | 1555 | + | 10.6881 | 8.61E-05 | 0.81 | GCACGTGGC |
| 6866 | mamo-s | *BGIBMGA008179* | up | 1529 | 1537 | - | 11.3303 | 4.96E-05 | 0.615 | GTGGGTGGC |
| 6867 | mamo-s | *BGIBMGA008180* | up | 152 | 160 | - | 11.3303 | 4.96E-05 | 0.615 | GTGGGTGGC |
| 6868 | mamo-s | *BGIBMGA008181* | up | 1019 | 1027 | + | 11.3303 | 4.96E-05 | 0.615 | GTGGGTGGC |
| 6869 | mamo-s | *BGIBMGA008181* | up | 1327 | 1335 | - | 11.0826 | 6.84E-05 | 0.709 | GTACGTGGC |
| 6870 | mamo-s | *BGIBMGA008183* | up | 1328 | 1336 | - | 11.3303 | 4.96E-05 | 0.615 | GTGGGTGGC |
| 6871 | mamo-s | *BGIBMGA008188* | up | 1038 | 1046 | - | 11.3303 | 4.96E-05 | 0.615 | GTGGGTGGC |
| 6872 | mamo-s | *BGIBMGA008188* | up | 1825 | 1833 | + | 11.3303 | 4.96E-05 | 0.615 | GTGGGTGGC |
| 6873 | mamo-s | *BGIBMGA008188* | down | 69 | 77 | - | 10.6881 | 8.61E-05 | 0.81 | GCACGTGGC |
| 6874 | mamo-s | *BGIBMGA008189* | up | 1390 | 1398 | - | 11.3303 | 4.96E-05 | 0.615 | GTGGGTGGC |
| 6875 | mamo-s | *BGIBMGA008191* | up | 1302 | 1310 | + | 10.9358 | 7.30E-05 | 0.728 | GCGGGTGGC |
| 6876 | mamo-s | *BGIBMGA008192* | up | 1249 | 1257 | + | 11.3303 | 4.96E-05 | 0.615 | GTGGGTGGC |
| 6877 | mamo-s | *BGIBMGA008192* | down | 1960 | 1968 | - | 11.3303 | 4.96E-05 | 0.615 | GTGGGTGGC |
| 6878 | mamo-s | *BGIBMGA008193* | up | 301 | 309 | - | 11.3303 | 4.96E-05 | 0.615 | GTGGGTGGC |
| 6879 | mamo-s | *BGIBMGA008193* | down | 514 | 522 | - | 11.3303 | 4.96E-05 | 0.615 | GTGGGTGGC |
| 6880 | mamo-s | *BGIBMGA008195* | up | 1712 | 1720 | + | 11.3303 | 4.96E-05 | 0.615 | GTGGGTGGC |
| 6881 | mamo-s | *BGIBMGA008195* | up | 643 | 651 | - | 11.0826 | 6.84E-05 | 0.709 | GTACGTGGC |
| 6882 | mamo-s | *BGIBMGA008196* | up | 527 | 535 | - | 11.0826 | 6.84E-05 | 0.709 | GTACGTGGC |
| 6883 | mamo-s | *BGIBMGA008198* | up | 203 | 211 | + | 11.3303 | 4.96E-05 | 0.615 | GTGGGTGGC |
| 6884 | mamo-s | *BGIBMGA008198* | up | 848 | 856 | - | 11.3303 | 4.96E-05 | 0.615 | GTGGGTGGC |
| 6885 | mamo-s | *BGIBMGA008198* | up | 1773 | 1781 | - | 10.5413 | 9.42E-05 | 0.857 | CAGCGTGTC |
| 6886 | mamo-s | *BGIBMGA008198* | up | 1290 | 1298 | - | 10.5138 | 9.75E-05 | 0.865 | GTGCGTGTT |
| 6887 | mamo-s | *BGIBMGA008200* | up | 630 | 638 | + | 11.3303 | 4.96E-05 | 0.615 | GTGGGTGGC |
| 6888 | mamo-s | *BGIBMGA008200* | down | 450 | 458 | - | 11.1743 | 5.90E-05 | 0.684 | CCGCGTGTC |
| 6889 | mamo-s | *BGIBMGA008206* | up | 964 | 972 | + | 11.3303 | 4.96E-05 | 0.615 | GTGGGTGGC |
| 6890 | mamo-s | *BGIBMGA008207* | up | 583 | 591 | - | 11.3303 | 4.96E-05 | 0.615 | GTGGGTGGC |
| 6891 | mamo-s | *BGIBMGA008207* | down | 1073 | 1081 | - | 10.7523 | 8.38E-05 | 0.797 | AAGCGTGGC |
| 6892 | mamo-s | *BGIBMGA008207* | up | 1514 | 1522 | - | 10.6881 | 8.61E-05 | 0.81 | GCACGTGGC |
| 6893 | mamo-s | *BGIBMGA008208* | down | 1496 | 1504 | + | 11.2844 | 5.18E-05 | 0.63 | CTGCGTGCC |
| 6894 | mamo-s | *BGIBMGA008209* | down | 1020 | 1028 | + | 11.5321 | 4.06E-05 | 0.615 | TTGCGTGGC |
| 6895 | mamo-s | *BGIBMGA008212* | up | 311 | 319 | + | 11.0826 | 6.84E-05 | 0.709 | GTACGTGGC |
| 6896 | mamo-s | *BGIBMGA008213* | down | 1937 | 1945 | - | 11.3303 | 4.96E-05 | 0.615 | GTGGGTGGC |
| 6897 | mamo-s | *BGIBMGA008214* | up | 988 | 996 | + | 11.3303 | 4.96E-05 | 0.615 | GTGGGTGGC |
| 6898 | mamo-s | *BGIBMGA008215* | down | 1538 | 1546 | + | 11.3303 | 4.96E-05 | 0.615 | GTGGGTGGC |
| 6899 | mamo-s | *BGIBMGA008215* | up | 561 | 569 | + | 11.1376 | 6.35E-05 | 0.709 | TCGCGTGGC |
| 6900 | mamo-s | *BGIBMGA008216* | down | 430 | 438 | + | 11.3303 | 4.96E-05 | 0.615 | GTGGGTGGC |
| 6901 | mamo-s | *BGIBMGA008216* | up | 1820 | 1828 | + | 11.3303 | 4.96E-05 | 0.615 | GTGGGTGGC |
| 6902 | mamo-s | *BGIBMGA008217* | up | 7 | 15 | - | 11.3303 | 4.96E-05 | 0.615 | GTGGGTGGC |
| 6903 | mamo-s | *BGIBMGA008217* | down | 1794 | 1802 | + | 11.3303 | 4.96E-05 | 0.615 | GTGGGTGGC |
| 6904 | mamo-s | *BGIBMGA008218* | down | 1547 | 1555 | - | 12.1009 | 2.43E-05 | 0.615 | GTGCGTGGT |
| 6905 | mamo-s | *BGIBMGA008222* | down | 461 | 469 | + | 11.3303 | 4.96E-05 | 0.615 | GTGGGTGGC |
| 6906 | mamo-s | *BGIBMGA008222* | up | 1472 | 1480 | - | 11.3303 | 4.96E-05 | 0.615 | GTGGGTGGC |
| 6907 | mamo-s | *BGIBMGA008222* | down | 1741 | 1749 | - | 11.3303 | 4.96E-05 | 0.615 | GTGGGTGGC |
| 6908 | mamo-s | *BGIBMGA008223* | up | 870 | 878 | + | 11.3303 | 4.96E-05 | 0.615 | GTGGGTGGC |
| 6909 | mamo-s | *BGIBMGA008224* | down | 1594 | 1602 | + | 12.4679 | 1.31E-05 | 0.615 | GTGCGTGTC |
| 6910 | mamo-s | *BGIBMGA008225* | down | 61 | 69 | + | 11.1743 | 5.90E-05 | 0.684 | CCGCGTGTC |
| 6911 | mamo-s | *BGIBMGA008227* | down | 1504 | 1512 | - | 11.7339 | 3.29E-05 | 0.615 | GCGCGAGGC |
| 6912 | mamo-s | *BGIBMGA008227* | up | 1477 | 1485 | + | 11.3303 | 4.96E-05 | 0.615 | GTGGGTGGC |
| 6913 | mamo-s | *BGIBMGA008227* | down | 1849 | 1857 | + | 10.6881 | 8.61E-05 | 0.81 | GCACGTGGC |
| 6914 | mamo-s | *BGIBMGA008229* | up | 132 | 140 | + | 11.3303 | 4.96E-05 | 0.615 | GTGGGTGGC |
| 6915 | mamo-s | *BGIBMGA008229* | down | 960 | 968 | + | 11.1743 | 5.90E-05 | 0.684 | CCGCGTGTC |
| 6916 | mamo-s | *BGIBMGA008229* | up | 1813 | 1821 | - | 10.5138 | 9.75E-05 | 0.865 | GTGCGTGTT |
| 6917 | mamo-s | *BGIBMGA008230* | up | 720 | 728 | - | 11.3303 | 4.96E-05 | 0.615 | GTGGGTGGC |
| 6918 | mamo-s | *BGIBMGA008230* | down | 944 | 952 | - | 11.3303 | 4.96E-05 | 0.615 | GTGGGTGGC |
| 6919 | mamo-s | *BGIBMGA008231* | down | 1463 | 1471 | - | 11.2844 | 5.18E-05 | 0.63 | CTGCGTGCC |
| 6920 | mamo-s | *BGIBMGA008232* | down | 180 | 188 | + | 11.3303 | 4.96E-05 | 0.615 | GTGGGTGGC |
| 6921 | mamo-s | *BGIBMGA008233* | up | 671 | 679 | - | 11.3303 | 4.96E-05 | 0.615 | GTGGGTGGC |
| 6922 | mamo-s | *BGIBMGA008234* | up | 708 | 716 | + | 11.3303 | 4.96E-05 | 0.615 | GTGGGTGGC |
| 6923 | mamo-s | *BGIBMGA008234* | down | 1494 | 1502 | + | 11.3303 | 4.96E-05 | 0.615 | GTGGGTGGC |
| 6924 | mamo-s | *BGIBMGA008235* | down | 1426 | 1434 | - | 10.9358 | 7.30E-05 | 0.728 | GCGGGTGGC |
| 6925 | mamo-s | *BGIBMGA008236* | down | 324 | 332 | - | 11.3303 | 4.96E-05 | 0.615 | GTGGGTGGC |
| 6926 | mamo-s | *BGIBMGA008236* | down | 1549 | 1557 | + | 11.3303 | 4.96E-05 | 0.615 | GTGGGTGGC |
| 6927 | mamo-s | *BGIBMGA008236* | down | 602 | 610 | - | 10.7523 | 8.38E-05 | 0.797 | AAGCGTGGC |
| 6928 | mamo-s | *BGIBMGA008237* | up | 274 | 282 | + | 11.3303 | 4.96E-05 | 0.615 | GTGGGTGGC |
| 6929 | mamo-s | *BGIBMGA008237* | down | 1912 | 1920 | - | 11.3303 | 4.96E-05 | 0.615 | GTGGGTGGC |
| 6930 | mamo-s | *BGIBMGA008238* | down | 587 | 595 | - | 12.1284 | 2.16E-05 | 0.615 | GTGCGAGGC |
| 6931 | mamo-s | *BGIBMGA008240* | up | 205 | 213 | + | 11.7064 | 3.52E-05 | 0.615 | GCGCGTGGT |
| 6932 | mamo-s | *BGIBMGA008240* | down | 1971 | 1979 | + | 10.5138 | 9.75E-05 | 0.865 | GTGCGTGTT |
| 6933 | mamo-s | *BGIBMGA008241* | up | 530 | 538 | + | 10.5138 | 9.75E-05 | 0.865 | GTGCGTGTT |
| 6934 | mamo-s | *BGIBMGA008242* | up | 829 | 837 | + | 11.3303 | 4.96E-05 | 0.615 | GTGGGTGGC |
| 6935 | mamo-s | *BGIBMGA008243* | down | 762 | 770 | - | 11.7798 | 3.11E-05 | 0.615 | ATGCGTGGC |
| 6936 | mamo-s | *BGIBMGA008244* | up | 175 | 183 | + | 11.3303 | 4.96E-05 | 0.615 | GTGGGTGGC |
| 6937 | mamo-s | *BGIBMGA008244* | up | 1108 | 1116 | + | 11.3303 | 4.96E-05 | 0.615 | GTGGGTGGC |
| 6938 | mamo-s | *BGIBMGA008247* | down | 393 | 401 | - | 11.3303 | 4.96E-05 | 0.615 | GTGGGTGGC |
| 6939 | mamo-s | *BGIBMGA008248* | up | 1674 | 1682 | - | 11.3303 | 4.96E-05 | 0.615 | GTGGGTGGC |
| 6940 | mamo-s | *BGIBMGA008248* | down | 1905 | 1913 | + | 11.0826 | 6.84E-05 | 0.709 | GTACGTGGC |
| 6941 | mamo-s | *BGIBMGA008249* | up | 83 | 91 | + | 13.6606 | 4.06E-06 | 0.615 | GCGCGTGGC |
| 6942 | mamo-s | *BGIBMGA008249* | down | 1745 | 1753 | - | 11.3303 | 4.96E-05 | 0.615 | GTGGGTGGC |
| 6943 | mamo-s | *BGIBMGA008250* | down | 1084 | 1092 | - | 13.6606 | 4.06E-06 | 0.615 | GCGCGTGGC |
| 6944 | mamo-s | *BGIBMGA008251* | down | 587 | 595 | + | 10.8073 | 7.89E-05 | 0.766 | CCGCGTGGT |
| 6945 | mamo-s | *BGIBMGA008255* | down | 851 | 859 | - | 11.4404 | 4.51E-05 | 0.615 | CGGCGTGGC |

| | | | | | | | | | | |
|---|---|---|---|---|---|---|---|---|---|---|
| 6946 | mamo-s | *BGIBMGA008255* | down | 513 | 521 | - | 11.3303 | 4.96E-05 | 0.615 | GTGGGTGGC |
| 6947 | mamo-s | *BGIBMGA008256* | down | 830 | 838 | + | 11.3303 | 4.96E-05 | 0.615 | GTGGGTGGC |
| 6948 | mamo-s | *BGIBMGA008257* | up | 1574 | 1582 | + | 11.3303 | 4.96E-05 | 0.615 | GTGGGTGGC |
| 6949 | mamo-s | *BGIBMGA008259* | down | 1740 | 1748 | + | 11.3303 | 4.96E-05 | 0.615 | GTGGGTGGC |
| 6950 | mamo-s | *BGIBMGA008260* | up | 1502 | 1510 | - | 11.2294 | 5.41E-05 | 0.65 | CTGCGAGGC |
| 6951 | mamo-s | *BGIBMGA008261* | down | 222 | 230 | + | 11.3303 | 4.96E-05 | 0.615 | GTGGGTGGC |
| 6952 | mamo-s | *BGIBMGA008262* | down | 1666 | 1674 | + | 11.4404 | 4.51E-05 | 0.615 | CGGCGTGGC |
| 6953 | mamo-s | *BGIBMGA008262* | up | 500 | 508 | - | 11.3303 | 4.96E-05 | 0.615 | GTGGGTGGC |
| 6954 | mamo-s | *BGIBMGA008262* | up | 1479 | 1487 | - | 11.3303 | 4.96E-05 | 0.615 | GTGGGTGGC |
| 6955 | mamo-s | *BGIBMGA008263* | down | 1325 | 1333 | + | 11.3303 | 4.96E-05 | 0.615 | GTGGGTGGC |
| 6956 | mamo-s | *BGIBMGA008264* | down | 1420 | 1428 | - | 11.3303 | 4.96E-05 | 0.615 | GTGGGTGGC |
| 6957 | mamo-s | *BGIBMGA008265* | down | 1682 | 1690 | + | 11.3303 | 4.96E-05 | 0.615 | GTGGGTGGC |
| 6958 | mamo-s | *BGIBMGA008268* | up | 1385 | 1393 | - | 11.3303 | 4.96E-05 | 0.615 | GTGGGTGGC |
| 6959 | mamo-s | *BGIBMGA008269* | down | 1141 | 1149 | - | 11.0826 | 6.84E-05 | 0.709 | GTACGTGGC |
| 6960 | mamo-s | *BGIBMGA008270* | up | 813 | 821 | - | 11.3303 | 4.96E-05 | 0.615 | GTGGGTGGC |
| 6961 | mamo-s | *BGIBMGA008271* | up | 1497 | 1505 | - | 11.3303 | 4.96E-05 | 0.615 | GTGGGTGGC |
| 6962 | mamo-s | *BGIBMGA008272* | up | 1736 | 1744 | - | 11.3303 | 4.96E-05 | 0.615 | GTGGGTGGC |
| 6963 | mamo-s | *BGIBMGA008273* | down | 1661 | 1669 | + | 11.3303 | 4.96E-05 | 0.615 | GTGGGTGGC |
| 6964 | mamo-s | *BGIBMGA008273* | down | 929 | 937 | + | 10.5688 | 8.88E-05 | 0.824 | GTGCGTGAC |
| 6965 | mamo-s | *BGIBMGA008274* | up | 255 | 263 | - | 11.3303 | 4.96E-05 | 0.615 | GTGGGTGGC |
| 6966 | mamo-s | *BGIBMGA008274* | down | 1043 | 1051 | - | 11.3303 | 4.96E-05 | 0.615 | GTGGGTGGC |
| 6967 | mamo-s | *BGIBMGA008275* | up | 1436 | 1444 | + | 10.7523 | 8.38E-05 | 0.797 | AAGCGTGGC |
| 6968 | mamo-s | *BGIBMGA008276* | down | 571 | 579 | - | 11.789 | 2.84E-05 | 0.615 | GCGCGTGCC |
| 6969 | mamo-s | *BGIBMGA008276* | down | 1835 | 1843 | + | 11.3303 | 4.96E-05 | 0.615 | GTGGGTGGC |
| 6970 | mamo-s | *BGIBMGA008276* | down | 565 | 573 | + | 11.0826 | 6.84E-05 | 0.709 | GTACGTGGC |
| 6971 | mamo-s | *BGIBMGA008276* | down | 888 | 896 | + | 10.8899 | 7.48E-05 | 0.739 | CCGCGTGCC |
| 6972 | mamo-s | *BGIBMGA008276* | up | 1973 | 1981 | + | 10.8899 | 7.48E-05 | 0.739 | CCGCGTGCC |
| 6973 | mamo-s | *BGIBMGA008277* | up | 1737 | 1745 | - | 11.789 | 2.84E-05 | 0.615 | GCGCGTGCC |
| 6974 | mamo-s | *BGIBMGA008277* | up | 997 | 1005 | - | 11.7064 | 3.52E-05 | 0.615 | GCGCGTGGT |
| 6975 | mamo-s | *BGIBMGA008277* | up | 793 | 801 | - | 11.3303 | 4.96E-05 | 0.615 | GTGGGTGGC |
| 6976 | mamo-s | *BGIBMGA008277* | up | 1731 | 1739 | + | 11.0826 | 6.84E-05 | 0.709 | GTACGTGGC |
| 6977 | mamo-s | *BGIBMGA008278* | up | 1267 | 1275 | - | 11.1376 | 6.35E-05 | 0.709 | TCGCGTGGC |
| 6978 | mamo-s | *BGIBMGA008280* | down | 1356 | 1364 | + | 13.0275 | 8.53E-06 | 0.615 | GAGCGTGGC |
| 6979 | mamo-s | *BGIBMGA008281* | up | 1393 | 1401 | + | 11.3303 | 4.96E-05 | 0.615 | GTGGGTGGC |
| 6980 | mamo-s | *BGIBMGA008282* | down | 570 | 578 | - | 11.3303 | 4.96E-05 | 0.615 | GTGGGTGGC |
| 6981 | mamo-s | *BGIBMGA008282* | down | 1163 | 1171 | + | 11.3303 | 4.96E-05 | 0.615 | GTGGGTGGC |
| 6982 | mamo-s | *BGIBMGA008282* | down | 1592 | 1600 | + | 11.3303 | 4.96E-05 | 0.615 | GTGGGTGGC |
| 6983 | mamo-s | *BGIBMGA008284* | up | 1564 | 1572 | - | 11.5321 | 4.06E-05 | 0.615 | TTGCGTGGC |
| 6984 | mamo-s | *BGIBMGA008285* | down | 468 | 476 | - | 11.2294 | 5.41E-05 | 0.65 | CTGCGAGGC |
| 6985 | mamo-s | *BGIBMGA008287* | down | 568 | 576 | + | 11.3303 | 4.96E-05 | 0.615 | GTGGGTGGC |
| 6986 | mamo-s | *BGIBMGA008287* | up | 1676 | 1684 | + | 11.3303 | 4.96E-05 | 0.615 | GTGGGTGGC |
| 6987 | mamo-s | *BGIBMGA008288* | down | 1029 | 1037 | + | 11.1009 | 6.57E-05 | 0.709 | GAGCGAGGC |
| 6988 | mamo-s | *BGIBMGA008288* | down | 38 | 46 | + | 10.9358 | 7.30E-05 | 0.728 | GCGGGTGGC |
| 6989 | mamo-s | *BGIBMGA008289* | up | 929 | 937 | + | 12.3394 | 1.49E-05 | 0.615 | GGGCGTGGC |
| 6990 | mamo-s | *BGIBMGA008289* | down | 1802 | 1810 | + | 11.3303 | 4.96E-05 | 0.615 | GTGGGTGGC |
| 6991 | mamo-s | *BGIBMGA008289* | up | 953 | 961 | + | 10.9358 | 7.30E-05 | 0.728 | GCGGGTGGC |
| 6992 | mamo-s | *BGIBMGA008289* | up | 1096 | 1104 | + | 10.9358 | 7.30E-05 | 0.728 | GCGGGTGGC |
| 6993 | mamo-s | *BGIBMGA008294* | down | 381 | 389 | + | 12.7615 | 1.04E-05 | 0.615 | CCGCGTGGC |
| 6994 | mamo-s | *BGIBMGA008294* | down | 1960 | 1968 | + | 11.4404 | 4.51E-05 | 0.615 | CGGCGTGGC |
| 6995 | mamo-s | *BGIBMGA008296* | down | 412 | 420 | + | 12.1009 | 2.43E-05 | 0.615 | GTGCGTGGT |
| 6996 | mamo-s | *BGIBMGA008297* | up | 160 | 168 | - | 11.3303 | 4.96E-05 | 0.615 | GTGGGTGGC |
| 6997 | mamo-s | *BGIBMGA008299* | up | 926 | 934 | - | 11.2294 | 5.41E-05 | 0.65 | CTGCGAGGC |
| 6998 | mamo-s | *BGIBMGA008299* | down | 915 | 923 | + | 10.7523 | 8.38E-05 | 0.797 | GGGCGTGTC |
| 6999 | mamo-s | *BGIBMGA008300* | down | 51 | 59 | - | 11.1743 | 5.90E-05 | 0.684 | CCGCGTGTC |
| 7000 | mamo-s | *BGIBMGA008303* | down | 945 | 953 | + | 10.8349 | 7.66E-05 | 0.751 | CCGCGAGGC |
| 7001 | mamo-s | *BGIBMGA008304* | up | 1767 | 1775 | + | 10.8349 | 7.66E-05 | 0.751 | CCGCGAGGC |
| 7002 | mamo-s | *BGIBMGA008305* | down | 1141 | 1149 | - | 10.8899 | 7.48E-05 | 0.739 | CCGCGTGCC |
| 7003 | mamo-s | *BGIBMGA008305* | down | 377 | 385 | - | 10.8073 | 7.89E-05 | 0.766 | CCGCGTGGT |
| 7004 | mamo-s | *BGIBMGA008305* | up | 401 | 409 | + | 10.5138 | 9.75E-05 | 0.865 | GTGCGTGTT |
| 7005 | mamo-s | *BGIBMGA008306* | up | 1013 | 1021 | + | 11.7798 | 3.11E-05 | 0.615 | ATGCGTGGC |
| 7006 | mamo-s | *BGIBMGA008308* | up | 402 | 410 | - | 12.4679 | 1.31E-05 | 0.615 | GTGCGTGTC |
| 7007 | mamo-s | *BGIBMGA008308* | down | 392 | 400 | + | 11.3303 | 4.96E-05 | 0.615 | GTGGGTGGC |
| 7008 | mamo-s | *BGIBMGA008309* | up | 223 | 231 | - | 11.3303 | 4.96E-05 | 0.615 | GTGGGTGGC |
| 7009 | mamo-s | *BGIBMGA008309* | up | 1245 | 1253 | + | 11.3303 | 4.96E-05 | 0.615 | GTGGGTGGC |
| 7010 | mamo-s | *BGIBMGA008309* | down | 1571 | 1579 | - | 11.3303 | 4.96E-05 | 0.615 | GTGGGTGGC |
| 7011 | mamo-s | *BGIBMGA008311* | down | 891 | 899 | + | 10.5138 | 9.75E-05 | 0.865 | GTGCGTGTT |
| 7012 | mamo-s | *BGIBMGA008316* | up | 1574 | 1582 | + | 11.3303 | 4.96E-05 | 0.615 | GTGGGTGGC |
| 7013 | mamo-s | *BGIBMGA008318* | down | 1822 | 1830 | - | 12.3394 | 1.49E-05 | 0.615 | GGGCGTGGC |
| 7014 | mamo-s | *BGIBMGA008321* | up | 1968 | 1976 | + | 13.0275 | 8.53E-06 | 0.615 | GAGCGTGGC |
| 7015 | mamo-s | *BGIBMGA008321* | up | 1223 | 1231 | - | 11.3303 | 4.96E-05 | 0.615 | GTGGGTGGC |
| 7016 | mamo-s | *BGIBMGA008326* | up | 427 | 435 | - | 12.3394 | 1.49E-05 | 0.615 | GGGCGTGGC |
| 7017 | mamo-s | *BGIBMGA008326* | up | 209 | 217 | + | 11.3303 | 4.96E-05 | 0.615 | GTGGGTGGC |
| 7018 | mamo-s | *BGIBMGA008327* | down | 1846 | 1854 | + | 11.3303 | 4.96E-05 | 0.615 | GTGGGTGGC |
| 7019 | mamo-s | *BGIBMGA008327* | up | 10 | 18 | + | 10.7523 | 8.38E-05 | 0.797 | AAGCGTGGC |
| 7020 | mamo-s | *BGIBMGA008329* | up | 1563 | 1571 | - | 10.8349 | 7.66E-05 | 0.751 | CCGCGAGGC |
| 7021 | mamo-s | *BGIBMGA008330* | up | 915 | 923 | + | 11.3303 | 4.96E-05 | 0.615 | GTGGGTGGC |
| 7022 | mamo-s | *BGIBMGA008332* | up | 1032 | 1040 | - | 11.3303 | 4.96E-05 | 0.615 | GTGGGTGGC |
| 7023 | mamo-s | *BGIBMGA008333* | down | 817 | 825 | - | 12.1284 | 2.16E-05 | 0.615 | CAGCGTGGC |
| 7024 | mamo-s | *BGIBMGA008333* | down | 103 | 111 | - | 12.1009 | 2.43E-05 | 0.615 | GTGCGTGGT |
| 7025 | mamo-s | *BGIBMGA008336* | down | 109 | 117 | + | 11.3303 | 4.96E-05 | 0.615 | GTGGGTGGC |
| 7026 | mamo-s | *BGIBMGA008337* | down | 282 | 290 | - | 12.1835 | 1.71E-05 | 0.615 | GTGCGTGCC |
| 7027 | mamo-s | *BGIBMGA008337* | down | 116 | 124 | - | 11.7339 | 3.29E-05 | 0.615 | GCGCGAGGC |
| 7028 | mamo-s | *BGIBMGA008339* | up | 1150 | 1158 | - | 11.3303 | 4.96E-05 | 0.615 | GTGGGTGGC |
| 7029 | mamo-s | *BGIBMGA008343* | down | 860 | 868 | - | 13.6606 | 4.06E-06 | 0.615 | GCGCGTGGC |
| 7030 | mamo-s | *BGIBMGA008346* | down | 1075 | 1083 | + | 11.7798 | 3.11E-05 | 0.615 | ATGCGTGGC |
| 7031 | mamo-s | *BGIBMGA008349* | up | 1291 | 1299 | + | 11.3303 | 4.96E-05 | 0.615 | GTGGGTGGC |
| 7032 | mamo-s | *BGIBMGA008349* | down | 1417 | 1425 | - | 11.3303 | 4.96E-05 | 0.615 | GTGGGTGGC |
| 7033 | mamo-s | *BGIBMGA008349* | up | 967 | 975 | - | 11.0826 | 6.84E-05 | 0.709 | GTACGTGGC |

| | | | | | | | | | | |
|---|---|---|---|---|---|---|---|---|---|---|
| 7034 | mamo-s | *BGIBMGA008350* | up | 1306 | 1314 | - | 11.1376 | 6.35E-05 | 0.709 | TCGCGTGGC |
| 7035 | mamo-s | *BGIBMGA008351* | up | 843 | 851 | - | 11.0826 | 6.84E-05 | 0.709 | GTACGTGGC |
| 7036 | mamo-s | *BGIBMGA008352* | up | 1573 | 1581 | + | 11.0826 | 6.84E-05 | 0.709 | GTACGTGGC |
| 7037 | mamo-s | *BGIBMGA008353* | up | 341 | 349 | + | 11.3303 | 4.96E-05 | 0.615 | GTGGGTGGC |
| 7038 | mamo-s | *BGIBMGA008353* | down | 1338 | 1346 | - | 11.3303 | 4.96E-05 | 0.615 | GTGGGTGGC |
| 7039 | mamo-s | *BGIBMGA008354* | up | 442 | 450 | - | 11.3303 | 4.96E-05 | 0.615 | GTGGGTGGC |
| 7040 | mamo-s | *BGIBMGA008354* | down | 625 | 633 | - | 11.3303 | 4.96E-05 | 0.615 | GTGGGTGGC |
| 7041 | mamo-s | *BGIBMGA008355* | up | 800 | 808 | + | 11.3303 | 4.96E-05 | 0.615 | GTGGGTGGC |
| 7042 | mamo-s | *BGIBMGA008356* | up | 1742 | 1750 | - | 10.5138 | 9.75E-05 | 0.865 | GTGCGTGTT |
| 7043 | mamo-s | *BGIBMGA008359* | up | 736 | 744 | + | 14.055 | 2.23E-06 | 0.615 | GTGCGTGGC |
| 7044 | mamo-s | *BGIBMGA008359* | down | 1929 | 1937 | + | 11.3303 | 4.96E-05 | 0.615 | GTGGGTGGC |
| 7045 | mamo-s | *BGIBMGA008360* | down | 1154 | 1162 | - | 11.3303 | 4.96E-05 | 0.615 | GTGGGTGGC |
| 7046 | mamo-s | *BGIBMGA008361* | up | 1747 | 1755 | + | 11.3303 | 4.96E-05 | 0.615 | GTGGGTGGC |
| 7047 | mamo-s | *BGIBMGA008361* | down | 795 | 803 | - | 11.0826 | 6.84E-05 | 0.709 | GTACGTGGC |
| 7048 | mamo-s | *BGIBMGA008362* | down | 728 | 736 | - | 10.5413 | 9.42E-05 | 0.857 | GTGCGAGTC |
| 7049 | mamo-s | *BGIBMGA008365* | up | 461 | 469 | + | 11.3303 | 4.96E-05 | 0.615 | GTGGGTGGC |
| 7050 | mamo-s | *BGIBMGA008366* | up | 1779 | 1787 | - | 11.3303 | 4.96E-05 | 0.615 | GTGGGTGGC |
| 7051 | mamo-s | *BGIBMGA008368* | down | 1949 | 1957 | - | 11.3303 | 4.96E-05 | 0.615 | GTGGGTGGC |
| 7052 | mamo-s | *BGIBMGA008368* | up | 1237 | 1245 | + | 11.0734 | 7.12E-05 | 0.726 | GAGCGTGGT |
| 7053 | mamo-s | *BGIBMGA008369* | down | 1413 | 1421 | + | 11.2294 | 5.41E-05 | 0.65 | CTGCGAGGC |
| 7054 | mamo-s | *BGIBMGA008369* | up | 1452 | 1460 | - | 11.1743 | 5.90E-05 | 0.684 | CCGCGTGTC |
| 7055 | mamo-s | *BGIBMGA008371* | down | 700 | 708 | + | 11.3303 | 4.96E-05 | 0.615 | GTGGGTGGC |
| 7056 | mamo-s | *BGIBMGA008373* | down | 375 | 383 | + | 14.055 | 2.23E-06 | 0.615 | GTGCGTGGC |
| 7057 | mamo-s | *BGIBMGA008374* | up | 177 | 185 | - | 12.1009 | 2.43E-05 | 0.615 | GTGCGTGGT |
| 7058 | mamo-s | *BGIBMGA008375* | down | 1353 | 1361 | + | 11.3303 | 4.96E-05 | 0.615 | GTGGGTGGC |
| 7059 | mamo-s | *BGIBMGA008375* | up | 101 | 109 | - | 11.0826 | 6.84E-05 | 0.709 | GTACGTGGC |
| 7060 | mamo-s | *BGIBMGA008376* | up | 1178 | 1186 | - | 12.0734 | 2.66E-05 | 0.615 | GCGCGTGTC |
| 7061 | mamo-s | *BGIBMGA008376* | up | 1299 | 1307 | - | 11.3853 | 4.74E-05 | 0.615 | ACGCGTGGC |
| 7062 | mamo-s | *BGIBMGA008379* | up | 1346 | 1354 | - | 12.4679 | 1.31E-05 | 0.615 | GTGCGTGTC |
| 7063 | mamo-s | *BGIBMGA008379* | up | 718 | 726 | + | 11.3303 | 4.96E-05 | 0.615 | GTGGGTGGC |
| 7064 | mamo-s | *BGIBMGA008380* | down | 1130 | 1138 | - | 11.3303 | 4.96E-05 | 0.615 | GTGGGTGGC |
| 7065 | mamo-s | *BGIBMGA008381* | up | 407 | 415 | - | 11.3303 | 4.96E-05 | 0.615 | GTGGGTGGC |
| 7066 | mamo-s | *BGIBMGA008381* | down | 654 | 662 | + | 11.0826 | 6.84E-05 | 0.709 | GTACGTGGC |
| 7067 | mamo-s | *BGIBMGA008382* | down | 75 | 83 | + | 11.5321 | 4.06E-05 | 0.615 | TTGCGTGGC |
| 7068 | mamo-s | *BGIBMGA008383* | up | 1191 | 1199 | + | 11.3303 | 4.96E-05 | 0.615 | GTGGGTGGC |
| 7069 | mamo-s | *BGIBMGA008384* | down | 375 | 383 | + | 11.0826 | 6.84E-05 | 0.709 | GTACGTGGC |
| 7070 | mamo-s | *BGIBMGA008384* | up | 1579 | 1587 | - | 10.8349 | 7.66E-05 | 0.751 | CCGCGAGGC |
| 7071 | mamo-s | *BGIBMGA008387* | down | 761 | 769 | + | 11.3303 | 4.96E-05 | 0.615 | GTGGGTGGC |
| 7072 | mamo-s | *BGIBMGA008387* | up | 87 | 95 | - | 11.1743 | 5.90E-05 | 0.684 | CCGCGTGTC |
| 7073 | mamo-s | *BGIBMGA008389* | up | 1167 | 1175 | - | 11.5688 | 3.79E-05 | 0.615 | CTGCGTGTC |
| 7074 | mamo-s | *BGIBMGA008390* | up | 1727 | 1735 | - | 11.3303 | 4.96E-05 | 0.615 | GTGGGTGGC |
| 7075 | mamo-s | *BGIBMGA008392* | down | 1439 | 1447 | - | 12.1284 | 2.16E-05 | 0.615 | GTGCGAGGC |
| 7076 | mamo-s | *BGIBMGA008394* | up | 76 | 84 | - | 11.3303 | 4.96E-05 | 0.615 | GTGGGTGGC |
| 7077 | mamo-s | *BGIBMGA008394* | up | 1752 | 1760 | + | 11.0734 | 7.12E-05 | 0.726 | GAGCGTGGT |
| 7078 | mamo-s | *BGIBMGA008396* | down | 554 | 562 | + | 14.055 | 2.23E-06 | 0.615 | GTGCGTGGC |
| 7079 | mamo-s | *BGIBMGA008396* | up | 684 | 692 | + | 11.3303 | 4.96E-05 | 0.615 | GTGGGTGGC |
| 7080 | mamo-s | *BGIBMGA008396* | up | 182 | 190 | + | 10.5138 | 9.75E-05 | 0.865 | GTGCGTGTT |
| 7081 | mamo-s | *BGIBMGA008397* | up | 997 | 1005 | - | 11.0826 | 6.84E-05 | 0.709 | GTACGTGGC |
| 7082 | mamo-s | *BGIBMGA008398* | up | 1510 | 1518 | + | 11.0826 | 6.84E-05 | 0.709 | GTACGTGGC |
| 7083 | mamo-s | *BGIBMGA008400* | down | 646 | 654 | - | 11.3303 | 4.96E-05 | 0.615 | GTGGGTGGC |
| 7084 | mamo-s | *BGIBMGA008400* | up | 1749 | 1757 | - | 11.3303 | 4.96E-05 | 0.615 | GTGGGTGGC |
| 7085 | mamo-s | *BGIBMGA008404* | up | 296 | 304 | - | 11.3303 | 4.96E-05 | 0.615 | GTGGGTGGC |
| 7086 | mamo-s | *BGIBMGA008404* | up | 834 | 842 | + | 10.5688 | 8.88E-05 | 0.824 | GTGCGTGAC |
| 7087 | mamo-s | *BGIBMGA008406* | up | 1208 | 1216 | - | 11.3303 | 4.96E-05 | 0.615 | GTGGGTGGC |
| 7088 | mamo-s | *BGIBMGA008408* | up | 428 | 436 | + | 11.7798 | 3.11E-05 | 0.615 | ATGCGTGGC |
| 7089 | mamo-s | *BGIBMGA008408* | down | 513 | 521 | + | 11.3303 | 4.96E-05 | 0.615 | GTGGGTGGC |
| 7090 | mamo-s | *BGIBMGA008409* | down | 1615 | 1623 | - | 11.3303 | 4.96E-05 | 0.615 | GTGGGTGGC |
| 7091 | mamo-s | *BGIBMGA008411* | up | 1746 | 1754 | - | 11.3303 | 4.96E-05 | 0.615 | GTGGGTGGC |
| 7092 | mamo-s | *BGIBMGA008412* | down | 1034 | 1042 | + | 11.3303 | 4.96E-05 | 0.615 | GTGGGTGGC |
| 7093 | mamo-s | *BGIBMGA008413* | down | 9 | 17 | + | 11.1743 | 5.90E-05 | 0.684 | CCGCGTGTC |
| 7094 | mamo-s | *BGIBMGA008417* | up | 1749 | 1757 | + | 11.3303 | 4.96E-05 | 0.615 | GTGGGTGGC |
| 7095 | mamo-s | *BGIBMGA008418* | down | 631 | 639 | + | 11.7798 | 3.11E-05 | 0.615 | ATGCGTGGC |
| 7096 | mamo-s | *BGIBMGA008418* | down | 150 | 158 | + | 10.5138 | 9.75E-05 | 0.865 | GTGCGTGTT |
| 7097 | mamo-s | *BGIBMGA008419* | down | 355 | 363 | + | 11.3303 | 4.96E-05 | 0.615 | GTGGGTGGC |
| 7098 | mamo-s | *BGIBMGA008420* | down | 1766 | 1774 | + | 11.3303 | 4.96E-05 | 0.615 | GTGGGTGGC |
| 7099 | mamo-s | *BGIBMGA008423* | down | 1874 | 1882 | - | 14.055 | 2.23E-06 | 0.615 | GTGCGTGGC |
| 7100 | mamo-s | *BGIBMGA008423* | up | 141 | 149 | + | 11.3303 | 4.96E-05 | 0.615 | GTGGGTGGC |
| 7101 | mamo-s | *BGIBMGA008426* | down | 1776 | 1784 | + | 11.0826 | 6.84E-05 | 0.709 | GTACGTGGC |
| 7102 | mamo-s | *BGIBMGA008427* | up | 1094 | 1102 | + | 11.0826 | 6.84E-05 | 0.709 | GTACGTGGC |
| 7103 | mamo-s | *BGIBMGA008428* | down | 244 | 252 | - | 11.3303 | 4.96E-05 | 0.615 | GTGGGTGGC |
| 7104 | mamo-s | *BGIBMGA008430* | up | 1242 | 1250 | - | 10.7523 | 8.38E-05 | 0.797 | GGGCGTGTC |
| 7105 | mamo-s | *BGIBMGA008431* | down | 1127 | 1135 | - | 11.5321 | 4.06E-05 | 0.615 | TTGCGTGGC |
| 7106 | mamo-s | *BGIBMGA008431* | down | 1104 | 1112 | + | 11.1009 | 6.57E-05 | 0.709 | GAGCGAGGC |
| 7107 | mamo-s | *BGIBMGA008431* | down | 1569 | 1577 | - | 11.0826 | 6.84E-05 | 0.709 | GTACGTGGC |
| 7108 | mamo-s | *BGIBMGA008432* | down | 267 | 275 | - | 11.2018 | 5.68E-05 | 0.674 | CTGCGTGGT |
| 7109 | mamo-s | *BGIBMGA008434* | up | 1414 | 1422 | - | 11.3303 | 4.96E-05 | 0.615 | GTGGGTGGC |
| 7110 | mamo-s | *BGIBMGA008435* | up | 1342 | 1350 | + | 11.3303 | 4.96E-05 | 0.615 | GTGGGTGGC |
| 7111 | mamo-s | *BGIBMGA008437* | up | 618 | 626 | - | 11.3303 | 4.96E-05 | 0.615 | GTGGGTGGC |
| 7112 | mamo-s | *BGIBMGA008437* | down | 1435 | 1443 | - | 11.0826 | 6.84E-05 | 0.709 | GTACGTGGC |
| 7113 | mamo-s | *BGIBMGA008437* | down | 1951 | 1959 | + | 10.5138 | 9.75E-05 | 0.865 | GTGCGTGTT |
| 7114 | mamo-s | *BGIBMGA008438* | up | 617 | 625 | + | 11.3303 | 4.96E-05 | 0.615 | GTGGGTGGC |
| 7115 | mamo-s | *BGIBMGA008439* | up | 1040 | 1048 | - | 12.7615 | 1.04E-05 | 0.615 | CCGCGTGGC |
| 7116 | mamo-s | *BGIBMGA008439* | down | 1284 | 1292 | - | 11.3303 | 4.96E-05 | 0.615 | GTGGGTGGC |
| 7117 | mamo-s | *BGIBMGA008441* | up | 527 | 535 | + | 12.3394 | 1.49E-05 | 0.615 | GGGCGTGGC |
| 7118 | mamo-s | *BGIBMGA008442* | up | 859 | 867 | + | 11.3303 | 4.96E-05 | 0.615 | GTGGGTGGC |
| 7119 | mamo-s | *BGIBMGA008442* | up | 1061 | 1069 | - | 11.3303 | 4.96E-05 | 0.615 | GTGGGTGGC |
| 7120 | mamo-s | *BGIBMGA008443* | up | 898 | 906 | - | 11.4404 | 4.51E-05 | 0.615 | GAGCGTGTC |
| 7121 | mamo-s | *BGIBMGA008444* | down | 1824 | 1832 | + | 12.4679 | 1.31E-05 | 0.615 | GTGCGTGTC |

| | | | | | | | | | | |
|---|---|---|---|---|---|---|---|---|---|---|
| 7122 | mamo-s | *BGIBMGA008444* | up | 529 | 537 | - | 11.3303 | 4.96E-05 | 0.615 | GTGGGTGGC |
| 7123 | mamo-s | *BGIBMGA008446* | up | 1170 | 1178 | + | 11.5321 | 4.06E-05 | 0.615 | TTGCGTGGC |
| 7124 | mamo-s | *BGIBMGA008446* | down | 1595 | 1603 | + | 11.3303 | 4.96E-05 | 0.615 | GTGGGTGGC |
| 7125 | mamo-s | *BGIBMGA008446* | up | 411 | 419 | - | 10.5138 | 9.75E-05 | 0.865 | GTGCGTGTT |
| 7126 | mamo-s | *BGIBMGA008447* | up | 443 | 451 | + | 11.3303 | 4.96E-05 | 0.615 | GTGGGTGGC |
| 7127 | mamo-s | *BGIBMGA008449* | up | 1611 | 1619 | - | 11.3303 | 4.96E-05 | 0.615 | GTGGGTGGC |
| 7128 | mamo-s | *BGIBMGA008450* | up | 1494 | 1502 | - | 14.055 | 2.23E-06 | 0.615 | GTGCGTGGC |
| 7129 | mamo-s | *BGIBMGA008450* | up | 130 | 138 | + | 11.3303 | 4.96E-05 | 0.615 | GTGGGTGGC |
| 7130 | mamo-s | *BGIBMGA008451* | up | 236 | 244 | - | 11.3303 | 4.96E-05 | 0.615 | GTGGGTGGC |
| 7131 | mamo-s | *BGIBMGA008452* | up | 1791 | 1799 | + | 11.7064 | 3.52E-05 | 0.615 | GCGCGTGGT |
| 7132 | mamo-s | *BGIBMGA008453* | up | 1094 | 1102 | - | 11.3303 | 4.96E-05 | 0.615 | GTGGGTGGC |
| 7133 | mamo-s | *BGIBMGA008456* | down | 273 | 281 | + | 11.3303 | 4.96E-05 | 0.615 | GTGGGTGGC |
| 7134 | mamo-s | *BGIBMGA008458* | up | 26 | 34 | - | 11.3303 | 4.96E-05 | 0.615 | GTGGGTGGC |
| 7135 | mamo-s | *BGIBMGA008458* | up | 447 | 455 | - | 11.0826 | 6.84E-05 | 0.709 | GTACGTGGC |
| 7136 | mamo-s | *BGIBMGA008461* | up | 1991 | 1999 | + | 14.055 | 2.23E-06 | 0.615 | GTGCGTGGC |
| 7137 | mamo-s | *BGIBMGA008461* | up | 1764 | 1772 | + | 11.3303 | 4.96E-05 | 0.615 | GTGGGTGGC |
| 7138 | mamo-s | *BGIBMGA008465* | up | 656 | 664 | + | 11.3303 | 4.96E-05 | 0.615 | GTGGGTGGC |
| 7139 | mamo-s | *BGIBMGA008465* | down | 1258 | 1266 | + | 11.3303 | 4.96E-05 | 0.615 | GTGGGTGGC |
| 7140 | mamo-s | *BGIBMGA008465* | down | 1704 | 1712 | - | 11.3303 | 4.96E-05 | 0.615 | GTGGGTGGC |
| 7141 | mamo-s | *BGIBMGA008466* | up | 710 | 718 | + | 11.3303 | 4.96E-05 | 0.615 | GTGGGTGGC |
| 7142 | mamo-s | *BGIBMGA008466* | down | 1576 | 1584 | - | 10.8349 | 7.66E-05 | 0.751 | CCGCGAGGC |
| 7143 | mamo-s | *BGIBMGA008467* | down | 1668 | 1676 | - | 11.3303 | 4.96E-05 | 0.615 | GTGGGTGGC |
| 7144 | mamo-s | *BGIBMGA008468* | down | 943 | 951 | + | 11.3303 | 4.96E-05 | 0.615 | GTGGGTGGC |
| 7145 | mamo-s | *BGIBMGA008468* | up | 1687 | 1695 | - | 11.3303 | 4.96E-05 | 0.615 | GTGGGTGGC |
| 7146 | mamo-s | *BGIBMGA008468* | down | 1202 | 1210 | - | 11.1743 | 5.90E-05 | 0.684 | CCGCGTGTC |
| 7147 | mamo-s | *BGIBMGA008469* | down | 664 | 672 | + | 11.3303 | 4.96E-05 | 0.615 | GTGGGTGGC |
| 7148 | mamo-s | *BGIBMGA008469* | down | 1287 | 1295 | + | 11.3303 | 4.96E-05 | 0.615 | GTGGGTGGC |
| 7149 | mamo-s | *BGIBMGA008470* | down | 1051 | 1059 | + | 12.3394 | 1.49E-05 | 0.615 | GGGCGTGGC |
| 7150 | mamo-s | *BGIBMGA008470* | up | 1126 | 1134 | + | 11.3303 | 4.96E-05 | 0.615 | GTGGGTGGC |
| 7151 | mamo-s | *BGIBMGA008470* | up | 1749 | 1757 | + | 11.3303 | 4.96E-05 | 0.615 | GTGGGTGGC |
| 7152 | mamo-s | *BGIBMGA008471* | up | 1034 | 1042 | + | 11.3303 | 4.96E-05 | 0.615 | GTGGGTGGC |
| 7153 | mamo-s | *BGIBMGA008471* | down | 1816 | 1824 | - | 11.2844 | 5.18E-05 | 0.63 | CTGCGTGCC |
| 7154 | mamo-s | *BGIBMGA008474* | down | 802 | 810 | + | 11.3303 | 4.96E-05 | 0.615 | GTGGGTGGC |
| 7155 | mamo-s | *BGIBMGA008474* | up | 556 | 564 | - | 10.8349 | 7.66E-05 | 0.751 | CCGCGAGGC |
| 7156 | mamo-s | *BGIBMGA008474* | up | 1262 | 1270 | + | 10.7523 | 8.38E-05 | 0.797 | AAGCGTGGC |
| 7157 | mamo-s | *BGIBMGA008479* | down | 398 | 406 | - | 11.7798 | 3.11E-05 | 0.615 | ATGCGTGGC |
| 7158 | mamo-s | *BGIBMGA008479* | up | 588 | 596 | + | 11.3303 | 4.96E-05 | 0.615 | GTGGGTGGC |
| 7159 | mamo-s | *BGIBMGA008481* | up | 442 | 450 | + | 12.4679 | 1.31E-05 | 0.615 | GTGCGTGTC |
| 7160 | mamo-s | *BGIBMGA008481* | up | 1084 | 1092 | + | 12.3394 | 1.49E-05 | 0.615 | GGGCGTGGC |
| 7161 | mamo-s | *BGIBMGA008484* | down | 1601 | 1609 | - | 11.3303 | 4.96E-05 | 0.615 | GTGGGTGGC |
| 7162 | mamo-s | *BGIBMGA008486* | up | 85 | 93 | + | 12.0734 | 2.66E-05 | 0.615 | GCGCGTGTC |
| 7163 | mamo-s | *BGIBMGA008486* | up | 1222 | 1230 | - | 11.3303 | 4.96E-05 | 0.615 | GTGGGTGGC |
| 7164 | mamo-s | *BGIBMGA008486* | up | 1756 | 1764 | - | 11.3303 | 4.96E-05 | 0.615 | GTGGGTGGC |
| 7165 | mamo-s | *BGIBMGA008488* | down | 252 | 260 | + | 12.1009 | 2.43E-05 | 0.615 | GTGCGTGGT |
| 7166 | mamo-s | *BGIBMGA008488* | down | 1576 | 1584 | + | 11.3303 | 4.96E-05 | 0.615 | GTGGGTGGC |
| 7167 | mamo-s | *BGIBMGA008489* | up | 1720 | 1728 | - | 11.3303 | 4.96E-05 | 0.615 | GTGGGTGGC |
| 7168 | mamo-s | *BGIBMGA008490* | up | 1796 | 1804 | + | 11.3303 | 4.96E-05 | 0.615 | GTGGGTGGC |
| 7169 | mamo-s | *BGIBMGA008496* | down | 1861 | 1869 | + | 10.5138 | 9.75E-05 | 0.865 | GTGCGTGTT |
| 7170 | mamo-s | *BGIBMGA008499* | up | 1852 | 1860 | + | 10.9358 | 7.30E-05 | 0.728 | GCGGGTGGC |
| 7171 | mamo-s | *BGIBMGA008500* | up | 49 | 57 | + | 11.3303 | 4.96E-05 | 0.615 | GTGGGTGGC |
| 7172 | mamo-s | *BGIBMGA008502* | up | 1 | 9 | + | 12.0734 | 2.66E-05 | 0.615 | GCGCGTGTC |
| 7173 | mamo-s | *BGIBMGA008502* | down | 1562 | 1570 | + | 11.3303 | 4.96E-05 | 0.615 | GTGGGTGGC |
| 7174 | mamo-s | *BGIBMGA008506* | down | 1053 | 1061 | + | 11.3303 | 4.96E-05 | 0.615 | GTGGGTGGC |
| 7175 | mamo-s | *BGIBMGA008507* | up | 565 | 573 | + | 11.3303 | 4.96E-05 | 0.615 | GTGGGTGGC |
| 7176 | mamo-s | *BGIBMGA008507* | down | 1715 | 1723 | + | 10.8349 | 7.66E-05 | 0.751 | CCGCGAGGC |
| 7177 | mamo-s | *BGIBMGA008508* | up | 1901 | 1909 | - | 13.156 | 6.30E-06 | 0.615 | CTGCGTGGC |
| 7178 | mamo-s | *BGIBMGA008508* | down | 440 | 448 | + | 11.3303 | 4.96E-05 | 0.615 | GTGGGTGGC |
| 7179 | mamo-s | *BGIBMGA008509* | down | 877 | 885 | - | 11.3303 | 4.96E-05 | 0.615 | GTGGGTGGC |
| 7180 | mamo-s | *BGIBMGA008510* | up | 1301 | 1309 | + | 11.1009 | 6.57E-05 | 0.709 | GAGCGAGGC |
| 7181 | mamo-s | *BGIBMGA008511* | down | 732 | 740 | - | 11.3303 | 4.96E-05 | 0.615 | GTGGGTGGC |
| 7182 | mamo-s | *BGIBMGA008512* | up | 774 | 782 | - | 11.3303 | 4.96E-05 | 0.615 | GTGGGTGGC |
| 7183 | mamo-s | *BGIBMGA008513* | up | 1036 | 1044 | + | 12.1284 | 2.16E-05 | 0.615 | GTGCGAGGC |
| 7184 | mamo-s | *BGIBMGA008513* | down | 993 | 1001 | - | 11.3303 | 4.96E-05 | 0.615 | GTGGGTGGC |
| 7185 | mamo-s | *BGIBMGA008513* | up | 1605 | 1613 | + | 11.2844 | 5.18E-05 | 0.63 | CTGCGTGCC |
| 7186 | mamo-s | *BGIBMGA008515* | down | 1714 | 1722 | + | 12.3394 | 1.49E-05 | 0.615 | GGGCGTGGC |
| 7187 | mamo-s | *BGIBMGA008519* | up | 980 | 988 | - | 13.6606 | 4.06E-06 | 0.615 | GCGCGTGGC |
| 7188 | mamo-s | *BGIBMGA008520* | up | 641 | 649 | - | 10.7523 | 8.38E-05 | 0.797 | AAGCGTGGC |
| 7189 | mamo-s | *BGIBMGA008521* | up | 1728 | 1736 | + | 11.5321 | 4.06E-05 | 0.615 | TTGCGTGGC |
| 7190 | mamo-s | *BGIBMGA008525* | up | 610 | 618 | - | 10.7523 | 8.38E-05 | 0.797 | AAGCGTGGC |
| 7191 | mamo-s | *BGIBMGA008526* | down | 946 | 954 | - | 11.3303 | 4.96E-05 | 0.615 | GTGGGTGGC |
| 7192 | mamo-s | *BGIBMGA008527* | up | 312 | 320 | - | 11.3303 | 4.96E-05 | 0.615 | GTGGGTGGC |
| 7193 | mamo-s | *BGIBMGA008528* | down | 1940 | 1948 | - | 11.3303 | 4.96E-05 | 0.615 | GTGGGTGGC |
| 7194 | mamo-s | *BGIBMGA008530* | down | 589 | 597 | - | 12.4679 | 1.31E-05 | 0.615 | GTGCGTGTC |
| 7195 | mamo-s | *BGIBMGA008530* | up | 1980 | 1988 | - | 12.4679 | 1.31E-05 | 0.615 | GTGCGTGTC |
| 7196 | mamo-s | *BGIBMGA008531* | up | 819 | 827 | + | 11.3303 | 4.96E-05 | 0.615 | GTGGGTGGC |
| 7197 | mamo-s | *BGIBMGA008532* | up | 1305 | 1313 | + | 11.3303 | 4.96E-05 | 0.615 | GTGGGTGGC |
| 7198 | mamo-s | *BGIBMGA008533* | down | 1951 | 1959 | - | 11.4404 | 4.51E-05 | 0.615 | CGGCGTGGC |
| 7199 | mamo-s | *BGIBMGA008533* | down | 552 | 560 | + | 11.3303 | 4.96E-05 | 0.615 | GTGGGTGGC |
| 7200 | mamo-s | *BGIBMGA008533* | up | 1608 | 1616 | + | 11.2844 | 5.18E-05 | 0.63 | CTGCGTGCC |
| 7201 | mamo-s | *BGIBMGA008537* | down | 515 | 523 | + | 11.3303 | 4.96E-05 | 0.615 | GTGGGTGGC |
| 7202 | mamo-s | *BGIBMGA008537* | up | 1303 | 1311 | + | 11.1743 | 5.90E-05 | 0.684 | CCGCGTGTC |
| 7203 | mamo-s | *BGIBMGA008538* | up | 381 | 389 | - | 11.5688 | 3.79E-05 | 0.615 | CTGCGTGTC |
| 7204 | mamo-s | *BGIBMGA008538* | down | 357 | 365 | + | 11.3303 | 4.96E-05 | 0.615 | GTGGGTGGC |
| 7205 | mamo-s | *BGIBMGA008539* | up | 1306 | 1314 | - | 11.3303 | 4.96E-05 | 0.615 | GTGGGTGGC |
| 7206 | mamo-s | *BGIBMGA008540* | down | 284 | 292 | + | 14.055 | 2.23E-06 | 0.615 | GTGCGTGGC |
| 7207 | mamo-s | *BGIBMGA008541* | up | 859 | 867 | + | 11.3303 | 4.96E-05 | 0.615 | GTGGGTGGC |
| 7208 | mamo-s | *BGIBMGA008542* | down | 673 | 681 | + | 12.3394 | 1.49E-05 | 0.615 | GGGCGTGGC |
| 7209 | mamo-s | *BGIBMGA008542* | down | 350 | 358 | - | 11.7798 | 3.11E-05 | 0.615 | ATGCGTGGC |

| | | | | | | | | | | |
|---|---|---|---|---|---|---|---|---|---|---|
| 7210 | mamo-s | *BGIBMGA008543* | up | 1751 | 1759 | + | 11.2844 | 5.18E-05 | 0.63 | CTGCGTGCC |
| 7211 | mamo-s | *BGIBMGA008544* | down | 609 | 617 | - | 11.3303 | 4.96E-05 | 0.615 | GTGGGTGGC |
| 7212 | mamo-s | *BGIBMGA008548* | down | 175 | 183 | - | 11.3303 | 4.96E-05 | 0.615 | GTGGGTGGC |
| 7213 | mamo-s | *BGIBMGA008549* | up | 1510 | 1518 | - | 11.3303 | 4.96E-05 | 0.615 | GTGGGTGGC |
| 7214 | mamo-s | *BGIBMGA008551* | down | 1296 | 1304 | - | 11.3303 | 4.96E-05 | 0.615 | GTGGGTGGC |
| 7215 | mamo-s | *BGIBMGA008551* | down | 1582 | 1590 | + | 11.3303 | 4.96E-05 | 0.615 | GTGGGTGGC |
| 7216 | mamo-s | *BGIBMGA008552* | up | 292 | 300 | + | 11.3303 | 4.96E-05 | 0.615 | GTGGGTGGC |
| 7217 | mamo-s | *BGIBMGA008554* | down | 125 | 133 | - | 11.3303 | 4.96E-05 | 0.615 | GTGGGTGGC |
| 7218 | mamo-s | *BGIBMGA008559* | down | 843 | 851 | - | 10.5138 | 9.75E-05 | 0.865 | GTGCGTGTT |
| 7219 | mamo-s | *BGIBMGA008560* | down | 1028 | 1036 | - | 11.2844 | 5.18E-05 | 0.63 | CTGCGTGCC |
| 7220 | mamo-s | *BGIBMGA008560* | up | 831 | 839 | - | 10.5138 | 9.75E-05 | 0.865 | GTGCGTGTT |
| 7221 | mamo-s | *BGIBMGA008561* | down | 1689 | 1697 | + | 11.3303 | 4.96E-05 | 0.615 | GTGGGTGGC |
| 7222 | mamo-s | *BGIBMGA008562* | up | 1968 | 1976 | + | 11.7339 | 3.29E-05 | 0.615 | GCGCGAGGC |
| 7223 | mamo-s | *BGIBMGA008562* | down | 1293 | 1301 | + | 11.3303 | 4.96E-05 | 0.615 | GTGGGTGGC |
| 7224 | mamo-s | *BGIBMGA008562* | up | 1546 | 1554 | - | 11.1376 | 6.35E-05 | 0.709 | TCGCGTGGC |
| 7225 | mamo-s | *BGIBMGA008563* | up | 60 | 68 | + | 11.0734 | 7.12E-05 | 0.726 | GAGCGTGGT |
| 7226 | mamo-s | *BGIBMGA008564* | down | 219 | 227 | - | 11.3303 | 4.96E-05 | 0.615 | GTGGGTGGC |
| 7227 | mamo-s | *BGIBMGA008564* | down | 1000 | 1008 | + | 11.3303 | 4.96E-05 | 0.615 | GTGGGTGGC |
| 7228 | mamo-s | *BGIBMGA008565* | up | 1459 | 1467 | + | 13.156 | 6.30E-06 | 0.615 | CTGCGTGGC |
| 7229 | mamo-s | *BGIBMGA008569* | down | 537 | 545 | - | 12.1284 | 2.16E-05 | 0.615 | GTGCGAGGC |
| 7230 | mamo-s | *BGIBMGA008569* | down | 514 | 522 | + | 11.2018 | 5.68E-05 | 0.674 | CTGCGTGGT |
| 7231 | mamo-s | *BGIBMGA008570* | up | 757 | 765 | - | 11.3303 | 4.96E-05 | 0.615 | GTGGGTGGC |
| 7232 | mamo-s | *BGIBMGA008572* | down | 1120 | 1128 | - | 11.4404 | 4.51E-05 | 0.615 | GAGCGTGTC |
| 7233 | mamo-s | *BGIBMGA008575* | down | 851 | 859 | - | 11.3303 | 4.96E-05 | 0.615 | GTGGGTGGC |
| 7234 | mamo-s | *BGIBMGA008576* | down | 1443 | 1451 | - | 12.1284 | 2.16E-05 | 0.615 | CAGCGTGGC |
| 7235 | mamo-s | *BGIBMGA008576* | up | 1158 | 1166 | - | 11.3303 | 4.96E-05 | 0.615 | GTGGGTGGC |
| 7236 | mamo-s | *BGIBMGA008576* | up | 405 | 413 | - | 11.2018 | 5.68E-05 | 0.674 | CTGCGTGGT |
| 7237 | mamo-s | *BGIBMGA008578* | down | 1354 | 1362 | + | 11.3303 | 4.96E-05 | 0.615 | GTGGGTGGC |
| 7238 | mamo-s | *BGIBMGA008580* | up | 1618 | 1626 | + | 11.3303 | 4.96E-05 | 0.615 | GTGGGTGGC |
| 7239 | mamo-s | *BGIBMGA008580* | up | 900 | 908 | - | 11.0826 | 6.84E-05 | 0.709 | GTACGTGGC |
| 7240 | mamo-s | *BGIBMGA008580* | up | 253 | 261 | + | 11.0734 | 7.12E-05 | 0.726 | GAGCGTGGT |
| 7241 | mamo-s | *BGIBMGA008581* | down | 454 | 462 | + | 11.3303 | 4.96E-05 | 0.615 | GTGGGTGGC |
| 7242 | mamo-s | *BGIBMGA008584* | up | 1526 | 1534 | + | 11.1743 | 5.90E-05 | 0.684 | CCGCGTGTC |
| 7243 | mamo-s | *BGIBMGA008585* | down | 1597 | 1605 | - | 10.7523 | 8.38E-05 | 0.797 | GGGCGTGTC |
| 7244 | mamo-s | *BGIBMGA008586* | down | 399 | 407 | + | 11.3303 | 4.96E-05 | 0.615 | GTGGGTGGC |
| 7245 | mamo-s | *BGIBMGA008586* | up | 466 | 474 | - | 11.3303 | 4.96E-05 | 0.615 | GTGGGTGGC |
| 7246 | mamo-s | *BGIBMGA008589* | down | 1290 | 1298 | - | 10.5413 | 9.42E-05 | 0.857 | CAGCGTGTC |
| 7247 | mamo-s | *BGIBMGA008591* | down | 765 | 773 | + | 10.6881 | 8.61E-05 | 0.81 | GCACGTGGC |
| 7248 | mamo-s | *BGIBMGA008592* | down | 1925 | 1933 | - | 11.3303 | 4.96E-05 | 0.615 | GTGGGTGGC |
| 7249 | mamo-s | *BGIBMGA008593* | down | 1843 | 1851 | - | 12.0734 | 2.66E-05 | 0.615 | GCGCGTGTC |
| 7250 | mamo-s | *BGIBMGA008593* | up | 508 | 516 | + | 11.3303 | 4.96E-05 | 0.615 | GTGGGTGGC |
| 7251 | mamo-s | *BGIBMGA008593* | up | 1733 | 1741 | - | 10.5138 | 9.75E-05 | 0.865 | GTGCGTGTT |
| 7252 | mamo-s | *BGIBMGA008597* | down | 1638 | 1646 | - | 13.156 | 6.30E-06 | 0.615 | CTGCGTGGC |
| 7253 | mamo-s | *BGIBMGA008597* | up | 1788 | 1796 | + | 11.3303 | 4.96E-05 | 0.615 | GTGGGTGGC |
| 7254 | mamo-s | *BGIBMGA008597* | down | 1625 | 1633 | + | 11.2018 | 5.68E-05 | 0.674 | CTGCGTGGT |
| 7255 | mamo-s | *BGIBMGA008598* | down | 1094 | 1102 | - | 14.055 | 2.23E-06 | 0.615 | GTGCGTGGC |
| 7256 | mamo-s | *BGIBMGA008599* | down | 1181 | 1189 | + | 10.9358 | 7.30E-05 | 0.728 | GCGGGTGGC |
| 7257 | mamo-s | *BGIBMGA008600* | up | 1029 | 1037 | + | 11.3303 | 4.96E-05 | 0.615 | GTGGGTGGC |
| 7258 | mamo-s | *BGIBMGA008601* | up | 419 | 427 | + | 11.3303 | 4.96E-05 | 0.615 | GTGGGTGGC |
| 7259 | mamo-s | *BGIBMGA008601* | down | 759 | 767 | - | 11.3303 | 4.96E-05 | 0.615 | GTGGGTGGC |
| 7260 | mamo-s | *BGIBMGA008605* | down | 548 | 556 | + | 10.6881 | 8.61E-05 | 0.81 | GCACGTGGC |
| 7261 | mamo-s | *BGIBMGA008606* | up | 557 | 565 | + | 11.3303 | 4.96E-05 | 0.615 | GTGGGTGGC |
| 7262 | mamo-s | *BGIBMGA008606* | down | 912 | 920 | + | 11.3303 | 4.96E-05 | 0.615 | GTGGGTGGC |
| 7263 | mamo-s | *BGIBMGA008606* | down | 1312 | 1320 | - | 11.3303 | 4.96E-05 | 0.615 | GTGGGTGGC |
| 7264 | mamo-s | *BGIBMGA008607* | up | 4 | 12 | - | 11.3303 | 4.96E-05 | 0.615 | GTGGGTGGC |
| 7265 | mamo-s | *BGIBMGA008609* | down | 1287 | 1295 | - | 11.3303 | 4.96E-05 | 0.615 | GTGGGTGGC |
| 7266 | mamo-s | *BGIBMGA008610* | up | 943 | 951 | - | 11.3303 | 4.96E-05 | 0.615 | GTGGGTGGC |
| 7267 | mamo-s | *BGIBMGA008610* | up | 668 | 676 | + | 11.0826 | 6.84E-05 | 0.709 | GTACGTGGC |
| 7268 | mamo-s | *BGIBMGA008611* | down | 1227 | 1235 | + | 11.3303 | 4.96E-05 | 0.615 | GTGGGTGGC |
| 7269 | mamo-s | *BGIBMGA008612* | up | 796 | 804 | + | 11.3303 | 4.96E-05 | 0.615 | GTGGGTGGC |
| 7270 | mamo-s | *BGIBMGA008613* | down | 1457 | 1465 | - | 12.0734 | 2.66E-05 | 0.615 | GCGCGTGTC |
| 7271 | mamo-s | *BGIBMGA008613* | up | 780 | 788 | - | 11.3303 | 4.96E-05 | 0.615 | GTGGGTGGC |
| 7272 | mamo-s | *BGIBMGA008613* | up | 616 | 624 | - | 10.5413 | 9.42E-05 | 0.857 | GTGCGAGTC |
| 7273 | mamo-s | *BGIBMGA008615* | down | 412 | 420 | - | 13.6606 | 4.06E-06 | 0.615 | GCGCGTGGC |
| 7274 | mamo-s | *BGIBMGA008616* | down | 48 | 56 | + | 11.7064 | 3.52E-05 | 0.615 | GCGCGTGGT |
| 7275 | mamo-s | *BGIBMGA008616* | up | 747 | 755 | - | 11.4404 | 4.51E-05 | 0.615 | CGGCGTGGC |
| 7276 | mamo-s | *BGIBMGA008617* | down | 1666 | 1674 | - | 10.7523 | 8.38E-05 | 0.797 | GGGCGTGTC |
| 7277 | mamo-s | *BGIBMGA008618* | up | 579 | 587 | - | 11.7064 | 3.52E-05 | 0.615 | GCGCGTGGT |
| 7278 | mamo-s | *BGIBMGA008618* | up | 1810 | 1818 | - | 11.3303 | 4.96E-05 | 0.615 | GTGGGTGGC |
| 7279 | mamo-s | *BGIBMGA008622* | down | 777 | 785 | + | 12.1835 | 1.71E-05 | 0.615 | GTGCGTGCC |
| 7280 | mamo-s | *BGIBMGA008623* | up | 1417 | 1425 | + | 11.3303 | 4.96E-05 | 0.615 | GTGGGTGGC |
| 7281 | mamo-s | *BGIBMGA008624* | up | 560 | 568 | - | 11.3853 | 4.74E-05 | 0.615 | ACGCGTGGC |
| 7282 | mamo-s | *BGIBMGA008628* | up | 1325 | 1333 | + | 12.3394 | 1.49E-05 | 0.615 | GGGCGTGGC |
| 7283 | mamo-s | *BGIBMGA008629* | down | 1229 | 1237 | + | 12.3394 | 1.49E-05 | 0.615 | GGGCGTGGC |
| 7284 | mamo-s | *BGIBMGA008633* | down | 848 | 856 | + | 11.2844 | 5.18E-05 | 0.63 | CTGCGTGCC |
| 7285 | mamo-s | *BGIBMGA008634* | down | 669 | 677 | - | 14.055 | 2.23E-06 | 0.615 | GTGCGTGGC |
| 7286 | mamo-s | *BGIBMGA008636* | up | 1526 | 1534 | - | 11.3303 | 4.96E-05 | 0.615 | GTGGGTGGC |
| 7287 | mamo-s | *BGIBMGA008636* | down | 1684 | 1692 | + | 10.6881 | 8.61E-05 | 0.81 | GCACGTGGC |
| 7288 | mamo-s | *BGIBMGA008637* | up | 269 | 277 | - | 11.1743 | 5.90E-05 | 0.684 | CCGCGTGTC |
| 7289 | mamo-s | *BGIBMGA008637* | up | 279 | 287 | - | 10.8899 | 7.48E-05 | 0.739 | CCGCGTGCC |
| 7290 | mamo-s | *BGIBMGA008638* | up | 132 | 140 | - | 10.8349 | 7.66E-05 | 0.751 | CCGCGAGGC |
| 7291 | mamo-s | *BGIBMGA008639* | up | 671 | 679 | + | 10.6881 | 8.61E-05 | 0.81 | GCACGTGGC |
| 7292 | mamo-s | *BGIBMGA008644* | down | 1022 | 1030 | - | 11.3303 | 4.96E-05 | 0.615 | GTGGGTGGC |
| 7293 | mamo-s | *BGIBMGA008645* | up | 2 | 10 | - | 11.0734 | 7.12E-05 | 0.726 | GAGCGTGGT |
| 7294 | mamo-s | *BGIBMGA008646* | up | 624 | 632 | + | 11.3303 | 4.96E-05 | 0.615 | GTGGGTGGC |
| 7295 | mamo-s | *BGIBMGA008646* | up | 1479 | 1487 | - | 11.0826 | 6.84E-05 | 0.709 | GTACGTGGC |
| 7296 | mamo-s | *BGIBMGA008647* | down | 654 | 662 | - | 12.4679 | 1.31E-05 | 0.615 | GTGCGTGTC |
| 7297 | mamo-s | *BGIBMGA008648* | up | 1047 | 1055 | + | 11.3303 | 4.96E-05 | 0.615 | GTGGGTGGC |

| | | | | | | | | | | |
|---|---|---|---|---|---|---|---|---|---|---|
| 7298 | mamo-s | *BGIBMGA008651* | up | 945 | 953 | - | 11.3303 | 4.96E-05 | 0.615 | GTGGGTGGC |
| 7299 | mamo-s | *BGIBMGA008652* | down | 718 | 726 | - | 11.3303 | 4.96E-05 | 0.615 | GTGGGTGGC |
| 7300 | mamo-s | *BGIBMGA008653* | down | 1323 | 1331 | + | 11.3303 | 4.96E-05 | 0.615 | GTGGGTGGC |
| 7301 | mamo-s | *BGIBMGA008656* | down | 605 | 613 | + | 11.3303 | 4.96E-05 | 0.615 | GTGGGTGGC |
| 7302 | mamo-s | *BGIBMGA008656* | down | 87 | 95 | - | 10.7523 | 8.38E-05 | 0.797 | GGGCGTGTC |
| 7303 | mamo-s | *BGIBMGA008657* | up | 93 | 101 | + | 11.3303 | 4.96E-05 | 0.615 | GTGGGTGGC |
| 7304 | mamo-s | *BGIBMGA008659* | down | 1610 | 1618 | - | 11.1376 | 6.35E-05 | 0.709 | TCGCGTGGC |
| 7305 | mamo-s | *BGIBMGA008660* | up | 1162 | 1170 | - | 13.6606 | 4.06E-06 | 0.615 | GCGCGTGGC |
| 7306 | mamo-s | *BGIBMGA008661* | down | 1496 | 1504 | + | 10.7523 | 8.38E-05 | 0.797 | GGGCGTGTC |
| 7307 | mamo-s | *BGIBMGA008661* | down | 668 | 676 | - | 10.5413 | 9.42E-05 | 0.857 | GTGCGAGTC |
| 7308 | mamo-s | *BGIBMGA008663* | down | 1808 | 1816 | - | 11.3303 | 4.96E-05 | 0.615 | GTGGGTGGC |
| 7309 | mamo-s | *BGIBMGA008667* | up | 415 | 423 | - | 11.3303 | 4.96E-05 | 0.615 | GTGGGTGGC |
| 7310 | mamo-s | *BGIBMGA008667* | up | 1137 | 1145 | - | 11.3303 | 4.96E-05 | 0.615 | GTGGGTGGC |
| 7311 | mamo-s | *BGIBMGA008668* | up | 1358 | 1366 | - | 11.0826 | 6.84E-05 | 0.709 | GTACGTGGC |
| 7312 | mamo-s | *BGIBMGA008669* | down | 219 | 227 | + | 11.3303 | 4.96E-05 | 0.615 | GTGGGTGGC |
| 7313 | mamo-s | *BGIBMGA008669* | down | 1756 | 1764 | + | 11.3303 | 4.96E-05 | 0.615 | GTGGGTGGC |
| 7314 | mamo-s | *BGIBMGA008672* | up | 122 | 130 | - | 11.3303 | 4.96E-05 | 0.615 | GTGGGTGGC |
| 7315 | mamo-s | *BGIBMGA008672* | up | 708 | 716 | + | 11.3303 | 4.96E-05 | 0.615 | GTGGGTGGC |
| 7316 | mamo-s | *BGIBMGA008672* | down | 1732 | 1740 | - | 11.0826 | 6.84E-05 | 0.709 | GTACGTGGC |
| 7317 | mamo-s | *BGIBMGA008673* | up | 1057 | 1065 | - | 11.3303 | 4.96E-05 | 0.615 | GTGGGTGGC |
| 7318 | mamo-s | *BGIBMGA008680* | down | 30 | 38 | + | 10.5688 | 8.88E-05 | 0.824 | GTGCGTGAC |
| 7319 | mamo-s | *BGIBMGA008681* | down | 47 | 55 | + | 12.7615 | 1.04E-05 | 0.615 | CCGCGTGGC |
| 7320 | mamo-s | *BGIBMGA008681* | up | 1235 | 1243 | + | 12.1009 | 2.43E-05 | 0.615 | GTGCGTGGT |
| 7321 | mamo-s | *BGIBMGA008682* | down | 806 | 814 | - | 11.3303 | 4.96E-05 | 0.615 | GTGGGTGGC |
| 7322 | mamo-s | *BGIBMGA008682* | up | 165 | 173 | - | 10.9358 | 7.30E-05 | 0.728 | GCGGGTGGC |
| 7323 | mamo-s | *BGIBMGA008683* | up | 950 | 958 | + | 10.6881 | 8.61E-05 | 0.81 | GCACGTGGC |
| 7324 | mamo-s | *BGIBMGA008686* | down | 940 | 948 | + | 11.3303 | 4.96E-05 | 0.615 | GTGGGTGGC |
| 7325 | mamo-s | *BGIBMGA008687* | down | 1742 | 1750 | + | 12.7615 | 1.04E-05 | 0.615 | CCGCGTGGC |
| 7326 | mamo-s | *BGIBMGA008687* | up | 1315 | 1323 | + | 11.3303 | 4.96E-05 | 0.615 | GTGGGTGGC |
| 7327 | mamo-s | *BGIBMGA008687* | down | 1525 | 1533 | + | 11.3303 | 4.96E-05 | 0.615 | GTGGGTGGC |
| 7328 | mamo-s | *BGIBMGA008688* | up | 1301 | 1309 | + | 11.3303 | 4.96E-05 | 0.615 | GTGGGTGGC |
| 7329 | mamo-s | *BGIBMGA008693* | up | 890 | 898 | + | 11.3303 | 4.96E-05 | 0.615 | GTGGGTGGC |
| 7330 | mamo-s | *BGIBMGA008694* | down | 1003 | 1011 | - | 11.156 | 6.13E-05 | 0.705 | GAGCGTGCC |
| 7331 | mamo-s | *BGIBMGA008695* | up | 33 | 41 | + | 11.4404 | 4.51E-05 | 0.615 | CGGCGTGGC |
| 7332 | mamo-s | *BGIBMGA008696* | down | 707 | 715 | + | 11.2294 | 5.41E-05 | 0.65 | CTGCGAGGC |
| 7333 | mamo-s | *BGIBMGA008697* | up | 1152 | 1160 | + | 11.2294 | 5.41E-05 | 0.65 | CTGCGAGGC |
| 7334 | mamo-s | *BGIBMGA008697* | down | 948 | 956 | - | 11.2018 | 5.68E-05 | 0.674 | CTGCGTGGT |
| 7335 | mamo-s | *BGIBMGA008698* | down | 712 | 720 | + | 12.7615 | 1.04E-05 | 0.615 | CCGCGTGGC |
| 7336 | mamo-s | *BGIBMGA008698* | up | 1332 | 1340 | - | 11.2018 | 5.68E-05 | 0.674 | CTGCGTGGT |
| 7337 | mamo-s | *BGIBMGA008699* | up | 933 | 941 | - | 12.1284 | 2.16E-05 | 0.615 | GTGCGAGGC |
| 7338 | mamo-s | *BGIBMGA008699* | down | 1351 | 1359 | + | 11.3303 | 4.96E-05 | 0.615 | GTGGGTGGC |
| 7339 | mamo-s | *BGIBMGA008701* | up | 484 | 492 | - | 10.8073 | 7.89E-05 | 0.766 | CCGCGTGGT |
| 7340 | mamo-s | *BGIBMGA008702* | down | 172 | 180 | - | 12.3394 | 1.49E-05 | 0.615 | GGGCGTGGC |
| 7341 | mamo-s | *BGIBMGA008702* | up | 497 | 505 | + | 12.1284 | 2.16E-05 | 0.615 | GTGCGAGGC |
| 7342 | mamo-s | *BGIBMGA008702* | down | 334 | 342 | + | 11.3853 | 4.74E-05 | 0.615 | ACGCGTGGC |
| 7343 | mamo-s | *BGIBMGA008702* | down | 1375 | 1383 | + | 11.1743 | 5.90E-05 | 0.684 | CCGCGTGTC |
| 7344 | mamo-s | *BGIBMGA008703* | down | 1165 | 1173 | + | 11.4404 | 4.51E-05 | 0.615 | CGGCGTGGC |
| 7345 | mamo-s | *BGIBMGA008703* | up | 746 | 754 | + | 11.1009 | 6.57E-05 | 0.709 | GAGCGAGGC |
| 7346 | mamo-s | *BGIBMGA008704* | up | 576 | 584 | + | 11.4404 | 4.51E-05 | 0.615 | CGGCGTGGC |
| 7347 | mamo-s | *BGIBMGA008704* | up | 1713 | 1721 | - | 11.3303 | 4.96E-05 | 0.615 | GTGGGTGGC |
| 7348 | mamo-s | *BGIBMGA008706* | up | 31 | 39 | - | 11.3303 | 4.96E-05 | 0.615 | GTGGGTGGC |
| 7349 | mamo-s | *BGIBMGA008706* | down | 510 | 518 | + | 11.3303 | 4.96E-05 | 0.615 | GTGGGTGGC |
| 7350 | mamo-s | *BGIBMGA008707* | up | 149 | 157 | + | 11.3303 | 4.96E-05 | 0.615 | GTGGGTGGC |
| 7351 | mamo-s | *BGIBMGA008708* | up | 315 | 323 | - | 11.3303 | 4.96E-05 | 0.615 | GTGGGTGGC |
| 7352 | mamo-s | *BGIBMGA008708* | down | 1671 | 1679 | - | 11.0826 | 6.84E-05 | 0.709 | GTACGTGGC |
| 7353 | mamo-s | *BGIBMGA008709* | down | 947 | 955 | + | 11.3303 | 4.96E-05 | 0.615 | GTGGGTGGC |
| 7354 | mamo-s | *BGIBMGA008709* | down | 1400 | 1408 | - | 11.3303 | 4.96E-05 | 0.615 | GTGGGTGGC |
| 7355 | mamo-s | *BGIBMGA008711* | up | 937 | 945 | - | 11.3303 | 4.96E-05 | 0.615 | GTGGGTGGC |
| 7356 | mamo-s | *BGIBMGA008715* | down | 716 | 724 | + | 11.3303 | 4.96E-05 | 0.615 | GTGGGTGGC |
| 7357 | mamo-s | *BGIBMGA008716* | down | 1076 | 1084 | + | 12.1835 | 1.71E-05 | 0.615 | GTGCGTGCC |
| 7358 | mamo-s | *BGIBMGA008716* | up | 278 | 286 | + | 11.789 | 2.84E-05 | 0.615 | GCGCGTGCC |
| 7359 | mamo-s | *BGIBMGA008716* | down | 318 | 326 | + | 11.3303 | 4.96E-05 | 0.615 | GTGGGTGGC |
| 7360 | mamo-s | *BGIBMGA008716* | up | 349 | 357 | - | 10.9358 | 7.30E-05 | 0.728 | GCGGGTGGC |
| 7361 | mamo-s | *BGIBMGA008719* | down | 198 | 206 | + | 10.5138 | 9.75E-05 | 0.865 | GTGCGTGTT |
| 7362 | mamo-s | *BGIBMGA008722* | up | 1381 | 1389 | - | 11.3303 | 4.96E-05 | 0.615 | GTGGGTGGC |
| 7363 | mamo-s | *BGIBMGA008723* | down | 145 | 153 | + | 11.3303 | 4.96E-05 | 0.615 | GTGGGTGGC |
| 7364 | mamo-s | *BGIBMGA008724* | down | 1720 | 1728 | - | 11.3303 | 4.96E-05 | 0.615 | GTGGGTGGC |
| 7365 | mamo-s | *BGIBMGA008725* | down | 1342 | 1350 | - | 11.3303 | 4.96E-05 | 0.615 | GTGGGTGGC |
| 7366 | mamo-s | *BGIBMGA008727* | up | 1886 | 1894 | + | 11.3303 | 4.96E-05 | 0.615 | GTGGGTGGC |
| 7367 | mamo-s | *BGIBMGA008727* | down | 522 | 530 | + | 11.2844 | 5.18E-05 | 0.63 | CTGCGTGCC |
| 7368 | mamo-s | *BGIBMGA008731* | down | 181 | 189 | - | 11.3303 | 4.96E-05 | 0.615 | GTGGGTGGC |
| 7369 | mamo-s | *BGIBMGA008731* | up | 1422 | 1430 | + | 11.3303 | 4.96E-05 | 0.615 | GTGGGTGGC |
| 7370 | mamo-s | *BGIBMGA008732* | up | 594 | 602 | + | 11.3303 | 4.96E-05 | 0.615 | GTGGGTGGC |
| 7371 | mamo-s | *BGIBMGA008733* | up | 426 | 434 | + | 11.3303 | 4.96E-05 | 0.615 | GTGGGTGGC |
| 7372 | mamo-s | *BGIBMGA008734* | up | 220 | 228 | - | 11.3303 | 4.96E-05 | 0.615 | GTGGGTGGC |
| 7373 | mamo-s | *BGIBMGA008734* | down | 401 | 409 | + | 11.3303 | 4.96E-05 | 0.615 | GTGGGTGGC |
| 7374 | mamo-s | *BGIBMGA008734* | down | 979 | 987 | - | 11.0826 | 6.84E-05 | 0.709 | GTACGTGGC |
| 7375 | mamo-s | *BGIBMGA008736* | down | 686 | 694 | + | 11.3303 | 4.96E-05 | 0.615 | GTGGGTGGC |
| 7376 | mamo-s | *BGIBMGA008737* | up | 1508 | 1516 | - | 12.3394 | 1.49E-05 | 0.615 | GGGCGTGGC |
| 7377 | mamo-s | *BGIBMGA008738* | down | 667 | 675 | + | 12.7615 | 1.04E-05 | 0.615 | CCGCGTGGC |
| 7378 | mamo-s | *BGIBMGA008738* | up | 1543 | 1551 | - | 10.8899 | 7.48E-05 | 0.739 | CCGCGTGCC |
| 7379 | mamo-s | *BGIBMGA008739* | up | 756 | 764 | - | 10.8899 | 7.48E-05 | 0.739 | CCGCGTGCC |
| 7380 | mamo-s | *BGIBMGA008740* | up | 118 | 126 | + | 12.7615 | 1.04E-05 | 0.615 | CCGCGTGGC |
| 7381 | mamo-s | *BGIBMGA008740* | down | 148 | 156 | + | 11.3303 | 4.96E-05 | 0.615 | GTGGGTGGC |
| 7382 | mamo-s | *BGIBMGA008741* | down | 1845 | 1853 | - | 11.3303 | 4.96E-05 | 0.615 | GTGGGTGGC |
| 7383 | mamo-s | *BGIBMGA008742* | up | 27 | 35 | - | 11.3303 | 4.96E-05 | 0.615 | GTGGGTGGC |
| 7384 | mamo-s | *BGIBMGA008742* | down | 1916 | 1924 | - | 11.3303 | 4.96E-05 | 0.615 | GTGGGTGGC |
| 7385 | mamo-s | *BGIBMGA008742* | down | 589 | 597 | - | 11.0826 | 6.84E-05 | 0.709 | GTACGTGGC |

| | | | | | | | | | | |
|---|---|---|---|---|---|---|---|---|---|---|
| 7386 | mamo-s | *BGIBMGA008743* | up | 491 | 499 | - | 11.3303 | 4.96E-05 | 0.615 | GTGGGTGGC |
| 7387 | mamo-s | *BGIBMGA008743* | down | 340 | 348 | - | 10.5413 | 9.42E-05 | 0.857 | GTGCGAGTC |
| 7388 | mamo-s | *BGIBMGA008744* | down | 337 | 345 | - | 11.3303 | 4.96E-05 | 0.615 | GTGGGTGGC |
| 7389 | mamo-s | *BGIBMGA008744* | up | 594 | 602 | - | 10.5413 | 9.42E-05 | 0.857 | GTGCGAGTC |
| 7390 | mamo-s | *BGIBMGA008745* | down | 902 | 910 | + | 14.055 | 2.23E-06 | 0.615 | GTGCGTGGC |
| 7391 | mamo-s | *BGIBMGA008745* | down | 1873 | 1881 | + | 11.3303 | 4.96E-05 | 0.615 | GTGGGTGGC |
| 7392 | mamo-s | *BGIBMGA008747* | up | 339 | 347 | - | 11.3303 | 4.96E-05 | 0.615 | GTGGGTGGC |
| 7393 | mamo-s | *BGIBMGA008749* | down | 1478 | 1486 | - | 11.3303 | 4.96E-05 | 0.615 | GTGGGTGGC |
| 7394 | mamo-s | *BGIBMGA008750* | down | 257 | 265 | - | 11.1376 | 6.35E-05 | 0.709 | TCGCGTGGC |
| 7395 | mamo-s | *BGIBMGA008753* | up | 1132 | 1140 | - | 11.3303 | 4.96E-05 | 0.615 | GTGGGTGGC |
| 7396 | mamo-s | *BGIBMGA008753* | down | 1854 | 1862 | + | 11.0826 | 6.84E-05 | 0.709 | GTACGTGGC |
| 7397 | mamo-s | *BGIBMGA008754* | up | 193 | 201 | - | 11.3303 | 4.96E-05 | 0.615 | GTGGGTGGC |
| 7398 | mamo-s | *BGIBMGA008754* | up | 85 | 93 | + | 10.5138 | 9.75E-05 | 0.865 | GTGCGTGTT |
| 7399 | mamo-s | *BGIBMGA008759* | up | 137 | 145 | - | 11.0734 | 7.12E-05 | 0.726 | GAGCGTGGT |
| 7400 | mamo-s | *BGIBMGA008760* | up | 1707 | 1715 | - | 11.0826 | 6.84E-05 | 0.709 | GTACGTGGC |
| 7401 | mamo-s | *BGIBMGA008761* | down | 1232 | 1240 | + | 10.6881 | 8.61E-05 | 0.81 | GCACGTGGC |
| 7402 | mamo-s | *BGIBMGA008763* | down | 1465 | 1473 | + | 12.7615 | 1.04E-05 | 0.615 | CCGCGTGGC |
| 7403 | mamo-s | *BGIBMGA008766* | down | 413 | 421 | - | 11.3303 | 4.96E-05 | 0.615 | GTGGGTGGC |
| 7404 | mamo-s | *BGIBMGA008766* | up | 276 | 284 | - | 11.0826 | 6.84E-05 | 0.709 | GTACGTGGC |
| 7405 | mamo-s | *BGIBMGA008769* | down | 737 | 745 | + | 11.7339 | 3.29E-05 | 0.615 | GCGCGAGGC |
| 7406 | mamo-s | *BGIBMGA008769* | down | 471 | 479 | + | 11.3303 | 4.96E-05 | 0.615 | GTGGGTGGC |
| 7407 | mamo-s | *BGIBMGA008770* | up | 2 | 10 | + | 11.7339 | 3.29E-05 | 0.615 | GCGCGAGGC |
| 7408 | mamo-s | *BGIBMGA008773* | down | 148 | 156 | + | 11.3303 | 4.96E-05 | 0.615 | GTGGGTGGC |
| 7409 | mamo-s | *BGIBMGA008773* | up | 1579 | 1587 | - | 11.0734 | 7.12E-05 | 0.726 | GAGCGTGGT |
| 7410 | mamo-s | *BGIBMGA008775* | up | 1598 | 1606 | + | 10.9358 | 7.30E-05 | 0.728 | GCGGGTGGC |
| 7411 | mamo-s | *BGIBMGA008776* | down | 1429 | 1437 | - | 11.3853 | 4.74E-05 | 0.615 | ACGCGTGGC |
| 7412 | mamo-s | *BGIBMGA008777* | down | 406 | 414 | - | 11.5321 | 4.06E-05 | 0.615 | TTGCGTGGC |
| 7413 | mamo-s | *BGIBMGA008777* | up | 1890 | 1898 | - | 11.3853 | 4.74E-05 | 0.615 | ACGCGTGGC |
| 7414 | mamo-s | *BGIBMGA008778* | down | 550 | 558 | - | 11.1009 | 6.57E-05 | 0.709 | GAGCGAGGC |
| 7415 | mamo-s | *BGIBMGA008780* | up | 907 | 915 | - | 11.3303 | 4.96E-05 | 0.615 | GTGGGTGGC |
| 7416 | mamo-s | *BGIBMGA008781* | down | 311 | 319 | + | 14.055 | 2.23E-06 | 0.615 | GTGCGTGGC |
| 7417 | mamo-s | *BGIBMGA008781* | down | 531 | 539 | - | 12.1284 | 2.16E-05 | 0.615 | GTGCGAGGC |
| 7418 | mamo-s | *BGIBMGA008781* | down | 1888 | 1896 | + | 11.3303 | 4.96E-05 | 0.615 | GTGGGTGGC |
| 7419 | mamo-s | *BGIBMGA008782* | down | 1131 | 1139 | - | 11.7798 | 3.11E-05 | 0.615 | ATGCGTGGC |
| 7420 | mamo-s | *BGIBMGA008783* | down | 1024 | 1032 | + | 11.1743 | 5.90E-05 | 0.684 | CCGCGTGTC |
| 7421 | mamo-s | *BGIBMGA008784* | up | 1294 | 1302 | + | 11.1743 | 5.90E-05 | 0.684 | CCGCGTGTC |
| 7422 | mamo-s | *BGIBMGA008784* | down | 1322 | 1330 | - | 11.0826 | 6.84E-05 | 0.709 | GTACGTGGC |
| 7423 | mamo-s | *BGIBMGA008785* | down | 474 | 482 | + | 11.3303 | 4.96E-05 | 0.615 | GTGGGTGGC |
| 7424 | mamo-s | *BGIBMGA008786* | down | 1096 | 1104 | - | 11.3853 | 4.74E-05 | 0.615 | ACGCGTGGC |
| 7425 | mamo-s | *BGIBMGA008787* | up | 1516 | 1524 | - | 11.3303 | 4.96E-05 | 0.615 | GTGGGTGGC |
| 7426 | mamo-s | *BGIBMGA008788* | down | 1431 | 1439 | - | 10.8899 | 7.48E-05 | 0.739 | CCGCGTGCC |
| 7427 | mamo-s | *BGIBMGA008789* | up | 1321 | 1329 | - | 11.2018 | 5.68E-05 | 0.674 | CTGCGTGGT |
| 7428 | mamo-s | *BGIBMGA008789* | down | 707 | 715 | + | 10.8349 | 7.66E-05 | 0.751 | CCGCGAGGC |
| 7429 | mamo-s | *BGIBMGA008790* | down | 821 | 829 | - | 11.3303 | 4.96E-05 | 0.615 | GTGGGTGGC |
| 7430 | mamo-s | *BGIBMGA008791* | down | 1961 | 1969 | + | 11.3303 | 4.96E-05 | 0.615 | GTGGGTGGC |
| 7431 | mamo-s | *BGIBMGA008791* | up | 340 | 348 | + | 11.0826 | 6.84E-05 | 0.709 | GTACGTGGC |
| 7432 | mamo-s | *BGIBMGA008791* | up | 1386 | 1394 | - | 11.0826 | 6.84E-05 | 0.709 | GTACGTGGC |
| 7433 | mamo-s | *BGIBMGA008792* | down | 1595 | 1603 | + | 11.3303 | 4.96E-05 | 0.615 | GTGGGTGGC |
| 7434 | mamo-s | *BGIBMGA008793* | up | 1297 | 1305 | + | 11.3303 | 4.96E-05 | 0.615 | GTGGGTGGC |
| 7435 | mamo-s | *BGIBMGA008794* | up | 469 | 477 | + | 11.3853 | 4.74E-05 | 0.615 | ACGCGTGGC |
| 7436 | mamo-s | *BGIBMGA008795* | down | 1183 | 1191 | + | 11.3303 | 4.96E-05 | 0.615 | GTGGGTGGC |
| 7437 | mamo-s | *BGIBMGA008796* | up | 789 | 797 | + | 11.3303 | 4.96E-05 | 0.615 | GTGGGTGGC |
| 7438 | mamo-s | *BGIBMGA008799* | up | 1426 | 1434 | + | 11.3303 | 4.96E-05 | 0.615 | GTGGGTGGC |
| 7439 | mamo-s | *BGIBMGA008800* | up | 424 | 432 | + | 11.3303 | 4.96E-05 | 0.615 | GTGGGTGGC |
| 7440 | mamo-s | *BGIBMGA008801* | up | 197 | 205 | - | 11.4404 | 4.51E-05 | 0.615 | CGGCGTGGC |
| 7441 | mamo-s | *BGIBMGA008801* | down | 1166 | 1174 | + | 11.3303 | 4.96E-05 | 0.615 | GTGGGTGGC |
| 7442 | mamo-s | *BGIBMGA008801* | up | 126 | 134 | - | 10.9358 | 7.30E-05 | 0.728 | GCGGGTGGC |
| 7443 | mamo-s | *BGIBMGA008804* | up | 1624 | 1632 | - | 11.3303 | 4.96E-05 | 0.615 | GTGGGTGGC |
| 7444 | mamo-s | *BGIBMGA008804* | up | 1129 | 1137 | - | 11.0826 | 6.84E-05 | 0.709 | GTACGTGGC |
| 7445 | mamo-s | *BGIBMGA008807* | down | 1973 | 1981 | - | 11.0826 | 6.84E-05 | 0.709 | GTACGTGGC |
| 7446 | mamo-s | *BGIBMGA008813* | down | 1970 | 1978 | + | 10.5688 | 8.88E-05 | 0.824 | GTGCGTGAC |
| 7447 | mamo-s | *BGIBMGA008814* | up | 452 | 460 | + | 10.5688 | 8.88E-05 | 0.824 | GTGCGTGAC |
| 7448 | mamo-s | *BGIBMGA008815* | up | 1209 | 1217 | + | 11.3303 | 4.96E-05 | 0.615 | GTGGGTGGC |
| 7449 | mamo-s | *BGIBMGA008816* | down | 1399 | 1407 | + | 11.3303 | 4.96E-05 | 0.615 | GTGGGTGGC |
| 7450 | mamo-s | *BGIBMGA008816* | down | 517 | 525 | + | 11.0826 | 6.84E-05 | 0.709 | GTACGTGGC |
| 7451 | mamo-s | *BGIBMGA008818* | down | 1543 | 1551 | - | 11.3303 | 4.96E-05 | 0.615 | GTGGGTGGC |
| 7452 | mamo-s | *BGIBMGA008819* | down | 261 | 269 | - | 11.3303 | 4.96E-05 | 0.615 | GTGGGTGGC |
| 7453 | mamo-s | *BGIBMGA008820* | up | 424 | 432 | - | 11.3303 | 4.96E-05 | 0.615 | GTGGGTGGC |
| 7454 | mamo-s | *BGIBMGA008820* | up | 1991 | 1999 | + | 11.0734 | 7.12E-05 | 0.726 | GAGCGTGGT |
| 7455 | mamo-s | *BGIBMGA008821* | up | 1145 | 1153 | + | 11.5321 | 4.06E-05 | 0.615 | TTGCGTGGC |
| 7456 | mamo-s | *BGIBMGA008821* | up | 1385 | 1393 | - | 11.3303 | 4.96E-05 | 0.615 | GTGGGTGGC |
| 7457 | mamo-s | *BGIBMGA008822* | down | 1007 | 1015 | + | 11.3303 | 4.96E-05 | 0.615 | GTGGGTGGC |
| 7458 | mamo-s | *BGIBMGA008822* | down | 169 | 177 | + | 11.2018 | 5.68E-05 | 0.674 | CTGCGTGGT |
| 7459 | mamo-s | *BGIBMGA008822* | down | 542 | 550 | + | 10.9358 | 7.30E-05 | 0.728 | GCGGGTGGC |
| 7460 | mamo-s | *BGIBMGA008823* | up | 564 | 572 | + | 14.055 | 2.23E-06 | 0.615 | GTGCGTGGC |
| 7461 | mamo-s | *BGIBMGA008824* | up | 1612 | 1620 | - | 11.0826 | 6.84E-05 | 0.709 | GTACGTGGC |
| 7462 | mamo-s | *BGIBMGA008830* | down | 643 | 651 | + | 12.1009 | 2.43E-05 | 0.615 | GTGCGTGGT |
| 7463 | mamo-s | *BGIBMGA008832* | down | 1628 | 1636 | - | 11.3303 | 4.96E-05 | 0.615 | GTGGGTGGC |
| 7464 | mamo-s | *BGIBMGA008833* | down | 597 | 605 | + | 11.3303 | 4.96E-05 | 0.615 | GTGGGTGGC |
| 7465 | mamo-s | *BGIBMGA008834* | down | 71 | 79 | + | 10.5138 | 9.75E-05 | 0.865 | GTGCGTGTT |
| 7466 | mamo-s | *BGIBMGA008837* | down | 1562 | 1570 | + | 11.3303 | 4.96E-05 | 0.615 | GTGGGTGGC |
| 7467 | mamo-s | *BGIBMGA008838* | up | 372 | 380 | + | 11.2844 | 5.18E-05 | 0.63 | CTGCGTGCC |
| 7468 | mamo-s | *BGIBMGA008839* | down | 773 | 781 | - | 11.3303 | 4.96E-05 | 0.615 | GTGGGTGGC |
| 7469 | mamo-s | *BGIBMGA008839* | down | 1817 | 1825 | + | 11.3303 | 4.96E-05 | 0.615 | GTGGGTGGC |
| 7470 | mamo-s | *BGIBMGA008840* | up | 673 | 681 | + | 11.3303 | 4.96E-05 | 0.615 | GTGGGTGGC |
| 7471 | mamo-s | *BGIBMGA008840* | down | 1008 | 1016 | - | 11.3303 | 4.96E-05 | 0.615 | GTGGGTGGC |
| 7472 | mamo-s | *BGIBMGA008843* | up | 1420 | 1428 | + | 11.3303 | 4.96E-05 | 0.615 | GTGGGTGGC |
| 7473 | mamo-s | *BGIBMGA008845* | up | 1540 | 1548 | - | 12.4679 | 1.31E-05 | 0.615 | GTGCGTGTC |

| | | | | | | | | | | |
|---|---|---|---|---|---|---|---|---|---|---|
| 7474 | mamo-s | *BGIBMGA008845* | down | 1347 | 1355 | + | 11.5321 | 4.06E-05 | 0.615 | TTGCGTGGC |
| 7475 | mamo-s | *BGIBMGA008846* | up | 1014 | 1022 | + | 11.5688 | 3.79E-05 | 0.615 | CTGCGTGTC |
| 7476 | mamo-s | *BGIBMGA008847* | down | 1227 | 1235 | - | 11.3303 | 4.96E-05 | 0.615 | GTGGGTGGC |
| 7477 | mamo-s | *BGIBMGA008847* | up | 1404 | 1412 | - | 11.3303 | 4.96E-05 | 0.615 | GTGGGTGGC |
| 7478 | mamo-s | *BGIBMGA008847* | down | 1967 | 1975 | + | 11.0826 | 6.84E-05 | 0.709 | GTACGTGGC |
| 7479 | mamo-s | *BGIBMGA008849* | down | 1082 | 1090 | - | 11.3303 | 4.96E-05 | 0.615 | GTGGGTGGC |
| 7480 | mamo-s | *BGIBMGA008850* | up | 47 | 55 | + | 11.3303 | 4.96E-05 | 0.615 | GTGGGTGGC |
| 7481 | mamo-s | *BGIBMGA008850* | down | 1149 | 1157 | + | 11.3303 | 4.96E-05 | 0.615 | GTGGGTGGC |
| 7482 | mamo-s | *BGIBMGA008851* | down | 442 | 450 | + | 11.3853 | 4.74E-05 | 0.615 | ACGCGTGGC |
| 7483 | mamo-s | *BGIBMGA008854* | up | 1755 | 1763 | - | 12.1835 | 1.71E-05 | 0.615 | GTGCGTGCC |
| 7484 | mamo-s | *BGIBMGA008854* | up | 803 | 811 | + | 11.3303 | 4.96E-05 | 0.615 | GTGGGTGGC |
| 7485 | mamo-s | *BGIBMGA008854* | down | 1214 | 1222 | + | 11.3303 | 4.96E-05 | 0.615 | GTGGGTGGC |
| 7486 | mamo-s | *BGIBMGA008855* | down | 308 | 316 | - | 11.3303 | 4.96E-05 | 0.615 | GTGGGTGGC |
| 7487 | mamo-s | *BGIBMGA008855* | up | 546 | 554 | - | 11.3303 | 4.96E-05 | 0.615 | GTGGGTGGC |
| 7488 | mamo-s | *BGIBMGA008858* | up | 1782 | 1790 | + | 11.3303 | 4.96E-05 | 0.615 | GTGGGTGGC |
| 7489 | mamo-s | *BGIBMGA008861* | up | 161 | 169 | - | 11.3303 | 4.96E-05 | 0.615 | GTGGGTGGC |
| 7490 | mamo-s | *BGIBMGA008861* | down | 816 | 824 | + | 11.3303 | 4.96E-05 | 0.615 | GTGGGTGGC |
| 7491 | mamo-s | *BGIBMGA008861* | down | 1939 | 1947 | + | 11.3303 | 4.96E-05 | 0.615 | GTGGGTGGC |
| 7492 | mamo-s | *BGIBMGA008862* | up | 492 | 500 | + | 11.3303 | 4.96E-05 | 0.615 | GTGGGTGGC |
| 7493 | mamo-s | *BGIBMGA008862* | up | 1615 | 1623 | + | 11.3303 | 4.96E-05 | 0.615 | GTGGGTGGC |
| 7494 | mamo-s | *BGIBMGA008866* | up | 1049 | 1057 | + | 11.5688 | 3.79E-05 | 0.615 | CTGCGTGTC |
| 7495 | mamo-s | *BGIBMGA008866* | up | 1220 | 1228 | - | 10.8349 | 7.66E-05 | 0.751 | CCGCGAGGC |
| 7496 | mamo-s | *BGIBMGA008882* | down | 1648 | 1656 | + | 11.3853 | 4.74E-05 | 0.615 | ACGCGTGGC |
| 7497 | mamo-s | *BGIBMGA008883* | down | 237 | 245 | + | 11.4404 | 4.51E-05 | 0.615 | GAGCGTGTC |
| 7498 | mamo-s | *BGIBMGA008886* | up | 335 | 343 | - | 13.0275 | 8.53E-06 | 0.615 | GAGCGTGGC |
| 7499 | mamo-s | *BGIBMGA008888* | up | 1759 | 1767 | + | 10.5138 | 9.75E-05 | 0.865 | GTGCGTGTT |
| 7500 | mamo-s | *BGIBMGA008890* | up | 297 | 305 | + | 10.6881 | 8.61E-05 | 0.81 | GCACGTGGC |
| 7501 | mamo-s | *BGIBMGA008892* | up | 670 | 678 | + | 10.7523 | 8.38E-05 | 0.797 | GGGCGTGTC |
| 7502 | mamo-s | *BGIBMGA008894* | up | 1802 | 1810 | + | 12.4679 | 1.31E-05 | 0.615 | GTGCGTGTC |
| 7503 | mamo-s | *BGIBMGA008895* | down | 1443 | 1451 | + | 11.3303 | 4.96E-05 | 0.615 | GTGGGTGGC |
| 7504 | mamo-s | *BGIBMGA008896* | up | 1497 | 1505 | + | 11.3303 | 4.96E-05 | 0.615 | GTGGGTGGC |
| 7505 | mamo-s | *BGIBMGA008896* | down | 1170 | 1178 | + | 10.5688 | 8.88E-05 | 0.824 | GTGCGTGAC |
| 7506 | mamo-s | *BGIBMGA008898* | up | 268 | 276 | - | 13.6606 | 4.06E-06 | 0.615 | GCGCGTGGC |
| 7507 | mamo-s | *BGIBMGA008901* | up | 1213 | 1221 | + | 11.2294 | 5.41E-05 | 0.65 | CTGCGAGGC |
| 7508 | mamo-s | *BGIBMGA008902* | down | 37 | 45 | - | 13.156 | 6.30E-06 | 0.615 | CTGCGTGGC |
| 7509 | mamo-s | *BGIBMGA008904* | up | 1649 | 1657 | + | 11.0826 | 6.84E-05 | 0.709 | GTACGTGGC |
| 7510 | mamo-s | *BGIBMGA008905* | up | 649 | 657 | - | 11.4404 | 4.51E-05 | 0.615 | CGGCGTGGC |
| 7511 | mamo-s | *BGIBMGA008905* | down | 1091 | 1099 | - | 11.3303 | 4.96E-05 | 0.615 | GTGGGTGGC |
| 7512 | mamo-s | *BGIBMGA008907* | up | 1306 | 1314 | + | 11.3303 | 4.96E-05 | 0.615 | GTGGGTGGC |
| 7513 | mamo-s | *BGIBMGA008915* | down | 40 | 48 | - | 12.1284 | 2.16E-05 | 0.615 | GTGCGAGGC |
| 7514 | mamo-s | *BGIBMGA008918* | up | 126 | 134 | + | 11.7339 | 3.29E-05 | 0.615 | GCGCGAGGC |
| 7515 | mamo-s | *BGIBMGA008918* | down | 1392 | 1400 | - | 11.3303 | 4.96E-05 | 0.615 | GTGGGTGGC |
| 7516 | mamo-s | *BGIBMGA008920* | down | 1870 | 1878 | - | 12.1835 | 1.71E-05 | 0.615 | GTGCGTGCC |
| 7517 | mamo-s | *BGIBMGA008921* | down | 1293 | 1301 | - | 14.055 | 2.23E-06 | 0.615 | GTGCGTGGC |
| 7518 | mamo-s | *BGIBMGA008921* | up | 501 | 509 | - | 10.7523 | 8.38E-05 | 0.797 | AAGCGTGGC |
| 7519 | mamo-s | *BGIBMGA008923* | down | 1025 | 1033 | - | 12.1009 | 2.43E-05 | 0.615 | GTGCGTGGT |
| 7520 | mamo-s | *BGIBMGA008923* | up | 201 | 209 | - | 11.3303 | 4.96E-05 | 0.615 | GTGGGTGGC |
| 7521 | mamo-s | *BGIBMGA008923* | up | 797 | 805 | + | 11.2018 | 5.68E-05 | 0.674 | CTGCGTGGT |
| 7522 | mamo-s | *BGIBMGA008924* | up | 804 | 812 | - | 11.3303 | 4.96E-05 | 0.615 | GTGGGTGGC |
| 7523 | mamo-s | *BGIBMGA008924* | down | 1613 | 1621 | - | 10.5138 | 9.75E-05 | 0.865 | GTGCGTGTT |
| 7524 | mamo-s | *BGIBMGA008925* | down | 1939 | 1947 | - | 11.156 | 6.13E-05 | 0.705 | GAGCGTGCC |
| 7525 | mamo-s | *BGIBMGA008927* | down | 127 | 135 | - | 11.3303 | 4.96E-05 | 0.615 | GTGGGTGGC |
| 7526 | mamo-s | *BGIBMGA008930* | up | 739 | 747 | - | 11.1376 | 6.35E-05 | 0.709 | TCGCGTGGC |
| 7527 | mamo-s | *BGIBMGA008930* | down | 591 | 599 | + | 10.5413 | 9.42E-05 | 0.857 | GTGCGAGTC |
| 7528 | mamo-s | *BGIBMGA008930* | up | 720 | 728 | - | 10.5413 | 9.42E-05 | 0.857 | CAGCGTGTC |
| 7529 | mamo-s | *BGIBMGA008932* | up | 583 | 591 | + | 12.1009 | 2.43E-05 | 0.615 | GTGCGTGGT |
| 7530 | mamo-s | *BGIBMGA008933* | up | 275 | 283 | + | 11.3303 | 4.96E-05 | 0.615 | GTGGGTGGC |
| 7531 | mamo-s | *BGIBMGA008933* | down | 874 | 882 | + | 11.0826 | 6.84E-05 | 0.709 | GTACGTGGC |
| 7532 | mamo-s | *BGIBMGA008935* | down | 1191 | 1199 | - | 11.3303 | 4.96E-05 | 0.615 | GTGGGTGGC |
| 7533 | mamo-s | *BGIBMGA008935* | up | 1743 | 1751 | + | 11.3303 | 4.96E-05 | 0.615 | GTGGGTGGC |
| 7534 | mamo-s | *BGIBMGA008935* | down | 735 | 743 | - | 11.0826 | 6.84E-05 | 0.709 | GTACGTGGC |
| 7535 | mamo-s | *BGIBMGA008935* | down | 471 | 479 | + | 11.0734 | 7.12E-05 | 0.726 | GAGCGTGGT |
| 7536 | mamo-s | *BGIBMGA008937* | down | 1290 | 1298 | + | 11.3303 | 4.96E-05 | 0.615 | GTGGGTGGC |
| 7537 | mamo-s | *BGIBMGA008937* | down | 707 | 715 | + | 10.8073 | 7.89E-05 | 0.766 | CCGCGTGGT |
| 7538 | mamo-s | *BGIBMGA008939* | down | 331 | 339 | - | 11.0826 | 6.84E-05 | 0.709 | GTACGTGGC |
| 7539 | mamo-s | *BGIBMGA008940* | down | 376 | 384 | - | 10.9358 | 7.30E-05 | 0.728 | GCGGGTGGC |
| 7540 | mamo-s | *BGIBMGA008941* | up | 1922 | 1930 | - | 12.3394 | 1.49E-05 | 0.615 | GGGCGTGGC |
| 7541 | mamo-s | *BGIBMGA008942* | up | 1136 | 1144 | + | 11.3303 | 4.96E-05 | 0.615 | GTGGGTGGC |
| 7542 | mamo-s | *BGIBMGA008942* | down | 1496 | 1504 | + | 11.3303 | 4.96E-05 | 0.615 | GTGGGTGGC |
| 7543 | mamo-s | *BGIBMGA008943* | up | 678 | 686 | - | 11.3303 | 4.96E-05 | 0.615 | GTGGGTGGC |
| 7544 | mamo-s | *BGIBMGA008945* | up | 611 | 619 | + | 11.3303 | 4.96E-05 | 0.615 | GTGGGTGGC |
| 7545 | mamo-s | *BGIBMGA008945* | down | 1574 | 1582 | + | 11.3303 | 4.96E-05 | 0.615 | GTGGGTGGC |
| 7546 | mamo-s | *BGIBMGA008945* | up | 1839 | 1847 | + | 11.3303 | 4.96E-05 | 0.615 | GTGGGTGGC |
| 7547 | mamo-s | *BGIBMGA008946* | down | 1013 | 1021 | + | 11.0826 | 6.84E-05 | 0.709 | GTACGTGGC |
| 7548 | mamo-s | *BGIBMGA008947* | down | 244 | 252 | + | 11.7798 | 3.11E-05 | 0.615 | ATGCGTGGC |
| 7549 | mamo-s | *BGIBMGA008948* | down | 1844 | 1852 | + | 11.3303 | 4.96E-05 | 0.615 | GTGGGTGGC |
| 7550 | mamo-s | *BGIBMGA008949* | up | 1852 | 1860 | + | 11.3303 | 4.96E-05 | 0.615 | GTGGGTGGC |
| 7551 | mamo-s | *BGIBMGA008950* | down | 267 | 275 | - | 11.3303 | 4.96E-05 | 0.615 | GTGGGTGGC |
| 7552 | mamo-s | *BGIBMGA008952* | up | 801 | 809 | + | 11.3303 | 4.96E-05 | 0.615 | GTGGGTGGC |
| 7553 | mamo-s | *BGIBMGA008954* | down | 232 | 240 | + | 12.1009 | 2.43E-05 | 0.615 | GTGCGTGGT |
| 7554 | mamo-s | *BGIBMGA008955* | up | 168 | 176 | - | 11.3303 | 4.96E-05 | 0.615 | GTGGGTGGC |
| 7555 | mamo-s | *BGIBMGA008955* | down | 1898 | 1906 | + | 11.3303 | 4.96E-05 | 0.615 | GTGGGTGGC |
| 7556 | mamo-s | *BGIBMGA008958* | up | 648 | 656 | - | 10.5413 | 9.42E-05 | 0.857 | GTGCGAGTC |
| 7557 | mamo-s | *BGIBMGA008959* | up | 141 | 149 | + | 11.3303 | 4.96E-05 | 0.615 | GTGGGTGGC |
| 7558 | mamo-s | *BGIBMGA008959* | down | 1397 | 1405 | + | 11.3303 | 4.96E-05 | 0.615 | GTGGGTGGC |
| 7559 | mamo-s | *BGIBMGA008961* | up | 1893 | 1901 | - | 11.2294 | 5.41E-05 | 0.65 | CTGCGAGGC |
| 7560 | mamo-s | *BGIBMGA008963* | up | 527 | 535 | + | 12.4679 | 1.31E-05 | 0.615 | GTGCGTGTC |
| 7561 | mamo-s | *BGIBMGA008964* | down | 1162 | 1170 | + | 11.3303 | 4.96E-05 | 0.615 | GTGGGTGGC |

| | | | | | | | | | | |
|---|---|---|---|---|---|---|---|---|---|---|
| 7562 | mamo-s | *BGIBMGA008965* | up | 1806 | 1814 | + | 12.0734 | 2.66E-05 | 0.615 | GCGCGTGTC |
| 7563 | mamo-s | *BGIBMGA008968* | down | 1893 | 1901 | - | 11.5321 | 4.06E-05 | 0.615 | TTGCGTGGC |
| 7564 | mamo-s | *BGIBMGA008969* | down | 695 | 703 | - | 13.6606 | 4.06E-06 | 0.615 | GCGCGTGGC |
| 7565 | mamo-s | *BGIBMGA008970* | up | 1771 | 1779 | - | 13.6606 | 4.06E-06 | 0.615 | GCGCGTGGC |
| 7566 | mamo-s | *BGIBMGA008970* | up | 238 | 246 | + | 11.1743 | 5.90E-05 | 0.684 | CCGCGTGTC |
| 7567 | mamo-s | *BGIBMGA008976* | down | 1938 | 1946 | + | 11.3303 | 4.96E-05 | 0.615 | GTGGGTGGC |
| 7568 | mamo-s | *BGIBMGA008977* | up | 11 | 19 | - | 11.2294 | 5.41E-05 | 0.65 | CTGCGAGGC |
| 7569 | mamo-s | *BGIBMGA008980* | down | 348 | 356 | - | 11.5688 | 3.79E-05 | 0.615 | CTGCGTGTC |
| 7570 | mamo-s | *BGIBMGA008983* | down | 1287 | 1295 | + | 11.789 | 2.84E-05 | 0.615 | GCGCGTGCC |
| 7571 | mamo-s | *BGIBMGA008983* | down | 407 | 415 | + | 11.3303 | 4.96E-05 | 0.615 | GTGGGTGGC |
| 7572 | mamo-s | *BGIBMGA008984* | up | 333 | 341 | - | 11.3303 | 4.96E-05 | 0.615 | GTGGGTGGC |
| 7573 | mamo-s | *BGIBMGA008984* | up | 1759 | 1767 | + | 11.3303 | 4.96E-05 | 0.615 | GTGGGTGGC |
| 7574 | mamo-s | *BGIBMGA008991* | down | 205 | 213 | + | 14.055 | 2.23E-06 | 0.615 | GTGCGTGGC |
| 7575 | mamo-s | *BGIBMGA008992* | down | 1667 | 1675 | + | 10.5413 | 9.42E-05 | 0.857 | GTGCGAGTC |
| 7576 | mamo-s | *BGIBMGA008993* | down | 783 | 791 | - | 10.5688 | 8.88E-05 | 0.824 | GTGCGTGAC |
| 7577 | mamo-s | *BGIBMGA008994* | up | 684 | 692 | + | 11.3303 | 4.96E-05 | 0.615 | GTGGGTGGC |
| 7578 | mamo-s | *BGIBMGA008996* | up | 417 | 425 | - | 10.9358 | 7.30E-05 | 0.728 | GCGGGTGGC |
| 7579 | mamo-s | *BGIBMGA008999* | up | 1228 | 1236 | + | 14.055 | 2.23E-06 | 0.615 | GTGCGTGGC |
| 7580 | mamo-s | *BGIBMGA008999* | up | 1610 | 1618 | - | 11.3303 | 4.96E-05 | 0.615 | GTGGGTGGC |
| 7581 | mamo-s | *BGIBMGA009003* | up | 1457 | 1465 | + | 10.5688 | 8.88E-05 | 0.824 | GTGCGTGAC |
| 7582 | mamo-s | *BGIBMGA009004* | up | 1449 | 1457 | + | 11.2018 | 5.68E-05 | 0.674 | CTGCGTGGT |
| 7583 | mamo-s | *BGIBMGA009005* | up | 23 | 31 | - | 12.3394 | 1.49E-05 | 0.615 | GGGCGTGGC |
| 7584 | mamo-s | *BGIBMGA009005* | up | 16 | 24 | - | 12.0734 | 2.66E-05 | 0.615 | GCGCGTGTC |
| 7585 | mamo-s | *BGIBMGA009005* | up | 1653 | 1661 | + | 11.789 | 2.84E-05 | 0.615 | GCGCGTGCC |
| 7586 | mamo-s | *BGIBMGA009008* | down | 1533 | 1541 | - | 12.1835 | 1.71E-05 | 0.615 | GTGCGTGCC |
| 7587 | mamo-s | *BGIBMGA009009* | up | 1283 | 1291 | - | 11.156 | 6.13E-05 | 0.705 | GAGCGTGCC |
| 7588 | mamo-s | *BGIBMGA009016* | down | 1391 | 1399 | + | 11.2844 | 5.18E-05 | 0.63 | CTGCGTGCC |
| 7589 | mamo-s | *BGIBMGA009016* | down | 262 | 270 | - | 10.5138 | 9.75E-05 | 0.865 | GTGCGTGTT |
| 7590 | mamo-s | *BGIBMGA009017* | up | 1397 | 1405 | - | 12.1284 | 2.16E-05 | 0.615 | GTGCGAGGC |
| 7591 | mamo-s | *BGIBMGA009018* | down | 628 | 636 | - | 11.3303 | 4.96E-05 | 0.615 | GTGGGTGGC |
| 7592 | mamo-s | *BGIBMGA009022* | up | 142 | 150 | + | 11.3303 | 4.96E-05 | 0.615 | GTGGGTGGC |
| 7593 | mamo-s | *BGIBMGA009023* | up | 168 | 176 | + | 11.3303 | 4.96E-05 | 0.615 | GTGGGTGGC |
| 7594 | mamo-s | *BGIBMGA009024* | down | 1161 | 1169 | - | 11.3303 | 4.96E-05 | 0.615 | GTGGGTGGC |
| 7595 | mamo-s | *BGIBMGA009024* | up | 1842 | 1850 | + | 11.3303 | 4.96E-05 | 0.615 | GTGGGTGGC |
| 7596 | mamo-s | *BGIBMGA009029* | down | 168 | 176 | + | 11.3303 | 4.96E-05 | 0.615 | GTGGGTGGC |
| 7597 | mamo-s | *BGIBMGA009031* | up | 1551 | 1559 | - | 11.3303 | 4.96E-05 | 0.615 | GTGGGTGGC |
| 7598 | mamo-s | *BGIBMGA009036* | up | 947 | 955 | + | 11.3303 | 4.96E-05 | 0.615 | GTGGGTGGC |
| 7599 | mamo-s | *BGIBMGA009036* | up | 1697 | 1705 | + | 11.3303 | 4.96E-05 | 0.615 | GTGGGTGGC |
| 7600 | mamo-s | *BGIBMGA009037* | up | 945 | 953 | - | 11.5688 | 3.79E-05 | 0.615 | CTGCGTGTC |
| 7601 | mamo-s | *BGIBMGA009038* | up | 1160 | 1168 | + | 11.3303 | 4.96E-05 | 0.615 | GTGGGTGGC |
| 7602 | mamo-s | *BGIBMGA009039* | up | 58 | 66 | - | 11.0826 | 6.84E-05 | 0.709 | GTACGTGGC |
| 7603 | mamo-s | *BGIBMGA009040* | up | 960 | 968 | - | 11.4404 | 4.51E-05 | 0.615 | GAGCGTGTC |
| 7604 | mamo-s | *BGIBMGA009040* | up | 1078 | 1086 | - | 11.3303 | 4.96E-05 | 0.615 | GTGGGTGGC |
| 7605 | mamo-s | *BGIBMGA009040* | down | 1487 | 1495 | - | 11.3303 | 4.96E-05 | 0.615 | GTGGGTGGC |
| 7606 | mamo-s | *BGIBMGA009041* | down | 1792 | 1800 | - | 11.789 | 2.84E-05 | 0.615 | GCGCGTGCC |
| 7607 | mamo-s | *BGIBMGA009041* | down | 1595 | 1603 | - | 11.2294 | 5.41E-05 | 0.65 | CTGCGAGGC |
| 7608 | mamo-s | *BGIBMGA009042* | up | 406 | 414 | - | 11.2018 | 5.68E-05 | 0.674 | CTGCGTGGT |
| 7609 | mamo-s | *BGIBMGA009047* | up | 1539 | 1547 | - | 11.3303 | 4.96E-05 | 0.615 | GTGGGTGGC |
| 7610 | mamo-s | *BGIBMGA009049* | up | 530 | 538 | - | 11.4404 | 4.51E-05 | 0.615 | GAGCGTGTC |
| 7611 | mamo-s | *BGIBMGA009050* | down | 881 | 889 | - | 11.7339 | 3.29E-05 | 0.615 | GCGCGAGGC |
| 7612 | mamo-s | *BGIBMGA009050* | up | 580 | 588 | - | 11.3303 | 4.96E-05 | 0.615 | GTGGGTGGC |
| 7613 | mamo-s | *BGIBMGA009051* | up | 250 | 258 | - | 11.7339 | 3.29E-05 | 0.615 | GCGCGAGGC |
| 7614 | mamo-s | *BGIBMGA009051* | down | 1422 | 1430 | - | 11.0826 | 6.84E-05 | 0.709 | GTACGTGGC |
| 7615 | mamo-s | *BGIBMGA009052* | up | 689 | 697 | - | 11.0826 | 6.84E-05 | 0.709 | GTACGTGGC |
| 7616 | mamo-s | *BGIBMGA009053* | up | 96 | 104 | - | 11.4404 | 4.51E-05 | 0.615 | GAGCGTGTC |
| 7617 | mamo-s | *BGIBMGA009056* | down | 1795 | 1803 | - | 11.4404 | 4.51E-05 | 0.615 | GAGCGTGTC |
| 7618 | mamo-s | *BGIBMGA009056* | down | 1886 | 1894 | + | 10.5138 | 9.75E-05 | 0.865 | GTGCGTGTT |
| 7619 | mamo-s | *BGIBMGA009057* | up | 119 | 127 | - | 11.3303 | 4.96E-05 | 0.615 | GTGGGTGGC |
| 7620 | mamo-s | *BGIBMGA009060* | up | 1800 | 1808 | - | 11.4404 | 4.51E-05 | 0.615 | CGGCGTGGC |
| 7621 | mamo-s | *BGIBMGA009064* | down | 304 | 312 | - | 11.3303 | 4.96E-05 | 0.615 | GTGGGTGGC |
| 7622 | mamo-s | *BGIBMGA009065* | up | 131 | 139 | + | 11.3303 | 4.96E-05 | 0.615 | GTGGGTGGC |
| 7623 | mamo-s | *BGIBMGA009067* | down | 725 | 733 | - | 14.055 | 2.23E-06 | 0.615 | GTGCGTGGC |
| 7624 | mamo-s | *BGIBMGA009067* | up | 1569 | 1577 | + | 11.3303 | 4.96E-05 | 0.615 | GTGGGTGGC |
| 7625 | mamo-s | *BGIBMGA009067* | down | 1770 | 1778 | - | 11.3303 | 4.96E-05 | 0.615 | GTGGGTGGC |
| 7626 | mamo-s | *BGIBMGA009069* | up | 1871 | 1879 | + | 11.3303 | 4.96E-05 | 0.615 | GTGGGTGGC |
| 7627 | mamo-s | *BGIBMGA009070* | down | 480 | 488 | + | 11.3303 | 4.96E-05 | 0.615 | GTGGGTGGC |
| 7628 | mamo-s | *BGIBMGA009070* | up | 1788 | 1796 | + | 11.3303 | 4.96E-05 | 0.615 | GTGGGTGGC |
| 7629 | mamo-s | *BGIBMGA009071* | down | 1231 | 1239 | + | 11.3303 | 4.96E-05 | 0.615 | GTGGGTGGC |
| 7630 | mamo-s | *BGIBMGA009071* | up | 1861 | 1869 | + | 11.3303 | 4.96E-05 | 0.615 | GTGGGTGGC |
| 7631 | mamo-s | *BGIBMGA009071* | up | 699 | 707 | - | 10.8073 | 7.89E-05 | 0.766 | CCGCGTGGT |
| 7632 | mamo-s | *BGIBMGA009073* | up | 1110 | 1118 | - | 11.3303 | 4.96E-05 | 0.615 | GTGGGTGGC |
| 7633 | mamo-s | *BGIBMGA009073* | up | 1468 | 1476 | - | 11.3303 | 4.96E-05 | 0.615 | GTGGGTGGC |
| 7634 | mamo-s | *BGIBMGA009074* | up | 1867 | 1875 | - | 12.0734 | 2.66E-05 | 0.615 | GCGCGTGTC |
| 7635 | mamo-s | *BGIBMGA009075* | up | 1618 | 1626 | + | 11.2844 | 5.18E-05 | 0.63 | CTGCGTGCC |
| 7636 | mamo-s | *BGIBMGA009076* | down | 1691 | 1699 | - | 14.055 | 2.23E-06 | 0.615 | GTGCGTGGC |
| 7637 | mamo-s | *BGIBMGA009077* | up | 1425 | 1433 | + | 12.1009 | 2.43E-05 | 0.615 | GTGCGTGGT |
| 7638 | mamo-s | *BGIBMGA009077* | up | 1513 | 1521 | - | 11.0826 | 6.84E-05 | 0.709 | GTACGTGGC |
| 7639 | mamo-s | *BGIBMGA009078* | up | 1001 | 1009 | - | 11.3303 | 4.96E-05 | 0.615 | GTGGGTGGC |
| 7640 | mamo-s | *BGIBMGA009078* | up | 1448 | 1456 | - | 11.3303 | 4.96E-05 | 0.615 | GTGGGTGGC |
| 7641 | mamo-s | *BGIBMGA009080* | up | 161 | 169 | - | 11.3303 | 4.96E-05 | 0.615 | GTGGGTGGC |
| 7642 | mamo-s | *BGIBMGA009080* | up | 940 | 948 | + | 11.3303 | 4.96E-05 | 0.615 | GTGGGTGGC |
| 7643 | mamo-s | *BGIBMGA009082* | down | 139 | 147 | - | 11.3303 | 4.96E-05 | 0.615 | GTGGGTGGC |
| 7644 | mamo-s | *BGIBMGA009082* | up | 416 | 424 | + | 11.3303 | 4.96E-05 | 0.615 | GTGGGTGGC |
| 7645 | mamo-s | *BGIBMGA009082* | up | 725 | 733 | - | 11.3303 | 4.96E-05 | 0.615 | GTGGGTGGC |
| 7646 | mamo-s | *BGIBMGA009082* | up | 1165 | 1173 | - | 11.3303 | 4.96E-05 | 0.615 | GTGGGTGGC |
| 7647 | mamo-s | *BGIBMGA009082* | down | 1576 | 1584 | - | 11.3303 | 4.96E-05 | 0.615 | GTGGGTGGC |
| 7648 | mamo-s | *BGIBMGA009083* | down | 1219 | 1227 | + | 11.3303 | 4.96E-05 | 0.615 | GTGGGTGGC |
| 7649 | mamo-s | *BGIBMGA009084* | down | 1098 | 1106 | + | 11.3303 | 4.96E-05 | 0.615 | GTGGGTGGC |

| | | | | | | | | | | |
|---|---|---|---|---|---|---|---|---|---|---|
| 7650 | mamo-s | *BGIBMGA009084* | up | 1843 | 1851 | + | 11.3303 | 4.96E-05 | 0.615 | GTGGGTGGC |
| 7651 | mamo-s | *BGIBMGA009085* | up | 1512 | 1520 | + | 11.3303 | 4.96E-05 | 0.615 | GTGGGTGGC |
| 7652 | mamo-s | *BGIBMGA009087* | up | 1034 | 1042 | + | 11.3303 | 4.96E-05 | 0.615 | GTGGGTGGC |
| 7653 | mamo-s | *BGIBMGA009087* | up | 1641 | 1649 | - | 11.3303 | 4.96E-05 | 0.615 | GTGGGTGGC |
| 7654 | mamo-s | *BGIBMGA009087* | down | 1855 | 1863 | + | 11.3303 | 4.96E-05 | 0.615 | GTGGGTGGC |
| 7655 | mamo-s | *BGIBMGA009090* | down | 141 | 149 | - | 11.4404 | 4.51E-05 | 0.615 | GAGCGTGTC |
| 7656 | mamo-s | *BGIBMGA009091* | up | 421 | 429 | - | 12.1284 | 2.16E-05 | 0.615 | CAGCGTGGC |
| 7657 | mamo-s | *BGIBMGA009094* | up | 1265 | 1273 | - | 11.2844 | 5.18E-05 | 0.63 | CTGCGTGCC |
| 7658 | mamo-s | *BGIBMGA009095* | down | 1147 | 1155 | + | 12.1009 | 2.43E-05 | 0.615 | GTGCGTGGT |
| 7659 | mamo-s | *BGIBMGA009095* | up | 530 | 538 | - | 11.3303 | 4.96E-05 | 0.615 | GTGGGTGGC |
| 7660 | mamo-s | *BGIBMGA009095* | up | 1023 | 1031 | + | 10.8349 | 7.66E-05 | 0.751 | CCGCGAGGC |
| 7661 | mamo-s | *BGIBMGA009100* | down | 661 | 669 | - | 11.3303 | 4.96E-05 | 0.615 | GTGGGTGGC |
| 7662 | mamo-s | *BGIBMGA009100* | down | 1092 | 1100 | - | 11.3303 | 4.96E-05 | 0.615 | GTGGGTGGC |
| 7663 | mamo-s | *BGIBMGA009103* | down | 1070 | 1078 | + | 11.3303 | 4.96E-05 | 0.615 | GTGGGTGGC |
| 7664 | mamo-s | *BGIBMGA009104* | up | 1312 | 1320 | - | 10.8899 | 7.48E-05 | 0.739 | CCGCGTGCC |
| 7665 | mamo-s | *BGIBMGA009105* | up | 1409 | 1417 | + | 11.3303 | 4.96E-05 | 0.615 | GTGGGTGGC |
| 7666 | mamo-s | *BGIBMGA009107* | down | 698 | 706 | - | 11.3853 | 4.74E-05 | 0.615 | ACGCGTGGC |
| 7667 | mamo-s | *BGIBMGA009110* | up | 783 | 791 | - | 11.3303 | 4.96E-05 | 0.615 | GTGGGTGGC |
| 7668 | mamo-s | *BGIBMGA009112* | up | 1671 | 1679 | + | 11.2294 | 5.41E-05 | 0.65 | CTGCGAGGC |
| 7669 | mamo-s | *BGIBMGA009116* | down | 959 | 967 | + | 11.0826 | 6.84E-05 | 0.709 | GTACGTGGC |
| 7670 | mamo-s | *BGIBMGA009117* | up | 852 | 860 | - | 11.4404 | 4.51E-05 | 0.615 | GAGCGTGTC |
| 7671 | mamo-s | *BGIBMGA009118* | down | 437 | 445 | + | 11.3303 | 4.96E-05 | 0.615 | GTGGGTGGC |
| 7672 | mamo-s | *BGIBMGA009118* | down | 1350 | 1358 | - | 11.3303 | 4.96E-05 | 0.615 | GTGGGTGGC |
| 7673 | mamo-s | *BGIBMGA009119* | down | 1783 | 1791 | + | 10.5138 | 9.75E-05 | 0.865 | GTGCGTGTT |
| 7674 | mamo-s | *BGIBMGA009121* | up | 1287 | 1295 | - | 11.3303 | 4.96E-05 | 0.615 | GTGGGTGGC |
| 7675 | mamo-s | *BGIBMGA009123* | up | 379 | 387 | + | 11.3303 | 4.96E-05 | 0.615 | GTGGGTGGC |
| 7676 | mamo-s | *BGIBMGA009123* | down | 857 | 865 | - | 11.3303 | 4.96E-05 | 0.615 | GTGGGTGGC |
| 7677 | mamo-s | *BGIBMGA009123* | up | 1175 | 1183 | - | 11.3303 | 4.96E-05 | 0.615 | GTGGGTGGC |
| 7678 | mamo-s | *BGIBMGA009124* | up | 1396 | 1404 | - | 11.3303 | 4.96E-05 | 0.615 | GTGGGTGGC |
| 7679 | mamo-s | *BGIBMGA009126* | down | 191 | 199 | - | 11.3303 | 4.96E-05 | 0.615 | GTGGGTGGC |
| 7680 | mamo-s | *BGIBMGA009126* | down | 1966 | 1974 | - | 11.3303 | 4.96E-05 | 0.615 | GTGGGTGGC |
| 7681 | mamo-s | *BGIBMGA009126* | down | 607 | 615 | - | 10.8899 | 7.48E-05 | 0.739 | CCGCGTGCC |
| 7682 | mamo-s | *BGIBMGA009127* | up | 1755 | 1763 | - | 11.3303 | 4.96E-05 | 0.615 | GTGGGTGGC |
| 7683 | mamo-s | *BGIBMGA009129* | up | 136 | 144 | + | 11.3303 | 4.96E-05 | 0.615 | GTGGGTGGC |
| 7684 | mamo-s | *BGIBMGA009129* | down | 685 | 693 | + | 11.0826 | 6.84E-05 | 0.709 | GTACGTGGC |
| 7685 | mamo-s | *BGIBMGA009130* | up | 1194 | 1202 | + | 11.0826 | 6.84E-05 | 0.709 | GTACGTGGC |
| 7686 | mamo-s | *BGIBMGA009131* | up | 659 | 667 | - | 11.3303 | 4.96E-05 | 0.615 | GTGGGTGGC |
| 7687 | mamo-s | *BGIBMGA009131* | up | 1856 | 1864 | + | 11.1009 | 6.57E-05 | 0.709 | GAGCGAGGC |
| 7688 | mamo-s | *BGIBMGA009132* | down | 529 | 537 | - | 12.1284 | 2.16E-05 | 0.615 | CAGCGTGGC |
| 7689 | mamo-s | *BGIBMGA009132* | up | 1526 | 1534 | - | 11.7339 | 3.29E-05 | 0.615 | GCGCGAGGC |
| 7690 | mamo-s | *BGIBMGA009133* | up | 795 | 803 | + | 11.3303 | 4.96E-05 | 0.615 | GTGGGTGGC |
| 7691 | mamo-s | *BGIBMGA009133* | down | 925 | 933 | - | 11.0734 | 7.12E-05 | 0.726 | GAGCGTGGT |
| 7692 | mamo-s | *BGIBMGA009137* | down | 1275 | 1283 | - | 11.3303 | 4.96E-05 | 0.615 | GTGGGTGGC |
| 7693 | mamo-s | *BGIBMGA009138* | down | 1647 | 1655 | - | 12.1284 | 2.16E-05 | 0.615 | CAGCGTGGC |
| 7694 | mamo-s | *BGIBMGA009139* | down | 1151 | 1159 | - | 11.2844 | 5.18E-05 | 0.63 | CTGCGTGCC |
| 7695 | mamo-s | *BGIBMGA009141* | up | 573 | 581 | + | 11.4404 | 4.51E-05 | 0.615 | GAGCGTGTC |
| 7696 | mamo-s | *BGIBMGA009141* | down | 446 | 454 | + | 11.2844 | 5.18E-05 | 0.63 | CTGCGTGCC |
| 7697 | mamo-s | *BGIBMGA009142* | up | 275 | 283 | - | 11.3303 | 4.96E-05 | 0.615 | GTGGGTGGC |
| 7698 | mamo-s | *BGIBMGA009143* | down | 870 | 878 | + | 11.2844 | 5.18E-05 | 0.63 | CTGCGTGCC |
| 7699 | mamo-s | *BGIBMGA009145* | down | 1014 | 1022 | - | 11.3303 | 4.96E-05 | 0.615 | GTGGGTGGC |
| 7700 | mamo-s | *BGIBMGA009145* | up | 1814 | 1822 | + | 11.3303 | 4.96E-05 | 0.615 | GTGGGTGGC |
| 7701 | mamo-s | *BGIBMGA009145* | down | 1938 | 1946 | + | 11.0826 | 6.84E-05 | 0.709 | GTACGTGGC |
| 7702 | mamo-s | *BGIBMGA009146* | down | 51 | 59 | + | 13.6606 | 4.06E-06 | 0.615 | GCGCGTGGC |
| 7703 | mamo-s | *BGIBMGA009146* | up | 1188 | 1196 | + | 11.3303 | 4.96E-05 | 0.615 | GTGGGTGGC |
| 7704 | mamo-s | *BGIBMGA009147* | up | 1751 | 1759 | - | 11.3303 | 4.96E-05 | 0.615 | GTGGGTGGC |
| 7705 | mamo-s | *BGIBMGA009148* | down | 581 | 589 | + | 11.3303 | 4.96E-05 | 0.615 | GTGGGTGGC |
| 7706 | mamo-s | *BGIBMGA009148* | up | 1167 | 1175 | + | 11.3303 | 4.96E-05 | 0.615 | GTGGGTGGC |
| 7707 | mamo-s | *BGIBMGA009148* | down | 1254 | 1262 | - | 11.3303 | 4.96E-05 | 0.615 | GTGGGTGGC |
| 7708 | mamo-s | *BGIBMGA009149* | down | 1241 | 1249 | - | 11.5688 | 3.79E-05 | 0.615 | CTGCGTGTC |
| 7709 | mamo-s | *BGIBMGA009150* | up | 342 | 350 | + | 11.3303 | 4.96E-05 | 0.615 | GTGGGTGGC |
| 7710 | mamo-s | *BGIBMGA009151* | up | 622 | 630 | - | 11.3303 | 4.96E-05 | 0.615 | GTGGGTGGC |
| 7711 | mamo-s | *BGIBMGA009152* | up | 1417 | 1425 | - | 10.7523 | 8.38E-05 | 0.797 | GGGCGTGTC |
| 7712 | mamo-s | *BGIBMGA009153* | up | 563 | 571 | - | 11.5688 | 3.79E-05 | 0.615 | CTGCGTGTC |
| 7713 | mamo-s | *BGIBMGA009153* | down | 107 | 115 | + | 11.0734 | 7.12E-05 | 0.726 | GAGCGTGGT |
| 7714 | mamo-s | *BGIBMGA009154* | up | 431 | 439 | + | 11.3303 | 4.96E-05 | 0.615 | GTGGGTGGC |
| 7715 | mamo-s | *BGIBMGA009154* | up | 1439 | 1447 | - | 11.0826 | 6.84E-05 | 0.709 | GTACGTGGC |
| 7716 | mamo-s | *BGIBMGA009155* | up | 122 | 130 | - | 11.3303 | 4.96E-05 | 0.615 | GTGGGTGGC |
| 7717 | mamo-s | *BGIBMGA009155* | down | 1168 | 1176 | - | 11.3303 | 4.96E-05 | 0.615 | GTGGGTGGC |
| 7718 | mamo-s | *BGIBMGA009156* | up | 1754 | 1762 | + | 11.3303 | 4.96E-05 | 0.615 | GTGGGTGGC |
| 7719 | mamo-s | *BGIBMGA009158* | up | 602 | 610 | + | 10.7523 | 8.38E-05 | 0.797 | AAGCGTGGC |
| 7720 | mamo-s | *BGIBMGA009159* | up | 1807 | 1815 | + | 11.3303 | 4.96E-05 | 0.615 | GTGGGTGGC |
| 7721 | mamo-s | *BGIBMGA009161* | up | 1859 | 1867 | + | 11.3303 | 4.96E-05 | 0.615 | GTGGGTGGC |
| 7722 | mamo-s | *BGIBMGA009161* | up | 859 | 867 | + | 10.9358 | 7.30E-05 | 0.728 | GCGGGTGGC |
| 7723 | mamo-s | *BGIBMGA009162* | up | 240 | 248 | - | 10.5138 | 9.75E-05 | 0.865 | GTGCGTGTT |
| 7724 | mamo-s | *BGIBMGA009163* | up | 776 | 784 | - | 11.5321 | 4.06E-05 | 0.615 | TTGCGTGGC |
| 7725 | mamo-s | *BGIBMGA009163* | down | 457 | 465 | - | 11.1376 | 6.35E-05 | 0.709 | TCGCGTGGC |
| 7726 | mamo-s | *BGIBMGA009163* | up | 1330 | 1338 | + | 11.0826 | 6.84E-05 | 0.709 | GTACGTGGC |
| 7727 | mamo-s | *BGIBMGA009165* | down | 120 | 128 | - | 11.3303 | 4.96E-05 | 0.615 | GTGGGTGGC |
| 7728 | mamo-s | *BGIBMGA009167* | up | 837 | 845 | - | 11.3303 | 4.96E-05 | 0.615 | GTGGGTGGC |
| 7729 | mamo-s | *BGIBMGA009169* | up | 421 | 429 | + | 11.7339 | 3.29E-05 | 0.615 | GCGCGAGGC |
| 7730 | mamo-s | *BGIBMGA009169* | up | 1590 | 1598 | + | 11.4404 | 4.51E-05 | 0.615 | CGGCGTGGC |
| 7731 | mamo-s | *BGIBMGA009169* | up | 650 | 658 | + | 11.1009 | 6.57E-05 | 0.709 | GAGCGAGGC |
| 7732 | mamo-s | *BGIBMGA009172* | down | 1765 | 1773 | - | 11.3303 | 4.96E-05 | 0.615 | GTGGGTGGC |
| 7733 | mamo-s | *BGIBMGA009172* | down | 1916 | 1924 | - | 10.7523 | 8.38E-05 | 0.797 | GGGCGTGTC |
| 7734 | mamo-s | *BGIBMGA009175* | down | 1469 | 1477 | + | 11.3303 | 4.96E-05 | 0.615 | GTGGGTGGC |
| 7735 | mamo-s | *BGIBMGA009175* | down | 648 | 656 | + | 11.1743 | 5.90E-05 | 0.684 | CCGCGTGTC |
| 7736 | mamo-s | *BGIBMGA009175* | up | 174 | 182 | + | 11.0826 | 6.84E-05 | 0.709 | GTACGTGGC |
| 7737 | mamo-s | *BGIBMGA009175* | up | 1475 | 1483 | + | 10.7523 | 8.38E-05 | 0.797 | GGGCGTGTC |

| | | | | | | | | | | |
|---|---|---|---|---|---|---|---|---|---|---|
| 7738 | mamo-s | *BGIBMGA009176* | up | 237 | 245 | - | 14.055 | 2.23E-06 | 0.615 | GTGCGTGGC |
| 7739 | mamo-s | *BGIBMGA009177* | down | 1238 | 1246 | + | 12.1835 | 1.71E-05 | 0.615 | GTGCGTGCC |
| 7740 | mamo-s | *BGIBMGA009177* | down | 560 | 568 | + | 11.3303 | 4.96E-05 | 0.615 | GTGGGTGGC |
| 7741 | mamo-s | *BGIBMGA009178* | down | 102 | 110 | + | 13.0275 | 8.53E-06 | 0.615 | GAGCGTGGC |
| 7742 | mamo-s | *BGIBMGA009178* | down | 1806 | 1814 | - | 11.3303 | 4.96E-05 | 0.615 | GTGGGTGGC |
| 7743 | mamo-s | *BGIBMGA009179* | down | 1474 | 1482 | - | 11.3303 | 4.96E-05 | 0.615 | GTGGGTGGC |
| 7744 | mamo-s | *BGIBMGA009180* | down | 1611 | 1619 | - | 13.156 | 6.30E-06 | 0.615 | CTGCGTGGC |
| 7745 | mamo-s | *BGIBMGA009180* | down | 1422 | 1430 | + | 11.3303 | 4.96E-05 | 0.615 | GTGGGTGGC |
| 7746 | mamo-s | *BGIBMGA009182* | down | 1648 | 1656 | - | 12.7615 | 1.04E-05 | 0.615 | CCGCGTGGC |
| 7747 | mamo-s | *BGIBMGA009182* | down | 1971 | 1979 | + | 12.0734 | 2.66E-05 | 0.615 | GCGCGTGTC |
| 7748 | mamo-s | *BGIBMGA009182* | up | 545 | 553 | - | 11.3303 | 4.96E-05 | 0.615 | GTGGGTGGC |
| 7749 | mamo-s | *BGIBMGA009182* | down | 1512 | 1520 | + | 11.156 | 6.13E-05 | 0.705 | GAGCGTGCC |
| 7750 | mamo-s | *BGIBMGA009184* | down | 1173 | 1181 | + | 11.2844 | 5.18E-05 | 0.63 | CTGCGTGCC |
| 7751 | mamo-s | *BGIBMGA009185* | up | 1910 | 1918 | - | 12.7615 | 1.04E-05 | 0.615 | CCGCGTGGC |
| 7752 | mamo-s | *BGIBMGA009185* | down | 443 | 451 | - | 10.9358 | 7.30E-05 | 0.728 | GCGGGTGGC |
| 7753 | mamo-s | *BGIBMGA009186* | down | 330 | 338 | - | 11.3303 | 4.96E-05 | 0.615 | GTGGGTGGC |
| 7754 | mamo-s | *BGIBMGA009186* | down | 951 | 959 | - | 11.3303 | 4.96E-05 | 0.615 | GTGGGTGGC |
| 7755 | mamo-s | *BGIBMGA009186* | down | 1331 | 1339 | + | 11.3303 | 4.96E-05 | 0.615 | GTGGGTGGC |
| 7756 | mamo-s | *BGIBMGA009190* | up | 1568 | 1576 | + | 14.055 | 2.23E-06 | 0.615 | GTGCGTGGC |
| 7757 | mamo-s | *BGIBMGA009190* | down | 9 | 17 | - | 11.156 | 6.13E-05 | 0.705 | GAGCGTGCC |
| 7758 | mamo-s | *BGIBMGA009191* | up | 1360 | 1368 | - | 11.3303 | 4.96E-05 | 0.615 | GTGGGTGGC |
| 7759 | mamo-s | *BGIBMGA009192* | down | 423 | 431 | - | 11.1009 | 6.57E-05 | 0.709 | GAGCGAGGC |
| 7760 | mamo-s | *BGIBMGA009194* | down | 1916 | 1924 | - | 12.4679 | 1.31E-05 | 0.615 | GTGCGTGTC |
| 7761 | mamo-s | *BGIBMGA009196* | up | 1386 | 1394 | + | 11.1743 | 5.90E-05 | 0.684 | CCGCGTGTC |
| 7762 | mamo-s | *BGIBMGA009196* | up | 496 | 504 | + | 10.8899 | 7.48E-05 | 0.739 | CCGCGTGCC |
| 7763 | mamo-s | *BGIBMGA009197* | down | 1031 | 1039 | - | 10.9358 | 7.30E-05 | 0.728 | GCGGGTGGC |
| 7764 | mamo-s | *BGIBMGA009202* | up | 1067 | 1075 | - | 11.3303 | 4.96E-05 | 0.615 | GTGGGTGGC |
| 7765 | mamo-s | *BGIBMGA009202* | down | 1268 | 1276 | + | 10.9358 | 7.30E-05 | 0.728 | GCGGGTGGC |
| 7766 | mamo-s | *BGIBMGA009203* | up | 634 | 642 | + | 11.3303 | 4.96E-05 | 0.615 | GTGGGTGGC |
| 7767 | mamo-s | *BGIBMGA009206* | down | 371 | 379 | - | 11.3303 | 4.96E-05 | 0.615 | GTGGGTGGC |
| 7768 | mamo-s | *BGIBMGA009207* | down | 1032 | 1040 | - | 11.3303 | 4.96E-05 | 0.615 | GTGGGTGGC |
| 7769 | mamo-s | *BGIBMGA009209* | down | 1359 | 1367 | + | 11.3303 | 4.96E-05 | 0.615 | GTGGGTGGC |
| 7770 | mamo-s | *BGIBMGA009210* | up | 1821 | 1829 | + | 11.3303 | 4.96E-05 | 0.615 | GTGGGTGGC |
| 7771 | mamo-s | *BGIBMGA009211* | up | 690 | 698 | + | 11.3303 | 4.96E-05 | 0.615 | GTGGGTGGC |
| 7772 | mamo-s | *BGIBMGA009215* | down | 1705 | 1713 | - | 11.2844 | 5.18E-05 | 0.63 | CTGCGTGCC |
| 7773 | mamo-s | *BGIBMGA009221* | down | 333 | 341 | - | 11.3303 | 4.96E-05 | 0.615 | GTGGGTGGC |
| 7774 | mamo-s | *BGIBMGA009221* | down | 963 | 971 | - | 11.3303 | 4.96E-05 | 0.615 | GTGGGTGGC |
| 7775 | mamo-s | *BGIBMGA009225* | up | 1503 | 1511 | - | 11.3303 | 4.96E-05 | 0.615 | GTGGGTGGC |
| 7776 | mamo-s | *BGIBMGA009226* | down | 1043 | 1051 | + | 11.3303 | 4.96E-05 | 0.615 | GTGGGTGGC |
| 7777 | mamo-s | *BGIBMGA009226* | down | 1914 | 1922 | - | 11.3303 | 4.96E-05 | 0.615 | GTGGGTGGC |
| 7778 | mamo-s | *BGIBMGA009227* | up | 828 | 836 | - | 11.3303 | 4.96E-05 | 0.615 | GTGGGTGGC |
| 7779 | mamo-s | *BGIBMGA009227* | down | 1827 | 1835 | - | 11.3303 | 4.96E-05 | 0.615 | GTGGGTGGC |
| 7780 | mamo-s | *BGIBMGA009228* | down | 1314 | 1322 | - | 11.7064 | 3.52E-05 | 0.615 | GCGCGTGGT |
| 7781 | mamo-s | *BGIBMGA009230* | down | 740 | 748 | + | 11.1743 | 5.90E-05 | 0.684 | CCGCGTGTC |
| 7782 | mamo-s | *BGIBMGA009231* | up | 145 | 153 | + | 11.1743 | 5.90E-05 | 0.684 | CCGCGTGTC |
| 7783 | mamo-s | *BGIBMGA009234* | down | 83 | 91 | - | 11.3303 | 4.96E-05 | 0.615 | GTGGGTGGC |
| 7784 | mamo-s | *BGIBMGA009236* | down | 820 | 828 | + | 11.3303 | 4.96E-05 | 0.615 | GTGGGTGGC |
| 7785 | mamo-s | *BGIBMGA009237* | up | 1698 | 1706 | + | 14.055 | 2.23E-06 | 0.615 | GTGCGTGGC |
| 7786 | mamo-s | *BGIBMGA009241* | down | 95 | 103 | - | 11.5688 | 3.79E-05 | 0.615 | CTGCGTGTC |
| 7787 | mamo-s | *BGIBMGA009241* | up | 1127 | 1135 | - | 11.3303 | 4.96E-05 | 0.615 | GTGGGTGGC |
| 7788 | mamo-s | *BGIBMGA009242* | up | 368 | 376 | + | 10.8073 | 7.89E-05 | 0.766 | CCGCGTGGT |
| 7789 | mamo-s | *BGIBMGA009245* | up | 1696 | 1704 | + | 11.3303 | 4.96E-05 | 0.615 | GTGGGTGGC |
| 7790 | mamo-s | *BGIBMGA009246* | down | 1285 | 1293 | - | 11.3303 | 4.96E-05 | 0.615 | GTGGGTGGC |
| 7791 | mamo-s | *BGIBMGA009247* | up | 123 | 131 | + | 11.3303 | 4.96E-05 | 0.615 | GTGGGTGGC |
| 7792 | mamo-s | *BGIBMGA009247* | up | 786 | 794 | + | 11.0826 | 6.84E-05 | 0.709 | GTACGTGGC |
| 7793 | mamo-s | *BGIBMGA009253* | down | 1239 | 1247 | - | 11.0826 | 6.84E-05 | 0.709 | GTACGTGGC |
| 7794 | mamo-s | *BGIBMGA009254* | down | 918 | 926 | + | 12.1835 | 1.71E-05 | 0.615 | GTGCGTGCC |
| 7795 | mamo-s | *BGIBMGA009254* | down | 512 | 520 | - | 11.3303 | 4.96E-05 | 0.615 | GTGGGTGGC |
| 7796 | mamo-s | *BGIBMGA009255* | up | 81 | 89 | - | 11.3303 | 4.96E-05 | 0.615 | GTGGGTGGC |
| 7797 | mamo-s | *BGIBMGA009255* | up | 1014 | 1022 | - | 11.3303 | 4.96E-05 | 0.615 | GTGGGTGGC |
| 7798 | mamo-s | *BGIBMGA009255* | up | 1829 | 1837 | - | 11.3303 | 4.96E-05 | 0.615 | GTGGGTGGC |
| 7799 | mamo-s | *BGIBMGA009257* | down | 196 | 204 | + | 11.3303 | 4.96E-05 | 0.615 | GTGGGTGGC |
| 7800 | mamo-s | *BGIBMGA009258* | down | 261 | 269 | + | 11.3303 | 4.96E-05 | 0.615 | GTGGGTGGC |
| 7801 | mamo-s | *BGIBMGA009258* | down | 839 | 847 | - | 11.3303 | 4.96E-05 | 0.615 | GTGGGTGGC |
| 7802 | mamo-s | *BGIBMGA009259* | down | 1314 | 1322 | + | 14.055 | 2.23E-06 | 0.615 | GTGCGTGGC |
| 7803 | mamo-s | *BGIBMGA009265* | down | 1047 | 1055 | - | 11.3303 | 4.96E-05 | 0.615 | GTGGGTGGC |
| 7804 | mamo-s | *BGIBMGA009266* | up | 803 | 811 | - | 11.3303 | 4.96E-05 | 0.615 | GTGGGTGGC |
| 7805 | mamo-s | *BGIBMGA009267* | up | 1862 | 1870 | + | 12.1284 | 2.16E-05 | 0.615 | CAGCGTGGC |
| 7806 | mamo-s | *BGIBMGA009271* | down | 1542 | 1550 | - | 13.6606 | 4.06E-06 | 0.615 | GCGCGTGGC |
| 7807 | mamo-s | *BGIBMGA009271* | down | 1074 | 1082 | - | 11.3303 | 4.96E-05 | 0.615 | GTGGGTGGC |
| 7808 | mamo-s | *BGIBMGA009273* | up | 451 | 459 | - | 12.1835 | 1.71E-05 | 0.615 | GTGCGTGCC |
| 7809 | mamo-s | *BGIBMGA009277* | down | 1586 | 1594 | + | 11.7064 | 3.52E-05 | 0.615 | GCGCGTGGT |
| 7810 | mamo-s | *BGIBMGA009278* | up | 1913 | 1921 | + | 11.5321 | 4.06E-05 | 0.615 | TTGCGTGGC |
| 7811 | mamo-s | *BGIBMGA009281* | up | 1135 | 1143 | + | 11.1009 | 6.57E-05 | 0.709 | GAGCGAGGC |
| 7812 | mamo-s | *BGIBMGA009283* | down | 1295 | 1303 | + | 11.3303 | 4.96E-05 | 0.615 | GTGGGTGGC |
| 7813 | mamo-s | *BGIBMGA009283* | down | 1637 | 1645 | - | 11.1376 | 6.35E-05 | 0.709 | TCGCGTGGC |
| 7814 | mamo-s | *BGIBMGA009284* | up | 860 | 868 | + | 11.3303 | 4.96E-05 | 0.615 | GTGGGTGGC |
| 7815 | mamo-s | *BGIBMGA009284* | up | 1202 | 1210 | - | 11.1376 | 6.35E-05 | 0.709 | TCGCGTGGC |
| 7816 | mamo-s | *BGIBMGA009286* | down | 1245 | 1253 | + | 11.7339 | 3.29E-05 | 0.615 | GCGCGAGGC |
| 7817 | mamo-s | *BGIBMGA009288* | up | 1388 | 1396 | - | 13.156 | 6.30E-06 | 0.615 | CTGCGTGGC |
| 7818 | mamo-s | *BGIBMGA009288* | down | 632 | 640 | - | 10.9358 | 7.30E-05 | 0.728 | GCGGGTGGC |
| 7819 | mamo-s | *BGIBMGA009290* | down | 426 | 434 | + | 10.5688 | 8.88E-05 | 0.824 | GTGCGTGAC |
| 7820 | mamo-s | *BGIBMGA009291* | down | 140 | 148 | - | 11.3303 | 4.96E-05 | 0.615 | GTGGGTGGC |
| 7821 | mamo-s | *BGIBMGA009294* | down | 239 | 247 | + | 11.3303 | 4.96E-05 | 0.615 | GTGGGTGGC |
| 7822 | mamo-s | *BGIBMGA009295* | down | 624 | 632 | + | 12.0734 | 2.66E-05 | 0.615 | GCGCGTGTC |
| 7823 | mamo-s | *BGIBMGA009295* | down | 1074 | 1082 | - | 11.3303 | 4.96E-05 | 0.615 | GTGGGTGGC |
| 7824 | mamo-s | *BGIBMGA009296* | up | 67 | 75 | - | 10.5138 | 9.75E-05 | 0.865 | GTGCGTGTT |
| 7825 | mamo-s | *BGIBMGA009297* | up | 893 | 901 | + | 11.3303 | 4.96E-05 | 0.615 | GTGGGTGGC |

| | | | | | | | | | | |
|---|---|---|---|---|---|---|---|---|---|---|
| 7826 | mamo-s | *BGIBMGA009297* | down | 103 | 111 | + | 10.7523 | 8.38E-05 | 0.797 | AAGCGTGGC |
| 7827 | mamo-s | *BGIBMGA009297* | down | 167 | 175 | + | 10.7523 | 8.38E-05 | 0.797 | AAGCGTGGC |
| 7828 | mamo-s | *BGIBMGA009298* | down | 490 | 498 | - | 11.3303 | 4.96E-05 | 0.615 | GTGGGTGGC |
| 7829 | mamo-s | *BGIBMGA009298* | down | 1691 | 1699 | + | 11.3303 | 4.96E-05 | 0.615 | GTGGGTGGC |
| 7830 | mamo-s | *BGIBMGA009300* | up | 299 | 307 | + | 11.1376 | 6.35E-05 | 0.709 | TCGCGTGGC |
| 7831 | mamo-s | *BGIBMGA009300* | up | 1067 | 1075 | + | 11.1009 | 6.57E-05 | 0.709 | GAGCGAGGC |
| 7832 | mamo-s | *BGIBMGA009304* | down | 1468 | 1476 | + | 12.1284 | 2.16E-05 | 0.615 | GTGCGAGGC |
| 7833 | mamo-s | *BGIBMGA009304* | up | 1630 | 1638 | - | 12.1284 | 2.16E-05 | 0.615 | GTGCGAGGC |
| 7834 | mamo-s | *BGIBMGA009305* | up | 1246 | 1254 | + | 13.6606 | 4.06E-06 | 0.615 | GCGCGTGGC |
| 7835 | mamo-s | *BGIBMGA009305* | down | 1068 | 1076 | + | 11.0826 | 6.84E-05 | 0.709 | GTACGTGGC |
| 7836 | mamo-s | *BGIBMGA009306* | up | 1037 | 1045 | + | 11.5688 | 3.79E-05 | 0.615 | CTGCGTGTC |
| 7837 | mamo-s | *BGIBMGA009306* | down | 971 | 979 | - | 11.3303 | 4.96E-05 | 0.615 | GTGGGTGGC |
| 7838 | mamo-s | *BGIBMGA009306* | up | 1104 | 1112 | + | 11.2844 | 5.18E-05 | 0.63 | CTGCGTGCC |
| 7839 | mamo-s | *BGIBMGA009307* | up | 1211 | 1219 | - | 11.2294 | 5.41E-05 | 0.65 | CTGCGAGGC |
| 7840 | mamo-s | *BGIBMGA009308* | down | 1719 | 1727 | + | 11.0734 | 7.12E-05 | 0.726 | GAGCGTGGT |
| 7841 | mamo-s | *BGIBMGA009309* | up | 252 | 260 | + | 11.3303 | 4.96E-05 | 0.615 | GTGGGTGGC |
| 7842 | mamo-s | *BGIBMGA009309* | down | 1394 | 1402 | - | 11.3303 | 4.96E-05 | 0.615 | GTGGGTGGC |
| 7843 | mamo-s | *BGIBMGA009313* | up | 709 | 717 | + | 11.3303 | 4.96E-05 | 0.615 | GTGGGTGGC |
| 7844 | mamo-s | *BGIBMGA009315* | down | 317 | 325 | - | 11.3303 | 4.96E-05 | 0.615 | GTGGGTGGC |
| 7845 | mamo-s | *BGIBMGA009317* | down | 1375 | 1383 | - | 12.1284 | 2.16E-05 | 0.615 | CAGCGTGGC |
| 7846 | mamo-s | *BGIBMGA009317* | down | 1299 | 1307 | + | 12.1009 | 2.43E-05 | 0.615 | GTGCGTGGT |
| 7847 | mamo-s | *BGIBMGA009317* | down | 1834 | 1842 | + | 11.5688 | 3.79E-05 | 0.615 | CTGCGTGTC |
| 7848 | mamo-s | *BGIBMGA009317* | up | 191 | 199 | + | 11.3303 | 4.96E-05 | 0.615 | GTGGGTGGC |
| 7849 | mamo-s | *BGIBMGA009317* | up | 1050 | 1058 | + | 11.3303 | 4.96E-05 | 0.615 | GTGGGTGGC |
| 7850 | mamo-s | *BGIBMGA009317* | up | 1203 | 1211 | + | 11.3303 | 4.96E-05 | 0.615 | GTGGGTGGC |
| 7851 | mamo-s | *BGIBMGA009319* | up | 1614 | 1622 | + | 11.3303 | 4.96E-05 | 0.615 | GTGGGTGGC |
| 7852 | mamo-s | *BGIBMGA009320* | down | 32 | 40 | + | 13.6606 | 4.06E-06 | 0.615 | GCGCGTGGC |
| 7853 | mamo-s | *BGIBMGA009320* | up | 1989 | 1997 | + | 13.6606 | 4.06E-06 | 0.615 | GCGCGTGGC |
| 7854 | mamo-s | *BGIBMGA009321* | down | 116 | 124 | + | 11.3303 | 4.96E-05 | 0.615 | GTGGGTGGC |
| 7855 | mamo-s | *BGIBMGA009322* | down | 1142 | 1150 | - | 11.3303 | 4.96E-05 | 0.615 | GTGGGTGGC |
| 7856 | mamo-s | *BGIBMGA009323* | up | 1865 | 1873 | - | 10.5688 | 8.88E-05 | 0.824 | GTGCGTGAC |
| 7857 | mamo-s | *BGIBMGA009324* | down | 413 | 421 | - | 11.3303 | 4.96E-05 | 0.615 | GTGGGTGGC |
| 7858 | mamo-s | *BGIBMGA009324* | up | 1559 | 1567 | - | 11.3303 | 4.96E-05 | 0.615 | GTGGGTGGC |
| 7859 | mamo-s | *BGIBMGA009325* | down | 489 | 497 | + | 11.3303 | 4.96E-05 | 0.615 | GTGGGTGGC |
| 7860 | mamo-s | *BGIBMGA009326* | up | 1005 | 1013 | + | 11.0826 | 6.84E-05 | 0.709 | GTACGTGGC |
| 7861 | mamo-s | *BGIBMGA009328* | down | 1605 | 1613 | - | 11.5688 | 3.79E-05 | 0.615 | CTGCGTGTC |
| 7862 | mamo-s | *BGIBMGA009328* | down | 996 | 1004 | + | 11.3303 | 4.96E-05 | 0.615 | GTGGGTGGC |
| 7863 | mamo-s | *BGIBMGA009331* | up | 857 | 865 | + | 11.3303 | 4.96E-05 | 0.615 | GTGGGTGGC |
| 7864 | mamo-s | *BGIBMGA009333* | down | 750 | 758 | + | 10.5138 | 9.75E-05 | 0.865 | GTGCGTGTT |
| 7865 | mamo-s | *BGIBMGA009334* | down | 918 | 926 | + | 10.9358 | 7.30E-05 | 0.728 | GCGGGTGGC |
| 7866 | mamo-s | *BGIBMGA009335* | up | 1983 | 1991 | + | 13.6606 | 4.06E-06 | 0.615 | GCGCGTGGC |
| 7867 | mamo-s | *BGIBMGA009336* | up | 409 | 417 | + | 11.3303 | 4.96E-05 | 0.615 | GTGGGTGGC |
| 7868 | mamo-s | *BGIBMGA009336* | down | 1642 | 1650 | - | 11.3303 | 4.96E-05 | 0.615 | GTGGGTGGC |
| 7869 | mamo-s | *BGIBMGA009337* | down | 546 | 554 | - | 13.0275 | 8.53E-06 | 0.615 | GAGCGTGGC |
| 7870 | mamo-s | *BGIBMGA009337* | down | 1783 | 1791 | - | 12.1009 | 2.43E-05 | 0.615 | GTGCGTGGT |
| 7871 | mamo-s | *BGIBMGA009337* | up | 862 | 870 | - | 10.9358 | 7.30E-05 | 0.728 | GCGGGTGGC |
| 7872 | mamo-s | *BGIBMGA009338* | down | 266 | 274 | + | 11.3303 | 4.96E-05 | 0.615 | GTGGGTGGC |
| 7873 | mamo-s | *BGIBMGA009338* | up | 1311 | 1319 | + | 11.3303 | 4.96E-05 | 0.615 | GTGGGTGGC |
| 7874 | mamo-s | *BGIBMGA009338* | down | 1367 | 1375 | + | 11.3303 | 4.96E-05 | 0.615 | GTGGGTGGC |
| 7875 | mamo-s | *BGIBMGA009339* | down | 697 | 705 | - | 11.3303 | 4.96E-05 | 0.615 | GTGGGTGGC |
| 7876 | mamo-s | *BGIBMGA009339* | up | 1588 | 1596 | + | 11.3303 | 4.96E-05 | 0.615 | GTGGGTGGC |
| 7877 | mamo-s | *BGIBMGA009341* | down | 163 | 171 | - | 11.5321 | 4.06E-05 | 0.615 | TTGCGTGGC |
| 7878 | mamo-s | *BGIBMGA009342* | up | 231 | 239 | - | 14.055 | 2.23E-06 | 0.615 | GTGCGTGGC |
| 7879 | mamo-s | *BGIBMGA009343* | up | 1296 | 1304 | - | 11.3303 | 4.96E-05 | 0.615 | GTGGGTGGC |
| 7880 | mamo-s | *BGIBMGA009344* | up | 805 | 813 | + | 11.2018 | 5.68E-05 | 0.674 | CTGCGTGGT |
| 7881 | mamo-s | *BGIBMGA009345* | up | 638 | 646 | - | 11.3303 | 4.96E-05 | 0.615 | GTGGGTGGC |
| 7882 | mamo-s | *BGIBMGA009345* | up | 1549 | 1557 | + | 11.3303 | 4.96E-05 | 0.615 | GTGGGTGGC |
| 7883 | mamo-s | *BGIBMGA009346* | down | 1749 | 1757 | - | 12.1835 | 1.71E-05 | 0.615 | GTGCGTGCC |
| 7884 | mamo-s | *BGIBMGA009347* | up | 676 | 684 | - | 13.6606 | 4.06E-06 | 0.615 | GCGCGTGGC |
| 7885 | mamo-s | *BGIBMGA009349* | down | 206 | 214 | + | 11.3303 | 4.96E-05 | 0.615 | GTGGGTGGC |
| 7886 | mamo-s | *BGIBMGA009351* | down | 1514 | 1522 | + | 11.3303 | 4.96E-05 | 0.615 | GTGGGTGGC |
| 7887 | mamo-s | *BGIBMGA009352* | up | 1618 | 1626 | - | 11.3303 | 4.96E-05 | 0.615 | GTGGGTGGC |
| 7888 | mamo-s | *BGIBMGA009352* | down | 1033 | 1041 | - | 10.5138 | 9.75E-05 | 0.865 | GTGCGTGTT |
| 7889 | mamo-s | *BGIBMGA009353* | down | 1818 | 1826 | - | 11.3303 | 4.96E-05 | 0.615 | GTGGGTGGC |
| 7890 | mamo-s | *BGIBMGA009354* | up | 1824 | 1832 | - | 11.3303 | 4.96E-05 | 0.615 | GTGGGTGGC |
| 7891 | mamo-s | *BGIBMGA009356* | up | 820 | 828 | - | 11.2018 | 5.68E-05 | 0.674 | CTGCGTGGT |
| 7892 | mamo-s | *BGIBMGA009357* | down | 1955 | 1963 | - | 13.156 | 6.30E-06 | 0.615 | CTGCGTGGC |
| 7893 | mamo-s | *BGIBMGA009358* | down | 22 | 30 | + | 11.7798 | 3.11E-05 | 0.615 | ATGCGTGGC |
| 7894 | mamo-s | *BGIBMGA009358* | up | 224 | 232 | - | 11.0734 | 7.12E-05 | 0.726 | GAGCGTGGT |
| 7895 | mamo-s | *BGIBMGA009361* | up | 1467 | 1475 | - | 11.3303 | 4.96E-05 | 0.615 | GTGGGTGGC |
| 7896 | mamo-s | *BGIBMGA009362* | up | 393 | 401 | - | 11.5688 | 3.79E-05 | 0.615 | CTGCGTGTC |
| 7897 | mamo-s | *BGIBMGA009362* | up | 907 | 915 | - | 10.6881 | 8.61E-05 | 0.81 | GCACGTGGC |
| 7898 | mamo-s | *BGIBMGA009364* | down | 1206 | 1214 | + | 12.1009 | 2.43E-05 | 0.615 | GTGCGTGGT |
| 7899 | mamo-s | *BGIBMGA009364* | down | 1093 | 1101 | + | 11.2294 | 5.41E-05 | 0.65 | CTGCGAGGC |
| 7900 | mamo-s | *BGIBMGA009366* | down | 1621 | 1629 | + | 11.3303 | 4.96E-05 | 0.615 | GTGGGTGGC |
| 7901 | mamo-s | *BGIBMGA009367* | up | 428 | 436 | + | 11.3303 | 4.96E-05 | 0.615 | GTGGGTGGC |
| 7902 | mamo-s | *BGIBMGA009368* | up | 131 | 139 | + | 11.3303 | 4.96E-05 | 0.615 | GTGGGTGGC |
| 7903 | mamo-s | *BGIBMGA009368* | down | 1117 | 1125 | + | 11.3303 | 4.96E-05 | 0.615 | GTGGGTGGC |
| 7904 | mamo-s | *BGIBMGA009368* | down | 1598 | 1606 | + | 11.3303 | 4.96E-05 | 0.615 | GTGGGTGGC |
| 7905 | mamo-s | *BGIBMGA009368* | down | 1449 | 1457 | + | 11.1009 | 6.57E-05 | 0.709 | GAGCGAGGC |
| 7906 | mamo-s | *BGIBMGA009369* | up | 381 | 389 | - | 11.3303 | 4.96E-05 | 0.615 | GTGGGTGGC |
| 7907 | mamo-s | *BGIBMGA009369* | down | 444 | 452 | - | 11.3303 | 4.96E-05 | 0.615 | GTGGGTGGC |
| 7908 | mamo-s | *BGIBMGA009371* | down | 396 | 404 | + | 11.3303 | 4.96E-05 | 0.615 | GTGGGTGGC |
| 7909 | mamo-s | *BGIBMGA009372* | down | 1558 | 1566 | - | 11.4404 | 4.51E-05 | 0.615 | GAGCGTGTC |
| 7910 | mamo-s | *BGIBMGA009372* | down | 850 | 858 | + | 11.3303 | 4.96E-05 | 0.615 | GTGGGTGGC |
| 7911 | mamo-s | *BGIBMGA009373* | up | 143 | 151 | + | 12.1284 | 2.16E-05 | 0.615 | CAGCGTGGC |
| 7912 | mamo-s | *BGIBMGA009373* | down | 1960 | 1968 | - | 11.3303 | 4.96E-05 | 0.615 | GTGGGTGGC |
| 7913 | mamo-s | *BGIBMGA009373* | up | 372 | 380 | - | 10.5688 | 8.88E-05 | 0.824 | GTGCGTGAC |

| | | | | | | | | | | |
|---|---|---|---|---|---|---|---|---|---|---|
| 7914 | mamo-s | *BGIBMGA009374* | down | 1475 | 1483 | - | 13.6606 | 4.06E-06 | 0.615 | GCGCGTGGC |
| 7915 | mamo-s | *BGIBMGA009376* | up | 934 | 942 | + | 14.055 | 2.23E-06 | 0.615 | GTGCGTGGC |
| 7916 | mamo-s | *BGIBMGA009376* | up | 1623 | 1631 | + | 11.3303 | 4.96E-05 | 0.615 | GTGGGTGGC |
| 7917 | mamo-s | *BGIBMGA009378* | up | 100 | 108 | - | 11.3303 | 4.96E-05 | 0.615 | GTGGGTGGC |
| 7918 | mamo-s | *BGIBMGA009379* | up | 340 | 348 | + | 12.0734 | 2.66E-05 | 0.615 | GCGCGTGTC |
| 7919 | mamo-s | *BGIBMGA009380* | up | 662 | 670 | - | 11.3303 | 4.96E-05 | 0.615 | GTGGGTGGC |
| 7920 | mamo-s | *BGIBMGA009381* | down | 865 | 873 | + | 11.3303 | 4.96E-05 | 0.615 | GTGGGTGGC |
| 7921 | mamo-s | *BGIBMGA009383* | down | 1513 | 1521 | - | 11.0826 | 6.84E-05 | 0.709 | GTACGTGGC |
| 7922 | mamo-s | *BGIBMGA009385* | up | 520 | 528 | - | 11.7064 | 3.52E-05 | 0.615 | GCGCGTGGT |
| 7923 | mamo-s | *BGIBMGA009385* | up | 799 | 807 | + | 11.1376 | 6.35E-05 | 0.709 | TCGCGTGGC |
| 7924 | mamo-s | *BGIBMGA009385* | up | 770 | 778 | + | 11.0734 | 7.12E-05 | 0.726 | GAGCGTGGT |
| 7925 | mamo-s | *BGIBMGA009386* | up | 110 | 118 | - | 11.5688 | 3.79E-05 | 0.615 | CTGCGTGTC |
| 7926 | mamo-s | *BGIBMGA009386* | up | 43 | 51 | - | 11.2844 | 5.18E-05 | 0.63 | CTGCGTGCC |
| 7927 | mamo-s | *BGIBMGA009386* | up | 1771 | 1779 | + | 11.1743 | 5.90E-05 | 0.684 | CCGCGTGTC |
| 7928 | mamo-s | *BGIBMGA009387* | up | 1761 | 1769 | - | 11.789 | 2.84E-05 | 0.615 | GCGCGTGCC |
| 7929 | mamo-s | *BGIBMGA009387* | up | 357 | 365 | - | 11.3303 | 4.96E-05 | 0.615 | GTGGGTGGC |
| 7930 | mamo-s | *BGIBMGA009388* | up | 610 | 618 | - | 11.0826 | 6.84E-05 | 0.709 | GTACGTGGC |
| 7931 | mamo-s | *BGIBMGA009390* | up | 762 | 770 | + | 12.1284 | 2.16E-05 | 0.615 | CAGCGTGGC |
| 7932 | mamo-s | *BGIBMGA009392* | up | 1412 | 1420 | - | 11.3303 | 4.96E-05 | 0.615 | GTGGGTGGC |
| 7933 | mamo-s | *BGIBMGA009392* | down | 1209 | 1217 | - | 11.0826 | 6.84E-05 | 0.709 | GTACGTGGC |
| 7934 | mamo-s | *BGIBMGA009393* | down | 988 | 996 | - | 11.3303 | 4.96E-05 | 0.615 | GTGGGTGGC |
| 7935 | mamo-s | *BGIBMGA009394* | down | 148 | 156 | - | 11.3303 | 4.96E-05 | 0.615 | GTGGGTGGC |
| 7936 | mamo-s | *BGIBMGA009395* | down | 1120 | 1128 | + | 11.3303 | 4.96E-05 | 0.615 | GTGGGTGGC |
| 7937 | mamo-s | *BGIBMGA009397* | down | 1120 | 1128 | - | 13.6606 | 4.06E-06 | 0.615 | GCGCGTGGC |
| 7938 | mamo-s | *BGIBMGA009397* | down | 1963 | 1971 | + | 11.3303 | 4.96E-05 | 0.615 | GTGGGTGGC |
| 7939 | mamo-s | *BGIBMGA009399* | up | 1702 | 1710 | - | 11.4404 | 4.51E-05 | 0.615 | CGGCGTGGC |
| 7940 | mamo-s | *BGIBMGA009400* | up | 157 | 165 | + | 11.3303 | 4.96E-05 | 0.615 | GTGGGTGGC |
| 7941 | mamo-s | *BGIBMGA009402* | up | 259 | 267 | - | 11.3303 | 4.96E-05 | 0.615 | GTGGGTGGC |
| 7942 | mamo-s | *BGIBMGA009402* | up | 1069 | 1077 | + | 11.3303 | 4.96E-05 | 0.615 | GTGGGTGGC |
| 7943 | mamo-s | *BGIBMGA009403* | up | 242 | 250 | + | 12.3394 | 1.49E-05 | 0.615 | GGGCGTGGC |
| 7944 | mamo-s | *BGIBMGA009406* | down | 1452 | 1460 | - | 11.4404 | 4.51E-05 | 0.615 | GAGCGTGTC |
| 7945 | mamo-s | *BGIBMGA009407* | up | 886 | 894 | + | 12.3394 | 1.49E-05 | 0.615 | GGGCGTGGC |
| 7946 | mamo-s | *BGIBMGA009407* | up | 576 | 584 | + | 11.3303 | 4.96E-05 | 0.615 | GTGGGTGGC |
| 7947 | mamo-s | *BGIBMGA009410* | down | 1316 | 1324 | + | 11.3303 | 4.96E-05 | 0.615 | GTGGGTGGC |
| 7948 | mamo-s | *BGIBMGA009411* | up | 923 | 931 | + | 11.3303 | 4.96E-05 | 0.615 | GTGGGTGGC |
| 7949 | mamo-s | *BGIBMGA009411* | down | 1320 | 1328 | - | 11.3303 | 4.96E-05 | 0.615 | GTGGGTGGC |
| 7950 | mamo-s | *BGIBMGA009412* | up | 1056 | 1064 | + | 11.3303 | 4.96E-05 | 0.615 | GTGGGTGGC |
| 7951 | mamo-s | *BGIBMGA009413* | up | 1564 | 1572 | - | 11.3303 | 4.96E-05 | 0.615 | GTGGGTGGC |
| 7952 | mamo-s | *BGIBMGA009414* | down | 406 | 414 | + | 11.3303 | 4.96E-05 | 0.615 | GTGGGTGGC |
| 7953 | mamo-s | *BGIBMGA009419* | up | 1934 | 1942 | - | 12.1284 | 2.16E-05 | 0.615 | GTGCGAGGC |
| 7954 | mamo-s | *BGIBMGA009421* | up | 999 | 1007 | + | 14.055 | 2.23E-06 | 0.615 | GTGCGTGGC |
| 7955 | mamo-s | *BGIBMGA009421* | down | 884 | 892 | + | 11.3303 | 4.96E-05 | 0.615 | GTGGGTGGC |
| 7956 | mamo-s | *BGIBMGA009421* | up | 1523 | 1531 | - | 11.0734 | 7.12E-05 | 0.726 | GAGCGTGGT |
| 7957 | mamo-s | *BGIBMGA009423* | up | 957 | 965 | - | 11.3303 | 4.96E-05 | 0.615 | GTGGGTGGC |
| 7958 | mamo-s | *BGIBMGA009425* | down | 92 | 100 | + | 12.0734 | 2.66E-05 | 0.615 | GCGCGTGTC |
| 7959 | mamo-s | *BGIBMGA009425* | down | 1747 | 1755 | - | 11.4404 | 4.51E-05 | 0.615 | CGGCGTGGC |
| 7960 | mamo-s | *BGIBMGA009425* | up | 132 | 140 | - | 11.3303 | 4.96E-05 | 0.615 | GTGGGTGGC |
| 7961 | mamo-s | *BGIBMGA009426* | up | 1107 | 1115 | + | 11.1743 | 5.90E-05 | 0.684 | CCGCGTGTC |
| 7962 | mamo-s | *BGIBMGA009428* | up | 526 | 534 | - | 11.3303 | 4.96E-05 | 0.615 | GTGGGTGGC |
| 7963 | mamo-s | *BGIBMGA009432* | down | 252 | 260 | + | 11.3303 | 4.96E-05 | 0.615 | GTGGGTGGC |
| 7964 | mamo-s | *BGIBMGA009432* | down | 686 | 694 | + | 11.3303 | 4.96E-05 | 0.615 | GTGGGTGGC |
| 7965 | mamo-s | *BGIBMGA009432* | up | 1442 | 1450 | - | 11.3303 | 4.96E-05 | 0.615 | GTGGGTGGC |
| 7966 | mamo-s | *BGIBMGA009433* | down | 1185 | 1193 | + | 11.2294 | 5.41E-05 | 0.65 | CTGCGAGGC |
| 7967 | mamo-s | *BGIBMGA009434* | up | 1633 | 1641 | + | 11.3303 | 4.96E-05 | 0.615 | GTGGGTGGC |
| 7968 | mamo-s | *BGIBMGA009436* | down | 970 | 978 | + | 11.3303 | 4.96E-05 | 0.615 | GTGGGTGGC |
| 7969 | mamo-s | *BGIBMGA009436* | down | 1888 | 1896 | + | 11.3303 | 4.96E-05 | 0.615 | GTGGGTGGC |
| 7970 | mamo-s | *BGIBMGA009439* | down | 1109 | 1117 | + | 11.5321 | 4.06E-05 | 0.615 | TTGCGTGGC |
| 7971 | mamo-s | *BGIBMGA009439* | up | 1175 | 1183 | - | 11.3303 | 4.96E-05 | 0.615 | GTGGGTGGC |
| 7972 | mamo-s | *BGIBMGA009439* | up | 1302 | 1310 | + | 10.8349 | 7.66E-05 | 0.751 | CCGCGAGGC |
| 7973 | mamo-s | *BGIBMGA009440* | up | 307 | 315 | + | 10.5413 | 9.42E-05 | 0.857 | GTGCGAGTC |
| 7974 | mamo-s | *BGIBMGA009441* | up | 445 | 453 | + | 11.3303 | 4.96E-05 | 0.615 | GTGGGTGGC |
| 7975 | mamo-s | *BGIBMGA009442* | down | 503 | 511 | - | 11.3303 | 4.96E-05 | 0.615 | GTGGGTGGC |
| 7976 | mamo-s | *BGIBMGA009446* | up | 1013 | 1021 | - | 11.3303 | 4.96E-05 | 0.615 | GTGGGTGGC |
| 7977 | mamo-s | *BGIBMGA009447* | up | 258 | 266 | + | 11.4404 | 4.51E-05 | 0.615 | GAGCGTGTC |
| 7978 | mamo-s | *BGIBMGA009447* | up | 378 | 386 | + | 11.4404 | 4.51E-05 | 0.615 | CGGCGTGGC |
| 7979 | mamo-s | *BGIBMGA009449* | down | 681 | 689 | - | 11.0734 | 7.12E-05 | 0.726 | GAGCGTGGT |
| 7980 | mamo-s | *BGIBMGA009450* | down | 714 | 722 | + | 11.3303 | 4.96E-05 | 0.615 | GTGGGTGGC |
| 7981 | mamo-s | *BGIBMGA009451* | up | 1032 | 1040 | + | 11.5321 | 4.06E-05 | 0.615 | TTGCGTGGC |
| 7982 | mamo-s | *BGIBMGA009451* | down | 665 | 673 | - | 11.3303 | 4.96E-05 | 0.615 | GTGGGTGGC |
| 7983 | mamo-s | *BGIBMGA009451* | down | 942 | 950 | + | 11.3303 | 4.96E-05 | 0.615 | GTGGGTGGC |
| 7984 | mamo-s | *BGIBMGA009452* | up | 174 | 182 | + | 10.5413 | 9.42E-05 | 0.857 | CAGCGTGTC |
| 7985 | mamo-s | *BGIBMGA009454* | up | 931 | 939 | - | 12.1284 | 2.16E-05 | 0.615 | CAGCGTGGC |
| 7986 | mamo-s | *BGIBMGA009455* | up | 600 | 608 | + | 11.2844 | 5.18E-05 | 0.63 | CTGCGTGCC |
| 7987 | mamo-s | *BGIBMGA009457* | up | 788 | 796 | - | 10.6881 | 8.61E-05 | 0.81 | GCACGTGGC |
| 7988 | mamo-s | *BGIBMGA009459* | down | 517 | 525 | + | 11.3303 | 4.96E-05 | 0.615 | GTGGGTGGC |
| 7989 | mamo-s | *BGIBMGA009460* | up | 1810 | 1818 | + | 11.0826 | 6.84E-05 | 0.709 | GTACGTGGC |
| 7990 | mamo-s | *BGIBMGA009461* | up | 447 | 455 | + | 11.3303 | 4.96E-05 | 0.615 | GTGGGTGGC |
| 7991 | mamo-s | *BGIBMGA009461* | down | 873 | 881 | + | 11.3303 | 4.96E-05 | 0.615 | GTGGGTGGC |
| 7992 | mamo-s | *BGIBMGA009461* | up | 1155 | 1163 | + | 11.0826 | 6.84E-05 | 0.709 | GTACGTGGC |
| 7993 | mamo-s | *BGIBMGA009464* | down | 641 | 649 | - | 10.5688 | 8.88E-05 | 0.824 | GTGCGTGAC |
| 7994 | mamo-s | *BGIBMGA009465* | down | 517 | 525 | + | 12.7615 | 1.04E-05 | 0.615 | CCGCGTGGC |
| 7995 | mamo-s | *BGIBMGA009467* | up | 1861 | 1869 | + | 11.3303 | 4.96E-05 | 0.615 | GTGGGTGGC |
| 7996 | mamo-s | *BGIBMGA009468* | down | 1219 | 1227 | + | 11.3303 | 4.96E-05 | 0.615 | GTGGGTGGC |
| 7997 | mamo-s | *BGIBMGA009468* | down | 278 | 286 | - | 11.2294 | 5.41E-05 | 0.65 | CTGCGAGGC |
| 7998 | mamo-s | *BGIBMGA009469* | down | 1890 | 1898 | - | 10.8349 | 7.66E-05 | 0.751 | CCGCGAGGC |
| 7999 | mamo-s | *BGIBMGA009470* | down | 1879 | 1887 | + | 11.3303 | 4.96E-05 | 0.615 | GTGGGTGGC |
| 8000 | mamo-s | *BGIBMGA009470* | down | 599 | 607 | + | 11.0826 | 6.84E-05 | 0.709 | GTACGTGGC |
| 8001 | mamo-s | *BGIBMGA009472* | down | 177 | 185 | + | 11.3303 | 4.96E-05 | 0.615 | GTGGGTGGC |

| | | | | | | | | | | |
|---|---|---|---|---|---|---|---|---|---|---|
| 8002 | mamo-s | *BGIBMGA009472* | up | 85 | 93 | + | 11.0826 | 6.84E-05 | 0.709 | GTACGTGGC |
| 8003 | mamo-s | *BGIBMGA009473* | down | 1105 | 1113 | + | 11.3303 | 4.96E-05 | 0.615 | GTGGGTGGC |
| 8004 | mamo-s | *BGIBMGA009474* | up | 1573 | 1581 | + | 11.3303 | 4.96E-05 | 0.615 | GTGGGTGGC |
| 8005 | mamo-s | *BGIBMGA009475* | up | 1757 | 1765 | - | 11.789 | 2.84E-05 | 0.615 | GCGCGTGCC |
| 8006 | mamo-s | *BGIBMGA009477* | up | 475 | 483 | - | 11.3303 | 4.96E-05 | 0.615 | GTGGGTGGC |
| 8007 | mamo-s | *BGIBMGA009477* | up | 1699 | 1707 | + | 11.3303 | 4.96E-05 | 0.615 | GTGGGTGGC |
| 8008 | mamo-s | *BGIBMGA009478* | up | 670 | 678 | - | 14.055 | 2.23E-06 | 0.615 | GTGCGTGGC |
| 8009 | mamo-s | *BGIBMGA009479* | down | 1195 | 1203 | - | 11.3303 | 4.96E-05 | 0.615 | GTGGGTGGC |
| 8010 | mamo-s | *BGIBMGA009479* | down | 1708 | 1716 | + | 11.1376 | 6.35E-05 | 0.709 | TCGCGTGGC |
| 8011 | mamo-s | *BGIBMGA009480* | down | 727 | 735 | + | 11.3303 | 4.96E-05 | 0.615 | GTGGGTGGC |
| 8012 | mamo-s | *BGIBMGA009480* | up | 1730 | 1738 | - | 11.0826 | 6.84E-05 | 0.709 | GTACGTGGC |
| 8013 | mamo-s | *BGIBMGA009483* | up | 1643 | 1651 | - | 11.3303 | 4.96E-05 | 0.615 | GTGGGTGGC |
| 8014 | mamo-s | *BGIBMGA009484* | up | 1245 | 1253 | + | 11.3303 | 4.96E-05 | 0.615 | GTGGGTGGC |
| 8015 | mamo-s | *BGIBMGA009484* | up | 1970 | 1978 | - | 10.8899 | 7.48E-05 | 0.739 | CCGCGTGCC |
| 8016 | mamo-s | *BGIBMGA009485* | down | 738 | 746 | - | 11.3303 | 4.96E-05 | 0.615 | GTGGGTGGC |
| 8017 | mamo-s | *BGIBMGA009485* | up | 760 | 768 | + | 11.3303 | 4.96E-05 | 0.615 | GTGGGTGGC |
| 8018 | mamo-s | *BGIBMGA009485* | up | 1485 | 1493 | - | 10.8899 | 7.48E-05 | 0.739 | CCGCGTGCC |
| 8019 | mamo-s | *BGIBMGA009486* | up | 55 | 63 | + | 11.3303 | 4.96E-05 | 0.615 | GTGGGTGGC |
| 8020 | mamo-s | *BGIBMGA009486* | up | 1076 | 1084 | - | 11.3303 | 4.96E-05 | 0.615 | GTGGGTGGC |
| 8021 | mamo-s | *BGIBMGA009486* | up | 1878 | 1886 | - | 11.3303 | 4.96E-05 | 0.615 | GTGGGTGGC |
| 8022 | mamo-s | *BGIBMGA009490* | down | 814 | 822 | + | 13.0275 | 8.53E-06 | 0.615 | GAGCGTGGC |
| 8023 | mamo-s | *BGIBMGA009490* | down | 1089 | 1097 | + | 11.3303 | 4.96E-05 | 0.615 | GTGGGTGGC |
| 8024 | mamo-s | *BGIBMGA009491* | up | 678 | 686 | + | 13.0275 | 8.53E-06 | 0.615 | GAGCGTGGC |
| 8025 | mamo-s | *BGIBMGA009491* | up | 953 | 961 | + | 11.3303 | 4.96E-05 | 0.615 | GTGGGTGGC |
| 8026 | mamo-s | *BGIBMGA009492* | down | 1118 | 1126 | - | 12.0734 | 2.66E-05 | 0.615 | GCGCGTGTC |
| 8027 | mamo-s | *BGIBMGA009492* | up | 1912 | 1920 | + | 11.0826 | 6.84E-05 | 0.709 | GTACGTGGC |
| 8028 | mamo-s | *BGIBMGA009494* | up | 277 | 285 | + | 11.1743 | 5.90E-05 | 0.684 | CCGCGTGTC |
| 8029 | mamo-s | *BGIBMGA009494* | up | 81 | 89 | - | 11.156 | 6.13E-05 | 0.705 | GAGCGTGCC |
| 8030 | mamo-s | *BGIBMGA009495* | down | 565 | 573 | + | 10.7523 | 8.38E-05 | 0.797 | AAGCGTGGC |
| 8031 | mamo-s | *BGIBMGA009497* | up | 48 | 56 | - | 11.3303 | 4.96E-05 | 0.615 | GTGGGTGGC |
| 8032 | mamo-s | *BGIBMGA009497* | down | 1786 | 1794 | + | 11.3303 | 4.96E-05 | 0.615 | GTGGGTGGC |
| 8033 | mamo-s | *BGIBMGA009498* | down | 1075 | 1083 | + | 11.3303 | 4.96E-05 | 0.615 | GTGGGTGGC |
| 8034 | mamo-s | *BGIBMGA009499* | up | 719 | 727 | - | 12.1284 | 2.16E-05 | 0.615 | GTGCGAGGC |
| 8035 | mamo-s | *BGIBMGA009499* | down | 659 | 667 | + | 11.3303 | 4.96E-05 | 0.615 | GTGGGTGGC |
| 8036 | mamo-s | *BGIBMGA009500* | down | 335 | 343 | - | 11.4404 | 4.51E-05 | 0.615 | CGGCGTGGC |
| 8037 | mamo-s | *BGIBMGA009500* | up | 503 | 511 | + | 11.3303 | 4.96E-05 | 0.615 | GTGGGTGGC |
| 8038 | mamo-s | *BGIBMGA009501* | up | 541 | 549 | + | 11.0826 | 6.84E-05 | 0.709 | GTACGTGGC |
| 8039 | mamo-s | *BGIBMGA009502* | down | 1758 | 1766 | + | 14.055 | 2.23E-06 | 0.615 | GTGCGTGGC |
| 8040 | mamo-s | *BGIBMGA009503* | up | 299 | 307 | + | 11.3303 | 4.96E-05 | 0.615 | GTGGGTGGC |
| 8041 | mamo-s | *BGIBMGA009504* | down | 1365 | 1373 | - | 12.1835 | 1.71E-05 | 0.615 | GTGCGTGCC |
| 8042 | mamo-s | *BGIBMGA009505* | up | 86 | 94 | + | 11.3303 | 4.96E-05 | 0.615 | GTGGGTGGC |
| 8043 | mamo-s | *BGIBMGA009507* | up | 880 | 888 | - | 11.3303 | 4.96E-05 | 0.615 | GTGGGTGGC |
| 8044 | mamo-s | *BGIBMGA009514* | up | 1248 | 1256 | + | 12.1284 | 2.16E-05 | 0.615 | CAGCGTGGC |
| 8045 | mamo-s | *BGIBMGA009514* | down | 1091 | 1099 | - | 11.3303 | 4.96E-05 | 0.615 | GTGGGTGGC |
| 8046 | mamo-s | *BGIBMGA009515* | up | 177 | 185 | + | 12.1284 | 2.16E-05 | 0.615 | GTGCGAGGC |
| 8047 | mamo-s | *BGIBMGA009516* | up | 1395 | 1403 | - | 11.3303 | 4.96E-05 | 0.615 | GTGGGTGGC |
| 8048 | mamo-s | *BGIBMGA009516* | up | 823 | 831 | - | 11.0826 | 6.84E-05 | 0.709 | GTACGTGGC |
| 8049 | mamo-s | *BGIBMGA009519* | up | 1191 | 1199 | - | 10.8899 | 7.48E-05 | 0.739 | CCGCGTGCC |
| 8050 | mamo-s | *BGIBMGA009523* | up | 97 | 105 | - | 11.3303 | 4.96E-05 | 0.615 | GTGGGTGGC |
| 8051 | mamo-s | *BGIBMGA009525* | up | 911 | 919 | + | 11.3303 | 4.96E-05 | 0.615 | GTGGGTGGC |
| 8052 | mamo-s | *BGIBMGA009526* | down | 1530 | 1538 | + | 11.3303 | 4.96E-05 | 0.615 | GTGGGTGGC |
| 8053 | mamo-s | *BGIBMGA009526* | up | 1576 | 1584 | - | 11.3303 | 4.96E-05 | 0.615 | GTGGGTGGC |
| 8054 | mamo-s | *BGIBMGA009527* | up | 533 | 541 | - | 10.8899 | 7.48E-05 | 0.739 | CCGCGTGCC |
| 8055 | mamo-s | *BGIBMGA009528* | up | 1192 | 1200 | - | 11.3303 | 4.96E-05 | 0.615 | GTGGGTGGC |
| 8056 | mamo-s | *BGIBMGA009528* | down | 1225 | 1233 | - | 11.3303 | 4.96E-05 | 0.615 | GTGGGTGGC |
| 8057 | mamo-s | *BGIBMGA009531* | up | 1121 | 1129 | + | 11.3303 | 4.96E-05 | 0.615 | GTGGGTGGC |
| 8058 | mamo-s | *BGIBMGA009533* | down | 730 | 738 | + | 11.4404 | 4.51E-05 | 0.615 | CGGCGTGGC |
| 8059 | mamo-s | *BGIBMGA009534* | down | 16 | 24 | - | 10.8073 | 7.89E-05 | 0.766 | CCGCGTGGT |
| 8060 | mamo-s | *BGIBMGA009535* | up | 1348 | 1356 | + | 10.7523 | 8.38E-05 | 0.797 | AAGCGTGGC |
| 8061 | mamo-s | *BGIBMGA009535* | up | 1075 | 1083 | + | 10.5138 | 9.75E-05 | 0.865 | GTGCGTGTT |
| 8062 | mamo-s | *BGIBMGA009538* | up | 1756 | 1764 | - | 12.1009 | 2.43E-05 | 0.615 | GTGCGTGGT |
| 8063 | mamo-s | *BGIBMGA009539* | up | 414 | 422 | - | 14.055 | 2.23E-06 | 0.615 | GTGCGTGGC |
| 8064 | mamo-s | *BGIBMGA009540* | down | 60 | 68 | - | 12.1009 | 2.43E-05 | 0.615 | GTGCGTGGT |
| 8065 | mamo-s | *BGIBMGA009542* | down | 706 | 714 | - | 12.3394 | 1.49E-05 | 0.615 | GGGCGTGGC |
| 8066 | mamo-s | *BGIBMGA009543* | down | 1783 | 1791 | + | 11.3303 | 4.96E-05 | 0.615 | GTGGGTGGC |
| 8067 | mamo-s | *BGIBMGA009544* | down | 625 | 633 | + | 11.0826 | 6.84E-05 | 0.709 | GTACGTGGC |
| 8068 | mamo-s | *BGIBMGA009544* | up | 962 | 970 | - | 10.8073 | 7.89E-05 | 0.766 | CCGCGTGGT |
| 8069 | mamo-s | *BGIBMGA009545* | up | 1567 | 1575 | - | 11.3303 | 4.96E-05 | 0.615 | GTGGGTGGC |
| 8070 | mamo-s | *BGIBMGA009548* | down | 753 | 761 | + | 11.3303 | 4.96E-05 | 0.615 | GTGGGTGGC |
| 8071 | mamo-s | *BGIBMGA009548* | down | 1443 | 1451 | - | 11.3303 | 4.96E-05 | 0.615 | GTGGGTGGC |
| 8072 | mamo-s | *BGIBMGA009548* | down | 1879 | 1887 | + | 11.3303 | 4.96E-05 | 0.615 | GTGGGTGGC |
| 8073 | mamo-s | *BGIBMGA009548* | up | 1589 | 1597 | - | 10.8349 | 7.66E-05 | 0.751 | CCGCGAGGC |
| 8074 | mamo-s | *BGIBMGA009550* | down | 1585 | 1593 | + | 11.3303 | 4.96E-05 | 0.615 | GTGGGTGGC |
| 8075 | mamo-s | *BGIBMGA009550* | up | 1139 | 1147 | - | 11.0826 | 6.84E-05 | 0.709 | GTACGTGGC |
| 8076 | mamo-s | *BGIBMGA009553* | up | 701 | 709 | - | 11.0826 | 6.84E-05 | 0.709 | GTACGTGGC |
| 8077 | mamo-s | *BGIBMGA009554* | down | 997 | 1005 | + | 11.3303 | 4.96E-05 | 0.615 | GTGGGTGGC |
| 8078 | mamo-s | *BGIBMGA009555* | down | 1066 | 1074 | - | 11.3303 | 4.96E-05 | 0.615 | GTGGGTGGC |
| 8079 | mamo-s | *BGIBMGA009555* | up | 735 | 743 | - | 11.2018 | 5.68E-05 | 0.674 | CTGCGTGGT |
| 8080 | mamo-s | *BGIBMGA009555* | down | 22 | 30 | - | 11.1009 | 6.57E-05 | 0.709 | GAGCGAGGC |
| 8081 | mamo-s | *BGIBMGA009556* | down | 1223 | 1231 | + | 11.3303 | 4.96E-05 | 0.615 | GTGGGTGGC |
| 8082 | mamo-s | *BGIBMGA009556* | down | 1973 | 1981 | + | 11.3303 | 4.96E-05 | 0.615 | GTGGGTGGC |
| 8083 | mamo-s | *BGIBMGA009558* | down | 187 | 195 | - | 11.3303 | 4.96E-05 | 0.615 | GTGGGTGGC |
| 8084 | mamo-s | *BGIBMGA009559* | down | 1468 | 1476 | - | 11.3303 | 4.96E-05 | 0.615 | GTGGGTGGC |
| 8085 | mamo-s | *BGIBMGA009563* | up | 55 | 63 | - | 11.0826 | 6.84E-05 | 0.709 | GTACGTGGC |
| 8086 | mamo-s | *BGIBMGA009563* | down | 1858 | 1866 | - | 11.0826 | 6.84E-05 | 0.709 | GTACGTGGC |
| 8087 | mamo-s | *BGIBMGA009564* | down | 450 | 458 | + | 12.1009 | 2.43E-05 | 0.615 | GTGCGTGGT |
| 8088 | mamo-s | *BGIBMGA009564* | down | 1057 | 1065 | - | 10.5138 | 9.75E-05 | 0.865 | GTGCGTGTT |
| 8089 | mamo-s | *BGIBMGA009569* | up | 1351 | 1359 | + | 12.0734 | 2.66E-05 | 0.615 | GCGCGTGTC |

| | | | | | | | | | | |
|---|---|---|---|---|---|---|---|---|---|---|
| 8090 | mamo-s | *BGIBMGA009569* | up | 1346 | 1354 | - | 11.7339 | 3.29E-05 | 0.615 | GCGCGAGGC |
| 8091 | mamo-s | *BGIBMGA009569* | down | 612 | 620 | - | 11.3303 | 4.96E-05 | 0.615 | GTGGGTGGC |
| 8092 | mamo-s | *BGIBMGA009570* | down | 408 | 416 | + | 13.6606 | 4.06E-06 | 0.615 | GCGCGTGGC |
| 8093 | mamo-s | *BGIBMGA009570* | up | 306 | 314 | + | 11.3303 | 4.96E-05 | 0.615 | GTGGGTGGC |
| 8094 | mamo-s | *BGIBMGA009573* | up | 160 | 168 | - | 11.3303 | 4.96E-05 | 0.615 | GTGGGTGGC |
| 8095 | mamo-s | *BGIBMGA009573* | down | 266 | 274 | - | 11.3303 | 4.96E-05 | 0.615 | GTGGGTGGC |
| 8096 | mamo-s | *BGIBMGA009577* | down | 904 | 912 | + | 13.0275 | 8.53E-06 | 0.615 | GAGCGTGGC |
| 8097 | mamo-s | *BGIBMGA009577* | up | 911 | 919 | + | 11.3303 | 4.96E-05 | 0.615 | GTGGGTGGC |
| 8098 | mamo-s | *BGIBMGA009579* | down | 780 | 788 | - | 12.3394 | 1.49E-05 | 0.615 | GGGCGTGGC |
| 8099 | mamo-s | *BGIBMGA009580* | up | 280 | 288 | + | 11.3303 | 4.96E-05 | 0.615 | GTGGGTGGC |
| 8100 | mamo-s | *BGIBMGA009580* | up | 1571 | 1579 | + | 10.9358 | 7.30E-05 | 0.728 | GCGGGTGGC |
| 8101 | mamo-s | *BGIBMGA009581* | down | 1644 | 1652 | + | 11.3303 | 4.96E-05 | 0.615 | GTGGGTGGC |
| 8102 | mamo-s | *BGIBMGA009582* | down | 891 | 899 | + | 13.0275 | 8.53E-06 | 0.615 | GAGCGTGGC |
| 8103 | mamo-s | *BGIBMGA009582* | down | 1075 | 1083 | + | 12.1835 | 1.71E-05 | 0.615 | GTGCGTGCC |
| 8104 | mamo-s | *BGIBMGA009582* | down | 531 | 539 | + | 11.3303 | 4.96E-05 | 0.615 | GTGGGTGGC |
| 8105 | mamo-s | *BGIBMGA009582* | up | 1663 | 1671 | - | 11.3303 | 4.96E-05 | 0.615 | GTGGGTGGC |
| 8106 | mamo-s | *BGIBMGA009583* | up | 1177 | 1185 | + | 11.2844 | 5.18E-05 | 0.63 | CTGCGTGCC |
| 8107 | mamo-s | *BGIBMGA009584* | down | 1866 | 1874 | - | 11.3303 | 4.96E-05 | 0.615 | GTGGGTGGC |
| 8108 | mamo-s | *BGIBMGA009585* | up | 172 | 180 | - | 11.3303 | 4.96E-05 | 0.615 | GTGGGTGGC |
| 8109 | mamo-s | *BGIBMGA009585* | down | 928 | 936 | - | 11.3303 | 4.96E-05 | 0.615 | GTGGGTGGC |
| 8110 | mamo-s | *BGIBMGA009586* | down | 689 | 697 | - | 14.055 | 2.23E-06 | 0.615 | GTGCGTGGC |
| 8111 | mamo-s | *BGIBMGA009586* | up | 250 | 258 | - | 11.3303 | 4.96E-05 | 0.615 | GTGGGTGGC |
| 8112 | mamo-s | *BGIBMGA009586* | down | 926 | 934 | + | 11.3303 | 4.96E-05 | 0.615 | GTGGGTGGC |
| 8113 | mamo-s | *BGIBMGA009587* | up | 711 | 719 | - | 14.055 | 2.23E-06 | 0.615 | GTGCGTGGC |
| 8114 | mamo-s | *BGIBMGA009587* | up | 948 | 956 | + | 11.3303 | 4.96E-05 | 0.615 | GTGGGTGGC |
| 8115 | mamo-s | *BGIBMGA009589* | up | 1136 | 1144 | + | 11.3303 | 4.96E-05 | 0.615 | GTGGGTGGC |
| 8116 | mamo-s | *BGIBMGA009589* | down | 1658 | 1666 | - | 11.3303 | 4.96E-05 | 0.615 | GTGGGTGGC |
| 8117 | mamo-s | *BGIBMGA009590* | down | 640 | 648 | - | 10.7523 | 8.38E-05 | 0.797 | AAGCGTGGC |
| 8118 | mamo-s | *BGIBMGA009592* | up | 512 | 520 | + | 11.0826 | 6.84E-05 | 0.709 | GTACGTGGC |
| 8119 | mamo-s | *BGIBMGA009593* | up | 157 | 165 | + | 13.6606 | 4.06E-06 | 0.615 | GCGCGTGGC |
| 8120 | mamo-s | *BGIBMGA009593* | up | 1781 | 1789 | - | 11.3853 | 4.74E-05 | 0.615 | ACGCGTGGC |
| 8121 | mamo-s | *BGIBMGA009593* | down | 482 | 490 | + | 11.3303 | 4.96E-05 | 0.615 | GTGGGTGGC |
| 8122 | mamo-s | *BGIBMGA009593* | up | 1324 | 1332 | - | 11.3303 | 4.96E-05 | 0.615 | GTGGGTGGC |
| 8123 | mamo-s | *BGIBMGA009594* | up | 295 | 303 | + | 10.5413 | 9.42E-05 | 0.857 | CAGCGTGTC |
| 8124 | mamo-s | *BGIBMGA009595* | down | 707 | 715 | - | 11.3303 | 4.96E-05 | 0.615 | GTGGGTGGC |
| 8125 | mamo-s | *BGIBMGA009595* | up | 1550 | 1558 | + | 10.9358 | 7.30E-05 | 0.728 | GCGGGTGGC |
| 8126 | mamo-s | *BGIBMGA009596* | down | 772 | 780 | - | 14.055 | 2.23E-06 | 0.615 | GTGCGTGGC |
| 8127 | mamo-s | *BGIBMGA009596* | down | 1402 | 1410 | + | 10.6881 | 8.61E-05 | 0.81 | GCACGTGGC |
| 8128 | mamo-s | *BGIBMGA009597* | down | 258 | 266 | + | 11.3303 | 4.96E-05 | 0.615 | GTGGGTGGC |
| 8129 | mamo-s | *BGIBMGA009597* | up | 1293 | 1301 | - | 11.3303 | 4.96E-05 | 0.615 | GTGGGTGGC |
| 8130 | mamo-s | *BGIBMGA009598* | down | 463 | 471 | + | 14.055 | 2.23E-06 | 0.615 | GTGCGTGGC |
| 8131 | mamo-s | *BGIBMGA009598* | down | 1424 | 1432 | - | 10.9358 | 7.30E-05 | 0.728 | GCGGGTGGC |
| 8132 | mamo-s | *BGIBMGA009599* | down | 936 | 944 | - | 11.3303 | 4.96E-05 | 0.615 | GTGGGTGGC |
| 8133 | mamo-s | *BGIBMGA009599* | down | 1378 | 1386 | - | 11.0826 | 6.84E-05 | 0.709 | GTACGTGGC |
| 8134 | mamo-s | *BGIBMGA009600* | down | 1372 | 1380 | + | 11.3303 | 4.96E-05 | 0.615 | GTGGGTGGC |
| 8135 | mamo-s | *BGIBMGA009601* | up | 1907 | 1915 | + | 11.4404 | 4.51E-05 | 0.615 | GAGCGTGTC |
| 8136 | mamo-s | *BGIBMGA009602* | up | 390 | 398 | - | 11.0826 | 6.84E-05 | 0.709 | GTACGTGGC |
| 8137 | mamo-s | *BGIBMGA009603* | down | 1265 | 1273 | - | 11.3303 | 4.96E-05 | 0.615 | GTGGGTGGC |
| 8138 | mamo-s | *BGIBMGA009604* | down | 437 | 445 | + | 11.3303 | 4.96E-05 | 0.615 | GTGGGTGGC |
| 8139 | mamo-s | *BGIBMGA009604* | up | 1686 | 1694 | - | 11.3303 | 4.96E-05 | 0.615 | GTGGGTGGC |
| 8140 | mamo-s | *BGIBMGA009605* | up | 1733 | 1741 | - | 11.3303 | 4.96E-05 | 0.615 | GTGGGTGGC |
| 8141 | mamo-s | *BGIBMGA009610* | up | 1581 | 1589 | - | 11.3303 | 4.96E-05 | 0.615 | GTGGGTGGC |
| 8142 | mamo-s | *BGIBMGA009611* | up | 10 | 18 | + | 11.3303 | 4.96E-05 | 0.615 | GTGGGTGGC |
| 8143 | mamo-s | *BGIBMGA009611* | up | 227 | 235 | + | 11.3303 | 4.96E-05 | 0.615 | GTGGGTGGC |
| 8144 | mamo-s | *BGIBMGA009612* | up | 1558 | 1566 | + | 11.2844 | 5.18E-05 | 0.63 | CTGCGTGCC |
| 8145 | mamo-s | *BGIBMGA009614* | up | 1366 | 1374 | + | 11.3303 | 4.96E-05 | 0.615 | GTGGGTGGC |
| 8146 | mamo-s | *BGIBMGA009615* | down | 135 | 143 | - | 11.3303 | 4.96E-05 | 0.615 | GTGGGTGGC |
| 8147 | mamo-s | *BGIBMGA009615* | down | 1714 | 1722 | + | 11.3303 | 4.96E-05 | 0.615 | GTGGGTGGC |
| 8148 | mamo-s | *BGIBMGA009616* | down | 1164 | 1172 | + | 12.3394 | 1.49E-05 | 0.615 | GGGCGTGGC |
| 8149 | mamo-s | *BGIBMGA009616* | down | 240 | 248 | - | 11.3303 | 4.96E-05 | 0.615 | GTGGGTGGC |
| 8150 | mamo-s | *BGIBMGA009616* | down | 1713 | 1721 | - | 11.0826 | 6.84E-05 | 0.709 | GTACGTGGC |
| 8151 | mamo-s | *BGIBMGA009617* | up | 315 | 323 | - | 11.3303 | 4.96E-05 | 0.615 | GTGGGTGGC |
| 8152 | mamo-s | *BGIBMGA009619* | down | 1354 | 1362 | + | 12.1835 | 1.71E-05 | 0.615 | GTGCGTGCC |
| 8153 | mamo-s | *BGIBMGA009621* | up | 3 | 11 | + | 10.5413 | 9.42E-05 | 0.857 | GTGCGAGTC |
| 8154 | mamo-s | *BGIBMGA009622* | down | 1018 | 1026 | - | 11.3303 | 4.96E-05 | 0.615 | GTGGGTGGC |
| 8155 | mamo-s | *BGIBMGA009623* | down | 471 | 479 | - | 12.1284 | 2.16E-05 | 0.615 | GTGCGAGGC |
| 8156 | mamo-s | *BGIBMGA009623* | up | 308 | 316 | - | 11.7064 | 3.52E-05 | 0.615 | GCGCGTGGT |
| 8157 | mamo-s | *BGIBMGA009623* | up | 1020 | 1028 | - | 11.3303 | 4.96E-05 | 0.615 | GTGGGTGGC |
| 8158 | mamo-s | *BGIBMGA009623* | up | 1353 | 1361 | - | 11.3303 | 4.96E-05 | 0.615 | GTGGGTGGC |
| 8159 | mamo-s | *BGIBMGA009626* | up | 239 | 247 | - | 11.3303 | 4.96E-05 | 0.615 | GTGGGTGGC |
| 8160 | mamo-s | *BGIBMGA009626* | down | 620 | 628 | - | 10.6881 | 8.61E-05 | 0.81 | GCACGTGGC |
| 8161 | mamo-s | *BGIBMGA009627* | up | 1795 | 1803 | + | 13.156 | 6.30E-06 | 0.615 | CTGCGTGGC |
| 8162 | mamo-s | *BGIBMGA009627* | down | 1566 | 1574 | + | 11.3303 | 4.96E-05 | 0.615 | GTGGGTGGC |
| 8163 | mamo-s | *BGIBMGA009627* | down | 1737 | 1745 | + | 11.3303 | 4.96E-05 | 0.615 | GTGGGTGGC |
| 8164 | mamo-s | *BGIBMGA009628* | down | 1464 | 1472 | - | 11.3303 | 4.96E-05 | 0.615 | GTGGGTGGC |
| 8165 | mamo-s | *BGIBMGA009632* | down | 337 | 345 | + | 11.3303 | 4.96E-05 | 0.615 | GTGGGTGGC |
| 8166 | mamo-s | *BGIBMGA009633* | up | 1404 | 1412 | - | 10.8349 | 7.66E-05 | 0.751 | CCGCGAGGC |
| 8167 | mamo-s | *BGIBMGA009634* | down | 1235 | 1243 | - | 11.3303 | 4.96E-05 | 0.615 | GTGGGTGGC |
| 8168 | mamo-s | *BGIBMGA009635* | down | 412 | 420 | + | 11.3303 | 4.96E-05 | 0.615 | GTGGGTGGC |
| 8169 | mamo-s | *BGIBMGA009635* | up | 636 | 644 | - | 11.3303 | 4.96E-05 | 0.615 | GTGGGTGGC |
| 8170 | mamo-s | *BGIBMGA009635* | down | 1731 | 1739 | - | 11.3303 | 4.96E-05 | 0.615 | GTGGGTGGC |
| 8171 | mamo-s | *BGIBMGA009637* | down | 1515 | 1523 | - | 11.3303 | 4.96E-05 | 0.615 | GTGGGTGGC |
| 8172 | mamo-s | *BGIBMGA009642* | down | 356 | 364 | + | 11.3303 | 4.96E-05 | 0.615 | GTGGGTGGC |
| 8173 | mamo-s | *BGIBMGA009642* | down | 274 | 282 | + | 11.2294 | 5.41E-05 | 0.65 | CTGCGAGGC |
| 8174 | mamo-s | *BGIBMGA009643* | down | 1620 | 1628 | - | 12.0734 | 2.66E-05 | 0.615 | GCGCGTGTC |
| 8175 | mamo-s | *BGIBMGA009643* | down | 428 | 436 | - | 11.1743 | 5.90E-05 | 0.684 | CCGCGTGTC |
| 8176 | mamo-s | *BGIBMGA009643* | down | 1158 | 1166 | - | 11.0826 | 6.84E-05 | 0.709 | GTACGTGGC |
| 8177 | mamo-s | *BGIBMGA009644* | down | 227 | 235 | + | 11.3303 | 4.96E-05 | 0.615 | GTGGGTGGC |

| | | | | | | | | | | |
|---|---|---|---|---|---|---|---|---|---|---|
| 8178 | mamo-s | *BGIBMGA009644* | up | 651 | 659 | - | 11.3303 | 4.96E-05 | 0.615 | GTGGGTGGC |
| 8179 | mamo-s | *BGIBMGA009644* | down | 1800 | 1808 | - | 11.3303 | 4.96E-05 | 0.615 | GTGGGTGGC |
| 8180 | mamo-s | *BGIBMGA009645* | up | 1672 | 1680 | - | 11.3303 | 4.96E-05 | 0.615 | GTGGGTGGC |
| 8181 | mamo-s | *BGIBMGA009645* | up | 1317 | 1325 | + | 10.5138 | 9.75E-05 | 0.865 | GTGCGTGTT |
| 8182 | mamo-s | *BGIBMGA009646* | down | 593 | 601 | + | 11.3303 | 4.96E-05 | 0.615 | GTGGGTGGC |
| 8183 | mamo-s | *BGIBMGA009650* | up | 266 | 274 | - | 11.0826 | 6.84E-05 | 0.709 | GTACGTGGC |
| 8184 | mamo-s | *BGIBMGA009651* | down | 388 | 396 | + | 11.3303 | 4.96E-05 | 0.615 | GTGGGTGGC |
| 8185 | mamo-s | *BGIBMGA009651* | down | 1475 | 1483 | - | 11.156 | 6.13E-05 | 0.705 | GAGCGTGCC |
| 8186 | mamo-s | *BGIBMGA009652* | up | 462 | 470 | - | 11.3303 | 4.96E-05 | 0.615 | GTGGGTGGC |
| 8187 | mamo-s | *BGIBMGA009653* | up | 1805 | 1813 | + | 11.4404 | 4.51E-05 | 0.615 | GAGCGTGTC |
| 8188 | mamo-s | *BGIBMGA009654* | up | 602 | 610 | + | 11.3303 | 4.96E-05 | 0.615 | GTGGGTGGC |
| 8189 | mamo-s | *BGIBMGA009655* | down | 363 | 371 | - | 13.6606 | 4.06E-06 | 0.615 | GCGCGTGGC |
| 8190 | mamo-s | *BGIBMGA009656* | down | 1514 | 1522 | - | 11.3303 | 4.96E-05 | 0.615 | GTGGGTGGC |
| 8191 | mamo-s | *BGIBMGA009656* | down | 1297 | 1305 | + | 11.0826 | 6.84E-05 | 0.709 | GTACGTGGC |
| 8192 | mamo-s | *BGIBMGA009657* | down | 1398 | 1406 | + | 11.3303 | 4.96E-05 | 0.615 | GTGGGTGGC |
| 8193 | mamo-s | *BGIBMGA009659* | up | 1033 | 1041 | + | 11.3303 | 4.96E-05 | 0.615 | GTGGGTGGC |
| 8194 | mamo-s | *BGIBMGA009660* | up | 650 | 658 | - | 12.7615 | 1.04E-05 | 0.615 | CCGCGTGGC |
| 8195 | mamo-s | *BGIBMGA009660* | down | 1046 | 1054 | + | 11.3303 | 4.96E-05 | 0.615 | GTGGGTGGC |
| 8196 | mamo-s | *BGIBMGA009661* | down | 883 | 891 | - | 14.055 | 2.23E-06 | 0.615 | GTGCGTGGC |
| 8197 | mamo-s | *BGIBMGA009661* | up | 1008 | 1016 | - | 12.7615 | 1.04E-05 | 0.615 | CCGCGTGGC |
| 8198 | mamo-s | *BGIBMGA009661* | up | 150 | 158 | + | 10.9358 | 7.30E-05 | 0.728 | GCGGGTGGC |
| 8199 | mamo-s | *BGIBMGA009661* | up | 326 | 334 | - | 10.8349 | 7.66E-05 | 0.751 | CCGCGAGGC |
| 8200 | mamo-s | *BGIBMGA009662* | down | 1394 | 1402 | + | 12.1284 | 2.16E-05 | 0.615 | GTGCGAGGC |
| 8201 | mamo-s | *BGIBMGA009662* | down | 1810 | 1818 | - | 11.3303 | 4.96E-05 | 0.615 | GTGGGTGGC |
| 8202 | mamo-s | *BGIBMGA009666* | up | 1570 | 1578 | - | 14.055 | 2.23E-06 | 0.615 | GTGCGTGGC |
| 8203 | mamo-s | *BGIBMGA009666* | down | 393 | 401 | - | 11.3303 | 4.96E-05 | 0.615 | GTGGGTGGC |
| 8204 | mamo-s | *BGIBMGA009666* | down | 974 | 982 | - | 11.3303 | 4.96E-05 | 0.615 | GTGGGTGGC |
| 8205 | mamo-s | *BGIBMGA009666* | up | 1373 | 1381 | - | 11.3303 | 4.96E-05 | 0.615 | GTGGGTGGC |
| 8206 | mamo-s | *BGIBMGA009668* | down | 582 | 590 | + | 11.3303 | 4.96E-05 | 0.615 | GTGGGTGGC |
| 8207 | mamo-s | *BGIBMGA009668* | up | 780 | 788 | - | 11.3303 | 4.96E-05 | 0.615 | GTGGGTGGC |
| 8208 | mamo-s | *BGIBMGA009669* | down | 1424 | 1432 | - | 11.0826 | 6.84E-05 | 0.709 | GTACGTGGC |
| 8209 | mamo-s | *BGIBMGA009670* | down | 64 | 72 | - | 13.156 | 6.30E-06 | 0.615 | CTGCGTGGC |
| 8210 | mamo-s | *BGIBMGA009672* | up | 278 | 286 | + | 13.0275 | 8.53E-06 | 0.615 | GAGCGTGGC |
| 8211 | mamo-s | *BGIBMGA009672* | up | 1138 | 1146 | + | 11.3303 | 4.96E-05 | 0.615 | GTGGGTGGC |
| 8212 | mamo-s | *BGIBMGA009674* | down | 499 | 507 | + | 11.0826 | 6.84E-05 | 0.709 | GTACGTGGC |
| 8213 | mamo-s | *BGIBMGA009676* | down | 934 | 942 | - | 11.3303 | 4.96E-05 | 0.615 | GTGGGTGGC |
| 8214 | mamo-s | *BGIBMGA009679* | down | 352 | 360 | - | 11.3303 | 4.96E-05 | 0.615 | GTGGGTGGC |
| 8215 | mamo-s | *BGIBMGA009679* | down | 1960 | 1968 | - | 11.3303 | 4.96E-05 | 0.615 | GTGGGTGGC |
| 8216 | mamo-s | *BGIBMGA009680* | up | 1804 | 1812 | - | 11.3303 | 4.96E-05 | 0.615 | GTGGGTGGC |
| 8217 | mamo-s | *BGIBMGA009681* | up | 1424 | 1432 | - | 12.0734 | 2.66E-05 | 0.615 | GCGCGTGTC |
| 8218 | mamo-s | *BGIBMGA009683* | down | 1414 | 1422 | - | 10.5413 | 9.42E-05 | 0.857 | CAGCGTGTC |
| 8219 | mamo-s | *BGIBMGA009686* | down | 1068 | 1076 | + | 11.3303 | 4.96E-05 | 0.615 | GTGGGTGGC |
| 8220 | mamo-s | *BGIBMGA009686* | down | 1803 | 1811 | + | 11.3303 | 4.96E-05 | 0.615 | GTGGGTGGC |
| 8221 | mamo-s | *BGIBMGA009687* | down | 97 | 105 | + | 10.5413 | 9.42E-05 | 0.857 | CAGCGTGTC |
| 8222 | mamo-s | *BGIBMGA009688* | up | 873 | 881 | + | 11.4404 | 4.51E-05 | 0.615 | GAGCGTGTC |
| 8223 | mamo-s | *BGIBMGA009688* | up | 1896 | 1904 | - | 11.2294 | 5.41E-05 | 0.65 | CTGCGAGGC |
| 8224 | mamo-s | *BGIBMGA009689* | up | 308 | 316 | - | 11.3303 | 4.96E-05 | 0.615 | GTGGGTGGC |
| 8225 | mamo-s | *BGIBMGA009689* | up | 874 | 882 | + | 11.3303 | 4.96E-05 | 0.615 | GTGGGTGGC |
| 8226 | mamo-s | *BGIBMGA009690* | up | 838 | 846 | - | 11.3303 | 4.96E-05 | 0.615 | GTGGGTGGC |
| 8227 | mamo-s | *BGIBMGA009690* | down | 1059 | 1067 | + | 10.8899 | 7.48E-05 | 0.739 | CCGCGTGCC |
| 8228 | mamo-s | *BGIBMGA009691* | down | 1881 | 1889 | + | 11.7064 | 3.52E-05 | 0.615 | GCGCGTGGT |
| 8229 | mamo-s | *BGIBMGA009691* | down | 1149 | 1157 | - | 11.5321 | 4.06E-05 | 0.615 | TTGCGTGGC |
| 8230 | mamo-s | *BGIBMGA009691* | down | 326 | 334 | + | 11.2844 | 5.18E-05 | 0.63 | CTGCGTGCC |
| 8231 | mamo-s | *BGIBMGA009694* | down | 426 | 434 | + | 11.3303 | 4.96E-05 | 0.615 | GTGGGTGGC |
| 8232 | mamo-s | *BGIBMGA009694* | up | 1098 | 1106 | + | 11.2844 | 5.18E-05 | 0.63 | CTGCGTGCC |
| 8233 | mamo-s | *BGIBMGA009695* | up | 1809 | 1817 | + | 11.3303 | 4.96E-05 | 0.615 | GTGGGTGGC |
| 8234 | mamo-s | *BGIBMGA009697* | up | 1504 | 1512 | + | 11.3303 | 4.96E-05 | 0.615 | GTGGGTGGC |
| 8235 | mamo-s | *BGIBMGA009700* | down | 295 | 303 | - | 11.3303 | 4.96E-05 | 0.615 | GTGGGTGGC |
| 8236 | mamo-s | *BGIBMGA009700* | down | 1867 | 1875 | - | 11.0826 | 6.84E-05 | 0.709 | GTACGTGGC |
| 8237 | mamo-s | *BGIBMGA009700* | up | 1443 | 1451 | + | 10.7523 | 8.38E-05 | 0.797 | AAGCGTGGC |
| 8238 | mamo-s | *BGIBMGA009701* | up | 951 | 959 | - | 11.3303 | 4.96E-05 | 0.615 | GTGGGTGGC |
| 8239 | mamo-s | *BGIBMGA009702* | down | 1527 | 1535 | + | 11.3303 | 4.96E-05 | 0.615 | GTGGGTGGC |
| 8240 | mamo-s | *BGIBMGA009703* | down | 582 | 590 | + | 11.7798 | 3.11E-05 | 0.615 | ATGCGTGGC |
| 8241 | mamo-s | *BGIBMGA009703* | up | 51 | 59 | + | 11.3303 | 4.96E-05 | 0.615 | GTGGGTGGC |
| 8242 | mamo-s | *BGIBMGA009703* | up | 201 | 209 | + | 11.3303 | 4.96E-05 | 0.615 | GTGGGTGGC |
| 8243 | mamo-s | *BGIBMGA009705* | up | 123 | 131 | + | 11.3303 | 4.96E-05 | 0.615 | GTGGGTGGC |
| 8244 | mamo-s | *BGIBMGA009705* | up | 1657 | 1665 | + | 11.3303 | 4.96E-05 | 0.615 | GTGGGTGGC |
| 8245 | mamo-s | *BGIBMGA009709* | down | 1071 | 1079 | - | 11.3303 | 4.96E-05 | 0.615 | GTGGGTGGC |
| 8246 | mamo-s | *BGIBMGA009710* | up | 1497 | 1505 | + | 11.3303 | 4.96E-05 | 0.615 | GTGGGTGGC |
| 8247 | mamo-s | *BGIBMGA009711* | up | 1156 | 1164 | + | 11.2294 | 5.41E-05 | 0.65 | CTGCGAGGC |
| 8248 | mamo-s | *BGIBMGA009712* | up | 882 | 890 | - | 11.3303 | 4.96E-05 | 0.615 | GTGGGTGGC |
| 8249 | mamo-s | *BGIBMGA009715* | up | 720 | 728 | - | 12.1009 | 2.43E-05 | 0.615 | GTGCGTGGT |
| 8250 | mamo-s | *BGIBMGA009715* | up | 650 | 658 | - | 10.8899 | 7.48E-05 | 0.739 | CCGCGTGCC |
| 8251 | mamo-s | *BGIBMGA009716* | up | 1677 | 1685 | - | 12.7615 | 1.04E-05 | 0.615 | CCGCGTGGC |
| 8252 | mamo-s | *BGIBMGA009716* | down | 1602 | 1610 | - | 11.3303 | 4.96E-05 | 0.615 | GTGGGTGGC |
| 8253 | mamo-s | *BGIBMGA009717* | down | 1316 | 1324 | + | 11.4404 | 4.51E-05 | 0.615 | CGGCGTGGC |
| 8254 | mamo-s | *BGIBMGA009717* | down | 1484 | 1492 | + | 11.4404 | 4.51E-05 | 0.615 | CGGCGTGGC |
| 8255 | mamo-s | *BGIBMGA009718* | up | 1220 | 1228 | + | 11.3303 | 4.96E-05 | 0.615 | GTGGGTGGC |
| 8256 | mamo-s | *BGIBMGA009719* | down | 641 | 649 | + | 14.055 | 2.23E-06 | 0.615 | GTGCGTGGC |
| 8257 | mamo-s | *BGIBMGA009719* | down | 1207 | 1215 | - | 11.3303 | 4.96E-05 | 0.615 | GTGGGTGGC |
| 8258 | mamo-s | *BGIBMGA009721* | down | 553 | 561 | - | 12.1284 | 2.16E-05 | 0.615 | GTGCGAGGC |
| 8259 | mamo-s | *BGIBMGA009721* | up | 1007 | 1015 | - | 11.3303 | 4.96E-05 | 0.615 | GTGGGTGGC |
| 8260 | mamo-s | *BGIBMGA009722* | down | 292 | 300 | + | 11.3303 | 4.96E-05 | 0.615 | GTGGGTGGC |
| 8261 | mamo-s | *BGIBMGA009722* | down | 1580 | 1588 | + | 11.3303 | 4.96E-05 | 0.615 | GTGGGTGGC |
| 8262 | mamo-s | *BGIBMGA009723* | down | 1019 | 1027 | - | 11.3303 | 4.96E-05 | 0.615 | GTGGGTGGC |
| 8263 | mamo-s | *BGIBMGA009724* | down | 536 | 544 | + | 10.5138 | 9.75E-05 | 0.865 | GTGCGTGTT |
| 8264 | mamo-s | *BGIBMGA009726* | down | 1250 | 1258 | + | 14.055 | 2.23E-06 | 0.615 | GTGCGTGGC |
| 8265 | mamo-s | *BGIBMGA009728* | up | 1525 | 1533 | + | 11.3303 | 4.96E-05 | 0.615 | GTGGGTGGC |

| | | | | | | | | | | |
|---|---|---|---|---|---|---|---|---|---|---|
| 8266 | mamo-s | *BGIBMGA009729* | up | 303 | 311 | - | 11.3303 | 4.96E-05 | 0.615 | GTGGGTGGC |
| 8267 | mamo-s | *BGIBMGA009730* | up | 1194 | 1202 | - | 11.3303 | 4.96E-05 | 0.615 | GTGGGTGGC |
| 8268 | mamo-s | *BGIBMGA009731* | down | 1918 | 1926 | - | 11.3303 | 4.96E-05 | 0.615 | GTGGGTGGC |
| 8269 | mamo-s | *BGIBMGA009731* | up | 1867 | 1875 | - | 10.7523 | 8.38E-05 | 0.797 | AAGCGTGGC |
| 8270 | mamo-s | *BGIBMGA009734* | down | 44 | 52 | - | 10.5413 | 9.42E-05 | 0.857 | GTGCGAGTC |
| 8271 | mamo-s | *BGIBMGA009740* | down | 82 | 90 | - | 11.5321 | 4.06E-05 | 0.615 | TTGCGTGGC |
| 8272 | mamo-s | *BGIBMGA009743* | up | 1733 | 1741 | + | 11.3303 | 4.96E-05 | 0.615 | GTGGGTGGC |
| 8273 | mamo-s | *BGIBMGA009746* | up | 1107 | 1115 | + | 11.3303 | 4.96E-05 | 0.615 | GTGGGTGGC |
| 8274 | mamo-s | *BGIBMGA009746* | down | 1173 | 1181 | + | 11.3303 | 4.96E-05 | 0.615 | GTGGGTGGC |
| 8275 | mamo-s | *BGIBMGA009747* | down | 1654 | 1662 | - | 11.7064 | 3.52E-05 | 0.615 | GCGCGTGGT |
| 8276 | mamo-s | *BGIBMGA009747* | up | 194 | 202 | - | 11.3303 | 4.96E-05 | 0.615 | GTGGGTGGC |
| 8277 | mamo-s | *BGIBMGA009747* | down | 385 | 393 | + | 11.3303 | 4.96E-05 | 0.615 | GTGGGTGGC |
| 8278 | mamo-s | *BGIBMGA009748* | down | 1174 | 1182 | - | 14.055 | 2.23E-06 | 0.615 | GTGCGTGGC |
| 8279 | mamo-s | *BGIBMGA009748* | up | 725 | 733 | + | 11.5321 | 4.06E-05 | 0.615 | TTGCGTGGC |
| 8280 | mamo-s | *BGIBMGA009748* | down | 136 | 144 | - | 11.3303 | 4.96E-05 | 0.615 | GTGGGTGGC |
| 8281 | mamo-s | *BGIBMGA009748* | up | 361 | 369 | + | 11.3303 | 4.96E-05 | 0.615 | GTGGGTGGC |
| 8282 | mamo-s | *BGIBMGA009748* | down | 1930 | 1938 | - | 11.0826 | 6.84E-05 | 0.709 | GTACGTGGC |
| 8283 | mamo-s | *BGIBMGA009749* | down | 271 | 279 | + | 11.3303 | 4.96E-05 | 0.615 | GTGGGTGGC |
| 8284 | mamo-s | *BGIBMGA009749* | up | 45 | 53 | - | 11.1009 | 6.57E-05 | 0.709 | GAGCGAGGC |
| 8285 | mamo-s | *BGIBMGA009749* | up | 1328 | 1336 | - | 10.7523 | 8.38E-05 | 0.797 | GGGCGTGTC |
| 8286 | mamo-s | *BGIBMGA009750* | down | 1871 | 1879 | + | 11.3303 | 4.96E-05 | 0.615 | GTGGGTGGC |
| 8287 | mamo-s | *BGIBMGA009750* | up | 949 | 957 | - | 10.9358 | 7.30E-05 | 0.728 | GCGGGTGGC |
| 8288 | mamo-s | *BGIBMGA009751* | down | 224 | 232 | - | 11.3303 | 4.96E-05 | 0.615 | GTGGGTGGC |
| 8289 | mamo-s | *BGIBMGA009752* | up | 616 | 624 | + | 12.7615 | 1.04E-05 | 0.615 | CCGCGTGGC |
| 8290 | mamo-s | *BGIBMGA009753* | down | 186 | 194 | - | 11.789 | 2.84E-05 | 0.615 | GCGCGTGCC |
| 8291 | mamo-s | *BGIBMGA009753* | down | 1415 | 1423 | + | 11.3303 | 4.96E-05 | 0.615 | GTGGGTGGC |
| 8292 | mamo-s | *BGIBMGA009754* | down | 458 | 466 | - | 11.3303 | 4.96E-05 | 0.615 | GTGGGTGGC |
| 8293 | mamo-s | *BGIBMGA009755* | up | 1527 | 1535 | - | 11.3303 | 4.96E-05 | 0.615 | GTGGGTGGC |
| 8294 | mamo-s | *BGIBMGA009755* | down | 1466 | 1474 | + | 11.1009 | 6.57E-05 | 0.709 | GAGCGAGGC |
| 8295 | mamo-s | *BGIBMGA009759* | up | 1540 | 1548 | - | 11.3303 | 4.96E-05 | 0.615 | GTGGGTGGC |
| 8296 | mamo-s | *BGIBMGA009761* | up | 934 | 942 | - | 11.3303 | 4.96E-05 | 0.615 | GTGGGTGGC |
| 8297 | mamo-s | *BGIBMGA009761* | up | 586 | 594 | - | 11.0826 | 6.84E-05 | 0.709 | GTACGTGGC |
| 8298 | mamo-s | *BGIBMGA009761* | down | 1390 | 1398 | + | 11.0826 | 6.84E-05 | 0.709 | GTACGTGGC |
| 8299 | mamo-s | *BGIBMGA009765* | down | 1238 | 1246 | - | 11.3303 | 4.96E-05 | 0.615 | GTGGGTGGC |
| 8300 | mamo-s | *BGIBMGA009767* | down | 1744 | 1752 | - | 11.3303 | 4.96E-05 | 0.615 | GTGGGTGGC |
| 8301 | mamo-s | *BGIBMGA009769* | up | 45 | 53 | + | 14.055 | 2.23E-06 | 0.615 | GTGCGTGGC |
| 8302 | mamo-s | *BGIBMGA009769* | up | 242 | 250 | + | 11.3303 | 4.96E-05 | 0.615 | GTGGGTGGC |
| 8303 | mamo-s | *BGIBMGA009769* | up | 1798 | 1806 | + | 11.3303 | 4.96E-05 | 0.615 | GTGGGTGGC |
| 8304 | mamo-s | *BGIBMGA009775* | up | 1437 | 1445 | - | 11.3303 | 4.96E-05 | 0.615 | GTGGGTGGC |
| 8305 | mamo-s | *BGIBMGA009776* | up | 1010 | 1018 | + | 11.3303 | 4.96E-05 | 0.615 | GTGGGTGGC |
| 8306 | mamo-s | *BGIBMGA009779* | up | 885 | 893 | - | 11.3303 | 4.96E-05 | 0.615 | GTGGGTGGC |
| 8307 | mamo-s | *BGIBMGA009783* | up | 1403 | 1411 | + | 11.0826 | 6.84E-05 | 0.709 | GTACGTGGC |
| 8308 | mamo-s | *BGIBMGA009784* | up | 1719 | 1727 | + | 13.6606 | 4.06E-06 | 0.615 | GCGCGTGGC |
| 8309 | mamo-s | *BGIBMGA009784* | down | 1079 | 1087 | - | 11.3303 | 4.96E-05 | 0.615 | GTGGGTGGC |
| 8310 | mamo-s | *BGIBMGA009784* | up | 1601 | 1609 | + | 11.3303 | 4.96E-05 | 0.615 | GTGGGTGGC |
| 8311 | mamo-s | *BGIBMGA009786* | down | 283 | 291 | - | 10.7523 | 8.38E-05 | 0.797 | AAGCGTGGC |
| 8312 | mamo-s | *BGIBMGA009787* | up | 437 | 445 | + | 11.3303 | 4.96E-05 | 0.615 | GTGGGTGGC |
| 8313 | mamo-s | *BGIBMGA009788* | down | 1846 | 1854 | - | 11.3303 | 4.96E-05 | 0.615 | GTGGGTGGC |
| 8314 | mamo-s | *BGIBMGA009788* | up | 1553 | 1561 | - | 11.0826 | 6.84E-05 | 0.709 | GTACGTGGC |
| 8315 | mamo-s | *BGIBMGA009789* | up | 1552 | 1560 | + | 11.789 | 2.84E-05 | 0.615 | GCGCGTGCC |
| 8316 | mamo-s | *BGIBMGA009789* | up | 666 | 674 | - | 11.3303 | 4.96E-05 | 0.615 | GTGGGTGGC |
| 8317 | mamo-s | *BGIBMGA009789* | down | 1250 | 1258 | + | 11.3303 | 4.96E-05 | 0.615 | GTGGGTGGC |
| 8318 | mamo-s | *BGIBMGA009789* | down | 1559 | 1567 | + | 11.3303 | 4.96E-05 | 0.615 | GTGGGTGGC |
| 8319 | mamo-s | *BGIBMGA009790* | down | 935 | 943 | - | 11.3303 | 4.96E-05 | 0.615 | GTGGGTGGC |
| 8320 | mamo-s | *BGIBMGA009790* | up | 1699 | 1707 | + | 11.3303 | 4.96E-05 | 0.615 | GTGGGTGGC |
| 8321 | mamo-s | *BGIBMGA009791* | down | 192 | 200 | + | 11.3303 | 4.96E-05 | 0.615 | GTGGGTGGC |
| 8322 | mamo-s | *BGIBMGA009791* | down | 1711 | 1719 | + | 11.3303 | 4.96E-05 | 0.615 | GTGGGTGGC |
| 8323 | mamo-s | *BGIBMGA009791* | up | 262 | 270 | + | 11.0826 | 6.84E-05 | 0.709 | GTACGTGGC |
| 8324 | mamo-s | *BGIBMGA009791* | up | 1888 | 1896 | + | 10.7523 | 8.38E-05 | 0.797 | GGGCGTGTC |
| 8325 | mamo-s | *BGIBMGA009791* | up | 1279 | 1287 | + | 10.5138 | 9.75E-05 | 0.865 | GTGCGTGTT |
| 8326 | mamo-s | *BGIBMGA009792* | up | 16 | 24 | - | 11.3303 | 4.96E-05 | 0.615 | GTGGGTGGC |
| 8327 | mamo-s | *BGIBMGA009792* | up | 1121 | 1129 | + | 11.3303 | 4.96E-05 | 0.615 | GTGGGTGGC |
| 8328 | mamo-s | *BGIBMGA009792* | down | 1669 | 1677 | + | 11.3303 | 4.96E-05 | 0.615 | GTGGGTGGC |
| 8329 | mamo-s | *BGIBMGA009794* | down | 769 | 777 | + | 11.3303 | 4.96E-05 | 0.615 | GTGGGTGGC |
| 8330 | mamo-s | *BGIBMGA009795* | up | 1142 | 1150 | + | 11.3303 | 4.96E-05 | 0.615 | GTGGGTGGC |
| 8331 | mamo-s | *BGIBMGA009797* | down | 1676 | 1684 | + | 11.3303 | 4.96E-05 | 0.615 | GTGGGTGGC |
| 8332 | mamo-s | *BGIBMGA009797* | up | 1965 | 1973 | + | 10.9358 | 7.30E-05 | 0.728 | GCGGGTGGC |
| 8333 | mamo-s | *BGIBMGA009797* | up | 1570 | 1578 | + | 10.7523 | 8.38E-05 | 0.797 | GGGCGTGTC |
| 8334 | mamo-s | *BGIBMGA009801* | up | 55 | 63 | - | 11.3303 | 4.96E-05 | 0.615 | GTGGGTGGC |
| 8335 | mamo-s | *BGIBMGA009801* | down | 1354 | 1362 | + | 11.3303 | 4.96E-05 | 0.615 | GTGGGTGGC |
| 8336 | mamo-s | *BGIBMGA009801* | up | 1470 | 1478 | - | 11.3303 | 4.96E-05 | 0.615 | GTGGGTGGC |
| 8337 | mamo-s | *BGIBMGA009801* | up | 660 | 668 | + | 10.8073 | 7.89E-05 | 0.766 | CCGCGTGGT |
| 8338 | mamo-s | *BGIBMGA009802* | up | 752 | 760 | + | 11.3303 | 4.96E-05 | 0.615 | GTGGGTGGC |
| 8339 | mamo-s | *BGIBMGA009803* | down | 771 | 779 | + | 11.3303 | 4.96E-05 | 0.615 | GTGGGTGGC |
| 8340 | mamo-s | *BGIBMGA009803* | down | 1264 | 1272 | - | 11.3303 | 4.96E-05 | 0.615 | GTGGGTGGC |
| 8341 | mamo-s | *BGIBMGA009803* | up | 792 | 800 | - | 11.156 | 6.13E-05 | 0.705 | GAGCGTGCC |
| 8342 | mamo-s | *BGIBMGA009804* | up | 812 | 820 | - | 11.3303 | 4.96E-05 | 0.615 | GTGGGTGGC |
| 8343 | mamo-s | *BGIBMGA009804* | down | 1185 | 1193 | - | 11.3303 | 4.96E-05 | 0.615 | GTGGGTGGC |
| 8344 | mamo-s | *BGIBMGA009807* | down | 1633 | 1641 | - | 12.1284 | 2.16E-05 | 0.615 | CAGCGTGGC |
| 8345 | mamo-s | *BGIBMGA009807* | down | 1748 | 1756 | + | 11.2018 | 5.68E-05 | 0.674 | CTGCGTGGT |
| 8346 | mamo-s | *BGIBMGA009808* | down | 580 | 588 | - | 11.3303 | 4.96E-05 | 0.615 | GTGGGTGGC |
| 8347 | mamo-s | *BGIBMGA009808* | up | 1698 | 1706 | - | 11.3303 | 4.96E-05 | 0.615 | GTGGGTGGC |
| 8348 | mamo-s | *BGIBMGA009809* | up | 1691 | 1699 | - | 14.055 | 2.23E-06 | 0.615 | GTGCGTGGC |
| 8349 | mamo-s | *BGIBMGA009809* | down | 389 | 397 | + | 11.7064 | 3.52E-05 | 0.615 | GCGCGTGGT |
| 8350 | mamo-s | *BGIBMGA009809* | down | 257 | 265 | + | 11.3303 | 4.96E-05 | 0.615 | GTGGGTGGC |
| 8351 | mamo-s | *BGIBMGA009813* | up | 1186 | 1194 | + | 11.3303 | 4.96E-05 | 0.615 | GTGGGTGGC |
| 8352 | mamo-s | *BGIBMGA009815* | up | 1803 | 1811 | + | 11.4404 | 4.51E-05 | 0.615 | GAGCGTGTC |
| 8353 | mamo-s | *BGIBMGA009815* | up | 1250 | 1258 | + | 11.0826 | 6.84E-05 | 0.709 | GTACGTGGC |

| | | | | | | | | | | |
|---|---|---|---|---|---|---|---|---|---|---|
| 8354 | mamo-s | *BGIBMGA009815* | up | 1009 | 1017 | + | 10.5138 | 9.75E-05 | 0.865 | GTGCGTGTT |
| 8355 | mamo-s | *BGIBMGA009816* | down | 1853 | 1861 | + | 11.3303 | 4.96E-05 | 0.615 | GTGGGTGGC |
| 8356 | mamo-s | *BGIBMGA009817* | up | 808 | 816 | + | 11.3303 | 4.96E-05 | 0.615 | GTGGGTGGC |
| 8357 | mamo-s | *BGIBMGA009818* | down | 762 | 770 | + | 11.7798 | 3.11E-05 | 0.615 | ATGCGTGGC |
| 8358 | mamo-s | *BGIBMGA009820* | down | 983 | 991 | - | 11.0826 | 6.84E-05 | 0.709 | GTACGTGGC |
| 8359 | mamo-s | *BGIBMGA009820* | up | 935 | 943 | + | 10.6881 | 8.61E-05 | 0.81 | GCACGTGGC |
| 8360 | mamo-s | *BGIBMGA009821* | up | 1824 | 1832 | + | 14.055 | 2.23E-06 | 0.615 | GTGCGTGGC |
| 8361 | mamo-s | *BGIBMGA009821* | up | 1799 | 1807 | + | 12.4679 | 1.31E-05 | 0.615 | GTGCGTGTC |
| 8362 | mamo-s | *BGIBMGA009821* | up | 66 | 74 | + | 12.1835 | 1.71E-05 | 0.615 | GTGCGTGCC |
| 8363 | mamo-s | *BGIBMGA009821* | down | 236 | 244 | - | 10.5688 | 8.88E-05 | 0.824 | GTGCGTGAC |
| 8364 | mamo-s | *BGIBMGA009822* | down | 308 | 316 | - | 10.9358 | 7.30E-05 | 0.728 | GCGGGTGGC |
| 8365 | mamo-s | *BGIBMGA009822* | down | 71 | 79 | - | 10.5138 | 9.75E-05 | 0.865 | GTGCGTGTT |
| 8366 | mamo-s | *BGIBMGA009824* | down | 444 | 452 | - | 10.8899 | 7.48E-05 | 0.739 | CCGCGTGCC |
| 8367 | mamo-s | *BGIBMGA009824* | down | 374 | 382 | + | 10.8073 | 7.89E-05 | 0.766 | CCGCGTGGT |
| 8368 | mamo-s | *BGIBMGA009824* | down | 349 | 357 | + | 10.5688 | 8.88E-05 | 0.824 | GTGCGTGAC |
| 8369 | mamo-s | *BGIBMGA009825* | down | 191 | 199 | - | 11.4404 | 4.51E-05 | 0.615 | CGGCGTGGC |
| 8370 | mamo-s | *BGIBMGA009825* | up | 1578 | 1586 | - | 11.4404 | 4.51E-05 | 0.615 | CGGCGTGGC |
| 8371 | mamo-s | *BGIBMGA009826* | down | 1314 | 1322 | + | 11.3303 | 4.96E-05 | 0.615 | GTGGGTGGC |
| 8372 | mamo-s | *BGIBMGA009827* | up | 1378 | 1386 | + | 11.3303 | 4.96E-05 | 0.615 | GTGGGTGGC |
| 8373 | mamo-s | *BGIBMGA009828* | up | 1697 | 1705 | + | 11.0826 | 6.84E-05 | 0.709 | GTACGTGGC |
| 8374 | mamo-s | *BGIBMGA009830* | up | 1703 | 1711 | + | 11.7339 | 3.29E-05 | 0.615 | GCGCGAGGC |
| 8375 | mamo-s | *BGIBMGA009831* | up | 1123 | 1131 | - | 11.1743 | 5.90E-05 | 0.684 | CCGCGTGTC |
| 8376 | mamo-s | *BGIBMGA009833* | down | 340 | 348 | - | 11.3303 | 4.96E-05 | 0.615 | GTGGGTGGC |
| 8377 | mamo-s | *BGIBMGA009835* | down | 1275 | 1283 | - | 11.789 | 2.84E-05 | 0.615 | GCGCGTGCC |
| 8378 | mamo-s | *BGIBMGA009836* | up | 1283 | 1291 | - | 11.3303 | 4.96E-05 | 0.615 | GTGGGTGGC |
| 8379 | mamo-s | *BGIBMGA009836* | down | 1304 | 1312 | + | 10.5138 | 9.75E-05 | 0.865 | GTGCGTGTT |
| 8380 | mamo-s | *BGIBMGA009837* | up | 1238 | 1246 | - | 11.3303 | 4.96E-05 | 0.615 | GTGGGTGGC |
| 8381 | mamo-s | *BGIBMGA009838* | up | 1825 | 1833 | - | 11.7798 | 3.11E-05 | 0.615 | ATGCGTGGC |
| 8382 | mamo-s | *BGIBMGA009839* | down | 917 | 925 | - | 14.055 | 2.23E-06 | 0.615 | GTGCGTGGC |
| 8383 | mamo-s | *BGIBMGA009839* | up | 1667 | 1675 | + | 11.3303 | 4.96E-05 | 0.615 | GTGGGTGGC |
| 8384 | mamo-s | *BGIBMGA009841* | down | 1605 | 1613 | + | 11.2294 | 5.41E-05 | 0.65 | CTGCGAGGC |
| 8385 | mamo-s | *BGIBMGA009846* | down | 363 | 371 | - | 11.3303 | 4.96E-05 | 0.615 | GTGGGTGGC |
| 8386 | mamo-s | *BGIBMGA009846* | up | 1482 | 1490 | - | 11.0826 | 6.84E-05 | 0.709 | GTACGTGGC |
| 8387 | mamo-s | *BGIBMGA009847* | down | 58 | 66 | + | 10.5138 | 9.75E-05 | 0.865 | GTGCGTGTT |
| 8388 | mamo-s | *BGIBMGA009849* | down | 191 | 199 | + | 11.3303 | 4.96E-05 | 0.615 | GTGGGTGGC |
| 8389 | mamo-s | *BGIBMGA009850* | up | 566 | 574 | - | 11.3303 | 4.96E-05 | 0.615 | GTGGGTGGC |
| 8390 | mamo-s | *BGIBMGA009851* | up | 1848 | 1856 | - | 11.2018 | 5.68E-05 | 0.674 | CTGCGTGGT |
| 8391 | mamo-s | *BGIBMGA009852* | down | 419 | 427 | + | 11.1743 | 5.90E-05 | 0.684 | CCGCGTGTC |
| 8392 | mamo-s | *BGIBMGA009853* | down | 473 | 481 | + | 11.3303 | 4.96E-05 | 0.615 | GTGGGTGGC |
| 8393 | mamo-s | *BGIBMGA009854* | up | 689 | 697 | + | 11.3303 | 4.96E-05 | 0.615 | GTGGGTGGC |
| 8394 | mamo-s | *BGIBMGA009855* | up | 559 | 567 | - | 11.2294 | 5.41E-05 | 0.65 | CTGCGAGGC |
| 8395 | mamo-s | *BGIBMGA009856* | down | 491 | 499 | - | 12.0734 | 2.66E-05 | 0.615 | GCGCGTGTC |
| 8396 | mamo-s | *BGIBMGA009857* | up | 92 | 100 | - | 11.2294 | 5.41E-05 | 0.65 | CTGCGAGGC |
| 8397 | mamo-s | *BGIBMGA009858* | up | 1946 | 1954 | + | 10.9358 | 7.30E-05 | 0.728 | GCGGGTGGC |
| 8398 | mamo-s | *BGIBMGA009859* | up | 1218 | 1226 | - | 11.4404 | 4.51E-05 | 0.615 | CGGCGTGGC |
| 8399 | mamo-s | *BGIBMGA009859* | down | 1154 | 1162 | + | 11.3303 | 4.96E-05 | 0.615 | GTGGGTGGC |
| 8400 | mamo-s | *BGIBMGA009859* | down | 1812 | 1820 | - | 11.3303 | 4.96E-05 | 0.615 | GTGGGTGGC |
| 8401 | mamo-s | *BGIBMGA009860* | down | 677 | 685 | - | 11.3303 | 4.96E-05 | 0.615 | GTGGGTGGC |
| 8402 | mamo-s | *BGIBMGA009861* | down | 1404 | 1412 | + | 11.0826 | 6.84E-05 | 0.709 | GTACGTGGC |
| 8403 | mamo-s | *BGIBMGA009862* | up | 1503 | 1511 | + | 11.0826 | 6.84E-05 | 0.709 | GTACGTGGC |
| 8404 | mamo-s | *BGIBMGA009863* | down | 646 | 654 | + | 11.3303 | 4.96E-05 | 0.615 | GTGGGTGGC |
| 8405 | mamo-s | *BGIBMGA009865* | up | 1065 | 1073 | + | 11.3303 | 4.96E-05 | 0.615 | GTGGGTGGC |
| 8406 | mamo-s | *BGIBMGA009865* | up | 1947 | 1955 | - | 11.1376 | 6.35E-05 | 0.709 | TCGCGTGGC |
| 8407 | mamo-s | *BGIBMGA009866* | up | 485 | 493 | + | 11.3303 | 4.96E-05 | 0.615 | GTGGGTGGC |
| 8408 | mamo-s | *BGIBMGA009866* | down | 1125 | 1133 | + | 11.3303 | 4.96E-05 | 0.615 | GTGGGTGGC |
| 8409 | mamo-s | *BGIBMGA009868* | up | 791 | 799 | - | 11.3303 | 4.96E-05 | 0.615 | GTGGGTGGC |
| 8410 | mamo-s | *BGIBMGA009868* | down | 1880 | 1888 | + | 11.3303 | 4.96E-05 | 0.615 | GTGGGTGGC |
| 8411 | mamo-s | *BGIBMGA009868* | up | 310 | 318 | - | 11.0826 | 6.84E-05 | 0.709 | GTACGTGGC |
| 8412 | mamo-s | *BGIBMGA009868* | down | 1594 | 1602 | - | 10.5688 | 8.88E-05 | 0.824 | GTGCGTGAC |
| 8413 | mamo-s | *BGIBMGA009870* | down | 102 | 110 | - | 11.3303 | 4.96E-05 | 0.615 | GTGGGTGGC |
| 8414 | mamo-s | *BGIBMGA009870* | up | 1076 | 1084 | - | 11.3303 | 4.96E-05 | 0.615 | GTGGGTGGC |
| 8415 | mamo-s | *BGIBMGA009872* | up | 160 | 168 | + | 10.8899 | 7.48E-05 | 0.739 | CCGCGTGCC |
| 8416 | mamo-s | *BGIBMGA009873* | up | 236 | 244 | - | 11.0734 | 7.12E-05 | 0.726 | GAGCGTGGT |
| 8417 | mamo-s | *BGIBMGA009875* | down | 1532 | 1540 | - | 11.3303 | 4.96E-05 | 0.615 | GTGGGTGGC |
| 8418 | mamo-s | *BGIBMGA009875* | down | 68 | 76 | - | 11.0734 | 7.12E-05 | 0.726 | GAGCGTGGT |
| 8419 | mamo-s | *BGIBMGA009878* | up | 1909 | 1917 | - | 12.1835 | 1.71E-05 | 0.615 | GTGCGTGCC |
| 8420 | mamo-s | *BGIBMGA009880* | down | 1020 | 1028 | + | 11.7064 | 3.52E-05 | 0.615 | GCGCGTGGT |
| 8421 | mamo-s | *BGIBMGA009882* | up | 572 | 580 | + | 11.4404 | 4.51E-05 | 0.615 | CGGCGTGGC |
| 8422 | mamo-s | *BGIBMGA009883* | down | 545 | 553 | + | 11.2844 | 5.18E-05 | 0.63 | CTGCGTGCC |
| 8423 | mamo-s | *BGIBMGA009884* | up | 399 | 407 | - | 11.3303 | 4.96E-05 | 0.615 | GTGGGTGGC |
| 8424 | mamo-s | *BGIBMGA009887* | down | 1600 | 1608 | - | 11.1743 | 5.90E-05 | 0.684 | CCGCGTGTC |
| 8425 | mamo-s | *BGIBMGA009889* | down | 910 | 918 | + | 11.0826 | 6.84E-05 | 0.709 | GTACGTGGC |
| 8426 | mamo-s | *BGIBMGA009890* | down | 1852 | 1860 | + | 11.3303 | 4.96E-05 | 0.615 | GTGGGTGGC |
| 8427 | mamo-s | *BGIBMGA009890* | up | 1358 | 1366 | - | 10.9358 | 7.30E-05 | 0.728 | GCGGGTGGC |
| 8428 | mamo-s | *BGIBMGA009891* | up | 1825 | 1833 | + | 11.3303 | 4.96E-05 | 0.615 | GTGGGTGGC |
| 8429 | mamo-s | *BGIBMGA009892* | up | 625 | 633 | - | 11.3303 | 4.96E-05 | 0.615 | GTGGGTGGC |
| 8430 | mamo-s | *BGIBMGA009895* | down | 1496 | 1504 | - | 12.4679 | 1.31E-05 | 0.615 | GTGCGTGTC |
| 8431 | mamo-s | *BGIBMGA009895* | up | 328 | 336 | - | 11.3303 | 4.96E-05 | 0.615 | GTGGGTGGC |
| 8432 | mamo-s | *BGIBMGA009898* | down | 1875 | 1883 | + | 11.0826 | 6.84E-05 | 0.709 | GTACGTGGC |
| 8433 | mamo-s | *BGIBMGA009901* | down | 399 | 407 | + | 11.5321 | 4.06E-05 | 0.615 | TTGCGTGGC |
| 8434 | mamo-s | *BGIBMGA009902* | up | 206 | 214 | - | 12.7615 | 1.04E-05 | 0.615 | CCGCGTGGC |
| 8435 | mamo-s | *BGIBMGA009903* | down | 859 | 867 | + | 12.1284 | 2.16E-05 | 0.615 | CAGCGTGGC |
| 8436 | mamo-s | *BGIBMGA009903* | down | 241 | 249 | - | 11.3303 | 4.96E-05 | 0.615 | GTGGGTGGC |
| 8437 | mamo-s | *BGIBMGA009903* | down | 66 | 74 | + | 10.6881 | 8.61E-05 | 0.81 | GCACGTGGC |
| 8438 | mamo-s | *BGIBMGA009904* | up | 377 | 385 | + | 11.3303 | 4.96E-05 | 0.615 | GTGGGTGGC |
| 8439 | mamo-s | *BGIBMGA009905* | up | 435 | 443 | - | 11.4404 | 4.51E-05 | 0.615 | GAGCGTGTC |
| 8440 | mamo-s | *BGIBMGA009905* | up | 76 | 84 | - | 11.3303 | 4.96E-05 | 0.615 | GTGGGTGGC |
| 8441 | mamo-s | *BGIBMGA009906* | up | 700 | 708 | - | 11.3303 | 4.96E-05 | 0.615 | GTGGGTGGC |

| | | | | | | | | | | |
|---|---|---|---|---|---|---|---|---|---|---|
| 8442 | mamo-s | *BGIBMGA009907* | up | 1881 | 1889 | - | 11.3303 | 4.96E-05 | 0.615 | GTGGGTGGC |
| 8443 | mamo-s | *BGIBMGA009909* | down | 68 | 76 | - | 12.4679 | 1.31E-05 | 0.615 | GTGCGTGTC |
| 8444 | mamo-s | *BGIBMGA009909* | down | 43 | 51 | + | 11.4404 | 4.51E-05 | 0.615 | GAGCGTGTC |
| 8445 | mamo-s | *BGIBMGA009910* | down | 154 | 162 | + | 10.8349 | 7.66E-05 | 0.751 | CCGCGAGGC |
| 8446 | mamo-s | *BGIBMGA009913* | up | 287 | 295 | - | 11.3303 | 4.96E-05 | 0.615 | GTGGGTGGC |
| 8447 | mamo-s | *BGIBMGA009913* | up | 1274 | 1282 | + | 11.3303 | 4.96E-05 | 0.615 | GTGGGTGGC |
| 8448 | mamo-s | *BGIBMGA009914* | down | 291 | 299 | + | 10.7523 | 8.38E-05 | 0.797 | AAGCGTGGC |
| 8449 | mamo-s | *BGIBMGA009916* | up | 1206 | 1214 | - | 11.3303 | 4.96E-05 | 0.615 | GTGGGTGGC |
| 8450 | mamo-s | *BGIBMGA009917* | down | 350 | 358 | - | 11.4404 | 4.51E-05 | 0.615 | CGGCGTGGC |
| 8451 | mamo-s | *BGIBMGA009917* | down | 360 | 368 | - | 11.4404 | 4.51E-05 | 0.615 | CGGCGTGGC |
| 8452 | mamo-s | *BGIBMGA009917* | down | 1922 | 1930 | + | 10.8349 | 7.66E-05 | 0.751 | CCGCGAGGC |
| 8453 | mamo-s | *BGIBMGA009917* | down | 335 | 343 | - | 10.7523 | 8.38E-05 | 0.797 | GGGCGTGTC |
| 8454 | mamo-s | *BGIBMGA009917* | down | 1457 | 1465 | - | 10.5413 | 9.42E-05 | 0.857 | GTGCGAGTC |
| 8455 | mamo-s | *BGIBMGA009918* | up | 1578 | 1586 | - | 10.5688 | 8.88E-05 | 0.824 | GTGCGTGAC |
| 8456 | mamo-s | *BGIBMGA009919* | up | 536 | 544 | - | 11.0826 | 6.84E-05 | 0.709 | GTACGTGGC |
| 8457 | mamo-s | *BGIBMGA009920* | down | 1590 | 1598 | + | 12.1284 | 2.16E-05 | 0.615 | CAGCGTGGC |
| 8458 | mamo-s | *BGIBMGA009920* | up | 1981 | 1989 | - | 11.0826 | 6.84E-05 | 0.709 | GTACGTGGC |
| 8459 | mamo-s | *BGIBMGA009921* | up | 1009 | 1017 | - | 11.1009 | 6.57E-05 | 0.709 | GAGCGAGGC |
| 8460 | mamo-s | *BGIBMGA009921* | up | 1724 | 1732 | - | 11.0826 | 6.84E-05 | 0.709 | GTACGTGGC |
| 8461 | mamo-s | *BGIBMGA009922* | down | 1590 | 1598 | + | 11.3303 | 4.96E-05 | 0.615 | GTGGGTGGC |
| 8462 | mamo-s | *BGIBMGA009924* | down | 515 | 523 | + | 11.3303 | 4.96E-05 | 0.615 | GTGGGTGGC |
| 8463 | mamo-s | *BGIBMGA009925* | down | 1375 | 1383 | - | 11.0826 | 6.84E-05 | 0.709 | GTACGTGGC |
| 8464 | mamo-s | *BGIBMGA009927* | down | 1904 | 1912 | - | 14.055 | 2.23E-06 | 0.615 | GTGCGTGGC |
| 8465 | mamo-s | *BGIBMGA009927* | up | 1989 | 1997 | - | 13.6606 | 4.06E-06 | 0.615 | GCGCGTGGC |
| 8466 | mamo-s | *BGIBMGA009927* | down | 1929 | 1937 | - | 12.4679 | 1.31E-05 | 0.615 | GTGCGTGTC |
| 8467 | mamo-s | *BGIBMGA009927* | down | 430 | 438 | + | 10.5688 | 8.88E-05 | 0.824 | GTGCGTGAC |
| 8468 | mamo-s | *BGIBMGA009930* | down | 1501 | 1509 | - | 11.3303 | 4.96E-05 | 0.615 | GTGGGTGGC |
| 8469 | mamo-s | *BGIBMGA009931* | up | 1564 | 1572 | - | 11.3303 | 4.96E-05 | 0.615 | GTGGGTGGC |
| 8470 | mamo-s | *BGIBMGA009932* | down | 629 | 637 | + | 11.3303 | 4.96E-05 | 0.615 | GTGGGTGGC |
| 8471 | mamo-s | *BGIBMGA009933* | up | 1960 | 1968 | + | 11.0826 | 6.84E-05 | 0.709 | GTACGTGGC |
| 8472 | mamo-s | *BGIBMGA009934* | up | 1218 | 1226 | - | 11.3303 | 4.96E-05 | 0.615 | GTGGGTGGC |
| 8473 | mamo-s | *BGIBMGA009935* | down | 1416 | 1424 | - | 11.0826 | 6.84E-05 | 0.709 | GTACGTGGC |
| 8474 | mamo-s | *BGIBMGA009937* | down | 203 | 211 | - | 11.3303 | 4.96E-05 | 0.615 | GTGGGTGGC |
| 8475 | mamo-s | *BGIBMGA009937* | up | 702 | 710 | + | 11.3303 | 4.96E-05 | 0.615 | GTGGGTGGC |
| 8476 | mamo-s | *BGIBMGA009938* | down | 1244 | 1252 | + | 11.3303 | 4.96E-05 | 0.615 | GTGGGTGGC |
| 8477 | mamo-s | *BGIBMGA009938* | up | 1556 | 1564 | + | 11.3303 | 4.96E-05 | 0.615 | GTGGGTGGC |
| 8478 | mamo-s | *BGIBMGA009938* | up | 177 | 185 | + | 10.7523 | 8.38E-05 | 0.797 | AAGCGTGGC |
| 8479 | mamo-s | *BGIBMGA009939* | up | 749 | 757 | + | 12.1284 | 2.16E-05 | 0.615 | CAGCGTGGC |
| 8480 | mamo-s | *BGIBMGA009943* | down | 1585 | 1593 | + | 11.3303 | 4.96E-05 | 0.615 | GTGGGTGGC |
| 8481 | mamo-s | *BGIBMGA009944* | down | 365 | 373 | + | 12.1835 | 1.71E-05 | 0.615 | GTGCGTGCC |
| 8482 | mamo-s | *BGIBMGA009944* | down | 1773 | 1781 | - | 11.3303 | 4.96E-05 | 0.615 | GTGGGTGGC |
| 8483 | mamo-s | *BGIBMGA009946* | up | 653 | 661 | - | 11.3303 | 4.96E-05 | 0.615 | GTGGGTGGC |
| 8484 | mamo-s | *BGIBMGA009946* | down | 1849 | 1857 | - | 11.3303 | 4.96E-05 | 0.615 | GTGGGTGGC |
| 8485 | mamo-s | *BGIBMGA009947* | up | 796 | 804 | - | 11.3303 | 4.96E-05 | 0.615 | GTGGGTGGC |
| 8486 | mamo-s | *BGIBMGA009948* | down | 883 | 891 | - | 12.1009 | 2.43E-05 | 0.615 | GTGCGTGGT |
| 8487 | mamo-s | *BGIBMGA009948* | up | 1669 | 1677 | + | 11.3303 | 4.96E-05 | 0.615 | GTGGGTGGC |
| 8488 | mamo-s | *BGIBMGA009949* | down | 890 | 898 | + | 11.3303 | 4.96E-05 | 0.615 | GTGGGTGGC |
| 8489 | mamo-s | *BGIBMGA009949* | down | 1277 | 1285 | - | 11.0826 | 6.84E-05 | 0.709 | GTACGTGGC |
| 8490 | mamo-s | *BGIBMGA009949* | down | 1846 | 1854 | - | 10.6881 | 8.61E-05 | 0.81 | GCACGTGGC |
| 8491 | mamo-s | *BGIBMGA009950* | down | 1270 | 1278 | - | 11.3303 | 4.96E-05 | 0.615 | GTGGGTGGC |
| 8492 | mamo-s | *BGIBMGA009951* | up | 1144 | 1152 | + | 11.3303 | 4.96E-05 | 0.615 | GTGGGTGGC |
| 8493 | mamo-s | *BGIBMGA009951* | down | 1283 | 1291 | - | 11.3303 | 4.96E-05 | 0.615 | GTGGGTGGC |
| 8494 | mamo-s | *BGIBMGA009952* | down | 295 | 303 | + | 11.3303 | 4.96E-05 | 0.615 | GTGGGTGGC |
| 8495 | mamo-s | *BGIBMGA009952* | up | 834 | 842 | + | 11.2844 | 5.18E-05 | 0.63 | CTGCGTGCC |
| 8496 | mamo-s | *BGIBMGA009957* | down | 887 | 895 | - | 11.3303 | 4.96E-05 | 0.615 | GTGGGTGGC |
| 8497 | mamo-s | *BGIBMGA009958* | up | 89 | 97 | + | 11.3303 | 4.96E-05 | 0.615 | GTGGGTGGC |
| 8498 | mamo-s | *BGIBMGA009958* | down | 482 | 490 | + | 10.8073 | 7.89E-05 | 0.766 | CCGCGTGGT |
| 8499 | mamo-s | *BGIBMGA009959* | down | 937 | 945 | - | 11.3303 | 4.96E-05 | 0.615 | GTGGGTGGC |
| 8500 | mamo-s | *BGIBMGA009959* | up | 1595 | 1603 | + | 10.8073 | 7.89E-05 | 0.766 | CCGCGTGGT |
| 8501 | mamo-s | *BGIBMGA009960* | down | 803 | 811 | - | 12.1009 | 2.43E-05 | 0.615 | GTGCGTGGT |
| 8502 | mamo-s | *BGIBMGA009960* | up | 2 | 10 | + | 10.8349 | 7.66E-05 | 0.751 | CCGCGAGGC |
| 8503 | mamo-s | *BGIBMGA009961* | down | 675 | 683 | + | 13.6606 | 4.06E-06 | 0.615 | GCGCGTGGC |
| 8504 | mamo-s | *BGIBMGA009961* | down | 786 | 794 | + | 11.0734 | 7.12E-05 | 0.726 | GAGCGTGGT |
| 8505 | mamo-s | *BGIBMGA009962* | down | 272 | 280 | + | 12.1835 | 1.71E-05 | 0.615 | GTGCGTGCC |
| 8506 | mamo-s | *BGIBMGA009963* | down | 1312 | 1320 | - | 11.3303 | 4.96E-05 | 0.615 | GTGGGTGGC |
| 8507 | mamo-s | *BGIBMGA009966* | up | 1841 | 1849 | + | 10.5688 | 8.88E-05 | 0.824 | GTGCGTGAC |
| 8508 | mamo-s | *BGIBMGA009968* | down | 524 | 532 | - | 11.5688 | 3.79E-05 | 0.615 | CTGCGTGTC |
| 8509 | mamo-s | *BGIBMGA009970* | down | 1097 | 1105 | - | 11.0826 | 6.84E-05 | 0.709 | GTACGTGGC |
| 8510 | mamo-s | *BGIBMGA009971* | up | 1120 | 1128 | + | 11.5688 | 3.79E-05 | 0.615 | CTGCGTGTC |
| 8511 | mamo-s | *BGIBMGA009972* | up | 1002 | 1010 | + | 12.1284 | 2.16E-05 | 0.615 | CAGCGTGGC |
| 8512 | mamo-s | *BGIBMGA009972* | up | 1260 | 1268 | - | 11.3303 | 4.96E-05 | 0.615 | GTGGGTGGC |
| 8513 | mamo-s | *BGIBMGA009973* | up | 488 | 496 | - | 11.3303 | 4.96E-05 | 0.615 | GTGGGTGGC |
| 8514 | mamo-s | *BGIBMGA009974* | up | 618 | 626 | - | 11.1743 | 5.90E-05 | 0.684 | CCGCGTGTC |
| 8515 | mamo-s | *BGIBMGA009975* | up | 1816 | 1824 | + | 11.3303 | 4.96E-05 | 0.615 | GTGGGTGGC |
| 8516 | mamo-s | *BGIBMGA009978* | up | 1033 | 1041 | - | 11.5688 | 3.79E-05 | 0.615 | CTGCGTGTC |
| 8517 | mamo-s | *BGIBMGA009978* | down | 652 | 660 | + | 11.1376 | 6.35E-05 | 0.709 | TCGCGTGGC |
| 8518 | mamo-s | *BGIBMGA009984* | down | 657 | 665 | + | 11.3303 | 4.96E-05 | 0.615 | GTGGGTGGC |
| 8519 | mamo-s | *BGIBMGA009985* | down | 1032 | 1040 | + | 11.3303 | 4.96E-05 | 0.615 | GTGGGTGGC |
| 8520 | mamo-s | *BGIBMGA009989* | down | 585 | 593 | + | 11.0826 | 6.84E-05 | 0.709 | GTACGTGGC |
| 8521 | mamo-s | *BGIBMGA009993* | down | 718 | 726 | - | 11.7339 | 3.29E-05 | 0.615 | GCGCGAGGC |
| 8522 | mamo-s | *BGIBMGA009993* | up | 425 | 433 | - | 11.3303 | 4.96E-05 | 0.615 | GTGGGTGGC |
| 8523 | mamo-s | *BGIBMGA009993* | up | 1280 | 1288 | - | 11.2844 | 5.18E-05 | 0.63 | CTGCGTGCC |
| 8524 | mamo-s | *BGIBMGA009994* | up | 534 | 542 | - | 11.7339 | 3.29E-05 | 0.615 | GCGCGAGGC |
| 8525 | mamo-s | *BGIBMGA009995* | down | 391 | 399 | - | 14.055 | 2.23E-06 | 0.615 | GTGCGTGGC |
| 8526 | mamo-s | *BGIBMGA009996* | up | 1457 | 1465 | + | 11.0734 | 7.12E-05 | 0.726 | GAGCGTGGT |
| 8527 | mamo-s | *BGIBMGA009998* | down | 762 | 770 | - | 13.0275 | 8.53E-06 | 0.615 | GAGCGTGGC |
| 8528 | mamo-s | *BGIBMGA009998* | down | 772 | 780 | - | 13.0275 | 8.53E-06 | 0.615 | GAGCGTGGC |
| 8529 | mamo-s | *BGIBMGA009998* | down | 782 | 790 | - | 13.0275 | 8.53E-06 | 0.615 | GAGCGTGGC |

| | | | | | | | | | | |
|---|---|---|---|---|---|---|---|---|---|---|
| 8530 | mamo-s | *BGIBMGA009998* | down | 792 | 800 | - | 13.0275 | 8.53E-06 | 0.615 | GAGCGTGGC |
| 8531 | mamo-s | *BGIBMGA009998* | down | 802 | 810 | - | 13.0275 | 8.53E-06 | 0.615 | GAGCGTGGC |
| 8532 | mamo-s | *BGIBMGA009998* | down | 1348 | 1356 | + | 11.3303 | 4.96E-05 | 0.615 | GTGGGTGGC |
| 8533 | mamo-s | *BGIBMGA009999* | up | 22 | 30 | - | 12.0734 | 2.66E-05 | 0.615 | GCGCGTGTC |
| 8534 | mamo-s | *BGIBMGA009999* | up | 42 | 50 | + | 10.8899 | 7.48E-05 | 0.739 | CCGCGTGCC |
| 8535 | mamo-s | *BGIBMGA010000* | up | 85 | 93 | + | 13.6606 | 4.06E-06 | 0.615 | GCGCGTGGC |
| 8536 | mamo-s | *BGIBMGA010000* | down | 1150 | 1158 | - | 12.4679 | 1.31E-05 | 0.615 | GTGCGTGTC |
| 8537 | mamo-s | *BGIBMGA010000* | down | 981 | 989 | - | 10.8899 | 7.48E-05 | 0.739 | CCGCGTGCC |
| 8538 | mamo-s | *BGIBMGA010000* | down | 942 | 950 | + | 10.5413 | 9.42E-05 | 0.857 | CAGCGTGTC |
| 8539 | mamo-s | *BGIBMGA010001* | up | 1254 | 1262 | - | 12.7615 | 1.04E-05 | 0.615 | CCGCGTGGC |
| 8540 | mamo-s | *BGIBMGA010001* | up | 1529 | 1537 | + | 12.0734 | 2.66E-05 | 0.615 | GCGCGTGTC |
| 8541 | mamo-s | *BGIBMGA010003* | up | 436 | 444 | - | 11.3303 | 4.96E-05 | 0.615 | GTGGGTGGC |
| 8542 | mamo-s | *BGIBMGA010008* | up | 524 | 532 | + | 11.3303 | 4.96E-05 | 0.615 | GTGGGTGGC |
| 8543 | mamo-s | *BGIBMGA010008* | up | 901 | 909 | - | 11.3303 | 4.96E-05 | 0.615 | GTGGGTGGC |
| 8544 | mamo-s | *BGIBMGA010010* | down | 1202 | 1210 | - | 12.7615 | 1.04E-05 | 0.615 | CCGCGTGGC |
| 8545 | mamo-s | *BGIBMGA010010* | up | 879 | 887 | + | 11.0826 | 6.84E-05 | 0.709 | GTACGTGGC |
| 8546 | mamo-s | *BGIBMGA010011* | up | 1360 | 1368 | - | 12.7615 | 1.04E-05 | 0.615 | CCGCGTGGC |
| 8547 | mamo-s | *BGIBMGA010012* | down | 744 | 752 | + | 11.3303 | 4.96E-05 | 0.615 | GTGGGTGGC |
| 8548 | mamo-s | *BGIBMGA010012* | down | 968 | 976 | - | 11.3303 | 4.96E-05 | 0.615 | GTGGGTGGC |
| 8549 | mamo-s | *BGIBMGA010012* | down | 1444 | 1452 | - | 11.3303 | 4.96E-05 | 0.615 | GTGGGTGGC |
| 8550 | mamo-s | *BGIBMGA010013* | down | 1892 | 1900 | + | 11.5321 | 4.06E-05 | 0.615 | TTGCGTGGC |
| 8551 | mamo-s | *BGIBMGA010014* | down | 387 | 395 | + | 11.3303 | 4.96E-05 | 0.615 | GTGGGTGGC |
| 8552 | mamo-s | *BGIBMGA010014* | up | 1598 | 1606 | - | 11.3303 | 4.96E-05 | 0.615 | GTGGGTGGC |
| 8553 | mamo-s | *BGIBMGA010015* | up | 870 | 878 | - | 11.3303 | 4.96E-05 | 0.615 | GTGGGTGGC |
| 8554 | mamo-s | *BGIBMGA010015* | up | 1544 | 1552 | + | 11.0826 | 6.84E-05 | 0.709 | GTACGTGGC |
| 8555 | mamo-s | *BGIBMGA010017* | up | 741 | 749 | - | 12.1009 | 2.43E-05 | 0.615 | GTGCGTGGT |
| 8556 | mamo-s | *BGIBMGA010017* | up | 197 | 205 | + | 11.3303 | 4.96E-05 | 0.615 | GTGGGTGGC |
| 8557 | mamo-s | *BGIBMGA010017* | up | 1212 | 1220 | + | 11.3303 | 4.96E-05 | 0.615 | GTGGGTGGC |
| 8558 | mamo-s | *BGIBMGA010017* | up | 97 | 105 | + | 11.0826 | 6.84E-05 | 0.709 | GTACGTGGC |
| 8559 | mamo-s | *BGIBMGA010021* | up | 1946 | 1954 | + | 11.5321 | 4.06E-05 | 0.615 | TTGCGTGGC |
| 8560 | mamo-s | *BGIBMGA010021* | up | 1468 | 1476 | - | 10.7523 | 8.38E-05 | 0.797 | AAGCGTGGC |
| 8561 | mamo-s | *BGIBMGA010022* | down | 1428 | 1436 | - | 11.3303 | 4.96E-05 | 0.615 | GTGGGTGGC |
| 8562 | mamo-s | *BGIBMGA010023* | up | 1251 | 1259 | - | 11.3303 | 4.96E-05 | 0.615 | GTGGGTGGC |
| 8563 | mamo-s | *BGIBMGA010024* | up | 215 | 223 | - | 11.3303 | 4.96E-05 | 0.615 | GTGGGTGGC |
| 8564 | mamo-s | *BGIBMGA010024* | down | 682 | 690 | + | 11.3303 | 4.96E-05 | 0.615 | GTGGGTGGC |
| 8565 | mamo-s | *BGIBMGA010024* | up | 934 | 942 | - | 11.3303 | 4.96E-05 | 0.615 | GTGGGTGGC |
| 8566 | mamo-s | *BGIBMGA010024* | up | 147 | 155 | - | 10.7523 | 8.38E-05 | 0.797 | GGGCGTGTC |
| 8567 | mamo-s | *BGIBMGA010024* | down | 817 | 825 | - | 10.5138 | 9.75E-05 | 0.865 | GTGCGTGTT |
| 8568 | mamo-s | *BGIBMGA010025* | down | 1476 | 1484 | - | 11.3303 | 4.96E-05 | 0.615 | GTGGGTGGC |
| 8569 | mamo-s | *BGIBMGA010027* | down | 623 | 631 | + | 11.3303 | 4.96E-05 | 0.615 | GTGGGTGGC |
| 8570 | mamo-s | *BGIBMGA010028* | down | 1839 | 1847 | + | 11.3303 | 4.96E-05 | 0.615 | GTGGGTGGC |
| 8571 | mamo-s | *BGIBMGA010033* | down | 1668 | 1676 | + | 11.3303 | 4.96E-05 | 0.615 | GTGGGTGGC |
| 8572 | mamo-s | *BGIBMGA010034* | down | 1851 | 1859 | - | 11.1376 | 6.35E-05 | 0.709 | TCGCGTGGC |
| 8573 | mamo-s | *BGIBMGA010036* | up | 71 | 79 | - | 11.3303 | 4.96E-05 | 0.615 | GTGGGTGGC |
| 8574 | mamo-s | *BGIBMGA010037* | up | 1337 | 1345 | - | 11.0826 | 6.84E-05 | 0.709 | GTACGTGGC |
| 8575 | mamo-s | *BGIBMGA010038* | down | 1619 | 1627 | + | 12.0734 | 2.66E-05 | 0.615 | GCGCGTGTC |
| 8576 | mamo-s | *BGIBMGA010038* | down | 20 | 28 | + | 10.8899 | 7.48E-05 | 0.739 | CCGCGTGCC |
| 8577 | mamo-s | *BGIBMGA010039* | up | 1991 | 1999 | - | 11.1376 | 6.35E-05 | 0.709 | TCGCGTGGC |
| 8578 | mamo-s | *BGIBMGA010041* | up | 1096 | 1104 | + | 13.156 | 6.30E-06 | 0.615 | CTGCGTGGC |
| 8579 | mamo-s | *BGIBMGA010043* | up | 1886 | 1894 | - | 12.7615 | 1.04E-05 | 0.615 | CCGCGTGGC |
| 8580 | mamo-s | *BGIBMGA010048* | down | 500 | 508 | - | 11.3303 | 4.96E-05 | 0.615 | GTGGGTGGC |
| 8581 | mamo-s | *BGIBMGA010049* | up | 507 | 515 | - | 11.3303 | 4.96E-05 | 0.615 | GTGGGTGGC |
| 8582 | mamo-s | *BGIBMGA010050* | up | 1828 | 1836 | - | 11.4404 | 4.51E-05 | 0.615 | CGGCGTGGC |
| 8583 | mamo-s | *BGIBMGA010052* | down | 731 | 739 | + | 11.3303 | 4.96E-05 | 0.615 | GTGGGTGGC |
| 8584 | mamo-s | *BGIBMGA010052* | up | 1302 | 1310 | + | 11.3303 | 4.96E-05 | 0.615 | GTGGGTGGC |
| 8585 | mamo-s | *BGIBMGA010052* | down | 1362 | 1370 | + | 11.3303 | 4.96E-05 | 0.615 | GTGGGTGGC |
| 8586 | mamo-s | *BGIBMGA010053* | down | 659 | 667 | + | 11.3303 | 4.96E-05 | 0.615 | GTGGGTGGC |
| 8587 | mamo-s | *BGIBMGA010057* | up | 1667 | 1675 | - | 11.5321 | 4.06E-05 | 0.615 | TTGCGTGGC |
| 8588 | mamo-s | *BGIBMGA010060* | up | 183 | 191 | + | 12.1835 | 1.71E-05 | 0.615 | GTGCGTGCC |
| 8589 | mamo-s | *BGIBMGA010061* | up | 1913 | 1921 | + | 11.3303 | 4.96E-05 | 0.615 | GTGGGTGGC |
| 8590 | mamo-s | *BGIBMGA010063* | up | 1056 | 1064 | - | 11.7339 | 3.29E-05 | 0.615 | GCGCGAGGC |
| 8591 | mamo-s | *BGIBMGA010063* | down | 1957 | 1965 | + | 10.5138 | 9.75E-05 | 0.865 | GTGCGTGTT |
| 8592 | mamo-s | *BGIBMGA010064* | up | 580 | 588 | - | 11.3303 | 4.96E-05 | 0.615 | GTGGGTGGC |
| 8593 | mamo-s | *BGIBMGA010064* | up | 1495 | 1503 | - | 11.3303 | 4.96E-05 | 0.615 | GTGGGTGGC |
| 8594 | mamo-s | *BGIBMGA010065* | down | 449 | 457 | + | 12.1009 | 2.43E-05 | 0.615 | GTGCGTGGT |
| 8595 | mamo-s | *BGIBMGA010065* | up | 749 | 757 | - | 10.9358 | 7.30E-05 | 0.728 | GCGGGTGGC |
| 8596 | mamo-s | *BGIBMGA010066* | down | 582 | 590 | - | 13.0275 | 8.53E-06 | 0.615 | GAGCGTGGC |
| 8597 | mamo-s | *BGIBMGA010067* | up | 647 | 655 | - | 11.3303 | 4.96E-05 | 0.615 | GTGGGTGGC |
| 8598 | mamo-s | *BGIBMGA010069* | up | 1387 | 1395 | - | 12.7615 | 1.04E-05 | 0.615 | CCGCGTGGC |
| 8599 | mamo-s | *BGIBMGA010069* | up | 69 | 77 | - | 11.3303 | 4.96E-05 | 0.615 | GTGGGTGGC |
| 8600 | mamo-s | *BGIBMGA010073* | up | 1218 | 1226 | - | 11.3303 | 4.96E-05 | 0.615 | GTGGGTGGC |
| 8601 | mamo-s | *BGIBMGA010073* | down | 1748 | 1756 | - | 11.3303 | 4.96E-05 | 0.615 | GTGGGTGGC |
| 8602 | mamo-s | *BGIBMGA010076* | down | 1431 | 1439 | + | 11.3303 | 4.96E-05 | 0.615 | GTGGGTGGC |
| 8603 | mamo-s | *BGIBMGA010077* | down | 992 | 1000 | - | 11.3303 | 4.96E-05 | 0.615 | GTGGGTGGC |
| 8604 | mamo-s | *BGIBMGA010077* | up | 935 | 943 | - | 11.0826 | 6.84E-05 | 0.709 | GTACGTGGC |
| 8605 | mamo-s | *BGIBMGA010078* | down | 1177 | 1185 | - | 11.3303 | 4.96E-05 | 0.615 | GTGGGTGGC |
| 8606 | mamo-s | *BGIBMGA010079* | up | 1671 | 1679 | + | 10.5413 | 9.42E-05 | 0.857 | CAGCGTGTC |
| 8607 | mamo-s | *BGIBMGA010081* | down | 1459 | 1467 | - | 12.1284 | 2.16E-05 | 0.615 | GTGCGAGGC |
| 8608 | mamo-s | *BGIBMGA010081* | down | 118 | 126 | - | 11.3303 | 4.96E-05 | 0.615 | GTGGGTGGC |
| 8609 | mamo-s | *BGIBMGA010082* | down | 596 | 604 | + | 11.3303 | 4.96E-05 | 0.615 | GTGGGTGGC |
| 8610 | mamo-s | *BGIBMGA010082* | down | 1365 | 1373 | + | 10.7523 | 8.38E-05 | 0.797 | GGGCGTGTC |
| 8611 | mamo-s | *BGIBMGA010083* | down | 1833 | 1841 | - | 11.3303 | 4.96E-05 | 0.615 | GTGGGTGGC |
| 8612 | mamo-s | *BGIBMGA010084* | up | 877 | 885 | - | 11.4404 | 4.51E-05 | 0.615 | CGGCGTGGC |
| 8613 | mamo-s | *BGIBMGA010084* | up | 642 | 650 | - | 11.3303 | 4.96E-05 | 0.615 | GTGGGTGGC |
| 8614 | mamo-s | *BGIBMGA010084* | up | 1290 | 1298 | - | 11.3303 | 4.96E-05 | 0.615 | GTGGGTGGC |
| 8615 | mamo-s | *BGIBMGA010087* | down | 1270 | 1278 | - | 11.3303 | 4.96E-05 | 0.615 | GTGGGTGGC |
| 8616 | mamo-s | *BGIBMGA010088* | down | 739 | 747 | - | 11.3303 | 4.96E-05 | 0.615 | GTGGGTGGC |
| 8617 | mamo-s | *BGIBMGA010088* | down | 961 | 969 | + | 11.3303 | 4.96E-05 | 0.615 | GTGGGTGGC |

| | | | | | | | | | | |
|---|---|---|---|---|---|---|---|---|---|---|
| 8618 | mamo-s | *BGIBMGA010090* | up | 1058 | 1066 | + | 11.3303 | 4.96E-05 | 0.615 | GTGGGTGGC |
| 8619 | mamo-s | *BGIBMGA010090* | up | 1261 | 1269 | + | 11.3303 | 4.96E-05 | 0.615 | GTGGGTGGC |
| 8620 | mamo-s | *BGIBMGA010090* | up | 1799 | 1807 | + | 11.1376 | 6.35E-05 | 0.709 | TCGCGTGGC |
| 8621 | mamo-s | *BGIBMGA010090* | up | 407 | 415 | - | 11.0826 | 6.84E-05 | 0.709 | GTACGTGGC |
| 8622 | mamo-s | *BGIBMGA010091* | down | 554 | 562 | + | 11.3303 | 4.96E-05 | 0.615 | GTGGGTGGC |
| 8623 | mamo-s | *BGIBMGA010091* | up | 797 | 805 | + | 11.3303 | 4.96E-05 | 0.615 | GTGGGTGGC |
| 8624 | mamo-s | *BGIBMGA010091* | down | 929 | 937 | + | 11.3303 | 4.96E-05 | 0.615 | GTGGGTGGC |
| 8625 | mamo-s | *BGIBMGA010092* | down | 100 | 108 | - | 11.3303 | 4.96E-05 | 0.615 | GTGGGTGGC |
| 8626 | mamo-s | *BGIBMGA010093* | down | 396 | 404 | + | 11.3303 | 4.96E-05 | 0.615 | GTGGGTGGC |
| 8627 | mamo-s | *BGIBMGA010093* | up | 1571 | 1579 | + | 10.9358 | 7.30E-05 | 0.728 | GCGGGTGGC |
| 8628 | mamo-s | *BGIBMGA010094* | up | 449 | 457 | + | 10.6881 | 8.61E-05 | 0.81 | GCACGTGGC |
| 8629 | mamo-s | *BGIBMGA010095* | down | 22 | 30 | + | 11.7798 | 3.11E-05 | 0.615 | ATGCGTGGC |
| 8630 | mamo-s | *BGIBMGA010095* | down | 360 | 368 | + | 11.0734 | 7.12E-05 | 0.726 | GAGCGTGGT |
| 8631 | mamo-s | *BGIBMGA010100* | down | 1545 | 1553 | + | 11.0826 | 6.84E-05 | 0.709 | GTACGTGGC |
| 8632 | mamo-s | *BGIBMGA010101* | down | 1844 | 1852 | + | 10.5688 | 8.88E-05 | 0.824 | GTGCGTGAC |
| 8633 | mamo-s | *BGIBMGA010102* | up | 1326 | 1334 | - | 11.5688 | 3.79E-05 | 0.615 | CTGCGTGTC |
| 8634 | mamo-s | *BGIBMGA010102* | down | 390 | 398 | + | 11.2844 | 5.18E-05 | 0.63 | CTGCGTGCC |
| 8635 | mamo-s | *BGIBMGA010106* | down | 436 | 444 | - | 11.3303 | 4.96E-05 | 0.615 | GTGGGTGGC |
| 8636 | mamo-s | *BGIBMGA010108* | down | 1729 | 1737 | + | 13.6606 | 4.06E-06 | 0.615 | GCGCGTGGC |
| 8637 | mamo-s | *BGIBMGA010110* | down | 987 | 995 | + | 11.3303 | 4.96E-05 | 0.615 | GTGGGTGGC |
| 8638 | mamo-s | *BGIBMGA010112* | up | 1763 | 1771 | - | 11.5688 | 3.79E-05 | 0.615 | CTGCGTGTC |
| 8639 | mamo-s | *BGIBMGA010112* | up | 648 | 656 | + | 10.5688 | 8.88E-05 | 0.824 | GTGCGTGAC |
| 8640 | mamo-s | *BGIBMGA010117* | down | 346 | 354 | + | 11.3303 | 4.96E-05 | 0.615 | GTGGGTGGC |
| 8641 | mamo-s | *BGIBMGA010120* | up | 1081 | 1089 | - | 11.3853 | 4.74E-05 | 0.615 | ACGCGTGGC |
| 8642 | mamo-s | *BGIBMGA010121* | down | 1 | 9 | - | 11.7339 | 3.29E-05 | 0.615 | GCGCGAGGC |
| 8643 | mamo-s | *BGIBMGA010126* | up | 1681 | 1689 | - | 11.3303 | 4.96E-05 | 0.615 | GTGGGTGGC |
| 8644 | mamo-s | *BGIBMGA010127* | down | 945 | 953 | - | 12.4679 | 1.31E-05 | 0.615 | GTGCGTGTC |
| 8645 | mamo-s | *BGIBMGA010127* | up | 930 | 938 | - | 10.5138 | 9.75E-05 | 0.865 | GTGCGTGTT |
| 8646 | mamo-s | *BGIBMGA010129* | up | 69 | 77 | + | 11.5688 | 3.79E-05 | 0.615 | CTGCGTGTC |
| 8647 | mamo-s | *BGIBMGA010131* | up | 1838 | 1846 | + | 12.1284 | 2.16E-05 | 0.615 | CAGCGTGGC |
| 8648 | mamo-s | *BGIBMGA010131* | down | 1347 | 1355 | - | 11.3303 | 4.96E-05 | 0.615 | GTGGGTGGC |
| 8649 | mamo-s | *BGIBMGA010131* | down | 1697 | 1705 | + | 11.3303 | 4.96E-05 | 0.615 | GTGGGTGGC |
| 8650 | mamo-s | *BGIBMGA010134* | up | 1683 | 1691 | + | 11.3303 | 4.96E-05 | 0.615 | GTGGGTGGC |
| 8651 | mamo-s | *BGIBMGA010135* | up | 1568 | 1576 | - | 12.1835 | 1.71E-05 | 0.615 | GTGCGTGCC |
| 8652 | mamo-s | *BGIBMGA010135* | up | 1737 | 1745 | - | 11.3303 | 4.96E-05 | 0.615 | GTGGGTGGC |
| 8653 | mamo-s | *BGIBMGA010136* | down | 1891 | 1899 | - | 11.3303 | 4.96E-05 | 0.615 | GTGGGTGGC |
| 8654 | mamo-s | *BGIBMGA010137* | down | 416 | 424 | - | 11.3303 | 4.96E-05 | 0.615 | GTGGGTGGC |
| 8655 | mamo-s | *BGIBMGA010140* | down | 1060 | 1068 | + | 12.4679 | 1.31E-05 | 0.615 | GTGCGTGTC |
| 8656 | mamo-s | *BGIBMGA010140* | up | 1446 | 1454 | + | 12.0734 | 2.66E-05 | 0.615 | GCGCGTGTC |
| 8657 | mamo-s | *BGIBMGA010141* | down | 589 | 597 | + | 11.3303 | 4.96E-05 | 0.615 | GTGGGTGGC |
| 8658 | mamo-s | *BGIBMGA010142* | up | 3 | 11 | - | 12.1284 | 2.16E-05 | 0.615 | GTGCGAGGC |
| 8659 | mamo-s | *BGIBMGA010142* | down | 1734 | 1742 | - | 11.3303 | 4.96E-05 | 0.615 | GTGGGTGGC |
| 8660 | mamo-s | *BGIBMGA010143* | up | 729 | 737 | - | 11.3303 | 4.96E-05 | 0.615 | GTGGGTGGC |
| 8661 | mamo-s | *BGIBMGA010146* | up | 1773 | 1781 | + | 10.7523 | 8.38E-05 | 0.797 | GGGCGTGTC |
| 8662 | mamo-s | *BGIBMGA010147* | down | 396 | 404 | + | 11.3303 | 4.96E-05 | 0.615 | GTGGGTGGC |
| 8663 | mamo-s | *BGIBMGA010148* | up | 904 | 912 | - | 11.0826 | 6.84E-05 | 0.709 | GTACGTGGC |
| 8664 | mamo-s | *BGIBMGA010151* | up | 614 | 622 | - | 11.1376 | 6.35E-05 | 0.709 | TCGCGTGGC |
| 8665 | mamo-s | *BGIBMGA010151* | up | 595 | 603 | - | 10.5413 | 9.42E-05 | 0.857 | CAGCGTGTC |
| 8666 | mamo-s | *BGIBMGA010152* | up | 109 | 117 | + | 11.3303 | 4.96E-05 | 0.615 | GTGGGTGGC |
| 8667 | mamo-s | *BGIBMGA010159* | down | 164 | 172 | + | 11.2018 | 5.68E-05 | 0.674 | CTGCGTGGT |
| 8668 | mamo-s | *BGIBMGA010160* | up | 628 | 636 | - | 13.0275 | 8.53E-06 | 0.615 | GAGCGTGGC |
| 8669 | mamo-s | *BGIBMGA010160* | up | 1563 | 1571 | - | 11.5688 | 3.79E-05 | 0.615 | CTGCGTGTC |
| 8670 | mamo-s | *BGIBMGA010160* | down | 1301 | 1309 | + | 11.3303 | 4.96E-05 | 0.615 | GTGGGTGGC |
| 8671 | mamo-s | *BGIBMGA010160* | down | 1825 | 1833 | + | 10.9358 | 7.30E-05 | 0.728 | GCGGGTGGC |
| 8672 | mamo-s | *BGIBMGA010162* | up | 1794 | 1802 | - | 10.9358 | 7.30E-05 | 0.728 | GCGGGTGGC |
| 8673 | mamo-s | *BGIBMGA010166* | down | 1578 | 1586 | + | 11.3303 | 4.96E-05 | 0.615 | GTGGGTGGC |
| 8674 | mamo-s | *BGIBMGA010168* | up | 264 | 272 | - | 10.6881 | 8.61E-05 | 0.81 | GCACGTGGC |
| 8675 | mamo-s | *BGIBMGA010168* | up | 269 | 277 | + | 10.5413 | 9.42E-05 | 0.857 | GTGCGAGTC |
| 8676 | mamo-s | *BGIBMGA010168* | down | 225 | 233 | - | 10.5138 | 9.75E-05 | 0.865 | GTGCGTGTT |
| 8677 | mamo-s | *BGIBMGA010169* | down | 980 | 988 | + | 11.0826 | 6.84E-05 | 0.709 | GTACGTGGC |
| 8678 | mamo-s | *BGIBMGA010174* | down | 1076 | 1084 | - | 11.3303 | 4.96E-05 | 0.615 | GTGGGTGGC |
| 8679 | mamo-s | *BGIBMGA010175* | up | 935 | 943 | - | 13.6606 | 4.06E-06 | 0.615 | GCGCGTGGC |
| 8680 | mamo-s | *BGIBMGA010176* | down | 526 | 534 | - | 11.1743 | 5.90E-05 | 0.684 | CCGCGTGTC |
| 8681 | mamo-s | *BGIBMGA010176* | up | 1108 | 1116 | - | 10.7523 | 8.38E-05 | 0.797 | AAGCGTGGC |
| 8682 | mamo-s | *BGIBMGA010177* | up | 1586 | 1594 | - | 11.3303 | 4.96E-05 | 0.615 | GTGGGTGGC |
| 8683 | mamo-s | *BGIBMGA010179* | down | 410 | 418 | - | 11.2018 | 5.68E-05 | 0.674 | CTGCGTGGT |
| 8684 | mamo-s | *BGIBMGA010180* | down | 456 | 464 | - | 11.3303 | 4.96E-05 | 0.615 | GTGGGTGGC |
| 8685 | mamo-s | *BGIBMGA010182* | down | 623 | 631 | - | 12.4679 | 1.31E-05 | 0.615 | GTGCGTGTC |
| 8686 | mamo-s | *BGIBMGA010182* | down | 1004 | 1012 | - | 11.3303 | 4.96E-05 | 0.615 | GTGGGTGGC |
| 8687 | mamo-s | *BGIBMGA010182* | up | 1065 | 1073 | - | 11.3303 | 4.96E-05 | 0.615 | GTGGGTGGC |
| 8688 | mamo-s | *BGIBMGA010182* | up | 1929 | 1937 | - | 11.3303 | 4.96E-05 | 0.615 | GTGGGTGGC |
| 8689 | mamo-s | *BGIBMGA010183* | up | 1318 | 1326 | - | 11.3303 | 4.96E-05 | 0.615 | GTGGGTGGC |
| 8690 | mamo-s | *BGIBMGA010184* | down | 128 | 136 | - | 11.3303 | 4.96E-05 | 0.615 | GTGGGTGGC |
| 8691 | mamo-s | *BGIBMGA010184* | down | 1116 | 1124 | + | 11.3303 | 4.96E-05 | 0.615 | GTGGGTGGC |
| 8692 | mamo-s | *BGIBMGA010185* | down | 1355 | 1363 | + | 10.5138 | 9.75E-05 | 0.865 | GTGCGTGTT |
| 8693 | mamo-s | *BGIBMGA010186* | up | 1739 | 1747 | - | 11.3303 | 4.96E-05 | 0.615 | GTGGGTGGC |
| 8694 | mamo-s | *BGIBMGA010187* | up | 1107 | 1115 | - | 11.4404 | 4.51E-05 | 0.615 | GAGCGTGTC |
| 8695 | mamo-s | *BGIBMGA010188* | down | 388 | 396 | + | 11.3303 | 4.96E-05 | 0.615 | GTGGGTGGC |
| 8696 | mamo-s | *BGIBMGA010188* | up | 1732 | 1740 | - | 10.7523 | 8.38E-05 | 0.797 | AAGCGTGGC |
| 8697 | mamo-s | *BGIBMGA010191* | up | 1886 | 1894 | - | 11.3303 | 4.96E-05 | 0.615 | GTGGGTGGC |
| 8698 | mamo-s | *BGIBMGA010191* | up | 1876 | 1884 | - | 10.5688 | 8.88E-05 | 0.824 | GTGCGTGAC |
| 8699 | mamo-s | *BGIBMGA010192* | up | 987 | 995 | + | 10.5138 | 9.75E-05 | 0.865 | GTGCGTGTT |
| 8700 | mamo-s | *BGIBMGA010193* | down | 948 | 956 | - | 11.789 | 2.84E-05 | 0.615 | GCGCGTGCC |
| 8701 | mamo-s | *BGIBMGA010193* | up | 1291 | 1299 | + | 11.5688 | 3.79E-05 | 0.615 | CTGCGTGTC |
| 8702 | mamo-s | *BGIBMGA010193* | up | 220 | 228 | - | 11.3303 | 4.96E-05 | 0.615 | GTGGGTGGC |
| 8703 | mamo-s | *BGIBMGA010193* | up | 867 | 875 | - | 11.3303 | 4.96E-05 | 0.615 | GTGGGTGGC |
| 8704 | mamo-s | *BGIBMGA010194* | up | 1546 | 1554 | - | 11.4404 | 4.51E-05 | 0.615 | CGGCGTGGC |
| 8705 | mamo-s | *BGIBMGA010194* | up | 409 | 417 | + | 10.6881 | 8.61E-05 | 0.81 | GCACGTGGC |

| | | | | | | | | | | |
|---|---|---|---|---|---|---|---|---|---|---|
| 8706 | mamo-s | *BGIBMGA010195* | down | 295 | 303 | - | 13.6606 | 4.06E-06 | 0.615 | GCGCGTGGC |
| 8707 | mamo-s | *BGIBMGA010195* | down | 325 | 333 | - | 12.4679 | 1.31E-05 | 0.615 | GTGCGTGTC |
| 8708 | mamo-s | *BGIBMGA010195* | down | 568 | 576 | - | 10.5413 | 9.42E-05 | 0.857 | GTGCGAGTC |
| 8709 | mamo-s | *BGIBMGA010202* | down | 1573 | 1581 | - | 11.0826 | 6.84E-05 | 0.709 | GTACGTGGC |
| 8710 | mamo-s | *BGIBMGA010204* | down | 1470 | 1478 | + | 11.3853 | 4.74E-05 | 0.615 | ACGCGTGGC |
| 8711 | mamo-s | *BGIBMGA010205* | up | 1090 | 1098 | - | 11.3303 | 4.96E-05 | 0.615 | GTGGGTGGC |
| 8712 | mamo-s | *BGIBMGA010206* | down | 1282 | 1290 | - | 11.3303 | 4.96E-05 | 0.615 | GTGGGTGGC |
| 8713 | mamo-s | *BGIBMGA010212* | up | 1118 | 1126 | + | 11.1009 | 6.57E-05 | 0.709 | GAGCGAGGC |
| 8714 | mamo-s | *BGIBMGA010213* | up | 1594 | 1602 | - | 11.0826 | 6.84E-05 | 0.709 | GTACGTGGC |
| 8715 | mamo-s | *BGIBMGA010214* | up | 1386 | 1394 | - | 11.3303 | 4.96E-05 | 0.615 | GTGGGTGGC |
| 8716 | mamo-s | *BGIBMGA010214* | down | 383 | 391 | + | 10.5138 | 9.75E-05 | 0.865 | GTGCGTGTT |
| 8717 | mamo-s | *BGIBMGA010215* | down | 655 | 663 | - | 10.9358 | 7.30E-05 | 0.728 | GCGGGTGGC |
| 8718 | mamo-s | *BGIBMGA010215* | up | 523 | 531 | + | 10.5138 | 9.75E-05 | 0.865 | GTGCGTGTT |
| 8719 | mamo-s | *BGIBMGA010216* | up | 801 | 809 | + | 11.3303 | 4.96E-05 | 0.615 | GTGGGTGGC |
| 8720 | mamo-s | *BGIBMGA010216* | up | 1125 | 1133 | - | 10.5688 | 8.88E-05 | 0.824 | GTGCGTGAC |
| 8721 | mamo-s | *BGIBMGA010217* | up | 877 | 885 | + | 11.3303 | 4.96E-05 | 0.615 | GTGGGTGGC |
| 8722 | mamo-s | *BGIBMGA010218* | up | 867 | 875 | + | 11.3303 | 4.96E-05 | 0.615 | GTGGGTGGC |
| 8723 | mamo-s | *BGIBMGA010219* | up | 1097 | 1105 | + | 11.3303 | 4.96E-05 | 0.615 | GTGGGTGGC |
| 8724 | mamo-s | *BGIBMGA010221* | up | 1495 | 1503 | - | 11.7339 | 3.29E-05 | 0.615 | GCGCGAGGC |
| 8725 | mamo-s | *BGIBMGA010221* | up | 1315 | 1323 | + | 10.5688 | 8.88E-05 | 0.824 | GTGCGTGAC |
| 8726 | mamo-s | *BGIBMGA010227* | up | 1858 | 1866 | + | 12.0734 | 2.66E-05 | 0.615 | GCGCGTGTC |
| 8727 | mamo-s | *BGIBMGA010228* | up | 1536 | 1544 | - | 11.0826 | 6.84E-05 | 0.709 | GTACGTGGC |
| 8728 | mamo-s | *BGIBMGA010231* | up | 273 | 281 | - | 11.7064 | 3.52E-05 | 0.615 | GCGCGTGGT |
| 8729 | mamo-s | *BGIBMGA010231* | down | 285 | 293 | - | 11.3303 | 4.96E-05 | 0.615 | GTGGGTGGC |
| 8730 | mamo-s | *BGIBMGA010232* | up | 1612 | 1620 | - | 11.7798 | 3.11E-05 | 0.615 | ATGCGTGGC |
| 8731 | mamo-s | *BGIBMGA010233* | down | 786 | 794 | - | 10.6881 | 8.61E-05 | 0.81 | GCACGTGGC |
| 8732 | mamo-s | *BGIBMGA010234* | up | 217 | 225 | + | 10.5688 | 8.88E-05 | 0.824 | GTGCGTGAC |
| 8733 | mamo-s | *BGIBMGA010235* | down | 974 | 982 | - | 11.7798 | 3.11E-05 | 0.615 | ATGCGTGGC |
| 8734 | mamo-s | *BGIBMGA010239* | up | 1398 | 1406 | + | 11.1743 | 5.90E-05 | 0.684 | CCGCGTGTC |
| 8735 | mamo-s | *BGIBMGA010241* | up | 612 | 620 | + | 11.2018 | 5.68E-05 | 0.674 | CTGCGTGGT |
| 8736 | mamo-s | *BGIBMGA010248* | up | 111 | 119 | - | 11.3303 | 4.96E-05 | 0.615 | GTGGGTGGC |
| 8737 | mamo-s | *BGIBMGA010249* | down | 1139 | 1147 | + | 11.3303 | 4.96E-05 | 0.615 | GTGGGTGGC |
| 8738 | mamo-s | *BGIBMGA010249* | down | 1846 | 1854 | - | 11.0826 | 6.84E-05 | 0.709 | GTACGTGGC |
| 8739 | mamo-s | *BGIBMGA010256* | up | 1186 | 1194 | + | 11.3303 | 4.96E-05 | 0.615 | GTGGGTGGC |
| 8740 | mamo-s | *BGIBMGA010257* | down | 1669 | 1677 | - | 11.3303 | 4.96E-05 | 0.615 | GTGGGTGGC |
| 8741 | mamo-s | *BGIBMGA010264* | down | 94 | 102 | + | 11.4404 | 4.51E-05 | 0.615 | CGGCGTGGC |
| 8742 | mamo-s | *BGIBMGA010264* | up | 1600 | 1608 | - | 11.3303 | 4.96E-05 | 0.615 | GTGGGTGGC |
| 8743 | mamo-s | *BGIBMGA010267* | up | 131 | 139 | + | 11.4404 | 4.51E-05 | 0.615 | CGGCGTGGC |
| 8744 | mamo-s | *BGIBMGA010267* | up | 477 | 485 | - | 11.3303 | 4.96E-05 | 0.615 | GTGGGTGGC |
| 8745 | mamo-s | *BGIBMGA010267* | down | 2 | 10 | - | 10.7523 | 8.38E-05 | 0.797 | AAGCGTGGC |
| 8746 | mamo-s | *BGIBMGA010268* | down | 925 | 933 | + | 10.9358 | 7.30E-05 | 0.728 | GCGGGTGGC |
| 8747 | mamo-s | *BGIBMGA010268* | down | 676 | 684 | - | 10.8073 | 7.89E-05 | 0.766 | CCGCGTGGT |
| 8748 | mamo-s | *BGIBMGA010269* | up | 521 | 529 | - | 12.1284 | 2.16E-05 | 0.615 | CAGCGTGGC |
| 8749 | mamo-s | *BGIBMGA010270* | up | 1584 | 1592 | + | 11.3303 | 4.96E-05 | 0.615 | GTGGGTGGC |
| 8750 | mamo-s | *BGIBMGA010271* | down | 724 | 732 | + | 12.4679 | 1.31E-05 | 0.615 | GTGCGTGTC |
| 8751 | mamo-s | *BGIBMGA010273* | down | 1472 | 1480 | - | 11.3303 | 4.96E-05 | 0.615 | GTGGGTGGC |
| 8752 | mamo-s | *BGIBMGA010274* | up | 840 | 848 | + | 10.5138 | 9.75E-05 | 0.865 | GTGCGTGTT |
| 8753 | mamo-s | *BGIBMGA010276* | up | 1556 | 1564 | - | 12.1009 | 2.43E-05 | 0.615 | GTGCGTGGT |
| 8754 | mamo-s | *BGIBMGA010276* | up | 1740 | 1748 | + | 12.1009 | 2.43E-05 | 0.615 | GTGCGTGGT |
| 8755 | mamo-s | *BGIBMGA010276* | down | 854 | 862 | - | 11.0826 | 6.84E-05 | 0.709 | GTACGTGGC |
| 8756 | mamo-s | *BGIBMGA010277* | up | 188 | 196 | - | 11.2018 | 5.68E-05 | 0.674 | CTGCGTGGT |
| 8757 | mamo-s | *BGIBMGA010277* | down | 1798 | 1806 | - | 11.1743 | 5.90E-05 | 0.684 | CCGCGTGTC |
| 8758 | mamo-s | *BGIBMGA010278* | up | 890 | 898 | - | 11.5321 | 4.06E-05 | 0.615 | TTGCGTGGC |
| 8759 | mamo-s | *BGIBMGA010279* | down | 1889 | 1897 | + | 12.7615 | 1.04E-05 | 0.615 | CCGCGTGGC |
| 8760 | mamo-s | *BGIBMGA010279* | down | 1515 | 1523 | - | 11.3303 | 4.96E-05 | 0.615 | GTGGGTGGC |
| 8761 | mamo-s | *BGIBMGA010280* | up | 135 | 143 | + | 11.1009 | 6.57E-05 | 0.709 | GAGCGAGGC |
| 8762 | mamo-s | *BGIBMGA010284* | up | 1809 | 1817 | + | 11.5688 | 3.79E-05 | 0.615 | CTGCGTGTC |
| 8763 | mamo-s | *BGIBMGA010284* | up | 568 | 576 | + | 10.8899 | 7.48E-05 | 0.739 | CCGCGTGCC |
| 8764 | mamo-s | *BGIBMGA010284* | down | 1666 | 1674 | - | 10.5688 | 8.88E-05 | 0.824 | GTGCGTGAC |
| 8765 | mamo-s | *BGIBMGA010286* | down | 1882 | 1890 | + | 11.3303 | 4.96E-05 | 0.615 | GTGGGTGGC |
| 8766 | mamo-s | *BGIBMGA010287* | down | 1579 | 1587 | + | 11.3303 | 4.96E-05 | 0.615 | GTGGGTGGC |
| 8767 | mamo-s | *BGIBMGA010288* | up | 356 | 364 | - | 10.5138 | 9.75E-05 | 0.865 | GTGCGTGTT |
| 8768 | mamo-s | *BGIBMGA010289* | down | 1460 | 1468 | - | 11.0826 | 6.84E-05 | 0.709 | GTACGTGGC |
| 8769 | mamo-s | *BGIBMGA010290* | up | 1674 | 1682 | + | 11.3303 | 4.96E-05 | 0.615 | GTGGGTGGC |
| 8770 | mamo-s | *BGIBMGA010290* | down | 901 | 909 | + | 10.7523 | 8.38E-05 | 0.797 | GGGCGTGTC |
| 8771 | mamo-s | *BGIBMGA010291* | up | 1693 | 1701 | + | 11.3303 | 4.96E-05 | 0.615 | GTGGGTGGC |
| 8772 | mamo-s | *BGIBMGA010291* | down | 1764 | 1772 | - | 11.3303 | 4.96E-05 | 0.615 | GTGGGTGGC |
| 8773 | mamo-s | *BGIBMGA010291* | down | 1044 | 1052 | - | 11.0826 | 6.84E-05 | 0.709 | GTACGTGGC |
| 8774 | mamo-s | *BGIBMGA010292* | up | 74 | 82 | - | 13.6606 | 4.06E-06 | 0.615 | GCGCGTGGC |
| 8775 | mamo-s | *BGIBMGA010292* | down | 366 | 374 | - | 11.3303 | 4.96E-05 | 0.615 | GTGGGTGGC |
| 8776 | mamo-s | *BGIBMGA010293* | down | 1015 | 1023 | - | 13.156 | 6.30E-06 | 0.615 | CTGCGTGGC |
| 8777 | mamo-s | *BGIBMGA010294* | down | 891 | 899 | - | 11.3303 | 4.96E-05 | 0.615 | GTGGGTGGC |
| 8778 | mamo-s | *BGIBMGA010294* | down | 1984 | 1992 | - | 11.3303 | 4.96E-05 | 0.615 | GTGGGTGGC |
| 8779 | mamo-s | *BGIBMGA010294* | up | 1639 | 1647 | - | 11.0826 | 6.84E-05 | 0.709 | GTACGTGGC |
| 8780 | mamo-s | *BGIBMGA010296* | down | 354 | 362 | + | 11.789 | 2.84E-05 | 0.615 | GCGCGTGCC |
| 8781 | mamo-s | *BGIBMGA010297* | down | 1292 | 1300 | - | 11.789 | 2.84E-05 | 0.615 | GCGCGTGCC |
| 8782 | mamo-s | *BGIBMGA010297* | up | 1849 | 1857 | - | 10.8349 | 7.66E-05 | 0.751 | CCGCGAGGC |
| 8783 | mamo-s | *BGIBMGA010298* | up | 395 | 403 | - | 11.3303 | 4.96E-05 | 0.615 | GTGGGTGGC |
| 8784 | mamo-s | *BGIBMGA010298* | down | 808 | 816 | - | 10.5688 | 8.88E-05 | 0.824 | GTGCGTGAC |
| 8785 | mamo-s | *BGIBMGA010299* | up | 629 | 637 | - | 11.789 | 2.84E-05 | 0.615 | GCGCGTGCC |
| 8786 | mamo-s | *BGIBMGA010301* | up | 165 | 173 | + | 12.3394 | 1.49E-05 | 0.615 | GGGCGTGGC |
| 8787 | mamo-s | *BGIBMGA010301* | down | 748 | 756 | - | 11.3303 | 4.96E-05 | 0.615 | GTGGGTGGC |
| 8788 | mamo-s | *BGIBMGA010301* | down | 1197 | 1205 | + | 10.8073 | 7.89E-05 | 0.766 | CCGCGTGGT |
| 8789 | mamo-s | *BGIBMGA010301* | down | 566 | 574 | + | 10.5688 | 8.88E-05 | 0.824 | GTGCGTGAC |
| 8790 | mamo-s | *BGIBMGA010302* | down | 1258 | 1266 | + | 12.0734 | 2.66E-05 | 0.615 | GCGCGTGTC |
| 8791 | mamo-s | *BGIBMGA010302* | down | 936 | 944 | + | 10.7523 | 8.38E-05 | 0.797 | AAGCGTGGC |
| 8792 | mamo-s | *BGIBMGA010304* | up | 1830 | 1838 | - | 12.1009 | 2.43E-05 | 0.615 | GTGCGTGGT |
| 8793 | mamo-s | *BGIBMGA010305* | down | 1718 | 1726 | + | 11.5321 | 4.06E-05 | 0.615 | TTGCGTGGC |

| | | | | | | | | | | |
|---|---|---|---|---|---|---|---|---|---|---|
| 8794 | mamo-s | *BGIBMGA010308* | up | 931 | 939 | + | 12.0734 | 2.66E-05 | 0.615 | GCGCGTGTC |
| 8795 | mamo-s | *BGIBMGA010308* | up | 1624 | 1632 | - | 11.3303 | 4.96E-05 | 0.615 | GTGGGTGGC |
| 8796 | mamo-s | *BGIBMGA010308* | up | 1782 | 1790 | + | 10.5413 | 9.42E-05 | 0.857 | CAGCGTGTC |
| 8797 | mamo-s | *BGIBMGA010311* | up | 1824 | 1832 | + | 11.1009 | 6.57E-05 | 0.709 | GAGCGAGGC |
| 8798 | mamo-s | *BGIBMGA010312* | up | 1118 | 1126 | + | 11.0826 | 6.84E-05 | 0.709 | GTACGTGGC |
| 8799 | mamo-s | *BGIBMGA010314* | up | 42 | 50 | + | 11.3303 | 4.96E-05 | 0.615 | GTGGGTGGC |
| 8800 | mamo-s | *BGIBMGA010315* | down | 1860 | 1868 | + | 10.8349 | 7.66E-05 | 0.751 | CCGCGAGGC |
| 8801 | mamo-s | *BGIBMGA010320* | up | 1292 | 1300 | + | 11.3303 | 4.96E-05 | 0.615 | GTGGGTGGC |
| 8802 | mamo-s | *BGIBMGA010321* | down | 1665 | 1673 | + | 11.3303 | 4.96E-05 | 0.615 | GTGGGTGGC |
| 8803 | mamo-s | *BGIBMGA010321* | up | 1759 | 1767 | - | 11.3303 | 4.96E-05 | 0.615 | GTGGGTGGC |
| 8804 | mamo-s | *BGIBMGA010323* | down | 1631 | 1639 | + | 11.3303 | 4.96E-05 | 0.615 | GTGGGTGGC |
| 8805 | mamo-s | *BGIBMGA010325* | down | 1408 | 1416 | + | 12.1009 | 2.43E-05 | 0.615 | GTGCGTGGT |
| 8806 | mamo-s | *BGIBMGA010327* | up | 555 | 563 | + | 10.7523 | 8.38E-05 | 0.797 | GGGCGTGTC |
| 8807 | mamo-s | *BGIBMGA010331* | up | 826 | 834 | + | 11.3303 | 4.96E-05 | 0.615 | GTGGGTGGC |
| 8808 | mamo-s | *BGIBMGA010334* | down | 418 | 426 | - | 11.3303 | 4.96E-05 | 0.615 | GTGGGTGGC |
| 8809 | mamo-s | *BGIBMGA010334* | up | 1135 | 1143 | + | 10.7523 | 8.38E-05 | 0.797 | AAGCGTGGC |
| 8810 | mamo-s | *BGIBMGA010335* | up | 1317 | 1325 | + | 11.7064 | 3.52E-05 | 0.615 | GCGCGTGGT |
| 8811 | mamo-s | *BGIBMGA010335* | up | 1371 | 1379 | + | 11.0734 | 7.12E-05 | 0.726 | GAGCGTGGT |
| 8812 | mamo-s | *BGIBMGA010337* | down | 1665 | 1673 | + | 11.5321 | 4.06E-05 | 0.615 | TTGCGTGGC |
| 8813 | mamo-s | *BGIBMGA010338* | down | 574 | 582 | - | 11.3303 | 4.96E-05 | 0.615 | GTGGGTGGC |
| 8814 | mamo-s | *BGIBMGA010340* | up | 147 | 155 | + | 11.3303 | 4.96E-05 | 0.615 | GTGGGTGGC |
| 8815 | mamo-s | *BGIBMGA010342* | down | 1238 | 1246 | + | 11.3303 | 4.96E-05 | 0.615 | GTGGGTGGC |
| 8816 | mamo-s | *BGIBMGA010343* | up | 1233 | 1241 | + | 11.3303 | 4.96E-05 | 0.615 | GTGGGTGGC |
| 8817 | mamo-s | *BGIBMGA010343* | down | 1274 | 1282 | - | 11.3303 | 4.96E-05 | 0.615 | GTGGGTGGC |
| 8818 | mamo-s | *BGIBMGA010345* | down | 1658 | 1666 | + | 12.1009 | 2.43E-05 | 0.615 | GTGCGTGGT |
| 8819 | mamo-s | *BGIBMGA010348* | down | 794 | 802 | - | 10.5138 | 9.75E-05 | 0.865 | GTGCGTGTT |
| 8820 | mamo-s | *BGIBMGA010349* | up | 491 | 499 | - | 10.5138 | 9.75E-05 | 0.865 | GTGCGTGTT |
| 8821 | mamo-s | *BGIBMGA010352* | up | 1181 | 1189 | + | 11.4404 | 4.51E-05 | 0.615 | CGGCGTGGC |
| 8822 | mamo-s | *BGIBMGA010352* | down | 721 | 729 | + | 11.3303 | 4.96E-05 | 0.615 | GTGGGTGGC |
| 8823 | mamo-s | *BGIBMGA010352* | up | 977 | 985 | + | 11.3303 | 4.96E-05 | 0.615 | GTGGGTGGC |
| 8824 | mamo-s | *BGIBMGA010352* | up | 1139 | 1147 | - | 11.1376 | 6.35E-05 | 0.709 | TCGCGTGGC |
| 8825 | mamo-s | *BGIBMGA010357* | down | 255 | 263 | + | 11.3303 | 4.96E-05 | 0.615 | GTGGGTGGC |
| 8826 | mamo-s | *BGIBMGA010359* | up | 1190 | 1198 | - | 10.5138 | 9.75E-05 | 0.865 | GTGCGTGTT |
| 8827 | mamo-s | *BGIBMGA010361* | down | 1705 | 1713 | - | 11.3303 | 4.96E-05 | 0.615 | GTGGGTGGC |
| 8828 | mamo-s | *BGIBMGA010364* | down | 1594 | 1602 | - | 11.3303 | 4.96E-05 | 0.615 | GTGGGTGGC |
| 8829 | mamo-s | *BGIBMGA010364* | down | 1306 | 1314 | - | 10.8899 | 7.48E-05 | 0.739 | CCGCGTGCC |
| 8830 | mamo-s | *BGIBMGA010369* | up | 537 | 545 | - | 12.0734 | 2.66E-05 | 0.615 | GCGCGTGTC |
| 8831 | mamo-s | *BGIBMGA010369* | down | 1450 | 1458 | + | 11.3303 | 4.96E-05 | 0.615 | GTGGGTGGC |
| 8832 | mamo-s | *BGIBMGA010369* | down | 125 | 133 | + | 10.8073 | 7.89E-05 | 0.766 | CCGCGTGGT |
| 8833 | mamo-s | *BGIBMGA010372* | up | 1854 | 1862 | + | 11.3303 | 4.96E-05 | 0.615 | GTGGGTGGC |
| 8834 | mamo-s | *BGIBMGA010373* | down | 163 | 171 | + | 10.5138 | 9.75E-05 | 0.865 | GTGCGTGTT |
| 8835 | mamo-s | *BGIBMGA010382* | up | 150 | 158 | + | 12.4679 | 1.31E-05 | 0.615 | GTGCGTGTC |
| 8836 | mamo-s | *BGIBMGA010384* | down | 1390 | 1398 | + | 11.3303 | 4.96E-05 | 0.615 | GTGGGTGGC |
| 8837 | mamo-s | *BGIBMGA010384* | down | 1915 | 1923 | - | 11.3303 | 4.96E-05 | 0.615 | GTGGGTGGC |
| 8838 | mamo-s | *BGIBMGA010386* | up | 1081 | 1089 | + | 11.0826 | 6.84E-05 | 0.709 | GTACGTGGC |
| 8839 | mamo-s | *BGIBMGA010388* | down | 434 | 442 | + | 11.3303 | 4.96E-05 | 0.615 | GTGGGTGGC |
| 8840 | mamo-s | *BGIBMGA010388* | down | 1290 | 1298 | + | 11.3303 | 4.96E-05 | 0.615 | GTGGGTGGC |
| 8841 | mamo-s | *BGIBMGA010392* | down | 1112 | 1120 | + | 11.3303 | 4.96E-05 | 0.615 | GTGGGTGGC |
| 8842 | mamo-s | *BGIBMGA010392* | up | 1991 | 1999 | + | 10.5413 | 9.42E-05 | 0.857 | CAGCGTGTC |
| 8843 | mamo-s | *BGIBMGA010393* | down | 1946 | 1954 | - | 13.6606 | 4.06E-06 | 0.615 | GCGCGTGGC |
| 8844 | mamo-s | *BGIBMGA010393* | down | 429 | 437 | - | 11.3303 | 4.96E-05 | 0.615 | GTGGGTGGC |
| 8845 | mamo-s | *BGIBMGA010400* | down | 710 | 718 | + | 11.3303 | 4.96E-05 | 0.615 | GTGGGTGGC |
| 8846 | mamo-s | *BGIBMGA010401* | up | 1297 | 1305 | + | 11.0826 | 6.84E-05 | 0.709 | GTACGTGGC |
| 8847 | mamo-s | *BGIBMGA010407* | up | 923 | 931 | - | 11.3303 | 4.96E-05 | 0.615 | GTGGGTGGC |
| 8848 | mamo-s | *BGIBMGA010408* | down | 1853 | 1861 | - | 11.3303 | 4.96E-05 | 0.615 | GTGGGTGGC |
| 8849 | mamo-s | *BGIBMGA010409* | down | 660 | 668 | - | 12.1009 | 2.43E-05 | 0.615 | GTGCGTGGT |
| 8850 | mamo-s | *BGIBMGA010411* | down | 229 | 237 | + | 10.7523 | 8.38E-05 | 0.797 | AAGCGTGGC |
| 8851 | mamo-s | *BGIBMGA010412* | down | 786 | 794 | - | 11.3303 | 4.96E-05 | 0.615 | GTGGGTGGC |
| 8852 | mamo-s | *BGIBMGA010413* | up | 1424 | 1432 | + | 11.4404 | 4.51E-05 | 0.615 | GAGCGTGTC |
| 8853 | mamo-s | *BGIBMGA010417* | up | 319 | 327 | + | 11.3303 | 4.96E-05 | 0.615 | GTGGGTGGC |
| 8854 | mamo-s | *BGIBMGA010418* | up | 19 | 27 | + | 10.5413 | 9.42E-05 | 0.857 | GTGCGAGTC |
| 8855 | mamo-s | *BGIBMGA010419* | down | 67 | 75 | + | 11.3303 | 4.96E-05 | 0.615 | GTGGGTGGC |
| 8856 | mamo-s | *BGIBMGA010419* | down | 1490 | 1498 | + | 10.9358 | 7.30E-05 | 0.728 | GCGGGTGGC |
| 8857 | mamo-s | *BGIBMGA010420* | down | 1 | 9 | - | 11.4404 | 4.51E-05 | 0.615 | CGGCGTGGC |
| 8858 | mamo-s | *BGIBMGA010423* | down | 838 | 846 | - | 14.055 | 2.23E-06 | 0.615 | GTGCGTGGC |
| 8859 | mamo-s | *BGIBMGA010424* | up | 492 | 500 | - | 11.3303 | 4.96E-05 | 0.615 | GTGGGTGGC |
| 8860 | mamo-s | *BGIBMGA010424* | up | 1500 | 1508 | - | 11.3303 | 4.96E-05 | 0.615 | GTGGGTGGC |
| 8861 | mamo-s | *BGIBMGA010425* | up | 990 | 998 | - | 11.3303 | 4.96E-05 | 0.615 | GTGGGTGGC |
| 8862 | mamo-s | *BGIBMGA010426* | up | 195 | 203 | + | 10.5688 | 8.88E-05 | 0.824 | GTGCGTGAC |
| 8863 | mamo-s | *BGIBMGA010426* | up | 520 | 528 | - | 10.5413 | 9.42E-05 | 0.857 | CAGCGTGTC |
| 8864 | mamo-s | *BGIBMGA010428* | down | 464 | 472 | + | 11.3303 | 4.96E-05 | 0.615 | GTGGGTGGC |
| 8865 | mamo-s | *BGIBMGA010428* | down | 1363 | 1371 | + | 11.3303 | 4.96E-05 | 0.615 | GTGGGTGGC |
| 8866 | mamo-s | *BGIBMGA010429* | up | 1298 | 1306 | - | 11.3303 | 4.96E-05 | 0.615 | GTGGGTGGC |
| 8867 | mamo-s | *BGIBMGA010430* | up | 1249 | 1257 | - | 12.0734 | 2.66E-05 | 0.615 | GCGCGTGTC |
| 8868 | mamo-s | *BGIBMGA010432* | up | 468 | 476 | + | 11.2844 | 5.18E-05 | 0.63 | CTGCGTGCC |
| 8869 | mamo-s | *BGIBMGA010434* | up | 843 | 851 | + | 11.3303 | 4.96E-05 | 0.615 | GTGGGTGGC |
| 8870 | mamo-s | *BGIBMGA010435* | up | 1147 | 1155 | + | 11.3303 | 4.96E-05 | 0.615 | GTGGGTGGC |
| 8871 | mamo-s | *BGIBMGA010439* | up | 648 | 656 | + | 11.3303 | 4.96E-05 | 0.615 | GTGGGTGGC |
| 8872 | mamo-s | *BGIBMGA010440* | down | 1830 | 1838 | + | 12.4679 | 1.31E-05 | 0.615 | GTGCGTGTC |
| 8873 | mamo-s | *BGIBMGA010441* | down | 1569 | 1577 | + | 11.3303 | 4.96E-05 | 0.615 | GTGGGTGGC |
| 8874 | mamo-s | *BGIBMGA010442* | up | 445 | 453 | - | 11.3303 | 4.96E-05 | 0.615 | GTGGGTGGC |
| 8875 | mamo-s | *BGIBMGA010442* | down | 1872 | 1880 | - | 11.0826 | 6.84E-05 | 0.709 | GTACGTGGC |
| 8876 | mamo-s | *BGIBMGA010443* | down | 686 | 694 | + | 11.3303 | 4.96E-05 | 0.615 | GTGGGTGGC |
| 8877 | mamo-s | *BGIBMGA010443* | up | 650 | 658 | - | 11.0826 | 6.84E-05 | 0.709 | GTACGTGGC |
| 8878 | mamo-s | *BGIBMGA010445* | down | 398 | 406 | + | 11.3303 | 4.96E-05 | 0.615 | GTGGGTGGC |
| 8879 | mamo-s | *BGIBMGA010446* | up | 1042 | 1050 | - | 11.3303 | 4.96E-05 | 0.615 | GTGGGTGGC |
| 8880 | mamo-s | *BGIBMGA010448* | up | 410 | 418 | + | 11.3303 | 4.96E-05 | 0.615 | GTGGGTGGC |
| 8881 | mamo-s | *BGIBMGA010448* | down | 616 | 624 | + | 11.2018 | 5.68E-05 | 0.674 | CTGCGTGGT |

| | | | | | | | | | | |
|---|---|---|---|---|---|---|---|---|---|---|
| 8882 | mamo-s | *BGIBMGA010448* | down | 639 | 647 | - | 10.8349 | 7.66E-05 | 0.751 | CCGCGAGGC |
| 8883 | mamo-s | *BGIBMGA010449* | up | 241 | 249 | - | 11.3303 | 4.96E-05 | 0.615 | GTGGGTGGC |
| 8884 | mamo-s | *BGIBMGA010450* | down | 1927 | 1935 | - | 11.5688 | 3.79E-05 | 0.615 | CTGCGTGTC |
| 8885 | mamo-s | *BGIBMGA010453* | down | 1015 | 1023 | - | 11.3303 | 4.96E-05 | 0.615 | GTGGGTGGC |
| 8886 | mamo-s | *BGIBMGA010453* | up | 1159 | 1167 | + | 11.3303 | 4.96E-05 | 0.615 | GTGGGTGGC |
| 8887 | mamo-s | *BGIBMGA010454* | down | 904 | 912 | + | 11.2844 | 5.18E-05 | 0.63 | CTGCGTGCC |
| 8888 | mamo-s | *BGIBMGA010454* | up | 699 | 707 | - | 11.0826 | 6.84E-05 | 0.709 | GTACGTGGC |
| 8889 | mamo-s | *BGIBMGA010455* | up | 471 | 479 | - | 12.1009 | 2.43E-05 | 0.615 | GTGCGTGGT |
| 8890 | mamo-s | *BGIBMGA010457* | down | 331 | 339 | - | 11.3303 | 4.96E-05 | 0.615 | GTGGGTGGC |
| 8891 | mamo-s | *BGIBMGA010458* | down | 784 | 792 | - | 11.4404 | 4.51E-05 | 0.615 | GAGCGTGTC |
| 8892 | mamo-s | *BGIBMGA010459* | up | 177 | 185 | + | 10.5413 | 9.42E-05 | 0.857 | GTGCGAGTC |
| 8893 | mamo-s | *BGIBMGA010461* | down | 952 | 960 | + | 13.156 | 6.30E-06 | 0.615 | CTGCGTGGC |
| 8894 | mamo-s | *BGIBMGA010461* | up | 1823 | 1831 | - | 11.3303 | 4.96E-05 | 0.615 | GTGGGTGGC |
| 8895 | mamo-s | *BGIBMGA010464* | down | 1735 | 1743 | + | 11.4404 | 4.51E-05 | 0.615 | CGGCGTGGC |
| 8896 | mamo-s | *BGIBMGA010464* | down | 1702 | 1710 | + | 11.3853 | 4.74E-05 | 0.615 | ACGCGTGGC |
| 8897 | mamo-s | *BGIBMGA010465* | down | 1750 | 1758 | - | 11.3303 | 4.96E-05 | 0.615 | GTGGGTGGC |
| 8898 | mamo-s | *BGIBMGA010467* | down | 1331 | 1339 | - | 11.3303 | 4.96E-05 | 0.615 | GTGGGTGGC |
| 8899 | mamo-s | *BGIBMGA010467* | up | 1368 | 1376 | - | 11.3303 | 4.96E-05 | 0.615 | GTGGGTGGC |
| 8900 | mamo-s | *BGIBMGA010468* | down | 1412 | 1420 | - | 12.7615 | 1.04E-05 | 0.615 | CCGCGTGGC |
| 8901 | mamo-s | *BGIBMGA010468* | up | 897 | 905 | + | 11.1009 | 6.57E-05 | 0.709 | GAGCGAGGC |
| 8902 | mamo-s | *BGIBMGA010469* | down | 1942 | 1950 | + | 11.3303 | 4.96E-05 | 0.615 | GTGGGTGGC |
| 8903 | mamo-s | *BGIBMGA010470* | up | 247 | 255 | + | 11.3303 | 4.96E-05 | 0.615 | GTGGGTGGC |
| 8904 | mamo-s | *BGIBMGA010470* | up | 1484 | 1492 | + | 11.3303 | 4.96E-05 | 0.615 | GTGGGTGGC |
| 8905 | mamo-s | *BGIBMGA010471* | down | 1002 | 1010 | - | 11.3303 | 4.96E-05 | 0.615 | GTGGGTGGC |
| 8906 | mamo-s | *BGIBMGA010472* | down | 22 | 30 | - | 10.5138 | 9.75E-05 | 0.865 | GTGCGTGTT |
| 8907 | mamo-s | *BGIBMGA010473* | down | 110 | 118 | - | 11.7064 | 3.52E-05 | 0.615 | GCGCGTGGT |
| 8908 | mamo-s | *BGIBMGA010473* | up | 1671 | 1679 | + | 11.3303 | 4.96E-05 | 0.615 | GTGGGTGGC |
| 8909 | mamo-s | *BGIBMGA010474* | up | 235 | 243 | - | 11.7064 | 3.52E-05 | 0.615 | GCGCGTGGT |
| 8910 | mamo-s | *BGIBMGA010474* | down | 1169 | 1177 | - | 11.3303 | 4.96E-05 | 0.615 | GTGGGTGGC |
| 8911 | mamo-s | *BGIBMGA010476* | up | 570 | 578 | + | 11.3303 | 4.96E-05 | 0.615 | GTGGGTGGC |
| 8912 | mamo-s | *BGIBMGA010479* | down | 734 | 742 | - | 11.3303 | 4.96E-05 | 0.615 | GTGGGTGGC |
| 8913 | mamo-s | *BGIBMGA010482* | down | 273 | 281 | + | 10.8073 | 7.89E-05 | 0.766 | CCGCGTGGT |
| 8914 | mamo-s | *BGIBMGA010484* | down | 1280 | 1288 | + | 11.3303 | 4.96E-05 | 0.615 | GTGGGTGGC |
| 8915 | mamo-s | *BGIBMGA010484* | down | 1095 | 1103 | + | 11.0826 | 6.84E-05 | 0.709 | GTACGTGGC |
| 8916 | mamo-s | *BGIBMGA010490* | up | 54 | 62 | + | 10.7523 | 8.38E-05 | 0.797 | GGGCGTGTC |
| 8917 | mamo-s | *BGIBMGA010490* | up | 1052 | 1060 | - | 10.5413 | 9.42E-05 | 0.857 | CAGCGTGTC |
| 8918 | mamo-s | *BGIBMGA010491* | down | 617 | 625 | + | 11.3303 | 4.96E-05 | 0.615 | GTGGGTGGC |
| 8919 | mamo-s | *BGIBMGA010491* | up | 1667 | 1675 | + | 11.3303 | 4.96E-05 | 0.615 | GTGGGTGGC |
| 8920 | mamo-s | *BGIBMGA010491* | up | 1260 | 1268 | - | 11.1376 | 6.35E-05 | 0.709 | TCGCGTGGC |
| 8921 | mamo-s | *BGIBMGA010492* | up | 1837 | 1845 | - | 11.3853 | 4.74E-05 | 0.615 | ACGCGTGGC |
| 8922 | mamo-s | *BGIBMGA010493* | down | 1907 | 1915 | + | 12.1009 | 2.43E-05 | 0.615 | GTGCGTGGT |
| 8923 | mamo-s | *BGIBMGA010493* | up | 701 | 709 | - | 11.3853 | 4.74E-05 | 0.615 | ACGCGTGGC |
| 8924 | mamo-s | *BGIBMGA010495* | down | 1051 | 1059 | - | 11.3303 | 4.96E-05 | 0.615 | GTGGGTGGC |
| 8925 | mamo-s | *BGIBMGA010495* | down | 1898 | 1906 | - | 10.5138 | 9.75E-05 | 0.865 | GTGCGTGTT |
| 8926 | mamo-s | *BGIBMGA010498* | up | 1632 | 1640 | + | 11.3303 | 4.96E-05 | 0.615 | GTGGGTGGC |
| 8927 | mamo-s | *BGIBMGA010499* | down | 1110 | 1118 | - | 11.3303 | 4.96E-05 | 0.615 | GTGGGTGGC |
| 8928 | mamo-s | *BGIBMGA010499* | down | 1295 | 1303 | - | 11.0826 | 6.84E-05 | 0.709 | GTACGTGGC |
| 8929 | mamo-s | *BGIBMGA010500* | down | 240 | 248 | + | 12.4679 | 1.31E-05 | 0.615 | GTGCGTGTC |
| 8930 | mamo-s | *BGIBMGA010500* | up | 475 | 483 | - | 11.3303 | 4.96E-05 | 0.615 | GTGGGTGGC |
| 8931 | mamo-s | *BGIBMGA010501* | up | 355 | 363 | - | 11.0826 | 6.84E-05 | 0.709 | GTACGTGGC |
| 8932 | mamo-s | *BGIBMGA010503* | up | 560 | 568 | - | 11.3853 | 4.74E-05 | 0.615 | ACGCGTGGC |
| 8933 | mamo-s | *BGIBMGA010503* | down | 1224 | 1232 | - | 11.3303 | 4.96E-05 | 0.615 | GTGGGTGGC |
| 8934 | mamo-s | *BGIBMGA010504* | up | 55 | 63 | - | 11.3303 | 4.96E-05 | 0.615 | GTGGGTGGC |
| 8935 | mamo-s | *BGIBMGA010504* | down | 202 | 210 | - | 11.3303 | 4.96E-05 | 0.615 | GTGGGTGGC |
| 8936 | mamo-s | *BGIBMGA010505* | up | 390 | 398 | - | 12.1009 | 2.43E-05 | 0.615 | GTGCGTGGT |
| 8937 | mamo-s | *BGIBMGA010505* | up | 676 | 684 | - | 11.3303 | 4.96E-05 | 0.615 | GTGGGTGGC |
| 8938 | mamo-s | *BGIBMGA010505* | up | 1539 | 1547 | + | 11.3303 | 4.96E-05 | 0.615 | GTGGGTGGC |
| 8939 | mamo-s | *BGIBMGA010505* | down | 1006 | 1014 | - | 11.1743 | 5.90E-05 | 0.684 | CCGCGTGTC |
| 8940 | mamo-s | *BGIBMGA010506* | up | 59 | 67 | - | 11.3303 | 4.96E-05 | 0.615 | GTGGGTGGC |
| 8941 | mamo-s | *BGIBMGA010507* | down | 1473 | 1481 | + | 11.3303 | 4.96E-05 | 0.615 | GTGGGTGGC |
| 8942 | mamo-s | *BGIBMGA010507* | down | 1260 | 1268 | - | 11.0826 | 6.84E-05 | 0.709 | GTACGTGGC |
| 8943 | mamo-s | *BGIBMGA010509* | down | 154 | 162 | + | 11.3303 | 4.96E-05 | 0.615 | GTGGGTGGC |
| 8944 | mamo-s | *BGIBMGA010512* | up | 340 | 348 | + | 11.3303 | 4.96E-05 | 0.615 | GTGGGTGGC |
| 8945 | mamo-s | *BGIBMGA010513* | up | 922 | 930 | - | 11.3303 | 4.96E-05 | 0.615 | GTGGGTGGC |
| 8946 | mamo-s | *BGIBMGA010516* | down | 1623 | 1631 | - | 12.1009 | 2.43E-05 | 0.615 | GTGCGTGGT |
| 8947 | mamo-s | *BGIBMGA010517* | up | 553 | 561 | - | 11.3303 | 4.96E-05 | 0.615 | GTGGGTGGC |
| 8948 | mamo-s | *BGIBMGA010518* | down | 1130 | 1138 | - | 11.3303 | 4.96E-05 | 0.615 | GTGGGTGGC |
| 8949 | mamo-s | *BGIBMGA010518* | down | 1853 | 1861 | - | 11.3303 | 4.96E-05 | 0.615 | GTGGGTGGC |
| 8950 | mamo-s | *BGIBMGA010519* | up | 1826 | 1834 | - | 11.3303 | 4.96E-05 | 0.615 | GTGGGTGGC |
| 8951 | mamo-s | *BGIBMGA010519* | up | 72 | 80 | - | 10.5413 | 9.42E-05 | 0.857 | GTGCGAGTC |
| 8952 | mamo-s | *BGIBMGA010520* | up | 149 | 157 | + | 11.7798 | 3.11E-05 | 0.615 | ATGCGTGGC |
| 8953 | mamo-s | *BGIBMGA010522* | down | 729 | 737 | - | 11.3303 | 4.96E-05 | 0.615 | GTGGGTGGC |
| 8954 | mamo-s | *BGIBMGA010529* | up | 771 | 779 | + | 11.3303 | 4.96E-05 | 0.615 | GTGGGTGGC |
| 8955 | mamo-s | *BGIBMGA010530* | down | 968 | 976 | + | 11.4404 | 4.51E-05 | 0.615 | CGGCGTGGC |
| 8956 | mamo-s | *BGIBMGA010530* | down | 1136 | 1144 | + | 11.4404 | 4.51E-05 | 0.615 | CGGCGTGGC |
| 8957 | mamo-s | *BGIBMGA010530* | up | 1140 | 1148 | + | 11.3853 | 4.74E-05 | 0.615 | ACGCGTGGC |
| 8958 | mamo-s | *BGIBMGA010530* | up | 995 | 1003 | + | 11.3303 | 4.96E-05 | 0.615 | GTGGGTGGC |
| 8959 | mamo-s | *BGIBMGA010531* | up | 1514 | 1522 | - | 11.3303 | 4.96E-05 | 0.615 | GTGGGTGGC |
| 8960 | mamo-s | *BGIBMGA010533* | up | 986 | 994 | - | 11.3303 | 4.96E-05 | 0.615 | GTGGGTGGC |
| 8961 | mamo-s | *BGIBMGA010534* | up | 1103 | 1111 | + | 11.3303 | 4.96E-05 | 0.615 | GTGGGTGGC |
| 8962 | mamo-s | *BGIBMGA010535* | up | 412 | 420 | + | 11.3303 | 4.96E-05 | 0.615 | GTGGGTGGC |
| 8963 | mamo-s | *BGIBMGA010535* | up | 704 | 712 | - | 10.5413 | 9.42E-05 | 0.857 | CAGCGTGTC |
| 8964 | mamo-s | *BGIBMGA010536* | up | 1855 | 1863 | - | 11.0826 | 6.84E-05 | 0.709 | GTACGTGGC |
| 8965 | mamo-s | *BGIBMGA010539* | down | 1252 | 1260 | - | 11.2844 | 5.18E-05 | 0.63 | CTGCGTGCC |
| 8966 | mamo-s | *BGIBMGA010539* | up | 651 | 659 | - | 11.2294 | 5.41E-05 | 0.65 | CTGCGAGGC |
| 8967 | mamo-s | *BGIBMGA010540* | up | 589 | 597 | + | 11.3303 | 4.96E-05 | 0.615 | GTGGGTGGC |
| 8968 | mamo-s | *BGIBMGA010545* | up | 746 | 754 | - | 10.5138 | 9.75E-05 | 0.865 | GTGCGTGTT |
| 8969 | mamo-s | *BGIBMGA010548* | down | 1988 | 1996 | + | 11.3303 | 4.96E-05 | 0.615 | GTGGGTGGC |

| | | | | | | | | | | |
|---|---|---|---|---|---|---|---|---|---|---|
| 8970 | mamo-s | *BGIBMGA010549* | down | 628 | 636 | + | 11.3303 | 4.96E-05 | 0.615 | GTGGGTGGC |
| 8971 | mamo-s | *BGIBMGA010550* | down | 243 | 251 | + | 11.3303 | 4.96E-05 | 0.615 | GTGGGTGGC |
| 8972 | mamo-s | *BGIBMGA010550* | down | 1411 | 1419 | + | 11.1743 | 5.90E-05 | 0.684 | CCGCGTGTC |
| 8973 | mamo-s | *BGIBMGA010552* | down | 1012 | 1020 | - | 11.3303 | 4.96E-05 | 0.615 | GTGGGTGGC |
| 8974 | mamo-s | *BGIBMGA010555* | up | 1806 | 1814 | + | 11.3303 | 4.96E-05 | 0.615 | GTGGGTGGC |
| 8975 | mamo-s | *BGIBMGA010556* | down | 1059 | 1067 | - | 11.3303 | 4.96E-05 | 0.615 | GTGGGTGGC |
| 8976 | mamo-s | *BGIBMGA010556* | down | 1302 | 1310 | + | 11.1376 | 6.35E-05 | 0.709 | TCGCGTGGC |
| 8977 | mamo-s | *BGIBMGA010556* | up | 1948 | 1956 | + | 11.0826 | 6.84E-05 | 0.709 | GTACGTGGC |
| 8978 | mamo-s | *BGIBMGA010558* | up | 1530 | 1538 | - | 11.3303 | 4.96E-05 | 0.615 | GTGGGTGGC |
| 8979 | mamo-s | *BGIBMGA010559* | down | 1017 | 1025 | + | 11.7339 | 3.29E-05 | 0.615 | GCGCGAGGC |
| 8980 | mamo-s | *BGIBMGA010563* | down | 305 | 313 | + | 11.0826 | 6.84E-05 | 0.709 | GTACGTGGC |
| 8981 | mamo-s | *BGIBMGA010565* | down | 473 | 481 | - | 11.3303 | 4.96E-05 | 0.615 | GTGGGTGGC |
| 8982 | mamo-s | *BGIBMGA010565* | down | 1329 | 1337 | - | 11.3303 | 4.96E-05 | 0.615 | GTGGGTGGC |
| 8983 | mamo-s | *BGIBMGA010569* | down | 420 | 428 | + | 11.0826 | 6.84E-05 | 0.709 | GTACGTGGC |
| 8984 | mamo-s | *BGIBMGA010574* | down | 1031 | 1039 | + | 11.4404 | 4.51E-05 | 0.615 | CGGCGTGGC |
| 8985 | mamo-s | *BGIBMGA010574* | down | 952 | 960 | - | 10.5413 | 9.42E-05 | 0.857 | CAGCGTGTC |
| 8986 | mamo-s | *BGIBMGA010583* | down | 895 | 903 | - | 13.156 | 6.30E-06 | 0.615 | CTGCGTGGC |
| 8987 | mamo-s | *BGIBMGA010583* | up | 1965 | 1973 | + | 12.1009 | 2.43E-05 | 0.615 | GTGCGTGGT |
| 8988 | mamo-s | *BGIBMGA010583* | up | 1802 | 1810 | + | 11.3303 | 4.96E-05 | 0.615 | GTGGGTGGC |
| 8989 | mamo-s | *BGIBMGA010585* | up | 327 | 335 | + | 11.3303 | 4.96E-05 | 0.615 | GTGGGTGGC |
| 8990 | mamo-s | *BGIBMGA010585* | down | 1625 | 1633 | + | 11.3303 | 4.96E-05 | 0.615 | GTGGGTGGC |
| 8991 | mamo-s | *BGIBMGA010592* | up | 1335 | 1343 | + | 11.7064 | 3.52E-05 | 0.615 | GCGCGTGGT |
| 8992 | mamo-s | *BGIBMGA010596* | up | 126 | 134 | + | 11.3303 | 4.96E-05 | 0.615 | GTGGGTGGC |
| 8993 | mamo-s | *BGIBMGA010597* | up | 200 | 208 | + | 11.3303 | 4.96E-05 | 0.615 | GTGGGTGGC |
| 8994 | mamo-s | *BGIBMGA010601* | down | 345 | 353 | + | 10.8899 | 7.48E-05 | 0.739 | CCGCGTGCC |
| 8995 | mamo-s | *BGIBMGA010602* | down | 1427 | 1435 | + | 11.0826 | 6.84E-05 | 0.709 | GTACGTGGC |
| 8996 | mamo-s | *BGIBMGA010603* | up | 212 | 220 | + | 11.0734 | 7.12E-05 | 0.726 | GAGCGTGGT |
| 8997 | mamo-s | *BGIBMGA010604* | up | 837 | 845 | - | 11.0826 | 6.84E-05 | 0.709 | GTACGTGGC |
| 8998 | mamo-s | *BGIBMGA010605* | up | 528 | 536 | - | 11.3303 | 4.96E-05 | 0.615 | GTGGGTGGC |
| 8999 | mamo-s | *BGIBMGA010606* | down | 1430 | 1438 | - | 11.5688 | 3.79E-05 | 0.615 | CTGCGTGTC |
| 9000 | mamo-s | *BGIBMGA010606* | up | 472 | 480 | - | 11.3303 | 4.96E-05 | 0.615 | GTGGGTGGC |
| 9001 | mamo-s | *BGIBMGA010606* | up | 1110 | 1118 | + | 11.3303 | 4.96E-05 | 0.615 | GTGGGTGGC |
| 9002 | mamo-s | *BGIBMGA010607* | down | 1559 | 1567 | + | 11.0734 | 7.12E-05 | 0.726 | GAGCGTGGT |
| 9003 | mamo-s | *BGIBMGA010609* | down | 1073 | 1081 | + | 11.3303 | 4.96E-05 | 0.615 | GTGGGTGGC |
| 9004 | mamo-s | *BGIBMGA010610* | down | 990 | 998 | + | 12.4679 | 1.31E-05 | 0.615 | GTGCGTGTC |
| 9005 | mamo-s | *BGIBMGA010612* | up | 463 | 471 | - | 11.3303 | 4.96E-05 | 0.615 | GTGGGTGGC |
| 9006 | mamo-s | *BGIBMGA010614* | down | 932 | 940 | + | 11.3303 | 4.96E-05 | 0.615 | GTGGGTGGC |
| 9007 | mamo-s | *BGIBMGA010618* | down | 740 | 748 | - | 11.3303 | 4.96E-05 | 0.615 | GTGGGTGGC |
| 9008 | mamo-s | *BGIBMGA010620* | up | 1199 | 1207 | - | 11.7064 | 3.52E-05 | 0.615 | GCGCGTGGT |
| 9009 | mamo-s | *BGIBMGA010621* | down | 1651 | 1659 | + | 11.3303 | 4.96E-05 | 0.615 | GTGGGTGGC |
| 9010 | mamo-s | *BGIBMGA010624* | down | 1150 | 1158 | + | 11.5321 | 4.06E-05 | 0.615 | TTGCGTGGC |
| 9011 | mamo-s | *BGIBMGA010626* | up | 1411 | 1419 | + | 11.3303 | 4.96E-05 | 0.615 | GTGGGTGGC |
| 9012 | mamo-s | *BGIBMGA010626* | down | 945 | 953 | - | 11.1009 | 6.57E-05 | 0.709 | GAGCGAGGC |
| 9013 | mamo-s | *BGIBMGA010629* | up | 1460 | 1468 | + | 11.1743 | 5.90E-05 | 0.684 | CCGCGTGTC |
| 9014 | mamo-s | *BGIBMGA010631* | up | 758 | 766 | + | 12.1835 | 1.71E-05 | 0.615 | GTGCGTGCC |
| 9015 | mamo-s | *BGIBMGA010631* | up | 24 | 32 | + | 11.789 | 2.84E-05 | 0.615 | GCGCGTGCC |
| 9016 | mamo-s | *BGIBMGA010632* | down | 926 | 934 | - | 11.3303 | 4.96E-05 | 0.615 | GTGGGTGGC |
| 9017 | mamo-s | *BGIBMGA010632* | up | 999 | 1007 | + | 11.3303 | 4.96E-05 | 0.615 | GTGGGTGGC |
| 9018 | mamo-s | *BGIBMGA010632* | down | 48 | 56 | - | 10.6881 | 8.61E-05 | 0.81 | GCACGTGGC |
| 9019 | mamo-s | *BGIBMGA010633* | up | 445 | 453 | - | 11.3303 | 4.96E-05 | 0.615 | GTGGGTGGC |
| 9020 | mamo-s | *BGIBMGA010633* | up | 578 | 586 | - | 11.3303 | 4.96E-05 | 0.615 | GTGGGTGGC |
| 9021 | mamo-s | *BGIBMGA010636* | down | 1128 | 1136 | - | 14.055 | 2.23E-06 | 0.615 | GTGCGTGGC |
| 9022 | mamo-s | *BGIBMGA010636* | up | 1046 | 1054 | - | 13.0275 | 8.53E-06 | 0.615 | GAGCGTGGC |
| 9023 | mamo-s | *BGIBMGA010636* | down | 1054 | 1062 | + | 11.3303 | 4.96E-05 | 0.615 | GTGGGTGGC |
| 9024 | mamo-s | *BGIBMGA010637* | down | 673 | 681 | + | 11.3303 | 4.96E-05 | 0.615 | GTGGGTGGC |
| 9025 | mamo-s | *BGIBMGA010638* | down | 475 | 483 | + | 11.1009 | 6.57E-05 | 0.709 | GAGCGAGGC |
| 9026 | mamo-s | *BGIBMGA010639* | down | 1641 | 1649 | + | 12.0734 | 2.66E-05 | 0.615 | GCGCGTGTC |
| 9027 | mamo-s | *BGIBMGA010639* | down | 318 | 326 | - | 11.3303 | 4.96E-05 | 0.615 | GTGGGTGGC |
| 9028 | mamo-s | *BGIBMGA010641* | up | 61 | 69 | + | 11.1743 | 5.90E-05 | 0.684 | CCGCGTGTC |
| 9029 | mamo-s | *BGIBMGA010642* | up | 1424 | 1432 | - | 11.3303 | 4.96E-05 | 0.615 | GTGGGTGGC |
| 9030 | mamo-s | *BGIBMGA010643* | down | 1730 | 1738 | - | 11.0826 | 6.84E-05 | 0.709 | GTACGTGGC |
| 9031 | mamo-s | *BGIBMGA010645* | up | 1116 | 1124 | - | 11.3303 | 4.96E-05 | 0.615 | GTGGGTGGC |
| 9032 | mamo-s | *BGIBMGA010645* | down | 1251 | 1259 | - | 11.0826 | 6.84E-05 | 0.709 | GTACGTGGC |
| 9033 | mamo-s | *BGIBMGA010646* | down | 1270 | 1278 | - | 11.3303 | 4.96E-05 | 0.615 | GTGGGTGGC |
| 9034 | mamo-s | *BGIBMGA010646* | up | 1656 | 1664 | - | 11.0826 | 6.84E-05 | 0.709 | GTACGTGGC |
| 9035 | mamo-s | *BGIBMGA010649* | up | 151 | 159 | + | 14.055 | 2.23E-06 | 0.615 | GTGCGTGGC |
| 9036 | mamo-s | *BGIBMGA010651* | up | 867 | 875 | + | 13.0275 | 8.53E-06 | 0.615 | GAGCGTGGC |
| 9037 | mamo-s | *BGIBMGA010651* | down | 1399 | 1407 | + | 11.3303 | 4.96E-05 | 0.615 | GTGGGTGGC |
| 9038 | mamo-s | *BGIBMGA010654* | up | 623 | 631 | - | 13.0275 | 8.53E-06 | 0.615 | GAGCGTGGC |
| 9039 | mamo-s | *BGIBMGA010655* | down | 1022 | 1030 | + | 11.3303 | 4.96E-05 | 0.615 | GTGGGTGGC |
| 9040 | mamo-s | *BGIBMGA010662* | down | 577 | 585 | + | 12.1284 | 2.16E-05 | 0.615 | CAGCGTGGC |
| 9041 | mamo-s | *BGIBMGA010662* | up | 557 | 565 | + | 11.3303 | 4.96E-05 | 0.615 | GTGGGTGGC |
| 9042 | mamo-s | *BGIBMGA010667* | up | 944 | 952 | - | 11.3303 | 4.96E-05 | 0.615 | GTGGGTGGC |
| 9043 | mamo-s | *BGIBMGA010668* | down | 1931 | 1939 | + | 11.0826 | 6.84E-05 | 0.709 | GTACGTGGC |
| 9044 | mamo-s | *BGIBMGA010669* | up | 1635 | 1643 | + | 11.7064 | 3.52E-05 | 0.615 | GCGCGTGGT |
| 9045 | mamo-s | *BGIBMGA010671* | down | 325 | 333 | - | 11.3303 | 4.96E-05 | 0.615 | GTGGGTGGC |
| 9046 | mamo-s | *BGIBMGA010672* | down | 571 | 579 | - | 11.3853 | 4.74E-05 | 0.615 | ACGCGTGGC |
| 9047 | mamo-s | *BGIBMGA010672* | down | 554 | 562 | + | 11.3303 | 4.96E-05 | 0.615 | GTGGGTGGC |
| 9048 | mamo-s | *BGIBMGA010675* | up | 1637 | 1645 | + | 10.5413 | 9.42E-05 | 0.857 | GTGCGAGTC |
| 9049 | mamo-s | *BGIBMGA010676* | up | 1466 | 1474 | - | 11.3303 | 4.96E-05 | 0.615 | GTGGGTGGC |
| 9050 | mamo-s | *BGIBMGA010681* | down | 1400 | 1408 | + | 14.055 | 2.23E-06 | 0.615 | GTGCGTGGC |
| 9051 | mamo-s | *BGIBMGA010681* | up | 122 | 130 | + | 11.3303 | 4.96E-05 | 0.615 | GTGGGTGGC |
| 9052 | mamo-s | *BGIBMGA010681* | up | 1365 | 1373 | - | 11.3303 | 4.96E-05 | 0.615 | GTGGGTGGC |
| 9053 | mamo-s | *BGIBMGA010685* | down | 1693 | 1701 | - | 11.3303 | 4.96E-05 | 0.615 | GTGGGTGGC |
| 9054 | mamo-s | *BGIBMGA010687* | down | 40 | 48 | + | 12.0734 | 2.66E-05 | 0.615 | GCGCGTGTC |
| 9055 | mamo-s | *BGIBMGA010688* | down | 225 | 233 | + | 10.5688 | 8.88E-05 | 0.824 | GTGCGTGAC |
| 9056 | mamo-s | *BGIBMGA010689* | down | 568 | 576 | + | 11.1009 | 6.57E-05 | 0.709 | GAGCGAGGC |
| 9057 | mamo-s | *BGIBMGA010689* | up | 1381 | 1389 | - | 11.0734 | 7.12E-05 | 0.726 | GAGCGTGGT |

| | | | | | | | | | | |
|---|---|---|---|---|---|---|---|---|---|---|
| 9058 | mamo-s | *BGIBMGA010690* | down | 124 | 132 | - | 11.3303 | 4.96E-05 | 0.615 | GTGGGTGGC |
| 9059 | mamo-s | *BGIBMGA010690* | down | 450 | 458 | - | 11.3303 | 4.96E-05 | 0.615 | GTGGGTGGC |
| 9060 | mamo-s | *BGIBMGA010690* | up | 781 | 789 | - | 11.0734 | 7.12E-05 | 0.726 | GAGCGTGGT |
| 9061 | mamo-s | *BGIBMGA010690* | up | 608 | 616 | - | 10.8073 | 7.89E-05 | 0.766 | CCGCGTGGT |
| 9062 | mamo-s | *BGIBMGA010691* | up | 546 | 554 | + | 11.7339 | 3.29E-05 | 0.615 | GCGCGAGGC |
| 9063 | mamo-s | *BGIBMGA010691* | down | 798 | 806 | + | 11.3303 | 4.96E-05 | 0.615 | GTGGGTGGC |
| 9064 | mamo-s | *BGIBMGA010691* | up | 569 | 577 | - | 11.2018 | 5.68E-05 | 0.674 | CTGCGTGGT |
| 9065 | mamo-s | *BGIBMGA010692* | up | 1265 | 1273 | - | 11.3303 | 4.96E-05 | 0.615 | GTGGGTGGC |
| 9066 | mamo-s | *BGIBMGA010693* | up | 1856 | 1864 | - | 12.3394 | 1.49E-05 | 0.615 | GGGCGTGGC |
| 9067 | mamo-s | *BGIBMGA010694* | up | 531 | 539 | + | 11.3303 | 4.96E-05 | 0.615 | GTGGGTGGC |
| 9068 | mamo-s | *BGIBMGA010698* | up | 403 | 411 | - | 11.5321 | 4.06E-05 | 0.615 | TTGCGTGGC |
| 9069 | mamo-s | *BGIBMGA010698* | down | 452 | 460 | - | 11.3303 | 4.96E-05 | 0.615 | GTGGGTGGC |
| 9070 | mamo-s | *BGIBMGA010700* | down | 897 | 905 | - | 12.0734 | 2.66E-05 | 0.615 | GCGCGTGTC |
| 9071 | mamo-s | *BGIBMGA010700* | down | 1140 | 1148 | + | 11.3303 | 4.96E-05 | 0.615 | GTGGGTGGC |
| 9072 | mamo-s | *BGIBMGA010702* | down | 1607 | 1615 | + | 11.0826 | 6.84E-05 | 0.709 | GTACGTGGC |
| 9073 | mamo-s | *BGIBMGA010703* | up | 488 | 496 | - | 11.3303 | 4.96E-05 | 0.615 | GTGGGTGGC |
| 9074 | mamo-s | *BGIBMGA010703* | up | 1340 | 1348 | + | 11.3303 | 4.96E-05 | 0.615 | GTGGGTGGC |
| 9075 | mamo-s | *BGIBMGA010704* | down | 787 | 795 | + | 11.5688 | 3.79E-05 | 0.615 | CTGCGTGTC |
| 9076 | mamo-s | *BGIBMGA010704* | up | 1094 | 1102 | - | 11.0734 | 7.12E-05 | 0.726 | GAGCGTGGT |
| 9077 | mamo-s | *BGIBMGA010705* | down | 120 | 128 | + | 11.3303 | 4.96E-05 | 0.615 | GTGGGTGGC |
| 9078 | mamo-s | *BGIBMGA010705* | down | 1524 | 1532 | + | 11.3303 | 4.96E-05 | 0.615 | GTGGGTGGC |
| 9079 | mamo-s | *BGIBMGA010706* | down | 683 | 691 | + | 11.3303 | 4.96E-05 | 0.615 | GTGGGTGGC |
| 9080 | mamo-s | *BGIBMGA010706* | down | 1316 | 1324 | - | 11.0826 | 6.84E-05 | 0.709 | GTACGTGGC |
| 9081 | mamo-s | *BGIBMGA010707* | up | 87 | 95 | - | 10.9358 | 7.30E-05 | 0.728 | GCGGGTGGC |
| 9082 | mamo-s | *BGIBMGA010708* | up | 1600 | 1608 | + | 13.0275 | 8.53E-06 | 0.615 | GAGCGTGGC |
| 9083 | mamo-s | *BGIBMGA010710* | down | 1553 | 1561 | - | 11.3303 | 4.96E-05 | 0.615 | GTGGGTGGC |
| 9084 | mamo-s | *BGIBMGA010711* | down | 315 | 323 | + | 11.3303 | 4.96E-05 | 0.615 | GTGGGTGGC |
| 9085 | mamo-s | *BGIBMGA010712* | up | 1424 | 1432 | + | 11.3303 | 4.96E-05 | 0.615 | GTGGGTGGC |
| 9086 | mamo-s | *BGIBMGA010713* | down | 386 | 394 | - | 10.7523 | 8.38E-05 | 0.797 | AAGCGTGGC |
| 9087 | mamo-s | *BGIBMGA010713* | up | 853 | 861 | + | 10.5138 | 9.75E-05 | 0.865 | GTGCGTGTT |
| 9088 | mamo-s | *BGIBMGA010714* | down | 542 | 550 | + | 11.3303 | 4.96E-05 | 0.615 | GTGGGTGGC |
| 9089 | mamo-s | *BGIBMGA010715* | up | 874 | 882 | + | 11.3303 | 4.96E-05 | 0.615 | GTGGGTGGC |
| 9090 | mamo-s | *BGIBMGA010717* | down | 377 | 385 | - | 11.3303 | 4.96E-05 | 0.615 | GTGGGTGGC |
| 9091 | mamo-s | *BGIBMGA010717* | down | 1988 | 1996 | + | 11.1743 | 5.90E-05 | 0.684 | CCGCGTGTC |
| 9092 | mamo-s | *BGIBMGA010717* | up | 1869 | 1877 | + | 10.8899 | 7.48E-05 | 0.739 | CCGCGTGCC |
| 9093 | mamo-s | *BGIBMGA010721* | up | 1223 | 1231 | - | 13.0275 | 8.53E-06 | 0.615 | GAGCGTGGC |
| 9094 | mamo-s | *BGIBMGA010722* | down | 1179 | 1187 | + | 10.5413 | 9.42E-05 | 0.857 | CAGCGTGTC |
| 9095 | mamo-s | *BGIBMGA010723* | up | 1988 | 1996 | - | 13.0275 | 8.53E-06 | 0.615 | GAGCGTGGC |
| 9096 | mamo-s | *BGIBMGA010723* | down | 1263 | 1271 | + | 11.3303 | 4.96E-05 | 0.615 | GTGGGTGGC |
| 9097 | mamo-s | *BGIBMGA010725* | down | 1902 | 1910 | - | 11.156 | 6.13E-05 | 0.705 | GAGCGTGCC |
| 9098 | mamo-s | *BGIBMGA010726* | up | 1839 | 1847 | + | 11.3303 | 4.96E-05 | 0.615 | GTGGGTGGC |
| 9099 | mamo-s | *BGIBMGA010727* | up | 876 | 884 | + | 11.3303 | 4.96E-05 | 0.615 | GTGGGTGGC |
| 9100 | mamo-s | *BGIBMGA010727* | up | 1379 | 1387 | + | 11.3303 | 4.96E-05 | 0.615 | GTGGGTGGC |
| 9101 | mamo-s | *BGIBMGA010729* | down | 32 | 40 | + | 13.6606 | 4.06E-06 | 0.615 | GCGCGTGGC |
| 9102 | mamo-s | *BGIBMGA010729* | down | 1683 | 1691 | + | 11.3303 | 4.96E-05 | 0.615 | GTGGGTGGC |
| 9103 | mamo-s | *BGIBMGA010729* | down | 1066 | 1074 | + | 11.2844 | 5.18E-05 | 0.63 | CTGCGTGCC |
| 9104 | mamo-s | *BGIBMGA010731* | up | 699 | 707 | - | 11.3303 | 4.96E-05 | 0.615 | GTGGGTGGC |
| 9105 | mamo-s | *BGIBMGA010732* | up | 41 | 49 | + | 11.3303 | 4.96E-05 | 0.615 | GTGGGTGGC |
| 9106 | mamo-s | *BGIBMGA010732* | down | 275 | 283 | - | 11.3303 | 4.96E-05 | 0.615 | GTGGGTGGC |
| 9107 | mamo-s | *BGIBMGA010733* | down | 1467 | 1475 | + | 11.3303 | 4.96E-05 | 0.615 | GTGGGTGGC |
| 9108 | mamo-s | *BGIBMGA010734* | down | 1443 | 1451 | + | 11.3303 | 4.96E-05 | 0.615 | GTGGGTGGC |
| 9109 | mamo-s | *BGIBMGA010735* | up | 1723 | 1731 | + | 11.2844 | 5.18E-05 | 0.63 | CTGCGTGCC |
| 9110 | mamo-s | *BGIBMGA010736* | up | 937 | 945 | - | 11.3303 | 4.96E-05 | 0.615 | GTGGGTGGC |
| 9111 | mamo-s | *BGIBMGA010736* | down | 1642 | 1650 | - | 11.3303 | 4.96E-05 | 0.615 | GTGGGTGGC |
| 9112 | mamo-s | *BGIBMGA010738* | up | 4 | 12 | - | 11.3303 | 4.96E-05 | 0.615 | GTGGGTGGC |
| 9113 | mamo-s | *BGIBMGA010739* | down | 1699 | 1707 | - | 12.1284 | 2.16E-05 | 0.615 | CAGCGTGGC |
| 9114 | mamo-s | *BGIBMGA010739* | down | 1992 | 2000 | + | 11.3303 | 4.96E-05 | 0.615 | GTGGGTGGC |
| 9115 | mamo-s | *BGIBMGA010740* | down | 789 | 797 | + | 14.055 | 2.23E-06 | 0.615 | GTGCGTGGC |
| 9116 | mamo-s | *BGIBMGA010740* | up | 817 | 825 | + | 10.8349 | 7.66E-05 | 0.751 | CCGCGAGGC |
| 9117 | mamo-s | *BGIBMGA010742* | up | 1439 | 1447 | + | 11.3303 | 4.96E-05 | 0.615 | GTGGGTGGC |
| 9118 | mamo-s | *BGIBMGA010742* | down | 1727 | 1735 | + | 11.0826 | 6.84E-05 | 0.709 | GTACGTGGC |
| 9119 | mamo-s | *BGIBMGA010742* | down | 1748 | 1756 | + | 11.0826 | 6.84E-05 | 0.709 | GTACGTGGC |
| 9120 | mamo-s | *BGIBMGA010742* | down | 1770 | 1778 | + | 11.0826 | 6.84E-05 | 0.709 | GTACGTGGC |
| 9121 | mamo-s | *BGIBMGA010742* | down | 1792 | 1800 | + | 11.0826 | 6.84E-05 | 0.709 | GTACGTGGC |
| 9122 | mamo-s | *BGIBMGA010743* | up | 507 | 515 | + | 11.3303 | 4.96E-05 | 0.615 | GTGGGTGGC |
| 9123 | mamo-s | *BGIBMGA010744* | up | 525 | 533 | - | 13.156 | 6.30E-06 | 0.615 | CTGCGTGGC |
| 9124 | mamo-s | *BGIBMGA010747* | down | 1591 | 1599 | - | 10.5688 | 8.88E-05 | 0.824 | GTGCGTGAC |
| 9125 | mamo-s | *BGIBMGA010749* | down | 118 | 126 | + | 11.3303 | 4.96E-05 | 0.615 | GTGGGTGGC |
| 9126 | mamo-s | *BGIBMGA010749* | down | 628 | 636 | + | 10.5413 | 9.42E-05 | 0.857 | GTGCGAGTC |
| 9127 | mamo-s | *BGIBMGA010751* | up | 475 | 483 | - | 11.3303 | 4.96E-05 | 0.615 | GTGGGTGGC |
| 9128 | mamo-s | *BGIBMGA010751* | down | 1783 | 1791 | + | 11.3303 | 4.96E-05 | 0.615 | GTGGGTGGC |
| 9129 | mamo-s | *BGIBMGA010752* | up | 61 | 69 | + | 11.3303 | 4.96E-05 | 0.615 | GTGGGTGGC |
| 9130 | mamo-s | *BGIBMGA010753* | down | 1123 | 1131 | - | 12.0734 | 2.66E-05 | 0.615 | GCGCGTGTC |
| 9131 | mamo-s | *BGIBMGA010757* | up | 1904 | 1912 | - | 12.0734 | 2.66E-05 | 0.615 | GCGCGTGTC |
| 9132 | mamo-s | *BGIBMGA010757* | up | 813 | 821 | + | 11.3303 | 4.96E-05 | 0.615 | GTGGGTGGC |
| 9133 | mamo-s | *BGIBMGA010758* | down | 287 | 295 | - | 11.3303 | 4.96E-05 | 0.615 | GTGGGTGGC |
| 9134 | mamo-s | *BGIBMGA010759* | up | 837 | 845 | + | 11.3303 | 4.96E-05 | 0.615 | GTGGGTGGC |
| 9135 | mamo-s | *BGIBMGA010761* | up | 1568 | 1576 | - | 11.3303 | 4.96E-05 | 0.615 | GTGGGTGGC |
| 9136 | mamo-s | *BGIBMGA010762* | up | 886 | 894 | - | 11.3303 | 4.96E-05 | 0.615 | GTGGGTGGC |
| 9137 | mamo-s | *BGIBMGA010762* | down | 383 | 391 | - | 10.7523 | 8.38E-05 | 0.797 | GGGCGTGTC |
| 9138 | mamo-s | *BGIBMGA010763* | down | 884 | 892 | + | 13.6606 | 4.06E-06 | 0.615 | GCGCGTGGC |
| 9139 | mamo-s | *BGIBMGA010763* | down | 657 | 665 | - | 11.3303 | 4.96E-05 | 0.615 | GTGGGTGGC |
| 9140 | mamo-s | *BGIBMGA010763* | up | 1857 | 1865 | + | 11.3303 | 4.96E-05 | 0.615 | GTGGGTGGC |
| 9141 | mamo-s | *BGIBMGA010764* | up | 1833 | 1841 | - | 11.3303 | 4.96E-05 | 0.615 | GTGGGTGGC |
| 9142 | mamo-s | *BGIBMGA010765* | up | 556 | 564 | - | 10.9358 | 7.30E-05 | 0.728 | GCGGGTGGC |
| 9143 | mamo-s | *BGIBMGA010768* | down | 1687 | 1695 | - | 11.3303 | 4.96E-05 | 0.615 | GTGGGTGGC |
| 9144 | mamo-s | *BGIBMGA010769* | down | 1686 | 1694 | + | 11.3303 | 4.96E-05 | 0.615 | GTGGGTGGC |
| 9145 | mamo-s | *BGIBMGA010771* | up | 1846 | 1854 | + | 11.3303 | 4.96E-05 | 0.615 | GTGGGTGGC |

| | | | | | | | | | | |
|---|---|---|---|---|---|---|---|---|---|---|
| 9146 | mamo-s | *BGIBMGA010773* | up | 82 | 90 | - | 11.3303 | 4.96E-05 | 0.615 | GTGGGTGGC |
| 9147 | mamo-s | *BGIBMGA010776* | down | 1146 | 1154 | - | 10.5413 | 9.42E-05 | 0.857 | CAGCGTGTC |
| 9148 | mamo-s | *BGIBMGA010777* | down | 729 | 737 | + | 12.4679 | 1.31E-05 | 0.615 | GTGCGTGTC |
| 9149 | mamo-s | *BGIBMGA010777* | down | 862 | 870 | - | 11.789 | 2.84E-05 | 0.615 | GCGCGTGCC |
| 9150 | mamo-s | *BGIBMGA010777* | down | 894 | 902 | + | 10.6881 | 8.61E-05 | 0.81 | GCACGTGGC |
| 9151 | mamo-s | *BGIBMGA010777* | down | 712 | 720 | - | 10.5413 | 9.42E-05 | 0.857 | CAGCGTGTC |
| 9152 | mamo-s | *BGIBMGA010778* | up | 880 | 888 | - | 12.1284 | 2.16E-05 | 0.615 | CAGCGTGGC |
| 9153 | mamo-s | *BGIBMGA010778* | up | 828 | 836 | + | 12.0734 | 2.66E-05 | 0.615 | GCGCGTGTC |
| 9154 | mamo-s | *BGIBMGA010778* | down | 1755 | 1763 | + | 11.789 | 2.84E-05 | 0.615 | GCGCGTGCC |
| 9155 | mamo-s | *BGIBMGA010778* | up | 922 | 930 | + | 11.4404 | 4.51E-05 | 0.615 | CGGCGTGGC |
| 9156 | mamo-s | *BGIBMGA010779* | up | 1236 | 1244 | - | 12.4679 | 1.31E-05 | 0.615 | GTGCGTGTC |
| 9157 | mamo-s | *BGIBMGA010779* | up | 1245 | 1253 | - | 10.5688 | 8.88E-05 | 0.824 | GTGCGTGAC |
| 9158 | mamo-s | *BGIBMGA010780* | down | 323 | 331 | + | 11.3853 | 4.74E-05 | 0.615 | ACGCGTGGC |
| 9159 | mamo-s | *BGIBMGA010780* | up | 1708 | 1716 | + | 11.3303 | 4.96E-05 | 0.615 | GTGGGTGGC |
| 9160 | mamo-s | *BGIBMGA010783* | up | 27 | 35 | - | 11.3303 | 4.96E-05 | 0.615 | GTGGGTGGC |
| 9161 | mamo-s | *BGIBMGA010784* | down | 1770 | 1778 | - | 11.2294 | 5.41E-05 | 0.65 | CTGCGAGGC |
| 9162 | mamo-s | *BGIBMGA010787* | up | 151 | 159 | - | 13.6606 | 4.06E-06 | 0.615 | GCGCGTGGC |
| 9163 | mamo-s | *BGIBMGA010788* | up | 1261 | 1269 | - | 11.3303 | 4.96E-05 | 0.615 | GTGGGTGGC |
| 9164 | mamo-s | *BGIBMGA010788* | down | 1754 | 1762 | + | 11.3303 | 4.96E-05 | 0.615 | GTGGGTGGC |
| 9165 | mamo-s | *BGIBMGA010788* | down | 1607 | 1615 | + | 11.0826 | 6.84E-05 | 0.709 | GTACGTGGC |
| 9166 | mamo-s | *BGIBMGA010790* | up | 1511 | 1519 | - | 11.3303 | 4.96E-05 | 0.615 | GTGGGTGGC |
| 9167 | mamo-s | *BGIBMGA010791* | down | 768 | 776 | - | 10.6881 | 8.61E-05 | 0.81 | GCACGTGGC |
| 9168 | mamo-s | *BGIBMGA010792* | down | 296 | 304 | + | 11.3303 | 4.96E-05 | 0.615 | GTGGGTGGC |
| 9169 | mamo-s | *BGIBMGA010793* | up | 207 | 215 | - | 11.0826 | 6.84E-05 | 0.709 | GTACGTGGC |
| 9170 | mamo-s | *BGIBMGA010796* | down | 212 | 220 | - | 11.3303 | 4.96E-05 | 0.615 | GTGGGTGGC |
| 9171 | mamo-s | *BGIBMGA010796* | up | 316 | 324 | - | 11.3303 | 4.96E-05 | 0.615 | GTGGGTGGC |
| 9172 | mamo-s | *BGIBMGA010797* | down | 15 | 23 | - | 11.1376 | 6.35E-05 | 0.709 | TCGCGTGGC |
| 9173 | mamo-s | *BGIBMGA010798* | down | 1962 | 1970 | - | 11.2018 | 5.68E-05 | 0.674 | CTGCGTGGT |
| 9174 | mamo-s | *BGIBMGA010799* | up | 1330 | 1338 | - | 11.4404 | 4.51E-05 | 0.615 | GAGCGTGTC |
| 9175 | mamo-s | *BGIBMGA010802* | down | 686 | 694 | + | 11.3303 | 4.96E-05 | 0.615 | GTGGGTGGC |
| 9176 | mamo-s | *BGIBMGA010802* | up | 316 | 324 | - | 10.7523 | 8.38E-05 | 0.797 | AAGCGTGGC |
| 9177 | mamo-s | *BGIBMGA010803* | up | 798 | 806 | + | 11.3303 | 4.96E-05 | 0.615 | GTGGGTGGC |
| 9178 | mamo-s | *BGIBMGA010804* | up | 930 | 938 | - | 12.7615 | 1.04E-05 | 0.615 | CCGCGTGGC |
| 9179 | mamo-s | *BGIBMGA010805* | up | 44 | 52 | - | 12.1284 | 2.16E-05 | 0.615 | GTGCGAGGC |
| 9180 | mamo-s | *BGIBMGA010807* | up | 1086 | 1094 | + | 11.3303 | 4.96E-05 | 0.615 | GTGGGTGGC |
| 9181 | mamo-s | *BGIBMGA010808* | up | 1239 | 1247 | + | 11.3303 | 4.96E-05 | 0.615 | GTGGGTGGC |
| 9182 | mamo-s | *BGIBMGA010809* | down | 1978 | 1986 | + | 12.1284 | 2.16E-05 | 0.615 | CAGCGTGGC |
| 9183 | mamo-s | *BGIBMGA010810* | down | 1491 | 1499 | - | 11.789 | 2.84E-05 | 0.615 | GCGCGTGCC |
| 9184 | mamo-s | *BGIBMGA010810* | up | 954 | 962 | + | 11.3303 | 4.96E-05 | 0.615 | GTGGGTGGC |
| 9185 | mamo-s | *BGIBMGA010811* | down | 1580 | 1588 | - | 11.3303 | 4.96E-05 | 0.615 | GTGGGTGGC |
| 9186 | mamo-s | *BGIBMGA010816* | down | 1446 | 1454 | - | 11.7798 | 3.11E-05 | 0.615 | ATGCGTGGC |
| 9187 | mamo-s | *BGIBMGA010816* | up | 1392 | 1400 | + | 11.0826 | 6.84E-05 | 0.709 | GTACGTGGC |
| 9188 | mamo-s | *BGIBMGA010817* | up | 1734 | 1742 | + | 11.3303 | 4.96E-05 | 0.615 | GTGGGTGGC |
| 9189 | mamo-s | *BGIBMGA010818* | down | 345 | 353 | + | 11.3303 | 4.96E-05 | 0.615 | GTGGGTGGC |
| 9190 | mamo-s | *BGIBMGA010818* | up | 1149 | 1157 | + | 11.3303 | 4.96E-05 | 0.615 | GTGGGTGGC |
| 9191 | mamo-s | *BGIBMGA010819* | down | 1839 | 1847 | - | 13.6606 | 4.06E-06 | 0.615 | GCGCGTGGC |
| 9192 | mamo-s | *BGIBMGA010820* | up | 1554 | 1562 | + | 11.3303 | 4.96E-05 | 0.615 | GTGGGTGGC |
| 9193 | mamo-s | *BGIBMGA010821* | up | 1617 | 1625 | - | 11.3303 | 4.96E-05 | 0.615 | GTGGGTGGC |
| 9194 | mamo-s | *BGIBMGA010822* | up | 1475 | 1483 | + | 11.0826 | 6.84E-05 | 0.709 | GTACGTGGC |
| 9195 | mamo-s | *BGIBMGA010824* | down | 65 | 73 | + | 11.5321 | 4.06E-05 | 0.615 | TTGCGTGGC |
| 9196 | mamo-s | *BGIBMGA010825* | up | 1643 | 1651 | + | 11.3303 | 4.96E-05 | 0.615 | GTGGGTGGC |
| 9197 | mamo-s | *BGIBMGA010825* | down | 1828 | 1836 | - | 11.3303 | 4.96E-05 | 0.615 | GTGGGTGGC |
| 9198 | mamo-s | *BGIBMGA010827* | down | 1194 | 1202 | - | 11.3303 | 4.96E-05 | 0.615 | GTGGGTGGC |
| 9199 | mamo-s | *BGIBMGA010829* | up | 951 | 959 | - | 11.3303 | 4.96E-05 | 0.615 | GTGGGTGGC |
| 9200 | mamo-s | *BGIBMGA010829* | down | 1130 | 1138 | - | 11.3303 | 4.96E-05 | 0.615 | GTGGGTGGC |
| 9201 | mamo-s | *BGIBMGA010830* | up | 1153 | 1161 | + | 11.0826 | 6.84E-05 | 0.709 | GTACGTGGC |
| 9202 | mamo-s | *BGIBMGA010832* | down | 743 | 751 | + | 11.3303 | 4.96E-05 | 0.615 | GTGGGTGGC |
| 9203 | mamo-s | *BGIBMGA010834* | up | 477 | 485 | - | 11.3303 | 4.96E-05 | 0.615 | GTGGGTGGC |
| 9204 | mamo-s | *BGIBMGA010834* | up | 52 | 60 | + | 10.7523 | 8.38E-05 | 0.797 | AAGCGTGGC |
| 9205 | mamo-s | *BGIBMGA010835* | down | 1691 | 1699 | + | 12.0734 | 2.66E-05 | 0.615 | GCGCGTGTC |
| 9206 | mamo-s | *BGIBMGA010836* | up | 1108 | 1116 | + | 12.0734 | 2.66E-05 | 0.615 | GCGCGTGTC |
| 9207 | mamo-s | *BGIBMGA010836* | up | 1760 | 1768 | - | 11.3303 | 4.96E-05 | 0.615 | GTGGGTGGC |
| 9208 | mamo-s | *BGIBMGA010838* | up | 1438 | 1446 | + | 11.3303 | 4.96E-05 | 0.615 | GTGGGTGGC |
| 9209 | mamo-s | *BGIBMGA010839* | down | 1403 | 1411 | + | 11.3303 | 4.96E-05 | 0.615 | GTGGGTGGC |
| 9210 | mamo-s | *BGIBMGA010840* | down | 1890 | 1898 | - | 11.3303 | 4.96E-05 | 0.615 | GTGGGTGGC |
| 9211 | mamo-s | *BGIBMGA010841* | down | 1726 | 1734 | + | 11.3303 | 4.96E-05 | 0.615 | GTGGGTGGC |
| 9212 | mamo-s | *BGIBMGA010843* | down | 1463 | 1471 | + | 12.1835 | 1.71E-05 | 0.615 | GTGCGTGCC |
| 9213 | mamo-s | *BGIBMGA010844* | up | 1563 | 1571 | + | 12.1835 | 1.71E-05 | 0.615 | GTGCGTGCC |
| 9214 | mamo-s | *BGIBMGA010845* | down | 1397 | 1405 | - | 11.3303 | 4.96E-05 | 0.615 | GTGGGTGGC |
| 9215 | mamo-s | *BGIBMGA010845* | down | 1648 | 1656 | + | 10.5688 | 8.88E-05 | 0.824 | GTGCGTGAC |
| 9216 | mamo-s | *BGIBMGA010848* | down | 1525 | 1533 | + | 11.3303 | 4.96E-05 | 0.615 | GTGGGTGGC |
| 9217 | mamo-s | *BGIBMGA010849* | up | 425 | 433 | + | 11.3303 | 4.96E-05 | 0.615 | GTGGGTGGC |
| 9218 | mamo-s | *BGIBMGA010849* | up | 1729 | 1737 | + | 11.3303 | 4.96E-05 | 0.615 | GTGGGTGGC |
| 9219 | mamo-s | *BGIBMGA010849* | down | 957 | 965 | + | 11.0826 | 6.84E-05 | 0.709 | GTACGTGGC |
| 9220 | mamo-s | *BGIBMGA010850* | up | 1300 | 1308 | + | 13.6606 | 4.06E-06 | 0.615 | GCGCGTGGC |
| 9221 | mamo-s | *BGIBMGA010850* | down | 261 | 269 | - | 11.3303 | 4.96E-05 | 0.615 | GTGGGTGGC |
| 9222 | mamo-s | *BGIBMGA010850* | up | 722 | 730 | - | 11.3303 | 4.96E-05 | 0.615 | GTGGGTGGC |
| 9223 | mamo-s | *BGIBMGA010850* | up | 1059 | 1067 | - | 11.3303 | 4.96E-05 | 0.615 | GTGGGTGGC |
| 9224 | mamo-s | *BGIBMGA010853* | up | 1820 | 1828 | + | 11.3303 | 4.96E-05 | 0.615 | GTGGGTGGC |
| 9225 | mamo-s | *BGIBMGA010853* | down | 1882 | 1890 | - | 11.2844 | 5.18E-05 | 0.63 | CTGCGTGCC |
| 9226 | mamo-s | *BGIBMGA010854* | down | 686 | 694 | + | 11.3303 | 4.96E-05 | 0.615 | GTGGGTGGC |
| 9227 | mamo-s | *BGIBMGA010854* | down | 1827 | 1835 | - | 11.3303 | 4.96E-05 | 0.615 | GTGGGTGGC |
| 9228 | mamo-s | *BGIBMGA010855* | down | 1335 | 1343 | - | 12.4679 | 1.31E-05 | 0.615 | GTGCGTGTC |
| 9229 | mamo-s | *BGIBMGA010855* | up | 1278 | 1286 | - | 11.3303 | 4.96E-05 | 0.615 | GTGGGTGGC |
| 9230 | mamo-s | *BGIBMGA010856* | down | 604 | 612 | + | 11.3303 | 4.96E-05 | 0.615 | GTGGGTGGC |
| 9231 | mamo-s | *BGIBMGA010859* | down | 222 | 230 | + | 11.2844 | 5.18E-05 | 0.63 | CTGCGTGCC |
| 9232 | mamo-s | *BGIBMGA010860* | down | 520 | 528 | - | 11.3303 | 4.96E-05 | 0.615 | GTGGGTGGC |
| 9233 | mamo-s | *BGIBMGA010860* | up | 593 | 601 | + | 11.3303 | 4.96E-05 | 0.615 | GTGGGTGGC |

| | | | | | | | | | | |
|---|---|---|---|---|---|---|---|---|---|---|
| 9234 | mamo-s | *BGIBMGA010861* | up | 1669 | 1677 | - | 11.3303 | 4.96E-05 | 0.615 | GTGGGTGGC |
| 9235 | mamo-s | *BGIBMGA010862* | up | 585 | 593 | - | 11.3303 | 4.96E-05 | 0.615 | GTGGGTGGC |
| 9236 | mamo-s | *BGIBMGA010862* | down | 1347 | 1355 | - | 10.5413 | 9.42E-05 | 0.857 | CAGCGTGTC |
| 9237 | mamo-s | *BGIBMGA010863* | down | 1142 | 1150 | - | 11.2294 | 5.41E-05 | 0.65 | CTGCGAGGC |
| 9238 | mamo-s | *BGIBMGA010865* | down | 300 | 308 | + | 11.3303 | 4.96E-05 | 0.615 | GTGGGTGGC |
| 9239 | mamo-s | *BGIBMGA010867* | up | 1950 | 1958 | + | 11.7339 | 3.29E-05 | 0.615 | GCGCGAGGC |
| 9240 | mamo-s | *BGIBMGA010868* | down | 836 | 844 | - | 11.3853 | 4.74E-05 | 0.615 | ACGCGTGGC |
| 9241 | mamo-s | *BGIBMGA010868* | up | 27 | 35 | + | 11.3303 | 4.96E-05 | 0.615 | GTGGGTGGC |
| 9242 | mamo-s | *BGIBMGA010869* | down | 895 | 903 | - | 11.3303 | 4.96E-05 | 0.615 | GTGGGTGGC |
| 9243 | mamo-s | *BGIBMGA010870* | down | 1097 | 1105 | + | 11.3303 | 4.96E-05 | 0.615 | GTGGGTGGC |
| 9244 | mamo-s | *BGIBMGA010872* | up | 1428 | 1436 | + | 11.0826 | 6.84E-05 | 0.709 | GTACGTGGC |
| 9245 | mamo-s | *BGIBMGA010873* | down | 1196 | 1204 | + | 14.055 | 2.23E-06 | 0.615 | GTGCGTGGC |
| 9246 | mamo-s | *BGIBMGA010873* | up | 1550 | 1558 | + | 11.3303 | 4.96E-05 | 0.615 | GTGGGTGGC |
| 9247 | mamo-s | *BGIBMGA010875* | down | 1964 | 1972 | + | 12.1284 | 2.16E-05 | 0.615 | CAGCGTGGC |
| 9248 | mamo-s | *BGIBMGA010878* | down | 1887 | 1895 | - | 11.3853 | 4.74E-05 | 0.615 | ACGCGTGGC |
| 9249 | mamo-s | *BGIBMGA010879* | up | 1412 | 1420 | - | 11.4404 | 4.51E-05 | 0.615 | CGGCGTGGC |
| 9250 | mamo-s | *BGIBMGA010879* | up | 1883 | 1891 | + | 11.3303 | 4.96E-05 | 0.615 | GTGGGTGGC |
| 9251 | mamo-s | *BGIBMGA010881* | down | 551 | 559 | - | 11.3303 | 4.96E-05 | 0.615 | GTGGGTGGC |
| 9252 | mamo-s | *BGIBMGA010881* | up | 771 | 779 | - | 11.0826 | 6.84E-05 | 0.709 | GTACGTGGC |
| 9253 | mamo-s | *BGIBMGA010881* | down | 1767 | 1775 | + | 10.9358 | 7.30E-05 | 0.728 | GCGGGTGGC |
| 9254 | mamo-s | *BGIBMGA010884* | down | 474 | 482 | - | 11.3303 | 4.96E-05 | 0.615 | GTGGGTGGC |
| 9255 | mamo-s | *BGIBMGA010886* | up | 1446 | 1454 | - | 11.3303 | 4.96E-05 | 0.615 | GTGGGTGGC |
| 9256 | mamo-s | *BGIBMGA010889* | down | 1750 | 1758 | - | 11.3303 | 4.96E-05 | 0.615 | GTGGGTGGC |
| 9257 | mamo-s | *BGIBMGA010890* | up | 947 | 955 | - | 11.1743 | 5.90E-05 | 0.684 | CCGCGTGTC |
| 9258 | mamo-s | *BGIBMGA010891* | down | 735 | 743 | + | 11.3303 | 4.96E-05 | 0.615 | GTGGGTGGC |
| 9259 | mamo-s | *BGIBMGA010893* | up | 619 | 627 | + | 11.3303 | 4.96E-05 | 0.615 | GTGGGTGGC |
| 9260 | mamo-s | *BGIBMGA010893* | up | 254 | 262 | + | 11.2018 | 5.68E-05 | 0.674 | CTGCGTGGT |
| 9261 | mamo-s | *BGIBMGA010894* | down | 1039 | 1047 | - | 13.0275 | 8.53E-06 | 0.615 | GAGCGTGGC |
| 9262 | mamo-s | *BGIBMGA010895* | down | 999 | 1007 | + | 11.3303 | 4.96E-05 | 0.615 | GTGGGTGGC |
| 9263 | mamo-s | *BGIBMGA010896* | up | 950 | 958 | + | 11.3303 | 4.96E-05 | 0.615 | GTGGGTGGC |
| 9264 | mamo-s | *BGIBMGA010897* | down | 799 | 807 | - | 11.3303 | 4.96E-05 | 0.615 | GTGGGTGGC |
| 9265 | mamo-s | *BGIBMGA010900* | down | 378 | 386 | + | 11.3303 | 4.96E-05 | 0.615 | GTGGGTGGC |
| 9266 | mamo-s | *BGIBMGA010900* | down | 761 | 769 | + | 11.3303 | 4.96E-05 | 0.615 | GTGGGTGGC |
| 9267 | mamo-s | *BGIBMGA010901* | down | 1935 | 1943 | - | 13.6606 | 4.06E-06 | 0.615 | GCGCGTGGC |
| 9268 | mamo-s | *BGIBMGA010901* | up | 41 | 49 | + | 11.3303 | 4.96E-05 | 0.615 | GTGGGTGGC |
| 9269 | mamo-s | *BGIBMGA010901* | down | 292 | 300 | + | 11.3303 | 4.96E-05 | 0.615 | GTGGGTGGC |
| 9270 | mamo-s | *BGIBMGA010901* | up | 1657 | 1665 | - | 11.3303 | 4.96E-05 | 0.615 | GTGGGTGGC |
| 9271 | mamo-s | *BGIBMGA010902* | down | 213 | 221 | - | 11.3303 | 4.96E-05 | 0.615 | GTGGGTGGC |
| 9272 | mamo-s | *BGIBMGA010902* | up | 1334 | 1342 | - | 11.3303 | 4.96E-05 | 0.615 | GTGGGTGGC |
| 9273 | mamo-s | *BGIBMGA010903* | up | 982 | 990 | + | 11.3303 | 4.96E-05 | 0.615 | GTGGGTGGC |
| 9274 | mamo-s | *BGIBMGA010905* | up | 109 | 117 | - | 11.2018 | 5.68E-05 | 0.674 | CTGCGTGGT |
| 9275 | mamo-s | *BGIBMGA010907* | up | 966 | 974 | + | 11.0826 | 6.84E-05 | 0.709 | GTACGTGGC |
| 9276 | mamo-s | *BGIBMGA010908* | up | 266 | 274 | - | 11.3853 | 4.74E-05 | 0.615 | ACGCGTGGC |
| 9277 | mamo-s | *BGIBMGA010910* | up | 1088 | 1096 | - | 11.3303 | 4.96E-05 | 0.615 | GTGGGTGGC |
| 9278 | mamo-s | *BGIBMGA010910* | up | 1614 | 1622 | - | 10.5413 | 9.42E-05 | 0.857 | GTGCGAGTC |
| 9279 | mamo-s | *BGIBMGA010913* | up | 733 | 741 | - | 11.3303 | 4.96E-05 | 0.615 | GTGGGTGGC |
| 9280 | mamo-s | *BGIBMGA010913* | down | 77 | 85 | + | 11.2018 | 5.68E-05 | 0.674 | CTGCGTGGT |
| 9281 | mamo-s | *BGIBMGA010915* | up | 271 | 279 | - | 11.3303 | 4.96E-05 | 0.615 | GTGGGTGGC |
| 9282 | mamo-s | *BGIBMGA010915* | up | 947 | 955 | - | 11.1743 | 5.90E-05 | 0.684 | CCGCGTGTC |
| 9283 | mamo-s | *BGIBMGA010916* | up | 1156 | 1164 | - | 10.5138 | 9.75E-05 | 0.865 | GTGCGTGTT |
| 9284 | mamo-s | *BGIBMGA010918* | up | 1849 | 1857 | - | 11.3303 | 4.96E-05 | 0.615 | GTGGGTGGC |
| 9285 | mamo-s | *BGIBMGA010919* | up | 515 | 523 | + | 11.3303 | 4.96E-05 | 0.615 | GTGGGTGGC |
| 9286 | mamo-s | *BGIBMGA010920* | down | 1896 | 1904 | - | 12.0734 | 2.66E-05 | 0.615 | GCGCGTGTC |
| 9287 | mamo-s | *BGIBMGA010920* | down | 1025 | 1033 | - | 11.3303 | 4.96E-05 | 0.615 | GTGGGTGGC |
| 9288 | mamo-s | *BGIBMGA010922* | up | 1720 | 1728 | + | 11.0826 | 6.84E-05 | 0.709 | GTACGTGGC |
| 9289 | mamo-s | *BGIBMGA010923* | down | 1432 | 1440 | + | 11.1376 | 6.35E-05 | 0.709 | TCGCGTGGC |
| 9290 | mamo-s | *BGIBMGA010925* | up | 991 | 999 | + | 13.6606 | 4.06E-06 | 0.615 | GCGCGTGGC |
| 9291 | mamo-s | *BGIBMGA010927* | up | 1028 | 1036 | + | 11.3303 | 4.96E-05 | 0.615 | GTGGGTGGC |
| 9292 | mamo-s | *BGIBMGA010927* | down | 1955 | 1963 | + | 11.3303 | 4.96E-05 | 0.615 | GTGGGTGGC |
| 9293 | mamo-s | *BGIBMGA010927* | down | 1794 | 1802 | - | 10.9358 | 7.30E-05 | 0.728 | GCGGGTGGC |
| 9294 | mamo-s | *BGIBMGA010928* | up | 515 | 523 | + | 11.3303 | 4.96E-05 | 0.615 | GTGGGTGGC |
| 9295 | mamo-s | *BGIBMGA010928* | up | 354 | 362 | - | 10.9358 | 7.30E-05 | 0.728 | GCGGGTGGC |
| 9296 | mamo-s | *BGIBMGA010928* | up | 1225 | 1233 | - | 10.8073 | 7.89E-05 | 0.766 | CCGCGTGGT |
| 9297 | mamo-s | *BGIBMGA010928* | down | 18 | 26 | - | 10.5688 | 8.88E-05 | 0.824 | GTGCGTGAC |
| 9298 | mamo-s | *BGIBMGA010929* | up | 1062 | 1070 | - | 11.3303 | 4.96E-05 | 0.615 | GTGGGTGGC |
| 9299 | mamo-s | *BGIBMGA010929* | up | 846 | 854 | + | 11.1376 | 6.35E-05 | 0.709 | TCGCGTGGC |
| 9300 | mamo-s | *BGIBMGA010931* | down | 226 | 234 | + | 11.3303 | 4.96E-05 | 0.615 | GTGGGTGGC |
| 9301 | mamo-s | *BGIBMGA010932* | up | 450 | 458 | + | 11.3303 | 4.96E-05 | 0.615 | GTGGGTGGC |
| 9302 | mamo-s | *BGIBMGA010932* | up | 156 | 164 | + | 10.9358 | 7.30E-05 | 0.728 | GCGGGTGGC |
| 9303 | mamo-s | *BGIBMGA010933* | up | 456 | 464 | + | 11.3303 | 4.96E-05 | 0.615 | GTGGGTGGC |
| 9304 | mamo-s | *BGIBMGA010933* | down | 1200 | 1208 | + | 11.1743 | 5.90E-05 | 0.684 | CCGCGTGTC |
| 9305 | mamo-s | *BGIBMGA010936* | down | 766 | 774 | + | 12.1284 | 2.16E-05 | 0.615 | CAGCGTGGC |
| 9306 | mamo-s | *BGIBMGA010936* | down | 342 | 350 | + | 11.4404 | 4.51E-05 | 0.615 | CGGCGTGGC |
| 9307 | mamo-s | *BGIBMGA010936* | up | 184 | 192 | - | 11.3303 | 4.96E-05 | 0.615 | GTGGGTGGC |
| 9308 | mamo-s | *BGIBMGA010936* | down | 169 | 177 | + | 10.5413 | 9.42E-05 | 0.857 | CAGCGTGTC |
| 9309 | mamo-s | *BGIBMGA010937* | down | 1606 | 1614 | - | 11.0826 | 6.84E-05 | 0.709 | GTACGTGGC |
| 9310 | mamo-s | *BGIBMGA010938* | down | 1906 | 1914 | + | 11.3303 | 4.96E-05 | 0.615 | GTGGGTGGC |
| 9311 | mamo-s | *BGIBMGA010939* | down | 395 | 403 | + | 11.3303 | 4.96E-05 | 0.615 | GTGGGTGGC |
| 9312 | mamo-s | *BGIBMGA010940* | down | 1948 | 1956 | + | 11.3303 | 4.96E-05 | 0.615 | GTGGGTGGC |
| 9313 | mamo-s | *BGIBMGA010944* | down | 1392 | 1400 | + | 11.3303 | 4.96E-05 | 0.615 | GTGGGTGGC |
| 9314 | mamo-s | *BGIBMGA010947* | up | 324 | 332 | + | 14.055 | 2.23E-06 | 0.615 | GTGCGTGGC |
| 9315 | mamo-s | *BGIBMGA010949* | up | 657 | 665 | + | 12.0734 | 2.66E-05 | 0.615 | GCGCGTGTC |
| 9316 | mamo-s | *BGIBMGA010950* | down | 1213 | 1221 | + | 11.3303 | 4.96E-05 | 0.615 | GTGGGTGGC |
| 9317 | mamo-s | *BGIBMGA010951* | down | 92 | 100 | + | 11.3303 | 4.96E-05 | 0.615 | GTGGGTGGC |
| 9318 | mamo-s | *BGIBMGA010955* | up | 533 | 541 | - | 11.3303 | 4.96E-05 | 0.615 | GTGGGTGGC |
| 9319 | mamo-s | *BGIBMGA010956* | down | 445 | 453 | - | 13.6606 | 4.06E-06 | 0.615 | GCGCGTGGC |
| 9320 | mamo-s | *BGIBMGA010956* | down | 1016 | 1024 | + | 11.3303 | 4.96E-05 | 0.615 | GTGGGTGGC |
| 9321 | mamo-s | *BGIBMGA010958* | down | 1457 | 1465 | + | 11.3303 | 4.96E-05 | 0.615 | GTGGGTGGC |

| | | | | | | | | | | |
|---|---|---|---|---|---|---|---|---|---|---|
| 9322 | mamo-s | *BGIBMGA010958* | down | 1888 | 1896 | - | 11.3303 | 4.96E-05 | 0.615 | GTGGGTGGC |
| 9323 | mamo-s | *BGIBMGA010958* | up | 1384 | 1392 | + | 11.1743 | 5.90E-05 | 0.684 | CCGCGTGTC |
| 9324 | mamo-s | *BGIBMGA010959* | up | 363 | 371 | - | 11.3303 | 4.96E-05 | 0.615 | GTGGGTGGC |
| 9325 | mamo-s | *BGIBMGA010960* | up | 431 | 439 | + | 11.2294 | 5.41E-05 | 0.65 | CTGCGAGGC |
| 9326 | mamo-s | *BGIBMGA010960* | up | 1869 | 1877 | - | 10.6881 | 8.61E-05 | 0.81 | GCACGTGGC |
| 9327 | mamo-s | *BGIBMGA010961* | up | 1229 | 1237 | + | 11.3303 | 4.96E-05 | 0.615 | GTGGGTGGC |
| 9328 | mamo-s | *BGIBMGA010961* | down | 1102 | 1110 | - | 11.0826 | 6.84E-05 | 0.709 | GTACGTGGC |
| 9329 | mamo-s | *BGIBMGA010961* | down | 510 | 518 | + | 11.0734 | 7.12E-05 | 0.726 | GAGCGTGGT |
| 9330 | mamo-s | *BGIBMGA010962* | down | 1272 | 1280 | - | 11.3303 | 4.96E-05 | 0.615 | GTGGGTGGC |
| 9331 | mamo-s | *BGIBMGA010962* | down | 911 | 919 | - | 10.9358 | 7.30E-05 | 0.728 | GCGGGTGGC |
| 9332 | mamo-s | *BGIBMGA010964* | up | 1612 | 1620 | + | 11.3303 | 4.96E-05 | 0.615 | GTGGGTGGC |
| 9333 | mamo-s | *BGIBMGA010967* | up | 324 | 332 | - | 11.3303 | 4.96E-05 | 0.615 | GTGGGTGGC |
| 9334 | mamo-s | *BGIBMGA010968* | up | 14 | 22 | - | 11.3303 | 4.96E-05 | 0.615 | GTGGGTGGC |
| 9335 | mamo-s | *BGIBMGA010968* | up | 591 | 599 | + | 11.3303 | 4.96E-05 | 0.615 | GTGGGTGGC |
| 9336 | mamo-s | *BGIBMGA010969* | down | 331 | 339 | + | 11.3303 | 4.96E-05 | 0.615 | GTGGGTGGC |
| 9337 | mamo-s | *BGIBMGA010972* | up | 1613 | 1621 | + | 13.6606 | 4.06E-06 | 0.615 | GCGCGTGGC |
| 9338 | mamo-s | *BGIBMGA010975* | down | 347 | 355 | - | 10.5413 | 9.42E-05 | 0.857 | GTGCGAGTC |
| 9339 | mamo-s | *BGIBMGA010977* | down | 736 | 744 | + | 11.3303 | 4.96E-05 | 0.615 | GTGGGTGGC |
| 9340 | mamo-s | *BGIBMGA010977* | down | 1705 | 1713 | + | 11.0826 | 6.84E-05 | 0.709 | GTACGTGGC |
| 9341 | mamo-s | *BGIBMGA010977* | up | 1571 | 1579 | + | 10.9358 | 7.30E-05 | 0.728 | GCGGGTGGC |
| 9342 | mamo-s | *BGIBMGA010979* | down | 440 | 448 | - | 11.3303 | 4.96E-05 | 0.615 | GTGGGTGGC |
| 9343 | mamo-s | *BGIBMGA010980* | up | 583 | 591 | - | 11.3303 | 4.96E-05 | 0.615 | GTGGGTGGC |
| 9344 | mamo-s | *BGIBMGA010980* | up | 1990 | 1998 | - | 11.1376 | 6.35E-05 | 0.709 | TCGCGTGGC |
| 9345 | mamo-s | *BGIBMGA010981* | down | 444 | 452 | + | 10.8899 | 7.48E-05 | 0.739 | CCGCGTGCC |
| 9346 | mamo-s | *BGIBMGA010983* | up | 1039 | 1047 | - | 10.5413 | 9.42E-05 | 0.857 | CAGCGTGTC |
| 9347 | mamo-s | *BGIBMGA010984* | down | 50 | 58 | + | 11.4404 | 4.51E-05 | 0.615 | CGGCGTGGC |
| 9348 | mamo-s | *BGIBMGA010985* | down | 589 | 597 | + | 11.3303 | 4.96E-05 | 0.615 | GTGGGTGGC |
| 9349 | mamo-s | *BGIBMGA010987* | up | 1283 | 1291 | - | 11.3303 | 4.96E-05 | 0.615 | GTGGGTGGC |
| 9350 | mamo-s | *BGIBMGA010987* | down | 1335 | 1343 | - | 11.3303 | 4.96E-05 | 0.615 | GTGGGTGGC |
| 9351 | mamo-s | *BGIBMGA010988* | down | 763 | 771 | + | 11.3303 | 4.96E-05 | 0.615 | GTGGGTGGC |
| 9352 | mamo-s | *BGIBMGA010988* | up | 1102 | 1110 | - | 11.3303 | 4.96E-05 | 0.615 | GTGGGTGGC |
| 9353 | mamo-s | *BGIBMGA010989* | up | 289 | 297 | + | 11.7798 | 3.11E-05 | 0.615 | ATGCGTGGC |
| 9354 | mamo-s | *BGIBMGA010989* | down | 106 | 114 | - | 11.3303 | 4.96E-05 | 0.615 | GTGGGTGGC |
| 9355 | mamo-s | *BGIBMGA010989* | up | 140 | 148 | + | 11.3303 | 4.96E-05 | 0.615 | GTGGGTGGC |
| 9356 | mamo-s | *BGIBMGA010989* | down | 323 | 331 | - | 11.3303 | 4.96E-05 | 0.615 | GTGGGTGGC |
| 9357 | mamo-s | *BGIBMGA010989* | down | 1132 | 1140 | + | 11.3303 | 4.96E-05 | 0.615 | GTGGGTGGC |
| 9358 | mamo-s | *BGIBMGA010989* | up | 1804 | 1812 | + | 11.3303 | 4.96E-05 | 0.615 | GTGGGTGGC |
| 9359 | mamo-s | *BGIBMGA010990* | down | 3 | 11 | + | 11.2294 | 5.41E-05 | 0.65 | CTGCGAGGC |
| 9360 | mamo-s | *BGIBMGA010991* | down | 1728 | 1736 | - | 11.3303 | 4.96E-05 | 0.615 | GTGGGTGGC |
| 9361 | mamo-s | *BGIBMGA010994* | up | 909 | 917 | + | 11.3303 | 4.96E-05 | 0.615 | GTGGGTGGC |
| 9362 | mamo-s | *BGIBMGA010994* | down | 1385 | 1393 | - | 11.3303 | 4.96E-05 | 0.615 | GTGGGTGGC |
| 9363 | mamo-s | *BGIBMGA010995* | down | 1385 | 1393 | + | 12.3394 | 1.49E-05 | 0.615 | GGGCGTGGC |
| 9364 | mamo-s | *BGIBMGA010997* | up | 703 | 711 | + | 11.3303 | 4.96E-05 | 0.615 | GTGGGTGGC |
| 9365 | mamo-s | *BGIBMGA010997* | down | 1070 | 1078 | + | 11.3303 | 4.96E-05 | 0.615 | GTGGGTGGC |
| 9366 | mamo-s | *BGIBMGA011000* | up | 141 | 149 | - | 11.1743 | 5.90E-05 | 0.684 | CCGCGTGTC |
| 9367 | mamo-s | *BGIBMGA011000* | down | 470 | 478 | - | 11.0734 | 7.12E-05 | 0.726 | GAGCGTGGT |
| 9368 | mamo-s | *BGIBMGA011001* | up | 1459 | 1467 | - | 12.0734 | 2.66E-05 | 0.615 | GCGCGTGTC |
| 9369 | mamo-s | *BGIBMGA011001* | down | 1385 | 1393 | + | 11.1743 | 5.90E-05 | 0.684 | CCGCGTGTC |
| 9370 | mamo-s | *BGIBMGA011002* | up | 591 | 599 | + | 11.1743 | 5.90E-05 | 0.684 | CCGCGTGTC |
| 9371 | mamo-s | *BGIBMGA011002* | down | 701 | 709 | - | 11.0734 | 7.12E-05 | 0.726 | GAGCGTGGT |
| 9372 | mamo-s | *BGIBMGA011004* | down | 965 | 973 | - | 11.1376 | 6.35E-05 | 0.709 | TCGCGTGGC |
| 9373 | mamo-s | *BGIBMGA011007* | down | 383 | 391 | - | 11.0826 | 6.84E-05 | 0.709 | GTACGTGGC |
| 9374 | mamo-s | *BGIBMGA011011* | up | 679 | 687 | + | 11.3303 | 4.96E-05 | 0.615 | GTGGGTGGC |
| 9375 | mamo-s | *BGIBMGA011012* | up | 1821 | 1829 | + | 11.3303 | 4.96E-05 | 0.615 | GTGGGTGGC |
| 9376 | mamo-s | *BGIBMGA011015* | down | 1817 | 1825 | - | 10.7523 | 8.38E-05 | 0.797 | AAGCGTGGC |
| 9377 | mamo-s | *BGIBMGA011016* | down | 842 | 850 | + | 11.3303 | 4.96E-05 | 0.615 | GTGGGTGGC |
| 9378 | mamo-s | *BGIBMGA011017* | down | 1191 | 1199 | + | 11.3303 | 4.96E-05 | 0.615 | GTGGGTGGC |
| 9379 | mamo-s | *BGIBMGA011018* | up | 1394 | 1402 | + | 11.3303 | 4.96E-05 | 0.615 | GTGGGTGGC |
| 9380 | mamo-s | *BGIBMGA011022* | up | 1046 | 1054 | - | 11.3303 | 4.96E-05 | 0.615 | GTGGGTGGC |
| 9381 | mamo-s | *BGIBMGA011023* | down | 102 | 110 | - | 11.2844 | 5.18E-05 | 0.63 | CTGCGTGCC |
| 9382 | mamo-s | *BGIBMGA011026* | up | 1910 | 1918 | + | 11.0826 | 6.84E-05 | 0.709 | GTACGTGGC |
| 9383 | mamo-s | *BGIBMGA011029* | down | 1877 | 1885 | + | 11.3303 | 4.96E-05 | 0.615 | GTGGGTGGC |
| 9384 | mamo-s | *BGIBMGA011030* | up | 1460 | 1468 | + | 11.3303 | 4.96E-05 | 0.615 | GTGGGTGGC |
| 9385 | mamo-s | *BGIBMGA011035* | up | 432 | 440 | - | 11.3303 | 4.96E-05 | 0.615 | GTGGGTGGC |
| 9386 | mamo-s | *BGIBMGA011035* | down | 622 | 630 | - | 11.3303 | 4.96E-05 | 0.615 | GTGGGTGGC |
| 9387 | mamo-s | *BGIBMGA011035* | up | 1053 | 1061 | + | 11.3303 | 4.96E-05 | 0.615 | GTGGGTGGC |
| 9388 | mamo-s | *BGIBMGA011036* | up | 1912 | 1920 | - | 11.7798 | 3.11E-05 | 0.615 | ATGCGTGGC |
| 9389 | mamo-s | *BGIBMGA011036* | down | 135 | 143 | - | 11.3303 | 4.96E-05 | 0.615 | GTGGGTGGC |
| 9390 | mamo-s | *BGIBMGA011036* | up | 150 | 158 | - | 11.3303 | 4.96E-05 | 0.615 | GTGGGTGGC |
| 9391 | mamo-s | *BGIBMGA011036* | down | 510 | 518 | + | 11.3303 | 4.96E-05 | 0.615 | GTGGGTGGC |
| 9392 | mamo-s | *BGIBMGA011036* | down | 1564 | 1572 | - | 11.3303 | 4.96E-05 | 0.615 | GTGGGTGGC |
| 9393 | mamo-s | *BGIBMGA011038* | up | 1861 | 1869 | - | 11.3853 | 4.74E-05 | 0.615 | ACGCGTGGC |
| 9394 | mamo-s | *BGIBMGA011039* | down | 1294 | 1302 | + | 12.0734 | 2.66E-05 | 0.615 | GCGCGTGTC |
| 9395 | mamo-s | *BGIBMGA011040* | up | 964 | 972 | + | 11.3303 | 4.96E-05 | 0.615 | GTGGGTGGC |
| 9396 | mamo-s | *BGIBMGA011040* | down | 1083 | 1091 | + | 10.5688 | 8.88E-05 | 0.824 | GTGCGTGAC |
| 9397 | mamo-s | *BGIBMGA011044* | down | 820 | 828 | + | 11.3303 | 4.96E-05 | 0.615 | GTGGGTGGC |
| 9398 | mamo-s | *BGIBMGA011045* | up | 1458 | 1466 | + | 11.5321 | 4.06E-05 | 0.615 | TTGCGTGGC |
| 9399 | mamo-s | *BGIBMGA011045* | down | 418 | 426 | - | 11.3303 | 4.96E-05 | 0.615 | GTGGGTGGC |
| 9400 | mamo-s | *BGIBMGA011046* | up | 775 | 783 | - | 11.3303 | 4.96E-05 | 0.615 | GTGGGTGGC |
| 9401 | mamo-s | *BGIBMGA011046* | down | 1812 | 1820 | + | 11.3303 | 4.96E-05 | 0.615 | GTGGGTGGC |
| 9402 | mamo-s | *BGIBMGA011047* | up | 139 | 147 | - | 11.3303 | 4.96E-05 | 0.615 | GTGGGTGGC |
| 9403 | mamo-s | *BGIBMGA011053* | down | 729 | 737 | + | 11.0826 | 6.84E-05 | 0.709 | GTACGTGGC |
| 9404 | mamo-s | *BGIBMGA011054* | up | 685 | 693 | + | 11.3303 | 4.96E-05 | 0.615 | GTGGGTGGC |
| 9405 | mamo-s | *BGIBMGA011054* | up | 71 | 79 | + | 11.0826 | 6.84E-05 | 0.709 | GTACGTGGC |
| 9406 | mamo-s | *BGIBMGA011055* | down | 198 | 206 | - | 11.3303 | 4.96E-05 | 0.615 | GTGGGTGGC |
| 9407 | mamo-s | *BGIBMGA011055* | up | 834 | 842 | - | 11.3303 | 4.96E-05 | 0.615 | GTGGGTGGC |
| 9408 | mamo-s | *BGIBMGA011055* | up | 1756 | 1764 | - | 11.3303 | 4.96E-05 | 0.615 | GTGGGTGGC |
| 9409 | mamo-s | *BGIBMGA011058* | down | 1609 | 1617 | - | 11.3853 | 4.74E-05 | 0.615 | ACGCGTGGC |

| | | | | | | | | | | |
|---|---|---|---|---|---|---|---|---|---|---|
| 9410 | mamo-s | *BGIBMGA011059* | down | 358 | 366 | - | 11.4404 | 4.51E-05 | 0.615 | GAGCGTGTC |
| 9411 | mamo-s | *BGIBMGA011059* | down | 1498 | 1506 | - | 11.1376 | 6.35E-05 | 0.709 | TCGCGTGGC |
| 9412 | mamo-s | *BGIBMGA011060* | up | 50 | 58 | - | 11.3303 | 4.96E-05 | 0.615 | GTGGGTGGC |
| 9413 | mamo-s | *BGIBMGA011064* | down | 265 | 273 | - | 14.055 | 2.23E-06 | 0.615 | GTGCGTGGC |
| 9414 | mamo-s | *BGIBMGA011064* | down | 981 | 989 | + | 11.3303 | 4.96E-05 | 0.615 | GTGGGTGGC |
| 9415 | mamo-s | *BGIBMGA011069* | down | 352 | 360 | - | 11.3303 | 4.96E-05 | 0.615 | GTGGGTGGC |
| 9416 | mamo-s | *BGIBMGA011069* | down | 1068 | 1076 | - | 10.7523 | 8.38E-05 | 0.797 | GGGCGTGTC |
| 9417 | mamo-s | *BGIBMGA011070* | down | 413 | 421 | + | 11.3303 | 4.96E-05 | 0.615 | GTGGGTGGC |
| 9418 | mamo-s | *BGIBMGA011071* | up | 647 | 655 | + | 11.3303 | 4.96E-05 | 0.615 | GTGGGTGGC |
| 9419 | mamo-s | *BGIBMGA011072* | up | 591 | 599 | - | 11.0826 | 6.84E-05 | 0.709 | GTACGTGGC |
| 9420 | mamo-s | *BGIBMGA011074* | up | 1539 | 1547 | + | 11.3303 | 4.96E-05 | 0.615 | GTGGGTGGC |
| 9421 | mamo-s | *BGIBMGA011076* | down | 1568 | 1576 | + | 11.1743 | 5.90E-05 | 0.684 | CCGCGTGTC |
| 9422 | mamo-s | *BGIBMGA011078* | down | 1809 | 1817 | + | 11.3303 | 4.96E-05 | 0.615 | GTGGGTGGC |
| 9423 | mamo-s | *BGIBMGA011079* | down | 1141 | 1149 | + | 14.055 | 2.23E-06 | 0.615 | GTGCGTGGC |
| 9424 | mamo-s | *BGIBMGA011085* | up | 928 | 936 | + | 12.0734 | 2.66E-05 | 0.615 | GCGCGTGTC |
| 9425 | mamo-s | *BGIBMGA011085* | up | 1780 | 1788 | + | 11.2018 | 5.68E-05 | 0.674 | CTGCGTGGT |
| 9426 | mamo-s | *BGIBMGA011086* | up | 557 | 565 | + | 11.3303 | 4.96E-05 | 0.615 | GTGGGTGGC |
| 9427 | mamo-s | *BGIBMGA011086* | down | 243 | 251 | - | 11.0826 | 6.84E-05 | 0.709 | GTACGTGGC |
| 9428 | mamo-s | *BGIBMGA011093* | up | 1885 | 1893 | + | 11.3303 | 4.96E-05 | 0.615 | GTGGGTGGC |
| 9429 | mamo-s | *BGIBMGA011093* | down | 1953 | 1961 | - | 11.0826 | 6.84E-05 | 0.709 | GTACGTGGC |
| 9430 | mamo-s | *BGIBMGA011097* | down | 1295 | 1303 | + | 11.5321 | 4.06E-05 | 0.615 | TTGCGTGGC |
| 9431 | mamo-s | *BGIBMGA011097* | down | 206 | 214 | + | 11.3303 | 4.96E-05 | 0.615 | GTGGGTGGC |
| 9432 | mamo-s | *BGIBMGA011098* | up | 472 | 480 | + | 14.055 | 2.23E-06 | 0.615 | GTGCGTGGC |
| 9433 | mamo-s | *BGIBMGA011101* | up | 1624 | 1632 | - | 11.3303 | 4.96E-05 | 0.615 | GTGGGTGGC |
| 9434 | mamo-s | *BGIBMGA011101* | down | 1797 | 1805 | - | 11.3303 | 4.96E-05 | 0.615 | GTGGGTGGC |
| 9435 | mamo-s | *BGIBMGA011101* | down | 1455 | 1463 | - | 11.1376 | 6.35E-05 | 0.709 | TCGCGTGGC |
| 9436 | mamo-s | *BGIBMGA011101* | down | 1178 | 1186 | - | 10.9358 | 7.30E-05 | 0.728 | GCGGGTGGC |
| 9437 | mamo-s | *BGIBMGA011102* | down | 635 | 643 | - | 12.1835 | 1.71E-05 | 0.615 | GTGCGTGCC |
| 9438 | mamo-s | *BGIBMGA011102* | down | 453 | 461 | - | 12.0734 | 2.66E-05 | 0.615 | GCGCGTGTC |
| 9439 | mamo-s | *BGIBMGA011102* | up | 11 | 19 | + | 11.3303 | 4.96E-05 | 0.615 | GTGGGTGGC |
| 9440 | mamo-s | *BGIBMGA011102* | down | 1957 | 1965 | + | 11.3303 | 4.96E-05 | 0.615 | GTGGGTGGC |
| 9441 | mamo-s | *BGIBMGA011102* | down | 7 | 15 | + | 11.156 | 6.13E-05 | 0.705 | GAGCGTGCC |
| 9442 | mamo-s | *BGIBMGA011102* | up | 207 | 215 | + | 11.0826 | 6.84E-05 | 0.709 | GTACGTGGC |
| 9443 | mamo-s | *BGIBMGA011103* | up | 938 | 946 | + | 13.156 | 6.30E-06 | 0.615 | CTGCGTGGC |
| 9444 | mamo-s | *BGIBMGA011103* | up | 1451 | 1459 | - | 11.3303 | 4.96E-05 | 0.615 | GTGGGTGGC |
| 9445 | mamo-s | *BGIBMGA011105* | down | 605 | 613 | - | 11.3303 | 4.96E-05 | 0.615 | GTGGGTGGC |
| 9446 | mamo-s | *BGIBMGA011105* | down | 1327 | 1335 | - | 11.0826 | 6.84E-05 | 0.709 | GTACGTGGC |
| 9447 | mamo-s | *BGIBMGA011108* | down | 953 | 961 | + | 11.0826 | 6.84E-05 | 0.709 | GTACGTGGC |
| 9448 | mamo-s | *BGIBMGA011109* | up | 1577 | 1585 | + | 11.3303 | 4.96E-05 | 0.615 | GTGGGTGGC |
| 9449 | mamo-s | *BGIBMGA011109* | up | 944 | 952 | + | 10.7523 | 8.38E-05 | 0.797 | GGGCGTGTC |
| 9450 | mamo-s | *BGIBMGA011110* | down | 227 | 235 | + | 11.3303 | 4.96E-05 | 0.615 | GTGGGTGGC |
| 9451 | mamo-s | *BGIBMGA011111* | down | 510 | 518 | - | 12.1009 | 2.43E-05 | 0.615 | GTGCGTGGT |
| 9452 | mamo-s | *BGIBMGA011111* | down | 27 | 35 | + | 11.156 | 6.13E-05 | 0.705 | GAGCGTGCC |
| 9453 | mamo-s | *BGIBMGA011113* | up | 1365 | 1373 | + | 12.4679 | 1.31E-05 | 0.615 | GTGCGTGTC |
| 9454 | mamo-s | *BGIBMGA011113* | up | 228 | 236 | + | 11.3303 | 4.96E-05 | 0.615 | GTGGGTGGC |
| 9455 | mamo-s | *BGIBMGA011114* | up | 305 | 313 | - | 11.3303 | 4.96E-05 | 0.615 | GTGGGTGGC |
| 9456 | mamo-s | *BGIBMGA011115* | down | 258 | 266 | + | 11.3303 | 4.96E-05 | 0.615 | GTGGGTGGC |
| 9457 | mamo-s | *BGIBMGA011115* | down | 1001 | 1009 | - | 11.3303 | 4.96E-05 | 0.615 | GTGGGTGGC |
| 9458 | mamo-s | *BGIBMGA011115* | up | 1823 | 1831 | - | 11.3303 | 4.96E-05 | 0.615 | GTGGGTGGC |
| 9459 | mamo-s | *BGIBMGA011116* | up | 1913 | 1921 | + | 13.156 | 6.30E-06 | 0.615 | CTGCGTGGC |
| 9460 | mamo-s | *BGIBMGA011116* | up | 1794 | 1802 | + | 11.3303 | 4.96E-05 | 0.615 | GTGGGTGGC |
| 9461 | mamo-s | *BGIBMGA011117* | up | 380 | 388 | + | 11.3303 | 4.96E-05 | 0.615 | GTGGGTGGC |
| 9462 | mamo-s | *BGIBMGA011117* | up | 504 | 512 | - | 11.3303 | 4.96E-05 | 0.615 | GTGGGTGGC |
| 9463 | mamo-s | *BGIBMGA011118* | down | 250 | 258 | - | 11.3303 | 4.96E-05 | 0.615 | GTGGGTGGC |
| 9464 | mamo-s | *BGIBMGA011118* | up | 1326 | 1334 | + | 10.8899 | 7.48E-05 | 0.739 | CCGCGTGCC |
| 9465 | mamo-s | *BGIBMGA011119* | down | 1934 | 1942 | + | 11.7064 | 3.52E-05 | 0.615 | GCGCGTGGT |
| 9466 | mamo-s | *BGIBMGA011120* | up | 625 | 633 | + | 11.7064 | 3.52E-05 | 0.615 | GCGCGTGGT |
| 9467 | mamo-s | *BGIBMGA011120* | up | 828 | 836 | + | 11.5688 | 3.79E-05 | 0.615 | CTGCGTGTC |
| 9468 | mamo-s | *BGIBMGA011121* | up | 1289 | 1297 | - | 11.3303 | 4.96E-05 | 0.615 | GTGGGTGGC |
| 9469 | mamo-s | *BGIBMGA011123* | down | 1586 | 1594 | + | 11.7798 | 3.11E-05 | 0.615 | ATGCGTGGC |
| 9470 | mamo-s | *BGIBMGA011123* | up | 741 | 749 | + | 11.2844 | 5.18E-05 | 0.63 | CTGCGTGCC |
| 9471 | mamo-s | *BGIBMGA011124* | down | 1626 | 1634 | + | 12.3394 | 1.49E-05 | 0.615 | GGGCGTGGC |
| 9472 | mamo-s | *BGIBMGA011125* | up | 739 | 747 | + | 11.3303 | 4.96E-05 | 0.615 | GTGGGTGGC |
| 9473 | mamo-s | *BGIBMGA011126* | up | 1396 | 1404 | + | 11.3303 | 4.96E-05 | 0.615 | GTGGGTGGC |
| 9474 | mamo-s | *BGIBMGA011126* | up | 1780 | 1788 | + | 11.1743 | 5.90E-05 | 0.684 | CCGCGTGTC |
| 9475 | mamo-s | *BGIBMGA011127* | down | 1139 | 1147 | - | 11.3303 | 4.96E-05 | 0.615 | GTGGGTGGC |
| 9476 | mamo-s | *BGIBMGA011132* | up | 473 | 481 | - | 11.3303 | 4.96E-05 | 0.615 | GTGGGTGGC |
| 9477 | mamo-s | *BGIBMGA011133* | down | 1842 | 1850 | + | 11.0734 | 7.12E-05 | 0.726 | GAGCGTGGT |
| 9478 | mamo-s | *BGIBMGA011134* | down | 11 | 19 | + | 12.4679 | 1.31E-05 | 0.615 | GTGCGTGTC |
| 9479 | mamo-s | *BGIBMGA011136* | up | 1302 | 1310 | + | 11.4404 | 4.51E-05 | 0.615 | GAGCGTGTC |
| 9480 | mamo-s | *BGIBMGA011136* | up | 121 | 129 | - | 11.0826 | 6.84E-05 | 0.709 | GTACGTGGC |
| 9481 | mamo-s | *BGIBMGA011137* | down | 569 | 577 | - | 11.3303 | 4.96E-05 | 0.615 | GTGGGTGGC |
| 9482 | mamo-s | *BGIBMGA011138* | down | 1708 | 1716 | - | 10.5138 | 9.75E-05 | 0.865 | GTGCGTGTT |
| 9483 | mamo-s | *BGIBMGA011139* | down | 367 | 375 | - | 11.3303 | 4.96E-05 | 0.615 | GTGGGTGGC |
| 9484 | mamo-s | *BGIBMGA011140* | down | 348 | 356 | + | 14.055 | 2.23E-06 | 0.615 | GTGCGTGGC |
| 9485 | mamo-s | *BGIBMGA011140* | up | 195 | 203 | + | 12.3394 | 1.49E-05 | 0.615 | GGGCGTGGC |
| 9486 | mamo-s | *BGIBMGA011143* | down | 1522 | 1530 | - | 11.3303 | 4.96E-05 | 0.615 | GTGGGTGGC |
| 9487 | mamo-s | *BGIBMGA011145* | up | 895 | 903 | + | 11.3303 | 4.96E-05 | 0.615 | GTGGGTGGC |
| 9488 | mamo-s | *BGIBMGA011145* | up | 1768 | 1776 | - | 11.2844 | 5.18E-05 | 0.63 | CTGCGTGCC |
| 9489 | mamo-s | *BGIBMGA011146* | up | 1056 | 1064 | + | 11.3303 | 4.96E-05 | 0.615 | GTGGGTGGC |
| 9490 | mamo-s | *BGIBMGA011146* | up | 1180 | 1188 | - | 11.3303 | 4.96E-05 | 0.615 | GTGGGTGGC |
| 9491 | mamo-s | *BGIBMGA011147* | down | 262 | 270 | - | 11.3303 | 4.96E-05 | 0.615 | GTGGGTGGC |
| 9492 | mamo-s | *BGIBMGA011147* | up | 771 | 779 | + | 11.3303 | 4.96E-05 | 0.615 | GTGGGTGGC |
| 9493 | mamo-s | *BGIBMGA011151* | up | 733 | 741 | + | 11.3303 | 4.96E-05 | 0.615 | GTGGGTGGC |
| 9494 | mamo-s | *BGIBMGA011151* | down | 1541 | 1549 | - | 10.8073 | 7.89E-05 | 0.766 | CCGCGTGGT |
| 9495 | mamo-s | *BGIBMGA011151* | up | 355 | 363 | - | 10.5413 | 9.42E-05 | 0.857 | GTGCGAGTC |
| 9496 | mamo-s | *BGIBMGA011154* | down | 1298 | 1306 | + | 11.1009 | 6.57E-05 | 0.709 | GAGCGAGGC |
| 9497 | mamo-s | *BGIBMGA011158* | up | 224 | 232 | + | 11.156 | 6.13E-05 | 0.705 | GAGCGTGCC |

| | | | | | | | | | | |
|---|---|---|---|---|---|---|---|---|---|---|
| 9498 | mamo-s | *BGIBMGA011159* | up | 1138 | 1146 | + | 13.6606 | 4.06E-06 | 0.615 | GCGCGTGGC |
| 9499 | mamo-s | *BGIBMGA011159* | down | 1780 | 1788 | + | 10.9358 | 7.30E-05 | 0.728 | GCGGGTGGC |
| 9500 | mamo-s | *BGIBMGA011159* | down | 1064 | 1072 | + | 10.5688 | 8.88E-05 | 0.824 | GTGCGTGAC |
| 9501 | mamo-s | *BGIBMGA011160* | down | 941 | 949 | + | 12.0734 | 2.66E-05 | 0.615 | GCGCGTGTC |
| 9502 | mamo-s | *BGIBMGA011160* | down | 1935 | 1943 | - | 11.3303 | 4.96E-05 | 0.615 | GTGGGTGGC |
| 9503 | mamo-s | *BGIBMGA011160* | down | 462 | 470 | - | 10.8899 | 7.48E-05 | 0.739 | CCGCGTGCC |
| 9504 | mamo-s | *BGIBMGA011160* | down | 1701 | 1709 | + | 10.7523 | 8.38E-05 | 0.797 | GGGCGTGTC |
| 9505 | mamo-s | *BGIBMGA011160* | up | 1706 | 1714 | + | 10.5688 | 8.88E-05 | 0.824 | GTGCGTGAC |
| 9506 | mamo-s | *BGIBMGA011161* | up | 1457 | 1465 | - | 11.156 | 6.13E-05 | 0.705 | GAGCGTGCC |
| 9507 | mamo-s | *BGIBMGA011163* | down | 956 | 964 | - | 11.3303 | 4.96E-05 | 0.615 | GTGGGTGGC |
| 9508 | mamo-s | *BGIBMGA011163* | up | 600 | 608 | - | 11.0826 | 6.84E-05 | 0.709 | GTACGTGGC |
| 9509 | mamo-s | *BGIBMGA011163* | down | 164 | 172 | + | 10.6881 | 8.61E-05 | 0.81 | GCACGTGGC |
| 9510 | mamo-s | *BGIBMGA011163* | up | 6 | 14 | + | 10.5413 | 9.42E-05 | 0.857 | GTGCGAGTC |
| 9511 | mamo-s | *BGIBMGA011166* | up | 1154 | 1162 | + | 12.1009 | 2.43E-05 | 0.615 | GTGCGTGGT |
| 9512 | mamo-s | *BGIBMGA011169* | up | 866 | 874 | - | 11.3303 | 4.96E-05 | 0.615 | GTGGGTGGC |
| 9513 | mamo-s | *BGIBMGA011169* | up | 721 | 729 | + | 11.2294 | 5.41E-05 | 0.65 | CTGCGAGGC |
| 9514 | mamo-s | *BGIBMGA011170* | down | 1659 | 1667 | - | 11.2294 | 5.41E-05 | 0.65 | CTGCGAGGC |
| 9515 | mamo-s | *BGIBMGA011171* | down | 1246 | 1254 | - | 11.3303 | 4.96E-05 | 0.615 | GTGGGTGGC |
| 9516 | mamo-s | *BGIBMGA011171* | down | 288 | 296 | - | 11.0826 | 6.84E-05 | 0.709 | GTACGTGGC |
| 9517 | mamo-s | *BGIBMGA011171* | down | 1638 | 1646 | + | 10.9358 | 7.30E-05 | 0.728 | GCGGGTGGC |
| 9518 | mamo-s | *BGIBMGA011172* | up | 1453 | 1461 | + | 11.3303 | 4.96E-05 | 0.615 | GTGGGTGGC |
| 9519 | mamo-s | *BGIBMGA011173* | down | 1736 | 1744 | - | 11.2018 | 5.68E-05 | 0.674 | CTGCGTGGT |
| 9520 | mamo-s | *BGIBMGA011174* | up | 122 | 130 | - | 11.2018 | 5.68E-05 | 0.674 | CTGCGTGGT |
| 9521 | mamo-s | *BGIBMGA011175* | up | 1838 | 1846 | + | 12.1835 | 1.71E-05 | 0.615 | GTGCGTGCC |
| 9522 | mamo-s | *BGIBMGA011176* | down | 190 | 198 | + | 11.3303 | 4.96E-05 | 0.615 | GTGGGTGGC |
| 9523 | mamo-s | *BGIBMGA011177* | down | 1910 | 1918 | + | 11.2844 | 5.18E-05 | 0.63 | CTGCGTGCC |
| 9524 | mamo-s | *BGIBMGA011177* | up | 1818 | 1826 | - | 10.9358 | 7.30E-05 | 0.728 | GCGGGTGGC |
| 9525 | mamo-s | *BGIBMGA011177* | up | 26 | 34 | + | 10.5413 | 9.42E-05 | 0.857 | CAGCGTGTC |
| 9526 | mamo-s | *BGIBMGA011179* | down | 1922 | 1930 | + | 11.2018 | 5.68E-05 | 0.674 | CTGCGTGGT |
| 9527 | mamo-s | *BGIBMGA011180* | down | 1922 | 1930 | + | 11.3303 | 4.96E-05 | 0.615 | GTGGGTGGC |
| 9528 | mamo-s | *BGIBMGA011182* | down | 170 | 178 | + | 11.3303 | 4.96E-05 | 0.615 | GTGGGTGGC |
| 9529 | mamo-s | *BGIBMGA011182* | down | 1384 | 1392 | + | 11.3303 | 4.96E-05 | 0.615 | GTGGGTGGC |
| 9530 | mamo-s | *BGIBMGA011182* | down | 1580 | 1588 | - | 11.2294 | 5.41E-05 | 0.65 | CTGCGAGGC |
| 9531 | mamo-s | *BGIBMGA011182* | up | 1612 | 1620 | - | 11.0826 | 6.84E-05 | 0.709 | GTACGTGGC |
| 9532 | mamo-s | *BGIBMGA011183* | down | 318 | 326 | - | 11.3303 | 4.96E-05 | 0.615 | GTGGGTGGC |
| 9533 | mamo-s | *BGIBMGA011183* | up | 654 | 662 | + | 11.3303 | 4.96E-05 | 0.615 | GTGGGTGGC |
| 9534 | mamo-s | *BGIBMGA011183* | up | 1868 | 1876 | + | 11.3303 | 4.96E-05 | 0.615 | GTGGGTGGC |
| 9535 | mamo-s | *BGIBMGA011185* | down | 1942 | 1950 | + | 11.3303 | 4.96E-05 | 0.615 | GTGGGTGGC |
| 9536 | mamo-s | *BGIBMGA011186* | down | 1022 | 1030 | + | 11.3303 | 4.96E-05 | 0.615 | GTGGGTGGC |
| 9537 | mamo-s | *BGIBMGA011186* | down | 371 | 379 | + | 10.8073 | 7.89E-05 | 0.766 | CCGCGTGGT |
| 9538 | mamo-s | *BGIBMGA011187* | up | 1156 | 1164 | + | 13.156 | 6.30E-06 | 0.615 | CTGCGTGGC |
| 9539 | mamo-s | *BGIBMGA011187* | up | 1796 | 1804 | + | 11.3303 | 4.96E-05 | 0.615 | GTGGGTGGC |
| 9540 | mamo-s | *BGIBMGA011188* | up | 325 | 333 | + | 11.4404 | 4.51E-05 | 0.615 | CGGCGTGGC |
| 9541 | mamo-s | *BGIBMGA011190* | up | 936 | 944 | + | 11.7798 | 3.11E-05 | 0.615 | ATGCGTGGC |
| 9542 | mamo-s | *BGIBMGA011190* | up | 449 | 457 | + | 11.3303 | 4.96E-05 | 0.615 | GTGGGTGGC |
| 9543 | mamo-s | *BGIBMGA011192* | up | 301 | 309 | - | 11.3303 | 4.96E-05 | 0.615 | GTGGGTGGC |
| 9544 | mamo-s | *BGIBMGA011192* | down | 1413 | 1421 | + | 11.3303 | 4.96E-05 | 0.615 | GTGGGTGGC |
| 9545 | mamo-s | *BGIBMGA011192* | down | 1724 | 1732 | - | 11.3303 | 4.96E-05 | 0.615 | GTGGGTGGC |
| 9546 | mamo-s | *BGIBMGA011194* | up | 1728 | 1736 | + | 13.6606 | 4.06E-06 | 0.615 | GCGCGTGGC |
| 9547 | mamo-s | *BGIBMGA011195* | down | 1708 | 1716 | - | 11.3303 | 4.96E-05 | 0.615 | GTGGGTGGC |
| 9548 | mamo-s | *BGIBMGA011197* | up | 229 | 237 | + | 11.3303 | 4.96E-05 | 0.615 | GTGGGTGGC |
| 9549 | mamo-s | *BGIBMGA011199* | down | 1405 | 1413 | - | 13.156 | 6.30E-06 | 0.615 | CTGCGTGGC |
| 9550 | mamo-s | *BGIBMGA011199* | down | 1139 | 1147 | - | 11.3303 | 4.96E-05 | 0.615 | GTGGGTGGC |
| 9551 | mamo-s | *BGIBMGA011199* | down | 1382 | 1390 | + | 11.3303 | 4.96E-05 | 0.615 | GTGGGTGGC |
| 9552 | mamo-s | *BGIBMGA011200* | up | 1808 | 1816 | - | 13.156 | 6.30E-06 | 0.615 | CTGCGTGGC |
| 9553 | mamo-s | *BGIBMGA011200* | up | 1542 | 1550 | - | 11.3303 | 4.96E-05 | 0.615 | GTGGGTGGC |
| 9554 | mamo-s | *BGIBMGA011200* | up | 1785 | 1793 | + | 11.3303 | 4.96E-05 | 0.615 | GTGGGTGGC |
| 9555 | mamo-s | *BGIBMGA011201* | down | 1786 | 1794 | - | 11.3303 | 4.96E-05 | 0.615 | GTGGGTGGC |
| 9556 | mamo-s | *BGIBMGA011201* | down | 1254 | 1262 | + | 11.2844 | 5.18E-05 | 0.63 | CTGCGTGCC |
| 9557 | mamo-s | *BGIBMGA011202* | down | 974 | 982 | - | 13.156 | 6.30E-06 | 0.615 | CTGCGTGGC |
| 9558 | mamo-s | *BGIBMGA011203* | down | 1753 | 1761 | - | 10.8899 | 7.48E-05 | 0.739 | CCGCGTGCC |
| 9559 | mamo-s | *BGIBMGA011204* | down | 450 | 458 | - | 11.3303 | 4.96E-05 | 0.615 | GTGGGTGGC |
| 9560 | mamo-s | *BGIBMGA011204* | down | 934 | 942 | - | 11.3303 | 4.96E-05 | 0.615 | GTGGGTGGC |
| 9561 | mamo-s | *BGIBMGA011204* | up | 925 | 933 | - | 10.8899 | 7.48E-05 | 0.739 | CCGCGTGCC |
| 9562 | mamo-s | *BGIBMGA011205* | down | 1072 | 1080 | - | 11.3303 | 4.96E-05 | 0.615 | GTGGGTGGC |
| 9563 | mamo-s | *BGIBMGA011206* | up | 1324 | 1332 | - | 11.3303 | 4.96E-05 | 0.615 | GTGGGTGGC |
| 9564 | mamo-s | *BGIBMGA011207* | down | 754 | 762 | + | 11.2294 | 5.41E-05 | 0.65 | CTGCGAGGC |
| 9565 | mamo-s | *BGIBMGA011208* | up | 334 | 342 | + | 11.2294 | 5.41E-05 | 0.65 | CTGCGAGGC |
| 9566 | mamo-s | *BGIBMGA011210* | down | 203 | 211 | + | 11.3303 | 4.96E-05 | 0.615 | GTGGGTGGC |
| 9567 | mamo-s | *BGIBMGA011211* | down | 845 | 853 | + | 11.3303 | 4.96E-05 | 0.615 | GTGGGTGGC |
| 9568 | mamo-s | *BGIBMGA011211* | down | 1583 | 1591 | - | 10.9358 | 7.30E-05 | 0.728 | GCGGGTGGC |
| 9569 | mamo-s | *BGIBMGA011212* | down | 1610 | 1618 | - | 11.3303 | 4.96E-05 | 0.615 | GTGGGTGGC |
| 9570 | mamo-s | *BGIBMGA011212* | up | 1737 | 1745 | + | 10.5138 | 9.75E-05 | 0.865 | GTGCGTGTT |
| 9571 | mamo-s | *BGIBMGA011213* | up | 953 | 961 | - | 11.3303 | 4.96E-05 | 0.615 | GTGGGTGGC |
| 9572 | mamo-s | *BGIBMGA011214* | down | 778 | 786 | + | 12.1284 | 2.16E-05 | 0.615 | CAGCGTGGC |
| 9573 | mamo-s | *BGIBMGA011217* | down | 279 | 287 | + | 11.4404 | 4.51E-05 | 0.615 | GAGCGTGTC |
| 9574 | mamo-s | *BGIBMGA011217* | up | 261 | 269 | + | 11.3303 | 4.96E-05 | 0.615 | GTGGGTGGC |
| 9575 | mamo-s | *BGIBMGA011217* | up | 1305 | 1313 | - | 11.3303 | 4.96E-05 | 0.615 | GTGGGTGGC |
| 9576 | mamo-s | *BGIBMGA011217* | down | 909 | 917 | + | 11.2844 | 5.18E-05 | 0.63 | CTGCGTGCC |
| 9577 | mamo-s | *BGIBMGA011218* | up | 1891 | 1899 | + | 11.3303 | 4.96E-05 | 0.615 | GTGGGTGGC |
| 9578 | mamo-s | *BGIBMGA011221* | up | 1607 | 1615 | + | 11.3303 | 4.96E-05 | 0.615 | GTGGGTGGC |
| 9579 | mamo-s | *BGIBMGA011221* | down | 1611 | 1619 | - | 11.2294 | 5.41E-05 | 0.65 | CTGCGAGGC |
| 9580 | mamo-s | *BGIBMGA011221* | down | 1233 | 1241 | - | 11.0826 | 6.84E-05 | 0.709 | GTACGTGGC |
| 9581 | mamo-s | *BGIBMGA011224* | up | 267 | 275 | - | 11.3303 | 4.96E-05 | 0.615 | GTGGGTGGC |
| 9582 | mamo-s | *BGIBMGA011224* | up | 479 | 487 | - | 11.3303 | 4.96E-05 | 0.615 | GTGGGTGGC |
| 9583 | mamo-s | *BGIBMGA011225* | up | 127 | 135 | + | 11.3303 | 4.96E-05 | 0.615 | GTGGGTGGC |
| 9584 | mamo-s | *BGIBMGA011225* | up | 495 | 503 | + | 11.3303 | 4.96E-05 | 0.615 | GTGGGTGGC |
| 9585 | mamo-s | *BGIBMGA011226* | up | 585 | 593 | - | 11.3303 | 4.96E-05 | 0.615 | GTGGGTGGC |

| | | | | | | | | | | |
|---|---|---|---|---|---|---|---|---|---|---|
| 9586 | mamo-s | *BGIBMGA011227* | up | 873 | 881 | - | 11.1743 | 5.90E-05 | 0.684 | CCGCGTGTC |
| 9587 | mamo-s | *BGIBMGA011228* | up | 712 | 720 | - | 11.3303 | 4.96E-05 | 0.615 | GTGGGTGGC |
| 9588 | mamo-s | *BGIBMGA011228* | up | 1215 | 1223 | - | 10.9358 | 7.30E-05 | 0.728 | GCGGGTGGC |
| 9589 | mamo-s | *BGIBMGA011235* | up | 1554 | 1562 | + | 10.5413 | 9.42E-05 | 0.857 | CAGCGTGTC |
| 9590 | mamo-s | *BGIBMGA011238* | up | 871 | 879 | + | 11.3303 | 4.96E-05 | 0.615 | GTGGGTGGC |
| 9591 | mamo-s | *BGIBMGA011239* | up | 416 | 424 | - | 13.6606 | 4.06E-06 | 0.615 | GCGCGTGGC |
| 9592 | mamo-s | *BGIBMGA011241* | up | 1477 | 1485 | + | 11.3303 | 4.96E-05 | 0.615 | GTGGGTGGC |
| 9593 | mamo-s | *BGIBMGA011242* | down | 1041 | 1049 | - | 12.1284 | 2.16E-05 | 0.615 | GTGCGAGGC |
| 9594 | mamo-s | *BGIBMGA011242* | down | 1034 | 1042 | + | 10.9358 | 7.30E-05 | 0.728 | GCGGGTGGC |
| 9595 | mamo-s | *BGIBMGA011242* | down | 541 | 549 | + | 10.6881 | 8.61E-05 | 0.81 | GCACGTGGC |
| 9596 | mamo-s | *BGIBMGA011243* | up | 1551 | 1559 | + | 10.6881 | 8.61E-05 | 0.81 | GCACGTGGC |
| 9597 | mamo-s | *BGIBMGA011244* | up | 80 | 88 | + | 11.3303 | 4.96E-05 | 0.615 | GTGGGTGGC |
| 9598 | mamo-s | *BGIBMGA011244* | down | 195 | 203 | + | 11.3303 | 4.96E-05 | 0.615 | GTGGGTGGC |
| 9599 | mamo-s | *BGIBMGA011244* | up | 495 | 503 | - | 11.3303 | 4.96E-05 | 0.615 | GTGGGTGGC |
| 9600 | mamo-s | *BGIBMGA011244* | down | 1233 | 1241 | + | 11.0826 | 6.84E-05 | 0.709 | GTACGTGGC |
| 9601 | mamo-s | *BGIBMGA011244* | up | 1468 | 1476 | - | 11.0826 | 6.84E-05 | 0.709 | GTACGTGGC |
| 9602 | mamo-s | *BGIBMGA011245* | down | 1171 | 1179 | - | 11.3303 | 4.96E-05 | 0.615 | GTGGGTGGC |
| 9603 | mamo-s | *BGIBMGA011245* | up | 1575 | 1583 | + | 11.3303 | 4.96E-05 | 0.615 | GTGGGTGGC |
| 9604 | mamo-s | *BGIBMGA011247* | down | 169 | 177 | + | 11.0826 | 6.84E-05 | 0.709 | GTACGTGGC |
| 9605 | mamo-s | *BGIBMGA011249* | down | 875 | 883 | + | 11.3303 | 4.96E-05 | 0.615 | GTGGGTGGC |
| 9606 | mamo-s | *BGIBMGA011249* | down | 1249 | 1257 | + | 11.0826 | 6.84E-05 | 0.709 | GTACGTGGC |
| 9607 | mamo-s | *BGIBMGA011251* | down | 1031 | 1039 | - | 11.3303 | 4.96E-05 | 0.615 | GTGGGTGGC |
| 9608 | mamo-s | *BGIBMGA011251* | down | 1806 | 1814 | + | 11.3303 | 4.96E-05 | 0.615 | GTGGGTGGC |
| 9609 | mamo-s | *BGIBMGA011252* | up | 1718 | 1726 | - | 11.3303 | 4.96E-05 | 0.615 | GTGGGTGGC |
| 9610 | mamo-s | *BGIBMGA011254* | down | 1140 | 1148 | + | 10.8349 | 7.66E-05 | 0.751 | CCGCGAGGC |
| 9611 | mamo-s | *BGIBMGA011254* | down | 316 | 324 | + | 10.5688 | 8.88E-05 | 0.824 | GTGCGTGAC |
| 9612 | mamo-s | *BGIBMGA011255* | up | 1965 | 1973 | + | 10.8349 | 7.66E-05 | 0.751 | CCGCGAGGC |
| 9613 | mamo-s | *BGIBMGA011255* | up | 1141 | 1149 | + | 10.5688 | 8.88E-05 | 0.824 | GTGCGTGAC |
| 9614 | mamo-s | *BGIBMGA011256* | down | 665 | 673 | - | 11.789 | 2.84E-05 | 0.615 | GCGCGTGCC |
| 9615 | mamo-s | *BGIBMGA011256* | up | 1392 | 1400 | - | 11.3303 | 4.96E-05 | 0.615 | GTGGGTGGC |
| 9616 | mamo-s | *BGIBMGA011256* | down | 1906 | 1914 | + | 11.3303 | 4.96E-05 | 0.615 | GTGGGTGGC |
| 9617 | mamo-s | *BGIBMGA011258* | up | 12 | 20 | - | 12.1284 | 2.16E-05 | 0.615 | GTGCGAGGC |
| 9618 | mamo-s | *BGIBMGA011258* | down | 218 | 226 | + | 11.3303 | 4.96E-05 | 0.615 | GTGGGTGGC |
| 9619 | mamo-s | *BGIBMGA011259* | down | 657 | 665 | + | 12.3394 | 1.49E-05 | 0.615 | GGGCGTGGC |
| 9620 | mamo-s | *BGIBMGA011259* | down | 1506 | 1514 | - | 11.4404 | 4.51E-05 | 0.615 | GAGCGTGTC |
| 9621 | mamo-s | *BGIBMGA011259* | up | 710 | 718 | + | 11.3303 | 4.96E-05 | 0.615 | GTGGGTGGC |
| 9622 | mamo-s | *BGIBMGA011260* | up | 792 | 800 | + | 12.4679 | 1.31E-05 | 0.615 | GTGCGTGTC |
| 9623 | mamo-s | *BGIBMGA011260* | up | 1434 | 1442 | + | 12.0734 | 2.66E-05 | 0.615 | GCGCGTGTC |
| 9624 | mamo-s | *BGIBMGA011260* | down | 1346 | 1354 | - | 10.7523 | 8.38E-05 | 0.797 | GGGCGTGTC |
| 9625 | mamo-s | *BGIBMGA011260* | up | 639 | 647 | + | 10.5138 | 9.75E-05 | 0.865 | GTGCGTGTT |
| 9626 | mamo-s | *BGIBMGA011261* | up | 1881 | 1889 | + | 11.789 | 2.84E-05 | 0.615 | GCGCGTGCC |
| 9627 | mamo-s | *BGIBMGA011262* | down | 1533 | 1541 | - | 12.1009 | 2.43E-05 | 0.615 | GTGCGTGGT |
| 9628 | mamo-s | *BGIBMGA011262* | down | 952 | 960 | + | 11.3303 | 4.96E-05 | 0.615 | GTGGGTGGC |
| 9629 | mamo-s | *BGIBMGA011262* | up | 842 | 850 | - | 10.8073 | 7.89E-05 | 0.766 | CCGCGTGGT |
| 9630 | mamo-s | *BGIBMGA011262* | up | 1143 | 1151 | + | 10.5688 | 8.88E-05 | 0.824 | GTGCGTGAC |
| 9631 | mamo-s | *BGIBMGA011266* | down | 1952 | 1960 | - | 14.055 | 2.23E-06 | 0.615 | GTGCGTGGC |
| 9632 | mamo-s | *BGIBMGA011266* | down | 1188 | 1196 | - | 11.3303 | 4.96E-05 | 0.615 | GTGGGTGGC |
| 9633 | mamo-s | *BGIBMGA011267* | up | 49 | 57 | + | 11.3303 | 4.96E-05 | 0.615 | GTGGGTGGC |
| 9634 | mamo-s | *BGIBMGA011267* | down | 293 | 301 | - | 11.3303 | 4.96E-05 | 0.615 | GTGGGTGGC |
| 9635 | mamo-s | *BGIBMGA011268* | up | 732 | 740 | - | 12.0734 | 2.66E-05 | 0.615 | GCGCGTGTC |
| 9636 | mamo-s | *BGIBMGA011268* | up | 743 | 751 | - | 11.4404 | 4.51E-05 | 0.615 | CGGCGTGGC |
| 9637 | mamo-s | *BGIBMGA011268* | down | 1160 | 1168 | - | 11.3303 | 4.96E-05 | 0.615 | GTGGGTGGC |
| 9638 | mamo-s | *BGIBMGA011268* | up | 347 | 355 | + | 11.0734 | 7.12E-05 | 0.726 | GAGCGTGGT |
| 9639 | mamo-s | *BGIBMGA011269* | down | 579 | 587 | + | 11.5321 | 4.06E-05 | 0.615 | TTGCGTGGC |
| 9640 | mamo-s | *BGIBMGA011269* | up | 1404 | 1412 | - | 11.3303 | 4.96E-05 | 0.615 | GTGGGTGGC |
| 9641 | mamo-s | *BGIBMGA011270* | down | 1786 | 1794 | - | 12.7615 | 1.04E-05 | 0.615 | CCGCGTGGC |
| 9642 | mamo-s | *BGIBMGA011270* | down | 1404 | 1412 | + | 11.0826 | 6.84E-05 | 0.709 | GTACGTGGC |
| 9643 | mamo-s | *BGIBMGA011271* | up | 90 | 98 | - | 10.9358 | 7.30E-05 | 0.728 | GCGGGTGGC |
| 9644 | mamo-s | *BGIBMGA011273* | down | 1857 | 1865 | + | 13.156 | 6.30E-06 | 0.615 | CTGCGTGGC |
| 9645 | mamo-s | *BGIBMGA011273* | down | 1349 | 1357 | + | 11.3303 | 4.96E-05 | 0.615 | GTGGGTGGC |
| 9646 | mamo-s | *BGIBMGA011276* | down | 919 | 927 | + | 11.3303 | 4.96E-05 | 0.615 | GTGGGTGGC |
| 9647 | mamo-s | *BGIBMGA011276* | down | 1835 | 1843 | - | 11.3303 | 4.96E-05 | 0.615 | GTGGGTGGC |
| 9648 | mamo-s | *BGIBMGA011279* | down | 166 | 174 | - | 11.3303 | 4.96E-05 | 0.615 | GTGGGTGGC |
| 9649 | mamo-s | *BGIBMGA011280* | up | 1154 | 1162 | + | 13.156 | 6.30E-06 | 0.615 | CTGCGTGGC |
| 9650 | mamo-s | *BGIBMGA011280* | up | 1787 | 1795 | + | 10.7523 | 8.38E-05 | 0.797 | GGGCGTGTC |
| 9651 | mamo-s | *BGIBMGA011281* | down | 1148 | 1156 | - | 10.8073 | 7.89E-05 | 0.766 | CCGCGTGGT |
| 9652 | mamo-s | *BGIBMGA011283* | up | 1532 | 1540 | - | 10.5413 | 9.42E-05 | 0.857 | CAGCGTGTC |
| 9653 | mamo-s | *BGIBMGA011284* | up | 971 | 979 | - | 10.5138 | 9.75E-05 | 0.865 | GTGCGTGTT |
| 9654 | mamo-s | *BGIBMGA011285* | up | 158 | 166 | + | 11.3303 | 4.96E-05 | 0.615 | GTGGGTGGC |
| 9655 | mamo-s | *BGIBMGA011301* | down | 1529 | 1537 | + | 11.3303 | 4.96E-05 | 0.615 | GTGGGTGGC |
| 9656 | mamo-s | *BGIBMGA011302* | up | 500 | 508 | + | 11.3303 | 4.96E-05 | 0.615 | GTGGGTGGC |
| 9657 | mamo-s | *BGIBMGA011302* | up | 1881 | 1889 | + | 11.3303 | 4.96E-05 | 0.615 | GTGGGTGGC |
| 9658 | mamo-s | *BGIBMGA011303* | up | 288 | 296 | - | 13.0275 | 8.53E-06 | 0.615 | GAGCGTGGC |
| 9659 | mamo-s | *BGIBMGA011306* | down | 382 | 390 | - | 11.3303 | 4.96E-05 | 0.615 | GTGGGTGGC |
| 9660 | mamo-s | *BGIBMGA011309* | up | 1111 | 1119 | + | 11.1743 | 5.90E-05 | 0.684 | CCGCGTGTC |
| 9661 | mamo-s | *BGIBMGA011312* | up | 1593 | 1601 | - | 10.8349 | 7.66E-05 | 0.751 | CCGCGAGGC |
| 9662 | mamo-s | *BGIBMGA011313* | down | 1191 | 1199 | - | 11.5688 | 3.79E-05 | 0.615 | CTGCGTGTC |
| 9663 | mamo-s | *BGIBMGA011316* | up | 1540 | 1548 | + | 11.4404 | 4.51E-05 | 0.615 | GAGCGTGTC |
| 9664 | mamo-s | *BGIBMGA011317* | down | 482 | 490 | + | 11.3303 | 4.96E-05 | 0.615 | GTGGGTGGC |
| 9665 | mamo-s | *BGIBMGA011318* | down | 269 | 277 | + | 11.1743 | 5.90E-05 | 0.684 | CCGCGTGTC |
| 9666 | mamo-s | *BGIBMGA011319* | down | 1983 | 1991 | - | 10.7523 | 8.38E-05 | 0.797 | GGGCGTGTC |
| 9667 | mamo-s | *BGIBMGA011320* | up | 1350 | 1358 | + | 11.3303 | 4.96E-05 | 0.615 | GTGGGTGGC |
| 9668 | mamo-s | *BGIBMGA011321* | up | 1096 | 1104 | + | 10.5138 | 9.75E-05 | 0.865 | GTGCGTGTT |
| 9669 | mamo-s | *BGIBMGA011322* | up | 1437 | 1445 | - | 12.1284 | 2.16E-05 | 0.615 | CAGCGTGGC |
| 9670 | mamo-s | *BGIBMGA011326* | up | 270 | 278 | - | 11.7798 | 3.11E-05 | 0.615 | ATGCGTGGC |
| 9671 | mamo-s | *BGIBMGA011327* | down | 886 | 894 | - | 12.4679 | 1.31E-05 | 0.615 | GTGCGTGTC |
| 9672 | mamo-s | *BGIBMGA011327* | down | 419 | 427 | - | 11.3303 | 4.96E-05 | 0.615 | GTGGGTGGC |
| 9673 | mamo-s | *BGIBMGA011327* | up | 909 | 917 | + | 10.9358 | 7.30E-05 | 0.728 | GCGGGTGGC |

| | | | | | | | | | | |
|---|---|---|---|---|---|---|---|---|---|---|
| 9674 | mamo-s | *BGIBMGA011328* | down | 652 | 660 | + | 11.3303 | 4.96E-05 | 0.615 | GTGGGTGGC |
| 9675 | mamo-s | *BGIBMGA011328* | down | 1128 | 1136 | - | 11.3303 | 4.96E-05 | 0.615 | GTGGGTGGC |
| 9676 | mamo-s | *BGIBMGA011330* | up | 1203 | 1211 | + | 10.6881 | 8.61E-05 | 0.81 | GCACGTGGC |
| 9677 | mamo-s | *BGIBMGA011331* | down | 914 | 922 | - | 12.3394 | 1.49E-05 | 0.615 | GGGCGTGGC |
| 9678 | mamo-s | *BGIBMGA011331* | down | 954 | 962 | - | 12.3394 | 1.49E-05 | 0.615 | GGGCGTGGC |
| 9679 | mamo-s | *BGIBMGA011333* | up | 469 | 477 | + | 11.1743 | 5.90E-05 | 0.684 | CCGCGTGTC |
| 9680 | mamo-s | *BGIBMGA011334* | down | 1652 | 1660 | + | 11.3303 | 4.96E-05 | 0.615 | GTGGGTGGC |
| 9681 | mamo-s | *BGIBMGA011336* | down | 460 | 468 | + | 10.8899 | 7.48E-05 | 0.739 | CCGCGTGCC |
| 9682 | mamo-s | *BGIBMGA011338* | down | 1517 | 1525 | - | 11.5321 | 4.06E-05 | 0.615 | TTGCGTGGC |
| 9683 | mamo-s | *BGIBMGA011338* | up | 1655 | 1663 | + | 11.3303 | 4.96E-05 | 0.615 | GTGGGTGGC |
| 9684 | mamo-s | *BGIBMGA011340* | down | 1074 | 1082 | - | 11.7064 | 3.52E-05 | 0.615 | GCGCGTGGT |
| 9685 | mamo-s | *BGIBMGA011340* | down | 951 | 959 | + | 10.9358 | 7.30E-05 | 0.728 | GCGGGTGGC |
| 9686 | mamo-s | *BGIBMGA011341* | up | 1114 | 1122 | - | 11.0734 | 7.12E-05 | 0.726 | GAGCGTGGT |
| 9687 | mamo-s | *BGIBMGA011342* | down | 300 | 308 | + | 11.0826 | 6.84E-05 | 0.709 | GTACGTGGC |
| 9688 | mamo-s | *BGIBMGA011346* | down | 190 | 198 | + | 11.3303 | 4.96E-05 | 0.615 | GTGGGTGGC |
| 9689 | mamo-s | *BGIBMGA011346* | down | 810 | 818 | + | 11.1743 | 5.90E-05 | 0.684 | CCGCGTGTC |
| 9690 | mamo-s | *BGIBMGA011347* | down | 1106 | 1114 | - | 11.3303 | 4.96E-05 | 0.615 | GTGGGTGGC |
| 9691 | mamo-s | *BGIBMGA011348* | down | 737 | 745 | + | 11.4404 | 4.51E-05 | 0.615 | GAGCGTGTC |
| 9692 | mamo-s | *BGIBMGA011348* | down | 1766 | 1774 | - | 10.9358 | 7.30E-05 | 0.728 | GCGGGTGGC |
| 9693 | mamo-s | *BGIBMGA011349* | up | 7 | 15 | + | 11.4404 | 4.51E-05 | 0.615 | CGGCGTGGC |
| 9694 | mamo-s | *BGIBMGA011350* | down | 968 | 976 | - | 12.3394 | 1.49E-05 | 0.615 | GGGCGTGGC |
| 9695 | mamo-s | *BGIBMGA011350* | down | 1978 | 1986 | + | 11.3303 | 4.96E-05 | 0.615 | GTGGGTGGC |
| 9696 | mamo-s | *BGIBMGA011352* | up | 1151 | 1159 | + | 11.3303 | 4.96E-05 | 0.615 | GTGGGTGGC |
| 9697 | mamo-s | *BGIBMGA011353* | up | 610 | 618 | + | 10.5138 | 9.75E-05 | 0.865 | GTGCGTGTT |
| 9698 | mamo-s | *BGIBMGA011355* | down | 1965 | 1973 | - | 11.3303 | 4.96E-05 | 0.615 | GTGGGTGGC |
| 9699 | mamo-s | *BGIBMGA011356* | up | 957 | 965 | - | 11.7339 | 3.29E-05 | 0.615 | GCGCGAGGC |
| 9700 | mamo-s | *BGIBMGA011356* | down | 1298 | 1306 | - | 11.3303 | 4.96E-05 | 0.615 | GTGGGTGGC |
| 9701 | mamo-s | *BGIBMGA011356* | down | 924 | 932 | - | 11.0826 | 6.84E-05 | 0.709 | GTACGTGGC |
| 9702 | mamo-s | *BGIBMGA011359* | down | 345 | 353 | - | 10.8073 | 7.89E-05 | 0.766 | CCGCGTGGT |
| 9703 | mamo-s | *BGIBMGA011360* | up | 1553 | 1561 | + | 12.1009 | 2.43E-05 | 0.615 | GTGCGTGGT |
| 9704 | mamo-s | *BGIBMGA011362* | down | 204 | 212 | + | 11.3303 | 4.96E-05 | 0.615 | GTGGGTGGC |
| 9705 | mamo-s | *BGIBMGA011362* | up | 1517 | 1525 | + | 11.3303 | 4.96E-05 | 0.615 | GTGGGTGGC |
| 9706 | mamo-s | *BGIBMGA011364* | up | 490 | 498 | - | 11.3303 | 4.96E-05 | 0.615 | GTGGGTGGC |
| 9707 | mamo-s | *BGIBMGA011365* | up | 586 | 594 | + | 11.0826 | 6.84E-05 | 0.709 | GTACGTGGC |
| 9708 | mamo-s | *BGIBMGA011368* | up | 869 | 877 | - | 11.3303 | 4.96E-05 | 0.615 | GTGGGTGGC |
| 9709 | mamo-s | *BGIBMGA011369* | down | 1481 | 1489 | - | 11.3303 | 4.96E-05 | 0.615 | GTGGGTGGC |
| 9710 | mamo-s | *BGIBMGA011369* | up | 874 | 882 | + | 11.1009 | 6.57E-05 | 0.709 | GAGCGAGGC |
| 9711 | mamo-s | *BGIBMGA011370* | up | 1689 | 1697 | - | 11.3303 | 4.96E-05 | 0.615 | GTGGGTGGC |
| 9712 | mamo-s | *BGIBMGA011370* | up | 1331 | 1339 | + | 11.2844 | 5.18E-05 | 0.63 | CTGCGTGCC |
| 9713 | mamo-s | *BGIBMGA011370* | up | 754 | 762 | + | 10.7523 | 8.38E-05 | 0.797 | AAGCGTGGC |
| 9714 | mamo-s | *BGIBMGA011371* | down | 113 | 121 | - | 11.3303 | 4.96E-05 | 0.615 | GTGGGTGGC |
| 9715 | mamo-s | *BGIBMGA011371* | down | 1577 | 1585 | - | 11.3303 | 4.96E-05 | 0.615 | GTGGGTGGC |
| 9716 | mamo-s | *BGIBMGA011371* | down | 1927 | 1935 | + | 11.3303 | 4.96E-05 | 0.615 | GTGGGTGGC |
| 9717 | mamo-s | *BGIBMGA011373* | down | 19 | 27 | - | 11.7064 | 3.52E-05 | 0.615 | GCGCGTGGT |
| 9718 | mamo-s | *BGIBMGA011373* | down | 1938 | 1946 | + | 11.3853 | 4.74E-05 | 0.615 | ACGCGTGGC |
| 9719 | mamo-s | *BGIBMGA011373* | down | 252 | 260 | - | 11.3303 | 4.96E-05 | 0.615 | GTGGGTGGC |
| 9720 | mamo-s | *BGIBMGA011373* | up | 1503 | 1511 | - | 10.9358 | 7.30E-05 | 0.728 | GCGGGTGGC |
| 9721 | mamo-s | *BGIBMGA011374* | up | 1763 | 1771 | - | 11.3303 | 4.96E-05 | 0.615 | GTGGGTGGC |
| 9722 | mamo-s | *BGIBMGA011375* | down | 891 | 899 | - | 12.1284 | 2.16E-05 | 0.615 | CAGCGTGGC |
| 9723 | mamo-s | *BGIBMGA011375* | down | 930 | 938 | - | 12.0734 | 2.66E-05 | 0.615 | GCGCGTGTC |
| 9724 | mamo-s | *BGIBMGA011375* | down | 377 | 385 | + | 11.3303 | 4.96E-05 | 0.615 | GTGGGTGGC |
| 9725 | mamo-s | *BGIBMGA011375* | down | 1091 | 1099 | - | 11.3303 | 4.96E-05 | 0.615 | GTGGGTGGC |
| 9726 | mamo-s | *BGIBMGA011376* | up | 510 | 518 | - | 14.055 | 2.23E-06 | 0.615 | GTGCGTGGC |
| 9727 | mamo-s | *BGIBMGA011376* | down | 195 | 203 | - | 12.1284 | 2.16E-05 | 0.615 | GTGCGAGGC |
| 9728 | mamo-s | *BGIBMGA011377* | down | 948 | 956 | + | 11.4404 | 4.51E-05 | 0.615 | CGGCGTGGC |
| 9729 | mamo-s | *BGIBMGA011378* | down | 1390 | 1398 | + | 14.055 | 2.23E-06 | 0.615 | GTGCGTGGC |
| 9730 | mamo-s | *BGIBMGA011379* | down | 1667 | 1675 | + | 12.1284 | 2.16E-05 | 0.615 | CAGCGTGGC |
| 9731 | mamo-s | *BGIBMGA011379* | down | 1891 | 1899 | - | 12.1009 | 2.43E-05 | 0.615 | GTGCGTGGT |
| 9732 | mamo-s | *BGIBMGA011383* | down | 337 | 345 | + | 11.3303 | 4.96E-05 | 0.615 | GTGGGTGGC |
| 9733 | mamo-s | *BGIBMGA011383* | up | 971 | 979 | - | 11.3303 | 4.96E-05 | 0.615 | GTGGGTGGC |
| 9734 | mamo-s | *BGIBMGA011383* | down | 701 | 709 | - | 11.0826 | 6.84E-05 | 0.709 | GTACGTGGC |
| 9735 | mamo-s | *BGIBMGA011387* | down | 212 | 220 | + | 11.3303 | 4.96E-05 | 0.615 | GTGGGTGGC |
| 9736 | mamo-s | *BGIBMGA011389* | up | 102 | 110 | - | 11.3303 | 4.96E-05 | 0.615 | GTGGGTGGC |
| 9737 | mamo-s | *BGIBMGA011389* | up | 215 | 223 | - | 11.3303 | 4.96E-05 | 0.615 | GTGGGTGGC |
| 9738 | mamo-s | *BGIBMGA011390* | up | 969 | 977 | + | 11.5321 | 4.06E-05 | 0.615 | TTGCGTGGC |
| 9739 | mamo-s | *BGIBMGA011390* | up | 437 | 445 | - | 11.1743 | 5.90E-05 | 0.684 | CCGCGTGTC |
| 9740 | mamo-s | *BGIBMGA011393* | up | 615 | 623 | + | 11.0826 | 6.84E-05 | 0.709 | GTACGTGGC |
| 9741 | mamo-s | *BGIBMGA011394* | down | 272 | 280 | - | 11.3303 | 4.96E-05 | 0.615 | GTGGGTGGC |
| 9742 | mamo-s | *BGIBMGA011394* | down | 1039 | 1047 | - | 11.3303 | 4.96E-05 | 0.615 | GTGGGTGGC |
| 9743 | mamo-s | *BGIBMGA011398* | down | 166 | 174 | - | 11.3303 | 4.96E-05 | 0.615 | GTGGGTGGC |
| 9744 | mamo-s | *BGIBMGA011399* | up | 1509 | 1517 | - | 12.4679 | 1.31E-05 | 0.615 | GTGCGTGTC |
| 9745 | mamo-s | *BGIBMGA011399* | down | 1791 | 1799 | + | 11.3303 | 4.96E-05 | 0.615 | GTGGGTGGC |
| 9746 | mamo-s | *BGIBMGA011400* | up | 536 | 544 | + | 11.3303 | 4.96E-05 | 0.615 | GTGGGTGGC |
| 9747 | mamo-s | *BGIBMGA011400* | up | 1837 | 1845 | + | 11.3303 | 4.96E-05 | 0.615 | GTGGGTGGC |
| 9748 | mamo-s | *BGIBMGA011401* | down | 1200 | 1208 | - | 12.7615 | 1.04E-05 | 0.615 | CCGCGTGGC |
| 9749 | mamo-s | *BGIBMGA011402* | up | 582 | 590 | - | 11.3303 | 4.96E-05 | 0.615 | GTGGGTGGC |
| 9750 | mamo-s | *BGIBMGA011403* | down | 990 | 998 | - | 14.055 | 2.23E-06 | 0.615 | GTGCGTGGC |
| 9751 | mamo-s | *BGIBMGA011403* | up | 1494 | 1502 | - | 11.4404 | 4.51E-05 | 0.615 | CGGCGTGGC |
| 9752 | mamo-s | *BGIBMGA011405* | up | 579 | 587 | + | 11.3303 | 4.96E-05 | 0.615 | GTGGGTGGC |
| 9753 | mamo-s | *BGIBMGA011405* | down | 1823 | 1831 | - | 11.3303 | 4.96E-05 | 0.615 | GTGGGTGGC |
| 9754 | mamo-s | *BGIBMGA011405* | down | 1448 | 1456 | + | 10.5413 | 9.42E-05 | 0.857 | CAGCGTGTC |
| 9755 | mamo-s | *BGIBMGA011406* | down | 1648 | 1656 | + | 11.3303 | 4.96E-05 | 0.615 | GTGGGTGGC |
| 9756 | mamo-s | *BGIBMGA011408* | down | 835 | 843 | - | 11.3303 | 4.96E-05 | 0.615 | GTGGGTGGC |
| 9757 | mamo-s | *BGIBMGA011408* | up | 1381 | 1389 | + | 11.3303 | 4.96E-05 | 0.615 | GTGGGTGGC |
| 9758 | mamo-s | *BGIBMGA011408* | up | 1704 | 1712 | - | 10.9358 | 7.30E-05 | 0.728 | GCGGGTGGC |
| 9759 | mamo-s | *BGIBMGA011409* | up | 1801 | 1809 | - | 11.3303 | 4.96E-05 | 0.615 | GTGGGTGGC |
| 9760 | mamo-s | *BGIBMGA011410* | down | 1729 | 1737 | - | 11.4404 | 4.51E-05 | 0.615 | GAGCGTGTC |
| 9761 | mamo-s | *BGIBMGA011410* | down | 1150 | 1158 | + | 11.3303 | 4.96E-05 | 0.615 | GTGGGTGGC |

| | | | | | | | | | | |
|---|---|---|---|---|---|---|---|---|---|---|
| 9762 | mamo-s | *BGIBMGA011411* | down | 1652 | 1660 | - | 11.3303 | 4.96E-05 | 0.615 | GTGGGTGGC |
| 9763 | mamo-s | *BGIBMGA011412* | down | 166 | 174 | + | 11.3303 | 4.96E-05 | 0.615 | GTGGGTGGC |
| 9764 | mamo-s | *BGIBMGA011412* | up | 1267 | 1275 | - | 11.3303 | 4.96E-05 | 0.615 | GTGGGTGGC |
| 9765 | mamo-s | *BGIBMGA011414* | down | 703 | 711 | - | 11.3303 | 4.96E-05 | 0.615 | GTGGGTGGC |
| 9766 | mamo-s | *BGIBMGA011414* | up | 1345 | 1353 | + | 11.0826 | 6.84E-05 | 0.709 | GTACGTGGC |
| 9767 | mamo-s | *BGIBMGA011414* | up | 1425 | 1433 | + | 10.9358 | 7.30E-05 | 0.728 | GCGGGTGGC |
| 9768 | mamo-s | *BGIBMGA011416* | up | 66 | 74 | - | 11.3303 | 4.96E-05 | 0.615 | GTGGGTGGC |
| 9769 | mamo-s | *BGIBMGA011416* | down | 494 | 502 | + | 11.3303 | 4.96E-05 | 0.615 | GTGGGTGGC |
| 9770 | mamo-s | *BGIBMGA011418* | up | 1555 | 1563 | - | 11.3303 | 4.96E-05 | 0.615 | GTGGGTGGC |
| 9771 | mamo-s | *BGIBMGA011418* | down | 1877 | 1885 | + | 11.3303 | 4.96E-05 | 0.615 | GTGGGTGGC |
| 9772 | mamo-s | *BGIBMGA011419* | up | 1556 | 1564 | - | 11.3303 | 4.96E-05 | 0.615 | GTGGGTGGC |
| 9773 | mamo-s | *BGIBMGA011419* | down | 67 | 75 | + | 10.5138 | 9.75E-05 | 0.865 | GTGCGTGTT |
| 9774 | mamo-s | *BGIBMGA011420* | down | 1407 | 1415 | - | 12.1835 | 1.71E-05 | 0.615 | GTGCGTGCC |
| 9775 | mamo-s | *BGIBMGA011420* | up | 535 | 543 | - | 10.9358 | 7.30E-05 | 0.728 | GCGGGTGGC |
| 9776 | mamo-s | *BGIBMGA011421* | down | 1131 | 1139 | + | 11.3303 | 4.96E-05 | 0.615 | GTGGGTGGC |
| 9777 | mamo-s | *BGIBMGA011422* | down | 878 | 886 | + | 11.3303 | 4.96E-05 | 0.615 | GTGGGTGGC |
| 9778 | mamo-s | *BGIBMGA011424* | up | 937 | 945 | - | 11.3303 | 4.96E-05 | 0.615 | GTGGGTGGC |
| 9779 | mamo-s | *BGIBMGA011425* | up | 198 | 206 | - | 11.3303 | 4.96E-05 | 0.615 | GTGGGTGGC |
| 9780 | mamo-s | *BGIBMGA011425* | down | 955 | 963 | + | 11.3303 | 4.96E-05 | 0.615 | GTGGGTGGC |
| 9781 | mamo-s | *BGIBMGA011425* | down | 1592 | 1600 | + | 11.3303 | 4.96E-05 | 0.615 | GTGGGTGGC |
| 9782 | mamo-s | *BGIBMGA011430* | up | 351 | 359 | - | 11.3303 | 4.96E-05 | 0.615 | GTGGGTGGC |
| 9783 | mamo-s | *BGIBMGA011432* | up | 730 | 738 | + | 11.3303 | 4.96E-05 | 0.615 | GTGGGTGGC |
| 9784 | mamo-s | *BGIBMGA011434* | up | 250 | 258 | + | 13.6606 | 4.06E-06 | 0.615 | GCGCGTGGC |
| 9785 | mamo-s | *BGIBMGA011437* | down | 816 | 824 | + | 11.3303 | 4.96E-05 | 0.615 | GTGGGTGGC |
| 9786 | mamo-s | *BGIBMGA011439* | up | 744 | 752 | + | 11.7798 | 3.11E-05 | 0.615 | ATGCGTGGC |
| 9787 | mamo-s | *BGIBMGA011440* | up | 1501 | 1509 | + | 11.789 | 2.84E-05 | 0.615 | GCGCGTGCC |
| 9788 | mamo-s | *BGIBMGA011441* | up | 1958 | 1966 | - | 10.5413 | 9.42E-05 | 0.857 | GTGCGAGTC |
| 9789 | mamo-s | *BGIBMGA011442* | down | 900 | 908 | - | 11.0826 | 6.84E-05 | 0.709 | GTACGTGGC |
| 9790 | mamo-s | *BGIBMGA011443* | down | 1225 | 1233 | + | 14.055 | 2.23E-06 | 0.615 | GTGCGTGGC |
| 9791 | mamo-s | *BGIBMGA011444* | up | 1868 | 1876 | + | 14.055 | 2.23E-06 | 0.615 | GTGCGTGGC |
| 9792 | mamo-s | *BGIBMGA011446* | down | 1278 | 1286 | + | 11.3303 | 4.96E-05 | 0.615 | GTGGGTGGC |
| 9793 | mamo-s | *BGIBMGA011451* | up | 566 | 574 | - | 10.7523 | 8.38E-05 | 0.797 | GGGCGTGTC |
| 9794 | mamo-s | *BGIBMGA011451* | down | 1316 | 1324 | - | 10.5138 | 9.75E-05 | 0.865 | GTGCGTGTT |
| 9795 | mamo-s | *BGIBMGA011452* | down | 459 | 467 | - | 11.3303 | 4.96E-05 | 0.615 | GTGGGTGGC |
| 9796 | mamo-s | *BGIBMGA011453* | down | 1243 | 1251 | + | 11.3303 | 4.96E-05 | 0.615 | GTGGGTGGC |
| 9797 | mamo-s | *BGIBMGA011454* | down | 1065 | 1073 | + | 11.3303 | 4.96E-05 | 0.615 | GTGGGTGGC |
| 9798 | mamo-s | *BGIBMGA011458* | up | 456 | 464 | - | 12.1284 | 2.16E-05 | 0.615 | CAGCGTGGC |
| 9799 | mamo-s | *BGIBMGA011460* | up | 1080 | 1088 | + | 11.3303 | 4.96E-05 | 0.615 | GTGGGTGGC |
| 9800 | mamo-s | *BGIBMGA011462* | down | 585 | 593 | + | 11.3303 | 4.96E-05 | 0.615 | GTGGGTGGC |
| 9801 | mamo-s | *BGIBMGA011465* | up | 1056 | 1064 | - | 10.5413 | 9.42E-05 | 0.857 | GTGCGAGTC |
| 9802 | mamo-s | *BGIBMGA011466* | down | 1123 | 1131 | - | 11.3303 | 4.96E-05 | 0.615 | GTGGGTGGC |
| 9803 | mamo-s | *BGIBMGA011466* | up | 1299 | 1307 | - | 11.0826 | 6.84E-05 | 0.709 | GTACGTGGC |
| 9804 | mamo-s | *BGIBMGA011467* | up | 1174 | 1182 | + | 11.3303 | 4.96E-05 | 0.615 | GTGGGTGGC |
| 9805 | mamo-s | *BGIBMGA011468* | down | 1929 | 1937 | + | 11.3303 | 4.96E-05 | 0.615 | GTGGGTGGC |
| 9806 | mamo-s | *BGIBMGA011470* | down | 1326 | 1334 | + | 11.3303 | 4.96E-05 | 0.615 | GTGGGTGGC |
| 9807 | mamo-s | *BGIBMGA011472* | up | 503 | 511 | + | 11.3303 | 4.96E-05 | 0.615 | GTGGGTGGC |
| 9808 | mamo-s | *BGIBMGA011475* | down | 880 | 888 | - | 11.3303 | 4.96E-05 | 0.615 | GTGGGTGGC |
| 9809 | mamo-s | *BGIBMGA011476* | down | 1337 | 1345 | - | 11.3303 | 4.96E-05 | 0.615 | GTGGGTGGC |
| 9810 | mamo-s | *BGIBMGA011477* | up | 1341 | 1349 | + | 13.6606 | 4.06E-06 | 0.615 | GCGCGTGGC |
| 9811 | mamo-s | *BGIBMGA011477* | up | 838 | 846 | - | 11.3303 | 4.96E-05 | 0.615 | GTGGGTGGC |
| 9812 | mamo-s | *BGIBMGA011478* | down | 135 | 143 | + | 11.3303 | 4.96E-05 | 0.615 | GTGGGTGGC |
| 9813 | mamo-s | *BGIBMGA011483* | up | 1377 | 1385 | - | 11.3303 | 4.96E-05 | 0.615 | GTGGGTGGC |
| 9814 | mamo-s | *BGIBMGA011483* | down | 824 | 832 | + | 11.156 | 6.13E-05 | 0.705 | GAGCGTGCC |
| 9815 | mamo-s | *BGIBMGA011487* | up | 1793 | 1801 | - | 11.3853 | 4.74E-05 | 0.615 | ACGCGTGGC |
| 9816 | mamo-s | *BGIBMGA011488* | up | 1723 | 1731 | + | 11.3303 | 4.96E-05 | 0.615 | GTGGGTGGC |
| 9817 | mamo-s | *BGIBMGA011490* | down | 977 | 985 | + | 12.3394 | 1.49E-05 | 0.615 | GGGCGTGGC |
| 9818 | mamo-s | *BGIBMGA011491* | down | 1757 | 1765 | - | 11.2018 | 5.68E-05 | 0.674 | CTGCGTGGT |
| 9819 | mamo-s | *BGIBMGA011491* | up | 7 | 15 | + | 10.7523 | 8.38E-05 | 0.797 | AAGCGTGGC |
| 9820 | mamo-s | *BGIBMGA011492* | up | 1346 | 1354 | + | 11.3303 | 4.96E-05 | 0.615 | GTGGGTGGC |
| 9821 | mamo-s | *BGIBMGA011494* | down | 829 | 837 | - | 11.2018 | 5.68E-05 | 0.674 | CTGCGTGGT |
| 9822 | mamo-s | *BGIBMGA011495* | down | 922 | 930 | + | 11.3303 | 4.96E-05 | 0.615 | GTGGGTGGC |
| 9823 | mamo-s | *BGIBMGA011499* | up | 1127 | 1135 | - | 10.5413 | 9.42E-05 | 0.857 | CAGCGTGTC |
| 9824 | mamo-s | *BGIBMGA011502* | up | 197 | 205 | - | 13.6606 | 4.06E-06 | 0.615 | GCGCGTGGC |
| 9825 | mamo-s | *BGIBMGA011502* | up | 283 | 291 | - | 11.789 | 2.84E-05 | 0.615 | GCGCGTGCC |
| 9826 | mamo-s | *BGIBMGA011503* | up | 970 | 978 | + | 11.3303 | 4.96E-05 | 0.615 | GTGGGTGGC |
| 9827 | mamo-s | *BGIBMGA011506* | down | 1245 | 1253 | + | 11.3303 | 4.96E-05 | 0.615 | GTGGGTGGC |
| 9828 | mamo-s | *BGIBMGA011510* | up | 888 | 896 | - | 11.3303 | 4.96E-05 | 0.615 | GTGGGTGGC |
| 9829 | mamo-s | *BGIBMGA011511* | down | 893 | 901 | - | 10.9358 | 7.30E-05 | 0.728 | GCGGGTGGC |
| 9830 | mamo-s | *BGIBMGA011514* | up | 554 | 562 | - | 11.3303 | 4.96E-05 | 0.615 | GTGGGTGGC |
| 9831 | mamo-s | *BGIBMGA011515* | down | 1040 | 1048 | - | 13.6606 | 4.06E-06 | 0.615 | GCGCGTGGC |
| 9832 | mamo-s | *BGIBMGA011516* | up | 1106 | 1114 | - | 11.3303 | 4.96E-05 | 0.615 | GTGGGTGGC |
| 9833 | mamo-s | *BGIBMGA011519* | down | 1906 | 1914 | + | 13.156 | 6.30E-06 | 0.615 | CTGCGTGGC |
| 9834 | mamo-s | *BGIBMGA011522* | down | 97 | 105 | + | 11.2018 | 5.68E-05 | 0.674 | CTGCGTGGT |
| 9835 | mamo-s | *BGIBMGA011523* | up | 1676 | 1684 | - | 10.7523 | 8.38E-05 | 0.797 | AAGCGTGGC |
| 9836 | mamo-s | *BGIBMGA011525* | up | 769 | 777 | - | 12.0734 | 2.66E-05 | 0.615 | GCGCGTGTC |
| 9837 | mamo-s | *BGIBMGA011526* | up | 1431 | 1439 | + | 11.1009 | 6.57E-05 | 0.709 | GAGCGAGGC |
| 9838 | mamo-s | *BGIBMGA011531* | down | 945 | 953 | + | 11.7339 | 3.29E-05 | 0.615 | GCGCGAGGC |
| 9839 | mamo-s | *BGIBMGA011531* | up | 343 | 351 | + | 11.3303 | 4.96E-05 | 0.615 | GTGGGTGGC |
| 9840 | mamo-s | *BGIBMGA011531* | up | 882 | 890 | + | 11.0826 | 6.84E-05 | 0.709 | GTACGTGGC |
| 9841 | mamo-s | *BGIBMGA011534* | down | 228 | 236 | - | 11.3303 | 4.96E-05 | 0.615 | GTGGGTGGC |
| 9842 | mamo-s | *BGIBMGA011535* | down | 199 | 207 | + | 11.3303 | 4.96E-05 | 0.615 | GTGGGTGGC |
| 9843 | mamo-s | *BGIBMGA011535* | up | 1199 | 1207 | - | 11.3303 | 4.96E-05 | 0.615 | GTGGGTGGC |
| 9844 | mamo-s | *BGIBMGA011536* | down | 579 | 587 | - | 11.3303 | 4.96E-05 | 0.615 | GTGGGTGGC |
| 9845 | mamo-s | *BGIBMGA011537* | up | 626 | 634 | - | 11.3303 | 4.96E-05 | 0.615 | GTGGGTGGC |
| 9846 | mamo-s | *BGIBMGA011537* | down | 1737 | 1745 | + | 11.3303 | 4.96E-05 | 0.615 | GTGGGTGGC |
| 9847 | mamo-s | *BGIBMGA011539* | down | 1315 | 1323 | - | 11.4404 | 4.51E-05 | 0.615 | CGGCGTGGC |
| 9848 | mamo-s | *BGIBMGA011540* | down | 1203 | 1211 | - | 11.3303 | 4.96E-05 | 0.615 | GTGGGTGGC |
| 9849 | mamo-s | *BGIBMGA011542* | down | 1058 | 1066 | - | 10.9358 | 7.30E-05 | 0.728 | GCGGGTGGC |

| | | | | | | | | | | |
|---|---|---|---|---|---|---|---|---|---|---|
| 9850 | mamo-s | *BGIBMGA011544* | up | 763 | 771 | - | 11.3303 | 4.96E-05 | 0.615 | GTGGGTGGC |
| 9851 | mamo-s | *BGIBMGA011546* | down | 93 | 101 | - | 11.3303 | 4.96E-05 | 0.615 | GTGGGTGGC |
| 9852 | mamo-s | *BGIBMGA011548* | up | 1111 | 1119 | + | 11.3303 | 4.96E-05 | 0.615 | GTGGGTGGC |
| 9853 | mamo-s | *BGIBMGA011550* | up | 1210 | 1218 | - | 11.3303 | 4.96E-05 | 0.615 | GTGGGTGGC |
| 9854 | mamo-s | *BGIBMGA011552* | down | 1304 | 1312 | - | 11.3303 | 4.96E-05 | 0.615 | GTGGGTGGC |
| 9855 | mamo-s | *BGIBMGA011554* | up | 1490 | 1498 | + | 11.3303 | 4.96E-05 | 0.615 | GTGGGTGGC |
| 9856 | mamo-s | *BGIBMGA011555* | down | 397 | 405 | + | 11.3303 | 4.96E-05 | 0.615 | GTGGGTGGC |
| 9857 | mamo-s | *BGIBMGA011556* | down | 1019 | 1027 | - | 11.0826 | 6.84E-05 | 0.709 | GTACGTGGC |
| 9858 | mamo-s | *BGIBMGA011558* | up | 1374 | 1382 | + | 12.4679 | 1.31E-05 | 0.615 | GTGCGTGTC |
| 9859 | mamo-s | *BGIBMGA011559* | up | 802 | 810 | + | 11.3303 | 4.96E-05 | 0.615 | GTGGGTGGC |
| 9860 | mamo-s | *BGIBMGA011564* | up | 942 | 950 | + | 11.3303 | 4.96E-05 | 0.615 | GTGGGTGGC |
| 9861 | mamo-s | *BGIBMGA011564* | up | 1159 | 1167 | + | 11.1743 | 5.90E-05 | 0.684 | CCGCGTGTC |
| 9862 | mamo-s | *BGIBMGA011565* | up | 1765 | 1773 | - | 11.0826 | 6.84E-05 | 0.709 | GTACGTGGC |
| 9863 | mamo-s | *BGIBMGA011568* | up | 1274 | 1282 | + | 11.3303 | 4.96E-05 | 0.615 | GTGGGTGGC |
| 9864 | mamo-s | *BGIBMGA011571* | up | 671 | 679 | + | 12.0734 | 2.66E-05 | 0.615 | GCGCGTGTC |
| 9865 | mamo-s | *BGIBMGA011572* | down | 1833 | 1841 | + | 10.8899 | 7.48E-05 | 0.739 | CCGCGTGCC |
| 9866 | mamo-s | *BGIBMGA011573* | up | 82 | 90 | - | 11.3303 | 4.96E-05 | 0.615 | GTGGGTGGC |
| 9867 | mamo-s | *BGIBMGA011573* | up | 860 | 868 | + | 11.3303 | 4.96E-05 | 0.615 | GTGGGTGGC |
| 9868 | mamo-s | *BGIBMGA011573* | up | 1437 | 1445 | + | 11.3303 | 4.96E-05 | 0.615 | GTGGGTGGC |
| 9869 | mamo-s | *BGIBMGA011574* | up | 324 | 332 | - | 11.3303 | 4.96E-05 | 0.615 | GTGGGTGGC |
| 9870 | mamo-s | *BGIBMGA011576* | down | 1174 | 1182 | + | 12.1009 | 2.43E-05 | 0.615 | GTGCGTGGT |
| 9871 | mamo-s | *BGIBMGA011579* | up | 1057 | 1065 | - | 14.055 | 2.23E-06 | 0.615 | GTGCGTGGC |
| 9872 | mamo-s | *BGIBMGA011580* | up | 1575 | 1583 | + | 11.3853 | 4.74E-05 | 0.615 | ACGCGTGGC |
| 9873 | mamo-s | *BGIBMGA011580* | down | 1738 | 1746 | - | 11.3303 | 4.96E-05 | 0.615 | GTGGGTGGC |
| 9874 | mamo-s | *BGIBMGA011580* | down | 1120 | 1128 | + | 11.0826 | 6.84E-05 | 0.709 | GTACGTGGC |
| 9875 | mamo-s | *BGIBMGA011584* | down | 769 | 777 | - | 11.3303 | 4.96E-05 | 0.615 | GTGGGTGGC |
| 9876 | mamo-s | *BGIBMGA011585* | down | 904 | 912 | - | 11.3303 | 4.96E-05 | 0.615 | GTGGGTGGC |
| 9877 | mamo-s | *BGIBMGA011588* | up | 1010 | 1018 | - | 11.3303 | 4.96E-05 | 0.615 | GTGGGTGGC |
| 9878 | mamo-s | *BGIBMGA011589* | up | 1834 | 1842 | + | 12.4679 | 1.31E-05 | 0.615 | GTGCGTGTC |
| 9879 | mamo-s | *BGIBMGA011589* | up | 211 | 219 | - | 11.3303 | 4.96E-05 | 0.615 | GTGGGTGGC |
| 9880 | mamo-s | *BGIBMGA011589* | up | 601 | 609 | + | 11.3303 | 4.96E-05 | 0.615 | GTGGGTGGC |
| 9881 | mamo-s | *BGIBMGA011593* | down | 1246 | 1254 | + | 11.0826 | 6.84E-05 | 0.709 | GTACGTGGC |
| 9882 | mamo-s | *BGIBMGA011594* | up | 359 | 367 | - | 11.3303 | 4.96E-05 | 0.615 | GTGGGTGGC |
| 9883 | mamo-s | *BGIBMGA011596* | down | 285 | 293 | - | 11.3303 | 4.96E-05 | 0.615 | GTGGGTGGC |
| 9884 | mamo-s | *BGIBMGA011597* | up | 849 | 857 | - | 11.3303 | 4.96E-05 | 0.615 | GTGGGTGGC |
| 9885 | mamo-s | *BGIBMGA011597* | up | 1467 | 1475 | + | 11.3303 | 4.96E-05 | 0.615 | GTGGGTGGC |
| 9886 | mamo-s | *BGIBMGA011597* | up | 445 | 453 | - | 11.0826 | 6.84E-05 | 0.709 | GTACGTGGC |
| 9887 | mamo-s | *BGIBMGA011597* | down | 1955 | 1963 | + | 11.0826 | 6.84E-05 | 0.709 | GTACGTGGC |
| 9888 | mamo-s | *BGIBMGA011598* | down | 736 | 744 | - | 11.3303 | 4.96E-05 | 0.615 | GTGGGTGGC |
| 9889 | mamo-s | *BGIBMGA011598* | down | 1712 | 1720 | + | 11.3303 | 4.96E-05 | 0.615 | GTGGGTGGC |
| 9890 | mamo-s | *BGIBMGA011600* | up | 983 | 991 | - | 11.3303 | 4.96E-05 | 0.615 | GTGGGTGGC |
| 9891 | mamo-s | *BGIBMGA011601* | up | 458 | 466 | + | 11.0826 | 6.84E-05 | 0.709 | GTACGTGGC |
| 9892 | mamo-s | *BGIBMGA011601* | up | 1073 | 1081 | + | 10.7523 | 8.38E-05 | 0.797 | GGGCGTGTC |
| 9893 | mamo-s | *BGIBMGA011602* | down | 1708 | 1716 | + | 11.3303 | 4.96E-05 | 0.615 | GTGGGTGGC |
| 9894 | mamo-s | *BGIBMGA011603* | down | 183 | 191 | - | 11.3303 | 4.96E-05 | 0.615 | GTGGGTGGC |
| 9895 | mamo-s | *BGIBMGA011603* | up | 202 | 210 | - | 11.3303 | 4.96E-05 | 0.615 | GTGGGTGGC |
| 9896 | mamo-s | *BGIBMGA011603* | up | 1436 | 1444 | - | 11.3303 | 4.96E-05 | 0.615 | GTGGGTGGC |
| 9897 | mamo-s | *BGIBMGA011604* | up | 398 | 406 | - | 11.2844 | 5.18E-05 | 0.63 | CTGCGTGCC |
| 9898 | mamo-s | *BGIBMGA011605* | up | 815 | 823 | + | 11.3303 | 4.96E-05 | 0.615 | GTGGGTGGC |
| 9899 | mamo-s | *BGIBMGA011606* | down | 716 | 724 | - | 11.3303 | 4.96E-05 | 0.615 | GTGGGTGGC |
| 9900 | mamo-s | *BGIBMGA011607* | down | 1536 | 1544 | - | 11.0826 | 6.84E-05 | 0.709 | GTACGTGGC |
| 9901 | mamo-s | *BGIBMGA011608* | up | 81 | 89 | - | 11.0826 | 6.84E-05 | 0.709 | GTACGTGGC |
| 9902 | mamo-s | *BGIBMGA011609* | up | 1863 | 1871 | + | 11.3303 | 4.96E-05 | 0.615 | GTGGGTGGC |
| 9903 | mamo-s | *BGIBMGA011610* | down | 1914 | 1922 | - | 13.6606 | 4.06E-06 | 0.615 | GCGCGTGGC |
| 9904 | mamo-s | *BGIBMGA011611* | down | 228 | 236 | + | 10.5138 | 9.75E-05 | 0.865 | GTGCGTGTT |
| 9905 | mamo-s | *BGIBMGA011612* | up | 663 | 671 | + | 11.3303 | 4.96E-05 | 0.615 | GTGGGTGGC |
| 9906 | mamo-s | *BGIBMGA011612* | up | 1809 | 1817 | + | 11.3303 | 4.96E-05 | 0.615 | GTGGGTGGC |
| 9907 | mamo-s | *BGIBMGA011612* | down | 1654 | 1662 | + | 11.2294 | 5.41E-05 | 0.65 | CTGCGAGGC |
| 9908 | mamo-s | *BGIBMGA011613* | down | 839 | 847 | + | 11.3303 | 4.96E-05 | 0.615 | GTGGGTGGC |
| 9909 | mamo-s | *BGIBMGA011616* | down | 1419 | 1427 | - | 13.156 | 6.30E-06 | 0.615 | CTGCGTGGC |
| 9910 | mamo-s | *BGIBMGA011617* | up | 1895 | 1903 | - | 11.3303 | 4.96E-05 | 0.615 | GTGGGTGGC |
| 9911 | mamo-s | *BGIBMGA011617* | up | 792 | 800 | - | 11.1743 | 5.90E-05 | 0.684 | CCGCGTGTC |
| 9912 | mamo-s | *BGIBMGA011618* | down | 473 | 481 | - | 11.3303 | 4.96E-05 | 0.615 | GTGGGTGGC |
| 9913 | mamo-s | *BGIBMGA011618* | down | 1531 | 1539 | - | 11.0826 | 6.84E-05 | 0.709 | GTACGTGGC |
| 9914 | mamo-s | *BGIBMGA011620* | up | 943 | 951 | - | 12.4679 | 1.31E-05 | 0.615 | GTGCGTGTC |
| 9915 | mamo-s | *BGIBMGA011621* | up | 1197 | 1205 | + | 13.6606 | 4.06E-06 | 0.615 | GCGCGTGGC |
| 9916 | mamo-s | *BGIBMGA011624* | up | 1728 | 1736 | + | 11.3303 | 4.96E-05 | 0.615 | GTGGGTGGC |
| 9917 | mamo-s | *BGIBMGA011628* | down | 1506 | 1514 | - | 12.7615 | 1.04E-05 | 0.615 | CCGCGTGGC |
| 9918 | mamo-s | *BGIBMGA011628* | up | 1003 | 1011 | + | 11.3303 | 4.96E-05 | 0.615 | GTGGGTGGC |
| 9919 | mamo-s | *BGIBMGA011630* | up | 1363 | 1371 | - | 11.7798 | 3.11E-05 | 0.615 | ATGCGTGGC |
| 9920 | mamo-s | *BGIBMGA011633* | up | 1477 | 1485 | + | 11.3303 | 4.96E-05 | 0.615 | GTGGGTGGC |
| 9921 | mamo-s | *BGIBMGA011633* | up | 156 | 164 | + | 11.1376 | 6.35E-05 | 0.709 | TCGCGTGGC |
| 9922 | mamo-s | *BGIBMGA011634* | up | 93 | 101 | - | 14.055 | 2.23E-06 | 0.615 | GTGCGTGGC |
| 9923 | mamo-s | *BGIBMGA011636* | up | 1868 | 1876 | + | 12.0734 | 2.66E-05 | 0.615 | GCGCGTGTC |
| 9924 | mamo-s | *BGIBMGA011636* | up | 1312 | 1320 | + | 10.5413 | 9.42E-05 | 0.857 | CAGCGTGTC |
| 9925 | mamo-s | *BGIBMGA011638* | down | 1676 | 1684 | - | 11.3303 | 4.96E-05 | 0.615 | GTGGGTGGC |
| 9926 | mamo-s | *BGIBMGA011639* | down | 246 | 254 | - | 11.0826 | 6.84E-05 | 0.709 | GTACGTGGC |
| 9927 | mamo-s | *BGIBMGA011640* | up | 1702 | 1710 | - | 11.0826 | 6.84E-05 | 0.709 | GTACGTGGC |
| 9928 | mamo-s | *BGIBMGA011641* | down | 574 | 582 | - | 13.6606 | 4.06E-06 | 0.615 | GCGCGTGGC |
| 9929 | mamo-s | *BGIBMGA011641* | down | 500 | 508 | + | 13.6606 | 4.06E-06 | 0.615 | GCGCGTGGC |
| 9930 | mamo-s | *BGIBMGA011641* | down | 170 | 178 | + | 11.3303 | 4.96E-05 | 0.615 | GTGGGTGGC |
| 9931 | mamo-s | *BGIBMGA011641* | down | 462 | 470 | + | 11.2844 | 5.18E-05 | 0.63 | CTGCGTGCC |
| 9932 | mamo-s | *BGIBMGA011641* | down | 456 | 464 | - | 10.5413 | 9.42E-05 | 0.857 | CAGCGTGTC |
| 9933 | mamo-s | *BGIBMGA011642* | up | 206 | 214 | - | 13.6606 | 4.06E-06 | 0.615 | GCGCGTGGC |
| 9934 | mamo-s | *BGIBMGA011642* | up | 132 | 140 | + | 13.6606 | 4.06E-06 | 0.615 | GCGCGTGGC |
| 9935 | mamo-s | *BGIBMGA011642* | up | 94 | 102 | + | 11.2844 | 5.18E-05 | 0.63 | CTGCGTGCC |
| 9936 | mamo-s | *BGIBMGA011642* | up | 88 | 96 | - | 10.5413 | 9.42E-05 | 0.857 | CAGCGTGTC |
| 9937 | mamo-s | *BGIBMGA011644* | down | 1260 | 1268 | - | 11.4404 | 4.51E-05 | 0.615 | GAGCGTGTC |

| | | | | | | | | | | |
|---|---|---|---|---|---|---|---|---|---|---|
| 9938 | mamo-s | *BGIBMGA011644* | up | 1241 | 1249 | - | 11.3303 | 4.96E-05 | 0.615 | GTGGGTGGC |
| 9939 | mamo-s | *BGIBMGA011645* | down | 1284 | 1292 | + | 13.0275 | 8.53E-06 | 0.615 | GAGCGTGGC |
| 9940 | mamo-s | *BGIBMGA011646* | down | 288 | 296 | - | 11.7798 | 3.11E-05 | 0.615 | ATGCGTGGC |
| 9941 | mamo-s | *BGIBMGA011647* | up | 1146 | 1154 | + | 11.3303 | 4.96E-05 | 0.615 | GTGGGTGGC |
| 9942 | mamo-s | *BGIBMGA011649* | up | 1825 | 1833 | + | 11.2018 | 5.68E-05 | 0.674 | CTGCGTGGT |
| 9943 | mamo-s | *BGIBMGA011651* | down | 310 | 318 | + | 11.3303 | 4.96E-05 | 0.615 | GTGGGTGGC |
| 9944 | mamo-s | *BGIBMGA011655* | down | 1987 | 1995 | + | 12.3394 | 1.49E-05 | 0.615 | GGGCGTGGC |
| 9945 | mamo-s | *BGIBMGA011657* | down | 1390 | 1398 | + | 13.6606 | 4.06E-06 | 0.615 | GCGCGTGGC |
| 9946 | mamo-s | *BGIBMGA011659* | down | 1781 | 1789 | + | 11.3303 | 4.96E-05 | 0.615 | GTGGGTGGC |
| 9947 | mamo-s | *BGIBMGA011662* | down | 1335 | 1343 | - | 11.3303 | 4.96E-05 | 0.615 | GTGGGTGGC |
| 9948 | mamo-s | *BGIBMGA011663* | up | 1389 | 1397 | + | 12.4679 | 1.31E-05 | 0.615 | GTGCGTGTC |
| 9949 | mamo-s | *BGIBMGA011663* | up | 286 | 294 | - | 11.3303 | 4.96E-05 | 0.615 | GTGGGTGGC |
| 9950 | mamo-s | *BGIBMGA011664* | down | 371 | 379 | + | 11.789 | 2.84E-05 | 0.615 | GCGCGTGCC |
| 9951 | mamo-s | *BGIBMGA011664* | down | 719 | 727 | + | 11.3303 | 4.96E-05 | 0.615 | GTGGGTGGC |
| 9952 | mamo-s | *BGIBMGA011664* | up | 1986 | 1994 | - | 10.8899 | 7.48E-05 | 0.739 | CCGCGTGCC |
| 9953 | mamo-s | *BGIBMGA011664* | down | 642 | 650 | - | 10.5138 | 9.75E-05 | 0.865 | GTGCGTGTT |
| 9954 | mamo-s | *BGIBMGA011665* | down | 1447 | 1455 | + | 11.3303 | 4.96E-05 | 0.615 | GTGGGTGGC |
| 9955 | mamo-s | *BGIBMGA011665* | up | 1527 | 1535 | + | 10.9358 | 7.30E-05 | 0.728 | GCGGGTGGC |
| 9956 | mamo-s | *BGIBMGA011672* | up | 1277 | 1285 | + | 12.1284 | 2.16E-05 | 0.615 | CAGCGTGGC |
| 9957 | mamo-s | *BGIBMGA011674* | down | 36 | 44 | - | 12.1009 | 2.43E-05 | 0.615 | GTGCGTGGT |
| 9958 | mamo-s | *BGIBMGA011675* | down | 1686 | 1694 | - | 11.3303 | 4.96E-05 | 0.615 | GTGGGTGGC |
| 9959 | mamo-s | *BGIBMGA011677* | up | 210 | 218 | + | 11.5321 | 4.06E-05 | 0.615 | TTGCGTGGC |
| 9960 | mamo-s | *BGIBMGA011677* | up | 969 | 977 | - | 11.1376 | 6.35E-05 | 0.709 | TCGCGTGGC |
| 9961 | mamo-s | *BGIBMGA011679* | down | 1327 | 1335 | - | 10.6881 | 8.61E-05 | 0.81 | GCACGTGGC |
| 9962 | mamo-s | *BGIBMGA011679* | down | 262 | 270 | - | 10.5413 | 9.42E-05 | 0.857 | GTGCGAGTC |
| 9963 | mamo-s | *BGIBMGA011680* | down | 1566 | 1574 | - | 11.3303 | 4.96E-05 | 0.615 | GTGGGTGGC |
| 9964 | mamo-s | *BGIBMGA011682* | down | 1175 | 1183 | - | 11.3303 | 4.96E-05 | 0.615 | GTGGGTGGC |
| 9965 | mamo-s | *BGIBMGA011683* | up | 1709 | 1717 | + | 10.7523 | 8.38E-05 | 0.797 | AAGCGTGGC |
| 9966 | mamo-s | *BGIBMGA011685* | up | 29 | 37 | - | 11.0734 | 7.12E-05 | 0.726 | GAGCGTGGT |
| 9967 | mamo-s | *BGIBMGA011686* | up | 1232 | 1240 | + | 10.7523 | 8.38E-05 | 0.797 | AAGCGTGGC |
| 9968 | mamo-s | *BGIBMGA011690* | up | 863 | 871 | - | 10.6881 | 8.61E-05 | 0.81 | GCACGTGGC |
| 9969 | mamo-s | *BGIBMGA011691* | up | 430 | 438 | + | 12.3394 | 1.49E-05 | 0.615 | GGGCGTGGC |
| 9970 | mamo-s | *BGIBMGA011691* | up | 418 | 426 | + | 11.5321 | 4.06E-05 | 0.615 | TTGCGTGGC |
| 9971 | mamo-s | *BGIBMGA011691* | down | 866 | 874 | - | 11.3303 | 4.96E-05 | 0.615 | GTGGGTGGC |
| 9972 | mamo-s | *BGIBMGA011691* | down | 1612 | 1620 | + | 11.3303 | 4.96E-05 | 0.615 | GTGGGTGGC |
| 9973 | mamo-s | *BGIBMGA011692* | down | 1301 | 1309 | + | 10.7523 | 8.38E-05 | 0.797 | GGGCGTGTC |
| 9974 | mamo-s | *BGIBMGA011693* | down | 303 | 311 | - | 13.0275 | 8.53E-06 | 0.615 | GAGCGTGGC |
| 9975 | mamo-s | *BGIBMGA011694* | up | 1272 | 1280 | - | 11.3303 | 4.96E-05 | 0.615 | GTGGGTGGC |
| 9976 | mamo-s | *BGIBMGA011695* | up | 703 | 711 | - | 11.3303 | 4.96E-05 | 0.615 | GTGGGTGGC |
| 9977 | mamo-s | *BGIBMGA011695* | down | 921 | 929 | - | 10.7523 | 8.38E-05 | 0.797 | GGGCGTGTC |
| 9978 | mamo-s | *BGIBMGA011698* | up | 968 | 976 | + | 11.3303 | 4.96E-05 | 0.615 | GTGGGTGGC |
| 9979 | mamo-s | *BGIBMGA011699* | up | 1958 | 1966 | - | 13.6606 | 4.06E-06 | 0.615 | GCGCGTGGC |
| 9980 | mamo-s | *BGIBMGA011699* | down | 857 | 865 | - | 11.3303 | 4.96E-05 | 0.615 | GTGGGTGGC |
| 9981 | mamo-s | *BGIBMGA011701* | down | 1544 | 1552 | - | 11.3303 | 4.96E-05 | 0.615 | GTGGGTGGC |
| 9982 | mamo-s | *BGIBMGA011702* | up | 450 | 458 | - | 11.0826 | 6.84E-05 | 0.709 | GTACGTGGC |
| 9983 | mamo-s | *BGIBMGA011704* | up | 1661 | 1669 | - | 11.3303 | 4.96E-05 | 0.615 | GTGGGTGGC |
| 9984 | mamo-s | *BGIBMGA011704* | down | 1081 | 1089 | + | 11.1009 | 6.57E-05 | 0.709 | GAGCGAGGC |
| 9985 | mamo-s | *BGIBMGA011705* | up | 697 | 705 | + | 11.3303 | 4.96E-05 | 0.615 | GTGGGTGGC |
| 9986 | mamo-s | *BGIBMGA011714* | down | 358 | 366 | + | 10.9358 | 7.30E-05 | 0.728 | GCGGGTGGC |
| 9987 | mamo-s | *BGIBMGA011717* | up | 1269 | 1277 | - | 11.3303 | 4.96E-05 | 0.615 | GTGGGTGGC |
| 9988 | mamo-s | *BGIBMGA011721* | down | 1492 | 1500 | - | 11.3303 | 4.96E-05 | 0.615 | GTGGGTGGC |
| 9989 | mamo-s | *BGIBMGA011722* | up | 292 | 300 | + | 11.3303 | 4.96E-05 | 0.615 | GTGGGTGGC |
| 9990 | mamo-s | *BGIBMGA011725* | up | 609 | 617 | + | 11.3303 | 4.96E-05 | 0.615 | GTGGGTGGC |
| 9991 | mamo-s | *BGIBMGA011728* | up | 1144 | 1152 | + | 12.1284 | 2.16E-05 | 0.615 | GTGCGAGGC |
| 9992 | mamo-s | *BGIBMGA011728* | up | 1489 | 1497 | - | 11.3303 | 4.96E-05 | 0.615 | GTGGGTGGC |
| 9993 | mamo-s | *BGIBMGA011729* | up | 931 | 939 | - | 10.7523 | 8.38E-05 | 0.797 | GGGCGTGTC |
| 9994 | mamo-s | *BGIBMGA011730* | up | 1117 | 1125 | + | 11.3303 | 4.96E-05 | 0.615 | GTGGGTGGC |
| 9995 | mamo-s | *BGIBMGA011731* | up | 1780 | 1788 | - | 14.055 | 2.23E-06 | 0.615 | GTGCGTGGC |
| 9996 | mamo-s | *BGIBMGA011731* | up | 450 | 458 | + | 11.1743 | 5.90E-05 | 0.684 | CCGCGTGTC |
| 9997 | mamo-s | *BGIBMGA011732* | down | 1653 | 1661 | + | 11.3303 | 4.96E-05 | 0.615 | GTGGGTGGC |
| 9998 | mamo-s | *BGIBMGA011737* | down | 1222 | 1230 | - | 11.3303 | 4.96E-05 | 0.615 | GTGGGTGGC |
| 9999 | mamo-s | *BGIBMGA011738* | down | 735 | 743 | - | 11.3303 | 4.96E-05 | 0.615 | GTGGGTGGC |
| 10000 | mamo-s | *BGIBMGA011739* | down | 1536 | 1544 | - | 11.789 | 2.84E-05 | 0.615 | GCGCGTGCC |
| 10001 | mamo-s | *BGIBMGA011739* | down | 1530 | 1538 | + | 11.1376 | 6.35E-05 | 0.709 | TCGCGTGGC |
| 10002 | mamo-s | *BGIBMGA011740* | up | 1931 | 1939 | - | 11.789 | 2.84E-05 | 0.615 | GCGCGTGCC |
| 10003 | mamo-s | *BGIBMGA011740* | up | 1925 | 1933 | + | 11.1376 | 6.35E-05 | 0.709 | TCGCGTGGC |
| 10004 | mamo-s | *BGIBMGA011744* | down | 282 | 290 | + | 11.3303 | 4.96E-05 | 0.615 | GTGGGTGGC |
| 10005 | mamo-s | *BGIBMGA011745* | up | 1431 | 1439 | + | 11.3303 | 4.96E-05 | 0.615 | GTGGGTGGC |
| 10006 | mamo-s | *BGIBMGA011745* | up | 1123 | 1131 | - | 10.5413 | 9.42E-05 | 0.857 | GTGCGAGTC |
| 10007 | mamo-s | *BGIBMGA011745* | up | 1138 | 1146 | - | 10.5413 | 9.42E-05 | 0.857 | CAGCGTGTC |
| 10008 | mamo-s | *BGIBMGA011747* | up | 678 | 686 | + | 11.3303 | 4.96E-05 | 0.615 | GTGGGTGGC |
| 10009 | mamo-s | *BGIBMGA011749* | down | 1323 | 1331 | + | 11.1743 | 5.90E-05 | 0.684 | CCGCGTGTC |
| 10010 | mamo-s | *BGIBMGA011750* | up | 1271 | 1279 | - | 11.5688 | 3.79E-05 | 0.615 | CTGCGTGTC |
| 10011 | mamo-s | *BGIBMGA011752* | up | 678 | 686 | - | 11.5321 | 4.06E-05 | 0.615 | TTGCGTGGC |
| 10012 | mamo-s | *BGIBMGA011752* | down | 1199 | 1207 | + | 11.1743 | 5.90E-05 | 0.684 | CCGCGTGTC |
| 10013 | mamo-s | *BGIBMGA011752* | up | 1069 | 1077 | - | 11.0826 | 6.84E-05 | 0.709 | GTACGTGGC |
| 10014 | mamo-s | *BGIBMGA011753* | down | 277 | 285 | - | 11.3303 | 4.96E-05 | 0.615 | GTGGGTGGC |
| 10015 | mamo-s | *BGIBMGA011756* | down | 402 | 410 | + | 11.5321 | 4.06E-05 | 0.615 | TTGCGTGGC |
| 10016 | mamo-s | *BGIBMGA011759* | up | 1057 | 1065 | - | 11.3303 | 4.96E-05 | 0.615 | GTGGGTGGC |
| 10017 | mamo-s | *BGIBMGA011760* | up | 1893 | 1901 | - | 12.1284 | 2.16E-05 | 0.615 | GTGCGAGGC |
| 10018 | mamo-s | *BGIBMGA011760* | up | 1548 | 1556 | + | 11.3303 | 4.96E-05 | 0.615 | GTGGGTGGC |
| 10019 | mamo-s | *BGIBMGA011762* | down | 1533 | 1541 | - | 11.3303 | 4.96E-05 | 0.615 | GTGGGTGGC |
| 10020 | mamo-s | *BGIBMGA011765* | down | 1667 | 1675 | - | 11.3303 | 4.96E-05 | 0.615 | GTGGGTGGC |
| 10021 | mamo-s | *BGIBMGA011766* | down | 902 | 910 | + | 11.3303 | 4.96E-05 | 0.615 | GTGGGTGGC |
| 10022 | mamo-s | *BGIBMGA011766* | down | 1904 | 1912 | - | 11.1743 | 5.90E-05 | 0.684 | CCGCGTGTC |
| 10023 | mamo-s | *BGIBMGA011769* | up | 170 | 178 | - | 11.3303 | 4.96E-05 | 0.615 | GTGGGTGGC |
| 10024 | mamo-s | *BGIBMGA011773* | down | 1038 | 1046 | - | 11.0826 | 6.84E-05 | 0.709 | GTACGTGGC |
| 10025 | mamo-s | *BGIBMGA011774* | down | 1403 | 1411 | - | 11.0826 | 6.84E-05 | 0.709 | GTACGTGGC |

| | | | | | | | | | | |
|---|---|---|---|---|---|---|---|---|---|---|
| 10026 | mamo-s | *BGIBMGA011777* | down | 1783 | 1791 | + | 11.3303 | 4.96E-05 | 0.615 | GTGGGTGGC |
| 10027 | mamo-s | *BGIBMGA011778* | up | 908 | 916 | + | 11.3303 | 4.96E-05 | 0.615 | GTGGGTGGC |
| 10028 | mamo-s | *BGIBMGA011778* | down | 1267 | 1275 | + | 11.3303 | 4.96E-05 | 0.615 | GTGGGTGGC |
| 10029 | mamo-s | *BGIBMGA011780* | down | 1549 | 1557 | - | 11.0734 | 7.12E-05 | 0.726 | GAGCGTGGT |
| 10030 | mamo-s | *BGIBMGA011781* | up | 1212 | 1220 | - | 11.0734 | 7.12E-05 | 0.726 | GAGCGTGGT |
| 10031 | mamo-s | *BGIBMGA011782* | down | 629 | 637 | + | 11.3303 | 4.96E-05 | 0.615 | GTGGGTGGC |
| 10032 | mamo-s | *BGIBMGA011785* | down | 800 | 808 | + | 11.0734 | 7.12E-05 | 0.726 | GAGCGTGGT |
| 10033 | mamo-s | *BGIBMGA011786* | up | 429 | 437 | + | 10.6881 | 8.61E-05 | 0.81 | GCACGTGGC |
| 10034 | mamo-s | *BGIBMGA011788* | up | 757 | 765 | - | 12.1835 | 1.71E-05 | 0.615 | GTGCGTGCC |
| 10035 | mamo-s | *BGIBMGA011792* | up | 289 | 297 | - | 10.9358 | 7.30E-05 | 0.728 | GCGGGTGGC |
| 10036 | mamo-s | *BGIBMGA011792* | down | 1474 | 1482 | - | 10.5138 | 9.75E-05 | 0.865 | GTGCGTGTT |
| 10037 | mamo-s | *BGIBMGA011793* | down | 1176 | 1184 | + | 11.3303 | 4.96E-05 | 0.615 | GTGGGTGGC |
| 10038 | mamo-s | *BGIBMGA011800* | down | 794 | 802 | + | 11.3303 | 4.96E-05 | 0.615 | GTGGGTGGC |
| 10039 | mamo-s | *BGIBMGA011801* | up | 1350 | 1358 | - | 11.3303 | 4.96E-05 | 0.615 | GTGGGTGGC |
| 10040 | mamo-s | *BGIBMGA011801* | up | 1557 | 1565 | - | 11.3303 | 4.96E-05 | 0.615 | GTGGGTGGC |
| 10041 | mamo-s | *BGIBMGA011802* | up | 1468 | 1476 | - | 11.0826 | 6.84E-05 | 0.709 | GTACGTGGC |
| 10042 | mamo-s | *BGIBMGA011806* | down | 820 | 828 | - | 11.3303 | 4.96E-05 | 0.615 | GTGGGTGGC |
| 10043 | mamo-s | *BGIBMGA011807* | down | 1870 | 1878 | - | 13.0275 | 8.53E-06 | 0.615 | GAGCGTGGC |
| 10044 | mamo-s | *BGIBMGA011807* | up | 1787 | 1795 | + | 11.3303 | 4.96E-05 | 0.615 | GTGGGTGGC |
| 10045 | mamo-s | *BGIBMGA011809* | up | 1455 | 1463 | - | 11.7339 | 3.29E-05 | 0.615 | GCGCGAGGC |
| 10046 | mamo-s | *BGIBMGA011809* | up | 625 | 633 | + | 11.2294 | 5.41E-05 | 0.65 | CTGCGAGGC |
| 10047 | mamo-s | *BGIBMGA011809* | down | 1498 | 1506 | + | 11.1376 | 6.35E-05 | 0.709 | TCGCGTGGC |
| 10048 | mamo-s | *BGIBMGA011810* | up | 726 | 734 | - | 11.3303 | 4.96E-05 | 0.615 | GTGGGTGGC |
| 10049 | mamo-s | *BGIBMGA011811* | down | 1328 | 1336 | + | 11.3303 | 4.96E-05 | 0.615 | GTGGGTGGC |
| 10050 | mamo-s | *BGIBMGA011812* | up | 701 | 709 | - | 11.3303 | 4.96E-05 | 0.615 | GTGGGTGGC |
| 10051 | mamo-s | *BGIBMGA011814* | up | 189 | 197 | + | 12.0734 | 2.66E-05 | 0.615 | GCGCGTGTC |
| 10052 | mamo-s | *BGIBMGA011816* | down | 1044 | 1052 | - | 14.055 | 2.23E-06 | 0.615 | GTGCGTGGC |
| 10053 | mamo-s | *BGIBMGA011823* | down | 1000 | 1008 | - | 12.3394 | 1.49E-05 | 0.615 | GGGCGTGGC |
| 10054 | mamo-s | *BGIBMGA011832* | down | 1918 | 1926 | - | 12.4679 | 1.31E-05 | 0.615 | GTGCGTGTC |
| 10055 | mamo-s | *BGIBMGA011832* | down | 722 | 730 | + | 11.7064 | 3.52E-05 | 0.615 | GCGCGTGGT |
| 10056 | mamo-s | *BGIBMGA011834* | down | 1493 | 1501 | + | 11.3303 | 4.96E-05 | 0.615 | GTGGGTGGC |
| 10057 | mamo-s | *BGIBMGA011835* | up | 889 | 897 | + | 11.3303 | 4.96E-05 | 0.615 | GTGGGTGGC |
| 10058 | mamo-s | *BGIBMGA011837* | down | 1988 | 1996 | + | 11.7798 | 3.11E-05 | 0.615 | ATGCGTGGC |
| 10059 | mamo-s | *BGIBMGA011837* | up | 638 | 646 | - | 10.6881 | 8.61E-05 | 0.81 | GCACGTGGC |
| 10060 | mamo-s | *BGIBMGA011840* | down | 866 | 874 | + | 11.4404 | 4.51E-05 | 0.615 | GAGCGTGTC |
| 10061 | mamo-s | *BGIBMGA011840* | up | 285 | 293 | - | 10.7523 | 8.38E-05 | 0.797 | AAGCGTGGC |
| 10062 | mamo-s | *BGIBMGA011841* | down | 408 | 416 | + | 11.3303 | 4.96E-05 | 0.615 | GTGGGTGGC |
| 10063 | mamo-s | *BGIBMGA011841* | up | 1179 | 1187 | + | 11.3303 | 4.96E-05 | 0.615 | GTGGGTGGC |
| 10064 | mamo-s | *BGIBMGA011843* | down | 976 | 984 | + | 11.3303 | 4.96E-05 | 0.615 | GTGGGTGGC |
| 10065 | mamo-s | *BGIBMGA011845* | down | 1878 | 1886 | + | 11.0826 | 6.84E-05 | 0.709 | GTACGTGGC |
| 10066 | mamo-s | *BGIBMGA011846* | down | 544 | 552 | + | 11.3303 | 4.96E-05 | 0.615 | GTGGGTGGC |
| 10067 | mamo-s | *BGIBMGA011847* | down | 1667 | 1675 | - | 11.3303 | 4.96E-05 | 0.615 | GTGGGTGGC |
| 10068 | mamo-s | *BGIBMGA011847* | up | 1009 | 1017 | + | 11.0826 | 6.84E-05 | 0.709 | GTACGTGGC |
| 10069 | mamo-s | *BGIBMGA011848* | up | 680 | 688 | - | 11.3303 | 4.96E-05 | 0.615 | GTGGGTGGC |
| 10070 | mamo-s | *BGIBMGA011849* | up | 610 | 618 | + | 11.0734 | 7.12E-05 | 0.726 | GAGCGTGGT |
| 10071 | mamo-s | *BGIBMGA011850* | down | 1583 | 1591 | - | 10.7523 | 8.38E-05 | 0.797 | AAGCGTGGC |
| 10072 | mamo-s | *BGIBMGA011851* | down | 377 | 385 | + | 11.0826 | 6.84E-05 | 0.709 | GTACGTGGC |
| 10073 | mamo-s | *BGIBMGA011851* | down | 705 | 713 | - | 10.5138 | 9.75E-05 | 0.865 | GTGCGTGTT |
| 10074 | mamo-s | *BGIBMGA011854* | up | 1701 | 1709 | + | 11.3303 | 4.96E-05 | 0.615 | GTGGGTGGC |
| 10075 | mamo-s | *BGIBMGA011855* | down | 1484 | 1492 | - | 10.8073 | 7.89E-05 | 0.766 | CCGCGTGGT |
| 10076 | mamo-s | *BGIBMGA011861* | down | 956 | 964 | + | 11.0826 | 6.84E-05 | 0.709 | GTACGTGGC |
| 10077 | mamo-s | *BGIBMGA011864* | down | 677 | 685 | + | 11.3303 | 4.96E-05 | 0.615 | GTGGGTGGC |
| 10078 | mamo-s | *BGIBMGA011867* | down | 1155 | 1163 | - | 10.5413 | 9.42E-05 | 0.857 | GTGCGAGTC |
| 10079 | mamo-s | *BGIBMGA011869* | down | 1104 | 1112 | - | 14.055 | 2.23E-06 | 0.615 | GTGCGTGGC |
| 10080 | mamo-s | *BGIBMGA011870* | up | 189 | 197 | + | 11.3303 | 4.96E-05 | 0.615 | GTGGGTGGC |
| 10081 | mamo-s | *BGIBMGA011872* | up | 1806 | 1814 | + | 11.3303 | 4.96E-05 | 0.615 | GTGGGTGGC |
| 10082 | mamo-s | *BGIBMGA011873* | up | 1355 | 1363 | + | 11.3303 | 4.96E-05 | 0.615 | GTGGGTGGC |
| 10083 | mamo-s | *BGIBMGA011874* | up | 464 | 472 | - | 11.3303 | 4.96E-05 | 0.615 | GTGGGTGGC |
| 10084 | mamo-s | *BGIBMGA011876* | up | 459 | 467 | - | 11.3303 | 4.96E-05 | 0.615 | GTGGGTGGC |
| 10085 | mamo-s | *BGIBMGA011876* | down | 954 | 962 | + | 11.3303 | 4.96E-05 | 0.615 | GTGGGTGGC |
| 10086 | mamo-s | *BGIBMGA011877* | down | 1331 | 1339 | - | 12.1284 | 2.16E-05 | 0.615 | CAGCGTGGC |
| 10087 | mamo-s | *BGIBMGA011878* | down | 1312 | 1320 | - | 11.789 | 2.84E-05 | 0.615 | GCGCGTGCC |
| 10088 | mamo-s | *BGIBMGA011878* | down | 558 | 566 | + | 11.3303 | 4.96E-05 | 0.615 | GTGGGTGGC |
| 10089 | mamo-s | *BGIBMGA011879* | up | 1212 | 1220 | - | 11.789 | 2.84E-05 | 0.615 | GCGCGTGCC |
| 10090 | mamo-s | *BGIBMGA011879* | up | 458 | 466 | + | 11.3303 | 4.96E-05 | 0.615 | GTGGGTGGC |
| 10091 | mamo-s | *BGIBMGA011881* | down | 1459 | 1467 | + | 12.0734 | 2.66E-05 | 0.615 | GCGCGTGTC |
| 10092 | mamo-s | *BGIBMGA011884* | down | 1775 | 1783 | + | 11.3303 | 4.96E-05 | 0.615 | GTGGGTGGC |
| 10093 | mamo-s | *BGIBMGA011884* | down | 1550 | 1558 | + | 10.8073 | 7.89E-05 | 0.766 | CCGCGTGGT |
| 10094 | mamo-s | *BGIBMGA011886* | up | 1177 | 1185 | + | 11.3303 | 4.96E-05 | 0.615 | GTGGGTGGC |
| 10095 | mamo-s | *BGIBMGA011887* | up | 1680 | 1688 | + | 11.3853 | 4.74E-05 | 0.615 | ACGCGTGGC |
| 10096 | mamo-s | *BGIBMGA011888* | down | 904 | 912 | + | 13.156 | 6.30E-06 | 0.615 | CTGCGTGGC |
| 10097 | mamo-s | *BGIBMGA011888* | down | 1597 | 1605 | - | 11.3303 | 4.96E-05 | 0.615 | GTGGGTGGC |
| 10098 | mamo-s | *BGIBMGA011889* | up | 1467 | 1475 | + | 10.9358 | 7.30E-05 | 0.728 | GCGGGTGGC |
| 10099 | mamo-s | *BGIBMGA011889* | up | 435 | 443 | + | 10.6881 | 8.61E-05 | 0.81 | GCACGTGGC |
| 10100 | mamo-s | *BGIBMGA011890* | down | 954 | 962 | + | 12.1835 | 1.71E-05 | 0.615 | GTGCGTGCC |
| 10101 | mamo-s | *BGIBMGA011892* | down | 7 | 15 | - | 13.6606 | 4.06E-06 | 0.615 | GCGCGTGGC |
| 10102 | mamo-s | *BGIBMGA011892* | down | 912 | 920 | - | 12.1009 | 2.43E-05 | 0.615 | GTGCGTGGT |
| 10103 | mamo-s | *BGIBMGA011892* | up | 254 | 262 | - | 11.3303 | 4.96E-05 | 0.615 | GTGGGTGGC |
| 10104 | mamo-s | *BGIBMGA011894* | up | 1850 | 1858 | + | 11.3303 | 4.96E-05 | 0.615 | GTGGGTGGC |
| 10105 | mamo-s | *BGIBMGA011896* | down | 784 | 792 | + | 12.1835 | 1.71E-05 | 0.615 | GTGCGTGCC |
| 10106 | mamo-s | *BGIBMGA011900* | down | 322 | 330 | - | 11.3303 | 4.96E-05 | 0.615 | GTGGGTGGC |
| 10107 | mamo-s | *BGIBMGA011900* | down | 1403 | 1411 | + | 10.5413 | 9.42E-05 | 0.857 | CAGCGTGTC |
| 10108 | mamo-s | *BGIBMGA011901* | down | 397 | 405 | - | 11.7798 | 3.11E-05 | 0.615 | ATGCGTGGC |
| 10109 | mamo-s | *BGIBMGA011901* | up | 159 | 167 | - | 11.3303 | 4.96E-05 | 0.615 | GTGGGTGGC |
| 10110 | mamo-s | *BGIBMGA011901* | up | 1240 | 1248 | + | 10.5413 | 9.42E-05 | 0.857 | CAGCGTGTC |
| 10111 | mamo-s | *BGIBMGA011904* | up | 315 | 323 | + | 11.3303 | 4.96E-05 | 0.615 | GTGGGTGGC |
| 10112 | mamo-s | *BGIBMGA011904* | down | 1082 | 1090 | - | 11.3303 | 4.96E-05 | 0.615 | GTGGGTGGC |
| 10113 | mamo-s | *BGIBMGA011907* | down | 225 | 233 | + | 11.7064 | 3.52E-05 | 0.615 | GCGCGTGGT |

| | | | | | | | | | | |
|---|---|---|---|---|---|---|---|---|---|---|
| 10114 | mamo-s | *BGIBMGA011907* | up | 1267 | 1275 | + | 11.3303 | 4.96E-05 | 0.615 | GTGGGTGGC |
| 10115 | mamo-s | *BGIBMGA011907* | down | 1905 | 1913 | + | 10.6881 | 8.61E-05 | 0.81 | GCACGTGGC |
| 10116 | mamo-s | *BGIBMGA011910* | up | 645 | 653 | - | 13.156 | 6.30E-06 | 0.615 | CTGCGTGGC |
| 10117 | mamo-s | *BGIBMGA011911* | down | 780 | 788 | - | 11.3303 | 4.96E-05 | 0.615 | GTGGGTGGC |
| 10118 | mamo-s | *BGIBMGA011913* | down | 1811 | 1819 | - | 11.0826 | 6.84E-05 | 0.709 | GTACGTGGC |
| 10119 | mamo-s | *BGIBMGA011914* | down | 918 | 926 | + | 11.789 | 2.84E-05 | 0.615 | GCGCGTGCC |
| 10120 | mamo-s | *BGIBMGA011914* | down | 1680 | 1688 | - | 11.3303 | 4.96E-05 | 0.615 | GTGGGTGGC |
| 10121 | mamo-s | *BGIBMGA011914* | down | 1704 | 1712 | - | 11.3303 | 4.96E-05 | 0.615 | GTGGGTGGC |
| 10122 | mamo-s | *BGIBMGA011915* | up | 87 | 95 | - | 13.6606 | 4.06E-06 | 0.615 | GCGCGTGGC |
| 10123 | mamo-s | *BGIBMGA011915* | up | 1877 | 1885 | + | 11.1743 | 5.90E-05 | 0.684 | CCGCGTGTC |
| 10124 | mamo-s | *BGIBMGA011917* | down | 228 | 236 | + | 12.0734 | 2.66E-05 | 0.615 | GCGCGTGTC |
| 10125 | mamo-s | *BGIBMGA011918* | up | 1412 | 1420 | - | 10.5138 | 9.75E-05 | 0.865 | GTGCGTGTT |
| 10126 | mamo-s | *BGIBMGA011920* | down | 1381 | 1389 | + | 12.0734 | 2.66E-05 | 0.615 | GCGCGTGTC |
| 10127 | mamo-s | *BGIBMGA011922* | down | 1419 | 1427 | - | 12.1284 | 2.16E-05 | 0.615 | CAGCGTGGC |
| 10128 | mamo-s | *BGIBMGA011925* | down | 79 | 87 | + | 11.3303 | 4.96E-05 | 0.615 | GTGGGTGGC |
| 10129 | mamo-s | *BGIBMGA011925* | down | 1065 | 1073 | - | 11.3303 | 4.96E-05 | 0.615 | GTGGGTGGC |
| 10130 | mamo-s | *BGIBMGA011926* | down | 1098 | 1106 | - | 11.3303 | 4.96E-05 | 0.615 | GTGGGTGGC |
| 10131 | mamo-s | *BGIBMGA011926* | up | 1272 | 1280 | + | 11.3303 | 4.96E-05 | 0.615 | GTGGGTGGC |
| 10132 | mamo-s | *BGIBMGA011927* | down | 1105 | 1113 | + | 12.7615 | 1.04E-05 | 0.615 | CCGCGTGGC |
| 10133 | mamo-s | *BGIBMGA011927* | up | 1895 | 1903 | - | 12.4679 | 1.31E-05 | 0.615 | GTGCGTGTC |
| 10134 | mamo-s | *BGIBMGA011927* | up | 1980 | 1988 | + | 11.4404 | 4.51E-05 | 0.615 | CGGCGTGGC |
| 10135 | mamo-s | *BGIBMGA011928* | up | 753 | 761 | - | 12.4679 | 1.31E-05 | 0.615 | GTGCGTGTC |
| 10136 | mamo-s | *BGIBMGA011928* | up | 872 | 880 | - | 12.4679 | 1.31E-05 | 0.615 | GTGCGTGTC |
| 10137 | mamo-s | *BGIBMGA011928* | up | 838 | 846 | + | 11.4404 | 4.51E-05 | 0.615 | CGGCGTGGC |
| 10138 | mamo-s | *BGIBMGA011928* | down | 1174 | 1182 | - | 11.156 | 6.13E-05 | 0.705 | GAGCGTGCC |
| 10139 | mamo-s | *BGIBMGA011930* | down | 387 | 395 | - | 12.4679 | 1.31E-05 | 0.615 | GTGCGTGTC |
| 10140 | mamo-s | *BGIBMGA011930* | down | 1632 | 1640 | - | 11.3303 | 4.96E-05 | 0.615 | GTGGGTGGC |
| 10141 | mamo-s | *BGIBMGA011931* | up | 1975 | 1983 | - | 13.0275 | 8.53E-06 | 0.615 | GAGCGTGGC |
| 10142 | mamo-s | *BGIBMGA011931* | up | 286 | 294 | - | 11.1743 | 5.90E-05 | 0.684 | CCGCGTGTC |
| 10143 | mamo-s | *BGIBMGA011932* | down | 848 | 856 | + | 11.3303 | 4.96E-05 | 0.615 | GTGGGTGGC |
| 10144 | mamo-s | *BGIBMGA011932* | up | 1975 | 1983 | + | 11.1009 | 6.57E-05 | 0.709 | GAGCGAGGC |
| 10145 | mamo-s | *BGIBMGA011934* | down | 1852 | 1860 | + | 11.3303 | 4.96E-05 | 0.615 | GTGGGTGGC |
| 10146 | mamo-s | *BGIBMGA011935* | down | 1320 | 1328 | - | 11.3303 | 4.96E-05 | 0.615 | GTGGGTGGC |
| 10147 | mamo-s | *BGIBMGA011939* | down | 473 | 481 | - | 11.3303 | 4.96E-05 | 0.615 | GTGGGTGGC |
| 10148 | mamo-s | *BGIBMGA011940* | down | 371 | 379 | - | 12.3394 | 1.49E-05 | 0.615 | GGGCGTGGC |
| 10149 | mamo-s | *BGIBMGA011942* | up | 41 | 49 | - | 12.1009 | 2.43E-05 | 0.615 | GTGCGTGGT |
| 10150 | mamo-s | *BGIBMGA011942* | up | 884 | 892 | + | 11.3303 | 4.96E-05 | 0.615 | GTGGGTGGC |
| 10151 | mamo-s | *BGIBMGA011942* | up | 1077 | 1085 | - | 11.3303 | 4.96E-05 | 0.615 | GTGGGTGGC |
| 10152 | mamo-s | *BGIBMGA011945* | down | 175 | 183 | - | 11.3303 | 4.96E-05 | 0.615 | GTGGGTGGC |
| 10153 | mamo-s | *BGIBMGA011946* | up | 336 | 344 | - | 11.3303 | 4.96E-05 | 0.615 | GTGGGTGGC |
| 10154 | mamo-s | *BGIBMGA011947* | up | 1449 | 1457 | - | 11.3303 | 4.96E-05 | 0.615 | GTGGGTGGC |
| 10155 | mamo-s | *BGIBMGA011948* | down | 1696 | 1704 | + | 11.0826 | 6.84E-05 | 0.709 | GTACGTGGC |
| 10156 | mamo-s | *BGIBMGA011949* | down | 1271 | 1279 | - | 11.3303 | 4.96E-05 | 0.615 | GTGGGTGGC |
| 10157 | mamo-s | *BGIBMGA011954* | up | 489 | 497 | - | 11.3303 | 4.96E-05 | 0.615 | GTGGGTGGC |
| 10158 | mamo-s | *BGIBMGA011955* | down | 864 | 872 | + | 14.055 | 2.23E-06 | 0.615 | GTGCGTGGC |
| 10159 | mamo-s | *BGIBMGA011956* | down | 162 | 170 | - | 11.4404 | 4.51E-05 | 0.615 | GAGCGTGTC |
| 10160 | mamo-s | *BGIBMGA011959* | up | 1552 | 1560 | + | 13.6606 | 4.06E-06 | 0.615 | GCGCGTGGC |
| 10161 | mamo-s | *BGIBMGA011959* | down | 45 | 53 | - | 11.0734 | 7.12E-05 | 0.726 | GAGCGTGGT |
| 10162 | mamo-s | *BGIBMGA011960* | up | 1078 | 1086 | - | 11.7064 | 3.52E-05 | 0.615 | GCGCGTGGT |
| 10163 | mamo-s | *BGIBMGA011962* | up | 1865 | 1873 | + | 11.3303 | 4.96E-05 | 0.615 | GTGGGTGGC |
| 10164 | mamo-s | *BGIBMGA011962* | down | 1648 | 1656 | - | 11.1743 | 5.90E-05 | 0.684 | CCGCGTGTC |
| 10165 | mamo-s | *BGIBMGA011966* | down | 1260 | 1268 | + | 10.5413 | 9.42E-05 | 0.857 | CAGCGTGTC |
| 10166 | mamo-s | *BGIBMGA011968* | down | 1491 | 1499 | + | 12.4679 | 1.31E-05 | 0.615 | GTGCGTGTC |
| 10167 | mamo-s | *BGIBMGA011968* | down | 246 | 254 | + | 11.3303 | 4.96E-05 | 0.615 | GTGGGTGGC |
| 10168 | mamo-s | *BGIBMGA011969* | up | 865 | 873 | + | 12.4679 | 1.31E-05 | 0.615 | GTGCGTGTC |
| 10169 | mamo-s | *BGIBMGA011970* | up | 1274 | 1282 | - | 12.7615 | 1.04E-05 | 0.615 | CCGCGTGGC |
| 10170 | mamo-s | *BGIBMGA011971* | up | 1153 | 1161 | + | 12.4679 | 1.31E-05 | 0.615 | GTGCGTGTC |
| 10171 | mamo-s | *BGIBMGA011971* | up | 1272 | 1280 | + | 12.4679 | 1.31E-05 | 0.615 | GTGCGTGTC |
| 10172 | mamo-s | *BGIBMGA011971* | up | 1187 | 1195 | - | 11.4404 | 4.51E-05 | 0.615 | CGGCGTGGC |
| 10173 | mamo-s | *BGIBMGA011972* | down | 1930 | 1938 | + | 11.3303 | 4.96E-05 | 0.615 | GTGGGTGGC |
| 10174 | mamo-s | *BGIBMGA011973* | down | 1801 | 1809 | + | 11.3303 | 4.96E-05 | 0.615 | GTGGGTGGC |
| 10175 | mamo-s | *BGIBMGA011976* | up | 699 | 707 | - | 11.156 | 6.13E-05 | 0.705 | GAGCGTGCC |
| 10176 | mamo-s | *BGIBMGA011979* | up | 1044 | 1052 | - | 12.0734 | 2.66E-05 | 0.615 | GCGCGTGTC |
| 10177 | mamo-s | *BGIBMGA011979* | up | 699 | 707 | + | 11.3303 | 4.96E-05 | 0.615 | GTGGGTGGC |
| 10178 | mamo-s | *BGIBMGA011981* | down | 1962 | 1970 | - | 12.4679 | 1.31E-05 | 0.615 | GTGCGTGTC |
| 10179 | mamo-s | *BGIBMGA011983* | down | 442 | 450 | + | 11.1743 | 5.90E-05 | 0.684 | CCGCGTGTC |
| 10180 | mamo-s | *BGIBMGA011984* | up | 1775 | 1783 | + | 11.1376 | 6.35E-05 | 0.709 | TCGCGTGGC |
| 10181 | mamo-s | *BGIBMGA011985* | up | 536 | 544 | + | 11.4404 | 4.51E-05 | 0.615 | CGGCGTGGC |
| 10182 | mamo-s | *BGIBMGA011985* | down | 710 | 718 | + | 11.3303 | 4.96E-05 | 0.615 | GTGGGTGGC |
| 10183 | mamo-s | *BGIBMGA011985* | up | 654 | 662 | + | 11.2294 | 5.41E-05 | 0.65 | CTGCGAGGC |
| 10184 | mamo-s | *BGIBMGA011986* | up | 1210 | 1218 | - | 11.0826 | 6.84E-05 | 0.709 | GTACGTGGC |
| 10185 | mamo-s | *BGIBMGA011989* | down | 839 | 847 | - | 11.7064 | 3.52E-05 | 0.615 | GCGCGTGGT |
| 10186 | mamo-s | *BGIBMGA011989* | down | 1415 | 1423 | - | 11.4404 | 4.51E-05 | 0.615 | GAGCGTGTC |
| 10187 | mamo-s | *BGIBMGA011997* | up | 619 | 627 | - | 11.3303 | 4.96E-05 | 0.615 | GTGGGTGGC |
| 10188 | mamo-s | *BGIBMGA011997* | down | 1883 | 1891 | - | 11.3303 | 4.96E-05 | 0.615 | GTGGGTGGC |
| 10189 | mamo-s | *BGIBMGA011998* | down | 1434 | 1442 | - | 12.1009 | 2.43E-05 | 0.615 | GTGCGTGGT |
| 10190 | mamo-s | *BGIBMGA011999* | down | 1016 | 1024 | + | 11.3303 | 4.96E-05 | 0.615 | GTGGGTGGC |
| 10191 | mamo-s | *BGIBMGA011999* | up | 1365 | 1373 | + | 11.3303 | 4.96E-05 | 0.615 | GTGGGTGGC |
| 10192 | mamo-s | *BGIBMGA012002* | down | 1990 | 1998 | - | 11.3303 | 4.96E-05 | 0.615 | GTGGGTGGC |
| 10193 | mamo-s | *BGIBMGA012003* | up | 1663 | 1671 | + | 11.3303 | 4.96E-05 | 0.615 | GTGGGTGGC |
| 10194 | mamo-s | *BGIBMGA012003* | up | 1141 | 1149 | - | 11.0826 | 6.84E-05 | 0.709 | GTACGTGGC |
| 10195 | mamo-s | *BGIBMGA012003* | down | 622 | 630 | - | 10.8899 | 7.48E-05 | 0.739 | CCGCGTGCC |
| 10196 | mamo-s | *BGIBMGA012005* | down | 1338 | 1346 | - | 12.1835 | 1.71E-05 | 0.615 | GTGCGTGCC |
| 10197 | mamo-s | *BGIBMGA012007* | down | 1439 | 1447 | + | 12.1284 | 2.16E-05 | 0.615 | CAGCGTGGC |
| 10198 | mamo-s | *BGIBMGA012009* | up | 293 | 301 | + | 11.7339 | 3.29E-05 | 0.615 | GCGCGAGGC |
| 10199 | mamo-s | *BGIBMGA012009* | up | 32 | 40 | - | 11.3303 | 4.96E-05 | 0.615 | GTGGGTGGC |
| 10200 | mamo-s | *BGIBMGA012009* | up | 1453 | 1461 | + | 10.5138 | 9.75E-05 | 0.865 | GTGCGTGTT |
| 10201 | mamo-s | *BGIBMGA012010* | up | 105 | 113 | - | 11.7339 | 3.29E-05 | 0.615 | GCGCGAGGC |

| | | | | | | | | | | |
|---|---|---|---|---|---|---|---|---|---|---|
| 10202 | mamo-s | *BGIBMGA012010* | up | 366 | 374 | + | 11.3303 | 4.96E-05 | 0.615 | GTGGGTGGC |
| 10203 | mamo-s | *BGIBMGA012011* | down | 404 | 412 | - | 12.1284 | 2.16E-05 | 0.615 | CAGCGTGGC |
| 10204 | mamo-s | *BGIBMGA012012* | down | 1020 | 1028 | - | 11.3303 | 4.96E-05 | 0.615 | GTGGGTGGC |
| 10205 | mamo-s | *BGIBMGA012015* | down | 337 | 345 | + | 11.3303 | 4.96E-05 | 0.615 | GTGGGTGGC |
| 10206 | mamo-s | *BGIBMGA012016* | up | 801 | 809 | + | 11.3303 | 4.96E-05 | 0.615 | GTGGGTGGC |
| 10207 | mamo-s | *BGIBMGA012017* | up | 252 | 260 | - | 11.0826 | 6.84E-05 | 0.709 | GTACGTGGC |
| 10208 | mamo-s | *BGIBMGA012018* | up | 920 | 928 | + | 11.3303 | 4.96E-05 | 0.615 | GTGGGTGGC |
| 10209 | mamo-s | *BGIBMGA012019* | up | 1587 | 1595 | - | 11.3303 | 4.96E-05 | 0.615 | GTGGGTGGC |
| 10210 | mamo-s | *BGIBMGA012020* | up | 421 | 429 | + | 12.1835 | 1.71E-05 | 0.615 | GTGCGTGCC |
| 10211 | mamo-s | *BGIBMGA012021* | up | 660 | 668 | + | 11.3303 | 4.96E-05 | 0.615 | GTGGGTGGC |
| 10212 | mamo-s | *BGIBMGA012022* | down | 418 | 426 | - | 11.1743 | 5.90E-05 | 0.684 | CCGCGTGTC |
| 10213 | mamo-s | *BGIBMGA012023* | down | 632 | 640 | + | 11.3853 | 4.74E-05 | 0.615 | ACGCGTGGC |
| 10214 | mamo-s | *BGIBMGA012023* | up | 313 | 321 | - | 11.3303 | 4.96E-05 | 0.615 | GTGGGTGGC |
| 10215 | mamo-s | *BGIBMGA012028* | down | 1417 | 1425 | - | 10.9358 | 7.30E-05 | 0.728 | GCGGGTGGC |
| 10216 | mamo-s | *BGIBMGA012030* | up | 596 | 604 | + | 11.0826 | 6.84E-05 | 0.709 | GTACGTGGC |
| 10217 | mamo-s | *BGIBMGA012034* | down | 908 | 916 | - | 10.6881 | 8.61E-05 | 0.81 | GCACGTGGC |
| 10218 | mamo-s | *BGIBMGA012035* | up | 1640 | 1648 | - | 11.3303 | 4.96E-05 | 0.615 | GTGGGTGGC |
| 10219 | mamo-s | *BGIBMGA012040* | down | 754 | 762 | + | 11.3303 | 4.96E-05 | 0.615 | GTGGGTGGC |
| 10220 | mamo-s | *BGIBMGA012041* | up | 611 | 619 | + | 11.3303 | 4.96E-05 | 0.615 | GTGGGTGGC |
| 10221 | mamo-s | *BGIBMGA012042* | down | 1764 | 1772 | + | 11.3303 | 4.96E-05 | 0.615 | GTGGGTGGC |
| 10222 | mamo-s | *BGIBMGA012047* | up | 54 | 62 | - | 11.3853 | 4.74E-05 | 0.615 | ACGCGTGGC |
| 10223 | mamo-s | *BGIBMGA012047* | down | 1880 | 1888 | + | 11.3303 | 4.96E-05 | 0.615 | GTGGGTGGC |
| 10224 | mamo-s | *BGIBMGA012050* | down | 1725 | 1733 | + | 11.3303 | 4.96E-05 | 0.615 | GTGGGTGGC |
| 10225 | mamo-s | *BGIBMGA012054* | down | 1077 | 1085 | + | 11.4404 | 4.51E-05 | 0.615 | CGGCGTGGC |
| 10226 | mamo-s | *BGIBMGA012054* | up | 1113 | 1121 | + | 11.3303 | 4.96E-05 | 0.615 | GTGGGTGGC |
| 10227 | mamo-s | *BGIBMGA012058* | down | 201 | 209 | + | 10.5138 | 9.75E-05 | 0.865 | GTGCGTGTT |
| 10228 | mamo-s | *BGIBMGA012060* | up | 856 | 864 | - | 11.3303 | 4.96E-05 | 0.615 | GTGGGTGGC |
| 10229 | mamo-s | *BGIBMGA012061* | down | 1765 | 1773 | + | 11.3303 | 4.96E-05 | 0.615 | GTGGGTGGC |
| 10230 | mamo-s | *BGIBMGA012064* | down | 1780 | 1788 | - | 11.3303 | 4.96E-05 | 0.615 | GTGGGTGGC |
| 10231 | mamo-s | *BGIBMGA012065* | down | 1654 | 1662 | + | 11.3303 | 4.96E-05 | 0.615 | GTGGGTGGC |
| 10232 | mamo-s | *BGIBMGA012067* | up | 115 | 123 | - | 11.3303 | 4.96E-05 | 0.615 | GTGGGTGGC |
| 10233 | mamo-s | *BGIBMGA012069* | down | 745 | 753 | - | 12.1284 | 2.16E-05 | 0.615 | CAGCGTGGC |
| 10234 | mamo-s | *BGIBMGA012070* | up | 1296 | 1304 | - | 11.3303 | 4.96E-05 | 0.615 | GTGGGTGGC |
| 10235 | mamo-s | *BGIBMGA012072* | down | 779 | 787 | - | 11.3303 | 4.96E-05 | 0.615 | GTGGGTGGC |
| 10236 | mamo-s | *BGIBMGA012079* | up | 281 | 289 | + | 11.3303 | 4.96E-05 | 0.615 | GTGGGTGGC |
| 10237 | mamo-s | *BGIBMGA012079* | down | 1688 | 1696 | - | 10.9358 | 7.30E-05 | 0.728 | GCGGGTGGC |
| 10238 | mamo-s | *BGIBMGA012080* | down | 1001 | 1009 | + | 11.0734 | 7.12E-05 | 0.726 | GAGCGTGGT |
| 10239 | mamo-s | *BGIBMGA012082* | up | 1403 | 1411 | - | 11.3303 | 4.96E-05 | 0.615 | GTGGGTGGC |
| 10240 | mamo-s | *BGIBMGA012082* | down | 729 | 737 | - | 10.5688 | 8.88E-05 | 0.824 | GTGCGTGAC |
| 10241 | mamo-s | *BGIBMGA012083* | down | 442 | 450 | + | 10.8349 | 7.66E-05 | 0.751 | CCGCGAGGC |
| 10242 | mamo-s | *BGIBMGA012084* | down | 876 | 884 | - | 11.3303 | 4.96E-05 | 0.615 | GTGGGTGGC |
| 10243 | mamo-s | *BGIBMGA012084* | up | 1954 | 1962 | + | 11.2018 | 5.68E-05 | 0.674 | CTGCGTGGT |
| 10244 | mamo-s | *BGIBMGA012085* | up | 1290 | 1298 | + | 11.3303 | 4.96E-05 | 0.615 | GTGGGTGGC |
| 10245 | mamo-s | *BGIBMGA012086* | up | 1830 | 1838 | + | 11.3303 | 4.96E-05 | 0.615 | GTGGGTGGC |
| 10246 | mamo-s | *BGIBMGA012089* | down | 1487 | 1495 | - | 11.1743 | 5.90E-05 | 0.684 | CCGCGTGTC |
| 10247 | mamo-s | *BGIBMGA012090* | down | 705 | 713 | - | 11.3303 | 4.96E-05 | 0.615 | GTGGGTGGC |
| 10248 | mamo-s | *BGIBMGA012090* | up | 1056 | 1064 | + | 11.0826 | 6.84E-05 | 0.709 | GTACGTGGC |
| 10249 | mamo-s | *BGIBMGA012094* | up | 1663 | 1671 | + | 11.3303 | 4.96E-05 | 0.615 | GTGGGTGGC |
| 10250 | mamo-s | *BGIBMGA012094* | up | 1691 | 1699 | + | 11.3303 | 4.96E-05 | 0.615 | GTGGGTGGC |
| 10251 | mamo-s | *BGIBMGA012097* | up | 264 | 272 | - | 11.3303 | 4.96E-05 | 0.615 | GTGGGTGGC |
| 10252 | mamo-s | *BGIBMGA012097* | up | 1841 | 1849 | + | 11.3303 | 4.96E-05 | 0.615 | GTGGGTGGC |
| 10253 | mamo-s | *BGIBMGA012098* | up | 934 | 942 | - | 11.3303 | 4.96E-05 | 0.615 | GTGGGTGGC |
| 10254 | mamo-s | *BGIBMGA012099* | down | 1313 | 1321 | - | 11.0826 | 6.84E-05 | 0.709 | GTACGTGGC |
| 10255 | mamo-s | *BGIBMGA012103* | down | 1734 | 1742 | - | 11.1743 | 5.90E-05 | 0.684 | CCGCGTGTC |
| 10256 | mamo-s | *BGIBMGA012104* | up | 1801 | 1809 | - | 12.4679 | 1.31E-05 | 0.615 | GTGCGTGTC |
| 10257 | mamo-s | *BGIBMGA012105* | up | 1555 | 1563 | + | 11.3303 | 4.96E-05 | 0.615 | GTGGGTGGC |
| 10258 | mamo-s | *BGIBMGA012105* | up | 359 | 367 | - | 10.5413 | 9.42E-05 | 0.857 | GTGCGAGTC |
| 10259 | mamo-s | *BGIBMGA012107* | down | 1015 | 1023 | + | 10.6881 | 8.61E-05 | 0.81 | GCACGTGGC |
| 10260 | mamo-s | *BGIBMGA012109* | down | 1801 | 1809 | + | 11.0826 | 6.84E-05 | 0.709 | GTACGTGGC |
| 10261 | mamo-s | *BGIBMGA012111* | down | 397 | 405 | + | 11.3303 | 4.96E-05 | 0.615 | GTGGGTGGC |
| 10262 | mamo-s | *BGIBMGA012112* | down | 1798 | 1806 | - | 11.3303 | 4.96E-05 | 0.615 | GTGGGTGGC |
| 10263 | mamo-s | *BGIBMGA012112* | up | 201 | 209 | - | 11.1009 | 6.57E-05 | 0.709 | GAGCGAGGC |
| 10264 | mamo-s | *BGIBMGA012115* | down | 912 | 920 | - | 13.6606 | 4.06E-06 | 0.615 | GCGCGTGGC |
| 10265 | mamo-s | *BGIBMGA012115* | up | 631 | 639 | - | 11.3303 | 4.96E-05 | 0.615 | GTGGGTGGC |
| 10266 | mamo-s | *BGIBMGA012126* | down | 1363 | 1371 | - | 11.2018 | 5.68E-05 | 0.674 | CTGCGTGGT |
| 10267 | mamo-s | *BGIBMGA012129* | up | 969 | 977 | - | 11.5688 | 3.79E-05 | 0.615 | CTGCGTGTC |
| 10268 | mamo-s | *BGIBMGA012129* | up | 764 | 772 | + | 11.3303 | 4.96E-05 | 0.615 | GTGGGTGGC |
| 10269 | mamo-s | *BGIBMGA012136* | up | 1979 | 1987 | - | 13.156 | 6.30E-06 | 0.615 | CTGCGTGGC |
| 10270 | mamo-s | *BGIBMGA012137* | down | 980 | 988 | + | 11.3303 | 4.96E-05 | 0.615 | GTGGGTGGC |
| 10271 | mamo-s | *BGIBMGA012138* | down | 885 | 893 | - | 11.3303 | 4.96E-05 | 0.615 | GTGGGTGGC |
| 10272 | mamo-s | *BGIBMGA012138* | up | 1633 | 1641 | + | 11.0826 | 6.84E-05 | 0.709 | GTACGTGGC |
| 10273 | mamo-s | *BGIBMGA012139* | up | 1138 | 1146 | - | 14.055 | 2.23E-06 | 0.615 | GTGCGTGGC |
| 10274 | mamo-s | *BGIBMGA012141* | down | 1012 | 1020 | - | 12.1009 | 2.43E-05 | 0.615 | GTGCGTGGT |
| 10275 | mamo-s | *BGIBMGA012141* | up | 724 | 732 | - | 11.3853 | 4.74E-05 | 0.615 | ACGCGTGGC |
| 10276 | mamo-s | *BGIBMGA012143* | up | 608 | 616 | - | 11.1009 | 6.57E-05 | 0.709 | GAGCGAGGC |
| 10277 | mamo-s | *BGIBMGA012145* | up | 1756 | 1764 | + | 11.3303 | 4.96E-05 | 0.615 | GTGGGTGGC |
| 10278 | mamo-s | *BGIBMGA012146* | up | 1349 | 1357 | + | 11.3303 | 4.96E-05 | 0.615 | GTGGGTGGC |
| 10279 | mamo-s | *BGIBMGA012147* | down | 268 | 276 | - | 11.3303 | 4.96E-05 | 0.615 | GTGGGTGGC |
| 10280 | mamo-s | *BGIBMGA012147* | down | 1624 | 1632 | - | 11.3303 | 4.96E-05 | 0.615 | GTGGGTGGC |
| 10281 | mamo-s | *BGIBMGA012147* | down | 1208 | 1216 | - | 11.0826 | 6.84E-05 | 0.709 | GTACGTGGC |
| 10282 | mamo-s | *BGIBMGA012149* | up | 529 | 537 | - | 11.3303 | 4.96E-05 | 0.615 | GTGGGTGGC |
| 10283 | mamo-s | *BGIBMGA012150* | down | 337 | 345 | + | 11.3303 | 4.96E-05 | 0.615 | GTGGGTGGC |
| 10284 | mamo-s | *BGIBMGA012152* | down | 1371 | 1379 | + | 11.3303 | 4.96E-05 | 0.615 | GTGGGTGGC |
| 10285 | mamo-s | *BGIBMGA012153* | down | 225 | 233 | - | 11.3303 | 4.96E-05 | 0.615 | GTGGGTGGC |
| 10286 | mamo-s | *BGIBMGA012154* | down | 1597 | 1605 | + | 11.3303 | 4.96E-05 | 0.615 | GTGGGTGGC |
| 10287 | mamo-s | *BGIBMGA012155* | up | 1427 | 1435 | - | 11.3303 | 4.96E-05 | 0.615 | GTGGGTGGC |
| 10288 | mamo-s | *BGIBMGA012156* | down | 80 | 88 | + | 11.4404 | 4.51E-05 | 0.615 | GAGCGTGTC |
| 10289 | mamo-s | *BGIBMGA012157* | up | 27 | 35 | + | 13.156 | 6.30E-06 | 0.615 | CTGCGTGGC |

| | | | | | | | | | | |
|---|---|---|---|---|---|---|---|---|---|---|
| 10290 | mamo-s | *BGIBMGA012157* | down | 1549 | 1557 | - | 11.3303 | 4.96E-05 | 0.615 | GTGGGTGGC |
| 10291 | mamo-s | *BGIBMGA012158* | down | 1341 | 1349 | + | 11.3303 | 4.96E-05 | 0.615 | GTGGGTGGC |
| 10292 | mamo-s | *BGIBMGA012161* | down | 894 | 902 | - | 11.3303 | 4.96E-05 | 0.615 | GTGGGTGGC |
| 10293 | mamo-s | *BGIBMGA012161* | up | 1801 | 1809 | + | 11.2294 | 5.41E-05 | 0.65 | CTGCGAGGC |
| 10294 | mamo-s | *BGIBMGA012161* | down | 594 | 602 | - | 10.5138 | 9.75E-05 | 0.865 | GTGCGTGTT |
| 10295 | mamo-s | *BGIBMGA012162* | up | 686 | 694 | - | 11.3303 | 4.96E-05 | 0.615 | GTGGGTGGC |
| 10296 | mamo-s | *BGIBMGA012162* | up | 386 | 394 | - | 10.5138 | 9.75E-05 | 0.865 | GTGCGTGTT |
| 10297 | mamo-s | *BGIBMGA012164* | up | 1368 | 1376 | + | 11.1009 | 6.57E-05 | 0.709 | GAGCGAGGC |
| 10298 | mamo-s | *BGIBMGA012165* | down | 370 | 378 | - | 11.3303 | 4.96E-05 | 0.615 | GTGGGTGGC |
| 10299 | mamo-s | *BGIBMGA012169* | down | 1000 | 1008 | + | 12.7615 | 1.04E-05 | 0.615 | CCGCGTGGC |
| 10300 | mamo-s | *BGIBMGA012169* | up | 1342 | 1350 | + | 11.3303 | 4.96E-05 | 0.615 | GTGGGTGGC |
| 10301 | mamo-s | *BGIBMGA012169* | up | 394 | 402 | - | 10.7523 | 8.38E-05 | 0.797 | AAGCGTGGC |
| 10302 | mamo-s | *BGIBMGA012173* | down | 1449 | 1457 | + | 11.3303 | 4.96E-05 | 0.615 | GTGGGTGGC |
| 10303 | mamo-s | *BGIBMGA012174* | down | 164 | 172 | + | 11.3303 | 4.96E-05 | 0.615 | GTGGGTGGC |
| 10304 | mamo-s | *BGIBMGA012175* | up | 276 | 284 | - | 12.4679 | 1.31E-05 | 0.615 | GTGCGTGTC |
| 10305 | mamo-s | *BGIBMGA012175* | up | 440 | 448 | + | 11.3303 | 4.96E-05 | 0.615 | GTGGGTGGC |
| 10306 | mamo-s | *BGIBMGA012175* | down | 1882 | 1890 | + | 11.3303 | 4.96E-05 | 0.615 | GTGGGTGGC |
| 10307 | mamo-s | *BGIBMGA012176* | up | 1366 | 1374 | + | 11.3303 | 4.96E-05 | 0.615 | GTGGGTGGC |
| 10308 | mamo-s | *BGIBMGA012176* | down | 1277 | 1285 | + | 10.5138 | 9.75E-05 | 0.865 | GTGCGTGTT |
| 10309 | mamo-s | *BGIBMGA012177* | up | 1252 | 1260 | - | 10.5138 | 9.75E-05 | 0.865 | GTGCGTGTT |
| 10310 | mamo-s | *BGIBMGA012177* | down | 1327 | 1335 | - | 10.5138 | 9.75E-05 | 0.865 | GTGCGTGTT |
| 10311 | mamo-s | *BGIBMGA012180* | down | 646 | 654 | - | 11.2294 | 5.41E-05 | 0.65 | CTGCGAGGC |
| 10312 | mamo-s | *BGIBMGA012181* | up | 1111 | 1119 | - | 11.0826 | 6.84E-05 | 0.709 | GTACGTGGC |
| 10313 | mamo-s | *BGIBMGA012183* | down | 1684 | 1692 | + | 10.5138 | 9.75E-05 | 0.865 | GTGCGTGTT |
| 10314 | mamo-s | *BGIBMGA012183* | down | 1828 | 1836 | - | 10.5138 | 9.75E-05 | 0.865 | GTGCGTGTT |
| 10315 | mamo-s | *BGIBMGA012184* | up | 690 | 698 | + | 10.5138 | 9.75E-05 | 0.865 | GTGCGTGTT |
| 10316 | mamo-s | *BGIBMGA012184* | up | 834 | 842 | - | 10.5138 | 9.75E-05 | 0.865 | GTGCGTGTT |
| 10317 | mamo-s | *BGIBMGA012186* | up | 1392 | 1400 | + | 11.3303 | 4.96E-05 | 0.615 | GTGGGTGGC |
| 10318 | mamo-s | *BGIBMGA012186* | up | 1483 | 1491 | + | 11.3303 | 4.96E-05 | 0.615 | GTGGGTGGC |
| 10319 | mamo-s | *BGIBMGA012186* | up | 1574 | 1582 | + | 11.3303 | 4.96E-05 | 0.615 | GTGGGTGGC |
| 10320 | mamo-s | *BGIBMGA012186* | down | 630 | 638 | - | 10.9358 | 7.30E-05 | 0.728 | GCGGGTGGC |
| 10321 | mamo-s | *BGIBMGA012186* | up | 1665 | 1673 | + | 10.9358 | 7.30E-05 | 0.728 | GCGGGTGGC |
| 10322 | mamo-s | *BGIBMGA012187* | down | 338 | 346 | - | 11.3303 | 4.96E-05 | 0.615 | GTGGGTGGC |
| 10323 | mamo-s | *BGIBMGA012188* | up | 1968 | 1976 | + | 13.0275 | 8.53E-06 | 0.615 | GAGCGTGGC |
| 10324 | mamo-s | *BGIBMGA012188* | down | 125 | 133 | - | 11.3303 | 4.96E-05 | 0.615 | GTGGGTGGC |
| 10325 | mamo-s | *BGIBMGA012189* | up | 1107 | 1115 | - | 11.1009 | 6.57E-05 | 0.709 | GAGCGAGGC |
| 10326 | mamo-s | *BGIBMGA012190* | down | 1570 | 1578 | - | 11.3303 | 4.96E-05 | 0.615 | GTGGGTGGC |
| 10327 | mamo-s | *BGIBMGA012191* | up | 483 | 491 | + | 11.3303 | 4.96E-05 | 0.615 | GTGGGTGGC |
| 10328 | mamo-s | *BGIBMGA012191* | down | 411 | 419 | - | 10.6881 | 8.61E-05 | 0.81 | GCACGTGGC |
| 10329 | mamo-s | *BGIBMGA012192* | down | 520 | 528 | - | 11.2844 | 5.18E-05 | 0.63 | CTGCGTGCC |
| 10330 | mamo-s | *BGIBMGA012193* | up | 24 | 32 | - | 11.3303 | 4.96E-05 | 0.615 | GTGGGTGGC |
| 10331 | mamo-s | *BGIBMGA012193* | down | 1830 | 1838 | + | 11.3303 | 4.96E-05 | 0.615 | GTGGGTGGC |
| 10332 | mamo-s | *BGIBMGA012194* | down | 1663 | 1671 | + | 12.7615 | 1.04E-05 | 0.615 | CCGCGTGGC |
| 10333 | mamo-s | *BGIBMGA012195* | down | 141 | 149 | + | 12.3394 | 1.49E-05 | 0.615 | GGGCGTGGC |
| 10334 | mamo-s | *BGIBMGA012195* | down | 1095 | 1103 | + | 11.5321 | 4.06E-05 | 0.615 | TTGCGTGGC |
| 10335 | mamo-s | *BGIBMGA012195* | down | 905 | 913 | + | 11.3853 | 4.74E-05 | 0.615 | ACGCGTGGC |
| 10336 | mamo-s | *BGIBMGA012195* | down | 757 | 765 | - | 11.3303 | 4.96E-05 | 0.615 | GTGGGTGGC |
| 10337 | mamo-s | *BGIBMGA012196* | up | 1723 | 1731 | - | 11.3303 | 4.96E-05 | 0.615 | GTGGGTGGC |
| 10338 | mamo-s | *BGIBMGA012199* | down | 1638 | 1646 | - | 11.3303 | 4.96E-05 | 0.615 | GTGGGTGGC |
| 10339 | mamo-s | *BGIBMGA012200* | up | 411 | 419 | - | 11.3303 | 4.96E-05 | 0.615 | GTGGGTGGC |
| 10340 | mamo-s | *BGIBMGA012200* | down | 1218 | 1226 | + | 11.3303 | 4.96E-05 | 0.615 | GTGGGTGGC |
| 10341 | mamo-s | *BGIBMGA012201* | up | 948 | 956 | - | 11.3303 | 4.96E-05 | 0.615 | GTGGGTGGC |
| 10342 | mamo-s | *BGIBMGA012201* | down | 1716 | 1724 | - | 11.0826 | 6.84E-05 | 0.709 | GTACGTGGC |
| 10343 | mamo-s | *BGIBMGA012202* | up | 352 | 360 | + | 14.055 | 2.23E-06 | 0.615 | GTGCGTGGC |
| 10344 | mamo-s | *BGIBMGA012202* | up | 950 | 958 | + | 12.1835 | 1.71E-05 | 0.615 | GTGCGTGCC |
| 10345 | mamo-s | *BGIBMGA012202* | up | 425 | 433 | - | 11.7339 | 3.29E-05 | 0.615 | GCGCGAGGC |
| 10346 | mamo-s | *BGIBMGA012203* | up | 638 | 646 | + | 11.2844 | 5.18E-05 | 0.63 | CTGCGTGCC |
| 10347 | mamo-s | *BGIBMGA012204* | down | 1517 | 1525 | + | 11.3303 | 4.96E-05 | 0.615 | GTGGGTGGC |
| 10348 | mamo-s | *BGIBMGA012205* | up | 1541 | 1549 | + | 11.3303 | 4.96E-05 | 0.615 | GTGGGTGGC |
| 10349 | mamo-s | *BGIBMGA012207* | up | 268 | 276 | + | 11.3303 | 4.96E-05 | 0.615 | GTGGGTGGC |
| 10350 | mamo-s | *BGIBMGA012207* | up | 1373 | 1381 | + | 11.0826 | 6.84E-05 | 0.709 | GTACGTGGC |
| 10351 | mamo-s | *BGIBMGA012209* | up | 380 | 388 | + | 11.3303 | 4.96E-05 | 0.615 | GTGGGTGGC |
| 10352 | mamo-s | *BGIBMGA012209* | down | 556 | 564 | - | 11.3303 | 4.96E-05 | 0.615 | GTGGGTGGC |
| 10353 | mamo-s | *BGIBMGA012210* | down | 303 | 311 | + | 11.3303 | 4.96E-05 | 0.615 | GTGGGTGGC |
| 10354 | mamo-s | *BGIBMGA012210* | down | 667 | 675 | + | 10.5138 | 9.75E-05 | 0.865 | GTGCGTGTT |
| 10355 | mamo-s | *BGIBMGA012211* | down | 6 | 14 | - | 12.0734 | 2.66E-05 | 0.615 | GCGCGTGTC |
| 10356 | mamo-s | *BGIBMGA012211* | down | 945 | 953 | - | 11.3303 | 4.96E-05 | 0.615 | GTGGGTGGC |
| 10357 | mamo-s | *BGIBMGA012211* | up | 1320 | 1328 | - | 11.0826 | 6.84E-05 | 0.709 | GTACGTGGC |
| 10358 | mamo-s | *BGIBMGA012214* | down | 775 | 783 | - | 11.3303 | 4.96E-05 | 0.615 | GTGGGTGGC |
| 10359 | mamo-s | *BGIBMGA012215* | down | 134 | 142 | + | 11.3303 | 4.96E-05 | 0.615 | GTGGGTGGC |
| 10360 | mamo-s | *BGIBMGA012216* | up | 1711 | 1719 | - | 14.055 | 2.23E-06 | 0.615 | GTGCGTGGC |
| 10361 | mamo-s | *BGIBMGA012217* | down | 53 | 61 | + | 11.7064 | 3.52E-05 | 0.615 | GCGCGTGGT |
| 10362 | mamo-s | *BGIBMGA012218* | down | 491 | 499 | + | 12.1284 | 2.16E-05 | 0.615 | GTGCGAGGC |
| 10363 | mamo-s | *BGIBMGA012218* | up | 681 | 689 | + | 11.5321 | 4.06E-05 | 0.615 | TTGCGTGGC |
| 10364 | mamo-s | *BGIBMGA012218* | down | 1751 | 1759 | - | 11.3303 | 4.96E-05 | 0.615 | GTGGGTGGC |
| 10365 | mamo-s | *BGIBMGA012219* | up | 1237 | 1245 | + | 11.3303 | 4.96E-05 | 0.615 | GTGGGTGGC |
| 10366 | mamo-s | *BGIBMGA012221* | up | 1489 | 1497 | + | 11.3303 | 4.96E-05 | 0.615 | GTGGGTGGC |
| 10367 | mamo-s | *BGIBMGA012221* | down | 1780 | 1788 | + | 11.3303 | 4.96E-05 | 0.615 | GTGGGTGGC |
| 10368 | mamo-s | *BGIBMGA012222* | up | 1314 | 1322 | + | 11.3303 | 4.96E-05 | 0.615 | GTGGGTGGC |
| 10369 | mamo-s | *BGIBMGA012222* | down | 1601 | 1609 | + | 11.3303 | 4.96E-05 | 0.615 | GTGGGTGGC |
| 10370 | mamo-s | *BGIBMGA012223* | up | 1000 | 1008 | + | 11.3303 | 4.96E-05 | 0.615 | GTGGGTGGC |
| 10371 | mamo-s | *BGIBMGA012225* | down | 1471 | 1479 | + | 11.3303 | 4.96E-05 | 0.615 | GTGGGTGGC |
| 10372 | mamo-s | *BGIBMGA012226* | down | 427 | 435 | - | 11.5321 | 4.06E-05 | 0.615 | TTGCGTGGC |
| 10373 | mamo-s | *BGIBMGA012227* | up | 1047 | 1055 | - | 11.2844 | 5.18E-05 | 0.63 | CTGCGTGCC |
| 10374 | mamo-s | *BGIBMGA012229* | up | 523 | 531 | - | 11.3303 | 4.96E-05 | 0.615 | GTGGGTGGC |
| 10375 | mamo-s | *BGIBMGA012229* | up | 1311 | 1319 | - | 11.3303 | 4.96E-05 | 0.615 | GTGGGTGGC |
| 10376 | mamo-s | *BGIBMGA012231* | down | 1242 | 1250 | + | 11.3303 | 4.96E-05 | 0.615 | GTGGGTGGC |
| 10377 | mamo-s | *BGIBMGA012234* | down | 25 | 33 | + | 11.4404 | 4.51E-05 | 0.615 | CGGCGTGGC |

| | | | | | | | | | | |
|---|---|---|---|---|---|---|---|---|---|---|
| 10378 | mamo-s | *BGIBMGA012234* | up | 1335 | 1343 | - | 10.7523 | 8.38E-05 | 0.797 | GGGCGTGTC |
| 10379 | mamo-s | *BGIBMGA012238* | down | 538 | 546 | + | 11.3303 | 4.96E-05 | 0.615 | GTGGGTGGC |
| 10380 | mamo-s | *BGIBMGA012238* | up | 1445 | 1453 | - | 11.3303 | 4.96E-05 | 0.615 | GTGGGTGGC |
| 10381 | mamo-s | *BGIBMGA012242* | up | 299 | 307 | - | 13.0275 | 8.53E-06 | 0.615 | GAGCGTGGC |
| 10382 | mamo-s | *BGIBMGA012243* | up | 1628 | 1636 | - | 11.7064 | 3.52E-05 | 0.615 | GCGCGTGGT |
| 10383 | mamo-s | *BGIBMGA012243* | down | 359 | 367 | + | 11.5321 | 4.06E-05 | 0.615 | TTGCGTGGC |
| 10384 | mamo-s | *BGIBMGA012244* | up | 285 | 293 | + | 11.3303 | 4.96E-05 | 0.615 | GTGGGTGGC |
| 10385 | mamo-s | *BGIBMGA012244* | down | 1547 | 1555 | + | 10.9358 | 7.30E-05 | 0.728 | GCGGGTGGC |
| 10386 | mamo-s | *BGIBMGA012246* | down | 1096 | 1104 | + | 11.7064 | 3.52E-05 | 0.615 | GCGCGTGGT |
| 10387 | mamo-s | *BGIBMGA012252* | down | 1037 | 1045 | + | 11.3303 | 4.96E-05 | 0.615 | GTGGGTGGC |
| 10388 | mamo-s | *BGIBMGA012252* | down | 1410 | 1418 | + | 10.7523 | 8.38E-05 | 0.797 | GGGCGTGTC |
| 10389 | mamo-s | *BGIBMGA012253* | up | 156 | 164 | - | 11.3303 | 4.96E-05 | 0.615 | GTGGGTGGC |
| 10390 | mamo-s | *BGIBMGA012256* | up | 1088 | 1096 | - | 11.3303 | 4.96E-05 | 0.615 | GTGGGTGGC |
| 10391 | mamo-s | *BGIBMGA012257* | up | 753 | 761 | - | 11.3303 | 4.96E-05 | 0.615 | GTGGGTGGC |
| 10392 | mamo-s | *BGIBMGA012258* | up | 1373 | 1381 | - | 12.1009 | 2.43E-05 | 0.615 | GTGCGTGGT |
| 10393 | mamo-s | *BGIBMGA012258* | down | 701 | 709 | - | 11.4404 | 4.51E-05 | 0.615 | CGGCGTGGC |
| 10394 | mamo-s | *BGIBMGA012258* | up | 522 | 530 | + | 10.8073 | 7.89E-05 | 0.766 | CCGCGTGGT |
| 10395 | mamo-s | *BGIBMGA012262* | down | 1646 | 1654 | + | 11.0826 | 6.84E-05 | 0.709 | GTACGTGGC |
| 10396 | mamo-s | *BGIBMGA012264* | up | 1925 | 1933 | + | 13.0275 | 8.53E-06 | 0.615 | GAGCGTGGC |
| 10397 | mamo-s | *BGIBMGA012265* | up | 816 | 824 | + | 10.5688 | 8.88E-05 | 0.824 | GTGCGTGAC |
| 10398 | mamo-s | *BGIBMGA012266* | down | 453 | 461 | - | 11.0826 | 6.84E-05 | 0.709 | GTACGTGGC |
| 10399 | mamo-s | *BGIBMGA012269* | down | 1491 | 1499 | - | 11.3303 | 4.96E-05 | 0.615 | GTGGGTGGC |
| 10400 | mamo-s | *BGIBMGA012270* | down | 1828 | 1836 | - | 11.3303 | 4.96E-05 | 0.615 | GTGGGTGGC |
| 10401 | mamo-s | *BGIBMGA012272* | up | 1516 | 1524 | - | 11.3303 | 4.96E-05 | 0.615 | GTGGGTGGC |
| 10402 | mamo-s | *BGIBMGA012273* | down | 1867 | 1875 | - | 11.3303 | 4.96E-05 | 0.615 | GTGGGTGGC |
| 10403 | mamo-s | *BGIBMGA012274* | up | 1391 | 1399 | - | 11.3303 | 4.96E-05 | 0.615 | GTGGGTGGC |
| 10404 | mamo-s | *BGIBMGA012276* | up | 1973 | 1981 | - | 12.0734 | 2.66E-05 | 0.615 | GCGCGTGTC |
| 10405 | mamo-s | *BGIBMGA012276* | down | 1671 | 1679 | + | 11.3303 | 4.96E-05 | 0.615 | GTGGGTGGC |
| 10406 | mamo-s | *BGIBMGA012276* | up | 389 | 397 | - | 10.5138 | 9.75E-05 | 0.865 | GTGCGTGTT |
| 10407 | mamo-s | *BGIBMGA012281* | down | 34 | 42 | - | 11.3303 | 4.96E-05 | 0.615 | GTGGGTGGC |
| 10408 | mamo-s | *BGIBMGA012283* | up | 1240 | 1248 | - | 12.1284 | 2.16E-05 | 0.615 | GTGCGAGGC |
| 10409 | mamo-s | *BGIBMGA012285* | up | 1611 | 1619 | + | 11.3303 | 4.96E-05 | 0.615 | GTGGGTGGC |
| 10410 | mamo-s | *BGIBMGA012286* | down | 835 | 843 | + | 11.3303 | 4.96E-05 | 0.615 | GTGGGTGGC |
| 10411 | mamo-s | *BGIBMGA012287* | up | 509 | 517 | + | 11.3303 | 4.96E-05 | 0.615 | GTGGGTGGC |
| 10412 | mamo-s | *BGIBMGA012288* | up | 33 | 41 | - | 11.3303 | 4.96E-05 | 0.615 | GTGGGTGGC |
| 10413 | mamo-s | *BGIBMGA012293* | up | 488 | 496 | + | 11.3303 | 4.96E-05 | 0.615 | GTGGGTGGC |
| 10414 | mamo-s | *BGIBMGA012293* | down | 1310 | 1318 | + | 11.3303 | 4.96E-05 | 0.615 | GTGGGTGGC |
| 10415 | mamo-s | *BGIBMGA012294* | up | 15 | 23 | + | 11.3303 | 4.96E-05 | 0.615 | GTGGGTGGC |
| 10416 | mamo-s | *BGIBMGA012296* | up | 1735 | 1743 | + | 11.2294 | 5.41E-05 | 0.65 | CTGCGAGGC |
| 10417 | mamo-s | *BGIBMGA012299* | up | 449 | 457 | + | 11.3303 | 4.96E-05 | 0.615 | GTGGGTGGC |
| 10418 | mamo-s | *BGIBMGA012301* | up | 870 | 878 | + | 12.1835 | 1.71E-05 | 0.615 | GTGCGTGCC |
| 10419 | mamo-s | *BGIBMGA012302* | down | 539 | 547 | - | 11.3303 | 4.96E-05 | 0.615 | GTGGGTGGC |
| 10420 | mamo-s | *BGIBMGA012302* | up | 1741 | 1749 | + | 10.5138 | 9.75E-05 | 0.865 | GTGCGTGTT |
| 10421 | mamo-s | *BGIBMGA012304* | down | 1892 | 1900 | - | 12.1284 | 2.16E-05 | 0.615 | GTGCGAGGC |
| 10422 | mamo-s | *BGIBMGA012304* | down | 951 | 959 | - | 10.5138 | 9.75E-05 | 0.865 | GTGCGTGTT |
| 10423 | mamo-s | *BGIBMGA012305* | down | 573 | 581 | - | 11.3303 | 4.96E-05 | 0.615 | GTGGGTGGC |
| 10424 | mamo-s | *BGIBMGA012306* | down | 1742 | 1750 | + | 11.3853 | 4.74E-05 | 0.615 | ACGCGTGGC |
| 10425 | mamo-s | *BGIBMGA012306* | up | 1504 | 1512 | - | 11.2844 | 5.18E-05 | 0.63 | CTGCGTGCC |
| 10426 | mamo-s | *BGIBMGA012309* | up | 1793 | 1801 | + | 12.1835 | 1.71E-05 | 0.615 | GTGCGTGCC |
| 10427 | mamo-s | *BGIBMGA012310* | down | 1594 | 1602 | - | 11.3303 | 4.96E-05 | 0.615 | GTGGGTGGC |
| 10428 | mamo-s | *BGIBMGA012314* | down | 1403 | 1411 | - | 11.3303 | 4.96E-05 | 0.615 | GTGGGTGGC |
| 10429 | mamo-s | *BGIBMGA012314* | up | 829 | 837 | - | 11.1743 | 5.90E-05 | 0.684 | CCGCGTGTC |
| 10430 | mamo-s | *BGIBMGA012316* | down | 797 | 805 | + | 10.9358 | 7.30E-05 | 0.728 | GCGGGTGGC |
| 10431 | mamo-s | *BGIBMGA012317* | down | 1707 | 1715 | + | 10.5413 | 9.42E-05 | 0.857 | GTGCGAGTC |
| 10432 | mamo-s | *BGIBMGA012318* | down | 289 | 297 | + | 11.3303 | 4.96E-05 | 0.615 | GTGGGTGGC |
| 10433 | mamo-s | *BGIBMGA012319* | up | 286 | 294 | + | 11.3303 | 4.96E-05 | 0.615 | GTGGGTGGC |
| 10434 | mamo-s | *BGIBMGA012319* | down | 1079 | 1087 | - | 11.3303 | 4.96E-05 | 0.615 | GTGGGTGGC |
| 10435 | mamo-s | *BGIBMGA012322* | up | 930 | 938 | + | 12.1835 | 1.71E-05 | 0.615 | GTGCGTGCC |
| 10436 | mamo-s | *BGIBMGA012322* | up | 465 | 473 | - | 11.2294 | 5.41E-05 | 0.65 | CTGCGAGGC |
| 10437 | mamo-s | *BGIBMGA012323* | up | 1423 | 1431 | - | 12.0734 | 2.66E-05 | 0.615 | GCGCGTGTC |
| 10438 | mamo-s | *BGIBMGA012323* | down | 137 | 145 | + | 10.5138 | 9.75E-05 | 0.865 | GTGCGTGTT |
| 10439 | mamo-s | *BGIBMGA012329* | down | 1986 | 1994 | + | 11.4404 | 4.51E-05 | 0.615 | GAGCGTGTC |
| 10440 | mamo-s | *BGIBMGA012335* | up | 1301 | 1309 | + | 11.5321 | 4.06E-05 | 0.615 | TTGCGTGGC |
| 10441 | mamo-s | *BGIBMGA012338* | up | 786 | 794 | + | 11.3303 | 4.96E-05 | 0.615 | GTGGGTGGC |
| 10442 | mamo-s | *BGIBMGA012344* | down | 762 | 770 | - | 11.3303 | 4.96E-05 | 0.615 | GTGGGTGGC |
| 10443 | mamo-s | *BGIBMGA012345* | down | 996 | 1004 | - | 11.7064 | 3.52E-05 | 0.615 | GCGCGTGGT |
| 10444 | mamo-s | *BGIBMGA012346* | up | 1499 | 1507 | + | 11.3303 | 4.96E-05 | 0.615 | GTGGGTGGC |
| 10445 | mamo-s | *BGIBMGA012347* | down | 558 | 566 | - | 10.5413 | 9.42E-05 | 0.857 | CAGCGTGTC |
| 10446 | mamo-s | *BGIBMGA012349* | up | 869 | 877 | - | 11.7798 | 3.11E-05 | 0.615 | ATGCGTGGC |
| 10447 | mamo-s | *BGIBMGA012350* | down | 843 | 851 | + | 11.5688 | 3.79E-05 | 0.615 | CTGCGTGTC |
| 10448 | mamo-s | *BGIBMGA012350* | down | 363 | 371 | + | 11.4404 | 4.51E-05 | 0.615 | GAGCGTGTC |
| 10449 | mamo-s | *BGIBMGA012350* | down | 938 | 946 | + | 11.3303 | 4.96E-05 | 0.615 | GTGGGTGGC |
| 10450 | mamo-s | *BGIBMGA012355* | down | 88 | 96 | - | 12.0734 | 2.66E-05 | 0.615 | GCGCGTGTC |
| 10451 | mamo-s | *BGIBMGA012356* | down | 429 | 437 | - | 11.3303 | 4.96E-05 | 0.615 | GTGGGTGGC |
| 10452 | mamo-s | *BGIBMGA012361* | down | 443 | 451 | + | 11.2018 | 5.68E-05 | 0.674 | CTGCGTGGT |
| 10453 | mamo-s | *BGIBMGA012363* | up | 1204 | 1212 | + | 11.3303 | 4.96E-05 | 0.615 | GTGGGTGGC |
| 10454 | mamo-s | *BGIBMGA012365* | up | 102 | 110 | + | 11.3303 | 4.96E-05 | 0.615 | GTGGGTGGC |
| 10455 | mamo-s | *BGIBMGA012365* | down | 580 | 588 | - | 11.3303 | 4.96E-05 | 0.615 | GTGGGTGGC |
| 10456 | mamo-s | *BGIBMGA012366* | down | 1433 | 1441 | + | 11.3303 | 4.96E-05 | 0.615 | GTGGGTGGC |
| 10457 | mamo-s | *BGIBMGA012369* | down | 595 | 603 | - | 11.3303 | 4.96E-05 | 0.615 | GTGGGTGGC |
| 10458 | mamo-s | *BGIBMGA012370* | up | 956 | 964 | + | 11.3303 | 4.96E-05 | 0.615 | GTGGGTGGC |
| 10459 | mamo-s | *BGIBMGA012372* | down | 1303 | 1311 | - | 11.3303 | 4.96E-05 | 0.615 | GTGGGTGGC |
| 10460 | mamo-s | *BGIBMGA012373* | down | 1548 | 1556 | - | 11.2294 | 5.41E-05 | 0.65 | CTGCGAGGC |
| 10461 | mamo-s | *BGIBMGA012374* | up | 539 | 547 | + | 11.3303 | 4.96E-05 | 0.615 | GTGGGTGGC |
| 10462 | mamo-s | *BGIBMGA012374* | up | 1468 | 1476 | - | 11.3303 | 4.96E-05 | 0.615 | GTGGGTGGC |
| 10463 | mamo-s | *BGIBMGA012374* | up | 11 | 19 | - | 11.2294 | 5.41E-05 | 0.65 | CTGCGAGGC |
| 10464 | mamo-s | *BGIBMGA012375* | down | 418 | 426 | - | 11.3303 | 4.96E-05 | 0.615 | GTGGGTGGC |
| 10465 | mamo-s | *BGIBMGA012375* | up | 401 | 409 | + | 11.0826 | 6.84E-05 | 0.709 | GTACGTGGC |

| | | | | | | | | | | |
|---|---|---|---|---|---|---|---|---|---|---|
| 10466 | mamo-s | *BGIBMGA012377* | down | 786 | 794 | - | 12.4679 | 1.31E-05 | 0.615 | GTGCGTGTC |
| 10467 | mamo-s | *BGIBMGA012377* | down | 430 | 438 | + | 11.1376 | 6.35E-05 | 0.709 | TCGCGTGGC |
| 10468 | mamo-s | *BGIBMGA012379* | down | 1387 | 1395 | + | 11.3303 | 4.96E-05 | 0.615 | GTGGGTGGC |
| 10469 | mamo-s | *BGIBMGA012380* | up | 1486 | 1494 | + | 11.3303 | 4.96E-05 | 0.615 | GTGGGTGGC |
| 10470 | mamo-s | *BGIBMGA012381* | down | 720 | 728 | + | 11.0826 | 6.84E-05 | 0.709 | GTACGTGGC |
| 10471 | mamo-s | *BGIBMGA012382* | down | 1566 | 1574 | + | 13.6606 | 4.06E-06 | 0.615 | GCGCGTGGC |
| 10472 | mamo-s | *BGIBMGA012383* | down | 1182 | 1190 | + | 12.3394 | 1.49E-05 | 0.615 | GGGCGTGGC |
| 10473 | mamo-s | *BGIBMGA012383* | up | 155 | 163 | + | 11.3303 | 4.96E-05 | 0.615 | GTGGGTGGC |
| 10474 | mamo-s | *BGIBMGA012384* | down | 1223 | 1231 | + | 11.2018 | 5.68E-05 | 0.674 | CTGCGTGGT |
| 10475 | mamo-s | *BGIBMGA012385* | down | 885 | 893 | + | 11.0826 | 6.84E-05 | 0.709 | GTACGTGGC |
| 10476 | mamo-s | *BGIBMGA012386* | down | 203 | 211 | + | 11.0826 | 6.84E-05 | 0.709 | GTACGTGGC |
| 10477 | mamo-s | *BGIBMGA012387* | down | 377 | 385 | + | 11.0826 | 6.84E-05 | 0.709 | GTACGTGGC |
| 10478 | mamo-s | *BGIBMGA012390* | up | 98 | 106 | + | 12.7615 | 1.04E-05 | 0.615 | CCGCGTGGC |
| 10479 | mamo-s | *BGIBMGA012392* | up | 267 | 275 | - | 11.3303 | 4.96E-05 | 0.615 | GTGGGTGGC |
| 10480 | mamo-s | *BGIBMGA012393* | up | 1456 | 1464 | - | 11.0826 | 6.84E-05 | 0.709 | GTACGTGGC |
| 10481 | mamo-s | *BGIBMGA012394* | up | 339 | 347 | - | 11.7064 | 3.52E-05 | 0.615 | GCGCGTGGT |
| 10482 | mamo-s | *BGIBMGA012394* | up | 881 | 889 | + | 11.3303 | 4.96E-05 | 0.615 | GTGGGTGGC |
| 10483 | mamo-s | *BGIBMGA012395* | up | 721 | 729 | + | 11.3303 | 4.96E-05 | 0.615 | GTGGGTGGC |
| 10484 | mamo-s | *BGIBMGA012395* | down | 1017 | 1025 | + | 11.3303 | 4.96E-05 | 0.615 | GTGGGTGGC |
| 10485 | mamo-s | *BGIBMGA012395* | down | 1278 | 1286 | - | 11.3303 | 4.96E-05 | 0.615 | GTGGGTGGC |
| 10486 | mamo-s | *BGIBMGA012397* | down | 426 | 434 | - | 11.156 | 6.13E-05 | 0.705 | GAGCGTGCC |
| 10487 | mamo-s | *BGIBMGA012398* | down | 1228 | 1236 | + | 11.3303 | 4.96E-05 | 0.615 | GTGGGTGGC |
| 10488 | mamo-s | *BGIBMGA012400* | up | 1679 | 1687 | + | 11.3303 | 4.96E-05 | 0.615 | GTGGGTGGC |
| 10489 | mamo-s | *BGIBMGA012401* | down | 354 | 362 | - | 13.6606 | 4.06E-06 | 0.615 | GCGCGTGGC |
| 10490 | mamo-s | *BGIBMGA012401* | down | 1203 | 1211 | + | 12.4679 | 1.31E-05 | 0.615 | GTGCGTGTC |
| 10491 | mamo-s | *BGIBMGA012401* | up | 248 | 256 | + | 11.3303 | 4.96E-05 | 0.615 | GTGGGTGGC |
| 10492 | mamo-s | *BGIBMGA012402* | up | 1193 | 1201 | - | 13.6606 | 4.06E-06 | 0.615 | GCGCGTGGC |
| 10493 | mamo-s | *BGIBMGA012404* | up | 1249 | 1257 | + | 11.7339 | 3.29E-05 | 0.615 | GCGCGAGGC |
| 10494 | mamo-s | *BGIBMGA012404* | up | 55 | 63 | + | 11.2844 | 5.18E-05 | 0.63 | CTGCGTGCC |
| 10495 | mamo-s | *BGIBMGA012404* | up | 1049 | 1057 | + | 10.6881 | 8.61E-05 | 0.81 | GCACGTGGC |
| 10496 | mamo-s | *BGIBMGA012405* | down | 251 | 259 | + | 11.3303 | 4.96E-05 | 0.615 | GTGGGTGGC |
| 10497 | mamo-s | *BGIBMGA012405* | down | 1951 | 1959 | - | 10.5138 | 9.75E-05 | 0.865 | GTGCGTGTT |
| 10498 | mamo-s | *BGIBMGA012406* | down | 171 | 179 | - | 11.3303 | 4.96E-05 | 0.615 | GTGGGTGGC |
| 10499 | mamo-s | *BGIBMGA012410* | up | 1140 | 1148 | + | 11.3303 | 4.96E-05 | 0.615 | GTGGGTGGC |
| 10500 | mamo-s | *BGIBMGA012411* | down | 806 | 814 | - | 11.7064 | 3.52E-05 | 0.615 | GCGCGTGGT |
| 10501 | mamo-s | *BGIBMGA012411* | down | 580 | 588 | + | 11.3303 | 4.96E-05 | 0.615 | GTGGGTGGC |
| 10502 | mamo-s | *BGIBMGA012412* | up | 1987 | 1995 | - | 11.7064 | 3.52E-05 | 0.615 | GCGCGTGGT |
| 10503 | mamo-s | *BGIBMGA012412* | down | 1447 | 1455 | + | 11.3303 | 4.96E-05 | 0.615 | GTGGGTGGC |
| 10504 | mamo-s | *BGIBMGA012412* | up | 1761 | 1769 | + | 11.3303 | 4.96E-05 | 0.615 | GTGGGTGGC |
| 10505 | mamo-s | *BGIBMGA012414* | up | 980 | 988 | + | 12.7615 | 1.04E-05 | 0.615 | CCGCGTGGC |
| 10506 | mamo-s | *BGIBMGA012414* | down | 352 | 360 | - | 11.3303 | 4.96E-05 | 0.615 | GTGGGTGGC |
| 10507 | mamo-s | *BGIBMGA012417* | down | 1918 | 1926 | + | 11.3303 | 4.96E-05 | 0.615 | GTGGGTGGC |
| 10508 | mamo-s | *BGIBMGA012419* | up | 511 | 519 | + | 11.3303 | 4.96E-05 | 0.615 | GTGGGTGGC |
| 10509 | mamo-s | *BGIBMGA012421* | down | 1589 | 1597 | - | 11.3303 | 4.96E-05 | 0.615 | GTGGGTGGC |
| 10510 | mamo-s | *BGIBMGA012422* | down | 1609 | 1617 | + | 11.4404 | 4.51E-05 | 0.615 | GAGCGTGTC |
| 10511 | mamo-s | *BGIBMGA012422* | down | 873 | 881 | + | 11.0826 | 6.84E-05 | 0.709 | GTACGTGGC |
| 10512 | mamo-s | *BGIBMGA012423* | down | 1149 | 1157 | - | 11.0826 | 6.84E-05 | 0.709 | GTACGTGGC |
| 10513 | mamo-s | *BGIBMGA012423* | up | 1646 | 1654 | - | 11.0734 | 7.12E-05 | 0.726 | GAGCGTGGT |
| 10514 | mamo-s | *BGIBMGA012424* | up | 1734 | 1742 | + | 11.3303 | 4.96E-05 | 0.615 | GTGGGTGGC |
| 10515 | mamo-s | *BGIBMGA012427* | up | 704 | 712 | - | 11.3303 | 4.96E-05 | 0.615 | GTGGGTGGC |
| 10516 | mamo-s | *BGIBMGA012434* | up | 1806 | 1814 | - | 12.1009 | 2.43E-05 | 0.615 | GTGCGTGGT |
| 10517 | mamo-s | *BGIBMGA012434* | up | 238 | 246 | - | 11.3303 | 4.96E-05 | 0.615 | GTGGGTGGC |
| 10518 | mamo-s | *BGIBMGA012434* | up | 647 | 655 | + | 11.0826 | 6.84E-05 | 0.709 | GTACGTGGC |
| 10519 | mamo-s | *BGIBMGA012436* | down | 6 | 14 | + | 10.5413 | 9.42E-05 | 0.857 | GTGCGAGTC |
| 10520 | mamo-s | *BGIBMGA012438* | down | 798 | 806 | + | 11.3303 | 4.96E-05 | 0.615 | GTGGGTGGC |
| 10521 | mamo-s | *BGIBMGA012441* | down | 1928 | 1936 | - | 11.3303 | 4.96E-05 | 0.615 | GTGGGTGGC |
| 10522 | mamo-s | *BGIBMGA012442* | down | 813 | 821 | + | 11.3303 | 4.96E-05 | 0.615 | GTGGGTGGC |
| 10523 | mamo-s | *BGIBMGA012443* | up | 1784 | 1792 | - | 11.3303 | 4.96E-05 | 0.615 | GTGGGTGGC |
| 10524 | mamo-s | *BGIBMGA012446* | up | 830 | 838 | - | 11.3303 | 4.96E-05 | 0.615 | GTGGGTGGC |
| 10525 | mamo-s | *BGIBMGA012446* | down | 20 | 28 | - | 10.9358 | 7.30E-05 | 0.728 | GCGGGTGGC |
| 10526 | mamo-s | *BGIBMGA012450* | up | 869 | 877 | + | 11.3303 | 4.96E-05 | 0.615 | GTGGGTGGC |
| 10527 | mamo-s | *BGIBMGA012451* | down | 1534 | 1542 | - | 11.5688 | 3.79E-05 | 0.615 | CTGCGTGTC |
| 10528 | mamo-s | *BGIBMGA012452* | up | 26 | 34 | - | 13.6606 | 4.06E-06 | 0.615 | GCGCGTGGC |
| 10529 | mamo-s | *BGIBMGA012453* | up | 775 | 783 | - | 11.0826 | 6.84E-05 | 0.709 | GTACGTGGC |
| 10530 | mamo-s | *BGIBMGA012453* | up | 1901 | 1909 | + | 10.7523 | 8.38E-05 | 0.797 | AAGCGTGGC |
| 10531 | mamo-s | *BGIBMGA012453* | up | 1222 | 1230 | - | 10.6881 | 8.61E-05 | 0.81 | GCACGTGGC |
| 10532 | mamo-s | *BGIBMGA012455* | up | 1407 | 1415 | - | 12.1284 | 2.16E-05 | 0.615 | CAGCGTGGC |
| 10533 | mamo-s | *BGIBMGA012456* | down | 1957 | 1965 | + | 11.3303 | 4.96E-05 | 0.615 | GTGGGTGGC |
| 10534 | mamo-s | *BGIBMGA012460* | up | 594 | 602 | + | 12.4679 | 1.31E-05 | 0.615 | GTGCGTGTC |
| 10535 | mamo-s | *BGIBMGA012464* | up | 1683 | 1691 | + | 11.3303 | 4.96E-05 | 0.615 | GTGGGTGGC |
| 10536 | mamo-s | *BGIBMGA012465* | up | 589 | 597 | + | 11.3303 | 4.96E-05 | 0.615 | GTGGGTGGC |
| 10537 | mamo-s | *BGIBMGA012467* | down | 168 | 176 | - | 11.3303 | 4.96E-05 | 0.615 | GTGGGTGGC |
| 10538 | mamo-s | *BGIBMGA012471* | down | 246 | 254 | + | 11.3303 | 4.96E-05 | 0.615 | GTGGGTGGC |
| 10539 | mamo-s | *BGIBMGA012471* | down | 1897 | 1905 | - | 11.3303 | 4.96E-05 | 0.615 | GTGGGTGGC |
| 10540 | mamo-s | *BGIBMGA012473* | up | 678 | 686 | + | 11.1009 | 6.57E-05 | 0.709 | GAGCGAGGC |
| 10541 | mamo-s | *BGIBMGA012476* | down | 421 | 429 | - | 11.3303 | 4.96E-05 | 0.615 | GTGGGTGGC |
| 10542 | mamo-s | *BGIBMGA012481* | up | 681 | 689 | + | 11.0826 | 6.84E-05 | 0.709 | GTACGTGGC |
| 10543 | mamo-s | *BGIBMGA012486* | up | 254 | 262 | - | 11.3303 | 4.96E-05 | 0.615 | GTGGGTGGC |
| 10544 | mamo-s | *BGIBMGA012487* | up | 1886 | 1894 | + | 11.156 | 6.13E-05 | 0.705 | GAGCGTGCC |
| 10545 | mamo-s | *BGIBMGA012487* | up | 1501 | 1509 | - | 11.0826 | 6.84E-05 | 0.709 | GTACGTGGC |
| 10546 | mamo-s | *BGIBMGA012491* | up | 1486 | 1494 | - | 11.3303 | 4.96E-05 | 0.615 | GTGGGTGGC |
| 10547 | mamo-s | *BGIBMGA012491* | up | 1789 | 1797 | - | 11.3303 | 4.96E-05 | 0.615 | GTGGGTGGC |
| 10548 | mamo-s | *BGIBMGA012492* | down | 651 | 659 | - | 11.7064 | 3.52E-05 | 0.615 | GCGCGTGGT |
| 10549 | mamo-s | *BGIBMGA012493* | up | 1588 | 1596 | - | 11.3303 | 4.96E-05 | 0.615 | GTGGGTGGC |
| 10550 | mamo-s | *BGIBMGA012496* | up | 2 | 10 | + | 11.3303 | 4.96E-05 | 0.615 | GTGGGTGGC |
| 10551 | mamo-s | *BGIBMGA012497* | up | 1737 | 1745 | - | 11.3303 | 4.96E-05 | 0.615 | GTGGGTGGC |
| 10552 | mamo-s | *BGIBMGA012498* | up | 1245 | 1253 | - | 11.3303 | 4.96E-05 | 0.615 | GTGGGTGGC |
| 10553 | mamo-s | *BGIBMGA012499* | down | 1978 | 1986 | + | 11.3303 | 4.96E-05 | 0.615 | GTGGGTGGC |

| | | | | | | | | | | |
|---|---|---|---|---|---|---|---|---|---|---|
| 10554 | mamo-s | *BGIBMGA012500* | up | 1188 | 1196 | - | 11.3303 | 4.96E-05 | 0.615 | GTGGGTGGC |
| 10555 | mamo-s | *BGIBMGA012501* | down | 818 | 826 | + | 11.3303 | 4.96E-05 | 0.615 | GTGGGTGGC |
| 10556 | mamo-s | *BGIBMGA012502* | up | 384 | 392 | - | 11.3303 | 4.96E-05 | 0.615 | GTGGGTGGC |
| 10557 | mamo-s | *BGIBMGA012503* | down | 7 | 15 | + | 11.7339 | 3.29E-05 | 0.615 | GCGCGAGGC |
| 10558 | mamo-s | *BGIBMGA012507* | down | 429 | 437 | - | 11.3303 | 4.96E-05 | 0.615 | GTGGGTGGC |
| 10559 | mamo-s | *BGIBMGA012510* | down | 579 | 587 | - | 11.3303 | 4.96E-05 | 0.615 | GTGGGTGGC |
| 10560 | mamo-s | *BGIBMGA012511* | up | 978 | 986 | + | 11.3303 | 4.96E-05 | 0.615 | GTGGGTGGC |
| 10561 | mamo-s | *BGIBMGA012511* | down | 1871 | 1879 | + | 11.3303 | 4.96E-05 | 0.615 | GTGGGTGGC |
| 10562 | mamo-s | *BGIBMGA012512* | up | 75 | 83 | + | 11.3303 | 4.96E-05 | 0.615 | GTGGGTGGC |
| 10563 | mamo-s | *BGIBMGA012512* | up | 745 | 753 | - | 11.1743 | 5.90E-05 | 0.684 | CCGCGTGTC |
| 10564 | mamo-s | *BGIBMGA012523* | up | 588 | 596 | + | 11.3303 | 4.96E-05 | 0.615 | GTGGGTGGC |
| 10565 | mamo-s | *BGIBMGA012523* | up | 1832 | 1840 | + | 11.3303 | 4.96E-05 | 0.615 | GTGGGTGGC |
| 10566 | mamo-s | *BGIBMGA012525* | up | 984 | 992 | - | 11.3303 | 4.96E-05 | 0.615 | GTGGGTGGC |
| 10567 | mamo-s | *BGIBMGA012533* | down | 1510 | 1518 | + | 12.1284 | 2.16E-05 | 0.615 | CAGCGTGGC |
| 10568 | mamo-s | *BGIBMGA012533* | down | 179 | 187 | - | 11.789 | 2.84E-05 | 0.615 | GCGCGTGCC |
| 10569 | mamo-s | *BGIBMGA012535* | up | 1545 | 1553 | - | 11.2018 | 5.68E-05 | 0.674 | CTGCGTGGT |
| 10570 | mamo-s | *BGIBMGA012536* | up | 119 | 127 | + | 10.5138 | 9.75E-05 | 0.865 | GTGCGTGTT |
| 10571 | mamo-s | *BGIBMGA012538* | up | 728 | 736 | - | 13.6606 | 4.06E-06 | 0.615 | GCGCGTGGC |
| 10572 | mamo-s | *BGIBMGA012542* | down | 700 | 708 | - | 14.055 | 2.23E-06 | 0.615 | GTGCGTGGC |
| 10573 | mamo-s | *BGIBMGA012544* | down | 1539 | 1547 | - | 11.1009 | 6.57E-05 | 0.709 | GAGCGAGGC |
| 10574 | mamo-s | *BGIBMGA012546* | down | 683 | 691 | + | 11.3303 | 4.96E-05 | 0.615 | GTGGGTGGC |
| 10575 | mamo-s | *BGIBMGA012548* | up | 1930 | 1938 | + | 10.5413 | 9.42E-05 | 0.857 | CAGCGTGTC |
| 10576 | mamo-s | *BGIBMGA012550* | down | 1237 | 1245 | - | 10.7523 | 8.38E-05 | 0.797 | GGGCGTGTC |
| 10577 | mamo-s | *BGIBMGA012551* | down | 553 | 561 | - | 12.1835 | 1.71E-05 | 0.615 | GTGCGTGCC |
| 10578 | mamo-s | *BGIBMGA012551* | down | 270 | 278 | + | 10.8899 | 7.48E-05 | 0.739 | CCGCGTGCC |
| 10579 | mamo-s | *BGIBMGA012552* | up | 1869 | 1877 | - | 11.5688 | 3.79E-05 | 0.615 | CTGCGTGTC |
| 10580 | mamo-s | *BGIBMGA012553* | down | 575 | 583 | + | 11.0826 | 6.84E-05 | 0.709 | GTACGTGGC |
| 10581 | mamo-s | *BGIBMGA012553* | down | 1254 | 1262 | - | 11.0734 | 7.12E-05 | 0.726 | GAGCGTGGT |
| 10582 | mamo-s | *BGIBMGA012554* | down | 888 | 896 | + | 11.3303 | 4.96E-05 | 0.615 | GTGGGTGGC |
| 10583 | mamo-s | *BGIBMGA012555* | down | 1237 | 1245 | - | 11.3303 | 4.96E-05 | 0.615 | GTGGGTGGC |
| 10584 | mamo-s | *BGIBMGA012559* | up | 201 | 209 | - | 12.3394 | 1.49E-05 | 0.615 | GGGCGTGGC |
| 10585 | mamo-s | *BGIBMGA012559* | up | 171 | 179 | - | 11.0826 | 6.84E-05 | 0.709 | GTACGTGGC |
| 10586 | mamo-s | *BGIBMGA012561* | down | 1923 | 1931 | - | 11.3303 | 4.96E-05 | 0.615 | GTGGGTGGC |
| 10587 | mamo-s | *BGIBMGA012562* | down | 1482 | 1490 | + | 11.3303 | 4.96E-05 | 0.615 | GTGGGTGGC |
| 10588 | mamo-s | *BGIBMGA012563* | down | 1731 | 1739 | - | 11.3303 | 4.96E-05 | 0.615 | GTGGGTGGC |
| 10589 | mamo-s | *BGIBMGA012564* | up | 1156 | 1164 | + | 11.3303 | 4.96E-05 | 0.615 | GTGGGTGGC |
| 10590 | mamo-s | *BGIBMGA012566* | down | 396 | 404 | + | 12.1835 | 1.71E-05 | 0.615 | GTGCGTGCC |
| 10591 | mamo-s | *BGIBMGA012567* | down | 1684 | 1692 | - | 11.3303 | 4.96E-05 | 0.615 | GTGGGTGGC |
| 10592 | mamo-s | *BGIBMGA012568* | down | 571 | 579 | - | 12.7615 | 1.04E-05 | 0.615 | CCGCGTGGC |
| 10593 | mamo-s | *BGIBMGA012568* | up | 777 | 785 | - | 10.5138 | 9.75E-05 | 0.865 | GTGCGTGTT |
| 10594 | mamo-s | *BGIBMGA012569* | up | 186 | 194 | + | 12.7615 | 1.04E-05 | 0.615 | CCGCGTGGC |
| 10595 | mamo-s | *BGIBMGA012571* | down | 1520 | 1528 | - | 12.0734 | 2.66E-05 | 0.615 | GCGCGTGTC |
| 10596 | mamo-s | *BGIBMGA012572* | down | 324 | 332 | - | 11.3303 | 4.96E-05 | 0.615 | GTGGGTGGC |
| 10597 | mamo-s | *BGIBMGA012572* | up | 888 | 896 | - | 11.3303 | 4.96E-05 | 0.615 | GTGGGTGGC |
| 10598 | mamo-s | *BGIBMGA012572* | down | 26 | 34 | + | 10.5413 | 9.42E-05 | 0.857 | GTGCGAGTC |
| 10599 | mamo-s | *BGIBMGA012574* | up | 1029 | 1037 | - | 11.3303 | 4.96E-05 | 0.615 | GTGGGTGGC |
| 10600 | mamo-s | *BGIBMGA012575* | up | 261 | 269 | - | 11.3303 | 4.96E-05 | 0.615 | GTGGGTGGC |
| 10601 | mamo-s | *BGIBMGA012577* | up | 161 | 169 | - | 11.3303 | 4.96E-05 | 0.615 | GTGGGTGGC |
| 10602 | mamo-s | *BGIBMGA012577* | up | 1337 | 1345 | + | 11.3303 | 4.96E-05 | 0.615 | GTGGGTGGC |
| 10603 | mamo-s | *BGIBMGA012578* | up | 1231 | 1239 | - | 14.055 | 2.23E-06 | 0.615 | GTGCGTGGC |
| 10604 | mamo-s | *BGIBMGA012578* | up | 8 | 16 | - | 11.7064 | 3.52E-05 | 0.615 | GCGCGTGGT |
| 10605 | mamo-s | *BGIBMGA012578* | up | 280 | 288 | - | 11.4404 | 4.51E-05 | 0.615 | CGGCGTGGC |
| 10606 | mamo-s | *BGIBMGA012578* | up | 1867 | 1875 | - | 11.2294 | 5.41E-05 | 0.65 | CTGCGAGGC |
| 10607 | mamo-s | *BGIBMGA012579* | down | 238 | 246 | - | 11.3303 | 4.96E-05 | 0.615 | GTGGGTGGC |
| 10608 | mamo-s | *BGIBMGA012582* | down | 573 | 581 | + | 11.5321 | 4.06E-05 | 0.615 | TTGCGTGGC |
| 10609 | mamo-s | *BGIBMGA012583* | up | 720 | 728 | - | 11.0826 | 6.84E-05 | 0.709 | GTACGTGGC |
| 10610 | mamo-s | *BGIBMGA012585* | up | 839 | 847 | - | 12.4679 | 1.31E-05 | 0.615 | GTGCGTGTC |
| 10611 | mamo-s | *BGIBMGA012585* | up | 1759 | 1767 | + | 11.3303 | 4.96E-05 | 0.615 | GTGGGTGGC |
| 10612 | mamo-s | *BGIBMGA012585* | down | 426 | 434 | + | 10.9358 | 7.30E-05 | 0.728 | GCGGGTGGC |
| 10613 | mamo-s | *BGIBMGA012586* | up | 142 | 150 | + | 11.3303 | 4.96E-05 | 0.615 | GTGGGTGGC |
| 10614 | mamo-s | *BGIBMGA012587* | up | 1351 | 1359 | + | 12.1009 | 2.43E-05 | 0.615 | GTGCGTGGT |
| 10615 | mamo-s | *BGIBMGA012587* | down | 122 | 130 | - | 11.3303 | 4.96E-05 | 0.615 | GTGGGTGGC |
| 10616 | mamo-s | *BGIBMGA012588* | down | 836 | 844 | - | 12.1284 | 2.16E-05 | 0.615 | CAGCGTGGC |
| 10617 | mamo-s | *BGIBMGA012591* | down | 665 | 673 | - | 12.4679 | 1.31E-05 | 0.615 | GTGCGTGTC |
| 10618 | mamo-s | *BGIBMGA012592* | down | 555 | 563 | + | 11.3303 | 4.96E-05 | 0.615 | GTGGGTGGC |
| 10619 | mamo-s | *BGIBMGA012592* | up | 800 | 808 | + | 11.0826 | 6.84E-05 | 0.709 | GTACGTGGC |
| 10620 | mamo-s | *BGIBMGA012594* | up | 36 | 44 | - | 11.3303 | 4.96E-05 | 0.615 | GTGGGTGGC |
| 10621 | mamo-s | *BGIBMGA012599* | up | 491 | 499 | - | 11.1376 | 6.35E-05 | 0.709 | TCGCGTGGC |
| 10622 | mamo-s | *BGIBMGA012602* | up | 853 | 861 | - | 12.3394 | 1.49E-05 | 0.615 | GGGCGTGGC |
| 10623 | mamo-s | *BGIBMGA012602* | up | 1091 | 1099 | - | 12.1009 | 2.43E-05 | 0.615 | GTGCGTGGT |
| 10624 | mamo-s | *BGIBMGA012602* | up | 39 | 47 | - | 11.3303 | 4.96E-05 | 0.615 | GTGGGTGGC |
| 10625 | mamo-s | *BGIBMGA012602* | up | 1468 | 1476 | + | 11.0826 | 6.84E-05 | 0.709 | GTACGTGGC |
| 10626 | mamo-s | *BGIBMGA012603* | up | 1262 | 1270 | + | 11.3303 | 4.96E-05 | 0.615 | GTGGGTGGC |
| 10627 | mamo-s | *BGIBMGA012605* | down | 1509 | 1517 | + | 11.3303 | 4.96E-05 | 0.615 | GTGGGTGGC |
| 10628 | mamo-s | *BGIBMGA012607* | down | 1010 | 1018 | + | 13.0275 | 8.53E-06 | 0.615 | GAGCGTGGC |
| 10629 | mamo-s | *BGIBMGA012609* | down | 1504 | 1512 | + | 12.4679 | 1.31E-05 | 0.615 | GTGCGTGTC |
| 10630 | mamo-s | *BGIBMGA012609* | up | 1022 | 1030 | - | 11.0826 | 6.84E-05 | 0.709 | GTACGTGGC |
| 10631 | mamo-s | *BGIBMGA012610* | down | 127 | 135 | - | 11.5688 | 3.79E-05 | 0.615 | CTGCGTGTC |
| 10632 | mamo-s | *BGIBMGA012610* | down | 1354 | 1362 | - | 11.3303 | 4.96E-05 | 0.615 | GTGGGTGGC |
| 10633 | mamo-s | *BGIBMGA012611* | up | 914 | 922 | + | 11.3303 | 4.96E-05 | 0.615 | GTGGGTGGC |
| 10634 | mamo-s | *BGIBMGA012611* | down | 1270 | 1278 | + | 11.3303 | 4.96E-05 | 0.615 | GTGGGTGGC |
| 10635 | mamo-s | *BGIBMGA012611* | up | 69 | 77 | - | 11.0826 | 6.84E-05 | 0.709 | GTACGTGGC |
| 10636 | mamo-s | *BGIBMGA012614* | down | 979 | 987 | + | 11.3303 | 4.96E-05 | 0.615 | GTGGGTGGC |
| 10637 | mamo-s | *BGIBMGA012618* | up | 60 | 68 | + | 11.3303 | 4.96E-05 | 0.615 | GTGGGTGGC |
| 10638 | mamo-s | *BGIBMGA012618* | up | 1560 | 1568 | - | 11.0826 | 6.84E-05 | 0.709 | GTACGTGGC |
| 10639 | mamo-s | *BGIBMGA012619* | down | 582 | 590 | + | 11.3303 | 4.96E-05 | 0.615 | GTGGGTGGC |
| 10640 | mamo-s | *BGIBMGA012619* | down | 752 | 760 | - | 11.3303 | 4.96E-05 | 0.615 | GTGGGTGGC |
| 10641 | mamo-s | *BGIBMGA012620* | down | 658 | 666 | - | 12.1009 | 2.43E-05 | 0.615 | GTGCGTGGT |

| | | | | | | | | | | |
|---|---|---|---|---|---|---|---|---|---|---|
| 10642 | mamo-s | *BGIBMGA012620* | up | 1595 | 1603 | + | 11.2018 | 5.68E-05 | 0.674 | CTGCGTGGT |
| 10643 | mamo-s | *BGIBMGA012621* | down | 525 | 533 | + | 13.156 | 6.30E-06 | 0.615 | CTGCGTGGC |
| 10644 | mamo-s | *BGIBMGA012621* | up | 204 | 212 | - | 11.3303 | 4.96E-05 | 0.615 | GTGGGTGGC |
| 10645 | mamo-s | *BGIBMGA012621* | down | 971 | 979 | - | 11.3303 | 4.96E-05 | 0.615 | GTGGGTGGC |
| 10646 | mamo-s | *BGIBMGA012621* | up | 1621 | 1629 | - | 10.9358 | 7.30E-05 | 0.728 | GCGGGTGGC |
| 10647 | mamo-s | *BGIBMGA012624* | down | 1149 | 1157 | + | 12.1835 | 1.71E-05 | 0.615 | GTGCGTGCC |
| 10648 | mamo-s | *BGIBMGA012624* | down | 869 | 877 | - | 11.0826 | 6.84E-05 | 0.709 | GTACGTGGC |
| 10649 | mamo-s | *BGIBMGA012624* | up | 1631 | 1639 | + | 11.0826 | 6.84E-05 | 0.709 | GTACGTGGC |
| 10650 | mamo-s | *BGIBMGA012625* | down | 503 | 511 | - | 11.3303 | 4.96E-05 | 0.615 | GTGGGTGGC |
| 10651 | mamo-s | *BGIBMGA012626* | down | 649 | 657 | + | 11.3303 | 4.96E-05 | 0.615 | GTGGGTGGC |
| 10652 | mamo-s | *BGIBMGA012626* | down | 850 | 858 | + | 10.6881 | 8.61E-05 | 0.81 | GCACGTGGC |
| 10653 | mamo-s | *BGIBMGA012627* | down | 1006 | 1014 | + | 11.3303 | 4.96E-05 | 0.615 | GTGGGTGGC |
| 10654 | mamo-s | *BGIBMGA012627* | up | 833 | 841 | + | 11.0826 | 6.84E-05 | 0.709 | GTACGTGGC |
| 10655 | mamo-s | *BGIBMGA012628* | down | 1639 | 1647 | - | 12.3394 | 1.49E-05 | 0.615 | GGGCGTGGC |
| 10656 | mamo-s | *BGIBMGA012628* | down | 1840 | 1848 | - | 12.3394 | 1.49E-05 | 0.615 | GGGCGTGGC |
| 10657 | mamo-s | *BGIBMGA012629* | down | 1299 | 1307 | - | 13.6606 | 4.06E-06 | 0.615 | GCGCGTGGC |
| 10658 | mamo-s | *BGIBMGA012629* | down | 1289 | 1297 | - | 11.2018 | 5.68E-05 | 0.674 | CTGCGTGGT |
| 10659 | mamo-s | *BGIBMGA012629* | up | 345 | 353 | + | 11.1009 | 6.57E-05 | 0.709 | GAGCGAGGC |
| 10660 | mamo-s | *BGIBMGA012630* | up | 1346 | 1354 | + | 14.055 | 2.23E-06 | 0.615 | GTGCGTGGC |
| 10661 | mamo-s | *BGIBMGA012630* | up | 710 | 718 | + | 11.2294 | 5.41E-05 | 0.65 | CTGCGAGGC |
| 10662 | mamo-s | *BGIBMGA012632* | down | 392 | 400 | - | 11.3303 | 4.96E-05 | 0.615 | GTGGGTGGC |
| 10663 | mamo-s | *BGIBMGA012632* | up | 480 | 488 | - | 11.0826 | 6.84E-05 | 0.709 | GTACGTGGC |
| 10664 | mamo-s | *BGIBMGA012633* | up | 443 | 451 | + | 14.055 | 2.23E-06 | 0.615 | GTGCGTGGC |
| 10665 | mamo-s | *BGIBMGA012634* | down | 510 | 518 | - | 11.3303 | 4.96E-05 | 0.615 | GTGGGTGGC |
| 10666 | mamo-s | *BGIBMGA012634* | down | 1874 | 1882 | - | 11.3303 | 4.96E-05 | 0.615 | GTGGGTGGC |
| 10667 | mamo-s | *BGIBMGA012638* | up | 457 | 465 | + | 11.3853 | 4.74E-05 | 0.615 | ACGCGTGGC |
| 10668 | mamo-s | *BGIBMGA012638* | down | 146 | 154 | + | 11.3303 | 4.96E-05 | 0.615 | GTGGGTGGC |
| 10669 | mamo-s | *BGIBMGA012640* | down | 659 | 667 | + | 11.3303 | 4.96E-05 | 0.615 | GTGGGTGGC |
| 10670 | mamo-s | *BGIBMGA012640* | up | 660 | 668 | - | 11.3303 | 4.96E-05 | 0.615 | GTGGGTGGC |
| 10671 | mamo-s | *BGIBMGA012640* | down | 1908 | 1916 | - | 11.3303 | 4.96E-05 | 0.615 | GTGGGTGGC |
| 10672 | mamo-s | *BGIBMGA012641* | down | 692 | 700 | + | 11.789 | 2.84E-05 | 0.615 | GCGCGTGCC |
| 10673 | mamo-s | *BGIBMGA012641* | up | 259 | 267 | + | 11.4404 | 4.51E-05 | 0.615 | CGGCGTGGC |
| 10674 | mamo-s | *BGIBMGA012641* | up | 1800 | 1808 | + | 11.3303 | 4.96E-05 | 0.615 | GTGGGTGGC |
| 10675 | mamo-s | *BGIBMGA012644* | up | 585 | 593 | + | 11.3303 | 4.96E-05 | 0.615 | GTGGGTGGC |
| 10676 | mamo-s | *BGIBMGA012644* | down | 817 | 825 | - | 11.2844 | 5.18E-05 | 0.63 | CTGCGTGCC |
| 10677 | mamo-s | *BGIBMGA012646* | up | 1763 | 1771 | + | 11.0826 | 6.84E-05 | 0.709 | GTACGTGGC |
| 10678 | mamo-s | *BGIBMGA012647* | down | 382 | 390 | - | 11.3303 | 4.96E-05 | 0.615 | GTGGGTGGC |
| 10679 | mamo-s | *BGIBMGA012647* | up | 747 | 755 | - | 11.3303 | 4.96E-05 | 0.615 | GTGGGTGGC |
| 10680 | mamo-s | *BGIBMGA012652* | down | 292 | 300 | + | 11.2844 | 5.18E-05 | 0.63 | CTGCGTGCC |
| 10681 | mamo-s | *BGIBMGA012654* | up | 1291 | 1299 | + | 11.3303 | 4.96E-05 | 0.615 | GTGGGTGGC |
| 10682 | mamo-s | *BGIBMGA012656* | down | 326 | 334 | - | 11.3303 | 4.96E-05 | 0.615 | GTGGGTGGC |
| 10683 | mamo-s | *BGIBMGA012657* | up | 575 | 583 | - | 11.3303 | 4.96E-05 | 0.615 | GTGGGTGGC |
| 10684 | mamo-s | *BGIBMGA012658* | up | 40 | 48 | + | 11.3303 | 4.96E-05 | 0.615 | GTGGGTGGC |
| 10685 | mamo-s | *BGIBMGA012659* | up | 1660 | 1668 | + | 12.1835 | 1.71E-05 | 0.615 | GTGCGTGCC |
| 10686 | mamo-s | *BGIBMGA012659* | up | 1422 | 1430 | + | 11.4404 | 4.51E-05 | 0.615 | CGGCGTGGC |
| 10687 | mamo-s | *BGIBMGA012659* | up | 165 | 173 | - | 11.3303 | 4.96E-05 | 0.615 | GTGGGTGGC |
| 10688 | mamo-s | *BGIBMGA012659* | up | 1897 | 1905 | + | 11.2844 | 5.18E-05 | 0.63 | CTGCGTGCC |
| 10689 | mamo-s | *BGIBMGA012660* | down | 704 | 712 | + | 11.3303 | 4.96E-05 | 0.615 | GTGGGTGGC |
| 10690 | mamo-s | *BGIBMGA012661* | up | 970 | 978 | + | 11.3303 | 4.96E-05 | 0.615 | GTGGGTGGC |
| 10691 | mamo-s | *BGIBMGA012661* | down | 1279 | 1287 | - | 11.0826 | 6.84E-05 | 0.709 | GTACGTGGC |
| 10692 | mamo-s | *BGIBMGA012662* | down | 1298 | 1306 | + | 11.3303 | 4.96E-05 | 0.615 | GTGGGTGGC |
| 10693 | mamo-s | *BGIBMGA012665* | down | 1955 | 1963 | + | 12.4679 | 1.31E-05 | 0.615 | GTGCGTGTC |
| 10694 | mamo-s | *BGIBMGA012666* | down | 942 | 950 | - | 11.3303 | 4.96E-05 | 0.615 | GTGGGTGGC |
| 10695 | mamo-s | *BGIBMGA012666* | up | 1886 | 1894 | + | 11.3303 | 4.96E-05 | 0.615 | GTGGGTGGC |
| 10696 | mamo-s | *BGIBMGA012668* | down | 1750 | 1758 | - | 11.3303 | 4.96E-05 | 0.615 | GTGGGTGGC |
| 10697 | mamo-s | *BGIBMGA012670* | up | 126 | 134 | - | 11.3303 | 4.96E-05 | 0.615 | GTGGGTGGC |
| 10698 | mamo-s | *BGIBMGA012671* | up | 88 | 96 | - | 10.5413 | 9.42E-05 | 0.857 | CAGCGTGTC |
| 10699 | mamo-s | *BGIBMGA012672* | up | 1405 | 1413 | - | 12.1284 | 2.16E-05 | 0.615 | GTGCGAGGC |
| 10700 | mamo-s | *BGIBMGA012675* | down | 254 | 262 | + | 11.3303 | 4.96E-05 | 0.615 | GTGGGTGGC |
| 10701 | mamo-s | *BGIBMGA012675* | down | 1411 | 1419 | + | 11.0734 | 7.12E-05 | 0.726 | GAGCGTGGT |
| 10702 | mamo-s | *BGIBMGA012676* | up | 337 | 345 | - | 11.3303 | 4.96E-05 | 0.615 | GTGGGTGGC |
| 10703 | mamo-s | *BGIBMGA012679* | down | 1508 | 1516 | - | 11.3303 | 4.96E-05 | 0.615 | GTGGGTGGC |
| 10704 | mamo-s | *BGIBMGA012680* | down | 246 | 254 | - | 11.3303 | 4.96E-05 | 0.615 | GTGGGTGGC |
| 10705 | mamo-s | *BGIBMGA012683* | up | 1702 | 1710 | + | 11.3303 | 4.96E-05 | 0.615 | GTGGGTGGC |
| 10706 | mamo-s | *BGIBMGA012684* | up | 473 | 481 | - | 11.3303 | 4.96E-05 | 0.615 | GTGGGTGGC |
| 10707 | mamo-s | *BGIBMGA012685* | up | 372 | 380 | + | 11.3303 | 4.96E-05 | 0.615 | GTGGGTGGC |
| 10708 | mamo-s | *BGIBMGA012685* | up | 595 | 603 | + | 11.2018 | 5.68E-05 | 0.674 | CTGCGTGGT |
| 10709 | mamo-s | *BGIBMGA012687* | down | 206 | 214 | + | 14.055 | 2.23E-06 | 0.615 | GTGCGTGGC |
| 10710 | mamo-s | *BGIBMGA012687* | up | 704 | 712 | + | 11.3303 | 4.96E-05 | 0.615 | GTGGGTGGC |
| 10711 | mamo-s | *BGIBMGA012689* | up | 17 | 25 | + | 11.0826 | 6.84E-05 | 0.709 | GTACGTGGC |
| 10712 | mamo-s | *BGIBMGA012689* | down | 629 | 637 | - | 10.5138 | 9.75E-05 | 0.865 | GTGCGTGTT |
| 10713 | mamo-s | *BGIBMGA012690* | up | 609 | 617 | - | 11.3303 | 4.96E-05 | 0.615 | GTGGGTGGC |
| 10714 | mamo-s | *BGIBMGA012692* | up | 584 | 592 | - | 10.7523 | 8.38E-05 | 0.797 | AAGCGTGGC |
| 10715 | mamo-s | *BGIBMGA012694* | up | 1296 | 1304 | + | 11.3303 | 4.96E-05 | 0.615 | GTGGGTGGC |
| 10716 | mamo-s | *BGIBMGA012695* | down | 456 | 464 | + | 11.3303 | 4.96E-05 | 0.615 | GTGGGTGGC |
| 10717 | mamo-s | *BGIBMGA012697* | up | 1537 | 1545 | + | 11.3303 | 4.96E-05 | 0.615 | GTGGGTGGC |
| 10718 | mamo-s | *BGIBMGA012697* | down | 755 | 763 | - | 10.5138 | 9.75E-05 | 0.865 | GTGCGTGTT |
| 10719 | mamo-s | *BGIBMGA012699* | up | 1343 | 1351 | + | 11.3303 | 4.96E-05 | 0.615 | GTGGGTGGC |
| 10720 | mamo-s | *BGIBMGA012700* | down | 531 | 539 | + | 11.3303 | 4.96E-05 | 0.615 | GTGGGTGGC |
| 10721 | mamo-s | *BGIBMGA012701* | down | 44 | 52 | - | 11.2294 | 5.41E-05 | 0.65 | CTGCGAGGC |
| 10722 | mamo-s | *BGIBMGA012701* | up | 264 | 272 | - | 10.7523 | 8.38E-05 | 0.797 | AAGCGTGGC |
| 10723 | mamo-s | *BGIBMGA012704* | up | 1308 | 1316 | - | 11.3303 | 4.96E-05 | 0.615 | GTGGGTGGC |
| 10724 | mamo-s | *BGIBMGA012706* | down | 315 | 323 | - | 11.3303 | 4.96E-05 | 0.615 | GTGGGTGGC |
| 10725 | mamo-s | *BGIBMGA012709* | down | 427 | 435 | - | 11.7798 | 3.11E-05 | 0.615 | ATGCGTGGC |
| 10726 | mamo-s | *BGIBMGA012709* | down | 1351 | 1359 | + | 11.1009 | 6.57E-05 | 0.709 | GAGCGAGGC |
| 10727 | mamo-s | *BGIBMGA012710* | down | 1557 | 1565 | + | 11.3303 | 4.96E-05 | 0.615 | GTGGGTGGC |
| 10728 | mamo-s | *BGIBMGA012712* | down | 645 | 653 | - | 11.3303 | 4.96E-05 | 0.615 | GTGGGTGGC |
| 10729 | mamo-s | *BGIBMGA012712* | up | 20 | 28 | + | 10.5413 | 9.42E-05 | 0.857 | CAGCGTGTC |

| | | | | | | | | | | |
|---|---|---|---|---|---|---|---|---|---|---|
| 10730 | mamo-s | *BGIBMGA012713* | up | 1547 | 1555 | - | 11.0826 | 6.84E-05 | 0.709 | GTACGTGGC |
| 10731 | mamo-s | *BGIBMGA012714* | up | 793 | 801 | + | 11.3303 | 4.96E-05 | 0.615 | GTGGGTGGC |
| 10732 | mamo-s | *BGIBMGA012715* | up | 131 | 139 | - | 11.3303 | 4.96E-05 | 0.615 | GTGGGTGGC |
| 10733 | mamo-s | *BGIBMGA012716* | up | 572 | 580 | - | 11.3303 | 4.96E-05 | 0.615 | GTGGGTGGC |
| 10734 | mamo-s | *BGIBMGA012722* | up | 1419 | 1427 | + | 10.5138 | 9.75E-05 | 0.865 | GTGCGTGTT |
| 10735 | mamo-s | *BGIBMGA012725* | up | 1219 | 1227 | + | 12.3394 | 1.49E-05 | 0.615 | GGGCGTGGC |
| 10736 | mamo-s | *BGIBMGA012725* | up | 1288 | 1296 | + | 12.0734 | 2.66E-05 | 0.615 | GCGCGTGTC |
| 10737 | mamo-s | *BGIBMGA012725* | down | 1354 | 1362 | + | 10.7523 | 8.38E-05 | 0.797 | AAGCGTGGC |
| 10738 | mamo-s | *BGIBMGA012727* | down | 348 | 356 | + | 11.3303 | 4.96E-05 | 0.615 | GTGGGTGGC |
| 10739 | mamo-s | *BGIBMGA012729* | up | 862 | 870 | - | 11.3303 | 4.96E-05 | 0.615 | GTGGGTGGC |
| 10740 | mamo-s | *BGIBMGA012729* | down | 980 | 988 | + | 11.3303 | 4.96E-05 | 0.615 | GTGGGTGGC |
| 10741 | mamo-s | *BGIBMGA012729* | up | 1464 | 1472 | + | 11.3303 | 4.96E-05 | 0.615 | GTGGGTGGC |
| 10742 | mamo-s | *BGIBMGA012730* | up | 490 | 498 | + | 14.055 | 2.23E-06 | 0.615 | GTGCGTGGC |
| 10743 | mamo-s | *BGIBMGA012730* | up | 1270 | 1278 | + | 11.3303 | 4.96E-05 | 0.615 | GTGGGTGGC |
| 10744 | mamo-s | *BGIBMGA012730* | up | 663 | 671 | - | 10.6881 | 8.61E-05 | 0.81 | GCACGTGGC |
| 10745 | mamo-s | *BGIBMGA012732* | down | 521 | 529 | - | 11.3303 | 4.96E-05 | 0.615 | GTGGGTGGC |
| 10746 | mamo-s | *BGIBMGA012733* | up | 1771 | 1779 | + | 11.3303 | 4.96E-05 | 0.615 | GTGGGTGGC |
| 10747 | mamo-s | *BGIBMGA012735* | up | 406 | 414 | + | 11.3303 | 4.96E-05 | 0.615 | GTGGGTGGC |
| 10748 | mamo-s | *BGIBMGA012735* | up | 544 | 552 | + | 11.3303 | 4.96E-05 | 0.615 | GTGGGTGGC |
| 10749 | mamo-s | *BGIBMGA012736* | down | 1556 | 1564 | - | 11.1743 | 5.90E-05 | 0.684 | CCGCGTGTC |
| 10750 | mamo-s | *BGIBMGA012737* | down | 243 | 251 | + | 11.3303 | 4.96E-05 | 0.615 | GTGGGTGGC |
| 10751 | mamo-s | *BGIBMGA012741* | down | 1621 | 1629 | + | 12.1284 | 2.16E-05 | 0.615 | GTGCGAGGC |
| 10752 | mamo-s | *BGIBMGA012741* | down | 1957 | 1965 | - | 11.3303 | 4.96E-05 | 0.615 | GTGGGTGGC |
| 10753 | mamo-s | *BGIBMGA012742* | down | 1362 | 1370 | + | 11.7798 | 3.11E-05 | 0.615 | ATGCGTGGC |
| 10754 | mamo-s | *BGIBMGA012742* | up | 1201 | 1209 | + | 11.0826 | 6.84E-05 | 0.709 | GTACGTGGC |
| 10755 | mamo-s | *BGIBMGA012743* | down | 950 | 958 | - | 10.7523 | 8.38E-05 | 0.797 | GGGCGTGTC |
| 10756 | mamo-s | *BGIBMGA012745* | down | 1050 | 1058 | - | 11.3303 | 4.96E-05 | 0.615 | GTGGGTGGC |
| 10757 | mamo-s | *BGIBMGA012746* | down | 926 | 934 | - | 11.3303 | 4.96E-05 | 0.615 | GTGGGTGGC |
| 10758 | mamo-s | *BGIBMGA012750* | up | 601 | 609 | - | 11.3303 | 4.96E-05 | 0.615 | GTGGGTGGC |
| 10759 | mamo-s | *BGIBMGA012750* | up | 1152 | 1160 | + | 11.3303 | 4.96E-05 | 0.615 | GTGGGTGGC |
| 10760 | mamo-s | *BGIBMGA012752* | down | 966 | 974 | - | 12.1835 | 1.71E-05 | 0.615 | GTGCGTGCC |
| 10761 | mamo-s | *BGIBMGA012752* | down | 603 | 611 | - | 11.1376 | 6.35E-05 | 0.709 | TCGCGTGGC |
| 10762 | mamo-s | *BGIBMGA012754* | up | 948 | 956 | - | 13.6606 | 4.06E-06 | 0.615 | GCGCGTGGC |
| 10763 | mamo-s | *BGIBMGA012754* | down | 436 | 444 | + | 13.0275 | 8.53E-06 | 0.615 | GAGCGTGGC |
| 10764 | mamo-s | *BGIBMGA012755* | up | 1337 | 1345 | - | 12.1284 | 2.16E-05 | 0.615 | GTGCGAGGC |
| 10765 | mamo-s | *BGIBMGA012755* | up | 608 | 616 | - | 12.1009 | 2.43E-05 | 0.615 | GTGCGTGGT |
| 10766 | mamo-s | *BGIBMGA012757* | down | 1838 | 1846 | + | 11.0734 | 7.12E-05 | 0.726 | GAGCGTGGT |
| 10767 | mamo-s | *BGIBMGA012758* | up | 1429 | 1437 | + | 11.3303 | 4.96E-05 | 0.615 | GTGGGTGGC |
| 10768 | mamo-s | *BGIBMGA012759* | down | 253 | 261 | - | 14.055 | 2.23E-06 | 0.615 | GTGCGTGGC |
| 10769 | mamo-s | *BGIBMGA012759* | up | 690 | 698 | - | 11.3303 | 4.96E-05 | 0.615 | GTGGGTGGC |
| 10770 | mamo-s | *BGIBMGA012761* | down | 264 | 272 | - | 14.055 | 2.23E-06 | 0.615 | GTGCGTGGC |
| 10771 | mamo-s | *BGIBMGA012761* | up | 110 | 118 | - | 11.3303 | 4.96E-05 | 0.615 | GTGGGTGGC |
| 10772 | mamo-s | *BGIBMGA012761* | down | 137 | 145 | - | 11.1376 | 6.35E-05 | 0.709 | TCGCGTGGC |
| 10773 | mamo-s | *BGIBMGA012763* | up | 61 | 69 | - | 14.055 | 2.23E-06 | 0.615 | GTGCGTGGC |
| 10774 | mamo-s | *BGIBMGA012763* | up | 224 | 232 | + | 11.3303 | 4.96E-05 | 0.615 | GTGGGTGGC |
| 10775 | mamo-s | *BGIBMGA012764* | up | 783 | 791 | - | 11.3303 | 4.96E-05 | 0.615 | GTGGGTGGC |
| 10776 | mamo-s | *BGIBMGA012764* | down | 798 | 806 | + | 11.3303 | 4.96E-05 | 0.615 | GTGGGTGGC |
| 10777 | mamo-s | *BGIBMGA012765* | up | 1270 | 1278 | + | 11.5688 | 3.79E-05 | 0.615 | CTGCGTGTC |
| 10778 | mamo-s | *BGIBMGA012765* | down | 1613 | 1621 | + | 11.1743 | 5.90E-05 | 0.684 | CCGCGTGTC |
| 10779 | mamo-s | *BGIBMGA012766* | down | 828 | 836 | - | 11.3303 | 4.96E-05 | 0.615 | GTGGGTGGC |
| 10780 | mamo-s | *BGIBMGA012768* | up | 318 | 326 | + | 10.9358 | 7.30E-05 | 0.728 | GCGGGTGGC |
| 10781 | mamo-s | *BGIBMGA012769* | down | 1846 | 1854 | - | 11.3303 | 4.96E-05 | 0.615 | GTGGGTGGC |
| 10782 | mamo-s | *BGIBMGA012769* | down | 1275 | 1283 | + | 11.0826 | 6.84E-05 | 0.709 | GTACGTGGC |
| 10783 | mamo-s | *BGIBMGA012770* | up | 850 | 858 | - | 11.1009 | 6.57E-05 | 0.709 | GAGCGAGGC |
| 10784 | mamo-s | *BGIBMGA012772* | up | 85 | 93 | - | 12.3394 | 1.49E-05 | 0.615 | GGGCGTGGC |
| 10785 | mamo-s | *BGIBMGA012772* | down | 976 | 984 | + | 11.3303 | 4.96E-05 | 0.615 | GTGGGTGGC |
| 10786 | mamo-s | *BGIBMGA012773* | down | 877 | 885 | - | 11.3303 | 4.96E-05 | 0.615 | GTGGGTGGC |
| 10787 | mamo-s | *BGIBMGA012773* | down | 1322 | 1330 | + | 11.0826 | 6.84E-05 | 0.709 | GTACGTGGC |
| 10788 | mamo-s | *BGIBMGA012778* | down | 1330 | 1338 | - | 11.4404 | 4.51E-05 | 0.615 | GAGCGTGTC |
| 10789 | mamo-s | *BGIBMGA012778* | up | 1655 | 1663 | - | 11.3303 | 4.96E-05 | 0.615 | GTGGGTGGC |
| 10790 | mamo-s | *BGIBMGA012778* | down | 1965 | 1973 | - | 10.8899 | 7.48E-05 | 0.739 | CCGCGTGCC |
| 10791 | mamo-s | *BGIBMGA012778* | down | 438 | 446 | - | 10.8073 | 7.89E-05 | 0.766 | CCGCGTGGT |
| 10792 | mamo-s | *BGIBMGA012782* | up | 244 | 252 | + | 10.5138 | 9.75E-05 | 0.865 | GTGCGTGTT |
| 10793 | mamo-s | *BGIBMGA012787* | down | 836 | 844 | + | 11.2844 | 5.18E-05 | 0.63 | CTGCGTGCC |
| 10794 | mamo-s | *BGIBMGA012788* | down | 449 | 457 | + | 11.3303 | 4.96E-05 | 0.615 | GTGGGTGGC |
| 10795 | mamo-s | *BGIBMGA012788* | up | 1267 | 1275 | + | 11.3303 | 4.96E-05 | 0.615 | GTGGGTGGC |
| 10796 | mamo-s | *BGIBMGA012789* | up | 70 | 78 | - | 11.3303 | 4.96E-05 | 0.615 | GTGGGTGGC |
| 10797 | mamo-s | *BGIBMGA012790* | down | 224 | 232 | + | 11.3303 | 4.96E-05 | 0.615 | GTGGGTGGC |
| 10798 | mamo-s | *BGIBMGA012790* | up | 1493 | 1501 | + | 11.1009 | 6.57E-05 | 0.709 | GAGCGAGGC |
| 10799 | mamo-s | *BGIBMGA012791* | up | 966 | 974 | + | 11.3303 | 4.96E-05 | 0.615 | GTGGGTGGC |
| 10800 | mamo-s | *BGIBMGA012793* | down | 133 | 141 | - | 11.3303 | 4.96E-05 | 0.615 | GTGGGTGGC |
| 10801 | mamo-s | *BGIBMGA012793* | up | 785 | 793 | + | 11.3303 | 4.96E-05 | 0.615 | GTGGGTGGC |
| 10802 | mamo-s | *BGIBMGA012794* | down | 1837 | 1845 | + | 11.3303 | 4.96E-05 | 0.615 | GTGGGTGGC |
| 10803 | mamo-s | *BGIBMGA012795* | up | 1186 | 1194 | - | 11.3303 | 4.96E-05 | 0.615 | GTGGGTGGC |
| 10804 | mamo-s | *BGIBMGA012796* | down | 1601 | 1609 | - | 12.1009 | 2.43E-05 | 0.615 | GTGCGTGGT |
| 10805 | mamo-s | *BGIBMGA012796* | down | 769 | 777 | + | 11.156 | 6.13E-05 | 0.705 | GAGCGTGCC |
| 10806 | mamo-s | *BGIBMGA012797* | down | 1314 | 1322 | - | 11.3303 | 4.96E-05 | 0.615 | GTGGGTGGC |
| 10807 | mamo-s | *BGIBMGA012799* | up | 1611 | 1619 | + | 10.5138 | 9.75E-05 | 0.865 | GTGCGTGTT |
| 10808 | mamo-s | *BGIBMGA012800* | down | 1525 | 1533 | + | 11.2844 | 5.18E-05 | 0.63 | CTGCGTGCC |
| 10809 | mamo-s | *BGIBMGA012803* | up | 382 | 390 | - | 11.3303 | 4.96E-05 | 0.615 | GTGGGTGGC |
| 10810 | mamo-s | *BGIBMGA012806* | down | 1869 | 1877 | - | 11.2018 | 5.68E-05 | 0.674 | CTGCGTGGT |
| 10811 | mamo-s | *BGIBMGA012809* | down | 475 | 483 | - | 11.3303 | 4.96E-05 | 0.615 | GTGGGTGGC |
| 10812 | mamo-s | *BGIBMGA012811* | down | 839 | 847 | + | 11.3303 | 4.96E-05 | 0.615 | GTGGGTGGC |
| 10813 | mamo-s | *BGIBMGA012811* | down | 1920 | 1928 | - | 11.3303 | 4.96E-05 | 0.615 | GTGGGTGGC |
| 10814 | mamo-s | *BGIBMGA012813* | down | 1856 | 1864 | + | 11.3303 | 4.96E-05 | 0.615 | GTGGGTGGC |
| 10815 | mamo-s | *BGIBMGA012814* | down | 1886 | 1894 | - | 11.3303 | 4.96E-05 | 0.615 | GTGGGTGGC |
| 10816 | mamo-s | *BGIBMGA012815* | up | 1934 | 1942 | + | 11.3303 | 4.96E-05 | 0.615 | GTGGGTGGC |
| 10817 | mamo-s | *BGIBMGA012815* | down | 1979 | 1987 | + | 11.3303 | 4.96E-05 | 0.615 | GTGGGTGGC |

| | | | | | | | | | | |
|---|---|---|---|---|---|---|---|---|---|---|
| 10818 | mamo-s | *BGIBMGA012816* | up | 158 | 166 | - | 11.3303 | 4.96E-05 | 0.615 | GTGGGTGGC |
| 10819 | mamo-s | *BGIBMGA012816* | up | 736 | 744 | + | 11.3303 | 4.96E-05 | 0.615 | GTGGGTGGC |
| 10820 | mamo-s | *BGIBMGA012816* | down | 1411 | 1419 | - | 11.3303 | 4.96E-05 | 0.615 | GTGGGTGGC |
| 10821 | mamo-s | *BGIBMGA012817* | up | 1294 | 1302 | - | 11.0826 | 6.84E-05 | 0.709 | GTACGTGGC |
| 10822 | mamo-s | *BGIBMGA012818* | up | 290 | 298 | - | 11.3303 | 4.96E-05 | 0.615 | GTGGGTGGC |
| 10823 | mamo-s | *BGIBMGA012818* | up | 970 | 978 | - | 11.3303 | 4.96E-05 | 0.615 | GTGGGTGGC |
| 10824 | mamo-s | *BGIBMGA012818* | up | 1615 | 1623 | + | 11.3303 | 4.96E-05 | 0.615 | GTGGGTGGC |
| 10825 | mamo-s | *BGIBMGA012822* | down | 1577 | 1585 | - | 14.055 | 2.23E-06 | 0.615 | GTGCGTGGC |
| 10826 | mamo-s | *BGIBMGA012826* | down | 451 | 459 | + | 11.3303 | 4.96E-05 | 0.615 | GTGGGTGGC |
| 10827 | mamo-s | *BGIBMGA012827* | down | 1478 | 1486 | - | 11.3303 | 4.96E-05 | 0.615 | GTGGGTGGC |
| 10828 | mamo-s | *BGIBMGA012827* | down | 1678 | 1686 | + | 10.6881 | 8.61E-05 | 0.81 | GCACGTGGC |
| 10829 | mamo-s | *BGIBMGA012828* | down | 359 | 367 | + | 13.6606 | 4.06E-06 | 0.615 | GCGCGTGGC |
| 10830 | mamo-s | *BGIBMGA012828* | down | 148 | 156 | + | 11.3303 | 4.96E-05 | 0.615 | GTGGGTGGC |
| 10831 | mamo-s | *BGIBMGA012828* | up | 1003 | 1011 | + | 11.3303 | 4.96E-05 | 0.615 | GTGGGTGGC |
| 10832 | mamo-s | *BGIBMGA012829* | down | 126 | 134 | + | 11.1009 | 6.57E-05 | 0.709 | GAGCGAGGC |
| 10833 | mamo-s | *BGIBMGA012833* | down | 882 | 890 | - | 11.5321 | 4.06E-05 | 0.615 | TTGCGTGGC |
| 10834 | mamo-s | *BGIBMGA012835* | down | 369 | 377 | + | 11.3303 | 4.96E-05 | 0.615 | GTGGGTGGC |
| 10835 | mamo-s | *BGIBMGA012835* | down | 1365 | 1373 | - | 11.3303 | 4.96E-05 | 0.615 | GTGGGTGGC |
| 10836 | mamo-s | *BGIBMGA012835* | up | 508 | 516 | - | 10.5688 | 8.88E-05 | 0.824 | GTGCGTGAC |
| 10837 | mamo-s | *BGIBMGA012836* | up | 525 | 533 | - | 11.3303 | 4.96E-05 | 0.615 | GTGGGTGGC |
| 10838 | mamo-s | *BGIBMGA012838* | up | 163 | 171 | - | 11.3303 | 4.96E-05 | 0.615 | GTGGGTGGC |
| 10839 | mamo-s | *BGIBMGA012841* | up | 245 | 253 | + | 11.3303 | 4.96E-05 | 0.615 | GTGGGTGGC |
| 10840 | mamo-s | *BGIBMGA012844* | up | 212 | 220 | + | 11.3303 | 4.96E-05 | 0.615 | GTGGGTGGC |
| 10841 | mamo-s | *BGIBMGA012849* | up | 684 | 692 | - | 11.5321 | 4.06E-05 | 0.615 | TTGCGTGGC |
| 10842 | mamo-s | *BGIBMGA012851* | down | 1395 | 1403 | + | 11.3853 | 4.74E-05 | 0.615 | ACGCGTGGC |
| 10843 | mamo-s | *BGIBMGA012854* | up | 1605 | 1613 | + | 10.5413 | 9.42E-05 | 0.857 | GTGCGAGTC |
| 10844 | mamo-s | *BGIBMGA012855* | down | 162 | 170 | + | 11.5688 | 3.79E-05 | 0.615 | CTGCGTGTC |
| 10845 | mamo-s | *BGIBMGA012855* | down | 1855 | 1863 | - | 11.3303 | 4.96E-05 | 0.615 | GTGGGTGGC |
| 10846 | mamo-s | *BGIBMGA012855* | down | 385 | 393 | + | 10.5413 | 9.42E-05 | 0.857 | GTGCGAGTC |
| 10847 | mamo-s | *BGIBMGA012858* | up | 1515 | 1523 | - | 11.3303 | 4.96E-05 | 0.615 | GTGGGTGGC |
| 10848 | mamo-s | *BGIBMGA012861* | down | 1474 | 1482 | - | 11.3303 | 4.96E-05 | 0.615 | GTGGGTGGC |
| 10849 | mamo-s | *BGIBMGA012862* | up | 226 | 234 | - | 11.3303 | 4.96E-05 | 0.615 | GTGGGTGGC |
| 10850 | mamo-s | *BGIBMGA012862* | up | 1025 | 1033 | - | 11.3303 | 4.96E-05 | 0.615 | GTGGGTGGC |
| 10851 | mamo-s | *BGIBMGA012863* | up | 1088 | 1096 | - | 11.3303 | 4.96E-05 | 0.615 | GTGGGTGGC |
| 10852 | mamo-s | *BGIBMGA012865* | down | 1202 | 1210 | + | 11.7339 | 3.29E-05 | 0.615 | GCGCGAGGC |
| 10853 | mamo-s | *BGIBMGA012865* | up | 954 | 962 | - | 11.3303 | 4.96E-05 | 0.615 | GTGGGTGGC |
| 10854 | mamo-s | *BGIBMGA012866* | up | 18 | 26 | + | 11.3303 | 4.96E-05 | 0.615 | GTGGGTGGC |
| 10855 | mamo-s | *BGIBMGA012866* | down | 593 | 601 | + | 11.3303 | 4.96E-05 | 0.615 | GTGGGTGGC |
| 10856 | mamo-s | *BGIBMGA012866* | up | 1690 | 1698 | + | 11.3303 | 4.96E-05 | 0.615 | GTGGGTGGC |
| 10857 | mamo-s | *BGIBMGA012868* | up | 1896 | 1904 | + | 11.3303 | 4.96E-05 | 0.615 | GTGGGTGGC |
| 10858 | mamo-s | *BGIBMGA012869* | down | 158 | 166 | - | 11.3303 | 4.96E-05 | 0.615 | GTGGGTGGC |
| 10859 | mamo-s | *BGIBMGA012869* | down | 1579 | 1587 | + | 11.3303 | 4.96E-05 | 0.615 | GTGGGTGGC |
| 10860 | mamo-s | *BGIBMGA012870* | down | 1016 | 1024 | + | 11.3303 | 4.96E-05 | 0.615 | GTGGGTGGC |
| 10861 | mamo-s | *BGIBMGA012876* | up | 1811 | 1819 | + | 11.3303 | 4.96E-05 | 0.615 | GTGGGTGGC |
| 10862 | mamo-s | *BGIBMGA012876* | down | 490 | 498 | + | 11.1376 | 6.35E-05 | 0.709 | TCGCGTGGC |
| 10863 | mamo-s | *BGIBMGA012876* | down | 593 | 601 | + | 10.5138 | 9.75E-05 | 0.865 | GTGCGTGTT |
| 10864 | mamo-s | *BGIBMGA012879* | up | 806 | 814 | - | 11.7064 | 3.52E-05 | 0.615 | GCGCGTGGT |
| 10865 | mamo-s | *BGIBMGA012881* | up | 1128 | 1136 | + | 10.8073 | 7.89E-05 | 0.766 | CCGCGTGGT |
| 10866 | mamo-s | *BGIBMGA012882* | down | 1437 | 1445 | - | 11.3303 | 4.96E-05 | 0.615 | GTGGGTGGC |
| 10867 | mamo-s | *BGIBMGA012884* | down | 239 | 247 | - | 10.5138 | 9.75E-05 | 0.865 | GTGCGTGTT |
| 10868 | mamo-s | *BGIBMGA012887* | down | 1295 | 1303 | + | 11.7339 | 3.29E-05 | 0.615 | GCGCGAGGC |
| 10869 | mamo-s | *BGIBMGA012888* | down | 1475 | 1483 | + | 11.3303 | 4.96E-05 | 0.615 | GTGGGTGGC |
| 10870 | mamo-s | *BGIBMGA012889* | up | 1305 | 1313 | + | 11.0734 | 7.12E-05 | 0.726 | GAGCGTGGT |
| 10871 | mamo-s | *BGIBMGA012892* | down | 1339 | 1347 | + | 11.2844 | 5.18E-05 | 0.63 | CTGCGTGCC |
| 10872 | mamo-s | *BGIBMGA012894* | down | 1246 | 1254 | + | 11.3303 | 4.96E-05 | 0.615 | GTGGGTGGC |
| 10873 | mamo-s | *BGIBMGA012894* | down | 1578 | 1586 | - | 11.3303 | 4.96E-05 | 0.615 | GTGGGTGGC |
| 10874 | mamo-s | *BGIBMGA012895* | down | 1621 | 1629 | + | 11.3303 | 4.96E-05 | 0.615 | GTGGGTGGC |
| 10875 | mamo-s | *BGIBMGA012897* | down | 924 | 932 | + | 11.7064 | 3.52E-05 | 0.615 | GCGCGTGGT |
| 10876 | mamo-s | *BGIBMGA012899* | down | 963 | 971 | - | 11.0826 | 6.84E-05 | 0.709 | GTACGTGGC |
| 10877 | mamo-s | *BGIBMGA012900* | down | 1012 | 1020 | + | 11.3303 | 4.96E-05 | 0.615 | GTGGGTGGC |
| 10878 | mamo-s | *BGIBMGA012901* | down | 1606 | 1614 | - | 14.055 | 2.23E-06 | 0.615 | GTGCGTGGC |
| 10879 | mamo-s | *BGIBMGA012901* | up | 253 | 261 | - | 11.3303 | 4.96E-05 | 0.615 | GTGGGTGGC |
| 10880 | mamo-s | *BGIBMGA012901* | up | 1847 | 1855 | + | 11.3303 | 4.96E-05 | 0.615 | GTGGGTGGC |
| 10881 | mamo-s | *BGIBMGA012907* | up | 927 | 935 | + | 11.3303 | 4.96E-05 | 0.615 | GTGGGTGGC |
| 10882 | mamo-s | *BGIBMGA012910* | up | 764 | 772 | + | 11.3303 | 4.96E-05 | 0.615 | GTGGGTGGC |
| 10883 | mamo-s | *BGIBMGA012910* | up | 1852 | 1860 | + | 11.3303 | 4.96E-05 | 0.615 | GTGGGTGGC |
| 10884 | mamo-s | *BGIBMGA012912* | down | 334 | 342 | + | 11.2018 | 5.68E-05 | 0.674 | CTGCGTGGT |
| 10885 | mamo-s | *BGIBMGA012913* | up | 1200 | 1208 | - | 12.1835 | 1.71E-05 | 0.615 | GTGCGTGCC |
| 10886 | mamo-s | *BGIBMGA012913* | down | 878 | 886 | + | 11.3303 | 4.96E-05 | 0.615 | GTGGGTGGC |
| 10887 | mamo-s | *BGIBMGA012915* | up | 662 | 670 | + | 11.3303 | 4.96E-05 | 0.615 | GTGGGTGGC |
| 10888 | mamo-s | *BGIBMGA012916* | down | 1777 | 1785 | + | 12.3394 | 1.49E-05 | 0.615 | GGGCGTGGC |
| 10889 | mamo-s | *BGIBMGA012916* | up | 156 | 164 | - | 11.3303 | 4.96E-05 | 0.615 | GTGGGTGGC |
| 10890 | mamo-s | *BGIBMGA012916* | down | 1491 | 1499 | + | 11.3303 | 4.96E-05 | 0.615 | GTGGGTGGC |
| 10891 | mamo-s | *BGIBMGA012917* | up | 1150 | 1158 | - | 11.3303 | 4.96E-05 | 0.615 | GTGGGTGGC |
| 10892 | mamo-s | *BGIBMGA012917* | up | 1784 | 1792 | - | 11.3303 | 4.96E-05 | 0.615 | GTGGGTGGC |
| 10893 | mamo-s | *BGIBMGA012918* | up | 45 | 53 | + | 11.3303 | 4.96E-05 | 0.615 | GTGGGTGGC |
| 10894 | mamo-s | *BGIBMGA012920* | up | 1131 | 1139 | + | 11.7064 | 3.52E-05 | 0.615 | GCGCGTGGT |
| 10895 | mamo-s | *BGIBMGA012920* | up | 641 | 649 | - | 11.3853 | 4.74E-05 | 0.615 | ACGCGTGGC |
| 10896 | mamo-s | *BGIBMGA012920* | down | 1112 | 1120 | - | 11.0826 | 6.84E-05 | 0.709 | GTACGTGGC |
| 10897 | mamo-s | *BGIBMGA012921* | up | 1413 | 1421 | + | 11.3303 | 4.96E-05 | 0.615 | GTGGGTGGC |
| 10898 | mamo-s | *BGIBMGA012923* | down | 987 | 995 | - | 11.3303 | 4.96E-05 | 0.615 | GTGGGTGGC |
| 10899 | mamo-s | *BGIBMGA012926* | up | 1032 | 1040 | + | 12.1835 | 1.71E-05 | 0.615 | GTGCGTGCC |
| 10900 | mamo-s | *BGIBMGA012926* | down | 937 | 945 | + | 11.0826 | 6.84E-05 | 0.709 | GTACGTGGC |
| 10901 | mamo-s | *BGIBMGA012927* | down | 1737 | 1745 | - | 11.2018 | 5.68E-05 | 0.674 | CTGCGTGGT |
| 10902 | mamo-s | *BGIBMGA012927* | up | 1530 | 1538 | + | 11.0826 | 6.84E-05 | 0.709 | GTACGTGGC |
| 10903 | mamo-s | *BGIBMGA012931* | up | 664 | 672 | + | 11.5688 | 3.79E-05 | 0.615 | CTGCGTGTC |
| 10904 | mamo-s | *BGIBMGA012933* | up | 362 | 370 | - | 11.3303 | 4.96E-05 | 0.615 | GTGGGTGGC |
| 10905 | mamo-s | *BGIBMGA012935* | down | 7 | 15 | - | 10.9358 | 7.30E-05 | 0.728 | GCGGGTGGC |

| | | | | | | | | | | |
|---|---|---|---|---|---|---|---|---|---|---|
| 10906 | mamo-s | *BGIBMGA012936* | up | 695 | 703 | + | 11.3303 | 4.96E-05 | 0.615 | GTGGGTGGC |
| 10907 | mamo-s | *BGIBMGA012938* | down | 261 | 269 | + | 11.3303 | 4.96E-05 | 0.615 | GTGGGTGGC |
| 10908 | mamo-s | *BGIBMGA012938* | up | 788 | 796 | - | 11.3303 | 4.96E-05 | 0.615 | GTGGGTGGC |
| 10909 | mamo-s | *BGIBMGA012940* | down | 1383 | 1391 | - | 11.3303 | 4.96E-05 | 0.615 | GTGGGTGGC |
| 10910 | mamo-s | *BGIBMGA012949* | up | 527 | 535 | - | 11.3303 | 4.96E-05 | 0.615 | GTGGGTGGC |
| 10911 | mamo-s | *BGIBMGA012950* | down | 1597 | 1605 | + | 11.0826 | 6.84E-05 | 0.709 | GTACGTGGC |
| 10912 | mamo-s | *BGIBMGA012951* | up | 95 | 103 | + | 11.3303 | 4.96E-05 | 0.615 | GTGGGTGGC |
| 10913 | mamo-s | *BGIBMGA012953* | up | 1563 | 1571 | - | 13.6606 | 4.06E-06 | 0.615 | GCGCGTGGC |
| 10914 | mamo-s | *BGIBMGA012953* | down | 349 | 357 | + | 11.3303 | 4.96E-05 | 0.615 | GTGGGTGGC |
| 10915 | mamo-s | *BGIBMGA012953* | up | 898 | 906 | + | 11.3303 | 4.96E-05 | 0.615 | GTGGGTGGC |
| 10916 | mamo-s | *BGIBMGA012955* | up | 1405 | 1413 | - | 11.3853 | 4.74E-05 | 0.615 | ACGCGTGGC |
| 10917 | mamo-s | *BGIBMGA012955* | up | 625 | 633 | - | 11.1743 | 5.90E-05 | 0.684 | CCGCGTGTC |
| 10918 | mamo-s | *BGIBMGA012956* | up | 722 | 730 | + | 11.3303 | 4.96E-05 | 0.615 | GTGGGTGGC |
| 10919 | mamo-s | *BGIBMGA012957* | up | 1658 | 1666 | - | 11.3303 | 4.96E-05 | 0.615 | GTGGGTGGC |
| 10920 | mamo-s | *BGIBMGA012960* | up | 1101 | 1109 | - | 11.3303 | 4.96E-05 | 0.615 | GTGGGTGGC |
| 10921 | mamo-s | *BGIBMGA012961* | down | 1297 | 1305 | - | 11.1009 | 6.57E-05 | 0.709 | GAGCGAGGC |
| 10922 | mamo-s | *BGIBMGA012962* | up | 697 | 705 | - | 11.3303 | 4.96E-05 | 0.615 | GTGGGTGGC |
| 10923 | mamo-s | *BGIBMGA012964* | up | 1886 | 1894 | - | 12.1835 | 1.71E-05 | 0.615 | GTGCGTGCC |
| 10924 | mamo-s | *BGIBMGA012968* | down | 942 | 950 | + | 11.3303 | 4.96E-05 | 0.615 | GTGGGTGGC |
| 10925 | mamo-s | *BGIBMGA012968* | down | 1732 | 1740 | + | 11.3303 | 4.96E-05 | 0.615 | GTGGGTGGC |
| 10926 | mamo-s | *BGIBMGA012968* | down | 386 | 394 | - | 11.0826 | 6.84E-05 | 0.709 | GTACGTGGC |
| 10927 | mamo-s | *BGIBMGA012969* | up | 207 | 215 | + | 11.3303 | 4.96E-05 | 0.615 | GTGGGTGGC |
| 10928 | mamo-s | *BGIBMGA012969* | up | 997 | 1005 | + | 11.3303 | 4.96E-05 | 0.615 | GTGGGTGGC |
| 10929 | mamo-s | *BGIBMGA012970* | up | 1427 | 1435 | - | 11.3303 | 4.96E-05 | 0.615 | GTGGGTGGC |
| 10930 | mamo-s | *BGIBMGA012971* | up | 596 | 604 | + | 11.3303 | 4.96E-05 | 0.615 | GTGGGTGGC |
| 10931 | mamo-s | *BGIBMGA012976* | up | 1287 | 1295 | - | 11.0734 | 7.12E-05 | 0.726 | GAGCGTGGT |
| 10932 | mamo-s | *BGIBMGA012978* | down | 288 | 296 | + | 11.3303 | 4.96E-05 | 0.615 | GTGGGTGGC |
| 10933 | mamo-s | *BGIBMGA012979* | up | 1129 | 1137 | + | 12.3394 | 1.49E-05 | 0.615 | GGGCGTGGC |
| 10934 | mamo-s | *BGIBMGA012979* | up | 719 | 727 | + | 11.4404 | 4.51E-05 | 0.615 | GAGCGTGTC |
| 10935 | mamo-s | *BGIBMGA012979* | up | 700 | 708 | - | 11.3853 | 4.74E-05 | 0.615 | ACGCGTGGC |
| 10936 | mamo-s | *BGIBMGA012979* | up | 1375 | 1383 | - | 11.3303 | 4.96E-05 | 0.615 | GTGGGTGGC |
| 10937 | mamo-s | *BGIBMGA012979* | down | 767 | 775 | + | 10.7523 | 8.38E-05 | 0.797 | AAGCGTGGC |
| 10938 | mamo-s | *BGIBMGA012981* | down | 758 | 766 | + | 11.3303 | 4.96E-05 | 0.615 | GTGGGTGGC |
| 10939 | mamo-s | *BGIBMGA012982* | down | 1037 | 1045 | - | 10.5413 | 9.42E-05 | 0.857 | GTGCGAGTC |
| 10940 | mamo-s | *BGIBMGA012983* | down | 1749 | 1757 | - | 11.3303 | 4.96E-05 | 0.615 | GTGGGTGGC |
| 10941 | mamo-s | *BGIBMGA012984* | down | 1620 | 1628 | + | 11.0826 | 6.84E-05 | 0.709 | GTACGTGGC |
| 10942 | mamo-s | *BGIBMGA012987* | up | 1321 | 1329 | + | 11.3303 | 4.96E-05 | 0.615 | GTGGGTGGC |
| 10943 | mamo-s | *BGIBMGA012988* | down | 1090 | 1098 | + | 11.3303 | 4.96E-05 | 0.615 | GTGGGTGGC |
| 10944 | mamo-s | *BGIBMGA012988* | down | 1965 | 1973 | + | 11.3303 | 4.96E-05 | 0.615 | GTGGGTGGC |
| 10945 | mamo-s | *BGIBMGA012992* | down | 123 | 131 | - | 11.3303 | 4.96E-05 | 0.615 | GTGGGTGGC |
| 10946 | mamo-s | *BGIBMGA012995* | up | 1313 | 1321 | + | 11.3303 | 4.96E-05 | 0.615 | GTGGGTGGC |
| 10947 | mamo-s | *BGIBMGA012995* | down | 1450 | 1458 | - | 11.3303 | 4.96E-05 | 0.615 | GTGGGTGGC |
| 10948 | mamo-s | *BGIBMGA012995* | up | 719 | 727 | - | 11.0826 | 6.84E-05 | 0.709 | GTACGTGGC |
| 10949 | mamo-s | *BGIBMGA012996* | up | 1180 | 1188 | + | 11.3303 | 4.96E-05 | 0.615 | GTGGGTGGC |
| 10950 | mamo-s | *BGIBMGA012997* | down | 948 | 956 | - | 11.7339 | 3.29E-05 | 0.615 | GCGCGAGGC |
| 10951 | mamo-s | *BGIBMGA012998* | down | 869 | 877 | - | 11.3303 | 4.96E-05 | 0.615 | GTGGGTGGC |
| 10952 | mamo-s | *BGIBMGA012999* | up | 498 | 506 | + | 11.3303 | 4.96E-05 | 0.615 | GTGGGTGGC |
| 10953 | mamo-s | *BGIBMGA012999* | down | 1980 | 1988 | + | 11.3303 | 4.96E-05 | 0.615 | GTGGGTGGC |
| 10954 | mamo-s | *BGIBMGA013000* | up | 1296 | 1304 | - | 10.8349 | 7.66E-05 | 0.751 | CCGCGAGGC |
| 10955 | mamo-s | *BGIBMGA013003* | up | 1201 | 1209 | - | 11.3303 | 4.96E-05 | 0.615 | GTGGGTGGC |
| 10956 | mamo-s | *BGIBMGA013006* | down | 143 | 151 | - | 11.3303 | 4.96E-05 | 0.615 | GTGGGTGGC |
| 10957 | mamo-s | *BGIBMGA013006* | up | 1642 | 1650 | - | 11.3303 | 4.96E-05 | 0.615 | GTGGGTGGC |
| 10958 | mamo-s | *BGIBMGA013007* | down | 1705 | 1713 | + | 14.055 | 2.23E-06 | 0.615 | GTGCGTGGC |
| 10959 | mamo-s | *BGIBMGA013007* | up | 994 | 1002 | + | 11.2844 | 5.18E-05 | 0.63 | CTGCGTGCC |
| 10960 | mamo-s | *BGIBMGA013008* | up | 415 | 423 | + | 14.055 | 2.23E-06 | 0.615 | GTGCGTGGC |
| 10961 | mamo-s | *BGIBMGA013009* | down | 1328 | 1336 | - | 10.6881 | 8.61E-05 | 0.81 | GCACGTGGC |
| 10962 | mamo-s | *BGIBMGA013014* | down | 1325 | 1333 | + | 11.3303 | 4.96E-05 | 0.615 | GTGGGTGGC |
| 10963 | mamo-s | *BGIBMGA013015* | down | 1893 | 1901 | - | 11.3303 | 4.96E-05 | 0.615 | GTGGGTGGC |
| 10964 | mamo-s | *BGIBMGA013016* | up | 123 | 131 | + | 11.3303 | 4.96E-05 | 0.615 | GTGGGTGGC |
| 10965 | mamo-s | *BGIBMGA013016* | up | 1516 | 1524 | - | 11.3303 | 4.96E-05 | 0.615 | GTGGGTGGC |
| 10966 | mamo-s | *BGIBMGA013017* | up | 20 | 28 | - | 11.3303 | 4.96E-05 | 0.615 | GTGGGTGGC |
| 10967 | mamo-s | *BGIBMGA013017* | down | 1703 | 1711 | + | 11.3303 | 4.96E-05 | 0.615 | GTGGGTGGC |
| 10968 | mamo-s | *BGIBMGA013019* | down | 405 | 413 | - | 11.1009 | 6.57E-05 | 0.709 | GAGCGAGGC |
| 10969 | mamo-s | *BGIBMGA013020* | up | 911 | 919 | - | 11.3303 | 4.96E-05 | 0.615 | GTGGGTGGC |
| 10970 | mamo-s | *BGIBMGA013022* | down | 397 | 405 | - | 11.3303 | 4.96E-05 | 0.615 | GTGGGTGGC |
| 10971 | mamo-s | *BGIBMGA013023* | down | 1618 | 1626 | - | 11.3303 | 4.96E-05 | 0.615 | GTGGGTGGC |
| 10972 | mamo-s | *BGIBMGA013023* | down | 512 | 520 | + | 10.9358 | 7.30E-05 | 0.728 | GCGGGTGGC |
| 10973 | mamo-s | *BGIBMGA013024* | up | 753 | 761 | - | 11.3303 | 4.96E-05 | 0.615 | GTGGGTGGC |
| 10974 | mamo-s | *BGIBMGA013024* | down | 1679 | 1687 | - | 11.3303 | 4.96E-05 | 0.615 | GTGGGTGGC |
| 10975 | mamo-s | *BGIBMGA013025* | down | 1280 | 1288 | - | 11.1376 | 6.35E-05 | 0.709 | TCGCGTGGC |
| 10976 | mamo-s | *BGIBMGA013025* | up | 84 | 92 | + | 10.9358 | 7.30E-05 | 0.728 | GCGGGTGGC |
| 10977 | mamo-s | *BGIBMGA013027* | down | 1293 | 1301 | + | 11.3303 | 4.96E-05 | 0.615 | GTGGGTGGC |
| 10978 | mamo-s | *BGIBMGA013027* | up | 1359 | 1367 | + | 11.0826 | 6.84E-05 | 0.709 | GTACGTGGC |
| 10979 | mamo-s | *BGIBMGA013028* | up | 1789 | 1797 | + | 11.3303 | 4.96E-05 | 0.615 | GTGGGTGGC |
| 10980 | mamo-s | *BGIBMGA013028* | up | 409 | 417 | + | 11.2844 | 5.18E-05 | 0.63 | CTGCGTGCC |
| 10981 | mamo-s | *BGIBMGA013029* | down | 689 | 697 | + | 11.3303 | 4.96E-05 | 0.615 | GTGGGTGGC |
| 10982 | mamo-s | *BGIBMGA013029* | down | 980 | 988 | - | 11.2844 | 5.18E-05 | 0.63 | CTGCGTGCC |
| 10983 | mamo-s | *BGIBMGA013029* | down | 1087 | 1095 | - | 11.2018 | 5.68E-05 | 0.674 | CTGCGTGGT |
| 10984 | mamo-s | *BGIBMGA013029* | down | 401 | 409 | - | 11.0826 | 6.84E-05 | 0.709 | GTACGTGGC |
| 10985 | mamo-s | *BGIBMGA013030* | up | 377 | 385 | + | 11.3303 | 4.96E-05 | 0.615 | GTGGGTGGC |
| 10986 | mamo-s | *BGIBMGA013030* | up | 668 | 676 | - | 11.2844 | 5.18E-05 | 0.63 | CTGCGTGCC |
| 10987 | mamo-s | *BGIBMGA013030* | up | 775 | 783 | - | 11.2018 | 5.68E-05 | 0.674 | CTGCGTGGT |
| 10988 | mamo-s | *BGIBMGA013030* | up | 89 | 97 | - | 11.0826 | 6.84E-05 | 0.709 | GTACGTGGC |
| 10989 | mamo-s | *BGIBMGA013031* | up | 798 | 806 | + | 11.3303 | 4.96E-05 | 0.615 | GTGGGTGGC |
| 10990 | mamo-s | *BGIBMGA013032* | up | 1171 | 1179 | - | 11.3303 | 4.96E-05 | 0.615 | GTGGGTGGC |
| 10991 | mamo-s | *BGIBMGA013032* | up | 958 | 966 | + | 10.6881 | 8.61E-05 | 0.81 | GCACGTGGC |
| 10992 | mamo-s | *BGIBMGA013032* | up | 1340 | 1348 | - | 10.6881 | 8.61E-05 | 0.81 | GCACGTGGC |
| 10993 | mamo-s | *BGIBMGA013035* | down | 169 | 177 | + | 11.3303 | 4.96E-05 | 0.615 | GTGGGTGGC |

| | | | | | | | | | | |
|---|---|---|---|---|---|---|---|---|---|---|
| 10994 | mamo-s | *BGIBMGA013035* | up | 1318 | 1326 | - | 11.3303 | 4.96E-05 | 0.615 | GTGGGTGGC |
| 10995 | mamo-s | *BGIBMGA013035* | down | 837 | 845 | - | 11.1743 | 5.90E-05 | 0.684 | CCGCGTGTC |
| 10996 | mamo-s | *BGIBMGA013036* | up | 1084 | 1092 | + | 11.7339 | 3.29E-05 | 0.615 | GCGCGAGGC |
| 10997 | mamo-s | *BGIBMGA013036* | up | 1373 | 1381 | + | 11.3303 | 4.96E-05 | 0.615 | GTGGGTGGC |
| 10998 | mamo-s | *BGIBMGA013036* | down | 1620 | 1628 | + | 11.3303 | 4.96E-05 | 0.615 | GTGGGTGGC |
| 10999 | mamo-s | *BGIBMGA013038* | up | 421 | 429 | + | 11.3303 | 4.96E-05 | 0.615 | GTGGGTGGC |
| 11000 | mamo-s | *BGIBMGA013038* | down | 1210 | 1218 | + | 11.3303 | 4.96E-05 | 0.615 | GTGGGTGGC |
| 11001 | mamo-s | *BGIBMGA013039* | up | 767 | 775 | + | 11.3303 | 4.96E-05 | 0.615 | GTGGGTGGC |
| 11002 | mamo-s | *BGIBMGA013040* | up | 540 | 548 | - | 11.7798 | 3.11E-05 | 0.615 | ATGCGTGGC |
| 11003 | mamo-s | *BGIBMGA013041* | down | 1131 | 1139 | + | 11.3853 | 4.74E-05 | 0.615 | ACGCGTGGC |
| 11004 | mamo-s | *BGIBMGA013042* | up | 268 | 276 | + | 12.1009 | 2.43E-05 | 0.615 | GTGCGTGGT |
| 11005 | mamo-s | *BGIBMGA013042* | down | 1781 | 1789 | - | 11.3303 | 4.96E-05 | 0.615 | GTGGGTGGC |
| 11006 | mamo-s | *BGIBMGA013043* | up | 1481 | 1489 | + | 12.7615 | 1.04E-05 | 0.615 | CCGCGTGGC |
| 11007 | mamo-s | *BGIBMGA013045* | down | 1850 | 1858 | + | 11.3303 | 4.96E-05 | 0.615 | GTGGGTGGC |
| 11008 | mamo-s | *BGIBMGA013046* | down | 372 | 380 | - | 14.055 | 2.23E-06 | 0.615 | GTGCGTGGC |
| 11009 | mamo-s | *BGIBMGA013051* | down | 844 | 852 | + | 11.3303 | 4.96E-05 | 0.615 | GTGGGTGGC |
| 11010 | mamo-s | *BGIBMGA013053* | down | 819 | 827 | + | 11.1743 | 5.90E-05 | 0.684 | CCGCGTGTC |
| 11011 | mamo-s | *BGIBMGA013053* | up | 1183 | 1191 | + | 10.5413 | 9.42E-05 | 0.857 | CAGCGTGTC |
| 11012 | mamo-s | *BGIBMGA013055* | up | 59 | 67 | + | 11.3303 | 4.96E-05 | 0.615 | GTGGGTGGC |
| 11013 | mamo-s | *BGIBMGA013055* | down | 1779 | 1787 | + | 11.1376 | 6.35E-05 | 0.709 | TCGCGTGGC |
| 11014 | mamo-s | *BGIBMGA013056* | up | 635 | 643 | + | 11.3303 | 4.96E-05 | 0.615 | GTGGGTGGC |
| 11015 | mamo-s | *BGIBMGA013056* | down | 1550 | 1558 | - | 11.3303 | 4.96E-05 | 0.615 | GTGGGTGGC |
| 11016 | mamo-s | *BGIBMGA013056* | up | 1818 | 1826 | + | 11.3303 | 4.96E-05 | 0.615 | GTGGGTGGC |
| 11017 | mamo-s | *BGIBMGA013058* | down | 767 | 775 | - | 11.3303 | 4.96E-05 | 0.615 | GTGGGTGGC |
| 11018 | mamo-s | *BGIBMGA013058* | up | 1645 | 1653 | - | 11.3303 | 4.96E-05 | 0.615 | GTGGGTGGC |
| 11019 | mamo-s | *BGIBMGA013060* | down | 521 | 529 | + | 11.3303 | 4.96E-05 | 0.615 | GTGGGTGGC |
| 11020 | mamo-s | *BGIBMGA013061* | up | 1771 | 1779 | + | 11.0826 | 6.84E-05 | 0.709 | GTACGTGGC |
| 11021 | mamo-s | *BGIBMGA013062* | down | 1449 | 1457 | - | 11.3303 | 4.96E-05 | 0.615 | GTGGGTGGC |
| 11022 | mamo-s | *BGIBMGA013064* | down | 1598 | 1606 | - | 11.1009 | 6.57E-05 | 0.709 | GAGCGAGGC |
| 11023 | mamo-s | *BGIBMGA013067* | down | 1527 | 1535 | + | 11.3303 | 4.96E-05 | 0.615 | GTGGGTGGC |
| 11024 | mamo-s | *BGIBMGA013069* | up | 377 | 385 | - | 11.3303 | 4.96E-05 | 0.615 | GTGGGTGGC |
| 11025 | mamo-s | *BGIBMGA013069* | up | 1842 | 1850 | + | 11.2294 | 5.41E-05 | 0.65 | CTGCGAGGC |
| 11026 | mamo-s | *BGIBMGA013071* | up | 390 | 398 | + | 12.3394 | 1.49E-05 | 0.615 | GGGCGTGGC |
| 11027 | mamo-s | *BGIBMGA013073* | up | 1067 | 1075 | - | 11.2018 | 5.68E-05 | 0.674 | CTGCGTGGT |
| 11028 | mamo-s | *BGIBMGA013074* | down | 1611 | 1619 | + | 11.3303 | 4.96E-05 | 0.615 | GTGGGTGGC |
| 11029 | mamo-s | *BGIBMGA013075* | up | 1169 | 1177 | - | 11.3303 | 4.96E-05 | 0.615 | GTGGGTGGC |
| 11030 | mamo-s | *BGIBMGA013078* | up | 1762 | 1770 | - | 11.3303 | 4.96E-05 | 0.615 | GTGGGTGGC |
| 11031 | mamo-s | *BGIBMGA013079* | down | 322 | 330 | - | 11.3303 | 4.96E-05 | 0.615 | GTGGGTGGC |
| 11032 | mamo-s | *BGIBMGA013079* | up | 626 | 634 | + | 11.3303 | 4.96E-05 | 0.615 | GTGGGTGGC |
| 11033 | mamo-s | *BGIBMGA013082* | down | 1457 | 1465 | + | 12.0734 | 2.66E-05 | 0.615 | GCGCGTGTC |
| 11034 | mamo-s | *BGIBMGA013084* | down | 115 | 123 | - | 11.3303 | 4.96E-05 | 0.615 | GTGGGTGGC |
| 11035 | mamo-s | *BGIBMGA013084* | up | 361 | 369 | + | 10.5138 | 9.75E-05 | 0.865 | GTGCGTGTT |
| 11036 | mamo-s | *BGIBMGA013085* | up | 155 | 163 | + | 11.3303 | 4.96E-05 | 0.615 | GTGGGTGGC |
| 11037 | mamo-s | *BGIBMGA013085* | down | 615 | 623 | - | 11.3303 | 4.96E-05 | 0.615 | GTGGGTGGC |
| 11038 | mamo-s | *BGIBMGA013086* | down | 848 | 856 | - | 11.3303 | 4.96E-05 | 0.615 | GTGGGTGGC |
| 11039 | mamo-s | *BGIBMGA013088* | up | 1455 | 1463 | + | 10.6881 | 8.61E-05 | 0.81 | GCACGTGGC |
| 11040 | mamo-s | *BGIBMGA013089* | down | 352 | 360 | - | 11.3303 | 4.96E-05 | 0.615 | GTGGGTGGC |
| 11041 | mamo-s | *BGIBMGA013090* | down | 552 | 560 | + | 11.3303 | 4.96E-05 | 0.615 | GTGGGTGGC |
| 11042 | mamo-s | *BGIBMGA013090* | down | 1038 | 1046 | + | 11.0826 | 6.84E-05 | 0.709 | GTACGTGGC |
| 11043 | mamo-s | *BGIBMGA013091* | up | 207 | 215 | - | 11.3303 | 4.96E-05 | 0.615 | GTGGGTGGC |
| 11044 | mamo-s | *BGIBMGA013091* | down | 1260 | 1268 | + | 11.3303 | 4.96E-05 | 0.615 | GTGGGTGGC |
| 11045 | mamo-s | *BGIBMGA013091* | down | 474 | 482 | - | 10.9358 | 7.30E-05 | 0.728 | GCGGGTGGC |
| 11046 | mamo-s | *BGIBMGA013092* | up | 1672 | 1680 | + | 12.1284 | 2.16E-05 | 0.615 | CAGCGTGGC |
| 11047 | mamo-s | *BGIBMGA013092* | up | 1454 | 1462 | - | 11.3303 | 4.96E-05 | 0.615 | GTGGGTGGC |
| 11048 | mamo-s | *BGIBMGA013092* | down | 1844 | 1852 | - | 11.3303 | 4.96E-05 | 0.615 | GTGGGTGGC |
| 11049 | mamo-s | *BGIBMGA013093* | down | 529 | 537 | + | 11.3303 | 4.96E-05 | 0.615 | GTGGGTGGC |
| 11050 | mamo-s | *BGIBMGA013095* | down | 587 | 595 | + | 11.3303 | 4.96E-05 | 0.615 | GTGGGTGGC |
| 11051 | mamo-s | *BGIBMGA013095* | up | 1577 | 1585 | - | 11.156 | 6.13E-05 | 0.705 | GAGCGTGCC |
| 11052 | mamo-s | *BGIBMGA013096* | up | 560 | 568 | - | 12.1835 | 1.71E-05 | 0.615 | GTGCGTGCC |
| 11053 | mamo-s | *BGIBMGA013096* | down | 1963 | 1971 | - | 12.0734 | 2.66E-05 | 0.615 | GCGCGTGTC |
| 11054 | mamo-s | *BGIBMGA013097* | up | 1669 | 1677 | + | 11.3303 | 4.96E-05 | 0.615 | GTGGGTGGC |
| 11055 | mamo-s | *BGIBMGA013099* | down | 943 | 951 | - | 10.8899 | 7.48E-05 | 0.739 | CCGCGTGCC |
| 11056 | mamo-s | *BGIBMGA013100* | down | 946 | 954 | - | 11.5688 | 3.79E-05 | 0.615 | CTGCGTGTC |
| 11057 | mamo-s | *BGIBMGA013102* | down | 685 | 693 | + | 10.9358 | 7.30E-05 | 0.728 | GCGGGTGGC |
| 11058 | mamo-s | *BGIBMGA013103* | up | 315 | 323 | + | 12.3394 | 1.49E-05 | 0.615 | GGGCGTGGC |
| 11059 | mamo-s | *BGIBMGA013104* | down | 1694 | 1702 | + | 11.7339 | 3.29E-05 | 0.615 | GCGCGAGGC |
| 11060 | mamo-s | *BGIBMGA013106* | down | 990 | 998 | + | 12.1284 | 2.16E-05 | 0.615 | GTGCGAGGC |
| 11061 | mamo-s | *BGIBMGA013106* | down | 1719 | 1727 | + | 12.1009 | 2.43E-05 | 0.615 | GTGCGTGGT |
| 11062 | mamo-s | *BGIBMGA013107* | down | 1295 | 1303 | - | 11.0734 | 7.12E-05 | 0.726 | GAGCGTGGT |
| 11063 | mamo-s | *BGIBMGA013108* | up | 1923 | 1931 | + | 14.055 | 2.23E-06 | 0.615 | GTGCGTGGC |
| 11064 | mamo-s | *BGIBMGA013108* | up | 1932 | 1940 | + | 14.055 | 2.23E-06 | 0.615 | GTGCGTGGC |
| 11065 | mamo-s | *BGIBMGA013108* | down | 1541 | 1549 | - | 11.3303 | 4.96E-05 | 0.615 | GTGGGTGGC |
| 11066 | mamo-s | *BGIBMGA013110* | down | 698 | 706 | + | 11.3303 | 4.96E-05 | 0.615 | GTGGGTGGC |
| 11067 | mamo-s | *BGIBMGA013114* | down | 1125 | 1133 | + | 13.156 | 6.30E-06 | 0.615 | CTGCGTGGC |
| 11068 | mamo-s | *BGIBMGA013117* | up | 1982 | 1990 | + | 11.7798 | 3.11E-05 | 0.615 | ATGCGTGGC |
| 11069 | mamo-s | *BGIBMGA013118* | down | 1970 | 1978 | - | 11.3303 | 4.96E-05 | 0.615 | GTGGGTGGC |
| 11070 | mamo-s | *BGIBMGA013118* | up | 251 | 259 | + | 11.2018 | 5.68E-05 | 0.674 | CTGCGTGGT |
| 11071 | mamo-s | *BGIBMGA013119* | down | 104 | 112 | + | 10.5413 | 9.42E-05 | 0.857 | GTGCGAGTC |
| 11072 | mamo-s | *BGIBMGA013122* | down | 219 | 227 | - | 12.4679 | 1.31E-05 | 0.615 | GTGCGTGTC |
| 11073 | mamo-s | *BGIBMGA013123* | up | 931 | 939 | + | 10.9358 | 7.30E-05 | 0.728 | GCGGGTGGC |
| 11074 | mamo-s | *BGIBMGA013124* | down | 1238 | 1246 | - | 10.7523 | 8.38E-05 | 0.797 | AAGCGTGGC |
| 11075 | mamo-s | *BGIBMGA013125* | down | 1358 | 1366 | + | 11.7339 | 3.29E-05 | 0.615 | GCGCGAGGC |
| 11076 | mamo-s | *BGIBMGA013126* | up | 1133 | 1141 | + | 11.7339 | 3.29E-05 | 0.615 | GCGCGAGGC |
| 11077 | mamo-s | *BGIBMGA013129* | down | 1156 | 1164 | - | 11.3303 | 4.96E-05 | 0.615 | GTGGGTGGC |
| 11078 | mamo-s | *BGIBMGA013130* | down | 415 | 423 | - | 11.3303 | 4.96E-05 | 0.615 | GTGGGTGGC |
| 11079 | mamo-s | *BGIBMGA013133* | up | 125 | 133 | - | 12.1284 | 2.16E-05 | 0.615 | GTGCGAGGC |
| 11080 | mamo-s | *BGIBMGA013136* | down | 1934 | 1942 | + | 10.7523 | 8.38E-05 | 0.797 | AAGCGTGGC |
| 11081 | mamo-s | *BGIBMGA013138* | down | 747 | 755 | + | 11.2844 | 5.18E-05 | 0.63 | CTGCGTGCC |

| | | | | | | | | | | |
|---|---|---|---|---|---|---|---|---|---|---|
| 11082 | mamo-s | *BGIBMGA013144* | down | 1423 | 1431 | - | 11.3303 | 4.96E-05 | 0.615 | GTGGGTGGC |
| 11083 | mamo-s | *BGIBMGA013148* | up | 1914 | 1922 | + | 11.789 | 2.84E-05 | 0.615 | GCGCGTGCC |
| 11084 | mamo-s | *BGIBMGA013148* | up | 620 | 628 | + | 11.3303 | 4.96E-05 | 0.615 | GTGGGTGGC |
| 11085 | mamo-s | *BGIBMGA013151* | down | 325 | 333 | - | 11.3303 | 4.96E-05 | 0.615 | GTGGGTGGC |
| 11086 | mamo-s | *BGIBMGA013152* | down | 891 | 899 | - | 11.3303 | 4.96E-05 | 0.615 | GTGGGTGGC |
| 11087 | mamo-s | *BGIBMGA013158* | up | 908 | 916 | + | 11.2844 | 5.18E-05 | 0.63 | CTGCGTGCC |
| 11088 | mamo-s | *BGIBMGA013158* | down | 359 | 367 | - | 10.8349 | 7.66E-05 | 0.751 | CCGCGAGGC |
| 11089 | mamo-s | *BGIBMGA013159* | up | 93 | 101 | - | 11.3303 | 4.96E-05 | 0.615 | GTGGGTGGC |
| 11090 | mamo-s | *BGIBMGA013160* | down | 1137 | 1145 | + | 11.3303 | 4.96E-05 | 0.615 | GTGGGTGGC |
| 11091 | mamo-s | *BGIBMGA013160* | up | 630 | 638 | - | 11.156 | 6.13E-05 | 0.705 | GAGCGTGCC |
| 11092 | mamo-s | *BGIBMGA013161* | up | 424 | 432 | - | 11.2294 | 5.41E-05 | 0.65 | CTGCGAGGC |
| 11093 | mamo-s | *BGIBMGA013170* | down | 1750 | 1758 | + | 11.3303 | 4.96E-05 | 0.615 | GTGGGTGGC |
| 11094 | mamo-s | *BGIBMGA013173* | up | 1433 | 1441 | - | 11.789 | 2.84E-05 | 0.615 | GCGCGTGCC |
| 11095 | mamo-s | *BGIBMGA013173* | down | 1456 | 1464 | + | 11.0734 | 7.12E-05 | 0.726 | GAGCGTGGT |
| 11096 | mamo-s | *BGIBMGA013174* | up | 569 | 577 | + | 11.156 | 6.13E-05 | 0.705 | GAGCGTGCC |
| 11097 | mamo-s | *BGIBMGA013175* | up | 1289 | 1297 | + | 13.156 | 6.30E-06 | 0.615 | CTGCGTGGC |
| 11098 | mamo-s | *BGIBMGA013175* | down | 567 | 575 | + | 11.3303 | 4.96E-05 | 0.615 | GTGGGTGGC |
| 11099 | mamo-s | *BGIBMGA013175* | up | 593 | 601 | - | 11.3303 | 4.96E-05 | 0.615 | GTGGGTGGC |
| 11100 | mamo-s | *BGIBMGA013176* | down | 1108 | 1116 | - | 11.5321 | 4.06E-05 | 0.615 | TTGCGTGGC |
| 11101 | mamo-s | *BGIBMGA013176* | down | 952 | 960 | - | 11.3303 | 4.96E-05 | 0.615 | GTGGGTGGC |
| 11102 | mamo-s | *BGIBMGA013176* | up | 1528 | 1536 | + | 11.2844 | 5.18E-05 | 0.63 | CTGCGTGCC |
| 11103 | mamo-s | *BGIBMGA013176* | down | 1793 | 1801 | - | 10.8073 | 7.89E-05 | 0.766 | CCGCGTGGT |
| 11104 | mamo-s | *BGIBMGA013178* | down | 77 | 85 | + | 11.3303 | 4.96E-05 | 0.615 | GTGGGTGGC |
| 11105 | mamo-s | *BGIBMGA013182* | down | 1120 | 1128 | + | 11.3303 | 4.96E-05 | 0.615 | GTGGGTGGC |
| 11106 | mamo-s | *BGIBMGA013182* | up | 1147 | 1155 | + | 11.3303 | 4.96E-05 | 0.615 | GTGGGTGGC |
| 11107 | mamo-s | *BGIBMGA013184* | down | 235 | 243 | - | 11.7339 | 3.29E-05 | 0.615 | GCGCGAGGC |
| 11108 | mamo-s | *BGIBMGA013185* | down | 1459 | 1467 | - | 11.1743 | 5.90E-05 | 0.684 | CCGCGTGTC |
| 11109 | mamo-s | *BGIBMGA013187* | down | 111 | 119 | - | 11.3303 | 4.96E-05 | 0.615 | GTGGGTGGC |
| 11110 | mamo-s | *BGIBMGA013187* | down | 1759 | 1767 | - | 11.3303 | 4.96E-05 | 0.615 | GTGGGTGGC |
| 11111 | mamo-s | *BGIBMGA013188* | up | 85 | 93 | - | 11.5321 | 4.06E-05 | 0.615 | TTGCGTGGC |
| 11112 | mamo-s | *BGIBMGA013189* | up | 1354 | 1362 | - | 11.3303 | 4.96E-05 | 0.615 | GTGGGTGGC |
| 11113 | mamo-s | *BGIBMGA013189* | down | 1389 | 1397 | - | 11.1009 | 6.57E-05 | 0.709 | GAGCGAGGC |
| 11114 | mamo-s | *BGIBMGA013191* | down | 272 | 280 | + | 11.3303 | 4.96E-05 | 0.615 | GTGGGTGGC |
| 11115 | mamo-s | *BGIBMGA013191* | down | 58 | 66 | + | 11.1743 | 5.90E-05 | 0.684 | CCGCGTGTC |
| 11116 | mamo-s | *BGIBMGA013192* | down | 512 | 520 | + | 14.055 | 2.23E-06 | 0.615 | GTGCGTGGC |
| 11117 | mamo-s | *BGIBMGA013193* | up | 914 | 922 | + | 14.055 | 2.23E-06 | 0.615 | GTGCGTGGC |
| 11118 | mamo-s | *BGIBMGA013194* | down | 868 | 876 | + | 11.3303 | 4.96E-05 | 0.615 | GTGGGTGGC |
| 11119 | mamo-s | *BGIBMGA013194* | down | 1306 | 1314 | + | 10.5413 | 9.42E-05 | 0.857 | GTGCGAGTC |
| 11120 | mamo-s | *BGIBMGA013195* | down | 50 | 58 | + | 12.4679 | 1.31E-05 | 0.615 | GTGCGTGTC |
| 11121 | mamo-s | *BGIBMGA013197* | up | 1683 | 1691 | - | 10.9358 | 7.30E-05 | 0.728 | GCGGGTGGC |
| 11122 | mamo-s | *BGIBMGA013198* | down | 1127 | 1135 | + | 11.5688 | 3.79E-05 | 0.615 | CTGCGTGTC |
| 11123 | mamo-s | *BGIBMGA013198* | down | 1783 | 1791 | + | 11.3303 | 4.96E-05 | 0.615 | GTGGGTGGC |
| 11124 | mamo-s | *BGIBMGA013199* | down | 647 | 655 | - | 11.1376 | 6.35E-05 | 0.709 | TCGCGTGGC |
| 11125 | mamo-s | *BGIBMGA013199* | down | 629 | 637 | + | 11.1009 | 6.57E-05 | 0.709 | GAGCGAGGC |
| 11126 | mamo-s | *BGIBMGA013200* | down | 1188 | 1196 | - | 11.3303 | 4.96E-05 | 0.615 | GTGGGTGGC |
| 11127 | mamo-s | *BGIBMGA013203* | up | 1214 | 1222 | - | 12.3394 | 1.49E-05 | 0.615 | GGGCGTGGC |
| 11128 | mamo-s | *BGIBMGA013203* | up | 1146 | 1154 | - | 12.0734 | 2.66E-05 | 0.615 | GCGCGTGTC |
| 11129 | mamo-s | *BGIBMGA013204* | down | 416 | 424 | - | 11.789 | 2.84E-05 | 0.615 | GCGCGTGCC |
| 11130 | mamo-s | *BGIBMGA013206* | up | 84 | 92 | - | 10.7523 | 8.38E-05 | 0.797 | GGGCGTGTC |
| 11131 | mamo-s | *BGIBMGA013210* | up | 332 | 340 | - | 11.3303 | 4.96E-05 | 0.615 | GTGGGTGGC |
| 11132 | mamo-s | *BGIBMGA013210* | down | 744 | 752 | + | 11.2844 | 5.18E-05 | 0.63 | CTGCGTGCC |
| 11133 | mamo-s | *BGIBMGA013211* | up | 29 | 37 | + | 11.7339 | 3.29E-05 | 0.615 | GCGCGAGGC |
| 11134 | mamo-s | *BGIBMGA013213* | down | 1726 | 1734 | + | 10.7523 | 8.38E-05 | 0.797 | GGGCGTGTC |
| 11135 | mamo-s | *BGIBMGA013214* | up | 773 | 781 | + | 12.1009 | 2.43E-05 | 0.615 | GTGCGTGGT |
| 11136 | mamo-s | *BGIBMGA013214* | up | 1967 | 1975 | - | 10.5688 | 8.88E-05 | 0.824 | GTGCGTGAC |
| 11137 | mamo-s | *BGIBMGA013215* | up | 1727 | 1735 | - | 11.3303 | 4.96E-05 | 0.615 | GTGGGTGGC |
| 11138 | mamo-s | *BGIBMGA013215* | up | 1188 | 1196 | - | 11.0826 | 6.84E-05 | 0.709 | GTACGTGGC |
| 11139 | mamo-s | *BGIBMGA013216* | down | 1890 | 1898 | + | 12.3394 | 1.49E-05 | 0.615 | GGGCGTGGC |
| 11140 | mamo-s | *BGIBMGA013216* | down | 621 | 629 | + | 11.789 | 2.84E-05 | 0.615 | GCGCGTGCC |
| 11141 | mamo-s | *BGIBMGA013217* | down | 447 | 455 | - | 11.3303 | 4.96E-05 | 0.615 | GTGGGTGGC |
| 11142 | mamo-s | *BGIBMGA013218* | up | 1616 | 1624 | + | 11.3303 | 4.96E-05 | 0.615 | GTGGGTGGC |
| 11143 | mamo-s | *BGIBMGA013218* | up | 578 | 586 | - | 10.7523 | 8.38E-05 | 0.797 | AAGCGTGGC |
| 11144 | mamo-s | *BGIBMGA013223* | up | 497 | 505 | + | 12.1835 | 1.71E-05 | 0.615 | GTGCGTGCC |
| 11145 | mamo-s | *BGIBMGA013223* | down | 1408 | 1416 | + | 11.3303 | 4.96E-05 | 0.615 | GTGGGTGGC |
| 11146 | mamo-s | *BGIBMGA013223* | down | 934 | 942 | + | 10.5688 | 8.88E-05 | 0.824 | GTGCGTGAC |
| 11147 | mamo-s | *BGIBMGA013224* | down | 1919 | 1927 | - | 12.1835 | 1.71E-05 | 0.615 | GTGCGTGCC |
| 11148 | mamo-s | *BGIBMGA013224* | down | 1075 | 1083 | - | 11.7339 | 3.29E-05 | 0.615 | GCGCGAGGC |
| 11149 | mamo-s | *BGIBMGA013224* | up | 967 | 975 | + | 11.3303 | 4.96E-05 | 0.615 | GTGGGTGGC |
| 11150 | mamo-s | *BGIBMGA013224* | up | 1730 | 1738 | - | 11.3303 | 4.96E-05 | 0.615 | GTGGGTGGC |
| 11151 | mamo-s | *BGIBMGA013224* | down | 234 | 242 | + | 11.2294 | 5.41E-05 | 0.65 | CTGCGAGGC |
| 11152 | mamo-s | *BGIBMGA013224* | down | 957 | 965 | + | 11.1009 | 6.57E-05 | 0.709 | GAGCGAGGC |
| 11153 | mamo-s | *BGIBMGA013224* | down | 694 | 702 | - | 11.0826 | 6.84E-05 | 0.709 | GTACGTGGC |
| 11154 | mamo-s | *BGIBMGA013224* | up | 493 | 501 | + | 10.5688 | 8.88E-05 | 0.824 | GTGCGTGAC |
| 11155 | mamo-s | *BGIBMGA013225* | up | 1454 | 1462 | - | 12.1835 | 1.71E-05 | 0.615 | GTGCGTGCC |
| 11156 | mamo-s | *BGIBMGA013225* | up | 610 | 618 | - | 11.7339 | 3.29E-05 | 0.615 | GCGCGAGGC |
| 11157 | mamo-s | *BGIBMGA013225* | up | 492 | 500 | + | 11.1009 | 6.57E-05 | 0.709 | GAGCGAGGC |
| 11158 | mamo-s | *BGIBMGA013225* | up | 229 | 237 | - | 11.0826 | 6.84E-05 | 0.709 | GTACGTGGC |
| 11159 | mamo-s | *BGIBMGA013225* | down | 552 | 560 | + | 10.5138 | 9.75E-05 | 0.865 | GTGCGTGTT |
| 11160 | mamo-s | *BGIBMGA013227* | down | 920 | 928 | + | 11.1009 | 6.57E-05 | 0.709 | GAGCGAGGC |
| 11161 | mamo-s | *BGIBMGA013229* | down | 1813 | 1821 | - | 11.789 | 2.84E-05 | 0.615 | GCGCGTGCC |
| 11162 | mamo-s | *BGIBMGA013229* | up | 776 | 784 | - | 11.3303 | 4.96E-05 | 0.615 | GTGGGTGGC |
| 11163 | mamo-s | *BGIBMGA013229* | up | 1216 | 1224 | + | 10.5138 | 9.75E-05 | 0.865 | GTGCGTGTT |
| 11164 | mamo-s | *BGIBMGA013230* | up | 816 | 824 | - | 11.789 | 2.84E-05 | 0.615 | GCGCGTGCC |
| 11165 | mamo-s | *BGIBMGA013233* | up | 1959 | 1967 | + | 12.4679 | 1.31E-05 | 0.615 | GTGCGTGTC |
| 11166 | mamo-s | *BGIBMGA013233* | up | 572 | 580 | + | 10.8899 | 7.48E-05 | 0.739 | CCGCGTGCC |
| 11167 | mamo-s | *BGIBMGA013234* | down | 192 | 200 | + | 11.1743 | 5.90E-05 | 0.684 | CCGCGTGTC |
| 11168 | mamo-s | *BGIBMGA013235* | up | 1259 | 1267 | - | 11.3303 | 4.96E-05 | 0.615 | GTGGGTGGC |
| 11169 | mamo-s | *BGIBMGA013237* | down | 159 | 167 | + | 11.3303 | 4.96E-05 | 0.615 | GTGGGTGGC |

| | | | | | | | | | | |
|---|---|---|---|---|---|---|---|---|---|---|
| 11170 | mamo-s | *BGIBMGA013238* | down | 450 | 458 | + | 11.3303 | 4.96E-05 | 0.615 | GTGGGTGGC |
| 11171 | mamo-s | *BGIBMGA013238* | up | 1783 | 1791 | - | 11.3303 | 4.96E-05 | 0.615 | GTGGGTGGC |
| 11172 | mamo-s | *BGIBMGA013239* | down | 1402 | 1410 | - | 11.5688 | 3.79E-05 | 0.615 | CTGCGTGTC |
| 11173 | mamo-s | *BGIBMGA013239* | up | 1344 | 1352 | - | 11.0734 | 7.12E-05 | 0.726 | GAGCGTGGT |
| 11174 | mamo-s | *BGIBMGA013239* | up | 1155 | 1163 | - | 10.7523 | 8.38E-05 | 0.797 | AAGCGTGGC |
| 11175 | mamo-s | *BGIBMGA013239* | down | 1298 | 1306 | - | 10.7523 | 8.38E-05 | 0.797 | AAGCGTGGC |
| 11176 | mamo-s | *BGIBMGA013240* | up | 592 | 600 | - | 11.3303 | 4.96E-05 | 0.615 | GTGGGTGGC |
| 11177 | mamo-s | *BGIBMGA013241* | up | 1516 | 1524 | + | 11.4404 | 4.51E-05 | 0.615 | CGGCGTGGC |
| 11178 | mamo-s | *BGIBMGA013242* | up | 255 | 263 | + | 11.3303 | 4.96E-05 | 0.615 | GTGGGTGGC |
| 11179 | mamo-s | *BGIBMGA013242* | up | 1066 | 1074 | - | 11.3303 | 4.96E-05 | 0.615 | GTGGGTGGC |
| 11180 | mamo-s | *BGIBMGA013243* | down | 682 | 690 | - | 11.3303 | 4.96E-05 | 0.615 | GTGGGTGGC |
| 11181 | mamo-s | *BGIBMGA013246* | down | 159 | 167 | - | 11.3303 | 4.96E-05 | 0.615 | GTGGGTGGC |
| 11182 | mamo-s | *BGIBMGA013246* | up | 213 | 221 | - | 11.3303 | 4.96E-05 | 0.615 | GTGGGTGGC |
| 11183 | mamo-s | *BGIBMGA013246* | up | 1117 | 1125 | - | 11.0826 | 6.84E-05 | 0.709 | GTACGTGGC |
| 11184 | mamo-s | *BGIBMGA013247* | down | 253 | 261 | + | 11.3303 | 4.96E-05 | 0.615 | GTGGGTGGC |
| 11185 | mamo-s | *BGIBMGA013248* | down | 176 | 184 | + | 11.0826 | 6.84E-05 | 0.709 | GTACGTGGC |
| 11186 | mamo-s | *BGIBMGA013249* | down | 350 | 358 | + | 10.5138 | 9.75E-05 | 0.865 | GTGCGTGTT |
| 11187 | mamo-s | *BGIBMGA013250* | down | 813 | 821 | - | 11.7339 | 3.29E-05 | 0.615 | GCGCGAGGC |
| 11188 | mamo-s | *BGIBMGA013251* | up | 463 | 471 | + | 11.156 | 6.13E-05 | 0.705 | GAGCGTGCC |
| 11189 | mamo-s | *BGIBMGA013253* | down | 1389 | 1397 | + | 11.3303 | 4.96E-05 | 0.615 | GTGGGTGGC |
| 11190 | mamo-s | *BGIBMGA013254* | up | 1542 | 1550 | + | 11.3303 | 4.96E-05 | 0.615 | GTGGGTGGC |
| 11191 | mamo-s | *BGIBMGA013254* | up | 812 | 820 | + | 10.7523 | 8.38E-05 | 0.797 | GGGCGTGTC |
| 11192 | mamo-s | *BGIBMGA013260* | down | 915 | 923 | - | 11.3303 | 4.96E-05 | 0.615 | GTGGGTGGC |
| 11193 | mamo-s | *BGIBMGA013261* | up | 497 | 505 | - | 11.3303 | 4.96E-05 | 0.615 | GTGGGTGGC |
| 11194 | mamo-s | *BGIBMGA013261* | down | 704 | 712 | + | 11.0826 | 6.84E-05 | 0.709 | GTACGTGGC |
| 11195 | mamo-s | *BGIBMGA013261* | up | 1460 | 1468 | + | 10.9358 | 7.30E-05 | 0.728 | GCGGGTGGC |
| 11196 | mamo-s | *BGIBMGA013264* | up | 1564 | 1572 | + | 11.0826 | 6.84E-05 | 0.709 | GTACGTGGC |
| 11197 | mamo-s | *BGIBMGA013264* | down | 1642 | 1650 | - | 11.0826 | 6.84E-05 | 0.709 | GTACGTGGC |
| 11198 | mamo-s | *BGIBMGA013265* | down | 318 | 326 | + | 11.3303 | 4.96E-05 | 0.615 | GTGGGTGGC |
| 11199 | mamo-s | *BGIBMGA013266* | up | 1684 | 1692 | - | 11.3303 | 4.96E-05 | 0.615 | GTGGGTGGC |
| 11200 | mamo-s | *BGIBMGA013268* | down | 731 | 739 | - | 11.3303 | 4.96E-05 | 0.615 | GTGGGTGGC |
| 11201 | mamo-s | *BGIBMGA013268* | up | 846 | 854 | - | 11.3303 | 4.96E-05 | 0.615 | GTGGGTGGC |
| 11202 | mamo-s | *BGIBMGA013268* | up | 1107 | 1115 | - | 10.8073 | 7.89E-05 | 0.766 | CCGCGTGGT |
| 11203 | mamo-s | *BGIBMGA013269* | up | 1737 | 1745 | - | 11.3303 | 4.96E-05 | 0.615 | GTGGGTGGC |
| 11204 | mamo-s | *BGIBMGA013269* | up | 408 | 416 | + | 10.5138 | 9.75E-05 | 0.865 | GTGCGTGTT |
| 11205 | mamo-s | *BGIBMGA013270* | down | 192 | 200 | + | 12.4679 | 1.31E-05 | 0.615 | GTGCGTGTC |
| 11206 | mamo-s | *BGIBMGA013270* | down | 1378 | 1386 | + | 10.9358 | 7.30E-05 | 0.728 | GCGGGTGGC |
| 11207 | mamo-s | *BGIBMGA013271* | up | 1851 | 1859 | - | 12.1009 | 2.43E-05 | 0.615 | GTGCGTGGT |
| 11208 | mamo-s | *BGIBMGA013271* | up | 348 | 356 | - | 11.3303 | 4.96E-05 | 0.615 | GTGGGTGGC |
| 11209 | mamo-s | *BGIBMGA013271* | up | 935 | 943 | + | 11.3303 | 4.96E-05 | 0.615 | GTGGGTGGC |
| 11210 | mamo-s | *BGIBMGA013272* | up | 1336 | 1344 | + | 11.3303 | 4.96E-05 | 0.615 | GTGGGTGGC |
| 11211 | mamo-s | *BGIBMGA013274* | down | 1290 | 1298 | - | 12.3394 | 1.49E-05 | 0.615 | GGGCGTGGC |
| 11212 | mamo-s | *BGIBMGA013274* | down | 1238 | 1246 | - | 11.3303 | 4.96E-05 | 0.615 | GTGGGTGGC |
| 11213 | mamo-s | *BGIBMGA013274* | down | 1633 | 1641 | + | 11.3303 | 4.96E-05 | 0.615 | GTGGGTGGC |
| 11214 | mamo-s | *BGIBMGA013276* | up | 1382 | 1390 | + | 11.3303 | 4.96E-05 | 0.615 | GTGGGTGGC |
| 11215 | mamo-s | *BGIBMGA013278* | up | 1883 | 1891 | - | 10.9358 | 7.30E-05 | 0.728 | GCGGGTGGC |
| 11216 | mamo-s | *BGIBMGA013282* | down | 1649 | 1657 | - | 11.3303 | 4.96E-05 | 0.615 | GTGGGTGGC |
| 11217 | mamo-s | *BGIBMGA013282* | down | 491 | 499 | + | 11.0826 | 6.84E-05 | 0.709 | GTACGTGGC |
| 11218 | mamo-s | *BGIBMGA013283* | down | 220 | 228 | - | 11.3303 | 4.96E-05 | 0.615 | GTGGGTGGC |
| 11219 | mamo-s | *BGIBMGA013283* | down | 423 | 431 | - | 11.3303 | 4.96E-05 | 0.615 | GTGGGTGGC |
| 11220 | mamo-s | *BGIBMGA013283* | up | 1598 | 1606 | + | 11.3303 | 4.96E-05 | 0.615 | GTGGGTGGC |
| 11221 | mamo-s | *BGIBMGA013285* | down | 1811 | 1819 | - | 11.3303 | 4.96E-05 | 0.615 | GTGGGTGGC |
| 11222 | mamo-s | *BGIBMGA013286* | up | 1291 | 1299 | - | 11.5321 | 4.06E-05 | 0.615 | TTGCGTGGC |
| 11223 | mamo-s | *BGIBMGA013286* | up | 1096 | 1104 | + | 11.0826 | 6.84E-05 | 0.709 | GTACGTGGC |
| 11224 | mamo-s | *BGIBMGA013290* | up | 1089 | 1097 | - | 11.3303 | 4.96E-05 | 0.615 | GTGGGTGGC |
| 11225 | mamo-s | *BGIBMGA013290* | down | 1115 | 1123 | - | 11.3303 | 4.96E-05 | 0.615 | GTGGGTGGC |
| 11226 | mamo-s | *BGIBMGA013292* | down | 1895 | 1903 | + | 12.4679 | 1.31E-05 | 0.615 | GTGCGTGTC |
| 11227 | mamo-s | *BGIBMGA013292* | up | 1005 | 1013 | - | 11.3303 | 4.96E-05 | 0.615 | GTGGGTGGC |
| 11228 | mamo-s | *BGIBMGA013293* | down | 128 | 136 | - | 11.3303 | 4.96E-05 | 0.615 | GTGGGTGGC |
| 11229 | mamo-s | *BGIBMGA013293* | up | 638 | 646 | + | 11.0826 | 6.84E-05 | 0.709 | GTACGTGGC |
| 11230 | mamo-s | *BGIBMGA013298* | up | 817 | 825 | - | 11.3303 | 4.96E-05 | 0.615 | GTGGGTGGC |
| 11231 | mamo-s | *BGIBMGA013299* | up | 1838 | 1846 | - | 11.7064 | 3.52E-05 | 0.615 | GCGCGTGGT |
| 11232 | mamo-s | *BGIBMGA013299* | down | 1558 | 1566 | + | 11.2844 | 5.18E-05 | 0.63 | CTGCGTGCC |
| 11233 | mamo-s | *BGIBMGA013301* | up | 1495 | 1503 | + | 11.3303 | 4.96E-05 | 0.615 | GTGGGTGGC |
| 11234 | mamo-s | *BGIBMGA013302* | down | 1459 | 1467 | - | 11.3303 | 4.96E-05 | 0.615 | GTGGGTGGC |
| 11235 | mamo-s | *BGIBMGA013303* | down | 386 | 394 | - | 11.3303 | 4.96E-05 | 0.615 | GTGGGTGGC |
| 11236 | mamo-s | *BGIBMGA013303* | up | 1062 | 1070 | + | 11.3303 | 4.96E-05 | 0.615 | GTGGGTGGC |
| 11237 | mamo-s | *BGIBMGA013303* | up | 1596 | 1604 | - | 11.3303 | 4.96E-05 | 0.615 | GTGGGTGGC |
| 11238 | mamo-s | *BGIBMGA013304* | up | 1915 | 1923 | + | 11.3303 | 4.96E-05 | 0.615 | GTGGGTGGC |
| 11239 | mamo-s | *BGIBMGA013305* | down | 507 | 515 | + | 11.3303 | 4.96E-05 | 0.615 | GTGGGTGGC |
| 11240 | mamo-s | *BGIBMGA013305* | down | 1382 | 1390 | + | 11.0826 | 6.84E-05 | 0.709 | GTACGTGGC |
| 11241 | mamo-s | *BGIBMGA013306* | up | 94 | 102 | + | 11.3303 | 4.96E-05 | 0.615 | GTGGGTGGC |
| 11242 | mamo-s | *BGIBMGA013308* | down | 1399 | 1407 | + | 11.3303 | 4.96E-05 | 0.615 | GTGGGTGGC |
| 11243 | mamo-s | *BGIBMGA013309* | up | 1337 | 1345 | + | 11.3303 | 4.96E-05 | 0.615 | GTGGGTGGC |
| 11244 | mamo-s | *BGIBMGA013310* | down | 1402 | 1410 | + | 10.8899 | 7.48E-05 | 0.739 | CCGCGTGCC |
| 11245 | mamo-s | *BGIBMGA013313* | up | 1584 | 1592 | - | 11.3303 | 4.96E-05 | 0.615 | GTGGGTGGC |
| 11246 | mamo-s | *BGIBMGA013314* | up | 1393 | 1401 | - | 11.3303 | 4.96E-05 | 0.615 | GTGGGTGGC |
| 11247 | mamo-s | *BGIBMGA013315* | up | 482 | 490 | - | 11.3303 | 4.96E-05 | 0.615 | GTGGGTGGC |
| 11248 | mamo-s | *BGIBMGA013315* | up | 1578 | 1586 | - | 11.3303 | 4.96E-05 | 0.615 | GTGGGTGGC |
| 11249 | mamo-s | *BGIBMGA013315* | down | 1464 | 1472 | - | 11.0826 | 6.84E-05 | 0.709 | GTACGTGGC |
| 11250 | mamo-s | *BGIBMGA013325* | down | 1008 | 1016 | + | 11.7798 | 3.11E-05 | 0.615 | ATGCGTGGC |
| 11251 | mamo-s | *BGIBMGA013325* | down | 1029 | 1037 | + | 11.7064 | 3.52E-05 | 0.615 | GCGCGTGGT |
| 11252 | mamo-s | *BGIBMGA013326* | down | 103 | 111 | - | 14.055 | 2.23E-06 | 0.615 | GTGCGTGGC |
| 11253 | mamo-s | *BGIBMGA013329* | up | 1912 | 1920 | + | 11.5321 | 4.06E-05 | 0.615 | TTGCGTGGC |
| 11254 | mamo-s | *BGIBMGA013333* | down | 1947 | 1955 | - | 11.1743 | 5.90E-05 | 0.684 | CCGCGTGTC |
| 11255 | mamo-s | *BGIBMGA013334* | up | 472 | 480 | - | 10.7523 | 8.38E-05 | 0.797 | AAGCGTGGC |
| 11256 | mamo-s | *BGIBMGA013335* | up | 269 | 277 | - | 11.0826 | 6.84E-05 | 0.709 | GTACGTGGC |
| 11257 | mamo-s | *BGIBMGA013335* | up | 1645 | 1653 | - | 11.0734 | 7.12E-05 | 0.726 | GAGCGTGGT |

| | | | | | | | | | | |
|---|---|---|---|---|---|---|---|---|---|---|
| 11258 | mamo-s | *BGIBMGA013336* | down | 1657 | 1665 | - | 11.3303 | 4.96E-05 | 0.615 | GTGGGTGGC |
| 11259 | mamo-s | *BGIBMGA013336* | down | 1906 | 1914 | - | 11.3303 | 4.96E-05 | 0.615 | GTGGGTGGC |
| 11260 | mamo-s | *BGIBMGA013338* | up | 180 | 188 | - | 11.3303 | 4.96E-05 | 0.615 | GTGGGTGGC |
| 11261 | mamo-s | *BGIBMGA013341* | down | 465 | 473 | - | 11.3303 | 4.96E-05 | 0.615 | GTGGGTGGC |
| 11262 | mamo-s | *BGIBMGA013342* | up | 1791 | 1799 | - | 11.3303 | 4.96E-05 | 0.615 | GTGGGTGGC |
| 11263 | mamo-s | *BGIBMGA013343* | down | 1764 | 1772 | - | 11.3303 | 4.96E-05 | 0.615 | GTGGGTGGC |
| 11264 | mamo-s | *BGIBMGA013344* | up | 1344 | 1352 | - | 11.4404 | 4.51E-05 | 0.615 | CGGCGTGGC |
| 11265 | mamo-s | *BGIBMGA013346* | down | 940 | 948 | + | 12.0734 | 2.66E-05 | 0.615 | GCGCGTGTC |
| 11266 | mamo-s | *BGIBMGA013347* | down | 542 | 550 | - | 12.7615 | 1.04E-05 | 0.615 | CCGCGTGGC |
| 11267 | mamo-s | *BGIBMGA013347* | down | 188 | 196 | - | 11.3303 | 4.96E-05 | 0.615 | GTGGGTGGC |
| 11268 | mamo-s | *BGIBMGA013352* | up | 448 | 456 | - | 11.0826 | 6.84E-05 | 0.709 | GTACGTGGC |
| 11269 | mamo-s | *BGIBMGA013353* | down | 1126 | 1134 | - | 11.3303 | 4.96E-05 | 0.615 | GTGGGTGGC |
| 11270 | mamo-s | *BGIBMGA013353* | down | 1891 | 1899 | - | 11.3303 | 4.96E-05 | 0.615 | GTGGGTGGC |
| 11271 | mamo-s | *BGIBMGA013353* | down | 1702 | 1710 | + | 10.9358 | 7.30E-05 | 0.728 | GCGGGTGGC |
| 11272 | mamo-s | *BGIBMGA013353* | up | 1992 | 2000 | + | 10.8899 | 7.48E-05 | 0.739 | CCGCGTGCC |
| 11273 | mamo-s | *BGIBMGA013354* | down | 1950 | 1958 | - | 11.2018 | 5.68E-05 | 0.674 | CTGCGTGGT |
| 11274 | mamo-s | *BGIBMGA013356* | up | 1883 | 1891 | + | 11.4404 | 4.51E-05 | 0.615 | GAGCGTGTC |
| 11275 | mamo-s | *BGIBMGA013356* | down | 805 | 813 | + | 11.1009 | 6.57E-05 | 0.709 | GAGCGAGGC |
| 11276 | mamo-s | *BGIBMGA013358* | up | 1716 | 1724 | - | 12.1009 | 2.43E-05 | 0.615 | GTGCGTGGT |
| 11277 | mamo-s | *BGIBMGA013358* | down | 1785 | 1793 | - | 11.1743 | 5.90E-05 | 0.684 | CCGCGTGTC |
| 11278 | mamo-s | *BGIBMGA013359* | up | 1219 | 1227 | - | 12.1284 | 2.16E-05 | 0.615 | GTGCGAGGC |
| 11279 | mamo-s | *BGIBMGA013359* | up | 287 | 295 | - | 11.3303 | 4.96E-05 | 0.615 | GTGGGTGGC |
| 11280 | mamo-s | *BGIBMGA013360* | down | 999 | 1007 | - | 11.1743 | 5.90E-05 | 0.684 | CCGCGTGTC |
| 11281 | mamo-s | *BGIBMGA013362* | down | 4 | 12 | + | 10.8899 | 7.48E-05 | 0.739 | CCGCGTGCC |
| 11282 | mamo-s | *BGIBMGA013362* | up | 1600 | 1608 | - | 10.8899 | 7.48E-05 | 0.739 | CCGCGTGCC |
| 11283 | mamo-s | *BGIBMGA013365* | down | 26 | 34 | + | 11.7339 | 3.29E-05 | 0.615 | GCGCGAGGC |
| 11284 | mamo-s | *BGIBMGA013365* | down | 1741 | 1749 | + | 10.5413 | 9.42E-05 | 0.857 | GTGCGAGTC |
| 11285 | mamo-s | *BGIBMGA013368* | down | 385 | 393 | - | 11.3303 | 4.96E-05 | 0.615 | GTGGGTGGC |
| 11286 | mamo-s | *BGIBMGA013368* | down | 1669 | 1677 | + | 11.3303 | 4.96E-05 | 0.615 | GTGGGTGGC |
| 11287 | mamo-s | *BGIBMGA013370* | up | 332 | 340 | - | 11.3303 | 4.96E-05 | 0.615 | GTGGGTGGC |
| 11288 | mamo-s | *BGIBMGA013370* | up | 1023 | 1031 | - | 11.3303 | 4.96E-05 | 0.615 | GTGGGTGGC |
| 11289 | mamo-s | *BGIBMGA013373* | down | 574 | 582 | + | 11.3303 | 4.96E-05 | 0.615 | GTGGGTGGC |
| 11290 | mamo-s | *BGIBMGA013373* | up | 1669 | 1677 | - | 11.0826 | 6.84E-05 | 0.709 | GTACGTGGC |
| 11291 | mamo-s | *BGIBMGA013374* | down | 356 | 364 | + | 11.3303 | 4.96E-05 | 0.615 | GTGGGTGGC |
| 11292 | mamo-s | *BGIBMGA013375* | up | 1808 | 1816 | + | 12.7615 | 1.04E-05 | 0.615 | CCGCGTGGC |
| 11293 | mamo-s | *BGIBMGA013376* | down | 139 | 147 | + | 11.3303 | 4.96E-05 | 0.615 | GTGGGTGGC |
| 11294 | mamo-s | *BGIBMGA013376* | down | 585 | 593 | - | 11.3303 | 4.96E-05 | 0.615 | GTGGGTGGC |
| 11295 | mamo-s | *BGIBMGA013377* | up | 7 | 15 | - | 11.3303 | 4.96E-05 | 0.615 | GTGGGTGGC |
| 11296 | mamo-s | *BGIBMGA013379* | down | 1749 | 1757 | + | 11.3303 | 4.96E-05 | 0.615 | GTGGGTGGC |
| 11297 | mamo-s | *BGIBMGA013381* | up | 601 | 609 | + | 11.3303 | 4.96E-05 | 0.615 | GTGGGTGGC |
| 11298 | mamo-s | *BGIBMGA013383* | down | 803 | 811 | + | 11.3303 | 4.96E-05 | 0.615 | GTGGGTGGC |
| 11299 | mamo-s | *BGIBMGA013384* | up | 872 | 880 | + | 14.055 | 2.23E-06 | 0.615 | GTGCGTGGC |
| 11300 | mamo-s | *BGIBMGA013384* | up | 1551 | 1559 | + | 11.3303 | 4.96E-05 | 0.615 | GTGGGTGGC |
| 11301 | mamo-s | *BGIBMGA013385* | up | 318 | 326 | + | 10.5138 | 9.75E-05 | 0.865 | GTGCGTGTT |
| 11302 | mamo-s | *BGIBMGA013387* | up | 91 | 99 | - | 11.0826 | 6.84E-05 | 0.709 | GTACGTGGC |
| 11303 | mamo-s | *BGIBMGA013391* | down | 332 | 340 | - | 11.0826 | 6.84E-05 | 0.709 | GTACGTGGC |
| 11304 | mamo-s | *BGIBMGA013393* | up | 444 | 452 | - | 11.3303 | 4.96E-05 | 0.615 | GTGGGTGGC |
| 11305 | mamo-s | *BGIBMGA013393* | down | 1411 | 1419 | - | 11.3303 | 4.96E-05 | 0.615 | GTGGGTGGC |
| 11306 | mamo-s | *BGIBMGA013395* | down | 1947 | 1955 | + | 11.789 | 2.84E-05 | 0.615 | GCGCGTGCC |
| 11307 | mamo-s | *BGIBMGA013395* | up | 729 | 737 | + | 11.3303 | 4.96E-05 | 0.615 | GTGGGTGGC |
| 11308 | mamo-s | *BGIBMGA013397* | down | 1308 | 1316 | + | 11.3303 | 4.96E-05 | 0.615 | GTGGGTGGC |
| 11309 | mamo-s | *BGIBMGA013399* | down | 323 | 331 | - | 11.3303 | 4.96E-05 | 0.615 | GTGGGTGGC |
| 11310 | mamo-s | *BGIBMGA013403* | down | 1047 | 1055 | + | 11.3303 | 4.96E-05 | 0.615 | GTGGGTGGC |
| 11311 | mamo-s | *BGIBMGA013404* | up | 1871 | 1879 | + | 11.3303 | 4.96E-05 | 0.615 | GTGGGTGGC |
| 11312 | mamo-s | *BGIBMGA013405* | down | 766 | 774 | + | 11.3303 | 4.96E-05 | 0.615 | GTGGGTGGC |
| 11313 | mamo-s | *BGIBMGA013405* | up | 742 | 750 | + | 11.0826 | 6.84E-05 | 0.709 | GTACGTGGC |
| 11314 | mamo-s | *BGIBMGA013407* | up | 400 | 408 | - | 12.1835 | 1.71E-05 | 0.615 | GTGCGTGCC |
| 11315 | mamo-s | *BGIBMGA013407* | down | 133 | 141 | - | 11.3303 | 4.96E-05 | 0.615 | GTGGGTGGC |
| 11316 | mamo-s | *BGIBMGA013407* | down | 1471 | 1479 | + | 11.3303 | 4.96E-05 | 0.615 | GTGGGTGGC |
| 11317 | mamo-s | *BGIBMGA013407* | up | 1080 | 1088 | - | 11.0826 | 6.84E-05 | 0.709 | GTACGTGGC |
| 11318 | mamo-s | *BGIBMGA013408* | up | 1662 | 1670 | + | 12.1284 | 2.16E-05 | 0.615 | CAGCGTGGC |
| 11319 | mamo-s | *BGIBMGA013408* | up | 1346 | 1354 | - | 11.3303 | 4.96E-05 | 0.615 | GTGGGTGGC |
| 11320 | mamo-s | *BGIBMGA013409* | up | 73 | 81 | + | 12.1284 | 2.16E-05 | 0.615 | CAGCGTGGC |
| 11321 | mamo-s | *BGIBMGA013409* | down | 1806 | 1814 | + | 11.3303 | 4.96E-05 | 0.615 | GTGGGTGGC |
| 11322 | mamo-s | *BGIBMGA013412* | up | 64 | 72 | - | 11.3303 | 4.96E-05 | 0.615 | GTGGGTGGC |
| 11323 | mamo-s | *BGIBMGA013412* | up | 478 | 486 | - | 10.5688 | 8.88E-05 | 0.824 | GTGCGTGAC |
| 11324 | mamo-s | *BGIBMGA013412* | down | 66 | 74 | + | 10.5413 | 9.42E-05 | 0.857 | CAGCGTGTC |
| 11325 | mamo-s | *BGIBMGA013413* | down | 658 | 666 | - | 11.3303 | 4.96E-05 | 0.615 | GTGGGTGGC |
| 11326 | mamo-s | *BGIBMGA013417* | down | 929 | 937 | - | 11.3303 | 4.96E-05 | 0.615 | GTGGGTGGC |
| 11327 | mamo-s | *BGIBMGA013418* | up | 1230 | 1238 | + | 11.0826 | 6.84E-05 | 0.709 | GTACGTGGC |
| 11328 | mamo-s | *BGIBMGA013419* | up | 1594 | 1602 | + | 11.3303 | 4.96E-05 | 0.615 | GTGGGTGGC |
| 11329 | mamo-s | *BGIBMGA013421* | down | 1286 | 1294 | - | 11.3303 | 4.96E-05 | 0.615 | GTGGGTGGC |
| 11330 | mamo-s | *BGIBMGA013423* | down | 1550 | 1558 | - | 11.3303 | 4.96E-05 | 0.615 | GTGGGTGGC |
| 11331 | mamo-s | *BGIBMGA013424* | down | 163 | 171 | - | 11.3303 | 4.96E-05 | 0.615 | GTGGGTGGC |
| 11332 | mamo-s | *BGIBMGA013425* | up | 1126 | 1134 | - | 12.7615 | 1.04E-05 | 0.615 | CCGCGTGGC |
| 11333 | mamo-s | *BGIBMGA013426* | up | 927 | 935 | - | 11.1376 | 6.35E-05 | 0.709 | TCGCGTGGC |
| 11334 | mamo-s | *BGIBMGA013426* | down | 1858 | 1866 | - | 10.7523 | 8.38E-05 | 0.797 | AAGCGTGGC |
| 11335 | mamo-s | *BGIBMGA013427* | up | 506 | 514 | - | 11.3303 | 4.96E-05 | 0.615 | GTGGGTGGC |
| 11336 | mamo-s | *BGIBMGA013428* | down | 419 | 427 | + | 11.3303 | 4.96E-05 | 0.615 | GTGGGTGGC |
| 11337 | mamo-s | *BGIBMGA013428* | up | 132 | 140 | - | 11.1009 | 6.57E-05 | 0.709 | GAGCGAGGC |
| 11338 | mamo-s | *BGIBMGA013429* | up | 1514 | 1522 | - | 13.6606 | 4.06E-06 | 0.615 | GCGCGTGGC |
| 11339 | mamo-s | *BGIBMGA013429* | up | 1521 | 1529 | - | 13.6606 | 4.06E-06 | 0.615 | GCGCGTGGC |
| 11340 | mamo-s | *BGIBMGA013429* | up | 1528 | 1536 | - | 13.0275 | 8.53E-06 | 0.615 | GAGCGTGGC |
| 11341 | mamo-s | *BGIBMGA013432* | up | 307 | 315 | - | 11.3303 | 4.96E-05 | 0.615 | GTGGGTGGC |
| 11342 | mamo-s | *BGIBMGA013432* | up | 1332 | 1340 | - | 11.3303 | 4.96E-05 | 0.615 | GTGGGTGGC |
| 11343 | mamo-s | *BGIBMGA013432* | up | 1710 | 1718 | + | 11.3303 | 4.96E-05 | 0.615 | GTGGGTGGC |
| 11344 | mamo-s | *BGIBMGA013432* | up | 1207 | 1215 | - | 10.5413 | 9.42E-05 | 0.857 | GTGCGAGTC |
| 11345 | mamo-s | *BGIBMGA013433* | up | 1526 | 1534 | + | 11.3303 | 4.96E-05 | 0.615 | GTGGGTGGC |

| | | | | | | | | | | |
|---|---|---|---|---|---|---|---|---|---|---|
| 11346 | mamo-s | *BGIBMGA013436* | down | 127 | 135 | - | 11.3303 | 4.96E-05 | 0.615 | GTGGGTGGC |
| 11347 | mamo-s | *BGIBMGA013436* | up | 147 | 155 | - | 11.3303 | 4.96E-05 | 0.615 | GTGGGTGGC |
| 11348 | mamo-s | *BGIBMGA013437* | down | 573 | 581 | - | 11.3303 | 4.96E-05 | 0.615 | GTGGGTGGC |
| 11349 | mamo-s | *BGIBMGA013438* | down | 883 | 891 | + | 11.3303 | 4.96E-05 | 0.615 | GTGGGTGGC |
| 11350 | mamo-s | *BGIBMGA013439* | up | 1039 | 1047 | + | 12.1284 | 2.16E-05 | 0.615 | CAGCGTGGC |
| 11351 | mamo-s | *BGIBMGA013441* | up | 415 | 423 | + | 11.2844 | 5.18E-05 | 0.63 | CTGCGTGCC |
| 11352 | mamo-s | *BGIBMGA013442* | down | 735 | 743 | + | 13.156 | 6.30E-06 | 0.615 | CTGCGTGGC |
| 11353 | mamo-s | *BGIBMGA013442* | down | 386 | 394 | + | 11.4404 | 4.51E-05 | 0.615 | GAGCGTGTC |
| 11354 | mamo-s | *BGIBMGA013442* | down | 893 | 901 | - | 11.3303 | 4.96E-05 | 0.615 | GTGGGTGGC |
| 11355 | mamo-s | *BGIBMGA013442* | down | 1001 | 1009 | - | 11.3303 | 4.96E-05 | 0.615 | GTGGGTGGC |
| 11356 | mamo-s | *BGIBMGA013442* | down | 706 | 714 | - | 10.5413 | 9.42E-05 | 0.857 | CAGCGTGTC |
| 11357 | mamo-s | *BGIBMGA013443* | up | 103 | 111 | + | 12.0734 | 2.66E-05 | 0.615 | GCGCGTGTC |
| 11358 | mamo-s | *BGIBMGA013443* | up | 1979 | 1987 | + | 11.4404 | 4.51E-05 | 0.615 | GAGCGTGTC |
| 11359 | mamo-s | *BGIBMGA013443* | down | 739 | 747 | - | 11.3303 | 4.96E-05 | 0.615 | GTGGGTGGC |
| 11360 | mamo-s | *BGIBMGA013444* | down | 770 | 778 | - | 11.3303 | 4.96E-05 | 0.615 | GTGGGTGGC |
| 11361 | mamo-s | *BGIBMGA013444* | down | 1665 | 1673 | + | 11.3303 | 4.96E-05 | 0.615 | GTGGGTGGC |
| 11362 | mamo-s | *BGIBMGA013445* | down | 815 | 823 | - | 11.3303 | 4.96E-05 | 0.615 | GTGGGTGGC |
| 11363 | mamo-s | *BGIBMGA013445* | up | 1598 | 1606 | - | 10.5413 | 9.42E-05 | 0.857 | CAGCGTGTC |
| 11364 | mamo-s | *BGIBMGA013445* | up | 591 | 599 | - | 10.5138 | 9.75E-05 | 0.865 | GTGCGTGTT |
| 11365 | mamo-s | *BGIBMGA013446* | up | 1246 | 1254 | + | 11.3303 | 4.96E-05 | 0.615 | GTGGGTGGC |
| 11366 | mamo-s | *BGIBMGA013449* | up | 1421 | 1429 | + | 11.3303 | 4.96E-05 | 0.615 | GTGGGTGGC |
| 11367 | mamo-s | *BGIBMGA013450* | down | 816 | 824 | + | 11.0826 | 6.84E-05 | 0.709 | GTACGTGGC |
| 11368 | mamo-s | *BGIBMGA013451* | up | 417 | 425 | + | 11.2844 | 5.18E-05 | 0.63 | CTGCGTGCC |
| 11369 | mamo-s | *BGIBMGA013453* | up | 1849 | 1857 | + | 11.5321 | 4.06E-05 | 0.615 | TTGCGTGGC |
| 11370 | mamo-s | *BGIBMGA013454* | up | 1427 | 1435 | + | 11.3303 | 4.96E-05 | 0.615 | GTGGGTGGC |
| 11371 | mamo-s | *BGIBMGA013455* | down | 1541 | 1549 | + | 11.5321 | 4.06E-05 | 0.615 | TTGCGTGGC |
| 11372 | mamo-s | *BGIBMGA013455* | down | 531 | 539 | + | 11.3303 | 4.96E-05 | 0.615 | GTGGGTGGC |
| 11373 | mamo-s | *BGIBMGA013456* | up | 1756 | 1764 | + | 11.5321 | 4.06E-05 | 0.615 | TTGCGTGGC |
| 11374 | mamo-s | *BGIBMGA013456* | down | 582 | 590 | + | 11.3303 | 4.96E-05 | 0.615 | GTGGGTGGC |
| 11375 | mamo-s | *BGIBMGA013456* | up | 746 | 754 | + | 11.3303 | 4.96E-05 | 0.615 | GTGGGTGGC |
| 11376 | mamo-s | *BGIBMGA013457* | down | 375 | 383 | - | 13.0275 | 8.53E-06 | 0.615 | GAGCGTGGC |
| 11377 | mamo-s | *BGIBMGA013457* | down | 1901 | 1909 | - | 11.3303 | 4.96E-05 | 0.615 | GTGGGTGGC |
| 11378 | mamo-s | *BGIBMGA013459* | up | 197 | 205 | - | 11.3303 | 4.96E-05 | 0.615 | GTGGGTGGC |
| 11379 | mamo-s | *BGIBMGA013459* | down | 981 | 989 | - | 11.3303 | 4.96E-05 | 0.615 | GTGGGTGGC |
| 11380 | mamo-s | *BGIBMGA013460* | up | 900 | 908 | + | 12.1835 | 1.71E-05 | 0.615 | GTGCGTGCC |
| 11381 | mamo-s | *BGIBMGA013464* | up | 729 | 737 | - | 11.7798 | 3.11E-05 | 0.615 | ATGCGTGGC |
| 11382 | mamo-s | *BGIBMGA013465* | up | 378 | 386 | - | 10.7523 | 8.38E-05 | 0.797 | AAGCGTGGC |
| 11383 | mamo-s | *BGIBMGA013467* | up | 435 | 443 | - | 11.3303 | 4.96E-05 | 0.615 | GTGGGTGGC |
| 11384 | mamo-s | *BGIBMGA013467* | up | 1771 | 1779 | + | 10.7523 | 8.38E-05 | 0.797 | AAGCGTGGC |
| 11385 | mamo-s | *BGIBMGA013469* | up | 1179 | 1187 | + | 11.789 | 2.84E-05 | 0.615 | GCGCGTGCC |
| 11386 | mamo-s | *BGIBMGA013469* | down | 1587 | 1595 | + | 11.3303 | 4.96E-05 | 0.615 | GTGGGTGGC |
| 11387 | mamo-s | *BGIBMGA013469* | down | 548 | 556 | + | 10.9358 | 7.30E-05 | 0.728 | GCGGGTGGC |
| 11388 | mamo-s | *BGIBMGA013470* | up | 1642 | 1650 | - | 14.055 | 2.23E-06 | 0.615 | GTGCGTGGC |
| 11389 | mamo-s | *BGIBMGA013470* | up | 810 | 818 | + | 11.3303 | 4.96E-05 | 0.615 | GTGGGTGGC |
| 11390 | mamo-s | *BGIBMGA013475* | down | 167 | 175 | + | 13.6606 | 4.06E-06 | 0.615 | GCGCGTGGC |
| 11391 | mamo-s | *BGIBMGA013475* | up | 372 | 380 | + | 13.0275 | 8.53E-06 | 0.615 | GAGCGTGGC |
| 11392 | mamo-s | *BGIBMGA013475* | down | 157 | 165 | + | 12.1284 | 2.16E-05 | 0.615 | CAGCGTGGC |
| 11393 | mamo-s | *BGIBMGA013476* | up | 1331 | 1339 | + | 12.1009 | 2.43E-05 | 0.615 | GTGCGTGGT |
| 11394 | mamo-s | *BGIBMGA013476* | down | 10 | 18 | + | 10.5138 | 9.75E-05 | 0.865 | GTGCGTGTT |
| 11395 | mamo-s | *BGIBMGA013477* | up | 509 | 517 | - | 11.3303 | 4.96E-05 | 0.615 | GTGGGTGGC |
| 11396 | mamo-s | *BGIBMGA013477* | down | 1764 | 1772 | + | 11.3303 | 4.96E-05 | 0.615 | GTGGGTGGC |
| 11397 | mamo-s | *BGIBMGA013478* | down | 1040 | 1048 | - | 10.5688 | 8.88E-05 | 0.824 | GTGCGTGAC |
| 11398 | mamo-s | *BGIBMGA013480* | up | 202 | 210 | + | 11.3303 | 4.96E-05 | 0.615 | GTGGGTGGC |
| 11399 | mamo-s | *BGIBMGA013482* | down | 1517 | 1525 | + | 12.1009 | 2.43E-05 | 0.615 | GTGCGTGGT |
| 11400 | mamo-s | *BGIBMGA013482* | down | 315 | 323 | + | 11.0734 | 7.12E-05 | 0.726 | GAGCGTGGT |
| 11401 | mamo-s | *BGIBMGA013483* | up | 882 | 890 | + | 12.1009 | 2.43E-05 | 0.615 | GTGCGTGGT |
| 11402 | mamo-s | *BGIBMGA013483* | up | 96 | 104 | - | 11.4404 | 4.51E-05 | 0.615 | GAGCGTGTC |
| 11403 | mamo-s | *BGIBMGA013483* | up | 1649 | 1657 | + | 11.0734 | 7.12E-05 | 0.726 | GAGCGTGGT |
| 11404 | mamo-s | *BGIBMGA013484* | up | 1923 | 1931 | + | 11.789 | 2.84E-05 | 0.615 | GCGCGTGCC |
| 11405 | mamo-s | *BGIBMGA013485* | up | 1822 | 1830 | - | 11.3303 | 4.96E-05 | 0.615 | GTGGGTGGC |
| 11406 | mamo-s | *BGIBMGA013486* | up | 1076 | 1084 | + | 12.3394 | 1.49E-05 | 0.615 | GGGCGTGGC |
| 11407 | mamo-s | *BGIBMGA013488* | down | 1502 | 1510 | + | 11.3303 | 4.96E-05 | 0.615 | GTGGGTGGC |
| 11408 | mamo-s | *BGIBMGA013490* | down | 1613 | 1621 | + | 11.3303 | 4.96E-05 | 0.615 | GTGGGTGGC |
| 11409 | mamo-s | *BGIBMGA013492* | up | 1303 | 1311 | - | 11.0826 | 6.84E-05 | 0.709 | GTACGTGGC |
| 11410 | mamo-s | *BGIBMGA013492* | down | 643 | 651 | - | 10.5688 | 8.88E-05 | 0.824 | GTGCGTGAC |
| 11411 | mamo-s | *BGIBMGA013493* | down | 210 | 218 | - | 11.5688 | 3.79E-05 | 0.615 | CTGCGTGTC |
| 11412 | mamo-s | *BGIBMGA013495* | up | 985 | 993 | - | 11.3303 | 4.96E-05 | 0.615 | GTGGGTGGC |
| 11413 | mamo-s | *BGIBMGA013497* | down | 1402 | 1410 | + | 11.4404 | 4.51E-05 | 0.615 | CGGCGTGGC |
| 11414 | mamo-s | *BGIBMGA013498* | down | 1147 | 1155 | - | 11.3303 | 4.96E-05 | 0.615 | GTGGGTGGC |
| 11415 | mamo-s | *BGIBMGA013498* | down | 1587 | 1595 | - | 11.3303 | 4.96E-05 | 0.615 | GTGGGTGGC |
| 11416 | mamo-s | *BGIBMGA013499* | down | 1283 | 1291 | + | 11.3303 | 4.96E-05 | 0.615 | GTGGGTGGC |
| 11417 | mamo-s | *BGIBMGA013499* | down | 1504 | 1512 | + | 10.6881 | 8.61E-05 | 0.81 | GCACGTGGC |
| 11418 | mamo-s | *BGIBMGA013500* | up | 1617 | 1625 | + | 10.5138 | 9.75E-05 | 0.865 | GTGCGTGTT |
| 11419 | mamo-s | *BGIBMGA013502* | down | 14 | 22 | - | 10.5688 | 8.88E-05 | 0.824 | GTGCGTGAC |
| 11420 | mamo-s | *BGIBMGA013503* | down | 1478 | 1486 | - | 11.3303 | 4.96E-05 | 0.615 | GTGGGTGGC |
| 11421 | mamo-s | *BGIBMGA013504* | down | 792 | 800 | + | 11.3303 | 4.96E-05 | 0.615 | GTGGGTGGC |
| 11422 | mamo-s | *BGIBMGA013504* | down | 1270 | 1278 | + | 11.0826 | 6.84E-05 | 0.709 | GTACGTGGC |
| 11423 | mamo-s | *BGIBMGA013505* | up | 518 | 526 | - | 13.156 | 6.30E-06 | 0.615 | CTGCGTGGC |
| 11424 | mamo-s | *BGIBMGA013505* | down | 1403 | 1411 | + | 11.7064 | 3.52E-05 | 0.615 | GCGCGTGGT |
| 11425 | mamo-s | *BGIBMGA013505* | up | 1560 | 1568 | + | 11.3303 | 4.96E-05 | 0.615 | GTGGGTGGC |
| 11426 | mamo-s | *BGIBMGA013506* | up | 657 | 665 | + | 11.7064 | 3.52E-05 | 0.615 | GCGCGTGGT |
| 11427 | mamo-s | *BGIBMGA013506* | down | 1365 | 1373 | + | 10.5688 | 8.88E-05 | 0.824 | GTGCGTGAC |
| 11428 | mamo-s | *BGIBMGA013509* | up | 894 | 902 | - | 10.5138 | 9.75E-05 | 0.865 | GTGCGTGTT |
| 11429 | mamo-s | *BGIBMGA013511* | up | 1591 | 1599 | - | 11.4404 | 4.51E-05 | 0.615 | CGGCGTGGC |
| 11430 | mamo-s | *BGIBMGA013513* | up | 49 | 57 | - | 11.1376 | 6.35E-05 | 0.709 | TCGCGTGGC |
| 11431 | mamo-s | *BGIBMGA013514* | up | 1247 | 1255 | + | 11.3303 | 4.96E-05 | 0.615 | GTGGGTGGC |
| 11432 | mamo-s | *BGIBMGA013517* | down | 867 | 875 | - | 11.3303 | 4.96E-05 | 0.615 | GTGGGTGGC |
| 11433 | mamo-s | *BGIBMGA013517* | down | 1869 | 1877 | - | 10.8899 | 7.48E-05 | 0.739 | CCGCGTGCC |

| | | | | | | | | | | |
|---|---|---|---|---|---|---|---|---|---|---|
| 11434 | mamo-s | *BGIBMGA013518* | up | 820 | 828 | + | 11.2018 | 5.68E-05 | 0.674 | CTGCGTGGT |
| 11435 | mamo-s | *BGIBMGA013518* | up | 1185 | 1193 | - | 11.0826 | 6.84E-05 | 0.709 | GTACGTGGC |
| 11436 | mamo-s | *BGIBMGA013520* | up | 969 | 977 | + | 11.0826 | 6.84E-05 | 0.709 | GTACGTGGC |
| 11437 | mamo-s | *BGIBMGA013522* | down | 829 | 837 | - | 11.3303 | 4.96E-05 | 0.615 | GTGGGTGGC |
| 11438 | mamo-s | *BGIBMGA013524* | down | 732 | 740 | - | 11.3303 | 4.96E-05 | 0.615 | GTGGGTGGC |
| 11439 | mamo-s | *BGIBMGA013525* | up | 412 | 420 | + | 11.3303 | 4.96E-05 | 0.615 | GTGGGTGGC |
| 11440 | mamo-s | *BGIBMGA013526* | down | 1575 | 1583 | - | 13.6606 | 4.06E-06 | 0.615 | GCGCGTGGC |
| 11441 | mamo-s | *BGIBMGA013526* | up | 1664 | 1672 | + | 11.3303 | 4.96E-05 | 0.615 | GTGGGTGGC |
| 11442 | mamo-s | *BGIBMGA013526* | up | 1884 | 1892 | + | 11.3303 | 4.96E-05 | 0.615 | GTGGGTGGC |
| 11443 | mamo-s | *BGIBMGA013526* | up | 1045 | 1053 | - | 10.6881 | 8.61E-05 | 0.81 | GCACGTGGC |
| 11444 | mamo-s | *BGIBMGA013529* | down | 1179 | 1187 | - | 10.9358 | 7.30E-05 | 0.728 | GCGGGTGGC |
| 11445 | mamo-s | *BGIBMGA013530* | up | 1542 | 1550 | + | 11.3303 | 4.96E-05 | 0.615 | GTGGGTGGC |
| 11446 | mamo-s | *BGIBMGA013538* | up | 675 | 683 | - | 11.3303 | 4.96E-05 | 0.615 | GTGGGTGGC |
| 11447 | mamo-s | *BGIBMGA013538* | down | 784 | 792 | - | 11.3303 | 4.96E-05 | 0.615 | GTGGGTGGC |
| 11448 | mamo-s | *BGIBMGA013539* | up | 1705 | 1713 | - | 12.1835 | 1.71E-05 | 0.615 | GTGCGTGCC |
| 11449 | mamo-s | *BGIBMGA013539* | up | 857 | 865 | - | 12.0734 | 2.66E-05 | 0.615 | GCGCGTGTC |
| 11450 | mamo-s | *BGIBMGA013539* | down | 184 | 192 | + | 11.3303 | 4.96E-05 | 0.615 | GTGGGTGGC |
| 11451 | mamo-s | *BGIBMGA013539* | up | 1698 | 1706 | - | 10.8899 | 7.48E-05 | 0.739 | CCGCGTGCC |
| 11452 | mamo-s | *BGIBMGA013540* | up | 1436 | 1444 | + | 11.3303 | 4.96E-05 | 0.615 | GTGGGTGGC |
| 11453 | mamo-s | *BGIBMGA013541* | down | 828 | 836 | + | 14.055 | 2.23E-06 | 0.615 | GTGCGTGGC |
| 11454 | mamo-s | *BGIBMGA013541* | down | 1179 | 1187 | - | 11.3303 | 4.96E-05 | 0.615 | GTGGGTGGC |
| 11455 | mamo-s | *BGIBMGA013544* | down | 134 | 142 | + | 11.3303 | 4.96E-05 | 0.615 | GTGGGTGGC |
| 11456 | mamo-s | *BGIBMGA013545* | down | 359 | 367 | + | 11.3303 | 4.96E-05 | 0.615 | GTGGGTGGC |
| 11457 | mamo-s | *BGIBMGA013545* | down | 1883 | 1891 | + | 11.1009 | 6.57E-05 | 0.709 | GAGCGAGGC |
| 11458 | mamo-s | *BGIBMGA013546* | up | 1062 | 1070 | + | 11.1009 | 6.57E-05 | 0.709 | GAGCGAGGC |
| 11459 | mamo-s | *BGIBMGA013547* | up | 1812 | 1820 | + | 11.3303 | 4.96E-05 | 0.615 | GTGGGTGGC |
| 11460 | mamo-s | *BGIBMGA013547* | up | 947 | 955 | + | 10.5138 | 9.75E-05 | 0.865 | GTGCGTGTT |
| 11461 | mamo-s | *BGIBMGA013553* | up | 744 | 752 | + | 11.3303 | 4.96E-05 | 0.615 | GTGGGTGGC |
| 11462 | mamo-s | *BGIBMGA013555* | up | 1311 | 1319 | - | 12.3394 | 1.49E-05 | 0.615 | GGGCGTGGC |
| 11463 | mamo-s | *BGIBMGA013555* | down | 1320 | 1328 | - | 12.0734 | 2.66E-05 | 0.615 | GCGCGTGTC |
| 11464 | mamo-s | *BGIBMGA013555* | down | 286 | 294 | - | 11.3303 | 4.96E-05 | 0.615 | GTGGGTGGC |
| 11465 | mamo-s | *BGIBMGA013557* | up | 136 | 144 | - | 11.0826 | 6.84E-05 | 0.709 | GTACGTGGC |
| 11466 | mamo-s | *BGIBMGA013557* | up | 262 | 270 | + | 10.7523 | 8.38E-05 | 0.797 | AAGCGTGGC |
| 11467 | mamo-s | *BGIBMGA013558* | up | 480 | 488 | + | 11.5321 | 4.06E-05 | 0.615 | TTGCGTGGC |
| 11468 | mamo-s | *BGIBMGA013558* | up | 204 | 212 | - | 11.3303 | 4.96E-05 | 0.615 | GTGGGTGGC |
| 11469 | mamo-s | *BGIBMGA013560* | down | 530 | 538 | - | 11.3303 | 4.96E-05 | 0.615 | GTGGGTGGC |
| 11470 | mamo-s | *BGIBMGA013561* | up | 282 | 290 | - | 11.3303 | 4.96E-05 | 0.615 | GTGGGTGGC |
| 11471 | mamo-s | *BGIBMGA013562* | up | 128 | 136 | + | 12.4679 | 1.31E-05 | 0.615 | GTGCGTGTC |
| 11472 | mamo-s | *BGIBMGA013562* | up | 373 | 381 | - | 11.3303 | 4.96E-05 | 0.615 | GTGGGTGGC |
| 11473 | mamo-s | *BGIBMGA013562* | up | 1678 | 1686 | + | 11.3303 | 4.96E-05 | 0.615 | GTGGGTGGC |
| 11474 | mamo-s | *BGIBMGA013563* | up | 1572 | 1580 | - | 11.3303 | 4.96E-05 | 0.615 | GTGGGTGGC |
| 11475 | mamo-s | *BGIBMGA013564* | down | 1711 | 1719 | + | 11.3303 | 4.96E-05 | 0.615 | GTGGGTGGC |
| 11476 | mamo-s | *BGIBMGA013565* | down | 377 | 385 | + | 10.5138 | 9.75E-05 | 0.865 | GTGCGTGTT |
| 11477 | mamo-s | *BGIBMGA013570* | down | 524 | 532 | - | 11.3303 | 4.96E-05 | 0.615 | GTGGGTGGC |
| 11478 | mamo-s | *BGIBMGA013571* | up | 230 | 238 | - | 11.3303 | 4.96E-05 | 0.615 | GTGGGTGGC |
| 11479 | mamo-s | *BGIBMGA013574* | up | 428 | 436 | - | 10.5138 | 9.75E-05 | 0.865 | GTGCGTGTT |
| 11480 | mamo-s | *BGIBMGA013575* | down | 482 | 490 | + | 11.3303 | 4.96E-05 | 0.615 | GTGGGTGGC |
| 11481 | mamo-s | *BGIBMGA013575* | down | 1038 | 1046 | - | 11.3303 | 4.96E-05 | 0.615 | GTGGGTGGC |
| 11482 | mamo-s | *BGIBMGA013575* | down | 1848 | 1856 | + | 11.2018 | 5.68E-05 | 0.674 | CTGCGTGGT |
| 11483 | mamo-s | *BGIBMGA013576* | down | 438 | 446 | + | 10.5138 | 9.75E-05 | 0.865 | GTGCGTGTT |
| 11484 | mamo-s | *BGIBMGA013577* | down | 1766 | 1774 | + | 11.3303 | 4.96E-05 | 0.615 | GTGGGTGGC |
| 11485 | mamo-s | *BGIBMGA013577* | up | 1768 | 1776 | - | 11.3303 | 4.96E-05 | 0.615 | GTGGGTGGC |
| 11486 | mamo-s | *BGIBMGA013577* | up | 526 | 534 | - | 10.5413 | 9.42E-05 | 0.857 | CAGCGTGTC |
| 11487 | mamo-s | *BGIBMGA013578* | up | 86 | 94 | + | 11.3303 | 4.96E-05 | 0.615 | GTGGGTGGC |
| 11488 | mamo-s | *BGIBMGA013578* | up | 1581 | 1589 | - | 11.3303 | 4.96E-05 | 0.615 | GTGGGTGGC |
| 11489 | mamo-s | *BGIBMGA013579* | up | 1800 | 1808 | - | 11.3303 | 4.96E-05 | 0.615 | GTGGGTGGC |
| 11490 | mamo-s | *BGIBMGA013579* | up | 371 | 379 | + | 10.5688 | 8.88E-05 | 0.824 | GTGCGTGAC |
| 11491 | mamo-s | *BGIBMGA013584* | up | 906 | 914 | + | 12.1284 | 2.16E-05 | 0.615 | GTGCGAGGC |
| 11492 | mamo-s | *BGIBMGA013584* | down | 236 | 244 | - | 11.3303 | 4.96E-05 | 0.615 | GTGGGTGGC |
| 11493 | mamo-s | *BGIBMGA013585* | up | 553 | 561 | - | 12.3394 | 1.49E-05 | 0.615 | GGGCGTGGC |
| 11494 | mamo-s | *BGIBMGA013585* | up | 546 | 554 | - | 12.0734 | 2.66E-05 | 0.615 | GCGCGTGTC |
| 11495 | mamo-s | *BGIBMGA013587* | down | 936 | 944 | + | 12.7615 | 1.04E-05 | 0.615 | CCGCGTGGC |
| 11496 | mamo-s | *BGIBMGA013588* | up | 1633 | 1641 | + | 11.0826 | 6.84E-05 | 0.709 | GTACGTGGC |
| 11497 | mamo-s | *BGIBMGA013589* | up | 1272 | 1280 | + | 12.1009 | 2.43E-05 | 0.615 | GTGCGTGGT |
| 11498 | mamo-s | *BGIBMGA013589* | down | 1376 | 1384 | + | 11.0826 | 6.84E-05 | 0.709 | GTACGTGGC |
| 11499 | mamo-s | *BGIBMGA013592* | up | 934 | 942 | - | 11.2844 | 5.18E-05 | 0.63 | CTGCGTGCC |
| 11500 | mamo-s | *BGIBMGA013593* | up | 543 | 551 | + | 11.3303 | 4.96E-05 | 0.615 | GTGGGTGGC |
| 11501 | mamo-s | *BGIBMGA013593* | up | 722 | 730 | + | 11.3303 | 4.96E-05 | 0.615 | GTGGGTGGC |
| 11502 | mamo-s | *BGIBMGA013595* | down | 1255 | 1263 | - | 11.0826 | 6.84E-05 | 0.709 | GTACGTGGC |
| 11503 | mamo-s | *BGIBMGA013595* | up | 307 | 315 | - | 10.9358 | 7.30E-05 | 0.728 | GCGGGTGGC |
| 11504 | mamo-s | *BGIBMGA013596* | down | 81 | 89 | + | 14.055 | 2.23E-06 | 0.615 | GTGCGTGGC |
| 11505 | mamo-s | *BGIBMGA013596* | down | 23 | 31 | - | 10.7523 | 8.38E-05 | 0.797 | GGGCGTGTC |
| 11506 | mamo-s | *BGIBMGA013597* | up | 128 | 136 | + | 14.055 | 2.23E-06 | 0.615 | GTGCGTGGC |
| 11507 | mamo-s | *BGIBMGA013597* | up | 70 | 78 | - | 10.7523 | 8.38E-05 | 0.797 | GGGCGTGTC |
| 11508 | mamo-s | *BGIBMGA013599* | up | 724 | 732 | + | 11.3303 | 4.96E-05 | 0.615 | GTGGGTGGC |
| 11509 | mamo-s | *BGIBMGA013600* | down | 1481 | 1489 | - | 13.6606 | 4.06E-06 | 0.615 | GCGCGTGGC |
| 11510 | mamo-s | *BGIBMGA013601* | up | 1758 | 1766 | - | 13.6606 | 4.06E-06 | 0.615 | GCGCGTGGC |
| 11511 | mamo-s | *BGIBMGA013602* | up | 960 | 968 | + | 11.3303 | 4.96E-05 | 0.615 | GTGGGTGGC |
| 11512 | mamo-s | *BGIBMGA013605* | down | 1689 | 1697 | + | 11.3303 | 4.96E-05 | 0.615 | GTGGGTGGC |
| 11513 | mamo-s | *BGIBMGA013606* | up | 1007 | 1015 | - | 11.3303 | 4.96E-05 | 0.615 | GTGGGTGGC |
| 11514 | mamo-s | *BGIBMGA013606* | down | 1736 | 1744 | - | 11.3303 | 4.96E-05 | 0.615 | GTGGGTGGC |
| 11515 | mamo-s | *BGIBMGA013607* | up | 1567 | 1575 | + | 11.3303 | 4.96E-05 | 0.615 | GTGGGTGGC |
| 11516 | mamo-s | *BGIBMGA013608* | up | 451 | 459 | + | 11.3303 | 4.96E-05 | 0.615 | GTGGGTGGC |
| 11517 | mamo-s | *BGIBMGA013609* | down | 511 | 519 | - | 13.156 | 6.30E-06 | 0.615 | CTGCGTGGC |
| 11518 | mamo-s | *BGIBMGA013609* | down | 715 | 723 | - | 11.3303 | 4.96E-05 | 0.615 | GTGGGTGGC |
| 11519 | mamo-s | *BGIBMGA013610* | up | 54 | 62 | - | 11.3303 | 4.96E-05 | 0.615 | GTGGGTGGC |
| 11520 | mamo-s | *BGIBMGA013612* | up | 323 | 331 | - | 11.3303 | 4.96E-05 | 0.615 | GTGGGTGGC |
| 11521 | mamo-s | *BGIBMGA013613* | down | 1373 | 1381 | + | 11.3303 | 4.96E-05 | 0.615 | GTGGGTGGC |

| | | | | | | | | | | |
|---|---|---|---|---|---|---|---|---|---|---|
| 11522 | mamo-s | *BGIBMGA013614* | down | 351 | 359 | + | 11.7798 | 3.11E-05 | 0.615 | ATGCGTGGC |
| 11523 | mamo-s | *BGIBMGA013614* | down | 210 | 218 | - | 10.5688 | 8.88E-05 | 0.824 | GTGCGTGAC |
| 11524 | mamo-s | *BGIBMGA013615* | up | 547 | 555 | + | 11.7798 | 3.11E-05 | 0.615 | ATGCGTGGC |
| 11525 | mamo-s | *BGIBMGA013615* | up | 406 | 414 | - | 10.5688 | 8.88E-05 | 0.824 | GTGCGTGAC |
| 11526 | mamo-s | *BGIBMGA013617* | down | 354 | 362 | + | 11.1009 | 6.57E-05 | 0.709 | GAGCGAGGC |
| 11527 | mamo-s | *BGIBMGA013619* | down | 1179 | 1187 | - | 11.3303 | 4.96E-05 | 0.615 | GTGGGTGGC |
| 11528 | mamo-s | *BGIBMGA013620* | up | 447 | 455 | + | 10.6881 | 8.61E-05 | 0.81 | GCACGTGGC |
| 11529 | mamo-s | *BGIBMGA013621* | up | 15 | 23 | - | 10.5688 | 8.88E-05 | 0.824 | GTGCGTGAC |
| 11530 | mamo-s | *BGIBMGA013623* | down | 1634 | 1642 | + | 11.3303 | 4.96E-05 | 0.615 | GTGGGTGGC |
| 11531 | mamo-s | *BGIBMGA013624* | down | 1685 | 1693 | + | 11.3303 | 4.96E-05 | 0.615 | GTGGGTGGC |
| 11532 | mamo-s | *BGIBMGA013624* | down | 1385 | 1393 | + | 11.0826 | 6.84E-05 | 0.709 | GTACGTGGC |
| 11533 | mamo-s | *BGIBMGA013626* | up | 1704 | 1712 | - | 12.1009 | 2.43E-05 | 0.615 | GTGCGTGGT |
| 11534 | mamo-s | *BGIBMGA013626* | down | 722 | 730 | + | 10.5688 | 8.88E-05 | 0.824 | GTGCGTGAC |
| 11535 | mamo-s | *BGIBMGA013627* | down | 763 | 771 | + | 11.3303 | 4.96E-05 | 0.615 | GTGGGTGGC |
| 11536 | mamo-s | *BGIBMGA013627* | up | 817 | 825 | + | 11.3303 | 4.96E-05 | 0.615 | GTGGGTGGC |
| 11537 | mamo-s | *BGIBMGA013630* | up | 1695 | 1703 | - | 11.3303 | 4.96E-05 | 0.615 | GTGGGTGGC |
| 11538 | mamo-s | *BGIBMGA013631* | up | 1033 | 1041 | - | 11.3303 | 4.96E-05 | 0.615 | GTGGGTGGC |
| 11539 | mamo-s | *BGIBMGA013632* | up | 585 | 593 | + | 11.3303 | 4.96E-05 | 0.615 | GTGGGTGGC |
| 11540 | mamo-s | *BGIBMGA013633* | down | 1257 | 1265 | + | 10.5138 | 9.75E-05 | 0.865 | GTGCGTGTT |
| 11541 | mamo-s | *BGIBMGA013636* | down | 982 | 990 | + | 11.3303 | 4.96E-05 | 0.615 | GTGGGTGGC |
| 11542 | mamo-s | *BGIBMGA013639* | up | 270 | 278 | - | 11.3303 | 4.96E-05 | 0.615 | GTGGGTGGC |
| 11543 | mamo-s | *BGIBMGA013639* | down | 779 | 787 | - | 11.3303 | 4.96E-05 | 0.615 | GTGGGTGGC |
| 11544 | mamo-s | *BGIBMGA013640* | up | 1633 | 1641 | + | 11.3303 | 4.96E-05 | 0.615 | GTGGGTGGC |
| 11545 | mamo-s | *BGIBMGA013641* | up | 189 | 197 | - | 11.3303 | 4.96E-05 | 0.615 | GTGGGTGGC |
| 11546 | mamo-s | *BGIBMGA013641* | down | 1038 | 1046 | + | 11.3303 | 4.96E-05 | 0.615 | GTGGGTGGC |
| 11547 | mamo-s | *BGIBMGA013641* | down | 1710 | 1718 | + | 11.3303 | 4.96E-05 | 0.615 | GTGGGTGGC |
| 11548 | mamo-s | *BGIBMGA013643* | down | 1059 | 1067 | - | 12.1284 | 2.16E-05 | 0.615 | GTGCGAGGC |
| 11549 | mamo-s | *BGIBMGA013643* | up | 1262 | 1270 | - | 11.3303 | 4.96E-05 | 0.615 | GTGGGTGGC |
| 11550 | mamo-s | *BGIBMGA013644* | up | 1705 | 1713 | + | 11.3303 | 4.96E-05 | 0.615 | GTGGGTGGC |
| 11551 | mamo-s | *BGIBMGA013644* | up | 901 | 909 | + | 10.8073 | 7.89E-05 | 0.766 | CCGCGTGGT |
| 11552 | mamo-s | *BGIBMGA013645* | down | 1748 | 1756 | + | 13.6606 | 4.06E-06 | 0.615 | GCGCGTGGC |
| 11553 | mamo-s | *BGIBMGA013645* | down | 163 | 171 | - | 11.3303 | 4.96E-05 | 0.615 | GTGGGTGGC |
| 11554 | mamo-s | *BGIBMGA013646* | up | 689 | 697 | + | 11.3303 | 4.96E-05 | 0.615 | GTGGGTGGC |
| 11555 | mamo-s | *BGIBMGA013646* | up | 1317 | 1325 | + | 11.3303 | 4.96E-05 | 0.615 | GTGGGTGGC |
| 11556 | mamo-s | *BGIBMGA013646* | down | 1440 | 1448 | + | 11.3303 | 4.96E-05 | 0.615 | GTGGGTGGC |
| 11557 | mamo-s | *BGIBMGA013648* | up | 118 | 126 | + | 11.5688 | 3.79E-05 | 0.615 | CTGCGTGTC |
| 11558 | mamo-s | *BGIBMGA013649* | down | 1119 | 1127 | - | 11.0734 | 7.12E-05 | 0.726 | GAGCGTGGT |
| 11559 | mamo-s | *BGIBMGA013649* | down | 331 | 339 | - | 10.5413 | 9.42E-05 | 0.857 | CAGCGTGTC |
| 11560 | mamo-s | *BGIBMGA013649* | down | 438 | 446 | + | 10.5413 | 9.42E-05 | 0.857 | CAGCGTGTC |
| 11561 | mamo-s | *BGIBMGA013650* | down | 1249 | 1257 | + | 14.055 | 2.23E-06 | 0.615 | GTGCGTGGC |
| 11562 | mamo-s | *BGIBMGA013651* | up | 146 | 154 | + | 14.055 | 2.23E-06 | 0.615 | GTGCGTGGC |
| 11563 | mamo-s | *BGIBMGA013655* | down | 1168 | 1176 | + | 11.3303 | 4.96E-05 | 0.615 | GTGGGTGGC |
| 11564 | mamo-s | *BGIBMGA013655* | up | 1687 | 1695 | + | 10.8899 | 7.48E-05 | 0.739 | CCGCGTGCC |
| 11565 | mamo-s | *BGIBMGA013656* | up | 1637 | 1645 | + | 10.6881 | 8.61E-05 | 0.81 | GCACGTGGC |
| 11566 | mamo-s | *BGIBMGA013659* | down | 1216 | 1224 | + | 10.5688 | 8.88E-05 | 0.824 | GTGCGTGAC |
| 11567 | mamo-s | *BGIBMGA013660* | down | 1968 | 1976 | + | 11.3303 | 4.96E-05 | 0.615 | GTGGGTGGC |
| 11568 | mamo-s | *BGIBMGA013660* | down | 1698 | 1706 | - | 11.0826 | 6.84E-05 | 0.709 | GTACGTGGC |
| 11569 | mamo-s | *BGIBMGA013660* | up | 20 | 28 | + | 10.6881 | 8.61E-05 | 0.81 | GCACGTGGC |
| 11570 | mamo-s | *BGIBMGA013663* | down | 1282 | 1290 | + | 10.9358 | 7.30E-05 | 0.728 | GCGGGTGGC |
| 11571 | mamo-s | *BGIBMGA013664* | up | 1312 | 1320 | - | 13.6606 | 4.06E-06 | 0.615 | GCGCGTGGC |
| 11572 | mamo-s | *BGIBMGA013664* | up | 143 | 151 | - | 11.3303 | 4.96E-05 | 0.615 | GTGGGTGGC |
| 11573 | mamo-s | *BGIBMGA013665* | down | 1286 | 1294 | + | 11.3303 | 4.96E-05 | 0.615 | GTGGGTGGC |
| 11574 | mamo-s | *BGIBMGA013665* | down | 1171 | 1179 | + | 10.9358 | 7.30E-05 | 0.728 | GCGGGTGGC |
| 11575 | mamo-s | *BGIBMGA013667* | up | 1172 | 1180 | + | 11.3303 | 4.96E-05 | 0.615 | GTGGGTGGC |
| 11576 | mamo-s | *BGIBMGA013669* | up | 195 | 203 | - | 11.789 | 2.84E-05 | 0.615 | GCGCGTGCC |
| 11577 | mamo-s | *BGIBMGA013669* | down | 1441 | 1449 | - | 11.3303 | 4.96E-05 | 0.615 | GTGGGTGGC |
| 11578 | mamo-s | *BGIBMGA013669* | up | 1925 | 1933 | + | 11.3303 | 4.96E-05 | 0.615 | GTGGGTGGC |
| 11579 | mamo-s | *BGIBMGA013672* | down | 1811 | 1819 | - | 11.3303 | 4.96E-05 | 0.615 | GTGGGTGGC |
| 11580 | mamo-s | *BGIBMGA013672* | down | 1989 | 1997 | - | 11.1743 | 5.90E-05 | 0.684 | CCGCGTGTC |
| 11581 | mamo-s | *BGIBMGA013673* | down | 1283 | 1291 | + | 11.3303 | 4.96E-05 | 0.615 | GTGGGTGGC |
| 11582 | mamo-s | *BGIBMGA013674* | up | 1257 | 1265 | + | 13.0275 | 8.53E-06 | 0.615 | GAGCGTGGC |
| 11583 | mamo-s | *BGIBMGA013675* | down | 984 | 992 | + | 11.0826 | 6.84E-05 | 0.709 | GTACGTGGC |
| 11584 | mamo-s | *BGIBMGA013678* | down | 1832 | 1840 | - | 11.3303 | 4.96E-05 | 0.615 | GTGGGTGGC |
| 11585 | mamo-s | *BGIBMGA013681* | up | 1519 | 1527 | - | 14.055 | 2.23E-06 | 0.615 | GTGCGTGGC |
| 11586 | mamo-s | *BGIBMGA013681* | down | 713 | 721 | - | 11.3303 | 4.96E-05 | 0.615 | GTGGGTGGC |
| 11587 | mamo-s | *BGIBMGA013681* | up | 1064 | 1072 | + | 11.3303 | 4.96E-05 | 0.615 | GTGGGTGGC |
| 11588 | mamo-s | *BGIBMGA013683* | up | 669 | 677 | + | 11.3303 | 4.96E-05 | 0.615 | GTGGGTGGC |
| 11589 | mamo-s | *BGIBMGA013685* | down | 1626 | 1634 | + | 11.3303 | 4.96E-05 | 0.615 | GTGGGTGGC |
| 11590 | mamo-s | *BGIBMGA013687* | up | 193 | 201 | - | 11.3303 | 4.96E-05 | 0.615 | GTGGGTGGC |
| 11591 | mamo-s | *BGIBMGA013687* | up | 1183 | 1191 | + | 11.2018 | 5.68E-05 | 0.674 | CTGCGTGGT |
| 11592 | mamo-s | *BGIBMGA013687* | up | 1699 | 1707 | + | 10.8073 | 7.89E-05 | 0.766 | CCGCGTGGT |
| 11593 | mamo-s | *BGIBMGA013687* | down | 15 | 23 | + | 10.5138 | 9.75E-05 | 0.865 | GTGCGTGTT |
| 11594 | mamo-s | *BGIBMGA013688* | down | 1500 | 1508 | + | 11.3303 | 4.96E-05 | 0.615 | GTGGGTGGC |
| 11595 | mamo-s | *BGIBMGA013689* | down | 1204 | 1212 | + | 11.3303 | 4.96E-05 | 0.615 | GTGGGTGGC |
| 11596 | mamo-s | *BGIBMGA013689* | up | 1663 | 1671 | + | 11.3303 | 4.96E-05 | 0.615 | GTGGGTGGC |
| 11597 | mamo-s | *BGIBMGA013692* | up | 1450 | 1458 | + | 11.1743 | 5.90E-05 | 0.684 | CCGCGTGTC |
| 11598 | mamo-s | *BGIBMGA013692* | down | 1080 | 1088 | - | 10.7523 | 8.38E-05 | 0.797 | AAGCGTGGC |
| 11599 | mamo-s | *BGIBMGA013693* | up | 61 | 69 | + | 10.8073 | 7.89E-05 | 0.766 | CCGCGTGGT |
| 11600 | mamo-s | *BGIBMGA013694* | up | 631 | 639 | + | 11.3853 | 4.74E-05 | 0.615 | ACGCGTGGC |
| 11601 | mamo-s | *BGIBMGA013695* | up | 329 | 337 | - | 11.3303 | 4.96E-05 | 0.615 | GTGGGTGGC |
| 11602 | mamo-s | *BGIBMGA013695* | down | 1719 | 1727 | + | 11.3303 | 4.96E-05 | 0.615 | GTGGGTGGC |
| 11603 | mamo-s | *BGIBMGA013700* | down | 310 | 318 | + | 11.3303 | 4.96E-05 | 0.615 | GTGGGTGGC |
| 11604 | mamo-s | *BGIBMGA013700* | down | 1515 | 1523 | + | 10.8073 | 7.89E-05 | 0.766 | CCGCGTGGT |
| 11605 | mamo-s | *BGIBMGA013701* | down | 563 | 571 | + | 11.3303 | 4.96E-05 | 0.615 | GTGGGTGGC |
| 11606 | mamo-s | *BGIBMGA013701* | down | 721 | 729 | + | 10.9358 | 7.30E-05 | 0.728 | GCGGGTGGC |
| 11607 | mamo-s | *BGIBMGA013702* | down | 1924 | 1932 | + | 13.6606 | 4.06E-06 | 0.615 | GCGCGTGGC |
| 11608 | mamo-s | *BGIBMGA013702* | up | 371 | 379 | - | 12.7615 | 1.04E-05 | 0.615 | CCGCGTGGC |
| 11609 | mamo-s | *BGIBMGA013704* | up | 1227 | 1235 | + | 11.4404 | 4.51E-05 | 0.615 | CGGCGTGGC |

| | | | | | | | | | | |
|---|---|---|---|---|---|---|---|---|---|---|
| 11610 | mamo-s | *BGIBMGA013704* | down | 765 | 773 | + | 11.3303 | 4.96E-05 | 0.615 | GTGGGTGGC |
| 11611 | mamo-s | *BGIBMGA013704* | up | 811 | 819 | + | 11.1009 | 6.57E-05 | 0.709 | GAGCGAGGC |
| 11612 | mamo-s | *BGIBMGA013705* | down | 1572 | 1580 | - | 11.1009 | 6.57E-05 | 0.709 | GAGCGAGGC |
| 11613 | mamo-s | *BGIBMGA013705* | down | 452 | 460 | - | 10.5688 | 8.88E-05 | 0.824 | GTGCGTGAC |
| 11614 | mamo-s | *BGIBMGA013706* | up | 239 | 247 | + | 10.5413 | 9.42E-05 | 0.857 | GTGCGAGTC |
| 11615 | mamo-s | *BGIBMGA013708* | up | 383 | 391 | - | 11.3303 | 4.96E-05 | 0.615 | GTGGGTGGC |
| 11616 | mamo-s | *BGIBMGA013709* | up | 1572 | 1580 | - | 12.0734 | 2.66E-05 | 0.615 | GCGCGTGTC |
| 11617 | mamo-s | *BGIBMGA013711* | down | 1121 | 1129 | - | 11.5321 | 4.06E-05 | 0.615 | TTGCGTGGC |
| 11618 | mamo-s | *BGIBMGA013711* | up | 1428 | 1436 | + | 11.3303 | 4.96E-05 | 0.615 | GTGGGTGGC |
| 11619 | mamo-s | *BGIBMGA013712* | down | 408 | 416 | - | 11.3303 | 4.96E-05 | 0.615 | GTGGGTGGC |
| 11620 | mamo-s | *BGIBMGA013714* | up | 474 | 482 | + | 11.3303 | 4.96E-05 | 0.615 | GTGGGTGGC |
| 11621 | mamo-s | *BGIBMGA013715* | down | 816 | 824 | - | 11.3303 | 4.96E-05 | 0.615 | GTGGGTGGC |
| 11622 | mamo-s | *BGIBMGA013715* | down | 658 | 666 | - | 10.9358 | 7.30E-05 | 0.728 | GCGGGTGGC |
| 11623 | mamo-s | *BGIBMGA013716* | down | 160 | 168 | - | 11.3303 | 4.96E-05 | 0.615 | GTGGGTGGC |
| 11624 | mamo-s | *BGIBMGA013716* | down | 2 | 10 | - | 10.9358 | 7.30E-05 | 0.728 | GCGGGTGGC |
| 11625 | mamo-s | *BGIBMGA013717* | down | 1146 | 1154 | - | 10.8073 | 7.89E-05 | 0.766 | CCGCGTGGT |
| 11626 | mamo-s | *BGIBMGA013719* | up | 188 | 196 | + | 14.055 | 2.23E-06 | 0.615 | GTGCGTGGC |
| 11627 | mamo-s | *BGIBMGA013719* | down | 1086 | 1094 | - | 11.3853 | 4.74E-05 | 0.615 | ACGCGTGGC |
| 11628 | mamo-s | *BGIBMGA013719* | up | 899 | 907 | - | 10.6881 | 8.61E-05 | 0.81 | GCACGTGGC |
| 11629 | mamo-s | *BGIBMGA013720* | up | 1404 | 1412 | + | 11.3303 | 4.96E-05 | 0.615 | GTGGGTGGC |
| 11630 | mamo-s | *BGIBMGA013722* | down | 1557 | 1565 | - | 11.3303 | 4.96E-05 | 0.615 | GTGGGTGGC |
| 11631 | mamo-s | *BGIBMGA013725* | up | 695 | 703 | - | 11.4404 | 4.51E-05 | 0.615 | GAGCGTGTC |
| 11632 | mamo-s | *BGIBMGA013726* | down | 1587 | 1595 | - | 11.1743 | 5.90E-05 | 0.684 | CCGCGTGTC |
| 11633 | mamo-s | *BGIBMGA013727* | up | 95 | 103 | - | 10.9358 | 7.30E-05 | 0.728 | GCGGGTGGC |
| 11634 | mamo-s | *BGIBMGA013729* | up | 780 | 788 | + | 10.5413 | 9.42E-05 | 0.857 | CAGCGTGTC |
| 11635 | mamo-s | *BGIBMGA013731* | down | 982 | 990 | + | 11.3303 | 4.96E-05 | 0.615 | GTGGGTGGC |
| 11636 | mamo-s | *BGIBMGA013731* | up | 275 | 283 | - | 11.1009 | 6.57E-05 | 0.709 | GAGCGAGGC |
| 11637 | mamo-s | *BGIBMGA013731* | up | 650 | 658 | - | 10.5413 | 9.42E-05 | 0.857 | CAGCGTGTC |
| 11638 | mamo-s | *BGIBMGA013732* | down | 1263 | 1271 | - | 13.0275 | 8.53E-06 | 0.615 | GAGCGTGGC |
| 11639 | mamo-s | *BGIBMGA013733* | up | 217 | 225 | - | 13.0275 | 8.53E-06 | 0.615 | GAGCGTGGC |
| 11640 | mamo-s | *BGIBMGA013734* | up | 1755 | 1763 | - | 11.789 | 2.84E-05 | 0.615 | GCGCGTGCC |
| 11641 | mamo-s | *BGIBMGA013734* | down | 565 | 573 | - | 11.1009 | 6.57E-05 | 0.709 | GAGCGAGGC |
| 11642 | mamo-s | *BGIBMGA013735* | up | 733 | 741 | - | 11.789 | 2.84E-05 | 0.615 | GCGCGTGCC |
| 11643 | mamo-s | *BGIBMGA013735* | down | 491 | 499 | - | 10.5413 | 9.42E-05 | 0.857 | CAGCGTGTC |
| 11644 | mamo-s | *BGIBMGA013736* | up | 336 | 344 | + | 11.2844 | 5.18E-05 | 0.63 | CTGCGTGCC |
| 11645 | mamo-s | *BGIBMGA013737* | down | 1548 | 1556 | - | 13.6606 | 4.06E-06 | 0.615 | GCGCGTGGC |
| 11646 | mamo-s | *BGIBMGA013737* | up | 3 | 11 | + | 11.3303 | 4.96E-05 | 0.615 | GTGGGTGGC |
| 11647 | mamo-s | *BGIBMGA013738* | up | 565 | 573 | + | 11.3303 | 4.96E-05 | 0.615 | GTGGGTGGC |
| 11648 | mamo-s | *BGIBMGA013738* | down | 1851 | 1859 | + | 11.3303 | 4.96E-05 | 0.615 | GTGGGTGGC |
| 11649 | mamo-s | *BGIBMGA013738* | down | 1673 | 1681 | + | 10.9358 | 7.30E-05 | 0.728 | GCGGGTGGC |
| 11650 | mamo-s | *BGIBMGA013739* | up | 1019 | 1027 | + | 12.3394 | 1.49E-05 | 0.615 | GGGCGTGGC |
| 11651 | mamo-s | *BGIBMGA013740* | down | 1856 | 1864 | + | 11.7339 | 3.29E-05 | 0.615 | GCGCGAGGC |
| 11652 | mamo-s | *BGIBMGA013740* | down | 472 | 480 | + | 11.3303 | 4.96E-05 | 0.615 | GTGGGTGGC |
| 11653 | mamo-s | *BGIBMGA013740* | down | 1879 | 1887 | - | 11.2018 | 5.68E-05 | 0.674 | CTGCGTGGT |
| 11654 | mamo-s | *BGIBMGA013742* | down | 271 | 279 | - | 11.3303 | 4.96E-05 | 0.615 | GTGGGTGGC |
| 11655 | mamo-s | *BGIBMGA013743* | down | 216 | 224 | + | 13.0275 | 8.53E-06 | 0.615 | GAGCGTGGC |
| 11656 | mamo-s | *BGIBMGA013743* | up | 1608 | 1616 | + | 11.3303 | 4.96E-05 | 0.615 | GTGGGTGGC |
| 11657 | mamo-s | *BGIBMGA013744* | down | 204 | 212 | + | 11.7798 | 3.11E-05 | 0.615 | ATGCGTGGC |
| 11658 | mamo-s | *BGIBMGA013744* | down | 1794 | 1802 | - | 11.3303 | 4.96E-05 | 0.615 | GTGGGTGGC |
| 11659 | mamo-s | *BGIBMGA013744* | up | 1545 | 1553 | - | 11.1743 | 5.90E-05 | 0.684 | CCGCGTGTC |
| 11660 | mamo-s | *BGIBMGA013745* | up | 747 | 755 | + | 11.0826 | 6.84E-05 | 0.709 | GTACGTGGC |
| 11661 | mamo-s | *BGIBMGA013747* | down | 709 | 717 | + | 11.3303 | 4.96E-05 | 0.615 | GTGGGTGGC |
| 11662 | mamo-s | *BGIBMGA013748* | down | 952 | 960 | + | 11.2018 | 5.68E-05 | 0.674 | CTGCGTGGT |
| 11663 | mamo-s | *BGIBMGA013749* | up | 226 | 234 | + | 11.3303 | 4.96E-05 | 0.615 | GTGGGTGGC |
| 11664 | mamo-s | *BGIBMGA013749* | down | 50 | 58 | + | 11.2018 | 5.68E-05 | 0.674 | CTGCGTGGT |
| 11665 | mamo-s | *BGIBMGA013750* | down | 1334 | 1342 | + | 11.3303 | 4.96E-05 | 0.615 | GTGGGTGGC |
| 11666 | mamo-s | *BGIBMGA013752* | down | 485 | 493 | - | 11.7798 | 3.11E-05 | 0.615 | ATGCGTGGC |
| 11667 | mamo-s | *BGIBMGA013754* | up | 1326 | 1334 | + | 11.3303 | 4.96E-05 | 0.615 | GTGGGTGGC |
| 11668 | mamo-s | *BGIBMGA013755* | up | 906 | 914 | + | 11.3303 | 4.96E-05 | 0.615 | GTGGGTGGC |
| 11669 | mamo-s | *BGIBMGA013755* | down | 1973 | 1981 | + | 11.3303 | 4.96E-05 | 0.615 | GTGGGTGGC |
| 11670 | mamo-s | *BGIBMGA013756* | up | 612 | 620 | + | 11.3303 | 4.96E-05 | 0.615 | GTGGGTGGC |
| 11671 | mamo-s | *BGIBMGA013756* | up | 775 | 783 | - | 11.0826 | 6.84E-05 | 0.709 | GTACGTGGC |
| 11672 | mamo-s | *BGIBMGA013758* | down | 1024 | 1032 | - | 11.3303 | 4.96E-05 | 0.615 | GTGGGTGGC |
| 11673 | mamo-s | *BGIBMGA013758* | up | 1479 | 1487 | - | 11.0826 | 6.84E-05 | 0.709 | GTACGTGGC |
| 11674 | mamo-s | *BGIBMGA013763* | up | 617 | 625 | - | 12.1009 | 2.43E-05 | 0.615 | GTGCGTGGT |
| 11675 | mamo-s | *BGIBMGA013764* | up | 1629 | 1637 | - | 12.7615 | 1.04E-05 | 0.615 | CCGCGTGGC |
| 11676 | mamo-s | *BGIBMGA013764* | up | 686 | 694 | + | 11.3303 | 4.96E-05 | 0.615 | GTGGGTGGC |
| 11677 | mamo-s | *BGIBMGA013764* | up | 1117 | 1125 | + | 11.156 | 6.13E-05 | 0.705 | GAGCGTGCC |
| 11678 | mamo-s | *BGIBMGA013765* | up | 152 | 160 | - | 14.055 | 2.23E-06 | 0.615 | GTGCGTGGC |
| 11679 | mamo-s | *BGIBMGA013766* | down | 1396 | 1404 | - | 11.3853 | 4.74E-05 | 0.615 | ACGCGTGGC |
| 11680 | mamo-s | *BGIBMGA013766* | down | 837 | 845 | - | 11.3303 | 4.96E-05 | 0.615 | GTGGGTGGC |
| 11681 | mamo-s | *BGIBMGA013767* | up | 28 | 36 | + | 11.3303 | 4.96E-05 | 0.615 | GTGGGTGGC |
| 11682 | mamo-s | *BGIBMGA013767* | up | 1833 | 1841 | + | 10.5688 | 8.88E-05 | 0.824 | GTGCGTGAC |
| 11683 | mamo-s | *BGIBMGA013767* | up | 1964 | 1972 | + | 10.5688 | 8.88E-05 | 0.824 | GTGCGTGAC |
| 11684 | mamo-s | *BGIBMGA013768* | up | 1308 | 1316 | + | 10.5138 | 9.75E-05 | 0.865 | GTGCGTGTT |
| 11685 | mamo-s | *BGIBMGA013770* | up | 864 | 872 | + | 11.3303 | 4.96E-05 | 0.615 | GTGGGTGGC |
| 11686 | mamo-s | *BGIBMGA013770* | down | 1332 | 1340 | - | 10.9358 | 7.30E-05 | 0.728 | GCGGGTGGC |
| 11687 | mamo-s | *BGIBMGA013772* | up | 204 | 212 | - | 11.3303 | 4.96E-05 | 0.615 | GTGGGTGGC |
| 11688 | mamo-s | *BGIBMGA013775* | down | 871 | 879 | - | 11.3303 | 4.96E-05 | 0.615 | GTGGGTGGC |
| 11689 | mamo-s | *BGIBMGA013777* | down | 1383 | 1391 | + | 12.7615 | 1.04E-05 | 0.615 | CCGCGTGGC |
| 11690 | mamo-s | *BGIBMGA013777* | down | 1603 | 1611 | + | 12.7615 | 1.04E-05 | 0.615 | CCGCGTGGC |
| 11691 | mamo-s | *BGIBMGA013778* | up | 1402 | 1410 | + | 14.055 | 2.23E-06 | 0.615 | GTGCGTGGC |
| 11692 | mamo-s | *BGIBMGA013778* | down | 115 | 123 | - | 11.3303 | 4.96E-05 | 0.615 | GTGGGTGGC |
| 11693 | mamo-s | *BGIBMGA013778* | up | 136 | 144 | + | 11.3303 | 4.96E-05 | 0.615 | GTGGGTGGC |
| 11694 | mamo-s | *BGIBMGA013778* | down | 600 | 608 | + | 11.3303 | 4.96E-05 | 0.615 | GTGGGTGGC |
| 11695 | mamo-s | *BGIBMGA013778* | up | 1659 | 1667 | - | 11.3303 | 4.96E-05 | 0.615 | GTGGGTGGC |
| 11696 | mamo-s | *BGIBMGA013779* | down | 252 | 260 | + | 11.3303 | 4.96E-05 | 0.615 | GTGGGTGGC |
| 11697 | mamo-s | *BGIBMGA013779* | up | 1849 | 1857 | - | 11.3303 | 4.96E-05 | 0.615 | GTGGGTGGC |

| | | | | | | | | | | |
|---|---|---|---|---|---|---|---|---|---|---|
| 11698 | mamo-s | *BGIBMGA013780* | up | 501 | 509 | + | 11.3303 | 4.96E-05 | 0.615 | GTGGGTGGC |
| 11699 | mamo-s | *BGIBMGA013781* | up | 1300 | 1308 | - | 12.1284 | 2.16E-05 | 0.615 | GTGCGAGGC |
| 11700 | mamo-s | *BGIBMGA013782* | up | 957 | 965 | + | 13.0275 | 8.53E-06 | 0.615 | GAGCGTGGC |
| 11701 | mamo-s | *BGIBMGA013783* | up | 1533 | 1541 | + | 11.0826 | 6.84E-05 | 0.709 | GTACGTGGC |
| 11702 | mamo-s | *BGIBMGA013784* | up | 1870 | 1878 | + | 11.2018 | 5.68E-05 | 0.674 | CTGCGTGGT |
| 11703 | mamo-s | *BGIBMGA013786* | down | 1979 | 1987 | - | 11.3303 | 4.96E-05 | 0.615 | GTGGGTGGC |
| 11704 | mamo-s | *BGIBMGA013787* | down | 562 | 570 | - | 11.0826 | 6.84E-05 | 0.709 | GTACGTGGC |
| 11705 | mamo-s | *BGIBMGA013787* | down | 1697 | 1705 | - | 11.0826 | 6.84E-05 | 0.709 | GTACGTGGC |
| 11706 | mamo-s | *BGIBMGA013788* | up | 1592 | 1600 | - | 11.3303 | 4.96E-05 | 0.615 | GTGGGTGGC |
| 11707 | mamo-s | *BGIBMGA013788* | up | 1971 | 1979 | + | 10.8899 | 7.48E-05 | 0.739 | CCGCGTGCC |
| 11708 | mamo-s | *BGIBMGA013790* | up | 882 | 890 | - | 11.3303 | 4.96E-05 | 0.615 | GTGGGTGGC |
| 11709 | mamo-s | *BGIBMGA013790* | down | 1384 | 1392 | + | 11.3303 | 4.96E-05 | 0.615 | GTGGGTGGC |
| 11710 | mamo-s | *BGIBMGA013792* | down | 946 | 954 | + | 10.7523 | 8.38E-05 | 0.797 | AAGCGTGGC |
| 11711 | mamo-s | *BGIBMGA013793* | up | 522 | 530 | + | 11.3303 | 4.96E-05 | 0.615 | GTGGGTGGC |
| 11712 | mamo-s | *BGIBMGA013794* | up | 47 | 55 | - | 11.3303 | 4.96E-05 | 0.615 | GTGGGTGGC |
| 11713 | mamo-s | *BGIBMGA013795* | down | 359 | 367 | - | 11.3303 | 4.96E-05 | 0.615 | GTGGGTGGC |
| 11714 | mamo-s | *BGIBMGA013796* | down | 1146 | 1154 | - | 11.2018 | 5.68E-05 | 0.674 | CTGCGTGGT |
| 11715 | mamo-s | *BGIBMGA013797* | up | 309 | 317 | - | 11.3303 | 4.96E-05 | 0.615 | GTGGGTGGC |
| 11716 | mamo-s | *BGIBMGA013797* | down | 1950 | 1958 | + | 11.3303 | 4.96E-05 | 0.615 | GTGGGTGGC |
| 11717 | mamo-s | *BGIBMGA013798* | down | 1504 | 1512 | - | 11.3303 | 4.96E-05 | 0.615 | GTGGGTGGC |
| 11718 | mamo-s | *BGIBMGA013799* | down | 1281 | 1289 | - | 11.7798 | 3.11E-05 | 0.615 | ATGCGTGGC |
| 11719 | mamo-s | *BGIBMGA013800* | down | 517 | 525 | - | 13.0275 | 8.53E-06 | 0.615 | GAGCGTGGC |
| 11720 | mamo-s | *BGIBMGA013800* | down | 1694 | 1702 | - | 11.3303 | 4.96E-05 | 0.615 | GTGGGTGGC |
| 11721 | mamo-s | *BGIBMGA013800* | up | 907 | 915 | + | 11.1743 | 5.90E-05 | 0.684 | CCGCGTGTC |
| 11722 | mamo-s | *BGIBMGA013801* | down | 952 | 960 | - | 11.3303 | 4.96E-05 | 0.615 | GTGGGTGGC |
| 11723 | mamo-s | *BGIBMGA013801* | up | 1367 | 1375 | - | 11.3303 | 4.96E-05 | 0.615 | GTGGGTGGC |
| 11724 | mamo-s | *BGIBMGA013803* | up | 558 | 566 | + | 11.3303 | 4.96E-05 | 0.615 | GTGGGTGGC |
| 11725 | mamo-s | *BGIBMGA013803* | down | 1601 | 1609 | + | 11.3303 | 4.96E-05 | 0.615 | GTGGGTGGC |
| 11726 | mamo-s | *BGIBMGA013803* | up | 231 | 239 | + | 11.1743 | 5.90E-05 | 0.684 | CCGCGTGTC |
| 11727 | mamo-s | *BGIBMGA013803* | down | 696 | 704 | + | 10.5413 | 9.42E-05 | 0.857 | GTGCGAGTC |
| 11728 | mamo-s | *BGIBMGA013806* | down | 403 | 411 | + | 13.156 | 6.30E-06 | 0.615 | CTGCGTGGC |
| 11729 | mamo-s | *BGIBMGA013807* | down | 230 | 238 | + | 11.3303 | 4.96E-05 | 0.615 | GTGGGTGGC |
| 11730 | mamo-s | *BGIBMGA013807* | up | 448 | 456 | + | 11.3303 | 4.96E-05 | 0.615 | GTGGGTGGC |
| 11731 | mamo-s | *BGIBMGA013808* | down | 282 | 290 | - | 11.1009 | 6.57E-05 | 0.709 | GAGCGAGGC |
| 11732 | mamo-s | *BGIBMGA013809* | up | 643 | 651 | + | 13.156 | 6.30E-06 | 0.615 | CTGCGTGGC |
| 11733 | mamo-s | *BGIBMGA013810* | down | 1482 | 1490 | - | 12.4679 | 1.31E-05 | 0.615 | GTGCGTGTC |
| 11734 | mamo-s | *BGIBMGA013810* | down | 262 | 270 | + | 11.3303 | 4.96E-05 | 0.615 | GTGGGTGGC |
| 11735 | mamo-s | *BGIBMGA013810* | up | 488 | 496 | - | 10.5413 | 9.42E-05 | 0.857 | CAGCGTGTC |
| 11736 | mamo-s | *BGIBMGA013811* | down | 1673 | 1681 | + | 12.1284 | 2.16E-05 | 0.615 | CAGCGTGGC |
| 11737 | mamo-s | *BGIBMGA013811* | up | 192 | 200 | + | 11.3303 | 4.96E-05 | 0.615 | GTGGGTGGC |
| 11738 | mamo-s | *BGIBMGA013813* | down | 1452 | 1460 | + | 12.4679 | 1.31E-05 | 0.615 | GTGCGTGTC |
| 11739 | mamo-s | *BGIBMGA013813* | up | 469 | 477 | - | 11.4404 | 4.51E-05 | 0.615 | GAGCGTGTC |
| 11740 | mamo-s | *BGIBMGA013813* | down | 1740 | 1748 | + | 11.3303 | 4.96E-05 | 0.615 | GTGGGTGGC |
| 11741 | mamo-s | *BGIBMGA013814* | up | 1915 | 1923 | - | 11.7798 | 3.11E-05 | 0.615 | ATGCGTGGC |
| 11742 | mamo-s | *BGIBMGA013814* | up | 1432 | 1440 | - | 11.3303 | 4.96E-05 | 0.615 | GTGGGTGGC |
| 11743 | mamo-s | *BGIBMGA013815* | up | 993 | 1001 | + | 11.3303 | 4.96E-05 | 0.615 | GTGGGTGGC |
| 11744 | mamo-s | *BGIBMGA013815* | up | 253 | 261 | - | 10.8349 | 7.66E-05 | 0.751 | CCGCGAGGC |
| 11745 | mamo-s | *BGIBMGA013815* | down | 848 | 856 | - | 10.5138 | 9.75E-05 | 0.865 | GTGCGTGTT |
| 11746 | mamo-s | *BGIBMGA013818* | up | 816 | 824 | - | 12.4679 | 1.31E-05 | 0.615 | GTGCGTGTC |
| 11747 | mamo-s | *BGIBMGA013819* | down | 1590 | 1598 | + | 11.3303 | 4.96E-05 | 0.615 | GTGGGTGGC |
| 11748 | mamo-s | *BGIBMGA013820* | down | 353 | 361 | - | 11.3303 | 4.96E-05 | 0.615 | GTGGGTGGC |
| 11749 | mamo-s | *BGIBMGA013821* | down | 1348 | 1356 | - | 11.3303 | 4.96E-05 | 0.615 | GTGGGTGGC |
| 11750 | mamo-s | *BGIBMGA013822* | up | 1548 | 1556 | - | 14.055 | 2.23E-06 | 0.615 | GTGCGTGGC |
| 11751 | mamo-s | *BGIBMGA013822* | down | 1337 | 1345 | + | 11.3303 | 4.96E-05 | 0.615 | GTGGGTGGC |
| 11752 | mamo-s | *BGIBMGA013823* | up | 1371 | 1379 | + | 14.055 | 2.23E-06 | 0.615 | GTGCGTGGC |
| 11753 | mamo-s | *BGIBMGA013823* | up | 310 | 318 | - | 12.1835 | 1.71E-05 | 0.615 | GTGCGTGCC |
| 11754 | mamo-s | *BGIBMGA013823* | up | 1107 | 1115 | - | 11.3303 | 4.96E-05 | 0.615 | GTGGGTGGC |
| 11755 | mamo-s | *BGIBMGA013823* | up | 1929 | 1937 | - | 11.0734 | 7.12E-05 | 0.726 | GAGCGTGGT |
| 11756 | mamo-s | *BGIBMGA013826* | up | 206 | 214 | - | 11.3303 | 4.96E-05 | 0.615 | GTGGGTGGC |
| 11757 | mamo-s | *BGIBMGA013826* | up | 966 | 974 | + | 11.3303 | 4.96E-05 | 0.615 | GTGGGTGGC |
| 11758 | mamo-s | *BGIBMGA013826* | up | 1857 | 1865 | + | 11.3303 | 4.96E-05 | 0.615 | GTGGGTGGC |
| 11759 | mamo-s | *BGIBMGA013827* | up | 469 | 477 | + | 11.3303 | 4.96E-05 | 0.615 | GTGGGTGGC |
| 11760 | mamo-s | *BGIBMGA013827* | down | 1657 | 1665 | - | 11.3303 | 4.96E-05 | 0.615 | GTGGGTGGC |
| 11761 | mamo-s | *BGIBMGA013829* | down | 772 | 780 | + | 11.3303 | 4.96E-05 | 0.615 | GTGGGTGGC |
| 11762 | mamo-s | *BGIBMGA013829* | up | 1208 | 1216 | + | 11.3303 | 4.96E-05 | 0.615 | GTGGGTGGC |
| 11763 | mamo-s | *BGIBMGA013830* | up | 1178 | 1186 | + | 11.3303 | 4.96E-05 | 0.615 | GTGGGTGGC |
| 11764 | mamo-s | *BGIBMGA013831* | down | 694 | 702 | + | 12.1284 | 2.16E-05 | 0.615 | GTGCGAGGC |
| 11765 | mamo-s | *BGIBMGA013831* | down | 772 | 780 | + | 11.0734 | 7.12E-05 | 0.726 | GAGCGTGGT |
| 11766 | mamo-s | *BGIBMGA013832* | up | 112 | 120 | + | 11.0734 | 7.12E-05 | 0.726 | GAGCGTGGT |
| 11767 | mamo-s | *BGIBMGA013833* | up | 1484 | 1492 | - | 10.9358 | 7.30E-05 | 0.728 | GCGGGTGGC |
| 11768 | mamo-s | *BGIBMGA013834* | up | 978 | 986 | - | 12.3394 | 1.49E-05 | 0.615 | GGGCGTGGC |
| 11769 | mamo-s | *BGIBMGA013834* | up | 1164 | 1172 | - | 11.5688 | 3.79E-05 | 0.615 | CTGCGTGTC |
| 11770 | mamo-s | *BGIBMGA013834* | down | 1123 | 1131 | + | 11.3303 | 4.96E-05 | 0.615 | GTGGGTGGC |
| 11771 | mamo-s | *BGIBMGA013835* | down | 1128 | 1136 | + | 12.4679 | 1.31E-05 | 0.615 | GTGCGTGTC |
| 11772 | mamo-s | *BGIBMGA013835* | down | 271 | 279 | - | 11.3303 | 4.96E-05 | 0.615 | GTGGGTGGC |
| 11773 | mamo-s | *BGIBMGA013836* | up | 1564 | 1572 | - | 11.0826 | 6.84E-05 | 0.709 | GTACGTGGC |
| 11774 | mamo-s | *BGIBMGA013837* | up | 256 | 264 | + | 11.3303 | 4.96E-05 | 0.615 | GTGGGTGGC |
| 11775 | mamo-s | *BGIBMGA013838* | up | 1023 | 1031 | - | 11.3303 | 4.96E-05 | 0.615 | GTGGGTGGC |
| 11776 | mamo-s | *BGIBMGA013839* | down | 830 | 838 | - | 11.3303 | 4.96E-05 | 0.615 | GTGGGTGGC |
| 11777 | mamo-s | *BGIBMGA013839* | up | 1362 | 1370 | - | 10.5688 | 8.88E-05 | 0.824 | GTGCGTGAC |
| 11778 | mamo-s | *BGIBMGA013840* | up | 1337 | 1345 | + | 11.1376 | 6.35E-05 | 0.709 | TCGCGTGGC |
| 11779 | mamo-s | *BGIBMGA013842* | down | 250 | 258 | + | 11.3303 | 4.96E-05 | 0.615 | GTGGGTGGC |
| 11780 | mamo-s | *BGIBMGA013843* | up | 1278 | 1286 | + | 11.3303 | 4.96E-05 | 0.615 | GTGGGTGGC |
| 11781 | mamo-s | *BGIBMGA013844* | up | 940 | 948 | - | 11.3303 | 4.96E-05 | 0.615 | GTGGGTGGC |
| 11782 | mamo-s | *BGIBMGA013844* | up | 1203 | 1211 | - | 11.3303 | 4.96E-05 | 0.615 | GTGGGTGGC |
| 11783 | mamo-s | *BGIBMGA013848* | up | 549 | 557 | + | 12.4679 | 1.31E-05 | 0.615 | GTGCGTGTC |
| 11784 | mamo-s | *BGIBMGA013848* | up | 584 | 592 | - | 11.4404 | 4.51E-05 | 0.615 | CGGCGTGGC |
| 11785 | mamo-s | *BGIBMGA013848* | up | 1637 | 1645 | + | 10.5688 | 8.88E-05 | 0.824 | GTGCGTGAC |

| | | | | | | | | | | |
|---|---|---|---|---|---|---|---|---|---|---|
| 11786 | mamo-s | *BGIBMGA013849* | up | 353 | 361 | + | 11.3303 | 4.96E-05 | 0.615 | GTGGGTGGC |
| 11787 | mamo-s | *BGIBMGA013853* | up | 392 | 400 | - | 11.0734 | 7.12E-05 | 0.726 | GAGCGTGGT |
| 11788 | mamo-s | *BGIBMGA013854* | down | 1449 | 1457 | - | 12.4679 | 1.31E-05 | 0.615 | GTGCGTGTC |
| 11789 | mamo-s | *BGIBMGA013854* | down | 293 | 301 | + | 11.3303 | 4.96E-05 | 0.615 | GTGGGTGGC |
| 11790 | mamo-s | *BGIBMGA013856* | up | 1738 | 1746 | + | 11.3303 | 4.96E-05 | 0.615 | GTGGGTGGC |
| 11791 | mamo-s | *BGIBMGA013856* | down | 1135 | 1143 | + | 11.0826 | 6.84E-05 | 0.709 | GTACGTGGC |
| 11792 | mamo-s | *BGIBMGA013857* | up | 515 | 523 | - | 11.3303 | 4.96E-05 | 0.615 | GTGGGTGGC |
| 11793 | mamo-s | *BGIBMGA013860* | up | 905 | 913 | - | 14.055 | 2.23E-06 | 0.615 | GTGCGTGGC |
| 11794 | mamo-s | *BGIBMGA013862* | down | 604 | 612 | - | 11.3303 | 4.96E-05 | 0.615 | GTGGGTGGC |
| 11795 | mamo-s | *BGIBMGA013862* | up | 394 | 402 | - | 10.8073 | 7.89E-05 | 0.766 | CCGCGTGGT |
| 11796 | mamo-s | *BGIBMGA013862* | up | 1510 | 1518 | - | 10.8073 | 7.89E-05 | 0.766 | CCGCGTGGT |
| 11797 | mamo-s | *BGIBMGA013863* | up | 322 | 330 | + | 11.3303 | 4.96E-05 | 0.615 | GTGGGTGGC |
| 11798 | mamo-s | *BGIBMGA013863* | up | 1284 | 1292 | + | 11.3303 | 4.96E-05 | 0.615 | GTGGGTGGC |
| 11799 | mamo-s | *BGIBMGA013864* | up | 743 | 751 | - | 11.3303 | 4.96E-05 | 0.615 | GTGGGTGGC |
| 11800 | mamo-s | *BGIBMGA013867* | down | 1570 | 1578 | + | 11.3303 | 4.96E-05 | 0.615 | GTGGGTGGC |
| 11801 | mamo-s | *BGIBMGA013867* | up | 1769 | 1777 | - | 11.3303 | 4.96E-05 | 0.615 | GTGGGTGGC |
| 11802 | mamo-s | *BGIBMGA013868* | down | 394 | 402 | - | 11.7064 | 3.52E-05 | 0.615 | GCGCGTGGT |
| 11803 | mamo-s | *BGIBMGA013868* | up | 822 | 830 | - | 11.3303 | 4.96E-05 | 0.615 | GTGGGTGGC |
| 11804 | mamo-s | *BGIBMGA013869* | up | 177 | 185 | - | 11.7064 | 3.52E-05 | 0.615 | GCGCGTGGT |
| 11805 | mamo-s | *BGIBMGA013872* | up | 162 | 170 | + | 11.3303 | 4.96E-05 | 0.615 | GTGGGTGGC |
| 11806 | mamo-s | *BGIBMGA013872* | down | 1493 | 1501 | + | 10.6881 | 8.61E-05 | 0.81 | GCACGTGGC |
| 11807 | mamo-s | *BGIBMGA013873* | down | 661 | 669 | - | 11.3303 | 4.96E-05 | 0.615 | GTGGGTGGC |
| 11808 | mamo-s | *BGIBMGA013873* | up | 1604 | 1612 | + | 11.3303 | 4.96E-05 | 0.615 | GTGGGTGGC |
| 11809 | mamo-s | *BGIBMGA013874* | down | 589 | 597 | + | 12.1009 | 2.43E-05 | 0.615 | GTGCGTGGT |
| 11810 | mamo-s | *BGIBMGA013874* | up | 933 | 941 | - | 11.3303 | 4.96E-05 | 0.615 | GTGGGTGGC |
| 11811 | mamo-s | *BGIBMGA013874* | down | 1846 | 1854 | + | 11.2844 | 5.18E-05 | 0.63 | CTGCGTGCC |
| 11812 | mamo-s | *BGIBMGA013875* | down | 1917 | 1925 | + | 11.3303 | 4.96E-05 | 0.615 | GTGGGTGGC |
| 11813 | mamo-s | *BGIBMGA013876* | up | 859 | 867 | + | 11.3303 | 4.96E-05 | 0.615 | GTGGGTGGC |
| 11814 | mamo-s | *BGIBMGA013878* | up | 1363 | 1371 | + | 11.3303 | 4.96E-05 | 0.615 | GTGGGTGGC |
| 11815 | mamo-s | *BGIBMGA013879* | down | 1639 | 1647 | + | 11.0826 | 6.84E-05 | 0.709 | GTACGTGGC |
| 11816 | mamo-s | *BGIBMGA013880* | up | 938 | 946 | + | 11.3303 | 4.96E-05 | 0.615 | GTGGGTGGC |
| 11817 | mamo-s | *BGIBMGA013881* | down | 1888 | 1896 | + | 12.4679 | 1.31E-05 | 0.615 | GTGCGTGTC |
| 11818 | mamo-s | *BGIBMGA013883* | up | 413 | 421 | + | 11.3303 | 4.96E-05 | 0.615 | GTGGGTGGC |
| 11819 | mamo-s | *BGIBMGA013884* | up | 519 | 527 | - | 13.156 | 6.30E-06 | 0.615 | CTGCGTGGC |
| 11820 | mamo-s | *BGIBMGA013884* | down | 1324 | 1332 | + | 11.3303 | 4.96E-05 | 0.615 | GTGGGTGGC |
| 11821 | mamo-s | *BGIBMGA013884* | down | 701 | 709 | + | 11.1009 | 6.57E-05 | 0.709 | GAGCGAGGC |
| 11822 | mamo-s | *BGIBMGA013884* | up | 1331 | 1339 | + | 10.5138 | 9.75E-05 | 0.865 | GTGCGTGTT |
| 11823 | mamo-s | *BGIBMGA013886* | up | 896 | 904 | + | 11.5321 | 4.06E-05 | 0.615 | TTGCGTGGC |
| 11824 | mamo-s | *BGIBMGA013887* | up | 1401 | 1409 | - | 11.3303 | 4.96E-05 | 0.615 | GTGGGTGGC |
| 11825 | mamo-s | *BGIBMGA013891* | down | 1961 | 1969 | - | 11.3303 | 4.96E-05 | 0.615 | GTGGGTGGC |
| 11826 | mamo-s | *BGIBMGA013892* | up | 1063 | 1071 | + | 11.3303 | 4.96E-05 | 0.615 | GTGGGTGGC |
| 11827 | mamo-s | *BGIBMGA013893* | down | 618 | 626 | + | 13.0275 | 8.53E-06 | 0.615 | GAGCGTGGC |
| 11828 | mamo-s | *BGIBMGA013894* | up | 1143 | 1151 | + | 13.0275 | 8.53E-06 | 0.615 | GAGCGTGGC |
| 11829 | mamo-s | *BGIBMGA013897* | up | 870 | 878 | + | 11.0734 | 7.12E-05 | 0.726 | GAGCGTGGT |
| 11830 | mamo-s | *BGIBMGA013899* | down | 1324 | 1332 | - | 11.5688 | 3.79E-05 | 0.615 | CTGCGTGTC |
| 11831 | mamo-s | *BGIBMGA013900* | up | 1895 | 1903 | - | 11.3303 | 4.96E-05 | 0.615 | GTGGGTGGC |
| 11832 | mamo-s | *BGIBMGA013901* | down | 1643 | 1651 | + | 11.0826 | 6.84E-05 | 0.709 | GTACGTGGC |
| 11833 | mamo-s | *BGIBMGA013901* | up | 1209 | 1217 | + | 10.9358 | 7.30E-05 | 0.728 | GCGGGTGGC |
| 11834 | mamo-s | *BGIBMGA013903* | down | 314 | 322 | + | 11.3303 | 4.96E-05 | 0.615 | GTGGGTGGC |
| 11835 | mamo-s | *BGIBMGA013903* | down | 1109 | 1117 | + | 11.3303 | 4.96E-05 | 0.615 | GTGGGTGGC |
| 11836 | mamo-s | *BGIBMGA013904* | up | 119 | 127 | + | 11.3303 | 4.96E-05 | 0.615 | GTGGGTGGC |
| 11837 | mamo-s | *BGIBMGA013904* | up | 914 | 922 | + | 11.3303 | 4.96E-05 | 0.615 | GTGGGTGGC |
| 11838 | mamo-s | *BGIBMGA013905* | up | 1702 | 1710 | - | 13.0275 | 8.53E-06 | 0.615 | GAGCGTGGC |
| 11839 | mamo-s | *BGIBMGA013906* | up | 642 | 650 | + | 11.3303 | 4.96E-05 | 0.615 | GTGGGTGGC |
| 11840 | mamo-s | *BGIBMGA013906* | down | 1035 | 1043 | - | 11.3303 | 4.96E-05 | 0.615 | GTGGGTGGC |
| 11841 | mamo-s | *BGIBMGA013908* | down | 612 | 620 | + | 11.3303 | 4.96E-05 | 0.615 | GTGGGTGGC |
| 11842 | mamo-s | *BGIBMGA013911* | up | 1488 | 1496 | + | 11.3303 | 4.96E-05 | 0.615 | GTGGGTGGC |
| 11843 | mamo-s | *BGIBMGA013912* | up | 1328 | 1336 | - | 11.3303 | 4.96E-05 | 0.615 | GTGGGTGGC |
| 11844 | mamo-s | *BGIBMGA013914* | up | 1446 | 1454 | - | 13.6606 | 4.06E-06 | 0.615 | GCGCGTGGC |
| 11845 | mamo-s | *BGIBMGA013914* | up | 147 | 155 | + | 11.3303 | 4.96E-05 | 0.615 | GTGGGTGGC |
| 11846 | mamo-s | *BGIBMGA013914* | up | 732 | 740 | - | 11.0826 | 6.84E-05 | 0.709 | GTACGTGGC |
| 11847 | mamo-s | *BGIBMGA013916* | down | 1153 | 1161 | - | 10.9358 | 7.30E-05 | 0.728 | GCGGGTGGC |
| 11848 | mamo-s | *BGIBMGA013918* | up | 766 | 774 | - | 11.5688 | 3.79E-05 | 0.615 | CTGCGTGTC |
| 11849 | mamo-s | *BGIBMGA013919* | down | 1188 | 1196 | + | 11.0826 | 6.84E-05 | 0.709 | GTACGTGGC |
| 11850 | mamo-s | *BGIBMGA013920* | up | 1934 | 1942 | - | 11.5321 | 4.06E-05 | 0.615 | TTGCGTGGC |
| 11851 | mamo-s | *BGIBMGA013920* | down | 307 | 315 | - | 11.3303 | 4.96E-05 | 0.615 | GTGGGTGGC |
| 11852 | mamo-s | *BGIBMGA013921* | up | 60 | 68 | + | 11.3303 | 4.96E-05 | 0.615 | GTGGGTGGC |
| 11853 | mamo-s | *BGIBMGA013924* | up | 16 | 24 | - | 11.3303 | 4.96E-05 | 0.615 | GTGGGTGGC |
| 11854 | mamo-s | *BGIBMGA013925* | up | 1794 | 1802 | + | 11.3303 | 4.96E-05 | 0.615 | GTGGGTGGC |
| 11855 | mamo-s | *BGIBMGA013926* | up | 737 | 745 | + | 11.3303 | 4.96E-05 | 0.615 | GTGGGTGGC |
| 11856 | mamo-s | *BGIBMGA013926* | up | 1054 | 1062 | - | 10.7523 | 8.38E-05 | 0.797 | GGGCGTGTC |
| 11857 | mamo-s | *BGIBMGA013927* | down | 1271 | 1279 | + | 11.3303 | 4.96E-05 | 0.615 | GTGGGTGGC |
| 11858 | mamo-s | *BGIBMGA013928* | up | 1897 | 1905 | + | 11.0826 | 6.84E-05 | 0.709 | GTACGTGGC |
| 11859 | mamo-s | *BGIBMGA013929* | down | 882 | 890 | + | 11.3303 | 4.96E-05 | 0.615 | GTGGGTGGC |
| 11860 | mamo-s | *BGIBMGA013929* | down | 1336 | 1344 | + | 11.3303 | 4.96E-05 | 0.615 | GTGGGTGGC |
| 11861 | mamo-s | *BGIBMGA013929* | down | 212 | 220 | + | 11.0826 | 6.84E-05 | 0.709 | GTACGTGGC |
| 11862 | mamo-s | *BGIBMGA013930* | down | 862 | 870 | - | 13.6606 | 4.06E-06 | 0.615 | GCGCGTGGC |
| 11863 | mamo-s | *BGIBMGA013931* | up | 1128 | 1136 | - | 11.3303 | 4.96E-05 | 0.615 | GTGGGTGGC |
| 11864 | mamo-s | *BGIBMGA013931* | up | 1833 | 1841 | - | 11.3303 | 4.96E-05 | 0.615 | GTGGGTGGC |
| 11865 | mamo-s | *BGIBMGA013931* | up | 1027 | 1035 | - | 10.5138 | 9.75E-05 | 0.865 | GTGCGTGTT |
| 11866 | mamo-s | *BGIBMGA013932* | up | 1793 | 1801 | - | 11.4404 | 4.51E-05 | 0.615 | GAGCGTGTC |
| 11867 | mamo-s | *BGIBMGA013934* | up | 622 | 630 | + | 11.7798 | 3.11E-05 | 0.615 | ATGCGTGGC |
| 11868 | mamo-s | *BGIBMGA013936* | down | 329 | 337 | + | 11.3303 | 4.96E-05 | 0.615 | GTGGGTGGC |
| 11869 | mamo-s | *BGIBMGA013936* | down | 981 | 989 | - | 11.3303 | 4.96E-05 | 0.615 | GTGGGTGGC |
| 11870 | mamo-s | *BGIBMGA013938* | up | 1531 | 1539 | - | 11.3853 | 4.74E-05 | 0.615 | ACGCGTGGC |
| 11871 | mamo-s | *BGIBMGA013939* | down | 1324 | 1332 | - | 11.0826 | 6.84E-05 | 0.709 | GTACGTGGC |
| 11872 | mamo-s | *BGIBMGA013939* | down | 1717 | 1725 | - | 10.7523 | 8.38E-05 | 0.797 | AAGCGTGGC |
| 11873 | mamo-s | *BGIBMGA013940* | up | 633 | 641 | + | 11.3303 | 4.96E-05 | 0.615 | GTGGGTGGC |

| | | | | | | | | | | |
|---|---|---|---|---|---|---|---|---|---|---|
| 11874 | mamo-s | *BGIBMGA013941* | up | 757 | 765 | - | 11.3303 | 4.96E-05 | 0.615 | GTGGGTGGC |
| 11875 | mamo-s | *BGIBMGA013941* | up | 153 | 161 | + | 11.0826 | 6.84E-05 | 0.709 | GTACGTGGC |
| 11876 | mamo-s | *BGIBMGA013941* | up | 502 | 510 | + | 10.6881 | 8.61E-05 | 0.81 | GCACGTGGC |
| 11877 | mamo-s | *BGIBMGA013942* | down | 1678 | 1686 | - | 10.8899 | 7.48E-05 | 0.739 | CCGCGTGCC |
| 11878 | mamo-s | *BGIBMGA013943* | up | 1251 | 1259 | - | 11.156 | 6.13E-05 | 0.705 | GAGCGTGCC |
| 11879 | mamo-s | *BGIBMGA013943* | up | 1625 | 1633 | - | 11.0826 | 6.84E-05 | 0.709 | GTACGTGGC |
| 11880 | mamo-s | *BGIBMGA013944* | down | 1045 | 1053 | - | 11.5688 | 3.79E-05 | 0.615 | CTGCGTGTC |
| 11881 | mamo-s | *BGIBMGA013946* | up | 470 | 478 | - | 11.4404 | 4.51E-05 | 0.615 | CGGCGTGGC |
| 11882 | mamo-s | *BGIBMGA013946* | up | 1515 | 1523 | - | 10.8899 | 7.48E-05 | 0.739 | CCGCGTGCC |
| 11883 | mamo-s | *BGIBMGA013947* | up | 1774 | 1782 | + | 11.3303 | 4.96E-05 | 0.615 | GTGGGTGGC |
| 11884 | mamo-s | *BGIBMGA013948* | down | 683 | 691 | + | 12.1835 | 1.71E-05 | 0.615 | GTGCGTGCC |
| 11885 | mamo-s | *BGIBMGA013948* | up | 1636 | 1644 | + | 11.1743 | 5.90E-05 | 0.684 | CCGCGTGTC |
| 11886 | mamo-s | *BGIBMGA013948* | up | 108 | 116 | - | 10.5688 | 8.88E-05 | 0.824 | GTGCGTGAC |
| 11887 | mamo-s | *BGIBMGA013950* | down | 1943 | 1951 | - | 11.3303 | 4.96E-05 | 0.615 | GTGGGTGGC |
| 11888 | mamo-s | *BGIBMGA013951* | up | 964 | 972 | - | 11.3303 | 4.96E-05 | 0.615 | GTGGGTGGC |
| 11889 | mamo-s | *BGIBMGA013952* | up | 1664 | 1672 | + | 11.7798 | 3.11E-05 | 0.615 | ATGCGTGGC |
| 11890 | mamo-s | *BGIBMGA013952* | up | 1793 | 1801 | + | 11.7064 | 3.52E-05 | 0.615 | GCGCGTGGT |
| 11891 | mamo-s | *BGIBMGA013952* | up | 47 | 55 | + | 11.3303 | 4.96E-05 | 0.615 | GTGGGTGGC |
| 11892 | mamo-s | *BGIBMGA013952* | down | 1280 | 1288 | - | 11.3303 | 4.96E-05 | 0.615 | GTGGGTGGC |
| 11893 | mamo-s | *BGIBMGA013953* | up | 408 | 416 | + | 11.3303 | 4.96E-05 | 0.615 | GTGGGTGGC |
| 11894 | mamo-s | *BGIBMGA013953* | up | 1187 | 1195 | + | 11.3303 | 4.96E-05 | 0.615 | GTGGGTGGC |
| 11895 | mamo-s | *BGIBMGA013954* | up | 1968 | 1976 | + | 12.1284 | 2.16E-05 | 0.615 | GTGCGAGGC |
| 11896 | mamo-s | *BGIBMGA013954* | up | 1978 | 1986 | + | 12.1284 | 2.16E-05 | 0.615 | GTGCGAGGC |
| 11897 | mamo-s | *BGIBMGA013954* | down | 1734 | 1742 | + | 11.3303 | 4.96E-05 | 0.615 | GTGGGTGGC |
| 11898 | mamo-s | *BGIBMGA013955* | up | 243 | 251 | - | 11.1009 | 6.57E-05 | 0.709 | GAGCGAGGC |
| 11899 | mamo-s | *BGIBMGA013956* | up | 1385 | 1393 | - | 11.0826 | 6.84E-05 | 0.709 | GTACGTGGC |
| 11900 | mamo-s | *BGIBMGA013957* | down | 534 | 542 | + | 11.3303 | 4.96E-05 | 0.615 | GTGGGTGGC |
| 11901 | mamo-s | *BGIBMGA013957* | down | 1467 | 1475 | - | 11.3303 | 4.96E-05 | 0.615 | GTGGGTGGC |
| 11902 | mamo-s | *BGIBMGA013961* | down | 31 | 39 | + | 11.3303 | 4.96E-05 | 0.615 | GTGGGTGGC |
| 11903 | mamo-s | *BGIBMGA013961* | up | 503 | 511 | - | 11.3303 | 4.96E-05 | 0.615 | GTGGGTGGC |
| 11904 | mamo-s | *BGIBMGA013961* | down | 1523 | 1531 | + | 10.8073 | 7.89E-05 | 0.766 | CCGCGTGGT |
| 11905 | mamo-s | *BGIBMGA013962* | down | 154 | 162 | + | 11.3303 | 4.96E-05 | 0.615 | GTGGGTGGC |
| 11906 | mamo-s | *BGIBMGA013962* | up | 956 | 964 | - | 11.3303 | 4.96E-05 | 0.615 | GTGGGTGGC |
| 11907 | mamo-s | *BGIBMGA013962* | down | 1102 | 1110 | - | 11.3303 | 4.96E-05 | 0.615 | GTGGGTGGC |
| 11908 | mamo-s | *BGIBMGA013962* | up | 1653 | 1661 | + | 11.3303 | 4.96E-05 | 0.615 | GTGGGTGGC |
| 11909 | mamo-s | *BGIBMGA013963* | up | 1922 | 1930 | + | 12.1835 | 1.71E-05 | 0.615 | GTGCGTGCC |
| 11910 | mamo-s | *BGIBMGA013963* | up | 1749 | 1757 | + | 10.5688 | 8.88E-05 | 0.824 | GTGCGTGAC |
| 11911 | mamo-s | *BGIBMGA013964* | down | 1204 | 1212 | - | 11.3303 | 4.96E-05 | 0.615 | GTGGGTGGC |
| 11912 | mamo-s | *BGIBMGA013964* | up | 1808 | 1816 | + | 11.3303 | 4.96E-05 | 0.615 | GTGGGTGGC |
| 11913 | mamo-s | *BGIBMGA013965* | down | 737 | 745 | + | 14.055 | 2.23E-06 | 0.615 | GTGCGTGGC |
| 11914 | mamo-s | *BGIBMGA013965* | up | 547 | 555 | - | 11.3303 | 4.96E-05 | 0.615 | GTGGGTGGC |
| 11915 | mamo-s | *BGIBMGA013965* | up | 1560 | 1568 | + | 11.3303 | 4.96E-05 | 0.615 | GTGGGTGGC |
| 11916 | mamo-s | *BGIBMGA013966* | up | 1286 | 1294 | - | 11.3303 | 4.96E-05 | 0.615 | GTGGGTGGC |
| 11917 | mamo-s | *BGIBMGA013967* | down | 296 | 304 | + | 11.3303 | 4.96E-05 | 0.615 | GTGGGTGGC |
| 11918 | mamo-s | *BGIBMGA013967* | up | 846 | 854 | - | 11.3303 | 4.96E-05 | 0.615 | GTGGGTGGC |
| 11919 | mamo-s | *BGIBMGA013968* | up | 57 | 65 | - | 11.3303 | 4.96E-05 | 0.615 | GTGGGTGGC |
| 11920 | mamo-s | *BGIBMGA013968* | down | 1725 | 1733 | + | 11.3303 | 4.96E-05 | 0.615 | GTGGGTGGC |
| 11921 | mamo-s | *BGIBMGA013968* | up | 802 | 810 | + | 10.9358 | 7.30E-05 | 0.728 | GCGGGTGGC |
| 11922 | mamo-s | *BGIBMGA013971* | up | 118 | 126 | + | 11.3303 | 4.96E-05 | 0.615 | GTGGGTGGC |
| 11923 | mamo-s | *BGIBMGA013975* | up | 1168 | 1176 | + | 11.3303 | 4.96E-05 | 0.615 | GTGGGTGGC |
| 11924 | mamo-s | *BGIBMGA013976* | up | 733 | 741 | - | 11.3303 | 4.96E-05 | 0.615 | GTGGGTGGC |
| 11925 | mamo-s | *BGIBMGA013977* | up | 547 | 555 | + | 10.8073 | 7.89E-05 | 0.766 | CCGCGTGGT |
| 11926 | mamo-s | *BGIBMGA013978* | up | 1233 | 1241 | - | 11.7798 | 3.11E-05 | 0.615 | ATGCGTGGC |
| 11927 | mamo-s | *BGIBMGA013978* | up | 1529 | 1537 | - | 11.7064 | 3.52E-05 | 0.615 | GCGCGTGGT |
| 11928 | mamo-s | *BGIBMGA013980* | up | 1447 | 1455 | + | 11.3303 | 4.96E-05 | 0.615 | GTGGGTGGC |
| 11929 | mamo-s | *BGIBMGA013980* | up | 179 | 187 | - | 11.2844 | 5.18E-05 | 0.63 | CTGCGTGCC |
| 11930 | mamo-s | *BGIBMGA013982* | up | 583 | 591 | + | 11.3303 | 4.96E-05 | 0.615 | GTGGGTGGC |
| 11931 | mamo-s | *BGIBMGA013984* | up | 721 | 729 | - | 11.3303 | 4.96E-05 | 0.615 | GTGGGTGGC |
| 11932 | mamo-s | *BGIBMGA013984* | up | 1734 | 1742 | + | 11.3303 | 4.96E-05 | 0.615 | GTGGGTGGC |
| 11933 | mamo-s | *BGIBMGA013985* | up | 345 | 353 | - | 11.3303 | 4.96E-05 | 0.615 | GTGGGTGGC |
| 11934 | mamo-s | *BGIBMGA013985* | down | 1536 | 1544 | + | 11.1743 | 5.90E-05 | 0.684 | CCGCGTGTC |
| 11935 | mamo-s | *BGIBMGA013987* | down | 906 | 914 | - | 11.3303 | 4.96E-05 | 0.615 | GTGGGTGGC |
| 11936 | mamo-s | *BGIBMGA013988* | up | 605 | 613 | + | 10.7523 | 8.38E-05 | 0.797 | AAGCGTGGC |
| 11937 | mamo-s | *BGIBMGA013989* | up | 1921 | 1929 | + | 11.7064 | 3.52E-05 | 0.615 | GCGCGTGGT |
| 11938 | mamo-s | *BGIBMGA013989* | up | 1904 | 1912 | + | 11.4404 | 4.51E-05 | 0.615 | CGGCGTGGC |
| 11939 | mamo-s | *BGIBMGA013990* | down | 1418 | 1426 | - | 10.5688 | 8.88E-05 | 0.824 | GTGCGTGAC |
| 11940 | mamo-s | *BGIBMGA013991* | up | 346 | 354 | - | 11.3303 | 4.96E-05 | 0.615 | GTGGGTGGC |
| 11941 | mamo-s | *BGIBMGA013992* | down | 195 | 203 | + | 11.3303 | 4.96E-05 | 0.615 | GTGGGTGGC |
| 11942 | mamo-s | *BGIBMGA013994* | up | 672 | 680 | + | 11.3303 | 4.96E-05 | 0.615 | GTGGGTGGC |
| 11943 | mamo-s | *BGIBMGA013994* | up | 1479 | 1487 | - | 11.2844 | 5.18E-05 | 0.63 | CTGCGTGCC |
| 11944 | mamo-s | *BGIBMGA013995* | up | 1079 | 1087 | - | 11.3303 | 4.96E-05 | 0.615 | GTGGGTGGC |
| 11945 | mamo-s | *BGIBMGA014000* | up | 1803 | 1811 | + | 11.3303 | 4.96E-05 | 0.615 | GTGGGTGGC |
| 11946 | mamo-s | *BGIBMGA014000* | down | 633 | 641 | - | 10.5138 | 9.75E-05 | 0.865 | GTGCGTGTT |
| 11947 | mamo-s | *BGIBMGA014001* | up | 65 | 73 | - | 11.3303 | 4.96E-05 | 0.615 | GTGGGTGGC |
| 11948 | mamo-s | *BGIBMGA014001* | down | 539 | 547 | + | 11.3303 | 4.96E-05 | 0.615 | GTGGGTGGC |
| 11949 | mamo-s | *BGIBMGA014001* | up | 1905 | 1913 | - | 10.5138 | 9.75E-05 | 0.865 | GTGCGTGTT |
| 11950 | mamo-s | *BGIBMGA014002* | up | 1738 | 1746 | + | 11.3303 | 4.96E-05 | 0.615 | GTGGGTGGC |
| 11951 | mamo-s | *BGIBMGA014002* | down | 1766 | 1774 | - | 11.3303 | 4.96E-05 | 0.615 | GTGGGTGGC |
| 11952 | mamo-s | *BGIBMGA014003* | down | 1338 | 1346 | + | 11.2294 | 5.41E-05 | 0.65 | CTGCGAGGC |
| 11953 | mamo-s | *BGIBMGA014003* | down | 495 | 503 | + | 11.0826 | 6.84E-05 | 0.709 | GTACGTGGC |
| 11954 | mamo-s | *BGIBMGA014003* | down | 826 | 834 | - | 10.8899 | 7.48E-05 | 0.739 | CCGCGTGCC |
| 11955 | mamo-s | *BGIBMGA014005* | down | 117 | 125 | - | 11.3303 | 4.96E-05 | 0.615 | GTGGGTGGC |
| 11956 | mamo-s | *BGIBMGA014005* | down | 696 | 704 | + | 11.3303 | 4.96E-05 | 0.615 | GTGGGTGGC |
| 11957 | mamo-s | *BGIBMGA014006* | down | 404 | 412 | - | 11.7339 | 3.29E-05 | 0.615 | GCGCGAGGC |
| 11958 | mamo-s | *BGIBMGA014006* | up | 530 | 538 | + | 11.3303 | 4.96E-05 | 0.615 | GTGGGTGGC |
| 11959 | mamo-s | *BGIBMGA014006* | down | 1571 | 1579 | - | 10.5138 | 9.75E-05 | 0.865 | GTGCGTGTT |
| 11960 | mamo-s | *BGIBMGA014007* | down | 1521 | 1529 | + | 13.6606 | 4.06E-06 | 0.615 | GCGCGTGGC |
| 11961 | mamo-s | *BGIBMGA014007* | up | 987 | 995 | - | 11.7798 | 3.11E-05 | 0.615 | ATGCGTGGC |

| | | | | | | | | | | |
|---|---|---|---|---|---|---|---|---|---|---|
| 11962 | mamo-s | *BGIBMGA014008* | down | 1695 | 1703 | - | 11.3303 | 4.96E-05 | 0.615 | GTGGGTGGC |
| 11963 | mamo-s | *BGIBMGA014009* | down | 271 | 279 | + | 11.3303 | 4.96E-05 | 0.615 | GTGGGTGGC |
| 11964 | mamo-s | *BGIBMGA014010* | up | 906 | 914 | - | 11.3303 | 4.96E-05 | 0.615 | GTGGGTGGC |
| 11965 | mamo-s | *BGIBMGA014011* | up | 315 | 323 | - | 11.789 | 2.84E-05 | 0.615 | GCGCGTGCC |
| 11966 | mamo-s | *BGIBMGA014011* | up | 392 | 400 | - | 11.0826 | 6.84E-05 | 0.709 | GTACGTGGC |
| 11967 | mamo-s | *BGIBMGA014013* | down | 566 | 574 | + | 11.3303 | 4.96E-05 | 0.615 | GTGGGTGGC |
| 11968 | mamo-s | *BGIBMGA014014* | up | 1793 | 1801 | + | 10.5688 | 8.88E-05 | 0.824 | GTGCGTGAC |
| 11969 | mamo-s | *BGIBMGA014016* | down | 1343 | 1351 | - | 11.3303 | 4.96E-05 | 0.615 | GTGGGTGGC |
| 11970 | mamo-s | *BGIBMGA014018* | down | 1552 | 1560 | + | 11.3303 | 4.96E-05 | 0.615 | GTGGGTGGC |
| 11971 | mamo-s | *BGIBMGA014019* | up | 382 | 390 | + | 11.3853 | 4.74E-05 | 0.615 | ACGCGTGGC |
| 11972 | mamo-s | *BGIBMGA014019* | up | 1310 | 1318 | + | 11.3303 | 4.96E-05 | 0.615 | GTGGGTGGC |
| 11973 | mamo-s | *BGIBMGA014020* | up | 767 | 775 | - | 12.3394 | 1.49E-05 | 0.615 | GGGCGTGGC |
| 11974 | mamo-s | *BGIBMGA014021* | down | 1811 | 1819 | + | 12.3394 | 1.49E-05 | 0.615 | GGGCGTGGC |
| 11975 | mamo-s | *BGIBMGA014021* | down | 1832 | 1840 | - | 12.0734 | 2.66E-05 | 0.615 | GCGCGTGTC |
| 11976 | mamo-s | *BGIBMGA014022* | down | 492 | 500 | + | 11.3303 | 4.96E-05 | 0.615 | GTGGGTGGC |
| 11977 | mamo-s | *BGIBMGA014023* | down | 513 | 521 | - | 13.6606 | 4.06E-06 | 0.615 | GCGCGTGGC |
| 11978 | mamo-s | *BGIBMGA014023* | down | 950 | 958 | - | 11.3303 | 4.96E-05 | 0.615 | GTGGGTGGC |
| 11979 | mamo-s | *BGIBMGA014024* | down | 1830 | 1838 | - | 11.7798 | 3.11E-05 | 0.615 | ATGCGTGGC |
| 11980 | mamo-s | *BGIBMGA014025* | down | 1180 | 1188 | - | 12.3394 | 1.49E-05 | 0.615 | GGGCGTGGC |
| 11981 | mamo-s | *BGIBMGA014025* | up | 965 | 973 | - | 11.1743 | 5.90E-05 | 0.684 | CCGCGTGTC |
| 11982 | mamo-s | *BGIBMGA014026* | down | 734 | 742 | + | 11.5321 | 4.06E-05 | 0.615 | TTGCGTGGC |
| 11983 | mamo-s | *BGIBMGA014026* | up | 1485 | 1493 | - | 11.3303 | 4.96E-05 | 0.615 | GTGGGTGGC |
| 11984 | mamo-s | *BGIBMGA014029* | down | 925 | 933 | + | 12.1009 | 2.43E-05 | 0.615 | GTGCGTGGT |
| 11985 | mamo-s | *BGIBMGA014030* | up | 1187 | 1195 | + | 11.3303 | 4.96E-05 | 0.615 | GTGGGTGGC |
| 11986 | mamo-s | *BGIBMGA014031* | up | 1190 | 1198 | + | 11.3303 | 4.96E-05 | 0.615 | GTGGGTGGC |
| 11987 | mamo-s | *BGIBMGA014033* | down | 1983 | 1991 | - | 12.0734 | 2.66E-05 | 0.615 | GCGCGTGTC |
| 11988 | mamo-s | *BGIBMGA014033* | up | 57 | 65 | - | 11.3303 | 4.96E-05 | 0.615 | GTGGGTGGC |
| 11989 | mamo-s | *BGIBMGA014034* | down | 420 | 428 | + | 12.1009 | 2.43E-05 | 0.615 | GTGCGTGGT |
| 11990 | mamo-s | *BGIBMGA014035* | up | 1359 | 1367 | + | 10.5138 | 9.75E-05 | 0.865 | GTGCGTGTT |
| 11991 | mamo-s | *BGIBMGA014037* | down | 282 | 290 | + | 11.3303 | 4.96E-05 | 0.615 | GTGGGTGGC |
| 11992 | mamo-s | *BGIBMGA014038* | up | 1113 | 1121 | - | 11.3303 | 4.96E-05 | 0.615 | GTGGGTGGC |
| 11993 | mamo-s | *BGIBMGA014038* | down | 1327 | 1335 | + | 10.5138 | 9.75E-05 | 0.865 | GTGCGTGTT |
| 11994 | mamo-s | *BGIBMGA014039* | up | 325 | 333 | + | 10.9358 | 7.30E-05 | 0.728 | GCGGGTGGC |
| 11995 | mamo-s | *BGIBMGA014040* | up | 972 | 980 | - | 14.055 | 2.23E-06 | 0.615 | GTGCGTGGC |
| 11996 | mamo-s | *BGIBMGA014046* | up | 1021 | 1029 | - | 11.3303 | 4.96E-05 | 0.615 | GTGGGTGGC |
| 11997 | mamo-s | *BGIBMGA014049* | up | 245 | 253 | + | 11.3303 | 4.96E-05 | 0.615 | GTGGGTGGC |
| 11998 | mamo-s | *BGIBMGA014049* | up | 912 | 920 | - | 11.3303 | 4.96E-05 | 0.615 | GTGGGTGGC |
| 11999 | mamo-s | *BGIBMGA014049* | up | 1490 | 1498 | - | 11.3303 | 4.96E-05 | 0.615 | GTGGGTGGC |
| 12000 | mamo-s | *BGIBMGA014052* | down | 1521 | 1529 | - | 11.3303 | 4.96E-05 | 0.615 | GTGGGTGGC |
| 12001 | mamo-s | *BGIBMGA014060* | up | 367 | 375 | - | 14.055 | 2.23E-06 | 0.615 | GTGCGTGGC |
| 12002 | mamo-s | *BGIBMGA014065* | up | 1028 | 1036 | + | 11.3303 | 4.96E-05 | 0.615 | GTGGGTGGC |
| 12003 | mamo-s | *BGIBMGA014066* | up | 1605 | 1613 | - | 11.3303 | 4.96E-05 | 0.615 | GTGGGTGGC |
| 12004 | mamo-s | *BGIBMGA014069* | up | 1926 | 1934 | + | 12.1009 | 2.43E-05 | 0.615 | GTGCGTGGT |
| 12005 | mamo-s | *BGIBMGA014069* | down | 435 | 443 | + | 11.3303 | 4.96E-05 | 0.615 | GTGGGTGGC |
| 12006 | mamo-s | *BGIBMGA014069* | down | 1627 | 1635 | + | 11.3303 | 4.96E-05 | 0.615 | GTGGGTGGC |
| 12007 | mamo-s | *BGIBMGA014069* | down | 21 | 29 | - | 11.1743 | 5.90E-05 | 0.684 | CCGCGTGTC |
| 12008 | mamo-s | *BGIBMGA014069* | down | 139 | 147 | - | 10.7523 | 8.38E-05 | 0.797 | GGGCGTGTC |
| 12009 | mamo-s | *BGIBMGA014074* | down | 259 | 267 | + | 11.3303 | 4.96E-05 | 0.615 | GTGGGTGGC |
| 12010 | mamo-s | *BGIBMGA014075* | down | 363 | 371 | - | 11.3303 | 4.96E-05 | 0.615 | GTGGGTGGC |
| 12011 | mamo-s | *BGIBMGA014075* | up | 527 | 535 | + | 11.3303 | 4.96E-05 | 0.615 | GTGGGTGGC |
| 12012 | mamo-s | *BGIBMGA014076* | up | 274 | 282 | + | 11.3303 | 4.96E-05 | 0.615 | GTGGGTGGC |
| 12013 | mamo-s | *BGIBMGA014076* | up | 1526 | 1534 | - | 10.7523 | 8.38E-05 | 0.797 | AAGCGTGGC |
| 12014 | mamo-s | *BGIBMGA014077* | up | 1903 | 1911 | + | 11.3303 | 4.96E-05 | 0.615 | GTGGGTGGC |
| 12015 | mamo-s | *BGIBMGA014077* | up | 1644 | 1652 | - | 11.0826 | 6.84E-05 | 0.709 | GTACGTGGC |
| 12016 | mamo-s | *BGIBMGA014079* | up | 45 | 53 | + | 11.3303 | 4.96E-05 | 0.615 | GTGGGTGGC |
| 12017 | mamo-s | *BGIBMGA014081* | up | 174 | 182 | + | 11.1376 | 6.35E-05 | 0.709 | TCGCGTGGC |
| 12018 | mamo-s | *BGIBMGA014082* | down | 485 | 493 | + | 11.0826 | 6.84E-05 | 0.709 | GTACGTGGC |
| 12019 | mamo-s | *BGIBMGA014084* | down | 509 | 517 | - | 14.055 | 2.23E-06 | 0.615 | GTGCGTGGC |
| 12020 | mamo-s | *BGIBMGA014084* | up | 1014 | 1022 | - | 11.3303 | 4.96E-05 | 0.615 | GTGGGTGGC |
| 12021 | mamo-s | *BGIBMGA014088* | down | 112 | 120 | + | 11.1376 | 6.35E-05 | 0.709 | TCGCGTGGC |
| 12022 | mamo-s | *BGIBMGA014088* | down | 210 | 218 | + | 11.1376 | 6.35E-05 | 0.709 | TCGCGTGGC |
| 12023 | mamo-s | *BGIBMGA014088* | down | 308 | 316 | + | 11.1376 | 6.35E-05 | 0.709 | TCGCGTGGC |
| 12024 | mamo-s | *BGIBMGA014088* | down | 406 | 414 | + | 11.1376 | 6.35E-05 | 0.709 | TCGCGTGGC |
| 12025 | mamo-s | *BGIBMGA014088* | down | 504 | 512 | + | 11.1376 | 6.35E-05 | 0.709 | TCGCGTGGC |
| 12026 | mamo-s | *BGIBMGA014089* | up | 84 | 92 | - | 11.3303 | 4.96E-05 | 0.615 | GTGGGTGGC |
| 12027 | mamo-s | *BGIBMGA014089* | up | 1933 | 1941 | + | 10.7523 | 8.38E-05 | 0.797 | AAGCGTGGC |
| 12028 | mamo-s | *BGIBMGA014092* | up | 898 | 906 | - | 11.3303 | 4.96E-05 | 0.615 | GTGGGTGGC |
| 12029 | mamo-s | *BGIBMGA014095* | up | 519 | 527 | + | 11.3303 | 4.96E-05 | 0.615 | GTGGGTGGC |
| 12030 | mamo-s | *BGIBMGA014096* | down | 878 | 886 | - | 14.055 | 2.23E-06 | 0.615 | GTGCGTGGC |
| 12031 | mamo-s | *BGIBMGA014096* | down | 1717 | 1725 | + | 11.3303 | 4.96E-05 | 0.615 | GTGGGTGGC |
| 12032 | mamo-s | *BGIBMGA014097* | up | 821 | 829 | - | 12.1284 | 2.16E-05 | 0.615 | GTGCGAGGC |
| 12033 | mamo-s | *BGIBMGA014097* | up | 300 | 308 | + | 11.3303 | 4.96E-05 | 0.615 | GTGGGTGGC |
| 12034 | mamo-s | *BGIBMGA014097* | up | 488 | 496 | - | 11.0734 | 7.12E-05 | 0.726 | GAGCGTGGT |
| 12035 | mamo-s | *BGIBMGA014097* | down | 1037 | 1045 | + | 10.8899 | 7.48E-05 | 0.739 | CCGCGTGCC |
| 12036 | mamo-s | *BGIBMGA014098* | up | 1455 | 1463 | - | 11.3303 | 4.96E-05 | 0.615 | GTGGGTGGC |
| 12037 | mamo-s | *BGIBMGA014100* | down | 10 | 18 | - | 11.1376 | 6.35E-05 | 0.709 | TCGCGTGGC |
| 12038 | mamo-s | *BGIBMGA014101* | down | 160 | 168 | - | 11.3303 | 4.96E-05 | 0.615 | GTGGGTGGC |
| 12039 | mamo-s | *BGIBMGA014105* | up | 910 | 918 | - | 11.3303 | 4.96E-05 | 0.615 | GTGGGTGGC |
| 12040 | mamo-s | *BGIBMGA014107* | up | 931 | 939 | + | 11.3303 | 4.96E-05 | 0.615 | GTGGGTGGC |
| 12041 | mamo-s | *BGIBMGA014108* | down | 1837 | 1845 | - | 11.3303 | 4.96E-05 | 0.615 | GTGGGTGGC |
| 12042 | mamo-s | *BGIBMGA014112* | down | 175 | 183 | + | 11.3303 | 4.96E-05 | 0.615 | GTGGGTGGC |
| 12043 | mamo-s | *BGIBMGA014113* | up | 878 | 886 | - | 11.1009 | 6.57E-05 | 0.709 | GAGCGAGGC |
| 12044 | mamo-s | *BGIBMGA014114* | down | 240 | 248 | - | 12.7615 | 1.04E-05 | 0.615 | CCGCGTGGC |
| 12045 | mamo-s | *BGIBMGA014114* | up | 1782 | 1790 | - | 12.1009 | 2.43E-05 | 0.615 | GTGCGTGGT |
| 12046 | mamo-s | *BGIBMGA014114* | down | 531 | 539 | + | 11.3303 | 4.96E-05 | 0.615 | GTGGGTGGC |
| 12047 | mamo-s | *BGIBMGA014118* | up | 1758 | 1766 | + | 12.0734 | 2.66E-05 | 0.615 | GCGCGTGTC |
| 12048 | mamo-s | *BGIBMGA014121* | up | 21 | 29 | - | 12.0734 | 2.66E-05 | 0.615 | GCGCGTGTC |
| 12049 | mamo-s | *BGIBMGA014122* | down | 300 | 308 | + | 11.3303 | 4.96E-05 | 0.615 | GTGGGTGGC |

| | | | | | | | | | | |
|---|---|---|---|---|---|---|---|---|---|---|
| 12050 | mamo-s | *BGIBMGA014122* | up | 1727 | 1735 | + | 10.7523 | 8.38E-05 | 0.797 | GGGCGTGTC |
| 12051 | mamo-s | *BGIBMGA014127* | down | 727 | 735 | - | 12.3394 | 1.49E-05 | 0.615 | GGGCGTGGC |
| 12052 | mamo-s | *BGIBMGA014128* | down | 885 | 893 | - | 11.3303 | 4.96E-05 | 0.615 | GTGGGTGGC |
| 12053 | mamo-s | *BGIBMGA014129* | down | 1844 | 1852 | + | 11.5688 | 3.79E-05 | 0.615 | CTGCGTGTC |
| 12054 | mamo-s | *BGIBMGA014129* | down | 694 | 702 | + | 11.3303 | 4.96E-05 | 0.615 | GTGGGTGGC |
| 12055 | mamo-s | *BGIBMGA014133* | down | 264 | 272 | - | 10.6881 | 8.61E-05 | 0.81 | GCACGTGGC |
| 12056 | mamo-s | *BGIBMGA014138* | up | 1172 | 1180 | + | 11.156 | 6.13E-05 | 0.705 | GAGCGTGCC |
| 12057 | mamo-s | *BGIBMGA014138* | up | 1314 | 1322 | + | 10.5138 | 9.75E-05 | 0.865 | GTGCGTGTT |
| 12058 | mamo-s | *BGIBMGA014139* | down | 110 | 118 | + | 12.1009 | 2.43E-05 | 0.615 | GTGCGTGGT |
| 12059 | mamo-s | *BGIBMGA014140* | down | 1974 | 1982 | + | 13.6606 | 4.06E-06 | 0.615 | GCGCGTGGC |
| 12060 | mamo-s | *BGIBMGA014140* | up | 1602 | 1610 | + | 11.3303 | 4.96E-05 | 0.615 | GTGGGTGGC |
| 12061 | mamo-s | *BGIBMGA014141* | down | 833 | 841 | - | 11.0826 | 6.84E-05 | 0.709 | GTACGTGGC |
| 12062 | mamo-s | *BGIBMGA014142* | up | 661 | 669 | - | 11.0826 | 6.84E-05 | 0.709 | GTACGTGGC |
| 12063 | mamo-s | *BGIBMGA014143* | up | 226 | 234 | - | 12.7615 | 1.04E-05 | 0.615 | CCGCGTGGC |
| 12064 | mamo-s | *BGIBMGA014143* | down | 845 | 853 | + | 11.3303 | 4.96E-05 | 0.615 | GTGGGTGGC |
| 12065 | mamo-s | *BGIBMGA014144* | up | 1362 | 1370 | + | 11.3303 | 4.96E-05 | 0.615 | GTGGGTGGC |
| 12066 | mamo-s | *BGIBMGA014146* | up | 1614 | 1622 | + | 11.3303 | 4.96E-05 | 0.615 | GTGGGTGGC |
| 12067 | mamo-s | *BGIBMGA014147* | down | 1621 | 1629 | + | 11.3303 | 4.96E-05 | 0.615 | GTGGGTGGC |
| 12068 | mamo-s | *BGIBMGA014148* | up | 102 | 110 | + | 11.3303 | 4.96E-05 | 0.615 | GTGGGTGGC |
| 12069 | mamo-s | *BGIBMGA014148* | down | 424 | 432 | + | 10.6881 | 8.61E-05 | 0.81 | GCACGTGGC |
| 12070 | mamo-s | *BGIBMGA014149* | up | 1539 | 1547 | - | 11.3303 | 4.96E-05 | 0.615 | GTGGGTGGC |
| 12071 | mamo-s | *BGIBMGA014149* | up | 1733 | 1741 | + | 11.3303 | 4.96E-05 | 0.615 | GTGGGTGGC |
| 12072 | mamo-s | *BGIBMGA014150* | down | 1146 | 1154 | + | 11.3303 | 4.96E-05 | 0.615 | GTGGGTGGC |
| 12073 | mamo-s | *BGIBMGA014151* | up | 586 | 594 | + | 12.1009 | 2.43E-05 | 0.615 | GTGCGTGGT |
| 12074 | mamo-s | *BGIBMGA014151* | down | 1011 | 1019 | - | 11.3303 | 4.96E-05 | 0.615 | GTGGGTGGC |
| 12075 | mamo-s | *BGIBMGA014151* | up | 1516 | 1524 | + | 11.3303 | 4.96E-05 | 0.615 | GTGGGTGGC |
| 12076 | mamo-s | *BGIBMGA014151* | down | 1917 | 1925 | - | 11.3303 | 4.96E-05 | 0.615 | GTGGGTGGC |
| 12077 | mamo-s | *BGIBMGA014152* | up | 1436 | 1444 | + | 11.3303 | 4.96E-05 | 0.615 | GTGGGTGGC |
| 12078 | mamo-s | *BGIBMGA014153* | up | 446 | 454 | + | 11.0826 | 6.84E-05 | 0.709 | GTACGTGGC |
| 12079 | mamo-s | *BGIBMGA014156* | up | 47 | 55 | - | 12.3394 | 1.49E-05 | 0.615 | GGGCGTGGC |
| 12080 | mamo-s | *BGIBMGA014159* | up | 578 | 586 | + | 12.4679 | 1.31E-05 | 0.615 | GTGCGTGTC |
| 12081 | mamo-s | *BGIBMGA014159* | down | 1144 | 1152 | - | 12.1835 | 1.71E-05 | 0.615 | GTGCGTGCC |
| 12082 | mamo-s | *BGIBMGA014159* | down | 1719 | 1727 | - | 11.3303 | 4.96E-05 | 0.615 | GTGGGTGGC |
| 12083 | mamo-s | *BGIBMGA014160* | up | 830 | 838 | - | 11.3303 | 4.96E-05 | 0.615 | GTGGGTGGC |
| 12084 | mamo-s | *BGIBMGA014161* | up | 1581 | 1589 | - | 11.3303 | 4.96E-05 | 0.615 | GTGGGTGGC |
| 12085 | mamo-s | *BGIBMGA014162* | down | 112 | 120 | + | 13.156 | 6.30E-06 | 0.615 | CTGCGTGGC |
| 12086 | mamo-s | *BGIBMGA014162* | down | 297 | 305 | + | 13.156 | 6.30E-06 | 0.615 | CTGCGTGGC |
| 12087 | mamo-s | *BGIBMGA014162* | down | 632 | 640 | - | 11.7339 | 3.29E-05 | 0.615 | GCGCGAGGC |
| 12088 | mamo-s | *BGIBMGA014162* | down | 1296 | 1304 | + | 11.5688 | 3.79E-05 | 0.615 | CTGCGTGTC |
| 12089 | mamo-s | *BGIBMGA014162* | down | 941 | 949 | - | 11.3303 | 4.96E-05 | 0.615 | GTGGGTGGC |
| 12090 | mamo-s | *BGIBMGA014164* | down | 992 | 1000 | + | 11.3303 | 4.96E-05 | 0.615 | GTGGGTGGC |
| 12091 | mamo-s | *BGIBMGA014164* | down | 1651 | 1659 | - | 11.0826 | 6.84E-05 | 0.709 | GTACGTGGC |
| 12092 | mamo-s | *BGIBMGA014165* | down | 1638 | 1646 | + | 11.3303 | 4.96E-05 | 0.615 | GTGGGTGGC |
| 12093 | mamo-s | *BGIBMGA014167* | down | 1041 | 1049 | - | 11.0826 | 6.84E-05 | 0.709 | GTACGTGGC |
| 12094 | mamo-s | *BGIBMGA014168* | down | 1844 | 1852 | - | 11.3303 | 4.96E-05 | 0.615 | GTGGGTGGC |
| 12095 | mamo-s | *BGIBMGA014169* | down | 204 | 212 | + | 11.3303 | 4.96E-05 | 0.615 | GTGGGTGGC |
| 12096 | mamo-s | *BGIBMGA014169* | up | 856 | 864 | - | 11.3303 | 4.96E-05 | 0.615 | GTGGGTGGC |
| 12097 | mamo-s | *BGIBMGA014172* | down | 1905 | 1913 | + | 11.3303 | 4.96E-05 | 0.615 | GTGGGTGGC |
| 12098 | mamo-s | *BGIBMGA014173* | up | 1514 | 1522 | + | 11.789 | 2.84E-05 | 0.615 | GCGCGTGCC |
| 12099 | mamo-s | *BGIBMGA014173* | down | 620 | 628 | + | 11.2294 | 5.41E-05 | 0.65 | CTGCGAGGC |
| 12100 | mamo-s | *BGIBMGA014173* | up | 1168 | 1176 | + | 10.5413 | 9.42E-05 | 0.857 | GTGCGAGTC |
| 12101 | mamo-s | *BGIBMGA014175* | down | 1116 | 1124 | + | 10.8073 | 7.89E-05 | 0.766 | CCGCGTGGT |
| 12102 | mamo-s | *BGIBMGA014177* | up | 607 | 615 | + | 11.3303 | 4.96E-05 | 0.615 | GTGGGTGGC |
| 12103 | mamo-s | *BGIBMGA014177* | down | 1703 | 1711 | + | 11.0826 | 6.84E-05 | 0.709 | GTACGTGGC |
| 12104 | mamo-s | *BGIBMGA014180* | down | 876 | 884 | + | 11.0826 | 6.84E-05 | 0.709 | GTACGTGGC |
| 12105 | mamo-s | *BGIBMGA014182* | down | 662 | 670 | - | 11.3303 | 4.96E-05 | 0.615 | GTGGGTGGC |
| 12106 | mamo-s | *BGIBMGA014183* | up | 294 | 302 | - | 11.3303 | 4.96E-05 | 0.615 | GTGGGTGGC |
| 12107 | mamo-s | *BGIBMGA014184* | down | 213 | 221 | - | 10.5688 | 8.88E-05 | 0.824 | GTGCGTGAC |
| 12108 | mamo-s | *BGIBMGA014187* | down | 1668 | 1676 | - | 11.3303 | 4.96E-05 | 0.615 | GTGGGTGGC |
| 12109 | mamo-s | *BGIBMGA014189* | up | 18 | 26 | + | 11.3303 | 4.96E-05 | 0.615 | GTGGGTGGC |
| 12110 | mamo-s | *BGIBMGA014191* | down | 118 | 126 | + | 12.1284 | 2.16E-05 | 0.615 | CAGCGTGGC |
| 12111 | mamo-s | *BGIBMGA014191* | up | 375 | 383 | + | 11.3303 | 4.96E-05 | 0.615 | GTGGGTGGC |
| 12112 | mamo-s | *BGIBMGA014192* | up | 32 | 40 | + | 11.3303 | 4.96E-05 | 0.615 | GTGGGTGGC |
| 12113 | mamo-s | *BGIBMGA014192* | down | 873 | 881 | - | 11.3303 | 4.96E-05 | 0.615 | GTGGGTGGC |
| 12114 | mamo-s | *BGIBMGA014194* | down | 443 | 451 | - | 13.6606 | 4.06E-06 | 0.615 | GCGCGTGGC |
| 12115 | mamo-s | *BGIBMGA014195* | up | 1371 | 1379 | + | 12.7615 | 1.04E-05 | 0.615 | CCGCGTGGC |
| 12116 | mamo-s | *BGIBMGA014195* | up | 247 | 255 | - | 11.3303 | 4.96E-05 | 0.615 | GTGGGTGGC |
| 12117 | mamo-s | *BGIBMGA014197* | up | 449 | 457 | + | 11.3303 | 4.96E-05 | 0.615 | GTGGGTGGC |
| 12118 | mamo-s | *BGIBMGA014197* | down | 1975 | 1983 | - | 11.3303 | 4.96E-05 | 0.615 | GTGGGTGGC |
| 12119 | mamo-s | *BGIBMGA014197* | down | 536 | 544 | - | 10.8899 | 7.48E-05 | 0.739 | CCGCGTGCC |
| 12120 | mamo-s | *BGIBMGA014198* | up | 68 | 76 | - | 12.3394 | 1.49E-05 | 0.615 | GGGCGTGGC |
| 12121 | mamo-s | *BGIBMGA014198* | down | 1961 | 1969 | - | 11.3303 | 4.96E-05 | 0.615 | GTGGGTGGC |
| 12122 | mamo-s | *BGIBMGA014199* | up | 93 | 101 | + | 11.3303 | 4.96E-05 | 0.615 | GTGGGTGGC |
| 12123 | mamo-s | *BGIBMGA014199* | up | 432 | 440 | + | 10.7523 | 8.38E-05 | 0.797 | AAGCGTGGC |
| 12124 | mamo-s | *BGIBMGA014201* | up | 1552 | 1560 | - | 11.3303 | 4.96E-05 | 0.615 | GTGGGTGGC |
| 12125 | mamo-s | *BGIBMGA014201* | down | 915 | 923 | + | 10.8073 | 7.89E-05 | 0.766 | CCGCGTGGT |
| 12126 | mamo-s | *BGIBMGA014202* | up | 648 | 656 | + | 10.8073 | 7.89E-05 | 0.766 | CCGCGTGGT |
| 12127 | mamo-s | *BGIBMGA014203* | down | 1375 | 1383 | + | 11.3303 | 4.96E-05 | 0.615 | GTGGGTGGC |
| 12128 | mamo-s | *BGIBMGA014204* | down | 521 | 529 | - | 11.3303 | 4.96E-05 | 0.615 | GTGGGTGGC |
| 12129 | mamo-s | *BGIBMGA014204* | up | 1323 | 1331 | - | 11.3303 | 4.96E-05 | 0.615 | GTGGGTGGC |
| 12130 | mamo-s | *BGIBMGA014206* | up | 1886 | 1894 | + | 11.1009 | 6.57E-05 | 0.709 | GAGCGAGGC |
| 12131 | mamo-s | *BGIBMGA014206* | down | 1480 | 1488 | - | 10.5413 | 9.42E-05 | 0.857 | GTGCGAGTC |
| 12132 | mamo-s | *BGIBMGA014207* | up | 645 | 653 | - | 11.3303 | 4.96E-05 | 0.615 | GTGGGTGGC |
| 12133 | mamo-s | *BGIBMGA014210* | down | 853 | 861 | + | 11.3303 | 4.96E-05 | 0.615 | GTGGGTGGC |
| 12134 | mamo-s | *BGIBMGA014212* | up | 373 | 381 | + | 12.0734 | 2.66E-05 | 0.615 | GCGCGTGTC |
| 12135 | mamo-s | *BGIBMGA014212* | up | 1751 | 1759 | + | 11.789 | 2.84E-05 | 0.615 | GCGCGTGCC |
| 12136 | mamo-s | *BGIBMGA014212* | down | 1965 | 1973 | + | 11.3303 | 4.96E-05 | 0.615 | GTGGGTGGC |
| 12137 | mamo-s | *BGIBMGA014212* | up | 1708 | 1716 | - | 11.156 | 6.13E-05 | 0.705 | GAGCGTGCC |

| | | | | | | | | | | |
|---|---|---|---|---|---|---|---|---|---|---|
| 12138 | mamo-s | *BGIBMGA014213* | up | 375 | 383 | - | 11.3303 | 4.96E-05 | 0.615 | GTGGGTGGC |
| 12139 | mamo-s | *BGIBMGA014214* | up | 49 | 57 | - | 11.3303 | 4.96E-05 | 0.615 | GTGGGTGGC |
| 12140 | mamo-s | *BGIBMGA014214* | up | 562 | 570 | + | 11.3303 | 4.96E-05 | 0.615 | GTGGGTGGC |
| 12141 | mamo-s | *BGIBMGA014215* | down | 1191 | 1199 | + | 11.0826 | 6.84E-05 | 0.709 | GTACGTGGC |
| 12142 | mamo-s | *BGIBMGA014217* | up | 1974 | 1982 | - | 11.4404 | 4.51E-05 | 0.615 | CGGCGTGGC |
| 12143 | mamo-s | *BGIBMGA014217* | down | 1543 | 1551 | - | 11.3303 | 4.96E-05 | 0.615 | GTGGGTGGC |
| 12144 | mamo-s | *BGIBMGA014217* | down | 1874 | 1882 | + | 11.0826 | 6.84E-05 | 0.709 | GTACGTGGC |
| 12145 | mamo-s | *BGIBMGA014218* | down | 812 | 820 | - | 11.3303 | 4.96E-05 | 0.615 | GTGGGTGGC |
| 12146 | mamo-s | *BGIBMGA014218* | up | 1795 | 1803 | - | 11.3303 | 4.96E-05 | 0.615 | GTGGGTGGC |
| 12147 | mamo-s | *BGIBMGA014222* | down | 695 | 703 | + | 11.3303 | 4.96E-05 | 0.615 | GTGGGTGGC |
| 12148 | mamo-s | *BGIBMGA014223* | down | 308 | 316 | + | 11.0826 | 6.84E-05 | 0.709 | GTACGTGGC |
| 12149 | mamo-s | *BGIBMGA014224* | up | 1809 | 1817 | + | 11.3303 | 4.96E-05 | 0.615 | GTGGGTGGC |
| 12150 | mamo-s | *BGIBMGA014225* | up | 1535 | 1543 | - | 11.3303 | 4.96E-05 | 0.615 | GTGGGTGGC |
| 12151 | mamo-s | *BGIBMGA014226* | down | 1676 | 1684 | + | 11.7339 | 3.29E-05 | 0.615 | GCGCGAGGC |
| 12152 | mamo-s | *BGIBMGA014228* | down | 551 | 559 | - | 12.1009 | 2.43E-05 | 0.615 | GTGCGTGGT |
| 12153 | mamo-s | *BGIBMGA014228* | up | 1767 | 1775 | - | 12.1009 | 2.43E-05 | 0.615 | GTGCGTGGT |
| 12154 | mamo-s | *BGIBMGA014231* | down | 135 | 143 | - | 11.3303 | 4.96E-05 | 0.615 | GTGGGTGGC |
| 12155 | mamo-s | *BGIBMGA014231* | up | 1270 | 1278 | + | 10.7523 | 8.38E-05 | 0.797 | GGGCGTGTC |
| 12156 | mamo-s | *BGIBMGA014232* | up | 944 | 952 | - | 11.3303 | 4.96E-05 | 0.615 | GTGGGTGGC |
| 12157 | mamo-s | *BGIBMGA014232* | up | 1673 | 1681 | - | 11.3303 | 4.96E-05 | 0.615 | GTGGGTGGC |
| 12158 | mamo-s | *BGIBMGA014232* | down | 1607 | 1615 | - | 10.8349 | 7.66E-05 | 0.751 | CCGCGAGGC |
| 12159 | mamo-s | *BGIBMGA014235* | down | 45 | 53 | + | 11.5321 | 4.06E-05 | 0.615 | TTGCGTGGC |
| 12160 | mamo-s | *BGIBMGA014235* | down | 1254 | 1262 | + | 11.1743 | 5.90E-05 | 0.684 | CCGCGTGTC |
| 12161 | mamo-s | *BGIBMGA014236* | up | 1947 | 1955 | - | 12.0734 | 2.66E-05 | 0.615 | GCGCGTGTC |
| 12162 | mamo-s | *BGIBMGA014236* | up | 709 | 717 | - | 11.7798 | 3.11E-05 | 0.615 | ATGCGTGGC |
| 12163 | mamo-s | *BGIBMGA014238* | up | 677 | 685 | - | 11.3303 | 4.96E-05 | 0.615 | GTGGGTGGC |
| 12164 | mamo-s | *BGIBMGA014239* | up | 1044 | 1052 | + | 11.3303 | 4.96E-05 | 0.615 | GTGGGTGGC |
| 12165 | mamo-s | *BGIBMGA014240* | down | 1261 | 1269 | + | 11.3303 | 4.96E-05 | 0.615 | GTGGGTGGC |
| 12166 | mamo-s | *BGIBMGA014240* | up | 1402 | 1410 | + | 11.3303 | 4.96E-05 | 0.615 | GTGGGTGGC |
| 12167 | mamo-s | *BGIBMGA014241* | up | 372 | 380 | + | 11.7798 | 3.11E-05 | 0.615 | ATGCGTGGC |
| 12168 | mamo-s | *BGIBMGA014241* | up | 708 | 716 | + | 11.3303 | 4.96E-05 | 0.615 | GTGGGTGGC |
| 12169 | mamo-s | *BGIBMGA014242* | up | 500 | 508 | + | 11.1743 | 5.90E-05 | 0.684 | CCGCGTGTC |
| 12170 | mamo-s | *BGIBMGA014243* | down | 859 | 867 | + | 14.055 | 2.23E-06 | 0.615 | GTGCGTGGC |
| 12171 | mamo-s | *BGIBMGA014244* | down | 712 | 720 | - | 13.0275 | 8.53E-06 | 0.615 | GAGCGTGGC |
| 12172 | mamo-s | *BGIBMGA014244* | down | 589 | 597 | + | 10.9358 | 7.30E-05 | 0.728 | GCGGGTGGC |
| 12173 | mamo-s | *BGIBMGA014245* | down | 537 | 545 | - | 13.0275 | 8.53E-06 | 0.615 | GAGCGTGGC |
| 12174 | mamo-s | *BGIBMGA014247* | down | 424 | 432 | - | 11.3303 | 4.96E-05 | 0.615 | GTGGGTGGC |
| 12175 | mamo-s | *BGIBMGA014248* | up | 452 | 460 | + | 12.0734 | 2.66E-05 | 0.615 | GCGCGTGTC |
| 12176 | mamo-s | *BGIBMGA014248* | up | 1114 | 1122 | - | 11.3303 | 4.96E-05 | 0.615 | GTGGGTGGC |
| 12177 | mamo-s | *BGIBMGA014249* | down | 1558 | 1566 | - | 11.5688 | 3.79E-05 | 0.615 | CTGCGTGTC |
| 12178 | mamo-s | *BGIBMGA014250* | up | 813 | 821 | - | 11.5688 | 3.79E-05 | 0.615 | CTGCGTGTC |
| 12179 | mamo-s | *BGIBMGA014251* | down | 175 | 183 | + | 11.3303 | 4.96E-05 | 0.615 | GTGGGTGGC |
| 12180 | mamo-s | *BGIBMGA014251* | up | 1070 | 1078 | - | 11.3303 | 4.96E-05 | 0.615 | GTGGGTGGC |
| 12181 | mamo-s | *BGIBMGA014252* | up | 224 | 232 | - | 11.3303 | 4.96E-05 | 0.615 | GTGGGTGGC |
| 12182 | mamo-s | *BGIBMGA014253* | up | 332 | 340 | - | 11.3303 | 4.96E-05 | 0.615 | GTGGGTGGC |
| 12183 | mamo-s | *BGIBMGA014254* | up | 222 | 230 | + | 11.3303 | 4.96E-05 | 0.615 | GTGGGTGGC |
| 12184 | mamo-s | *BGIBMGA014255* | down | 117 | 125 | + | 12.1284 | 2.16E-05 | 0.615 | GTGCGAGGC |
| 12185 | mamo-s | *BGIBMGA014256* | down | 1040 | 1048 | - | 13.0275 | 8.53E-06 | 0.615 | GAGCGTGGC |
| 12186 | mamo-s | *BGIBMGA014256* | up | 1497 | 1505 | + | 12.1284 | 2.16E-05 | 0.615 | GTGCGAGGC |
| 12187 | mamo-s | *BGIBMGA014256* | down | 1653 | 1661 | - | 12.1009 | 2.43E-05 | 0.615 | GTGCGTGGT |
| 12188 | mamo-s | *BGIBMGA014256* | down | 30 | 38 | + | 10.8899 | 7.48E-05 | 0.739 | CCGCGTGCC |
| 12189 | mamo-s | *BGIBMGA014258* | down | 292 | 300 | + | 11.3303 | 4.96E-05 | 0.615 | GTGGGTGGC |
| 12190 | mamo-s | *BGIBMGA014258* | down | 1189 | 1197 | + | 11.2844 | 5.18E-05 | 0.63 | CTGCGTGCC |
| 12191 | mamo-s | *BGIBMGA014259* | up | 1853 | 1861 | - | 10.5138 | 9.75E-05 | 0.865 | GTGCGTGTT |
| 12192 | mamo-s | *BGIBMGA014260* | down | 1195 | 1203 | + | 11.3303 | 4.96E-05 | 0.615 | GTGGGTGGC |
| 12193 | mamo-s | *BGIBMGA014260* | down | 1660 | 1668 | - | 11.3303 | 4.96E-05 | 0.615 | GTGGGTGGC |
| 12194 | mamo-s | *BGIBMGA014261* | down | 1105 | 1113 | - | 10.5688 | 8.88E-05 | 0.824 | GTGCGTGAC |
| 12195 | mamo-s | *BGIBMGA014261* | up | 459 | 467 | - | 10.5138 | 9.75E-05 | 0.865 | GTGCGTGTT |
| 12196 | mamo-s | *BGIBMGA014262* | up | 1798 | 1806 | - | 11.3303 | 4.96E-05 | 0.615 | GTGGGTGGC |
| 12197 | mamo-s | *BGIBMGA014263* | up | 686 | 694 | - | 12.1009 | 2.43E-05 | 0.615 | GTGCGTGGT |
| 12198 | mamo-s | *BGIBMGA014264* | up | 1124 | 1132 | - | 13.6606 | 4.06E-06 | 0.615 | GCGCGTGGC |
| 12199 | mamo-s | *BGIBMGA014267* | up | 519 | 527 | + | 11.3303 | 4.96E-05 | 0.615 | GTGGGTGGC |
| 12200 | mamo-s | *BGIBMGA014267* | up | 1102 | 1110 | - | 11.3303 | 4.96E-05 | 0.615 | GTGGGTGGC |
| 12201 | mamo-s | *BGIBMGA014268* | down | 1416 | 1424 | - | 10.8349 | 7.66E-05 | 0.751 | CCGCGAGGC |
| 12202 | mamo-s | *BGIBMGA014270* | up | 471 | 479 | - | 11.7064 | 3.52E-05 | 0.615 | GCGCGTGGT |
| 12203 | mamo-s | *BGIBMGA014272* | down | 1941 | 1949 | + | 11.3303 | 4.96E-05 | 0.615 | GTGGGTGGC |
| 12204 | mamo-s | *BGIBMGA014274* | down | 1892 | 1900 | + | 11.3303 | 4.96E-05 | 0.615 | GTGGGTGGC |
| 12205 | mamo-s | *BGIBMGA014275* | up | 822 | 830 | - | 11.3303 | 4.96E-05 | 0.615 | GTGGGTGGC |
| 12206 | mamo-s | *BGIBMGA014278* | up | 1773 | 1781 | + | 11.3303 | 4.96E-05 | 0.615 | GTGGGTGGC |
| 12207 | mamo-s | *BGIBMGA014282* | up | 1705 | 1713 | + | 11.3303 | 4.96E-05 | 0.615 | GTGGGTGGC |
| 12208 | mamo-s | *BGIBMGA014284* | down | 448 | 456 | + | 11.3303 | 4.96E-05 | 0.615 | GTGGGTGGC |
| 12209 | mamo-s | *BGIBMGA014285* | up | 617 | 625 | + | 11.3303 | 4.96E-05 | 0.615 | GTGGGTGGC |
| 12210 | mamo-s | *BGIBMGA014288* | down | 488 | 496 | + | 12.0734 | 2.66E-05 | 0.615 | GCGCGTGTC |
| 12211 | mamo-s | *BGIBMGA014288* | down | 1045 | 1053 | + | 11.3303 | 4.96E-05 | 0.615 | GTGGGTGGC |
| 12212 | mamo-s | *BGIBMGA014289* | down | 1599 | 1607 | - | 11.3303 | 4.96E-05 | 0.615 | GTGGGTGGC |
| 12213 | mamo-s | *BGIBMGA014290* | up | 1805 | 1813 | - | 11.3303 | 4.96E-05 | 0.615 | GTGGGTGGC |
| 12214 | mamo-s | *BGIBMGA014292* | down | 1099 | 1107 | - | 11.0826 | 6.84E-05 | 0.709 | GTACGTGGC |
| 12215 | mamo-s | *BGIBMGA014294* | down | 709 | 717 | - | 10.5413 | 9.42E-05 | 0.857 | GTGCGAGTC |
| 12216 | mamo-s | *BGIBMGA014295* | up | 769 | 777 | - | 14.055 | 2.23E-06 | 0.615 | GTGCGTGGC |
| 12217 | mamo-s | *BGIBMGA014296* | down | 1673 | 1681 | + | 11.7339 | 3.29E-05 | 0.615 | GCGCGAGGC |
| 12218 | mamo-s | *BGIBMGA014297* | down | 226 | 234 | + | 11.3303 | 4.96E-05 | 0.615 | GTGGGTGGC |
| 12219 | mamo-s | *BGIBMGA014297* | up | 1215 | 1223 | + | 11.3303 | 4.96E-05 | 0.615 | GTGGGTGGC |
| 12220 | mamo-s | *BGIBMGA014298* | down | 1112 | 1120 | + | 11.3303 | 4.96E-05 | 0.615 | GTGGGTGGC |
| 12221 | mamo-s | *BGIBMGA014303* | up | 126 | 134 | + | 10.9358 | 7.30E-05 | 0.728 | GCGGGTGGC |
| 12222 | mamo-s | *BGIBMGA014305* | up | 1263 | 1271 | + | 11.3303 | 4.96E-05 | 0.615 | GTGGGTGGC |
| 12223 | mamo-s | *BGIBMGA014306* | down | 193 | 201 | + | 11.3303 | 4.96E-05 | 0.615 | GTGGGTGGC |
| 12224 | mamo-s | *BGIBMGA014307* | down | 724 | 732 | + | 13.6606 | 4.06E-06 | 0.615 | GCGCGTGGC |
| 12225 | mamo-s | *BGIBMGA014307* | up | 392 | 400 | - | 11.3303 | 4.96E-05 | 0.615 | GTGGGTGGC |

| | | | | | | | | | | |
|---|---|---|---|---|---|---|---|---|---|---|
| 12226 | mamo-s | *BGIBMGA014308* | down | 970 | 978 | + | 11.3303 | 4.96E-05 | 0.615 | GTGGGTGGC |
| 12227 | mamo-s | *BGIBMGA014308* | down | 1782 | 1790 | - | 11.0826 | 6.84E-05 | 0.709 | GTACGTGGC |
| 12228 | mamo-s | *BGIBMGA014308* | up | 386 | 394 | - | 10.5688 | 8.88E-05 | 0.824 | GTGCGTGAC |
| 12229 | mamo-s | *BGIBMGA014309* | down | 364 | 372 | + | 11.3303 | 4.96E-05 | 0.615 | GTGGGTGGC |
| 12230 | mamo-s | *BGIBMGA014311* | down | 163 | 171 | - | 11.3303 | 4.96E-05 | 0.615 | GTGGGTGGC |
| 12231 | mamo-s | *BGIBMGA014311* | down | 1286 | 1294 | + | 11.3303 | 4.96E-05 | 0.615 | GTGGGTGGC |
| 12232 | mamo-s | *BGIBMGA014312* | down | 162 | 170 | - | 10.8349 | 7.66E-05 | 0.751 | CCGCGAGGC |
| 12233 | mamo-s | *BGIBMGA014313* | down | 743 | 751 | + | 11.3303 | 4.96E-05 | 0.615 | GTGGGTGGC |
| 12234 | mamo-s | *BGIBMGA014313* | down | 1570 | 1578 | - | 11.3303 | 4.96E-05 | 0.615 | GTGGGTGGC |
| 12235 | mamo-s | *BGIBMGA014313* | up | 368 | 376 | + | 10.7523 | 8.38E-05 | 0.797 | GGGCGTGTC |
| 12236 | mamo-s | *BGIBMGA014316* | up | 1961 | 1969 | + | 12.1009 | 2.43E-05 | 0.615 | GTGCGTGGT |
| 12237 | mamo-s | *BGIBMGA014316* | down | 1018 | 1026 | + | 11.3303 | 4.96E-05 | 0.615 | GTGGGTGGC |
| 12238 | mamo-s | *BGIBMGA014316* | down | 1777 | 1785 | + | 11.3303 | 4.96E-05 | 0.615 | GTGGGTGGC |
| 12239 | mamo-s | *BGIBMGA014319* | down | 1024 | 1032 | + | 11.1376 | 6.35E-05 | 0.709 | TCGCGTGGC |
| 12240 | mamo-s | *BGIBMGA014321* | down | 236 | 244 | - | 11.3303 | 4.96E-05 | 0.615 | GTGGGTGGC |
| 12241 | mamo-s | *BGIBMGA014324* | up | 1564 | 1572 | - | 11.3303 | 4.96E-05 | 0.615 | GTGGGTGGC |
| 12242 | mamo-s | *BGIBMGA014325* | up | 106 | 114 | - | 11.3303 | 4.96E-05 | 0.615 | GTGGGTGGC |
| 12243 | mamo-s | *BGIBMGA014325* | up | 678 | 686 | - | 10.5138 | 9.75E-05 | 0.865 | GTGCGTGTT |
| 12244 | mamo-s | *BGIBMGA014327* | up | 1060 | 1068 | + | 14.055 | 2.23E-06 | 0.615 | GTGCGTGGC |
| 12245 | mamo-s | *BGIBMGA014327* | up | 1459 | 1467 | + | 11.3303 | 4.96E-05 | 0.615 | GTGGGTGGC |
| 12246 | mamo-s | *BGIBMGA014327* | up | 152 | 160 | + | 11.0826 | 6.84E-05 | 0.709 | GTACGTGGC |
| 12247 | mamo-s | *BGIBMGA014328* | up | 664 | 672 | + | 10.5138 | 9.75E-05 | 0.865 | GTGCGTGTT |
| 12248 | mamo-s | *BGIBMGA014330* | up | 189 | 197 | + | 13.0275 | 8.53E-06 | 0.615 | GAGCGTGGC |
| 12249 | mamo-s | *BGIBMGA014331* | down | 39 | 47 | + | 11.7798 | 3.11E-05 | 0.615 | ATGCGTGGC |
| 12250 | mamo-s | *BGIBMGA014332* | down | 197 | 205 | + | 11.3303 | 4.96E-05 | 0.615 | GTGGGTGGC |
| 12251 | mamo-s | *BGIBMGA014333* | down | 925 | 933 | + | 11.3303 | 4.96E-05 | 0.615 | GTGGGTGGC |
| 12252 | mamo-s | *BGIBMGA014339* | down | 907 | 915 | - | 11.3303 | 4.96E-05 | 0.615 | GTGGGTGGC |
| 12253 | mamo-s | *BGIBMGA014341* | up | 1003 | 1011 | + | 11.3303 | 4.96E-05 | 0.615 | GTGGGTGGC |
| 12254 | mamo-s | *BGIBMGA014342* | down | 647 | 655 | + | 11.3303 | 4.96E-05 | 0.615 | GTGGGTGGC |
| 12255 | mamo-s | *BGIBMGA014345* | down | 652 | 660 | + | 11.3303 | 4.96E-05 | 0.615 | GTGGGTGGC |
| 12256 | mamo-s | *BGIBMGA014345* | down | 1359 | 1367 | + | 11.3303 | 4.96E-05 | 0.615 | GTGGGTGGC |
| 12257 | mamo-s | *BGIBMGA014348* | down | 738 | 746 | + | 14.055 | 2.23E-06 | 0.615 | GTGCGTGGC |
| 12258 | mamo-s | *BGIBMGA014349* | up | 212 | 220 | + | 11.3303 | 4.96E-05 | 0.615 | GTGGGTGGC |
| 12259 | mamo-s | *BGIBMGA014351* | down | 1508 | 1516 | + | 11.7798 | 3.11E-05 | 0.615 | ATGCGTGGC |
| 12260 | mamo-s | *BGIBMGA014355* | down | 333 | 341 | + | 11.3303 | 4.96E-05 | 0.615 | GTGGGTGGC |
| 12261 | mamo-s | *BGIBMGA014357* | up | 169 | 177 | + | 11.7064 | 3.52E-05 | 0.615 | GCGCGTGGT |
| 12262 | mamo-s | *BGIBMGA014358* | down | 432 | 440 | + | 10.9358 | 7.30E-05 | 0.728 | GCGGGTGGC |
| 12263 | mamo-s | *BGIBMGA014358* | down | 1635 | 1643 | - | 10.5413 | 9.42E-05 | 0.857 | GTGCGAGTC |
| 12264 | mamo-s | *BGIBMGA014359* | down | 1699 | 1707 | - | 11.0826 | 6.84E-05 | 0.709 | GTACGTGGC |
| 12265 | mamo-s | *BGIBMGA014360* | up | 1249 | 1257 | - | 12.1284 | 2.16E-05 | 0.615 | GTGCGAGGC |
| 12266 | mamo-s | *BGIBMGA014360* | up | 1269 | 1277 | - | 11.4404 | 4.51E-05 | 0.615 | CGGCGTGGC |
| 12267 | mamo-s | *BGIBMGA014360* | up | 1521 | 1529 | - | 10.8073 | 7.89E-05 | 0.766 | CCGCGTGGT |
| 12268 | mamo-s | *BGIBMGA014362* | down | 755 | 763 | + | 10.7523 | 8.38E-05 | 0.797 | AAGCGTGGC |
| 12269 | mamo-s | *BGIBMGA014363* | down | 1980 | 1988 | - | 10.5413 | 9.42E-05 | 0.857 | CAGCGTGTC |
| 12270 | mamo-s | *BGIBMGA014366* | up | 1938 | 1946 | + | 13.0275 | 8.53E-06 | 0.615 | GAGCGTGGC |
| 12271 | mamo-s | *BGIBMGA014366* | down | 1494 | 1502 | + | 11.0734 | 7.12E-05 | 0.726 | GAGCGTGGT |
| 12272 | mamo-s | *BGIBMGA014368* | up | 1172 | 1180 | + | 12.0734 | 2.66E-05 | 0.615 | GCGCGTGTC |
| 12273 | mamo-s | *BGIBMGA014368* | up | 1344 | 1352 | - | 11.4404 | 4.51E-05 | 0.615 | GAGCGTGTC |
| 12274 | mamo-s | *BGIBMGA014370* | up | 1176 | 1184 | - | 12.1009 | 2.43E-05 | 0.615 | GTGCGTGGT |
| 12275 | mamo-s | *BGIBMGA014370* | up | 1836 | 1844 | - | 11.1376 | 6.35E-05 | 0.709 | TCGCGTGGC |
| 12276 | mamo-s | *BGIBMGA014371* | up | 1866 | 1874 | - | 11.3303 | 4.96E-05 | 0.615 | GTGGGTGGC |
| 12277 | mamo-s | *BGIBMGA014372* | up | 342 | 350 | + | 11.5688 | 3.79E-05 | 0.615 | CTGCGTGTC |
| 12278 | mamo-s | *BGIBMGA014374* | up | 1797 | 1805 | + | 10.7523 | 8.38E-05 | 0.797 | AAGCGTGGC |
| 12279 | mamo-s | *BGIBMGA014375* | up | 15 | 23 | + | 12.0734 | 2.66E-05 | 0.615 | GCGCGTGTC |
| 12280 | mamo-s | *BGIBMGA014375* | down | 509 | 517 | + | 11.3303 | 4.96E-05 | 0.615 | GTGGGTGGC |
| 12281 | mamo-s | *BGIBMGA014377* | up | 1289 | 1297 | - | 13.6606 | 4.06E-06 | 0.615 | GCGCGTGGC |
| 12282 | mamo-s | *BGIBMGA014377* | up | 1254 | 1262 | - | 11.5688 | 3.79E-05 | 0.615 | CTGCGTGTC |
| 12283 | mamo-s | *BGIBMGA014377* | down | 1700 | 1708 | + | 11.5321 | 4.06E-05 | 0.615 | TTGCGTGGC |
| 12284 | mamo-s | *BGIBMGA014377* | down | 941 | 949 | + | 11.3303 | 4.96E-05 | 0.615 | GTGGGTGGC |
| 12285 | mamo-s | *BGIBMGA014378* | up | 919 | 927 | - | 10.5138 | 9.75E-05 | 0.865 | GTGCGTGTT |
| 12286 | mamo-s | *BGIBMGA014382* | down | 296 | 304 | - | 11.3303 | 4.96E-05 | 0.615 | GTGGGTGGC |
| 12287 | mamo-s | *BGIBMGA014385* | down | 1185 | 1193 | + | 11.2018 | 5.68E-05 | 0.674 | CTGCGTGGT |
| 12288 | mamo-s | *BGIBMGA014391* | down | 27 | 35 | + | 11.3853 | 4.74E-05 | 0.615 | ACGCGTGGC |
| 12289 | mamo-s | *BGIBMGA014391* | up | 1193 | 1201 | - | 10.5138 | 9.75E-05 | 0.865 | GTGCGTGTT |
| 12290 | mamo-s | *BGIBMGA014392* | down | 702 | 710 | + | 13.6606 | 4.06E-06 | 0.615 | GCGCGTGGC |
| 12291 | mamo-s | *BGIBMGA014392* | up | 1940 | 1948 | - | 10.9358 | 7.30E-05 | 0.728 | GCGGGTGGC |
| 12292 | mamo-s | *BGIBMGA014393* | up | 1687 | 1695 | - | 11.1743 | 5.90E-05 | 0.684 | CCGCGTGTC |
| 12293 | mamo-s | *BGIBMGA014393* | up | 1892 | 1900 | + | 11.0826 | 6.84E-05 | 0.709 | GTACGTGGC |
| 12294 | mamo-s | *BGIBMGA014395* | down | 397 | 405 | + | 11.789 | 2.84E-05 | 0.615 | GCGCGTGCC |
| 12295 | mamo-s | *BGIBMGA014397* | down | 1726 | 1734 | + | 11.3303 | 4.96E-05 | 0.615 | GTGGGTGGC |
| 12296 | mamo-s | *BGIBMGA014399* | down | 1779 | 1787 | - | 10.5413 | 9.42E-05 | 0.857 | GTGCGAGTC |
| 12297 | mamo-s | *BGIBMGA014400* | up | 1083 | 1091 | + | 11.0826 | 6.84E-05 | 0.709 | GTACGTGGC |
| 12298 | mamo-s | *BGIBMGA014402* | up | 1465 | 1473 | - | 11.3303 | 4.96E-05 | 0.615 | GTGGGTGGC |
| 12299 | mamo-s | *BGIBMGA014402* | up | 476 | 484 | + | 11.2844 | 5.18E-05 | 0.63 | CTGCGTGCC |
| 12300 | mamo-s | *BGIBMGA014407* | down | 795 | 803 | + | 11.3303 | 4.96E-05 | 0.615 | GTGGGTGGC |
| 12301 | mamo-s | *BGIBMGA014409* | up | 1223 | 1231 | - | 13.156 | 6.30E-06 | 0.615 | CTGCGTGGC |
| 12302 | mamo-s | *BGIBMGA014410* | up | 764 | 772 | - | 13.156 | 6.30E-06 | 0.615 | CTGCGTGGC |
| 12303 | mamo-s | *BGIBMGA014411* | up | 1831 | 1839 | - | 12.4679 | 1.31E-05 | 0.615 | GTGCGTGTC |
| 12304 | mamo-s | *BGIBMGA014411* | up | 579 | 587 | - | 12.3394 | 1.49E-05 | 0.615 | GGGCGTGGC |
| 12305 | mamo-s | *BGIBMGA014413* | up | 1943 | 1951 | + | 11.789 | 2.84E-05 | 0.615 | GCGCGTGCC |
| 12306 | mamo-s | *BGIBMGA014413* | down | 404 | 412 | + | 10.8899 | 7.48E-05 | 0.739 | CCGCGTGCC |
| 12307 | mamo-s | *BGIBMGA014414* | down | 460 | 468 | + | 11.7339 | 3.29E-05 | 0.615 | GCGCGAGGC |
| 12308 | mamo-s | *BGIBMGA014415* | down | 217 | 225 | - | 12.3394 | 1.49E-05 | 0.615 | GGGCGTGGC |
| 12309 | mamo-s | *BGIBMGA014415* | down | 569 | 577 | + | 11.4404 | 4.51E-05 | 0.615 | CGGCGTGGC |
| 12310 | mamo-s | *BGIBMGA014415* | down | 379 | 387 | + | 11.3853 | 4.74E-05 | 0.615 | ACGCGTGGC |
| 12311 | mamo-s | *BGIBMGA014415* | down | 870 | 878 | + | 11.3303 | 4.96E-05 | 0.615 | GTGGGTGGC |
| 12312 | mamo-s | *BGIBMGA014415* | down | 579 | 587 | + | 11.1743 | 5.90E-05 | 0.684 | CCGCGTGTC |
| 12313 | mamo-s | *BGIBMGA014416* | down | 1460 | 1468 | - | 12.7615 | 1.04E-05 | 0.615 | CCGCGTGGC |

| | | | | | | | | | | |
|---|---|---|---|---|---|---|---|---|---|---|
| 12314 | mamo-s | *BGIBMGA014416* | down | 1576 | 1584 | + | 12.1284 | 2.16E-05 | 0.615 | CAGCGTGGC |
| 12315 | mamo-s | *BGIBMGA014416* | down | 879 | 887 | - | 10.8899 | 7.48E-05 | 0.739 | CCGCGTGCC |
| 12316 | mamo-s | *BGIBMGA014416* | down | 1886 | 1894 | - | 10.7523 | 8.38E-05 | 0.797 | GGGCGTGTC |
| 12317 | mamo-s | *BGIBMGA014417* | down | 693 | 701 | - | 11.3303 | 4.96E-05 | 0.615 | GTGGGTGGC |
| 12318 | mamo-s | *BGIBMGA014419* | up | 1914 | 1922 | - | 14.055 | 2.23E-06 | 0.615 | GTGCGTGGC |
| 12319 | mamo-s | *BGIBMGA014419* | up | 6 | 14 | + | 11.3303 | 4.96E-05 | 0.615 | GTGGGTGGC |
| 12320 | mamo-s | *BGIBMGA014419* | down | 602 | 610 | + | 10.5138 | 9.75E-05 | 0.865 | GTGCGTGTT |
| 12321 | mamo-s | *BGIBMGA014420* | down | 1207 | 1215 | + | 11.7064 | 3.52E-05 | 0.615 | GCGCGTGGT |
| 12322 | mamo-s | *BGIBMGA014420* | down | 1792 | 1800 | + | 11.4404 | 4.51E-05 | 0.615 | CGGCGTGGC |
| 12323 | mamo-s | *BGIBMGA014420* | down | 835 | 843 | - | 11.3303 | 4.96E-05 | 0.615 | GTGGGTGGC |
| 12324 | mamo-s | *BGIBMGA014420* | down | 1435 | 1443 | + | 11.2844 | 5.18E-05 | 0.63 | CTGCGTGCC |
| 12325 | mamo-s | *BGIBMGA014420* | up | 254 | 262 | + | 11.1009 | 6.57E-05 | 0.709 | GAGCGAGGC |
| 12326 | mamo-s | *BGIBMGA014420* | up | 111 | 119 | - | 10.5688 | 8.88E-05 | 0.824 | GTGCGTGAC |
| 12327 | mamo-s | *BGIBMGA014421* | up | 75 | 83 | + | 11.4404 | 4.51E-05 | 0.615 | CGGCGTGGC |
| 12328 | mamo-s | *BGIBMGA014422* | up | 990 | 998 | + | 11.3303 | 4.96E-05 | 0.615 | GTGGGTGGC |
| 12329 | mamo-s | *BGIBMGA014422* | down | 709 | 717 | + | 11.0734 | 7.12E-05 | 0.726 | GAGCGTGGT |
| 12330 | mamo-s | *BGIBMGA014422* | up | 1401 | 1409 | + | 11.0734 | 7.12E-05 | 0.726 | GAGCGTGGT |
| 12331 | mamo-s | *BGIBMGA014422* | up | 533 | 541 | + | 10.5138 | 9.75E-05 | 0.865 | GTGCGTGTT |
| 12332 | mamo-s | *BGIBMGA014423* | down | 230 | 238 | + | 11.4404 | 4.51E-05 | 0.615 | CGGCGTGGC |
| 12333 | mamo-s | *BGIBMGA014423* | down | 1492 | 1500 | + | 11.3303 | 4.96E-05 | 0.615 | GTGGGTGGC |
| 12334 | mamo-s | *BGIBMGA014423* | down | 436 | 444 | + | 10.8073 | 7.89E-05 | 0.766 | CCGCGTGGT |
| 12335 | mamo-s | *BGIBMGA014424* | down | 748 | 756 | - | 11.7064 | 3.52E-05 | 0.615 | GCGCGTGGT |
| 12336 | mamo-s | *BGIBMGA014424* | down | 703 | 711 | - | 11.4404 | 4.51E-05 | 0.615 | CGGCGTGGC |
| 12337 | mamo-s | *BGIBMGA014424* | up | 400 | 408 | - | 11.3303 | 4.96E-05 | 0.615 | GTGGGTGGC |
| 12338 | mamo-s | *BGIBMGA014425* | down | 1109 | 1117 | - | 13.0275 | 8.53E-06 | 0.615 | GAGCGTGGC |
| 12339 | mamo-s | *BGIBMGA014426* | down | 657 | 665 | - | 11.0826 | 6.84E-05 | 0.709 | GTACGTGGC |
| 12340 | mamo-s | *BGIBMGA014429* | up | 454 | 462 | - | 12.3394 | 1.49E-05 | 0.615 | GGGCGTGGC |
| 12341 | mamo-s | *BGIBMGA014429* | up | 879 | 887 | + | 11.2018 | 5.68E-05 | 0.674 | CTGCGTGGT |
| 12342 | mamo-s | *BGIBMGA014429* | up | 1738 | 1746 | - | 11.1743 | 5.90E-05 | 0.684 | CCGCGTGTC |
| 12343 | mamo-s | *BGIBMGA014431* | up | 187 | 195 | - | 12.1284 | 2.16E-05 | 0.615 | CAGCGTGGC |
| 12344 | mamo-s | *BGIBMGA014431* | down | 456 | 464 | + | 11.1743 | 5.90E-05 | 0.684 | CCGCGTGTC |
| 12345 | mamo-s | *BGIBMGA014431* | down | 1466 | 1474 | + | 11.0826 | 6.84E-05 | 0.709 | GTACGTGGC |
| 12346 | mamo-s | *BGIBMGA014432* | up | 928 | 936 | - | 11.3853 | 4.74E-05 | 0.615 | ACGCGTGGC |
| 12347 | mamo-s | *BGIBMGA014432* | down | 823 | 831 | - | 11.2844 | 5.18E-05 | 0.63 | CTGCGTGCC |
| 12348 | mamo-s | *BGIBMGA014432* | down | 1444 | 1452 | + | 11.2844 | 5.18E-05 | 0.63 | CTGCGTGCC |
| 12349 | mamo-s | *BGIBMGA014433* | up | 489 | 497 | - | 11.2844 | 5.18E-05 | 0.63 | CTGCGTGCC |
| 12350 | mamo-s | *BGIBMGA014433* | up | 1110 | 1118 | + | 11.2844 | 5.18E-05 | 0.63 | CTGCGTGCC |
| 12351 | mamo-s | *BGIBMGA014434* | up | 1542 | 1550 | + | 11.156 | 6.13E-05 | 0.705 | GAGCGTGCC |
| 12352 | mamo-s | *BGIBMGA014436* | down | 861 | 869 | - | 12.7615 | 1.04E-05 | 0.615 | CCGCGTGGC |
| 12353 | mamo-s | *BGIBMGA014436* | down | 593 | 601 | + | 11.1743 | 5.90E-05 | 0.684 | CCGCGTGTC |
| 12354 | mamo-s | *BGIBMGA014437* | down | 1792 | 1800 | - | 11.1009 | 6.57E-05 | 0.709 | GAGCGAGGC |
| 12355 | mamo-s | *BGIBMGA014438* | down | 1741 | 1749 | + | 12.0734 | 2.66E-05 | 0.615 | GCGCGTGTC |
| 12356 | mamo-s | *BGIBMGA014438* | up | 1294 | 1302 | + | 10.7523 | 8.38E-05 | 0.797 | GGGCGTGTC |
| 12357 | mamo-s | *BGIBMGA014440* | up | 650 | 658 | - | 13.6606 | 4.06E-06 | 0.615 | GCGCGTGGC |
| 12358 | mamo-s | *BGIBMGA014444* | up | 1186 | 1194 | - | 11.3303 | 4.96E-05 | 0.615 | GTGGGTGGC |
| 12359 | mamo-s | *BGIBMGA014445* | up | 909 | 917 | + | 11.3303 | 4.96E-05 | 0.615 | GTGGGTGGC |
| 12360 | mamo-s | *BGIBMGA014446* | down | 415 | 423 | + | 12.1284 | 2.16E-05 | 0.615 | CAGCGTGGC |
| 12361 | mamo-s | *BGIBMGA014446* | down | 400 | 408 | + | 10.5138 | 9.75E-05 | 0.865 | GTGCGTGTT |
| 12362 | mamo-s | *BGIBMGA014450* | up | 874 | 882 | + | 11.4404 | 4.51E-05 | 0.615 | GAGCGTGTC |
| 12363 | mamo-s | *BGIBMGA014450* | down | 1903 | 1911 | + | 11.3303 | 4.96E-05 | 0.615 | GTGGGTGGC |
| 12364 | mamo-s | *BGIBMGA014451* | up | 1002 | 1010 | + | 11.3303 | 4.96E-05 | 0.615 | GTGGGTGGC |
| 12365 | mamo-s | *BGIBMGA014451* | up | 1319 | 1327 | + | 11.3303 | 4.96E-05 | 0.615 | GTGGGTGGC |
| 12366 | mamo-s | *BGIBMGA014452* | down | 1754 | 1762 | - | 11.3303 | 4.96E-05 | 0.615 | GTGGGTGGC |
| 12367 | mamo-s | *BGIBMGA014452* | up | 594 | 602 | - | 11.1743 | 5.90E-05 | 0.684 | CCGCGTGTC |
| 12368 | mamo-s | *BGIBMGA014452* | up | 1585 | 1593 | - | 10.5413 | 9.42E-05 | 0.857 | CAGCGTGTC |
| 12369 | mamo-s | *BGIBMGA014453* | down | 1189 | 1197 | - | 12.0734 | 2.66E-05 | 0.615 | GCGCGTGTC |
| 12370 | mamo-s | *BGIBMGA014453* | down | 230 | 238 | - | 11.3303 | 4.96E-05 | 0.615 | GTGGGTGGC |
| 12371 | mamo-s | *BGIBMGA014454* | up | 790 | 798 | - | 12.0734 | 2.66E-05 | 0.615 | GCGCGTGTC |
| 12372 | mamo-s | *BGIBMGA014455* | down | 305 | 313 | + | 11.3303 | 4.96E-05 | 0.615 | GTGGGTGGC |
| 12373 | mamo-s | *BGIBMGA014456* | down | 876 | 884 | + | 13.6606 | 4.06E-06 | 0.615 | GCGCGTGGC |
| 12374 | mamo-s | *BGIBMGA014456* | down | 1012 | 1020 | - | 11.789 | 2.84E-05 | 0.615 | GCGCGTGCC |
| 12375 | mamo-s | *BGIBMGA014456* | down | 161 | 169 | - | 10.8349 | 7.66E-05 | 0.751 | CCGCGAGGC |
| 12376 | mamo-s | *BGIBMGA014457* | down | 1214 | 1222 | + | 11.3303 | 4.96E-05 | 0.615 | GTGGGTGGC |
| 12377 | mamo-s | *BGIBMGA014459* | up | 606 | 614 | - | 11.3303 | 4.96E-05 | 0.615 | GTGGGTGGC |
| 12378 | mamo-s | *BGIBMGA014459* | up | 991 | 999 | + | 11.3303 | 4.96E-05 | 0.615 | GTGGGTGGC |
| 12379 | mamo-s | *BGIBMGA014460* | up | 1192 | 1200 | - | 12.3394 | 1.49E-05 | 0.615 | GGGCGTGGC |
| 12380 | mamo-s | *BGIBMGA014460* | up | 1123 | 1131 | - | 12.0734 | 2.66E-05 | 0.615 | GCGCGTGTC |
| 12381 | mamo-s | *BGIBMGA014461* | up | 1052 | 1060 | - | 10.6881 | 8.61E-05 | 0.81 | GCACGTGGC |
| 12382 | mamo-s | *BGIBMGA014463* | down | 1232 | 1240 | - | 11.3303 | 4.96E-05 | 0.615 | GTGGGTGGC |
| 12383 | mamo-s | *BGIBMGA014463* | down | 1910 | 1918 | + | 11.1376 | 6.35E-05 | 0.709 | TCGCGTGGC |
| 12384 | mamo-s | *BGIBMGA014463* | down | 1346 | 1354 | - | 10.9358 | 7.30E-05 | 0.728 | GCGGGTGGC |
| 12385 | mamo-s | *BGIBMGA014463* | down | 208 | 216 | - | 10.5688 | 8.88E-05 | 0.824 | GTGCGTGAC |
| 12386 | mamo-s | *BGIBMGA014467* | up | 854 | 862 | + | 11.3303 | 4.96E-05 | 0.615 | GTGGGTGGC |
| 12387 | mamo-s | *BGIBMGA014468* | down | 1175 | 1183 | + | 11.3303 | 4.96E-05 | 0.615 | GTGGGTGGC |
| 12388 | mamo-s | *BGIBMGA014469* | up | 1280 | 1288 | - | 12.0734 | 2.66E-05 | 0.615 | GCGCGTGTC |
| 12389 | mamo-s | *BGIBMGA014469* | up | 541 | 549 | + | 11.4404 | 4.51E-05 | 0.615 | GAGCGTGTC |
| 12390 | mamo-s | *BGIBMGA014469* | down | 609 | 617 | - | 11.3303 | 4.96E-05 | 0.615 | GTGGGTGGC |
| 12391 | mamo-s | *BGIBMGA014469* | down | 1185 | 1193 | - | 11.0826 | 6.84E-05 | 0.709 | GTACGTGGC |
| 12392 | mamo-s | *BGIBMGA014470* | up | 1244 | 1252 | + | 11.3303 | 4.96E-05 | 0.615 | GTGGGTGGC |
| 12393 | mamo-s | *BGIBMGA014470* | down | 1723 | 1731 | - | 10.9358 | 7.30E-05 | 0.728 | GCGGGTGGC |
| 12394 | mamo-s | *BGIBMGA014471* | up | 969 | 977 | - | 11.3303 | 4.96E-05 | 0.615 | GTGGGTGGC |
| 12395 | mamo-s | *BGIBMGA014471* | down | 1616 | 1624 | + | 11.0826 | 6.84E-05 | 0.709 | GTACGTGGC |
| 12396 | mamo-s | *BGIBMGA014471* | up | 496 | 504 | - | 10.9358 | 7.30E-05 | 0.728 | GCGGGTGGC |
| 12397 | mamo-s | *BGIBMGA014472* | down | 207 | 215 | - | 11.3303 | 4.96E-05 | 0.615 | GTGGGTGGC |
| 12398 | mamo-s | *BGIBMGA014472* | up | 197 | 205 | + | 11.0826 | 6.84E-05 | 0.709 | GTACGTGGC |
| 12399 | mamo-s | *BGIBMGA014472* | up | 1980 | 1988 | + | 10.9358 | 7.30E-05 | 0.728 | GCGGGTGGC |
| 12400 | mamo-s | *BGIBMGA014473* | down | 1634 | 1642 | + | 12.1835 | 1.71E-05 | 0.615 | GTGCGTGCC |
| 12401 | mamo-s | *BGIBMGA014473* | down | 1571 | 1579 | + | 12.1284 | 2.16E-05 | 0.615 | CAGCGTGGC |

| | | | | | | | | | | |
|---|---|---|---|---|---|---|---|---|---|---|
| 12402 | mamo-s | *BGIBMGA014474* | down | 1276 | 1284 | - | 11.7339 | 3.29E-05 | 0.615 | GCGCGAGGC |
| 12403 | mamo-s | *BGIBMGA014474* | down | 1431 | 1439 | - | 11.5321 | 4.06E-05 | 0.615 | TTGCGTGGC |
| 12404 | mamo-s | *BGIBMGA014478* | down | 1924 | 1932 | + | 11.3303 | 4.96E-05 | 0.615 | GTGGGTGGC |
| 12405 | mamo-s | *BGIBMGA014478* | down | 557 | 565 | - | 11.0826 | 6.84E-05 | 0.709 | GTACGTGGC |
| 12406 | mamo-s | *BGIBMGA014480* | up | 456 | 464 | - | 12.1009 | 2.43E-05 | 0.615 | GTGCGTGGT |
| 12407 | mamo-s | *BGIBMGA014480* | down | 1224 | 1232 | - | 11.3303 | 4.96E-05 | 0.615 | GTGGGTGGC |
| 12408 | mamo-s | *BGIBMGA014481* | down | 1910 | 1918 | + | 11.3303 | 4.96E-05 | 0.615 | GTGGGTGGC |
| 12409 | mamo-s | *BGIBMGA014482* | down | 1866 | 1874 | - | 11.789 | 2.84E-05 | 0.615 | GCGCGTGCC |
| 12410 | mamo-s | *BGIBMGA014482* | down | 1299 | 1307 | + | 11.156 | 6.13E-05 | 0.705 | GAGCGTGCC |
| 12411 | mamo-s | *BGIBMGA014482* | up | 554 | 562 | + | 11.1376 | 6.35E-05 | 0.709 | TCGCGTGGC |
| 12412 | mamo-s | *BGIBMGA014483* | up | 1585 | 1593 | + | 11.3303 | 4.96E-05 | 0.615 | GTGGGTGGC |
| 12413 | mamo-s | *BGIBMGA014484* | down | 846 | 854 | + | 11.3303 | 4.96E-05 | 0.615 | GTGGGTGGC |
| 12414 | mamo-s | *BGIBMGA014484* | up | 1091 | 1099 | - | 11.3303 | 4.96E-05 | 0.615 | GTGGGTGGC |
| 12415 | mamo-s | *BGIBMGA014484* | down | 559 | 567 | - | 10.7523 | 8.38E-05 | 0.797 | GGGCGTGTC |
| 12416 | mamo-s | *BGIBMGA014484* | down | 1435 | 1443 | + | 10.5688 | 8.88E-05 | 0.824 | GTGCGTGAC |
| 12417 | mamo-s | *BGIBMGA014485* | down | 50 | 58 | + | 11.789 | 2.84E-05 | 0.615 | GCGCGTGCC |
| 12418 | mamo-s | *BGIBMGA014485* | down | 617 | 625 | - | 11.156 | 6.13E-05 | 0.705 | GAGCGTGCC |
| 12419 | mamo-s | *BGIBMGA014487* | up | 1923 | 1931 | + | 10.7523 | 8.38E-05 | 0.797 | AAGCGTGGC |
| 12420 | mamo-s | *BGIBMGA014488* | down | 1236 | 1244 | + | 11.3303 | 4.96E-05 | 0.615 | GTGGGTGGC |
| 12421 | mamo-s | *BGIBMGA014489* | down | 1715 | 1723 | + | 11.3303 | 4.96E-05 | 0.615 | GTGGGTGGC |
| 12422 | mamo-s | *BGIBMGA014492* | up | 1509 | 1517 | + | 13.6606 | 4.06E-06 | 0.615 | GCGCGTGGC |
| 12423 | mamo-s | *BGIBMGA014493* | down | 1368 | 1376 | + | 12.1835 | 1.71E-05 | 0.615 | GTGCGTGCC |
| 12424 | mamo-s | *BGIBMGA014494* | up | 221 | 229 | - | 11.3303 | 4.96E-05 | 0.615 | GTGGGTGGC |
| 12425 | mamo-s | *BGIBMGA014494* | down | 255 | 263 | - | 11.3303 | 4.96E-05 | 0.615 | GTGGGTGGC |
| 12426 | mamo-s | *BGIBMGA014494* | up | 1249 | 1257 | - | 10.7523 | 8.38E-05 | 0.797 | AAGCGTGGC |
| 12427 | mamo-s | *BGIBMGA014495* | up | 492 | 500 | - | 11.789 | 2.84E-05 | 0.615 | GCGCGTGCC |
| 12428 | mamo-s | *BGIBMGA014495* | down | 258 | 266 | - | 11.3303 | 4.96E-05 | 0.615 | GTGGGTGGC |
| 12429 | mamo-s | *BGIBMGA014496* | up | 136 | 144 | - | 11.1009 | 6.57E-05 | 0.709 | GAGCGAGGC |
| 12430 | mamo-s | *BGIBMGA014497* | up | 1274 | 1282 | - | 12.7615 | 1.04E-05 | 0.615 | CCGCGTGGC |
| 12431 | mamo-s | *BGIBMGA014497* | down | 999 | 1007 | + | 11.1009 | 6.57E-05 | 0.709 | GAGCGAGGC |
| 12432 | mamo-s | *BGIBMGA014498* | up | 750 | 758 | - | 12.7615 | 1.04E-05 | 0.615 | CCGCGTGGC |
| 12433 | mamo-s | *BGIBMGA014498* | up | 401 | 409 | + | 11.3303 | 4.96E-05 | 0.615 | GTGGGTGGC |
| 12434 | mamo-s | *BGIBMGA014498* | down | 432 | 440 | + | 11.3303 | 4.96E-05 | 0.615 | GTGGGTGGC |
| 12435 | mamo-s | *BGIBMGA014498* | up | 1677 | 1685 | + | 10.7523 | 8.38E-05 | 0.797 | AAGCGTGGC |
| 12436 | mamo-s | *BGIBMGA014500* | down | 493 | 501 | + | 13.156 | 6.30E-06 | 0.615 | CTGCGTGGC |
| 12437 | mamo-s | *BGIBMGA014500* | up | 395 | 403 | - | 11.3303 | 4.96E-05 | 0.615 | GTGGGTGGC |
| 12438 | mamo-s | *BGIBMGA014500* | up | 1179 | 1187 | + | 11.3303 | 4.96E-05 | 0.615 | GTGGGTGGC |
| 12439 | mamo-s | *BGIBMGA014500* | up | 746 | 754 | - | 10.5138 | 9.75E-05 | 0.865 | GTGCGTGTT |
| 12440 | mamo-s | *BGIBMGA014501* | up | 863 | 871 | - | 11.3303 | 4.96E-05 | 0.615 | GTGGGTGGC |
| 12441 | mamo-s | *BGIBMGA014505* | down | 136 | 144 | - | 11.3303 | 4.96E-05 | 0.615 | GTGGGTGGC |
| 12442 | mamo-s | *BGIBMGA014506* | up | 529 | 537 | + | 11.0826 | 6.84E-05 | 0.709 | GTACGTGGC |
| 12443 | mamo-s | *BGIBMGA014507* | down | 1938 | 1946 | + | 11.7798 | 3.11E-05 | 0.615 | ATGCGTGGC |
| 12444 | mamo-s | *BGIBMGA014508* | down | 307 | 315 | - | 11.3303 | 4.96E-05 | 0.615 | GTGGGTGGC |
| 12445 | mamo-s | *BGIBMGA014508* | up | 538 | 546 | - | 11.3303 | 4.96E-05 | 0.615 | GTGGGTGGC |
| 12446 | mamo-s | *BGIBMGA014509* | down | 272 | 280 | + | 11.3303 | 4.96E-05 | 0.615 | GTGGGTGGC |
| 12447 | mamo-s | *BGIBMGA014512* | down | 1473 | 1481 | - | 11.0734 | 7.12E-05 | 0.726 | GAGCGTGGT |
| 12448 | mamo-s | *BGIBMGA014513* | up | 1041 | 1049 | - | 12.4679 | 1.31E-05 | 0.615 | GTGCGTGTC |
| 12449 | mamo-s | *BGIBMGA014514* | down | 1374 | 1382 | - | 12.3394 | 1.49E-05 | 0.615 | GGGCGTGGC |
| 12450 | mamo-s | *BGIBMGA014518* | down | 466 | 474 | - | 13.6606 | 4.06E-06 | 0.615 | GCGCGTGGC |
| 12451 | mamo-s | *BGIBMGA014518* | down | 455 | 463 | - | 11.789 | 2.84E-05 | 0.615 | GCGCGTGCC |
| 12452 | mamo-s | *BGIBMGA014518* | down | 818 | 826 | - | 11.3303 | 4.96E-05 | 0.615 | GTGGGTGGC |
| 12453 | mamo-s | *BGIBMGA014518* | down | 1973 | 1981 | - | 11.156 | 6.13E-05 | 0.705 | GAGCGTGCC |
| 12454 | mamo-s | *BGIBMGA014519* | up | 1229 | 1237 | + | 11.3303 | 4.96E-05 | 0.615 | GTGGGTGGC |
| 12455 | mamo-s | *BGIBMGA014520* | down | 1077 | 1085 | - | 11.3303 | 4.96E-05 | 0.615 | GTGGGTGGC |
| 12456 | mamo-s | *BGIBMGA014522* | up | 1367 | 1375 | - | 12.3394 | 1.49E-05 | 0.615 | GGGCGTGGC |
| 12457 | mamo-s | *BGIBMGA014522* | down | 566 | 574 | + | 11.3303 | 4.96E-05 | 0.615 | GTGGGTGGC |
| 12458 | mamo-s | *BGIBMGA014524* | up | 1512 | 1520 | - | 11.4404 | 4.51E-05 | 0.615 | CGGCGTGGC |
| 12459 | mamo-s | *BGIBMGA014524* | up | 928 | 936 | - | 11.3853 | 4.74E-05 | 0.615 | ACGCGTGGC |
| 12460 | mamo-s | *BGIBMGA014524* | up | 1745 | 1753 | - | 11.1743 | 5.90E-05 | 0.684 | CCGCGTGTC |
| 12461 | mamo-s | *BGIBMGA014524* | up | 1561 | 1569 | - | 11.1376 | 6.35E-05 | 0.709 | TCGCGTGGC |
| 12462 | mamo-s | *BGIBMGA014527* | down | 169 | 177 | - | 10.5138 | 9.75E-05 | 0.865 | GTGCGTGTT |
| 12463 | mamo-s | *BGIBMGA014527* | up | 398 | 406 | - | 10.5138 | 9.75E-05 | 0.865 | GTGCGTGTT |
| 12464 | mamo-s | *BGIBMGA014528* | down | 338 | 346 | - | 10.5138 | 9.75E-05 | 0.865 | GTGCGTGTT |
| 12465 | mamo-s | *BGIBMGA014529* | up | 1229 | 1237 | - | 11.4404 | 4.51E-05 | 0.615 | CGGCGTGGC |
| 12466 | mamo-s | *BGIBMGA014529* | down | 380 | 388 | + | 11.3303 | 4.96E-05 | 0.615 | GTGGGTGGC |
| 12467 | mamo-s | *BGIBMGA014530* | up | 1844 | 1852 | - | 11.5688 | 3.79E-05 | 0.615 | CTGCGTGTC |
| 12468 | mamo-s | *BGIBMGA014530* | up | 376 | 384 | + | 11.0734 | 7.12E-05 | 0.726 | GAGCGTGGT |
| 12469 | mamo-s | *BGIBMGA014531* | down | 886 | 894 | + | 11.3303 | 4.96E-05 | 0.615 | GTGGGTGGC |
| 12470 | mamo-s | *BGIBMGA014531* | up | 1440 | 1448 | + | 11.3303 | 4.96E-05 | 0.615 | GTGGGTGGC |
| 12471 | mamo-s | *BGIBMGA014531* | down | 240 | 248 | + | 11.0826 | 6.84E-05 | 0.709 | GTACGTGGC |
| 12472 | mamo-s | *BGIBMGA014532* | up | 996 | 1004 | + | 11.3303 | 4.96E-05 | 0.615 | GTGGGTGGC |
| 12473 | mamo-s | *BGIBMGA014532* | up | 350 | 358 | + | 11.0826 | 6.84E-05 | 0.709 | GTACGTGGC |
| 12474 | mamo-s | *BGIBMGA014533* | down | 468 | 476 | + | 11.4404 | 4.51E-05 | 0.615 | CGGCGTGGC |
| 12475 | mamo-s | *BGIBMGA014535* | up | 1378 | 1386 | + | 11.3303 | 4.96E-05 | 0.615 | GTGGGTGGC |
| 12476 | mamo-s | *BGIBMGA014537* | down | 1403 | 1411 | - | 11.1743 | 5.90E-05 | 0.684 | CCGCGTGTC |
| 12477 | mamo-s | *BGIBMGA014538* | down | 559 | 567 | + | 11.3303 | 4.96E-05 | 0.615 | GTGGGTGGC |
| 12478 | mamo-s | *BGIBMGA014540* | down | 1419 | 1427 | - | 11.3303 | 4.96E-05 | 0.615 | GTGGGTGGC |
| 12479 | mamo-s | *BGIBMGA014540* | up | 1583 | 1591 | + | 11.3303 | 4.96E-05 | 0.615 | GTGGGTGGC |
| 12480 | mamo-s | *BGIBMGA014541* | up | 1813 | 1821 | - | 11.3303 | 4.96E-05 | 0.615 | GTGGGTGGC |
| 12481 | mamo-s | *BGIBMGA014542* | up | 1822 | 1830 | + | 11.3303 | 4.96E-05 | 0.615 | GTGGGTGGC |
| 12482 | mamo-s | *BGIBMGA014542* | down | 352 | 360 | - | 10.6881 | 8.61E-05 | 0.81 | GCACGTGGC |
| 12483 | mamo-s | *BGIBMGA014544* | up | 461 | 469 | - | 11.789 | 2.84E-05 | 0.615 | GCGCGTGCC |
| 12484 | mamo-s | *BGIBMGA014544* | up | 1246 | 1254 | - | 11.3303 | 4.96E-05 | 0.615 | GTGGGTGGC |
| 12485 | mamo-s | *BGIBMGA014545* | down | 1938 | 1946 | + | 11.3303 | 4.96E-05 | 0.615 | GTGGGTGGC |
| 12486 | mamo-s | *BGIBMGA014546* | down | 194 | 202 | + | 11.3303 | 4.96E-05 | 0.615 | GTGGGTGGC |
| 12487 | mamo-s | *BGIBMGA014546* | down | 954 | 962 | + | 11.3303 | 4.96E-05 | 0.615 | GTGGGTGGC |
| 12488 | mamo-s | *BGIBMGA014546* | up | 567 | 575 | + | 10.7523 | 8.38E-05 | 0.797 | GGGCGTGTC |
| 12489 | mamo-s | *BGIBMGA014548* | down | 181 | 189 | - | 10.9358 | 7.30E-05 | 0.728 | GCGGGTGGC |

| | | | | | | | | | | |
|---|---|---|---|---|---|---|---|---|---|---|
| 12490 | mamo-s | *BGIBMGA014549* | up | 533 | 541 | - | 11.3303 | 4.96E-05 | 0.615 | GTGGGTGGC |
| 12491 | mamo-s | *BGIBMGA014552* | down | 320 | 328 | + | 11.3303 | 4.96E-05 | 0.615 | GTGGGTGGC |
| 12492 | mamo-s | *BGIBMGA014555* | up | 1688 | 1696 | - | 11.156 | 6.13E-05 | 0.705 | GAGCGTGCC |
| 12493 | mamo-s | *BGIBMGA014556* | up | 801 | 809 | - | 11.7798 | 3.11E-05 | 0.615 | ATGCGTGGC |
| 12494 | mamo-s | *BGIBMGA014557* | down | 1503 | 1511 | - | 11.2844 | 5.18E-05 | 0.63 | CTGCGTGCC |
| 12495 | mamo-s | *BGIBMGA014558* | down | 602 | 610 | + | 11.3303 | 4.96E-05 | 0.615 | GTGGGTGGC |
| 12496 | mamo-s | *BGIBMGA014558* | down | 831 | 839 | + | 11.3303 | 4.96E-05 | 0.615 | GTGGGTGGC |
| 12497 | mamo-s | *BGIBMGA014559* | up | 384 | 392 | + | 13.6606 | 4.06E-06 | 0.615 | GCGCGTGGC |
| 12498 | mamo-s | *BGIBMGA014566* | up | 1189 | 1197 | - | 12.4679 | 1.31E-05 | 0.615 | GTGCGTGTC |
| 12499 | mamo-s | *BGIBMGA014566* | up | 1154 | 1162 | + | 11.4404 | 4.51E-05 | 0.615 | CGGCGTGGC |
| 12500 | mamo-s | *BGIBMGA014566* | down | 1818 | 1826 | + | 11.3303 | 4.96E-05 | 0.615 | GTGGGTGGC |
| 12501 | mamo-s | *BGIBMGA014567* | up | 1526 | 1534 | + | 11.3303 | 4.96E-05 | 0.615 | GTGGGTGGC |
| 12502 | mamo-s | *BGIBMGA014568* | up | 431 | 439 | + | 11.3303 | 4.96E-05 | 0.615 | GTGGGTGGC |
| 12503 | mamo-s | *BGIBMGA014568* | down | 1534 | 1542 | - | 11.3303 | 4.96E-05 | 0.615 | GTGGGTGGC |
| 12504 | mamo-s | *BGIBMGA014568* | up | 1497 | 1505 | + | 11.2844 | 5.18E-05 | 0.63 | CTGCGTGCC |
| 12505 | mamo-s | *BGIBMGA014570* | down | 583 | 591 | - | 11.3303 | 4.96E-05 | 0.615 | GTGGGTGGC |
| 12506 | mamo-s | *BGIBMGA014571* | down | 1173 | 1181 | + | 11.0826 | 6.84E-05 | 0.709 | GTACGTGGC |
| 12507 | mamo-s | *BGIBMGA014572* | down | 499 | 507 | + | 11.3303 | 4.96E-05 | 0.615 | GTGGGTGGC |
| 12508 | mamo-s | *BGIBMGA014572* | down | 1138 | 1146 | + | 11.3303 | 4.96E-05 | 0.615 | GTGGGTGGC |
| 12509 | mamo-s | *BGIBMGA014573* | down | 1356 | 1364 | + | 11.3303 | 4.96E-05 | 0.615 | GTGGGTGGC |
| 12510 | mamo-s | *BGIBMGA014573* | up | 52 | 60 | - | 11.0734 | 7.12E-05 | 0.726 | GAGCGTGGT |
| 12511 | mamo-s | *BGIBMGA014574* | down | 442 | 450 | + | 11.3303 | 4.96E-05 | 0.615 | GTGGGTGGC |
| 12512 | mamo-s | *BGIBMGA014574* | down | 1626 | 1634 | - | 11.3303 | 4.96E-05 | 0.615 | GTGGGTGGC |
| 12513 | mamo-s | *BGIBMGA014575* | down | 468 | 476 | - | 14.055 | 2.23E-06 | 0.615 | GTGCGTGGC |
| 12514 | mamo-s | *BGIBMGA014575* | up | 896 | 904 | - | 11.3303 | 4.96E-05 | 0.615 | GTGGGTGGC |
| 12515 | mamo-s | *BGIBMGA014575* | down | 1799 | 1807 | - | 11.3303 | 4.96E-05 | 0.615 | GTGGGTGGC |
| 12516 | mamo-s | *BGIBMGA014586* | up | 774 | 782 | + | 11.3303 | 4.96E-05 | 0.615 | GTGGGTGGC |
| 12517 | mamo-s | *BGIBMGA014586* | down | 1899 | 1907 | - | 11.3303 | 4.96E-05 | 0.615 | GTGGGTGGC |
| 12518 | mamo-s | *BGIBMGA014586* | up | 352 | 360 | - | 11.2844 | 5.18E-05 | 0.63 | CTGCGTGCC |
| 12519 | mamo-s | *BGIBMGA014587* | down | 1867 | 1875 | + | 11.7798 | 3.11E-05 | 0.615 | ATGCGTGGC |
| 12520 | mamo-s | *BGIBMGA014587* | down | 636 | 644 | - | 11.3303 | 4.96E-05 | 0.615 | GTGGGTGGC |
| 12521 | mamo-s | *BGIBMGA014587* | up | 801 | 809 | - | 11.3303 | 4.96E-05 | 0.615 | GTGGGTGGC |
| 12522 | mamo-s | *BGIBMGA014588* | up | 1003 | 1011 | - | 10.5413 | 9.42E-05 | 0.857 | GTGCGAGTC |
| 12523 | mamo-s | *BGIBMGA014589* | up | 1725 | 1733 | - | 11.3303 | 4.96E-05 | 0.615 | GTGGGTGGC |
| 12524 | mamo-s | *BGIBMGA014590* | down | 1197 | 1205 | - | 11.2018 | 5.68E-05 | 0.674 | CTGCGTGGT |
| 12525 | mamo-s | *BGIBMGA014591* | up | 1550 | 1558 | + | 11.3303 | 4.96E-05 | 0.615 | GTGGGTGGC |
| 12526 | mamo-s | *BGIBMGA014591* | down | 1785 | 1793 | + | 10.5138 | 9.75E-05 | 0.865 | GTGCGTGTT |
| 12527 | mamo-s | *BGIBMGA014593* | down | 1282 | 1290 | + | 11.3303 | 4.96E-05 | 0.615 | GTGGGTGGC |
| 12528 | mamo-s | *BGIBMGA014594* | down | 1545 | 1553 | + | 11.3303 | 4.96E-05 | 0.615 | GTGGGTGGC |
| 12529 | mamo-s | *BGIBMGA014595* | up | 609 | 617 | + | 11.3303 | 4.96E-05 | 0.615 | GTGGGTGGC |
| 12530 | mamo-s | *BGIBMGA014595* | up | 1125 | 1133 | - | 11.3303 | 4.96E-05 | 0.615 | GTGGGTGGC |
| 12531 | mamo-s | *BGIBMGA014596* | up | 1252 | 1260 | - | 11.0734 | 7.12E-05 | 0.726 | GAGCGTGGT |
| 12532 | mamo-s | *BGIBMGA014597* | down | 764 | 772 | + | 11.3303 | 4.96E-05 | 0.615 | GTGGGTGGC |
| 12533 | mamo-s | *BGIBMGA014598* | down | 1030 | 1038 | + | 13.6606 | 4.06E-06 | 0.615 | GCGCGTGGC |
| 12534 | mamo-s | *BGIBMGA014598* | down | 1730 | 1738 | + | 12.3394 | 1.49E-05 | 0.615 | GGGCGTGGC |
| 12535 | mamo-s | *BGIBMGA014601* | up | 765 | 773 | + | 11.3303 | 4.96E-05 | 0.615 | GTGGGTGGC |
| 12536 | mamo-s | *BGIBMGA014601* | up | 1210 | 1218 | - | 11.2018 | 5.68E-05 | 0.674 | CTGCGTGGT |
| 12537 | mamo-s | *BGIBMGA014603* | down | 142 | 150 | - | 11.3303 | 4.96E-05 | 0.615 | GTGGGTGGC |
| 12538 | mamo-s | *BGIBMGA014604* | up | 1282 | 1290 | - | 11.2844 | 5.18E-05 | 0.63 | CTGCGTGCC |
| 12539 | mamo-s | *BGIBMGA014604* | down | 639 | 647 | - | 11.1743 | 5.90E-05 | 0.684 | CCGCGTGTC |
| 12540 | mamo-s | *BGIBMGA014604* | up | 862 | 870 | - | 11.0826 | 6.84E-05 | 0.709 | GTACGTGGC |
| 12541 | mamo-s | *BGIBMGA014605* | down | 973 | 981 | + | 11.3303 | 4.96E-05 | 0.615 | GTGGGTGGC |
| 12542 | mamo-s | *BGIBMGA014607* | down | 574 | 582 | - | 11.3303 | 4.96E-05 | 0.615 | GTGGGTGGC |
| 12543 | mamo-s | *BGIBMGA014607* | up | 880 | 888 | - | 11.3303 | 4.96E-05 | 0.615 | GTGGGTGGC |
| 12544 | mamo-s | *BGIBMGA014609* | down | 157 | 165 | - | 11.3303 | 4.96E-05 | 0.615 | GTGGGTGGC |
| 12545 | mamo-s | *BGIBMGA014610* | up | 276 | 284 | + | 11.3303 | 4.96E-05 | 0.615 | GTGGGTGGC |
| 12546 | mamo-s | *BGIBMGA014613* | down | 453 | 461 | - | 11.0826 | 6.84E-05 | 0.709 | GTACGTGGC |
| 12547 | mamo-s | *BGIBMGA014614* | up | 828 | 836 | - | 11.0826 | 6.84E-05 | 0.709 | GTACGTGGC |
| 12548 | mamo-s | *BGIBMGA014617* | up | 522 | 530 | + | 12.3394 | 1.49E-05 | 0.615 | GGGCGTGGC |
| 12549 | mamo-s | *BGIBMGA014619* | down | 1363 | 1371 | + | 11.3303 | 4.96E-05 | 0.615 | GTGGGTGGC |
| 12550 | mamo-s | *BGIBMGA014623* | down | 958 | 966 | - | 11.3303 | 4.96E-05 | 0.615 | GTGGGTGGC |
| 12551 | mamo-s | *BGIBMGA014623* | down | 1199 | 1207 | - | 11.1743 | 5.90E-05 | 0.684 | CCGCGTGTC |

**Supplementary Table S5 Differentially expressed genes between *bd* /*bd* and +/*bd* individulas**

| Number | classification | Gene_id | readcount_*bd* /*bd* | readcount_+ /*bd* | log2.Fold_change. | pvalue | qvalue | Signature | Blast against nr database | |
|---|---|---|---|---|---|---|---|---|---|---|
| | | | | | | | | | E-value | Best hit |
| 1 | Cuticular protein | *BGIBMGA000324* | 348.7065101 | 40.54875149 | 3.1043 | 1.1629E-52 | 3.7028E-50 | TRUE | 7E-65 | ref\|NP_001037001.1\| cuticular protein RR-1 motif 46 precursor [Bombyx mori] |
| 2 | | *BGIBMGA000326* | 4.383227074 | 29.30633681 | -2.7411 | 7.4631E-07 | 0.000039881 | TRUE | 1E-69 | ref\|NP_001166710.1\| cuticular protein RR-1 motif 44 precursor [Bombyx mori] |
| 3 | | *BGIBMGA000328* | 388.1875481 | 29.94808141 | 3.6962 | 4.8524E-69 | 2.0043E-66 | TRUE | 3E-97 | ref\|NP_001166712.1\| cuticular protein RR-1 motif 42 precursor [Bombyx mori] |
| 4 | | *BGIBMGA000329* | 2149.253007 | 458.7523169 | 2.228 | 1.7962E-205 | 1.5251E-202 | TRUE | 1E-126 | tpd\|FAA00544.1\| TPA: putative cuticle protein [Bombyx mori] |
| 5 | | *BGIBMGA000333* | 2378.364606 | 363.9642624 | 2.7081 | 1.7692E-296 | 2.7039E-293 | TRUE | 1E-108 | ref\|NP_001036828.1\| larval cuticle protein LCP-22 precursor [Bombyx mori] |
| 6 | | *BGIBMGA000341* | 342.8835369 | 97.21242298 | 1.8185 | 2.6736E-25 | 4.4902E-23 | TRUE | 2E-46 | ref\|NP_001166722.1\| cuticular protein RR-1 motif 28 precursor [Bombyx mori] |
| 7 | | *BGIBMGA000520* | 72.21126647 | 25.4558692 | 1.5042 | 0.000056876 | 0.0020996 | TRUE | 2E-58 | ref\|NP_001036950.1\| cuticular protein tweedle motif 2 precursor [Bombyx mori] |
| 8 | | *BGIBMGA002384* | 1017.068653 | 438.7393926 | 1.213 | 3.2473E-34 | 6.99E-32 | TRUE | 6E-93 | ref\|NP_001166790.1\| cuticular protein glycine-rich 13 precursor [Bombyx mori] |
| 9 | | *BGIBMGA002548* | 1005.51869 | 112.3528419 | 3.1618 | 2.7804E-151 | 1.7706E-148 | TRUE | 5E-50 | ref\|NP_001036835.1\| cuticular protein RR-1 motif 4 precursor [Bombyx mori] |
| 10 | | *BGIBMGA002549* | 1671.129318 | 243.6252654 | 2.7781 | 1.3266E-215 | 1.1926E-212 | TRUE | 3E-61 | ref\|NP_001166743.1\| cuticular protein RR-1 motif 5 precursor [Bombyx mori] |
| 11 | | *BGIBMGA005259* | 274.6715652 | 55.42771892 | 2.309 | 3.2753E-29 | 6.0309E-27 | TRUE | 4E-35 | ref\|NP_001036951.1\| cuticular protein tweedle motif 3 [Bombyx mori] |
| 12 | | *BGIBMGA005277* | 2831.276741 | 69.49856352 | 5.3483 | 0 | 0 | TRUE | 3E-87 | tpd\|FAA00504.1\| TPA: putative cuticle protein [Bombyx mori] |
| 13 | | *BGIBMGA005278* | 369.6308204 | 53.66886335 | 2.7839 | 2.8627E-49 | 8.4136E-47 | TRUE | 3E-51 | ref\|NP_001166744.1\| cuticular protein RR-1 motif 3 precursor [Bombyx mori] |
| 14 | | *BGIBMGA008255* | 2642.030112 | 421.2696785 | 2.6488 | 0 | 0 | TRUE | / | No hits |
| 15 | | *BGIBMGA011761* | 90.41605629 | 37.22118689 | 1.2805 | 0.00012032 | 0.0041321 | TRUE | 1E-16 | ref\|NP_001166758.1\| cuticular protein hypothetical 19 precursor [Bombyx mori] |
| 16 | | *BGIBMGA011765* | 1788.996533 | 63.27126405 | 4.8215 | 0 | 0 | TRUE | 0.003 | ref\|NP_001166753.1\| cuticular protein hypothetical 24 precursor [Bombyx mori] |
| 17 | | *BGIBMGA011766* | 1440.897916 | 301.3347429 | 2.2575 | 4.7063E-141 | 2.6639E-138 | TRUE | 0.0000001 | ref\|XP_045887395.1\| flocculation protein FLO11-like [Micropterus dolomieu] |
| 18 | | *BGIBMGA012601* | 381.9166539 | 1253.184597 | -1.7143 | 1.9537E-142 | 1.1484E-139 | TRUE | 1E-168 | ref\|NP_001166737.1\| cuticular protein RR-1 motif 11 precursor [Bombyx mori] |
| 19 | | *Novel01073* | 221.1769983 | 61.96400653 | 1.8357 | 3.9885E-17 | 4.549E-15 | TRUE | 1E-56 | ref\|NP_001157379.1\|cuticular protein CPFL family 1 precursor [Bombyx mori] |
| 20 | Antibacterial peptide | *BGIBMGA000021* | 221.7209024 | 69.83131998 | 1.6668 | 9.2957E-15 | 9.2853E-13 | TRUE | 8E-37 | ref\|NP_001037460.1\| cecropin-B precursor [Bombyx mori] |
| 21 | | *BGIBMGA000023* | 82.48145546 | 25.69355238 | 1.6827 | 1.8819E-06 | 0.000095552 | TRUE | 8E-37 | ref\|NP_001037460.1\| cecropin-B precursor [Bombyx mori] |
| 22 | | *BGIBMGA000024* | 56.88596889 | 18.08769044 | 1.6531 | 0.000097881 | 0.0034628 | TRUE | 8E-37 | ref\|NP_001037460.1\| cecropin-B precursor [Bombyx mori] |
| 23 | | *BGIBMGA000039* | 25.21155427 | 3.589016105 | 2.8124 | 0.00010558 | 0.0037181 | TRUE | 2E-20 | ref\|NP_001091844.1\| antibacterial peptide enbocin 2 precursor [Bombyx mori] |
| 24 | | *BGIBMGA002739* | 188.1268263 | 47.13257574 | 1.9969 | 9.2826E-17 | 1.0355E-14 | TRUE | 7E-142 | ref\|XP_004926758.1\| attacin-like [Bombyx mori] |
| 25 | | *BGIBMGA002747* | 127.0815908 | 32.18230335 | 1.9814 | 1.1187E-11 | 9.4461E-10 | TRUE | 1E-142 | ref\|NP_001037006.1\| attacin precursor [Bombyx mori] |
| 26 | | *BGIBMGA005658* | 98.51062892 | 11.50386619 | 3.0982 | 5.1287E-16 | 5.4432E-14 | TRUE | 1E-125 | ref\|NP_001037683.1\| gloverin 2 precursor [Bombyx mori] |
| 27 | | *BGIBMGA005948* | 267.6648008 | 67.28810989 | 1.992 | 4.2779E-23 | 6.2865E-21 | TRUE | 1E-103 | ref\|NP_001165397.1\| C-type lectin 4 precursor [Bombyx mori] |
| 28 | | *BGIBMGA006280* | 224.6003946 | 88.63205997 | 1.3415 | 2.1084E-10 | 1.5952E-08 | TRUE | 1E-22 | ref\|NP_001037462.1\| cecropin-A precursor [Bombyx mori] |
| 29 | | *BGIBMGA006694* | 25.78745271 | 65.83824246 | -1.3523 | 4.2345E-07 | 0.000023113 | TRUE | 0 | ref\|NP_001104817.1\| hemocytin [Bombyx mori] |

| | | | | | | | | | |
|---|---|---|---|---|---|---|---|---|---|
| 30 | *BGIBMGA006775* | 184.9593848 | 39.43164052 | 2.2298 | 2.8565E-19 | 3.7312E-17 | TRUE | 1E-98 | ref\|NP_001119732.1\| lebocin-3 precursor [Bombyx mori] |
| 31 | *BGIBMGA011607* | 101.2301494 | 271.3866615 | -1.4227 | 3.2869E-26 | 5.6442E-24 | TRUE | 0 | ref\|NP_001128672.1\| beta-1,3-glucan recognition protein 3 precursor [Bombyx mori] |
| 32 | *BGIBMGA013803* | 273.7437288 | 77.93631661 | 1.8125 | 2.0494E-20 | 2.7474E-18 | TRUE | 1E-125 | ref\|NP_001093312.1\| gloverin 3 precursor [Bombyx mori] |
| 33 | *BGIBMGA013863* | 226.9999715 | 32.65766972 | 2.7972 | 4.2967E-31 | 8.6403E-29 | TRUE | 1E-118 | ref\|NP_001036930.1\| gloverin 1 precursor [Bombyx mori] |
| 34 | *BGIBMGA014285* | 112.2362086 | 31.51679043 | 1.8323 | 2.1796E-09 | 1.5073E-07 | TRUE | 1E-22 | ref\|NP_001037462.1\| cecropin-A precursor [Bombyx mori] |
| 35 | *Novel01661* | 191.6782 | 94.02746829 | 1.0275 | 0.000010292 | 0.00044558 | TRUE | 0 | ref\|NP_001036879.1\|nimrod B precursor [Bombyx mori] |
| 36 | *Novel02143* | 103.5337432 | 16.56651805 | 2.6438 | 4.1069E-14 | 3.9788E-12 | TRUE | 9E-27 | ref\|NP_001037392.1\|cecropin precursor [Bombyx mori] |
| 37 | *BGIBMGA000291* | 5.663001403 | 24.90919787 | -2.137 | 0.000052612 | 0.0019564 | TRUE | 0 | ref\|XP_004931946.1\| elongation of very long chain fatty acids protein AAEL008004 [Bombyx mori] |
| 38 | *BGIBMGA000301* | 109.6766599 | 26.12138212 | 2.07 | 6.5663E-11 | 5.2267E-09 | TRUE | 0 | ref\|XP_021207482.2\| pancreatic triacylglycerol lipase isoform X2 [Bombyx mori] |
| 39 | *BGIBMGA000402* | 19.19661492 | 52.48044742 | -1.4509 | 2.4489E-06 | 0.00012231 | TRUE | 0 | ref\|KOB74522.1\| putative multiple inositol polyphosphate phosphatase [Operophtera brumata] |
| 40 | *BGIBMGA000424* | 22.4280451 | 62.15415308 | -1.4705 | 2.3438E-07 | 0.000013169 | TRUE | 3E-131 | ref\|XP_028030207.1\| elongation of very long chain fatty acids protein AAEL008004-like [Bombyx mandarina] |
| 41 | *BGIBMGA001333* | 178.4645301 | 65.38664441 | 1.4486 | 1.0234E-09 | 7.309E-08 | TRUE | 0 | ref\|NP_001037382.1\| uricase [Bombyx mori] |
| 42 | *BGIBMGA001891* | 132.6166148 | 25.09934442 | 2.4015 | 1.0866E-15 | 1.1375E-13 | TRUE | 0 | ref\|NP_001153673.1\| carboxypeptidase B-like precursor [Bombyx mori] |
| 43 | *BGIBMGA001968* | 118.2191536 | 20.36944902 | 2.537 | 3.8622E-15 | 3.9351E-13 | TRUE | 2E-95 | ref\|XP_021208460.1\| A disintegrin and metalloproteinase with thrombospondin motifs adt-2 isoform X2 [Bombyx mori] |
| 44 | *BGIBMGA002432* | 23.09992662 | 77.55602352 | -1.7474 | 1.6719E-10 | 1.284E-08 | TRUE | 0 | ref\|XP_004931128.1\| V-type proton ATPase 116 kDa subunit a1 [Bombyx mori] |
| 45 | *BGIBMGA002457* | 9.246369522 | 32.94288955 | -1.833 | 0.000018845 | 0.00075791 | TRUE | 0 | ref\|XP_004931115.1\| retinal dehydrogenase 1 [Bombyx mori] |
| 46 | *BGIBMGA002592* | 79.18603656 | 160.2460038 | -1.017 | 3.6915E-11 | 2.985E-09 | TRUE | 9E-131 | gb\|AWK60411.1\| prolyl oligopeptidase variant 1 [Bombyx mori] |
| 47 | *BGIBMGA003461* | 923.7731045 | 352.222713 | 1.3911 | 1.051E-40 | 2.5098E-38 | TRUE | 0 | ref\|XP_021206214.2\| proclotting enzyme [Bombyx mori] |
| 48 | *BGIBMGA004176* | 38.16926934 | 6.916580706 | 2.4643 | 0.000011954 | 0.0005075 | TRUE | 0 | ref\|XP_037873557.1\| alpha-protein kinase 1 [Bombyx mori] |
| 49 | *BGIBMGA004421* | 180.1922254 | 86.65928953 | 1.0561 | 0.000010219 | 0.00044367 | TRUE | 9E-173 | ref\|XP_028029511.1\| uncharacterized peptidase C1-like protein F26E4.3 [Bombyx mandarina] |
| 50 | *BGIBMGA004655* | 21.0522877 | 48.70128476 | -1.21 | 0.00004834 | 0.0018107 | TRUE | 0 | ref\|XP_037870080.1\| fatty acid synthase [Bombyx mori] |
| 51 | *BGIBMGA005052* | 105.8053426 | 46.79981928 | 1.1768 | 0.00013985 | 0.0047496 | TRUE | 1E-166 | ref\|NP_001036900.1\| ecdysteroid-phosphate phosphatase [Bombyx mori] |
| 52 | *BGIBMGA005320* | 136.7118926 | 34.98696495 | 1.9662 | 2.568E-12 | 2.2818E-10 | TRUE | 0 | ref\|XP_028039390.1\| carboxypeptidase D-like isoform X2 [Bombyx mandarina] |
| 53 | *BGIBMGA005321* | 53.27060641 | 13.97577132 | 1.9304 | 0.000016738 | 0.00068517 | TRUE | 3E-31 | ref\|XP_037868240.1\| carboxypeptidase D isoform X1 [Bombyx mori] |
| 54 | *BGIBMGA005695* | 135.944028 | 65.1251929 | 1.0617 | 0.00011545 | 0.0039921 | TRUE | 0 | gb\|AGN90922.1\| acid lipase-1 [Bombyx mori] |
| 55 | *BGIBMGA005785* | 169.9220364 | 70.18784476 | 1.2756 | 1.5103E-07 | 8.8437E-06 | TRUE | 0 | ref\|XP_004923866.1\| myrosinase 1 [Bombyx mori] |
| 56 | *BGIBMGA006456* | 19.6765303 | 51.98131273 | -1.4015 | 4.3695E-06 | 0.00020921 | TRUE | 0 | ref\|NP_001040411.1\| carboxylesterase clade H, member 1 precursor [Bombyx mori] |
| 57 | *BGIBMGA006745* | 79.9219068 | 27.1196515 | 1.5593 | 0.000011919 | 0.0005075 | TRUE | 0 | dbj\|BAG70891.1\| laccase 2A [Bombyx mori] |
| 58 | *BGIBMGA006816* | 165.8267586 | 339.2689794 | -1.0328 | 2.4718E-22 | 3.4979E-20 | TRUE | 0 | ref\|XP_028041682.1\| CAD protein isoform X1 [Bombyx mandarina] |
| 59 | *BGIBMGA006874* | 1491.704957 | 3830.478454 | -1.3606 | 0 | 0 | TRUE | 0 | ref\|XP_028042303.1\|probable chitinase 10 [Bombyx mandarina] |
| 60 | *BGIBMGA006885* | 104.5255683 | 43.13949822 | 1.2768 | 0.000037478 | 0.0014428 | TRUE | 0 | ref\|XP_028031881.1\| atrial natriuretic peptide-converting enzyme [Bombyx mandarina] |

| | | | | | | | | | | |
|---|---|---|---|---|---|---|---|---|---|---|
| 61 | | *BGIBMGA008029* | 41.46468824 | 91.93585626 | -1.1487 | 6.4296E-08 | 3.9305E-06 | TRUE | 1E-157 | ref\|XP_037868821.1\| allantoicase [Bombyx mori] |
| 62 | | *BGIBMGA008709* | 8184.892699 | 2064.444847 | 1.9872 | 0 | 0 | TRUE | 0 | ref\|NP_001108407.1\| chitinase precursor [Bombyx mori] |
| 63 | Enzyme | *BGIBMGA008815* | 6.270894208 | 25.40833256 | -2.0186 | 0.000074315 | 0.0026914 | TRUE | 7E-124 | pdb\|6KTH_A\| Chain A, Juvenile hormone diol kinase [Bombyx mori] |
| 64 | | *BGIBMGA009184* | 214.5221718 | 57.51933096 | 1.899 | 1.5553E-17 | 1.8285E-15 | TRUE | 0 | gb\|ADA67926.1\| putative peptidase [Bombyx mori] |
| 65 | | *BGIBMGA009518* | 329.7338557 | 662.9459417 | -1.0076 | 7.7729E-41 | 1.8856E-38 | TRUE | 5E-77 | ref\|XP_012552702.2\| alpha-N-acetylgalactosaminidase isoform X2 [Bombyx mori] |
| 66 | | *BGIBMGA009614* | 361.4402647 | 177.3829615 | 1.0269 | 1.4208E-09 | 1.0006E-07 | TRUE | 5E-154 | ref\|XP_012545939.1\| lysosomal thiol reductase IP30 isoform X2 [Bombyx mori] |
| 67 | | *BGIBMGA009743* | 251.2197006 | 58.01846565 | 2.1144 | 9.4284E-24 | 1.4703E-21 | TRUE | 0 | ref\|XP_004933269.3\| serine protease filzig [Bombyx mori] |
| 68 | | *BGIBMGA009744* | 122.3144314 | 33.25187769 | 1.8791 | 1.7751E-10 | 1.3565E-08 | TRUE | 0 | ref\|XP_004933271.1\| decaprenyl-diphosphate synthase subunit 2 [Bombyx mori] |
| 69 | | *BGIBMGA009750* | 484.3305945 | 198.4892284 | 1.2869 | 3.6417E-19 | 4.677E-17 | TRUE | 0 | ref\|XP_012551872.2\| serine proteinase stubble [Bombyx mori] |
| 70 | | *BGIBMGA010214* | 6243.091111 | 2291.907655 | 1.4457 | 1.5412E-285 | 2.1413E-282 | TRUE | 0 | ref\|NP_001139703.1\| serine protease inhibitor 10 precursor [Bombyx mori] |
| 71 | | *BGIBMGA010240* | 2602.069159 | 706.6321113 | 1.8806 | 8.3194E-191 | 6.3572E-188 | TRUE | 0 | dbj\|BAB13481.1\| chitinase precursor [Bombyx mori] |
| 72 | | *BGIBMGA010439* | 284.2698727 | 66.31360883 | 2.0999 | 2.1254E-26 | 3.6913E-24 | TRUE | 5E-99 | ref\|NP_001037448.1\| lysozyme precursor [Bombyx mori] |
| 73 | | *BGIBMGA010681* | 12.92572072 | 40.71512972 | -1.6553 | 7.2255E-06 | 0.00032671 | TRUE | 0 | ref\|NP_001037018.1\| acyl-CoA desaturase [Bombyx mori] |
| 74 | | *BGIBMGA010763* | 306.4099685 | 692.2998152 | -1.1759 | 3.1653E-51 | 9.8724E-49 | TRUE | 0 | gb\|AFK85028.1\| aminopeptidase N-12 [Bombyx mori] |
| 75 | | *BGIBMGA010764* | 199.356846 | 422.2204112 | -1.0826 | 5.8525E-29 | 1.0648E-26 | TRUE | 0 | gb\|AFK85028.1\| aminopeptidase N-12 [Bombyx mori] |
| 76 | | *BGIBMGA011342* | 571.8351642 | 267.1796691 | 1.0978 | 2.2171E-16 | 2.4203E-14 | TRUE | 0 | ref\|NP_001037464.2\| fibroinase precursor [Bombyx mori] |
| 77 | | *BGIBMGA012526* | 156.1644624 | 72.01800529 | 1.1166 | 0.000012369 | 0.00052076 | TRUE | 0 | ref\|XP_004925227.1\| alanine--glyoxylate aminotransferase 2-like [Bombyx mori] |
| 78 | | *BGIBMGA012831* | 766.616817 | 244.3145467 | 1.6498 | 2.9649E-46 | 8.2388E-44 | TRUE | 0 | ref\|XP_004928930.1\| aldo-keto reductase AKR2E4 isoform X1 [Bombyx mori] |
| 79 | | *BGIBMGA012862* | 21.18026513 | 57.28164777 | -1.4354 | 1.0015E-06 | 0.000052777 | TRUE | 2E-27 | ref\|XP_037872352.1\| glucose dehydrogenase [FAD, quinone] [Bombyx mori] |
| 80 | | *BGIBMGA012863* | 87.92049635 | 215.3885029 | -1.2927 | 5.543E-19 | 7.0011E-17 | TRUE | 0 | ref\|XP_037872352.1\| glucose dehydrogenase [FAD, quinone] [Bombyx mori] |
| 81 | | *BGIBMGA012997* | 58.32571501 | 1.49740407 | 5.2836 | 4.2545E-14 | 4.0894E-12 | TRUE | 0 | ref\|XP_004928920.1\| glucose dehydrogenase [FAD, quinone]-like isoform X1 [Bombyx mori] |
| 82 | | *BGIBMGA013009* | 202.2363382 | 510.9475444 | -1.3371 | 1.6347E-44 | 4.383E-42 | TRUE | 0 | ref\|XP_028043358.1\| glucose dehydrogenase [FAD, quinone]-like [Bombyx mandarina] |
| 83 | | *BGIBMGA013737* | 29.01888289 | 101.7521718 | -1.81 | 8.2883E-14 | 7.8192E-12 | TRUE | 0 | ref\|XP_028030680.1\| CLIP domain-containing serine protease 2-like [Bombyx mandarina] |
| 84 | | *BGIBMGA013816* | 80.81774883 | 295.3451266 | -1.8697 | 2.2523E-38 | 5.1377E-36 | TRUE | 0 | ref\|XP_028035390.1\| hydroxylysine kinase isoform X1 [Bombyx mandarina] |
| 85 | | *BGIBMGA014198* | 416.8864874 | 199.1072047 | 1.0661 | 1.17E-11 | 9.8251E-10 | TRUE | 0 | ref\|XP_004927708.1\| apyrase [Bombyx mori] |
| 86 | | *BGIBMGA014292* | 3013.100678 | 589.3354591 | 2.3541 | 0 | 0 | TRUE | 1E-13 | ref\|KAG1657830.1\| Heparan-alpha-glucosaminide N-acetyltransferase [Nymphon striatum] |
| 87 | | *BGIBMGA014453* | 35.00182788 | 165.5701072 | -2.2419 | 1.1664E-26 | 2.0489E-24 | TRUE | 0 | gb\|ADQ89807.1\| aldo-keto reductase [Bombyx mori] |
| 88 | | *Novel00038* | 53.04664591 | 108.5974476 | -1.0337 | 3.7559E-08 | 2.3918E-06 | TRUE | 9E-34 | gi\|AWK60411.1\| prolyl oligopeptidase variant 1 [Bombyx mori] |
| 89 | | *Novel00078* | 23.16391534 | 49.62824919 | -1.0993 | 0.00011113 | 0.0038688 | TRUE | 1E-65 | ref\|XP_037868821.1\|allantoicase [Bombyx mori] |
| 90 | | *BGIBMGA000319* | 27.03523268 | 110.0710833 | -2.0255 | 1.4335E-16 | 1.5875E-14 | TRUE | 0 | ref\|XP_021207484.2\| PR domain zinc finger protein 1 isoform X2 [Bombyx mori] |
| 91 | | *BGIBMGA000987* | 9.310358238 | 40.4299099 | -2.1185 | 2.9248E-07 | 0.000016137 | TRUE | 9E-14 | ref\|AFI61405.1\| OVO protein [Bombyx mori] |

| | | | | | | | | | | |
|---|---|---|---|---|---|---|---|---|---|---|
| 92 | Transcription factor | *BGIBMGA000988* | 60.66130316 | 262.7112252 | -2.1146 | 5.5393E-39 | 1.3024E-36 | TRUE | 0 | gb\|XP_021204666.2\| protein ovo isoform X1 [Bombyx mori] |
| 93 | | *BGIBMGA003874* | 62.13304364 | 20.77351044 | 1.5806 | 0.000092415 | 0.0032922 | TRUE | 2E-142 | ref\|XP_012544801.1\| hepatic leukemia factor isoform X11 [Bombyx mori] |
| 94 | | *BGIBMGA007065* | 1602.821363 | 410.9542282 | 1.9636 | 9.415E-127 | 4.9617E-124 | TRUE | 1E-144 | ref\|XP_012551266.1\| zinc finger protein hangover isoform X2 [Bombyx mori] |
| 95 | Transporter protein | *BGIBMGA002430* | 7.102747522 | 33.91739061 | -2.2556 | 1.2223E-06 | 0.000063325 | TRUE | 0 | ref\|XP_021206944.1\| synaptic vesicle glycoprotein 2B isoform X1 [Bombyx mori] |
| 96 | | *BGIBMGA013242* | 19.6765303 | 1.140879292 | 4.1083 | 0.000037626 | 0.0014448 | TRUE | 0 | ref\|NP_001243989.1\| major facilitator superfamily protein [Bombyx mori] |
| 97 | | *BGIBMGA013572* | 6.398871641 | 28.07038424 | -2.1332 | 0.000017984 | 0.00072711 | TRUE | 0 | ref\|XP_004925280.1\| lysosome membrane protein 2 isoform X2 [Bombyx mori] |
| 98 | | *BGIBMGA013576* | 152.2931451 | 70.35422299 | 1.1141 | 0.000016634 | 0.00068337 | TRUE | 0 | ref\|XP_004925273.3\| organic cation/carnitine transporter 7 [Bombyx mori] |
| 99 | | *BGIBMGA013656* | 151.749241 | 59.68224795 | 1.3463 | 1.5976E-07 | 9.3193E-06 | TRUE | 0 | ref\|XP_037874017.1\| solute carrier family 2, facilitated glucose transporter member 6 [Bombyx mori] |
| 100 | | *BGIBMGA011016* | 91.47187011 | 231.0042882 | -1.3365 | 5.0693E-21 | 6.9173E-19 | TRUE | 0 | ref\|XP_004921746.1\| organic cation transporter protein [Bombyx mori] |
| 101 | Cytochrome | *BGIBMGA001162* | 117.6752495 | 466.5483253 | -1.9872 | 1.3205E-63 | 4.9221E-61 | TRUE | 0 | dbj\|BAM73799.1\| cytochrome P450 [Bombyx mori] |
| 102 | | *BGIBMGA001753* | 7.166736238 | 96.64198333 | -3.7533 | 6.0528E-24 | 9.5366E-22 | TRUE | 0 | dbj\|BAH47267.1\| cytochrome P450 [Bombyx mori] |
| 103 | | *BGIBMGA006916* | 173.7933538 | 56.99642795 | 1.6084 | 3.0564E-11 | 2.4979E-09 | TRUE | 0 | ref\|NP_001077078.1\| cytochrome P450 18a1 [Bombyx mori] |
| 104 | | *BGIBMGA009522* | 163.811114 | 67.59709803 | 1.277 | 2.4448E-07 | 0.000013686 | TRUE | 0 | dbj\|BAM73845.1\| cytochrome P450, partial [Bombyx mori] |
| 105 | | *BGIBMGA009523* | 203.5481069 | 60.15761432 | 1.7586 | 7.1932E-15 | 7.2325E-13 | TRUE | 0 | ref\|NP_001274759.1\| probable cytochrome P450 49a1-like [Bombyx mori] |
| 106 | | *BGIBMGA000010* | 591.2237453 | 148.4331495 | 1.9939 | 4.6733E-49 | 1.3476E-46 | TRUE | 1E-131 | ref\|NP_001129360.1\| osiris 9 precursor [Bombyx mori] |
| 107 | | *BGIBMGA000012* | 1814.815981 | 263.9709461 | 2.7814 | 2.5999E-234 | 2.8382E-231 | TRUE | 1E-136 | ref\|XP_004926115.1\| uncharacterized protein LOC101741151 [Bombyx mori] |
| 108 | | *BGIBMGA000013* | 57.42987298 | 10.81458495 | 2.4088 | 1.246E-07 | 7.3526E-06 | TRUE | 1E-96 | gi\|389610551\|dbj\|BAM18887.1\| osiris 9A [Papilio polytes] |
| 109 | | *BGIBMGA000025* | 74.2589054 | 13.14388017 | 2.4982 | 7.0749E-10 | 5.1489E-08 | TRUE | 0 | ref\|XP_004926131.1\| uncharacterized protein LOC101743499 [Bombyx mori] |
| 110 | | *BGIBMGA000045* | 1239.397448 | 157.6077205 | 2.9752 | 6.1495E-174 | 4.4754E-171 | TRUE | 1E-111 | ref\|XP_004926113.1\| uncharacterized protein LOC101740865 [Bombyx mori] |
| 111 | | *BGIBMGA000590* | 368.7669727 | 179.7597934 | 1.0366 | 6.3829E-10 | 4.6675E-08 | TRUE | 6E-66 | ref\|NP_001108476.1\| pleiotrophin-like protein precursor [Bombyx mori |
| 112 | | *BGIBMGA000691* | 2567.739212 | 1262.929608 | 1.0237 | 3.0423E-58 | 1.0108E-55 | TRUE | 0 | ref\|XP_012549077.2\| collagen alpha-2(I) chain [Bombyx mori] |
| 113 | | *BGIBMGA000884* | 209.723018 | 78.19776812 | 1.4233 | 7.6423E-11 | 6.0205E-09 | TRUE | 1E-94 | ref\|XP_037870846.1\| lipopolysaccharide-induced tumor necrosis factor-alpha factor homolog isoform X1 [Bombyx mori] |
| 114 | | *BGIBMGA001325* | 34.9058448 | 82.7850536 | -1.2459 | 6.8733E-08 | 4.1684E-06 | TRUE | 1E-148 | ref\|NP_001036945.1\| juvenile hormone binding protein an-0128 precursor [Bombyx mori] |
| 115 | | *BGIBMGA001452* | 54.42240331 | 8.913119466 | 2.6102 | 5.5129E-08 | 3.3973E-06 | TRUE | 1E-129 | ref\|XP_012548820.1\| flocculation protein FLO11 isoform X2 [Bombyx mori] |
| 116 | | *BGIBMGA001713* | 214.9061041 | 104.74698 | 1.0368 | 2.3717E-06 | 0.00011884 | TRUE | 0 | ref\|XP_037870105.1\| protein toll isoform X3 [Bombyx mori] |
| 117 | | *BGIBMGA001899* | 333.0292746 | 110.4513764 | 1.5922 | 8.0081E-20 | 1.0642E-17 | TRUE | 0 | ref\|XP_012552045.1\| uncharacterized protein LOC101743429 [Bombyx mori] |
| 118 | | *BGIBMGA002168* | 233.1428882 | 96.28545856 | 1.2758 | 7.6301E-10 | 5.5266E-08 | TRUE | 0 | ref\|XP_004924462.1\| ichor [Bombyx mori] |
| 119 | | *BGIBMGA002561* | 13.30965301 | 68.42898918 | -2.3621 | 1.7491E-12 | 1.5724E-10 | TRUE | / | No hits |
| 120 | | *BGIBMGA002562* | 28.02705779 | 167.1150479 | -2.5759 | 1.4777E-30 | 2.8587E-28 | TRUE | 3E-43 | ref\|XP_037867253.1\| spidroin-1 isoform X2 [Bombyx mori] |
| 121 | | *BGIBMGA002566* | 21.11627642 | 47.06127078 | -1.1562 | 0.00010293 | 0.0036329 | TRUE | 9E-17 | ref\|XP_037867173.1\| glycine-rich cell wall structural protein [Bombyx mori] |
| 122 | | *BGIBMGA002567* | 35.41775453 | 127.2555777 | -1.8452 | 3.1763E-17 | 3.6775E-15 | TRUE | / | No hits |

| | | | | | | | | | | |
|---|---|---|---|---|---|---|---|---|---|---|
| 123 | | *BGIBMGA002624* | 394.5864198 | 78.31660971 | 2.333 | 8.1707E-42 | 2.0471E-39 | TRUE | 0 | ref\|KAI8438262.1\| hypothetical protein MSG28_010859 [Choristoneura fumiferana] |
| 124 | | *BGIBMGA002668* | 155.5565696 | 20.79727875 | 2.903 | 9.5222E-23 | 1.386E-20 | TRUE | 4E-114 | ref\|XP_004932345.1\| uncharacterized protein LOC101736422 [Bombyx mori] |
| 125 | | *BGIBMGA002696* | 28.4749788 | 58.77905184 | -1.0456 | 0.000045642 | 0.0017223 | TRUE | 0 | ref\|XP_037866855.1\| adhesive plaque matrix protein isoform X2 [Bombyx mori] |
| 126 | | *BGIBMGA003122* | 149.9575569 | 45.82531822 | 1.7103 | 7.0774E-11 | 5.6043E-09 | TRUE | 0 | ref\|XP_012550007.2\| leucine-rich repeat and fibronectin type III domain-containing protein 1-like protein [Bombyx mori] |
| 127 | | *BGIBMGA003274* | 129.4811677 | 33.63217079 | 1.9448 | 1.4603E-11 | 1.2196E-09 | TRUE | 4E-127 | ref\|XP_037873326.1\| uncharacterized protein LOC101741103 [Bombyx mori] |
| 128 | | *BGIBMGA003481* | 84.62507746 | 242.0090198 | -1.5159 | 2.8013E-25 | 4.6535E-23 | TRUE | 0 | ref\|NP_001036961.1\| wing disc-specific protein [Bombyx mori] |
| 129 | | *BGIBMGA003510* | 271.0242084 | 73.30149449 | 1.8865 | 1.6054E-21 | 2.2305E-19 | TRUE | 0 | ref\|XP_012549213.2\| uncharacterized protein LOC101739696 [Bombyx mori] |
| 130 | | *BGIBMGA003773* | 100.0463581 | 8.081228316 | 3.63 | 9.1918E-19 | 1.1329E-16 | TRUE | 8E-97 | ref\|CAH0726225.1\| unnamed protein product [Brenthis ino] |
| 131 | | *BGIBMGA004035* | 384.156259 | 785.2339408 | -1.0314 | 2.0219E-49 | 6.059E-47 | TRUE | 1E-78 | ref\|NP_001037066.1\| chemosensory protein precursor [Bombyx mori] |
| 132 | | *BGIBMGA004040* | 37.75334268 | 217.5038833 | -2.5264 | 1.3947E-38 | 3.2296E-36 | TRUE | 2E-88 | ref\|NP_001037067.1\| chemosensory protein 8 precursor [Bombyx mori] |
| 133 | | *BGIBMGA004045* | 66.03635534 | 165.7127171 | -1.3274 | 2.113E-15 | 2.1819E-13 | TRUE | 7E-87 | ref\|NP_001037052.1\| chemosensory protein 4 precursor [Bombyx mori] |
| 134 | | *BGIBMGA004063* | 0 | 16.28129823 | -7.9912 | 0.000043176 | 0.0016333 | TRUE | 0 | ref\|XP_028032517.1\| uncharacterized protein LOC114244806 [Bombyx mandarina] |
| 135 | | *BGIBMGA004073* | 16.50908883 | 80.71720989 | -2.2896 | 4.6267E-14 | 4.4193E-12 | TRUE | / | No hits |
| 136 | | *BGIBMGA004378* | 108.4608743 | 26.95327327 | 2.0086 | 2.2692E-10 | 0.000000017 | TRUE | 0 | ref\|XP_004922581.2\| retinaldehyde-binding protein 1 isoform X1 [Bombyx mori] |
| 137 | | *BGIBMGA004397* | 28.69893931 | 81.43025944 | -1.5046 | 1.988E-09 | 1.381E-07 | TRUE | 0 | ref\|NP_001266309.1\| low molecular mass 30 kDa lipoprotein 19G1-like precursor [Bombyx mori] |
| 138 | | *BGIBMGA004398* | 24.41169531 | 52.19522759 | -1.0963 | 0.000075894 | 0.0027356 | TRUE | 1E-178 | ref\|NP_001266375.1\| low molecular 30 kDa lipoprotein PBMHPC-19-like precursor [Bombyx mori] |
| 139 | | *BGIBMGA004613* | 174.3692522 | 56.85381804 | 1.6168 | 2.2895E-11 | 1.8913E-09 | TRUE | 1E-174 | ref\|XP_026180192.1\| heat shock 70 kDa protein 1-like [Mastacembelus armatus] |
| 140 | | *BGIBMGA004614* | 210.970798 | 63.50894724 | 1.732 | 5.3689E-15 | 5.434E-13 | TRUE | 0 | ref\|OWR50327.1\| HSP70 protein [Danaus plexippus plexippus] |
| 141 | | *BGIBMGA004906* | 503.7191756 | 203.3379654 | 1.3087 | 1.6647E-20 | 2.2514E-18 | TRUE | 0 | ref\|CAH4036867.1\| unnamed protein product [Pieris brassicae] |
| 142 | | *BGIBMGA005119* | 266.5449982 | 96.35676351 | 1.4679 | 4.1134E-14 | 3.9788E-12 | TRUE | 0 | ref\|XP_012545066.1\| balbiani ring protein 3 isoform X2 [Bombyx mori] |
| 143 | | *BGIBMGA005823* | 258.0344989 | 67.35941485 | 1.9376 | 2.0412E-21 | 2.8104E-19 | TRUE | 1E-114 | ref\|NP_001276564.1\| alpha-crystallin B chain-like precursor [Bombyx mori] |
| 144 | | *BGIBMGA006288* | 737.5339454 | 178.4049992 | 2.0476 | 2.7417E-63 | 9.9766E-61 | TRUE | 0 | ref\|XP_012553311.1\| uncharacterized protein LOC101739351 [Bombyx mori] |
| 145 | | *BGIBMGA006315* | 131.3048461 | 61.01327379 | 1.1057 | 0.000073439 | 0.0026659 | TRUE | 0 | ref\|XP_004923549.1\| uncharacterized protein LOC101740768 [Bombyx mori] |
| 146 | | *BGIBMGA006730* | 3698.70778 | 1159.085824 | 1.674 | 6.3426E-222 | 6.0584E-219 | TRUE | 1E-79 | ref\|XP_004930398.1\| uncharacterized protein LOC101746207 [Bombyx mori] |
| 147 | | *BGIBMGA006731* | 1789.348471 | 465.0509213 | 1.944 | 4.8437E-139 | 2.6438E-136 | TRUE | 1E-83 | ref\|XP_004930399.1\| uncharacterized protein LOC101746349 [Bombyx mori] |
| 148 | other | *BGIBMGA006742* | 109.8366317 | 37.07857698 | 1.5667 | 2.5194E-07 | 0.000014052 | TRUE | 5E-95 | ref\|XP_028038519.1\| troponin C-like [Bombyx mandarina] |
| 149 | | *BGIBMGA007040* | 37.01747244 | 7.106727255 | 2.3809 | 0.000025564 | 0.001007 | TRUE | 1E-82 | ref\|XP_004932476.1\| uncharacterized protein LOC101738364 [Bombyx mori] |
| 150 | | *BGIBMGA007250* | 7.422691104 | 37.55394335 | -2.339 | 1.9979E-07 | 0.000011351 | TRUE | 0 | ref\|XP_037872693.1\| mucin-22 [Bombyx mori] |
| 151 | | *BGIBMGA007388* | 46.48780247 | 152.5213003 | -1.7141 | 7.8986E-19 | 9.8142E-17 | TRUE | 0 | ref\|XP_012543959.1\| Ig-like and fibronectin type-III domain-containing protein 1 isoform X1 [Bombyx mori] |
| 152 | | *BGIBMGA007827* | 152.5171056 | 65.98085237 | 1.2089 | 2.5422E-06 | 0.00012614 | TRUE | 3E-23 | ref\|KAF9795210.1\| hypothetical protein SFRURICE_004582 [Spodoptera frugiperda] |
| 153 | | *BGIBMGA007828* | 417.3664028 | 177.9058645 | 1.2302 | 2.2313E-15 | 2.2887E-13 | TRUE | 1E-151 | ref\|XP_004928543.1\| uncharacterized protein LOC101740597 [Bombyx mori] |

| | | | | | | | | | |
|---|---|---|---|---|---|---|---|---|---|
| 154 | *BGIBMGA007829* | 853.4175108 | 359.3056719 | 1.248 | 1.1866E-30 | 2.325E-28 | TRUE | 0 | ref\|XP_004928543.1\| uncharacterized protein LOC101740597 [Bombyx mori] |
| 155 | *BGIBMGA008091* | 138.1516387 | 60.11007768 | 1.2006 | 0.000008778 | 0.00038773 | TRUE | 3E-178 | ref\|XP_004925664.1\| uncharacterized protein LOC101736398 [Bombyx mori] |
| 156 | *BGIBMGA008346* | 782.8059622 | 189.9326337 | 2.0432 | 7.6618E-67 | 3.0024E-64 | TRUE | 1E-146 | ref\|XP_004928246.1\| PREDICTED: uncharacterized protein LOC101745570 isoform X1 [Bombyx mori] |
| 157 | *BGIBMGA008575* | 1023.147581 | 510.2820315 | 1.0036 | 3.3788E-23 | 5.0626E-21 | TRUE | 0 | ref\|XP_037873351.1\| uncharacterized protein LOC101738605 [Bombyx mori] |
| 158 | *BGIBMGA008841* | 171.2977938 | 47.51286884 | 1.8501 | 9.1436E-14 | 8.5731E-12 | TRUE | 0 | ref\|NP_001243947.1\| spatzle-like protein [Bombyx mori] |
| 159 | *BGIBMGA009108* | 0 | 19.63263114 | -8.2612 | 9.5948E-06 | 0.00042017 | TRUE | 3E-10 | emb\|CAG4956167.1\| unnamed protein product [Parnassius apollo] |
| 160 | *BGIBMGA010029* | 4.703170656 | 29.52025167 | -2.65 | 9.963E-07 | 0.000052687 | TRUE | 0 | ref\|XP_004923319.2\| mucin-5AC [Bombyx mori] |
| 161 | *BGIBMGA010716* | 56.98195197 | 13.97577132 | 2.0276 | 3.6382E-06 | 0.00017764 | TRUE | 0 | ref\|NP_001129362.1\| osiris 19 precursor [Bombyx mori] |
| 162 | *BGIBMGA010779* | 927.0685234 | 434.5324002 | 1.0932 | 2.3252E-25 | 3.9485E-23 | TRUE | 0 | ref\|XP_012551662.2\| protein Skeletor, isoforms D/E isoform X1 [Bombyx mori] |
| 163 | *BGIBMGA011094* | 514.3413025 | 224.4680006 | 1.1962 | 1.2819E-17 | 1.5187E-15 | TRUE | 0 | ref\|XP_004921750.1\| proline-rich extensin-like protein EPR1 [Bombyx mori] |
| 164 | *BGIBMGA011298* | 222.7767162 | 110.3087665 | 1.0141 | 2.8186E-06 | 0.00013941 | TRUE | 0 | ref\|XP_004930684.1\| uncharacterized protein LOC101740799 [Bombyx mori] |
| 165 | *BGIBMGA011399* | 237.0781943 | 80.76474652 | 1.5536 | 5.6014E-14 | 5.3172E-12 | TRUE | 2E-51 | ref\|XP_004930780.1\| uncharacterized protein LOC101741978 [Bombyx mori] |
| 166 | *BGIBMGA011666* | 67.63607325 | 20.41698566 | 1.728 | 9.9824E-06 | 0.00043465 | TRUE | 0 | ref\|XP_037870070.1\|uncharacterized protein LOC105842372 isoform X5 [Bombyx mori] |
| 167 | *BGIBMGA011901* | 21.18026513 | 0 | 7.4775 | 0.000007857 | 0.00035317 | TRUE | 0 | ref\|NP_001166287.1\| sericin 2 isoform 1 precursor [Bombyx mori] |
| 168 | *BGIBMGA011904* | 247.0604341 | 29.54401999 | 3.0639 | 2.664E-37 | 5.9874E-35 | TRUE | 0 | ref\|XP_037869958.1\|extensin-2 isoform X2 [Bombyx mori] |
| 169 | *BGIBMGA012419* | 5.151091671 | 27.7613961 | -2.4301 | 5.2321E-06 | 0.00024528 | TRUE | 2E-129 | ref\|XP_021203323.1\| uncharacterized protein LOC110385097 [Bombyx mori] |
| 170 | *BGIBMGA012957* | 69.71570653 | 24.88542955 | 1.4862 | 0.000092128 | 0.0032897 | TRUE | 9E-77 | ref\|CAH2093933.1\| unnamed protein product [Euphydryas editha] |
| 171 | *BGIBMGA012958* | 241.2694552 | 85.23319042 | 1.5012 | 2.0695E-13 | 1.9169E-11 | TRUE | 3E-127 | ref\|XP_028035292.1\| uncharacterized protein LOC114246796 isoform X1 [Bombyx mandarina] |
| 172 | *BGIBMGA012968* | 405.4005128 | 156.3955362 | 1.3741 | 2.2789E-18 | 2.7642E-16 | TRUE | 0 | ref\|XP_028035271.1\| neurogenic locus notch homolog protein 1 [Bombyx mandarina] |
| 173 | *BGIBMGA013017* | 109.2927276 | 46.56213609 | 1.231 | 0.000049268 | 0.001841 | TRUE | 0 | ref\|NP_001040355.1\| cellular retinaldehyde-binding protein [Bombyx mori] |
| 174 | *BGIBMGA013026* | 61.97307185 | 12.24068407 | 2.34 | 7.4406E-08 | 4.4947E-06 | TRUE | 0 | ref\|XP_012548192.1\| uncharacterized protein LOC105842064 isoform X2 [Bombyx mori] |
| 175 | *BGIBMGA013041* | 109.7086543 | 48.24968671 | 1.1851 | 0.000093425 | 0.0033205 | TRUE | 5E-47 | ref\|XP_004928949.1\| somatomedin-B and thrombospondin type-1 domain-containing protein [Bombyx mori] |
| 176 | *BGIBMGA013042* | 489.3217144 | 217.7891031 | 1.1679 | 4.5038E-16 | 4.8134E-14 | TRUE | 8E-93 | ref\|XP_004928949.1\| somatomedin-B and thrombospondin type-1 domain-containing protein [Bombyx mori] |
| 177 | *BGIBMGA013078* | 13.21366994 | 58.61267361 | -2.1492 | 4.9279E-10 | 3.6383E-08 | TRUE | 1E-119 | ref\|XP_021205411.1\| uncharacterized protein LOC101739604 isoform X2 [Bombyx mori] |
| 178 | *BGIBMGA013376* | 434.8993111 | 124.2607695 | 1.8073 | 2.47E-31 | 5.0333E-29 | TRUE | 0 | ref\|XP_037876760.1\|uncharacterized protein LOC101743303 isoform X1 [Bombyx mori] |
| 179 | *BGIBMGA013377* | 78.99407041 | 19.25233805 | 2.0367 | 4.4481E-08 | 2.7747E-06 | TRUE | 6E-131 | ref\|XP_037876763.1\| uncharacterized protein LOC101743303 isoform X4 [Bombyx mori] |
| 180 | *BGIBMGA013448* | 1256.770385 | 530.2949558 | 1.2449 | 4.0823E-44 | 1.0757E-41 | TRUE | 0 | ref\|NP_001040421.1\| SPARC precursor [Bombyx mori] |
| 181 | *BGIBMGA013655* | 705.6355702 | 220.0708617 | 1.681 | 4.2129E-44 | 1.0913E-41 | TRUE | 0 | ref\|XP_004927378.1\| agrin [Bombyx mori] |
| 182 | *BGIBMGA013881* | 103.9816642 | 36.93596707 | 1.4932 | 1.6091E-06 | 0.000082247 | TRUE | 2E-128 | ref\|XP_028035368.1\| uncharacterized protein LOC114246851 [Bombyx mandarina] |
| 183 | *BGIBMGA013949* | 142.6628432 | 14.80766247 | 3.2682 | 1.0108E-23 | 1.5604E-21 | TRUE | 0 | ref\|XP_004933585.1\| PREDICTED: flocculation protein FLO11-like [Bombyx mori] |
| 184 | *BGIBMGA014039* | 3559.404345 | 1639.966445 | 1.118 | 3.5713E-97 | 1.7057E-94 | TRUE | 0 | ref\|XP_037877659.1\| collagen, type IV isoform X1 [Bombyx mori] |

| | | | | | | | | | |
|---|---|---|---|---|---|---|---|---|---|
| 185 | *BGIBMGA014040* | 2639.278597 | 1247.90803 | 1.0806 | 2.216E-67 | 8.9122E-65 | TRUE | 0 | ref\|XP_028043402.1\| collagen alpha-2(IV) chain [Bombyx mandarina] |
| 186 | *BGIBMGA014559* | 67.79604504 | 18.6818984 | 1.8596 | 2.4961E-06 | 0.00012426 | TRUE | 0 | ref\|XP_037867691.1\| protein PFC0760c [Bombyx mori] |
| 187 | *BGIBMGA014618* | 116.5554469 | 32.65766972 | 1.8355 | 1.0166E-09 | 7.2943E-08 | TRUE | 1E-173 | ref\|XP_015193124.1\| PREDICTED: heat shock cognate 70 kDa protein [Lepisosteus oculatus] |
| 188 | *Novel00049* | 793.5240722 | 2131.661652 | -1.4256 | 2.6543E-194 | 2.135E-191 | TRUE | 1.6 | ref\|XP_010237988.1\|PREDICTED: nucleoporin nup189-like [Brachypodium distachyon] |
| 189 | *Novel01771* | 595.0310739 | 168.8501352 | 1.8172 | 1.2731E-42 | 3.2428E-40 | TRUE | 0 | ref\|XP_004929514.1\|glucose dehydrogenase [FAD, quinone] [Bombyx mori] |
| 190 | *Novel01918* | 8.478504925 | 33.06173114 | -1.9633 | 8.5091E-06 | 0.00037914 | TRUE | / | No hits |
| 191 | *Novel02144* | 139.6873679 | 39.17018901 | 1.8344 | 2.359E-11 | 1.9383E-09 | TRUE | 0.0001 | gi\|238623776\|dbj\|BAH66362.1\|olfactory receptor [Bombyx mori] |

Supplementary Table S6. The ortholog of 19 cuticular protein genes.

| Name | Family | Gene in SilkDB | Gene in Silkbase | Gene in *D.melanogaster* | Gene in *Danaus plexippus* | Accession number of *Papilio xuthus* |
|---|---|---|---|---|---|---|
| *BmorCPR2* | RR-1 | *BGIBMGA005277* | *KWMTBOMO12562* | *Cuticular protein 67Fb* | *DPOGS204934* | NP_001299352.1 |
| *BmorCPR3* | RR-1 | *BGIBMGA005278* | *KWMTBOMO04385* | *Cuticular protein 65Ec* | *DPOGS204935* | NP_001299389.1 |
| *BmorCPR4* | RR-1 | *BGIBMGA002548* | *KWMTBOMO05376* | *Cuticular protein 65Av* | *DPOGS210226* | NP_001299425.1 |
| *BmorCPR5* | RR-1 | *BGIBMGA002549* | *KWMTBOMO05377* | *Cuticular protein 65Av* | *DPOGS21022* | NP_001299125.1 |
| *BmorCPR11* | RR-1 | *BGIBMGA012601* | *KWMTBOMO11214* | *Cuticular protein 65Eb* | *DPOGS209413* | XP_013163146.1 |
| *BmorCPR28* | RR-1 | *BGIBMGA000341* | *KWMTBOMO13158* | *Adult cuticle protein 65Aa* | *DPOGS207987* | NP_001299546.1 |
| *BmorCPR38* | RR-1 | *BGIBMGA000333* | *KWMTBOMO13146* | *Cuticular protein 67Fb* | *DPOGS207981* | NP_001299095.1 |
| *BmorCPR41* | RR-1 | *BGIBMGA000329* | *KWMTBOMO13141* | *Cuticular protein 49Ab* | *DPOGS207978* | NP_001299345.1 |
| *BmorCPR42* | RR-1 | *BGIBMGA000328* | *KWMTBOMO13140* | *Cuticular protein 67Fb* | *DPOGS207976* | NP_001299352.1 |
| *BmorCPR44* | RR-1 | *BGIBMGA000326* | *KWMTBOMO13138* | *Cuticular protein 49Aa* | *DPOGS207976* | KPI97901.1 |
| *BmorCPR46* | RR-1 | *BGIBMGA000324* | *KWMTBOMO13135* | *Cuticular protein 11B* | *DPOGS207973* | NP_001298836.1 |
| *BmorCPH19* | CPH | *BGIBMGA011761* | *KWMTBOMO06685* | | *DPOGS204868* | NP_001298992.1 |
| *BmorCPH24* | CPH | *BGIBMGA011765* | *KWMTBOMO06675* | | | |
| *BmorCPH25* | CPH | *BGIBMGA011766* | *KWMTBOMO06674* | | | |
| *BmorCPT2* | CPT | *BGIBMGA000520* | *KWMTBOMO00354* | *TweedleE* | *DPOGS206032* | KPI91969.1 |
| *BmorCPT3* | CPT | *BGIBMGA005259* | *KWMTBOMO04356* | *TweedleE* | *DPOGS206331* | KPI91969.1 |
| *BmorCPG4* | CPG | *BGIBMGA008255* | *KWMTBOMO10672* | *CG32564* | | BAG30773.1 |
| *BmorCPG13* | CPG | *BGIBMGA002384* | *KWMTBOMO05344* | *Vajk4* | *DPOGS210253* | KPJ05455.1 |
| *BmorCPFL1* | CPFL | *Novel01073* | *KWMTBOMO00895* | | | |

Supplementary Table S7. The cuticular protein genes with high expression in the black epidermal region of *papilio xuthu* caterpillar

| Name of *Bombyx mori* | Family | Gene number in *SilkDB* | Name of *Papilio xuthu* | Accession number of *Papilio xuthu* | Gene in *Danaus plexippus* | Gene in *Drosophila melanogaster* |
|---|---|---|---|---|---|---|
| *BmorCPR67* | RR-2 | *BGIBMGA010145* | *PxutCPR67Bc* | AB264688 | *DPGLEAN07212* | *Ccp84Ad* |
| *BmorCPR71* | RR-2 | *BGIBMGA010142* | *PxutCPR71* | AB264671 | *DPGLEAN07212* | *Ccp84Ad* |
| *BmorCPR76* | RR-2 | *BGIBMGA001486* | *PxutCPR76b* | AK401331 | *DPGLEAN07395* | *Ccp84Ad* |
| *BmorCPR79* | RR-2 | *BGIBMGA000285* | *PxutCPR79* | AK401617 | *DPGLEAN16380* | *Cuticular protein 76Bd* |
| *BmorCPR99* | RR-2 | *BGIBMGA000265* | *PxutCPR97Bb* | AK401509 | *DPGLEAN08085* | *CG34461* |
| *BmorCPR107* | RR-2 | *BGIBMGA000259* | *PxutCPR107* | AK401841 | *DPGLEAN16484* | *Cuticular protein 62Bc* |
| *BmorCPR129* | RR-2 | *BGIBMGA000246* | *PxutCPR129* | AK401839 | *DPGLEAN07718* | *CG34461* |
| *BmorCPG4* | CPG | *BGIBMGA008255* | *PxutCPG4* | AB264698 | *DPGLEAN07899* | *CG1368* |
| *BmorCPG27* | CPG | *BGIBMGA008213* | *PxutCPG3* | AB264657 | *DPGLEAN07900* | *CG15597* |
| *BmorCPG38* | CPG | *BGIBMGA008396* | *PxutCPG38* | AK403802 | *DPGLEAN10638* | *CG16885* |
| *BmorCPT4* | CPT | *BGIBMGA006826* | *PxutCPT4* | AK401206 | *DPGLEAN11147* | *TweedleE* |
| *BmorCPFL4* | CPFL | *BGIBMGA011455* | *PxutCPFL6Ba* | AK402093 | *DPGLEAN19242* | *CG32603* |
| *BmorCPH5* | CPH | *BGIBMGA005397* | *PxutCPH5* | AK401958 | *DPGLEAN19902* | |

Futahashi R,et al. Comprehensive microarray-based analysis for stage-specific larval camouflage pattern-associated genes in the swallowtail butterfly, *Papilio xuthus* . *BMC Biol* . 2012;10:46. Published 2012 May 31. doi:10.1186/1741-7007-10-46

**Supplementary Table S8. The temporal expression of *mamo* gene in *Drosophila melannogaster***

| Library Collection ID | Parent Library Name | Gene Symbol | FBgn ID | Category ID | Category Descriptive Name | RPKM Expression Value | Expression Value BinID |
|---|---|---|---|---|---|---|---|
| FBlc0000086 | modENCODE_mRNA-Seq_development | *mamo* | FBgn0267033 | 40 | mE_mRNA_em0-2hr | 0 | 1 |
| FBlc0000087 | modENCODE_mRNA-Seq_development | *mamo* | FBgn0267033 | 170 | mE_mRNA_em2-4hr | 0 | 1 |
| FBlc0000088 | modENCODE_mRNA-Seq_development | *mamo* | FBgn0267033 | 280 | mE_mRNA_em4-6hr | 0 | 1 |
| FBlc0000089 | modENCODE_mRNA-Seq_development | *mamo* | FBgn0267033 | 310 | mE_mRNA_em6-8hr | 0 | 1 |
| FBlc0000090 | modENCODE_mRNA-Seq_development | *mamo* | FBgn0267033 | 320 | mE_mRNA_em8-10hr | 0 | 1 |
| FBlc0000091 | modENCODE_mRNA-Seq_development | *mamo* | FBgn0267033 | 340 | mE_mRNA_em10-12hr | 0 | 1 |
| FBlc0000092 | modENCODE_mRNA-Seq_development | *mamo* | FBgn0267033 | 360 | mE_mRNA_em12-14hr | 0 | 1 |
| FBlc0000093 | modENCODE_mRNA-Seq_development | *mamo* | FBgn0267033 | 380 | mE_mRNA_em14-16hr | 0 | 1 |
| FBlc0000094 | modENCODE_mRNA-Seq_development | *mamo* | FBgn0267033 | 391 | mE_mRNA_em16-18hr | 0 | 1 |
| FBlc0000095 | modENCODE_mRNA-Seq_development | *mamo* | FBgn0267033 | 392 | mE_mRNA_em18-20hr | 0 | 1 |
| FBlc0000096 | modENCODE_mRNA-Seq_development | *mamo* | FBgn0267033 | 393 | mE_mRNA_em20-22hr | 0 | 1 |
| FBlc0000097 | modENCODE_mRNA-Seq_development | *mamo* | FBgn0267033 | 394 | mE_mRNA_em22-24hr | 0 | 1 |
| FBlc0000098 | modENCODE_mRNA-Seq_development | *mamo* | FBgn0267033 | 410 | mE_mRNA_L1 | 0 | 1 |
| FBlc0000099 | modENCODE_mRNA-Seq_development | *mamo* | FBgn0267033 | 420 | mE_mRNA_L2 | 0 | 1 |
| FBlc0000100 | modENCODE_mRNA-Seq_development | *mamo* | FBgn0267033 | 440 | mE_mRNA_L3_12hr | 2 | 2 |
| FBlc0000101 | modENCODE_mRNA-Seq_development | *mamo* | FBgn0267033 | 462 | mE_mRNA_L3_PS1-2 | 1 | 2 |
| FBlc0000102 | modENCODE_mRNA-Seq_development | *mamo* | FBgn0267033 | 465 | mE_mRNA_L3_PS3-6 | 1 | 2 |
| FBlc0000103 | modENCODE_mRNA-Seq_development | *mamo* | FBgn0267033 | 467 | mE_mRNA_L3_PS7-9 | 2 | 2 |
| FBlc0000104 | modENCODE_mRNA-Seq_development | *mamo* | FBgn0267033 | 510 | mE_mRNA_WPP | 2 | 2 |
| FBlc0000105 | modENCODE_mRNA-Seq_development | *mamo* | FBgn0267033 | 580 | mE_mRNA_P5 | 2 | 2 |
| FBlc0000106 | modENCODE_mRNA-Seq_development | *mamo* | FBgn0267033 | 610 | mE_mRNA_P6 | 3 | 2 |
| FBlc0000107 | modENCODE_mRNA-Seq_development | *mamo* | FBgn0267033 | 630 | mE_mRNA_P8 | 3 | 2 |
| FBlc0000108 | modENCODE_mRNA-Seq_development | *mamo* | FBgn0267033 | 640 | mE_mRNA_P9-10 | 1 | 2 |
| FBlc0000109 | modENCODE_mRNA-Seq_development | *mamo* | FBgn0267033 | 740 | mE_mRNA_P15 | 4 | 3 |
| FBlc0000113 | modENCODE_mRNA-Seq_development | *mamo* | FBgn0267033 | 810 | mE_mRNA_AdM_Ecl_1days | 4 | 3 |
| FBlc0000114 | modENCODE_mRNA-Seq_development | *mamo* | FBgn0267033 | 850 | mE_mRNA_AdM_Ecl_5days | 4 | 3 |
| FBlc0000115 | modENCODE_mRNA-Seq_development | *mamo* | FBgn0267033 | 1100 | mE_mRNA_AdM_Ecl_30days | 4 | 3 |
| FBlc0000110 | modENCODE_mRNA-Seq_development | *mamo* | FBgn0267033 | 810 | mE_mRNA_AdF_Ecl_1days | 4 | 3 |
| FBlc0000111 | modENCODE_mRNA-Seq_development | *mamo* | FBgn0267033 | 850 | mE_mRNA_AdF_Ecl_5days | 2 | 2 |
| FBlc0000112 | modENCODE_mRNA-Seq_development | *mamo* | FBgn0267033 | 1100 | mE_mRNA_AdF_Ecl_30days | 2 | 2 |

Brown JB, Boley N, Eisman R, et al. Diversity and dynamics of the *Drosophila* transcriptome. Nature. 2014;512(7515):393-399. doi:10.1038/nature12962

Supplementary Table S9. The information of Silkworm Strains

| Number | Sample Id | Name | Phenotype | Group | Location |
|---|---|---|---|---|---|
| 1 | BomW3 | | | Wild | Shaanxi, China |
| 2 | BomW15 | | | Wild | Shaanxi, China |
| 3 | BomW16 | | | Wild | Shaanxi, China |
| 4 | BomW17 | | | Wild | Shaanxi, China |
| 5 | BomW37 | | | Wild | Beijing, China |
| 6 | BomW40 | | | Wild | Hebei, China |
| 7 | BomW43 | | | Wild | Anhui, China |
| 8 | BomW45 | | | Wild | Zhejiang, China |
| 9 | BomW46 | | | Wild | Shanghai, China |
| 10 | BomW47 | | | Wild | Shanghai, China |
| 11 | BomW49 | | | Wild | Jiangsu, China |
| 12 | BomW51 | | | Wild | Jiangsu, China |
| 13 | BomL10 | Tu No.1 | p3 | Local | Shanxi, China |
| 14 | BomL100 | Xupu | p3 | Local | Hunan, China |
| 15 | BomL193 | C108N | p | Local | China |
| 16 | Dazao | Dazao | p3 | Local | China |
| 17 | BomL112 | N4 | p | Local | India |
| 18 | BomL113 | Huangban | - | Local | Guangdong, China |
| 19 | BomL122 | Lao | p | Local | Laos |
| 20 | BomL123 | Fang 4 | p | Local | Guangxi, China |
| 21 | BomL125 | H2-2 | p3 | Local | Vietnam |
| 22 | BomL138 | Gaohua | p3 | Local | Guangdong, China |
| 23 | BomL165 | Cambodia | p | Local | Cambodia |
| 24 | BomL211 | 5G | - | Local | unknown |

The p indicates larva has a pale body color and have no markings. The p3 indicates larva has a pale body color and have eye spots, lunar spots, and star spots. The hyphen indicates not available.

Supplementary Table S10. Primers for this research

| Number | Name | F | R | |
|---|---|---|---|---|
| 1 | *BmorCPR2* | CGCCCCAGTGGTCAAATC | CGATGGCGGGGTATTCTG | |
| 2 | *BmorCPR3* | TTACATCTTCCTCGCTTTGGTC | CTGATTCCGTTGCTGGTCTG | |
| 3 | *BmorCPR4* | TGTCGCTCTCTTCGTCGCC | CCTGAGCCTTGATTCCGTTG | |
| 4 | *BmorCPR5* | GAGGGTTACGGCTTTGGCT | ACGTCGTCTCGTGGACCTTT | |
| 5 | *BmorCPR11* | CGAAGCGAAACCCGAAAGA | GCCACGCCATTTTCTCCG | |
| 6 | *BmorCPR27* | GCCAATCTGATGGGACCGT | GCCGGGACCTTGCTCTATTT | |
| 7 | *BmorCPR28* | GTTATTAGCATTGTGTTTGGTGG | CCTGCTCAAATACGGTCCCA | |
| 8 | *BmorCPR38* | ACTCCTACTCCGTTTCTGCCTATC | CGATACCGTTGTTGGTTTCGT | |
| 9 | *BmorCPR41* | CTACGTCGGAACCAAGGAGG | TGATGCCGTATTTGTAGTCGTAGC | |
| 10 | *BmorCPR42* | TCTTGGTTTTGGCTTTATGCG | GCTTCACTGCCGCTGACTACT | |
| 11 | *BmorCPR44* | GTGTTGGTTTCTATGCTGGTCG | GGTATGACCCGTCGCCGTT | |
| 12 | *BmorCPR46* | TCTCGCCCTTGTGGCTACC | ACGCACGATGCCGTTGGA | |
| 13 | *BmorCPR67* | CACACCCTCAATACAGTTTCGC | CGTTGAAGCCGTTGTGGG | |
| 14 | *BmorCPR76* | GTTCACGACGAGCAAAGCG | TAGCGTTAAATCCGTTTACTTTGT | |
| 15 | *BmorCPR79* | TCATCAAGGACCTGTAGTATCGC | GGAATCTGTAGGGCGGGTAA | |
| 16 | *BmorCPR71* | GGAATACGACGCTCATCCTCAG | GGGTCAACGACGGAGTAGGA | |
| 17 | *BmorCPR99* | ACTTCTCCACTCCGCTACTCCT | ACAGGAGCGGCGTATTGG | |
| 18 | *BmorCPR107* | CGTCGTGAAGGGCGTGTA | ACATGGCGAGCATGACCC | |
| 19 | *BmorCPH5* | ACGAAGTCGGTGACGCTGAA | TACCGTCAGGAAGAAGCACTCG | |
| 20 | *BmorCPH19* | GCCAGCAGGGCATTGAGTC | GGGAGAGTGTAGGGAGCCG | |
| 21 | *BmorCPH24* | TCAGAGTCCTTGTCGCATTCG | GCGGTGTAGGCGTAGGTGG | |
| 22 | *BmorCPH25* | CCCGCTCTCACCACCTCC | CTACGACGGGAGCGGTGA | Primers for qRT-PCR |
| 23 | *BmorCPH34* | CGCACCAAATCCACACACC | GGGCTGCGTAAGTTCGGG | |
| 24 | *BmorCPT2* | TGAACCTCCAGTGGGATACAGC | GGAAATGCTGCCTCCAGAAAG | |
| 25 | *BmorCPT3* | CACCCATCATCCCCGCTC | TGGGTGGGAGCAGCAGTG | |
| 26 | *BmorCPT4* | GCCACCGCCAGATAACGA | CGGTCGGTGTTGGAGGTGT | |
| 27 | *BmorCPG4* | CGCCAAGACAGAAGACAAGG | GCTGTAAGACGAGGGAGAGGA | |
| 28 | *BmorCPG13* | CTACATCGGACACGGACAAGG | GTATGTCCACCGAAGTTGAGGC | |
| 29 | *BmorCPFL4* | AGTCGGCTGTCACATATTCTGC | ACCGCTGAAGGAGATGTGAGAC | |
| 30 | *RPL3* | CGGTGTTGTTGGATACATTGAG | GCTCATCCTGCCATTTCTTACT | |
| 31 | *yellow* | TCGCTTACTCGTGGGAACAGA | ATAGGAGACGCCGTGATGCC | |
| 32 | *ddc* | AGCCTTGGACTGCGGTGAT | ATAGCGGGATACGAGTTAGCG | |
| 33 | *ebony* | GACGCCCAGGTCAAGATTCG | CGTAGCATAGCACCACGCACT | |
| 34 | *TH* | TTGATGCCCAAACACGC | TCGCAGGGTAAAGCCAGT | |
| 35 | *black* | GTGGGCGGATGTCCTCAAA | GCAAGCCTTCCCGATTACG | |
| 36 | *Laccase2* | CATCTCCCAGCCATCCCACT | CGAACCATCATAAACTCCCGT | |
| 37 | *tan* | CCACGGTTCTCGTCTTTATGCTA | AGGAGGAGTGTTTCCAGGCTTT | |
| 38 | *iAANAT* | TGAATCTCGCCGTCAATCTG | GAAACTCCATCGCTCAAGGTAG | |
| 39 | *Bm-mamo* | TGCCTAAAGATGAACCACCTGAC | TCTAGGGCTTGAAAGTGATGGAA | |
| 40 | *BGIBMGA012516* | ACTCCTCTCAAAGTTAGCAGTC | GTTCATCTATGCTGTTGCTCTG | |
| 41 | *BGIBMGA012517* | AATCTCGCGATCTTAGGGAAAT | TGGAATCTCTAACGTCTCGAAG | |
| 42 | *BGIBMGA012518* | GTACTGACAGCAAAAACGAGAG | GCTCTGCCACAAAATAGACAAA | |
| 43 | *BGIBMGA012519* | GTAATGTGATGCTGTTCGGTAC | GAATCGTCCCTGTTTCCAATTC | |
| 44 | *BGIBMGA014089* | GCGACTGAACATGTACTCATTC | TACAACTTCTGTTTCACGCAAG | |
| 45 | *KWMTBOMO05082* | CCATTGGTATGGTGGTTGTGC | TGGTGAGGGCCAATCATCAC | |
| 46 | *KWMTBOMO05084* | AATACTTGCCGACGCTGGTG | CTCGTCAGGCTTGTGAGGTTTT | |
| 47 | *KWMTBOMO05085* | CTGGCGGTGTCTCTTGACAT | AAATAGGACCCAACCAGCCG | |
| 48 | a | CAGGAACAGGGATGTGGACTTG | TTGCTATCGGAGCTCTGGGA | |
| 49 | b | GGAGGGCTCGCTAAGTCG | GCTCATTGGGCACGGTC | |
| 50 | c | AATGTGAATCTCGGAAACGG | CATTTAAGCCTTTAGGGATTGG | |
| 51 | d | GCTCTATTTATTTCCCAACTTCCAC | TTACTTTCGCTGTCATATCACCA | |
| 52 | e | GGCAGAAATAGGCAGGGTG | CTGCTTTATCTGGTGGTGGG | |
| 53 | f | ATAGTTTTTCGCATTAGGGTCAT | AGTCAGATAGTTGGGCTTTCTTAAT | |
| 54 | g | GCCGTAGCCTAAATGATAAGACC | TGTTCATCACAAGCCGTAGGA | Primers for mapping |
| 55 | h | AGTGTAGTTTGAAGGGTAGGG | AACCGATCCTACTCTTCTGC | |
| 56 | i | GCGTATTTAGTCGTCCGTGAC | ACCGCTAATACCCGTAGACACT | |
| 57 | j | CTTTCCTCACTTTACTCCTCCAC | CTTTTTCCGCTTTCTTCCTTT | |
| 58 | k | CATGCTTACTTTTCTGCGATCT | TTGTGGCCCAACTTTTGTAGTA | |
| 59 | l | GAGTACAATGCCAGTAGCCAAGT | GCATAAGCCATCTAATCCCTATT | |
| 60 | m | GCAATAGGTGTTCCCGAAAGAG | GAAAGTGATTGATGCCTGGTGT | |
| 61 | *BGIBMGA00012517* | ATGGGAAGTGAGCACTATTGC | TCATTGATGTGGACTCTCCTCTT | Primers for colning |
| 62 | *BGIBMGA00012517/18* | ATGGGAAGTGAGCACTATTGC | TCATCGAGTGTTATGTACGTGTG | |
